# A stress response that allows highly mutated eukaryotic cells to survive and proliferate

**DOI:** 10.1101/515460

**Authors:** Rebecca A. Zabinsky, Jonathan Mares, Richard She, Michelle K. Zeman, Thomas R. Silvers, Daniel F. Jarosz

## Abstract

Rapid mutation fuels the evolution of many cancers and pathogens. Much of the ensuing genetic variation is detrimental, but cells can survive by limiting the cost of accumulating mutation burden. We investigated this behavior by propagating hypermutating yeast lineages to create independent populations harboring thousands of distinct genetic variants. Mutation rate and spectrum remained unchanged throughout the experiment, yet lesions that arose early were more deleterious than those that arose later. Although the lineages shared no mutations in common, each mounted a similar transcriptional response to mutation burden. The proteins involved in this response formed a highly connected network that has not previously been identified. Inhibiting this response increased the cost of accumulated mutations, selectively killing highly mutated cells. A similar gene expression program exists in hypermutating human cancers and is linked to survival. Our data thus define a conserved stress response that buffers the cost of accumulating genetic lesions and further suggest that this network could be targeted therapeutically.

## INTRODUCTION

A multitude of DNA repair factors promote faithful transmission of genetic information from one generation to the next (Friedberg et al., 2014). However, in response to oncogenic transformation, infection, and environmental stress, many cells become hypermutators (Alexandrov et al., 2013; Bielas et al., 2006; Cairns and Foster, 1991; Galhardo et al., 2007a, b; Lawrence et al., 2013; Loeb, 2016; Oliver et al., 2000). Human cancers (e.g., melanoma, lung cancer, and colorectal cancer; Lawrence et al., 2013) often have mutation rates that are three orders of magnitude higher than those of normal tissue. Although mutations can occasionally be adaptive, theoretical and experimental studies in a wide range of organisms suggest that they are on average slightly deleterious (Bloom et al., 2004; Eyre-Walker and Keightley, 2007; Firnberg et al., 2014; Pakula and Sauer, 1989; Tokuriki and Tawfik, 2009). Therefore, although hypermutation can fuel rapid evolution, it is also associated with fitness costs. In the simplest scenarios, random mutations can disrupt catalysis by an enzyme, decrease the strength of protein–protein interactions, or perturb baseline gene expression levels. In addition, empirical studies have shown that the majority of mutations in coding regions destabilize protein structure (Fersht, 1998), a property that may have slowed evolution of highly expressed genes (Drummond et al., 2005; Drummond and Wilke, 2008). Thus, as mutations accumulate, fitness is expected to decline. Synergistic epistasis among accumulating mutations could in principle accelerate this effect (de Visser et al., 2011; Elena and Lenski, 1997).

Despite the toxic consequences of mutation, many cells, ranging from pathogenic bacteria and fungi (Fares et al., 2002b; Maisnier-Patin et al., 2005; Oliver et al., 2000) to human tumor cells (Andor et al., 2017; Lawrence et al., 2013) can survive considerable mutation burdens. High mutation rates arising from loss of DNA mismatch repair genes occur in >20% of pathogenic *Pseudomonas aeruginosa* colonizing the lungs of cystic fibrosis patients (Oliver et al., 2000). Extreme mutators can also be found among *Helicobacter pylori* and *Neisseria meningitidis* isolated from human patients (Hall and Henderson-Begg, 2006). In humans, mutations in DNA mismatch repair genes, including *MSH2* and *MSH6*, cause Lynch Syndrome, a disease characterized by an elevated risk of colorectal and other cancers (Lynch et al., 2015; Sijmons and Hofstra, 2016). Many human cancers that do not arise from hereditary mutations in DNA mismatch repair genes also exhibit elevated mutation burden (Campbell et al., 2017; Lawrence et al., 2013; Yousif et al., 2018).

Each of these systems has a mutation rate that approaches the theoretical limits for a ‘error catastrophe,’ in which the fidelity of information transfer from one generation to the next is too low to support viability (Eigen, 2002; McFarland et al., 2017; Sole and Deisboeck, 2004). As these cells accumulate more mutations, however, they do not die, but instead remain capable of surviving, proliferating, and evolving new traits with devastating clinical consequences (e.g., resistance to antibiotics or chemotherapeutics). In bacteria, the capacity of cells to buffer the cost of mutations has been linked to the protein homeostasis network (Aguilar-Rodriguez et al., 2016; Fares et al., 2002b; Maisnier-Patin et al., 2005; Moran, 1996; Sabater-Munoz et al., 2015). It remains unclear, however, whether analogous pathways exist in eukaryotes to dampen the effects of deleterious mutations.

One potential modulator of mutation cost in eukaryotes, <u>h</u>eat <u>s</u>hock <u>p</u>rotein 90 (Hsp90), has been proposed to influence the phenotypic manifestation of natural genetic variation (Burga et al., 2011; Cowen and Lindquist, 2005; Jarosz and Lindquist, 2010; Queitsch et al., 2002; Rohner et al., 2013; Rutherford and Lindquist, 1998). Hsp90 is a highly conserved molecular chaperone that functions with dozens of co-chaperones (Taipale et al., 2010; Taipale et al., 2014) to fold hundreds of client proteins, most of which are key regulators of growth and development. From yeast to humans, Hsp90 can strongly influence the phenotypic effects of genetic and epigenetic variation that naturally arises within populations (Burga et al., 2011; Cowen and Lindquist, 2005; Jarosz, 2016; Jarosz and Lindquist, 2010; Karras et al., 2017; Queitsch et al., 2002; Rohner et al., 2013; Rutherford and Lindquist, 1998; Sangster et al., 2004). Although Hsp90 has been shown to enhance phenotypes derived from some recently accumulated genetic variants (Geiler-Samerotte et al., 2016; Mason et al., 2018), the full effects of this chaperone on recently accumulated mutations have yet to be fully characterized.

In this study, we harnessed the power of a simple model eukaryote, *Saccharomyces cerevisiae*, in which a wealth of genomic tools enabled us to investigate the interplay between *de novo* mutations and stress response pathways. We created obligate hypermutator lineages by disrupting DNA polymerase proofreading and/or mismatch repair, mimicking the situation in Lynch Syndrome patients and many hypermutating tumors. We then evolved the lineages in parallel using a mutation accumulation (MA) framework that mirrors the population bottlenecks that characterize infectious pathogens, solid tumors, and metastatic or refractory disease. These experiments revealed a strong bifurcation in the cost of accumulating mutations: lesions that arose in early generations had a much stronger impact on fitness than those that arose in later generations. Although no single mutation was shared among all lineages, mRNA sequencing revealed a shared response to mutation burden, which we term the Eukaryotic Mutation Burden Response (EMBR). EMBR is distinct from previously characterized stress responses and its components form a highly coherent network of protein–protein interactions in yeast, as do their homologs in humans. Targeting the EMBR network revealed that highly mutated yeast and colon cancer cells depend on EMBR gene functions to buffer the cost of accumulating mutations, representing an addiction that could be exploited therapeutically.

## RESULTS

### A sequenced collection of mutation accumulation lineages

Cancer cells on the brink of error catastrophe are apparently able to buffer the fitness consequences of accumulating mutation burden, but the molecular origins of this capacity remain unclear. We explored this question using mutation accumulation (MA) experiments, which have served as valuable tools in studies of a wide range of biological processes, from adaptive fitness trajectories in evolution to DNA polymerase usage at the replication fork (Denver et al., 2009; Huang et al., 2016; Lujan et al., 2014; Ossowski et al., 2010; Uchimura et al., 2015; Zhu et al., 2014b). By propagating independent lineages through single-cell bottlenecks, we mimicked the genetic drift that can occur in the proliferation of solid tumors and in metastatic events (Barrick and Lenski, 2013; Sun et al., 2017).

Specifically, we constructed *de novo* MA lineages in the budding yeast *S. cerevisiae* (Figure 1A). Due to its concise genome and faithful recapitulation of fundamental eukaryotic biology, *S. cerevisiae* has been widely used to model the fundamental biology of cancer and stress responses (Hartwell, 2004; Khurana and Lindquist, 2010). To reproduce mutation rates that correspond to human tumors, we made use of extensive data about patterns of mutagenesis amassed in both *S. cerevisiae* and human patients (Figure 1A-B; Lawrence et al., 2013; Lujan et al., 2014; Roberts and Gordenin, 2014; Serero et al., 2014; Supek and Lehner, 2015; Wielgoss et al., 2013). We constructed mutator yeast strains by deleting the mismatch repair gene *MSH6*, and generated hypermutator strains by combining the *msh6Δ* allele with a point mutation in DNA polymerase δ that reduces its proofreading activity (*pol3-L612M*). Combining these alleles amplifies mutation frequency while preserving the mutation spectrum inherent to loss of *MSH6* function (base pair substitutions; Lujan et al., 2014; Nick McElhinny et al., 2008). Indeed, mutation or misregulation of human *MSH6* co-occurs with mutation in pol δ in patients with Lynch syndrome (Jansen et al., 2016). Sequencing of multiple mutation accumulation clones (see below) revealed that the *msh6Δ* mutator exhibited a 47-fold increase in mutation rate (^~^0.08 mutations per cell division) whereas the *pol3-L612M msh6Δ* double mutant exhibited a ^~^1,100-fold increase in mutation rate (^~^1.5 mutations per cell division; Figure 1B). Similar values have previously been reported using canavanine mutagenesis assays (Herr et al., 2011). Hereafter, we refer to these strains as “mutator” (*msh6Δ*) and “hypermutator” (*pol3-L612M msh6Δ*). This range encompasses the relative mutation rates in rapidly mutating human tumors (e.g., melanomas, glioblastomas, and colorectal cancers (Lawrence et al., 2013); Figure 1B; see SI for additional discussion).

**Figure 1.**
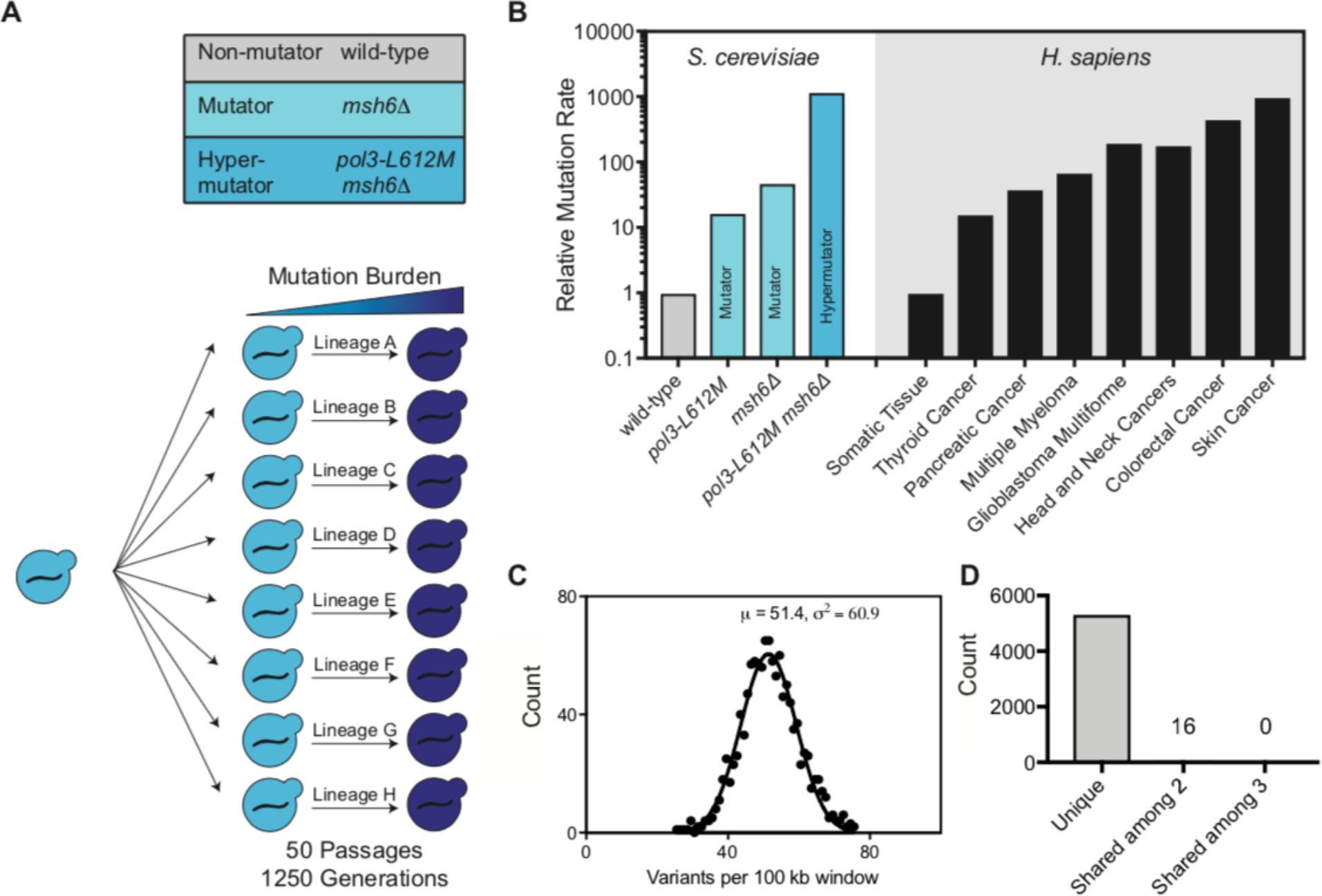
Yeast MA lines as a model for accumulating mutation burden. (**A**) Three distinct genotypes were used to generate MA lines; non-mutator, mutator, and hypermutator. Eight independent lineages (arrows labeled A–H) of each ancestral strain was passaged, resulting in unique mutations in each lineage. The wild-type ancestor accumulated mutations at a negligible rate, the *msh6Δ* mutator strains at a slow rate (^~^0.08 mutations/generation), and the *pol3-L612M msh6Δ* hypermutator strains at a rapid rate (^~^1.5 mutations/generation). The dark blue hue represents increasing mutation burden. After 50 passages (^~^1250 generations), the mutator lineages had accumulated ^~^100 mutations each, and the hypermutator lineages had accumulated ^~^2,000 mutations each. (**B**) Relative mutation rates of our yeast strains and human cancers. Shown are *H. sapiens* cancer types with enhanced mutation rates, estimated from comparisons of tumors versus normal tissue (Lawrence et al., 2013). (**C**) The number of variants observed within 100-kb windows fits a normal Gaussian distribution, as expected for randomly generated mutations. (**D**) Over 5,000 unique mutations arose in sequencing of three parallel hypermutator lineages. Only 16 mutations were shared between two lineages, and none were shared among all three.

### The mutations in these lineages are random

We passaged sixteen parallel lineages of each MA genotype for over 1,000 generations to create independent descendants with distinct mutation spectra (Figure 1A). To promote random mutational trajectories and minimize the effects of selection, we employed single-cell bottlenecks, picking average-sized colonies every ~25 generations. Whole-genome sequencing (Table S1) established that the lineages accumulated around 100 (for mutators) or 2,000 (for hypermutators) unique mutations over the course of passaging. Lineages from the same generation did not share more mutations than expected by chance (Figure 1C-D), in stark contrast to the stereotyped parallelism observed in chemostat and other evolution experiments in large population sizes (Hope et al., 2017; Venkataram et al., 2016).

Several lines of evidence demonstrate that our MA experiment preserved a wide range of naturally occurring mutations, with the exception of lethal or near-lethal mutations: 1) In whole-genome sequencing data from multiple parallel lineages after 500 and 1,250 generations of propagation, we observed no systematic bias in the physical locations in which mutations arose; the number of mutations in 100 kilobase windows conformed closely to random expectation (Figure 1C); 2) Estimates of mutation rate from whole-genome sequencing data established that the hypermutator phenotype was maintained throughout passaging (Figure 1 – figure supplement 1A, Table S1); 3) The mutation spectrum was dominated by base substitutions and enriched for transitions over transversions both early (passages 1 – 20) and late (passages 20 – 50) in the experiment, also consistent with the neutral expectation (Figure 1 – figure supplement 1B; Lujan et al., 2014); 4) Mutations accumulated independently of the local sequence context (Figure 1 – figure supplement 1C); 5) Given the number of cell divisions that occurred during the experiment and the elevated mutation rate, each lineage could have explored mutations covering virtually every base pair in the genome (see SI for further discussion). If specific mutations were selected for their fitness benefit, they would be expected to appear in multiple independent lineages, as is the case with cancer driver mutations (Sidow and Spies, 2015). Very few individual mutations occurred in more than one independent lineage, and none were shared in more than two lineages (Figure 1D); 6) Several genes were mutated (at different locations) in three lineages, but fewer genes with missense mutations were shared and no genes with severe mutations (stop codon gains or frameshifts) were shared (Figure 1 – figure supplement 1D). The number of mutations in a given ORF is primarily driven by gene length as expected from minimal selection (Figure 1 – figure supplement 1E); 7) Deleterious mutations predicted by the SIFT algorithm (Kumar et al., 2009) occurred at a constant rate throughout passaging (Figure 1 – figure supplement 1F). We further compared the fraction of deleterious variants in our strains relative to natural genetic variants from sequenced wild yeast strains that have experienced extensive selective pressure (Bergstrom et al., 2014; Liti et al., 2009). The fraction of predicted deleterious variants in our lineages was much larger than the fraction in natural variants from wild yeasts (Figure 1 – figure supplement 1G). We therefore conclude that the MA lines were largely free from selective pressures that could bias their evolution toward convergent genotypes. Taken together, these results support our use of these MA lines as a model for investigating the accumulation of random mutations and the resulting mutation burden.

### The impact of accumulated mutations on fitness

To assess the phenotypic impact of accumulated mutation burden, we measured the doubling times of the various mutated lineages relative to their ancestors. As a critical control, we passaged eight independent lineages of wild-type cells for the same number of generations and under the same conditions. These lineages exhibited little change in genotype (they accumulated one or zero mutations) or fitness over the entire experiment when grown on glucose or non-fermentable glycerol (Figure 2A, Figure 2 – figure supplement 1A, Table S1), providing further evidence that our propagation scheme did not impose undue selection. Likewise, in the mutator lineages (*msh6Δ*) we observed no fitness decline in multiple growth environments (Figure 2A, Figure 2 – figure supplement 1A).

The fitness of the hypermutator lines (*pol3-L612M msh6Δ*) did decline. Moreover, the cost of each successive mutation decreased over the course of the experiment (Figure 2B, Figure 2 – figure supplement 1B), in striking contrast to the expectation had the effects of the accumulating mutations been independent (i.e., additive). In a generic regression model 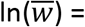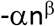, a β value of 1 would correspond to a linear, purely additive model (Lenski et al., 1999; Maisnier-Patin et al., 2005). However, our data were best fit by β = 0.460 ± 0.106 (for growth on glucose; *p* < 0.0001), and β = 0.575 ± 0.119 (for growth on glycerol; *p* < 0.0001 by extra-sum-of-squares F-test). These values are indicative of antagonistic epistasis (Figure 2B, Figure 2 – figure supplement 1B). That is, the apparent fitness cost per mutation was much higher for mutations that arose early in these lineages than for those that arose later (Figure 2C, Figure 2 – figure supplement 1C). This effect is remarkable considering that there was no discernable difference between the types of mutations that accumulated in the earliest and latest passages.

**Figure 2.**
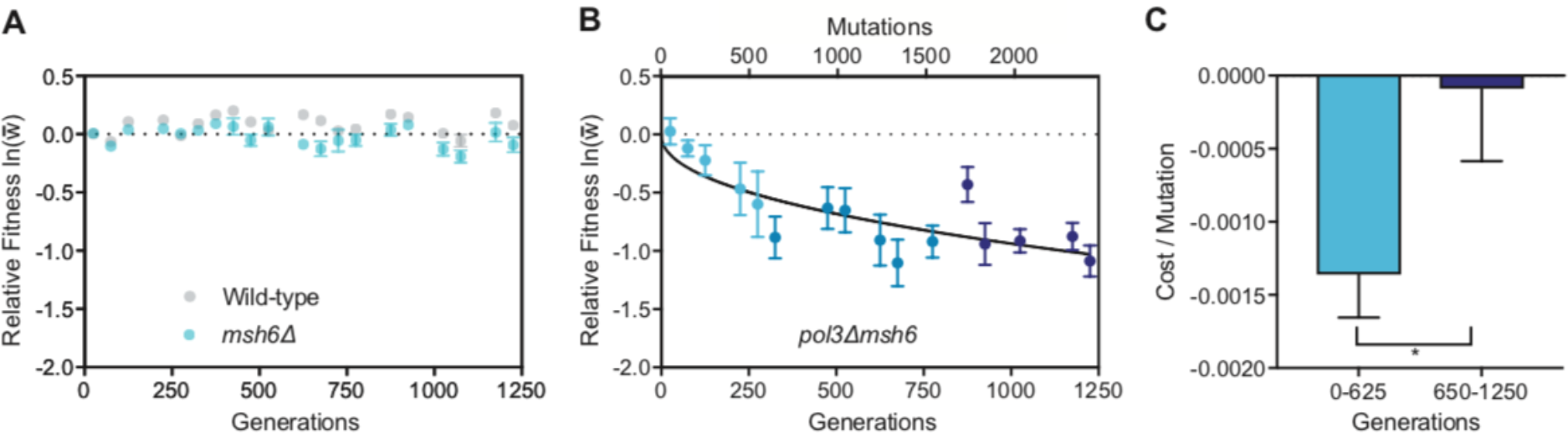
Fitness cost declines with increasing mutation burden. (**A**) Mean relative fitness of eight independently passaged lineages of the control wild-type strain BY4741 and its *msh6Δ* derivative. Relative fitness 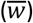 calculated based on doubling time relative to wild-type at passage 1. Error bars represent SEM from eight biological replicates. (**B**) Fitness trajectories of five hypermutator lineages passaged on glucose media. Fits are based on a regression model for detecting epistasis: 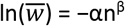 (α = 0.039  0.028, β = 0.460 ± 0.106; Maisnier-Patin et al., 2005). If β = 1, the regression model is linear, and no epistasis among mutations is detectable. β > 1 indicates synergistic epistasis and 0 < β < 1 indicates antagonistic epistasis. Parameter estimates for the model were fitted with a least squares regression. The goodness-of-fit of this model was superior to the simpler additive model (β = 1), p < 0.0001, extra-sum-of-squares F-test. Error bars represent SEM from five biological replicates. (**C**) Fitness cost per mutation across the first 625 generations and last 600 generations, calculated by linear regression, in passaged lineages. Error bars represent SE of best-fit values. * p < 0.05, extra-sum-of-squares F-test.

A simple explanation for such apparent epistasis would be an inability to observe colonies with fitness below a certain threshold (because they would not grow to a sufficient size before the next round of propagation). This would inadvertently select for progressively less deleterious mutations over the course of the experiment. However, populations with much lower fitness than we observed are capable of growing into sizeable colonies within our propagation schema (see SI for additional discussion). As described below, we could also measure even slower growth of the MA strains we generated after chemical inhibition (Figure 3B). Therefore, we conclude that the fitness trajectory of accumulating mutations is indeed characterized by apparent antagonistic epistasis.

In an evolution experiment where selection had dominated, the likeliest origin of antagonistic epistasis would be genetic suppression. That is, the fitness costs of an ancestral mutation could be compensated by newly arising variants. However, in our schema the variants accumulated randomly. Thus, for genetic suppression to entirely explain the effect, the number of antagonistic genetic interactions would need to be greater than the number of synergistic genetic interactions. However, data from systematic double deletion libraries have revealed significantly higher numbers of synergistic interactions than antagonistic interactions (^~^1.5-fold; Costanzo et al., 2016). The effect sizes of synergistic interactions also tend to be larger in magnitude than those of antagonistic interactions. We therefore searched for a cellular response that might buffer the cost of mutation load.

### Protein folding is a key contributor to the cost of accumulating mutations

The antagonistic epistasis we observed was strikingly reminiscent of mutation accumulation experiments in *Salmonella typhimurium* (Maisnier-Patin et al., 2005) and *Escherichia coli* (Fares et al., 2002b). In these systems, upregulation of the bacterial protein chaperone GroEL buffers the fitness cost of accumulating mutation burden. Hence, we investigated the influence of molecular chaperones. The eukaryotic GroEL homolog Hsp60 has an exclusively mitochondrial function (Zeilstra-Ryalls et al., 1991), but the Hsp90 chaperone can exert a strong influence on the capacity of natural genetic variants to produce phenotypes (Burga et al., 2011; Cowen and Lindquist, 2005; Jarosz and Lindquist, 2010; Queitsch et al., 2002; Rohner et al., 2013; Rutherford and Lindquist, 1998). Whether this chaperone buffers the phenotypic outcome of *de novo* mutations remains controversial (Geiler-Samerotte et al., 2016; Mason et al., 2018). Therefore, we asked whether the behavior of our MA lineages would be influenced by inhibition of Hsp90.

We first exposed the mutated lineages to radicicol, a potent inhibitor of Hsp90 function. The mutated cells were much more sensitive to the drug than their unmutated ancestors (Figure 3A). Indeed, the fitness cost per mutation was amplified by this treatment. That is, the alpha value of this generic regression model decreased (untreated α = 0.006 ± 0.004, treated α = 0.010 ± 0.007; *p* < 0.05). Hsp90 inhibition also resulted in a more additive relationship between accumulating mutations and fitness (untreated β = 0.699 ± 0.096, treated β = 0.749 ± 0.095; Figure 3B-C). When α was constrained to 0.01 (the fitness cost per mutation in the presence of Hsp90 inhibitor), the β value of the untreated fit decreased further to 0.612. These data suggest that the apparent antagonistic epistasis we observed is in part driven by an active cellular response involving Hsp90. Importantly, both mutated and ancestral cells were equally sensitive to other, unrelated stressors such as hydrogen peroxide and the antifungal fluconazole (Figure 3D-E). Because no mutation was shared by all of the lineages, we conclude that Hsp90 has a broad impact on the phenotypic outcome of multiple new mutations.

**Figure 3.**
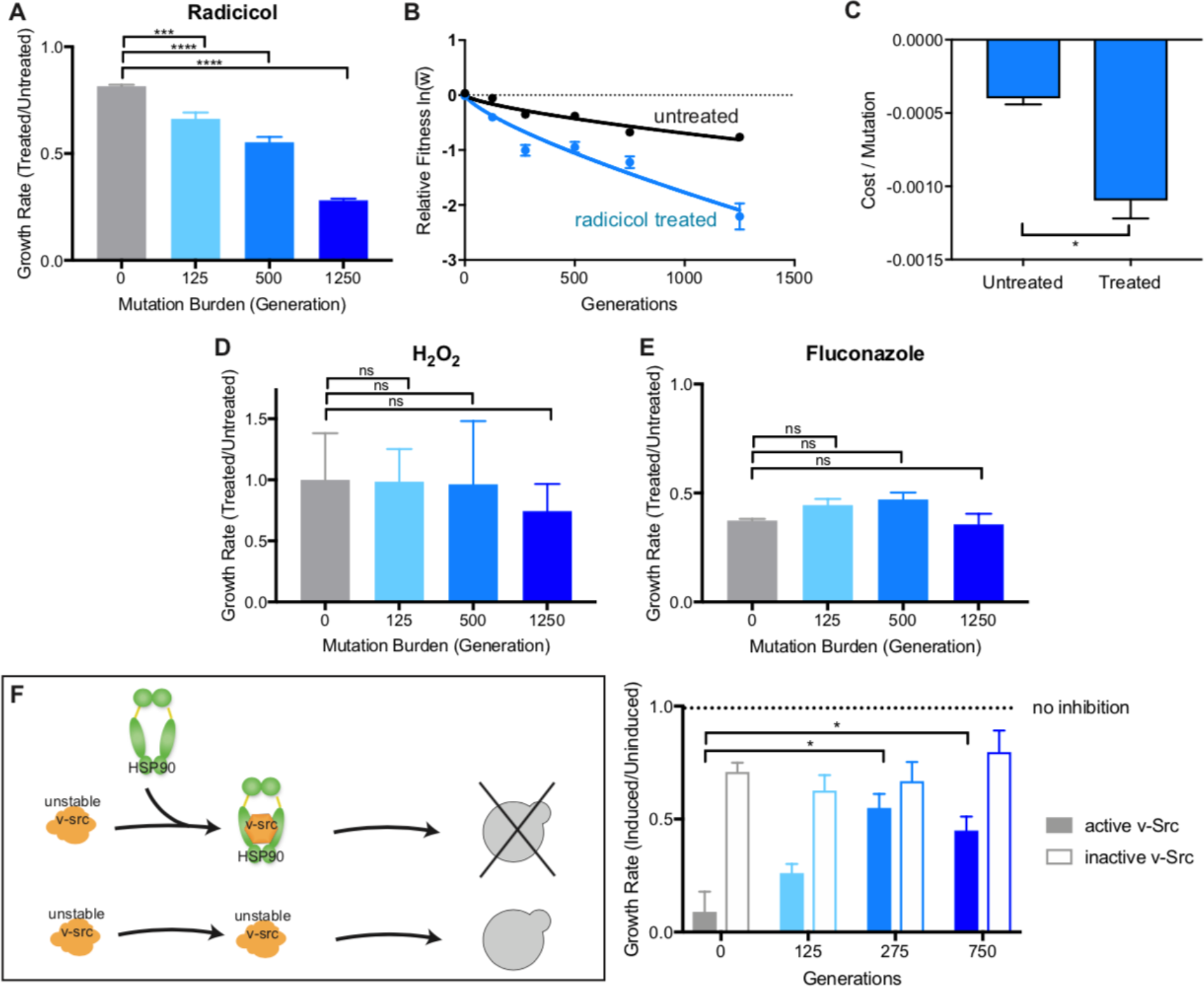
Protein misfolding is a source of toxicity associated with increasing mutation burden. (**A**) Strains were grown in the presence or absence of 100µM radicicol, an Hsp90 inhibitor, in rich medium. Error bars represent SD from three biological replicates. *** p < 0.001, **** p < 0.0001, student’s t-test. (**B**) Fitness trajectories of hypermutator lineages grown in rich media with and without 100uM radicicol. Fits are based on a regression model for detecting epistasis: 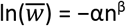 (untreated α = 0.006 ± 0.004, β = 0.699 ± 0.096, treated α = 0.010 ± 0.007, β = 0.749 ± 0.095) described in Figure 2 (Maisnier-Patin et al., 2005). The goodness of fit of this model was superior to the simpler additive model (β = 1), p < 0.0001, extra-sum-of-squares F-test. Error bars represent SEM from six-eight biological replicates. (**C**) Fitness cost per mutation across the untreated and treated samples, calculated by linear regression, in passaged lineages. Error bars represent SE of best-fit values. * p < 0.05, extra-sum-of-squares F-test. (**D**) Strains were growth in the presence or absence of 4.4mM H_2_O_2_ in rich medium. Error bars represent SD from three biological replicates. (**E**) Strains were growth in the presence or absence of 50µM fluconazole in rich medium. Error bars represent SD from three biological replicates. (**F**) Expression of the active v-Src kinase is toxic in yeast. However, this is dependent on HSP90 activity and v-Src activity. Inactivating v-Src with a point mutation eliminates the toxicity. Toxicity of v-Src expression in MA lines was measured by comparing the growth rate of cells expressing active v-Src (solid bars) and cells expressing an inactive v-Src control (open bars). Error bars represent SEM from three-twenty biological replicates. * p < 0.05, student’s t-test.

To assess Hsp90 activity in the MA lineages, we measured the activity of the Hsp90-dependent tyrosine kinase v-Src from the transforming Rous Sarcoma Virus (Xu and Lindquist, 1993; Zabinsky et al., 2018). This widely used assay relies on the fact that v-Src kinase inhibits yeast growth. However, because v-Src is an obligate Hsp90 client, its toxicity requires abundant and available Hsp90 to chaperone the folding of the kinase (Brugge et al., 1987; Falsone et al., 2004; Xu and Lindquist, 1993). We reasoned that if Hsp90 activity were consumed by demand from mutated proteins in the MA lineages, the toxic effect of v-Src would be mitigated. Indeed, v-Src toxicity was markedly reduced in the MA lineages compared to ancestral lines (Figure 3E– F). Collectively, these data suggest that the Hsp90 chaperone plays a central role in buffering the costs of accumulating mutations.

### The lineages mount a shared stress response

Each MA lineage harbored a unique set of mutations (Figure 1E, Figure 1 – figure supplement 1D) but all lineages we examined shared a physiological response. This led us to investigate whether independent lineages might engage a common gene expression response to mutation burden. To address this question, we performed mRNA-sequencing (RNA-seq) to examine gene expression across multiple hypermutator lineages, choosing time points corresponding to both the ‘high-cost’ and ‘low-cost’ phases of their fitness trajectories. We first tested whether continuous passage alone had an impact on gene expression, examining the transcriptome of wild-type lineages passaged for 1,250 generations. Gene expression patterns in these cells were virtually unchanged relative to the wild-type ancestor, and the eight genes that were differentially expressed were not enriched for any shared function or protein–protein interactions (Table S2). Thus, passaging itself had a minimal impact on gene expression.

Increasing mutation burden in the hypermutator lineages, by contrast, was associated with marked changes in gene expression. As expected, transcriptional profiles diverged with increasing genetic distance between each passaged MA lineage and its ancestor (represented by gray arrow in Figure 4A, and gray points in Figure 4B). We also compared sister lineages, whose genetic distance from each other was twice that of a single lineage to its own ancestor (orange arrows and points). In these cases, the increased genetic distance was not associated with a concomitant transcriptional divergence (Figure 4B). Thus, despite having accumulated completely different sets of mutations, the congruent gene expression profiles of the highly mutated lineages point to a shared response to mutation burden.

**Figure 4.**
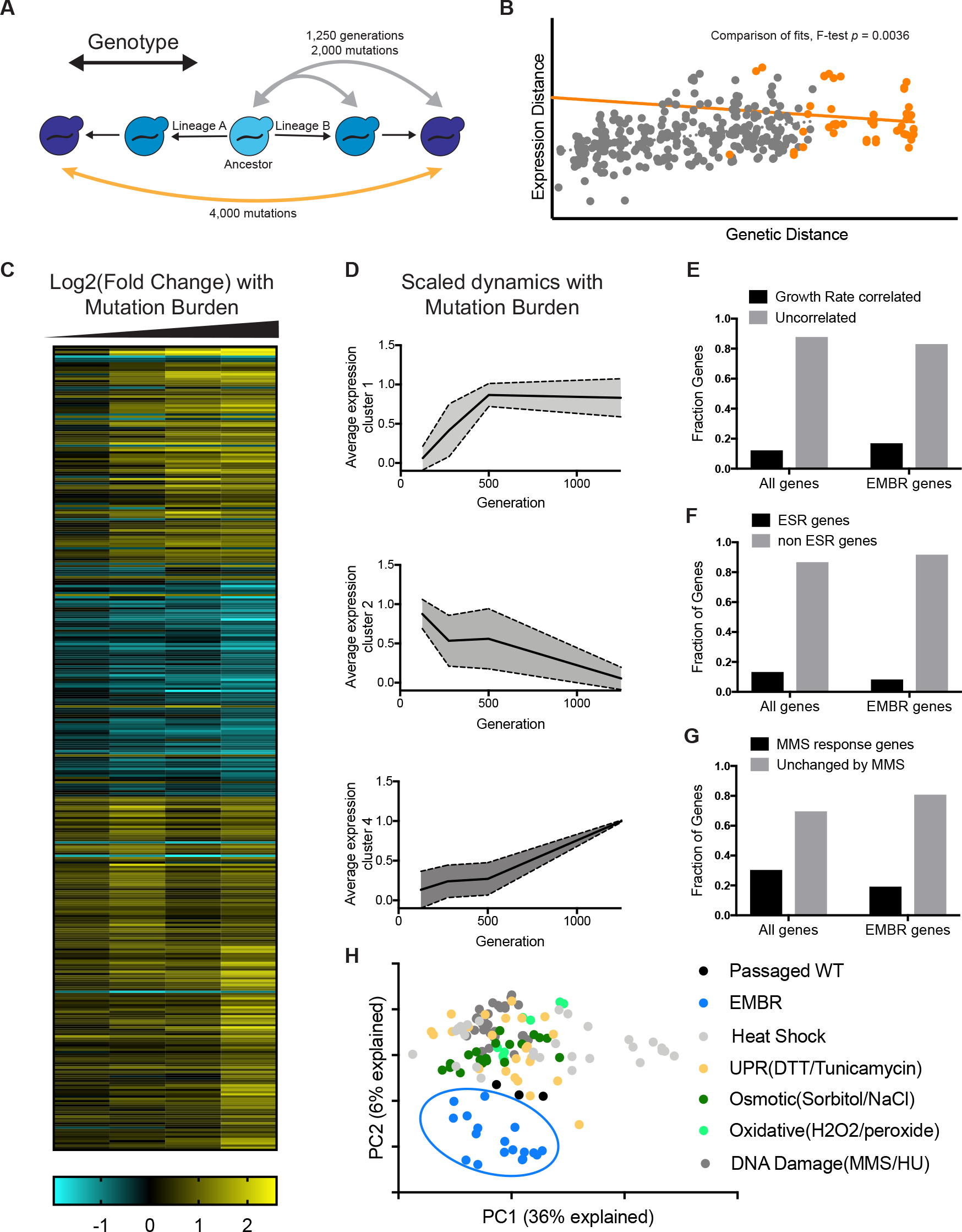
A shared transcriptional response to mutation burden. (**A**) Cartoon representation of the genotypic distance between sister MA lineages (orange) and between a passaged lineage and its ancestor (gray). (**B**) Relationship between genetic distance (measured by number of variant calls) and expression distance (measured by Euclidean distance between gene expression vectors). (**C**) Gene expression changes (at generations 125, 275, 500, 1250) were scaled per gene (row), then clustered by k-means for heat map. Color represents Log2 fold change in expression relative to the ancestor. (**D**) Average scaled expression of each cluster. Dashed lines represent standard deviation of the mean of nearly 100 genes in each cluster. (**E**) Comparison between EMBR genes and genes differentially expressed during slow growth (Brauer et al., 2008). (**F**) Comparison between EMBR genes and genes differentially expressed during the environmental stress response (ESR; Gasch et al., 2000). (**G**) Comparison between EMBR genes and genes differentially expressed during the DNA damage response (DDR; Caba et al., 2005). (**H**) Principal component analysis comparing MA gene expression data to published gene expression responses from the SPELL database (https://spell.yeastgenome.org/).

Further examination of our data revealed shared changes in gene expression among the independent lineages, which we term the Eukaryotic Mutation Burden Response (EMBR). After ^~^1,250 generations in the hypermutator lineages, expression of this cohort of 360 genes (FDR *Q* < 0.05, Figure 4C-D, Table S2-7) changed robustly relative to the ancestor. EMBR was not dominated by the effects of one individual lineage, nor was the response simply due to the slower growth rate of lineages with high mutation burden, as we observed only a slight enrichment in genes whose expression is correlated with growth rate (Figure 4E; Brauer et al., 2008). We therefore conclude that, despite the diversity of genetic lesions they accumulated, the hypermutating lineages each mounted a similar stress response.

### EMBR is distinct from previously reported stress responses

To further characterize the transcriptional response to mutation burden, we analyzed the shared pathways and functions of differentially expressed genes. The only Gene Ontology (GO) term (Szklarczyk et al., 2015) enriched among EMBR genes that increased in expression was ‘protein refolding,’ consistent with our prior observations of sensitivity to Hsp90 chaperone inhibition. However, this enrichment was very weak (FDR = 0.00295), and largely centered on Hsp90. Down regulated genes showed even less enrichment; ‘regulation of transcription from RNA polymerase II promoter’ was the only GO term that emerged (FDR = 0.0117).

We next compared these gene expression changes to other stress conditions that have been transcriptionally profiled in yeast. EMBR was not enriched for environmental stress response genes (Gasch et al., 2000) that characterize many such gene expression profiles (Figure 4F) or the DNA damage response to MMS, bleomycin, and cisplatin (Caba et al., 2005; Figure 4G). Although direct comparisons of gene expression datasets can be complicated by batch effects, a principal component analysis grouped EMBR as a separate response from well-studied responses to heat shock, unfolded protein, osmotic stress, oxidative stress, and DNA damage (Figure 4H). Comparison to all of the 2,400 gene expression profiles that have previously been reported in yeast (Hibbs et al., 2007) did not reveal stronger overlaps with other known stress responses.

To examine the network connectivity amongst proteins involved in EMBR, we took advantage of the protein–protein interaction networks that have been systematically mapped in *S. cerevisiae* (Costanzo et al., 2010; Szklarczyk et al., 2015). In contrast to the weak GO term enrichment, we observed an extremely strong enrichment for protein–protein interactions among EMBR components (Figure 5A; *p* < 10^−16^ for components that increase with mutation burden; *p* < 10^−7^ for those that decrease with mutation burden; *p* < 10^−15^ for all components). This high degree of connectivity suggests that EMBR is a concerted physiological response.

**Figure 5.**
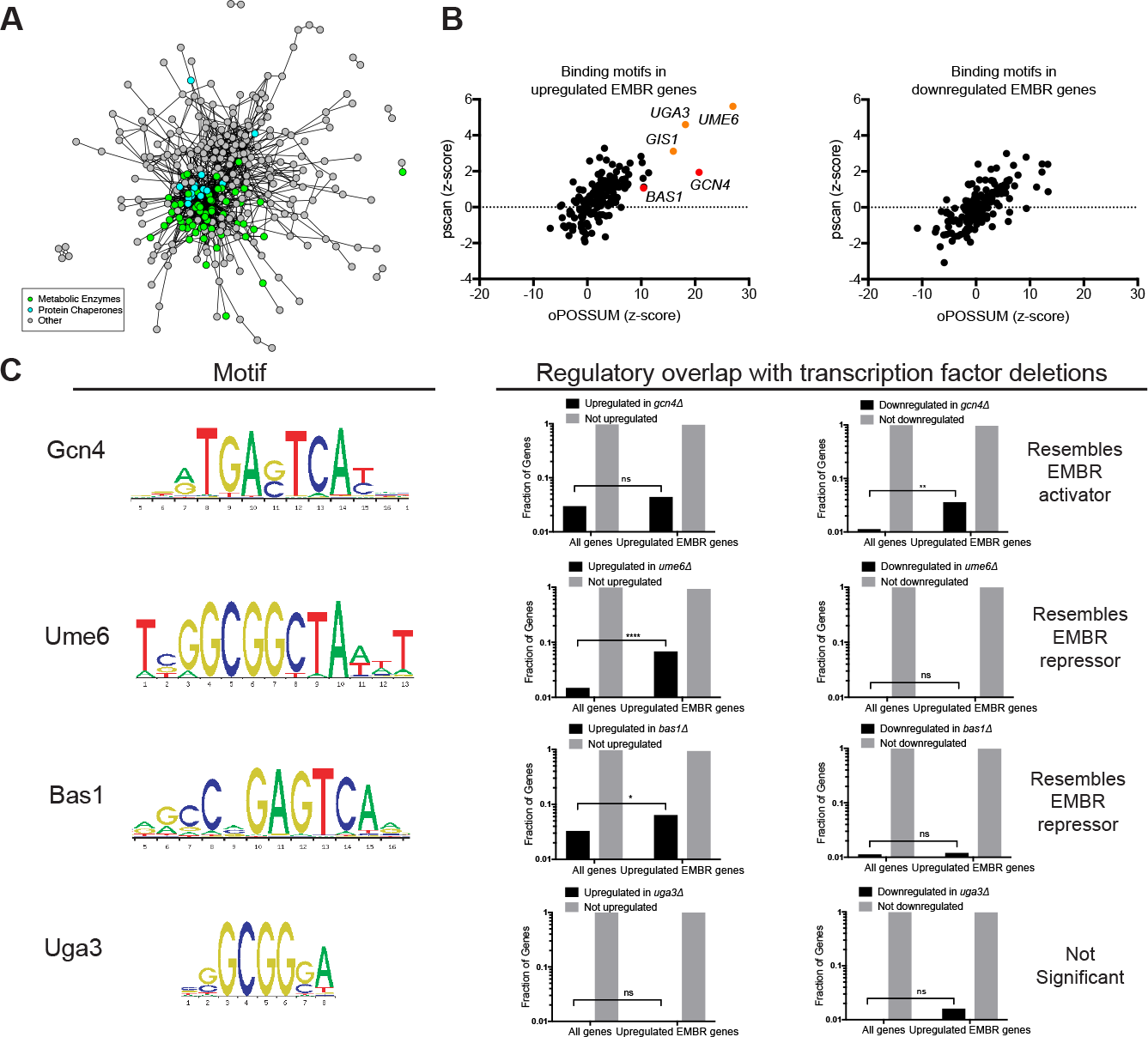
Transcription factors control EMBR genes. (**A**) Network of EMBR genes; edges represent protein-protein interactions and genetic interactions from the STRING database. Metabolic enzymes are highlighted in green, protein chaperones/co-chaperones are highlighted in blue. (**B**) Transcription factor binding sites enriched in upregulated and downregulated EMBR gene promoters. Ranking by Pscan (Zambelli et al., 2009) on the y axis and oPOSSUM (Kwon et al., 2012) on the x axis. In red are genes that were also identified by gene expression comparisons to the mutant via SPELL. (**C**) Transcription factor motifs calculated by JASPER (Khan et al., 2018). Bar graphs represent enrichment of upregulated EMBR genes *upregulated* or *downregulated* in transcription factor mutants. * p < 0.05, ** p < 0.01, **** p < 0.0001, Fisher’s exact test.

We next sought to identify factors that might coordinate EMBR. We searched for enrichment of transcription factor binding sites among the promoters of EMBR genes using two complementary algorithms: Pscan (Zambelli et al., 2009) and oPOSSUM (Kwon et al., 2012). Among the genes that were upregulated, binding sites for four transcription factors were clearly enriched: Ume6, Gcn4, Uga3, and Gis1 (Figure 5B). By contrast, the targets of transcription factors critical for mounting other stress responses, such as Hsf1 or Msn2/4, were not enriched in the mutated lineages (*p* = 0.5 by Pscan).

Ume6 represses meiosis-specific promoters by binding to upstream repressive sequences and recruiting the histone deacetylase complex Rpd3–Sin3 (Hahn and Young, 2011). Ume6 is also part of a signaling cascade that regulates autophagy via repression of *ATG8* (Bartholomew et al., 2012) and its repression and activation by Sin3 and Ime3 is analogous to the stepwise activation of human Myc/Max/Mad axis (Washburn and Esposito, 2001). Gcn4 is a central regulator of metabolism, notably amino acid biosynthesis, and several stress responses (Natarajan et al., 2001). Uga3 is a transcriptional activator for γ-aminobutyrate (GABA)-responsive genes (Andre, 1990). Gis1 is a histone demethylase and transcription factor that regulates genes during nutrient limitation (Tu et al., 2007) and hypoxia (Dastidar et al., 2012). We also compared EMBR to gene sets that were differentially regulated in strains in which transcription factor genes had been deleted, using the SPELL database (Hibbs et al., 2007). This analysis revealed overlap between EMBR and two mutant datasets, *gcn4∆* and *bas1∆* (Fendt et al., 2010). Both of these genes were also identified by Pscan and oPOSSUM (highlighted in red in Figure 5B). Bas1 regulates the expression of enzymes in the histidine, purine, and pyrimidine biosynthetic pathways (Arndt et al., 1987; Daignan-Fornier and Fink, 1992; Denis et al., 1998; Denis and Daignan-Fornier, 1998) and is homologous to the myb proto-oncogene family (Tice-Baldwin et al., 1989).

The overlap of genes in EMBR with those that are differentially regulated in *gcn4∆* and *bas1∆* mutants suggests that these transcription factors are likely important regulators of EMBR. In total, 33% of EMBR genes are experimentally validated targets of at least one of these five transcription factors (Cameroni et al., 2004; Cherry et al., 2012), whereas the validated targets of all five compose 22% of all genes in the genome. We reasoned that if the transcription factors acted predominantly as an EMBR activator, its deletion should reduce expression of upregulated EMBR genes and conversely if a transcription factor acted predominantly as an EMBR repressor, its deletion should increase expression of upregulated EMBR genes. Genes that were downregulated in the *gcn4∆* mutant were significantly enriched in upregulated EMBR genes, suggesting Gcn4 primarily functions as an EMBR activator (Figure 5C; *p* < 0.01, Fisher’s exact test). Genes upregulated in the *ume6∆* and *bas1∆* mutants were enriched in upregulated EMBR genes suggesting Ume6 and Bas1 function as EMBR repressors (Figure 5C; *p* < 0.05, Fisher’s exact test). Very few EMBR genes changed expression in the *uga3∆* mutant (Hu et al., 2007). Understanding the full role of these transcription factors in EMBR activation and repression, and the regulation of the remaining 67% of EMBR genes, stands as a goalpost for future studies. However, the strong connectivity that we observe among EMBR genes led us to investigate whether the response had adaptive value.

### EMBR components are required for the survival of mutated cells

To determine the importance of EMBR genes in buffering mutation burden, we perturbed the activity of several proteins that were: i) upregulated with accumulating mutations; ii) engaged in several physical and genetic interactions with other genes involved in the same biological processes; iii) amenable to pharmacological inhibition; and iv) homologous to well-described human proteins. First, we observed upregulation of several autophagy-related genes that are crucial for phagophore initiation (e.g., *ATG1*, *ATG2*, *ATG13*), expansion of the autophagosome (*ATG7*), and fusion of the autophagosome with the vacuole (*VAM6*, *MON1*). Autophagy not only serves as a way to recycle macromolecules in the face of nutrient limitation, but also as a mechanism to clear damaged organelles and misfolded protein aggregates. We wondered whether the mutation burden, and consequent protein misfolding, in our mutated cells might require autophagy to clear. To investigate, we exposed the MA lineages to bafilomycin, a well-characterized inhibitor of autophagy (Yoshimori et al., 1991). Indeed, sensitivity to this compound increased with the mutation burden (Figure 6A-B), establishing the protective effect of autophagy for mutation burden.

**Figure 6.**
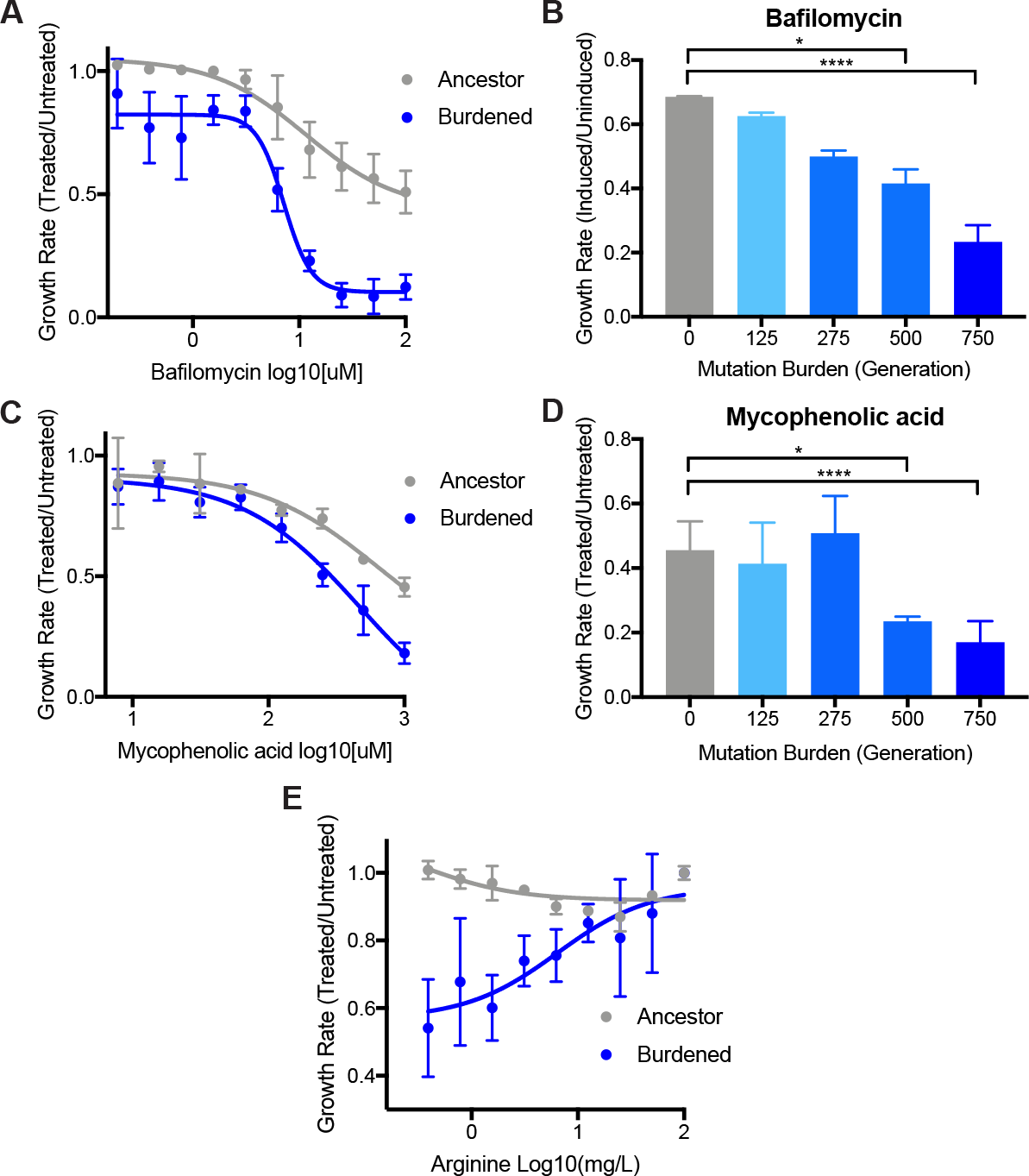
EMBR components are required for the proliferation of highly mutated cells. (**A**) Dose response to bafilomycin of MA lineages and ancestor control. Error bars represent SEM from three biological replicates. (**B**) Normalized growth rate after treatment with 12.5µM bafilomycin as a function of mutation burden. Error bars represent SD from three biological replicates. * p < 0.05, **** p < 0.0001, student’s t-test. (**C**) Dose response to mycophenolic acid of MA lineages and ancestor control. Error bars represent SEM from three biological replicates. (**D**) Normalized growth rate after treatment with 1mM mycophenolic acid as a function of mutation burden. Error bars represent SD from three biological replicates. * p < 0.05, **** p < 0.0001, student’s t-test. (**E**) Dose response to addition of arginine to synthetic media without arginine of MA lineages and ancestor control. Error bars represent SEM from three biological replicates.

We also observed upregulation of the inosine-5’-monophosphate (IMP) dehydrogenases, *IMD1/2. IMD2* is an important but non-essential gene in yeast; it catalyzes the rate-limiting step in GTP synthesis and is regulated by Ume6. Heterozygous mutations in one of its human homologs, *IMPDH1*, are associated with retinitis pigmentosa (McKusick, 2007), and expression of its other human homolog, *IMPDH2,* is correlated with poor prognosis of nasopharyngeal carcinoma (Xu et al., 2017), subtypes of which have high mutation load (Zhang et al., 2017). The encoded protein can be inhibited *in vivo* by mycophenolic acid (Fleming et al., 1996), which is currently used to limit transplant rejection and autoimmune disease and has recently been introduced as an anti-cancer agent in clinical trials (Chen and Pankiewicz, 2007). The requirement for *IMPDH2* in cancer is not completely understood (Xu et al., 2017). The mutated lineages were much more sensitive to treatment with this inhibitor than ancestral controls (Figure 6C-D), establishing the importance of this EMBR component for withstanding accumulating mutation burden.

We further observed upregulation of several arginine biosynthesis genes: *ARG1, ARG2, ARG5/6*. To test if the mutated lineages became unusually dependent on arginine for proliferation, we decreased levels of arginine in the growth medium and measured the effect on growth. Whereas the ancestral strain was tolerant to arginine removal, the burdened lineages were strikingly sensitive to decreased arginine availability (Figure 6E). The precise mechanisms by which these and other factors limit the cost of mutation burden demand future study. However, these data clearly establish that EMBR components can be critical for the survival of hypermutating cells.

### Conservation of EMBR in humans

We wondered whether a response resembling EMBR might be conserved in humans. To investigate, we identified 568 human homologs of the *S. cerevisiae* EMBR genes through deltaBLAST searches (Boratyn et al., 2012). The human homologs of EMBR genes (hEMBR) form a highly enriched protein–protein interaction network (Figure 7A; *p* < 1.1 × 10^−16^, hypergeometric test using physical and genetic interactions from BIOGRID; www.thebiogrid.org). However, because they are involved in deeply rooted biological processes, random sets of genes conserved from yeast to humans also encode proteins that are enriched in interactions. To test whether hEMBR was enriched in protein–protein interactions beyond what would be expected for conserved proteins, we sampled the protein–protein interactions of 500 random sets of proteins conserved from yeast to humans to define a null distribution. This bootstrapping analysis revealed that hEMBR was significantly enriched in protein–protein interactions relative to random expectation (*P* = 7.0×10^−4^), suggesting that the underlying structure of EMBR is conserved in humans.

**Figure 7.**
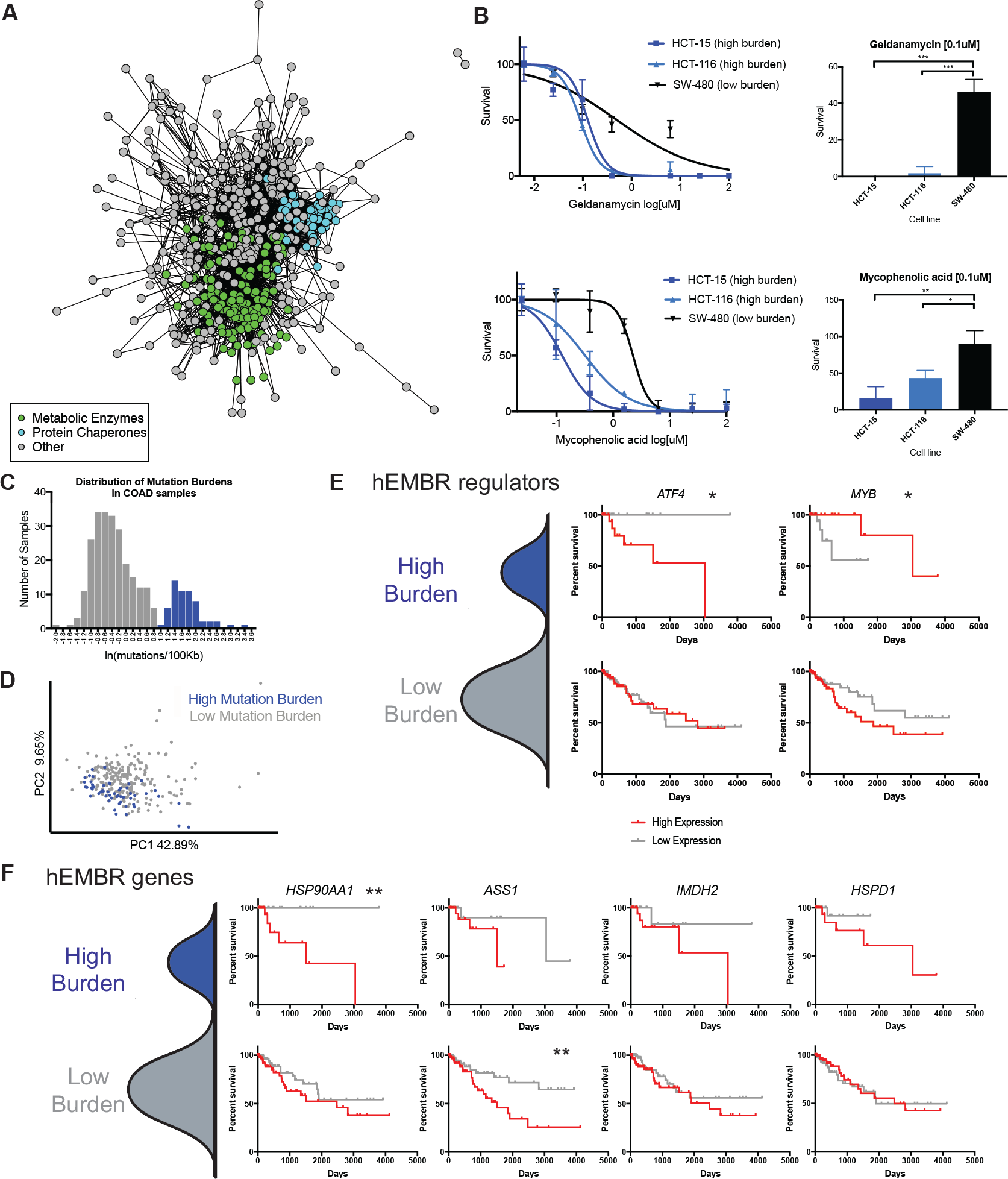
EMBR homologs in human cell lines and tumors. (**A**) Human homologs of EMBR genes (hEMBR) are highly enriched for protein–protein interactions. Network of hEMBR genes; edges represent protein-protein interactions and genetic interactions. Metabolic enzymes are highlighted in green, protein chaperones/co-chaperones are highlighted in blue. (**B**) Cell line survival upon inhibition with geldanamycin and mycophenolic acid. Bar graphs are representative of treatment with 0.1µM geldanamycin and 0.1µM mycophenolic acid. Error bars represent SD from three replicates. * = p < 0.05, ** = p < 0.01, *** p < 0.001 unpaired student’s t-test. (**C**) Bimodal distribution of mutation burden across colon cancer tumors (data from TCGA). (**D**) Principal component analysis of gene expression data separates colon cancer cell lines with high (blue) and low (gray) mutation burdens. (**E and F**) Patient survival data in samples with high and low expression of the genes of interest in all colon cancer tumors with high mutation burden (top row), or low mutation burden (bottom row). * = p < 0.05, ** = p < 0.01, Log-rank (Mantel-Cox) test.

If hEMBR were important for cancers with a high mutation burden, we would predict that cell lines derived from such tumors would be sensitive to inhibition of hEMBR genes. We used an inhibitor of Hsp90, geldanamycin, and an inhibitor of *IMPDH2*, mycophenolic acid, to investigate. We used two colon cancer cell lines that are confirmed mutators with thousands of variants measured, HCT-116 and HCT-15 (Barretina et al., 2012; Vilar and Gruber, 2010), and a non-mutator colon cancer cell line SW480 (Barretina et al., 2012). Indeed, HCT-116 and HCT-15 were far more sensitive to both inhibitors than SW480 was (^~^5-fold for geldanamycin, ~10-fold for mycophenolic acid; Figure 7B). These same cell lines are equivalently sensitive to multiple other drugs (Iorio et al., 2016). This result is further supported by anti-cancer activity of mycophenolic acid and another IMP-dehydrogenase inhibitor, Tiazofurin (Chen and Pankiewicz, 2007; Malek et al., 2004).

Can yeast EMBR further inform our understanding of the response to mutation burden in human cancer cells? Autophagy was important for yeast cells to withstand mutation burden, and this mechanism also provides a promising target for cancer therapies (Levy et al., 2017). Arginine was also important for yeast cells to withstand mutation burden, and the human homolog of the EMBR gene *ARG1, ASS1,* is activated in colon cancers, where it is thought to contribute to pathogenicity (Bateman et al., 2017). Tumors have varying mutation burdens, as well as diverse genotypes and mechanisms of oncogenesis. They thus constitute an incredibly complex dataset, with many confounding variables. Although these analyses are fundamentally under-powered and are limited to genes conserved from yeast to humans, we nevertheless searched for an hEMBR signature within curated datasets from The Cancer Genome Atlas (TCGA). Colon adenocarcinomas exhibited a bimodal distribution of mutation burdens, allowing us to separate tumors with high and low mutation burdens into distinct groups (Figure 7C). To narrow our scope and identify which hEMBR components drive the largest biological differences among colon cancer tumors, we performed a principal component analysis on the transcriptomes of the tumor samples. High mutation burden tumors roughly clustered together along the first two principal components (Figure 7D). We then asked what hEMBR factors contributed the most to the distinction between high and low mutation burden in the principal component analysis. We found *HSP90AA1* (a human homolog of Hsp90), *ASS1* (the human homolog of *ARG1*), *IMPDH2* (the human homolog of *IMD2*), and *HSPD1* (a human homolog of Hsp60; Figure 7 – figure supplement 1).

We next asked whether the expression of homologs of the EMBR regulators we identified were associated with survival of patients with tumors harboring high and low mutation burden. We hypothesized that activation of hEMBR due to differential expression of these hEMBR regulators would be beneficial for cancer cells with high mutation burdens but detrimental to patient survival. *GCN4* is homologous to the transcriptional activator *ATF4* (Murguia and Serrano, 2012), and *BAS1* is homologous to *MYB* oncogene (Tice-Baldwin et al., 1989). Indeed, although there was no distinction between survival curves in low mutation burden cancers regarding *ATF4* expression (Figure 7E), in high mutation burden cancers, high *ATF4* was associated with significantly decreased survival (Figure 7E). This is consistent with Gcn4 functioning to activate EMBR in yeast. *ATF4* is known to promote survival under stress while carefully balancing induction of apoptosis (Wortel et al., 2017). Based on these observations we propose that an additional role could be activation of hEMBR in high mutation burden cancers. We next performed a similar analysis for the *BAS1* homolog *MYB*. There was no distinction between survival curves for patients with low mutation burden cancers with respect to MYB expression (Figure 7E). But in cancers with high mutation burdens, high *MYB* expression was associated with increased survival (Figure 7E). This is consistent with Bas1 functioning to repress EMBR in yeast. *MYB* mostly acts as a transcriptional activator balancing cell differentiation, is linked to several leukemias, and can be activated in both colon and breast cancers (Ramsay and Gonda, 2008). Our results suggest that regulation of hEMBR by both *ATF4* and *MYB* could be protective against high mutation burden in cancers and thus lead to poor patient prognosis.

Although *HSP90* expression has been implicated in cancer survival in several contexts (Jaeger and Whitesell, 2019; Whitesell and Lindquist, 2005; Whitesell et al., 2014), the ability to compare expression to patient survival data is limited by the low number of patients, particularly when stratifying samples. Despite these limitations we observed a striking correlation in colon cancer patient data (Chandrashekar et al., 2017) between increased *HSP90* expression and decreased patient survival that was only evident in cancers with high mutation burden (Figure 7E). We thus propose that mutational buffering could offer an explanation for the strong correlation of *HSP90* expression and tumor growth (Ciocca and Calderwood, 2005; Jaeger and Whitesell, 2019; Santagata et al., 2011). The other hEMBR genes that we identified above (Figure 7 – figure supplement 1) appear to show a similar trend in our analysis (Figure 7E). Further study will be required to fully evaluate the effects of hEMBR genes on how tumors tolerate high mutation burdens and how this impacts patient survival. Nonetheless, these analyses suggest that hEMBR may have an important role in hypermutating human cancers.

## DISCUSSION

Many pathogenic bacteria and cancer cells exhibit elevated mutation rates, which can confer a wide variety of benefits such as drug resistance and rapid escape from immune surveillance (Davoli et al., 2017; Oliver et al., 2000). However, most mutations are predicted to be deleterious. Accordingly, to optimize fitness, such hypermutator phenotypes should be transient and revert once a beneficial mutation has been assimilated (Sniegowski et al., 1997; Sniegowski and Murphy, 2006). Indeed, this behavior has recently been observed in experimental evolution in *E. coli* (Swings et al., 2017). However, in a surprisingly large number of biological scenarios, including many human cancers, hypermutator phenotypes are stable, and cells remain capable of surviving, proliferating, and evolving new traits without succumbing to error catastrophe. Our observation that eukaryotic cells possess an intrinsic robustness against accumulating mutations provides a potential resolution for this paradox.

By generating a set of mutation accumulation (MA) lineages with defined mutation rate and spectrum, we were able to systematically examine the relationship between fitness and mutation burden. Rapid acquisition of non-overlapping sets of mutations led to a consistent, non-linear relationship between mutation burden and fitness: mutations acquired early were on average costlier than mutations acquired later. Remarkably, this behavior was consistent across lineages, despite the fact that no single mutation was universal. Because the MA lineages maintained their hypermutator phenotypes over the course of the experiment, we conclude that the cells induced a protective response that limited the fitness cost of mutations that accumulated later in the buffered phase of the experiment.

Similar behavior has been observed in bacteria, in which buffering is mediated by the chaperonin GroEL in several species. For example, GroEL suppresses the phenotypes of temperature-sensitive mutations in the replication protein DnaA (Jenkins et al., 1986) and some phage proteins (Van Dyk et al., 1989). Highly mutated *S. typhimurium* express high levels of GroEL, and artificial overexpression of the chaperonin further improves fitness (Maisnier-Patin et al., 2005). Bacteria that naturally sustain high mutation burdens, such as the aphid endosymbiont *Buchnera*, also strongly express GroEL, and this has been linked to their capacity to withstand the extremely high levels of genetic drift caused by their small effective population size (Fares et al., 2002a). However, because the eukaryotic GroEL homolog Hsp60 has limited mitochondrial function, it has remained unclear whether and how buffering might be mediated in eukaryotes.

The stress response that we report here differs from any that have previously been described. A key component of EMBR is the Hsp90 chaperone and several of its accessory co-chaperones. Data from flies (Rutherford and Lindquist, 1998), plants (Queitsch et al., 2002; Sangster et al., 2007; Sangster et al., 2008), yeast (Cowen and Lindquist, 2005; Jarosz and Lindquist, 2010), worms (Burga et al., 2011), Mexican cavefish (Rohner et al., 2013), and most recently humans (Karras et al., 2017) suggests that this chaperone plays a key role in the phenotypic manifestation of genetic variation. In mechanistic terms, Hsp90 can assist in the folding of unstable gain-of-function protein variants, thereby potentiating their immediate phenotypic effect. It can also buffer the impact of other genetic variants, silencing their phenotypic impact. Here we found that inhibiting Hsp90 amplified the fitness defects later in the MA lineages. Our data thus establish that Hsp90 can buffer the cost of *de novo* mutations. A prior study that did not reach the same conclusion used a much smaller number of mutations (roughly four mutations per lineage in 94 separate lineages; Geiler-Samerotte et al., 2016). In that regime, the biochemical effects of specific mutations themselves might have dominated the fitness measurements. In our experiment, in which a larger number of polymorphisms were examined, the effects of Hsp90 were strongest when mutation burden was sufficient to affect a much larger swath of the proteome.

We identified hundreds of other EMBR components. These were strongly enriched for protein–protein interactions. By contrast, gene ontology enrichments were sparse, and surprisingly weak where they existed. Thus, EMBR is very different from other stress responses that have previously been reported. The strong enrichment for interacting proteins, and the fact that inhibition of EMBR components impaired the fitness of highly mutated cells, suggest that this stress response is likely to be adaptive.

In pathogens hypermutator phenotypes can facilitate antibiotic resistance, immune evasion, or morphological innovation such as filamentous growth. Although the clinical use of antibiotics is a relatively recent phenomenon, host–pathogen conflict is ancient. Conservation of a response that allows pathogens to adopt hypermutator states with minimal fitness cost could be beneficial for evolution over short and long timescales. Indeed, many microbial pathogens, and even wild fungal populations, frequently adopt a hypermutator state (Bui et al., 2017; Guo et al., 2018; Raghavan et al., 2018). The alleles that govern this behavior have been retained across a considerable evolutionary distance (Bui et al., 2017), suggesting that it confers an adaptive advantage.

Hypermutating cancers are evolutionary experts, capable of buffering the cost of accumulating mutations while simultaneously harnessing the potential of this raw genetic material to rapidly evolve new traits. This evolutionary virtuosity can have devastating consequences for human health, such as the emergence of resistance to chemotherapies. Here, we show that eukaryotic cells can broadly limit the cost of accumulating mutations by mounting a conserved stress response that we term EMBR. We posit that highly mutated cells are addicted to EMBR, analogous to oncogene addiction but in this case driven by a gene expression network rather than a single driver mutation. Our data suggest that this addiction may expose multiple Achilles’ heels with potential for therapeutic intervention.

## METHODS

### Strain construction and propagation

BY4741 haploid yeast were transformed to replace the *MSH6* gene with an antibiotic resistance marker (hygromycin) or to replace the *POL3* gene with a variant that lacks proofreading activity, *pol3-L612M* (Nick McElhinny et al., 2007). Double mutants were constructed by performing both types of transformations in series, *pol3-L612M* followed by *msh6*∆. Transformants were grown up from single colonies and frozen as passage 0. We constructed our accelerated mutation accumulation (MA) lines in haploid yeast so that we could study the effects of all polymorphisms, including recessive alleles.

Passage 0 strains (parent BY, *msh6∆*, *pol3-L612M*, and double mutant *pol3-L612M msh6∆)* were streaked on both YPD (yeast extract-peptone-dextrose medium) and YPG (yeast extract-peptone-glycerol medium) plates and grown at 25°C for at least 96 hours. Individual colonies of average size were selected, resuspended in a 96-well plate, and then streaked onto a fresh YPD or YPG plate for another round of growth (passage 1). Because *S. cerevisiae* can lose their ability to respire on fermentable carbon sources, we propagated one set of eight lineages on media containing glucose as a carbon source, and another set of eight lineages on media containing glycerol as a carbon source, so that potential loss of respiration would not confound our findings. At odd-numbered passages, small volumes of the stock plates were then transferred to fresh liquid YPD, and growth in solution was measured every ^~^5 minutes for 84 hours. At the end of this period, the cultures were diluted and spotted onto YPD, YPG, phosphonoacetic acid (Li et al., 2005), and canavanine plates to assess general growth, growth on glycerol (YPD strains only), maintenance of the *pol3-L612M* variant (which confers resistance to PPA), and mutation rate. Any loss-of-function mutation in the *CAN1* arginine permease gene gives rise to canavanine resistance, providing a readout of mutagenesis. This process was repeated for 50 passages. During the course of the experiment on YPD, one hypermutator lineage lost its mutator phenotype, one went extinct. During the course of the experiment on YPG, one hypermutator lineage lost its mutator phenotype early on, one lost its mutator phenotype halfway through, and three went extinct. The frequency of these events is consistent with expectation from other mutation accumulation experiments in bacteria (Aguilar-Rodriguez et al., 2016). To study the biological consequences of cells surviving despite mutation burden, we focused our phenotypic analysis on lineages that survived for all 50 passages, or approximately 1,250 generations, while maintaining the mutator phenotype.

### Growth measurements

At odd-numbered passages, surviving strains were inoculated in liquid culture to measure OD_600_ on a Synergy Multi-Mode Microplate Reader. Doubling time was extrapolated using a Bayesian curve-fitting model and converted to fitness (w) relative to the first passage on the same media: 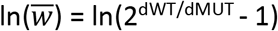. WT = wild-type, MUT = hypermutator.

### Drug treatment

Hyperutator and wildtype yeast strains were grown to saturation in YPD media in 96-well plates. Dilutions of drugs obtained from Selleckchem, MicroSource, and Sigma were prepared in DMSO and added to YPD; 10-100µM radicicol; 4.4mM H_2_O_2_; 10µM fluconazole; 12.5µM bafilomycin, 1mM mycophenolic acid. Media lacking arginine and uracil was obtained from Sunrise Science.

Yeast were diluted, and OD measurements were taken using a Synergy Multi-Mode Microplate Reader (Biotek). Mean and standard error were calculated for three technical replicates of each lineage, and then across independent mutated lineages. Growth rates were compared to extract a dose-response curve and inhibitory concentrations.

### Sequencing, variant calling, and gene expression analysis

Variant calls were made using the GATK pipeline and annotations were made with SnpEff (Cingolani et al., 2012; McKenna et al., 2010). Only variants called in multiple biological replicates were considered for comparison between lineages (Fig 1E). Excluding variants present in the shared ancestor, variant calls unique to each lineage, shared between any two lineages, or shared across lineages were counted. The predicted phenotypic impact of these variants was characterized using SIFT (Kumar et al., 2009). Genes in which variants occurred within ORFs were also categorized as essential (as defined by the Stanford Deletion Project) or non-essential.

For gene expression analysis, mutator and wildtype strains were collected during log-phase growth. Pellets were flash frozen and stored at −80°C. Yeast were lysed, RNA extracted, and RNA quality was assessed on a Bioanalyzer. A sequencing library was prepared using the TruSeq RNA Library Prep Kit v2. Reads were filtered based on quality scores and aligned to the genome using Bowtie2 (Langmead and Salzberg, 2012) with a current reference genome (Engel et al., 2014). HTSeq was used to quantify counts (Anders et al., 2015). Normalization, batch effect correction, and differential expression testing was performed using the DESeq2 software suite (Love et al., 2014). For downstream graphing purposes, limma was used for batch effect correction. Cluster 3.0 was used for k-means clustering of gene expression data after row normalization (Figure 4C-D; de Hoon et al., 2004). Motifs and lists of EMBR genes containing transcription factor motifs were obtained from JASPAR (Khan et al., 2018). Gene expression data from deletions were collected from Hu et al. and Cameroni et al. (Cameroni et al., 2004; Hu et al., 2007).

### v-Src assay

The plasmid carrying the v-Src gene for expression in yeast, Yep-src (Brugge et al., 1987), was a gift from the Lindquist laboratory. Hypermutator and wild-type yeast were transformed via electroporation, and the plasmid was maintained by growth on SD-Ura. Expression of v-Src was induced by diluting saturated yeast (grown in 2% raffinose media) into 2% galactose media. Growth was monitored by measuring OD_600_.

### Principal component analysis

Yeast expression datasets for environmental stress responses, DNA damage responses, and the unfolded protein response (Fry et al., 2003; Gasch et al., 2001; Gasch et al., 2000; Travers et al., 2000; Travesa et al., 2012) were obtained from the SPELL database (spell.yeastgenome.org; Hibbs et al., 2007). RNA expression data from the mutation accumulation lines were normalized against those of their un-passaged parent and log_2_-normalized. Expression data from passaged WT lines were normalized to their WT parent. All other expression data were normalized to their internal untreated controls. These expression data were collectively visualized through principal component analysis (PCA), a technique that reduces high-dimensional expression data (Jolliffe and Cadima, 2016).

### EMBR human homolog protein–protein interaction enrichment

Human homologs of yeast EMBR genes were identified through deltaBLAST searches using default settings. Only hits with e-values ≤ 1 × 10^−10^ were collected as homologs. Hits were then mapped to unique Entrez IDs. A total of 568 human homologs of yeast EMBR genes (hEMBR) were identified. The protein–protein interactions of the hEMBR network were mapped using the BIOGRID database. The protein–protein interactions of 500 equally sized sets of random human genes and 500 equally sized random sets of annotated human homologs of all yeast genes (ENSEMBL) were also mapped. A hypergeometric test was computed for the protein–protein interactions in the hEMBR set and the median interactions in the random human gene and human homolog gene sets. To compute the hypergeometric test, key parameters were determined by mapping the total interactions between all annotated (HUGO) human genes in BIOGRID. The amount of total possible interactions was defined as (n*(n-1))/2 for n nodes in an undirected network. The protein–protein interactions of the 500 random sets of human homologs were fit to a gamma distribution. From this null distribution, 10,000 random samples were drawn to compute a p-value against the null hypothesis that the number of protein–protein interactions in the hEMBR is not larger than expected from the null distribution.

### Cell lines

Colon cancer cell lines HCT-116, HCT-15, and SW480 were purchased from ATCC and cultured in RPMI supplemented with 10% FBS or recommended medium. Cell viability was measured after 48 hours of growth in drug or DMSO control with PrestoBlue (Thermo Fisher Scientific). Background was subtracted from OD measurements, outliers removed, and normalized to untreated controls. Each condition was repeated in triplicate. IC50 values were calculated using a nonlinear regression (GraphPad Prism v7).

### EMBR signature in Colon Cancer

Whole exome sequencing, RNA sequencing, and patient clinical data were retrieved from TCGA (Cancer Genome Atlas, 2012) and assembled using the TCGA-Assembler R package (Zhu et al., 2014a). Tumor mutation burden of each sample was defined as the sum of total variant calls (relative to normal tissue) for each sample from whole exome sequencing and inferring mutations per 100kb based on the approximate length of genomic sequence captured by the exome sequencing preparations used, ^~^44Mbs, and assuming equal distribution of mutations across the entire genome. A histogram of tumor mutation burdens for the colon adenocarcinoma (COAD) dataset revealed a bimodal distribution with low mutation burden (< 2.55 mutations/100kb) and high mutation burden cohorts (> 2.55 mutations/100kb). For comparison with EMBR, only genes with annotated yeast homologs were kept for subsequent analysis. Expression levels of a given gene were defined by the scaled estimate of counts as reported in TCGA. A principal component analysis was performed on the expression data across all tumor samples to identify any clustering among the high mutation burden cohort. The norm vectors of the coefficients for principal components 1 and 2 (explaining 52.5% of the variance) were rank ordered and the top 8 EMBR homologs were chosen for further analysis on their impact on clinical outcomes. For the top 8 EMBR homologs, expression levels across the high mutation burden tumor samples, or across low mutation burden tumor samples in a parallel analysis, were grouped into low (less than median expression of the given gene in the dataset) and high expression (greater than median expression of the given gene in the dataset) cohorts. Kaplan-Meier survival plots for the high and low expression cohorts were created and compared using the Log-rank (Mantel-Cox) test.

## ACKNOWLEDGEMENTS

We thank K. Cimprich, A. Fire, S. Lujan, and members of the Jarosz laboratory for materials, helpful discussions, and comments on the manuscript. This work was supported by an NIH Pathway to Independence Award (R00-GM098600), a Searle Scholar Award (14-SSP-210), and a Kimmel Scholar Award (SKF-15-154) to DFJ. DFJ is also a Science and Engineering Fellow of the David and Lucile Packard Foundation. RS was supported by a Stanford Graduate Fellowship and a Gerald J. Lieberman Fellowship. RAZ and MKZ were supported by a Stanford Dean’s Postdoctoral Fellowship. MKZ was also supported by a postdoctoral fellowship from the Walter and Idun Berry Family foundation. JM was supported by NIH training grant T32HG000044 and TS was supported by NIH training grant T32-CA009302. Results shown here are in part based upon data generated by the TCGA Research Network: http://cancergenome.nih.gov/.

## AUTHOR CONTRIBUTIONS

RAZ, JM, RS, MKZ, and DFJ designed the research; RAZ, JM, RS, MKZ, and TRS performed the research and analyzed the data. MKZ passaged the mutator yeast lineages. RAZ, JM, RS, and DFJ wrote the paper. DFJ supervised all aspects of the work.

## Supplementary Tables

**Table S1.** Mutations called from DNA sequencing.

**Table S2.** Differential expression of passaged wild-type compared to wild-type ancestor.

**Table S3.** Differential expression of hypermutator ancestor compared to wild-type ancestor.

**Table S4.** Differential expression of hypermutator after 125 generations compared to hypermutator ancestor.

**Table S5.** Differential expression of hypermutator after 275 generations compared to hypermutator ancestor.

**Table S6.** Differential expression of hypermutator after 500 generations compared to hypermutator ancestor.

**Table S7.** Differential expression of hypermutator after 1250 generations compared to hypermutator ancestor.

### Supplementary Information

#### Comparison of mutation rates in yeast lineages and human tumors

We calculated the yeast mutation rate as the number of sequenced mutations relative to the parent strain, divided by the number of generations (assuming 25 generations per passage). This was normalized against the *S. cerevisiae* mutation rate as calculated by Zhu et al. (Zhu et al., 2014b), 1.67 × 10^−10^ per base per generation. Multiple estimates of the human somatic cell mutation rate have been reported. We used the average of reported somatic cell mutation rate estimates, 0.77 × 10^−9^, as a reference (Lynch, 2010). We converted the reported frequency of mutations in different tumor types (Lawrence et al., 2013) to mutation rate by assuming that a 1-cm^3^ tumor sample contains 1 × 10^9^ cells, which would require 30 divisions to grow from a single cell (Del Monte, 2009).

#### Distribution of mutations

Using a Poisson distribution, we calculated the probability that any given base pair would be mutated over the course of our accelerated evolution experiment. At each generation, around two new mutations arise, satisfying the assumption of the Poisson distribution that the probability of any given base being mutated is small 2/(12 × 10^6^). However, the mean number of mutations that arise in the population between the expansion of a single cell into a visible colony is roughly equal to twice the number of cells in the final colony. Given our assumption of 25 generations per bottleneck, we expect to observe 2^21^ (^~^2 × 10^6^) mutations arising in the population per 20 generations. Integrated over 1000 generations, we have 10^8^ mutations arising in each replicate of our experiment, or 8.7 mutations per base pair in the genome. The probability that a base pair is never mutated, assuming an equal chance of any base pair being mutated, is thus e^−8.7^ (^~^1.67 × 10^−4^). If we assume 25 generations per bottleneck, then this larger effective population size implies that ^~^280 mutations arise for each base pair across the experiment.

#### Expected target size of epistatic interactions

Approximately 1,000 of the ^~^6,000 genes in *S. cerevisiae* are essential (Giaever et al., 2002). Digenic interactions cause synthetic lethality of many more combinations of genetic knockouts; 10,000 interactions involved over 3,000 genes. That is ^~^3% of all gene pairs display a negative genetic interaction or synergistic epistasis (Costanzo et al., 2016). Positive genetic interactions or antagonistic epistasis was far less common (1.9%). This multitude of synthetic lethal interactions predicts that accumulation of random mutations will likely result in accelerated fitness decline. Fewer available trajectories or increasing number of positive genetic interactions does not explain the decreased fitness cost observed (Figure 2, Figure 2 – figure supplement 1).

#### Limit of detection

We examined the three smallest reasonably manipulatable colony of a highly passaged strain and directly measured the area from a microscope image. From this area (^~^0.25mm), we estimated the total number of yeast cells in the colony (assuming a half-sphere for volume and assuming an average haploid cell volume from BioNumbers, 37 µm^3^ (Milo et al., 2010)). This allowed us to estimate the average growth rate based on its appearance on the plate after 4 days 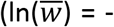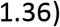. The growth rate was well below the minimum measured for the MA lines (Figure 2, Figure 2 – figure supplement 1) suggesting that we were not limited by failing to detect small colonies.

**Figure 1 – figure supplement 1.**
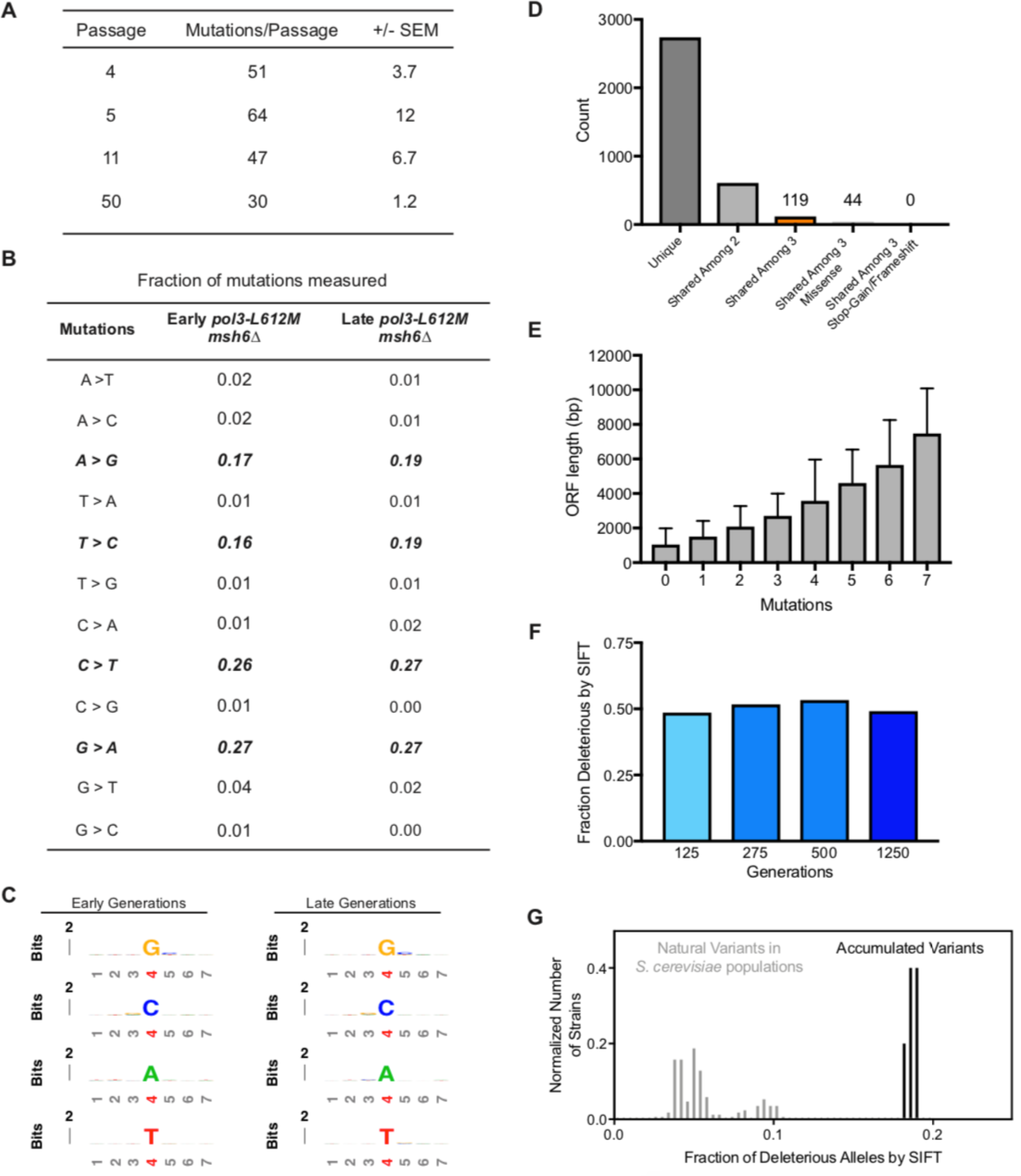
MA lines accumulate mutations at a constant rate with expected bias. (**A**) Samples were collected for sequencing before passaging, after 20 passages (500 generations), and after 50 passages (1250 generations). Values represent the mean mutation rate calculated based on variants called from RNA-seq after accounting for generations from three independently passaged lineages ± SEM. (**B**) Mutation bias favored transitions (italicized, bold) over transversions and was consistent between early passages (1–20) and late passages (20–50). (**C**) No sequence signature surrounds mutations in each mutated nucleotide, in either early or late generations, across all data collected. Sequence alignments were visualized with kpLogo (Wu and Bartel, 2017). (**D**) Counts of uniquely mutated ORFs, mutated ORFs shared among two lineages, mutated ORFs shared across three lineages, missense mutated ORFs shared across three lineages, and counts of stop-gain/frameshift mutated ORFs shared across three lineages. (**E**) Length of ORFs with different numbers of mutations mapped across three lineages. (**F**) Fraction of deleterious mutations in data collected at generation 125, 275, 500, and 1250 called by SIFT (Kumar et al., 2009). (**G**) Based on deleterious SIFT scores, a large fraction of MA lines (y axis) contain a large fraction of deleterious alleles (x axis) relative to the natural variation observed in strains catalogued by the Saccharomyces Genome Resequencing Project (SGRP, http://www.sanger.ac.uk/research/projects/genomeinformatics/sgrp.html). Increased accumulation of deleterious alleles confirms the limited selection imposed upon these lines.

**Figure 2 – figure supplement 1.**
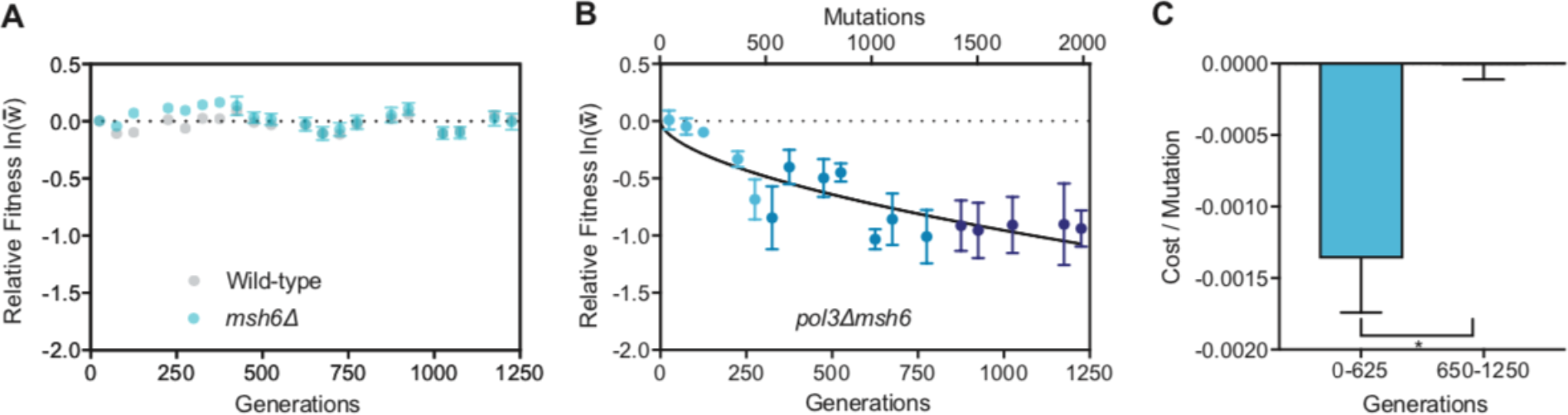
Fitness cost declines with increasing mutation burden on glycerol. (**A**) Mean relative fitness of eight independently passaged lineages of the control wild-type strain BY4741 and its *msh6Δ* derivative. Relative fitness 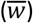 calculated based on doubling time relative to wild-type at passage 1. Error bars represent SEM from eight biological replicates. (**B**) Fitness trajectories of four hypermutator lineages passaged on glycerol medium. Fits are based on a regression model for detecting epistasis: 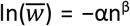 (α = 0.018 ± 0.014, β = 0.575 ± 0.119) described in Figure 2 (Maisnier-Patin et al., 2005). The goodness of fit of this model was superior to the simpler additive model (β = 1), p < 0.0001, extra-sum-of-squares F-test. Error bars represent SEM from three-four biological replicates. (**C**) Fitness cost per mutation across the first 625 generations and last 600 generations, calculated by linear regression, in passaged lineages. Error bars represent SE of best-fit values. * p < 0.05, extra-sum-of-squares F-test.

**Figure 7 – figure supplement 1.**
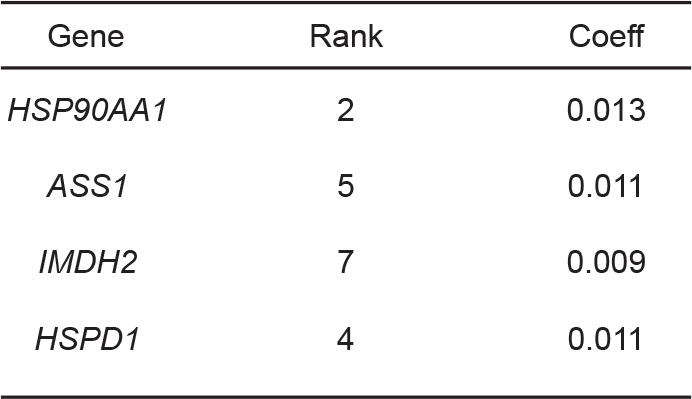
EMBR genes are expressed in human tumors with mutation burden. Based on the principal component analysis in Figure 7D, each gene was graphed based on rank and coefficient. The top ranking EMBR homologs are highlighted in this table.

## REFERENCES

Aguilar-Rodriguez, J., Sabater-Munoz, B., Montagud-Martinez, R., Berlanga, V., Alvarez-Ponce, D., Wagner, A., and Fares, M.A. (2016). The Molecular Chaperone DnaK Is a Source of Mutational Robustness. Genome Biol Evol 8, 2979–2991.

Alexandrov, L.B., Nik-Zainal, S., Wedge, D.C., Aparicio, S.A., Behjati, S., Biankin, A.V., Bignell, G.R., Bolli, N., Borg, A., Borresen-Dale, A.L., et al. (2013). Signatures of mutational processes in human cancer. Nature 500, 415–421.

Anders, S., Pyl, P.T., and Huber, W. (2015). HTSeq--a Python framework to work with high-throughput sequencing data. Bioinformatics 31, 166–169.

Andor, N., Maley, C.C., and Ji, H.P. (2017). Genomic Instability in Cancer: Teetering on the Limit of Tolerance. Cancer Res 77, 2179–2185.

Andre, B. (1990). The UGA3 gene regulating the GABA catabolic pathway in Saccharomyces cerevisiae codes for a putative zinc-finger protein acting on RNA amount. Mol Gen Genet 220, 269–276.

Arndt, K.T., Styles, C., and Fink, G.R. (1987). Multiple global regulators control HIS4 transcription in yeast. Science 237, 874–880.

Barretina, J., Caponigro, G., Stransky, N., Venkatesan, K., Margolin, A.A., Kim, S., Wilson, C.J., Lehar, J., Kryukov, G.V., Sonkin, D., et al. (2012). The Cancer Cell Line Encyclopedia enables predictive modelling of anticancer drug sensitivity. Nature 483, 603–607.

Barrick, J.E., and Lenski, R.E. (2013). Genome dynamics during experimental evolution. Nat Rev Genet 14, 827–839.

Bartholomew, C.R., Suzuki, T., Du, Z., Backues, S.K., Jin, M., Lynch-Day, M.A., Umekawa, M., Kamath, A., Zhao, M., Xie, Z., et al. (2012). Ume6 transcription factor is part of a signaling cascade that regulates autophagy. Proc Natl Acad Sci U S A 109, 11206–11210.

Bateman, L.A., Ku, W.M., Heslin, M.J., Contreras, C.M., Skibola, C.F., and Nomura, D.K. (2017). Argininosuccinate Synthase 1 is a Metabolic Regulator of Colorectal Cancer Pathogenicity. ACS Chem Biol 12, 905–911.

Bergstrom, A., Simpson, J.T., Salinas, F., Barre, B., Parts, L., Zia, A., Nguyen Ba, A.N., Moses, A.M., Louis, E.J., Mustonen, V., et al. (2014). A high-definition view of functional genetic variation from natural yeast genomes. Mol Biol Evol 31, 872–888.

Bielas, J.H., Loeb, K.R., Rubin, B.P., True, L.D., and Loeb, L.A. (2006). Human cancers express a mutator phenotype. Proc Natl Acad Sci USA 103, 18238–18242.

Bloom, J.D., Wilke, C.O., Arnold, F.H., and Adami, C. (2004). Stability and the evolvability of function in a model protein. Biophys J 86, 2758–2764.

Boratyn, G.M., Schaffer, A.A., Agarwala, R., Altschul, S.F., Lipman, D.J., and Madden, T.L. (2012). Domain enhanced lookup time accelerated BLAST. Biol Direct 7, 12.

Brauer, M.J., Huttenhower, C., Airoldi, E.M., Rosenstein, R., Matese, J.C., Gresham, D., Boer, V.M., Troyanskaya, O.G., and Botstein, D. (2008). Coordination of growth rate, cell cycle, stress response, and metabolic activity in yeast. Mol Biol Cell 19, 352–367.

Brugge, J.S., Jarosik, G., Andersen, J., Queral-Lustig, A., Fedor-Chaiken, M., and Broach, J.R. (1987). Expression of Rous Sarcoma Virus Transforming Protein pp6oV-src in Saccharomyces cerevisiae Cells. Molecular and Cellular Biology 7, 2180–2187.

Bui, D.T., Friedrich, A., Al-Sweel, N., Liti, G., Schacherer, J., Aquadro, C.F., and Alani, E. (2017). Mismatch Repair Incompatibilities in Diverse Yeast Populations. Genetics 205, 1459–1471.

Burga, A., Casanueva, M.O., and Lehner, B. (2011). Predicting mutation outcome from early stochastic variation in genetic interaction partners. Nature 480, 250–253.

Caba, E., Dickinson, D.A., Warnes, G.R., and Aubrecht, J. (2005). Differentiating mechanisms of toxicity using global gene expression analysis in Saccharomyces cerevisiae. Mutat Res 575, 34–46.

Cairns, J., and Foster, P.L. (1991). Adaptive reversion of a frameshift mutation in Escherichia coli. Genetics 128, 695–701.

Cameroni, E., Hulo, N., Roosen, J., Winderickx, J., and De Virgilio, C. (2004). The novel yeast PAS kinase Rim 15 orchestrates G0-associated antioxidant defense mechanisms. Cell Cycle 3, 462–468.

Campbell, B.B., Light, N., Fabrizio, D., Zatzman, M., Fuligni, F., de Borja, R., Davidson, S., Edwards, M., Elvin, J.A., Hodel, K.P., et al. (2017). Comprehensive Analysis of Hypermutation in Human Cancer. Cell 171, 1042–1056 e1010.

Cancer Genome Atlas, N. (2012). Comprehensive molecular characterization of human colon and rectal cancer. Nature 487, 330–337.

Chandrashekar, D.S., Bashel, B., Balasubramanya, S.A.H., Creighton, C.J., Ponce-Rodriguez, I., Chakravarthi, B., and Varambally, S. (2017). UALCAN: A Portal for Facilitating Tumor Subgroup Gene Expression and Survival Analyses. Neoplasia 19, 649–658.

Chen, L., and Pankiewicz, K.W. (2007). Recent development of IMP dehydrogenase inhibitors for the treatment of cancer. Curr Opin Drug Discov Devel 10, 403–412.

Cherry, J.M., Hong, E.L., Amundsen, C., Balakrishnan, R., Binkley, G., Chan, E.T., Christie, K.R., Costanzo, M.C., Dwight, S.S., Engel, S.R., et al. (2012). Saccharomyces Genome Database: the genomics resource of budding yeast. Nucleic Acids Res 40, D700–705.

Cingolani, P., Platts, A., Wang le, L., Coon, M., Nguyen, T., Wang, L., Land, S.J., Lu, X., and Ruden, D.M. (2012). A program for annotating and predicting the effects of single nucleotide polymorphisms, SnpEff: SNPs in the genome of Drosophila melanogaster strain w1118; iso-2; iso-3. Fly (Austin) 6, 80–92.

Ciocca, D.R., and Calderwood, S.K. (2005). Heat shock proteins in cancer: diagnostic, prognostic, predictive, and treatment implications. Cell Stress Chaperones 10, 86–103.

Costanzo, M., Baryshnikova, A., Bellay, J., Kim, Y., Spear, E.D., Sevier, C.S., Ding, H., Koh, J.L., Toufighi, K., Mostafavi, S., et al. (2010). The genetic landscape of a cell. Science 327, 425–431.

Costanzo, M., VanderSluis, B., Koch, E.N., Baryshnikova, A., Pons, C., Tan, G., Wang, W., Usaj, M., Hanchard, J., Lee, S.D., et al. (2016). A global genetic interaction network maps a wiring diagram of cellular function. Science 353.

Cowen, L.E., and Lindquist, S. (2005). Hsp90 potentiates the rapid evolution of new traits: drug resistance in diverse fungi. Science 309, 2185–2189.

Daignan-Fornier, B., and Fink, G.R. (1992). Coregulation of purine and histidine biosynthesis by the transcriptional activators BAS1 and BAS2. Proc Natl Acad Sci U S A 89, 6746–6750.

Dastidar, R.G., Hooda, J., Shah, A., Cao, T.M., Henke, R.M., and Zhang, L. (2012). The nuclear localization of SWI/SNF proteins is subjected to oxygen regulation. Cell & Bioscience 2, 30.

Davoli, T., Uno, H., Wooten, E.C., and Elledge, S.J. (2017). Tumor aneuploidy correlates with markers of immune evasion and with reduced response to immunotherapy. Science 355.

de Hoon, M.J., Imoto, S., Nolan, J., and Miyano, S. (2004). Open source clustering software. Bioinformatics 20, 1453–1454.

de Visser, J.A., Cooper, T.F., and Elena, S.F. (2011). The causes of epistasis. Proc Biol Sci 278, 3617–3624.

Del Monte, U. (2009). Does the cell number 10(9) still really fit one gram of tumor tissue? Cell Cycle 8, 505–506.

Denis, V., Boucherie, H., Monribot, C., and Daignan-Fornier, B. (1998). Role of the myb-like protein bas1p in Saccharomyces cerevisiae: a proteome analysis. Mol Microbiol 30, 557–566.

Denis, V., and Daignan-Fornier, B. (1998). Synthesis of glutamine, glycine and 10-formyl tetrahydrofolate is coregulated with purine biosynthesis in Saccharomyces cerevisiae. Mol Gen Genet 259, 246–255.

Denver, D.R., Dolan, P.C., Wilhelm, L.J., Sung, W., Lucas-Lledo, J.I., Howe, D.K., Lewis, S.C., Okamoto, K., Thomas, W.K., Lynch, M., et al. (2009). A genome-wide view of Caenorhabditis elegans base-substitution mutation processes. Proc Natl Acad Sci U S A 106, 16310–16314.

Drummond, D.A., Bloom, J.D., Adami, C., Wilke, C.O., and Arnold, F.H. (2005). Why highly expressed proteins evolve slowly. Proc Natl Acad Sci U S A 102, 14338–14343.

Drummond, D.A., and Wilke, C.O. (2008). Mistranslation-induced protein misfolding as a dominant constraint on coding-sequence evolution. Cell 134, 341–352.

Eigen, M. (2002). Error catastrophe and antiviral strategy. Proc Natl Acad Sci U S A 99, 13374–13376.

Elena, S.F., and Lenski, R.E. (1997). Test of synergistic interactions among deleterious mutations in bacteria. Nature 390, 395–398.

Engel, S.R., Dietrich, F.S., Fisk, D.G., Binkley, G., Balakrishnan, R., Costanzo, M.C., Dwight, S.S., Hitz, B.C., Karra, K., Nash, R.S., et al. (2014). The reference genome sequence of Saccharomyces cerevisiae: then and now. G3 (Bethesda) 4, 389–398.

Eyre-Walker, A., and Keightley, P.D. (2007). The distribution of fitness effects of new mutations. Nat Rev Genet 8, 610–618.

Falsone, S.F., Leptihn, S., Osterauer, A., Haslbeck, M., and Buchner, J. (2004). Oncogenic mutations reduce the stability of SRC kinase. J Mol Biol 344, 281–291.

Fares, M.A., Barrio, E., Sabater-Munoz, B., and Moya, A. (2002a). The evolution of the heat-shock protein GroEL from Buchnera, the primary endosymbiont of aphids, is governed by positive selection. Mol Biol Evol 19, 1162–1170.

Fares, M.A., Ruiz-Gonzalez, M.X., Moya, A., Elena, S.F., and Barrio, E. (2002b). Endosymbiotic bacteria: groEL buffers against deleterious mutations. Nature 417, 398.

Fendt, S.M., Oliveira, A.P., Christen, S., Picotti, P., Dechant, R.C., and Sauer, U. (2010). Unraveling condition-dependent networks of transcription factors that control metabolic pathway activity in yeast. Mol Syst Biol 6, 432.

Fersht, A.R. (1998). Structure and Mechanism in Protein Science: A Guide to Enzyme Catalysis and Protein Folding, 1st edn (W.H. Freeman).

Firnberg, E., Labonte, J.W., Gray, J.J., and Ostermeier, M. (2014). A comprehensive, high-resolution map of a gene's fitness landscape. Mol Biol Evol 31, 1581–1592.

Fleming, M.A., Chambers, S.P., Connelly, P.R., Nimmesgern, E., Fox, T., Bruzzese, F.J., Hoe, S.T., Fulghum, J.R., Livingston, D.J., Stuver, C.M., et al. (1996). Inhibition of IMPDH by mycophenolic acid: dissection of forward and reverse pathways using capillary electrophoresis. Biochemistry 35, 6990–6997.

Friedberg, E.C., Elledge, S.J., Lehmann, A.R., LIndahl, T., and Muzi-Falconi, M. (2014). DNA Repair, Mutagenesis, and Other Responses to DNA Damage, 1st edn (Cold Spring Harbor, New York: Cold Spring Harbor Laboratory press).

Fry, R.C., Sambandan, T.G., and Rha, C. (2003). DNA damage and stress transcripts in Saccharomyces cerevisiae mutant sgs1. Mech Ageing Dev 124, 839–846.

Galhardo, R.S., Hastings, P.J., and Rosenberg, S.M. (2007a). Mutation as a stress response and the regulation of evolvability. Crit Rev Biochem Mol Biol 42, 399–435.

Galhardo, R.S., Hastings, P.J., and Rosenberg, S.M. (2007b). Mutation as a stress response and the regulation of evolvability. Critical reviews in biochemistry and molecular biology 42, 399–435.

Gasch, A.P., Huang, M., Metzner, S., Botstein, D., Elledge, S.J., and Brown, P.O. (2001). Genomic expression responses to DNA-damaging agents and the regulatory role of the yeast ATR homolog Mec1p. Mol Biol Cell 12, 2987–3003.

Gasch, A.P., Spellman, P.T., Kao, C.M., Carmel-Harel, O., Eisen, M.B., Storz, G., Botstein, D., and Brown, P.O. (2000). Genomic expression programs in the response of yeast cells to environmental changes. Mol Biol Cell 11, 4241–4257.

Geiler-Samerotte, K.A., Zhu, Y.O., Goulet, B.E., Hall, D.W., and Siegal, M.L. (2016). Selection Transforms the Landscape of Genetic Variation Interacting with Hsp90. PLoS Biol 14, e2000465.

Giaever, G., Chu, A.M., Ni, L., Connelly, C., Riles, L., Veronneau, S., Dow, S., Lucau-Danila, A., Anderson, K., Andre, B., et al. (2002). Functional profiling of the Saccharomyces cerevisiae genome. Nature 418, 387–391.

Guo, L., Bloom, J.S., and Kruglyak, L. (2018). The Genetic Basis of Mutation Rate Variation in Yeast. Genetics.

Hahn, S., and Young, E.T. (2011). Transcriptional regulation in Saccharomyces cerevisiae: transcription factor regulation and function, mechanisms of initiation, and roles of activators and coactivators. Genetics 189, 705–736.

Hall, L.M., and Henderson-Begg, S.K. (2006). Hypermutable bacteria isolated from humans--a critical analysis. Microbiology 152, 2505–2514.

Hartwell, L.H. (2004). Yeast and cancer. Bioscience reports 24, 523–544.

Herr, A.J., Ogawa, M., Lawrence, N.A., Williams, L.N., Eggington, J.M., Singh, M., Smith, R.A., and Preston, B.D. (2011). Mutator suppression and escape from replication error-induced extinction in yeast. PLoS Genet 7, e1002282.

Hibbs, M.A., Hess, D.C., Myers, C.L., Huttenhower, C., Li, K., and Troyanskaya, O.G. (2007). Exploring the functional landscape of gene expression: directed search of large microarray compendia. Bioinformatics 23, 2692–2699.

Hope, E.A., Amorosi, C.J., Miller, A.W., Dang, K., Heil, C.S., and Dunham, M.J. (2017). Experimental Evolution Reveals Favored Adaptive Routes to Cell Aggregation in Yeast. Genetics 206, 1153–1167.

Hu, Z., Killion, P.J., and Iyer, V.R. (2007). Genetic reconstruction of a functional transcriptional regulatory network. Nat Genet 39, 683–687.

Huang, W., Lyman, R.F., Lyman, R.A., Carbone, M.A., Harbison, S.T., Magwire, M.M., and Mackay, T.F. (2016). Spontaneous mutations and the origin and maintenance of quantitative genetic variation. Elife 5.

Iorio, F., Knijnenburg, T.A., Vis, D.J., Bignell, G.R., Menden, M.P., Schubert, M., Aben, N., Goncalves, E., Barthorpe, S., Lightfoot, H., et al. (2016). A Landscape of Pharmacogenomic Interactions in Cancer. Cell 166, 740–754.

Jaeger, A.M., and Whitesell, L. (2019). HSP90: Enabler of Cancer Adaptation. Annual Review of Cancer Biology.

Jansen, A.M., van Wezel, T., van den Akker, B.E., Ventayol Garcia, M., Ruano, D., Tops, C.M., Wagner, A., Letteboer, T.G., Gomez-Garcia, E.B., Devilee, P., et al. (2016). Combined mismatch repair and POLE/POLD1 defects explain unresolved suspected Lynch syndrome cancers. Eur J Hum Genet 24, 1089–1092.

Jarosz, D. (2016). Hsp90: A Global Regulator of the Genotype-to-Phenotype Map in Cancers. Adv Cancer Res 129, 225–247.

Jarosz, D.F., and Lindquist, S. (2010). Hsp90 and environmental stress transform the adaptive value of natural genetic variation. Science 330, 1820–1824.

Jenkins, A.J., March, J.B., Oliver, I.R., and Masters, M. (1986). A DNA fragment containing the groE genes can suppress mutations in the Escherichia coli dnaA gene. Mol Gen Genet 202, 446–454.

Jolliffe, I.T., and Cadima, J. (2016). Principal component analysis: a review and recent developments. Philos Trans A Math Phys Eng Sci 374, 20150202.

Karras, G.I., Yi, S., Sahni, N., Fischer, M., Xie, J., Vidal, M., D’Andrea, A.D., Whitesell, L., and Lindquist, S. (2017). HSP90 Shapes the Consequences of Human Genetic Variation. Cell 168, 856–866 e812.

Khan, A., Fornes, O., Stigliani, A., Gheorghe, M., Castro-Mondragon, J.A., van der Lee, R., Bessy, A., Cheneby, J., Kulkarni, S.R., Tan, G., et al. (2018). JASPAR 2018: update of the open-access database of transcription factor binding profiles and its web framework. Nucleic Acids Res 46, D260–D266.

Khurana, V., and Lindquist, S. (2010). Modelling neurodegeneration in Saccharomyces cerevisiae: why cook with baker’s yeast? Nat Rev Neurosci 11, 436–449.

Kumar, P., Henikoff, S., and Ng, P.C. (2009). Predicting the effects of coding non-synonymous variants on protein function using the SIFT algorithm. Nat Protoc 4, 1073–1081.

Kwon, A.T., Arenillas, D.J., Worsley Hunt, R., and Wasserman, W.W. (2012). oPOSSUM-3: advanced analysis of regulatory motif over-representation across genes or ChIP-Seq datasets. G3 (Bethesda) 2, 987–1002.

Langmead, B., and Salzberg, S.L. (2012). Fast gapped-read alignment with Bowtie 2. Nat Methods 9, 357–359.

Lawrence, M.S., Stojanov, P., Polak, P., Kryukov, G.V., Cibulskis, K., Sivachenko, A., Carter, S.L., Stewart, C., Mermel, C.H., Roberts, S.A., et al. (2013). Mutational heterogeneity in cancer and the search for new cancer-associated genes. Nature 499, 214–218.

Lenski, R.E., Ofria, C., Collier, T.C., and Adami, C. (1999). Genome complexity, robustness and genetic interactions in digital organisms. Nature 400, 661–664.

Levy, J.M.M., Towers, C.G., and Thorburn, A. (2017). Targeting autophagy in cancer. Nat Rev Cancer 17, 528–542.

Li, L., Murphy, K.M., Kanevets, U., and Reha-Krantz, L.J. (2005). Sensitivity to phosphonoacetic acid: a new phenotype to probe DNA polymerase delta in Saccharomyces cerevisiae. Genetics 170, 569–580.

Liti, G., Carter, D.M., Moses, A.M., Warringer, J., Parts, L., James, S.A., Davey, R.P., Roberts, I.N., Burt, A., Koufopanou, V., et al. (2009). Population genomics of domestic and wild yeasts. Nature 458, 337–341.

Loeb, L.A. (2016). Human Cancers Express a Mutator Phenotype: Hypothesis, Origin, and Consequences. Cancer Res 76, 2057–2059.

Love, M.I., Huber, W., and Anders, S. (2014). Moderated estimation of fold change and dispersion for RNA-seq data with DESeq2. Genome Biol 15, 550.

Lujan, S.A., Clausen, A.R., Clark, A.B., MacAlpine, H.K., MacAlpine, D.M., Malc, E.P., Mieczkowski, P.A., Burkholder, A.B., Fargo, D.C., Gordenin, D.A., et al. (2014). Heterogeneous polymerase fidelity and mismatch repair bias genome variation and composition. Genome Research 24, 1751–1764.

Lynch, H.T., Snyder, C.L., Shaw, T.G., Heinen, C.D., and Hitchins, M.P. (2015). Milestones of Lynch syndrome: 1895-2015. Nat Rev Cancer 15, 181–194.

Lynch, M. (2010). Rate, molecular spectrum, and consequences of human mutation. Proc Natl Acad Sci U S A 107, 961–968.

Maisnier-Patin, S., Roth, J.R., Fredriksson, A., Nystrom, T., Berg, O.G., and Andersson, D.I. (2005). Genomic buffering mitigates the effects of deleterious mutations in bacteria. Nat Genet 37, 1376–1379.

Malek, K., Boosalis, M.S., Waraska, K., Mitchell, B.S., and Wright, D.G. (2004). Effects of the IMP-dehydrogenase inhibitor, Tiazofurin, in bcr-abl positive acute myelogenous leukemia. Part I. In vivo studies. Leuk Res 28, 1125–1136.

Mason, G.A., Carlson, K.D., Press, M.O., Bubb, K.L., and Queitsch, C. (2018). HSP90 buffers newly induced mutations in massively mutated plant lines. bioRxiv.

McFarland, C.D., Yaglom, J.A., Wojtkowiak, J.W., Scott, J.G., Morse, D.L., Sherman, M.Y., and Mirny, L.A. (2017). The Damaging Effect of Passenger Mutations on Cancer Progression. Cancer Res 77, 4763–4772.

McKenna, A., Hanna, M., Banks, E., Sivachenko, A., Cibulskis, K., Kernytsky, A., Garimella, K., Altshuler, D., Gabriel, S., Daly, M., et al. (2010). The Genome Analysis Toolkit: a MapReduce framework for analyzing next-generation DNA sequencing data. Genome Research 20, 1297–1303.

McKusick, V.A. (2007). Mendelian Inheritance in Man and its online version, OMIM. Am J Hum Genet 80, 588–604.

Milo, R., Jorgensen, P., Moran, U., Weber, G., and Springer, M. (2010). BioNumbers--the database of key numbers in molecular and cell biology. Nucleic Acids Res 38, D750–753.

Moran, N.A. (1996). Accelerated evolution and Muller's rachet in endosymbiotic bacteria. Proc Natl Acad Sci U S A 93, 2873–2878.

Murguia, J.R., and Serrano, R. (2012). New functions of protein kinase Gcn2 in yeast and mammals. IUBMB Life 64, 971–974.

Natarajan, K., Meyer, M.R., Jackson, B.M., Slade, D., Roberts, C., Hinnebusch, A.G., and Marton, M.J. (2001). Transcriptional profiling shows that Gcn4p is a master regulator of gene expression during amino acid starvation in yeast. Mol Cell Biol 21, 4347–4368.

Nick McElhinny, S.A., Gordenin, D.A., Stith, C.M., Burgers, P.M., and Kunkel, T.A. (2008). Division of labor at the eukaryotic replication fork. Mol Cell 30, 137–144.

Nick McElhinny, S.A., Stith, C.M., Burgers, P.M., and Kunkel, T.A. (2007). Inefficient proofreading and biased error rates during inaccurate DNA synthesis by a mutant derivative of Saccharomyces cerevisiae DNA polymerase delta. J Biol Chem 282, 2324–2332.

Oliver, A., Canton, R., Campo, P., Baquero, F., and Blazquez, J. (2000). High frequency of hypermutable Pseudomonas aeruginosa in cystic fibrosis lung infection. Science 288, 1251–1254.

Ossowski, S., Schneeberger, K., Lucas-Lledo, J.I., Warthmann, N., Clark, R.M., Shaw, R.G., Weigel, D., and Lynch, M. (2010). The rate and molecular spectrum of spontaneous mutations in Arabidopsis thaliana. Science 327, 92–94.

Pakula, A.A., and Sauer, R.T. (1989). Genetic analysis of protein stability and function. Annu Rev Genet 23, 289–310.

Queitsch, C., Sangster, T.A., and Lindquist, S. (2002). Hsp90 as a capacitor of phenotypic variation. Nature 417, 618.

Raghavan, V., Bui, D.T., Al-Sweel, N., Friedrich, A., Schacherer, J., Aquadro, C.F., and Alani, E. (2018). Incompatibilities in Mismatch Repair Genes MLH1-PMS1 Contribute to a Wide Range of Mutation Rates in Human Isolates of Baker’s Yeast. Genetics 210, 1253–1266.

Ramsay, R.G., and Gonda, T.J. (2008). MYB function in normal and cancer cells. Nat Rev Cancer 8, 523–534.

Roberts, S.A., and Gordenin, D.A. (2014). Hypermutation in human cancer genomes: footprints and mechanisms. Nat Rev Cancer 14, 786–800.

Rohner, N., Jarosz, D.F., Kowalko, J.E., Yoshizawa, M., Jeffery, W.R., Borowsky, R.L., Lindquist, S., and Tabin, C.J. (2013). Cryptic variation in morphological evolution: HSP90 as a capacitor for loss of eyes in cavefish. Science 342, 1372–1375.

Rutherford, S.L., and Lindquist, S. (1998). Hsp90 as a capacitor for morphological evolution. Nature 396, 336–342.

Sabater-Munoz, B., Prats-Escriche, M., Montagud-Martinez, R., Lopez-Cerdan, A., Toft, C., Aguilar-Rodriguez, J., Wagner, A., and Fares, M.A. (2015). Fitness Trade-Offs Determine the Role of the Molecular Chaperonin GroEL in Buffering Mutations. Mol Biol Evol 32, 2681–2693.

Sangster, T.A., Bahrami, A., Wilczek, A., Watanabe, E., Schellenberg, K., McLellan, C., Kelley, A., Kong, S.W., Queitsch, C., and Lindquist, S. (2007). Phenotypic diversity and altered environmental plasticity in Arabidopsis thaliana with reduced Hsp90 levels. PLoS One 2, e648.

Sangster, T.A., Lindquist, S., and Queitsch, C. (2004). Under cover: causes, effects and implications of Hsp90-mediated genetic capacitance. Bioessays 26, 348–362.

Sangster, T.A., Salathia, N., Lee, H.N., Watanabe, E., Schellenberg, K., Morneau, K., Wang, H., Undurraga, S., Queitsch, C., and Lindquist, S. (2008). HSP90-buffered genetic variation is common in Arabidopsis thaliana. Proc Natl Acad Sci U S A 105, 2969–2974.

Santagata, S., Hu, R., Lin, N.U., Mendillo, M.L., Collins, L.C., Hankinson, S.E., Schnitt, S.J., Whitesell, L., Tamimi, R.M., Lindquist, S., et al. (2011). High levels of nuclear heat-shock factor 1 (HSF1) are associated with poor prognosis in breast cancer. Proc Natl Acad Sci U S A 108, 18378–18383.

Serero, A., Jubin, C., Loeillet, S., Legoix-Ne, P., and Nicolas, A.G. (2014). Mutational landscape of yeast mutator strains. Proc Natl Acad Sci U S A 111, 1897–1902.

Sidow, A., and Spies, N. (2015). Concepts in solid tumor evolution. Trends Genet 31, 208–214.

Sijmons, R.H., and Hofstra, R.M. (2016). Review: Clinical aspects of hereditary DNA Mismatch repair gene mutations. DNA repair 38, 155–162.

Sniegowski, P.D., Gerrish, P.J., and Lenski, R.E. (1997). Evolution of high mutation rates in experimental populations of E. coli. Nature 387, 703–705.

Sniegowski, P.D., and Murphy, H.A. (2006). Evolvability. Curr Biol 16, R831–834.

Sole, R.V., and Deisboeck, T.S. (2004). An error catastrophe in cancer? J Theor Biol 228, 47–54.

Sun, R., Hu, Z., Sottoriva, A., Graham, T.A., Harpak, A., Ma, Z., Fischer, J.M., Shibata, D., and Curtis, C. (2017). Between-region genetic divergence reflects the mode and tempo of tumor evolution. Nat Genet 49, 1015–1024.

Supek, F., and Lehner, B. (2015). Differential DNA mismatch repair underlies mutation rate variation across the human genome. Nature 521, 81–84.

Swings, T., Van den Bergh, B., Wuyts, S., Oeyen, E., Voordeckers, K., Verstrepen, K.J., Fauvart, M., Verstraeten, N., and Michiels, J. (2017). Adaptive tuning of mutation rates allows fast response to lethal stress in Escherichia coli. Elife 6.

Szklarczyk, D., Franceschini, A., Wyder, S., Forslund, K., Heller, D., Huerta-Cepas, J., Simonovic, M., Roth, A., Santos, A., Tsafou, K.P., et al. (2015). STRING v10: protein-protein interaction networks, integrated over the tree of life. Nucleic Acids Res 43, D447–452.

Taipale, M., Jarosz, D.F., and Lindquist, S. (2010). HSP90 at the hub of protein homeostasis: emerging mechanistic insights. Nat Rev Mol Cell Biol 11, 515–528.

Taipale, M., Tucker, G., Peng, J., Krykbaeva, I., Lin, Z.Y., Larsen, B., Choi, H., Berger, B., Gingras, A.C., and Lindquist, S. (2014). A quantitative chaperone interaction network reveals the architecture of cellular protein homeostasis pathways. Cell 158, 434–448.

Tice-Baldwin, K., Fink, G.R., and Arndt, K.T. (1989). BAS1 has a Myb motif and activates HIS4 transcription only in combination with BAS2. Science 246, 931–935.

Tokuriki, N., and Tawfik, D.S. (2009). Stability effects of mutations and protein evolvability. Curr Opin Struct Biol 19, 596–604.

Travers, K.J., Patil, C.K., Wodicka, L., Lockhart, D.J., Weissman, J.S., and Walter, P. (2000). Functional and genomic analyses reveal an essential coordination between the unfolded protein response and ER-associated degradation. Cell 101, 249–258.

Travesa, A., Kuo, D., de Bruin, R.A., Kalashnikova, T.I., Guaderrama, M., Thai, K., Aslanian, A., Smolka, M.B., Yates, J.R., 3rd, Ideker, T., et al. (2012). DNA replication stress differentially regulates G1/S genes via Rad53-dependent inactivation of Nrm1. EMBO J 31, 1811–1822.

Tu, S., Bulloch, E.M., Yang, L., Ren, C., Huang, W.C., Hsu, P.H., Chen, C.H., Liao, C.L., Yu, H.M., Lo, W.S., et al. (2007). Identification of histone demethylases in Saccharomyces cerevisiae. J Biol Chem 282, 14262–14271.

Uchimura, A., Higuchi, M., Minakuchi, Y., Ohno, M., Toyoda, A., Fujiyama, A., Miura, I., Wakana, S., Nishino, J., and Yagi, T. (2015). Germline mutation rates and the long-term phenotypic effects of mutation accumulation in wild-type laboratory mice and mutator mice. Genome Research 25, 1125–1134.

Van Dyk, T.K., Gatenby, A.A., and LaRossa, R.A. (1989). Demonstration by genetic suppression of interaction of GroE products with many proteins. Nature 342, 451–453.

Venkataram, S., Dunn, B., Li, Y., Agarwala, A., Chang, J., Ebel, E.R., Geiler-Samerotte, K., Herissant, L., Blundell, J.R., Levy, S.F., et al. (2016). Development of a Comprehensive Genotype-to-Fitness Map of Adaptation-Driving Mutations in Yeast. Cell 166, 1585–1596 e1522.

Vilar, E., and Gruber, S.B. (2010). Microsatellite instability in colorectal cancer-the stable evidence. Nat Rev Clin Oncol 7, 153–162.

Washburn, B.K., and Esposito, R.E. (2001). Identification of the Sin3-binding site in Ume6 defines a two-step process for conversion of Ume6 from a transcriptional repressor to an activator in yeast. Mol Cell Biol 21, 2057–2069.

Whitesell, L., and Lindquist, S.L. (2005). HSP90 and the chaperoning of cancer. Nat Rev Cancer 5, 761–772.

Whitesell, L., Santagata, S., Mendillo, M.L., Lin, N.U., Proia, D.A., and Lindquist, S. (2014). HSP90 empowers evolution of resistance to hormonal therapy in human breast cancer models. Proc Natl Acad Sci U S A 111, 18297–18302.

Wielgoss, S., Barrick, J.E., Tenaillon, O., Wiser, M.J., Dittmar, W.J., Cruveiller, S., Chane-Woon-Ming, B., Medigue, C., Lenski, R.E., and Schneider, D. (2013). Mutation rate dynamics in a bacterial population reflect tension between adaptation and genetic load. Proc Natl Acad Sci U S A 110, 222–227.

Wortel, I.M.N., van der Meer, L.T., Kilberg, M.S., and van Leeuwen, F.N. (2017). Surviving Stress: Modulation of ATF4-Mediated Stress Responses in Normal and Malignant Cells. Trends Endocrinol Metab 28, 794–806.

Wu, X., and Bartel, D.P. (2017). kpLogo: positional k-mer analysis reveals hidden specificity in biological sequences. Nucleic Acids Res 45, W534–W538.

Xu, Y., and Lindquist, S. (1993). Heat-shock protein hsp90 governs the activity of pp60v-src kinase. Proc Natl Acad Sci U S A 90, 7074–7078.

Xu, Y., Zheng, Z., Gao, Y., Duan, S., Chen, C., Rong, J., Wang, K., Yun, M., Weng, H., Ye, S., et al. (2017). High expression of IMPDH2 is associated with aggressive features and poor prognosis of primary nasopharyngeal carcinoma. Sci Rep 7, 745.

Yoshimori, T., Yamamoto, A., Moriyama, Y., Futai, M., and Tashiro, Y. (1991). Bafilomycin A1, a specific inhibitor of vacuolar-type H(+)-ATPase, inhibits acidification and protein degradation in lysosomes of cultured cells. J Biol Chem 266, 17707–17712.

Yousif, F., Prokopec, S., Sun, R.X., Fan, F., Lalansingh, C.M., Drysdale, E., Park, D.H., Szyca, L., and Boutros, P.C. (2018). The Origins and Consequences of Localized Global Somatic Hypermutation. bioRxiv 287839.

Zabinsky, R.A., Mason, G.A., Queitsch, C., and Jarosz, D.F. (2018). It’s not magic - Hsp90 and its effects on genetic and epigenetic variation. Seminars in Cell and Developmental Biology.

Zambelli, F., Pesole, G., and Pavesi, G. (2009). Pscan: finding over-represented transcription factor binding site motifs in sequences from co-regulated or co-expressed genes. Nucleic Acids Res 37, W247–252.

Zeilstra-Ryalls, J., Fayet, O., and Georgopoulos, C. (1991). The universally conserved GroE (Hsp60) chaperonins. Annu Rev Microbiol 45, 301–325.

Zhang, L., MacIsaac, K.D., Zhou, T., Huang, P.Y., Xin, C., Dobson, J.R., Yu, K., Chiang, D.Y., Fan, Y., Pelletier, M., et al. (2017). Genomic Analysis of Nasopharyngeal Carcinoma Reveals TME-Based Subtypes. Mol Cancer Res 15, 1722–1732.

Zhu, Y., Qiu, P., and Ji, Y. (2014a). TCGA-assembler: open-source software for retrieving and processing TCGA data. Nat Methods 11, 599–600.

Zhu, Y.O., Siegal, M.L., Hall, D.W., and Petrov, D.A. (2014b). Precise estimates of mutation rate and spectrum in yeast. Proc Natl Acad Sci U S A 111, E2310–2318.

