## Supplementary material for "A stress response that allows highly mutated eukaryotic cells to survive and proliferate": Table S1

| Sample Genotype | Media | Lineage | Passage | Batch | # Of Mutations | Read Count | Coverage |
|---|---|---|---|---|---|---|---|
| WT | YPD | ancestor | 0 | 2 | 161 | 17937284 | 56.05 |
| WT | YPG | ancestor | 0 | 2 | 196 | 22566316 | 70.52 |
| msh6 | YPD | ancestor | 0 | 7 | 170 | 21524176 | 67.26 |
| msh6 | YPG | ancestor | 0 | 7 | 175 | 18751500 | 58.60 |
| WT | YPD | 1 | 50 | 1 | 162 | 15390964 | 48.10 |
| msh6 | YPD | 1 | 50 | 3 | 225 | 16252244 | 50.79 |
| msh6 | YPD | 2 | 50 | 3 | 248 | 14867144 | 46.46 |
| msh6 | YPD | 3 | 50 | 3 | 220 | 16585316 | 51.83 |
| msh6 | YPD | 4 | 50 | 3 | 229 | 17984036 | 56.20 |
| msh6 | YPD | 5 | 50 | 3 | 373 | 27243016 | 85.13 |
| msh6 | YPD | 6 | 50 | 3 | 1 | 10324 | 0.03 |
| msh6 | YPD | 7 | 50 | 3 | 481 | 19957896 | 62.37 |
| msh6 | YPD | 8 | 50 | 3 | 249 | 26962880 | 84.26 |
| msh6 | YPG | 1 | 50 | 3 | 295 | 21163244 | 66.14 |
| msh6 | YPG | 2 | 50 | 3 | 257 | 12902696 | 40.32 |
| msh6 | YPG | 3 | 50 | 3 | 270 | 17084368 | 53.39 |
| msh6 | YPG | 4 | 50 | 3 | 211 | 17648264 | 55.15 |
| msh6 | YPG | 5 | 50 | 3 | 265 | 17439804 | 54.50 |
| msh6 | YPG | 6 | 50 | 3 | 261 | 16459292 | 51.44 |
| msh6 | YPG | 7 | 50 | 3 | 283 | 18055476 | 56.42 |
| msh6 | YPG | 8 | 50 | 3 | 262 | 17834796 | 55.73 |
| pol3 | YPD | ancestor | 0 | 7 | 168 | 13891124 | 43.41 |
| pol3 | YPG | ancestor | 0 | 7 | 181 | 20767212 | 64.90 |
| pol3 | YPD | 1 | 50 | 4 | 203 | 22543708 | 70.45 |
| pol3 | YPD | 2 | 50 | 4 | 199 | 16194556 | 50.61 |
| pol3 | YPD | 3 | 50 | 4 | 188 | 16684364 | 52.14 |
| pol3 | YPD | 4 | 50 | 4 | 225 | 17487788 | 54.65 |
| pol3 | YPD | 5 | 50 | 4 | 243 | 17904232 | 55.95 |
| pol3 | YPD | 6 | 50 | 4 | 189 | 18981108 | 59.32 |
| pol3 | YPD | 7 | 50 | 4 | 219 | 18647192 | 58.27 |
| pol3 | YPD | 8 | 50 | 4 | 213 | 18967628 | 59.27 |
| pol3 | YPG | 1 | 50 | 4 | 243 | 19096972 | 59.68 |
| pol3 | YPG | 2 | 50 | 4 | 229 | 21298724 | 66.56 |
| pol3 | YPG | 3 | 50 | 4 | 224 | 16314140 | 50.98 |
| pol3 | YPG | 4 | 50 | 4 | 204 | 16048952 | 50.15 |
| pol3 | YPG | 5 | 50 | 4 | 254 | 22360524 | 69.88 |
| pol3 | YPG | 6 | 50 | 4 | 214 | 12167536 | 38.02 |
| pol3 | YPG | 7 | 50 | 4 | 242 | 17751272 | 55.47 |
| pol3 | YPG | 8 | 50 | 4 | 217 | 17543296 | 54.82 |
| pol3 msh6 | YPD | ancestor | 0 | 6 | 324 | 16944764 | 52.95 |
| pol3 msh6 | YPG | ancestor | 0 | 6 | 366 | 24194772 | 75.61 |
| pol3 msh6 | YPD | 1 | 20 | 6 | 1572 | 16437324 | 51.37 |
| pol3 msh6 | YPD | 2 | 20 | 6 | 1287 | 16558372 | 51.74 |
| pol3 msh6 | YPD | 3 | 20 | 6 | 1475 | 19525588 | 61.02 |
| pol3 msh6 | YPD | 5 | 20 | 6 | 263 | 15379976 | 48.06 |
| pol3 msh6 | YPD | 6 | 20 | 6 | 1628 | 15761000 | 49.25 |
| pol3 msh6 | YPD | 7 | 20 | 6 | 1538 | 16297104 | 50.93 |
| pol3 msh6 | YPD | 8 | 20 | 6 | 1521 | 21497952 | 67.18 |

| | | | | | | | |
|---|---|---|---|---|---|---|---|
| pol3 msh6 | YPG | 1 | 20 | 6 | 192 | 14985848 | 46.83 |
| pol3 msh6 | YPG | 2 | 20 | 6 | 1389 | 23312184 | 72.85 |
| pol3 msh6 | YPG | 3 | 20 | 6 | 1240 | 35518504 | 111.00 |
| pol3 msh6 | YPG | 4 | 20 | 6 | 1232 | 13085548 | 40.89 |
| pol3 msh6 | YPG | 5 | 20 | 6 | 95 | 124716 | 0.39 |
| pol3 msh6 | YPG | 6 | 20 | 6 | 1144 | 12782528 | 39.95 |
| pol3 msh6 | YPG | 7 | 20 | 6 | 1322 | 17370696 | 54.28 |
| pol3 msh6 | YPG | 8 | 20 | 6 | 1764 | 15660776 | 48.94 |
| pol3 msh6 | YPD | 1 | 50 | 7 | 2214 | 21214060 | 66.29 |
| pol3 msh6 | YPD | 2 | 50 | 7 | 2906 | 13492612 | 42.16 |
| pol3 msh6 | YPD | 3 | 50 | 7 | 2205 | 18109888 | 56.59 |
| pol3 msh6 | YPD | 5 | 50 | 7 | 364 | 19254104 | 60.17 |
| pol3 msh6 | YPD | 6 | 50 | 7 | 2569 | 17827708 | 55.71 |
| pol3 msh6 | YPD | 7 | 50 | 7 | 2289 | 18015076 | 56.30 |
| pol3 msh6 | YPD | 8 | 50 | 7 | 2617 | 18986976 | 59.33 |
| pol3 msh6 | YPG | 1 | 50 | 7 | 216 | 19857200 | 62.05 |
| pol3 msh6 | YPG | 2 | 50 | 7 | 2592 | 21770556 | 68.03 |
| pol3 msh6 | YPG | 3 | 50 | 7 | 2095 | 15075252 | 47.11 |
| pol3 msh6 | YPG | 4 | 50 | 7 | 1396 | 20519512 | 64.12 |
| pol3 msh6 | YPG | 6 | 50 | 7 | 2073 | 20090360 | 62.78 |
