## Supplementary material for "A stress response that allows highly mutated eukaryotic cells to survive and proliferate": Table S2

| Gene | baseMean | log2FoldChange | pvalue | padj |
|---|---|---|---|---|
| YJL016W | 1543.218497 | 1.372318649 | 3.94E-06 | 0.023 |
| YBL091C | 4543.955358 | -0.891270827 | 3.30E-05 | 0.052 |
| YFL039C | 29152.57267 | -0.833555657 | 3.55E-05 | 0.052 |
| YLR154C | 335.1264893 | -1.069152558 | 1.99E-05 | 0.052 |
| YDL121C | 1113.601796 | -1.352192677 | 5.32E-05 | 0.062 |
| YFL025C | 800.9686963 | 0.687275268 | 0.000123261 | 0.098 |
| YFR035C | 184.0960805 | -1.005149981 | 0.000133479 | 0.098 |
| YLR183C | 714.9768804 | -1.125225034 | 0.00010447 | 0.098 |
| YDL213C | 1003.203847 | -1.347957626 | 0.000197206 | 0.129 |
| YBR118W | 10015.00281 | -1.171750367 | 0.000254533 | 0.132 |
| YBR263W | 3971.725103 | -0.850326838 | 0.000273934 | 0.132 |
| YER120W | 4329.489139 | -0.812221086 | 0.000338574 | 0.132 |
| YNL312W | 897.7476795 | -0.989104642 | 0.000239087 | 0.132 |
| YNR017W | 1847.028777 | -0.880242394 | 0.000360175 | 0.132 |
| YOR160W | 1175.925728 | 0.616834362 | 0.000346282 | 0.132 |
| YOR252W | 480.3993858 | -1.193472989 | 0.000340442 | 0.132 |
| YBR206W | 255.7378889 | -1.149100748 | 0.000426994 | 0.147 |
| YKR015C | 299.9934309 | 0.912166509 | 0.00056348 | 0.184 |
| YBR127C | 10020.63379 | -0.832916186 | 0.000728268 | 0.214 |
| YDR010C | 32.06483583 | 1.087163539 | 0.000697233 | 0.214 |
| YBL077W | 139.1714064 | -1.486944808 | 0.000924087 | 0.217 |
| YGR127W | 732.3981957 | 1.173209511 | 0.000833217 | 0.217 |
| YLR335W | 2838.169755 | -0.702445599 | 0.000903394 | 0.217 |
| YPL063W | 2649.060944 | -0.68303572 | 0.000882149 | 0.217 |
| YPR070W | 926.8338353 | 0.587046599 | 0.00084744 | 0.217 |
| YLR107W | 1035.729202 | 0.905332231 | 0.001010703 | 0.228 |
| YOR197W | 2558.031802 | -0.716242382 | 0.001053033 | 0.229 |
| YBR109C | 2942.937927 | -0.753900785 | 0.001154791 | 0.234 |
| YNL150W | 83.29807569 | -1.220861678 | 0.001137122 | 0.234 |
| YHR049C-A | 129.3206508 | -0.734922845 | 0.001250869 | 0.245 |
| YGR262C | 876.7108592 | -0.612948296 | 0.001389365 | 0.263 |
| YER006W | 1977.06385 | -1.153794749 | 0.001504525 | 0.270 |
| YPR174C | 411.2973194 | -0.83440577 | 0.001516029 | 0.270 |
| YBR155W | 833.2201083 | -1.156916052 | 0.001758253 | 0.279 |
| YDR027C | 1560.996774 | 0.512764937 | 0.00173178 | 0.279 |
| YER087C-B | 1311.571506 | -1.047781751 | 0.001711891 | 0.279 |
| YJR072C | 1860.139893 | -0.933619337 | 0.001639753 | 0.279 |
| YHR155W | 1225.572051 | 0.690863585 | 0.001818411 | 0.281 |
| YDR179C | 161.4536724 | -0.812012391 | 0.001916447 | 0.281 |
| YHR139C | 204.3743283 | 1.226205547 | 0.001879338 | 0.281 |

| | | | | |
|---|---|---|---|---|
| YHR088W | 1005.281896 | -1.173678223 | 0.002053639 | 0.287 |
| YPL218W | 2797.046919 | -0.84872508 | 0.002039429 | 0.287 |
| YBR091C | 248.9757925 | -0.928364938 | 0.002154611 | 0.287 |
| YML085C | 4155.33263 | -0.786113446 | 0.002148636 | 0.287 |
| YAL042W | 3676.592816 | -0.666534156 | 0.003534343 | 0.289 |
| YBL028C | 410.9135571 | -1.106791174 | 0.00271034 | 0.289 |
| YDL133W | 835.1418998 | 0.570411883 | 0.003471702 | 0.289 |
| YDR017C | 2063.432291 | 0.591126286 | 0.003292655 | 0.289 |
| YDR137W | 962.3953612 | 0.571150421 | 0.002807266 | 0.289 |
| YDR321W | 1332.488079 | -0.953753499 | 0.003105211 | 0.289 |
| YDR484W | 738.7697722 | 0.571367398 | 0.003061804 | 0.289 |
| YER072W | 2000.095671 | -1.02893023 | 0.003115239 | 0.289 |
| YER176W | 1918.703807 | -0.870543679 | 0.002850241 | 0.289 |
| YFR057W | 37.29733796 | 0.855818141 | 0.003132772 | 0.289 |
| YGL130W | 854.7206722 | -0.464375777 | 0.003587751 | 0.289 |
| YGR003W | 648.3625673 | 0.705807999 | 0.00263124 | 0.289 |
| YGR007W | 1236.749371 | -0.71093511 | 0.003360203 | 0.289 |
| YHR063W-A | 174.3401986 | -1.266777571 | 0.002993883 | 0.289 |
| YIL073C | 530.2219401 | 0.698791593 | 0.002623222 | 0.289 |
| YKL191W | 1251.863612 | -0.993857247 | 0.002789453 | 0.289 |
| YLR027C | 7716.105415 | -0.888443729 | 0.002943965 | 0.289 |
| YLR221C | 955.7495307 | -1.023086821 | 0.002512333 | 0.289 |
| YLR273C | 457.5126435 | 0.871340062 | 0.003280487 | 0.289 |
| YLR446W | 527.2109038 | 0.911547996 | 0.002517324 | 0.289 |
| YMR232W | 290.2765579 | 1.383442667 | 0.002898996 | 0.289 |
| YMR292W | 429.0322087 | -0.947089741 | 0.00233533 | 0.289 |
| YNL016W | 4774.57143 | -0.809902343 | 0.003355386 | 0.289 |
| YOL142W | 655.5671288 | -0.873708572 | 0.003592743 | 0.289 |
| YOL146W | 364.144711 | -0.928318607 | 0.003102601 | 0.289 |
| YOR004W | 494.9946687 | -0.774984286 | 0.003435969 | 0.289 |
| YOR021C | 1473.922183 | -1.004847417 | 0.002672789 | 0.289 |
| YOR095C | 614.382855 | -1.089930106 | 0.00344271 | 0.289 |
| YPL234C | 1583.101568 | -0.614423644 | 0.002854231 | 0.289 |
| YCL059C | 1691.420964 | -1.103878824 | 0.004028929 | 0.292 |
| YDR454C | 2624.226849 | -0.884829663 | 0.003899635 | 0.292 |
| YKR070W | 801.1916976 | -0.506827299 | 0.003741735 | 0.292 |
| YLL022C | 865.5611491 | -0.878943279 | 0.003842072 | 0.292 |
| YLR116W | 1156.147687 | 0.626227973 | 0.003819537 | 0.292 |
| YLR192C | 3457.976113 | -0.737084378 | 0.003977418 | 0.292 |
| YLR354C | 11943.28845 | -0.757240535 | 0.003760622 | 0.292 |
| YMR128W | 2322.124433 | -1.126059312 | 0.004018798 | 0.292 |
| YIL096C | 413.8812975 | -1.008635303 | 0.00414965 | 0.294 |
| YML060W | 354.1411795 | -0.909654586 | 0.004157078 | 0.294 |

| | | | | |
|---|---|---|---|---|
| YGL124C | 833.4516769 | 0.71098479 | 0.004367432 | 0.302 |
| YOR249C | 647.5824875 | 0.776534502 | 0.004355453 | 0.302 |
| YHR049W | 2093.457795 | -0.641211031 | 0.004448136 | 0.302 |
| YOR286W | 1162.614528 | -0.715221044 | 0.004471752 | 0.302 |
| YJL173C | 671.1948643 | -0.911215947 | 0.004563801 | 0.302 |
| YPR143W | 537.9356283 | -0.946253723 | 0.004581394 | 0.302 |
| YCR028C-A | 1716.279601 | -0.847737191 | 0.005039283 | 0.308 |
| YDR263C | 317.4342443 | 0.962755466 | 0.004823427 | 0.308 |
| YEL026W | 1802.083858 | -1.025322546 | 0.004800451 | 0.308 |
| YGR245C | 1499.530458 | -1.052113029 | 0.005042158 | 0.308 |
| YKR081C | 1261.655674 | -1.094284326 | 0.005043601 | 0.308 |
| YNR039C | 749.4014615 | 0.58606622 | 0.004948184 | 0.308 |
| YOR210W | 598.8398742 | -0.930578653 | 0.004978945 | 0.308 |
| YOR296W | 873.0215241 | 0.713291517 | 0.005145643 | 0.311 |
| YLR301W | 1464.12998 | -0.861081237 | 0.005229463 | 0.313 |
| YBL018C | 137.4735368 | -0.974052098 | 0.010692031 | 0.314 |
| YBL026W | 682.7010545 | -0.599463937 | 0.00894359 | 0.314 |
| YBL092W | 15431.27049 | -0.885601748 | 0.006109856 | 0.314 |
| YBR048W | 5960.519296 | -0.795351232 | 0.010617315 | 0.314 |
| YBR088C | 178.1843606 | -0.949240888 | 0.006488232 | 0.314 |
| YBR089C-A | 560.6070076 | -0.778474582 | 0.010620159 | 0.314 |
| YBR191W | 11850.59787 | -0.666098119 | 0.010018627 | 0.314 |
| YBR245C | 1580.844401 | -0.582021058 | 0.010554252 | 0.314 |
| YBR247C | 781.1952554 | -0.899610405 | 0.008620653 | 0.314 |
| YCR016W | 907.9551315 | -1.050891271 | 0.006882954 | 0.314 |
| YCR024C | 641.2724541 | 0.581892294 | 0.009583792 | 0.314 |
| YCR038C | 411.3006768 | 0.72126616 | 0.009715822 | 0.314 |
| YCR047C | 558.0855162 | -0.927353249 | 0.006312748 | 0.314 |
| YDL055C | 11798.61054 | -0.601090476 | 0.008763997 | 0.314 |
| YDL081C | 10855.42483 | -0.858089609 | 0.007818522 | 0.314 |
| YDL153C | 1719.824839 | -1.077112805 | 0.006778295 | 0.314 |
| YDL192W | 4665.318464 | -0.936637666 | 0.008020448 | 0.314 |
| YDL212W | 1994.74877 | -0.780381868 | 0.008738388 | 0.314 |
| YDL219W | 330.3585764 | -0.993150251 | 0.007182349 | 0.314 |
| YDR064W | 9582.184184 | -0.671514923 | 0.008845805 | 0.314 |
| YDR084C | 664.1695601 | -0.678966732 | 0.005786765 | 0.314 |
| YDR098C | 1711.271644 | -0.60117603 | 0.007957664 | 0.314 |
| YDR120C | 1240.636645 | -0.985366956 | 0.008604571 | 0.314 |
| YDR327W | 614.5550695 | -0.686162818 | 0.007769413 | 0.314 |
| YDR378C | 714.4468599 | -0.665886952 | 0.007067659 | 0.314 |
| YDR382W | 7559.11107 | -0.984787935 | 0.005934779 | 0.314 |
| YDR398W | 1282.093858 | -0.919314557 | 0.007725184 | 0.314 |
| YER002W | 1431.744689 | -0.904391482 | 0.008708567 | 0.314 |

| | | | | |
|---|---|---|---|---|
| YER007C-A | 951.757768 | -0.809998353 | 0.010167989 | 0.314 |
| YER009W | 3909.301197 | -0.933277972 | 0.006816098 | 0.314 |
| YER110C | 3689.619393 | -0.932702847 | 0.008515332 | 0.314 |
| YER145C | 1496.957868 | -0.841960264 | 0.010001899 | 0.314 |
| YER146W | 140.8937845 | -0.867964696 | 0.008560238 | 0.314 |
| YER165W | 11478.83612 | -0.702908814 | 0.009809496 | 0.314 |
| YFR001W | 1018.480753 | -0.907347082 | 0.007813364 | 0.314 |
| YFR031C-A | 6438.897656 | -0.748173319 | 0.010528761 | 0.314 |
| YGL029W | 383.3919755 | -1.013754443 | 0.006202118 | 0.314 |
| YGL030W | 7624.667323 | -0.70017979 | 0.005949863 | 0.314 |
| YGL070C | 462.601344 | -0.85408424 | 0.007124078 | 0.314 |
| YGL137W | 4577.146804 | -0.645557622 | 0.010140816 | 0.314 |
| YGL147C | 5508.177604 | -0.58120958 | 0.00983241 | 0.314 |
| YGL160W | 1159.891272 | 0.60691083 | 0.008818263 | 0.314 |
| YGR034W | 5016.73135 | -0.823238398 | 0.007869228 | 0.314 |
| YGR103W | 3122.539574 | -1.019329593 | 0.008927193 | 0.314 |
| YGR155W | 6507.68049 | -0.799908533 | 0.005671169 | 0.314 |
| YGR214W | 11003.56684 | -0.793552399 | 0.008633744 | 0.314 |
| YGR271C-A | 454.6880839 | -0.978479702 | 0.006408278 | 0.314 |
| YHL034W-A | 585.1049194 | -0.879893394 | 0.005719118 | 0.314 |
| YHR021W-A | 41.37134005 | 1.002782872 | 0.005758616 | 0.314 |
| YHR022C | 158.1795629 | 0.884979721 | 0.008206345 | 0.314 |
| YHR072W-A | 1183.992345 | -1.042574871 | 0.006942671 | 0.314 |
| YHR143W-A | 554.7529102 | -0.991378667 | 0.007677014 | 0.314 |
| YIL008W | 681.0099622 | -1.115046299 | 0.009146952 | 0.314 |
| YIL016W | 483.5722155 | -0.918772111 | 0.009773459 | 0.314 |
| YIL127C | 639.7542142 | -1.081624489 | 0.006587915 | 0.314 |
| YIL153W | 970.2297401 | -0.469993886 | 0.005508001 | 0.314 |
| YJL011C | 110.249814 | -1.022502624 | 0.008874875 | 0.314 |
| YJR067C | 196.0036818 | -0.692977575 | 0.009980744 | 0.314 |
| YKL006W | 2404.725348 | -0.74860571 | 0.005938175 | 0.314 |
| YKL056C | 11023.03919 | -0.993660533 | 0.005591177 | 0.314 |
| YKL143W | 1110.759937 | -0.997521389 | 0.007378714 | 0.314 |
| YKR059W | 1551.844916 | -0.823734093 | 0.006727235 | 0.314 |
| YLR017W | 1271.451959 | -0.889971468 | 0.010167539 | 0.314 |
| YLR106C | 6573.73141 | -0.759231741 | 0.007465157 | 0.314 |
| YLR172C | 1976.093187 | -0.862543262 | 0.010674078 | 0.314 |
| YLR195C | 1211.827498 | -0.693077172 | 0.010225336 | 0.314 |
| YLR297W | 487.8704755 | 0.930499818 | 0.008333784 | 0.314 |
| YLR323C | 363.5088652 | -0.823776196 | 0.009745631 | 0.314 |
| YML022W | 1077.904293 | -0.946236927 | 0.006765194 | 0.314 |
| YML054C | 1101.015099 | 1.129218398 | 0.009531222 | 0.314 |
| YMR154C | 501.7366665 | 0.910266704 | 0.005599768 | 0.314 |

| | | | | |
|---|---|---|---|---|
| YMR194W | 1484.319594 | -0.905923988 | 0.007576938 | 0.314 |
| YMR202W | 2009.003075 | -0.67107794 | 0.007480415 | 0.314 |
| YMR210W | 1188.233193 | 0.692287607 | 0.009905101 | 0.314 |
| YMR260C | 3410.062096 | -0.891662239 | 0.008593697 | 0.314 |
| YNL049C | 1288.390098 | -0.695566772 | 0.008639552 | 0.314 |
| YNL056W | 768.1896527 | -0.667195457 | 0.01049463 | 0.314 |
| YNL075W | 678.2006525 | -0.920059305 | 0.006930592 | 0.314 |
| YNL102W | 1324.90783 | -0.636773686 | 0.006691252 | 0.314 |
| YNL110C | 899.4305976 | -0.931286447 | 0.005925457 | 0.314 |
| YNL302C | 5878.096109 | -0.81815353 | 0.010616396 | 0.314 |
| YOL120C | 9488.699457 | -0.650911464 | 0.005838777 | 0.314 |
| YOL124C | 564.6964356 | -0.831962748 | 0.008993465 | 0.314 |
| YOR107W | 180.9819764 | 0.776935106 | 0.00855389 | 0.314 |
| YOR133W | 1425.340865 | -0.873679215 | 0.009183892 | 0.314 |
| YOR164C | 1461.403289 | -0.72951368 | 0.006081933 | 0.314 |
| YOR182C | 1767.616754 | -0.959834027 | 0.006831652 | 0.314 |
| YOR219C | 1764.610148 | 0.656936317 | 0.010533124 | 0.314 |
| YOR236W | 448.7599801 | -0.882684536 | 0.009196313 | 0.314 |
| YOR248W | 35.3813631 | -0.643132154 | 0.01017189 | 0.314 |
| YOR265W | 445.8131326 | -0.796819073 | 0.007921978 | 0.314 |
| YPL043W | 1057.06526 | -0.955053089 | 0.007103694 | 0.314 |
| YPL076W | 380.2952376 | 0.855494575 | 0.009453018 | 0.314 |
| YPL106C | 19467.56403 | -0.773572261 | 0.010260232 | 0.314 |
| YPL127C | 2242.895143 | -0.717204816 | 0.01067752 | 0.314 |
| YPL146C | 923.2028372 | -0.960546255 | 0.006937673 | 0.314 |
| YPL237W | 2463.364966 | -0.834215856 | 0.010660727 | 0.314 |
| YPL266W | 1132.936461 | -0.983834886 | 0.006674174 | 0.314 |
| YPR064W | 39.20156943 | 1.137553963 | 0.006416687 | 0.314 |
| YPR106W | 760.0851492 | 0.69851213 | 0.008296459 | 0.314 |
| YPR170W-B | 216.8408172 | -0.826835985 | 0.009173915 | 0.314 |
| YPR183W | 3494.823853 | -0.710992434 | 0.008987655 | 0.314 |
| YBR187W | 950.8054053 | -0.64226583 | 0.010760412 | 0.314 |
| YDR470C | 683.2878102 | 0.576065453 | 0.010918746 | 0.316 |
| YKR057W | 2516.350053 | -1.061839689 | 0.010997071 | 0.316 |
| YMR290C | 2185.404412 | -0.956509305 | 0.010974522 | 0.316 |
| YBR205W | 1374.246383 | -0.642797164 | 0.011368504 | 0.317 |
| YIL064W | 505.7570311 | -1.065615165 | 0.011090319 | 0.317 |
| YJL094C | 1236.455255 | 0.641736348 | 0.011555822 | 0.317 |
| YML024W | 8930.152891 | -0.740245265 | 0.011542151 | 0.317 |
| YNL002C | 1677.703778 | -0.917931606 | 0.011446878 | 0.317 |
| YOR066W | 1115.721778 | 0.901958299 | 0.011161281 | 0.317 |
| YOR340C | 775.5259752 | -0.784487929 | 0.011272705 | 0.317 |
| YPL177C | 1454.555337 | 0.651815991 | 0.011518073 | 0.317 |

| | | | | |
|---|---|---|---|---|
| YPL178W | 1119.186631 | -0.528118281 | 0.011312557 | 0.317 |
| YPR123C | 99.99461053 | -0.670880794 | 0.011293363 | 0.317 |
| YLR185W | 4884.991766 | -0.808145468 | 0.011724042 | 0.319 |
| YOL022C | 1058.507084 | -0.911769992 | 0.01168835 | 0.319 |
| YBL027W | 10124.8249 | -0.854588693 | 0.013278888 | 0.320 |
| YBL039C | 1528.083632 | -0.895464687 | 0.013113584 | 0.320 |
| YBR113W | 115.7153994 | -0.727542945 | 0.014053976 | 0.320 |
| YBR121C | 7616.904588 | -0.757769543 | 0.013616179 | 0.320 |
| YBR213W | 228.053134 | -0.685735195 | 0.013334655 | 0.320 |
| YDL083C | 6219.514303 | -0.651512385 | 0.014180696 | 0.320 |
| YDL208W | 1913.168285 | -0.862784877 | 0.013543714 | 0.320 |
| YDR167W | 803.2507476 | -0.669569299 | 0.013601219 | 0.320 |
| YDR353W | 5059.752159 | -0.837299811 | 0.013450102 | 0.320 |
| YDR429C | 3118.563913 | -0.807251466 | 0.012875109 | 0.320 |
| YDR465C | 1088.193625 | -1.002161461 | 0.012325089 | 0.320 |
| YDR538W | 133.1462476 | -0.715596577 | 0.013025367 | 0.320 |
| YEL027W | 5744.186516 | -0.656187565 | 0.012359797 | 0.320 |
| YFL037W | 5301.265498 | -0.689703648 | 0.012568299 | 0.320 |
| YGL189C | 20518.43042 | -0.733285955 | 0.012954581 | 0.320 |
| YGL190C | 1598.926776 | 0.690772517 | 0.013195438 | 0.320 |
| YGL225W | 1789.038412 | -0.617835214 | 0.012945301 | 0.320 |
| YGR030C | 291.5946813 | -0.937625165 | 0.012636789 | 0.320 |
| YGR083C | 2563.158798 | -0.858493887 | 0.013913541 | 0.320 |
| YGR280C | 1635.461549 | -1.091963827 | 0.012520311 | 0.320 |
| YGR285C | 10708.64158 | -0.859415955 | 0.013663415 | 0.320 |
| YHR107C | 1509.6288 | -0.645739776 | 0.013696386 | 0.320 |
| YJL138C | 1955.737437 | -0.7637282 | 0.012502094 | 0.320 |
| YJL158C | 11506.92019 | -0.695415781 | 0.012458112 | 0.320 |
| YKL005C | 1108.015733 | 0.522131496 | 0.013989096 | 0.320 |
| YKL081W | 14463.37631 | -0.627536241 | 0.014152433 | 0.320 |
| YKL113C | 707.9229549 | -0.83871246 | 0.011968656 | 0.320 |
| YKR043C | 2568.82772 | -0.890335045 | 0.012799562 | 0.320 |
| YKR092C | 1945.135383 | -1.031142032 | 0.013393492 | 0.320 |
| YLL008W | 1923.071209 | -0.999138394 | 0.014050443 | 0.320 |
| YLL018C | 8408.142457 | -0.628112974 | 0.014078402 | 0.320 |
| YLR146C | 511.7510501 | -0.719736439 | 0.011856377 | 0.320 |
| YLR149C-A | 61.1192566 | -0.676259402 | 0.013978902 | 0.320 |
| YLR229C | 1318.158779 | -0.658070021 | 0.012091765 | 0.320 |
| YMR071C | 1073.602658 | -0.391939996 | 0.012412303 | 0.320 |
| YMR131C | 2416.896733 | -0.892557425 | 0.012383558 | 0.320 |
| YOL012C | 810.102208 | -0.662945987 | 0.01408294 | 0.320 |
| YOR005C | 988.1912317 | 0.746987942 | 0.0128742 | 0.320 |
| YOR293W | 11411.33478 | -0.636576212 | 0.012585872 | 0.320 |

| | | | | |
|---|---|---|---|---|
| YOR294W | 771.7312013 | -1.099535416 | 0.012087117 | 0.320 |
| YOR332W | 2877.284969 | -0.62143349 | 0.013991382 | 0.320 |
| YOR361C | 4132.636454 | -0.787750647 | 0.013879004 | 0.320 |
| YPL211W | 668.7255746 | -1.027469233 | 0.011999536 | 0.320 |
| YPR133W-A | 1114.918643 | -0.791921114 | 0.013811082 | 0.320 |
| YAR002C-A | 2600.594621 | -0.725557938 | 0.016727732 | 0.322 |
| YBL057C | 768.8630819 | -0.702296085 | 0.016124201 | 0.322 |
| YBL068W | 791.3128306 | -0.815293955 | 0.014379847 | 0.322 |
| YBR181C | 1794.107741 | -0.744963102 | 0.014963892 | 0.322 |
| YBR189W | 11760.64407 | -0.61836926 | 0.015043056 | 0.322 |
| YBR243C | 997.8241569 | -0.695244728 | 0.016113426 | 0.322 |
| YBR257W | 391.5498488 | -0.642515779 | 0.015937513 | 0.322 |
| YDR161W | 1294.242267 | -0.912582722 | 0.014453591 | 0.322 |
| YDR292C | 1617.849006 | -0.592165798 | 0.016462763 | 0.322 |
| YDR412W | 170.5135109 | -0.759712465 | 0.016741218 | 0.322 |
| YDR424C | 383.6752581 | -0.592693058 | 0.01671913 | 0.322 |
| YDR447C | 4088.999597 | -0.726908658 | 0.015651978 | 0.322 |
| YER003C | 3520.842344 | -0.671114806 | 0.0152787 | 0.322 |
| YGL102C | 355.1928609 | -0.884720897 | 0.016092381 | 0.322 |
| YGL123W | 15975.90289 | -0.773424389 | 0.016760901 | 0.322 |
| YGL148W | 4268.613078 | -0.9260788 | 0.016315652 | 0.322 |
| YGL234W | 9210.041986 | -0.650123302 | 0.016735992 | 0.322 |
| YGR110W | 407.9203872 | 0.674925692 | 0.015218344 | 0.322 |
| YHL015W | 17106.50707 | -0.743724268 | 0.016428479 | 0.322 |
| YHR021C | 5458.86145 | -0.738992714 | 0.016973426 | 0.322 |
| YIL019W | 543.1334136 | -1.20617193 | 0.015460712 | 0.322 |
| YIL071C | 381.3119395 | 0.574779201 | 0.016068738 | 0.322 |
| YJL124C | 1576.189687 | -0.613230918 | 0.014670721 | 0.322 |
| YJR043C | 619.0166524 | -0.722688246 | 0.015557511 | 0.322 |
| YJR097W | 96.20405896 | -0.89776598 | 0.015383953 | 0.322 |
| YJR105W | 3472.841224 | -0.840418155 | 0.015186623 | 0.322 |
| YJR135W-A | 435.3473743 | -0.775997934 | 0.015011335 | 0.322 |
| YKL121W | 1073.818565 | 0.767961852 | 0.014830843 | 0.322 |
| YKL172W | 2863.993194 | -1.045762523 | 0.014600013 | 0.322 |
| YKL213C | 1964.500982 | 0.521854799 | 0.014827166 | 0.322 |
| YKR025W | 782.5030894 | -0.92212706 | 0.015045889 | 0.322 |
| YLR047C | 709.1954371 | 0.506779569 | 0.015753158 | 0.322 |
| YLR293C | 8683.145693 | -0.821863716 | 0.015966123 | 0.322 |
| YML012W | 3525.480868 | -0.681224905 | 0.016856165 | 0.322 |
| YMR116C | 12111.33345 | -0.646677656 | 0.015259697 | 0.322 |
| YMR181C | 1531.918567 | 0.847042926 | 0.014761372 | 0.322 |
| YMR318C | 2412.871237 | -0.711537055 | 0.015891732 | 0.322 |
| YNL113W | 221.4093861 | -0.814188263 | 0.016557613 | 0.322 |

| | | | | |
|---|---|---|---|---|
| YNL175C | 613.6716753 | -1.019542165 | 0.01690842 | 0.322 |
| YNL290W | 854.0353401 | -0.672052252 | 0.016733526 | 0.322 |
| YNR003C | 554.144982 | -0.826518747 | 0.016963936 | 0.322 |
| YNR058W | 282.8560691 | 0.691283804 | 0.014470245 | 0.322 |
| YOL085C | 30.97884527 | 1.142971079 | 0.016774111 | 0.322 |
| YOL116W | 722.658756 | 0.700207978 | 0.016374 | 0.322 |
| YOR074C | 848.568103 | -0.688765575 | 0.016193874 | 0.322 |
| YOR239W | 2726.169468 | -0.657671151 | 0.016967026 | 0.322 |
| YOR339C | 48.09399516 | 0.872326706 | 0.016218356 | 0.322 |
| YPL010W | 2111.463364 | -0.669356532 | 0.016142368 | 0.322 |
| YPR163C | 5059.100182 | -0.929289292 | 0.014901492 | 0.322 |
| YAL059W | 208.6556716 | -0.868598857 | 0.017132338 | 0.324 |
| YDR500C | 6683.941473 | -0.89759128 | 0.017316305 | 0.325 |
| YLR384C | 2550.492331 | -0.579264498 | 0.017371602 | 0.325 |
| YNR015W | 1147.190779 | -0.696232265 | 0.017244918 | 0.325 |
| YPR062W | 2367.986615 | -0.592184597 | 0.017354798 | 0.325 |
| YDR433W | 87.14790436 | -0.995900528 | 0.017516312 | 0.326 |
| YBR061C | 630.0720407 | -0.729076202 | 0.018124789 | 0.328 |
| YDR399W | 3185.764567 | -0.828462959 | 0.018320633 | 0.328 |
| YER131W | 9395.024759 | -0.599408919 | 0.018314651 | 0.328 |
| YGL229C | 708.5937653 | 0.742619605 | 0.018360174 | 0.328 |
| YGR027C | 9077.452786 | -0.841980854 | 0.017844765 | 0.328 |
| YHR056C | 832.3445362 | 0.68973598 | 0.017740863 | 0.328 |
| YJL015C | 72.86729941 | 0.609111522 | 0.017993402 | 0.328 |
| YJL104W | 974.9529404 | -0.718913732 | 0.017883347 | 0.328 |
| YJR014W | 1708.927082 | -0.561487028 | 0.018374868 | 0.328 |
| YJR069C | 644.3818806 | -0.773766293 | 0.017940983 | 0.328 |
| YML094W | 513.2141562 | -0.711438166 | 0.018057519 | 0.328 |
| YMR242C | 8203.831599 | -0.734882833 | 0.017884135 | 0.328 |
| YOL077C | 1854.977032 | -0.880683188 | 0.017796137 | 0.328 |
| YOR190W | 351.1788756 | 0.763011838 | 0.018356952 | 0.328 |
| YDR018C | 446.8147304 | 1.000074874 | 0.018576987 | 0.329 |
| YPR092W | 28.34006108 | 0.794871587 | 0.018531468 | 0.329 |
| YDR323C | 314.788297 | 0.499939072 | 0.018763904 | 0.330 |
| YJR070C | 1785.660344 | -0.688831502 | 0.018729351 | 0.330 |
| YPR124W | 1891.733227 | -0.816228591 | 0.018676664 | 0.330 |
| YMR269W | 489.6101925 | -0.953153518 | 0.018888161 | 0.331 |
| YNL135C | 5166.62026 | -0.772252091 | 0.019101482 | 0.331 |
| YOR275C | 1566.16532 | 0.535695613 | 0.019083461 | 0.331 |
| YOR326W | 4223.653773 | -0.588376118 | 0.019069314 | 0.331 |
| YPL140C | 588.07308 | 0.700426409 | 0.018981045 | 0.331 |
| YBL087C | 3378.344673 | -0.82000114 | 0.021724043 | 0.331 |
| YBR090C | 79.12993298 | -0.880451237 | 0.019572679 | 0.331 |

| | | | | |
|---|---|---|---|---|
| YBR249C | 4275.177673 | -0.619662797 | 0.020857957 | 0.331 |
| YBR270C | 464.1113888 | 0.714443732 | 0.020130491 | 0.331 |
| YCL054W | 1476.642208 | -0.988704126 | 0.021493967 | 0.331 |
| YDL075W | 5187.384084 | -0.795904379 | 0.020585699 | 0.331 |
| YDL206W | 597.9682559 | 0.670513822 | 0.021796618 | 0.331 |
| YDR014W | 486.9454278 | 0.627919977 | 0.020645681 | 0.331 |
| YDR083W | 1132.402572 | -0.85684185 | 0.021888171 | 0.331 |
| YDR228C | 901.0372572 | 0.573009995 | 0.019709083 | 0.331 |
| YDR450W | 8491.535949 | -0.723504228 | 0.020619135 | 0.331 |
| YER074W | 4108.232098 | -0.840273349 | 0.022227162 | 0.331 |
| YER093C | 1449.072568 | 0.568555917 | 0.019364358 | 0.331 |
| YER102W | 2473.647383 | -0.71031488 | 0.020184337 | 0.331 |
| YER118C | 889.8446464 | -0.707931708 | 0.02077373 | 0.331 |
| YGL176C | 596.0143813 | 0.544902333 | 0.019634033 | 0.331 |
| YGR148C | 7673.362198 | -0.693802794 | 0.020213148 | 0.331 |
| YHL013C | 503.0447213 | -0.756831689 | 0.021237152 | 0.331 |
| YHR061C | 329.4509042 | 0.68663678 | 0.022380742 | 0.331 |
| YHR154W | 1331.473724 | -0.591783405 | 0.021852749 | 0.331 |
| YIL052C | 4784.674042 | -0.753202929 | 0.022215213 | 0.331 |
| YIL078W | 8337.626041 | -0.733070262 | 0.022108294 | 0.331 |
| YJL115W | 931.9259287 | -0.729444428 | 0.021108959 | 0.331 |
| YJL121C | 2601.821139 | -0.588833811 | 0.021399803 | 0.331 |
| YJL132W | 534.7548177 | 0.81959617 | 0.022170421 | 0.331 |
| YJL136C | 5415.03448 | -0.752212455 | 0.021381223 | 0.331 |
| YJR057W | 461.680688 | -0.577020833 | 0.022299326 | 0.331 |
| YJR061W | 732.5988419 | 0.694588453 | 0.020598232 | 0.331 |
| YKL009W | 1409.022835 | -0.886531365 | 0.020557068 | 0.331 |
| YKL045W | 918.6422418 | -0.536913087 | 0.02136364 | 0.331 |
| YKL214C | 653.6592896 | -0.615109094 | 0.019335949 | 0.331 |
| YKR047W | 143.8726915 | -0.656647778 | 0.021928653 | 0.331 |
| YKR085C | 837.2824572 | -0.662235588 | 0.0216128 | 0.331 |
| YLL045C | 7512.686429 | -0.638498772 | 0.021619648 | 0.331 |
| YLL050C | 6100.834604 | -0.6948629 | 0.019799294 | 0.331 |
| YLR167W | 28914.13243 | -0.633733901 | 0.021095151 | 0.331 |
| YLR287C-A | 5234.314487 | -0.819663346 | 0.021466535 | 0.331 |
| YLR314C | 3042.070083 | -0.523137456 | 0.021302249 | 0.331 |
| YLR340W | 21021.60532 | -0.734887087 | 0.021728762 | 0.331 |
| YLR378C | 3906.764345 | -0.584259108 | 0.021599039 | 0.331 |
| YLR449W | 2574.095007 | -0.985547149 | 0.020192085 | 0.331 |
| YMR049C | 1525.516535 | -0.870243955 | 0.021098851 | 0.331 |
| YMR142C | 11187.93902 | -0.747625039 | 0.022237401 | 0.331 |
| YMR298W | 1278.824067 | -0.619133676 | 0.021116107 | 0.331 |
| YNL255C | 4025.75008 | -0.885348665 | 0.022399008 | 0.331 |

| | | | | |
|---|---|---|---|---|
| YOL039W | 5643.898973 | -0.834478043 | 0.022140416 | 0.331 |
| YOL127W | 9841.72384 | -0.78127016 | 0.0195567 | 0.331 |
| YOR167C | 4868.170026 | -0.994504328 | 0.021879917 | 0.331 |
| YOR232W | 2857.566582 | -0.65527425 | 0.020769482 | 0.331 |
| YOR251C | 1296.455018 | -0.497525909 | 0.019626951 | 0.331 |
| YOR272W | 1204.374523 | -0.906906877 | 0.019532315 | 0.331 |
| YOR369C | 15497.1337 | -0.884798349 | 0.020344333 | 0.331 |
| YPL054W | 345.0757807 | 0.988456529 | 0.020710587 | 0.331 |
| YPL101W | 1017.044188 | -0.686807275 | 0.020743244 | 0.331 |
| YPL117C | 2245.25569 | -0.637285434 | 0.021982732 | 0.331 |
| YPL246C | 2119.845638 | -0.570197235 | 0.019716362 | 0.331 |
| YPL263C | 1770.940205 | -0.743023131 | 0.02092408 | 0.331 |
| snR49 | 77.03202196 | -0.244883368 | 0.021415699 | 0.331 |
| YBR261C | 744.8069304 | -0.67270572 | 0.023385355 | 0.332 |
| YCL048W-A | 89.40578665 | 1.188818082 | 0.022936736 | 0.332 |
| YDL228C | 55.92892077 | -0.80026086 | 0.0234399 | 0.332 |
| YDR131C | 626.2979697 | 0.589149031 | 0.022669931 | 0.332 |
| YER039C | 101.9948572 | 0.769908733 | 0.023210184 | 0.332 |
| YER154W | 1666.340326 | -0.482094189 | 0.023289811 | 0.332 |
| YGL076C | 4229.319259 | -0.589971487 | 0.022926891 | 0.332 |
| YGR089W | 1598.86435 | 0.597516729 | 0.02254997 | 0.332 |
| YHR010W | 11885.41138 | -0.614773903 | 0.023043838 | 0.332 |
| YLL024C | 20764.11389 | -0.519222635 | 0.023281483 | 0.332 |
| YLR247C | 1785.222725 | 0.59556867 | 0.0231077 | 0.332 |
| YML026C | 7646.727007 | -0.570205087 | 0.023227727 | 0.332 |
| YML047C | 45.4588878 | 1.345181164 | 0.023477092 | 0.332 |
| YNL149C | 291.2466735 | -0.915976789 | 0.023081741 | 0.332 |
| YOL080C | 199.8794327 | -0.820244146 | 0.022946295 | 0.332 |
| YOR063W | 34774.24031 | -0.638810405 | 0.023087946 | 0.332 |
| YOR145C | 791.624005 | -0.809190939 | 0.022576586 | 0.332 |
| YPL198W | 414.9903406 | -0.713525309 | 0.023336723 | 0.332 |
| YJR129C | 222.0216409 | -0.709799101 | 0.023543018 | 0.332 |
| YBR162W-A | 580.1318829 | -0.862770473 | 0.024005179 | 0.333 |
| YDL084W | 7170.004042 | -0.645902582 | 0.023967106 | 0.333 |
| YDL166C | 434.2604045 | -0.876777085 | 0.023804461 | 0.333 |
| YFR032C-A | 2429.038295 | -0.792875425 | 0.024444542 | 0.333 |
| YGR145W | 1225.275886 | -0.891409069 | 0.024295162 | 0.333 |
| YHR025W | 4807.328396 | -0.589999042 | 0.024241019 | 0.333 |
| YJR155W | 885.0501737 | 0.874514549 | 0.02430444 | 0.333 |
| YLR150W | 21689.73235 | -0.725982058 | 0.024278685 | 0.333 |
| YLR227C | 512.1356286 | 0.483706659 | 0.024241996 | 0.333 |
| YOL025W | 668.044846 | 0.760105977 | 0.02441612 | 0.333 |
| YOL132W | 124.9358227 | 0.976943285 | 0.023918984 | 0.333 |

| | | | | |
|---|---|---|---|---|
| YOR253W | 809.4249218 | -0.769660217 | 0.024111283 | 0.333 |
| YPR102C | 3154.540675 | -0.827749312 | 0.02415074 | 0.333 |
| YPR112C | 1587.350101 | -0.98398857 | 0.023710762 | 0.333 |
| YPR192W | 327.8980894 | 0.720062153 | 0.024407345 | 0.333 |
| YBL020W | 997.2826168 | -0.56338437 | 0.027735263 | 0.334 |
| YBL021C | 336.3286264 | -0.655446454 | 0.025653906 | 0.334 |
| YBR058C-A | 528.5056217 | -0.705633629 | 0.025085285 | 0.334 |
| YBR070C | 438.6999546 | -0.618578414 | 0.027377808 | 0.334 |
| YBR105C | 2095.906659 | 0.811243632 | 0.026447289 | 0.334 |
| YBR221C | 5794.723277 | -0.563196823 | 0.026679992 | 0.334 |
| YCR031C | 11369.4013 | -0.736108342 | 0.02573112 | 0.334 |
| YDL031W | 1680.703139 | -0.916959422 | 0.025498491 | 0.334 |
| YDL064W | 884.0529801 | -0.692528658 | 0.026680549 | 0.334 |
| YDL102W | 928.5717968 | -0.479106157 | 0.02765995 | 0.334 |
| YDR043C | 825.8596949 | 0.734779245 | 0.026410348 | 0.334 |
| YDR487C | 2644.625402 | -0.579594124 | 0.026675764 | 0.334 |
| YEL054C | 3213.78471 | -0.497606966 | 0.026515561 | 0.334 |
| YER005W | 999.8103274 | 0.42014669 | 0.026588108 | 0.334 |
| YER019C-A | 1962.324665 | -0.643624667 | 0.027483287 | 0.334 |
| YER101C | 634.792482 | 0.786847358 | 0.026783567 | 0.334 |
| YFL004W | 2271.622342 | -0.476772461 | 0.027286453 | 0.334 |
| YGR082W | 1782.261898 | -0.602030076 | 0.026338408 | 0.334 |
| YGR216C | 759.5937639 | 0.503638082 | 0.026511237 | 0.334 |
| YGR218W | 3998.195199 | 0.411069315 | 0.027216902 | 0.334 |
| YHL011C | 1690.318772 | -0.563158089 | 0.026655618 | 0.334 |
| YHR068W | 2922.590435 | -0.664981436 | 0.024570823 | 0.334 |
| YHR091C | 596.7962742 | -0.475670088 | 0.025237092 | 0.334 |
| YIL107C | 1780.529737 | 0.780821653 | 0.026202118 | 0.334 |
| YIL148W | 9852.705743 | -0.645246448 | 0.025305165 | 0.334 |
| YJL050W | 1800.303824 | -0.834306348 | 0.027333392 | 0.334 |
| YJL074C | 945.199767 | -0.569005032 | 0.025583565 | 0.334 |
| YJR002W | 1689.001321 | -0.711373973 | 0.025890301 | 0.334 |
| YJR094W-A | 1689.898779 | -0.851133061 | 0.02769988 | 0.334 |
| YKL124W | 623.7006372 | 0.632650719 | 0.025188999 | 0.334 |
| YKL192C | 3222.400243 | -0.540039973 | 0.026802908 | 0.334 |
| YLR003C | 782.5858822 | -0.838164588 | 0.025726144 | 0.334 |
| YLR135W | 729.3825255 | 0.629211985 | 0.025948199 | 0.334 |
| YLR142W | 1292.784521 | 0.951435244 | 0.024852543 | 0.334 |
| YLR153C | 3246.572959 | -0.621694981 | 0.024791229 | 0.334 |
| YLR188W | 1233.172467 | -0.536355721 | 0.025604095 | 0.334 |
| YLR288C | 449.0939426 | 0.435242322 | 0.026032516 | 0.334 |
| YLR388W | 1672.166035 | -0.755753593 | 0.025154225 | 0.334 |
| YML063W | 16224.08176 | -0.567988394 | 0.026302017 | 0.334 |

| | | | | |
|---|---|---|---|---|
| YMR138W | 253.6379214 | -0.676846295 | 0.027451571 | 0.334 |
| YNL012W | 592.4635068 | 0.804488492 | 0.02670956 | 0.334 |
| YNL061W | 2817.950295 | -0.78481533 | 0.026854944 | 0.334 |
| YNL162W-A | 54.12294867 | -1.228236548 | 0.026744081 | 0.334 |
| YNL178W | 12909.44004 | -0.681174929 | 0.025454108 | 0.334 |
| YNL181W | 1100.958937 | -0.530169819 | 0.027076372 | 0.334 |
| YNL313C | 1104.908095 | -0.680246431 | 0.025082819 | 0.334 |
| YOL139C | 5554.572319 | -0.840871938 | 0.025920638 | 0.334 |
| YOR143C | 346.0541178 | -0.679406035 | 0.02774847 | 0.334 |
| YOR271C | 1520.089436 | -0.792997494 | 0.025723483 | 0.334 |
| YOR351C | 100.6577504 | 0.875014572 | 0.027486436 | 0.334 |
| YPL036W | 288.1590263 | 0.760300652 | 0.027160408 | 0.334 |
| YPL084W | 2075.944666 | 0.474434506 | 0.02470917 | 0.334 |
| YPL090C | 2392.913651 | -0.745795572 | 0.025736324 | 0.334 |
| YPL163C | 3377.565521 | -0.547934964 | 0.0273572 | 0.334 |
| YPL217C | 2396.710824 | -0.727764253 | 0.026898101 | 0.334 |
| YPL231W | 11637.98709 | -0.623413381 | 0.027526212 | 0.334 |
| YPR058W | 588.0067957 | -0.82026499 | 0.026668839 | 0.334 |
| YLR196W | 1358.156963 | -0.917768868 | 0.027805851 | 0.334 |
| YBR164C | 1099.977428 | -0.725249139 | 0.027907829 | 0.334 |
| YLR209C | 881.8739647 | -0.653187165 | 0.028037574 | 0.334 |
| YLR264W | 525.6829336 | -0.865257781 | 0.028025455 | 0.334 |
| YMR230W | 6828.386976 | -0.708080087 | 0.028046175 | 0.334 |
| YGL031C | 8332.907395 | -0.591791747 | 0.028243515 | 0.334 |
| YJL148W | 1042.119748 | -0.829600113 | 0.028208859 | 0.334 |
| YPL079W | 6102.629794 | -0.640706235 | 0.028201992 | 0.334 |
| YER043C | 7073.679738 | -0.557918456 | 0.028429328 | 0.334 |
| YGR085C | 1835.900279 | -0.975254679 | 0.028552367 | 0.334 |
| YGR196C | 1285.850548 | 0.527091021 | 0.028552621 | 0.334 |
| YJL080C | 5735.124683 | -0.531296101 | 0.028454058 | 0.334 |
| YML055W | 811.1131816 | -0.596297232 | 0.028509276 | 0.334 |
| YDL051W | 1178.81834 | -0.878220971 | 0.02867346 | 0.335 |
| YML073C | 8568.253747 | -0.694578792 | 0.028622207 | 0.335 |
| YDR299W | 1150.958917 | -0.963259677 | 0.028804788 | 0.335 |
| YPR151C | 43.40930441 | 0.926896371 | 0.028826098 | 0.335 |
| YGL226C-A | 628.3645694 | -0.678268194 | 0.029059845 | 0.335 |
| YKL156W | 2081.100672 | -0.941795648 | 0.028980975 | 0.335 |
| YLR001C | 1116.57839 | 0.612419435 | 0.028958332 | 0.335 |
| YLR372W | 2301.005993 | -0.3703759 | 0.029108154 | 0.335 |
| YNL070W | 685.2481589 | -0.628992303 | 0.029228479 | 0.335 |
| YOR225W | 54.88254174 | -0.986517071 | 0.029209684 | 0.335 |
| YOR312C | 9424.205996 | -0.623890192 | 0.02917269 | 0.335 |
| YBL040C | 1584.786506 | -0.587431122 | 0.029326587 | 0.335 |

| | | | | |
|---|---|---|---|---|
| YJL090C | 629.2586035 | 0.564307674 | 0.029383522 | 0.335 |
| YNL162W | 2350.306083 | -0.761138971 | 0.029855932 | 0.340 |
| YBR188C | 339.0880427 | -0.549617198 | 0.029940435 | 0.340 |
| YAL035W | 9252.51472 | -0.901214243 | 0.030093081 | 0.341 |
| YLR285C-A | 41.10839909 | -0.914998771 | 0.03004707 | 0.341 |
| YLR009W | 1043.475365 | -0.802652787 | 0.030272528 | 0.342 |
| YMR038C | 1941.251831 | -0.612682241 | 0.030278119 | 0.342 |
| YLR043C | 5741.406917 | -0.629487465 | 0.030448316 | 0.342 |
| YOR045W | 622.8710581 | -0.709940169 | 0.030441877 | 0.342 |
| YKL162C | 452.2195831 | 0.632773839 | 0.030517653 | 0.342 |
| YNL010W | 2912.199227 | -0.684482166 | 0.030603503 | 0.343 |
| YJL009W | 66.31151028 | -0.788048367 | 0.030807082 | 0.344 |
| YAL036C | 1045.610431 | -0.782416301 | 0.031066308 | 0.345 |
| YAR030C | 64.62074774 | 0.726705264 | 0.032292558 | 0.345 |
| YBR084C-A | 8707.666036 | -0.76869951 | 0.031123552 | 0.345 |
| YBR284W | 530.3934343 | 0.73984007 | 0.032664235 | 0.345 |
| YDR037W | 9604.757078 | -0.710276847 | 0.032452117 | 0.345 |
| YDR170C | 4884.61556 | -0.528694034 | 0.031527246 | 0.345 |
| YDR308C | 370.9057891 | -0.570780632 | 0.031875736 | 0.345 |
| YER034W | 308.6813768 | -0.819913144 | 0.032241233 | 0.345 |
| YER117W | 4528.496655 | -0.549008224 | 0.031938838 | 0.345 |
| YFL040W | 216.9489399 | 0.990322842 | 0.031373692 | 0.345 |
| YGL022W | 3864.561612 | -0.592996264 | 0.031063272 | 0.345 |
| YGR001C | 885.4111524 | -0.445007738 | 0.031879626 | 0.345 |
| YGR225W | 197.8168855 | 0.926045556 | 0.032530109 | 0.345 |
| YGR269W | 48.40957868 | 0.512497221 | 0.032407558 | 0.345 |
| YHR089C | 2056.029582 | -0.831819027 | 0.032110156 | 0.345 |
| YHR127W | 498.1945386 | -0.670368219 | 0.032048972 | 0.345 |
| YHR193C | 8396.042296 | -0.698498682 | 0.031453172 | 0.345 |
| YIL133C | 6678.70118 | -0.60422475 | 0.032649516 | 0.345 |
| YJL122W | 643.9700151 | -0.77070956 | 0.03266873 | 0.345 |
| YJR119C | 341.555846 | 0.629025748 | 0.032606162 | 0.345 |
| YKL033W-A | 1409.695613 | -0.545497796 | 0.032140502 | 0.345 |
| YKL082C | 1057.576571 | -0.823209581 | 0.031662713 | 0.345 |
| YKL090W | 556.2185917 | 0.662524482 | 0.0314262 | 0.345 |
| YLR008C | 578.2897755 | -0.638341479 | 0.031337456 | 0.345 |
| YLR363W-A | 269.4630453 | -0.738211405 | 0.032127015 | 0.345 |
| YNL099C | 699.7662248 | -0.617115294 | 0.031952744 | 0.345 |
| YNL132W | 1681.161762 | -0.697216457 | 0.031711678 | 0.345 |
| YNL147W | 569.2824244 | -0.593776809 | 0.031344391 | 0.345 |
| YOL097C | 2168.680575 | -0.636440782 | 0.032000098 | 0.345 |
| YOR091W | 1558.309002 | -0.687818244 | 0.032316106 | 0.345 |
| YOR309C | 407.1266153 | -0.692889003 | 0.03247555 | 0.345 |

| | | | | |
|---|---|---|---|---|
| YPR181C | 4442.448271 | -0.51066022 | 0.03273869 | 0.345 |
| YPL238C | 56.97812647 | -0.970144327 | 0.032820367 | 0.345 |
| YDL082W | 7404.050186 | -0.746138123 | 0.032975253 | 0.346 |
| YOR012W | 60.32598792 | 0.61820024 | 0.033106265 | 0.347 |
| YDR297W | 2184.395638 | -0.598936845 | 0.033391199 | 0.349 |
| YMR065W | 380.856771 | 0.523477281 | 0.033420208 | 0.349 |
| YAL002W | 1569.483592 | 0.505760122 | 0.035313822 | 0.350 |
| YAL010C | 552.0969494 | 0.466181835 | 0.034371139 | 0.350 |
| YBL032W | 1731.697141 | -0.676948227 | 0.033717869 | 0.350 |
| YBR031W | 10790.74196 | -0.573096244 | 0.035272759 | 0.350 |
| YBR084W | 2864.241089 | -0.691161686 | 0.035664644 | 0.350 |
| YDL061C | 2048.312712 | -0.777636431 | 0.035176692 | 0.350 |
| YDL184C | 118.3516335 | -1.048896231 | 0.033771604 | 0.350 |
| YDR086C | 1629.638329 | -0.70276349 | 0.035126839 | 0.350 |
| YDR267C | 660.9899359 | -0.495955712 | 0.035102733 | 0.350 |
| YGL059W | 1140.532396 | 0.699749998 | 0.033868473 | 0.350 |
| YGR106C | 3546.453004 | -0.510256947 | 0.034520009 | 0.350 |
| YGR208W | 739.3899853 | -0.611214063 | 0.035599911 | 0.350 |
| YHL012W | 237.0696554 | 0.595840104 | 0.035129885 | 0.350 |
| YIL003W | 534.8841649 | -0.611328239 | 0.0345521 | 0.350 |
| YIR026C | 772.7201823 | -0.764692959 | 0.034242213 | 0.350 |
| YJL177W | 7142.560719 | -0.609800457 | 0.034392762 | 0.350 |
| YKL183W | 567.8673175 | -0.568101015 | 0.034965271 | 0.350 |
| YKR094C | 5698.150477 | -0.622497044 | 0.034949131 | 0.350 |
| YLR029C | 12797.37208 | -0.585270921 | 0.035684457 | 0.350 |
| YLR180W | 3585.379162 | -0.768978445 | 0.035119251 | 0.350 |
| YLR198C | 85.68816582 | -1.019277799 | 0.034868059 | 0.350 |
| YLR200W | 774.013828 | -0.625102347 | 0.035323937 | 0.350 |
| YLR358C | 39.08246637 | -0.646961429 | 0.034423716 | 0.350 |
| YMR108W | 5770.966467 | -0.586364047 | 0.034532894 | 0.350 |
| YMR143W | 7020.519511 | -0.754269133 | 0.03463243 | 0.350 |
| YMR217W | 5540.397432 | -0.747590814 | 0.034826825 | 0.350 |
| YNL151C | 742.0050839 | -0.840997347 | 0.034545674 | 0.350 |
| YNR057C | 196.7239761 | 0.556505814 | 0.035483791 | 0.350 |
| YOL061W | 1754.163485 | -0.607760438 | 0.035271659 | 0.350 |
| YOL103W-B | 37.1755698 | 0.767751278 | 0.034363055 | 0.350 |
| YOR102W | 49.45521107 | -0.780710769 | 0.035542379 | 0.350 |
| YOR198C | 6733.130979 | -0.661706896 | 0.035386148 | 0.350 |
| YPR010C | 3009.126793 | -0.679699908 | 0.033831245 | 0.350 |
| YPR041W | 3343.881383 | -0.685134825 | 0.033995813 | 0.350 |
| YPR052C | 667.2448711 | -0.750252034 | 0.035649266 | 0.350 |
| YPR187W | 1425.65006 | -0.710704279 | 0.034452083 | 0.350 |
| YAR008W | 281.6693003 | -0.704270297 | 0.036674345 | 0.350 |

| | | | | |
|---|---|---|---|---|
| YBR299W | 41.66988027 | 0.759554073 | 0.036867593 | 0.350 |
| YCR084C | 3633.028548 | -0.583785047 | 0.037245034 | 0.350 |
| YDR016C | 195.2031569 | -0.634601347 | 0.037331235 | 0.350 |
| YDR085C | 1259.372351 | 0.77507466 | 0.036184666 | 0.350 |
| YDR152W | 1737.468978 | -0.664350328 | 0.037027884 | 0.350 |
| YDR418W | 2788.265019 | -0.707733366 | 0.036165087 | 0.350 |
| YER077C | 789.2511196 | 0.453491393 | 0.036414782 | 0.350 |
| YGL135W | 2224.916712 | -0.803129856 | 0.037357709 | 0.350 |
| YGL200C | 3717.842992 | -0.503556716 | 0.036322541 | 0.350 |
| YGR159C | 2044.642627 | -0.777533872 | 0.036501056 | 0.350 |
| YHR064C | 6332.657586 | -0.726736488 | 0.037497584 | 0.350 |
| YHR179W | 8288.962491 | -0.662695328 | 0.036444261 | 0.350 |
| YJR108W | 40.18860279 | 0.778517652 | 0.036165852 | 0.350 |
| YJR123W | 26166.00582 | -0.565746518 | 0.036238233 | 0.350 |
| YJR139C | 4827.24467 | -0.641448835 | 0.036634022 | 0.350 |
| YLR110C | 106387.3494 | -0.632109638 | 0.037392446 | 0.350 |
| YLR406C | 2407.447144 | -0.549931797 | 0.036100714 | 0.350 |
| YML059C | 1781.099171 | 0.625682507 | 0.036335349 | 0.350 |
| YMR003W | 306.5250482 | -0.635948507 | 0.037449062 | 0.350 |
| YMR014W | 1014.309235 | -0.767624928 | 0.037254694 | 0.350 |
| YMR023C | 334.2712538 | 0.633152226 | 0.037416169 | 0.350 |
| YNL262W | 2257.387419 | -0.515643487 | 0.036491175 | 0.350 |
| YOL040C | 16073.93418 | -0.652827525 | 0.037118519 | 0.350 |
| YOR048C | 2248.629889 | -0.554313148 | 0.037409988 | 0.350 |
| YPL212C | 1188.64058 | -0.742308968 | 0.035806536 | 0.350 |
| YPL220W | 1761.369957 | -0.805174505 | 0.036883199 | 0.350 |
| YPR013C | 1083.106605 | 0.729732098 | 0.036957351 | 0.350 |
| YPR110C | 2091.025042 | -0.731321065 | 0.036319642 | 0.350 |
| YPR138C | 1408.826707 | 0.811462461 | 0.037033833 | 0.350 |
| YAL003W | 12374.8526 | -0.557025869 | 0.03842432 | 0.351 |
| YDR002W | 3692.839207 | -0.730796583 | 0.037654431 | 0.351 |
| YDR012W | 4314.996695 | -0.531628524 | 0.037997959 | 0.351 |
| YDR156W | 897.9312179 | -0.67884845 | 0.03873456 | 0.351 |
| YDR302W | 389.0867763 | -0.590653043 | 0.03874932 | 0.351 |
| YDR347W | 1409.321423 | -0.374829325 | 0.037785094 | 0.351 |
| YEL058W | 1683.65634 | -0.662887537 | 0.037926966 | 0.351 |
| YER073W | 2130.104252 | -0.910372923 | 0.038475957 | 0.351 |
| YGL245W | 8715.959502 | -0.592387262 | 0.038422874 | 0.351 |
| YHL038C | 821.5884324 | 0.555739474 | 0.038708941 | 0.351 |
| YHL043W | 209.5139873 | 0.787502522 | 0.038310122 | 0.351 |
| YHR096C | 1194.376628 | 0.776821966 | 0.038487178 | 0.351 |
| YKL196C | 2255.642273 | -0.544881591 | 0.038673855 | 0.351 |
| YLR268W | 967.406712 | -0.509444911 | 0.038312686 | 0.351 |

| | | | | |
|---|---|---|---|---|
| YNL209W | 13780.76239 | -0.689844723 | 0.038237721 | 0.351 |
| YOL048C | 1220.015423 | 0.740579979 | 0.038133411 | 0.351 |
| YOR214C | 33.33902204 | 0.849054125 | 0.038636563 | 0.351 |
| YOR234C | 2998.326335 | -0.668867053 | 0.038438996 | 0.351 |
| YPL202C | 824.3595251 | 0.470367341 | 0.037755593 | 0.351 |
| YPL208W | 646.0074609 | -0.558818889 | 0.038690225 | 0.351 |
| YBR233W-A | 253.0169158 | -0.730852842 | 0.038892722 | 0.352 |
| YCL038C | 682.2522644 | 0.598188332 | 0.039224518 | 0.352 |
| YER036C | 4277.526152 | -0.595419811 | 0.039112473 | 0.352 |
| YGL104C | 947.9922368 | 0.708300347 | 0.039386909 | 0.352 |
| YKL083W | 46.77827339 | -1.089581202 | 0.039083904 | 0.352 |
| YLR435W | 440.7011003 | -0.776719009 | 0.039361483 | 0.352 |
| YOR096W | 6750.76705 | -0.696422096 | 0.03930105 | 0.352 |
| YOR224C | 1759.04057 | -0.821498521 | 0.039314055 | 0.352 |
| YPL086C | 1089.007189 | -0.677388035 | 0.039139697 | 0.352 |
| YLR325C | 5760.035975 | -0.78323584 | 0.039635144 | 0.353 |
| YGL008C | 26806.32689 | -0.297089194 | 0.039977232 | 0.356 |
| YGR143W | 1625.66982 | 0.651450605 | 0.040254352 | 0.356 |
| YJL084C | 1615.45903 | 0.622645231 | 0.040106039 | 0.356 |
| YKR069W | 646.4230382 | 0.515844912 | 0.040239375 | 0.356 |
| YMR153W | 881.8064044 | -0.42613984 | 0.04023267 | 0.356 |
| YPR132W | 5266.120182 | -0.753469562 | 0.040367 | 0.357 |
| YGL180W | 843.8043135 | 0.626355054 | 0.040459776 | 0.357 |
| YDL020C | 3590.678769 | 0.688048653 | 0.041272511 | 0.358 |
| YER074W-A | 546.9637993 | -0.50581332 | 0.041371532 | 0.358 |
| YER088C-A | 302.6604055 | -0.649138643 | 0.040833119 | 0.358 |
| YGR105W | 477.947228 | -0.672643854 | 0.041046261 | 0.358 |
| YHR042W | 4638.852766 | -0.536107642 | 0.04084844 | 0.358 |
| YHR197W | 861.9145076 | -0.731691414 | 0.040876063 | 0.358 |
| YJL179W | 307.4969877 | -0.773160236 | 0.04063104 | 0.358 |
| YJL188C | 893.4798181 | -0.851432808 | 0.040811242 | 0.358 |
| YKR005C | 237.9372028 | 0.602155216 | 0.041050144 | 0.358 |
| YKR023W | 810.8865639 | -0.560598478 | 0.041085842 | 0.358 |
| YKR101W | 347.6961828 | 0.610043444 | 0.04125945 | 0.358 |
| YLR044C | 136554.4679 | -0.54563955 | 0.041314461 | 0.358 |
| YML127W | 993.038483 | -0.591329754 | 0.04137388 | 0.358 |
| YOR338W | 133.0538926 | 0.532709721 | 0.041396773 | 0.358 |
| YAL040C | 2404.494026 | 0.699522197 | 0.041826432 | 0.358 |
| YDR364C | 596.6987029 | 0.435398031 | 0.041667264 | 0.358 |
| YFL045C | 5242.345974 | -0.589673923 | 0.041686955 | 0.358 |
| YGR124W | 7850.205704 | -0.884627416 | 0.041614319 | 0.358 |
| YJL144W | 785.7166274 | 0.889548607 | 0.041557687 | 0.358 |
| YLR175W | 4387.69014 | -0.632527056 | 0.041770322 | 0.358 |

| | | | | |
|---|---|---|---|---|
| YNL160W | 19462.39163 | 1.008496069 | 0.041773768 | 0.358 |
| YMR186W | 31490.05511 | -0.637481719 | 0.041944499 | 0.358 |
| YKL179C | 1404.902826 | 0.496890524 | 0.042068756 | 0.358 |
| YNL308C | 3277.042014 | -0.856794314 | 0.042020516 | 0.358 |
| YJL143W | 1167.282332 | -0.577394335 | 0.04221368 | 0.359 |
| YNR027W | 549.891736 | -0.646379485 | 0.042312307 | 0.359 |
| YDR280W | 709.9216811 | -0.68763187 | 0.042374006 | 0.359 |
| YKL189W | 702.1789039 | 0.733758818 | 0.042515516 | 0.360 |
| YLR002C | 907.6259007 | -0.534275706 | 0.042658309 | 0.361 |
| YFL022C | 4340.34362 | -0.605629054 | 0.042937864 | 0.361 |
| YGL184C | 275.5662791 | 0.496764527 | 0.042963623 | 0.361 |
| YGR052W | 642.70953 | 0.760895587 | 0.042943809 | 0.361 |
| YIL021W | 814.8146206 | -0.516360514 | 0.042862529 | 0.361 |
| YNL279W | 510.6368292 | 1.022728611 | 0.042971947 | 0.361 |
| YDL014W | 4306.735257 | -0.69987914 | 0.043355977 | 0.361 |
| YDR071C | 2792.908431 | -0.619448389 | 0.043373571 | 0.361 |
| YLR066W | 1030.577166 | -0.586642412 | 0.043289515 | 0.361 |
| YML014W | 592.0947476 | -0.759980907 | 0.043347166 | 0.361 |
| YNL243W | 3599.331124 | -0.461838926 | 0.043151379 | 0.361 |
| snR63 | 94.54093227 | -0.524724935 | 0.043372794 | 0.361 |
| YDR025W | 5331.251911 | -0.610713147 | 0.043550954 | 0.362 |
| YHL033C | 4723.149428 | -0.544174981 | 0.043679929 | 0.363 |
| YHR121W | 1103.128777 | -0.450595677 | 0.043921172 | 0.363 |
| YMR182C | 564.4635256 | 0.597349262 | 0.043813294 | 0.363 |
| YNL210W | 106.0752581 | 0.663345785 | 0.043872817 | 0.363 |
| YGL120C | 2147.315673 | -0.733721292 | 0.044030584 | 0.363 |
| YNL096C | 3035.774305 | -0.628734215 | 0.04415327 | 0.364 |
| YER127W | 556.9285849 | -0.812026317 | 0.044221693 | 0.364 |
| YAL046C | 313.7043981 | -0.611543526 | 0.044296739 | 0.364 |
| YBR283C | 3928.577615 | -0.547924566 | 0.044696318 | 0.365 |
| YHR066W | 786.1240577 | -0.761269983 | 0.044547268 | 0.365 |
| YIL018W | 10226.88893 | -0.598220318 | 0.04457368 | 0.365 |
| YLR051C | 783.2029443 | -0.781939482 | 0.044698292 | 0.365 |
| YOR341W | 3809.897668 | -0.582540934 | 0.044620867 | 0.365 |
| YBR242W | 549.7068913 | -0.571901005 | 0.044892989 | 0.365 |
| YBR288C | 774.8863305 | -0.556230251 | 0.044862385 | 0.365 |
| YDR069C | 1204.898386 | 0.630390665 | 0.044984082 | 0.366 |
| YHR164C | 1299.336952 | 0.533996042 | 0.045331919 | 0.366 |
| YIL092W | 426.021742 | 0.445540319 | 0.045253096 | 0.366 |
| YMR184W | 674.607137 | -0.551056721 | 0.045217061 | 0.366 |
| YPR037C | 567.1835844 | -0.615851005 | 0.045281098 | 0.366 |
| YLR212C | 601.9431586 | -0.665165234 | 0.045453122 | 0.367 |
| YDR139C | 266.4093081 | -0.62141833 | 0.045546328 | 0.367 |

| | | | | |
|---|---|---|---|---|
| YKL127W | 1973.346678 | -0.62019371 | 0.045636048 | 0.367 |
| YLR263W | 707.0718116 | 0.717541585 | 0.04567574 | 0.367 |
| YDL137W | 4443.238326 | -0.546103435 | 0.046221141 | 0.367 |
| YDR158W | 6048.209084 | -0.640644176 | 0.046018534 | 0.367 |
| YIL022W | 2268.335855 | -0.349366134 | 0.045928525 | 0.367 |
| YIL159W | 678.524918 | -0.545510931 | 0.045789884 | 0.367 |
| YJL057C | 1203.837592 | 0.826501374 | 0.046186382 | 0.367 |
| YKL046C | 1814.036225 | -0.474694302 | 0.046280643 | 0.367 |
| YKL188C | 680.4138758 | 0.635226672 | 0.04604168 | 0.367 |
| YNL187W | 311.4857358 | 0.577141275 | 0.046247237 | 0.367 |
| YOL084W | 492.0065357 | 0.695765111 | 0.046190115 | 0.367 |
| YMR133W | 502.5352221 | 0.772794189 | 0.046583491 | 0.369 |
| YNL069C | 13320.02635 | -0.513572546 | 0.046540329 | 0.369 |
| YDR524W-C | 1641.998262 | -0.791251501 | 0.046762079 | 0.370 |
| YEL062W | 560.1717097 | 0.527747812 | 0.046987941 | 0.371 |
| YKL069W | 336.0140703 | -0.485166038 | 0.047009166 | 0.371 |
| YAL023C | 4478.973811 | -0.484249093 | 0.048023595 | 0.372 |
| YBR067C | 11273.4348 | 0.666349945 | 0.048153795 | 0.372 |
| YBR143C | 4538.418741 | -0.694082067 | 0.047994811 | 0.372 |
| YBR148W | 178.9686946 | 0.811420404 | 0.047414071 | 0.372 |
| YDL130W | 4709.707857 | -0.754282039 | 0.047943423 | 0.372 |
| YDL157C | 952.2291983 | -0.731967142 | 0.047473933 | 0.372 |
| YGR195W | 892.1333078 | -0.509171875 | 0.047939594 | 0.372 |
| YJR063W | 331.564801 | -0.754324288 | 0.047664786 | 0.372 |
| YKR088C | 850.0044261 | -0.49634481 | 0.048162055 | 0.372 |
| YLL049W | 333.5985502 | -0.59853123 | 0.048226395 | 0.372 |
| YLR448W | 5493.030487 | -0.651467255 | 0.048032278 | 0.372 |
| YPL059W | 1008.866988 | -0.453295598 | 0.04800259 | 0.372 |
| YPL108W | 159.4425969 | -0.718638485 | 0.048186193 | 0.372 |
| YPL216W | 730.7653507 | 0.605037744 | 0.048208277 | 0.372 |
| YPL261C | 48.62531076 | 0.717735401 | 0.047661681 | 0.372 |
| YPR019W | 1285.937535 | -0.386662334 | 0.048024119 | 0.372 |
| YEL051W | 3229.319305 | -0.583842777 | 0.048312815 | 0.373 |
| YMR041C | 591.2121868 | 0.792687757 | 0.048420621 | 0.373 |
| YBL076C | 7280.098127 | -0.583722184 | 0.048756965 | 0.374 |
| YDL077C | 1087.970908 | 0.388919697 | 0.048647317 | 0.374 |
| YDR524C-B | 18745.499 | -0.750151245 | 0.049210508 | 0.374 |
| YGR020C | 3100.566353 | -0.647089451 | 0.048861997 | 0.374 |
| YHR101C | 345.7524508 | 0.578530361 | 0.049122603 | 0.374 |
| YIL118W | 1012.394304 | -0.631755006 | 0.049178295 | 0.374 |
| YMR237W | 2047.326568 | 0.352103736 | 0.048868867 | 0.374 |
| YOL138C | 1055.538589 | 0.492395033 | 0.049124978 | 0.374 |
| YOR122C | 4814.102226 | -0.604709022 | 0.04918347 | 0.374 |

| | | | | |
|---|---|---|---|---|
| YPL143W | 740.5431297 | -0.838796295 | 0.049176284 | 0.374 |
| YHR131C | 966.4131265 | 0.651243014 | 0.049373195 | 0.374 |
| YJL133C-A | 1683.932792 | 0.650169341 | 0.049422253 | 0.374 |
| YJR144W | 1299.865299 | -0.478678407 | 0.049496062 | 0.374 |
| YML008C | 4220.321447 | -0.642181838 | 0.049504624 | 0.374 |
| YGR234W | 5131.339655 | -0.462414223 | 0.049625918 | 0.375 |
| YOL126C | 3179.903333 | 0.92863109 | 0.049724134 | 0.375 |
| YEL042W | 1714.856815 | -0.59530829 | 0.049814144 | 0.375 |
| YAL034C | 1899.332638 | 0.69730939 | 0.049972293 | 0.375 |
| YHL016C | 664.2747126 | 0.719486452 | 0.049908627 | 0.375 |
| YAL028W | 836.6833612 | 0.651308444 | 0.050322902 | 0.377 |
| YFL017C | 717.9144755 | -0.502447356 | 0.05027467 | 0.377 |
| YPL226W | 2793.678288 | -0.607780802 | 0.050311748 | 0.377 |
| YGL092W | 1819.13534 | -0.522283824 | 0.050482472 | 0.377 |
| YJL183W | 732.387825 | -0.640113365 | 0.050432744 | 0.377 |
| YBL072C | 3997.128198 | -0.766720294 | 0.050715613 | 0.377 |
| YLR240W | 1128.041716 | 0.533856517 | 0.050697873 | 0.377 |
| YPL027W | 50.11860961 | 0.746203191 | 0.050642778 | 0.377 |
| YBR159W | 2299.591935 | -0.608832393 | 0.051009627 | 0.378 |
| YGL259W | 86.59077684 | 1.264212776 | 0.051084533 | 0.378 |
| YNR034W | 660.948473 | 0.603879112 | 0.051088853 | 0.378 |
| YOR093C | 1072.002934 | 0.437078047 | 0.050998737 | 0.378 |
| YPR044C | 114.0311076 | -0.664380466 | 0.050889391 | 0.378 |
| YBR142W | 1284.846746 | -0.705192087 | 0.051181746 | 0.378 |
| YCR051W | 1038.832374 | -0.773406227 | 0.051717509 | 0.378 |
| YDL183C | 401.7373652 | 0.624443564 | 0.051523422 | 0.378 |
| YER112W | 1280.704105 | -0.549223744 | 0.051994766 | 0.378 |
| YGL251C | 608.1790507 | 0.666612117 | 0.051656948 | 0.378 |
| YHR141C | 3868.981705 | -0.634971289 | 0.052146842 | 0.378 |
| YHR175W-A | 167.202498 | -0.658028051 | 0.052066514 | 0.378 |
| YLR075W | 9637.869009 | -0.666376252 | 0.051564899 | 0.378 |
| YMR093W | 713.9051908 | -0.642273891 | 0.051866818 | 0.378 |
| YMR120C | 11760.55672 | -0.21125639 | 0.051784144 | 0.378 |
| YMR160W | 946.3755314 | 0.64013067 | 0.051489391 | 0.378 |
| YMR284W | 845.7287906 | 0.574959548 | 0.052172402 | 0.378 |
| YPL058C | 2629.343095 | 0.900101962 | 0.051974326 | 0.378 |
| YPL066W | 1565.740738 | 0.418542399 | 0.051449672 | 0.378 |
| YPR068C | 284.1750342 | 0.435681884 | 0.051434148 | 0.378 |
| snR4 | 89.67640266 | -0.534530361 | 0.05197888 | 0.378 |
| YIL072W | 537.0665215 | 0.57046815 | 0.052494711 | 0.379 |
| YJL145W | 1081.572718 | -0.636887548 | 0.052544783 | 0.379 |
| YLR048W | 13755.73587 | -0.61962663 | 0.052564502 | 0.379 |
| YNR021W | 3396.726535 | -0.566650955 | 0.052708868 | 0.380 |

| | | | | |
|---|---|---|---|---|
| YEL005C | 730.8016186 | 0.623403843 | 0.053133445 | 0.381 |
| YEL018W | 225.091547 | -0.499222668 | 0.053545579 | 0.381 |
| YER086W | 3514.638296 | -0.305170041 | 0.053046676 | 0.381 |
| YGR077C | 1152.926191 | 0.486241409 | 0.053305177 | 0.381 |
| YHR026W | 2431.689374 | -0.503171086 | 0.053335592 | 0.381 |
| YIR010W | 867.0379525 | 0.489661446 | 0.052990804 | 0.381 |
| YLL013C | 908.878738 | -0.424293262 | 0.053481295 | 0.381 |
| YLR176C | 916.0927001 | 0.549916181 | 0.053013796 | 0.381 |
| YLR438C-A | 659.2030513 | -0.741971642 | 0.053394544 | 0.381 |
| YMR105W-A | 43.59841519 | 0.539997052 | 0.053562894 | 0.381 |
| YPR114W | 2316.846378 | -0.404310234 | 0.053257348 | 0.381 |
| YBR045C | 255.946774 | 0.51469671 | 0.05440596 | 0.382 |
| YBR254C | 311.1708692 | -0.701556124 | 0.055580098 | 0.382 |
| YCL009C | 7647.273312 | -0.59925222 | 0.055450557 | 0.382 |
| YCL017C | 1588.391961 | -0.447721407 | 0.054572848 | 0.382 |
| YCL050C | 4005.943189 | -0.55699608 | 0.053951721 | 0.382 |
| YDL134C | 1962.281647 | 0.260453622 | 0.055949088 | 0.382 |
| YDR092W | 1261.21405 | -0.493859618 | 0.055282324 | 0.382 |
| YDR396W | 125.4408117 | -0.696769395 | 0.055504632 | 0.382 |
| YDR519W | 900.2252274 | -0.638391989 | 0.053859146 | 0.382 |
| YEL023C | 623.1672116 | 0.604882903 | 0.054885435 | 0.382 |
| YER055C | 2945.986692 | -0.496244597 | 0.055544747 | 0.382 |
| YGL112C | 1298.501075 | -0.36680832 | 0.055328935 | 0.382 |
| YGR051C | 51.37808205 | 0.59558998 | 0.055095056 | 0.382 |
| YGR207C | 1419.844599 | -0.456034116 | 0.055091083 | 0.382 |
| YHR104W | 3547.736943 | 0.664804612 | 0.055724324 | 0.382 |
| YIL158W | 651.6543059 | -0.77825014 | 0.055651204 | 0.382 |
| YJL192C | 2669.267366 | -0.538264171 | 0.05546806 | 0.382 |
| YJR136C | 423.3917741 | 0.393798606 | 0.05461719 | 0.382 |
| YJR145C | 6502.704468 | -0.609665022 | 0.055908752 | 0.382 |
| YKL060C | 152874.8661 | -0.548648801 | 0.05572631 | 0.382 |
| YKL122C | 1563.613747 | -0.644017028 | 0.053832052 | 0.382 |
| YLL014W | 750.2093462 | -0.526071846 | 0.054358652 | 0.382 |
| YLL062C | 209.8685733 | 0.840718429 | 0.055547548 | 0.382 |
| YLR151C | 483.8723388 | 0.548703895 | 0.055415351 | 0.382 |
| YLR431C | 452.0758432 | 0.51668487 | 0.054047435 | 0.382 |
| YNL244C | 2167.186944 | -0.745116642 | 0.055328213 | 0.382 |
| YNL247W | 4374.768503 | -0.709553791 | 0.055257473 | 0.382 |
| YOL047C | 302.1448912 | 0.783997749 | 0.055783809 | 0.382 |
| YOL144W | 766.01586 | -0.83585289 | 0.054449045 | 0.382 |
| YOR276W | 906.6874329 | -0.587519422 | 0.055677427 | 0.382 |
| YOR310C | 3221.373678 | -0.590547327 | 0.054933004 | 0.382 |
| YPL001W | 324.2471356 | -0.715045963 | 0.055892383 | 0.382 |

| | | | | |
|---|---|---|---|---|
| YPL160W | 4849.742262 | -0.56902922 | 0.054616252 | 0.382 |
| YPL249C-A | 77.54661876 | -1.00175895 | 0.05560197 | 0.382 |
| YPL272C | 344.5733609 | 0.596292828 | 0.054763483 | 0.382 |
| YER039C-A | 49.35745349 | 0.673198307 | 0.056057089 | 0.382 |
| YBL004W | 2589.356792 | -0.570741888 | 0.05631445 | 0.383 |
| YER049W | 1494.465582 | -0.623493198 | 0.056420448 | 0.384 |
| YIL035C | 1690.415511 | -0.421022733 | 0.056653651 | 0.385 |
| YEL003W | 487.0478849 | -0.716250018 | 0.056786342 | 0.385 |
| YDR121W | 548.7039891 | -0.481857453 | 0.056857801 | 0.385 |
| YEL013W | 1708.200753 | 0.510899755 | 0.057345799 | 0.386 |
| YER075C | 845.204575 | 0.460370278 | 0.057107061 | 0.386 |
| YER162C | 931.7095211 | 0.574546812 | 0.05714542 | 0.386 |
| YGL195W | 6234.838225 | -0.442872945 | 0.057355774 | 0.386 |
| YGR275W | 659.6729657 | -0.592953697 | 0.057450279 | 0.386 |
| YML106W | 2946.051592 | -0.540135902 | 0.057438934 | 0.386 |
| YNR064C | 215.5475844 | 0.543612676 | 0.057331517 | 0.386 |
| YPL142C | 228.9766843 | -0.571430834 | 0.057218283 | 0.386 |
| YEL037C | 3155.723536 | -0.525270576 | 0.057585978 | 0.386 |
| YNR016C | 6951.886561 | -0.384345924 | 0.057642612 | 0.386 |
| YKL168C | 972.1957599 | 0.57505355 | 0.057811276 | 0.386 |
| YPL124W | 247.3933842 | -0.576595038 | 0.057798266 | 0.386 |
| YBR244W | 574.3609399 | -0.738546733 | 0.058031685 | 0.387 |
| YGL171W | 777.7803509 | -0.641175423 | 0.058048951 | 0.387 |
| snR34 | 37.83791355 | -0.654410887 | 0.058045952 | 0.387 |
| YHR203C | 5825.930229 | -0.517884803 | 0.058232706 | 0.387 |
| YIL162W | 3263.600398 | 0.79390306 | 0.058391911 | 0.387 |
| YMR123W | 1008.572929 | -0.636812992 | 0.058396101 | 0.387 |
| YOR039W | 1072.032394 | -0.484796931 | 0.058406336 | 0.387 |
| YBR252W | 608.885071 | -0.698004869 | 0.058667155 | 0.389 |
| YBR096W | 835.4279583 | -0.656233458 | 0.059310946 | 0.390 |
| YCL001W | 1214.350549 | -0.519134396 | 0.05928028 | 0.390 |
| YGR054W | 2529.613046 | -0.578725949 | 0.059204691 | 0.390 |
| YIL013C | 704.2538412 | 0.527811357 | 0.059262711 | 0.390 |
| YKL078W | 299.5473506 | -0.749237619 | 0.059218797 | 0.390 |
| YKL180W | 12036.25542 | -0.598498889 | 0.058940729 | 0.390 |
| YOL104C | 616.0912562 | 1.004860107 | 0.059353247 | 0.390 |
| YPR007C | 168.1873056 | 0.776786446 | 0.059061321 | 0.390 |
| YNL195C | 538.1405229 | 0.70423977 | 0.05947662 | 0.390 |
| YDR262W | 1927.726265 | 0.753079022 | 0.059726877 | 0.390 |
| YER056C-A | 4887.303568 | -0.579430507 | 0.059809599 | 0.390 |
| YFL057C | 160.9643278 | 0.48659563 | 0.059747726 | 0.390 |
| YJL033W | 1448.230029 | -0.80597018 | 0.059701481 | 0.390 |
| YMR223W | 516.9298345 | 0.401584051 | 0.059577207 | 0.390 |

| | | | | |
|---|---|---|---|---|
| YBL106C | 800.8986155 | 0.495378666 | 0.060032459 | 0.391 |
| YFL049W | 806.3019635 | 0.424352539 | 0.06030426 | 0.392 |
| YKL099C | 686.8097655 | -0.699691592 | 0.060502523 | 0.393 |
| YPL040C | 952.0787614 | 0.679934536 | 0.060614679 | 0.393 |
| YBL056W | 3616.602261 | -0.36289132 | 0.061165988 | 0.394 |
| YBR002C | 607.1395928 | -0.670025421 | 0.061044085 | 0.394 |
| YBR271W | 387.9475512 | -0.568156518 | 0.060947996 | 0.394 |
| YER012W | 3318.681234 | -0.487138715 | 0.061441568 | 0.394 |
| YGL003C | 856.5289623 | 0.507726738 | 0.061500703 | 0.394 |
| YHR005C-A | 2208.832297 | -0.508714774 | 0.06118255 | 0.394 |
| YIL142W | 3130.184557 | -0.554943714 | 0.060820706 | 0.394 |
| YJL206C | 539.1174838 | 0.510948199 | 0.061441436 | 0.394 |
| YKL055C | 140.4891808 | 0.672754848 | 0.060855429 | 0.394 |
| YLR056W | 3964.313149 | -0.517079888 | 0.061212285 | 0.394 |
| YLR068W | 534.5964698 | -0.818702443 | 0.061466134 | 0.394 |
| YLR453C | 293.5520271 | 0.501869054 | 0.061347697 | 0.394 |
| YOR206W | 2506.527701 | -0.666813748 | 0.061526731 | 0.394 |
| YOL102C | 532.3526812 | -0.404021749 | 0.06170559 | 0.394 |
| YPL184C | 1810.247002 | -0.596608183 | 0.061680081 | 0.394 |
| YPR118W | 1223.179848 | -0.693736676 | 0.062037064 | 0.396 |
| YBR239C | 441.6078035 | 0.473820961 | 0.062338311 | 0.397 |
| YCR034W | 1456.119271 | -0.518964686 | 0.063441556 | 0.397 |
| YDL100C | 5446.231089 | -0.524980742 | 0.063168542 | 0.397 |
| YDR332W | 604.027567 | 0.392564364 | 0.063559025 | 0.397 |
| YER156C | 1503.86508 | -0.592227525 | 0.062375528 | 0.397 |
| YGL035C | 1849.649908 | 0.523141448 | 0.062676382 | 0.397 |
| YGL107C | 935.2339762 | 0.285186835 | 0.063605604 | 0.397 |
| YGR192C | 181337.4741 | -0.486842886 | 0.063210281 | 0.397 |
| YHR039C-A | 2505.228571 | -0.51469513 | 0.06324936 | 0.397 |
| YHR062C | 561.4502383 | -0.596843684 | 0.062443701 | 0.397 |
| YIR031C | 471.7138035 | 0.692779597 | 0.063218561 | 0.397 |
| YJR032W | 585.2468444 | -0.636338842 | 0.062737107 | 0.397 |
| YJR087W | 189.3064917 | -0.661466052 | 0.062924865 | 0.397 |
| YNL300W | 1681.600577 | -0.599240156 | 0.062983802 | 0.397 |
| YNL301C | 1146.542523 | -0.471660213 | 0.063055565 | 0.397 |
| YNR054C | 724.980203 | -0.629093427 | 0.063627442 | 0.397 |
| YOL045W | 1132.896142 | 0.372023638 | 0.062544283 | 0.397 |
| YOL100W | 1483.492555 | 0.621127498 | 0.062560876 | 0.397 |
| YOR165W | 1637.432086 | -0.446244973 | 0.063437282 | 0.397 |
| YOR363C | 1212.557429 | 0.602163915 | 0.063275379 | 0.397 |
| YPL264C | 197.1729292 | 0.496394727 | 0.063603001 | 0.397 |
| YDR183C-A | 47.62983085 | -0.278224686 | 0.063771887 | 0.397 |
| YNL295W | 664.526791 | 0.48565834 | 0.063885137 | 0.398 |

| | | | | |
|---|---|---|---|---|
| YAL025C | 983.4578298 | -0.723243542 | 0.064170375 | 0.398 |
| YLR230W | 287.1818195 | -0.530429746 | 0.064119073 | 0.398 |
| YLR451W | 1124.900564 | 0.460639568 | 0.064240611 | 0.398 |
| YCL005W-A | 247.3743167 | -0.504925485 | 0.064501667 | 0.399 |
| YHR182C-A | 130.5541577 | -0.553498759 | 0.064535225 | 0.399 |
| YLR080W | 639.2701305 | 0.818348822 | 0.064642927 | 0.399 |
| YNL025C | 730.0152463 | 0.508408648 | 0.064630367 | 0.399 |
| YOL147C | 576.9213691 | -0.631821939 | 0.064725937 | 0.399 |
| YPL113C | 241.6280316 | 0.56396184 | 0.064706147 | 0.399 |
| YBR060C | 829.7516444 | 0.416424722 | 0.065006541 | 0.399 |
| YDR522C | 159.2048735 | 0.659033503 | 0.06494209 | 0.399 |
| YJL134W | 603.1655535 | -0.562615549 | 0.065021864 | 0.399 |
| YPL012W | 1836.869245 | -0.673331003 | 0.065042076 | 0.399 |
| YNL249C | 367.0811931 | 0.442149763 | 0.065162842 | 0.400 |
| YAL012W | 6420.264953 | -0.685434014 | 0.065317874 | 0.400 |
| YML009C-A | 52.04514359 | -0.671074713 | 0.065272611 | 0.400 |
| YOL105C | 760.4552602 | 0.61564266 | 0.06541697 | 0.400 |
| YMR146C | 5181.578577 | -0.55859609 | 0.065787449 | 0.401 |
| YMR235C | 2269.617315 | -0.570171508 | 0.065743786 | 0.401 |
| YCR007C | 318.2333147 | 0.570161202 | 0.066050672 | 0.402 |
| YEL063C | 1734.961887 | 0.658047874 | 0.066181625 | 0.402 |
| YJL218W | 146.9624687 | -0.59122801 | 0.065965816 | 0.402 |
| YJR007W | 3429.418011 | -0.547999904 | 0.066315653 | 0.402 |
| YKR074W | 1111.118511 | -0.481615451 | 0.066236997 | 0.402 |
| YLR076C | 95.93637703 | -0.491743295 | 0.06625477 | 0.402 |
| YLR244C | 3158.461036 | -0.642209789 | 0.066212642 | 0.402 |
| YMR137C | 598.7111524 | 0.520537806 | 0.066661384 | 0.402 |
| YOR186W | 27.95959597 | 0.811797536 | 0.066593004 | 0.402 |
| YPL169C | 1689.790258 | -0.463942994 | 0.06660613 | 0.402 |
| YDR471W | 4845.236011 | -0.619638977 | 0.067133934 | 0.404 |
| YIL166C | 439.1240138 | 0.644251132 | 0.067122361 | 0.404 |
| YKL220C | 368.1832027 | 0.691749654 | 0.067053423 | 0.404 |
| YLR065C | 843.2361186 | -0.733635759 | 0.067132294 | 0.404 |
| YHR019C | 9735.850466 | -0.627137235 | 0.067360224 | 0.405 |
| YCL027W | 675.5367401 | 1.003687821 | 0.067447903 | 0.405 |
| YGL042C | 343.2123261 | -0.683546145 | 0.06757269 | 0.405 |
| YGL099W | 1111.20672 | -0.620770256 | 0.067700182 | 0.405 |
| YPL131W | 24700.79965 | -0.635468433 | 0.067727409 | 0.405 |
| YBR154C | 1262.041062 | -0.642549094 | 0.067999555 | 0.406 |
| YJL014W | 2604.88629 | -0.588591896 | 0.06801876 | 0.406 |
| YAR015W | 7474.541729 | -0.416167261 | 0.068526131 | 0.407 |
| YBR162C | 7471.337212 | -0.656026878 | 0.068508366 | 0.407 |
| YDR254W | 574.0903535 | 0.455291788 | 0.068333775 | 0.407 |

| | | | | |
|---|---|---|---|---|
| YMR229C | 3400.401838 | -0.651432169 | 0.06838761 | 0.407 |
| YOL082W | 2476.571718 | 0.588875512 | 0.068511411 | 0.407 |
| YJR143C | 2572.640819 | -0.480249309 | 0.068609589 | 0.407 |
| YLR367W | 2289.066484 | -0.453180969 | 0.068759125 | 0.408 |
| YLR126C | 468.0585451 | -0.655774732 | 0.068906839 | 0.408 |
| YHR052W | 1127.946092 | -0.558199978 | 0.069188737 | 0.409 |
| YNL223W | 533.8185201 | 0.52504957 | 0.069142494 | 0.409 |
| YFL017W-A | 221.0675394 | -0.598661674 | 0.069475861 | 0.409 |
| YHL024W | 910.8922734 | 0.512416177 | 0.069359183 | 0.409 |
| YJR112W | 374.389265 | -0.58215445 | 0.069402132 | 0.409 |
| YMR214W | 1048.466602 | -0.571994752 | 0.069453071 | 0.409 |
| YDL136W | 915.4579307 | -0.7063365 | 0.06968503 | 0.410 |
| YML004C | 3324.450509 | 0.6168542 | 0.069761372 | 0.410 |
| YBR034C | 1111.628509 | -0.638698017 | 0.070274512 | 0.411 |
| YGR111W | 1711.371366 | 0.602132468 | 0.070284212 | 0.411 |
| YHR075C | 548.2435732 | 0.478539256 | 0.070126092 | 0.411 |
| YKL163W | 2089.664663 | 0.818056525 | 0.07003724 | 0.411 |
| YOL013C | 1216.146773 | 0.405358212 | 0.070323595 | 0.411 |
| YPR169W-A | 32.89680766 | -0.735565084 | 0.07027674 | 0.411 |
| YPL273W | 2617.65817 | -0.738294786 | 0.07043621 | 0.411 |
| YGR037C | 2287.491924 | -0.68719573 | 0.07064323 | 0.411 |
| YPR084W | 689.7868138 | 0.307773498 | 0.070685212 | 0.411 |
| YBR171W | 1056.70053 | -0.600109782 | 0.07108553 | 0.412 |
| YDR060W | 3331.110803 | -0.602938609 | 0.070896427 | 0.412 |
| YIL076W | 2351.989968 | -0.545410526 | 0.07101812 | 0.412 |
| YLR441C | 16750.1152 | -0.632212428 | 0.07108429 | 0.412 |
| YML010W | 2999.463402 | -0.436991663 | 0.071136479 | 0.412 |
| YPR017C | 239.3606668 | -0.566004745 | 0.070921397 | 0.412 |
| YHR175W | 775.2689431 | -0.504896952 | 0.071645135 | 0.414 |
| YLR264C-A | 117.4733387 | -0.692624342 | 0.071670962 | 0.414 |
| YBR282W | 512.5901703 | -0.482530824 | 0.071814064 | 0.414 |
| YBL009W | 764.0807255 | -0.498541478 | 0.07190342 | 0.414 |
| YDL226C | 1711.631594 | -0.542782895 | 0.07202112 | 0.414 |
| YGR172C | 884.3306623 | -0.444424907 | 0.072171176 | 0.414 |
| YNL042W | 593.1711026 | 0.597102666 | 0.072094583 | 0.414 |
| YNL131W | 2877.827714 | -0.487818931 | 0.072121414 | 0.414 |
| YPL126W | 2185.981195 | -0.652538655 | 0.072280198 | 0.415 |
| YDL158C | 89.30551169 | -0.62524345 | 0.07242738 | 0.415 |
| YEL028W | 113.7795172 | 0.552420176 | 0.072504055 | 0.415 |
| YGR104C | 540.5492389 | 0.344831006 | 0.072500214 | 0.415 |
| YDR483W | 4802.755253 | -0.367629901 | 0.072641365 | 0.415 |
| YGL131C | 1060.332301 | 0.75821124 | 0.07292554 | 0.416 |
| YGR002C | 912.0190954 | 0.495116474 | 0.072959988 | 0.416 |

| | | | | |
|---|---|---|---|---|
| YLR307C-A | 38.91718273 | 0.510943256 | 0.072827892 | 0.416 |
| YDR023W | 6264.678536 | -0.540285842 | 0.0730547 | 0.416 |
| YDR056C | 1611.517655 | -0.439464579 | 0.073200823 | 0.416 |
| YLR276C | 2051.953288 | -0.598907658 | 0.073311747 | 0.416 |
| YOL032W | 1063.876828 | 0.676511293 | 0.073286482 | 0.416 |
| YNL093W | 262.8484398 | 0.785061912 | 0.073620474 | 0.417 |
| YNL251C | 2085.43933 | 0.714480241 | 0.073696403 | 0.417 |
| YNL303W | 97.83657889 | -0.783998545 | 0.073986854 | 0.419 |
| YDR045C | 241.2309895 | -0.649887452 | 0.07407409 | 0.419 |
| YBR267W | 453.4917639 | -0.680914063 | 0.074192311 | 0.419 |
| YHR108W | 3358.554888 | -0.477708289 | 0.074316834 | 0.419 |
| YNL112W | 1857.470789 | -0.669597507 | 0.074377839 | 0.419 |
| YDR051C | 1050.532227 | 0.389154763 | 0.074595722 | 0.420 |
| YPL129W | 1819.666295 | -0.429534184 | 0.074597373 | 0.420 |
| YLL003W | 928.3757705 | 0.362456578 | 0.074950587 | 0.421 |
| YOL041C | 1321.58671 | -0.66740137 | 0.075010971 | 0.421 |
| YPR061C | 393.0462601 | 0.609234083 | 0.075126081 | 0.421 |
| YDR153C | 1161.100144 | 0.492758134 | 0.075531095 | 0.421 |
| YDR188W | 2974.432832 | -0.472019292 | 0.075988544 | 0.421 |
| YER007W | 271.3998551 | 0.598134606 | 0.075588814 | 0.421 |
| YER031C | 2004.561338 | -0.582041484 | 0.075935702 | 0.421 |
| YGL066W | 1160.200958 | 0.482863682 | 0.075827752 | 0.421 |
| YGL210W | 815.3164128 | -0.454801426 | 0.075538168 | 0.421 |
| YGR171C | 566.9877726 | 0.454859801 | 0.075904278 | 0.421 |
| YHR160C | 281.5696466 | 0.62540973 | 0.075410361 | 0.421 |
| YJR038C | 33.2004855 | 0.809259966 | 0.075839748 | 0.421 |
| YLR179C | 2935.995773 | -0.518680258 | 0.075231578 | 0.421 |
| YMR320W | 113.0593369 | 0.683667701 | 0.075298779 | 0.421 |
| YOL119C | 1253.305663 | 0.642182575 | 0.075888279 | 0.421 |
| YIL002C | 1259.267656 | 0.462483115 | 0.076101042 | 0.422 |
| YGR261C | 1931.303256 | -0.450819357 | 0.076281516 | 0.422 |
| YNL292W | 950.1678617 | -0.544475308 | 0.076373162 | 0.422 |
| YOR035C | 1268.519702 | 0.732045721 | 0.076363734 | 0.422 |
| YDR061W | 1142.905546 | 0.402324288 | 0.076569848 | 0.423 |
| YPR190C | 965.2019513 | -0.555024022 | 0.076621384 | 0.423 |
| YMR091C | 998.5083437 | -0.509741295 | 0.07690889 | 0.424 |
| YGL162W | 395.3975497 | 0.463516912 | 0.077289272 | 0.425 |
| YDR344C | 46.73109222 | -0.671556694 | 0.077562515 | 0.427 |
| YER126C | 1138.972805 | -0.660003227 | 0.077683317 | 0.427 |
| NME1 | 51.73748825 | -0.549957481 | 0.078175554 | 0.427 |
| YBR038W | 1800.515437 | -0.460909187 | 0.079175383 | 0.427 |
| YBR169C | 4813.961053 | 0.924760258 | 0.078026382 | 0.427 |
| YDL050C | 34.4743764 | -0.817537168 | 0.079041346 | 0.427 |

| | | | | |
|---|---|---|---|---|
| YDR087C | 817.7593103 | -0.783794184 | 0.078977266 | 0.427 |
| YDR165W | 1014.540812 | -0.612419037 | 0.078629942 | 0.427 |
| YER150W | 3467.414047 | 0.833529831 | 0.078527781 | 0.427 |
| YER157W | 974.6953246 | 0.396562263 | 0.077853275 | 0.427 |
| YFR025C | 947.4315897 | -0.340931852 | 0.078244548 | 0.427 |
| YGR094W | 6966.605497 | -0.566442481 | 0.079135838 | 0.427 |
| YGR248W | 2366.128573 | 0.874011488 | 0.078386308 | 0.427 |
| YHR202W | 670.4759746 | 0.609682175 | 0.078285698 | 0.427 |
| YJL117W | 1366.561839 | -0.54444627 | 0.078937285 | 0.427 |
| YML001W | 1453.0376 | -0.437839485 | 0.078473215 | 0.427 |
| YML082W | 766.1872079 | -0.511075227 | 0.07817888 | 0.427 |
| YNL207W | 833.7029402 | -0.581830711 | 0.078980931 | 0.427 |
| YNL326C | 366.0651518 | 0.363192423 | 0.079031598 | 0.427 |
| YOL064C | 847.1293742 | -0.612235496 | 0.078222946 | 0.427 |
| YOL121C | 3681.856963 | -0.632990578 | 0.079085988 | 0.427 |
| YPR018W | 922.6100673 | -0.591269404 | 0.078868863 | 0.427 |
| YBL082C | 673.6132807 | -0.559089175 | 0.079542922 | 0.427 |
| YDL140C | 8072.034708 | -0.466526116 | 0.079558043 | 0.427 |
| YDL198C | 1156.426648 | -0.456610126 | 0.07946459 | 0.427 |
| YMR243C | 2759.15272 | -0.524232453 | 0.079603927 | 0.427 |
| YNL141W | 1283.346108 | -0.748100362 | 0.079319748 | 0.427 |
| YLR004C | 257.8975095 | 0.673890713 | 0.079691145 | 0.427 |
| YIL165C | 180.1950386 | 0.656809224 | 0.079768454 | 0.427 |
| YDR425W | 1030.779588 | 0.526140645 | 0.079902691 | 0.428 |
| YFR032C | 47.36854765 | 0.361313859 | 0.079923172 | 0.428 |
| YCR100C | 233.2545617 | 0.757088525 | 0.080111902 | 0.428 |
| YDR097C | 218.1671058 | -0.492925424 | 0.080129727 | 0.428 |
| YOL131W | 75.70252965 | 0.521823548 | 0.080252443 | 0.428 |
| YKL036C | 247.9148773 | 0.701400298 | 0.080467777 | 0.428 |
| YML023C | 695.7405584 | 0.359210466 | 0.080405672 | 0.428 |
| YDR385W | 1851.31364 | -0.634569643 | 0.081119333 | 0.431 |
| YGL153W | 279.9517565 | 0.542663515 | 0.081129868 | 0.431 |
| YDR242W | 619.518258 | 0.551015814 | 0.081402943 | 0.431 |
| YJR035W | 878.6193365 | 0.522937108 | 0.081732071 | 0.431 |
| YKL080W | 5080.565803 | -0.413143636 | 0.081605195 | 0.431 |
| YLR262C-A | 800.3537696 | -0.600330214 | 0.081666195 | 0.431 |
| YLR272C | 941.972981 | -0.421813246 | 0.081632706 | 0.431 |
| YML019W | 694.8842248 | -0.420755947 | 0.081604257 | 0.431 |
| YMR129W | 1909.721365 | -0.372526219 | 0.08160461 | 0.431 |
| YNL125C | 2090.525445 | 0.55490258 | 0.081247725 | 0.431 |
| YDL201W | 813.4402377 | -0.599950547 | 0.082001443 | 0.432 |
| YDR233C | 7397.691198 | -0.512362453 | 0.082097532 | 0.432 |
| YDR312W | 598.3889984 | -0.641984132 | 0.082255296 | 0.432 |

| | | | | |
|---|---|---|---|---|
| YGL105W | 8369.251023 | -0.560584919 | 0.081954238 | 0.432 |
| YHR013C | 1158.208101 | -0.51146455 | 0.082432683 | 0.432 |
| YKL207W | 2006.977142 | -0.487071817 | 0.082376896 | 0.432 |
| YLR243W | 616.8482711 | -0.540746447 | 0.08191579 | 0.432 |
| YNL067W | 1749.269832 | -0.557555588 | 0.082413223 | 0.432 |
| snR30 | 275.4218461 | -0.330787219 | 0.082223167 | 0.432 |
| YDR317W | 303.4699364 | 0.580247654 | 0.082659217 | 0.432 |
| YDL060W | 1265.050122 | -0.57003824 | 0.082959085 | 0.432 |
| YGL044C | 659.773802 | 0.326135867 | 0.082982988 | 0.432 |
| YLR318W | 479.1884801 | 0.448655612 | 0.082778486 | 0.432 |
| YPL123C | 1037.088707 | 0.719439602 | 0.082881088 | 0.432 |
| YBL097W | 438.5923101 | -0.481638699 | 0.083586475 | 0.434 |
| YDR361C | 744.7914399 | -0.654081738 | 0.083665538 | 0.434 |
| YMR155W | 707.9678984 | 0.558233151 | 0.083605875 | 0.434 |
| YMR187C | 357.2988229 | 0.623635839 | 0.083471633 | 0.434 |
| YAL037C-B | 133.1022568 | -0.464244281 | 0.084498509 | 0.435 |
| YBL036C | 1223.509384 | -0.475098735 | 0.084779726 | 0.435 |
| YDR003W-A | 67.78719135 | 0.471585508 | 0.084611489 | 0.435 |
| YDR480W | 507.0349407 | -0.324717552 | 0.084500762 | 0.435 |
| YER023W | 2339.829047 | -0.546254627 | 0.084398274 | 0.435 |
| YGL019W | 1587.591155 | -0.344275637 | 0.084750147 | 0.435 |
| YHR100C | 468.2098929 | -0.454128282 | 0.08443623 | 0.435 |
| YHR128W | 1328.56215 | -0.547078464 | 0.084108776 | 0.435 |
| YJL167W | 5562.553131 | -0.458961949 | 0.084099741 | 0.435 |
| YLR344W | 3310.960859 | -0.674385216 | 0.084296092 | 0.435 |
| YML092C | 2758.585269 | -0.401862825 | 0.08455786 | 0.435 |
| YMR224C | 500.4037572 | 0.396806665 | 0.084460593 | 0.435 |
| YNR032C-A | 183.6670278 | -0.57069955 | 0.084713701 | 0.435 |
| YPL239W | 1334.632299 | -0.620789697 | 0.084755807 | 0.435 |
| YAL018C | 88.41004538 | 0.761564753 | 0.08490053 | 0.435 |
| YDL229W | 7732.532873 | -0.423501202 | 0.085694912 | 0.438 |
| YNL231C | 1567.48489 | -0.417061575 | 0.085726932 | 0.438 |
| YOR247W | 5496.630424 | -0.43103794 | 0.085541766 | 0.438 |
| YOR254C | 3232.98471 | -0.567093834 | 0.08559296 | 0.438 |
| YOL091W | 152.6270626 | 0.412127126 | 0.085802268 | 0.438 |
| YPL267W | 353.8623989 | -0.515678283 | 0.085894304 | 0.438 |
| YCL007C | 102.0392471 | -0.589001773 | 0.085984548 | 0.438 |
| YFL051C | 87.84562174 | 0.873424329 | 0.086115997 | 0.438 |
| YBR014C | 1010.158039 | -0.404512963 | 0.086433644 | 0.439 |
| YBR104W | 350.9392427 | -0.829691216 | 0.086858893 | 0.439 |
| YCR061W | 1066.525845 | 0.717792453 | 0.086580822 | 0.439 |
| YDR034W-B | 127.2105401 | 0.462088596 | 0.086467656 | 0.439 |
| YGR256W | 1242.341744 | 0.969304356 | 0.086848777 | 0.439 |

| | | | | |
|---|---|---|---|---|
| YJL092W | 1243.946562 | -0.441090986 | 0.086829058 | 0.439 |
| YJL161W | 501.2537696 | 0.750682405 | 0.086374078 | 0.439 |
| YJL190C | 9254.873132 | -0.515530521 | 0.086653029 | 0.439 |
| YLL034C | 960.4010793 | -0.71184786 | 0.086504202 | 0.439 |
| YBR030W | 579.6269936 | -0.571898137 | 0.087131071 | 0.440 |
| YDL063C | 706.2977789 | -0.711261807 | 0.087764942 | 0.442 |
| YDL149W | 944.7875532 | 0.597802503 | 0.087641542 | 0.442 |
| YML126C | 4399.479194 | -0.501766472 | 0.08774264 | 0.442 |
| YNL185C | 474.6914722 | -0.464403582 | 0.087918277 | 0.442 |
| YDR338C | 695.8448786 | 0.502763977 | 0.08822843 | 0.443 |
| YLR208W | 2849.872811 | -0.442635886 | 0.088273578 | 0.443 |
| YDR367W | 606.3521729 | -0.468365492 | 0.088456885 | 0.443 |
| YIR012W | 1025.35668 | -0.692428659 | 0.088449702 | 0.443 |
| YKL125W | 592.2613366 | 0.466745922 | 0.088679403 | 0.444 |
| YOR069W | 1035.064237 | 0.363301091 | 0.088986338 | 0.445 |
| YMR072W | 3717.086972 | -0.428023829 | 0.089189156 | 0.446 |
| YBR141C | 286.9036203 | -0.569262965 | 0.089935164 | 0.446 |
| YDL145C | 6294.266606 | -0.40235711 | 0.089778929 | 0.446 |
| YDL148C | 2147.255757 | -0.724539843 | 0.089910902 | 0.446 |
| YDR288W | 555.4180316 | 0.488347196 | 0.08998163 | 0.446 |
| YIL043C | 4110.226673 | -0.449041878 | 0.089429016 | 0.446 |
| YMR122W-A | 5293.670712 | -0.66990401 | 0.089544139 | 0.446 |
| YOR287C | 297.2565187 | -0.678523755 | 0.089604571 | 0.446 |
| YPR026W | 1343.846593 | 0.630569258 | 0.089749622 | 0.446 |
| snR3 | 32.48912912 | -0.170890821 | 0.089545633 | 0.446 |
| YOR120W | 2918.156498 | 0.753125877 | 0.090071461 | 0.446 |
| YAL034W-A | 187.40184 | -0.596657602 | 0.090351138 | 0.447 |
| YHR093W | 35.27687991 | 0.789093018 | 0.090635472 | 0.448 |
| YBR173C | 2339.401983 | -0.456553865 | 0.091311319 | 0.449 |
| YDL154W | 661.9950458 | 0.494244969 | 0.091648063 | 0.449 |
| YDR223W | 358.2338356 | 0.479570693 | 0.091341573 | 0.449 |
| YEL033W | 1403.07415 | -0.620581344 | 0.091746082 | 0.449 |
| YER169W | 1395.485471 | 0.445607562 | 0.091696663 | 0.449 |
| YGL045W | 904.8478281 | 0.600646429 | 0.091111154 | 0.449 |
| YGL133W | 1877.257774 | 0.442305035 | 0.091541368 | 0.449 |
| YGR264C | 2825.266453 | -0.556312847 | 0.090991684 | 0.449 |
| YJL097W | 1713.191405 | -0.51475884 | 0.09116407 | 0.449 |
| YLR034C | 1366.422953 | 0.599901717 | 0.091054106 | 0.449 |
| YML035C | 2451.342109 | 0.567183075 | 0.09150093 | 0.449 |
| YNL273W | 713.7141054 | -0.438975008 | 0.09127011 | 0.449 |
| YPR103W | 3867.741987 | -0.56040929 | 0.091157978 | 0.449 |
| YPR153W | 81.58409703 | 0.588210184 | 0.091590866 | 0.449 |
| YCL029C | 771.878883 | -0.422967204 | 0.091874873 | 0.449 |

| | | | | |
|---|---|---|---|---|
| YDR122W | 2792.779664 | -0.467039865 | 0.092285087 | 0.450 |
| YJL047C | 709.4046862 | 0.507554678 | 0.092159531 | 0.450 |
| YOR019W | 709.5102111 | 0.577134285 | 0.092203218 | 0.450 |
| YPR165W | 4589.151851 | -0.386296873 | 0.092308302 | 0.450 |
| YPL219W | 632.4863798 | 0.451545337 | 0.092429495 | 0.450 |
| YMR279C | 707.3104958 | 0.44057133 | 0.092555918 | 0.450 |
| YPR166C | 431.6523763 | -0.479488001 | 0.092652677 | 0.450 |
| YDR436W | 1255.862426 | 0.533399149 | 0.092750707 | 0.450 |
| YDR226W | 6443.35181 | -0.520127497 | 0.092858469 | 0.450 |
| YCR020C-A | 266.9746924 | -0.4906952 | 0.093412147 | 0.453 |
| YFR037C | 1666.566511 | -0.4532546 | 0.09349607 | 0.453 |
| YGL017W | 502.0325199 | 0.366853043 | 0.093600403 | 0.453 |
| YIL059C | 89.463829 | 0.606946254 | 0.093636411 | 0.453 |
| YNL074C | 2355.063484 | 0.479921583 | 0.094230758 | 0.455 |
| YPL165C | 810.9716846 | 0.510091754 | 0.094201819 | 0.455 |
| YLR292C | 1010.798537 | -0.548631501 | 0.094352406 | 0.455 |
| YPR154W | 2486.651934 | 0.508639019 | 0.094522393 | 0.455 |
| YLR343W | 1249.632226 | 0.68799819 | 0.094701531 | 0.456 |
| YLL059C | 81.93358505 | 0.513614389 | 0.094958467 | 0.457 |
| YLR423C | 531.1083054 | 0.550627434 | 0.095050205 | 0.457 |
| YEL002C | 3293.830024 | -0.36215016 | 0.095214472 | 0.457 |
| YBR008C | 906.2949136 | 0.430874083 | 0.096521723 | 0.457 |
| YBR011C | 18392.14922 | -0.559088258 | 0.095551057 | 0.457 |
| YDR142C | 534.1730573 | 0.533394017 | 0.09573583 | 0.457 |
| YDR523C | 105.2296251 | 0.463081663 | 0.096626115 | 0.457 |
| YER015W | 725.609463 | 0.439893771 | 0.09631454 | 0.457 |
| YGL097W | 1307.008078 | -0.674815254 | 0.096034957 | 0.457 |
| YGR123C | 655.6704403 | -0.467045594 | 0.09554349 | 0.457 |
| YGR199W | 855.4590705 | 0.538171831 | 0.09628885 | 0.457 |
| YGR224W | 693.0347479 | 0.449748686 | 0.096140162 | 0.457 |
| YHR020W | 3665.040866 | -0.610913904 | 0.095909013 | 0.457 |
| YIL069C | 2018.584495 | -0.734094547 | 0.096228698 | 0.457 |
| YLR219W | 2484.562083 | 0.469622139 | 0.096360036 | 0.457 |
| YMR007W | 90.19118303 | 0.353873763 | 0.095747961 | 0.457 |
| YMR059W | 205.0702465 | -0.355676996 | 0.096593909 | 0.457 |
| YNL071W | 6818.92242 | -0.383760981 | 0.096497298 | 0.457 |
| YNR046W | 947.7260883 | -0.659095003 | 0.096277844 | 0.457 |
| YOL149W | 642.6006688 | -0.528854373 | 0.096636933 | 0.457 |
| YOR059C | 1230.819833 | 0.463041793 | 0.095639309 | 0.457 |
| YBR106W | 4289.033597 | -0.541075315 | 0.096882202 | 0.457 |
| YLR333C | 2119.808061 | -0.802297156 | 0.096785471 | 0.457 |
| YOR349W | 572.160655 | 0.446784703 | 0.096847454 | 0.457 |
| YBL071W-A | 297.3219208 | -0.786659408 | 0.097483768 | 0.458 |

| | | | | |
|---|---|---|---|---|
| YCL025C | 3652.132376 | 0.64126587 | 0.097685292 | 0.458 |
| YDR238C | 4318.542325 | -0.374302819 | 0.098581718 | 0.458 |
| YDR341C | 4249.5314 | -0.483500116 | 0.098310789 | 0.458 |
| YHR176W | 373.1735066 | 0.500109774 | 0.097559707 | 0.458 |
| YIR025W | 303.8110514 | 0.391433518 | 0.098105226 | 0.458 |
| YJL100W | 1049.188324 | 0.473900198 | 0.098530099 | 0.458 |
| YJL155C | 1149.698916 | 0.618504263 | 0.098331484 | 0.458 |
| YKR034W | 49.93480154 | 0.682172336 | 0.098269468 | 0.458 |
| YKR039W | 765.789828 | 0.744702238 | 0.098276701 | 0.458 |
| YKR064W | 1141.244604 | 0.412085485 | 0.098218664 | 0.458 |
| YLL021W | 2378.696108 | -0.528278208 | 0.097182989 | 0.458 |
| YLR165C | 405.6673301 | 0.464591613 | 0.097600026 | 0.458 |
| YLR259C | 10469.29817 | -0.350539864 | 0.098066375 | 0.458 |
| YML083C | 177.180785 | 0.621041848 | 0.098446632 | 0.458 |
| YMR258C | 733.537512 | 0.511165569 | 0.098679864 | 0.458 |
| YMR283C | 413.1862547 | 0.406124055 | 0.09859238 | 0.458 |
| YNL159C | 572.5140174 | 0.510905217 | 0.097117985 | 0.458 |
| YNR061C | 596.8560089 | -0.522331289 | 0.097836591 | 0.458 |
| YPL067C | 285.3555196 | -0.642189822 | 0.097881251 | 0.458 |
| YPL144W | 173.5181567 | -0.551029175 | 0.098674572 | 0.458 |
| YPR155C | 883.1515984 | 0.533690384 | 0.097931613 | 0.458 |
| YNL118C | 3319.845542 | 0.372513359 | 0.098764309 | 0.458 |
| YLR171W | 25.60461968 | 0.351579188 | 0.098903891 | 0.458 |
| YPL250W-A | 88.80290187 | -0.721354922 | 0.098990448 | 0.458 |
| YPR036W | 6660.698271 | -0.424759253 | 0.099049732 | 0.458 |
| YBR234C | 4120.481034 | -0.396678996 | 0.099226182 | 0.458 |
| YMR085W | 377.7619403 | 0.384628594 | 0.099163231 | 0.458 |
| YIL160C | 460.4774311 | 0.66843016 | 0.099378009 | 0.458 |
| YDR034C | 1497.715383 | 0.609038913 | 0.099456773 | 0.458 |
| YJR033C | 1583.049905 | 0.446285383 | 0.099708526 | 0.459 |
| YPR131C | 480.7629468 | -0.44928655 | 0.099805379 | 0.459 |
| YJL191W | 879.6997619 | -0.7934035 | 0.099919391 | 0.460 |
| YDL190C | 2381.415921 | 0.389424677 | 0.100257956 | 0.461 |
| YAL001C | 1631.198822 | 0.493140324 | 0.10079042 | 0.462 |
| YNL064C | 7494.727656 | -0.393961797 | 0.100722646 | 0.462 |
| YHR153C | 73.13873436 | -0.382162249 | 0.101200364 | 0.464 |
| YPL037C | 6009.036088 | -0.546868202 | 0.10115348 | 0.464 |
| YDR201W | 235.4443499 | -0.591217096 | 0.101325772 | 0.464 |
| YER140W | 473.4616837 | 0.343334893 | 0.101362594 | 0.464 |
| YMR240C | 592.2234425 | 0.539248725 | 0.10149403 | 0.464 |
| YAR023C | 164.0948317 | 0.488095904 | 0.1016433 | 0.464 |
| YIL014W | 862.4746572 | 0.532762121 | 0.101888239 | 0.465 |
| YJR076C | 1668.359247 | -0.327781116 | 0.101867616 | 0.465 |

| | | | | |
|---|---|---|---|---|
| YMR125W | 2280.761082 | -0.385604839 | 0.101986328 | 0.465 |
| YOR136W | 7814.887983 | -0.407791058 | 0.102091712 | 0.465 |
| YAL007C | 1549.491558 | -0.486125575 | 0.102512852 | 0.465 |
| YCR021C | 10522.50846 | 0.832601694 | 0.102355567 | 0.465 |
| YHR187W | 633.7803075 | -0.384619027 | 0.102404682 | 0.465 |
| YNR051C | 2926.448335 | -0.544858434 | 0.10248353 | 0.465 |
| YJL208C | 631.1153592 | -0.623629652 | 0.102596651 | 0.465 |
| YBL031W | 539.4437355 | -0.576439283 | 0.102911129 | 0.466 |
| YCR066W | 426.4825775 | 0.440086564 | 0.102938539 | 0.466 |
| YHR195W | 1250.379878 | 0.502782347 | 0.102978474 | 0.466 |
| YEL057C | 119.2283707 | 0.713270465 | 0.103218975 | 0.466 |
| YJR055W | 115.4214659 | -0.536840135 | 0.103168686 | 0.466 |
| YMR126C | 404.3389603 | 0.49377243 | 0.103249943 | 0.466 |
| YNL114C | 62.61749004 | -0.815034034 | 0.10334444 | 0.466 |
| YDL150W | 525.2641532 | -0.521582251 | 0.103971088 | 0.466 |
| YDL235C | 741.3905909 | -0.480962026 | 0.104012214 | 0.466 |
| YER098W | 930.5704049 | 0.535744895 | 0.103537755 | 0.466 |
| YJL163C | 815.8405692 | 0.596244386 | 0.103853278 | 0.466 |
| YLR164W | 251.3796178 | 0.552657799 | 0.104050883 | 0.466 |
| YML075C | 1371.306293 | 0.413991341 | 0.104061523 | 0.466 |
| YMR206W | 572.161077 | 0.680581604 | 0.10375247 | 0.466 |
| YOR052C | 2749.089296 | 0.615437904 | 0.103950658 | 0.466 |
| YER145C-A | 50.64924556 | -0.475946973 | 0.104283609 | 0.467 |
| YOL092W | 1398.16665 | -0.491165049 | 0.10435763 | 0.467 |
| YPR074C | 10075.28978 | -0.422773709 | 0.10438744 | 0.467 |
| YKR058W | 2130.69981 | 0.591892397 | 0.104618065 | 0.468 |
| YCL028W | 2867.256977 | -0.423806073 | 0.105105625 | 0.468 |
| YDL191W | 1342.250708 | -0.631706126 | 0.105469792 | 0.468 |
| YDR076W | 307.3025521 | 0.481784266 | 0.105495619 | 0.468 |
| YER053C-A | 179.222376 | 0.873325144 | 0.105747357 | 0.468 |
| YFL056C | 232.3744781 | 0.494374658 | 0.105058948 | 0.468 |
| YHR085W | 335.931842 | -0.622339051 | 0.105423693 | 0.468 |
| YJR075W | 2038.161429 | -0.421295371 | 0.105670689 | 0.468 |
| YKL140W | 1755.744554 | 0.362207758 | 0.105795332 | 0.468 |
| YLR408C | 156.1371802 | -0.509144401 | 0.104943348 | 0.468 |
| YLR452C | 1814.469836 | 0.866723358 | 0.105759572 | 0.468 |
| YML097C | 708.2199714 | -0.42778445 | 0.105320561 | 0.468 |
| YMR165C | 1579.930507 | 0.633615967 | 0.104925732 | 0.468 |
| YNL038W | 473.5438124 | -0.602036953 | 0.105258086 | 0.468 |
| YBR025C | 6377.054239 | -0.575101667 | 0.106044296 | 0.469 |
| YBL003C | 4258.595501 | -0.722067713 | 0.106364224 | 0.469 |
| YBL049W | 377.5575063 | 0.623673618 | 0.106314995 | 0.469 |
| YDR195W | 649.2195359 | 0.369699868 | 0.106330954 | 0.469 |

| | | | | |
|---|---|---|---|---|
| YGR099W | 582.6782539 | -0.403822643 | 0.106721735 | 0.470 |
| YIL002W-A | 495.5817191 | -0.440680476 | 0.106833728 | 0.470 |
| YKL018C-A | 261.9017664 | -0.4339885 | 0.106787174 | 0.470 |
| YIL048W | 1213.90232 | 0.394407807 | 0.107026738 | 0.470 |
| YNL227C | 372.4616162 | -0.485442753 | 0.107029802 | 0.470 |
| YFR048W | 703.4879681 | 0.446738132 | 0.107117299 | 0.470 |
| YOR077W | 225.6560466 | 0.476205159 | 0.107576795 | 0.472 |
| YDR057W | 882.7471612 | 0.433230452 | 0.107851926 | 0.472 |
| YHR170W | 1876.524363 | -0.643731608 | 0.107806635 | 0.472 |
| YMR081C | 3745.685883 | 0.662839009 | 0.107962146 | 0.472 |
| YPL190C | 2551.586572 | -0.443096096 | 0.10795714 | 0.472 |
| YPL189C-A | 341.9617668 | 0.590953915 | 0.108089083 | 0.473 |
| YDR315C | 362.5235632 | -0.468000154 | 0.108203968 | 0.473 |
| YIL037C | 352.6358083 | 0.805140383 | 0.108430434 | 0.473 |
| YKL154W | 669.8187345 | -0.530740993 | 0.108333336 | 0.473 |
| YLR213C | 172.0132438 | 0.568822921 | 0.108382387 | 0.473 |
| YGL163C | 996.8369865 | 0.50245944 | 0.108847697 | 0.474 |
| YJR024C | 696.9415683 | -0.474550997 | 0.108827991 | 0.474 |
| YJR115W | 567.164948 | 0.489720527 | 0.10887365 | 0.474 |
| YDL079C | 1497.675817 | 0.557942837 | 0.109221784 | 0.474 |
| YDL133C-A | 72.78703027 | -1.138750109 | 0.109291882 | 0.474 |
| YDL170W | 672.20876 | 0.468802805 | 0.109334021 | 0.474 |
| YKL033W | 666.8103831 | 0.448155607 | 0.109046247 | 0.474 |
| YMR079W | 1633.407315 | -0.307744681 | 0.109282736 | 0.474 |
| YEL001C | 2451.407888 | -0.490196418 | 0.109428792 | 0.474 |
| YBR117C | 1336.758461 | 0.662746102 | 0.109546283 | 0.474 |
| YJL116C | 947.324176 | 0.589579216 | 0.109676011 | 0.474 |
| YOR020C | 3482.406455 | -0.373544859 | 0.109786352 | 0.474 |
| YKL023C-A | 532.0907152 | -0.481285884 | 0.110426445 | 0.477 |
| YDR273W | 249.707739 | 0.45681547 | 0.110718494 | 0.478 |
| YGL111W | 1214.214479 | -0.514829154 | 0.110761797 | 0.478 |
| YJR052W | 627.3978725 | 0.383237662 | 0.110974725 | 0.478 |
| YML105C | 1352.843987 | -0.414068193 | 0.110984621 | 0.478 |
| YDL202W | 808.8072189 | -0.408146859 | 0.111099731 | 0.478 |
| YDR464W | 2373.668914 | -0.565898482 | 0.111239315 | 0.478 |
| YER091C | 3130.444513 | 0.545785141 | 0.111269472 | 0.478 |
| YPL203W | 1453.358861 | 0.539076258 | 0.111413453 | 0.478 |
| snR17b | 27.76593566 | 0.051183013 | 0.11211843 | 0.481 |
| YBR191W-A | 25.03964351 | -0.373472926 | 0.113518902 | 0.482 |
| YDL143W | 3248.97051 | -0.505029893 | 0.113430305 | 0.482 |
| YDR205W | 692.1640308 | 0.409657554 | 0.113429504 | 0.482 |
| YER097W | 28.63993449 | 0.915545262 | 0.112637884 | 0.482 |
| YGL230C | 33.65395104 | 0.630413594 | 0.113384952 | 0.482 |

| | | | | |
|---|---|---|---|---|
| YGL242C | 665.893361 | -0.577429324 | 0.112749793 | 0.482 |
| YGL262W | 77.76796003 | 0.662277202 | 0.113246211 | 0.482 |
| YGR088W | 3462.726793 | 0.630514458 | 0.112966826 | 0.482 |
| YJR068W | 1198.277592 | -0.496177921 | 0.113424008 | 0.482 |
| YKL052C | 501.3271381 | -0.458852162 | 0.113450863 | 0.482 |
| YLR062C | 194.5995134 | -0.331275456 | 0.113583862 | 0.482 |
| YLR285W | 889.692098 | -0.48415091 | 0.113463219 | 0.482 |
| YMR017W | 57.51133406 | 0.542510627 | 0.113410851 | 0.482 |
| YPL018W | 471.7341201 | 0.464487207 | 0.113272847 | 0.482 |
| YPR016C | 1938.157861 | -0.456057115 | 0.112475988 | 0.482 |
| YFR022W | 849.2747048 | 0.569911046 | 0.11380264 | 0.482 |
| YKL062W | 2465.244192 | 0.49375768 | 0.113784171 | 0.482 |
| YBR275C | 1768.794249 | -0.401743512 | 0.114500812 | 0.483 |
| YGL079W | 448.9297125 | -0.378263838 | 0.114622668 | 0.483 |
| YGR060W | 3114.502526 | -0.57854367 | 0.11435982 | 0.483 |
| YJL103C | 874.1846601 | 0.508609599 | 0.114379834 | 0.483 |
| YJL147C | 286.6419753 | 0.405572323 | 0.114185345 | 0.483 |
| YKL144C | 415.786387 | -0.592949074 | 0.114627301 | 0.483 |
| YNL248C | 1500.899157 | -0.516069417 | 0.114591365 | 0.483 |
| YPL070W | 1079.494033 | 0.459613964 | 0.114380599 | 0.483 |
| YDL239C | 522.0657782 | 0.442194641 | 0.114734482 | 0.483 |
| YDL085W | 757.5796414 | 0.747932135 | 0.114924783 | 0.483 |
| YNL325C | 453.4130958 | 0.541749746 | 0.114861364 | 0.483 |
| YIL146C | 613.496777 | 0.414607975 | 0.115321644 | 0.484 |
| YKL101W | 3903.982421 | -0.469673768 | 0.115322748 | 0.484 |
| YNL022C | 938.4563852 | -0.494352768 | 0.115238267 | 0.484 |
| YOR371C | 940.0880721 | -0.436287555 | 0.115379621 | 0.484 |
| YBR147W | 531.1286909 | 0.681881991 | 0.115507675 | 0.484 |
| YKL135C | 1473.22037 | 0.421591549 | 0.115606086 | 0.484 |
| YHR017W | 1256.836076 | 0.503197304 | 0.115850576 | 0.485 |
| YDL092W | 1065.77792 | -0.456512803 | 0.11604853 | 0.485 |
| YLR269C | 113.3738155 | 0.52121197 | 0.116011667 | 0.485 |
| YPL109C | 614.5552418 | 0.664228679 | 0.116289101 | 0.485 |
| YPR015C | 179.9077993 | 0.66205249 | 0.116321652 | 0.485 |
| YHR174W | 94635.8071 | -0.347781048 | 0.116867896 | 0.487 |
| YDL091C | 1187.448267 | 0.590806027 | 0.117185332 | 0.487 |
| YDL111C | 757.760422 | -0.46282161 | 0.117089534 | 0.487 |
| YJL203W | 345.939173 | -0.353460335 | 0.117133839 | 0.487 |
| YDL182W | 2531.684902 | -0.866858426 | 0.117292579 | 0.487 |
| YNL328C | 68.99382563 | -0.421651835 | 0.117485815 | 0.488 |
| YOR389W | 203.2493311 | 0.673359688 | 0.117444111 | 0.488 |
| YOR207C | 1745.865963 | -0.477937968 | 0.117778539 | 0.488 |
| YML028W | 27162.20889 | -0.444884495 | 0.117863974 | 0.488 |

| | | | | |
|---|---|---|---|---|
| YBL010C | 437.6652565 | -0.351306327 | 0.118353238 | 0.489 |
| YOL010W | 524.4137956 | -0.607241674 | 0.118282729 | 0.489 |
| YPL017C | 762.3634934 | 0.461045299 | 0.118290783 | 0.489 |
| YPL166W | 366.5892241 | 0.50539963 | 0.118105244 | 0.489 |
| YDR354W | 871.1284672 | -0.602334179 | 0.118770182 | 0.490 |
| YHR041C | 584.7828437 | -0.454536037 | 0.11920059 | 0.492 |
| YPL096W | 660.7866918 | 0.373144718 | 0.119145458 | 0.492 |
| YGL020C | 1129.205349 | -0.438816907 | 0.119413462 | 0.492 |
| YLR023C | 1130.122098 | 0.399505101 | 0.11935451 | 0.492 |
| YGR200C | 1330.804216 | -0.439237665 | 0.119771135 | 0.493 |
| YGR135W | 3309.689244 | -0.454363833 | 0.120085383 | 0.493 |
| YJR017C | 953.2894932 | -0.444539532 | 0.119989439 | 0.493 |
| YKL123W | 31.46802918 | 0.231710256 | 0.120122618 | 0.493 |
| YHR037W | 3383.544 | 0.536615937 | 0.120328279 | 0.493 |
| YPL227C | 650.9518323 | -0.505901483 | 0.120331616 | 0.493 |
| YBR116C | 35.41794033 | 0.70855258 | 0.12070171 | 0.494 |
| YGL149W | 44.44871907 | -0.717542347 | 0.12055894 | 0.494 |
| YJL216C | 203.2730397 | 0.523269883 | 0.120480598 | 0.494 |
| YJR071W | 37.86358004 | -0.441983053 | 0.12070873 | 0.494 |
| YGR081C | 443.1057485 | -0.679630071 | 0.120828728 | 0.494 |
| YPL048W | 7812.579467 | -0.388260084 | 0.120931368 | 0.494 |
| YBR079C | 5290.065687 | -0.507139053 | 0.121404905 | 0.494 |
| YDR062W | 2339.714713 | -0.427252118 | 0.121161069 | 0.494 |
| YDR496C | 1668.42234 | -0.504163391 | 0.121543812 | 0.494 |
| YDR507C | 1606.54523 | -0.434418514 | 0.121710136 | 0.494 |
| YGR118W | 4349.282861 | -0.617838491 | 0.121466645 | 0.494 |
| YHR148W | 337.7936716 | -0.666575363 | 0.121769406 | 0.494 |
| YJL060W | 1402.078784 | -0.406962113 | 0.121548627 | 0.494 |
| YKL112W | 917.1897974 | -0.354036295 | 0.121741142 | 0.494 |
| YML034W | 2522.763719 | 0.368909448 | 0.121797379 | 0.494 |
| YBR272C | 606.5333168 | 0.438068266 | 0.122311679 | 0.495 |
| YER054C | 1167.402398 | 0.595083728 | 0.122384589 | 0.495 |
| YIL061C | 381.0352818 | -0.438373082 | 0.122315155 | 0.495 |
| YLR447C | 5880.017489 | -0.364981752 | 0.122136386 | 0.495 |
| YKR096W | 1782.315311 | 0.521043034 | 0.122560946 | 0.495 |
| YLR424W | 404.0091344 | 0.34094958 | 0.122607704 | 0.495 |
| YDL027C | 1559.088359 | 0.538992429 | 0.12284049 | 0.496 |
| SRG1 | 1084.659985 | 0.560270858 | 0.12297475 | 0.496 |
| YDL236W | 1309.137891 | -0.499849434 | 0.123168412 | 0.497 |
| YOR031W | 276.3450169 | 0.634356815 | 0.123385496 | 0.497 |
| YOR135C | 405.7747912 | -0.338701382 | 0.12373223 | 0.498 |
| YML020W | 864.3038212 | 0.381023904 | 0.123860072 | 0.498 |
| YER087W | 372.1981021 | 0.552710054 | 0.1240649 | 0.499 |

| | | | | |
|---|---|---|---|---|
| YGL078C | 1925.6458 | -0.556646222 | 0.12403564 | 0.499 |
| YGL106W | 2381.873892 | -0.437066392 | 0.124157816 | 0.499 |
| YOR108W | 1787.909268 | -0.51994505 | 0.124420472 | 0.499 |
| YKR065C | 1072.834618 | -0.414473305 | 0.124678355 | 0.500 |
| YHR117W | 2228.445201 | -0.399977495 | 0.124893489 | 0.501 |
| YPL028W | 3100.321144 | -0.424324882 | 0.1249884 | 0.501 |
| YHR004C | 921.2706765 | 0.455552584 | 0.125278383 | 0.501 |
| YKL096W-A | 39924.89359 | -0.446278853 | 0.125152852 | 0.501 |
| YMR025W | 229.9295307 | 0.680219384 | 0.12522009 | 0.501 |
| YDR126W | 326.9726421 | 0.517640096 | 0.125628479 | 0.501 |
| YER041W | 647.7988303 | 0.454740792 | 0.125831442 | 0.501 |
| YLR383W | 888.8933966 | -0.362726701 | 0.125944809 | 0.501 |
| YNL232W | 718.0441816 | -0.460009815 | 0.12572353 | 0.501 |
| YPL249C | 1511.229449 | 0.397270595 | 0.125535473 | 0.501 |
| YPR128C | 787.6455183 | -0.384543498 | 0.125938445 | 0.501 |
| YDL095W | 3378.459132 | -0.418832004 | 0.126212171 | 0.502 |
| YFR051C | 3059.620652 | -0.448158416 | 0.126321974 | 0.502 |
| YKL164C | 7734.898504 | -0.52023024 | 0.126146735 | 0.502 |
| YKL190W | 813.5201171 | -0.506236892 | 0.12625591 | 0.502 |
| YIL045W | 1955.748951 | 0.718494803 | 0.126428355 | 0.502 |
| YGL098W | 531.3017666 | 0.474298117 | 0.126623149 | 0.502 |
| YKR017C | 571.9042365 | 0.445458578 | 0.126690342 | 0.502 |
| YMR070W | 1232.669923 | -0.665490819 | 0.126840871 | 0.502 |
| YBR160W | 885.0084099 | -0.443175182 | 0.127729006 | 0.504 |
| YDL216C | 462.7023205 | 0.52076364 | 0.127885145 | 0.504 |
| YDR151C | 1113.580859 | 0.4132865 | 0.127494935 | 0.504 |
| YGL258W-A | 56.62039045 | 0.818376774 | 0.127928162 | 0.504 |
| YGR041W | 794.2535473 | 0.404289059 | 0.127873526 | 0.504 |
| YIL058W | 79.78173189 | 0.518195681 | 0.127674935 | 0.504 |
| YKL093W | 994.1827608 | 0.506456944 | 0.127644354 | 0.504 |
| YLR053C | 106.3162181 | 0.691173612 | 0.12754107 | 0.504 |
| YBR029C | 1375.442278 | -0.447049453 | 0.128785182 | 0.504 |
| YDL195W | 7573.167842 | -0.359784605 | 0.129100622 | 0.504 |
| YDR009W | 246.880862 | 0.68328544 | 0.129214019 | 0.504 |
| YDR417C | 435.6659374 | -0.778151755 | 0.129076714 | 0.504 |
| YER035W | 1131.143378 | 0.607869895 | 0.129213398 | 0.504 |
| YFR017C | 3258.918594 | 0.619131186 | 0.128422266 | 0.504 |
| YJL164C | 1399.537088 | 0.510254968 | 0.129235524 | 0.504 |
| YLR173W | 824.1208493 | 0.561923343 | 0.12871084 | 0.504 |
| YNL142W | 1026.084342 | 0.821781852 | 0.128766414 | 0.504 |
| YOR267C | 3236.118357 | 0.487362811 | 0.128739762 | 0.504 |
| YPL260W | 2621.937685 | 0.473158585 | 0.128700593 | 0.504 |
| YPR078C | 88.79558885 | 0.471949698 | 0.128834434 | 0.504 |

| | | | | |
|---|---|---|---|---|
| YPR121W | 333.11981 | 0.478402098 | 0.128804748 | 0.504 |
| YPR182W | 302.4381877 | -0.441989904 | 0.128934778 | 0.504 |
| YMR314W | 2980.188475 | -0.343591 | 0.129433171 | 0.505 |
| YCL001W-A | 26.77980078 | 0.920473352 | 0.130043194 | 0.506 |
| YJR010W | 677.4384961 | 0.446441344 | 0.130011203 | 0.506 |
| YKL211C | 1647.230083 | -0.570138906 | 0.130017093 | 0.506 |
| YBR080C | 2198.347068 | -0.389129855 | 0.130209614 | 0.506 |
| YPR080W | 5311.40989 | -0.432033441 | 0.130254019 | 0.506 |
| YHR080C | 2045.316793 | 0.431039577 | 0.130571038 | 0.507 |
| YBR003W | 1468.050788 | 0.385483395 | 0.130969835 | 0.508 |
| YDL089W | 963.5832446 | 0.612574063 | 0.132000408 | 0.508 |
| YDR463W | 1963.287278 | 0.42827202 | 0.131417883 | 0.508 |
| YFL021W | 926.9442787 | 0.486847207 | 0.130896053 | 0.508 |
| YGR279C | 11712.04252 | -0.285234307 | 0.132170679 | 0.508 |
| YIL094C | 3348.951104 | -0.439662178 | 0.131686723 | 0.508 |
| YKR012C | 142.5434976 | -0.518738787 | 0.132089298 | 0.508 |
| YKR041W | 32.67705338 | -0.54276596 | 0.131922768 | 0.508 |
| YLR100W | 1453.024103 | -0.499430103 | 0.131996851 | 0.508 |
| YLR339C | 501.9621213 | -0.663181212 | 0.131729008 | 0.508 |
| YLR422W | 2135.341687 | 0.443054003 | 0.130787157 | 0.508 |
| YML074C | 7957.336564 | -0.637162143 | 0.131053519 | 0.508 |
| YML093W | 1683.648354 | -0.608511633 | 0.13198741 | 0.508 |
| YMR198W | 512.8655409 | 0.4144632 | 0.131983838 | 0.508 |
| YNL233W | 992.805205 | -0.327349026 | 0.131878696 | 0.508 |
| YOL095C | 382.7305205 | 0.422816277 | 0.131863469 | 0.508 |
| YOR237W | 107.6435638 | 0.461821846 | 0.132122155 | 0.508 |
| YBR131W | 285.0666359 | -0.485491096 | 0.132401027 | 0.508 |
| YOR058C | 1144.395841 | 0.435286689 | 0.132442258 | 0.508 |
| YMR225C | 419.9415237 | -0.435330733 | 0.132795904 | 0.509 |
| YPL005W | 369.5032375 | 0.492060836 | 0.132767306 | 0.509 |
| YHR152W | 1107.927047 | -0.356175436 | 0.133049016 | 0.509 |
| YCR057C | 1417.95789 | -0.528112888 | 0.133310615 | 0.510 |
| YGR146C | 1344.611896 | 0.416593932 | 0.133309726 | 0.510 |
| YKL077W | 3042.957164 | -0.341304336 | 0.133373561 | 0.510 |
| YHR043C | 1297.179518 | -0.428016655 | 0.133584066 | 0.510 |
| TLC1 | 1417.425959 | -0.255189861 | 0.133852415 | 0.511 |
| YNL259C | 262.1284302 | -0.46103913 | 0.134036486 | 0.511 |
| YDR255C | 870.1597032 | 0.589733744 | 0.134986724 | 0.512 |
| YEL034C-A | 2748.462234 | -0.459155026 | 0.134712098 | 0.512 |
| YGL005C | 924.1279481 | 0.456288342 | 0.134867414 | 0.512 |
| YHR191C | 210.2719774 | 0.314395679 | 0.134954649 | 0.512 |
| YIL020C-A | 191.6015626 | -0.494387648 | 0.134686327 | 0.512 |
| YML098W | 504.9087228 | -0.471852 | 0.134604189 | 0.512 |

| | | | | |
|---|---|---|---|---|
| YNL267W | 1939.373479 | 0.305167094 | 0.134544186 | 0.512 |
| YPL100W | 911.5930892 | 0.389679323 | 0.134783501 | 0.512 |
| YPL159C | 943.1509361 | 0.508955417 | 0.134538542 | 0.512 |
| YHL034C | 6448.755806 | -0.333083227 | 0.135224905 | 0.512 |
| YOR323C | 2339.246441 | -0.415249812 | 0.135320877 | 0.512 |
| YER027C | 1561.298312 | -0.34571151 | 0.135612376 | 0.513 |
| YDR306C | 1159.558737 | 0.379790717 | 0.135786216 | 0.513 |
| YPR091C | 1815.236332 | 0.369750259 | 0.136279833 | 0.515 |
| YBL048W | 125.8109055 | 0.537328427 | 0.136664362 | 0.515 |
| YDL211C | 175.1726895 | -0.607728765 | 0.137183421 | 0.515 |
| YGR144W | 97.38216331 | 0.577288521 | 0.136709446 | 0.515 |
| YGR173W | 798.4212433 | -0.640873896 | 0.136975811 | 0.515 |
| YGR219W | 32.06784529 | -0.666544158 | 0.137116292 | 0.515 |
| YLR098C | 706.1029207 | 0.456302806 | 0.137073147 | 0.515 |
| YLR342W-A | 51.47657195 | 0.287431739 | 0.136538282 | 0.515 |
| YML078W | 2283.677343 | -0.427774496 | 0.136781469 | 0.515 |
| YMR300C | 1920.32889 | -0.452119433 | 0.137187734 | 0.515 |
| YOR245C | 966.5341375 | 0.527412738 | 0.137126481 | 0.515 |
| YGR053C | 339.3237027 | 0.446022588 | 0.13732177 | 0.515 |
| YIL041W | 4311.545468 | -0.426040399 | 0.137575946 | 0.515 |
| YJR039W | 935.4373703 | 0.458752635 | 0.137402328 | 0.515 |
| YLR102C | 476.3043851 | 0.51944674 | 0.137538158 | 0.515 |
| YNR055C | 1151.83484 | 0.415101504 | 0.137706108 | 0.515 |
| YCR052W | 1334.780982 | -0.379617962 | 0.138036873 | 0.516 |
| YDR351W | 1313.626306 | 0.411605615 | 0.137985244 | 0.516 |
| YFL001W | 478.2892235 | -0.446055232 | 0.138146012 | 0.516 |
| YFL005W | 2091.331245 | -0.420867004 | 0.137927715 | 0.516 |
| YFL055W | 265.1999324 | 0.725640379 | 0.138198611 | 0.516 |
| YER149C | 245.5366344 | -0.431782658 | 0.138803538 | 0.516 |
| YJL008C | 2286.818099 | -0.432116537 | 0.138770946 | 0.516 |
| YJL111W | 2828.454028 | -0.483218311 | 0.138845247 | 0.516 |
| YMR271C | 754.1723733 | 0.736643348 | 0.138843766 | 0.516 |
| YNL037C | 5393.729596 | -0.458848569 | 0.138677645 | 0.516 |
| YBR253W | 396.1883397 | -0.38983428 | 0.139548681 | 0.517 |
| YDL232W | 292.6096052 | -0.503001885 | 0.13960667 | 0.517 |
| YGR120C | 376.5121986 | -0.213579072 | 0.139513197 | 0.517 |
| YLR083C | 1368.675892 | -0.30387384 | 0.139114624 | 0.517 |
| YNL126W | 705.9981899 | -0.395639809 | 0.139277486 | 0.517 |
| YNL170W | 85.88966089 | -0.611667335 | 0.139411265 | 0.517 |
| YOR123C | 2187.626634 | -0.27660876 | 0.13953678 | 0.517 |
| YOR243C | 1479.808571 | -0.293076375 | 0.139426132 | 0.517 |
| YAR020C | 43.79066228 | 0.51954131 | 0.13993878 | 0.517 |
| YDR469W | 524.7044195 | -0.515682568 | 0.139862894 | 0.517 |

| | | | | |
|---|---|---|---|---|
| YGR158C | 631.124363 | -0.564055646 | 0.14020154 | 0.517 |
| YGR187C | 552.0431833 | -0.43179347 | 0.139886849 | 0.517 |
| YIL170W | 172.5743318 | 0.694095586 | 0.140196972 | 0.517 |
| YLR361C | 559.6543534 | 0.402003366 | 0.140174663 | 0.517 |
| YER045C | 814.7735686 | 0.472587031 | 0.140355819 | 0.517 |
| YHR081W | 846.2639082 | -0.597145799 | 0.140435116 | 0.517 |
| YBR294W | 590.8243932 | 0.502629485 | 0.140571048 | 0.517 |
| YEL038W | 943.0798183 | -0.42842617 | 0.140996268 | 0.518 |
| YJR118C | 868.4456577 | -0.367919256 | 0.14092644 | 0.518 |
| YMR312W | 719.0027121 | -0.459345263 | 0.141081819 | 0.518 |
| YOR090C | 1602.791656 | 0.32806882 | 0.141039597 | 0.518 |
| YGL060W | 769.6330407 | 0.420792314 | 0.141389519 | 0.518 |
| YLR409C | 1307.908763 | -0.480476362 | 0.141356813 | 0.518 |
| YIL060W | 54.27024793 | 0.648162183 | 0.141671826 | 0.519 |
| YPL110C | 931.3902179 | 0.474665786 | 0.141626009 | 0.519 |
| YER179W | 293.0579118 | 0.560853659 | 0.142190779 | 0.519 |
| YGR071C | 709.1837053 | 0.368438148 | 0.142111625 | 0.519 |
| YGR181W | 835.8007421 | -0.400522817 | 0.142094017 | 0.519 |
| YJR149W | 627.5970928 | 0.529627164 | 0.142093001 | 0.519 |
| YOR018W | 939.5138966 | 0.313719498 | 0.142132008 | 0.519 |
| YLR046C | 345.7378309 | 0.267735211 | 0.14231401 | 0.519 |
| YDL248W | 224.0356228 | 0.749660908 | 0.142461419 | 0.519 |
| YGR211W | 3320.437529 | -0.518453492 | 0.142737159 | 0.519 |
| YJR046W | 1243.286799 | -0.333682379 | 0.142652552 | 0.519 |
| YJR101W | 534.1436387 | -0.374457717 | 0.142613132 | 0.519 |
| YER158C | 1028.126375 | 0.603517216 | 0.142880591 | 0.519 |
| YDR066C | 173.6197188 | 0.548440738 | 0.143235659 | 0.520 |
| YGL173C | 8071.132916 | -0.276896859 | 0.143198155 | 0.520 |
| YLR186W | 830.8214037 | -0.465964927 | 0.1431236 | 0.520 |
| YDR313C | 770.3934735 | 0.502883613 | 0.143624507 | 0.520 |
| YGR281W | 2050.830805 | 0.404352293 | 0.14356604 | 0.520 |
| YHR183W | 10217.98699 | -0.422799163 | 0.143685036 | 0.520 |
| YDR277C | 2772.925083 | 0.4293065 | 0.144092789 | 0.521 |
| YKL050C | 526.5764301 | 0.361565185 | 0.144079821 | 0.521 |
| YLR092W | 690.3329885 | 0.487116859 | 0.144150005 | 0.521 |
| YOR387C | 76.98828773 | 0.620180676 | 0.144109365 | 0.521 |
| YPR111W | 665.5579675 | 0.496362591 | 0.143818103 | 0.521 |
| YHR060W | 732.5771452 | -0.437647347 | 0.144357687 | 0.521 |
| YDR358W | 1790.698239 | 0.514653706 | 0.145101421 | 0.523 |
| YFR023W | 96.03552642 | 0.678212534 | 0.14495005 | 0.523 |
| YHR163W | 2218.862273 | -0.564466286 | 0.145093484 | 0.523 |
| YPR175W | 704.7198488 | -0.422162214 | 0.145155271 | 0.523 |
| YGR128C | 1414.526792 | -0.400659743 | 0.145314167 | 0.523 |

| | | | | |
|---|---|---|---|---|
| YPL230W | 2214.598115 | 0.471526335 | 0.145344903 | 0.523 |
| YAR031W | 151.5423779 | 0.533050923 | 0.145536277 | 0.523 |
| YLR440C | 889.0207433 | 0.558137718 | 0.145485726 | 0.523 |
| YKR078W | 765.3811944 | 0.392710978 | 0.145701999 | 0.523 |
| YJR152W | 334.6737416 | 0.451737138 | 0.145827768 | 0.523 |
| YGL227W | 1564.861142 | 0.539732394 | 0.145940243 | 0.523 |
| YAR027W | 1481.56887 | 0.394653026 | 0.146546194 | 0.523 |
| YGR161C-D | 41.89517477 | 0.753370023 | 0.146452635 | 0.523 |
| YJL204C | 683.8445536 | 0.407411574 | 0.14653375 | 0.523 |
| YML064C | 587.9701312 | 0.294708522 | 0.146339828 | 0.523 |
| YMR118C | 295.0484477 | 0.370344852 | 0.146573787 | 0.523 |
| YMR169C | 812.630576 | 0.645667002 | 0.146606074 | 0.523 |
| YPL151C | 670.1565001 | 0.27786135 | 0.146550639 | 0.523 |
| YBR134W | 28.00305569 | -0.906931099 | 0.146739514 | 0.523 |
| YJR008W | 1161.824512 | 0.550963829 | 0.147062663 | 0.524 |
| YER183C | 378.9572978 | -0.392685197 | 0.147192148 | 0.524 |
| YKL017C | 627.7815414 | 0.423756966 | 0.147550323 | 0.524 |
| YLR312C | 417.3618401 | 0.610225374 | 0.14741839 | 0.524 |
| YML002W | 853.1962719 | 0.384666239 | 0.147542699 | 0.524 |
| YPR021C | 1682.395818 | 0.490113956 | 0.147441159 | 0.524 |
| YFL020C | 87.69189529 | 0.438311112 | 0.147770339 | 0.525 |
| YNL044W | 495.5352853 | -0.465044196 | 0.148074704 | 0.526 |
| YIL007C | 541.263808 | 0.437457758 | 0.14816539 | 0.526 |
| LSR1 | 186.1998782 | -0.094824792 | 0.148429484 | 0.526 |
| YPR081C | 985.9306388 | 0.491240147 | 0.148450243 | 0.526 |
| YDL193W | 1212.895326 | 0.347201146 | 0.148741452 | 0.527 |
| YOR025W | 737.1839475 | -0.493894291 | 0.148796888 | 0.527 |
| YEL049W | 54.33605347 | 0.4713487 | 0.149041924 | 0.527 |
| YBR279W | 1844.253736 | -0.337525351 | 0.14951926 | 0.528 |
| YHR122W | 579.3180386 | -0.442092275 | 0.149826925 | 0.528 |
| YKL181W | 2705.955695 | -0.489661533 | 0.149329931 | 0.528 |
| YLR205C | 412.8411079 | 0.344245134 | 0.149797368 | 0.528 |
| YNL079C | 8141.583459 | -0.484793419 | 0.149472209 | 0.528 |
| YNR044W | 3311.34449 | 0.861247034 | 0.149697318 | 0.528 |
| YPR134W | 542.0304792 | 0.368006116 | 0.149726155 | 0.528 |
| YEL004W | 410.7288824 | 0.466552443 | 0.149939765 | 0.528 |
| YPL075W | 2226.650529 | 0.461344928 | 0.150081825 | 0.528 |
| YCR091W | 1488.499833 | 0.533227429 | 0.150202414 | 0.528 |
| YLR074C | 1095.319455 | -0.648109737 | 0.150323518 | 0.528 |
| YGL040C | 1261.243607 | -0.325784928 | 0.150771907 | 0.529 |
| YJL129C | 2269.469864 | 0.309323004 | 0.150696473 | 0.529 |
| YMR031C | 3707.880046 | 0.433264537 | 0.150540839 | 0.529 |
| YPR060C | 619.4282416 | -0.55029109 | 0.150769208 | 0.529 |

| | | | | |
|---|---|---|---|---|
| YEL048C | 96.72112286 | -0.830357503 | 0.151214392 | 0.529 |
| YER084W | 124.9336603 | 0.480845838 | 0.151024678 | 0.529 |
| YHR180W | 212.5255364 | 0.570654264 | 0.151350348 | 0.529 |
| YMR098C | 840.7673515 | 0.531523618 | 0.151205634 | 0.529 |
| YNL107W | 513.6532475 | -0.317924687 | 0.151458347 | 0.529 |
| YPL056C | 172.8020928 | 0.412420796 | 0.151432392 | 0.529 |
| YMR162C | 1643.610674 | 0.496306533 | 0.151558561 | 0.529 |
| YNL072W | 456.0809998 | -0.384271464 | 0.151813897 | 0.529 |
| YPL026C | 1074.82514 | 0.57934733 | 0.151828683 | 0.529 |
| YJL120W | 37.94521827 | -0.644348554 | 0.151989632 | 0.530 |
| YGL012W | 2257.199778 | -0.246192012 | 0.152443324 | 0.531 |
| YMR270C | 708.6444493 | -0.409316435 | 0.152495446 | 0.531 |
| YCR018C | 861.2432726 | 0.520210709 | 0.153574514 | 0.533 |
| YDL024C | 228.1924469 | 0.80972462 | 0.153961136 | 0.533 |
| YDL131W | 1846.278027 | -0.502437547 | 0.153213931 | 0.533 |
| YDR324C | 1092.059972 | -0.455671275 | 0.153895998 | 0.533 |
| YGL090W | 485.8897575 | 0.338602463 | 0.153974158 | 0.533 |
| YGL125W | 958.6893776 | 0.424983421 | 0.154006669 | 0.533 |
| YKR004C | 281.3110563 | 0.405755114 | 0.153797728 | 0.533 |
| YKR029C | 884.4269483 | 0.379445068 | 0.153907855 | 0.533 |
| YMR018W | 66.38637523 | 0.632314422 | 0.153428693 | 0.533 |
| YMR194C-B | 137.2124333 | 0.69088097 | 0.154157951 | 0.533 |
| YNR031C | 1352.088361 | -0.367405968 | 0.153634552 | 0.533 |
| YOR177C | 142.866635 | 0.813949865 | 0.154100015 | 0.533 |
| YER178W | 9418.417677 | -0.32457216 | 0.154493372 | 0.533 |
| YGR078C | 839.5108181 | -0.461376155 | 0.154508172 | 0.533 |
| YIL097W | 1123.344615 | 0.518360996 | 0.154613226 | 0.533 |
| YJL118W | 178.5211681 | -0.529298358 | 0.15469899 | 0.533 |
| YLR216C | 5399.303942 | -0.376484539 | 0.154935815 | 0.533 |
| YNL153C | 847.6509295 | -0.524769374 | 0.154817597 | 0.533 |
| YNL299W | 424.5188557 | -0.589549969 | 0.154924078 | 0.533 |
| YOL009C | 380.9976744 | 0.383372092 | 0.154758255 | 0.533 |
| YDR119W-A | 458.2326718 | 0.691356086 | 0.155303169 | 0.534 |
| YDR197W | 858.0497536 | 0.448737276 | 0.155249675 | 0.534 |
| YDL040C | 3238.410502 | -0.301655283 | 0.155489214 | 0.534 |
| YDL214C | 464.7760152 | 0.284490384 | 0.155415562 | 0.534 |
| YGR095C | 969.1047325 | -0.485879087 | 0.156895765 | 0.535 |
| YHL018W | 70.12879735 | 0.435813852 | 0.156615 | 0.535 |
| YIL046W | 1246.958862 | 0.325333302 | 0.156112909 | 0.535 |
| YKL160W | 1725.224125 | -0.404195691 | 0.156649095 | 0.535 |
| YKR002W | 1434.178622 | 0.387236716 | 0.156563498 | 0.535 |
| YKR076W | 1274.281879 | 0.676160056 | 0.156581527 | 0.535 |
| YLR287C | 617.6178514 | -0.375261735 | 0.156939814 | 0.535 |

| | | | | |
|---|---|---|---|---|
| YOR089C | 2098.945136 | 0.412816231 | 0.15618697 | 0.535 |
| YOR159C | 320.0573697 | -0.472094411 | 0.156832566 | 0.535 |
| YPL093W | 1935.182804 | -0.482234358 | 0.156362542 | 0.535 |
| YPL150W | 1067.29309 | 0.387296358 | 0.156960064 | 0.535 |
| YDR101C | 1247.120699 | -0.614156957 | 0.15709384 | 0.535 |
| YPR196W | 243.4727794 | 0.538792608 | 0.157233558 | 0.535 |
| YDL142C | 615.3529618 | 0.392535658 | 0.157791864 | 0.536 |
| YIL055C | 666.1215044 | 0.356450224 | 0.157681632 | 0.536 |
| YJL201W | 1030.90288 | 0.311460814 | 0.157510589 | 0.536 |
| YML117W | 3482.921508 | 0.398153735 | 0.157488003 | 0.536 |
| YML125C | 2139.072841 | -0.568390796 | 0.157797071 | 0.536 |
| YPL003W | 547.4702758 | 0.396903329 | 0.157629525 | 0.536 |
| YIL017C | 1438.735299 | 0.462806261 | 0.158078369 | 0.536 |
| YJR015W | 1958.669387 | -0.409320266 | 0.158082933 | 0.536 |
| YJR141W | 396.4509931 | -0.444408217 | 0.158198396 | 0.536 |
| YLR125W | 49.15160341 | 0.511316828 | 0.158255594 | 0.536 |
| YOL083W | 724.7720388 | 0.595936754 | 0.157999573 | 0.536 |
| YBR072W | 9138.655064 | 1.174571227 | 0.158479846 | 0.536 |
| YCR001W | 33.60447039 | 0.730042 | 0.158686376 | 0.536 |
| YHR181W | 756.8304441 | -0.377789993 | 0.158783662 | 0.536 |
| YBR085W | 162.0814879 | -0.696332443 | 0.159086737 | 0.537 |
| YBR250W | 80.81966717 | 0.632292867 | 0.15944541 | 0.537 |
| YCR010C | 273.3869127 | 0.577127873 | 0.159452082 | 0.537 |
| YDR390C | 1683.184066 | -0.305374873 | 0.15940092 | 0.537 |
| YJL041W | 2583.756357 | -0.305757935 | 0.159414378 | 0.537 |
| YDL139C | 356.8731646 | 0.303044174 | 0.160271741 | 0.537 |
| YDR446W | 60.61919013 | 0.246404935 | 0.159739211 | 0.537 |
| YDR458C | 1309.381626 | 0.405382371 | 0.159767576 | 0.537 |
| YFL003C | 165.623347 | 0.561181897 | 0.160197728 | 0.537 |
| YGL027C | 1397.054052 | -0.333502424 | 0.160302473 | 0.537 |
| YJL127C | 657.9035431 | 0.315847194 | 0.160204731 | 0.537 |
| YKR095W-A | 172.6357846 | -0.605394179 | 0.160261642 | 0.537 |
| YML017W | 751.7187367 | 0.481177022 | 0.15986309 | 0.537 |
| YNL194C | 937.042565 | 0.495683141 | 0.160211689 | 0.537 |
| YCR059C | 1130.721478 | -0.489652565 | 0.160754879 | 0.538 |
| YFL054C | 1511.844206 | 0.527745183 | 0.160601198 | 0.538 |
| YLR022C | 541.692103 | -0.451658247 | 0.160687683 | 0.538 |
| YMR201C | 569.2749285 | 0.434446034 | 0.160852939 | 0.538 |
| YOR305W | 732.3331193 | -0.511411447 | 0.16091645 | 0.538 |
| YDR109C | 536.0726963 | 0.27374339 | 0.161251571 | 0.538 |
| YJL149W | 1033.010855 | 0.323979054 | 0.161174069 | 0.538 |
| YOR061W | 1662.759146 | -0.37478468 | 0.16113551 | 0.538 |
| YDR104C | 711.7535955 | 0.434881653 | 0.161388079 | 0.538 |

| | | | | |
|---|---|---|---|---|
| YLR037C | 167.7921303 | 0.345957228 | 0.161438479 | 0.538 |
| YMR031W-A | 40.58383151 | 0.573390974 | 0.161562384 | 0.538 |
| YEL031W | 4279.639928 | -0.368959883 | 0.162103839 | 0.539 |
| YJR064W | 2718.261132 | -0.406956613 | 0.162183975 | 0.539 |
| YKL182W | 10979.47839 | -0.255025172 | 0.16208844 | 0.539 |
| YKL119C | 426.0044347 | -0.439081749 | 0.162326729 | 0.539 |
| YGL206C | 5144.593261 | 0.402006159 | 0.162520207 | 0.540 |
| YJL187C | 1084.409994 | -0.388887505 | 0.162763072 | 0.540 |
| YMR148W | 862.6307015 | 0.42100027 | 0.162796605 | 0.540 |
| YOR080W | 677.2163658 | 0.254983065 | 0.16270331 | 0.540 |
| YOL005C | 1554.462517 | -0.553941744 | 0.163142634 | 0.541 |
| YHL021C | 7761.479525 | 0.571372968 | 0.163428505 | 0.541 |
| YNL331C | 347.7391832 | 0.418465915 | 0.16343108 | 0.541 |
| YBR246W | 994.4891912 | -0.328051566 | 0.163758825 | 0.541 |
| YML009C | 743.0693468 | -0.355022396 | 0.163591779 | 0.541 |
| YNL091W | 4224.416881 | -0.388483956 | 0.163668449 | 0.541 |
| YDR395W | 2175.578678 | -0.404401519 | 0.164551581 | 0.541 |
| YER078C | 919.3267409 | 0.349129139 | 0.16403195 | 0.541 |
| YGR061C | 5651.499061 | -0.292869515 | 0.16418646 | 0.541 |
| YKR001C | 4012.227533 | -0.323624582 | 0.163958607 | 0.541 |
| YMR161W | 724.2845416 | -0.328327553 | 0.164271053 | 0.541 |
| YOR092W | 648.6032701 | 0.393834179 | 0.16441566 | 0.541 |
| YPR082C | 431.92377 | -0.445442301 | 0.164515218 | 0.541 |
| YPR137W | 608.8911311 | -0.507771824 | 0.164403115 | 0.541 |
| YER063W | 1431.45469 | -0.309902791 | 0.164663155 | 0.541 |
| YJR077C | 4794.775519 | -0.297530309 | 0.164800732 | 0.541 |
| YMR301C | 1249.636815 | -0.378043634 | 0.164838379 | 0.541 |
| YBR200W | 1077.368125 | -0.308982494 | 0.165702105 | 0.542 |
| YGL061C | 446.8633151 | -0.377495038 | 0.165364984 | 0.542 |
| YHR069C | 771.2357851 | -0.486924532 | 0.165211789 | 0.542 |
| YIR022W | 1186.448374 | -0.381676399 | 0.165700001 | 0.542 |
| YJL199C | 163.0335414 | 0.294529674 | 0.165379632 | 0.542 |
| YMR043W | 2143.509763 | 0.42266012 | 0.165393668 | 0.542 |
| YMR087W | 259.2299237 | 0.391304931 | 0.16566991 | 0.542 |
| YMR317W | 752.1354442 | 0.457097461 | 0.165687801 | 0.542 |
| YBL023C | 762.982123 | -0.376159961 | 0.166314332 | 0.542 |
| YDL164C | 704.8057939 | -0.324939752 | 0.167971053 | 0.542 |
| YDR200C | 868.5642679 | 0.295177362 | 0.166611078 | 0.542 |
| YDR202C | 884.8533036 | 0.446108999 | 0.166369065 | 0.542 |
| YDR225W | 3245.605018 | -0.626488621 | 0.166710994 | 0.542 |
| YDR241W | 32.77525977 | 0.426241872 | 0.167408613 | 0.542 |
| YDR422C | 1006.650102 | 0.397125954 | 0.167446993 | 0.542 |
| YGR180C | 8240.520071 | -0.440406727 | 0.16678733 | 0.542 |

| | | | | |
|---|---|---|---|---|
| YHR138C | 1476.126547 | 0.33873596 | 0.167517522 | 0.542 |
| YIL050W | 1083.890647 | 0.388566471 | 0.166237338 | 0.542 |
| YKL063C | 828.820505 | -0.40377787 | 0.167843507 | 0.542 |
| YKL068W-A | 172.0547206 | -0.52358451 | 0.166378323 | 0.542 |
| YKL129C | 2233.401318 | 0.38475344 | 0.167997453 | 0.542 |
| YLR336C | 993.7017255 | -0.600035216 | 0.167742217 | 0.542 |
| YLR430W | 3065.658429 | -0.325707875 | 0.16793181 | 0.542 |
| YLR460C | 187.4228264 | -0.522106866 | 0.167588218 | 0.542 |
| YMR075W | 761.6920474 | 0.376160433 | 0.167987783 | 0.542 |
| YMR083W | 8478.832219 | -0.451337801 | 0.166265521 | 0.542 |
| YMR112C | 344.1352482 | -0.435616678 | 0.16662603 | 0.542 |
| YNL108C | 290.0576123 | -0.41546021 | 0.16685951 | 0.542 |
| YNL200C | 2258.279724 | 0.729405971 | 0.166482801 | 0.542 |
| YNL202W | 289.1619632 | 0.494336864 | 0.167182517 | 0.542 |
| YOR129C | 585.8837465 | 0.349053172 | 0.167242883 | 0.542 |
| YPL255W | 357.750249 | -0.469388769 | 0.167542192 | 0.542 |
| YCL073C | 26.5180597 | 0.349779871 | 0.168550697 | 0.542 |
| YDR115W | 505.144241 | -0.411604178 | 0.168306435 | 0.542 |
| YJL079C | 9145.850933 | 0.702134752 | 0.168753718 | 0.542 |
| YJL169W | 25.36216702 | -0.582744288 | 0.168840054 | 0.542 |
| YJR096W | 2414.849421 | 0.61552894 | 0.168160815 | 0.542 |
| YMR208W | 1260.14289 | -0.367308723 | 0.168352983 | 0.542 |
| YNL216W | 1187.634305 | -0.34661952 | 0.168739905 | 0.542 |
| YOR156C | 466.9299753 | 0.400210412 | 0.168652066 | 0.542 |
| YOR357C | 469.3315493 | -0.378709055 | 0.168750725 | 0.542 |
| YER057C | 2325.694126 | -0.451245461 | 0.168965625 | 0.542 |
| YBR087W | 809.749781 | -0.430664722 | 0.16930021 | 0.543 |
| YML104C | 923.4299097 | 0.409921796 | 0.169451179 | 0.543 |
| YPR109W | 650.4688515 | 0.38191163 | 0.169487281 | 0.543 |
| YCR005C | 6272.799933 | 0.379598513 | 0.169646473 | 0.543 |
| YDR030C | 444.8820441 | 0.35645179 | 0.169821418 | 0.543 |
| YGR043C | 1676.63942 | 0.705382213 | 0.170273452 | 0.543 |
| YHL008C | 1005.082799 | 0.413998237 | 0.16995332 | 0.543 |
| YHR115C | 896.8338505 | 0.430621844 | 0.170350665 | 0.543 |
| YJR153W | 431.5896283 | 0.733482005 | 0.170373104 | 0.543 |
| YKL018W | 496.5039326 | -0.368413369 | 0.170316017 | 0.543 |
| YKL035W | 11294.21789 | 0.447347796 | 0.170392616 | 0.543 |
| YKL134C | 542.9022847 | 0.366594351 | 0.170259042 | 0.543 |
| YGL194C-A | 142.847988 | 0.525024056 | 0.170735158 | 0.544 |
| YPR032W | 901.7856576 | 0.388946414 | 0.170768712 | 0.544 |
| YAR019C | 904.0673159 | 0.352079489 | 0.171085686 | 0.544 |
| YBL019W | 515.3822708 | 0.432207552 | 0.171181041 | 0.544 |
| YHR087W | 3070.022032 | 0.58848898 | 0.171122584 | 0.544 |

| | | | | |
|---|---|---|---|---|
| YBL002W | 4494.23128 | -0.484533849 | 0.171947503 | 0.545 |
| YBR135W | 639.2382675 | -0.544445912 | 0.17187754 | 0.545 |
| YDL238C | 479.1219346 | 0.445981115 | 0.171838591 | 0.545 |
| YDR449C | 532.8954421 | -0.503507179 | 0.171912629 | 0.545 |
| YNL030W | 3372.305745 | -0.540564319 | 0.171906515 | 0.545 |
| YOR078W | 697.5265362 | -0.495718411 | 0.17175668 | 0.545 |
| YDR492W | 689.8420252 | -0.537627204 | 0.172076829 | 0.545 |
| YJL096W | 663.1647142 | -0.358121538 | 0.172206555 | 0.545 |
| YBL042C | 1258.710363 | 0.671233287 | 0.173261754 | 0.545 |
| YER025W | 3287.005691 | -0.317872288 | 0.172891906 | 0.545 |
| YFR020W | 385.8178556 | 0.368053869 | 0.173007334 | 0.545 |
| YIL040W | 372.5621769 | -0.417164735 | 0.172660097 | 0.545 |
| YKL034W | 1217.582813 | 0.575500718 | 0.172895272 | 0.545 |
| YKL059C | 865.3974506 | 0.277155031 | 0.173131075 | 0.545 |
| YLR249W | 32716.59454 | -0.337368497 | 0.173103368 | 0.545 |
| YLR291C | 1399.72151 | -0.263374237 | 0.172977027 | 0.545 |
| YOR246C | 1262.599493 | -0.520056461 | 0.1731335 | 0.545 |
| YPR030W | 2156.768509 | 0.536882972 | 0.173169978 | 0.545 |
| YCR002C | 1893.656785 | -0.364149974 | 0.173622486 | 0.546 |
| YLR289W | 462.7410856 | 0.396912558 | 0.173660638 | 0.546 |
| YDR173C | 369.7980333 | 0.356975632 | 0.173797287 | 0.546 |
| YGL053W | 527.1405514 | 0.685544283 | 0.174302916 | 0.548 |
| YER121W | 184.2483 | 0.535349968 | 0.174480588 | 0.548 |
| YLR359W | 9224.21068 | -0.366026753 | 0.174819202 | 0.549 |
| YBR082C | 6782.616874 | -0.486718536 | 0.175171974 | 0.549 |
| YCR043C | 438.4206899 | -0.442206929 | 0.175101115 | 0.549 |
| YMR004W | 1474.539468 | 0.345615001 | 0.175166169 | 0.549 |
| YJL036W | 1275.214013 | 0.368243366 | 0.175267044 | 0.549 |
| YDR283C | 1675.592568 | 0.423654309 | 0.1758566 | 0.549 |
| YFR044C | 5702.058987 | -0.414546118 | 0.175784598 | 0.549 |
| YHR133C | 977.6515666 | -0.365181932 | 0.175542167 | 0.549 |
| YJL185C | 564.2478549 | 0.5017829 | 0.175758435 | 0.549 |
| YNL050C | 614.5421658 | -0.339480484 | 0.175818001 | 0.549 |
| YDR143C | 1247.877225 | 0.386473321 | 0.176237134 | 0.549 |
| YDR293C | 6400.531719 | 0.415387645 | 0.176210846 | 0.549 |
| YML080W | 296.6855777 | -0.557838715 | 0.176269165 | 0.549 |
| YPL196W | 943.9259177 | 0.522336548 | 0.176216595 | 0.549 |
| YDR236C | 550.9804022 | -0.377531835 | 0.176558349 | 0.550 |
| YMR076C | 1386.678863 | -0.297737087 | 0.176569175 | 0.550 |
| YHR139C-A | 36.30751378 | 0.538891646 | 0.177420572 | 0.551 |
| YIL128W | 1273.394888 | 0.418613687 | 0.177551014 | 0.551 |
| YJR088C | 954.591927 | -0.341297587 | 0.177477581 | 0.551 |
| YLR197W | 4160.668892 | -0.520703784 | 0.177357959 | 0.551 |

| | | | | |
|---|---|---|---|---|
| YNR008W | 899.8873602 | -0.397408456 | 0.177174968 | 0.551 |
| YOR304W | 1385.923767 | 0.26248554 | 0.177290391 | 0.551 |
| YBR001C | 1318.791969 | 0.403257464 | 0.17769291 | 0.551 |
| YFR028C | 1133.072845 | -0.308967432 | 0.178038789 | 0.552 |
| YGL141W | 1274.03239 | 0.319856555 | 0.17799212 | 0.552 |
| YIL068C | 981.7929067 | 0.258960413 | 0.178199596 | 0.552 |
| YGR267C | 1681.678228 | -0.279090519 | 0.17836917 | 0.552 |
| YDR245W | 1164.324708 | -0.409954609 | 0.178542341 | 0.552 |
| YDR279W | 575.5796481 | -0.32748016 | 0.17916991 | 0.552 |
| YGL258W | 52.00397703 | 0.575027134 | 0.179057175 | 0.552 |
| YKL019W | 1109.764572 | -0.367023274 | 0.179066372 | 0.552 |
| YLL007C | 554.0715465 | 0.431226636 | 0.178896929 | 0.552 |
| YLR060W | 5384.459239 | -0.398980266 | 0.178948194 | 0.552 |
| YOR098C | 2501.292329 | -0.338527488 | 0.179155951 | 0.552 |
| YKL222C | 519.5767688 | 0.536057049 | 0.179371542 | 0.553 |
| YLR019W | 1293.150392 | 0.349564449 | 0.179359074 | 0.553 |
| YCR014C | 618.1633306 | 0.328777969 | 0.180126413 | 0.553 |
| YDL199C | 559.7723367 | 0.406089145 | 0.180434896 | 0.553 |
| YDR007W | 490.6292629 | -0.29737264 | 0.179818947 | 0.553 |
| YER107W-A | 76.0626659 | -0.454781326 | 0.180404491 | 0.553 |
| YHR028C | 2661.078301 | 0.445094529 | 0.180052266 | 0.553 |
| YIL109C | 4372.511551 | -0.271162294 | 0.180393528 | 0.553 |
| YIL122W | 600.3807645 | 0.38172824 | 0.180345952 | 0.553 |
| YJL063C | 928.728548 | -0.318097922 | 0.179739894 | 0.553 |
| YJL184W | 578.763074 | -0.406814139 | 0.179713361 | 0.553 |
| YKL212W | 2578.339284 | -0.334849801 | 0.180350582 | 0.553 |
| YNL217W | 1768.453008 | -0.570262318 | 0.180229775 | 0.553 |
| YML029W | 1034.498713 | 0.362322478 | 0.180612552 | 0.553 |
| YMR100W | 811.7962257 | 0.449333424 | 0.180749276 | 0.553 |
| YDR160W | 879.6911624 | 0.269256156 | 0.181420369 | 0.553 |
| YGL170C | 69.87350446 | 0.40240046 | 0.180909905 | 0.553 |
| YKL126W | 2370.590453 | -0.324404063 | 0.181388947 | 0.553 |
| YOR308C | 716.1626853 | -0.485106064 | 0.181234523 | 0.553 |
| YOR331C | 254.0815524 | 0.233630964 | 0.181303394 | 0.553 |
| YPL068C | 273.8997414 | -0.575635318 | 0.181115441 | 0.553 |
| YPL174C | 736.6858257 | 0.297218602 | 0.180988072 | 0.553 |
| YCR063W | 60.70870613 | -0.462684863 | 0.181657296 | 0.553 |
| YHR070W | 688.4026548 | -0.366831733 | 0.181866303 | 0.554 |
| YJL214W | 204.5476536 | 0.506450283 | 0.182177025 | 0.554 |
| YOR157C | 2318.284571 | -0.293176075 | 0.182189632 | 0.554 |
| YPL186C | 1114.795126 | 0.59611208 | 0.182269275 | 0.554 |
| YPL236C | 570.2921643 | 0.451705046 | 0.18209924 | 0.554 |
| YPL244C | 876.1384791 | -0.391520761 | 0.181906667 | 0.554 |

| | | | | |
|---|---|---|---|---|
| YNR060W | 291.930447 | 0.408013118 | 0.182494058 | 0.554 |
| YBR274W | 529.7886878 | -0.360086536 | 0.182778737 | 0.555 |
| YDR300C | 1204.265207 | -0.424288139 | 0.183298996 | 0.556 |
| YER087C-A | 36.6678806 | -0.631225206 | 0.183315923 | 0.556 |
| YAL032C | 524.818116 | 0.259726411 | 0.183788822 | 0.556 |
| YBR185C | 732.2746685 | -0.306141498 | 0.184110282 | 0.556 |
| YCL008C | 1030.586387 | 0.318840927 | 0.18414184 | 0.556 |
| YGR038W | 610.0071877 | -0.300858867 | 0.184080313 | 0.556 |
| YLR368W | 767.6389144 | 0.405572643 | 0.183854023 | 0.556 |
| YOR266W | 469.4838324 | 0.300832551 | 0.183628192 | 0.556 |
| YPL111W | 2984.765722 | 0.296321533 | 0.183928575 | 0.556 |
| YPL252C | 462.6644596 | -0.455611695 | 0.183664936 | 0.556 |
| YAL033W | 350.8440347 | -0.579133384 | 0.184729866 | 0.556 |
| YBL074C | 232.3789197 | 0.394598832 | 0.184855207 | 0.556 |
| YCR042C | 1593.772478 | 0.289898696 | 0.184747384 | 0.556 |
| YDR379C-A | 462.1957351 | 0.419934092 | 0.184464299 | 0.556 |
| YGR212W | 361.1538773 | 0.370781759 | 0.18491585 | 0.556 |
| YLR058C | 11510.40084 | -0.348326558 | 0.184797451 | 0.556 |
| YOR213C | 459.655666 | -0.451471409 | 0.184983871 | 0.556 |
| YCR036W | 1075.104784 | 0.270388328 | 0.185210232 | 0.557 |
| YMR307W | 11748.41017 | -0.230515316 | 0.185250504 | 0.557 |
| YDR490C | 1094.352116 | 0.409145033 | 0.185796673 | 0.557 |
| YGR152C | 296.8399677 | -0.413906842 | 0.185608124 | 0.557 |
| YKL178C | 114.5867776 | 0.886094019 | 0.185741321 | 0.557 |
| YLR134W | 556.7123022 | 0.323368592 | 0.185635137 | 0.557 |
| YOL096C | 679.2678424 | 0.380217932 | 0.185901006 | 0.557 |
| YPL147W | 471.9776317 | 0.283155295 | 0.185906994 | 0.557 |
| YDR212W | 2389.189098 | -0.39488598 | 0.186765721 | 0.557 |
| YGL010W | 399.2167766 | 0.379691831 | 0.186194246 | 0.557 |
| YGR185C | 4359.206315 | -0.455445941 | 0.186408996 | 0.557 |
| YHL036W | 545.2189465 | 0.311966624 | 0.186700534 | 0.557 |
| YIL131C | 677.2016628 | -0.377375416 | 0.186668035 | 0.557 |
| YJR090C | 1653.355955 | 0.326633865 | 0.186814042 | 0.557 |
| YMR153C-A | 60.08017136 | -0.641382709 | 0.186407669 | 0.557 |
| YPL222W | 848.2140163 | 0.557641515 | 0.186626743 | 0.557 |
| YPR048W | 424.7867171 | -0.459968188 | 0.186411024 | 0.557 |
| YDR004W | 628.5806399 | 0.258267899 | 0.186917216 | 0.557 |
| YGL054C | 1677.23256 | -0.369865206 | 0.187096444 | 0.557 |
| YGR201C | 974.6666707 | 0.533154536 | 0.187224859 | 0.557 |
| YAL019W | 1321.957689 | -0.487637564 | 0.187781771 | 0.558 |
| YBR280C | 1033.06933 | 0.439922988 | 0.188155005 | 0.558 |
| YHR070C-A | 129.6976848 | -0.442963626 | 0.188340982 | 0.558 |
| YJL180C | 680.1821358 | -0.333009016 | 0.188087237 | 0.558 |

| | | | | |
|---|---|---|---|---|
| YML068W | 512.1804017 | 0.286035703 | 0.187653742 | 0.558 |
| YMR276W | 5216.693498 | -0.399378244 | 0.187861685 | 0.558 |
| YNR053C | 1005.221474 | -0.402265184 | 0.188236614 | 0.558 |
| YPL050C | 1693.46287 | -0.373010221 | 0.188295404 | 0.558 |
| YPL258C | 548.5976694 | 0.436154033 | 0.188261689 | 0.558 |
| YKL096W | 1798.684379 | -0.413789252 | 0.188635933 | 0.558 |
| YPL006W | 1523.461434 | 0.548081284 | 0.188615642 | 0.558 |
| YAL031C | 1097.389337 | 0.37781942 | 0.189233328 | 0.559 |
| YDR239C | 796.3576703 | 0.330256013 | 0.188857445 | 0.559 |
| YLR437C | 329.593239 | -0.43020679 | 0.189015225 | 0.559 |
| YMR084W | 174.3110976 | 0.35041037 | 0.189271603 | 0.559 |
| YNL310C | 605.4678235 | -0.318025822 | 0.189082876 | 0.559 |
| YOL011W | 1428.931177 | 0.350208399 | 0.189350426 | 0.559 |
| YOR223W | 687.6095973 | 0.356463659 | 0.189202496 | 0.559 |
| YDR363W-A | 1308.889566 | -0.42339515 | 0.190210203 | 0.559 |
| YEL059W | 50.35325287 | 0.275055294 | 0.189593084 | 0.559 |
| YFL024C | 1322.826078 | 0.305574034 | 0.189978054 | 0.559 |
| YGL101W | 448.9089719 | -0.41135523 | 0.189849387 | 0.559 |
| YGL145W | 1156.588747 | 0.30882725 | 0.19047224 | 0.559 |
| YGL154C | 461.1131948 | 0.300000435 | 0.19052713 | 0.559 |
| YJR161C | 124.4036715 | 0.699495488 | 0.189863271 | 0.559 |
| YML088W | 784.353378 | -0.428619694 | 0.189829149 | 0.559 |
| YMR215W | 1997.531105 | -0.427318124 | 0.190034855 | 0.559 |
| YNL087W | 1914.774752 | -0.34129742 | 0.190098833 | 0.559 |
| YOL042W | 569.9826336 | 0.444540785 | 0.190379376 | 0.559 |
| YPL277C | 206.5316036 | 0.537562494 | 0.190373872 | 0.559 |
| YAL022C | 845.7051212 | -0.286929266 | 0.191952434 | 0.559 |
| YBR112C | 3373.883429 | -0.388433204 | 0.191770187 | 0.559 |
| YDR334W | 1700.440041 | 0.34895643 | 0.191357795 | 0.559 |
| YDR339C | 1137.925374 | -0.542059718 | 0.190844745 | 0.559 |
| YGL096W | 785.2336716 | 0.542771297 | 0.191200248 | 0.559 |
| YHR144C | 383.3281151 | -0.311531932 | 0.191292932 | 0.559 |
| YIL164C | 260.9748713 | 0.452007032 | 0.191552863 | 0.559 |
| YJL108C | 258.931357 | 0.386800446 | 0.191316549 | 0.559 |
| YNL166C | 1009.247872 | -0.4441688 | 0.191634405 | 0.559 |
| YNL321W | 1495.615211 | 0.364794816 | 0.191782891 | 0.559 |
| YOL093W | 662.3279101 | -0.418158547 | 0.191258757 | 0.559 |
| YOR051C | 2007.894821 | -0.48347452 | 0.191834473 | 0.559 |
| YOR321W | 1083.064286 | -0.328910769 | 0.191857922 | 0.559 |
| YPL199C | 1017.89655 | -0.44374035 | 0.192154219 | 0.560 |
| YBR042C | 577.29725 | -0.355645459 | 0.192682946 | 0.560 |
| YDR219C | 850.7025719 | 0.345606223 | 0.192612271 | 0.560 |
| YKR056W | 1143.695694 | -0.471701654 | 0.192781754 | 0.560 |

| | | | | |
|---|---|---|---|---|
| YML069W | 2256.742432 | -0.437777702 | 0.192323878 | 0.560 |
| YMR250W | 7553.858966 | 0.731001931 | 0.192430284 | 0.560 |
| YPL247C | 2827.171119 | 0.416656079 | 0.192758916 | 0.560 |
| YLR220W | 930.48531 | 0.368010952 | 0.193088169 | 0.560 |
| YML108W | 173.1265371 | -0.626578648 | 0.193035005 | 0.560 |
| YOR386W | 897.2436472 | 0.619392008 | 0.192997527 | 0.560 |
| YHR031C | 752.7001486 | -0.432309061 | 0.193370089 | 0.560 |
| YMR176W | 1256.155137 | 0.430954466 | 0.193501196 | 0.560 |
| YKL037W | 43.69720214 | -0.746666282 | 0.193754607 | 0.561 |
| YGR093W | 815.5848449 | 0.299799657 | 0.193932247 | 0.561 |
| YKL084W | 479.3447424 | -0.582878158 | 0.19436896 | 0.562 |
| YHR169W | 496.2218428 | -0.435841548 | 0.194650641 | 0.563 |
| YDR091C | 1728.536947 | -0.387932008 | 0.194818697 | 0.563 |
| YDR510W | 2598.34956 | -0.352451351 | 0.194844701 | 0.563 |
| YDL048C | 3537.203448 | 0.512047931 | 0.195727037 | 0.564 |
| YGR046W | 906.1577966 | 0.431230025 | 0.195711948 | 0.564 |
| YGR059W | 83.10299922 | 0.648903958 | 0.195487181 | 0.564 |
| YHR106W | 1096.925363 | 0.333432525 | 0.195562926 | 0.564 |
| YMR304W | 1821.67097 | 0.40888236 | 0.195435149 | 0.564 |
| YPR063C | 1556.762503 | -0.370613746 | 0.195625614 | 0.564 |
| YOR298W | 171.3347166 | 0.37690479 | 0.195966698 | 0.564 |
| YAL063C | 272.6184868 | 0.387324227 | 0.199607235 | 0.565 |
| YBL041W | 3024.581572 | -0.264072913 | 0.196628765 | 0.565 |
| YBR094W | 1036.817856 | -0.332991532 | 0.198406027 | 0.565 |
| YBR183W | 786.7022482 | 0.605374217 | 0.197763887 | 0.565 |
| YBR268W | 567.5701433 | -0.343918708 | 0.199642385 | 0.565 |
| YCL024W | 493.5504233 | -0.302099933 | 0.196392573 | 0.565 |
| YDL052C | 1575.894255 | -0.313130252 | 0.196998285 | 0.565 |
| YDL222C | 906.6701758 | 0.544835718 | 0.199360713 | 0.565 |
| YDR065W | 277.2481422 | 0.504081244 | 0.199541342 | 0.565 |
| YDR106W | 46.31624816 | 0.507184603 | 0.2002525 | 0.565 |
| YDR124W | 300.8040212 | 0.394879037 | 0.199231567 | 0.565 |
| YDR132C | 453.7495244 | 0.513758758 | 0.199120497 | 0.565 |
| YDR171W | 12610.16854 | 0.536777493 | 0.198131993 | 0.565 |
| YDR274C | 62.88697688 | 0.602299939 | 0.200386468 | 0.565 |
| YDR421W | 1031.149921 | 0.316367357 | 0.19866465 | 0.565 |
| YEL022W | 2372.854744 | -0.33585 | 0.198847992 | 0.565 |
| YEL036C | 1490.088365 | -0.484283227 | 0.197685897 | 0.565 |
| YER016W | 838.7636501 | -0.370046099 | 0.199408142 | 0.565 |
| YGL128C | 505.7413235 | 0.299118109 | 0.198748383 | 0.565 |
| YGL215W | 3518.182479 | -0.302992778 | 0.197149179 | 0.565 |
| YHR058C | 391.0666567 | -0.348011539 | 0.198832433 | 0.565 |
| YHR097C | 2968.055756 | 0.476290422 | 0.199356578 | 0.565 |

| | | | | |
|---|---|---|---|---|
| YIL110W | 498.0390037 | -0.441804344 | 0.199369893 | 0.565 |
| YIL113W | 384.3761033 | 0.278633019 | 0.19684241 | 0.565 |
| YJL058C | 326.9260026 | -0.473463308 | 0.199946461 | 0.565 |
| YJL070C | 972.7876692 | 0.390679079 | 0.197695043 | 0.565 |
| YJR100C | 772.6621126 | 0.320026399 | 0.199376065 | 0.565 |
| YKL051W | 1194.148337 | 0.432271171 | 0.199720563 | 0.565 |
| YKR049C | 1528.555977 | 0.477312146 | 0.200265462 | 0.565 |
| YKR066C | 4134.117453 | 0.472441178 | 0.197042192 | 0.565 |
| YLR045C | 1304.494239 | -0.291605496 | 0.200509798 | 0.565 |
| YLR073C | 322.9361924 | -0.447309684 | 0.196444786 | 0.565 |
| YLR147C | 302.8969457 | -0.387709364 | 0.199585738 | 0.565 |
| YLR178C | 3271.315417 | 0.640888003 | 0.200518865 | 0.565 |
| YLR187W | 889.7815093 | -0.327964357 | 0.198719169 | 0.565 |
| YLR425W | 1120.132672 | 0.335300278 | 0.198007449 | 0.565 |
| YLR426W | 420.438408 | -0.372471005 | 0.197706562 | 0.565 |
| YLR437C-A | 26.88165292 | 0.638483171 | 0.198820411 | 0.565 |
| YNR043W | 2170.763038 | -0.368483184 | 0.196717742 | 0.565 |
| YOL057W | 2160.171024 | -0.362933225 | 0.200481677 | 0.565 |
| YOL153C | 1352.172864 | 0.410530875 | 0.20016301 | 0.565 |
| YOR304C-A | 201.0708852 | -0.530604307 | 0.19819856 | 0.565 |
| YOR380W | 392.7551247 | 0.340821005 | 0.197993652 | 0.565 |
| YPL077C | 211.278667 | 0.334665333 | 0.199658959 | 0.565 |
| YPL265W | 2999.757451 | 0.555953788 | 0.200282856 | 0.565 |
| YDR144C | 842.4657292 | -0.291022839 | 0.200872296 | 0.565 |
| YHR167W | 161.5282471 | -0.49928462 | 0.200862208 | 0.565 |
| YKL091C | 918.9800837 | 0.516379627 | 0.200932522 | 0.565 |
| YLR201C | 1199.611196 | 0.429635557 | 0.200784988 | 0.565 |
| YBL086C | 1904.650929 | 0.274938514 | 0.202515782 | 0.565 |
| YCL051W | 1104.388423 | 0.31311615 | 0.202214532 | 0.565 |
| YCR099C | 126.2405817 | 0.5519447 | 0.202195373 | 0.565 |
| YDL165W | 697.5260403 | -0.284169246 | 0.202001992 | 0.565 |
| YDR438W | 373.0593006 | 0.324973892 | 0.20252277 | 0.565 |
| YGL108C | 328.3815974 | 0.357302007 | 0.202518452 | 0.565 |
| YGL121C | 888.9061681 | 0.859860911 | 0.201469959 | 0.565 |
| YIL084C | 434.165782 | 0.402238328 | 0.20150965 | 0.565 |
| YJL099W | 747.8275315 | 0.336913146 | 0.202189474 | 0.565 |
| YJR021C | 884.0147888 | 0.329530268 | 0.202346515 | 0.565 |
| YNR024W | 280.6447987 | -0.435173628 | 0.202276528 | 0.565 |
| YOL130W | 1532.382846 | 0.254749194 | 0.202098087 | 0.565 |
| YOR173W | 2098.419186 | 0.629684803 | 0.202285303 | 0.565 |
| YOR319W | 365.6759573 | -0.355834589 | 0.201304778 | 0.565 |
| YPR164W | 1248.455329 | -0.261542038 | 0.201224718 | 0.565 |
| YCR013C | 2806.322738 | -0.319919672 | 0.202694547 | 0.565 |

| | | | | |
|---|---|---|---|---|
| YNL023C | 950.809325 | -0.360418946 | 0.202799015 | 0.565 |
| YDR440W | 548.6066441 | -0.352344937 | 0.203167853 | 0.566 |
| YLL053C | 96.70396409 | 0.285662563 | 0.203212908 | 0.566 |
| YMR140W | 1470.388235 | 0.356484609 | 0.203262431 | 0.566 |
| YIL075C | 7548.161259 | -0.3987683 | 0.203477358 | 0.566 |
| YAL005C | 15020.34614 | -0.28968046 | 0.2036556 | 0.566 |
| YBR054W | 4277.398072 | 0.822126485 | 0.206364477 | 0.567 |
| YBR216C | 879.1909045 | 0.323930988 | 0.205361742 | 0.567 |
| YBR248C | 1552.941582 | -0.456245193 | 0.203882972 | 0.567 |
| YCL052C | 1614.200073 | 0.362068947 | 0.206392597 | 0.567 |
| YDL099W | 2397.811912 | -0.431291688 | 0.206370891 | 0.567 |
| YEL039C | 1783.57122 | 0.518926712 | 0.205432746 | 0.567 |
| YGL058W | 714.9977444 | -0.432681051 | 0.206515326 | 0.567 |
| YGR066C | 223.0827395 | 0.300267033 | 0.20391589 | 0.567 |
| YGR154C | 333.0792836 | 0.326635551 | 0.206174668 | 0.567 |
| YGR215W | 453.6817551 | -0.412050063 | 0.204785472 | 0.567 |
| YIL039W | 1826.424519 | -0.284161272 | 0.206631749 | 0.567 |
| YJL178C | 1325.547528 | -0.296084543 | 0.204903494 | 0.567 |
| YKL138C-A | 204.185803 | -0.406002784 | 0.205193082 | 0.567 |
| YKR106W | 110.7787445 | 0.915752724 | 0.206038089 | 0.567 |
| YLL060C | 550.1729711 | 0.349755614 | 0.205161235 | 0.567 |
| YLR254C | 554.1871051 | 0.310632013 | 0.2054106 | 0.567 |
| YLR411W | 132.197405 | 0.369429463 | 0.206647106 | 0.567 |
| YLR450W | 2180.849651 | 0.448390982 | 0.206378899 | 0.567 |
| YML095C | 255.548255 | 0.403967927 | 0.204911314 | 0.567 |
| YMR002W | 4776.428936 | -0.430720051 | 0.206696858 | 0.567 |
| YMR204C | 437.305049 | 0.395718618 | 0.206416702 | 0.567 |
| YMR293C | 413.7964048 | 0.385761275 | 0.204738094 | 0.567 |
| YNL119W | 513.2444289 | -0.515040087 | 0.204465918 | 0.567 |
| YNL139C | 1577.578713 | 0.345887121 | 0.204212166 | 0.567 |
| YNL182C | 900.9513347 | -0.505642338 | 0.206256791 | 0.567 |
| YNL269W | 50.67416069 | 0.212336258 | 0.205132977 | 0.567 |
| YNR007C | 864.4677932 | 0.49991664 | 0.205534379 | 0.567 |
| YOL108C | 360.9401457 | 0.396606497 | 0.20635234 | 0.567 |
| YOR350C | 702.3901449 | 0.440211994 | 0.204730114 | 0.567 |
| YPL167C | 937.9139202 | 0.339008693 | 0.205557007 | 0.567 |
| YPL248C | 793.6864415 | 0.364147397 | 0.206707908 | 0.567 |
| YDL056W | 1411.798857 | 0.352861503 | 0.207416708 | 0.567 |
| YEL029C | 331.9088854 | -0.454639001 | 0.20750493 | 0.567 |
| YER048W-A | 659.0895735 | -0.232739191 | 0.206987683 | 0.567 |
| YGL006W | 3158.91505 | 0.354527002 | 0.207349465 | 0.567 |
| YGR238C | 880.0595867 | 0.395523132 | 0.207250231 | 0.567 |
| YGR239C | 410.924928 | -0.425653324 | 0.207325445 | 0.567 |

| | | | | |
|---|---|---|---|---|
| YHR140W | 190.2878906 | 0.978552409 | 0.207190979 | 0.567 |
| YJL037W | 69.21169324 | 0.515427151 | 0.207537709 | 0.567 |
| YMR097C | 670.8866515 | 0.288933093 | 0.207674767 | 0.567 |
| YDR531W | 969.2146497 | -0.425315924 | 0.208473114 | 0.567 |
| YGL156W | 2401.006767 | 0.636216986 | 0.208626346 | 0.567 |
| YKL021C | 1160.236373 | -0.445539162 | 0.208707816 | 0.567 |
| YKL166C | 506.1904776 | -0.426321226 | 0.208311923 | 0.567 |
| YLL015W | 2174.31995 | 0.353699297 | 0.208215144 | 0.567 |
| YNL283C | 1549.072925 | -0.316440759 | 0.208515717 | 0.567 |
| YOL089C | 836.6158144 | 0.314852515 | 0.208472534 | 0.567 |
| YPL094C | 2895.773005 | -0.294857748 | 0.208703622 | 0.567 |
| YPR073C | 916.4024804 | -0.344306578 | 0.207858851 | 0.567 |
| YPR146C | 99.73914934 | -0.273635508 | 0.208003808 | 0.567 |
| YPL098C | 678.1300721 | -0.32220916 | 0.208811163 | 0.567 |
| YMR219W | 3001.660673 | 0.469163334 | 0.209275605 | 0.568 |
| YPR156C | 1539.255789 | 0.556013871 | 0.209844253 | 0.569 |
| YGR284C | 2427.894256 | -0.325854473 | 0.209994067 | 0.569 |
| YBR056W | 2794.886987 | 0.412005691 | 0.210293339 | 0.570 |
| YMR158W | 693.5786925 | -0.314949382 | 0.210425309 | 0.570 |
| YER066W | 204.0645964 | 0.259527255 | 0.210687926 | 0.570 |
| YGL100W | 1500.808318 | -0.250157423 | 0.210677691 | 0.570 |
| YGR149W | 1082.04674 | 0.411690167 | 0.210943217 | 0.571 |
| YOR226C | 454.5609099 | -0.432891253 | 0.210996079 | 0.571 |
| YPL096C-A | 145.3313639 | -0.449368274 | 0.211299946 | 0.571 |
| YJL001W | 1978.350649 | -0.326534862 | 0.211397599 | 0.571 |
| YBR128C | 336.3131952 | 0.419123535 | 0.211727562 | 0.572 |
| YOR148C | 393.9247418 | -0.355993142 | 0.21185705 | 0.572 |
| YGR292W | 46.2187461 | 0.495068249 | 0.21229119 | 0.573 |
| RPR1 | 36.63381865 | -0.266955022 | 0.213518214 | 0.574 |
| YBL033C | 1274.437806 | 0.280567981 | 0.213518843 | 0.574 |
| YDR125C | 356.5508843 | 0.524468955 | 0.213464381 | 0.574 |
| YDR403W | 327.0898139 | 0.736882498 | 0.213562202 | 0.574 |
| YHL039W | 1028.011858 | -0.361715017 | 0.213611259 | 0.574 |
| YKL007W | 2212.362326 | -0.219849715 | 0.213629597 | 0.574 |
| YLR412W | 993.3823084 | -0.365446424 | 0.213688357 | 0.574 |
| YOR056C | 1339.644393 | -0.37377683 | 0.212967503 | 0.574 |
| YPL189W | 119.6862802 | -0.363732873 | 0.213203464 | 0.574 |
| YGR116W | 3744.798928 | -0.322311113 | 0.213895108 | 0.574 |
| YHR109W | 446.2718996 | 0.51690392 | 0.213845972 | 0.574 |
| YCL030C | 8176.65539 | -0.379047421 | 0.214140927 | 0.575 |
| YBR055C | 925.8039594 | -0.247148054 | 0.214270468 | 0.575 |
| YJL012C | 2507.923216 | -0.40293896 | 0.214366935 | 0.575 |
| YNL128W | 87.91000712 | 0.674899861 | 0.214454359 | 0.575 |

| | | | | |
|---|---|---|---|---|
| YBR018C | 184.0387102 | 0.353217585 | 0.21496182 | 0.575 |
| YBR040W | 134.4656267 | 0.727871192 | 0.215139961 | 0.575 |
| YCL048W | 92.73705911 | 0.635478071 | 0.215086114 | 0.575 |
| YDL180W | 735.4223348 | 0.415648323 | 0.215318929 | 0.575 |
| YDR537C | 28.61472288 | -0.490448812 | 0.214843175 | 0.575 |
| YER070W | 2200.790509 | -0.092682298 | 0.215488699 | 0.575 |
| YJL023C | 307.217395 | 0.269400731 | 0.215438823 | 0.575 |
| YJR045C | 25492.38268 | -0.225311415 | 0.215568424 | 0.575 |
| YJR131W | 581.4205166 | 0.46573049 | 0.215780179 | 0.575 |
| YLR099W-A | 174.4936248 | -0.399571192 | 0.215614152 | 0.575 |
| YMR180C | 510.7680392 | 0.290653979 | 0.215764476 | 0.575 |
| YOR023C | 2117.560133 | 0.319292016 | 0.21566811 | 0.575 |
| YOR046C | 2190.495993 | -0.395313852 | 0.215775868 | 0.575 |
| YHR110W | 1458.33201 | -0.251392009 | 0.215985251 | 0.575 |
| YDL210W | 142.1653441 | 0.66718455 | 0.216291741 | 0.576 |
| YBR139W | 2470.205803 | 0.44290182 | 0.216495416 | 0.576 |
| YOL086C | 59663.42444 | -0.315738041 | 0.216585643 | 0.576 |
| YJL067W | 49.69168928 | 0.700000664 | 0.216701158 | 0.576 |
| YGL036W | 2417.923989 | 0.286828401 | 0.21712777 | 0.576 |
| YGL220W | 745.8780305 | -0.404238191 | 0.217193187 | 0.576 |
| YKL029C | 3291.164423 | -0.317141341 | 0.217062776 | 0.576 |
| YBL059C-A | 504.0749236 | -0.371791289 | 0.217309873 | 0.576 |
| YNL237W | 756.7694566 | 0.350796877 | 0.217555467 | 0.577 |
| YMR012W | 7425.571889 | -0.218771373 | 0.217714175 | 0.577 |
| YOL114C | 133.5426332 | 0.462684575 | 0.217852757 | 0.577 |
| YJL213W | 560.8105438 | 0.450209102 | 0.218195015 | 0.578 |
| YLL017W | 60.73191178 | 0.480465383 | 0.218156397 | 0.578 |
| YJL093C | 1063.991787 | 0.272685247 | 0.2183152 | 0.578 |
| YCR089W | 1832.043276 | 0.504132957 | 0.218608534 | 0.578 |
| YOR221C | 1523.763907 | 0.355773219 | 0.21874027 | 0.578 |
| YCL037C | 1123.648124 | -0.174318421 | 0.218951569 | 0.578 |
| YLR309C | 2329.969598 | -0.480507728 | 0.218976357 | 0.578 |
| YDR291W | 877.4001219 | 0.279906858 | 0.219172233 | 0.579 |
| YGR009C | 1451.464581 | 0.282372188 | 0.219356124 | 0.579 |
| YHR112C | 1466.272889 | 0.434675359 | 0.21933413 | 0.579 |
| YJR137C | 1214.839856 | 0.379491727 | 0.219580773 | 0.579 |
| YGL011C | 3486.685044 | -0.32025069 | 0.219752084 | 0.579 |
| YOR228C | 505.0466268 | 0.444789371 | 0.219803556 | 0.579 |
| YOL001W | 429.8453508 | 0.370808446 | 0.220154576 | 0.580 |
| YHR028W-A | 38.81881491 | 0.659334673 | 0.22028029 | 0.580 |
| YIL103W | 637.74726 | -0.366808531 | 0.220433246 | 0.580 |
| YKL107W | 249.2824796 | 0.602414501 | 0.220750993 | 0.580 |
| YOR072W | 29.04315607 | 0.355014662 | 0.220765826 | 0.580 |

| | | | | |
|---|---|---|---|---|
| YNL009W | 277.6331838 | 0.500382232 | 0.220973502 | 0.580 |
| YOR101W | 962.4280426 | -0.38953955 | 0.221010133 | 0.580 |
| YDL058W | 1998.717774 | -0.337795225 | 0.221731955 | 0.581 |
| YKL054C | 4561.844374 | -0.381394042 | 0.221643598 | 0.581 |
| YLR362W | 1083.342817 | 0.405643923 | 0.221610293 | 0.581 |
| YNL263C | 1345.46904 | -0.337263092 | 0.221664132 | 0.581 |
| YDR363W | 641.2311125 | 0.413337297 | 0.222333042 | 0.582 |
| YNL311C | 782.0547648 | 0.295956129 | 0.222253972 | 0.582 |
| YPR005C | 426.8209175 | 0.262874815 | 0.222454906 | 0.582 |
| YLR374C | 34.93928504 | 0.332347665 | 0.222671262 | 0.583 |
| YLR223C | 2252.47739 | -0.546549584 | 0.222919909 | 0.583 |
| YLR403W | 956.7749718 | 0.338246963 | 0.222971637 | 0.583 |
| YNL024C-A | 249.9046551 | -0.354919209 | 0.223172295 | 0.583 |
| YNL264C | 1013.510724 | -0.392385515 | 0.223105456 | 0.583 |
| YGL207W | 4342.654723 | -0.293944564 | 0.223531923 | 0.583 |
| YCR020W-B | 152.1239298 | -0.714939998 | 0.223756315 | 0.584 |
| YOR127W | 1893.705383 | 0.313806238 | 0.223813103 | 0.584 |
| YCR090C | 636.078747 | -0.39026722 | 0.224096583 | 0.584 |
| YKL115C | 42.82706794 | -0.388641365 | 0.224062041 | 0.584 |
| YDR505C | 1755.081331 | 0.472578432 | 0.224315019 | 0.584 |
| YPL225W | 2696.542417 | -0.339103604 | 0.224600544 | 0.585 |
| YBL029W | 662.4144794 | 0.429877492 | 0.22750012 | 0.585 |
| YBL055C | 719.3024768 | -0.389956442 | 0.226106959 | 0.585 |
| YCL045C | 3038.105977 | -0.375810744 | 0.225099242 | 0.585 |
| YCR068W | 666.4091136 | 0.345194029 | 0.227142268 | 0.585 |
| YDL009C | 106.1081082 | -0.406042603 | 0.226707428 | 0.585 |
| YDL110C | 2473.385616 | 0.387609938 | 0.225556546 | 0.585 |
| YDR209C | 344.9577194 | -0.587752592 | 0.224965183 | 0.585 |
| YDR295C | 798.8722941 | 0.251552423 | 0.227453657 | 0.585 |
| YDR402C | 76.67558535 | 0.539148298 | 0.226779964 | 0.585 |
| YEL040W | 3165.37713 | -0.367637842 | 0.225664146 | 0.585 |
| YEL053C | 1328.366975 | 0.223886748 | 0.227511148 | 0.585 |
| YER053C | 3873.936022 | 0.511696015 | 0.227111622 | 0.585 |
| YGL081W | 335.8828397 | 0.417014831 | 0.22645649 | 0.585 |
| YGL158W | 260.9463165 | 0.318712198 | 0.226268165 | 0.585 |
| YGR028W | 820.509086 | 0.39185996 | 0.22642087 | 0.585 |
| YGR162W | 4487.780728 | -0.326026086 | 0.226759452 | 0.585 |
| YIR039C | 1029.181658 | 0.781379111 | 0.226738274 | 0.585 |
| YKL100C | 2362.710996 | 0.355559262 | 0.225630891 | 0.585 |
| YKR098C | 1187.222635 | 0.474085889 | 0.226852487 | 0.585 |
| YLR311C | 30.19399839 | 0.616963408 | 0.226459558 | 0.585 |
| YNL058C | 1080.922928 | -0.36261206 | 0.225359364 | 0.585 |
| YOR041C | 37.3273586 | 0.632769242 | 0.227242542 | 0.585 |

| | | | | |
|---|---|---|---|---|
| YOR087W | 1019.099344 | 0.422989419 | 0.226249888 | 0.585 |
| YOR174W | 544.9117735 | -0.308624331 | 0.225966714 | 0.585 |
| YOR269W | 302.0905548 | 0.311896351 | 0.22684269 | 0.585 |
| YOR292C | 645.9873627 | 0.280293132 | 0.226440321 | 0.585 |
| YPR186C | 328.1330267 | -0.405047487 | 0.227140699 | 0.585 |
| YLR457C | 335.6948937 | -0.28636648 | 0.227890446 | 0.586 |
| YDR408C | 1967.993247 | -0.444499629 | 0.229955772 | 0.587 |
| YER030W | 1421.194205 | -0.432520913 | 0.229891898 | 0.587 |
| YER123W | 1029.367215 | 0.325311233 | 0.229011218 | 0.587 |
| YGR102C | 796.5538343 | 0.391540385 | 0.229362088 | 0.587 |
| YHR120W | 606.7255558 | 0.242505526 | 0.228723771 | 0.587 |
| YIL027C | 793.9722774 | -0.356794519 | 0.229266979 | 0.587 |
| YIL087C | 960.1717362 | 0.582714596 | 0.229197045 | 0.587 |
| YJL005W | 4176.860231 | 0.482282101 | 0.229737939 | 0.587 |
| YJL048C | 1114.208522 | 0.449971889 | 0.229442376 | 0.587 |
| YKL026C | 613.714189 | 0.5168421 | 0.228930235 | 0.587 |
| YKR061W | 573.6088479 | 0.434222086 | 0.228932324 | 0.587 |
| YLR021W | 410.9321978 | -0.287180447 | 0.228979398 | 0.587 |
| YLR026C | 1116.422754 | -0.348281241 | 0.230010753 | 0.587 |
| YML118W | 395.2746742 | 0.590429093 | 0.22853755 | 0.587 |
| YNL073W | 1409.044777 | 0.397667 | 0.228754784 | 0.587 |
| YOL049W | 2360.13263 | -0.334582191 | 0.229948184 | 0.587 |
| YOR017W | 870.202076 | 0.278452722 | 0.228371232 | 0.587 |
| YOR373W | 745.8071516 | -0.370016311 | 0.229511697 | 0.587 |
| YFL011W | 70.61598103 | 0.472946187 | 0.230221368 | 0.587 |
| YIL001W | 471.4395924 | 0.474946536 | 0.230159673 | 0.587 |
| YBR083W | 2339.410841 | 0.418331194 | 0.230518413 | 0.587 |
| YOR110W | 712.9241656 | 0.287881495 | 0.2304903 | 0.587 |
| YAL011W | 648.981818 | -0.425237575 | 0.230867326 | 0.588 |
| YIL144W | 1230.130031 | 0.363998511 | 0.231043376 | 0.588 |
| YOL002C | 621.7558878 | -0.298102208 | 0.231378626 | 0.588 |
| YOL007C | 839.2247362 | -0.369243838 | 0.231388367 | 0.588 |
| YBR021W | 496.9097626 | 0.490537129 | 0.232422575 | 0.589 |
| YDL021W | 1151.606846 | 0.886318739 | 0.232754042 | 0.589 |
| YDR005C | 1131.574288 | 0.327865308 | 0.2322783 | 0.589 |
| YGL231C | 1251.65783 | -0.294373327 | 0.23276491 | 0.589 |
| YHR077C | 2925.324128 | 0.302532872 | 0.23247442 | 0.589 |
| YHR215W | 110.5268911 | -0.690008467 | 0.23247738 | 0.589 |
| YLR284C | 336.5089725 | 0.299379341 | 0.232749146 | 0.589 |
| YLR332W | 1136.948721 | 0.356222482 | 0.232242179 | 0.589 |
| YNR041C | 337.6551047 | -0.324431617 | 0.231871709 | 0.589 |
| YOR085W | 1806.530962 | -0.379811431 | 0.23266124 | 0.589 |
| YOR315W | 1158.774212 | -0.137787983 | 0.232394537 | 0.589 |

| | | | | |
|---|---|---|---|---|
| YPL250C | 749.2294185 | 0.576098378 | 0.232261749 | 0.589 |
| YCR072C | 412.6699919 | -0.503688036 | 0.233915155 | 0.589 |
| YDR008C | 37.32753147 | -0.336584449 | 0.233899364 | 0.589 |
| YDR534C | 224.6689044 | -0.866313997 | 0.233012041 | 0.589 |
| YIR027C | 227.1514133 | 0.550819125 | 0.233129708 | 0.589 |
| YLR132C | 337.4595812 | 0.287177329 | 0.233730341 | 0.589 |
| YLR266C | 1095.178781 | 0.328524169 | 0.23353133 | 0.589 |
| YML096W | 652.0319638 | -0.304119716 | 0.234040071 | 0.589 |
| YMR151W | 69.13225197 | 0.311722348 | 0.233793799 | 0.589 |
| YMR246W | 3290.570155 | -0.320297655 | 0.233108387 | 0.589 |
| YOL021C | 1670.215668 | -0.284970496 | 0.233547836 | 0.589 |
| YOR081C | 1945.1418 | 0.387191562 | 0.233247968 | 0.589 |
| YOR353C | 984.6133479 | 0.363788829 | 0.233731498 | 0.589 |
| YPL074W | 944.5738824 | 0.31876551 | 0.23408684 | 0.589 |
| YBL024W | 2012.558866 | -0.386344678 | 0.234417213 | 0.589 |
| YGL068W | 385.5514446 | -0.417949463 | 0.234350391 | 0.589 |
| YIL020C | 394.2399705 | -0.517489338 | 0.234226617 | 0.589 |
| YGL243W | 434.8625344 | -0.217761221 | 0.234709559 | 0.589 |
| YHR083W | 1675.31411 | -0.280659288 | 0.234704895 | 0.589 |
| YOR134W | 351.2595047 | 0.37626463 | 0.235339708 | 0.590 |
| YOR368W | 442.6825001 | 0.273018444 | 0.235523256 | 0.591 |
| YLR138W | 1645.971988 | -0.46263894 | 0.235650384 | 0.591 |
| YHR014W | 138.7802733 | 0.170016843 | 0.235856147 | 0.591 |
| YPL023C | 1340.132366 | -0.281253235 | 0.235756663 | 0.591 |
| YLR031W | 139.756576 | 0.227996008 | 0.236075923 | 0.591 |
| YPL119C-A | 58.40963327 | 0.353889831 | 0.236058371 | 0.591 |
| YOL163W | 57.10674557 | 0.461563852 | 0.23647555 | 0.591 |
| YLR030W | 185.5800298 | 0.24687682 | 0.236665564 | 0.592 |
| YLR206W | 3965.73873 | 0.288186758 | 0.236847556 | 0.592 |
| YLR427W | 1359.694674 | 0.171173495 | 0.237121792 | 0.592 |
| YNL277W | 546.4771573 | 0.322903865 | 0.237042637 | 0.592 |
| snR37 | 153.9250937 | -0.167660205 | 0.236976662 | 0.592 |
| YAL037W | 195.7634153 | 0.578425168 | 0.238112679 | 0.592 |
| YDL234C | 2388.462086 | 0.310640071 | 0.237953297 | 0.592 |
| YGR288W | 284.2488249 | 0.434647919 | 0.238216747 | 0.592 |
| YLR248W | 3268.331309 | 0.436422577 | 0.237336227 | 0.592 |
| YMR302C | 2126.727676 | 0.440646793 | 0.237618747 | 0.592 |
| YNR014W | 1455.118637 | 0.261654932 | 0.237659184 | 0.592 |
| YOL055C | 533.3864453 | 0.336675634 | 0.238105023 | 0.592 |
| YOL137W | 547.8586694 | 0.286482975 | 0.238222765 | 0.592 |
| YOR038C | 676.9008791 | 0.284647241 | 0.238158505 | 0.592 |
| YPL180W | 1077.43255 | 0.248378109 | 0.237820357 | 0.592 |
| YGL221C | 1756.035958 | -0.329606222 | 0.238425174 | 0.592 |

| | | | | |
|---|---|---|---|---|
| YDL112W | 1910.821894 | -0.354313118 | 0.238553359 | 0.592 |
| YOR343C | 63.14207102 | 0.615061729 | 0.238628772 | 0.592 |
| YAL041W | 1345.73957 | 0.37331363 | 0.238901339 | 0.593 |
| YDR070C | 1811.421734 | 0.629104662 | 0.239082345 | 0.593 |
| YKL074C | 747.5415477 | 0.240817148 | 0.239051254 | 0.593 |
| YDR090C | 473.3492687 | 0.316326206 | 0.239736813 | 0.593 |
| YGR035W-A | 93.40510398 | -0.397101799 | 0.239478406 | 0.593 |
| YJL010C | 755.184634 | -0.350225379 | 0.239675116 | 0.593 |
| YLR241W | 1618.382874 | 0.305023563 | 0.23966991 | 0.593 |
| YPL060W | 360.6791713 | 0.507976743 | 0.239587196 | 0.593 |
| YAR075W | 64.72959559 | -0.043815853 | 0.240017249 | 0.593 |
| YDR328C | 1849.787592 | -0.289041126 | 0.240047104 | 0.593 |
| YMR094W | 306.3491696 | -0.408978065 | 0.240213275 | 0.594 |
| YGR029W | 703.0611046 | -0.304448621 | 0.240595019 | 0.594 |
| YKR026C | 956.8364878 | -0.333697293 | 0.241238587 | 0.596 |
| YER106W | 34.98872113 | 0.624340748 | 0.241369682 | 0.596 |
| YNR025C | 48.20698063 | -0.859356064 | 0.24165879 | 0.596 |
| YLR342W | 12036.59934 | -0.174650085 | 0.241871999 | 0.596 |
| YPR179C | 773.0608325 | 0.311995329 | 0.242068133 | 0.597 |
| YPL039W | 426.2421359 | 0.323605368 | 0.24219584 | 0.597 |
| YJR016C | 6106.441237 | -0.43346871 | 0.242304434 | 0.597 |
| YFL052W | 300.3969613 | 0.552999186 | 0.242445173 | 0.597 |
| YDR172W | 3599.759088 | -0.240557215 | 0.242764062 | 0.597 |
| YDR187C | 25.29702409 | -0.581766904 | 0.242742358 | 0.597 |
| YNR037C | 584.4173485 | -0.373929452 | 0.242601362 | 0.597 |
| YIR007W | 898.4409432 | 0.333315619 | 0.242974724 | 0.597 |
| YJL098W | 1058.267184 | -0.382989754 | 0.24305099 | 0.597 |
| YAL044C | 2900.555908 | -0.463390201 | 0.243246574 | 0.597 |
| YLR345W | 2235.165084 | 0.361077868 | 0.243397796 | 0.597 |
| YDL071C | 49.14928453 | -0.452493175 | 0.244112431 | 0.599 |
| YDR252W | 88.93914678 | 0.532131588 | 0.244332417 | 0.599 |
| YEL017W | 1022.681916 | -0.232496097 | 0.244354063 | 0.599 |
| YOL052C | 1181.165007 | -0.264969384 | 0.244270227 | 0.599 |
| YOR242C | 110.2984083 | 0.073253825 | 0.24440761 | 0.599 |
| YBL107C | 641.7007619 | -0.335348159 | 0.245799347 | 0.599 |
| YBR076W | 114.1570412 | 0.368822537 | 0.245781063 | 0.599 |
| YDL209C | 238.6569979 | -0.497611727 | 0.245678298 | 0.599 |
| YGR204W | 10459.91176 | -0.349355824 | 0.245888209 | 0.599 |
| YGR217W | 881.6554263 | -0.294145059 | 0.246063563 | 0.599 |
| YHL001W | 1027.533504 | -0.544695401 | 0.245308035 | 0.599 |
| YIL126W | 2338.60398 | -0.300746192 | 0.245481726 | 0.599 |
| YKL218C | 773.1891416 | 0.50615358 | 0.245434082 | 0.599 |
| YLR024C | 1454.81679 | 0.517346809 | 0.245531005 | 0.599 |

| | | | | |
|---|---|---|---|---|
| YMR185W | 803.0326457 | -0.441335942 | 0.245034549 | 0.599 |
| YMR304C-A | 599.6439516 | -0.492089098 | 0.246078144 | 0.599 |
| YNL138W-A | 114.4265169 | -0.419205977 | 0.245086087 | 0.599 |
| YOL063C | 983.1121387 | 0.281686082 | 0.245669851 | 0.599 |
| YOR281C | 1009.23062 | -0.403698813 | 0.245607369 | 0.599 |
| YOR372C | 1526.129326 | 0.355909705 | 0.245438436 | 0.599 |
| YLR270W | 2664.450215 | 0.525716928 | 0.246381202 | 0.599 |
| YDL080C | 732.479931 | 0.247218878 | 0.246531528 | 0.599 |
| YGL063W | 321.9330079 | -0.503968128 | 0.246603747 | 0.599 |
| YCL058C | 103.80183 | -0.234448852 | 0.246847352 | 0.600 |
| YOR086C | 3224.270398 | 0.313505636 | 0.246858199 | 0.600 |
| YIL151C | 1650.033227 | 0.333933958 | 0.24715916 | 0.600 |
| YJL193W | 438.7222894 | 0.356851997 | 0.247291609 | 0.600 |
| YLR094C | 1048.349221 | 0.27169621 | 0.247226119 | 0.600 |
| YPR033C | 3601.656962 | -0.344095215 | 0.247129567 | 0.600 |
| YMR170C | 596.7981642 | 0.478937581 | 0.24773774 | 0.600 |
| YDR282C | 383.6210181 | 0.416553267 | 0.248424131 | 0.601 |
| YGR174W-A | 47.91809201 | 0.432194943 | 0.248622595 | 0.601 |
| YML077W | 412.7293887 | -0.365912065 | 0.248592448 | 0.601 |
| YMR167W | 578.0579077 | 0.357948274 | 0.248602265 | 0.601 |
| YPL157W | 499.4077603 | -0.401658033 | 0.248234936 | 0.601 |
| YNL201C | 1076.736749 | 0.265851804 | 0.248725351 | 0.601 |
| YBR175W | 264.1838842 | -0.595935611 | 0.249077495 | 0.602 |
| YOL087C | 2320.349339 | 0.317338707 | 0.249014168 | 0.602 |
| YHR156C | 433.8648918 | 0.336629975 | 0.249603706 | 0.602 |
| YLR307W | 82.37745124 | 0.574612451 | 0.249583768 | 0.602 |
| YLR351C | 1673.534289 | -0.28252832 | 0.249503765 | 0.602 |
| YBR012W-B | 321.6022163 | 0.553010265 | 0.251562388 | 0.606 |
| YDR089W | 994.7212521 | 0.397838859 | 0.251659169 | 0.606 |
| YDR373W | 760.4394828 | -0.311730007 | 0.251913657 | 0.606 |
| YDR381W | 4059.585724 | -0.325273275 | 0.25203072 | 0.606 |
| YFR014C | 1739.196956 | 0.329887129 | 0.251884854 | 0.606 |
| YHR149C | 909.9713222 | -0.382036557 | 0.252076097 | 0.606 |
| YIR018C-A | 31.48971955 | 0.430385143 | 0.251643341 | 0.606 |
| YJR062C | 945.4060688 | 0.310179512 | 0.251534396 | 0.606 |
| YNL138W | 4814.982276 | -0.287655132 | 0.252001254 | 0.606 |
| YOR175C | 2021.823765 | -0.28185474 | 0.251770776 | 0.606 |
| YBR225W | 1153.176358 | 0.34771768 | 0.2523578 | 0.606 |
| YLL057C | 252.3608873 | 0.373796963 | 0.252755636 | 0.607 |
| YDR520C | 591.1970741 | 0.485736845 | 0.25339293 | 0.607 |
| YFL062W | 38.28792094 | 0.994615011 | 0.253050099 | 0.607 |
| YGL113W | 616.1660547 | 0.293507352 | 0.253540008 | 0.607 |
| YHR216W | 3639.691774 | -0.207620522 | 0.253526435 | 0.607 |

| | | | | |
|---|---|---|---|---|
| YNR034W-A | 4345.183948 | 0.538087448 | 0.253213634 | 0.607 |
| YOR003W | 499.5835225 | 0.236243167 | 0.253533213 | 0.607 |
| YCL002C | 469.4119103 | -0.42691319 | 0.254005316 | 0.608 |
| YER092W | 378.6245953 | -0.316203598 | 0.253910121 | 0.608 |
| YIL114C | 411.0869055 | -0.397527713 | 0.254248066 | 0.608 |
| YLR104W | 635.3306236 | -0.381662817 | 0.254186991 | 0.608 |
| YOR336W | 1255.740018 | 0.363568356 | 0.254849923 | 0.609 |
| YKL171W | 1421.131993 | 0.362813434 | 0.254971018 | 0.609 |
| YHR071C-A | 39.05421358 | -0.467074434 | 0.255155974 | 0.609 |
| YGR087C | 287.4531423 | 0.449944933 | 0.255486711 | 0.610 |
| YHR098C | 2961.484376 | -0.219868713 | 0.255552942 | 0.610 |
| YLL010C | 1089.845187 | 0.269179543 | 0.255496621 | 0.610 |
| YBR241C | 1365.325299 | 0.484697634 | 0.256220415 | 0.610 |
| YDL013W | 806.2108951 | 0.320367571 | 0.256250283 | 0.610 |
| YDL204W | 5424.611742 | 0.554845032 | 0.255996947 | 0.610 |
| YML128C | 9230.440998 | 0.561781092 | 0.256140205 | 0.610 |
| YNL129W | 332.9229067 | -0.251020303 | 0.256315183 | 0.610 |
| YNL220W | 4979.508941 | -0.321989193 | 0.256249005 | 0.610 |
| YFL050C | 563.7473526 | 0.343170952 | 0.256820991 | 0.610 |
| YJL198W | 970.0448734 | -0.461093835 | 0.256796897 | 0.610 |
| YLL016W | 1012.667191 | 0.487358914 | 0.256737929 | 0.610 |
| YKL151C | 2678.459393 | 0.482530355 | 0.257280499 | 0.611 |
| YMR010W | 733.3907547 | -0.321741928 | 0.257395623 | 0.611 |
| YOL156W | 95.75918972 | 0.685779135 | 0.257442961 | 0.611 |
| YOR191W | 2372.954185 | 0.351561435 | 0.257172682 | 0.611 |
| YBR032W | 28.5083245 | 0.242394601 | 0.257883882 | 0.611 |
| YBR156C | 807.2285578 | 0.249503548 | 0.259802514 | 0.611 |
| YDR105C | 933.9584198 | 0.311760216 | 0.259545383 | 0.611 |
| YER184C | 397.7514223 | 0.368449429 | 0.259955321 | 0.611 |
| YGR070W | 2198.55855 | 0.391864156 | 0.259397503 | 0.611 |
| YHR065C | 1260.588152 | -0.370023428 | 0.258190415 | 0.611 |
| YHR123W | 699.5030842 | -0.221441655 | 0.25863679 | 0.611 |
| YIL099W | 500.7923991 | 0.233979944 | 0.259832257 | 0.611 |
| YIL123W | 4293.520238 | -0.22891253 | 0.259612873 | 0.611 |
| YIR005W | 167.3355793 | 0.253731996 | 0.2592039 | 0.611 |
| YJL045W | 286.1990705 | 0.461120464 | 0.259063133 | 0.611 |
| YJL083W | 610.9975272 | 0.272280072 | 0.259882758 | 0.611 |
| YJL110C | 684.2542418 | 0.319586454 | 0.259987546 | 0.611 |
| YJL196C | 1508.45467 | -0.248296887 | 0.259140261 | 0.611 |
| YLR117C | 1230.224284 | 0.260102687 | 0.258013095 | 0.611 |
| YLR238W | 614.7184945 | 0.290130331 | 0.259563525 | 0.611 |
| YLR262C | 283.7623055 | -0.327739554 | 0.257727926 | 0.611 |
| YOR118W | 1462.615579 | 0.224679636 | 0.25795094 | 0.611 |

| | | | | |
|---|---|---|---|---|
| YOR178C | 2730.240932 | 0.472812675 | 0.2588822 | 0.611 |
| YOR211C | 1075.422511 | 0.301708165 | 0.258973909 | 0.611 |
| YOR365C | 309.6078398 | 0.356366415 | 0.258800618 | 0.611 |
| YPL164C | 886.5830673 | 0.276179466 | 0.259149091 | 0.611 |
| YPL182C | 43.60254699 | 0.547879911 | 0.259931096 | 0.611 |
| YLR059C | 1264.300383 | -0.268325286 | 0.260236409 | 0.611 |
| YPL112C | 806.8455711 | -0.283192818 | 0.260224415 | 0.611 |
| YLR350W | 3571.832696 | 0.281974789 | 0.260382831 | 0.611 |
| YDL227C | 308.6751335 | -0.262940877 | 0.260658002 | 0.612 |
| YDR477W | 1886.671826 | 0.25492883 | 0.260857869 | 0.612 |
| YLR324W | 1737.806437 | 0.309323856 | 0.261018408 | 0.612 |
| YMR099C | 2686.030784 | -0.222736692 | 0.261135111 | 0.612 |
| YJL026W | 4324.794509 | -0.30260106 | 0.261366782 | 0.612 |
| YLR039C | 1333.277955 | 0.300794208 | 0.261274928 | 0.612 |
| YNL256W | 1084.449905 | -0.374135095 | 0.261807038 | 0.613 |
| YJL066C | 1041.928086 | 0.625415181 | 0.262041773 | 0.613 |
| YNL291C | 628.873147 | -0.284502044 | 0.261939781 | 0.613 |
| YJL131C | 712.8647158 | 0.377589791 | 0.262246969 | 0.613 |
| SCR1 | 181.7417356 | 0.334633485 | 0.262511882 | 0.613 |
| YAL016W | 2950.826603 | 0.300360058 | 0.263126191 | 0.613 |
| YIL006W | 374.9591791 | 0.277064196 | 0.262984924 | 0.613 |
| YKR048C | 4547.539769 | -0.35783772 | 0.263021791 | 0.613 |
| YML048W | 4451.705992 | -0.271170954 | 0.262643718 | 0.613 |
| YNL189W | 2792.970706 | -0.259219494 | 0.263084681 | 0.613 |
| YNL265C | 447.0625511 | 0.277750477 | 0.262781209 | 0.613 |
| YOL135C | 544.3922283 | -0.294097754 | 0.262893119 | 0.613 |
| YGR130C | 4694.98638 | 0.350069146 | 0.263523534 | 0.614 |
| YIL056W | 1630.203256 | 0.284882522 | 0.263421574 | 0.614 |
| YKL095W | 323.4844945 | 0.260102108 | 0.263652262 | 0.614 |
| YDR392W | 654.5457347 | 0.226848019 | 0.263855738 | 0.614 |
| YDL176W | 672.0622412 | 0.324625547 | 0.264275189 | 0.615 |
| YFR045W | 698.908615 | 0.435568617 | 0.264283522 | 0.615 |
| YCR009C | 3474.642891 | -0.267871137 | 0.264448133 | 0.615 |
| YER047C | 911.6547256 | 0.301640714 | 0.264579799 | 0.615 |
| YAL038W | 41041.91188 | -0.205523158 | 0.264927317 | 0.615 |
| YKL040C | 858.7842501 | -0.317484927 | 0.265194573 | 0.616 |
| YLR145W | 262.3006502 | -0.356308911 | 0.265199862 | 0.616 |
| YGL222C | 1086.796722 | 0.520964022 | 0.265450691 | 0.616 |
| YMR144W | 347.0360461 | -0.322601756 | 0.265375758 | 0.616 |
| YJR124C | 485.3455454 | -0.491048529 | 0.266361265 | 0.618 |
| YAL008W | 865.7638687 | 0.482686262 | 0.267368656 | 0.618 |
| YAL044W-A | 228.0761832 | -0.578318746 | 0.268660064 | 0.618 |
| YAL051W | 1222.388192 | 0.289691926 | 0.269317451 | 0.618 |

| | | | | |
|---|---|---|---|---|
| YAL067C | 87.22318015 | 1.230394172 | 0.268262418 | 0.618 |
| YDL029W | 3675.857405 | -0.174546947 | 0.268030393 | 0.618 |
| YDL169C | 664.258116 | 0.292342007 | 0.268851871 | 0.618 |
| YDL186W | 120.8658792 | 0.503994858 | 0.267764662 | 0.618 |
| YDR114C | 36.82268978 | 0.214553285 | 0.269241417 | 0.618 |
| YDR432W | 4551.301879 | -0.233700658 | 0.26923928 | 0.618 |
| YFL038C | 5600.505944 | -0.276100248 | 0.269567822 | 0.618 |
| YGL257C | 1193.728393 | -0.292620852 | 0.268223525 | 0.618 |
| YHL005C | 24.92643688 | 0.700311354 | 0.268778885 | 0.618 |
| YIL111W | 1481.52504 | 0.469664174 | 0.269275655 | 0.618 |
| YIR021W | 261.891268 | -0.323944884 | 0.267841127 | 0.618 |
| YLL052C | 77.86079202 | 0.268557549 | 0.267732854 | 0.618 |
| YLR239C | 753.9388732 | 0.351849592 | 0.268586695 | 0.618 |
| YLR267W | 1321.159087 | 0.232626503 | 0.26820239 | 0.618 |
| YLR330W | 3759.858385 | -0.211605194 | 0.267200061 | 0.618 |
| YMR134W | 453.7137928 | -0.327971282 | 0.269631103 | 0.618 |
| YNL062C | 805.5436558 | -0.381662371 | 0.26705396 | 0.618 |
| YNL080C | 1115.372394 | -0.333834398 | 0.269541872 | 0.618 |
| YNL130C | 1326.028586 | -0.248365279 | 0.268393205 | 0.618 |
| YOR026W | 568.5531387 | -0.45356745 | 0.267779448 | 0.618 |
| YOR169C | 39.59953477 | -1.012983382 | 0.267118466 | 0.618 |
| YOR220W | 2249.332399 | 0.370142052 | 0.269036096 | 0.618 |
| YPL119C | 673.8655489 | 0.348218272 | 0.26696325 | 0.618 |
| YPL268W | 929.1963896 | 0.423334408 | 0.267697898 | 0.618 |
| YPR069C | 2243.3628 | -0.438988953 | 0.268203357 | 0.618 |
| YPR125W | 636.6611003 | -0.305381507 | 0.26957938 | 0.618 |
| YNL115C | 1894.716258 | 0.401391911 | 0.269970427 | 0.619 |
| YNL103W | 1585.703492 | 0.325641132 | 0.270372707 | 0.619 |
| YLL063C | 302.5298246 | 0.455746542 | 0.27079004 | 0.619 |
| YMR228W | 316.010038 | -0.288729026 | 0.270561308 | 0.619 |
| YNL063W | 522.2571621 | 0.252860863 | 0.270633539 | 0.619 |
| YOR044W | 320.4821165 | 0.28340714 | 0.270805488 | 0.619 |
| YNL095C | 702.5523068 | 0.299009179 | 0.271298828 | 0.620 |
| YBL030C | 9599.31549 | 0.022698242 | 0.271771484 | 0.621 |
| YMR030W | 574.6177195 | 0.35411288 | 0.271851696 | 0.621 |
| YHL042W | 88.3151764 | 0.497016057 | 0.272140594 | 0.621 |
| YPL007C | 583.714763 | -0.200556216 | 0.272556673 | 0.622 |
| YLR054C | 138.716349 | 0.410703027 | 0.272900367 | 0.622 |
| YNL123W | 2060.867856 | -0.333544288 | 0.272870102 | 0.622 |
| YER119C-A | 190.6802462 | -0.348418419 | 0.273301032 | 0.623 |
| YJR107W | 474.7713781 | 0.394226481 | 0.273094858 | 0.623 |
| YKR075C | 2465.019565 | 0.161669046 | 0.273262506 | 0.623 |
| YGR210C | 1863.079648 | -0.286046238 | 0.273717569 | 0.623 |

| | | | | |
|---|---|---|---|---|
| YPL259C | 962.6624104 | 0.231964461 | 0.273637642 | 0.623 |
| YDR423C | 1205.3164 | 0.381039654 | 0.273871217 | 0.623 |
| YMR033W | 1176.392622 | -0.283182133 | 0.273986978 | 0.623 |
| YEL030W | 524.4916329 | 0.291414276 | 0.274408226 | 0.624 |
| YLR337C | 2433.363455 | -0.324438677 | 0.274506333 | 0.624 |
| YBL099W | 11787.99535 | -0.200280099 | 0.274737248 | 0.624 |
| YEL056W | 977.5251167 | 0.23852117 | 0.275522174 | 0.625 |
| YGR189C | 5490.403188 | -0.436165722 | 0.275575608 | 0.625 |
| YOR291W | 1661.347699 | 0.25911026 | 0.275398598 | 0.625 |
| YOR376W | 25.04102901 | 0.506164801 | 0.275285978 | 0.625 |
| YPR051W | 72.40075259 | -0.410434517 | 0.27533837 | 0.625 |
| YBR069C | 2910.073987 | -0.429702622 | 0.275745518 | 0.625 |
| YDL123W | 682.3334241 | 0.165434658 | 0.275802034 | 0.625 |
| YPL197C | 174.3224877 | -0.32955577 | 0.27600968 | 0.625 |
| YLR246W | 470.2308665 | 0.47398767 | 0.276148884 | 0.625 |
| YDR350C | 1056.007335 | 0.256534971 | 0.276713134 | 0.626 |
| snR17a | 59.62016603 | -0.213787764 | 0.276719923 | 0.626 |
| YGR024C | 991.4360363 | -0.228450205 | 0.277754093 | 0.626 |
| YHR200W | 3347.025252 | -0.299794598 | 0.277185251 | 0.626 |
| YKL079W | 964.5499269 | 0.22731261 | 0.277449189 | 0.626 |
| YLR067C | 640.6258307 | 0.281626717 | 0.277457752 | 0.626 |
| YMR245W | 80.78367086 | -0.635847555 | 0.277721125 | 0.626 |
| YMR278W | 1574.862371 | 0.360882247 | 0.277406203 | 0.626 |
| YOR097C | 623.6386998 | 0.278823462 | 0.277706098 | 0.626 |
| YPL116W | 1071.716343 | 0.292655056 | 0.277562013 | 0.626 |
| YEL053W-A | 186.8810681 | -0.244896521 | 0.277904357 | 0.626 |
| YKL010C | 2290.07323 | 0.226993499 | 0.278130022 | 0.627 |
| YBL081W | 976.0547384 | -0.444863199 | 0.278310284 | 0.627 |
| YCL026C-B | 269.8485152 | 0.527131347 | 0.278659426 | 0.627 |
| YCL032W | 909.5801687 | 0.258967371 | 0.278752328 | 0.627 |
| YKL174C | 903.8221117 | 0.196185778 | 0.278732359 | 0.627 |
| snR19 | 122.3490857 | -0.234866239 | 0.278616953 | 0.627 |
| YGL183C | 72.4162606 | 0.229620846 | 0.278867532 | 0.627 |
| YDR184C | 548.4562077 | -0.426369424 | 0.279286911 | 0.627 |
| YJL062W | 1004.118516 | -0.213807978 | 0.27922881 | 0.627 |
| YGL065C | 1014.84416 | -0.234821719 | 0.27975133 | 0.628 |
| YDR006C | 2310.266671 | 0.256751254 | 0.279986715 | 0.629 |
| YFL023W | 930.5339775 | -0.474374633 | 0.280913956 | 0.630 |
| YCL021W-A | 277.7575158 | -0.347078487 | 0.281473736 | 0.631 |
| YGR057C | 395.64537 | -0.327966546 | 0.281505751 | 0.631 |
| YDR393W | 653.7265915 | 0.289888962 | 0.281862876 | 0.632 |
| YGL237C | 638.8350245 | 0.331155805 | 0.282025574 | 0.632 |
| YGR268C | 875.2811265 | 0.24789811 | 0.281985776 | 0.632 |

| | | | | |
|---|---|---|---|---|
| YLR245C | 188.3206886 | -0.457643791 | 0.282114214 | 0.632 |
| YCR020C | 245.5464449 | 0.234511422 | 0.282426276 | 0.632 |
| YGL095C | 1579.460918 | 0.268668613 | 0.283005673 | 0.633 |
| YNL101W | 1154.001663 | 0.333665177 | 0.282985376 | 0.633 |
| YCR053W | 6107.267981 | -0.354017972 | 0.283642128 | 0.634 |
| YHR192W | 784.2571123 | -0.283982902 | 0.28351457 | 0.634 |
| YNL274C | 1846.518968 | 0.431453802 | 0.283704654 | 0.634 |
| YDL245C | 34.45281204 | 0.791743878 | 0.283962257 | 0.634 |
| YDR155C | 17689.29075 | -0.400453094 | 0.283868873 | 0.634 |
| YBR013C | 207.4427581 | 0.351006331 | 0.287271465 | 0.634 |
| YBR130C | 1160.408881 | -0.329436111 | 0.287576889 | 0.634 |
| YBR196C | 23554.20363 | -0.275200129 | 0.287836251 | 0.634 |
| YBR198C | 1683.847961 | -0.292108383 | 0.286479372 | 0.634 |
| YBR265W | 1130.650771 | -0.243736717 | 0.285241256 | 0.634 |
| YCR028C | 527.4192161 | 0.237906353 | 0.284524126 | 0.634 |
| YDR040C | 132.5886651 | 0.215671261 | 0.284833401 | 0.634 |
| YDR058C | 448.3672811 | 0.423529112 | 0.286465522 | 0.634 |
| YDR186C | 2571.215729 | 0.330052632 | 0.286234388 | 0.634 |
| YDR271C | 28.54979964 | -0.450794883 | 0.286883199 | 0.634 |
| YDR391C | 1742.594469 | 0.370875927 | 0.285538157 | 0.634 |
| YEL006W | 705.1819844 | 0.335400774 | 0.287690424 | 0.634 |
| YFR012W | 37.52047909 | 0.489771822 | 0.285461773 | 0.634 |
| YFR053C | 31755.34613 | 0.596136725 | 0.287555395 | 0.634 |
| YHL048W | 1666.986201 | 0.533351504 | 0.28543953 | 0.634 |
| YHR047C | 1885.311487 | -0.320879193 | 0.287196928 | 0.634 |
| YHR199C-A | 146.5882374 | 0.325452715 | 0.287650879 | 0.634 |
| YIL155C | 2791.794832 | 0.355177475 | 0.28647761 | 0.634 |
| YIR019C | 622.3080892 | 0.365655391 | 0.287352903 | 0.634 |
| YKL014C | 1967.775598 | -0.341291155 | 0.286007481 | 0.634 |
| YKR037C | 412.1583252 | -0.241961194 | 0.285329767 | 0.634 |
| YKR077W | 759.0676767 | 0.343771282 | 0.28778089 | 0.634 |
| YLR144C | 964.8658379 | 0.357871538 | 0.284954019 | 0.634 |
| YML090W | 156.4475378 | -0.31419433 | 0.286703367 | 0.634 |
| YMR102C | 1653.341009 | -0.54685107 | 0.28457705 | 0.634 |
| YMR114C | 1303.792032 | 0.265801746 | 0.28743922 | 0.634 |
| YMR141C | 167.2172498 | 0.209724888 | 0.286880328 | 0.634 |
| YNL011C | 614.0144377 | 0.337624905 | 0.287145176 | 0.634 |
| YNL305C | 2021.497779 | 0.43470333 | 0.285222868 | 0.634 |
| YOL052C-A | 1535.771374 | 0.494693785 | 0.287477297 | 0.634 |
| YOR258W | 218.0314211 | 0.375004031 | 0.286995255 | 0.634 |
| YPR003C | 637.4553595 | 0.341179611 | 0.286516845 | 0.634 |
| YPR173C | 1842.223717 | -0.318803598 | 0.287157921 | 0.634 |
| YBL071C | 215.271228 | -0.638467526 | 0.289075612 | 0.636 |

| | | | | |
|---|---|---|---|---|
| YBR201C-A | 201.3285883 | 0.289995837 | 0.289152426 | 0.636 |
| YDR118W | 391.9679351 | 0.224589353 | 0.288808761 | 0.636 |
| YFL034C-A | 576.1615287 | -0.665027383 | 0.289203081 | 0.636 |
| YGR133W | 486.059603 | -0.346571039 | 0.288971524 | 0.636 |
| YMR105C | 9001.650714 | 0.472973001 | 0.289203431 | 0.636 |
| YMR254C | 26.76419027 | 0.312493823 | 0.288860166 | 0.636 |
| YCR075W-A | 129.8943621 | -0.458593698 | 0.2896326 | 0.637 |
| YDL125C | 2310.617952 | -0.393402127 | 0.29024285 | 0.638 |
| YDR028C | 3835.45418 | 0.335830378 | 0.290603652 | 0.638 |
| YEL046C | 2157.479841 | -0.322365895 | 0.290684066 | 0.638 |
| YOR007C | 8060.679187 | -0.280327859 | 0.291221355 | 0.639 |
| YKL133C | 601.5672696 | 0.303685303 | 0.291689155 | 0.640 |
| YNL206C | 942.6169972 | -0.232076452 | 0.29172092 | 0.640 |
| YGL057C | 252.5789153 | 0.259656828 | 0.292008581 | 0.640 |
| YOL015W | 325.5388955 | 0.262480671 | 0.292097934 | 0.640 |
| YLR194C | 2445.708189 | -0.623897472 | 0.292264324 | 0.640 |
| YCR032W | 1688.564969 | 0.392722889 | 0.292914995 | 0.640 |
| YGR194C | 2007.303758 | 0.369204534 | 0.293058866 | 0.640 |
| YIL077C | 964.8137743 | 0.371387033 | 0.292863565 | 0.640 |
| YJR010C-A | 212.1436636 | -0.359140456 | 0.292920793 | 0.640 |
| YOL141W | 189.2930206 | -0.398773584 | 0.293065941 | 0.640 |
| YPL185W | 39.28495564 | 0.249849954 | 0.292681923 | 0.640 |
| YPR089W | 1983.427567 | 0.246989828 | 0.292418602 | 0.640 |
| YDR386W | 496.5773156 | 0.282492579 | 0.293261303 | 0.640 |
| YGR230W | 330.0627222 | -0.344132544 | 0.293718015 | 0.641 |
| YHL050C | 143.7644195 | 0.436723081 | 0.293572706 | 0.641 |
| YLL041C | 3315.150059 | 0.605143412 | 0.293763091 | 0.641 |
| YIL053W | 6767.756759 | -0.533466865 | 0.29396778 | 0.641 |
| YLR070C | 346.2836033 | 0.277101549 | 0.294007536 | 0.641 |
| YCR012W | 43096.09714 | -0.151677263 | 0.294262467 | 0.641 |
| YBL063W | 882.9945164 | -0.292307826 | 0.295074789 | 0.641 |
| YDR210W-B | 26.10421514 | 0.299532197 | 0.29452289 | 0.641 |
| YDR326C | 1698.894045 | 0.265224483 | 0.295082844 | 0.641 |
| YJL153C | 833.1459195 | 0.042667065 | 0.295171644 | 0.641 |
| YLR413W | 2093.12908 | -0.105886024 | 0.294908115 | 0.641 |
| YML056C | 4012.776938 | -0.432598726 | 0.294912289 | 0.641 |
| YMR090W | 1879.223335 | 0.462900088 | 0.294686113 | 0.641 |
| YPL030W | 552.0003609 | -0.342063864 | 0.295138265 | 0.641 |
| YML076C | 1190.303539 | 0.281587265 | 0.29533175 | 0.641 |
| YPR160W | 12134.47846 | 0.57609698 | 0.295526185 | 0.642 |
| YLR432W | 4140.896048 | -0.4110157 | 0.295902907 | 0.642 |
| YHR002W | 437.310458 | 0.336894838 | 0.296211985 | 0.642 |
| YOR076C | 709.8237452 | 0.264680988 | 0.296228541 | 0.642 |

| | | | | |
|---|---|---|---|---|
| YBR009C | 3036.286227 | -0.590643407 | 0.296785881 | 0.643 |
| YJL215C | 44.17616213 | 0.541282347 | 0.296842555 | 0.643 |
| YJR013W | 645.2124988 | 0.268311551 | 0.296530452 | 0.643 |
| YLR214W | 1682.159024 | -0.510965496 | 0.296622047 | 0.643 |
| YMR101C | 112.86694 | 0.407885309 | 0.29678267 | 0.643 |
| YAL064C-A | 47.35030167 | 0.764973451 | 0.29718743 | 0.643 |
| YOR022C | 608.0869392 | 0.399700719 | 0.297189332 | 0.643 |
| YFR042W | 519.19235 | -0.309359308 | 0.297617544 | 0.643 |
| YKL023W | 1129.71016 | 0.369495972 | 0.297536495 | 0.643 |
| YKL170W | 81.53735369 | -0.264824963 | 0.297670071 | 0.643 |
| YDR217C | 884.5738376 | -0.360070037 | 0.298807703 | 0.644 |
| YER148W | 733.3492834 | -0.352299223 | 0.29868527 | 0.644 |
| YFR038W | 1186.649674 | -0.34171832 | 0.298405577 | 0.644 |
| YGL047W | 625.4320388 | -0.193215459 | 0.298535364 | 0.644 |
| YGR063C | 116.1850697 | -0.311147602 | 0.298279088 | 0.644 |
| YML124C | 2648.714801 | -0.310489231 | 0.29846366 | 0.644 |
| YPL224C | 949.2587858 | 0.367333774 | 0.298735931 | 0.644 |
| YBL065W | 30.86397396 | -0.00784949 | 0.299161864 | 0.644 |
| YIL010W | 1043.363564 | -0.311051471 | 0.299142156 | 0.644 |
| YDR001C | 5262.491506 | 0.351241688 | 0.299305949 | 0.644 |
| YBR240C | 118.1647641 | 0.37482905 | 0.299446708 | 0.644 |
| YDL093W | 807.8112275 | -0.247896908 | 0.300013592 | 0.645 |
| YDR281C | 42.74964081 | -0.6140644 | 0.300701567 | 0.645 |
| YDR374C | 71.50374083 | 0.15599361 | 0.300641419 | 0.645 |
| YFL018C | 6022.957872 | -0.211803988 | 0.300532556 | 0.645 |
| YJL109C | 2576.133571 | -0.274788406 | 0.300644036 | 0.645 |
| YKL024C | 1589.092419 | -0.308080354 | 0.300108298 | 0.645 |
| YLL025W | 92.59299699 | 0.20903106 | 0.300330654 | 0.645 |
| YMR080C | 1907.455726 | -0.313588798 | 0.300196567 | 0.645 |
| YMR139W | 1446.676083 | 0.366077616 | 0.30001838 | 0.645 |
| YNL018C | 116.9826504 | 0.376059074 | 0.300243174 | 0.645 |
| YDR235W | 670.6497022 | 0.239976945 | 0.300823846 | 0.645 |
| YBL035C | 596.37002 | -0.180098091 | 0.301312817 | 0.645 |
| YBR015C | 2379.48263 | -0.257425378 | 0.30121425 | 0.645 |
| YBR052C | 2651.004465 | -0.294768894 | 0.301361342 | 0.645 |
| YDR453C | 889.6622872 | 0.407899933 | 0.30144639 | 0.645 |
| YBR065C | 528.4307138 | 0.334331293 | 0.301756057 | 0.645 |
| YBR071W | 990.0527368 | -0.453847177 | 0.301845182 | 0.645 |
| YPR162C | 653.0734249 | 0.4705725 | 0.301855465 | 0.645 |
| YBL051C | 3664.33556 | -0.334112694 | 0.302293253 | 0.646 |
| YDR224C | 8028.945616 | -0.433006442 | 0.302650941 | 0.646 |
| YDR439W | 511.333794 | 0.353397798 | 0.302550919 | 0.646 |
| YOR126C | 423.6858153 | -0.221506503 | 0.30286041 | 0.646 |

| | | | | |
|---|---|---|---|---|
| YOL117W | 709.0122211 | 0.382682878 | 0.303152316 | 0.647 |
| YDR410C | 388.474414 | -0.352320368 | 0.303492158 | 0.647 |
| YKR013W | 4305.106637 | -0.341041792 | 0.303665442 | 0.647 |
| YLR393W | 270.1892054 | 0.324042343 | 0.303603359 | 0.647 |
| YMR247C | 1642.766432 | 0.333579133 | 0.30343093 | 0.647 |
| YPR076W | 41.22373968 | 0.57368574 | 0.304082602 | 0.648 |
| YPR188C | 534.7061592 | -0.309936649 | 0.304204527 | 0.648 |
| YBR285W | 449.6014678 | 0.501249539 | 0.3047886 | 0.648 |
| YDL185W | 10910.49268 | -0.220773055 | 0.305054052 | 0.648 |
| YDR441C | 376.4209723 | -0.360815546 | 0.305461705 | 0.648 |
| YDR513W | 3464.363476 | 0.431026721 | 0.305459262 | 0.648 |
| YER151C | 2013.443986 | -0.248791476 | 0.305563284 | 0.648 |
| YGL073W | 2411.314838 | 0.271932647 | 0.305717015 | 0.648 |
| YJR122W | 745.1766826 | 0.362049101 | 0.305748753 | 0.648 |
| YJR130C | 1173.075557 | 0.30627335 | 0.305174461 | 0.648 |
| YKL139W | 677.7577592 | 0.269031144 | 0.305434659 | 0.648 |
| YLR327C | 4824.371416 | 0.368608995 | 0.305635079 | 0.648 |
| YMR013C | 348.9768631 | 0.327718265 | 0.305733931 | 0.648 |
| YNL111C | 246.1352445 | -0.318995312 | 0.305350815 | 0.648 |
| YPR147C | 688.0952487 | -0.207834189 | 0.30565302 | 0.648 |
| YLR321C | 729.7163579 | -0.314933182 | 0.306242197 | 0.649 |
| YCR023C | 1565.302118 | 0.377594859 | 0.307131438 | 0.649 |
| YDR405W | 1203.736123 | -0.262804381 | 0.306681981 | 0.649 |
| YER128W | 281.9899223 | 0.298927858 | 0.306710431 | 0.649 |
| YGL048C | 4170.423196 | -0.218791383 | 0.306928005 | 0.649 |
| YKL011C | 416.7502341 | 0.292600734 | 0.307034352 | 0.649 |
| YOL003C | 636.2491343 | -0.236455337 | 0.306581679 | 0.649 |
| YOR283W | 529.1178643 | -0.311428378 | 0.306922401 | 0.649 |
| YDR077W | 88090.2349 | 0.368142258 | 0.307955212 | 0.649 |
| YER038C | 327.709464 | 0.382621921 | 0.307974548 | 0.649 |
| YFL019C | 43.77288423 | 0.365011265 | 0.30785508 | 0.649 |
| YGR174C | 1048.598299 | 0.352545246 | 0.308071166 | 0.649 |
| YIR016W | 1147.104809 | 0.293357352 | 0.307501173 | 0.649 |
| YNL076W | 1058.446405 | 0.231070332 | 0.307743999 | 0.649 |
| YPL009C | 4156.098331 | -0.418589949 | 0.308086545 | 0.649 |
| YNL281W | 1354.281332 | -0.25538398 | 0.308320401 | 0.649 |
| YPL088W | 468.228703 | 0.179471396 | 0.308211608 | 0.649 |
| YDL057W | 396.4937822 | 0.406733532 | 0.309246969 | 0.651 |
| YDR198C | 539.1037114 | 0.248744615 | 0.309350629 | 0.651 |
| YER132C | 1149.8348 | -0.309546547 | 0.309361773 | 0.651 |
| YNR035C | 3358.997637 | -0.336390896 | 0.309232742 | 0.651 |
| YGL232W | 728.6792639 | -0.223903314 | 0.309622874 | 0.651 |
| YPR169W | 1659.207747 | -0.37207829 | 0.309518403 | 0.651 |

| | | | | |
|---|---|---|---|---|
| YDL023C | 219.5343501 | 0.379256831 | 0.310088907 | 0.651 |
| YDL126C | 11516.69474 | -0.24415791 | 0.310345467 | 0.651 |
| YGL089C | 132.0787924 | 0.245926937 | 0.309891096 | 0.651 |
| YGR008C | 4582.699628 | 0.401927172 | 0.310371867 | 0.651 |
| YHR001W | 1515.116914 | 0.278716139 | 0.310036906 | 0.651 |
| YIL071W-A | 73.5379669 | 0.124764148 | 0.309946682 | 0.651 |
| YJR094C | 253.8151399 | 0.545201861 | 0.310414795 | 0.651 |
| YLR299W | 1242.489768 | 0.28667917 | 0.310539144 | 0.651 |
| YDL088C | 1480.552376 | 0.241585836 | 0.310904125 | 0.651 |
| YHL044W | 369.968591 | 0.525060745 | 0.310961694 | 0.651 |
| YJR089W | 953.5011996 | 0.237686605 | 0.310820294 | 0.651 |
| YOR262W | 658.3672371 | -0.424447149 | 0.311255531 | 0.651 |
| YDR384C | 561.1890906 | -0.485150044 | 0.311549138 | 0.652 |
| YJL038C | 194.4642451 | 0.642390863 | 0.31176049 | 0.652 |
| YLR407W | 355.7060391 | 0.259771141 | 0.311926558 | 0.652 |
| YGL146C | 201.790122 | 0.388544314 | 0.313044945 | 0.654 |
| YHR033W | 1072.035586 | 0.094035117 | 0.313374208 | 0.654 |
| YIR011C | 1166.213259 | 0.243792028 | 0.313488658 | 0.654 |
| YIR043C | 49.74477082 | 0.186382543 | 0.313441557 | 0.654 |
| YKL150W | 7424.00365 | 0.474561474 | 0.313622759 | 0.654 |
| YLL032C | 878.0650576 | 0.307415369 | 0.313513437 | 0.654 |
| YLR005W | 967.4119109 | 0.223997281 | 0.313132421 | 0.654 |
| YLR129W | 1609.244371 | -0.368164412 | 0.313596477 | 0.654 |
| YOR062C | 585.4618256 | 0.12827008 | 0.313605389 | 0.654 |
| YGL249W | 273.3583474 | 0.245563681 | 0.314176606 | 0.654 |
| YPR157W | 448.7836668 | 0.431565304 | 0.314239087 | 0.654 |
| YEL032W | 1185.048516 | 0.247961033 | 0.314373131 | 0.654 |
| YBR220C | 693.632069 | -0.212927637 | 0.316351213 | 0.654 |
| YDR021W | 377.9993278 | -0.554063039 | 0.316089122 | 0.654 |
| YEL052W | 1984.785259 | 0.394311946 | 0.315742863 | 0.654 |
| YER141W | 2454.288383 | 0.47208148 | 0.314728348 | 0.654 |
| YFL012W | 27.05345673 | 0.429963412 | 0.315220325 | 0.654 |
| YFR019W | 2589.385757 | 0.30725263 | 0.31673509 | 0.654 |
| YGR113W | 643.4472036 | -0.183108299 | 0.315369418 | 0.654 |
| YGR209C | 5181.82162 | -0.27174145 | 0.315890372 | 0.654 |
| YHR177W | 301.5515693 | 0.421929706 | 0.31532207 | 0.654 |
| YIR004W | 1446.338254 | -0.263249234 | 0.314899253 | 0.654 |
| YIR029W | 388.4137592 | 0.492181525 | 0.316388186 | 0.654 |
| YKR080W | 1412.607481 | -0.410310833 | 0.315513301 | 0.654 |
| YMR029C | 1228.193843 | 0.331168622 | 0.315711479 | 0.654 |
| YMR199W | 1528.60034 | -0.323078037 | 0.316626109 | 0.654 |
| YNL041C | 1185.16925 | 0.284643099 | 0.315707017 | 0.654 |
| YNL199C | 1154.934737 | 0.153336995 | 0.316458173 | 0.654 |

| | | | | |
|---|---|---|---|---|
| YNR056C | 522.7352776 | 0.30896466 | 0.31673749 | 0.654 |
| YOR132W | 1718.595531 | 0.378642364 | 0.314612764 | 0.654 |
| YOR158W | 1525.323441 | -0.266842523 | 0.31570913 | 0.654 |
| YOR274W | 529.1848909 | 0.301994643 | 0.315659852 | 0.654 |
| YPR171W | 883.9863961 | 0.266587054 | 0.31604496 | 0.654 |
| YGR222W | 726.7683442 | 0.252562506 | 0.316989077 | 0.655 |
| YMR035W | 535.4792987 | -0.341260616 | 0.316937596 | 0.655 |
| YDL103C | 1449.48201 | -0.23444091 | 0.317746542 | 0.655 |
| YDR199W | 75.09684199 | 0.309341929 | 0.317630481 | 0.655 |
| YHR038W | 426.523879 | -0.25268243 | 0.317361698 | 0.655 |
| YIL079C | 348.2789788 | -0.384901765 | 0.317513526 | 0.655 |
| YIR035C | 1381.941223 | -0.29612194 | 0.317373918 | 0.655 |
| YKR027W | 720.4128865 | 0.235441181 | 0.317635281 | 0.655 |
| YMR296C | 3004.03015 | -0.351643598 | 0.318090223 | 0.655 |
| YPL183C | 913.1902093 | -0.322205403 | 0.317983623 | 0.655 |
| YER100W | 1553.364761 | -0.285673946 | 0.318546293 | 0.655 |
| YLL056C | 287.6766171 | 0.215841448 | 0.318707312 | 0.655 |
| YMR174C | 615.2184477 | 0.465652356 | 0.318827201 | 0.655 |
| YMR257C | 582.2395624 | 0.340488344 | 0.318860644 | 0.655 |
| YPL104W | 889.4598617 | 0.413651187 | 0.318548834 | 0.655 |
| YCR101C | 105.2670668 | 0.403990423 | 0.319243431 | 0.656 |
| YDL053C | 1309.962591 | -0.221685576 | 0.319824389 | 0.656 |
| YER113C | 743.4950149 | 0.276717263 | 0.319636054 | 0.656 |
| YGL142C | 1131.213732 | 0.33946524 | 0.319651365 | 0.656 |
| YLR057W | 1092.547525 | 0.271784223 | 0.319500419 | 0.656 |
| YOR352W | 729.6147343 | 0.241423744 | 0.319772448 | 0.656 |
| YCR026C | 1309.803177 | 0.334715229 | 0.320017958 | 0.656 |
| YJL168C | 610.0870071 | -0.30556023 | 0.320091246 | 0.656 |
| YCR094W | 731.6068684 | -0.220542218 | 0.321146797 | 0.657 |
| YDL174C | 2966.298062 | 0.324756829 | 0.321561937 | 0.657 |
| YDR154C | 3531.276985 | -0.486642318 | 0.321320938 | 0.657 |
| YEL055C | 1803.113225 | -0.228539154 | 0.320908248 | 0.657 |
| YHL003C | 1333.653462 | -0.204455118 | 0.320509127 | 0.657 |
| YHL010C | 843.3803368 | 0.41165755 | 0.3212716 | 0.657 |
| YHR007C-A | 61.56889406 | 0.149474255 | 0.320851752 | 0.657 |
| YKL153W | 3671.352718 | -0.204721725 | 0.321534069 | 0.657 |
| YLR109W | 26967.32056 | -0.281253332 | 0.321351738 | 0.657 |
| YNR068C | 378.9501574 | 0.3350443 | 0.320853811 | 0.657 |
| YPL175W | 580.2634042 | -0.248919524 | 0.3214461 | 0.657 |
| RUF20 | 86.17514557 | -0.45220785 | 0.322387968 | 0.657 |
| YBR276C | 1528.649976 | -0.183060384 | 0.32288844 | 0.657 |
| YFL027C | 673.7008422 | 0.365081077 | 0.32360382 | 0.657 |
| YFL042C | 1495.188388 | 0.412240156 | 0.322527218 | 0.657 |

| | | | | |
|---|---|---|---|---|
| YGR205W | 695.7192252 | 0.385150501 | 0.322282592 | 0.657 |
| YHL004W | 1819.11881 | -0.13688147 | 0.323309227 | 0.657 |
| YHR182W | 713.7981983 | 0.398996029 | 0.32369373 | 0.657 |
| YJL030W | 384.5980723 | -0.272955208 | 0.323013872 | 0.657 |
| YJR098C | 316.068717 | 0.321623126 | 0.323427899 | 0.657 |
| YLL018C-A | 305.6957299 | 0.404551352 | 0.323188257 | 0.657 |
| YLL020C | 30.70630028 | 0.247826375 | 0.323197531 | 0.657 |
| YLR035C | 528.4128752 | 0.355554327 | 0.323771198 | 0.657 |
| YML109W | 760.9435844 | -0.309651432 | 0.322861917 | 0.657 |
| YMR020W | 992.3354434 | 0.417789253 | 0.323666143 | 0.657 |
| YOL101C | 182.685157 | 0.396863156 | 0.323598838 | 0.657 |
| YOR049C | 767.2844506 | 0.311892486 | 0.323536348 | 0.657 |
| YDL045C | 1046.513314 | 0.182038304 | 0.323963189 | 0.658 |
| YGL043W | 1337.950161 | -0.294953454 | 0.324610779 | 0.658 |
| YKR031C | 1778.007412 | 0.290368117 | 0.324923453 | 0.658 |
| YOL038W | 4309.883987 | -0.296008935 | 0.324829575 | 0.658 |
| YOR036W | 1460.876014 | 0.230430409 | 0.324559002 | 0.658 |
| YOR099W | 2318.668874 | -0.208466941 | 0.324940187 | 0.658 |
| YOR256C | 1260.844527 | 0.273729058 | 0.324705705 | 0.658 |
| YCR077C | 3337.529518 | -0.223184811 | 0.325263281 | 0.659 |
| YCR035C | 836.2083583 | -0.214747131 | 0.325645621 | 0.659 |
| YIL163C | 43.51788641 | 0.367393015 | 0.325862106 | 0.659 |
| YGR198W | 1689.551002 | -0.323041023 | 0.326391562 | 0.660 |
| YLR380W | 1647.260273 | -0.305463696 | 0.326407485 | 0.660 |
| YNL158W | 434.8285056 | 0.191055316 | 0.326488541 | 0.660 |
| YAR007C | 1627.673812 | -0.241181002 | 0.326933777 | 0.661 |
| YGL122C | 2999.105342 | -0.398820279 | 0.327169768 | 0.661 |
| YLR455W | 572.5909218 | -0.284969322 | 0.327685094 | 0.662 |
| YBR102C | 889.0122882 | -0.181022334 | 0.328033973 | 0.662 |
| YCR105W | 210.4429239 | 0.677530749 | 0.327979075 | 0.662 |
| YBR050C | 396.3143748 | 0.394149867 | 0.328225652 | 0.662 |
| YDL167C | 1031.301724 | -0.315698155 | 0.32839643 | 0.662 |
| YGL178W | 2073.150568 | 0.312382797 | 0.328528347 | 0.662 |
| YMR110C | 3066.365125 | 0.396025321 | 0.328741136 | 0.662 |
| YPR028W | 4646.798863 | -0.274763462 | 0.329024796 | 0.663 |
| YCL001W-B | 30.32607662 | 0.455103302 | 0.329288778 | 0.663 |
| YKL128C | 1025.460861 | -0.31602266 | 0.32931376 | 0.663 |
| YLR379W | 289.3293491 | -0.083494652 | 0.329429502 | 0.663 |
| YEL044W | 582.0177494 | -0.307015055 | 0.329900594 | 0.663 |
| YMR305C | 6521.89708 | -0.303337196 | 0.329685996 | 0.663 |
| YPL274W | 1036.210642 | 0.41209898 | 0.329837076 | 0.663 |
| YER052C | 2274.895414 | -0.350855261 | 0.330563484 | 0.664 |
| YNL176C | 1925.34034 | 0.239197645 | 0.330479673 | 0.664 |

| | | | | |
|---|---|---|---|---|
| YOL086W-A | 318.8786531 | -0.555969708 | 0.330355243 | 0.664 |
| YGL023C | 1541.23519 | -0.307601264 | 0.33069734 | 0.664 |
| YNL088W | 2851.433881 | -0.286213321 | 0.331276333 | 0.664 |
| YNL121C | 5779.129433 | -0.204524837 | 0.331323004 | 0.664 |
| YOR199W | 28.95971662 | 0.013604522 | 0.33140451 | 0.664 |
| YOR335C | 3110.388093 | -0.209800432 | 0.331390269 | 0.664 |
| YOR189W | 408.1511083 | -0.269387139 | 0.331793634 | 0.665 |
| YAL056W | 1700.26388 | 0.354772017 | 0.33313523 | 0.665 |
| YDL177C | 218.4776378 | -0.463811152 | 0.333080781 | 0.665 |
| YDL218W | 253.6505308 | 0.220705345 | 0.332605798 | 0.665 |
| YFR009W | 2637.309837 | -0.197630707 | 0.332661496 | 0.665 |
| YJL095W | 1317.430087 | -0.207183282 | 0.333113133 | 0.665 |
| YLL023C | 2955.471664 | 0.348198557 | 0.332360184 | 0.665 |
| YLR290C | 1190.940137 | 0.285092261 | 0.332508839 | 0.665 |
| YNL031C | 9825.730059 | -0.370355212 | 0.333156611 | 0.665 |
| YNL196C | 204.2397635 | 0.195496559 | 0.332376815 | 0.665 |
| YOR141C | 1550.614472 | -0.270859036 | 0.332341741 | 0.665 |
| YOR342C | 1114.452608 | -0.250596438 | 0.332907814 | 0.665 |
| YPR012W | 66.76164095 | 0.275933468 | 0.331936985 | 0.665 |
| YDR190C | 2078.525809 | -0.273248193 | 0.3334215 | 0.665 |
| YER059W | 910.2356733 | 0.322573841 | 0.333395287 | 0.665 |
| YMR130W | 646.8299959 | -0.352681362 | 0.333543753 | 0.665 |
| YDL220C | 416.2502353 | -0.018145595 | 0.334358186 | 0.665 |
| YGL150C | 2988.328671 | 0.195070372 | 0.333866434 | 0.665 |
| YGL192W | 1154.271063 | 0.261243098 | 0.334107784 | 0.665 |
| YHR198C | 872.0543676 | 0.411873806 | 0.334226417 | 0.665 |
| YKL058W | 636.5138347 | -0.259807141 | 0.333766634 | 0.665 |
| YKL088W | 1161.516822 | 0.318911294 | 0.334262181 | 0.665 |
| YNL146W | 85.60940575 | 0.192929683 | 0.334373055 | 0.665 |
| YMR111C | 424.0046487 | 0.208144201 | 0.334661727 | 0.665 |
| YBR208C | 3562.373289 | 0.472917497 | 0.335435214 | 0.667 |
| YER107C | 624.5620743 | -0.316147085 | 0.335533264 | 0.667 |
| YHR086W | 910.6374675 | 0.424518604 | 0.335379474 | 0.667 |
| YDL054C | 853.6605887 | 0.242685345 | 0.33580198 | 0.667 |
| YJL031C | 712.299831 | 0.289819266 | 0.335929572 | 0.667 |
| YDR368W | 2244.977391 | -0.247432555 | 0.336227063 | 0.667 |
| YFL067W | 57.17560859 | 0.516604314 | 0.336145597 | 0.667 |
| YLR399C | 2272.649731 | 0.234141271 | 0.3363726 | 0.667 |
| YHR157W | 84.3613017 | 0.228805313 | 0.336610805 | 0.667 |
| YHR208W | 7460.933205 | -0.275818502 | 0.336589982 | 0.667 |
| YDL115C | 894.0269756 | 0.205611999 | 0.336945921 | 0.667 |
| YOR124C | 2791.881139 | 0.318942297 | 0.336977372 | 0.667 |
| YAL048C | 583.0194511 | 0.304085625 | 0.337577593 | 0.668 |

| | | | | |
|---|---|---|---|---|
| YMR163C | 451.0209425 | 0.221664546 | 0.337894948 | 0.669 |
| YOR193W | 437.421385 | 0.24268225 | 0.338048663 | 0.669 |
| YGL250W | 555.5711852 | 0.255947822 | 0.338795512 | 0.670 |
| YLR143W | 501.3258454 | -0.235492113 | 0.339139596 | 0.670 |
| YPL125W | 1168.744838 | 0.400365319 | 0.339100838 | 0.670 |
| YPL194W | 451.5443271 | 0.286050478 | 0.339390649 | 0.671 |
| YBR211C | 479.4755282 | 0.206233267 | 0.339594863 | 0.671 |
| YCR086W | 338.9363584 | -0.329489413 | 0.340002773 | 0.671 |
| YER042W | 653.285425 | -0.579398866 | 0.34005448 | 0.671 |
| YKL142W | 4281.67462 | 0.365065638 | 0.340150638 | 0.671 |
| YNL054W | 1269.220328 | -0.256888502 | 0.340022989 | 0.671 |
| YLR041W | 27.97790779 | 0.705597542 | 0.340406207 | 0.671 |
| YBR033W | 734.5856264 | 0.388713379 | 0.340745474 | 0.672 |
| YMR196W | 3241.463398 | 0.465750182 | 0.340956375 | 0.672 |
| YGR223C | 1134.541576 | 0.222838621 | 0.341285909 | 0.672 |
| YJL170C | 341.173457 | 0.595180958 | 0.341887877 | 0.673 |
| YER082C | 1253.635463 | -0.325293868 | 0.342720791 | 0.674 |
| YLR207W | 1228.065049 | 0.253550614 | 0.34243217 | 0.674 |
| YOR070C | 1064.02202 | 0.185225137 | 0.342647883 | 0.674 |
| YPR055W | 1160.769461 | 0.286649412 | 0.342771399 | 0.674 |
| YLR072W | 910.1580571 | 0.262685263 | 0.342950409 | 0.674 |
| YMR277W | 1413.809425 | -0.271701228 | 0.343060623 | 0.674 |
| YNL332W | 25.85198322 | 0.070684006 | 0.343180975 | 0.674 |
| YJL140W | 1600.235301 | -0.217559924 | 0.343779212 | 0.674 |
| YJL141C | 4385.516839 | 0.213713711 | 0.343761898 | 0.674 |
| YOL023W | 636.5704075 | 0.310312826 | 0.343583083 | 0.674 |
| YOL054W | 494.8918395 | -0.291449461 | 0.343778894 | 0.674 |
| YIL143C | 1830.368647 | 0.277139258 | 0.344532196 | 0.676 |
| YGR232W | 824.5178442 | -0.283726366 | 0.344732064 | 0.676 |
| YNL167C | 1564.476742 | 0.256522079 | 0.344881511 | 0.676 |
| YGL216W | 962.5040676 | 0.221094769 | 0.34568829 | 0.677 |
| YIL108W | 1452.736233 | 0.349096478 | 0.345895784 | 0.677 |
| YLR118C | 939.2371521 | -0.226420223 | 0.345843012 | 0.677 |
| YMR273C | 1360.501919 | 0.315424895 | 0.346110615 | 0.677 |
| YGL211W | 636.0371021 | -0.280256724 | 0.346369735 | 0.678 |
| YGR108W | 622.4932134 | -0.224167399 | 0.347509341 | 0.680 |
| YBR010W | 4241.31261 | -0.558765402 | 0.348292871 | 0.680 |
| YDR127W | 3620.75176 | -0.217425954 | 0.347997523 | 0.680 |
| YGL197W | 2610.651509 | 0.363610617 | 0.348533452 | 0.680 |
| YJL186W | 1797.326564 | -0.219591 | 0.348160166 | 0.680 |
| YML067C | 1114.441541 | -0.166939074 | 0.348497403 | 0.680 |
| YML101C | 335.3248863 | -0.171341905 | 0.348524886 | 0.680 |
| YOL090W | 1084.862315 | -0.206224165 | 0.348650418 | 0.680 |

| | | | | |
|---|---|---|---|---|
| YER177W | 16691.55629 | -0.184799339 | 0.348789452 | 0.680 |
| YIL167W | 108.465152 | 0.344871548 | 0.349092744 | 0.681 |
| YBR019C | 339.0837633 | 0.293140834 | 0.349543545 | 0.681 |
| YER175C | 1510.549619 | 0.338594985 | 0.349751967 | 0.681 |
| YGL037C | 14451.53815 | 0.336714727 | 0.349718841 | 0.681 |
| YER079W | 1934.057175 | 0.379331545 | 0.349897648 | 0.681 |
| YBL079W | 2598.015595 | -0.216648159 | 0.35044739 | 0.681 |
| YDR362C | 764.8048646 | 0.230440535 | 0.350707798 | 0.681 |
| YGL077C | 1372.265624 | -0.199111773 | 0.350614965 | 0.681 |
| YHL046W-A | 47.85481112 | 0.642733742 | 0.350656914 | 0.681 |
| YOR032C | 566.9607469 | 0.286261595 | 0.350157175 | 0.681 |
| YPL168W | 515.986609 | 0.3492809 | 0.350726802 | 0.681 |
| YPR199C | 796.7118888 | 0.405548339 | 0.350315486 | 0.681 |
| YCL042W | 285.5413665 | 0.29024357 | 0.351219602 | 0.682 |
| YCR048W | 995.8304266 | -0.323242363 | 0.352421742 | 0.682 |
| YDL016C | 38.72923011 | -0.346690483 | 0.351924448 | 0.682 |
| YDL124W | 12433.87204 | 0.381055604 | 0.352840177 | 0.682 |
| YDR208W | 1907.44225 | 0.237859162 | 0.353179283 | 0.682 |
| YDR311W | 727.3776002 | 0.229380082 | 0.351621528 | 0.682 |
| YDR475C | 1828.530627 | 0.355771303 | 0.352763572 | 0.682 |
| YFL034W | 949.1269789 | -0.2232845 | 0.351984365 | 0.682 |
| YGL219C | 1273.719314 | 0.32462022 | 0.353042321 | 0.682 |
| YGL226W | 346.3222245 | -0.255904189 | 0.352121678 | 0.682 |
| YGR084C | 1041.679379 | -0.179614049 | 0.352362196 | 0.682 |
| YHL019C | 544.6940331 | 0.346274435 | 0.351146986 | 0.682 |
| YKL110C | 846.8769824 | -0.327840472 | 0.352764503 | 0.682 |
| YLL061W | 447.0050952 | 0.357057969 | 0.353181448 | 0.682 |
| YLR281C | 33.92288358 | -0.07140139 | 0.352570108 | 0.682 |
| YLR421C | 1264.683888 | -0.300796224 | 0.352676899 | 0.682 |
| YML036W | 292.5583071 | -0.244311369 | 0.352335973 | 0.682 |
| YMR220W | 1203.063636 | 0.262213036 | 0.352829089 | 0.682 |
| YOL056W | 561.7509431 | -0.355705806 | 0.351827586 | 0.682 |
| YOR261C | 3484.142585 | -0.264389343 | 0.352074243 | 0.682 |
| YOR311C | 1031.860298 | -0.233981732 | 0.351037567 | 0.682 |
| YER094C | 2268.208556 | -0.243409092 | 0.353419905 | 0.682 |
| YMR145C | 7155.424672 | -0.16104228 | 0.353495236 | 0.682 |
| YDR140W | 525.20568 | -0.247390882 | 0.353624955 | 0.682 |
| YBR007C | 740.5673067 | -0.298619185 | 0.354172689 | 0.682 |
| YDR251W | 2664.249522 | 0.269577135 | 0.354333363 | 0.682 |
| YKL152C | 30648.47356 | -0.233595018 | 0.354345073 | 0.682 |
| YKR099W | 800.367483 | -0.324045496 | 0.353906886 | 0.682 |
| YOR054C | 904.8796156 | 0.266852386 | 0.354310096 | 0.682 |
| YOR348C | 282.9627385 | 0.457337223 | 0.354180683 | 0.682 |

| | | | | |
|---|---|---|---|---|
| YBR258C | 250.7365151 | -0.260042298 | 0.355565161 | 0.682 |
| YDR013W | 271.9851003 | -0.356376347 | 0.355232733 | 0.682 |
| YHR050W | 1110.031456 | -0.261456203 | 0.354935966 | 0.682 |
| YJL056C | 734.4930024 | 0.164240095 | 0.355443343 | 0.682 |
| YKR019C | 546.7471303 | 0.207148269 | 0.355183334 | 0.682 |
| YLL012W | 268.5425901 | 0.542084143 | 0.354964146 | 0.682 |
| YLR225C | 1476.977135 | 0.243057442 | 0.354727009 | 0.682 |
| YLR275W | 260.6216284 | -0.363218832 | 0.354711646 | 0.682 |
| YMR122C | 59.11544635 | 0.297303567 | 0.355599592 | 0.682 |
| YMR310C | 757.4056331 | -0.410624993 | 0.354564457 | 0.682 |
| YOR354C | 1688.06875 | 0.234598656 | 0.355639501 | 0.682 |
| YNL320W | 323.6388355 | -0.292859841 | 0.355909199 | 0.682 |
| YDR427W | 2923.459081 | -0.252546301 | 0.356291322 | 0.682 |
| YHR027C | 5786.339997 | -0.313670427 | 0.356380908 | 0.682 |
| YIL130W | 1279.051362 | 0.323939386 | 0.356356754 | 0.682 |
| YLR397C | 907.763381 | -0.210523808 | 0.356133583 | 0.682 |
| YKR100C | 1038.230895 | 0.234874788 | 0.35669509 | 0.682 |
| YLR320W | 753.4878726 | 0.238869511 | 0.356700665 | 0.682 |
| YDR285W | 510.8247622 | 0.029016312 | 0.35697147 | 0.683 |
| YGR161C | 2432.867838 | 0.205047653 | 0.357111887 | 0.683 |
| YMR019W | 977.1459301 | 0.330669675 | 0.357421973 | 0.683 |
| YMR066W | 755.7979974 | 0.25772561 | 0.357493378 | 0.683 |
| YNL168C | 1117.770385 | -0.176070148 | 0.357760913 | 0.683 |
| YJR048W | 1046.168345 | -0.140944265 | 0.357972368 | 0.683 |
| YJL002C | 4648.101511 | -0.216716594 | 0.358749369 | 0.685 |
| YGL244W | 2087.284965 | -0.279187561 | 0.359021377 | 0.685 |
| YDR174W | 3004.282379 | -0.223289371 | 0.359416165 | 0.685 |
| YNL327W | 5207.566398 | -0.167309681 | 0.359422867 | 0.685 |
| snR86 | 894.0383097 | -0.307038005 | 0.359281247 | 0.685 |
| YML031W | 1341.560963 | -0.292410495 | 0.359861484 | 0.685 |
| YMR113W | 461.667001 | 0.226710427 | 0.359771046 | 0.685 |
| YFR052C-A | 3217.393186 | 0.468368456 | 0.360082258 | 0.686 |
| YLR124W | 42.32116711 | 0.32168349 | 0.36032187 | 0.686 |
| YDR014W-A | 36.16776295 | 0.185023444 | 0.360627201 | 0.686 |
| YGR220C | 677.1113476 | -0.191689996 | 0.360670636 | 0.686 |
| YDR343C | 868.7911596 | 0.325786603 | 0.360815443 | 0.686 |
| YGR122W | 624.9209214 | 0.292049547 | 0.360969104 | 0.686 |
| YGR257C | 921.5740821 | -0.267236128 | 0.361249442 | 0.686 |
| YML043C | 371.5090553 | -0.565799058 | 0.361270301 | 0.686 |
| YGR250C | 3005.303955 | 0.342964606 | 0.361548772 | 0.687 |
| YKR089C | 3634.661722 | 0.266685721 | 0.361967752 | 0.687 |
| YBL061C | 962.4150781 | -0.260661246 | 0.362447032 | 0.688 |
| YDR222W | 466.2824915 | 0.27441461 | 0.362831164 | 0.688 |

| | | | | |
|---|---|---|---|---|
| YHL035C | 3064.413674 | 0.434867639 | 0.362835649 | 0.688 |
| YOR390W | 45.23234325 | 0.728137328 | 0.363087463 | 0.688 |
| YER105C | 2233.871003 | -0.149809975 | 0.363213306 | 0.688 |
| YMR306W | 518.3037802 | 0.25812558 | 0.363710806 | 0.689 |
| YDR179W-A | 321.5877637 | -0.259331014 | 0.364141067 | 0.690 |
| YOR152C | 552.1803625 | 0.368824036 | 0.364231841 | 0.690 |
| YLL033W | 166.5445731 | 0.226974907 | 0.364532594 | 0.690 |
| YNR040W | 422.1360755 | -0.276996031 | 0.365268117 | 0.691 |
| YCR092C | 872.6618833 | 0.22988195 | 0.365646153 | 0.691 |
| YGL185C | 473.3669461 | 0.338474778 | 0.365561259 | 0.691 |
| YKL108W | 391.5450461 | -0.326297962 | 0.365492488 | 0.691 |
| YIL145C | 1126.99064 | -0.246210896 | 0.365929642 | 0.691 |
| YPL243W | 2085.694288 | -0.247452126 | 0.36586137 | 0.691 |
| YLR149C | 1780.729906 | 0.244990001 | 0.366208373 | 0.692 |
| YOL094C | 437.8167325 | -0.271375752 | 0.366583921 | 0.692 |
| YBR297W | 465.5986861 | 0.443077742 | 0.36675412 | 0.692 |
| YDL036C | 422.8116952 | -0.196980345 | 0.367639247 | 0.693 |
| YGR249W | 613.9099062 | 0.19639237 | 0.367420384 | 0.693 |
| YIL024C | 346.2449471 | 0.378322046 | 0.367537693 | 0.693 |
| YJL054W | 1689.096692 | -0.263269557 | 0.367051443 | 0.693 |
| YML021C | 560.3287148 | -0.234577958 | 0.36732435 | 0.693 |
| YOR071C | 651.6605909 | 0.409797173 | 0.36724468 | 0.693 |
| YDR415C | 529.3228874 | 0.366685404 | 0.368002115 | 0.693 |
| YJR099W | 741.8529597 | -0.237276589 | 0.367905807 | 0.693 |
| YGL203C | 1571.604512 | 0.329628722 | 0.368247485 | 0.693 |
| YDR372C | 1904.188965 | -0.235183179 | 0.368375517 | 0.693 |
| YMR294W-A | 229.7802387 | -0.30389267 | 0.368628128 | 0.693 |
| YOR279C | 315.7050157 | 0.486839974 | 0.368518911 | 0.693 |
| YHR184W | 81.12732022 | 0.59540032 | 0.36882532 | 0.693 |
| YOR205C | 770.9234478 | 0.270789313 | 0.36888569 | 0.693 |
| YKL130C | 628.0827477 | -0.323508813 | 0.369289296 | 0.694 |
| YGL175C | 162.4549988 | 0.225476652 | 0.37023223 | 0.695 |
| YPL139C | 637.4930321 | 0.192835613 | 0.370165577 | 0.695 |
| YGL056C | 2031.538502 | 0.251209356 | 0.370511335 | 0.695 |
| YNL163C | 1921.119184 | -0.310340607 | 0.370625048 | 0.695 |
| YJR127C | 3525.789326 | 0.262855366 | 0.370913348 | 0.696 |
| YLR356W | 2015.024039 | 0.337886948 | 0.371006978 | 0.696 |
| YDR216W | 2843.606015 | 0.293947013 | 0.37161405 | 0.696 |
| YGL134W | 814.0702966 | 0.246626211 | 0.371496381 | 0.696 |
| YJL217W | 2088.445581 | -0.455987758 | 0.371461661 | 0.696 |
| YNL253W | 477.8813502 | 0.181003216 | 0.372035625 | 0.696 |
| YPR108W | 2749.924825 | -0.217408321 | 0.371923367 | 0.696 |
| YPL071C | 441.9177148 | -0.346506114 | 0.372195003 | 0.696 |

| | | | | |
|---|---|---|---|---|
| YBL029C-A | 1079.485322 | 0.260084661 | 0.372341308 | 0.696 |
| YBR157C | 589.2630079 | 0.194966816 | 0.373365712 | 0.697 |
| YDR068W | 1258.879245 | -0.157269101 | 0.373447827 | 0.697 |
| YDR369C | 596.1613966 | 0.332854337 | 0.373178423 | 0.697 |
| YGL052W | 83.8653604 | 0.351768964 | 0.373526069 | 0.697 |
| YLR152C | 1272.570191 | 0.279847876 | 0.372859395 | 0.697 |
| YLR405W | 359.0345633 | -0.335865042 | 0.372924747 | 0.697 |
| YMR135C | 1809.513775 | 0.421744577 | 0.373032656 | 0.697 |
| YMR315W | 4197.096296 | -0.284618297 | 0.373282119 | 0.697 |
| snR32 | 25.20381189 | -0.333393291 | 0.373442421 | 0.697 |
| YEL069C | 125.5249506 | 0.525172785 | 0.373990598 | 0.697 |
| YJL146W | 600.4909491 | 0.184450307 | 0.373934685 | 0.697 |
| YHR099W | 4737.26123 | -0.159538618 | 0.374388511 | 0.698 |
| YDL012C | 999.0124699 | -0.32862226 | 0.375078047 | 0.698 |
| YDL072C | 3778.461007 | -0.205682874 | 0.375103574 | 0.698 |
| YJR058C | 464.9848205 | -0.235563878 | 0.375281068 | 0.698 |
| YOR130C | 372.7954083 | -0.482573729 | 0.375369672 | 0.698 |
| YPL206C | 1410.718758 | -0.242279981 | 0.375367892 | 0.698 |
| YDR141C | 2153.367361 | 0.225583785 | 0.375621287 | 0.698 |
| YNL218W | 370.2566133 | -0.220255887 | 0.375649686 | 0.698 |
| YPR198W | 724.9139692 | -0.34106516 | 0.375755887 | 0.698 |
| YBL067C | 1161.166699 | -0.197946563 | 0.376189671 | 0.699 |
| YPL240C | 19482.15572 | -0.212832188 | 0.376285487 | 0.699 |
| YCL047C | 434.4156263 | -0.275432598 | 0.376573178 | 0.699 |
| YPR152C | 381.9563274 | 0.23410517 | 0.376558561 | 0.699 |
| YER159C | 1569.026083 | -0.30502917 | 0.377195261 | 0.699 |
| YGL039W | 719.9626894 | -0.334631855 | 0.377461475 | 0.699 |
| YGL085W | 385.7388603 | 0.214242652 | 0.377454532 | 0.699 |
| YHR159W | 783.3955676 | 0.253697046 | 0.377326979 | 0.699 |
| YML119W | 1027.81525 | 0.241547657 | 0.377286179 | 0.699 |
| YPL279C | 37.92327787 | 0.458085123 | 0.377360164 | 0.699 |
| YDL114W | 43.38451628 | 0.333409664 | 0.377628244 | 0.699 |
| YKL104C | 3816.518475 | -0.208037009 | 0.37776294 | 0.699 |
| YDL130W-A | 2765.774643 | 0.4480781 | 0.378677054 | 0.700 |
| YDR456W | 1671.501856 | 0.231640358 | 0.37829999 | 0.700 |
| YGL157W | 1718.848717 | -0.263416062 | 0.378825637 | 0.700 |
| YGR167W | 3020.873713 | -0.184983463 | 0.378729334 | 0.700 |
| YIL036W | 2051.229049 | 0.294301229 | 0.378194551 | 0.700 |
| YJL078C | 3272.163203 | -0.361941847 | 0.378722816 | 0.700 |
| YOL043C | 577.8464057 | 0.187102118 | 0.378562982 | 0.700 |
| YPR201W | 177.3808268 | 0.606786059 | 0.378249862 | 0.700 |
| YEL064C | 340.0873494 | 0.306443754 | 0.379079046 | 0.700 |
| YHR040W | 740.6643017 | -0.577093384 | 0.379015962 | 0.700 |

| | | | | |
|---|---|---|---|---|
| YBR210W | 464.9591183 | -0.246761132 | 0.379328699 | 0.700 |
| YLR105C | 592.2672821 | 0.293308548 | 0.379693869 | 0.700 |
| YDR388W | 5322.26629 | -0.176232672 | 0.379823808 | 0.700 |
| YML114C | 955.0918143 | -0.234436891 | 0.380126488 | 0.701 |
| YIL121W | 2357.398488 | 0.478691078 | 0.380650649 | 0.701 |
| YOR384W | 1087.084219 | 0.251073506 | 0.380706112 | 0.701 |
| YDR533C | 1411.920932 | 0.575522567 | 0.381077245 | 0.702 |
| YDL022W | 5632.112705 | 0.247468016 | 0.38175963 | 0.702 |
| YER029C | 871.4448752 | -0.208260648 | 0.382038148 | 0.702 |
| YGR112W | 838.4700607 | 0.25963121 | 0.381602794 | 0.702 |
| YGR243W | 1286.002734 | 0.285590687 | 0.381506831 | 0.702 |
| YJR117W | 3603.291074 | -0.266281715 | 0.381983835 | 0.702 |
| YJR150C | 147.0716932 | -0.059423633 | 0.381864425 | 0.702 |
| YKL215C | 1029.488394 | 0.290433973 | 0.381485407 | 0.702 |
| YPR009W | 346.2208437 | -0.4703723 | 0.382226262 | 0.702 |
| YDR272W | 1625.208653 | 0.354304151 | 0.382658996 | 0.703 |
| YGR119C | 2276.598641 | -0.209759132 | 0.382740933 | 0.703 |
| YOR128C | 2990.654925 | -0.233421425 | 0.382604022 | 0.703 |
| YDR204W | 1297.475345 | 0.411818924 | 0.382879239 | 0.703 |
| YNL184C | 77.22136504 | -0.109537744 | 0.38304151 | 0.703 |
| YGR163W | 840.8307527 | 0.285785254 | 0.383348998 | 0.703 |
| YKL004W | 2100.85667 | -0.201631626 | 0.38350485 | 0.703 |
| YLR120C | 2959.418798 | 0.259789869 | 0.383583041 | 0.703 |
| YBR026C | 2145.982583 | 0.337276215 | 0.383758387 | 0.703 |
| YDL109C | 380.7977413 | 0.318130294 | 0.383981446 | 0.703 |
| YGL241W | 933.7531382 | -0.195974274 | 0.38386615 | 0.703 |
| YJR003C | 586.7430541 | -0.266869275 | 0.384066891 | 0.703 |
| YER187W | 132.2577032 | 0.634565254 | 0.384265481 | 0.703 |
| YGR062C | 462.3300151 | 0.312384496 | 0.38449338 | 0.703 |
| YOL070C | 1742.264613 | -0.332015447 | 0.384387326 | 0.703 |
| YIL070C | 1550.574512 | -0.155521101 | 0.385047941 | 0.703 |
| YLR361C-A | 262.7999201 | -0.286674517 | 0.385086644 | 0.703 |
| YNR030W | 1357.008714 | -0.272025829 | 0.384826952 | 0.703 |
| YOR195W | 1326.708475 | -0.245335787 | 0.385115521 | 0.703 |
| YDL107W | 383.3863117 | 0.251998033 | 0.385365961 | 0.704 |
| YBR295W | 678.020174 | 0.244922525 | 0.385819215 | 0.704 |
| YDR022C | 413.59294 | 0.378469859 | 0.387140803 | 0.704 |
| YER104W | 241.2969985 | 0.247572452 | 0.386021362 | 0.704 |
| YER139C | 313.6167797 | 0.217938124 | 0.387293617 | 0.704 |
| YGR065C | 698.7091847 | 0.33977134 | 0.387485107 | 0.704 |
| YIR028W | 189.55325 | 0.597544224 | 0.38687476 | 0.704 |
| YJR006W | 1054.347239 | -0.236729086 | 0.387337405 | 0.704 |
| YJR025C | 1368.606225 | 0.122935547 | 0.387171422 | 0.704 |

| | | | | |
|---|---|---|---|---|
| YKR084C | 1262.417524 | -0.242070922 | 0.386974356 | 0.704 |
| YLL046C | 28.77526537 | 0.316046877 | 0.386360198 | 0.704 |
| YML041C | 413.9344954 | 0.174066514 | 0.387246986 | 0.704 |
| YNL144C | 1767.380134 | 0.206143736 | 0.386497257 | 0.704 |
| YNL190W | 6079.357238 | 0.344710932 | 0.38752384 | 0.704 |
| YNR069C | 436.7652166 | 0.275015457 | 0.386761505 | 0.704 |
| YPR049C | 968.2983267 | 0.286445332 | 0.386226586 | 0.704 |
| YPR184W | 5293.583109 | 0.472306593 | 0.386269705 | 0.704 |
| YJR156C | 47.49998049 | 0.582229255 | 0.387725264 | 0.704 |
| YOR161C | 928.1721785 | 0.02082513 | 0.387890291 | 0.704 |
| YDL194W | 1083.690954 | 0.208829609 | 0.388277373 | 0.705 |
| YDL120W | 712.7094342 | -0.248552009 | 0.388624188 | 0.705 |
| YML100W | 14497.66976 | 0.318530869 | 0.388674124 | 0.705 |
| YBR259W | 488.2154125 | -0.192056049 | 0.390871391 | 0.708 |
| YCL043C | 17735.82228 | -0.271040242 | 0.390599617 | 0.708 |
| YEL012W | 1045.847779 | 0.216715256 | 0.390858402 | 0.708 |
| YEL020C | 1045.827861 | 0.192368506 | 0.390685562 | 0.708 |
| YJR004C | 1252.603429 | 0.157710242 | 0.390608494 | 0.708 |
| YNL215W | 1103.384408 | 0.222613033 | 0.39084327 | 0.708 |
| YBR158W | 2964.217702 | -0.336270509 | 0.391049661 | 0.708 |
| YER051W | 512.8476233 | 0.320693586 | 0.391166326 | 0.708 |
| YKL094W | 2161.93051 | 0.169398369 | 0.391508462 | 0.708 |
| YJL089W | 344.4116839 | 0.077802046 | 0.391921766 | 0.709 |
| YGR117C | 888.1364455 | 0.281524139 | 0.392255201 | 0.709 |
| YNL124W | 1308.097286 | -0.401766716 | 0.392130844 | 0.709 |
| YNR050C | 3301.317883 | -0.451402631 | 0.392490515 | 0.709 |
| YOR346W | 878.2111125 | -0.243876181 | 0.392491092 | 0.709 |
| YDR304C | 4614.533404 | -0.207561964 | 0.393313695 | 0.709 |
| YHL032C | 2561.780761 | 0.218374792 | 0.393074746 | 0.709 |
| YKL109W | 7924.096762 | 0.276723164 | 0.393245274 | 0.709 |
| YNL152W | 687.0558678 | 0.169166751 | 0.392979274 | 0.709 |
| YPL052W | 344.0806128 | -0.415282187 | 0.393164272 | 0.709 |
| YDR286C | 156.0831168 | -0.288419491 | 0.393469145 | 0.709 |
| YPR035W | 16997.33407 | 0.362773931 | 0.39430795 | 0.711 |
| YCR067C | 1807.391426 | -0.215339155 | 0.394429688 | 0.711 |
| YGR255C | 1753.194799 | 0.306533696 | 0.394812703 | 0.711 |
| YML086C | 1768.316639 | -0.191547307 | 0.395007437 | 0.711 |
| YPR115W | 2825.752527 | 0.267354244 | 0.394919482 | 0.711 |
| YBR115C | 2897.179739 | -0.383493108 | 0.395739114 | 0.711 |
| YCL057W | 3573.183746 | -0.201346072 | 0.396779267 | 0.711 |
| YDR214W | 2305.632085 | -0.085063465 | 0.396737595 | 0.711 |
| YDR381C-A | 448.1284196 | 0.386358262 | 0.395861838 | 0.711 |
| YJR005W | 1677.202028 | 0.190805383 | 0.396132789 | 0.711 |

| | | | | |
|---|---|---|---|---|
| YKL187C | 740.5340162 | 0.262778786 | 0.396046758 | 0.711 |
| YMR264W | 1111.60862 | -0.227937014 | 0.395641744 | 0.711 |
| YMR303C | 445.8492587 | -0.029510544 | 0.396318756 | 0.711 |
| YNL270C | 240.3573834 | 0.080921394 | 0.39556041 | 0.711 |
| YNL309W | 553.1266537 | -0.228634182 | 0.396628928 | 0.711 |
| YOL129W | 2548.890951 | -0.192961181 | 0.396703292 | 0.711 |
| YOR215C | 2322.968103 | 0.408988201 | 0.396568283 | 0.711 |
| YPR148C | 2535.151148 | -0.19140881 | 0.396733572 | 0.711 |
| YBR194W | 226.3319567 | 0.323685856 | 0.397539949 | 0.712 |
| YDR181C | 744.5427058 | 0.271957759 | 0.397485093 | 0.712 |
| YDR504C | 165.3464621 | 0.230555351 | 0.397416792 | 0.712 |
| YDR207C | 1436.193266 | -0.336261083 | 0.397731218 | 0.712 |
| YPL152W | 506.4565509 | 0.18430855 | 0.398037631 | 0.713 |
| YGL136C | 674.1887477 | 0.190604049 | 0.398497594 | 0.713 |
| YPL016W | 1163.813997 | 0.32783279 | 0.398393502 | 0.713 |
| YEL070W | 298.2975483 | 0.274662393 | 0.398953802 | 0.713 |
| YGL049C | 2260.025541 | 0.173342877 | 0.398703377 | 0.713 |
| YJR151C | 560.5480031 | 0.328132008 | 0.399048334 | 0.713 |
| YMR192W | 859.8324449 | -0.198034057 | 0.399066524 | 0.713 |
| YDL003W | 760.5875817 | -0.155607555 | 0.399434044 | 0.713 |
| YJR009C | 33615.59982 | -0.100937289 | 0.399313754 | 0.713 |
| YOR016C | 992.9033585 | -0.213107762 | 0.399544468 | 0.713 |
| YBR168W | 396.8925406 | 0.180522443 | 0.400248711 | 0.713 |
| YDR305C | 387.9870014 | -0.235290502 | 0.400809283 | 0.713 |
| YEL073C | 122.6223904 | 0.666275425 | 0.400491767 | 0.713 |
| YGR032W | 3378.619384 | 0.261824046 | 0.400872774 | 0.713 |
| YJL076W | 2985.809725 | -0.212396293 | 0.3998553 | 0.713 |
| YJL091C | 524.9582628 | 0.308139356 | 0.400446578 | 0.713 |
| YJR060W | 664.6110196 | -0.187976915 | 0.39996391 | 0.713 |
| YML102W | 604.0898534 | -0.205040343 | 0.40073565 | 0.713 |
| YNL188W | 592.8677499 | -0.331119349 | 0.400915001 | 0.713 |
| YNR070W | 728.5997471 | 0.37445061 | 0.400885753 | 0.713 |
| YPR022C | 1532.191812 | 0.196385182 | 0.399967536 | 0.713 |
| YIL062C | 1798.738473 | -0.252400076 | 0.401302435 | 0.714 |
| YDR387C | 888.6103683 | 0.19696509 | 0.402238639 | 0.715 |
| YGL172W | 1277.929348 | -0.28989513 | 0.402197896 | 0.715 |
| YNR065C | 953.8955562 | 0.192012268 | 0.402251681 | 0.715 |
| ICR1 | 346.589135 | 0.328685661 | 0.403773418 | 0.715 |
| YDR371W | 631.8329655 | 0.270624199 | 0.404026566 | 0.715 |
| YDR400W | 652.8362371 | -0.172981664 | 0.403799753 | 0.715 |
| YDR506C | 448.3958033 | 0.218207017 | 0.40345183 | 0.715 |
| YGR100W | 1479.985263 | 0.151363086 | 0.404001155 | 0.715 |
| YIR030C | 90.29914152 | 0.618845418 | 0.40347549 | 0.715 |

| | | | | |
|---|---|---|---|---|
| YJL046W | 632.8219176 | 0.247243049 | 0.403332151 | 0.715 |
| YJL061W | 1142.158213 | 0.258767053 | 0.402922331 | 0.715 |
| YKL068W | 1722.980976 | 0.274876568 | 0.402700658 | 0.715 |
| YLR352W | 926.4882915 | 0.191693627 | 0.403525225 | 0.715 |
| YML018C | 644.4081194 | -0.347238608 | 0.403520642 | 0.715 |
| YNR022C | 907.2227542 | -0.188244264 | 0.402838491 | 0.715 |
| YOR034C | 667.8994742 | 0.330303281 | 0.402683514 | 0.715 |
| YPL229W | 514.8866138 | 0.250599022 | 0.403591099 | 0.715 |
| YDL047W | 1531.758576 | -0.247870838 | 0.404320249 | 0.715 |
| YMR239C | 477.7892356 | -0.318736343 | 0.404716838 | 0.716 |
| YOR115C | 581.5147795 | -0.240908059 | 0.404735197 | 0.716 |
| YGL126W | 1063.33427 | -0.242653011 | 0.405104155 | 0.716 |
| YLL001W | 1443.400677 | 0.333946925 | 0.404994127 | 0.716 |
| YBR068C | 7312.593911 | 0.221758986 | 0.405293473 | 0.716 |
| YDL042C | 548.3117902 | -0.11955608 | 0.407138449 | 0.717 |
| YDL241W | 250.4921648 | -0.434293709 | 0.407066378 | 0.717 |
| YDR237W | 832.9477874 | -0.173404192 | 0.406453421 | 0.717 |
| YGR056W | 1165.037029 | -0.307695096 | 0.407068598 | 0.717 |
| YIL120W | 363.836718 | 0.376680756 | 0.406750374 | 0.717 |
| YNL039W | 1798.324816 | 0.170234965 | 0.407136143 | 0.717 |
| YOL006C | 1492.750425 | 0.195182141 | 0.406606729 | 0.717 |
| YOL128C | 161.4112011 | 0.176001707 | 0.406966488 | 0.717 |
| YAL017W | 3309.438838 | 0.304143234 | 0.408252518 | 0.718 |
| YCR081W | 992.4265821 | 0.228026991 | 0.40816415 | 0.718 |
| YDR451C | 513.5586606 | 0.306927752 | 0.408269878 | 0.718 |
| YER089C | 1927.936186 | -0.256977494 | 0.408072235 | 0.718 |
| YGR290W | 27.11482068 | 0.059068481 | 0.408314337 | 0.718 |
| YKL221W | 211.2597824 | 0.454582111 | 0.407879553 | 0.718 |
| YOL111C | 1821.468977 | -0.16021925 | 0.408487292 | 0.718 |
| YOR150W | 702.6825687 | -0.259654605 | 0.408408683 | 0.718 |
| YPL133C | 580.4116075 | 0.243391568 | 0.407990012 | 0.718 |
| YKR051W | 666.9246654 | 0.246583311 | 0.408875607 | 0.718 |
| YML113W | 567.9041791 | -0.40807795 | 0.409196727 | 0.718 |
| YNR047W | 1084.837265 | 0.266489821 | 0.409128819 | 0.718 |
| YPL019C | 1922.931958 | -0.368319891 | 0.409282307 | 0.718 |
| YBL100C | 649.5552869 | -0.007246521 | 0.410091006 | 0.719 |
| YBR126C | 9994.323143 | 0.211718162 | 0.409802018 | 0.719 |
| YDR227W | 1551.753891 | -0.336177333 | 0.409865345 | 0.719 |
| YDR491C | 102.3610769 | -0.099012741 | 0.410063678 | 0.719 |
| YMR056C | 319.9319845 | 0.315849924 | 0.409710598 | 0.719 |
| YDL175C | 355.6103486 | 0.375797828 | 0.410261005 | 0.719 |
| YNR020C | 315.5485889 | -0.237329228 | 0.410379522 | 0.719 |
| YBR046C | 1727.421703 | 0.369775406 | 0.410782127 | 0.719 |

| | | | | |
|---|---|---|---|---|
| YIR018W | 398.3013208 | 0.308257376 | 0.41068732 | 0.719 |
| YIL023C | 838.1130985 | 0.265358427 | 0.411225801 | 0.719 |
| YPL188W | 716.8167679 | 0.23786091 | 0.41110739 | 0.719 |
| YDR479C | 1259.888261 | 0.249816739 | 0.411456449 | 0.720 |
| YBR076C-A | 84.52994015 | 0.222325366 | 0.413492445 | 0.720 |
| YBR161W | 517.5371153 | 0.240382314 | 0.412988995 | 0.720 |
| YCL075W | 37.39387639 | 0.15289128 | 0.412609647 | 0.720 |
| YCR079W | 984.6018828 | 0.163892151 | 0.412130781 | 0.720 |
| YDR356W | 1114.540575 | -0.236280928 | 0.412527913 | 0.720 |
| YER020W | 2824.873338 | 0.230772884 | 0.413095612 | 0.720 |
| YGL205W | 247.6080273 | 0.421368146 | 0.413300358 | 0.720 |
| YJL106W | 294.2742704 | 0.054420772 | 0.412868685 | 0.720 |
| YLR089C | 5710.768483 | 0.338604734 | 0.411700958 | 0.720 |
| YML005W | 433.1547412 | -0.165882155 | 0.412034558 | 0.720 |
| YMR055C | 275.6053073 | -0.249215568 | 0.413296106 | 0.720 |
| YMR103C | 375.8522499 | 0.151280134 | 0.412945286 | 0.720 |
| YNL236W | 1176.860728 | -0.204369113 | 0.413493819 | 0.720 |
| YOR360C | 883.5535106 | 0.342516031 | 0.412463496 | 0.720 |
| YPL156C | 858.4762505 | 0.269127128 | 0.413328149 | 0.720 |
| YPR054W | 265.2938746 | 0.391713127 | 0.413530405 | 0.720 |
| YAR002W | 1776.649309 | -0.200764201 | 0.414033857 | 0.720 |
| YJR083C | 550.0152446 | 0.224110455 | 0.413919568 | 0.720 |
| YCR011C | 2808.118139 | 0.237902271 | 0.414908617 | 0.721 |
| YDR342C | 1706.769269 | 0.141939832 | 0.414831828 | 0.721 |
| YOR301W | 501.5102259 | -0.157839591 | 0.414926366 | 0.721 |
| YGR289C | 671.1201555 | 0.128076804 | 0.415396837 | 0.722 |
| YLR369W | 1148.442381 | 0.275695385 | 0.415689765 | 0.722 |
| YML131W | 1116.004425 | 0.452248146 | 0.41547954 | 0.722 |
| YOR289W | 937.4069291 | 0.389995952 | 0.415570725 | 0.722 |
| YNL053W | 544.6626736 | 0.219950941 | 0.416005843 | 0.722 |
| YMR024W | 882.2534561 | -0.081820552 | 0.41633824 | 0.722 |
| YDR258C | 9048.121163 | 0.440267547 | 0.416839888 | 0.723 |
| YLR234W | 283.065063 | -0.329901486 | 0.416696197 | 0.723 |
| YML057C-A | 86.62699701 | 0.244337984 | 0.416800293 | 0.723 |
| YKL208W | 299.0743362 | 0.249860207 | 0.417384667 | 0.723 |
| YKL031W | 35.11063328 | 0.474838492 | 0.417704871 | 0.724 |
| YOR125C | 859.4168859 | 0.344837451 | 0.417742726 | 0.724 |
| YIL149C | 3262.267006 | -0.3588067 | 0.417966655 | 0.724 |
| YPL081W | 1016.551966 | -0.16600146 | 0.418194998 | 0.724 |
| YCL031C | 1450.481764 | -0.242394312 | 0.419217014 | 0.724 |
| YFL036W | 2229.86904 | 0.295627422 | 0.4190929 | 0.724 |
| YGR170W | 2142.624103 | 0.237376383 | 0.418967639 | 0.724 |
| YJL003W | 223.2586603 | 0.236617281 | 0.419347343 | 0.724 |

| | | | | |
|---|---|---|---|---|
| YJL152W | 449.6754094 | 0.243209564 | 0.418572157 | 0.724 |
| YKL205W | 1310.174229 | -0.213170486 | 0.418870383 | 0.724 |
| YLR394W | 730.9999237 | 0.182611146 | 0.418827284 | 0.724 |
| YNR018W | 1764.18714 | -0.151577978 | 0.418797495 | 0.724 |
| YPR100W | 188.3881606 | -0.396013342 | 0.419226999 | 0.724 |
| YJL165C | 2120.172359 | 0.095825443 | 0.419550305 | 0.724 |
| YDR497C | 3420.957422 | -0.186451899 | 0.420034569 | 0.725 |
| YHR076W | 1124.266555 | -0.178941336 | 0.420466086 | 0.725 |
| YJL062W-A | 1034.508423 | -0.184910923 | 0.420386602 | 0.725 |
| YOR322C | 1313.48176 | 0.210689849 | 0.420684015 | 0.725 |
| YOR362C | 3907.49208 | -0.198171487 | 0.42095246 | 0.725 |
| YBL090W | 640.7121751 | -0.235125462 | 0.424282478 | 0.726 |
| YBR022W | 363.9709831 | -0.295376078 | 0.424303933 | 0.726 |
| YCL055W | 639.3184492 | 0.33891322 | 0.423612457 | 0.726 |
| YDL141W | 1031.602218 | -0.158321031 | 0.422679091 | 0.726 |
| YDL144C | 951.4980585 | -0.215243577 | 0.423418853 | 0.726 |
| YDL168W | 1463.24926 | -0.308227895 | 0.425776004 | 0.726 |
| YDR244W | 923.2552739 | 0.180742981 | 0.422923868 | 0.726 |
| YER033C | 1095.361309 | 0.289559575 | 0.425071708 | 0.726 |
| YER093C-A | 366.8172657 | -0.216737166 | 0.42220789 | 0.726 |
| YFL009W | 860.3502379 | 0.159556344 | 0.423425923 | 0.726 |
| YGL004C | 1031.613443 | 0.289465397 | 0.424513331 | 0.726 |
| YGL224C | 544.3672231 | -0.242916165 | 0.422476297 | 0.726 |
| YGR004W | 331.5972983 | 0.203313082 | 0.422713034 | 0.726 |
| YGR013W | 916.8674279 | 0.190597903 | 0.424722768 | 0.726 |
| YIL150C | 295.6540585 | -0.143807616 | 0.424435907 | 0.726 |
| YJL101C | 2271.091311 | 0.213250098 | 0.42369074 | 0.726 |
| YKL075C | 477.0577168 | 0.215598174 | 0.423160226 | 0.726 |
| YKL216W | 7973.975155 | 0.443609872 | 0.423774577 | 0.726 |
| YLR300W | 3505.549383 | -0.280226235 | 0.422821971 | 0.726 |
| YLR312W-A | 784.781583 | -0.180413353 | 0.425534405 | 0.726 |
| YMR149W | 2739.886665 | -0.192483497 | 0.425521102 | 0.726 |
| YMR274C | 249.323644 | -0.188688587 | 0.424046331 | 0.726 |
| YNL008C | 1006.822611 | 0.191298083 | 0.424955131 | 0.726 |
| YNL027W | 1180.449634 | 0.200129618 | 0.421692996 | 0.726 |
| YNL066W | 1969.929236 | -0.189666079 | 0.424864654 | 0.726 |
| YNL094W | 1768.042353 | 0.187697997 | 0.425502556 | 0.726 |
| YNL289W | 446.6126721 | -0.269278656 | 0.423536403 | 0.726 |
| YOL077W-A | 1163.041858 | -0.285073228 | 0.422845224 | 0.726 |
| YOR002W | 2112.418166 | 0.235582018 | 0.425149258 | 0.726 |
| YOR006C | 758.4019018 | 0.132312546 | 0.425338529 | 0.726 |
| YOR057W | 859.8202576 | -0.294685029 | 0.424827824 | 0.726 |
| YOR188W | 1393.160444 | -0.166712028 | 0.422841424 | 0.726 |

| | | | | |
|---|---|---|---|---|
| YOR328W | 830.771978 | 0.096172292 | 0.423375964 | 0.726 |
| YPR016W-A | 37.91128357 | -0.616893061 | 0.423939895 | 0.726 |
| YPR079W | 754.8054031 | 0.158790449 | 0.425685751 | 0.726 |
| YPR116W | 103.378319 | -0.236942041 | 0.421685502 | 0.726 |
| YCR045C | 398.9252367 | 0.208244306 | 0.426431133 | 0.726 |
| YDR088C | 645.3535973 | 0.258890701 | 0.426185213 | 0.726 |
| YDR307W | 749.860008 | 0.273061179 | 0.42615663 | 0.726 |
| YKR009C | 961.8765586 | 0.166371625 | 0.426189943 | 0.726 |
| snR31 | 65.06005548 | -0.156665329 | 0.426315388 | 0.726 |
| YBR140C | 4380.796749 | -0.144319236 | 0.427514289 | 0.726 |
| YDL085C-A | 569.0392384 | -0.313454129 | 0.428325678 | 0.726 |
| YDR166C | 1344.376251 | 0.232162521 | 0.428222721 | 0.726 |
| YDR330W | 1850.284876 | 0.161073753 | 0.426834833 | 0.726 |
| YER103W | 3269.463849 | 0.610971586 | 0.428318804 | 0.726 |
| YFR050C | 3854.503578 | -0.231434083 | 0.427111956 | 0.726 |
| YGL075C | 312.3284424 | 0.195555614 | 0.428010859 | 0.726 |
| YHR194W | 859.058911 | 0.269108479 | 0.428296447 | 0.726 |
| YIL105C | 1531.713028 | 0.130867259 | 0.427891061 | 0.726 |
| YJR110W | 661.1509666 | 0.156834773 | 0.427401793 | 0.726 |
| YMR032W | 652.0530589 | -0.218868258 | 0.428388555 | 0.726 |
| YMR052C-A | 61.89756679 | -0.363979207 | 0.427538669 | 0.726 |
| YOR079C | 215.4299264 | -0.486626727 | 0.426725088 | 0.726 |
| YOR377W | 997.8606777 | 0.140393354 | 0.427164803 | 0.726 |
| YPR178W | 455.7725824 | 0.186415798 | 0.426997736 | 0.726 |
| YAL016C-A | 61.31300348 | 0.17045875 | 0.428523745 | 0.726 |
| YJL013C | 787.2498623 | 0.174975838 | 0.4288583 | 0.727 |
| YBR269C | 1256.661555 | 0.274316566 | 0.430309782 | 0.728 |
| YKR104W | 184.9350151 | 0.260397417 | 0.43002731 | 0.728 |
| YLR127C | 955.0961502 | 0.190257846 | 0.430237433 | 0.728 |
| YOL148C | 713.6397629 | -0.199382222 | 0.43019166 | 0.728 |
| YLR242C | 302.599813 | -0.243929695 | 0.430538276 | 0.728 |
| YKL193C | 1757.604095 | 0.287326763 | 0.430740745 | 0.729 |
| YGR258C | 1398.526849 | 0.214356722 | 0.431193481 | 0.729 |
| YMR203W | 3020.852994 | -0.157012114 | 0.431322469 | 0.729 |
| YOL162W | 65.72006374 | 0.274920372 | 0.431357602 | 0.729 |
| YOR381W | 508.0002714 | 0.243615327 | 0.431078547 | 0.729 |
| YDR003W | 807.4785109 | 0.164639522 | 0.431541631 | 0.729 |
| YKR046C | 2535.087216 | 0.268167053 | 0.431928428 | 0.729 |
| YJR138W | 1461.069987 | 0.308981537 | 0.432716045 | 0.730 |
| YOR337W | 623.7332196 | -0.25870157 | 0.432595999 | 0.730 |
| YER026C | 2306.614486 | -0.218367243 | 0.433070405 | 0.730 |
| YJL072C | 353.6349072 | -0.175698791 | 0.433091598 | 0.730 |
| YNL293W | 734.6319527 | 0.202142855 | 0.432959647 | 0.730 |

| | | | | |
|---|---|---|---|---|
| YOR121C | 64.33732395 | 0.396080802 | 0.433219851 | 0.730 |
| YBL022C | 3958.243828 | -0.205357533 | 0.433898003 | 0.730 |
| YDL230W | 846.3743425 | 0.191996208 | 0.434549624 | 0.730 |
| YER090W | 2828.394196 | -0.364593614 | 0.434449378 | 0.730 |
| YER182W | 1071.612284 | 0.301738841 | 0.434298443 | 0.730 |
| YHL007C | 2278.179389 | -0.225948481 | 0.434386628 | 0.730 |
| YIL116W | 2678.75766 | -0.263201684 | 0.433959094 | 0.730 |
| YLR170C | 116.6208147 | -0.302977526 | 0.434219442 | 0.730 |
| YMR252C | 396.7363367 | -0.322216801 | 0.433905509 | 0.730 |
| YOL109W | 9848.185932 | -0.387922748 | 0.433891446 | 0.730 |
| YOR149C | 790.0054864 | 0.266599652 | 0.43397664 | 0.730 |
| YKL176C | 1422.060464 | 0.287138571 | 0.434709201 | 0.730 |
| YGL228W | 2378.82392 | 0.225675666 | 0.434950652 | 0.731 |
| YLL043W | 1838.000712 | -0.248176625 | 0.435321593 | 0.731 |
| YKL027W | 701.3136707 | 0.328671676 | 0.435711204 | 0.731 |
| YPL051W | 362.5719952 | -0.264286469 | 0.435809286 | 0.731 |
| YPL145C | 2994.997329 | -0.147677983 | 0.435837315 | 0.731 |
| YPR129W | 1545.384555 | -0.202685777 | 0.435661981 | 0.731 |
| YLR166C | 1318.264285 | 0.190270739 | 0.436335955 | 0.732 |
| YOL059W | 2587.338376 | 0.255330076 | 0.436424071 | 0.732 |
| YER019W | 1037.358398 | -0.300199517 | 0.436897233 | 0.732 |
| YJL156C | 869.0736001 | 0.207353039 | 0.437283515 | 0.733 |
| YGL212W | 286.2266484 | -0.215922747 | 0.438055574 | 0.734 |
| YLR443W | 472.8026492 | -0.319610475 | 0.438318918 | 0.734 |
| YPL024W | 252.5020697 | -0.31961195 | 0.438424443 | 0.734 |
| YOR094W | 410.9080628 | -0.25589651 | 0.438908801 | 0.735 |
| YFL007W | 4593.396937 | -0.189764028 | 0.439416468 | 0.735 |
| YGL248W | 1428.88377 | 0.284098429 | 0.439394192 | 0.735 |
| YPR158W-B | 29.96033516 | 0.384709536 | 0.439215844 | 0.735 |
| YMR291W | 2090.324495 | 0.148433379 | 0.439670453 | 0.735 |
| YOR343W-B | 107.9180972 | 0.349415757 | 0.439609875 | 0.735 |
| YPL038W | 299.6200602 | -0.37489992 | 0.439932029 | 0.735 |
| YGR253C | 3517.823624 | -0.177426415 | 0.440248854 | 0.735 |
| YHR006W | 1327.030831 | 0.184508715 | 0.440430679 | 0.735 |
| YDL207W | 1207.359156 | -0.149901449 | 0.44071906 | 0.736 |
| YDL215C | 3694.89417 | 0.299240904 | 0.440839243 | 0.736 |
| YDR129C | 5762.337606 | 0.185437675 | 0.440724609 | 0.736 |
| YGL091C | 545.7319922 | -0.172061054 | 0.441169201 | 0.736 |
| YEL050W-A | 71.35641565 | -0.315226538 | 0.441723011 | 0.736 |
| YJR134C | 1708.030399 | -0.244999629 | 0.441975206 | 0.736 |
| YKL072W | 554.6822525 | 0.166930383 | 0.441768013 | 0.736 |
| YKL194C | 593.5236151 | 0.23590547 | 0.441671839 | 0.736 |
| YLR189C | 2093.552884 | 0.171468682 | 0.441970463 | 0.736 |

| | | | | |
|---|---|---|---|---|
| YOR202W | 1479.782603 | -0.380182679 | 0.441827235 | 0.736 |
| YBR182C | 159.913284 | 0.145793499 | 0.442143127 | 0.736 |
| YML107C | 389.7043417 | 0.139707818 | 0.442247391 | 0.736 |
| YDR049W | 886.770398 | 0.146593839 | 0.442448604 | 0.736 |
| YER067W | 2559.902963 | 0.179114838 | 0.44292287 | 0.736 |
| YGR025W | 100.9215658 | -0.071909244 | 0.442650549 | 0.736 |
| YIL067C | 437.2526192 | 0.157645107 | 0.442775986 | 0.736 |
| YIL101C | 1402.735777 | -0.005431419 | 0.442942393 | 0.736 |
| YDL007W | 5356.140616 | -0.219372258 | 0.443880532 | 0.737 |
| YDR524C | 904.6436686 | 0.270718755 | 0.444144224 | 0.737 |
| YGL246C | 646.4635665 | -0.213664795 | 0.443920154 | 0.737 |
| YIL136W | 7495.331143 | 0.360574977 | 0.44400456 | 0.737 |
| YOL103W | 1490.877623 | -0.152653495 | 0.444202987 | 0.737 |
| YOR227W | 3483.244777 | 0.28212592 | 0.44367646 | 0.737 |
| YHR207C | 677.6706609 | -0.188101531 | 0.444643144 | 0.737 |
| YMR021C | 731.4049333 | 0.109752699 | 0.444705354 | 0.737 |
| YOR176W | 2270.643704 | 0.372385167 | 0.444625678 | 0.737 |
| YER022W | 948.246119 | 0.163190751 | 0.444933956 | 0.737 |
| YNL156C | 1413.410846 | 0.184839571 | 0.445289744 | 0.738 |
| YOL037C | 71.18780393 | 0.349804381 | 0.445343923 | 0.738 |
| YIL140W | 1095.766571 | -0.206241024 | 0.445867813 | 0.738 |
| YPR034W | 1306.898874 | -0.217425062 | 0.445990098 | 0.738 |
| YKL025C | 1634.326653 | 0.189728777 | 0.446201853 | 0.738 |
| YDR194C | 2378.698014 | -0.125077488 | 0.446461011 | 0.739 |
| YDR289C | 776.5077999 | 0.262553737 | 0.446653593 | 0.739 |
| YEL011W | 8924.284313 | 0.244045788 | 0.44657928 | 0.739 |
| YGL055W | 3060.50732 | -0.1962164 | 0.448217509 | 0.740 |
| YKR006C | 1505.723079 | -0.134122528 | 0.448331359 | 0.740 |
| YKR010C | 516.9970849 | -0.154220629 | 0.448132983 | 0.740 |
| YOR027W | 11708.63489 | -0.088565903 | 0.448294355 | 0.740 |
| YOR040W | 503.3947844 | 0.332675142 | 0.448016306 | 0.740 |
| YPR088C | 2257.094394 | -0.19815913 | 0.447727078 | 0.740 |
| YBR132C | 1205.603555 | 0.165447037 | 0.448696126 | 0.740 |
| YAR018C | 738.6960091 | 0.198116813 | 0.449877214 | 0.742 |
| YGL109W | 79.04302724 | 0.195358347 | 0.450053458 | 0.742 |
| YPL149W | 607.7809907 | -0.456415104 | 0.450122679 | 0.742 |
| YLR040C | 271.3228626 | 0.652873434 | 0.450902959 | 0.743 |
| YOR374W | 6364.239227 | 0.17077357 | 0.450860792 | 0.743 |
| YBL064C | 4712.365868 | 0.331421797 | 0.45160489 | 0.743 |
| YBR223C | 483.3688068 | 0.20240738 | 0.452464636 | 0.743 |
| YDR183W | 267.9198317 | -0.180715727 | 0.452326398 | 0.743 |
| YER001W | 1455.504127 | -0.228381563 | 0.451112191 | 0.743 |
| YGL253W | 16247.71261 | -0.064163302 | 0.451739692 | 0.743 |

| | | | | |
|---|---|---|---|---|
| YGR076C | 617.1306078 | -0.240200953 | 0.45279643 | 0.743 |
| YHR189W | 309.5122663 | 0.17165412 | 0.452404729 | 0.743 |
| YHR199C | 2007.264009 | 0.123606415 | 0.451453458 | 0.743 |
| YJL160C | 84.43696653 | 0.452660263 | 0.453170399 | 0.743 |
| YJL174W | 2482.371966 | -0.150605399 | 0.453301952 | 0.743 |
| YLL042C | 71.42312394 | -0.314177834 | 0.45335803 | 0.743 |
| YLR258W | 9832.7404 | 0.197368741 | 0.452193652 | 0.743 |
| YLR414C | 1706.526088 | 0.113140692 | 0.45345928 | 0.743 |
| YML003W | 222.6670877 | 0.21876759 | 0.453028671 | 0.743 |
| YML016C | 1375.407219 | 0.150062749 | 0.452653227 | 0.743 |
| YOL071W | 1035.627926 | 0.293927847 | 0.453202402 | 0.743 |
| YOR008C | 1383.478149 | 0.252871255 | 0.452346434 | 0.743 |
| YOR043W | 1570.483617 | 0.281167008 | 0.451469212 | 0.743 |
| YOR260W | 2245.68858 | -0.198568472 | 0.451364824 | 0.743 |
| YPL138C | 483.5208101 | 0.195155959 | 0.452694057 | 0.743 |
| YLR402W | 27.81687827 | 0.238800706 | 0.453974123 | 0.744 |
| YOR284W | 207.2496443 | -0.209635084 | 0.454340237 | 0.744 |
| YGL025C | 784.0082065 | 0.20084564 | 0.454787862 | 0.744 |
| YPL053C | 1363.897708 | -0.265997293 | 0.454673113 | 0.744 |
| YGR270W | 2956.664275 | 0.177876149 | 0.454931291 | 0.744 |
| YER116C | 578.3983863 | 0.203901323 | 0.455349283 | 0.745 |
| YJL071W | 590.1206135 | 0.167861217 | 0.455789278 | 0.745 |
| YMR259C | 1208.426766 | -0.213962716 | 0.455790113 | 0.745 |
| YER164W | 2428.238913 | -0.194184408 | 0.456277535 | 0.746 |
| YMR227C | 799.7644018 | -0.27094195 | 0.456327732 | 0.746 |
| YDL179W | 455.9668158 | -0.162258137 | 0.456562252 | 0.746 |
| YDR093W | 1759.127414 | -0.125867082 | 0.457631715 | 0.747 |
| YLR016C | 414.1257784 | -0.249075527 | 0.457624262 | 0.747 |
| YLR392C | 638.9786435 | 0.162973286 | 0.457522244 | 0.747 |
| YOR140W | 905.6577941 | -0.201126238 | 0.457241344 | 0.747 |
| YPL061W | 6576.454349 | -0.548498392 | 0.457642413 | 0.747 |
| YDR539W | 882.7410932 | 0.603156881 | 0.457950101 | 0.747 |
| YLR050C | 545.8709868 | -0.216284892 | 0.458050802 | 0.747 |
| YNL083W | 670.395816 | 0.240071607 | 0.457995866 | 0.747 |
| YNL161W | 1329.930942 | -0.154792775 | 0.458419984 | 0.747 |
| YBL052C | 898.9203912 | 0.199699633 | 0.458708063 | 0.747 |
| YBL080C | 486.2802333 | 0.167352352 | 0.459726842 | 0.747 |
| YBR298C | 1312.084765 | 0.057661416 | 0.45922738 | 0.747 |
| YGR175C | 3323.313876 | -0.146759088 | 0.459657389 | 0.747 |
| YKL048C | 583.1360838 | 0.200785743 | 0.45954442 | 0.747 |
| YLR148W | 974.658438 | 0.212321885 | 0.458800077 | 0.747 |
| YLR279W | 36.37996098 | -0.174980806 | 0.459115859 | 0.747 |
| YNL180C | 2386.54101 | 0.300326295 | 0.459643168 | 0.747 |

| | | | | |
|---|---|---|---|---|
| YOR216C | 1165.7555 | -0.201613673 | 0.458969587 | 0.747 |
| YPL210C | 1902.73736 | -0.208937794 | 0.459342212 | 0.747 |
| YOL075C | 712.6275688 | 0.259698799 | 0.46000005 | 0.747 |
| YOR307C | 1226.108407 | 0.182080352 | 0.459956807 | 0.747 |
| YGR017W | 1237.157084 | -0.168401224 | 0.460561379 | 0.747 |
| YKL073W | 1776.826393 | -0.152363927 | 0.460608323 | 0.747 |
| YPL245W | 615.8162164 | -0.457618433 | 0.460497431 | 0.747 |
| YBL005W | 1420.668611 | 0.248136939 | 0.461518526 | 0.748 |
| YCL004W | 588.1019431 | 0.196473537 | 0.461170704 | 0.748 |
| YER173W | 470.1702433 | 0.14680134 | 0.461387671 | 0.748 |
| YGL208W | 855.2756461 | 0.361572286 | 0.461606816 | 0.748 |
| YOR320C | 1070.658594 | -0.12865546 | 0.461373917 | 0.748 |
| YBL084C | 817.2451769 | 0.137841775 | 0.461938831 | 0.748 |
| YLR093C | 1657.689066 | 0.273296068 | 0.462016687 | 0.748 |
| YEL007W | 3320.15412 | 0.272589311 | 0.462193551 | 0.748 |
| YDL231C | 1653.703222 | 0.13265883 | 0.462848808 | 0.749 |
| YIL080W | 141.3218365 | 0.234520957 | 0.462730986 | 0.749 |
| YIL026C | 1582.807889 | -0.166815818 | 0.463354405 | 0.749 |
| YKL120W | 1575.003705 | -0.161990219 | 0.463296268 | 0.749 |
| YOR181W | 1611.589546 | -0.310528586 | 0.463257002 | 0.749 |
| YER165C-A | 28.00543952 | -0.049287579 | 0.464166779 | 0.749 |
| YGR227W | 1061.56859 | 0.346783424 | 0.464023457 | 0.749 |
| YLR355C | 9452.460146 | 0.312410474 | 0.463780534 | 0.749 |
| YNL100W | 1262.153408 | 0.331642143 | 0.464086852 | 0.749 |
| YPL008W | 976.9490792 | 0.219805447 | 0.463811122 | 0.749 |
| YPR085C | 667.2581144 | 0.12027614 | 0.463549752 | 0.749 |
| YGL174W | 430.9199773 | -0.193749656 | 0.464356266 | 0.749 |
| YCR046C | 1534.125343 | -0.137844862 | 0.464668423 | 0.750 |
| YDL119C | 290.4231748 | 0.203855698 | 0.464895065 | 0.750 |
| YFR011C | 642.0102271 | -0.141632757 | 0.46514508 | 0.750 |
| YPL270W | 1768.066094 | 0.263871509 | 0.465031657 | 0.750 |
| YBR255W | 917.9047892 | 0.152739502 | 0.465594902 | 0.750 |
| YBR290W | 1203.208406 | 0.130750449 | 0.467693713 | 0.750 |
| YDL002C | 564.8870678 | 0.15361198 | 0.467882667 | 0.750 |
| YDR530C | 1226.853067 | 0.297303572 | 0.467818873 | 0.750 |
| YER004W | 2233.162604 | -0.276896057 | 0.466762218 | 0.750 |
| YGL067W | 999.1477018 | -0.163540186 | 0.46800554 | 0.750 |
| YGL213C | 596.0286848 | -0.148874796 | 0.466543766 | 0.750 |
| YHR119W | 1485.712969 | -0.16609799 | 0.466011709 | 0.750 |
| YIL029C | 174.946717 | 0.119376187 | 0.4671186 | 0.750 |
| YIL066C | 409.4135175 | 0.303547916 | 0.467354269 | 0.750 |
| YLR416C | 28.8196825 | -0.140541898 | 0.467701082 | 0.750 |
| YMR287C | 795.4088763 | 0.20554175 | 0.466502687 | 0.750 |

| | | | | |
|---|---|---|---|---|
| YNL078W | 1260.908163 | -0.225282913 | 0.466582914 | 0.750 |
| YOL024W | 52.91060507 | 0.423386833 | 0.466072179 | 0.750 |
| YOR014W | 3387.711726 | 0.231906373 | 0.467319634 | 0.750 |
| YOR192C | 247.398385 | 0.352805815 | 0.467318748 | 0.750 |
| YOR295W | 256.0382172 | -0.1792464 | 0.46624222 | 0.750 |
| YPL121C | 25.34339254 | 0.316760477 | 0.467826176 | 0.750 |
| YPL191C | 278.5044925 | 0.285982876 | 0.465791465 | 0.750 |
| YPR127W | 1734.545853 | 0.209244438 | 0.467251976 | 0.750 |
| YPR168W | 170.9858215 | 0.146571582 | 0.46581043 | 0.750 |
| YDR229W | 1110.14149 | 0.165342479 | 0.468301 | 0.750 |
| YGR263C | 454.4543224 | 0.208981245 | 0.468439771 | 0.750 |
| YDR377W | 1422.817631 | -0.168778976 | 0.468986716 | 0.751 |
| YAR029W | 75.14481749 | 0.140389065 | 0.469474052 | 0.751 |
| YIR003W | 1806.796543 | 0.201971011 | 0.469547215 | 0.751 |
| YKL041W | 833.3256523 | 0.181229962 | 0.469598176 | 0.751 |
| YGL026C | 4379.972897 | -0.281415002 | 0.470283761 | 0.752 |
| YHR007C | 4279.432771 | -0.101619015 | 0.470294162 | 0.752 |
| YHR039C | 2261.246675 | -0.107368873 | 0.470287331 | 0.752 |
| YDR020C | 268.9612697 | -0.241410609 | 0.470679261 | 0.752 |
| YKL105C | 1423.737383 | 0.194885028 | 0.470630923 | 0.752 |
| YDL200C | 293.2384509 | -0.217251368 | 0.47098201 | 0.752 |
| YER147C-A | 275.4373645 | -0.203212209 | 0.471791894 | 0.753 |
| YJR036C | 438.6482321 | 0.149302579 | 0.471785603 | 0.753 |
| YNR009W | 344.9832981 | -0.164746917 | 0.471724116 | 0.753 |
| YOR015W | 345.4012917 | 0.162099983 | 0.47215207 | 0.753 |
| YBL054W | 675.0753311 | -0.528342912 | 0.472439641 | 0.754 |
| YHR032W | 1019.136565 | -0.269589373 | 0.473070227 | 0.754 |
| YML025C | 1270.924858 | 0.13855725 | 0.472981894 | 0.754 |
| YIL134W | 238.1456546 | -0.198849502 | 0.473311808 | 0.755 |
| YER066C-A | 1483.59707 | 0.175014225 | 0.473501491 | 0.755 |
| YDL066W | 3505.644872 | -0.076448301 | 0.474157441 | 0.755 |
| YGL201C | 994.5170532 | -0.137600016 | 0.474154439 | 0.755 |
| YOR010C | 455.6940822 | -0.038863131 | 0.474407369 | 0.755 |
| YOR084W | 822.8839971 | 0.195409118 | 0.474434502 | 0.755 |
| YOR231W | 881.0751627 | 0.209304883 | 0.475155543 | 0.756 |
| YDR035W | 3632.420651 | -0.280995792 | 0.475374002 | 0.756 |
| YEL074W | 35.0602757 | 0.144655581 | 0.475623749 | 0.757 |
| YBR149W | 7039.888751 | 0.244667101 | 0.47647467 | 0.757 |
| YBR166C | 859.3257193 | -0.181363001 | 0.47796497 | 0.757 |
| YDL001W | 483.443293 | 0.177288454 | 0.477369753 | 0.757 |
| YDL017W | 421.2237274 | 0.122033474 | 0.477674386 | 0.757 |
| YDL225W | 1750.506735 | -0.201866232 | 0.478100743 | 0.757 |
| YDR253C | 229.9334863 | 0.236553222 | 0.476769457 | 0.757 |

| | | | | |
|---|---|---|---|---|
| YHR015W | 104.6987227 | 0.566301639 | 0.477649808 | 0.757 |
| YHR063C | 1432.676855 | -0.198507844 | 0.47787104 | 0.757 |
| YJR049C | 1397.771565 | 0.291341579 | 0.477934808 | 0.757 |
| YKL071W | 436.0298233 | 0.135051195 | 0.47680241 | 0.757 |
| YLR271W | 587.0390943 | 0.257628184 | 0.476703137 | 0.757 |
| YMR106C | 546.3577506 | 0.216367622 | 0.476310478 | 0.757 |
| YNL230C | 466.9526022 | 0.269712604 | 0.477481168 | 0.757 |
| YNR045W | 861.3548214 | 0.288912296 | 0.477863479 | 0.757 |
| YOL029C | 366.4284423 | -0.202243192 | 0.476299212 | 0.757 |
| YOR030W | 929.3153858 | 0.16351479 | 0.477981974 | 0.757 |
| snR44 | 201.2193719 | -0.218347952 | 0.477236136 | 0.757 |
| YGR129W | 435.9932211 | 0.244716143 | 0.4783882 | 0.757 |
| YJL102W | 772.7229577 | -0.096844352 | 0.478642906 | 0.757 |
| YDL106C | 1181.738765 | 0.19385266 | 0.479582663 | 0.758 |
| YDR159W | 1698.043068 | 0.165391595 | 0.479497848 | 0.758 |
| YKL053C-A | 473.2870725 | -0.213388985 | 0.479553281 | 0.758 |
| YMR117C | 204.4706004 | -0.281222287 | 0.479528612 | 0.758 |
| YPL034W | 171.3658521 | -0.312896644 | 0.479412513 | 0.758 |
| YGL202W | 6879.756501 | -0.153417568 | 0.480080704 | 0.759 |
| YBR017C | 2405.834993 | -0.143981209 | 0.480226933 | 0.759 |
| YHL017W | 1099.085857 | 0.148012822 | 0.480826818 | 0.759 |
| YIL138C | 1495.862581 | -0.275711123 | 0.480698787 | 0.759 |
| YOR163W | 1216.735785 | -0.158149016 | 0.481303369 | 0.760 |
| YCL061C | 1120.832529 | -0.146001308 | 0.482442644 | 0.760 |
| YEL066W | 476.9220074 | -0.23475046 | 0.48258441 | 0.760 |
| YER076C | 386.5202214 | 0.07031362 | 0.48212849 | 0.760 |
| YIL051C | 4289.573013 | -0.251469481 | 0.481768698 | 0.760 |
| YJL207C | 2307.188321 | -0.197645387 | 0.482014097 | 0.760 |
| YNL179C | 130.2526057 | 0.225634086 | 0.482306063 | 0.760 |
| YNL306W | 1534.453742 | -0.165668745 | 0.481766646 | 0.760 |
| YNL336W | 467.9927857 | 0.412265041 | 0.482498922 | 0.760 |
| YOR147W | 1293.879256 | 0.213426025 | 0.482262941 | 0.760 |
| YAL039C | 1453.711163 | 0.156634941 | 0.483973463 | 0.761 |
| YBR073W | 1901.278512 | -0.345920377 | 0.484817738 | 0.761 |
| YBR222C | 4562.01386 | 0.190873172 | 0.48386952 | 0.761 |
| YDR430C | 1380.899649 | 0.132061396 | 0.484327551 | 0.761 |
| YFL041W | 1108.963199 | 0.22728174 | 0.484012024 | 0.761 |
| YGR147C | 943.7636147 | -0.179368685 | 0.48490237 | 0.761 |
| YLR064W | 1412.390781 | 0.193259613 | 0.484919967 | 0.761 |
| YLR329W | 110.7708514 | 0.151836848 | 0.48401761 | 0.761 |
| YOR162C | 795.3084144 | 0.11976792 | 0.484517335 | 0.761 |
| YOR359W | 584.0345426 | 0.263185604 | 0.484309276 | 0.761 |
| YPR053C | 166.5759994 | 0.223866527 | 0.484533303 | 0.761 |

| | | | | |
|---|---|---|---|---|
| YLR373C | 1362.586121 | 0.153883414 | 0.485081308 | 0.761 |
| YBR095C | 539.8071954 | 0.160863842 | 0.485491559 | 0.761 |
| YDR266C | 1529.276069 | 0.224739382 | 0.486169083 | 0.761 |
| YDR493W | 345.6632951 | -0.162874685 | 0.485791116 | 0.761 |
| YEL020C-B | 34.39724677 | 0.256389126 | 0.485979402 | 0.761 |
| YHR094C | 525.1336673 | -0.772643194 | 0.48583781 | 0.761 |
| YIL042C | 814.6115743 | 0.157325879 | 0.486167309 | 0.761 |
| YLR077W | 986.4922801 | 0.152079718 | 0.486013365 | 0.761 |
| YNL254C | 501.4634557 | 0.153564774 | 0.485535913 | 0.761 |
| YDR320C | 1271.658982 | 0.196100991 | 0.486489281 | 0.762 |
| YPL107W | 480.1557675 | 0.244689548 | 0.486793435 | 0.762 |
| YLR319C | 843.3891881 | 0.138858116 | 0.487071987 | 0.762 |
| YPR172W | 1219.274092 | 0.259455314 | 0.487029789 | 0.762 |
| YDR284C | 1216.7009 | 0.132968393 | 0.487983883 | 0.763 |
| YDR448W | 426.123254 | -0.154366189 | 0.487857003 | 0.763 |
| YJL053W | 1232.219717 | 0.17866612 | 0.487841265 | 0.763 |
| YDL037C | 92.66763176 | -0.683929002 | 0.488139639 | 0.763 |
| YAR071W | 43.79477313 | -0.439410379 | 0.488982765 | 0.763 |
| YDR133C | 1765.716231 | -0.39011751 | 0.48939975 | 0.763 |
| YDR257C | 688.7119517 | -0.240118672 | 0.488980669 | 0.763 |
| YER099C | 790.1735302 | 0.130174497 | 0.488800344 | 0.763 |
| YJR031C | 1083.680948 | -0.079793787 | 0.48957561 | 0.763 |
| YLR006C | 1262.134466 | -0.276693954 | 0.489153682 | 0.763 |
| YMR095C | 600.2590168 | -0.432030459 | 0.489394211 | 0.763 |
| YOL053W | 775.6100204 | 0.273758889 | 0.489610718 | 0.763 |
| YDL217C | 416.6172424 | -0.144336921 | 0.490570724 | 0.765 |
| YDR213W | 568.9307813 | -0.270176865 | 0.490847508 | 0.765 |
| YJR050W | 528.5017956 | 0.190876393 | 0.491065379 | 0.765 |
| YML050W | 378.8334393 | 0.197225817 | 0.490815615 | 0.765 |
| YPL233W | 252.806959 | -0.173246645 | 0.491042498 | 0.765 |
| YDL004W | 2874.441994 | -0.249016746 | 0.49195999 | 0.765 |
| YDR047W | 1762.459456 | -0.147572136 | 0.492028792 | 0.765 |
| YJL052W | 23370.62399 | 0.768951056 | 0.491942052 | 0.765 |
| YNL122C | 325.2385534 | -0.174961848 | 0.491821472 | 0.765 |
| YPR144C | 1036.348245 | -0.172633393 | 0.491830556 | 0.765 |
| YDR270W | 829.6044968 | 0.224442068 | 0.492684927 | 0.766 |
| YHR137W | 2665.655739 | -0.244445268 | 0.49249391 | 0.766 |
| YPL057C | 1620.500312 | 0.097487047 | 0.492621691 | 0.766 |
| YDR380W | 4921.676121 | -0.291375391 | 0.493058578 | 0.766 |
| YGL080W | 877.4500948 | -0.266292844 | 0.493402932 | 0.766 |
| YIR032C | 113.4974223 | 0.137422948 | 0.492914035 | 0.766 |
| YKR103W | 526.4204375 | 0.338049511 | 0.493360846 | 0.766 |
| YMR121C | 804.8314095 | -0.279415671 | 0.493283578 | 0.766 |

| | | | | |
|---|---|---|---|---|
| YBL034C | 918.1202298 | -0.170310127 | 0.493549299 | 0.766 |
| YDL015C | 3069.696513 | -0.125520272 | 0.494199689 | 0.766 |
| YDL242W | 58.94192665 | -0.314451108 | 0.494431877 | 0.766 |
| YFL047W | 1142.35423 | 0.27913569 | 0.494861751 | 0.766 |
| YGR169C | 437.5350343 | -0.186412866 | 0.494511 | 0.766 |
| YGR251W | 285.9824856 | -0.219090997 | 0.494775304 | 0.766 |
| YHR171W | 640.7534835 | 0.202659726 | 0.495286304 | 0.766 |
| YIL028W | 59.74955175 | 0.051531247 | 0.494381741 | 0.766 |
| YIL139C | 280.0724876 | -0.154811079 | 0.49521406 | 0.766 |
| YLR082C | 386.5514271 | 0.188979736 | 0.495294431 | 0.766 |
| YLR146W-A | 192.5995673 | -0.277204845 | 0.495087578 | 0.766 |
| YMR064W | 1294.977576 | 0.177402364 | 0.495413824 | 0.766 |
| YNL006W | 936.4455341 | 0.298855597 | 0.495023286 | 0.766 |
| YOL154W | 197.7584736 | 0.114648742 | 0.494467105 | 0.766 |
| snR70 | 64.50681846 | 0.218731699 | 0.49435686 | 0.766 |
| YDR082W | 550.4664737 | 0.101651284 | 0.495969067 | 0.766 |
| YNL146C-A | 29.52790111 | 0.169710935 | 0.496263794 | 0.767 |
| YOL036W | 1888.124512 | 0.258931979 | 0.496384987 | 0.767 |
| YDL097C | 4494.471702 | -0.188377752 | 0.496765934 | 0.767 |
| YDR247W | 1026.990415 | 0.249446214 | 0.496829489 | 0.767 |
| YDR478W | 183.0152712 | -0.180814336 | 0.496910084 | 0.767 |
| YBR296C | 209.9913106 | -0.751595319 | 0.497524141 | 0.768 |
| YBL104C | 644.4185242 | -0.152249069 | 0.497766507 | 0.768 |
| RUF21 | 43.48350584 | -0.083528602 | 0.498130661 | 0.768 |
| YML099C | 1070.053648 | 0.201011462 | 0.498078977 | 0.768 |
| YER058W | 286.3082478 | -0.209083074 | 0.498408429 | 0.768 |
| YLR283W | 409.1057668 | -0.171943701 | 0.498386161 | 0.768 |
| YEL060C | 10805.49764 | 0.226551469 | 0.498596006 | 0.768 |
| YER096W | 439.4636005 | -0.134464671 | 0.498769787 | 0.768 |
| YHR172W | 514.8081565 | -0.050873534 | 0.498950002 | 0.768 |
| YBR233W | 474.558741 | -0.235539818 | 0.499787739 | 0.769 |
| YKR091W | 1094.538594 | 0.193619385 | 0.499803603 | 0.769 |
| YEL017C-A | 672.6729484 | -0.11245193 | 0.500157479 | 0.769 |
| YNL261W | 688.6478875 | 0.243396084 | 0.500172924 | 0.769 |
| YOR028C | 706.5725488 | -0.047355039 | 0.500321284 | 0.769 |
| YGL110C | 882.4868812 | 0.1588614 | 0.500661371 | 0.770 |
| YLR434C | 48.33837913 | -0.166397914 | 0.501279092 | 0.770 |
| YLR182W | 1421.620302 | 0.185227137 | 0.501510628 | 0.771 |
| YIL171W | 27.73883289 | 0.131001154 | 0.501802288 | 0.771 |
| YJL082W | 1623.02873 | 0.195398749 | 0.501924659 | 0.771 |
| YOR209C | 3422.936842 | -0.111537046 | 0.501704886 | 0.771 |
| YCL035C | 4330.054767 | 0.200817839 | 0.503818739 | 0.772 |
| YDR113C | 648.1289584 | -0.122003397 | 0.503545867 | 0.772 |

| | | | | |
|---|---|---|---|---|
| YKR093W | 2668.023949 | 0.204670504 | 0.503326046 | 0.772 |
| YLR177W | 3573.62477 | 0.061708317 | 0.50350471 | 0.772 |
| YMR299C | 640.5800002 | -0.232199595 | 0.503703995 | 0.772 |
| YNL084C | 2582.808264 | -0.12775483 | 0.503735823 | 0.772 |
| YDR033W | 12974.70679 | -0.042014864 | 0.504302419 | 0.773 |
| YLR445W | 32.03943686 | -0.138267193 | 0.504412998 | 0.773 |
| YNR038W | 1115.811097 | -0.244610616 | 0.504446007 | 0.773 |
| YGL168W | 276.6572721 | -0.373682503 | 0.504717615 | 0.773 |
| YNL051W | 867.7540206 | 0.192151018 | 0.504609667 | 0.773 |
| YJR091C | 3359.251305 | 0.13870043 | 0.504964737 | 0.773 |
| YMR068W | 420.8665046 | 0.12242708 | 0.505272374 | 0.773 |
| YKR008W | 1399.761894 | -0.19325096 | 0.505966923 | 0.774 |
| YNL090W | 89.81803356 | -0.151526101 | 0.506115626 | 0.774 |
| YIR014W | 256.3884009 | -0.373924179 | 0.506510579 | 0.774 |
| YJL049W | 674.8429005 | 0.163457228 | 0.506821289 | 0.775 |
| YNL077W | 1205.161408 | 0.235095966 | 0.506872627 | 0.775 |
| YGL256W | 1304.621904 | 0.174658431 | 0.507062165 | 0.775 |
| YPR140W | 900.9897118 | 0.37265539 | 0.507500625 | 0.775 |
| YGR283C | 529.6205161 | -0.274811731 | 0.507660853 | 0.775 |
| YLR222C | 1088.378581 | -0.199389226 | 0.508183005 | 0.776 |
| YCL036W | 402.9926525 | -0.575519516 | 0.508906479 | 0.776 |
| YDR163W | 579.7588355 | -0.208220924 | 0.508611924 | 0.776 |
| YDR459C | 457.0405938 | 0.093757645 | 0.508713156 | 0.776 |
| YLL055W | 836.4946306 | 0.188552141 | 0.508511915 | 0.776 |
| YLR103C | 462.3096359 | -0.162546571 | 0.508962533 | 0.776 |
| YBR035C | 2483.374886 | -0.192301241 | 0.509554774 | 0.776 |
| YBR201W | 456.7810734 | -0.206138476 | 0.509717956 | 0.776 |
| YLR036C | 360.058471 | 0.233501604 | 0.50932133 | 0.776 |
| YOL030W | 6136.814993 | 0.23022725 | 0.510062623 | 0.776 |
| YPL004C | 14429.78474 | 0.196573298 | 0.510063284 | 0.776 |
| YPL029W | 470.8569792 | -0.144537996 | 0.510101435 | 0.776 |
| YPR042C | 1338.934548 | 0.178340931 | 0.509888921 | 0.776 |
| YBL059W | 273.8243146 | -0.138295166 | 0.510572357 | 0.776 |
| YFR024C-A | 2497.30533 | -0.197098703 | 0.510320578 | 0.776 |
| YNL040W | 832.5546055 | 0.190987439 | 0.510531006 | 0.776 |
| YGL001C | 1350.851203 | -0.162805821 | 0.511607408 | 0.777 |
| YGL018C | 470.8004765 | 0.219853247 | 0.511433729 | 0.777 |
| YIL065C | 779.4474724 | 0.165906337 | 0.511407464 | 0.777 |
| YLL011W | 676.0042616 | -0.237654487 | 0.511477495 | 0.777 |
| YOL157C | 78.40010831 | 0.238210068 | 0.511389689 | 0.777 |
| YBL113C | 286.5211358 | 0.294974839 | 0.512183696 | 0.777 |
| YGR240C | 9640.703605 | -0.113029816 | 0.511991532 | 0.777 |
| YLR396C | 522.7183585 | 0.147095477 | 0.512306051 | 0.777 |

| | | | | |
|---|---|---|---|---|
| YOR273C | 1760.091485 | -0.440886112 | 0.512128645 | 0.777 |
| YGL144C | 301.1617021 | 0.162718266 | 0.512863694 | 0.777 |
| YIR037W | 2878.72569 | -0.194423904 | 0.512843847 | 0.777 |
| YLR090W | 931.4547969 | 0.191504589 | 0.512786697 | 0.777 |
| YML057W | 1931.217961 | 0.165819062 | 0.513500153 | 0.778 |
| YJL051W | 1072.541155 | 0.171065289 | 0.514254132 | 0.779 |
| YLR193C | 745.5545212 | 0.28677127 | 0.514008054 | 0.779 |
| YPL103C | 590.7071612 | 0.207990147 | 0.514127126 | 0.779 |
| YDL221W | 36.90750404 | 0.39698578 | 0.51466019 | 0.779 |
| YJR135C | 188.8236464 | -0.187174385 | 0.515104339 | 0.780 |
| YOL159C | 274.8863973 | 0.448857044 | 0.5157223 | 0.780 |
| YOR103C | 111.7299261 | -0.194425557 | 0.516229213 | 0.781 |
| YDL159W | 513.5387781 | 0.136990741 | 0.517455402 | 0.782 |
| YGR101W | 624.5372953 | 0.168210057 | 0.51768688 | 0.782 |
| YIL044C | 893.1546159 | -0.142854308 | 0.517667724 | 0.782 |
| YKL039W | 2424.281879 | 0.196172402 | 0.517930372 | 0.782 |
| YKL138C | 823.724232 | -0.186992917 | 0.517928691 | 0.782 |
| YNR004W | 166.7322391 | -0.258196698 | 0.517639653 | 0.782 |
| YOR330C | 1527.333333 | -0.203462699 | 0.517398768 | 0.782 |
| YIL011W | 777.5395764 | -0.620503658 | 0.518638299 | 0.783 |
| YBL007C | 3937.826759 | -0.177591888 | 0.520409557 | 0.783 |
| YBR287W | 2396.597397 | 0.089262918 | 0.520212404 | 0.783 |
| YER119C | 1204.859566 | 0.286492271 | 0.519580672 | 0.783 |
| YHL025W | 1272.013263 | 0.223784743 | 0.519335084 | 0.783 |
| YJR147W | 698.6444516 | -0.223096793 | 0.519730065 | 0.783 |
| YLL002W | 366.0264394 | -0.123652962 | 0.520089048 | 0.783 |
| YLL051C | 1170.125266 | -0.204142077 | 0.520288153 | 0.783 |
| YMR241W | 1393.972884 | -0.14111405 | 0.520451974 | 0.783 |
| YNL197C | 1704.51539 | -0.156209166 | 0.520564773 | 0.783 |
| YNL252C | 974.4576049 | -0.118200733 | 0.519544986 | 0.783 |
| YNL280C | 1498.81009 | -0.174479245 | 0.519092153 | 0.783 |
| YNL317W | 817.8553177 | 0.132495371 | 0.519644206 | 0.783 |
| YPL195W | 2059.660784 | -0.110891747 | 0.520637398 | 0.783 |
| YPR067W | 664.4484323 | -0.140525897 | 0.519321479 | 0.783 |
| YAR050W | 236.1195337 | 0.415572338 | 0.521202267 | 0.783 |
| YBR228W | 415.0546527 | 0.111893808 | 0.521608518 | 0.783 |
| YDR489W | 358.8647918 | -0.237962905 | 0.521244042 | 0.783 |
| YIL031W | 1396.422579 | 0.137870139 | 0.521055039 | 0.783 |
| YIL135C | 1336.175722 | 0.179722519 | 0.521799438 | 0.783 |
| YKR055W | 476.0228776 | 0.122630693 | 0.521449651 | 0.783 |
| YNL026W | 973.5528021 | -0.122349917 | 0.52179443 | 0.783 |
| YNL268W | 2190.621688 | 0.224539579 | 0.521766675 | 0.783 |
| YIR006C | 5095.375019 | -0.133272017 | 0.52208495 | 0.783 |

| | | | | |
|---|---|---|---|---|
| YKR024C | 515.1058632 | -0.248965299 | 0.522043814 | 0.783 |
| YAL053W | 2097.963607 | -0.345858433 | 0.522717121 | 0.784 |
| YGL164C | 861.3475827 | 0.169541824 | 0.522748614 | 0.784 |
| YNL001W | 629.9634099 | -0.159387795 | 0.522718576 | 0.784 |
| YBR133C | 1160.137007 | -0.256676883 | 0.523090821 | 0.784 |
| YIL091C | 989.7593991 | -0.285032485 | 0.523506464 | 0.784 |
| YLL005C | 615.6111971 | 0.316582495 | 0.523528488 | 0.784 |
| YPR036W-A | 8058.010483 | 0.205175405 | 0.524108176 | 0.785 |
| YPL204W | 2957.690828 | 0.153672162 | 0.524381329 | 0.785 |
| YJR005C-A | 81.403139 | -0.925663928 | 0.524717372 | 0.785 |
| YDR473C | 858.9200777 | 0.149323688 | 0.52485628 | 0.785 |
| YBR215W | 1081.67593 | 0.158673033 | 0.525878015 | 0.787 |
| YBL060W | 624.3164774 | 0.127111577 | 0.526750871 | 0.787 |
| YBR111W-A | 229.152125 | -0.20737535 | 0.527686411 | 0.787 |
| YBR146W | 1432.145995 | -0.077651524 | 0.5269401 | 0.787 |
| YBR217W | 353.7676501 | -0.23579515 | 0.527455345 | 0.787 |
| YDR264C | 2392.197288 | -0.126033604 | 0.526971411 | 0.787 |
| YGL139W | 1235.197549 | 0.131704583 | 0.527197264 | 0.787 |
| YHR012W | 963.21973 | -0.177512598 | 0.527171174 | 0.787 |
| YIL104C | 354.0212252 | 0.123549861 | 0.527881819 | 0.787 |
| YJR047C | 407.0063415 | -0.480746238 | 0.526440099 | 0.787 |
| YML015C | 859.349761 | -0.28977966 | 0.527923384 | 0.787 |
| YMR238W | 935.1001777 | -0.152126449 | 0.52753371 | 0.787 |
| YNL014W | 784.7565686 | 0.106047234 | 0.528244027 | 0.787 |
| YNL021W | 893.5874562 | -0.10122326 | 0.527461953 | 0.787 |
| YOR285W | 4227.371531 | 0.292092946 | 0.528112604 | 0.787 |
| YPL041C | 160.4382575 | -0.147950043 | 0.528214143 | 0.787 |
| YBR203W | 994.1971392 | 0.038361432 | 0.529094629 | 0.787 |
| YDR031W | 342.8572195 | 0.275224661 | 0.52885744 | 0.787 |
| YDR536W | 315.8429448 | 0.318973219 | 0.52861844 | 0.787 |
| YJL123C | 2650.72374 | -0.20333927 | 0.528664426 | 0.787 |
| YMR037C | 1272.708272 | -0.189908743 | 0.529023676 | 0.787 |
| YPL078C | 4809.066893 | -0.116830043 | 0.528767677 | 0.787 |
| YGL151W | 1728.404115 | 0.175901935 | 0.529779172 | 0.788 |
| YIL129C | 2649.636452 | 0.218788954 | 0.530605143 | 0.789 |
| YOL076W | 856.9107843 | -0.173222096 | 0.530518638 | 0.789 |
| YGR010W | 527.974638 | 0.201024182 | 0.531079068 | 0.790 |
| YBR238C | 642.2716869 | -0.14306527 | 0.531268307 | 0.790 |
| YHR103W | 1865.030837 | 0.183139788 | 0.533003388 | 0.791 |
| YKR079C | 916.529157 | -0.210961755 | 0.532991897 | 0.791 |
| YLL027W | 1273.25217 | 0.136676827 | 0.532841035 | 0.791 |
| YMR178W | 1157.445412 | 0.264625497 | 0.532885497 | 0.791 |
| YBR167C | 275.1433522 | -0.257077181 | 0.533193322 | 0.791 |

| | | | | |
|---|---|---|---|---|
| YPL082C | 1537.0483 | -0.146161285 | 0.53349203 | 0.792 |
| YJL219W | 39.81053365 | 0.49287148 | 0.534293905 | 0.792 |
| YNL065W | 531.9531231 | 0.402395327 | 0.534153176 | 0.792 |
| YOR073W | 590.4176603 | -0.150461019 | 0.534284597 | 0.792 |
| YER044C-A | 102.3682402 | 0.094499098 | 0.534790113 | 0.793 |
| YKR060W | 417.425685 | -0.256187211 | 0.534659284 | 0.793 |
| YMR268C | 351.0386447 | 0.112106267 | 0.534913054 | 0.793 |
| YDR011W | 4201.778813 | -0.196457102 | 0.535176922 | 0.793 |
| YDR294C | 2749.492975 | 0.16526311 | 0.535776979 | 0.793 |
| YDR376W | 985.9130091 | 0.177709068 | 0.536131887 | 0.793 |
| YGR277C | 611.2932905 | -0.170507484 | 0.53603035 | 0.793 |
| YHR114W | 1802.212821 | 0.149760036 | 0.536260768 | 0.793 |
| YJL151C | 3281.2426 | 0.23114867 | 0.53579583 | 0.793 |
| YKL137W | 420.0820122 | -0.077928717 | 0.536197978 | 0.793 |
| YMR244C-A | 108.6393324 | 0.189100544 | 0.535370936 | 0.793 |
| YPL085W | 4050.180373 | -0.137854742 | 0.535940018 | 0.793 |
| YJR085C | 2130.783694 | -0.179398557 | 0.536952614 | 0.794 |
| YHR147C | 857.7185719 | -0.102860317 | 0.537715871 | 0.794 |
| YHR161C | 2280.636973 | 0.160182354 | 0.537603276 | 0.794 |
| YIL063C | 1081.563362 | -0.184361861 | 0.537615452 | 0.794 |
| YLR168C | 702.4056037 | -0.200651613 | 0.537519711 | 0.794 |
| YDR411C | 1171.452043 | -0.127930353 | 0.538038985 | 0.794 |
| YLR095C | 1759.144757 | -0.15277961 | 0.538200874 | 0.794 |
| YMR195W | 1764.107553 | 0.106564519 | 0.538074539 | 0.794 |
| YPL262W | 5873.810031 | -0.041240533 | 0.538409912 | 0.794 |
| YLR251W | 493.6892026 | 0.125939568 | 0.538936951 | 0.795 |
| YLR081W | 221.380283 | 0.315387839 | 0.539307538 | 0.795 |
| YKR020W | 476.7599628 | -0.237480116 | 0.539620158 | 0.795 |
| YMR053C | 712.958308 | 0.129960772 | 0.539698109 | 0.795 |
| YJL200C | 1688.489143 | -0.250045073 | 0.540019624 | 0.795 |
| YKL173W | 1269.160496 | 0.118613616 | 0.540161239 | 0.795 |
| YOL020W | 976.6345917 | 0.232704788 | 0.540048843 | 0.795 |
| YGL166W | 257.6413584 | 0.086234313 | 0.540372237 | 0.795 |
| YPL091W | 3000.977941 | -0.108517447 | 0.540487395 | 0.795 |
| YDR044W | 1839.127613 | -0.169717349 | 0.541246316 | 0.796 |
| YBL037W | 1072.022099 | 0.166808359 | 0.54297754 | 0.797 |
| YBR180W | 89.72716224 | 0.600538774 | 0.542942274 | 0.797 |
| YDL113C | 1161.019898 | 0.088655001 | 0.543151996 | 0.797 |
| YDL128W | 1754.772543 | 0.169955462 | 0.543358591 | 0.797 |
| YEL009C | 21167.59097 | -0.239911792 | 0.542501186 | 0.797 |
| YGL177W | 36.28800868 | -0.228074433 | 0.542965247 | 0.797 |
| YGL198W | 494.0719347 | -0.297314003 | 0.543393643 | 0.797 |
| YGR019W | 2682.062979 | 0.196440046 | 0.542336774 | 0.797 |

| | | | | |
|---|---|---|---|---|
| YGR161W-C | 303.4318056 | 0.150577805 | 0.541478638 | 0.797 |
| YHR001W-A | 1152.222923 | 0.465184223 | 0.541971135 | 0.797 |
| YHR209W | 694.2958547 | 0.266333982 | 0.542787249 | 0.797 |
| YKL013C | 2002.473422 | -0.146430881 | 0.542550546 | 0.797 |
| YLR357W | 954.9234442 | -0.186177231 | 0.543506229 | 0.797 |
| YLR363C | 329.7816062 | 0.067058767 | 0.542487537 | 0.797 |
| YPL087W | 1694.761164 | 0.270354863 | 0.542521656 | 0.797 |
| snR41 | 28.2332558 | -0.093183254 | 0.542349061 | 0.797 |
| YGR137W | 236.2341776 | -0.262352196 | 0.543717873 | 0.797 |
| YDR075W | 328.511722 | -0.21067344 | 0.543990785 | 0.797 |
| YJR059W | 3492.702056 | 0.135327704 | 0.544058548 | 0.797 |
| YOR306C | 1215.578289 | -0.365030213 | 0.544470593 | 0.797 |
| YIL141W | 35.62085203 | -0.168844079 | 0.544840034 | 0.797 |
| YCL049C | 935.682406 | 0.356838082 | 0.546163991 | 0.798 |
| YDR319C | 485.2377065 | 0.184673859 | 0.545467643 | 0.798 |
| YFL010W-A | 308.9446486 | -0.256782754 | 0.546046579 | 0.798 |
| YGR096W | 275.6515308 | 0.105340679 | 0.546284545 | 0.798 |
| YJL042W | 3416.022292 | 0.176903263 | 0.54586652 | 0.798 |
| YKL076C | 34.14810955 | 0.073462863 | 0.545492402 | 0.798 |
| YLR113W | 3162.32221 | -0.171594015 | 0.545262378 | 0.798 |
| YMR086C-A | 82.59944884 | 0.104465973 | 0.546017723 | 0.798 |
| YMR309C | 2879.75717 | -0.049957461 | 0.545930863 | 0.798 |
| YJL142C | 106.3041783 | -0.044715088 | 0.546632132 | 0.798 |
| YAL009W | 404.8562545 | -0.166386141 | 0.54704645 | 0.798 |
| YDR046C | 2206.66233 | 0.134617724 | 0.546892618 | 0.798 |
| YLR277C | 732.8482737 | -0.171102811 | 0.546926777 | 0.798 |
| YBR224W | 27.9935508 | 0.383495933 | 0.547683545 | 0.798 |
| YDR510C-A | 439.1253795 | -0.379157274 | 0.547458386 | 0.798 |
| YFL031C-A | 101.8569422 | -0.102843255 | 0.547466803 | 0.798 |
| YGL028C | 1677.067671 | 0.222437692 | 0.547798834 | 0.798 |
| YIL005W | 1687.731259 | 0.202656 | 0.547607995 | 0.798 |
| YBR145W | 1370.985162 | 0.111680682 | 0.548585718 | 0.798 |
| YGR091W | 441.73971 | 0.184951972 | 0.548397409 | 0.798 |
| YLR130C | 1253.100793 | 0.18927059 | 0.548571928 | 0.798 |
| YLR348C | 661.9197696 | -0.38894992 | 0.548588015 | 0.798 |
| YOR204W | 2305.73227 | -0.041679354 | 0.548680216 | 0.798 |
| snR35 | 41.6109753 | -0.360420315 | 0.548146168 | 0.798 |
| YDR434W | 1180.144828 | 0.229811393 | 0.549118787 | 0.799 |
| YMR221C | 1062.231053 | -0.155762384 | 0.549002851 | 0.799 |
| YGL117W | 750.1030158 | 0.060413186 | 0.549364257 | 0.799 |
| YHL002W | 1516.40559 | 0.160557489 | 0.549469748 | 0.799 |
| YJR133W | 618.7266674 | -0.198101373 | 0.549726007 | 0.799 |
| YNL271C | 3740.617123 | -0.159479015 | 0.549896492 | 0.799 |

| | | | | |
|---|---|---|---|---|
| YOL073C | 1132.754224 | 0.177465026 | 0.549838099 | 0.799 |
| YDR346C | 2820.625886 | -0.21460614 | 0.550270876 | 0.799 |
| YCR069W | 2249.27644 | 0.166486582 | 0.550718858 | 0.799 |
| YOL027C | 2880.865523 | -0.133573634 | 0.55071656 | 0.799 |
| YCL014W | 1364.401777 | -0.162007072 | 0.55097511 | 0.799 |
| YGL188C | 26.0866713 | 0.315652002 | 0.551301056 | 0.800 |
| YML046W | 728.0597624 | 0.198067981 | 0.551390817 | 0.800 |
| YJR148W | 6070.611654 | 0.259943811 | 0.551865146 | 0.800 |
| YDL189W | 1070.015735 | -0.188010619 | 0.553069204 | 0.800 |
| YDR042C | 81.10570396 | -0.016698843 | 0.553006997 | 0.800 |
| YDR215C | 67.50004526 | 0.332834136 | 0.552624922 | 0.800 |
| YIL152W | 734.4896336 | 0.121555967 | 0.552559447 | 0.800 |
| YKR053C | 486.0346558 | 0.088591921 | 0.55277724 | 0.800 |
| YMR009W | 601.1334204 | 0.099724113 | 0.552780851 | 0.800 |
| YOL122C | 1128.161192 | -0.104566087 | 0.553149883 | 0.800 |
| YPL172C | 484.663762 | -0.116090298 | 0.553042931 | 0.800 |
| YAL029C | 2871.619875 | 0.141512106 | 0.553365646 | 0.801 |
| YPR045C | 1121.270528 | 0.186966244 | 0.553451484 | 0.801 |
| YLR420W | 2792.570329 | -0.01748987 | 0.553735259 | 0.801 |
| YJL209W | 626.0794008 | 0.133464265 | 0.554212103 | 0.801 |
| YKR090W | 850.9383093 | -0.100631069 | 0.554338254 | 0.801 |
| YNL241C | 6779.137631 | -0.214812768 | 0.554609259 | 0.801 |
| YOL164W | 729.2266525 | 0.119284245 | 0.554600907 | 0.801 |
| YER068W | 1672.188885 | 0.19538287 | 0.55499104 | 0.802 |
| YLR313C | 272.7990365 | -0.235793565 | 0.555171225 | 0.802 |
| YCL057C-A | 1206.33424 | -0.158839456 | 0.555657298 | 0.802 |
| YDR080W | 1103.850689 | 0.189908861 | 0.555660528 | 0.802 |
| YER188W | 461.4895098 | 0.619585006 | 0.555721896 | 0.802 |
| YDR054C | 1985.938955 | 0.206506073 | 0.556294733 | 0.802 |
| YKL219W | 500.3272007 | 0.244787015 | 0.556661113 | 0.803 |
| YOR367W | 630.0179766 | -0.193107468 | 0.556762494 | 0.803 |
| YER081W | 9451.152045 | -0.083990829 | 0.556938071 | 0.803 |
| YOR050C | 67.3008437 | -0.291740434 | 0.557140574 | 0.803 |
| YPL254W | 446.984746 | 0.163641668 | 0.557315591 | 0.803 |
| YMR282C | 677.7599955 | 0.168638997 | 0.557607022 | 0.803 |
| YEL024W | 1858.219132 | 0.355487975 | 0.558535233 | 0.804 |
| YJR034W | 337.4607648 | -0.198551793 | 0.558527428 | 0.804 |
| YKL049C | 577.3829424 | -0.153287751 | 0.558271347 | 0.804 |
| YLR087C | 2042.517235 | -0.08859266 | 0.558215786 | 0.804 |
| YIL093C | 871.5653388 | -0.077780547 | 0.5587028 | 0.804 |
| YBR195C | 288.8600834 | -0.172506336 | 0.559958755 | 0.804 |
| YDR352W | 489.6613604 | -0.06368546 | 0.560037045 | 0.804 |
| YEL068C | 304.3910786 | -0.223086291 | 0.559924845 | 0.804 |

| | | | | |
|---|---|---|---|---|
| YER060W-A | 584.2480367 | -0.106093689 | 0.559190984 | 0.804 |
| YJR111C | 500.5453341 | 0.173727168 | 0.559529074 | 0.804 |
| YKR071C | 2094.595777 | -0.138188008 | 0.559736042 | 0.804 |
| YOL068C | 1498.234041 | -0.166490427 | 0.559175204 | 0.804 |
| YOR116C | 1548.881592 | -0.166502327 | 0.55933546 | 0.804 |
| YOR192C-B | 24.91802453 | -0.315313813 | 0.559918936 | 0.804 |
| YFL016C | 3333.34381 | -0.149023352 | 0.560816573 | 0.805 |
| YDR221W | 936.1837264 | -0.09904303 | 0.561134011 | 0.805 |
| YCR083W | 821.7942376 | 0.250579468 | 0.56160054 | 0.805 |
| YDR540C | 110.2480797 | 0.579624915 | 0.561606425 | 0.805 |
| YMR211W | 431.1277672 | 0.102463745 | 0.561665206 | 0.805 |
| YDR146C | 1456.032171 | -0.091997163 | 0.561833271 | 0.805 |
| YOR185C | 2527.056266 | 0.206024279 | 0.562097546 | 0.805 |
| YDL233W | 547.8994161 | -0.242551776 | 0.562585234 | 0.806 |
| YER050C | 1168.128624 | -0.165930708 | 0.562486149 | 0.806 |
| YHL023C | 1253.971871 | 0.126145304 | 0.562634711 | 0.806 |
| YKL022C | 1018.942477 | 0.147164476 | 0.5626941 | 0.806 |
| YMR063W | 168.4962103 | 0.124293097 | 0.563357946 | 0.806 |
| YNR010W | 212.8810604 | 0.168364069 | 0.563452268 | 0.806 |
| YOL159C-A | 58.15453861 | 0.374964075 | 0.563318786 | 0.806 |
| YER069W | 925.6063303 | 0.107162644 | 0.564024933 | 0.807 |
| YHL028W | 1137.369035 | 0.377275393 | 0.564464511 | 0.807 |
| YGR114C | 36.93060097 | -0.745199562 | 0.564611577 | 0.807 |
| YKR105C | 96.82105819 | 0.256076346 | 0.565199958 | 0.807 |
| YLR326W | 402.567239 | -0.182559989 | 0.565166258 | 0.807 |
| YNL307C | 2792.714562 | -0.111390906 | 0.565151495 | 0.807 |
| YPL161C | 652.0526344 | 0.116137605 | 0.565115648 | 0.807 |
| YIL033C | 4130.82095 | 0.23666105 | 0.565478238 | 0.807 |
| YPR185W | 1716.783017 | 0.148092479 | 0.565575387 | 0.807 |
| YBR226C | 66.35832606 | 0.057227814 | 0.566840697 | 0.808 |
| YBR229C | 1792.230472 | -0.102296963 | 0.566974255 | 0.808 |
| YBR260C | 1246.430976 | -0.170821264 | 0.566584591 | 0.808 |
| YMR015C | 1137.039851 | 0.386244844 | 0.566216597 | 0.808 |
| YNL157W | 1475.809734 | -0.164469501 | 0.566498844 | 0.808 |
| YOR184W | 6279.813274 | -0.244833902 | 0.566648318 | 0.808 |
| YPR029C | 1930.500091 | -0.105000236 | 0.567000991 | 0.808 |
| YGR055W | 1342.856307 | 0.13643061 | 0.567968231 | 0.809 |
| YKL165C | 1154.053703 | 0.074583721 | 0.567845144 | 0.809 |
| YAL060W | 4666.556071 | 0.422419099 | 0.570052753 | 0.810 |
| YCL026C-A | 44.13033287 | 0.209545909 | 0.569520219 | 0.810 |
| YDR231C | 627.5516275 | 0.128962255 | 0.569833316 | 0.810 |
| YFR003C | 1017.431306 | 0.122916075 | 0.570007903 | 0.810 |
| YKR050W | 1256.621968 | 0.084559666 | 0.569210735 | 0.810 |

| | | | | |
|---|---|---|---|---|
| YLL035W | 581.4295973 | -0.210193204 | 0.569645176 | 0.810 |
| YLR139C | 783.7450537 | 0.151177267 | 0.569368764 | 0.810 |
| YML049C | 1684.51308 | -0.187964776 | 0.569891751 | 0.810 |
| YOL044W | 473.6470695 | 0.137445258 | 0.569230375 | 0.810 |
| YPL181W | 1430.008347 | 0.187094831 | 0.569901891 | 0.810 |
| YPL251W | 76.6529722 | -0.210387328 | 0.57019786 | 0.810 |
| YNL304W | 349.9176759 | -0.180948267 | 0.570533899 | 0.810 |
| YMR078C | 885.7332334 | -0.118534035 | 0.571067768 | 0.811 |
| YPL269W | 614.929882 | 0.132969242 | 0.571051469 | 0.811 |
| YJL081C | 1547.994145 | -0.070107295 | 0.571344376 | 0.811 |
| YNL286W | 504.2093629 | 0.182601975 | 0.571389567 | 0.811 |
| YJL159W | 37900.6137 | 0.272509327 | 0.571875565 | 0.811 |
| YNL143C | 26.87962177 | -0.139755005 | 0.572121259 | 0.811 |
| YDL073W | 971.6916673 | -0.161516946 | 0.572985757 | 0.812 |
| YDR059C | 828.9203049 | 0.175882243 | 0.573334682 | 0.812 |
| YGR109W-B | 857.2496031 | 0.195244755 | 0.573307354 | 0.812 |
| YIR008C | 729.6399623 | 0.129717663 | 0.573221895 | 0.812 |
| YMR311C | 3644.500712 | 0.157614464 | 0.573288128 | 0.812 |
| YOR217W | 1207.361787 | 0.184637229 | 0.573148736 | 0.812 |
| YPL241C | 283.7799123 | -0.077614989 | 0.573988761 | 0.813 |
| YLR028C | 4846.630925 | -0.075363066 | 0.574601913 | 0.813 |
| YMR115W | 1106.490843 | 0.155917443 | 0.574704315 | 0.813 |
| YHL022C | 136.8231823 | 0.167119173 | 0.575583327 | 0.814 |
| YBL093C | 1069.915355 | -0.176536885 | 0.575737383 | 0.814 |
| YPL153C | 363.2997459 | -0.164207474 | 0.576047402 | 0.815 |
| YDR394W | 3757.628989 | -0.132179239 | 0.576581674 | 0.815 |
| YKL030W | 149.8142583 | -0.639656813 | 0.57646457 | 0.815 |
| YDR414C | 448.7530762 | -0.176704075 | 0.577768454 | 0.816 |
| YFR052W | 2758.803837 | -0.22298284 | 0.578441142 | 0.817 |
| YGR125W | 1519.849762 | 0.144958854 | 0.578181287 | 0.817 |
| YIL147C | 2000.977172 | 0.138978353 | 0.578055334 | 0.817 |
| YMR156C | 369.432093 | 0.229549978 | 0.578330324 | 0.817 |
| YCR017C | 2986.294618 | -0.13564025 | 0.579107116 | 0.817 |
| YDL034W | 29.64111763 | 0.390747789 | 0.579480592 | 0.817 |
| YFL044C | 776.7355212 | 0.214682517 | 0.579936291 | 0.817 |
| YHR059W | 401.0845698 | -0.113331388 | 0.579148374 | 0.817 |
| YHR079C | 1259.061897 | 0.111874758 | 0.578692738 | 0.817 |
| YLR025W | 2034.136821 | 0.133859722 | 0.579443315 | 0.817 |
| YLR211C | 375.8938446 | 0.064115517 | 0.578859647 | 0.817 |
| YLR286C | 14573.14708 | -0.144840004 | 0.579048473 | 0.817 |
| YML070W | 2763.480937 | 0.231678938 | 0.579787267 | 0.817 |
| YML099W-A | 34.73959572 | 0.052403911 | 0.580376474 | 0.817 |
| YML115C | 1520.660585 | 0.131239848 | 0.580157308 | 0.817 |

| | | | | |
|---|---|---|---|---|
| YNL034W | 79.93693595 | 0.321732851 | 0.580266667 | 0.817 |
| YOR034C-A | 45.66152085 | -0.290603413 | 0.580296244 | 0.817 |
| YOR117W | 3352.309698 | -0.177387832 | 0.5803992 | 0.817 |
| YDR192C | 476.3597916 | 0.092898066 | 0.581907497 | 0.817 |
| YER125W | 2425.853452 | 0.173149403 | 0.581979268 | 0.817 |
| YGR121C | 1176.230558 | 0.091552649 | 0.58172608 | 0.817 |
| YIL095W | 775.5717487 | 0.135772229 | 0.581478528 | 0.817 |
| YJL034W | 12765.00653 | 0.042035243 | 0.582150955 | 0.817 |
| YJR154W | 284.604362 | 0.202167 | 0.581889811 | 0.817 |
| YKL006C-A | 282.6893408 | -0.156443404 | 0.582237743 | 0.817 |
| YKL201C | 1146.760044 | 0.097432448 | 0.581328658 | 0.817 |
| YNL137C | 1635.261502 | 0.193660645 | 0.581218914 | 0.817 |
| YCL039W | 1768.904925 | 0.190857625 | 0.582560198 | 0.817 |
| YDL224C | 1328.137391 | -0.154927694 | 0.582440543 | 0.817 |
| YLR032W | 823.8658356 | -0.124435597 | 0.582808689 | 0.817 |
| YMR272C | 4343.78123 | 0.250536556 | 0.582936682 | 0.817 |
| YNL116W | 793.4753006 | 0.175998602 | 0.582899202 | 0.817 |
| YFR030W | 1464.99151 | 0.080356185 | 0.584581266 | 0.818 |
| YHR034C | 511.3439371 | -0.187119267 | 0.583872266 | 0.818 |
| YHR124W | 347.8549107 | 0.254416201 | 0.584335723 | 0.818 |
| YLR417W | 736.9680806 | 0.11060359 | 0.584641765 | 0.818 |
| YMR200W | 1797.595559 | -0.128652817 | 0.583921152 | 0.818 |
| YNL221C | 1802.455751 | -0.167716727 | 0.584091856 | 0.818 |
| YOR298C-A | 6204.241243 | 0.219158268 | 0.584273963 | 0.818 |
| YPR004C | 1678.278628 | -0.050138569 | 0.584480463 | 0.818 |
| YFL029C | 966.6378114 | 0.171838574 | 0.585491757 | 0.819 |
| YDR099W | 8428.443895 | 0.140992224 | 0.585996461 | 0.819 |
| YER174C | 586.1669315 | 0.174274214 | 0.586207026 | 0.819 |
| YML123C | 2594.845401 | -0.195645378 | 0.585860416 | 0.819 |
| YMR191W | 2004.986398 | 0.189827242 | 0.585823523 | 0.819 |
| YOL098C | 2415.997601 | -0.094732243 | 0.586166393 | 0.819 |
| YDR333C | 1359.98522 | 0.13952974 | 0.586505669 | 0.819 |
| YBL098W | 1032.627619 | 0.144595997 | 0.586919028 | 0.820 |
| YBR028C | 772.1128092 | -0.142494768 | 0.587635613 | 0.820 |
| YBR199W | 2706.736809 | 0.07710937 | 0.58786664 | 0.820 |
| YCL011C | 2495.29652 | -0.220757635 | 0.587581543 | 0.820 |
| YDL138W | 666.9826314 | 0.128589653 | 0.588035887 | 0.820 |
| YDL160C-A | 95.22130843 | -0.27196617 | 0.588900027 | 0.820 |
| YDR029W | 118.6548652 | 0.253628049 | 0.587048843 | 0.820 |
| YDR050C | 39610.42901 | -0.180116666 | 0.588881915 | 0.820 |
| YDR169C | 1729.231302 | 0.174434975 | 0.5882806 | 0.820 |
| YDR287W | 832.7406735 | 0.258022139 | 0.587286227 | 0.820 |
| YIL115C | 2357.200299 | 0.109406448 | 0.58751361 | 0.820 |

| | | | | |
|---|---|---|---|---|
| YKL186C | 1093.769583 | -0.225726942 | 0.588984845 | 0.820 |
| YLR341W | 104.3766662 | 0.2537537 | 0.588640679 | 0.820 |
| YLR398C | 2110.607765 | -0.113630872 | 0.588534965 | 0.820 |
| YNL284C | 1102.406706 | 0.216113774 | 0.587961437 | 0.820 |
| YOR020W-A | 1169.212022 | 0.123884437 | 0.588465416 | 0.820 |
| YOR112W | 1049.770792 | 0.133612948 | 0.587946257 | 0.820 |
| YLR035C-A | 40.57446476 | 0.031314689 | 0.589142752 | 0.820 |
| YBL006C | 611.4550235 | -0.180600186 | 0.589859784 | 0.820 |
| YLR265C | 441.14354 | -0.294201763 | 0.589873303 | 0.820 |
| YNL148C | 367.0903378 | -0.065346154 | 0.589852122 | 0.820 |
| YOL016C | 3937.741808 | 0.075657683 | 0.58944932 | 0.820 |
| YOR166C | 666.3506234 | 0.124903115 | 0.589566869 | 0.820 |
| YER018C | 488.1395433 | -0.157173112 | 0.590285951 | 0.820 |
| YER163C | 538.8237534 | -0.142144473 | 0.590138379 | 0.820 |
| YKR087C | 773.1699867 | 0.135635823 | 0.590336261 | 0.820 |
| YFR034C | 832.1624053 | -0.154453831 | 0.590857169 | 0.820 |
| YLR454W | 3660.437031 | 0.226898999 | 0.590956368 | 0.820 |
| YFL002C | 1100.586944 | 0.104543162 | 0.591111506 | 0.820 |
| YBL102W | 1291.754797 | -0.169336265 | 0.592023384 | 0.821 |
| YBR063C | 508.4120941 | 0.068164703 | 0.592059212 | 0.821 |
| YBR214W | 5062.049562 | 0.099007421 | 0.591776173 | 0.821 |
| YGR202C | 1624.722457 | -0.110832695 | 0.591941601 | 0.821 |
| YPR180W | 730.2139766 | 0.149137576 | 0.592483626 | 0.821 |
| YGL013C | 1490.03588 | -0.153318555 | 0.592728992 | 0.821 |
| YKL145W | 5482.552156 | -0.174177961 | 0.593064525 | 0.822 |
| YOL018C | 882.7418897 | 0.092137895 | 0.593503468 | 0.822 |
| YAL030W | 729.4057845 | -0.160701573 | 0.594666141 | 0.823 |
| YER180C-A | 68.54860649 | 0.115798524 | 0.595781154 | 0.824 |
| YHR044C | 186.9132602 | 0.276356666 | 0.595575241 | 0.824 |
| YHR136C | 132.1966559 | -0.585750026 | 0.595841826 | 0.824 |
| YAR009C | 236.1235724 | 0.380486372 | 0.596177877 | 0.825 |
| YER129W | 2291.22394 | 0.098514727 | 0.596100216 | 0.825 |
| YDL237W | 1508.528864 | -0.062716331 | 0.59696266 | 0.825 |
| YDR337W | 923.5124951 | -0.06931228 | 0.597344062 | 0.825 |
| YIL168W | 92.19596636 | 0.244260388 | 0.597371377 | 0.825 |
| YPL011C | 726.714861 | 0.101612879 | 0.597388817 | 0.825 |
| snR9 | 75.73439823 | -0.037114234 | 0.597796148 | 0.826 |
| YKR007W | 738.2939867 | -0.14326953 | 0.597942882 | 0.826 |
| YPR072W | 2174.009813 | -0.08110186 | 0.598152011 | 0.826 |
| YJR012C | 286.1622087 | 0.119418956 | 0.598397641 | 0.826 |
| YKL161C | 387.0663817 | -0.137202204 | 0.59857982 | 0.826 |
| YNR012W | 597.0846222 | -0.261219171 | 0.598508499 | 0.826 |
| YDR512C | 741.4198181 | 0.127334361 | 0.598760503 | 0.826 |

| | | | | |
|---|---|---|---|---|
| YBR262C | 557.8168642 | -0.2411956 | 0.598983693 | 0.826 |
| YBR023C | 1741.008098 | -0.148569978 | 0.599786303 | 0.827 |
| YHR113W | 1875.547688 | -0.114139791 | 0.599768151 | 0.827 |
| YCL063W | 606.1625178 | -0.090315906 | 0.600374713 | 0.827 |
| YGR080W | 1465.883709 | -0.149755017 | 0.600735971 | 0.827 |
| YJL065C | 199.2698224 | -0.155289091 | 0.600721216 | 0.827 |
| YPR020W | 1129.530709 | 0.005816928 | 0.60080308 | 0.827 |
| YDR365W-B | 142.2689351 | 0.581495595 | 0.600962178 | 0.827 |
| YBR078W | 17879.15973 | -0.083464602 | 0.601347768 | 0.828 |
| YER062C | 1876.234052 | 0.141016534 | 0.601469354 | 0.828 |
| YPL042C | 568.6329785 | 0.106012562 | 0.601194804 | 0.828 |
| YBR122C | 1316.133892 | -0.118332687 | 0.601976953 | 0.828 |
| YLR459W | 960.9364848 | 0.193779339 | 0.602016151 | 0.828 |
| YDR178W | 2932.392626 | 0.372808106 | 0.602498929 | 0.828 |
| YJL202C | 29.15207009 | -0.199581883 | 0.602481928 | 0.828 |
| YNR001C | 10779.43573 | 0.31564652 | 0.603346532 | 0.829 |
| YDR096W | 1946.105843 | 0.06869474 | 0.603718288 | 0.829 |
| YGR193C | 2486.614356 | 0.162233531 | 0.603930141 | 0.829 |
| YOL123W | 4114.119417 | -0.245953246 | 0.603765516 | 0.829 |
| snR83 | 177.6034874 | -0.100468529 | 0.603797883 | 0.829 |
| YAL026C-A | 170.5922701 | 0.065341104 | 0.604531013 | 0.829 |
| YEL061C | 1022.169501 | -0.112816116 | 0.604647451 | 0.829 |
| YGR247W | 484.9641046 | 0.139468472 | 0.605120604 | 0.829 |
| YKR036C | 843.7972906 | 0.094328272 | 0.605078264 | 0.829 |
| YLL058W | 677.8123693 | 0.115981589 | 0.604195365 | 0.829 |
| YLR386W | 1077.114184 | 0.123749724 | 0.605052636 | 0.829 |
| YNR059W | 692.5145797 | 0.082601956 | 0.604529484 | 0.829 |
| snR8 | 32.91757815 | -0.134359818 | 0.604707614 | 0.829 |
| YEL050C | 1046.082243 | -0.082954474 | 0.606273425 | 0.830 |
| YFR015C | 6189.813754 | -0.042378302 | 0.605709458 | 0.830 |
| YGL009C | 9382.784081 | -0.173237087 | 0.605738717 | 0.830 |
| YIL038C | 1659.427705 | -0.180581577 | 0.606158826 | 0.830 |
| YMR323W | 206.5265565 | 0.134581535 | 0.606127358 | 0.830 |
| YPL137C | 1258.265301 | -0.264547278 | 0.606191631 | 0.830 |
| YBR193C | 556.4922631 | 0.138337416 | 0.607049734 | 0.830 |
| YDR038C | 125.3516014 | -0.247986896 | 0.60718339 | 0.830 |
| YER147C | 529.8378852 | 0.066566969 | 0.607125096 | 0.830 |
| YGL074C | 37.00723578 | -0.020679382 | 0.607236601 | 0.830 |
| YKL197C | 902.4571586 | 0.186181459 | 0.607287714 | 0.830 |
| YMR183C | 2217.146681 | -0.133792996 | 0.607007038 | 0.830 |
| YMR251W | 257.8128007 | 0.029401767 | 0.606575759 | 0.830 |
| YJR056C | 431.2081935 | 0.12269457 | 0.607519751 | 0.830 |
| YER158W-A | 247.6290209 | -0.216424521 | 0.607740651 | 0.830 |

| | | | | |
|---|---|---|---|---|
| YGR178C | 3642.360299 | 0.184994435 | 0.607939588 | 0.830 |
| YPR057W | 656.961666 | -0.171601071 | 0.608155992 | 0.830 |
| YBR230C | 2523.139558 | 0.033413307 | 0.609173806 | 0.831 |
| YDR034C-D | 94.28631679 | -0.207043025 | 0.608860335 | 0.831 |
| YJR082C | 608.3949667 | -0.234771942 | 0.609204228 | 0.831 |
| YML081C-A | 1028.969006 | -0.245759929 | 0.609229233 | 0.831 |
| YOR104W | 252.8227861 | 0.137255477 | 0.609362141 | 0.831 |
| YKL043W | 3024.896012 | -0.159783935 | 0.60960095 | 0.831 |
| YML013W | 1108.615798 | 0.22376636 | 0.609866218 | 0.831 |
| YGR132C | 2816.110199 | -0.113556 | 0.610469465 | 0.832 |
| YKL198C | 1074.404315 | 0.066325653 | 0.610642889 | 0.832 |
| YOR075W | 634.1618193 | -0.235184367 | 0.610613761 | 0.832 |
| YDR359C | 1438.122379 | 0.249022419 | 0.610823802 | 0.832 |
| YDL039C | 373.0125142 | -0.51034874 | 0.611351975 | 0.832 |
| YKR068C | 742.7822083 | -0.148705007 | 0.611998085 | 0.833 |
| YPL022W | 1488.077225 | 0.114191943 | 0.611994136 | 0.833 |
| YER109C | 1065.439283 | -0.171232388 | 0.612293559 | 0.833 |
| YJL029C | 1512.843835 | 0.087949858 | 0.612519853 | 0.833 |
| YLL036C | 873.0418955 | 0.098844126 | 0.612626434 | 0.833 |
| YPL120W | 588.9261154 | 0.12237303 | 0.61284842 | 0.833 |
| YPL162C | 476.9994766 | 0.125745192 | 0.612714392 | 0.833 |
| YDR379W | 1429.073535 | 0.100381083 | 0.613275201 | 0.833 |
| YBL103C | 858.4740089 | 0.086979019 | 0.614115572 | 0.834 |
| YFL053W | 516.855343 | 0.520275229 | 0.614318008 | 0.834 |
| YKL020C | 2087.802283 | 0.100834631 | 0.61438113 | 0.834 |
| YMR124W | 1653.834225 | 0.131649293 | 0.614273716 | 0.834 |
| YMR280C | 1028.334214 | 0.096934358 | 0.614295213 | 0.834 |
| YOR082C | 31.57473045 | 0.170640092 | 0.6142683 | 0.834 |
| YDR444W | 770.8185864 | 0.142067745 | 0.614682344 | 0.834 |
| YDR296W | 1076.454873 | -0.071488912 | 0.615104674 | 0.834 |
| YGL119W | 689.6648811 | -0.102211784 | 0.615164515 | 0.834 |
| YHL031C | 431.2428898 | -0.113695601 | 0.615428674 | 0.834 |
| YBR066C | 407.0918605 | -0.441403613 | 0.615801026 | 0.834 |
| YJL125C | 686.7482617 | -0.2019323 | 0.61576875 | 0.834 |
| YDR502C | 1141.466361 | -0.215211981 | 0.616973507 | 0.835 |
| YEL020W-A | 192.2562057 | -0.139559834 | 0.616590521 | 0.835 |
| YFL013C | 826.2401076 | 0.171971768 | 0.617491635 | 0.835 |
| YGL223C | 914.5737412 | 0.14020535 | 0.617382156 | 0.835 |
| YHR166C | 498.330332 | 0.157066723 | 0.617095025 | 0.835 |
| YIL089W | 155.6273305 | 0.153117611 | 0.617154326 | 0.835 |
| YKR014C | 1917.457601 | 0.171117204 | 0.617001875 | 0.835 |
| YLR204W | 619.6469206 | -0.149400512 | 0.617436195 | 0.835 |
| YMR016C | 1109.247117 | -0.150000682 | 0.61633782 | 0.835 |

| | | | | |
|---|---|---|---|---|
| YPR145W | 8044.845508 | -0.28031735 | 0.617583365 | 0.835 |
| YMR119W | 1094.314263 | 0.181318486 | 0.618606471 | 0.836 |
| YBL058W | 3576.371774 | -0.161480332 | 0.619527132 | 0.837 |
| YKR054C | 2620.574744 | -0.001610957 | 0.619806335 | 0.837 |
| YNL186W | 1470.462386 | -0.147245137 | 0.620067408 | 0.837 |
| YHR016C | 1896.688185 | 0.183751029 | 0.620858556 | 0.838 |
| YLR360W | 653.1816733 | 0.134928422 | 0.620839121 | 0.838 |
| YIL074C | 1391.44586 | -0.164174098 | 0.621045015 | 0.838 |
| YGR295C | 1429.22247 | 0.369881857 | 0.621523386 | 0.839 |
| YDR019C | 2663.732875 | -0.080318841 | 0.622644407 | 0.839 |
| YDR435C | 760.2512519 | 0.110997905 | 0.622520137 | 0.839 |
| YGR109C | 138.0694558 | -0.200466751 | 0.622374294 | 0.839 |
| YHR074W | 2090.525023 | -0.073085913 | 0.622626632 | 0.839 |
| YGR140W | 791.394714 | -0.089360905 | 0.622796415 | 0.839 |
| YMR262W | 628.7777309 | 0.08140928 | 0.623196686 | 0.840 |
| YBL095W | 182.0968065 | -0.010342732 | 0.623456538 | 0.840 |
| YKL057C | 1611.669894 | 0.208132194 | 0.623765513 | 0.840 |
| YBL008W | 564.4162656 | -0.18506806 | 0.62425369 | 0.840 |
| YDL044C | 428.1351393 | 0.128965283 | 0.624500296 | 0.840 |
| YGR033C | 1013.438168 | -0.105618249 | 0.624638483 | 0.840 |
| YLR020C | 866.2619741 | -0.089759774 | 0.624494016 | 0.840 |
| YOR119C | 842.963497 | 0.119266568 | 0.624394541 | 0.840 |
| YER136W | 4119.832209 | 0.123215705 | 0.624823428 | 0.840 |
| YNL173C | 2504.496471 | 0.132906383 | 0.62493815 | 0.840 |
| YBR024W | 889.4330761 | 0.188454565 | 0.625159195 | 0.840 |
| YER180C | 405.292426 | 0.093449542 | 0.6255726 | 0.841 |
| YGR031W | 376.7830976 | 0.172140976 | 0.625683904 | 0.841 |
| YOR144C | 421.6026059 | -0.094577957 | 0.626414028 | 0.842 |
| YPR149W | 11719.12039 | 0.123896055 | 0.626572766 | 0.842 |
| YER064C | 1674.781352 | -0.285132806 | 0.627273365 | 0.842 |
| YFR056C | 41.82065454 | 0.166934066 | 0.626932462 | 0.842 |
| YOR327C | 881.0062218 | -0.211180206 | 0.627077715 | 0.842 |
| YPR098C | 1032.023923 | 0.235762265 | 0.62716237 | 0.842 |
| YAL061W | 2752.646997 | 0.02919709 | 0.627907287 | 0.842 |
| YBR041W | 1541.15066 | -0.104973829 | 0.628341761 | 0.842 |
| YML037C | 139.9825756 | -0.141835338 | 0.628342329 | 0.842 |
| YNL136W | 1295.54364 | -0.233949985 | 0.628109807 | 0.842 |
| YJL171C | 1558.602902 | 0.179313075 | 0.629060789 | 0.843 |
| YLR400W | 49.34542517 | 0.03145364 | 0.629188289 | 0.843 |
| YPR117W | 2597.357181 | 0.147479685 | 0.62900522 | 0.843 |
| YFR010W | 3072.848472 | 0.132174409 | 0.629748029 | 0.843 |
| YGR229C | 1916.022007 | -0.069196331 | 0.629884994 | 0.843 |
| YDR481C | 2453.405315 | 0.057004409 | 0.630374194 | 0.844 |

| | | | | |
|---|---|---|---|---|
| YJR074W | 487.5630441 | -0.178878506 | 0.630289696 | 0.844 |
| YER172C | 2101.126018 | 0.131086126 | 0.630657401 | 0.844 |
| YOR033C | 730.7503196 | 0.104409026 | 0.630759617 | 0.844 |
| YCR076C | 593.9023385 | 0.18442374 | 0.630924498 | 0.844 |
| YDL203C | 667.0945544 | -0.099197803 | 0.631216858 | 0.844 |
| YGL247W | 197.7711302 | -0.214680339 | 0.631570002 | 0.844 |
| YNR026C | 756.2847135 | -0.123971935 | 0.631738513 | 0.844 |
| YBL050W | 1823.454847 | -0.120799387 | 0.632291845 | 0.845 |
| YLR099C | 1018.600053 | -0.258270776 | 0.632193832 | 0.845 |
| YPL275W | 31.2658679 | 0.026394508 | 0.632534154 | 0.845 |
| YDL160C | 1362.561057 | 0.292653691 | 0.632954086 | 0.845 |
| YDL185C-A | 100.139643 | 0.206979025 | 0.633238314 | 0.845 |
| YGR050C | 100.6325721 | 0.164137015 | 0.633214735 | 0.845 |
| YBR150C | 848.9681448 | -0.215068225 | 0.633703711 | 0.846 |
| YNL225C | 446.0966712 | -0.144854362 | 0.633827733 | 0.846 |
| YPL148C | 142.8618352 | -0.192417641 | 0.633968081 | 0.846 |
| YLR227W-B | 558.9698813 | -0.053539943 | 0.634266703 | 0.846 |
| YJL166W | 2486.295448 | 0.189441474 | 0.634713357 | 0.846 |
| YPL089C | 1858.715154 | 0.078930843 | 0.634604651 | 0.846 |
| YKR102W | 639.1097035 | 0.27444421 | 0.634858375 | 0.846 |
| YDR210C-D | 43.7176532 | 0.045136925 | 0.635265581 | 0.846 |
| YFL032W | 146.5638958 | 0.049624655 | 0.635413834 | 0.846 |
| YJL006C | 336.1143852 | -0.188624398 | 0.635646929 | 0.847 |
| YMR316W | 898.4662527 | 0.002777507 | 0.635892844 | 0.847 |
| YDL018C | 497.6776418 | -0.020103811 | 0.636772089 | 0.848 |
| YDR218C | 78.04916244 | 0.204516742 | 0.636791392 | 0.848 |
| YGL062W | 7360.866755 | 0.104655908 | 0.636985683 | 0.848 |
| YBR085C-A | 1789.206511 | 0.022937057 | 0.637301172 | 0.848 |
| YMR152W | 1727.69017 | 0.121392999 | 0.637206402 | 0.848 |
| YCL005W | 658.6999475 | -0.099668054 | 0.638770172 | 0.849 |
| YCR030C | 2439.320353 | 0.150754121 | 0.639755826 | 0.849 |
| YDR541C | 514.5384389 | 0.503042711 | 0.639069188 | 0.849 |
| YFR018C | 1513.157132 | -0.127060131 | 0.640267096 | 0.849 |
| YGL196W | 1262.908167 | 0.205359372 | 0.639768712 | 0.849 |
| YHR168W | 379.0957302 | -0.147291277 | 0.639991561 | 0.849 |
| YJR106W | 683.5325213 | 0.178159615 | 0.639269416 | 0.849 |
| YLR119W | 415.3763546 | -0.153923672 | 0.638867585 | 0.849 |
| YLR347C | 1405.641169 | -0.054029389 | 0.639563705 | 0.849 |
| YLR429W | 3551.790145 | -0.050997181 | 0.639694664 | 0.849 |
| YNL085W | 3478.261348 | -0.047456167 | 0.63993752 | 0.849 |
| YOL065C | 517.0688786 | 0.107027882 | 0.638547849 | 0.849 |
| YOL140W | 502.3006349 | -0.220634558 | 0.638578955 | 0.849 |
| YOR180C | 257.2908648 | 0.05138132 | 0.640215603 | 0.849 |

| | | | | |
|---|---|---|---|---|
| YOR278W | 477.7903224 | -0.024808942 | 0.638243381 | 0.849 |
| YLR163C | 1121.057782 | 0.124988063 | 0.640568126 | 0.849 |
| YHR162W | 4274.252821 | -0.172607383 | 0.641121453 | 0.849 |
| YLR133W | 1071.862832 | -0.053164954 | 0.64091506 | 0.849 |
| YMR197C | 923.7331716 | -0.157228101 | 0.641055889 | 0.849 |
| YDL108W | 550.3870487 | 0.082974934 | 0.641587225 | 0.849 |
| YKR045C | 368.478154 | -0.058296855 | 0.641532223 | 0.849 |
| YPL069C | 455.5840403 | -0.174371282 | 0.641387702 | 0.849 |
| YAL054C | 1109.295849 | 3.37E-05 | 0.642728588 | 0.849 |
| YBR043C | 1241.74171 | 0.025108522 | 0.643262215 | 0.849 |
| YIL117C | 1107.127061 | 0.204974803 | 0.642229069 | 0.849 |
| YLR199C | 824.459216 | 0.108694526 | 0.643240275 | 0.849 |
| YLR203C | 2343.4384 | 0.171597342 | 0.642520931 | 0.849 |
| YMR212C | 1054.541147 | -0.148636085 | 0.642984061 | 0.849 |
| YNL052W | 2172.615322 | -0.074005726 | 0.643398151 | 0.849 |
| YNL155W | 1378.201426 | -0.218652053 | 0.642916715 | 0.849 |
| YNL193W | 690.484771 | -0.208145622 | 0.642484468 | 0.849 |
| YOR194C | 1465.560671 | 0.118225632 | 0.642851259 | 0.849 |
| YPL155C | 690.291389 | 0.136134882 | 0.642050993 | 0.849 |
| YBL011W | 1645.010666 | -0.117962673 | 0.643772451 | 0.850 |
| YCL034W | 1930.992154 | 0.099930309 | 0.645214105 | 0.850 |
| YEL008W | 30.07335462 | 0.159684262 | 0.644261258 | 0.850 |
| YFL008W | 1609.672053 | -0.058474234 | 0.644671762 | 0.850 |
| YGL014W | 2156.619039 | -0.209261236 | 0.644518182 | 0.850 |
| YGR098C | 1335.545417 | -0.075628405 | 0.645227518 | 0.850 |
| YKL117W | 7112.851481 | -0.177171366 | 0.64514328 | 0.850 |
| YLR078C | 672.0208317 | -0.166520655 | 0.644889768 | 0.850 |
| YLR252W | 254.6818923 | -0.035617385 | 0.644019155 | 0.850 |
| YNL169C | 1167.217459 | 0.105967622 | 0.644884112 | 0.850 |
| YNR011C | 902.2961481 | -0.143157183 | 0.644534099 | 0.850 |
| YLR011W | 284.831779 | -0.148614594 | 0.645537866 | 0.850 |
| YOR358W | 499.4768456 | 0.102128314 | 0.645456449 | 0.850 |
| YCL016C | 442.1886812 | 0.079206245 | 0.646493284 | 0.851 |
| YHR132C | 1760.575956 | -0.11776481 | 0.646342919 | 0.851 |
| YOL125W | 710.1287675 | -0.134346556 | 0.646593807 | 0.851 |
| YNR075W | 186.6264191 | 0.382247712 | 0.647296989 | 0.851 |
| YDR331W | 571.3069512 | 0.178785496 | 0.647521753 | 0.851 |
| YLR274W | 1064.782127 | 0.158417209 | 0.6477168 | 0.851 |
| YMR036C | 631.5049431 | 0.050700366 | 0.647806708 | 0.851 |
| YBR103W | 849.3179685 | 0.109490248 | 0.648873032 | 0.853 |
| YDR119W | 1442.902962 | -0.046507463 | 0.649067517 | 0.853 |
| YDR349C | 1807.872356 | -0.076860687 | 0.649330107 | 0.853 |
| YDR508C | 2566.963439 | 0.187525165 | 0.649262781 | 0.853 |

| | | | | |
|---|---|---|---|---|
| YFR047C | 2554.161875 | 0.418194305 | 0.649668927 | 0.853 |
| YML087C | 330.7889367 | 0.274507087 | 0.649627302 | 0.853 |
| YKR067W | 2101.266331 | 0.105660052 | 0.650180516 | 0.853 |
| YPL122C | 549.6598792 | -0.104843324 | 0.650082641 | 0.853 |
| YAL026C | 1397.824394 | -0.109245941 | 0.652524278 | 0.853 |
| YBR057C | 866.0815496 | 0.124372125 | 0.65202107 | 0.853 |
| YDR249C | 239.973621 | 0.059199619 | 0.652706433 | 0.853 |
| YGL002W | 690.1637106 | 0.122190183 | 0.651592036 | 0.853 |
| YHR072W | 1833.218722 | -0.118264646 | 0.652727107 | 0.853 |
| YIL119C | 838.0211886 | 0.075095085 | 0.652448457 | 0.853 |
| YJL073W | 1063.600671 | -0.053591353 | 0.652569804 | 0.853 |
| YLR018C | 838.7588705 | -0.12892167 | 0.652303818 | 0.853 |
| YLR141W | 304.974896 | 0.105154073 | 0.652011111 | 0.853 |
| YNL222W | 196.3405424 | -0.093670926 | 0.652217031 | 0.853 |
| YOL151W | 926.9123911 | -0.132009633 | 0.651252953 | 0.853 |
| YOR083W | 334.3141069 | 0.10346852 | 0.651039879 | 0.853 |
| YOR280C | 559.6900755 | -0.192717011 | 0.651197638 | 0.853 |
| YPR122W | 1503.146832 | 0.238523419 | 0.65158353 | 0.853 |
| snR33 | 120.9149393 | -0.162733559 | 0.651972611 | 0.853 |
| YKR038C | 1110.135443 | -0.134434115 | 0.653193305 | 0.854 |
| YLR071C | 1557.164849 | 0.112506461 | 0.653419318 | 0.854 |
| YER088C | 4317.842613 | -0.118403801 | 0.654171931 | 0.855 |
| YGR179C | 503.8137475 | -0.081589211 | 0.654399444 | 0.855 |
| YOL034W | 735.3581814 | 0.093763663 | 0.654515558 | 0.855 |
| YPL097W | 440.8818332 | 0.161920585 | 0.654885803 | 0.855 |
| YOR302W | 66.2485987 | -0.044159785 | 0.655167328 | 0.855 |
| YHR046C | 620.1635411 | -0.155024587 | 0.655661272 | 0.856 |
| YOL072W | 594.5699607 | 0.071720904 | 0.655673867 | 0.856 |
| YMR171C | 1518.290743 | 0.170333627 | 0.655862924 | 0.856 |
| YDL101C | 860.843029 | -0.117863522 | 0.656739897 | 0.856 |
| YDR467C | 75.26126934 | 0.032893549 | 0.656468844 | 0.856 |
| YKL015W | 1286.742474 | -0.269815413 | 0.656593269 | 0.856 |
| YNL234W | 710.7866789 | -0.069110916 | 0.65681098 | 0.856 |
| YOR196C | 1243.855488 | -0.173410538 | 0.656582859 | 0.856 |
| YDR488C | 747.4804028 | -0.202932455 | 0.658030684 | 0.857 |
| YJR121W | 23130.92602 | 0.188551294 | 0.657950726 | 0.857 |
| YEL065W | 2278.709245 | 0.244394339 | 0.658758773 | 0.858 |
| YKL175W | 2025.203634 | 0.118500247 | 0.658922154 | 0.858 |
| YLR202C | 55.2829566 | -0.035929297 | 0.658836201 | 0.858 |
| YNL204C | 121.2325338 | 0.012168618 | 0.658643604 | 0.858 |
| YML007W | 2781.196668 | 0.111756253 | 0.659253785 | 0.858 |
| YGL041W-A | 345.8516558 | 0.102334368 | 0.659475287 | 0.858 |
| YBR152W | 271.4958287 | 0.054924185 | 0.659977057 | 0.858 |

| | | | | |
|---|---|---|---|---|
| YFR043C | 412.3285976 | 0.076434758 | 0.66016648 | 0.858 |
| YAL055W | 312.0534701 | 0.128780904 | 0.660547677 | 0.858 |
| YDL156W | 331.6691228 | -0.052764614 | 0.660645256 | 0.858 |
| YPL049C | 2306.704815 | 0.113537843 | 0.660429541 | 0.858 |
| YLR052W | 1071.581443 | 0.056749766 | 0.661133784 | 0.858 |
| YLR304C | 13028.11852 | 0.069473113 | 0.661052353 | 0.858 |
| YNR048W | 344.510222 | 0.119970831 | 0.660915773 | 0.858 |
| YDR275W | 363.6731352 | 0.073356414 | 0.661552665 | 0.859 |
| YER115C | 529.525795 | -0.00036438 | 0.661738083 | 0.859 |
| YDR529C | 3824.487001 | 0.320991937 | 0.662411151 | 0.859 |
| YMR205C | 7427.558626 | 0.015523048 | 0.662383564 | 0.859 |
| YOL112W | 963.5678133 | -0.137110695 | 0.662136072 | 0.859 |
| YBL039W-B | 151.3574507 | 0.125694139 | 0.662821301 | 0.859 |
| YER186C | 678.5275475 | -0.119380192 | 0.662678016 | 0.859 |
| YHR071W | 3724.97766 | -0.085589577 | 0.663133582 | 0.860 |
| YDL076C | 828.6133083 | -0.10248916 | 0.663280804 | 0.860 |
| YBR062C | 819.8607412 | 0.110622806 | 0.664765785 | 0.861 |
| YHR011W | 509.988716 | 0.26393194 | 0.6646841 | 0.861 |
| YLR096W | 1948.609356 | -0.165886096 | 0.664975603 | 0.861 |
| YOR383C | 13192.95676 | 0.168313149 | 0.664849781 | 0.861 |
| YNL015W | 3387.233502 | 0.19225272 | 0.665190866 | 0.861 |
| YBR092C | 2232.790112 | -0.426716438 | 0.666346637 | 0.862 |
| YCL069W | 36.22888945 | 0.011245151 | 0.667365094 | 0.862 |
| YCR027C | 255.2084978 | -0.04219663 | 0.666471914 | 0.862 |
| YDR081C | 1262.178022 | 0.099771998 | 0.666947721 | 0.862 |
| YDR389W | 894.8507584 | -0.16565892 | 0.667095593 | 0.862 |
| YGR278W | 442.5587334 | 0.121057756 | 0.667186975 | 0.862 |
| YKL003C | 652.5154668 | -0.12216515 | 0.666622116 | 0.862 |
| YNL097C | 784.7174827 | 0.116673712 | 0.667099491 | 0.862 |
| YNR049C | 561.0637018 | 0.109461006 | 0.667409027 | 0.862 |
| YPL257W | 181.2677801 | 0.228355398 | 0.666816838 | 0.862 |
| YIL034C | 3381.997418 | -0.068174687 | 0.667564449 | 0.862 |
| YBR176W | 422.8364361 | -0.160510449 | 0.668136263 | 0.862 |
| snR11 | 261.7635615 | 0.228532386 | 0.668000495 | 0.862 |
| YBR036C | 2605.132077 | 0.11646229 | 0.66932676 | 0.864 |
| YPR200C | 70.27684883 | 0.412079142 | 0.669347092 | 0.864 |
| YLR253W | 1177.596684 | 0.249023872 | 0.669614933 | 0.864 |
| YOL019W | 928.0908204 | -0.155036221 | 0.669679999 | 0.864 |
| YDR485C | 1410.914463 | -0.1982143 | 0.670187053 | 0.864 |
| YMR164C | 711.3389073 | -0.080663947 | 0.670128852 | 0.864 |
| YBL101C | 1814.259973 | 0.081414466 | 0.670562463 | 0.864 |
| YDR164C | 1143.654091 | -0.038928457 | 0.67056819 | 0.864 |
| YGL233W | 1565.055775 | 0.098176172 | 0.671250066 | 0.864 |

| | | | | |
|---|---|---|---|---|
| YIR015W | 351.8671415 | -0.220932612 | 0.671184199 | 0.864 |
| YPR014C | 81.20498688 | 0.140518648 | 0.671100256 | 0.864 |
| YCR037C | 1044.612368 | 0.133835102 | 0.671815662 | 0.865 |
| YGL240W | 171.0354676 | -0.133044334 | 0.672624361 | 0.866 |
| YCR024C-A | 1033.453894 | -0.147190244 | 0.672989754 | 0.866 |
| YKL047W | 711.1080868 | 0.125469027 | 0.673457636 | 0.866 |
| YNL092W | 338.5129162 | 0.141412708 | 0.673547658 | 0.866 |
| YOR151C | 4196.260493 | 0.183308015 | 0.673961124 | 0.867 |
| YDR494W | 1564.77137 | 0.17137194 | 0.67498811 | 0.868 |
| YMR189W | 2797.782123 | -0.025594936 | 0.675118162 | 0.868 |
| YPL170W | 622.4061116 | -0.123887726 | 0.674840974 | 0.868 |
| YOR329C | 1982.861977 | 0.162012446 | 0.675526164 | 0.868 |
| YNL172W | 1164.569131 | 0.119018117 | 0.67571833 | 0.868 |
| YBR086C | 4766.880566 | 0.114005236 | 0.676349614 | 0.868 |
| YBR236C | 870.5665803 | 0.147307962 | 0.676410584 | 0.868 |
| YDR079W | 403.891688 | 0.214939047 | 0.676009966 | 0.868 |
| YOR047C | 1007.814108 | 0.011458854 | 0.676334381 | 0.868 |
| YER083C | 1868.758699 | -0.099583832 | 0.676559742 | 0.868 |
| YGR274C | 1344.663557 | 0.067888346 | 0.676832086 | 0.868 |
| YFR008W | 307.6442666 | 0.074114331 | 0.677658798 | 0.869 |
| YDR443C | 1163.053889 | 0.193025892 | 0.677923864 | 0.869 |
| YGR186W | 2388.772284 | 0.146756475 | 0.678166581 | 0.869 |
| YHR102W | 1800.525655 | 0.120622525 | 0.678406222 | 0.869 |
| YMR213W | 660.1636091 | 0.064076239 | 0.678052215 | 0.869 |
| YNL005C | 1578.642517 | 0.145974265 | 0.678703767 | 0.869 |
| YNR067C | 2460.572138 | -0.06097802 | 0.678528309 | 0.869 |
| YPL228W | 971.5333391 | 0.113451847 | 0.678670785 | 0.869 |
| YBR108W | 1821.037124 | 0.120011398 | 0.679352756 | 0.870 |
| YGR074W | 145.6756142 | -0.184342308 | 0.679364172 | 0.870 |
| YCL010C | 597.9803619 | 0.038544281 | 0.679654589 | 0.870 |
| YNL238W | 1373.585003 | -0.144459941 | 0.680021492 | 0.870 |
| YOR037W | 696.733578 | 0.105662561 | 0.679883859 | 0.870 |
| YDL035C | 1472.135402 | 0.09036656 | 0.680246088 | 0.870 |
| YHL030W | 2779.309717 | 0.134878101 | 0.680553852 | 0.870 |
| YER171W | 948.7525289 | -0.10426794 | 0.680928713 | 0.870 |
| YFR031C | 1205.359626 | 0.066527586 | 0.681029515 | 0.870 |
| YOR187W | 4877.755588 | 0.004836436 | 0.681116339 | 0.870 |
| YDR189W | 2199.458913 | -0.107431464 | 0.682350445 | 0.871 |
| YDR191W | 146.592055 | 0.137135946 | 0.682091544 | 0.871 |
| YGL038C | 1572.160144 | 0.171122742 | 0.682067166 | 0.871 |
| YLL019C | 1900.382246 | 0.076870398 | 0.682257984 | 0.871 |
| YML112W | 572.6104301 | -0.113555446 | 0.682224369 | 0.871 |
| YOL158C | 1769.752458 | -0.276594868 | 0.681794144 | 0.871 |

| | | | | |
|---|---|---|---|---|
| YAL024C | 799.2694521 | 0.173605624 | 0.68481217 | 0.871 |
| YBR273C | 1305.311458 | -0.11187887 | 0.684874373 | 0.871 |
| YDR041W | 1214.356247 | 0.153605079 | 0.683887116 | 0.871 |
| YDR276C | 2763.13579 | -0.097534422 | 0.684686543 | 0.871 |
| YDR468C | 586.9211769 | 0.117941449 | 0.683348872 | 0.871 |
| YDR525W-A | 275.582834 | 0.155292659 | 0.683251656 | 0.871 |
| YEL072W | 207.4067694 | 0.046569463 | 0.684212532 | 0.871 |
| YER076W-A | 76.7457419 | -0.000241974 | 0.682864476 | 0.871 |
| YGL127C | 488.4323072 | 0.055612162 | 0.683618815 | 0.871 |
| YGR035C | 329.4527926 | -0.329886862 | 0.684391226 | 0.871 |
| YKR083C | 244.3380659 | -0.293219795 | 0.683601999 | 0.871 |
| YLL028W | 724.5266873 | 0.010728904 | 0.683808067 | 0.871 |
| YLR122C | 30.05120318 | 0.124460867 | 0.682548762 | 0.871 |
| YML030W | 762.1544554 | 0.160479874 | 0.684786204 | 0.871 |
| YML091C | 3874.618724 | -0.043520319 | 0.683846687 | 0.871 |
| YMR319C | 2635.007516 | 0.064560775 | 0.68345032 | 0.871 |
| YNR072W | 156.9410464 | -0.157275721 | 0.684089284 | 0.871 |
| YLR410W | 2881.535918 | 0.112945771 | 0.68544543 | 0.871 |
| YCL012C | 422.2313505 | -0.267984633 | 0.688362798 | 0.872 |
| YCR054C | 374.3747854 | 0.114643122 | 0.685852221 | 0.872 |
| YDR240C | 401.4217535 | -0.072350669 | 0.688100153 | 0.872 |
| YDR336W | 301.9017741 | 0.027036363 | 0.688067901 | 0.872 |
| YDR461C-A | 326.4764294 | -0.020565292 | 0.686150651 | 0.872 |
| YGL161C | 1505.046156 | -0.082087152 | 0.687177382 | 0.872 |
| YIL015W | 2424.316267 | 0.357157053 | 0.687372927 | 0.872 |
| YIR009W | 172.3613335 | -0.278349531 | 0.688223573 | 0.872 |
| YJL004C | 587.5114449 | -0.120890468 | 0.686135012 | 0.872 |
| YJL020C | 4552.045062 | 0.187893922 | 0.688535134 | 0.872 |
| YJR051W | 1291.045072 | 0.296937553 | 0.686815064 | 0.872 |
| YJR066W | 2254.068141 | 0.172086564 | 0.687515169 | 0.872 |
| YJR125C | 2124.757426 | -0.116756085 | 0.6883407 | 0.872 |
| YLR294C | 1804.172401 | -0.089745292 | 0.688513138 | 0.872 |
| YML103C | 3218.889825 | 0.164996797 | 0.686313252 | 0.872 |
| YML110C | 4679.227772 | -0.028898942 | 0.68658621 | 0.872 |
| YMR073C | 579.5582083 | 0.178847273 | 0.687823377 | 0.872 |
| YMR109W | 3069.730753 | -0.099841934 | 0.687513911 | 0.872 |
| YNL164C | 278.0976276 | -0.181836638 | 0.686233512 | 0.872 |
| snR51 | 25.09375178 | -0.18918501 | 0.686654295 | 0.872 |
| YJR001W | 1763.85175 | 0.138701289 | 0.689075152 | 0.872 |
| YOR299W | 1060.38632 | -0.150352379 | 0.689405984 | 0.872 |
| YPR024W | 1909.485964 | -0.128351855 | 0.689430494 | 0.872 |
| YDL244W | 51.51006224 | 0.225911078 | 0.690231753 | 0.872 |
| YKR022C | 681.4384392 | -0.154114233 | 0.689906025 | 0.872 |

| | | | | |
|---|---|---|---|---|
| YLR401C | 1341.53932 | -0.062639476 | 0.690025885 | 0.872 |
| YMR244W | 46.42719777 | 0.27883131 | 0.690088165 | 0.872 |
| YDL173W | 1092.267621 | -0.18599592 | 0.690439461 | 0.872 |
| YDR374W-A | 251.03932 | 0.142172445 | 0.690608257 | 0.872 |
| YDR335W | 3040.69958 | 0.09609443 | 0.691289609 | 0.873 |
| YGL155W | 739.9393467 | -0.057007326 | 0.692055218 | 0.874 |
| YGL209W | 1219.680335 | -0.313559021 | 0.692470837 | 0.874 |
| YJR018W | 57.57815418 | -0.192308027 | 0.692297086 | 0.874 |
| YNL297C | 2104.946506 | -0.094496036 | 0.692391541 | 0.874 |
| YPL047W | 255.7983457 | 0.025616749 | 0.691920497 | 0.874 |
| YGL193C | 193.4216907 | -0.029007432 | 0.692823603 | 0.874 |
| YML116W-A | 166.4577303 | -0.16408561 | 0.692809946 | 0.874 |
| YDR248C | 767.0999761 | 0.103452881 | 0.693305983 | 0.874 |
| YDR314C | 384.5141242 | 0.168037832 | 0.693047998 | 0.874 |
| YLR101C | 65.86852303 | -0.265655711 | 0.693299655 | 0.874 |
| YPL064C | 883.912951 | 0.061476901 | 0.693452266 | 0.874 |
| YOR344C | 1985.74596 | 0.355597555 | 0.694271827 | 0.875 |
| YLR436C | 2590.336596 | 0.114708064 | 0.694469163 | 0.875 |
| YDR230W | 158.0472236 | 0.063839836 | 0.694751495 | 0.875 |
| YFL033C | 2297.741868 | -0.075552665 | 0.695222832 | 0.875 |
| YHR111W | 861.1772037 | -0.095457555 | 0.695326209 | 0.875 |
| YJL069C | 705.1093909 | -0.210406903 | 0.696624031 | 0.876 |
| YJL162C | 484.7743599 | -0.144302817 | 0.696847905 | 0.876 |
| YJR104C | 12449.61979 | -0.172809087 | 0.696816431 | 0.876 |
| YKL085W | 9612.778538 | 0.459534286 | 0.696808891 | 0.876 |
| YPL256C | 1702.304083 | 0.010006849 | 0.696987535 | 0.876 |
| YBR278W | 245.0159717 | -0.20553797 | 0.697892875 | 0.877 |
| YDL067C | 1308.526792 | -0.01615265 | 0.698118188 | 0.877 |
| YFR013W | 1397.161166 | 0.103502618 | 0.698137162 | 0.877 |
| YGL051W | 80.44841429 | 0.163892604 | 0.697532738 | 0.877 |
| YJL035C | 163.2101994 | -0.128452589 | 0.698129146 | 0.877 |
| YLL006W | 812.6766975 | 0.177186043 | 0.698208152 | 0.877 |
| YMR086W | 2080.250939 | 0.088530019 | 0.697921445 | 0.877 |
| YBR281C | 961.503364 | -0.105536626 | 0.698549924 | 0.877 |
| YMR028W | 753.5245359 | -0.099514574 | 0.699100285 | 0.877 |
| YGR197C | 588.1649926 | 0.087429508 | 0.699400798 | 0.877 |
| YNL250W | 994.2437698 | 0.142704681 | 0.699376157 | 0.877 |
| YER175W-A | 27.41241786 | -0.063871791 | 0.700559333 | 0.878 |
| YHL047C | 1687.543568 | 0.241392909 | 0.700464907 | 0.878 |
| YOR114W | 311.2740858 | 0.103401932 | 0.700445629 | 0.878 |
| YKL044W | 194.2159883 | -0.122971738 | 0.701112298 | 0.879 |
| YPR027C | 342.9171324 | -0.070477049 | 0.701086801 | 0.879 |
| YNL260C | 268.8863879 | -0.084279733 | 0.701308175 | 0.879 |

| | | | | |
|---|---|---|---|---|
| YDR170W-A | 128.0923816 | 0.031271259 | 0.701776493 | 0.879 |
| YDR026C | 840.5603663 | -0.100009234 | 0.70315734 | 0.880 |
| YEL019C | 187.0813824 | -0.136645487 | 0.70253032 | 0.880 |
| YFL021C-A | 41.5780796 | 0.320532712 | 0.703098032 | 0.880 |
| YJR030C | 708.7385852 | 0.007187897 | 0.703115465 | 0.880 |
| YMR005W | 1198.923005 | -0.160086185 | 0.702657536 | 0.880 |
| YPR001W | 533.0217534 | 0.022144686 | 0.702803094 | 0.880 |
| YDR462W | 726.8986569 | -0.046759033 | 0.703739033 | 0.880 |
| YHR146W | 3577.393266 | -0.223610851 | 0.703681864 | 0.880 |
| YNL211C | 132.5864716 | -0.121673488 | 0.703620764 | 0.880 |
| YFL014W | 10835.63862 | 0.193959982 | 0.704147888 | 0.880 |
| YAL015C | 634.3594781 | 0.088192153 | 0.70450042 | 0.880 |
| YBR227C | 856.5044097 | -0.121133789 | 0.70459038 | 0.880 |
| YOR288C | 898.6918954 | 0.183026126 | 0.704633278 | 0.880 |
| YBR192W | 512.8528961 | -0.078167649 | 0.705222754 | 0.880 |
| YDR482C | 359.2978323 | -0.125196317 | 0.705145776 | 0.880 |
| YMR132C | 213.3288234 | -0.045008842 | 0.704933773 | 0.880 |
| YAR033W | 105.2011719 | 0.171111568 | 0.705425834 | 0.880 |
| YPL013C | 664.0145708 | -0.185348941 | 0.706664183 | 0.882 |
| YML051W | 1749.382621 | -0.155305381 | 0.707302391 | 0.882 |
| YBR179C | 1264.938407 | 0.161709685 | 0.707711509 | 0.883 |
| YDL116W | 1341.682086 | 0.107632488 | 0.708224216 | 0.883 |
| YMR147W | 251.4438843 | 0.274206946 | 0.708651939 | 0.883 |
| YNL043C | 66.28583242 | -0.187444853 | 0.708392068 | 0.883 |
| YOR370C | 2847.263179 | -0.09293488 | 0.708526552 | 0.883 |
| YJR065C | 3820.877578 | -0.065408393 | 0.709043199 | 0.883 |
| YJR086W | 331.1910032 | -0.111651691 | 0.708909213 | 0.883 |
| YNL020C | 852.0700517 | 0.034700455 | 0.709228038 | 0.883 |
| YPR047W | 774.0936392 | 0.165273266 | 0.709711993 | 0.884 |
| YKL042W | 561.3749184 | -0.065028761 | 0.709981801 | 0.884 |
| YGR131W | 100.6509022 | -0.330734874 | 0.710404439 | 0.884 |
| YPL173W | 902.2046084 | -0.076277096 | 0.710702674 | 0.884 |
| YCL040W | 21911.49524 | 0.285474535 | 0.71086504 | 0.884 |
| YBL038W | 1166.095027 | -0.105331104 | 0.712015932 | 0.885 |
| YDL028C | 1026.396259 | 0.114560376 | 0.712249156 | 0.885 |
| YER067C-A | 258.7943359 | 0.221443892 | 0.712263013 | 0.885 |
| YHL027W | 1629.795881 | 0.049686319 | 0.711870361 | 0.885 |
| YKR018C | 3562.456922 | 0.186100819 | 0.711832364 | 0.885 |
| YNL106C | 963.8876185 | 0.10230126 | 0.711700629 | 0.885 |
| YBR020W | 310.6456705 | -0.001987322 | 0.712950594 | 0.885 |
| YIR023W | 1089.743487 | -0.098577754 | 0.712650001 | 0.885 |
| YKL118W | 29.53687639 | 0.015561269 | 0.713065783 | 0.885 |
| YMR166C | 428.7049362 | 0.06087678 | 0.713059148 | 0.885 |

| | | | | |
|---|---|---|---|---|
| YKR072C | 2109.757496 | 0.053685236 | 0.713996894 | 0.886 |
| YMR265C | 688.687566 | 0.065879406 | 0.713950265 | 0.886 |
| YOR222W | 1007.50754 | -0.116103483 | 0.714074933 | 0.886 |
| YEL071W | 3890.356415 | -0.020955297 | 0.714561832 | 0.886 |
| YIR024C | 366.6225047 | 0.046376859 | 0.714785994 | 0.886 |
| YOR131C | 796.6694001 | 0.097374928 | 0.714707823 | 0.886 |
| YIL098C | 444.8949209 | -0.12636377 | 0.715291858 | 0.887 |
| YBR081C | 1882.249025 | 0.064961175 | 0.715643462 | 0.887 |
| YOL028C | 419.6545084 | -0.108822882 | 0.715936443 | 0.887 |
| YOR250C | 655.6893916 | -0.081005102 | 0.715963235 | 0.887 |
| YDR063W | 1183.071975 | 0.072216888 | 0.716878592 | 0.887 |
| YHR092C | 9528.876224 | 0.076058649 | 0.716837874 | 0.887 |
| YJL156W-A | 83.97878785 | 0.025910054 | 0.716905551 | 0.887 |
| YPL032C | 2772.776631 | 0.120674838 | 0.716834587 | 0.887 |
| YPL132W | 679.0950944 | -0.119881566 | 0.716871536 | 0.887 |
| YPR158C-D | 130.4698158 | -0.185948284 | 0.717014914 | 0.887 |
| YHL006C | 116.1902464 | 0.1272233 | 0.71726253 | 0.887 |
| YOR060C | 313.4302218 | 0.129481644 | 0.71745487 | 0.887 |
| YDL090C | 724.6090506 | -0.095686285 | 0.717722753 | 0.887 |
| YJL112W | 1088.839683 | 0.103209928 | 0.717839559 | 0.887 |
| YDL147W | 5209.098548 | 0.107573719 | 0.718992383 | 0.888 |
| YDR320C-A | 373.904445 | -0.224201726 | 0.71910873 | 0.888 |
| YER011W | 1426.306813 | -0.497852573 | 0.719072808 | 0.888 |
| YGR048W | 875.7732177 | -0.152521985 | 0.718996875 | 0.888 |
| YMR316C-A | 33.48519679 | -0.210780483 | 0.718638545 | 0.888 |
| YPL193W | 620.6543173 | -0.122715695 | 0.718863036 | 0.888 |
| YML032C | 1432.723324 | 0.014084461 | 0.719279891 | 0.888 |
| YEL067C | 149.0124198 | -0.300984961 | 0.719683612 | 0.888 |
| YDR404C | 728.6666751 | -0.070165098 | 0.720126039 | 0.888 |
| YPR031W | 763.0479333 | 0.132079649 | 0.720019309 | 0.888 |
| YBR097W | 952.5745358 | 0.116236718 | 0.720296452 | 0.888 |
| YJL059W | 446.5285115 | 0.098546329 | 0.721092023 | 0.889 |
| YLR121C | 1254.397824 | 0.054612694 | 0.72098821 | 0.889 |
| YNL219C | 1817.162227 | 0.097834613 | 0.721039918 | 0.889 |
| YIR001C | 869.6360964 | -0.045649529 | 0.721265207 | 0.889 |
| YDR452W | 4294.15743 | 0.075023916 | 0.72208739 | 0.889 |
| YHR003C | 820.5090626 | 0.174210157 | 0.72214715 | 0.889 |
| YKL136W | 25.19505527 | 0.130249497 | 0.721757588 | 0.889 |
| YLR298C | 806.4703735 | -0.180808133 | 0.721921487 | 0.889 |
| YPL055C | 1072.396721 | -0.122698043 | 0.722475575 | 0.889 |
| YPL095C | 828.2325169 | 0.301512249 | 0.722547031 | 0.889 |
| YPR176C | 535.2741493 | -0.060964627 | 0.722341668 | 0.889 |
| YDL006W | 361.6658456 | 0.055371337 | 0.722711215 | 0.889 |

| | | | | |
|---|---|---|---|---|
| YLR128W | 336.2576961 | 0.151139564 | 0.722866412 | 0.889 |
| YNL258C | 964.2763945 | 0.119854913 | 0.723013537 | 0.889 |
| YDL135C | 948.2549942 | -0.10087415 | 0.723840685 | 0.889 |
| YDR234W | 1972.622196 | 0.064660467 | 0.72349052 | 0.889 |
| YDR370C | 533.8393634 | -0.106281427 | 0.723692374 | 0.889 |
| YGR235C | 1048.42614 | -0.117625223 | 0.723926048 | 0.889 |
| YKL204W | 2110.527656 | -0.102195776 | 0.723285628 | 0.889 |
| YOL078W | 796.3214789 | 0.114478774 | 0.723652505 | 0.889 |
| YDL030W | 676.7024249 | 0.082540085 | 0.724105562 | 0.889 |
| YMR289W | 967.6114857 | -0.073583734 | 0.724725028 | 0.890 |
| YKR063C | 686.126014 | 0.087193762 | 0.725334082 | 0.890 |
| YKR082W | 1772.94275 | 0.1018182 | 0.725242701 | 0.890 |
| YNL287W | 4413.325689 | -0.033334692 | 0.725396897 | 0.890 |
| YBR053C | 1693.976608 | 0.149586453 | 0.725596337 | 0.890 |
| YDR261W-B | 58.90647804 | 0.236133361 | 0.727948772 | 0.892 |
| YFR046C | 527.2575594 | 0.087920536 | 0.728043434 | 0.892 |
| YHL029C | 889.4266622 | 0.039102473 | 0.727932557 | 0.892 |
| YML061C | 1035.299962 | 0.093567322 | 0.727652585 | 0.892 |
| YGR126W | 190.2789593 | 0.12007738 | 0.728330131 | 0.892 |
| YKR021W | 1328.600543 | 0.036616957 | 0.728950331 | 0.893 |
| YPL232W | 2081.428239 | -0.113179628 | 0.729870469 | 0.894 |
| YAR003W | 588.5199738 | 0.120119172 | 0.730078437 | 0.894 |
| YBR129C | 610.3882792 | -0.128681631 | 0.730437886 | 0.894 |
| YBR153W | 413.971492 | 0.035006032 | 0.730798836 | 0.894 |
| YKL210W | 8749.432253 | -0.07489545 | 0.731026811 | 0.894 |
| YOR233W | 853.5809378 | 0.079577525 | 0.730888796 | 0.894 |
| YBL078C | 809.4078099 | 0.025153727 | 0.731352433 | 0.895 |
| YJL157C | 1263.741636 | 0.063133541 | 0.732012913 | 0.895 |
| YBL089W | 493.6798455 | 0.127200813 | 0.732422312 | 0.895 |
| YKL209C | 3091.85579 | 0.252104754 | 0.732379163 | 0.895 |
| YDR246W | 916.4165129 | -0.09519752 | 0.733031467 | 0.896 |
| YJL068C | 1815.839599 | 0.111619383 | 0.733052468 | 0.896 |
| YKL061W | 244.3100746 | 0.048541956 | 0.733096946 | 0.896 |
| YGR183C | 999.4364709 | 0.243843717 | 0.733759824 | 0.896 |
| YIR036C | 938.6857022 | 0.099926545 | 0.734050754 | 0.896 |
| YOL152W | 484.6917609 | -0.108130489 | 0.733923459 | 0.896 |
| YAR035W | 1245.571751 | 0.080828471 | 0.734259741 | 0.896 |
| YFL034C-B | 921.0599209 | -0.144519744 | 0.735399355 | 0.897 |
| YOR170W | 97.11015677 | -0.165552803 | 0.735158945 | 0.897 |
| YPR065W | 1458.659767 | 0.07118624 | 0.735443468 | 0.897 |
| YDL155W | 763.3718322 | -0.084393109 | 0.736156431 | 0.898 |
| YMR026C | 666.726482 | -0.086234091 | 0.736198492 | 0.898 |
| YPL235W | 1507.98677 | -0.086602434 | 0.736329229 | 0.898 |

| | | | | |
|---|---|---|---|---|
| YOR113W | 1039.201076 | 0.102469909 | 0.736863307 | 0.898 |
| YOR259C | 3766.746436 | -0.138841096 | 0.736783172 | 0.898 |
| YFL048C | 1163.307711 | -0.100955466 | 0.737664868 | 0.899 |
| YLL039C | 7097.91074 | 0.064650997 | 0.738105055 | 0.899 |
| YDL033C | 356.3492838 | 0.063211309 | 0.738598254 | 0.900 |
| YPR093C | 386.3642488 | 0.288525513 | 0.738821466 | 0.900 |
| YML130C | 2617.920044 | 0.072217761 | 0.739106968 | 0.900 |
| YIL106W | 893.3105346 | -0.069486135 | 0.739520511 | 0.900 |
| YCR098C | 141.8503964 | 0.274275692 | 0.739934623 | 0.900 |
| YLR433C | 1164.793625 | 0.147456717 | 0.740446433 | 0.901 |
| YPL213W | 340.9140453 | -0.131019911 | 0.7404351 | 0.901 |
| YDR176W | 1688.228212 | 0.084922553 | 0.7407241 | 0.901 |
| YIR034C | 2228.546885 | -0.097079592 | 0.741505102 | 0.901 |
| YJL113W | 119.706187 | 0.202797584 | 0.741413703 | 0.901 |
| YMR039C | 1898.602952 | -0.138085159 | 0.742073208 | 0.902 |
| YPR194C | 1121.813284 | 0.275199047 | 0.742196141 | 0.902 |
| YGL186C | 1156.394032 | -0.051546706 | 0.742524391 | 0.902 |
| YHR206W | 1203.868698 | 0.109910054 | 0.74250418 | 0.902 |
| YAR028W | 513.5910617 | 0.181630442 | 0.744343143 | 0.903 |
| YFR049W | 788.1728678 | 0.244377957 | 0.744149981 | 0.903 |
| YGR254W | 35005.12746 | 0.261266787 | 0.743984722 | 0.903 |
| YJR140C | 1086.888351 | 0.193974317 | 0.744360286 | 0.903 |
| YLR169W | 310.0591441 | -0.12529708 | 0.744083175 | 0.903 |
| YOL062C | 1303.409819 | -0.115379317 | 0.743866661 | 0.903 |
| YOR208W | 1224.294967 | -0.064731346 | 0.743655859 | 0.903 |
| YPR133C | 1652.526675 | -0.079018703 | 0.743820549 | 0.903 |
| YDR232W | 2313.644723 | -0.06913567 | 0.744870297 | 0.903 |
| YJL127C-B | 263.7714489 | -0.128383716 | 0.744815065 | 0.903 |
| YLR257W | 6468.084537 | -0.02914778 | 0.745143528 | 0.903 |
| YFR041C | 978.3498425 | -0.047570037 | 0.745492005 | 0.903 |
| YPL015C | 1643.336309 | -0.138311262 | 0.745494862 | 0.903 |
| YBR137W | 861.5111114 | -0.126086279 | 0.746239765 | 0.904 |
| YER122C | 4064.222569 | -0.064867875 | 0.746555769 | 0.904 |
| YER080W | 3267.440935 | 0.148624374 | 0.747115714 | 0.904 |
| YGR011W | 95.88242328 | 0.150611042 | 0.747383208 | 0.905 |
| YIL049W | 383.4643737 | 0.03797333 | 0.747479398 | 0.905 |
| YCR088W | 5938.374856 | -0.07413416 | 0.748014003 | 0.905 |
| YGR221C | 838.7959631 | 0.136413443 | 0.748114048 | 0.905 |
| YNL298W | 1195.295385 | -0.121086223 | 0.747836978 | 0.905 |
| YOR355W | 1652.733408 | 0.125766417 | 0.747862919 | 0.905 |
| YHL045W | 26.03413018 | 0.16140396 | 0.748479066 | 0.905 |
| YNR013C | 2464.582186 | -0.073648224 | 0.748498233 | 0.905 |
| YMR022W | 1239.988654 | -0.100905008 | 0.749338682 | 0.905 |

| | | | | |
|---|---|---|---|---|
| YPL215W | 1126.607522 | -0.038316754 | 0.749263941 | 0.905 |
| YER133W-A | 95.44590047 | -0.167224149 | 0.749708318 | 0.906 |
| YDR457W | 5588.916733 | 0.013876393 | 0.749952245 | 0.906 |
| YDL127W | 422.9534044 | -0.064634808 | 0.750220071 | 0.906 |
| YDR503C | 368.4756431 | -0.083699062 | 0.750580453 | 0.906 |
| YIL137C | 2189.193159 | -0.038496835 | 0.750444593 | 0.906 |
| YBR151W | 3044.846839 | -0.116462208 | 0.751021219 | 0.906 |
| YDL122W | 1760.896516 | 0.086279488 | 0.751140118 | 0.906 |
| YGL181W | 1793.449442 | -0.10729037 | 0.751160359 | 0.906 |
| YOL115W | 646.4767705 | -0.122004154 | 0.751255168 | 0.906 |
| YNR032W | 607.7463894 | 0.100307571 | 0.751455296 | 0.906 |
| YDL074C | 1505.723255 | -0.075410741 | 0.751682802 | 0.906 |
| YBR123C | 731.156432 | 0.075104666 | 0.752085307 | 0.906 |
| YGL021W | 1221.378254 | -0.091131129 | 0.752888035 | 0.907 |
| YGR006W | 239.4957069 | -0.083098854 | 0.75274957 | 0.907 |
| YJR084W | 669.2500078 | -0.051520407 | 0.752655058 | 0.907 |
| YCR006C | 58.71736424 | 0.235149162 | 0.753375922 | 0.907 |
| YPL183W-A | 354.0512636 | -0.072447601 | 0.753618126 | 0.907 |
| YDR128W | 2008.877758 | 0.077396943 | 0.75411645 | 0.908 |
| YDR516C | 6053.645479 | 0.058856385 | 0.754319953 | 0.908 |
| YMR088C | 861.7053416 | -0.067721532 | 0.754384558 | 0.908 |
| YGL115W | 1145.902181 | 0.05942389 | 0.75515875 | 0.908 |
| YGL194C | 571.5770023 | -0.083336225 | 0.755039557 | 0.908 |
| YML122C | 47.85383085 | 0.11807304 | 0.755453892 | 0.908 |
| YDL205C | 1039.950483 | 0.042697224 | 0.755920206 | 0.909 |
| YNR063W | 178.9442947 | 0.009905351 | 0.755973127 | 0.909 |
| YKR011C | 1595.285427 | 0.067077798 | 0.756474101 | 0.909 |
| YDR419W | 525.8031882 | 0.111732564 | 0.756932742 | 0.909 |
| YMR011W | 5664.342762 | -0.647172701 | 0.75704635 | 0.909 |
| YOR109W | 2024.801836 | 0.099013469 | 0.75690724 | 0.909 |
| YKL067W | 3440.072656 | 0.250417785 | 0.757316626 | 0.909 |
| YKL103C | 3936.941911 | 0.245834915 | 0.757501853 | 0.909 |
| YPL209C | 162.5627938 | -0.138350909 | 0.757705056 | 0.909 |
| YER061C | 958.2654695 | 0.150426724 | 0.758793528 | 0.910 |
| YJL197W | 1228.673265 | -0.048804894 | 0.758347102 | 0.910 |
| YKL086W | 218.7788586 | -0.261558173 | 0.758662764 | 0.910 |
| YNL086W | 191.101467 | 0.035126287 | 0.758276785 | 0.910 |
| YOL088C | 665.7794028 | -0.071113037 | 0.75866214 | 0.910 |
| YGR079W | 359.5947801 | -0.230545966 | 0.759107364 | 0.910 |
| YMR295C | 1636.767669 | -0.079048566 | 0.759012222 | 0.910 |
| YHR142W | 1336.189865 | -0.088285384 | 0.759865579 | 0.910 |
| YMR096W | 4030.130556 | -0.474021932 | 0.75960354 | 0.910 |
| YPR097W | 1353.173051 | -0.067211038 | 0.759861988 | 0.910 |

| | | | | |
|---|---|---|---|---|
| YIR002C | 1205.196083 | 0.037243208 | 0.760038986 | 0.910 |
| YDR486C | 1425.158924 | 0.055782873 | 0.761000563 | 0.911 |
| YKR086W | 1098.768049 | 0.049706995 | 0.761268107 | 0.911 |
| YOR263C | 29.84403306 | 0.07371104 | 0.761570771 | 0.911 |
| YOR376W-A | 83.93269889 | -0.055677582 | 0.761857668 | 0.912 |
| YGR233C | 1147.708039 | -0.105911823 | 0.762084356 | 0.912 |
| YBL005W-B | 450.3577253 | -0.220441934 | 0.762406175 | 0.912 |
| YDR130C | 501.1168326 | 0.159720695 | 0.762281915 | 0.912 |
| YEL025C | 1231.625482 | 0.044041723 | 0.762789225 | 0.912 |
| YLL031C | 1740.005527 | -0.05562863 | 0.76288563 | 0.912 |
| YNL330C | 1017.919377 | 0.23444114 | 0.763222839 | 0.912 |
| YBR136W | 1309.50165 | 0.098500722 | 0.763751076 | 0.912 |
| YCR003W | 578.0986481 | -0.104990452 | 0.763753481 | 0.912 |
| YMR234W | 552.7610201 | -0.066285956 | 0.763715308 | 0.912 |
| YGR014W | 3080.238337 | 0.09798774 | 0.763978129 | 0.912 |
| YGL169W | 682.8701947 | -0.141662973 | 0.764169034 | 0.912 |
| YCR102C | 144.1370507 | -0.137609688 | 0.764708037 | 0.912 |
| YDR420W | 1513.909661 | -0.110631216 | 0.764891798 | 0.912 |
| YFR021W | 746.0129726 | -0.089236953 | 0.764518742 | 0.912 |
| YLR250W | 1395.702651 | -0.07823329 | 0.764946442 | 0.912 |
| YLR306W | 266.7883268 | -0.061392313 | 0.764938526 | 0.912 |
| YBL091C-A | 508.1746239 | 0.069422473 | 0.76537529 | 0.912 |
| YMR233W | 360.9564304 | 0.184462513 | 0.765281344 | 0.912 |
| YMR231W | 864.2847521 | -0.044221528 | 0.765824876 | 0.913 |
| YPL253C | 481.3229549 | 0.124671119 | 0.766072459 | 0.913 |
| YOR238W | 549.7060682 | 0.085636008 | 0.766424291 | 0.913 |
| YER040W | 900.0166243 | -0.184129896 | 0.766756839 | 0.913 |
| YLR410W-B | 66.3723325 | -0.209850796 | 0.76667244 | 0.913 |
| YGR138C | 1416.124697 | -0.035045973 | 0.76708119 | 0.913 |
| YKR062W | 891.4921763 | 0.081068475 | 0.767414374 | 0.914 |
| YDR182W | 1361.286881 | 0.084000726 | 0.767665604 | 0.914 |
| YBR289W | 1372.875089 | -0.183213617 | 0.768058756 | 0.914 |
| YMR173W | 743.4251563 | 0.236537849 | 0.769098957 | 0.915 |
| YNL239W | 3388.853667 | -0.045397493 | 0.768959667 | 0.915 |
| YNL315C | 1636.297254 | -0.029796764 | 0.769188828 | 0.915 |
| YLR419W | 1247.341194 | -0.056045539 | 0.770160715 | 0.916 |
| YMR062C | 1501.153431 | -0.113154059 | 0.770095327 | 0.916 |
| YJL019W | 605.7582629 | -0.061805269 | 0.770859653 | 0.916 |
| YMR104C | 1639.396102 | -0.088249967 | 0.771017117 | 0.916 |
| YPR023C | 1837.667458 | -0.042579172 | 0.771080363 | 0.916 |
| YPR086W | 1622.357285 | 0.061736017 | 0.77073196 | 0.916 |
| YDR256C | 561.4619469 | -0.041713263 | 0.771762697 | 0.917 |
| YLL044W | 94.44590477 | -0.345313088 | 0.772011123 | 0.917 |

| | | | | |
|---|---|---|---|---|
| YMR048W | 313.0835867 | -0.016170578 | 0.772058515 | 0.917 |
| YDL065C | 1387.029558 | -0.089093459 | 0.772775266 | 0.917 |
| YHR165C | 3249.517075 | -0.04352207 | 0.772857895 | 0.917 |
| YPL014W | 2476.516022 | -0.039462767 | 0.773500859 | 0.918 |
| YKL167C | 608.6144606 | -0.053582528 | 0.774663736 | 0.919 |
| YLR375W | 1937.904522 | 0.059669636 | 0.775127409 | 0.919 |
| YNL203C | 26.22039237 | -0.076283254 | 0.775013379 | 0.919 |
| YCL064C | 3239.199963 | 0.308722253 | 0.776396875 | 0.919 |
| YDR428C | 292.2817321 | -0.016331933 | 0.775701003 | 0.919 |
| YER111C | 1455.121069 | 0.018713292 | 0.776682027 | 0.919 |
| YGL093W | 1066.84026 | 0.054644319 | 0.776397773 | 0.919 |
| YGR244C | 5502.307672 | 0.253565909 | 0.776663126 | 0.919 |
| YHR178W | 530.4654676 | -0.074530766 | 0.776291426 | 0.919 |
| YLR382C | 967.5198513 | 0.072802124 | 0.776680697 | 0.919 |
| YNL213C | 587.1355028 | -0.143808642 | 0.775870274 | 0.919 |
| YNR023W | 727.2388585 | -0.062384187 | 0.775904286 | 0.919 |
| YIL154C | 2075.807869 | 0.057395844 | 0.776851155 | 0.919 |
| YCR004C | 3730.565062 | -0.087525022 | 0.777289569 | 0.919 |
| YKL001C | 376.5252013 | -0.336457427 | 0.777281278 | 0.919 |
| YMR222C | 712.4011542 | -0.073731269 | 0.77781363 | 0.919 |
| YNL191W | 591.9956262 | -0.018080047 | 0.777600169 | 0.919 |
| YOR201C | 630.2633868 | -0.036716024 | 0.777784661 | 0.919 |
| YGR188C | 503.2494794 | 0.070620672 | 0.778128083 | 0.920 |
| YJL181W | 405.181886 | -0.088829646 | 0.778600714 | 0.920 |
| YGR213C | 279.0368426 | -0.024111034 | 0.779385235 | 0.920 |
| YMR042W | 272.8014796 | -0.156826121 | 0.77924632 | 0.920 |
| YPL171C | 308.1636844 | 0.116014527 | 0.779115678 | 0.920 |
| YPL141C | 1338.030401 | 0.065345345 | 0.779779642 | 0.921 |
| YDL129W | 428.0062351 | 0.016773917 | 0.779957554 | 0.921 |
| YBL014C | 840.8039898 | -0.223772479 | 0.780377044 | 0.921 |
| YGR282C | 11912.38121 | -0.009231871 | 0.780451651 | 0.921 |
| YLR063W | 280.3060801 | -0.21757604 | 0.78073475 | 0.921 |
| YML027W | 523.9520569 | -0.124861726 | 0.780579298 | 0.921 |
| YGR168C | 490.2041357 | -0.1191191 | 0.781203138 | 0.921 |
| YJL077W-B | 26.1550574 | -0.192927598 | 0.781311961 | 0.921 |
| YOR067C | 1025.694017 | 0.187965008 | 0.781453534 | 0.921 |
| YFR016C | 3711.04018 | 0.1850447 | 0.781777229 | 0.921 |
| YHR186C | 1649.917336 | 0.118847875 | 0.782028954 | 0.921 |
| YLR049C | 719.3459985 | -0.038118811 | 0.782083634 | 0.921 |
| YDL008W | 381.8332185 | -0.035750103 | 0.782478401 | 0.922 |
| YGR182C | 629.4422912 | -0.158452367 | 0.783682353 | 0.923 |
| YLL029W | 3427.507563 | 0.07311968 | 0.783649577 | 0.923 |
| YNL314W | 384.9311948 | -0.108737243 | 0.783922285 | 0.923 |

| | | | | |
|---|---|---|---|---|
| YBR218C | 7147.332039 | 0.107717091 | 0.785089968 | 0.924 |
| YCL044C | 1238.471005 | -0.154294644 | 0.78510186 | 0.924 |
| YNR006W | 1506.041195 | 0.042308236 | 0.785427869 | 0.924 |
| YOR303W | 3621.710527 | -0.280423652 | 0.785416157 | 0.924 |
| YPL083C | 495.9825937 | -0.01406337 | 0.785541784 | 0.924 |
| YPR071W | 302.2135475 | 0.073285343 | 0.785131663 | 0.924 |
| YBR197C | 458.4082334 | -0.108129747 | 0.786069974 | 0.924 |
| YCR044C | 1029.447304 | -0.040115365 | 0.786173148 | 0.924 |
| YPL105C | 3260.005512 | -0.082199937 | 0.785874784 | 0.924 |
| YOL014W | 428.9922641 | -0.289174355 | 0.786564918 | 0.924 |
| YML058W | 2577.833646 | 0.042118951 | 0.78692341 | 0.924 |
| YGR068C | 535.7602898 | 0.062152966 | 0.787651793 | 0.925 |
| YHR008C | 8653.583569 | 0.22226362 | 0.787461839 | 0.925 |
| YMR006C | 894.5004784 | -0.024356613 | 0.787618443 | 0.925 |
| YBR044C | 837.2738234 | 0.120217695 | 0.787810454 | 0.925 |
| YPR056W | 512.7372419 | -0.039435457 | 0.788360247 | 0.925 |
| YER170W | 376.8018239 | -0.063688847 | 0.789317561 | 0.925 |
| YKL012W | 1327.340355 | 0.038249943 | 0.789051138 | 0.925 |
| YLR140W | 51.21550356 | 0.059221001 | 0.788927193 | 0.925 |
| YMR054W | 1855.817154 | -0.021225876 | 0.789104707 | 0.925 |
| YOR179C | 265.678804 | 0.195762726 | 0.789275571 | 0.925 |
| YAL043C | 1560.889418 | 0.100884136 | 0.789764112 | 0.925 |
| YJL137C | 923.3034038 | 0.269147792 | 0.789669241 | 0.925 |
| YOL107W | 336.081692 | 0.217845314 | 0.790127995 | 0.926 |
| YHR048W | 458.0188382 | -0.008296137 | 0.790614265 | 0.926 |
| YNL257C | 1662.623362 | 0.066075204 | 0.790551687 | 0.926 |
| YGR184C | 3421.297226 | -0.025532867 | 0.791486026 | 0.926 |
| YKL169C | 32.06083426 | -0.130064538 | 0.791336825 | 0.926 |
| YLR295C | 2976.605613 | -0.008617317 | 0.791167744 | 0.926 |
| YNL329C | 1184.177512 | 0.091164929 | 0.79134187 | 0.926 |
| YAR042W | 3250.319067 | 0.098848661 | 0.791940909 | 0.926 |
| YDR180W | 980.8547337 | -0.077843752 | 0.791820618 | 0.926 |
| YFR055W | 975.2716539 | -0.194981569 | 0.792004779 | 0.926 |
| YDR148C | 4531.818706 | 0.296627233 | 0.792325114 | 0.926 |
| YPR025C | 1080.062057 | 0.034301041 | 0.792210807 | 0.926 |
| YBR202W | 984.15721 | -0.080292707 | 0.792881157 | 0.927 |
| YLR233C | 453.3027113 | 0.070564627 | 0.79377196 | 0.927 |
| YLR438W | 1335.790008 | 0.188608425 | 0.793944946 | 0.927 |
| YMR218C | 1133.912405 | 0.064161375 | 0.793947057 | 0.927 |
| YGL143C | 536.4770627 | 0.176851561 | 0.794456691 | 0.927 |
| YHR005C | 1534.146963 | 0.244945345 | 0.794768666 | 0.927 |
| YJR078W | 516.4472306 | 0.231437265 | 0.794362976 | 0.927 |
| YLR228C | 906.7831503 | -0.216659897 | 0.794633789 | 0.927 |

| | | | | |
|---|---|---|---|---|
| YPR159W | 2743.676738 | 0.147584095 | 0.794282097 | 0.927 |
| YDR268W | 586.9385642 | -0.003011045 | 0.795321746 | 0.927 |
| YLR097C | 748.1202732 | 0.030736655 | 0.795314474 | 0.927 |
| YLR191W | 472.4930163 | -0.132926708 | 0.795117089 | 0.927 |
| YLR370C | 2370.21826 | 0.048754659 | 0.795519099 | 0.927 |
| YDR476C | 1082.582617 | 0.037431395 | 0.795765935 | 0.928 |
| YNL007C | 10095.98848 | 0.022435546 | 0.796231994 | 0.928 |
| YHR210C | 320.0019552 | 0.390429844 | 0.796608012 | 0.928 |
| YBL013W | 160.0267849 | -0.10605201 | 0.797545132 | 0.928 |
| YBR110W | 1287.287964 | -0.075636835 | 0.797963555 | 0.928 |
| YBR165W | 515.6336536 | 0.023991252 | 0.797745314 | 0.928 |
| YDL043C | 369.3400281 | -0.079249729 | 0.79738616 | 0.928 |
| YFL059W | 70.79713485 | 0.432452144 | 0.797742956 | 0.928 |
| YGR203W | 222.1887092 | 0.122961007 | 0.797967219 | 0.928 |
| YML121W | 941.5006143 | -0.076538188 | 0.796903429 | 0.928 |
| YOR334W | 302.0782196 | 0.049654175 | 0.798002736 | 0.928 |
| YNR066C | 265.0332103 | -0.06568349 | 0.798287036 | 0.929 |
| YDR515W | 694.9239372 | 0.112785066 | 0.798525625 | 0.929 |
| YIR038C | 2847.18512 | 0.067482508 | 0.798611783 | 0.929 |
| YJL154C | 2489.813399 | 0.048905475 | 0.799229945 | 0.929 |
| YOL133W | 270.3290721 | -0.222964327 | 0.799080897 | 0.929 |
| YNL284C-B | 190.6577576 | 0.052163074 | 0.800255623 | 0.930 |
| YDR511W | 614.1187303 | 0.233305276 | 0.80150278 | 0.931 |
| YHL009C | 603.1387723 | 0.2426863 | 0.801520084 | 0.931 |
| YJL055W | 2135.334979 | 0.08300995 | 0.801545711 | 0.931 |
| YOR241W | 933.790879 | -0.050112292 | 0.801455266 | 0.931 |
| YPR107C | 422.2754774 | 0.014731778 | 0.801870716 | 0.931 |
| YPR141C | 623.3342075 | 0.122036331 | 0.802076023 | 0.931 |
| YGR058W | 565.2067235 | 0.095972105 | 0.802588054 | 0.931 |
| YPR104C | 970.815797 | 0.076600596 | 0.802645785 | 0.931 |
| YDR055W | 3654.499745 | 0.123721324 | 0.802902888 | 0.931 |
| YMR077C | 308.7109529 | -0.024175586 | 0.803027371 | 0.931 |
| YGR157W | 3036.942102 | 0.108812064 | 0.803614562 | 0.931 |
| YKL002W | 1601.062288 | 0.072967548 | 0.803287558 | 0.931 |
| YKL064W | 2390.234747 | 0.083329666 | 0.803676483 | 0.931 |
| YOR171C | 1509.079417 | 0.070603317 | 0.803462376 | 0.931 |
| YOR347C | 1046.083791 | 0.155708787 | 0.804045853 | 0.932 |
| YER048C | 2682.830499 | -0.068214417 | 0.804652762 | 0.932 |
| YJR132W | 1306.62171 | 0.102467265 | 0.804576599 | 0.932 |
| YOR264W | 932.8104963 | 0.102776984 | 0.804443544 | 0.932 |
| YBR012C | 184.3218002 | -0.257747886 | 0.806282717 | 0.932 |
| YBR101C | 3697.675555 | 0.040137419 | 0.805715964 | 0.932 |
| YDL045W-A | 694.1101578 | 0.012094415 | 0.805907137 | 0.932 |

| | | | | |
|---|---|---|---|---|
| YDR067C | 673.9208846 | -0.063071133 | 0.805490292 | 0.932 |
| YDR206W | 1789.144398 | -0.086500343 | 0.805753017 | 0.932 |
| YGL263W | 253.3051631 | 0.743316243 | 0.806047215 | 0.932 |
| YIL169C | 2177.35422 | 0.201581463 | 0.805357956 | 0.932 |
| YJL107C | 129.4785502 | -0.423004826 | 0.80561883 | 0.932 |
| YLR055C | 1018.744824 | -0.170572771 | 0.806240982 | 0.932 |
| YLR364W | 131.7044529 | -0.133369265 | 0.804932859 | 0.932 |
| YAL031W-A | 36.5218107 | 0.030234852 | 0.807305892 | 0.933 |
| YMR313C | 988.4819991 | -0.066389959 | 0.807327215 | 0.933 |
| YDL049C | 168.7509482 | -0.084805028 | 0.809466124 | 0.934 |
| YDR072C | 1794.903187 | 0.013820595 | 0.809148464 | 0.934 |
| YDR168W | 2674.002145 | -0.071439107 | 0.810492426 | 0.934 |
| YDR261C | 1216.328788 | -0.039644894 | 0.810237255 | 0.934 |
| YGR026W | 4521.168167 | -0.050032644 | 0.810446198 | 0.934 |
| YGR141W | 841.9095851 | 0.210999136 | 0.80961344 | 0.934 |
| YIL009W | 578.8586392 | -0.063382708 | 0.810367957 | 0.934 |
| YIR033W | 2182.959679 | -0.059421256 | 0.809719036 | 0.934 |
| YJR041C | 1150.543526 | -0.048752995 | 0.810543286 | 0.934 |
| YLR131C | 1246.543234 | 0.034604002 | 0.809767369 | 0.934 |
| YMR261C | 3042.037261 | 0.034447072 | 0.809358142 | 0.934 |
| YNR052C | 1154.692673 | -0.058357819 | 0.810080979 | 0.934 |
| YOR042W | 2668.884695 | 0.082386517 | 0.810227432 | 0.934 |
| YPL154C | 14185.44316 | 0.133469281 | 0.810442469 | 0.934 |
| YDR185C | 264.7477216 | 0.019717684 | 0.811140483 | 0.934 |
| YGR023W | 927.8650533 | -0.239939046 | 0.811396836 | 0.934 |
| YDR134C | 8310.682574 | -0.351680242 | 0.811707338 | 0.934 |
| YJL211C | 56.87482788 | 0.021487426 | 0.811995185 | 0.935 |
| YKL217W | 2399.793467 | 0.057529041 | 0.812186968 | 0.935 |
| YKL070W | 302.0799748 | -0.114723614 | 0.812530572 | 0.935 |
| YMR288W | 774.5062547 | -0.093116846 | 0.812545523 | 0.935 |
| YDL025C | 1555.623411 | 0.029885219 | 0.81282126 | 0.935 |
| YBR058C | 1360.304304 | -0.032748827 | 0.813426016 | 0.935 |
| YNR028W | 918.8142146 | 0.006042371 | 0.813302338 | 0.935 |
| YER166W | 2859.051695 | -0.043121698 | 0.815025959 | 0.936 |
| YGR237C | 1848.787726 | -0.098790604 | 0.814293258 | 0.936 |
| YIL132C | 143.1249549 | -0.102344372 | 0.814721278 | 0.936 |
| YKL028W | 1511.073698 | -0.109001243 | 0.814218461 | 0.936 |
| YKL155C | 791.2689341 | 0.074022548 | 0.814407283 | 0.936 |
| YKL195W | 3025.520768 | 0.102663831 | 0.815066457 | 0.936 |
| YLR215C | 433.3667117 | 0.056090524 | 0.814832855 | 0.936 |
| YPL035C | 100.9354993 | 0.034718104 | 0.814784663 | 0.936 |
| YAL047C | 589.7842721 | -0.057745608 | 0.815991959 | 0.936 |
| YBR111C | 3318.167647 | -0.083722861 | 0.816755885 | 0.936 |

| | | | | |
|---|---|---|---|---|
| YBR126W-A | 695.852559 | 0.133069078 | 0.816517563 | 0.936 |
| YDR145W | 1845.179932 | -0.096472644 | 0.815706328 | 0.936 |
| YGL254W | 331.7870674 | 0.119626182 | 0.816490796 | 0.936 |
| YGR036C | 221.213258 | -0.025293918 | 0.816019122 | 0.936 |
| YHR105W | 228.2746463 | 0.05351043 | 0.816426822 | 0.936 |
| YMR294W | 447.832337 | 0.040218411 | 0.816621508 | 0.936 |
| YPR066W | 302.1036782 | 0.087231921 | 0.815853709 | 0.936 |
| YCR093W | 4708.644101 | -0.019576458 | 0.817155123 | 0.936 |
| YNL214W | 315.125881 | 0.034047439 | 0.817132681 | 0.936 |
| YGL165C | 78.38274117 | -0.233149357 | 0.81772911 | 0.936 |
| YOL081W | 5088.832739 | 0.10535777 | 0.817842154 | 0.936 |
| YBL088C | 1866.759984 | 0.074475014 | 0.818906972 | 0.936 |
| YBR207W | 1493.475156 | -0.082210578 | 0.818387299 | 0.936 |
| YCR008W | 1187.11323 | -0.106062269 | 0.819614188 | 0.936 |
| YDR074W | 10482.83127 | -0.088820118 | 0.81905615 | 0.936 |
| YFL031W | 3441.588262 | 0.011692035 | 0.819637391 | 0.936 |
| YHR201C | 1092.809897 | 0.005032929 | 0.818696763 | 0.936 |
| YJL043W | 47.06314645 | 0.054747714 | 0.819006694 | 0.936 |
| YJL210W | 543.5019911 | 0.027686638 | 0.819187223 | 0.936 |
| YKR095W | 4316.14506 | 0.090222222 | 0.819554374 | 0.936 |
| YNL316C | 479.8242776 | -0.050189559 | 0.818894571 | 0.936 |
| YOR203W | 129.138005 | -0.366010323 | 0.818387762 | 0.936 |
| YDR325W | 750.6903469 | -0.02259678 | 0.820018166 | 0.937 |
| YDL178W | 1397.749906 | 0.075758783 | 0.820675677 | 0.937 |
| YDR514C | 275.4543514 | -0.043945881 | 0.82072473 | 0.937 |
| YMR216C | 1336.629718 | 0.089553425 | 0.820559117 | 0.937 |
| YDR157W | 53.23769391 | -0.38754957 | 0.821050186 | 0.937 |
| YOR001W | 1523.051345 | -0.042969391 | 0.820927197 | 0.937 |
| YBR286W | 17567.33584 | 0.071816363 | 0.821483482 | 0.937 |
| YER071C | 272.1086234 | -0.097957386 | 0.822538742 | 0.937 |
| YFR007W | 462.3779448 | 0.113158725 | 0.822364525 | 0.937 |
| YGL086W | 1388.801297 | 0.05823484 | 0.822149269 | 0.937 |
| YIR042C | 196.4981605 | -0.024735586 | 0.821618896 | 0.937 |
| YJL024C | 668.6768051 | 0.053945432 | 0.822578355 | 0.937 |
| YJR103W | 3899.480126 | 0.148241418 | 0.822720033 | 0.937 |
| YLR404W | 245.17757 | -0.041038347 | 0.821929697 | 0.937 |
| YNL192W | 2879.150646 | -0.160840896 | 0.821376036 | 0.937 |
| YOR385W | 901.596547 | -0.010346429 | 0.822716909 | 0.937 |
| YMR207C | 1385.212155 | 0.082754907 | 0.822902233 | 0.937 |
| YEL041W | 348.1575551 | 0.013403012 | 0.823373526 | 0.937 |
| YGR265W | 63.64327138 | 0.048619876 | 0.823612404 | 0.937 |
| YHR129C | 426.0946941 | -0.069588165 | 0.82363461 | 0.937 |
| YPL031C | 1442.118487 | -0.010013567 | 0.823397994 | 0.937 |

| | | | | |
|---|---|---|---|---|
| YDR498C | 1014.743074 | 0.035122011 | 0.823875935 | 0.937 |
| YBR204C | 1064.479878 | 0.125083836 | 0.824692947 | 0.937 |
| YDR357C | 553.215464 | 0.1501216 | 0.825157391 | 0.937 |
| YFR039C | 736.7528646 | 0.07280306 | 0.824517051 | 0.937 |
| YGL167C | 2492.369981 | 0.07332915 | 0.825131673 | 0.937 |
| YGR142W | 4162.125638 | 0.317107397 | 0.825249579 | 0.937 |
| YJR092W | 2324.987782 | -0.014203445 | 0.825140294 | 0.937 |
| YLR231C | 929.429053 | 0.478586835 | 0.825208735 | 0.937 |
| YOL136C | 1016.324902 | 0.133072047 | 0.824805953 | 0.937 |
| YLR042C | 282.794653 | 0.300505957 | 0.825572458 | 0.938 |
| YGR049W | 549.8511767 | 0.006445347 | 0.826114352 | 0.938 |
| YNL032W | 670.5607595 | -0.09663681 | 0.826138695 | 0.938 |
| YLR115W | 1204.082371 | 0.056980858 | 0.827603882 | 0.939 |
| YDR466W | 1549.812513 | 0.034451212 | 0.828424405 | 0.940 |
| YKL141W | 4100.397187 | 0.233275009 | 0.828665419 | 0.940 |
| YKR052C | 558.5950233 | -0.077636789 | 0.828657426 | 0.940 |
| YNL059C | 1845.140766 | 0.060853272 | 0.829069571 | 0.940 |
| YDR135C | 3272.34562 | -0.046311266 | 0.829383638 | 0.940 |
| YGL179C | 645.0803373 | 0.002427929 | 0.829682575 | 0.941 |
| YJR040W | 752.4605261 | 0.103779206 | 0.829895379 | 0.941 |
| YER143W | 1597.506506 | 0.000361871 | 0.830128438 | 0.941 |
| YMR008C | 3679.477965 | -0.04442809 | 0.830300188 | 0.941 |
| YBR037C | 852.7234834 | 0.069703071 | 0.831077059 | 0.941 |
| YER037W | 745.1407721 | -0.126484708 | 0.831237671 | 0.941 |
| YOR138C | 1254.624127 | -0.086122381 | 0.831536685 | 0.941 |
| YOR142W | 3533.236265 | 0.016566018 | 0.831215081 | 0.941 |
| snR82 | 454.6100305 | -0.191914793 | 0.83143429 | 0.941 |
| YER133W | 3950.455243 | -0.004253229 | 0.832015032 | 0.942 |
| YHR134W | 365.9132551 | 0.000381285 | 0.832079917 | 0.942 |
| YGR005C | 1346.509283 | -0.076586838 | 0.832661102 | 0.942 |
| YNR029C | 598.402815 | -0.031885044 | 0.832554144 | 0.942 |
| YLR260W | 983.6383575 | 0.016681838 | 0.833015887 | 0.942 |
| YCL056C | 401.4808141 | -0.097487125 | 0.833987259 | 0.943 |
| YPL201C | 234.758662 | -0.070819555 | 0.833840156 | 0.943 |
| YNL208W | 6496.983749 | -0.136672717 | 0.834713121 | 0.943 |
| YJR113C | 864.1046061 | -0.036730364 | 0.835127709 | 0.944 |
| YDR310C | 1697.719061 | -0.003435782 | 0.835482367 | 0.944 |
| YDL104C | 329.3472486 | -0.062963184 | 0.835816328 | 0.944 |
| YJL176C | 1289.4859 | 0.044103449 | 0.835772167 | 0.944 |
| YHR196W | 929.0161741 | -0.071213893 | 0.836194143 | 0.944 |
| YFL030W | 1073.611575 | 0.014214914 | 0.836543907 | 0.944 |
| YNL081C | 921.3541155 | -0.063671273 | 0.83643965 | 0.944 |
| YAL049C | 1497.895567 | 0.030901425 | 0.837141777 | 0.944 |

| | | | | |
|---|---|---|---|---|
| YGR156W | 932.64115 | -0.059032092 | 0.837277813 | 0.944 |
| YJL126W | 376.2312005 | -0.067718947 | 0.836966643 | 0.944 |
| YML066C | 127.9180195 | -0.169931501 | 0.837241271 | 0.944 |
| YDR472W | 675.6541934 | -0.038245847 | 0.837820488 | 0.945 |
| YHR032W-A | 60.96537533 | -0.110574361 | 0.8379225 | 0.945 |
| YLR007W | 741.9239686 | -0.049963152 | 0.837975369 | 0.945 |
| YBR047W | 453.8190728 | -0.164273838 | 0.83817704 | 0.945 |
| YER032W | 1018.880889 | -0.017870221 | 0.838301418 | 0.945 |
| YLR085C | 664.5290334 | -0.067280419 | 0.838653701 | 0.945 |
| YOR155C | 1051.816257 | 0.081746177 | 0.838593187 | 0.945 |
| YBR005W | 1310.131508 | -0.160627188 | 0.839304555 | 0.945 |
| YMR060C | 493.2206546 | -0.081652569 | 0.839169648 | 0.945 |
| YNL323W | 681.0384119 | -0.026395056 | 0.839355395 | 0.945 |
| YDL087C | 335.3146294 | 0.005263242 | 0.839945713 | 0.945 |
| YLR381W | 607.7067419 | 0.047802896 | 0.84007687 | 0.945 |
| YOR229W | 494.4703852 | -0.043612602 | 0.840267813 | 0.945 |
| YMR308C | 3347.564301 | 0.055069371 | 0.840571903 | 0.945 |
| YLR328W | 869.5602715 | -0.080233948 | 0.840756089 | 0.945 |
| YKR044W | 684.1987114 | 0.021374242 | 0.841135423 | 0.946 |
| YOR111W | 390.0644929 | 0.034997002 | 0.841397475 | 0.946 |
| YGR042W | 764.6829972 | -0.148297603 | 0.842210974 | 0.946 |
| YGR191W | 1267.875828 | 0.087711951 | 0.842398088 | 0.946 |
| YJL205C | 132.3580317 | -0.062943331 | 0.842363096 | 0.946 |
| YNL212W | 1282.46661 | 0.108014818 | 0.842298997 | 0.946 |
| YPR075C | 1855.120278 | -0.025493504 | 0.841806372 | 0.946 |
| YAL027W | 394.2818606 | -0.089276037 | 0.84357695 | 0.946 |
| YDR116C | 1042.960963 | -0.019524855 | 0.843361022 | 0.946 |
| YDR309C | 2613.4939 | 0.042330678 | 0.84327924 | 0.946 |
| YJL075C | 78.82663296 | -0.239575848 | 0.84277091 | 0.946 |
| YMR172W | 685.1558079 | -0.048061072 | 0.8428722 | 0.946 |
| YOR375C | 17984.54886 | -0.047149736 | 0.843185882 | 0.946 |
| YPR170C | 83.5281119 | 0.044564559 | 0.843600007 | 0.946 |
| YHL040C | 1041.983492 | 0.203181329 | 0.844293429 | 0.946 |
| YHR073W | 1977.66425 | -0.036174726 | 0.844282136 | 0.946 |
| YHR118C | 526.5681517 | -0.028134337 | 0.844274723 | 0.946 |
| YNL133C | 297.07226 | -0.132413377 | 0.844457942 | 0.946 |
| snR42 | 83.98583809 | -0.035176868 | 0.845293513 | 0.947 |
| YBR098W | 547.1579177 | -0.048475152 | 0.846198171 | 0.947 |
| YGR012W | 677.5451845 | -0.066007924 | 0.845920871 | 0.947 |
| YGR072W | 838.1301736 | -0.142016486 | 0.846084616 | 0.947 |
| YHR023W | 2927.24121 | -0.035632214 | 0.84614839 | 0.947 |
| YMR157C | 556.2957769 | 0.102212182 | 0.846351603 | 0.947 |
| YPR161C | 1370.124428 | -0.06840524 | 0.84581522 | 0.947 |

| | | | | |
|---|---|---|---|---|
| YDR196C | 880.6195121 | 0.025592715 | 0.847809355 | 0.947 |
| YDR517W | 2232.436932 | 0.109279095 | 0.847711912 | 0.947 |
| YEL043W | 2331.741844 | 0.046372257 | 0.847163679 | 0.947 |
| YKL016C | 3252.389125 | 0.008563988 | 0.84773906 | 0.947 |
| YLL026W | 27987.51895 | 0.182574287 | 0.847508176 | 0.947 |
| YLL040C | 5253.851757 | 0.276609498 | 0.847186994 | 0.947 |
| YLR136C | 655.8429852 | 0.204978482 | 0.847705074 | 0.947 |
| YLR346C | 131.9806369 | -0.01839384 | 0.84739533 | 0.947 |
| YCR073C | 984.1918819 | -0.10736862 | 0.848035042 | 0.947 |
| YDL117W | 1322.092729 | 0.032486063 | 0.848424429 | 0.948 |
| YNL127W | 1075.477959 | 0.131368329 | 0.848383787 | 0.948 |
| YDR461W | 1830.189511 | 0.167742309 | 0.84872439 | 0.948 |
| YDL078C | 3642.136284 | 0.032445033 | 0.849048926 | 0.948 |
| YDL098C | 246.2120054 | 0.041987968 | 0.850086156 | 0.948 |
| YDR211W | 2365.993285 | -0.014740463 | 0.849175784 | 0.948 |
| YHR132W-A | 721.2397547 | 0.076264253 | 0.850298791 | 0.948 |
| YLR038C | 1207.825369 | -0.005410068 | 0.849969337 | 0.948 |
| YLR112W | 96.84022753 | 0.006728153 | 0.850172304 | 0.948 |
| YLR442C | 902.5050218 | 0.019300257 | 0.850084847 | 0.948 |
| YNL278W | 749.3245241 | 0.033414891 | 0.849539538 | 0.948 |
| YPL192C | 128.9945052 | -0.049126269 | 0.849589324 | 0.948 |
| YJR053W | 291.1479634 | 0.019259363 | 0.851032585 | 0.948 |
| YMR266W | 1578.35636 | -0.016877186 | 0.850901684 | 0.948 |
| YNL104C | 8329.192681 | -0.04635627 | 0.851045263 | 0.948 |
| YOR317W | 4625.356852 | 0.116275828 | 0.85098402 | 0.948 |
| YDL146W | 626.6868048 | 0.127473984 | 0.851765657 | 0.948 |
| YER137C | 65.6651259 | -0.24942664 | 0.851708844 | 0.948 |
| YBL100W-B | 175.5232464 | 0.068877852 | 0.852069937 | 0.948 |
| YGL033W | 39.3922468 | -0.029543961 | 0.852156831 | 0.948 |
| YNR033W | 1803.332991 | -0.027618026 | 0.85244679 | 0.949 |
| YBR235W | 1364.419973 | -0.083369462 | 0.852882164 | 0.949 |
| YOR153W | 11636.99259 | 0.087054092 | 0.85300299 | 0.949 |
| YDL059C | 182.996654 | -0.003926292 | 0.853367283 | 0.949 |
| YFR026C | 243.3675063 | -0.181362661 | 0.853913101 | 0.949 |
| YKR042W | 18766.32248 | -0.007523048 | 0.853951529 | 0.949 |
| YPR193C | 57.30247136 | -0.051127328 | 0.853880078 | 0.949 |
| YDL086W | 1034.431113 | -0.060576649 | 0.855127653 | 0.949 |
| YDR100W | 731.7492012 | -0.054922637 | 0.85471874 | 0.949 |
| YDR123C | 353.8570592 | -0.156240116 | 0.854338951 | 0.949 |
| YEL016C | 826.6367047 | -0.056748296 | 0.854953456 | 0.949 |
| YHR135C | 3300.939762 | -0.025713404 | 0.855349426 | 0.949 |
| YHR205W | 2582.464034 | -0.098870954 | 0.855481108 | 0.949 |
| YMR047C | 2343.921258 | -0.05378979 | 0.854198667 | 0.949 |

| | | | | |
|---|---|---|---|---|
| YMR074C | 1117.334751 | -0.125708975 | 0.854987195 | 0.949 |
| YNL275W | 352.1195966 | -0.091396446 | 0.855571389 | 0.949 |
| YOL033W | 423.9709748 | 0.091617095 | 0.855200773 | 0.949 |
| YBR138C | 569.4593536 | 0.017540931 | 0.856587975 | 0.950 |
| YGR039W | 40.79718114 | 0.064299418 | 0.857529777 | 0.950 |
| YIL083C | 1169.719343 | 0.045920206 | 0.857075245 | 0.950 |
| YKL087C | 723.7436146 | -0.052848097 | 0.857444868 | 0.950 |
| YLR310C | 3300.526123 | 0.099095866 | 0.857108837 | 0.950 |
| YML072C | 6041.463861 | 0.111084339 | 0.856951372 | 0.950 |
| YNL246W | 1195.836863 | 0.028239661 | 0.856223905 | 0.950 |
| YOL110W | 433.4687434 | -0.034214237 | 0.857561252 | 0.950 |
| YOR011W | 696.3198627 | 0.02247069 | 0.85683182 | 0.950 |
| YPL099C | 334.8201937 | 0.134643303 | 0.857423494 | 0.950 |
| YBR039W | 6878.817001 | 0.125991629 | 0.858472539 | 0.950 |
| YDR243C | 512.1645164 | 0.010809638 | 0.858407007 | 0.950 |
| YDR495C | 1040.925179 | 0.103250262 | 0.858337919 | 0.950 |
| YER114C | 1610.317658 | -0.041762927 | 0.858233614 | 0.950 |
| YNL098C | 3291.418535 | 0.03369294 | 0.857935303 | 0.950 |
| YHR045W | 983.5331682 | 0.018735327 | 0.858645756 | 0.950 |
| YLR376C | 277.9237935 | -0.105592362 | 0.859308555 | 0.950 |
| YHL020C | 961.5750146 | -0.036664423 | 0.860237636 | 0.951 |
| YMR044W | 903.5982709 | 0.018273946 | 0.860251412 | 0.951 |
| YNL145W | 1813.203416 | -0.038322474 | 0.860060591 | 0.951 |
| YKR003W | 1162.972466 | 0.014491915 | 0.860512768 | 0.951 |
| YKL065C | 4326.57925 | 0.047234314 | 0.860944843 | 0.951 |
| YPL115C | 1185.518438 | 0.006227988 | 0.860918914 | 0.951 |
| YLR224W | 1026.056679 | -0.164052368 | 0.861214651 | 0.951 |
| YHR204W | 1257.868137 | -0.066544696 | 0.861802546 | 0.951 |
| YJL194W | 154.6134221 | -0.144156304 | 0.861676609 | 0.951 |
| YCR071C | 461.1120902 | -0.067758689 | 0.862196788 | 0.952 |
| YML081W | 1706.956949 | -0.044615392 | 0.86218318 | 0.952 |
| YOR270C | 5846.210473 | -0.006901895 | 0.862655547 | 0.952 |
| YBR255C-A | 311.2384417 | -0.052012087 | 0.863652309 | 0.952 |
| YGL238W | 2547.939909 | -0.036888799 | 0.86321339 | 0.952 |
| YHR024C | 1377.744705 | -0.032046089 | 0.863465116 | 0.952 |
| YIL161W | 488.7537738 | -0.013054788 | 0.863595192 | 0.952 |
| YMR092C | 2509.767947 | 0.04970337 | 0.863517254 | 0.952 |
| YNL003C | 715.0467326 | 0.093568325 | 0.863604053 | 0.952 |
| YCR075C | 329.3656461 | 0.071207975 | 0.864800824 | 0.952 |
| YGR252W | 827.8896718 | -0.021932869 | 0.864542749 | 0.952 |
| YHR009C | 3439.835058 | 0.055356945 | 0.864574895 | 0.952 |
| YHR082C | 4569.103984 | -0.021354961 | 0.865081607 | 0.952 |
| YKL149C | 355.2699607 | -0.09962897 | 0.864133523 | 0.952 |

| | | | | |
|---|---|---|---|---|
| YML052W | 1751.911847 | -0.095204369 | 0.864998561 | 0.952 |
| YOR378W | 266.6242043 | -0.004339896 | 0.864787916 | 0.952 |
| YLR079W | 2171.764341 | 0.006945992 | 0.865292647 | 0.952 |
| YCR106W | 394.726015 | 0.210252958 | 0.865587583 | 0.952 |
| YGL191W | 2937.202908 | -0.007505676 | 0.865917554 | 0.953 |
| YLL054C | 624.0988471 | 0.129051907 | 0.866233499 | 0.953 |
| YDL161W | 1895.847844 | 0.018689095 | 0.867811813 | 0.953 |
| YEL077C | 280.3488731 | 0.327866312 | 0.867563852 | 0.953 |
| YER095W | 1267.382435 | 0.025757724 | 0.867940154 | 0.953 |
| YGL050W | 416.7826537 | 0.102423722 | 0.866880922 | 0.953 |
| YJR044C | 1233.447551 | -0.026131753 | 0.867520988 | 0.953 |
| YJR116W | 691.1206507 | -0.11830754 | 0.867312159 | 0.953 |
| YMR275C | 1521.663704 | -0.035595507 | 0.868003993 | 0.953 |
| YPR010C-A | 733.8429902 | -0.009569024 | 0.867956565 | 0.953 |
| YCR073W-A | 1340.952291 | -0.108153025 | 0.868742805 | 0.954 |
| YMR150C | 511.1258272 | 0.064116464 | 0.868829171 | 0.954 |
| YGR271W | 1597.427923 | 0.113299984 | 0.870525714 | 0.956 |
| YBR231C | 448.4132605 | -0.164043204 | 0.872010891 | 0.956 |
| YDL243C | 368.4938618 | -0.118522645 | 0.872272875 | 0.956 |
| YDR375C | 907.9536977 | 0.053794344 | 0.872213553 | 0.956 |
| YDR416W | 716.6573712 | -0.067829182 | 0.872374774 | 0.956 |
| YER161C | 1183.000093 | 0.026744967 | 0.872402174 | 0.956 |
| YML111W | 1306.672498 | -0.007379645 | 0.872107927 | 0.956 |
| YMR285C | 827.213202 | 0.039362757 | 0.871798637 | 0.956 |
| YPR011C | 560.157333 | 0.124082235 | 0.871827087 | 0.956 |
| YPR083W | 614.8326597 | -0.099930012 | 0.871981101 | 0.956 |
| YPR095C | 1088.661894 | 0.017504581 | 0.872636765 | 0.956 |
| YNR074C | 411.2345917 | 0.054338204 | 0.872813853 | 0.956 |
| YEL047C | 2163.496037 | -0.002584109 | 0.872983942 | 0.956 |
| YMR136W | 2477.221542 | -0.255344109 | 0.873682087 | 0.957 |
| snR5 | 32.14065526 | 0.182308198 | 0.873947548 | 0.957 |
| YBL045C | 7514.298009 | 0.313903993 | 0.874523798 | 0.957 |
| YDR162C | 515.4142839 | -0.060765051 | 0.875019293 | 0.957 |
| YFL028C | 788.0732688 | -0.054075853 | 0.874638302 | 0.957 |
| YHR051W | 2680.129991 | 0.187843736 | 0.87500478 | 0.957 |
| YMR236W | 1041.170124 | -0.047857618 | 0.87510266 | 0.957 |
| YPR008W | 1796.059535 | -0.003835172 | 0.87491355 | 0.957 |
| YNL048W | 1078.322808 | -0.017077856 | 0.876026448 | 0.958 |
| YHR057C | 1425.81956 | -0.12005935 | 0.876620287 | 0.958 |
| YBR107C | 343.2589721 | -0.008330059 | 0.881568823 | 0.959 |
| YCR015C | 255.0577346 | 0.077192734 | 0.879000111 | 0.959 |
| YCR065W | 756.8632472 | 0.153525596 | 0.881145691 | 0.959 |
| YDL069C | 126.889784 | 0.042333646 | 0.882052529 | 0.959 |

| | | | | |
|---|---|---|---|---|
| YDL240W | 734.1749038 | 0.058230824 | 0.881120994 | 0.959 |
| YDR032C | 4060.720659 | -0.099564345 | 0.877864155 | 0.959 |
| YDR177W | 1727.041157 | -0.017028698 | 0.881064845 | 0.959 |
| YDR345C | 12971.54144 | -0.096419783 | 0.881162183 | 0.959 |
| YDR532C | 597.6785104 | -0.004672572 | 0.882273803 | 0.959 |
| YER168C | 714.524667 | 0.029426682 | 0.877239256 | 0.959 |
| YER185W | 79.91884135 | -0.039923296 | 0.879497717 | 0.959 |
| YFR002W | 1555.318988 | -0.029037814 | 0.877434796 | 0.959 |
| YGL204C | 78.35386912 | 0.216713198 | 0.881381407 | 0.959 |
| YGR090W | 2037.446504 | 0.001427875 | 0.878182365 | 0.959 |
| YGR169C-A | 750.5850996 | -0.085795798 | 0.88079783 | 0.959 |
| YGR177C | 234.3513182 | -0.022785747 | 0.87808571 | 0.959 |
| YHR188C | 2845.286709 | 0.033639597 | 0.877631173 | 0.959 |
| YIL009C-A | 425.8589138 | -0.080635086 | 0.88070989 | 0.959 |
| YIL014C-A | 253.291496 | 0.047432174 | 0.878388308 | 0.959 |
| YJR073C | 2980.954697 | 0.015081485 | 0.882030641 | 0.959 |
| YJR080C | 1270.85858 | 0.245028466 | 0.882134099 | 0.959 |
| YKL089W | 415.1913401 | 0.043795976 | 0.879670862 | 0.959 |
| YKL114C | 901.3872837 | -0.059983465 | 0.880912298 | 0.959 |
| YKL146W | 1758.494435 | 0.010684876 | 0.880378991 | 0.959 |
| YLR371W | 1780.890302 | 0.011433565 | 0.881741939 | 0.959 |
| YMR052W | 219.8800281 | -0.093161802 | 0.882052926 | 0.959 |
| YMR286W | 654.5341281 | 0.027701185 | 0.879104754 | 0.959 |
| YMR297W | 8256.874804 | -0.026263141 | 0.881444646 | 0.959 |
| YNL154C | 2670.283788 | 0.040791871 | 0.879023658 | 0.959 |
| YNL165W | 500.9806009 | -0.176426448 | 0.880005831 | 0.959 |
| YNL294C | 1109.535055 | -0.031670106 | 0.879864274 | 0.959 |
| YPR191W | 6094.519828 | 0.25378464 | 0.880874603 | 0.959 |
| YER152C | 1264.809581 | -0.050046699 | 0.882704374 | 0.959 |
| YLR091W | 340.1364791 | -0.058050708 | 0.883009042 | 0.959 |
| YOL066C | 908.4756316 | -0.134526733 | 0.883258772 | 0.959 |
| YHR029C | 965.1862947 | -0.027153768 | 0.883757626 | 0.960 |
| YHL026C | 386.086378 | -0.1804595 | 0.884342144 | 0.960 |
| YBR237W | 937.0201345 | 0.001747636 | 0.884801909 | 0.960 |
| YFL026W | 6326.346816 | 0.261956292 | 0.884731013 | 0.960 |
| YCR033W | 2347.054478 | -0.079606381 | 0.885062012 | 0.960 |
| YDR111C | 552.8400061 | 0.069293689 | 0.885419398 | 0.960 |
| YMR089C | 2367.558674 | 0.008564033 | 0.885267486 | 0.960 |
| YOR356W | 1275.917226 | 0.060490815 | 0.885885624 | 0.961 |
| YBR251W | 1030.356421 | -0.033728938 | 0.88675622 | 0.961 |
| YIL102C-A | 112.0491515 | 0.022992725 | 0.886791173 | 0.961 |
| YBR291C | 379.9577466 | -0.204772881 | 0.887939975 | 0.962 |
| YDR108W | 495.5770673 | -0.073852669 | 0.887852376 | 0.962 |

| | | | | |
|---|---|---|---|---|
| YMR251W-A | 5878.547147 | -0.10593724 | 0.887817742 | 0.962 |
| YDL005C | 1599.265031 | 0.024077957 | 0.888225408 | 0.962 |
| YKR030W | 649.1186189 | -0.045326092 | 0.888785625 | 0.963 |
| YOR137C | 579.6398816 | -0.149882014 | 0.888915246 | 0.963 |
| YGR150C | 1213.396537 | 0.078257408 | 0.889267842 | 0.963 |
| YOR290C | 3907.230484 | 0.058332235 | 0.889603141 | 0.963 |
| YBR114W | 883.2430403 | -0.0613946 | 0.890542982 | 0.964 |
| YDR303C | 1032.483515 | 0.033377584 | 0.89039045 | 0.964 |
| YMR263W | 652.3046851 | -0.164880105 | 0.891327398 | 0.964 |
| YPL205C | 28.51967725 | 0.060227303 | 0.891385905 | 0.964 |
| YLR226W | 384.869465 | 0.02702741 | 0.891556997 | 0.964 |
| YBL047C | 5698.969568 | 0.041480509 | 0.892060978 | 0.964 |
| YER056C | 2408.22934 | 0.052949595 | 0.89190307 | 0.964 |
| YKL098W | 807.0315194 | -0.055292906 | 0.892522368 | 0.964 |
| YML053C | 803.1994724 | 0.035802945 | 0.892520967 | 0.964 |
| YOR316C | 1558.064909 | 0.014354092 | 0.892475281 | 0.964 |
| YNL288W | 932.0705303 | -0.045581885 | 0.892704167 | 0.964 |
| YMR034C | 377.1167506 | -0.199752069 | 0.893680863 | 0.965 |
| YDR298C | 3635.360191 | 0.114412191 | 0.893971714 | 0.965 |
| YFR033C | 3330.696407 | 0.132220222 | 0.894641358 | 0.965 |
| YJR114W | 33.45969567 | -0.064670102 | 0.894674957 | 0.965 |
| YLR086W | 1588.915329 | 0.064000308 | 0.894658514 | 0.965 |
| YMR001C | 1386.236499 | 0.054589215 | 0.894316708 | 0.965 |
| YMR255W | 1118.071772 | -0.097900878 | 0.894313845 | 0.965 |
| YJL085W | 484.172969 | 0.157952373 | 0.894985877 | 0.965 |
| YGL082W | 1508.755452 | -0.027281931 | 0.895980534 | 0.966 |
| YHL006W-A | 55.86611926 | 0.063719854 | 0.895756861 | 0.966 |
| YHR090C | 634.1320105 | -0.038796206 | 0.895980207 | 0.966 |
| YLR278C | 1940.968689 | 0.04115899 | 0.896019907 | 0.966 |
| YNL082W | 573.887894 | 0.125083203 | 0.895436668 | 0.966 |
| YBR264C | 359.5103772 | -0.005822478 | 0.896595641 | 0.966 |
| YCR024C-B | 2025.586967 | 0.001858351 | 0.896688597 | 0.966 |
| YNL282W | 259.9576363 | 0.001026167 | 0.896674816 | 0.966 |
| YPR101W | 539.0655902 | -0.095231436 | 0.896713209 | 0.966 |
| YBL025W | 212.6114748 | -0.00334269 | 0.897250094 | 0.966 |
| YDR117C | 1080.281382 | 0.061179959 | 0.897478581 | 0.966 |
| YER152W-A | 37.9606133 | -0.25126256 | 0.897113517 | 0.966 |
| YJL133W | 754.0202192 | 0.000417237 | 0.897448948 | 0.966 |
| YDL046W | 2942.234491 | 0.049189188 | 0.899566113 | 0.966 |
| YDR499W | 868.5785284 | -0.148907331 | 0.898361421 | 0.966 |
| YER134C | 346.3610386 | -0.021713709 | 0.899318174 | 0.966 |
| YFR005C | 788.6190865 | -0.038311978 | 0.898120207 | 0.966 |
| YGL094C | 1570.488055 | 0.047825662 | 0.897812052 | 0.966 |

| | | | | |
|---|---|---|---|---|
| YGR134W | 745.1524745 | 0.062182149 | 0.898930687 | 0.966 |
| YGR190C | 89.92851229 | 0.011980383 | 0.89950524 | 0.966 |
| YJR102C | 338.1234787 | 0.021554731 | 0.898579834 | 0.966 |
| YKL038W | 1738.382543 | 0.033591802 | 0.898850029 | 0.966 |
| YKL159C | 432.357669 | -0.011451082 | 0.899116126 | 0.966 |
| YNL183C | 1035.511832 | 0.002180179 | 0.898141522 | 0.966 |
| YNR073C | 195.2436335 | 0.031048078 | 0.899117431 | 0.966 |
| YGL140C | 1656.222341 | 0.021204655 | 0.89999043 | 0.966 |
| YIL047C | 1300.508261 | 0.03839659 | 0.900061318 | 0.966 |
| YMR193C-A | 123.8016037 | -0.122410685 | 0.900022022 | 0.966 |
| YER013W | 829.6032387 | -0.013208043 | 0.900943803 | 0.966 |
| YGL252C | 1693.94272 | 0.031258726 | 0.900501957 | 0.966 |
| YML120C | 4283.544511 | 0.221110273 | 0.900696982 | 0.966 |
| YNL177C | 1301.893277 | 0.012874374 | 0.900942966 | 0.966 |
| YCR082W | 1363.300313 | -0.06461285 | 0.902278303 | 0.966 |
| YEL059C-A | 318.8027126 | -0.047039363 | 0.901559792 | 0.966 |
| YER144C | 1434.76784 | -0.116272685 | 0.901798257 | 0.966 |
| YGR015C | 438.1021585 | -0.003016062 | 0.901405121 | 0.966 |
| YGR241C | 1428.572307 | -0.012326981 | 0.90202663 | 0.966 |
| YLR256W | 4392.10169 | 0.147933095 | 0.902331841 | 0.966 |
| YMR119W-A | 41.58140006 | 0.012350472 | 0.901979554 | 0.966 |
| YPR195C | 54.58354851 | 0.092680185 | 0.901286869 | 0.966 |
| YDR383C | 354.9421834 | -0.085606988 | 0.902627297 | 0.966 |
| YKL148C | 4538.261302 | 0.145409032 | 0.903104164 | 0.967 |
| YDR138W | 884.1934286 | 0.047000259 | 0.903437881 | 0.967 |
| YJL139C | 556.1081958 | 0.068939841 | 0.903589309 | 0.967 |
| YBR230W-A | 211.0787097 | -0.153658988 | 0.904611297 | 0.968 |
| YDL105W | 423.3140946 | -0.001105327 | 0.90458048 | 0.968 |
| YNL224C | 704.2866123 | 0.020486596 | 0.904900821 | 0.968 |
| YOL155C | 4950.84233 | -0.1056142 | 0.905505128 | 0.968 |
| YCL041C | 119.8839828 | -0.01194712 | 0.906117812 | 0.969 |
| YMR179W | 627.9998767 | 0.043431705 | 0.906078482 | 0.969 |
| YBL066C | 1237.560937 | -0.016101244 | 0.906381738 | 0.969 |
| YGR166W | 818.3285643 | -0.064666371 | 0.906637699 | 0.969 |
| YOR244W | 816.0935801 | 0.060251404 | 0.906737915 | 0.969 |
| YHR211W | 534.044984 | 0.052928973 | 0.907058049 | 0.969 |
| YDL019C | 2417.237505 | -0.058691933 | 0.907442022 | 0.969 |
| YFR004W | 2781.166019 | 0.014646916 | 0.907446667 | 0.969 |
| YDL181W | 1061.423493 | 0.259429269 | 0.907872231 | 0.969 |
| YBR049C | 1277.509775 | 0.087894965 | 0.90907599 | 0.969 |
| YDL132W | 2917.981172 | 0.0189735 | 0.910249626 | 0.969 |
| YDR036C | 1560.530814 | 0.074803688 | 0.909917608 | 0.969 |
| YDR107C | 540.536576 | 0.051907639 | 0.910126934 | 0.969 |

| | | | | |
|---|---|---|---|---|
| YDR261C-D | 88.68167414 | -0.10562774 | 0.909764881 | 0.969 |
| YGR092W | 957.140587 | 0.109077581 | 0.909296505 | 0.969 |
| YGR260W | 2070.398504 | 0.196995061 | 0.910414375 | 0.969 |
| YHR050W-A | 204.2549426 | 0.061306925 | 0.909382843 | 0.969 |
| YJR093C | 790.9035242 | -0.046492679 | 0.909507403 | 0.969 |
| YLL038C | 326.4334207 | -0.096882595 | 0.909897587 | 0.969 |
| YLR137W | 268.6657732 | -0.039061335 | 0.90864954 | 0.969 |
| YNL054W-B | 451.4424074 | -0.134755107 | 0.910392971 | 0.969 |
| YNL134C | 7013.757129 | 0.009154591 | 0.909808811 | 0.969 |
| YPL214C | 1002.445323 | -0.002817169 | 0.908991241 | 0.969 |
| YPR158W | 1022.854173 | 0.013018979 | 0.90941886 | 0.969 |
| YGR136W | 2019.817506 | -0.022764322 | 0.911332888 | 0.970 |
| YPL179W | 1527.082249 | 0.051247166 | 0.911190965 | 0.970 |
| YPL128C | 628.8955352 | 0.048521984 | 0.911748017 | 0.970 |
| YBR059C | 2706.219813 | 0.004643856 | 0.91259165 | 0.970 |
| YHR035W | 231.8124454 | 0.067101349 | 0.912376759 | 0.970 |
| YKL008C | 1304.945966 | 0.05833536 | 0.912807479 | 0.970 |
| YKL157W | 4702.190162 | 0.146832171 | 0.912832711 | 0.970 |
| YLR456W | 255.5161749 | -0.008123132 | 0.912407187 | 0.970 |
| YNL240C | 1001.196601 | -0.093351479 | 0.912789853 | 0.970 |
| YAL013W | 1274.881408 | -0.023151537 | 0.913032226 | 0.970 |
| YER124C | 2039.69695 | 0.14006124 | 0.913233664 | 0.970 |
| YDR316W | 792.8873428 | 0.102532267 | 0.914232988 | 0.971 |
| YLR387C | 1423.423086 | 0.027069623 | 0.914086339 | 0.971 |
| YLR084C | 1027.133398 | -0.011256105 | 0.914560047 | 0.971 |
| YMR141W-A | 54.52612508 | -0.229916808 | 0.914757452 | 0.971 |
| YLR218C | 291.942794 | -0.134900658 | 0.914995699 | 0.971 |
| YJR042W | 1389.72183 | 0.032221743 | 0.915702085 | 0.972 |
| YMR127C | 400.3588714 | -0.016398165 | 0.91573359 | 0.972 |
| YNL029C | 467.6043522 | -0.078755071 | 0.915588776 | 0.972 |
| YGL236C | 488.164276 | -0.043676018 | 0.915962516 | 0.972 |
| YCR095C | 483.8306058 | 0.02469799 | 0.916295425 | 0.972 |
| YAL020C | 521.9507996 | -0.09501134 | 0.919964871 | 0.972 |
| YBR077C | 472.5529414 | -0.059440335 | 0.920396048 | 0.972 |
| YBR125C | 746.1231571 | -0.04604736 | 0.921343428 | 0.972 |
| YBR170C | 1144.125576 | 0.02541627 | 0.920517347 | 0.972 |
| YBR177C | 2530.709586 | 0.098282194 | 0.916933422 | 0.972 |
| YDR078C | 147.0952169 | -0.16629189 | 0.921181004 | 0.972 |
| YER021W | 5518.8989 | 0.040258731 | 0.916956461 | 0.972 |
| YGL087C | 1504.49763 | 0.029936711 | 0.919321666 | 0.972 |
| YGL114W | 1125.03715 | -0.033769034 | 0.92111959 | 0.972 |
| YGR086C | 9907.562538 | 0.03740605 | 0.920954308 | 0.972 |
| YHR151C | 595.1830514 | -0.006120569 | 0.921357425 | 0.972 |

| | | | | |
|---|---|---|---|---|
| YHR173C | 88.05682976 | -0.091952419 | 0.919078047 | 0.972 |
| YHR214C-B | 235.6005636 | 0.23634659 | 0.917968171 | 0.972 |
| YKL184W | 1354.204344 | 0.03732006 | 0.919750139 | 0.972 |
| YKL185W | 2577.028628 | 0.103105528 | 0.921447483 | 0.972 |
| YLL009C | 581.7236454 | -0.00098126 | 0.921540945 | 0.972 |
| YLR315W | 96.87354306 | -0.101505925 | 0.921239801 | 0.972 |
| YLR353W | 907.9621406 | 0.058603651 | 0.920674919 | 0.972 |
| YLR385C | 52.50375543 | 0.006302574 | 0.91993738 | 0.972 |
| YML038C | 1118.066731 | -0.096883459 | 0.921333755 | 0.972 |
| YML065W | 967.920824 | -0.03275602 | 0.920873076 | 0.972 |
| YMR209C | 810.1729969 | 0.038889996 | 0.918382135 | 0.972 |
| YNL046W | 536.2842104 | -0.094078606 | 0.918960926 | 0.972 |
| YNR019W | 1395.465767 | 0.044748588 | 0.918084603 | 0.972 |
| YNR036C | 1583.864307 | 0.081599015 | 0.918538013 | 0.972 |
| YOL118C | 79.69627109 | -0.083549084 | 0.919288483 | 0.972 |
| YOR382W | 2866.519088 | 0.027195585 | 0.919078607 | 0.972 |
| YPR002W | 1192.998192 | -0.006840482 | 0.917309727 | 0.972 |
| YPR006C | 675.2062996 | -0.011055158 | 0.920540319 | 0.972 |
| YPR167C | 421.1211 | -0.060681441 | 0.919516329 | 0.972 |
| YGR153W | 193.310946 | -0.040328505 | 0.922274462 | 0.972 |
| YKL206C | 563.6605725 | -0.031760209 | 0.922214264 | 0.972 |
| YKL203C | 2057.606082 | 0.030739446 | 0.922964088 | 0.973 |
| YLR390W-A | 12426.05045 | 0.032944417 | 0.923740659 | 0.974 |
| YNL045W | 3993.621371 | 0.271456419 | 0.924311201 | 0.974 |
| YPR135W | 846.2041741 | 0.08275274 | 0.924594056 | 0.974 |
| YPL176C | 1288.392675 | -0.032135554 | 0.924772372 | 0.974 |
| YBR163W | 719.9181119 | -0.072899342 | 0.925282571 | 0.974 |
| YCR019W | 382.8804523 | 0.013958775 | 0.925607483 | 0.974 |
| YDR365C | 4849.022737 | 0.013206725 | 0.925803565 | 0.974 |
| YJR142W | 1091.921242 | -0.017068163 | 0.925857916 | 0.974 |
| YOR106W | 543.2030625 | -0.026624089 | 0.925984157 | 0.974 |
| YPL158C | 1350.795148 | -0.046049507 | 0.92590065 | 0.974 |
| YAL014C | 833.2543927 | -0.122939781 | 0.926393222 | 0.975 |
| YGL084C | 1084.484417 | 0.051583864 | 0.926561513 | 0.975 |
| YOR257W | 743.3095021 | -0.029104549 | 0.926915858 | 0.975 |
| YKL066W | 1252.501288 | 0.075401437 | 0.927515848 | 0.975 |
| YML071C | 1055.622411 | 0.00143894 | 0.928678849 | 0.976 |
| YMR267W | 919.8795189 | 0.077716263 | 0.92863005 | 0.976 |
| YDR110W | 523.1803163 | -0.08775809 | 0.929036623 | 0.976 |
| YFR036W | 228.7837864 | 0.032658432 | 0.929399397 | 0.977 |
| YPL221W | 1548.403188 | -0.100386906 | 0.929695621 | 0.977 |
| YAL058W | 573.3781549 | 0.108737239 | 0.930894405 | 0.977 |
| YBL012C | 41.44037077 | -0.030748937 | 0.930313442 | 0.977 |

| | | | | |
|---|---|---|---|---|
| YCR097W | 1013.079937 | 0.198208701 | 0.930637388 | 0.977 |
| YDL171C | 4109.615185 | 0.125562499 | 0.931320725 | 0.977 |
| YGL159W | 407.2189918 | 0.081179035 | 0.930883416 | 0.977 |
| YIL030C | 2092.978365 | 0.01715064 | 0.931559398 | 0.977 |
| YJL088W | 187.9199165 | -0.313458704 | 0.931499766 | 0.977 |
| YKR028W | 1923.731148 | 0.026279179 | 0.931542137 | 0.977 |
| YLR316C | 329.0837142 | -0.015979496 | 0.93112252 | 0.977 |
| snR84 | 162.4042194 | 0.093146856 | 0.930376034 | 0.977 |
| YHR078W | 1137.159604 | 0.049630151 | 0.931733611 | 0.977 |
| YCR025C | 44.00460202 | -0.226890068 | 0.933586183 | 0.979 |
| YGL071W | 1392.981634 | -0.004720032 | 0.933750835 | 0.979 |
| YJL105W | 247.8507967 | -0.099111799 | 0.933905056 | 0.979 |
| YOR297C | 445.9780063 | 0.044463151 | 0.93351564 | 0.979 |
| YPR113W | 3675.90658 | 0.033528267 | 0.934248332 | 0.979 |
| YOL060C | 2188.941543 | -0.006228938 | 0.934613031 | 0.979 |
| YER142C | 720.8941802 | 0.000536095 | 0.935017212 | 0.979 |
| YJR109C | 3577.314205 | -0.1824776 | 0.935139331 | 0.979 |
| YPR126C | 82.68269767 | -0.158973836 | 0.93490053 | 0.979 |
| YBR184W | 226.9887473 | 0.008183254 | 0.935974352 | 0.979 |
| YCL074W | 140.2978941 | 0.191186251 | 0.937098641 | 0.979 |
| YFR029W | 834.7381381 | -0.154441862 | 0.936766387 | 0.979 |
| YLR010C | 127.8600135 | -0.098360637 | 0.936346013 | 0.979 |
| YNL035C | 765.5856918 | 0.026064048 | 0.936566542 | 0.979 |
| YOL069W | 510.0668666 | 0.108113803 | 0.936805082 | 0.979 |
| YOL143C | 5652.711754 | 0.122360932 | 0.935889717 | 0.979 |
| YPL045W | 887.7248714 | 0.017694361 | 0.93614666 | 0.979 |
| YPL207W | 1844.153383 | -0.093371888 | 0.937012763 | 0.979 |
| YGR206W | 264.0711736 | -0.000433928 | 0.937439333 | 0.980 |
| YOR324C | 520.2886814 | 0.026120792 | 0.938323259 | 0.980 |
| YAR014C | 1593.739874 | 0.04413511 | 0.939031812 | 0.981 |
| YDR409W | 969.523209 | 0.004473631 | 0.939291052 | 0.981 |
| YER008C | 2582.281632 | -0.086372339 | 0.939303504 | 0.981 |
| YPR189W | 1450.192608 | 0.041909498 | 0.939415781 | 0.981 |
| YDR147W | 507.4257469 | 0.046517837 | 0.940351146 | 0.981 |
| YDR407C | 1886.152895 | 0.011234202 | 0.940212495 | 0.981 |
| YDR528W | 366.4782046 | -0.061557886 | 0.940244387 | 0.981 |
| YNL245C | 408.0242862 | -0.03211005 | 0.940224087 | 0.981 |
| YBR256C | 1957.968457 | -0.0879274 | 0.940719116 | 0.981 |
| YLR389C | 2863.752537 | 0.026284721 | 0.940755061 | 0.981 |
| YGL032C | 1976.542723 | 0.064456591 | 0.941085235 | 0.981 |
| YPL092W | 1197.47636 | -0.08502543 | 0.941104777 | 0.981 |
| YGR040W | 494.1182396 | 0.1550474 | 0.941503188 | 0.981 |
| YDR103W | 991.2634766 | -0.151156353 | 0.942270298 | 0.982 |

| | | | | |
|---|---|---|---|---|
| YMR188C | 1203.74754 | -0.046986702 | 0.94214935 | 0.982 |
| YFR027W | 221.6537022 | -0.041769572 | 0.942981005 | 0.982 |
| YJR112W-A | 356.5088081 | -0.008625936 | 0.943573269 | 0.983 |
| YMR193W | 776.035786 | 0.097234963 | 0.943609578 | 0.983 |
| YPR119W | 682.8278436 | 0.132008173 | 0.943386804 | 0.983 |
| YDR329C | 1759.988143 | -0.044134278 | 0.944614744 | 0.983 |
| YGR242W | 37.21832003 | 0.040770724 | 0.944539735 | 0.983 |
| YJL087C | 474.7532502 | 0.094134199 | 0.943888394 | 0.983 |
| YMR253C | 606.2113411 | -0.008879719 | 0.944236194 | 0.983 |
| YMR281W | 530.4325943 | -0.003195679 | 0.944641094 | 0.983 |
| YOL113W | 1261.758159 | -0.220395136 | 0.944661931 | 0.983 |
| YNL229C | 1349.306513 | -0.007811105 | 0.94490323 | 0.983 |
| YDR348C | 1537.308452 | 0.016070611 | 0.945586895 | 0.983 |
| YPR046W | 225.601515 | -0.123839756 | 0.94545597 | 0.983 |
| YDR518W | 1466.448011 | 0.105590864 | 0.946537845 | 0.984 |
| YMR061W | 1751.530821 | 0.026467008 | 0.947016685 | 0.984 |
| YMR190C | 1250.870637 | -0.005942291 | 0.946960012 | 0.984 |
| YPL135W | 1241.708085 | 0.032363961 | 0.947098732 | 0.984 |
| YKL092C | 887.6693114 | 0.025803591 | 0.947424503 | 0.984 |
| YDR079C-A | 102.409036 | -0.0711268 | 0.947646589 | 0.984 |
| YJL130C | 21427.36188 | 0.450031774 | 0.948702483 | 0.985 |
| YNL024C | 148.4866774 | -0.0777572 | 0.948551529 | 0.985 |
| YHR084W | 1211.807628 | 0.102944927 | 0.94932687 | 0.985 |
| YML006C | 696.4182875 | -0.02161148 | 0.949273421 | 0.985 |
| YOL145C | 2673.103928 | -0.028870065 | 0.949441687 | 0.985 |
| YML079W | 1250.560699 | -0.004988714 | 0.949650403 | 0.985 |
| YER044C | 1096.772117 | 0.098115271 | 0.949925171 | 0.985 |
| YIL085C | 1498.284283 | -0.03588218 | 0.95000432 | 0.985 |
| YHR036W | 340.3963184 | 0.025126688 | 0.950511617 | 0.986 |
| YDR360W | 36.19514521 | 0.255285252 | 0.951015071 | 0.986 |
| YMR040W | 194.5584091 | 0.02073702 | 0.951197991 | 0.986 |
| YBR004C | 976.5395701 | -0.060317102 | 0.952058495 | 0.986 |
| YCR060W | 437.7609902 | 0.035557167 | 0.953251299 | 0.986 |
| YDL010W | 362.0027109 | -0.015222693 | 0.953025496 | 0.986 |
| YDR260C | 314.9401976 | -0.061194216 | 0.952480711 | 0.986 |
| YGR246C | 1106.975461 | 0.044422347 | 0.9525067 | 0.986 |
| YHR067W | 339.2689826 | 0.092710343 | 0.952795209 | 0.986 |
| YLL004W | 381.8327061 | 0.031805006 | 0.952202087 | 0.986 |
| YLR033W | 1049.258607 | 0.035330707 | 0.952225079 | 0.986 |
| YMR168C | 528.3657431 | 0.041867776 | 0.953268833 | 0.986 |
| YPL046C | 139.4253961 | -0.016983671 | 0.952618426 | 0.986 |
| YBR119W | 605.7970902 | -0.015798265 | 0.953618187 | 0.987 |
| YLR114C | 1321.213644 | 0.092163387 | 0.953956779 | 0.987 |

| | | | | |
|---|---|---|---|---|
| YBL043W | 324.5838638 | -0.050064293 | 0.95414063 | 0.987 |
| YGR165W | 998.5420975 | 0.030489221 | 0.955366042 | 0.988 |
| YPR040W | 901.3829077 | -0.055307969 | 0.955410976 | 0.988 |
| YER153C | 351.5393272 | -0.163769664 | 0.955767456 | 0.988 |
| YPR120C | 354.4464034 | -0.019121994 | 0.955788467 | 0.988 |
| YLR181C | 881.938971 | -0.007965424 | 0.956292591 | 0.988 |
| YNL242W | 1398.740797 | 0.023194875 | 0.956379845 | 0.988 |
| YBR212W | 2837.682435 | -0.103323274 | 0.957576487 | 0.988 |
| YJL172W | 1932.176519 | 0.198046493 | 0.957405532 | 0.988 |
| YLR305C | 2169.127998 | 0.076531539 | 0.957677878 | 0.988 |
| YMR067C | 1220.387108 | 0.020641383 | 0.957475317 | 0.988 |
| YMR069W | 79.44065746 | 0.168384683 | 0.957729475 | 0.988 |
| YNL322C | 3831.740608 | 0.026629169 | 0.957141629 | 0.988 |
| YPL065W | 569.9929052 | -0.01377404 | 0.957275996 | 0.988 |
| YKL132C | 214.3347933 | -0.022089892 | 0.958457592 | 0.989 |
| YLR015W | 562.8104697 | 0.035058601 | 0.958358016 | 0.989 |
| YMR173W-A | 260.3952806 | 0.058369172 | 0.958545639 | 0.989 |
| YBR172C | 2003.113678 | -0.012992465 | 0.960622046 | 0.989 |
| YDR175C | 1042.356621 | 0.049081908 | 0.95984419 | 0.989 |
| YER028C | 507.0764562 | -0.24266014 | 0.96057619 | 0.989 |
| YGL069C | 85.63194987 | -0.021183636 | 0.960710288 | 0.989 |
| YGL116W | 933.3944118 | -0.007480467 | 0.959728968 | 0.989 |
| YLL048C | 4130.734576 | 0.027132145 | 0.960012328 | 0.989 |
| YLR069C | 1475.235001 | 0.062182254 | 0.960054972 | 0.989 |
| YMR177W | 436.2483597 | 0.020635179 | 0.959780357 | 0.989 |
| YOR064C | 481.1143371 | -0.03559587 | 0.959267978 | 0.989 |
| YPL134C | 1579.391088 | -0.00947235 | 0.960695031 | 0.989 |
| YLR418C | 850.5544851 | 0.054594861 | 0.961120092 | 0.989 |
| YBR074W | 1213.234406 | 0.004767679 | 0.961594764 | 0.989 |
| YJR146W | 99.77471099 | 0.037741713 | 0.961526725 | 0.989 |
| YHL014C | 453.0111576 | 0.060415905 | 0.962147692 | 0.990 |
| YMR256C | 1012.905586 | -0.001853627 | 0.962550892 | 0.990 |
| YOR172W | 824.7713828 | 0.013903463 | 0.962455854 | 0.990 |
| YDR460W | 574.0402083 | -0.022608977 | 0.962976757 | 0.990 |
| YIL088C | 2503.905976 | 0.090023277 | 0.962960765 | 0.990 |
| YBR093C | 949.3588178 | -0.384277335 | 0.963623476 | 0.990 |
| YDR322W | 1031.616879 | 0.119689823 | 0.963707874 | 0.990 |
| YNL103W-A | 104.3621644 | 0.482234987 | 0.963334293 | 0.990 |
| YOL035C | 58.4740954 | -0.2274574 | 0.9637897 | 0.990 |
| YPR105C | 1243.404818 | -0.029719974 | 0.963992686 | 0.990 |
| YPR204W | 106.9528255 | 0.291263532 | 0.964074416 | 0.990 |
| YHR150W | 670.7977171 | -0.117649547 | 0.964831866 | 0.990 |
| YLR395C | 1418.063046 | 0.152298474 | 0.964735076 | 0.990 |

| | | | | |
|---|---|---|---|---|
| YHR030C | 2133.377408 | -0.076031873 | 0.965079201 | 0.991 |
| YLR256W-A | 92.61969338 | 0.168931912 | 0.965426749 | 0.991 |
| YCR107W | 236.9080811 | 0.095007049 | 0.965988031 | 0.991 |
| YGR075C | 302.7071594 | -0.012789758 | 0.966148554 | 0.991 |
| YNL047C | 720.3103825 | -0.135529035 | 0.965859095 | 0.991 |
| HRA1 | 34.22771229 | 0.208944226 | 0.971769881 | 0.991 |
| YBL075C | 2650.065607 | -0.181346951 | 0.970751893 | 0.991 |
| YBL105C | 2392.516684 | 0.052693791 | 0.971424762 | 0.991 |
| YBR006W | 1760.927773 | -0.104401666 | 0.967230301 | 0.991 |
| YCL033C | 876.4314652 | -0.10303194 | 0.970399648 | 0.991 |
| YDL188C | 1068.696328 | 0.070409394 | 0.967873714 | 0.991 |
| YDR073W | 640.4950618 | -0.117887444 | 0.971699255 | 0.991 |
| YDR259C | 452.5952789 | -0.027102249 | 0.967840329 | 0.991 |
| YDR301W | 1341.743029 | 0.045094495 | 0.969429557 | 0.991 |
| YDR318W | 508.7245087 | 0.012890528 | 0.968247815 | 0.991 |
| YDR501W | 295.5069374 | 0.005946287 | 0.971283196 | 0.991 |
| YEL015W | 1435.206957 | 0.004669436 | 0.967450912 | 0.991 |
| YER046W | 177.0920564 | -0.119778298 | 0.971794971 | 0.991 |
| YER130C | 1003.132192 | 0.116352063 | 0.968625881 | 0.991 |
| YER155C | 4972.789594 | 0.083879678 | 0.971036586 | 0.991 |
| YGL016W | 794.4716524 | 0.12966152 | 0.969598529 | 0.991 |
| YGR097W | 2251.701342 | -0.102635752 | 0.969040694 | 0.991 |
| YGR231C | 3688.450165 | -0.000498097 | 0.970521047 | 0.991 |
| YHR158C | 1689.928736 | 0.011575983 | 0.970760628 | 0.991 |
| YIL030W-A | 74.85518125 | -0.211660842 | 0.968257262 | 0.991 |
| YJL025W | 374.9847479 | -0.134951132 | 0.969653645 | 0.991 |
| YJL128C | 1506.583064 | 0.001141188 | 0.968097149 | 0.991 |
| YJR019C | 596.1541169 | -0.078700939 | 0.968820792 | 0.991 |
| YJR027W | 207.2279874 | 0.235499315 | 0.966896286 | 0.991 |
| YJR126C | 1827.592541 | -0.019596113 | 0.969787728 | 0.991 |
| YLR088W | 986.2933263 | -0.001708599 | 0.971033474 | 0.991 |
| YLR190W | 1584.336567 | 0.022864507 | 0.9669084 | 0.991 |
| YML129C | 536.8329539 | 0.016196677 | 0.969332429 | 0.991 |
| YNL333W | 85.09307581 | 0.054785577 | 0.96702375 | 0.991 |
| YNR062C | 190.9739726 | 0.011831959 | 0.969024323 | 0.991 |
| YOR009W | 307.6360112 | -0.355872489 | 0.970385974 | 0.991 |
| YOR065W | 3707.159237 | 0.25078321 | 0.971079295 | 0.991 |
| YPR139C | 1065.385559 | 0.059915445 | 0.971112289 | 0.991 |
| YIL156W-B | 291.2591429 | -0.021163774 | 0.972332575 | 0.991 |
| YBL017C | 3980.2423 | -0.015006865 | 0.972533226 | 0.991 |
| YBR056W-A | 43.21007303 | -0.315734873 | 0.973171305 | 0.992 |
| YDR150W | 4371.450407 | 0.048103148 | 0.97330182 | 0.992 |
| YLR444C | 25.68564702 | -0.540653616 | 0.973229743 | 0.992 |

| | | | | |
|---|---|---|---|---|
| YOL051W | 2569.23447 | 0.00271827 | 0.973308318 | 0.992 |
| YIL125W | 5872.779859 | 0.228552563 | 0.973828197 | 0.992 |
| YIL112W | 2132.63932 | -0.017046483 | 0.974153859 | 0.992 |
| YOR013W | 86.58484926 | 0.048328058 | 0.974527319 | 0.992 |
| YOR313C | 174.7411276 | 0.089837353 | 0.974580241 | 0.992 |
| YJL212C | 1268.474729 | -0.207003336 | 0.974920281 | 0.992 |
| YOL026C | 840.8767019 | -0.081808831 | 0.974785803 | 0.992 |
| YGL129C | 1219.243345 | 0.047972978 | 0.975581609 | 0.992 |
| YNL004W | 1614.267096 | 0.016161932 | 0.975449948 | 0.992 |
| YOL067C | 283.6975055 | -0.044717254 | 0.975530662 | 0.992 |
| YMR226C | 4108.834639 | -0.030957902 | 0.975762179 | 0.992 |
| YHR018C | 3547.596925 | -0.004650971 | 0.976189723 | 0.992 |
| YOR212W | 1199.500024 | 0.044893157 | 0.976173755 | 0.992 |
| YPL242C | 2035.672722 | 0.019618649 | 0.976340816 | 0.992 |
| YBL069W | 640.7564823 | -0.035990883 | 0.977401167 | 0.993 |
| YBR186W | 272.8183513 | 0.054499895 | 0.97695851 | 0.993 |
| YDR052C | 1279.720929 | 0.03703996 | 0.977640498 | 0.993 |
| YML011C | 267.8459983 | -0.01644138 | 0.977440034 | 0.993 |
| YOL008W | 381.7473344 | 0.009320928 | 0.977675914 | 0.993 |
| YPL002C | 492.3419627 | -0.041121522 | 0.977071312 | 0.993 |
| YKR016W | 2416.547829 | 0.160735241 | 0.97803504 | 0.993 |
| YML058W-A | 99.58508334 | -0.313655763 | 0.978512269 | 0.993 |
| YLR237W | 1061.722449 | -0.207059511 | 0.978958677 | 0.994 |
| YLR390W | 418.8392228 | 0.013281712 | 0.979365553 | 0.994 |
| YDR437W | 89.57715872 | 0.135997077 | 0.980423125 | 0.994 |
| YGR021W | 818.8357092 | -0.016027331 | 0.980143922 | 0.994 |
| YPL271W | 1376.293801 | 0.050858174 | 0.980458029 | 0.994 |
| YLR014C | 353.4875684 | -0.03724175 | 0.981163679 | 0.995 |
| YBL015W | 5485.33881 | 0.088033148 | 0.981747667 | 0.995 |
| YGR287C | 287.959068 | -0.146817147 | 0.981839997 | 0.995 |
| YDR265W | 518.8074155 | -0.037917869 | 0.982896507 | 0.995 |
| YGR266W | 1152.085768 | -0.062196221 | 0.982315837 | 0.995 |
| YLR108C | 407.9615838 | -0.187141109 | 0.982567636 | 0.995 |
| YNL272C | 1304.645862 | 0.031810585 | 0.982531337 | 0.995 |
| YPL020C | 1208.520388 | 0.00541833 | 0.982879232 | 0.995 |
| YHR079C-A | 27.69347479 | -0.191667784 | 0.983168049 | 0.996 |
| YBL016W | 1090.490568 | 0.105496037 | 0.98360424 | 0.996 |
| YBR126W-B | 156.4452467 | 0.002975369 | 0.98361246 | 0.996 |
| YER014W | 690.0676914 | 0.015556833 | 0.98504665 | 0.996 |
| YFR040W | 3140.971961 | 0.054351816 | 0.985085976 | 0.996 |
| YGL083W | 940.5046339 | 0.025443575 | 0.985790682 | 0.996 |
| YGL255W | 1201.748656 | -0.404320138 | 0.983894522 | 0.996 |
| YGR044C | 1106.237884 | 0.037466764 | 0.98596782 | 0.996 |

| | | | | |
|---|---|---|---|---|
| YGR047C | 884.2267464 | -0.03110579 | 0.984312499 | 0.996 |
| YGR276C | 1041.106244 | 0.000168983 | 0.985053044 | 0.996 |
| YGR286C | 924.5538048 | -0.069071054 | 0.984787067 | 0.996 |
| YJL044C | 1379.126824 | -0.001719594 | 0.984743214 | 0.996 |
| YJR054W | 539.7820263 | 0.016861634 | 0.986105098 | 0.996 |
| YKL116C | 778.7224786 | -0.00455623 | 0.985592868 | 0.996 |
| YLR439W | 1150.159115 | 0.068433287 | 0.984750088 | 0.996 |
| YMR058W | 5542.022572 | 0.147309024 | 0.985894317 | 0.996 |
| YOR230W | 8762.727513 | 0.07481696 | 0.985565202 | 0.996 |
| YPR094W | 154.1332011 | -0.032892367 | 0.985737238 | 0.996 |
| YAR073W | 171.304609 | -0.027607464 | 0.986466247 | 0.996 |
| YBR120C | 395.0224857 | 0.071229505 | 0.986325087 | 0.996 |
| YFR006W | 3007.667423 | 0.046324223 | 0.986631346 | 0.996 |
| YJL039C | 2015.018256 | 0.109353278 | 0.987057824 | 0.996 |
| YOR318C | 25.32067042 | 0.072043645 | 0.986987927 | 0.996 |
| YDL070W | 4583.373904 | -0.112315632 | 0.987825682 | 0.996 |
| YDR406W | 3009.969096 | 0.023822922 | 0.987877531 | 0.996 |
| YDR527W | 984.8264926 | -0.055068756 | 0.987527049 | 0.996 |
| YKL106W | 462.66436 | -0.040315256 | 0.98771277 | 0.996 |
| YFL046W | 596.7158239 | 0.021555181 | 0.98845595 | 0.996 |
| YLR210W | 386.6683639 | 0.013231255 | 0.988377787 | 0.996 |
| YPL072W | 410.1680408 | 0.064351523 | 0.98841869 | 0.996 |
| YBR016W | 1677.943304 | -0.084621528 | 0.989750374 | 0.996 |
| YBR293W | 445.7090525 | -0.210076196 | 0.989826718 | 0.996 |
| YIL004C | 332.265247 | -0.020754792 | 0.989093172 | 0.996 |
| YIL157C | 795.6414589 | 0.058682582 | 0.989565458 | 0.996 |
| YJL077C | 162.6090211 | -0.160650467 | 0.989302226 | 0.996 |
| YML062C | 782.2227983 | -0.062505167 | 0.989689495 | 0.996 |
| YOR154W | 667.1567549 | -0.016023765 | 0.989675849 | 0.996 |
| YBL001C | 1539.933777 | -0.078869928 | 0.990392216 | 0.996 |
| YNL055C | 20028.30046 | 0.083487407 | 0.990467567 | 0.996 |
| YOL004W | 4094.946984 | 0.017955981 | 0.990576012 | 0.996 |
| YBL046W | 691.3216368 | -0.063319646 | 0.99108339 | 0.997 |
| YFL013W-A | 84.24870204 | -0.119720373 | 0.991307725 | 0.997 |
| YGR016W | 101.0221921 | -0.044691905 | 0.991754706 | 0.997 |
| YOR168W | 3054.619284 | 0.048021655 | 0.991686432 | 0.997 |
| YHR143W | 3407.641358 | 0.105789093 | 0.992301159 | 0.997 |
| YPL118W | 2044.672532 | 0.047325335 | 0.992655622 | 0.997 |
| YGR240C-A | 46.68747277 | 0.053308576 | 0.993114538 | 0.998 |
| YIL156W | 1109.423201 | 0.017789807 | 0.993206362 | 0.998 |
| YER010C | 1015.031336 | 0.037697229 | 0.993785057 | 0.998 |
| YER060W | 750.1023347 | 0.002215568 | 0.993710798 | 0.998 |
| YMR045C | 326.5494461 | -0.050534266 | 0.994074216 | 0.998 |

| | | | | |
|---|---|---|---|---|
| YOL031C | 1089.984363 | 0.003140059 | 0.99399765 | 0.998 |
| YPR145C-A | 45.3897147 | -0.339405081 | 0.993401632 | 0.998 |
| YER167W | 1421.029082 | -0.031112631 | 0.995109411 | 0.999 |
| YIL090W | 898.4488003 | -0.011932824 | 0.995520182 | 0.999 |
| YNL068C | 1421.244951 | -0.011105884 | 0.995656896 | 0.999 |
| YDL197C | 997.8504474 | 0.02893161 | 0.996254999 | 0.999 |
| YDR322C-A | 892.3853165 | -0.019391376 | 0.996405061 | 0.999 |
| YFL010C | 2890.139798 | 0.002206209 | 0.996470242 | 0.999 |
| YHR116W | 348.981213 | -0.034949157 | 0.996851362 | 0.999 |
| YOL058W | 2645.481694 | -0.439495247 | 0.996876889 | 0.999 |
| YML116W | 562.6593031 | -0.160383765 | 0.997257257 | 0.999 |
| YGL187C | 1692.180783 | 0.223471312 | 0.997473441 | 0.999 |
| YBL085W | 2143.648048 | -0.060617484 | 0.99778376 | 0.999 |
| YAL021C | 1595.906288 | -0.006046143 | 0.998674477 | 1.000 |
| YHR190W | 5071.299505 | -0.045688146 | 0.99820102 | 1.000 |
| YIL124W | 2615.425395 | 0.010003404 | 0.998816522 | 1.000 |
| YJR011C | 218.4017674 | -0.009993776 | 0.998596526 | 1.000 |
| YKL032C | 3133.578964 | -0.031314027 | 0.998284632 | 1.000 |
| YNL171C | 59.8874962 | -0.008578763 | 0.998829024 | 1.000 |
| YGL064C | 383.7385393 | 0.029127112 | 0.999141463 | 1.000 |
| YOL017W | 430.0355777 | 0.048713049 | 0.999276146 | 1.000 |
| YMR027W | 2670.4492 | 0.057819395 | 0.999595156 | 1.000 |
| YER017C | 1876.346183 | -0.011889773 | 0.99993368 | 1.000 |
| RNA170 | 10.42132142 | -0.351051747 | 0.72638896 | NA |
| RUF22 | 1.126113591 | 0.043173889 | 0.994983957 | NA |
| RUF23 | 8.975541829 | -0.185574537 | 0.788634424 | NA |
| YAL016C-B | 15.45888685 | -0.253729888 | 0.597579293 | NA |
| YAL019W-A | 5.164115052 | -0.982911729 | 0.058314117 | NA |
| YAL034C-B | 9.086766243 | -0.615531881 | 0.209275992 | NA |
| YAL037C-A | 1.195205434 | -0.163850184 | 0.615791904 | NA |
| YAL042C-A | 8.764234173 | -1.181418071 | 0.361834947 | NA |
| YAL045C | 8.89025023 | -0.642175907 | 0.884039015 | NA |
| YAL047W-A | 1.181246705 | 0.35656455 | 0.994454939 | NA |
| YAL056C-A | 23.07572137 | -0.415367818 | 0.196641362 | NA |
| YAL062W | 2248.194645 | 0.176343181 | NA | NA |
| YAL063C-A | 8.265236292 | 0.967065837 | 0.084549805 | NA |
| YAL064W | 3.131576444 | 0.326562581 | 0.254653625 | NA |
| YAL064W-B | 18.20372016 | 0.145821251 | 0.900723589 | NA |
| YAL065C | 13.48444795 | 0.475363514 | 0.921586979 | NA |
| YAL066W | 0.770254238 | 0.069358591 | 0.837505731 | NA |
| YAL067W-A | 0.291097411 | 0.009560724 | 0.872979907 | NA |
| YAL068C | 5.759798843 | 0.315362792 | 0.45204467 | NA |
| YAL068W-A | 0.241654238 | 0.011896906 | 0.894258047 | NA |

| | | | | |
|---|---|---|---|---|
| YAL069W | 0.918104495 | -0.001996195 | 0.782721806 | NA |
| YAR019W-A | 5.564052438 | -0.372453506 | 0.720035989 | NA |
| YAR035C-A | 1.313184395 | 0.735780154 | 0.044613738 | NA |
| YAR047C | 21.66468268 | -0.119740274 | 0.407763216 | NA |
| YAR053W | 9.523462605 | 0.321295193 | 0.321138403 | NA |
| YAR064W | 14.8431874 | 0.802068641 | 0.62703459 | NA |
| YAR066W | 12.65917974 | 0.683252707 | 0.590819835 | NA |
| YAR068W | 4.842260677 | -0.178077836 | 0.97166005 | NA |
| YAR070C | 0.925768202 | 0.587677396 | 0.275875956 | NA |
| YBL008W-A | 2.748531795 | 0.016756912 | 0.947789422 | NA |
| YBL044W | 3.828780377 | -0.243226065 | 0.977884037 | NA |
| YBL053W | 19.02715806 | -0.354948166 | 0.660252532 | NA |
| YBL062W | 0.924462825 | -0.138634173 | 0.606791315 | NA |
| YBL070C | 21.83445099 | -0.744011329 | 0.192294598 | NA |
| YBL071C-B | 1.645198606 | -0.531675187 | 0.496779132 | NA |
| YBL073W | 6.010181019 | 0.752415817 | 0.394835684 | NA |
| YBL083C | 7.154873991 | -0.608734831 | 0.38849853 | NA |
| YBL094C | 5.123149382 | -0.149832084 | 0.90986303 | NA |
| YBL096C | 11.76381799 | 0.057426408 | 0.921930143 | NA |
| YBL100W-C | 6.925265405 | 0.7639059 | 0.1021964 | NA |
| YBL108W | 0.561379936 | 0.280528893 | 0.410443352 | NA |
| YBL109W | 16.60491535 | 0.561479175 | 0.116624093 | NA |
| YBR027C | 8.649310737 | -0.093833207 | 0.357063603 | NA |
| YBR051W | 12.61851356 | 0.580970669 | 0.187750532 | NA |
| YBR064W | 3.158811985 | -0.324887015 | 0.621819477 | NA |
| YBR072C-A | 2.769886284 | -0.174438409 | 0.39057898 | NA |
| YBR089W | 6.735755394 | -0.001758136 | 0.980943245 | NA |
| YBR103C-A | 1.570441558 | -0.24227972 | 0.356768432 | NA |
| YBR109W-A | 20.49364363 | 0.33031392 | 0.538234844 | NA |
| YBR124W | 16.74512171 | 0.500479058 | 0.329996184 | NA |
| YBR144C | 8.772859821 | -0.430937543 | 0.862849427 | NA |
| YBR174C | 5.135026563 | -0.770506045 | 0.087698029 | NA |
| YBR178W | 20.74423611 | -0.116206463 | 0.496038866 | NA |
| YBR182C-A | 7.795727576 | -0.140775483 | 0.93846029 | NA |
| YBR190W | 3.930160671 | 0.02547174 | 0.87801163 | NA |
| YBR196C-A | 7.414882286 | -0.540384043 | 0.895615452 | NA |
| YBR196C-B | 11.09747944 | -0.280117695 | 0.585408016 | NA |
| YBR200W-A | 19.77475914 | 0.277972656 | 0.780501749 | NA |
| YBR209W | 5.517408049 | -0.072597358 | 0.6985991 | NA |
| YBR219C | 1.059885032 | -0.225840353 | 0.347126308 | NA |
| YBR221W-A | 0.837044541 | 0.210968947 | 0.365906315 | NA |
| YBR232C | 24.08316402 | 0.053342811 | 0.844592879 | NA |
| YBR277C | 4.283921599 | -0.15690423 | 0.345141983 | NA |

| | | | | |
|---|---|---|---|---|
| YBR292C | 15.20409235 | -0.035535319 | 0.838407512 | NA |
| YBR296C-A | 3.00001637 | -0.222161284 | 0.865001179 | NA |
| YBR298C-A | 3.08929577 | 0.747412687 | 0.162945458 | NA |
| YBR300C | 5.482970764 | -1.150576961 | 0.013139404 | NA |
| YBR301W | 22.99326035 | 0.15343423 | 0.604600494 | NA |
| YCL023C | 8.286348364 | 0.362385114 | 0.355567343 | NA |
| YCL046W | 22.38513549 | -0.033942238 | 0.471016753 | NA |
| YCL065W | 1.135078196 | 0.000887478 | 0.821572313 | NA |
| YCL068C | 4.465179414 | 0.000533375 | 0.91092476 | NA |
| YCL076W | 24.7210401 | 0.133696957 | 0.653095295 | NA |
| YCR018C-A | 8.72999001 | -0.938359559 | 0.243972315 | NA |
| YCR022C | 20.80731429 | 0.037142699 | 0.508922321 | NA |
| YCR041W | 16.4079087 | -0.244488427 | 0.380200553 | NA |
| YCR049C | 7.558405616 | -0.522414948 | 0.623077472 | NA |
| YCR050C | 17.22525728 | -0.219653402 | 0.988477738 | NA |
| YCR064C | 12.51518247 | -0.453535337 | 0.171676895 | NA |
| YCR085W | 15.93952821 | -0.603139396 | 0.081454888 | NA |
| YCR087W | 24.15940244 | -1.017500037 | 0.139946374 | NA |
| YCR095W-A | 20.07196858 | 1.154443528 | 0.008356112 | NA |
| YCR097W-A | 1.60235674 | 0.012184964 | 0.833766356 | NA |
| YCR102W-A | 9.4716423 | -0.104071409 | 0.686027403 | NA |
| YCR104W | 19.10661275 | -0.691160066 | 0.343327571 | NA |
| YCR108C | 1.912873177 | 0.000446067 | 0.868075289 | NA |
| YDL007C-A | 3.853033258 | -0.068308981 | 0.708270338 | NA |
| YDL011C | 7.393723597 | 1.038540698 | 0.13703201 | NA |
| YDL022C-A | 6.814086928 | 0.241884113 | 0.491284016 | NA |
| YDL026W | 16.56831607 | 0.955627625 | 0.021081638 | NA |
| YDL032W | 4.179935405 | -0.602399175 | 0.696505094 | NA |
| YDL041W | 0.97089065 | -0.139453189 | 0.596071182 | NA |
| YDL062W | 21.94852935 | -0.070146811 | 0.895380637 | NA |
| YDL068W | 4.863206233 | 0.425218125 | 0.558430507 | NA |
| YDL094C | 21.86703582 | -0.599748268 | 0.590377532 | NA |
| YDL096C | 9.504570723 | 0.407116168 | 0.516588168 | NA |
| YDL114W-A | 24.54000287 | 0.201432926 | 0.457869898 | NA |
| YDL118W | 21.28319356 | 1.040057136 | 0.004537264 | NA |
| YDL151C | 4.616503935 | -0.621503394 | 0.310308082 | NA |
| YDL152W | 5.390111654 | -0.614779283 | 0.974769252 | NA |
| YDL159W-A | 1.037341025 | 0.113820084 | 0.416684652 | NA |
| YDL162C | 2.875843937 | 0.277590207 | 0.911113773 | NA |
| YDL163W | 0.057163039 | 0.012332478 | 0.977259767 | NA |
| YDL172C | 10.58915184 | -0.162798182 | 0.907223804 | NA |
| YDL187C | 3.68310749 | 0.213627873 | 0.394004317 | NA |
| YDL196W | 15.17669232 | 0.179338755 | 0.622955801 | NA |

| | | | | |
|---|---|---|---|---|
| YDL223C | 1796.784019 | 0.625386276 | NA | NA |
| YDL240C-A | 1.077122984 | 0.130373864 | 0.652797222 | NA |
| YDL246C | 6.191078768 | -0.21855744 | 0.910838025 | NA |
| YDL247W | 3.022100866 | 0.695253269 | 0.719406883 | NA |
| YDR024W | 7.360211428 | -0.508081464 | 0.903092215 | NA |
| YDR034C-A | 4.06626248 | 0.13548567 | 0.605221537 | NA |
| YDR048C | 13.8983324 | 0.078510129 | 0.420785728 | NA |
| YDR053W | 7.370760006 | 0.123530956 | 0.367353955 | NA |
| YDR094W | 4.116771726 | 0.177025401 | 0.964594942 | NA |
| YDR095C | 5.026425328 | -0.441181746 | 0.290892611 | NA |
| YDR098C-B | 19.81780983 | 0.382897173 | 0.4482975 | NA |
| YDR102C | 12.68433027 | 0.182798555 | 0.735453834 | NA |
| YDR112W | 22.56100124 | 0.404904103 | 0.97036811 | NA |
| YDR118W-A | 11.72360647 | -0.160488886 | 0.891120056 | NA |
| YDR136C | 21.36116146 | -0.130825568 | 0.915517447 | NA |
| YDR149C | 14.02290454 | 0.004962876 | 0.517798404 | NA |
| YDR169C-A | 13.43403719 | 0.053012744 | 0.774820582 | NA |
| YDR182W-A | 24.64411227 | 0.696765787 | 0.010105867 | NA |
| YDR193W | 12.98139175 | 0.496303903 | 0.519960919 | NA |
| YDR194W-A | 0.453289942 | -0.042098606 | 0.994444644 | NA |
| YDR203W | 15.81601128 | 0.782574137 | 0.05858258 | NA |
| YDR220C | 3.958958383 | 0.169730418 | 0.51398218 | NA |
| YDR246W-A | 23.46436155 | 0.141039467 | 0.886043815 | NA |
| YDR250C | 6.490017442 | 0.387951109 | 0.078070709 | NA |
| YDR269C | 1.063098833 | 0.206225358 | 0.849644173 | NA |
| YDR278C | 0.649229666 | -0.080868879 | 0.952217671 | NA |
| YDR290W | 8.55083412 | 0.109143324 | 0.730402881 | NA |
| YDR320W-B | 5.22193975 | -0.145862477 | 0.482023187 | NA |
| YDR340W | 4.60165433 | -0.009600347 | 0.639084678 | NA |
| YDR355C | 4.46006282 | -0.17820694 | 0.770725904 | NA |
| YDR366C | 7.819864358 | 0.208270819 | 0.809945689 | NA |
| YDR401W | 0.227003213 | 0.031169213 | 0.696997516 | NA |
| YDR406W-A | 12.68467301 | 0.082932647 | 0.583499518 | NA |
| YDR413C | 10.51150758 | -0.726219763 | 0.101749266 | NA |
| YDR426C | 18.4381422 | 0.302492135 | 0.208997275 | NA |
| YDR431W | 11.62652329 | 0.203274589 | 0.747640857 | NA |
| YDR442W | 17.52253569 | -0.224369326 | 0.510033486 | NA |
| YDR445C | 3.142353253 | -0.1880747 | 0.494940317 | NA |
| YDR455C | 13.39180516 | 0.108496846 | 0.818098288 | NA |
| YDR509W | 13.68760112 | 0.508500769 | 0.198399597 | NA |
| YDR521W | 13.62209314 | -0.488183986 | 0.369465553 | NA |
| YDR524C-A | 2.188681507 | 0.229006453 | 0.401597807 | NA |
| YDR525W | 4.560372998 | 0.215243436 | 0.173265425 | NA |

| | | | | |
|---|---|---|---|---|
| YDR526C | 4.767823074 | -0.99016275 | 0.246175842 | NA |
| YDR535C | 20.65811424 | 0.634863223 | 0.219520622 | NA |
| YDR542W | 1.676832288 | 0.379293014 | 0.149151115 | NA |
| YDR543C | 0.805732287 | 0.010685686 | 0.675855466 | NA |
| YDR544C | 2.336721419 | 0.1397703 | 0.656192438 | NA |
| YEL009C-A | 8.998175135 | 0.501041015 | 0.093565312 | NA |
| YEL010W | 4.427753031 | 0.325652516 | 0.330491539 | NA |
| YEL014C | 21.68196635 | -0.641800547 | 0.739326343 | NA |
| YEL018C-A | 0.416927005 | -0.183416035 | 0.687171162 | NA |
| YEL030C-A | 7.262884205 | 0.124487483 | 0.44604558 | NA |
| YEL035C | 12.59177912 | 0.472088771 | 0.107611884 | NA |
| YEL045C | 16.86688846 | -1.275231972 | 0.050390178 | NA |
| YER006C-A | 2.225353622 | 0.142501258 | 0.770835548 | NA |
| YER024W | 2782.885664 | -0.012897782 | NA | NA |
| YER038W-A | 0.198141795 | 0.01374763 | 0.988055869 | NA |
| YER046W-A | 13.59530817 | 0.999819125 | 0.028604466 | NA |
| YER065C | 3480.324226 | -0.143630424 | NA | NA |
| YER068C-A | 10.30512685 | 0.60661853 | 0.026134495 | NA |
| YER078W-A | 4.281084499 | 0.429937303 | 0.083966648 | NA |
| YER079C-A | 18.60445932 | 0.762413155 | 0.039290345 | NA |
| YER084W-A | 0.719422121 | 0.018374304 | 0.854140015 | NA |
| YER085C | 1.090623589 | -0.204886886 | 0.770108259 | NA |
| YER091C-A | 0.485272411 | 0.042117181 | 0.990334092 | NA |
| YER135C | 23.51565939 | 0.385422077 | 0.67516585 | NA |
| YER137W-A | 3.086112355 | 0.127269582 | 0.200033593 | NA |
| YER138W-A | 1.112909077 | -0.346142637 | 0.21654471 | NA |
| YER148W-A | 0.781507553 | 0.014432279 | 0.73666515 | NA |
| YER172C-A | 0.861078059 | -0.110977014 | 0.650958519 | NA |
| YER181C | 6.5610832 | 0.14281479 | 0.726379652 | NA |
| YER188C-A | 23.15774642 | 0.607607242 | 0.03734245 | NA |
| YER189W | 0.915658928 | -0.288297376 | 0.533207627 | NA |
| YER190W | 15.31140442 | -0.080775828 | 0.459337995 | NA |
| YFL012W-A | 14.51855676 | 0.574636108 | 0.375561481 | NA |
| YFL015C | 19.00869044 | 0.215135892 | 0.495531574 | NA |
| YFL015W-A | 1.143841678 | -0.010386173 | 0.841287364 | NA |
| YFL041W-A | 23.17885335 | 0.258070724 | 0.309922137 | NA |
| YFL058W | 7.345596516 | 0.914966182 | 0.043904402 | NA |
| YFL060C | 16.02228195 | 0.763985819 | 0.714639895 | NA |
| YFL061W | 11.58614878 | 0.8532854 | 0.338766136 | NA |
| YFL063W | 4.881871603 | 0.23252141 | 0.471135285 | NA |
| YFL064C | 3.015892609 | 0.543875338 | 0.275316909 | NA |
| YFL065C | 1.354222162 | 0.38571114 | 0.209510351 | NA |
| YFL066C | 2.343434624 | 0.426419856 | 0.357375143 | NA |

| | | | | |
|---|---|---|---|---|
| YFR010W-A | 16.18554247 | -0.239519262 | 0.320798717 | NA |
| YFR012W-A | 2.137477958 | 0.310367686 | 0.623475563 | NA |
| YFR032C-B | 19.14700684 | -0.420300331 | 0.285452877 | NA |
| YFR034W-A | 2.98827912 | -0.30449628 | 0.503716317 | NA |
| YFR036W-A | 15.15766896 | -0.60842551 | 0.027922697 | NA |
| YFR054C | 20.51319414 | 1.167231074 | 0.025317184 | NA |
| YGL006W-A | 3.943531239 | -0.96747569 | 0.358019687 | NA |
| YGL007C-A | 2.085025417 | 0.143072858 | 0.673160629 | NA |
| YGL007W | 19.49434054 | 0.120968571 | 0.50460731 | NA |
| YGL015C | 22.18286877 | 0.030597086 | 0.313008749 | NA |
| YGL024W | 7.765530916 | 0.217879817 | 0.986984258 | NA |
| YGL034C | 9.87258369 | -0.010664593 | 0.682970696 | NA |
| YGL041C-B | 21.05072236 | -0.16166793 | 0.920118519 | NA |
| YGL072C | 15.1439202 | 0.294197197 | 0.05212701 | NA |
| YGL088W | 0.432313012 | 0.141391504 | 0.560175635 | NA |
| YGL103W | 19.59323296 | -0.927468665 | 0.155525313 | NA |
| YGL118C | 2.533670207 | 0.130648747 | 0.528251691 | NA |
| YGL132W | 16.53764806 | 0.553680977 | 0.394419153 | NA |
| YGL138C | 22.53163453 | 0.767996326 | 0.027967365 | NA |
| YGL152C | 7.084888852 | -0.030464572 | 0.570948345 | NA |
| YGL182C | 10.33516089 | 0.157579821 | 0.816770176 | NA |
| YGL188C-A | 18.77666255 | -0.149613666 | 0.553705592 | NA |
| YGL199C | 2.062524245 | -1.690511473 | 0.039585701 | NA |
| YGL214W | 9.34196156 | 0.236922154 | 0.925799884 | NA |
| YGL217C | 8.839690893 | 0.745150531 | 0.425263255 | NA |
| YGL218W | 11.00510591 | 0.399691319 | 0.243947078 | NA |
| YGL235W | 1.903973228 | -0.239982696 | 0.830620707 | NA |
| YGL239C | 18.18533761 | -0.144725013 | 0.491129048 | NA |
| YGL260W | 1.415207671 | 0.169068652 | 0.710049632 | NA |
| YGL261C | 11.97177178 | 0.435195497 | 0.610819177 | NA |
| YGR018C | 8.799849222 | 0.205302205 | 0.36370494 | NA |
| YGR022C | 1.81368062 | 0.492535873 | 0.306170082 | NA |
| YGR045C | 23.80619598 | 0.444130372 | 0.230636151 | NA |
| YGR064W | 14.24950712 | -0.69028799 | 0.359617723 | NA |
| YGR067C | 601.4253985 | 0.065791412 | NA | NA |
| YGR069W | 7.64600337 | 0.157127663 | 0.840604505 | NA |
| YGR073C | 7.437514438 | 0.383330101 | 0.563869432 | NA |
| YGR107W | 13.44440994 | -0.11358575 | 0.906105933 | NA |
| YGR121W-A | 8.749518959 | -0.124236143 | 0.82388171 | NA |
| YGR122C-A | 15.60192201 | 0.006954853 | 0.938473069 | NA |
| YGR139W | 13.99976418 | 0.18159878 | 0.743884521 | NA |
| YGR146C-A | 13.60921533 | -0.329025756 | 0.74251276 | NA |
| YGR151C | 2.6142374 | -0.795826724 | 0.016689442 | NA |

| | | | | |
|---|---|---|---|---|
| YGR160W | 19.24237613 | -0.618841294 | 0.077976441 | NA |
| YGR164W | 4.051111594 | -0.622143549 | 0.305060263 | NA |
| YGR176W | 11.59700922 | 0.037385493 | 0.622413897 | NA |
| YGR204C-A | 13.54650687 | -0.510219612 | 0.341331359 | NA |
| YGR228W | 14.43502374 | -0.581942226 | 0.39830664 | NA |
| YGR236C | 237.8810767 | 0.891803383 | NA | NA |
| YGR259C | 0.780911468 | -0.145827279 | 0.674462793 | NA |
| YGR273C | 7.812625483 | 0.177853822 | 0.726527942 | NA |
| YGR291C | 1.566792632 | -0.153283775 | 0.54377383 | NA |
| YGR293C | 7.546648827 | -0.100188738 | 0.453825823 | NA |
| YGR294W | 10.95232902 | 0.21136663 | 0.317175041 | NA |
| YHL002C-A | 8.009981089 | 0.586547854 | 0.090733335 | NA |
| YHL009W-B | 5.78416984 | 0.059972427 | 0.689897284 | NA |
| YHL015W-A | 4.315791202 | 0.286966694 | 0.71765704 | NA |
| YHL019W-A | 10.01596588 | 0.072996057 | 0.587556253 | NA |
| YHL030W-A | 22.0345387 | 0.657387049 | 0.071893789 | NA |
| YHL037C | 20.29589679 | -0.395379347 | 0.906133178 | NA |
| YHL041W | 3.288023996 | -0.5924286 | 0.177098646 | NA |
| YHL046C | 9.174623277 | 0.099910699 | 0.669285476 | NA |
| YHL048C-A | 2.76683096 | 0.617848788 | 0.247391742 | NA |
| YHR022C-A | 3.937421936 | 0.511277289 | 0.163431113 | NA |
| YHR032C-A | 12.55755435 | -0.353083481 | 0.61900261 | NA |
| YHR056W-A | 0.867587117 | -0.090869098 | 0.79153962 | NA |
| YHR069C-A | 0.116121444 | 0.012337369 | 0.951740623 | NA |
| YHR086W-A | 3.107825184 | -1.045616996 | 0.283571934 | NA |
| YHR095W | 5.929557488 | 0.243651101 | 0.441140073 | NA |
| YHR125W | 4.057342841 | 0.955355034 | 0.080102742 | NA |
| YHR126C | 18.42016663 | 0.525262673 | 0.244752828 | NA |
| YHR130C | 24.10330458 | 0.411054366 | 0.136620132 | NA |
| YHR131W-A | 19.588359 | 0.313401345 | 0.455302173 | NA |
| YHR145C | 0.415497853 | -0.013768706 | 0.879705402 | NA |
| YHR165W-A | 6.420816 | -0.005741269 | 0.324425841 | NA |
| YHR180W-A | 0.36648756 | -0.019968868 | 0.937505608 | NA |
| YHR185C | 7.642313868 | 0.208427029 | 0.91651642 | NA |
| YHR193C-A | 1.288094432 | 0.352385652 | 0.532275919 | NA |
| YHR213W-B | 12.16949824 | 0.499520875 | 0.67931094 | NA |
| YHR214C-E | 1.813646714 | 0.056966222 | 0.534209748 | NA |
| YHR214W | 18.69123236 | 0.565750127 | 0.229581115 | NA |
| YHR214W-A | 22.9859353 | 0.125975921 | 0.554470781 | NA |
| YHR217C | 1.225581631 | 0.10235325 | 0.991054085 | NA |
| YHR219W | 0.159331785 | 0.041035636 | 0.924536993 | NA |
| YIL012W | 10.18010829 | 0.948148149 | 0.050703044 | NA |
| YIL025C | 3.736035911 | -0.195285454 | 0.668718384 | NA |

| | | | | |
|---|---|---|---|---|
| YIL029W-A | 8.108002149 | -0.130089732 | 0.607054026 | NA |
| YIL032C | 13.12625335 | 0.485849866 | 0.185198342 | NA |
| YIL046W-A | 7.695578865 | 0.650250092 | 0.208116445 | NA |
| YIL047C-A | 10.41380776 | -0.266831442 | 0.473971474 | NA |
| YIL054W | 8.069700908 | 0.399208838 | 0.524781394 | NA |
| YIL057C | 383.2825996 | 0.221807221 | NA | NA |
| YIL066W-A | 3.308676015 | -0.373332084 | 0.967806633 | NA |
| YIL068W-A | 5.981971104 | -0.439199388 | 0.80288022 | NA |
| YIL086C | 23.45932813 | 0.619602307 | 0.054328784 | NA |
| YIL100C-A | 6.139255363 | -0.371028032 | 0.54286201 | NA |
| YIL100W | 3.722972412 | 0.356865966 | 0.546472838 | NA |
| YIL102C | 13.70446277 | 0.072766904 | 0.872986089 | NA |
| YIL115W-A | 23.70554908 | -1.026612703 | 0.059764769 | NA |
| YIL134C-A | 3.121789303 | -0.080513854 | 0.718206539 | NA |
| YIL142C-A | 7.771319082 | -0.34433048 | 0.751379718 | NA |
| YIL156W-A | 6.740450198 | -1.004274568 | 0.331149485 | NA |
| YIL173W | 14.66085042 | 1.088383182 | 0.064320433 | NA |
| YIL177C | 0.203940714 | -0.049614875 | 0.82048517 | NA |
| YIR013C | 17.07044727 | 0.13554645 | 0.258809327 | NA |
| YIR017C | 6.4006803 | 0.151630188 | 0.140101593 | NA |
| YIR017W-A | 3.982442675 | 0.943445993 | 0.087581497 | NA |
| YIR020C-B | 5.673146071 | 0.134222516 | 0.393657959 | NA |
| YIR021W-A | 4.66148073 | -0.121337941 | 0.41718263 | NA |
| YIR023C-A | 13.57792364 | -0.831582458 | 0.126874309 | NA |
| YIR030W-A | 17.57088172 | -0.271788151 | 0.592882921 | NA |
| YIR036W-A | 10.01589679 | 0.393201969 | 0.054468486 | NA |
| YIR040C | 0.167590483 | 0.01071366 | 0.923022794 | NA |
| YIR041W | 5.637300496 | -0.118371749 | 0.582808309 | NA |
| YIR044C | 6.331727114 | 0.475637881 | 0.615524669 | NA |
| YJL007C | 3.350774349 | -0.157880834 | 0.540420905 | NA |
| YJL022W | 2.660856745 | 0.077425531 | 0.660941002 | NA |
| YJL027C | 10.13200317 | -0.188529271 | 0.928452259 | NA |
| YJL028W | 3.906046468 | -0.176624765 | 0.589183941 | NA |
| YJL032W | 6.67566349 | 0.268817551 | 0.154922438 | NA |
| YJL047C-A | 19.06572924 | 0.017592133 | 0.90921493 | NA |
| YJL052C-A | 1.688849253 | -0.277479358 | 0.625641761 | NA |
| YJL064W | 3.253557565 | 0.018226571 | 0.937821183 | NA |
| YJL086C | 9.129343542 | 0.366896383 | 0.927779489 | NA |
| YJL114W | 3.593194288 | 0.419332414 | 0.946537831 | NA |
| YJL119C | 7.109202711 | -0.006923328 | 0.861004545 | NA |
| YJL127W-A | 12.80997666 | -0.694611384 | 0.089710078 | NA |
| YJL135W | 5.319281498 | -0.226251388 | 0.986682609 | NA |
| YJL136W-A | 0.34890375 | -0.095539092 | 0.848961325 | NA |

| | | | | |
|---|---|---|---|---|
| YJL150W | 10.09973158 | 0.58539611 | 0.304541366 | NA |
| YJL175W | 4.732791699 | 0.578850338 | 0.171793694 | NA |
| YJL182C | 1.793622545 | -0.461279484 | 0.602508133 | NA |
| YJL195C | 5.108398903 | -0.764563012 | 0.971944015 | NA |
| YJL222W | 13.76304146 | 0.923872444 | 0.08403007 | NA |
| YJL225C | 0.280381926 | 0.052282816 | 0.955482724 | NA |
| YJR020W | 8.221696255 | -0.875501541 | 0.218916074 | NA |
| YJR022W | 18.26801566 | -0.199700678 | 0.834342951 | NA |
| YJR023C | 6.716930152 | 0.051831925 | 0.9705865 | NA |
| YJR037W | 10.84309032 | 0.654843351 | 0.342323364 | NA |
| YJR079W | 11.62148612 | -0.010227853 | 0.590593071 | NA |
| YJR095W | 1024.664601 | -0.048085273 | NA | NA |
| YJR120W | 18.09619799 | 0.405995376 | 0.17192775 | NA |
| YJR128W | 14.84064674 | -0.048757589 | 0.918816958 | NA |
| YJR157W | 1.945110995 | 0.790743971 | 0.12379969 | NA |
| YJR159W | 6.187748904 | 0.304869234 | 0.644606805 | NA |
| YJR160C | 11.90922489 | 0.74546292 | 0.233587056 | NA |
| YKL053W | 17.58730316 | -0.095398521 | 0.807596804 | NA |
| YKL065W-A | 60.35011298 | 0.692714687 | NA | NA |
| YKL096C-B | 8.469479675 | 0.263919974 | 0.549187394 | NA |
| YKL097C | 10.27058523 | 0.432303725 | 0.390811688 | NA |
| YKL102C | 18.56354003 | -0.271231037 | 0.513708337 | NA |
| YKL106C-A | 1.684117879 | 0.384757768 | 0.807002671 | NA |
| YKL111C | 22.34564795 | -1.001818927 | 0.02067959 | NA |
| YKL131W | 17.12686382 | 0.329014116 | 0.426200688 | NA |
| YKL147C | 17.64954721 | 0.107880416 | 0.247015045 | NA |
| YKL162C-A | 16.50216676 | -0.162652525 | 0.899022675 | NA |
| YKL165C-A | 6.96473103 | -0.386219822 | 0.958494502 | NA |
| YKL177W | 2.5279419 | -0.523102122 | 0.724978001 | NA |
| YKL183C-A | 2.603283005 | 0.55667731 | 0.519396791 | NA |
| YKL202W | 21.88607995 | -0.294673285 | 0.859341602 | NA |
| YKL223W | 0.064032033 | 0.012318354 | 0.967041033 | NA |
| YKL224C | 0.174623486 | -0.013833002 | 0.986381552 | NA |
| YKL225W | 0.626369128 | 0.011145338 | 0.837893542 | NA |
| YKR032W | 0.979223699 | -0.067092635 | 0.886003563 | NA |
| YKR033C | 0.132511194 | 0.041834605 | 0.72881702 | NA |
| YKR035C | 23.50975344 | -0.343867214 | 0.891693168 | NA |
| YKR040C | 4.454389 | -0.112123848 | 0.379174427 | NA |
| YKR073C | 18.81130727 | 0.242831084 | 0.98234766 | NA |
| YKR097W | 1618.765281 | -0.018388451 | NA | NA |
| YLL006W-A | 2.814822079 | 0.008646406 | 0.904667441 | NA |
| YLL030C | 4.099886808 | -0.322657041 | 0.821130542 | NA |
| YLL037W | 13.52665894 | 0.071640848 | 0.573854086 | NA |

| | | | | |
|---|---|---|---|---|
| YLL047W | 4.30351014 | -0.259722993 | 0.964040311 | NA |
| YLL065W | 0.583562943 | 0.017031261 | 0.788593299 | NA |
| YLL066C | 1.397171575 | 0.463235724 | 0.613361997 | NA |
| YLL066W-B | 2.220126499 | 0.66048317 | 0.919922155 | NA |
| YLL067C | 0.999643856 | -0.112929542 | 0.987724132 | NA |
| YLR012C | 11.76101632 | 0.25564728 | 0.283158014 | NA |
| YLR013W | 3.60279815 | 0.409663219 | 0.351919091 | NA |
| YLR111W | 15.73315875 | 0.327953735 | 0.469119626 | NA |
| YLR120W-A | 22.81140064 | -0.682045686 | 0.110220825 | NA |
| YLR123C | 17.47407493 | 0.713411477 | 0.408816743 | NA |
| YLR157C-B | 0.750636836 | 0.528278158 | 0.880105035 | NA |
| YLR162W | 2.387178616 | 0.274395946 | 0.670955627 | NA |
| YLR162W-A | 8.884728875 | -0.378432442 | 0.021670404 | NA |
| YLR174W | 1214.876474 | -0.029065923 | NA | NA |
| YLR184W | 0.707088895 | 0.007333184 | 0.76880067 | NA |
| YLR217W | 16.95655225 | 0.178414248 | 0.495018619 | NA |
| YLR222C-A | 13.05158283 | 0.217009264 | 0.83881039 | NA |
| YLR232W | 2.842598345 | -0.298231637 | 0.240629408 | NA |
| YLR235C | 18.73304548 | -0.624831659 | 0.205310218 | NA |
| YLR236C | 21.91002832 | -0.141525041 | 0.968838189 | NA |
| YLR255C | 23.50063941 | 0.076452243 | 0.668200019 | NA |
| YLR261C | 10.17984332 | -0.855941187 | 0.205213902 | NA |
| YLR282C | 4.588773458 | -1.115996249 | 0.206096801 | NA |
| YLR296W | 4.726955448 | -0.07864084 | 0.978462612 | NA |
| YLR302C | 0.305761563 | 0.016295918 | 0.955277182 | NA |
| YLR308W | 11.06339218 | 0.354066274 | 0.179658921 | NA |
| YLR317W | 2.198595472 | -0.005390863 | 0.17526044 | NA |
| YLR322W | 7.310944976 | -0.75809505 | 0.119092018 | NA |
| YLR331C | 3.488696978 | -0.360938106 | 0.91817345 | NA |
| YLR334C | 3.854398197 | -0.379175023 | 0.300843205 | NA |
| YLR338W | 19.58277131 | 0.125058728 | 0.697631787 | NA |
| YLR349W | 23.19539208 | -0.51246648 | 0.674679631 | NA |
| YLR364C-A | 0.303554837 | 0.048938536 | 0.806860643 | NA |
| YLR365W | 4.223598425 | 0.346449762 | 0.345689887 | NA |
| YLR366W | 1.593075245 | 0.89929917 | 0.047797007 | NA |
| YLR377C | 564.6863452 | -0.020063553 | NA | NA |
| YLR406C-A | 2.560257022 | -0.058625574 | 0.930795117 | NA |
| YLR412C-A | 1.32995427 | 0.39712522 | 0.874607766 | NA |
| YLR415C | 24.41748512 | -0.15577775 | 0.682420381 | NA |
| YLR428C | 15.96009415 | -0.147281476 | 0.93588121 | NA |
| YLR458W | 2.467406493 | -0.614209103 | 0.839685301 | NA |
| YLR461W | 7.039838185 | 0.652655966 | 0.31964606 | NA |
| YLR462W | 4.00598291 | 0.449683799 | 0.220487069 | NA |

| | | | | |
|---|---|---|---|---|
| YLR466C-B | 12.5768575 | 0.715393713 | 0.825911214 | NA |
| YML007C-A | 2.984197591 | 0.532766184 | 0.47079286 | NA |
| YML012C-A | 4.439298629 | -0.149775858 | 0.789436796 | NA |
| YML031C-A | 2.97142876 | 0.385289759 | 0.16124826 | NA |
| YML034C-A | 23.30372078 | -0.044452358 | 0.790537356 | NA |
| YML039W | 1.131094598 | -0.037164342 | 0.557220623 | NA |
| YML042W | 2686.768376 | 0.133181624 | NA | NA |
| YML047W-A | 2.39499993 | 1.4736185 | 0.001223136 | NA |
| YML054C-A | 11.9894805 | -0.119564259 | 0.913866248 | NA |
| YML084W | 2.063062303 | -0.750013419 | 0.470481877 | NA |
| YML089C | 19.45677984 | 0.233663056 | 0.461884149 | NA |
| YML094C-A | 15.34009962 | 0.058729002 | 0.591766278 | NA |
| YML100W-A | 6.289539175 | 0.083838926 | 0.454061957 | NA |
| YML101C-A | 7.327177589 | -0.9563866 | 0.150608667 | NA |
| YML133C | 9.588165535 | -0.37479144 | 0.530347804 | NA |
| YMR001C-A | 16.08451551 | -0.420338361 | 0.083316382 | NA |
| YMR030W-A | 15.82228062 | -0.183199656 | 0.967618682 | NA |
| YMR057C | 13.35741525 | -0.127551692 | 0.440913262 | NA |
| YMR075C-A | 15.14864191 | -0.220270242 | 0.486114032 | NA |
| YMR082C | 5.748276588 | 0.830115723 | 0.276244718 | NA |
| YMR107W | 2022.649302 | 0.550912083 | NA | NA |
| YMR135W-A | 18.48988281 | -0.415925411 | 0.448738768 | NA |
| YMR158C-A | 22.93483608 | 0.61679155 | 0.081765506 | NA |
| YMR158W-B | 1.160282167 | 0.297495401 | 0.240181202 | NA |
| YMR159C | 22.87110149 | 0.585073716 | 0.100621806 | NA |
| YMR172C-A | 11.37396813 | -0.319614459 | 0.98368359 | NA |
| YMR175W | 966.6429644 | 0.10833617 | NA | NA |
| YMR175W-A | 25.9663659 | 0.064581615 | NA | NA |
| YMR182W-A | 4.590688512 | 0.53349473 | 0.125706533 | NA |
| YMR194C-A | 22.19182237 | 0.504114371 | 0.628749479 | NA |
| YMR230W-A | 4.038623613 | 0.065145392 | 0.451306631 | NA |
| YMR242W-A | 1.825788145 | 0.05326134 | 0.353400362 | NA |
| YMR247W-A | 22.89498367 | 0.607366179 | 0.143723479 | NA |
| YMR272W-B | 3.438828838 | -0.035042654 | 0.82174184 | NA |
| YMR290W-A | 11.59226843 | -1.429682558 | 0.172501301 | NA |
| YMR306C-A | 3.882802162 | -0.568118036 | 0.247401858 | NA |
| YMR315W-A | 6.002010905 | 0.650325889 | 0.182614085 | NA |
| YMR316C-B | 10.98411933 | 0.273929009 | 0.347549366 | NA |
| YMR321C | 24.07796373 | 1.317814279 | 0.038809291 | NA |
| YMR322C | 22.28920933 | 0.369375735 | 0.25626187 | NA |
| YMR324C | 2.573506675 | 0.325684519 | 0.796665624 | NA |
| YMR325W | 4.52283349 | 0.714079595 | 0.109047244 | NA |
| YMR326C | 0.406623722 | 0.034305927 | 0.861054214 | NA |

| | | | | |
|---|---|---|---|---|
| YNL013C | 5.629906296 | 0.031249766 | 0.504248928 | NA |
| YNL017C | 1.615564791 | -0.078876373 | 0.541405776 | NA |
| YNL019C | 10.29601328 | 0.786165911 | 0.108304785 | NA |
| YNL028W | 6.323808974 | 0.733898766 | 0.245180529 | NA |
| YNL033W | 5.453513622 | -0.41790912 | 0.480448083 | NA |
| YNL036W | 2481.742596 | -0.312615658 | NA | NA |
| YNL042W-B | 14.4556208 | -0.250507724 | 0.536888803 | NA |
| YNL057W | 19.3681916 | -0.907353074 | 0.093407182 | NA |
| YNL067W-A | 0.709210999 | 0.532941311 | 0.258417348 | NA |
| YNL067W-B | 0.339190409 | 0.026079614 | 0.851421563 | NA |
| YNL089C | 16.65282477 | -0.362972352 | 0.489687531 | NA |
| YNL097C-B | 13.11125706 | 0.383231641 | 0.397442463 | NA |
| YNL105W | 8.716167534 | 0.309688349 | 0.452025827 | NA |
| YNL109W | 19.36367514 | 0.227930384 | 0.751799226 | NA |
| YNL117W | 3801.736785 | -0.002617496 | NA | NA |
| YNL120C | 0.227028385 | 0.065970484 | 0.711377041 | NA |
| YNL130C-A | 3.921452687 | 0.887922168 | 0.132767291 | NA |
| YNL140C | 6.604161499 | 0.905795276 | 0.016182714 | NA |
| YNL174W | 7.421723425 | -0.795618806 | 0.220532546 | NA |
| YNL198C | 23.75937545 | -0.132740214 | 0.689092745 | NA |
| YNL205C | 11.3162081 | 0.088799963 | 0.504969841 | NA |
| YNL226W | 19.21780824 | -0.034019698 | 0.79719032 | NA |
| YNL228W | 12.1493579 | 0.284442093 | 0.371347386 | NA |
| YNL235C | 7.928094321 | -0.508403199 | 0.610254327 | NA |
| YNL266W | 15.42456766 | 0.564714551 | 0.260438957 | NA |
| YNL276C | 5.892936072 | -0.439313793 | 0.312309896 | NA |
| YNL277W-A | 6.085182075 | 0.081324207 | 0.50771391 | NA |
| YNL285W | 3.926788582 | 0.221442128 | 0.324563915 | NA |
| YNL296W | 0.17572267 | 0.011561405 | 0.960616976 | NA |
| YNL318C | 22.2967731 | 0.448421307 | 0.195357295 | NA |
| YNL319W | 5.59444275 | -0.082648008 | 0.387369908 | NA |
| YNL324W | 2.168671185 | 1.001185843 | 0.037428372 | NA |
| YNL334C | 19.40823403 | 0.548931468 | 0.584386753 | NA |
| YNL335W | 13.42162756 | 0.520263165 | 0.52467487 | NA |
| YNL337W | 0.659062417 | 0.08462256 | 0.472706855 | NA |
| YNL338W | 0.154202725 | 0.067433572 | 0.948884979 | NA |
| YNR001W-A | 2.398336405 | 0.811269388 | 0.093485429 | NA |
| YNR002C | 907.6027094 | 0.534119973 | NA | NA |
| YNR005C | 1.56429782 | -0.428160228 | 0.939473133 | NA |
| YNR042W | 0.825755426 | -0.125743598 | 0.896752299 | NA |
| YNR071C | 317.5831201 | 0.252429787 | NA | NA |
| YNR075C-A | 12.77963038 | -0.202805727 | 0.180087619 | NA |
| YNR077C | 0.24124529 | 0.014217215 | 0.931254855 | NA |

| | | | | |
|---|---|---|---|---|
| YOL013W-A | 5.152321446 | 0.564623196 | 0.727862769 | NA |
| YOL013W-B | 4.383672181 | 0.973097639 | 0.430309681 | NA |
| YOL019W-A | 0.720433626 | 0.018980231 | 0.763959535 | NA |
| YOL038C-A | 23.65291474 | -0.04697753 | 0.824478326 | NA |
| YOL046C | 5.940874887 | 0.499060423 | 0.138714308 | NA |
| YOL050C | 16.33482394 | -0.377350388 | 0.386353548 | NA |
| YOL079W | 9.64455915 | -0.444832755 | 0.480829571 | NA |
| YOL085W-A | 20.71787704 | 1.098802714 | 0.056599495 | NA |
| YOL097W-A | 14.8332561 | -0.420177374 | 0.247374162 | NA |
| YOL099C | 11.41692104 | 0.078749935 | 0.819553986 | NA |
| YOL106W | 14.51135789 | 0.1383512 | 0.485088895 | NA |
| YOL134C | 23.62819378 | -0.276851314 | 0.886218899 | NA |
| YOL150C | 3.831416512 | 0.875896193 | 0.166922263 | NA |
| YOL160W | 15.67971505 | 0.293138574 | 0.417215228 | NA |
| YOL161C | 3.517453047 | 0.296906462 | 0.136627269 | NA |
| YOL164W-A | 24.89009234 | 0.408001138 | 0.310687495 | NA |
| YOL165C | 22.45477283 | 0.284692275 | 0.705530168 | NA |
| YOL166C | 2.808993483 | -0.062378762 | 0.878900238 | NA |
| YOL166W-A | 0.347622843 | 0.007454854 | 0.8774579 | NA |
| YOR008C-A | 9.788861467 | 0.383469427 | 0.60225176 | NA |
| YOR008W-B | 4.890160442 | -0.161500841 | 0.530167109 | NA |
| YOR011W-A | 3.164669454 | 0.424297232 | 0.33903154 | NA |
| YOR024W | 4.641545194 | 1.103479354 | 0.02068185 | NA |
| YOR029W | 6.24277143 | 0.015322151 | 0.569298209 | NA |
| YOR032W-A | 11.5403846 | -0.037868663 | 0.38510991 | NA |
| YOR053W | 4.40231234 | -0.86785626 | 0.548091035 | NA |
| YOR055W | 14.54954616 | -0.22538126 | 0.548572092 | NA |
| YOR072W-A | 14.05662092 | -0.034431476 | 0.523430401 | NA |
| YOR072W-B | 0.082611248 | 0.065114227 | 0.981932033 | NA |
| YOR073W-A | 11.62724158 | -0.432898096 | 0.467877652 | NA |
| YOR100C | 795.4961629 | 0.179337145 | NA | NA |
| YOR105W | 21.94253227 | 0.544293161 | 0.139183934 | NA |
| YOR139C | 5.37072037 | -0.391179059 | 0.695537324 | NA |
| YOR146W | 6.962391281 | -0.968468075 | 0.801410038 | NA |
| YOR161C-C | 9.014220677 | 0.192002872 | 0.244695517 | NA |
| YOR183W | 10.65070105 | -0.135060677 | 0.696505897 | NA |
| YOR192C-C | 17.72254902 | 0.01544557 | 0.577745386 | NA |
| YOR200W | 9.969440735 | 0.809540736 | 0.016368542 | NA |
| YOR218C | 5.229072963 | 0.183771533 | 0.60273832 | NA |
| YOR235W | 3.373424746 | 0.428799772 | 0.213287293 | NA |
| YOR255W | 23.76826284 | 0.929282689 | 0.03096473 | NA |
| YOR268C | 11.87888145 | 0.290177323 | 0.783662194 | NA |
| YOR277C | 6.042620409 | -0.630832523 | 0.019592796 | NA |

| | | | | |
|---|---|---|---|---|
| YOR282W | 14.73409586 | -0.690593794 | 0.461925023 | NA |
| YOR293C-A | 2.608736587 | 0.142239673 | 0.430183894 | NA |
| YOR300W | 17.40293291 | 0.239942017 | 0.314412056 | NA |
| YOR314W | 17.17022364 | -0.080953435 | 0.311881917 | NA |
| YOR314W-A | 0.578252409 | 0.241785073 | 0.828341135 | NA |
| YOR316C-A | 8.072827245 | 0.014489154 | 0.715051782 | NA |
| YOR325W | 4.631536169 | -0.351300005 | 0.331201004 | NA |
| YOR329W-A | 20.44917844 | 0.428742263 | 0.42380039 | NA |
| YOR333C | 3.992636415 | 0.271480773 | 0.514239022 | NA |
| YOR345C | 1.957434997 | 0.225970612 | 0.284121246 | NA |
| YOR364W | 7.913433737 | 0.542659056 | 0.385570998 | NA |
| YOR366W | 3.244594662 | -0.442299045 | 0.955721704 | NA |
| YOR379C | 0.107512111 | 0.010022914 | 0.997169526 | NA |
| YOR381W-A | 9.096748957 | 0.439126937 | 0.240472527 | NA |
| YOR388C | 10.58733181 | 0.549740073 | 0.166065702 | NA |
| YOR391C | 10.43191586 | 1.1059614 | 0.065563943 | NA |
| YOR392W | 4.708188968 | 0.932736983 | 0.056113535 | NA |
| YOR394C-A | 4.135778786 | 0.237058786 | 0.958500024 | NA |
| YPL021W | 14.06563719 | 0.239058344 | 0.572791865 | NA |
| YPL025C | 15.86269313 | 0.006434493 | 0.589167466 | NA |
| YPL033C | 16.58126869 | 0.637518641 | 0.175739155 | NA |
| YPL038W-A | 14.34188488 | 0.056052851 | 0.359726271 | NA |
| YPL044C | 1.74310561 | -0.325159921 | 0.557476067 | NA |
| YPL060C-A | 3.51919416 | -0.066741381 | 0.712242095 | NA |
| YPL062W | 4.463478236 | 0.151184196 | 0.536507558 | NA |
| YPL073C | 1.14139142 | -0.113217929 | 0.79219688 | NA |
| YPL080C | 11.15479793 | -0.633293472 | 0.140884597 | NA |
| YPL102C | 13.71806551 | 0.373727894 | 0.233783024 | NA |
| YPL114W | 1.969765016 | -0.220899624 | 0.979486675 | NA |
| YPL130W | 24.21083011 | -0.084799824 | 0.759883841 | NA |
| YPL136W | 0.97000745 | -0.249915941 | 0.156081421 | NA |
| YPL152W-A | 1.958413515 | -0.273617988 | 0.184480213 | NA |
| YPL187W | 17.17000227 | 0.505620573 | 0.26618231 | NA |
| YPL200W | 21.68345646 | -0.042527769 | 0.704105082 | NA |
| YPL223C | 915.9171589 | 0.608750106 | NA | NA |
| YPL276W | 15.28614474 | 0.0806062 | 0.553202715 | NA |
| YPL278C | 8.613341367 | 0.328229844 | 0.255653777 | NA |
| YPR002C-A | 1.918449357 | -0.506225503 | 0.639000092 | NA |
| YPR038W | 18.2580769 | -0.036480846 | 0.944376905 | NA |
| YPR039W | 9.707681236 | -0.589651226 | 0.580217751 | NA |
| YPR050C | 1.567602133 | -0.017714416 | 0.469396244 | NA |
| YPR059C | 12.62828581 | -1.310260066 | 0.006587152 | NA |
| YPR074W-A | 0.407755058 | 0.017287709 | 0.889792046 | NA |

| | | | | |
|---|---|---|---|---|
| YPR077C | 1.201353204 | 0.209082067 | 0.248101881 | NA |
| YPR087W | 20.87255534 | -0.443167321 | 0.753822959 | NA |
| YPR096C | 16.87382422 | 0.110103195 | 0.801742737 | NA |
| YPR099C | 24.60634511 | -0.747289027 | 0.516488788 | NA |
| YPR108W-A | 1.090591333 | 0.276514631 | 0.235170659 | NA |
| YPR130C | 10.79124991 | -0.285643026 | 0.540670308 | NA |
| YPR136C | 12.04995248 | -0.402896479 | 0.597577944 | NA |
| YPR142C | 12.52047692 | -1.078250448 | 0.025426499 | NA |
| YPR150W | 6.435730199 | 0.078193204 | 0.362164218 | NA |
| YPR159C-A | 2.271891006 | -0.348505163 | 0.887968415 | NA |
| YPR170W-A | 19.17302177 | -0.45942596 | 0.156411658 | NA |
| YPR177C | 1.844393858 | 0.117896915 | 0.639139552 | NA |
| YPR197C | 5.599447647 | 0.031879656 | 0.763973294 | NA |
| YPR202W | 17.57397274 | 0.005071875 | 0.25278223 | NA |
| snR10 | 11.52677515 | -0.368048613 | 0.317924032 | NA |
| snR128 | 13.17491061 | -0.553791809 | 0.014519068 | NA |
| snR13 | 24.86462889 | -0.53609593 | 0.178932836 | NA |
| snR14 | 19.85838705 | -0.331119736 | 0.824234836 | NA |
| snR161 | 8.988952116 | 0.058770164 | 0.630745621 | NA |
| snR18 | 14.34494847 | -0.65298207 | 0.128735231 | NA |
| snR189 | 9.691874789 | 0.179160934 | 0.62889945 | NA |
| snR190 | 17.01701234 | -0.181346309 | 0.659926631 | NA |
| snR191 | 16.29529748 | -0.028911759 | 0.108209381 | NA |
| snR24 | 8.183960631 | -0.789126673 | 0.223251258 | NA |
| snR36 | 18.91914442 | 0.594857552 | 0.345427625 | NA |
| snR38 | 6.274197792 | -0.366007684 | 0.127533939 | NA |
| snR39 | 11.68210812 | -0.509592549 | 0.427616208 | NA |
| snR39B | 13.58422976 | -0.592407103 | 0.27299092 | NA |
| snR40 | 6.538551087 | 0.379018102 | 0.702697804 | NA |
| snR43 | 20.09245082 | 0.180535262 | 0.55555546 | NA |
| snR45 | 13.49639451 | 0.007356193 | 0.655012351 | NA |
| snR46 | 23.07413489 | -0.342811733 | 0.135375787 | NA |
| snR47 | 5.450594875 | 0.318994175 | 0.350832027 | NA |
| snR48 | 2.926793808 | -0.171500587 | 0.529053473 | NA |
| snR50 | 8.290616263 | 0.540395334 | 0.224373982 | NA |
| snR52 | 0.834128967 | -0.051561734 | 0.821675104 | NA |
| snR53 | 0.80815136 | -0.085580454 | 0.477105362 | NA |
| snR54 | 8.052861878 | -0.429928954 | 0.60412079 | NA |
| snR55 | 9.080532067 | 0.089458155 | 0.877116814 | NA |
| snR56 | 6.041353852 | 0.205637745 | 0.131675578 | NA |
| snR57 | 17.27289396 | 0.169767298 | 0.696225664 | NA |
| snR58 | 5.413360595 | 0.928461565 | 0.102118072 | NA |
| snR59 | 2.356669998 | -0.302587604 | 0.329648553 | NA |

| | | | | |
|---|---|---|---|---|
| snR6 | 0.308944371 | 0.117996738 | 0.902285653 | NA |
| snR60 | 11.58332067 | 0.597705822 | 0.126205814 | NA |
| snR61 | 12.13748594 | -0.211188084 | 0.336541297 | NA |
| snR62 | 6.46452847 | 0.020961958 | 0.765106069 | NA |
| snR64 | 4.034418541 | -0.014757476 | 0.455126252 | NA |
| snR65 | 5.68201134 | -0.583587528 | 0.452869854 | NA |
| snR66 | 7.293652703 | 0.405835462 | 0.904977455 | NA |
| snR67 | 6.800863812 | -0.888433927 | 0.316781972 | NA |
| snR68 | 10.29817386 | -0.059681859 | 0.991377393 | NA |
| snR69 | 9.747852558 | 1.20856365 | 0.008760016 | NA |
| snR7-L | 6.909111604 | -0.006726345 | 0.547221624 | NA |
| snR71 | 12.58626851 | -0.464818594 | 0.852922952 | NA |
| snR72 | 8.104041704 | -0.28056318 | 0.668233464 | NA |
| snR73 | 0.982818557 | 0.123693121 | 0.778400912 | NA |
| snR74 | 1.214578707 | 0.170439415 | 0.66280949 | NA |
| snR75 | 1.574604411 | -0.289375794 | 0.397816273 | NA |
| snR76 | 8.640963018 | 0.029798223 | 0.72167786 | NA |
| snR77 | 9.325617253 | 0.018355053 | 0.502983121 | NA |
| snR78 | 1.974544426 | -0.319975729 | 0.804045109 | NA |
| snR79 | 6.23839996 | -0.230628804 | 0.249249941 | NA |
| snR80 | 3.471418478 | 0.032741698 | 0.214644021 | NA |
| snR81 | 9.309990788 | -0.239913977 | 0.132390097 | NA |
| snR85 | 18.57666508 | -0.307664009 | 0.529382645 | NA |
| snR87 | 6.424473744 | -0.188670563 | 0.624320938 | NA |
| tD(GUC)N | 0.865673422 | 0.414106036 | 0.540148647 | NA |
| tG(CCC)O | 0.202424979 | 0.01375506 | 0.953677546 | NA |
| tL(GAG)G | 0.599720226 | -0.031138459 | 0.971042573 | NA |
| tQ(UUG)B | 0.084186605 | 0.009981901 | 0.96140879 | NA |
| tQ(UUG)E2 | 0.083088204 | 0.013091469 | 0.942373725 | NA |
| tR(ACG)J | 0.2026169 | 0.01283314 | 0.992986744 | NA |
| tR(CCG)L | 0.085991782 | 0.013119717 | 0.965280491 | NA |
| tR(CCU)J | 0.0941958 | 0.011564254 | 0.977266963 | NA |
| tT(CGU)K | 0.429968112 | -0.30360795 | 0.694794847 | NA |
