## Supplementary material for "A stress response that allows highly mutated eukaryotic cells to survive and proliferate": Table S3

| Gene | baseMean | log2FoldChange | pvalue | padj |
|---|---|---|---|---|
| YDR097C | 218.1671058 | -7.746082354 | 1.56E-58 | 0.00 |
| YKR070W | 801.1916976 | -0.80421126 | 5.01E-10 | 0.00 |
| YFL025C | 800.9686963 | 0.723241248 | 5.99E-09 | 0.00 |
| YGL198W | 494.0719347 | -1.494699509 | 1.83E-08 | 0.00 |
| YKL033W | 666.8103831 | 1.004791152 | 1.71E-08 | 0.00 |
| YMR071C | 1073.602658 | -0.596073972 | 4.03E-08 | 0.00 |
| YLR451W | 1124.900564 | 0.931960825 | 1.92E-07 | 0.00 |
| YBL091C | 4543.955358 | -0.783676565 | 8.15E-07 | 0.00 |
| YBR109C | 2942.937927 | -0.815129443 | 2.26E-06 | 0.00 |
| YKL068W | 1722.980976 | 0.989466293 | 5.80E-06 | 0.00 |
| YOR129C | 585.8837465 | 0.715615034 | 1.44E-05 | 0.01 |
| YLR047C | 709.1954371 | 0.622703427 | 2.07E-05 | 0.01 |
| YIL002W-A | 495.5817191 | -0.854234406 | 2.37E-05 | 0.01 |
| YOR332W | 2877.284969 | -0.86511483 | 2.68E-05 | 0.01 |
| YML017W | 751.7187367 | 0.851486615 | 2.99E-05 | 0.01 |
| YNL071W | 6818.92242 | -0.852867578 | 3.36E-05 | 0.01 |
| YDR139C | 266.4093081 | -1.015715401 | 4.06E-05 | 0.01 |
| YML001W | 1453.0376 | -0.780723424 | 3.91E-05 | 0.01 |
| YBR052C | 2651.004465 | -0.841153252 | 5.29E-05 | 0.01 |
| YHR155W | 1225.572051 | 0.633569063 | 5.40E-05 | 0.01 |
| YPR070W | 926.8338353 | 0.488182759 | 5.22E-05 | 0.01 |
| YMR314W | 2980.188475 | -0.689407298 | 6.71E-05 | 0.02 |
| YDL137W | 4443.238326 | -0.895595881 | 7.55E-05 | 0.02 |
| YER129W | 2291.22394 | 0.50145491 | 7.64E-05 | 0.02 |
| YBR035C | 2483.374886 | -0.84391825 | 9.26E-05 | 0.02 |
| YJL127C | 657.9035431 | 0.595573757 | 9.37E-05 | 0.02 |
| YMR197C | 923.7331716 | -0.721419896 | 9.40E-05 | 0.02 |
| YNL053W | 544.6626736 | 0.800332337 | 8.35E-05 | 0.02 |
| YOR331C | 254.0815524 | -0.909691241 | 9.58E-05 | 0.02 |
| YGR003W | 648.3625673 | 0.674880415 | 0.000102739 | 0.02 |
| YDR017C | 2063.432291 | 0.585947765 | 0.000112598 | 0.02 |
| YDR065W | 277.2481422 | 1.042686271 | 0.000124792 | 0.02 |
| YEL038W | 943.0798183 | -0.832808562 | 0.000127861 | 0.02 |
| YGL107C | 935.2339762 | 0.405355471 | 0.000122103 | 0.02 |
| YOL129W | 2548.890951 | -0.643323902 | 0.000120345 | 0.02 |
| YDL029W | 3675.857405 | -0.470336118 | 0.000148164 | 0.02 |
| YGR041W | 794.2535473 | 0.809551571 | 0.000151894 | 0.02 |
| YGR216C | 759.5937639 | 0.621229736 | 0.000155859 | 0.02 |
| YHL034C | 6448.755806 | -0.759420547 | 0.000153738 | 0.02 |
| YHL034W-A | 585.1049194 | -0.878502891 | 0.000145435 | 0.02 |

| | | | | |
|---|---|---|---|---|
| YDR118W | 391.9679351 | 0.526225484 | 0.000170933 | 0.02 |
| YBR234C | 4120.481034 | -0.650005542 | 0.000176999 | 0.02 |
| YDL072C | 3778.461007 | -0.62805072 | 0.000179608 | 0.02 |
| YBL041W | 3024.581572 | -0.534094513 | 0.000189285 | 0.02 |
| YDR363W-A | 1308.889566 | -0.840141348 | 0.00018849 | 0.02 |
| YBR253W | 396.1883397 | -0.651275484 | 0.000196251 | 0.02 |
| YER074W-A | 546.9637993 | -0.696267866 | 0.000235026 | 0.03 |
| YOR107W | 180.9819764 | 0.947752217 | 0.000255657 | 0.03 |
| YIL115C | 2357.200299 | 0.443154935 | 0.000269648 | 0.03 |
| YDR378C | 714.4468599 | -0.651310296 | 0.000288756 | 0.03 |
| YER060W | 750.1023347 | 1.067478944 | 0.000277881 | 0.03 |
| YER075C | 845.204575 | 0.68346028 | 0.000302236 | 0.03 |
| YGR135W | 3309.689244 | -0.835570627 | 0.000318616 | 0.03 |
| YIL092W | 426.021742 | 0.572422266 | 0.000330405 | 0.03 |
| YLL050C | 6100.834604 | -0.929902626 | 0.000328216 | 0.03 |
| YML092C | 2758.585269 | -0.630006699 | 0.00031308 | 0.03 |
| YMR072W | 3717.086972 | -0.689505483 | 0.000312421 | 0.03 |
| YNR032C-A | 183.6670278 | -0.891754966 | 0.000324088 | 0.03 |
| YPL059W | 1008.866988 | -0.679875411 | 0.000313218 | 0.03 |
| YPL082C | 1537.0483 | 0.889193319 | 0.000323569 | 0.03 |
| YDR359C | 1438.122379 | 0.885015738 | 0.000353949 | 0.03 |
| YJR061W | 732.5988419 | 0.872006402 | 0.000342614 | 0.03 |
| YMR276W | 5216.693498 | -0.744970816 | 0.00034622 | 0.03 |
| YOR265W | 445.8131326 | -0.845302878 | 0.000350998 | 0.03 |
| YML085C | 4155.33263 | -0.781801156 | 0.000372825 | 0.03 |
| YOR357C | 469.3315493 | -0.65677677 | 0.000407355 | 0.03 |
| YGL011C | 3486.685044 | -0.646354888 | 0.000430575 | 0.04 |
| YML059C | 1781.099171 | 0.782498311 | 0.000454876 | 0.04 |
| YDR424C | 383.6752581 | -0.628752968 | 0.000510745 | 0.04 |
| YGL106W | 2381.873892 | -0.703005921 | 0.000500326 | 0.04 |
| YMR295C | 1636.767669 | -0.546260947 | 0.000514664 | 0.04 |
| YGR076C | 617.1306078 | -0.812360674 | 0.000551874 | 0.04 |
| YDR263C | 317.4342443 | 1.046711418 | 0.000597388 | 0.04 |
| YER048W-A | 659.0895735 | -0.423125608 | 0.00058412 | 0.04 |
| YGL200C | 3717.842992 | -0.667525474 | 0.000599404 | 0.04 |
| YER057C | 2325.694126 | -0.775580775 | 0.000656892 | 0.05 |
| YKR015C | 299.9934309 | 0.774834485 | 0.000655294 | 0.05 |
| YDR328C | 1849.787592 | -0.617792533 | 0.000682701 | 0.05 |
| YER093C | 1449.072568 | 0.603766964 | 0.00069095 | 0.05 |
| YGR108W | 622.4932134 | 1.076695976 | 0.000698453 | 0.05 |
| YLR216C | 5399.303942 | -0.929326729 | 0.000704233 | 0.05 |
| YDR089W | 994.7212521 | 0.892622993 | 0.000749007 | 0.05 |
| YER056C | 2408.22934 | 1.122608048 | 0.000745343 | 0.05 |

| | | | | |
|---|---|---|---|---|
| YGR209C | 5181.82162 | -0.734297512 | 0.000737571 | 0.05 |
| YML104C | 923.4299097 | 0.700540648 | 0.000761298 | 0.05 |
| YOL026C | 840.8767019 | -0.697944128 | 0.000770998 | 0.05 |
| YER012W | 3318.681234 | -0.643967316 | 0.000804025 | 0.05 |
| YMR154C | 501.7366665 | 0.804469621 | 0.000804515 | 0.05 |
| YBR058C-A | 528.5056217 | -0.80107603 | 0.000830528 | 0.05 |
| YKL176C | 1422.060464 | 0.777460751 | 0.000825234 | 0.05 |
| YMR167W | 578.0579077 | 0.733177244 | 0.000840636 | 0.05 |
| YBR173C | 2339.401983 | -0.667882159 | 0.000895428 | 0.05 |
| YHR043C | 1297.179518 | -0.840013181 | 0.000941177 | 0.05 |
| YLR214W | 1682.159024 | 1.22993653 | 0.000921762 | 0.05 |
| YOR164C | 1461.403289 | -0.75940349 | 0.000940076 | 0.05 |
| YOR251C | 1296.455018 | -0.530919351 | 0.000921584 | 0.05 |
| YDR084C | 664.1695601 | -0.679294902 | 0.000967457 | 0.05 |
| YDR071C | 2792.908431 | -0.822675506 | 0.001076913 | 0.06 |
| YDR137W | 962.3953612 | 0.454495739 | 0.001124928 | 0.06 |
| YGR253C | 3517.823624 | -0.616172059 | 0.001065758 | 0.06 |
| YJR017C | 953.2894932 | -0.754518718 | 0.001074579 | 0.06 |
| YJR087W | 189.3064917 | -0.756366107 | 0.001122658 | 0.06 |
| YNR039C | 749.4014615 | 0.503345749 | 0.001074275 | 0.06 |
| YOL104C | 616.0912562 | 1.252179458 | 0.001123759 | 0.06 |
| YOR109W | 2024.801836 | 0.587577058 | 0.001022747 | 0.06 |
| YPL036W | 288.1590263 | 1.024811149 | 0.001048494 | 0.06 |
| YPL234C | 1583.101568 | -0.522999926 | 0.001087462 | 0.06 |
| YPR188C | 534.7061592 | -0.700367968 | 0.001123676 | 0.06 |
| YPR037C | 567.1835844 | -0.7810983 | 0.001139687 | 0.06 |
| YGR218W | 3998.195199 | 0.40543457 | 0.001161093 | 0.06 |
| YLR262C-A | 800.3537696 | -0.9240632 | 0.001267697 | 0.06 |
| YAL030W | 729.4057845 | -0.713036236 | 0.001349793 | 0.06 |
| YCR004C | 3730.565062 | -0.684139466 | 0.001339959 | 0.06 |
| YDR141C | 2153.367361 | 0.494767262 | 0.001339107 | 0.06 |
| YOR189W | 408.1511083 | -0.704817762 | 0.001339926 | 0.06 |
| YMR298W | 1278.824067 | -0.762549865 | 0.001399593 | 0.07 |
| YDR056C | 1611.517655 | -0.617135225 | 0.001458239 | 0.07 |
| YGR232W | 824.5178442 | -0.704381025 | 0.001459359 | 0.07 |
| YGR080W | 1465.883709 | -0.684491136 | 0.001473062 | 0.07 |
| YBR127C | 10020.63379 | -0.63541912 | 0.001529199 | 0.07 |
| YKR023W | 810.8865639 | -0.634849976 | 0.001520199 | 0.07 |
| YLR309C | 2329.969598 | -1.005229145 | 0.001504979 | 0.07 |
| YHR146W | 3577.393266 | -0.808381629 | 0.001552134 | 0.07 |
| YIL041W | 4311.545468 | -0.714298636 | 0.001586352 | 0.07 |
| YHR060W | 732.5771452 | -0.757414429 | 0.00164978 | 0.07 |
| YNL139C | 1577.578713 | 0.539069731 | 0.001702612 | 0.07 |

| | | | | |
|---|---|---|---|---|
| YBR273C | 1305.311458 | -0.564706031 | 0.001720999 | 0.07 |
| YJL121C | 2601.821139 | -0.641166054 | 0.00173488 | 0.07 |
| YDR373W | 760.4394828 | -0.574776222 | 0.001806852 | 0.08 |
| YLR027C | 7716.105415 | -0.763530696 | 0.001807125 | 0.08 |
| YBR014C | 1010.158039 | -0.599182119 | 0.001821552 | 0.08 |
| YDR327W | 614.5550695 | -0.588287581 | 0.00192143 | 0.08 |
| YGR133W | 486.059603 | -0.616442558 | 0.00192699 | 0.08 |
| YMR279C | 707.3104958 | 0.603193075 | 0.001922648 | 0.08 |
| YJL129C | 2269.469864 | 0.496066484 | 0.00196469 | 0.08 |
| YLR353W | 907.9621406 | 0.524803305 | 0.00195326 | 0.08 |
| YDR179C | 161.4536724 | -0.568724792 | 0.002050373 | 0.08 |
| YHL012W | 237.0696554 | 1.109252047 | 0.002040998 | 0.08 |
| YDL099W | 2397.811912 | -0.73656733 | 0.002107978 | 0.08 |
| YJL090C | 629.2586035 | 0.587009256 | 0.002138311 | 0.08 |
| YGL082W | 1508.755452 | -0.458996973 | 0.002231627 | 0.09 |
| YFR035C | 184.0960805 | -0.645264053 | 0.00228985 | 0.09 |
| YJL001W | 1978.350649 | -0.558723265 | 0.002318458 | 0.09 |
| YJL039C | 2015.018256 | 0.715203757 | 0.002330603 | 0.09 |
| YPL240C | 19482.15572 | -1.096558416 | 0.002329735 | 0.09 |
| YDR092W | 1261.21405 | -0.616533569 | 0.002347548 | 0.09 |
| YER178W | 9418.417677 | -0.587744722 | 0.002407765 | 0.09 |
| YFL038C | 5600.505944 | -0.570539598 | 0.002492716 | 0.09 |
| YHR115C | 896.8338505 | 0.767743984 | 0.002507573 | 0.09 |
| YIR037W | 2878.72569 | -0.80478165 | 0.002465643 | 0.09 |
| YJR135W-A | 435.3473743 | -0.817611263 | 0.002455409 | 0.09 |
| YNL230C | 466.9526022 | 0.866910566 | 0.002435986 | 0.09 |
| YPR103W | 3867.741987 | -0.787711405 | 0.002506829 | 0.09 |
| YOR359W | 584.0345426 | 1.038219288 | 0.002526876 | 0.09 |
| YIL062C | 1798.738473 | -0.627523882 | 0.002559606 | 0.09 |
| YER114C | 1610.317658 | 0.443470705 | 0.002660152 | 0.09 |
| YDR143C | 1247.877225 | 0.627675314 | 0.0027241 | 0.09 |
| YER023W | 2339.829047 | -0.803836084 | 0.002792324 | 0.09 |
| YIL030C | 2092.978365 | 0.768426932 | 0.002779592 | 0.09 |
| YIR014W | 256.3884009 | -1.006991191 | 0.00274307 | 0.09 |
| YKL065C | 4326.57925 | -0.613181059 | 0.002802475 | 0.09 |
| YKR065C | 1072.834618 | -0.672789524 | 0.002738745 | 0.09 |
| YLR066W | 1030.577166 | -0.69507739 | 0.002761789 | 0.09 |
| YNL283C | 1549.072925 | 0.61992867 | 0.002683544 | 0.09 |
| YJL061W | 1142.158213 | 0.542225312 | 0.002820704 | 0.09 |
| YBL058W | 3576.371774 | -0.589971613 | 0.002876962 | 0.09 |
| YPL170W | 622.4061116 | -0.535828203 | 0.002865029 | 0.09 |
| YDR388W | 5322.26629 | -0.477265271 | 0.002929725 | 0.10 |
| YBL001C | 1539.933777 | -0.91221212 | 0.003067557 | 0.10 |

| | | | | |
|---|---|---|---|---|
| YFL005W | 2091.331245 | -0.641977301 | 0.003054992 | 0.10 |
| YGL077C | 1372.265624 | 0.626776562 | 0.003042929 | 0.10 |
| YIL153W | 970.2297401 | -0.346662728 | 0.003044065 | 0.10 |
| YLR116W | 1156.147687 | 0.485005688 | 0.002986608 | 0.10 |
| YML075C | 1371.306293 | 0.564821544 | 0.003064718 | 0.10 |
| YPL048W | 7812.579467 | -0.591124183 | 0.002992617 | 0.10 |
| YLR408C | 156.1371802 | -0.658531936 | 0.003112434 | 0.10 |
| YDR368W | 2244.977391 | -0.775220219 | 0.003186568 | 0.10 |
| YOR296W | 873.0215241 | 0.521175594 | 0.003172117 | 0.10 |
| YEL020W-A | 192.2562057 | -0.735760175 | 0.003251701 | 0.10 |
| YMR099C | 2686.030784 | -0.523498844 | 0.00326521 | 0.10 |
| YDR093W | 1759.127414 | 0.61723399 | 0.003300994 | 0.10 |
| YOR160W | 1175.925728 | 0.359537647 | 0.003436692 | 0.10 |
| YJR125C | 2124.757426 | -0.576088867 | 0.003457699 | 0.10 |
| YOL138C | 1055.538589 | 0.570712336 | 0.003479688 | 0.10 |
| YCL017C | 1588.391961 | -0.624802978 | 0.00356989 | 0.10 |
| YKL207W | 2006.977142 | -0.689131013 | 0.003551375 | 0.10 |
| YPL225W | 2696.542417 | -0.617499438 | 0.003541652 | 0.10 |
| YER083C | 1868.758699 | -0.469800598 | 0.003623939 | 0.11 |
| YER072W | 2000.095671 | -1.014816426 | 0.003676697 | 0.11 |
| YHR039C-A | 2505.228571 | -0.59851972 | 0.003705716 | 0.11 |
| YKL165C | 1154.053703 | 0.86271446 | 0.003667616 | 0.11 |
| YLR403W | 956.7749718 | 0.729158611 | 0.003695839 | 0.11 |
| YBR164C | 1099.977428 | -0.796333943 | 0.003766603 | 0.11 |
| YDR304C | 4614.533404 | -0.686930796 | 0.003809764 | 0.11 |
| YNL221C | 1802.455751 | 0.692692571 | 0.003815189 | 0.11 |
| YNR035C | 3358.997637 | -0.784841083 | 0.003861791 | 0.11 |
| YIL027C | 793.9722774 | -0.677769094 | 0.003904004 | 0.11 |
| YOR033C | 730.7503196 | 0.478257797 | 0.003901496 | 0.11 |
| YER030W | 1421.194205 | -0.7246138 | 0.003957533 | 0.11 |
| YHR057C | 1425.81956 | -0.730488433 | 0.003981778 | 0.11 |
| YJL178C | 1325.547528 | -0.516278186 | 0.003968446 | 0.11 |
| YOR043W | 1570.483617 | 0.674155616 | 0.004016154 | 0.11 |
| YBL036C | 1223.509384 | -0.564155459 | 0.00416487 | 0.11 |
| YER004W | 2233.162604 | -0.716884039 | 0.004167372 | 0.11 |
| YJL091C | 524.9582628 | 0.583232154 | 0.004187374 | 0.11 |
| YKR020W | 476.7599628 | -0.639835015 | 0.004136115 | 0.11 |
| YMR223W | 516.9298345 | 0.440448487 | 0.004214756 | 0.11 |
| YNL249C | 367.0811931 | 0.546560796 | 0.004218517 | 0.11 |
| YOR261C | 3484.142585 | -0.772632486 | 0.004171754 | 0.11 |
| YPR073C | 916.4024804 | -0.658361489 | 0.004205801 | 0.11 |
| YCR009C | 3474.642891 | -0.601944826 | 0.004241115 | 0.11 |
| YFR050C | 3854.503578 | -0.633658081 | 0.004334007 | 0.11 |

| | | | | |
|---|---|---|---|---|
| YLR118C | 939.2371521 | -0.591864462 | 0.004331391 | 0.11 |
| YER060W-A | 584.2480367 | 0.975862586 | 0.004366664 | 0.11 |
| YGL100W | 1500.808318 | -0.444053877 | 0.004390684 | 0.11 |
| YKL214C | 653.6592896 | -0.556150534 | 0.004413101 | 0.11 |
| YLR119W | 415.3763546 | -0.574528486 | 0.004507178 | 0.11 |
| YOL038W | 4309.883987 | -0.626740262 | 0.004478671 | 0.11 |
| YPL128C | 628.8955352 | 0.678858509 | 0.004494983 | 0.11 |
| YPR104C | 970.815797 | 0.550523859 | 0.004584436 | 0.11 |
| YPL009C | 4156.098331 | -0.898747475 | 0.004620215 | 0.11 |
| YDR523C | 105.2296251 | 1.25942995 | 0.004660943 | 0.11 |
| YFL039C | 29152.57267 | -0.459055808 | 0.004736198 | 0.11 |
| YKL182W | 10979.47839 | 0.671531336 | 0.004715047 | 0.11 |
| YLR024C | 1454.81679 | 0.741036103 | 0.004728108 | 0.11 |
| YPL032C | 2772.776631 | 0.802810755 | 0.004753918 | 0.11 |
| YPR156C | 1539.255789 | 0.881691719 | 0.004724841 | 0.11 |
| YGL091C | 545.7319922 | -0.389005673 | 0.004918512 | 0.12 |
| YJR014W | 1708.927082 | -0.507109444 | 0.005018912 | 0.12 |
| YNL135C | 5166.62026 | -0.809717154 | 0.005287371 | 0.13 |
| YAL035W | 9252.51472 | -1.008066175 | 0.005382965 | 0.13 |
| YOR020C | 3482.406455 | -0.818826804 | 0.005392627 | 0.13 |
| YOR253W | 809.4249218 | -0.815771304 | 0.005381145 | 0.13 |
| YPR183W | 3494.823853 | -0.678891115 | 0.005326891 | 0.13 |
| YHR186C | 1649.917336 | 0.515592289 | 0.005594882 | 0.13 |
| YOR034C | 667.8994742 | 0.745455008 | 0.005625175 | 0.13 |
| YPL140C | 588.07308 | 0.661608081 | 0.005608481 | 0.13 |
| YKL013C | 2002.473422 | -0.548320411 | 0.005752755 | 0.13 |
| YDR239C | 796.3576703 | 0.493954234 | 0.005811175 | 0.13 |
| YER140W | 473.4616837 | 0.425915919 | 0.005837929 | 0.13 |
| YPL125W | 1168.744838 | 0.797099449 | 0.005852576 | 0.13 |
| YKR063C | 686.126014 | 0.454834585 | 0.005942786 | 0.13 |
| YCR028C-A | 1716.279601 | -0.757110311 | 0.006020127 | 0.13 |
| YLL014W | 750.2093462 | -0.678159704 | 0.005986025 | 0.13 |
| YLR347C | 1405.641169 | 0.517969167 | 0.006021357 | 0.13 |
| YMR001C | 1386.236499 | 0.675913332 | 0.00605767 | 0.14 |
| YNL267W | 1939.373479 | 0.379495669 | 0.006090417 | 0.14 |
| YOR149C | 790.0054864 | 0.532753875 | 0.006149943 | 0.14 |
| YOR233W | 853.5809378 | 0.713304874 | 0.006190522 | 0.14 |
| YIR022W | 1186.448374 | -0.616254656 | 0.006238202 | 0.14 |
| YLR169W | 310.0591441 | -0.557864001 | 0.006296145 | 0.14 |
| YML074C | 7957.336564 | -0.898123479 | 0.006292486 | 0.14 |
| YAL005C | 15020.34614 | -0.736971099 | 0.006405283 | 0.14 |
| YLR043C | 5741.406917 | -0.802249683 | 0.006381193 | 0.14 |
| YDL007W | 5356.140616 | -0.60720176 | 0.006562558 | 0.14 |

| | | | | |
|---|---|---|---|---|
| YDR422C | 1006.650102 | 0.601992703 | 0.006561951 | 0.14 |
| YLR440C | 889.0207433 | 0.729312302 | 0.006495423 | 0.14 |
| YOR038C | 676.9008791 | 0.435520013 | 0.00655633 | 0.14 |
| YBR060C | 829.7516444 | 0.464722481 | 0.006853276 | 0.15 |
| YMR129W | 1909.721365 | 0.444795488 | 0.006927845 | 0.15 |
| YNL199C | 1154.934737 | 0.29343971 | 0.006940236 | 0.15 |
| YGR167W | 3020.873713 | -0.470062226 | 0.007010294 | 0.15 |
| YLR046C | 345.7378309 | 0.420690698 | 0.007002376 | 0.15 |
| YGL231C | 1251.65783 | -0.49251935 | 0.007085292 | 0.15 |
| YHR026W | 2431.689374 | -0.66641058 | 0.007148233 | 0.15 |
| YIL043C | 4110.226673 | -0.60248716 | 0.007138428 | 0.15 |
| YPR063C | 1556.762503 | -0.586215024 | 0.007377779 | 0.15 |
| YIL047C | 1300.508261 | 0.627310148 | 0.007519528 | 0.15 |
| YJL030W | 384.5980723 | -0.6257913 | 0.007494062 | 0.15 |
| YNL181W | 1100.958937 | -0.499672704 | 0.007571668 | 0.15 |
| YGR180C | 8240.520071 | -0.738276266 | 0.007707745 | 0.16 |
| YHR110W | 1458.33201 | -0.504614463 | 0.007667652 | 0.16 |
| YPL016W | 1163.813997 | 0.750392883 | 0.007703217 | 0.16 |
| YKR048C | 4547.539769 | -0.656431899 | 0.00778968 | 0.16 |
| YDL097C | 4494.471702 | -0.531911245 | 0.008021911 | 0.16 |
| YIL003W | 534.8841649 | -0.509447198 | 0.00797901 | 0.16 |
| YLR343W | 1249.632226 | 1.143606939 | 0.007893741 | 0.16 |
| YNL329C | 1184.177512 | 0.533126303 | 0.007936466 | 0.16 |
| YPL178W | 1119.186631 | -0.404054365 | 0.008031106 | 0.16 |
| YPR036W | 6660.698271 | -0.633065695 | 0.007959484 | 0.16 |
| YBR011C | 18392.14922 | -0.755369344 | 0.008109521 | 0.16 |
| YDL165W | 697.5260403 | -0.428375993 | 0.008141332 | 0.16 |
| YGR132C | 2816.110199 | -0.610743384 | 0.008146625 | 0.16 |
| YMR074C | 1117.334751 | -0.724152922 | 0.008171559 | 0.16 |
| YDR364C | 596.6987029 | 0.435017165 | 0.008266568 | 0.16 |
| YER007W | 271.3998551 | 0.683621545 | 0.008281529 | 0.16 |
| YER120W | 4329.489139 | -0.474826019 | 0.008236229 | 0.16 |
| YDR291W | 877.4001219 | 0.412704913 | 0.008353366 | 0.16 |
| YOL111C | 1821.468977 | -0.396872637 | 0.008564363 | 0.16 |
| YER071C | 272.1086234 | -0.635938116 | 0.008618078 | 0.16 |
| YGR090W | 2037.446504 | 0.99043499 | 0.008679341 | 0.16 |
| YJL184W | 578.763074 | -0.5637583 | 0.008937034 | 0.17 |
| YHR147C | 857.7185719 | -0.487212465 | 0.00898215 | 0.17 |
| YML023C | 695.7405584 | 0.474442652 | 0.009039494 | 0.17 |
| YMR168C | 528.3657431 | 0.534895382 | 0.009169017 | 0.17 |
| YOR163W | 1216.735785 | -0.518385338 | 0.00915954 | 0.17 |
| YIR036C | 938.6857022 | -0.647149098 | 0.009242461 | 0.17 |
| YKR044W | 684.1987114 | 0.51254019 | 0.009278299 | 0.17 |

| | | | | |
|---|---|---|---|---|
| YDR374W-A | 251.03932 | -0.704549629 | 0.009356121 | 0.17 |
| YEL066W | 476.9220074 | -0.886098089 | 0.009562202 | 0.17 |
| YOL105C | 760.4552602 | 0.890185969 | 0.0095911 | 0.17 |
| YDL232W | 292.6096052 | -0.757474856 | 0.009632633 | 0.18 |
| YAR002C-A | 2600.594621 | -0.70903294 | 0.009856918 | 0.18 |
| YBL042C | 1258.710363 | 0.878948167 | 0.009793362 | 0.18 |
| YKL117W | 7112.851481 | -0.626930365 | 0.009930769 | 0.18 |
| YML078W | 2283.677343 | -0.786409677 | 0.00993095 | 0.18 |
| YNL079C | 8141.583459 | -0.702462314 | 0.009942045 | 0.18 |
| YOR327C | 881.0062218 | -0.612696449 | 0.009910337 | 0.18 |
| YOR349W | 572.160655 | 0.612888268 | 0.009937505 | 0.18 |
| YBL026W | 682.7010545 | -0.43006262 | 0.010144299 | 0.18 |
| YDR098C | 1711.271644 | -0.444415408 | 0.010137682 | 0.18 |
| YJR140C | 1086.888351 | 0.671870049 | 0.010205786 | 0.18 |
| YPL058C | 2629.343095 | 0.830727405 | 0.010178189 | 0.18 |
| YDL121C | 1113.601796 | -0.716833043 | 0.010250237 | 0.18 |
| YEL051W | 3229.319305 | -0.697191813 | 0.01032516 | 0.18 |
| YML081W | 1706.956949 | 0.692372763 | 0.010575776 | 0.18 |
| YIL075C | 7548.161259 | -0.639822854 | 0.010664798 | 0.18 |
| YPR148C | 2535.151148 | -0.456615477 | 0.010679856 | 0.18 |
| YBL056W | 3616.602261 | -0.37752382 | 0.010797983 | 0.19 |
| YAL029C | 2871.619875 | 0.408676467 | 0.010955415 | 0.19 |
| YBL052C | 898.9203912 | 0.490050387 | 0.011066654 | 0.19 |
| YDR155C | 17689.29075 | -0.88952566 | 0.011079104 | 0.19 |
| YKL058W | 636.5138347 | -0.527871619 | 0.011071049 | 0.19 |
| YLR073C | 322.9361924 | 0.976044857 | 0.010972935 | 0.19 |
| YMR216C | 1336.629718 | 0.544489077 | 0.010923033 | 0.19 |
| YOR355W | 1652.733408 | 0.903109378 | 0.011053628 | 0.19 |
| YCR002C | 1893.656785 | -0.629866487 | 0.011211317 | 0.19 |
| YDR524C | 904.6436686 | 0.552749003 | 0.011146983 | 0.19 |
| YEL037C | 3155.723536 | -0.59985648 | 0.011229834 | 0.19 |
| YJR088C | 954.591927 | -0.58115759 | 0.011254899 | 0.19 |
| YOR093C | 1072.002934 | 0.395117762 | 0.011171792 | 0.19 |
| YNL024C-A | 249.9046551 | -0.547167437 | 0.01135502 | 0.19 |
| YDR362C | 764.8048646 | 0.430412339 | 0.011523907 | 0.19 |
| YLR034C | 1366.422953 | 0.702505547 | 0.011500425 | 0.19 |
| YLR397C | 907.763381 | 0.631908216 | 0.011474516 | 0.19 |
| YMR186W | 31490.05511 | -1.002237059 | 0.011429113 | 0.19 |
| YOR156C | 466.9299753 | 0.566882444 | 0.011783127 | 0.19 |
| YKR027W | 720.4128865 | 0.443049045 | 0.011987956 | 0.19 |
| YNL084C | 2582.808264 | -0.360191094 | 0.01199301 | 0.19 |
| YDR381W | 4059.585724 | -0.716229887 | 0.012091078 | 0.19 |
| YLR421C | 1264.683888 | -0.518372987 | 0.012056085 | 0.19 |

| | | | | |
|---|---|---|---|---|
| YIL013C | 704.2538412 | 0.619545778 | 0.012199531 | 0.20 |
| YLR438C-A | 659.2030513 | -0.696659249 | 0.012215853 | 0.20 |
| YBL106C | 800.8986155 | 0.687866381 | 0.012501341 | 0.20 |
| YBR089C-A | 560.6070076 | -0.653699603 | 0.01269207 | 0.20 |
| YBR205W | 1374.246383 | -0.522053299 | 0.012629145 | 0.20 |
| YBR279W | 1844.253736 | -0.446843827 | 0.012433941 | 0.20 |
| YDL220C | 416.2502353 | 0.854150897 | 0.012652191 | 0.20 |
| YGR089W | 1598.86435 | 0.503928301 | 0.012684503 | 0.20 |
| YIL051C | 4289.573013 | -0.849853229 | 0.012656596 | 0.20 |
| YJL117W | 1366.561839 | -0.580603131 | 0.012530357 | 0.20 |
| YOR188W | 1393.160444 | 0.57574967 | 0.012509271 | 0.20 |
| YPL098C | 678.1300721 | -0.502014207 | 0.012625758 | 0.20 |
| YFL028C | 788.0732688 | -0.510370634 | 0.012935047 | 0.20 |
| YGL020C | 1129.205349 | -0.599034844 | 0.012900003 | 0.20 |
| YJL124C | 1576.189687 | -0.515930645 | 0.012796847 | 0.20 |
| YJR144W | 1299.865299 | -0.444162486 | 0.012896047 | 0.20 |
| YMR264W | 1111.60862 | -0.514992465 | 0.012840579 | 0.20 |
| YMR294W-A | 229.7802387 | -0.468468212 | 0.012952496 | 0.20 |
| YDR016C | 195.2031569 | -0.598322248 | 0.013115955 | 0.20 |
| YDL212W | 1994.74877 | -0.690821812 | 0.013163522 | 0.20 |
| YBR221C | 5794.723277 | -0.571156946 | 0.013350177 | 0.20 |
| YDL080C | 732.479931 | 0.415485563 | 0.013357092 | 0.20 |
| YLR355C | 9452.460146 | 0.836557505 | 0.013292783 | 0.20 |
| YHR100C | 468.2098929 | -0.501395831 | 0.01346403 | 0.20 |
| YLR023C | 1130.122098 | 0.500052171 | 0.013468291 | 0.20 |
| YDR167W | 803.2507476 | -0.49331878 | 0.01359314 | 0.20 |
| YLR170C | 116.6208147 | -0.760851912 | 0.013675975 | 0.20 |
| YMR226C | 4108.834639 | -0.615505751 | 0.013768482 | 0.20 |
| YGR282C | 11912.38121 | -0.47970611 | 0.013811038 | 0.20 |
| YDR063W | 1183.071975 | -0.400711802 | 0.013880002 | 0.21 |
| YPL127C | 2242.895143 | -0.614222787 | 0.013939218 | 0.21 |
| YDR325W | 750.6903469 | 0.608794242 | 0.014078073 | 0.21 |
| YGL216W | 962.5040676 | 0.444324262 | 0.01414811 | 0.21 |
| YPL254W | 446.984746 | 0.55095887 | 0.014184472 | 0.21 |
| YDR152W | 1737.468978 | -0.561562707 | 0.014270108 | 0.21 |
| YIL034C | 3381.997418 | -0.349306623 | 0.01424611 | 0.21 |
| YDR228C | 901.0372572 | 0.505244786 | 0.014342665 | 0.21 |
| YDR469W | 524.7044195 | -0.641228823 | 0.01454812 | 0.21 |
| YER044C | 1096.772117 | -0.700236663 | 0.014631642 | 0.21 |
| YGR040W | 494.1182396 | 0.758033657 | 0.014720755 | 0.21 |
| YIL124W | 2615.425395 | -0.679022672 | 0.014706859 | 0.21 |
| YFL016C | 3333.34381 | -0.668336171 | 0.014777931 | 0.21 |
| YOR127W | 1893.705383 | 0.498804742 | 0.014820825 | 0.21 |

| | | | | |
|---|---|---|---|---|
| YMR022W | 1239.988654 | -0.46742266 | 0.014932 | 0.21 |
| YML055W | 811.1131816 | -0.51181646 | 0.015059588 | 0.21 |
| YBR282W | 512.5901703 | -0.59247258 | 0.01517039 | 0.22 |
| YML006C | 696.4182875 | 0.448089066 | 0.015235095 | 0.22 |
| YBR206W | 255.7378889 | -0.57108263 | 0.015446369 | 0.22 |
| YLR154C | 335.1264893 | -0.454853905 | 0.015421926 | 0.22 |
| YNL138W | 4814.982276 | -0.508760189 | 0.015441582 | 0.22 |
| YJL068C | 1815.839599 | -0.648591752 | 0.015643144 | 0.22 |
| YMR308C | 3347.564301 | 0.750169915 | 0.015671844 | 0.22 |
| YOR074C | 848.568103 | -0.713342384 | 0.015646245 | 0.22 |
| YGL028C | 1677.067671 | 0.737004417 | 0.015756519 | 0.22 |
| YDR038C | 125.3516014 | 0.601725265 | 0.015900498 | 0.22 |
| YEL013W | 1708.200753 | 0.450614769 | 0.016332342 | 0.23 |
| YGR026W | 4521.168167 | -0.51807838 | 0.0165196 | 0.23 |
| YKR053C | 486.0346558 | 0.625685361 | 0.016525348 | 0.23 |
| YGR275W | 659.6729657 | -0.558020661 | 0.016579019 | 0.23 |
| YJR132W | 1306.62171 | 0.540159712 | 0.016700923 | 0.23 |
| YGR092W | 957.140587 | 0.71739848 | 0.016799204 | 0.23 |
| YGR083C | 2563.158798 | -0.741790306 | 0.017036142 | 0.23 |
| YLR351C | 1673.534289 | -0.465843474 | 0.016992708 | 0.23 |
| YMR184W | 674.607137 | -0.520886549 | 0.017051628 | 0.23 |
| YNL186W | 1470.462386 | 0.514194538 | 0.017028459 | 0.23 |
| YFR052W | 2758.803837 | -0.619411725 | 0.017362975 | 0.23 |
| YNL268W | 2190.621688 | 0.565333995 | 0.017398203 | 0.23 |
| YER070W | 2200.790509 | 1.007611332 | 0.017497504 | 0.23 |
| YPR162C | 653.0734249 | 0.625923464 | 0.01754849 | 0.24 |
| YBR091C | 248.9757925 | -0.572195443 | 0.017664551 | 0.24 |
| YJL098W | 1058.267184 | 0.747133664 | 0.017833605 | 0.24 |
| YHR119W | 1485.712969 | 0.542302315 | 0.017903175 | 0.24 |
| YLR460C | 187.4228264 | -0.705694562 | 0.018289093 | 0.24 |
| YHR109W | 446.2718996 | 0.55485273 | 0.018348364 | 0.24 |
| YDR086C | 1629.638329 | -0.707981235 | 0.018551878 | 0.24 |
| YDL180W | 735.4223348 | 0.55036978 | 0.018661277 | 0.25 |
| YJR082C | 608.3949667 | -0.541586128 | 0.01867326 | 0.25 |
| YER113C | 743.4950149 | 0.455972027 | 0.018799272 | 0.25 |
| YBR200W | 1077.368125 | -0.397461641 | 0.018974067 | 0.25 |
| YCR024C | 641.2724541 | 0.373035505 | 0.018965341 | 0.25 |
| YDR007W | 490.6292629 | -0.401123268 | 0.018901887 | 0.25 |
| YPL256C | 1702.304083 | 0.838538578 | 0.019156471 | 0.25 |
| YLR230W | 287.1818195 | -0.604246032 | 0.01924314 | 0.25 |
| YER105C | 2233.871003 | 0.358581785 | 0.019332102 | 0.25 |
| YJL092W | 1243.946562 | 0.56593651 | 0.019440926 | 0.25 |
| YJL197W | 1228.673265 | 0.402822396 | 0.019428416 | 0.25 |

| | | | | |
|---|---|---|---|---|
| YNL099C | 699.7662248 | -0.516340518 | 0.019435213 | 0.25 |
| YOR027W | 11708.63489 | -0.711328383 | 0.01952101 | 0.25 |
| YDR068W | 1258.879245 | -0.267295613 | 0.019621939 | 0.25 |
| YGR206W | 264.0711736 | -0.525314182 | 0.019682054 | 0.25 |
| YJR099W | 741.8529597 | -0.488551361 | 0.019861912 | 0.25 |
| YLR430W | 3065.658429 | 0.49702486 | 0.019870949 | 0.25 |
| YOL006C | 1492.750425 | 0.477346831 | 0.01984344 | 0.25 |
| YKR007W | 738.2939867 | -0.44936358 | 0.01999871 | 0.25 |
| YKR010C | 516.9970849 | 0.663725664 | 0.019955211 | 0.25 |
| YGR271W | 1597.427923 | 0.459967899 | 0.020051197 | 0.25 |
| YBR239C | 441.6078035 | 0.47510716 | 0.020428942 | 0.25 |
| YDL122W | 1760.896516 | 0.416221156 | 0.020396581 | 0.25 |
| YKR047W | 143.8726915 | -0.448098044 | 0.02045761 | 0.25 |
| YNL010W | 2912.199227 | -0.772494212 | 0.020466099 | 0.25 |
| YOR116C | 1548.881592 | 0.628663951 | 0.020411125 | 0.25 |
| YFR024C-A | 2497.30533 | -0.500109212 | 0.02053667 | 0.25 |
| YBR263W | 3971.725103 | -0.429191035 | 0.020615723 | 0.25 |
| YJR031C | 1083.680948 | 0.56548359 | 0.02066527 | 0.25 |
| YOR298W | 171.3347166 | 0.726907209 | 0.020711899 | 0.25 |
| YML012W | 3525.480868 | -0.578657533 | 0.020850403 | 0.26 |
| YNL133C | 297.07226 | -0.633278044 | 0.020981721 | 0.26 |
| YOR198C | 6733.130979 | -0.66395507 | 0.020927653 | 0.26 |
| YOR210W | 598.8398742 | -0.762572594 | 0.020961109 | 0.26 |
| YOR067C | 1025.694017 | 0.493276266 | 0.021051914 | 0.26 |
| YDR339C | 1137.925374 | -0.763709157 | 0.021290941 | 0.26 |
| YGL131C | 1060.332301 | 0.738926512 | 0.021278572 | 0.26 |
| YMR152W | 1727.69017 | -0.446483311 | 0.02122832 | 0.26 |
| YDR154C | 3531.276985 | -0.804581814 | 0.021494445 | 0.26 |
| YLR259C | 10469.29817 | -0.515323524 | 0.021494915 | 0.26 |
| YML121W | 941.5006143 | -0.409311702 | 0.021476848 | 0.26 |
| YCR028C | 527.4192161 | 0.369996176 | 0.021946226 | 0.26 |
| YCR086W | 338.9363584 | -0.653145515 | 0.022319478 | 0.26 |
| YDR242W | 619.518258 | 0.61188 | 0.022176419 | 0.26 |
| YDR482C | 359.2978323 | -0.381693905 | 0.021884346 | 0.26 |
| YDR486C | 1425.158924 | -0.508884043 | 0.022117763 | 0.26 |
| YER180C | 405.292426 | 0.458205345 | 0.021652878 | 0.26 |
| YGR200C | 1330.804216 | 0.697032835 | 0.022068097 | 0.26 |
| YHL031C | 431.2428898 | -0.435219157 | 0.021661433 | 0.26 |
| YIL010W | 1043.363564 | -0.560140435 | 0.02232665 | 0.26 |
| YKL192C | 3222.400243 | -0.452286192 | 0.021999691 | 0.26 |
| YNL155W | 1378.201426 | -0.615519863 | 0.021816529 | 0.26 |
| YOL146W | 364.144711 | -0.575889859 | 0.021771754 | 0.26 |
| YOL156W | 95.75918972 | 1.029404562 | 0.021747957 | 0.26 |

| | | | | |
|---|---|---|---|---|
| YOR154W | 667.1567549 | 0.423616116 | 0.022052507 | 0.26 |
| YOR286W | 1162.614528 | -0.498072476 | 0.021934525 | 0.26 |
| YOR362C | 3907.49208 | -0.428213732 | 0.022329198 | 0.26 |
| YPL106C | 19467.56403 | -0.738109464 | 0.0223151 | 0.26 |
| YEL015W | 1435.206957 | 0.376389863 | 0.022852456 | 0.26 |
| YKL054C | 4561.844374 | 0.641921422 | 0.022721456 | 0.26 |
| YKR011C | 1595.285427 | -0.722727791 | 0.022815027 | 0.26 |
| YMR059W | 205.0702465 | -0.378678337 | 0.022723911 | 0.26 |
| YOR117W | 3352.309698 | -0.509492592 | 0.022861243 | 0.26 |
| YNL278W | 749.3245241 | 0.499525065 | 0.022967308 | 0.26 |
| YLR218C | 291.942794 | -0.607638262 | 0.023112106 | 0.26 |
| YGR284C | 2427.894256 | -0.610441991 | 0.023194443 | 0.26 |
| YBR137W | 861.5111114 | -0.84222234 | 0.023455621 | 0.27 |
| YMR150C | 511.1258272 | -0.48703444 | 0.023531698 | 0.27 |
| YDL157C | 952.2291983 | -0.850816051 | 0.023639785 | 0.27 |
| YKL145W | 5482.552156 | -0.583498111 | 0.023627493 | 0.27 |
| YDR032C | 4060.720659 | -0.690159904 | 0.023803912 | 0.27 |
| YJL026W | 4324.794509 | -0.555889677 | 0.023722681 | 0.27 |
| YLR200W | 774.013828 | -0.50582375 | 0.023778181 | 0.27 |
| YPL272C | 344.5733609 | 0.771670373 | 0.02384816 | 0.27 |
| YGR267C | 1681.678228 | -0.357110343 | 0.024011543 | 0.27 |
| YML110C | 4679.227772 | -0.706586076 | 0.023999385 | 0.27 |
| YER048C | 2682.830499 | -0.51518581 | 0.024319871 | 0.27 |
| YOR157C | 2318.284571 | -0.381836526 | 0.024508156 | 0.27 |
| YPL216W | 730.7653507 | 0.504327495 | 0.024498203 | 0.27 |
| YBR062C | 819.8607412 | -0.61512939 | 0.024600117 | 0.27 |
| YER009W | 3909.301197 | -0.794186756 | 0.024651436 | 0.27 |
| YCR075W-A | 129.8943621 | -0.649257932 | 0.024727442 | 0.27 |
| YGR199W | 855.4590705 | 0.534996117 | 0.024753095 | 0.27 |
| YNR063W | 178.9442947 | 0.678769536 | 0.024995884 | 0.27 |
| YDR246W | 916.4165129 | -0.383913816 | 0.02523952 | 0.28 |
| YGL122C | 2999.105342 | -0.644694862 | 0.02533235 | 0.28 |
| YIL156W-B | 291.2591429 | -0.487630984 | 0.025338309 | 0.28 |
| YMR013C | 348.9768631 | 0.646835081 | 0.025286623 | 0.28 |
| YOR021C | 1473.922183 | -0.61097812 | 0.025388944 | 0.28 |
| YOR150W | 702.6825687 | -0.619461986 | 0.025408921 | 0.28 |
| YCR020C-A | 266.9746924 | -0.61299128 | 0.025502938 | 0.28 |
| YGL220W | 745.8780305 | -0.519811305 | 0.0256357 | 0.28 |
| YGR262C | 876.7108592 | -0.303950686 | 0.025646661 | 0.28 |
| YNL095C | 702.5523068 | 0.633284928 | 0.025606106 | 0.28 |
| YDL200C | 293.2384509 | -0.508226734 | 0.025767795 | 0.28 |
| YAL002W | 1569.483592 | 0.386919999 | 0.026118833 | 0.28 |
| YER142C | 720.8941802 | -0.685906324 | 0.02607074 | 0.28 |

| | | | | |
|---|---|---|---|---|
| YNL107W | 513.6532475 | -0.342242256 | 0.026053475 | 0.28 |
| YNL310C | 605.4678235 | -0.392321346 | 0.025975926 | 0.28 |
| YDR248C | 767.0999761 | -0.636408536 | 0.026261215 | 0.28 |
| YFR044C | 5702.058987 | -0.582023545 | 0.026212309 | 0.28 |
| YKL119C | 426.0044347 | -0.543569722 | 0.02630066 | 0.28 |
| YFR003C | 1017.431306 | -0.592588038 | 0.026657954 | 0.28 |
| YOR269W | 302.0905548 | 0.409526305 | 0.026689905 | 0.28 |
| YER172C | 2101.126018 | 0.323684777 | 0.026801017 | 0.28 |
| YIL076W | 2351.989968 | -0.643179363 | 0.026940237 | 0.28 |
| YGL066W | 1160.200958 | 0.467904317 | 0.027028346 | 0.28 |
| YGR024C | 991.4360363 | -0.335518639 | 0.027084548 | 0.28 |
| YKL125W | 592.2613366 | 0.551191189 | 0.027122466 | 0.28 |
| YGL130W | 854.7206722 | -0.243211317 | 0.027340003 | 0.29 |
| YER051W | 512.8476233 | 0.689989041 | 0.027451763 | 0.29 |
| YHR113W | 1875.547688 | -0.523205118 | 0.027511994 | 0.29 |
| YMR028W | 753.5245359 | -0.329248572 | 0.027779131 | 0.29 |
| YOL008W | 381.7473344 | -0.381292941 | 0.028121859 | 0.29 |
| YNL208W | 6496.983749 | -0.543948974 | 0.02839181 | 0.29 |
| YBR159W | 2299.591935 | -0.670088931 | 0.028510271 | 0.29 |
| YBR264C | 359.5103772 | -0.404887498 | 0.028664109 | 0.29 |
| YHR164C | 1299.336952 | 0.435765312 | 0.028644612 | 0.29 |
| YGR191W | 1267.875828 | 0.5856101 | 0.028770829 | 0.30 |
| YPL183C | 913.1902093 | 0.709548108 | 0.028845702 | 0.30 |
| YAL044W-A | 228.0761832 | -0.786644779 | 0.029070791 | 0.30 |
| YER100W | 1553.364761 | -0.473285711 | 0.029243534 | 0.30 |
| YGR120C | 376.5121986 | -0.229877397 | 0.029370459 | 0.30 |
| YGR157W | 3036.942102 | 0.759218107 | 0.029006901 | 0.30 |
| YJL016W | 1543.218497 | 0.54011779 | 0.029282174 | 0.30 |
| YKL063C | 828.820505 | -0.471786036 | 0.029408225 | 0.30 |
| YKL196C | 2255.642273 | -0.433465365 | 0.029392452 | 0.30 |
| YLL026W | 27987.51895 | -0.694678317 | 0.029092938 | 0.30 |
| YML048W | 4451.705992 | -0.472378135 | 0.029304253 | 0.30 |
| YPL065W | 569.9929052 | -0.52843749 | 0.02986805 | 0.30 |
| YDR347W | 1409.321423 | -0.300165771 | 0.030102492 | 0.30 |
| YJR119C | 341.555846 | 0.575318094 | 0.03008682 | 0.30 |
| YER184C | 397.7514223 | 0.590768801 | 0.030382807 | 0.30 |
| YIL008W | 681.0099622 | -0.746143038 | 0.03036772 | 0.30 |
| YKL057C | 1611.669894 | 0.548958769 | 0.030315306 | 0.30 |
| YLR301W | 1464.12998 | -0.867128627 | 0.030317632 | 0.30 |
| YOR337W | 623.7332196 | 0.540360863 | 0.030536648 | 0.30 |
| YBL059C-A | 504.0749236 | -0.78065297 | 0.0308477 | 0.30 |
| YER010C | 1015.031336 | -0.413252048 | 0.030859663 | 0.30 |
| YKR054C | 2620.574744 | 0.47038213 | 0.030972898 | 0.30 |

| | | | | |
|---|---|---|---|---|
| YLR059C | 1264.300383 | -0.513439405 | 0.031010283 | 0.30 |
| YLR401C | 1341.53932 | 0.501563058 | 0.031081818 | 0.30 |
| YMR005W | 1198.923005 | -0.536358691 | 0.030765822 | 0.30 |
| YMR244C-A | 108.6393324 | -0.571798237 | 0.030909882 | 0.30 |
| YOR204W | 2305.73227 | 0.6032888 | 0.031090748 | 0.30 |
| YDR487C | 2644.625402 | -0.413255172 | 0.031325415 | 0.30 |
| YEL058W | 1683.65634 | -0.635249037 | 0.031356575 | 0.30 |
| YFL040W | 216.9489399 | 0.856563918 | 0.031404135 | 0.30 |
| YNR055C | 1151.83484 | 0.50619478 | 0.031408846 | 0.30 |
| YPR108W | 2749.924825 | -0.446124771 | 0.0312791 | 0.30 |
| YOR126C | 423.6858153 | -0.331826527 | 0.031481663 | 0.30 |
| YCL028W | 2867.256977 | -0.459996949 | 0.031804805 | 0.31 |
| YMR100W | 811.7962257 | 0.616968299 | 0.031934912 | 0.31 |
| YOR020W-A | 1169.212022 | -0.774476259 | 0.032282294 | 0.31 |
| YLL033W | 166.5445731 | 0.420108485 | 0.032466251 | 0.31 |
| YOR217W | 1207.361787 | 0.393149256 | 0.032465865 | 0.31 |
| YPR028W | 4646.798863 | -0.570791539 | 0.032597636 | 0.31 |
| YEL053C | 1328.366975 | 0.296407267 | 0.032709145 | 0.31 |
| YOR166C | 666.3506234 | 0.342538972 | 0.032751651 | 0.31 |
| YDL103C | 1449.48201 | -0.464602972 | 0.032933164 | 0.31 |
| YNL312W | 897.7476795 | -0.480520991 | 0.032932463 | 0.31 |
| YER031C | 2004.561338 | -0.622966692 | 0.032996966 | 0.31 |
| YDL015C | 3069.696513 | -0.34340403 | 0.033353153 | 0.32 |
| YKR062W | 891.4921763 | -0.340616218 | 0.03347075 | 0.32 |
| YGR212W | 361.1538773 | 0.411667087 | 0.033621871 | 0.32 |
| YBR251W | 1030.356421 | -0.522005654 | 0.033906784 | 0.32 |
| YMR236W | 1041.170124 | -0.319630444 | 0.033836066 | 0.32 |
| YOR252W | 480.3993858 | -0.600038746 | 0.033944555 | 0.32 |
| YHR059W | 401.0845698 | -0.490083319 | 0.034032998 | 0.32 |
| YGR005C | 1346.509283 | -0.356335372 | 0.034346472 | 0.32 |
| YOR259C | 3766.746436 | -0.549802474 | 0.03428144 | 0.32 |
| YPR107C | 422.2754774 | -0.365710375 | 0.034341615 | 0.32 |
| YKL160W | 1725.224125 | -0.444867338 | 0.034414302 | 0.32 |
| YCR038C | 411.3006768 | 0.521337498 | 0.034599896 | 0.32 |
| YNL201C | 1076.736749 | 0.338225294 | 0.03462459 | 0.32 |
| YIL095W | 775.5717487 | 0.360207978 | 0.034729288 | 0.32 |
| YNL048W | 1078.322808 | 0.323210829 | 0.034909805 | 0.32 |
| YOL045W | 1132.896142 | 0.326475511 | 0.034994723 | 0.32 |
| YDR026C | 840.5603663 | 0.353767135 | 0.035269832 | 0.32 |
| YKR028W | 1923.731148 | 0.446748468 | 0.03522368 | 0.32 |
| YGR020C | 3100.566353 | -0.663471028 | 0.035383836 | 0.33 |
| YGR230W | 330.0627222 | -0.439935566 | 0.035524881 | 0.33 |
| YLL039C | 7097.91074 | -0.688383761 | 0.035683563 | 0.33 |

| | | | | |
|---|---|---|---|---|
| YHR102W | 1800.525655 | 0.400782229 | 0.035871487 | 0.33 |
| YLR146W-A | 192.5995673 | -0.609259633 | 0.036015029 | 0.33 |
| YMR222C | 712.4011542 | -0.379760026 | 0.035902407 | 0.33 |
| YPL013C | 664.0145708 | -0.606318114 | 0.035957419 | 0.33 |
| YJL133W | 754.0202192 | 0.536165832 | 0.036113561 | 0.33 |
| YJR034W | 337.4607648 | -0.62769956 | 0.036208304 | 0.33 |
| YIL090W | 898.4488003 | 0.389030321 | 0.036297377 | 0.33 |
| YGL048C | 4170.423196 | -0.380010371 | 0.036662307 | 0.33 |
| YER005W | 999.8103274 | 0.294036469 | 0.036797327 | 0.33 |
| YGR071C | 709.1837053 | 0.430749166 | 0.036876496 | 0.33 |
| YIL073C | 530.2219401 | 0.407452503 | 0.037091179 | 0.33 |
| YLR361C | 559.6543534 | 0.432374346 | 0.037125519 | 0.33 |
| YJL051W | 1072.541155 | 0.341164055 | 0.037195673 | 0.33 |
| YEL047C | 2163.496037 | -0.5393282 | 0.037366979 | 0.33 |
| YCR073W-A | 1340.952291 | -0.648646336 | 0.038014495 | 0.34 |
| YDR046C | 2206.66233 | 0.602858674 | 0.038032651 | 0.34 |
| YER077C | 789.2511196 | 0.348338653 | 0.037939383 | 0.34 |
| YKL190W | 813.5201171 | -0.550280858 | 0.03791963 | 0.34 |
| YBR122C | 1316.133892 | -0.517340605 | 0.038303578 | 0.34 |
| YFR021W | 746.0129726 | -0.311429292 | 0.038300969 | 0.34 |
| YLR456W | 255.5161749 | -0.501123052 | 0.03850887 | 0.34 |
| YMR198W | 512.8655409 | 0.402283335 | 0.038676584 | 0.34 |
| YMR266W | 1578.35636 | 0.505920832 | 0.03875194 | 0.34 |
| YJR085C | 2130.783694 | -0.537335632 | 0.039019321 | 0.34 |
| YBR156C | 807.2285578 | 0.354135589 | 0.039223257 | 0.34 |
| YKL003C | 652.5154668 | -0.534298475 | 0.039197909 | 0.34 |
| YBL102W | 1291.754797 | -0.650691818 | 0.039578468 | 0.35 |
| YNL149C | 291.2466735 | -0.701711325 | 0.039455928 | 0.35 |
| YPL081W | 1016.551966 | 0.936630115 | 0.039559461 | 0.35 |
| YCR059C | 1130.721478 | -0.494186067 | 0.039653477 | 0.35 |
| YBL050W | 1823.454847 | -0.391586523 | 0.04020442 | 0.35 |
| YDL100C | 5446.231089 | -0.57274298 | 0.040178314 | 0.35 |
| YDR405W | 1203.736123 | -0.416296664 | 0.040088469 | 0.35 |
| YOR304W | 1385.923767 | 0.29177898 | 0.040312949 | 0.35 |
| YEL068C | 304.3910786 | -0.618723866 | 0.04044695 | 0.35 |
| YJL201W | 1030.90288 | 0.365888996 | 0.040504267 | 0.35 |
| YGL016W | 794.4716524 | 0.76120034 | 0.040594048 | 0.35 |
| YLR320W | 753.4878726 | 0.409341308 | 0.040638529 | 0.35 |
| YBR069C | 2910.073987 | 0.669545645 | 0.041081646 | 0.35 |
| YER019C-A | 1962.324665 | -0.455171935 | 0.041015133 | 0.35 |
| YLR067C | 640.6258307 | 0.423629046 | 0.041115818 | 0.35 |
| YML016C | 1375.407219 | 0.474856211 | 0.041211454 | 0.35 |
| YEL027W | 5744.186516 | -0.460777782 | 0.041308822 | 0.35 |

| | | | | |
|---|---|---|---|---|
| YPL179W | 1527.082249 | 0.520121605 | 0.041762883 | 0.36 |
| YFR043C | 412.3285976 | -0.397643876 | 0.041851991 | 0.36 |
| YJL205C | 132.3580317 | -0.642524292 | 0.042226611 | 0.36 |
| YJR011C | 218.4017674 | -0.379087298 | 0.042313491 | 0.36 |
| YDR002W | 3692.839207 | -0.740995134 | 0.042716518 | 0.36 |
| YEL032W | 1185.048516 | 0.362770712 | 0.043161518 | 0.36 |
| YER058W | 286.3082478 | -0.507053163 | 0.043058007 | 0.36 |
| YHR061C | 329.4509042 | 0.466560301 | 0.042992058 | 0.36 |
| YLR370C | 2370.21826 | -0.484107479 | 0.043109762 | 0.36 |
| YOR280C | 559.6900755 | -0.50772995 | 0.043119583 | 0.36 |
| YPL117C | 2245.25569 | -0.454684435 | 0.043008685 | 0.36 |
| YJL024C | 668.6768051 | -0.363423953 | 0.043347787 | 0.36 |
| YCL034W | 1930.992154 | -0.517380031 | 0.043552256 | 0.37 |
| YDR454C | 2624.226849 | -0.656923108 | 0.043863956 | 0.37 |
| YPL177C | 1454.555337 | 0.423506092 | 0.043821825 | 0.37 |
| YAL040C | 2404.494026 | 0.68635345 | 0.043933231 | 0.37 |
| YDR195W | 649.2195359 | 0.410310247 | 0.044389242 | 0.37 |
| YMR306W | 518.3037802 | 0.610497313 | 0.0443868 | 0.37 |
| YDL117W | 1322.092729 | 0.375392165 | 0.044874885 | 0.37 |
| YLR305C | 2169.127998 | 0.517838474 | 0.045211476 | 0.38 |
| YHR056C | 832.3445362 | 0.422473439 | 0.045414904 | 0.38 |
| YPR082C | 431.92377 | -0.521146373 | 0.045349525 | 0.38 |
| YHR182W | 713.7981983 | 0.55278741 | 0.04563471 | 0.38 |
| YKL053C-A | 473.2870725 | -0.627045338 | 0.045966522 | 0.38 |
| YDR027C | 1560.996774 | 0.234146738 | 0.046360205 | 0.38 |
| YJR147W | 698.6444516 | 0.807987481 | 0.046676808 | 0.38 |
| YNL218W | 370.2566133 | 0.478704389 | 0.046660303 | 0.38 |
| YNL131W | 2877.827714 | -0.473004082 | 0.047135113 | 0.39 |
| YIL002C | 1259.267656 | 0.376125853 | 0.047416222 | 0.39 |
| YML041C | 413.9344954 | -0.433787648 | 0.047403616 | 0.39 |
| YMR232W | 290.2765579 | 0.864810485 | 0.047343956 | 0.39 |
| YPL037C | 6009.036088 | -0.606144732 | 0.047519856 | 0.39 |
| YKR093W | 2668.023949 | 0.758178953 | 0.047736678 | 0.39 |
| YPL237W | 2463.364966 | -0.582059869 | 0.047678498 | 0.39 |
| YER093C-A | 366.8172657 | -0.506576125 | 0.047933457 | 0.39 |
| YBR230W-A | 211.0787097 | -0.955320601 | 0.048046729 | 0.39 |
| YJL126W | 376.2312005 | -0.330109461 | 0.04828248 | 0.39 |
| YCR024C-B | 2025.586967 | 0.591930759 | 0.048822971 | 0.39 |
| YGL226C-A | 628.3645694 | -0.548270915 | 0.04895404 | 0.39 |
| YKL002W | 1601.062288 | -0.406747596 | 0.048953095 | 0.39 |
| YLR105C | 592.2672821 | 0.441812232 | 0.048849301 | 0.39 |
| YPL087W | 1694.761164 | -0.591085289 | 0.048892644 | 0.39 |
| YPR159W | 2743.676738 | 0.55223724 | 0.048761848 | 0.39 |

| | | | | |
|---|---|---|---|---|
| YAL039C | 1453.711163 | -0.513220993 | 0.050422894 | 0.39 |
| YBL004W | 2589.356792 | 0.719383644 | 0.049703871 | 0.39 |
| YCR082W | 1363.300313 | -0.599680968 | 0.050037197 | 0.39 |
| YDL047W | 1531.758576 | -0.42114079 | 0.050230136 | 0.39 |
| YDR163W | 579.7588355 | -0.459701829 | 0.049673518 | 0.39 |
| YDR427W | 2923.459081 | -0.458253688 | 0.050097663 | 0.39 |
| YER034W | 308.6813768 | -0.586694524 | 0.050477307 | 0.39 |
| YER112W | 1280.704105 | -0.431154177 | 0.049262797 | 0.39 |
| YFR025C | 947.4315897 | -0.310578748 | 0.050165362 | 0.39 |
| YGL047W | 625.4320388 | -0.395607893 | 0.049538368 | 0.39 |
| YGL144C | 301.1617021 | 0.531234587 | 0.050183453 | 0.39 |
| YGR207C | 1419.844599 | -0.382132844 | 0.049997064 | 0.39 |
| YKL138C-A | 204.185803 | -0.447953406 | 0.049243887 | 0.39 |
| YKR083C | 244.3380659 | -0.651247534 | 0.050466697 | 0.39 |
| YLR025W | 2034.136821 | -0.476730945 | 0.049564236 | 0.39 |
| YLR229C | 1318.158779 | -0.440573456 | 0.0501947 | 0.39 |
| YLR272C | 941.972981 | 0.449357388 | 0.050425112 | 0.39 |
| YOL133W | 270.3290721 | -0.565731697 | 0.050027902 | 0.39 |
| YOR042W | 2668.884695 | -0.528277828 | 0.050462524 | 0.39 |
| YOR367W | 630.0179766 | -0.464797368 | 0.04970984 | 0.39 |
| YLR204W | 619.6469206 | -0.551247518 | 0.050595377 | 0.39 |
| YER171W | 948.7525289 | 0.370027835 | 0.051072492 | 0.40 |
| YKL173W | 1269.160496 | 0.274300681 | 0.05129448 | 0.40 |
| YHR200W | 3347.025252 | -0.431360953 | 0.051408493 | 0.40 |
| YOR230W | 8762.727513 | -0.560667222 | 0.051570862 | 0.40 |
| YDL147W | 5209.098548 | -0.374374707 | 0.051889221 | 0.40 |
| YGR037C | 2287.491924 | -0.623289478 | 0.051839229 | 0.40 |
| YBL078C | 809.4078099 | -0.84199386 | 0.052761466 | 0.40 |
| YDR519W | 900.2252274 | -0.672086712 | 0.052541757 | 0.40 |
| YER001W | 1455.504127 | 0.641499913 | 0.052307764 | 0.40 |
| YGL054C | 1677.23256 | -0.423482365 | 0.05236094 | 0.40 |
| YGR140W | 791.394714 | 0.559495652 | 0.052651587 | 0.40 |
| YJL055W | 2135.334979 | -0.450841379 | 0.052643419 | 0.40 |
| YJR072C | 1860.139893 | -0.483066322 | 0.052627316 | 0.40 |
| YKR061W | 573.6088479 | 0.775761025 | 0.052732137 | 0.40 |
| YLL018C | 8408.142457 | -0.432599605 | 0.052203415 | 0.40 |
| YLR215C | 433.3667117 | 0.352172089 | 0.052474199 | 0.40 |
| YPR196W | 243.4727794 | 0.612964252 | 0.052417789 | 0.40 |
| YGL087C | 1504.49763 | -0.566558859 | 0.053010965 | 0.40 |
| YJL192C | 2669.267366 | -0.56962228 | 0.052958522 | 0.40 |
| YMR320W | 113.0593369 | 0.804209747 | 0.053077711 | 0.40 |
| YNR060W | 291.930447 | 0.544678844 | 0.053086274 | 0.40 |
| YAR033W | 105.2011719 | -0.41069089 | 0.05387334 | 0.40 |

| | | | | |
|---|---|---|---|---|
| YBR149W | 7039.888751 | -0.465320368 | 0.053943649 | 0.40 |
| YDL184C | 118.3516335 | -0.743903774 | 0.053666631 | 0.40 |
| YGL197W | 2610.651509 | 0.539502917 | 0.053783421 | 0.40 |
| YOR155C | 1051.816257 | -0.40395809 | 0.053905662 | 0.40 |
| snR44 | 201.2193719 | 0.914414474 | 0.05376426 | 0.40 |
| YIR005W | 167.3355793 | -0.380804812 | 0.054330118 | 0.41 |
| YJR069C | 644.3818806 | -0.800572566 | 0.054446099 | 0.41 |
| YMR193C-A | 123.8016037 | 0.796709717 | 0.054545775 | 0.41 |
| YDR224C | 8028.945616 | -0.608177097 | 0.054651983 | 0.41 |
| YDL086W | 1034.431113 | -0.612501205 | 0.054950271 | 0.41 |
| YGR221C | 838.7959631 | 0.499033879 | 0.054973739 | 0.41 |
| YNL239W | 3388.853667 | -0.451872834 | 0.055064566 | 0.41 |
| YMR188C | 1203.74754 | -0.482477245 | 0.055429639 | 0.41 |
| YPR113W | 3675.90658 | -0.452631356 | 0.055359411 | 0.41 |
| YDR497C | 3420.957422 | 0.361117118 | 0.055514861 | 0.41 |
| YOR162C | 795.3084144 | 0.264210505 | 0.055638964 | 0.41 |
| YGR231C | 3688.450165 | -0.540955744 | 0.055750728 | 0.41 |
| YGR268C | 875.2811265 | -0.456763883 | 0.055940713 | 0.41 |
| YPR149W | 11719.12039 | -0.627209869 | 0.055904223 | 0.41 |
| YCR036W | 1075.104784 | -0.377360171 | 0.056223102 | 0.41 |
| YER076C | 386.5202214 | -0.353697917 | 0.056413095 | 0.41 |
| YIL048W | 1213.90232 | 0.331728901 | 0.056224741 | 0.41 |
| YOL034W | 735.3581814 | 0.471232553 | 0.056386286 | 0.41 |
| YOL109W | 9848.185932 | -0.593345438 | 0.056479319 | 0.41 |
| YOR158W | 1525.323441 | -0.450296708 | 0.056416543 | 0.41 |
| YGL155W | 739.9393467 | 0.453653782 | 0.056738497 | 0.41 |
| YJL218W | 146.9624687 | -0.76137554 | 0.057000315 | 0.41 |
| YDR205W | 692.1640308 | 0.37500361 | 0.057182728 | 0.42 |
| YGR202C | 1624.722457 | -0.338857438 | 0.057291579 | 0.42 |
| YHR050W-A | 204.2549426 | -0.535830982 | 0.057532198 | 0.42 |
| YOR131C | 796.6694001 | -0.434110764 | 0.057478685 | 0.42 |
| YOL003C | 636.2491343 | -0.373431555 | 0.057664823 | 0.42 |
| YBL064C | 4712.365868 | -0.74455061 | 0.057831492 | 0.42 |
| YMR176W | 1256.155137 | 0.462769147 | 0.057765295 | 0.42 |
| YDR223W | 358.2338356 | 0.595337067 | 0.058030098 | 0.42 |
| YBR185C | 732.2746685 | -0.412265217 | 0.058156668 | 0.42 |
| YAL042W | 3676.592816 | -0.326215628 | 0.058256651 | 0.42 |
| YKL069W | 336.0140703 | -0.527295166 | 0.058470673 | 0.42 |
| YML015C | 859.349761 | -0.551007479 | 0.058441299 | 0.42 |
| YPL075W | 2226.650529 | 0.502714002 | 0.058324455 | 0.42 |
| YOL139C | 5554.572319 | -0.661622019 | 0.058791497 | 0.42 |
| YDR286C | 156.0831168 | -0.524743116 | 0.059001249 | 0.42 |
| YFL034C-A | 576.1615287 | 0.870949207 | 0.058991186 | 0.42 |

| | | | | |
|---|---|---|---|---|
| YGR048W | 875.7732177 | -0.422810521 | 0.059117656 | 0.42 |
| YOR322C | 1313.48176 | 0.390380724 | 0.059071129 | 0.42 |
| YKL172W | 2863.993194 | -0.72422908 | 0.059522559 | 0.42 |
| YBL039W-B | 151.3574507 | -0.603304368 | 0.059765379 | 0.42 |
| YGR235C | 1048.42614 | -0.5416081 | 0.059636999 | 0.42 |
| YJL152W | 449.6754094 | -0.626873507 | 0.059709663 | 0.42 |
| YBR133C | 1160.137007 | 0.717574547 | 0.060460335 | 0.42 |
| YDL219W | 330.3585764 | -0.696735924 | 0.060714363 | 0.42 |
| YGR143W | 1625.66982 | 0.534418885 | 0.060503773 | 0.42 |
| YGR179C | 503.8137475 | 0.301903313 | 0.060322356 | 0.42 |
| YJL173C | 671.1948643 | -0.524049146 | 0.060604774 | 0.42 |
| YKL055C | 140.4891808 | 0.536902077 | 0.060714766 | 0.42 |
| YLR011W | 284.831779 | -0.421872078 | 0.060564338 | 0.42 |
| YLR360W | 653.1816733 | 0.318129255 | 0.059878425 | 0.42 |
| YMR039C | 1898.602952 | -0.543391446 | 0.06057572 | 0.42 |
| YMR162C | 1643.610674 | 0.462483721 | 0.060676126 | 0.42 |
| YMR203W | 3020.852994 | -0.345800364 | 0.059964582 | 0.42 |
| YOR329C | 1982.861977 | 0.452620265 | 0.060668819 | 0.42 |
| YOR007C | 8060.679187 | -0.487627658 | 0.060859929 | 0.42 |
| YER124C | 2039.69695 | 0.504026368 | 0.060948156 | 0.42 |
| YAL024C | 799.2694521 | 0.487086031 | 0.061344299 | 0.42 |
| YDL069C | 126.889784 | -0.488323181 | 0.061366071 | 0.42 |
| YFL010W-A | 308.9446486 | -0.505213812 | 0.061294053 | 0.42 |
| YJL019W | 605.7582629 | 0.476118806 | 0.061441692 | 0.42 |
| YJL167W | 5562.553131 | -0.466665316 | 0.061314852 | 0.42 |
| YHR029C | 965.1862947 | -0.646480651 | 0.061953947 | 0.43 |
| YHR077C | 2925.324128 | 0.351962618 | 0.061984191 | 0.43 |
| YNL065W | 531.9531231 | 0.831480774 | 0.061968315 | 0.43 |
| YMR187C | 357.2988229 | 0.483518559 | 0.062462842 | 0.43 |
| YNL068C | 1421.244951 | 0.473360064 | 0.062401472 | 0.43 |
| YHR117W | 2228.445201 | -0.464907391 | 0.062602327 | 0.43 |
| YDL048C | 3537.203448 | 0.777282448 | 0.062949637 | 0.43 |
| YJR068W | 1198.277592 | -0.458067192 | 0.062841571 | 0.43 |
| YLR115W | 1204.082371 | 0.276909094 | 0.062893448 | 0.43 |
| YDR307W | 749.860008 | 0.373001018 | 0.063031751 | 0.43 |
| YHR063W-A | 174.3401986 | -0.686895294 | 0.063308952 | 0.43 |
| YKL122C | 1563.613747 | -0.571459447 | 0.063532958 | 0.43 |
| YEL059C-A | 318.8027126 | -0.655814586 | 0.064208436 | 0.43 |
| YGR184C | 3421.297226 | 0.298271511 | 0.064214846 | 0.43 |
| YNL190W | 6079.357238 | 0.457054692 | 0.064222106 | 0.43 |
| YBR111C | 3318.167647 | -0.711320381 | 0.064496499 | 0.44 |
| YMR237W | 2047.326568 | 0.213912142 | 0.064430491 | 0.44 |
| YNL176C | 1925.34034 | 0.436677397 | 0.06463078 | 0.44 |

| | | | | |
|---|---|---|---|---|
| YOR291W | 1661.347699 | 0.316569735 | 0.064614137 | 0.44 |
| YLR298C | 806.4703735 | -0.444859794 | 0.064948332 | 0.44 |
| YKL033W-A | 1409.695613 | -0.395849961 | 0.065037054 | 0.44 |
| YNL327W | 5207.566398 | 0.753655242 | 0.065464889 | 0.44 |
| YGR215W | 453.6817551 | -0.564902216 | 0.065717744 | 0.44 |
| YKL019W | 1109.764572 | -0.377015213 | 0.065727459 | 0.44 |
| YLR346C | 131.9806369 | -0.876021101 | 0.066202663 | 0.44 |
| YLL003W | 928.3757705 | 0.280345796 | 0.066302262 | 0.44 |
| YOR066W | 1115.721778 | 0.546686535 | 0.066464828 | 0.44 |
| YOL069W | 510.0668666 | 0.413772813 | 0.066565618 | 0.44 |
| YLR425W | 1120.132672 | 0.367037615 | 0.066808229 | 0.44 |
| YDR221W | 936.1837264 | 0.346210503 | 0.066985386 | 0.44 |
| YML082W | 766.1872079 | 0.424241744 | 0.066934877 | 0.44 |
| YCR042C | 1593.772478 | 0.284833687 | 0.067492404 | 0.45 |
| YFL037W | 5301.265498 | -0.471422328 | 0.0675224 | 0.45 |
| YGR281W | 2050.830805 | 0.340712386 | 0.067613777 | 0.45 |
| YHR204W | 1257.868137 | 0.482253062 | 0.06743063 | 0.45 |
| YKL040C | 858.7842501 | -0.417848816 | 0.067368097 | 0.45 |
| YLL054C | 624.0988471 | 0.3753668 | 0.067844576 | 0.45 |
| YLR195C | 1211.827498 | -0.433601843 | 0.067854009 | 0.45 |
| YLR287C-A | 5234.314487 | -0.640668365 | 0.067305221 | 0.45 |
| YOR308C | 716.1626853 | -0.548297329 | 0.067689601 | 0.45 |
| YPR062W | 2367.986615 | -0.341476863 | 0.067708506 | 0.45 |
| YNL256W | 1084.449905 | 0.520961704 | 0.06799914 | 0.45 |
| YJL104W | 974.9529404 | -0.486235593 | 0.068107053 | 0.45 |
| YLR342W | 12036.59934 | 0.550607434 | 0.068263046 | 0.45 |
| YOR336W | 1255.740018 | 0.393106136 | 0.068238964 | 0.45 |
| YML009C | 743.0693468 | -0.554159329 | 0.068799047 | 0.45 |
| YDR041W | 1214.356247 | -0.436945292 | 0.069423959 | 0.45 |
| YGL150C | 2988.328671 | 0.296704792 | 0.069577718 | 0.45 |
| YKL184W | 1354.204344 | 0.405163662 | 0.069561296 | 0.45 |
| YKL018C-A | 261.9017664 | -0.326969146 | 0.070105064 | 0.45 |
| YKR101W | 347.6961828 | 0.422821796 | 0.07003016 | 0.45 |
| YBR111W-A | 229.152125 | -0.501346645 | 0.070481683 | 0.46 |
| YFR042W | 519.19235 | -0.444954389 | 0.070854907 | 0.46 |
| YGL008C | 26806.32689 | 0.702404428 | 0.0708278 | 0.46 |
| YGL043W | 1337.950161 | -0.436039329 | 0.070905531 | 0.46 |
| YIL170W | 172.5743318 | 0.763462529 | 0.070794007 | 0.46 |
| YNL233W | 992.805205 | 0.328473489 | 0.070873076 | 0.46 |
| YPL002C | 492.3419627 | -0.339010032 | 0.070709765 | 0.46 |
| YBR217W | 353.7676501 | -0.378913956 | 0.071106986 | 0.46 |
| YHR079C | 1259.061897 | 0.34279314 | 0.07107303 | 0.46 |
| YPL010W | 2111.463364 | -0.427011699 | 0.071158622 | 0.46 |

| | | | | |
|---|---|---|---|---|
| YDR127W | 3620.75176 | 0.605869562 | 0.071277665 | 0.46 |
| YNL301C | 1146.542523 | 0.74648906 | 0.071542348 | 0.46 |
| YIL151C | 1650.033227 | 0.409270634 | 0.071771602 | 0.46 |
| YPR131C | 480.7629468 | -0.364565613 | 0.071981236 | 0.46 |
| YJL109C | 2576.133571 | 0.631786279 | 0.072172593 | 0.46 |
| YCR003W | 578.0986481 | -0.499218744 | 0.072559564 | 0.46 |
| YEL002C | 3293.830024 | -0.342838362 | 0.072484044 | 0.46 |
| YMR112C | 344.1352482 | -0.454435869 | 0.072764037 | 0.46 |
| YBL090W | 640.7121751 | -0.522428317 | 0.073031669 | 0.46 |
| YCL012C | 422.2313505 | -0.587357968 | 0.07306672 | 0.46 |
| YDL139C | 356.8731646 | -0.369093328 | 0.073708539 | 0.46 |
| YDL177C | 218.4776378 | -0.512693343 | 0.073585984 | 0.46 |
| YHR139C | 204.3743283 | 0.718944598 | 0.073398222 | 0.46 |
| YKL179C | 1404.902826 | 0.33141575 | 0.073484723 | 0.46 |
| YNL173C | 2504.496471 | -0.532715835 | 0.073746275 | 0.46 |
| YNR016C | 6951.886561 | 0.525319842 | 0.073581552 | 0.46 |
| YNR040W | 422.1360755 | -0.471627449 | 0.073731897 | 0.46 |
| YNR067C | 2460.572138 | 0.585029854 | 0.073280397 | 0.46 |
| YGL137W | 4577.146804 | -0.367483005 | 0.073896659 | 0.46 |
| YER157W | 974.6953246 | 0.281339505 | 0.074039242 | 0.46 |
| YGL037C | 14451.53815 | -0.57372153 | 0.074424265 | 0.47 |
| YKL130C | 628.0827477 | -0.480745298 | 0.074669668 | 0.47 |
| YNL168C | 1117.770385 | -0.513074016 | 0.074683934 | 0.47 |
| YOR224C | 1759.04057 | -0.687861275 | 0.074809854 | 0.47 |
| YCL033C | 876.4314652 | -0.710949737 | 0.074981889 | 0.47 |
| YIR015W | 351.8671415 | -0.502159383 | 0.075093306 | 0.47 |
| YMR038C | 1941.251831 | -0.469712477 | 0.075193007 | 0.47 |
| YDL077C | 1087.970908 | 0.260272491 | 0.075682321 | 0.47 |
| YJL085W | 484.172969 | 0.54402135 | 0.075600172 | 0.47 |
| YMR161W | 724.2845416 | -0.290935919 | 0.075511951 | 0.47 |
| YMR273C | 1360.501919 | 0.455522151 | 0.075648372 | 0.47 |
| YDR168W | 2674.002145 | -0.562630844 | 0.075984867 | 0.47 |
| YPR018W | 922.6100673 | -0.516540689 | 0.076060056 | 0.47 |
| YKR074W | 1111.118511 | -0.415091406 | 0.076273263 | 0.47 |
| YLR099W-A | 174.4936248 | -0.453454605 | 0.076359307 | 0.47 |
| YBR077C | 472.5529414 | -0.402254464 | 0.076806808 | 0.47 |
| YDL189W | 1070.015735 | 0.467179879 | 0.07684425 | 0.47 |
| YDR283C | 1675.592568 | 0.383545342 | 0.076765924 | 0.47 |
| YER050C | 1168.128624 | -0.548142814 | 0.076775732 | 0.47 |
| YDR407C | 1886.152895 | 0.363729654 | 0.07702549 | 0.47 |
| YBR118W | 10015.00281 | -0.629353894 | 0.077350448 | 0.47 |
| YBR202W | 984.15721 | 0.46008012 | 0.07746035 | 0.47 |
| YDR052C | 1279.720929 | 0.350973454 | 0.077715669 | 0.47 |

| | | | | |
|---|---|---|---|---|
| YNL076W | 1058.446405 | 0.325290875 | 0.077646998 | 0.47 |
| YNL255C | 4025.75008 | -0.565153608 | 0.077611055 | 0.47 |
| YNL275W | 352.1195966 | 0.349674185 | 0.077583578 | 0.47 |
| YGL105W | 8369.251023 | -0.577845786 | 0.078098376 | 0.47 |
| YPR138C | 1408.826707 | 0.571438704 | 0.078182431 | 0.47 |
| YKL004W | 2100.85667 | 0.511928118 | 0.078275248 | 0.47 |
| YBL021C | 336.3286264 | -0.381441392 | 0.078690371 | 0.47 |
| YER169W | 1395.485471 | 0.451904382 | 0.078918451 | 0.47 |
| YGL221C | 1756.035958 | -0.422464566 | 0.078759036 | 0.47 |
| YLR208W | 2849.872811 | -0.414702278 | 0.078822068 | 0.47 |
| YML094W | 513.2141562 | -0.423560052 | 0.078878899 | 0.47 |
| YOL021C | 1670.215668 | 0.371548313 | 0.078555656 | 0.47 |
| YPL218W | 2797.046919 | -0.441678088 | 0.078400723 | 0.47 |
| YDR514C | 275.4543514 | 0.545460713 | 0.079092582 | 0.48 |
| YGR150C | 1213.396537 | 0.318005357 | 0.079506734 | 0.48 |
| YOL011W | 1428.931177 | 0.538065789 | 0.079846879 | 0.48 |
| YPL194W | 451.5443271 | 0.353154103 | 0.079699628 | 0.48 |
| YPR129W | 1545.384555 | -0.385665171 | 0.079859771 | 0.48 |
| YJR057W | 461.680688 | -0.337457356 | 0.079951269 | 0.48 |
| YHR016C | 1896.688185 | -0.474441672 | 0.080299929 | 0.48 |
| YKL005C | 1108.015733 | 0.315179615 | 0.080240493 | 0.48 |
| YBL040C | 1584.786506 | -0.455829768 | 0.080878647 | 0.48 |
| YLR057W | 1092.547525 | 0.389440875 | 0.081243648 | 0.48 |
| YMR182C | 564.4635256 | 0.496468087 | 0.081445441 | 0.48 |
| YMR311C | 3644.500712 | -0.521322135 | 0.081383837 | 0.48 |
| YKL153W | 3671.352718 | -0.707161124 | 0.082005569 | 0.49 |
| YMR128W | 2322.124433 | -0.697759917 | 0.081937904 | 0.49 |
| YPL204W | 2957.690828 | 0.297377236 | 0.082026021 | 0.49 |
| YPL253C | 481.3229549 | 0.439684731 | 0.082148314 | 0.49 |
| YDR394W | 3757.628989 | -0.336273191 | 0.083197737 | 0.49 |
| YHR076W | 1124.266555 | -0.403504812 | 0.082951672 | 0.49 |
| YJL151C | 3281.2426 | -0.628906007 | 0.08308342 | 0.49 |
| YJR030C | 708.7385852 | 0.42506424 | 0.083235751 | 0.49 |
| YMR043W | 2143.509763 | 0.466204717 | 0.083206189 | 0.49 |
| YOL096C | 679.2678424 | -0.43578715 | 0.083204978 | 0.49 |
| YGL186C | 1156.394032 | 0.495892425 | 0.083353085 | 0.49 |
| YHR038W | 426.523879 | -0.410360569 | 0.083414812 | 0.49 |
| YDR201W | 235.4443499 | -0.474159155 | 0.083848181 | 0.49 |
| YLR384C | 2550.492331 | 0.335546276 | 0.083870504 | 0.49 |
| YML079W | 1250.560699 | -0.559798301 | 0.083777977 | 0.49 |
| YOR167C | 4868.170026 | -0.709196118 | 0.083790917 | 0.49 |
| YIR038C | 2847.18512 | -0.630827966 | 0.084123455 | 0.49 |
| YML029W | 1034.498713 | 0.322319775 | 0.08426872 | 0.49 |

| | | | | |
|---|---|---|---|---|
| YJR116W | 691.1206507 | -0.497990085 | 0.084485644 | 0.49 |
| YKL152C | 30648.47356 | -0.706141229 | 0.0846893 | 0.49 |
| YKR030W | 649.1186189 | -0.298525647 | 0.084675612 | 0.49 |
| YMR259C | 1208.426766 | 0.509187727 | 0.084419068 | 0.49 |
| YOR373W | 745.8071516 | 0.489764696 | 0.084734608 | 0.49 |
| YGR098C | 1335.545417 | 0.433460707 | 0.084900933 | 0.49 |
| YJL060W | 1402.078784 | -0.424024205 | 0.084925577 | 0.49 |
| YBR170C | 1144.125576 | -0.28953034 | 0.085489048 | 0.49 |
| YDR206W | 1789.144398 | 0.34993792 | 0.08542004 | 0.49 |
| YLR147C | 302.8969457 | -0.339571064 | 0.085407898 | 0.49 |
| YML068W | 512.1804017 | 0.325924182 | 0.085346976 | 0.49 |
| YPR069C | 2243.3628 | -0.703518077 | 0.085329084 | 0.49 |
| YGL058W | 714.9977444 | -0.504630824 | 0.085702407 | 0.49 |
| YMR297W | 8256.874804 | -0.44766687 | 0.085683768 | 0.49 |
| YDR135C | 3272.34562 | 0.317484998 | 0.085934417 | 0.49 |
| YMR138W | 253.6379214 | -0.377329895 | 0.086099992 | 0.49 |
| YDR484W | 738.7697722 | 0.23672017 | 0.086271203 | 0.49 |
| YLR407W | 355.7060391 | 0.399553421 | 0.086529243 | 0.49 |
| YGR105W | 477.947228 | -0.497944239 | 0.086633775 | 0.49 |
| YOL020W | 976.6345917 | 0.547168891 | 0.087178251 | 0.50 |
| YOL086W-A | 318.8786531 | -0.63214137 | 0.087116386 | 0.50 |
| YDR066C | 173.6197188 | 0.528417624 | 0.087515397 | 0.50 |
| YGL251C | 608.1790507 | 0.425989149 | 0.087554407 | 0.50 |
| YNL021W | 893.5874562 | 0.250462748 | 0.087466341 | 0.50 |
| YHR103W | 1865.030837 | 0.412484682 | 0.087691361 | 0.50 |
| YER090W | 2828.394196 | -0.614192619 | 0.087855486 | 0.50 |
| YDR179W-A | 321.5877637 | 0.494398185 | 0.088093255 | 0.50 |
| YCL050C | 4005.943189 | -0.731172565 | 0.088356388 | 0.50 |
| YLR288C | 449.0939426 | 0.269049093 | 0.088357583 | 0.50 |
| YLR405W | 359.0345633 | 0.501768009 | 0.088459057 | 0.50 |
| YBR186W | 272.8183513 | 0.724280044 | 0.088570939 | 0.50 |
| YPL076W | 380.2952376 | 0.439168527 | 0.088893503 | 0.50 |
| YBR024W | 889.4330761 | -0.557010938 | 0.089302934 | 0.50 |
| YBR163W | 719.9181119 | 0.455997563 | 0.089271793 | 0.50 |
| YCR045C | 398.9252367 | 0.431740648 | 0.089607359 | 0.50 |
| YDL144C | 951.4980585 | -0.395672723 | 0.089500947 | 0.50 |
| YML096W | 652.0319638 | 0.395525718 | 0.089110677 | 0.50 |
| YOL005C | 1554.462517 | -0.504610163 | 0.089630888 | 0.50 |
| YPL246C | 2119.845638 | -0.31598782 | 0.089456022 | 0.50 |
| YJL217W | 2088.445581 | -0.681196111 | 0.089889498 | 0.50 |
| YDR353W | 5059.752159 | -0.504549855 | 0.090079123 | 0.50 |
| YJR106W | 683.5325213 | 0.404175126 | 0.090231018 | 0.50 |
| YJL097W | 1713.191405 | -0.545321186 | 0.090327746 | 0.50 |

| | | | | |
|---|---|---|---|---|
| YAR014C | 1593.739874 | 0.443166967 | 0.090456282 | 0.50 |
| YLR419W | 1247.341194 | 0.423258358 | 0.090514647 | 0.50 |
| YGL139W | 1235.197549 | 0.311850141 | 0.09063148 | 0.50 |
| YBR162W-A | 580.1318829 | -0.50845317 | 0.09084363 | 0.50 |
| YLR283W | 409.1057668 | -0.282138605 | 0.091032872 | 0.50 |
| YBR231C | 448.4132605 | -0.487933893 | 0.091204849 | 0.50 |
| YAL011W | 648.981818 | -0.3872936 | 0.091373491 | 0.51 |
| YER132C | 1149.8348 | 0.457854375 | 0.091796615 | 0.51 |
| YIL154C | 2075.807869 | -0.47467798 | 0.091854114 | 0.51 |
| YJL132W | 534.7548177 | 0.585328334 | 0.091716474 | 0.51 |
| YJL003W | 223.2586603 | -0.417612737 | 0.092086162 | 0.51 |
| YDL074C | 1505.723255 | 0.439837851 | 0.092777967 | 0.51 |
| YJR142W | 1091.921242 | -0.258921559 | 0.092700445 | 0.51 |
| YKR088C | 850.0044261 | -0.32044339 | 0.092829748 | 0.51 |
| YNR021W | 3396.726535 | -0.527340155 | 0.092867803 | 0.51 |
| YCR088W | 5938.374856 | -0.429354856 | 0.093286891 | 0.51 |
| YJR127C | 3525.789326 | 0.391386866 | 0.093658882 | 0.51 |
| YLR045C | 1304.494239 | 0.396311642 | 0.09357707 | 0.51 |
| YNL169C | 1167.217459 | 0.239838798 | 0.09348203 | 0.51 |
| YDR099W | 8428.443895 | -0.261796046 | 0.093861107 | 0.51 |
| YPL242C | 2035.672722 | 0.384250637 | 0.094120548 | 0.51 |
| YLR172C | 1976.093187 | -0.539425006 | 0.094284141 | 0.51 |
| YGR192C | 181337.4741 | -0.659897743 | 0.094384383 | 0.51 |
| YDR510W | 2598.34956 | -0.359155423 | 0.094622863 | 0.51 |
| YHR172W | 514.8081565 | 0.594617554 | 0.094644602 | 0.51 |
| YLR389C | 2863.752537 | 0.295660829 | 0.094733298 | 0.51 |
| YDR020C | 268.9612697 | 0.570592939 | 0.094993361 | 0.51 |
| YOL102C | 532.3526812 | -0.295401224 | 0.095170389 | 0.52 |
| YGL001C | 1350.851203 | -0.552129928 | 0.095403918 | 0.52 |
| YPL029W | 470.8569792 | 0.304899072 | 0.095500008 | 0.52 |
| YLR051C | 783.2029443 | -0.526154446 | 0.096053404 | 0.52 |
| YOR219C | 1764.610148 | 0.330380777 | 0.096010487 | 0.52 |
| YPL001W | 324.2471356 | -0.566186121 | 0.09622992 | 0.52 |
| YBR023C | 1741.008098 | 0.420056837 | 0.096418146 | 0.52 |
| YLR103C | 462.3096359 | 0.452248772 | 0.096977637 | 0.52 |
| YDR081C | 1262.178022 | 0.343534733 | 0.097079441 | 0.52 |
| YBR074W | 1213.234406 | 0.270270563 | 0.097508394 | 0.52 |
| YIL014W | 862.4746572 | 0.398950383 | 0.097473985 | 0.52 |
| YJL147C | 286.6419753 | 0.318878076 | 0.097618511 | 0.52 |
| YJL193W | 438.7222894 | 0.374046497 | 0.09759652 | 0.52 |
| YLR452C | 1814.469836 | 0.664169075 | 0.097597106 | 0.52 |
| YER021W | 5518.8989 | -0.389509064 | 0.097775566 | 0.52 |
| YDR461C-A | 326.4764294 | -0.618070892 | 0.09853901 | 0.52 |

| | | | | |
|---|---|---|---|---|
| YER003C | 3520.842344 | -0.57053655 | 0.098250871 | 0.52 |
| YGL041W-A | 345.8516558 | -0.404116434 | 0.098302037 | 0.52 |
| YJR090C | 1653.355955 | 0.313883994 | 0.098545934 | 0.52 |
| YMR260C | 3410.062096 | -0.525763466 | 0.098625752 | 0.52 |
| YOR122C | 4814.102226 | -0.477313894 | 0.098074373 | 0.52 |
| YPL197C | 174.3224877 | 0.788419739 | 0.098579812 | 0.52 |
| YER022W | 948.246119 | 0.223823302 | 0.098825931 | 0.52 |
| YKL185W | 2577.028628 | 0.49392904 | 0.098875214 | 0.52 |
| YJR012C | 286.1622087 | -0.320755008 | 0.099148968 | 0.52 |
| snR83 | 177.6034874 | 0.786322941 | 0.099135503 | 0.52 |
| YNR051C | 2926.448335 | -0.551851302 | 0.099780098 | 0.53 |
| YER143W | 1597.506506 | -0.387023601 | 0.100043979 | 0.53 |
| YLR278C | 1940.968689 | 0.260054453 | 0.100000265 | 0.53 |
| YBR082C | 6782.616874 | -0.540838661 | 0.100342582 | 0.53 |
| YGR077C | 1152.926191 | 0.311720211 | 0.100366399 | 0.53 |
| YLR035C | 528.4128752 | 0.427140162 | 0.100252478 | 0.53 |
| YAL009W | 404.8562545 | -0.318442084 | 0.100507855 | 0.53 |
| YBR053C | 1693.976608 | -0.581696663 | 0.100562787 | 0.53 |
| YCL035C | 4330.054767 | -0.642656094 | 0.100975161 | 0.53 |
| YDR159W | 1698.043068 | 0.284614466 | 0.100787087 | 0.53 |
| YKL056C | 11023.03919 | -0.627886148 | 0.100970011 | 0.53 |
| YKR084C | 1262.417524 | -0.374735429 | 0.101081986 | 0.53 |
| YNL122C | 325.2385534 | -0.477074213 | 0.101010436 | 0.53 |
| YJR107W | 474.7713781 | 0.52396444 | 0.101461597 | 0.53 |
| YLL010C | 1089.845187 | 0.2790588 | 0.101450455 | 0.53 |
| YOR111W | 390.0644929 | -0.362605186 | 0.101350757 | 0.53 |
| YAL049C | 1497.895567 | -0.443229376 | 0.101613014 | 0.53 |
| YGR078C | 839.5108181 | -0.480074864 | 0.101983072 | 0.53 |
| YCR024C-A | 1033.453894 | 0.504903905 | 0.102512063 | 0.53 |
| YNL212W | 1282.46661 | 0.367825661 | 0.102483714 | 0.53 |
| YOR133W | 1425.340865 | -0.522164095 | 0.102497604 | 0.53 |
| YOR285W | 4227.371531 | -0.58662571 | 0.102373576 | 0.53 |
| snR82 | 454.6100305 | -0.797593313 | 0.102582474 | 0.53 |
| YOR363C | 1212.557429 | 0.438112974 | 0.102706534 | 0.53 |
| YDR330W | 1850.284876 | -0.53071375 | 0.103712749 | 0.53 |
| YDR399W | 3185.764567 | 0.695969125 | 0.103730482 | 0.53 |
| YGR099W | 582.6782539 | 0.374083339 | 0.103621932 | 0.53 |
| snR63 | 94.54093227 | 0.693938324 | 0.103590182 | 0.53 |
| YDR416W | 716.6573712 | 0.325467834 | 0.104038862 | 0.53 |
| YHR193C | 8396.042296 | -0.505403989 | 0.103946603 | 0.53 |
| YJR065C | 3820.877578 | -0.263086461 | 0.104096234 | 0.53 |
| YHR064C | 6332.657586 | -0.585155492 | 0.104227207 | 0.53 |
| YPL175W | 580.2634042 | 0.333086421 | 0.10477056 | 0.54 |

| | | | | |
|---|---|---|---|---|
| YGL083W | 940.5046339 | 0.315136408 | 0.104989092 | 0.54 |
| YNL029C | 467.6043522 | 0.411756648 | 0.104922917 | 0.54 |
| YLR014C | 353.4875684 | 0.434917165 | 0.105244835 | 0.54 |
| YOR242C | 110.2984083 | 0.555520936 | 0.105426733 | 0.54 |
| YMR173W | 743.4251563 | -0.51688793 | 0.105709431 | 0.54 |
| YJL123C | 2650.72374 | -0.361276705 | 0.106279232 | 0.54 |
| YAL027W | 394.2818606 | -0.3071099 | 0.106622145 | 0.54 |
| YNR053C | 1005.221474 | 0.656818645 | 0.10667776 | 0.54 |
| YOR174W | 544.9117735 | -0.306179665 | 0.106596364 | 0.54 |
| YPR006C | 675.2062996 | -0.672224667 | 0.106527608 | 0.54 |
| YCL005W-A | 247.3743167 | -0.366627405 | 0.107538451 | 0.54 |
| YDR448W | 426.123254 | 0.444408733 | 0.107511318 | 0.54 |
| YFL018C | 6022.957872 | -0.428155093 | 0.10747118 | 0.54 |
| YGR252W | 827.8896718 | 0.257664517 | 0.107114029 | 0.54 |
| YLL061W | 447.0050952 | 0.513993844 | 0.107351896 | 0.54 |
| YLR361C-A | 262.7999201 | -0.384449615 | 0.107364647 | 0.54 |
| YNL037C | 5393.729596 | -0.471529516 | 0.107117733 | 0.54 |
| YHR120W | 606.7255558 | 0.262932618 | 0.107678515 | 0.54 |
| YOR192C | 247.398385 | 0.636019279 | 0.107894776 | 0.54 |
| YDR326C | 1698.894045 | 0.347320142 | 0.108010478 | 0.54 |
| YDR452W | 4294.15743 | -0.437014109 | 0.108249501 | 0.54 |
| YFR015C | 6189.813754 | -0.740324211 | 0.108412266 | 0.54 |
| YFR031C | 1205.359626 | 0.278478632 | 0.108418578 | 0.54 |
| YMR300C | 1920.32889 | 0.774100494 | 0.108400599 | 0.54 |
| YNL147W | 569.2824244 | -0.373769315 | 0.108498963 | 0.54 |
| YML129C | 536.8329539 | -0.487208143 | 0.108675859 | 0.54 |
| YCR016W | 907.9551315 | -0.595072798 | 0.110432202 | 0.54 |
| YDL012C | 999.0124699 | -0.421052545 | 0.109143481 | 0.54 |
| YDR258C | 9048.121163 | -0.613683591 | 0.110162223 | 0.54 |
| YFL017C | 717.9144755 | -0.323513518 | 0.109207899 | 0.54 |
| YGL113W | 616.1660547 | 0.379486068 | 0.109228768 | 0.54 |
| YGL163C | 996.8369865 | 0.400589512 | 0.109202355 | 0.54 |
| YKL186C | 1093.769583 | -0.39495802 | 0.109123424 | 0.54 |
| YKR001C | 4012.227533 | -0.34292701 | 0.109957081 | 0.54 |
| YLL005C | 615.6111971 | 0.605451706 | 0.110314477 | 0.54 |
| YLL009C | 581.7236454 | -0.728999706 | 0.110302933 | 0.54 |
| YLR273C | 457.5126435 | 0.399307446 | 0.110596144 | 0.54 |
| YML028W | 27162.20889 | -0.387233079 | 0.109766759 | 0.54 |
| YMR009W | 601.1334204 | -0.406994854 | 0.110512241 | 0.54 |
| YMR054W | 1855.817154 | 0.385958007 | 0.109669616 | 0.54 |
| YMR231W | 864.2847521 | 0.319788457 | 0.109707601 | 0.54 |
| YNL082W | 573.887894 | 0.392883973 | 0.109765699 | 0.54 |
| YOL070C | 1742.264613 | -0.444423567 | 0.110475407 | 0.54 |

| | | | | |
|---|---|---|---|---|
| YPL030W | 552.0003609 | 0.478720345 | 0.110564776 | 0.54 |
| YPR022C | 1532.191812 | 0.287534501 | 0.109875891 | 0.54 |
| YOR304C-A | 201.0708852 | -0.471213316 | 0.110966352 | 0.54 |
| YPR171W | 883.9863961 | 0.313890124 | 0.11090058 | 0.54 |
| YER134C | 346.3610386 | -0.528374063 | 0.111493333 | 0.55 |
| YLR109W | 26967.32056 | -0.438262473 | 0.111439587 | 0.55 |
| YCL009C | 7647.273312 | -0.436390923 | 0.113188066 | 0.55 |
| YDL046W | 2942.234491 | -0.403109924 | 0.113781796 | 0.55 |
| YDR031W | 342.8572195 | -0.582711482 | 0.112017656 | 0.55 |
| YDR034C | 1497.715383 | 0.487684989 | 0.112427444 | 0.55 |
| YDR122W | 2792.779664 | -0.332669414 | 0.112435358 | 0.55 |
| YDR477W | 1886.671826 | 0.246743538 | 0.113552382 | 0.55 |
| YGL140C | 1656.222341 | 0.244359698 | 0.113841734 | 0.55 |
| YHR118C | 526.5681517 | 0.217934787 | 0.112367145 | 0.55 |
| YIL098C | 444.8949209 | -0.456504748 | 0.113118191 | 0.55 |
| YIL103W | 637.74726 | 0.428280112 | 0.113188794 | 0.55 |
| YJL087C | 474.7532502 | 0.469905048 | 0.112205623 | 0.55 |
| YKL041W | 833.3256523 | -0.373124658 | 0.113762511 | 0.55 |
| YKL132C | 214.3347933 | 0.423926971 | 0.112825346 | 0.55 |
| YKR037C | 412.1583252 | -0.245256455 | 0.113839525 | 0.55 |
| YKR055W | 476.0228776 | -0.385391304 | 0.113391761 | 0.55 |
| YLR223C | 2252.47739 | -0.659211527 | 0.113855265 | 0.55 |
| YMR109W | 3069.730753 | 0.352373709 | 0.113142343 | 0.55 |
| YNR064C | 215.5475844 | 0.699605262 | 0.113239386 | 0.55 |
| YOR197W | 2558.031802 | -0.266078706 | 0.113288633 | 0.55 |
| YOR301W | 501.5102259 | 0.572149331 | 0.112973615 | 0.55 |
| YPL259C | 962.6624104 | 0.268659912 | 0.113112279 | 0.55 |
| YLR145W | 262.3006502 | -0.352502893 | 0.114150967 | 0.55 |
| YBL091C-A | 508.1746239 | -0.439911854 | 0.114498691 | 0.55 |
| YER042W | 653.285425 | -0.58632048 | 0.114903425 | 0.55 |
| YER128W | 281.9899223 | -0.389558018 | 0.114813209 | 0.55 |
| YHL011C | 1690.318772 | -0.588525971 | 0.114639386 | 0.55 |
| YKL135C | 1473.22037 | 0.296972831 | 0.114613236 | 0.55 |
| YOR190W | 351.1788756 | 0.48850284 | 0.114763507 | 0.55 |
| YIL138C | 1495.862581 | -0.45755311 | 0.115253327 | 0.55 |
| snR33 | 120.9149393 | -0.454307786 | 0.11521518 | 0.55 |
| YCL044C | 1238.471005 | -0.442324094 | 0.115529435 | 0.55 |
| YJR045C | 25492.38268 | -0.531424429 | 0.115502474 | 0.55 |
| YMR189W | 2797.782123 | 0.77452692 | 0.115799657 | 0.55 |
| YGR172C | 884.3306623 | -0.331901389 | 0.116405556 | 0.55 |
| YHL004W | 1819.11881 | 0.291979649 | 0.116419697 | 0.55 |
| YDL004W | 2874.441994 | -0.677871087 | 0.117034008 | 0.55 |
| YDL085C-A | 569.0392384 | -0.518735925 | 0.11711545 | 0.55 |

| | | | | |
|---|---|---|---|---|
| YLL051C | 1170.125266 | 0.377432438 | 0.117168153 | 0.55 |
| YNL252C | 974.4576049 | -0.454558468 | 0.11712988 | 0.55 |
| YIL067C | 437.2526192 | 0.274526187 | 0.11750249 | 0.56 |
| YER182W | 1071.612284 | -0.577705422 | 0.117805734 | 0.56 |
| YKL060C | 152874.8661 | -0.653222195 | 0.118187504 | 0.56 |
| YNL157W | 1475.809734 | -0.262874715 | 0.118654619 | 0.56 |
| YPL040C | 952.0787614 | 0.410997317 | 0.118764815 | 0.56 |
| YAR050W | 236.1195337 | 0.697237807 | 0.119621398 | 0.56 |
| YBR016W | 1677.943304 | -0.297698253 | 0.120210611 | 0.56 |
| YDL176W | 672.0622412 | 0.365805166 | 0.119609854 | 0.56 |
| YDL237W | 1508.528864 | -0.40459915 | 0.120124666 | 0.56 |
| YDR014W | 486.9454278 | 0.309071196 | 0.121396919 | 0.56 |
| YDR107C | 540.536576 | 0.327818655 | 0.119258719 | 0.56 |
| YDR180W | 980.8547337 | 0.434787298 | 0.11997454 | 0.56 |
| YDR501W | 295.5069374 | 0.326097268 | 0.121068949 | 0.56 |
| YEL003W | 487.0478849 | -0.495517506 | 0.121468853 | 0.56 |
| YEL006W | 705.1819844 | 0.341463925 | 0.119508595 | 0.56 |
| YFR001W | 1018.480753 | -0.45043979 | 0.12041425 | 0.56 |
| YGL042C | 343.2123261 | -0.418509167 | 0.119865495 | 0.56 |
| YGL201C | 994.5170532 | 0.411587821 | 0.12129469 | 0.56 |
| YGR285C | 10708.64158 | -0.642892181 | 0.120357571 | 0.56 |
| YHR131C | 966.4131265 | 0.430790347 | 0.119165973 | 0.56 |
| YJR139C | 4827.24467 | -0.514719715 | 0.12034404 | 0.56 |
| YKL045W | 918.6422418 | -0.325543281 | 0.12132037 | 0.56 |
| YLL012W | 268.5425901 | 0.668895008 | 0.120691954 | 0.56 |
| YML067C | 1114.441541 | -0.247945083 | 0.120213491 | 0.56 |
| YML069W | 2256.742432 | -0.44919053 | 0.121432172 | 0.56 |
| YMR126C | 404.3389603 | 0.386157475 | 0.120986322 | 0.56 |
| YMR261C | 3042.037261 | -0.383569172 | 0.119033373 | 0.56 |
| YNL156C | 1413.410846 | -0.320715291 | 0.119015982 | 0.56 |
| YNL251C | 2085.43933 | 0.576982617 | 0.120696045 | 0.56 |
| YNL277W | 546.4771573 | 0.525288576 | 0.120487579 | 0.56 |
| YNR037C | 584.4173485 | -0.494261159 | 0.12130771 | 0.56 |
| YKR045C | 368.478154 | 0.367850099 | 0.121672795 | 0.56 |
| YOR144C | 421.6026059 | 0.390297506 | 0.121762396 | 0.56 |
| YHR143W-A | 554.7529102 | -0.493783265 | 0.12207653 | 0.56 |
| YKR068C | 742.7822083 | -0.315153577 | 0.12221751 | 0.56 |
| YOR123C | 2187.626634 | -0.214376389 | 0.122133471 | 0.56 |
| YOR368W | 442.6825001 | 0.280434893 | 0.12203014 | 0.56 |
| YDR115W | 505.144241 | -0.359673248 | 0.12249807 | 0.56 |
| YFR010W | 3072.848472 | -0.364465108 | 0.122590694 | 0.56 |
| YMR239C | 477.7892356 | 0.520302547 | 0.122560089 | 0.56 |
| YGR061C | 5651.499061 | 0.642480921 | 0.123215666 | 0.56 |

| | | | | |
|---|---|---|---|---|
| YMR251W-A | 5878.547147 | -0.632041016 | 0.123174986 | 0.56 |
| YMR012W | 7425.571889 | 0.368983798 | 0.123527753 | 0.56 |
| YML116W | 562.6593031 | 0.6169735 | 0.123663789 | 0.56 |
| YLR341W | 104.3766662 | 0.556829492 | 0.123827453 | 0.56 |
| YLR042C | 282.794653 | 0.754071093 | 0.124416359 | 0.57 |
| YDL213C | 1003.203847 | -0.618367236 | 0.125190667 | 0.57 |
| YDR483W | 4802.755253 | -0.260001699 | 0.125131483 | 0.57 |
| YMR006C | 894.5004784 | 0.403930559 | 0.125079374 | 0.57 |
| YHR075C | 548.2435732 | -0.326508125 | 0.125446624 | 0.57 |
| YGR068C | 535.7602898 | 0.299771955 | 0.125904418 | 0.57 |
| YPR144C | 1036.348245 | 0.544315551 | 0.125930458 | 0.57 |
| YBR068C | 7312.593911 | 0.579384005 | 0.126460536 | 0.57 |
| YCL057W | 3573.183746 | -0.337792872 | 0.126726733 | 0.57 |
| YDR209C | 344.9577194 | -0.517227329 | 0.126737483 | 0.57 |
| YGL070C | 462.601344 | -0.429391758 | 0.12736443 | 0.57 |
| YGL161C | 1505.046156 | -0.205552518 | 0.127306635 | 0.57 |
| YGR181W | 835.8007421 | -0.341590265 | 0.127228716 | 0.57 |
| YGR188C | 503.2494794 | 0.418354191 | 0.127644568 | 0.57 |
| YIR009W | 172.3613335 | -0.392174021 | 0.126195404 | 0.57 |
| YJR105W | 3472.841224 | -0.507583202 | 0.126826269 | 0.57 |
| YJR117W | 3603.291074 | -0.542331677 | 0.127509694 | 0.57 |
| YKL046C | 1814.036225 | -0.280228441 | 0.126411915 | 0.57 |
| YLR078C | 672.0208317 | -0.300287441 | 0.127919896 | 0.57 |
| YLR250W | 1395.702651 | -0.34894628 | 0.127364431 | 0.57 |
| YNL088W | 2851.433881 | -0.359498155 | 0.127606607 | 0.57 |
| YOL084W | 492.0065357 | 0.643454062 | 0.127182957 | 0.57 |
| YOR098C | 2501.292329 | 0.342180856 | 0.127960967 | 0.57 |
| YOR295W | 256.0382172 | -0.345839272 | 0.127788406 | 0.57 |
| YPL004C | 14429.78474 | -0.406244531 | 0.126891252 | 0.57 |
| YPL116W | 1071.716343 | 0.347305948 | 0.127727404 | 0.57 |
| YAL015C | 634.3594781 | -0.439907758 | 0.128802954 | 0.57 |
| YBR006W | 1760.927773 | -0.575688862 | 0.128710893 | 0.57 |
| YBR171W | 1056.70053 | -0.515759558 | 0.128933118 | 0.57 |
| YDR429C | 3118.563913 | -0.465798978 | 0.128764243 | 0.57 |
| YGL226W | 346.3222245 | -0.325617294 | 0.129103462 | 0.57 |
| YLR330W | 3759.858385 | -0.237965188 | 0.128734144 | 0.57 |
| YML103C | 3218.889825 | 0.378103931 | 0.129004603 | 0.57 |
| YMR255W | 1118.071772 | -0.459354951 | 0.128941018 | 0.57 |
| YGL050W | 416.7826537 | 0.4622813 | 0.129296503 | 0.57 |
| YNL187W | 311.4857358 | 0.363017627 | 0.129319055 | 0.57 |
| YCR037C | 1044.612368 | 0.325868148 | 0.130467061 | 0.57 |
| YGL019W | 1587.591155 | -0.218908636 | 0.129926738 | 0.57 |
| YGL185C | 473.3669461 | -0.461483866 | 0.130240291 | 0.57 |

| | | | | |
|---|---|---|---|---|
| YJL010C | 755.184634 | 0.568198995 | 0.1304638 | 0.57 |
| YML056C | 4012.776938 | 0.595370448 | 0.130104855 | 0.57 |
| YMR023C | 334.2712538 | 0.376080191 | 0.130456459 | 0.57 |
| YMR228W | 316.010038 | -0.316330582 | 0.130299029 | 0.57 |
| YOR057W | 859.8202576 | -0.386304494 | 0.130148473 | 0.57 |
| YER144C | 1434.76784 | 0.384476056 | 0.130977904 | 0.58 |
| YDR444W | 770.8185864 | 0.3492541 | 0.131194676 | 0.58 |
| YPL052W | 344.0806128 | -0.417350139 | 0.131141918 | 0.58 |
| YER014W | 690.0676914 | -0.320719906 | 0.131398664 | 0.58 |
| YIL159W | 678.524918 | 0.382059584 | 0.131512182 | 0.58 |
| YPL173W | 902.2046084 | -0.349761706 | 0.131470364 | 0.58 |
| YNL264C | 1013.510724 | -0.515052878 | 0.131759478 | 0.58 |
| YOR352W | 729.6147343 | -0.303032871 | 0.132018781 | 0.58 |
| YNL023C | 950.809325 | 0.299264547 | 0.132161218 | 0.58 |
| YBR085C-A | 1789.206511 | -0.582245604 | 0.132564154 | 0.58 |
| YGL205W | 247.6080273 | 0.607666206 | 0.132535095 | 0.58 |
| YDR253C | 229.9334863 | 0.385047146 | 0.132762371 | 0.58 |
| YIL068C | 981.7929067 | 0.196565432 | 0.133158803 | 0.58 |
| YMR208W | 1260.14289 | -0.368998635 | 0.133061519 | 0.58 |
| YNL259C | 262.1284302 | -0.566717119 | 0.13309493 | 0.58 |
| YCR051W | 1038.832374 | -0.458096092 | 0.133272926 | 0.58 |
| YDL192W | 4665.318464 | -0.620382855 | 0.134010141 | 0.58 |
| YDR363W | 641.2311125 | 0.370475661 | 0.134005331 | 0.58 |
| YML005W | 433.1547412 | 0.215455917 | 0.133979698 | 0.58 |
| YML065W | 967.920824 | 0.420116451 | 0.134023439 | 0.58 |
| YMR133W | 502.5352221 | 0.565237316 | 0.13445788 | 0.58 |
| YGL236C | 488.164276 | 0.46257933 | 0.134612424 | 0.58 |
| YDR320C-A | 373.904445 | -0.388793011 | 0.134881344 | 0.58 |
| YDR516C | 6053.645479 | -0.646706114 | 0.134833274 | 0.58 |
| YIR024C | 366.6225047 | -0.292871277 | 0.134941619 | 0.58 |
| YBL037W | 1072.022099 | 0.298435111 | 0.135083582 | 0.58 |
| YER133W-A | 95.44590047 | -0.532961844 | 0.135487854 | 0.58 |
| YFL010C | 2890.139798 | -0.437395704 | 0.135753361 | 0.58 |
| YKL084W | 479.3447424 | -0.493494772 | 0.135666271 | 0.58 |
| YLR314C | 3042.070083 | -0.269976279 | 0.135403589 | 0.58 |
| YPL055C | 1072.396721 | -0.307187821 | 0.135264703 | 0.58 |
| YPR100W | 188.3881606 | -0.453081577 | 0.13566846 | 0.58 |
| YBL057C | 768.8630819 | -0.345730348 | 0.135978666 | 0.58 |
| YNL244C | 2167.186944 | -0.457676438 | 0.135878276 | 0.58 |
| YHR111W | 861.1772037 | -0.31534883 | 0.136199714 | 0.58 |
| YDL076C | 828.6133083 | -0.250865371 | 0.136513018 | 0.58 |
| YOR249C | 647.5824875 | 0.310600158 | 0.136459251 | 0.58 |
| YER041W | 647.7988303 | 0.369564613 | 0.136795108 | 0.58 |

| | | | | |
|---|---|---|---|---|
| YKL210W | 8749.432253 | -0.288756435 | 0.136942144 | 0.58 |
| YMR002W | 4776.428936 | -0.378271047 | 0.137482673 | 0.59 |
| YMR024W | 882.2534561 | 0.321629288 | 0.137463552 | 0.59 |
| YNL273W | 713.7141054 | 0.380490734 | 0.137600427 | 0.59 |
| YER155C | 4972.789594 | 0.31767868 | 0.137897334 | 0.59 |
| YMR083W | 8478.832219 | -0.399589644 | 0.137952901 | 0.59 |
| YOL017W | 430.0355777 | 0.326399627 | 0.13792814 | 0.59 |
| YDR230W | 158.0472236 | -0.530125595 | 0.138164887 | 0.59 |
| YOR232W | 2857.566582 | -0.38025749 | 0.138206844 | 0.59 |
| YFR028C | 1133.072845 | 0.291875757 | 0.13836102 | 0.59 |
| YBR204C | 1064.479878 | -0.395965878 | 0.140887876 | 0.59 |
| YCR035C | 836.2083583 | 0.257858593 | 0.140946888 | 0.59 |
| YDR226W | 6443.35181 | -0.44996801 | 0.139247714 | 0.59 |
| YDR508C | 2566.963439 | 0.641358157 | 0.139320362 | 0.59 |
| YEL063C | 1734.961887 | 0.447021753 | 0.140943631 | 0.59 |
| YER091C | 3130.444513 | 0.459487142 | 0.139942106 | 0.59 |
| YHR190W | 5071.299505 | -0.374159352 | 0.139488879 | 0.59 |
| YIL016W | 483.5722155 | -0.49764349 | 0.139000507 | 0.59 |
| YIR001C | 869.6360964 | -0.271169599 | 0.139850012 | 0.59 |
| YKL220C | 368.1832027 | 0.583516288 | 0.140629044 | 0.59 |
| YLR114C | 1321.213644 | 0.323654889 | 0.139602606 | 0.59 |
| YLR245C | 188.3206886 | -0.455736945 | 0.140490673 | 0.59 |
| YLR268W | 967.406712 | -0.308465445 | 0.140564555 | 0.59 |
| YLR381W | 607.7067419 | 0.290330833 | 0.140640377 | 0.59 |
| YNL004W | 1614.267096 | 0.259260156 | 0.140058672 | 0.59 |
| YNL042W | 593.1711026 | 0.474388433 | 0.140103643 | 0.59 |
| YNL291C | 628.873147 | 0.362542585 | 0.139865105 | 0.59 |
| YOL082W | 2476.571718 | -0.414936641 | 0.139551216 | 0.59 |
| YOR276W | 906.6874329 | -0.542192498 | 0.140211539 | 0.59 |
| YPL206C | 1410.718758 | -0.43826452 | 0.138821607 | 0.59 |
| YPR067W | 664.4484323 | -0.239354271 | 0.138746178 | 0.59 |
| YPR180W | 730.2139766 | -0.3196212 | 0.140112036 | 0.59 |
| YJR054W | 539.7820263 | 0.334920969 | 0.141340341 | 0.59 |
| YHR036W | 340.3963184 | 0.43039718 | 0.141776853 | 0.59 |
| YIL021W | 814.8146206 | -0.330646241 | 0.1416581 | 0.59 |
| YLR222C | 1088.378581 | 0.589170266 | 0.141784653 | 0.59 |
| YNL159C | 572.5140174 | 0.37393126 | 0.141953747 | 0.59 |
| YBL038W | 1166.095027 | -0.512876881 | 0.142170561 | 0.59 |
| YDR462W | 726.8986569 | -0.340870647 | 0.142079274 | 0.59 |
| YJL199C | 163.0335414 | -0.325347934 | 0.142462542 | 0.59 |
| YMR016C | 1109.247117 | 0.473229472 | 0.142370096 | 0.59 |
| YDR045C | 241.2309895 | -0.438417369 | 0.14271328 | 0.59 |
| YDR367W | 606.3521729 | -0.373297161 | 0.142755799 | 0.59 |

| | | | | |
|---|---|---|---|---|
| YOL012C | 810.102208 | -0.343186598 | 0.142807703 | 0.59 |
| YGL148W | 4268.613078 | -0.552494947 | 0.142974881 | 0.59 |
| YGR106C | 3546.453004 | -0.348148935 | 0.143247219 | 0.59 |
| YHR011W | 509.988716 | 0.515484597 | 0.143298125 | 0.59 |
| YMR052W | 219.8800281 | -0.449116044 | 0.143440626 | 0.59 |
| YPL215W | 1126.607522 | -0.319624503 | 0.143943496 | 0.59 |
| YDL168W | 1463.24926 | -0.462517707 | 0.144186763 | 0.59 |
| YOR334W | 302.0782196 | 0.304613935 | 0.144091095 | 0.59 |
| YMR004W | 1474.539468 | -0.300143386 | 0.144472993 | 0.59 |
| YDR183W | 267.9198317 | -0.288543961 | 0.144947482 | 0.59 |
| YGL129C | 1219.243345 | -0.31830775 | 0.145027646 | 0.59 |
| YLL062C | 209.8685733 | 0.48217796 | 0.144836066 | 0.59 |
| YOR275C | 1566.16532 | 0.256471377 | 0.145030371 | 0.59 |
| YGL190C | 1598.926776 | 0.351338117 | 0.145429786 | 0.59 |
| YMR312W | 719.0027121 | -0.344367892 | 0.145676979 | 0.59 |
| YMR145C | 7155.424672 | 0.489499799 | 0.146301938 | 0.60 |
| YLR356W | 2015.024039 | -0.577579024 | 0.146574969 | 0.60 |
| YLR446W | 527.2109038 | 0.418161887 | 0.146582595 | 0.60 |
| YBL035C | 596.37002 | 0.451794046 | 0.147480403 | 0.60 |
| YBR071W | 990.0527368 | -0.508364677 | 0.147533438 | 0.60 |
| YFL004W | 2271.622342 | -0.272233602 | 0.147385083 | 0.60 |
| YFR037C | 1666.566511 | -0.349460224 | 0.147624652 | 0.60 |
| YFR048W | 703.4879681 | 0.338406836 | 0.147686457 | 0.60 |
| YIR032C | 113.4974223 | -0.686558286 | 0.146829257 | 0.60 |
| YLR084C | 1027.133398 | 0.481834054 | 0.147619435 | 0.60 |
| YOR264W | 932.8104963 | 0.361821904 | 0.14718197 | 0.60 |
| YPR166C | 431.6523763 | -0.41090795 | 0.147657601 | 0.60 |
| YBR131W | 285.0666359 | -0.341213012 | 0.147971612 | 0.60 |
| YPR095C | 1088.661894 | 0.268875079 | 0.148899007 | 0.60 |
| YDR419W | 525.8031882 | 0.37855853 | 0.149230593 | 0.60 |
| YGR237C | 1848.787726 | -0.336378751 | 0.149270575 | 0.60 |
| YHR138C | 1476.126547 | -0.492082509 | 0.149447257 | 0.60 |
| YOL152W | 484.6917609 | 0.692107634 | 0.149773691 | 0.60 |
| snR84 | 162.4042194 | 0.396108072 | 0.149711159 | 0.60 |
| YGL176C | 596.0143813 | 0.285052416 | 0.150121506 | 0.60 |
| YNL308C | 3277.042014 | -0.494941053 | 0.150572768 | 0.61 |
| YCR073C | 984.1918819 | 0.254796314 | 0.151071049 | 0.61 |
| YHR049W | 2093.457795 | -0.578489257 | 0.150896988 | 0.61 |
| YLR019W | 1293.150392 | 0.294259195 | 0.151073284 | 0.61 |
| YPR010C | 3009.126793 | 0.610008883 | 0.1510393 | 0.61 |
| YDR272W | 1625.208653 | -0.459092602 | 0.151210436 | 0.61 |
| YBR086C | 4766.880566 | 0.274788552 | 0.152469006 | 0.61 |
| YBR188C | 339.0880427 | -0.273694887 | 0.151825602 | 0.61 |

| | | | | |
|---|---|---|---|---|
| YCR081W | 992.4265821 | 0.272248923 | 0.152352377 | 0.61 |
| YGL094C | 1570.488055 | 0.343488031 | 0.151671805 | 0.61 |
| YGR270W | 2956.664275 | 0.24750215 | 0.15277224 | 0.61 |
| YHR008C | 8653.583569 | -0.553396124 | 0.152009887 | 0.61 |
| YIL079C | 348.2789788 | 0.594920382 | 0.151492714 | 0.61 |
| YJL073W | 1063.600671 | 0.441253785 | 0.152508345 | 0.61 |
| YJR134C | 1708.030399 | -0.351363121 | 0.151922901 | 0.61 |
| YLL006W | 812.6766975 | -0.534857716 | 0.152214123 | 0.61 |
| YLR150W | 21689.73235 | -0.554087757 | 0.15280141 | 0.61 |
| YML081C-A | 1028.969006 | -0.621778927 | 0.152478289 | 0.61 |
| YOR135C | 405.7747912 | -0.380493487 | 0.152739394 | 0.61 |
| YML095C | 255.548255 | 0.32054364 | 0.153146316 | 0.61 |
| YNL161W | 1329.930942 | 0.228256273 | 0.153137594 | 0.61 |
| YNL247W | 4374.768503 | -0.509278544 | 0.152965328 | 0.61 |
| YEL007W | 3320.15412 | 0.404829161 | 0.153897544 | 0.61 |
| YKL139W | 677.7577592 | 0.288741277 | 0.154050422 | 0.61 |
| YDL153C | 1719.824839 | -0.502882892 | 0.15489093 | 0.61 |
| YGR177C | 234.3513182 | 0.628074982 | 0.154867375 | 0.61 |
| YBR270C | 464.1113888 | 0.392829475 | 0.155229699 | 0.61 |
| YOL029C | 366.4284423 | 0.253046356 | 0.155212568 | 0.61 |
| YOR103C | 111.7299261 | -0.372682777 | 0.155012292 | 0.61 |
| YHR112C | 1466.272889 | -0.40582708 | 0.155689993 | 0.61 |
| YLL024C | 20764.11389 | -0.621880742 | 0.155710328 | 0.61 |
| YMR078C | 885.7332334 | 0.273846805 | 0.155526915 | 0.61 |
| YGR014W | 3080.238337 | 0.590619745 | 0.155844147 | 0.61 |
| YDR049W | 886.770398 | 0.193829293 | 0.155976874 | 0.61 |
| YPL083C | 495.9825937 | 0.248276795 | 0.156085487 | 0.61 |
| YPR157W | 448.7836668 | 0.65217109 | 0.156269194 | 0.61 |
| YHR005C-A | 2208.832297 | -0.413181284 | 0.156443063 | 0.61 |
| YBR021W | 496.9097626 | 0.400782672 | 0.156810589 | 0.61 |
| YDL001W | 483.443293 | 0.259569739 | 0.156873153 | 0.61 |
| YDR315C | 362.5235632 | -0.355063555 | 0.15674049 | 0.61 |
| YGR224W | 693.0347479 | 0.380200077 | 0.15720139 | 0.61 |
| YLR275W | 260.6216284 | -0.371124789 | 0.157212006 | 0.61 |
| YMR269W | 489.6101925 | -0.463567158 | 0.157370905 | 0.61 |
| YOR371C | 940.0880721 | -0.356831251 | 0.157300408 | 0.61 |
| YDL235C | 741.3905909 | -0.34637305 | 0.15750469 | 0.61 |
| YDR415C | 529.3228874 | 0.387494586 | 0.157924807 | 0.61 |
| YLR130C | 1253.100793 | 0.375096113 | 0.157722962 | 0.61 |
| YPL008W | 976.9490792 | 0.290447955 | 0.157883575 | 0.61 |
| YHR192W | 784.2571123 | -0.306168814 | 0.15837103 | 0.61 |
| YHR206W | 1203.868698 | 0.412981153 | 0.158513473 | 0.61 |
| YKL092C | 887.6693114 | 0.329044776 | 0.158462407 | 0.61 |

| | | | | |
|---|---|---|---|---|
| YNR024W | 280.6447987 | -0.359911285 | 0.158679869 | 0.61 |
| YPR047W | 774.0936392 | -0.291351442 | 0.158792399 | 0.61 |
| YJL179W | 307.4969877 | -0.498951886 | 0.159078017 | 0.62 |
| YOR058C | 1144.395841 | 0.324082535 | 0.159128974 | 0.62 |
| YBR037C | 852.7234834 | -0.352069587 | 0.159563304 | 0.62 |
| YGL075C | 312.3284424 | 0.279204343 | 0.159697911 | 0.62 |
| YDR054C | 1985.938955 | -0.354228438 | 0.160406926 | 0.62 |
| YGR287C | 287.959068 | -0.439511814 | 0.160374289 | 0.62 |
| YJL006C | 336.1143852 | -0.345957545 | 0.160182965 | 0.62 |
| YJL062W-A | 1034.508423 | -0.425435654 | 0.16054669 | 0.62 |
| YKL103C | 3936.941911 | -0.588423934 | 0.159987809 | 0.62 |
| YOL058W | 2645.481694 | 0.634042597 | 0.160286046 | 0.62 |
| YOR196C | 1243.855488 | -0.344569588 | 0.160562322 | 0.62 |
| YBL093C | 1069.915355 | -0.24854409 | 0.160826664 | 0.62 |
| YDR236C | 550.9804022 | -0.319681552 | 0.160883226 | 0.62 |
| YKL028W | 1511.073698 | -0.33582223 | 0.161175492 | 0.62 |
| YBR259W | 488.2154125 | 0.222881848 | 0.161544753 | 0.62 |
| YDL090C | 724.6090506 | -0.328688232 | 0.161493539 | 0.62 |
| YGL174W | 430.9199773 | -0.286339698 | 0.161657654 | 0.62 |
| YDL065C | 1387.029558 | -0.381802441 | 0.162400604 | 0.62 |
| YEL046C | 2157.479841 | 0.552954217 | 0.162365299 | 0.62 |
| YGL026C | 4379.972897 | -0.463916401 | 0.162026208 | 0.62 |
| YGL244W | 2087.284965 | -0.356798551 | 0.161969848 | 0.62 |
| YLR050C | 545.8709868 | -0.477293491 | 0.162427109 | 0.62 |
| YML105C | 1352.843987 | -0.289999868 | 0.162454591 | 0.62 |
| YOR351C | 100.6577504 | 0.505433679 | 0.162050592 | 0.62 |
| YGR286C | 924.5538048 | -0.610301204 | 0.162624238 | 0.62 |
| YDR346C | 2820.625886 | 0.344165928 | 0.162927849 | 0.62 |
| YDL173W | 1092.267621 | -0.599467186 | 0.163319232 | 0.62 |
| YKL077W | 3042.957164 | -0.280873029 | 0.163252056 | 0.62 |
| YPL213W | 340.9140453 | -0.340477939 | 0.163195503 | 0.62 |
| YFL022C | 4340.34362 | -0.456919798 | 0.16369557 | 0.62 |
| YLR380W | 1647.260273 | -0.4249657 | 0.163680192 | 0.62 |
| YOL151W | 926.9123911 | -0.443622354 | 0.163482806 | 0.62 |
| YNL016W | 4774.57143 | -0.343644179 | 0.164035012 | 0.62 |
| YPL070W | 1079.494033 | 0.307053154 | 0.163945296 | 0.62 |
| YLR015W | 562.8104697 | 0.332545631 | 0.164220524 | 0.62 |
| YJR075W | 2038.161429 | -0.343237445 | 0.165191646 | 0.62 |
| YCR057C | 1417.95789 | 0.501181896 | 0.165346024 | 0.62 |
| YBR160W | 885.0084099 | -0.339660236 | 0.165702066 | 0.62 |
| YKL035W | 11294.21789 | -0.39091639 | 0.165778893 | 0.62 |
| YMR307W | 11748.41017 | 0.578039403 | 0.165603579 | 0.62 |
| YGL038C | 1572.160144 | 0.350870539 | 0.165893419 | 0.62 |

| | | | | |
|---|---|---|---|---|
| YGL055W | 3060.50732 | 0.460376111 | 0.166284697 | 0.63 |
| YDL022W | 5632.112705 | -0.508172033 | 0.166536973 | 0.63 |
| YNL231C | 1567.48489 | -0.318948512 | 0.166596065 | 0.63 |
| YIR008C | 729.6399623 | -0.234546049 | 0.167120108 | 0.63 |
| YER063W | 1431.45469 | -0.324607111 | 0.167342838 | 0.63 |
| YPL091W | 3000.977941 | -0.296014689 | 0.167349275 | 0.63 |
| YAL007C | 1549.491558 | -0.518686254 | 0.168445176 | 0.63 |
| YBR126W-B | 156.4452467 | -0.546048838 | 0.168560152 | 0.63 |
| YDR464W | 2373.668914 | -0.431130112 | 0.168143984 | 0.63 |
| YEL001C | 2451.407888 | -0.476238844 | 0.168648095 | 0.63 |
| YGL092W | 1819.13534 | 0.334325997 | 0.168405666 | 0.63 |
| YHR196W | 929.0161741 | 0.560268134 | 0.167794877 | 0.63 |
| YLR247C | 1785.222725 | 0.29231985 | 0.168532756 | 0.63 |
| YLR263W | 707.0718116 | 0.574877857 | 0.168021206 | 0.63 |
| YNL011C | 614.0144377 | 0.38902853 | 0.167757954 | 0.63 |
| YNL306W | 1534.453742 | -0.275753875 | 0.168219202 | 0.63 |
| YPL202C | 824.3595251 | 0.263670293 | 0.16852647 | 0.63 |
| YDL175C | 355.6103486 | 0.425954983 | 0.169301269 | 0.63 |
| YJL145W | 1081.572718 | -0.43430831 | 0.169104044 | 0.63 |
| YLR004C | 257.8975095 | 0.427133126 | 0.169290233 | 0.63 |
| YML054C | 1101.015099 | 0.648910274 | 0.169191532 | 0.63 |
| YDR059C | 828.9203049 | -0.40309409 | 0.169741436 | 0.63 |
| YDR198C | 539.1037114 | 0.316442623 | 0.169631582 | 0.63 |
| YOR372C | 1526.129326 | 0.313200034 | 0.169760805 | 0.63 |
| YLR306W | 266.7883268 | -0.419570144 | 0.169876074 | 0.63 |
| YLL048C | 4130.734576 | 0.303167679 | 0.170011059 | 0.63 |
| YMR277W | 1413.809425 | 0.306795747 | 0.170664453 | 0.63 |
| YGR015C | 438.1021585 | -0.234733136 | 0.171230191 | 0.63 |
| YGR161W-C | 303.4318056 | -0.515633699 | 0.171515017 | 0.63 |
| YLL032C | 878.0650576 | 0.290086747 | 0.17160466 | 0.63 |
| YNL265C | 447.0625511 | -0.242922515 | 0.171714922 | 0.63 |
| YPL115C | 1185.518438 | 0.340405381 | 0.171681934 | 0.63 |
| YPL180W | 1077.43255 | 0.21169196 | 0.17132771 | 0.63 |
| YLR335W | 2838.169755 | -0.230784812 | 0.171928558 | 0.63 |
| YMR288W | 774.5062547 | 0.284758265 | 0.172215496 | 0.63 |
| YNL007C | 10095.98848 | -0.36749655 | 0.172274418 | 0.63 |
| YER170W | 376.8018239 | -0.283857907 | 0.172472009 | 0.63 |
| YLR097C | 748.1202732 | -0.321562885 | 0.172819759 | 0.63 |
| YNL214W | 315.125881 | -0.209319412 | 0.172763213 | 0.63 |
| YAL046C | 313.7043981 | -0.405892246 | 0.173045912 | 0.63 |
| YDR050C | 39610.42901 | -0.491593942 | 0.173368843 | 0.63 |
| YHL028W | 1137.369035 | 0.53987778 | 0.173364394 | 0.63 |
| YDR079W | 403.891688 | -0.466820402 | 0.173529759 | 0.63 |

| | | | | |
|---|---|---|---|---|
| YDL108W | 550.3870487 | -0.227166885 | 0.17370455 | 0.63 |
| YIL009C-A | 425.8589138 | -0.350846681 | 0.173890698 | 0.63 |
| YBR212W | 2837.682435 | -0.407313982 | 0.174955818 | 0.64 |
| YDL188C | 1068.696328 | 0.347827639 | 0.175273552 | 0.64 |
| YDR123C | 353.8570592 | 0.424731254 | 0.175204406 | 0.64 |
| YDR365W-B | 142.2689351 | -0.69906593 | 0.174623283 | 0.64 |
| YEL028W | 113.7795172 | 0.396646871 | 0.175096032 | 0.64 |
| YER029C | 871.4448752 | -0.23246132 | 0.174269947 | 0.64 |
| YGR004W | 331.5972983 | 0.311609753 | 0.174833452 | 0.64 |
| YGR021W | 818.8357092 | -0.337802186 | 0.174694329 | 0.64 |
| YIL065C | 779.4474724 | -0.430501857 | 0.174463298 | 0.64 |
| YIL150C | 295.6540585 | 0.322901179 | 0.174870923 | 0.64 |
| YJR053W | 291.1479634 | 0.329397342 | 0.174973236 | 0.64 |
| YKL051W | 1194.148337 | 0.458891497 | 0.175599179 | 0.64 |
| YLR028C | 4846.630925 | -0.139168236 | 0.175540795 | 0.64 |
| YBL099W | 11787.99535 | -0.606727155 | 0.176175629 | 0.64 |
| YBL105C | 2392.516684 | 0.314943855 | 0.176289485 | 0.64 |
| YCL041C | 119.8839828 | -0.435074701 | 0.176031854 | 0.64 |
| YGR271C-A | 454.6880839 | -0.567154856 | 0.176311332 | 0.64 |
| YLR323C | 363.5088652 | -0.317867816 | 0.176264267 | 0.64 |
| YER088C-A | 302.6604055 | 0.610700164 | 0.17672879 | 0.64 |
| YIL160C | 460.4774311 | 0.633921851 | 0.176720331 | 0.64 |
| YBR271W | 387.9475512 | 0.343493267 | 0.176996751 | 0.64 |
| YJR092W | 2324.987782 | 0.410867608 | 0.177362698 | 0.64 |
| YDR111C | 552.8400061 | 0.524214573 | 0.177639087 | 0.64 |
| YJR041C | 1150.543526 | 0.461544896 | 0.177705934 | 0.64 |
| YLR290C | 1190.940137 | -0.361416497 | 0.178019755 | 0.64 |
| YLR413W | 2093.12908 | 0.669737192 | 0.178289005 | 0.64 |
| YPR173C | 1842.223717 | -0.369753286 | 0.178229754 | 0.64 |
| YHR098C | 2961.484376 | 0.314991432 | 0.178514592 | 0.64 |
| YDR503C | 368.4756431 | -0.318540234 | 0.178689605 | 0.64 |
| YAL012W | 6420.264953 | -0.515387961 | 0.180636842 | 0.64 |
| YBR056W | 2794.886987 | -0.4527614 | 0.180484398 | 0.64 |
| YCL024W | 493.5504233 | 0.389503402 | 0.180118631 | 0.64 |
| YER130C | 1003.132192 | 0.54634383 | 0.180439285 | 0.64 |
| YIL166C | 439.1240138 | 0.468381702 | 0.18059924 | 0.64 |
| YIR006C | 5095.375019 | 0.231209558 | 0.179205498 | 0.64 |
| YJR047C | 407.0063415 | -0.631430085 | 0.179598949 | 0.64 |
| YNL325C | 453.4130958 | 0.351763601 | 0.180471423 | 0.64 |
| YOL027C | 2880.865523 | -0.358097833 | 0.179870808 | 0.64 |
| YOL095C | 382.7305205 | 0.30540857 | 0.179860161 | 0.64 |
| YOR108W | 1787.909268 | 0.539522595 | 0.180272895 | 0.64 |
| YOR257W | 743.3095021 | -0.493462826 | 0.179386438 | 0.64 |

| | | | | |
|---|---|---|---|---|
| YPL078C | 4809.066893 | -0.503165372 | 0.180237312 | 0.64 |
| YPL110C | 931.3902179 | 0.330309844 | 0.180126719 | 0.64 |
| YPL154C | 14185.44316 | -0.484812671 | 0.180608957 | 0.64 |
| YPR189W | 1450.192608 | 0.307637935 | 0.180111361 | 0.64 |
| YGL085W | 385.7388603 | -0.294702143 | 0.181291478 | 0.64 |
| YAL001C | 1631.198822 | 0.284242801 | 0.181546948 | 0.64 |
| YMR027W | 2670.4492 | -0.378964356 | 0.181657228 | 0.64 |
| YMR160W | 946.3755314 | 0.352307196 | 0.181632442 | 0.64 |
| YDL171C | 4109.615185 | 0.399978439 | 0.182448486 | 0.64 |
| YDR208W | 1907.44225 | 0.324606615 | 0.182481498 | 0.64 |
| YDR231C | 627.5516275 | -0.474741502 | 0.182696636 | 0.64 |
| YDR276C | 2763.13579 | -0.354030208 | 0.182411549 | 0.64 |
| YHR001W | 1515.116914 | -0.289759907 | 0.182135404 | 0.64 |
| YIL128W | 1273.394888 | 0.343482016 | 0.182574552 | 0.64 |
| YJL096W | 663.1647142 | -0.340998333 | 0.18272295 | 0.64 |
| YOR073W | 590.4176603 | 0.402225406 | 0.182134281 | 0.64 |
| YPR020W | 1129.530709 | -0.592166221 | 0.18283928 | 0.64 |
| YPR088C | 2257.094394 | -0.29821729 | 0.182549361 | 0.64 |
| YKR043C | 2568.82772 | -0.584017857 | 0.183061366 | 0.64 |
| YJR013W | 645.2124988 | 0.215063623 | 0.18333275 | 0.64 |
| YOR185C | 2527.056266 | -0.511117885 | 0.183331052 | 0.64 |
| YJL095W | 1317.430087 | 0.226290673 | 0.183576984 | 0.64 |
| YDR132C | 453.7495244 | 0.361149367 | 0.183910798 | 0.64 |
| YDR349C | 1807.872356 | 0.367131672 | 0.184019864 | 0.64 |
| YKR090W | 850.9383093 | 0.32215416 | 0.184043381 | 0.64 |
| YDR170C | 4884.61556 | -0.279870066 | 0.184491379 | 0.64 |
| YGR193C | 2486.614356 | -0.25372038 | 0.184425353 | 0.64 |
| YOL028C | 419.6545084 | 0.266079121 | 0.184593928 | 0.64 |
| YAL008W | 865.7638687 | -0.457991255 | 0.184884636 | 0.64 |
| YER067C-A | 258.7943359 | -0.681087804 | 0.185264177 | 0.65 |
| YGR082W | 1782.261898 | -0.325608683 | 0.185190314 | 0.65 |
| YBR049C | 1277.509775 | 0.327631343 | 0.186904818 | 0.65 |
| YER062C | 1876.234052 | -0.447514409 | 0.186783733 | 0.65 |
| YFR055W | 975.2716539 | 0.600289772 | 0.186507748 | 0.65 |
| YGL004C | 1031.613443 | -0.331093457 | 0.186450035 | 0.65 |
| YGL206C | 5144.593261 | 0.257655646 | 0.186051826 | 0.65 |
| YJL128C | 1506.583064 | 0.224508579 | 0.186586884 | 0.65 |
| YJL214W | 204.5476536 | 0.384339912 | 0.186021926 | 0.65 |
| YKL080W | 5080.565803 | -0.353801874 | 0.187191999 | 0.65 |
| YKL167C | 608.6144606 | -0.328584037 | 0.185764158 | 0.65 |
| YLL035W | 581.4295973 | 0.450894969 | 0.187176714 | 0.65 |
| YLR262C | 283.7623055 | -0.316680373 | 0.187189446 | 0.65 |
| YML064C | 587.9701312 | 0.207074662 | 0.185850114 | 0.65 |

| | | | | |
|---|---|---|---|---|
| YMR047C | 2343.921258 | 0.278647543 | 0.186656058 | 0.65 |
| YNL055C | 20028.30046 | -0.572584756 | 0.186691745 | 0.65 |
| YOR378W | 266.6242043 | 0.463182068 | 0.185969754 | 0.65 |
| YPL152W | 506.4565509 | -0.242305104 | 0.186752723 | 0.65 |
| YPR133W-A | 1114.918643 | -0.467571363 | 0.187483004 | 0.65 |
| YDL109C | 380.7977413 | 0.379415989 | 0.188141903 | 0.65 |
| YOR324C | 520.2886814 | 0.328666407 | 0.188027564 | 0.65 |
| YJR040W | 752.4605261 | 0.298858295 | 0.188519146 | 0.65 |
| YDR296W | 1076.454873 | -0.342468851 | 0.188969159 | 0.65 |
| YJL004C | 587.5114449 | -0.257157995 | 0.188918844 | 0.65 |
| YNL315C | 1636.297254 | -0.318186256 | 0.189461392 | 0.65 |
| YDL102W | 928.5717968 | 0.229787365 | 0.189597573 | 0.65 |
| YOR009W | 307.6360112 | 0.545133865 | 0.189860485 | 0.65 |
| YPL094C | 2895.773005 | -0.293190551 | 0.18993717 | 0.65 |
| YBR042C | 577.29725 | -0.292602982 | 0.190472902 | 0.65 |
| YDR121W | 548.7039891 | -0.253640256 | 0.190683548 | 0.65 |
| YIL022W | 2268.335855 | -0.169904874 | 0.190718264 | 0.65 |
| YNL261W | 688.6478875 | 0.340646 | 0.190507454 | 0.65 |
| YOL136C | 1016.324902 | 0.490992388 | 0.190794681 | 0.65 |
| YBL088C | 1866.759984 | 0.315294896 | 0.191309213 | 0.65 |
| YHL043W | 209.5139873 | 0.457608326 | 0.191349607 | 0.65 |
| YDR511W | 614.1187303 | -0.458666913 | 0.191470593 | 0.65 |
| YGL245W | 8715.959502 | -0.396629928 | 0.1917991 | 0.65 |
| YKL049C | 577.3829424 | -0.296054059 | 0.191908379 | 0.65 |
| YOL122C | 1128.161192 | 0.428736648 | 0.191912629 | 0.65 |
| YNR017W | 1847.028777 | -0.283275812 | 0.19223947 | 0.65 |
| YER123W | 1029.367215 | 0.258617352 | 0.192676361 | 0.66 |
| YPR143W | 537.9356283 | -0.39298778 | 0.19300363 | 0.66 |
| YDL240W | 734.1749038 | 0.440257397 | 0.193387037 | 0.66 |
| YGL084C | 1084.484417 | 0.231440967 | 0.193709491 | 0.66 |
| YIL111W | 1481.52504 | -0.460084605 | 0.193919657 | 0.66 |
| YML077W | 412.7293887 | -0.386409538 | 0.193794608 | 0.66 |
| YNL263C | 1345.46904 | -0.35224552 | 0.193941336 | 0.66 |
| YPL141C | 1338.030401 | 0.381082846 | 0.194016403 | 0.66 |
| YNL102W | 1324.90783 | 0.343814736 | 0.194292297 | 0.66 |
| YNL225C | 446.0966712 | -0.233640762 | 0.19421371 | 0.66 |
| YKL061W | 244.3100746 | -0.420963872 | 0.19471175 | 0.66 |
| YOR326W | 4223.653773 | -0.328275581 | 0.194735359 | 0.66 |
| YJL008C | 2286.818099 | -0.333225026 | 0.194891776 | 0.66 |
| YLR409C | 1307.908763 | 0.525012154 | 0.195023737 | 0.66 |
| YBL009W | 764.0807255 | 0.380368786 | 0.195437373 | 0.66 |
| YDR481C | 2453.405315 | -0.407530888 | 0.195577132 | 0.66 |
| YLL036C | 873.0418955 | 0.178354737 | 0.196202564 | 0.66 |

| | | | | |
|---|---|---|---|---|
| YHR140W | 190.2878906 | -0.567282283 | 0.196344261 | 0.66 |
| YLR348C | 661.9197696 | -0.541684027 | 0.196526538 | 0.66 |
| YIL105C | 1531.713028 | -0.34565535 | 0.196663739 | 0.66 |
| YDR047W | 1762.459456 | -0.275345295 | 0.196953907 | 0.66 |
| YDR439W | 511.333794 | 0.29002464 | 0.197197128 | 0.66 |
| YML125C | 2139.072841 | -0.449447755 | 0.19723587 | 0.66 |
| YOL018C | 882.7418897 | -0.164153353 | 0.197609685 | 0.66 |
| YPR052C | 667.2448711 | -0.431874982 | 0.197790894 | 0.66 |
| YPR025C | 1080.062057 | -0.334369544 | 0.198122945 | 0.66 |
| YBR010W | 4241.31261 | -0.524129952 | 0.198383854 | 0.66 |
| YPL184C | 1810.247002 | -0.371961729 | 0.198370956 | 0.66 |
| YOR191W | 2372.954185 | 0.288991821 | 0.198567658 | 0.66 |
| YDR470C | 683.2878102 | 0.217428562 | 0.198885197 | 0.66 |
| YGR162W | 4487.780728 | 0.432487069 | 0.199170497 | 0.66 |
| YER045C | 814.7735686 | 0.420160807 | 0.199883544 | 0.67 |
| YGR163W | 840.8307527 | 0.266974527 | 0.200176044 | 0.67 |
| YJR019C | 596.1541169 | -0.346382154 | 0.199704258 | 0.67 |
| YLR427W | 1359.694674 | 0.126511537 | 0.200272807 | 0.67 |
| YOR159C | 320.0573697 | -0.317136358 | 0.200039479 | 0.67 |
| YPR058W | 588.0067957 | -0.411045305 | 0.20024888 | 0.67 |
| YKL163W | 2089.664663 | 0.577509047 | 0.200538254 | 0.67 |
| YGR052W | 642.70953 | 0.635647666 | 0.201193417 | 0.67 |
| YBR199W | 2706.736809 | -0.14462101 | 0.202178242 | 0.67 |
| YBR210W | 464.9591183 | -0.326944973 | 0.201985846 | 0.67 |
| YDR384C | 561.1890906 | 0.619222203 | 0.20225521 | 0.67 |
| YJL180C | 680.1821358 | -0.265312149 | 0.20223413 | 0.67 |
| YCL047C | 434.4156263 | -0.359668319 | 0.203676161 | 0.67 |
| YDR534C | 224.6689044 | -0.563109106 | 0.203073547 | 0.67 |
| YGR225W | 197.8168855 | 0.581111906 | 0.203212131 | 0.67 |
| YJL198W | 970.0448734 | 0.413757367 | 0.203812941 | 0.67 |
| YKL211C | 1647.230083 | -0.475562912 | 0.203106419 | 0.67 |
| YOL128C | 161.4112011 | 0.345077839 | 0.203792422 | 0.67 |
| YOR092W | 648.6032701 | 0.429593526 | 0.203648469 | 0.67 |
| YOR115C | 581.5147795 | -0.295334965 | 0.20371121 | 0.67 |
| YPL248C | 793.6864415 | 0.38090531 | 0.203061882 | 0.67 |
| YPR098C | 1032.023923 | -0.466551371 | 0.203411651 | 0.67 |
| YBL079W | 2598.015595 | 0.21406758 | 0.20430554 | 0.67 |
| YBR257W | 391.5498488 | -0.245581777 | 0.204694582 | 0.67 |
| YDL045C | 1046.513314 | -0.182900514 | 0.204955485 | 0.67 |
| YDL197C | 997.8504474 | 0.315168208 | 0.204860121 | 0.67 |
| YDR119W | 1442.902962 | 0.464382864 | 0.204808674 | 0.67 |
| YLR422W | 2135.341687 | 0.289877201 | 0.204461443 | 0.67 |
| YOR002W | 2112.418166 | 0.245209597 | 0.204795111 | 0.67 |

| | | | | |
|---|---|---|---|---|
| YOR243C | 1479.808571 | 0.515348345 | 0.204342211 | 0.67 |
| YDL006W | 361.6658456 | -0.203525312 | 0.207271084 | 0.67 |
| YDL110C | 2473.385616 | -0.46814318 | 0.205930961 | 0.67 |
| YDR164C | 1143.654091 | 0.208155812 | 0.20606205 | 0.67 |
| YDR406W | 3009.969096 | -0.585770928 | 0.20734708 | 0.67 |
| YDR435C | 760.2512519 | -0.359054156 | 0.206731539 | 0.67 |
| YEL017C-A | 672.6729484 | 0.40669721 | 0.207004059 | 0.67 |
| YGL191W | 2937.202908 | -0.577693996 | 0.205519852 | 0.67 |
| YHR136C | 132.1966559 | -0.670194603 | 0.206395933 | 0.67 |
| YHR182C-A | 130.5541577 | -0.611694395 | 0.206595512 | 0.67 |
| YLR065C | 843.2361186 | -0.436911884 | 0.205776822 | 0.67 |
| YML098W | 504.9087228 | -0.335517857 | 0.207093351 | 0.67 |
| YNL081C | 921.3541155 | -0.23834198 | 0.206701914 | 0.67 |
| YNL154C | 2670.283788 | 0.348033153 | 0.207329069 | 0.67 |
| YOL078W | 796.3214789 | 0.33438679 | 0.206698093 | 0.67 |
| YOR267C | 3236.118357 | 0.33930294 | 0.207160965 | 0.67 |
| YOR380W | 392.7551247 | 0.306303642 | 0.205906909 | 0.67 |
| YPL195W | 2059.660784 | -0.203891011 | 0.20546066 | 0.67 |
| YEL053W-A | 186.8810681 | 0.622647178 | 0.207612675 | 0.67 |
| YHR175W | 775.2689431 | -0.361460076 | 0.20761008 | 0.67 |
| YDR323C | 314.788297 | 0.202577563 | 0.207955703 | 0.67 |
| YOR305W | 732.3331193 | -0.331694141 | 0.207916137 | 0.67 |
| YBR287W | 2396.597397 | -0.330980215 | 0.208479695 | 0.67 |
| YNR043W | 2170.763038 | -0.366714702 | 0.208494285 | 0.67 |
| YOR060C | 313.4302218 | 0.322648122 | 0.208396091 | 0.67 |
| YPR084W | 689.7868138 | 0.158694446 | 0.209198974 | 0.68 |
| YGR125W | 1519.849762 | 0.224283136 | 0.209561877 | 0.68 |
| YDR144C | 842.4657292 | 0.614698408 | 0.209729398 | 0.68 |
| YGR277C | 611.2932905 | -0.31894347 | 0.209937188 | 0.68 |
| YCR013C | 2806.322738 | -0.533376918 | 0.210971937 | 0.68 |
| YCR092C | 872.6618833 | 0.248221842 | 0.21010757 | 0.68 |
| YDL133W | 835.1418998 | 0.193493808 | 0.211390308 | 0.68 |
| YER109C | 1065.439283 | 0.254412476 | 0.210469818 | 0.68 |
| YFL009W | 860.3502379 | 0.191506032 | 0.210461799 | 0.68 |
| YGL194C | 571.5770023 | -0.235061562 | 0.211573184 | 0.68 |
| YGR137W | 236.2341776 | -0.362939455 | 0.210593124 | 0.68 |
| YHR086W | 910.6374675 | 0.306325773 | 0.211195191 | 0.68 |
| YIR033W | 2182.959679 | 0.280416931 | 0.211114685 | 0.68 |
| YJR104C | 12449.61979 | -0.32555666 | 0.210935908 | 0.68 |
| YNL087W | 1914.774752 | 0.299384184 | 0.21151034 | 0.68 |
| YNL274C | 1846.518968 | -0.471686274 | 0.210525147 | 0.68 |
| YOR119C | 842.963497 | 0.282263394 | 0.211479867 | 0.68 |
| YDL202W | 808.8072189 | -0.325450178 | 0.211760326 | 0.68 |

| | | | | |
|---|---|---|---|---|
| YAL051W | 1222.388192 | 0.256381397 | 0.213929596 | 0.68 |
| YBR172C | 2003.113678 | 0.331570517 | 0.213303991 | 0.68 |
| YBR177C | 2530.709586 | -0.362835146 | 0.2128963 | 0.68 |
| YDL155W | 763.3718322 | 0.256685257 | 0.213475137 | 0.68 |
| YDR075W | 328.511722 | 0.381374733 | 0.213561605 | 0.68 |
| YDR254W | 574.0903535 | 0.240598859 | 0.212599739 | 0.68 |
| YGR116W | 3744.798928 | -0.271163078 | 0.212170959 | 0.68 |
| YGR280C | 1635.461549 | -0.479338641 | 0.212244567 | 0.68 |
| YHR047C | 1885.311487 | -0.402752368 | 0.213821182 | 0.68 |
| YHR081W | 846.2639082 | -0.338584929 | 0.214007895 | 0.68 |
| YIL020C | 394.2399705 | -0.404289077 | 0.213449337 | 0.68 |
| YJL153C | 833.1459195 | 0.512650509 | 0.213331251 | 0.68 |
| YJR124C | 485.3455454 | 0.473049573 | 0.213706109 | 0.68 |
| YLR373C | 1362.586121 | 0.232012367 | 0.212022341 | 0.68 |
| YMR258C | 733.537512 | 0.309062827 | 0.213094185 | 0.68 |
| YOL130W | 1532.382846 | 0.245390428 | 0.21352156 | 0.68 |
| YPL056C | 172.8020928 | 0.501405718 | 0.213591589 | 0.68 |
| YPL103C | 590.7071612 | 0.282711483 | 0.213495476 | 0.68 |
| YNL279W | 510.6368292 | 0.512424243 | 0.214278185 | 0.68 |
| YPR085C | 667.2581144 | 0.241786987 | 0.214300194 | 0.68 |
| YBR274W | 529.7886878 | -0.23705287 | 0.216612661 | 0.68 |
| YDL023C | 219.5343501 | -0.461972281 | 0.214964281 | 0.68 |
| YDL164C | 704.8057939 | 0.245280617 | 0.215356424 | 0.68 |
| YDL231C | 1653.703222 | 0.351981323 | 0.215422222 | 0.68 |
| YDR130C | 501.1168326 | 0.303102621 | 0.215792082 | 0.68 |
| YDR438W | 373.0593006 | 0.325598583 | 0.215223937 | 0.68 |
| YER176W | 1918.703807 | -0.280201814 | 0.215751674 | 0.68 |
| YFR016C | 3711.04018 | -0.379791817 | 0.216146072 | 0.68 |
| YGR025W | 100.9215658 | -0.381259454 | 0.216160863 | 0.68 |
| YGR244C | 5502.307672 | -0.429067098 | 0.216265857 | 0.68 |
| YLR256W | 4392.10169 | 0.388888086 | 0.216481311 | 0.68 |
| YMR120C | 11760.55672 | 0.575936839 | 0.216202686 | 0.68 |
| YMR183C | 2217.146681 | -0.319226567 | 0.216631959 | 0.68 |
| YNL119W | 513.2444289 | 0.407022719 | 0.214709662 | 0.68 |
| YOR315W | 1158.774212 | 0.597865214 | 0.215761275 | 0.68 |
| YPR031W | 763.0479333 | 0.334305447 | 0.21606989 | 0.68 |
| YPR119W | 682.8278436 | 0.515791839 | 0.216281623 | 0.68 |
| YHL010C | 843.3803368 | 0.34874185 | 0.217105329 | 0.68 |
| YMR218C | 1133.912405 | 0.224457671 | 0.217144368 | 0.68 |
| YGL057C | 252.5789153 | 0.205509711 | 0.217543248 | 0.68 |
| YHL019C | 544.6940331 | 0.352692736 | 0.217676457 | 0.68 |
| YJR004C | 1252.603429 | 0.479328249 | 0.217588531 | 0.68 |
| YKL034W | 1217.582813 | 0.352022877 | 0.217718184 | 0.68 |

| | | | | |
|---|---|---|---|---|
| YBL051C | 3664.33556 | -0.469710868 | 0.218092218 | 0.68 |
| YJL209W | 626.0794008 | 0.23880622 | 0.217939854 | 0.68 |
| YOL137W | 547.8586694 | 0.211742649 | 0.217982548 | 0.68 |
| YKR099W | 800.367483 | 0.38671518 | 0.218253476 | 0.68 |
| YOL114C | 133.5426332 | -0.387437269 | 0.218453633 | 0.68 |
| YDR430C | 1380.899649 | 0.151289246 | 0.219515413 | 0.68 |
| YLR062C | 194.5995134 | 0.560396729 | 0.219972384 | 0.68 |
| YKL193C | 1757.604095 | -0.413704372 | 0.220207792 | 0.68 |
| YFL046W | 596.7158239 | -0.241617325 | 0.220399421 | 0.68 |
| YPL258C | 548.5976694 | 0.426100116 | 0.221215708 | 0.69 |
| YOR071C | 651.6605909 | 0.416792677 | 0.221638025 | 0.69 |
| YDL236W | 1309.137891 | -0.407539166 | 0.222607549 | 0.69 |
| YHR082C | 4569.103984 | -0.36163811 | 0.222571026 | 0.69 |
| YIL063C | 1081.563362 | -0.373955553 | 0.222656958 | 0.69 |
| YNL070W | 685.2481589 | -0.318010951 | 0.222299142 | 0.69 |
| YPL155C | 690.291389 | 0.302282279 | 0.222661794 | 0.69 |
| YBR088C | 178.1843606 | -0.442171123 | 0.22295734 | 0.69 |
| YCL011C | 2495.29652 | 0.313222397 | 0.222888773 | 0.69 |
| YIR031C | 471.7138035 | 0.43579084 | 0.223188982 | 0.69 |
| YBL054W | 675.0753311 | 0.496177619 | 0.223980479 | 0.69 |
| YBR080C | 2198.347068 | -0.291809229 | 0.22382582 | 0.69 |
| YDR485C | 1410.914463 | -0.324579693 | 0.223921403 | 0.69 |
| YGR249W | 613.9099062 | 0.587765185 | 0.223703695 | 0.69 |
| YHR088W | 1005.281896 | -0.440109643 | 0.224138976 | 0.69 |
| YJR007W | 3429.418011 | -0.400304088 | 0.223655441 | 0.69 |
| YOL071W | 1035.627926 | -0.437243515 | 0.224056954 | 0.69 |
| YDR459C | 457.0405938 | 0.199712552 | 0.224486161 | 0.69 |
| YPR182W | 302.4381877 | -0.27041927 | 0.224774738 | 0.69 |
| YGL233W | 1565.055775 | 0.185669445 | 0.224945958 | 0.69 |
| YIL129C | 2649.636452 | 0.25158864 | 0.225077016 | 0.69 |
| YBL046W | 691.3216368 | -0.224812379 | 0.226228293 | 0.69 |
| YCR060W | 437.7609902 | -0.335819323 | 0.225388922 | 0.69 |
| YCR079W | 984.6018828 | -0.379449514 | 0.226048894 | 0.69 |
| YDR043C | 825.8596949 | 0.470607728 | 0.226140023 | 0.69 |
| YKL205W | 1310.174229 | 0.383244594 | 0.226229674 | 0.69 |
| YNL164C | 278.0976276 | 0.336762036 | 0.225596576 | 0.69 |
| YNL241C | 6779.137631 | -0.399508785 | 0.22614603 | 0.69 |
| YNL290W | 854.0353401 | -0.317737633 | 0.22553703 | 0.69 |
| YOR335C | 3110.388093 | 0.343048301 | 0.225688435 | 0.69 |
| YKL114C | 901.3872837 | -0.298617168 | 0.226538221 | 0.69 |
| YOR350C | 702.3901449 | 0.323120619 | 0.226485878 | 0.69 |
| YBR005W | 1310.131508 | -0.480240846 | 0.227173896 | 0.69 |
| YNL086W | 191.101467 | -0.32508756 | 0.227196647 | 0.69 |

| | | | | |
|---|---|---|---|---|
| YJR118C | 868.4456577 | -0.339112032 | 0.227791287 | 0.69 |
| YJR056C | 431.2081935 | 0.232523734 | 0.227918537 | 0.69 |
| YDR108W | 495.5770673 | 0.226696521 | 0.228103413 | 0.69 |
| YLR447C | 5880.017489 | -0.266284464 | 0.228201807 | 0.69 |
| YGR276C | 1041.106244 | -0.171230096 | 0.228576164 | 0.69 |
| YOL075C | 712.6275688 | 0.326153931 | 0.228921618 | 0.69 |
| YOR017W | 870.202076 | 0.220173478 | 0.228891096 | 0.69 |
| YBR256C | 1957.968457 | -0.367292257 | 0.229131826 | 0.69 |
| YKL007W | 2212.362326 | -0.167464023 | 0.229192359 | 0.69 |
| YKL086W | 218.7788586 | -0.481076658 | 0.229358286 | 0.69 |
| YDL078C | 3642.136284 | -0.396704719 | 0.229918434 | 0.70 |
| YDR153C | 1161.100144 | 0.258354639 | 0.230565601 | 0.70 |
| YGL207W | 4342.654723 | -0.274741529 | 0.230138439 | 0.70 |
| YHR024C | 1377.744705 | -0.205398056 | 0.230273115 | 0.70 |
| YHR143W | 3407.641358 | 0.417519301 | 0.230426292 | 0.70 |
| YLR332W | 1136.948721 | 0.373942528 | 0.230342849 | 0.70 |
| YNL321W | 1495.615211 | 0.257473368 | 0.230151823 | 0.70 |
| YOR140W | 905.6577941 | 0.283556922 | 0.230499222 | 0.70 |
| YPR124W | 1891.733227 | 0.47712522 | 0.230692129 | 0.70 |
| YPR147C | 688.0952487 | -0.239398608 | 0.230991 | 0.70 |
| YLR173W | 824.1208493 | 0.42709223 | 0.231415954 | 0.70 |
| YPR033C | 3601.656962 | -0.349579832 | 0.231466937 | 0.70 |
| YBR286W | 17567.33584 | -0.350469485 | 0.231662528 | 0.70 |
| YNL166C | 1009.247872 | -0.39728253 | 0.232018088 | 0.70 |
| YLR270W | 2664.450215 | -0.443252843 | 0.232511349 | 0.70 |
| YDR287W | 832.7406735 | -0.395785939 | 0.232780553 | 0.70 |
| YJR102C | 338.1234787 | -0.285158545 | 0.232784869 | 0.70 |
| YDR100W | 731.7492012 | -0.460166094 | 0.233896133 | 0.70 |
| YER087C-B | 1311.571506 | -0.430775836 | 0.233944301 | 0.70 |
| YGL108C | 328.3815974 | -0.286950101 | 0.234108359 | 0.70 |
| YHR178W | 530.4654676 | 0.374428872 | 0.23415184 | 0.70 |
| YJL035C | 163.2101994 | 0.418301447 | 0.233193857 | 0.70 |
| YKL024C | 1589.092419 | -0.330340602 | 0.233435459 | 0.70 |
| YKL106W | 462.66436 | 0.356378892 | 0.233760871 | 0.70 |
| YLR026C | 1116.422754 | -0.30569502 | 0.233622085 | 0.70 |
| YLR038C | 1207.825369 | -0.544136876 | 0.233991472 | 0.70 |
| YPL026C | 1074.82514 | 0.481424093 | 0.233490803 | 0.70 |
| YDR267C | 660.9899359 | -0.210376157 | 0.234424285 | 0.70 |
| YCR089W | 1832.043276 | 0.426950526 | 0.23464509 | 0.70 |
| YPL220W | 1761.369957 | -0.513402556 | 0.23489684 | 0.70 |
| YJL084C | 1615.45903 | 0.30065148 | 0.235114461 | 0.70 |
| YPR121W | 333.11981 | 0.359606167 | 0.235175885 | 0.70 |
| YEL044W | 582.0177494 | -0.265268621 | 0.236272522 | 0.70 |

| | | | | |
|---|---|---|---|---|
| YGL097W | 1307.008078 | -0.485022207 | 0.236229387 | 0.70 |
| YGR060W | 3114.502526 | -0.494081102 | 0.236300258 | 0.70 |
| YCR014C | 618.1633306 | 0.297106574 | 0.237255892 | 0.70 |
| YDL093W | 807.8112275 | 0.258924726 | 0.236833293 | 0.70 |
| YFL049W | 806.3019635 | 0.201580956 | 0.23676213 | 0.70 |
| YHR122W | 579.3180386 | -0.324499363 | 0.237619045 | 0.70 |
| YJR133W | 618.7266674 | -0.377509594 | 0.236630674 | 0.70 |
| YKL094W | 2161.93051 | -0.260750077 | 0.237224514 | 0.70 |
| YLR260W | 983.6383575 | -0.343831402 | 0.237910766 | 0.70 |
| YMR212C | 1054.541147 | 0.355848312 | 0.237900238 | 0.70 |
| YMR263W | 652.3046851 | -0.319437296 | 0.237836603 | 0.70 |
| YNL058C | 1080.922928 | -0.445581116 | 0.237068616 | 0.70 |
| YNR038W | 1115.811097 | 0.340753751 | 0.237632037 | 0.70 |
| YOR338W | 133.0538926 | 0.37663586 | 0.237422027 | 0.70 |
| YPL192C | 128.9945052 | 0.406882608 | 0.238277908 | 0.70 |
| YCL025C | 3652.132376 | 0.504953332 | 0.239033111 | 0.70 |
| YJR001W | 1763.85175 | 0.210372286 | 0.238926941 | 0.70 |
| YMR229C | 3400.401838 | 0.449925023 | 0.23880111 | 0.70 |
| YOR059C | 1230.819833 | 0.262517052 | 0.238883783 | 0.70 |
| YPL267W | 353.8623989 | -0.38946072 | 0.238588679 | 0.70 |
| YKL222C | 519.5767688 | 0.344474266 | 0.239454576 | 0.70 |
| YGR030C | 291.5946813 | -0.35522462 | 0.239921431 | 0.70 |
| YGR251W | 285.9824856 | 0.311577153 | 0.239921703 | 0.70 |
| YLR264W | 525.6829336 | 0.481329556 | 0.239800585 | 0.70 |
| YCL057C-A | 1206.33424 | -0.443417989 | 0.241020326 | 0.70 |
| YDL020C | 3590.678769 | 0.415792357 | 0.241537663 | 0.70 |
| YDR227W | 1551.753891 | -0.372686985 | 0.24073873 | 0.70 |
| YFR004W | 2781.166019 | -0.209566965 | 0.24067837 | 0.70 |
| YGR134W | 745.1524745 | 0.256909091 | 0.241323946 | 0.70 |
| YHL014C | 453.0111576 | 0.298248129 | 0.240571192 | 0.70 |
| YHR009C | 3439.835058 | -0.241644531 | 0.24038077 | 0.70 |
| YJL012C | 2507.923216 | -0.436182429 | 0.24169152 | 0.70 |
| YLR264C-A | 117.4733387 | 0.568176853 | 0.241457085 | 0.70 |
| YML126C | 4399.479194 | -0.374633372 | 0.240378772 | 0.70 |
| YNL066W | 1969.929236 | 0.36895944 | 0.240393501 | 0.70 |
| YOR077W | 225.6560466 | 0.294421731 | 0.241549533 | 0.70 |
| YPL122C | 549.6598792 | 0.190813528 | 0.241014999 | 0.70 |
| YPR054W | 265.2938746 | 0.447102582 | 0.241745538 | 0.70 |
| YHR068W | 2922.590435 | -0.434195599 | 0.242259202 | 0.70 |
| YKL194C | 593.5236151 | -0.259070725 | 0.242067786 | 0.70 |
| YNL054W | 1269.220328 | 0.258622189 | 0.241970577 | 0.70 |
| YPR128C | 787.6455183 | -0.221853725 | 0.24225938 | 0.70 |
| YGR211W | 3320.437529 | -0.445774054 | 0.242410076 | 0.70 |

| | | | | |
|---|---|---|---|---|
| YNL136W | 1295.54364 | -0.388332218 | 0.242631938 | 0.70 |
| YPL024W | 252.5020697 | 0.443898418 | 0.242800957 | 0.71 |
| YAL043C | 1560.889418 | 0.256360284 | 0.243830451 | 0.71 |
| YGL133W | 1877.257774 | 0.242953332 | 0.244092642 | 0.71 |
| YHL016C | 664.2747126 | 0.51227564 | 0.244337508 | 0.71 |
| YJL031C | 712.299831 | -0.273105308 | 0.243822345 | 0.71 |
| YLR072W | 910.1580571 | 0.237391789 | 0.244093723 | 0.71 |
| YLR254C | 554.1871051 | -0.288948904 | 0.244468683 | 0.71 |
| YMR067C | 1220.387108 | -0.221624464 | 0.244340092 | 0.71 |
| YOL143C | 5652.711754 | -0.421979436 | 0.244443726 | 0.71 |
| YOR125C | 859.4168859 | -0.373209414 | 0.244324867 | 0.71 |
| YOR215C | 2322.968103 | -0.441939812 | 0.243791282 | 0.71 |
| YPR186C | 328.1330267 | -0.280128945 | 0.244755079 | 0.71 |
| YKL023W | 1129.71016 | -0.387524289 | 0.244925117 | 0.71 |
| YDL120W | 712.7094342 | -0.265585759 | 0.246565646 | 0.71 |
| YDR476C | 1082.582617 | -0.28786062 | 0.245906462 | 0.71 |
| YEL050C | 1046.082243 | -0.237313162 | 0.246089253 | 0.71 |
| YGL210W | 815.3164128 | -0.241898782 | 0.246173721 | 0.71 |
| YLR081W | 221.380283 | 0.472612194 | 0.246805055 | 0.71 |
| YLR106C | 6573.73141 | 0.372283714 | 0.245905051 | 0.71 |
| YLR357W | 954.9234442 | 0.296355043 | 0.24655548 | 0.71 |
| YMR163C | 451.0209425 | 0.211238656 | 0.246042309 | 0.71 |
| YMR247C | 1642.766432 | 0.253945049 | 0.246575816 | 0.71 |
| YMR257C | 582.2395624 | 0.349312723 | 0.246883697 | 0.71 |
| YNL121C | 5779.129433 | -0.30046717 | 0.246908194 | 0.71 |
| YOR016C | 992.9033585 | -0.236343764 | 0.245754633 | 0.71 |
| YMR075W | 761.6920474 | 0.215271048 | 0.24759525 | 0.71 |
| YBL100C | 649.5552869 | -0.518000412 | 0.248030316 | 0.71 |
| YIR016W | 1147.104809 | -0.383539063 | 0.248632235 | 0.71 |
| YLR131C | 1246.543234 | 0.258519942 | 0.24882657 | 0.71 |
| YMR199W | 1528.60034 | 0.43222277 | 0.248746234 | 0.71 |
| YMR224C | 500.4037572 | 0.198730692 | 0.248860561 | 0.71 |
| YNL197C | 1704.51539 | 0.312278032 | 0.248538706 | 0.71 |
| YPL093W | 1935.182804 | 0.487462624 | 0.248363922 | 0.71 |
| YOL098C | 2415.997601 | 0.24531698 | 0.249275457 | 0.71 |
| YDR409W | 969.523209 | 0.248669443 | 0.249752095 | 0.71 |
| YOR080W | 677.2163658 | 0.171512464 | 0.249746603 | 0.71 |
| YMR214W | 1048.466602 | -0.329958002 | 0.249911474 | 0.71 |
| YCL043C | 17735.82228 | -0.368696346 | 0.250811814 | 0.71 |
| YKL195W | 3025.520768 | -0.218452695 | 0.250842147 | 0.71 |
| YML111W | 1306.672498 | 0.240114707 | 0.250960495 | 0.71 |
| YOR205C | 770.9234478 | 0.266021385 | 0.250536382 | 0.71 |
| YPL183W-A | 354.0512636 | -0.278368236 | 0.25089141 | 0.71 |

| | | | | |
|---|---|---|---|---|
| YML010W | 2999.463402 | -0.264978446 | 0.251561643 | 0.71 |
| YOR045W | 622.8710581 | -0.380190195 | 0.251570161 | 0.71 |
| YER080W | 3267.440935 | -0.335884563 | 0.251720088 | 0.71 |
| YBR262C | 557.8168642 | -0.414265275 | 0.252474884 | 0.71 |
| YDR480W | 507.0349407 | -0.150101825 | 0.252768011 | 0.71 |
| YKR067W | 2101.266331 | -0.382124356 | 0.252431087 | 0.71 |
| YLR227C | 512.1356286 | 0.198853076 | 0.252807386 | 0.71 |
| YNL126W | 705.9981899 | 0.323334945 | 0.25283564 | 0.71 |
| YOR081C | 1945.1418 | 0.285739164 | 0.252511299 | 0.71 |
| YOR118W | 1462.615579 | 0.179546354 | 0.25255682 | 0.71 |
| YBL067C | 1161.166699 | 0.161889565 | 0.253891194 | 0.72 |
| YEL017W | 1022.681916 | 0.458020471 | 0.25389297 | 0.72 |
| YHR160C | 281.5696466 | 0.395690979 | 0.253884818 | 0.72 |
| YBL089W | 493.6798455 | 0.242206555 | 0.255336019 | 0.72 |
| YBR276C | 1528.649976 | 0.138144167 | 0.254977936 | 0.72 |
| YDR249C | 239.973621 | 0.331080086 | 0.255363771 | 0.72 |
| YGR079W | 359.5947801 | 0.460337488 | 0.25535544 | 0.72 |
| YHR091C | 596.7962742 | -0.17015939 | 0.255357685 | 0.72 |
| YHR149C | 909.9713222 | 0.391981753 | 0.255409423 | 0.72 |
| YLR098C | 706.1029207 | 0.294290164 | 0.255046175 | 0.72 |
| YNL152W | 687.0558678 | 0.173321599 | 0.254566706 | 0.72 |
| YOR001W | 1523.051345 | 0.264115204 | 0.25519676 | 0.72 |
| YOR075W | 634.1618193 | -0.376154204 | 0.254365399 | 0.72 |
| YDR158W | 6048.209084 | -0.354391217 | 0.255545649 | 0.72 |
| YDR194C | 2378.698014 | 0.274687035 | 0.255901559 | 0.72 |
| YDR076W | 307.3025521 | 0.291534346 | 0.256456651 | 0.72 |
| YDR266C | 1529.276069 | 0.261173853 | 0.256652822 | 0.72 |
| YOL123W | 4114.119417 | -0.365037154 | 0.256690758 | 0.72 |
| YOR151C | 4196.260493 | 0.287281483 | 0.256617786 | 0.72 |
| YPL071C | 441.9177148 | -0.311057676 | 0.256599128 | 0.72 |
| YDR520C | 591.1970741 | 0.358425988 | 0.257051846 | 0.72 |
| YFR022W | 849.2747048 | 0.382456887 | 0.257673146 | 0.72 |
| YML071C | 1055.622411 | -0.269037642 | 0.257817756 | 0.72 |
| YMR124W | 1653.834225 | 0.251286771 | 0.257763457 | 0.72 |
| YOL019W | 928.0908204 | 0.314955902 | 0.257993153 | 0.72 |
| YJL156C | 869.0736001 | 0.219582845 | 0.258513555 | 0.72 |
| YKR095W-A | 172.6357846 | -0.373944128 | 0.258505413 | 0.72 |
| YLL023C | 2955.471664 | -0.356273371 | 0.258863857 | 0.72 |
| YOR317W | 4625.356852 | -0.406536783 | 0.259276193 | 0.72 |
| YDL064W | 884.0529801 | -0.326045476 | 0.259678129 | 0.72 |
| YLL058W | 677.8123693 | -0.279895738 | 0.259877978 | 0.72 |
| YMR177W | 436.2483597 | 0.292136659 | 0.260043092 | 0.72 |
| YNL236W | 1176.860728 | 0.231423582 | 0.259963652 | 0.72 |

| | | | | |
|---|---|---|---|---|
| YBR255C-A | 311.2384417 | -0.348302595 | 0.260326792 | 0.72 |
| YML070W | 2763.480937 | -0.317827855 | 0.260580359 | 0.72 |
| YMR092C | 2509.767947 | -0.250305378 | 0.260519167 | 0.72 |
| YML062C | 782.2227983 | -0.223511121 | 0.26099747 | 0.72 |
| YNR033W | 1803.332991 | -0.262048711 | 0.261215 | 0.72 |
| YNR049C | 561.0637018 | -0.20192311 | 0.261128189 | 0.72 |
| YGR124W | 7850.205704 | -0.523401066 | 0.261484827 | 0.72 |
| YPR106W | 760.0851492 | 0.245262652 | 0.261795088 | 0.72 |
| YMR207C | 1385.212155 | 0.254663833 | 0.262084186 | 0.73 |
| YDR352W | 489.6613604 | 0.355716549 | 0.262575811 | 0.73 |
| YLR450W | 2180.849651 | 0.339970221 | 0.263281784 | 0.73 |
| YDR457W | 5588.916733 | 0.252076554 | 0.263682421 | 0.73 |
| YLR058C | 11510.40084 | 0.501434971 | 0.263691251 | 0.73 |
| YMR055C | 275.6053073 | 0.266989882 | 0.263681756 | 0.73 |
| YJL062W | 1004.118516 | 0.205190674 | 0.264027167 | 0.73 |
| YMR135C | 1809.513775 | 0.478353037 | 0.264394519 | 0.73 |
| YOR202W | 1479.782603 | -0.380090262 | 0.264867138 | 0.73 |
| YHR163W | 2218.862273 | -0.488672731 | 0.265341214 | 0.73 |
| YCR032W | 1688.564969 | 0.311145415 | 0.265883637 | 0.73 |
| YCR102C | 144.1370507 | -0.439620481 | 0.265675336 | 0.73 |
| YNL094W | 1768.042353 | 0.224938403 | 0.265800545 | 0.73 |
| YDR116C | 1042.960963 | -0.241026896 | 0.266404231 | 0.73 |
| YJR146W | 99.77471099 | 0.497991123 | 0.266625478 | 0.73 |
| YLR192C | 3457.976113 | -0.269778981 | 0.266762555 | 0.73 |
| YNL040W | 832.5546055 | -0.346395061 | 0.266329772 | 0.73 |
| YPL051W | 362.5719952 | -0.263020678 | 0.266789979 | 0.73 |
| YPL074W | 944.5738824 | 0.276737437 | 0.266205537 | 0.73 |
| YHR012W | 963.21973 | -0.280249833 | 0.267088686 | 0.73 |
| YPL096W | 660.7866918 | -0.213888198 | 0.267099539 | 0.73 |
| YBR155W | 833.2201083 | -0.474509964 | 0.267657417 | 0.73 |
| YDR138W | 884.1934286 | 0.234627021 | 0.268685246 | 0.73 |
| YDR146C | 1456.032171 | 0.339262602 | 0.268121715 | 0.73 |
| YGR154C | 333.0792836 | 0.324111358 | 0.268754751 | 0.73 |
| YKR022C | 681.4384392 | -0.220784325 | 0.268599044 | 0.73 |
| YKR079C | 916.529157 | 0.293097955 | 0.268436174 | 0.73 |
| YPR117W | 2597.357181 | 0.216324513 | 0.268442302 | 0.73 |
| YLL002W | 366.0264394 | 0.307323343 | 0.269082754 | 0.73 |
| YGR217W | 881.6554263 | 0.281955321 | 0.269430641 | 0.73 |
| YBR162C | 7471.337212 | -0.435000427 | 0.270020366 | 0.74 |
| YBR165W | 515.6336536 | -0.213531285 | 0.270274354 | 0.74 |
| YDR420W | 1513.909661 | 0.351853255 | 0.270066651 | 0.74 |
| YGL089C | 132.0787924 | 0.22290706 | 0.270148567 | 0.74 |
| YHR101C | 345.7524508 | 0.232606838 | 0.270421367 | 0.74 |

| | | | | |
|---|---|---|---|---|
| YDR379C-A | 462.1957351 | -0.359521379 | 0.271234585 | 0.74 |
| YDR432W | 4551.301879 | -0.351439158 | 0.271124781 | 0.74 |
| YGR035C | 329.4527926 | -0.493847096 | 0.271001895 | 0.74 |
| YLR390W-A | 12426.05045 | -0.222220337 | 0.271019245 | 0.74 |
| YNL238W | 1373.585003 | 0.240233856 | 0.271223051 | 0.74 |
| YPL038W | 299.6200602 | -0.337994227 | 0.270590408 | 0.74 |
| YHR007C | 4279.432771 | 0.439434536 | 0.271628617 | 0.74 |
| YIL078W | 8337.626041 | -0.41458439 | 0.271511017 | 0.74 |
| YPL221W | 1548.403188 | 0.310695741 | 0.271655897 | 0.74 |
| YBR235W | 1364.419973 | 0.349101242 | 0.272295022 | 0.74 |
| YKL121W | 1073.818565 | 0.279754608 | 0.272235746 | 0.74 |
| YOL148C | 713.6397629 | -0.22063905 | 0.272625633 | 0.74 |
| YDR161W | 1294.242267 | -0.360495954 | 0.272853448 | 0.74 |
| YCL005W | 658.6999475 | -0.150717703 | 0.27316632 | 0.74 |
| YBR029C | 1375.442278 | -0.292498526 | 0.274147642 | 0.74 |
| YBR096W | 835.4279583 | -0.399615352 | 0.273977302 | 0.74 |
| YGL164C | 861.3475827 | -0.261734519 | 0.273807658 | 0.74 |
| YIR021W | 261.891268 | -0.23929444 | 0.274186887 | 0.74 |
| YLR354C | 11943.28845 | -0.287414416 | 0.273826617 | 0.74 |
| YPL108W | 159.4425969 | -0.290759279 | 0.274093603 | 0.74 |
| YEL069C | 125.5249506 | 0.555030406 | 0.274387733 | 0.74 |
| YIL104C | 354.0212252 | 0.276367258 | 0.274638216 | 0.74 |
| YLR457C | 335.6948937 | -0.192399813 | 0.274851627 | 0.74 |
| YBR213W | 228.053134 | -0.252135776 | 0.275504279 | 0.74 |
| YLL018C-A | 305.6957299 | 0.413856967 | 0.27541987 | 0.74 |
| YNL061W | 2817.950295 | -0.375569634 | 0.275723232 | 0.74 |
| YNL113W | 221.4093861 | -0.443036423 | 0.275715677 | 0.74 |
| YOR287C | 297.2565187 | -0.434710551 | 0.275854625 | 0.74 |
| YPR068C | 284.1750342 | 0.174938248 | 0.275772239 | 0.74 |
| YMR220W | 1203.063636 | 0.200096979 | 0.276332542 | 0.74 |
| YKL138C | 823.724232 | -0.311400516 | 0.2769171 | 0.74 |
| YLR376C | 277.9237935 | -0.315563913 | 0.276860158 | 0.74 |
| YML076C | 1190.303539 | 0.257798289 | 0.276858112 | 0.74 |
| YLR199C | 824.459216 | 0.161199856 | 0.277168002 | 0.74 |
| YNL101W | 1154.001663 | 0.284241188 | 0.277295861 | 0.74 |
| YGL119W | 689.6648811 | -0.188543677 | 0.27753499 | 0.74 |
| YIL113W | 384.3761033 | -0.43743181 | 0.277771666 | 0.74 |
| YOL103W | 1490.877623 | 0.176395921 | 0.277928033 | 0.74 |
| YPL151C | 670.1565001 | 0.166676831 | 0.277991712 | 0.74 |
| YNL245C | 408.0242862 | 0.26801357 | 0.278250251 | 0.74 |
| YER008C | 2582.281632 | -0.333109539 | 0.278687261 | 0.74 |
| YLR269C | 113.3738155 | -0.247143625 | 0.278678362 | 0.74 |
| YPL274W | 1036.210642 | 0.45163125 | 0.278539412 | 0.74 |

| | | | | |
|---|---|---|---|---|
| YMR058W | 5542.022572 | 0.483290915 | 0.278868321 | 0.74 |
| YIL001W | 471.4395924 | 0.348632495 | 0.279147173 | 0.74 |
| YNL305C | 2021.497779 | -0.398044502 | 0.279133307 | 0.74 |
| YDL182W | 2531.684902 | 0.461305459 | 0.279583747 | 0.74 |
| YDL185W | 10910.49268 | -0.197045849 | 0.279649027 | 0.74 |
| YLR252W | 254.6818923 | -0.402192296 | 0.279624425 | 0.74 |
| YJL036W | 1275.214013 | -0.286843646 | 0.279855603 | 0.74 |
| YBR214W | 5062.049562 | -0.390369699 | 0.280118019 | 0.74 |
| YDR001C | 5262.491506 | -0.33036974 | 0.280410413 | 0.74 |
| YOR128C | 2990.654925 | 0.429373257 | 0.280936417 | 0.74 |
| YDR512C | 741.4198181 | -0.244548006 | 0.281513525 | 0.74 |
| YLR246W | 470.2308665 | 0.348025642 | 0.281148138 | 0.74 |
| YMR178W | 1157.445412 | -0.261091633 | 0.281260556 | 0.74 |
| YNL298W | 1195.295385 | 0.326671655 | 0.281500107 | 0.74 |
| YPL109C | 614.5552418 | 0.424334099 | 0.28194582 | 0.75 |
| YDR240C | 401.4217535 | 0.177818362 | 0.282399284 | 0.75 |
| YGL018C | 470.8004765 | -0.26759175 | 0.282365967 | 0.75 |
| YNL138W-A | 114.4265169 | -0.250782374 | 0.282147671 | 0.75 |
| YAR027W | 1481.56887 | -0.238739015 | 0.283628588 | 0.75 |
| YBR008C | 906.2949136 | 0.293766678 | 0.28400403 | 0.75 |
| YFR036W | 228.7837864 | 0.248456916 | 0.283242998 | 0.75 |
| YGL209W | 1219.680335 | 0.373921988 | 0.283995888 | 0.75 |
| YHL050C | 143.7644195 | 0.488412368 | 0.283734726 | 0.75 |
| YHR035W | 231.8124454 | 0.315505296 | 0.283332424 | 0.75 |
| YKL157W | 4702.190162 | -0.327561617 | 0.283382712 | 0.75 |
| YLR197W | 4160.668892 | -0.435236006 | 0.283885757 | 0.75 |
| YER007C-A | 951.757768 | -0.326264446 | 0.284513882 | 0.75 |
| YPL257W | 181.2677801 | 0.469511269 | 0.284842443 | 0.75 |
| YHR073W | 1977.66425 | 0.256264053 | 0.285353073 | 0.75 |
| YKL137W | 420.0820122 | -0.335304646 | 0.285440616 | 0.75 |
| YPR040W | 901.3829077 | -0.29213095 | 0.285229487 | 0.75 |
| YNL091W | 4224.416881 | -0.250061619 | 0.286163266 | 0.75 |
| YKR014C | 1917.457601 | -0.287283953 | 0.286404099 | 0.75 |
| YNL292W | 950.1678617 | -0.285463393 | 0.286580736 | 0.75 |
| YER111C | 1455.121069 | 0.260119344 | 0.286728592 | 0.75 |
| YGL241W | 933.7531382 | 0.194373351 | 0.287052827 | 0.75 |
| YJR150C | 147.0716932 | 0.510236486 | 0.287350889 | 0.75 |
| YLR040C | 271.3228626 | 0.498453536 | 0.287317687 | 0.75 |
| YAR003W | 588.5199738 | 0.31382777 | 0.290983032 | 0.75 |
| YBR106W | 4289.033597 | -0.574039958 | 0.289392964 | 0.75 |
| YCR027C | 255.2084978 | 0.253424583 | 0.288760407 | 0.75 |
| YDL186W | 120.8658792 | 0.397867078 | 0.289764258 | 0.75 |
| YDR067C | 673.9208846 | -0.203829619 | 0.290391305 | 0.75 |

| | | | | |
|---|---|---|---|---|
| YDR324C | 1092.059972 | 0.463812264 | 0.287863644 | 0.75 |
| YDR390C | 1683.184066 | -0.175898479 | 0.291618319 | 0.75 |
| YEL054C | 3213.78471 | 0.466024493 | 0.289473065 | 0.75 |
| YEL077C | 280.3488731 | 0.444593378 | 0.290198094 | 0.75 |
| YFL002C | 1100.586944 | 0.151608247 | 0.288599605 | 0.75 |
| YFR005C | 788.6190865 | 0.250195682 | 0.290387435 | 0.75 |
| YGR158C | 631.124363 | -0.338312455 | 0.290699325 | 0.75 |
| YIL011W | 777.5395764 | -0.527126894 | 0.288485095 | 0.75 |
| YJR042W | 1389.72183 | 0.224925128 | 0.289100127 | 0.75 |
| YKL029C | 3291.164423 | 0.479667981 | 0.291589261 | 0.75 |
| YKL078W | 299.5473506 | 0.473405209 | 0.29164218 | 0.75 |
| YKL142W | 4281.67462 | -0.364390197 | 0.290790628 | 0.75 |
| YLR017W | 1271.451959 | -0.327994384 | 0.290872585 | 0.75 |
| YLR071C | 1557.164849 | 0.246111833 | 0.289613497 | 0.75 |
| YLR126C | 468.0585451 | -0.371658958 | 0.288635218 | 0.75 |
| YLR213C | 172.0132438 | 0.336421329 | 0.290456591 | 0.75 |
| YLR312W-A | 784.781583 | -0.257028279 | 0.288300353 | 0.75 |
| YML018C | 644.4081194 | 0.370240557 | 0.289536978 | 0.75 |
| YMR252C | 396.7363367 | -0.267089279 | 0.290872813 | 0.75 |
| YOR313C | 174.7411276 | 0.541898717 | 0.291341333 | 0.75 |
| YPL063W | 2649.060944 | -0.180946788 | 0.291304158 | 0.75 |
| YPR120C | 354.4464034 | 0.272608353 | 0.28943261 | 0.75 |
| YPR145W | 8044.845508 | 0.478563794 | 0.291420133 | 0.75 |
| YPR164W | 1248.455329 | 0.190674103 | 0.290053419 | 0.75 |
| YKL203C | 2057.606082 | 0.21600398 | 0.292151608 | 0.75 |
| YDR527W | 984.8264926 | 0.30765445 | 0.292367672 | 0.75 |
| YIL114C | 411.0869055 | -0.356379392 | 0.292447127 | 0.75 |
| YBR105C | 2095.906659 | 0.388137414 | 0.293216509 | 0.75 |
| YDL092W | 1065.77792 | -0.248560469 | 0.292843152 | 0.75 |
| YDR451C | 513.5586606 | 0.34331505 | 0.293413462 | 0.75 |
| YEL055C | 1803.113225 | 0.289708479 | 0.293382908 | 0.75 |
| YGL158W | 260.9463165 | 0.57532264 | 0.292884417 | 0.75 |
| YLR318W | 479.1884801 | 0.241656988 | 0.293304716 | 0.75 |
| YOL126C | 3179.903333 | 0.476513923 | 0.293548434 | 0.75 |
| YOR231W | 881.0751627 | 0.266264402 | 0.294212063 | 0.76 |
| YGR122W | 624.9209214 | 0.290347444 | 0.294629045 | 0.76 |
| YOR124C | 2791.881139 | -0.266117649 | 0.2947217 | 0.76 |
| YMR032W | 652.0530589 | 0.287412892 | 0.294932523 | 0.76 |
| YJR059W | 3492.702056 | -0.289467003 | 0.295288025 | 0.76 |
| YCR061W | 1066.525845 | 0.336357345 | 0.295427855 | 0.76 |
| YJR153W | 431.5896283 | 0.496845814 | 0.295875495 | 0.76 |
| YNL118C | 3319.845542 | 0.176095569 | 0.295913802 | 0.76 |
| YOL057W | 2160.171024 | -0.293895564 | 0.29577556 | 0.76 |

| | | | | |
|---|---|---|---|---|
| YDL230W | 846.3743425 | -0.268934962 | 0.296133092 | 0.76 |
| YDR261C | 1216.328788 | 0.35293183 | 0.296430742 | 0.76 |
| YER133W | 3950.455243 | -0.211523101 | 0.296671065 | 0.76 |
| YIL157C | 795.6414589 | -0.312426866 | 0.296536384 | 0.76 |
| YOR035C | 1268.519702 | 0.335678478 | 0.297012793 | 0.76 |
| YFL003C | 165.623347 | 0.406326066 | 0.297204853 | 0.76 |
| YGL141W | 1274.03239 | -0.196120519 | 0.297578582 | 0.76 |
| YNR061C | 596.8560089 | -0.305770856 | 0.297690856 | 0.76 |
| YPR163C | 5059.100182 | -0.395167613 | 0.297724063 | 0.76 |
| YBL033C | 1274.437806 | -0.195494419 | 0.299033533 | 0.76 |
| YBR296C | 209.9913106 | -0.515242611 | 0.298992113 | 0.76 |
| YDR171W | 12610.16854 | -0.401621176 | 0.299442763 | 0.76 |
| YEL023C | 623.1672116 | 0.267537556 | 0.299470768 | 0.76 |
| YEL031W | 4279.639928 | 0.355585778 | 0.298324487 | 0.76 |
| YGL160W | 1159.891272 | 0.17444296 | 0.298854057 | 0.76 |
| YHR049C-A | 129.3206508 | -0.380716286 | 0.299161012 | 0.76 |
| YIL069C | 2018.584495 | 0.424185155 | 0.299525611 | 0.76 |
| YLR210W | 386.6683639 | 0.240961892 | 0.299362102 | 0.76 |
| YML127W | 993.038483 | -0.235257319 | 0.298451201 | 0.76 |
| YPL092W | 1197.47636 | 0.419371263 | 0.299110094 | 0.76 |
| snR30 | 275.4218461 | 0.419822572 | 0.29834821 | 0.76 |
| YBR192W | 512.8528961 | 0.191558535 | 0.299755819 | 0.76 |
| YGR127W | 732.3981957 | 0.316458096 | 0.300105776 | 0.76 |
| YCR033W | 2347.054478 | 0.309947471 | 0.300897615 | 0.76 |
| YHR107C | 1509.6288 | -0.259345067 | 0.300556896 | 0.76 |
| YIL132C | 143.1249549 | 0.367765841 | 0.300722613 | 0.76 |
| YKL050C | 526.5764301 | 0.335123826 | 0.30077328 | 0.76 |
| YPR032W | 901.7856576 | 0.214477125 | 0.300782235 | 0.76 |
| YBR040W | 134.4656267 | 0.42254415 | 0.301770098 | 0.76 |
| YER079W | 1934.057175 | -0.46533286 | 0.302000311 | 0.76 |
| YIR035C | 1381.941223 | -0.262041629 | 0.301863231 | 0.76 |
| YKL001C | 376.5252013 | -0.373538349 | 0.301338756 | 0.76 |
| YLR319C | 843.3891881 | 0.14776528 | 0.301648297 | 0.76 |
| YPR176C | 535.2741493 | 0.23947636 | 0.301950921 | 0.76 |
| YLR424W | 404.0091344 | 0.170461451 | 0.302287722 | 0.76 |
| YBR026C | 2145.982583 | -0.291963723 | 0.302675609 | 0.76 |
| YDR012W | 4314.996695 | 0.443742347 | 0.303407577 | 0.76 |
| YHR042W | 4638.852766 | -0.242078747 | 0.303347749 | 0.76 |
| YLR091W | 340.1364791 | 0.196590992 | 0.303072614 | 0.76 |
| YPL157W | 499.4077603 | 0.363685846 | 0.303332985 | 0.76 |
| YAL037C-B | 133.1022568 | -0.440310594 | 0.30374996 | 0.76 |
| YHR121W | 1103.128777 | -0.213321145 | 0.303678969 | 0.76 |
| YDR312W | 598.3889984 | -0.347086315 | 0.304323489 | 0.76 |

| | | | | |
|---|---|---|---|---|
| YPL022W | 1488.077225 | 0.268706473 | 0.304399877 | 0.76 |
| YDL084W | 7170.004042 | -0.334422695 | 0.304554881 | 0.76 |
| YCR012W | 43096.09714 | -0.4758717 | 0.304833087 | 0.76 |
| YOL051W | 2569.23447 | 0.246799596 | 0.304936548 | 0.76 |
| YCR046C | 1534.125343 | -0.252638719 | 0.305258993 | 0.76 |
| YDR306C | 1159.558737 | -0.259103451 | 0.305604251 | 0.76 |
| YFR014C | 1739.196956 | -0.335833797 | 0.305422562 | 0.76 |
| YLR233C | 453.3027113 | 0.332601432 | 0.30562879 | 0.76 |
| YMR283C | 413.1862547 | 0.189321311 | 0.305711212 | 0.76 |
| YMR035W | 535.4792987 | -0.282458855 | 0.30649346 | 0.76 |
| YBR278W | 245.0159717 | -0.314926546 | 0.306894535 | 0.77 |
| YKR080W | 1412.607481 | 0.426644955 | 0.307044808 | 0.77 |
| YLR244C | 3158.461036 | -0.314406518 | 0.306819754 | 0.77 |
| YML086C | 1768.316639 | -0.186461092 | 0.307153315 | 0.77 |
| YCL040W | 21911.49524 | -0.409435399 | 0.307766302 | 0.77 |
| YDR072C | 1794.903187 | 0.326379908 | 0.308405813 | 0.77 |
| YGR095C | 969.1047325 | -0.280779973 | 0.308328338 | 0.77 |
| YLL029W | 3427.507563 | -0.311283967 | 0.308506634 | 0.77 |
| YML008C | 4220.321447 | -0.373313142 | 0.308491067 | 0.77 |
| YNL160W | 19462.39163 | 0.419348537 | 0.308524355 | 0.77 |
| YKL067W | 3440.072656 | -0.441459176 | 0.309181277 | 0.77 |
| YNR048W | 344.510222 | 0.220676129 | 0.309699824 | 0.77 |
| YOL077W-A | 1163.041858 | -0.417749723 | 0.309923738 | 0.77 |
| YHR083W | 1675.31411 | -0.207465023 | 0.310749632 | 0.77 |
| YIR003W | 1806.796543 | -0.252176365 | 0.310714278 | 0.77 |
| YMR272C | 4343.78123 | -0.241668436 | 0.310738108 | 0.77 |
| YKR059W | 1551.844916 | -0.404825971 | 0.311081376 | 0.77 |
| YDL135C | 948.2549942 | -0.239171227 | 0.311282652 | 0.77 |
| YGL169W | 682.8701947 | 0.307203815 | 0.311609859 | 0.77 |
| YML004C | 3324.450509 | -0.29723658 | 0.311583662 | 0.77 |
| YDR389W | 894.8507584 | 0.233214812 | 0.311792236 | 0.77 |
| YOR028C | 706.5725488 | 0.50896888 | 0.312107612 | 0.77 |
| YBR117C | 1336.758461 | 0.482906536 | 0.31270483 | 0.77 |
| YDR414C | 448.7530762 | 0.349327265 | 0.312617292 | 0.77 |
| YGL093W | 1066.84026 | 0.143562935 | 0.312627411 | 0.77 |
| YPR133C | 1652.526675 | -0.204964773 | 0.312632866 | 0.77 |
| YCR053W | 6107.267981 | -0.357607361 | 0.314725454 | 0.77 |
| YDL116W | 1341.682086 | 0.189855305 | 0.314437772 | 0.77 |
| YDL227C | 308.6751335 | 0.536399996 | 0.314804973 | 0.77 |
| YDR022C | 413.59294 | -0.405472865 | 0.314670746 | 0.77 |
| YDR310C | 1697.719061 | 0.190435862 | 0.314036984 | 0.77 |
| YDR525W-A | 275.582834 | -0.380227997 | 0.313858021 | 0.77 |
| YER044C-A | 102.3682402 | -0.420686476 | 0.313806307 | 0.77 |

| | | | | |
|---|---|---|---|---|
| YHR034C | 511.3439371 | -0.205113393 | 0.314647212 | 0.77 |
| YHR116W | 348.981213 | -0.282778715 | 0.313448286 | 0.77 |
| YJL159W | 37900.6137 | 0.32525787 | 0.313424058 | 0.77 |
| YJR076C | 1668.359247 | -0.16474772 | 0.314940474 | 0.77 |
| YLR293C | 8683.145693 | -0.364532197 | 0.314319031 | 0.77 |
| YNL322C | 3831.740608 | -0.319924921 | 0.314338016 | 0.77 |
| YNL103W | 1585.703492 | 0.27996607 | 0.315172616 | 0.77 |
| YOL073C | 1132.754224 | -0.340028653 | 0.315245011 | 0.77 |
| YCR019W | 382.8804523 | -0.181496253 | 0.315529957 | 0.77 |
| YBR107C | 343.2589721 | 0.28520911 | 0.316136936 | 0.77 |
| YDR105C | 933.9584198 | 0.194521608 | 0.316137048 | 0.77 |
| YER139C | 313.6167797 | -0.210721767 | 0.315968309 | 0.77 |
| YKL159C | 432.357669 | -0.293692161 | 0.315759477 | 0.77 |
| YNL189W | 2792.970706 | -0.237072342 | 0.316361178 | 0.77 |
| YOL004W | 4094.946984 | 0.185470237 | 0.31639994 | 0.77 |
| YDR383C | 354.9421834 | -0.261186312 | 0.316944484 | 0.77 |
| YBR061C | 630.0720407 | -0.247050746 | 0.318985671 | 0.77 |
| YBR098W | 547.1579177 | 0.19580212 | 0.317505156 | 0.77 |
| YBR125C | 746.1231571 | 0.29406466 | 0.319261017 | 0.77 |
| YBR238C | 642.2716869 | 0.392755864 | 0.317343708 | 0.77 |
| YDR311W | 727.3776002 | 0.22805537 | 0.318588363 | 0.77 |
| YDR473C | 858.9200777 | -0.194767011 | 0.31790615 | 0.77 |
| YHR014W | 138.7802733 | 0.396676647 | 0.318653222 | 0.77 |
| YLR143W | 501.3258454 | 0.264451639 | 0.31773445 | 0.77 |
| YLR176C | 916.0927001 | 0.234675338 | 0.318118498 | 0.77 |
| YLR189C | 2093.552884 | -0.260711769 | 0.319065861 | 0.77 |
| YMR309C | 2879.75717 | 0.34688627 | 0.317305936 | 0.77 |
| YOR051C | 2007.894821 | -0.474480995 | 0.318909997 | 0.77 |
| YPL268W | 929.1963896 | 0.313416628 | 0.319292998 | 0.77 |
| YPR010C-A | 733.8429902 | -0.509130016 | 0.318961863 | 0.77 |
| YPR158W | 1022.854173 | -0.392176577 | 0.319309897 | 0.77 |
| YPR167C | 421.1211 | -0.321317421 | 0.318529253 | 0.77 |
| YGR032W | 3378.619384 | 0.407708455 | 0.320175098 | 0.78 |
| YOR281C | 1009.23062 | -0.274287019 | 0.320180008 | 0.78 |
| YAL036C | 1045.610431 | -0.354859891 | 0.320852396 | 0.78 |
| YER039C | 101.9948572 | 0.332586116 | 0.320748165 | 0.78 |
| YJL139C | 556.1081958 | 0.326287007 | 0.320839988 | 0.78 |
| YPL207W | 1844.153383 | 0.300376956 | 0.320700935 | 0.78 |
| YAR007C | 1627.673812 | -0.237381113 | 0.321724352 | 0.78 |
| YBR126C | 9994.323143 | -0.37408199 | 0.321467947 | 0.78 |
| YBR258C | 250.7365151 | -0.185980363 | 0.321485159 | 0.78 |
| YIL091C | 989.7593991 | 0.380039018 | 0.321126874 | 0.78 |
| YLR139C | 783.7450537 | 0.272709548 | 0.321585222 | 0.78 |

| | | | | |
|---|---|---|---|---|
| YLR396C | 522.7183585 | 0.159275342 | 0.321609721 | 0.78 |
| YBR288C | 774.8863305 | -0.213835064 | 0.324201801 | 0.78 |
| YDR185C | 264.7477216 | 0.336161841 | 0.323600074 | 0.78 |
| YDR222W | 466.2824915 | 0.414098596 | 0.322322304 | 0.78 |
| YDR496C | 1668.42234 | 0.388780735 | 0.32252301 | 0.78 |
| YHR090C | 634.1320105 | -0.292782396 | 0.322396991 | 0.78 |
| YIL149C | 3262.267006 | -0.32578852 | 0.322700913 | 0.78 |
| YJL088W | 187.9199165 | 0.477574538 | 0.323388965 | 0.78 |
| YJL127C-B | 263.7714489 | -0.230735077 | 0.32379858 | 0.78 |
| YKL164C | 7734.898504 | 0.403664176 | 0.324337387 | 0.78 |
| YLR052W | 1071.581443 | 0.099677865 | 0.324252115 | 0.78 |
| YLR201C | 1199.611196 | -0.25945992 | 0.32345065 | 0.78 |
| YLR386W | 1077.114184 | 0.160595365 | 0.324040253 | 0.78 |
| YMR010W | 733.3907547 | 0.321685587 | 0.323695562 | 0.78 |
| YMR278W | 1574.862371 | -0.28676214 | 0.3238503 | 0.78 |
| YOL086C | 59663.42444 | -0.458285869 | 0.324312769 | 0.78 |
| YOL116W | 722.658756 | 0.248443635 | 0.324414428 | 0.78 |
| YPR035W | 16997.33407 | 0.432095532 | 0.323591395 | 0.78 |
| YDL145C | 6294.266606 | 0.220664586 | 0.324740744 | 0.78 |
| YMR225C | 419.9415237 | -0.284227873 | 0.324692016 | 0.78 |
| YHR152W | 1107.927047 | -0.218324892 | 0.325192607 | 0.78 |
| YLR001C | 1116.57839 | 0.200963027 | 0.325156013 | 0.78 |
| YML112W | 572.6104301 | -0.204172991 | 0.325045221 | 0.78 |
| YDL193W | 1212.895326 | -0.184007679 | 0.326581698 | 0.78 |
| YDL209C | 238.6569979 | -0.300621426 | 0.327081852 | 0.78 |
| YER059W | 910.2356733 | 0.262752191 | 0.326397628 | 0.78 |
| YJL047C | 709.4046862 | 0.223872153 | 0.327351644 | 0.78 |
| YJR046W | 1243.286799 | -0.189378355 | 0.326648816 | 0.78 |
| YJR058C | 464.9848205 | -0.205171599 | 0.326942809 | 0.78 |
| YJR074W | 487.5630441 | -0.342192007 | 0.326948098 | 0.78 |
| YLR432W | 4140.896048 | 0.414117685 | 0.327349708 | 0.78 |
| YPR065W | 1458.659767 | 0.398313908 | 0.327174148 | 0.78 |
| YPR185W | 1716.783017 | -0.210857846 | 0.326179911 | 0.78 |
| YOR136W | 7814.887983 | -0.268170138 | 0.3275856 | 0.78 |
| YBR167C | 275.1433522 | -0.244187853 | 0.32824204 | 0.78 |
| YBR291C | 379.9577466 | 0.312770663 | 0.328106559 | 0.78 |
| YDR162C | 515.4142839 | -0.189024044 | 0.327908935 | 0.78 |
| YKR006C | 1505.723079 | -0.230990492 | 0.3284532 | 0.78 |
| YOR085W | 1806.530962 | -0.301896014 | 0.328341139 | 0.78 |
| YDR256C | 561.4619469 | 0.38989698 | 0.329674083 | 0.78 |
| YOR341W | 3809.897668 | 0.380720835 | 0.329695687 | 0.78 |
| YAL041W | 1345.73957 | 0.26612162 | 0.330942821 | 0.78 |
| YBR022W | 363.9709831 | -0.323348627 | 0.331648931 | 0.78 |

| | | | | |
|---|---|---|---|---|
| YCL007C | 102.0392471 | -0.364205973 | 0.330819226 | 0.78 |
| YDR074W | 10482.83127 | -0.425602568 | 0.330822185 | 0.78 |
| YGL145W | 1156.588747 | 0.166563428 | 0.331196878 | 0.78 |
| YJL131C | 712.8647158 | -0.275116398 | 0.33174966 | 0.78 |
| YJR003C | 586.7430541 | 0.33869052 | 0.331614353 | 0.78 |
| YKR103W | 526.4204375 | 0.387712396 | 0.331547381 | 0.78 |
| YOL049W | 2360.13263 | -0.253350347 | 0.331766969 | 0.78 |
| YDR317W | 303.4699364 | 0.286597986 | 0.332483087 | 0.78 |
| YGL021W | 1221.378254 | 0.210339675 | 0.332853872 | 0.78 |
| YHR179W | 8288.962491 | -0.288525323 | 0.332937539 | 0.78 |
| YKR050W | 1256.621968 | 0.316859174 | 0.332848989 | 0.78 |
| YOR171C | 1509.079417 | -0.222734515 | 0.332996011 | 0.78 |
| YPL064C | 883.912951 | -0.147964388 | 0.332863339 | 0.78 |
| YNL281W | 1354.281332 | -0.376055117 | 0.333384267 | 0.78 |
| YHR169W | 496.2218428 | 0.427520826 | 0.333536285 | 0.78 |
| YCL059C | 1691.420964 | -0.406570488 | 0.334170856 | 0.79 |
| YKR024C | 515.1058632 | 0.332263563 | 0.334127389 | 0.79 |
| YNL015W | 3387.233502 | -0.450177689 | 0.333961605 | 0.79 |
| YBL086C | 1904.650929 | -0.266822712 | 0.33698257 | 0.79 |
| YDR338C | 695.8448786 | 0.22792478 | 0.337143896 | 0.79 |
| YDR531W | 969.2146497 | -0.289960884 | 0.335303724 | 0.79 |
| YIL106W | 893.3105346 | 0.191329524 | 0.335488713 | 0.79 |
| YJR073C | 2980.954697 | -0.434286885 | 0.336282107 | 0.79 |
| YLL022C | 865.5611491 | -0.233542728 | 0.336127901 | 0.79 |
| YLR187W | 889.7815093 | 0.180209493 | 0.336056716 | 0.79 |
| YLR328W | 869.5602715 | 0.283450663 | 0.335840745 | 0.79 |
| YLR382C | 967.5198513 | 0.185074649 | 0.337015837 | 0.79 |
| YMR289W | 967.6114857 | -0.190646356 | 0.337115791 | 0.79 |
| YNL331C | 347.7391832 | 0.315789322 | 0.335571788 | 0.79 |
| YOL059W | 2587.338376 | -0.312709735 | 0.336866949 | 0.79 |
| YOL065C | 517.0688786 | -0.226809438 | 0.336931623 | 0.79 |
| YOR342C | 1114.452608 | 0.255381472 | 0.336850131 | 0.79 |
| YPR046W | 225.601515 | -0.263996012 | 0.337181929 | 0.79 |
| YOR344C | 1985.74596 | 0.389014479 | 0.337437535 | 0.79 |
| YFL001W | 478.2892235 | 0.265656602 | 0.337716978 | 0.79 |
| YIL037C | 352.6358083 | 0.418197495 | 0.338256211 | 0.79 |
| YKL082C | 1057.576571 | -0.324633116 | 0.33821879 | 0.79 |
| YOL159C | 274.8863973 | -0.351025251 | 0.33851225 | 0.79 |
| YKL110C | 846.8769824 | -0.240342308 | 0.339011578 | 0.79 |
| YLR079W | 2171.764341 | -0.269855519 | 0.338877889 | 0.79 |
| YBL087C | 3378.344673 | -0.4652483 | 0.340649163 | 0.79 |
| YBR252W | 608.885071 | -0.416100529 | 0.340241725 | 0.79 |
| YDL150W | 525.2641532 | 0.291522926 | 0.340631297 | 0.79 |

| | | | | |
|---|---|---|---|---|
| YEL025C | 1231.625482 | 0.22609529 | 0.340838211 | 0.79 |
| YGR091W | 441.73971 | 0.253378913 | 0.340294538 | 0.79 |
| YJL110C | 684.2542418 | 0.215768077 | 0.340909493 | 0.79 |
| YJR015W | 1958.669387 | -0.223041239 | 0.34024246 | 0.79 |
| YJR136C | 423.3917741 | 0.150869318 | 0.340559928 | 0.79 |
| YKL027W | 701.3136707 | 0.313238242 | 0.341154512 | 0.79 |
| YLR010C | 127.8600135 | -0.261200788 | 0.341340873 | 0.79 |
| YLR193C | 745.5545212 | -0.272024532 | 0.341235617 | 0.79 |
| YLR321C | 729.7163579 | -0.250220526 | 0.340892954 | 0.79 |
| YCL042W | 285.5413665 | -0.41154034 | 0.34161329 | 0.79 |
| YGL180W | 843.8043135 | 0.26096857 | 0.341915821 | 0.79 |
| YKL017C | 627.7815414 | 0.217912447 | 0.341897145 | 0.79 |
| YNL257C | 1662.623362 | 0.158549586 | 0.342041701 | 0.79 |
| YFL029C | 966.6378114 | -0.386275402 | 0.343254543 | 0.79 |
| YGL079W | 448.9297125 | -0.170608822 | 0.34308921 | 0.79 |
| YGR036C | 221.213258 | 0.256800808 | 0.343115835 | 0.79 |
| YHR032W | 1019.136565 | 0.334160859 | 0.343243425 | 0.79 |
| YIL137C | 2189.193159 | -0.287850934 | 0.343178359 | 0.79 |
| YDR069C | 1204.898386 | 0.242410618 | 0.344080219 | 0.79 |
| YDR351W | 1313.626306 | 0.239378929 | 0.343860147 | 0.79 |
| YLR211C | 375.8938446 | -0.310653384 | 0.343661889 | 0.79 |
| YNR059W | 692.5145797 | 0.250066566 | 0.344035323 | 0.79 |
| YGL234W | 9210.041986 | 0.41357809 | 0.344294455 | 0.79 |
| YDL143W | 3248.97051 | -0.258722181 | 0.345025637 | 0.79 |
| YDR337W | 923.5124951 | -0.195783845 | 0.345367788 | 0.79 |
| YDR441C | 376.4209723 | -0.199246181 | 0.344814877 | 0.79 |
| YFR008W | 307.6442666 | -0.230042663 | 0.344769762 | 0.79 |
| YGL098W | 531.3017666 | 0.253885891 | 0.345532183 | 0.79 |
| YJR098C | 316.068717 | 0.233841405 | 0.345049735 | 0.79 |
| YOR271C | 1520.089436 | -0.402661214 | 0.345403652 | 0.79 |
| YPL266W | 1132.936461 | -0.369122199 | 0.345136094 | 0.79 |
| YPL273W | 2617.65817 | -0.42742621 | 0.345746243 | 0.79 |
| YER006W | 1977.06385 | -0.378632858 | 0.345977043 | 0.79 |
| YER148W | 733.3492834 | -0.254448609 | 0.346288525 | 0.79 |
| YLR064W | 1412.390781 | -0.213044771 | 0.346448346 | 0.79 |
| YPL134C | 1579.391088 | -0.446619923 | 0.346548317 | 0.79 |
| YER126C | 1138.972805 | -0.379088865 | 0.346856189 | 0.79 |
| YPL132W | 679.0950944 | -0.240279546 | 0.346760381 | 0.79 |
| YDL013W | 806.2108951 | 0.259108383 | 0.347106451 | 0.79 |
| YMR115W | 1106.490843 | -0.226512504 | 0.347638394 | 0.79 |
| YDR006C | 2310.266671 | 0.23266072 | 0.350441773 | 0.79 |
| YDR299W | 1150.958917 | -0.380189917 | 0.350587404 | 0.79 |
| YEL043W | 2331.741844 | 0.237934182 | 0.349642556 | 0.79 |

| | | | | |
|---|---|---|---|---|
| YER047C | 911.6547256 | 0.216954823 | 0.348806766 | 0.79 |
| YGL127C | 488.4323072 | -0.252056366 | 0.349773777 | 0.79 |
| YGL195W | 6234.838225 | 0.261636433 | 0.348749445 | 0.79 |
| YGL242C | 665.893361 | -0.330416227 | 0.348656156 | 0.79 |
| YGR104C | 540.5492389 | -0.139379398 | 0.348693383 | 0.79 |
| YGR117C | 888.1364455 | 0.214301773 | 0.350431865 | 0.79 |
| YGR173W | 798.4212433 | -0.314126224 | 0.348159492 | 0.79 |
| YJR080C | 1270.85858 | -0.332553112 | 0.350196082 | 0.79 |
| YLR271W | 587.0390943 | -0.35595053 | 0.350054615 | 0.79 |
| YLR453C | 293.5520271 | 0.205021377 | 0.349458684 | 0.79 |
| YML025C | 1270.924858 | -0.129730298 | 0.350549047 | 0.79 |
| YMR181C | 1531.918567 | 0.290386569 | 0.349135595 | 0.79 |
| YMR284W | 845.7287906 | -0.250584626 | 0.350312892 | 0.79 |
| YNL106C | 963.8876185 | 0.167185722 | 0.349812662 | 0.79 |
| YOL100W | 1483.492555 | 0.253927829 | 0.348855396 | 0.79 |
| YOR239W | 2726.169468 | -0.244251364 | 0.349149108 | 0.79 |
| YPR041W | 3343.881383 | -0.30165802 | 0.348433709 | 0.79 |
| YMR281W | 530.4325943 | -0.227841152 | 0.350833877 | 0.80 |
| YMR122W-A | 5293.670712 | -0.300920197 | 0.351263528 | 0.80 |
| YOL087C | 2320.349339 | 0.23206073 | 0.351396956 | 0.80 |
| YGR027C | 9077.452786 | -0.449938036 | 0.352342166 | 0.80 |
| YGR254W | 35005.12746 | -0.477941169 | 0.352349417 | 0.80 |
| YLL027W | 1273.25217 | -0.259153115 | 0.352269485 | 0.80 |
| YPL146C | 923.2028372 | -0.318743932 | 0.351932832 | 0.80 |
| YBR267W | 453.4917639 | 0.288325205 | 0.352641324 | 0.80 |
| YLR053C | 106.3162181 | -0.41525244 | 0.352548366 | 0.80 |
| YDL010W | 362.0027109 | -0.192539908 | 0.352900664 | 0.80 |
| YMR014W | 1014.309235 | -0.324877157 | 0.353393542 | 0.80 |
| YMR268C | 351.0386447 | 0.174556321 | 0.353361525 | 0.80 |
| YOL025W | 668.044846 | 0.255858023 | 0.353385522 | 0.80 |
| YEL067C | 149.0124198 | -0.30314644 | 0.353702298 | 0.80 |
| YMR319C | 2635.007516 | 0.339284408 | 0.353799037 | 0.80 |
| YLR324W | 1737.806437 | -0.278592809 | 0.354426384 | 0.80 |
| YOL119C | 1253.305663 | 0.361406169 | 0.354303538 | 0.80 |
| YGL172W | 1277.929348 | -0.217140093 | 0.35550902 | 0.80 |
| YHL035C | 3064.413674 | 0.388083264 | 0.355647834 | 0.80 |
| YDL031W | 1680.703139 | -0.30855592 | 0.356628371 | 0.80 |
| YER166W | 2859.051695 | 0.209387993 | 0.355966645 | 0.80 |
| YGL002W | 690.1637106 | -0.218448452 | 0.356573956 | 0.80 |
| YGL124C | 833.4516769 | 0.192110991 | 0.356615293 | 0.80 |
| YGL151W | 1728.404115 | 0.196653958 | 0.356442703 | 0.80 |
| YLR083C | 1368.675892 | 0.353644832 | 0.356339975 | 0.80 |
| YGL003C | 856.5289623 | 0.213573208 | 0.356786891 | 0.80 |

| | | | | |
|---|---|---|---|---|
| YAL048C | 583.0194511 | 0.236123657 | 0.35761234 | 0.80 |
| YCR083W | 821.7942376 | -0.357235691 | 0.357970134 | 0.80 |
| YFR047C | 2554.161875 | -0.398670642 | 0.357796583 | 0.80 |
| YGL116W | 933.3944118 | 0.225292813 | 0.357732583 | 0.80 |
| YGR072W | 838.1301736 | -0.223534402 | 0.35803518 | 0.80 |
| YLR039C | 1333.277955 | 0.180188614 | 0.358070209 | 0.80 |
| YNL295W | 664.526791 | 0.215734059 | 0.357808253 | 0.80 |
| YAL038W | 41041.91188 | -0.378232148 | 0.364738258 | 0.80 |
| YBL010C | 437.6652565 | -0.153460755 | 0.365001716 | 0.80 |
| YBL072C | 3997.128198 | -0.444144005 | 0.361142519 | 0.80 |
| YBL077W | 139.1714064 | -0.377154682 | 0.366928371 | 0.80 |
| YBL081W | 976.0547384 | 0.366572613 | 0.36280747 | 0.80 |
| YDL060W | 1265.050122 | 0.308639063 | 0.366373553 | 0.80 |
| YDL087C | 335.3146294 | -0.250263256 | 0.364684853 | 0.80 |
| YDL115C | 894.0269756 | -0.169535637 | 0.360066049 | 0.80 |
| YDL225W | 1750.506735 | -0.219447184 | 0.360633996 | 0.80 |
| YDR058C | 448.3672811 | -0.305352891 | 0.36339011 | 0.80 |
| YDR090C | 473.3492687 | 0.196828738 | 0.362244144 | 0.80 |
| YDR151C | 1113.580859 | 0.175139337 | 0.362779064 | 0.80 |
| YDR302W | 389.0867763 | -0.265753681 | 0.364380961 | 0.80 |
| YER069W | 925.6063303 | 0.304909752 | 0.366019446 | 0.80 |
| YER119C-A | 190.6802462 | -0.265045251 | 0.359194369 | 0.80 |
| YER145C | 1496.957868 | 0.293896497 | 0.359616765 | 0.80 |
| YER151C | 2013.443986 | 0.186949703 | 0.364623198 | 0.80 |
| YER174C | 586.1669315 | -0.238590297 | 0.366445319 | 0.80 |
| YFR006W | 3007.667423 | -0.15981659 | 0.360135757 | 0.80 |
| YGL080W | 877.4500948 | -0.249854979 | 0.360827385 | 0.80 |
| YGL250W | 555.5711852 | -0.310046871 | 0.359445272 | 0.80 |
| YGL263W | 253.3051631 | 0.417611254 | 0.366394116 | 0.80 |
| YGR029W | 703.0611046 | -0.200166874 | 0.365558001 | 0.80 |
| YGR093W | 815.5848449 | 0.209870269 | 0.361430379 | 0.80 |
| YGR178C | 3642.360299 | 0.185591447 | 0.366069992 | 0.80 |
| YGR187C | 552.0431833 | 0.378936693 | 0.36255433 | 0.80 |
| YGR196C | 1285.850548 | 0.166937041 | 0.360400226 | 0.80 |
| YGR197C | 588.1649926 | 0.342496677 | 0.361619511 | 0.80 |
| YHR018C | 3547.596925 | -0.406704489 | 0.366667096 | 0.80 |
| YHR027C | 5786.339997 | -0.298474684 | 0.358587548 | 0.80 |
| YHR114W | 1802.212821 | -0.137167279 | 0.366922618 | 0.80 |
| YIL112W | 2132.63932 | -0.300817968 | 0.364659503 | 0.80 |
| YIR034C | 2228.546885 | -0.411995997 | 0.359894299 | 0.80 |
| YJL099W | 747.8275315 | 0.206959408 | 0.36484895 | 0.80 |
| YJR024C | 696.9415683 | -0.289721086 | 0.360738765 | 0.80 |
| YKL018W | 496.5039326 | -0.176900706 | 0.363555198 | 0.80 |

| | | | | |
|---|---|---|---|---|
| YKL096W | 1798.684379 | 0.405322974 | 0.364516078 | 0.80 |
| YKL141W | 4100.397187 | -0.41412619 | 0.36280276 | 0.80 |
| YLL004W | 381.8327061 | 0.248572093 | 0.366672419 | 0.80 |
| YLR206W | 3965.73873 | 0.177246462 | 0.366548524 | 0.80 |
| YLR442C | 902.5050218 | 0.156490995 | 0.366842542 | 0.80 |
| YML124C | 2648.714801 | -0.281382214 | 0.360803316 | 0.80 |
| YMR019W | 977.1459301 | 0.268101552 | 0.366271546 | 0.80 |
| YMR080C | 1907.455726 | 0.222441049 | 0.366194514 | 0.80 |
| YMR147W | 251.4438843 | 0.354080048 | 0.366207777 | 0.80 |
| YMR158W | 693.5786925 | -0.27551844 | 0.363332079 | 0.80 |
| YMR315W | 4197.096296 | -0.338781579 | 0.365461282 | 0.80 |
| YNL056W | 768.1896527 | -0.179673848 | 0.359087139 | 0.80 |
| YNR022C | 907.2227542 | -0.314966928 | 0.36117465 | 0.80 |
| YNR070W | 728.5997471 | 0.293880587 | 0.360901509 | 0.80 |
| YOL066C | 908.4756316 | -0.241826751 | 0.366077255 | 0.80 |
| YOL149W | 642.6006688 | -0.274493857 | 0.364384118 | 0.80 |
| YOR134W | 351.2595047 | 0.40452195 | 0.360419675 | 0.80 |
| YOR213C | 459.655666 | 0.208093992 | 0.364682145 | 0.80 |
| YOR358W | 499.4768456 | -0.239234909 | 0.363396638 | 0.80 |
| YOR389W | 203.2493311 | 0.340547503 | 0.365632467 | 0.80 |
| YPL005W | 369.5032375 | 0.260298027 | 0.362602232 | 0.80 |
| YPL077C | 211.278667 | 0.198160376 | 0.365474299 | 0.80 |
| YPR057W | 656.961666 | 0.180593614 | 0.362909944 | 0.80 |
| YPR111W | 665.5579675 | 0.254751538 | 0.366527094 | 0.80 |
| YBR045C | 255.946774 | 0.27462933 | 0.369759075 | 0.80 |
| YBR166C | 859.3257193 | -0.209347493 | 0.369332852 | 0.80 |
| YDL051W | 1178.81834 | -0.367615683 | 0.367510806 | 0.80 |
| YDL169C | 664.258116 | -0.439341499 | 0.369152833 | 0.80 |
| YDR285W | 510.8247622 | -0.339516929 | 0.370606192 | 0.80 |
| YDR288W | 555.4180316 | 0.222604644 | 0.368598758 | 0.80 |
| YDR386W | 496.5773156 | 0.234198335 | 0.368402235 | 0.80 |
| YDR411C | 1171.452043 | 0.315696039 | 0.37162762 | 0.80 |
| YER094C | 2268.208556 | -0.176435502 | 0.369506185 | 0.80 |
| YGR007W | 1236.749371 | -0.17345535 | 0.37114927 | 0.80 |
| YGR170W | 2142.624103 | 0.200011391 | 0.370191682 | 0.80 |
| YGR266W | 1152.085768 | -0.249214037 | 0.367750986 | 0.80 |
| YIR019C | 622.3080892 | 0.34326436 | 0.370263583 | 0.80 |
| YJL191W | 879.6997619 | 0.402453879 | 0.368567433 | 0.80 |
| YKL023C-A | 532.0907152 | -0.299381663 | 0.369612927 | 0.80 |
| YKL030W | 149.8142583 | 0.420086285 | 0.367250057 | 0.80 |
| YKL039W | 2424.281879 | 0.166999436 | 0.371429725 | 0.80 |
| YKL066W | 1252.501288 | -0.373339924 | 0.370541351 | 0.80 |
| YKL134C | 542.9022847 | 0.195986854 | 0.36857267 | 0.80 |

| | | | | |
|---|---|---|---|---|
| YLR292C | 1010.798537 | -0.316448235 | 0.370255653 | 0.80 |
| YMR200W | 1797.595559 | -0.198371737 | 0.371392773 | 0.80 |
| YMR234W | 552.7610201 | 0.159970343 | 0.369976009 | 0.80 |
| YNL051W | 867.7540206 | -0.177761734 | 0.371276165 | 0.80 |
| YNL204C | 121.2325338 | 0.361331908 | 0.371512427 | 0.80 |
| YNL243W | 3599.331124 | -0.15455568 | 0.371197624 | 0.80 |
| YOR237W | 107.6435638 | 0.414180235 | 0.370446708 | 0.80 |
| YOR319W | 365.6759573 | -0.182130291 | 0.371038702 | 0.80 |
| YOR356W | 1275.917226 | -0.235119552 | 0.370473238 | 0.80 |
| YPL133C | 580.4116075 | 0.256970892 | 0.369298807 | 0.80 |
| YPR060C | 619.4282416 | -0.364779363 | 0.371468631 | 0.80 |
| YPR122W | 1503.146832 | 0.445326353 | 0.368618905 | 0.80 |
| YDL129W | 428.0062351 | 0.231218979 | 0.37185883 | 0.80 |
| YDR147W | 507.4257469 | 0.262711993 | 0.372092321 | 0.80 |
| YGL112C | 1298.501075 | 0.133672842 | 0.372093262 | 0.80 |
| YNL323W | 681.0384119 | 0.170079623 | 0.372245658 | 0.80 |
| YHR094C | 525.1336673 | 0.442605815 | 0.372924378 | 0.80 |
| YIL152W | 734.4896336 | -0.318664966 | 0.372974761 | 0.80 |
| YBR269C | 1256.661555 | -0.327497179 | 0.373284222 | 0.80 |
| YNL134C | 7013.757129 | -0.396052116 | 0.373156176 | 0.80 |
| YBR063C | 508.4120941 | -0.191779959 | 0.37472425 | 0.80 |
| YBR294W | 590.8243932 | 0.269383189 | 0.373627839 | 0.80 |
| YGR042W | 764.6829972 | -0.241558736 | 0.375042399 | 0.80 |
| YIL139C | 280.0724876 | -0.160794946 | 0.374358111 | 0.80 |
| YLR285W | 889.692098 | -0.288299631 | 0.375415964 | 0.80 |
| YML090W | 156.4475378 | -0.446302555 | 0.375307669 | 0.80 |
| YMR246W | 3290.570155 | 0.36788516 | 0.374905419 | 0.80 |
| YMR290C | 2185.404412 | -0.421175495 | 0.374068728 | 0.80 |
| YOR207C | 1745.865963 | 0.256991714 | 0.374291673 | 0.80 |
| YOR370C | 2847.263179 | 0.200098302 | 0.374784539 | 0.80 |
| YPL012W | 1836.869245 | 0.385445958 | 0.375000623 | 0.80 |
| YPL118W | 2044.672532 | -0.210756408 | 0.375355295 | 0.80 |
| YPL262W | 5873.810031 | -0.419531257 | 0.374566905 | 0.80 |
| YPR165W | 4589.151851 | -0.232880323 | 0.374336875 | 0.80 |
| YNL299W | 424.5188557 | 0.3433665 | 0.376017746 | 0.80 |
| YPL198W | 414.9903406 | 0.296602477 | 0.376237252 | 0.80 |
| YLR406C | 2407.447144 | 0.337488844 | 0.37666253 | 0.80 |
| YBL097W | 438.5923101 | 0.24674403 | 0.377467167 | 0.80 |
| YDL130W | 4709.707857 | -0.414072227 | 0.377924599 | 0.80 |
| YDR524C-B | 18745.499 | -0.391973995 | 0.378628236 | 0.80 |
| YER002W | 1431.744689 | -0.24889658 | 0.378714663 | 0.80 |
| YHR174W | 94635.8071 | -0.481515218 | 0.378317123 | 0.80 |
| YJL118W | 178.5211681 | 0.317360056 | 0.378296091 | 0.80 |

| | | | | |
|---|---|---|---|---|
| YML117W | 3482.921508 | 0.193288067 | 0.378477917 | 0.80 |
| YOR201C | 630.2633868 | 0.186957345 | 0.377658325 | 0.80 |
| YOR227W | 3483.244777 | 0.264208111 | 0.377443179 | 0.80 |
| YOR383C | 13192.95676 | -0.455798982 | 0.378541812 | 0.80 |
| YPL162C | 476.9994766 | 0.231742175 | 0.378370624 | 0.80 |
| YPR021C | 1682.395818 | 0.240909043 | 0.377194091 | 0.80 |
| YPR101W | 539.0655902 | -0.309263441 | 0.377564927 | 0.80 |
| YPR127W | 1734.545853 | -0.330050133 | 0.378575576 | 0.80 |
| YDR019C | 2663.732875 | 0.430096807 | 0.379214976 | 0.80 |
| YLR390W | 418.8392228 | -0.162674567 | 0.379189977 | 0.80 |
| YDR202C | 884.8533036 | -0.290119366 | 0.379757865 | 0.81 |
| YML061C | 1035.299962 | 0.23349127 | 0.379744792 | 0.81 |
| YPL263C | 1770.940205 | -0.255839706 | 0.379498379 | 0.81 |
| YLR032W | 823.8658356 | 0.21229314 | 0.380230714 | 0.81 |
| YJL108C | 258.931357 | 0.284240963 | 0.381068714 | 0.81 |
| YNL112W | 1857.470789 | 0.42445693 | 0.381040204 | 0.81 |
| YIL133C | 6678.70118 | 0.342915766 | 0.381301211 | 0.81 |
| YMR256C | 1012.905586 | -0.422123602 | 0.381716466 | 0.81 |
| YBR047W | 453.8190728 | -0.342556739 | 0.381996597 | 0.81 |
| YBR236C | 870.5665803 | -0.205507888 | 0.382507457 | 0.81 |
| YCR071C | 461.1120902 | -0.286784658 | 0.382507725 | 0.81 |
| YGL167C | 2492.369981 | 0.234280336 | 0.382331474 | 0.81 |
| YKL079W | 964.5499269 | 0.137770463 | 0.383018235 | 0.81 |
| YHR080C | 2045.316793 | 0.224063726 | 0.38363264 | 0.81 |
| YLR005W | 967.4119109 | -0.160296208 | 0.383396982 | 0.81 |
| YOR011W | 696.3198627 | 0.259531598 | 0.383517945 | 0.81 |
| YFR051C | 3059.620652 | -0.264824538 | 0.383917369 | 0.81 |
| YDR133C | 1765.716231 | -0.343875064 | 0.385713619 | 0.81 |
| YDR329C | 1759.988143 | -0.270868718 | 0.385183713 | 0.81 |
| YKL022C | 1018.942477 | 0.184573084 | 0.385572782 | 0.81 |
| YKR082W | 1772.94275 | 0.205049781 | 0.385727264 | 0.81 |
| YNL242W | 1398.740797 | -0.212747733 | 0.385367409 | 0.81 |
| YDR348C | 1537.308452 | 0.110549033 | 0.38622078 | 0.81 |
| YJL216C | 203.2730397 | 0.322225583 | 0.386149472 | 0.81 |
| YPL019C | 1922.931958 | -0.371774772 | 0.386176794 | 0.81 |
| YIL088C | 2503.905976 | -0.232047233 | 0.386452092 | 0.81 |
| YAL034C | 1899.332638 | 0.342763981 | 0.386639039 | 0.81 |
| YBL104C | 644.4185242 | 0.143962903 | 0.386786068 | 0.81 |
| YDR035W | 3632.420651 | -0.270563143 | 0.387468652 | 0.81 |
| YER066C-A | 1483.59707 | -0.445194826 | 0.387528276 | 0.81 |
| YHL007C | 2278.179389 | 0.215611913 | 0.387334652 | 0.81 |
| YLR182W | 1421.620302 | 0.220627736 | 0.387282737 | 0.81 |
| YNL129W | 332.9229067 | -0.356574264 | 0.387180149 | 0.81 |

| | | | | |
|---|---|---|---|---|
| YDR513W | 3464.363476 | -0.357131696 | 0.388042141 | 0.81 |
| YLR345W | 2235.165084 | -0.274302655 | 0.387965055 | 0.81 |
| YBR025C | 6377.054239 | -0.430869056 | 0.388735862 | 0.81 |
| YDR517W | 2232.436932 | -0.253093554 | 0.388641312 | 0.81 |
| YJL078C | 3272.163203 | 0.251600518 | 0.388695879 | 0.81 |
| YJR089W | 953.5011996 | 0.167391014 | 0.38872729 | 0.81 |
| YAL016W | 2950.826603 | -0.180082918 | 0.390242855 | 0.81 |
| YAL022C | 845.7051212 | 0.246715443 | 0.391588894 | 0.81 |
| YBR197C | 458.4082334 | 0.260459915 | 0.39057771 | 0.81 |
| YDL183C | 401.7373652 | 0.283234171 | 0.390652168 | 0.81 |
| YDR077W | 88090.2349 | 0.316425525 | 0.389798824 | 0.81 |
| YDR396W | 125.4408117 | -0.24670365 | 0.3915199 | 0.81 |
| YER146W | 140.8937845 | -0.315393864 | 0.391015564 | 0.81 |
| YGR019W | 2682.062979 | -0.363445782 | 0.391449357 | 0.81 |
| YGR084C | 1041.679379 | -0.169633493 | 0.391308692 | 0.81 |
| YGR239C | 410.924928 | 0.224130208 | 0.391188152 | 0.81 |
| YGR247W | 484.9641046 | -0.259975444 | 0.389338165 | 0.81 |
| YLL015W | 2174.31995 | 0.21500762 | 0.39175042 | 0.81 |
| YLR339C | 501.9621213 | -0.357563656 | 0.39181213 | 0.81 |
| YOL052C | 1181.165007 | 0.163703787 | 0.391649764 | 0.81 |
| YOR132W | 1718.595531 | 0.221138688 | 0.390464468 | 0.81 |
| YOR168W | 3054.619284 | 0.243421311 | 0.391347834 | 0.81 |
| YOR292C | 645.9873627 | -0.269812122 | 0.391578743 | 0.81 |
| YOR299W | 1060.38632 | 0.2389628 | 0.390643277 | 0.81 |
| YPL249C | 1511.229449 | 0.148792288 | 0.389240808 | 0.81 |
| YLR459W | 960.9364848 | 0.203966602 | 0.392440298 | 0.81 |
| YPR094W | 154.1332011 | -0.226901844 | 0.392421078 | 0.81 |
| YPR172W | 1219.274092 | -0.258416046 | 0.392403364 | 0.81 |
| YPL208W | 646.0074609 | -0.236136532 | 0.392723577 | 0.81 |
| YLR266C | 1095.178781 | 0.232886217 | 0.392908841 | 0.81 |
| YGR002C | 912.0190954 | 0.174778231 | 0.393303328 | 0.81 |
| YKL198C | 1074.404315 | 0.254981531 | 0.393454846 | 0.81 |
| YDR273W | 249.707739 | 0.29165436 | 0.394048397 | 0.82 |
| YCR072C | 412.6699919 | 0.368352844 | 0.39602906 | 0.82 |
| YCR093W | 4708.644101 | 0.228737311 | 0.395823196 | 0.82 |
| YDR331W | 571.3069512 | 0.175291769 | 0.394399264 | 0.82 |
| YDR532C | 597.6785104 | -0.288645894 | 0.396173025 | 0.82 |
| YER163C | 538.8237534 | -0.256618687 | 0.394459716 | 0.82 |
| YGR062C | 462.3300151 | -0.209430187 | 0.395475577 | 0.82 |
| YIL024C | 346.2449471 | 0.31498761 | 0.394964257 | 0.82 |
| YJL053W | 1232.219717 | -0.186003946 | 0.396288224 | 0.82 |
| YML037C | 139.9825756 | 0.222516464 | 0.394644487 | 0.82 |
| YMR157C | 556.2957769 | -0.226135346 | 0.394796984 | 0.82 |

| | | | | |
|---|---|---|---|---|
| YNL216W | 1187.634305 | 0.187127702 | 0.395964593 | 0.82 |
| YNL253W | 477.8813502 | 0.117381854 | 0.396229833 | 0.82 |
| YOL091W | 152.6270626 | 0.258346111 | 0.395587912 | 0.82 |
| YOL124C | 564.6964356 | 0.385480354 | 0.395647699 | 0.82 |
| YOR003W | 499.5835225 | -0.236487003 | 0.396064028 | 0.82 |
| YCR054C | 374.3747854 | 0.231976075 | 0.396973 | 0.82 |
| YCR065W | 756.8632472 | 0.283519875 | 0.396689793 | 0.82 |
| YKL047W | 711.1080868 | 0.168996034 | 0.396959099 | 0.82 |
| YNR044W | 3311.34449 | 0.375981619 | 0.396928946 | 0.82 |
| YAL003W | 12374.8526 | -0.427425986 | 0.397665918 | 0.82 |
| YDR493W | 345.6632951 | -0.237087315 | 0.397916229 | 0.82 |
| YGL181W | 1793.449442 | -0.172940445 | 0.39793001 | 0.82 |
| YGL223C | 914.5737412 | 0.140401408 | 0.397454732 | 0.82 |
| YLL011W | 676.0042616 | 0.28196143 | 0.39723737 | 0.82 |
| YLR092W | 690.3329885 | 0.287392584 | 0.397630337 | 0.82 |
| YCL014W | 1364.401777 | 0.218767238 | 0.398373148 | 0.82 |
| YGL192W | 1154.271063 | 0.319903117 | 0.39856865 | 0.82 |
| YAL061W | 2752.646997 | -0.444005052 | 0.399403718 | 0.82 |
| YGR175C | 3323.313876 | -0.277295321 | 0.399834968 | 0.82 |
| YML099C | 1070.053648 | 0.254854076 | 0.399767947 | 0.82 |
| YGR165W | 998.5420975 | -0.188559352 | 0.400121178 | 0.82 |
| YHL029C | 889.4266622 | 0.310275674 | 0.400171677 | 0.82 |
| YDL132W | 2917.981172 | -0.144267227 | 0.400485846 | 0.82 |
| YNL287W | 4413.325689 | 0.123813483 | 0.400660275 | 0.82 |
| YCR090C | 636.078747 | -0.300141705 | 0.401255752 | 0.82 |
| YOL001W | 429.8453508 | 0.264487597 | 0.401335779 | 0.82 |
| YBL003C | 4258.595501 | -0.438633445 | 0.4039112 | 0.82 |
| YCL074W | 140.2978941 | 0.215852992 | 0.40374956 | 0.82 |
| YDR009W | 246.880862 | 0.330151519 | 0.403698316 | 0.82 |
| YDR023W | 6264.678536 | -0.251482851 | 0.402599256 | 0.82 |
| YDR382W | 7559.11107 | -0.375476456 | 0.402280469 | 0.82 |
| YER177W | 16691.55629 | -0.227160049 | 0.4026733 | 0.82 |
| YER183C | 378.9572978 | -0.187968019 | 0.404156916 | 0.82 |
| YGL229C | 708.5937653 | 0.195871417 | 0.40301376 | 0.82 |
| YGR086C | 9907.562538 | -0.168224974 | 0.404155478 | 0.82 |
| YJR005W | 1677.202028 | 0.169852816 | 0.402058711 | 0.82 |
| YJR113C | 864.1046061 | -0.217485231 | 0.402219101 | 0.82 |
| YKL140W | 1755.744554 | -0.132995934 | 0.403321328 | 0.82 |
| YLR294C | 1804.172401 | -0.323010695 | 0.404205865 | 0.82 |
| YLR295C | 2976.605613 | -0.319780399 | 0.403825864 | 0.82 |
| YML102W | 604.0898534 | 0.149738645 | 0.404002295 | 0.82 |
| YMR166C | 428.7049362 | -0.125507493 | 0.403535325 | 0.82 |
| YLL007C | 554.0715465 | 0.227229129 | 0.404891182 | 0.82 |

| | | | | |
|---|---|---|---|---|
| YPL034W | 171.3658521 | -0.200453287 | 0.404636359 | 0.82 |
| YPR152C | 381.9563274 | 0.19523753 | 0.40483996 | 0.82 |
| YPL099C | 334.8201937 | -0.217983115 | 0.40516927 | 0.82 |
| YJL137C | 923.3034038 | 0.38095949 | 0.405413114 | 0.82 |
| YOL031C | 1089.984363 | 0.320833192 | 0.405781343 | 0.82 |
| YJL186W | 1797.326564 | 0.166463455 | 0.40626377 | 0.82 |
| YFL041W | 1108.963199 | 0.206736246 | 0.407185373 | 0.82 |
| YJL023C | 307.217395 | -0.191279399 | 0.407153554 | 0.82 |
| YKR005C | 237.9372028 | 0.269229899 | 0.407226384 | 0.82 |
| YIL147C | 2000.977172 | 0.216339947 | 0.407577521 | 0.82 |
| YJR149W | 627.5970928 | 0.354592783 | 0.407547981 | 0.82 |
| YNR009W | 344.9832981 | 0.303678983 | 0.40773365 | 0.82 |
| YNL172W | 1164.569131 | 0.171922528 | 0.407993761 | 0.82 |
| YDR358W | 1790.698239 | -0.275215032 | 0.408305583 | 0.82 |
| YJR044C | 1233.447551 | -0.186341861 | 0.40874003 | 0.82 |
| YKR019C | 546.7471303 | 0.207691201 | 0.408777631 | 0.82 |
| YOL054W | 494.8918395 | -0.168450825 | 0.408762649 | 0.82 |
| YEL039C | 1783.57122 | -0.393970478 | 0.408929357 | 0.82 |
| YFL044C | 776.7355212 | -0.252554845 | 0.409194762 | 0.82 |
| YGL073W | 2411.314838 | 0.181025782 | 0.409479296 | 0.82 |
| YOR360C | 883.5535106 | 0.242350552 | 0.409607285 | 0.82 |
| YBR085W | 162.0814879 | -0.418965658 | 0.410511597 | 0.83 |
| YDL211C | 175.1726895 | 0.27770666 | 0.41031097 | 0.83 |
| YLR449W | 2574.095007 | -0.40963545 | 0.410493874 | 0.83 |
| YNR031C | 1352.088361 | 0.209306768 | 0.410451355 | 0.83 |
| YNR062C | 190.9739726 | 0.302978684 | 0.410141521 | 0.83 |
| YDL238C | 479.1219346 | -0.265522764 | 0.41071115 | 0.83 |
| YGR171C | 566.9877726 | 0.144109059 | 0.411041603 | 0.83 |
| YCL055W | 639.3184492 | -0.284991589 | 0.411398056 | 0.83 |
| YDR186C | 2571.215729 | 0.263248865 | 0.411552605 | 0.83 |
| YPR192W | 327.8980894 | 0.313389519 | 0.411946417 | 0.83 |
| YAR073W | 171.304609 | 0.104395579 | 0.412664455 | 0.83 |
| YDR421W | 1031.149921 | 0.186471197 | 0.412855872 | 0.83 |
| YEL011W | 8924.284313 | -0.393266966 | 0.412861916 | 0.83 |
| YNL309W | 553.1266537 | 0.187412055 | 0.412582623 | 0.83 |
| YPR097W | 1353.173051 | 0.138146039 | 0.412122961 | 0.83 |
| YPR139C | 1065.385559 | -0.230617017 | 0.412300274 | 0.83 |
| YER158C | 1028.126375 | 0.294393449 | 0.413257718 | 0.83 |
| YOL016C | 3937.741808 | -0.335201226 | 0.413119084 | 0.83 |
| YGL045W | 904.8478281 | 0.303968335 | 0.413467205 | 0.83 |
| YBL061C | 962.4150781 | -0.190328944 | 0.417699135 | 0.83 |
| YDL033C | 356.3492838 | 0.258414006 | 0.416353851 | 0.83 |
| YDL045W-A | 694.1101578 | -0.234534044 | 0.416864891 | 0.83 |

| | | | | |
|---|---|---|---|---|
| YDL112W | 1910.821894 | 0.268496958 | 0.416653226 | 0.83 |
| YDR232W | 2313.644723 | 0.227022663 | 0.415344553 | 0.83 |
| YDR372C | 1904.188965 | -0.222371664 | 0.417573484 | 0.83 |
| YEL012W | 1045.847779 | -0.303010683 | 0.416599419 | 0.83 |
| YGR195W | 892.1333078 | -0.170506691 | 0.416353284 | 0.83 |
| YHR166C | 498.330332 | 0.218989875 | 0.416662304 | 0.83 |
| YJL196C | 1508.45467 | -0.311680445 | 0.415966406 | 0.83 |
| YLR129W | 1609.244371 | 0.316979253 | 0.415084751 | 0.83 |
| YLR258W | 9832.7404 | -0.348217014 | 0.416365715 | 0.83 |
| YLR274W | 1064.782127 | 0.197291299 | 0.41643553 | 0.83 |
| YLR277C | 732.8482737 | 0.175580862 | 0.417813122 | 0.83 |
| YML011C | 267.8459983 | -0.148047228 | 0.417623682 | 0.83 |
| YMR169C | 812.630576 | 0.43375825 | 0.41613013 | 0.83 |
| YNL026W | 973.5528021 | -0.119967119 | 0.416643955 | 0.83 |
| YNL035C | 765.5856918 | 0.189870596 | 0.417639149 | 0.83 |
| YNR012W | 597.0846222 | 0.261336842 | 0.417815681 | 0.83 |
| YNR027W | 549.891736 | -0.214829781 | 0.415131689 | 0.83 |
| YOR347C | 1046.083791 | -0.394084994 | 0.415136932 | 0.83 |
| YPL174C | 736.6858257 | 0.141198662 | 0.417525637 | 0.83 |
| YDR491C | 102.3610769 | 0.341680063 | 0.418196934 | 0.83 |
| YIL042C | 814.6115743 | -0.299404063 | 0.41819161 | 0.83 |
| YML130C | 2617.920044 | 0.291151549 | 0.41848085 | 0.83 |
| YOR018W | 939.5138966 | -0.18457439 | 0.41853772 | 0.83 |
| YOR172W | 824.7713828 | 0.15408741 | 0.418881088 | 0.83 |
| YDL059C | 182.996654 | -0.218549518 | 0.419377838 | 0.83 |
| YLR375W | 1937.904522 | 0.149576436 | 0.419444214 | 0.83 |
| YOR238W | 549.7060682 | -0.204834447 | 0.419236075 | 0.83 |
| YDR321W | 1332.488079 | -0.368106437 | 0.41992074 | 0.83 |
| YNL307C | 2792.714562 | -0.183806349 | 0.419960478 | 0.83 |
| YBR083W | 2339.410841 | 0.295146484 | 0.420139092 | 0.83 |
| YGR012W | 677.5451845 | -0.16079409 | 0.420803801 | 0.83 |
| YKL171W | 1421.131993 | 0.283041136 | 0.421424798 | 0.83 |
| YPL072W | 410.1680408 | 0.18747316 | 0.421329374 | 0.83 |
| YCL036W | 402.9926525 | 0.347194877 | 0.422397525 | 0.83 |
| YDR148C | 4531.818706 | -0.359545325 | 0.422573605 | 0.83 |
| YER046W | 177.0920564 | -0.321211237 | 0.422318784 | 0.83 |
| YKL116C | 778.7224786 | 0.107159427 | 0.421965056 | 0.83 |
| YOR049C | 767.2844506 | -0.295397205 | 0.422498524 | 0.83 |
| YPL101W | 1017.044188 | -0.184983332 | 0.421889661 | 0.83 |
| YPL190C | 2551.586572 | -0.189342457 | 0.422266226 | 0.83 |
| YBR158W | 2964.217702 | 0.266043019 | 0.425510662 | 0.83 |
| YCL054W | 1476.642208 | -0.342942368 | 0.424707582 | 0.83 |
| YCL064C | 3239.199963 | -0.315191786 | 0.425290742 | 0.83 |

| | | | | |
|---|---|---|---|---|
| YCR020W-B | 152.1239298 | -0.291719673 | 0.425938922 | 0.83 |
| YDL052C | 1575.894255 | 0.288189439 | 0.425207532 | 0.83 |
| YDL066W | 3505.644872 | -0.379801679 | 0.423297005 | 0.83 |
| YDL130W-A | 2765.774643 | -0.380268828 | 0.424877577 | 0.83 |
| YDR061W | 1142.905546 | 0.130188804 | 0.425628521 | 0.83 |
| YER159C | 1569.026083 | -0.275093887 | 0.425394501 | 0.83 |
| YER164W | 2428.238913 | 0.150514848 | 0.424455833 | 0.83 |
| YGR112W | 838.4700607 | -0.267446071 | 0.423089262 | 0.83 |
| YIL033C | 4130.82095 | -0.216093698 | 0.425804011 | 0.83 |
| YIL094C | 3348.951104 | -0.194579468 | 0.425345455 | 0.83 |
| YKL072W | 554.6822525 | 0.158189809 | 0.423669784 | 0.83 |
| YLR379W | 289.3293491 | 0.229535687 | 0.424644544 | 0.83 |
| YMR021C | 731.4049333 | -0.17864868 | 0.423709937 | 0.83 |
| YMR105C | 9001.650714 | -0.388408295 | 0.425800243 | 0.83 |
| YMR287C | 795.4088763 | 0.160241408 | 0.422988089 | 0.83 |
| YNL024C | 148.4866774 | 0.408179212 | 0.424097486 | 0.83 |
| YNL108C | 290.0576123 | -0.211874885 | 0.425789817 | 0.83 |
| YOL132W | 124.9358227 | 0.308227195 | 0.424878415 | 0.83 |
| YOL142W | 655.5671288 | -0.195090168 | 0.423403067 | 0.83 |
| YDR303C | 1032.483515 | 0.236915747 | 0.426455246 | 0.83 |
| YPL232W | 2081.428239 | -0.164516144 | 0.426562961 | 0.83 |
| YBR230C | 2523.139558 | -0.349497915 | 0.428689281 | 0.83 |
| YER038C | 327.709464 | 0.267811592 | 0.428601693 | 0.83 |
| YFL017W-A | 221.0675394 | -0.231807333 | 0.427251331 | 0.83 |
| YIL061C | 381.0352818 | 0.176372727 | 0.428942784 | 0.83 |
| YJR032W | 585.2468444 | -0.281257532 | 0.428208092 | 0.83 |
| YLR086W | 1588.915329 | 0.185451209 | 0.428447506 | 0.83 |
| YLR181C | 881.938971 | -0.158088934 | 0.427776502 | 0.83 |
| YLR310C | 3300.526123 | 0.195853753 | 0.428618684 | 0.83 |
| YLR368W | 767.6389144 | 0.220348143 | 0.42793835 | 0.83 |
| YMR030W | 574.6177195 | -0.30949332 | 0.428721471 | 0.83 |
| YMR114C | 1303.792032 | -0.329087175 | 0.428643588 | 0.83 |
| YOR195W | 1326.708475 | -0.24010297 | 0.428923356 | 0.83 |
| YOR323C | 2339.246441 | -0.223399751 | 0.428244936 | 0.83 |
| YPL214C | 1002.445323 | -0.178483345 | 0.428013315 | 0.83 |
| YPL250C | 749.2294185 | 0.329058529 | 0.427832133 | 0.83 |
| YML021C | 560.3287148 | -0.185611246 | 0.429128416 | 0.83 |
| YDR003W | 807.4785109 | -0.229210864 | 0.429397256 | 0.83 |
| YFL045C | 5242.345974 | -0.324692113 | 0.42955572 | 0.83 |
| YKL161C | 387.0663817 | 0.300475271 | 0.429665266 | 0.83 |
| YDL067C | 1308.526792 | -0.379007519 | 0.430431613 | 0.83 |
| YDR124W | 300.8040212 | 0.288513635 | 0.430700325 | 0.83 |
| YGR057C | 395.64537 | -0.152194941 | 0.431017734 | 0.83 |

| | | | | |
|---|---|---|---|---|
| YKL133C | 601.5672696 | -0.285320253 | 0.431111441 | 0.83 |
| YKR100C | 1038.230895 | -0.169978928 | 0.431041636 | 0.83 |
| YPR141C | 623.3342075 | 0.220939699 | 0.431083901 | 0.83 |
| YPL211W | 668.7255746 | -0.404113007 | 0.431534433 | 0.83 |
| YML014W | 592.0947476 | -0.261790165 | 0.431842029 | 0.83 |
| YBL047C | 5698.969568 | 0.151055416 | 0.432397752 | 0.84 |
| YDL206W | 597.9682559 | 0.196576021 | 0.432147043 | 0.84 |
| YKR066C | 4134.117453 | -0.28805091 | 0.432422468 | 0.84 |
| YOR094W | 410.9080628 | -0.191976769 | 0.433029688 | 0.84 |
| YOR114W | 311.2740858 | -0.269455672 | 0.433002049 | 0.84 |
| YDL170W | 672.20876 | 0.240461667 | 0.434161991 | 0.84 |
| YGL006W | 3158.91505 | -0.201710235 | 0.434162203 | 0.84 |
| YGR033C | 1013.438168 | 0.136443283 | 0.433972177 | 0.84 |
| YLR149C | 1780.729906 | -0.349872398 | 0.434019603 | 0.84 |
| YOR137C | 579.6398816 | -0.251105197 | 0.433540476 | 0.84 |
| YOR346W | 878.2111125 | 0.227464568 | 0.433812812 | 0.84 |
| YFL056C | 232.3744781 | 0.280484802 | 0.434875308 | 0.84 |
| YBR121C | 7616.904588 | -0.293729961 | 0.435196278 | 0.84 |
| YDL056W | 1411.798857 | 0.205325956 | 0.436476306 | 0.84 |
| YDR083W | 1132.402572 | -0.258685009 | 0.43744951 | 0.84 |
| YGL114W | 1125.03715 | 0.186568623 | 0.437515691 | 0.84 |
| YIL035C | 1690.415511 | -0.149810619 | 0.437074635 | 0.84 |
| YJL013C | 787.2498623 | 0.13149319 | 0.436542681 | 0.84 |
| YJL052W | 23370.62399 | -0.415639826 | 0.436573041 | 0.84 |
| YJL187C | 1084.409994 | 0.198886662 | 0.436808666 | 0.84 |
| YJR126C | 1827.592541 | -0.122173159 | 0.437536152 | 0.84 |
| YKL090W | 556.2185917 | 0.221780408 | 0.437521662 | 0.84 |
| YKR012C | 142.5434976 | -0.267881752 | 0.437128389 | 0.84 |
| YLR188W | 1233.172467 | -0.150438714 | 0.436307009 | 0.84 |
| YOL053W | 775.6100204 | -0.272343368 | 0.437117312 | 0.84 |
| YDL205C | 1039.950483 | 0.278466198 | 0.438640958 | 0.84 |
| YDR188W | 2974.432832 | -0.154668945 | 0.438087404 | 0.84 |
| YHR211W | 534.044984 | 0.340966233 | 0.438630911 | 0.84 |
| YOL022C | 1058.507084 | -0.25908686 | 0.438474365 | 0.84 |
| YOR065W | 3707.159237 | -0.353443907 | 0.438231023 | 0.84 |
| YOR343W-B | 107.9180972 | 0.320191608 | 0.43855581 | 0.84 |
| YBR152W | 271.4958287 | 0.142744323 | 0.439348842 | 0.84 |
| YGR213C | 279.0368426 | 0.264277334 | 0.440077303 | 0.84 |
| YGR241C | 1428.572307 | 0.161845826 | 0.439981882 | 0.84 |
| YHR132C | 1760.575956 | -0.192302315 | 0.440039279 | 0.84 |
| YKR016W | 2416.547829 | -0.304035249 | 0.439466465 | 0.84 |
| YNL282W | 259.9576363 | 0.185861875 | 0.438956668 | 0.84 |
| YNR074C | 411.2345917 | -0.328484494 | 0.439546414 | 0.84 |

| | | | | |
|---|---|---|---|---|
| YOL089C | 836.6158144 | 0.167146665 | 0.43977538 | 0.84 |
| YOR046C | 2190.495993 | -0.251709792 | 0.439026178 | 0.84 |
| YDR025W | 5331.251911 | 0.318479468 | 0.440816812 | 0.84 |
| YER032W | 1018.880889 | 0.240658392 | 0.44129686 | 0.84 |
| YJL140W | 1600.235301 | -0.139412348 | 0.441095209 | 0.84 |
| YPL150W | 1067.29309 | 0.181767158 | 0.441330794 | 0.84 |
| YDL138W | 666.9826314 | 0.182411319 | 0.441987108 | 0.84 |
| YER067W | 2559.902963 | -0.38492413 | 0.44256747 | 0.84 |
| YHR177W | 301.5515693 | 0.238253564 | 0.442447649 | 0.84 |
| YNR058W | 282.8560691 | 0.222621238 | 0.442449662 | 0.84 |
| YBR126W-A | 695.852559 | -0.310631778 | 0.442822652 | 0.84 |
| YHR051W | 2680.129991 | -0.353705675 | 0.443082233 | 0.84 |
| YIL131C | 677.2016628 | 0.271567158 | 0.443166889 | 0.84 |
| YLR316C | 329.0837142 | 0.137739166 | 0.443584426 | 0.84 |
| YBR246W | 994.4891912 | -0.128639658 | 0.443991483 | 0.84 |
| YPL138C | 483.5208101 | 0.162342723 | 0.444519345 | 0.84 |
| YPR175W | 704.7198488 | 0.235820354 | 0.444588777 | 0.84 |
| YDL185C-A | 100.139643 | -0.215210826 | 0.444871913 | 0.84 |
| YDR013W | 271.9851003 | -0.215107934 | 0.445832053 | 0.84 |
| YGR138C | 1416.124697 | 0.350741178 | 0.445548461 | 0.84 |
| YHR181W | 756.8304441 | -0.287966458 | 0.445612627 | 0.84 |
| YJR141W | 396.4509931 | 0.177452989 | 0.44582471 | 0.84 |
| YOR278W | 477.7903224 | 0.170265106 | 0.446101395 | 0.84 |
| YKL016C | 3252.389125 | -0.28960785 | 0.446954114 | 0.85 |
| YDR377W | 1422.817631 | -0.327113947 | 0.447323116 | 0.85 |
| YCR067C | 1807.391426 | 0.16112971 | 0.447593766 | 0.85 |
| YGL071W | 1392.981634 | 0.182357443 | 0.447864339 | 0.85 |
| YER088C | 4317.842613 | 0.242560578 | 0.448259679 | 0.85 |
| YJR122W | 745.1766826 | 0.201622339 | 0.448492843 | 0.85 |
| YPL239W | 1334.632299 | -0.272135731 | 0.448457059 | 0.85 |
| YPL120W | 588.9261154 | -0.189781461 | 0.448661696 | 0.85 |
| YML026C | 7646.727007 | 0.288444024 | 0.449108557 | 0.85 |
| YML051W | 1749.382621 | -0.231183507 | 0.44951392 | 0.85 |
| YHR108W | 3358.554888 | -0.221852105 | 0.450272094 | 0.85 |
| YOL064C | 847.1293742 | -0.248860907 | 0.450498266 | 0.85 |
| YHR063C | 1432.676855 | -0.267527848 | 0.451484852 | 0.85 |
| YPR089W | 1983.427567 | 0.134141724 | 0.451356721 | 0.85 |
| YDR044W | 1839.127613 | -0.270148931 | 0.451970379 | 0.85 |
| YGL023C | 1541.23519 | -0.168237649 | 0.452162472 | 0.85 |
| YIL014C-A | 253.291496 | -0.415065306 | 0.452248642 | 0.85 |
| YAL047C | 589.7842721 | -0.142090963 | 0.453978471 | 0.85 |
| YBL018C | 137.4735368 | -0.398827842 | 0.459203643 | 0.85 |
| YBL022C | 3958.243828 | -0.185391515 | 0.453281893 | 0.85 |

| | | | | |
|---|---|---|---|---|
| YBR013C | 207.4427581 | -0.26446677 | 0.458099007 | 0.85 |
| YBR261C | 744.8069304 | -0.278954537 | 0.457172494 | 0.85 |
| YDL159W | 513.5387781 | -0.205688305 | 0.45933848 | 0.85 |
| YDL160C | 1362.561057 | 0.22887833 | 0.455514313 | 0.85 |
| YDL179W | 455.9668158 | 0.151219996 | 0.458238153 | 0.85 |
| YDR104C | 711.7535955 | 0.191987819 | 0.457799564 | 0.85 |
| YDR189W | 2199.458913 | -0.160861498 | 0.457502883 | 0.85 |
| YDR255C | 870.1597032 | 0.236949907 | 0.456323588 | 0.85 |
| YDR524W-C | 1641.998262 | -0.344668947 | 0.455927007 | 0.85 |
| YEL060C | 10805.49764 | -0.296359383 | 0.458913524 | 0.85 |
| YFL024C | 1322.826078 | 0.163677426 | 0.457085708 | 0.85 |
| YFL055W | 265.1999324 | 0.320358799 | 0.454274897 | 0.85 |
| YGR142W | 4162.125638 | -0.292661157 | 0.454521542 | 0.85 |
| YGR194C | 2007.303758 | -0.257803494 | 0.454152769 | 0.85 |
| YHR019C | 9735.850466 | -0.288892184 | 0.454786768 | 0.85 |
| YHR022C | 158.1795629 | 0.264698386 | 0.45787488 | 0.85 |
| YHR104W | 3547.736943 | -0.194139792 | 0.459416838 | 0.85 |
| YHR175W-A | 167.202498 | 0.310013814 | 0.455876677 | 0.85 |
| YIL122W | 600.3807645 | 0.251769462 | 0.452839249 | 0.85 |
| YIL169C | 2177.35422 | 0.278715947 | 0.457185195 | 0.85 |
| YJL056C | 734.4930024 | 0.227763335 | 0.458634402 | 0.85 |
| YJL157C | 1263.741636 | 0.377654501 | 0.455028766 | 0.85 |
| YJL174W | 2482.371966 | -0.122374286 | 0.456031115 | 0.85 |
| YJR010C-A | 212.1436636 | -0.259863225 | 0.456655098 | 0.85 |
| YJR010W | 677.4384961 | 0.234196751 | 0.458459142 | 0.85 |
| YJR151C | 560.5480031 | 0.276106251 | 0.456259816 | 0.85 |
| YKR038C | 1110.135443 | 0.13428199 | 0.45636602 | 0.85 |
| YKR057W | 2516.350053 | -0.413510403 | 0.458537987 | 0.85 |
| YLR068W | 534.5964698 | -0.24803587 | 0.45761044 | 0.85 |
| YLR207W | 1228.065049 | 0.152650221 | 0.454307918 | 0.85 |
| YMR036C | 631.5049431 | 0.173278733 | 0.457346984 | 0.85 |
| YMR060C | 493.2206546 | 0.161055592 | 0.457771747 | 0.85 |
| YMR064W | 1294.977576 | 0.172880774 | 0.457871363 | 0.85 |
| YMR241W | 1393.972884 | 0.235777397 | 0.454758585 | 0.85 |
| YNL127W | 1075.477959 | 0.207614605 | 0.456411525 | 0.85 |
| YNL167C | 1564.476742 | 0.183583915 | 0.459009093 | 0.85 |
| YNL220W | 4979.508941 | 0.251173779 | 0.458945953 | 0.85 |
| YNR003C | 554.144982 | -0.2986761 | 0.458130238 | 0.85 |
| YOL044W | 473.6470695 | -0.21794465 | 0.45539987 | 0.85 |
| YOR086C | 3224.270398 | 0.146940196 | 0.453315286 | 0.85 |
| YOR260W | 2245.68858 | 0.255490881 | 0.454912661 | 0.85 |
| YPL089C | 1858.715154 | 0.205020301 | 0.452915789 | 0.85 |
| YPL145C | 2994.997329 | 0.213124232 | 0.459054351 | 0.85 |

| | | | | |
|---|---|---|---|---|
| YBR003W | 1468.050788 | -0.135657621 | 0.462491345 | 0.85 |
| YBR222C | 4562.01386 | -0.260898435 | 0.46186941 | 0.85 |
| YDL224C | 1328.137391 | 0.172779232 | 0.461155807 | 0.85 |
| YEL062W | 560.1717097 | 0.152224779 | 0.463106072 | 0.85 |
| YFL013C | 826.2401076 | -0.169431045 | 0.46272596 | 0.85 |
| YGL134W | 814.0702966 | 0.207880652 | 0.462682821 | 0.85 |
| YGL224C | 544.3672231 | -0.232382863 | 0.460089498 | 0.85 |
| YGR008C | 4582.699628 | -0.357586176 | 0.463281973 | 0.85 |
| YHR030C | 2133.377408 | 0.339631191 | 0.462447567 | 0.85 |
| YHR207C | 677.6706609 | -0.201167414 | 0.461590732 | 0.85 |
| YIL055C | 666.1215044 | 0.262423981 | 0.461001301 | 0.85 |
| YIL118W | 1012.394304 | -0.25169433 | 0.46336765 | 0.85 |
| YKL174C | 903.8221117 | 0.103895612 | 0.460563005 | 0.85 |
| YLL053C | 96.70396409 | 0.324611267 | 0.461190088 | 0.85 |
| YLR135W | 729.3825255 | 0.160917017 | 0.462239133 | 0.85 |
| YLR209C | 881.8739647 | -0.210126614 | 0.462879382 | 0.85 |
| YLR412W | 993.3823084 | -0.177303892 | 0.46281274 | 0.85 |
| YML034W | 2522.763719 | 0.141559412 | 0.46364071 | 0.85 |
| YMR063W | 168.4962103 | 0.202373558 | 0.462377856 | 0.85 |
| YMR121C | 804.8314095 | -0.227339345 | 0.461551236 | 0.85 |
| YNL041C | 1185.16925 | 0.134731177 | 0.460642074 | 0.85 |
| YNL165W | 500.9806009 | -0.196440967 | 0.46266792 | 0.85 |
| YNL209W | 13780.76239 | -0.309815187 | 0.463542247 | 0.85 |
| YPL042C | 568.6329785 | 0.213596782 | 0.460397529 | 0.85 |
| YPR045C | 1121.270528 | -0.1802246 | 0.461181196 | 0.85 |
| YBL113C | 286.5211358 | 0.252808369 | 0.464342798 | 0.85 |
| YDR365C | 4849.022737 | 0.197096716 | 0.464144598 | 0.85 |
| YLR431C | 452.0758432 | 0.152701728 | 0.464598204 | 0.85 |
| YML100W | 14497.66976 | -0.307312289 | 0.464207106 | 0.85 |
| YMR037C | 1272.708272 | 0.242469274 | 0.46464438 | 0.85 |
| YCL016C | 442.1886812 | -0.171868395 | 0.465016455 | 0.85 |
| YGR063C | 116.1850697 | -0.140603501 | 0.464983196 | 0.85 |
| YBR143C | 4538.418741 | -0.257281891 | 0.465356595 | 0.85 |
| YCL056C | 401.4808141 | -0.116643717 | 0.465309671 | 0.85 |
| YBR182C | 159.913284 | 0.305743927 | 0.466211232 | 0.85 |
| YNL210W | 106.0752581 | 0.23017107 | 0.466246919 | 0.85 |
| YBL028C | 410.9135571 | -0.319986346 | 0.46679799 | 0.85 |
| YGR256W | 1242.341744 | 0.292989762 | 0.467142182 | 0.85 |
| YJL183W | 732.387825 | -0.243297669 | 0.467138753 | 0.85 |
| YER188W | 461.4895098 | -0.386294746 | 0.467347705 | 0.85 |
| YIL039W | 1826.424519 | -0.173577705 | 0.467497132 | 0.85 |
| YLR033W | 1049.258607 | 0.127286103 | 0.468211444 | 0.86 |
| YBR036C | 2605.132077 | -0.178246079 | 0.468532532 | 0.86 |

| | | | | |
|---|---|---|---|---|
| YAL017W | 3309.438838 | -0.257308108 | 0.468731192 | 0.86 |
| YML020W | 864.3038212 | 0.159352207 | 0.469014347 | 0.86 |
| YKR085C | 837.2824572 | -0.194861895 | 0.469365916 | 0.86 |
| YOL117W | 709.0122211 | -0.258749335 | 0.469256688 | 0.86 |
| YKR049C | 1528.555977 | -0.346911427 | 0.469762516 | 0.86 |
| YPL199C | 1017.89655 | -0.233274702 | 0.469779083 | 0.86 |
| YAL063C | 272.6184868 | 0.256002996 | 0.47005116 | 0.86 |
| YNL052W | 2172.615322 | -0.343861608 | 0.470195492 | 0.86 |
| YHR015W | 104.6987227 | 0.347615273 | 0.470581705 | 0.86 |
| YOL097C | 2168.680575 | -0.309207474 | 0.470471029 | 0.86 |
| YPL053C | 1363.897708 | -0.227625435 | 0.470694694 | 0.86 |
| YDR197W | 858.0497536 | 0.185921338 | 0.471181162 | 0.86 |
| YHL033C | 4723.149428 | 0.225282726 | 0.471079977 | 0.86 |
| YLR177W | 3573.62477 | -0.261388607 | 0.471548442 | 0.86 |
| YOR112W | 1049.770792 | 0.118708419 | 0.471473453 | 0.86 |
| YDR080W | 1103.850689 | -0.192251716 | 0.472761435 | 0.86 |
| YDR334W | 1700.440041 | 0.180873536 | 0.473044472 | 0.86 |
| YDR478W | 183.0152712 | -0.158746879 | 0.472824103 | 0.86 |
| YER086W | 3514.638296 | 0.309158606 | 0.473096336 | 0.86 |
| YFL047W | 1142.35423 | 0.196369339 | 0.474437537 | 0.86 |
| YGR066C | 223.0827395 | -0.314434359 | 0.473494872 | 0.86 |
| YHR158C | 1689.928736 | 0.168578006 | 0.473778565 | 0.86 |
| YKR064W | 1141.244604 | 0.146061614 | 0.473351007 | 0.86 |
| YLR029C | 12797.37208 | 0.254010248 | 0.474042651 | 0.86 |
| YLR226W | 384.869465 | 0.126932796 | 0.474520617 | 0.86 |
| YMR091C | 998.5083437 | -0.191681762 | 0.472300379 | 0.86 |
| YNL031C | 9825.730059 | -0.256525835 | 0.474442606 | 0.86 |
| YNL096C | 3035.774305 | 0.251345191 | 0.474359562 | 0.86 |
| YNL182C | 900.9513347 | 0.301194181 | 0.473061559 | 0.86 |
| YOR220W | 2249.332399 | -0.250442371 | 0.473732218 | 0.86 |
| YOR320C | 1070.658594 | 0.158886024 | 0.474075338 | 0.86 |
| YPR204W | 106.9528255 | 0.336139703 | 0.474148613 | 0.86 |
| YOR106W | 543.2030625 | 0.111711383 | 0.474901168 | 0.86 |
| YJL116C | 947.324176 | 0.31808202 | 0.475240803 | 0.86 |
| YLR383W | 888.8933966 | 0.154914219 | 0.475392428 | 0.86 |
| YGR174C | 1048.598299 | -0.267232781 | 0.475816568 | 0.86 |
| YMR211W | 431.1277672 | 0.11976761 | 0.47572058 | 0.86 |
| YDR505C | 1755.081331 | 0.219993901 | 0.476183905 | 0.86 |
| YFL034C-B | 921.0599209 | -0.226262818 | 0.47608019 | 0.86 |
| YML131W | 1116.004425 | -0.31553068 | 0.476383695 | 0.86 |
| YFL050C | 563.7473526 | 0.228424987 | 0.476616064 | 0.86 |
| YOR036W | 1460.876014 | -0.21078893 | 0.476929968 | 0.86 |
| YDR471W | 4845.236011 | 0.289591436 | 0.477408756 | 0.86 |

| | | | | |
|---|---|---|---|---|
| YDR200C | 868.5642679 | 0.11380333 | 0.4793193 | 0.86 |
| YDR289C | 776.5077999 | -0.200724133 | 0.479407546 | 0.86 |
| YDR510C-A | 439.1253795 | -0.194719809 | 0.478519048 | 0.86 |
| YDR518W | 1466.448011 | -0.239127194 | 0.479010038 | 0.86 |
| YER147C | 529.8378852 | 0.171179083 | 0.479135687 | 0.86 |
| YLR108C | 407.9615838 | -0.239157999 | 0.479399985 | 0.86 |
| YLR221C | 955.7495307 | -0.224269748 | 0.479372104 | 0.86 |
| YNL115C | 1894.716258 | -0.307387685 | 0.479265163 | 0.86 |
| YPR135W | 846.2041741 | 0.147688887 | 0.47930651 | 0.86 |
| YDR322W | 1031.616879 | -0.210396447 | 0.479733813 | 0.86 |
| YHR072W-A | 1183.992345 | -0.271197433 | 0.480238497 | 0.86 |
| YAL023C | 4478.973811 | 0.19711094 | 0.480594813 | 0.86 |
| YCR076C | 593.9023385 | -0.217480188 | 0.482883428 | 0.86 |
| YDR361C | 744.7914399 | 0.268401209 | 0.482225053 | 0.86 |
| YGL063W | 321.9330079 | 0.242868869 | 0.480975188 | 0.86 |
| YGR017W | 1237.157084 | -0.125696381 | 0.482582667 | 0.86 |
| YHL009C | 603.1387723 | 0.189401776 | 0.481918456 | 0.86 |
| YHR165C | 3249.517075 | 0.213650351 | 0.483019085 | 0.86 |
| YIL130W | 1279.051362 | 0.22625114 | 0.482970247 | 0.86 |
| YJL100W | 1049.188324 | 0.183344008 | 0.482568063 | 0.86 |
| YJR115W | 567.164948 | 0.372142186 | 0.482602027 | 0.86 |
| YLL031C | 1740.005527 | 0.155509897 | 0.48226373 | 0.86 |
| YLR194C | 2445.708189 | 0.285361649 | 0.482779843 | 0.86 |
| YML030W | 762.1544554 | -0.257267437 | 0.481373188 | 0.86 |
| YNL006W | 936.4455341 | 0.183660772 | 0.480989724 | 0.86 |
| YNR011C | 902.2961481 | 0.165206114 | 0.48193439 | 0.86 |
| YOR223W | 687.6095973 | -0.193046535 | 0.481232212 | 0.86 |
| YPR013C | 1083.106605 | 0.211466731 | 0.482803773 | 0.86 |
| YBR066C | 407.0918605 | -0.328794281 | 0.48324359 | 0.86 |
| YBR110W | 1287.287964 | -0.157619759 | 0.483948751 | 0.86 |
| YDL239C | 522.0657782 | 0.173326782 | 0.484233064 | 0.86 |
| YDR033W | 12974.70679 | 0.249655469 | 0.484342036 | 0.86 |
| YHL036W | 545.2189465 | 0.274278648 | 0.484060531 | 0.86 |
| YGL179C | 645.0803373 | 0.327316215 | 0.485609468 | 0.86 |
| YJL105W | 247.8507967 | -0.232230137 | 0.485530812 | 0.86 |
| YBL107C | 641.7007619 | -0.278723159 | 0.486225838 | 0.86 |
| YDR466W | 1549.812513 | 0.13464668 | 0.486113649 | 0.86 |
| YML115C | 1520.660585 | -0.12448503 | 0.48624598 | 0.86 |
| YMR065W | 380.856771 | 0.143144099 | 0.486058853 | 0.86 |
| YBR101C | 3697.675555 | -0.199676748 | 0.486936699 | 0.87 |
| YOL015W | 325.5388955 | 0.267426339 | 0.487251128 | 0.87 |
| YDR087C | 817.7593103 | -0.268373931 | 0.48756584 | 0.87 |
| YBR120C | 395.0224857 | -0.181610007 | 0.489825374 | 0.87 |

| | | | | |
|---|---|---|---|---|
| YBR247C | 781.1952554 | 0.259700992 | 0.489123723 | 0.87 |
| YDR192C | 476.3597916 | -0.146621253 | 0.48945055 | 0.87 |
| YDR260C | 314.9401976 | -0.159951249 | 0.489450055 | 0.87 |
| YER064C | 1674.781352 | 0.229367048 | 0.488243742 | 0.87 |
| YIL019W | 543.1334136 | -0.280509948 | 0.488618074 | 0.87 |
| YJL164C | 1399.537088 | -0.182440282 | 0.489326576 | 0.87 |
| YLR448W | 5493.030487 | 0.302561003 | 0.489843144 | 0.87 |
| YNR030W | 1357.008714 | -0.2431713 | 0.489511457 | 0.87 |
| YOR037W | 696.733578 | -0.150024409 | 0.488037089 | 0.87 |
| YOR170W | 97.11015677 | -0.236262555 | 0.487871839 | 0.87 |
| YOR236W | 448.7599801 | -0.281451634 | 0.488563993 | 0.87 |
| YOR361C | 4132.636454 | -0.214370232 | 0.489258301 | 0.87 |
| YPL147W | 471.9776317 | -0.273889754 | 0.489687071 | 0.87 |
| YNR075W | 186.6264191 | 0.28574058 | 0.490043635 | 0.87 |
| YDL073W | 971.6916673 | 0.168138052 | 0.491275833 | 0.87 |
| YDL146W | 626.6868048 | 0.246538961 | 0.490886603 | 0.87 |
| YER053C-A | 179.222376 | 0.351300215 | 0.491400797 | 0.87 |
| YGR220C | 677.1113476 | -0.162334263 | 0.491433541 | 0.87 |
| YIL155C | 2791.794832 | -0.339385357 | 0.491189836 | 0.87 |
| YKL129C | 2233.401318 | 0.158394815 | 0.490904534 | 0.87 |
| YML114C | 955.0918143 | 0.126038219 | 0.491431881 | 0.87 |
| YMR089C | 2367.558674 | -0.214242264 | 0.490619945 | 0.87 |
| YGR250C | 3005.303955 | 0.200506014 | 0.492407442 | 0.87 |
| YLR237W | 1061.722449 | -0.289517026 | 0.492800865 | 0.87 |
| YNL085W | 3478.261348 | 0.181058825 | 0.493372103 | 0.87 |
| YPL228W | 971.5333391 | -0.161659397 | 0.493546879 | 0.87 |
| YLR087C | 2042.517235 | 0.166225855 | 0.493855712 | 0.87 |
| YMR251W | 257.8128007 | -0.313231161 | 0.493960054 | 0.87 |
| YJL093C | 1063.991787 | 0.147952976 | 0.494193105 | 0.87 |
| YFL031W | 3441.588262 | 0.254579 | 0.494479443 | 0.87 |
| YOR234C | 2998.326335 | 0.262931135 | 0.494614138 | 0.87 |
| YPL124W | 247.3933842 | -0.181360003 | 0.495033585 | 0.87 |
| YJL101C | 2271.091311 | 0.150031659 | 0.495199262 | 0.87 |
| YLR166C | 1318.264285 | 0.106174639 | 0.495953509 | 0.87 |
| YDR178W | 2932.392626 | -0.320672404 | 0.496335803 | 0.87 |
| YDR196C | 880.6195121 | -0.182287969 | 0.496454804 | 0.87 |
| YNL175C | 613.6716753 | -0.299159088 | 0.497057076 | 0.87 |
| YDR113C | 648.1289584 | -0.141559215 | 0.497423264 | 0.87 |
| YPR019W | 1285.937535 | 0.099952148 | 0.497613946 | 0.87 |
| YBR195C | 288.8600834 | 0.173922042 | 0.498445227 | 0.87 |
| YPR072W | 2174.009813 | -0.134864112 | 0.49872413 | 0.87 |
| YDL039C | 373.0125142 | 0.340448187 | 0.499308745 | 0.87 |
| YJL200C | 1688.489143 | 0.247513224 | 0.499708915 | 0.87 |

| | | | | |
|---|---|---|---|---|
| YMR205C | 7427.558626 | 0.247114993 | 0.500053165 | 0.88 |
| YPL144W | 173.5181567 | -0.250518372 | 0.500111098 | 0.88 |
| YAR031W | 151.5423779 | 0.224154622 | 0.501355152 | 0.88 |
| YBR081C | 1882.249025 | -0.173419091 | 0.504336522 | 0.88 |
| YBR095C | 539.8071954 | 0.133403896 | 0.503019098 | 0.88 |
| YBR290W | 1203.208406 | -0.161204231 | 0.506131655 | 0.88 |
| YCL051W | 1104.388423 | 0.159740143 | 0.502407515 | 0.88 |
| YDL203C | 667.0945544 | 0.097734496 | 0.503960669 | 0.88 |
| YDR037W | 9604.757078 | -0.314650433 | 0.502378319 | 0.88 |
| YDR078C | 147.0952169 | -0.232222784 | 0.501278273 | 0.88 |
| YDR336W | 301.9017741 | -0.161093717 | 0.505014133 | 0.88 |
| YDR536W | 315.8429448 | 0.290697024 | 0.50478493 | 0.88 |
| YEL029C | 331.9088854 | -0.226260446 | 0.501516309 | 0.88 |
| YER068W | 1672.188885 | 0.171558027 | 0.506947359 | 0.88 |
| YER173W | 470.1702433 | 0.153702591 | 0.501356366 | 0.88 |
| YFR041C | 978.3498425 | -0.17943767 | 0.505727097 | 0.88 |
| YGL159W | 407.2189918 | -0.206935382 | 0.503409959 | 0.88 |
| YGL162W | 395.3975497 | 0.291304187 | 0.504654079 | 0.88 |
| YGR103W | 3122.539574 | -0.315848687 | 0.505100472 | 0.88 |
| YIL031W | 1396.422579 | 0.152902972 | 0.504822936 | 0.88 |
| YIL070C | 1550.574512 | -0.238082325 | 0.500858654 | 0.88 |
| YIL126W | 2338.60398 | 0.207435224 | 0.506289506 | 0.88 |
| YIL127C | 639.7542142 | -0.251371282 | 0.506813249 | 0.88 |
| YJL083W | 610.9975272 | 0.166976984 | 0.504927779 | 0.88 |
| YJL094C | 1236.455255 | 0.161053997 | 0.503181575 | 0.88 |
| YJL166W | 2486.295448 | -0.300508737 | 0.506753735 | 0.88 |
| YJL203W | 345.939173 | 0.120010324 | 0.507082389 | 0.88 |
| YKL148C | 4538.261302 | -0.32542904 | 0.505702774 | 0.88 |
| YKL209C | 3091.85579 | 0.355331733 | 0.503545579 | 0.88 |
| YKR086W | 1098.768049 | 0.144311689 | 0.504099099 | 0.88 |
| YKR105C | 96.82105819 | 0.269679255 | 0.503068804 | 0.88 |
| YLR132C | 337.4595812 | 0.118656232 | 0.506877323 | 0.88 |
| YLR378C | 3906.764345 | 0.180952226 | 0.502222402 | 0.88 |
| YML097C | 708.2199714 | 0.161164236 | 0.505370779 | 0.88 |
| YMR149W | 2739.886665 | -0.171428628 | 0.501094388 | 0.88 |
| YMR179W | 627.9998767 | 0.236050983 | 0.506062859 | 0.88 |
| YMR209C | 810.1729969 | 0.187779017 | 0.503096433 | 0.88 |
| YNL059C | 1845.140766 | -0.146215099 | 0.505145923 | 0.88 |
| YOL088C | 665.7794028 | -0.1664557 | 0.505897715 | 0.88 |
| YOR023C | 2117.560133 | 0.180187741 | 0.505198745 | 0.88 |
| YOR194C | 1465.560671 | -0.201614571 | 0.503917703 | 0.88 |
| YOR211C | 1075.422511 | 0.137369429 | 0.502076909 | 0.88 |
| YPL244C | 876.1384791 | -0.180758066 | 0.50172545 | 0.88 |

| | | | | |
|---|---|---|---|---|
| YPR112C | 1587.350101 | -0.262150748 | 0.502568571 | 0.88 |
| YPR118W | 1223.179848 | -0.290443912 | 0.502467144 | 0.88 |
| YFR011C | 642.0102271 | -0.250419147 | 0.508397376 | 0.88 |
| YGL068W | 385.5514446 | -0.23018341 | 0.508614749 | 0.88 |
| YJR094C | 253.8151399 | 0.264387539 | 0.508457098 | 0.88 |
| YKR025W | 782.5030894 | -0.19112114 | 0.508564063 | 0.88 |
| YOR063W | 34774.24031 | 0.218237154 | 0.508195251 | 0.88 |
| YPR071W | 302.2135475 | 0.244593296 | 0.5081892 | 0.88 |
| YBR031W | 10790.74196 | 0.248589178 | 0.510282627 | 0.88 |
| YDR011W | 4201.778813 | 0.231423661 | 0.512044129 | 0.88 |
| YDR142C | 534.1730573 | 0.178143542 | 0.510910798 | 0.88 |
| YDR500C | 6683.941473 | -0.291051097 | 0.511505382 | 0.88 |
| YER020W | 2824.873338 | -0.246196415 | 0.511654782 | 0.88 |
| YER035W | 1131.143378 | -0.245217889 | 0.511980505 | 0.88 |
| YFR019W | 2589.385757 | 0.146643191 | 0.510609398 | 0.88 |
| YGL143C | 536.4770627 | 0.208743965 | 0.512188562 | 0.88 |
| YHL008C | 1005.082799 | 0.226608136 | 0.510296528 | 0.88 |
| YHL040C | 1041.983492 | 0.27230766 | 0.511190824 | 0.88 |
| YIR010W | 867.0379525 | 0.144252144 | 0.512116125 | 0.88 |
| YJL025W | 374.9847479 | 0.241431274 | 0.511581756 | 0.88 |
| YJL059W | 446.5285115 | -0.171603233 | 0.511781812 | 0.88 |
| YLR096W | 1948.609356 | 0.149223131 | 0.512021861 | 0.88 |
| YML031W | 1341.560963 | 0.154318892 | 0.511358019 | 0.88 |
| YMR081C | 3745.685883 | 0.272977915 | 0.511429071 | 0.88 |
| YMR110C | 3066.365125 | -0.265178647 | 0.51228617 | 0.88 |
| YNL254C | 501.4634557 | 0.186418448 | 0.511393474 | 0.88 |
| YNR019W | 1395.465767 | -0.195340883 | 0.511040403 | 0.88 |
| YOL007C | 839.2247362 | 0.249646817 | 0.509746611 | 0.88 |
| YOR010C | 455.6940822 | 0.286927884 | 0.50978095 | 0.88 |
| YOR374W | 6364.239227 | -0.312503888 | 0.510073437 | 0.88 |
| YGL128C | 505.7413235 | -0.16654719 | 0.512653911 | 0.88 |
| YNR018W | 1764.18714 | -0.209138877 | 0.513302015 | 0.88 |
| YBR059C | 2706.219813 | -0.166362954 | 0.514013581 | 0.88 |
| YBR220C | 693.632069 | 0.18915578 | 0.513527128 | 0.88 |
| YCR095C | 483.8306058 | -0.150293348 | 0.514679772 | 0.88 |
| YDR156W | 897.9312179 | -0.168601614 | 0.514032474 | 0.88 |
| YDR370C | 533.8393634 | 0.135204279 | 0.514598225 | 0.88 |
| YDR410C | 388.474414 | -0.322557949 | 0.514905515 | 0.88 |
| YGR126W | 190.2789593 | 0.189452927 | 0.514605325 | 0.88 |
| YMR108W | 5770.966467 | 0.176451621 | 0.514919765 | 0.88 |
| YNL177C | 1301.893277 | -0.094335655 | 0.51465598 | 0.88 |
| YNR065C | 953.8955562 | 0.170451345 | 0.514212485 | 0.88 |
| YBR092C | 2232.790112 | 0.31596654 | 0.517126882 | 0.88 |

| | | | | |
|---|---|---|---|---|
| YDR085C | 1259.372351 | 0.244086765 | 0.516448227 | 0.88 |
| YDR295C | 798.8722941 | 0.111062182 | 0.515718462 | 0.88 |
| YER027C | 1561.298312 | -0.139356961 | 0.516454455 | 0.88 |
| YJR112W-A | 356.5088081 | -0.154834819 | 0.516617624 | 0.88 |
| YML060W | 354.1411795 | -0.238835978 | 0.516702879 | 0.88 |
| YMR195W | 1764.107553 | -0.312461571 | 0.516765709 | 0.88 |
| YNL185C | 474.6914722 | -0.176133831 | 0.51603061 | 0.88 |
| YNL303W | 97.83657889 | 0.31193093 | 0.516928924 | 0.88 |
| YNR068C | 378.9501574 | -0.270640498 | 0.516167325 | 0.88 |
| YOR354C | 1688.06875 | 0.103452373 | 0.517096483 | 0.88 |
| YPL191C | 278.5044925 | 0.197827879 | 0.516559824 | 0.88 |
| YPR198W | 724.9139692 | -0.22726822 | 0.516264337 | 0.88 |
| YDR468C | 586.9211769 | 0.148922167 | 0.517564594 | 0.88 |
| YLR363W-A | 269.4630453 | -0.249472628 | 0.517457566 | 0.88 |
| YMR262W | 628.7777309 | -0.203043816 | 0.517783798 | 0.88 |
| YMR286W | 654.5341281 | -0.182855181 | 0.517877432 | 0.88 |
| YDR166C | 1344.376251 | 0.133942608 | 0.518372564 | 0.88 |
| YHL044W | 369.968591 | -0.290922666 | 0.518874845 | 0.88 |
| YJR103W | 3899.480126 | -0.322525463 | 0.519158359 | 0.88 |
| YLR063W | 280.3060801 | 0.256664764 | 0.51954975 | 0.88 |
| YMR294W | 447.832337 | -0.185102349 | 0.520037441 | 0.88 |
| YBR138C | 569.4593536 | -0.144179888 | 0.520221498 | 0.88 |
| YJR100C | 772.6621126 | -0.12104781 | 0.520754801 | 0.88 |
| YMR293C | 413.7964048 | -0.153734524 | 0.521017768 | 0.88 |
| YER152C | 1264.809581 | -0.20985001 | 0.521269706 | 0.88 |
| YNL032W | 670.5607595 | 0.127159024 | 0.521393372 | 0.88 |
| snR37 | 153.9250937 | 0.269827895 | 0.521698314 | 0.88 |
| YBL029C-A | 1079.485322 | -0.278896886 | 0.522343596 | 0.88 |
| YJL212C | 1268.474729 | 0.295369896 | 0.522055262 | 0.88 |
| YMR139W | 1446.676083 | -0.206693267 | 0.522326562 | 0.88 |
| YER103W | 3269.463849 | -0.287744257 | 0.522755726 | 0.88 |
| YCR008W | 1187.11323 | 0.197741542 | 0.524690508 | 0.88 |
| YDL218W | 253.6505308 | 0.238422148 | 0.5243513 | 0.88 |
| YDR109C | 536.0726963 | 0.105560842 | 0.52502171 | 0.88 |
| YDR301W | 1341.743029 | 0.171168956 | 0.525031782 | 0.88 |
| YER115C | 529.525795 | -0.176567827 | 0.524306099 | 0.88 |
| YGL049C | 2260.025541 | 0.090145866 | 0.523631611 | 0.88 |
| YGL194C-A | 142.847988 | 0.215614722 | 0.523799555 | 0.88 |
| YGR038W | 610.0071877 | 0.172015654 | 0.524717794 | 0.88 |
| YGR182C | 629.4422912 | -0.222034564 | 0.524161107 | 0.88 |
| YJL194W | 154.6134221 | -0.274846692 | 0.524259334 | 0.88 |
| YMR191W | 2004.986398 | -0.300157479 | 0.524660897 | 0.88 |
| YPR134W | 542.0304792 | 0.119026647 | 0.524483828 | 0.88 |

| | | | | |
|---|---|---|---|---|
| YNR045W | 861.3548214 | 0.24220759 | 0.525567174 | 0.88 |
| YGR110W | 407.9203872 | 0.204738503 | 0.526238713 | 0.88 |
| YKL180W | 12036.25542 | 0.249913545 | 0.526642198 | 0.88 |
| YKR096W | 1782.315311 | 0.170520686 | 0.526893502 | 0.88 |
| YMR265C | 688.687566 | 0.17956187 | 0.526803912 | 0.88 |
| YNL137C | 1635.261502 | 0.115081887 | 0.527018628 | 0.88 |
| YNL316C | 479.8242776 | -0.134950829 | 0.526553163 | 0.88 |
| YOR097C | 623.6386998 | -0.13653815 | 0.526839084 | 0.88 |
| YBR193C | 556.4922631 | -0.136431297 | 0.527733459 | 0.88 |
| YDL057W | 396.4937822 | -0.241816134 | 0.52774149 | 0.88 |
| YDR034W-B | 127.2105401 | 0.26840079 | 0.528032317 | 0.89 |
| YGL243W | 434.8625344 | 0.08606131 | 0.528725369 | 0.89 |
| YKL014C | 1967.775598 | 0.239620451 | 0.528685714 | 0.89 |
| YLR102C | 476.3043851 | -0.194070364 | 0.528768703 | 0.89 |
| YNL038W | 473.5438124 | -0.191970914 | 0.528569518 | 0.89 |
| YOR047C | 1007.814108 | 0.262876058 | 0.529227455 | 0.89 |
| YGR149W | 1082.04674 | 0.173194085 | 0.529836225 | 0.89 |
| YKR081C | 1261.655674 | -0.254753652 | 0.529520434 | 0.89 |
| YNL125C | 2090.525445 | 0.181529309 | 0.529834565 | 0.89 |
| YJL106W | 294.2742704 | 0.21630641 | 0.5300312 | 0.89 |
| YCR069W | 2249.27644 | -0.198611358 | 0.530733539 | 0.89 |
| YHR162W | 4274.252821 | -0.217776461 | 0.530862991 | 0.89 |
| YLR225C | 1476.977135 | -0.230717061 | 0.531117773 | 0.89 |
| TLC1 | 1417.425959 | 0.267081851 | 0.533563972 | 0.89 |
| YBR017C | 2405.834993 | -0.130485303 | 0.533672193 | 0.89 |
| YDR318W | 508.7245087 | 0.141508135 | 0.533243673 | 0.89 |
| YDR479C | 1259.888261 | 0.170064862 | 0.531900777 | 0.89 |
| YFR027W | 221.6537022 | -0.154435724 | 0.53270086 | 0.89 |
| YFR049W | 788.1728678 | -0.247849074 | 0.534967892 | 0.89 |
| YFR053C | 31755.34613 | -0.315355475 | 0.534768208 | 0.89 |
| YGL228W | 2378.82392 | -0.176355419 | 0.534806561 | 0.89 |
| YGR085C | 1835.900279 | 0.239831484 | 0.532867258 | 0.89 |
| YKL099C | 686.8097655 | -0.234450986 | 0.53396093 | 0.89 |
| YML063W | 16224.08176 | 0.239816535 | 0.535044826 | 0.89 |
| YMR008C | 3679.477965 | 0.185747603 | 0.531813311 | 0.89 |
| YMR084W | 174.3110976 | 0.247499497 | 0.534568453 | 0.89 |
| YMR238W | 935.1001777 | 0.219335676 | 0.534121096 | 0.89 |
| YMR285C | 827.213202 | 0.100447632 | 0.533384058 | 0.89 |
| YMR323W | 206.5265565 | 0.055126537 | 0.534857494 | 0.89 |
| YNL073W | 1409.044777 | 0.163806944 | 0.534064924 | 0.89 |
| YNR015W | 1147.190779 | -0.140656127 | 0.53380929 | 0.89 |
| YOR006C | 758.4019018 | -0.126200303 | 0.534393422 | 0.89 |
| YOR176W | 2270.643704 | -0.196635108 | 0.53427483 | 0.89 |

| | | | | |
|---|---|---|---|---|
| YOR274W | 529.1848909 | 0.159610771 | 0.533233719 | 0.89 |
| YPR123C | 99.99461053 | 0.228546754 | 0.533514415 | 0.89 |
| YDR262W | 1927.726265 | 0.235456513 | 0.535451037 | 0.89 |
| YEL061C | 1022.169501 | 0.194077841 | 0.535691634 | 0.89 |
| YGL255W | 1201.748656 | 0.290875788 | 0.535742316 | 0.89 |
| YLR082C | 386.5514271 | -0.136922067 | 0.535371043 | 0.89 |
| YBR123C | 731.156432 | 0.108987095 | 0.536797174 | 0.89 |
| YKL011C | 416.7502341 | -0.174998419 | 0.536886805 | 0.89 |
| YKL219W | 500.3272007 | -0.191815973 | 0.536259251 | 0.89 |
| YNL246W | 1195.836863 | 0.093927709 | 0.536473418 | 0.89 |
| YNR073C | 195.2436335 | -0.25283753 | 0.535946787 | 0.89 |
| YPL169C | 1689.790258 | -0.131839936 | 0.536699917 | 0.89 |
| YPR194C | 1121.813284 | 0.309480691 | 0.536375089 | 0.89 |
| YFR052C-A | 3217.393186 | -0.287332575 | 0.538104841 | 0.89 |
| YGR088W | 3462.726793 | 0.287582093 | 0.53764752 | 0.89 |
| YJL115W | 931.9259287 | -0.191641299 | 0.538465941 | 0.89 |
| YKL073W | 1776.826393 | 0.216482358 | 0.53788729 | 0.89 |
| YLR030W | 185.5800298 | 0.273115755 | 0.53851603 | 0.89 |
| YLR191W | 472.4930163 | -0.138035809 | 0.537906969 | 0.89 |
| YPR007C | 168.1873056 | 0.257118141 | 0.538363492 | 0.89 |
| YBR135W | 639.2382675 | -0.227646202 | 0.538713423 | 0.89 |
| YIL144W | 1230.130031 | 0.157375436 | 0.539384349 | 0.89 |
| YLR344W | 3310.960859 | 0.226513394 | 0.5392755 | 0.89 |
| YDR251W | 2664.249522 | 0.131139451 | 0.539634992 | 0.89 |
| YDR506C | 448.3958033 | -0.119949841 | 0.539889618 | 0.89 |
| YLR287C | 617.6178514 | 0.226280874 | 0.539941303 | 0.89 |
| YBR203W | 994.1971392 | -0.274600323 | 0.54012117 | 0.89 |
| YBR242W | 549.7068913 | -0.133770038 | 0.54079755 | 0.89 |
| YOL141W | 189.2930206 | 0.209129774 | 0.540718496 | 0.89 |
| YBR227C | 856.5044097 | -0.166268336 | 0.541616128 | 0.89 |
| YDR129C | 5762.337606 | -0.138643606 | 0.541912415 | 0.89 |
| YHR106W | 1096.925363 | -0.202075742 | 0.541822516 | 0.89 |
| YDL005C | 1599.265031 | -0.107328943 | 0.542377528 | 0.89 |
| YDL063C | 706.2977789 | 0.275884583 | 0.54345669 | 0.89 |
| YGL064C | 383.7385393 | 0.134485608 | 0.543275638 | 0.89 |
| YGL166W | 257.6413584 | -0.15560761 | 0.54228537 | 0.89 |
| YKL154W | 669.8187345 | -0.246511148 | 0.543641744 | 0.89 |
| YKL156W | 2081.100672 | 0.245470244 | 0.543063367 | 0.89 |
| YKL221W | 211.2597824 | -0.34466374 | 0.543018428 | 0.89 |
| YLR336C | 993.7017255 | -0.197598301 | 0.543105153 | 0.89 |
| YLR437C | 329.593239 | -0.179501272 | 0.543626592 | 0.89 |
| YOL101C | 182.685157 | 0.215097054 | 0.542497446 | 0.89 |
| YOR179C | 265.678804 | 0.186246711 | 0.543698774 | 0.89 |

| | | | | |
|---|---|---|---|---|
| YBL005W | 1420.668611 | 0.148904339 | 0.545114042 | 0.89 |
| YBR033W | 734.5856264 | 0.285475502 | 0.546991716 | 0.89 |
| YBR054W | 4277.398072 | 0.266420392 | 0.549475595 | 0.89 |
| YBR139W | 2470.205803 | -0.19147637 | 0.54524938 | 0.89 |
| YDL002C | 564.8870678 | -0.099240851 | 0.550234996 | 0.89 |
| YDL008W | 381.8332185 | -0.171978994 | 0.549205084 | 0.89 |
| YDL217C | 416.6172424 | -0.133187091 | 0.54930726 | 0.89 |
| YDR322C-A | 892.3853165 | -0.260070842 | 0.54457423 | 0.89 |
| YDR393W | 653.7265915 | 0.114732976 | 0.544979119 | 0.89 |
| YFL027C | 673.7008422 | 0.207875558 | 0.545148683 | 0.89 |
| YFR002W | 1555.318988 | 0.152953006 | 0.548994222 | 0.89 |
| YGL111W | 1214.214479 | 0.170333392 | 0.546112837 | 0.89 |
| YHR135C | 3300.939762 | 0.112885327 | 0.547344922 | 0.89 |
| YIR004W | 1446.338254 | -0.122205528 | 0.544974307 | 0.89 |
| YJL066C | 1041.928086 | -0.25238859 | 0.548581872 | 0.89 |
| YJL069C | 705.1093909 | 0.251603487 | 0.54938052 | 0.89 |
| YJL076W | 2985.809725 | 0.143007014 | 0.546847168 | 0.89 |
| YJL168C | 610.0870071 | 0.204795857 | 0.54998432 | 0.89 |
| YKL093W | 994.1827608 | 0.258029552 | 0.545701288 | 0.89 |
| YKR104W | 184.9350151 | 0.222685983 | 0.550182736 | 0.89 |
| YLR060W | 5384.459239 | -0.235675527 | 0.545959725 | 0.89 |
| YLR455W | 572.5909218 | 0.148106271 | 0.550069047 | 0.89 |
| YML107C | 389.7043417 | 0.152746296 | 0.549123566 | 0.89 |
| YMR101C | 112.86694 | 0.224363501 | 0.547989421 | 0.89 |
| YMR148W | 862.6307015 | -0.155259678 | 0.544744664 | 0.89 |
| YMR193W | 776.035786 | -0.191623049 | 0.548809792 | 0.89 |
| YNL215W | 1103.384408 | -0.171676602 | 0.544438502 | 0.89 |
| YNL330C | 1017.919377 | -0.223610712 | 0.546187816 | 0.89 |
| YOL144W | 766.01586 | -0.218977601 | 0.546958977 | 0.89 |
| YOL147C | 576.9213691 | -0.194840855 | 0.549279058 | 0.89 |
| YOR187W | 4877.755588 | -0.139742399 | 0.549530146 | 0.89 |
| YOR216C | 1165.7555 | -0.190015428 | 0.547991939 | 0.89 |
| YOR254C | 3232.98471 | -0.214334429 | 0.548183919 | 0.89 |
| YPL060W | 360.6791713 | 0.199699431 | 0.546979112 | 0.89 |
| YPL086C | 1089.007189 | -0.202224446 | 0.549136645 | 0.89 |
| YPL100W | 911.5930892 | -0.152167356 | 0.550088692 | 0.89 |
| YPL123C | 1037.088707 | -0.226269874 | 0.548635589 | 0.89 |
| YPL135W | 1241.708085 | 0.275213185 | 0.548522624 | 0.89 |
| YPR053C | 166.5759994 | 0.200102826 | 0.547175189 | 0.89 |
| YBR241C | 1365.325299 | -0.283486123 | 0.550643326 | 0.89 |
| YPL006W | 1523.461434 | 0.172812483 | 0.550907385 | 0.89 |
| YIL050W | 1083.890647 | -0.180494439 | 0.551239954 | 0.89 |
| YNL020C | 852.0700517 | 0.162334922 | 0.551456695 | 0.89 |

| | | | | |
|---|---|---|---|---|
| YPL049C | 2306.704815 | 0.126756903 | 0.551456901 | 0.89 |
| YBR019C | 339.0837633 | 0.245171438 | 0.551962074 | 0.89 |
| YLR009W | 1043.475365 | 0.240926197 | 0.551864396 | 0.89 |
| YBL006C | 611.4550235 | -0.173429894 | 0.556870877 | 0.89 |
| YBR055C | 925.8039594 | 0.093266308 | 0.557055453 | 0.89 |
| YBR187W | 950.8054053 | 0.190058668 | 0.556144038 | 0.89 |
| YBR225W | 1153.176358 | 0.172673715 | 0.554469166 | 0.89 |
| YBR255W | 917.9047892 | -0.122874377 | 0.555349915 | 0.89 |
| YBR260C | 1246.430976 | -0.151617153 | 0.557118461 | 0.89 |
| YCL031C | 1450.481764 | -0.153923835 | 0.55652 | 0.89 |
| YCL032W | 909.5801687 | -0.139878535 | 0.55275409 | 0.89 |
| YDL128W | 1754.772543 | 0.112229957 | 0.555578619 | 0.89 |
| YDL167C | 1031.301724 | 0.259088794 | 0.556014773 | 0.89 |
| YDR234W | 1972.622196 | 0.210389556 | 0.558141485 | 0.89 |
| YER025W | 3287.005691 | 0.162829668 | 0.554289749 | 0.89 |
| YER053C | 3873.936022 | -0.263954666 | 0.55301553 | 0.89 |
| YGL027C | 1397.054052 | 0.140376743 | 0.555692076 | 0.89 |
| YGL247W | 197.7711302 | 0.155087269 | 0.557633775 | 0.89 |
| YGR028W | 820.509086 | -0.153837241 | 0.558388109 | 0.89 |
| YGR146C | 1344.611896 | 0.227265693 | 0.558396058 | 0.89 |
| YGR227W | 1061.56859 | 0.163738301 | 0.558107003 | 0.89 |
| YGR246C | 1106.975461 | -0.140686033 | 0.553202384 | 0.89 |
| YHR187W | 633.7803075 | -0.102323294 | 0.555654619 | 0.89 |
| YIR023W | 1089.743487 | 0.146946834 | 0.558061387 | 0.89 |
| YJL143W | 1167.282332 | -0.148220194 | 0.55619759 | 0.89 |
| YJR009C | 33615.59982 | -0.326759398 | 0.558093297 | 0.89 |
| YJR077C | 4794.775519 | -0.221805319 | 0.553969573 | 0.89 |
| YKL036C | 247.9148773 | -0.199238075 | 0.556425248 | 0.89 |
| YKR069W | 646.4230382 | 0.157279824 | 0.558556412 | 0.89 |
| YKR077W | 759.0676767 | 0.186385331 | 0.558696237 | 0.89 |
| YKR102W | 639.1097035 | 0.25413028 | 0.556960802 | 0.89 |
| YLR008C | 578.2897755 | -0.18873429 | 0.552439322 | 0.89 |
| YLR205C | 412.8411079 | 0.243998706 | 0.553384302 | 0.89 |
| YLR369W | 1148.442381 | 0.187453748 | 0.554883881 | 0.89 |
| YMR062C | 1501.153431 | -0.227302768 | 0.553129261 | 0.89 |
| YMR215W | 1997.531105 | -0.222351529 | 0.558070588 | 0.89 |
| YNL005C | 1578.642517 | -0.11048693 | 0.554823027 | 0.89 |
| YNL153C | 847.6509295 | -0.192112624 | 0.557277881 | 0.89 |
| YOL107W | 336.081692 | 0.187280455 | 0.557218892 | 0.89 |
| YPL046C | 139.4253961 | -0.157089069 | 0.55650895 | 0.89 |
| YPL112C | 806.8455711 | -0.132196761 | 0.555850081 | 0.89 |
| YPL217C | 2396.710824 | 0.160903643 | 0.555051951 | 0.89 |
| YPL271W | 1376.293801 | -0.251809033 | 0.557763893 | 0.89 |

| | | | | |
|---|---|---|---|---|
| YIR028W | 189.55325 | 0.302160671 | 0.558861287 | 0.89 |
| YJR109C | 3577.314205 | 0.26759636 | 0.559301052 | 0.89 |
| YLR006C | 1262.134466 | -0.159632166 | 0.55939162 | 0.89 |
| YOR270C | 5846.210473 | 0.138726064 | 0.559133073 | 0.89 |
| YGR128C | 1414.526792 | 0.269773448 | 0.5595673 | 0.89 |
| YER011W | 1426.306813 | -0.291609414 | 0.55988849 | 0.89 |
| YER150W | 3467.414047 | 0.293367264 | 0.560543679 | 0.89 |
| YDL040C | 3238.410502 | 0.135736382 | 0.560954243 | 0.89 |
| YEL019C | 187.0813824 | 0.191444124 | 0.561392833 | 0.89 |
| YJL079C | 9145.850933 | 0.235761347 | 0.561923634 | 0.90 |
| YOR290C | 3907.230484 | 0.125007572 | 0.562149776 | 0.90 |
| YDR533C | 1411.920932 | -0.286782737 | 0.562995352 | 0.90 |
| YJL057C | 1203.837592 | 0.221483387 | 0.56266675 | 0.90 |
| YML101C | 335.3248863 | 0.164692322 | 0.563068941 | 0.90 |
| YOL041C | 1321.58671 | -0.247064934 | 0.562829885 | 0.90 |
| YOR328W | 830.771978 | 0.222085256 | 0.562938432 | 0.90 |
| YGR152C | 296.8399677 | -0.214756033 | 0.563638634 | 0.90 |
| YLR178C | 3271.315417 | -0.279826832 | 0.563538605 | 0.90 |
| YML022W | 1077.904293 | -0.246059234 | 0.56387613 | 0.90 |
| snR86 | 894.0383097 | 0.23091701 | 0.56396018 | 0.90 |
| YHR134W | 365.9132551 | 0.183572743 | 0.564282569 | 0.90 |
| YDR495C | 1040.925179 | -0.154880591 | 0.564489338 | 0.90 |
| YJR129C | 222.0216409 | -0.195986983 | 0.564702557 | 0.90 |
| YMR204C | 437.305049 | 0.153369305 | 0.564931267 | 0.90 |
| YCL027W | 675.5367401 | 0.253377319 | 0.565895272 | 0.90 |
| YDR313C | 770.3934735 | -0.194576427 | 0.566150222 | 0.90 |
| YDR404C | 728.6666751 | 0.129431963 | 0.56700645 | 0.90 |
| YDR428C | 292.2817321 | -0.22543112 | 0.567421981 | 0.90 |
| YER104W | 241.2969985 | 0.146373432 | 0.566585008 | 0.90 |
| YIL165C | 180.1950386 | 0.232548418 | 0.565527768 | 0.90 |
| YJL208C | 631.1153592 | -0.242426973 | 0.567590397 | 0.90 |
| YKR008W | 1399.761894 | 0.122749752 | 0.56580855 | 0.90 |
| YLL016W | 1012.667191 | 0.237749609 | 0.56554492 | 0.90 |
| YLL038C | 326.4334207 | 0.173584093 | 0.566318734 | 0.90 |
| YLR093C | 1657.689066 | -0.147183498 | 0.567275904 | 0.90 |
| YMR098C | 840.7673515 | 0.146597497 | 0.567521996 | 0.90 |
| YNL163C | 1921.119184 | 0.16119239 | 0.567292234 | 0.90 |
| YOL154W | 197.7584736 | 0.145296576 | 0.567075235 | 0.90 |
| YOR353C | 984.6133479 | 0.156395433 | 0.566710264 | 0.90 |
| YPR191W | 6094.519828 | -0.266037056 | 0.565430013 | 0.90 |
| YAL056W | 1700.26388 | 0.189769893 | 0.567963107 | 0.90 |
| YHR199C | 2007.264009 | -0.197702352 | 0.568178326 | 0.90 |
| YLR056W | 3964.313149 | 0.227321208 | 0.568397071 | 0.90 |

| | | | | |
|---|---|---|---|---|
| YEL004W | 410.7288824 | 0.120799529 | 0.568725464 | 0.90 |
| YBR039W | 6878.817001 | -0.258145688 | 0.569192893 | 0.90 |
| YBR237W | 937.0201345 | -0.164543944 | 0.570055655 | 0.90 |
| YDL036C | 422.8116952 | 0.134059959 | 0.5703035 | 0.90 |
| YGL065C | 1014.84416 | -0.100527313 | 0.5704271 | 0.90 |
| YGL146C | 201.790122 | 0.239909389 | 0.570480373 | 0.90 |
| YGR185C | 4359.206315 | -0.240412766 | 0.569671778 | 0.90 |
| YLR121C | 1254.397824 | 0.173526448 | 0.569082716 | 0.90 |
| YLR152C | 1272.570191 | 0.216013233 | 0.570408025 | 0.90 |
| YMR172W | 685.1558079 | 0.133606685 | 0.570108758 | 0.90 |
| YPL031C | 1442.118487 | -0.141135577 | 0.570373411 | 0.90 |
| YBR196C | 23554.20363 | -0.176132669 | 0.571054393 | 0.90 |
| YDL156W | 331.6691228 | 0.179052533 | 0.570944294 | 0.90 |
| YNR036C | 1583.864307 | -0.190138607 | 0.571053232 | 0.90 |
| YDR309C | 2613.4939 | -0.214582099 | 0.571377991 | 0.90 |
| YBL013W | 160.0267849 | 0.171163027 | 0.57213207 | 0.90 |
| YBR151W | 3044.846839 | -0.147341702 | 0.572104728 | 0.90 |
| YDL123W | 682.3334241 | -0.169421499 | 0.572027437 | 0.90 |
| YLR329W | 110.7708514 | 0.219487157 | 0.571776635 | 0.90 |
| YLR371W | 1780.890302 | 0.153035785 | 0.572510545 | 0.90 |
| YDL018C | 497.6776418 | 0.162805502 | 0.573750904 | 0.90 |
| YHR013C | 1158.208101 | -0.196196662 | 0.573774493 | 0.90 |
| YHR132W-A | 721.2397547 | -0.159282904 | 0.573191043 | 0.90 |
| YJL154C | 2489.813399 | 0.06384945 | 0.573550936 | 0.90 |
| YLR099C | 1018.600053 | -0.201442176 | 0.573849384 | 0.90 |
| YNL179C | 130.2526057 | -0.209964056 | 0.573760288 | 0.90 |
| YOR256C | 1260.844527 | 0.120626227 | 0.573808151 | 0.90 |
| YDR357C | 553.215464 | -0.177635702 | 0.574414258 | 0.90 |
| YBR154C | 1262.041062 | -0.180642539 | 0.574975685 | 0.90 |
| YOL093W | 662.3279101 | 0.17088092 | 0.57538892 | 0.90 |
| YPL168W | 515.986609 | 0.17579234 | 0.575306822 | 0.90 |
| YPR081C | 985.9306388 | 0.138043015 | 0.57578153 | 0.90 |
| YJR083C | 550.0152446 | -0.129972748 | 0.576099493 | 0.90 |
| YML109W | 760.9435844 | 0.13046437 | 0.576192451 | 0.90 |
| YDR489W | 358.8647918 | 0.137398805 | 0.57662206 | 0.90 |
| YDR538W | 133.1462476 | -0.127871146 | 0.576669127 | 0.90 |
| YKR017C | 571.9042365 | 0.184184058 | 0.576807784 | 0.90 |
| YML093W | 1683.648354 | 0.207574487 | 0.57702851 | 0.90 |
| YMR044W | 903.5982709 | -0.128968663 | 0.577039033 | 0.90 |
| YHR089C | 2056.029582 | -0.257518949 | 0.577240088 | 0.90 |
| YOR266W | 469.4838324 | 0.087622291 | 0.577443112 | 0.90 |
| YCR052W | 1334.780982 | -0.116074484 | 0.578702643 | 0.90 |
| YDL088C | 1480.552376 | 0.124975198 | 0.579152551 | 0.90 |

| | | | | |
|---|---|---|---|---|
| YDL107W | 383.3863117 | -0.140109373 | 0.579788941 | 0.90 |
| YDR244W | 923.2552739 | -0.107574991 | 0.580261594 | 0.90 |
| YER096W | 439.4636005 | -0.333793701 | 0.579339639 | 0.90 |
| YFL032W | 146.5638958 | 0.221496427 | 0.579873205 | 0.90 |
| YGR109C | 138.0694558 | -0.278239945 | 0.578486314 | 0.90 |
| YHR002W | 437.310458 | 0.197710641 | 0.580424973 | 0.90 |
| YKL089W | 415.1913401 | -0.164221996 | 0.579371684 | 0.90 |
| YKL181W | 2705.955695 | 0.205455008 | 0.580143683 | 0.90 |
| YLR196W | 1358.156963 | 0.226222374 | 0.580465716 | 0.90 |
| YLR367W | 2289.066484 | 0.179627588 | 0.579329537 | 0.90 |
| YMR003W | 306.5250482 | -0.179532686 | 0.57868354 | 0.90 |
| YPL167C | 937.9139202 | 0.141847273 | 0.578323419 | 0.90 |
| YPL203W | 1453.358861 | -0.172599803 | 0.579697304 | 0.90 |
| YGR119C | 2276.598641 | -0.097944434 | 0.580687622 | 0.90 |
| YJL161W | 501.2537696 | -0.246375449 | 0.581032288 | 0.90 |
| YLR304C | 13028.11852 | -0.21217328 | 0.581529073 | 0.90 |
| YPL139C | 637.4930321 | 0.084820635 | 0.581879217 | 0.90 |
| YDR028C | 3835.45418 | 0.132463438 | 0.58240022 | 0.90 |
| YOL060C | 2188.941543 | -0.141440651 | 0.582556282 | 0.90 |
| YDL210W | 142.1653441 | 0.251361088 | 0.583167357 | 0.90 |
| YNL188W | 592.8677499 | 0.174993134 | 0.583785389 | 0.90 |
| YPL043W | 1057.06526 | 0.212885647 | 0.583908698 | 0.90 |
| YDR342C | 1706.769269 | -0.259305446 | 0.585656239 | 0.90 |
| YGR161C | 2432.867838 | 0.23731758 | 0.585903941 | 0.90 |
| YHR189W | 309.5122663 | 0.189504883 | 0.584625768 | 0.90 |
| YJL176C | 1289.4859 | 0.136291274 | 0.585808764 | 0.90 |
| YKR075C | 2465.019565 | 0.275218675 | 0.584683832 | 0.90 |
| YLR163C | 1121.057782 | -0.110203753 | 0.584722061 | 0.90 |
| YNL142W | 1026.084342 | 0.238725843 | 0.585158479 | 0.90 |
| YNL314W | 384.9311948 | -0.12497178 | 0.585650157 | 0.90 |
| YOR262W | 658.3672371 | -0.152250419 | 0.58485282 | 0.90 |
| YPL189C-A | 341.9617668 | 0.210585274 | 0.585192963 | 0.90 |
| YPR075C | 1855.120278 | -0.138754429 | 0.585976372 | 0.90 |
| YCR010C | 273.3869127 | 0.273466852 | 0.586475627 | 0.90 |
| YGL252C | 1693.94272 | 0.073793666 | 0.586861218 | 0.90 |
| YGR006W | 239.4957069 | -0.114196751 | 0.586669645 | 0.90 |
| YHL021C | 7761.479525 | -0.246101856 | 0.586313544 | 0.90 |
| YJR110W | 661.1509666 | 0.085546977 | 0.58680999 | 0.90 |
| YGL249W | 273.3583474 | 0.228978813 | 0.587155168 | 0.91 |
| YBR268W | 567.5701433 | -0.150835658 | 0.587383021 | 0.91 |
| YAL020C | 521.9507996 | -0.142916653 | 0.587671273 | 0.91 |
| YJR137C | 1214.839856 | 0.147732613 | 0.587942956 | 0.91 |
| YPL260W | 2621.937685 | -0.144411808 | 0.587824498 | 0.91 |

| | | | | |
|---|---|---|---|---|
| YMR117C | 204.4706004 | 0.146623661 | 0.588171448 | 0.91 |
| YGL115W | 1145.902181 | -0.076280605 | 0.588629036 | 0.91 |
| YAR019C | 904.0673159 | 0.145675736 | 0.589169333 | 0.91 |
| YGR097W | 2251.701342 | -0.159540614 | 0.589037662 | 0.91 |
| YGR123C | 655.6704403 | 0.242077363 | 0.589937958 | 0.91 |
| YJL125C | 686.7482617 | 0.164385533 | 0.589883929 | 0.91 |
| YKL081W | 14463.37631 | -0.263359671 | 0.589792277 | 0.91 |
| YKR098C | 1187.222635 | 0.181279675 | 0.589864501 | 0.91 |
| YNL078W | 1260.908163 | 0.174174351 | 0.589368199 | 0.91 |
| YBR012W-B | 321.6022163 | 0.224619605 | 0.590385248 | 0.91 |
| YGR155W | 6507.68049 | 0.131609283 | 0.590442507 | 0.91 |
| YKL215C | 1029.488394 | 0.128385504 | 0.590501544 | 0.91 |
| YGL196W | 1262.908167 | 0.207311957 | 0.590833753 | 0.91 |
| YDL061C | 2048.312712 | -0.277115313 | 0.591316078 | 0.91 |
| YNL178W | 12909.44004 | 0.17522905 | 0.591559763 | 0.91 |
| YDR297W | 2184.395638 | -0.147223109 | 0.591777638 | 0.91 |
| YBR103W | 849.3179685 | -0.114841969 | 0.592669539 | 0.91 |
| YGL090W | 485.8897575 | 0.115924092 | 0.592566782 | 0.91 |
| YIL084C | 434.165782 | 0.13922551 | 0.592335648 | 0.91 |
| YMR165C | 1579.930507 | 0.217337413 | 0.592278344 | 0.91 |
| YPL201C | 234.758662 | 0.234067689 | 0.592382344 | 0.91 |
| YMR119W | 1094.314263 | 0.129460806 | 0.593208124 | 0.91 |
| YIL080W | 141.3218365 | 0.247579749 | 0.593719762 | 0.91 |
| YLR146C | 511.7510501 | -0.124411073 | 0.593398467 | 0.91 |
| YLR429W | 3551.790145 | -0.2301834 | 0.593658525 | 0.91 |
| YGL184C | 275.5662791 | 0.179718303 | 0.594189977 | 0.91 |
| YKL183W | 567.8673175 | -0.174463704 | 0.595004541 | 0.91 |
| YPL188W | 716.8167679 | -0.1392777 | 0.595233299 | 0.91 |
| YDL127W | 422.9534044 | 0.16095156 | 0.595711541 | 0.91 |
| YBR044C | 837.2738234 | 0.128036471 | 0.596229715 | 0.91 |
| YDR320C | 1271.658982 | 0.093577001 | 0.596272108 | 0.91 |
| YOR208W | 1224.294967 | -0.188515955 | 0.595987235 | 0.91 |
| YLR241W | 1618.382874 | 0.104782679 | 0.596614167 | 0.91 |
| YGL061C | 446.8633151 | -0.158732482 | 0.597293968 | 0.91 |
| YGR169C-A | 750.5850996 | -0.128299123 | 0.597064915 | 0.91 |
| YNL272C | 1304.645862 | -0.094749064 | 0.59714781 | 0.91 |
| YKR072C | 2109.757496 | 0.148069399 | 0.597552279 | 0.91 |
| YEL026W | 1802.083858 | -0.284039758 | 0.598749723 | 0.91 |
| YHR023W | 2927.24121 | 0.167998528 | 0.598646827 | 0.91 |
| YMR015C | 1137.039851 | 0.187707116 | 0.598490005 | 0.91 |
| YOR306C | 1215.578289 | 0.232190231 | 0.59853661 | 0.91 |
| YOR044W | 320.4821165 | 0.146225496 | 0.59902751 | 0.91 |
| YPR091C | 1815.236332 | 0.095471058 | 0.599282062 | 0.91 |

| | | | | |
|---|---|---|---|---|
| YBR043C | 1241.74171 | 0.217344103 | 0.600176264 | 0.91 |
| YBR176W | 422.8364361 | -0.131395171 | 0.600533221 | 0.91 |
| YDL208W | 1913.168285 | -0.211652375 | 0.601939363 | 0.91 |
| YDR259C | 452.5952789 | -0.208007707 | 0.602304139 | 0.91 |
| YDR381C-A | 448.1284196 | -0.188957786 | 0.601764092 | 0.91 |
| YEL072W | 207.4067694 | -0.224874167 | 0.600522636 | 0.91 |
| YFL057C | 160.9643278 | 0.165761804 | 0.600970182 | 0.91 |
| YGR043C | 1676.63942 | 0.260496183 | 0.602104095 | 0.91 |
| YHR092C | 9528.876224 | -0.296784682 | 0.601961054 | 0.91 |
| YHR159W | 783.3955676 | -0.147919116 | 0.601903069 | 0.91 |
| YKR106W | 110.7787445 | -0.272239713 | 0.602203229 | 0.91 |
| YLR142W | 1292.784521 | 0.24012148 | 0.600926222 | 0.91 |
| YLR423C | 531.1083054 | -0.169545256 | 0.600612552 | 0.91 |
| YML002W | 853.1962719 | 0.121261296 | 0.601581162 | 0.91 |
| YML058W | 2577.833646 | -0.163977792 | 0.600180069 | 0.91 |
| YNL064C | 7494.727656 | -0.225191647 | 0.60155748 | 0.91 |
| YBR070C | 438.6999546 | -0.199393148 | 0.603326837 | 0.91 |
| YBR132C | 1205.603555 | -0.222102555 | 0.603199202 | 0.91 |
| YDR270W | 829.6044968 | 0.151091984 | 0.602819563 | 0.91 |
| YFL034W | 949.1269789 | 0.086850676 | 0.60271743 | 0.91 |
| YGR295C | 1429.22247 | 0.206589269 | 0.603329379 | 0.91 |
| YPR161C | 1370.124428 | -0.126067231 | 0.603269412 | 0.91 |
| YDL079C | 1497.675817 | 0.257221694 | 0.603879953 | 0.91 |
| YGR233C | 1147.708039 | 0.155533091 | 0.604010672 | 0.91 |
| YDR064W | 9582.184184 | 0.165502747 | 0.604375517 | 0.91 |
| YHR001W-A | 1152.222923 | -0.239646951 | 0.604293627 | 0.91 |
| YER165W | 11478.83612 | -0.15408273 | 0.605051571 | 0.91 |
| YGR255C | 1753.194799 | -0.12168518 | 0.604944272 | 0.91 |
| YLR300W | 3505.549383 | 0.22083595 | 0.605116852 | 0.91 |
| YOL120C | 9488.699457 | 0.111062706 | 0.604856877 | 0.91 |
| YAR035W | 1245.571751 | -0.259309758 | 0.605707504 | 0.91 |
| YBR048W | 5960.519296 | -0.29159231 | 0.60618832 | 0.91 |
| YDR235W | 670.6497022 | 0.096283942 | 0.606149915 | 0.91 |
| YGL222C | 1086.796722 | -0.255898186 | 0.605541632 | 0.91 |
| YIL146C | 613.496777 | 0.179729319 | 0.60586495 | 0.91 |
| YGR113W | 643.4472036 | -0.067294414 | 0.606460699 | 0.91 |
| YLR203C | 2343.4384 | -0.10386257 | 0.606709473 | 0.91 |
| YMR093W | 713.9051908 | 0.201658517 | 0.606785645 | 0.91 |
| YMR073C | 579.5582083 | 0.140931708 | 0.60722362 | 0.91 |
| YDR110W | 523.1803163 | 0.1569558 | 0.608427704 | 0.92 |
| YGL010W | 399.2167766 | -0.182272414 | 0.608421523 | 0.92 |
| YOR382W | 2866.519088 | -0.312276874 | 0.608705114 | 0.92 |
| YDL226C | 1711.631594 | -0.166447756 | 0.610069209 | 0.92 |

| | | | | |
|---|---|---|---|---|
| YER019W | 1037.358398 | 0.190335048 | 0.609845178 | 0.92 |
| YGR034W | 5016.73135 | -0.293508694 | 0.609664502 | 0.92 |
| YMR153W | 881.8064044 | -0.084497711 | 0.609687936 | 0.92 |
| YNL002C | 1677.703778 | -0.210089261 | 0.60989444 | 0.92 |
| YNL080C | 1115.372394 | 0.124385408 | 0.610357206 | 0.92 |
| YOR173W | 2098.419186 | -0.244952844 | 0.61037165 | 0.92 |
| YGR081C | 443.1057485 | 0.213480243 | 0.610599562 | 0.92 |
| YGR130C | 4694.98638 | -0.167071868 | 0.611091312 | 0.92 |
| YNL191W | 591.9956262 | -0.128214791 | 0.611215141 | 0.92 |
| YDL160C-A | 95.22130843 | -0.296361551 | 0.611524609 | 0.92 |
| YPL265W | 2999.757451 | 0.245260709 | 0.611850312 | 0.92 |
| YGL256W | 1304.621904 | 0.124551661 | 0.612155438 | 0.92 |
| YBR211C | 479.4755282 | -0.131438953 | 0.612541825 | 0.92 |
| YHR085W | 335.931842 | 0.19552939 | 0.612445656 | 0.92 |
| YPR170W-B | 216.8408172 | -0.163909167 | 0.612730192 | 0.92 |
| YOR203W | 129.138005 | -0.194052031 | 0.612902422 | 0.92 |
| YBR129C | 610.3882792 | -0.138953671 | 0.613681555 | 0.92 |
| YMR313C | 988.4819991 | 0.108956978 | 0.614388544 | 0.92 |
| YGL153W | 279.9517565 | -0.122556464 | 0.614739304 | 0.92 |
| YMR123W | 1008.572929 | -0.200369332 | 0.614774041 | 0.92 |
| YKL128C | 1025.460861 | -0.159717327 | 0.615260213 | 0.92 |
| YNL202W | 289.1619632 | 0.210142342 | 0.615560031 | 0.92 |
| YOL010W | 524.4137956 | 0.19571199 | 0.61562968 | 0.92 |
| YPL226W | 2793.678288 | 0.194193493 | 0.615618768 | 0.92 |
| YAR042W | 3250.319067 | -0.156920076 | 0.61690462 | 0.92 |
| YOR222W | 1007.50754 | 0.172612487 | 0.616870316 | 0.92 |
| YNR056C | 522.7352776 | 0.237466522 | 0.617212631 | 0.92 |
| YCL052C | 1614.200073 | -0.121415733 | 0.618034967 | 0.92 |
| YGR264C | 2825.266453 | 0.209881295 | 0.617836095 | 0.92 |
| YLR153C | 3246.572959 | 0.131377548 | 0.618061948 | 0.92 |
| YOL047C | 302.1448912 | 0.226614236 | 0.617975902 | 0.92 |
| YEL048C | 96.72112286 | -0.191008301 | 0.618599156 | 0.92 |
| YDL125C | 2310.617952 | -0.147328312 | 0.618797254 | 0.92 |
| YER168C | 714.524667 | -0.079383598 | 0.619004305 | 0.92 |
| YPL269W | 614.929882 | 0.12717078 | 0.619276346 | 0.92 |
| YBR028C | 772.1128092 | 0.149155257 | 0.619873646 | 0.92 |
| YJR067C | 196.0036818 | -0.10315233 | 0.619742569 | 0.92 |
| YKL101W | 3903.982421 | 0.188616974 | 0.619868432 | 0.92 |
| YCR044C | 1029.447304 | -0.103482596 | 0.620251923 | 0.92 |
| YGR210C | 1863.079648 | 0.138253995 | 0.620460571 | 0.92 |
| YDR465C | 1088.193625 | -0.219845946 | 0.620950836 | 0.92 |
| YGR289C | 671.1201555 | -0.220725465 | 0.621193729 | 0.92 |
| YOL030W | 6136.814993 | -0.140868947 | 0.621152889 | 0.92 |

| | | | | |
|---|---|---|---|---|
| YIL108W | 1452.736233 | -0.187056891 | 0.62154598 | 0.92 |
| YDR212W | 2389.189098 | -0.132031856 | 0.621884907 | 0.92 |
| YNL223W | 533.8185201 | -0.139209367 | 0.621885474 | 0.92 |
| YDL141W | 1031.602218 | 0.089287703 | 0.622327578 | 0.92 |
| YBR087W | 809.749781 | 0.128625916 | 0.625286371 | 0.92 |
| YBR201W | 456.7810734 | -0.116958184 | 0.626140772 | 0.92 |
| YBR207W | 1493.475156 | -0.146559634 | 0.624756697 | 0.92 |
| YBR295W | 678.020174 | 0.157620262 | 0.625580087 | 0.92 |
| YCR100C | 233.2545617 | 0.195922747 | 0.623003426 | 0.92 |
| YDL215C | 3694.89417 | 0.220866719 | 0.62316401 | 0.92 |
| YDR040C | 132.5886651 | 0.204295393 | 0.62514594 | 0.92 |
| YDR528W | 366.4782046 | -0.231486559 | 0.625765627 | 0.92 |
| YGL060W | 769.6330407 | 0.104637218 | 0.62497788 | 0.92 |
| YGL202W | 6879.756501 | -0.135019734 | 0.624945168 | 0.92 |
| YHR065C | 1260.588152 | -0.171570436 | 0.623881052 | 0.92 |
| YIL023C | 838.1130985 | 0.121662169 | 0.626133886 | 0.92 |
| YIL089W | 155.6273305 | -0.214790497 | 0.623544546 | 0.92 |
| YJR093C | 790.9035242 | -0.068330233 | 0.624354262 | 0.92 |
| YJR138W | 1461.069987 | 0.13524176 | 0.623593687 | 0.92 |
| YLR134W | 556.7123022 | 0.156445985 | 0.624550037 | 0.92 |
| YML106W | 2946.051592 | -0.181623661 | 0.62503463 | 0.92 |
| YNL092W | 338.5129162 | -0.163759107 | 0.62587969 | 0.92 |
| YOR004W | 494.9946687 | 0.213595117 | 0.626251626 | 0.92 |
| YOR289W | 937.4069291 | -0.23137808 | 0.623082292 | 0.92 |
| YPL105C | 3260.005512 | -0.093423694 | 0.625018923 | 0.92 |
| YPR174C | 411.2973194 | -0.11792833 | 0.623473235 | 0.92 |
| YPR190C | 965.2019513 | 0.169611838 | 0.623614323 | 0.92 |
| YDR530C | 1226.853067 | -0.171421503 | 0.62673337 | 0.92 |
| YER061C | 958.2654695 | 0.126876178 | 0.626985167 | 0.92 |
| YLR036C | 360.058471 | -0.115107029 | 0.627084424 | 0.92 |
| YHR127W | 498.1945386 | -0.190799423 | 0.627511351 | 0.92 |
| YOR152C | 552.1803625 | -0.177787179 | 0.62759199 | 0.92 |
| YER037W | 745.1407721 | -0.246751888 | 0.628785775 | 0.93 |
| YFR007W | 462.3779448 | -0.1522015 | 0.62943451 | 0.93 |
| YKR018C | 3562.456922 | -0.163163864 | 0.629486924 | 0.93 |
| YMR180C | 510.7680392 | -0.116656677 | 0.628981049 | 0.93 |
| YOL083W | 724.7720388 | -0.200152501 | 0.628564412 | 0.93 |
| YOR283W | 529.1178643 | -0.133231561 | 0.62916898 | 0.93 |
| YPR146C | 99.73914934 | -0.127722785 | 0.629430591 | 0.93 |
| YBR119W | 605.7970902 | 0.161515398 | 0.629936777 | 0.93 |
| YFR045W | 698.908615 | -0.148228143 | 0.630393481 | 0.93 |
| YBL068W | 791.3128306 | -0.129272676 | 0.630798672 | 0.93 |
| YPL159C | 943.1509361 | -0.147903264 | 0.63075566 | 0.93 |

| | | | | |
|---|---|---|---|---|
| YBL045C | 7514.298009 | -0.209065305 | 0.63097057 | 0.93 |
| YLR020C | 866.2619741 | 0.124713976 | 0.631145421 | 0.93 |
| YDL035C | 1472.135402 | 0.118200108 | 0.631748824 | 0.93 |
| YJR154W | 284.604362 | 0.182884495 | 0.632031182 | 0.93 |
| YLR240W | 1128.041716 | 0.101364906 | 0.631894769 | 0.93 |
| YER017C | 1876.346183 | -0.086318892 | 0.632223272 | 0.93 |
| YBL024W | 2012.558866 | 0.159364109 | 0.632751641 | 0.93 |
| YBL031W | 539.4437355 | -0.148155219 | 0.636006489 | 0.93 |
| YDR170W-A | 128.0923816 | 0.24941849 | 0.633647263 | 0.93 |
| YDR213W | 568.9307813 | 0.15513123 | 0.635485401 | 0.93 |
| YGL013C | 1490.03588 | 0.099544167 | 0.635926651 | 0.93 |
| YGL032C | 1976.542723 | -0.268910703 | 0.63565008 | 0.93 |
| YGL059W | 1140.532396 | 0.146469708 | 0.635786898 | 0.93 |
| YGL147C | 5508.177604 | 0.128774533 | 0.634111064 | 0.93 |
| YGL157W | 1718.848717 | -0.256175424 | 0.634572236 | 0.93 |
| YGL211W | 636.0371021 | -0.086414704 | 0.633241832 | 0.93 |
| YIL123W | 4293.520238 | 0.163682478 | 0.635974628 | 0.93 |
| YKL206C | 563.6605725 | -0.113586314 | 0.634778072 | 0.93 |
| YLL034C | 960.4010793 | -0.196557153 | 0.634924447 | 0.93 |
| YLR120C | 2959.418798 | 0.194964521 | 0.634863837 | 0.93 |
| YLR127C | 955.0961502 | -0.108777088 | 0.63359757 | 0.93 |
| YMR227C | 799.7644018 | -0.136304969 | 0.633146815 | 0.93 |
| YMR291W | 2090.324495 | 0.225720801 | 0.635001466 | 0.93 |
| YNL286W | 504.2093629 | 0.142237541 | 0.633820792 | 0.93 |
| YOL125W | 710.1287675 | 0.128741198 | 0.633941422 | 0.93 |
| YPL023C | 1340.132366 | -0.089632776 | 0.634561381 | 0.93 |
| YPR155C | 883.1515984 | 0.146392361 | 0.634242323 | 0.93 |
| YIL121W | 2357.398488 | 0.223721295 | 0.636482754 | 0.93 |
| YLR297W | 487.8704755 | 0.166594877 | 0.636875111 | 0.93 |
| YPR105C | 1243.404818 | -0.090160264 | 0.636827286 | 0.93 |
| YPR049C | 968.2983267 | 0.14706671 | 0.637060594 | 0.93 |
| YPR137W | 608.8911311 | 0.169871882 | 0.637252738 | 0.93 |
| YMR155W | 707.9678984 | 0.130308636 | 0.637459837 | 0.93 |
| YBL100W-B | 175.5232464 | -0.227899709 | 0.638801479 | 0.93 |
| YBR215W | 1081.67593 | 0.078954329 | 0.638507484 | 0.93 |
| YDR499W | 868.5785284 | -0.15620719 | 0.639008719 | 0.93 |
| YER081W | 9451.152045 | 0.22212821 | 0.638554949 | 0.93 |
| YGL014W | 2156.619039 | -0.160123185 | 0.639033456 | 0.93 |
| YKL204W | 2110.527656 | -0.10762176 | 0.638653702 | 0.93 |
| YNL336W | 467.9927857 | -0.193068486 | 0.638225508 | 0.93 |
| YNL008C | 1006.822611 | -0.118575256 | 0.639307096 | 0.93 |
| YCL039W | 1768.904925 | -0.149721097 | 0.640200466 | 0.93 |
| YDR169C | 1729.231302 | 0.125837305 | 0.64062436 | 0.93 |

| | | | | |
|---|---|---|---|---|
| YJR111C | 500.5453341 | 0.148886632 | 0.639794253 | 0.93 |
| YKL112W | 917.1897974 | 0.12712043 | 0.640616991 | 0.93 |
| YOR244W | 816.0935801 | 0.107248739 | 0.640636816 | 0.93 |
| YPR086W | 1622.357285 | -0.119591685 | 0.640326277 | 0.93 |
| YCL037C | 1123.648124 | 0.196291332 | 0.64154972 | 0.93 |
| YHR154W | 1331.473724 | -0.160433826 | 0.641549 | 0.93 |
| YJL063C | 928.728548 | -0.137367383 | 0.640914256 | 0.93 |
| YLR088W | 986.2933263 | 0.081312397 | 0.641229182 | 0.93 |
| YNL069C | 13320.02635 | 0.180688532 | 0.641488321 | 0.93 |
| YLL019C | 1900.382246 | -0.190958783 | 0.641720384 | 0.93 |
| YDR494W | 1564.77137 | -0.103155363 | 0.642526809 | 0.93 |
| YJL206C | 539.1174838 | 0.128172098 | 0.642362722 | 0.93 |
| YLR251W | 493.6892026 | -0.151392203 | 0.642260857 | 0.93 |
| YMR086W | 2080.250939 | 0.095753264 | 0.643072664 | 0.93 |
| YGL044C | 659.773802 | 0.064931411 | 0.643422935 | 0.93 |
| YMR310C | 757.4056331 | 0.158404494 | 0.643684212 | 0.93 |
| YJL011C | 110.249814 | -0.219332532 | 0.644034997 | 0.93 |
| YNL083W | 670.395816 | -0.15531976 | 0.644346293 | 0.93 |
| YDR354W | 871.1284672 | -0.211742056 | 0.645025723 | 0.93 |
| YKR078W | 765.3811944 | 0.111127096 | 0.644902637 | 0.93 |
| YLR100W | 1453.024103 | -0.155303388 | 0.644869704 | 0.93 |
| YOL063C | 983.1121387 | 0.094286878 | 0.645098802 | 0.93 |
| YJL112W | 1088.839683 | 0.120794019 | 0.645279589 | 0.93 |
| YNL100W | 1262.153408 | -0.166145407 | 0.645454321 | 0.93 |
| YKR087C | 773.1699867 | 0.099220833 | 0.645639555 | 0.93 |
| YBR065C | 528.4307138 | -0.137564376 | 0.650138556 | 0.93 |
| YBR112C | 3373.883429 | 0.117647316 | 0.649933816 | 0.93 |
| YBR201C-A | 201.3285883 | -0.217587755 | 0.65012443 | 0.93 |
| YDL241W | 250.4921648 | -0.213406046 | 0.648953167 | 0.93 |
| YDL243C | 368.4938618 | -0.224803888 | 0.647777962 | 0.93 |
| YDR119W-A | 458.2326718 | 0.203494651 | 0.648804845 | 0.93 |
| YDR125C | 356.5508843 | 0.174195438 | 0.649917595 | 0.93 |
| YDR308C | 370.9057891 | -0.138423378 | 0.649046841 | 0.93 |
| YDR335W | 3040.69958 | -0.110723231 | 0.649838642 | 0.93 |
| YEL052W | 1984.785259 | 0.13335534 | 0.64982806 | 0.93 |
| YFR033C | 3330.696407 | -0.220089589 | 0.648049627 | 0.93 |
| YGL095C | 1579.460918 | 0.07371049 | 0.650224077 | 0.93 |
| YGL225W | 1789.038412 | -0.221768397 | 0.648422549 | 0.93 |
| YGR053C | 339.3237027 | -0.148102138 | 0.649264127 | 0.93 |
| YGR131W | 100.6509022 | -0.184485997 | 0.650136414 | 0.93 |
| YGR166W | 818.3285643 | 0.113463243 | 0.647831598 | 0.93 |
| YGR279C | 11712.04252 | -0.101527895 | 0.648974519 | 0.93 |
| YLR069C | 1475.235001 | -0.115022182 | 0.646013367 | 0.93 |

| | | | | |
|---|---|---|---|---|
| YLR426W | 420.438408 | 0.116055544 | 0.647615563 | 0.93 |
| YML066C | 127.9180195 | 0.224847414 | 0.646450439 | 0.93 |
| YMR164C | 711.3389073 | 0.098172718 | 0.648918347 | 0.93 |
| YMR196W | 3241.463398 | -0.196463388 | 0.647508428 | 0.93 |
| YMR210W | 1188.233193 | 0.09523046 | 0.64663514 | 0.93 |
| YOR297C | 445.9780063 | 0.09873872 | 0.649903367 | 0.93 |
| YPL020C | 1208.520388 | -0.067918957 | 0.647235359 | 0.93 |
| YPL066W | 1565.740738 | 0.08494253 | 0.650199165 | 0.93 |
| YPR066W | 302.1036782 | -0.138370721 | 0.647047729 | 0.93 |
| YHR031C | 752.7001486 | 0.142701001 | 0.651059009 | 0.93 |
| YJL054W | 1689.096692 | -0.112583904 | 0.651140715 | 0.93 |
| YKR089C | 3634.661722 | 0.115107328 | 0.651355861 | 0.93 |
| YML035C | 2451.342109 | 0.130562114 | 0.651364064 | 0.93 |
| YNL260C | 268.8863879 | 0.085724753 | 0.651594037 | 0.93 |
| YOR311C | 1031.860298 | -0.11135771 | 0.651486813 | 0.93 |
| YBL082C | 673.6132807 | -0.118290256 | 0.651921842 | 0.93 |
| YJL041W | 2583.756357 | -0.079801917 | 0.652053904 | 0.93 |
| LSR1 | 186.1998782 | -0.197609106 | 0.652504109 | 0.93 |
| YDL070W | 4583.373904 | -0.139216787 | 0.652912838 | 0.93 |
| YDL204W | 5424.611742 | 0.213105152 | 0.652648227 | 0.93 |
| YLR141W | 304.974896 | -0.122588204 | 0.652825338 | 0.93 |
| YNL195C | 538.1405229 | 0.193101837 | 0.652707014 | 0.93 |
| YPR024W | 1909.485964 | 0.090081997 | 0.653296673 | 0.93 |
| ICR1 | 346.589135 | 0.137770765 | 0.657568237 | 0.93 |
| YBR298C | 1312.084765 | -0.215571939 | 0.655336699 | 0.93 |
| YDL126C | 11516.69474 | -0.094486728 | 0.657389946 | 0.93 |
| YDR298C | 3635.360191 | -0.197508309 | 0.655202697 | 0.93 |
| YEL022W | 2372.854744 | 0.129521187 | 0.656915653 | 0.93 |
| YGL203C | 1571.604512 | 0.11130957 | 0.656654224 | 0.93 |
| YGL254W | 331.7870674 | 0.130647506 | 0.65712123 | 0.93 |
| YGR111W | 1711.371366 | -0.123774429 | 0.658743009 | 0.93 |
| YGR159C | 2044.642627 | 0.223772883 | 0.654391995 | 0.93 |
| YHL013C | 503.0447213 | -0.138385726 | 0.658904389 | 0.93 |
| YJR033C | 1583.049905 | 0.092058487 | 0.657359589 | 0.93 |
| YJR070C | 1785.660344 | 0.162430257 | 0.657345917 | 0.93 |
| YJR161C | 124.4036715 | -0.174641123 | 0.65767021 | 0.93 |
| YKL009W | 1409.022835 | -0.210296662 | 0.656003916 | 0.93 |
| YLR420W | 2792.570329 | -0.169926244 | 0.654701211 | 0.93 |
| YML046W | 728.0597624 | 0.112716045 | 0.654318188 | 0.93 |
| YML058W-A | 99.58508334 | -0.229520009 | 0.656902479 | 0.93 |
| YMR025W | 229.9295307 | -0.156696633 | 0.657611567 | 0.93 |
| YMR070W | 1232.669923 | 0.154232888 | 0.658001337 | 0.93 |
| YMR077C | 308.7109529 | -0.108651068 | 0.658500227 | 0.93 |

| | | | | |
|---|---|---|---|---|
| YMR127C | 400.3588714 | 0.111417162 | 0.656348935 | 0.93 |
| YNL074C | 2355.063484 | -0.110273491 | 0.655462678 | 0.93 |
| YNL304W | 349.9176759 | -0.097182095 | 0.65794985 | 0.93 |
| YNR007C | 864.4677932 | -0.18329439 | 0.653938018 | 0.93 |
| YOL080C | 199.8794327 | 0.208536795 | 0.656960795 | 0.93 |
| YOR062C | 585.4618256 | 0.189428504 | 0.65810445 | 0.93 |
| YOR247W | 5496.630424 | 0.187820269 | 0.655887409 | 0.93 |
| YOR307C | 1226.108407 | -0.103148632 | 0.658838093 | 0.93 |
| YPL090C | 2392.913651 | -0.262609678 | 0.654042012 | 0.93 |
| YPL095C | 828.2325169 | 0.126549029 | 0.658706629 | 0.93 |
| YPL171C | 308.1636844 | 0.217179502 | 0.658605427 | 0.93 |
| YBL008W | 564.4162656 | 0.122418802 | 0.659472017 | 0.93 |
| YDR117C | 1080.281382 | 0.073275079 | 0.660839078 | 0.93 |
| YDR184C | 548.4562077 | 0.166347191 | 0.659853711 | 0.93 |
| YER161C | 1183.000093 | 0.098424319 | 0.660164005 | 0.93 |
| YER186C | 678.5275475 | 0.129793644 | 0.659874999 | 0.93 |
| YGL135W | 2224.916712 | 0.140428115 | 0.660872648 | 0.93 |
| YLR183C | 714.9768804 | -0.154539329 | 0.660950702 | 0.93 |
| YLR387C | 1423.423086 | -0.133997934 | 0.660150151 | 0.93 |
| YNR050C | 3301.317883 | -0.194696037 | 0.660474449 | 0.93 |
| YOR241W | 933.790879 | 0.108339538 | 0.660879372 | 0.93 |
| YPL111W | 2984.765722 | 0.119179296 | 0.660074302 | 0.93 |
| YPR004C | 1678.278628 | -0.146096721 | 0.659559505 | 0.93 |
| YCL010C | 597.9803619 | 0.126146911 | 0.661280456 | 0.93 |
| YOR375C | 17984.54886 | -0.207614367 | 0.661308901 | 0.93 |
| YHR078W | 1137.159604 | 0.137555915 | 0.66150212 | 0.93 |
| YNL284C-B | 190.6577576 | 0.203456647 | 0.661882601 | 0.93 |
| YHR170W | 1876.524363 | 0.184431507 | 0.662071016 | 0.93 |
| YLR085C | 664.5290334 | -0.069187937 | 0.662250114 | 0.93 |
| YBR128C | 336.3131952 | -0.148929177 | 0.6626562 | 0.93 |
| YAL032C | 524.818116 | 0.07777793 | 0.662888819 | 0.93 |
| YIR002C | 1205.196083 | 0.121748076 | 0.663196091 | 0.93 |
| YDR134C | 8310.682574 | -0.162417907 | 0.663653861 | 0.93 |
| YMR049C | 1525.516535 | 0.189734339 | 0.663783636 | 0.93 |
| YBL015W | 5485.33881 | -0.198703473 | 0.666668683 | 0.93 |
| YDL222C | 906.6701758 | 0.233385127 | 0.664770863 | 0.93 |
| YDR160W | 879.6911624 | 0.06850228 | 0.666522928 | 0.93 |
| YDR333C | 1359.98522 | 0.078155463 | 0.666127211 | 0.93 |
| YDR440W | 548.6066441 | 0.11583643 | 0.666848279 | 0.93 |
| YDR540C | 110.2480797 | -0.202552706 | 0.665142308 | 0.93 |
| YER107C | 624.5620743 | 0.126246711 | 0.665607694 | 0.93 |
| YFR018C | 1513.157132 | -0.120870111 | 0.665609905 | 0.93 |
| YGL142C | 1131.213732 | 0.112881148 | 0.664838709 | 0.93 |

| | | | | |
|---|---|---|---|---|
| YIL097W | 1123.344615 | -0.171364664 | 0.664259125 | 0.93 |
| YIL116W | 2678.75766 | -0.154723723 | 0.666010176 | 0.93 |
| YLR276C | 2051.953288 | 0.159725865 | 0.666869459 | 0.93 |
| YMR026C | 666.726482 | 0.089037759 | 0.666823008 | 0.93 |
| YNL207W | 833.7029402 | 0.121375302 | 0.666171821 | 0.93 |
| YNR069C | 436.7652166 | -0.171716047 | 0.666066591 | 0.93 |
| YOL002C | 621.7558878 | -0.144337768 | 0.66488741 | 0.93 |
| YOL042W | 569.9826336 | 0.116272336 | 0.666914449 | 0.93 |
| YOR316C | 1558.064909 | -0.129633492 | 0.665350117 | 0.93 |
| YBR179C | 1264.938407 | 0.101038395 | 0.667217343 | 0.93 |
| YKR095W | 4316.14506 | 0.09356745 | 0.667443805 | 0.93 |
| YJR155W | 885.0501737 | 0.156391376 | 0.667773591 | 0.93 |
| YDL195W | 7573.167842 | 0.108375786 | 0.668342792 | 0.93 |
| YKL064W | 2390.234747 | 0.098448726 | 0.668265795 | 0.93 |
| YKR056W | 1143.695694 | 0.13859669 | 0.668184657 | 0.93 |
| YGR096W | 275.6515308 | 0.124296059 | 0.668614135 | 0.94 |
| YGR201C | 974.6666707 | -0.220826168 | 0.668884637 | 0.94 |
| YJL122W | 643.9700151 | 0.200573491 | 0.66880504 | 0.94 |
| YDR238C | 4318.542325 | -0.084956719 | 0.669964381 | 0.94 |
| YGL120C | 2147.315673 | -0.178618696 | 0.670029195 | 0.94 |
| YHR048W | 458.0188382 | 0.16880984 | 0.669681153 | 0.94 |
| YIR011C | 1166.213259 | 0.09561965 | 0.670053377 | 0.94 |
| YJR035W | 878.6193365 | 0.111000363 | 0.670252502 | 0.94 |
| YDR204W | 1297.475345 | -0.162624887 | 0.670612155 | 0.94 |
| YPR003C | 637.4553595 | 0.124007675 | 0.670462493 | 0.94 |
| YLL043W | 1838.000712 | -0.094406441 | 0.671119117 | 0.94 |
| YKL151C | 2678.459393 | -0.198993608 | 0.671631883 | 0.94 |
| YMR097C | 670.8866515 | -0.084859638 | 0.671504695 | 0.94 |
| YNL063W | 522.2571621 | 0.080178007 | 0.671554573 | 0.94 |
| YGR065C | 698.7091847 | 0.135233459 | 0.671874842 | 0.94 |
| YGR055W | 1342.856307 | 0.139825288 | 0.672362538 | 0.94 |
| YPR008W | 1796.059535 | 0.089084184 | 0.672433202 | 0.94 |
| YJL102W | 772.7229577 | 0.123122874 | 0.672710873 | 0.94 |
| YHR028C | 2661.078301 | 0.099998632 | 0.673127089 | 0.94 |
| YPR125W | 636.6611003 | -0.108953576 | 0.672998246 | 0.94 |
| YDL085W | 757.5796414 | 0.167800752 | 0.674553083 | 0.94 |
| YDL101C | 860.843029 | -0.113645497 | 0.674970996 | 0.94 |
| YDL140C | 8072.034708 | 0.112625426 | 0.674647953 | 0.94 |
| YGL005C | 924.1279481 | -0.098687313 | 0.674876539 | 0.94 |
| YHR209W | 694.2958547 | 0.165983965 | 0.675213867 | 0.94 |
| YKL188C | 680.4138758 | 0.136538177 | 0.674311296 | 0.94 |
| YML088W | 784.353378 | 0.113654807 | 0.673514234 | 0.94 |
| YOL092W | 1398.16665 | -0.130954461 | 0.673980113 | 0.94 |

| | | | | |
|---|---|---|---|---|
| YOL115W | 646.4767705 | 0.116871824 | 0.674438803 | 0.94 |
| YPL003W | 547.4702758 | 0.120503989 | 0.675057699 | 0.94 |
| YPL085W | 4050.180373 | 0.109328745 | 0.67507086 | 0.94 |
| YPL142C | 228.9766843 | 0.147718549 | 0.675548865 | 0.94 |
| YBL007C | 3937.826759 | 0.08107136 | 0.679114917 | 0.94 |
| YCR075C | 329.3656461 | -0.084116908 | 0.67833792 | 0.94 |
| YDR145W | 1845.179932 | -0.107016273 | 0.679158521 | 0.94 |
| YDR293C | 6400.531719 | 0.112342621 | 0.679123017 | 0.94 |
| YER056C-A | 4887.303568 | 0.155831799 | 0.677611346 | 0.94 |
| YER136W | 4119.832209 | -0.06598775 | 0.678452208 | 0.94 |
| YGL022W | 3864.561612 | -0.105008868 | 0.678751655 | 0.94 |
| YGL053W | 527.1405514 | 0.152363076 | 0.678628058 | 0.94 |
| YGL232W | 728.6792639 | -0.14427136 | 0.679130832 | 0.94 |
| YGR169C | 437.5350343 | 0.125472302 | 0.679210581 | 0.94 |
| YHL047C | 1687.543568 | -0.198666947 | 0.678830434 | 0.94 |
| YJL170C | 341.173457 | 0.228447421 | 0.678544805 | 0.94 |
| YKL126W | 2370.590453 | -0.082750536 | 0.678383316 | 0.94 |
| YKR046C | 2535.087216 | 0.131464295 | 0.676897112 | 0.94 |
| YLL049W | 333.5985502 | -0.126242459 | 0.678444058 | 0.94 |
| YMR029C | 1228.193843 | -0.101305327 | 0.678067334 | 0.94 |
| YMR171C | 1518.290743 | 0.165649207 | 0.678546446 | 0.94 |
| YNL193W | 690.484771 | -0.162957758 | 0.678599265 | 0.94 |
| YOR312C | 9424.205996 | 0.143622304 | 0.678466246 | 0.94 |
| YOR384W | 1087.084219 | 0.139238626 | 0.678740284 | 0.94 |
| YFR034C | 832.1624053 | 0.110318403 | 0.679603588 | 0.94 |
| YBL076C | 7280.098127 | -0.156562878 | 0.680473094 | 0.94 |
| YDR522C | 159.2048735 | 0.138727747 | 0.680865769 | 0.94 |
| YER179W | 293.0579118 | 0.161014273 | 0.680817868 | 0.94 |
| YOR310C | 3221.373678 | -0.188179408 | 0.680668662 | 0.94 |
| SRG1 | 1084.659985 | -0.218867955 | 0.681616077 | 0.94 |
| YBL074C | 232.3789197 | 0.117910989 | 0.681688709 | 0.94 |
| YGL178W | 2073.150568 | 0.106689515 | 0.681755346 | 0.94 |
| YMR270C | 708.6444493 | -0.087430453 | 0.681414999 | 0.94 |
| YOL094C | 437.8167325 | -0.125751243 | 0.681830783 | 0.94 |
| YBR265W | 1130.650771 | -0.085994861 | 0.684104342 | 0.94 |
| YCR023C | 1565.302118 | 0.117755644 | 0.683529438 | 0.94 |
| YDL142C | 615.3529618 | -0.103189328 | 0.684055 | 0.94 |
| YER089C | 1927.936186 | 0.142634438 | 0.682953296 | 0.94 |
| YGL036W | 2417.923989 | 0.080405666 | 0.682795952 | 0.94 |
| YGL039W | 719.9626894 | -0.133948466 | 0.684222428 | 0.94 |
| YHR142W | 1336.189865 | 0.133972927 | 0.682918302 | 0.94 |
| YKL048C | 583.1360838 | -0.138813519 | 0.683217968 | 0.94 |
| YLR186W | 830.8214037 | 0.147064029 | 0.683852944 | 0.94 |

| | | | | |
|---|---|---|---|---|
| YLR438W | 1335.790008 | -0.155505165 | 0.684201392 | 0.94 |
| YMR170C | 596.7981642 | -0.175520251 | 0.682673543 | 0.94 |
| YOL056W | 561.7509431 | -0.152265904 | 0.683976805 | 0.94 |
| YGL238W | 2547.939909 | 0.079327893 | 0.684569675 | 0.94 |
| YHR096C | 1194.376628 | 0.176716256 | 0.684624812 | 0.94 |
| YJR049C | 1397.771565 | 0.109540641 | 0.684991559 | 0.94 |
| YKR071C | 2094.595777 | -0.103932133 | 0.68543697 | 0.94 |
| YLR249W | 32716.59454 | 0.170197917 | 0.685309801 | 0.94 |
| YOR212W | 1199.500024 | 0.209997207 | 0.685372233 | 0.94 |
| YOR381W | 508.0002714 | 0.170464556 | 0.684965773 | 0.94 |
| YDL216C | 462.7023205 | 0.132424085 | 0.686186312 | 0.94 |
| YFR009W | 2637.309837 | 0.114573277 | 0.685719165 | 0.94 |
| YHR099W | 4737.26123 | 0.120117256 | 0.685983189 | 0.94 |
| YKL071W | 436.0298233 | 0.08756931 | 0.686038617 | 0.94 |
| YPR044C | 114.0311076 | 0.164884884 | 0.686347687 | 0.94 |
| YBR243C | 997.8241569 | -0.118482598 | 0.68706289 | 0.94 |
| YPR093C | 386.3642488 | 0.176883972 | 0.68729076 | 0.94 |
| YAL014C | 833.2543927 | -0.110313998 | 0.687546319 | 0.94 |
| YBR169C | 4813.961053 | 0.164530527 | 0.687742591 | 0.94 |
| YBR284W | 530.3934343 | 0.131945923 | 0.688848021 | 0.94 |
| YPL270W | 1768.066094 | 0.118736368 | 0.688895635 | 0.94 |
| YJL207C | 2307.188321 | -0.11383823 | 0.689682996 | 0.94 |
| YMR233W | 360.9564304 | 0.132524188 | 0.689828945 | 0.94 |
| YNR057C | 196.7239761 | 0.119122776 | 0.689780326 | 0.94 |
| YOR294W | 771.7312013 | -0.159902402 | 0.690097189 | 0.94 |
| YPL229W | 514.8866138 | -0.128015987 | 0.690543658 | 0.94 |
| YDR450W | 8491.535949 | -0.223436942 | 0.690973462 | 0.94 |
| YGR141W | 841.9095851 | 0.151990165 | 0.690978899 | 0.94 |
| YDR264C | 2392.197288 | 0.083295056 | 0.691229618 | 0.94 |
| YDL081C | 10855.42483 | -0.224264203 | 0.692027005 | 0.94 |
| YIL029C | 174.946717 | -0.148207554 | 0.692524981 | 0.94 |
| YIL066C | 409.4135175 | 0.162756233 | 0.691700416 | 0.94 |
| YLR044C | 136554.4679 | -0.203477876 | 0.692330676 | 0.94 |
| YMR102C | 1653.341009 | -0.174724546 | 0.692597957 | 0.94 |
| YNL180C | 2386.54101 | -0.147626343 | 0.692336829 | 0.94 |
| YNL196C | 204.2397635 | -0.220895813 | 0.692689275 | 0.94 |
| YPL196W | 943.9259177 | -0.154208451 | 0.692896446 | 0.94 |
| YNR046W | 947.7260883 | -0.152831478 | 0.693413027 | 0.94 |
| YLL044W | 94.44590477 | 0.291779985 | 0.693645021 | 0.94 |
| YER036C | 4277.526152 | 0.133451818 | 0.693956669 | 0.94 |
| YGR144W | 97.38216331 | 0.165738615 | 0.694184874 | 0.94 |
| YER013W | 829.6032387 | -0.084348099 | 0.694619589 | 0.94 |
| YLR433C | 1164.793625 | -0.110628593 | 0.69493709 | 0.94 |

| | | | | |
|---|---|---|---|---|
| YNL320W | 323.6388355 | -0.11842716 | 0.694734193 | 0.94 |
| YPL069C | 455.5840403 | -0.100884229 | 0.694957989 | 0.94 |
| YDR305C | 387.9870014 | -0.126039236 | 0.695759975 | 0.94 |
| YPR115W | 2825.752527 | 0.127729268 | 0.695595505 | 0.94 |
| YBL060W | 624.3164774 | 0.087014011 | 0.696482134 | 0.95 |
| YJL146W | 600.4909491 | 0.06386203 | 0.696557558 | 0.95 |
| YLR138W | 1645.971988 | 0.111862674 | 0.696358084 | 0.95 |
| YIR025W | 303.8110514 | 0.075446002 | 0.696939587 | 0.95 |
| YKR092C | 1945.135383 | -0.146527833 | 0.697141274 | 0.95 |
| YHR041C | 584.7828437 | 0.123454138 | 0.69767437 | 0.95 |
| YJR055W | 115.4214659 | 0.097752608 | 0.69768299 | 0.95 |
| YNR023W | 727.2388585 | 0.076785832 | 0.69747306 | 0.95 |
| YDR211W | 2365.993285 | 0.107971518 | 0.698146927 | 0.95 |
| YHL038C | 821.5884324 | 0.077366431 | 0.69871464 | 0.95 |
| YHR188C | 2845.286709 | -0.106742868 | 0.698032566 | 0.95 |
| YIL007C | 541.263808 | -0.114284497 | 0.698945755 | 0.95 |
| YJR131W | 581.4205166 | 0.102365917 | 0.699151139 | 0.95 |
| YKL201C | 1146.760044 | -0.115210453 | 0.698888317 | 0.95 |
| YML053C | 803.1994724 | -0.106883351 | 0.699035649 | 0.95 |
| YOR165W | 1637.432086 | -0.089802033 | 0.698499288 | 0.95 |
| YBR084C-A | 8707.666036 | -0.212844019 | 0.70091517 | 0.95 |
| YBR150C | 848.9681448 | -0.09424615 | 0.700714705 | 0.95 |
| YCL029C | 771.878883 | 0.073707272 | 0.700971403 | 0.95 |
| YDR332W | 604.027567 | 0.070562845 | 0.701413885 | 0.95 |
| YDR502C | 1141.466361 | -0.126127632 | 0.701172129 | 0.95 |
| YKL062W | 2465.244192 | 0.141051454 | 0.701574685 | 0.95 |
| YLR003C | 782.5858822 | -0.15616013 | 0.70142395 | 0.95 |
| YLR110C | 106387.3494 | -0.182360236 | 0.700585416 | 0.95 |
| YLR364W | 131.7044529 | -0.124323039 | 0.700234782 | 0.95 |
| YNR054C | 724.980203 | -0.144044904 | 0.700532703 | 0.95 |
| YOR091W | 1558.309002 | -0.123698395 | 0.700816999 | 0.95 |
| YPL035C | 100.9354993 | -0.109837815 | 0.700493516 | 0.95 |
| YPL047W | 255.7983457 | 0.084105698 | 0.699509898 | 0.95 |
| YHR156C | 433.8648918 | 0.103526538 | 0.701842974 | 0.95 |
| YBR046C | 1727.421703 | -0.137421332 | 0.702827852 | 0.95 |
| YBR233W-A | 253.0169158 | -0.139487937 | 0.70216856 | 0.95 |
| YER131W | 9395.024759 | 0.139338506 | 0.702473232 | 0.95 |
| YHR067W | 339.2689826 | 0.142194048 | 0.702674795 | 0.95 |
| YHR199C-A | 146.5882374 | 0.162887602 | 0.702654414 | 0.95 |
| YDL131W | 1846.278027 | 0.112710875 | 0.703202392 | 0.95 |
| YDR539W | 882.7410932 | -0.169092533 | 0.704170229 | 0.95 |
| YBL025W | 212.6114748 | 0.118549479 | 0.704692618 | 0.95 |
| YBL084C | 817.2451769 | 0.100096715 | 0.704921808 | 0.95 |

| | | | | |
|---|---|---|---|---|
| YMR031C | 3707.880046 | -0.138518971 | 0.705165471 | 0.95 |
| YHR046C | 620.1635411 | 0.129178593 | 0.705341792 | 0.95 |
| YOR228C | 505.0466268 | -0.13964619 | 0.70559122 | 0.95 |
| YGL175C | 162.4549988 | 0.082912839 | 0.706315103 | 0.95 |
| YOR147W | 1293.879256 | -0.143638182 | 0.706317879 | 0.95 |
| YGR056W | 1165.037029 | 0.097524864 | 0.706537268 | 0.95 |
| YDR101C | 1247.120699 | 0.181100763 | 0.707263703 | 0.95 |
| YIL046W | 1246.958862 | 0.106421641 | 0.706923666 | 0.95 |
| YKL213C | 1964.500982 | 0.06105422 | 0.707025597 | 0.95 |
| YNL194C | 937.042565 | 0.188901766 | 0.707192775 | 0.95 |
| YOR084W | 822.8839971 | 0.099790671 | 0.707503533 | 0.95 |
| YHL015W | 17106.50707 | -0.213675454 | 0.707757388 | 0.95 |
| YLR291C | 1399.72151 | -0.068681964 | 0.707893588 | 0.95 |
| YHR208W | 7460.933205 | -0.15299141 | 0.70826941 | 0.95 |
| YPL088W | 468.228703 | 0.144452616 | 0.70837045 | 0.95 |
| YAL055W | 312.0534701 | -0.110613983 | 0.709271567 | 0.95 |
| YDR498C | 1014.743074 | 0.04424569 | 0.709142681 | 0.95 |
| YGR205W | 695.7192252 | -0.132403781 | 0.709373986 | 0.95 |
| YLL028W | 724.5266873 | -0.116899608 | 0.709918237 | 0.95 |
| YOR026W | 568.5531387 | 0.109605222 | 0.709974843 | 0.95 |
| YOR120W | 2918.156498 | 0.166351565 | 0.709353403 | 0.95 |
| YOR221C | 1523.763907 | -0.088809298 | 0.709355236 | 0.95 |
| YPL096C-A | 145.3313639 | 0.111190821 | 0.710006842 | 0.95 |
| YPL212C | 1188.64058 | -0.144171382 | 0.709550684 | 0.95 |
| YAL037W | 195.7634153 | 0.131280444 | 0.710406493 | 0.95 |
| YGR109W-B | 857.2496031 | 0.152517664 | 0.710662061 | 0.95 |
| YNR010W | 212.8810604 | 0.091050671 | 0.710764786 | 0.95 |
| YKL074C | 747.5415477 | 0.06665959 | 0.711194563 | 0.95 |
| YDR268W | 586.9385642 | -0.088006704 | 0.712906499 | 0.95 |
| YDR392W | 654.5457347 | 0.065141757 | 0.713276883 | 0.95 |
| YDR456W | 1671.501856 | 0.072316505 | 0.712758305 | 0.95 |
| YGL025C | 784.0082065 | 0.073044902 | 0.713301593 | 0.95 |
| YIL072W | 537.0665215 | 0.10682328 | 0.713004039 | 0.95 |
| YLR394W | 730.9999237 | 0.083384754 | 0.712987361 | 0.95 |
| YNL062C | 805.5436558 | 0.154159354 | 0.712847261 | 0.95 |
| YNL317W | 817.8553177 | 0.051109858 | 0.712466379 | 0.95 |
| YPL041C | 160.4382575 | 0.128583283 | 0.713856705 | 0.95 |
| YDL042C | 548.3117902 | 0.153409093 | 0.715012733 | 0.95 |
| YDL148C | 2147.255757 | -0.134046927 | 0.71466133 | 0.95 |
| YDR460W | 574.0402083 | -0.048991979 | 0.715122942 | 0.95 |
| YER073W | 2130.104252 | 0.152767577 | 0.714193457 | 0.95 |
| YER153C | 351.5393272 | -0.12271006 | 0.714494671 | 0.95 |
| YGL173C | 8071.132916 | 0.061060341 | 0.714598197 | 0.95 |

| | | | | |
|---|---|---|---|---|
| YKL052C | 501.3271381 | -0.081209312 | 0.714895753 | 0.95 |
| YAL026C-A | 170.5922701 | -0.104492612 | 0.715445529 | 0.95 |
| YJL070C | 972.7876692 | -0.10675846 | 0.715728703 | 0.95 |
| YKL100C | 2362.710996 | -0.09277228 | 0.715815927 | 0.95 |
| YMR316W | 898.4662527 | 0.154464559 | 0.715938035 | 0.95 |
| YLR048W | 13755.73587 | 0.134476487 | 0.716160564 | 0.95 |
| YDL017W | 421.2237274 | 0.066668811 | 0.716807482 | 0.95 |
| YDL234C | 2388.462086 | -0.125808013 | 0.716927759 | 0.95 |
| YGR204W | 10459.91176 | -0.094911567 | 0.716436927 | 0.95 |
| YNL289W | 446.6126721 | -0.178302697 | 0.716793134 | 0.95 |
| YBL075C | 2650.065607 | -0.209842544 | 0.722502378 | 0.95 |
| YCL001W | 1214.350549 | 0.104027666 | 0.720170689 | 0.95 |
| YCR018C | 861.2432726 | 0.133783183 | 0.718025235 | 0.95 |
| YCR031C | 11369.4013 | -0.19453665 | 0.723372659 | 0.95 |
| YCR106W | 394.726015 | 0.173939882 | 0.723559312 | 0.95 |
| YDL009C | 106.1081082 | -0.09140327 | 0.723470841 | 0.95 |
| YDL149W | 944.7875532 | 0.109724863 | 0.722516671 | 0.95 |
| YDR341C | 4249.5314 | -0.148095294 | 0.721499471 | 0.95 |
| YDR398W | 1282.093858 | 0.14081458 | 0.719195297 | 0.95 |
| YDR449C | 532.8954421 | 0.17586613 | 0.7186923 | 0.95 |
| YER015W | 725.609463 | 0.111289435 | 0.720979075 | 0.95 |
| YGR074W | 145.6756142 | -0.085398312 | 0.720410263 | 0.95 |
| YGR075C | 302.7071594 | 0.078555825 | 0.719924123 | 0.95 |
| YGR229C | 1916.022007 | -0.094547149 | 0.722809322 | 0.95 |
| YHR025W | 4807.328396 | -0.163415169 | 0.72306394 | 0.95 |
| YHR210C | 320.0019552 | -0.153778248 | 0.722344676 | 0.95 |
| YIL026C | 1582.807889 | -0.091128662 | 0.718962321 | 0.95 |
| YIL083C | 1169.719343 | -0.125616389 | 0.722320358 | 0.95 |
| YIL093C | 871.5653388 | -0.086767391 | 0.723255475 | 0.95 |
| YJL181W | 405.181886 | 0.078044963 | 0.717658326 | 0.95 |
| YJR016C | 6106.441237 | -0.126844168 | 0.721994669 | 0.95 |
| YJR063W | 331.564801 | -0.172397857 | 0.723516283 | 0.95 |
| YJR086W | 331.1910032 | -0.237154206 | 0.717227992 | 0.95 |
| YKL021C | 1160.236373 | -0.118543179 | 0.721632076 | 0.95 |
| YLR037C | 167.7921303 | 0.113348442 | 0.722819156 | 0.95 |
| YLR070C | 346.2836033 | -0.141411472 | 0.717830412 | 0.95 |
| YLR076C | 95.93637703 | -0.283277819 | 0.720279693 | 0.95 |
| YLR220W | 930.48531 | -0.089743885 | 0.722957166 | 0.95 |
| YLR299W | 1242.489768 | 0.104543272 | 0.721846668 | 0.95 |
| YLR359W | 9224.21068 | 0.138704801 | 0.721404409 | 0.95 |
| YLR411W | 132.197405 | -0.104237762 | 0.720200195 | 0.95 |
| YNL025C | 730.0152463 | 0.088451625 | 0.721459952 | 0.95 |
| YNL270C | 240.3573834 | 0.175447638 | 0.720105891 | 0.95 |

| | | | | |
|---|---|---|---|---|
| YOR025W | 737.1839475 | 0.092186182 | 0.723444724 | 0.95 |
| YOR089C | 2098.945136 | -0.089775944 | 0.723406855 | 0.95 |
| YOR180C | 257.2908648 | 0.1243824 | 0.719163787 | 0.95 |
| YPL241C | 283.7799123 | 0.114855305 | 0.720944942 | 0.95 |
| YPR009W | 346.2208437 | -0.166892 | 0.723506182 | 0.95 |
| YBL069W | 640.7564823 | 0.113769287 | 0.726520813 | 0.95 |
| YBR189W | 11760.64407 | 0.083011138 | 0.724806319 | 0.95 |
| YBR280C | 1033.06933 | -0.14891519 | 0.727178928 | 0.95 |
| YCR026C | 1309.803177 | 0.096133947 | 0.72592 | 0.95 |
| YDL083C | 6219.514303 | 0.066744355 | 0.724709107 | 0.95 |
| YDR004W | 628.5806399 | 0.060864763 | 0.727416499 | 0.95 |
| YEL073C | 122.6223904 | 0.176753645 | 0.727123104 | 0.95 |
| YFR040W | 3140.971961 | 0.124746626 | 0.7245454 | 0.95 |
| YHL026C | 386.086378 | 0.114851465 | 0.724338664 | 0.95 |
| YIL004C | 332.265247 | -0.060094098 | 0.726303975 | 0.95 |
| YIL071C | 381.3119395 | 0.066278243 | 0.727474348 | 0.95 |
| YJL111W | 2828.454028 | -0.111239193 | 0.727032746 | 0.95 |
| YJR050W | 528.5017956 | -0.092759929 | 0.726873745 | 0.95 |
| YLL001W | 1443.400677 | 0.089059502 | 0.726614159 | 0.95 |
| YLR021W | 410.9321978 | -0.067462661 | 0.726308715 | 0.95 |
| YLR136C | 655.8429852 | -0.194442443 | 0.727219926 | 0.95 |
| YMR090W | 1879.223335 | -0.169376505 | 0.724229331 | 0.95 |
| YMR242C | 8203.831599 | 0.110709681 | 0.727469066 | 0.95 |
| YNL132W | 1681.161762 | 0.140991229 | 0.727507286 | 0.95 |
| YNL232W | 718.0441816 | -0.081314113 | 0.726895624 | 0.95 |
| YOL076W | 856.9107843 | 0.120139798 | 0.724929644 | 0.95 |
| YOR113W | 1039.201076 | -0.12830091 | 0.72471013 | 0.95 |
| YBR007C | 740.5673067 | 0.104627489 | 0.730555692 | 0.95 |
| YCR005C | 6272.799933 | 0.158282023 | 0.732057558 | 0.95 |
| YDL119C | 290.4231748 | -0.080461363 | 0.728934077 | 0.95 |
| YDR280W | 709.9216811 | 0.100469662 | 0.728394197 | 0.95 |
| YDR314C | 384.5141242 | 0.11536124 | 0.731169258 | 0.95 |
| YDR375C | 907.9536977 | 0.058149396 | 0.731345342 | 0.95 |
| YGR145W | 1225.275886 | -0.140216704 | 0.731122436 | 0.95 |
| YHR150W | 670.7977171 | -0.114986079 | 0.729753215 | 0.95 |
| YHR168W | 379.0957302 | 0.072459386 | 0.73121902 | 0.95 |
| YIR012W | 1025.35668 | -0.13791744 | 0.731713716 | 0.95 |
| YJR039W | 935.4373703 | 0.112967639 | 0.730325034 | 0.95 |
| YLL021W | 2378.696108 | -0.101240385 | 0.732672815 | 0.95 |
| YLR002C | 907.6259007 | 0.117524751 | 0.731418375 | 0.95 |
| YLR031W | 139.756576 | -0.155288488 | 0.731083346 | 0.95 |
| YLR313C | 272.7990365 | 0.106516741 | 0.729290526 | 0.95 |
| YLR372W | 2301.005993 | 0.140486819 | 0.730334729 | 0.95 |

| | | | | |
|---|---|---|---|---|
| YML019W | 694.8842248 | -0.070659659 | 0.729944411 | 0.95 |
| YMR202W | 2009.003075 | -0.194219564 | 0.732343527 | 0.95 |
| YMR206W | 572.161077 | 0.169362636 | 0.728543882 | 0.95 |
| YMR282C | 677.7599955 | 0.072069231 | 0.729711105 | 0.95 |
| YNL018C | 116.9826504 | 0.10533391 | 0.730863487 | 0.95 |
| YNL200C | 2258.279724 | -0.159835891 | 0.731835542 | 0.95 |
| YNL313C | 1104.908095 | 0.123568574 | 0.728930471 | 0.95 |
| YNR026C | 756.2847135 | 0.085886565 | 0.72880201 | 0.95 |
| YOR369C | 15497.1337 | -0.202630583 | 0.732457017 | 0.95 |
| YPL161C | 652.0526344 | 0.056663052 | 0.73266682 | 0.95 |
| YPR181C | 4442.448271 | -0.079727871 | 0.731833419 | 0.95 |
| YBR194W | 226.3319567 | 0.103363597 | 0.733065624 | 0.95 |
| YJL065C | 199.2698224 | -0.061871524 | 0.732909311 | 0.95 |
| YPL158C | 1350.795148 | 0.107774541 | 0.733320443 | 0.95 |
| YER117W | 4528.496655 | 0.070261056 | 0.733731573 | 0.95 |
| YOR070C | 1064.02202 | 0.072387165 | 0.733770554 | 0.95 |
| YMR103C | 375.8522499 | -0.138199258 | 0.734090164 | 0.95 |
| YOR130C | 372.7954083 | -0.128261215 | 0.734527734 | 0.95 |
| YPL164C | 886.5830673 | -0.09731651 | 0.734762242 | 0.95 |
| YCR107W | 236.9080811 | 0.165450568 | 0.735064375 | 0.95 |
| YNL302C | 5878.096109 | -0.219370253 | 0.735359092 | 0.95 |
| YHR133C | 977.6515666 | -0.151982133 | 0.735912003 | 0.95 |
| YLR399C | 2272.649731 | 0.073008925 | 0.735904697 | 0.95 |
| YDR475C | 1828.530627 | 0.117357446 | 0.736600204 | 0.96 |
| YGL035C | 1849.649908 | 0.104905017 | 0.737033218 | 0.96 |
| YKL113C | 707.9229549 | -0.106297774 | 0.737075632 | 0.96 |
| YDR091C | 1728.536947 | 0.138149945 | 0.737444959 | 0.96 |
| YAL044C | 2900.555908 | -0.169690678 | 0.748666832 | 0.96 |
| YBL030C | 9599.31549 | -0.154605044 | 0.739358274 | 0.96 |
| YBL034C | 918.1202298 | 0.081679522 | 0.753540755 | 0.96 |
| YBL063W | 882.9945164 | 0.075490923 | 0.745037492 | 0.96 |
| YBL092W | 15431.27049 | -0.239846245 | 0.74972428 | 0.96 |
| YBR058C | 1360.304304 | -0.04204875 | 0.751409429 | 0.96 |
| YBR078W | 17879.15973 | -0.12226144 | 0.743638797 | 0.96 |
| YBR115C | 2897.179739 | 0.107548673 | 0.74236145 | 0.96 |
| YBR136W | 1309.50165 | 0.085107136 | 0.752090973 | 0.96 |
| YCL021W-A | 277.7575158 | 0.143048026 | 0.751665299 | 0.96 |
| YCR020C | 245.5464449 | 0.125792361 | 0.748504451 | 0.96 |
| YDL053C | 1309.962591 | -0.068061441 | 0.751780977 | 0.96 |
| YDL106C | 1181.738765 | -0.092543255 | 0.741701891 | 0.96 |
| YDL191W | 1342.250708 | -0.168745789 | 0.741907967 | 0.96 |
| YDL233W | 547.8994161 | -0.082682285 | 0.742582276 | 0.96 |
| YDR057W | 882.7471612 | 0.067270334 | 0.747026936 | 0.96 |

| | | | | |
|---|---|---|---|---|
| YDR096W | 1946.105843 | -0.11601201 | 0.74656816 | 0.96 |
| YDR350C | 1056.007335 | 0.080705621 | 0.751300957 | 0.96 |
| YDR385W | 1851.31364 | 0.11511915 | 0.750842201 | 0.96 |
| YDR472W | 675.6541934 | 0.059886835 | 0.750243328 | 0.96 |
| YDR529C | 3824.487001 | -0.153238999 | 0.740224098 | 0.96 |
| YEL018W | 225.091547 | -0.057000919 | 0.753521601 | 0.96 |
| YEL020C | 1045.827861 | 0.104382913 | 0.751636114 | 0.96 |
| YEL064C | 340.0873494 | 0.089566475 | 0.750590574 | 0.96 |
| YER101C | 634.792482 | 0.096027844 | 0.748319329 | 0.96 |
| YFR039C | 736.7528646 | 0.084684827 | 0.746220346 | 0.96 |
| YGL067W | 999.1477018 | -0.081433763 | 0.739547248 | 0.96 |
| YGL193C | 193.4216907 | 0.18940016 | 0.742149113 | 0.96 |
| YGL237C | 638.8350245 | -0.096572118 | 0.751182551 | 0.96 |
| YGR121C | 1176.230558 | 0.095600691 | 0.752279212 | 0.96 |
| YGR274C | 1344.663557 | 0.089496791 | 0.737914822 | 0.96 |
| YHR003C | 820.5090626 | 0.078733118 | 0.74542032 | 0.96 |
| YHR072W | 1833.218722 | 0.109479554 | 0.743732269 | 0.96 |
| YHR161C | 2280.636973 | 0.122627565 | 0.740509699 | 0.96 |
| YHR202W | 670.4759746 | 0.0904558 | 0.753063257 | 0.96 |
| YIL005W | 1687.731259 | -0.06393273 | 0.738182185 | 0.96 |
| YIL045W | 1955.748951 | -0.163945184 | 0.741501021 | 0.96 |
| YIL099W | 500.7923991 | -0.208303476 | 0.741045578 | 0.96 |
| YIL135C | 1336.175722 | -0.089255189 | 0.74610921 | 0.96 |
| YIR007W | 898.4409432 | -0.073264499 | 0.738662963 | 0.96 |
| YJL034W | 12765.00653 | 0.122773664 | 0.749026928 | 0.96 |
| YJL042W | 3416.022292 | 0.080115983 | 0.750273957 | 0.96 |
| YJL044C | 1379.126824 | -0.078115658 | 0.747620691 | 0.96 |
| YJL162C | 484.7743599 | -0.064274969 | 0.743723224 | 0.96 |
| YJR006W | 1054.347239 | -0.073349861 | 0.747270445 | 0.96 |
| YJR043C | 619.0166524 | -0.141127631 | 0.748318421 | 0.96 |
| YKL010C | 2290.07323 | 0.050501469 | 0.738771987 | 0.96 |
| YKL032C | 3133.578964 | -0.123566087 | 0.73845457 | 0.96 |
| YKL043W | 3024.896012 | -0.157134954 | 0.753452633 | 0.96 |
| YKL088W | 1161.516822 | 0.091227644 | 0.747863178 | 0.96 |
| YKL098W | 807.0315194 | -0.07954336 | 0.751918609 | 0.96 |
| YKL144C | 415.786387 | -0.105643659 | 0.745264712 | 0.96 |
| YKL162C | 452.2195831 | -0.099941908 | 0.74924722 | 0.96 |
| YKR021W | 1328.600543 | 0.061593035 | 0.749235849 | 0.96 |
| YKR094C | 5698.150477 | 0.100465686 | 0.74512147 | 0.96 |
| YLR022C | 541.692103 | 0.086653135 | 0.750079016 | 0.96 |
| YLR074C | 1095.319455 | -0.102773998 | 0.753630461 | 0.96 |
| YLR077W | 986.4922801 | 0.049037855 | 0.750657443 | 0.96 |
| YLR104W | 635.3306236 | -0.099731457 | 0.746367979 | 0.96 |

| | | | | |
|---|---|---|---|---|
| YLR107W | 1035.729202 | 0.075525993 | 0.744725412 | 0.96 |
| YLR212C | 601.9431586 | 0.109326392 | 0.753655611 | 0.96 |
| YLR286C | 14573.14708 | 0.081951821 | 0.74502713 | 0.96 |
| YLR312C | 417.3618401 | 0.153674783 | 0.751558907 | 0.96 |
| YLR315W | 96.87354306 | -0.103880791 | 0.752287141 | 0.96 |
| YML024W | 8930.152891 | -0.20584038 | 0.752929699 | 0.96 |
| YML050W | 378.8334393 | -0.098461438 | 0.750062378 | 0.96 |
| YML091C | 3874.618724 | -0.155322161 | 0.741678999 | 0.96 |
| YML116W-A | 166.4577303 | -0.079464107 | 0.741993117 | 0.96 |
| YMR111C | 424.0046487 | -0.065782255 | 0.745877878 | 0.96 |
| YMR132C | 213.3288234 | -0.070790871 | 0.753255588 | 0.96 |
| YMR185W | 803.0326457 | 0.09549955 | 0.748972898 | 0.96 |
| YMR219W | 3001.660673 | 0.092014749 | 0.749105128 | 0.96 |
| YMR243C | 2759.15272 | 0.083010704 | 0.751662091 | 0.96 |
| YMR253C | 606.2113411 | 0.088411705 | 0.752188004 | 0.96 |
| YNL158W | 434.8285056 | 0.074945172 | 0.752778838 | 0.96 |
| YNL258C | 964.2763945 | 0.060257938 | 0.750579898 | 0.96 |
| YOL036W | 1888.124512 | 0.108012517 | 0.753577447 | 0.96 |
| YOL043C | 577.8464057 | -0.092453082 | 0.751559936 | 0.96 |
| YOR008C | 1383.478149 | 0.082279008 | 0.738158923 | 0.96 |
| YOR052C | 2749.089296 | 0.115123839 | 0.744493606 | 0.96 |
| YOR056C | 1339.644393 | 0.070643401 | 0.748874787 | 0.96 |
| YOR069W | 1035.064237 | 0.053180929 | 0.751877014 | 0.96 |
| YOR083W | 334.3141069 | -0.079535912 | 0.750971538 | 0.96 |
| YOR138C | 1254.624127 | -0.057289611 | 0.742310308 | 0.96 |
| YOR161C | 928.1721785 | -0.127776143 | 0.750792335 | 0.96 |
| YOR258W | 218.0314211 | 0.080582513 | 0.747988439 | 0.96 |
| YOR279C | 315.7050157 | 0.091551643 | 0.751047639 | 0.96 |
| YOR284W | 207.2496443 | 0.148397834 | 0.742448364 | 0.96 |
| YOR365C | 309.6078398 | -0.163405663 | 0.752331495 | 0.96 |
| YPL107W | 480.1557675 | -0.088061708 | 0.753209735 | 0.96 |
| YPL113C | 241.6280316 | 0.099406672 | 0.752998767 | 0.96 |
| YPR160W | 12134.47846 | -0.153391819 | 0.747816384 | 0.96 |
| snR19 | 122.3490857 | 0.135033213 | 0.752684126 | 0.96 |
| YGR257C | 921.5740821 | -0.076579427 | 0.754272461 | 0.96 |
| YMR143W | 7020.519511 | 0.090349521 | 0.75422548 | 0.96 |
| YAR023C | 164.0948317 | 0.113549445 | 0.755978125 | 0.96 |
| YDR387C | 888.6103683 | 0.080516342 | 0.755327824 | 0.96 |
| YEL057C | 119.2283707 | 0.157398777 | 0.755882028 | 0.96 |
| YGL056C | 2031.538502 | 0.088601656 | 0.755156859 | 0.96 |
| YJR052W | 627.3978725 | -0.063901237 | 0.754828842 | 0.96 |
| YKL175W | 2025.203634 | -0.068052274 | 0.754754127 | 0.96 |
| YLR392C | 638.9786435 | 0.102890734 | 0.755656635 | 0.96 |

| | | | | |
|---|---|---|---|---|
| YNL183C | 1035.511832 | -0.072350444 | 0.755172242 | 0.96 |
| YPL045W | 887.7248714 | -0.058531292 | 0.755964 | 0.96 |
| YAL021C | 1595.906288 | 0.056548131 | 0.757449162 | 0.96 |
| YBR140C | 4380.796749 | 0.074333696 | 0.756643726 | 0.96 |
| YGL215W | 3518.182479 | -0.082035943 | 0.757369101 | 0.96 |
| YKL208W | 299.0743362 | -0.070830177 | 0.757464579 | 0.96 |
| YMR095C | 600.2590168 | 0.170950771 | 0.757297227 | 0.96 |
| YNL075W | 678.2006525 | -0.134072281 | 0.757212672 | 0.96 |
| YMR146C | 5181.578577 | -0.084005543 | 0.757817832 | 0.96 |
| YMR104C | 1639.396102 | -0.128267512 | 0.758638178 | 0.96 |
| YOR022C | 608.0869392 | 0.08472897 | 0.758467981 | 0.96 |
| YLR164W | 251.3796178 | 0.132429507 | 0.759152386 | 0.96 |
| YBL005W-B | 450.3577253 | -0.153057015 | 0.764755264 | 0.96 |
| YBL049W | 377.5575063 | 0.13935224 | 0.762123227 | 0.96 |
| YBL098W | 1032.627619 | 0.153465804 | 0.762992722 | 0.96 |
| YBR050C | 396.3143748 | -0.147498958 | 0.759898617 | 0.96 |
| YDL014W | 4306.735257 | 0.121329785 | 0.761369148 | 0.96 |
| YDL027C | 1559.088359 | -0.086390534 | 0.763777854 | 0.96 |
| YDL161W | 1895.847844 | 0.069694095 | 0.764543023 | 0.96 |
| YDR214W | 2305.632085 | 0.098885544 | 0.765376906 | 0.96 |
| YDR507C | 1606.54523 | 0.121545447 | 0.762695827 | 0.96 |
| YER119C | 1204.859566 | -0.089521641 | 0.761064138 | 0.96 |
| YER125W | 2425.853452 | 0.056559131 | 0.762611167 | 0.96 |
| YER154W | 1666.340326 | 0.053040384 | 0.765307847 | 0.96 |
| YFR023W | 96.03552642 | 0.126689327 | 0.764370112 | 0.96 |
| YHL017W | 1099.085857 | 0.08334982 | 0.759828635 | 0.96 |
| YHR044C | 186.9132602 | -0.129665875 | 0.764062156 | 0.96 |
| YHR050W | 1110.031456 | 0.117983168 | 0.7645319 | 0.96 |
| YHR071W | 3724.97766 | -0.143148108 | 0.764872009 | 0.96 |
| YHR097C | 2968.055756 | -0.130458556 | 0.763527222 | 0.96 |
| YHR137W | 2665.655739 | -0.151345132 | 0.761986872 | 0.96 |
| YIL087C | 960.1717362 | -0.14165605 | 0.760347866 | 0.96 |
| YIL109C | 4372.511551 | -0.05313649 | 0.759647228 | 0.96 |
| YIL145C | 1126.99064 | 0.088027224 | 0.762662199 | 0.96 |
| YKL085W | 9612.778538 | -0.140536776 | 0.762571626 | 0.96 |
| YLR417W | 736.9680806 | 0.06360984 | 0.762982549 | 0.96 |
| YMR141C | 167.2172498 | -0.104004919 | 0.765007954 | 0.96 |
| YMR240C | 592.2234425 | 0.080606574 | 0.765234544 | 0.96 |
| YNL009W | 277.6331838 | -0.144654908 | 0.762516677 | 0.96 |
| YNL067W | 1749.269832 | -0.172501832 | 0.765474864 | 0.96 |
| YNL104C | 8329.192681 | -0.135866409 | 0.763843483 | 0.96 |
| YNL123W | 2060.867856 | -0.098034274 | 0.759635071 | 0.96 |
| YOL033W | 423.9709748 | -0.064059763 | 0.761257281 | 0.96 |

| | | | | |
|---|---|---|---|---|
| YOL135C | 544.3922283 | -0.072644834 | 0.761002696 | 0.96 |
| YOR087W | 1019.099344 | 0.075230829 | 0.764864763 | 0.96 |
| YOR090C | 1602.791656 | -0.043001318 | 0.76427122 | 0.96 |
| YOR206W | 2506.527701 | 0.114954036 | 0.764839928 | 0.96 |
| YPL057C | 1620.500312 | 0.117613159 | 0.765132434 | 0.96 |
| YOL039W | 5643.898973 | -0.175283317 | 0.765983625 | 0.96 |
| YNL162W | 2350.306083 | 0.084568478 | 0.766411246 | 0.96 |
| YHL025W | 1272.013263 | 0.093162195 | 0.767034158 | 0.96 |
| YKR060W | 417.425685 | 0.104806248 | 0.766858346 | 0.96 |
| YPL245W | 615.8162164 | -0.11257317 | 0.767139109 | 0.96 |
| YJR148W | 6070.611654 | -0.111764335 | 0.768021548 | 0.96 |
| YMR113W | 461.667001 | -0.069805509 | 0.76806415 | 0.96 |
| YOL113W | 1261.758159 | -0.113329431 | 0.767757096 | 0.96 |
| YLR234W | 283.065063 | 0.096398523 | 0.76861637 | 0.96 |
| YNL326C | 366.0651518 | 0.042949297 | 0.7686106 | 0.96 |
| YPR102C | 3154.540675 | -0.154757562 | 0.768361635 | 0.96 |
| YAL026C | 1397.824394 | 0.064400161 | 0.776564232 | 0.96 |
| YAR009C | 236.1235724 | 0.104081434 | 0.7709063 | 0.96 |
| YBL020W | 997.2826168 | -0.060560421 | 0.776334891 | 0.96 |
| YBR038W | 1800.515437 | 0.064639767 | 0.773737002 | 0.96 |
| YBR057C | 866.0815496 | -0.080632942 | 0.772303453 | 0.96 |
| YBR142W | 1284.846746 | -0.0966011 | 0.774315559 | 0.96 |
| YCR048W | 995.8304266 | -0.064200691 | 0.775600715 | 0.96 |
| YDL181W | 1061.423493 | -0.123698472 | 0.772109532 | 0.96 |
| YDL194W | 1083.690954 | -0.103803038 | 0.776497236 | 0.96 |
| YDR219C | 850.7025719 | -0.066932454 | 0.771904086 | 0.96 |
| YEL033W | 1403.07415 | -0.115012552 | 0.776269928 | 0.96 |
| YEL065W | 2278.709245 | 0.140559438 | 0.774094454 | 0.96 |
| YER175C | 1510.549619 | -0.1365879 | 0.774964388 | 0.96 |
| YFL021W | 926.9442787 | -0.133650465 | 0.772290698 | 0.96 |
| YGR011W | 95.88242328 | -0.060270882 | 0.771145673 | 0.96 |
| YGR156W | 932.64115 | 0.073560869 | 0.770129652 | 0.96 |
| YGR198W | 1689.551002 | 0.083211278 | 0.770186981 | 0.96 |
| YHR020W | 3665.040866 | 0.109716689 | 0.774770566 | 0.96 |
| YHR040W | 740.6643017 | -0.107277064 | 0.774303289 | 0.96 |
| YHR123W | 699.5030842 | 0.126404359 | 0.775157692 | 0.96 |
| YJL045W | 286.1990705 | 0.108524644 | 0.769963578 | 0.96 |
| YJR025C | 1368.606225 | -0.158738863 | 0.771089714 | 0.96 |
| YKL020C | 2087.802283 | 0.075294846 | 0.770596191 | 0.96 |
| YKL087C | 723.7436146 | 0.062406992 | 0.775894651 | 0.96 |
| YKL189W | 702.1789039 | 0.085445596 | 0.775967614 | 0.96 |
| YLL008W | 1923.071209 | -0.106763641 | 0.775201179 | 0.96 |
| YLR179C | 2935.995773 | -0.15753687 | 0.771991993 | 0.96 |

| | | | | |
|---|---|---|---|---|
| YLR190W | 1584.336567 | 0.09075601 | 0.768934542 | 0.96 |
| YLR289W | 462.7410856 | 0.084168868 | 0.77593285 | 0.96 |
| YML027W | 523.9520569 | -0.123705952 | 0.770341357 | 0.96 |
| YML123C | 2594.845401 | -0.182201495 | 0.776623654 | 0.96 |
| YMR011W | 5664.342762 | -0.263215506 | 0.770563311 | 0.96 |
| YMR087W | 259.2299237 | 0.074161294 | 0.776349222 | 0.96 |
| YMR317W | 752.1354442 | 0.07895785 | 0.774957574 | 0.96 |
| YNL003C | 715.0467326 | -0.08446227 | 0.775884558 | 0.96 |
| YNL151C | 742.0050839 | -0.105258117 | 0.775353902 | 0.96 |
| YNR014W | 1455.118637 | 0.146816065 | 0.770411068 | 0.96 |
| YNR032W | 607.7463894 | -0.062904102 | 0.771829332 | 0.96 |
| YOL061W | 1754.163485 | -0.079568676 | 0.773210071 | 0.96 |
| YOL067C | 283.6975055 | 0.078707822 | 0.775375318 | 0.96 |
| YPL011C | 726.714861 | 0.053880737 | 0.771608117 | 0.96 |
| YPL067C | 285.3555196 | -0.115576021 | 0.76937765 | 0.96 |
| YPL181W | 1430.008347 | 0.087495758 | 0.775360543 | 0.96 |
| YPL219W | 632.4863798 | -0.080752659 | 0.77141762 | 0.96 |
| YPR169W | 1659.207747 | 0.076549678 | 0.773108911 | 0.96 |
| YBR281C | 961.503364 | -0.06995695 | 0.77689362 | 0.96 |
| YER033C | 1095.361309 | 0.090290663 | 0.777088276 | 0.96 |
| YKR036C | 843.7972906 | 0.046437352 | 0.777220197 | 0.96 |
| YBR275C | 1768.794249 | 0.069800354 | 0.778029501 | 0.96 |
| YGR243W | 1286.002734 | 0.152341234 | 0.77842078 | 0.96 |
| YJR036C | 438.6482321 | -0.094978016 | 0.778359875 | 0.96 |
| YMR094W | 306.3491696 | 0.087511725 | 0.778301491 | 0.96 |
| YHR144C | 383.3281151 | 0.110292135 | 0.77868439 | 0.96 |
| YDL028C | 1026.396259 | 0.063586748 | 0.779753897 | 0.96 |
| YER049W | 1494.465582 | 0.095235595 | 0.779706981 | 0.96 |
| YKL070W | 302.0799748 | -0.061525217 | 0.780360476 | 0.96 |
| YML052W | 1751.911847 | 0.063541593 | 0.77997739 | 0.96 |
| YMR194W | 1484.319594 | 0.056117244 | 0.780344813 | 0.96 |
| YNL110C | 899.4305976 | -0.12471662 | 0.779720136 | 0.96 |
| YPR114W | 2316.846378 | -0.050184362 | 0.779864645 | 0.96 |
| YPR178W | 455.7725824 | -0.057631843 | 0.780310867 | 0.96 |
| YDL198C | 1156.426648 | -0.098032457 | 0.780730706 | 0.96 |
| YKL149C | 355.2699607 | -0.067178979 | 0.781116113 | 0.96 |
| YMR053C | 712.958308 | -0.085011167 | 0.781433454 | 0.96 |
| YMR140W | 1470.388235 | 0.078412118 | 0.781534504 | 0.96 |
| YBR289W | 1372.875089 | -0.078346388 | 0.782187537 | 0.96 |
| YKL168C | 972.1957599 | 0.076656265 | 0.782104467 | 0.96 |
| YOR005C | 988.1912317 | 0.075331513 | 0.781893016 | 0.96 |
| YPL243W | 2085.694288 | 0.082040085 | 0.782242287 | 0.96 |
| YAL010C | 552.0969494 | 0.044101798 | 0.791242915 | 0.96 |

| | | | | |
|---|---|---|---|---|
| YAL019W | 1321.957689 | 0.076722289 | 0.803581697 | 0.96 |
| YAL034W-A | 187.40184 | 0.073872731 | 0.790099666 | 0.96 |
| YAR002W | 1776.649309 | -0.066658066 | 0.795512896 | 0.96 |
| YBL017C | 3980.2423 | 0.071650665 | 0.80410134 | 0.96 |
| YBL103C | 858.4740089 | -0.066103387 | 0.800959393 | 0.96 |
| YBR041W | 1541.15066 | -0.062684576 | 0.787857367 | 0.96 |
| YBR076W | 114.1570412 | 0.126265942 | 0.790288396 | 0.96 |
| YBR084W | 2864.241089 | 0.099296555 | 0.799192017 | 0.96 |
| YBR141C | 286.9036203 | -0.062105353 | 0.803652959 | 0.96 |
| YBR146W | 1432.145995 | -0.05470718 | 0.798944266 | 0.96 |
| YBR168W | 396.8925406 | -0.060756058 | 0.789221821 | 0.96 |
| YBR175W | 264.1838842 | -0.087146916 | 0.795045813 | 0.96 |
| YBR191W | 11850.59787 | 0.044202066 | 0.798528061 | 0.96 |
| YBR229C | 1792.230472 | 0.040208457 | 0.802041191 | 0.96 |
| YBR254C | 311.1708692 | 0.074193247 | 0.802143791 | 0.96 |
| YBR285W | 449.6014678 | -0.127631134 | 0.803571824 | 0.96 |
| YCL038C | 682.2522644 | -0.060768325 | 0.801108632 | 0.96 |
| YCR015C | 255.0577346 | 0.051456553 | 0.792366692 | 0.96 |
| YCR068W | 666.4091136 | 0.073169544 | 0.801052587 | 0.96 |
| YDL030W | 676.7024249 | 0.050745441 | 0.788114303 | 0.96 |
| YDL043C | 369.3400281 | -0.051237303 | 0.803341675 | 0.96 |
| YDL055C | 11798.61054 | 0.045929162 | 0.799736688 | 0.96 |
| YDL089W | 963.5832446 | 0.083967977 | 0.796408719 | 0.96 |
| YDL113C | 1161.019898 | -0.075525703 | 0.783209964 | 0.96 |
| YDL154W | 661.9950458 | 0.075882872 | 0.798741145 | 0.96 |
| YDL248W | 224.0356228 | 0.096604831 | 0.803394692 | 0.96 |
| YDR062W | 2339.714713 | -0.065224592 | 0.793835502 | 0.96 |
| YDR177W | 1727.041157 | 0.060098612 | 0.802606595 | 0.96 |
| YDR233C | 7397.691198 | -0.093451146 | 0.792087033 | 0.96 |
| YDR245W | 1164.324708 | 0.078610799 | 0.789757616 | 0.96 |
| YDR279W | 575.5796481 | -0.053641879 | 0.79314795 | 0.96 |
| YDR400W | 652.8362371 | -0.065666139 | 0.788182039 | 0.96 |
| YDR403W | 327.0898139 | 0.112798405 | 0.790341457 | 0.96 |
| YDR412W | 170.5135109 | -0.083746549 | 0.801121707 | 0.96 |
| YDR488C | 747.4804028 | 0.076165516 | 0.798570134 | 0.96 |
| YDR490C | 1094.352116 | 0.062126517 | 0.795244723 | 0.96 |
| YEL040W | 3165.37713 | -0.08189057 | 0.797106876 | 0.96 |
| YEL071W | 3890.356415 | -0.115548398 | 0.800078461 | 0.96 |
| YER110C | 3689.619393 | -0.104787135 | 0.802298625 | 0.96 |
| YFL026W | 6326.346816 | 0.190097464 | 0.803101674 | 0.96 |
| YFL053W | 516.855343 | 0.129398895 | 0.793940113 | 0.96 |
| YFR020W | 385.8178556 | 0.058386582 | 0.794037577 | 0.96 |
| YFR026C | 243.3675063 | 0.105087893 | 0.791062369 | 0.96 |

| | | | | |
|---|---|---|---|---|
| YFR029W | 834.7381381 | -0.085810715 | 0.798728452 | 0.96 |
| YGL009C | 9382.784081 | -0.124964993 | 0.796764397 | 0.96 |
| YGL029W | 383.3919755 | -0.13395846 | 0.784333029 | 0.96 |
| YGL099W | 1111.20672 | 0.089439997 | 0.785841836 | 0.96 |
| YGL126W | 1063.33427 | 0.089173381 | 0.798290872 | 0.96 |
| YGR013W | 916.8674279 | -0.063031759 | 0.790410327 | 0.96 |
| YGR031W | 376.7830976 | -0.075634369 | 0.789313227 | 0.96 |
| YGR136W | 2019.817506 | -0.062424261 | 0.792876719 | 0.96 |
| YGR147C | 943.7636147 | -0.053148843 | 0.796093884 | 0.96 |
| YGR223C | 1134.541576 | -0.084825347 | 0.799503765 | 0.96 |
| YGR263C | 454.4543224 | -0.050412977 | 0.799729094 | 0.96 |
| YGR278W | 442.5587334 | 0.050212061 | 0.804182702 | 0.96 |
| YGR283C | 529.6205161 | 0.069069275 | 0.793335595 | 0.96 |
| YHL020C | 961.5750146 | 0.067449993 | 0.798235521 | 0.96 |
| YHR010W | 11885.41138 | 0.086319008 | 0.789634329 | 0.96 |
| YHR045W | 983.5331682 | 0.085719418 | 0.789515157 | 0.96 |
| YHR074W | 2090.525023 | -0.058419682 | 0.788409773 | 0.96 |
| YHR129C | 426.0946941 | 0.064870987 | 0.798584336 | 0.96 |
| YHR167W | 161.5282471 | -0.099650228 | 0.787280481 | 0.96 |
| YHR198C | 872.0543676 | -0.09793344 | 0.804132433 | 0.96 |
| YIL020C-A | 191.6015626 | -0.068949864 | 0.785355206 | 0.96 |
| YIL036W | 2051.229049 | 0.068160204 | 0.796707825 | 0.96 |
| YIL044C | 893.1546159 | -0.066559705 | 0.787469295 | 0.96 |
| YIL085C | 1498.284283 | 0.049787704 | 0.786308154 | 0.96 |
| YJL049W | 674.8429005 | -0.074366525 | 0.791052387 | 0.96 |
| YJL058C | 326.9260026 | -0.061111303 | 0.798155798 | 0.96 |
| YJL072C | 353.6349072 | -0.073854014 | 0.789883198 | 0.96 |
| YJL130C | 21427.36188 | -0.116473816 | 0.795476516 | 0.96 |
| YJL165C | 2120.172359 | -0.072424 | 0.795353099 | 0.96 |
| YJL210W | 543.5019911 | -0.080416648 | 0.795619404 | 0.96 |
| YJR060W | 664.6110196 | -0.045414949 | 0.789134233 | 0.96 |
| YJR066W | 2254.068141 | 0.061001688 | 0.801181625 | 0.96 |
| YJR112W | 374.389265 | -0.072516638 | 0.78694331 | 0.96 |
| YJR152W | 334.6737416 | 0.089785681 | 0.789226914 | 0.96 |
| YKR004C | 281.3110563 | -0.054493966 | 0.802647368 | 0.96 |
| YLL057C | 252.3608873 | 0.100805118 | 0.792721936 | 0.96 |
| YLR018C | 838.7588705 | -0.054080985 | 0.786982909 | 0.96 |
| YLR089C | 5710.768483 | -0.118331839 | 0.801175956 | 0.96 |
| YLR224W | 1026.056679 | -0.067503851 | 0.790625446 | 0.96 |
| YLR227W-B | 558.9698813 | -0.143110839 | 0.790514717 | 0.96 |
| YLR350W | 3571.832696 | 0.074033988 | 0.799806962 | 0.96 |
| YML043C | 371.5090553 | -0.107504701 | 0.785862576 | 0.96 |
| YML108W | 173.1265371 | -0.084285383 | 0.797465977 | 0.96 |

| | | | | |
|---|---|---|---|---|
| YML113W | 567.9041791 | 0.072628403 | 0.802260722 | 0.96 |
| YMR048W | 313.0835867 | 0.034798002 | 0.790872144 | 0.96 |
| YMR076C | 1386.678863 | 0.088022654 | 0.795085644 | 0.96 |
| YMR118C | 295.0484477 | 0.091439653 | 0.803132893 | 0.96 |
| YMR174C | 615.2184477 | -0.148826137 | 0.795552206 | 0.96 |
| YMR213W | 660.1636091 | 0.063196052 | 0.787533934 | 0.96 |
| YMR230W | 6828.386976 | -0.136960013 | 0.788441823 | 0.96 |
| YNL001W | 629.9634099 | -0.072594794 | 0.787009095 | 0.96 |
| YNL044W | 495.5352853 | -0.105200598 | 0.789363555 | 0.96 |
| YNL050C | 614.5421658 | 0.050322854 | 0.785364613 | 0.96 |
| YNL192W | 2879.150646 | 0.113284877 | 0.793135696 | 0.96 |
| YNL288W | 932.0705303 | -0.047617395 | 0.793167084 | 0.96 |
| YNL294C | 1109.535055 | 0.0809842 | 0.797076617 | 0.96 |
| YNR029C | 598.402815 | 0.054750188 | 0.801364969 | 0.96 |
| YNR034W | 660.948473 | -0.072076057 | 0.791005947 | 0.96 |
| YOL009C | 380.9976744 | -0.05638467 | 0.804183586 | 0.96 |
| YOL013C | 1216.146773 | 0.046245185 | 0.79397138 | 0.96 |
| YOL023W | 636.5704075 | 0.067584207 | 0.802314267 | 0.96 |
| YOL032W | 1063.876828 | -0.082405612 | 0.80414442 | 0.96 |
| YOL055C | 533.3864453 | 0.076943137 | 0.798410884 | 0.96 |
| YOL072W | 594.5699607 | 0.052557433 | 0.794970847 | 0.96 |
| YOL081W | 5088.832739 | 0.08671628 | 0.796924902 | 0.96 |
| YOL090W | 1084.862315 | -0.075553362 | 0.787376863 | 0.96 |
| YOR178C | 2730.240932 | 0.133069728 | 0.789632351 | 0.96 |
| YOR250C | 655.6893916 | 0.045018656 | 0.800697205 | 0.96 |
| YOR272W | 1204.374523 | -0.095148048 | 0.79583214 | 0.96 |
| YOR348C | 282.9627385 | 0.123467334 | 0.788578785 | 0.96 |
| YPL017C | 762.3634934 | 0.080872148 | 0.79556277 | 0.96 |
| YPL054W | 345.0757807 | -0.114336161 | 0.799702397 | 0.96 |
| YPL143W | 740.5431297 | -0.163499368 | 0.804066137 | 0.96 |
| YPL148C | 142.8618352 | 0.08521831 | 0.799438549 | 0.96 |
| YPL230W | 2214.598115 | 0.130913745 | 0.784518537 | 0.96 |
| YPL235W | 1507.98677 | -0.05260088 | 0.793688898 | 0.96 |
| YPL264C | 197.1729292 | 0.060989652 | 0.793427203 | 0.96 |
| YPR023C | 1837.667458 | -0.091090682 | 0.786559235 | 0.96 |
| YPR036W-A | 8058.010483 | -0.130318727 | 0.79299479 | 0.96 |
| YPR116W | 103.378319 | 0.089742096 | 0.788524065 | 0.96 |
| YAL058W | 573.3781549 | -0.058679786 | 0.805542045 | 0.96 |
| YBL011W | 1645.010666 | -0.050472166 | 0.806096905 | 0.96 |
| YBR181C | 1794.107741 | -0.227791167 | 0.805975517 | 0.96 |
| YDR181C | 744.5427058 | -0.076754861 | 0.805513501 | 0.96 |
| YIL056W | 1630.203256 | 0.078628532 | 0.805647283 | 0.96 |
| YIL164C | 260.9748713 | 0.094641544 | 0.805841755 | 0.96 |

| | | | | |
|---|---|---|---|---|
| YOR040W | 503.3947844 | -0.096778159 | 0.805568109 | 0.96 |
| YOR175C | 2021.823765 | -0.048313427 | 0.804983179 | 0.96 |
| YPL149W | 607.7809907 | 0.076196262 | 0.805604297 | 0.96 |
| YER127W | 556.9285849 | -0.111743606 | 0.806588712 | 0.96 |
| YHR105W | 228.2746463 | -0.076491496 | 0.806517894 | 0.96 |
| YHR128W | 1328.56215 | -0.128391511 | 0.807145455 | 0.96 |
| YPR048W | 424.7867171 | 0.075358122 | 0.80692378 | 0.96 |
| YPR110C | 2091.025042 | -0.09860089 | 0.807115042 | 0.96 |
| YDR073W | 640.4950618 | 0.079548141 | 0.80790432 | 0.96 |
| YMR040W | 194.5584091 | -0.097143058 | 0.807644309 | 0.96 |
| YMR041C | 591.2121868 | -0.100987667 | 0.807878361 | 0.96 |
| YHL024W | 910.8922734 | 0.074133761 | 0.808096995 | 0.96 |
| YOR177C | 142.866635 | -0.126284146 | 0.808316837 | 0.96 |
| YCR084C | 3633.028548 | 0.072556195 | 0.808519952 | 0.96 |
| YCL002C | 469.4119103 | 0.063912979 | 0.808906139 | 0.96 |
| YMR303C | 445.8492587 | 0.07477314 | 0.80900224 | 0.96 |
| YCL063W | 606.1625178 | 0.07054923 | 0.813524533 | 0.96 |
| YCR047C | 558.0855162 | -0.065003787 | 0.811501488 | 0.96 |
| YDL134C | 1962.281647 | 0.023027578 | 0.811139606 | 0.96 |
| YDR051C | 1050.532227 | -0.038487443 | 0.811211518 | 0.96 |
| YDR217C | 884.5738376 | 0.057839187 | 0.814568928 | 0.96 |
| YER084W | 124.9336603 | -0.210699938 | 0.812519403 | 0.96 |
| YER116C | 578.3983863 | 0.07306844 | 0.811028596 | 0.96 |
| YER149C | 245.5366344 | -0.0930844 | 0.813916658 | 0.96 |
| YGL154C | 461.1131948 | -0.055972753 | 0.814282724 | 0.96 |
| YGL208W | 855.2756461 | 0.101558171 | 0.812244167 | 0.96 |
| YGR258C | 1398.526849 | -0.057402419 | 0.811183959 | 0.96 |
| YHL023C | 1253.971871 | -0.041130601 | 0.812449908 | 0.96 |
| YIL101C | 1402.735777 | -0.112435717 | 0.811153938 | 0.96 |
| YIL125W | 5872.779859 | 0.111563022 | 0.812432932 | 0.96 |
| YIL162W | 3263.600398 | 0.088668441 | 0.811772844 | 0.96 |
| YIR018W | 398.3013208 | -0.092002616 | 0.81175627 | 0.96 |
| YJL002C | 4648.101511 | -0.066199945 | 0.812192121 | 0.96 |
| YKL095W | 323.4844945 | 0.065906422 | 0.812913265 | 0.96 |
| YLL045C | 7512.686429 | 0.054071485 | 0.812346512 | 0.96 |
| YML119W | 1027.81525 | 0.050036542 | 0.810055205 | 0.96 |
| YML120C | 4283.544511 | -0.10543724 | 0.813967647 | 0.96 |
| YMR056C | 319.9319845 | -0.10614652 | 0.813187206 | 0.96 |
| YMR275C | 1521.663704 | 0.054859975 | 0.810968843 | 0.96 |
| YNL141W | 1283.346108 | -0.121583296 | 0.814247969 | 0.96 |
| YNL144C | 1767.380134 | -0.111444698 | 0.813378871 | 0.96 |
| YOR145C | 791.624005 | 0.073336926 | 0.813660478 | 0.96 |
| YPL176C | 1288.392675 | 0.056204486 | 0.813215507 | 0.96 |

| | | | | |
|---|---|---|---|---|
| YPL210C | 1902.73736 | -0.069613026 | 0.814442491 | 0.96 |
| YPR056W | 512.7372419 | 0.03728524 | 0.812501521 | 0.96 |
| YBL071C | 215.271228 | 0.072561793 | 0.815183629 | 0.96 |
| YHL030W | 2779.309717 | -0.076072867 | 0.81519618 | 0.96 |
| YIL038C | 1659.427705 | -0.066130816 | 0.81520139 | 0.96 |
| YIL049W | 383.4643737 | -0.082006215 | 0.815806578 | 0.96 |
| YIL140W | 1095.766571 | 0.090245814 | 0.81578142 | 0.96 |
| YCR098C | 141.8503964 | 0.129994709 | 0.816616401 | 0.96 |
| YLR094C | 1048.349221 | 0.060755545 | 0.816870719 | 0.96 |
| YNL224C | 704.2866123 | -0.041159093 | 0.816823451 | 0.96 |
| YNR004W | 166.7322391 | 0.057133407 | 0.817012563 | 0.96 |
| YPL018W | 471.7341201 | 0.081335067 | 0.817233167 | 0.96 |
| YOL140W | 502.3006349 | 0.09765121 | 0.817443386 | 0.96 |
| YKL143W | 1110.759937 | -0.084639521 | 0.817732733 | 0.96 |
| YOR246C | 1262.599493 | 0.062910885 | 0.817881316 | 0.96 |
| YLR175W | 4387.69014 | 0.074375124 | 0.819406334 | 0.97 |
| YMR088C | 861.7053416 | 0.060796769 | 0.819550241 | 0.97 |
| YCR101C | 105.2670668 | 0.092763592 | 0.8200314 | 0.97 |
| YDR425W | 1030.779588 | -0.066768774 | 0.820030262 | 0.97 |
| YER087W | 372.1981021 | 0.067613277 | 0.82132611 | 0.97 |
| YER122C | 4064.222569 | -0.054440205 | 0.820910942 | 0.97 |
| YGR049W | 549.8511767 | -0.060993076 | 0.82070533 | 0.97 |
| YIL009W | 578.8586392 | 0.08293754 | 0.820121168 | 0.97 |
| YML003W | 222.6670877 | -0.069387051 | 0.820451568 | 0.97 |
| YNL103W-A | 104.3621644 | 0.099818407 | 0.821231614 | 0.97 |
| YNL300W | 1681.600577 | -0.11427817 | 0.821229084 | 0.97 |
| YOR377W | 997.8606777 | -0.062645764 | 0.820526141 | 0.97 |
| YER147C-A | 275.4373645 | -0.072587571 | 0.822107088 | 0.97 |
| YGR168C | 490.2041357 | -0.045265765 | 0.821977483 | 0.97 |
| YIL110W | 498.0390037 | -0.074180142 | 0.822288211 | 0.97 |
| YML073C | 8568.253747 | 0.064977233 | 0.822228364 | 0.97 |
| YKL105C | 1423.737383 | 0.056621856 | 0.822837465 | 0.97 |
| YGL168W | 276.6572721 | -0.065079694 | 0.823655339 | 0.97 |
| YGR044C | 1106.237884 | -0.126196704 | 0.823913237 | 0.97 |
| YHR033W | 1072.035586 | 0.06805512 | 0.824169371 | 0.97 |
| YKL150W | 7424.00365 | -0.102689442 | 0.823223917 | 0.97 |
| YLR443W | 472.8026492 | 0.056891931 | 0.823870848 | 0.97 |
| YNL072W | 456.0809998 | 0.047157438 | 0.823721001 | 0.97 |
| YPL163C | 3377.565521 | 0.088089576 | 0.824209796 | 0.97 |
| YBR198C | 1683.847961 | 0.045294803 | 0.82453167 | 0.97 |
| YDR417C | 435.6659374 | -0.09693297 | 0.824765186 | 0.97 |
| YJL144W | 785.7166274 | 0.086397651 | 0.824804035 | 0.97 |
| YBR161W | 517.5371153 | 0.076187389 | 0.826465498 | 0.97 |

| | | | | |
|---|---|---|---|---|
| YBR233W | 474.558741 | -0.066258954 | 0.825934896 | 0.97 |
| YCL004W | 588.1019431 | 0.051699262 | 0.827471349 | 0.97 |
| YCL045C | 3038.105977 | -0.063601406 | 0.826498878 | 0.97 |
| YCR105W | 210.4429239 | 0.107694784 | 0.826499934 | 0.97 |
| YDL166C | 434.2604045 | -0.078357554 | 0.826734169 | 0.97 |
| YFL008W | 1609.672053 | -0.062936364 | 0.827885554 | 0.97 |
| YHR180W | 212.5255364 | -0.090208974 | 0.825946688 | 0.97 |
| YIL156W | 1109.423201 | 0.053083053 | 0.826862492 | 0.97 |
| YJL148W | 1042.119748 | -0.109915602 | 0.82539551 | 0.97 |
| YKR026C | 956.8364878 | -0.065928383 | 0.827402212 | 0.97 |
| YKR051W | 666.9246654 | 0.049975238 | 0.82597531 | 0.97 |
| YLR242C | 302.599813 | 0.051870085 | 0.827676727 | 0.97 |
| YMR304C-A | 599.6439516 | -0.085279059 | 0.826157938 | 0.97 |
| YOR019W | 709.5102111 | -0.106210296 | 0.827027203 | 0.97 |
| YPR001W | 533.0217534 | -0.115698219 | 0.827732507 | 0.97 |
| YDR140W | 525.20568 | 0.050710523 | 0.828972021 | 0.97 |
| YIL134W | 238.1456546 | 0.085014006 | 0.828532577 | 0.97 |
| YKL124W | 623.7006372 | 0.054594435 | 0.829259803 | 0.97 |
| YLR436C | 2590.336596 | 0.055129872 | 0.829026951 | 0.97 |
| YML072C | 6041.463861 | 0.047710647 | 0.829203748 | 0.97 |
| YNL234W | 710.7866789 | 0.08673253 | 0.828915408 | 0.97 |
| YOR209C | 3422.936842 | -0.066342912 | 0.828895906 | 0.97 |
| YBR183W | 786.7022482 | -0.095426837 | 0.831215519 | 0.97 |
| YDR018C | 446.8147304 | 0.086596266 | 0.830531055 | 0.97 |
| YDR247W | 1026.990415 | 0.079331316 | 0.831062347 | 0.97 |
| YDR423C | 1205.3164 | 0.062673335 | 0.830543007 | 0.97 |
| YGL040C | 1261.243607 | -0.06165428 | 0.83047094 | 0.97 |
| YGR102C | 796.5538343 | 0.074663952 | 0.830517226 | 0.97 |
| YGR234W | 5131.339655 | 0.075269038 | 0.830674342 | 0.97 |
| YJL005W | 4176.860231 | 0.064898448 | 0.830917413 | 0.97 |
| YLR439W | 1150.159115 | 0.061541634 | 0.829965988 | 0.97 |
| YOL077C | 1854.977032 | 0.080867658 | 0.830566323 | 0.97 |
| YOR095C | 614.382855 | 0.062115875 | 0.830922402 | 0.97 |
| YBR130C | 1160.408881 | 0.050520465 | 0.831537129 | 0.97 |
| YDL098C | 246.2120054 | -0.050534012 | 0.833057899 | 0.97 |
| YDR034C-D | 94.28631679 | 0.079360328 | 0.833264204 | 0.97 |
| YGL212W | 286.2266484 | -0.047001632 | 0.832819081 | 0.97 |
| YGR261C | 1931.303256 | -0.049742205 | 0.832929277 | 0.97 |
| YHR037W | 3383.544 | -0.057270569 | 0.833302694 | 0.97 |
| YHR070C-A | 129.6976848 | 0.075192572 | 0.833338887 | 0.97 |
| YJL050W | 1800.303824 | 0.079363342 | 0.833180473 | 0.97 |
| YKL042W | 561.3749184 | -0.061330295 | 0.832337085 | 0.97 |
| YMR134W | 453.7137928 | -0.05647659 | 0.833343083 | 0.97 |

| | | | | |
|---|---|---|---|---|
| YOR054C | 904.8796156 | -0.064259854 | 0.83319332 | 0.97 |
| YPR179C | 773.0608325 | 0.043218807 | 0.833358479 | 0.97 |
| YIL102C-A | 112.0491515 | -0.059714162 | 0.833712181 | 0.97 |
| YAL028W | 836.6833612 | 0.063431707 | 0.835409337 | 0.97 |
| YDR029W | 118.6548652 | -0.072778877 | 0.836178038 | 0.97 |
| YDR126W | 326.9726421 | 0.054459184 | 0.834594578 | 0.97 |
| YDR229W | 1110.14149 | -0.042874746 | 0.834688206 | 0.97 |
| YFR031C-A | 6438.897656 | 0.013573068 | 0.835915294 | 0.97 |
| YGL219C | 1273.719314 | -0.067121694 | 0.834615829 | 0.97 |
| YHR062C | 561.4502383 | -0.062470386 | 0.835222443 | 0.97 |
| YJL074C | 945.199767 | 0.057310705 | 0.836197312 | 0.97 |
| YLR327C | 4824.371416 | -0.132182755 | 0.834738761 | 0.97 |
| YNL130C | 1326.028586 | -0.072532711 | 0.834834151 | 0.97 |
| YOR143C | 346.0541178 | -0.072576523 | 0.836056334 | 0.97 |
| YOR303W | 3621.710527 | 0.095170131 | 0.836010855 | 0.97 |
| YPR168W | 170.9858215 | 0.059679065 | 0.835340437 | 0.97 |
| YCL049C | 935.682406 | -0.083780533 | 0.836417706 | 0.97 |
| YJR084W | 669.2500078 | -0.032444905 | 0.836680161 | 0.97 |
| YNL148C | 367.0903378 | 0.080245557 | 0.837381742 | 0.97 |
| YPL007C | 583.714763 | 0.029454776 | 0.837896115 | 0.97 |
| YPL252C | 462.6644596 | -0.06941036 | 0.837567967 | 0.97 |
| YPR017C | 239.3606668 | -0.062123775 | 0.837843874 | 0.97 |
| YIL052C | 4784.674042 | -0.141633683 | 0.838213487 | 0.97 |
| YNL077W | 1205.161408 | 0.070262434 | 0.838793832 | 0.97 |
| YCR021C | 10522.50846 | 0.09898633 | 0.839199084 | 0.97 |
| YBL048W | 125.8109055 | 0.084250306 | 0.84135973 | 0.97 |
| YBR009C | 3036.286227 | -0.13770668 | 0.841209909 | 0.97 |
| YDR128W | 2008.877758 | 0.052017813 | 0.841154798 | 0.97 |
| YER074W | 4108.232098 | -0.180959864 | 0.839916802 | 0.97 |
| YIL018W | 10226.88893 | 0.014881885 | 0.841087266 | 0.97 |
| YLR148W | 974.658438 | -0.040405497 | 0.84106233 | 0.97 |
| YMR235C | 2269.617315 | -0.084279789 | 0.83994635 | 0.97 |
| YOR101W | 962.4280426 | 0.056669862 | 0.840187913 | 0.97 |
| YPL172C | 484.663762 | 0.039452516 | 0.840752515 | 0.97 |
| YPR034W | 1306.898874 | 0.046903141 | 0.841192883 | 0.97 |
| YPL014W | 2476.516022 | 0.08384151 | 0.841924481 | 0.97 |
| YJL014W | 2604.88629 | -0.064088107 | 0.842291957 | 0.97 |
| YNL219C | 1817.162227 | -0.042675243 | 0.842152236 | 0.97 |
| YAL025C | 983.4578298 | -0.07721391 | 0.843170568 | 0.97 |
| YGL110C | 882.4868812 | -0.037121892 | 0.843085749 | 0.97 |
| YMR280C | 1028.334214 | 0.090072554 | 0.843226561 | 0.97 |
| YNL012W | 592.4635068 | 0.059629586 | 0.842753871 | 0.97 |
| YNL027W | 1180.449634 | 0.046137981 | 0.842699239 | 0.97 |

| | | | | |
|---|---|---|---|---|
| YNL054W-B | 451.4424074 | -0.118419112 | 0.843565933 | 0.97 |
| YDR515W | 694.9239372 | -0.059175547 | 0.843802162 | 0.97 |
| YJR051W | 1291.045072 | -0.066704591 | 0.844094897 | 0.97 |
| YGR203W | 222.1887092 | 0.075456649 | 0.844578584 | 0.97 |
| YHL032C | 2561.780761 | 0.08998963 | 0.84443799 | 0.97 |
| YER043C | 7073.679738 | -0.077129598 | 0.844833002 | 0.97 |
| YAL013W | 1274.881408 | 0.042704693 | 0.847182453 | 0.97 |
| YAR008W | 281.6693003 | -0.071195642 | 0.846245287 | 0.97 |
| YBR030W | 579.6269936 | 0.053596218 | 0.847507925 | 0.97 |
| YBR223C | 483.3688068 | -0.053095998 | 0.84942027 | 0.97 |
| YBR272C | 606.5333168 | -0.039785769 | 0.850724694 | 0.97 |
| YCR030C | 2439.320353 | 0.045432507 | 0.850490932 | 0.97 |
| YDL190C | 2381.415921 | 0.032647306 | 0.849283644 | 0.97 |
| YDR182W | 1361.286881 | 0.033806228 | 0.849401533 | 0.97 |
| YEL016C | 826.6367047 | -0.031873513 | 0.849493613 | 0.97 |
| YER187W | 132.2577032 | -0.089324256 | 0.846788101 | 0.97 |
| YGR009C | 1451.464581 | -0.0345608 | 0.850496652 | 0.97 |
| YGR058W | 565.2067235 | 0.039251555 | 0.849046151 | 0.97 |
| YHR191C | 210.2719774 | 0.030716009 | 0.849225851 | 0.97 |
| YIL006W | 374.9591791 | -0.107140886 | 0.850690296 | 0.97 |
| YIL015W | 2424.316267 | 0.164066701 | 0.849827901 | 0.97 |
| YIR026C | 772.7201823 | 0.075707836 | 0.850844183 | 0.97 |
| YJL149W | 1033.010855 | 0.03751075 | 0.845863549 | 0.97 |
| YJR027W | 207.2279874 | 0.073414428 | 0.850226564 | 0.97 |
| YKL178C | 114.5867776 | 0.094146426 | 0.847912605 | 0.97 |
| YLR165C | 405.6673301 | 0.052358397 | 0.846423132 | 0.97 |
| YLR257W | 6468.084537 | -0.065226471 | 0.850545944 | 0.97 |
| YLR441C | 16750.1152 | -0.111825563 | 0.849833049 | 0.97 |
| YML038C | 1118.066731 | -0.044193509 | 0.848956616 | 0.97 |
| YMR045C | 326.5494461 | -0.091641282 | 0.848217723 | 0.97 |
| YMR299C | 640.5800002 | 0.052570553 | 0.848285667 | 0.97 |
| YMR304W | 1821.67097 | 0.045660524 | 0.845311931 | 0.97 |
| YMR318C | 2412.871237 | -0.094937392 | 0.849385455 | 0.97 |
| YNL097C | 784.7174827 | -0.051033979 | 0.846678985 | 0.97 |
| YOR048C | 2248.629889 | 0.044142956 | 0.849441862 | 0.97 |
| YOR061W | 1662.759146 | -0.044001263 | 0.848338227 | 0.97 |
| YOR298C-A | 6204.241243 | -0.038804103 | 0.850900872 | 0.97 |
| YOR321W | 1083.064286 | 0.044668853 | 0.846993143 | 0.97 |
| YPL247C | 2827.171119 | -0.089774998 | 0.849576408 | 0.97 |
| YPR199C | 796.7118888 | -0.058606897 | 0.846516928 | 0.97 |
| YCL061C | 1120.832529 | 0.047904978 | 0.851192007 | 0.97 |
| YDR174W | 3004.282379 | -0.04017187 | 0.851566559 | 0.97 |
| YKL015W | 1286.742474 | -0.069989343 | 0.851474125 | 0.97 |

| | | | | |
|---|---|---|---|---|
| YOL145C | 2673.103928 | -0.046245126 | 0.85177734 | 0.97 |
| YBR020W | 310.6456705 | 0.062091614 | 0.853140758 | 0.97 |
| YBR034C | 1111.628509 | 0.061521158 | 0.85643002 | 0.97 |
| YBR245C | 1580.844401 | 0.035135416 | 0.852866767 | 0.97 |
| YCR077C | 3337.529518 | -0.034113688 | 0.856084893 | 0.97 |
| YCR099C | 126.2405817 | -0.094587798 | 0.85605991 | 0.97 |
| YDL021W | 1151.606846 | -0.084181614 | 0.857001956 | 0.97 |
| YDL024C | 228.1924469 | 0.082940454 | 0.852879135 | 0.97 |
| YDL136W | 915.4579307 | -0.116328935 | 0.856199624 | 0.97 |
| YDR082W | 550.4664737 | 0.051371457 | 0.854775968 | 0.97 |
| YDR191W | 146.592055 | 0.057507869 | 0.853798106 | 0.97 |
| YDR282C | 383.6210181 | -0.056746285 | 0.853089061 | 0.97 |
| YDR319C | 485.2377065 | -0.046195665 | 0.856410238 | 0.97 |
| YDR447C | 4088.999597 | -0.17170288 | 0.852553501 | 0.97 |
| YER052C | 2274.895414 | 0.082598853 | 0.857067853 | 0.97 |
| YFR030W | 1464.99151 | -0.07393337 | 0.855297186 | 0.97 |
| YGL076C | 4229.319259 | -0.167033211 | 0.855114511 | 0.97 |
| YHR066W | 786.1240577 | -0.077624889 | 0.854240927 | 0.97 |
| YHR087W | 3070.022032 | -0.093877731 | 0.856740816 | 0.97 |
| YHR176W | 373.1735066 | -0.046215463 | 0.855261866 | 0.97 |
| YIL167W | 108.465152 | -0.05742264 | 0.855795118 | 0.97 |
| YJL029C | 1512.843835 | -0.02944545 | 0.8572175 | 0.97 |
| YJL134W | 603.1655535 | -0.058235289 | 0.852759689 | 0.97 |
| YLR231C | 929.429053 | -0.09335489 | 0.854877733 | 0.97 |
| YLR243W | 616.8482711 | -0.055545958 | 0.856311117 | 0.97 |
| YLR418C | 850.5544851 | -0.035928299 | 0.852989542 | 0.97 |
| YML013W | 1108.615798 | 0.0483773 | 0.857712656 | 0.97 |
| YML057W | 1931.217961 | -0.035357872 | 0.855087351 | 0.97 |
| YML083C | 177.180785 | -0.121549045 | 0.857276762 | 0.97 |
| YMR131C | 2416.896733 | 0.070018018 | 0.857872023 | 0.97 |
| YMR144W | 347.0360461 | 0.058326267 | 0.85768937 | 0.97 |
| YNL047C | 720.3103825 | 0.049284202 | 0.855826059 | 0.97 |
| YNR001C | 10779.43573 | -0.080669962 | 0.857383616 | 0.97 |
| YOL014W | 428.9922641 | -0.146067355 | 0.853369772 | 0.97 |
| YOR245C | 966.5341375 | 0.052113541 | 0.858011605 | 0.97 |
| YPR140W | 900.9897118 | -0.053088174 | 0.854125245 | 0.97 |
| YER040W | 900.0166243 | 0.051600387 | 0.858673375 | 0.97 |
| YER102W | 2473.647383 | -0.155690858 | 0.85881903 | 0.97 |
| YKL096W-A | 39924.89359 | 0.067075748 | 0.85852939 | 0.97 |
| YMR020W | 992.3354434 | 0.074891296 | 0.859484848 | 0.97 |
| YCL030C | 8176.65539 | -0.057940791 | 0.860448993 | 0.97 |
| YEL030W | 524.4916329 | 0.049418382 | 0.861285879 | 0.97 |
| YGL123W | 15975.90289 | 0.029095304 | 0.861334419 | 0.97 |

| | | | | |
|---|---|---|---|---|
| YGL125W | 958.6893776 | -0.067221488 | 0.861056873 | 0.97 |
| YJL113W | 119.706187 | 0.060964741 | 0.860961396 | 0.97 |
| YLR185W | 4884.991766 | 0.021496015 | 0.860954376 | 0.97 |
| YLR219W | 2484.562083 | 0.045969219 | 0.860247727 | 0.97 |
| YNL262W | 2257.387419 | -0.053152914 | 0.861483373 | 0.97 |
| YPL084W | 2075.944666 | -0.027166123 | 0.860507267 | 0.97 |
| snR11 | 261.7635615 | -0.056236638 | 0.861229104 | 0.97 |
| YCR066W | 426.4825775 | 0.039379488 | 0.861842683 | 0.97 |
| YGL102C | 355.1928609 | 0.046225369 | 0.8616912 | 0.97 |
| YBR148W | 178.9686946 | -0.066977228 | 0.862090575 | 0.97 |
| YAL031C | 1097.389337 | 0.043723544 | 0.868728981 | 0.97 |
| YAL054C | 1109.295849 | -0.072248654 | 0.871135433 | 0.97 |
| YBR004C | 976.5395701 | -0.05090153 | 0.869373053 | 0.97 |
| YBR108W | 1821.037124 | 0.035561052 | 0.865815834 | 0.97 |
| YCR017C | 2986.294618 | 0.047961042 | 0.87023171 | 0.97 |
| YDL054C | 853.6605887 | 0.046619128 | 0.864938495 | 0.97 |
| YDL095W | 3378.459132 | 0.04359576 | 0.868308545 | 0.97 |
| YDL214C | 464.7760152 | -0.15875583 | 0.865209514 | 0.97 |
| YDR005C | 1131.574288 | -0.036357563 | 0.86923904 | 0.97 |
| YDR265W | 518.8074155 | 0.037618645 | 0.870321997 | 0.97 |
| YDR343C | 868.7911596 | -0.074413333 | 0.862838494 | 0.97 |
| YDR356W | 1114.540575 | -0.044619743 | 0.863150368 | 0.97 |
| YDR418W | 2788.265019 | 0.020466441 | 0.870658654 | 0.97 |
| YDR504C | 165.3464621 | -0.0441728 | 0.863343527 | 0.97 |
| YEL042W | 1714.856815 | 0.049767812 | 0.869977894 | 0.97 |
| YFL048C | 1163.307711 | -0.042516526 | 0.866726091 | 0.97 |
| YGL101W | 448.9089719 | -0.063906946 | 0.86571311 | 0.97 |
| YGR214W | 11003.56684 | -0.125391864 | 0.865439697 | 0.97 |
| YHR215W | 110.5268911 | -0.151302561 | 0.86961246 | 0.97 |
| YIL117C | 1107.127061 | -0.065681809 | 0.870487859 | 0.97 |
| YIL143C | 1830.368647 | -0.054361178 | 0.868081728 | 0.97 |
| YJL171C | 1558.602902 | -0.04326685 | 0.87106325 | 0.97 |
| YJR064W | 2718.261132 | 0.046103375 | 0.87157355 | 0.97 |
| YJR097W | 96.20405896 | 0.076641222 | 0.864284028 | 0.97 |
| YJR101W | 534.1436387 | 0.027951255 | 0.871662983 | 0.97 |
| YKL008C | 1304.945966 | 0.051078468 | 0.870674694 | 0.97 |
| YKL107W | 249.2824796 | 0.054939668 | 0.871124953 | 0.97 |
| YKL155C | 791.2689341 | -0.03460514 | 0.865602503 | 0.97 |
| YKL166C | 506.1904776 | -0.045043043 | 0.865877336 | 0.97 |
| YKR042W | 18766.32248 | -0.052600451 | 0.869526865 | 0.97 |
| YKR058W | 2130.69981 | 0.069024667 | 0.871169676 | 0.97 |
| YLL041C | 3315.150059 | -0.082036749 | 0.869692566 | 0.97 |
| YLR090W | 931.4547969 | -0.048140524 | 0.867059591 | 0.97 |

| | | | | |
|---|---|---|---|---|
| YLR337C | 2433.363455 | -0.035041451 | 0.866455593 | 0.97 |
| YLR388W | 1672.166035 | 0.034362301 | 0.864779583 | 0.97 |
| YMR033W | 1176.392622 | 0.04594039 | 0.864179866 | 0.97 |
| YMR192W | 859.8324449 | 0.040013151 | 0.868459991 | 0.97 |
| YMR274C | 249.323644 | -0.055530603 | 0.863910931 | 0.97 |
| YNL039W | 1798.324816 | -0.034330401 | 0.867023176 | 0.97 |
| YNL045W | 3993.621371 | -0.081570788 | 0.865585095 | 0.97 |
| YNL111C | 246.1352445 | 0.071640116 | 0.868521063 | 0.97 |
| YNL211C | 132.5864716 | -0.054872535 | 0.864993357 | 0.97 |
| YNL271C | 3740.617123 | 0.035342658 | 0.869838193 | 0.97 |
| YNR006W | 1506.041195 | -0.030669677 | 0.868772732 | 0.97 |
| YOL048C | 1220.015423 | -0.054830881 | 0.869328238 | 0.97 |
| YOL158C | 1769.752458 | -0.087344607 | 0.870691507 | 0.97 |
| YOR031W | 276.3450169 | -0.092932519 | 0.870882498 | 0.97 |
| YOR064C | 481.1143371 | -0.063871591 | 0.863245452 | 0.97 |
| YOR330C | 1527.333333 | 0.032837436 | 0.865682145 | 0.97 |
| YPL028W | 3100.321144 | -0.072138916 | 0.871627482 | 0.97 |
| YPL137C | 1258.265301 | 0.050784928 | 0.867877554 | 0.97 |
| YPL277C | 206.5316036 | 0.066655661 | 0.864020467 | 0.97 |
| YPR015C | 179.9077993 | -0.072306989 | 0.866564367 | 0.97 |
| YPR030W | 2156.768509 | 0.081154877 | 0.869572528 | 0.97 |
| YAL060W | 4666.556071 | -0.072304855 | 0.874099668 | 0.97 |
| YBL039C | 1528.083632 | -0.074028933 | 0.872389224 | 0.97 |
| YBL043W | 324.5838638 | -0.090870367 | 0.873357847 | 0.97 |
| YBL101C | 1814.259973 | 0.045559452 | 0.872965711 | 0.97 |
| YBR079C | 5290.065687 | 0.054736096 | 0.872438432 | 0.97 |
| YCR097W | 1013.079937 | -0.063321807 | 0.874232648 | 0.97 |
| YDR176W | 1688.228212 | -0.034873382 | 0.872734574 | 0.97 |
| YDR376W | 985.9130091 | -0.034420965 | 0.874445433 | 0.97 |
| YDR380W | 4921.676121 | 0.082410936 | 0.87254807 | 0.97 |
| YER095W | 1267.382435 | 0.038805563 | 0.872041083 | 0.97 |
| YGR245C | 1499.530458 | -0.066012609 | 0.873713416 | 0.97 |
| YHR214C-B | 235.6005636 | 0.073154483 | 0.873560981 | 0.97 |
| YJR135C | 188.8236464 | 0.038935414 | 0.87425268 | 0.97 |
| YOR148C | 393.9247418 | -0.033205144 | 0.873184669 | 0.97 |
| YOR193W | 437.421385 | 0.041331832 | 0.87457003 | 0.97 |
| YPL039W | 426.2421359 | 0.046738015 | 0.873804225 | 0.97 |
| YIR027C | 227.1514133 | 0.064530819 | 0.874793319 | 0.97 |
| YBR145W | 1370.985162 | -0.069728613 | 0.875707965 | 0.97 |
| YKL187C | 740.5340162 | -0.076701404 | 0.875590566 | 0.97 |
| YNL237W | 756.7694566 | 0.044963225 | 0.875727494 | 0.97 |
| YOR182C | 1767.616754 | -0.102058701 | 0.876114579 | 0.97 |
| YDR434W | 1180.144828 | 0.048632941 | 0.876675074 | 0.97 |

| | | | | |
|---|---|---|---|---|
| YFR046C | 527.2575594 | -0.034359982 | 0.876917118 | 0.97 |
| YER141W | 2454.288383 | -0.044644437 | 0.877440083 | 0.97 |
| YNL206C | 942.6169972 | 0.02811936 | 0.877438749 | 0.97 |
| YHR171W | 640.7534835 | -0.046923827 | 0.877944951 | 0.97 |
| YIR029W | 388.4137592 | 0.06915864 | 0.878047974 | 0.97 |
| YDL049C | 168.7509482 | -0.045723162 | 0.8782687 | 0.97 |
| YPL126W | 2185.981195 | 0.070105918 | 0.878443436 | 0.97 |
| YJL071W | 590.1206135 | 0.032493656 | 0.878763243 | 0.97 |
| YIL120W | 363.836718 | -0.049636823 | 0.879093738 | 0.97 |
| YBR113W | 115.7153994 | -0.047371321 | 0.880029085 | 0.98 |
| YBR249C | 4275.177673 | 0.010976269 | 0.879964263 | 0.98 |
| YLL060C | 550.1729711 | -0.06401898 | 0.880157025 | 0.98 |
| YLR414C | 1706.526088 | 0.049830559 | 0.879803394 | 0.98 |
| YAL033W | 350.8440347 | -0.065340633 | 0.882328627 | 0.98 |
| YBL023C | 762.982123 | -0.030069244 | 0.888215359 | 0.98 |
| YBR073W | 1901.278512 | -0.059383098 | 0.887462796 | 0.98 |
| YDL044C | 428.1351393 | 0.029677091 | 0.882273675 | 0.98 |
| YDL082W | 7404.050186 | 0.02915523 | 0.884266127 | 0.98 |
| YDL178W | 1397.749906 | 0.023716459 | 0.884658449 | 0.98 |
| YDL199C | 559.7723367 | -0.064378062 | 0.881778707 | 0.98 |
| YDR131C | 626.2979697 | 0.028484878 | 0.882260715 | 0.98 |
| YDR175C | 1042.356621 | -0.037961991 | 0.88733174 | 0.98 |
| YDR190C | 2078.525809 | 0.038874394 | 0.887446818 | 0.98 |
| YEL056W | 977.5251167 | -0.023785555 | 0.883217897 | 0.98 |
| YER098W | 930.5704049 | 0.043385782 | 0.882306863 | 0.98 |
| YGL213C | 596.0286848 | -0.030667428 | 0.887211604 | 0.98 |
| YHL022C | 136.8231823 | -0.064949911 | 0.887740205 | 0.98 |
| YHL039W | 1028.011858 | 0.049938638 | 0.881177298 | 0.98 |
| YHR005C | 1534.146963 | 0.097568801 | 0.889043089 | 0.98 |
| YHR052W | 1127.946092 | 0.064803084 | 0.886473747 | 0.98 |
| YHR151C | 595.1830514 | -0.048749654 | 0.881877379 | 0.98 |
| YHR201C | 1092.809897 | 0.043687395 | 0.888781561 | 0.98 |
| YHR205W | 2582.464034 | -0.04259717 | 0.887866191 | 0.98 |
| YIL074C | 1391.44586 | 0.046694854 | 0.889105992 | 0.98 |
| YIL096C | 413.8812975 | -0.05230593 | 0.880805525 | 0.98 |
| YIR042C | 196.4981605 | -0.081750042 | 0.886170301 | 0.98 |
| YJL077C | 162.6090211 | -0.042559199 | 0.883756903 | 0.98 |
| YJL081C | 1547.994145 | 0.021918025 | 0.887536968 | 0.98 |
| YJR130C | 1173.075557 | 0.049567242 | 0.883148884 | 0.98 |
| YKL109W | 7924.096762 | 0.051757372 | 0.88816082 | 0.98 |
| YKR029C | 884.4269483 | 0.031665768 | 0.885619923 | 0.98 |
| YLR151C | 483.8723388 | 0.034068191 | 0.887002587 | 0.98 |
| YML032C | 1432.723324 | -0.048297573 | 0.886382958 | 0.98 |

| | | | | |
|---|---|---|---|---|
| YMR130W | 646.8299959 | -0.057924813 | 0.886356374 | 0.98 |
| YMR136W | 2477.221542 | 0.052381309 | 0.88419358 | 0.98 |
| YNL098C | 3291.418535 | -0.029493552 | 0.888936712 | 0.98 |
| YNL124W | 1308.097286 | 0.038857696 | 0.886803114 | 0.98 |
| YNL250W | 994.2437698 | 0.037131754 | 0.889404717 | 0.98 |
| YNR020C | 315.5485889 | -0.035259713 | 0.888663766 | 0.98 |
| YNR047W | 1084.837265 | 0.042519223 | 0.886097007 | 0.98 |
| YNR052C | 1154.692673 | -0.025788018 | 0.882725322 | 0.98 |
| YOL121C | 3681.856963 | 0.018722853 | 0.886281188 | 0.98 |
| YOL153C | 1352.172864 | -0.060361384 | 0.882776344 | 0.98 |
| YOL164W | 729.2266525 | -0.049123719 | 0.885210673 | 0.98 |
| YOR014W | 3387.711726 | 0.035723511 | 0.88504284 | 0.98 |
| YOR015W | 345.4012917 | 0.033101934 | 0.889244647 | 0.98 |
| YOR032C | 566.9607469 | -0.07572847 | 0.886584889 | 0.98 |
| YOR110W | 712.9241656 | -0.030960164 | 0.886358975 | 0.98 |
| YOR385W | 901.596547 | -0.060736161 | 0.883532892 | 0.98 |
| YOR386W | 897.2436472 | 0.060179156 | 0.884064236 | 0.98 |
| YPL131W | 24700.79965 | 0.040567591 | 0.888721908 | 0.98 |
| YPL156C | 858.4762505 | -0.063856857 | 0.888986157 | 0.98 |
| YPL160W | 4849.742262 | 0.051378079 | 0.880516251 | 0.98 |
| YPR074C | 10075.28978 | -0.054614999 | 0.885632708 | 0.98 |
| YPR201W | 177.3808268 | 0.069609501 | 0.887334698 | 0.98 |
| YIL148W | 9852.705743 | 0.011127345 | 0.889585313 | 0.98 |
| YFL030W | 1073.611575 | -0.073274057 | 0.889855478 | 0.98 |
| YIL107C | 1780.529737 | 0.037085461 | 0.890665808 | 0.98 |
| YPR002W | 1192.998192 | 0.066873756 | 0.891162722 | 0.98 |
| YBL014C | 840.8039898 | -0.045842744 | 0.89195371 | 0.98 |
| YBR147W | 531.1286909 | 0.06097571 | 0.8919649 | 0.98 |
| YER018C | 488.1395433 | 0.062880342 | 0.892254164 | 0.98 |
| YHR004C | 921.2706765 | -0.036428682 | 0.892126731 | 0.98 |
| YKR091W | 1094.538594 | -0.056654361 | 0.891817871 | 0.98 |
| YMR061W | 1751.530821 | 0.028681086 | 0.891901889 | 0.98 |
| YBL095W | 182.0968065 | 0.0480479 | 0.893257857 | 0.98 |
| YDR237W | 832.9477874 | -0.034253627 | 0.892802964 | 0.98 |
| YGL227W | 1564.861142 | -0.047563452 | 0.89334682 | 0.98 |
| YGR222W | 726.7683442 | 0.028990237 | 0.892880796 | 0.98 |
| YNL022C | 938.4563852 | -0.032181331 | 0.893247342 | 0.98 |
| YNL229C | 1349.306513 | 0.01792135 | 0.89277747 | 0.98 |
| YLR080W | 639.2701305 | -0.055383431 | 0.894062958 | 0.98 |
| YFL023W | 930.5339775 | 0.041257029 | 0.894707934 | 0.98 |
| YOL068C | 1498.234041 | 0.028428376 | 0.894832837 | 0.98 |
| YPR083W | 614.8326597 | -0.061570161 | 0.894613662 | 0.98 |
| YAR015W | 7474.541729 | 0.030520785 | 0.896211566 | 0.98 |

| | | | | |
|---|---|---|---|---|
| YBR093C | 949.3588178 | 0.064803924 | 0.896398462 | 0.98 |
| YBR114W | 883.2430403 | -0.031241139 | 0.896298288 | 0.98 |
| YDR369C | 596.1613966 | 0.035575618 | 0.895608956 | 0.98 |
| YHR058C | 391.0666567 | -0.029777139 | 0.89594129 | 0.98 |
| YLR363C | 329.7816062 | -0.035861816 | 0.896531764 | 0.98 |
| YMR079W | 1633.407315 | -0.043810926 | 0.896244693 | 0.98 |
| YOR229W | 494.4703852 | 0.028637463 | 0.896614442 | 0.98 |
| YML036W | 292.5583071 | -0.043273731 | 0.896930076 | 0.98 |
| YMR085W | 377.7619403 | 0.039947244 | 0.897306695 | 0.98 |
| YBR072W | 9138.655064 | 0.096547261 | 0.89858328 | 0.98 |
| YBR216C | 879.1909045 | -0.024539512 | 0.899045207 | 0.98 |
| YDR408C | 1967.993247 | 0.03997273 | 0.89844066 | 0.98 |
| YJR048W | 1046.168345 | -0.071290631 | 0.898428391 | 0.98 |
| YKL068W-A | 172.0547206 | 0.03935031 | 0.898396162 | 0.98 |
| YKL197C | 902.4571586 | 0.032325328 | 0.899427568 | 0.98 |
| YLR253W | 1177.596684 | -0.045583128 | 0.897892894 | 0.98 |
| YLR454W | 3660.437031 | 0.032588075 | 0.898550525 | 0.98 |
| YMR267W | 919.8795189 | -0.046377514 | 0.899115183 | 0.98 |
| YNL248C | 1500.899157 | 0.063236196 | 0.89932515 | 0.98 |
| YNL311C | 782.0547648 | 0.02635563 | 0.898962798 | 0.98 |
| YBR157C | 589.2630079 | 0.060869291 | 0.900546162 | 0.98 |
| YER158W-A | 247.6290209 | -0.041941319 | 0.900441663 | 0.98 |
| YHR148W | 337.7936716 | 0.057005307 | 0.901091755 | 0.98 |
| YHR197W | 861.9145076 | 0.048611599 | 0.899967913 | 0.98 |
| YMR096W | 4030.130556 | 0.081018828 | 0.90106786 | 0.98 |
| YMR296C | 3004.03015 | 0.038034491 | 0.900968364 | 0.98 |
| YNL145W | 1813.203416 | -0.095286845 | 0.900734469 | 0.98 |
| YPL079W | 6102.629794 | -0.108996378 | 0.90035923 | 0.98 |
| YGR238C | 880.0595867 | -0.034129152 | 0.902124207 | 0.98 |
| YJL046W | 632.8219176 | 0.024022417 | 0.901749814 | 0.98 |
| YJL133C-A | 1683.932792 | 0.054855876 | 0.902299533 | 0.98 |
| YJL185C | 564.2478549 | 0.05341051 | 0.90159639 | 0.98 |
| YLR144C | 964.8658379 | 0.029771252 | 0.90241654 | 0.98 |
| YMR156C | 369.432093 | 0.049562947 | 0.902209835 | 0.98 |
| YIL158W | 651.6543059 | 0.036826855 | 0.902878034 | 0.98 |
| YKR039W | 765.789828 | -0.13578205 | 0.903973438 | 0.98 |
| YLL063C | 302.5298246 | 0.03636098 | 0.904565304 | 0.98 |
| YMR301C | 1249.636815 | -0.02536035 | 0.90413131 | 0.98 |
| YNL046W | 536.2842104 | 0.03406405 | 0.90446776 | 0.98 |
| YOR226C | 454.5609099 | -0.050092954 | 0.903644304 | 0.98 |
| YPL186C | 1114.795126 | -0.059825804 | 0.9041727 | 0.98 |
| YPL224C | 949.2587858 | -0.035866244 | 0.904282386 | 0.98 |
| SCR1 | 181.7417356 | 0.044732947 | 0.906710461 | 0.98 |

| | | | | |
|---|---|---|---|---|
| YAR028W | 513.5910617 | 0.026034495 | 0.906810122 | 0.98 |
| YJL158C | 11506.92019 | 0.027912171 | 0.906404865 | 0.98 |
| YJL163C | 815.8405692 | 0.048583781 | 0.906481572 | 0.98 |
| YKL146W | 1758.494435 | -0.053722382 | 0.906567454 | 0.98 |
| YOR181W | 1611.589546 | 0.035431842 | 0.906335549 | 0.98 |
| YPL104W | 889.4598617 | 0.031694897 | 0.905872684 | 0.98 |
| YHR070W | 688.4026548 | 0.035170649 | 0.907129214 | 0.98 |
| YFL014W | 10835.63862 | -0.035643655 | 0.90758343 | 0.98 |
| YKL108W | 391.5450461 | 0.039447839 | 0.908195587 | 0.98 |
| YPR184W | 5293.583109 | -0.060017857 | 0.908114103 | 0.98 |
| YGL031C | 8332.907395 | -0.017467009 | 0.908720231 | 0.98 |
| YJL089W | 344.4116839 | -0.005633226 | 0.908719804 | 0.98 |
| YKL012W | 1327.340355 | -0.016490879 | 0.908956579 | 0.98 |
| YKL127W | 1973.346678 | 0.026670001 | 0.909365955 | 0.98 |
| YMR042W | 272.8014796 | 0.041543815 | 0.909515912 | 0.98 |
| YBL029W | 662.4144794 | -0.036741958 | 0.910008857 | 0.98 |
| YBR218C | 7147.332039 | 0.029235956 | 0.910680664 | 0.98 |
| YGL171W | 777.7803509 | 0.054579605 | 0.910642312 | 0.98 |
| YNL116W | 793.4753006 | 0.027294917 | 0.910522733 | 0.98 |
| YPL050C | 1693.46287 | 0.031884631 | 0.910683512 | 0.98 |
| YPL209C | 162.5627938 | -0.03405194 | 0.910700074 | 0.98 |
| YJL038C | 194.4642451 | -0.060274366 | 0.910982486 | 0.98 |
| YBR012C | 184.3218002 | -0.046985744 | 0.914805175 | 0.98 |
| YDL201W | 813.4402377 | 0.032814807 | 0.913972285 | 0.98 |
| YDR036C | 1560.530814 | 0.035468978 | 0.912840264 | 0.98 |
| YDR395W | 2175.578678 | -0.030889613 | 0.913694149 | 0.98 |
| YEL024W | 1858.219132 | -0.048511145 | 0.914558706 | 0.98 |
| YFR032C-A | 2429.038295 | -0.065269042 | 0.912977617 | 0.98 |
| YHR084W | 1211.807628 | -0.030813785 | 0.912110379 | 0.98 |
| YIL142W | 3130.184557 | -0.026700223 | 0.912367066 | 0.98 |
| YJL155C | 1149.698916 | 0.039547673 | 0.911707374 | 0.98 |
| YJL172W | 1932.176519 | 0.039453381 | 0.913178832 | 0.98 |
| YJL177W | 7142.560719 | 0.008677922 | 0.914163801 | 0.98 |
| YJR096W | 2414.849421 | 0.045113019 | 0.913420721 | 0.98 |
| YKL044W | 194.2159883 | -0.045518305 | 0.912077413 | 0.98 |
| YKL075C | 477.0577168 | -0.025347729 | 0.913458225 | 0.98 |
| YLR362W | 1083.342817 | 0.029126818 | 0.914722643 | 0.98 |
| YMR271C | 754.1723733 | -0.074744773 | 0.912156339 | 0.98 |
| YOR141C | 1550.614472 | -0.021942525 | 0.913957128 | 0.98 |
| YOR184W | 6279.813274 | -0.044002292 | 0.911345711 | 0.98 |
| YPL165C | 810.9716846 | 0.035261676 | 0.911472796 | 0.98 |
| YPL255W | 357.750249 | 0.035652615 | 0.913708587 | 0.98 |
| YPR061C | 393.0462601 | -0.052416063 | 0.914468103 | 0.98 |

| | | | | |
|---|---|---|---|---|
| YDL025C | 1555.623411 | 0.029681442 | 0.917168518 | 0.98 |
| YDR316W | 792.8873428 | -0.025868821 | 0.917638325 | 0.98 |
| YDR458C | 1309.381626 | 0.024895566 | 0.915342373 | 0.98 |
| YDR492W | 689.8420252 | 0.036208723 | 0.917468892 | 0.98 |
| YGL078C | 1925.6458 | 0.040920592 | 0.917795801 | 0.98 |
| YGL121C | 888.9061681 | -0.057560683 | 0.917465075 | 0.98 |
| YGR023W | 927.8650533 | -0.059724794 | 0.916469425 | 0.98 |
| YGR288W | 284.2488249 | 0.044827543 | 0.915922644 | 0.98 |
| YLR435W | 440.7011003 | -0.041279135 | 0.915324858 | 0.98 |
| YMR106C | 546.3577506 | 0.024764338 | 0.916936912 | 0.98 |
| YNR013C | 2464.582186 | 0.025403108 | 0.915457206 | 0.98 |
| YOL040C | 16073.93418 | 0.012553474 | 0.916481703 | 0.98 |
| YOL110W | 433.4687434 | 0.044778242 | 0.91584207 | 0.98 |
| YOR076C | 709.8237452 | 0.021323215 | 0.916856565 | 0.98 |
| YOR142W | 3533.236265 | 0.024414056 | 0.917174434 | 0.98 |
| YPL097W | 440.8818332 | 0.023370227 | 0.916640006 | 0.98 |
| YPR154W | 2486.651934 | -0.025598397 | 0.917958431 | 0.98 |
| YDL105W | 423.3140946 | 0.021502304 | 0.919547728 | 0.98 |
| YJR143C | 2572.640819 | 0.030321412 | 0.91933672 | 0.98 |
| YKL120W | 1575.003705 | 0.047190153 | 0.919480485 | 0.98 |
| YKL212W | 2578.339284 | 0.022131981 | 0.919299563 | 0.98 |
| YLR398C | 2110.607765 | -0.01986225 | 0.919401746 | 0.98 |
| YOR030W | 929.3153858 | 0.023540784 | 0.919580951 | 0.98 |
| YGL246C | 646.4635665 | 0.019012411 | 0.920005013 | 0.98 |
| YIL053W | 6767.756759 | 0.008184483 | 0.919894663 | 0.98 |
| YBL019W | 515.3822708 | -0.026469513 | 0.920513075 | 0.98 |
| YGR186W | 2388.772284 | 0.024372381 | 0.920414437 | 0.98 |
| YLR007W | 741.9239686 | -0.027530181 | 0.920843369 | 0.98 |
| YOL127W | 9841.72384 | -0.123982058 | 0.920994559 | 0.98 |
| YBR097W | 952.5745358 | 0.018902255 | 0.922186656 | 0.98 |
| YBR248C | 1552.941582 | -0.040680848 | 0.922870148 | 0.98 |
| YER167W | 1421.029082 | 0.028731664 | 0.922878147 | 0.98 |
| YHL003C | 1333.653462 | -0.046400777 | 0.922765867 | 0.98 |
| YJL082W | 1623.02873 | -0.033072171 | 0.921392462 | 0.98 |
| YLR238W | 614.7184945 | 0.020280099 | 0.92285333 | 0.98 |
| YLR326W | 402.567239 | -0.026376311 | 0.921329751 | 0.98 |
| YMR190C | 1250.870637 | 0.028641478 | 0.922576443 | 0.98 |
| YOR309C | 407.1266153 | -0.044351436 | 0.922974274 | 0.98 |
| YPL166W | 366.5892241 | -0.043179051 | 0.922581974 | 0.98 |
| YPL233W | 252.806959 | 0.018392621 | 0.922822898 | 0.98 |
| YBR094W | 1036.817856 | -0.019872558 | 0.924574602 | 0.98 |
| YCR091W | 1488.499833 | 0.035428731 | 0.923166142 | 0.98 |
| YER121W | 184.2483 | 0.049898588 | 0.924246707 | 0.98 |

| | | | | |
|---|---|---|---|---|
| YGR046W | 906.1577966 | 0.030189723 | 0.924804606 | 0.98 |
| YGR153W | 193.310946 | -0.030970598 | 0.924956229 | 0.98 |
| YGR208W | 739.3899853 | 0.025139773 | 0.924573207 | 0.98 |
| YKL218C | 773.1891416 | 0.038270761 | 0.923917198 | 0.98 |
| YLL056C | 287.6766171 | -0.044189244 | 0.924768276 | 0.98 |
| YMR292W | 429.0322087 | -0.049727069 | 0.924342473 | 0.98 |
| YOL108C | 360.9401457 | 0.034425475 | 0.923551149 | 0.98 |
| YPR042C | 1338.934548 | 0.022666464 | 0.923566728 | 0.98 |
| YBR015C | 2379.48263 | -0.019034937 | 0.925516182 | 0.98 |
| YBR067C | 11273.4348 | 0.01984148 | 0.926206057 | 0.98 |
| YAR018C | 738.6960091 | -0.019250992 | 0.934064829 | 0.98 |
| YBL066C | 1237.560937 | -0.039364066 | 0.929979054 | 0.98 |
| YBR001C | 1318.791969 | 0.027010313 | 0.927422114 | 0.98 |
| YBR018C | 184.0387102 | -0.051109805 | 0.934134519 | 0.98 |
| YBR208C | 3562.373289 | 0.056577606 | 0.934015282 | 0.98 |
| YDL058W | 1998.717774 | -0.017719462 | 0.933164264 | 0.98 |
| YDL111C | 757.760422 | -0.023724057 | 0.934171231 | 0.98 |
| YDL229W | 7732.532873 | -0.06775371 | 0.927229534 | 0.98 |
| YDR060W | 3331.110803 | -0.022969779 | 0.93370414 | 0.98 |
| YDR243C | 512.1645164 | 0.016074018 | 0.932210974 | 0.98 |
| YDR294C | 2749.492975 | -0.015095314 | 0.933904565 | 0.98 |
| YDR371W | 631.8329655 | -0.01991331 | 0.93300463 | 0.98 |
| YDR461W | 1830.189511 | -0.054077029 | 0.929527044 | 0.98 |
| YER026C | 2306.614486 | -0.035264876 | 0.930269461 | 0.98 |
| YER099C | 790.1735302 | 0.013042211 | 0.931019986 | 0.98 |
| YFL052W | 300.3969613 | 0.026038829 | 0.929911774 | 0.98 |
| YGL081W | 335.8828397 | 0.031018195 | 0.934076525 | 0.98 |
| YHL001W | 1027.533504 | 0.004311408 | 0.933035846 | 0.98 |
| YIL040W | 372.5621769 | -0.026414995 | 0.932547032 | 0.98 |
| YIR039C | 1029.181658 | -0.044343163 | 0.933132135 | 0.98 |
| YJL142C | 106.3041783 | -0.021727117 | 0.927598753 | 0.98 |
| YKL025C | 1634.326653 | -0.016874568 | 0.929978257 | 0.98 |
| YKL038W | 1738.382543 | 0.030743484 | 0.929842083 | 0.98 |
| YKR031C | 1778.007412 | 0.022329551 | 0.931797224 | 0.98 |
| YKR052C | 558.5950233 | 0.025522892 | 0.933139769 | 0.98 |
| YLL040C | 5253.851757 | 0.029873308 | 0.931694956 | 0.98 |
| YLR112W | 96.84022753 | 0.036959309 | 0.927781206 | 0.98 |
| YLR113W | 3162.32221 | -0.021147167 | 0.929443755 | 0.98 |
| YLR239C | 753.9388732 | 0.020220584 | 0.931714144 | 0.98 |
| YLR284C | 336.5089725 | -0.033164659 | 0.926626737 | 0.98 |
| YML007W | 2781.196668 | -0.023520883 | 0.933679158 | 0.98 |
| YMR142C | 11187.93902 | -0.030269987 | 0.929868024 | 0.98 |
| YMR201C | 569.2749285 | -0.028898931 | 0.930286226 | 0.98 |

| | | | | |
|---|---|---|---|---|
| YMR250W | 7553.858966 | -0.042813939 | 0.929767323 | 0.98 |
| YNL014W | 784.7565686 | -0.052426082 | 0.929289583 | 0.98 |
| YNL030W | 3372.305745 | -0.01469601 | 0.93395451 | 0.98 |
| YNR034W-A | 4345.183948 | 0.03529707 | 0.934204097 | 0.98 |
| YNR072W | 156.9410464 | -0.029746962 | 0.932566966 | 0.98 |
| YOL112W | 963.5678133 | 0.019760809 | 0.932132896 | 0.98 |
| YOL155C | 4950.84233 | 0.040083254 | 0.933856058 | 0.98 |
| YOR293W | 11411.33478 | 0.003266175 | 0.929749211 | 0.98 |
| YPR011C | 560.157333 | -0.030208863 | 0.929845322 | 0.98 |
| YPR029C | 1930.500091 | 0.013982339 | 0.93179337 | 0.98 |
| YPR187W | 1425.65006 | 0.014170024 | 0.932578558 | 0.98 |
| YAL059W | 208.6556716 | 0.025135912 | 0.935363999 | 0.98 |
| YDR030C | 444.8820441 | 0.026136857 | 0.936516665 | 0.98 |
| YDR173C | 369.7980333 | 0.024520264 | 0.936799374 | 0.98 |
| YDR225W | 3245.605018 | -0.030824924 | 0.93635396 | 0.98 |
| YER055C | 2945.986692 | 0.015389705 | 0.935479302 | 0.98 |
| YGL086W | 1388.801297 | -0.021841723 | 0.934966264 | 0.98 |
| YGL096W | 785.2336716 | -0.040754251 | 0.935772269 | 0.98 |
| YHR141C | 3868.981705 | -0.090365931 | 0.935986468 | 0.98 |
| YIL017C | 1438.735299 | -0.02967397 | 0.934496016 | 0.98 |
| YJL213W | 560.8105438 | -0.035647287 | 0.936019854 | 0.98 |
| YKR013W | 4305.106637 | 0.035205146 | 0.935024969 | 0.98 |
| YKR076W | 1274.281879 | -0.040212508 | 0.934706208 | 0.98 |
| YLR095C | 1759.144757 | -0.014845961 | 0.93666962 | 0.98 |
| YNL217W | 1768.453008 | 0.01966562 | 0.936384641 | 0.98 |
| YPR016C | 1938.157861 | -0.021435566 | 0.936981927 | 0.98 |
| YOR099W | 2318.668874 | 0.017045143 | 0.937166649 | 0.98 |
| YFL042C | 1495.188388 | -0.03198874 | 0.938278215 | 0.98 |
| YGR001C | 885.4111524 | -0.001862771 | 0.937527092 | 0.98 |
| YHL027W | 1629.795881 | -0.025829816 | 0.938126173 | 0.98 |
| YOR039W | 1072.032394 | -0.020074519 | 0.938312451 | 0.98 |
| YPL015C | 1643.336309 | 0.019265479 | 0.93791854 | 0.98 |
| YPL068C | 273.8997414 | 0.030166242 | 0.937804311 | 0.98 |
| YFL007W | 4593.396937 | 0.018384895 | 0.938606428 | 0.98 |
| YOR340C | 775.5259752 | 0.026706016 | 0.938674899 | 0.98 |
| YDR103W | 991.2634766 | 0.039922348 | 0.939714152 | 0.98 |
| YDR541C | 514.5384389 | 0.038523213 | 0.939639259 | 0.98 |
| YGL062W | 7360.866755 | -0.045417017 | 0.939425303 | 0.98 |
| YGL253W | 16247.71261 | -0.045786645 | 0.940139642 | 0.98 |
| YHR017W | 1256.836076 | 0.021298883 | 0.939489816 | 0.98 |
| YHR039C | 2261.246675 | 0.023645155 | 0.940311276 | 0.98 |
| YIL064W | 505.7570311 | -0.034163706 | 0.940083526 | 0.98 |
| YJR078W | 516.4472306 | 0.036211006 | 0.940117933 | 0.98 |

| | | | | |
|---|---|---|---|---|
| YLR180W | 3585.379162 | 0.007599239 | 0.939323809 | 0.98 |
| YKL059C | 865.3974506 | 0.011218449 | 0.940653955 | 0.98 |
| YDR463W | 1963.287278 | 0.014895104 | 0.941840655 | 0.98 |
| YLR049C | 719.3459985 | 0.023911149 | 0.941812189 | 0.98 |
| YMR194C-B | 137.2124333 | 0.036623078 | 0.940976679 | 0.98 |
| YNL049C | 1288.390098 | -0.015634347 | 0.941558613 | 0.98 |
| YOR153W | 11636.99259 | -0.066211604 | 0.941686049 | 0.98 |
| YPR132W | 5266.120182 | -0.109035967 | 0.941752399 | 0.98 |
| YCL058C | 103.80183 | -0.019332242 | 0.943220221 | 0.99 |
| YDR443C | 1163.053889 | 0.018812879 | 0.943580986 | 0.99 |
| YGL187C | 1692.180783 | 0.029545868 | 0.943073984 | 0.99 |
| YGR189C | 5490.403188 | 0.036434574 | 0.942786617 | 0.99 |
| YKL006C-A | 282.6893408 | 0.017646413 | 0.942455371 | 0.99 |
| YKL026C | 613.714189 | -0.039129322 | 0.942225845 | 0.99 |
| YLR352W | 926.4882915 | 0.013220706 | 0.942925997 | 0.99 |
| YPL119C | 673.8655489 | -0.035071164 | 0.943411513 | 0.99 |
| YPR079W | 754.8054031 | -0.012175004 | 0.943656695 | 0.99 |
| YCR034W | 1456.119271 | -0.038574737 | 0.943915871 | 0.99 |
| YNL297C | 2104.946506 | 0.013344622 | 0.944078815 | 0.99 |
| YDR172W | 3599.759088 | -0.012798947 | 0.944598025 | 0.99 |
| YGR047C | 884.2267464 | -0.013784502 | 0.94461827 | 0.99 |
| YJR002W | 1689.001321 | 0.020622262 | 0.946011401 | 0.99 |
| YCR011C | 2808.118139 | 0.014629769 | 0.946453131 | 0.99 |
| YOR078W | 697.5265362 | -0.019088319 | 0.94669506 | 0.99 |
| YDL207W | 1207.359156 | 0.00928588 | 0.947516317 | 0.99 |
| YEL005C | 730.8016186 | -0.025453127 | 0.947879132 | 0.99 |
| YGL189C | 20518.43042 | -0.051365057 | 0.946992931 | 0.99 |
| YGR101W | 624.5372953 | -0.011647164 | 0.947878176 | 0.99 |
| YKR003W | 1162.972466 | -0.017657949 | 0.947609827 | 0.99 |
| YLR325C | 5760.035975 | -0.067851829 | 0.94724331 | 0.99 |
| YCL026C-B | 269.8485152 | -0.024134663 | 0.94833935 | 0.99 |
| YGR100W | 1479.985263 | 0.008491827 | 0.948372919 | 0.99 |
| YGL030W | 7624.667323 | -0.102515366 | 0.949026737 | 0.99 |
| YKR009C | 961.8765586 | 0.026002029 | 0.94952644 | 0.99 |
| YNL293W | 734.6319527 | 0.018329391 | 0.949717611 | 0.99 |
| YPL231W | 11637.98709 | -0.022945744 | 0.950095258 | 0.99 |
| YPL227C | 650.9518323 | 0.02042559 | 0.95030536 | 0.99 |
| YLR137W | 268.6657732 | -0.018920806 | 0.950668807 | 0.99 |
| YBL055C | 719.3024768 | -0.014877711 | 0.951436722 | 0.99 |
| YBR283C | 3928.577615 | 0.006982759 | 0.953394722 | 0.99 |
| YDR055W | 3654.499745 | -0.014414781 | 0.952676017 | 0.99 |
| YDR391C | 1742.594469 | -0.02230735 | 0.953541296 | 0.99 |
| YEL036C | 1490.088365 | 0.020330988 | 0.951944288 | 0.99 |

| | | | | |
|---|---|---|---|---|
| YER078C | 919.3267409 | -0.012915425 | 0.952229753 | 0.99 |
| YHR069C | 771.2357851 | 0.018632454 | 0.953500811 | 0.99 |
| YJL107C | 129.4785502 | 0.024924336 | 0.953513279 | 0.99 |
| YJL138C | 1955.737437 | -0.019989668 | 0.953135854 | 0.99 |
| YJR062C | 945.4060688 | 0.012909801 | 0.95177879 | 0.99 |
| YKL217W | 2399.793467 | 0.024176469 | 0.953078767 | 0.99 |
| YPL193W | 620.6543173 | -0.011249235 | 0.952865198 | 0.99 |
| YPR080W | 5311.40989 | -0.052324138 | 0.954068775 | 0.99 |
| YBL080C | 486.2802333 | 0.013814785 | 0.955133364 | 0.99 |
| YMR068W | 420.8665046 | -0.017869427 | 0.955735916 | 0.99 |
| YDR257C | 688.7119517 | -0.016382191 | 0.95594908 | 0.99 |
| YBL085W | 2143.648048 | 0.013636946 | 0.956813488 | 0.99 |
| YCR007C | 318.2333147 | 0.009168411 | 0.95734013 | 0.99 |
| YER162C | 931.7095211 | 0.009058257 | 0.956567914 | 0.99 |
| YJL033W | 1448.230029 | 0.028951408 | 0.957094746 | 0.99 |
| YKL191W | 1251.863612 | 0.014084365 | 0.957406306 | 0.99 |
| YKR002W | 1434.178622 | 0.009758539 | 0.957151766 | 0.99 |
| YMR305C | 6521.89708 | -0.017286047 | 0.957454879 | 0.99 |
| YPL222W | 848.2140163 | 0.014071218 | 0.956768153 | 0.99 |
| YBR104W | 350.9392427 | 0.023104547 | 0.959172307 | 0.99 |
| YDR207C | 1436.193266 | -0.015450573 | 0.95851883 | 0.99 |
| YDR436W | 1255.862426 | 0.012424552 | 0.959383937 | 0.99 |
| YFL031C-A | 101.8569422 | 0.017004233 | 0.958990987 | 0.99 |
| YLR395C | 1418.063046 | -0.024876604 | 0.958732025 | 0.99 |
| YLR404W | 245.17757 | -0.013652085 | 0.958390187 | 0.99 |
| YPR055W | 1160.769461 | 0.01178964 | 0.959313534 | 0.99 |
| YPR158C-D | 130.4698158 | -0.048682555 | 0.958130478 | 0.99 |
| YLL055W | 836.4946306 | -0.025880994 | 0.959710171 | 0.99 |
| YBR293W | 445.7090525 | -0.016664124 | 0.962019278 | 0.99 |
| YCR094W | 731.6068684 | 0.010798632 | 0.961194983 | 0.99 |
| YDR284C | 1216.7009 | 0.00571349 | 0.96217778 | 0.99 |
| YEL034C-A | 2748.462234 | -0.023051657 | 0.962071227 | 0.99 |
| YEL041W | 348.1575551 | 0.017595363 | 0.96043999 | 0.99 |
| YLR265C | 441.14354 | 0.001004417 | 0.96205162 | 0.99 |
| YLR333C | 2119.808061 | -0.06632106 | 0.961447199 | 0.99 |
| YLR410W | 2881.535918 | 0.010250575 | 0.96075572 | 0.99 |
| YMR125W | 2280.761082 | 0.013443289 | 0.961096945 | 0.99 |
| YNL093W | 262.8484398 | 0.020488178 | 0.961682556 | 0.99 |
| YNL284C | 1102.406706 | -0.011361508 | 0.961510294 | 0.99 |
| YNR066C | 265.0332103 | -0.016582555 | 0.960362283 | 0.99 |
| YMR137C | 598.7111524 | 0.008695298 | 0.962389642 | 0.99 |
| YMR217W | 5540.397432 | 0.011522786 | 0.962593358 | 0.99 |
| YBL027W | 10124.8249 | -0.10896069 | 0.962935782 | 0.99 |

| | | | | |
|---|---|---|---|---|
| YBL002W | 4494.23128 | -0.043008286 | 0.969408139 | 0.99 |
| YBL016W | 1090.490568 | 0.027246889 | 0.967853655 | 0.99 |
| YBR228W | 415.0546527 | -0.010380308 | 0.967997644 | 0.99 |
| YCR043C | 438.4206899 | -0.017927763 | 0.965614683 | 0.99 |
| YDR088C | 645.3535973 | 0.009709503 | 0.969239951 | 0.99 |
| YEL009C | 21167.59097 | 0.013905735 | 0.969168355 | 0.99 |
| YER092W | 378.6245953 | 0.01127762 | 0.969209047 | 0.99 |
| YER156C | 1503.86508 | -0.024781762 | 0.963361676 | 0.99 |
| YFR017C | 3258.918594 | -0.020371833 | 0.966098337 | 0.99 |
| YGL156W | 2401.006767 | 0.014048038 | 0.965277304 | 0.99 |
| YGR070W | 2198.55855 | -0.026451984 | 0.968678579 | 0.99 |
| YGR087C | 287.4531423 | -0.028191733 | 0.964522161 | 0.99 |
| YGR183C | 999.4364709 | -0.023086995 | 0.963971016 | 0.99 |
| YHL006C | 116.1902464 | -0.014912372 | 0.967016836 | 0.99 |
| YHR021C | 5458.86145 | -0.021622007 | 0.96383928 | 0.99 |
| YHR124W | 347.8549107 | -0.01544041 | 0.964226554 | 0.99 |
| YIL119C | 838.0211886 | 0.02004311 | 0.969258149 | 0.99 |
| YIL136W | 7495.331143 | -0.015272882 | 0.969662052 | 0.99 |
| YJL048C | 1114.208522 | 0.012618615 | 0.965296185 | 0.99 |
| YJL188C | 893.4798181 | -0.093952499 | 0.965674446 | 0.99 |
| YJR008W | 1161.824512 | 0.01924957 | 0.967365574 | 0.99 |
| YJR091C | 3359.251305 | -0.013731502 | 0.968319834 | 0.99 |
| YKL091C | 918.9800837 | -0.014354069 | 0.968025587 | 0.99 |
| YKL216W | 7973.975155 | -0.026889263 | 0.967847257 | 0.99 |
| YLL013C | 908.878738 | -0.014447507 | 0.969192196 | 0.99 |
| YLR054C | 138.716349 | -0.022699951 | 0.965386624 | 0.99 |
| YLR167W | 28914.13243 | -0.05137437 | 0.966551674 | 0.99 |
| YLR267W | 1321.159087 | 0.014130985 | 0.967146096 | 0.99 |
| YLR393W | 270.1892054 | -0.013809441 | 0.969494206 | 0.99 |
| YML049C | 1684.51308 | -0.012435471 | 0.964475448 | 0.99 |
| YML080W | 296.6855777 | 0.0215595 | 0.966732024 | 0.99 |
| YML128C | 9230.440998 | 0.031344899 | 0.966987727 | 0.99 |
| YMR116C | 12111.33345 | -0.124567324 | 0.965505182 | 0.99 |
| YMR173W-A | 260.3952806 | -0.015388031 | 0.967082352 | 0.99 |
| YOR273C | 1760.091485 | 0.011283215 | 0.968704071 | 0.99 |
| YPL236C | 570.2921643 | -0.013297552 | 0.969527062 | 0.99 |
| YBR102C | 889.0122882 | 0.005395344 | 0.970030822 | 0.99 |
| YBR240C | 118.1647641 | 0.006669811 | 0.970934915 | 0.99 |
| YDR275W | 363.6731352 | 0.016377559 | 0.97141924 | 0.99 |
| YDR379W | 1429.073535 | -0.010264974 | 0.970944514 | 0.99 |
| YER028C | 507.0764562 | -0.00947818 | 0.971115115 | 0.99 |
| YGR118W | 4349.282861 | -0.053339464 | 0.9711736 | 0.99 |
| YLR133W | 1071.862832 | -0.007321041 | 0.970925802 | 0.99 |

| | | | | |
|---|---|---|---|---|
| YML087C | 330.7889367 | 0.015584492 | 0.971331806 | 0.99 |
| YMR034C | 377.1167506 | -0.012316947 | 0.970188819 | 0.99 |
| YDL003W | 760.5875817 | 0.025005998 | 0.972511536 | 0.99 |
| YDR277C | 2772.925083 | 0.018406127 | 0.973060764 | 0.99 |
| YDR345C | 12971.54144 | -0.022855844 | 0.972527262 | 0.99 |
| YER082C | 1253.635463 | -0.007753966 | 0.972595537 | 0.99 |
| YER118C | 889.8446464 | 0.011743283 | 0.972679658 | 0.99 |
| YGL240W | 171.0354676 | 0.007574014 | 0.972352698 | 0.99 |
| YGR010W | 527.974638 | 0.008347501 | 0.973306508 | 0.99 |
| YIL161W | 488.7537738 | 0.008368883 | 0.9729778 | 0.99 |
| YJL080C | 5735.124683 | -0.018094461 | 0.971829243 | 0.99 |
| YNL240C | 1001.196601 | 0.01021772 | 0.973300539 | 0.99 |
| YHL002W | 1516.40559 | 0.006922815 | 0.975123235 | 0.99 |
| YJR123W | 26166.00582 | -0.050568833 | 0.974850483 | 0.99 |
| YMR302C | 2126.727676 | -0.012661438 | 0.975086887 | 0.99 |
| YBL059W | 273.8243146 | 0.006329449 | 0.975532835 | 0.99 |
| YGR054W | 2529.613046 | 0.013911331 | 0.976076857 | 0.99 |
| YHR203C | 5825.930229 | -0.064752953 | 0.976163511 | 0.99 |
| YPR027C | 342.9171324 | 0.011328509 | 0.976289723 | 0.99 |
| YDL019C | 2417.237505 | -0.006033911 | 0.97725683 | 1.00 |
| YJR145C | 6502.704468 | -0.073587496 | 0.977477638 | 1.00 |
| YDR453C | 889.6622872 | -0.013055903 | 0.97798021 | 1.00 |
| YLR228C | 906.7831503 | -0.009918846 | 0.977899385 | 1.00 |
| YBL032W | 1731.697141 | -0.023631739 | 0.978294272 | 1.00 |
| YBR184W | 226.9887473 | -0.0077337 | 0.983470899 | 1.00 |
| YDL104C | 329.3472486 | -0.00619055 | 0.981609136 | 1.00 |
| YDL124W | 12433.87204 | -0.005696432 | 0.982690141 | 1.00 |
| YDR165W | 1014.540812 | 0.01074624 | 0.980035115 | 1.00 |
| YDR292C | 1617.849006 | -0.008487108 | 0.980873854 | 1.00 |
| YEL070W | 298.2975483 | -0.015187444 | 0.982233735 | 1.00 |
| YFL036W | 2229.86904 | 0.007316483 | 0.980786218 | 1.00 |
| YGL012W | 2257.199778 | -0.010498415 | 0.981437007 | 1.00 |
| YGL017W | 502.0325199 | 0.003666671 | 0.982902289 | 1.00 |
| YGL248W | 1428.88377 | -0.008522642 | 0.983122902 | 1.00 |
| YGR094W | 6966.605497 | -0.014266542 | 0.983225797 | 1.00 |
| YGR148C | 7673.362198 | -0.028346385 | 0.978974461 | 1.00 |
| YGR240C | 9640.703605 | 0.004693127 | 0.983084898 | 1.00 |
| YGR248W | 2366.128573 | -0.014546874 | 0.981347253 | 1.00 |
| YHR216W | 3639.691774 | 0.002226332 | 0.981962386 | 1.00 |
| YJL103C | 874.1846601 | 0.001854367 | 0.978778375 | 1.00 |
| YJL136C | 5415.03448 | -0.059311291 | 0.980002119 | 1.00 |
| YJL141C | 4385.516839 | -0.010987037 | 0.981732848 | 1.00 |
| YJL190C | 9254.873132 | -0.046728721 | 0.983517582 | 1.00 |

| | | | | |
|---|---|---|---|---|
| YJL204C | 683.8445536 | -0.005142429 | 0.981253655 | 1.00 |
| YJR021C | 884.0147888 | 0.006949035 | 0.979814441 | 1.00 |
| YLR055C | 1018.744824 | 0.007407345 | 0.979721821 | 1.00 |
| YMR221C | 1062.231053 | -0.006523388 | 0.981008378 | 1.00 |
| YNL213C | 587.1355028 | -0.011263489 | 0.981825705 | 1.00 |
| YNL227C | 372.4616162 | 0.006856049 | 0.981227283 | 1.00 |
| YNL280C | 1498.81009 | 0.005174971 | 0.983290095 | 1.00 |
| YNR028W | 918.8142146 | 0.0049508 | 0.979782806 | 1.00 |
| YOL052C-A | 1535.771374 | 0.012670526 | 0.982704408 | 1.00 |
| YOL062C | 1303.409819 | 0.004599859 | 0.980546307 | 1.00 |
| YBL071W-A | 297.3219208 | -0.005904244 | 0.984348326 | 1.00 |
| YBR002C | 607.1395928 | -0.009804244 | 0.984223547 | 1.00 |
| YDR070C | 1811.421734 | 0.004117192 | 0.98436829 | 1.00 |
| YMR066W | 755.7979974 | 0.003712368 | 0.984200327 | 1.00 |
| YKL104C | 3816.518475 | 0.013033904 | 0.984560464 | 1.00 |
| YHR006W | 1327.030831 | -0.012012575 | 0.985195562 | 1.00 |
| YIL077C | 964.8137743 | -0.007353485 | 0.985239784 | 1.00 |
| YDR150W | 4371.450407 | -0.009315987 | 0.985494115 | 1.00 |
| YOR288C | 898.6918954 | 0.004986349 | 0.985681733 | 1.00 |
| YFL033C | 2297.741868 | 0.003594834 | 0.986287231 | 1.00 |
| YGR050C | 100.6325721 | -0.00579537 | 0.986230464 | 1.00 |
| YNL222W | 196.3405424 | 0.008263255 | 0.986081889 | 1.00 |
| YAL053W | 2097.963607 | -0.005115813 | 0.988627796 | 1.00 |
| YBR153W | 413.971492 | -0.00169398 | 0.988941214 | 1.00 |
| YBR244W | 574.3609399 | -0.065027351 | 0.986929129 | 1.00 |
| YCL008C | 1030.586387 | -1.38E-05 | 0.988519826 | 1.00 |
| YER016W | 838.7636501 | -0.013834738 | 0.987214295 | 1.00 |
| YFL054C | 1511.844206 | -0.002497531 | 0.988325078 | 1.00 |
| YFR013W | 1397.161166 | -0.003806413 | 0.989045051 | 1.00 |
| YFR038W | 1186.649674 | -0.004843201 | 0.989157371 | 1.00 |
| YGL117W | 750.1030158 | 0.005422995 | 0.988814472 | 1.00 |
| YGR016W | 101.0221921 | 0.003204303 | 0.989233503 | 1.00 |
| YGR129W | 435.9932211 | 0.003322995 | 0.988906471 | 1.00 |
| YHL048W | 1666.986201 | -0.006245348 | 0.988787618 | 1.00 |
| YLR075W | 9637.869009 | -0.058109352 | 0.987860304 | 1.00 |
| YLR168C | 702.4056037 | 0.005966582 | 0.988262671 | 1.00 |
| YOR096W | 6750.76705 | -0.085317238 | 0.987180278 | 1.00 |
| YJR121W | 23130.92602 | -0.004895201 | 0.989958413 | 1.00 |
| YNR041C | 337.6551047 | 0.000706897 | 0.990585438 | 1.00 |
| YOR079C | 215.4299264 | -0.01102243 | 0.990405552 | 1.00 |
| YGL257C | 1193.728393 | -0.0026873 | 0.990784551 | 1.00 |
| YDL091C | 1187.448267 | -0.014750933 | 0.991007078 | 1.00 |
| YHR194W | 859.058911 | -0.003466346 | 0.991420754 | 1.00 |

| | | | | |
|---|---|---|---|---|
| YLR128W | 336.2576961 | 0.003476206 | 0.991353308 | 1.00 |
| YDR216W | 2843.606015 | -0.010787623 | 0.991764484 | 1.00 |
| YPR005C | 426.8209175 | -0.000786594 | 0.991989412 | 1.00 |
| YPR109W | 650.4688515 | -0.002527976 | 0.992058632 | 1.00 |
| YDR120C | 1240.636645 | -0.008260096 | 0.992286796 | 1.00 |
| YDR300C | 1204.265207 | -0.003158237 | 0.992644209 | 1.00 |
| YKL006W | 2404.725348 | -0.100888366 | 0.992870364 | 1.00 |
| YNR008W | 899.8873602 | -0.003094761 | 0.993063594 | 1.00 |
| YBR297W | 465.5986861 | 0.004612414 | 0.994717101 | 1.00 |
| YER054C | 1167.402398 | -0.004534814 | 0.994379672 | 1.00 |
| YGL104C | 947.9922368 | -0.001644366 | 0.993856177 | 1.00 |
| YGR260W | 2070.398504 | 0.002918997 | 0.994691111 | 1.00 |
| YHR195W | 1250.379878 | -0.003910908 | 0.994385642 | 1.00 |
| YLR117C | 1230.224284 | -0.002541191 | 0.994730492 | 1.00 |
| YLR340W | 21021.60532 | -0.018819811 | 0.993409386 | 1.00 |
| YPL061W | 6576.454349 | -0.092420523 | 0.994462511 | 1.00 |
| YPL153C | 363.2997459 | -0.002689044 | 0.993978851 | 1.00 |
| YGL136C | 674.1887477 | 0.001857114 | 0.995772687 | 1.00 |
| YJR094W-A | 1689.898779 | -0.045181129 | 0.995163468 | 1.00 |
| YLR016C | 414.1257784 | -0.000875773 | 0.995679318 | 1.00 |
| YLR248W | 3268.331309 | 9.65E-05 | 0.99602855 | 1.00 |
| YPL129W | 1819.666295 | -0.00532365 | 0.995907596 | 1.00 |
| YPR026W | 1343.846593 | -0.018911267 | 0.995698299 | 1.00 |
| YDL075W | 5187.384084 | -0.070850139 | 0.997213212 | 1.00 |
| YER066W | 204.0645964 | -0.007376689 | 0.996861112 | 1.00 |
| YJL020C | 4552.045062 | -0.000759223 | 0.996943562 | 1.00 |
| YOR104W | 252.8227861 | -0.000737403 | 0.997161955 | 1.00 |
| YDR021W | 377.9993278 | 0.008285141 | 0.997545804 | 1.00 |
| YDR079C-A | 102.409036 | -0.000314374 | 0.997770777 | 1.00 |
| YPL189W | 119.6862802 | -0.000677354 | 0.998166231 | 1.00 |
| YHR183W | 10217.98699 | -0.007503644 | 0.998808856 | 1.00 |
| YDL174C | 2966.298062 | 0.001764267 | 0.999268065 | 1.00 |
| YML118W | 395.2746742 | -0.005157518 | 0.999275975 | 1.00 |
| HRA1 | 34.22771229 | -0.308381106 | 0.389881714 | NA |
| NME1 | 51.73748825 | -0.237713511 | 0.643136768 | NA |
| RNA170 | 10.42132142 | 0.514724396 | 0.245876096 | NA |
| RPR1 | 36.63381865 | -0.159358555 | 0.792114392 | NA |
| RUF20 | 86.17514557 | -0.078428218 | 0.809364498 | NA |
| RUF21 | 43.48350584 | -0.151302106 | 0.669398311 | NA |
| RUF22 | 1.126113591 | 0.603802241 | 0.144807611 | NA |
| RUF23 | 8.975541829 | -0.434377033 | 0.412516604 | NA |
| YAL016C-A | 61.31300348 | -0.305318915 | 0.403424485 | NA |
| YAL016C-B | 15.45888685 | -0.446214534 | 0.295323232 | NA |

| | | | | |
|---|---|---|---|---|
| YAL018C | 88.41004538 | 0.426161693 | 0.24585708 | NA |
| YAL019W-A | 5.164115052 | 0.044494397 | 0.929307145 | NA |
| YAL031W-A | 36.5218107 | -0.23407371 | 0.452539611 | NA |
| YAL034C-B | 9.086766243 | 0.262183594 | 0.585951266 | NA |
| YAL037C-A | 1.195205434 | 0.270606459 | 0.542941396 | NA |
| YAL042C-A | 8.764234173 | -0.363480706 | 0.436966793 | NA |
| YAL045C | 8.89025023 | -0.526885836 | 0.327478497 | NA |
| YAL047W-A | 1.181246705 | 0.271440473 | 0.623089132 | NA |
| YAL056C-A | 23.07572137 | -0.579112922 | 0.097608986 | NA |
| YAL062W | 2248.194645 | -0.021168743 | NA | NA |
| YAL063C-A | 8.265236292 | 0.458084138 | 0.389669106 | NA |
| YAL064C-A | 47.35030167 | 0.338156779 | 0.468028818 | NA |
| YAL064W | 3.131576444 | 0.340760571 | 0.489633561 | NA |
| YAL064W-B | 18.20372016 | 0.176922138 | 0.712681224 | NA |
| YAL065C | 13.48444795 | -0.25379563 | 0.604556225 | NA |
| YAL066W | 0.770254238 | -0.067435311 | 0.757937571 | NA |
| YAL067C | 87.22318015 | 0.264535368 | 0.599025836 | NA |
| YAL067W-A | 0.291097411 | 0.047759145 | 0.792062275 | NA |
| YAL068C | 5.759798843 | -0.339559036 | 0.446250027 | NA |
| YAL068W-A | 0.241654238 | -0.004038728 | 0.99574098 | NA |
| YAL069W | 0.918104495 | 0.078640767 | 0.732462367 | NA |
| YAR019W-A | 5.564052438 | 0.153090276 | 0.741823134 | NA |
| YAR020C | 43.79066228 | -0.069408686 | 0.879886326 | NA |
| YAR029W | 75.14481749 | 0.330389174 | 0.383958606 | NA |
| YAR030C | 64.62074774 | 0.88378436 | 0.013640621 | NA |
| YAR035C-A | 1.313184395 | 0.33327247 | 0.387474433 | NA |
| YAR047C | 21.66468268 | -0.608493076 | 0.184944268 | NA |
| YAR053W | 9.523462605 | -0.610970039 | 0.19534555 | NA |
| YAR064W | 14.8431874 | 1.258493466 | 0.006013988 | NA |
| YAR066W | 12.65917974 | 0.879893381 | 0.06827778 | NA |
| YAR068W | 4.842260677 | 0.327265455 | 0.50181087 | NA |
| YAR070C | 0.925768202 | 0.045308478 | 0.998231323 | NA |
| YAR071W | 43.79477313 | -0.14580012 | 0.809590139 | NA |
| YAR075W | 64.72959559 | -0.001949427 | 0.982164238 | NA |
| YBL008W-A | 2.748531795 | 1.117758943 | 0.045414497 | NA |
| YBL012C | 41.44037077 | 0.148408112 | 0.698378343 | NA |
| YBL044W | 3.828780377 | 0.566302783 | 0.29109659 | NA |
| YBL053W | 19.02715806 | 0.385733248 | 0.452579683 | NA |
| YBL062W | 0.924462825 | -0.200149519 | 0.672669271 | NA |
| YBL065W | 30.86397396 | -0.34247293 | 0.499016075 | NA |
| YBL070C | 21.83445099 | 0.460750854 | 0.298210944 | NA |
| YBL071C-B | 1.645198606 | -0.03078981 | 0.941054033 | NA |
| YBL073W | 6.010181019 | 0.374256531 | 0.461952099 | NA |

| | | | | |
|---|---|---|---|---|
| YBL083C | 7.154873991 | 0.13474823 | 0.767865841 | NA |
| YBL094C | 5.123149382 | -0.182226268 | 0.725319501 | NA |
| YBL096C | 11.76381799 | 0.879427185 | 0.07103878 | NA |
| YBL100W-C | 6.925265405 | 0.451583136 | 0.347853066 | NA |
| YBL108W | 0.561379936 | 0.075551172 | 0.806523391 | NA |
| YBL109W | 16.60491535 | 0.36536576 | 0.404106864 | NA |
| YBR027C | 8.649310737 | -0.19074961 | 0.665184806 | NA |
| YBR032W | 28.5083245 | 0.655740058 | 0.17516342 | NA |
| YBR051W | 12.61851356 | 0.178790471 | 0.735068534 | NA |
| YBR056W-A | 43.21007303 | -0.094979779 | 0.880633612 | NA |
| YBR064W | 3.158811985 | -0.194863286 | 0.779442766 | NA |
| YBR072C-A | 2.769886284 | -0.125590697 | 0.840267607 | NA |
| YBR076C-A | 84.52994015 | 0.163308437 | 0.723396392 | NA |
| YBR089W | 6.735755394 | -0.562034198 | 0.272510629 | NA |
| YBR090C | 79.12993298 | -0.474407909 | 0.167944673 | NA |
| YBR103C-A | 1.570441558 | -0.030780342 | 0.961595321 | NA |
| YBR109W-A | 20.49364363 | -0.119221536 | 0.778979164 | NA |
| YBR116C | 35.41794033 | 0.675492163 | 0.182476877 | NA |
| YBR124W | 16.74512171 | 0.351245006 | 0.367079538 | NA |
| YBR134W | 28.00305569 | -0.117060574 | 0.83825855 | NA |
| YBR144C | 8.772859821 | -0.407508923 | 0.419374256 | NA |
| YBR174C | 5.135026563 | -0.320613572 | 0.518240053 | NA |
| YBR178W | 20.74423611 | -0.583087258 | 0.17553114 | NA |
| YBR180W | 89.72716224 | 0.569191477 | 0.231362349 | NA |
| YBR182C-A | 7.795727576 | 0.310439632 | 0.50648454 | NA |
| YBR190W | 3.930160671 | 0.4255122 | 0.512665781 | NA |
| YBR191W-A | 25.03964351 | -0.440720044 | 0.888230304 | NA |
| YBR196C-A | 7.414882286 | -0.106224183 | 0.856617396 | NA |
| YBR196C-B | 11.09747944 | 0.317790696 | 0.544823528 | NA |
| YBR200W-A | 19.77475914 | 0.309976268 | 0.47376813 | NA |
| YBR209W | 5.517408049 | 0.187922457 | 0.746757676 | NA |
| YBR219C | 1.059885032 | -0.543273634 | 0.465876209 | NA |
| YBR221W-A | 0.837044541 | 0.091804706 | 0.82144502 | NA |
| YBR224W | 27.9935508 | 0.045228709 | 0.913295956 | NA |
| YBR226C | 66.35832606 | -0.197479681 | 0.584969493 | NA |
| YBR232C | 24.08316402 | -0.138790461 | 0.734815985 | NA |
| YBR250W | 80.81966717 | 0.23349868 | 0.590542734 | NA |
| YBR277C | 4.283921599 | 0.011231797 | 0.99348537 | NA |
| YBR292C | 15.20409235 | -0.050838368 | 0.874897753 | NA |
| YBR296C-A | 3.00001637 | 0.316981471 | 0.515108678 | NA |
| YBR298C-A | 3.08929577 | 1.213174607 | 0.02728991 | NA |
| YBR299W | 41.66988027 | 0.019975159 | 0.945899964 | NA |
| YBR300C | 5.482970764 | -0.298841624 | 0.562474133 | NA |

| | | | | |
|---|---|---|---|---|
| YBR301W | 22.99326035 | -0.161496531 | 0.755528319 | NA |
| YCL001W-A | 26.77980078 | 0.365080485 | 0.417458386 | NA |
| YCL001W-B | 30.32607662 | 0.268094987 | 0.536707293 | NA |
| YCL023C | 8.286348364 | 0.133220481 | 0.755766124 | NA |
| YCL026C-A | 44.13033287 | -0.046795751 | 0.913081488 | NA |
| YCL046W | 22.38513549 | 0.172115782 | 0.67304893 | NA |
| YCL048W | 92.73705911 | 0.496520517 | 0.276135661 | NA |
| YCL048W-A | 89.40578665 | 0.485714751 | 0.28972566 | NA |
| YCL065W | 1.135078196 | -0.000327028 | 0.995033607 | NA |
| YCL068C | 4.465179414 | -0.000113452 | 0.994658042 | NA |
| YCL069W | 36.22888945 | 0.002132554 | 0.888475146 | NA |
| YCL073C | 26.5180597 | -0.003588295 | 0.774612163 | NA |
| YCL075W | 37.39387639 | 0.427571397 | 0.108067497 | NA |
| YCL076W | 24.7210401 | 0.271159418 | 0.286987037 | NA |
| YCR001W | 33.60447039 | 0.035600299 | 0.928040797 | NA |
| YCR006C | 58.71736424 | 0.406118594 | 0.31047463 | NA |
| YCR018C-A | 8.72999001 | -0.75030721 | 0.117572038 | NA |
| YCR022C | 20.80731429 | 0.236119 | 0.549637116 | NA |
| YCR025C | 44.00460202 | -0.094156148 | 0.836674747 | NA |
| YCR041W | 16.4079087 | -0.063302534 | 0.875032101 | NA |
| YCR049C | 7.558405616 | 0.226354143 | 0.581395619 | NA |
| YCR050C | 17.22525728 | 0.201535191 | 0.630391329 | NA |
| YCR063W | 60.70870613 | 0.196567772 | 0.443182142 | NA |
| YCR064C | 12.51518247 | 0.104728455 | 0.787982665 | NA |
| YCR085W | 15.93952821 | 0.624710918 | 0.184457755 | NA |
| YCR087W | 24.15940244 | -0.419756079 | 0.295261883 | NA |
| YCR095W-A | 20.07196858 | 0.345193897 | 0.389994463 | NA |
| YCR097W-A | 1.60235674 | -0.002566531 | 0.991389042 | NA |
| YCR102W-A | 9.4716423 | -0.504235354 | 0.339986571 | NA |
| YCR104W | 19.10661275 | -0.477133623 | 0.328564858 | NA |
| YCR108C | 1.912873177 | -0.000143669 | 0.99540095 | NA |
| YDL007C-A | 3.853033258 | 0.135078951 | 0.634655856 | NA |
| YDL011C | 7.393723597 | 0.487989599 | 0.33062759 | NA |
| YDL016C | 38.72923011 | -0.20853088 | 0.390578151 | NA |
| YDL022C-A | 6.814086928 | -0.364916409 | 0.474091772 | NA |
| YDL026W | 16.56831607 | 0.01704599 | 0.899176245 | NA |
| YDL032W | 4.179935405 | 0.254902957 | 0.633430874 | NA |
| YDL034W | 29.64111763 | 0.148452576 | 0.693531438 | NA |
| YDL037C | 92.66763176 | 0.230094396 | 0.649433443 | NA |
| YDL041W | 0.97089065 | -0.246152666 | 0.658731855 | NA |
| YDL050C | 34.4743764 | 0.162914721 | 0.721645996 | NA |
| YDL062W | 21.94852935 | -0.130351897 | 0.765694698 | NA |
| YDL068W | 4.863206233 | 0.480478769 | 0.393930107 | NA |

| | | | | |
|---|---|---|---|---|
| YDL071C | 49.14928453 | 0.08935831 | 0.783765455 | NA |
| YDL094C | 21.86703582 | 0.060200914 | 0.913674158 | NA |
| YDL096C | 9.504570723 | 0.558014765 | 0.247441426 | NA |
| YDL114W | 43.38451628 | 0.463026089 | 0.304826616 | NA |
| YDL114W-A | 24.54000287 | 0.234955718 | 0.607816049 | NA |
| YDL118W | 21.28319356 | 0.403853689 | 0.234270048 | NA |
| YDL133C-A | 72.78703027 | -0.674480558 | 0.151631778 | NA |
| YDL151C | 4.616503935 | -0.061398048 | 0.911845868 | NA |
| YDL152W | 5.390111654 | -0.443167895 | 0.495854835 | NA |
| YDL158C | 89.30551169 | -0.641362763 | 0.068591243 | NA |
| YDL159W-A | 1.037341025 | -0.095439177 | 0.721393176 | NA |
| YDL162C | 2.875843937 | 0.962679863 | 0.075809565 | NA |
| YDL163W | 0.057163039 | 0.026970009 | 0.883673227 | NA |
| YDL172C | 10.58915184 | -0.286670968 | 0.568768652 | NA |
| YDL187C | 3.68310749 | 0.699274441 | 0.166959273 | NA |
| YDL196W | 15.17669232 | 0.180909362 | 0.657359566 | NA |
| YDL221W | 36.90750404 | 0.881089784 | 0.035442139 | NA |
| YDL223C | 1796.784019 | 0.408099971 | NA | NA |
| YDL228C | 55.92892077 | 0.076436984 | 0.944681784 | NA |
| YDL240C-A | 1.077122984 | 0.357879657 | 0.562693958 | NA |
| YDL242W | 58.94192665 | -0.095714812 | 0.798617292 | NA |
| YDL244W | 51.51006224 | -0.10843545 | 0.839278328 | NA |
| YDL245C | 34.45281204 | 0.207797367 | 0.65748869 | NA |
| YDL246C | 6.191078768 | -0.069020415 | 0.843035218 | NA |
| YDL247W | 3.022100866 | 0.611114795 | 0.234141968 | NA |
| YDR003W-A | 67.78719135 | -0.04512367 | 0.886229958 | NA |
| YDR008C | 37.32753147 | -0.291389256 | 0.232973858 | NA |
| YDR010C | 32.06483583 | 1.08943468 | 0.000980714 | NA |
| YDR014W-A | 36.16776295 | -0.164997919 | 0.885063369 | NA |
| YDR024W | 7.360211428 | 0.136969829 | 0.774827927 | NA |
| YDR034C-A | 4.06626248 | 0.70922433 | 0.169265609 | NA |
| YDR042C | 81.10570396 | 0.424777549 | 0.341669778 | NA |
| YDR048C | 13.8983324 | 0.38597402 | 0.407864518 | NA |
| YDR053W | 7.370760006 | -0.053407198 | 0.910957864 | NA |
| YDR094W | 4.116771726 | 0.331785274 | 0.522433965 | NA |
| YDR095C | 5.026425328 | 1.338887781 | 0.010275318 | NA |
| YDR098C-B | 19.81780983 | 0.422797552 | 0.360827639 | NA |
| YDR102C | 12.68433027 | 0.087799851 | 0.863638835 | NA |
| YDR106W | 46.31624816 | 0.280275704 | 0.409339494 | NA |
| YDR112W | 22.56100124 | 1.133327427 | 0.014959427 | NA |
| YDR114C | 36.82268978 | -0.070782105 | 0.808944298 | NA |
| YDR118W-A | 11.72360647 | 0.256165848 | 0.531848906 | NA |
| YDR136C | 21.36116146 | 0.227682321 | 0.495074911 | NA |

| | | | | |
|---|---|---|---|---|
| YDR149C | 14.02290454 | -0.292169212 | 0.531048598 | NA |
| YDR157W | 53.23769391 | -0.207091822 | 0.608731316 | NA |
| YDR169C-A | 13.43403719 | 0.026054414 | 0.969057154 | NA |
| YDR182W-A | 24.64411227 | -0.322336557 | 0.320264743 | NA |
| YDR183C-A | 47.62983085 | 0.483788334 | 0.261275855 | NA |
| YDR187C | 25.29702409 | -0.352104003 | 0.360606198 | NA |
| YDR193W | 12.98139175 | 0.188694307 | 0.732966276 | NA |
| YDR194W-A | 0.453289942 | -0.064869334 | 0.803396391 | NA |
| YDR199W | 75.09684199 | 0.130699393 | 0.600734391 | NA |
| YDR203W | 15.81601128 | 0.637609347 | 0.106913331 | NA |
| YDR210C-D | 43.7176532 | 0.015053518 | 0.831714182 | NA |
| YDR210W-B | 26.10421514 | 0.052016376 | 0.891777885 | NA |
| YDR215C | 67.50004526 | 0.295547634 | 0.549111866 | NA |
| YDR218C | 78.04916244 | -0.053492766 | 0.927519538 | NA |
| YDR220C | 3.958958383 | 0.303889375 | 0.513048326 | NA |
| YDR241W | 32.77525977 | 0.076140818 | 0.829879876 | NA |
| YDR246W-A | 23.46436155 | 0.153745219 | 0.711419366 | NA |
| YDR250C | 6.490017442 | -0.363654028 | 0.47878626 | NA |
| YDR252W | 88.93914678 | 0.237403862 | 0.575517272 | NA |
| YDR261C-D | 88.68167414 | -0.034212816 | 0.977889057 | NA |
| YDR261W-B | 58.90647804 | 0.045855768 | 0.918422008 | NA |
| YDR269C | 1.063098833 | 0.299713401 | 0.498280857 | NA |
| YDR271C | 28.54979964 | -0.441139687 | 0.295053416 | NA |
| YDR274C | 62.88697688 | 0.593719171 | 0.162598719 | NA |
| YDR278C | 0.649229666 | -0.126675036 | 0.692817164 | NA |
| YDR281C | 42.74964081 | -0.707524695 | 0.155430636 | NA |
| YDR290W | 8.55083412 | -0.293568914 | 0.512649895 | NA |
| YDR320W-B | 5.22193975 | -0.55019635 | 0.245406836 | NA |
| YDR340W | 4.60165433 | 0.20217926 | 0.71667635 | NA |
| YDR344C | 46.73109222 | 0.103727214 | 0.817669005 | NA |
| YDR355C | 4.46006282 | -0.632041898 | 0.264726398 | NA |
| YDR360W | 36.19514521 | 0.668993677 | 0.057588782 | NA |
| YDR366C | 7.819864358 | 0.334875363 | 0.541929264 | NA |
| YDR374C | 71.50374083 | -0.279845568 | 0.64629488 | NA |
| YDR401W | 0.227003213 | 0.000720353 | 0.996711899 | NA |
| YDR402C | 76.67558535 | 0.548496606 | 0.210195978 | NA |
| YDR406W-A | 12.68467301 | -0.59224442 | 0.217862298 | NA |
| YDR413C | 10.51150758 | 0.178985593 | 0.698575441 | NA |
| YDR426C | 18.4381422 | 0.111174163 | 0.70555891 | NA |
| YDR431W | 11.62652329 | -0.007325408 | 0.981594711 | NA |
| YDR433W | 87.14790436 | -0.467638436 | 0.133744418 | NA |
| YDR437W | 89.57715872 | 0.664980438 | 0.040393364 | NA |
| YDR442W | 17.52253569 | 0.083903494 | 0.840087275 | NA |

| | | | | |
|---|---|---|---|---|
| YDR445C | 3.142353253 | 0.49880725 | 0.388278686 | NA |
| YDR446W | 60.61919013 | -0.168103979 | 0.902678542 | NA |
| YDR455C | 13.39180516 | 0.26371148 | 0.527978541 | NA |
| YDR467C | 75.26126934 | -0.020313274 | 0.942928024 | NA |
| YDR509W | 13.68760112 | 0.821964527 | 0.103317632 | NA |
| YDR521W | 13.62209314 | -0.241933108 | 0.595869399 | NA |
| YDR524C-A | 2.188681507 | 0.948762691 | 0.079694244 | NA |
| YDR525W | 4.560372998 | 0.339356049 | 0.411625518 | NA |
| YDR526C | 4.767823074 | -0.837001547 | 0.102124402 | NA |
| YDR535C | 20.65811424 | 0.254316103 | 0.542891246 | NA |
| YDR537C | 28.61472288 | -0.422226126 | 0.268933396 | NA |
| YDR542W | 1.676832288 | -0.2818877 | 0.612590734 | NA |
| YDR543C | 0.805732287 | 0.029212765 | 0.774544964 | NA |
| YDR544C | 2.336721419 | 0.219295828 | 0.513254308 | NA |
| YEL008W | 30.07335462 | -0.164459704 | 0.731495949 | NA |
| YEL009C-A | 8.998175135 | 0.531499435 | 0.230889406 | NA |
| YEL010W | 4.427753031 | -0.100747854 | 0.866809933 | NA |
| YEL014C | 21.68196635 | -0.292575495 | 0.527051147 | NA |
| YEL018C-A | 0.416927005 | -0.191469734 | 0.523283633 | NA |
| YEL020C-B | 34.39724677 | -0.520480322 | 0.105729234 | NA |
| YEL030C-A | 7.262884205 | -0.838684256 | 0.077192498 | NA |
| YEL035C | 12.59177912 | 0.263897851 | 0.523756463 | NA |
| YEL045C | 16.86688846 | -0.561540824 | 0.195929117 | NA |
| YEL049W | 54.33605347 | 0.095591318 | 0.793838796 | NA |
| YEL050W-A | 71.35641565 | -0.221409454 | 0.485981501 | NA |
| YEL059W | 50.35325287 | -0.105554524 | 0.817740279 | NA |
| YEL074W | 35.0602757 | 0.340424233 | 0.486960755 | NA |
| YER006C-A | 2.225353622 | 0.034055105 | 0.942567598 | NA |
| YER024W | 2782.885664 | 0.022437879 | NA | NA |
| YER038W-A | 0.198141795 | 0.082745364 | 0.688812737 | NA |
| YER039C-A | 49.35745349 | 0.534556598 | 0.158343488 | NA |
| YER046W-A | 13.59530817 | 0.714904798 | 0.102061109 | NA |
| YER065C | 3480.324226 | 0.04377344 | NA | NA |
| YER068C-A | 10.30512685 | 0.632427905 | 0.14575746 | NA |
| YER076W-A | 76.7457419 | -0.350309705 | 0.145976134 | NA |
| YER078W-A | 4.281084499 | -0.188004084 | 0.724154209 | NA |
| YER079C-A | 18.60445932 | -0.366432795 | 0.520642593 | NA |
| YER084W-A | 0.719422121 | 0.130910979 | 0.684920223 | NA |
| YER085C | 1.090623589 | -0.204834489 | 0.565240118 | NA |
| YER087C-A | 36.6678806 | -0.125245721 | 0.741846845 | NA |
| YER091C-A | 0.485272411 | 0.242068616 | 0.449593993 | NA |
| YER097W | 28.63993449 | 0.292691459 | 0.468497989 | NA |
| YER106W | 34.98872113 | 0.042088605 | 0.850793773 | NA |

| | | | | |
|---|---|---|---|---|
| YER107W-A | 76.0626659 | -0.087102687 | 0.843521758 | NA |
| YER135C | 23.51565939 | 0.006380665 | 0.987605901 | NA |
| YER137C | 65.6651259 | -0.296088818 | 0.428818007 | NA |
| YER137W-A | 3.086112355 | -0.20781832 | 0.73574382 | NA |
| YER138W-A | 1.112909077 | -0.608191993 | 0.173607503 | NA |
| YER145C-A | 50.64924556 | -0.114032479 | 0.793321711 | NA |
| YER148W-A | 0.781507553 | 0.617480308 | 0.280968281 | NA |
| YER152W-A | 37.9606133 | -0.020312234 | 0.965158274 | NA |
| YER165C-A | 28.00543952 | 0.194260426 | 0.554596864 | NA |
| YER172C-A | 0.861078059 | 0.270721435 | 0.565039657 | NA |
| YER175W-A | 27.41241786 | -0.173714208 | 0.603860781 | NA |
| YER180C-A | 68.54860649 | 0.160113884 | 0.539046376 | NA |
| YER181C | 6.5610832 | 0.011771253 | 0.977418548 | NA |
| YER185W | 79.91884135 | 0.365899401 | 0.333145491 | NA |
| YER188C-A | 23.15774642 | 0.990029326 | 0.027889219 | NA |
| YER189W | 0.915658928 | 0.243638304 | 0.655099852 | NA |
| YER190W | 15.31140442 | -0.093616434 | 0.863216818 | NA |
| YFL011W | 70.61598103 | -0.282526589 | 0.584127892 | NA |
| YFL012W | 27.05345673 | -0.353214427 | 0.589227058 | NA |
| YFL012W-A | 14.51855676 | 0.239031999 | 0.599489033 | NA |
| YFL013W-A | 84.24870204 | -0.615660834 | 0.022539662 | NA |
| YFL015C | 19.00869044 | -1.148321716 | 0.009031682 | NA |
| YFL015W-A | 1.143841678 | 0.119777608 | 0.73518269 | NA |
| YFL019C | 43.77288423 | -0.056572975 | 0.908837117 | NA |
| YFL020C | 87.69189529 | -0.339370091 | 0.596192514 | NA |
| YFL021C-A | 41.5780796 | 0.484938614 | 0.296302716 | NA |
| YFL041W-A | 23.17885335 | -0.066393825 | 0.872823319 | NA |
| YFL051C | 87.84562174 | 0.287788717 | 0.552746669 | NA |
| YFL058W | 7.345596516 | 0.599276024 | 0.131091748 | NA |
| YFL059W | 70.79713485 | -0.161916494 | 0.740024108 | NA |
| YFL060C | 16.02228195 | 0.406240661 | 0.408099006 | NA |
| YFL061W | 11.58614878 | 0.584279445 | 0.228451619 | NA |
| YFL062W | 38.28792094 | -0.145366073 | 0.739684652 | NA |
| YFL063W | 4.881871603 | 0.252480584 | 0.333887327 | NA |
| YFL064C | 3.015892609 | 0.359391693 | 0.229075828 | NA |
| YFL065C | 1.354222162 | 0.193175957 | 0.472542276 | NA |
| YFL066C | 2.343434624 | 0.494023681 | 0.18717849 | NA |
| YFL067W | 57.17560859 | 0.425495936 | 0.277096856 | NA |
| YFR010W-A | 16.18554247 | -0.604428745 | 0.215236903 | NA |
| YFR012W | 37.52047909 | 0.192162621 | 0.703295655 | NA |
| YFR012W-A | 2.137477958 | -0.051916758 | 0.855826707 | NA |
| YFR032C | 47.36854765 | 0.855085088 | 0.043988736 | NA |
| YFR032C-B | 19.14700684 | -0.241686152 | 0.562629687 | NA |

| | | | | |
|---|---|---|---|---|
| YFR034W-A | 2.98827912 | -0.283088782 | 0.564408825 | NA |
| YFR036W-A | 15.15766896 | -0.167049553 | 0.676875531 | NA |
| YFR054C | 20.51319414 | 0.593928553 | 0.174773905 | NA |
| YFR056C | 41.82065454 | 0.195517565 | 0.630231854 | NA |
| YFR057W | 37.29733796 | 0.009188474 | 0.853575761 | NA |
| YGL006W-A | 3.943531239 | -0.802152724 | 0.137492475 | NA |
| YGL007C-A | 2.085025417 | -0.73149161 | 0.196358292 | NA |
| YGL007W | 19.49434054 | -0.015724986 | 0.971107896 | NA |
| YGL015C | 22.18286877 | -0.688189868 | 0.170635841 | NA |
| YGL024W | 7.765530916 | 0.436865236 | 0.347771386 | NA |
| YGL033W | 39.3922468 | -0.753323834 | 0.124604358 | NA |
| YGL034C | 9.87258369 | -0.059093208 | 0.91109452 | NA |
| YGL041C-B | 21.05072236 | -0.065806487 | 0.870018564 | NA |
| YGL051W | 80.44841429 | -0.08574618 | 0.788683618 | NA |
| YGL052W | 83.8653604 | 0.258588901 | 0.479872429 | NA |
| YGL069C | 85.63194987 | -0.087337336 | 0.789617297 | NA |
| YGL072C | 15.1439202 | 0.084394408 | 0.788560618 | NA |
| YGL074C | 37.00723578 | 0.480402058 | 0.248164773 | NA |
| YGL088W | 0.432313012 | -0.012502482 | 0.991745176 | NA |
| YGL103W | 19.59323296 | 0.703893263 | 0.165697065 | NA |
| YGL109W | 79.04302724 | -0.221921186 | 0.376260249 | NA |
| YGL118C | 2.533670207 | 0.421464216 | 0.381126549 | NA |
| YGL132W | 16.53764806 | 0.893904668 | 0.055813021 | NA |
| YGL138C | 22.53163453 | 0.110825424 | 0.608291157 | NA |
| YGL149W | 44.44871907 | 0.138443491 | 0.695479671 | NA |
| YGL152C | 7.084888852 | -0.483786075 | 0.242461336 | NA |
| YGL165C | 78.38274117 | -0.079375568 | 0.74121433 | NA |
| YGL170C | 69.87350446 | 1.186591514 | 0.011957018 | NA |
| YGL177W | 36.28800868 | -0.345087809 | 0.297267586 | NA |
| YGL182C | 10.33516089 | 0.382881188 | 0.404335688 | NA |
| YGL183C | 72.4162606 | 0.340820549 | 0.464060925 | NA |
| YGL188C | 26.0866713 | 0.341661478 | 0.475343396 | NA |
| YGL188C-A | 18.77666255 | 0.514982832 | 0.311573801 | NA |
| YGL199C | 2.062524245 | -1.934238704 | 0.001552374 | NA |
| YGL204C | 78.35386912 | -0.184965676 | 0.629073758 | NA |
| YGL214W | 9.34196156 | 0.247210311 | 0.600229545 | NA |
| YGL217C | 8.839690893 | 0.948218484 | 0.040723624 | NA |
| YGL218W | 11.00510591 | 0.362431762 | 0.427188694 | NA |
| YGL230C | 33.65395104 | -0.196908914 | 0.686383159 | NA |
| YGL235W | 1.903973228 | 0.369804385 | 0.487374653 | NA |
| YGL239C | 18.18533761 | 0.11598737 | 0.766082227 | NA |
| YGL258W | 52.00397703 | 0.469102202 | 0.281848179 | NA |
| YGL258W-A | 56.62039045 | -0.012188534 | 0.991011839 | NA |

| | | | | |
|---|---|---|---|---|
| YGL259W | 86.59077684 | 0.288594682 | 0.518715227 | NA |
| YGL260W | 1.415207671 | 0.177383942 | 0.731345093 | NA |
| YGL261C | 11.97177178 | -0.270199942 | 0.592926633 | NA |
| YGL262W | 77.76796003 | 0.150676013 | 0.551509371 | NA |
| YGR018C | 8.799849222 | 0.078019085 | 0.88151222 | NA |
| YGR022C | 1.81368062 | -0.293655831 | 0.401693104 | NA |
| YGR035W-A | 93.40510398 | 0.333478042 | 0.346462579 | NA |
| YGR039W | 40.79718114 | 0.39355693 | 0.274184514 | NA |
| YGR045C | 23.80619598 | 0.185742264 | 0.633595305 | NA |
| YGR051C | 51.37808205 | 0.191059695 | 0.558307463 | NA |
| YGR059W | 83.10299922 | 0.245425704 | 0.596903434 | NA |
| YGR064W | 14.24950712 | -0.047760492 | 0.929288821 | NA |
| YGR067C | 601.4253985 | -0.078313477 | NA | NA |
| YGR069W | 7.64600337 | -0.191099877 | 0.697455187 | NA |
| YGR073C | 7.437514438 | 0.967371079 | 0.043951213 | NA |
| YGR107W | 13.44440994 | 0.061038569 | 0.902271822 | NA |
| YGR114C | 36.93060097 | -1.609111825 | 0.00059507 | NA |
| YGR121W-A | 8.749518959 | -0.335378742 | 0.510605941 | NA |
| YGR122C-A | 15.60192201 | -0.136436248 | 0.759041814 | NA |
| YGR139W | 13.99976418 | 0.060979408 | 0.891877889 | NA |
| YGR146C-A | 13.60921533 | 0.221831165 | 0.651862353 | NA |
| YGR151C | 2.6142374 | -0.562545123 | 0.367430724 | NA |
| YGR160W | 19.24237613 | -0.03271357 | 0.965760337 | NA |
| YGR161C-D | 41.89517477 | 0.700146844 | 0.105056632 | NA |
| YGR164W | 4.051111594 | 1.070424473 | 0.063098428 | NA |
| YGR174W-A | 47.91809201 | -0.039125963 | 0.951125529 | NA |
| YGR176W | 11.59700922 | 0.621708559 | 0.230270089 | NA |
| YGR190C | 89.92851229 | 0.332224445 | 0.289583476 | NA |
| YGR204C-A | 13.54650687 | -0.470201474 | 0.316582408 | NA |
| YGR219W | 32.06784529 | -0.472669591 | 0.132244601 | NA |
| YGR228W | 14.43502374 | -0.834785994 | 0.070196874 | NA |
| YGR236C | 237.8810767 | 0.250697487 | NA | NA |
| YGR240C-A | 46.68747277 | 0.430372206 | 0.335911548 | NA |
| YGR242W | 37.21832003 | 0.139276261 | 0.674258431 | NA |
| YGR259C | 0.780911468 | -0.16370459 | 0.520025213 | NA |
| YGR265W | 63.64327138 | 0.124948195 | 0.743601919 | NA |
| YGR269W | 48.40957868 | -0.003571122 | 0.972186111 | NA |
| YGR273C | 7.812625483 | -0.250402842 | 0.600489954 | NA |
| YGR290W | 27.11482068 | -0.135059379 | 0.760139853 | NA |
| YGR291C | 1.566792632 | 0.177947811 | 0.650541274 | NA |
| YGR292W | 46.2187461 | 0.072918607 | 0.891817968 | NA |
| YGR293C | 7.546648827 | 0.018488676 | 0.970965173 | NA |
| YGR294W | 10.95232902 | -0.095419795 | 0.877752686 | NA |

| | | | | |
|---|---|---|---|---|
| YHL002C-A | 8.009981089 | -0.313865362 | 0.572916136 | NA |
| YHL005C | 24.92643688 | 0.33217096 | 0.320547883 | NA |
| YHL006W-A | 55.86611926 | 0.849148038 | 0.016611855 | NA |
| YHL009W-B | 5.78416984 | -0.401258708 | 0.423269458 | NA |
| YHL015W-A | 4.315791202 | 0.187471045 | 0.728421447 | NA |
| YHL018W | 70.12879735 | -0.000779224 | 0.993172913 | NA |
| YHL019W-A | 10.01596588 | 0.641535425 | 0.163779172 | NA |
| YHL030W-A | 22.0345387 | -0.563763025 | 0.208904695 | NA |
| YHL037C | 20.29589679 | -0.178540868 | 0.718431514 | NA |
| YHL041W | 3.288023996 | 0.018881747 | 0.96518358 | NA |
| YHL042W | 88.3151764 | -0.112702015 | 0.746652677 | NA |
| YHL045W | 26.03413018 | -0.086991788 | 0.846886868 | NA |
| YHL046C | 9.174623277 | 0.082026141 | 0.845895783 | NA |
| YHL046W-A | 47.85481112 | 0.429272249 | 0.371211415 | NA |
| YHL048C-A | 2.76683096 | 0.419757279 | 0.163834306 | NA |
| YHR007C-A | 61.56889406 | 0.246712494 | 0.547650071 | NA |
| YHR021W-A | 41.37134005 | -0.042916086 | 0.946972482 | NA |
| YHR022C-A | 3.937421936 | 0.160826583 | 0.776814816 | NA |
| YHR028W-A | 38.81881491 | 0.089014834 | 0.779596756 | NA |
| YHR032C-A | 12.55755435 | 0.030457513 | 0.962780695 | NA |
| YHR032W-A | 60.96537533 | 0.410055355 | 0.284431506 | NA |
| YHR056W-A | 0.867587117 | -0.302141677 | 0.373733503 | NA |
| YHR069C-A | 0.116121444 | 0.026970009 | 0.883680447 | NA |
| YHR071C-A | 39.05421358 | -0.300287789 | 0.44234725 | NA |
| YHR079C-A | 27.69347479 | -0.044605278 | 0.936367386 | NA |
| YHR086W-A | 3.107825184 | -0.446312204 | 0.42291938 | NA |
| YHR093W | 35.27687991 | 0.566130887 | 0.143267825 | NA |
| YHR095W | 5.929557488 | 0.421273625 | 0.421012015 | NA |
| YHR125W | 4.057342841 | -0.042766591 | 0.949189407 | NA |
| YHR126C | 18.42016663 | 0.088152079 | 0.853031233 | NA |
| YHR130C | 24.10330458 | 0.562703179 | 0.193613973 | NA |
| YHR131W-A | 19.588359 | 0.436697103 | 0.275542521 | NA |
| YHR139C-A | 36.30751378 | -0.639788158 | 0.153665918 | NA |
| YHR145C | 0.415497853 | 0.034303853 | 0.910880705 | NA |
| YHR153C | 73.13873436 | -0.097579413 | 0.794343641 | NA |
| YHR157W | 84.3613017 | 0.006876374 | 0.971762942 | NA |
| YHR165W-A | 6.420816 | -0.087378835 | 0.876416486 | NA |
| YHR173C | 88.05682976 | 0.320660265 | 0.373568069 | NA |
| YHR180W-A | 0.36648756 | 0.02867792 | 0.917938843 | NA |
| YHR184W | 81.12732022 | 0.02582982 | 0.958609013 | NA |
| YHR185C | 7.642313868 | -0.034582081 | 0.941348155 | NA |
| YHR193C-A | 1.288094432 | 0.280769975 | 0.680140641 | NA |
| YHR213W-B | 12.16949824 | 0.954790365 | 0.040576243 | NA |

| | | | | |
|---|---|---|---|---|
| YHR214C-E | 1.813646714 | 0.408528615 | 0.344631253 | NA |
| YHR214W | 18.69123236 | 0.989295096 | 0.030919388 | NA |
| YHR214W-A | 22.9859353 | 0.642391032 | 0.169828166 | NA |
| YHR217C | 1.225581631 | 0.008864726 | 0.991333441 | NA |
| YHR219W | 0.159331785 | -0.000977006 | 0.995760437 | NA |
| YIL012W | 10.18010829 | 1.016312101 | 0.028302409 | NA |
| YIL025C | 3.736035911 | 0.250011608 | 0.614561787 | NA |
| YIL028W | 59.74955175 | -0.049725754 | 0.897506645 | NA |
| YIL029W-A | 8.108002149 | 0.366304869 | 0.426257407 | NA |
| YIL030W-A | 74.85518125 | -0.10268214 | 0.695035963 | NA |
| YIL032C | 13.12625335 | 0.782144703 | 0.039526603 | NA |
| YIL046W-A | 7.695578865 | 0.919179122 | 0.038626024 | NA |
| YIL047C-A | 10.41380776 | 0.012630952 | 0.977909875 | NA |
| YIL054W | 8.069700908 | 0.366310364 | 0.417968363 | NA |
| YIL057C | 383.2825996 | 0.044635409 | NA | NA |
| YIL058W | 79.78173189 | 0.581320957 | 0.071527517 | NA |
| YIL059C | 89.463829 | 0.139298496 | 0.766724785 | NA |
| YIL060W | 54.27024793 | -0.039260061 | 0.941733921 | NA |
| YIL066W-A | 3.308676015 | -0.174659572 | 0.741202345 | NA |
| YIL068W-A | 5.981971104 | -0.182405175 | 0.715310749 | NA |
| YIL071W-A | 73.5379669 | 0.059708681 | 0.773550309 | NA |
| YIL086C | 23.45932813 | 0.211054184 | 0.57043858 | NA |
| YIL100C-A | 6.139255363 | 0.297681733 | 0.582668928 | NA |
| YIL100W | 3.722972412 | 0.114847815 | 0.803546725 | NA |
| YIL102C | 13.70446277 | -0.352194835 | 0.494250785 | NA |
| YIL115W-A | 23.70554908 | 0.272124459 | 0.478222028 | NA |
| YIL134C-A | 3.121789303 | -0.436158814 | 0.393665255 | NA |
| YIL141W | 35.62085203 | -0.125453603 | 0.752036904 | NA |
| YIL142C-A | 7.771319082 | 0.17023801 | 0.682386079 | NA |
| YIL156W-A | 6.740450198 | -0.038100904 | 0.899972375 | NA |
| YIL163C | 43.51788641 | 0.351177305 | 0.324204085 | NA |
| YIL168W | 92.19596636 | 0.127951484 | 0.708645517 | NA |
| YIL171W | 27.73883289 | 0.279854013 | 0.546252541 | NA |
| YIL173W | 14.66085042 | 0.828182345 | 0.078556927 | NA |
| YIL177C | 0.203940714 | -0.034615421 | 0.833083491 | NA |
| YIR013C | 17.07044727 | 0.686395884 | 0.140827716 | NA |
| YIR017C | 6.4006803 | -0.487275548 | 0.344629693 | NA |
| YIR017W-A | 3.982442675 | 0.564400088 | 0.188353676 | NA |
| YIR018C-A | 31.48971955 | -0.046812443 | 0.887914909 | NA |
| YIR020C-B | 5.673146071 | -0.021937205 | 0.979076486 | NA |
| YIR021W-A | 4.66148073 | -0.269509 | 0.569989074 | NA |
| YIR023C-A | 13.57792364 | -0.080322671 | 0.886331031 | NA |
| YIR030C | 90.29914152 | -0.135055355 | 0.764164595 | NA |

| | | | | |
|---|---|---|---|---|
| YIR030W-A | 17.57088172 | -0.517290578 | 0.272591562 | NA |
| YIR036W-A | 10.01589679 | -0.957480337 | 0.038003173 | NA |
| YIR040C | 0.167590483 | -0.000977006 | 0.995771626 | NA |
| YIR041W | 5.637300496 | -0.606915538 | 0.228321697 | NA |
| YIR043C | 49.74477082 | 0.659451259 | 0.178763379 | NA |
| YIR044C | 6.331727114 | 0.742937862 | 0.132678665 | NA |
| YJL007C | 3.350774349 | 0.341855765 | 0.457125685 | NA |
| YJL009W | 66.31151028 | 0.12921167 | 0.689779367 | NA |
| YJL015C | 72.86729941 | 0.438982844 | 0.160235068 | NA |
| YJL022W | 2.660856745 | -0.405873616 | 0.447896853 | NA |
| YJL027C | 10.13200317 | -0.424299782 | 0.350968298 | NA |
| YJL028W | 3.906046468 | -0.719530372 | 0.146546688 | NA |
| YJL032W | 6.67566349 | -0.09211818 | 0.809675232 | NA |
| YJL037W | 69.21169324 | 0.116393514 | 0.75250806 | NA |
| YJL043W | 47.06314645 | -0.021052976 | 0.951224287 | NA |
| YJL047C-A | 19.06572924 | 1.044776688 | 0.024698921 | NA |
| YJL052C-A | 1.688849253 | 0.03607246 | 0.880049667 | NA |
| YJL064W | 3.253557565 | 0.364156223 | 0.463723439 | NA |
| YJL067W | 49.69168928 | 0.090517327 | 0.811830531 | NA |
| YJL075C | 78.82663296 | -0.122043795 | 0.663141151 | NA |
| YJL077W-B | 26.1550574 | 0.122207864 | 0.794694746 | NA |
| YJL086C | 9.129343542 | 0.109051222 | 0.812256475 | NA |
| YJL114W | 3.593194288 | -0.611099203 | 0.240746364 | NA |
| YJL119C | 7.109202711 | 0.191411641 | 0.689286398 | NA |
| YJL120W | 37.94521827 | -0.621865313 | 0.091665776 | NA |
| YJL127W-A | 12.80997666 | -0.454457978 | 0.254740343 | NA |
| YJL135W | 5.319281498 | 0.35340916 | 0.498712195 | NA |
| YJL136W-A | 0.34890375 | -0.07182556 | 0.735581118 | NA |
| YJL150W | 10.09973158 | 0.280884405 | 0.550657582 | NA |
| YJL156W-A | 83.97878785 | 0.067701217 | 0.792596043 | NA |
| YJL160C | 84.43696653 | 0.119000305 | 0.760155093 | NA |
| YJL169W | 25.36216702 | 0.098641403 | 0.831390256 | NA |
| YJL175W | 4.732791699 | 0.450626676 | 0.333838139 | NA |
| YJL182C | 1.793622545 | -0.277947653 | 0.636504418 | NA |
| YJL195C | 5.108398903 | -0.95706916 | 0.094811614 | NA |
| YJL202C | 29.15207009 | 0.110598828 | 0.745800353 | NA |
| YJL211C | 56.87482788 | 0.311802815 | 0.403427462 | NA |
| YJL215C | 44.17616213 | 0.533256132 | 0.174012675 | NA |
| YJL219W | 39.81053365 | -0.119111798 | 0.82128093 | NA |
| YJL222W | 13.76304146 | 0.789486765 | 0.075703012 | NA |
| YJL225C | 0.280381926 | -0.042304137 | 0.821594801 | NA |
| YJR005C-A | 81.403139 | -0.648777127 | 0.214013239 | NA |
| YJR018W | 57.57815418 | -0.832930869 | 0.004728082 | NA |

| | | | | |
|---|---|---|---|---|
| YJR020W | 8.221696255 | -0.635459224 | 0.155219398 | NA |
| YJR022W | 18.26801566 | -0.86689966 | 0.036566969 | NA |
| YJR023C | 6.716930152 | -0.175077935 | 0.708003819 | NA |
| YJR037W | 10.84309032 | 0.924204785 | 0.032332418 | NA |
| YJR038C | 33.2004855 | 0.535688315 | 0.201138798 | NA |
| YJR071W | 37.86358004 | 0.077004056 | 0.850897293 | NA |
| YJR079W | 11.62148612 | -0.347963797 | 0.461291198 | NA |
| YJR095W | 1024.664601 | -0.050899461 | NA | NA |
| YJR108W | 40.18860279 | 0.598579436 | 0.082979162 | NA |
| YJR114W | 33.45969567 | -0.045194651 | 0.907362196 | NA |
| YJR120W | 18.09619799 | 0.046035177 | 0.901132313 | NA |
| YJR128W | 14.84064674 | 0.618581089 | 0.165263222 | NA |
| YJR156C | 47.49998049 | -0.056684433 | 0.912128728 | NA |
| YJR157W | 1.945110995 | 1.265855104 | 0.006479002 | NA |
| YJR159W | 6.187748904 | 0.305780418 | 0.541256548 | NA |
| YJR160C | 11.90922489 | 0.609251306 | 0.18199551 | NA |
| YKL031W | 35.11063328 | 0.224981523 | 0.666789708 | NA |
| YKL037W | 43.69720214 | -0.200663665 | 0.599356146 | NA |
| YKL053W | 17.58730316 | -0.046661072 | 0.88697673 | NA |
| YKL065W-A | 60.35011298 | -0.04958657 | NA | NA |
| YKL076C | 34.14810955 | -0.000958285 | 0.997123081 | NA |
| YKL083W | 46.77827339 | -0.290398035 | 0.359289579 | NA |
| YKL096C-B | 8.469479675 | 0.926220039 | 0.049222803 | NA |
| YKL097C | 10.27058523 | 0.0118235 | 0.953306552 | NA |
| YKL102C | 18.56354003 | 0.233857718 | 0.669060643 | NA |
| YKL106C-A | 1.684117879 | 0.288646195 | 0.525087509 | NA |
| YKL111C | 22.34564795 | 0.228993103 | 0.478250299 | NA |
| YKL115C | 42.82706794 | 0.530915203 | 0.05386838 | NA |
| YKL118W | 29.53687639 | -0.719691978 | 0.009989611 | NA |
| YKL123W | 31.46802918 | -0.154647402 | 0.661433006 | NA |
| YKL131W | 17.12686382 | 1.583395945 | 0.0001051 | NA |
| YKL136W | 25.19505527 | 0.470868744 | 0.140852505 | NA |
| YKL147C | 17.64954721 | -0.049811711 | 0.955491279 | NA |
| YKL162C-A | 16.50216676 | -0.383417948 | 0.396845045 | NA |
| YKL165C-A | 6.96473103 | -0.320080689 | 0.485421891 | NA |
| YKL169C | 32.06083426 | -0.186014399 | 0.579642422 | NA |
| YKL170W | 81.53735369 | -0.088347686 | 0.786027672 | NA |
| YKL177W | 2.5279419 | -0.269088429 | 0.620487468 | NA |
| YKL183C-A | 2.603283005 | 1.05500299 | 0.046202059 | NA |
| YKL202W | 21.88607995 | -0.297605798 | 0.369107592 | NA |
| YKL223W | 0.064032033 | -0.000977006 | 0.995757316 | NA |
| YKL224C | 0.174623486 | -0.031378577 | 0.876367258 | NA |
| YKL225W | 0.626369128 | -0.007763315 | 0.99407335 | NA |

| | | | | |
|---|---|---|---|---|
| YKR032W | 0.979223699 | -0.616918949 | 0.122949979 | NA |
| YKR033C | 0.132511194 | -0.000977006 | 0.995726403 | NA |
| YKR034W | 49.93480154 | 0.219682324 | 0.469071193 | NA |
| YKR035C | 23.50975344 | -0.306637594 | 0.336342464 | NA |
| YKR040C | 4.454389 | -0.207055034 | 0.702052795 | NA |
| YKR041W | 32.67705338 | 0.331961985 | 0.430963999 | NA |
| YKR073C | 18.81130727 | -0.264115448 | 0.479436243 | NA |
| YKR097W | 1618.765281 | -0.010365864 | NA | NA |
| YLL006W-A | 2.814822079 | 0.204701076 | 0.650962367 | NA |
| YLL017W | 60.73191178 | 0.383884112 | 0.380197035 | NA |
| YLL020C | 30.70630028 | -0.092399927 | 0.856938946 | NA |
| YLL025W | 92.59299699 | -0.003124855 | 0.98685418 | NA |
| YLL030C | 4.099886808 | -0.386143412 | 0.437432961 | NA |
| YLL037W | 13.52665894 | -0.244834647 | 0.549175497 | NA |
| YLL042C | 71.42312394 | -0.06976548 | 0.8309259 | NA |
| YLL046C | 28.77526537 | -0.223004642 | 0.691544339 | NA |
| YLL047W | 4.30351014 | -0.090711617 | 0.889581263 | NA |
| YLL052C | 77.86079202 | 0.421733441 | 0.322152212 | NA |
| YLL059C | 81.93358505 | 0.33468691 | 0.273420386 | NA |
| YLL065W | 0.583562943 | 0.022051584 | 0.838678363 | NA |
| YLL066C | 1.397171575 | 3.76E-05 | 0.98770177 | NA |
| YLL066W-B | 2.220126499 | 0.141648909 | 0.675388694 | NA |
| YLL067C | 0.999643856 | -0.156779067 | 0.736515205 | NA |
| YLR012C | 11.76101632 | -0.002158591 | 0.928729612 | NA |
| YLR013W | 3.60279815 | -0.011430756 | 0.945199016 | NA |
| YLR035C-A | 40.57446476 | -0.03420345 | 0.812687297 | NA |
| YLR041W | 27.97790779 | 0.073991401 | 0.88585477 | NA |
| YLR101C | 65.86852303 | -0.301117171 | 0.297014284 | NA |
| YLR111W | 15.73315875 | 0.26552622 | 0.555244086 | NA |
| YLR120W-A | 22.81140064 | 0.197060661 | 0.627838618 | NA |
| YLR122C | 30.05120318 | 0.382520928 | 0.382311817 | NA |
| YLR123C | 17.47407493 | 0.393356745 | 0.407924561 | NA |
| YLR124W | 42.32116711 | 0.281533231 | 0.589635988 | NA |
| YLR125W | 49.15160341 | 0.270589821 | 0.496754023 | NA |
| YLR140W | 51.21550356 | -0.333506345 | 0.139025711 | NA |
| YLR149C-A | 61.1192566 | -0.546487203 | 0.279339988 | NA |
| YLR157C-B | 0.750636836 | 0.144448191 | 0.751445997 | NA |
| YLR162W | 2.387178616 | -0.054325287 | 0.876817159 | NA |
| YLR162W-A | 8.884728875 | -0.34973556 | 0.492873349 | NA |
| YLR171W | 25.60461968 | -0.831159192 | 0.023814801 | NA |
| YLR174W | 1214.876474 | 0.141312082 | NA | NA |
| YLR184W | 0.707088895 | 0.04804054 | 0.780457874 | NA |
| YLR198C | 85.68816582 | -0.490991205 | 0.297442576 | NA |

| | | | | |
|---|---|---|---|---|
| YLR202C | 55.2829566 | -0.093883693 | 0.776031178 | NA |
| YLR217W | 16.95655225 | -0.618791448 | 0.156135097 | NA |
| YLR222C-A | 13.05158283 | 0.41306889 | 0.397711245 | NA |
| YLR232W | 2.842598345 | 0.110253265 | 0.938927444 | NA |
| YLR235C | 18.73304548 | -0.012221567 | 0.975594068 | NA |
| YLR236C | 21.91002832 | 0.205112824 | 0.599488681 | NA |
| YLR255C | 23.50063941 | 0.116011879 | 0.773488787 | NA |
| YLR256W-A | 92.61969338 | -0.339506495 | 0.356993657 | NA |
| YLR261C | 10.17984332 | -0.272615076 | 0.518387943 | NA |
| YLR279W | 36.37996098 | -0.192219355 | 0.697608855 | NA |
| YLR281C | 33.92288358 | -0.173158177 | 0.720939925 | NA |
| YLR282C | 4.588773458 | -0.411062228 | 0.41933751 | NA |
| YLR285C-A | 41.10839909 | 0.150216438 | 0.691974142 | NA |
| YLR296W | 4.726955448 | -0.172395431 | 0.662999294 | NA |
| YLR302C | 0.305761563 | -0.004732254 | 0.994958619 | NA |
| YLR307C-A | 38.91718273 | 0.280889112 | 0.267063648 | NA |
| YLR307W | 82.37745124 | 0.152493571 | 0.743328308 | NA |
| YLR308W | 11.06339218 | 0.751058132 | 0.06935706 | NA |
| YLR311C | 30.19399839 | -0.157831114 | 0.75180555 | NA |
| YLR317W | 2.198595472 | 0.268992642 | 0.634956733 | NA |
| YLR322W | 7.310944976 | -0.065365861 | 0.945796399 | NA |
| YLR331C | 3.488696978 | -0.471494231 | 0.35088463 | NA |
| YLR334C | 3.854398197 | 0.331982621 | 0.490016655 | NA |
| YLR338W | 19.58277131 | -0.269397837 | 0.456410534 | NA |
| YLR342W-A | 51.47657195 | 0.885364689 | 0.030021372 | NA |
| YLR349W | 23.19539208 | -0.103522612 | 0.788530571 | NA |
| YLR358C | 39.08246637 | 0.158035562 | 0.503846425 | NA |
| YLR364C-A | 0.303554837 | 0.072493656 | 0.689108885 | NA |
| YLR365W | 4.223598425 | -0.221972147 | 0.674038124 | NA |
| YLR366W | 1.593075245 | -0.202679573 | 0.730740386 | NA |
| YLR374C | 34.93928504 | 0.319477725 | 0.344429045 | NA |
| YLR377C | 564.6863452 | 0.037133949 | NA | NA |
| YLR385C | 52.50375543 | 0.431206696 | 0.146047057 | NA |
| YLR400W | 49.34542517 | 1.281331342 | 0.005346832 | NA |
| YLR402W | 27.81687827 | 0.639384501 | 0.055755293 | NA |
| YLR406C-A | 2.560257022 | 0.924638057 | 0.074925371 | NA |
| YLR410W-B | 66.3723325 | 0.470747548 | 0.274035937 | NA |
| YLR412C-A | 1.32995427 | 0.50020777 | 0.356330138 | NA |
| YLR415C | 24.41748512 | 0.044396743 | 0.909612508 | NA |
| YLR416C | 28.8196825 | 0.007265282 | 0.956653613 | NA |
| YLR428C | 15.96009415 | -0.533995474 | 0.181997089 | NA |
| YLR434C | 48.33837913 | -0.06031468 | 0.899655807 | NA |
| YLR437C-A | 26.88165292 | -0.059265595 | 0.886099859 | NA |

| | | | | |
|---|---|---|---|---|
| YLR444C | 25.68564702 | -0.089326675 | 0.903984611 | NA |
| YLR445W | 32.03943686 | 0.073625835 | 0.873039382 | NA |
| YLR458W | 2.467406493 | -0.25356219 | 0.684489215 | NA |
| YLR461W | 7.039838185 | 0.036593958 | 0.911935066 | NA |
| YLR462W | 4.00598291 | 0.047029969 | 0.906194434 | NA |
| YLR466C-B | 12.5768575 | 0.489074523 | 0.279236432 | NA |
| YML007C-A | 2.984197591 | 0.232048874 | 0.685490251 | NA |
| YML009C-A | 52.04514359 | -0.922964101 | 0.009386917 | NA |
| YML012C-A | 4.439298629 | -0.151822759 | 0.795509656 | NA |
| YML031C-A | 2.97142876 | -0.138653507 | 0.770857615 | NA |
| YML034C-A | 23.30372078 | 0.043336295 | 0.906495369 | NA |
| YML039W | 1.131094598 | 0.232683893 | 0.623298492 | NA |
| YML042W | 2686.768376 | -0.071932236 | NA | NA |
| YML047C | 45.4588878 | 0.803847369 | 0.065839801 | NA |
| YML047W-A | 2.39499993 | 0.739144631 | 0.107859774 | NA |
| YML054C-A | 11.9894805 | -0.098243303 | 0.800376629 | NA |
| YML057C-A | 86.62699701 | -0.277455521 | 0.279225122 | NA |
| YML084W | 2.063062303 | -0.614037488 | 0.263993247 | NA |
| YML089C | 19.45677984 | -0.277796463 | 0.48516581 | NA |
| YML094C-A | 15.34009962 | 0.560770721 | 0.105261388 | NA |
| YML099W-A | 34.73959572 | 0.217087935 | 0.418291261 | NA |
| YML100W-A | 6.289539175 | -0.830997722 | 0.086799896 | NA |
| YML101C-A | 7.327177589 | -0.028750949 | 0.948767859 | NA |
| YML122C | 47.85383085 | 0.583312628 | 0.15753448 | NA |
| YML133C | 9.588165535 | -0.089136384 | 0.859729282 | NA |
| YMR001C-A | 16.08451551 | -0.403532993 | 0.334676733 | NA |
| YMR007W | 90.19118303 | 0.743457079 | 0.072879849 | NA |
| YMR017W | 57.51133406 | -0.107360318 | 0.829462758 | NA |
| YMR018W | 66.38637523 | 0.683024279 | 0.079221582 | NA |
| YMR030W-A | 15.82228062 | 0.155981865 | 0.726537348 | NA |
| YMR031W-A | 40.58383151 | 0.0268178 | 0.942620539 | NA |
| YMR052C-A | 61.89756679 | -0.070551269 | 0.808804102 | NA |
| YMR057C | 13.35741525 | 0.169415665 | 0.716039374 | NA |
| YMR069W | 79.44065746 | 0.562914108 | 0.235500008 | NA |
| YMR075C-A | 15.14864191 | -0.195341661 | 0.643708805 | NA |
| YMR082C | 5.748276588 | 0.424576222 | 0.350447508 | NA |
| YMR086C-A | 82.59944884 | 0.434717581 | 0.158481924 | NA |
| YMR105W-A | 43.59841519 | 0.414640943 | 0.118725391 | NA |
| YMR107W | 2022.649302 | 0.135098321 | NA | NA |
| YMR119W-A | 41.58140006 | 0.19651717 | 0.593004099 | NA |
| YMR122C | 59.11544635 | 0.002437916 | 0.982271882 | NA |
| YMR135W-A | 18.48988281 | 0.471054696 | 0.286913736 | NA |
| YMR141W-A | 54.52612508 | 0.965771923 | 0.03962493 | NA |

| | | | | |
|---|---|---|---|---|
| YMR151W | 69.13225197 | 0.034213345 | 0.907405137 | NA |
| YMR153C-A | 60.08017136 | -0.352494934 | 0.213025266 | NA |
| YMR158C-A | 22.93483608 | 0.139893324 | 0.714993172 | NA |
| YMR158W-B | 1.160282167 | 0.357568904 | 0.515491896 | NA |
| YMR159C | 22.87110149 | 0.170523985 | 0.557200932 | NA |
| YMR172C-A | 11.37396813 | -0.482057145 | 0.183005646 | NA |
| YMR175W | 966.6429644 | -0.035704122 | NA | NA |
| YMR175W-A | 25.9663659 | -0.042162168 | NA | NA |
| YMR182W-A | 4.590688512 | 0.298644502 | 0.531181856 | NA |
| YMR194C-A | 22.19182237 | 0.60762069 | 0.172274009 | NA |
| YMR230W-A | 4.038623613 | 0.113873986 | 0.892359775 | NA |
| YMR242W-A | 1.825788145 | 0.362278035 | 0.491587581 | NA |
| YMR244W | 46.42719777 | 0.629678509 | 0.193899813 | NA |
| YMR245W | 80.78367086 | -0.16418314 | 0.651239674 | NA |
| YMR247W-A | 22.89498367 | 0.538808161 | 0.250025903 | NA |
| YMR254C | 26.76419027 | 0.237220972 | 0.499359521 | NA |
| YMR272W-B | 3.438828838 | 1.02127796 | 0.048336457 | NA |
| YMR290W-A | 11.59226843 | -0.580976704 | 0.268864195 | NA |
| YMR306C-A | 3.882802162 | -0.854833456 | 0.093989855 | NA |
| YMR315W-A | 6.002010905 | 0.024947854 | 0.866279069 | NA |
| YMR316C-A | 33.48519679 | -0.018556319 | 0.906911254 | NA |
| YMR316C-B | 10.98411933 | -0.005993427 | 0.997936407 | NA |
| YMR321C | 24.07796373 | 0.772703658 | 0.091039755 | NA |
| YMR322C | 22.28920933 | 0.170525996 | 0.46508588 | NA |
| YMR324C | 2.573506675 | 0.323378942 | 0.429304528 | NA |
| YMR325W | 4.52283349 | 0.137628043 | 0.699709528 | NA |
| YMR326C | 0.406623722 | -0.005260013 | 0.995748254 | NA |
| YNL013C | 5.629906296 | -0.181663559 | 0.741581538 | NA |
| YNL017C | 1.615564791 | -0.120569858 | 0.820640153 | NA |
| YNL019C | 10.29601328 | 0.7325897 | 0.112281552 | NA |
| YNL028W | 6.323808974 | 0.826822348 | 0.082780305 | NA |
| YNL033W | 5.453513622 | 0.101156889 | 0.842758644 | NA |
| YNL034W | 79.93693595 | -0.20798432 | 0.651018656 | NA |
| YNL036W | 2481.742596 | -0.234381705 | NA | NA |
| YNL042W-B | 14.4556208 | -0.010607457 | 0.94681226 | NA |
| YNL043C | 66.28583242 | 0.860295857 | 0.024973817 | NA |
| YNL057W | 19.3681916 | -0.670201677 | 0.191878825 | NA |
| YNL067W-A | 0.709210999 | 0.307621023 | 0.360062047 | NA |
| YNL067W-B | 0.339190409 | -0.064565268 | 0.735631987 | NA |
| YNL089C | 16.65282477 | -0.059588673 | 0.914140482 | NA |
| YNL090W | 89.81803356 | -0.06087748 | 0.853197035 | NA |
| YNL097C-B | 13.11125706 | 0.679635874 | 0.165430081 | NA |
| YNL105W | 8.716167534 | 0.12285972 | 0.781137073 | NA |

| | | | | |
|---|---|---|---|---|
| YNL109W | 19.36367514 | -0.032506602 | 0.918190159 | NA |
| YNL114C | 62.61749004 | -0.554057057 | 0.142618113 | NA |
| YNL117W | 3801.736785 | -0.021013295 | NA | NA |
| YNL120C | 0.227028385 | 0.026970009 | 0.883709887 | NA |
| YNL128W | 87.91000712 | 0.220780244 | 0.607026596 | NA |
| YNL130C-A | 3.921452687 | -0.153877884 | 0.77226889 | NA |
| YNL140C | 6.604161499 | 0.671390696 | 0.154454218 | NA |
| YNL143C | 26.87962177 | 0.354181455 | 0.360309038 | NA |
| YNL146C-A | 29.52790111 | 0.549714314 | 0.301305074 | NA |
| YNL146W | 85.60940575 | -0.166881576 | 0.558594373 | NA |
| YNL150W | 83.29807569 | -0.863901025 | 0.01022434 | NA |
| YNL162W-A | 54.12294867 | 0.136404348 | 0.746671853 | NA |
| YNL170W | 85.88966089 | -0.185750558 | 0.579318984 | NA |
| YNL171C | 59.8874962 | -0.1956889 | 0.543416148 | NA |
| YNL174W | 7.421723425 | -0.532697202 | 0.35422643 | NA |
| YNL184C | 77.22136504 | 0.037217535 | 0.898052016 | NA |
| YNL198C | 23.75937545 | -0.161958563 | 0.670630747 | NA |
| YNL203C | 26.22039237 | 0.187699878 | 0.604880832 | NA |
| YNL205C | 11.3162081 | 0.137415795 | 0.790461443 | NA |
| YNL226W | 19.21780824 | -0.043833379 | 0.92244599 | NA |
| YNL228W | 12.1493579 | 0.472300881 | 0.290563684 | NA |
| YNL235C | 7.928094321 | -0.252330894 | 0.599449713 | NA |
| YNL266W | 15.42456766 | 0.064765015 | 0.841093342 | NA |
| YNL269W | 50.67416069 | 0.477316198 | 0.232547434 | NA |
| YNL276C | 5.892936072 | -0.0162936 | 0.976638918 | NA |
| YNL277W-A | 6.085182075 | 0.587021644 | 0.238231515 | NA |
| YNL285W | 3.926788582 | 0.294256831 | 0.573169663 | NA |
| YNL296W | 0.17572267 | -0.028924021 | 0.864229296 | NA |
| YNL318C | 22.2967731 | 0.71999391 | 0.06576246 | NA |
| YNL319W | 5.59444275 | 0.763031249 | 0.160522678 | NA |
| YNL324W | 2.168671185 | 0.612815509 | 0.137285024 | NA |
| YNL328C | 68.99382563 | 0.404936411 | 0.136103371 | NA |
| YNL332W | 25.85198322 | 0.231425653 | 0.628309153 | NA |
| YNL333W | 85.09307581 | -0.391749402 | 0.394915946 | NA |
| YNL334C | 19.40823403 | 0.143138334 | 0.787718446 | NA |
| YNL335W | 13.42162756 | 0.443453802 | 0.356789165 | NA |
| YNL337W | 0.659062417 | -0.010585082 | 0.992455555 | NA |
| YNL338W | 0.154202725 | -0.000977006 | 0.995772101 | NA |
| YNR001W-A | 2.398336405 | 0.52674787 | 0.306263424 | NA |
| YNR002C | 907.6027094 | 0.269680537 | NA | NA |
| YNR005C | 1.56429782 | -0.044028047 | 0.94162051 | NA |
| YNR025C | 48.20698063 | -0.341083649 | 0.31794275 | NA |
| YNR042W | 0.825755426 | 0.357599292 | 0.56534334 | NA |

| | | | | |
|---|---|---|---|---|
| YNR071C | 317.5831201 | 0.214552217 | NA | NA |
| YNR075C-A | 12.77963038 | 0.316222039 | 0.551359818 | NA |
| YNR077C | 0.24124529 | 0.067504483 | 0.75981506 | NA |
| YOL013W-A | 5.152321446 | 1.592466041 | 0.004662858 | NA |
| YOL013W-B | 4.383672181 | 0.330697894 | 0.519098038 | NA |
| YOL019W-A | 0.720433626 | -0.013724876 | 0.992869997 | NA |
| YOL024W | 52.91060507 | 0.433002948 | 0.335926419 | NA |
| YOL035C | 58.4740954 | 0.070158146 | 0.828067751 | NA |
| YOL037C | 71.18780393 | 0.054088724 | 0.861745194 | NA |
| YOL038C-A | 23.65291474 | -1.293384774 | 1.37E-05 | NA |
| YOL046C | 5.940874887 | 1.121609387 | 0.021825228 | NA |
| YOL050C | 16.33482394 | -0.100983436 | 0.79950367 | NA |
| YOL079W | 9.64455915 | 0.164993843 | 0.754100242 | NA |
| YOL085C | 30.97884527 | 0.625805796 | 0.170254672 | NA |
| YOL085W-A | 20.71787704 | 0.52468982 | 0.272656787 | NA |
| YOL097W-A | 14.8332561 | -0.150972491 | 0.66192235 | NA |
| YOL099C | 11.41692104 | -0.19793396 | 0.624456203 | NA |
| YOL103W-B | 37.1755698 | 0.437548348 | 0.289358479 | NA |
| YOL106W | 14.51135789 | 0.759733506 | 0.095121988 | NA |
| YOL118C | 79.69627109 | 0.042349875 | 0.930421032 | NA |
| YOL131W | 75.70252965 | 0.079003048 | 0.826474514 | NA |
| YOL134C | 23.62819378 | -0.866116108 | 0.002108858 | NA |
| YOL150C | 3.831416512 | 0.194916169 | 0.692362222 | NA |
| YOL157C | 78.40010831 | -0.142438577 | 0.762530983 | NA |
| YOL159C-A | 58.15453861 | -0.21358182 | 0.599428305 | NA |
| YOL160W | 15.67971505 | -0.202905524 | 0.604792612 | NA |
| YOL161C | 3.517453047 | -0.464849689 | 0.216616233 | NA |
| YOL162W | 65.72006374 | -0.140884093 | 0.782227347 | NA |
| YOL163W | 57.10674557 | 0.031615776 | 0.922575695 | NA |
| YOL164W-A | 24.89009234 | -0.11991543 | 0.790457552 | NA |
| YOL165C | 22.45477283 | -0.032595318 | 0.942208349 | NA |
| YOL166C | 2.808993483 | 0.190929146 | 0.61813776 | NA |
| YOL166W-A | 0.347622843 | -0.005361856 | 0.995681612 | NA |
| YOR008C-A | 9.788861467 | 0.323666747 | 0.505149536 | NA |
| YOR008W-B | 4.890160442 | 0.235158589 | 0.635829328 | NA |
| YOR011W-A | 3.164669454 | 0.124261301 | 0.75673904 | NA |
| YOR012W | 60.32598792 | 0.180802267 | 0.468073227 | NA |
| YOR013W | 86.58484926 | -0.174487868 | 0.547541999 | NA |
| YOR024W | 4.641545194 | 0.697535579 | 0.101970364 | NA |
| YOR029W | 6.24277143 | -0.091562862 | 0.863612546 | NA |
| YOR032W-A | 11.5403846 | 1.330644835 | 0.002366429 | NA |
| YOR034C-A | 45.66152085 | 0.024128736 | 0.956137119 | NA |
| YOR041C | 37.3273586 | -0.035859444 | 0.91136011 | NA |

| | | | | |
|---|---|---|---|---|
| YOR050C | 67.3008437 | 0.201358223 | 0.496652851 | NA |
| YOR053W | 4.40231234 | -0.423265537 | 0.460697781 | NA |
| YOR055W | 14.54954616 | -0.031498155 | 0.934304082 | NA |
| YOR072W | 29.04315607 | 0.426625596 | 0.245894805 | NA |
| YOR072W-A | 14.05662092 | 0.438397982 | 0.307056212 | NA |
| YOR072W-B | 0.082611248 | -0.000977006 | 0.995785278 | NA |
| YOR073W-A | 11.62724158 | -0.997976398 | 0.037440194 | NA |
| YOR082C | 31.57473045 | 0.029323849 | 0.919938806 | NA |
| YOR100C | 795.4961629 | -0.051881374 | NA | NA |
| YOR102W | 49.45521107 | -0.348142472 | 0.289599372 | NA |
| YOR105W | 21.94253227 | 0.527545815 | 0.113556454 | NA |
| YOR121C | 64.33732395 | 0.049100451 | 0.886083618 | NA |
| YOR139C | 5.37072037 | -0.429244598 | 0.400037119 | NA |
| YOR146W | 6.962391281 | 0.318492579 | 0.4868245 | NA |
| YOR161C-C | 9.014220677 | 0.325671357 | 0.495845821 | NA |
| YOR169C | 39.59953477 | 0.344728861 | 0.476363622 | NA |
| YOR183W | 10.65070105 | 0.124680511 | 0.786359839 | NA |
| YOR186W | 27.95959597 | 0.337332306 | 0.436992887 | NA |
| YOR192C-B | 24.91802453 | -0.268129078 | 0.621823059 | NA |
| YOR192C-C | 17.72254902 | -0.083096282 | 0.829113335 | NA |
| YOR199W | 28.95971662 | -0.03772095 | 0.930954376 | NA |
| YOR200W | 9.969440735 | 0.658013941 | 0.151739006 | NA |
| YOR214C | 33.33902204 | 0.391120738 | 0.343391376 | NA |
| YOR218C | 5.229072963 | 0.308878006 | 0.530920104 | NA |
| YOR225W | 54.88254174 | 0.147322513 | 0.745761154 | NA |
| YOR235W | 3.373424746 | 0.699945936 | 0.142816763 | NA |
| YOR248W | 35.3813631 | 0.290898192 | 0.485725337 | NA |
| YOR255W | 23.76826284 | 0.650370064 | 0.077684423 | NA |
| YOR263C | 29.84403306 | 0.03549564 | 0.917286698 | NA |
| YOR268C | 11.87888145 | -0.260264255 | 0.602191472 | NA |
| YOR277C | 6.042620409 | -0.492019773 | 0.328623439 | NA |
| YOR282W | 14.73409586 | -0.58832301 | 0.126604048 | NA |
| YOR293C-A | 2.608736587 | 0.786001979 | 0.150293139 | NA |
| YOR300W | 17.40293291 | 0.337661261 | 0.380366022 | NA |
| YOR302W | 66.2485987 | 0.416485771 | 0.383868743 | NA |
| YOR314W | 17.17022364 | 0.332373149 | 0.522018795 | NA |
| YOR314W-A | 0.578252409 | 0.04851277 | 0.785053474 | NA |
| YOR316C-A | 8.072827245 | 0.545126182 | 0.275392272 | NA |
| YOR318C | 25.32067042 | 0.176731726 | 0.656232495 | NA |
| YOR325W | 4.631536169 | 0.370234729 | 0.533523677 | NA |
| YOR329W-A | 20.44917844 | 0.574578273 | 0.109593315 | NA |
| YOR333C | 3.992636415 | 0.341062764 | 0.511436121 | NA |
| YOR339C | 48.09399516 | 0.111459657 | 0.739948442 | NA |

| | | | | |
|---|---|---|---|---|
| YOR343C | 63.14207102 | 0.313034357 | 0.503067884 | NA |
| YOR345C | 1.957434997 | 0.354707456 | 0.48014625 | NA |
| YOR364W | 7.913433737 | -0.725127579 | 0.149207202 | NA |
| YOR366W | 3.244594662 | -0.061663009 | 0.949482092 | NA |
| YOR376W | 25.04102901 | -0.137592377 | 0.814590704 | NA |
| YOR376W-A | 83.93269889 | 0.001700096 | 0.993348439 | NA |
| YOR379C | 0.107512111 | 0.026970009 | 0.883673227 | NA |
| YOR381W-A | 9.096748957 | 0.252215872 | 0.600040763 | NA |
| YOR387C | 76.98828773 | 0.459818841 | 0.265329226 | NA |
| YOR388C | 10.58733181 | 0.047727251 | 0.826964155 | NA |
| YOR390W | 45.23234325 | 0.506676126 | 0.212476693 | NA |
| YOR391C | 10.43191586 | 0.270824314 | 0.510098519 | NA |
| YOR392W | 4.708188968 | -0.006261869 | 0.996795617 | NA |
| YOR394C-A | 4.135778786 | 0.942893932 | 0.06633296 | NA |
| YPL021W | 14.06563719 | -0.053584228 | 0.88780672 | NA |
| YPL025C | 15.86269313 | 0.362996861 | 0.391820999 | NA |
| YPL027W | 50.11860961 | 0.585358002 | 0.17901349 | NA |
| YPL033C | 16.58126869 | 0.607743232 | 0.18769685 | NA |
| YPL038W-A | 14.34188488 | -0.036298654 | 0.925311982 | NA |
| YPL044C | 1.74310561 | 0.013067815 | 0.931444135 | NA |
| YPL060C-A | 3.51919416 | -0.700054808 | 0.178540849 | NA |
| YPL062W | 4.463478236 | 0.474694956 | 0.334121016 | NA |
| YPL073C | 1.14139142 | 0.002817053 | 0.996613272 | NA |
| YPL080C | 11.15479793 | 1.377216876 | 0.005441498 | NA |
| YPL102C | 13.71806551 | -0.2228219 | 0.620147524 | NA |
| YPL114W | 1.969765016 | -0.393311139 | 0.465725937 | NA |
| YPL119C-A | 58.40963327 | -0.264135017 | 0.625014066 | NA |
| YPL121C | 25.34339254 | -0.14822842 | 0.750628953 | NA |
| YPL130W | 24.21083011 | 0.717425267 | 0.064510132 | NA |
| YPL136W | 0.97000745 | -0.388142741 | 0.330752628 | NA |
| YPL152W-A | 1.958413515 | 0.495184932 | 0.384217205 | NA |
| YPL182C | 43.60254699 | 0.495842591 | 0.096938085 | NA |
| YPL185W | 39.28495564 | -0.334159345 | 0.392014389 | NA |
| YPL187W | 17.17000227 | 0.02203541 | 0.927021828 | NA |
| YPL200W | 21.68345646 | 0.578730093 | 0.24867125 | NA |
| YPL205C | 28.51967725 | -0.121692858 | 0.707237309 | NA |
| YPL223C | 915.9171589 | 0.396363446 | NA | NA |
| YPL238C | 56.97812647 | -0.999136921 | 0.000631671 | NA |
| YPL249C-A | 77.54661876 | -0.213067733 | 0.598595665 | NA |
| YPL250W-A | 88.80290187 | -0.348721579 | 0.366601375 | NA |
| YPL251W | 76.6529722 | 0.032558291 | 0.923439923 | NA |
| YPL261C | 48.62531076 | -0.286020138 | 0.42310955 | NA |
| YPL275W | 31.2658679 | 0.267044108 | 0.610956818 | NA |

| | | | | |
|---|---|---|---|---|
| YPL276W | 15.28614474 | 0.445954526 | 0.309903665 | NA |
| YPL278C | 8.613341367 | 0.789416533 | 0.081091799 | NA |
| YPL279C | 37.92327787 | 0.453736505 | 0.243403646 | NA |
| YPR002C-A | 1.918449357 | -0.477491603 | 0.343012925 | NA |
| YPR012W | 66.76164095 | -0.418203497 | 0.170569251 | NA |
| YPR014C | 81.20498688 | 0.207985224 | 0.572379958 | NA |
| YPR016W-A | 37.91128357 | 0.3639162 | 0.356501358 | NA |
| YPR038W | 18.2580769 | -0.515943881 | 0.119801966 | NA |
| YPR039W | 9.707681236 | -0.127122754 | 0.812877242 | NA |
| YPR050C | 1.567602133 | 0.573040828 | 0.384586675 | NA |
| YPR051W | 72.40075259 | 0.37381977 | 0.367642183 | NA |
| YPR059C | 12.62828581 | -0.250842709 | 0.554603286 | NA |
| YPR064W | 39.20156943 | 0.524814167 | 0.183325376 | NA |
| YPR074W-A | 0.407755058 | 0.164667705 | 0.583715854 | NA |
| YPR076W | 41.22373968 | -0.185657861 | 0.638436604 | NA |
| YPR077C | 1.201353204 | 0.456358076 | 0.277668362 | NA |
| YPR078C | 88.79558885 | 0.061426468 | 0.841205268 | NA |
| YPR087W | 20.87255534 | 0.027057224 | 0.932969004 | NA |
| YPR092W | 28.34006108 | 1.056398417 | 0.004796867 | NA |
| YPR096C | 16.87382422 | 0.203422167 | 0.639564499 | NA |
| YPR099C | 24.60634511 | -0.977629441 | 0.00162801 | NA |
| YPR108W-A | 1.090591333 | 0.459173189 | 0.154286254 | NA |
| YPR126C | 82.68269767 | 0.238930676 | 0.481683689 | NA |
| YPR130C | 10.79124991 | 0.190358593 | 0.671574282 | NA |
| YPR136C | 12.04995248 | 0.026465257 | 0.951297963 | NA |
| YPR142C | 12.52047692 | -0.615838979 | 0.140457876 | NA |
| YPR145C-A | 45.3897147 | -0.164810287 | 0.729820904 | NA |
| YPR150W | 6.435730199 | -0.368846741 | 0.507190081 | NA |
| YPR151C | 43.40930441 | 0.163744783 | 0.620753953 | NA |
| YPR153W | 81.58409703 | 0.424606091 | 0.151845196 | NA |
| YPR158W-B | 29.96033516 | 0.186228439 | 0.635352316 | NA |
| YPR159C-A | 2.271891006 | -0.572765847 | 0.231417899 | NA |
| YPR169W-A | 32.89680766 | -0.442346841 | 0.129304789 | NA |
| YPR170C | 83.5281119 | 0.042131867 | 0.876975176 | NA |
| YPR170W-A | 19.17302177 | -0.152288436 | 0.694799581 | NA |
| YPR177C | 1.844393858 | -0.6057735 | 0.266469881 | NA |
| YPR193C | 57.30247136 | -0.090976229 | 0.842603899 | NA |
| YPR195C | 54.58354851 | 0.502973588 | 0.242935104 | NA |
| YPR197C | 5.599447647 | -0.476086483 | 0.395007343 | NA |
| YPR200C | 70.27684883 | -0.372220265 | 0.356091424 | NA |
| YPR202W | 17.57397274 | 0.317375104 | 0.511617552 | NA |
| snR10 | 11.52677515 | 0.130470182 | 0.688238701 | NA |
| snR128 | 13.17491061 | 0.028875355 | 0.861660687 | NA |

| | | | | |
|---|---|---|---|---|
| snR13 | 24.86462889 | 0.384947327 | 0.454658883 | NA |
| snR14 | 19.85838705 | -0.95200976 | 0.021318899 | NA |
| snR161 | 8.988952116 | 0.714842707 | 0.101183506 | NA |
| snR17a | 59.62016603 | -0.610661461 | 0.198620785 | NA |
| snR17b | 27.76593566 | 0.158717284 | 0.741163031 | NA |
| snR18 | 14.34494847 | -0.152671533 | 0.728214633 | NA |
| snR189 | 9.691874789 | 0.251923745 | 0.630092353 | NA |
| snR190 | 17.01701234 | -0.724848547 | 0.14656405 | NA |
| snR191 | 16.29529748 | 0.993279836 | 0.022104875 | NA |
| snR24 | 8.183960631 | -0.161024626 | 0.780443075 | NA |
| snR3 | 32.48912912 | -0.011830858 | 0.990468659 | NA |
| snR31 | 65.06005548 | 0.558546689 | 0.176684903 | NA |
| snR32 | 25.20381189 | 0.224003989 | 0.629819289 | NA |
| snR34 | 37.83791355 | -0.022573533 | 0.88261605 | NA |
| snR35 | 41.6109753 | -0.158165695 | 0.702388992 | NA |
| snR36 | 18.91914442 | 0.439015213 | 0.298157175 | NA |
| snR38 | 6.274197792 | -0.019057075 | 0.964757365 | NA |
| snR39 | 11.68210812 | 0.611579161 | 0.245230322 | NA |
| snR39B | 13.58422976 | 0.439315323 | 0.391602436 | NA |
| snR4 | 89.67640266 | 0.172798901 | 0.69552018 | NA |
| snR40 | 6.538551087 | 0.263533012 | 0.597957561 | NA |
| snR41 | 28.2332558 | 0.593605426 | 0.152180115 | NA |
| snR42 | 83.98583809 | 0.479864913 | 0.257027519 | NA |
| snR43 | 20.09245082 | 0.4403841 | 0.25379802 | NA |
| snR45 | 13.49639451 | -0.174417634 | 0.734279215 | NA |
| snR46 | 23.07413489 | 0.379205429 | 0.391537741 | NA |
| snR47 | 5.450594875 | 0.959840771 | 0.06241062 | NA |
| snR48 | 2.926793808 | -0.184981511 | 0.750616923 | NA |
| snR49 | 77.03202196 | 0.31472163 | 0.455187382 | NA |
| snR5 | 32.14065526 | -0.360192522 | 0.38076412 | NA |
| snR50 | 8.290616263 | 0.633018472 | 0.183529355 | NA |
| snR51 | 25.09375178 | 0.425366457 | 0.374639205 | NA |
| snR52 | 0.834128967 | 0.295910285 | 0.427510276 | NA |
| snR53 | 0.80815136 | -0.036527368 | 0.883054137 | NA |
| snR54 | 8.052861878 | 0.230927086 | 0.643487452 | NA |
| snR55 | 9.080532067 | 0.097870569 | 0.814777457 | NA |
| snR56 | 6.041353852 | 0.143068114 | 0.772099174 | NA |
| snR57 | 17.27289396 | -0.038642041 | 0.946826632 | NA |
| snR58 | 5.413360595 | 0.972110319 | 0.052017385 | NA |
| snR59 | 2.356669998 | 0.443437059 | 0.382937238 | NA |
| snR6 | 0.308944371 | 0.045186629 | 0.838283991 | NA |
| snR60 | 11.58332067 | 0.933462116 | 0.051298877 | NA |
| snR61 | 12.13748594 | 0.350172706 | 0.499710332 | NA |

| | | | | |
|---|---|---|---|---|
| snR62 | 6.46452847 | 0.061735037 | 0.891350673 | NA |
| snR64 | 4.034418541 | -0.066373873 | 0.89929396 | NA |
| snR65 | 5.68201134 | 0.143787171 | 0.771531681 | NA |
| snR66 | 7.293652703 | 0.685859054 | 0.154411824 | NA |
| snR67 | 6.800863812 | -0.607589005 | 0.226033282 | NA |
| snR68 | 10.29817386 | -0.034396136 | 0.936941707 | NA |
| snR69 | 9.747852558 | 0.615727682 | 0.176395895 | NA |
| snR7-L | 6.909111604 | -0.05474669 | 0.915972613 | NA |
| snR70 | 64.50681846 | -0.076966905 | 0.850558431 | NA |
| snR71 | 12.58626851 | 0.018401505 | 0.962181999 | NA |
| snR72 | 8.104041704 | 0.507709071 | 0.301308143 | NA |
| snR73 | 0.982818557 | 0.699245957 | 0.099636783 | NA |
| snR74 | 1.214578707 | 0.273878279 | 0.359924721 | NA |
| snR75 | 1.574604411 | -0.024224001 | 0.947720823 | NA |
| snR76 | 8.640963018 | 0.632691989 | 0.199972051 | NA |
| snR77 | 9.325617253 | 0.597347291 | 0.213561501 | NA |
| snR78 | 1.974544426 | -0.335555263 | 0.466168981 | NA |
| snR79 | 6.23839996 | 0.406662872 | 0.415120041 | NA |
| snR8 | 32.91757815 | 0.232611676 | 0.565069169 | NA |
| snR80 | 3.471418478 | 0.799908703 | 0.116813885 | NA |
| snR81 | 9.309990788 | 0.706579206 | 0.132338351 | NA |
| snR85 | 18.57666508 | 0.494525636 | 0.31940795 | NA |
| snR87 | 6.424473744 | 0.103322868 | 0.830947899 | NA |
| snR9 | 75.73439823 | -0.144396737 | 0.70612319 | NA |
| tD(GUC)N | 0.865673422 | 0.261110031 | 0.65187296 | NA |
| tG(CCC)O | 0.202424979 | 0.054790145 | 0.801157015 | NA |
| tL(GAG)G | 0.599720226 | 0.200133441 | 0.623058457 | NA |
| tQ(UUG)B | 0.084186605 | -0.000977006 | 0.995767047 | NA |
| tQ(UUG)E2 | 0.083088204 | -0.000977006 | 0.995752613 | NA |
| tR(ACG)J | 0.2026169 | 0.110474323 | 0.645538688 | NA |
| tR(CCG)L | 0.085991782 | 0.054917023 | 0.79048973 | NA |
| tR(CCU)J | 0.0941958 | 0.026970009 | 0.883675634 | NA |
| tT(CGU)K | 0.429968112 | -0.299471989 | 0.455219846 | NA |
