## Supplementary material for "A stress response that allows highly mutated eukaryotic cells to survive and proliferate": Table S4

| Gene | baseMean | log2FoldChange | pvalue | padj |
|---|---|---|---|---|
| YFR035C | 184.0960805 | -1.379637273 | 4.03E-05 | 0.26 |
| YFL039C | 29152.57267 | -0.849291103 | 0.000347552 | 0.75 |
| YNR017W | 1847.028777 | -1.10685049 | 0.000236209 | 0.75 |
| YDL133W | 835.1418998 | 0.73734367 | 0.002138711 | 0.77 |
| YER154W | 1666.340326 | -0.869213686 | 0.001777997 | 0.77 |
| YGL072C | 15.1439202 | 0.950189449 | 0.001054905 | 0.77 |
| YGL165C | 78.38274117 | 1.097583794 | 0.000898789 | 0.77 |
| YGR269W | 48.40957868 | 1.098057773 | 0.001870906 | 0.77 |
| YHL004W | 1819.11881 | -0.872522721 | 0.002073251 | 0.77 |
| YHR049C-A | 129.3206508 | -0.714854837 | 0.001466476 | 0.77 |
| YIL073C | 530.2219401 | 0.881042981 | 0.001838998 | 0.77 |
| YJL016W | 1543.218497 | 1.06317431 | 0.000695364 | 0.77 |
| YJL094C | 1236.455255 | 0.982608186 | 0.002059063 | 0.77 |
| YKL005C | 1108.015733 | 0.889105561 | 0.001162799 | 0.77 |
| YNR064C | 215.5475844 | 1.488676998 | 0.001409809 | 0.77 |
| YPL063W | 2649.060944 | -0.848302703 | 0.000686188 | 0.77 |
| YPL111W | 2984.765722 | 1.258239698 | 0.001432795 | 0.77 |
| YPR146C | 99.73914934 | -1.233476533 | 0.001919147 | 0.77 |
| YGL124C | 833.4516769 | 0.933128047 | 0.002303862 | 0.79 |
| YNR057C | 196.7239761 | 1.120991064 | 0.002499941 | 0.81 |
| LSR1 | 186.1998782 | -0.213106166 | 0.100788562 | 0.84 |
| RPR1 | 36.63381865 | -0.432126791 | 0.154506737 | 0.84 |
| SCR1 | 181.7417356 | -0.184756417 | 0.077028913 | 0.84 |
| TLC1 | 1417.425959 | -0.675490315 | 0.048442914 | 0.84 |
| YAL003W | 12374.8526 | -0.170639368 | 0.125278677 | 0.84 |
| YAL007C | 1549.491558 | -0.467895383 | 0.138531318 | 0.84 |
| YAL010C | 552.0969494 | 0.336573252 | 0.171144555 | 0.84 |
| YAL022C | 845.7051212 | -0.576553909 | 0.072623704 | 0.84 |
| YAL023C | 4478.973811 | -0.516354295 | 0.057447855 | 0.84 |
| YAL025C | 983.4578298 | -0.413296226 | 0.202598314 | 0.84 |
| YAL028W | 836.6833612 | 0.921573166 | 0.023475728 | 0.84 |
| YAL032C | 524.818116 | 0.375066604 | 0.123190276 | 0.84 |
| YAL034C | 1899.332638 | 0.395383989 | 0.117244046 | 0.84 |
| YAL034W-A | 187.40184 | -0.548114604 | 0.160639587 | 0.84 |
| YAL037C-B | 133.1022568 | -0.501786919 | 0.08547624 | 0.84 |
| YAL040C | 2404.494026 | 0.495255302 | 0.121064437 | 0.84 |
| YAL042W | 3676.592816 | -0.352771835 | 0.146177969 | 0.84 |
| YAL045C | 8.89025023 | 0.264192043 | 0.184610699 | 0.84 |
| YAL046C | 313.7043981 | -0.854500085 | 0.024860837 | 0.84 |
| YAL059W | 208.6556716 | -0.428515665 | 0.150176169 | 0.84 |

| | | | | |
|---|---|---|---|---|
| YAL063C | 272.6184868 | 0.739150779 | 0.07523465 | 0.84 |
| YAL064W | 3.131576444 | 0.442333472 | 0.14605271 | 0.84 |
| YAL067C | 87.22318015 | -1.348097535 | 0.026610176 | 0.84 |
| YAR002C-A | 2600.594621 | -0.568557548 | 0.074579283 | 0.84 |
| YAR008W | 281.6693003 | -0.403248165 | 0.152908189 | 0.84 |
| YAR015W | 7474.541729 | 0.073648264 | 0.214964657 | 0.84 |
| YAR019C | 904.0673159 | 0.645094277 | 0.062370538 | 0.84 |
| YAR019W-A | 5.564052438 | 0.272447817 | 0.036287387 | 0.84 |
| YAR020C | 43.79066228 | 0.752698887 | 0.087948942 | 0.84 |
| YAR023C | 164.0948317 | 0.229106935 | 0.191078935 | 0.84 |
| YAR030C | 64.62074774 | 0.174836819 | 0.212493139 | 0.84 |
| YAR031W | 151.5423779 | 0.571638427 | 0.174930201 | 0.84 |
| YAR064W | 14.8431874 | -0.807736753 | 0.113555005 | 0.84 |
| YBL002W | 4494.23128 | -0.420541201 | 0.204596758 | 0.84 |
| YBL004W | 2589.356792 | -0.370796492 | 0.133211139 | 0.84 |
| YBL009W | 764.0807255 | -0.472681902 | 0.108394861 | 0.84 |
| YBL018C | 137.4735368 | -0.338739394 | 0.098222781 | 0.84 |
| YBL020W | 997.2826168 | -0.309070434 | 0.216765259 | 0.84 |
| YBL026W | 682.7010545 | -0.595547413 | 0.032340938 | 0.84 |
| YBL027W | 10124.8249 | -0.40590343 | 0.064915666 | 0.84 |
| YBL028C | 410.9135571 | -0.482338437 | 0.055692014 | 0.84 |
| YBL030C | 9599.31549 | -0.103502014 | 0.189582779 | 0.84 |
| YBL032W | 1731.697141 | -0.338194434 | 0.188285651 | 0.84 |
| YBL035C | 596.37002 | -0.685172983 | 0.056903339 | 0.84 |
| YBL039C | 1528.083632 | -0.260767782 | 0.136704243 | 0.84 |
| YBL040C | 1584.786506 | -0.4874355 | 0.080830945 | 0.84 |
| YBL042C | 1258.710363 | -0.909273125 | 0.157653474 | 0.84 |
| YBL046W | 691.3216368 | 0.398548514 | 0.148538318 | 0.84 |
| YBL048W | 125.8109055 | 0.512743467 | 0.151231738 | 0.84 |
| YBL049W | 377.5575063 | 0.325077062 | 0.185971293 | 0.84 |
| YBL056W | 3616.602261 | -0.358101153 | 0.121015214 | 0.84 |
| YBL057C | 768.8630819 | -0.506671493 | 0.11340914 | 0.84 |
| YBL059W | 273.8243146 | -0.551416371 | 0.162979565 | 0.84 |
| YBL060W | 624.3164774 | 0.473999405 | 0.157469 | 0.84 |
| YBL065W | 30.86397396 | 0.503261276 | 0.072927409 | 0.84 |
| YBL072C | 3997.128198 | -0.336907404 | 0.192598786 | 0.84 |
| YBL074C | 232.3789197 | 0.43107456 | 0.197686069 | 0.84 |
| YBL077W | 139.1714064 | -0.104816575 | 0.203166644 | 0.84 |
| YBL086C | 1904.650929 | 0.862108024 | 0.01580181 | 0.84 |
| YBL087C | 3378.344673 | -0.343084021 | 0.117838523 | 0.84 |
| YBL091C | 4543.955358 | -0.548266943 | 0.022873952 | 0.84 |
| YBL092W | 15431.27049 | -0.307113079 | 0.051604207 | 0.84 |
| YBL097W | 438.5923101 | -0.328652737 | 0.189308605 | 0.84 |

| | | | | |
|---|---|---|---|---|
| YBL106C | 800.8986155 | 0.682308946 | 0.03183796 | 0.84 |
| YBL107C | 641.7007619 | -0.557362272 | 0.149721565 | 0.84 |
| YBL109W | 16.60491535 | 0.938294781 | 0.071535353 | 0.84 |
| YBR001C | 1318.791969 | 0.489592305 | 0.154738451 | 0.84 |
| YBR006W | 1760.927773 | 0.842362567 | 0.140564285 | 0.84 |
| YBR008C | 906.2949136 | 0.531483987 | 0.076400684 | 0.84 |
| YBR012W-B | 321.6022163 | 0.818669175 | 0.211850683 | 0.84 |
| YBR027C | 8.649310737 | 0.430080881 | 0.036633413 | 0.84 |
| YBR029C | 1375.442278 | -0.520082158 | 0.140970805 | 0.84 |
| YBR031W | 10790.74196 | -0.424584571 | 0.067197758 | 0.84 |
| YBR032W | 28.5083245 | 0.356285774 | 0.125827131 | 0.84 |
| YBR043C | 1241.74171 | 0.703626076 | 0.144502545 | 0.84 |
| YBR045C | 255.946774 | 0.947658896 | 0.009271307 | 0.84 |
| YBR047W | 453.8190728 | 1.136244148 | 0.031464997 | 0.84 |
| YBR048W | 5960.519296 | -0.333408648 | 0.059809964 | 0.84 |
| YBR055C | 925.8039594 | -0.508351503 | 0.051908442 | 0.84 |
| YBR056W | 2794.886987 | 0.450596175 | 0.209160416 | 0.84 |
| YBR060C | 829.7516444 | 0.316005168 | 0.214373906 | 0.84 |
| YBR061C | 630.0720407 | -0.510779119 | 0.125101365 | 0.84 |
| YBR064W | 3.158811985 | 0.440421963 | 0.148540056 | 0.84 |
| YBR068C | 7312.593911 | 0.74240554 | 0.109919535 | 0.84 |
| YBR070C | 438.6999546 | -0.542649485 | 0.049882184 | 0.84 |
| YBR072C-A | 2.769886284 | -0.593422443 | 0.196372093 | 0.84 |
| YBR076W | 114.1570412 | 0.341769109 | 0.209944299 | 0.84 |
| YBR079C | 5290.065687 | -0.514196788 | 0.162432507 | 0.84 |
| YBR080C | 2198.347068 | -0.345537024 | 0.211690438 | 0.84 |
| YBR084C-A | 8707.666036 | -0.271051138 | 0.151259126 | 0.84 |
| YBR085C-A | 1789.206511 | 0.804327092 | 0.096389186 | 0.84 |
| YBR088C | 178.1843606 | -0.863898609 | 0.021061982 | 0.84 |
| YBR089C-A | 560.6070076 | -0.514599317 | 0.092325374 | 0.84 |
| YBR091C | 248.9757925 | -0.648936198 | 0.048090624 | 0.84 |
| YBR093C | 949.3588178 | 0.718706843 | 0.148339813 | 0.84 |
| YBR096W | 835.4279583 | -0.405342008 | 0.184204316 | 0.84 |
| YBR105C | 2095.906659 | 0.439912567 | 0.119632461 | 0.84 |
| YBR109C | 2942.937927 | -0.34992971 | 0.155687312 | 0.84 |
| YBR113W | 115.7153994 | -0.515181708 | 0.052818338 | 0.84 |
| YBR114W | 883.2430403 | 0.703812301 | 0.059417026 | 0.84 |
| YBR118W | 10015.00281 | -0.629808975 | 0.013882084 | 0.84 |
| YBR120C | 395.0224857 | -0.574175737 | 0.156385622 | 0.84 |
| YBR121C | 7616.904588 | -0.150864581 | 0.170689997 | 0.84 |
| YBR126C | 9994.323143 | 0.467639712 | 0.217272498 | 0.84 |
| YBR127C | 10020.63379 | -0.554203738 | 0.032262675 | 0.84 |
| YBR132C | 1205.603555 | 0.494519503 | 0.208535259 | 0.84 |

| | | | | |
|---|---|---|---|---|
| YBR139W | 2470.205803 | 0.532718348 | 0.208167017 | 0.84 |
| YBR140C | 4380.796749 | -0.361701576 | 0.207306921 | 0.84 |
| YBR145W | 1370.985162 | 0.652083971 | 0.167183571 | 0.84 |
| YBR146W | 1432.145995 | -0.837235525 | 0.008390704 | 0.84 |
| YBR147W | 531.1286909 | 0.679739021 | 0.095941607 | 0.84 |
| YBR152W | 271.4958287 | 0.436548437 | 0.123958563 | 0.84 |
| YBR154C | 1262.041062 | -0.453774108 | 0.203060707 | 0.84 |
| YBR155W | 833.2201083 | -0.167621377 | 0.122371792 | 0.84 |
| YBR160W | 885.0084099 | -0.406537573 | 0.216651292 | 0.84 |
| YBR164C | 1099.977428 | -0.456686489 | 0.175521461 | 0.84 |
| YBR171W | 1056.70053 | -0.54318184 | 0.1275616 | 0.84 |
| YBR181C | 1794.107741 | -0.20611106 | 0.078112827 | 0.84 |
| YBR185C | 732.2746685 | -0.420547417 | 0.140845586 | 0.84 |
| YBR187W | 950.8054053 | -0.436648967 | 0.025261176 | 0.84 |
| YBR188C | 339.0880427 | -0.348135678 | 0.204783864 | 0.84 |
| YBR189W | 11760.64407 | -0.276798515 | 0.048320754 | 0.84 |
| YBR191W | 11850.59787 | -0.31044162 | 0.037582308 | 0.84 |
| YBR191W-A | 25.03964351 | 0.217380715 | 0.205144586 | 0.84 |
| YBR196C-A | 7.414882286 | 0.397619643 | 0.128719815 | 0.84 |
| YBR203W | 994.1971392 | 0.804081893 | 0.085847157 | 0.84 |
| YBR205W | 1374.246383 | -0.277960349 | 0.217885069 | 0.84 |
| YBR206W | 255.7378889 | -0.48427155 | 0.0822388 | 0.84 |
| YBR211C | 479.4755282 | 0.404482691 | 0.178060193 | 0.84 |
| YBR212W | 2837.682435 | 0.497261982 | 0.216542745 | 0.84 |
| YBR213W | 228.053134 | -0.470943896 | 0.094722228 | 0.84 |
| YBR218C | 7147.332039 | 0.657566756 | 0.177921276 | 0.84 |
| YBR221C | 5794.723277 | -0.264520952 | 0.207917883 | 0.84 |
| YBR222C | 4562.01386 | 0.766712294 | 0.119299532 | 0.84 |
| YBR230C | 2523.139558 | 0.553923802 | 0.209014577 | 0.84 |
| YBR233W-A | 253.0169158 | -0.561027852 | 0.118712808 | 0.84 |
| YBR243C | 997.8241569 | -0.560872898 | 0.056564126 | 0.84 |
| YBR244W | 574.3609399 | -0.549163545 | 0.117989673 | 0.84 |
| YBR245C | 1580.844401 | -0.601195769 | 0.023624216 | 0.84 |
| YBR247C | 781.1952554 | -0.324670238 | 0.117990514 | 0.84 |
| YBR249C | 4275.177673 | 0.247712624 | 0.216821335 | 0.84 |
| YBR252W | 608.885071 | -0.409466068 | 0.17014784 | 0.84 |
| YBR255W | 917.9047892 | 0.493733972 | 0.121266458 | 0.84 |
| YBR256C | 1957.968457 | 0.621278854 | 0.213772719 | 0.84 |
| YBR258C | 250.7365151 | -0.457661419 | 0.197436316 | 0.84 |
| YBR261C | 744.8069304 | -0.579634719 | 0.048096207 | 0.84 |
| YBR263W | 3971.725103 | -0.456795549 | 0.05125565 | 0.84 |
| YBR265W | 1130.650771 | -0.599836188 | 0.060631558 | 0.84 |
| YBR268W | 567.5701433 | -0.606254412 | 0.087781973 | 0.84 |

| | | | | |
|---|---|---|---|---|
| YBR270C | 464.1113888 | 0.428013998 | 0.126258647 | 0.84 |
| YBR271W | 387.9475512 | -0.449523385 | 0.168247763 | 0.84 |
| YBR277C | 4.283921599 | 0.265883062 | 0.05638973 | 0.84 |
| YBR279W | 1844.253736 | -0.354596691 | 0.210574125 | 0.84 |
| YBR280C | 1033.06933 | 0.52754009 | 0.148375826 | 0.84 |
| YBR282W | 512.5901703 | -0.667149236 | 0.044740265 | 0.84 |
| YBR283C | 3928.577615 | -0.514092392 | 0.070463953 | 0.84 |
| YBR284W | 530.3934343 | 0.687863914 | 0.045564858 | 0.84 |
| YBR287W | 2396.597397 | 0.476934766 | 0.137811298 | 0.84 |
| YBR298C | 1312.084765 | 0.595131211 | 0.111052994 | 0.84 |
| YBR299W | 41.66988027 | 0.483323219 | 0.124649327 | 0.84 |
| YBR300C | 5.482970764 | -0.317469837 | 0.057288515 | 0.84 |
| YCL001W | 1214.350549 | -0.358461036 | 0.124023521 | 0.84 |
| YCL005W-A | 247.3743167 | -0.907496824 | 0.010879504 | 0.84 |
| YCL007C | 102.0392471 | -0.475543415 | 0.154388595 | 0.84 |
| YCL008C | 1030.586387 | 0.389350624 | 0.18051329 | 0.84 |
| YCL017C | 1588.391961 | -0.303700696 | 0.178871376 | 0.84 |
| YCL023C | 8.286348364 | 0.870363629 | 0.206802377 | 0.84 |
| YCL024W | 493.5504233 | -0.318692486 | 0.207095119 | 0.84 |
| YCL025C | 3652.132376 | 0.529477637 | 0.134429349 | 0.84 |
| YCL032W | 909.5801687 | 0.413931171 | 0.17880378 | 0.84 |
| YCL037C | 1123.648124 | -0.250928181 | 0.168588888 | 0.84 |
| YCL038C | 682.2522644 | 0.44773977 | 0.148521853 | 0.84 |
| YCL050C | 4005.943189 | -0.115078254 | 0.209893619 | 0.84 |
| YCL051W | 1104.388423 | 0.550753461 | 0.088583771 | 0.84 |
| YCL059C | 1691.420964 | -0.184998197 | 0.164024698 | 0.84 |
| YCR005C | 6272.799933 | 0.713656867 | 0.047583704 | 0.84 |
| YCR007C | 318.2333147 | 0.341169917 | 0.205651823 | 0.84 |
| YCR009C | 3474.642891 | -0.404359541 | 0.187082352 | 0.84 |
| YCR013C | 2806.322738 | -0.357875705 | 0.191712311 | 0.84 |
| YCR014C | 618.1633306 | 0.497726152 | 0.108743718 | 0.84 |
| YCR015C | 255.0577346 | -0.418178648 | 0.202094108 | 0.84 |
| YCR016W | 907.9551315 | -0.313372666 | 0.188326515 | 0.84 |
| YCR020C-A | 266.9746924 | -0.42722809 | 0.171076508 | 0.84 |
| YCR021C | 10522.50846 | 0.515824391 | 0.18721224 | 0.84 |
| YCR022C | 20.80731429 | 0.385361936 | 0.216423584 | 0.84 |
| YCR027C | 255.2084978 | -0.701896792 | 0.055307572 | 0.84 |
| YCR028C-A | 1716.279601 | -0.597408615 | 0.048770949 | 0.84 |
| YCR031C | 11369.4013 | -0.375954401 | 0.096053821 | 0.84 |
| YCR034W | 1456.119271 | -0.216659305 | 0.163087715 | 0.84 |
| YCR035C | 836.2083583 | -0.408045674 | 0.152741922 | 0.84 |
| YCR036W | 1075.104784 | 0.454049387 | 0.086024489 | 0.84 |
| YCR038C | 411.3006768 | 0.957340237 | 0.004623044 | 0.84 |

| | | | | |
|---|---|---|---|---|
| YCR046C | 1534.125343 | -0.587037902 | 0.089610579 | 0.84 |
| YCR061W | 1066.525845 | 0.600077928 | 0.21687049 | 0.84 |
| YCR064C | 12.51518247 | -0.483282317 | 0.164243927 | 0.84 |
| YCR066W | 426.4825775 | 0.361204185 | 0.205905641 | 0.84 |
| YCR068W | 666.4091136 | 0.719194294 | 0.075415617 | 0.84 |
| YCR073C | 984.1918819 | 0.554685105 | 0.055452777 | 0.84 |
| YCR077C | 3337.529518 | -0.404507167 | 0.171364432 | 0.84 |
| YCR084C | 3633.028548 | -0.478811481 | 0.08883268 | 0.84 |
| YCR085W | 15.93952821 | -0.741170094 | 0.064652012 | 0.84 |
| YCR091W | 1488.499833 | 0.381438026 | 0.205651068 | 0.84 |
| YDL003W | 760.5875817 | -0.586238345 | 0.145404171 | 0.84 |
| YDL014W | 4306.735257 | -0.232964403 | 0.177627643 | 0.84 |
| YDL017W | 421.2237274 | 0.619121179 | 0.035361178 | 0.84 |
| YDL020C | 3590.678769 | 0.480516549 | 0.099521773 | 0.84 |
| YDL023C | 219.5343501 | 0.567976737 | 0.199397959 | 0.84 |
| YDL026W | 16.56831607 | 0.543095087 | 0.080715344 | 0.84 |
| YDL036C | 422.8116952 | -0.449519694 | 0.160294782 | 0.84 |
| YDL040C | 3238.410502 | -0.375833585 | 0.136091155 | 0.84 |
| YDL045W-A | 694.1101578 | -0.654433534 | 0.126914196 | 0.84 |
| YDL052C | 1575.894255 | -0.504382713 | 0.122345314 | 0.84 |
| YDL054C | 853.6605887 | 0.739615454 | 0.065317601 | 0.84 |
| YDL055C | 11798.61054 | -0.681819318 | 0.00825446 | 0.84 |
| YDL061C | 2048.312712 | -0.458330285 | 0.114894316 | 0.84 |
| YDL064W | 884.0529801 | -0.439320486 | 0.13644823 | 0.84 |
| YDL075W | 5187.384084 | -0.390807134 | 0.081999484 | 0.84 |
| YDL077C | 1087.970908 | 0.52223761 | 0.032800194 | 0.84 |
| YDL079C | 1497.675817 | 0.45858762 | 0.077968528 | 0.84 |
| YDL081C | 10855.42483 | -0.524965365 | 0.037066895 | 0.84 |
| YDL082W | 7404.050186 | -0.313571363 | 0.13600574 | 0.84 |
| YDL083C | 6219.514303 | -0.413914914 | 0.033922257 | 0.84 |
| YDL084W | 7170.004042 | -0.329329178 | 0.123727335 | 0.84 |
| YDL102W | 928.5717968 | -0.369375641 | 0.115152156 | 0.84 |
| YDL110C | 2473.385616 | 0.447542301 | 0.201871995 | 0.84 |
| YDL111C | 757.760422 | -0.50329115 | 0.143951994 | 0.84 |
| YDL113C | 1161.019898 | 0.487131366 | 0.151770322 | 0.84 |
| YDL118W | 21.28319356 | 0.676057912 | 0.039790605 | 0.84 |
| YDL121C | 1113.601796 | -0.215395287 | 0.199284861 | 0.84 |
| YDL123W | 682.3334241 | 0.946458503 | 0.009489793 | 0.84 |
| YDL127W | 422.9534044 | -0.611743186 | 0.195753436 | 0.84 |
| YDL130W | 4709.707857 | -0.384317923 | 0.169825072 | 0.84 |
| YDL134C | 1962.281647 | 0.313182483 | 0.064581507 | 0.84 |
| YDL136W | 915.4579307 | -0.38095302 | 0.201938091 | 0.84 |
| YDL137W | 4443.238326 | -0.351350564 | 0.216163444 | 0.84 |

| | | | | |
|---|---|---|---|---|
| YDL139C | 356.8731646 | 0.880392858 | 0.005536236 | 0.84 |
| YDL140C | 8072.034708 | -0.570377985 | 0.071990351 | 0.84 |
| YDL142C | 615.3529618 | 0.460216165 | 0.158657489 | 0.84 |
| YDL145C | 6294.266606 | -0.357880899 | 0.155323973 | 0.84 |
| YDL149W | 944.7875532 | 0.558583685 | 0.141569402 | 0.84 |
| YDL150W | 525.2641532 | -0.507931603 | 0.150566841 | 0.84 |
| YDL154W | 661.9950458 | 0.518587606 | 0.098945944 | 0.84 |
| YDL164C | 704.8057939 | -0.340723831 | 0.21473575 | 0.84 |
| YDL166C | 434.2604045 | -0.459214064 | 0.212572104 | 0.84 |
| YDL169C | 664.258116 | 0.537109203 | 0.137533701 | 0.84 |
| YDL170W | 672.20876 | 0.581158771 | 0.089643556 | 0.84 |
| YDL179W | 455.9668158 | -0.597092911 | 0.070831167 | 0.84 |
| YDL183C | 401.7373652 | 0.629800726 | 0.063167746 | 0.84 |
| YDL190C | 2381.415921 | 0.462368788 | 0.113066766 | 0.84 |
| YDL192W | 4665.318464 | -0.383441305 | 0.095680262 | 0.84 |
| YDL194W | 1083.690954 | 0.864430513 | 0.062468862 | 0.84 |
| YDL195W | 7573.167842 | -0.321529733 | 0.186598623 | 0.84 |
| YDL198C | 1156.426648 | -0.300111791 | 0.166430114 | 0.84 |
| YDL199C | 559.7723367 | 0.554726682 | 0.121562941 | 0.84 |
| YDL201W | 813.4402377 | -0.396470349 | 0.211978033 | 0.84 |
| YDL202W | 808.8072189 | -0.852221673 | 0.018072385 | 0.84 |
| YDL206W | 597.9682559 | 0.563144984 | 0.072668701 | 0.84 |
| YDL208W | 1913.168285 | -0.53558973 | 0.07438578 | 0.84 |
| YDL212W | 1994.74877 | -0.629148815 | 0.044613011 | 0.84 |
| YDL213C | 1003.203847 | -0.356264555 | 0.046653524 | 0.84 |
| YDL214C | 464.7760152 | 0.396059925 | 0.084689588 | 0.84 |
| YDL217C | 416.6172424 | -0.391524552 | 0.217143178 | 0.84 |
| YDL218W | 253.6505308 | 0.545095957 | 0.178294517 | 0.84 |
| YDL219W | 330.3585764 | -0.277260222 | 0.178372705 | 0.84 |
| YDL220C | 416.2502353 | -0.477853307 | 0.075548038 | 0.84 |
| YDL226C | 1711.631594 | -0.406670974 | 0.161941302 | 0.84 |
| YDL227C | 308.6751335 | -0.847800217 | 0.093052378 | 0.84 |
| YDL228C | 55.92892077 | -0.445651314 | 0.06136503 | 0.84 |
| YDL230W | 846.3743425 | 0.553333928 | 0.138936404 | 0.84 |
| YDL231C | 1653.703222 | 0.443287828 | 0.185946539 | 0.84 |
| YDL234C | 2388.462086 | 0.653892009 | 0.082215057 | 0.84 |
| YDL236W | 1309.137891 | -0.420035796 | 0.211220724 | 0.84 |
| YDL238C | 479.1219346 | 0.507452052 | 0.174033819 | 0.84 |
| YDL239C | 522.0657782 | 0.512378735 | 0.120743143 | 0.84 |
| YDL245C | 34.45281204 | -0.869678661 | 0.209835034 | 0.84 |
| YDL247W | 3.022100866 | -0.772061962 | 0.043802641 | 0.84 |
| YDR003W | 807.4785109 | 0.444701086 | 0.187696751 | 0.84 |
| YDR003W-A | 67.78719135 | 0.516084577 | 0.088216376 | 0.84 |

| | | | | |
|---|---|---|---|---|
| YDR004W | 628.5806399 | 0.511306769 | 0.051593893 | 0.84 |
| YDR006C | 2310.266671 | 0.491389381 | 0.133313322 | 0.84 |
| YDR007W | 490.6292629 | -0.38256667 | 0.166901434 | 0.84 |
| YDR010C | 32.06483583 | 0.834024559 | 0.003891357 | 0.84 |
| YDR012W | 4314.996695 | -0.267261369 | 0.090513765 | 0.84 |
| YDR014W-A | 36.16776295 | 0.727249764 | 0.17221063 | 0.84 |
| YDR016C | 195.2031569 | -0.539696091 | 0.12402371 | 0.84 |
| YDR017C | 2063.432291 | 0.668068413 | 0.006229062 | 0.84 |
| YDR018C | 446.8147304 | 0.905954414 | 0.03112161 | 0.84 |
| YDR025W | 5331.251911 | -0.427288583 | 0.083678751 | 0.84 |
| YDR027C | 1560.996774 | 0.501493011 | 0.011691231 | 0.84 |
| YDR030C | 444.8820441 | 0.651671671 | 0.063089219 | 0.84 |
| YDR034C-A | 4.06626248 | -0.770706376 | 0.131806338 | 0.84 |
| YDR034W-B | 127.2105401 | 0.403060776 | 0.019987485 | 0.84 |
| YDR040C | 132.5886651 | 0.351431413 | 0.163183187 | 0.84 |
| YDR043C | 825.8596949 | 0.436411521 | 0.062655139 | 0.84 |
| YDR047W | 1762.459456 | -0.40209696 | 0.210557598 | 0.84 |
| YDR048C | 13.8983324 | 0.539096255 | 0.105281863 | 0.84 |
| YDR057W | 882.7471612 | 0.431747305 | 0.173242068 | 0.84 |
| YDR064W | 9582.184184 | -0.386072116 | 0.028271413 | 0.84 |
| YDR069C | 1204.898386 | 0.428247224 | 0.189382218 | 0.84 |
| YDR074W | 10482.83127 | 0.796794216 | 0.14365529 | 0.84 |
| YDR076W | 307.3025521 | 0.515757485 | 0.136144458 | 0.84 |
| YDR082W | 550.4664737 | 0.410104177 | 0.195426041 | 0.84 |
| YDR084C | 664.1695601 | -0.428665414 | 0.08703741 | 0.84 |
| YDR085C | 1259.372351 | 0.479670361 | 0.105775714 | 0.84 |
| YDR091C | 1728.536947 | -0.418552982 | 0.212690171 | 0.84 |
| YDR092W | 1261.21405 | -0.465117935 | 0.122193099 | 0.84 |
| YDR093W | 1759.127414 | -0.4793167 | 0.104895397 | 0.84 |
| YDR096W | 1946.105843 | 0.553811912 | 0.196707785 | 0.84 |
| YDR098C | 1711.271644 | -0.383788878 | 0.123552752 | 0.84 |
| YDR109C | 536.0726963 | 0.350539581 | 0.1419987 | 0.84 |
| YDR113C | 648.1289584 | -0.407967029 | 0.171256753 | 0.84 |
| YDR115W | 505.144241 | -0.633567155 | 0.095453699 | 0.84 |
| YDR116C | 1042.960963 | -0.506148897 | 0.156693423 | 0.84 |
| YDR121W | 548.7039891 | -0.493772095 | 0.102194539 | 0.84 |
| YDR124W | 300.8040212 | 0.498202017 | 0.163929373 | 0.84 |
| YDR131C | 626.2979697 | 0.367615726 | 0.174169359 | 0.84 |
| YDR136C | 21.36116146 | 0.516390357 | 0.185505489 | 0.84 |
| YDR137W | 962.3953612 | 0.335961233 | 0.118206711 | 0.84 |
| YDR164C | 1143.654091 | -0.376582048 | 0.123898861 | 0.84 |
| YDR172W | 3599.759088 | -0.547273041 | 0.074030514 | 0.84 |
| YDR173C | 369.7980333 | 0.479438361 | 0.1199943 | 0.84 |

| | | | | |
|---|---|---|---|---|
| YDR182W-A | 24.64411227 | 0.751759837 | 0.012218998 | 0.84 |
| YDR183C-A | 47.62983085 | -0.441504082 | 0.039038323 | 0.84 |
| YDR194C | 2378.698014 | -0.598397121 | 0.072899874 | 0.84 |
| YDR195W | 649.2195359 | 0.581015094 | 0.046385676 | 0.84 |
| YDR200C | 868.5642679 | 0.391235147 | 0.137722389 | 0.84 |
| YDR214W | 2305.632085 | -0.442990974 | 0.149427491 | 0.84 |
| YDR223W | 358.2338356 | 0.471100748 | 0.112805256 | 0.84 |
| YDR225W | 3245.605018 | -0.62097506 | 0.203720703 | 0.84 |
| YDR228C | 901.0372572 | 0.536982343 | 0.052006796 | 0.84 |
| YDR233C | 7397.691198 | -0.458621748 | 0.129482729 | 0.84 |
| YDR236C | 550.9804022 | -0.625729539 | 0.083008636 | 0.84 |
| YDR237W | 832.9477874 | -0.668260102 | 0.060162735 | 0.84 |
| YDR245W | 1164.324708 | -0.429770057 | 0.213592305 | 0.84 |
| YDR250C | 6.490017442 | 1.081143844 | 0.005410965 | 0.84 |
| YDR263C | 317.4342443 | 0.525154355 | 0.078183782 | 0.84 |
| YDR273W | 249.707739 | 0.399615808 | 0.142701543 | 0.84 |
| YDR277C | 2772.925083 | 0.691930614 | 0.048730804 | 0.84 |
| YDR280W | 709.9216811 | -0.536468574 | 0.122836804 | 0.84 |
| YDR285W | 510.8247622 | 0.790602382 | 0.02437393 | 0.84 |
| YDR292C | 1617.849006 | -0.611354721 | 0.030490139 | 0.84 |
| YDR293C | 6400.531719 | 0.55354319 | 0.140811862 | 0.84 |
| YDR296W | 1076.454873 | -0.477945155 | 0.175896456 | 0.84 |
| YDR297W | 2184.395638 | -0.48213631 | 0.094540398 | 0.84 |
| YDR302W | 389.0867763 | -0.611072888 | 0.056845466 | 0.84 |
| YDR306C | 1159.558737 | 0.565147167 | 0.076570605 | 0.84 |
| YDR308C | 370.9057891 | -0.678394686 | 0.028351698 | 0.84 |
| YDR320W-B | 5.22193975 | 0.344011153 | 0.10716182 | 0.84 |
| YDR321W | 1332.488079 | -0.168638381 | 0.08128877 | 0.84 |
| YDR323C | 314.788297 | 0.413617963 | 0.095603512 | 0.84 |
| YDR327W | 614.5550695 | -0.527840748 | 0.075694763 | 0.84 |
| YDR330W | 1850.284876 | 0.407141355 | 0.218620427 | 0.84 |
| YDR331W | 571.3069512 | -0.624135694 | 0.077033309 | 0.84 |
| YDR332W | 604.027567 | 0.635514845 | 0.019024052 | 0.84 |
| YDR336W | 301.9017741 | 0.447112255 | 0.166639329 | 0.84 |
| YDR337W | 923.5124951 | -0.508963701 | 0.092102642 | 0.84 |
| YDR342C | 1706.769269 | 0.358335946 | 0.214487681 | 0.84 |
| YDR347W | 1409.321423 | -0.557035899 | 0.014124154 | 0.84 |
| YDR350C | 1056.007335 | 0.524283334 | 0.11223587 | 0.84 |
| YDR353W | 5059.752159 | -0.343306087 | 0.213518567 | 0.84 |
| YDR358W | 1790.698239 | 0.545588026 | 0.167274429 | 0.84 |
| YDR364C | 596.6987029 | 0.381420979 | 0.122312695 | 0.84 |
| YDR367W | 606.3521729 | -0.596347442 | 0.07426698 | 0.84 |
| YDR374C | 71.50374083 | 0.297731084 | 0.144895356 | 0.84 |

| | | | | |
|---|---|---|---|---|
| YDR381W | 4059.585724 | -0.499648536 | 0.181977921 | 0.84 |
| YDR382W | 7559.11107 | -0.651080369 | 0.036580698 | 0.84 |
| YDR386W | 496.5773156 | 0.585463457 | 0.111587393 | 0.84 |
| YDR387C | 888.6103683 | 0.448435782 | 0.184042061 | 0.84 |
| YDR391C | 1742.594469 | 0.656671788 | 0.15331726 | 0.84 |
| YDR396W | 125.4408117 | -0.716397601 | 0.093589363 | 0.84 |
| YDR398W | 1282.093858 | -0.381387813 | 0.100459433 | 0.84 |
| YDR399W | 3185.764567 | -0.056779806 | 0.172368608 | 0.84 |
| YDR400W | 652.8362371 | -0.427380225 | 0.174320053 | 0.84 |
| YDR412W | 170.5135109 | -0.400843506 | 0.11617365 | 0.84 |
| YDR413C | 10.51150758 | -0.647277729 | 0.135711031 | 0.84 |
| YDR418W | 2788.265019 | -0.299792086 | 0.125653051 | 0.84 |
| YDR421W | 1031.149921 | 0.563489955 | 0.08379236 | 0.84 |
| YDR424C | 383.6752581 | -0.423367349 | 0.135215377 | 0.84 |
| YDR425W | 1030.779588 | 0.579903286 | 0.097604991 | 0.84 |
| YDR429C | 3118.563913 | -0.459306354 | 0.11492833 | 0.84 |
| YDR436W | 1255.862426 | 0.551601066 | 0.117786306 | 0.84 |
| YDR438W | 373.0593006 | 0.39633408 | 0.179846574 | 0.84 |
| YDR446W | 60.61919013 | 0.858882856 | 0.021120516 | 0.84 |
| YDR447C | 4088.999597 | -0.301254464 | 0.065823646 | 0.84 |
| YDR450W | 8491.535949 | -0.249971734 | 0.101209999 | 0.84 |
| YDR454C | 2624.226849 | -0.485873346 | 0.051697042 | 0.84 |
| YDR459C | 457.0405938 | 0.359120518 | 0.134185508 | 0.84 |
| YDR462W | 726.8986569 | -0.501241566 | 0.15006846 | 0.84 |
| YDR463W | 1963.287278 | 0.576938984 | 0.099471888 | 0.84 |
| YDR471W | 4845.236011 | -0.395851444 | 0.136902249 | 0.84 |
| YDR481C | 2453.405315 | 0.869668206 | 0.054752011 | 0.84 |
| YDR483W | 4802.755253 | -0.4687695 | 0.061595985 | 0.84 |
| YDR490C | 1094.352116 | 0.598186095 | 0.124524841 | 0.84 |
| YDR493W | 345.6632951 | -0.521161382 | 0.175261441 | 0.84 |
| YDR500C | 6683.941473 | -0.450755583 | 0.115261901 | 0.84 |
| YDR507C | 1606.54523 | -0.309782852 | 0.202482597 | 0.84 |
| YDR510W | 2598.34956 | -0.412099398 | 0.211178752 | 0.84 |
| YDR519W | 900.2252274 | -0.31681586 | 0.214663689 | 0.84 |
| YDR523C | 105.2296251 | 0.475504859 | 0.075027835 | 0.84 |
| YDR524C-A | 2.188681507 | -0.157449945 | 0.149560871 | 0.84 |
| YDR524C-B | 18745.499 | -0.615906197 | 0.104837038 | 0.84 |
| YDR524W-C | 1641.998262 | -0.752293059 | 0.080458285 | 0.84 |
| YDR525W | 4.560372998 | 0.37700986 | 0.066541447 | 0.84 |
| YDR538W | 133.1462476 | -0.576644379 | 0.064719707 | 0.84 |
| YDR540C | 110.2480797 | -1.102294089 | 0.105366857 | 0.84 |
| YDR541C | 514.5384389 | -0.904532782 | 0.205346174 | 0.84 |
| YDR542W | 1.676832288 | 0.410904265 | 0.069077627 | 0.84 |

| | | | | |
|---|---|---|---|---|
| YEL001C | 2451.407888 | -0.512837516 | 0.131924933 | 0.84 |
| YEL002C | 3293.830024 | -0.445606487 | 0.091078773 | 0.84 |
| YEL009C-A | 8.998175135 | 0.978730958 | 0.027286409 | 0.84 |
| YEL011W | 8924.284313 | 0.74367805 | 0.155981007 | 0.84 |
| YEL012W | 1045.847779 | 0.513949176 | 0.187085491 | 0.84 |
| YEL018W | 225.091547 | -0.475639889 | 0.123887074 | 0.84 |
| YEL020W-A | 192.2562057 | -0.525188604 | 0.207708478 | 0.84 |
| YEL026W | 1802.083858 | -0.390552764 | 0.061763456 | 0.84 |
| YEL027W | 5744.186516 | -0.503188714 | 0.072016819 | 0.84 |
| YEL030C-A | 7.262884205 | 1.257669165 | 0.028704485 | 0.84 |
| YEL030W | 524.4916329 | 0.740290557 | 0.058108632 | 0.84 |
| YEL031W | 4279.639928 | -0.389177009 | 0.173943902 | 0.84 |
| YEL033W | 1403.07415 | -0.870784695 | 0.058630705 | 0.84 |
| YEL034C-A | 2748.462234 | -0.633396461 | 0.097069967 | 0.84 |
| YEL035C | 12.59177912 | 0.321108034 | 0.096758044 | 0.84 |
| YEL037C | 3155.723536 | -0.38317994 | 0.188656772 | 0.84 |
| YEL039C | 1783.57122 | 0.420757901 | 0.18658836 | 0.84 |
| YEL042W | 1714.856815 | -0.232604056 | 0.203977061 | 0.84 |
| YEL050C | 1046.082243 | -0.44621263 | 0.14828763 | 0.84 |
| YEL053C | 1328.366975 | 0.350071875 | 0.134591752 | 0.84 |
| YEL054C | 3213.78471 | -0.29527212 | 0.05588853 | 0.84 |
| YEL057C | 119.2283707 | 0.217713176 | 0.199899039 | 0.84 |
| YEL058W | 1683.65634 | -0.404946603 | 0.17786463 | 0.84 |
| YER003C | 3520.842344 | -0.541945916 | 0.041493498 | 0.84 |
| YER005W | 999.8103274 | 0.437295275 | 0.057049536 | 0.84 |
| YER006W | 1977.06385 | -0.256866867 | 0.101237675 | 0.84 |
| YER007C-A | 951.757768 | -0.591558551 | 0.058746883 | 0.84 |
| YER009W | 3909.301197 | -0.523251152 | 0.073714534 | 0.84 |
| YER015W | 725.609463 | 0.776050307 | 0.027793337 | 0.84 |
| YER016W | 838.7636501 | -0.427355097 | 0.191582909 | 0.84 |
| YER025W | 3287.005691 | -0.321958236 | 0.19788783 | 0.84 |
| YER036C | 4277.526152 | -0.371242937 | 0.114501481 | 0.84 |
| YER039C | 101.9948572 | 0.369449618 | 0.142445875 | 0.84 |
| YER039C-A | 49.35745349 | 0.517016064 | 0.105869927 | 0.84 |
| YER043C | 7073.679738 | -0.447262786 | 0.057707442 | 0.84 |
| YER044C-A | 102.3682402 | 1.076464077 | 0.054391603 | 0.84 |
| YER054C | 1167.402398 | 0.55568899 | 0.106733145 | 0.84 |
| YER055C | 2945.986692 | -0.00876832 | 0.19515416 | 0.84 |
| YER056C-A | 4887.303568 | -0.350575295 | 0.123475542 | 0.84 |
| YER063W | 1431.45469 | -0.564105469 | 0.063067553 | 0.84 |
| YER066C-A | 1483.59707 | 0.870769988 | 0.109684254 | 0.84 |
| YER066W | 204.0645964 | 0.591490688 | 0.060920651 | 0.84 |
| YER067W | 2559.902963 | 0.77131448 | 0.114321222 | 0.84 |

| | | | | |
|---|---|---|---|---|
| YER068C-A | 10.30512685 | 0.627641663 | 0.01005846 | 0.84 |
| YER070W | 2200.790509 | -0.976720172 | 0.017099143 | 0.84 |
| YER072W | 2000.095671 | -0.552798891 | 0.056773363 | 0.84 |
| YER074W | 4108.232098 | -0.429470637 | 0.089112713 | 0.84 |
| YER074W-A | 546.9637993 | -0.34769317 | 0.205695908 | 0.84 |
| YER075C | 845.204575 | 0.332255364 | 0.209406484 | 0.84 |
| YER076C | 386.5202214 | 0.518624418 | 0.043852136 | 0.84 |
| YER078C | 919.3267409 | 0.507809924 | 0.105312239 | 0.84 |
| YER078W-A | 4.281084499 | 0.030368849 | 0.212016299 | 0.84 |
| YER079C-A | 18.60445932 | 0.373174192 | 0.134200535 | 0.84 |
| YER081W | 9451.152045 | 1.56841914 | 0.007636679 | 0.84 |
| YER086W | 3514.638296 | 0.118182368 | 0.166297711 | 0.84 |
| YER087C-B | 1311.571506 | -0.603123172 | 0.031208955 | 0.84 |
| YER088C-A | 302.6604055 | -0.064584458 | 0.217512381 | 0.84 |
| YER091C | 3130.444513 | 1.027650127 | 0.025940666 | 0.84 |
| YER093C | 1449.072568 | 0.404873009 | 0.149427778 | 0.84 |
| YER096W | 439.4636005 | 0.967826951 | 0.037381344 | 0.84 |
| YER098W | 930.5704049 | 0.572552735 | 0.125974353 | 0.84 |
| YER099C | 790.1735302 | 0.352871117 | 0.163047154 | 0.84 |
| YER101C | 634.792482 | 0.528512468 | 0.12598395 | 0.84 |
| YER102W | 2473.647383 | -0.056904393 | 0.134157472 | 0.84 |
| YER105C | 2233.871003 | -0.405721727 | 0.085710462 | 0.84 |
| YER110C | 3689.619393 | -0.067914793 | 0.2163174 | 0.84 |
| YER115C | 529.525795 | 0.382345022 | 0.204549512 | 0.84 |
| YER116C | 578.3983863 | 0.529874482 | 0.198196787 | 0.84 |
| YER117W | 4528.496655 | -0.325621091 | 0.062454796 | 0.84 |
| YER118C | 889.8446464 | -0.180415533 | 0.158109695 | 0.84 |
| YER120W | 4329.489139 | -0.639390377 | 0.011483522 | 0.84 |
| YER121W | 184.2483 | 0.29487952 | 0.210947926 | 0.84 |
| YER131W | 9395.024759 | -0.298823769 | 0.055013903 | 0.84 |
| YER137C | 65.6651259 | 0.476830815 | 0.142611904 | 0.84 |
| YER137W-A | 3.086112355 | 0.410845155 | 0.126404445 | 0.84 |
| YER145C | 1496.957868 | -0.804576893 | 0.020846243 | 0.84 |
| YER146W | 140.8937845 | -0.889911852 | 0.013758048 | 0.84 |
| YER147C | 529.8378852 | 0.466409017 | 0.151332344 | 0.84 |
| YER152W-A | 37.9606133 | 0.570481032 | 0.196824423 | 0.84 |
| YER153C | 351.5393272 | 0.563527777 | 0.180563834 | 0.84 |
| YER156C | 1503.86508 | -0.56642263 | 0.094846125 | 0.84 |
| YER162C | 931.7095211 | 0.598895899 | 0.069482342 | 0.84 |
| YER165W | 11478.83612 | -0.582862912 | 0.03318233 | 0.84 |
| YER169W | 1395.485471 | 0.826555999 | 0.019657538 | 0.84 |
| YER175C | 1510.549619 | 0.771705934 | 0.15110972 | 0.84 |
| YER175W-A | 27.41241786 | 0.660248441 | 0.104959159 | 0.84 |

| | | | | |
|---|---|---|---|---|
| YER177W | 16691.55629 | -0.38574552 | 0.188601974 | 0.84 |
| YER179W | 293.0579118 | 0.558735028 | 0.174991811 | 0.84 |
| YER183C | 378.9572978 | -0.429241687 | 0.184151976 | 0.84 |
| YER188C-A | 23.15774642 | 0.855979267 | 0.017830972 | 0.84 |
| YFL004W | 2271.622342 | -0.515123754 | 0.041108273 | 0.84 |
| YFL017C | 717.9144755 | -0.425876294 | 0.145401629 | 0.84 |
| YFL017W-A | 221.0675394 | -0.644537921 | 0.092610516 | 0.84 |
| YFL020C | 87.69189529 | 0.124013647 | 0.216207424 | 0.84 |
| YFL021W | 926.9442787 | 0.638545319 | 0.111249568 | 0.84 |
| YFL022C | 4340.34362 | -0.280279379 | 0.190960484 | 0.84 |
| YFL024C | 1322.826078 | 0.371238164 | 0.177801731 | 0.84 |
| YFL031C-A | 101.8569422 | -0.758583806 | 0.08066819 | 0.84 |
| YFL037W | 5301.265498 | -0.36277846 | 0.12805316 | 0.84 |
| YFL041W-A | 23.17885335 | 0.321000299 | 0.213480551 | 0.84 |
| YFL045C | 5242.345974 | -0.501278328 | 0.072461169 | 0.84 |
| YFL049W | 806.3019635 | 0.34175617 | 0.189646888 | 0.84 |
| YFL054C | 1511.844206 | 0.311001055 | 0.198755881 | 0.84 |
| YFL056C | 232.3744781 | 0.397378005 | 0.143699367 | 0.84 |
| YFL057C | 160.9643278 | 0.827806006 | 0.015105771 | 0.84 |
| YFL059W | 70.79713485 | -1.16957323 | 0.087546875 | 0.84 |
| YFL060C | 16.02228195 | -1.726216411 | 0.004348433 | 0.84 |
| YFL061W | 11.58614878 | -0.917641372 | 0.136776671 | 0.84 |
| YFR001W | 1018.480753 | -0.382976328 | 0.188236884 | 0.84 |
| YFR009W | 2637.309837 | -0.408143327 | 0.180997949 | 0.84 |
| YFR015C | 6189.813754 | 0.689245788 | 0.111534502 | 0.84 |
| YFR017C | 3258.918594 | 0.540308603 | 0.125149439 | 0.84 |
| YFR020W | 385.8178556 | 0.40302244 | 0.207554743 | 0.84 |
| YFR028C | 1133.072845 | -0.592510003 | 0.050789177 | 0.84 |
| YFR030W | 1464.99151 | 0.540354561 | 0.20256176 | 0.84 |
| YFR031C | 1205.359626 | 0.429703157 | 0.13178564 | 0.84 |
| YFR031C-A | 6438.897656 | -0.259566289 | 0.057779772 | 0.84 |
| YFR032C | 47.36854765 | 0.073706693 | 0.101859628 | 0.84 |
| YFR032C-A | 2429.038295 | -0.617892915 | 0.059166897 | 0.84 |
| YFR036W-A | 15.15766896 | -1.192216694 | 0.003267806 | 0.84 |
| YFR037C | 1666.566511 | -0.366314828 | 0.207815378 | 0.84 |
| YFR054C | 20.51319414 | -1.160458989 | 0.135567006 | 0.84 |
| YFR057W | 37.29733796 | -0.001615897 | 0.06832633 | 0.84 |
| YGL003C | 856.5289623 | 0.352393898 | 0.199656523 | 0.84 |
| YGL006W | 3158.91505 | 0.468586305 | 0.168284735 | 0.84 |
| YGL008C | 26806.32689 | -0.364108031 | 0.031195177 | 0.84 |
| YGL010W | 399.2167766 | 0.416134135 | 0.167582354 | 0.84 |
| YGL012W | 2257.199778 | -0.776308086 | 0.039111525 | 0.84 |
| YGL015C | 22.18286877 | 0.705507366 | 0.04296665 | 0.84 |

| | | | | |
|---|---|---|---|---|
| YGL017W | 502.0325199 | 0.364475626 | 0.156774245 | 0.84 |
| YGL019W | 1587.591155 | -0.302218341 | 0.207347944 | 0.84 |
| YGL020C | 1129.205349 | -0.431026213 | 0.189536739 | 0.84 |
| YGL022W | 3864.561612 | -0.297425967 | 0.202135444 | 0.84 |
| YGL029W | 383.3919755 | -0.303671143 | 0.102349345 | 0.84 |
| YGL030W | 7624.667323 | -0.468112019 | 0.015583375 | 0.84 |
| YGL031C | 8332.907395 | -0.109474921 | 0.110293544 | 0.84 |
| YGL035C | 1849.649908 | 0.643317604 | 0.041202541 | 0.84 |
| YGL036W | 2417.923989 | 0.537549048 | 0.080919639 | 0.84 |
| YGL040C | 1261.243607 | -0.629158836 | 0.04605218 | 0.84 |
| YGL044C | 659.773802 | 0.335446528 | 0.139232855 | 0.84 |
| YGL045W | 904.8478281 | 0.361214945 | 0.203244317 | 0.84 |
| YGL047W | 625.4320388 | -0.359127897 | 0.157949269 | 0.84 |
| YGL059W | 1140.532396 | 1.024097937 | 0.012039925 | 0.84 |
| YGL068W | 385.5514446 | -0.58520095 | 0.180236374 | 0.84 |
| YGL070C | 462.601344 | -0.409165468 | 0.134929525 | 0.84 |
| YGL076C | 4229.319259 | -0.20653572 | 0.075058926 | 0.84 |
| YGL077C | 1372.265624 | -0.412180589 | 0.163081419 | 0.84 |
| YGL081W | 335.8828397 | 0.969265441 | 0.06674885 | 0.84 |
| YGL090W | 485.8897575 | 0.447500826 | 0.121854859 | 0.84 |
| YGL092W | 1819.13534 | -0.503134949 | 0.089674294 | 0.84 |
| YGL102C | 355.1928609 | -0.373442292 | 0.098087174 | 0.84 |
| YGL104C | 947.9922368 | 0.473225652 | 0.131171089 | 0.84 |
| YGL112C | 1298.501075 | -0.450912259 | 0.053161568 | 0.84 |
| YGL113W | 616.1660547 | 0.446282344 | 0.176110012 | 0.84 |
| YGL117W | 750.1030158 | 0.87762782 | 0.100348471 | 0.84 |
| YGL123W | 15975.90289 | -0.472582188 | 0.057781507 | 0.84 |
| YGL125W | 958.6893776 | 0.797415256 | 0.050393226 | 0.84 |
| YGL128C | 505.7413235 | 0.45142917 | 0.127516567 | 0.84 |
| YGL130W | 854.7206722 | -0.263891179 | 0.15809024 | 0.84 |
| YGL137W | 4577.146804 | -0.351504531 | 0.151573729 | 0.84 |
| YGL138C | 22.53163453 | 0.192788726 | 0.069153831 | 0.84 |
| YGL141W | 1274.03239 | 0.450971578 | 0.128803721 | 0.84 |
| YGL147C | 5508.177604 | -0.243236731 | 0.03349282 | 0.84 |
| YGL150C | 2988.328671 | 0.36942788 | 0.161826296 | 0.84 |
| YGL154C | 461.1131948 | 0.457663223 | 0.114844104 | 0.84 |
| YGL158W | 260.9463165 | 0.574293 | 0.019427746 | 0.84 |
| YGL162W | 395.3975497 | 0.350580588 | 0.053189414 | 0.84 |
| YGL166W | 257.6413584 | 0.773776397 | 0.03739189 | 0.84 |
| YGL170C | 69.87350446 | 0.407553616 | 0.153369099 | 0.84 |
| YGL173C | 8071.132916 | -0.339119115 | 0.135059984 | 0.84 |
| YGL175C | 162.4549988 | 0.660343495 | 0.067600341 | 0.84 |
| YGL176C | 596.0143813 | 0.74911251 | 0.009641551 | 0.84 |

| | | | | |
|---|---|---|---|---|
| YGL180W | 843.8043135 | 0.694709227 | 0.048816543 | 0.84 |
| YGL183C | 72.4162606 | 0.975170777 | 0.044245357 | 0.84 |
| YGL184C | 275.5662791 | 1.018287788 | 0.003463639 | 0.84 |
| YGL189C | 20518.43042 | -0.395334997 | 0.054903682 | 0.84 |
| YGL190C | 1598.926776 | 0.443725202 | 0.099770159 | 0.84 |
| YGL195W | 6234.838225 | -0.553992885 | 0.045305155 | 0.84 |
| YGL204C | 78.35386912 | -0.993248488 | 0.042007746 | 0.84 |
| YGL210W | 815.3164128 | -0.453315227 | 0.12757248 | 0.84 |
| YGL215W | 3518.182479 | -0.325021053 | 0.209752854 | 0.84 |
| YGL225W | 1789.038412 | -0.475673237 | 0.029409491 | 0.84 |
| YGL226C-A | 628.3645694 | -0.950711975 | 0.014927531 | 0.84 |
| YGL229C | 708.5937653 | 0.495210811 | 0.163889812 | 0.84 |
| YGL230C | 33.65395104 | 0.248403209 | 0.20091056 | 0.84 |
| YGL232W | 728.6792639 | -0.377577848 | 0.206344351 | 0.84 |
| YGL234W | 9210.041986 | 0.108374662 | 0.171075527 | 0.84 |
| YGL241W | 933.7531382 | -0.405783018 | 0.173026882 | 0.84 |
| YGL243W | 434.8625344 | -0.436561731 | 0.067072265 | 0.84 |
| YGL245W | 8715.959502 | -0.226705027 | 0.215505582 | 0.84 |
| YGL255W | 1201.748656 | 1.096921474 | 0.04555062 | 0.84 |
| YGL263W | 253.3051631 | -0.769433133 | 0.085030261 | 0.84 |
| YGR001C | 885.4111524 | -0.175004898 | 0.076312597 | 0.84 |
| YGR003W | 648.3625673 | 0.53043409 | 0.050909117 | 0.84 |
| YGR007W | 1236.749371 | -0.644527528 | 0.020659972 | 0.84 |
| YGR020C | 3100.566353 | -0.399659068 | 0.18235415 | 0.84 |
| YGR023W | 927.8650533 | 0.505008606 | 0.171987028 | 0.84 |
| YGR025W | 100.9215658 | -0.393510956 | 0.177396586 | 0.84 |
| YGR027C | 9077.452786 | -0.348505275 | 0.107909246 | 0.84 |
| YGR034W | 5016.73135 | -0.394138692 | 0.044830424 | 0.84 |
| YGR035W-A | 93.40510398 | -0.708736217 | 0.118435715 | 0.84 |
| YGR038W | 610.0071877 | -0.520640101 | 0.091046856 | 0.84 |
| YGR039W | 40.79718114 | -0.643485913 | 0.209593138 | 0.84 |
| YGR051C | 51.37808205 | 0.610112376 | 0.063702611 | 0.84 |
| YGR052W | 642.70953 | 0.404330975 | 0.042049446 | 0.84 |
| YGR053C | 339.3237027 | 0.444199945 | 0.152829478 | 0.84 |
| YGR054W | 2529.613046 | -0.519836839 | 0.103682822 | 0.84 |
| YGR066C | 223.0827395 | 0.521297249 | 0.091400795 | 0.84 |
| YGR070W | 2198.55855 | 0.597959902 | 0.176510692 | 0.84 |
| YGR072W | 838.1301736 | 0.55836901 | 0.140433709 | 0.84 |
| YGR078C | 839.5108181 | -0.605573259 | 0.135705065 | 0.84 |
| YGR082W | 1782.261898 | -0.619680792 | 0.045696619 | 0.84 |
| YGR084C | 1041.679379 | -0.558969679 | 0.06115427 | 0.84 |
| YGR088W | 3462.726793 | 0.267291108 | 0.16529006 | 0.84 |
| YGR093W | 815.5848449 | 0.470104413 | 0.112332944 | 0.84 |

| | | | | |
|---|---|---|---|---|
| YGR096W | 275.6515308 | 0.545216592 | 0.140058807 | 0.84 |
| YGR097W | 2251.701342 | 0.581063985 | 0.161962039 | 0.84 |
| YGR099W | 582.6782539 | -0.47884009 | 0.104714869 | 0.84 |
| YGR104C | 540.5492389 | 0.445564189 | 0.061403745 | 0.84 |
| YGR105W | 477.947228 | -0.417216221 | 0.192025423 | 0.84 |
| YGR106C | 3546.453004 | -0.416109746 | 0.091408119 | 0.84 |
| YGR108W | 622.4932134 | -0.89254532 | 0.041549349 | 0.84 |
| YGR110W | 407.9203872 | 0.857969434 | 0.006018248 | 0.84 |
| YGR112W | 838.4700607 | 0.602516777 | 0.161270559 | 0.84 |
| YGR114C | 36.93060097 | 0.440993806 | 0.095071394 | 0.84 |
| YGR120C | 376.5121986 | -0.385975511 | 0.039931493 | 0.84 |
| YGR123C | 655.6704403 | -0.423472907 | 0.126382538 | 0.84 |
| YGR127W | 732.3981957 | 0.429463205 | 0.081995804 | 0.84 |
| YGR130C | 4694.98638 | 0.475041614 | 0.20983143 | 0.84 |
| YGR131W | 100.6509022 | 1.038260081 | 0.008206409 | 0.84 |
| YGR143W | 1625.66982 | 0.3839113 | 0.180711885 | 0.84 |
| YGR146C | 1344.611896 | 0.972017867 | 0.018622439 | 0.84 |
| YGR148C | 7673.362198 | -0.346392157 | 0.072234136 | 0.84 |
| YGR149W | 1082.04674 | 0.661250206 | 0.116441719 | 0.84 |
| YGR150C | 1213.396537 | -0.489383636 | 0.104010688 | 0.84 |
| YGR151C | 2.6142374 | -0.093479497 | 0.055302641 | 0.84 |
| YGR152C | 296.8399677 | -0.538690918 | 0.160977109 | 0.84 |
| YGR154C | 333.0792836 | 0.55925298 | 0.099137819 | 0.84 |
| YGR155W | 6507.68049 | -0.348072268 | 0.159981981 | 0.84 |
| YGR165W | 998.5420975 | -0.522076855 | 0.157340485 | 0.84 |
| YGR169C-A | 750.5850996 | 0.401566853 | 0.176052153 | 0.84 |
| YGR172C | 884.3306623 | -0.432152038 | 0.128731891 | 0.84 |
| YGR175C | 3323.313876 | -0.494499469 | 0.212829226 | 0.84 |
| YGR181W | 835.8007421 | -0.538653106 | 0.110968586 | 0.84 |
| YGR187C | 552.0431833 | -0.452107846 | 0.160418762 | 0.84 |
| YGR192C | 181337.4741 | -0.156131117 | 0.186281005 | 0.84 |
| YGR195W | 892.1333078 | -0.380680709 | 0.172953375 | 0.84 |
| YGR200C | 1330.804216 | -0.416319171 | 0.163783675 | 0.84 |
| YGR201C | 974.6666707 | 0.499677328 | 0.184959382 | 0.84 |
| YGR207C | 1419.844599 | -0.370894 | 0.158669874 | 0.84 |
| YGR208W | 739.3899853 | -0.254779691 | 0.114004563 | 0.84 |
| YGR214W | 11003.56684 | -0.283127123 | 0.060495946 | 0.84 |
| YGR215W | 453.6817551 | -0.515563488 | 0.190413181 | 0.84 |
| YGR216C | 759.5937639 | 0.333260518 | 0.204197123 | 0.84 |
| YGR219W | 32.06784529 | -0.703938389 | 0.198750473 | 0.84 |
| YGR220C | 677.1113476 | -0.60397551 | 0.072969215 | 0.84 |
| YGR223C | 1134.541576 | 0.420152777 | 0.200243822 | 0.84 |
| YGR224W | 693.0347479 | 0.338326869 | 0.187741448 | 0.84 |

| | | | | |
|---|---|---|---|---|
| YGR225W | 197.8168855 | 0.779668805 | 0.049581051 | 0.84 |
| YGR249W | 613.9099062 | 0.760640362 | 0.098443554 | 0.84 |
| YGR252W | 827.8896718 | -0.455415678 | 0.107077632 | 0.84 |
| YGR261C | 1931.303256 | -0.403862224 | 0.138886309 | 0.84 |
| YGR268C | 875.2811265 | 0.39829747 | 0.18122235 | 0.84 |
| YGR271C-A | 454.6880839 | -0.246723599 | 0.126984763 | 0.84 |
| YGR271W | 1597.427923 | -0.421971144 | 0.159446117 | 0.84 |
| YGR279C | 11712.04252 | -0.553549088 | 0.038525424 | 0.84 |
| YGR285C | 10708.64158 | -0.100317346 | 0.211843329 | 0.84 |
| YGR287C | 287.959068 | 0.628989272 | 0.132308264 | 0.84 |
| YGR290W | 27.11482068 | 0.420121556 | 0.105557315 | 0.84 |
| YGR293C | 7.546648827 | -0.148520493 | 0.133041953 | 0.84 |
| YHL002C-A | 8.009981089 | 0.678520877 | 0.065160455 | 0.84 |
| YHL008C | 1005.082799 | 0.496198298 | 0.143316399 | 0.84 |
| YHL011C | 1690.318772 | -0.335223557 | 0.0825938 | 0.84 |
| YHL012W | 237.0696554 | 0.621623726 | 0.027084502 | 0.84 |
| YHL013C | 503.0447213 | -0.458677132 | 0.125739724 | 0.84 |
| YHL015W | 17106.50707 | -0.221461167 | 0.099374931 | 0.84 |
| YHL016C | 664.2747126 | 0.780794377 | 0.03896339 | 0.84 |
| YHL017W | 1099.085857 | 0.435383538 | 0.212515658 | 0.84 |
| YHL018W | 70.12879735 | 0.467997814 | 0.178465337 | 0.84 |
| YHL021C | 7761.479525 | 0.737337172 | 0.101133293 | 0.84 |
| YHL022C | 136.8231823 | 0.804178153 | 0.185713454 | 0.84 |
| YHL024W | 910.8922734 | 0.57690385 | 0.054974226 | 0.84 |
| YHL028W | 1137.369035 | -1.035201859 | 0.133057801 | 0.84 |
| YHL030W-A | 22.0345387 | 0.611288968 | 0.096322014 | 0.84 |
| YHL033C | 4723.149428 | -0.283302003 | 0.081008516 | 0.84 |
| YHL036W | 545.2189465 | 1.026117201 | 0.015325551 | 0.84 |
| YHL037C | 20.29589679 | 0.93583235 | 0.048594931 | 0.84 |
| YHR004C | 921.2706765 | 0.536298881 | 0.124640467 | 0.84 |
| YHR005C-A | 2208.832297 | -0.620517119 | 0.053338191 | 0.84 |
| YHR010W | 11885.41138 | -0.087794589 | 0.116087669 | 0.84 |
| YHR013C | 1158.208101 | -0.377451507 | 0.162021758 | 0.84 |
| YHR014W | 138.7802733 | 0.882572026 | 0.021827601 | 0.84 |
| YHR017W | 1256.836076 | 0.4783021 | 0.179476881 | 0.84 |
| YHR021C | 5458.86145 | -0.488396424 | 0.049019769 | 0.84 |
| YHR021W-A | 41.37134005 | 0.362271253 | 0.043522586 | 0.84 |
| YHR022C | 158.1795629 | 0.564276671 | 0.035073611 | 0.84 |
| YHR025W | 4807.328396 | 0.047315707 | 0.181790306 | 0.84 |
| YHR026W | 2431.689374 | -0.588025585 | 0.058355136 | 0.84 |
| YHR033W | 1072.035586 | 0.365015563 | 0.125599524 | 0.84 |
| YHR039C-A | 2505.228571 | -0.448622976 | 0.173323912 | 0.84 |
| YHR041C | 584.7828437 | -1.021499009 | 0.018933553 | 0.84 |

| | | | | |
|---|---|---|---|---|
| YHR049W | 2093.457795 | -0.555758332 | 0.009946074 | 0.84 |
| YHR052W | 1127.946092 | -0.246735942 | 0.202982652 | 0.84 |
| YHR059W | 401.0845698 | -0.692064253 | 0.064180387 | 0.84 |
| YHR062C | 561.4502383 | -0.395958086 | 0.198345227 | 0.84 |
| YHR063W-A | 174.3401986 | -0.35818823 | 0.192713145 | 0.84 |
| YHR068W | 2922.590435 | -0.191879686 | 0.16824997 | 0.84 |
| YHR071W | 3724.97766 | 0.726833699 | 0.117664222 | 0.84 |
| YHR072W-A | 1183.992345 | -0.395394086 | 0.14215567 | 0.84 |
| YHR075C | 548.2435732 | 0.507230208 | 0.104944301 | 0.84 |
| YHR080C | 2045.316793 | 0.627848498 | 0.079229939 | 0.84 |
| YHR088W | 1005.281896 | -0.354352196 | 0.133993069 | 0.84 |
| YHR089C | 2056.029582 | -0.489816448 | 0.12935109 | 0.84 |
| YHR095W | 5.929557488 | -0.618340101 | 0.104122979 | 0.84 |
| YHR096C | 1194.376628 | 0.380624729 | 0.051001476 | 0.84 |
| YHR097C | 2968.055756 | 0.513956744 | 0.180349874 | 0.84 |
| YHR098C | 2961.484376 | -0.372216212 | 0.151459185 | 0.84 |
| YHR106W | 1096.925363 | 0.630472305 | 0.076475775 | 0.84 |
| YHR107C | 1509.6288 | -0.521713867 | 0.053644536 | 0.84 |
| YHR108W | 3358.554888 | -0.354198776 | 0.158575668 | 0.84 |
| YHR109W | 446.2718996 | -0.453770729 | 0.188378519 | 0.84 |
| YHR110W | 1458.33201 | -0.300582967 | 0.211729928 | 0.84 |
| YHR120W | 606.7255558 | 0.476490896 | 0.071840759 | 0.84 |
| YHR121W | 1103.128777 | -0.699241024 | 0.014644937 | 0.84 |
| YHR127W | 498.1945386 | -0.329728472 | 0.128183265 | 0.84 |
| YHR130C | 24.10330458 | 0.179186497 | 0.149468106 | 0.84 |
| YHR133C | 977.6515666 | -0.34603839 | 0.19992233 | 0.84 |
| YHR138C | 1476.126547 | 0.892299302 | 0.020352493 | 0.84 |
| YHR139C | 204.3743283 | 0.218274935 | 0.061230785 | 0.84 |
| YHR139C-A | 36.30751378 | 0.653222673 | 0.146108381 | 0.84 |
| YHR141C | 3868.981705 | -0.249299042 | 0.15361763 | 0.84 |
| YHR143W-A | 554.7529102 | -0.415818616 | 0.187788461 | 0.84 |
| YHR144C | 383.3281151 | -0.343932961 | 0.187706902 | 0.84 |
| YHR146W | 3577.393266 | 0.547485888 | 0.154340852 | 0.84 |
| YHR147C | 857.7185719 | -0.561842511 | 0.057863734 | 0.84 |
| YHR152W | 1107.927047 | -0.341764541 | 0.191086094 | 0.84 |
| YHR153C | 73.13873436 | -0.518221873 | 0.07280961 | 0.84 |
| YHR154W | 1331.473724 | -0.424872262 | 0.060354954 | 0.84 |
| YHR159W | 783.3955676 | 0.739095578 | 0.082763097 | 0.84 |
| YHR160C | 281.5696466 | 0.441754266 | 0.175145795 | 0.84 |
| YHR164C | 1299.336952 | 0.402416356 | 0.192043131 | 0.84 |
| YHR171W | 640.7534835 | 0.546531175 | 0.216399222 | 0.84 |
| YHR174W | 94635.8071 | -0.116702039 | 0.196249879 | 0.84 |
| YHR175W-A | 167.202498 | -0.160975825 | 0.21659151 | 0.84 |

| | | | | |
|---|---|---|---|---|
| YHR189W | 309.5122663 | 0.538938684 | 0.184797928 | 0.84 |
| YHR193C | 8396.042296 | -0.734460273 | 0.048814388 | 0.84 |
| YHR195W | 1250.379878 | 0.479662086 | 0.129264413 | 0.84 |
| YHR202W | 670.4759746 | 0.566250229 | 0.146609738 | 0.84 |
| YHR203C | 5825.930229 | -0.005425129 | 0.210608658 | 0.84 |
| YIL002W-A | 495.5817191 | -0.442001246 | 0.18601091 | 0.84 |
| YIL013C | 704.2538412 | 0.612089988 | 0.062684124 | 0.84 |
| YIL016W | 483.5722155 | -0.342468415 | 0.176500789 | 0.84 |
| YIL017C | 1438.735299 | 0.498691426 | 0.175174225 | 0.84 |
| YIL018W | 10226.88893 | -0.320994972 | 0.098606636 | 0.84 |
| YIL020C-A | 191.6015626 | -0.488971228 | 0.197067604 | 0.84 |
| YIL021W | 814.8146206 | -0.436922767 | 0.114656252 | 0.84 |
| YIL022W | 2268.335855 | -0.421713164 | 0.050825688 | 0.84 |
| YIL030W-A | 74.85518125 | 0.610996919 | 0.112061263 | 0.84 |
| YIL035C | 1690.415511 | -0.511409541 | 0.055020962 | 0.84 |
| YIL039W | 1826.424519 | -0.356014084 | 0.186766547 | 0.84 |
| YIL040W | 372.5621769 | -0.482394868 | 0.180200221 | 0.84 |
| YIL046W | 1246.958862 | 0.551660523 | 0.071856686 | 0.84 |
| YIL050W | 1083.890647 | 0.521501876 | 0.12228218 | 0.84 |
| YIL052C | 4784.674042 | -0.298160341 | 0.104472025 | 0.84 |
| YIL055C | 666.1215044 | 0.818715054 | 0.033539676 | 0.84 |
| YIL059C | 89.463829 | 0.280666555 | 0.143606695 | 0.84 |
| YIL069C | 2018.584495 | -0.470583591 | 0.210722214 | 0.84 |
| YIL070C | 1550.574512 | -0.450920618 | 0.177703653 | 0.84 |
| YIL071C | 381.3119395 | 0.562889679 | 0.044342411 | 0.84 |
| YIL071W-A | 73.5379669 | 0.531626924 | 0.046433286 | 0.84 |
| YIL072W | 537.0665215 | 1.052759431 | 0.008779601 | 0.84 |
| YIL076W | 2351.989968 | -0.404622816 | 0.183514305 | 0.84 |
| YIL086C | 23.45932813 | 0.234821864 | 0.163085083 | 0.84 |
| YIL093C | 871.5653388 | -0.635067136 | 0.065697668 | 0.84 |
| YIL096C | 413.8812975 | -0.464864135 | 0.075734689 | 0.84 |
| YIL099W | 500.7923991 | 0.511284837 | 0.120305514 | 0.84 |
| YIL100W | 3.722972412 | -0.770411803 | 0.1855687 | 0.84 |
| YIL101C | 1402.735777 | 0.605024648 | 0.078268122 | 0.84 |
| YIL105C | 1531.713028 | 0.458970348 | 0.137038585 | 0.84 |
| YIL107C | 1780.529737 | 0.475825748 | 0.161531766 | 0.84 |
| YIL109C | 4372.511551 | -0.307600523 | 0.200517606 | 0.84 |
| YIL113W | 384.3761033 | 0.698803069 | 0.043060688 | 0.84 |
| YIL118W | 1012.394304 | -0.482323788 | 0.122867283 | 0.84 |
| YIL123W | 4293.520238 | -0.554657268 | 0.104723731 | 0.84 |
| YIL126W | 2338.60398 | -0.379009667 | 0.218587602 | 0.84 |
| YIL127C | 639.7542142 | -0.274046704 | 0.211139391 | 0.84 |
| YIL133C | 6678.70118 | -0.214957778 | 0.096469986 | 0.84 |

| | | | | |
|---|---|---|---|---|
| YIL142W | 3130.184557 | -0.481834321 | 0.146859851 | 0.84 |
| YIL146C | 613.496777 | 0.745063547 | 0.036266747 | 0.84 |
| YIL148W | 9852.705743 | -0.241984189 | 0.093207612 | 0.84 |
| YIL155C | 2791.794832 | 0.538486662 | 0.187189461 | 0.84 |
| YIL158W | 651.6543059 | -0.400636035 | 0.213811663 | 0.84 |
| YIL159W | 678.524918 | -0.619757002 | 0.052165343 | 0.84 |
| YIL161W | 488.7537738 | -0.482541967 | 0.202125282 | 0.84 |
| YIL162W | 3263.600398 | 0.261015108 | 0.172832604 | 0.84 |
| YIL165C | 180.1950386 | 0.437134676 | 0.154959735 | 0.84 |
| YIL166C | 439.1240138 | 0.516433724 | 0.135915766 | 0.84 |
| YIR013C | 17.07044727 | 0.655774156 | 0.061914996 | 0.84 |
| YIR017C | 6.4006803 | 0.42599669 | 0.046137058 | 0.84 |
| YIR021W-A | 4.66148073 | -0.596523064 | 0.150469941 | 0.84 |
| YIR023C-A | 13.57792364 | -0.972621899 | 0.099384235 | 0.84 |
| YIR036W-A | 10.01589679 | 0.558653454 | 0.032741338 | 0.84 |
| YIR041W | 5.637300496 | 0.497425314 | 0.040322442 | 0.84 |
| YIR042C | 196.4981605 | -0.922411578 | 0.209292954 | 0.84 |
| YJL009W | 66.31151028 | -0.819891095 | 0.049521592 | 0.84 |
| YJL011C | 110.249814 | -0.365959583 | 0.117787368 | 0.84 |
| YJL015C | 72.86729941 | 0.559879235 | 0.021088046 | 0.84 |
| YJL023C | 307.217395 | 0.560017081 | 0.066133097 | 0.84 |
| YJL036W | 1275.214013 | 0.372248882 | 0.207203394 | 0.84 |
| YJL039C | 2015.018256 | -0.518374079 | 0.116935778 | 0.84 |
| YJL056C | 734.4930024 | 0.570963333 | 0.117829876 | 0.84 |
| YJL062W-A | 1034.508423 | -0.532363713 | 0.160444016 | 0.84 |
| YJL063C | 928.728548 | -0.832256469 | 0.022519017 | 0.84 |
| YJL070C | 972.7876692 | 0.505993241 | 0.167093741 | 0.84 |
| YJL071W | 590.1206135 | 0.66462264 | 0.072245945 | 0.84 |
| YJL074C | 945.199767 | -0.603984418 | 0.035908233 | 0.84 |
| YJL075C | 78.82663296 | 0.544116943 | 0.077201325 | 0.84 |
| YJL080C | 5735.124683 | -0.085894636 | 0.142292007 | 0.84 |
| YJL081C | 1547.994145 | -0.359800008 | 0.123296821 | 0.84 |
| YJL083W | 610.9975272 | 0.575156502 | 0.089771738 | 0.84 |
| YJL084C | 1615.45903 | 0.429959267 | 0.16886185 | 0.84 |
| YJL089W | 344.4116839 | 0.684778209 | 0.132027541 | 0.84 |
| YJL091C | 524.9582628 | -0.484151893 | 0.138741369 | 0.84 |
| YJL092W | 1243.946562 | -0.45952867 | 0.111753512 | 0.84 |
| YJL093C | 1063.991787 | 0.589228471 | 0.055913666 | 0.84 |
| YJL096W | 663.1647142 | -0.687697103 | 0.052878516 | 0.84 |
| YJL097W | 1713.191405 | -0.403152813 | 0.180397987 | 0.84 |
| YJL100W | 1049.188324 | 0.381467885 | 0.194288177 | 0.84 |
| YJL103C | 874.1846601 | 0.571177788 | 0.1059539 | 0.84 |
| YJL104W | 974.9529404 | -0.417261405 | 0.144016561 | 0.84 |

| | | | | |
|---|---|---|---|---|
| YJL106W | 294.2742704 | 0.966994984 | 0.017942525 | 0.84 |
| YJL107C | 129.4785502 | 0.677101325 | 0.173114693 | 0.84 |
| YJL108C | 258.931357 | 0.656191201 | 0.095222958 | 0.84 |
| YJL115W | 931.9259287 | -0.468345733 | 0.105600993 | 0.84 |
| YJL121C | 2601.821139 | -0.354439106 | 0.180399157 | 0.84 |
| YJL122W | 643.9700151 | -0.448325329 | 0.113843678 | 0.84 |
| YJL124C | 1576.189687 | -0.541482088 | 0.056570582 | 0.84 |
| YJL129C | 2269.469864 | 0.365685142 | 0.166157612 | 0.84 |
| YJL132W | 534.7548177 | 0.580596606 | 0.078681672 | 0.84 |
| YJL133C-A | 1683.932792 | 0.532267739 | 0.035652448 | 0.84 |
| YJL134W | 603.1655535 | -0.355235449 | 0.215949417 | 0.84 |
| YJL136C | 5415.03448 | -0.308210482 | 0.1010248 | 0.84 |
| YJL138C | 1955.737437 | -0.466584441 | 0.110699595 | 0.84 |
| YJL141C | 4385.516839 | 0.824078358 | 0.06335951 | 0.84 |
| YJL142C | 106.3041783 | 0.659608267 | 0.079250312 | 0.84 |
| YJL143W | 1167.282332 | -0.641009546 | 0.056250223 | 0.84 |
| YJL144W | 785.7166274 | 0.355530969 | 0.202038986 | 0.84 |
| YJL148W | 1042.119748 | -0.156747198 | 0.218791266 | 0.84 |
| YJL149W | 1033.010855 | 0.415159434 | 0.139469451 | 0.84 |
| YJL153C | 833.1459195 | 0.787681684 | 0.037811658 | 0.84 |
| YJL155C | 1149.698916 | 0.532600367 | 0.152838453 | 0.84 |
| YJL158C | 11506.92019 | -0.370987538 | 0.057684304 | 0.84 |
| YJL163C | 815.8405692 | 0.564486519 | 0.118283323 | 0.84 |
| YJL165C | 2120.172359 | 0.778756354 | 0.027925673 | 0.84 |
| YJL169W | 25.36216702 | -0.774038476 | 0.131327788 | 0.84 |
| YJL173C | 671.1948643 | -0.489098869 | 0.097041431 | 0.84 |
| YJL177W | 7142.560719 | -0.243997523 | 0.11014801 | 0.84 |
| YJL180C | 680.1821358 | -0.531889006 | 0.099907788 | 0.84 |
| YJL183W | 732.387825 | -0.483880308 | 0.134167037 | 0.84 |
| YJL188C | 893.4798181 | -0.56769072 | 0.110946648 | 0.84 |
| YJL195C | 5.108398903 | 0.623920933 | 0.085293934 | 0.84 |
| YJL196C | 1508.45467 | -0.486247258 | 0.154157404 | 0.84 |
| YJL199C | 163.0335414 | 0.537934086 | 0.060664076 | 0.84 |
| YJL201W | 1030.90288 | 0.414733014 | 0.12768186 | 0.84 |
| YJL203W | 345.939173 | -0.317777112 | 0.210151632 | 0.84 |
| YJL206C | 539.1174838 | 0.671423111 | 0.039846855 | 0.84 |
| YJL212C | 1268.474729 | 0.857470475 | 0.150607865 | 0.84 |
| YJR004C | 1252.603429 | 0.55774571 | 0.166720249 | 0.84 |
| YJR005C-A | 81.403139 | 0.886212565 | 0.065035534 | 0.84 |
| YJR007W | 3429.418011 | -0.46055305 | 0.127046169 | 0.84 |
| YJR008W | 1161.824512 | 0.505810831 | 0.140004834 | 0.84 |
| YJR010W | 677.4384961 | 0.320287426 | 0.213651488 | 0.84 |
| YJR014W | 1708.927082 | -0.536186828 | 0.059581258 | 0.84 |

| | | | | |
|---|---|---|---|---|
| YJR018W | 57.57815418 | 0.695528071 | 0.055657707 | 0.84 |
| YJR024C | 696.9415683 | -0.823562441 | 0.039099689 | 0.84 |
| YJR030C | 708.7385852 | -0.403939023 | 0.177425879 | 0.84 |
| YJR031C | 1083.680948 | -0.560989605 | 0.076183442 | 0.84 |
| YJR035W | 878.6193365 | 0.543227065 | 0.110767059 | 0.84 |
| YJR036C | 438.6482321 | 0.695309297 | 0.10663022 | 0.84 |
| YJR039W | 935.4373703 | 0.594120063 | 0.101857297 | 0.84 |
| YJR043C | 619.0166524 | -0.249964268 | 0.099430986 | 0.84 |
| YJR045C | 25492.38268 | -0.297278287 | 0.190995153 | 0.84 |
| YJR052W | 627.3978725 | 0.448904358 | 0.11987244 | 0.84 |
| YJR057W | 461.680688 | -0.426532728 | 0.132750845 | 0.84 |
| YJR063W | 331.564801 | -0.413961688 | 0.159608498 | 0.84 |
| YJR067C | 196.0036818 | -0.42128263 | 0.129316194 | 0.84 |
| YJR069C | 644.3818806 | -0.488366886 | 0.071838932 | 0.84 |
| YJR070C | 1785.660344 | -0.35893227 | 0.065508147 | 0.84 |
| YJR072C | 1860.139893 | -0.466721558 | 0.088924732 | 0.84 |
| YJR076C | 1668.359247 | -0.364776559 | 0.126836948 | 0.84 |
| YJR091C | 3359.251305 | 0.576324308 | 0.117049104 | 0.84 |
| YJR094W-A | 1689.898779 | -0.710656814 | 0.061625009 | 0.84 |
| YJR097W | 96.20405896 | -0.059320895 | 0.207454698 | 0.84 |
| YJR101W | 534.1436387 | -0.635843437 | 0.061240764 | 0.84 |
| YJR109C | 3577.314205 | 0.958763122 | 0.214699295 | 0.84 |
| YJR110W | 661.1509666 | 0.398630875 | 0.134907988 | 0.84 |
| YJR115W | 567.164948 | 0.295969415 | 0.045930878 | 0.84 |
| YJR118C | 868.4456577 | -0.584696131 | 0.075182736 | 0.84 |
| YJR120W | 18.09619799 | 0.531508268 | 0.12145738 | 0.84 |
| YJR123W | 26166.00582 | -0.125245295 | 0.122561993 | 0.84 |
| YJR129C | 222.0216409 | -0.225362333 | 0.184154547 | 0.84 |
| YJR130C | 1173.075557 | 0.463995957 | 0.212712707 | 0.84 |
| YJR135W-A | 435.3473743 | -0.666700447 | 0.062834156 | 0.84 |
| YJR136C | 423.3917741 | 0.409809513 | 0.094164214 | 0.84 |
| YJR143C | 2572.640819 | -0.275715247 | 0.171485779 | 0.84 |
| YJR145C | 6502.704468 | -0.294213818 | 0.138208585 | 0.84 |
| YJR149W | 627.5970928 | 0.701170637 | 0.104924325 | 0.84 |
| YJR150C | 147.0716932 | 0.966070372 | 0.009951868 | 0.84 |
| YJR152W | 334.6737416 | 0.290180997 | 0.215860707 | 0.84 |
| YJR155W | 885.0501737 | 0.465119067 | 0.062811604 | 0.84 |
| YKL006W | 2404.725348 | -0.312327435 | 0.031418683 | 0.84 |
| YKL007W | 2212.362326 | -0.373100823 | 0.099015458 | 0.84 |
| YKL009W | 1409.022835 | -0.430011605 | 0.118866962 | 0.84 |
| YKL035W | 11294.21789 | 0.552326355 | 0.155651025 | 0.84 |
| YKL045W | 918.6422418 | -0.634248355 | 0.023047454 | 0.84 |
| YKL050C | 526.5764301 | 0.815180771 | 0.032231497 | 0.84 |

| | | | | |
|---|---|---|---|---|
| YKL053C-A | 473.2870725 | -0.60996174 | 0.186795796 | 0.84 |
| YKL056C | 11023.03919 | -0.444486234 | 0.089736929 | 0.84 |
| YKL058W | 636.5138347 | -0.582428484 | 0.103106174 | 0.84 |
| YKL060C | 152874.8661 | -0.168677245 | 0.185258979 | 0.84 |
| YKL062W | 2465.244192 | 0.54910368 | 0.094647553 | 0.84 |
| YKL069W | 336.0140703 | -0.777189863 | 0.015908221 | 0.84 |
| YKL074C | 747.5415477 | 0.38520354 | 0.144838962 | 0.84 |
| YKL080W | 5080.565803 | -0.529579624 | 0.064384802 | 0.84 |
| YKL081W | 14463.37631 | -0.114329676 | 0.089466473 | 0.84 |
| YKL086W | 218.7788586 | 0.714476536 | 0.05595426 | 0.84 |
| YKL090W | 556.2185917 | 0.410100202 | 0.133720954 | 0.84 |
| YKL093W | 994.1827608 | 0.54712405 | 0.104509594 | 0.84 |
| YKL094W | 2161.93051 | 0.38538226 | 0.178886562 | 0.84 |
| YKL095W | 323.4844945 | 0.6107119 | 0.07989726 | 0.84 |
| YKL096W | 1798.684379 | -0.574727598 | 0.147696143 | 0.84 |
| YKL096W-A | 39924.89359 | -0.640962835 | 0.086890391 | 0.84 |
| YKL105C | 1423.737383 | 0.535426363 | 0.17268215 | 0.84 |
| YKL111C | 22.34564795 | -1.433063761 | 0.006872603 | 0.84 |
| YKL112W | 917.1897974 | -0.6153863 | 0.049393155 | 0.84 |
| YKL113C | 707.9229549 | -0.462550597 | 0.118129025 | 0.84 |
| YKL119C | 426.0044347 | -0.493348769 | 0.205354775 | 0.84 |
| YKL121W | 1073.818565 | 0.421542528 | 0.168903158 | 0.84 |
| YKL122C | 1563.613747 | -0.485210105 | 0.156125822 | 0.84 |
| YKL123W | 31.46802918 | 0.961393101 | 0.00452598 | 0.84 |
| YKL124W | 623.7006372 | 0.684707345 | 0.030929975 | 0.84 |
| YKL125W | 592.2613366 | 0.398353633 | 0.17132629 | 0.84 |
| YKL129C | 2233.401318 | 0.4747225 | 0.161825131 | 0.84 |
| YKL137W | 420.0820122 | -0.694549374 | 0.078678691 | 0.84 |
| YKL138C | 823.724232 | -0.553604156 | 0.199238891 | 0.84 |
| YKL143W | 1110.759937 | -0.214712736 | 0.18820822 | 0.84 |
| YKL147C | 17.64954721 | 0.286614849 | 0.093271183 | 0.84 |
| YKL153W | 3671.352718 | -0.427672406 | 0.19185856 | 0.84 |
| YKL156W | 2081.100672 | -0.525992352 | 0.143205785 | 0.84 |
| YKL162C | 452.2195831 | 0.809662224 | 0.019047156 | 0.84 |
| YKL165C | 1154.053703 | -0.676499682 | 0.039968294 | 0.84 |
| YKL168C | 972.1957599 | 0.52897061 | 0.101145009 | 0.84 |
| YKL170W | 81.53735369 | -0.804193865 | 0.046634539 | 0.84 |
| YKL174C | 903.8221117 | 0.424627682 | 0.07577319 | 0.84 |
| YKL180W | 12036.25542 | -0.369090202 | 0.123978238 | 0.84 |
| YKL182W | 10979.47839 | -0.576826958 | 0.038075636 | 0.84 |
| YKL183W | 567.8673175 | -0.64779967 | 0.034874466 | 0.84 |
| YKL188C | 680.4138758 | 0.871655495 | 0.02201425 | 0.84 |
| YKL191W | 1251.863612 | -0.294250417 | 0.192419994 | 0.84 |

| | | | | |
|---|---|---|---|---|
| YKL192C | 3222.400243 | -0.531782391 | 0.061619505 | 0.84 |
| YKL202W | 21.88607995 | 0.499714206 | 0.142449953 | 0.84 |
| YKL207W | 2006.977142 | -0.535075442 | 0.110844582 | 0.84 |
| YKL213C | 1964.500982 | 0.481397705 | 0.055216467 | 0.84 |
| YKR005C | 237.9372028 | 0.779497193 | 0.024807113 | 0.84 |
| YKR006C | 1505.723079 | -0.641159471 | 0.058578316 | 0.84 |
| YKR015C | 299.9934309 | 0.863304627 | 0.00346732 | 0.84 |
| YKR017C | 571.9042365 | 0.464611252 | 0.127151146 | 0.84 |
| YKR019C | 546.7471303 | 0.542848744 | 0.109995047 | 0.84 |
| YKR021W | 1328.600543 | 0.407761244 | 0.173773471 | 0.84 |
| YKR026C | 956.8364878 | -0.499757247 | 0.171074989 | 0.84 |
| YKR041W | 32.67705338 | -0.618570711 | 0.139135134 | 0.84 |
| YKR045C | 368.478154 | -0.515277835 | 0.130944143 | 0.84 |
| YKR049C | 1528.555977 | 0.40534636 | 0.215248929 | 0.84 |
| YKR050W | 1256.621968 | 0.519728492 | 0.193306997 | 0.84 |
| YKR054C | 2620.574744 | -0.547873828 | 0.055127363 | 0.84 |
| YKR055W | 476.0228776 | 0.400007434 | 0.216827243 | 0.84 |
| YKR057W | 2516.350053 | -0.537592804 | 0.082459327 | 0.84 |
| YKR058W | 2130.69981 | 0.658594702 | 0.08424944 | 0.84 |
| YKR059W | 1551.844916 | -0.381596325 | 0.057958712 | 0.84 |
| YKR065C | 1072.834618 | -0.464721147 | 0.151495312 | 0.84 |
| YKR069W | 646.4230382 | 0.676140131 | 0.022789985 | 0.84 |
| YKR070W | 801.1916976 | -0.336493286 | 0.095626758 | 0.84 |
| YKR074W | 1111.118511 | -0.369910584 | 0.174639118 | 0.84 |
| YKR075C | 2465.019565 | 0.488387859 | 0.057281669 | 0.84 |
| YKR078W | 765.3811944 | 0.516465249 | 0.117354052 | 0.84 |
| YKR081C | 1261.655674 | -0.336210898 | 0.13332228 | 0.84 |
| YKR085C | 837.2824572 | -0.7940548 | 0.021234742 | 0.84 |
| YKR094C | 5698.150477 | -0.238123593 | 0.108822681 | 0.84 |
| YKR098C | 1187.222635 | 0.590188359 | 0.213222884 | 0.84 |
| YLL013C | 908.878738 | -0.406398924 | 0.067194712 | 0.84 |
| YLL014W | 750.2093462 | -0.783813443 | 0.024771851 | 0.84 |
| YLL015W | 2174.31995 | 0.710149489 | 0.060340919 | 0.84 |
| YLL022C | 865.5611491 | -0.319909233 | 0.217316845 | 0.84 |
| YLL024C | 20764.11389 | 0.069720544 | 0.156255158 | 0.84 |
| YLL025W | 92.59299699 | 0.520539862 | 0.124417923 | 0.84 |
| YLL030C | 4.099886808 | 0.37477722 | 0.12569975 | 0.84 |
| YLL045C | 7512.686429 | -0.274583843 | 0.073476931 | 0.84 |
| YLL049W | 333.5985502 | -0.53053027 | 0.09903413 | 0.84 |
| YLL050C | 6100.834604 | -0.416999874 | 0.152024045 | 0.84 |
| YLL053C | 96.70396409 | 0.184946058 | 0.181725554 | 0.84 |
| YLL056C | 287.6766171 | 0.60232752 | 0.113056487 | 0.84 |
| YLL058W | 677.8123693 | 0.758294025 | 0.059581096 | 0.84 |

| | | | | |
|---|---|---|---|---|
| YLL059C | 81.93358505 | 0.699184229 | 0.063172053 | 0.84 |
| YLR002C | 907.6259007 | -0.743031661 | 0.026405549 | 0.84 |
| YLR003C | 782.5858822 | -0.411522898 | 0.149037469 | 0.84 |
| YLR008C | 578.2897755 | -0.703332686 | 0.039455853 | 0.84 |
| YLR009W | 1043.475365 | -0.424616803 | 0.139512123 | 0.84 |
| YLR023C | 1130.122098 | 0.390916223 | 0.193244246 | 0.84 |
| YLR026C | 1116.422754 | -0.456946961 | 0.202558112 | 0.84 |
| YLR029C | 12797.37208 | -0.366709505 | 0.067701598 | 0.84 |
| YLR030W | 185.5800298 | 0.575527203 | 0.075294397 | 0.84 |
| YLR031W | 139.756576 | 0.654265682 | 0.056693542 | 0.84 |
| YLR033W | 1049.258607 | -0.425725018 | 0.125761677 | 0.84 |
| YLR037C | 167.7921303 | 0.287873931 | 0.177157284 | 0.84 |
| YLR043C | 5741.406917 | -0.492196576 | 0.091145467 | 0.84 |
| YLR044C | 136554.4679 | -0.046333334 | 0.175313781 | 0.84 |
| YLR046C | 345.7378309 | 0.583817024 | 0.018037192 | 0.84 |
| YLR047C | 709.1954371 | 0.376777707 | 0.133037055 | 0.84 |
| YLR048W | 13755.73587 | -0.196085834 | 0.177458925 | 0.84 |
| YLR052W | 1071.581443 | 0.238819299 | 0.182929808 | 0.84 |
| YLR056W | 3964.313149 | -0.338296506 | 0.116720746 | 0.84 |
| YLR062C | 194.5995134 | -0.320116631 | 0.120165006 | 0.84 |
| YLR070C | 346.2836033 | 0.518256138 | 0.154759162 | 0.84 |
| YLR073C | 322.9361924 | -0.686285673 | 0.126037896 | 0.84 |
| YLR076C | 95.93637703 | -0.010041296 | 0.185832178 | 0.84 |
| YLR083C | 1368.675892 | -0.469706938 | 0.092715445 | 0.84 |
| YLR089C | 5710.768483 | 0.783756784 | 0.199355043 | 0.84 |
| YLR094C | 1048.349221 | 0.441319244 | 0.150254248 | 0.84 |
| YLR099W-A | 174.4936248 | -0.79064808 | 0.063074963 | 0.84 |
| YLR106C | 6573.73141 | -0.400036562 | 0.059500496 | 0.84 |
| YLR107W | 1035.729202 | 0.843651612 | 0.004117931 | 0.84 |
| YLR108C | 407.9615838 | 0.661813933 | 0.125424917 | 0.84 |
| YLR110C | 106387.3494 | -0.42522086 | 0.087022738 | 0.84 |
| YLR116W | 1156.147687 | 0.443877933 | 0.067263986 | 0.84 |
| YLR117C | 1230.224284 | 0.372840194 | 0.191257465 | 0.84 |
| YLR126C | 468.0585451 | -0.598490326 | 0.126387747 | 0.84 |
| YLR134W | 556.7123022 | 0.337788105 | 0.175794683 | 0.84 |
| YLR135W | 729.3825255 | 0.449033758 | 0.147889221 | 0.84 |
| YLR142W | 1292.784521 | 0.760500604 | 0.023180341 | 0.84 |
| YLR143W | 501.3258454 | -0.458452898 | 0.181657578 | 0.84 |
| YLR146C | 511.7510501 | -0.438589118 | 0.136071573 | 0.84 |
| YLR149C | 1780.729906 | 0.492640743 | 0.205685478 | 0.84 |
| YLR149C-A | 61.1192566 | -0.101093065 | 0.082866603 | 0.84 |
| YLR150W | 21689.73235 | -0.243500364 | 0.162048957 | 0.84 |
| YLR152C | 1272.570191 | 0.598697067 | 0.181368993 | 0.84 |

| | | | | |
|---|---|---|---|---|
| YLR153C | 3246.572959 | -0.341389029 | 0.146547809 | 0.84 |
| YLR154C | 335.1264893 | -0.826102488 | 0.003143705 | 0.84 |
| YLR162W-A | 8.884728875 | 0.042929252 | 0.041666967 | 0.84 |
| YLR164W | 251.3796178 | 0.277477264 | 0.152676604 | 0.84 |
| YLR165C | 405.6673301 | 0.443157776 | 0.131643432 | 0.84 |
| YLR167W | 28914.13243 | -0.133980416 | 0.09586916 | 0.84 |
| YLR171W | 25.60461968 | 0.763777759 | 0.029915173 | 0.84 |
| YLR173W | 824.1208493 | 0.558336909 | 0.168044479 | 0.84 |
| YLR175W | 4387.69014 | -0.453438785 | 0.114452475 | 0.84 |
| YLR176C | 916.0927001 | 0.575858069 | 0.082469238 | 0.84 |
| YLR177W | 3573.62477 | 0.756964072 | 0.06890697 | 0.84 |
| YLR179C | 2935.995773 | -0.245594114 | 0.165412701 | 0.84 |
| YLR183C | 714.9768804 | -0.739287533 | 0.003510341 | 0.84 |
| YLR185W | 4884.991766 | -0.319039543 | 0.067779151 | 0.84 |
| YLR188W | 1233.172467 | -0.337337756 | 0.173066636 | 0.84 |
| YLR189C | 2093.552884 | 0.719357744 | 0.062043613 | 0.84 |
| YLR192C | 3457.976113 | -0.548929352 | 0.032866969 | 0.84 |
| YLR195C | 1211.827498 | -0.550752432 | 0.054643063 | 0.84 |
| YLR205C | 412.8411079 | 0.899831727 | 0.018357007 | 0.84 |
| YLR206W | 3965.73873 | 0.436432671 | 0.157413051 | 0.84 |
| YLR208W | 2849.872811 | -0.344117128 | 0.201886019 | 0.84 |
| YLR211C | 375.8938446 | 0.499582769 | 0.181484626 | 0.84 |
| YLR212C | 601.9431586 | -0.356313585 | 0.178702076 | 0.84 |
| YLR219W | 2484.562083 | 0.886795857 | 0.02005338 | 0.84 |
| YLR221C | 955.7495307 | -0.463712852 | 0.091716691 | 0.84 |
| YLR227C | 512.1356286 | 0.554855725 | 0.031824132 | 0.84 |
| YLR228C | 906.7831503 | 0.644744047 | 0.171203007 | 0.84 |
| YLR229C | 1318.158779 | -0.675458665 | 0.026471595 | 0.84 |
| YLR232W | 2.842598345 | -0.589636929 | 0.160267274 | 0.84 |
| YLR237W | 1061.722449 | 0.678782344 | 0.212659193 | 0.84 |
| YLR240W | 1128.041716 | 0.505964433 | 0.111934575 | 0.84 |
| YLR247C | 1785.222725 | 0.576077698 | 0.056948982 | 0.84 |
| YLR252W | 254.6818923 | 0.411429325 | 0.209965172 | 0.84 |
| YLR257W | 6468.084537 | 0.610264577 | 0.19039174 | 0.84 |
| YLR258W | 9832.7404 | 0.721236606 | 0.140712353 | 0.84 |
| YLR259C | 10469.29817 | -0.532558666 | 0.052363905 | 0.84 |
| YLR262C-A | 800.3537696 | -0.506350461 | 0.201778674 | 0.84 |
| YLR263W | 707.0718116 | 0.770227093 | 0.045493251 | 0.84 |
| YLR264C-A | 117.4733387 | -0.736992364 | 0.090053704 | 0.84 |
| YLR264W | 525.6829336 | -0.843549163 | 0.042030383 | 0.84 |
| YLR267W | 1321.159087 | 0.579630993 | 0.08389308 | 0.84 |
| YLR268W | 967.406712 | -0.591674188 | 0.044891955 | 0.84 |
| YLR272C | 941.972981 | -0.47703341 | 0.087882304 | 0.84 |

| | | | | |
|---|---|---|---|---|
| YLR273C | 457.5126435 | 0.974068406 | 0.005168791 | 0.84 |
| YLR276C | 2051.953288 | -0.425995204 | 0.173513265 | 0.84 |
| YLR279W | 36.37996098 | -0.657666877 | 0.16692073 | 0.84 |
| YLR281C | 33.92288358 | -0.573220318 | 0.110722311 | 0.84 |
| YLR284C | 336.5089725 | 0.3228196 | 0.218163449 | 0.84 |
| YLR285C-A | 41.10839909 | -0.572481722 | 0.12652242 | 0.84 |
| YLR287C | 617.6178514 | -0.335358204 | 0.196091658 | 0.84 |
| YLR287C-A | 5234.314487 | -0.39531519 | 0.160949808 | 0.84 |
| YLR288C | 449.0939426 | 0.575565222 | 0.017107303 | 0.84 |
| YLR289W | 462.7410856 | 0.402135507 | 0.197371912 | 0.84 |
| YLR291C | 1399.72151 | -0.625546056 | 0.02343006 | 0.84 |
| YLR292C | 1010.798537 | -0.442575017 | 0.174165762 | 0.84 |
| YLR293C | 8683.145693 | -0.397424487 | 0.124946225 | 0.84 |
| YLR297W | 487.8704755 | 1.14515783 | 0.003886797 | 0.84 |
| YLR299W | 1242.489768 | 0.58036587 | 0.13789182 | 0.84 |
| YLR301W | 1464.12998 | -0.348281582 | 0.06434316 | 0.84 |
| YLR308W | 11.06339218 | 0.597091671 | 0.103155117 | 0.84 |
| YLR312C | 417.3618401 | 0.402461476 | 0.206181904 | 0.84 |
| YLR312W-A | 784.781583 | -0.508776457 | 0.142948899 | 0.84 |
| YLR317W | 2.198595472 | -0.067849959 | 0.13629063 | 0.84 |
| YLR324W | 1737.806437 | 0.679208962 | 0.084173352 | 0.84 |
| YLR325C | 5760.035975 | -0.368126929 | 0.163537759 | 0.84 |
| YLR329W | 110.7708514 | 0.51098094 | 0.209333386 | 0.84 |
| YLR331C | 3.488696978 | 1.031777993 | 0.056267906 | 0.84 |
| YLR335W | 2838.169755 | -0.472569294 | 0.035309158 | 0.84 |
| YLR340W | 21021.60532 | -0.374371 | 0.089459609 | 0.84 |
| YLR342W | 12036.59934 | -0.669772248 | 0.039122365 | 0.84 |
| YLR342W-A | 51.47657195 | 0.555300069 | 0.038426619 | 0.84 |
| YLR344W | 3310.960859 | -0.372622234 | 0.196975407 | 0.84 |
| YLR345W | 2235.165084 | 0.480271194 | 0.194171297 | 0.84 |
| YLR350W | 3571.832696 | 0.441394993 | 0.162551506 | 0.84 |
| YLR352W | 926.4882915 | 0.518556607 | 0.114050934 | 0.84 |
| YLR354C | 11943.28845 | -0.52475305 | 0.034401762 | 0.84 |
| YLR358C | 39.08246637 | -0.54573795 | 0.045430745 | 0.84 |
| YLR363C | 329.7816062 | 0.406821838 | 0.17612437 | 0.84 |
| YLR363W-A | 269.4630453 | -0.278035448 | 0.190363919 | 0.84 |
| YLR366W | 1.593075245 | 0.038218476 | 0.172067528 | 0.84 |
| YLR367W | 2289.066484 | -0.171080174 | 0.128664616 | 0.84 |
| YLR372W | 2301.005993 | -0.247362498 | 0.050130063 | 0.84 |
| YLR378C | 3906.764345 | -0.785720516 | 0.012275452 | 0.84 |
| YLR379W | 289.3293491 | -0.415835776 | 0.099513122 | 0.84 |
| YLR388W | 1672.166035 | -0.60806615 | 0.052267742 | 0.84 |
| YLR393W | 270.1892054 | 0.57144478 | 0.169175064 | 0.84 |

| | | | | |
|---|---|---|---|---|
| YLR394W | 730.9999237 | 0.399047339 | 0.216913046 | 0.84 |
| YLR406C | 2407.447144 | -0.294506539 | 0.072746167 | 0.84 |
| YLR411W | 132.197405 | 0.423773071 | 0.203728559 | 0.84 |
| YLR413W | 2093.12908 | -0.378547509 | 0.175073584 | 0.84 |
| YLR414C | 1706.526088 | 0.445451372 | 0.170458851 | 0.84 |
| YLR415C | 24.41748512 | 0.891386145 | 0.051696681 | 0.84 |
| YLR416C | 28.8196825 | 0.984282008 | 0.01914595 | 0.84 |
| YLR417W | 736.9680806 | 0.418574844 | 0.191130283 | 0.84 |
| YLR420W | 2792.570329 | -0.4402694 | 0.215981943 | 0.84 |
| YLR423C | 531.1083054 | 0.597505643 | 0.107381858 | 0.84 |
| YLR437C-A | 26.88165292 | 0.72698242 | 0.21856581 | 0.84 |
| YLR442C | 902.5050218 | 0.362383591 | 0.214445175 | 0.84 |
| YLR445W | 32.03943686 | 0.695957948 | 0.043586149 | 0.84 |
| YLR446W | 527.2109038 | 0.809737013 | 0.00904951 | 0.84 |
| YLR448W | 5493.030487 | -0.32265467 | 0.134216708 | 0.84 |
| YLR450W | 2180.849651 | 0.586886526 | 0.176100038 | 0.84 |
| YLR453C | 293.5520271 | 0.460200923 | 0.131644084 | 0.84 |
| YLR462W | 4.00598291 | 0.042747 | 0.097352078 | 0.84 |
| YLR466C-B | 12.5768575 | -0.57956996 | 0.154770347 | 0.84 |
| YML002W | 853.1962719 | 0.496519662 | 0.125012062 | 0.84 |
| YML008C | 4220.321447 | -0.335603614 | 0.181553081 | 0.84 |
| YML009C | 743.0693468 | -0.715171689 | 0.051448798 | 0.84 |
| YML010W | 2999.463402 | -0.329061819 | 0.173833455 | 0.84 |
| YML012W | 3525.480868 | -0.326922884 | 0.188405103 | 0.84 |
| YML016C | 1375.407219 | 0.380180189 | 0.212019103 | 0.84 |
| YML019W | 694.8842248 | -0.499037117 | 0.086108144 | 0.84 |
| YML022W | 1077.904293 | -0.358357143 | 0.098335907 | 0.84 |
| YML023C | 695.7405584 | 0.666677836 | 0.013850961 | 0.84 |
| YML024W | 8930.152891 | -0.34430691 | 0.051998314 | 0.84 |
| YML026C | 7646.727007 | -0.40517386 | 0.042869068 | 0.84 |
| YML031C-A | 2.97142876 | 0.349082888 | 0.173529623 | 0.84 |
| YML034C-A | 23.30372078 | 0.652123203 | 0.187824293 | 0.84 |
| YML034W | 2522.763719 | 0.750289884 | 0.016930808 | 0.84 |
| YML035C | 2451.342109 | 0.51030988 | 0.167398337 | 0.84 |
| YML036W | 292.5583071 | -0.573768098 | 0.155054675 | 0.84 |
| YML041C | 413.9344954 | 0.346753385 | 0.216150376 | 0.84 |
| YML047W-A | 2.39499993 | -0.020710535 | 0.146931036 | 0.84 |
| YML054C | 1101.015099 | 0.070013964 | 0.099319923 | 0.84 |
| YML057C-A | 86.62699701 | 0.508410641 | 0.201290459 | 0.84 |
| YML060W | 354.1411795 | -0.14798369 | 0.126366114 | 0.84 |
| YML063W | 16224.08176 | -0.208586438 | 0.081320499 | 0.84 |
| YML064C | 587.9701312 | 0.326045587 | 0.181343971 | 0.84 |
| YML067C | 1114.441541 | -0.43395506 | 0.084355975 | 0.84 |

| | | | | |
|---|---|---|---|---|
| YML073C | 8568.253747 | -0.398714197 | 0.083463412 | 0.84 |
| YML078W | 2283.677343 | -0.557277636 | 0.122425722 | 0.84 |
| YML085C | 4155.33263 | -0.402578953 | 0.087865314 | 0.84 |
| YML094W | 513.2141562 | -0.440667303 | 0.150233402 | 0.84 |
| YML097C | 708.2199714 | -0.442488518 | 0.140845329 | 0.84 |
| YML100W | 14497.66976 | 0.770232641 | 0.151656968 | 0.84 |
| YML100W-A | 6.289539175 | 1.043989765 | 0.052996364 | 0.84 |
| YML101C | 335.3248863 | -0.782843787 | 0.041762931 | 0.84 |
| YML106W | 2946.051592 | -0.433917314 | 0.105106172 | 0.84 |
| YML130C | 2617.920044 | -0.471077501 | 0.216280655 | 0.84 |
| YMR001C | 1386.236499 | -0.547556511 | 0.128520615 | 0.84 |
| YMR001C-A | 16.08451551 | -0.28212068 | 0.168264078 | 0.84 |
| YMR003W | 306.5250482 | -0.478920481 | 0.102078479 | 0.84 |
| YMR007W | 90.19118303 | 0.564470671 | 0.026238432 | 0.84 |
| YMR009W | 601.1334204 | 0.491416319 | 0.151658223 | 0.84 |
| YMR010W | 733.3907547 | -0.431585606 | 0.211714594 | 0.84 |
| YMR012W | 7425.571889 | -0.490620515 | 0.071577843 | 0.84 |
| YMR015C | 1137.039851 | -0.792971476 | 0.131120247 | 0.84 |
| YMR017W | 57.51133406 | 0.299970003 | 0.193662759 | 0.84 |
| YMR021C | 731.4049333 | 0.633308212 | 0.041173682 | 0.84 |
| YMR024W | 882.2534561 | -0.913321992 | 0.003840033 | 0.84 |
| YMR031C | 3707.880046 | 0.618678206 | 0.087641147 | 0.84 |
| YMR031W-A | 40.58383151 | 0.904328277 | 0.079138308 | 0.84 |
| YMR033W | 1176.392622 | -0.400470348 | 0.217698145 | 0.84 |
| YMR034C | 377.1167506 | 0.565221814 | 0.151405645 | 0.84 |
| YMR038C | 1941.251831 | -0.344419315 | 0.169683401 | 0.84 |
| YMR048W | 313.0835867 | -0.507512328 | 0.022527252 | 0.84 |
| YMR049C | 1525.516535 | -0.253586238 | 0.198596033 | 0.84 |
| YMR053C | 712.958308 | 0.573081853 | 0.163041286 | 0.84 |
| YMR057C | 13.35741525 | 0.367613037 | 0.042771544 | 0.84 |
| YMR062C | 1501.153431 | 1.098978313 | 0.029525566 | 0.84 |
| YMR068W | 420.8665046 | 0.412445848 | 0.218201575 | 0.84 |
| YMR076C | 1386.678863 | -0.386161722 | 0.147326631 | 0.84 |
| YMR079W | 1633.407315 | -0.632312326 | 0.035944518 | 0.84 |
| YMR081C | 3745.685883 | 0.431547507 | 0.110076023 | 0.84 |
| YMR084W | 174.3110976 | 0.731642685 | 0.061895553 | 0.84 |
| YMR085W | 377.7619403 | 0.853836492 | 0.01349827 | 0.84 |
| YMR091C | 998.5083437 | -0.362836224 | 0.204180062 | 0.84 |
| YMR093W | 713.9051908 | -0.377956754 | 0.16604549 | 0.84 |
| YMR096W | 4030.130556 | 1.178328832 | 0.050507178 | 0.84 |
| YMR097C | 670.8866515 | 0.349370209 | 0.202199458 | 0.84 |
| YMR099C | 2686.030784 | -0.335427637 | 0.182449055 | 0.84 |
| YMR103C | 375.8522499 | 0.449125622 | 0.190222593 | 0.84 |

| | | | | |
|---|---|---|---|---|
| YMR104C | 1639.396102 | 0.733261012 | 0.134729174 | 0.84 |
| YMR105W-A | 43.59841519 | 0.621541295 | 0.050852225 | 0.84 |
| YMR111C | 424.0046487 | 0.384046102 | 0.181142855 | 0.84 |
| YMR114C | 1303.792032 | 0.580367425 | 0.127769577 | 0.84 |
| YMR116C | 12111.33345 | -0.113986562 | 0.076037451 | 0.84 |
| YMR118C | 295.0484477 | 0.756441557 | 0.044402796 | 0.84 |
| YMR120C | 11760.55672 | -0.002290222 | 0.093339232 | 0.84 |
| YMR122W-A | 5293.670712 | -0.649938507 | 0.16374884 | 0.84 |
| YMR125W | 2280.761082 | -0.589305206 | 0.05664314 | 0.84 |
| YMR131C | 2416.896733 | -0.167684583 | 0.201351223 | 0.84 |
| YMR133W | 502.5352221 | 0.728879797 | 0.074699884 | 0.84 |
| YMR137C | 598.7111524 | 0.635399938 | 0.058127615 | 0.84 |
| YMR140W | 1470.388235 | 0.600594306 | 0.102013396 | 0.84 |
| YMR141C | 167.2172498 | 0.362885352 | 0.170697116 | 0.84 |
| YMR142C | 11187.93902 | -0.432274522 | 0.068459076 | 0.84 |
| YMR143W | 7020.519511 | -0.399058645 | 0.111300043 | 0.84 |
| YMR146C | 5181.578577 | -0.378344237 | 0.211795062 | 0.84 |
| YMR148W | 862.6307015 | 0.565686192 | 0.133568773 | 0.84 |
| YMR151W | 69.13225197 | 0.569453247 | 0.108479304 | 0.84 |
| YMR153W | 881.8064044 | -0.530423785 | 0.036528984 | 0.84 |
| YMR158C-A | 22.93483608 | 0.50138407 | 0.128868917 | 0.84 |
| YMR158W | 693.5786925 | -0.589121016 | 0.092463172 | 0.84 |
| YMR158W-B | 1.160282167 | -0.109982459 | 0.218396191 | 0.84 |
| YMR165C | 1579.930507 | 0.483014958 | 0.175431148 | 0.84 |
| YMR181C | 1531.918567 | 0.59865065 | 0.078499658 | 0.84 |
| YMR182C | 564.4635256 | 0.873114945 | 0.018452682 | 0.84 |
| YMR194W | 1484.319594 | -0.675413802 | 0.02584104 | 0.84 |
| YMR201C | 569.2749285 | 0.574412367 | 0.123722465 | 0.84 |
| YMR202W | 2009.003075 | -0.314787437 | 0.035268372 | 0.84 |
| YMR203W | 3020.852994 | -0.411453743 | 0.154412232 | 0.84 |
| YMR206W | 572.161077 | 0.209405955 | 0.166502091 | 0.84 |
| YMR214W | 1048.466602 | -0.431303087 | 0.183097653 | 0.84 |
| YMR225C | 419.9415237 | -0.717592651 | 0.057291813 | 0.84 |
| YMR230W | 6828.386976 | -0.188232359 | 0.159578664 | 0.84 |
| YMR232W | 290.2765579 | 0.38600437 | 0.110012722 | 0.84 |
| YMR235C | 2269.617315 | -0.371982446 | 0.157655156 | 0.84 |
| YMR242C | 8203.831599 | -0.400605447 | 0.062440431 | 0.84 |
| YMR242W-A | 1.825788145 | -0.461192121 | 0.122961904 | 0.84 |
| YMR260C | 3410.062096 | -0.426355099 | 0.127827147 | 0.84 |
| YMR284W | 845.7287906 | 0.72894216 | 0.040883378 | 0.84 |
| YMR290C | 2185.404412 | -0.028208533 | 0.212351017 | 0.84 |
| YMR292W | 429.0322087 | -0.738208257 | 0.014299838 | 0.84 |
| YMR298W | 1278.824067 | -0.277297617 | 0.211421789 | 0.84 |

| | | | | |
|---|---|---|---|---|
| YMR300C | 1920.32889 | -0.345608203 | 0.195405998 | 0.84 |
| YMR303C | 445.8492587 | 0.40550701 | 0.095136981 | 0.84 |
| YMR306W | 518.3037802 | 0.480496736 | 0.218800868 | 0.84 |
| YMR307W | 11748.41017 | -0.527930493 | 0.088654762 | 0.84 |
| YMR318C | 2412.871237 | -0.473176248 | 0.054885232 | 0.84 |
| YNL002C | 1677.703778 | -0.461858156 | 0.097345532 | 0.84 |
| YNL010W | 2912.199227 | -0.240227602 | 0.218844589 | 0.84 |
| YNL012W | 592.4635068 | 0.576393368 | 0.108957285 | 0.84 |
| YNL016W | 4774.57143 | -0.55439945 | 0.043970375 | 0.84 |
| YNL025C | 730.0152463 | 0.667619104 | 0.045015687 | 0.84 |
| YNL039W | 1798.324816 | 0.370453975 | 0.19443115 | 0.84 |
| YNL040W | 832.5546055 | 0.640244001 | 0.162304518 | 0.84 |
| YNL044W | 495.5352853 | -0.562376566 | 0.137888095 | 0.84 |
| YNL061W | 2817.950295 | -0.345033485 | 0.197760761 | 0.84 |
| YNL064C | 7494.727656 | -0.248946175 | 0.163316556 | 0.84 |
| YNL067W | 1749.269832 | -0.49395025 | 0.11210652 | 0.84 |
| YNL069C | 13320.02635 | -0.248895917 | 0.105336608 | 0.84 |
| YNL070W | 685.2481589 | -0.657118904 | 0.046144398 | 0.84 |
| YNL075W | 678.2006525 | -0.292210887 | 0.102301434 | 0.84 |
| YNL093W | 262.8484398 | 0.274550338 | 0.199128318 | 0.84 |
| YNL096C | 3035.774305 | -0.330828764 | 0.110629517 | 0.84 |
| YNL102W | 1324.90783 | -0.866710139 | 0.002974004 | 0.84 |
| YNL110C | 899.4305976 | -0.306560716 | 0.086198066 | 0.84 |
| YNL111C | 246.1352445 | -0.733744005 | 0.14344475 | 0.84 |
| YNL113W | 221.4093861 | -0.56248916 | 0.063394414 | 0.84 |
| YNL122C | 325.2385534 | -0.567928987 | 0.171280053 | 0.84 |
| YNL125C | 2090.525445 | 0.731782966 | 0.05982895 | 0.84 |
| YNL126W | 705.9981899 | -0.608924034 | 0.076661723 | 0.84 |
| YNL131W | 2877.827714 | -0.372451401 | 0.189474573 | 0.84 |
| YNL135C | 5166.62026 | -0.652778168 | 0.068901481 | 0.84 |
| YNL137C | 1635.261502 | -0.428063247 | 0.144260613 | 0.84 |
| YNL140C | 6.604161499 | 0.435182418 | 0.052120506 | 0.84 |
| YNL144C | 1767.380134 | 0.614444701 | 0.141755295 | 0.84 |
| YNL146W | 85.60940575 | 0.461292469 | 0.141216394 | 0.84 |
| YNL147W | 569.2824244 | -0.657125031 | 0.04479338 | 0.84 |
| YNL150W | 83.29807569 | -0.772364626 | 0.029101493 | 0.84 |
| YNL158W | 434.8285056 | 0.50541254 | 0.099872141 | 0.84 |
| YNL162W | 2350.306083 | -0.3598448 | 0.108134385 | 0.84 |
| YNL165W | 500.9806009 | 0.691166555 | 0.081991264 | 0.84 |
| YNL176C | 1925.34034 | 0.553468809 | 0.112362102 | 0.84 |
| YNL177C | 1301.893277 | -0.436201603 | 0.065680933 | 0.84 |
| YNL178W | 12909.44004 | -0.358225001 | 0.072410994 | 0.84 |
| YNL184C | 77.22136504 | -0.684583849 | 0.077020799 | 0.84 |

| | | | | |
|---|---|---|---|---|
| YNL185C | 474.6914722 | -0.79163568 | 0.026844002 | 0.84 |
| YNL187W | 311.4857358 | 0.371193898 | 0.206472644 | 0.84 |
| YNL191W | 591.9956262 | 0.68864267 | 0.062447054 | 0.84 |
| YNL194C | 937.042565 | 0.150288795 | 0.152591825 | 0.84 |
| YNL195C | 538.1405229 | 0.290008266 | 0.078859123 | 0.84 |
| YNL198C | 23.75937545 | 0.689866639 | 0.121975113 | 0.84 |
| YNL210W | 106.0752581 | 0.357243431 | 0.181550211 | 0.84 |
| YNL214W | 315.125881 | 0.341815596 | 0.196052109 | 0.84 |
| YNL216W | 1187.634305 | -0.414569075 | 0.168144405 | 0.84 |
| YNL223W | 533.8185201 | 0.661321975 | 0.053433785 | 0.84 |
| YNL226W | 19.21780824 | 0.871154873 | 0.14842071 | 0.84 |
| YNL231C | 1567.48489 | -0.764465366 | 0.018948343 | 0.84 |
| YNL233W | 992.805205 | -0.558836032 | 0.045708595 | 0.84 |
| YNL234W | 710.7866789 | 0.593660872 | 0.135421862 | 0.84 |
| YNL249C | 367.0811931 | 0.328323114 | 0.20835033 | 0.84 |
| YNL252C | 974.4576049 | -0.480591252 | 0.193220238 | 0.84 |
| YNL254C | 501.4634557 | 0.725296215 | 0.081106442 | 0.84 |
| YNL262W | 2257.387419 | -0.256905172 | 0.139489959 | 0.84 |
| YNL269W | 50.67416069 | 0.604268387 | 0.070740462 | 0.84 |
| YNL270C | 240.3573834 | 0.862942897 | 0.047598365 | 0.84 |
| YNL273W | 713.7141054 | -0.47217014 | 0.107398629 | 0.84 |
| YNL276C | 5.892936072 | -0.67694057 | 0.203113194 | 0.84 |
| YNL290W | 854.0353401 | -0.667633039 | 0.035390081 | 0.84 |
| YNL291C | 628.873147 | -0.402404833 | 0.202668272 | 0.84 |
| YNL295W | 664.526791 | 0.52144015 | 0.083064352 | 0.84 |
| YNL300W | 1681.600577 | -0.45752927 | 0.111590203 | 0.84 |
| YNL301C | 1146.542523 | -0.38895562 | 0.070136362 | 0.84 |
| YNL302C | 5878.096109 | -0.452676709 | 0.046460111 | 0.84 |
| YNL303W | 97.83657889 | -0.453052164 | 0.216787272 | 0.84 |
| YNL306W | 1534.453742 | -0.465257576 | 0.156496994 | 0.84 |
| YNL311C | 782.0547648 | 0.681053398 | 0.052246455 | 0.84 |
| YNL312W | 897.7476795 | -0.798109902 | 0.00682245 | 0.84 |
| YNL313C | 1104.908095 | -0.453699769 | 0.098056669 | 0.84 |
| YNL315C | 1636.297254 | -0.493152956 | 0.180165351 | 0.84 |
| YNL323W | 681.0384119 | -0.401070341 | 0.210003835 | 0.84 |
| YNL327W | 5207.566398 | -0.462316165 | 0.189778895 | 0.84 |
| YNL328C | 68.99382563 | -0.574029601 | 0.078567756 | 0.84 |
| YNL332W | 25.85198322 | 0.033097214 | 0.101831424 | 0.84 |
| YNL334C | 19.40823403 | -1.098631501 | 0.167813671 | 0.84 |
| YNL335W | 13.42162756 | -0.93634992 | 0.21191434 | 0.84 |
| YNR003C | 554.144982 | -0.452019275 | 0.094559821 | 0.84 |
| YNR014W | 1455.118637 | 0.664086921 | 0.05936187 | 0.84 |
| YNR016C | 6951.886561 | -0.450489788 | 0.050160698 | 0.84 |

| | | | | |
|---|---|---|---|---|
| YNR021W | 3396.726535 | -0.40158892 | 0.143303077 | 0.84 |
| YNR022C | 907.2227542 | -0.541104571 | 0.165162226 | 0.84 |
| YNR028W | 918.8142146 | -0.433402992 | 0.121308066 | 0.84 |
| YNR031C | 1352.088361 | -0.444418884 | 0.143580052 | 0.84 |
| YNR032C-A | 183.6670278 | -0.511900964 | 0.202895379 | 0.84 |
| YNR033W | 1803.332991 | 0.528120397 | 0.125144661 | 0.84 |
| YNR034W | 660.948473 | 0.483404241 | 0.138509911 | 0.84 |
| YNR037C | 584.4173485 | -0.543958827 | 0.175813274 | 0.84 |
| YNR039C | 749.4014615 | 0.356107781 | 0.12522747 | 0.84 |
| YNR041C | 337.6551047 | -0.442622282 | 0.184178733 | 0.84 |
| YNR053C | 1005.221474 | -0.582334616 | 0.1423073 | 0.84 |
| YNR054C | 724.980203 | -0.419464368 | 0.160890179 | 0.84 |
| YNR058W | 282.8560691 | 0.735327038 | 0.015960736 | 0.84 |
| YNR065C | 953.8955562 | 0.587851466 | 0.116487616 | 0.84 |
| YNR075C-A | 12.77963038 | -0.796500285 | 0.056669321 | 0.84 |
| YOL002C | 621.7558878 | -0.689826459 | 0.079602433 | 0.84 |
| YOL012C | 810.102208 | -0.719541462 | 0.023585373 | 0.84 |
| YOL013C | 1216.146773 | 0.413767526 | 0.116632671 | 0.84 |
| YOL013W-B | 4.383672181 | -0.650069772 | 0.164280826 | 0.84 |
| YOL015W | 325.5388955 | 0.414427654 | 0.188358633 | 0.84 |
| YOL017W | 430.0355777 | -0.522840125 | 0.14760928 | 0.84 |
| YOL025W | 668.044846 | 0.398620533 | 0.210288176 | 0.84 |
| YOL039W | 5643.898973 | -0.330769184 | 0.126226156 | 0.84 |
| YOL040C | 16073.93418 | -0.414460172 | 0.088145355 | 0.84 |
| YOL045W | 1132.896142 | 0.55792864 | 0.026362479 | 0.84 |
| YOL047C | 302.1448912 | 0.168614964 | 0.15758672 | 0.84 |
| YOL048C | 1220.015423 | 0.419969367 | 0.169592706 | 0.84 |
| YOL058W | 2645.481694 | 1.257841359 | 0.017337455 | 0.84 |
| YOL061W | 1754.163485 | -0.569259363 | 0.072650147 | 0.84 |
| YOL077C | 1854.977032 | -0.124161849 | 0.20471896 | 0.84 |
| YOL080C | 199.8794327 | -0.259189207 | 0.150014284 | 0.84 |
| YOL084W | 492.0065357 | 0.13519906 | 0.174721372 | 0.84 |
| YOL087C | 2320.349339 | 0.511905687 | 0.145829602 | 0.84 |
| YOL090W | 1084.862315 | -0.415374574 | 0.201275751 | 0.84 |
| YOL091W | 152.6270626 | 0.696746944 | 0.026759594 | 0.84 |
| YOL094C | 437.8167325 | -0.514645963 | 0.218234039 | 0.84 |
| YOL095C | 382.7305205 | 0.41965337 | 0.197844306 | 0.84 |
| YOL097C | 2168.680575 | -0.269456129 | 0.126556408 | 0.84 |
| YOL097W-A | 14.8332561 | -0.735526021 | 0.173122421 | 0.84 |
| YOL102C | 532.3526812 | -0.490323325 | 0.061318708 | 0.84 |
| YOL105C | 760.4552602 | 0.423425362 | 0.154568563 | 0.84 |
| YOL116W | 722.658756 | 0.585736816 | 0.060558011 | 0.84 |
| YOL119C | 1253.305663 | 0.790269067 | 0.056920255 | 0.84 |

| | | | | |
|---|---|---|---|---|
| YOL120C | 9488.699457 | -0.263191192 | 0.02371909 | 0.84 |
| YOL121C | 3681.856963 | -0.475765135 | 0.138663133 | 0.84 |
| YOL124C | 564.6964356 | -0.129048782 | 0.111163966 | 0.84 |
| YOL127W | 9841.72384 | -0.287379293 | 0.094349042 | 0.84 |
| YOL130W | 1532.382846 | 0.791442962 | 0.007960919 | 0.84 |
| YOL131W | 75.70252965 | 0.952345911 | 0.013971907 | 0.84 |
| YOL134C | 23.62819378 | 0.633723282 | 0.047675642 | 0.84 |
| YOL135C | 544.3922283 | -0.398887989 | 0.216717853 | 0.84 |
| YOL138C | 1055.538589 | 0.417798513 | 0.142978255 | 0.84 |
| YOL139C | 5554.572319 | -0.397550252 | 0.205093808 | 0.84 |
| YOL142W | 655.5671288 | -0.399043331 | 0.14967056 | 0.84 |
| YOL143C | 5652.711754 | -0.907097194 | 0.131765217 | 0.84 |
| YOL146W | 364.144711 | -0.750808722 | 0.033000359 | 0.84 |
| YOL149W | 642.6006688 | -0.499544655 | 0.152526652 | 0.84 |
| YOL153C | 1352.172864 | 0.403062728 | 0.190614019 | 0.84 |
| YOL159C | 274.8863973 | -0.752851987 | 0.191196502 | 0.84 |
| YOL161C | 3.517453047 | 0.403741871 | 0.062382468 | 0.84 |
| YOL165C | 22.45477283 | -0.375862055 | 0.046181943 | 0.84 |
| YOR003W | 499.5835225 | 0.544687671 | 0.0821041 | 0.84 |
| YOR004W | 494.9946687 | -0.312561511 | 0.028477295 | 0.84 |
| YOR005C | 988.1912317 | 0.917055118 | 0.009422715 | 0.84 |
| YOR006C | 758.4019018 | 0.499513955 | 0.081129406 | 0.84 |
| YOR008C-A | 9.788861467 | -0.381516748 | 0.147094609 | 0.84 |
| YOR008W-B | 4.890160442 | 0.399151867 | 0.064316851 | 0.84 |
| YOR009W | 307.6360112 | 0.560092717 | 0.148910809 | 0.84 |
| YOR018W | 939.5138966 | 0.714520629 | 0.019913146 | 0.84 |
| YOR019W | 709.5102111 | 0.376822608 | 0.132997138 | 0.84 |
| YOR020C | 3482.406455 | -0.625895492 | 0.049230399 | 0.84 |
| YOR023C | 2117.560133 | 0.440217936 | 0.169907879 | 0.84 |
| YOR028C | 706.5725488 | 0.221633638 | 0.192144039 | 0.84 |
| YOR029W | 6.24277143 | 0.452552144 | 0.207999696 | 0.84 |
| YOR031W | 276.3450169 | 0.436388405 | 0.144736834 | 0.84 |
| YOR039W | 1072.032394 | -0.478384848 | 0.102235316 | 0.84 |
| YOR044W | 320.4821165 | 0.476589018 | 0.153689609 | 0.84 |
| YOR045W | 622.8710581 | -1.024164212 | 0.013277609 | 0.84 |
| YOR052C | 2749.089296 | 0.540336269 | 0.164532247 | 0.84 |
| YOR061W | 1662.759146 | -0.466920502 | 0.152115594 | 0.84 |
| YOR062C | 585.4618256 | 0.250377994 | 0.155821643 | 0.84 |
| YOR063W | 34774.24031 | -0.348434437 | 0.060352147 | 0.84 |
| YOR067C | 1025.694017 | -0.378407359 | 0.188051411 | 0.84 |
| YOR070C | 1064.02202 | 0.382449946 | 0.162267842 | 0.84 |
| YOR072W | 29.04315607 | 0.798057881 | 0.071449944 | 0.84 |
| YOR072W-A | 14.05662092 | 0.415447798 | 0.147408021 | 0.84 |

| | | | | |
|---|---|---|---|---|
| YOR074C | 848.568103 | -0.468735225 | 0.078327788 | 0.84 |
| YOR077W | 225.6560466 | 0.659184692 | 0.073280849 | 0.84 |
| YOR080W | 677.2163658 | 0.277438201 | 0.198577155 | 0.84 |
| YOR091W | 1558.309002 | -0.591629057 | 0.077738871 | 0.84 |
| YOR095C | 614.382855 | -0.299350542 | 0.107344072 | 0.84 |
| YOR096W | 6750.76705 | -0.383536501 | 0.103692716 | 0.84 |
| YOR099W | 2318.668874 | -0.443376791 | 0.176334466 | 0.84 |
| YOR102W | 49.45521107 | -0.368214425 | 0.204476717 | 0.84 |
| YOR107W | 180.9819764 | 0.614805531 | 0.044868948 | 0.84 |
| YOR110W | 712.9241656 | 0.356205194 | 0.216120065 | 0.84 |
| YOR120W | 2918.156498 | 0.585395438 | 0.141931477 | 0.84 |
| YOR122C | 4814.102226 | -0.474706438 | 0.130252778 | 0.84 |
| YOR123C | 2187.626634 | -0.299789028 | 0.187961504 | 0.84 |
| YOR133W | 1425.340865 | -0.331719638 | 0.161252424 | 0.84 |
| YOR134W | 351.2595047 | 0.562607842 | 0.135671707 | 0.84 |
| YOR136W | 7814.887983 | -0.448718172 | 0.1126745 | 0.84 |
| YOR137C | 579.6398816 | 0.574496712 | 0.128895348 | 0.84 |
| YOR138C | 1254.624127 | 0.360780784 | 0.210484129 | 0.84 |
| YOR143C | 346.0541178 | -0.570494436 | 0.069621575 | 0.84 |
| YOR145C | 791.624005 | -0.636822323 | 0.083620664 | 0.84 |
| YOR146W | 6.962391281 | 0.139955807 | 0.12355372 | 0.84 |
| YOR150W | 702.6825687 | -0.586287805 | 0.177402708 | 0.84 |
| YOR153W | 11636.99259 | -0.927131482 | 0.14389289 | 0.84 |
| YOR157C | 2318.284571 | -0.345805037 | 0.200109945 | 0.84 |
| YOR159C | 320.0573697 | -0.511200453 | 0.211217921 | 0.84 |
| YOR160W | 1175.925728 | 0.324199116 | 0.100650269 | 0.84 |
| YOR161C | 928.1721785 | 0.547847717 | 0.064725375 | 0.84 |
| YOR161C-C | 9.014220677 | 0.238174074 | 0.149646831 | 0.84 |
| YOR164C | 1461.403289 | -0.371644938 | 0.125256255 | 0.84 |
| YOR165W | 1637.432086 | -0.744036582 | 0.018558916 | 0.84 |
| YOR182C | 1767.616754 | -0.630278928 | 0.036709787 | 0.84 |
| YOR186W | 27.95959597 | 0.556600968 | 0.156659928 | 0.84 |
| YOR187W | 4877.755588 | -0.455358029 | 0.124252136 | 0.84 |
| YOR190W | 351.1788756 | 0.448009246 | 0.102396058 | 0.84 |
| YOR197W | 2558.031802 | -0.363784748 | 0.102101449 | 0.84 |
| YOR200W | 9.969440735 | 0.080593854 | 0.111784266 | 0.84 |
| YOR203W | 129.138005 | 0.707770313 | 0.159392585 | 0.84 |
| YOR204W | 2305.73227 | -0.605675847 | 0.084177123 | 0.84 |
| YOR206W | 2506.527701 | -0.434099275 | 0.189064927 | 0.84 |
| YOR208W | 1224.294967 | 0.628220035 | 0.129660712 | 0.84 |
| YOR210W | 598.8398742 | -0.47546235 | 0.077794729 | 0.84 |
| YOR214C | 33.33902204 | 0.681929845 | 0.069695143 | 0.84 |
| YOR219C | 1764.610148 | 0.47918124 | 0.092700521 | 0.84 |

| | | | | |
|---|---|---|---|---|
| YOR232W | 2857.566582 | -0.653480542 | 0.042285695 | 0.84 |
| YOR234C | 2998.326335 | -0.280651427 | 0.118139601 | 0.84 |
| YOR236W | 448.7599801 | -0.543250431 | 0.054528695 | 0.84 |
| YOR237W | 107.6435638 | 0.567288806 | 0.090472791 | 0.84 |
| YOR239W | 2726.169468 | -0.441239924 | 0.093930982 | 0.84 |
| YOR242C | 110.2984083 | 0.574654889 | 0.030958779 | 0.84 |
| YOR243C | 1479.808571 | -0.259805128 | 0.157054773 | 0.84 |
| YOR247W | 5496.630424 | -0.36810598 | 0.114543167 | 0.84 |
| YOR248W | 35.3813631 | -1.111788614 | 0.002715351 | 0.84 |
| YOR252W | 480.3993858 | -0.322729402 | 0.153670001 | 0.84 |
| YOR265W | 445.8131326 | -0.499865153 | 0.113930945 | 0.84 |
| YOR275C | 1566.16532 | 0.471106484 | 0.074845959 | 0.84 |
| YOR276W | 906.6874329 | -0.227629613 | 0.216148557 | 0.84 |
| YOR277C | 6.042620409 | -0.272219105 | 0.057192796 | 0.84 |
| YOR278W | 477.7903224 | -0.632744811 | 0.049714663 | 0.84 |
| YOR279C | 315.7050157 | -0.636012652 | 0.136024824 | 0.84 |
| YOR286W | 1162.614528 | -0.792751172 | 0.007520931 | 0.84 |
| YOR292C | 645.9873627 | 0.807964388 | 0.032751146 | 0.84 |
| YOR293W | 11411.33478 | -0.171204054 | 0.06636247 | 0.84 |
| YOR298C-A | 6204.241243 | -0.515555024 | 0.114910889 | 0.84 |
| YOR301W | 501.5102259 | -0.455315949 | 0.204009008 | 0.84 |
| YOR302W | 66.2485987 | 0.688454263 | 0.170654077 | 0.84 |
| YOR309C | 407.1266153 | -0.267167508 | 0.126599059 | 0.84 |
| YOR310C | 3221.373678 | -0.178815938 | 0.199602401 | 0.84 |
| YOR312C | 9424.205996 | -0.23971926 | 0.097237542 | 0.84 |
| YOR314W | 17.17022364 | -0.609683115 | 0.117939751 | 0.84 |
| YOR315W | 1158.774212 | -0.742405949 | 0.064350366 | 0.84 |
| YOR320C | 1070.658594 | -0.355047967 | 0.209004091 | 0.84 |
| YOR326W | 4223.653773 | -0.355256073 | 0.106241027 | 0.84 |
| YOR328W | 830.771978 | 0.884371426 | 0.05230936 | 0.84 |
| YOR331C | 254.0815524 | 0.984768364 | 0.00413023 | 0.84 |
| YOR332W | 2877.284969 | -0.414904334 | 0.120068495 | 0.84 |
| YOR338W | 133.0538926 | 0.925864935 | 0.006410164 | 0.84 |
| YOR339C | 48.09399516 | 0.328286539 | 0.112641868 | 0.84 |
| YOR340C | 775.5259752 | -0.334604473 | 0.068228331 | 0.84 |
| YOR341W | 3809.897668 | -0.171412286 | 0.187535945 | 0.84 |
| YOR345C | 1.957434997 | 0.418576046 | 0.063422369 | 0.84 |
| YOR361C | 4132.636454 | -0.53288449 | 0.081638496 | 0.84 |
| YOR369C | 15497.1337 | -0.453433039 | 0.103614221 | 0.84 |
| YOR374W | 6364.239227 | 0.494557667 | 0.198280232 | 0.84 |
| YOR377W | 997.8606777 | 0.639415279 | 0.070864142 | 0.84 |
| YOR390W | 45.23234325 | -1.09861673 | 0.070005394 | 0.84 |
| YPL003W | 547.4702758 | 0.380126636 | 0.194868884 | 0.84 |

| | | | | |
|---|---|---|---|---|
| YPL007C | 583.714763 | -0.459671368 | 0.057916913 | 0.84 |
| YPL010W | 2111.463364 | -0.492424432 | 0.093157217 | 0.84 |
| YPL017C | 762.3634934 | 0.554434651 | 0.102628596 | 0.84 |
| YPL018W | 471.7341201 | 0.828887367 | 0.033252107 | 0.84 |
| YPL027W | 50.11860961 | 0.51598376 | 0.088532895 | 0.84 |
| YPL028W | 3100.321144 | -0.313281972 | 0.197392846 | 0.84 |
| YPL032C | 2772.776631 | -0.629795879 | 0.054067756 | 0.84 |
| YPL036W | 288.1590263 | 0.348447244 | 0.199155312 | 0.84 |
| YPL037C | 6009.036088 | -0.462045762 | 0.199468159 | 0.84 |
| YPL038W-A | 14.34188488 | 0.458904723 | 0.093279043 | 0.84 |
| YPL043W | 1057.06526 | -0.374916626 | 0.105129286 | 0.84 |
| YPL047W | 255.7983457 | 0.477516651 | 0.120857226 | 0.84 |
| YPL050C | 1693.46287 | -0.392690931 | 0.216097771 | 0.84 |
| YPL054W | 345.0757807 | 0.350157276 | 0.141271592 | 0.84 |
| YPL056C | 172.8020928 | 0.257027422 | 0.197684362 | 0.84 |
| YPL059W | 1008.866988 | -0.528214944 | 0.057023275 | 0.84 |
| YPL066W | 1565.740738 | 0.571262796 | 0.030511846 | 0.84 |
| YPL070W | 1079.494033 | 0.447548275 | 0.196061159 | 0.84 |
| YPL072W | 410.1680408 | -0.518071841 | 0.165409904 | 0.84 |
| YPL079W | 6102.629794 | -0.377428134 | 0.072054399 | 0.84 |
| YPL090C | 2392.913651 | -0.195329532 | 0.148487669 | 0.84 |
| YPL092W | 1197.47636 | 0.943702456 | 0.125931954 | 0.84 |
| YPL094C | 2895.773005 | -0.447259887 | 0.135363951 | 0.84 |
| YPL095C | 828.2325169 | -0.896671003 | 0.047580969 | 0.84 |
| YPL098C | 678.1300721 | -0.472897923 | 0.145491262 | 0.84 |
| YPL100W | 911.5930892 | 0.417684078 | 0.149446243 | 0.84 |
| YPL102C | 13.71806551 | 0.715035805 | 0.128497394 | 0.84 |
| YPL106C | 19467.56403 | -0.138985069 | 0.204753136 | 0.84 |
| YPL110C | 931.3902179 | 0.513839658 | 0.188458174 | 0.84 |
| YPL113C | 241.6280316 | 0.593396366 | 0.071576927 | 0.84 |
| YPL119C | 673.8655489 | 0.634503889 | 0.141526499 | 0.84 |
| YPL119C-A | 58.40963327 | 0.541824361 | 0.158986383 | 0.84 |
| YPL124W | 247.3933842 | -0.428503395 | 0.169534609 | 0.84 |
| YPL127C | 2242.895143 | -0.494451853 | 0.078632595 | 0.84 |
| YPL129W | 1819.666295 | -0.695069786 | 0.025042493 | 0.84 |
| YPL130W | 24.21083011 | 0.160297207 | 0.212035076 | 0.84 |
| YPL140C | 588.07308 | 0.358231931 | 0.214340755 | 0.84 |
| YPL142C | 228.9766843 | -0.470489901 | 0.083972577 | 0.84 |
| YPL143W | 740.5431297 | -0.352002885 | 0.215616929 | 0.84 |
| YPL144W | 173.5181567 | -0.417727521 | 0.183779435 | 0.84 |
| YPL146C | 923.2028372 | -0.323270238 | 0.164879012 | 0.84 |
| YPL147W | 471.9776317 | 0.533324346 | 0.071102544 | 0.84 |
| YPL150W | 1067.29309 | 0.490147385 | 0.141031294 | 0.84 |

| | | | | |
|---|---|---|---|---|
| YPL151C | 670.1565001 | 0.2861885 | 0.20481522 | 0.84 |
| YPL152W-A | 1.958413515 | -0.539766846 | 0.138038827 | 0.84 |
| YPL163C | 3377.565521 | -0.301278523 | 0.076258605 | 0.84 |
| YPL164C | 886.5830673 | 0.92369063 | 0.020729885 | 0.84 |
| YPL165C | 810.9716846 | 0.49390516 | 0.101526826 | 0.84 |
| YPL166W | 366.5892241 | 0.541533569 | 0.105333353 | 0.84 |
| YPL167C | 937.9139202 | 0.625416298 | 0.079766119 | 0.84 |
| YPL169C | 1689.790258 | -0.382494796 | 0.166050167 | 0.84 |
| YPL174C | 736.6858257 | 0.381712116 | 0.163682264 | 0.84 |
| YPL175W | 580.2634042 | -0.393890776 | 0.218817491 | 0.84 |
| YPL178W | 1119.186631 | -0.558128952 | 0.028586335 | 0.84 |
| YPL183W-A | 354.0512636 | -0.572435943 | 0.158562355 | 0.84 |
| YPL185W | 39.28495564 | 0.7492637 | 0.072594892 | 0.84 |
| YPL186C | 1114.795126 | 0.487741896 | 0.20352784 | 0.84 |
| YPL189C-A | 341.9617668 | 0.36178854 | 0.215773864 | 0.84 |
| YPL198W | 414.9903406 | 0.031119545 | 0.159660668 | 0.84 |
| YPL202C | 824.3595251 | 0.445082874 | 0.08138547 | 0.84 |
| YPL203W | 1453.358861 | 0.535269185 | 0.1485279 | 0.84 |
| YPL208W | 646.0074609 | -0.517925827 | 0.073299972 | 0.84 |
| YPL211W | 668.7255746 | -0.219008055 | 0.171000096 | 0.84 |
| YPL215W | 1126.607522 | -0.55545926 | 0.111538675 | 0.84 |
| YPL218W | 2797.046919 | -0.799807783 | 0.009060475 | 0.84 |
| YPL219W | 632.4863798 | 0.722520585 | 0.035337389 | 0.84 |
| YPL226W | 2793.678288 | -0.269367069 | 0.176405832 | 0.84 |
| YPL227C | 650.9518323 | -0.43777126 | 0.183456702 | 0.84 |
| YPL230W | 2214.598115 | 0.483529431 | 0.095674998 | 0.84 |
| YPL231W | 11637.98709 | -0.300679816 | 0.121220259 | 0.84 |
| YPL234C | 1583.101568 | -0.596841421 | 0.015060266 | 0.84 |
| YPL237W | 2463.364966 | -0.426264631 | 0.138376057 | 0.84 |
| YPL241C | 283.7799123 | -0.545763592 | 0.174564079 | 0.84 |
| YPL247C | 2827.171119 | 0.62293135 | 0.09744918 | 0.84 |
| YPL248C | 793.6864415 | 0.601892417 | 0.10535786 | 0.84 |
| YPL252C | 462.6644596 | -0.485564425 | 0.213479459 | 0.84 |
| YPL256C | 1702.304083 | -0.516956356 | 0.213889664 | 0.84 |
| YPL261C | 48.62531076 | 0.727735899 | 0.066787531 | 0.84 |
| YPL264C | 197.1729292 | 0.42137945 | 0.131703351 | 0.84 |
| YPL267W | 353.8623989 | -0.75713782 | 0.052704793 | 0.84 |
| YPL272C | 344.5733609 | 0.390965944 | 0.133274093 | 0.84 |
| YPR005C | 426.8209175 | 0.773033314 | 0.03395963 | 0.84 |
| YPR007C | 168.1873056 | 0.70515444 | 0.089407154 | 0.84 |
| YPR010C | 3009.126793 | -0.265519061 | 0.148880152 | 0.84 |
| YPR016C | 1938.157861 | -0.578840742 | 0.099755119 | 0.84 |
| YPR017C | 239.3606668 | -0.506765363 | 0.141318993 | 0.84 |

| | | | | |
|---|---|---|---|---|
| YPR019W | 1285.937535 | -0.425070587 | 0.071471333 | 0.84 |
| YPR026W | 1343.846593 | 0.466798548 | 0.173460586 | 0.84 |
| YPR041W | 3343.881383 | -0.477659483 | 0.123092923 | 0.84 |
| YPR044C | 114.0311076 | -0.697341466 | 0.059460537 | 0.84 |
| YPR050C | 1.567602133 | -0.828777834 | 0.160139218 | 0.84 |
| YPR052C | 667.2448711 | -0.856041277 | 0.042555948 | 0.84 |
| YPR056W | 512.7372419 | -0.340299194 | 0.20190243 | 0.84 |
| YPR059C | 12.62828581 | 0.121936289 | 0.055592946 | 0.84 |
| YPR061C | 393.0462601 | 0.570877952 | 0.081888219 | 0.84 |
| YPR064W | 39.20156943 | 0.528715664 | 0.089601688 | 0.84 |
| YPR072W | 2174.009813 | -0.406746857 | 0.168463276 | 0.84 |
| YPR074C | 10075.28978 | -0.220776032 | 0.212539523 | 0.84 |
| YPR077C | 1.201353204 | 0.231544384 | 0.154085776 | 0.84 |
| YPR080W | 5311.40989 | -0.625687415 | 0.087840701 | 0.84 |
| YPR085C | 667.2581144 | 0.485193116 | 0.088802607 | 0.84 |
| YPR091C | 1815.236332 | 0.405243981 | 0.181212346 | 0.84 |
| YPR092W | 28.34006108 | 0.660452591 | 0.037248464 | 0.84 |
| YPR102C | 3154.540675 | -0.416722412 | 0.117272871 | 0.84 |
| YPR106W | 760.0851492 | 0.562521656 | 0.050042361 | 0.84 |
| YPR110C | 2091.025042 | -0.342201062 | 0.170521399 | 0.84 |
| YPR111W | 665.5579675 | 0.493368246 | 0.213885806 | 0.84 |
| YPR114W | 2316.846378 | -0.647010851 | 0.015413286 | 0.84 |
| YPR123C | 99.99461053 | 0.041121064 | 0.107925721 | 0.84 |
| YPR124W | 1891.733227 | -0.164921904 | 0.179694369 | 0.84 |
| YPR132W | 5266.120182 | -0.404729522 | 0.121005085 | 0.84 |
| YPR133W-A | 1114.918643 | -0.571880703 | 0.055969979 | 0.84 |
| YPR142C | 12.52047692 | -0.686289101 | 0.038588801 | 0.84 |
| YPR143W | 537.9356283 | -0.375972655 | 0.140714308 | 0.84 |
| YPR145C-A | 45.3897147 | 0.565724728 | 0.204240511 | 0.84 |
| YPR147C | 688.0952487 | -0.366707428 | 0.172759816 | 0.84 |
| YPR150W | 6.435730199 | 0.441671703 | 0.145942925 | 0.84 |
| YPR151C | 43.40930441 | 0.247135156 | 0.127478578 | 0.84 |
| YPR155C | 883.1515984 | 0.596861745 | 0.102422379 | 0.84 |
| YPR159C-A | 2.271891006 | 0.142413995 | 0.20608724 | 0.84 |
| YPR163C | 5059.100182 | -0.306721909 | 0.20787756 | 0.84 |
| YPR165W | 4589.151851 | -0.607472969 | 0.042039334 | 0.84 |
| YPR166C | 431.6523763 | -1.067909498 | 0.007971101 | 0.84 |
| YPR168W | 170.9858215 | 0.450635215 | 0.186717196 | 0.84 |
| YPR170W-A | 19.17302177 | -0.892158116 | 0.049982455 | 0.84 |
| YPR170W-B | 216.8408172 | -0.496040805 | 0.09009513 | 0.84 |
| YPR174C | 411.2973194 | -0.656565565 | 0.015409882 | 0.84 |
| YPR181C | 4442.448271 | -0.515280763 | 0.053655449 | 0.84 |
| YPR182W | 302.4381877 | -0.443714441 | 0.19893906 | 0.84 |

| | | | | |
|---|---|---|---|---|
| YPR183W | 3494.823853 | -0.605751093 | 0.039541132 | 0.84 |
| YPR185W | 1716.783017 | 0.537307484 | 0.140658771 | 0.84 |
| YPR187W | 1425.65006 | -0.540794938 | 0.084684346 | 0.84 |
| YPR192W | 327.8980894 | 0.133706817 | 0.114675291 | 0.84 |
| YPR200C | 70.27684883 | -0.682529808 | 0.21546362 | 0.84 |
| snR128 | 13.17491061 | 0.417804862 | 0.103120711 | 0.84 |
| snR13 | 24.86462889 | -0.728237655 | 0.151713702 | 0.84 |
| snR17a | 59.62016603 | -0.488510889 | 0.159488962 | 0.84 |
| snR17b | 27.76593566 | -0.243410257 | 0.053263054 | 0.84 |
| snR189 | 9.691874789 | -0.927346965 | 0.074405096 | 0.84 |
| snR19 | 122.3490857 | -0.623754802 | 0.128556472 | 0.84 |
| snR191 | 16.29529748 | -0.481951358 | 0.030288184 | 0.84 |
| snR3 | 32.48912912 | -0.476402712 | 0.031010878 | 0.84 |
| snR30 | 275.4218461 | -0.367673199 | 0.072571312 | 0.84 |
| snR34 | 37.83791355 | -0.14246983 | 0.187049883 | 0.84 |
| snR37 | 153.9250937 | -0.551842738 | 0.095504718 | 0.84 |
| snR4 | 89.67640266 | -0.336814091 | 0.099504233 | 0.84 |
| snR46 | 23.07413489 | -0.495394932 | 0.095763929 | 0.84 |
| snR47 | 5.450594875 | -0.071066312 | 0.176482796 | 0.84 |
| snR49 | 77.03202196 | -0.289684306 | 0.019420269 | 0.84 |
| snR51 | 25.09375178 | 0.447306631 | 0.144292147 | 0.84 |
| snR56 | 6.041353852 | 0.294429704 | 0.124789205 | 0.84 |
| snR63 | 94.54093227 | 0.027601229 | 0.146641783 | 0.84 |
| snR76 | 8.640963018 | -1.060118362 | 0.117442048 | 0.84 |
| snR79 | 6.23839996 | -0.520229034 | 0.18888995 | 0.84 |
| snR80 | 3.471418478 | -0.550156024 | 0.071718497 | 0.84 |
| snR81 | 9.309990788 | -0.887108718 | 0.035626699 | 0.84 |
| snR84 | 162.4042194 | -0.601044883 | 0.183797445 | 0.84 |
| YMR309C | 2879.75717 | -0.43251919 | 0.219057163 | 0.84 |
| YAL019W-A | 5.164115052 | -0.392229213 | 0.222199826 | 0.84 |
| YBL003C | 4258.595501 | -0.497828648 | 0.219623423 | 0.84 |
| YBL100W-C | 6.925265405 | 0.498849123 | 0.224596883 | 0.84 |
| YBR108W | 1821.037124 | 0.471482936 | 0.224871257 | 0.84 |
| YBR159W | 2299.591935 | -0.320359184 | 0.224951484 | 0.84 |
| YBR254C | 311.1708692 | -0.402252801 | 0.221029185 | 0.84 |
| YDL129W | 428.0062351 | 0.450133324 | 0.222500658 | 0.84 |
| YDL216C | 462.7023205 | 0.422812145 | 0.219839126 | 0.84 |
| YDR160W | 879.6911624 | 0.295212864 | 0.221902045 | 0.84 |
| YEL049W | 54.33605347 | 0.306804787 | 0.225747134 | 0.84 |
| YEL063C | 1734.961887 | 0.433824165 | 0.226049825 | 0.84 |
| YER023W | 2339.829047 | -0.41489541 | 0.225272056 | 0.84 |
| YER027C | 1561.298312 | -0.297739804 | 0.224948663 | 0.84 |
| YER028C | 507.0764562 | 0.564062817 | 0.224577107 | 0.84 |

| | | | | |
|---|---|---|---|---|
| YGL061C | 446.8633151 | -0.339326546 | 0.221550286 | 0.84 |
| YGL134W | 814.0702966 | 0.451462989 | 0.2209207 | 0.84 |
| YGL227W | 1564.861142 | 0.490009963 | 0.223006634 | 0.84 |
| YGR060W | 3114.502526 | -0.435531791 | 0.22581659 | 0.84 |
| YGR085C | 1835.900279 | -0.338914685 | 0.226011794 | 0.84 |
| YGR128C | 1414.526792 | -0.272519839 | 0.224438258 | 0.84 |
| YGR139W | 13.99976418 | 1.00182326 | 0.225304208 | 0.84 |
| YGR245C | 1499.530458 | -0.030978327 | 0.219857542 | 0.84 |
| YHL038C | 821.5884324 | 0.354678412 | 0.226007466 | 0.84 |
| YHR038W | 426.523879 | -0.395208015 | 0.222970518 | 0.84 |
| YHR199C | 2007.264009 | 0.375561315 | 0.224301907 | 0.84 |
| YIL028W | 59.74955175 | 0.308624529 | 0.224317878 | 0.84 |
| YIL029C | 174.946717 | 0.38839071 | 0.223630153 | 0.84 |
| YJL190C | 9254.873132 | -0.101879819 | 0.220126765 | 0.84 |
| YJR071W | 37.86358004 | -0.168389147 | 0.221280847 | 0.84 |
| YJR075W | 2038.161429 | -0.319825908 | 0.222919672 | 0.84 |
| YKL011C | 416.7502341 | 0.438798715 | 0.220660283 | 0.84 |
| YKL201C | 1146.760044 | 0.452356473 | 0.22033587 | 0.84 |
| YKR043C | 2568.82772 | -0.044640608 | 0.221273327 | 0.84 |
| YKR090W | 850.9383093 | -0.388772084 | 0.221754165 | 0.84 |
| YKR095W-A | 172.6357846 | -0.63566547 | 0.223248509 | 0.84 |
| YLR053C | 106.3162181 | 0.587540656 | 0.22202564 | 0.84 |
| YLR243W | 616.8482711 | -0.331887011 | 0.223538933 | 0.84 |
| YLR332W | 1136.948721 | 0.40993272 | 0.224938 | 0.84 |
| YLR362W | 1083.342817 | 0.473778899 | 0.223902118 | 0.84 |
| YLR365W | 4.223598425 | 0.615020015 | 0.225069939 | 0.84 |
| YLR399C | 2272.649731 | 0.378913105 | 0.225756641 | 0.84 |
| YLR426W | 420.438408 | -0.405026808 | 0.223690013 | 0.84 |
| YML020W | 864.3038212 | 0.317680524 | 0.222625114 | 0.84 |
| YML096W | 652.0319638 | -0.373623661 | 0.222938438 | 0.84 |
| YML105C | 1352.843987 | -0.364351856 | 0.221861486 | 0.84 |
| YML116W | 562.6593031 | 0.649355571 | 0.22197635 | 0.84 |
| YMR041C | 591.2121868 | 0.25643761 | 0.225006024 | 0.84 |
| YMR199W | 1528.60034 | -0.519697215 | 0.22074376 | 0.84 |
| YNL066W | 1969.929236 | -0.457619764 | 0.221587413 | 0.84 |
| YNL173C | 2504.496471 | 0.543492246 | 0.223392086 | 0.84 |
| YNR068C | 378.9501574 | 0.538105232 | 0.223430883 | 0.84 |
| YOR118W | 1462.615579 | 0.299912295 | 0.223525088 | 0.84 |
| YPL039W | 426.2421359 | 0.386960514 | 0.221695094 | 0.84 |
| snR74 | 1.214578707 | -0.225877919 | 0.223165972 | 0.84 |
| YBR067C | 11273.4348 | 0.282936648 | 0.227926307 | 0.84 |
| YDL095W | 3378.459132 | -0.329186749 | 0.227152308 | 0.84 |
| YER049W | 1494.465582 | -0.30185374 | 0.226532347 | 0.84 |

| | | | | |
|---|---|---|---|---|
| YGL231C | 1251.65783 | -0.371470536 | 0.22657155 | 0.84 |
| YGR037C | 2287.491924 | -0.47981302 | 0.227873005 | 0.84 |
| YJL088W | 187.9199165 | 0.476779038 | 0.228181523 | 0.84 |
| YJL130C | 21427.36188 | -0.374203284 | 0.227698667 | 0.84 |
| YKL023C-A | 532.0907152 | -0.315720498 | 0.227411995 | 0.84 |
| YLL002W | 366.0264394 | -0.422891039 | 0.226464036 | 0.84 |
| YLR347C | 1405.641169 | -0.311593958 | 0.227030519 | 0.84 |
| YLR392C | 638.9786435 | 0.430620576 | 0.228018367 | 0.84 |
| YMR129W | 1909.721365 | -0.285772758 | 0.228087845 | 0.84 |
| YOR349W | 572.160655 | 0.332121739 | 0.227212466 | 0.84 |
| YPL244C | 876.1384791 | -0.403328931 | 0.228147295 | 0.84 |
| YPL265W | 2999.757451 | 0.468080583 | 0.227752451 | 0.84 |
| YPL266W | 1132.936461 | -0.092477727 | 0.227910869 | 0.84 |
| YDR322W | 1031.616879 | -0.475424018 | 0.228479296 | 0.84 |
| YDR053W | 7.370760006 | 0.195832427 | 0.22863326 | 0.84 |
| YJL014W | 2604.88629 | -0.339737026 | 0.22883329 | 0.84 |
| YNR020C | 315.5485889 | -0.459791584 | 0.22937022 | 0.84 |
| YHR149C | 909.9713222 | -0.488988684 | 0.229521836 | 0.84 |
| YPL001W | 324.2471356 | -0.417939135 | 0.229643254 | 0.84 |
| YDL193W | 1212.895326 | 0.340667363 | 0.22986594 | 0.84 |
| YDR299W | 1150.958917 | -0.416747717 | 0.229956601 | 0.84 |
| YNL048W | 1078.322808 | -0.316222347 | 0.230044693 | 0.84 |
| YGR019W | 2682.062979 | 0.656448454 | 0.230331477 | 0.84 |
| YHR157W | 84.3613017 | 0.379412291 | 0.231037155 | 0.84 |
| YLR021W | 410.9321978 | -0.349079785 | 0.230998672 | 0.84 |
| YBL101C | 1814.259973 | 0.477353691 | 0.233655129 | 0.84 |
| YBR144C | 8.772859821 | 0.650168496 | 0.231879729 | 0.84 |
| YBR223C | 483.3688068 | 0.45996022 | 0.231741554 | 0.84 |
| YBR288C | 774.8863305 | -0.355646624 | 0.234055688 | 0.84 |
| YCL042W | 285.5413665 | 0.547714524 | 0.232334926 | 0.84 |
| YCR079W | 984.6018828 | 0.362932379 | 0.235826069 | 0.84 |
| YDL029W | 3675.857405 | -0.233381976 | 0.235769948 | 0.84 |
| YDR295C | 798.8722941 | 0.296381425 | 0.235551303 | 0.84 |
| YDR457W | 5588.916733 | -0.319290768 | 0.235947931 | 0.84 |
| YEL051W | 3229.319305 | -0.308808898 | 0.234611276 | 0.84 |
| YER019C-A | 1962.324665 | -0.368661712 | 0.235798878 | 0.84 |
| YER144C | 1434.76784 | 0.460638672 | 0.231417978 | 0.84 |
| YFR023W | 96.03552642 | 0.49480953 | 0.232608577 | 0.84 |
| YGL171W | 777.7803509 | -0.23928071 | 0.232835449 | 0.84 |
| YGL201C | 994.5170532 | -0.37978357 | 0.234945887 | 0.84 |
| YGR159C | 2044.642627 | -0.112235179 | 0.233334325 | 0.84 |
| YGR234W | 5131.339655 | 0.145905472 | 0.23602322 | 0.84 |
| YGR289C | 671.1201555 | 0.263697892 | 0.235196717 | 0.84 |

| | | | | |
|---|---|---|---|---|
| YHR066W | 786.1240577 | -0.204867639 | 0.2355354 | 0.84 |
| YIR021W | 261.891268 | -0.435852296 | 0.232705045 | 0.84 |
| YJR059W | 3492.702056 | 0.438559227 | 0.236143329 | 0.84 |
| YJR139C | 4827.24467 | -0.216141763 | 0.232121428 | 0.84 |
| YKR067W | 2101.266331 | 0.559795537 | 0.233366461 | 0.84 |
| YKR072C | 2109.757496 | 0.477216853 | 0.236147021 | 0.84 |
| YLR390W | 418.8392228 | -0.405504038 | 0.232740654 | 0.84 |
| YLR437C | 329.593239 | -0.447928125 | 0.233772133 | 0.84 |
| YLR447C | 5880.017489 | -0.284863792 | 0.235442947 | 0.84 |
| YML028W | 27162.20889 | -0.371978621 | 0.234673937 | 0.84 |
| YML121W | 941.5006143 | 0.378184736 | 0.234742361 | 0.84 |
| YNL030W | 3372.305745 | -0.450162499 | 0.235404724 | 0.84 |
| YNL121C | 5779.129433 | -0.341808757 | 0.233271837 | 0.84 |
| YNL307C | 2792.714562 | -0.393696013 | 0.232620074 | 0.84 |
| YOL010W | 524.4137956 | -0.487908591 | 0.234547581 | 0.84 |
| YOL013W-A | 5.152321446 | -1.074546089 | 0.235885335 | 0.84 |
| YOL154W | 197.7584736 | 0.346687617 | 0.236141731 | 0.84 |
| YOR098C | 2501.292329 | -0.337060082 | 0.234183397 | 0.84 |
| YPL212C | 1188.64058 | -0.201898355 | 0.235697303 | 0.84 |
| snR32 | 25.20381189 | -0.641086149 | 0.235596841 | 0.84 |
| YCR044C | 1029.447304 | -0.39894695 | 0.236494217 | 0.84 |
| YNL237W | 756.7694566 | 0.372194534 | 0.236528517 | 0.84 |
| YEL022W | 2372.854744 | -0.33270964 | 0.237172481 | 0.84 |
| YJL187C | 1084.409994 | -0.358036454 | 0.237138442 | 0.84 |
| YAL031C | 1097.389337 | 0.35738402 | 0.243189792 | 0.84 |
| YAL051W | 1222.388192 | 0.375107736 | 0.245546544 | 0.84 |
| YBR106W | 4289.033597 | -0.161557562 | 0.239087158 | 0.84 |
| YBR242W | 549.7068913 | -0.362791899 | 0.241662069 | 0.84 |
| YCR020C | 245.5464449 | 0.225599827 | 0.241683911 | 0.84 |
| YCR052W | 1334.780982 | -0.336558724 | 0.244798973 | 0.84 |
| YCR095W-A | 20.07196858 | 0.233336679 | 0.245632498 | 0.84 |
| YDL042C | 548.3117902 | -0.361027446 | 0.240970615 | 0.84 |
| YDL045C | 1046.513314 | 0.273567702 | 0.240345783 | 0.84 |
| YDR171W | 12610.16854 | 0.513898213 | 0.244560316 | 0.84 |
| YDR210W-B | 26.10421514 | 0.386052702 | 0.239628072 | 0.84 |
| YDR222W | 466.2824915 | 0.458157571 | 0.24502152 | 0.84 |
| YDR340W | 4.60165433 | -0.656592899 | 0.24518973 | 0.84 |
| YER002W | 1431.744689 | -0.345975608 | 0.237782173 | 0.84 |
| YER069W | 925.6063303 | 0.484322153 | 0.237774738 | 0.84 |
| YER112W | 1280.704105 | -0.35606096 | 0.243556601 | 0.84 |
| YER145C-A | 50.64924556 | -0.164259426 | 0.238571062 | 0.84 |
| YER148W | 733.3492834 | -0.508157546 | 0.242891077 | 0.84 |
| YER187W | 132.2577032 | -0.997823518 | 0.23875647 | 0.84 |

| | | | | |
|---|---|---|---|---|
| YGL135W | 2224.916712 | -0.167859966 | 0.241150506 | 0.84 |
| YGL200C | 3717.842992 | -0.282202216 | 0.243438459 | 0.84 |
| YGR004W | 331.5972983 | 0.401024148 | 0.23948316 | 0.84 |
| YGR061C | 5651.499061 | -0.115646306 | 0.24576557 | 0.84 |
| YGR161C | 2432.867838 | 0.304540823 | 0.238685387 | 0.84 |
| YHL032C | 2561.780761 | 0.425607152 | 0.241693906 | 0.84 |
| YHR043C | 1297.179518 | -0.364959714 | 0.24201188 | 0.84 |
| YHR172W | 514.8081565 | -0.343209207 | 0.242793048 | 0.84 |
| YIL031W | 1396.422579 | 0.391274208 | 0.241722731 | 0.84 |
| YIL056W | 1630.203256 | 0.320313483 | 0.243616058 | 0.84 |
| YIL142C-A | 7.771319082 | 0.357698127 | 0.243833792 | 0.84 |
| YJL062W | 1004.118516 | -0.279157954 | 0.24366022 | 0.84 |
| YJL077C | 162.6090211 | 0.399108327 | 0.241364997 | 0.84 |
| YJL099W | 747.8275315 | 0.360432247 | 0.239018883 | 0.84 |
| YJL145W | 1081.572718 | -0.309202361 | 0.243970763 | 0.84 |
| YKL161C | 387.0663817 | 0.141913602 | 0.238025987 | 0.84 |
| YKL164C | 7734.898504 | -0.265128351 | 0.245346079 | 0.84 |
| YLL060C | 550.1729711 | 0.230573715 | 0.238764391 | 0.84 |
| YLR075W | 9637.869009 | -0.100465344 | 0.237679882 | 0.84 |
| YLR092W | 690.3329885 | 0.363122016 | 0.241543614 | 0.84 |
| YLR124W | 42.32116711 | 0.546260892 | 0.238258121 | 0.84 |
| YLR198C | 85.68816582 | -0.352523965 | 0.245212895 | 0.84 |
| YLR318W | 479.1884801 | 0.283502932 | 0.244782876 | 0.84 |
| YML048W | 4451.705992 | -0.348780615 | 0.240048074 | 0.84 |
| YML082W | 766.1872079 | -0.363063697 | 0.244034618 | 0.84 |
| YMR014W | 1014.309235 | -0.32962618 | 0.238517586 | 0.84 |
| YMR059W | 205.0702465 | -0.269074059 | 0.241643857 | 0.84 |
| YMR101C | 112.86694 | 0.692506127 | 0.244061735 | 0.84 |
| YMR279C | 707.3104958 | 0.355819455 | 0.238652986 | 0.84 |
| YNL014W | 784.7565686 | 0.54400814 | 0.241140742 | 0.84 |
| YNL074C | 2355.063484 | 0.353895814 | 0.242969278 | 0.84 |
| YNL095C | 702.5523068 | 0.365884701 | 0.245141754 | 0.84 |
| YNL175C | 613.6716753 | -0.238091328 | 0.245777279 | 0.84 |
| YNL244C | 2167.186944 | -0.491665154 | 0.243879305 | 0.84 |
| YNL259C | 262.1284302 | -0.284018629 | 0.243313974 | 0.84 |
| YNR009W | 344.9832981 | -0.474682134 | 0.243955093 | 0.84 |
| YOL011W | 1428.931177 | 0.292825078 | 0.243642173 | 0.84 |
| YOL021C | 1670.215668 | -0.332160911 | 0.240736526 | 0.84 |
| YOL055C | 533.3864453 | 0.363308341 | 0.243386572 | 0.84 |
| YOL082W | 2476.571718 | 0.36530317 | 0.239409222 | 0.84 |
| YOL086C | 59663.42444 | -0.253881869 | 0.240841285 | 0.84 |
| YOR032W-A | 11.5403846 | -0.42833965 | 0.244981192 | 0.84 |
| YOR036W | 1460.876014 | 0.332380075 | 0.242497507 | 0.84 |

| | | | | |
|---|---|---|---|---|
| YPL029W | 470.8569792 | -0.350906738 | 0.239444459 | 0.84 |
| YPL086C | 1089.007189 | -0.235841295 | 0.241861841 | 0.84 |
| YPL206C | 1410.718758 | -0.461079832 | 0.245465349 | 0.84 |
| YPL243W | 2085.694288 | -0.432552917 | 0.244621813 | 0.84 |
| YBR168W | 396.8925406 | 0.334545689 | 0.246675788 | 0.84 |
| YDR104C | 711.7535955 | 0.409795814 | 0.246756875 | 0.84 |
| YGR196C | 1285.850548 | 0.295224997 | 0.24626484 | 0.84 |
| YHR085W | 335.931842 | -0.403309911 | 0.246011697 | 0.84 |
| YJL140W | 1600.235301 | -0.337115856 | 0.246279202 | 0.84 |
| YKL051W | 1194.148337 | 0.402131324 | 0.246896254 | 0.84 |
| YKL091C | 918.9800837 | 0.457111199 | 0.246833344 | 0.84 |
| YKL127W | 1973.346678 | -0.050365424 | 0.247060539 | 0.84 |
| YLL018C | 8408.142457 | -0.221429143 | 0.247144961 | 0.84 |
| YLR322W | 7.310944976 | -0.105506693 | 0.247264055 | 0.84 |
| YMR215W | 1997.531105 | -0.394191086 | 0.246546945 | 0.84 |
| YPR157W | 448.7836668 | 0.642524664 | 0.247394802 | 0.84 |
| YBR065C | 528.4307138 | 0.461389738 | 0.250549909 | 0.84 |
| YBR220C | 693.632069 | -0.30682318 | 0.247719709 | 0.84 |
| YCR047C | 558.0855162 | -0.324022923 | 0.254019681 | 0.84 |
| YDL048C | 3537.203448 | 0.446040123 | 0.247914012 | 0.84 |
| YDL074C | 1505.723255 | -0.490183003 | 0.252756812 | 0.84 |
| YDR051C | 1050.532227 | 0.292314122 | 0.253312409 | 0.84 |
| YDR120C | 1240.636645 | -0.163508506 | 0.251224551 | 0.84 |
| YDR544C | 2.336721419 | 0.934722394 | 0.249657128 | 0.84 |
| YEL003W | 487.0478849 | -0.420000763 | 0.252483123 | 0.84 |
| YEL050W-A | 71.35641565 | -0.607070385 | 0.250010106 | 0.84 |
| YER042W | 653.285425 | 0.708886969 | 0.25292935 | 0.84 |
| YER087C-A | 36.6678806 | -0.565074209 | 0.252904327 | 0.84 |
| YFL013W-A | 84.24870204 | 0.459560039 | 0.247912163 | 0.84 |
| YFR051C | 3059.620652 | -0.30875608 | 0.253483223 | 0.84 |
| YGL133W | 1877.257774 | 0.329991464 | 0.251035206 | 0.84 |
| YGL250W | 555.5711852 | 0.3821765 | 0.253291589 | 0.84 |
| YGR288W | 284.2488249 | 0.423978776 | 0.248353288 | 0.84 |
| YIL044C | 893.1546159 | -0.396458639 | 0.251198526 | 0.84 |
| YIR009W | 172.3613335 | 0.357780357 | 0.2534398 | 0.84 |
| YIR026C | 772.7201823 | -0.100475132 | 0.250790017 | 0.84 |
| YJR096W | 2414.849421 | 0.420207057 | 0.251790804 | 0.84 |
| YJR126C | 1827.592541 | 0.319319284 | 0.253582946 | 0.84 |
| YKL109W | 7924.096762 | 0.490614118 | 0.248999481 | 0.84 |
| YKL181W | 2705.955695 | -0.35556411 | 0.25282898 | 0.84 |
| YKL220C | 368.1832027 | 0.251856928 | 0.250340933 | 0.84 |
| YKR010C | 516.9970849 | -0.369563101 | 0.249658303 | 0.84 |
| YKR064W | 1141.244604 | 0.303739726 | 0.252290598 | 0.84 |

| | | | | |
|---|---|---|---|---|
| YLR007W | 741.9239686 | 0.398467967 | 0.252171562 | 0.84 |
| YLR103C | 462.3096359 | -0.432320745 | 0.248830566 | 0.84 |
| YLR307C-A | 38.91718273 | 0.020086792 | 0.253845314 | 0.84 |
| YLR384C | 2550.492331 | -0.253170657 | 0.252328788 | 0.84 |
| YML068W | 512.1804017 | 0.279726088 | 0.250513454 | 0.84 |
| YML083C | 177.180785 | 0.24172749 | 0.250549324 | 0.84 |
| YMR030W | 574.6177195 | 0.388983307 | 0.252478546 | 0.84 |
| YMR291W | 2090.324495 | 0.3602961 | 0.248987691 | 0.84 |
| YMR320W | 113.0593369 | 0.239826242 | 0.251854701 | 0.84 |
| YNL101W | 1154.001663 | 0.437472158 | 0.253643581 | 0.84 |
| YNL289W | 446.6126721 | -0.634300829 | 0.250187312 | 0.84 |
| YNR005C | 1.56429782 | 0.231256015 | 0.25138945 | 0.84 |
| YNR060W | 291.930447 | 0.378354849 | 0.249720206 | 0.84 |
| YNR069C | 436.7652166 | 0.500577989 | 0.252051432 | 0.84 |
| YOL016C | 3937.741808 | 0.459352967 | 0.251624478 | 0.84 |
| YOL022C | 1058.507084 | -0.245000859 | 0.251422221 | 0.84 |
| YOL100W | 1483.492555 | 0.394383914 | 0.248108546 | 0.84 |
| YOL103W-B | 37.1755698 | 0.025946785 | 0.248135895 | 0.84 |
| YOL147C | 576.9213691 | -0.333769241 | 0.25401491 | 0.84 |
| YOR167C | 4868.170026 | -0.343491728 | 0.25075501 | 0.84 |
| YPL067C | 285.3555196 | -0.408124272 | 0.249018231 | 0.84 |
| YPL112C | 806.8455711 | -0.354984735 | 0.24977564 | 0.84 |
| YPL249C-A | 77.54661876 | -0.532791952 | 0.252862316 | 0.84 |
| snR31 | 65.06005548 | -0.409484597 | 0.251291384 | 0.84 |
| YBR034C | 1111.628509 | -0.285504724 | 0.25758512 | 0.84 |
| YBR162C | 7471.337212 | -0.288527732 | 0.25842779 | 0.84 |
| YBR162W-A | 580.1318829 | -0.427371036 | 0.257430711 | 0.84 |
| YBR214W | 5062.049562 | 0.463268448 | 0.259627929 | 0.84 |
| YCL063W | 606.1625178 | -0.419923781 | 0.258570215 | 0.84 |
| YCR043C | 438.4206899 | -0.387942323 | 0.257433805 | 0.84 |
| YDL050C | 34.4743764 | -0.364527782 | 0.258236037 | 0.84 |
| YDL070W | 4583.373904 | 0.491895161 | 0.25718442 | 0.84 |
| YDL235C | 741.3905909 | -0.362420598 | 0.259799781 | 0.84 |
| YDR202C | 884.8533036 | 0.33928147 | 0.25970926 | 0.84 |
| YDR509W | 13.68760112 | 0.396762211 | 0.255695028 | 0.84 |
| YEL017W | 1022.681916 | -0.172031019 | 0.258651807 | 0.84 |
| YER053C | 3873.936022 | 0.465491223 | 0.256625882 | 0.84 |
| YGR035C | 329.4527926 | -1.000664552 | 0.257965586 | 0.84 |
| YGR103W | 3122.539574 | 0.033910839 | 0.258859891 | 0.84 |
| YGR179C | 503.8137475 | -0.3036763 | 0.256758518 | 0.84 |
| YIL097W | 1123.344615 | 0.354460571 | 0.259474708 | 0.84 |
| YJL161W | 501.2537696 | 0.296295341 | 0.257352668 | 0.84 |
| YJL185C | 564.2478549 | 0.370888014 | 0.25846252 | 0.84 |

| | | | | |
|---|---|---|---|---|
| YJR032W | 585.2468444 | -0.284854748 | 0.254229274 | 0.84 |
| YJR038C | 33.2004855 | 0.437958411 | 0.259755882 | 0.84 |
| YJR108W | 40.18860279 | 0.268226471 | 0.2599298 | 0.84 |
| YKL046C | 1814.036225 | -0.292774257 | 0.257247735 | 0.84 |
| YKL140W | 1755.744554 | 0.305431821 | 0.25964709 | 0.84 |
| YKL212W | 2578.339284 | -0.306812027 | 0.256458365 | 0.84 |
| YLL023C | 2955.471664 | 0.510577094 | 0.255504022 | 0.84 |
| YLL028W | 724.5266873 | -0.451868192 | 0.256057607 | 0.84 |
| YLR045C | 1304.494239 | -0.268905672 | 0.256526467 | 0.84 |
| YLR151C | 483.8723388 | 0.299917019 | 0.260101859 | 0.84 |
| YLR172C | 1976.093187 | -0.181139006 | 0.259359692 | 0.84 |
| YLR230W | 287.1818195 | -0.286772257 | 0.259337322 | 0.84 |
| YML076C | 1190.303539 | 0.382516256 | 0.255539773 | 0.84 |
| YML077W | 412.7293887 | -0.424958762 | 0.255673444 | 0.84 |
| YMR230W-A | 4.038623613 | -0.123186547 | 0.259893557 | 0.84 |
| YMR316C-A | 33.48519679 | 0.176188125 | 0.258103128 | 0.84 |
| YNL042W | 593.1711026 | 0.284868795 | 0.258160793 | 0.84 |
| YNL189W | 2792.970706 | -0.316027691 | 0.254646801 | 0.84 |
| YNL209W | 13780.76239 | -0.081255609 | 0.259739697 | 0.84 |
| YOR012W | 60.32598792 | 0.269494312 | 0.257506453 | 0.84 |
| YOR066W | 1115.721778 | 0.29543939 | 0.255632077 | 0.84 |
| YOR069W | 1035.064237 | 0.261724508 | 0.260189141 | 0.84 |
| YOR267C | 3236.118357 | 0.397506601 | 0.255310044 | 0.84 |
| YOR293C-A | 2.608736587 | -0.253520915 | 0.254871733 | 0.84 |
| YPL081W | 1016.551966 | -0.377978784 | 0.255235361 | 0.84 |
| YPL184C | 1810.247002 | -0.324378383 | 0.2572816 | 0.84 |
| YPL220W | 1761.369957 | -0.199304854 | 0.256633055 | 0.84 |
| YPL251W | 76.6529722 | -0.573326846 | 0.258377223 | 0.84 |
| YKL036C | 247.9148773 | 0.437912675 | 0.260527538 | 0.84 |
| YNL277W | 546.4771573 | 0.29918235 | 0.260507223 | 0.84 |
| YAL002W | 1569.483592 | 0.307634698 | 0.263006296 | 0.85 |
| YBR102C | 889.0122882 | -0.259694166 | 0.262208196 | 0.85 |
| YDR242W | 619.518258 | 0.349610922 | 0.262815415 | 0.85 |
| YDR279W | 575.5796481 | -0.311889393 | 0.262472954 | 0.85 |
| YER149C | 245.5366344 | -0.232137258 | 0.263086112 | 0.85 |
| YGR238C | 880.0595867 | 0.407283439 | 0.26367253 | 0.85 |
| YIL147C | 2000.977172 | 0.440842906 | 0.26248352 | 0.85 |
| YIR044C | 6.331727114 | -0.551596945 | 0.263542795 | 0.85 |
| YKR035C | 23.50975344 | 0.143008425 | 0.263391094 | 0.85 |
| YMR004W | 1474.539468 | 0.327752128 | 0.263548019 | 0.85 |
| YMR123W | 1008.572929 | -0.18739787 | 0.262475063 | 0.85 |
| YMR155W | 707.9678984 | 0.347737462 | 0.262925647 | 0.85 |
| YOR223W | 687.6095973 | 0.305458829 | 0.263469083 | 0.85 |

| | | | | |
|---|---|---|---|---|
| YOR354C | 1688.06875 | -0.307882446 | 0.262747362 | 0.85 |
| YBL103C | 858.4740089 | 0.379639501 | 0.263844277 | 0.85 |
| YHR007C-A | 61.56889406 | 0.20250538 | 0.264174427 | 0.85 |
| YOR297C | 445.9780063 | -0.351868183 | 0.264086005 | 0.85 |
| YDR144C | 842.4657292 | -0.180769512 | 0.264716169 | 0.85 |
| YDR212W | 2389.189098 | -0.377405211 | 0.264819376 | 0.85 |
| YJR112W | 374.389265 | -0.35169033 | 0.264757178 | 0.85 |
| YPL131W | 24700.79965 | -0.124418884 | 0.264920419 | 0.85 |
| YDL114W-A | 24.54000287 | 0.465623366 | 0.265962928 | 0.85 |
| YFL009W | 860.3502379 | 0.284900459 | 0.265682195 | 0.85 |
| YGR194C | 2007.303758 | 0.458280118 | 0.265447159 | 0.85 |
| YHR099W | 4737.26123 | -0.28402004 | 0.266232785 | 0.85 |
| YJR137C | 1214.839856 | 0.392353432 | 0.266100278 | 0.85 |
| YKL072W | 554.6822525 | 0.323775193 | 0.265892955 | 0.85 |
| YKR001C | 4012.227533 | -0.27535081 | 0.265526295 | 0.85 |
| YMR228W | 316.010038 | -0.362262188 | 0.265383055 | 0.85 |
| YNL331C | 347.7391832 | 0.286026475 | 0.266193309 | 0.85 |
| YOR271C | 1520.089436 | -0.074552188 | 0.265164898 | 0.85 |
| YBL019W | 515.3822708 | 0.384216797 | 0.268127224 | 0.85 |
| YBR275C | 1768.794249 | -0.249856051 | 0.266502069 | 0.85 |
| YDL043C | 369.3400281 | -0.441355848 | 0.268010883 | 0.85 |
| YDR181C | 744.5427058 | 0.469647826 | 0.267541648 | 0.85 |
| YDR262W | 1927.726265 | 0.300081802 | 0.268144137 | 0.85 |
| YDR465C | 1088.193625 | -0.079348646 | 0.268912403 | 0.85 |
| YDR526C | 4.767823074 | 1.389689808 | 0.268519546 | 0.85 |
| YER077C | 789.2511196 | 0.24679934 | 0.266692457 | 0.85 |
| YFL005W | 2091.331245 | -0.370341291 | 0.268644425 | 0.85 |
| YFR014C | 1739.196956 | 0.31982485 | 0.267047005 | 0.85 |
| YHR042W | 4638.852766 | -0.230943576 | 0.268363108 | 0.85 |
| YIL106W | 893.3105346 | -0.367551036 | 0.268301502 | 0.85 |
| YJL157C | 1263.741636 | -0.61545835 | 0.268167542 | 0.85 |
| YLR022C | 541.692103 | -0.389652894 | 0.267095422 | 0.85 |
| YLR418C | 850.5544851 | -0.340414778 | 0.267608235 | 0.85 |
| YLR435W | 440.7011003 | -0.177551054 | 0.268821459 | 0.85 |
| YOL093W | 662.3279101 | -0.38388168 | 0.268887498 | 0.85 |
| YOR048C | 2248.629889 | -0.258189371 | 0.266833303 | 0.85 |
| YPL118W | 2044.672532 | -0.415574308 | 0.268843819 | 0.85 |
| YPR176C | 535.2741493 | -0.392876064 | 0.266629215 | 0.85 |
| YBR148W | 178.9686946 | 0.353261802 | 0.269457344 | 0.85 |
| YDR379W | 1429.073535 | 0.371270606 | 0.269382144 | 0.85 |
| YGL027C | 1397.054052 | -0.250072422 | 0.269628702 | 0.85 |
| YNL130C | 1326.028586 | -0.266409995 | 0.269799226 | 0.85 |
| YBR250W | 80.81966717 | 0.484193533 | 0.270463607 | 0.85 |

| | | | | |
|---|---|---|---|---|
| YCL028W | 2867.256977 | -0.309519983 | 0.270348126 | 0.85 |
| YDR090C | 473.3492687 | 0.356861418 | 0.270044115 | 0.85 |
| YNR061C | 596.8560089 | -0.331337955 | 0.270201831 | 0.85 |
| YOL089C | 836.6158144 | 0.314224873 | 0.270224517 | 0.85 |
| YCL029C | 771.878883 | -0.302910779 | 0.271910843 | 0.85 |
| YDR167W | 803.2507476 | -0.320217761 | 0.271114381 | 0.85 |
| YHR079C-A | 27.69347479 | 0.627268872 | 0.271146115 | 0.85 |
| YIR010W | 867.0379525 | 0.232605951 | 0.27159225 | 0.85 |
| YJL012C | 2507.923216 | -0.382817206 | 0.271324116 | 0.85 |
| YLL052C | 77.86079202 | 0.156453678 | 0.27172884 | 0.85 |
| YMR159C | 22.87110149 | 0.381841401 | 0.271829011 | 0.85 |
| YNL005C | 1578.642517 | -0.336350374 | 0.271250036 | 0.85 |
| YNL319W | 5.59444275 | -0.571795064 | 0.271150836 | 0.85 |
| YOL043C | 577.8464057 | 0.304470518 | 0.271597057 | 0.85 |
| YER092W | 378.6245953 | -0.322647238 | 0.272316609 | 0.85 |
| YHR190W | 5071.299505 | 0.446142429 | 0.272281693 | 0.85 |
| YJR077C | 4794.775519 | -0.116490065 | 0.272170624 | 0.85 |
| YAR007C | 1627.673812 | -0.343523881 | 0.273875877 | 0.85 |
| YBR201C-A | 201.3285883 | 0.288010121 | 0.273626187 | 0.85 |
| YBR267W | 453.4917639 | -0.432811834 | 0.274216347 | 0.85 |
| YDL016C | 38.72923011 | -0.575225341 | 0.273393924 | 0.85 |
| YGR160W | 19.24237613 | -0.035777236 | 0.27430823 | 0.85 |
| YGR282C | 11912.38121 | -0.304658628 | 0.273225844 | 0.85 |
| YHR094C | 525.1336673 | 0.463487 | 0.274352563 | 0.85 |
| YLL057C | 252.3608873 | 0.35154803 | 0.274303322 | 0.85 |
| YLR241W | 1618.382874 | 0.344411223 | 0.274068839 | 0.85 |
| YMR036C | 631.5049431 | 0.320575354 | 0.273894867 | 0.85 |
| YOR209C | 3422.936842 | -0.351097685 | 0.272768622 | 0.85 |
| YPL093W | 1935.182804 | -0.31953768 | 0.274394083 | 0.85 |
| YPL173W | 902.2046084 | -0.404140061 | 0.274435753 | 0.85 |
| YPR125W | 636.6611003 | -0.364727458 | 0.273497595 | 0.85 |
| YNL132W | 1681.161762 | -0.055087147 | 0.274605655 | 0.85 |
| YPL188W | 716.8167679 | 0.411300603 | 0.274739636 | 0.85 |
| YDL094C | 21.86703582 | 0.598859125 | 0.27545596 | 0.85 |
| YDR385W | 1851.31364 | -0.257075762 | 0.275303899 | 0.85 |
| YLR368W | 767.6389144 | 0.355423527 | 0.275488788 | 0.85 |
| YNL206C | 942.6169972 | -0.291024859 | 0.276697938 | 0.86 |
| NME1 | 51.73748825 | 0.018845265 | 0.2844926 | 0.86 |
| YCR041W | 16.4079087 | -0.410770264 | 0.278425774 | 0.86 |
| YCR094W | 731.6068684 | -0.291948149 | 0.278171034 | 0.86 |
| YDL022C-A | 6.814086928 | 0.565341498 | 0.279439321 | 0.86 |
| YDL049C | 168.7509482 | 0.326296565 | 0.280012281 | 0.86 |
| YDL232W | 292.6096052 | -0.402444537 | 0.282883594 | 0.86 |

| | | | | |
|---|---|---|---|---|
| YDR238C | 4318.542325 | -0.233565732 | 0.282342167 | 0.86 |
| YDR254W | 574.0903535 | 0.287201318 | 0.285012473 | 0.86 |
| YDR310C | 1697.719061 | -0.29753149 | 0.284347747 | 0.86 |
| YDR338C | 695.8448786 | 0.334672157 | 0.278105538 | 0.86 |
| YDR360W | 36.19514521 | -0.308090271 | 0.284654816 | 0.86 |
| YDR496C | 1668.42234 | -0.250593171 | 0.281206528 | 0.86 |
| YEL005C | 730.8016186 | 0.239156362 | 0.283857753 | 0.86 |
| YEL025C | 1231.625482 | 0.381624908 | 0.281708567 | 0.86 |
| YEL059W | 50.35325287 | -0.016274532 | 0.283115727 | 0.86 |
| YER020W | 2824.873338 | 0.396918307 | 0.282866993 | 0.86 |
| YER150W | 3467.414047 | 0.045740953 | 0.282832434 | 0.86 |
| YFL001W | 478.2892235 | -0.305447387 | 0.284205574 | 0.86 |
| YGL066W | 1160.200958 | 0.318994944 | 0.279825197 | 0.86 |
| YGL096W | 785.2336716 | 0.26570152 | 0.283869842 | 0.86 |
| YGR063C | 116.1850697 | -0.441679364 | 0.277894225 | 0.86 |
| YGR118W | 4349.282861 | -0.359067856 | 0.278163094 | 0.86 |
| YHL023C | 1253.971871 | 0.315866663 | 0.278701159 | 0.86 |
| YHR175W | 775.2689431 | -0.205325388 | 0.284800834 | 0.86 |
| YIL007C | 541.263808 | 0.306489655 | 0.280896481 | 0.86 |
| YIL045W | 1955.748951 | 0.246345568 | 0.282274987 | 0.86 |
| YIL065C | 779.4474724 | 0.418857003 | 0.28295849 | 0.86 |
| YJL034W | 12765.00653 | -0.274127014 | 0.281654222 | 0.86 |
| YJL109C | 2576.133571 | -0.330708363 | 0.283163492 | 0.86 |
| YJL192C | 2669.267366 | -0.175185301 | 0.282415636 | 0.86 |
| YJR017C | 953.2894932 | -0.345353759 | 0.278553045 | 0.86 |
| YJR033C | 1583.049905 | 0.324203014 | 0.283815363 | 0.86 |
| YJR098C | 316.068717 | 0.443977452 | 0.277364519 | 0.86 |
| YJR105W | 3472.841224 | -0.179475824 | 0.284947636 | 0.86 |
| YKL154W | 669.8187345 | -0.230003197 | 0.279750129 | 0.86 |
| YLL003W | 928.3757705 | 0.247618711 | 0.278819737 | 0.86 |
| YLL054C | 624.0988471 | -0.30068095 | 0.280605014 | 0.86 |
| YLR209C | 881.8739647 | -0.195332794 | 0.283917418 | 0.86 |
| YLR225C | 1476.977135 | 0.314339825 | 0.284875148 | 0.86 |
| YLR285W | 889.692098 | -0.271078083 | 0.278430451 | 0.86 |
| YLR343W | 1249.632226 | 0.317791708 | 0.2802703 | 0.86 |
| YMR149W | 2739.886665 | -0.360435891 | 0.282128263 | 0.86 |
| YMR186W | 31490.05511 | -0.070022611 | 0.279709109 | 0.86 |
| YMR217W | 5540.397432 | -0.006251073 | 0.284146047 | 0.86 |
| YMR263W | 652.3046851 | 0.377468704 | 0.283081975 | 0.86 |
| YNL007C | 10095.98848 | -0.3677446 | 0.280005247 | 0.86 |
| YNL151C | 742.0050839 | -0.320963945 | 0.282433013 | 0.86 |
| YNL248C | 1500.899157 | -0.182892098 | 0.279860615 | 0.86 |
| YNL293W | 734.6319527 | 0.376847934 | 0.282561368 | 0.86 |

| | | | | |
|---|---|---|---|---|
| YNR055C | 1151.83484 | 0.319196642 | 0.279363048 | 0.86 |
| YOL009C | 380.9976744 | 0.313570809 | 0.281425743 | 0.86 |
| YOL044W | 473.6470695 | 0.422198252 | 0.283848893 | 0.86 |
| YOL046C | 5.940874887 | 0.090745817 | 0.282040605 | 0.86 |
| YOL103W | 1490.877623 | -0.27982344 | 0.280048313 | 0.86 |
| YOR086C | 3224.270398 | 0.361116612 | 0.27883181 | 0.86 |
| YOR128C | 2990.654925 | -0.403614039 | 0.278480226 | 0.86 |
| YOR158W | 1525.323441 | -0.354866943 | 0.280103986 | 0.86 |
| YOR199W | 28.95971662 | -0.002444643 | 0.279657598 | 0.86 |
| YOR221C | 1523.763907 | 0.366910557 | 0.284276085 | 0.86 |
| YOR272W | 1204.374523 | -0.191059335 | 0.281689939 | 0.86 |
| YOR342C | 1114.452608 | -0.339593211 | 0.285070748 | 0.86 |
| YPL076W | 380.2952376 | 0.321905365 | 0.277618611 | 0.86 |
| YPR178W | 455.7725824 | 0.328106801 | 0.283699429 | 0.86 |
| YDR014W | 486.9454278 | 0.313760265 | 0.28529096 | 0.86 |
| YEL020C | 1045.827861 | 0.31556925 | 0.286166225 | 0.86 |
| YGR210C | 1863.079648 | -0.30919169 | 0.285988412 | 0.86 |
| YJR010C-A | 212.1436636 | -0.430890553 | 0.286009636 | 0.86 |
| YLR120W-A | 22.81140064 | -0.130801153 | 0.286056728 | 0.86 |
| YLR254C | 554.1871051 | 0.267932246 | 0.286088528 | 0.86 |
| YNL115C | 1894.716258 | 0.414014064 | 0.285664215 | 0.86 |
| YPL123C | 1037.088707 | 0.333005103 | 0.285587659 | 0.86 |
| YEL047C | 2163.496037 | 0.597045292 | 0.28647026 | 0.86 |
| YFR021W | 746.0129726 | 0.266834396 | 0.286475668 | 0.86 |
| YJL032W | 6.67566349 | -0.201645159 | 0.287114036 | 0.86 |
| YPR175W | 704.7198488 | -0.260158956 | 0.287173557 | 0.86 |
| YPR190C | 965.2019513 | -0.19932556 | 0.287170305 | 0.86 |
| YBR016W | 1677.943304 | 0.258626112 | 0.288821843 | 0.86 |
| YCR012W | 43096.09714 | -0.11219755 | 0.290425175 | 0.86 |
| YDL025C | 1555.623411 | 0.467910299 | 0.290252165 | 0.86 |
| YDR146C | 1456.032171 | -0.36246508 | 0.289792377 | 0.86 |
| YGR145W | 1225.275886 | -0.123500352 | 0.290486606 | 0.86 |
| YGR162W | 4487.780728 | -0.278093978 | 0.290151928 | 0.86 |
| YHR191C | 210.2719774 | 0.253060468 | 0.289550593 | 0.86 |
| YIL006W | 374.9591791 | 0.182896673 | 0.289914972 | 0.86 |
| YIR031C | 471.7138035 | 0.25336587 | 0.288127638 | 0.86 |
| YJL179W | 307.4969877 | -0.29060634 | 0.288039723 | 0.86 |
| YLR005W | 967.4119109 | 0.288368567 | 0.288240069 | 0.86 |
| YLR019W | 1293.150392 | 0.31866396 | 0.289013484 | 0.86 |
| YLR120C | 2959.418798 | 0.375083466 | 0.289064356 | 0.86 |
| YMR011W | 5664.342762 | 0.16988286 | 0.290466831 | 0.86 |
| YMR128W | 2322.124433 | -0.008144034 | 0.287398908 | 0.86 |
| YMR267W | 919.8795189 | -0.45266461 | 0.28878762 | 0.86 |

| | | | | |
|---|---|---|---|---|
| YNL215W | 1103.384408 | 0.357186036 | 0.289706143 | 0.86 |
| YNL222W | 196.3405424 | -0.477590975 | 0.290415394 | 0.86 |
| YOL159C-A | 58.15453861 | -0.732561701 | 0.287772551 | 0.86 |
| YOR127W | 1893.705383 | 0.325839308 | 0.28876167 | 0.86 |
| YOR162C | 795.3084144 | 0.237352664 | 0.289993223 | 0.86 |
| YOR178C | 2730.240932 | 0.293786282 | 0.289892574 | 0.86 |
| YOR351C | 100.6577504 | 0.255081479 | 0.287811083 | 0.86 |
| snR70 | 64.50681846 | 0.507288234 | 0.288128837 | 0.86 |
| snR86 | 894.0383097 | -0.424171501 | 0.289590269 | 0.86 |
| YBR199W | 2706.736809 | 0.201927644 | 0.291160431 | 0.86 |
| YJR127C | 3525.789326 | 0.392908765 | 0.290973816 | 0.86 |
| YLR441C | 16750.1152 | -0.115794379 | 0.291162561 | 0.86 |
| YBL010C | 437.6652565 | -0.266547555 | 0.292802097 | 0.86 |
| YBL071W-A | 297.3219208 | 0.49374252 | 0.293560698 | 0.86 |
| YCR090C | 636.078747 | -0.345828444 | 0.293577535 | 0.86 |
| YDR024W | 7.360211428 | 0.074388277 | 0.292434401 | 0.86 |
| YDR392W | 654.5457347 | 0.247003765 | 0.293380302 | 0.86 |
| YGL207W | 4342.654723 | -0.279742124 | 0.293107238 | 0.86 |
| YHR022C-A | 3.937421936 | -0.249172065 | 0.293556827 | 0.86 |
| YJR054W | 539.7820263 | -0.414068667 | 0.293020926 | 0.86 |
| YML069W | 2256.742432 | -0.403286992 | 0.29247257 | 0.86 |
| YNL008C | 1006.822611 | 0.335549361 | 0.292036147 | 0.86 |
| YOR030W | 929.3153858 | 0.353757828 | 0.292941252 | 0.86 |
| YPL005W | 369.5032375 | 0.344489667 | 0.292418 | 0.86 |
| snR58 | 5.413360595 | -1.385085533 | 0.293398733 | 0.86 |
| YDR448W | 426.123254 | -0.351425901 | 0.293751031 | 0.86 |
| YCR063W | 60.70870613 | -0.410245713 | 0.295032764 | 0.86 |
| YHR128W | 1328.56215 | -0.053162245 | 0.294238082 | 0.86 |
| YIL060W | 54.27024793 | 0.236812108 | 0.294240777 | 0.86 |
| YJL010C | 755.184634 | -0.308328233 | 0.29505293 | 0.86 |
| YKL006C-A | 282.6893408 | -0.431674643 | 0.294740996 | 0.86 |
| YKL172W | 2863.993194 | -0.284284062 | 0.294621002 | 0.86 |
| YKR036C | 843.7972906 | 0.277335472 | 0.294832633 | 0.86 |
| YMR105C | 9001.650714 | 0.427398744 | 0.294827401 | 0.86 |
| YNR007C | 864.4677932 | 0.388660541 | 0.295051999 | 0.86 |
| YMR272W-B | 3.438828838 | 0.266512813 | 0.295597707 | 0.86 |
| YPL246C | 2119.845638 | -0.257386702 | 0.295496967 | 0.86 |
| YAL056C-A | 23.07572137 | -0.195593527 | 0.298559628 | 0.86 |
| YBR259W | 488.2154125 | -0.292520721 | 0.29863868 | 0.86 |
| YCR057C | 1417.95789 | -0.321654534 | 0.295932045 | 0.86 |
| YDL191W | 1342.250708 | -0.270615736 | 0.299066383 | 0.86 |
| YDR046C | 2206.66233 | 0.371614367 | 0.296751131 | 0.86 |
| YDR165W | 1014.540812 | -0.222881339 | 0.297896585 | 0.86 |

| | | | | |
|---|---|---|---|---|
| YDR221W | 936.1837264 | -0.271164671 | 0.298292153 | 0.86 |
| YDR288W | 555.4180316 | 0.280580283 | 0.297090411 | 0.86 |
| YER038C | 327.709464 | 0.461516113 | 0.297888223 | 0.86 |
| YER087W | 372.1981021 | 0.385610783 | 0.29893697 | 0.86 |
| YER126C | 1138.972805 | -0.24423617 | 0.297032401 | 0.86 |
| YGR257C | 921.5740821 | -0.379664196 | 0.298753767 | 0.86 |
| YJL118W | 178.5211681 | -0.355183995 | 0.299136903 | 0.86 |
| YJR061W | 732.5988419 | 0.265995904 | 0.297566163 | 0.86 |
| YKL214C | 653.6592896 | -0.288061535 | 0.297704448 | 0.86 |
| YKR013W | 4305.106637 | -0.407661964 | 0.297905114 | 0.86 |
| YLR327C | 4824.371416 | 0.233102066 | 0.296868489 | 0.86 |
| YNL281W | 1354.281332 | -0.294681043 | 0.298760651 | 0.86 |
| YOL038C-A | 23.65291474 | 0.375358734 | 0.299050988 | 0.86 |
| YOL063C | 983.1121387 | 0.291291158 | 0.299022137 | 0.86 |
| YOL106W | 14.51135789 | 0.274612323 | 0.298430518 | 0.86 |
| YPL012W | 1836.869245 | -0.178844535 | 0.297672212 | 0.86 |
| YPL250C | 749.2294185 | 0.565705731 | 0.297353371 | 0.86 |
| YPR046W | 225.601515 | 0.309964654 | 0.297026855 | 0.86 |
| YPR164W | 1248.455329 | -0.22756015 | 0.299020258 | 0.86 |
| snR41 | 28.2332558 | -0.404183873 | 0.298376123 | 0.86 |
| YBR138C | 569.4593536 | 0.368333356 | 0.299856584 | 0.86 |
| YDR484W | 738.7697722 | 0.215795204 | 0.299857743 | 0.86 |
| YOR089C | 2098.945136 | 0.311036819 | 0.299656925 | 0.86 |
| YBR143C | 4538.418741 | -0.229493899 | 0.301180545 | 0.86 |
| YBR208C | 3562.373289 | 0.596206159 | 0.301845283 | 0.86 |
| YDL060W | 1265.050122 | -0.229921079 | 0.300891148 | 0.86 |
| YER155C | 4972.789594 | -0.305154367 | 0.301214154 | 0.86 |
| YFR022W | 849.2747048 | 0.321020699 | 0.302283174 | 0.86 |
| YHL015W-A | 4.315791202 | -0.816073875 | 0.301931114 | 0.86 |
| YIL078W | 8337.626041 | 0.009410199 | 0.302392299 | 0.86 |
| YJR089W | 953.5011996 | 0.294492873 | 0.302406722 | 0.86 |
| YKL012W | 1327.340355 | 0.2486263 | 0.301370721 | 0.86 |
| YKR088C | 850.0044261 | -0.27614909 | 0.302066653 | 0.86 |
| YML001W | 1453.0376 | -0.289069496 | 0.302149436 | 0.86 |
| YNL112W | 1857.470789 | -0.119474552 | 0.30027094 | 0.86 |
| YOL111C | 1821.468977 | -0.254733621 | 0.301761125 | 0.86 |
| YOR024W | 4.641545194 | 0.230294725 | 0.301776134 | 0.86 |
| YOR085W | 1806.530962 | -0.366644136 | 0.30035213 | 0.86 |
| YOR283W | 529.1178643 | -0.387027924 | 0.301585236 | 0.86 |
| YPL096W | 660.7866918 | 0.266665894 | 0.300539074 | 0.86 |
| YPL263C | 1770.940205 | -0.195996866 | 0.301070061 | 0.86 |
| YPR003C | 637.4553595 | 0.382652906 | 0.302071965 | 0.86 |
| YDR467C | 75.26126934 | 0.327589724 | 0.302659464 | 0.86 |

| | | | | |
|---|---|---|---|---|
| YIR001C | 869.6360964 | -0.277218172 | 0.302707542 | 0.86 |
| YKL101W | 3903.982421 | -0.147636842 | 0.303037161 | 0.86 |
| YGR264C | 2825.266453 | -0.129069505 | 0.303244243 | 0.87 |
| YPL128C | 628.8955352 | -0.329623146 | 0.303514288 | 0.87 |
| YOR198C | 6733.130979 | -0.226984041 | 0.303908925 | 0.87 |
| YDR126W | 326.9726421 | 0.34451299 | 0.305439497 | 0.87 |
| YHR105W | 228.2746463 | 0.51740933 | 0.305204302 | 0.87 |
| YIL153W | 970.2297401 | -0.197860672 | 0.304525154 | 0.87 |
| YKR086W | 1098.768049 | 0.353261868 | 0.304811694 | 0.87 |
| YLR266C | 1095.178781 | 0.289635075 | 0.305465387 | 0.87 |
| YLR290C | 1190.940137 | 0.366617311 | 0.304139785 | 0.87 |
| YMR261C | 3042.037261 | 0.385199964 | 0.305554057 | 0.87 |
| YNL011C | 614.0144377 | 0.375452762 | 0.304577926 | 0.87 |
| YOR298W | 171.3347166 | 0.255108698 | 0.305211175 | 0.87 |
| YPL236C | 570.2921643 | 0.357532198 | 0.30431629 | 0.87 |
| YPR030W | 2156.768509 | 0.091069899 | 0.305037131 | 0.87 |
| snR83 | 177.6034874 | -0.539117727 | 0.304727067 | 0.87 |
| YDL159W-A | 1.037341025 | 0.138030884 | 0.309453088 | 0.87 |
| YDL167C | 1031.301724 | -0.416753533 | 0.307498621 | 0.87 |
| YDR062W | 2339.714713 | -0.270227635 | 0.306297476 | 0.87 |
| YDR188W | 2974.432832 | -0.305060158 | 0.309021026 | 0.87 |
| YDR324C | 1092.059972 | -0.203635547 | 0.308886668 | 0.87 |
| YDR426C | 18.4381422 | 0.117327709 | 0.308735137 | 0.87 |
| YDR432W | 4551.301879 | -0.204369146 | 0.309208688 | 0.87 |
| YER067C-A | 258.7943359 | 0.805507789 | 0.309134917 | 0.87 |
| YER093C-A | 366.8172657 | -0.357349358 | 0.308085799 | 0.87 |
| YER095W | 1267.382435 | -0.330030336 | 0.309184371 | 0.87 |
| YER104W | 241.2969985 | 0.364602715 | 0.308281037 | 0.87 |
| YER107C | 624.5620743 | -0.4057692 | 0.308828354 | 0.87 |
| YGL079W | 448.9297125 | -0.275841447 | 0.309613536 | 0.87 |
| YHR156C | 433.8648918 | 0.325292048 | 0.309321507 | 0.87 |
| YIL104C | 354.0212252 | 0.319728651 | 0.305941227 | 0.87 |
| YIR002C | 1205.196083 | 0.399726628 | 0.307830792 | 0.87 |
| YJL116C | 947.324176 | 0.044440159 | 0.307732636 | 0.87 |
| YJR021C | 884.0147888 | 0.226603939 | 0.308295076 | 0.87 |
| YJR025C | 1368.606225 | 0.047867597 | 0.3081675 | 0.87 |
| YJR119C | 341.555846 | 0.183229801 | 0.309135047 | 0.87 |
| YJR148W | 6070.611654 | 0.613351649 | 0.309590386 | 0.87 |
| YLR409C | 1307.908763 | -0.222828772 | 0.308047043 | 0.87 |
| YMR136W | 2477.221542 | 0.572873886 | 0.306668025 | 0.87 |
| YMR319C | 2635.007516 | 0.450155797 | 0.307329519 | 0.87 |
| YOL007C | 839.2247362 | -0.311366367 | 0.307020359 | 0.87 |
| YOR224C | 1759.04057 | -0.255367125 | 0.309054187 | 0.87 |

| | | | | |
|---|---|---|---|---|
| YOR380W | 392.7551247 | 0.280273666 | 0.308082997 | 0.87 |
| YPL057C | 1620.500312 | 0.256902861 | 0.309328498 | 0.87 |
| YPR049C | 968.2983267 | 0.44477922 | 0.309189105 | 0.87 |
| YPR051W | 72.40075259 | -0.404999314 | 0.305794155 | 0.87 |
| YAL064W-B | 18.20372016 | -0.551950355 | 0.31090237 | 0.87 |
| YAR029W | 75.14481749 | 0.281659806 | 0.318790673 | 0.87 |
| YBL059C-A | 504.0749236 | -0.272612466 | 0.311834379 | 0.87 |
| YBL067C | 1161.166699 | 0.25446234 | 0.310787971 | 0.87 |
| YBL090W | 640.7121751 | -0.385480788 | 0.318968381 | 0.87 |
| YBR018C | 184.0387102 | 0.07512686 | 0.311278773 | 0.87 |
| YBR087W | 809.749781 | -0.331059625 | 0.313485756 | 0.87 |
| YCL058C | 103.80183 | -0.118120179 | 0.315616488 | 0.87 |
| YDL013W | 806.2108951 | 0.302399988 | 0.314067732 | 0.87 |
| YDL022W | 5632.112705 | 0.31566436 | 0.316549929 | 0.87 |
| YDL089W | 963.5832446 | 0.418440486 | 0.317350156 | 0.87 |
| YDR001C | 5262.491506 | 0.393493811 | 0.318064162 | 0.87 |
| YDR002W | 3692.839207 | -0.15153449 | 0.314926607 | 0.87 |
| YDR059C | 828.9203049 | 0.456428455 | 0.310425283 | 0.87 |
| YDR094W | 4.116771726 | -0.736489196 | 0.318252937 | 0.87 |
| YDR170C | 4884.61556 | -0.186254883 | 0.315353593 | 0.87 |
| YDR183W | 267.9198317 | -0.317212759 | 0.316115584 | 0.87 |
| YDR198C | 539.1037114 | 0.287597215 | 0.31458433 | 0.87 |
| YDR316W | 792.8873428 | -0.383748895 | 0.317193749 | 0.87 |
| YDR344C | 46.73109222 | -0.069893974 | 0.311736544 | 0.87 |
| YDR422C | 1006.650102 | 0.326904081 | 0.318758231 | 0.87 |
| YEL041W | 348.1575551 | 0.488430529 | 0.315232402 | 0.87 |
| YEL073C | 122.6223904 | -0.746549081 | 0.312283052 | 0.87 |
| YER037W | 745.1407721 | 0.467432551 | 0.317157804 | 0.87 |
| YER111C | 1455.121069 | -0.273319087 | 0.319010736 | 0.87 |
| YER158W-A | 247.6290209 | -0.552409204 | 0.315864021 | 0.87 |
| YER173W | 470.1702433 | 0.278015163 | 0.317100828 | 0.87 |
| YFL011W | 70.61598103 | 0.089043302 | 0.317310287 | 0.87 |
| YFL050C | 563.7473526 | 0.30481348 | 0.316728582 | 0.87 |
| YFR003C | 1017.431306 | 0.345625106 | 0.310781303 | 0.87 |
| YGL188C-A | 18.77666255 | -0.566719119 | 0.317795487 | 0.87 |
| YGL249W | 273.3583474 | 0.160383811 | 0.318407408 | 0.87 |
| YGR008C | 4582.699628 | 0.349287021 | 0.313785954 | 0.87 |
| YGR073C | 7.437514438 | -1.095497629 | 0.316033376 | 0.87 |
| YGR153W | 193.310946 | 0.439169539 | 0.316398212 | 0.87 |
| YHR006W | 1327.030831 | 0.33806258 | 0.312210113 | 0.87 |
| YHR007C | 4279.432771 | -0.2973878 | 0.316171864 | 0.87 |
| YHR039C | 2261.246675 | -0.262192788 | 0.318653934 | 0.87 |
| YHR079C | 1259.061897 | 0.28866415 | 0.31584191 | 0.87 |

| | | | | |
|---|---|---|---|---|
| YHR123W | 699.5030842 | -0.111160087 | 0.317289836 | 0.87 |
| YHR161C | 2280.636973 | 0.429744098 | 0.318846027 | 0.87 |
| YIR022W | 1186.448374 | -0.313911588 | 0.311097008 | 0.87 |
| YJL028W | 3.906046468 | 0.104964446 | 0.314256636 | 0.87 |
| YJL191W | 879.6997619 | -0.318804937 | 0.311440168 | 0.87 |
| YJL216C | 203.2730397 | 0.183021701 | 0.310247689 | 0.87 |
| YJR082C | 608.3949667 | 0.345365504 | 0.312539612 | 0.87 |
| YJR084W | 669.2500078 | -0.261344546 | 0.317000746 | 0.87 |
| YJR113C | 864.1046061 | -0.415419287 | 0.318611979 | 0.87 |
| YKL171W | 1421.131993 | 0.307522901 | 0.317150244 | 0.87 |
| YKL184W | 1354.204344 | -0.311273267 | 0.318819138 | 0.87 |
| YLR098C | 706.1029207 | 0.319922714 | 0.31283534 | 0.87 |
| YLR180W | 3585.379162 | 0.059660832 | 0.314064571 | 0.87 |
| YLR300W | 3505.549383 | -0.518414072 | 0.310089814 | 0.87 |
| YML127W | 993.038483 | -0.289784045 | 0.315570704 | 0.87 |
| YMR045C | 326.5494461 | 0.788901628 | 0.312990751 | 0.87 |
| YMR065W | 380.856771 | 0.197312664 | 0.313055098 | 0.87 |
| YMR144W | 347.0360461 | -0.294069498 | 0.31335955 | 0.87 |
| YMR194C-A | 22.19182237 | -0.565428462 | 0.315927861 | 0.87 |
| YMR247W-A | 22.89498367 | 0.171124064 | 0.319102648 | 0.87 |
| YMR251W | 257.8128007 | 0.322105117 | 0.31820282 | 0.87 |
| YMR262W | 628.7777309 | 0.35896401 | 0.318587547 | 0.87 |
| YNL037C | 5393.729596 | -0.298949665 | 0.311301231 | 0.87 |
| YNL181W | 1100.958937 | -0.228249882 | 0.318059577 | 0.87 |
| YNR006W | 1506.041195 | 0.296689479 | 0.314127168 | 0.87 |
| YOL049W | 2360.13263 | -0.301565588 | 0.319114985 | 0.87 |
| YOR114W | 311.2740858 | 0.51215128 | 0.316217981 | 0.87 |
| YPL089C | 1858.715154 | 0.372092259 | 0.313626265 | 0.87 |
| YPL096C-A | 145.3313639 | -0.395531425 | 0.311629674 | 0.87 |
| YPL255W | 357.750249 | -0.335913831 | 0.318368111 | 0.87 |
| YPL278C | 8.613341367 | 0.120526299 | 0.312275866 | 0.87 |
| YJL111W | 2828.454028 | -0.291503629 | 0.319612988 | 0.87 |
| YLR458W | 2.467406493 | 0.171651519 | 0.319994909 | 0.87 |
| YKL166C | 506.1904776 | -0.385447704 | 0.320194153 | 0.87 |
| YBR002C | 607.1395928 | -0.327480743 | 0.324397824 | 0.87 |
| YBR130C | 1160.408881 | -0.371176924 | 0.324384615 | 0.87 |
| YCR065W | 756.8632472 | -0.494000549 | 0.324487363 | 0.87 |
| YDL030W | 676.7024249 | 0.3236772 | 0.324160079 | 0.87 |
| YDL085W | 757.5796414 | 0.276318433 | 0.322425209 | 0.87 |
| YDL196W | 15.17669232 | 0.58767701 | 0.322243734 | 0.87 |
| YDR033W | 12974.70679 | -0.223021856 | 0.322435482 | 0.87 |
| YEL060C | 10805.49764 | 0.462799267 | 0.322035256 | 0.87 |
| YGL100W | 1500.808318 | -0.229622773 | 0.324552429 | 0.87 |

| | | | | |
|---|---|---|---|---|
| YGR071C | 709.1837053 | 0.260500079 | 0.321690148 | 0.87 |
| YHL009C | 603.1387723 | -0.286844803 | 0.324722343 | 0.87 |
| YHR140W | 190.2878906 | -0.610344144 | 0.321100874 | 0.87 |
| YHR201C | 1092.809897 | -0.416956435 | 0.322590777 | 0.87 |
| YJL082W | 1623.02873 | 0.403287039 | 0.320921896 | 0.87 |
| YJL164C | 1399.537088 | 0.359027795 | 0.324650767 | 0.87 |
| YJR002W | 1689.001321 | -0.215022101 | 0.323975186 | 0.87 |
| YJR088C | 954.591927 | -0.263296164 | 0.324072464 | 0.87 |
| YLL055W | 836.4946306 | 0.448102483 | 0.32386702 | 0.87 |
| YLR196W | 1358.156963 | -0.210907598 | 0.32342392 | 0.87 |
| YLR249W | 32716.59454 | -0.036833881 | 0.321338267 | 0.87 |
| YLR431C | 452.0758432 | 0.262670106 | 0.324752529 | 0.87 |
| YML004C | 3324.450509 | 0.287122086 | 0.323941891 | 0.87 |
| YMR254C | 26.76419027 | 0.269232321 | 0.322217655 | 0.87 |
| YNL108C | 290.0576123 | -0.297819276 | 0.323547434 | 0.87 |
| YNL263C | 1345.46904 | -0.28633806 | 0.323076909 | 0.87 |
| YNL284C | 1102.406706 | -0.32889588 | 0.323377599 | 0.87 |
| YOL117W | 709.0122211 | 0.425911998 | 0.322577614 | 0.87 |
| YOR235W | 3.373424746 | 0.067465829 | 0.324603476 | 0.87 |
| YPL210C | 1902.73736 | -0.396243968 | 0.320908853 | 0.87 |
| YPR036W | 6660.698271 | -0.218947529 | 0.321425533 | 0.87 |
| YPR084W | 689.7868138 | 0.185594942 | 0.324664173 | 0.87 |
| snR5 | 32.14065526 | -0.661441204 | 0.32378655 | 0.87 |
| YDR231C | 627.5516275 | 0.385827533 | 0.325581644 | 0.87 |
| YDR334W | 1700.440041 | 0.253222713 | 0.325087005 | 0.87 |
| YIL131C | 677.2016628 | -0.196629862 | 0.325595169 | 0.87 |
| YKL190W | 813.5201171 | -0.354493207 | 0.325140555 | 0.87 |
| YKR040C | 4.454389 | -0.180224595 | 0.325420359 | 0.87 |
| YNL168C | 1117.770385 | -0.219674642 | 0.32531752 | 0.87 |
| YLR186W | 830.8214037 | -0.215134435 | 0.325835239 | 0.87 |
| YER041W | 647.7988303 | 0.293108125 | 0.326414997 | 0.87 |
| YGR013W | 916.8674279 | 0.303101837 | 0.326190877 | 0.87 |
| YJL041W | 2583.756357 | -0.229930505 | 0.326500287 | 0.87 |
| YLR361C | 559.6543534 | 0.303373515 | 0.326441178 | 0.87 |
| YDR506C | 448.3958033 | 0.322991565 | 0.32685855 | 0.87 |
| RNA170 | 10.42132142 | 0.630548948 | 0.331132141 | 0.87 |
| YAL012W | 6420.264953 | -0.168501967 | 0.351908046 | 0.87 |
| YAL020C | 521.9507996 | 0.26918686 | 0.336156482 | 0.87 |
| YAL034C-B | 9.086766243 | -0.40548063 | 0.34299672 | 0.87 |
| YAL068C | 5.759798843 | 0.458619095 | 0.348276319 | 0.87 |
| YAR002W | 1776.649309 | -0.30118579 | 0.346540045 | 0.87 |
| YBL024W | 2012.558866 | -0.294064849 | 0.351035307 | 0.87 |
| YBL031W | 539.4437355 | -0.280002248 | 0.359256689 | 0.87 |

| | | | | |
|---|---|---|---|---|
| YBL033C | 1274.437806 | 0.23475181 | 0.341830299 | 0.87 |
| YBL081W | 976.0547384 | -0.405298215 | 0.349424815 | 0.87 |
| YBL095W | 182.0968065 | 0.209016798 | 0.329354517 | 0.87 |
| YBR005W | 1310.131508 | 0.266886616 | 0.329462568 | 0.87 |
| YBR025C | 6377.054239 | -0.084858617 | 0.338294356 | 0.87 |
| YBR028C | 772.1128092 | -0.386166426 | 0.349868367 | 0.87 |
| YBR046C | 1727.421703 | 0.526843445 | 0.344649717 | 0.87 |
| YBR082C | 6782.616874 | -0.316956263 | 0.355451582 | 0.87 |
| YBR192W | 512.8528961 | -0.293659952 | 0.337848074 | 0.87 |
| YBR219C | 1.059885032 | -0.29691268 | 0.359514298 | 0.87 |
| YBR290W | 1203.208406 | 0.232923731 | 0.337211723 | 0.87 |
| YCL010C | 597.9803619 | 0.296217946 | 0.341397678 | 0.87 |
| YCL016C | 442.1886812 | 0.291794795 | 0.346225839 | 0.87 |
| YCR018C | 861.2432726 | 0.210804123 | 0.351373755 | 0.87 |
| YCR072C | 412.6699919 | -0.401940928 | 0.356994886 | 0.87 |
| YCR089W | 1832.043276 | 0.441901732 | 0.342520447 | 0.87 |
| YDL008W | 381.8332185 | -0.397248388 | 0.333313902 | 0.87 |
| YDL015C | 3069.696513 | -0.228777691 | 0.350549715 | 0.87 |
| YDL018C | 497.6776418 | -0.260250192 | 0.334923001 | 0.87 |
| YDL053C | 1309.962591 | -0.247193259 | 0.333751268 | 0.87 |
| YDL085C-A | 569.0392384 | -0.502753995 | 0.331877161 | 0.87 |
| YDL092W | 1065.77792 | -0.289916463 | 0.335608805 | 0.87 |
| YDL153C | 1719.824839 | -0.151952171 | 0.340918105 | 0.87 |
| YDL159W | 513.5387781 | 0.285141106 | 0.351861259 | 0.87 |
| YDL215C | 3694.89417 | 0.460619659 | 0.349973917 | 0.87 |
| YDL229W | 7732.532873 | 0.230592914 | 0.34641774 | 0.87 |
| YDL237W | 1508.528864 | -0.254468515 | 0.349309129 | 0.87 |
| YDR023W | 6264.678536 | -0.185792401 | 0.332570596 | 0.87 |
| YDR037W | 9604.757078 | 0.154849843 | 0.336737189 | 0.87 |
| YDR083W | 1132.402572 | -0.212546929 | 0.331492744 | 0.87 |
| YDR119W-A | 458.2326718 | 0.024490547 | 0.338584291 | 0.87 |
| YDR129C | 5762.337606 | 0.297372383 | 0.329842478 | 0.87 |
| YDR130C | 501.1168326 | -0.360557161 | 0.341998164 | 0.87 |
| YDR149C | 14.02290454 | -0.188959191 | 0.355388394 | 0.87 |
| YDR161W | 1294.242267 | -0.183145738 | 0.350437726 | 0.87 |
| YDR190C | 2078.525809 | -0.315603671 | 0.346487918 | 0.87 |
| YDR249C | 239.973621 | 0.302893034 | 0.347537867 | 0.87 |
| YDR258C | 9048.121163 | 0.619277972 | 0.349868353 | 0.87 |
| YDR267C | 660.9899359 | -0.233557268 | 0.343206209 | 0.87 |
| YDR303C | 1032.483515 | -0.391886447 | 0.344980639 | 0.87 |
| YDR305C | 387.9870014 | -0.311527694 | 0.3548946 | 0.87 |
| YDR312W | 598.3889984 | -0.295412384 | 0.329761786 | 0.87 |
| YDR317W | 303.4699364 | 0.255184648 | 0.337817093 | 0.87 |

| | | | | |
|---|---|---|---|---|
| YDR326C | 1698.894045 | 0.286194152 | 0.333560674 | 0.87 |
| YDR341C | 4249.5314 | -0.017549302 | 0.346152049 | 0.87 |
| YDR351W | 1313.626306 | 0.232284699 | 0.338673181 | 0.87 |
| YDR395W | 2175.578678 | -0.205489287 | 0.343603792 | 0.87 |
| YDR402C | 76.67558535 | 0.487804572 | 0.342052564 | 0.87 |
| YDR404C | 728.6666751 | -0.331203582 | 0.340714323 | 0.87 |
| YDR405W | 1203.736123 | -0.283989467 | 0.357907759 | 0.87 |
| YDR407C | 1886.152895 | -0.316767214 | 0.337037945 | 0.87 |
| YDR411C | 1171.452043 | -0.342848787 | 0.344279095 | 0.87 |
| YDR453C | 889.6622872 | 0.274163043 | 0.342972076 | 0.87 |
| YDR475C | 1828.530627 | 0.413936841 | 0.357791424 | 0.87 |
| YDR480W | 507.0349407 | -0.216376038 | 0.329901959 | 0.87 |
| YDR511W | 614.1187303 | -0.465618159 | 0.353575524 | 0.87 |
| YEL028W | 113.7795172 | 0.172731579 | 0.342213185 | 0.87 |
| YEL040W | 3165.37713 | -0.28757485 | 0.350640112 | 0.87 |
| YEL061C | 1022.169501 | -0.366875023 | 0.346452068 | 0.87 |
| YEL067C | 149.0124198 | 0.395892885 | 0.346300847 | 0.87 |
| YER011W | 1426.306813 | 0.576734254 | 0.333924628 | 0.87 |
| YER031C | 2004.561338 | -0.258468348 | 0.341368147 | 0.87 |
| YER079W | 1934.057175 | 0.407606468 | 0.342889839 | 0.87 |
| YER127W | 556.9285849 | -0.038857679 | 0.358660481 | 0.87 |
| YER151C | 2013.443986 | -0.276224736 | 0.334433927 | 0.87 |
| YER184C | 397.7514223 | 0.342592458 | 0.351776494 | 0.87 |
| YER186C | 678.5275475 | -0.431970134 | 0.33522685 | 0.87 |
| YFL031W | 3441.588262 | -0.431618828 | 0.355021239 | 0.87 |
| YFL042C | 1495.188388 | 0.458188065 | 0.34302029 | 0.87 |
| YFR008W | 307.6442666 | 0.299194744 | 0.359095014 | 0.87 |
| YFR029W | 834.7381381 | 0.390232717 | 0.32777994 | 0.87 |
| YFR041C | 978.3498425 | -0.358760011 | 0.343006706 | 0.87 |
| YGL001C | 1350.851203 | -0.35454476 | 0.358635067 | 0.87 |
| YGL062W | 7360.866755 | 0.522818946 | 0.355944499 | 0.87 |
| YGL065C | 1014.84416 | -0.231011405 | 0.359428914 | 0.87 |
| YGL098W | 531.3017666 | 0.283148774 | 0.33307124 | 0.87 |
| YGL101W | 448.9089719 | -0.252982433 | 0.345110003 | 0.87 |
| YGL105W | 8369.251023 | -0.198068375 | 0.332177573 | 0.87 |
| YGL140C | 1656.222341 | 0.258188188 | 0.351775537 | 0.87 |
| YGL144C | 301.1617021 | 0.313673748 | 0.357703867 | 0.87 |
| YGL198W | 494.0719347 | 0.383609387 | 0.332893291 | 0.87 |
| YGL242C | 665.893361 | -0.261551802 | 0.356838964 | 0.87 |
| YGL257C | 1193.728393 | -0.279960745 | 0.343990063 | 0.87 |
| YGL258W | 52.00397703 | 0.34835808 | 0.342906475 | 0.87 |
| YGR045C | 23.80619598 | 0.302202109 | 0.346845531 | 0.87 |
| YGR101W | 624.5372953 | -0.319296348 | 0.336062787 | 0.87 |

| | | | | |
|---|---|---|---|---|
| YGR102C | 796.5538343 | 0.252913709 | 0.359545874 | 0.87 |
| YGR111W | 1711.371366 | 0.270629906 | 0.353673214 | 0.87 |
| YGR113W | 643.4472036 | -0.21242242 | 0.344333116 | 0.87 |
| YGR126W | 190.2789593 | 0.467006131 | 0.343013834 | 0.87 |
| YGR237C | 1848.787726 | 0.390952443 | 0.335619569 | 0.87 |
| YGR292W | 46.2187461 | 0.087711191 | 0.343800187 | 0.87 |
| YHL003C | 1333.653462 | -0.168207923 | 0.350049896 | 0.87 |
| YHL030W | 2779.309717 | 0.483474396 | 0.335471628 | 0.87 |
| YHR002W | 437.310458 | 0.307996778 | 0.331608967 | 0.87 |
| YHR018C | 3547.596925 | 0.740258045 | 0.348654584 | 0.87 |
| YHR037W | 3383.544 | 0.336616627 | 0.342783779 | 0.87 |
| YHR058C | 391.0666567 | -0.289788949 | 0.330779673 | 0.87 |
| YHR083W | 1675.31411 | -0.2374774 | 0.351958051 | 0.87 |
| YHR117W | 2228.445201 | -0.203879417 | 0.342341812 | 0.87 |
| YHR131C | 966.4131265 | 0.263057463 | 0.337321809 | 0.87 |
| YHR165W-A | 6.420816 | 0.112546059 | 0.342498703 | 0.87 |
| YHR169W | 496.2218428 | -0.212489274 | 0.348454104 | 0.87 |
| YHR197W | 861.9145076 | -0.167456512 | 0.34426513 | 0.87 |
| YIL026C | 1582.807889 | -0.299990582 | 0.33752882 | 0.87 |
| YIL043C | 4110.226673 | -0.227102388 | 0.349751129 | 0.87 |
| YIL051C | 4289.573013 | -0.487318092 | 0.334100351 | 0.87 |
| YIL058W | 79.78173189 | 0.240099215 | 0.350234781 | 0.87 |
| YIL061C | 381.0352818 | -0.279171919 | 0.346045074 | 0.87 |
| YIL092W | 426.021742 | 0.22663362 | 0.351037887 | 0.87 |
| YIL122W | 600.3807645 | 0.148422606 | 0.346879684 | 0.87 |
| YIL139C | 280.0724876 | -0.279218984 | 0.348450584 | 0.87 |
| YIL143C | 1830.368647 | 0.30486148 | 0.356985984 | 0.87 |
| YIR004W | 1446.338254 | -0.293741222 | 0.353725474 | 0.87 |
| YIR005W | 167.3355793 | 0.253507678 | 0.328324278 | 0.87 |
| YIR014W | 256.3884009 | 0.518219065 | 0.352388346 | 0.87 |
| YIR025W | 303.8110514 | 0.21685379 | 0.345682017 | 0.87 |
| YIR034C | 2228.546885 | 0.24231538 | 0.352335542 | 0.87 |
| YJL002C | 4648.101511 | -0.258787642 | 0.345632535 | 0.87 |
| YJL048C | 1114.208522 | 0.345825255 | 0.332757127 | 0.87 |
| YJL057C | 1203.837592 | 0.127916038 | 0.353578309 | 0.87 |
| YJL101C | 2271.091311 | 0.328057126 | 0.329394715 | 0.87 |
| YJL168C | 610.0870071 | -0.323045112 | 0.341264763 | 0.87 |
| YJR003C | 586.7430541 | -0.340194276 | 0.357617163 | 0.87 |
| YJR034W | 337.4607648 | -0.455106672 | 0.334527104 | 0.87 |
| YJR156C | 47.49998049 | -0.815531 | 0.350527373 | 0.87 |
| YKL042W | 561.3749184 | -0.399988982 | 0.33625538 | 0.87 |
| YKL098W | 807.0315194 | 0.28134098 | 0.350229257 | 0.87 |
| YKL133C | 601.5672696 | 0.218341422 | 0.356057656 | 0.87 |

| | | | | |
|---|---|---|---|---|
| YKL151C | 2678.459393 | 0.322202537 | 0.355467139 | 0.87 |
| YKL160W | 1725.224125 | -0.314038847 | 0.3466958 | 0.87 |
| YKL177W | 2.5279419 | 0.262193265 | 0.348114926 | 0.87 |
| YKL189W | 702.1789039 | 0.263501408 | 0.344469997 | 0.87 |
| YKL216W | 7973.975155 | -0.961261408 | 0.350147907 | 0.87 |
| YKR009C | 961.8765586 | 0.212502409 | 0.357629255 | 0.87 |
| YKR029C | 884.4269483 | 0.235366255 | 0.354112467 | 0.87 |
| YKR039W | 765.789828 | 0.18169415 | 0.339337318 | 0.87 |
| YKR100C | 1038.230895 | 0.290350804 | 0.34514125 | 0.87 |
| YKR104W | 184.9350151 | 0.403938532 | 0.354032201 | 0.87 |
| YLL021W | 2378.696108 | -0.21397252 | 0.357108412 | 0.87 |
| YLR017W | 1271.451959 | -0.154728586 | 0.328361914 | 0.87 |
| YLR054C | 138.716349 | 0.24248773 | 0.345428819 | 0.87 |
| YLR058C | 11510.40084 | 0.017983749 | 0.356136231 | 0.87 |
| YLR066W | 1030.577166 | -0.25686703 | 0.347643258 | 0.87 |
| YLR080W | 639.2701305 | 0.343198906 | 0.334532392 | 0.87 |
| YLR204W | 619.6469206 | -0.426214458 | 0.333352255 | 0.87 |
| YLR207W | 1228.065049 | 0.307330699 | 0.349563535 | 0.87 |
| YLR222C-A | 13.05158283 | -0.735717334 | 0.343741226 | 0.87 |
| YLR244C | 3158.461036 | -0.261170283 | 0.348410854 | 0.87 |
| YLR304C | 13028.11852 | -0.248840435 | 0.344100818 | 0.87 |
| YLR314C | 3042.070083 | -0.189584844 | 0.336062275 | 0.87 |
| YLR383W | 888.8933966 | -0.17899994 | 0.354010101 | 0.87 |
| YLR412W | 993.3823084 | -0.314889113 | 0.338507357 | 0.87 |
| YLR425W | 1120.132672 | 0.284622215 | 0.342462494 | 0.87 |
| YLR449W | 2574.095007 | 0.015676468 | 0.329502668 | 0.87 |
| YLR455W | 572.5909218 | -0.339849782 | 0.332228772 | 0.87 |
| YML055W | 811.1131816 | -0.256914671 | 0.33212905 | 0.87 |
| YML092C | 2758.585269 | -0.241407416 | 0.350561985 | 0.87 |
| YML094C-A | 15.34009962 | -0.175741922 | 0.341070465 | 0.87 |
| YML098W | 504.9087228 | -0.309787858 | 0.331769246 | 0.87 |
| YML104C | 923.4299097 | 0.334674481 | 0.357709014 | 0.87 |
| YMR086C-A | 82.59944884 | 0.298279928 | 0.348998086 | 0.87 |
| YMR086W | 2080.250939 | 0.314833394 | 0.350847028 | 0.87 |
| YMR145C | 7155.424672 | -0.155791658 | 0.352464136 | 0.87 |
| YMR160W | 946.3755314 | 0.275987559 | 0.353905087 | 0.87 |
| YMR170C | 596.7981642 | 0.381503758 | 0.359444965 | 0.87 |
| YMR191W | 2004.986398 | 0.57687478 | 0.331829818 | 0.87 |
| YMR195W | 1764.107553 | 0.276202533 | 0.35532809 | 0.87 |
| YMR222C | 712.4011542 | -0.322517861 | 0.336255105 | 0.87 |
| YMR229C | 3400.401838 | -0.142355358 | 0.347754074 | 0.87 |
| YMR304W | 1821.67097 | 0.341699895 | 0.355175523 | 0.87 |
| YNL021W | 893.5874562 | -0.207173552 | 0.33967482 | 0.87 |

| | | | | |
|---|---|---|---|---|
| YNL027W | 1180.449634 | 0.293726135 | 0.345664914 | 0.87 |
| YNL031C | 9825.730059 | -0.439967083 | 0.335293666 | 0.87 |
| YNL049C | 1288.390098 | -0.175184483 | 0.358101464 | 0.87 |
| YNL050C | 614.5421658 | -0.270286693 | 0.353649176 | 0.87 |
| YNL103W | 1585.703492 | 0.312039954 | 0.339403352 | 0.87 |
| YNL123W | 2060.867856 | -0.285384135 | 0.357181484 | 0.87 |
| YNL129W | 332.9229067 | -0.119856655 | 0.350048961 | 0.87 |
| YNL154C | 2670.283788 | -0.355280405 | 0.327314576 | 0.87 |
| YNL159C | 572.5140174 | 0.289069855 | 0.330619077 | 0.87 |
| YNL188W | 592.8677499 | -0.458370635 | 0.356379993 | 0.87 |
| YNL196C | 204.2397635 | 0.093057632 | 0.334328955 | 0.87 |
| YNL232W | 718.0441816 | -0.303530006 | 0.349876223 | 0.87 |
| YNL253W | 477.8813502 | -0.242267717 | 0.351757666 | 0.87 |
| YNL330C | 1017.919377 | -0.608842525 | 0.34093315 | 0.87 |
| YNL333W | 85.09307581 | -0.640027609 | 0.350847876 | 0.87 |
| YNR023W | 727.2388585 | -0.317609831 | 0.344030607 | 0.87 |
| YOL041C | 1321.58671 | -0.129458572 | 0.346459312 | 0.87 |
| YOL085C | 30.97884527 | 0.000697845 | 0.349864602 | 0.87 |
| YOR013W | 86.58484926 | -0.450450742 | 0.358306274 | 0.87 |
| YOR054C | 904.8796156 | 0.326600899 | 0.335826049 | 0.87 |
| YOR078W | 697.5265362 | -0.344487417 | 0.353489469 | 0.87 |
| YOR092W | 648.6032701 | 0.118788929 | 0.343848239 | 0.87 |
| YOR097C | 623.6386998 | 0.287388347 | 0.333899946 | 0.87 |
| YOR183W | 10.65070105 | 0.097564086 | 0.354574433 | 0.87 |
| YOR220W | 2249.332399 | 0.283856693 | 0.357737117 | 0.87 |
| YOR249C | 647.5824875 | 0.222364311 | 0.333265082 | 0.87 |
| YOR253W | 809.4249218 | -0.242533476 | 0.327364883 | 0.87 |
| YOR287C | 297.2565187 | -0.250503908 | 0.349030843 | 0.87 |
| YOR294W | 771.7312013 | -0.181383687 | 0.337754102 | 0.87 |
| YOR352W | 729.6147343 | 0.263527162 | 0.353857998 | 0.87 |
| YOR363C | 1212.557429 | 0.26746677 | 0.349084885 | 0.87 |
| YOR381W-A | 9.096748957 | 0.232583512 | 0.348318702 | 0.87 |
| YPL074W | 944.5738824 | 0.246452388 | 0.359057186 | 0.87 |
| YPL088W | 468.228703 | 0.050337076 | 0.330221 | 0.87 |
| YPL101W | 1017.044188 | -0.246411225 | 0.352059878 | 0.87 |
| YPL121C | 25.34339254 | 0.559286186 | 0.35539143 | 0.87 |
| YPL126W | 2185.981195 | -0.131436589 | 0.357741028 | 0.87 |
| YPL180W | 1077.43255 | 0.249010791 | 0.327495371 | 0.87 |
| YPL190C | 2551.586572 | -0.243929679 | 0.339315595 | 0.87 |
| YPL196W | 943.9259177 | 0.268214228 | 0.359398127 | 0.87 |
| YPL199C | 1017.89655 | -0.319390332 | 0.339683367 | 0.87 |
| YPL207W | 1844.153383 | 0.435498646 | 0.359165585 | 0.87 |
| YPL249C | 1511.229449 | 0.316075949 | 0.335573487 | 0.87 |

| | | | | |
|---|---|---|---|---|
| YPL250W-A | 88.80290187 | -0.356901507 | 0.331483164 | 0.87 |
| YPL258C | 548.5976694 | 0.264544683 | 0.342493918 | 0.87 |
| YPL279C | 37.92327787 | -0.78744416 | 0.346585254 | 0.87 |
| YPR002C-A | 1.918449357 | 0.573760905 | 0.338712763 | 0.87 |
| YPR015C | 179.9077993 | 0.269770276 | 0.338915655 | 0.87 |
| YPR065W | 1458.659767 | -0.352925724 | 0.333740375 | 0.87 |
| YPR078C | 88.79558885 | -0.090144011 | 0.333186953 | 0.87 |
| YPR089W | 1983.427567 | 0.278794151 | 0.329578066 | 0.87 |
| YPR112C | 1587.350101 | -0.224949703 | 0.333140981 | 0.87 |
| YPR115W | 2825.752527 | 0.359863449 | 0.35126743 | 0.87 |
| YPR118W | 1223.179848 | -0.114815361 | 0.336364032 | 0.87 |
| YPR137W | 608.8911311 | -0.3229643 | 0.342640204 | 0.87 |
| YPR138C | 1408.826707 | 0.28428454 | 0.345464934 | 0.87 |
| YPR160W | 12134.47846 | 0.511669129 | 0.359123259 | 0.87 |
| YPR189W | 1450.192608 | -0.318029991 | 0.329643149 | 0.87 |
| YPR202W | 17.57397274 | 0.358644997 | 0.357334648 | 0.87 |
| snR53 | 0.80815136 | -0.161942194 | 0.341491103 | 0.87 |
| snR64 | 4.034418541 | -0.195543986 | 0.336982735 | 0.87 |
| snR69 | 9.747852558 | -0.044456798 | 0.356639757 | 0.87 |
| snR77 | 9.325617253 | -0.258846382 | 0.343015261 | 0.87 |
| YIL032C | 13.12625335 | 0.503636023 | 0.360230747 | 0.87 |
| YJL167W | 5562.553131 | -0.17518258 | 0.360063511 | 0.87 |
| YJR064W | 2718.261132 | -0.20326015 | 0.359998004 | 0.87 |
| YKL150W | 7424.00365 | 0.43705453 | 0.360016517 | 0.87 |
| YKR092C | 1945.135383 | -0.13952291 | 0.360433663 | 0.87 |
| YLR065C | 843.2361186 | -0.312014534 | 0.360432456 | 0.87 |
| YBR166C | 859.3257193 | -0.30890224 | 0.360807148 | 0.87 |
| YGL037C | 14451.53815 | 0.378699391 | 0.361022843 | 0.87 |
| YBR084W | 2864.241089 | 0.059619256 | 0.361557463 | 0.87 |
| YHR176W | 373.1735066 | 0.256070361 | 0.361564215 | 0.87 |
| YOR119C | 842.963497 | 0.321109408 | 0.361463553 | 0.87 |
| YBR128C | 336.3131952 | 0.234697025 | 0.363896379 | 0.87 |
| YBR195C | 288.8600834 | -0.364859853 | 0.36311818 | 0.87 |
| YBR228W | 415.0546527 | 0.234343862 | 0.362255424 | 0.87 |
| YCL054W | 1476.642208 | -0.065533282 | 0.362457342 | 0.87 |
| YDL211C | 175.1726895 | -0.358743896 | 0.363001332 | 0.87 |
| YDR449C | 532.8954421 | -0.229180002 | 0.363554439 | 0.87 |
| YER010C | 1015.031336 | -0.265776594 | 0.363486675 | 0.87 |
| YKL023W | 1129.71016 | 0.366511413 | 0.36249092 | 0.87 |
| YKL079W | 964.5499269 | 0.22705609 | 0.363786902 | 0.87 |
| YMR250W | 7553.858966 | 0.49366338 | 0.363473505 | 0.87 |
| YNL202W | 289.1619632 | 0.111309293 | 0.363327745 | 0.87 |
| YOR007C | 8060.679187 | -0.258134647 | 0.363831095 | 0.87 |

| | | | | |
|---|---|---|---|---|
| YOR043W | 1570.483617 | -0.367890087 | 0.362708202 | 0.87 |
| YPL041C | 160.4382575 | -0.333195436 | 0.361964953 | 0.87 |
| YPL277C | 206.5316036 | 0.292683364 | 0.363375904 | 0.87 |
| YPR079W | 754.8054031 | 0.224955801 | 0.362969184 | 0.87 |
| snR50 | 8.290616263 | 0.355706209 | 0.362900046 | 0.87 |
| YBR014C | 1010.158039 | -0.205946433 | 0.364645036 | 0.87 |
| YDL132W | 2917.981172 | 0.273918349 | 0.36600527 | 0.87 |
| YDR080W | 1103.850689 | 0.387115326 | 0.365549772 | 0.87 |
| YDR086C | 1629.638329 | -0.179083024 | 0.367860922 | 0.87 |
| YDR307W | 749.860008 | -0.283008731 | 0.368143938 | 0.87 |
| YDR466W | 1549.812513 | 0.287200061 | 0.368386744 | 0.87 |
| YDR470C | 683.2878102 | 0.197206054 | 0.367616317 | 0.87 |
| YDR504C | 165.3464621 | 0.2968906 | 0.367794498 | 0.87 |
| YEL010W | 4.427753031 | 0.556704696 | 0.366280973 | 0.87 |
| YEL053W-A | 186.8810681 | -0.107006811 | 0.366587316 | 0.87 |
| YER114C | 1610.317658 | 0.24619138 | 0.367184679 | 0.87 |
| YER133W | 3950.455243 | -0.253788713 | 0.367155408 | 0.87 |
| YFL018C | 6022.957872 | -0.169499824 | 0.367568871 | 0.87 |
| YGL136C | 674.1887477 | 0.245729406 | 0.365171532 | 0.87 |
| YGL260W | 1.415207671 | -0.224245641 | 0.368517361 | 0.87 |
| YIR032C | 113.4974223 | 0.274216192 | 0.365890193 | 0.87 |
| YJL030W | 384.5980723 | -0.300585978 | 0.364476035 | 0.87 |
| YJL050W | 1800.303824 | -0.124489724 | 0.36528339 | 0.87 |
| YJR141W | 396.4509931 | -0.320854089 | 0.367652148 | 0.87 |
| YKL033W-A | 1409.695613 | -0.168079902 | 0.365046921 | 0.87 |
| YLL042C | 71.42312394 | -0.472050494 | 0.368465631 | 0.87 |
| YLR306W | 266.7883268 | -0.409427058 | 0.366143918 | 0.87 |
| YLR422W | 2135.341687 | 0.28409502 | 0.368029862 | 0.87 |
| YML117W | 3482.921508 | 0.277909286 | 0.367845524 | 0.87 |
| YMR161W | 724.2845416 | -0.26367355 | 0.365759749 | 0.87 |
| YNL287W | 4413.325689 | -0.184199562 | 0.366018989 | 0.87 |
| YOL057W | 2160.171024 | -0.230697894 | 0.36719986 | 0.87 |
| YOL092W | 1398.16665 | -0.164305429 | 0.365102554 | 0.87 |
| YOR010C | 455.6940822 | -0.023572792 | 0.366274944 | 0.87 |
| YOR032C | 566.9607469 | 0.178838796 | 0.368325699 | 0.87 |
| YPL080C | 11.15479793 | -1.001458149 | 0.365490716 | 0.87 |
| YPL082C | 1537.0483 | -0.295317538 | 0.36518165 | 0.87 |
| snR44 | 201.2193719 | -0.403668636 | 0.367240373 | 0.87 |
| YGR121C | 1176.230558 | 0.281296777 | 0.369011474 | 0.87 |
| YNL020C | 852.0700517 | 0.278463213 | 0.369036589 | 0.87 |
| YNL082W | 573.887894 | -0.297855105 | 0.36905695 | 0.87 |
| YDR060W | 3331.110803 | -0.114872728 | 0.369562147 | 0.87 |
| YDR445C | 3.142353253 | -0.531216412 | 0.369705658 | 0.87 |

| | | | | |
|---|---|---|---|---|
| YFL028C | 788.0732688 | 0.331613713 | 0.369901129 | 0.87 |
| YKL078W | 299.5473506 | -0.069890882 | 0.369727801 | 0.87 |
| YKR091W | 1094.538594 | 0.315514517 | 0.369876978 | 0.87 |
| YNL079C | 8141.583459 | -0.325959885 | 0.369433873 | 0.87 |
| YKR003W | 1162.972466 | -0.313431413 | 0.370100698 | 0.87 |
| YGR209C | 5181.82162 | -0.292812182 | 0.370645833 | 0.87 |
| YKL077W | 3042.957164 | -0.184562496 | 0.37033932 | 0.87 |
| YPR121W | 333.11981 | 0.194749613 | 0.370636288 | 0.87 |
| snR10 | 11.52677515 | -0.262408668 | 0.37043936 | 0.87 |
| YGR009C | 1451.464581 | 0.232599057 | 0.370889478 | 0.87 |
| YDL184C | 118.3516335 | -0.39976332 | 0.371237614 | 0.87 |
| YER152C | 1264.809581 | 0.372343987 | 0.371276374 | 0.87 |
| snR18 | 14.34494847 | -0.229097375 | 0.371342636 | 0.87 |
| YBL068W | 791.3128306 | -0.2246851 | 0.371538703 | 0.87 |
| YMR180C | 510.7680392 | 0.17998278 | 0.371625443 | 0.87 |
| YDR274C | 62.88697688 | 0.412296324 | 0.371983528 | 0.87 |
| YNL282W | 259.9576363 | 0.337997251 | 0.371963087 | 0.87 |
| YOR188W | 1393.160444 | -0.222202391 | 0.372047099 | 0.87 |
| YDL178W | 1397.749906 | -0.236712091 | 0.372384999 | 0.87 |
| YNL197C | 1704.51539 | -0.304118516 | 0.372422118 | 0.87 |
| SRG1 | 1084.659985 | 0.014245032 | 0.373567117 | 0.87 |
| YAL054C | 1109.295849 | 0.304395423 | 0.379264486 | 0.87 |
| YBR210W | 464.9591183 | -0.297530193 | 0.378480559 | 0.87 |
| YCL061C | 1120.832529 | -0.243342438 | 0.379203655 | 0.87 |
| YCR024C-B | 2025.586967 | -0.410720716 | 0.377342585 | 0.87 |
| YCR102W-A | 9.4716423 | 0.225148143 | 0.378173854 | 0.87 |
| YDR035W | 3632.420651 | 0.587912742 | 0.375862779 | 0.87 |
| YDR107C | 540.536576 | -0.217416287 | 0.378990418 | 0.87 |
| YDR522C | 159.2048735 | 0.086655061 | 0.378081833 | 0.87 |
| YER139C | 313.6167797 | 0.271226773 | 0.374962326 | 0.87 |
| YGL056C | 2031.538502 | 0.283254945 | 0.37929344 | 0.87 |
| YGL214W | 9.34196156 | -0.523872972 | 0.377705196 | 0.87 |
| YGR083C | 2563.158798 | -0.132186991 | 0.373950845 | 0.87 |
| YGR089W | 1598.86435 | 0.206234114 | 0.378298395 | 0.87 |
| YGR202C | 1624.722457 | -0.248618856 | 0.378980099 | 0.87 |
| YHL034C | 6448.755806 | -0.174027168 | 0.375594125 | 0.87 |
| YHR028W-A | 38.81881491 | 0.593330699 | 0.375075051 | 0.87 |
| YHR076W | 1124.266555 | -0.245530564 | 0.376163817 | 0.87 |
| YHR181W | 756.8304441 | -0.062442554 | 0.374374773 | 0.87 |
| YIL067C | 437.2526192 | 0.2288703 | 0.373782673 | 0.87 |
| YIL103W | 637.74726 | -0.263357529 | 0.373829329 | 0.87 |
| YIL164C | 260.9748713 | 0.160018384 | 0.378877773 | 0.87 |
| YIR012W | 1025.35668 | -0.224092994 | 0.377324162 | 0.87 |

| | | | | |
|---|---|---|---|---|
| YJL060W | 1402.078784 | -0.18079809 | 0.378397137 | 0.87 |
| YJL098W | 1058.267184 | -0.286838328 | 0.376553746 | 0.87 |
| YJL178C | 1325.547528 | -0.238637082 | 0.379137005 | 0.87 |
| YJR062C | 945.4060688 | 0.27060282 | 0.37891486 | 0.87 |
| YKL019W | 1109.764572 | -0.280292449 | 0.376008578 | 0.87 |
| YKL076C | 34.14810955 | 0.204478254 | 0.376606929 | 0.87 |
| YKL107W | 249.2824796 | 0.452600372 | 0.373027919 | 0.87 |
| YKL152C | 30648.47356 | -0.217199513 | 0.374690983 | 0.87 |
| YLL039C | 7097.91074 | 0.360434605 | 0.375447273 | 0.87 |
| YLR027C | 7716.105415 | -0.13986824 | 0.374439912 | 0.87 |
| YLR118C | 939.2371521 | -0.24971443 | 0.378277475 | 0.87 |
| YLR370C | 2370.21826 | 0.325191676 | 0.377832043 | 0.87 |
| YLR389C | 2863.752537 | 0.300577728 | 0.378542122 | 0.87 |
| YML061C | 1035.299962 | 0.375421833 | 0.378888422 | 0.87 |
| YMR210W | 1188.233193 | 0.187009347 | 0.374295942 | 0.87 |
| YNL240C | 1001.196601 | 0.370304909 | 0.378834707 | 0.87 |
| YNL283C | 1549.072925 | -0.233150637 | 0.377875822 | 0.87 |
| YNL318C | 22.2967731 | 0.157273859 | 0.378633489 | 0.87 |
| YOR185C | 2527.056266 | 0.4513115 | 0.373819983 | 0.87 |
| YOR207C | 1745.865963 | -0.20524141 | 0.374675931 | 0.87 |
| YOR254C | 3232.98471 | -0.11583993 | 0.373571514 | 0.87 |
| YPL077C | 211.278667 | 0.235578156 | 0.379261566 | 0.87 |
| YPL084W | 2075.944666 | 0.204187716 | 0.375134384 | 0.87 |
| YPR021C | 1682.395818 | 0.313612342 | 0.374084079 | 0.87 |
| YPR028W | 4646.798863 | -0.284599113 | 0.378040929 | 0.87 |
| snR60 | 11.58332067 | 0.083894473 | 0.377367187 | 0.87 |
| snR71 | 12.58626851 | 0.131186498 | 0.378979853 | 0.87 |
| YDR045C | 241.2309895 | -0.30034176 | 0.379581845 | 0.88 |
| YIL036W | 2051.229049 | 0.357852184 | 0.379839309 | 0.88 |
| YDR379C-A | 462.1957351 | 0.214442147 | 0.380167567 | 0.88 |
| YFR025C | 947.4315897 | -0.150688441 | 0.380324097 | 0.88 |
| YNL167C | 1564.476742 | 0.273757295 | 0.380276184 | 0.88 |
| YKL165C-A | 6.96473103 | 0.323580709 | 0.38051 | 0.88 |
| YDR135C | 3272.34562 | 0.320029905 | 0.380773742 | 0.88 |
| YBR180W | 89.72716224 | -0.502307554 | 0.381370907 | 0.88 |
| YPL083C | 495.9825937 | -0.202576741 | 0.381578591 | 0.88 |
| YBL085W | 2143.648048 | 0.332660307 | 0.382221145 | 0.88 |
| YDR152W | 1737.468978 | -0.31257936 | 0.382845042 | 0.88 |
| YEL023C | 623.1672116 | 0.224105643 | 0.383069985 | 0.88 |
| YFR032C-B | 19.14700684 | -0.285827669 | 0.382378353 | 0.88 |
| YGL006W-A | 3.943531239 | 0.641552575 | 0.382419184 | 0.88 |
| YKL041W | 833.3256523 | 0.278845602 | 0.381999272 | 0.88 |
| YMR192W | 859.8324449 | -0.243373832 | 0.382944368 | 0.88 |

| | | | | |
|---|---|---|---|---|
| YMR223W | 516.9298345 | 0.206225894 | 0.381894478 | 0.88 |
| YNL292W | 950.1678617 | -0.206238701 | 0.382652625 | 0.88 |
| YOL113W | 1261.758159 | 0.340831814 | 0.382571387 | 0.88 |
| YPL025C | 15.86269313 | 0.229233638 | 0.382855459 | 0.88 |
| YDR247W | 1026.990415 | 0.41082245 | 0.383465432 | 0.88 |
| YGL120C | 2147.315673 | 0.042078528 | 0.384352324 | 0.88 |
| YNL277W-A | 6.085182075 | -0.085027329 | 0.384123289 | 0.88 |
| YOL052C | 1181.165007 | -0.213273594 | 0.38402567 | 0.88 |
| YOR321W | 1083.064286 | -0.199644597 | 0.384264318 | 0.88 |
| YOR373W | 745.8071516 | -0.258062441 | 0.384041807 | 0.88 |
| YDL224C | 1328.137391 | -0.326269434 | 0.385029897 | 0.88 |
| YOR303W | 3621.710527 | 0.604274806 | 0.384910317 | 0.88 |
| YOR384W | 1087.084219 | 0.268611105 | 0.384824269 | 0.88 |
| YBL023C | 762.982123 | -0.239625212 | 0.392125199 | 0.88 |
| YBL093C | 1069.915355 | 0.238543427 | 0.390912326 | 0.88 |
| YBR090C | 79.12993298 | -0.090076505 | 0.391405263 | 0.88 |
| YBR225W | 1153.176358 | 0.24281389 | 0.39163581 | 0.88 |
| YBR295W | 678.020174 | 0.262751055 | 0.389308871 | 0.88 |
| YCR098C | 141.8503964 | -0.651408642 | 0.387162998 | 0.88 |
| YDL007C-A | 3.853033258 | -0.316970059 | 0.386227193 | 0.88 |
| YDL009C | 106.1081082 | -0.346646402 | 0.391445887 | 0.88 |
| YDR410C | 388.474414 | -0.289653645 | 0.388542012 | 0.88 |
| YDR534C | 224.6689044 | 0.627335559 | 0.38821056 | 0.88 |
| YEL071W | 3890.356415 | 0.288096758 | 0.390376786 | 0.88 |
| YEL077C | 280.3488731 | -0.336138773 | 0.392312036 | 0.88 |
| YER026C | 2306.614486 | -0.299115279 | 0.387188723 | 0.88 |
| YER058W | 286.3082478 | -0.339544104 | 0.390409787 | 0.88 |
| YER188W | 461.4895098 | -0.653303915 | 0.38985595 | 0.88 |
| YFL019C | 43.77288423 | 0.080970439 | 0.391136966 | 0.88 |
| YGL093W | 1066.84026 | 0.222710256 | 0.385455431 | 0.88 |
| YGL099W | 1111.20672 | -0.157509046 | 0.391942209 | 0.88 |
| YGL218W | 11.00510591 | 0.093148241 | 0.386366417 | 0.88 |
| YGL253W | 16247.71261 | -0.113973385 | 0.39196884 | 0.88 |
| YGR032W | 3378.619384 | 0.270107631 | 0.389627606 | 0.88 |
| YGR229C | 1916.022007 | -0.260611787 | 0.391825501 | 0.88 |
| YGR248W | 2366.128573 | -0.047456423 | 0.387129853 | 0.88 |
| YGR270W | 2956.664275 | 0.255204656 | 0.39251021 | 0.88 |
| YHL034W-A | 585.1049194 | -0.130062443 | 0.388717069 | 0.88 |
| YIL157C | 795.6414589 | -0.385820425 | 0.386540221 | 0.88 |
| YJL073W | 1063.600671 | -0.269380154 | 0.390400003 | 0.88 |
| YJL175W | 4.732791699 | 0.198715339 | 0.388782088 | 0.88 |
| YJR009C | 33615.59982 | -0.059094357 | 0.386339324 | 0.88 |
| YKL004W | 2100.85667 | -0.212941779 | 0.388174743 | 0.88 |

| | | | | |
|---|---|---|---|---|
| YKL100C | 2362.710996 | 0.257575855 | 0.38890861 | 0.88 |
| YKL187C | 740.5340162 | 0.291622916 | 0.391806053 | 0.88 |
| YKR052C | 558.5950233 | 0.259788001 | 0.388323267 | 0.88 |
| YKR066C | 4134.117453 | 0.261307308 | 0.385258863 | 0.88 |
| YLR216C | 5399.303942 | -0.161076558 | 0.389980338 | 0.88 |
| YLR333C | 2119.808061 | -0.261093097 | 0.387089157 | 0.88 |
| YLR369W | 1148.442381 | 0.347626798 | 0.391489573 | 0.88 |
| YLR400W | 49.34542517 | 0.271940048 | 0.385720209 | 0.88 |
| YLR407W | 355.7060391 | 0.241986836 | 0.387125157 | 0.88 |
| YML101C-A | 7.327177589 | -0.330084526 | 0.390435486 | 0.88 |
| YMR023C | 334.2712538 | 0.200507555 | 0.388431851 | 0.88 |
| YMR184W | 674.607137 | -0.207444073 | 0.389014138 | 0.88 |
| YMR272C | 4343.78123 | -0.303822638 | 0.389698616 | 0.88 |
| YMR305C | 6521.89708 | -0.253941352 | 0.389203366 | 0.88 |
| YNL062C | 805.5436558 | -0.254586248 | 0.392428116 | 0.88 |
| YNL321W | 1495.615211 | 0.272075654 | 0.386506985 | 0.88 |
| YNL326C | 366.0651518 | -0.2561799 | 0.392227964 | 0.88 |
| YOL003C | 636.2491343 | -0.228692959 | 0.385802883 | 0.88 |
| YOR189W | 408.1511083 | -0.292589341 | 0.391089098 | 0.88 |
| YOR365C | 309.6078398 | 0.149402891 | 0.38927796 | 0.88 |
| YPL136W | 0.97000745 | 0.11214555 | 0.391658339 | 0.88 |
| YPL183C | 913.1902093 | -0.291017259 | 0.390146464 | 0.88 |
| YPR029C | 1930.500091 | -0.211936913 | 0.390111313 | 0.88 |
| YPR107C | 422.2754774 | 0.201856289 | 0.389049664 | 0.88 |
| snR7-L | 6.909111604 | -0.219982351 | 0.392181301 | 0.88 |
| YDL158C | 89.30551169 | -0.09465369 | 0.392954967 | 0.88 |
| YGR243W | 1286.002734 | 0.028208506 | 0.392951233 | 0.88 |
| YBR017C | 2405.834993 | -0.226810733 | 0.393409284 | 0.88 |
| YLR178C | 3271.315417 | 0.277759491 | 0.39334221 | 0.88 |
| YDR372C | 1904.188965 | -0.252353536 | 0.393556519 | 0.88 |
| YCR023C | 1565.302118 | 0.371541383 | 0.393864958 | 0.88 |
| YDR139C | 266.4093081 | -0.234320709 | 0.393914657 | 0.88 |
| YKL082C | 1057.576571 | -0.170301263 | 0.39438126 | 0.88 |
| YJL047C | 709.4046862 | 0.270889241 | 0.394827649 | 0.88 |
| YMR280C | 1028.334214 | 0.348586236 | 0.394852133 | 0.88 |
| YAL016W | 2950.826603 | 0.254951323 | 0.403283205 | 0.88 |
| YAL039C | 1453.711163 | 0.24038072 | 0.398415083 | 0.88 |
| YBL082C | 673.6132807 | -0.240111802 | 0.397429519 | 0.88 |
| YBR010W | 4241.31261 | -0.633255435 | 0.401524014 | 0.88 |
| YBR112C | 3373.883429 | -0.236624968 | 0.402764835 | 0.88 |
| YBR115C | 2897.179739 | 0.554440465 | 0.396699217 | 0.88 |
| YDL006W | 361.6658456 | 0.210102062 | 0.399713072 | 0.88 |
| YDR029W | 118.6548652 | 0.506231358 | 0.400549525 | 0.88 |

| | | | | |
|---|---|---|---|---|
| YDR034C | 1497.715383 | 0.278929084 | 0.401579572 | 0.88 |
| YDR066C | 173.6197188 | 0.301592214 | 0.39914955 | 0.88 |
| YDR175C | 1042.356621 | -0.374566236 | 0.395185219 | 0.88 |
| YDR224C | 8028.945616 | -0.435715392 | 0.401537182 | 0.88 |
| YDR304C | 4614.533404 | -0.246113537 | 0.399537195 | 0.88 |
| YER124C | 2039.69695 | -0.27913701 | 0.400124725 | 0.88 |
| YER172C | 2101.126018 | -0.189488611 | 0.39969573 | 0.88 |
| YFR012W | 37.52047909 | 0.34620913 | 0.401214363 | 0.88 |
| YFR043C | 412.3285976 | 0.231764594 | 0.400748835 | 0.88 |
| YGL007C-A | 2.085025417 | 0.439513821 | 0.400830524 | 0.88 |
| YGL103W | 19.59323296 | -0.660189029 | 0.398889312 | 0.88 |
| YGL126W | 1063.33427 | -0.263409518 | 0.401974935 | 0.88 |
| YGL178W | 2073.150568 | 0.311770443 | 0.403198789 | 0.88 |
| YGR057C | 395.64537 | 0.274952861 | 0.397859258 | 0.88 |
| YGR135W | 3309.689244 | -0.255432802 | 0.400903002 | 0.88 |
| YGR169C | 437.5350343 | -0.29682613 | 0.400315892 | 0.88 |
| YGR185C | 4359.206315 | -0.15896205 | 0.400792127 | 0.88 |
| YGR253C | 3517.823624 | -0.242990361 | 0.402672049 | 0.88 |
| YHR064C | 6332.657586 | -0.042899495 | 0.400175907 | 0.88 |
| YHR092C | 9528.876224 | 0.49295815 | 0.403125386 | 0.88 |
| YIR011C | 1166.213259 | 0.214303517 | 0.403222112 | 0.88 |
| YKL099C | 686.8097655 | -0.098083244 | 0.398881613 | 0.88 |
| YKL205W | 1310.174229 | -0.259645791 | 0.402132189 | 0.88 |
| YLL026W | 27987.51895 | 0.618351618 | 0.403025716 | 0.88 |
| YLR060W | 5384.459239 | -0.095206094 | 0.402747283 | 0.88 |
| YLR251W | 493.6892026 | 0.23804594 | 0.40300622 | 0.88 |
| YLR321C | 729.7163579 | -0.293050402 | 0.401294271 | 0.88 |
| YLR330W | 3759.858385 | -0.180618913 | 0.396538955 | 0.88 |
| YLR457C | 335.6948937 | -0.229793492 | 0.396499022 | 0.88 |
| YML075C | 1371.306293 | 0.230564164 | 0.401873835 | 0.88 |
| YML102W | 604.0898534 | -0.254048482 | 0.398870648 | 0.88 |
| YMR032W | 652.0530589 | -0.279662425 | 0.40081753 | 0.88 |
| YMR095C | 600.2590168 | 0.800769987 | 0.396194953 | 0.88 |
| YMR132C | 213.3288234 | -0.238074699 | 0.395392237 | 0.88 |
| YMR304C-A | 599.6439516 | -0.349656763 | 0.402609717 | 0.88 |
| YNL019C | 10.29601328 | 0.280187992 | 0.396987175 | 0.88 |
| YNL056W | 768.1896527 | -0.204877239 | 0.396742869 | 0.88 |
| YNR027W | 549.891736 | -0.201241064 | 0.398978289 | 0.88 |
| YNR030W | 1357.008714 | -0.308388204 | 0.401083971 | 0.88 |
| YOL033W | 423.9709748 | -0.278406202 | 0.402707518 | 0.88 |
| YOL096C | 679.2678424 | 0.217842725 | 0.401550387 | 0.88 |
| YOR121C | 64.33732395 | 0.507010256 | 0.398108671 | 0.88 |
| YOR225W | 54.88254174 | 0.040595829 | 0.402764832 | 0.88 |

| | | | | |
|---|---|---|---|---|
| YOR245C | 966.5341375 | 0.278747525 | 0.398908302 | 0.88 |
| YOR300W | 17.40293291 | 0.07863472 | 0.402590301 | 0.88 |
| YOR376W | 25.04102901 | 0.34439945 | 0.401557421 | 0.88 |
| YPL030W | 552.0003609 | -0.286672859 | 0.401145148 | 0.88 |
| YPL099C | 334.8201937 | -0.310874711 | 0.397737637 | 0.88 |
| YPL108W | 159.4425969 | -0.328541871 | 0.398518089 | 0.88 |
| YPL189W | 119.6862802 | -0.199749245 | 0.399243016 | 0.88 |
| YPR013C | 1083.106605 | 0.154821872 | 0.403044616 | 0.88 |
| YPR060C | 619.4282416 | -0.208960208 | 0.395362941 | 0.88 |
| YPR103W | 3867.741987 | -0.292863891 | 0.395635954 | 0.88 |
| snR54 | 8.052861878 | 0.409638278 | 0.399343056 | 0.88 |
| YAR066W | 12.65917974 | -0.313596347 | 0.405512182 | 0.88 |
| YBR030W | 579.6269936 | -0.230218564 | 0.405679459 | 0.88 |
| YCL046W | 22.38513549 | -0.070906868 | 0.40547856 | 0.88 |
| YDL143W | 3248.97051 | -0.233978536 | 0.405094363 | 0.88 |
| YDR275W | 363.6731352 | 0.33161818 | 0.4048186 | 0.88 |
| YDR284C | 1216.7009 | -0.172092324 | 0.403940872 | 0.88 |
| YDR433W | 87.14790436 | -0.266851148 | 0.405598357 | 0.88 |
| YDR435C | 760.2512519 | 0.290729083 | 0.404171008 | 0.88 |
| YHR186C | 1649.917336 | -0.225332797 | 0.405331388 | 0.88 |
| YIL027C | 793.9722774 | -0.284106208 | 0.405142798 | 0.88 |
| YJR159W | 6.187748904 | -0.557648963 | 0.405033614 | 0.88 |
| YLR129W | 1609.244371 | -0.326657399 | 0.405185736 | 0.88 |
| YNL038W | 473.5438124 | -0.296616723 | 0.404182539 | 0.88 |
| YNL072W | 456.0809998 | -0.221821116 | 0.404555873 | 0.88 |
| YNL153C | 847.6509295 | -0.28178559 | 0.403793662 | 0.88 |
| YNL155W | 1378.201426 | 0.355584299 | 0.405376972 | 0.88 |
| YOR059C | 1230.819833 | 0.216923257 | 0.405676899 | 0.88 |
| YDR134C | 8310.682574 | 0.246844329 | 0.406472658 | 0.88 |
| YHR070C-A | 129.6976848 | -0.165189052 | 0.406522183 | 0.88 |
| YJR048W | 1046.168345 | -0.062262934 | 0.406336671 | 0.88 |
| YKL134C | 542.9022847 | 0.224246902 | 0.406758302 | 0.88 |
| YPR083W | 614.8326597 | 0.360368121 | 0.406678972 | 0.88 |
| YAL036C | 1045.610431 | -0.046849017 | 0.409813559 | 0.88 |
| YBL029C-A | 1079.485322 | 0.18030157 | 0.410004741 | 0.88 |
| YBR058C-A | 528.5056217 | -0.236710795 | 0.411225687 | 0.88 |
| YDL088C | 1480.552376 | 0.200042543 | 0.411486567 | 0.88 |
| YDL114W | 43.38451628 | 0.268033085 | 0.411932879 | 0.88 |
| YDL176W | 672.0622412 | 0.266608828 | 0.411276753 | 0.88 |
| YDR071C | 2792.908431 | -0.202928863 | 0.407885687 | 0.88 |
| YDR142C | 534.1730573 | 0.213508025 | 0.408662602 | 0.88 |
| YDR210C-D | 43.7176532 | 0.488782108 | 0.409772291 | 0.88 |
| YDR539W | 882.7410932 | -0.53970115 | 0.411570746 | 0.88 |

| | | | | |
|---|---|---|---|---|
| YER084W | 124.9336603 | -0.056985659 | 0.410582752 | 0.88 |
| YER172C-A | 0.861078059 | -0.425491448 | 0.411683401 | 0.88 |
| YGL055W | 3060.50732 | -0.251627954 | 0.41209035 | 0.88 |
| YGL142C | 1131.213732 | 0.337588587 | 0.410499074 | 0.88 |
| YGR022C | 1.81368062 | 0.299494425 | 0.40744129 | 0.88 |
| YGR258C | 1398.526849 | 0.274838566 | 0.411695104 | 0.88 |
| YHR087W | 3070.022032 | 0.00949627 | 0.408742905 | 0.88 |
| YHR199C-A | 146.5882374 | 0.0876617 | 0.408451974 | 0.88 |
| YIL152W | 734.4896336 | 0.265510769 | 0.411920257 | 0.88 |
| YJR058C | 464.9848205 | -0.261884586 | 0.412042098 | 0.88 |
| YKL108W | 391.5450461 | -0.336079292 | 0.409200805 | 0.88 |
| YLL047W | 4.30351014 | 0.403841729 | 0.408040163 | 0.88 |
| YLR028C | 4846.630925 | -0.142653674 | 0.410541286 | 0.88 |
| YLR238W | 614.7184945 | 0.219611392 | 0.41019626 | 0.88 |
| YLR359W | 9224.21068 | 0.097351028 | 0.410848384 | 0.88 |
| YLR424W | 404.0091344 | 0.201038823 | 0.408376355 | 0.88 |
| YMR002W | 4776.428936 | -0.32651158 | 0.411419475 | 0.88 |
| YMR072W | 3717.086972 | -0.219313397 | 0.410185337 | 0.88 |
| YMR216C | 1336.629718 | -0.300789504 | 0.411444363 | 0.88 |
| YMR240C | 592.2234425 | 0.209190407 | 0.411252015 | 0.88 |
| YMR265C | 688.687566 | 0.306936616 | 0.407118112 | 0.88 |
| YMR271C | 754.1723733 | 0.227735079 | 0.407540358 | 0.88 |
| YNL001W | 629.9634099 | -0.275919855 | 0.410148565 | 0.88 |
| YNL142W | 1026.084342 | 0.342614225 | 0.408307779 | 0.88 |
| YNL278W | 749.3245241 | 0.298092544 | 0.409840449 | 0.88 |
| YOL088C | 665.7794028 | -0.33337426 | 0.410482004 | 0.88 |
| YPL161C | 652.0526344 | 0.215956257 | 0.411746824 | 0.88 |
| YPR177C | 1.844393858 | 0.545134682 | 0.409319946 | 0.88 |
| YDR473C | 858.9200777 | 0.249277439 | 0.412245971 | 0.88 |
| YAL044C | 2900.555908 | -0.207152637 | 0.415062879 | 0.88 |
| YAL048C | 583.0194511 | 0.303799422 | 0.413589893 | 0.88 |
| YAR033W | 105.2011719 | -0.25158028 | 0.416409256 | 0.88 |
| YBL006C | 611.4550235 | -0.359754147 | 0.41293118 | 0.88 |
| YBL064C | 4712.365868 | 0.409323224 | 0.414776575 | 0.88 |
| YBR076C-A | 84.52994015 | 0.170225991 | 0.415446807 | 0.88 |
| YCL052C | 1614.200073 | 0.227825382 | 0.416718731 | 0.88 |
| YDL091C | 1187.448267 | 0.184250657 | 0.415894993 | 0.88 |
| YDR219C | 850.7025719 | 0.202205077 | 0.414055599 | 0.88 |
| YDR226W | 6443.35181 | -0.134327248 | 0.414037052 | 0.88 |
| YDR240C | 401.4217535 | -0.211583835 | 0.417545004 | 0.88 |
| YDR256C | 561.4619469 | 0.279857677 | 0.416595114 | 0.88 |
| YDR315C | 362.5235632 | -0.187734948 | 0.416365256 | 0.88 |
| YDR408C | 1967.993247 | -0.245662168 | 0.417112798 | 0.88 |

| | | | | |
|---|---|---|---|---|
| YEL046C | 2157.479841 | -0.177562245 | 0.416081281 | 0.88 |
| YFL015C | 19.00869044 | 0.416261967 | 0.414758452 | 0.88 |
| YFL029C | 966.6378114 | 0.405358637 | 0.417472182 | 0.88 |
| YHR001W | 1515.116914 | 0.258270324 | 0.41400973 | 0.88 |
| YIL098C | 444.8949209 | -0.402356195 | 0.416867404 | 0.88 |
| YIL110W | 498.0390037 | -0.24215058 | 0.416781386 | 0.88 |
| YIL140W | 1095.766571 | -0.269611649 | 0.417292487 | 0.88 |
| YJL022W | 2.660856745 | 0.357315526 | 0.415730911 | 0.88 |
| YJL077W-B | 26.1550574 | 0.699306826 | 0.415156167 | 0.88 |
| YJL146W | 600.4909491 | 0.201397367 | 0.417191366 | 0.88 |
| YJL150W | 10.09973158 | 0.530979265 | 0.412828611 | 0.88 |
| YKL014C | 1967.775598 | -0.216312719 | 0.416386048 | 0.88 |
| YKL115C | 42.82706794 | -0.204213844 | 0.415368041 | 0.88 |
| YLL008W | 1923.071209 | -0.026232732 | 0.41463541 | 0.88 |
| YLR132C | 337.4595812 | 0.226515788 | 0.417584549 | 0.88 |
| YLR217W | 16.95655225 | 0.359311754 | 0.41305531 | 0.88 |
| YMR205C | 7427.558626 | -0.199337513 | 0.416181034 | 0.88 |
| YMR314W | 2980.188475 | -0.199023495 | 0.417148669 | 0.88 |
| YMR317W | 752.1354442 | 0.25147963 | 0.413770905 | 0.88 |
| YOL085W-A | 20.71787704 | 0.283962369 | 0.412827008 | 0.88 |
| YOL151W | 926.9123911 | -0.376546288 | 0.414185688 | 0.88 |
| YOR346W | 878.2111125 | -0.262668606 | 0.416520992 | 0.88 |
| YPL109C | 614.5552418 | 0.208359145 | 0.412654744 | 0.88 |
| YPL157W | 499.4077603 | -0.243373776 | 0.415937864 | 0.88 |
| snR61 | 12.13748594 | -0.061508989 | 0.417466328 | 0.88 |
| YHR091C | 596.7962742 | -0.187439547 | 0.418028856 | 0.88 |
| YMR278W | 1574.862371 | 0.280282096 | 0.418013629 | 0.88 |
| YBL113C | 286.5211358 | 0.463998744 | 0.418617658 | 0.88 |
| YDL063C | 706.2977789 | -0.106679738 | 0.419026966 | 0.88 |
| YKL167C | 608.6144606 | -0.300648066 | 0.419317378 | 0.88 |
| YNL013C | 5.629906296 | 0.039238663 | 0.419174456 | 0.88 |
| YNL243W | 3599.331124 | -0.176577492 | 0.419383418 | 0.88 |
| YNL274C | 1846.518968 | 0.305886707 | 0.419475428 | 0.88 |
| YOR017W | 870.202076 | 0.203581029 | 0.418977553 | 0.88 |
| YAL026C-A | 170.5922701 | 0.22185118 | 0.42107729 | 0.88 |
| YBL034C | 918.1202298 | -0.253281822 | 0.420147836 | 0.88 |
| YBL076C | 7280.098127 | 0.12131642 | 0.420217224 | 0.88 |
| YBR063C | 508.4120941 | 0.175259058 | 0.421235672 | 0.88 |
| YBR157C | 589.2630079 | -0.076652026 | 0.422933415 | 0.88 |
| YBR161W | 517.5371153 | 0.237676695 | 0.422905875 | 0.88 |
| YCL057C-A | 1206.33424 | -0.32528151 | 0.421231703 | 0.88 |
| YCL064C | 3239.199963 | -0.734128652 | 0.422802771 | 0.88 |
| YDR230W | 158.0472236 | 0.328647226 | 0.421831217 | 0.88 |

| | | | | |
|---|---|---|---|---|
| YDR442W | 17.52253569 | -0.352544605 | 0.422363149 | 0.88 |
| YER050C | 1168.128624 | -0.325420999 | 0.421642634 | 0.88 |
| YFL002C | 1100.586944 | 0.20005809 | 0.421915612 | 0.88 |
| YGL069C | 85.63194987 | -0.502215126 | 0.421843136 | 0.88 |
| YGR065C | 698.7091847 | 0.38459282 | 0.421269823 | 0.88 |
| YJL208C | 631.1153592 | -0.084685191 | 0.422232909 | 0.88 |
| YJR050W | 528.5017956 | 0.279625889 | 0.422860984 | 0.88 |
| YLL011W | 676.0042616 | -0.384649151 | 0.420591195 | 0.88 |
| YLL027W | 1273.25217 | 0.237930358 | 0.420041917 | 0.88 |
| YLR012C | 11.76101632 | 0.060302009 | 0.421398789 | 0.88 |
| YLR191W | 472.4930163 | 0.265919295 | 0.421989529 | 0.88 |
| YMR211W | 431.1277672 | 0.188282277 | 0.422933855 | 0.88 |
| YMR316C-B | 10.98411933 | 0.142935437 | 0.422567171 | 0.88 |
| YNR036C | 1583.864307 | -0.443005088 | 0.421936562 | 0.88 |
| YNR056C | 522.7352776 | 0.098949969 | 0.421105729 | 0.88 |
| YPL260W | 2621.937685 | 0.205103046 | 0.422661025 | 0.88 |
| YBR038W | 1800.515437 | -0.167186723 | 0.424778062 | 0.88 |
| YCR006C | 58.71736424 | -0.52667963 | 0.424014592 | 0.88 |
| YDL112W | 1910.821894 | -0.176723071 | 0.424069401 | 0.88 |
| YDR329C | 1759.988143 | 0.34964254 | 0.424949381 | 0.88 |
| YDR352W | 489.6613604 | -0.167146098 | 0.425090774 | 0.88 |
| YGL149W | 44.44871907 | -0.275232382 | 0.424924603 | 0.88 |
| YHR167W | 161.5282471 | -0.26575963 | 0.425001602 | 0.88 |
| YLR059C | 1264.300383 | -0.169432392 | 0.42419771 | 0.88 |
| YLR097C | 748.1202732 | 0.260092851 | 0.423527772 | 0.88 |
| YML039W | 1.131094598 | -0.199443588 | 0.423624536 | 0.88 |
| YNL092W | 338.5129162 | 0.400658231 | 0.423415413 | 0.88 |
| YNR046W | 947.7260883 | -0.168324919 | 0.424332397 | 0.88 |
| YOL032W | 1063.876828 | 0.197643404 | 0.424015518 | 0.88 |
| YOR025W | 737.1839475 | -0.295563153 | 0.424725711 | 0.88 |
| YOR316C-A | 8.072827245 | -0.370576957 | 0.424172903 | 0.88 |
| YHR074W | 2090.525023 | -0.198762959 | 0.425387102 | 0.88 |
| YDR192C | 476.3597916 | 0.197620744 | 0.425791843 | 0.89 |
| YLR109W | 26967.32056 | -0.230574453 | 0.425923792 | 0.89 |
| YPR012W | 66.76164095 | 0.225698703 | 0.425985053 | 0.89 |
| YBR117C | 1336.758461 | -0.216400874 | 0.42614286 | 0.89 |
| YGR213C | 279.0368426 | 0.27332251 | 0.426971954 | 0.89 |
| YHR115C | 896.8338505 | 0.24931774 | 0.426869478 | 0.89 |
| YLR255C | 23.50063941 | 0.376868376 | 0.426841552 | 0.89 |
| YOR027W | 11708.63489 | -0.101027773 | 0.426678472 | 0.89 |
| YBR209W | 5.517408049 | -0.467918356 | 0.427146361 | 0.89 |
| YJL052W | 23370.62399 | -0.47875124 | 0.427327129 | 0.89 |
| YNL094W | 1768.042353 | 0.225920243 | 0.427971095 | 0.89 |

| | | | | |
|---|---|---|---|---|
| YOR051C | 2007.894821 | -0.122104583 | 0.427904748 | 0.89 |
| YBL080C | 486.2802333 | 0.209188015 | 0.428739971 | 0.89 |
| YBR217W | 353.7676501 | 0.252165886 | 0.428809011 | 0.89 |
| YDR075W | 328.511722 | -0.358331494 | 0.428512735 | 0.89 |
| YDR339C | 1137.925374 | -0.366670432 | 0.428627188 | 0.89 |
| YOL001W | 429.8453508 | 0.157202042 | 0.428560356 | 0.89 |
| YPR149W | 11719.12039 | 0.299453142 | 0.428329203 | 0.89 |
| YCL005W | 658.6999475 | -0.209430608 | 0.429089715 | 0.89 |
| YJR154W | 284.604362 | 0.421021982 | 0.429137427 | 0.89 |
| YAL033W | 350.8440347 | -0.301209801 | 0.431509915 | 0.89 |
| YBL071C | 215.271228 | 0.236489295 | 0.432057828 | 0.89 |
| YDL103C | 1449.48201 | -0.196050512 | 0.432063939 | 0.89 |
| YDL174C | 2966.298062 | 0.150569006 | 0.429451575 | 0.89 |
| YDL185C-A | 100.139643 | 0.420463832 | 0.430399999 | 0.89 |
| YDR050C | 39610.42901 | -0.370401968 | 0.431348905 | 0.89 |
| YDR077W | 88090.2349 | 0.270750245 | 0.430717454 | 0.89 |
| YDR244W | 923.2552739 | 0.210251313 | 0.431785344 | 0.89 |
| YDR300C | 1204.265207 | -0.21844195 | 0.42959916 | 0.89 |
| YDR311W | 727.3776002 | 0.208616048 | 0.430230036 | 0.89 |
| YDR378C | 714.4468599 | -0.188833375 | 0.431072955 | 0.89 |
| YER119C-A | 190.6802462 | -0.233087179 | 0.431719918 | 0.89 |
| YGL074C | 37.00723578 | -0.186796726 | 0.429465892 | 0.89 |
| YHR056C | 832.3445362 | 0.234309352 | 0.429965973 | 0.89 |
| YHR104W | 3547.736943 | 0.264452447 | 0.4306835 | 0.89 |
| YJR111C | 500.5453341 | 0.323469312 | 0.431452989 | 0.89 |
| YML009C-A | 52.04514359 | -0.077067665 | 0.430202357 | 0.89 |
| YMR197C | 923.7331716 | 0.22178637 | 0.430124179 | 0.89 |
| YNL054W-B | 451.4424074 | 0.745510269 | 0.430490292 | 0.89 |
| YOL083W | 724.7720388 | 0.207581343 | 0.432173767 | 0.89 |
| YOR021C | 1473.922183 | -0.104272144 | 0.431588973 | 0.89 |
| YPL160W | 4849.742262 | 0.08303196 | 0.430814726 | 0.89 |
| YDR118W-A | 11.72360647 | 0.29564046 | 0.43234409 | 0.89 |
| YBR241C | 1365.325299 | 0.214701777 | 0.433035399 | 0.89 |
| YCL056C | 401.4808141 | 0.171267397 | 0.432987803 | 0.89 |
| YGL106W | 2381.873892 | -0.268316818 | 0.432730726 | 0.89 |
| YGL155W | 739.9393467 | -0.225749283 | 0.43266555 | 0.89 |
| YNR026C | 756.2847135 | -0.286925743 | 0.432982888 | 0.89 |
| YCR002C | 1893.656785 | -0.174704817 | 0.434083073 | 0.89 |
| YCR011C | 2808.118139 | 0.281657817 | 0.434663822 | 0.89 |
| YCR071C | 461.1120902 | -0.451506573 | 0.43551751 | 0.89 |
| YDL068W | 4.863206233 | -0.76271176 | 0.435011441 | 0.89 |
| YDR028C | 3835.45418 | 0.292388463 | 0.434302847 | 0.89 |
| YDR044W | 1839.127613 | -0.318522555 | 0.433723192 | 0.89 |

| | | | | |
|---|---|---|---|---|
| YEL074W | 35.0602757 | 0.19306567 | 0.435109659 | 0.89 |
| YER076W-A | 76.7457419 | 0.139045245 | 0.433958291 | 0.89 |
| YFL038C | 5600.505944 | -0.22893793 | 0.435273299 | 0.89 |
| YGL160W | 1159.891272 | 0.176760344 | 0.43555519 | 0.89 |
| YGR157W | 3036.942102 | -0.161962855 | 0.434478601 | 0.89 |
| YHL043W | 209.5139873 | 0.067514145 | 0.435170681 | 0.89 |
| YHR119W | 1485.712969 | -0.211674146 | 0.433836958 | 0.89 |
| YMR243C | 2759.15272 | -0.143392612 | 0.435553653 | 0.89 |
| YNL138W | 4814.982276 | -0.210850859 | 0.433870335 | 0.89 |
| YNL141W | 1283.346108 | -0.044957465 | 0.434452838 | 0.89 |
| YOR251C | 1296.455018 | -0.151756466 | 0.43534249 | 0.89 |
| YPR042C | 1338.934548 | 0.267359919 | 0.434934265 | 0.89 |
| YGR030C | 291.5946813 | -0.176496399 | 0.435756347 | 0.89 |
| YNL320W | 323.6388355 | -0.266555382 | 0.43586512 | 0.89 |
| YFL051C | 87.84562174 | 0.093198527 | 0.436840711 | 0.89 |
| YJL127C | 657.9035431 | 0.213100483 | 0.436176397 | 0.89 |
| YLR269C | 113.3738155 | 0.302167527 | 0.436491865 | 0.89 |
| YLR380W | 1647.260273 | -0.252788547 | 0.436799922 | 0.89 |
| YNL043C | 66.28583242 | -0.536021446 | 0.436849574 | 0.89 |
| snR48 | 2.926793808 | -0.257097679 | 0.436735665 | 0.89 |
| YAL065C | 13.48444795 | -0.105632923 | 0.438043288 | 0.89 |
| YDR478W | 183.0152712 | -0.258823146 | 0.439162673 | 0.89 |
| YER089C | 1927.936186 | -0.249221254 | 0.439334416 | 0.89 |
| YER129W | 2291.22394 | 0.18093034 | 0.438612736 | 0.89 |
| YGL159W | 407.2189918 | -0.274092153 | 0.439038539 | 0.89 |
| YGL163C | 996.8369865 | 0.229119366 | 0.438805697 | 0.89 |
| YGR188C | 503.2494794 | 0.342061021 | 0.439284277 | 0.89 |
| YJL029C | 1512.843835 | 0.184501021 | 0.438789902 | 0.89 |
| YJR012C | 286.1622087 | 0.233968031 | 0.437787438 | 0.89 |
| YMR274C | 249.323644 | -0.192738743 | 0.438050124 | 0.89 |
| YOL108C | 360.9401457 | 0.075347706 | 0.438821976 | 0.89 |
| YOR291W | 1661.347699 | 0.220033484 | 0.43881312 | 0.89 |
| YPL022W | 1488.077225 | 0.256313847 | 0.438170104 | 0.89 |
| YPL053C | 1363.897708 | -0.352140657 | 0.438207396 | 0.89 |
| YPR173C | 1842.223717 | -0.243321153 | 0.437495664 | 0.89 |
| YIL066W-A | 3.308676015 | 0.37455919 | 0.43959674 | 0.89 |
| YMR078C | 885.7332334 | -0.213560448 | 0.439721097 | 0.89 |
| snR38 | 6.274197792 | 0.74290126 | 0.439876108 | 0.89 |
| YIR020C-B | 5.673146071 | -0.046447008 | 0.440296246 | 0.89 |
| YJL061W | 1142.158213 | -0.21154565 | 0.440042868 | 0.89 |
| YKL071W | 436.0298233 | 0.173482107 | 0.440184591 | 0.89 |
| YLR102C | 476.3043851 | 0.211990157 | 0.440490896 | 0.89 |
| YLL046C | 28.77526537 | 0.282017516 | 0.441190534 | 0.89 |

| | | | | |
|---|---|---|---|---|
| YLR224W | 1026.056679 | 0.220916825 | 0.441125486 | 0.89 |
| YNL258C | 964.2763945 | -0.240481197 | 0.441332605 | 0.89 |
| YAL038W | 41041.91188 | 0.067614058 | 0.442393887 | 0.89 |
| YBL069W | 640.7564823 | 0.431303697 | 0.44208996 | 0.89 |
| YBR024W | 889.4330761 | 0.391372024 | 0.44199095 | 0.89 |
| YBR251W | 1030.356421 | -0.334805259 | 0.442850448 | 0.89 |
| YCR105W | 210.4429239 | -0.731754241 | 0.442963794 | 0.89 |
| YJL037W | 69.21169324 | 0.124001778 | 0.442758891 | 0.89 |
| YKR032W | 0.979223699 | 0.404547388 | 0.443468178 | 0.89 |
| YML038C | 1118.066731 | 0.23306492 | 0.443487292 | 0.89 |
| YMR208W | 1260.14289 | -0.159538485 | 0.443279711 | 0.89 |
| YNL116W | 793.4753006 | 0.350388452 | 0.443464212 | 0.89 |
| YNL207W | 833.7029402 | -0.184189257 | 0.441888498 | 0.89 |
| YNL228W | 12.1493579 | 0.164417518 | 0.443204537 | 0.89 |
| YNL236W | 1176.860728 | -0.227727852 | 0.442775272 | 0.89 |
| YOR388C | 10.58733181 | 0.077160523 | 0.442047648 | 0.89 |
| snR66 | 7.293652703 | -0.122914471 | 0.443192058 | 0.89 |
| YDR052C | 1279.720929 | -0.240308502 | 0.443764979 | 0.89 |
| YBL014C | 840.8039898 | 0.2962113 | 0.445215513 | 0.89 |
| YBR156C | 807.2285578 | 0.188759093 | 0.444236426 | 0.89 |
| YBR296C-A | 3.00001637 | 0.257420803 | 0.445876834 | 0.89 |
| YDR264C | 2392.197288 | -0.196660454 | 0.445187139 | 0.89 |
| YDR505C | 1755.081331 | 0.315734935 | 0.445578602 | 0.89 |
| YER053C-A | 179.222376 | 0.143933391 | 0.444840496 | 0.89 |
| YHL039W | 1028.011858 | -0.149010298 | 0.444543401 | 0.89 |
| YIL002C | 1259.267656 | 0.216065035 | 0.44439421 | 0.89 |
| YJL131C | 712.8647158 | 0.26356127 | 0.445232705 | 0.89 |
| YMR122C | 59.11544635 | 0.203284923 | 0.445344443 | 0.89 |
| YNL174W | 7.421723425 | 0.098940784 | 0.444659484 | 0.89 |
| YNL255C | 4025.75008 | -0.180395115 | 0.44570636 | 0.89 |
| YNL275W | 352.1195966 | 0.233167399 | 0.445820608 | 0.89 |
| YOL069W | 510.0668666 | -0.201942083 | 0.444186818 | 0.89 |
| YPR198W | 724.9139692 | -0.333834345 | 0.444966328 | 0.89 |
| YGL060W | 769.6330407 | 0.233754355 | 0.446087468 | 0.89 |
| YLR085C | 664.5290334 | 0.197367782 | 0.446325405 | 0.89 |
| YLR157C-B | 0.750636836 | -0.194567333 | 0.446601889 | 0.89 |
| YOL076W | 856.9107843 | -0.28241497 | 0.446564216 | 0.89 |
| YOR218C | 5.229072963 | 0.457143409 | 0.446741787 | 0.89 |
| YDR335W | 3040.69958 | 0.275516866 | 0.447257482 | 0.89 |
| YFL008W | 1609.672053 | -0.214362161 | 0.44760656 | 0.89 |
| YGR240C | 9640.703605 | -0.169635868 | 0.448136189 | 0.89 |
| YHR019C | 9735.850466 | 0.02906322 | 0.448064756 | 0.89 |
| YJR055W | 115.4214659 | -0.250499049 | 0.448051814 | 0.89 |

| | | | | |
|---|---|---|---|---|
| YML099C | 1070.053648 | 0.283804295 | 0.448713112 | 0.89 |
| YOL018C | 882.7418897 | 0.16635132 | 0.4486009 | 0.89 |
| YBR050C | 396.3143748 | 0.10727181 | 0.449738728 | 0.89 |
| YCL076W | 24.7210401 | 0.082526129 | 0.449976303 | 0.89 |
| YDL100C | 5446.231089 | -0.05002029 | 0.451876728 | 0.89 |
| YDL108W | 550.3870487 | 0.187122581 | 0.452021054 | 0.89 |
| YDL133C-A | 72.78703027 | -0.577456516 | 0.449430888 | 0.89 |
| YDR011W | 4201.778813 | -0.315086186 | 0.452043451 | 0.89 |
| YDR185C | 264.7477216 | 0.328973109 | 0.449602783 | 0.89 |
| YDR452W | 4294.15743 | 0.274051744 | 0.451399697 | 0.89 |
| YER047C | 911.6547256 | 0.205952455 | 0.451174276 | 0.89 |
| YGL164C | 861.3475827 | 0.250885367 | 0.451172466 | 0.89 |
| YGL217C | 8.839690893 | -0.302614224 | 0.451872527 | 0.89 |
| YGR262C | 876.7108592 | -0.138317344 | 0.450830076 | 0.89 |
| YKR106W | 110.7787445 | -0.70693775 | 0.449765986 | 0.89 |
| YLL061W | 447.0050952 | 0.331401492 | 0.449579037 | 0.89 |
| YLR248W | 3268.331309 | 0.317480645 | 0.451711606 | 0.89 |
| YLR439W | 1150.159115 | -0.311410282 | 0.45069861 | 0.89 |
| YML017W | 751.7187367 | -0.209838983 | 0.449269383 | 0.89 |
| YML093W | 1683.648354 | -0.232743276 | 0.451728189 | 0.89 |
| YML126C | 4399.479194 | -0.023710044 | 0.45094654 | 0.89 |
| YMR139W | 1446.676083 | 0.261690201 | 0.451564208 | 0.89 |
| YNR059W | 692.5145797 | 0.185090501 | 0.451144902 | 0.89 |
| YOR046C | 2190.495993 | -0.177207708 | 0.450756059 | 0.89 |
| YOR173W | 2098.419186 | 0.188127468 | 0.450193206 | 0.89 |
| YPL222W | 848.2140163 | 0.13563976 | 0.449202571 | 0.89 |
| YHR020W | 3665.040866 | -0.035298349 | 0.452309433 | 0.89 |
| YHL006W-A | 55.86611926 | -0.310727707 | 0.452453114 | 0.89 |
| YFL034C-A | 576.1615287 | -0.53381794 | 0.452899662 | 0.89 |
| YLL066W-B | 2.220126499 | 0.050613885 | 0.452850735 | 0.89 |
| YNL047C | 720.3103825 | 0.226141397 | 0.452840241 | 0.89 |
| YKL024C | 1589.092419 | -0.219187225 | 0.453261832 | 0.89 |
| YKL163W | 2089.664663 | -0.135805071 | 0.453589963 | 0.89 |
| YNL299W | 424.5188557 | -0.213431395 | 0.453666095 | 0.89 |
| YOR366W | 3.244594662 | -0.127206115 | 0.453432397 | 0.89 |
| YBR021W | 496.9097626 | -0.407888452 | 0.454607342 | 0.89 |
| YDR365C | 4849.022737 | 0.347348124 | 0.454064238 | 0.89 |
| YLR315W | 96.87354306 | 0.224688307 | 0.45404691 | 0.89 |
| YNL042W-B | 14.4556208 | -0.358833674 | 0.454606926 | 0.89 |
| YPL085W | 4050.180373 | -0.210147434 | 0.454333155 | 0.89 |
| snR85 | 18.57666508 | -0.469322519 | 0.454281191 | 0.89 |
| YLR430W | 3065.658429 | -0.131541587 | 0.455114278 | 0.89 |
| YOR179C | 265.678804 | -0.336697868 | 0.45518374 | 0.89 |

| | | | | |
|---|---|---|---|---|
| YDR201W | 235.4443499 | -0.283528135 | 0.45542034 | 0.89 |
| YER159C | 1569.026083 | -0.278484115 | 0.455593023 | 0.89 |
| YNL018C | 116.9826504 | 0.231962697 | 0.455634851 | 0.89 |
| YGR094W | 6966.605497 | 0.016624693 | 0.45615298 | 0.89 |
| YNR074C | 411.2345917 | -0.404938396 | 0.456224504 | 0.89 |
| YPL004C | 14429.78474 | 0.272404191 | 0.456075108 | 0.89 |
| YPR139C | 1065.385559 | -0.285202645 | 0.456199011 | 0.89 |
| RUF21 | 43.48350584 | -0.091912205 | 0.467132168 | 0.89 |
| RUF22 | 1.126113591 | -0.545060214 | 0.472670152 | 0.89 |
| YAL037C-A | 1.195205434 | -0.386552868 | 0.489347546 | 0.89 |
| YAL049C | 1497.895567 | 0.296503864 | 0.462015686 | 0.89 |
| YBL038W | 1166.095027 | -0.331273775 | 0.489454515 | 0.89 |
| YBL084C | 817.2451769 | 0.121284229 | 0.479296741 | 0.89 |
| YBL104C | 644.4185242 | -0.194040926 | 0.482812636 | 0.89 |
| YBR009C | 3036.286227 | -0.394439984 | 0.489515316 | 0.89 |
| YBR012C | 184.3218002 | 0.52294391 | 0.47771059 | 0.89 |
| YBR033W | 734.5856264 | 0.233342786 | 0.457497221 | 0.89 |
| YBR078W | 17879.15973 | -0.226530736 | 0.458800896 | 0.89 |
| YBR083W | 2339.410841 | 0.126845708 | 0.476021669 | 0.89 |
| YBR116C | 35.41794033 | -0.172372359 | 0.482393466 | 0.89 |
| YBR122C | 1316.133892 | -0.226768731 | 0.471315534 | 0.89 |
| YBR136W | 1309.50165 | 0.327429986 | 0.476810233 | 0.89 |
| YBR153W | 413.971492 | 0.174767145 | 0.47594219 | 0.89 |
| YBR158W | 2964.217702 | -0.318490423 | 0.482828661 | 0.89 |
| YBR169C | 4813.961053 | 0.266866623 | 0.462643683 | 0.89 |
| YBR183W | 786.7022482 | 0.206900916 | 0.460182723 | 0.89 |
| YBR198C | 1683.847961 | -0.238182293 | 0.458138239 | 0.89 |
| YBR285W | 449.6014678 | 0.116413946 | 0.484827002 | 0.89 |
| YCL048W-A | 89.40578665 | -0.404534482 | 0.477283187 | 0.89 |
| YCR003W | 578.0986481 | -0.348192346 | 0.479588907 | 0.89 |
| YCR010C | 273.3869127 | -0.031498202 | 0.458462003 | 0.89 |
| YCR024C | 641.2724541 | 0.166865545 | 0.481035742 | 0.89 |
| YCR048W | 995.8304266 | 0.282674396 | 0.471691497 | 0.89 |
| YCR073W-A | 1340.952291 | 0.373159569 | 0.46724358 | 0.89 |
| YDL035C | 1472.135402 | 0.245527488 | 0.474649088 | 0.89 |
| YDL056W | 1411.798857 | 0.130102003 | 0.486914185 | 0.89 |
| YDL124W | 12433.87204 | 0.329437446 | 0.46091969 | 0.89 |
| YDL157C | 952.2291983 | -0.008586135 | 0.465118621 | 0.89 |
| YDL162C | 2.875843937 | -0.579299546 | 0.479945801 | 0.89 |
| YDL189W | 1070.015735 | -0.292845911 | 0.468238152 | 0.89 |
| YDL205C | 1039.950483 | -0.171881725 | 0.480185385 | 0.89 |
| YDL243C | 368.4938618 | 0.323734177 | 0.457387747 | 0.89 |
| YDR101C | 1247.120699 | -0.174982078 | 0.485165345 | 0.89 |

| | | | | |
|---|---|---|---|---|
| YDR119W | 1442.902962 | -0.183174644 | 0.488577579 | 0.89 |
| YDR125C | 356.5508843 | 0.220659786 | 0.489114435 | 0.89 |
| YDR156W | 897.9312179 | -0.156776179 | 0.489620742 | 0.89 |
| YDR186C | 2571.215729 | 0.175941476 | 0.465436011 | 0.89 |
| YDR205W | 692.1640308 | 0.176764699 | 0.488163729 | 0.89 |
| YDR216W | 2843.606015 | 0.12514794 | 0.485544876 | 0.89 |
| YDR234W | 1972.622196 | 0.29633298 | 0.486660683 | 0.89 |
| YDR282C | 383.6210181 | 0.232570854 | 0.466898436 | 0.89 |
| YDR291W | 877.4001219 | 0.205641424 | 0.460282023 | 0.89 |
| YDR390C | 1683.184066 | -0.161526781 | 0.467394326 | 0.89 |
| YDR427W | 2923.459081 | -0.22251801 | 0.472554829 | 0.89 |
| YDR482C | 359.2978323 | 0.173723306 | 0.488914054 | 0.89 |
| YDR494W | 1564.77137 | -0.237113199 | 0.485643072 | 0.89 |
| YDR502C | 1141.466361 | 0.464580728 | 0.481940555 | 0.89 |
| YDR510C-A | 439.1253795 | -0.537179582 | 0.481585822 | 0.89 |
| YDR516C | 6053.645479 | 0.334945454 | 0.476862276 | 0.89 |
| YDR524C | 904.6436686 | -0.249815697 | 0.482956991 | 0.89 |
| YDR531W | 969.2146497 | -0.221262565 | 0.46636075 | 0.89 |
| YEL015W | 1435.206957 | -0.193334749 | 0.486324034 | 0.89 |
| YEL036C | 1490.088365 | -0.24317485 | 0.461503576 | 0.89 |
| YER045C | 814.7735686 | 0.062841798 | 0.482278264 | 0.89 |
| YER046W-A | 13.59530817 | -0.119478826 | 0.478871684 | 0.89 |
| YER125W | 2425.853452 | -0.212801741 | 0.483014849 | 0.89 |
| YER128W | 281.9899223 | 0.23417324 | 0.46074116 | 0.89 |
| YER132C | 1149.8348 | -0.220921296 | 0.459396848 | 0.89 |
| YER138W-A | 1.112909077 | 0.093875599 | 0.477280846 | 0.89 |
| YER148W-A | 0.781507553 | -0.475366822 | 0.463906771 | 0.89 |
| YER176W | 1918.703807 | -0.107549375 | 0.472099966 | 0.89 |
| YFL012W | 27.05345673 | 0.174521638 | 0.47180172 | 0.89 |
| YFL047W | 1142.35423 | -0.360014636 | 0.477609101 | 0.89 |
| YFL052W | 300.3969613 | 0.113272639 | 0.478551462 | 0.89 |
| YFR011C | 642.0102271 | -0.142082994 | 0.463983024 | 0.89 |
| YGL007W | 19.49434054 | 0.056081952 | 0.485989893 | 0.89 |
| YGL033W | 39.3922468 | 0.462626848 | 0.470085094 | 0.89 |
| YGL048C | 4170.423196 | -0.156642 | 0.48426426 | 0.89 |
| YGL049C | 2260.025541 | 0.188790809 | 0.459829623 | 0.89 |
| YGL064C | 383.7385393 | -0.256599772 | 0.467860617 | 0.89 |
| YGL073W | 2411.314838 | 0.21277855 | 0.463136451 | 0.89 |
| YGL075C | 312.3284424 | 0.21967005 | 0.458506067 | 0.89 |
| YGL078C | 1925.6458 | -0.116058407 | 0.458124124 | 0.89 |
| YGL108C | 328.3815974 | 0.177810829 | 0.477931591 | 0.89 |
| YGL139W | 1235.197549 | 0.185894377 | 0.473514843 | 0.89 |
| YGL148W | 4268.613078 | 0.064504288 | 0.481469988 | 0.89 |

| | | | | |
|---|---|---|---|---|
| YGL157W | 1718.848717 | -0.132270098 | 0.468513452 | 0.89 |
| YGL213C | 596.0286848 | -0.174848601 | 0.462042065 | 0.89 |
| YGL219C | 1273.719314 | 0.278533025 | 0.466027427 | 0.89 |
| YGL237C | 638.8350245 | 0.191268364 | 0.46177256 | 0.89 |
| YGL252C | 1693.94272 | -0.174263627 | 0.469135058 | 0.89 |
| YGR040W | 494.1182396 | -0.242128739 | 0.469414398 | 0.89 |
| YGR043C | 1676.63942 | -0.006634076 | 0.463189921 | 0.89 |
| YGR046W | 906.1577966 | 0.117269963 | 0.479805276 | 0.89 |
| YGR059W | 83.10299922 | 0.135819205 | 0.489438486 | 0.89 |
| YGR076C | 617.1306078 | -0.297126941 | 0.468142381 | 0.89 |
| YGR092W | 957.140587 | -0.163251596 | 0.477860837 | 0.89 |
| YGR098C | 1335.545417 | -0.198022507 | 0.467051048 | 0.89 |
| YGR116W | 3744.798928 | -0.17320228 | 0.483326217 | 0.89 |
| YGR129W | 435.9932211 | 0.286900215 | 0.483174156 | 0.89 |
| YGR133W | 486.059603 | 0.2230117 | 0.462606496 | 0.89 |
| YGR161C-D | 41.89517477 | 0.358335553 | 0.485970775 | 0.89 |
| YGR167W | 3020.873713 | -0.174242682 | 0.469949193 | 0.89 |
| YGR211W | 3320.437529 | -0.071973496 | 0.486473674 | 0.89 |
| YGR217W | 881.6554263 | -0.116424415 | 0.484276346 | 0.89 |
| YGR266W | 1152.085768 | 0.291458463 | 0.469415298 | 0.89 |
| YGR276C | 1041.106244 | 0.179150473 | 0.485512241 | 0.89 |
| YGR294W | 10.95232902 | -0.229615572 | 0.472729387 | 0.89 |
| YHL019C | 544.6940331 | 0.318373555 | 0.463383004 | 0.89 |
| YHL027W | 1629.795881 | 0.231689631 | 0.485513876 | 0.89 |
| YHL046C | 9.174623277 | 0.359621927 | 0.465418542 | 0.89 |
| YHR029C | 965.1862947 | 0.515680834 | 0.481893672 | 0.89 |
| YHR032W-A | 60.96537533 | -0.498129542 | 0.476288965 | 0.89 |
| YHR040W | 740.6643017 | 0.334782868 | 0.485058052 | 0.89 |
| YHR148W | 337.7936716 | -0.12334216 | 0.483538684 | 0.89 |
| YHR170W | 1876.524363 | -0.063958007 | 0.469262172 | 0.89 |
| YHR183W | 10217.98699 | 0.153410343 | 0.469629508 | 0.89 |
| YHR192W | 784.2571123 | -0.194136161 | 0.487235773 | 0.89 |
| YIL024C | 346.2449471 | 0.32737218 | 0.48864561 | 0.89 |
| YIL033C | 4130.82095 | 0.364820242 | 0.477134009 | 0.89 |
| YIL042C | 814.6115743 | 0.140680144 | 0.484249704 | 0.89 |
| YIL064W | 505.7570311 | 0.053576163 | 0.476303295 | 0.89 |
| YIL089W | 155.6273305 | 0.313869476 | 0.477871591 | 0.89 |
| YIL114C | 411.0869055 | -0.195939877 | 0.487879446 | 0.89 |
| YIR007W | 898.4409432 | 0.226553014 | 0.482418654 | 0.89 |
| YIR019C | 622.3080892 | 0.157586615 | 0.486908168 | 0.89 |
| YJL027C | 10.13200317 | 0.455551717 | 0.459350665 | 0.89 |
| YJL033W | 1448.230029 | -0.124284061 | 0.459246371 | 0.89 |
| YJL127W-A | 12.80997666 | 0.088876226 | 0.468163951 | 0.89 |

| | | | | |
|---|---|---|---|---|
| YJL154C | 2489.813399 | 0.164318186 | 0.461455931 | 0.89 |
| YJL213W | 560.8105438 | 0.041592526 | 0.478684722 | 0.89 |
| YJL218W | 146.9624687 | 0.145178044 | 0.469010485 | 0.89 |
| YJL219W | 39.81053365 | -0.452329357 | 0.488046825 | 0.89 |
| YJR005W | 1677.202028 | 0.169598565 | 0.482176024 | 0.89 |
| YJR053W | 291.1479634 | -0.187883495 | 0.487389522 | 0.89 |
| YJR157W | 1.945110995 | -0.816690525 | 0.479362111 | 0.89 |
| YKL008C | 1304.945966 | -0.318373184 | 0.459109856 | 0.89 |
| YKL013C | 2002.473422 | -0.210871331 | 0.489284062 | 0.89 |
| YKL025C | 1634.326653 | 0.21180826 | 0.485806779 | 0.89 |
| YKL026C | 613.714189 | 0.018960219 | 0.462369044 | 0.89 |
| YKL052C | 501.3271381 | -0.197091091 | 0.484468405 | 0.89 |
| YKL053W | 17.58730316 | 0.1643535 | 0.473798742 | 0.89 |
| YKL055C | 140.4891808 | 0.212808599 | 0.45665525 | 0.89 |
| YKL073W | 1776.826393 | -0.151615619 | 0.459939383 | 0.89 |
| YKL104C | 3816.518475 | -0.119188101 | 0.482574316 | 0.89 |
| YKL195W | 3025.520768 | -0.197993235 | 0.479389159 | 0.89 |
| YKL198C | 1074.404315 | 0.150032586 | 0.483681106 | 0.89 |
| YKL203C | 2057.606082 | -0.188071322 | 0.465469532 | 0.89 |
| YKL204W | 2110.527656 | -0.282718865 | 0.470417319 | 0.89 |
| YKR011C | 1595.285427 | 0.281098671 | 0.468747417 | 0.89 |
| YKR012C | 142.5434976 | -0.050750437 | 0.473855497 | 0.89 |
| YKR076W | 1274.281879 | 0.135654663 | 0.466424991 | 0.89 |
| YKR096W | 1782.315311 | 0.22959419 | 0.462492804 | 0.89 |
| YLL012W | 268.5425901 | -0.624184816 | 0.484945395 | 0.89 |
| YLL018C-A | 305.6957299 | 0.259689958 | 0.486901656 | 0.89 |
| YLR050C | 545.8709868 | -0.234609611 | 0.467280371 | 0.89 |
| YLR069C | 1475.235001 | -0.287806068 | 0.457688846 | 0.89 |
| YLR104W | 635.3306236 | -0.196336773 | 0.488673343 | 0.89 |
| YLR121C | 1254.397824 | 0.235820458 | 0.485692985 | 0.89 |
| YLR125W | 49.15160341 | 0.025235528 | 0.477025145 | 0.89 |
| YLR127C | 955.0961502 | 0.197899266 | 0.464259613 | 0.89 |
| YLR200W | 774.013828 | -0.18236895 | 0.486384047 | 0.89 |
| YLR201C | 1199.611196 | 0.242328071 | 0.481940189 | 0.89 |
| YLR220W | 930.48531 | 0.168590327 | 0.469991612 | 0.89 |
| YLR222C | 1088.378581 | -0.262593596 | 0.477839291 | 0.89 |
| YLR298C | 806.4703735 | 0.239993066 | 0.476065904 | 0.89 |
| YLR356W | 2015.024039 | 0.284314317 | 0.459677018 | 0.89 |
| YLR373C | 1362.586121 | 0.187129678 | 0.483223549 | 0.89 |
| YLR397C | 907.763381 | -0.127861328 | 0.487236402 | 0.89 |
| YLR401C | 1341.53932 | -0.193565149 | 0.480453834 | 0.89 |
| YLR403W | 956.7749718 | 0.15702045 | 0.485067547 | 0.89 |
| YLR410W | 2881.535918 | 0.271992197 | 0.469325839 | 0.89 |

| | | | | |
|---|---|---|---|---|
| YLR429W | 3551.790145 | -0.237031591 | 0.470694707 | 0.89 |
| YLR459W | 960.9364848 | -0.292586661 | 0.488800972 | 0.89 |
| YML027W | 523.9520569 | -0.50661505 | 0.457194192 | 0.89 |
| YML089C | 19.45677984 | 0.270076171 | 0.470439393 | 0.89 |
| YML107C | 389.7043417 | 0.096159872 | 0.489164062 | 0.89 |
| YMR020W | 992.3354434 | 0.252943114 | 0.479866214 | 0.89 |
| YMR025W | 229.9295307 | 0.292540497 | 0.472505808 | 0.89 |
| YMR063W | 168.4962103 | 0.190931148 | 0.486065148 | 0.89 |
| YMR066W | 755.7979974 | 0.212526455 | 0.463983926 | 0.89 |
| YMR106C | 546.3577506 | 0.257036417 | 0.486595958 | 0.89 |
| YMR110C | 3066.365125 | 0.248481415 | 0.482132067 | 0.89 |
| YMR124W | 1653.834225 | 0.248784742 | 0.476091941 | 0.89 |
| YMR219W | 3001.660673 | 0.270947317 | 0.479567381 | 0.89 |
| YMR233W | 360.9564304 | -0.389406619 | 0.478608025 | 0.89 |
| YMR234W | 552.7610201 | -0.213777845 | 0.485294607 | 0.89 |
| YMR238W | 935.1001777 | -0.208473896 | 0.478193732 | 0.89 |
| YMR268C | 351.0386447 | 0.163071474 | 0.473974044 | 0.89 |
| YMR269W | 489.6101925 | -0.169274988 | 0.475676497 | 0.89 |
| YMR316W | 898.4662527 | 0.094479135 | 0.471856139 | 0.89 |
| YNL068C | 1421.244951 | 0.334703587 | 0.477223906 | 0.89 |
| YNL077W | 1205.161408 | 0.294128159 | 0.48074367 | 0.89 |
| YNL087W | 1914.774752 | -0.133043257 | 0.479477956 | 0.89 |
| YNL104C | 8329.192681 | 0.723786705 | 0.467589928 | 0.89 |
| YNL148C | 367.0903378 | -0.170833263 | 0.471685674 | 0.89 |
| YNL172W | 1164.569131 | 0.270548625 | 0.465806285 | 0.89 |
| YNL179C | 130.2526057 | 0.259393336 | 0.471711279 | 0.89 |
| YNL183C | 1035.511832 | 0.236356035 | 0.487340118 | 0.89 |
| YNL204C | 121.2325338 | 0.13972495 | 0.474540146 | 0.89 |
| YNR034W-A | 4345.183948 | 0.101702768 | 0.474208284 | 0.89 |
| YNR067C | 2460.572138 | -0.247470057 | 0.459265875 | 0.89 |
| YOL008W | 381.7473344 | -0.224716036 | 0.485551713 | 0.89 |
| YOL059W | 2587.338376 | 0.263240084 | 0.462174204 | 0.89 |
| YOL060C | 2188.941543 | 0.392262726 | 0.461654228 | 0.89 |
| YOL077W-A | 1163.041858 | -0.241878203 | 0.484344712 | 0.89 |
| YOL118C | 79.69627109 | 0.372954323 | 0.484587464 | 0.89 |
| YOR040W | 503.3947844 | 0.334593274 | 0.486859353 | 0.89 |
| YOR058C | 1144.395841 | 0.194976796 | 0.471427257 | 0.89 |
| YOR101W | 962.4280426 | -0.206127064 | 0.466290945 | 0.89 |
| YOR103C | 111.7299261 | -0.281798844 | 0.469773257 | 0.89 |
| YOR104W | 252.8227861 | 0.25615755 | 0.473600022 | 0.89 |
| YOR108W | 1787.909268 | 0.019293592 | 0.478901478 | 0.89 |
| YOR115C | 581.5147795 | -0.252957904 | 0.462175217 | 0.89 |
| YOR124C | 2791.881139 | 0.288624109 | 0.464946116 | 0.89 |

| | | | | |
|---|---|---|---|---|
| YOR152C | 552.1803625 | 0.308652783 | 0.475224306 | 0.89 |
| YOR156C | 466.9299753 | 0.212339981 | 0.471692822 | 0.89 |
| YOR163W | 1216.735785 | -0.194685403 | 0.470915267 | 0.89 |
| YOR228C | 505.0466268 | 0.149468188 | 0.464835543 | 0.89 |
| YOR256C | 1260.844527 | 0.219843996 | 0.48590623 | 0.89 |
| YOR307C | 1226.108407 | 0.198610714 | 0.479626866 | 0.89 |
| YOR313C | 174.7411276 | -0.510855474 | 0.477531152 | 0.89 |
| YOR323C | 2339.246441 | -0.09118718 | 0.468269369 | 0.89 |
| YOR325W | 4.631536169 | -0.208150678 | 0.459341438 | 0.89 |
| YOR353C | 984.6133479 | 0.188700279 | 0.471131371 | 0.89 |
| YOR381W | 508.0002714 | 0.165973164 | 0.483367072 | 0.89 |
| YOR394C-A | 4.135778786 | -0.5409082 | 0.487800789 | 0.89 |
| YPL033C | 16.58126869 | 0.171887093 | 0.465321597 | 0.89 |
| YPL035C | 100.9354993 | 0.271269411 | 0.486373547 | 0.89 |
| YPL064C | 883.912951 | 0.159773844 | 0.479655445 | 0.89 |
| YPL114W | 1.969765016 | 0.391100924 | 0.460645483 | 0.89 |
| YPL133C | 580.4116075 | 0.215622313 | 0.47807861 | 0.89 |
| YPL162C | 476.9994766 | 0.259779574 | 0.456785755 | 0.89 |
| YPL177C | 1454.555337 | 0.084742015 | 0.464899774 | 0.89 |
| YPL214C | 1002.445323 | 0.235110196 | 0.478413258 | 0.89 |
| YPL235W | 1507.98677 | -0.265570598 | 0.458173881 | 0.89 |
| YPL275W | 31.2658679 | 0.351220111 | 0.468637499 | 0.89 |
| YPR018W | 922.6100673 | -0.143166983 | 0.480356897 | 0.89 |
| YPR034W | 1306.898874 | -0.254537647 | 0.461276152 | 0.89 |
| YPR035W | 16997.33407 | 0.274747797 | 0.48865346 | 0.89 |
| YPR037C | 567.1835844 | -0.188519353 | 0.465290384 | 0.89 |
| YPR067W | 664.4484323 | -0.198779166 | 0.465463233 | 0.89 |
| YPR119W | 682.8278436 | -0.322958154 | 0.461228383 | 0.89 |
| YPR127W | 1734.545853 | 0.18773158 | 0.477626543 | 0.89 |
| YPR128C | 787.6455183 | -0.185077117 | 0.479595539 | 0.89 |
| YPR133C | 1652.526675 | -0.25255501 | 0.467031036 | 0.89 |
| YPR154W | 2486.651934 | 0.219281099 | 0.468970605 | 0.89 |
| YPR158W | 1022.854173 | 0.464205225 | 0.462542217 | 0.89 |
| snR161 | 8.988952116 | 0.247121032 | 0.470633219 | 0.89 |
| YGR087C | 287.4531423 | 0.01517405 | 0.490410727 | 0.90 |
| YGL111W | 1214.214479 | -0.141898232 | 0.491203151 | 0.90 |
| YGL208W | 855.2756461 | 0.399981202 | 0.491588989 | 0.90 |
| YER035W | 1131.143378 | 0.124704389 | 0.492187611 | 0.90 |
| YKL003C | 652.5154668 | -0.271562979 | 0.492025038 | 0.90 |
| YKR022C | 681.4384392 | 0.199853261 | 0.491940759 | 0.90 |
| YPR022C | 1532.191812 | 0.191253586 | 0.492232881 | 0.90 |
| YDL051W | 1178.81834 | 0.061438082 | 0.492508443 | 0.90 |
| ICR1 | 346.589135 | 0.312751479 | 0.495137063 | 0.90 |

| | | | | |
|---|---|---|---|---|
| YAL018C | 88.41004538 | 0.016273629 | 0.497823672 | 0.90 |
| YAR027W | 1481.56887 | 0.149112985 | 0.50303272 | 0.90 |
| YBR101C | 3697.675555 | -0.157467274 | 0.500822224 | 0.90 |
| YBR142W | 1284.846746 | -0.022080411 | 0.496417619 | 0.90 |
| YBR234C | 4120.481034 | -0.183603969 | 0.497937579 | 0.90 |
| YBR272C | 606.5333168 | 0.200918712 | 0.498207839 | 0.90 |
| YCL045C | 3038.105977 | -0.151535361 | 0.501132082 | 0.90 |
| YCR018C-A | 8.72999001 | 0.417102856 | 0.497794659 | 0.90 |
| YCR054C | 374.3747854 | 0.275187631 | 0.496484699 | 0.90 |
| YDL027C | 1559.088359 | 0.209184943 | 0.501867611 | 0.90 |
| YDL204W | 5424.611742 | 0.097639488 | 0.503066189 | 0.90 |
| YDL207W | 1207.359156 | -0.160504825 | 0.495862876 | 0.90 |
| YDR005C | 1131.574288 | 0.183246212 | 0.500917607 | 0.90 |
| YDR087C | 817.7593103 | -0.178503466 | 0.497470677 | 0.90 |
| YDR128W | 2008.877758 | 0.285324088 | 0.498410766 | 0.90 |
| YDR140W | 525.20568 | -0.181257844 | 0.498055877 | 0.90 |
| YDR174W | 3004.282379 | -0.179402895 | 0.499496432 | 0.90 |
| YDR204W | 1297.475345 | 0.362055314 | 0.497669619 | 0.90 |
| YDR208W | 1907.44225 | 0.164551515 | 0.498871946 | 0.90 |
| YDR361C | 744.7914399 | 0.026283763 | 0.501861168 | 0.90 |
| YDR461C-A | 326.4764294 | 0.104385565 | 0.494064219 | 0.90 |
| YER007W | 271.3998551 | 0.202659306 | 0.499732931 | 0.90 |
| YER014W | 690.0676914 | 0.29355979 | 0.502901669 | 0.90 |
| YER085C | 1.090623589 | 0.551783725 | 0.501515706 | 0.90 |
| YFR046C | 527.2575594 | 0.244410998 | 0.494725793 | 0.90 |
| YGL026C | 4379.972897 | 0.526111086 | 0.493582205 | 0.90 |
| YGL042C | 343.2123261 | -0.249804148 | 0.498717038 | 0.90 |
| YGR018C | 8.799849222 | -0.053192835 | 0.496826061 | 0.90 |
| YGR036C | 221.213258 | -0.247938485 | 0.494571247 | 0.90 |
| YGR081C | 443.1057485 | -0.130459195 | 0.494884292 | 0.90 |
| YGR197C | 588.1649926 | 0.289417761 | 0.497626353 | 0.90 |
| YGR277C | 611.2932905 | -0.233015146 | 0.497238057 | 0.90 |
| YGR280C | 1635.461549 | -0.019248622 | 0.503646736 | 0.90 |
| YGR284C | 2427.894256 | -0.096180825 | 0.500601628 | 0.90 |
| YHL031C | 431.2428898 | -0.200354562 | 0.495356173 | 0.90 |
| YHR065C | 1260.588152 | -0.129368004 | 0.503591627 | 0.90 |
| YHR086W | 910.6374675 | -0.197933051 | 0.496687474 | 0.90 |
| YHR151C | 595.1830514 | 0.3196465 | 0.496655035 | 0.90 |
| YHR200W | 3347.025252 | -0.194127604 | 0.503594917 | 0.90 |
| YHR214W | 18.69123236 | 0.232997802 | 0.499412942 | 0.90 |
| YIL037C | 352.6358083 | 0.203597413 | 0.497522905 | 0.90 |
| YIL079C | 348.2789788 | -0.189032893 | 0.502662372 | 0.90 |
| YIR016W | 1147.104809 | 0.101350698 | 0.495751924 | 0.90 |

| | | | | |
|---|---|---|---|---|
| YJR013W | 645.2124988 | -0.180125541 | 0.494093904 | 0.90 |
| YKL010C | 2290.07323 | 0.165151429 | 0.501330821 | 0.90 |
| YKL059C | 865.3974506 | 0.143491226 | 0.497584218 | 0.90 |
| YKL142W | 4281.67462 | 0.244934342 | 0.499934806 | 0.90 |
| YKR004C | 281.3110563 | 0.198117497 | 0.495935036 | 0.90 |
| YKR093W | 2668.023949 | 0.213919301 | 0.502425356 | 0.90 |
| YKR101W | 347.6961828 | 0.153281787 | 0.499759028 | 0.90 |
| YLL006W-A | 2.814822079 | 0.50442858 | 0.50009482 | 0.90 |
| YLL019C | 1900.382246 | 0.264151707 | 0.496671893 | 0.90 |
| YLL034C | 960.4010793 | -0.031162433 | 0.494676004 | 0.90 |
| YLR006C | 1262.134466 | 0.35925359 | 0.503640636 | 0.90 |
| YLR122C | 30.05120318 | 0.361987762 | 0.499146062 | 0.90 |
| YLR168C | 702.4056037 | -0.275472717 | 0.496523455 | 0.90 |
| YLR203C | 2343.4384 | -0.20118576 | 0.502524099 | 0.90 |
| YML032C | 1432.723324 | 0.165717241 | 0.50337612 | 0.90 |
| YMR130W | 646.8299959 | -0.205385112 | 0.499860994 | 0.90 |
| YNL106C | 963.8876185 | 0.230913757 | 0.503521616 | 0.90 |
| YNR015W | 1147.190779 | -0.128000407 | 0.498608067 | 0.90 |
| YOL006C | 1492.750425 | 0.172975942 | 0.503251568 | 0.90 |
| YOL026C | 840.8767019 | 0.158178493 | 0.49342267 | 0.90 |
| YOL081W | 5088.832739 | -0.095353507 | 0.502715914 | 0.90 |
| YOL098C | 2415.997601 | -0.157931352 | 0.494954011 | 0.90 |
| YOL132W | 124.9358227 | -0.157641589 | 0.503164314 | 0.90 |
| YOR035C | 1268.519702 | 0.240337322 | 0.498152988 | 0.90 |
| YOR076C | 709.8237452 | 0.175910045 | 0.497953167 | 0.90 |
| YOR135C | 405.7747912 | 0.038493894 | 0.498434936 | 0.90 |
| YOR250C | 655.6893916 | -0.201403766 | 0.501607801 | 0.90 |
| YOR260W | 2245.68858 | -0.174613213 | 0.494931214 | 0.90 |
| YOR282W | 14.73409586 | 0.493347816 | 0.495152895 | 0.90 |
| YPL172C | 484.663762 | -0.163302333 | 0.497787694 | 0.90 |
| YPL274W | 1036.210642 | 0.245361173 | 0.50096946 | 0.90 |
| YPR001W | 533.0217534 | 0.273729558 | 0.501768149 | 0.90 |
| YPR062W | 2367.986615 | -0.138564444 | 0.494076642 | 0.90 |
| YPR076W | 41.22373968 | 0.445104914 | 0.502504783 | 0.90 |
| YPR095C | 1088.661894 | 0.19794933 | 0.496950795 | 0.90 |
| YDR366C | 7.819864358 | -0.524240264 | 0.503935519 | 0.90 |
| YBR052C | 2651.004465 | -0.22791995 | 0.505650483 | 0.90 |
| YDL138W | 666.9826314 | 0.207053003 | 0.504672642 | 0.90 |
| YDR042C | 81.10570396 | -0.09180719 | 0.505205197 | 0.90 |
| YHR084W | 1211.807628 | -0.327863261 | 0.505718335 | 0.90 |
| YHR112C | 1466.272889 | 0.228023613 | 0.504496854 | 0.90 |
| YIL066C | 409.4135175 | 0.313451485 | 0.505309001 | 0.90 |
| YLR338W | 19.58277131 | 0.364878577 | 0.505024803 | 0.90 |

| | | | | |
|---|---|---|---|---|
| YMR173W-A | 260.3952806 | -0.422727811 | 0.50443681 | 0.90 |
| YNL149C | 291.2466735 | -0.092751987 | 0.505591791 | 0.90 |
| YNL284C-B | 190.6577576 | 0.518326939 | 0.5042376 | 0.90 |
| YOR289W | 937.4069291 | 0.295099028 | 0.505253164 | 0.90 |
| snR8 | 32.91757815 | -0.248616712 | 0.505550242 | 0.90 |
| YIL163C | 43.51788641 | 0.206778289 | 0.506122699 | 0.90 |
| YBR086C | 4766.880566 | 0.229702684 | 0.506841325 | 0.90 |
| YDL128W | 1754.772543 | -0.236686395 | 0.507634955 | 0.90 |
| YDL248W | 224.0356228 | 0.339712832 | 0.507085897 | 0.90 |
| YGL224C | 544.3672231 | -0.203034829 | 0.50679211 | 0.90 |
| YHR069C | 771.2357851 | -0.195401659 | 0.507619713 | 0.90 |
| YMR246W | 3290.570155 | 0.056853761 | 0.507050026 | 0.90 |
| YMR277W | 1413.809425 | -0.211335786 | 0.507632522 | 0.90 |
| YNR047W | 1084.837265 | 0.201078822 | 0.507413183 | 0.90 |
| YBL021C | 336.3286264 | -0.166730366 | 0.508298069 | 0.90 |
| YIR017W-A | 3.982442675 | 0.276915242 | 0.508328686 | 0.90 |
| YKL039W | 2424.281879 | -0.203901087 | 0.508344074 | 0.90 |
| YPR131C | 480.7629468 | -0.18902722 | 0.508211281 | 0.90 |
| YBR077C | 472.5529414 | 0.236805766 | 0.509577596 | 0.90 |
| YCL026C-B | 269.8485152 | 0.349486792 | 0.50897803 | 0.90 |
| YDR235W | 670.6497022 | 0.159005404 | 0.509761563 | 0.90 |
| YDR376W | 985.9130091 | 0.235742706 | 0.509403526 | 0.90 |
| YFR048W | 703.4879681 | 0.121512838 | 0.509842252 | 0.90 |
| YHR050W | 1110.031456 | -0.084547451 | 0.50969234 | 0.90 |
| YJL162C | 484.7743599 | 0.196095646 | 0.509644299 | 0.90 |
| YLR025W | 2034.136821 | 0.196621285 | 0.509794573 | 0.90 |
| YMR313C | 988.4819991 | 0.283215085 | 0.509424526 | 0.90 |
| YGL182C | 10.33516089 | 0.683888882 | 0.510004774 | 0.90 |
| YBL036C | 1223.509384 | -0.202512784 | 0.514269458 | 0.90 |
| YBL078C | 809.4078099 | 0.217342326 | 0.513047452 | 0.90 |
| YBR051W | 12.61851356 | -0.073272237 | 0.513096839 | 0.90 |
| YBR059C | 2706.219813 | 0.258866307 | 0.511183012 | 0.90 |
| YBR174C | 5.135026563 | 0.547443476 | 0.511005729 | 0.90 |
| YBR196C | 23554.20363 | -0.080913777 | 0.51036266 | 0.90 |
| YBR235W | 1364.419973 | 0.383499377 | 0.514247146 | 0.90 |
| YCL012C | 422.2313505 | 0.287870972 | 0.51412083 | 0.90 |
| YCR093W | 4708.644101 | -0.220295329 | 0.514007114 | 0.90 |
| YDL032W | 4.179935405 | -0.138868016 | 0.510533561 | 0.90 |
| YDR440W | 548.6066441 | -0.107459531 | 0.512518302 | 0.90 |
| YEL068C | 304.3910786 | -0.321893989 | 0.514295341 | 0.90 |
| YEL070W | 298.2975483 | 0.056265903 | 0.511803234 | 0.90 |
| YER017C | 1876.346183 | 0.202746979 | 0.511354743 | 0.90 |
| YER029C | 871.4448752 | -0.186059231 | 0.514510329 | 0.90 |

| | | | | |
|---|---|---|---|---|
| YER040W | 900.0166243 | 0.298392435 | 0.511825728 | 0.90 |
| YGR227W | 1061.56859 | -0.271344642 | 0.51459039 | 0.90 |
| YKL197C | 902.4571586 | 0.304016333 | 0.513119648 | 0.90 |
| YLL020C | 30.70630028 | -0.173731138 | 0.514379239 | 0.90 |
| YLR020C | 866.2619741 | -0.175324726 | 0.513918462 | 0.90 |
| YLR039C | 1333.277955 | 0.201471249 | 0.513135753 | 0.90 |
| YMR054W | 1855.817154 | -0.18664716 | 0.510457703 | 0.90 |
| YMR058W | 5542.022572 | -0.374130826 | 0.514045816 | 0.90 |
| YMR087W | 259.2299237 | 0.108191038 | 0.514094267 | 0.90 |
| YMR213W | 660.1636091 | 0.173401045 | 0.51230869 | 0.90 |
| YNL078W | 1260.908163 | -0.220362734 | 0.511221007 | 0.90 |
| YNR043W | 2170.763038 | -0.089200915 | 0.51291938 | 0.90 |
| YOL073C | 1132.754224 | 0.231021908 | 0.512976053 | 0.90 |
| YOR047C | 1007.814108 | 0.131049834 | 0.512996 | 0.90 |
| YOR093C | 1072.002934 | -0.209839744 | 0.512887901 | 0.90 |
| YOR193W | 437.421385 | 0.122808978 | 0.511575023 | 0.90 |
| YOR295W | 256.0382172 | -0.181457343 | 0.513346255 | 0.90 |
| YPR144C | 1036.348245 | -0.157633241 | 0.511392678 | 0.90 |
| YBL051C | 3664.33556 | -0.12622224 | 0.515185321 | 0.90 |
| YKL017C | 627.7815414 | 0.177020509 | 0.515276946 | 0.90 |
| YLR018C | 838.7588705 | -0.235962364 | 0.515264035 | 0.90 |
| YBR200W | 1077.368125 | -0.149306773 | 0.515507841 | 0.90 |
| YHR036W | 340.3963184 | -0.280151335 | 0.516468929 | 0.90 |
| YIL009W | 578.8586392 | -0.34473736 | 0.516517366 | 0.90 |
| YKL144C | 415.786387 | -0.158501096 | 0.516332074 | 0.90 |
| YKR007W | 738.2939867 | -0.223295866 | 0.516217875 | 0.90 |
| YMR224C | 500.4037572 | 0.143522134 | 0.516031941 | 0.90 |
| YNL054W | 1269.220328 | -0.182185855 | 0.516124601 | 0.90 |
| YPR039W | 9.707681236 | 0.571753925 | 0.516385208 | 0.90 |
| YDL151C | 4.616503935 | -0.451904686 | 0.516889359 | 0.90 |
| YNL242W | 1398.740797 | 0.361430922 | 0.516941598 | 0.90 |
| YAL005C | 15020.34614 | -0.011094541 | 0.51921079 | 0.90 |
| YCR087W | 24.15940244 | 0.400927065 | 0.519936573 | 0.90 |
| YDR108W | 495.5770673 | 0.17869489 | 0.517647668 | 0.90 |
| YER001W | 1455.504127 | -0.192828265 | 0.51877962 | 0.90 |
| YER034W | 308.6813768 | -0.166584163 | 0.517395662 | 0.90 |
| YHR216W | 3639.691774 | 0.218692233 | 0.519997872 | 0.90 |
| YIL047C-A | 10.41380776 | -0.221979113 | 0.51931801 | 0.90 |
| YIL115W-A | 23.70554908 | -0.208402246 | 0.519205585 | 0.90 |
| YIR015W | 351.8671415 | 0.282484608 | 0.518906077 | 0.90 |
| YJL013C | 787.2498623 | 0.167933274 | 0.519979307 | 0.90 |
| YKL015W | 1286.742474 | 0.392768167 | 0.51736846 | 0.90 |
| YKL215C | 1029.488394 | 0.260118242 | 0.518791727 | 0.90 |

| | | | | |
|---|---|---|---|---|
| YML086C | 1768.316639 | -0.164254376 | 0.519450013 | 0.90 |
| YML119W | 1027.81525 | 0.203617788 | 0.51949638 | 0.90 |
| YMR221C | 1062.231053 | -0.202219597 | 0.519521005 | 0.90 |
| YNL090W | 89.81803356 | -0.168946131 | 0.517773993 | 0.90 |
| YNL200C | 2258.279724 | 0.18357103 | 0.519327006 | 0.90 |
| YNR029C | 598.402815 | -0.207787506 | 0.519743524 | 0.90 |
| YOL156W | 95.75918972 | -0.616060199 | 0.519712666 | 0.90 |
| YPL068C | 273.8997414 | -0.155464342 | 0.518320215 | 0.90 |
| YPL181W | 1430.008347 | 0.269559964 | 0.518047511 | 0.90 |
| YPR025C | 1080.062057 | 0.213484283 | 0.519965746 | 0.90 |
| YGR140W | 791.394714 | -0.165821323 | 0.52043088 | 0.90 |
| YOL019W | 928.0908204 | -0.303915462 | 0.520512749 | 0.90 |
| YER107W-A | 76.0626659 | 0.007839292 | 0.521721227 | 0.90 |
| YIR024C | 366.6225047 | 0.143824814 | 0.521676544 | 0.90 |
| YJL194W | 154.6134221 | -0.629485771 | 0.521730741 | 0.90 |
| YKL210W | 8749.432253 | 0.264672007 | 0.521424698 | 0.90 |
| YMR112C | 344.1352482 | -0.177016736 | 0.521361482 | 0.90 |
| YOL128C | 161.4112011 | 0.082181271 | 0.521629612 | 0.90 |
| YKL092C | 887.6693114 | -0.20631374 | 0.521879886 | 0.90 |
| YAL029C | 2871.619875 | 0.191880675 | 0.523212757 | 0.90 |
| YBR004C | 976.5395701 | 0.230198228 | 0.524611314 | 0.90 |
| YBR182C | 159.913284 | -0.017213361 | 0.523330029 | 0.90 |
| YCL031C | 1450.481764 | -0.211578704 | 0.524127549 | 0.90 |
| YCL035C | 4330.054767 | 0.175112562 | 0.524521085 | 0.90 |
| YDR179W-A | 321.5877637 | -0.151649199 | 0.524299576 | 0.90 |
| YDR414C | 448.7530762 | -0.251465744 | 0.523455854 | 0.90 |
| YFL040W | 216.9489399 | 0.064158068 | 0.524417064 | 0.90 |
| YGR124W | 7850.205704 | 0.135023658 | 0.523329791 | 0.90 |
| YHR103W | 1865.030837 | 0.227710811 | 0.523924429 | 0.90 |
| YHR118C | 526.5681517 | -0.146621773 | 0.523758578 | 0.90 |
| YIL160C | 460.4774311 | -0.206615657 | 0.523727181 | 0.90 |
| YJR015W | 1958.669387 | -0.16025768 | 0.524806119 | 0.90 |
| YKL029C | 3291.164423 | 0.150239695 | 0.52309616 | 0.90 |
| YLR087C | 2042.517235 | -0.107262541 | 0.523300452 | 0.90 |
| YLR404W | 245.17757 | -0.23891708 | 0.523128713 | 0.90 |
| YMR006C | 894.5004784 | -0.192014198 | 0.524262161 | 0.90 |
| YMR308C | 3347.564301 | -0.152447475 | 0.523816239 | 0.90 |
| YNL156C | 1413.410846 | 0.173855458 | 0.522788877 | 0.90 |
| YPL217C | 2396.710824 | -0.0823939 | 0.524884592 | 0.90 |
| YPL239W | 1334.632299 | -0.009261842 | 0.522453392 | 0.90 |
| YBR054W | 4277.398072 | 0.452403403 | 0.527130242 | 0.90 |
| YCR051W | 1038.832374 | -0.20932304 | 0.528947671 | 0.90 |
| YDL120W | 712.7094342 | -0.205088934 | 0.525745158 | 0.90 |

| | | | | |
|---|---|---|---|---|
| YDR239C | 796.3576703 | 0.184444937 | 0.525581845 | 0.90 |
| YDR486C | 1425.158924 | 0.195932402 | 0.526784447 | 0.90 |
| YGL054C | 1677.23256 | -0.181646067 | 0.526037085 | 0.90 |
| YGL212W | 286.2266484 | -0.204356109 | 0.52814598 | 0.90 |
| YGL228W | 2378.82392 | 0.174717032 | 0.528365616 | 0.90 |
| YGR232W | 824.5178442 | -0.230439325 | 0.525916395 | 0.90 |
| YHL041W | 3.288023996 | 0.167817742 | 0.526029672 | 0.90 |
| YJR046W | 1243.286799 | -0.099269707 | 0.528775413 | 0.90 |
| YKL128C | 1025.460861 | -0.169953693 | 0.52783465 | 0.90 |
| YLR136C | 655.8429852 | -0.453135466 | 0.528036658 | 0.90 |
| YLR305C | 2169.127998 | -0.161373412 | 0.526829296 | 0.90 |
| YML080W | 296.6855777 | -0.190714656 | 0.527560188 | 0.90 |
| YMR019W | 977.1459301 | 0.248746864 | 0.526635262 | 0.90 |
| YMR090W | 1879.223335 | 0.059444568 | 0.527794435 | 0.90 |
| YMR126C | 404.3389603 | 0.133472581 | 0.528619685 | 0.90 |
| YMR193W | 776.035786 | -0.279354579 | 0.528795386 | 0.90 |
| YMR241W | 1393.972884 | -0.141251796 | 0.526355406 | 0.90 |
| YMR258C | 733.537512 | 0.149875979 | 0.526405378 | 0.90 |
| YMR322C | 22.28920933 | -0.273969731 | 0.528377981 | 0.90 |
| YNL004W | 1614.267096 | -0.195940664 | 0.525249552 | 0.90 |
| YNL081C | 921.3541155 | -0.230967171 | 0.528876744 | 0.90 |
| YNL166C | 1009.247872 | -0.112218104 | 0.526410042 | 0.90 |
| YNR051C | 2926.448335 | -0.029716982 | 0.527704361 | 0.90 |
| YOR020W-A | 1169.212022 | 0.17269554 | 0.525323242 | 0.90 |
| YOR144C | 421.6026059 | -0.16670626 | 0.527495642 | 0.90 |
| YPR027C | 342.9171324 | -0.020334507 | 0.527263272 | 0.90 |
| YBR246W | 994.4891912 | -0.167505258 | 0.529137281 | 0.90 |
| YAL015C | 634.3594781 | 0.224697202 | 0.52932779 | 0.90 |
| YOR375C | 17984.54886 | 0.708837953 | 0.529535906 | 0.90 |
| YBR231C | 448.4132605 | 0.192862405 | 0.530167194 | 0.90 |
| YDR369C | 596.1613966 | 0.283162803 | 0.530077467 | 0.90 |
| YJL008C | 2286.818099 | -0.116547537 | 0.530242197 | 0.90 |
| YGL248W | 1428.88377 | 0.206842668 | 0.530464451 | 0.90 |
| YLR004C | 257.8975095 | 0.17301029 | 0.530711625 | 0.90 |
| YDL130W-A | 2765.774643 | 0.317167783 | 0.531702077 | 0.90 |
| YGL187C | 1692.180783 | -0.234661427 | 0.531632362 | 0.90 |
| YHR185C | 7.642313868 | -0.335812796 | 0.5308788 | 0.90 |
| YLR316C | 329.0837142 | 0.151299441 | 0.531552523 | 0.90 |
| YOR284W | 207.2496443 | -0.097662958 | 0.531617805 | 0.90 |
| YPL191C | 278.5044925 | 0.289949762 | 0.531224084 | 0.90 |
| YPR004C | 1678.278628 | -0.072493523 | 0.531318515 | 0.90 |
| YGL196W | 1262.908167 | 0.354757249 | 0.532070224 | 0.90 |
| YMR077C | 308.7109529 | 0.108914214 | 0.532250781 | 0.90 |

| | | | | |
|---|---|---|---|---|
| YPL115C | 1185.518438 | -0.185568185 | 0.532155684 | 0.90 |
| YAL017W | 3309.438838 | 0.226768355 | 0.532679615 | 0.90 |
| YAR028W | 513.5910617 | -0.157836882 | 0.540643951 | 0.90 |
| YBL071C-B | 1.645198606 | 0.472249021 | 0.533706003 | 0.90 |
| YBL105C | 2392.516684 | -0.207014567 | 0.537187696 | 0.90 |
| YBR007C | 740.5673067 | -0.185949895 | 0.53297817 | 0.90 |
| YBR238C | 642.2716869 | -0.152984142 | 0.538672716 | 0.90 |
| YBR301W | 22.99326035 | 0.215889584 | 0.539828072 | 0.90 |
| YCL021W-A | 277.7575158 | 0.016634053 | 0.536653307 | 0.90 |
| YCR028C | 527.4192161 | 0.160085407 | 0.535945229 | 0.90 |
| YDL144C | 951.4980585 | -0.177654271 | 0.541184725 | 0.90 |
| YDL177C | 218.4776378 | 0.247328998 | 0.540845214 | 0.90 |
| YDR013W | 271.9851003 | -0.280443664 | 0.539699281 | 0.90 |
| YDR182W | 1361.286881 | -0.19549976 | 0.534607198 | 0.90 |
| YDR375C | 907.9536977 | -0.172577189 | 0.54150356 | 0.90 |
| YDR456W | 1671.501856 | 0.191637908 | 0.533733673 | 0.90 |
| YER171W | 948.7525289 | -0.200600755 | 0.540296783 | 0.90 |
| YGL114W | 1125.03715 | 0.27595626 | 0.541418667 | 0.90 |
| YGL226W | 346.3222245 | -0.187099118 | 0.536714011 | 0.90 |
| YGR029W | 703.0611046 | -0.134962874 | 0.540152918 | 0.90 |
| YGR049W | 549.8511767 | -0.169226219 | 0.538433715 | 0.90 |
| YGR158C | 631.124363 | -0.18073635 | 0.538339499 | 0.90 |
| YGR184C | 3421.297226 | -0.127848404 | 0.538886413 | 0.90 |
| YHR180W | 212.5255364 | 0.037014947 | 0.534097417 | 0.90 |
| YHR187W | 633.7803075 | -0.144070643 | 0.53657501 | 0.90 |
| YIL003W | 534.8841649 | 0.210942754 | 0.538391485 | 0.90 |
| YIL034C | 3381.997418 | -0.128662751 | 0.540827325 | 0.90 |
| YIL077C | 964.8137743 | 0.036757122 | 0.537988639 | 0.90 |
| YIL087C | 960.1717362 | 0.157372341 | 0.536056495 | 0.90 |
| YIL136W | 7495.331143 | 0.225978127 | 0.533125117 | 0.90 |
| YJL120W | 37.94521827 | -0.103747528 | 0.535513453 | 0.90 |
| YKL037W | 43.69720214 | -0.485926406 | 0.534533115 | 0.90 |
| YKL063C | 828.820505 | -0.193614852 | 0.538501923 | 0.90 |
| YKL132C | 214.3347933 | 0.245330699 | 0.533227641 | 0.90 |
| YKL194C | 593.5236151 | 0.228529375 | 0.538808538 | 0.90 |
| YKL218C | 773.1891416 | 0.181016987 | 0.532734237 | 0.90 |
| YKR031C | 1778.007412 | 0.156172542 | 0.5380855 | 0.90 |
| YKR053C | 486.0346558 | 0.081309587 | 0.538976162 | 0.90 |
| YLR072W | 910.1580571 | 0.200758341 | 0.5355968 | 0.90 |
| YLR114C | 1321.213644 | -0.147645431 | 0.532486296 | 0.90 |
| YLR131C | 1246.543234 | 0.197010561 | 0.537578452 | 0.90 |
| YLR261C | 10.17984332 | -0.006297086 | 0.535669469 | 0.90 |
| YLR277C | 732.8482737 | -0.21904354 | 0.535484505 | 0.90 |

| | | | | |
|---|---|---|---|---|
| YLR412C-A | 1.32995427 | -0.224572248 | 0.534820229 | 0.90 |
| YLR427W | 1359.694674 | 0.110005585 | 0.535762132 | 0.90 |
| YML014W | 592.0947476 | -0.041335133 | 0.536226252 | 0.90 |
| YML084W | 2.063062303 | 0.115254892 | 0.536649074 | 0.90 |
| YMR056C | 319.9319845 | 0.14352651 | 0.541377244 | 0.90 |
| YMR109W | 3069.730753 | -0.206157896 | 0.536539857 | 0.90 |
| YNL015W | 3387.233502 | 0.363824068 | 0.540750474 | 0.90 |
| YNL041C | 1185.16925 | -0.190113982 | 0.533269564 | 0.90 |
| YNL114C | 62.61749004 | -0.251588563 | 0.538228988 | 0.90 |
| YNL119W | 513.2444289 | -0.232367682 | 0.536860423 | 0.90 |
| YNL130C-A | 3.921452687 | 0.176502258 | 0.534067588 | 0.90 |
| YNL160W | 19462.39163 | 0.091634956 | 0.536095568 | 0.90 |
| YOL075C | 712.6275688 | 0.260635854 | 0.53676923 | 0.90 |
| YOR022C | 608.0869392 | 0.280229926 | 0.539637039 | 0.90 |
| YOR140W | 905.6577941 | -0.175658124 | 0.541321102 | 0.90 |
| YOR147W | 1293.879256 | 0.162972411 | 0.540195346 | 0.90 |
| YOR154W | 667.1567549 | 0.198971196 | 0.534006183 | 0.90 |
| YPL197C | 174.3224877 | 0.151268728 | 0.541233913 | 0.90 |
| YPL216W | 730.7653507 | 0.183255949 | 0.538114794 | 0.90 |
| YPR020W | 1129.530709 | -0.046972877 | 0.53836032 | 0.90 |
| YPR036W-A | 8058.010483 | 0.130682517 | 0.541577251 | 0.90 |
| YPR063C | 1556.762503 | -0.182915851 | 0.539910491 | 0.90 |
| YPR162C | 653.0734249 | -0.201259658 | 0.540209756 | 0.90 |
| YPR167C | 421.1211 | 0.214595591 | 0.533629693 | 0.90 |
| YPR172W | 1219.274092 | 0.271587379 | 0.53774478 | 0.90 |
| YCL039W | 1768.904925 | 0.268640818 | 0.542514936 | 0.91 |
| YDR117C | 1080.281382 | -0.146555013 | 0.54238345 | 0.91 |
| YDR169C-A | 13.43403719 | -0.175014942 | 0.542398563 | 0.91 |
| YOR385W | 901.596547 | 0.223968217 | 0.542330696 | 0.91 |
| YDR255C | 870.1597032 | 0.19020715 | 0.542960888 | 0.91 |
| YER178W | 9418.417677 | -0.09867522 | 0.543049968 | 0.91 |
| YHL007C | 2278.179389 | -0.1900138 | 0.54302942 | 0.91 |
| YAL042C-A | 8.764234173 | 0.031797438 | 0.543525294 | 0.91 |
| YPL023C | 1340.132366 | -0.140665661 | 0.543469823 | 0.91 |
| YPL073C | 1.14139142 | -0.455605526 | 0.543335225 | 0.91 |
| YCR032W | 1688.564969 | 0.257227724 | 0.543815703 | 0.91 |
| YMR013C | 348.9768631 | 0.158663041 | 0.543806836 | 0.91 |
| YMR220W | 1203.063636 | -0.193996434 | 0.544031713 | 0.91 |
| YAL027W | 394.2818606 | 0.163594631 | 0.546899558 | 0.91 |
| YBL005W-B | 450.3577253 | 0.590482368 | 0.54906708 | 0.91 |
| YBR107C | 343.2589721 | 0.208187029 | 0.547183772 | 0.91 |
| YCR025C | 44.00460202 | 0.045317239 | 0.548415699 | 0.91 |
| YDL116W | 1341.682086 | -0.207928501 | 0.546130111 | 0.91 |

| | | | | |
|---|---|---|---|---|
| YDL160C | 1362.561057 | -0.234352285 | 0.54673967 | 0.91 |
| YDL222C | 906.6701758 | -0.384562637 | 0.547692311 | 0.91 |
| YER122C | 4064.222569 | -0.197223712 | 0.545460329 | 0.91 |
| YGL063W | 321.9330079 | -0.253554638 | 0.546149533 | 0.91 |
| YGR274C | 1344.663557 | 0.1619307 | 0.546829739 | 0.91 |
| YHL019W-A | 10.01596588 | -0.043241463 | 0.549338659 | 0.91 |
| YHL042W | 88.3151764 | 0.2930784 | 0.545764737 | 0.91 |
| YHR126C | 18.42016663 | 3.08E-05 | 0.54664683 | 0.91 |
| YIL102C-A | 112.0491515 | -0.388254267 | 0.546836595 | 0.91 |
| YIL134W | 238.1456546 | -0.120578814 | 0.544950955 | 0.91 |
| YIL145C | 1126.99064 | -0.109821047 | 0.545636541 | 0.91 |
| YJL204C | 683.8445536 | 0.157810036 | 0.549119898 | 0.91 |
| YKL054C | 4561.844374 | -0.128619304 | 0.548075455 | 0.91 |
| YLR214W | 1682.159024 | -0.32709768 | 0.545218869 | 0.91 |
| YLR386W | 1077.114184 | 0.181878788 | 0.545878846 | 0.91 |
| YLR402W | 27.81687827 | -0.638150804 | 0.54563429 | 0.91 |
| YLR461W | 7.039838185 | 0.529431313 | 0.545035782 | 0.91 |
| YML122C | 47.85383085 | -0.02206264 | 0.54910105 | 0.91 |
| YML124C | 2648.714801 | -0.114832196 | 0.547674931 | 0.91 |
| YMR108W | 5770.966467 | 0.074343641 | 0.547773778 | 0.91 |
| YMR226C | 4108.834639 | 0.214929036 | 0.54834446 | 0.91 |
| YMR244C-A | 108.6393324 | 0.217567675 | 0.549220674 | 0.91 |
| YMR293C | 413.7964048 | 0.176716229 | 0.545256391 | 0.91 |
| YNL203C | 26.22039237 | -0.296592853 | 0.549309553 | 0.91 |
| YNL245C | 408.0242862 | 0.187749164 | 0.546685 | 0.91 |
| YNR073C | 195.2436335 | 0.39061281 | 0.548101246 | 0.91 |
| YOL101C | 182.685157 | 0.226091407 | 0.545786091 | 0.91 |
| YOR335C | 3110.388093 | -0.029563317 | 0.54928149 | 0.91 |
| YPL031C | 1442.118487 | -0.171062456 | 0.547170009 | 0.91 |
| YPL075W | 2226.650529 | 0.143744003 | 0.548133298 | 0.91 |
| YPL134C | 1579.391088 | 0.473862886 | 0.544641402 | 0.91 |
| YPL193W | 620.6543173 | 0.180520026 | 0.545079238 | 0.91 |
| YPR158W-B | 29.96033516 | 0.384484526 | 0.548362096 | 0.91 |
| YGR250C | 3005.303955 | 0.245520646 | 0.549861972 | 0.91 |
| YDR251W | 2664.249522 | 0.20179795 | 0.550624246 | 0.91 |
| YFR002W | 1555.318988 | -0.243634855 | 0.550481175 | 0.91 |
| YHR100C | 468.2098929 | -0.129909974 | 0.550289004 | 0.91 |
| YHR210C | 320.0019552 | -0.18903458 | 0.550556922 | 0.91 |
| YPR156C | 1539.255789 | 0.312155122 | 0.550075332 | 0.91 |
| YGL118C | 2.533670207 | -0.003406221 | 0.550793934 | 0.91 |
| YDL109C | 380.7977413 | 0.243384633 | 0.551516533 | 0.91 |
| YGL034C | 9.87258369 | 0.028073467 | 0.551647853 | 0.91 |
| YGR041W | 794.2535473 | 0.120686063 | 0.551584832 | 0.91 |

| | | | | |
|---|---|---|---|---|
| YIL094C | 3348.951104 | -0.126283554 | 0.551353432 | 0.91 |
| YPR101W | 539.0655902 | 0.13079357 | 0.551407688 | 0.91 |
| YDR537C | 28.61472288 | -0.094279097 | 0.552073629 | 0.91 |
| YDL200C | 293.2384509 | 0.285832372 | 0.553428247 | 0.91 |
| YDR151C | 1113.580859 | -0.240992649 | 0.553493388 | 0.91 |
| YGL153W | 279.9517565 | 0.160726342 | 0.553463617 | 0.91 |
| YGR005C | 1346.509283 | 0.150085654 | 0.553639648 | 0.91 |
| YGR091W | 441.73971 | 0.216601205 | 0.553372422 | 0.91 |
| YML125C | 2139.072841 | -0.1669947 | 0.553059986 | 0.91 |
| YNL024C-A | 249.9046551 | -0.19830491 | 0.552477418 | 0.91 |
| YNL057W | 19.3681916 | 0.052802429 | 0.553204093 | 0.91 |
| YNL182C | 900.9513347 | -0.079230895 | 0.553827439 | 0.91 |
| YOL052C-A | 1535.771374 | -0.149034416 | 0.553733237 | 0.91 |
| YPR140W | 900.9897118 | -0.182343697 | 0.553176712 | 0.91 |
| YDR243C | 512.1645164 | 0.156292697 | 0.554432692 | 0.91 |
| YJL085W | 484.172969 | -0.196057297 | 0.554334972 | 0.91 |
| YOL031C | 1089.984363 | -0.412256604 | 0.554494597 | 0.91 |
| YAR068W | 4.842260677 | 0.057875444 | 0.555738321 | 0.91 |
| YBR013C | 207.4427581 | 0.072865998 | 0.560109044 | 0.91 |
| YBR074W | 1213.234406 | -0.154083569 | 0.557403009 | 0.91 |
| YBR111W-A | 229.152125 | -0.228621215 | 0.559267166 | 0.91 |
| YDL080C | 732.479931 | 0.121955395 | 0.555367734 | 0.91 |
| YDL244W | 51.51006224 | -0.570797468 | 0.555691235 | 0.91 |
| YDR110W | 523.1803163 | 0.272305987 | 0.55710092 | 0.91 |
| YDR123C | 353.8570592 | 0.249732133 | 0.5548095 | 0.91 |
| YEL038W | 943.0798183 | -0.177716887 | 0.559307297 | 0.91 |
| YER051W | 512.8476233 | 0.226078559 | 0.55663874 | 0.91 |
| YGL097W | 1307.008078 | 0.01132714 | 0.559766512 | 0.91 |
| YGL233W | 1565.055775 | -0.163790028 | 0.559391743 | 0.91 |
| YGR189C | 5490.403188 | -0.127977585 | 0.559021573 | 0.91 |
| YHL035C | 3064.413674 | 0.26761978 | 0.560079001 | 0.91 |
| YIR027C | 227.1514133 | 0.005098522 | 0.558631988 | 0.91 |
| YJL026W | 4324.794509 | -0.119707288 | 0.559824482 | 0.91 |
| YJL046W | 632.8219176 | -0.202567593 | 0.559874096 | 0.91 |
| YKL001C | 376.5252013 | 0.086219912 | 0.557972107 | 0.91 |
| YKL020C | 2087.802283 | 0.14481991 | 0.55918377 | 0.91 |
| YKL030W | 149.8142583 | 0.026087908 | 0.558895573 | 0.91 |
| YLR042C | 282.794653 | -0.332378515 | 0.556794043 | 0.91 |
| YLR123C | 17.47407493 | -0.281509391 | 0.55581567 | 0.91 |
| YLR140W | 51.21550356 | -0.300051141 | 0.559601769 | 0.91 |
| YLR278C | 1940.968689 | -0.155407031 | 0.558671305 | 0.91 |
| YML003W | 222.6670877 | 0.154751124 | 0.560065313 | 0.91 |
| YML118W | 395.2746742 | 0.146344211 | 0.558587048 | 0.91 |

| | | | | |
|---|---|---|---|---|
| YMR127C | 400.3588714 | 0.18846574 | 0.558839153 | 0.91 |
| YMR196W | 3241.463398 | 0.251673127 | 0.558496767 | 0.91 |
| YNL026W | 973.5528021 | -0.135606477 | 0.557685749 | 0.91 |
| YNL071W | 6818.92242 | -0.05464694 | 0.556711839 | 0.91 |
| YNL251C | 2085.43933 | -0.002551653 | 0.5588101 | 0.91 |
| YNR001W-A | 2.398336405 | -0.226158706 | 0.55803788 | 0.91 |
| YOL064C | 847.1293742 | -0.020955043 | 0.558478415 | 0.91 |
| YOL114C | 133.5426332 | 0.183345531 | 0.556546609 | 0.91 |
| YOR227W | 3483.244777 | 0.247848373 | 0.555681904 | 0.91 |
| YOR281C | 1009.23062 | -0.204444399 | 0.55614259 | 0.91 |
| YOR386W | 897.2436472 | 0.169005549 | 0.557733386 | 0.91 |
| YPL048W | 7812.579467 | -0.118025283 | 0.556604908 | 0.91 |
| YPR040W | 901.3829077 | 0.187529329 | 0.555515138 | 0.91 |
| YDL019C | 2417.237505 | 0.214008391 | 0.560438313 | 0.91 |
| YBL050W | 1823.454847 | -0.185921641 | 0.562737286 | 0.91 |
| YBL089W | 493.6798455 | -0.202518079 | 0.561644595 | 0.91 |
| YGR064W | 14.24950712 | -0.571731559 | 0.56141885 | 0.91 |
| YJL001W | 1978.350649 | -0.179706879 | 0.56217726 | 0.91 |
| YJL005W | 4176.860231 | 0.231205729 | 0.5629722 | 0.91 |
| YJL054W | 1689.096692 | -0.165733367 | 0.562201348 | 0.91 |
| YJL207C | 2307.188321 | -0.167390624 | 0.561346384 | 0.91 |
| YJR011C | 218.4017674 | -0.219070051 | 0.561222449 | 0.91 |
| YKL178C | 114.5867776 | -0.617389638 | 0.561732638 | 0.91 |
| YLL004W | 381.8327061 | -0.290852059 | 0.562481184 | 0.91 |
| YLR181C | 881.938971 | 0.178177462 | 0.563042642 | 0.91 |
| YML128C | 9230.440998 | -0.026574462 | 0.562105903 | 0.91 |
| YNL294C | 1109.535055 | 0.187796413 | 0.56251065 | 0.91 |
| YNL305C | 2021.497779 | 0.168170078 | 0.561838158 | 0.91 |
| YNL329C | 1184.177512 | -0.196979298 | 0.562830079 | 0.91 |
| YPL055C | 1072.396721 | 0.203791752 | 0.562553388 | 0.91 |
| YPL145C | 2994.997329 | -0.041407162 | 0.56252563 | 0.91 |
| snR59 | 2.356669998 | -0.268721558 | 0.562041989 | 0.91 |
| YAR035C-A | 1.313184395 | -0.155604505 | 0.565299638 | 0.91 |
| YBR135W | 639.2382675 | -0.110952551 | 0.564594898 | 0.91 |
| YDR056C | 1611.517655 | -0.100174561 | 0.565086459 | 0.91 |
| YEL055C | 1803.113225 | -0.038015014 | 0.564144991 | 0.91 |
| YGL005C | 924.1279481 | 0.160651992 | 0.564466158 | 0.91 |
| YHR131W-A | 19.588359 | 0.225389262 | 0.564356326 | 0.91 |
| YHR165C | 3249.517075 | -0.194240418 | 0.564999399 | 0.91 |
| YIL169C | 2177.35422 | -0.312005712 | 0.563799262 | 0.91 |
| YLR079W | 2171.764341 | 0.205579752 | 0.564411516 | 0.91 |
| YML007C-A | 2.984197591 | -0.50017821 | 0.563940408 | 0.91 |
| YNL029C | 467.6043522 | 0.214073833 | 0.565053078 | 0.91 |

| | | | | |
|---|---|---|---|---|
| YOR171C | 1509.079417 | 0.226356008 | 0.565260677 | 0.91 |
| YPL273W | 2617.65817 | 0.150322526 | 0.564805592 | 0.91 |
| YPR108W | 2749.924825 | -0.144380227 | 0.564197279 | 0.91 |
| YNL218W | 370.2566133 | -0.113556371 | 0.565500176 | 0.91 |
| YCR024C-A | 1033.453894 | -0.269386603 | 0.566015901 | 0.91 |
| YPL117C | 2245.25569 | -0.06019056 | 0.566001854 | 0.91 |
| YIR003W | 1806.796543 | 0.171220446 | 0.56653117 | 0.91 |
| YLR111W | 15.73315875 | 0.247439981 | 0.566342877 | 0.91 |
| YLR313C | 272.7990365 | -0.281038794 | 0.56654401 | 0.91 |
| YJL051W | 1072.541155 | 0.193604885 | 0.566694848 | 0.91 |
| YGR156W | 932.64115 | -0.239520027 | 0.567093121 | 0.91 |
| YHR179W | 8288.962491 | 0.029057216 | 0.56728576 | 0.91 |
| YPR116W | 103.378319 | -0.099861275 | 0.567625126 | 0.91 |
| YKR002W | 1434.178622 | 0.161897784 | 0.567922082 | 0.91 |
| YIL084C | 434.165782 | 0.141534909 | 0.56835634 | 0.91 |
| YJL035C | 163.2101994 | -0.265921967 | 0.568246808 | 0.91 |
| YNL143C | 26.87962177 | -0.144166773 | 0.568178404 | 0.91 |
| YDR374W-A | 251.03932 | -0.331275766 | 0.568579707 | 0.91 |
| YPR082C | 431.92377 | -0.144411275 | 0.568767953 | 0.91 |
| YML111W | 1306.672498 | -0.158836938 | 0.56891231 | 0.91 |
| YLR307W | 82.37745124 | 0.044312997 | 0.569095564 | 0.91 |
| YDL148C | 2147.255757 | -0.062072429 | 0.570539591 | 0.91 |
| YDR406W | 3009.969096 | -0.455687507 | 0.570679271 | 0.91 |
| YEL064C | 340.0873494 | 0.21667031 | 0.570757755 | 0.91 |
| YER082C | 1253.635463 | -0.182590983 | 0.569543531 | 0.91 |
| YER097W | 28.63993449 | 0.397860449 | 0.571662436 | 0.91 |
| YER143W | 1597.506506 | 0.135658614 | 0.571183288 | 0.91 |
| YHR124W | 347.8549107 | 0.318507526 | 0.569820106 | 0.91 |
| YIL144W | 1230.130031 | 0.133060398 | 0.571659991 | 0.91 |
| YJL202C | 29.15207009 | -0.261889088 | 0.569393793 | 0.91 |
| YJR161C | 124.4036715 | 0.320834873 | 0.571289047 | 0.91 |
| YLR405W | 359.0345633 | -0.2312787 | 0.57169874 | 0.91 |
| YML114C | 955.0918143 | -0.184874049 | 0.570741343 | 0.91 |
| YNL099C | 699.7662248 | -0.108545359 | 0.570390105 | 0.91 |
| YNL171C | 59.8874962 | 0.353146728 | 0.571408159 | 0.91 |
| YNL285W | 3.926788582 | -0.417520275 | 0.570944809 | 0.91 |
| YPL026C | 1074.82514 | -0.002349338 | 0.571331516 | 0.91 |
| YPL224C | 949.2587858 | 0.174588673 | 0.571548687 | 0.91 |
| YPR169W-A | 32.89680766 | -0.10960392 | 0.570812629 | 0.91 |
| YBL052C | 898.9203912 | 0.184632308 | 0.573013155 | 0.91 |
| YBR072W | 9138.655064 | -0.522215018 | 0.572948898 | 0.91 |
| YDR170W-A | 128.0923816 | 0.112647776 | 0.572829002 | 0.91 |
| YER147C-A | 275.4373645 | -0.145981831 | 0.572301477 | 0.91 |

| | | | | |
|---|---|---|---|---|
| YER163C | 538.8237534 | -0.17036476 | 0.57283373 | 0.91 |
| YKL057C | 1611.669894 | -0.230391204 | 0.573532711 | 0.91 |
| YLR271W | 587.0390943 | 0.195685709 | 0.57237393 | 0.91 |
| YLR320W | 753.4878726 | 0.160248286 | 0.573526443 | 0.91 |
| YLR361C-A | 262.7999201 | -0.223353573 | 0.573265406 | 0.91 |
| YNL084C | 2582.808264 | -0.130193939 | 0.573520369 | 0.91 |
| YNR018W | 1764.18714 | -0.005956441 | 0.573651712 | 0.91 |
| YOR014W | 3387.711726 | 0.207737325 | 0.572627897 | 0.91 |
| YOR231W | 881.0751627 | 0.182091343 | 0.573441956 | 0.91 |
| YFL016C | 3333.34381 | -0.165772348 | 0.573906345 | 0.91 |
| YAL035W | 9252.51472 | -0.028667128 | 0.574411691 | 0.91 |
| YHR045W | 983.5331682 | -0.187760525 | 0.574281723 | 0.91 |
| YOR129C | 585.8837465 | -0.163427717 | 0.574143359 | 0.91 |
| YHR070W | 688.4026548 | 0.017173091 | 0.57456691 | 0.91 |
| YJL068C | 1815.839599 | 0.244431059 | 0.574917013 | 0.91 |
| YLR260W | 983.6383575 | 0.196861044 | 0.575050455 | 0.91 |
| YPL200W | 21.68345646 | -0.040746355 | 0.575033237 | 0.91 |
| YOR382W | 2866.519088 | -0.371798026 | 0.575348879 | 0.91 |
| YIL053W | 6767.756759 | -0.142559588 | 0.575611861 | 0.91 |
| YAL056W | 1700.26388 | 0.165355408 | 0.577344321 | 0.91 |
| YBL098W | 1032.627619 | 0.222107258 | 0.580268512 | 0.91 |
| YBR042C | 577.29725 | -0.122362672 | 0.580187224 | 0.91 |
| YBR097W | 952.5745358 | -0.178821619 | 0.576943839 | 0.91 |
| YBR150C | 848.9681448 | 0.187772355 | 0.579996995 | 0.91 |
| YCL043C | 17735.82228 | -0.137322056 | 0.578793824 | 0.91 |
| YCR083W | 821.7942376 | 0.277248489 | 0.579740071 | 0.91 |
| YDL105W | 423.3140946 | -0.166084732 | 0.581244928 | 0.91 |
| YDL225W | 1750.506735 | -0.179523051 | 0.580040114 | 0.91 |
| YDR008C | 37.32753147 | -0.047567556 | 0.575865866 | 0.91 |
| YDR301W | 1341.743029 | -0.232593477 | 0.580837473 | 0.91 |
| YEL014C | 21.68196635 | -0.247985482 | 0.578582421 | 0.91 |
| YEL044W | 582.0177494 | -0.207649648 | 0.581585985 | 0.91 |
| YFR018C | 1513.157132 | -0.186456 | 0.580335251 | 0.91 |
| YFR053C | 31755.34613 | 0.208812904 | 0.581420843 | 0.91 |
| YGL009C | 9382.784081 | 0.790642111 | 0.580549738 | 0.91 |
| YGL067W | 999.1477018 | -0.107979478 | 0.578755211 | 0.91 |
| YGR062C | 462.3300151 | -0.257625683 | 0.580201367 | 0.91 |
| YGR125W | 1519.849762 | 0.181594857 | 0.581681675 | 0.91 |
| YHR214C-E | 1.813646714 | -0.080347374 | 0.579003538 | 0.91 |
| YIL011W | 777.5395764 | 0.273327989 | 0.57657944 | 0.91 |
| YIL014C-A | 253.291496 | -0.319389562 | 0.576217867 | 0.91 |
| YIL054W | 8.069700908 | 0.481451226 | 0.579735197 | 0.91 |
| YJL152W | 449.6754094 | 0.129050808 | 0.578596889 | 0.91 |

| | | | | |
|---|---|---|---|---|
| YJR121W | 23130.92602 | 0.192215529 | 0.581954084 | 0.91 |
| YKL175W | 2025.203634 | 0.190884024 | 0.577379427 | 0.91 |
| YKL183C-A | 2.603283005 | -0.498759059 | 0.580565486 | 0.91 |
| YKR084C | 1262.417524 | -0.166198236 | 0.580899306 | 0.91 |
| YML079W | 1250.560699 | -0.302303486 | 0.581659841 | 0.91 |
| YMR037C | 1272.708272 | -0.170254018 | 0.57934079 | 0.91 |
| YMR075C-A | 15.14864191 | -0.00782282 | 0.580935855 | 0.91 |
| YMR311C | 3644.500712 | 0.17406093 | 0.581044549 | 0.91 |
| YNL009W | 277.6331838 | 0.050872625 | 0.580947782 | 0.91 |
| YNL211C | 132.5864716 | -0.242716374 | 0.581818646 | 0.91 |
| YNL260C | 268.8863879 | -0.159823226 | 0.58090753 | 0.91 |
| YNL337W | 0.659062417 | 0.018558143 | 0.578909176 | 0.91 |
| YOL107W | 336.081692 | -0.197221459 | 0.576424475 | 0.91 |
| YOL144W | 766.01586 | -0.080401982 | 0.577836687 | 0.91 |
| YOR343C | 63.14207102 | 0.220167996 | 0.578498555 | 0.91 |
| YOR368W | 442.6825001 | 0.117960656 | 0.581180019 | 0.91 |
| YOR387C | 76.98828773 | 0.02183671 | 0.580353167 | 0.91 |
| YPL065W | 569.9929052 | 0.200390702 | 0.579279547 | 0.91 |
| YPL159C | 943.1509361 | 0.06933671 | 0.576433829 | 0.91 |
| YPR014C | 81.20498688 | -0.528567869 | 0.580718978 | 0.91 |
| YPR045C | 1121.270528 | 0.215015157 | 0.579429342 | 0.91 |
| YIL090W | 898.4488003 | 0.17513994 | 0.582243627 | 0.91 |
| YOL133W | 270.3290721 | 0.119624594 | 0.582162536 | 0.91 |
| YGL091C | 545.7319922 | 0.120214913 | 0.582385538 | 0.91 |
| YGR198W | 1689.551002 | -0.164903919 | 0.582617001 | 0.91 |
| YMR028W | 753.5245359 | 0.132730147 | 0.582814388 | 0.91 |
| YNL107W | 513.6532475 | 0.175998833 | 0.582743865 | 0.91 |
| YBL061C | 962.4150781 | -0.161214903 | 0.583947211 | 0.91 |
| YDL059C | 182.996654 | 0.160492084 | 0.58465121 | 0.91 |
| YDR141C | 2153.367361 | -0.123572615 | 0.584468528 | 0.91 |
| YDR163W | 579.7588355 | -0.205731023 | 0.584607308 | 0.91 |
| YDR513W | 3464.363476 | 0.10224265 | 0.584041477 | 0.91 |
| YFR026C | 243.3675063 | 0.090195497 | 0.583357547 | 0.91 |
| YGR017W | 1237.157084 | -0.141523068 | 0.584133054 | 0.91 |
| YJR060W | 664.6110196 | -0.137405479 | 0.582994862 | 0.91 |
| YKL040C | 858.7842501 | -0.152633569 | 0.584218247 | 0.91 |
| YMR176W | 1256.155137 | 0.194043802 | 0.584428434 | 0.91 |
| YNL058C | 1080.922928 | 0.053992996 | 0.583546558 | 0.91 |
| YNL180C | 2386.54101 | 0.226879473 | 0.584625889 | 0.91 |
| YPR153W | 81.58409703 | 0.078200827 | 0.584099401 | 0.91 |
| YBR019C | 339.0837633 | -0.004550998 | 0.585626277 | 0.91 |
| YBR240C | 118.1647641 | -0.027129971 | 0.585218347 | 0.91 |
| YDL209C | 238.6569979 | -0.275263068 | 0.585872159 | 0.91 |

| | | | | |
|---|---|---|---|---|
| YDR318W | 508.7245087 | -0.234280064 | 0.585746921 | 0.91 |
| YHL026C | 386.086378 | 0.173987653 | 0.585491632 | 0.91 |
| YKL193C | 1757.604095 | 0.177379885 | 0.58555051 | 0.91 |
| YLL007C | 554.0715465 | 0.10733891 | 0.585356465 | 0.91 |
| YPL254W | 446.984746 | -0.279466494 | 0.585463292 | 0.91 |
| YIL041W | 4311.545468 | -0.131480891 | 0.586156077 | 0.91 |
| YNL097C | 784.7174827 | 0.192967762 | 0.586055351 | 0.91 |
| YBR089W | 6.735755394 | 0.469485013 | 0.586357553 | 0.91 |
| YMR169C | 812.630576 | -0.448189908 | 0.586502351 | 0.91 |
| YDR207C | 1436.193266 | 0.276705975 | 0.586856862 | 0.91 |
| YDR394W | 3757.628989 | -0.158445549 | 0.586790764 | 0.91 |
| YJL087C | 474.7532502 | -0.208554605 | 0.586929468 | 0.91 |
| YDL106C | 1181.738765 | 0.140029408 | 0.587607071 | 0.91 |
| YDR261C-D | 88.68167414 | -0.461541229 | 0.587406023 | 0.91 |
| YLR101C | 65.86852303 | 0.19870088 | 0.587500678 | 0.91 |
| YHR090C | 634.1320105 | 0.151024881 | 0.587787747 | 0.91 |
| YLR024C | 1454.81679 | -0.16477116 | 0.587951009 | 0.91 |
| YFL062W | 38.28792094 | -0.029860436 | 0.588728517 | 0.91 |
| YFR019W | 2589.385757 | 0.192113692 | 0.588801517 | 0.91 |
| YHR057C | 1425.81956 | 0.157463558 | 0.588667663 | 0.91 |
| YJL090C | 629.2586035 | 0.113278295 | 0.588320588 | 0.91 |
| YOR026W | 568.5531387 | -0.260258188 | 0.588788991 | 0.91 |
| YDL119C | 290.4231748 | 0.161805217 | 0.590204109 | 0.91 |
| YEL066W | 476.9220074 | -0.173044532 | 0.590385135 | 0.91 |
| YER032W | 1018.880889 | -0.203187067 | 0.59023054 | 0.91 |
| YGR047C | 884.2267464 | 0.157107581 | 0.590649178 | 0.91 |
| YGR221C | 838.7959631 | -0.239549605 | 0.590567386 | 0.91 |
| YHR114W | 1802.212821 | 0.168154636 | 0.590161783 | 0.91 |
| YLR144C | 964.8658379 | 0.191246859 | 0.589894979 | 0.91 |
| YOL005C | 1554.462517 | -0.216334713 | 0.589223706 | 0.91 |
| YOR255W | 23.76826284 | -0.200571216 | 0.590289903 | 0.91 |
| YOR273C | 1760.091485 | 0.318343648 | 0.589844566 | 0.91 |
| YPL138C | 483.5208101 | 0.153207001 | 0.590366755 | 0.91 |
| snR9 | 75.73439823 | -0.005522334 | 0.590650951 | 0.91 |
| YBR081C | 1882.249025 | 0.150362132 | 0.591379004 | 0.91 |
| YDL188C | 1068.696328 | -0.17355154 | 0.592395283 | 0.91 |
| YDR232W | 2313.644723 | -0.178331865 | 0.591178831 | 0.91 |
| YDR313C | 770.3934735 | 0.029557237 | 0.591592774 | 0.91 |
| YGR176W | 11.59700922 | -0.011901224 | 0.592002881 | 0.91 |
| YHR048W | 458.0188382 | 0.146986746 | 0.592103762 | 0.91 |
| YLR408C | 156.1371802 | 0.272135919 | 0.591192955 | 0.91 |
| YMR055C | 275.6053073 | -0.172973058 | 0.590927232 | 0.91 |
| YMR092C | 2509.767947 | 0.206393488 | 0.592180092 | 0.91 |

| | | | | |
|---|---|---|---|---|
| YPL042C | 568.6329785 | 0.111186203 | 0.592073588 | 0.91 |
| YPR048W | 424.7867171 | -0.10720695 | 0.592085485 | 0.91 |
| YPR105C | 1243.404818 | 0.164682433 | 0.592371248 | 0.91 |
| YLR112W | 96.84022753 | -0.229478033 | 0.592558936 | 0.91 |
| YLR353W | 907.9621406 | -0.119510264 | 0.592687085 | 0.91 |
| YKL173W | 1269.160496 | 0.126341079 | 0.592847885 | 0.91 |
| YLR270W | 2664.450215 | 0.176300852 | 0.593664672 | 0.91 |
| YML129C | 536.8329539 | -0.306009703 | 0.593731474 | 0.91 |
| snR33 | 120.9149393 | -0.252230442 | 0.593613251 | 0.91 |
| YBL058W | 3576.371774 | 0.16908562 | 0.595493594 | 0.91 |
| YBR026C | 2145.982583 | 0.235701639 | 0.596883076 | 0.91 |
| YBR092C | 2232.790112 | 0.293001395 | 0.594024636 | 0.91 |
| YBR176W | 422.8364361 | 0.239739923 | 0.595228081 | 0.91 |
| YDR009W | 246.880862 | 0.079246148 | 0.596050059 | 0.91 |
| YFL003C | 165.623347 | 0.004634309 | 0.596726349 | 0.91 |
| YGL011C | 3486.685044 | -0.167495698 | 0.595147801 | 0.91 |
| YGL156W | 2401.006767 | 0.150592508 | 0.596288866 | 0.91 |
| YGL192W | 1154.271063 | -0.06949749 | 0.596539207 | 0.91 |
| YGR228W | 14.43502374 | 0.49337105 | 0.596736184 | 0.91 |
| YHL001W | 1027.533504 | -0.13331731 | 0.5965047 | 0.91 |
| YIL012W | 10.18010829 | -0.121572814 | 0.594317203 | 0.91 |
| YJL045W | 286.1990705 | 0.029952932 | 0.595426461 | 0.91 |
| YJL058C | 326.9260026 | 0.217691586 | 0.59461701 | 0.91 |
| YJR001W | 1763.85175 | -0.128200623 | 0.595656786 | 0.91 |
| YKL061W | 244.3100746 | 0.139288141 | 0.596238623 | 0.91 |
| YKL135C | 1473.22037 | 0.160454133 | 0.596211209 | 0.91 |
| YMR302C | 2126.727676 | 0.153162258 | 0.594887929 | 0.91 |
| YNR008W | 899.8873602 | -0.067438458 | 0.596111606 | 0.91 |
| YPR008W | 1796.059535 | 0.130601447 | 0.595552144 | 0.91 |
| YPR087W | 20.87255534 | 0.043107028 | 0.596998225 | 0.91 |
| YPR113W | 3675.90658 | -0.232130075 | 0.595381985 | 0.91 |
| snR14 | 19.85838705 | 0.040845143 | 0.595257232 | 0.91 |
| YKR089C | 3634.661722 | 0.135186464 | 0.597256221 | 0.91 |
| YDR325W | 750.6903469 | -0.152726818 | 0.597968154 | 0.92 |
| YDR437W | 89.57715872 | -0.143033684 | 0.597835741 | 0.92 |
| YGR174C | 1048.598299 | 0.040455304 | 0.597852091 | 0.92 |
| YMR135C | 1809.513775 | 0.204273915 | 0.597770648 | 0.92 |
| YOR177C | 142.866635 | 0.151757917 | 0.598083974 | 0.92 |
| YDR428C | 292.2817321 | -0.160161761 | 0.5990848 | 0.92 |
| YGL186C | 1156.394032 | -0.151123435 | 0.598959128 | 0.92 |
| YGR161W-C | 303.4318056 | 0.06483365 | 0.598710797 | 0.92 |
| YLL037W | 13.52665894 | 0.014566189 | 0.598785157 | 0.92 |
| YLR055C | 1018.744824 | 0.189952118 | 0.598993849 | 0.92 |

| | | | | |
|---|---|---|---|---|
| YMR182W-A | 4.590688512 | -0.38815487 | 0.59830073 | 0.92 |
| YPL152W | 506.4565509 | 0.112783336 | 0.598920273 | 0.92 |
| YPL149W | 607.7809907 | 0.171434489 | 0.599267436 | 0.92 |
| YBL099W | 11787.99535 | 0.262684687 | 0.600502713 | 0.92 |
| YDR377W | 1422.817631 | -0.01662159 | 0.600054451 | 0.92 |
| YFL066C | 2.343434624 | -0.38108379 | 0.600393042 | 0.92 |
| YMR071C | 1073.602658 | -0.096408346 | 0.600034098 | 0.92 |
| YOR192C | 247.398385 | 0.289898638 | 0.600146034 | 0.92 |
| YOR324C | 520.2886814 | -0.247929629 | 0.600218644 | 0.92 |
| YPL232W | 2081.428239 | -0.214448131 | 0.600272265 | 0.92 |
| YDR406W-A | 12.68467301 | -0.058192516 | 0.600997495 | 0.92 |
| YLR199C | 824.459216 | -0.134373349 | 0.600976821 | 0.92 |
| YDR197W | 858.0497536 | 0.110233997 | 0.601515561 | 0.92 |
| YDR220C | 3.958958383 | 0.075417715 | 0.601327483 | 0.92 |
| YGL179C | 645.0803373 | 0.230602675 | 0.601703528 | 0.92 |
| YHR155W | 1225.572051 | 0.101046619 | 0.601915345 | 0.92 |
| YLR233C | 453.3027113 | 0.235076864 | 0.602042659 | 0.92 |
| YLR355C | 9452.460146 | 0.253112762 | 0.60182363 | 0.92 |
| YOR304C-A | 201.0708852 | -0.24758846 | 0.60200777 | 0.92 |
| YDR286C | 156.0831168 | -0.185582436 | 0.60261361 | 0.92 |
| YJL137C | 923.3034038 | 0.593511103 | 0.602784901 | 0.92 |
| YBR022W | 363.9709831 | -0.180723053 | 0.603301147 | 0.92 |
| YBR103W | 849.3179685 | 0.155321214 | 0.603220837 | 0.92 |
| YER060W-A | 584.2480367 | -0.056962426 | 0.603178343 | 0.92 |
| YLR010C | 127.8600135 | 0.163302245 | 0.603535381 | 0.92 |
| YAL011W | 648.981818 | 0.195899948 | 0.609498828 | 0.92 |
| YBL066C | 1237.560937 | 0.261886359 | 0.604108936 | 0.92 |
| YBR196C-B | 11.09747944 | -0.319542358 | 0.607550374 | 0.92 |
| YBR239C | 441.6078035 | 0.05748497 | 0.605850077 | 0.92 |
| YCL044C | 1238.471005 | 0.214299713 | 0.606392132 | 0.92 |
| YDL115C | 894.0269756 | 0.090556659 | 0.609471387 | 0.92 |
| YDL165W | 697.5260403 | -0.123656189 | 0.608564186 | 0.92 |
| YDR159W | 1698.043068 | 0.143569515 | 0.605928494 | 0.92 |
| YDR290W | 8.55083412 | 0.1897741 | 0.609511328 | 0.92 |
| YDR479C | 1259.888261 | 0.139325621 | 0.607827834 | 0.92 |
| YDR499W | 868.5785284 | 0.187606566 | 0.608156527 | 0.92 |
| YGL085W | 385.7388603 | 0.104650388 | 0.604345362 | 0.92 |
| YGR273C | 7.812625483 | -0.178506126 | 0.606747091 | 0.92 |
| YHL020C | 961.5750146 | 0.092918117 | 0.605789311 | 0.92 |
| YHR061C | 329.4509042 | 0.079975999 | 0.609769904 | 0.92 |
| YHR072W | 1833.218722 | -0.165640537 | 0.609701381 | 0.92 |
| YIL112W | 2132.63932 | 0.275739036 | 0.606332286 | 0.92 |
| YJL072C | 353.6349072 | -0.051890379 | 0.608340339 | 0.92 |

| | | | | |
|---|---|---|---|---|
| YJL159W | 37900.6137 | 0.312268637 | 0.607171768 | 0.92 |
| YJL172W | 1932.176519 | -0.008526786 | 0.606694356 | 0.92 |
| YJL197W | 1228.673265 | -0.118858169 | 0.60867955 | 0.92 |
| YJR087W | 189.3064917 | -0.226650775 | 0.604728913 | 0.92 |
| YJR131W | 581.4205166 | 0.22893198 | 0.606376307 | 0.92 |
| YKR018C | 3562.456922 | 0.32761372 | 0.609770156 | 0.92 |
| YLL040C | 5253.851757 | -0.070869556 | 0.608389068 | 0.92 |
| YLR419W | 1247.341194 | -0.157545533 | 0.60897818 | 0.92 |
| YLR456W | 255.5161749 | 0.163604386 | 0.607347391 | 0.92 |
| YML029W | 1034.498713 | 0.16750547 | 0.606171556 | 0.92 |
| YML062C | 782.2227983 | 0.101516988 | 0.607374802 | 0.92 |
| YMR207C | 1385.212155 | 0.231267083 | 0.606548176 | 0.92 |
| YMR259C | 1208.426766 | -0.118559177 | 0.605879894 | 0.92 |
| YNL017C | 1.615564791 | 0.050555759 | 0.608616557 | 0.92 |
| YNL246W | 1195.836863 | 0.128300095 | 0.60459109 | 0.92 |
| YOL068C | 1498.234041 | -0.17807323 | 0.605619703 | 0.92 |
| YOR174W | 544.9117735 | -0.134922085 | 0.609666516 | 0.92 |
| YOR180C | 257.2908648 | 0.055496882 | 0.607571241 | 0.92 |
| YOR192C-C | 17.72254902 | -0.078205018 | 0.605718108 | 0.92 |
| YOR350C | 702.3901449 | 0.164607647 | 0.608993349 | 0.92 |
| YOR358W | 499.4768456 | 0.135503519 | 0.606676853 | 0.92 |
| YPL097W | 440.8818332 | -0.214159581 | 0.605526203 | 0.92 |
| YPR081C | 985.9306388 | 0.179523459 | 0.608789046 | 0.92 |
| YPR193C | 57.30247136 | 0.176566444 | 0.604694388 | 0.92 |
| YBR201W | 456.7810734 | -0.187872988 | 0.610169526 | 0.92 |
| YOL065C | 517.0688786 | 0.136461482 | 0.610206086 | 0.92 |
| YOR111W | 390.0644929 | 0.162206148 | 0.610618829 | 0.92 |
| YMR283C | 413.1862547 | 0.109258411 | 0.610934099 | 0.92 |
| YMR187C | 357.2988229 | 0.190195753 | 0.611388015 | 0.92 |
| YFL065C | 1.354222162 | -0.053549007 | 0.611688841 | 0.92 |
| YHR093W | 35.27687991 | 0.087025642 | 0.611822161 | 0.92 |
| YOR264W | 932.8104963 | -0.218428466 | 0.611925248 | 0.92 |
| YBR292C | 15.20409235 | 0.180586374 | 0.612396725 | 0.92 |
| YPL141C | 1338.030401 | 0.207978306 | 0.612315891 | 0.92 |
| YPR126C | 82.68269767 | -0.471319509 | 0.612377944 | 0.92 |
| YJR117W | 3603.291074 | -0.074503928 | 0.612905693 | 0.92 |
| YMR162C | 1643.610674 | 0.194841019 | 0.612795736 | 0.92 |
| YNL265C | 447.0625511 | 0.143645576 | 0.612823904 | 0.92 |
| YER090W | 2828.394196 | 0.40574884 | 0.6133606 | 0.92 |
| YGL143C | 536.4770627 | -0.22343979 | 0.613784499 | 0.92 |
| YLL005C | 615.6111971 | 0.297302126 | 0.613783431 | 0.92 |
| YOR216C | 1165.7555 | -0.089213954 | 0.613603329 | 0.92 |
| YBR071W | 990.0527368 | -0.23027353 | 0.615325258 | 0.92 |

| | | | | |
|---|---|---|---|---|
| YBR104W | 350.9392427 | -0.04948858 | 0.614312579 | 0.92 |
| YBR193C | 556.4922631 | 0.163225841 | 0.614906315 | 0.92 |
| YCL033C | 876.4314652 | 0.190376432 | 0.615412537 | 0.92 |
| YCL074W | 140.2978941 | 0.108523664 | 0.615009922 | 0.92 |
| YGR058W | 565.2067235 | -0.162321837 | 0.614311749 | 0.92 |
| YHL029C | 889.4266622 | 0.139903562 | 0.614923857 | 0.92 |
| YHR135C | 3300.939762 | -0.137228122 | 0.615081118 | 0.92 |
| YLR202C | 55.2829566 | -0.048517203 | 0.614881462 | 0.92 |
| YOL037C | 71.18780393 | 0.246849163 | 0.614409346 | 0.92 |
| snR24 | 8.183960631 | 0.635334625 | 0.615328014 | 0.92 |
| YAR073W | 171.304609 | -0.141665906 | 0.616075693 | 0.92 |
| YNL109W | 19.36367514 | -0.256323604 | 0.616014662 | 0.92 |
| YIR035C | 1381.941223 | -0.124800285 | 0.616902593 | 0.92 |
| YKL118W | 29.53687639 | 0.090552513 | 0.616977341 | 0.92 |
| YLR001C | 1116.57839 | 0.118390444 | 0.61679545 | 0.92 |
| YDR319C | 485.2377065 | -0.286929176 | 0.618066291 | 0.92 |
| YGR180C | 8240.520071 | -0.078451623 | 0.618715332 | 0.92 |
| YMR030W-A | 15.82228062 | 0.082602 | 0.618660541 | 0.92 |
| YOR087W | 1019.099344 | 0.191726004 | 0.618833211 | 0.92 |
| YGR077C | 1152.926191 | 0.091566936 | 0.619504417 | 0.92 |
| YDR261W-B | 58.90647804 | 0.451591083 | 0.619810101 | 0.92 |
| YNL091W | 4224.416881 | -0.07575859 | 0.619979737 | 0.92 |
| YNL138W-A | 114.4265169 | -0.244269712 | 0.619861858 | 0.92 |
| YAL058W | 573.3781549 | -0.104794771 | 0.621968514 | 0.92 |
| YBL025W | 212.6114748 | 0.19057094 | 0.621113934 | 0.92 |
| YDL044C | 428.1351393 | -0.205742802 | 0.620914795 | 0.92 |
| YDR079W | 403.891688 | -0.321882734 | 0.622468085 | 0.92 |
| YDR458C | 1309.381626 | 0.061886966 | 0.622600022 | 0.92 |
| YER190W | 15.31140442 | 0.20128354 | 0.621422163 | 0.92 |
| YGR002C | 912.0190954 | 0.114307845 | 0.622351441 | 0.92 |
| YHR016C | 1896.688185 | 0.22364739 | 0.621737691 | 0.92 |
| YJL065C | 199.2698224 | -0.183823528 | 0.622908739 | 0.92 |
| YJL102W | 772.7229577 | 0.036295524 | 0.622239865 | 0.92 |
| YKL075C | 477.0577168 | 0.126099975 | 0.622152208 | 0.92 |
| YKL169C | 32.06083426 | -0.306472151 | 0.622899552 | 0.92 |
| YLR231C | 929.429053 | -0.042222743 | 0.621970279 | 0.92 |
| YML112W | 572.6104301 | -0.170803037 | 0.621977421 | 0.92 |
| YNL105W | 8.716167534 | 0.14522052 | 0.621563622 | 0.92 |
| YPL238C | 56.97812647 | 0.378136194 | 0.622999451 | 0.92 |
| snR57 | 17.27289396 | 0.181194405 | 0.622907284 | 0.92 |
| YBR207W | 1493.475156 | -0.24721657 | 0.624620584 | 0.92 |
| YDR184C | 548.4562077 | -0.092178967 | 0.624674012 | 0.92 |
| YER018C | 488.1395433 | -0.127371437 | 0.624036083 | 0.92 |

| | | | | |
|---|---|---|---|---|
| YGL258W-A | 56.62039045 | -0.642587178 | 0.624365365 | 0.92 |
| YGR100W | 1479.985263 | 0.102314853 | 0.625148172 | 0.92 |
| YJR006W | 1054.347239 | -0.126754069 | 0.62378254 | 0.92 |
| YJR116W | 691.1206507 | 0.18995932 | 0.623562241 | 0.92 |
| YKL126W | 2370.590453 | -0.081390792 | 0.625031331 | 0.92 |
| YKL136W | 25.19505527 | -0.38967778 | 0.62369363 | 0.92 |
| YML012C-A | 4.439298629 | -0.31450804 | 0.624803168 | 0.92 |
| YNL247W | 4374.768503 | 0.095836969 | 0.62382692 | 0.92 |
| YOL115W | 646.4767705 | -0.243823849 | 0.625065551 | 0.92 |
| YOR229W | 494.4703852 | 0.241109854 | 0.623894073 | 0.92 |
| YOR304W | 1385.923767 | 0.095824886 | 0.624406348 | 0.92 |
| YPR031W | 763.0479333 | 0.235525404 | 0.623258591 | 0.92 |
| RUF23 | 8.975541829 | -0.085546061 | 0.626203044 | 0.93 |
| YDR118W | 391.9679351 | -0.125419449 | 0.626086119 | 0.93 |
| YBR023C | 1741.008098 | -0.158788265 | 0.626678729 | 0.93 |
| YER064C | 1674.781352 | 0.227873603 | 0.626867677 | 0.93 |
| YGL235W | 1.903973228 | -0.681616305 | 0.626916075 | 0.93 |
| YBR011C | 18392.14922 | -0.055059635 | 0.628204437 | 0.93 |
| YDR444W | 770.8185864 | 0.163454719 | 0.627689376 | 0.93 |
| YGR235C | 1048.42614 | -0.219456216 | 0.627817849 | 0.93 |
| YKR105C | 96.82105819 | -0.483150082 | 0.628182333 | 0.93 |
| YLL016W | 1012.667191 | 0.070849139 | 0.627515246 | 0.93 |
| YLR100W | 1453.024103 | 0.027541909 | 0.627652117 | 0.93 |
| YLR242C | 302.599813 | -0.162993032 | 0.62751342 | 0.93 |
| YNR040W | 422.1360755 | -0.127078078 | 0.628102602 | 0.93 |
| YOR360C | 883.5535106 | -0.28547005 | 0.62811207 | 0.93 |
| YHL014C | 453.0111576 | -0.166723328 | 0.628698165 | 0.93 |
| YJR086W | 331.1910032 | -0.280749787 | 0.628765002 | 0.93 |
| YPR117W | 2597.357181 | 0.187608808 | 0.629111508 | 0.93 |
| YBL011W | 1645.010666 | -0.150080752 | 0.629311292 | 0.93 |
| YDL240C-A | 1.077122984 | -0.765751214 | 0.629846273 | 0.93 |
| YHR051W | 2680.129991 | -0.030756236 | 0.629855174 | 0.93 |
| YOR343W-B | 107.9180972 | 0.194594884 | 0.629799216 | 0.93 |
| YEL020C-B | 34.39724677 | -0.392961293 | 0.63013216 | 0.93 |
| YBR257W | 391.5498488 | -0.087047926 | 0.630745111 | 0.93 |
| YKR056W | 1143.695694 | -0.078707774 | 0.63050604 | 0.93 |
| YLR197W | 4160.668892 | 0.02059369 | 0.630654353 | 0.93 |
| YOL164W | 729.2266525 | 0.038610117 | 0.630366273 | 0.93 |
| YBR035C | 2483.374886 | 0.213217775 | 0.631861517 | 0.93 |
| YDR054C | 1985.938955 | 0.201501817 | 0.632604179 | 0.93 |
| YIL083C | 1169.719343 | 0.241364215 | 0.632376725 | 0.93 |
| YKL064W | 2390.234747 | 0.202880949 | 0.632775337 | 0.93 |
| YKL159C | 432.357669 | -0.202638505 | 0.632699578 | 0.93 |

| | | | | |
|---|---|---|---|---|
| YLL041C | 3315.150059 | 0.248245783 | 0.631416617 | 0.93 |
| YLR057W | 1092.547525 | 0.116642489 | 0.632372548 | 0.93 |
| YMR070W | 1232.669923 | -0.117187011 | 0.631620926 | 0.93 |
| YMR157C | 556.2957769 | -0.197014776 | 0.631534749 | 0.93 |
| YMR174C | 615.2184477 | -0.01046226 | 0.632634045 | 0.93 |
| YOR073W | 590.4176603 | -0.080081886 | 0.631653409 | 0.93 |
| YOR336W | 1255.740018 | 0.193523008 | 0.631976609 | 0.93 |
| YPR024W | 1909.485964 | 0.164658214 | 0.631345966 | 0.93 |
| YBR269C | 1256.661555 | 0.086109354 | 0.636109071 | 0.93 |
| YDL126C | 11516.69474 | -0.078156696 | 0.636142785 | 0.93 |
| YDR061W | 1142.905546 | 0.105338018 | 0.636108077 | 0.93 |
| YDR070C | 1811.421734 | -0.013568161 | 0.637872432 | 0.93 |
| YDR194W-A | 0.453289942 | 0.240406391 | 0.635243208 | 0.93 |
| YDR423C | 1205.3164 | 0.128294226 | 0.637739335 | 0.93 |
| YER181C | 6.5610832 | 0.250294239 | 0.636052381 | 0.93 |
| YFR027W | 221.6537022 | 0.120944035 | 0.636985566 | 0.93 |
| YGL107C | 935.2339762 | -0.104818377 | 0.635795932 | 0.93 |
| YGL129C | 1219.243345 | -0.140426479 | 0.637135899 | 0.93 |
| YGR075C | 302.7071594 | -0.236198307 | 0.636578723 | 0.93 |
| YGR086C | 9907.562538 | -0.168813915 | 0.637956778 | 0.93 |
| YGR090W | 2037.446504 | -0.208037983 | 0.633387429 | 0.93 |
| YGR199W | 855.4590705 | 0.138755345 | 0.636007435 | 0.93 |
| YHL040C | 1041.983492 | -0.271693559 | 0.637042889 | 0.93 |
| YIR006C | 5095.375019 | -0.105276164 | 0.637810309 | 0.93 |
| YJL042W | 3416.022292 | 0.155093373 | 0.634232889 | 0.93 |
| YJL117W | 1366.561839 | -0.12228049 | 0.635167241 | 0.93 |
| YJR027W | 207.2279874 | 0.011791141 | 0.636614281 | 0.93 |
| YJR151C | 560.5480031 | 0.132178792 | 0.637202022 | 0.93 |
| YKL031W | 35.11063328 | 0.307934068 | 0.637936724 | 0.93 |
| YKL068W-A | 172.0547206 | -0.039151414 | 0.634858493 | 0.93 |
| YKL176C | 1422.060464 | -0.178122564 | 0.633653578 | 0.93 |
| YKR020W | 476.7599628 | 0.142993243 | 0.633521017 | 0.93 |
| YKR024C | 515.1058632 | -0.19194626 | 0.634984713 | 0.93 |
| YKR068C | 742.7822083 | -0.173019215 | 0.637876218 | 0.93 |
| YLL017W | 60.73191178 | -0.078977393 | 0.633650787 | 0.93 |
| YLR035C-A | 40.57446476 | 0.066704565 | 0.635547095 | 0.93 |
| YNL034W | 79.93693595 | 0.361784492 | 0.636790821 | 0.93 |
| YNL065W | 531.9531231 | -0.359307576 | 0.635795285 | 0.93 |
| YNL086W | 191.101467 | 0.105965254 | 0.637665075 | 0.93 |
| YOR001W | 1523.051345 | -0.152907898 | 0.637588937 | 0.93 |
| YOR056C | 1339.644393 | -0.145275907 | 0.633794804 | 0.93 |
| YPL011C | 726.714861 | 0.100032246 | 0.63562831 | 0.93 |
| YPR130C | 10.79124991 | -0.18587371 | 0.634222649 | 0.93 |

| | | | | |
|---|---|---|---|---|
| YPR145W | 8044.845508 | 0.479953051 | 0.637046416 | 0.93 |
| YIL008W | 681.0099622 | 0.013423733 | 0.638716874 | 0.93 |
| YKR099W | 800.367483 | -0.114346717 | 0.638676453 | 0.93 |
| YNR010W | 212.8810604 | -0.279993338 | 0.638710345 | 0.93 |
| YOR184W | 6279.813274 | 0.468832126 | 0.638632373 | 0.93 |
| YPR179C | 773.0608325 | 0.111236111 | 0.638315941 | 0.93 |
| YCL036W | 402.9926525 | 0.183722639 | 0.639552097 | 0.93 |
| YER033C | 1095.361309 | 0.133601302 | 0.639714343 | 0.93 |
| YHR150W | 670.7977171 | 0.14571822 | 0.639388102 | 0.93 |
| YHR166C | 498.330332 | 0.171548995 | 0.639746253 | 0.93 |
| YML090W | 156.4475378 | 0.20622762 | 0.639692178 | 0.93 |
| RUF20 | 86.17514557 | -0.280478053 | 0.641602375 | 0.93 |
| YBR236C | 870.5665803 | -0.204556923 | 0.641333825 | 0.93 |
| YCR082W | 1363.300313 | 0.069254594 | 0.64102411 | 0.93 |
| YDL012C | 999.0124699 | -0.200218805 | 0.641043968 | 0.93 |
| YDL156W | 331.6691228 | -0.051881502 | 0.640928775 | 0.93 |
| YDR259C | 452.5952789 | 0.346205153 | 0.641171912 | 0.93 |
| YIL062C | 1798.738473 | -0.176710584 | 0.64108697 | 0.93 |
| YJL049W | 674.8429005 | 0.074344629 | 0.641485474 | 0.93 |
| YLR133W | 1071.862832 | -0.047565025 | 0.641531482 | 0.93 |
| YPR071W | 302.2135475 | 0.206498755 | 0.640771453 | 0.93 |
| YBR178W | 20.74423611 | 0.103953963 | 0.642047147 | 0.93 |
| YPR073C | 916.4024804 | -0.077808346 | 0.64225755 | 0.93 |
| YBR041W | 1541.15066 | -0.110582394 | 0.642722642 | 0.93 |
| YAL061W | 2752.646997 | -0.136246998 | 0.646855013 | 0.93 |
| YBL022C | 3958.243828 | -0.093986313 | 0.648988372 | 0.93 |
| YBR129C | 610.3882792 | -0.21458538 | 0.649941569 | 0.93 |
| YBR202W | 984.15721 | -0.192391737 | 0.647147353 | 0.93 |
| YCL026C-A | 44.13033287 | 0.151977418 | 0.649645054 | 0.93 |
| YCR104W | 19.10661275 | 0.380163786 | 0.650177331 | 0.93 |
| YDL066W | 3505.644872 | 0.180846392 | 0.643904463 | 0.93 |
| YDR097C | 218.1671058 | -0.13479396 | 0.645751838 | 0.93 |
| YDR416W | 716.6573712 | 0.139345315 | 0.649868225 | 0.93 |
| YDR514C | 275.4543514 | -0.180577576 | 0.644731793 | 0.93 |
| YDR528W | 366.4782046 | -0.424844824 | 0.64613925 | 0.93 |
| YDR543C | 0.805732287 | 0.133538432 | 0.649749119 | 0.93 |
| YFL015W-A | 1.143841678 | -0.450177789 | 0.650254391 | 0.93 |
| YFL048C | 1163.307711 | -0.192598533 | 0.644635658 | 0.93 |
| YGL082W | 1508.755452 | -0.127812271 | 0.645650105 | 0.93 |
| YGL223C | 914.5737412 | -0.117833255 | 0.64430923 | 0.93 |
| YGL259W | 86.59077684 | -0.539286422 | 0.647624102 | 0.93 |
| YGR042W | 764.6829972 | 0.129336431 | 0.646558335 | 0.93 |
| YGR147C | 943.7636147 | -0.13003407 | 0.64580227 | 0.93 |

| | | | | |
|---|---|---|---|---|
| YGR177C | 234.3513182 | 0.193197089 | 0.644878806 | 0.93 |
| YHL050C | 143.7644195 | -0.01763313 | 0.643710499 | 0.93 |
| YHR028C | 2661.078301 | 0.16278532 | 0.648775514 | 0.93 |
| YHR030C | 2133.377408 | 0.230520749 | 0.647834689 | 0.93 |
| YHR056W-A | 0.867587117 | 0.445992793 | 0.643148884 | 0.93 |
| YHR134W | 365.9132551 | 0.117446468 | 0.648634973 | 0.93 |
| YIR029W | 388.4137592 | 0.091367509 | 0.64671207 | 0.93 |
| YJL052C-A | 1.688849253 | -0.32577607 | 0.647039482 | 0.93 |
| YJR019C | 596.1541169 | 0.138683982 | 0.643310921 | 0.93 |
| YKL018W | 496.5039326 | -0.121818973 | 0.649694101 | 0.93 |
| YKL021C | 1160.236373 | -0.038946953 | 0.645024316 | 0.93 |
| YKL044W | 194.2159883 | -0.241988994 | 0.64339471 | 0.93 |
| YKR044W | 684.1987114 | 0.114807731 | 0.645608578 | 0.93 |
| YKR061W | 573.6088479 | -0.095958903 | 0.648743611 | 0.93 |
| YLR032W | 823.8658356 | -0.096648133 | 0.646537672 | 0.93 |
| YLR049C | 719.3459985 | -0.142034774 | 0.644544514 | 0.93 |
| YLR210W | 386.6683639 | -0.170555985 | 0.650059984 | 0.93 |
| YLR398C | 2110.607765 | -0.101134761 | 0.647785935 | 0.93 |
| YML072C | 6041.463861 | 0.266824666 | 0.643781039 | 0.93 |
| YMR052W | 219.8800281 | 0.217538097 | 0.648557686 | 0.93 |
| YMR113W | 461.667001 | 0.06160387 | 0.648882159 | 0.93 |
| YMR172C-A | 11.37396813 | -0.050661779 | 0.649331378 | 0.93 |
| YMR264W | 1111.60862 | -0.134021904 | 0.643807324 | 0.93 |
| YMR321C | 24.07796373 | -0.423173641 | 0.649833497 | 0.93 |
| YMR323W | 206.5265565 | -0.076228611 | 0.645771005 | 0.93 |
| YNL227C | 372.4616162 | 0.069365084 | 0.646062731 | 0.93 |
| YOL034W | 735.3581814 | 0.107594041 | 0.649983113 | 0.93 |
| YOL067C | 283.6975055 | 0.159000524 | 0.6474807 | 0.93 |
| YPL024W | 252.5020697 | -0.15229066 | 0.649087586 | 0.93 |
| YPL052W | 344.0806128 | 0.14329737 | 0.644250326 | 0.93 |
| YPR011C | 560.157333 | -0.205924932 | 0.64879809 | 0.93 |
| YPR184W | 5293.583109 | 0.154832164 | 0.6475181 | 0.93 |
| YPR204W | 106.9528255 | -0.006956879 | 0.644640897 | 0.93 |
| YJR134C | 1708.030399 | -0.158711712 | 0.650565453 | 0.93 |
| YML123C | 2594.845401 | -0.114963844 | 0.650508718 | 0.93 |
| YBL096C | 11.76381799 | -0.53705732 | 0.651698648 | 0.93 |
| YBR126W-B | 156.4452467 | -0.311050956 | 0.651039932 | 0.93 |
| YCL014W | 1364.401777 | -0.11476889 | 0.653665509 | 0.93 |
| YCR030C | 2439.320353 | 0.176577096 | 0.653794425 | 0.93 |
| YDL011C | 7.393723597 | -0.501785011 | 0.651913302 | 0.93 |
| YDR041W | 1214.356247 | -0.189731242 | 0.653705928 | 0.93 |
| YDR527W | 984.8264926 | 0.165271295 | 0.652913124 | 0.93 |
| YER013W | 829.6032387 | 0.102731233 | 0.652407774 | 0.93 |

| | | | | |
|---|---|---|---|---|
| YFR056C | 41.82065454 | 0.152969815 | 0.653023433 | 0.93 |
| YGL177W | 36.28800868 | 0.414338227 | 0.653195061 | 0.93 |
| YGL221C | 1756.035958 | -0.072298979 | 0.65213865 | 0.93 |
| YGR109C | 138.0694558 | -0.155976384 | 0.651248999 | 0.93 |
| YGR203W | 222.1887092 | -0.376354826 | 0.654219211 | 0.93 |
| YHL045W | 26.03413018 | 0.335655984 | 0.651831367 | 0.93 |
| YIL095W | 775.5717487 | -0.150036481 | 0.65235306 | 0.93 |
| YIL130W | 1279.051362 | 0.100272969 | 0.651642278 | 0.93 |
| YJR049C | 1397.771565 | 0.239928855 | 0.651013061 | 0.93 |
| YKL196C | 2255.642273 | -0.073708886 | 0.652695286 | 0.93 |
| YLR090W | 931.4547969 | 0.103084382 | 0.654078793 | 0.93 |
| YLR349W | 23.19539208 | -0.018799118 | 0.653627249 | 0.93 |
| YMR236W | 1041.170124 | -0.146960621 | 0.651252338 | 0.93 |
| YOR130C | 372.7954083 | 0.325604048 | 0.654336038 | 0.93 |
| YOR246C | 1262.599493 | -0.148619392 | 0.653114241 | 0.93 |
| YOR348C | 282.9627385 | 0.08631025 | 0.654226971 | 0.93 |
| YPR032W | 901.7856576 | 0.127382917 | 0.652628806 | 0.93 |
| snR39B | 13.58422976 | -0.058679471 | 0.653528055 | 0.93 |
| YMR042W | 272.8014796 | -0.317637955 | 0.654892629 | 0.93 |
| YEL062W | 560.1717097 | 0.042470418 | 0.655627628 | 0.93 |
| YER103W | 3269.463849 | -0.27575414 | 0.655625298 | 0.93 |
| YGL013C | 1490.03588 | -0.152240853 | 0.655770049 | 0.93 |
| YGL088W | 0.432313012 | 0.067642738 | 0.655575783 | 0.93 |
| YJR093C | 790.9035242 | 0.082798962 | 0.65566547 | 0.93 |
| YDL034W | 29.64111763 | 0.42077644 | 0.656727311 | 0.93 |
| YDR229W | 1110.14149 | 0.084497059 | 0.657734138 | 0.93 |
| YDR246W-A | 23.46436155 | -0.265234038 | 0.657033431 | 0.93 |
| YDR477W | 1886.671826 | 0.121364259 | 0.656756151 | 0.93 |
| YER167W | 1421.029082 | 0.214430726 | 0.657826591 | 0.93 |
| YGR171C | 566.9877726 | -0.150162952 | 0.657260227 | 0.93 |
| YJR047C | 407.0063415 | 0.323763018 | 0.656433284 | 0.93 |
| YLR374C | 34.93928504 | -0.162605715 | 0.656627891 | 0.93 |
| YNL139C | 1577.578713 | -0.108805318 | 0.657814605 | 0.93 |
| YNR019W | 1395.465767 | 0.244681513 | 0.657788188 | 0.93 |
| YOR132W | 1718.595531 | 0.207932527 | 0.657307458 | 0.93 |
| YOR202W | 1479.782603 | 0.300514476 | 0.657632867 | 0.93 |
| YOR241W | 933.790879 | -0.143651661 | 0.657924098 | 0.93 |
| YPL276W | 15.28614474 | 0.089970616 | 0.657541035 | 0.93 |
| YDR112W | 22.56100124 | 0.208051685 | 0.658444263 | 0.93 |
| YDR356W | 1114.540575 | -0.095839532 | 0.658559518 | 0.93 |
| YMR189W | 2797.782123 | -0.007307816 | 0.65889517 | 0.93 |
| YAL047W-A | 1.181246705 | -0.003537171 | 0.659499443 | 0.93 |
| YBR103C-A | 1.570441558 | 0.164048027 | 0.659629749 | 0.93 |

| | | | | |
|---|---|---|---|---|
| YBR177C | 2530.709586 | -0.171208901 | 0.659791635 | 0.93 |
| YCR081W | 992.4265821 | 0.148206138 | 0.661169567 | 0.93 |
| YDR294C | 2749.492975 | 0.14880632 | 0.66022553 | 0.93 |
| YGL119W | 689.6648811 | -0.103390882 | 0.660169644 | 0.93 |
| YGL181W | 1793.449442 | 0.151590118 | 0.659606324 | 0.93 |
| YGR170W | 2142.624103 | 0.140524012 | 0.661099225 | 0.93 |
| YJR079W | 11.62148612 | 0.175854745 | 0.660685385 | 0.93 |
| YLR067C | 640.6258307 | 0.092482918 | 0.660579795 | 0.93 |
| YNL297C | 2104.946506 | -0.13026906 | 0.660815063 | 0.93 |
| YOL160W | 15.67971505 | -0.52207337 | 0.660893306 | 0.93 |
| YPL045W | 887.7248714 | 0.146215301 | 0.660985729 | 0.93 |
| YPL120W | 588.9261154 | 0.110034239 | 0.659882583 | 0.93 |
| YPR086W | 1622.357285 | 0.136643911 | 0.661021882 | 0.93 |
| YBR226C | 66.35832606 | -0.050468174 | 0.661603681 | 0.93 |
| YCR042C | 1593.772478 | 0.102522275 | 0.661915527 | 0.93 |
| YMR204C | 437.305049 | 0.063584191 | 0.662153243 | 0.93 |
| YOR383C | 13192.95676 | 0.142535954 | 0.662012575 | 0.93 |
| YBL063W | 882.9945164 | -0.0849825 | 0.662583835 | 0.93 |
| YPL155C | 690.291389 | 0.151680933 | 0.662894433 | 0.93 |
| YGL239C | 18.18533761 | 0.044486424 | 0.663091138 | 0.93 |
| YBL073W | 6.010181019 | -0.260222948 | 0.664752379 | 0.93 |
| YBR190W | 3.930160671 | -0.605003677 | 0.66677297 | 0.93 |
| YDL240W | 734.1749038 | -0.301316 | 0.666660884 | 0.93 |
| YDR049W | 886.770398 | 0.09660061 | 0.664790022 | 0.93 |
| YDR191W | 146.592055 | -0.322352209 | 0.666417527 | 0.93 |
| YDR203W | 15.81601128 | -0.290384789 | 0.664550057 | 0.93 |
| YDR209C | 344.9577194 | -0.275607367 | 0.66381993 | 0.93 |
| YDR460W | 574.0402083 | 0.088538087 | 0.665431858 | 0.93 |
| YER100W | 1553.364761 | -0.127853808 | 0.663649608 | 0.93 |
| YGL244W | 2087.284965 | -0.116375101 | 0.665690327 | 0.93 |
| YGR163W | 840.8307527 | -0.140623765 | 0.66716189 | 0.93 |
| YHR071C-A | 39.05421358 | 0.031417561 | 0.667015619 | 0.93 |
| YHR136C | 132.1966559 | 0.175270989 | 0.664324842 | 0.93 |
| YJL174W | 2482.371966 | -0.087192946 | 0.66460674 | 0.93 |
| YKL033W | 666.8103831 | -0.111664015 | 0.664081324 | 0.93 |
| YLR013W | 3.60279815 | 0.24892065 | 0.667062609 | 0.93 |
| YLR041W | 27.97790779 | 0.392507758 | 0.664950858 | 0.93 |
| YML007W | 2781.196668 | 0.124171003 | 0.666468926 | 0.93 |
| YML103C | 3218.889825 | -0.176051032 | 0.666905391 | 0.93 |
| YNL118C | 3319.845542 | 0.078112166 | 0.663527739 | 0.93 |
| YNL304W | 349.9176759 | -0.173816598 | 0.664237142 | 0.93 |
| YOL137W | 547.8586694 | 0.118098194 | 0.666293738 | 0.93 |
| YOR131C | 796.6694001 | 0.143198805 | 0.666880272 | 0.93 |

| | | | | |
|---|---|---|---|---|
| YOR311C | 1031.860298 | -0.044973369 | 0.664564417 | 0.93 |
| YOR344C | 1985.74596 | -0.162146421 | 0.666813415 | 0.93 |
| YPR002W | 1192.998192 | 0.272213022 | 0.666584825 | 0.93 |
| YPR047W | 774.0936392 | -0.09296132 | 0.665108978 | 0.93 |
| YPR075C | 1855.120278 | -0.129182215 | 0.665224582 | 0.93 |
| YAR018C | 738.6960091 | 0.111156379 | 0.667535789 | 0.93 |
| YGL131C | 1060.332301 | 0.1215507 | 0.667751817 | 0.93 |
| YOR226C | 454.5609099 | 0.154100452 | 0.667485513 | 0.93 |
| YPL013C | 664.0145708 | -0.277348268 | 0.66771651 | 0.93 |
| YOR389W | 203.2493311 | 0.013197254 | 0.668035622 | 0.93 |
| YLR421C | 1264.683888 | 0.133459282 | 0.668531837 | 0.93 |
| YBL102W | 1291.754797 | -0.141102074 | 0.67006918 | 0.93 |
| YCL048W | 92.73705911 | 0.069668744 | 0.670013089 | 0.93 |
| YCR001W | 33.60447039 | -0.41808573 | 0.669418319 | 0.93 |
| YDR512C | 741.4198181 | 0.106757499 | 0.669936175 | 0.93 |
| YFR052C-A | 3217.393186 | 0.055527823 | 0.67009911 | 0.93 |
| YGR241C | 1428.572307 | 0.094885745 | 0.670230385 | 0.93 |
| YJR094C | 253.8151399 | 0.250141634 | 0.669346102 | 0.93 |
| YMR153C-A | 60.08017136 | -0.24304447 | 0.669517899 | 0.93 |
| YPL262W | 5873.810031 | 0.150639555 | 0.669758687 | 0.93 |
| YER091C-A | 0.485272411 | -0.189242576 | 0.670774927 | 0.94 |
| YER185W | 79.91884135 | 0.366694375 | 0.670699029 | 0.94 |
| YCR069W | 2249.27644 | 0.07632832 | 0.671017811 | 0.94 |
| YDL071C | 49.14928453 | -0.0514345 | 0.671251864 | 0.94 |
| YMR200W | 1797.595559 | -0.101226756 | 0.671169558 | 0.94 |
| YML050W | 378.8334393 | 0.06901431 | 0.671429134 | 0.94 |
| YDR199W | 75.09684199 | 0.071191158 | 0.67218477 | 0.94 |
| YDR485C | 1410.914463 | 0.168220977 | 0.671701253 | 0.94 |
| YGR024C | 991.4360363 | -0.099381941 | 0.672168485 | 0.94 |
| YKL179C | 1404.902826 | 0.055684321 | 0.672119643 | 0.94 |
| YPR135W | 846.2041741 | -0.086014353 | 0.67181643 | 0.94 |
| YDR187C | 25.29702409 | -0.165725456 | 0.674323118 | 0.94 |
| YDR271C | 28.54979964 | 0.14393381 | 0.673154532 | 0.94 |
| YER044C | 1096.772117 | -0.138147981 | 0.673526595 | 0.94 |
| YGL095C | 1579.460918 | -0.111736157 | 0.673432079 | 0.94 |
| YGL236C | 488.164276 | -0.233943053 | 0.674154534 | 0.94 |
| YIR023W | 1089.743487 | -0.138304242 | 0.673407059 | 0.94 |
| YJR092W | 2324.987782 | -0.115180167 | 0.674044959 | 0.94 |
| YKR048C | 4547.539769 | -0.120340446 | 0.67434897 | 0.94 |
| YLR234W | 283.065063 | 0.374601428 | 0.673761559 | 0.94 |
| YLR239C | 753.9388732 | 0.134873109 | 0.672843979 | 0.94 |
| YOL035C | 58.4740954 | 0.006080573 | 0.673489141 | 0.94 |
| YOR305W | 732.3331193 | -0.157688799 | 0.673890959 | 0.94 |

| | | | | |
|---|---|---|---|---|
| YPL078C | 4809.066893 | 0.020649225 | 0.673286886 | 0.94 |
| YPR136C | 12.04995248 | 0.397002213 | 0.673724617 | 0.94 |
| YBL083C | 7.154873991 | -0.407312757 | 0.67509848 | 0.94 |
| YFL030W | 1073.611575 | 0.215343458 | 0.674918854 | 0.94 |
| YHL046W-A | 47.85481112 | -0.400077478 | 0.675215732 | 0.94 |
| YMR270C | 708.6444493 | -0.107446117 | 0.674590615 | 0.94 |
| YNL145W | 1813.203416 | -0.347265697 | 0.674946422 | 0.94 |
| YPL270W | 1768.066094 | 0.150068691 | 0.6747399 | 0.94 |
| YOR055W | 14.54954616 | -0.10506557 | 0.675714846 | 0.94 |
| YBL037W | 1072.022099 | -0.174404306 | 0.676643584 | 0.94 |
| YCL075W | 37.39387639 | -0.063680181 | 0.677852972 | 0.94 |
| YDL005C | 1599.265031 | 0.117841141 | 0.677628757 | 0.94 |
| YDR020C | 268.9612697 | -0.078647437 | 0.679085094 | 0.94 |
| YDR354W | 871.1284672 | 0.232858564 | 0.67715496 | 0.94 |
| YEL013W | 1708.200753 | -0.169047899 | 0.678775752 | 0.94 |
| YGR240C-A | 46.68747277 | -0.278221577 | 0.678359493 | 0.94 |
| YIR043C | 49.74477082 | -0.603240584 | 0.676845257 | 0.94 |
| YJL193W | 438.7222894 | 0.129207386 | 0.677887248 | 0.94 |
| YJR107W | 474.7713781 | 0.062139809 | 0.678510727 | 0.94 |
| YKL043W | 3024.896012 | -0.106924989 | 0.678670053 | 0.94 |
| YKL049C | 577.3829424 | -0.111206841 | 0.677045534 | 0.94 |
| YKR008W | 1399.761894 | -0.140252648 | 0.677905374 | 0.94 |
| YKR073C | 18.81130727 | 0.6038653 | 0.678068859 | 0.94 |
| YLR337C | 2433.363455 | 0.170083042 | 0.678983187 | 0.94 |
| YML054C-A | 11.9894805 | 0.033576546 | 0.678908028 | 0.94 |
| YMR183C | 2217.146681 | -0.119788323 | 0.676557642 | 0.94 |
| YOR037W | 696.733578 | 0.131219314 | 0.677639121 | 0.94 |
| YOR201C | 630.2633868 | -0.087393176 | 0.67800533 | 0.94 |
| YPL153C | 363.2997459 | -0.116909081 | 0.678166127 | 0.94 |
| YPR023C | 1837.667458 | -0.109189213 | 0.678026883 | 0.94 |
| YDR401W | 0.227003213 | 0.028759609 | 0.681754077 | 0.94 |
| YDR430C | 1380.899649 | -0.088465692 | 0.681637171 | 0.94 |
| YER006C-A | 2.225353622 | 0.346273036 | 0.680591146 | 0.94 |
| YER170W | 376.8018239 | -0.134472394 | 0.680262762 | 0.94 |
| YGL028C | 1677.067671 | -0.305025977 | 0.680675544 | 0.94 |
| YGR144W | 97.38216331 | -0.216702424 | 0.680549512 | 0.94 |
| YHR163W | 2218.862273 | 0.178672947 | 0.681857329 | 0.94 |
| YHR194W | 859.058911 | 0.11035345 | 0.681471315 | 0.94 |
| YIL128W | 1273.394888 | 0.053878644 | 0.68196812 | 0.94 |
| YJR022W | 18.26801566 | 0.241210334 | 0.679879472 | 0.94 |
| YJR080C | 1270.85858 | -0.035011819 | 0.681512921 | 0.94 |
| YKL083W | 46.77827339 | -0.091155197 | 0.681473187 | 0.94 |
| YML059C | 1781.099171 | 0.073874405 | 0.681822359 | 0.94 |

| | | | | |
|---|---|---|---|---|
| YNL152W | 687.0558678 | -0.153762709 | 0.680197912 | 0.94 |
| YNL280C | 1498.81009 | -0.048688024 | 0.681677464 | 0.94 |
| YNR001C | 10779.43573 | 0.277703001 | 0.681608119 | 0.94 |
| YOR113W | 1039.201076 | 0.157948566 | 0.681084249 | 0.94 |
| snR75 | 1.574604411 | -0.033378421 | 0.680602326 | 0.94 |
| YGL222C | 1086.796722 | 0.076004122 | 0.682125804 | 0.94 |
| YDL241W | 250.4921648 | 0.573269993 | 0.68243214 | 0.94 |
| YLR063W | 280.3060801 | 0.241189815 | 0.682323118 | 0.94 |
| YDR068W | 1258.879245 | 0.080692741 | 0.68273082 | 0.94 |
| YOR116C | 1548.881592 | -0.105633983 | 0.682684501 | 0.94 |
| YCR075W-A | 129.8943621 | -0.23602501 | 0.683701252 | 0.94 |
| YDR019C | 2663.732875 | 0.036899763 | 0.683173591 | 0.94 |
| YJR124C | 485.3455454 | -0.096729627 | 0.68346018 | 0.94 |
| YMR044W | 903.5982709 | 0.108785568 | 0.683855859 | 0.94 |
| YNL286W | 504.2093629 | 0.126067393 | 0.683867952 | 0.94 |
| YOL050C | 16.33482394 | -0.024406777 | 0.683696969 | 0.94 |
| YDL062W | 21.94852935 | 0.426150426 | 0.68403931 | 0.94 |
| YBR293W | 445.7090525 | -0.028568661 | 0.686113922 | 0.94 |
| YDL186W | 120.8658792 | 0.12108017 | 0.684909343 | 0.94 |
| YDL203C | 667.0945544 | 0.119623137 | 0.68592213 | 0.94 |
| YER094C | 2268.208556 | -0.111156884 | 0.685368938 | 0.94 |
| YGL109W | 79.04302724 | 0.063649516 | 0.686150794 | 0.94 |
| YGL146C | 201.790122 | -0.062632077 | 0.684608634 | 0.94 |
| YGR164W | 4.051111594 | -0.645235325 | 0.685320519 | 0.94 |
| YHR142W | 1336.189865 | -0.163072416 | 0.685906537 | 0.94 |
| YJL078C | 3272.163203 | 0.24488873 | 0.685675956 | 0.94 |
| YLR113W | 3162.32221 | -0.131526223 | 0.68563499 | 0.94 |
| YMR075W | 761.6920474 | 0.135756944 | 0.684820424 | 0.94 |
| YNL309W | 553.1266537 | -0.072663738 | 0.684246589 | 0.94 |
| YPL062W | 4.463478236 | -0.130251586 | 0.686030918 | 0.94 |
| YPL116W | 1071.716343 | 0.061308536 | 0.685051847 | 0.94 |
| YPL257W | 181.2677801 | 0.263899935 | 0.686235606 | 0.94 |
| YPR199C | 796.7118888 | -0.246327254 | 0.686789955 | 0.94 |
| YLR311C | 30.19399839 | -0.008753016 | 0.686955761 | 0.94 |
| YAL016C-B | 15.45888685 | -0.192392953 | 0.689454504 | 0.94 |
| YCL041C | 119.8839828 | 0.316239325 | 0.68808982 | 0.94 |
| YDL057W | 396.4937822 | 0.043633561 | 0.689007523 | 0.94 |
| YDR417C | 435.6659374 | 0.039852733 | 0.689106603 | 0.94 |
| YDR476C | 1082.582617 | 0.09717239 | 0.688982139 | 0.94 |
| YGL199C | 2.062524245 | 0.883686499 | 0.688992321 | 0.94 |
| YGR031W | 376.7830976 | 0.162827732 | 0.687675662 | 0.94 |
| YJL095W | 1317.430087 | -0.066143008 | 0.688028818 | 0.94 |
| YJL222W | 13.76304146 | 0.003229101 | 0.689457036 | 0.94 |

| | | | | |
|---|---|---|---|---|
| YKL048C | 583.1360838 | -0.00718359 | 0.688619332 | 0.94 |
| YMR018W | 66.38637523 | 0.010349478 | 0.689540982 | 0.94 |
| YMR117C | 204.4706004 | 0.208839983 | 0.68847346 | 0.94 |
| YMR227C | 799.7644018 | -0.147396588 | 0.68846723 | 0.94 |
| YOL053W | 775.6100204 | 0.133120474 | 0.687320107 | 0.94 |
| YOR367W | 630.0179766 | -0.160962489 | 0.689518005 | 0.94 |
| YOR371C | 940.0880721 | 0.036261745 | 0.687947069 | 0.94 |
| snR52 | 0.834128967 | -0.187099692 | 0.689271835 | 0.94 |
| YCR020W-B | 152.1239298 | 0.17539578 | 0.691545916 | 0.94 |
| YDR268W | 586.9385642 | -0.045321234 | 0.692317458 | 0.94 |
| YDR365W-B | 142.2689351 | -0.096375441 | 0.692140631 | 0.94 |
| YDR472W | 675.6541934 | -0.111547151 | 0.691406403 | 0.94 |
| YEL004W | 410.7288824 | -0.127700459 | 0.692037976 | 0.94 |
| YEL017C-A | 672.6729484 | 0.072551411 | 0.690503289 | 0.94 |
| YGL194C-A | 142.847988 | 0.009769549 | 0.691719501 | 0.94 |
| YJL114W | 3.593194288 | 0.05768183 | 0.690982119 | 0.94 |
| YKR080W | 1412.607481 | 0.147156756 | 0.691525093 | 0.94 |
| YLR105C | 592.2672821 | -0.163825226 | 0.691268304 | 0.94 |
| YLR406C-A | 2.560257022 | -0.113423025 | 0.692342121 | 0.94 |
| YMR297W | 8256.874804 | -0.154688256 | 0.690920556 | 0.94 |
| YNL103W-A | 104.3621644 | 0.999005602 | 0.69085202 | 0.94 |
| YOR230W | 8762.727513 | -0.119740023 | 0.690386996 | 0.94 |
| YOR316C | 1558.064909 | -0.120063342 | 0.692263808 | 0.94 |
| YOR330C | 1527.333333 | 0.11575806 | 0.69195534 | 0.94 |
| YPL187W | 17.17000227 | -0.155323751 | 0.689969304 | 0.94 |
| YPL221W | 1548.403188 | 0.112462293 | 0.690403713 | 0.94 |
| YPR069C | 2243.3628 | 0.006488275 | 0.690016955 | 0.94 |
| YHR113W | 1875.547688 | -0.075777299 | 0.692628608 | 0.94 |
| YCR050C | 17.22525728 | 0.095299523 | 0.694319006 | 0.94 |
| YDL155W | 763.3718322 | -0.123233254 | 0.694248623 | 0.94 |
| YDR089W | 994.7212521 | 0.115941556 | 0.693554039 | 0.94 |
| YDR373W | 760.4394828 | 0.132695921 | 0.694087061 | 0.94 |
| YFL041W | 1108.963199 | 0.134593364 | 0.694671767 | 0.94 |
| YGL041C-B | 21.05072236 | 0.061679289 | 0.694772084 | 0.94 |
| YGR122C-A | 15.60192201 | -0.386361563 | 0.694065822 | 0.94 |
| YHR060W | 732.5771452 | -0.061921543 | 0.694327585 | 0.94 |
| YNR052C | 1154.692673 | -0.123298463 | 0.694534747 | 0.94 |
| YOR015W | 345.4012917 | 0.035721634 | 0.694585842 | 0.94 |
| YPL170W | 622.4061116 | 0.113087535 | 0.693393332 | 0.94 |
| YPR100W | 188.3881606 | -0.28187522 | 0.694237234 | 0.94 |
| YDL125C | 2310.617952 | -0.127454952 | 0.695697404 | 0.94 |
| YHR046C | 620.1635411 | -0.151025544 | 0.696340764 | 0.94 |
| YMR134W | 453.7137928 | -0.058215458 | 0.696151785 | 0.94 |

| | | | | |
|---|---|---|---|---|
| YMR190C | 1250.870637 | 0.203799922 | 0.696343911 | 0.94 |
| YDR388W | 5322.26629 | -0.069558419 | 0.697381559 | 0.94 |
| YEL009C | 21167.59097 | -0.140772054 | 0.696615753 | 0.94 |
| YGL050W | 416.7826537 | -0.181872122 | 0.696790666 | 0.94 |
| YGL174W | 430.9199773 | -0.108124763 | 0.696820369 | 0.94 |
| YMR022W | 1239.988654 | 0.122920738 | 0.697163818 | 0.94 |
| YMR115W | 1106.490843 | 0.10791045 | 0.697345663 | 0.94 |
| YMR296C | 3004.03015 | -0.071270794 | 0.697309591 | 0.94 |
| YBR163W | 719.9181119 | 0.139935695 | 0.697756675 | 0.94 |
| YKR046C | 2535.087216 | 0.08493111 | 0.697914906 | 0.94 |
| YNL157W | 1475.809734 | 0.091107523 | 0.698076043 | 0.94 |
| YOR215C | 2322.968103 | 0.18174415 | 0.698117558 | 0.94 |
| YPL046C | 139.4253961 | -0.18587363 | 0.697618542 | 0.94 |
| YLR166C | 1318.264285 | -0.096691899 | 0.698297708 | 0.94 |
| YDL010W | 362.0027109 | 0.152562025 | 0.698606095 | 0.94 |
| YLR326W | 402.567239 | 0.307275334 | 0.698898107 | 0.94 |
| YER048C | 2682.830499 | -0.150245942 | 0.699128397 | 0.94 |
| YOR244W | 816.0935801 | -0.134252557 | 0.699213489 | 0.94 |
| YBR262C | 557.8168642 | -0.205983611 | 0.700148613 | 0.94 |
| YFR055W | 975.2716539 | 0.304164248 | 0.700250636 | 0.94 |
| YJR065C | 3820.877578 | -0.082227365 | 0.700246262 | 0.94 |
| YMR172W | 685.1558079 | -0.135689992 | 0.700161885 | 0.94 |
| YPL156C | 858.4762505 | -0.109292015 | 0.700096985 | 0.94 |
| YIL150C | 295.6540585 | 0.036985615 | 0.70051338 | 0.94 |
| YFR052W | 2758.803837 | 0.161370237 | 0.700938525 | 0.94 |
| YGL023C | 1541.23519 | 0.182483447 | 0.701237344 | 0.94 |
| YLR246W | 470.2308665 | 0.165954327 | 0.701586277 | 0.94 |
| YDR153C | 1161.100144 | 0.042212878 | 0.702377153 | 0.94 |
| YGL086W | 1388.801297 | 0.143222198 | 0.702188974 | 0.94 |
| YIL129C | 2649.636452 | -0.106947122 | 0.70227001 | 0.94 |
| YLR436C | 2590.336596 | 0.134428593 | 0.702198717 | 0.94 |
| YBL047C | 5698.969568 | 0.140891497 | 0.702610682 | 0.94 |
| YKL208W | 299.0743362 | -0.209809435 | 0.702921689 | 0.94 |
| YAL013W | 1274.881408 | 0.071856452 | 0.703678192 | 0.94 |
| YBR015C | 2379.48263 | -0.078347833 | 0.704099088 | 0.94 |
| YER141W | 2454.288383 | -0.13631923 | 0.704535817 | 0.94 |
| YJR135C | 188.8236464 | -0.092208087 | 0.704123941 | 0.94 |
| YJR144W | 1299.865299 | -0.067721548 | 0.704533039 | 0.94 |
| YMR043W | 2143.509763 | 0.004844524 | 0.703696769 | 0.94 |
| YMR154C | 501.7366665 | 0.064706321 | 0.703825891 | 0.94 |
| YNL241C | 6779.137631 | 0.342633643 | 0.704284107 | 0.94 |
| YOR050C | 67.3008437 | -0.276827285 | 0.703254027 | 0.94 |
| YPL176C | 1288.392675 | 0.174546616 | 0.703748165 | 0.94 |

| | | | | |
|---|---|---|---|---|
| YGL024W | 7.765530916 | -0.010721247 | 0.704721533 | 0.94 |
| YDR488C | 747.4804028 | 0.20566671 | 0.705370794 | 0.94 |
| YJR106W | 683.5325213 | 0.186338163 | 0.705382928 | 0.94 |
| YDL024C | 228.1924469 | 0.035155183 | 0.706275376 | 0.94 |
| YGR275W | 659.6729657 | -0.068970562 | 0.706053992 | 0.94 |
| YLL067C | 0.999643856 | 0.28469095 | 0.706472095 | 0.94 |
| YLR428C | 15.96009415 | 0.012397672 | 0.705756822 | 0.94 |
| YMR100W | 811.7962257 | 0.025771078 | 0.706096404 | 0.94 |
| YMR168C | 528.3657431 | -0.109056878 | 0.706400469 | 0.94 |
| YAR047C | 21.66468268 | 0.336623045 | 0.709238025 | 0.94 |
| YBL013W | 160.0267849 | 0.227761447 | 0.70969816 | 0.94 |
| YBL029W | 662.4144794 | 0.073534599 | 0.709615055 | 0.94 |
| YBL070C | 21.83445099 | -0.100992264 | 0.710526349 | 0.94 |
| YBL079W | 2598.015595 | -0.090465214 | 0.708749378 | 0.94 |
| YBR020W | 310.6456705 | -0.031260648 | 0.708944603 | 0.94 |
| YDL098C | 246.2120054 | 0.129176074 | 0.710048576 | 0.94 |
| YDL160C-A | 95.22130843 | -0.135247358 | 0.707245018 | 0.94 |
| YEL072W | 207.4067694 | -0.041669046 | 0.706849058 | 0.94 |
| YER182W | 1071.612284 | 0.062348581 | 0.710532772 | 0.94 |
| YFL058W | 7.345596516 | -0.187265342 | 0.710804778 | 0.94 |
| YGL211W | 636.0371021 | 0.086380565 | 0.707611019 | 0.94 |
| YGR015C | 438.1021585 | 0.069744 | 0.707632961 | 0.94 |
| YHR035W | 231.8124454 | 0.236437695 | 0.707428228 | 0.94 |
| YJR042W | 1389.72183 | -0.151847192 | 0.7099137 | 0.94 |
| YKL097C | 10.27058523 | 0.154845567 | 0.708926058 | 0.94 |
| YKR071C | 2094.595777 | 0.300892197 | 0.708778321 | 0.94 |
| YLR226W | 384.869465 | 0.111277749 | 0.709990126 | 0.94 |
| YML095C | 255.548255 | -0.181866802 | 0.707803934 | 0.94 |
| YMR156C | 369.432093 | 0.13425438 | 0.707891387 | 0.94 |
| YMR285C | 827.213202 | -0.110650438 | 0.710890416 | 0.94 |
| YMR306C-A | 3.882802162 | 0.386644716 | 0.71076963 | 0.94 |
| YNL083W | 670.395816 | 0.035997377 | 0.708567599 | 0.94 |
| YNL308C | 3277.042014 | 0.018462167 | 0.70936243 | 0.94 |
| YOL004W | 4094.946984 | -0.095483116 | 0.710300126 | 0.94 |
| YOL122C | 1128.161192 | 0.043686705 | 0.708954438 | 0.94 |
| YOR317W | 4625.356852 | 0.259537746 | 0.709578444 | 0.94 |
| YPL229W | 514.8866138 | 0.011514419 | 0.707201065 | 0.94 |
| YPR096C | 16.87382422 | 0.224412873 | 0.709619469 | 0.94 |
| YPR197C | 5.599447647 | 0.153193163 | 0.710706491 | 0.94 |
| YBL005W | 1420.668611 | -0.199663707 | 0.711927491 | 0.94 |
| YBL075C | 2650.065607 | 0.102142907 | 0.713364723 | 0.94 |
| YDR241W | 32.77525977 | -0.23485993 | 0.712085908 | 0.94 |
| YDR359C | 1438.122379 | -0.077028114 | 0.713221522 | 0.94 |

| | | | | |
|---|---|---|---|---|
| YDR389W | 894.8507584 | -0.186134876 | 0.713127115 | 0.94 |
| YEL043W | 2331.741844 | 0.143231946 | 0.713702745 | 0.94 |
| YFL021C-A | 41.5780796 | -0.231419628 | 0.71207781 | 0.94 |
| YGR109W-B | 857.2496031 | 0.126359764 | 0.71252499 | 0.94 |
| YGR186W | 2388.772284 | 0.163266083 | 0.713666133 | 0.94 |
| YIL080W | 141.3218365 | -0.043713186 | 0.712485888 | 0.94 |
| YLR294C | 1804.172401 | -0.049599141 | 0.713543326 | 0.94 |
| YMR064W | 1294.977576 | 0.075899127 | 0.712950651 | 0.94 |
| YMR289W | 967.6114857 | -0.093172631 | 0.712329358 | 0.94 |
| YOL129W | 2548.890951 | -0.093062407 | 0.712443469 | 0.94 |
| YOL166C | 2.808993483 | -0.244161555 | 0.713350826 | 0.94 |
| YOR192C-B | 24.91802453 | -0.121218882 | 0.71193729 | 0.94 |
| YPL021W | 14.06563719 | 0.158762272 | 0.713189338 | 0.94 |
| YPL225W | 2696.542417 | -0.087113817 | 0.713126188 | 0.94 |
| YBR184W | 226.9887473 | 0.352740426 | 0.714223532 | 0.94 |
| YDL041W | 0.97089065 | 0.066095637 | 0.714134913 | 0.94 |
| YER019W | 1037.358398 | -0.069642144 | 0.714118507 | 0.94 |
| YNL316C | 479.8242776 | -0.113123948 | 0.714507202 | 0.94 |
| YPR054W | 265.2938746 | -0.363849869 | 0.714401002 | 0.94 |
| YMR141W-A | 54.52612508 | -0.013526526 | 0.714921854 | 0.94 |
| YOL051W | 2569.23447 | 0.187675012 | 0.714674294 | 0.94 |
| YOR262W | 658.3672371 | 0.178937562 | 0.714943771 | 0.94 |
| YAR075W | 64.72959559 | 0.019452612 | 0.717201574 | 0.94 |
| YBR056W-A | 43.21007303 | -0.169319986 | 0.71894829 | 0.94 |
| YBR057C | 866.0815496 | 0.09568458 | 0.715909057 | 0.94 |
| YCR086W | 338.9363584 | -0.094865325 | 0.718632556 | 0.94 |
| YDR218C | 78.04916244 | 0.123599641 | 0.719578543 | 0.94 |
| YDR498C | 1014.743074 | -0.086064514 | 0.718871957 | 0.94 |
| YFR004W | 2781.166019 | 0.093560345 | 0.719095807 | 0.94 |
| YGL216W | 962.5040676 | 0.074162602 | 0.716734355 | 0.94 |
| YHL002W | 1516.40559 | 0.065327711 | 0.719440386 | 0.94 |
| YHR005C | 1534.146963 | -0.390090549 | 0.719511334 | 0.94 |
| YHR101C | 345.7524508 | 0.088348798 | 0.719919971 | 0.94 |
| YHR182W | 713.7981983 | 0.182613234 | 0.716250195 | 0.94 |
| YIL111W | 1481.52504 | 0.072703162 | 0.718623792 | 0.94 |
| YJL020C | 4552.045062 | 0.218452322 | 0.718052195 | 0.94 |
| YJL123C | 2650.72374 | -0.1425434 | 0.719049358 | 0.94 |
| YJL166W | 2486.295448 | 0.083656569 | 0.716810164 | 0.94 |
| YKR077W | 759.0676767 | -0.044726124 | 0.7183642 | 0.94 |
| YLL029W | 3427.507563 | 0.131334223 | 0.716289585 | 0.94 |
| YLR095C | 1759.144757 | -0.09998534 | 0.718169871 | 0.94 |
| YLR357W | 954.9234442 | -0.10566746 | 0.716500329 | 0.94 |
| YLR443W | 472.8026492 | 0.125679664 | 0.717346734 | 0.94 |

| | | | | |
|---|---|---|---|---|
| YML091C | 3874.618724 | 0.006414074 | 0.719488673 | 0.94 |
| YMR281W | 530.4325943 | 0.118821323 | 0.719890925 | 0.94 |
| YMR310C | 757.4056331 | -0.136846479 | 0.718528435 | 0.94 |
| YNL120C | 0.227028385 | 0.038602857 | 0.71566168 | 0.94 |
| YNR066C | 265.0332103 | 0.304382104 | 0.717376297 | 0.94 |
| YOR081C | 1945.1418 | 0.089044018 | 0.717324785 | 0.94 |
| YOR105W | 21.94253227 | -0.118744346 | 0.716228224 | 0.94 |
| YOR169C | 39.59953477 | -0.20718642 | 0.71729985 | 0.94 |
| YOR319W | 365.6759573 | -0.092594122 | 0.717290283 | 0.94 |
| YPL122C | 549.6598792 | -0.101133 | 0.719676282 | 0.94 |
| YPR094W | 154.1332011 | -0.242952028 | 0.717668339 | 0.94 |
| YPR186C | 328.1330267 | -0.114609226 | 0.718521911 | 0.94 |
| snR68 | 10.29817386 | 0.26602203 | 0.720101822 | 0.94 |
| YBR179C | 1264.938407 | -0.108210894 | 0.721098215 | 0.95 |
| YHR173C | 88.05682976 | -0.292809666 | 0.721508925 | 0.95 |
| YHR182C-A | 130.5541577 | 0.646270857 | 0.721200288 | 0.95 |
| YPL195W | 2059.660784 | -0.04582825 | 0.721447644 | 0.95 |
| YPL233W | 252.806959 | -0.092120719 | 0.721525028 | 0.95 |
| YBR248C | 1552.941582 | 0.185587323 | 0.722042861 | 0.95 |
| YCR075C | 329.3656461 | -0.103565248 | 0.722293806 | 0.95 |
| YGL089C | 132.0787924 | 0.039723041 | 0.722206276 | 0.95 |
| YGL193C | 193.4216907 | -0.226916064 | 0.721885471 | 0.95 |
| YJL135W | 5.319281498 | -0.011262629 | 0.722690975 | 0.95 |
| YGL145W | 1156.588747 | 0.078254511 | 0.72297175 | 0.95 |
| YLR262C | 283.7623055 | -0.055163497 | 0.723284129 | 0.95 |
| YNL325C | 453.4130958 | 0.06937973 | 0.723215359 | 0.95 |
| YOR280C | 559.6900755 | 0.126232829 | 0.723493639 | 0.95 |
| YHR077C | 2925.324128 | 0.079677128 | 0.723994455 | 0.95 |
| YLR351C | 1673.534289 | -0.055128675 | 0.724047115 | 0.95 |
| YNL073W | 1409.044777 | 0.066117109 | 0.724276137 | 0.95 |
| YPR120C | 354.4464034 | -0.23035953 | 0.724340918 | 0.95 |
| YCL034W | 1930.992154 | 0.054012485 | 0.725097805 | 0.95 |
| YHR034C | 511.3439371 | 0.103236053 | 0.725171451 | 0.95 |
| YLR011W | 284.831779 | -0.142220891 | 0.725022745 | 0.95 |
| YOR288C | 898.6918954 | -0.165626434 | 0.724604556 | 0.95 |
| YPR194C | 1121.813284 | 0.382585844 | 0.72483753 | 0.95 |
| YER133W-A | 95.44590047 | -0.240428348 | 0.725730683 | 0.95 |
| YKR028W | 1923.731148 | -0.139569716 | 0.725630625 | 0.95 |
| YLR310C | 3300.526123 | -0.111469811 | 0.725652184 | 0.95 |
| YMR288W | 774.5062547 | 0.109896076 | 0.725869226 | 0.95 |
| YGR247W | 484.9641046 | 0.055889795 | 0.726203211 | 0.95 |
| YJL044C | 1379.126824 | 0.143050411 | 0.726374741 | 0.95 |
| YNL169C | 1167.217459 | -0.081093622 | 0.726447568 | 0.95 |

| | | | | |
|---|---|---|---|---|
| YNL134C | 7013.757129 | 0.401506274 | 0.726641594 | 0.95 |
| YHL009W-B | 5.78416984 | 0.154767161 | 0.727254708 | 0.95 |
| YLR282C | 4.588773458 | -0.117465081 | 0.727463496 | 0.95 |
| YMR067C | 1220.387108 | -0.1162321 | 0.727589043 | 0.95 |
| YNR004W | 166.7322391 | 0.134935537 | 0.727552978 | 0.95 |
| YPL168W | 515.986609 | 0.082313978 | 0.727012271 | 0.95 |
| YLL044W | 94.44590477 | -0.453627595 | 0.727885527 | 0.95 |
| YIL009C-A | 425.8589138 | 0.129088091 | 0.728452693 | 0.95 |
| YJL038C | 194.4642451 | -0.234213937 | 0.728353581 | 0.95 |
| snR82 | 454.6100305 | -0.488097975 | 0.728516408 | 0.95 |
| YAR042W | 3250.319067 | 0.175671139 | 0.732827529 | 0.95 |
| YBL039W-B | 151.3574507 | 0.085241777 | 0.732598007 | 0.95 |
| YBR126W-A | 695.852559 | -0.267098937 | 0.729032898 | 0.95 |
| YBR173C | 2339.401983 | -0.062482077 | 0.730805472 | 0.95 |
| YDL067C | 1308.526792 | 0.051158635 | 0.730949473 | 0.95 |
| YDR343C | 868.7911596 | -0.147332255 | 0.731525715 | 0.95 |
| YER008C | 2582.281632 | 0.123192113 | 0.731489932 | 0.95 |
| YER164W | 2428.238913 | 0.142485553 | 0.73182074 | 0.95 |
| YFR038W | 1186.649674 | 0.173785549 | 0.729804861 | 0.95 |
| YGL032C | 1976.542723 | -0.271547702 | 0.730147262 | 0.95 |
| YGR132C | 2816.110199 | -0.066739483 | 0.730257209 | 0.95 |
| YGR190C | 89.92851229 | 0.304376364 | 0.731896577 | 0.95 |
| YHR008C | 8653.583569 | 0.332493702 | 0.732113083 | 0.95 |
| YIR018W | 398.3013208 | 0.0193185 | 0.730745241 | 0.95 |
| YJL059W | 446.5285115 | 0.105223485 | 0.732191514 | 0.95 |
| YJR122W | 745.1766826 | 0.110823932 | 0.732466637 | 0.95 |
| YLR145W | 262.3006502 | -0.125317262 | 0.732798204 | 0.95 |
| YLR227W-B | 558.9698813 | 0.143531324 | 0.73312731 | 0.95 |
| YMR164C | 711.3389073 | -0.053440659 | 0.732199695 | 0.95 |
| YMR231W | 864.2847521 | -0.061871283 | 0.731250163 | 0.95 |
| YMR299C | 640.5800002 | -0.110883942 | 0.731955508 | 0.95 |
| YMR325W | 4.52283349 | 0.148336323 | 0.730496295 | 0.95 |
| YNL080C | 1115.372394 | -0.057682223 | 0.731864263 | 0.95 |
| YNL085W | 3478.261348 | 0.029064696 | 0.732856399 | 0.95 |
| YNL205C | 11.3162081 | -0.257249537 | 0.732402969 | 0.95 |
| YNL324W | 2.168671185 | -0.463023036 | 0.732857382 | 0.95 |
| YOL028C | 419.6545084 | 0.128691243 | 0.733066098 | 0.95 |
| YOR082C | 31.57473045 | 0.084773567 | 0.73173179 | 0.95 |
| YOR196C | 1243.855488 | 0.133560664 | 0.732751812 | 0.95 |
| YOR261C | 3484.142585 | -0.028864079 | 0.731126334 | 0.95 |
| YDR252W | 88.93914678 | -0.010652311 | 0.73335947 | 0.95 |
| YIL102C | 13.70446277 | -0.034879638 | 0.733714995 | 0.95 |
| YNR045W | 861.3548214 | 0.076620045 | 0.733460653 | 0.95 |

| | | | | |
|---|---|---|---|---|
| YPR108W-A | 1.090591333 | -0.234184839 | 0.733722101 | 0.95 |
| YDL007W | 5356.140616 | -0.096571006 | 0.734159306 | 0.95 |
| YOL070C | 1742.264613 | 0.212101418 | 0.734006428 | 0.95 |
| YOR290C | 3907.230484 | -0.113784114 | 0.734248257 | 0.95 |
| YBL094C | 5.123149382 | -0.371633883 | 0.73607916 | 0.95 |
| YCR017C | 2986.294618 | -0.029508599 | 0.736932956 | 0.95 |
| YDR533C | 1411.920932 | -0.449730481 | 0.736790083 | 0.95 |
| YGR107W | 13.44440994 | 0.288104589 | 0.735200886 | 0.95 |
| YGR206W | 264.0711736 | -0.183201346 | 0.735276483 | 0.95 |
| YHL047C | 1687.543568 | -0.311921552 | 0.734858141 | 0.95 |
| YIL019W | 543.1334136 | 0.088181013 | 0.736446289 | 0.95 |
| YIL135C | 1336.175722 | 0.05973543 | 0.736160366 | 0.95 |
| YJR085C | 2130.783694 | -0.079314003 | 0.736300077 | 0.95 |
| YKL070W | 302.0799748 | 0.084667151 | 0.736815727 | 0.95 |
| YKL088W | 1161.516822 | 0.043819655 | 0.735351298 | 0.95 |
| YLR346C | 131.9806369 | 0.057047205 | 0.73489524 | 0.95 |
| YLR434C | 48.33837913 | 0.150527798 | 0.736779325 | 0.95 |
| YMR290W-A | 11.59226843 | 0.224811421 | 0.73605429 | 0.95 |
| YNL076W | 1058.446405 | 0.045630082 | 0.736104146 | 0.95 |
| YPL071C | 441.9177148 | -0.151323513 | 0.734543749 | 0.95 |
| YPR180W | 730.2139766 | 0.098562922 | 0.736776122 | 0.95 |
| snR67 | 6.800863812 | 0.146133872 | 0.736641573 | 0.95 |
| YKL139W | 677.7577592 | -0.184153777 | 0.737562989 | 0.95 |
| YBR049C | 1277.509775 | -0.087256427 | 0.738183548 | 0.95 |
| YGR230W | 330.0627222 | 0.107243743 | 0.738320302 | 0.95 |
| YHR132W-A | 721.2397547 | 0.170191722 | 0.738259913 | 0.95 |
| YHR198C | 872.0543676 | 0.021063011 | 0.737814467 | 0.95 |
| YMR166C | 428.7049362 | -0.126821913 | 0.737899154 | 0.95 |
| YOR357C | 469.3315493 | 0.10956295 | 0.738443191 | 0.95 |
| YKR082W | 1772.94275 | 0.117926093 | 0.738655973 | 0.95 |
| YML056C | 4012.776938 | 0.011046059 | 0.738760261 | 0.95 |
| YAL008W | 865.7638687 | 0.074120606 | 0.739840794 | 0.95 |
| YBR133C | 1160.137007 | -0.073428682 | 0.741810655 | 0.95 |
| YBR137W | 861.5111114 | -0.167739594 | 0.741637824 | 0.95 |
| YCL047C | 434.4156263 | -0.034711032 | 0.74184521 | 0.95 |
| YDL181W | 1061.423493 | 0.003667653 | 0.739680726 | 0.95 |
| YDR193W | 12.98139175 | 0.311288439 | 0.742525009 | 0.95 |
| YDR270W | 829.6044968 | 0.072440886 | 0.739811538 | 0.95 |
| YER168C | 714.524667 | 0.086265191 | 0.74026581 | 0.95 |
| YGR168C | 490.2041357 | 0.070216471 | 0.742032081 | 0.95 |
| YGR204C-A | 13.54650687 | 0.006588433 | 0.740071407 | 0.95 |
| YHL044W | 369.968591 | 0.04239323 | 0.74086177 | 0.95 |
| YHR067W | 339.2689826 | 0.311613278 | 0.741605854 | 0.95 |

| | | | | |
|---|---|---|---|---|
| YIL085C | 1498.284283 | 0.086156188 | 0.739662905 | 0.95 |
| YJR100C | 772.6621126 | 0.059924946 | 0.742229543 | 0.95 |
| YKL102C | 18.56354003 | -0.042281288 | 0.742655803 | 0.95 |
| YKR025W | 782.5030894 | -0.008178544 | 0.739717946 | 0.95 |
| YLL009C | 581.7236454 | -0.197199155 | 0.742269246 | 0.95 |
| YLR213C | 172.0132438 | -0.084230358 | 0.741748552 | 0.95 |
| YML037C | 139.9825756 | -0.089109493 | 0.742371141 | 0.95 |
| YMR247C | 1642.766432 | 0.135981799 | 0.74126973 | 0.95 |
| YNL088W | 2851.433881 | 0.223704045 | 0.742265064 | 0.95 |
| YNL220W | 4979.508941 | 0.195249379 | 0.741451789 | 0.95 |
| YNL272C | 1304.645862 | -0.075919616 | 0.740763695 | 0.95 |
| YNR025C | 48.20698063 | -0.081376015 | 0.741081392 | 0.95 |
| YPR033C | 3601.656962 | 0.038106979 | 0.740860645 | 0.95 |
| YPR148C | 2535.151148 | -0.062801158 | 0.74240139 | 0.95 |
| YBR237W | 937.0201345 | 0.088230217 | 0.742998223 | 0.95 |
| YLR360W | 653.1816733 | -0.08794942 | 0.743142514 | 0.95 |
| YNR038W | 1115.811097 | -0.110061963 | 0.7430566 | 0.95 |
| YMR094W | 306.3491696 | 0.384577459 | 0.74353778 | 0.95 |
| YBR216C | 879.1909045 | 0.060725199 | 0.744026283 | 0.95 |
| YDL002C | 564.8870678 | -0.138729138 | 0.743936519 | 0.95 |
| YDR441C | 376.4209723 | 0.06543566 | 0.744111565 | 0.95 |
| YGL052W | 83.8653604 | 0.03637419 | 0.744667421 | 0.95 |
| YJR068W | 1198.277592 | -0.033697463 | 0.744618 | 0.95 |
| YOL036W | 1888.124512 | 0.068417334 | 0.744266186 | 0.95 |
| YOL072W | 594.5699607 | 0.03087773 | 0.744496882 | 0.95 |
| YBR230W-A | 211.0787097 | -0.580020958 | 0.746065726 | 0.95 |
| YBR276C | 1528.649976 | 0.068869056 | 0.745917981 | 0.95 |
| YCR097W-A | 1.60235674 | 0.08521213 | 0.746229245 | 0.95 |
| YEL024W | 1858.219132 | 0.210902567 | 0.746576537 | 0.95 |
| YER165C-A | 28.00543952 | 0.245281805 | 0.745786791 | 0.95 |
| YKR060W | 417.425685 | -0.113347146 | 0.746551993 | 0.95 |
| YLR077W | 986.4922801 | 0.083112789 | 0.74587916 | 0.95 |
| YLR451W | 1124.900564 | 0.061985892 | 0.746168006 | 0.95 |
| YMR026C | 666.726482 | -0.09910207 | 0.746559664 | 0.95 |
| YOR175C | 2021.823765 | -0.046013646 | 0.74607888 | 0.95 |
| YOR194C | 1465.560671 | 0.044764391 | 0.74591965 | 0.95 |
| YPR093C | 386.3642488 | -0.147185611 | 0.745217426 | 0.95 |
| YNL052W | 2172.615322 | 0.061391128 | 0.746923431 | 0.95 |
| YIL168W | 92.19596636 | 0.174157045 | 0.747134876 | 0.95 |
| YAL001C | 1631.198822 | 0.092910297 | 0.747848956 | 0.95 |
| YEL069C | 125.5249506 | 0.118143826 | 0.747594669 | 0.95 |
| YGR119C | 2276.598641 | -0.077504301 | 0.747875229 | 0.95 |
| YPR038W | 18.2580769 | -0.289664298 | 0.747850295 | 0.95 |

| | | | | |
|---|---|---|---|---|
| YCR101C | 105.2670668 | -0.152294262 | 0.749371188 | 0.95 |
| YDL172C | 10.58915184 | -0.433214807 | 0.75013799 | 0.95 |
| YDR102C | 12.68433027 | -0.36352289 | 0.748520395 | 0.95 |
| YFL063W | 4.881871603 | 0.166014977 | 0.750117578 | 0.95 |
| YFR034W-A | 2.98827912 | 0.061151271 | 0.749370345 | 0.95 |
| YGL203C | 1571.604512 | 0.143862071 | 0.74999531 | 0.95 |
| YGR055W | 1342.856307 | -0.029565555 | 0.750164112 | 0.95 |
| YGR138C | 1416.124697 | 0.538614894 | 0.75031008 | 0.95 |
| YJL024C | 668.6768051 | -0.115997457 | 0.749157978 | 0.95 |
| YJR073C | 2980.954697 | 0.138518072 | 0.748561252 | 0.95 |
| YKL186C | 1093.769583 | -0.193962295 | 0.748282995 | 0.95 |
| YLR265C | 441.14354 | -0.12752632 | 0.75007203 | 0.95 |
| YMR209C | 810.1729969 | -0.177189258 | 0.748911326 | 0.95 |
| YNL229C | 1349.306513 | -0.071180108 | 0.749719656 | 0.95 |
| YOL024W | 52.91060507 | 0.234027457 | 0.749263984 | 0.95 |
| YDR491C | 102.3610769 | 0.341783332 | 0.751026166 | 0.95 |
| YER142C | 720.8941802 | 0.120280659 | 0.750904913 | 0.95 |
| YLR071C | 1557.164849 | 0.077337762 | 0.750836722 | 0.95 |
| YGR260W | 2070.398504 | -0.086558425 | 0.751286136 | 0.95 |
| YML011C | 267.8459983 | 0.072746721 | 0.751424868 | 0.95 |
| YAL024C | 799.2694521 | -0.124338664 | 0.751817658 | 0.95 |
| YKR034W | 49.93480154 | -0.292792013 | 0.751705952 | 0.95 |
| YLR182W | 1421.620302 | 0.049263502 | 0.75217758 | 0.95 |
| YML025C | 1270.924858 | -0.095098594 | 0.752091628 | 0.95 |
| YDL210W | 142.1653441 | 0.061813825 | 0.753820201 | 0.95 |
| YDR487C | 2644.625402 | -0.038820293 | 0.753217082 | 0.95 |
| YEL032W | 1185.048516 | 0.076473455 | 0.752917094 | 0.95 |
| YGR281W | 2050.830805 | -0.106952465 | 0.753078124 | 0.95 |
| YIL138C | 1495.862581 | -0.133243966 | 0.753303436 | 0.95 |
| YJL066C | 1041.928086 | 0.117587849 | 0.753552748 | 0.95 |
| YKL209C | 3091.85579 | -0.369923535 | 0.75344236 | 0.95 |
| YKR103W | 526.4204375 | 0.162950718 | 0.753647103 | 0.95 |
| YML043C | 371.5090553 | -0.226671699 | 0.753935845 | 0.95 |
| YDR508C | 2566.963439 | 0.085090607 | 0.754654633 | 0.95 |
| YBR297W | 465.5986861 | 0.102314649 | 0.756836548 | 0.95 |
| YCL057W | 3573.183746 | -0.017752241 | 0.755982325 | 0.95 |
| YDL168W | 1463.24926 | 0.275029659 | 0.757790409 | 0.95 |
| YDL187C | 3.68310749 | -0.495643814 | 0.758205992 | 0.95 |
| YDR106W | 46.31624816 | -0.011410333 | 0.757357681 | 0.95 |
| YDR497C | 3420.957422 | -0.052924793 | 0.75545586 | 0.95 |
| YDR532C | 597.6785104 | 0.071598119 | 0.756490054 | 0.95 |
| YEL056W | 977.5251167 | 0.058313784 | 0.758180337 | 0.95 |
| YGL127C | 488.4323072 | -0.001108813 | 0.755313625 | 0.95 |

| | | | | |
|---|---|---|---|---|
| YHR011W | 509.988716 | -0.147907516 | 0.756313557 | 0.95 |
| YHR143W | 3407.641358 | -0.049324365 | 0.756488093 | 0.95 |
| YIL030C | 2092.978365 | -0.073413078 | 0.758064308 | 0.95 |
| YIR030W-A | 17.57088172 | -0.040745858 | 0.757195596 | 0.95 |
| YJL043W | 47.06314645 | -0.416867773 | 0.75718866 | 0.95 |
| YKL149C | 355.2699607 | 0.081248041 | 0.757656358 | 0.95 |
| YLR371W | 1780.890302 | 0.085994052 | 0.756646164 | 0.95 |
| YML006C | 696.4182875 | 0.102354391 | 0.755162746 | 0.95 |
| YMR266W | 1578.35636 | -0.0659592 | 0.756722461 | 0.95 |
| YOL029C | 366.4284423 | -0.118314174 | 0.75746588 | 0.95 |
| YOL150C | 3.831416512 | 0.002016696 | 0.756752665 | 0.95 |
| YOR233W | 853.5809378 | 0.076388767 | 0.757606772 | 0.95 |
| YPL125W | 1168.744838 | 0.15455438 | 0.757753151 | 0.95 |
| YPL179W | 1527.082249 | -0.133717908 | 0.7554675 | 0.95 |
| YPR134W | 542.0304792 | 0.063309781 | 0.757868429 | 0.95 |
| YBR094W | 1036.817856 | -0.024872153 | 0.759305732 | 0.95 |
| YBR221W-A | 0.837044541 | -0.132384807 | 0.759148847 | 0.95 |
| YHR162W | 4274.252821 | -0.106770502 | 0.759446992 | 0.95 |
| YHR207C | 677.6706609 | 0.016307823 | 0.759427082 | 0.95 |
| YLR381W | 607.7067419 | -0.129240283 | 0.758770671 | 0.95 |
| YNR035C | 3358.997637 | -0.067895825 | 0.759473244 | 0.95 |
| YPL201C | 234.758662 | -0.00629856 | 0.758750531 | 0.95 |
| YAL044W-A | 228.0761832 | -0.163279946 | 0.75993627 | 0.95 |
| YDL087C | 335.3146294 | 0.039599148 | 0.760589664 | 0.95 |
| YDR368W | 2244.977391 | 0.024761062 | 0.760566287 | 0.95 |
| YFL067W | 57.17560859 | -0.326232758 | 0.760555294 | 0.95 |
| YIL010W | 1043.363564 | -0.045604484 | 0.759783323 | 0.95 |
| YJR040W | 752.4605261 | -0.08379543 | 0.760360636 | 0.95 |
| YOR195W | 1326.708475 | 0.033388298 | 0.760173572 | 0.95 |
| YMR295C | 1636.767669 | 0.076417524 | 0.761025172 | 0.95 |
| YJL003W | 223.2586603 | -0.229645684 | 0.761603815 | 0.95 |
| YNL256W | 1084.449905 | -0.012044315 | 0.761499444 | 0.95 |
| YNL133C | 297.07226 | 0.077575642 | 0.761992884 | 0.95 |
| YOR327C | 881.0062218 | 0.064499529 | 0.762541817 | 0.95 |
| YJR056C | 431.2081935 | 0.073089443 | 0.762850196 | 0.96 |
| YML110C | 4679.227772 | 0.032408926 | 0.763265037 | 0.96 |
| YOL110W | 433.4687434 | 0.005287025 | 0.763246616 | 0.96 |
| YOR376W-A | 83.93269889 | -0.050474836 | 0.762982411 | 0.96 |
| YCR107W | 236.9080811 | -0.16546554 | 0.763993395 | 0.96 |
| YIL049W | 383.4643737 | -0.002989608 | 0.763930368 | 0.96 |
| YKL103C | 3936.941911 | 0.25427713 | 0.763920761 | 0.96 |
| YNL317W | 817.8553177 | -0.07929208 | 0.764017147 | 0.96 |
| YBR264C | 359.5103772 | 0.040313039 | 0.764456052 | 0.96 |

| | | | | |
|---|---|---|---|---|
| YFR044C | 5702.058987 | 0.003012521 | 0.764956017 | 0.96 |
| YHR177W | 301.5515693 | -0.230818718 | 0.764900452 | 0.96 |
| YPR055W | 1160.769461 | 0.045083438 | 0.764951167 | 0.96 |
| YNL098C | 3291.418535 | 0.081830018 | 0.765235554 | 0.96 |
| YDL031W | 1680.703139 | 0.096663845 | 0.766189126 | 0.96 |
| YAL019W | 1321.957689 | -0.019020307 | 0.766986747 | 0.96 |
| YIR033W | 2182.959679 | 0.158010578 | 0.767192959 | 0.96 |
| YJR104C | 12449.61979 | 0.105456532 | 0.76678302 | 0.96 |
| YLR382C | 967.5198513 | -0.119082789 | 0.766888467 | 0.96 |
| YOR314W-A | 0.578252409 | 0.031161933 | 0.767128048 | 0.96 |
| YBL007C | 3937.826759 | 0.121901429 | 0.768315884 | 0.96 |
| YBR125C | 746.1231571 | 0.15677157 | 0.769474464 | 0.96 |
| YBR255C-A | 311.2384417 | -0.030274475 | 0.769338797 | 0.96 |
| YDR022C | 413.59294 | 0.019021307 | 0.767547149 | 0.96 |
| YDR269C | 1.063098833 | -0.182456383 | 0.768989398 | 0.96 |
| YDR420W | 1513.909661 | -0.126831172 | 0.769169582 | 0.96 |
| YDR469W | 524.7044195 | -0.05833812 | 0.769082968 | 0.96 |
| YIL091C | 989.7593991 | -0.100321262 | 0.769517094 | 0.96 |
| YJL182C | 1.793622545 | 0.208278664 | 0.769457594 | 0.96 |
| YLR215C | 433.3667117 | -0.111481953 | 0.76775913 | 0.96 |
| YLR438C-A | 659.2030513 | -0.076653605 | 0.768045216 | 0.96 |
| YMR252C | 396.7363367 | 0.096146101 | 0.768294468 | 0.96 |
| YNL063W | 522.2571621 | 0.021198812 | 0.768197848 | 0.96 |
| YPL044C | 1.74310561 | 0.139130693 | 0.768501398 | 0.96 |
| YPR009W | 346.2208437 | -0.097043464 | 0.769432703 | 0.96 |
| YJL186W | 1797.326564 | -0.022487067 | 0.769733884 | 0.96 |
| YBL054W | 675.0753311 | 0.091129394 | 0.770153142 | 0.96 |
| YGR006W | 239.4957069 | -0.090050171 | 0.770262433 | 0.96 |
| YKL084W | 479.3447424 | 0.269420099 | 0.770226375 | 0.96 |
| YDL090C | 724.6090506 | -0.08729658 | 0.77042988 | 0.96 |
| YBL016W | 1090.490568 | -0.223457557 | 0.771290457 | 0.96 |
| YIL068W-A | 5.981971104 | -0.094783095 | 0.771519254 | 0.96 |
| YNL279W | 510.6368292 | -0.144299081 | 0.771311473 | 0.96 |
| YPL104W | 889.4598617 | -0.121754369 | 0.771582438 | 0.96 |
| YER130C | 1003.132192 | -0.091918561 | 0.771962198 | 0.96 |
| YMR193C-A | 123.8016037 | -0.105722957 | 0.772038164 | 0.96 |
| YOR329W-A | 20.44917844 | 0.176978752 | 0.772167199 | 0.96 |
| YPL271W | 1376.293801 | -0.164541366 | 0.772369084 | 0.96 |
| YAR035W | 1245.571751 | 0.091809113 | 0.772627902 | 0.96 |
| YLR339C | 501.9621213 | 0.144302484 | 0.772913149 | 0.96 |
| YDR333C | 1359.98522 | -0.106242486 | 0.773460902 | 0.96 |
| YDR383C | 354.9421834 | -0.208402249 | 0.773242716 | 0.96 |
| YFR050C | 3854.503578 | -0.078338312 | 0.773398259 | 0.96 |

| | | | | |
|---|---|---|---|---|
| YER059W | 910.2356733 | 0.052083288 | 0.774178726 | 0.96 |
| YIL134C-A | 3.121789303 | 0.702513467 | 0.774724898 | 0.96 |
| YLR375W | 1937.904522 | 0.071611967 | 0.774578219 | 0.96 |
| YOR238W | 549.7060682 | 0.098316968 | 0.774517049 | 0.96 |
| YOR372C | 1526.129326 | 0.068530246 | 0.774449397 | 0.96 |
| YHR158C | 1689.928736 | 0.160619538 | 0.774899525 | 0.96 |
| YCR099C | 126.2405817 | -0.22125976 | 0.776203846 | 0.96 |
| YDR189W | 2199.458913 | -0.0802218 | 0.776033869 | 0.96 |
| YGL121C | 888.9061681 | 0.047904633 | 0.776237085 | 0.96 |
| YKL028W | 1511.073698 | 0.086725507 | 0.775739664 | 0.96 |
| YLR051C | 783.2029443 | 0.062681652 | 0.776105825 | 0.96 |
| YMR273C | 1360.501919 | 0.07824545 | 0.776226642 | 0.96 |
| YDR088C | 645.3535973 | 0.067835388 | 0.776780615 | 0.96 |
| YKL018C-A | 261.9017664 | -0.08878215 | 0.776780425 | 0.96 |
| YOR269W | 302.0905548 | -0.132317215 | 0.776590839 | 0.96 |
| YLR319C | 843.3891881 | 0.062121849 | 0.777472685 | 0.96 |
| YMR005W | 1198.923005 | 0.095875266 | 0.777655471 | 0.96 |
| YNL170W | 85.88966089 | 0.108868498 | 0.777721864 | 0.96 |
| YOL140W | 502.3006349 | -0.11766386 | 0.777726025 | 0.96 |
| YOR391C | 10.43191586 | -0.26539498 | 0.777708332 | 0.96 |
| YBR204C | 1064.479878 | 0.19322377 | 0.777984742 | 0.96 |
| YDR155C | 17689.29075 | 0.031758905 | 0.778044777 | 0.96 |
| YLL038C | 326.4334207 | -0.230710751 | 0.778565422 | 0.96 |
| YAL053W | 2097.963607 | 0.244481999 | 0.779528308 | 0.96 |
| YER119C | 1204.859566 | 0.168151111 | 0.779420583 | 0.96 |
| YIL063C | 1081.563362 | -0.029434108 | 0.779420384 | 0.96 |
| YJL126W | 376.2312005 | 0.058443377 | 0.779617365 | 0.96 |
| YJR153W | 431.5896283 | -0.10309824 | 0.779630629 | 0.96 |
| YAL043C | 1560.889418 | -0.0898287 | 0.780817466 | 0.96 |
| YFL010W-A | 308.9446486 | -0.157759286 | 0.780293256 | 0.96 |
| YFL014W | 10835.63862 | -0.371066693 | 0.779887423 | 0.96 |
| YJL047C-A | 19.06572924 | -0.375352571 | 0.781032913 | 0.96 |
| YJL210W | 543.5019911 | 0.042695557 | 0.780324291 | 0.96 |
| YKL146W | 1758.494435 | 0.108957267 | 0.780562675 | 0.96 |
| YLR078C | 672.0208317 | 0.052743317 | 0.781155057 | 0.96 |
| YLR091W | 340.1364791 | 0.076056176 | 0.78064869 | 0.96 |
| YMR047C | 2343.921258 | -0.104498245 | 0.780388685 | 0.96 |
| YOL099C | 11.41692104 | 0.121523661 | 0.781070316 | 0.96 |
| YBR111C | 3318.167647 | 0.266563521 | 0.781795082 | 0.96 |
| YGL256W | 1304.621904 | 0.029010372 | 0.781936508 | 0.96 |
| YHR196W | 929.0161741 | -0.128469005 | 0.781673586 | 0.96 |
| YNL067W-B | 0.339190409 | -0.021696306 | 0.781863877 | 0.96 |
| YPL268W | 929.1963896 | 0.029043291 | 0.781657942 | 0.96 |

| | | | | |
|---|---|---|---|---|
| YML099W-A | 34.73959572 | -0.151681368 | 0.782237625 | 0.96 |
| YIL151C | 1650.033227 | 0.036132806 | 0.782481492 | 0.96 |
| YNL224C | 704.2866123 | 0.074642639 | 0.782676284 | 0.96 |
| YOR141C | 1550.614472 | -0.072173883 | 0.78273865 | 0.96 |
| YML088W | 784.353378 | 0.003857939 | 0.78308885 | 0.96 |
| YBR039W | 6878.817001 | 0.108872341 | 0.783396691 | 0.96 |
| YJR020W | 8.221696255 | 0.052246719 | 0.783365757 | 0.96 |
| YLR040C | 271.3228626 | -0.054649788 | 0.783602913 | 0.96 |
| YKR037C | 412.1583252 | 0.086178051 | 0.783889281 | 0.96 |
| YPL148C | 142.8618352 | -0.104491101 | 0.783980016 | 0.96 |
| YAL047C | 589.7842721 | 0.107715504 | 0.786681307 | 0.96 |
| YAL055W | 312.0534701 | 0.049283913 | 0.789445909 | 0.96 |
| YAR071W | 43.79477313 | -0.128285795 | 0.785218936 | 0.96 |
| YBL001C | 1539.933777 | 0.022835502 | 0.788198796 | 0.96 |
| YBR291C | 379.9577466 | -0.054674226 | 0.78674989 | 0.96 |
| YCR092C | 872.6618833 | 0.046692809 | 0.789941033 | 0.96 |
| YDR248C | 767.0999761 | 0.041850629 | 0.785873271 | 0.96 |
| YDR298C | 3635.360191 | 0.056229606 | 0.788300768 | 0.96 |
| YER158C | 1028.126375 | -0.098546819 | 0.789682349 | 0.96 |
| YFL032W | 146.5638958 | -0.127213868 | 0.786572801 | 0.96 |
| YFR042W | 519.19235 | -0.019672883 | 0.788482773 | 0.96 |
| YGL057C | 252.5789153 | -0.092331676 | 0.785150888 | 0.96 |
| YGR079W | 359.5947801 | 0.181456461 | 0.789006553 | 0.96 |
| YGR204W | 10459.91176 | 0.272999554 | 0.789467305 | 0.96 |
| YGR205W | 695.7192252 | -0.036109286 | 0.785799186 | 0.96 |
| YGR259C | 0.780911468 | 0.170250999 | 0.788162739 | 0.96 |
| YIL038C | 1659.427705 | -0.088295858 | 0.789936738 | 0.96 |
| YJL076W | 2985.809725 | 0.001142844 | 0.785756269 | 0.96 |
| YJL156W-A | 83.97878785 | -0.042970653 | 0.785702197 | 0.96 |
| YJR160C | 11.90922489 | -0.352758465 | 0.78950712 | 0.96 |
| YKL002W | 1601.062288 | -0.111066306 | 0.785786563 | 0.96 |
| YKL162C-A | 16.50216676 | -0.283221646 | 0.787581842 | 0.96 |
| YKL185W | 2577.028628 | -0.059668349 | 0.78784369 | 0.96 |
| YKR033C | 0.132511194 | 0.014458325 | 0.786946205 | 0.96 |
| YLR328W | 869.5602715 | -0.139099213 | 0.785753319 | 0.96 |
| YLR432W | 4140.896048 | 0.184246604 | 0.784309509 | 0.96 |
| YML018C | 644.4081194 | -0.056812998 | 0.786316278 | 0.96 |
| YMR171C | 1518.290743 | 0.056129215 | 0.786851492 | 0.96 |
| YMR188C | 1203.74754 | -0.160642323 | 0.784768555 | 0.96 |
| YNL035C | 765.5856918 | -0.038099481 | 0.787140657 | 0.96 |
| YNL199C | 1154.934737 | -0.068586905 | 0.789019013 | 0.96 |
| YNL271C | 3740.617123 | 0.155650689 | 0.789821418 | 0.96 |
| YNL298W | 1195.295385 | -0.109001029 | 0.788433926 | 0.96 |

| | | | | |
|---|---|---|---|---|
| YNL310C | 605.4678235 | -0.047164056 | 0.787564795 | 0.96 |
| YOL056W | 561.7509431 | 0.026989065 | 0.784438301 | 0.96 |
| YOR057W | 859.8202576 | 0.136098139 | 0.788095938 | 0.96 |
| YOR212W | 1199.500024 | -0.277323714 | 0.786304926 | 0.96 |
| YOR213C | 459.655666 | 0.121724268 | 0.785602446 | 0.96 |
| YOR274W | 529.1848909 | 0.008784163 | 0.787458316 | 0.96 |
| YPR016W-A | 37.91128357 | -0.024994836 | 0.789927877 | 0.96 |
| YBL015W | 5485.33881 | 0.398333481 | 0.790141755 | 0.96 |
| YDR403W | 327.0898139 | 0.077560367 | 0.790416679 | 0.96 |
| YNL235C | 7.928094321 | -0.407442045 | 0.790346256 | 0.96 |
| YIL173W | 14.66085042 | -0.497279896 | 0.790576315 | 0.96 |
| YFL025C | 800.9686963 | 0.027799534 | 0.791642422 | 0.96 |
| YGR242W | 37.21832003 | 0.259129022 | 0.791273281 | 0.96 |
| YHR031C | 752.7001486 | 0.102672439 | 0.791606941 | 0.96 |
| YHR214C-B | 235.6005636 | 0.222857055 | 0.791405337 | 0.96 |
| YIL115C | 2357.200299 | -0.043298978 | 0.791057183 | 0.96 |
| YKR027W | 720.4128865 | 0.047393303 | 0.7909149 | 0.96 |
| YMR119W | 1094.314263 | 0.12131387 | 0.79178742 | 0.96 |
| YOR333C | 3.992636415 | -0.093876956 | 0.79087948 | 0.96 |
| YMR121C | 804.8314095 | -0.100938021 | 0.791961392 | 0.96 |
| YCR106W | 394.726015 | -0.109178434 | 0.792476627 | 0.96 |
| YBL043W | 324.5838638 | 0.08677909 | 0.792742701 | 0.96 |
| YJL086C | 9.129343542 | 0.170863584 | 0.793309026 | 0.96 |
| YDR357C | 553.215464 | -0.084734735 | 0.793991604 | 0.96 |
| YGL058W | 714.9977444 | 0.021581783 | 0.793789465 | 0.96 |
| YOR355W | 1652.733408 | -0.237424245 | 0.793876694 | 0.96 |
| YGL220W | 745.8780305 | -0.082578803 | 0.794415461 | 0.96 |
| YDR461W | 1830.189511 | -0.360660272 | 0.794607978 | 0.96 |
| YDR078C | 147.0952169 | -0.01422592 | 0.794758057 | 0.96 |
| YKR030W | 649.1186189 | 0.067450503 | 0.795325683 | 0.96 |
| YML013W | 1108.615798 | 0.160562389 | 0.795267829 | 0.96 |
| YNL221C | 1802.455751 | -0.071783447 | 0.795083214 | 0.96 |
| YDL101C | 860.843029 | -0.039860717 | 0.795955489 | 0.96 |
| YGR122W | 624.9209214 | 0.008539541 | 0.795964687 | 0.96 |
| YMR245W | 80.78367086 | 0.128232185 | 0.795877576 | 0.96 |
| YPL015C | 1643.336309 | 0.086620721 | 0.796073205 | 0.96 |
| YAL016C-A | 61.31300348 | -0.19314031 | 0.796542273 | 0.96 |
| YLR138W | 1645.971988 | 0.199919924 | 0.796602268 | 0.96 |
| YDR127W | 3620.75176 | 0.202219621 | 0.7968094 | 0.96 |
| YAL026C | 1397.824394 | -0.047485021 | 0.798555632 | 0.96 |
| YAL041W | 1345.73957 | -0.02896399 | 0.803450226 | 0.96 |
| YBR149W | 7039.888751 | 0.096124703 | 0.803316664 | 0.96 |
| YBR224W | 27.9935508 | 0.206267546 | 0.804087478 | 0.96 |

| | | | | |
|---|---|---|---|---|
| YCR026C | 1309.803177 | 0.016524564 | 0.8005882 | 0.96 |
| YDL004W | 2874.441994 | 0.058162985 | 0.804780928 | 0.96 |
| YDL104C | 329.3472486 | -0.097716268 | 0.800962431 | 0.96 |
| YDL107W | 383.3863117 | 0.006821316 | 0.800885808 | 0.96 |
| YDL161W | 1895.847844 | 0.049027819 | 0.802709042 | 0.96 |
| YDR111C | 552.8400061 | 0.038689408 | 0.798692123 | 0.96 |
| YDR328C | 1849.787592 | -0.047260135 | 0.801377096 | 0.96 |
| YDR419W | 525.8031882 | 0.113853367 | 0.797378434 | 0.96 |
| YDR495C | 1040.925179 | -0.08211177 | 0.80250466 | 0.96 |
| YEL006W | 705.1819844 | -0.116638781 | 0.803176186 | 0.96 |
| YEL016C | 826.6367047 | 0.061253331 | 0.804221392 | 0.96 |
| YER030W | 1421.194205 | -0.100159209 | 0.802262426 | 0.96 |
| YER134C | 346.3610386 | -0.115664787 | 0.798939384 | 0.96 |
| YFL034C-B | 921.0599209 | -0.133105948 | 0.802565471 | 0.96 |
| YFL036W | 2229.86904 | 0.034824833 | 0.799466686 | 0.96 |
| YFR045W | 698.908615 | 0.040547263 | 0.804058992 | 0.96 |
| YGL025C | 784.0082065 | 0.065775175 | 0.800372296 | 0.96 |
| YGL152C | 7.084888852 | -0.306636783 | 0.799573273 | 0.96 |
| YGL251C | 608.1790507 | 0.044176563 | 0.800535879 | 0.96 |
| YGR256W | 1242.341744 | -0.242791256 | 0.803290254 | 0.96 |
| YHR032C-A | 12.55755435 | 0.23391096 | 0.803952393 | 0.96 |
| YHR129C | 426.0946941 | -0.102523834 | 0.79723235 | 0.96 |
| YHR193C-A | 1.288094432 | -0.451340831 | 0.799791964 | 0.96 |
| YHR208W | 7460.933205 | 0.628119331 | 0.801698788 | 0.96 |
| YIL015W | 2424.316267 | -0.444128146 | 0.80414513 | 0.96 |
| YIL088C | 2503.905976 | -0.021290442 | 0.797815283 | 0.96 |
| YIL108W | 1452.736233 | -0.069475317 | 0.803610346 | 0.96 |
| YJL053W | 1232.219717 | 0.035960328 | 0.79739503 | 0.96 |
| YJR066W | 2254.068141 | -0.068856843 | 0.804727002 | 0.96 |
| YKL016C | 3252.389125 | -0.013326588 | 0.798845186 | 0.96 |
| YKR087C | 773.1699867 | 0.046888235 | 0.802755075 | 0.96 |
| YLR096W | 1948.609356 | 0.085904763 | 0.803050945 | 0.96 |
| YLR170C | 116.6208147 | -0.088083965 | 0.801544773 | 0.96 |
| YLR223C | 2252.47739 | 0.095543861 | 0.803924901 | 0.96 |
| YLR410W-B | 66.3723325 | 0.19350093 | 0.800263096 | 0.96 |
| YLR454W | 3660.437031 | 0.15780448 | 0.803849666 | 0.96 |
| YML053C | 803.1994724 | -0.148949054 | 0.804558277 | 0.96 |
| YMR294W-A | 229.7802387 | 0.022819558 | 0.80060924 | 0.96 |
| YNL023C | 950.809325 | 0.12280588 | 0.801674827 | 0.96 |
| YNL053W | 544.6626736 | -0.133415333 | 0.804372586 | 0.96 |
| YNL124W | 1308.097286 | 0.181025019 | 0.797732017 | 0.96 |
| YNL219C | 1817.162227 | 0.075591986 | 0.799426772 | 0.96 |
| YNL257C | 1662.623362 | 0.078160039 | 0.804775855 | 0.96 |

| | | | | |
|---|---|---|---|---|
| YOL054W | 494.8918395 | 0.091081331 | 0.800900344 | 0.96 |
| YOR148C | 393.9247418 | -0.072526022 | 0.801670388 | 0.96 |
| YPL016W | 1163.813997 | 0.100323783 | 0.800000863 | 0.96 |
| YPL058C | 2629.343095 | -0.192020474 | 0.797601111 | 0.96 |
| YPL192C | 128.9945052 | -0.043544043 | 0.803488359 | 0.96 |
| YPL245W | 615.8162164 | 0.114293029 | 0.799072986 | 0.96 |
| tR(ACG)J | 0.2026169 | -0.098436367 | 0.804832955 | 0.96 |
| YGL084C | 1084.484417 | -0.042770036 | 0.805100934 | 0.97 |
| YAR003W | 588.5199738 | 0.08607139 | 0.807795385 | 0.97 |
| YDL039C | 373.0125142 | -0.464081351 | 0.808923362 | 0.97 |
| YDL233W | 547.8994161 | 0.07273909 | 0.805801914 | 0.97 |
| YDR063W | 1183.071975 | -0.097480961 | 0.806998848 | 0.97 |
| YDR143C | 1247.877225 | 0.047670715 | 0.807743897 | 0.97 |
| YDR283C | 1675.592568 | 0.083587536 | 0.809412876 | 0.97 |
| YDR381C-A | 448.1284196 | 0.05844401 | 0.809241512 | 0.97 |
| YDR464W | 2373.668914 | 0.109906969 | 0.807636534 | 0.97 |
| YER060W | 750.1023347 | -0.125901673 | 0.806561369 | 0.97 |
| YER113C | 743.4950149 | -0.101527471 | 0.810028676 | 0.97 |
| YFR036W | 228.7837864 | -0.09989559 | 0.808983034 | 0.97 |
| YGL188C | 26.0866713 | -0.366641107 | 0.808836931 | 0.97 |
| YGL209W | 1219.680335 | -0.011800053 | 0.806996421 | 0.97 |
| YGR074W | 145.6756142 | 0.042737741 | 0.808531796 | 0.97 |
| YGR233C | 1147.708039 | -0.093513018 | 0.807881023 | 0.97 |
| YGR286C | 924.5538048 | 0.090621188 | 0.805889422 | 0.97 |
| YHR027C | 5786.339997 | 0.03578041 | 0.807295953 | 0.97 |
| YHR078W | 1137.159604 | 0.162588779 | 0.80996421 | 0.97 |
| YKR014C | 1917.457601 | 0.073635317 | 0.807928668 | 0.97 |
| YKR079C | 916.529157 | -0.066561388 | 0.808674822 | 0.97 |
| YLR035C | 528.4128752 | 0.083409274 | 0.807742116 | 0.97 |
| YLR194C | 2445.708189 | 0.189172899 | 0.806300382 | 0.97 |
| YLR364W | 131.7044529 | -0.179205369 | 0.807766633 | 0.97 |
| YMR244W | 46.42719777 | -0.230645505 | 0.806573308 | 0.97 |
| YNL146C-A | 29.52790111 | -0.416627936 | 0.809773007 | 0.97 |
| YOL162W | 65.72006374 | -0.534983696 | 0.808784136 | 0.97 |
| YOR064C | 481.1143371 | 0.080751831 | 0.805979126 | 0.97 |
| YOR370C | 2847.263179 | -0.052744645 | 0.809464155 | 0.97 |
| YPL060W | 360.6791713 | 0.047614879 | 0.809260135 | 0.97 |
| YPL105C | 3260.005512 | 0.07791703 | 0.809764364 | 0.97 |
| YPL107W | 480.1557675 | 0.069677544 | 0.805878325 | 0.97 |
| YPR088C | 2257.094394 | -0.023707266 | 0.808876567 | 0.97 |
| YHR003C | 820.5090626 | -0.039891795 | 0.810674806 | 0.97 |
| YBR095C | 539.8071954 | 0.021778102 | 0.811220893 | 0.97 |
| YBR194W | 226.3319567 | 0.058582676 | 0.811563557 | 0.97 |

| | | | | |
|---|---|---|---|---|
| YNL163C | 1921.119184 | 0.221556117 | 0.811565056 | 0.97 |
| YLR088W | 986.2933263 | 0.088636198 | 0.811760528 | 0.97 |
| YDR515W | 694.9239372 | -0.142190679 | 0.812394355 | 0.97 |
| YLR148W | 974.658438 | -0.107689883 | 0.81296243 | 0.97 |
| YAL063C-A | 8.265236292 | -0.288301267 | 0.813485245 | 0.97 |
| YMR251W-A | 5878.547147 | -0.136641997 | 0.813382358 | 0.97 |
| YOL126C | 3179.903333 | -0.590347089 | 0.813440542 | 0.97 |
| HRA1 | 34.22771229 | 0.462354881 | 0.816349773 | 0.97 |
| YBL017C | 3980.2423 | 0.124184176 | 0.81624419 | 0.97 |
| YBR273C | 1305.311458 | 0.070024666 | 0.817064003 | 0.97 |
| YDL046W | 2942.234491 | -0.113949637 | 0.815789666 | 0.97 |
| YDL073W | 971.6916673 | -0.041857014 | 0.815967277 | 0.97 |
| YDL122W | 1760.896516 | -0.075214332 | 0.814729764 | 0.97 |
| YDR530C | 1226.853067 | 0.033988743 | 0.816705464 | 0.97 |
| YER038W-A | 0.198141795 | -0.06905409 | 0.814561408 | 0.97 |
| YER073W | 2130.104252 | 0.752380175 | 0.816707616 | 0.97 |
| YGR033C | 1013.438168 | 0.10656776 | 0.817168232 | 0.97 |
| YGR048W | 875.7732177 | -0.155849804 | 0.814675926 | 0.97 |
| YHL048W | 1666.986201 | -0.027750938 | 0.815351515 | 0.97 |
| YHR063C | 1432.676855 | 0.095026582 | 0.816303439 | 0.97 |
| YHR206W | 1203.868698 | 0.085545168 | 0.814911911 | 0.97 |
| YJR103W | 3899.480126 | 0.189463229 | 0.814699352 | 0.97 |
| YJR140C | 1086.888351 | -0.012466002 | 0.816974672 | 0.97 |
| YLL033W | 166.5445731 | 0.027487163 | 0.817006529 | 0.97 |
| YLR184W | 0.707088895 | 0.055133684 | 0.816919786 | 0.97 |
| YMR185W | 803.0326457 | -0.037738599 | 0.814605329 | 0.97 |
| YNL089C | 16.65282477 | -0.024133012 | 0.815117911 | 0.97 |
| YOL145C | 2673.103928 | -0.122225987 | 0.816346358 | 0.97 |
| YPR201W | 177.3808268 | -0.28397339 | 0.813935507 | 0.97 |
| snR42 | 83.98583809 | -0.07876732 | 0.816368502 | 0.97 |
| YOL019W-A | 0.720433626 | 0.031587993 | 0.817365323 | 0.97 |
| YBR175W | 264.1838842 | 0.093766519 | 0.820015057 | 0.97 |
| YCR067C | 1807.391426 | -0.011484265 | 0.820924512 | 0.97 |
| YDL147W | 5209.098548 | 0.088996054 | 0.818963849 | 0.97 |
| YDR026C | 840.5603663 | 0.065285139 | 0.82134391 | 0.97 |
| YDR079C-A | 102.409036 | 0.033277676 | 0.821147072 | 0.97 |
| YER083C | 1868.758699 | -0.069513622 | 0.819142667 | 0.97 |
| YFR006W | 3007.667423 | -0.015789363 | 0.818916352 | 0.97 |
| YGL116W | 933.3944118 | 0.120982909 | 0.81847419 | 0.97 |
| YGR137W | 236.2341776 | 0.10969935 | 0.821427972 | 0.97 |
| YGR146C-A | 13.60921533 | 0.057315831 | 0.820849423 | 0.97 |
| YGR222W | 726.7683442 | -0.011624352 | 0.818983903 | 0.97 |
| YHL048C-A | 2.76683096 | 0.342447825 | 0.818787895 | 0.97 |

| | | | | |
|---|---|---|---|---|
| YHR073W | 1977.66425 | -0.052720952 | 0.817986304 | 0.97 |
| YHR188C | 2845.286709 | 0.024469215 | 0.819946428 | 0.97 |
| YHR205W | 2582.464034 | 0.096914511 | 0.818895968 | 0.97 |
| YIL048W | 1213.90232 | 0.043657853 | 0.820465829 | 0.97 |
| YJR041C | 1150.543526 | -0.037032481 | 0.819827115 | 0.97 |
| YMR060C | 493.2206546 | -0.118324898 | 0.820529625 | 0.97 |
| YNL067W-A | 0.709210999 | -0.232415624 | 0.820990193 | 0.97 |
| YNL268W | 2190.621688 | -0.103363104 | 0.820551665 | 0.97 |
| YOL066C | 908.4756316 | -0.003956032 | 0.820346234 | 0.97 |
| YOR002W | 2112.418166 | -0.069703868 | 0.820893208 | 0.97 |
| YOR072W-B | 0.082611248 | 0.008775963 | 0.819450362 | 0.97 |
| YPL002C | 492.3419627 | 0.030189606 | 0.817724815 | 0.97 |
| YPL009C | 4156.098331 | 0.205072975 | 0.819714929 | 0.97 |
| YPL038W | 299.6200602 | -0.180892379 | 0.818160878 | 0.97 |
| YAL064C-A | 47.35030167 | -0.154056481 | 0.822592649 | 0.97 |
| YDR114C | 36.82268978 | -0.300849461 | 0.821728693 | 0.97 |
| YEL018C-A | 0.416927005 | 0.089355552 | 0.822873098 | 0.97 |
| YIL119C | 838.0211886 | -0.218706175 | 0.822950394 | 0.97 |
| YKL217W | 2399.793467 | 0.064593496 | 0.822563681 | 0.97 |
| YLL063C | 302.5298246 | -0.033681962 | 0.822621834 | 0.97 |
| YLR014C | 353.4875684 | 0.076445665 | 0.822349121 | 0.97 |
| YOL078W | 796.3214789 | 0.082802684 | 0.822056709 | 0.97 |
| YOR125C | 859.4168859 | 0.076631857 | 0.822692982 | 0.97 |
| YOR259C | 3766.746436 | -0.136609461 | 0.822953613 | 0.97 |
| YML046W | 728.0597624 | -0.140825423 | 0.823264689 | 0.97 |
| YOR060C | 313.4302218 | -0.119536399 | 0.823145277 | 0.97 |
| YCR100C | 233.2545617 | -0.399566645 | 0.825830009 | 0.97 |
| YDL097C | 4494.471702 | -0.06627854 | 0.826101527 | 0.97 |
| YDR246W | 916.4165129 | -0.081927955 | 0.826262315 | 0.97 |
| YFL064C | 3.015892609 | -0.069547953 | 0.826384043 | 0.97 |
| YHR012W | 963.21973 | -0.021034874 | 0.82516926 | 0.97 |
| YJL170C | 341.173457 | -0.11778787 | 0.826385252 | 0.97 |
| YJR023C | 6.716930152 | -0.067540293 | 0.826449467 | 0.97 |
| YKL038W | 1738.382543 | 0.127394002 | 0.825531763 | 0.97 |
| YKL087C | 723.7436146 | -0.081960977 | 0.824632925 | 0.97 |
| YKL120W | 1575.003705 | 0.306499915 | 0.824577612 | 0.97 |
| YKR063C | 686.126014 | -0.066098207 | 0.824966699 | 0.97 |
| YLL066C | 1.397171575 | 0.363469833 | 0.825451759 | 0.97 |
| YNL213C | 587.1355028 | -0.132719485 | 0.825862413 | 0.97 |
| YPL209C | 162.5627938 | 0.130238334 | 0.826358482 | 0.97 |
| snR62 | 6.46452847 | -0.104624433 | 0.825167241 | 0.97 |
| YBR124W | 16.74512171 | 0.052693875 | 0.827695103 | 0.97 |
| YBR286W | 17567.33584 | 0.084357228 | 0.829400815 | 0.97 |

| | | | | |
|---|---|---|---|---|
| YCL001W-B | 30.32607662 | -0.078977881 | 0.828770631 | 0.97 |
| YCR008W | 1187.11323 | 0.123132321 | 0.828396568 | 0.97 |
| YER021W | 5518.8989 | 0.096636322 | 0.829047877 | 0.97 |
| YFR034C | 832.1624053 | -0.019037177 | 0.828900506 | 0.97 |
| YIL029W-A | 8.108002149 | 0.291919035 | 0.829546884 | 0.97 |
| YIR037W | 2878.72569 | -0.021948356 | 0.828704639 | 0.97 |
| YKL131W | 17.12686382 | -0.112622588 | 0.829316227 | 0.97 |
| YKR042W | 18766.32248 | -0.015920833 | 0.826886122 | 0.97 |
| YLR256W-A | 92.61969338 | 0.115942839 | 0.829629117 | 0.97 |
| YML047C | 45.4588878 | -0.184187329 | 0.827587758 | 0.97 |
| YNL250W | 994.2437698 | 0.091205097 | 0.829509594 | 0.97 |
| YOL030W | 6136.814993 | 0.044238559 | 0.829723542 | 0.97 |
| YOL071W | 1035.627926 | -0.012062878 | 0.827640109 | 0.97 |
| YOL125W | 710.1287675 | -0.032758534 | 0.828060057 | 0.97 |
| YOR033C | 730.7503196 | 0.045662349 | 0.828526445 | 0.97 |
| YOR170W | 97.11015677 | -0.115231768 | 0.829033897 | 0.97 |
| YPL204W | 2957.690828 | -0.079242368 | 0.829810625 | 0.97 |
| YPL240C | 19482.15572 | 0.191342138 | 0.827421854 | 0.97 |
| YPR152C | 381.9563274 | -0.013435723 | 0.827016397 | 0.97 |
| YPR158C-D | 130.4698158 | -0.005753514 | 0.829199508 | 0.97 |
| YJL007C | 3.350774349 | -0.756020108 | 0.830160121 | 0.97 |
| YIL125W | 5872.779859 | 0.116780034 | 0.830489013 | 0.97 |
| YMR218C | 1133.912405 | 0.063266104 | 0.831015903 | 0.97 |
| YDR489W | 358.8647918 | 0.093319157 | 0.831512449 | 0.97 |
| YER061C | 958.2654695 | -0.055450093 | 0.831631581 | 0.97 |
| YJL217W | 2088.445581 | 0.263526263 | 0.831807561 | 0.97 |
| YDR103W | 991.2634766 | -0.015634479 | 0.832483379 | 0.97 |
| YIL137C | 2189.193159 | 0.031603147 | 0.832354785 | 0.97 |
| YKL089W | 415.1913401 | 0.088216812 | 0.832187031 | 0.97 |
| YLR286C | 14573.14708 | -0.004539208 | 0.8324396 | 0.97 |
| YCR053W | 6107.267981 | 0.153021842 | 0.833009537 | 0.97 |
| YGR263C | 454.4543224 | 0.071701525 | 0.833355108 | 0.97 |
| YHR009C | 3439.835058 | -0.072627431 | 0.833701354 | 0.97 |
| YML081C-A | 1028.969006 | -0.060286675 | 0.833334396 | 0.97 |
| YMR040W | 194.5584091 | 0.130411332 | 0.833488554 | 0.97 |
| YMR286W | 654.5341281 | -0.164321936 | 0.833470227 | 0.97 |
| YOR094W | 410.9080628 | -0.063037041 | 0.833169521 | 0.97 |
| snR40 | 6.538551087 | -0.087961838 | 0.833581338 | 0.97 |
| YLR334C | 3.854398197 | 0.263410026 | 0.833938955 | 0.97 |
| YKR051W | 666.9246654 | -0.136320457 | 0.83426665 | 0.97 |
| snR35 | 41.6109753 | 0.161093988 | 0.834279624 | 0.97 |
| YBR227C | 856.5044097 | -0.060787609 | 0.83625271 | 0.97 |
| YDR257C | 688.7119517 | -0.010619017 | 0.836180335 | 0.97 |

| | | | | |
|---|---|---|---|---|
| YDR362C | 764.8048646 | -0.073958932 | 0.835566141 | 0.97 |
| YFL026W | 6326.346816 | -0.205598193 | 0.83476466 | 0.97 |
| YGL051W | 80.44841429 | 0.099302488 | 0.835743931 | 0.97 |
| YGL110C | 882.4868812 | 0.032749574 | 0.83548634 | 0.97 |
| YIL171W | 27.73883289 | -0.381806197 | 0.834969372 | 0.97 |
| YJL067W | 49.69168928 | -0.231520352 | 0.835464392 | 0.97 |
| YLR193C | 745.5545212 | 0.124813466 | 0.835994015 | 0.97 |
| YLR295C | 2976.605613 | 0.051596865 | 0.835000528 | 0.97 |
| YOL109W | 9848.185932 | -0.182745646 | 0.835838911 | 0.97 |
| snR43 | 20.09245082 | -0.437622724 | 0.835615145 | 0.97 |
| snR78 | 1.974544426 | 0.157563121 | 0.835924764 | 0.97 |
| YAL031W-A | 36.5218107 | -0.230624343 | 0.837200625 | 0.97 |
| YCL073C | 26.5180597 | 0.022754023 | 0.837129339 | 0.97 |
| YDL021W | 1151.606846 | -0.310199442 | 0.838387673 | 0.97 |
| YDL180W | 735.4223348 | 0.07064371 | 0.837127138 | 0.97 |
| YDR345C | 12971.54144 | 0.216633919 | 0.838791554 | 0.97 |
| YGL167C | 2492.369981 | 0.080825889 | 0.836877914 | 0.97 |
| YGR173W | 798.4212433 | 0.009810355 | 0.838029367 | 0.97 |
| YGR278W | 442.5587334 | -0.090250117 | 0.838364912 | 0.97 |
| YKL022C | 1018.942477 | 0.030496225 | 0.837710127 | 0.97 |
| YKL065C | 4326.57925 | 0.065584769 | 0.838844919 | 0.97 |
| YLR235C | 18.73304548 | 0.140103672 | 0.838480372 | 0.97 |
| YLR438W | 1335.790008 | 0.21270388 | 0.837629365 | 0.97 |
| YMR178W | 1157.445412 | -0.03127039 | 0.837430697 | 0.97 |
| YMR315W-A | 6.002010905 | -0.251277787 | 0.837959669 | 0.97 |
| YOL038W | 4309.883987 | -0.051779196 | 0.836563586 | 0.97 |
| YOR083W | 334.3141069 | 0.000225564 | 0.838163899 | 0.97 |
| snR55 | 9.080532067 | 0.526706935 | 0.838769448 | 0.97 |
| YGR166W | 818.3285643 | 0.066147785 | 0.83945447 | 0.97 |
| YHL025W | 1272.013263 | 0.007752181 | 0.839657404 | 0.97 |
| YIL014W | 862.4746572 | 0.01697937 | 0.839925259 | 0.97 |
| YJR037W | 10.84309032 | -0.049365117 | 0.839983493 | 0.97 |
| YJR090C | 1653.355955 | 0.005185931 | 0.84003375 | 0.97 |
| YOR217W | 1207.361787 | -0.023639647 | 0.839662846 | 0.97 |
| YBR037C | 852.7234834 | -0.107866482 | 0.841094437 | 0.97 |
| YBR053C | 1693.976608 | 0.07216719 | 0.840966205 | 0.97 |
| YDR147W | 507.4257469 | -0.091075147 | 0.841056861 | 0.97 |
| YDR169C | 1729.231302 | 0.039944351 | 0.841380197 | 0.97 |
| YIL167W | 108.465152 | -0.022105543 | 0.841334287 | 0.97 |
| YKL206C | 563.6605725 | 0.091525045 | 0.840480888 | 0.97 |
| YMR179W | 627.9998767 | -0.15100367 | 0.841246229 | 0.97 |
| YOR362C | 3907.49208 | -0.035213623 | 0.841174975 | 0.97 |
| YBL008W-A | 2.748531795 | -0.17719094 | 0.843171106 | 0.97 |

| | | | | |
|---|---|---|---|---|
| YBR167C | 275.1433522 | -0.136307781 | 0.843179976 | 0.97 |
| YBR200W-A | 19.77475914 | -0.017337173 | 0.844647685 | 0.97 |
| YBR253W | 396.1883397 | -0.077790659 | 0.844469711 | 0.97 |
| YDR067C | 673.9208846 | 0.079342034 | 0.845949411 | 0.97 |
| YDR206W | 1789.144398 | 0.058820083 | 0.842000834 | 0.97 |
| YDR281C | 42.74964081 | 0.065023676 | 0.843410135 | 0.97 |
| YER068W | 1672.188885 | 0.050671105 | 0.845477689 | 0.97 |
| YFR010W-A | 16.18554247 | 0.32971549 | 0.845954307 | 0.97 |
| YFR016C | 3711.04018 | 0.206302408 | 0.845772001 | 0.97 |
| YGL004C | 1031.613443 | 0.080660199 | 0.845326437 | 0.97 |
| YGL043W | 1337.950161 | -0.007121111 | 0.845064688 | 0.97 |
| YGL161C | 1505.046156 | 0.045809136 | 0.845442587 | 0.97 |
| YJL004C | 587.5114449 | 0.062390914 | 0.846106804 | 0.97 |
| YKR062W | 891.4921763 | -0.043112638 | 0.844330891 | 0.97 |
| YLL035W | 581.4295973 | 0.003910859 | 0.844152708 | 0.97 |
| YML015C | 859.349761 | 0.073153265 | 0.844036789 | 0.97 |
| YML116W-A | 166.4577303 | -0.11189721 | 0.843603976 | 0.97 |
| YMR212C | 1054.541147 | -0.025219502 | 0.844563103 | 0.97 |
| YMR237W | 2047.326568 | -0.020260276 | 0.846108755 | 0.97 |
| YMR276W | 5216.693498 | -0.069636152 | 0.845193466 | 0.97 |
| YNL186W | 1470.462386 | -0.058932087 | 0.845575916 | 0.97 |
| YNL314W | 384.9311948 | 0.064312003 | 0.842610958 | 0.97 |
| YOL027C | 2880.865523 | 0.047971316 | 0.842306882 | 0.97 |
| YOL086W-A | 318.8786531 | 0.081522198 | 0.843282245 | 0.97 |
| YOR139C | 5.37072037 | 0.13717035 | 0.842955215 | 0.97 |
| YOR329C | 1982.861977 | -0.067560714 | 0.84189595 | 0.97 |
| YPL051W | 362.5719952 | -0.074864416 | 0.843605265 | 0.97 |
| YPL154C | 14185.44316 | 0.114467118 | 0.844510348 | 0.97 |
| snR6 | 0.308944371 | 0.010911725 | 0.844082334 | 0.97 |
| tD(GUC)N | 0.865673422 | 0.052556488 | 0.842741225 | 0.97 |
| YLR141W | 304.974896 | -0.017351314 | 0.846689687 | 0.97 |
| YML071C | 1055.622411 | 0.040581752 | 0.846590935 | 0.97 |
| YNR011C | 902.2961481 | 0.117849995 | 0.846791623 | 0.97 |
| YKL116C | 778.7224786 | -0.049544052 | 0.847942421 | 0.97 |
| YOR222W | 1007.50754 | 0.228584143 | 0.848190804 | 0.97 |
| YIR018C-A | 31.48971955 | -0.16174848 | 0.848648106 | 0.97 |
| YPL091W | 3000.977941 | 0.048308534 | 0.848572569 | 0.97 |
| YBL008W | 564.4162656 | 0.125636421 | 0.848905595 | 0.97 |
| YAL009W | 404.8562545 | -0.061656426 | 0.84928795 | 0.97 |
| YBL100W-B | 175.5232464 | 0.095108627 | 0.85029602 | 0.97 |
| YER109C | 1065.439283 | 0.049869669 | 0.850023069 | 0.97 |
| YFL013C | 826.2401076 | 0.077928853 | 0.849822678 | 0.97 |
| YFL033C | 2297.741868 | 0.141243468 | 0.849716035 | 0.97 |

| | | | | |
|---|---|---|---|---|
| YGR283C | 529.6205161 | 0.064278255 | 0.850014257 | 0.97 |
| YKL027W | 701.3136707 | 0.046521868 | 0.850507441 | 0.97 |
| YML058W-A | 99.58508334 | -0.612214599 | 0.850766631 | 0.97 |
| YML074C | 7957.336564 | 0.007372155 | 0.850813277 | 0.97 |
| YMR073C | 579.5582083 | -0.071269469 | 0.850390768 | 0.97 |
| YMR312W | 719.0027121 | 0.004293987 | 0.850180683 | 0.97 |
| YOR268C | 11.87888145 | -0.015127593 | 0.849097445 | 0.97 |
| YOR318C | 25.32067042 | -0.051357847 | 0.850870868 | 0.97 |
| YMR089C | 2367.558674 | 0.001095684 | 0.851285033 | 0.97 |
| YLR099C | 1018.600053 | -0.131450041 | 0.852119891 | 0.97 |
| YBR058C | 1360.304304 | -0.026077546 | 0.852338845 | 0.97 |
| YMR294W | 447.832337 | 0.009734917 | 0.852896581 | 0.98 |
| YLR081W | 221.380283 | -0.203338771 | 0.853097769 | 0.98 |
| YBR233W | 474.558741 | -0.008375132 | 0.853315411 | 0.98 |
| YJL064W | 3.253557565 | -0.088902512 | 0.853584473 | 0.98 |
| YDR073W | 640.4950618 | -0.267459042 | 0.853774656 | 0.98 |
| YBR296C | 209.9913106 | -0.206224982 | 0.854236074 | 0.98 |
| YGR191W | 1267.875828 | -0.078726193 | 0.854400038 | 0.98 |
| YAR070C | 0.925768202 | -0.012287877 | 0.85711792 | 0.98 |
| YBL108W | 0.561379936 | -0.06928422 | 0.855146627 | 0.98 |
| YDL099W | 2397.811912 | -0.060512875 | 0.856184573 | 0.98 |
| YDR138W | 884.1934286 | -0.058865008 | 0.857039813 | 0.98 |
| YDR349C | 1807.872356 | 0.067722167 | 0.856590623 | 0.98 |
| YDR355C | 4.46006282 | 0.076926374 | 0.856186268 | 0.98 |
| YDR529C | 3824.487001 | 0.195504473 | 0.857145744 | 0.98 |
| YGR016W | 101.0221921 | 0.021752703 | 0.857410451 | 0.98 |
| YIL001W | 471.4395924 | -0.033793217 | 0.855368823 | 0.98 |
| YJL069C | 705.1093909 | -0.105048545 | 0.856137335 | 0.98 |
| YJL225C | 0.280381926 | 0.017252384 | 0.857388041 | 0.98 |
| YJR083C | 550.0152446 | -0.002493871 | 0.856708416 | 0.98 |
| YKL047W | 711.1080868 | -0.066842912 | 0.856920523 | 0.98 |
| YKL222C | 519.5767688 | -0.166684776 | 0.854964437 | 0.98 |
| YLR038C | 1207.825369 | 0.002399616 | 0.85542762 | 0.98 |
| YLR162W | 2.387178616 | 0.666531689 | 0.854813269 | 0.98 |
| YLR444C | 25.68564702 | -0.380169064 | 0.856347496 | 0.98 |
| YML052W | 1751.911847 | -0.135397079 | 0.855753766 | 0.98 |
| YMR029C | 1228.193843 | 0.033506608 | 0.855560137 | 0.98 |
| YNR075W | 186.6264191 | 0.014836198 | 0.855257624 | 0.98 |
| YOR038C | 676.9008791 | -0.049642691 | 0.85769244 | 0.98 |
| YBR182C-A | 7.795727576 | -0.223771912 | 0.858411522 | 0.98 |
| YER071C | 272.1086234 | -0.114464399 | 0.858237335 | 0.98 |
| YFR005C | 788.6190865 | -0.071234593 | 0.858491487 | 0.98 |
| YER135C | 23.51565939 | 0.038824699 | 0.858872345 | 0.98 |

| | | | | |
|---|---|---|---|---|
| YML065W | 967.920824 | 0.107698707 | 0.858987855 | 0.98 |
| YGL039W | 719.9626894 | 0.01496842 | 0.85938206 | 0.98 |
| YGR183C | 999.4364709 | 0.159784941 | 0.859383903 | 0.98 |
| YAL037W | 195.7634153 | 0.066387967 | 0.861827817 | 0.98 |
| YBL041W | 3024.581572 | -0.035194688 | 0.861407364 | 0.98 |
| YBR229C | 1792.230472 | 0.102889138 | 0.862025403 | 0.98 |
| YCL065W | 1.135078196 | 0.001030904 | 0.861855668 | 0.98 |
| YDL242W | 58.94192665 | 0.229526343 | 0.861181629 | 0.98 |
| YDR038C | 125.3516014 | 0.016135531 | 0.862942885 | 0.98 |
| YDR289C | 776.5077999 | 0.010797019 | 0.860085296 | 0.98 |
| YDR468C | 586.9211769 | 0.042585756 | 0.861306357 | 0.98 |
| YGL053W | 527.1405514 | -0.20110045 | 0.861650619 | 0.98 |
| YGL083W | 940.5046339 | 0.094138526 | 0.859994516 | 0.98 |
| YGR134W | 745.1524745 | -0.039973565 | 0.860992324 | 0.98 |
| YGR174W-A | 47.91809201 | -0.5187078 | 0.862112489 | 0.98 |
| YGR178C | 3642.360299 | 0.109460415 | 0.862506609 | 0.98 |
| YHR081W | 846.2639082 | 0.084347217 | 0.861299064 | 0.98 |
| YIL100C-A | 6.139255363 | -0.050394185 | 0.86238456 | 0.98 |
| YIL156W | 1109.423201 | -0.044338471 | 0.86295624 | 0.98 |
| YLR068W | 534.5964698 | 0.057257456 | 0.862884519 | 0.98 |
| YLR256W | 4392.10169 | 0.034086862 | 0.861593153 | 0.98 |
| YML108W | 173.1265371 | 0.112554092 | 0.862076297 | 0.98 |
| YMR150C | 511.1258272 | -0.053339455 | 0.862814777 | 0.98 |
| YNL261W | 688.6478875 | 0.061861739 | 0.861059601 | 0.98 |
| YPR097W | 1353.173051 | 0.068607471 | 0.861806741 | 0.98 |
| YPR099C | 24.60634511 | -0.198496569 | 0.85976404 | 0.98 |
| YDR099W | 8428.443895 | 0.071843995 | 0.86358837 | 0.98 |
| YPR171W | 883.9863961 | 0.024750772 | 0.86351118 | 0.98 |
| YDL086W | 1034.431113 | 0.131805419 | 0.863951316 | 0.98 |
| YHR215W | 110.5268911 | 0.261557071 | 0.864598293 | 0.98 |
| YJL128C | 1506.583064 | -0.036903609 | 0.864734642 | 0.98 |
| YNR012W | 597.0846222 | 0.091313702 | 0.864861642 | 0.98 |
| YOR191W | 2372.954185 | 0.036154617 | 0.864696428 | 0.98 |
| YOR266W | 469.4838324 | -0.061482464 | 0.864331799 | 0.98 |
| YPL019C | 1922.931958 | 0.072356139 | 0.864202281 | 0.98 |
| YIR039C | 1029.181658 | -0.292513206 | 0.865261461 | 0.98 |
| YDR058C | 448.3672811 | -0.068067047 | 0.866499393 | 0.98 |
| YDR154C | 3531.276985 | -0.064718466 | 0.866514004 | 0.98 |
| YEL008W | 30.07335462 | -0.369593271 | 0.865836136 | 0.98 |
| YGR069W | 7.64600337 | -0.14557168 | 0.865543078 | 0.98 |
| YGR267C | 1681.678228 | 0.000350102 | 0.866203056 | 0.98 |
| YJR114W | 33.45969567 | -0.346977887 | 0.866164111 | 0.98 |
| YMR135W-A | 18.48988281 | 0.229257415 | 0.866513587 | 0.98 |

| | | | | |
|---|---|---|---|---|
| snR72 | 8.104041704 | 0.077321029 | 0.865782339 | 0.98 |
| YCR045C | 398.9252367 | -0.06046055 | 0.86723802 | 0.98 |
| YDR314C | 384.5141242 | 0.064570204 | 0.867262169 | 0.98 |
| YDR393W | 653.7265915 | -0.07671999 | 0.867035628 | 0.98 |
| YGL080W | 877.4500948 | 0.096766073 | 0.867167898 | 0.98 |
| YBR141C | 286.9036203 | 0.015857997 | 0.868840767 | 0.98 |
| YCL004W | 588.1019431 | -0.021742591 | 0.867543593 | 0.98 |
| YDL146W | 626.6868048 | 0.155800232 | 0.868135574 | 0.98 |
| YDR227W | 1551.753891 | -0.02720048 | 0.869257589 | 0.98 |
| YFL055W | 265.1999324 | -0.196963062 | 0.867943931 | 0.98 |
| YGL132W | 16.53764806 | -0.193246266 | 0.868439985 | 0.98 |
| YJR016C | 6106.441237 | 0.374720303 | 0.868921126 | 0.98 |
| YLR250W | 1395.702651 | -0.039379541 | 0.868527906 | 0.98 |
| YMR008C | 3679.477965 | 0.165807126 | 0.868986184 | 0.98 |
| YNL264C | 1013.510724 | 0.211601798 | 0.869094323 | 0.98 |
| YOR049C | 767.2844506 | -0.245941711 | 0.868454964 | 0.98 |
| YPR129W | 1545.384555 | 0.139744616 | 0.869243949 | 0.98 |
| YPR195C | 54.58354851 | -0.11727964 | 0.869204714 | 0.98 |
| YML021C | 560.3287148 | 0.046610756 | 0.869880728 | 0.98 |
| YNL208W | 6496.983749 | -0.019073375 | 0.869821772 | 0.98 |
| YER174C | 586.1669315 | 0.027736615 | 0.870215402 | 0.98 |
| YPR070W | 926.8338353 | 0.019782878 | 0.87019622 | 0.98 |
| YBR289W | 1372.875089 | -0.004724247 | 0.871133882 | 0.98 |
| YCL055W | 639.3184492 | -0.260532015 | 0.871151431 | 0.98 |
| YCR049C | 7.558405616 | -0.144370437 | 0.87102291 | 0.98 |
| YJL139C | 556.1081958 | 0.114677107 | 0.870552302 | 0.98 |
| YNL100W | 1262.153408 | 0.017110972 | 0.871066099 | 0.98 |
| YGR026W | 4521.168167 | 0.098048792 | 0.871815445 | 0.98 |
| YDL096C | 9.504570723 | -0.243343413 | 0.87245629 | 0.98 |
| YER046W | 177.0920564 | -0.031438536 | 0.873517485 | 0.98 |
| YFR012W-A | 2.137477958 | 0.122102617 | 0.873835019 | 0.98 |
| YGL151W | 1728.404115 | 0.030889425 | 0.872656736 | 0.98 |
| YIL156W-A | 6.740450198 | -0.349652389 | 0.8726143 | 0.98 |
| YNL338W | 0.154202725 | 0.010948126 | 0.873817634 | 0.98 |
| YNR049C | 561.0637018 | -0.060716242 | 0.873679058 | 0.98 |
| YNR072W | 156.9410464 | 0.342476324 | 0.873668684 | 0.98 |
| YOR053W | 4.40231234 | -0.289186513 | 0.873124659 | 0.98 |
| YOR109W | 2024.801836 | -0.028188415 | 0.873502921 | 0.98 |
| YPL171C | 308.1636844 | 0.02887354 | 0.8723559 | 0.98 |
| YPR098C | 1032.023923 | -0.146607688 | 0.873108806 | 0.98 |
| YPR109W | 650.4688515 | -0.062365196 | 0.87399344 | 0.98 |
| YOL141W | 189.2930206 | 0.339665579 | 0.874159917 | 0.98 |
| YDR168W | 2674.002145 | 0.143108546 | 0.874364011 | 0.98 |

| | | | | |
|---|---|---|---|---|
| YCL030C | 8176.65539 | 0.419488301 | 0.874685292 | 0.98 |
| YFL044C | 776.7355212 | 0.034084529 | 0.874969313 | 0.98 |
| YMR324C | 2.573506675 | 0.070980648 | 0.875102722 | 0.98 |
| YOR117W | 3352.309698 | -0.063826411 | 0.874907085 | 0.98 |
| YER012W | 3318.681234 | 0.006322024 | 0.876394126 | 0.98 |
| YIL117C | 1107.127061 | -0.161936761 | 0.87623734 | 0.98 |
| YJL110C | 684.2542418 | -0.154972251 | 0.87616321 | 0.98 |
| YJL214W | 204.5476536 | -0.018337489 | 0.875866416 | 0.98 |
| YKL225W | 0.626369128 | 0.021454586 | 0.875911628 | 0.98 |
| YLL010C | 1089.845187 | 0.023261208 | 0.876271325 | 0.98 |
| YLR364C-A | 0.303554837 | 0.012724113 | 0.87595102 | 0.98 |
| YNL033W | 5.453513622 | -0.041504276 | 0.875535597 | 0.98 |
| YBR073W | 1901.278512 | 0.039473754 | 0.879127355 | 0.98 |
| YBR186W | 272.8183513 | 0.174554873 | 0.877723661 | 0.98 |
| YCR059C | 1130.721478 | -0.056326552 | 0.88023035 | 0.98 |
| YDR501W | 295.5069374 | 0.086550113 | 0.879220087 | 0.98 |
| YER056C | 2408.22934 | 0.072341095 | 0.87931548 | 0.98 |
| YER106W | 34.98872113 | -0.339817767 | 0.878823817 | 0.98 |
| YGR056W | 1165.037029 | -0.05349685 | 0.878807555 | 0.98 |
| YHR132C | 1760.575956 | 0.001349986 | 0.880275781 | 0.98 |
| YJL055W | 2135.334979 | 0.056438249 | 0.880134053 | 0.98 |
| YJL136W-A | 0.34890375 | 0.049215501 | 0.880168644 | 0.98 |
| YJL211C | 56.87482788 | -0.058561142 | 0.878498393 | 0.98 |
| YLR348C | 661.9197696 | -0.15181618 | 0.879538608 | 0.98 |
| YMR119W-A | 41.58140006 | 0.025247905 | 0.87815331 | 0.98 |
| YMR138W | 253.6379214 | 0.016563247 | 0.877355925 | 0.98 |
| YMR194C-B | 137.2124333 | -0.387289246 | 0.877928702 | 0.98 |
| YNL006W | 936.4455341 | -0.018194332 | 0.88017835 | 0.98 |
| YNL288W | 932.0705303 | -0.065265979 | 0.878938551 | 0.98 |
| YOL042W | 569.9826336 | -0.037967989 | 0.879571199 | 0.98 |
| YOR016C | 992.9033585 | 0.002151858 | 0.879842572 | 0.98 |
| YOR270C | 5846.210473 | 0.006952963 | 0.877335322 | 0.98 |
| YOR306C | 1215.578289 | -0.087647199 | 0.879655454 | 0.98 |
| YPL069C | 455.5840403 | 0.081424497 | 0.878993187 | 0.98 |
| YFR047C | 2554.161875 | 0.051664289 | 0.880684924 | 0.98 |
| YAR009C | 236.1235724 | 0.241422134 | 0.88120571 | 0.98 |
| YDR260C | 314.9401976 | 0.005014428 | 0.881165389 | 0.98 |
| YGL206C | 5144.593261 | -0.053579759 | 0.88134123 | 0.98 |
| YLR395C | 1418.063046 | 0.131837855 | 0.881017838 | 0.98 |
| YBL091C-A | 508.1746239 | -0.005173568 | 0.882098709 | 0.98 |
| YDR518W | 1466.448011 | 0.208982669 | 0.882115675 | 0.98 |
| YKL211C | 1647.230083 | 0.267879144 | 0.881707068 | 0.98 |
| YOL155C | 4950.84233 | -0.162532887 | 0.881957757 | 0.98 |

| | | | | |
|---|---|---|---|---|
| YOR151C | 4196.260493 | -0.032468071 | 0.881792227 | 0.98 |
| YBR131W | 285.0666359 | 0.141618513 | 0.882332017 | 0.98 |
| YBR109W-A | 20.49364363 | 0.037732423 | 0.884580317 | 0.98 |
| YCL040W | 21911.49524 | 0.198703682 | 0.883190523 | 0.98 |
| YCR019W | 382.8804523 | 0.03288855 | 0.882807872 | 0.98 |
| YDR278C | 0.649229666 | 0.051455443 | 0.882981272 | 0.98 |
| YDR431W | 11.62652329 | 0.092152426 | 0.882546224 | 0.98 |
| YGL094C | 1570.488055 | -0.060223696 | 0.884584704 | 0.98 |
| YGR265W | 63.64327138 | 0.367635009 | 0.885283185 | 0.98 |
| YHR047C | 1885.311487 | 0.349102006 | 0.8856086 | 0.98 |
| YHR086W-A | 3.107825184 | -0.103747106 | 0.885262459 | 0.98 |
| YJL160C | 84.43696653 | 0.109801825 | 0.885772395 | 0.98 |
| YJR132W | 1306.62171 | -0.03031074 | 0.883649542 | 0.98 |
| YLR082C | 386.5514271 | -0.113351093 | 0.884261616 | 0.98 |
| YLR084C | 1027.133398 | -0.027065278 | 0.884759828 | 0.98 |
| YLR130C | 1253.100793 | 0.00277283 | 0.884954132 | 0.98 |
| YML081W | 1706.956949 | -0.033020466 | 0.883869531 | 0.98 |
| YMR016C | 1109.247117 | 0.048766315 | 0.885533885 | 0.98 |
| YMR083W | 8478.832219 | 0.07037657 | 0.883995778 | 0.98 |
| YMR315W | 4197.096296 | 0.194472919 | 0.885671543 | 0.98 |
| YNL230C | 466.9526022 | 0.029803549 | 0.883651845 | 0.98 |
| YNR024W | 280.6447987 | 0.140099097 | 0.882872594 | 0.98 |
| YPL253C | 481.3229549 | -0.047424205 | 0.885829985 | 0.98 |
| YPR068C | 284.1750342 | 0.002078087 | 0.884524165 | 0.98 |
| snR39 | 11.68210812 | -0.387543865 | 0.885560144 | 0.98 |
| YML057W | 1931.217961 | 0.020337345 | 0.886046048 | 0.98 |
| YCL009C | 7647.273312 | 0.335048688 | 0.88661101 | 0.98 |
| YHR214W-A | 22.9859353 | -0.303291773 | 0.886383776 | 0.98 |
| YML133C | 9.588165535 | -0.045488063 | 0.886628504 | 0.98 |
| YJR138W | 1461.069987 | 0.056063031 | 0.886821981 | 0.98 |
| YCL069W | 36.22888945 | 0.000893756 | 0.887205917 | 0.98 |
| YAL030W | 729.4057845 | 0.07690648 | 0.889257855 | 0.98 |
| YAR050W | 236.1195337 | 0.127884416 | 0.895492406 | 0.98 |
| YBL053W | 19.02715806 | -0.052102616 | 0.894263012 | 0.98 |
| YBL100C | 649.5552869 | 0.641885637 | 0.898937051 | 0.98 |
| YBR044C | 837.2738234 | -0.027966716 | 0.88914402 | 0.98 |
| YBR062C | 819.8607412 | -0.026287556 | 0.891602337 | 0.98 |
| YBR098W | 547.1579177 | 0.063017358 | 0.894793744 | 0.98 |
| YBR119W | 605.7970902 | 0.01884334 | 0.890476646 | 0.98 |
| YBR232C | 24.08316402 | 0.338684051 | 0.895385456 | 0.98 |
| YCL002C | 469.4119103 | -0.026915525 | 0.898194121 | 0.98 |
| YCR033W | 2347.054478 | -0.090959889 | 0.896529348 | 0.98 |
| YCR108C | 1.912873177 | 0.000548393 | 0.898900731 | 0.98 |

| | | | | |
|---|---|---|---|---|
| YDL076C | 828.6133083 | 0.051728158 | 0.892384912 | 0.98 |
| YDL117W | 1322.092729 | 0.019517714 | 0.891040888 | 0.98 |
| YDL141W | 1031.602218 | 0.040325341 | 0.893113641 | 0.98 |
| YDL152W | 5.390111654 | -0.372979983 | 0.897346721 | 0.98 |
| YDR095C | 5.026425328 | -0.326730239 | 0.894942179 | 0.98 |
| YDR176W | 1688.228212 | 0.024601469 | 0.896650763 | 0.98 |
| YDR253C | 229.9334863 | -0.112003101 | 0.897276237 | 0.98 |
| YDR370C | 533.8393634 | -0.049868737 | 0.893412142 | 0.98 |
| YDR409W | 969.523209 | 0.026117916 | 0.895330481 | 0.98 |
| YDR517W | 2232.436932 | 0.115001225 | 0.888393048 | 0.98 |
| YEL048C | 96.72112286 | -0.001625979 | 0.89718729 | 0.98 |
| YER062C | 1876.234052 | -0.112424867 | 0.892662125 | 0.98 |
| YER157W | 974.6953246 | 0.018342284 | 0.891004039 | 0.98 |
| YFL023W | 930.5339775 | 0.174518466 | 0.898224102 | 0.98 |
| YFR049W | 788.1728678 | -0.021737399 | 0.895816361 | 0.98 |
| YGL191W | 2937.202908 | 0.015898122 | 0.895630504 | 0.98 |
| YGL197W | 2610.651509 | 0.038576632 | 0.88988898 | 0.98 |
| YGL238W | 2547.939909 | -0.027643192 | 0.896204007 | 0.98 |
| YGL247W | 197.7711302 | 0.025220063 | 0.895482032 | 0.98 |
| YGL254W | 331.7870674 | 0.099638074 | 0.896331045 | 0.98 |
| YGL261C | 11.97177178 | 0.177747415 | 0.89547519 | 0.98 |
| YGR117C | 888.1364455 | 0.040243408 | 0.898525081 | 0.98 |
| YGR121W-A | 8.749518959 | 0.227712439 | 0.889032063 | 0.98 |
| YGR141W | 841.9095851 | 0.208739751 | 0.891399017 | 0.98 |
| YGR239C | 410.924928 | 0.03287822 | 0.894234725 | 0.98 |
| YHL006C | 116.1902464 | -0.18665091 | 0.895372613 | 0.98 |
| YHR168W | 379.0957302 | -0.052201217 | 0.898870739 | 0.98 |
| YHR204W | 1257.868137 | 0.070837081 | 0.894559826 | 0.98 |
| YIL004C | 332.265247 | 0.017347902 | 0.889575598 | 0.98 |
| YIL025C | 3.736035911 | 0.427789489 | 0.892362117 | 0.98 |
| YIL141W | 35.62085203 | 0.391723106 | 0.894542334 | 0.98 |
| YIR028W | 189.55325 | 0.044271049 | 0.896128159 | 0.98 |
| YJL019W | 605.7582629 | 0.133686706 | 0.897595191 | 0.98 |
| YJL031C | 712.299831 | -0.129592142 | 0.894803114 | 0.98 |
| YJL105W | 247.8507967 | -0.126317595 | 0.893241414 | 0.98 |
| YJL209W | 626.0794008 | -0.007425142 | 0.897029238 | 0.98 |
| YJR078W | 516.4472306 | 0.203453077 | 0.899366378 | 0.98 |
| YJR112W-A | 356.5088081 | 0.006087196 | 0.898648564 | 0.98 |
| YKL068W | 1722.980976 | -0.047429905 | 0.891743446 | 0.98 |
| YKL221W | 211.2597824 | -0.228598298 | 0.899260176 | 0.98 |
| YKR095W | 4316.14506 | 0.077893355 | 0.899259968 | 0.98 |
| YLL006W | 812.6766975 | 0.002581509 | 0.898693873 | 0.98 |
| YLL048C | 4130.734576 | 0.117886551 | 0.899254223 | 0.98 |

| | | | | |
|---|---|---|---|---|
| YLR036C | 360.058471 | -0.060590572 | 0.888225669 | 0.98 |
| YLR302C | 0.305761563 | 0.05990859 | 0.891822455 | 0.98 |
| YLR323C | 363.5088652 | 0.074394726 | 0.893454762 | 0.98 |
| YLR433C | 1164.793625 | 0.085462151 | 0.893791071 | 0.98 |
| YML049C | 1684.51308 | -0.003201658 | 0.890598878 | 0.98 |
| YML051W | 1749.382621 | -0.051006755 | 0.895664493 | 0.98 |
| YML087C | 330.7889367 | -0.176958289 | 0.887870203 | 0.98 |
| YMR039C | 1898.602952 | 0.06662655 | 0.895797637 | 0.98 |
| YMR098C | 840.7673515 | 0.045521254 | 0.89169179 | 0.98 |
| YMR282C | 677.7599955 | -0.069026478 | 0.896208422 | 0.98 |
| YNL128W | 87.91000712 | -0.069206191 | 0.894348841 | 0.98 |
| YNL164C | 278.0976276 | -0.095465889 | 0.887996279 | 0.98 |
| YNL238W | 1373.585003 | 0.038814194 | 0.89255126 | 0.98 |
| YNR044W | 3311.34449 | -0.131881844 | 0.88977462 | 0.98 |
| YOL112W | 963.5678133 | -0.040010666 | 0.896187528 | 0.98 |
| YOL148C | 713.6397629 | -0.000453874 | 0.887572197 | 0.98 |
| YOL164W-A | 24.89009234 | -0.404107017 | 0.892540146 | 0.98 |
| YOR090C | 1602.791656 | -0.013783296 | 0.891352844 | 0.98 |
| YOR155C | 1051.816257 | 0.080735074 | 0.888708099 | 0.98 |
| YOR392W | 4.708188968 | -0.123196609 | 0.898489822 | 0.98 |
| YPL006W | 1523.461434 | 0.052145799 | 0.892067473 | 0.98 |
| YPL040C | 952.0787614 | -0.005550179 | 0.890088025 | 0.98 |
| YPR141C | 623.3342075 | -0.043630053 | 0.891205823 | 0.98 |
| YPR188C | 534.7061592 | -0.029060546 | 0.899151753 | 0.98 |
| YPR191W | 6094.519828 | 0.08126309 | 0.896760857 | 0.98 |
| YNL127W | 1075.477959 | 0.013438335 | 0.899593076 | 0.98 |
| YAL060W | 4666.556071 | 0.156053289 | 0.900164956 | 0.98 |
| YGL202W | 6879.756501 | 0.259581098 | 0.899941887 | 0.98 |
| YIR008C | 729.6399623 | -0.057234786 | 0.900094301 | 0.98 |
| YBL012C | 41.44037077 | -0.027889759 | 0.900868648 | 0.98 |
| YBR003W | 1468.050788 | 0.002200083 | 0.901765643 | 0.98 |
| YER180C-A | 68.54860649 | -0.084487385 | 0.900932764 | 0.98 |
| YGL115W | 1145.902181 | 0.023330708 | 0.90166309 | 0.98 |
| YJL215C | 44.17616213 | -0.202430484 | 0.901429993 | 0.98 |
| YJR147W | 698.6444516 | 0.148595777 | 0.901042005 | 0.98 |
| YML030W | 762.1544554 | -0.005457072 | 0.90121795 | 0.98 |
| YMR061W | 1751.530821 | -0.041631741 | 0.901675232 | 0.98 |
| YOL158C | 1769.752458 | 0.071699339 | 0.901279085 | 0.98 |
| YPL014W | 2476.516022 | 0.345159207 | 0.900949659 | 0.98 |
| YCL027W | 675.5367401 | -0.307523824 | 0.90263329 | 0.98 |
| YCR037C | 1044.612368 | -0.052258803 | 0.902539515 | 0.98 |
| YMR052C-A | 61.89756679 | 0.060398357 | 0.902747083 | 0.98 |
| YMR198W | 512.8655409 | -0.074214079 | 0.902393413 | 0.98 |

| | | | | |
|---|---|---|---|---|
| YOR168W | 3054.619284 | 0.125487011 | 0.902775679 | 0.98 |
| YOR364W | 7.913433737 | 0.014435869 | 0.902729527 | 0.98 |
| YML070W | 2763.480937 | -0.08704339 | 0.904158795 | 0.98 |
| YML131W | 1116.004425 | -0.09094714 | 0.903923395 | 0.98 |
| YMR239C | 477.7892356 | 0.070984737 | 0.903736057 | 0.98 |
| YNR013C | 2464.582186 | 0.005423671 | 0.90424454 | 0.98 |
| YOR257W | 743.3095021 | -0.045330019 | 0.90407407 | 0.98 |
| YFL027C | 673.7008422 | -0.135832742 | 0.905024076 | 0.98 |
| YJR051W | 1291.045072 | 0.062668855 | 0.905014847 | 0.98 |
| YLR296W | 4.726955448 | -0.127306695 | 0.90477987 | 0.98 |
| YNL161W | 1329.930942 | -0.003064722 | 0.904656169 | 0.98 |
| YNL225C | 446.0966712 | 0.01333136 | 0.904770182 | 0.98 |
| YAL021C | 1595.906288 | 0.031345677 | 0.905817602 | 0.98 |
| YHR082C | 4569.103984 | 0.150500662 | 0.90559628 | 0.98 |
| YIL120W | 363.836718 | 0.023385356 | 0.906492982 | 0.98 |
| YJL113W | 119.706187 | -0.032959643 | 0.906200677 | 0.98 |
| YLR169W | 310.0591441 | 0.011165091 | 0.906417156 | 0.98 |
| YOL136C | 1016.324902 | -0.070955067 | 0.906009863 | 0.98 |
| YOL166W-A | 0.347622843 | 0.010927507 | 0.906138759 | 0.98 |
| YBL055C | 719.3024768 | -0.01032419 | 0.907258535 | 0.98 |
| YCL011C | 2495.29652 | -0.077684908 | 0.907361093 | 0.98 |
| YDR122W | 2792.779664 | 0.037986429 | 0.907892343 | 0.98 |
| YEL029C | 331.9088854 | 0.17469119 | 0.907631192 | 0.98 |
| YGL262W | 77.76796003 | -0.174146062 | 0.907772207 | 0.98 |
| YIR030C | 90.29914152 | 0.120538336 | 0.907353025 | 0.98 |
| YJL151C | 3281.2426 | -0.065874289 | 0.908018023 | 0.98 |
| YLR093C | 1657.689066 | -0.064445958 | 0.907887175 | 0.98 |
| YMR163C | 451.0209425 | -0.016627715 | 0.907843279 | 0.98 |
| tR(CCG)L | 0.085991782 | -0.042484115 | 0.907585829 | 0.98 |
| YDR055W | 3654.499745 | -0.027397927 | 0.908413488 | 0.98 |
| YGL002W | 690.1637106 | -0.003872267 | 0.908529314 | 0.98 |
| YGR212W | 361.1538773 | -0.057617938 | 0.90836472 | 0.98 |
| YOL123W | 4114.119417 | -0.08434546 | 0.908718925 | 0.98 |
| YJL171C | 1558.602902 | 0.033497214 | 0.909042971 | 0.98 |
| YHR125W | 4.057342841 | -0.128180094 | 0.9092558 | 0.98 |
| YNR070W | 728.5997471 | 0.035633796 | 0.909449663 | 0.98 |
| YDR455C | 13.39180516 | 0.026534557 | 0.909721376 | 0.98 |
| YKL067W | 3440.072656 | 0.157563765 | 0.91007976 | 0.98 |
| YNR063W | 178.9442947 | 0.15997243 | 0.910297732 | 0.98 |
| YOR042W | 2668.884695 | 0.035530844 | 0.910264331 | 0.98 |
| YDL221W | 36.90750404 | -0.105876035 | 0.910603087 | 0.98 |
| YAL066W | 0.770254238 | 0.13941688 | 0.923051342 | 0.98 |
| YAL068W-A | 0.241654238 | 0.014597415 | 0.919539007 | 0.98 |

| | | | | |
|---|---|---|---|---|
| YBR066C | 407.0918605 | -0.254116145 | 0.915519944 | 0.98 |
| YBR069C | 2910.073987 | 0.086830845 | 0.921797695 | 0.98 |
| YBR123C | 731.156432 | -0.043863854 | 0.921303504 | 0.98 |
| YBR134W | 28.00305569 | 0.331105518 | 0.922418668 | 0.98 |
| YBR165W | 515.6336536 | -0.039176669 | 0.922933816 | 0.98 |
| YBR294W | 590.8243932 | -0.188129086 | 0.92325019 | 0.98 |
| YCR076C | 593.9023385 | 0.007359803 | 0.915301793 | 0.98 |
| YCR097W | 1013.079937 | 0.117043204 | 0.92052683 | 0.98 |
| YDL131W | 1846.278027 | 0.129581489 | 0.912292291 | 0.98 |
| YDL173W | 1092.267621 | -0.005013608 | 0.911819101 | 0.98 |
| YDL185W | 10910.49268 | 0.043465404 | 0.913026427 | 0.98 |
| YDR021W | 377.9993278 | -0.025303141 | 0.914862654 | 0.98 |
| YDR081C | 1262.178022 | -0.005444796 | 0.916070489 | 0.98 |
| YDR105C | 933.9584198 | -0.057222022 | 0.912064051 | 0.98 |
| YDR145W | 1845.179932 | 0.039161011 | 0.916705006 | 0.98 |
| YDR177W | 1727.041157 | 0.103225078 | 0.918557612 | 0.98 |
| YDR178W | 2932.392626 | 0.103835981 | 0.919116166 | 0.98 |
| YDR346C | 2820.625886 | 0.050108523 | 0.919427508 | 0.98 |
| YDR371W | 631.8329655 | -0.046464555 | 0.91496987 | 0.98 |
| YDR384C | 561.1890906 | 0.273673275 | 0.917823972 | 0.98 |
| YDR451C | 513.5586606 | -0.072198735 | 0.920214151 | 0.98 |
| YER057C | 2325.694126 | -0.029650867 | 0.91708477 | 0.98 |
| YER123W | 1029.367215 | -0.003124284 | 0.916473423 | 0.98 |
| YER136W | 4119.832209 | 0.040509347 | 0.921836837 | 0.98 |
| YER161C | 1183.000093 | -0.084053932 | 0.91487516 | 0.98 |
| YGL041W-A | 345.8516558 | -0.038462094 | 0.915290939 | 0.98 |
| YGL087C | 1504.49763 | -0.069241881 | 0.918831896 | 0.98 |
| YGL122C | 2999.105342 | -0.034982808 | 0.923320959 | 0.98 |
| YGL168W | 276.6572721 | -0.004243715 | 0.922714853 | 0.98 |
| YGR014W | 3080.238337 | -0.201629307 | 0.919371841 | 0.98 |
| YGR142W | 4162.125638 | 0.191586668 | 0.920720865 | 0.98 |
| YHR015W | 104.6987227 | 0.002434639 | 0.922157522 | 0.98 |
| YHR024C | 1377.744705 | -0.01639005 | 0.922628017 | 0.98 |
| YHR111W | 861.1772037 | -0.001780642 | 0.918948035 | 0.98 |
| YHR122W | 579.3180386 | 0.260971181 | 0.918401439 | 0.98 |
| YHR180W-A | 0.36648756 | -0.011726511 | 0.922687656 | 0.98 |
| YHR184W | 81.12732022 | 0.013608126 | 0.915037797 | 0.98 |
| YHR211W | 534.044984 | 0.091520958 | 0.918190337 | 0.98 |
| YHR213W-B | 12.16949824 | 0.210396108 | 0.915439381 | 0.98 |
| YIL124W | 2615.425395 | -0.042275593 | 0.922706501 | 0.98 |
| YIL156W-B | 291.2591429 | -0.090086144 | 0.916656108 | 0.98 |
| YJL025W | 374.9847479 | -0.097491184 | 0.917073866 | 0.98 |
| YJL112W | 1088.839683 | -0.001468268 | 0.913885202 | 0.98 |

| | | | | |
|---|---|---|---|---|
| YJL184W | 578.763074 | -0.007606095 | 0.913558608 | 0.98 |
| YJL205C | 132.3580317 | -0.015519241 | 0.914361126 | 0.98 |
| YKL106C-A | 1.684117879 | 0.272465544 | 0.9189538 | 0.98 |
| YKL106W | 462.66436 | -0.121814454 | 0.920380034 | 0.98 |
| YKL145W | 5482.552156 | -0.01590132 | 0.91174882 | 0.98 |
| YKL219W | 500.3272007 | -0.098755108 | 0.918915183 | 0.98 |
| YKR016W | 2416.547829 | 0.111202072 | 0.911401174 | 0.98 |
| YKR047W | 143.8726915 | 0.043929455 | 0.911304586 | 0.98 |
| YKR102W | 639.1097035 | 0.08614463 | 0.91569854 | 0.98 |
| YLL032C | 878.0650576 | 0.031135181 | 0.911657938 | 0.98 |
| YLR064W | 1412.390781 | -0.016829147 | 0.915731991 | 0.98 |
| YLR115W | 1204.082371 | 0.043437526 | 0.912988401 | 0.98 |
| YLR253W | 1177.596684 | -0.010451028 | 0.919594327 | 0.98 |
| YLR274W | 1064.782127 | -0.055891517 | 0.91753277 | 0.98 |
| YMR080C | 1907.455726 | 0.051360505 | 0.921734268 | 0.98 |
| YMR173W | 743.4251563 | 0.16333414 | 0.915071202 | 0.98 |
| YMR275C | 1521.663704 | 0.076272632 | 0.920868127 | 0.98 |
| YMR287C | 795.4088763 | -0.051260413 | 0.922993011 | 0.98 |
| YNL022C | 938.4563852 | 0.14509099 | 0.922841021 | 0.98 |
| YNL055C | 20028.30046 | 0.168550985 | 0.913848981 | 0.98 |
| YNL336W | 467.9927857 | -0.112615292 | 0.923006556 | 0.98 |
| YNR032W | 607.7463894 | -0.04445052 | 0.916945625 | 0.98 |
| YOL014W | 428.9922641 | 0.010168055 | 0.920148401 | 0.98 |
| YOL020W | 976.6345917 | -0.152931151 | 0.914196542 | 0.98 |
| YOL104C | 616.0912562 | -0.019823968 | 0.917476724 | 0.98 |
| YOR008C | 1383.478149 | -0.076428007 | 0.92143665 | 0.98 |
| YOR011W-A | 3.164669454 | -0.24144835 | 0.921700007 | 0.98 |
| YOR065W | 3707.159237 | 0.169876156 | 0.919824405 | 0.98 |
| YOR084W | 822.8839971 | -0.070059383 | 0.911379472 | 0.98 |
| YOR149C | 790.0054864 | 0.020085622 | 0.912709166 | 0.98 |
| YOR285W | 4227.371531 | 0.018218232 | 0.911309278 | 0.98 |
| YOR334W | 302.0782196 | -0.088418849 | 0.911688203 | 0.98 |
| YPL008W | 976.9490792 | -0.042956163 | 0.918032763 | 0.98 |
| YPL034W | 171.3658521 | -0.14704573 | 0.916984995 | 0.98 |
| YPL213W | 340.9140453 | 0.026295564 | 0.919047806 | 0.98 |
| YPL228W | 971.5333391 | -0.000639538 | 0.913020799 | 0.98 |
| snR45 | 13.49639451 | 0.414780354 | 0.919664727 | 0.98 |
| tG(CCC)O | 0.202424979 | -0.04109887 | 0.922683272 | 0.98 |
| tL(GAG)G | 0.599720226 | -0.079770562 | 0.914749726 | 0.98 |
| YIL074C | 1391.44586 | 0.064861041 | 0.923638057 | 0.98 |
| YJL147C | 286.6419753 | -0.030627461 | 0.923712514 | 0.98 |
| YLR452C | 1814.469836 | -0.20985461 | 0.924262901 | 0.98 |
| YPL259C | 962.6624104 | -0.034659464 | 0.924432631 | 0.98 |

| | | | | |
|---|---|---|---|---|
| YML031W | 1341.560963 | -0.001360004 | 0.924820508 | 0.98 |
| YBR281C | 961.503364 | 0.095640907 | 0.926867204 | 0.99 |
| YDL033C | 356.3492838 | -0.085067722 | 0.926032988 | 0.99 |
| YDR148C | 4531.818706 | 0.088215899 | 0.925904649 | 0.99 |
| YGL014W | 2156.619039 | -0.025209857 | 0.927045138 | 0.99 |
| YHR178W | 530.4654676 | 0.034075546 | 0.926568875 | 0.99 |
| YKL114C | 901.3872837 | 0.028974909 | 0.926427925 | 0.99 |
| YKR083C | 244.3380659 | -0.087906048 | 0.926882081 | 0.99 |
| YLL036C | 873.0418955 | -0.031354062 | 0.925204026 | 0.99 |
| YLL065W | 0.583562943 | -0.006871712 | 0.925647543 | 0.99 |
| YOR034C-A | 45.66152085 | -0.014752545 | 0.926493095 | 0.99 |
| YOR041C | 37.3273586 | -0.102529901 | 0.926077443 | 0.99 |
| YOR075W | 634.1618193 | -0.039134769 | 0.926221342 | 0.99 |
| YOR112W | 1049.770792 | 0.029720911 | 0.927041433 | 0.99 |
| YPL269W | 614.929882 | -0.075941667 | 0.926105926 | 0.99 |
| YGR080W | 1465.883709 | -0.019369329 | 0.927250113 | 0.99 |
| YBR260C | 1246.430976 | 0.028736108 | 0.92787054 | 0.99 |
| YCR102C | 144.1370507 | 0.001517629 | 0.927859316 | 0.99 |
| YFL046W | 596.7158239 | -0.008507686 | 0.927812745 | 0.99 |
| YOR359W | 584.0345426 | -0.172942259 | 0.928011249 | 0.99 |
| snR11 | 261.7635615 | -0.146281186 | 0.928337422 | 0.99 |
| YIL154C | 2075.807869 | -0.041908098 | 0.928803107 | 0.99 |
| YKL117W | 7112.851481 | -0.064869765 | 0.928740961 | 0.99 |
| YOR142W | 3533.236265 | 0.192403806 | 0.928807087 | 0.99 |
| YBR110W | 1287.287964 | -0.022153487 | 0.933479815 | 0.99 |
| YCL068C | 4.465179414 | 0.000700238 | 0.93392923 | 0.99 |
| YCR060W | 437.7609902 | -0.017939318 | 0.935666056 | 0.99 |
| YDL028C | 1026.396259 | 0.021142612 | 0.933598592 | 0.99 |
| YDL093W | 807.8112275 | 0.096170104 | 0.931060802 | 0.99 |
| YDL171C | 4109.615185 | 0.17477493 | 0.935868586 | 0.99 |
| YDR036C | 1560.530814 | 0.083173528 | 0.934097119 | 0.99 |
| YDR100W | 731.7492012 | 0.009874843 | 0.936314332 | 0.99 |
| YDR150W | 4371.450407 | 0.051883814 | 0.929577674 | 0.99 |
| YDR322C-A | 892.3853165 | -0.097770001 | 0.935590907 | 0.99 |
| YDR443C | 1163.053889 | 0.077326291 | 0.929689213 | 0.99 |
| YDR521W | 13.62209314 | 0.400285858 | 0.933328649 | 0.99 |
| YEL045C | 16.86688846 | -0.104987131 | 0.931599469 | 0.99 |
| YER022W | 948.246119 | 0.034838744 | 0.930179753 | 0.99 |
| YFR010W | 3072.848472 | -0.024125823 | 0.93100253 | 0.99 |
| YGL021W | 1221.378254 | 0.050722388 | 0.935519113 | 0.99 |
| YGR021W | 818.8357092 | 0.026725613 | 0.935894476 | 0.99 |
| YGR068C | 535.7602898 | 0.008058055 | 0.932584847 | 0.99 |
| YGR193C | 2486.614356 | -0.007472142 | 0.932006322 | 0.99 |

| | | | | |
|---|---|---|---|---|
| YGR218W | 3998.195199 | 0.024667589 | 0.93030905 | 0.99 |
| YHR032W | 1019.136565 | 0.099858295 | 0.935673407 | 0.99 |
| YIR040C | 0.167590483 | 0.012467015 | 0.934605366 | 0.99 |
| YJL119C | 7.109202711 | 0.124951292 | 0.934053021 | 0.99 |
| YJL156C | 869.0736001 | -0.056100561 | 0.930595653 | 0.99 |
| YJR044C | 1233.447551 | 0.078422382 | 0.933993644 | 0.99 |
| YJR146W | 99.77471099 | -0.066068058 | 0.929994413 | 0.99 |
| YKL148C | 4538.261302 | 0.227952517 | 0.930091677 | 0.99 |
| YKR023W | 810.8865639 | 0.104928054 | 0.936155814 | 0.99 |
| YLL043W | 1838.000712 | 0.046240614 | 0.933869532 | 0.99 |
| YLR086W | 1588.915329 | 0.0647057 | 0.935282448 | 0.99 |
| YLR163C | 1121.057782 | 0.01121572 | 0.93381744 | 0.99 |
| YLR460C | 187.4228264 | 0.085084898 | 0.933058577 | 0.99 |
| YMR069W | 79.44065746 | 0.14559214 | 0.936177675 | 0.99 |
| YMR082C | 5.748276588 | -0.002425233 | 0.934569154 | 0.99 |
| YMR152W | 1727.69017 | 0.008301596 | 0.93236411 | 0.99 |
| YMR167W | 578.0579077 | 0.006282383 | 0.935734948 | 0.99 |
| YNL045W | 3993.621371 | 0.37315439 | 0.932251615 | 0.99 |
| YNL097C-B | 13.11125706 | -0.276581708 | 0.933787091 | 0.99 |
| YNL267W | 1939.373479 | -0.024569661 | 0.93358167 | 0.99 |
| YNL296W | 0.17572267 | 0.01098955 | 0.935569017 | 0.99 |
| YNR048W | 344.510222 | -0.067087564 | 0.930826882 | 0.99 |
| YOL079W | 9.64455915 | 0.160106823 | 0.934525703 | 0.99 |
| YOR322C | 1313.48176 | -0.020126273 | 0.936043674 | 0.99 |
| YOR337W | 623.7332196 | 0.015876933 | 0.930687567 | 0.99 |
| YPL060C-A | 3.51919416 | 0.452819367 | 0.931874636 | 0.99 |
| YPL205C | 28.51967725 | -0.058464399 | 0.933981365 | 0.99 |
| YPR159W | 2743.676738 | 0.101004611 | 0.935148155 | 0.99 |
| YPR161C | 1370.124428 | -0.032391495 | 0.935640935 | 0.99 |
| YDL065C | 1387.029558 | 0.079300238 | 0.936572428 | 0.99 |
| YDL001W | 483.443293 | 0.002948232 | 0.937066926 | 0.99 |
| YER084W-A | 0.719422121 | -0.09732772 | 0.937110598 | 0.99 |
| YKL141W | 4100.397187 | 0.182454132 | 0.936819284 | 0.99 |
| YDR272W | 1625.208653 | -0.037403664 | 0.938326451 | 0.99 |
| YDR503C | 368.4756431 | 0.006861974 | 0.939283925 | 0.99 |
| YFR024C-A | 2497.30533 | 0.048301432 | 0.93807476 | 0.99 |
| YGL016W | 794.4716524 | 0.109334767 | 0.93917812 | 0.99 |
| YHR050W-A | 204.2549426 | -0.018069332 | 0.938228576 | 0.99 |
| YIL177C | 0.203940714 | -0.014948216 | 0.939245048 | 0.99 |
| YJR128W | 14.84064674 | -0.038232781 | 0.937911967 | 0.99 |
| YLR128W | 336.2576961 | 0.019441897 | 0.937944138 | 0.99 |
| YML113W | 567.9041791 | -0.082248558 | 0.938748241 | 0.99 |
| YNL212W | 1282.46661 | 0.09337251 | 0.938388327 | 0.99 |

| | | | | |
|---|---|---|---|---|
| YOR073W-A | 11.62724158 | 0.266419431 | 0.938622723 | 0.99 |
| YOR106W | 543.2030625 | 0.01618395 | 0.939362213 | 0.99 |
| YOR181W | 1611.589546 | -0.016438722 | 0.939312886 | 0.99 |
| YDR520C | 591.1970741 | -0.024698606 | 0.939696682 | 0.99 |
| YLR034C | 1366.422953 | -0.10787556 | 0.940238289 | 0.99 |
| YNL032W | 670.5607595 | -0.019812883 | 0.940153317 | 0.99 |
| YNL201C | 1076.736749 | -0.034062052 | 0.940258507 | 0.99 |
| YAL014C | 833.2543927 | -0.093935759 | 0.94173963 | 0.99 |
| YER140W | 473.4616837 | -0.028839132 | 0.941524269 | 0.99 |
| YHR217C | 1.225581631 | 0.250696457 | 0.941763867 | 0.99 |
| YLR016C | 414.1257784 | -0.007055926 | 0.94171059 | 0.99 |
| YLR390W-A | 12426.05045 | -0.018713718 | 0.942592708 | 0.99 |
| YDR213W | 568.9307813 | 0.14001363 | 0.943407757 | 0.99 |
| YJL181W | 405.181886 | 0.036076497 | 0.943640035 | 0.99 |
| YDR196C | 880.6195121 | -0.110931739 | 0.944520125 | 0.99 |
| YLR336C | 993.7017255 | 0.143262231 | 0.944622188 | 0.99 |
| YML115C | 1520.660585 | 0.000230279 | 0.944611538 | 0.99 |
| YMR102C | 1653.341009 | 0.118315849 | 0.944334353 | 0.99 |
| YIL116W | 2678.75766 | 0.260747779 | 0.944936274 | 0.99 |
| YDR217C | 884.5738376 | 0.067466923 | 0.945314816 | 0.99 |
| YGL071W | 1392.981634 | -0.0141891 | 0.945418106 | 0.99 |
| YNL028W | 6.323808974 | -0.105142041 | 0.945507122 | 0.99 |
| YPL139C | 637.4930321 | -0.036161866 | 0.945706862 | 0.99 |
| tT(CGU)K | 0.429968112 | -0.005425503 | 0.94568141 | 0.99 |
| YGR136W | 2019.817506 | 0.052938768 | 0.946183917 | 0.99 |
| YLR275W | 260.6216284 | 0.033938648 | 0.946804056 | 0.99 |
| YOR166C | 666.3506234 | -0.026310959 | 0.946655508 | 0.99 |
| YEL007W | 3320.15412 | -0.084733312 | 0.94709526 | 0.99 |
| YBR170C | 1144.125576 | -0.020745711 | 0.94816422 | 0.99 |
| YBR172C | 2003.113678 | -0.010730185 | 0.948575779 | 0.99 |
| YDL163W | 0.057163039 | -0.015287262 | 0.948927766 | 0.99 |
| YER180C | 405.292426 | -0.103498307 | 0.94844964 | 0.99 |
| YHR044C | 186.9132602 | -0.143587467 | 0.948831292 | 0.99 |
| YIL068C | 981.7929067 | 0.013198764 | 0.948987276 | 0.99 |
| YFL007W | 4593.396937 | 0.153074655 | 0.94962295 | 0.99 |
| YFR033C | 3330.696407 | 0.035819628 | 0.949805344 | 0.99 |
| YGR244C | 5502.307672 | 0.122326399 | 0.949856795 | 0.99 |
| YDL047W | 1531.758576 | -0.003746323 | 0.951648177 | 0.99 |
| YDL078C | 3642.136284 | 0.146227911 | 0.953076354 | 0.99 |
| YDR098C-B | 19.81780983 | -0.134505126 | 0.950373648 | 0.99 |
| YDR179C | 161.4536724 | 0.084776352 | 0.95288405 | 0.99 |
| YFL053W | 516.855343 | 0.299625086 | 0.951443781 | 0.99 |
| YGR050C | 100.6325721 | -0.125440462 | 0.951749018 | 0.99 |

| | | | | |
|---|---|---|---|---|
| YGR254W | 35005.12746 | 0.101610617 | 0.952842514 | 0.99 |
| YHR137W | 2665.655739 | 0.315715017 | 0.951504822 | 0.99 |
| YKL138C-A | 204.185803 | 0.059861386 | 0.952254195 | 0.99 |
| YKL224C | 0.174623486 | 0.017060693 | 0.950570543 | 0.99 |
| YLL062C | 209.8685733 | -0.096739684 | 0.951925866 | 0.99 |
| YLR119W | 415.3763546 | -0.044109229 | 0.953116268 | 0.99 |
| YLR137W | 268.6657732 | -0.178095402 | 0.95312978 | 0.99 |
| YLR146W-A | 192.5995673 | 0.016269786 | 0.95309179 | 0.99 |
| YNR050C | 3301.317883 | 0.099166722 | 0.950923321 | 0.99 |
| YOL163W | 57.10674557 | -0.513550989 | 0.951824399 | 0.99 |
| YOR079C | 215.4299264 | -0.066173113 | 0.952320088 | 0.99 |
| YOR263C | 29.84403306 | -0.084426897 | 0.952167253 | 0.99 |
| YPR122W | 1503.146832 | -0.209713122 | 0.951343063 | 0.99 |
| snR73 | 0.982818557 | -0.162963667 | 0.951327039 | 0.99 |
| YDR157W | 53.23769391 | -0.249754281 | 0.953427107 | 0.99 |
| YDR158W | 6048.209084 | 0.424919326 | 0.953572155 | 0.99 |
| YMR257C | 582.2395624 | -0.122824016 | 0.953956487 | 0.99 |
| YDR065W | 277.2481422 | -0.049193572 | 0.954502154 | 0.99 |
| YGL240W | 171.0354676 | 0.034373536 | 0.954513053 | 0.99 |
| YBR278W | 245.0159717 | -0.004533574 | 0.955703159 | 0.99 |
| YDR536W | 315.8429448 | -0.06249398 | 0.955610427 | 0.99 |
| YGR028W | 820.509086 | -0.091297853 | 0.955317566 | 0.99 |
| YHR219W | 0.159331785 | 0.012876966 | 0.95594238 | 0.99 |
| YLR283W | 409.1057668 | -0.024125977 | 0.956023461 | 0.99 |
| YML058W | 2577.833646 | -0.177248632 | 0.955582392 | 0.99 |
| YMR177W | 436.2483597 | -0.027172234 | 0.955862819 | 0.99 |
| YNL051W | 867.7540206 | 0.012142746 | 0.955967129 | 0.99 |
| YPR074W-A | 0.407755058 | -0.07617203 | 0.95529651 | 0.99 |
| YDR380W | 4921.676121 | 0.274514586 | 0.956238568 | 0.99 |
| YDR415C | 529.3228874 | 0.026786224 | 0.957571539 | 0.99 |
| YER052C | 2274.895414 | 0.480634216 | 0.957739422 | 0.99 |
| YHR102W | 1800.525655 | 0.025985864 | 0.957886098 | 0.99 |
| YIL075C | 7548.161259 | 0.078210693 | 0.957259884 | 0.99 |
| YJL006C | 336.1143852 | -0.053605769 | 0.95675023 | 0.99 |
| YMR035W | 535.4792987 | 0.068802135 | 0.957423195 | 0.99 |
| YOL152W | 484.6917609 | 0.164220209 | 0.957313616 | 0.99 |
| YOR356W | 1275.917226 | 0.033504071 | 0.957046173 | 0.99 |
| YPL194W | 451.5443271 | -0.020739795 | 0.957196411 | 0.99 |
| tQ(UUG)E2 | 0.083088204 | 0.013357699 | 0.957991433 | 0.99 |
| YDR363W-A | 1308.889566 | 0.036399272 | 0.959211875 | 0.99 |
| YFR040W | 3140.971961 | 0.102605958 | 0.959519722 | 0.99 |
| YGR295C | 1429.22247 | 0.143238931 | 0.959502905 | 0.99 |
| YKL096C-B | 8.469479675 | -0.178435644 | 0.959367244 | 0.99 |

| | | | | |
|---|---|---|---|---|
| YLR147C | 302.8969457 | -0.024875718 | 0.95853114 | 0.99 |
| YLR218C | 291.942794 | -0.174152303 | 0.958313372 | 0.99 |
| YLR385C | 52.50375543 | 0.056933223 | 0.959164851 | 0.99 |
| YOL157C | 78.40010831 | -0.372745191 | 0.95910842 | 0.99 |
| YPL135W | 1241.708085 | 0.015814143 | 0.958912825 | 0.99 |
| YJL125C | 686.7482617 | 0.006393487 | 0.959800282 | 0.99 |
| YDR320C | 1271.658982 | 0.004141047 | 0.959985511 | 0.99 |
| YDL072C | 3778.461007 | 0.033108852 | 0.960808478 | 0.99 |
| YDR180W | 980.8547337 | 0.083338661 | 0.960829344 | 0.99 |
| YHR145C | 0.415497853 | -0.051526232 | 0.9605815 | 0.99 |
| YIR036C | 938.6857022 | -0.060068838 | 0.960165337 | 0.99 |
| YJR133W | 618.7266674 | 0.090599092 | 0.960913895 | 0.99 |
| YPR104C | 970.815797 | -0.010301854 | 0.960633504 | 0.99 |
| YDR032C | 4060.720659 | -0.093984717 | 0.961448236 | 0.99 |
| YIL047C | 1300.508261 | 0.012902921 | 0.961536567 | 0.99 |
| YDR265W | 518.8074155 | -0.048321448 | 0.962204957 | 0.99 |
| YER088C | 4317.842613 | 0.163872035 | 0.961957769 | 0.99 |
| YMR088C | 861.7053416 | 0.059461382 | 0.96224152 | 0.99 |
| YDR166C | 1344.376251 | 0.002009349 | 0.962798706 | 0.99 |
| YDR439W | 511.333794 | -0.004938277 | 0.962490456 | 0.99 |
| YOR034C | 667.8994742 | 0.019630901 | 0.962902841 | 0.99 |
| YPR053C | 166.5759994 | -0.198465972 | 0.962767539 | 0.99 |
| YBR197C | 458.4082334 | -0.006015039 | 0.964073946 | 0.99 |
| YER189W | 0.915658928 | 0.009252689 | 0.96387651 | 0.99 |
| YGL172W | 1277.929348 | 0.020581567 | 0.963488517 | 0.99 |
| YIL170W | 172.5743318 | -0.364361519 | 0.963836986 | 0.99 |
| YNL190W | 6079.357238 | 0.068591493 | 0.964022609 | 0.99 |
| YNL193W | 690.484771 | 0.12479902 | 0.963716283 | 0.99 |
| snR65 | 5.68201134 | -0.029652359 | 0.963209664 | 0.99 |
| YDL197C | 997.8504474 | 0.067388967 | 0.964285768 | 0.99 |
| YKL066W | 1252.501288 | 0.086526385 | 0.964880619 | 0.99 |
| YLR187W | 889.7815093 | 0.046238897 | 0.964670414 | 0.99 |
| YLR341W | 104.3766662 | -0.085594466 | 0.96458626 | 0.99 |
| YPL182C | 43.60254699 | 0.110496614 | 0.964931112 | 0.99 |
| tQ(UUG)B | 0.084186605 | 0.010307913 | 0.96524168 | 0.99 |
| YER166W | 2859.051695 | 0.072224177 | 0.965928813 | 0.99 |
| YML005W | 433.1547412 | 0.005178254 | 0.965656274 | 0.99 |
| YNR042W | 0.825755426 | -0.346875857 | 0.965776949 | 0.99 |
| YOR205C | 770.9234478 | -0.03547691 | 0.965811568 | 0.99 |
| YDR266C | 1529.276069 | 0.000572198 | 0.966479964 | 0.99 |
| YDR535C | 20.65811424 | -0.11536361 | 0.966378935 | 0.99 |
| YNL322C | 3831.740608 | 0.027042726 | 0.966423665 | 0.99 |
| YBL088C | 1866.759984 | 0.008875957 | 0.966921059 | 0.99 |

| | | | | |
|---|---|---|---|---|
| snR190 | 17.01701234 | 0.257359067 | 0.96703812 | 0.99 |
| YBL045C | 7514.298009 | 0.254136267 | 0.967374819 | 0.99 |
| YBR151W | 3044.846839 | 0.02637475 | 0.970545214 | 0.99 |
| YBR274W | 529.7886878 | 0.012518659 | 0.968654634 | 0.99 |
| YDL135C | 948.2549942 | 0.012526024 | 0.968222177 | 0.99 |
| YDR034C-D | 94.28631679 | 0.181889222 | 0.968684399 | 0.99 |
| YEL065W | 2278.709245 | -0.099229248 | 0.970452347 | 0.99 |
| YER048W-A | 659.0895735 | 0.006255652 | 0.969844363 | 0.99 |
| YFL034W | 949.1269789 | 0.025924904 | 0.968019009 | 0.99 |
| YFR013W | 1397.161166 | -0.017783644 | 0.969246534 | 0.99 |
| YGR251W | 285.9824856 | 0.098900651 | 0.970130178 | 0.99 |
| YHR069C-A | 0.116121444 | -0.015752678 | 0.969276281 | 0.99 |
| YIL046W-A | 7.695578865 | -0.272170549 | 0.968557076 | 0.99 |
| YIL121W | 2357.398488 | -0.229303043 | 0.971058646 | 0.99 |
| YIR038C | 2847.18512 | -0.163329623 | 0.967750392 | 0.99 |
| YJR074W | 487.5630441 | 0.077318684 | 0.967978182 | 0.99 |
| YLL001W | 1443.400677 | 0.047699224 | 0.971036932 | 0.99 |
| YLL051C | 1170.125266 | 0.023648515 | 0.970629073 | 0.99 |
| YLR139C | 783.7450537 | -0.098082804 | 0.970962699 | 0.99 |
| YLR190W | 1584.336567 | 0.032924385 | 0.969445064 | 0.99 |
| YLR245C | 188.3206886 | 0.066227953 | 0.969937033 | 0.99 |
| YLR309C | 2329.969598 | 0.154423568 | 0.96777142 | 0.99 |
| YNL239W | 3388.853667 | 0.096389337 | 0.969982539 | 0.99 |
| YPL061W | 6576.454349 | 0.008491609 | 0.968216824 | 0.99 |
| YPL103C | 590.7071612 | -0.002030467 | 0.970943716 | 0.99 |
| YPL132W | 679.0950944 | 0.005675977 | 0.970232863 | 0.99 |
| YPR196W | 243.4727794 | -0.159233211 | 0.969481973 | 0.99 |
| YBL062W | 0.924462825 | 0.3585921 | 0.97412925 | 0.99 |
| YBR085W | 162.0814879 | 0.436906036 | 0.975207408 | 0.99 |
| YCL001W-A | 26.77980078 | -0.142277752 | 0.971456176 | 0.99 |
| YCL049C | 935.682406 | -0.015763029 | 0.97255022 | 0.99 |
| YDL069C | 126.889784 | -0.012921487 | 0.978486362 | 0.99 |
| YDL175C | 355.6103486 | 0.013766817 | 0.978141638 | 0.99 |
| YDL246C | 6.191078768 | -0.195031037 | 0.976111404 | 0.99 |
| YDR072C | 1794.903187 | -0.173240204 | 0.976083598 | 0.99 |
| YDR133C | 1765.716231 | -0.020661699 | 0.977129431 | 0.99 |
| YDR309C | 2613.4939 | -0.216237882 | 0.974209688 | 0.99 |
| YDR348C | 1537.308452 | -0.002367183 | 0.976164676 | 0.99 |
| YDR363W | 641.2311125 | -0.035013203 | 0.975964461 | 0.99 |
| YDR492W | 689.8420252 | 0.20456484 | 0.978698852 | 0.99 |
| YFR039C | 736.7528646 | -0.038547044 | 0.978299259 | 0.99 |
| YGL205W | 247.6080273 | -0.110473842 | 0.97520687 | 0.99 |
| YGR011W | 95.88242328 | 0.071868809 | 0.977121752 | 0.99 |

| | | | | |
|---|---|---|---|---|
| YGR044C | 1106.237884 | 0.037773365 | 0.976707403 | 0.99 |
| YGR246C | 1106.975461 | 0.048397401 | 0.977853513 | 0.99 |
| YGR291C | 1.566792632 | 0.417195059 | 0.971389442 | 0.99 |
| YIL020C | 394.2399705 | 0.096710689 | 0.976728178 | 0.99 |
| YJL176C | 1289.4859 | -0.028939176 | 0.971603612 | 0.99 |
| YJL200C | 1688.489143 | 0.127461037 | 0.978255538 | 0.99 |
| YJR099W | 741.8529597 | 0.072947695 | 0.977222888 | 0.99 |
| YJR102C | 338.1234787 | -0.02404311 | 0.97712712 | 0.99 |
| YJR142W | 1091.921242 | -0.00332738 | 0.975959558 | 0.99 |
| YKL034W | 1217.582813 | 0.019236172 | 0.978815678 | 0.99 |
| YKL155C | 791.2689341 | -0.005052429 | 0.976433222 | 0.99 |
| YKL223W | 0.064032033 | 0.012633645 | 0.97768054 | 0.99 |
| YLL031C | 1740.005527 | 0.076260138 | 0.975914241 | 0.99 |
| YLR074C | 1095.319455 | 0.049486233 | 0.972685876 | 0.99 |
| YLR376C | 277.9237935 | -0.026778962 | 0.977809718 | 0.99 |
| YLR396C | 522.7183585 | -0.02367513 | 0.97573667 | 0.99 |
| YML120C | 4283.544511 | 0.201142585 | 0.977296361 | 0.99 |
| YMR027W | 2670.4492 | 0.065061562 | 0.977321259 | 0.99 |
| YMR074C | 1117.334751 | -0.102371478 | 0.978867948 | 0.99 |
| YMR147W | 251.4438843 | 0.11419992 | 0.975480669 | 0.99 |
| YMR256C | 1012.905586 | -0.106932429 | 0.972294697 | 0.99 |
| YMR301C | 1249.636815 | 0.071957019 | 0.974127518 | 0.99 |
| YNL024C | 148.4866774 | -0.020302273 | 0.974451871 | 0.99 |
| YNL059C | 1845.140766 | -0.004512746 | 0.976814348 | 0.99 |
| YNL162W-A | 54.12294867 | 0.297506722 | 0.97656542 | 0.99 |
| YNL217W | 1768.453008 | 0.273321009 | 0.973401561 | 0.99 |
| YOR071C | 651.6605909 | -0.042087083 | 0.975287007 | 0.99 |
| YOR126C | 423.6858153 | 0.013015715 | 0.976259716 | 0.99 |
| YOR299W | 1060.38632 | 0.005558029 | 0.976309125 | 0.99 |
| YOR308C | 716.1626853 | 0.144964057 | 0.975658134 | 0.99 |
| YOR378W | 266.6242043 | 0.162348961 | 0.974415152 | 0.99 |
| YPL087W | 1694.761164 | 0.001532396 | 0.974940075 | 0.99 |
| YPL242C | 2035.672722 | 0.031305354 | 0.972572633 | 0.99 |
| YPR169W | 1659.207747 | 0.054943559 | 0.977548887 | 0.99 |
| snR36 | 18.91914442 | 0.032645376 | 0.972256321 | 0.99 |
| YBR040W | 134.4656267 | -0.157053176 | 0.979605566 | 1.00 |
| YER004W | 2233.162604 | -0.008702998 | 0.979497834 | 1.00 |
| YIL132C | 143.1249549 | 0.051205039 | 0.979688231 | 1.00 |
| YDR261C | 1216.328788 | 0.101948825 | 0.980102867 | 1.00 |
| YKL032C | 3133.578964 | -0.020266014 | 0.980965612 | 1.00 |
| YKR038C | 1110.135443 | -0.029336695 | 0.981002382 | 1.00 |
| YMR253C | 606.2113411 | 0.017394906 | 0.981077141 | 1.00 |
| YNL136W | 1295.54364 | -0.011110573 | 0.981021075 | 1.00 |

| | | | | |
|---|---|---|---|---|
| YNL192W | 2879.150646 | -0.025546053 | 0.980968748 | 1.00 |
| YOR211C | 1075.422511 | -0.051602518 | 0.98083659 | 1.00 |
| YAL067W-A | 0.291097411 | -0.039575613 | 0.98254169 | 1.00 |
| YAL069W | 0.918104495 | -0.08449197 | 0.982575855 | 1.00 |
| YDR276C | 2763.13579 | 0.099657638 | 0.982312169 | 1.00 |
| YGL018C | 470.8004765 | -0.024088897 | 0.982122125 | 1.00 |
| YHL005C | 24.92643688 | 0.099860683 | 0.98190413 | 1.00 |
| YKL110C | 846.8769824 | 0.012753411 | 0.982211298 | 1.00 |
| YLR236C | 21.91002832 | -0.203003183 | 0.982425571 | 1.00 |
| YOR011W | 696.3198627 | 0.165846227 | 0.982082569 | 1.00 |
| YPL020C | 1208.520388 | 0.00751208 | 0.982229004 | 1.00 |
| YDR320C-A | 373.904445 | -0.110040345 | 0.98275877 | 1.00 |
| YKL130C | 628.0827477 | 0.033562946 | 0.982914888 | 1.00 |
| YGL185C | 473.3669461 | -0.121089101 | 0.983564259 | 1.00 |
| YAR014C | 1593.739874 | 0.032190892 | 0.988776052 | 1.00 |
| YBL044W | 3.828780377 | -0.79304508 | 0.986780571 | 1.00 |
| YBR036C | 2605.132077 | -0.054929064 | 0.988891958 | 1.00 |
| YBR215W | 1081.67593 | 0.00528119 | 0.988050879 | 1.00 |
| YBR298C-A | 3.08929577 | -0.444630773 | 0.986076227 | 1.00 |
| YCR004C | 3730.565062 | -0.010698849 | 0.986455603 | 1.00 |
| YCR095C | 483.8306058 | -0.025149819 | 0.987462774 | 1.00 |
| YDL037C | 92.66763176 | -0.314840786 | 0.989548903 | 1.00 |
| YDL058W | 1998.717774 | 0.062444319 | 0.985253937 | 1.00 |
| YDR031W | 342.8572195 | -0.103796139 | 0.988622849 | 1.00 |
| YDR132C | 453.7495244 | 0.005147777 | 0.987769562 | 1.00 |
| YDR434W | 1180.144828 | -0.058499986 | 0.988608551 | 1.00 |
| YEL052W | 1984.785259 | -0.068954231 | 0.988390412 | 1.00 |
| YEL059C-A | 318.8027126 | 0.010344784 | 0.984768597 | 1.00 |
| YER080W | 3267.440935 | 0.010790794 | 0.987498093 | 1.00 |
| YFL010C | 2890.139798 | 0.009996493 | 0.985513474 | 1.00 |
| YFL012W-A | 14.51855676 | -0.006186912 | 0.987057799 | 1.00 |
| YGL169W | 682.8701947 | -0.00042668 | 0.989691164 | 1.00 |
| YGL194C | 571.5770023 | 0.017378467 | 0.984892363 | 1.00 |
| YGL246C | 646.4635665 | -0.001792 | 0.990075775 | 1.00 |
| YGR010W | 527.974638 | -0.039620045 | 0.988371788 | 1.00 |
| YGR182C | 629.4422912 | -0.02544474 | 0.989996252 | 1.00 |
| YGR231C | 3688.450165 | 0.012429945 | 0.989730002 | 1.00 |
| YGR255C | 1753.194799 | 0.022794787 | 0.985156124 | 1.00 |
| YHR001W-A | 1152.222923 | 0.059493681 | 0.989972022 | 1.00 |
| YHR209W | 694.2958547 | -0.12957502 | 0.984577262 | 1.00 |
| YIL005W | 1687.731259 | 0.036455805 | 0.985285399 | 1.00 |
| YIL023C | 838.1130985 | -0.045353208 | 0.989211009 | 1.00 |
| YIL149C | 3262.267006 | 0.06171993 | 0.987857058 | 1.00 |

| | | | | |
|---|---|---|---|---|
| YKL085W | 9612.778538 | 0.212997558 | 0.985617123 | 1.00 |
| YLR015W | 562.8104697 | 0.073017261 | 0.983859331 | 1.00 |
| YLR387C | 1423.423086 | -0.026356919 | 0.988287578 | 1.00 |
| YNL046W | 536.2842104 | -0.06989638 | 0.987176476 | 1.00 |
| YOL023W | 636.5704075 | -0.093744345 | 0.985767358 | 1.00 |
| YOL062C | 1303.409819 | -0.040586211 | 0.989785892 | 1.00 |
| YOR176W | 2270.643704 | 0.006531467 | 0.984811439 | 1.00 |
| YPL137C | 1258.265301 | -0.048197073 | 0.986089419 | 1.00 |
| YPL158C | 1350.795148 | -0.007374201 | 0.987788675 | 1.00 |
| YPR057W | 656.961666 | -0.025646045 | 0.988922431 | 1.00 |
| YPR066W | 302.1036782 | -0.016140872 | 0.986166594 | 1.00 |
| snR87 | 6.424473744 | 0.340548472 | 0.986453294 | 1.00 |
| tR(CCU)J | 0.0941958 | 0.011470282 | 0.985839765 | 1.00 |
| YNR077C | 0.24124529 | -0.019913442 | 0.990334243 | 1.00 |
| YHL010C | 843.3803368 | 0.005022127 | 0.99086692 | 1.00 |
| YNL266W | 15.42456766 | -0.140882241 | 0.99056052 | 1.00 |
| YOR258W | 218.0314211 | -0.023282874 | 0.990793024 | 1.00 |
| YNR062C | 190.9739726 | 0.003407326 | 0.991045469 | 1.00 |
| YML109W | 760.9435844 | 0.062691675 | 0.991472567 | 1.00 |
| YFR007W | 462.3779448 | 0.018202218 | 0.992059672 | 1.00 |
| YDR215C | 67.50004526 | -0.124896917 | 0.992231537 | 1.00 |
| YOR347C | 1046.083791 | -0.037668352 | 0.992547322 | 1.00 |
| YAR053W | 9.523462605 | -0.264346258 | 0.992984517 | 1.00 |
| YDR162C | 515.4142839 | -0.021869509 | 0.993618258 | 1.00 |
| YGR095C | 969.1047325 | 0.136046044 | 0.993276248 | 1.00 |
| YJL198W | 970.0448734 | 0.131368255 | 0.993633993 | 1.00 |
| YJR125C | 2124.757426 | -0.002468796 | 0.993788865 | 1.00 |
| YMR326C | 0.406623722 | 0.010116559 | 0.993015173 | 1.00 |
| YOR296W | 873.0215241 | -0.026371305 | 0.993592147 | 1.00 |
| YOR379C | 0.107512111 | 0.009556758 | 0.993767073 | 1.00 |
| YLR440C | 889.0207433 | -0.023620285 | 0.994756176 | 1.00 |
| YOR172W | 824.7713828 | 0.004023809 | 0.994497414 | 1.00 |
| YPR006C | 675.2062996 | 0.082806996 | 0.994542789 | 1.00 |
| YPR170C | 83.5281119 | -0.042650581 | 0.994716527 | 1.00 |
| YJL079C | 9145.850933 | -0.204427675 | 0.995272526 | 1.00 |
| YPR010C-A | 733.8429902 | 0.173052773 | 0.995261922 | 1.00 |
| YDR525W-A | 275.582834 | -0.138305547 | 0.996031055 | 1.00 |
| YHR023W | 2927.24121 | 0.094020616 | 0.996034288 | 1.00 |
| YPL049C | 2306.704815 | -0.034534355 | 0.995852813 | 1.00 |
| YDL182W | 2531.684902 | 0.166871415 | 0.996796629 | 1.00 |
| YGR012W | 677.5451845 | -0.013271021 | 0.996840217 | 1.00 |
| YJL127C-B | 263.7714489 | -0.022448138 | 0.997194704 | 1.00 |
| YCR088W | 5938.374856 | 0.061899663 | 0.997658363 | 1.00 |

| | | | | |
|---|---|---|---|---|
| YKL157W | 4702.190162 | 0.144851759 | 0.998128698 | 1.00 |
| YML066C | 127.9180195 | -0.046845618 | 0.998094328 | 1.00 |
| YMR255W | 1118.071772 | -0.062278017 | 0.997911325 | 1.00 |
| YPR058W | 588.0067957 | 0.325430415 | 0.998062644 | 1.00 |
| YJL133W | 754.0202192 | -0.072866872 | 0.99865147 | 1.00 |
| YEL019C | 187.0813824 | 0.084673515 | 0.998856251 | 1.00 |
| YDR211W | 2365.993285 | 0.100866662 | 0.999943028 | 1.00 |
| YDR287W | 832.7406735 | -0.01103143 | 0.999528161 | 1.00 |
| YGL038C | 1572.160144 | 0.02673265 | 0.999892611 | 1.00 |
| YHR116W | 348.981213 | -0.046977126 | 0.999904459 | 1.00 |
| YNL003C | 715.0467326 | 0.026121596 | 0.999348488 | 1.00 |
| YAL062W | 2248.194645 | 0.114552851 | NA | NA |
| YDL223C | 1796.784019 | -0.108717494 | NA | NA |
| YER024W | 2782.885664 | 0.205499858 | NA | NA |
| YER065C | 3480.324226 | 0.114212919 | NA | NA |
| YGR067C | 601.4253985 | 0.470227846 | NA | NA |
| YGR236C | 237.8810767 | -0.235321009 | NA | NA |
| YIL057C | 383.2825996 | -0.075812136 | NA | NA |
| YJR095W | 1024.664601 | 0.174146923 | NA | NA |
| YKL065W-A | 60.35011298 | -0.376455716 | NA | NA |
| YKR097W | 1618.765281 | 0.093085488 | NA | NA |
| YLR174W | 1214.876474 | 0.191085389 | NA | NA |
| YLR377C | 564.6863452 | 0.034182904 | NA | NA |
| YML042W | 2686.768376 | 0.340618872 | NA | NA |
| YMR107W | 2022.649302 | -0.147890928 | NA | NA |
| YMR175W | 966.6429644 | -0.103283147 | NA | NA |
| YMR175W-A | 25.9663659 | 0.342254501 | NA | NA |
| YNL036W | 2481.742596 | 0.144164635 | NA | NA |
| YNL117W | 3801.736785 | 0.147265676 | NA | NA |
| YNR002C | 907.6027094 | 0.17716803 | NA | NA |
| YNR071C | 317.5831201 | 0.067202011 | NA | NA |
| YOR100C | 795.4961629 | 0.657565801 | NA | NA |
| YPL223C | 915.9171589 | -0.37297679 | NA | NA |
