## Supplementary material for "A stress response that allows highly mutated eukaryotic cells to survive and proliferate": Table S5

| Gene | baseMean | log2FoldChange | pvalue | padj |
|---|---|---|---|---|
| YGL184C | 275.5662791 | 1.963922348 | 2.71E-05 | 0.12 |
| YMR048W | 313.0835867 | -0.973064939 | 3.73E-05 | 0.12 |
| YHR049C-A | 129.3206508 | -1.329935708 | 8.96E-05 | 0.15 |
| YPL063W | 2649.060944 | -1.016756287 | 8.61E-05 | 0.15 |
| YFL039C | 29152.57267 | -0.935252654 | 0.0001122 | 0.15 |
| YBR045C | 255.946774 | 1.558598716 | 0.000279808 | 0.17 |
| YFL004W | 2271.622342 | -1.034112352 | 0.000415258 | 0.17 |
| YFR057W | 37.29733796 | 0.868382206 | 0.000449845 | 0.17 |
| YGL072C | 15.1439202 | 1.320430608 | 0.000238005 | 0.17 |
| YGL165C | 78.38274117 | 1.296209608 | 0.000208732 | 0.17 |
| YGR110W | 407.9203872 | 1.358282545 | 0.000382472 | 0.17 |
| YGR269W | 48.40957868 | 1.370448215 | 0.000376946 | 0.17 |
| YLR107W | 1035.729202 | 1.144466802 | 0.000462965 | 0.17 |
| YNR017W | 1847.028777 | -1.083914212 | 0.000295732 | 0.17 |
| YNR057C | 196.7239761 | 1.414045758 | 0.000432232 | 0.17 |
| YOL131W | 75.70252965 | 2.046826341 | 0.000369204 | 0.17 |
| YOR338W | 133.0538926 | 1.426570708 | 0.000406055 | 0.17 |
| YPL111W | 2984.765722 | 1.55129108 | 0.000167547 | 0.17 |
| YDR182W-A | 24.64411227 | 1.314647133 | 0.000520003 | 0.17 |
| YLR446W | 527.2109038 | 1.270093732 | 0.000528585 | 0.17 |
| YPL234C | 1583.101568 | -0.897812403 | 0.000569534 | 0.18 |
| YBL065W | 30.86397396 | 2.168301993 | 0.000626245 | 0.19 |
| YDL133W | 835.1418998 | 0.828593807 | 0.000708032 | 0.20 |
| YJL094C | 1236.455255 | 1.106662139 | 0.000785731 | 0.21 |
| YDL139C | 356.8731646 | 1.09080914 | 0.000883438 | 0.22 |
| YGL183C | 72.4162606 | 2.308442624 | 0.000909149 | 0.22 |
| YJL153C | 833.1459195 | 1.936121327 | 0.000897792 | 0.22 |
| YJL016W | 1543.218497 | 1.005681766 | 0.001087344 | 0.22 |
| YKL213C | 1964.500982 | 0.893698626 | 0.001088678 | 0.22 |
| YNR058W | 282.8560691 | 1.238794793 | 0.00106185 | 0.22 |
| YPL095C | 828.2325169 | -1.614997297 | 0.001051509 | 0.22 |
| YPL147W | 471.9776317 | 1.619782566 | 0.001042385 | 0.22 |
| YGR120C | 376.5121986 | -0.629616107 | 0.001177165 | 0.23 |
| YGL124C | 833.4516769 | 1.00350315 | 0.00125697 | 0.24 |
| YKL005C | 1108.015733 | 0.864263537 | 0.001506619 | 0.27 |
| YNL102W | 1324.90783 | -0.966511404 | 0.00151712 | 0.27 |
| YDR034W-B | 127.2105401 | 1.270543067 | 0.001642856 | 0.29 |
| YHL004W | 1819.11881 | -0.892521477 | 0.001707965 | 0.29 |
| YER145C | 1496.957868 | -1.336947372 | 0.001848388 | 0.31 |
| YHL024W | 910.8922734 | 1.38029606 | 0.00191717 | 0.31 |

| | | | | |
|---|---|---|---|---|
| YIL072W | 537.0665215 | 1.353238103 | 0.001994469 | 0.31 |
| YJL124C | 1576.189687 | -0.996754458 | 0.002006515 | 0.31 |
| YDR010C | 32.06483583 | 0.940245994 | 0.002317805 | 0.31 |
| YHR014W | 138.7802733 | 1.484432949 | 0.002183339 | 0.31 |
| YHR049W | 2093.457795 | -0.888414701 | 0.002352908 | 0.31 |
| YKL123W | 31.46802918 | 1.092184312 | 0.002371778 | 0.31 |
| YLR154C | 335.1264893 | -0.861388667 | 0.002289111 | 0.31 |
| YLR183C | 714.9768804 | -0.840141296 | 0.002091256 | 0.31 |
| YLR273C | 457.5126435 | 1.092922527 | 0.002289626 | 0.31 |
| YLR192C | 3457.976113 | -0.941059098 | 0.002568889 | 0.33 |
| YER015W | 725.609463 | 1.280951737 | 0.002641291 | 0.34 |
| YHL036W | 545.2189465 | 1.439736233 | 0.002877497 | 0.35 |
| YML034W | 2522.763719 | 0.965426796 | 0.002805983 | 0.35 |
| YOR248W | 35.3813631 | -1.098457306 | 0.002866533 | 0.35 |
| YER068C-A | 10.30512685 | 1.07038573 | 0.003347636 | 0.39 |
| YHR138C | 1476.126547 | 1.283399279 | 0.003359373 | 0.39 |
| YOL102C | 532.3526812 | -0.841018941 | 0.003378552 | 0.39 |
| YGL180W | 843.8043135 | 1.186209051 | 0.003513909 | 0.39 |
| YDR027C | 1560.996774 | 0.586074108 | 0.003632713 | 0.39 |
| YGL204C | 78.35386912 | -1.581762542 | 0.003747899 | 0.39 |
| YGR007W | 1236.749371 | -0.851882958 | 0.00381108 | 0.39 |
| YJL081C | 1547.994145 | -0.756619968 | 0.003737271 | 0.39 |
| YKR069W | 646.4230382 | 1.010908259 | 0.003827486 | 0.39 |
| YHR121W | 1103.128777 | -0.867947276 | 0.003961062 | 0.40 |
| YDR292C | 1617.849006 | -0.9019089 | 0.004087528 | 0.41 |
| YJL015C | 72.86729941 | 0.904841613 | 0.004242772 | 0.42 |
| YDL123W | 682.3334241 | 1.071274644 | 0.004752668 | 0.45 |
| YLR291C | 1399.72151 | -0.81705061 | 0.004829021 | 0.45 |
| YMR085W | 377.7619403 | 1.144416487 | 0.004828713 | 0.45 |
| YPL218W | 2797.046919 | -0.899554473 | 0.004684168 | 0.45 |
| YKR005C | 237.9372028 | 1.136845177 | 0.005016183 | 0.46 |
| YDR285W | 510.8247622 | 1.159061312 | 0.005309123 | 0.46 |
| YEL027W | 5744.186516 | -0.914575458 | 0.005351409 | 0.46 |
| YJL133C-A | 1683.932792 | 1.119757764 | 0.005281467 | 0.46 |
| YKL182W | 10979.47839 | -0.897191982 | 0.005166194 | 0.46 |
| YDL055C | 11798.61054 | -0.775893169 | 0.005743855 | 0.47 |
| YDR446W | 60.61919013 | 1.351395525 | 0.005755591 | 0.47 |
| YER120W | 4329.489139 | -0.718937538 | 0.005661322 | 0.47 |
| YKL050C | 526.5764301 | 1.222113614 | 0.005539039 | 0.47 |
| YDR030C | 444.8820441 | 1.165871021 | 0.005998085 | 0.48 |
| YDR332W | 604.027567 | 0.772158764 | 0.006162528 | 0.48 |
| YGL176C | 596.0143813 | 0.802286262 | 0.006313215 | 0.48 |
| YKL069W | 336.0140703 | -0.953228745 | 0.005901738 | 0.48 |

| | | | | |
|---|---|---|---|---|
| YKL111C | 22.34564795 | -1.453285989 | 0.006442344 | 0.48 |
| YMR007W | 90.19118303 | 0.968665382 | 0.006393599 | 0.48 |
| YNL270C | 240.3573834 | 1.560497236 | 0.006192307 | 0.48 |
| YPR114W | 2316.846378 | -0.749281365 | 0.006271155 | 0.48 |
| YPR146C | 99.73914934 | -1.045382648 | 0.006351146 | 0.48 |
| YLR445W | 32.03943686 | 1.378881859 | 0.006691859 | 0.49 |
| YER063W | 1431.45469 | -0.931253029 | 0.006810798 | 0.49 |
| YMR292W | 429.0322087 | -0.893020935 | 0.007062538 | 0.50 |
| YBL086C | 1904.650929 | 1.002678598 | 0.007163498 | 0.51 |
| YBR263W | 3971.725103 | -0.687802958 | 0.008792106 | 0.51 |
| YBR284W | 530.3934343 | 1.150416866 | 0.008294288 | 0.51 |
| YER078W-A | 4.281084499 | 1.331949429 | 0.008596801 | 0.51 |
| YER165W | 11478.83612 | -0.848526373 | 0.008668864 | 0.51 |
| YFL060C | 16.02228195 | -1.576551451 | 0.00810104 | 0.51 |
| YGR279C | 11712.04252 | -0.791230705 | 0.008193695 | 0.51 |
| YHR139C | 204.3743283 | 0.831327783 | 0.008809749 | 0.51 |
| YJL149W | 1033.010855 | 0.857594731 | 0.008701312 | 0.51 |
| YJL165C | 2120.172359 | 1.004396656 | 0.008646608 | 0.51 |
| YKL045W | 918.6422418 | -0.778854149 | 0.008318854 | 0.51 |
| YKL162C | 452.2195831 | 0.976469757 | 0.00809805 | 0.51 |
| YKR015C | 299.9934309 | 0.742869875 | 0.008273063 | 0.51 |
| YLR219W | 2484.562083 | 1.06263225 | 0.007742412 | 0.51 |
| YLR288C | 449.0939426 | 0.657160999 | 0.007809313 | 0.51 |
| YLR342W-A | 51.47657195 | 1.03047368 | 0.007565859 | 0.51 |
| YMR021C | 731.4049333 | 0.87640292 | 0.008541826 | 0.51 |
| YOL045W | 1132.896142 | 0.690563404 | 0.007333834 | 0.51 |
| YOR005C | 988.1912317 | 0.954954892 | 0.00760512 | 0.51 |
| YOR242C | 110.2984083 | 0.90194095 | 0.008363761 | 0.51 |
| YPL018W | 471.7341201 | 1.163026223 | 0.008184664 | 0.51 |
| YBL033C | 1274.437806 | 0.779213584 | 0.01201243 | 0.52 |
| YBL106C | 800.8986155 | 0.869338906 | 0.011952354 | 0.52 |
| YBR047W | 453.8190728 | 1.469980205 | 0.009548901 | 0.52 |
| YBR055C | 925.8039594 | -0.683263453 | 0.011384134 | 0.52 |
| YBR118W | 10015.00281 | -0.645317496 | 0.012957675 | 0.52 |
| YBR127C | 10020.63379 | -0.70322156 | 0.011002187 | 0.52 |
| YCR073C | 984.1918819 | 0.761621892 | 0.011140182 | 0.52 |
| YDL077C | 1087.970908 | 0.640631891 | 0.010191533 | 0.52 |
| YDL183C | 401.7373652 | 1.051076056 | 0.010983834 | 0.52 |
| YDR250C | 6.490017442 | 0.904573272 | 0.009338705 | 0.52 |
| YDR263C | 317.4342443 | 0.942485177 | 0.012369222 | 0.52 |
| YDR308C | 370.9057891 | -0.856226489 | 0.011426636 | 0.52 |
| YER070W | 2200.790509 | -1.121875245 | 0.010228528 | 0.52 |
| YER072W | 2000.095671 | -0.918354141 | 0.013014632 | 0.52 |

| | | | | |
|---|---|---|---|---|
| YER076C | 386.5202214 | 0.690160341 | 0.012140313 | 0.52 |
| YGL008C | 26806.32689 | -0.665265412 | 0.011171937 | 0.52 |
| YGL030W | 7624.667323 | -0.62773416 | 0.009392662 | 0.52 |
| YGL112C | 1298.501075 | -0.608674424 | 0.012573772 | 0.52 |
| YGL243W | 434.8625344 | -0.614350967 | 0.012058568 | 0.52 |
| YGR146C | 1344.611896 | 1.09548953 | 0.011709929 | 0.52 |
| YHL037C | 20.29589679 | 1.585209459 | 0.011424638 | 0.52 |
| YHR005C-A | 2208.832297 | -0.928085452 | 0.011919311 | 0.52 |
| YHR106W | 1096.925363 | 1.078871514 | 0.011915222 | 0.52 |
| YHR110W | 1458.33201 | -0.752487927 | 0.011342943 | 0.52 |
| YHR154W | 1331.473724 | -0.877041632 | 0.010059011 | 0.52 |
| YIL071W-A | 73.5379669 | 0.750399286 | 0.013000379 | 0.52 |
| YIL073C | 530.2219401 | 0.684707939 | 0.009841152 | 0.52 |
| YIL146C | 613.496777 | 1.063876475 | 0.010351053 | 0.52 |
| YIL162W | 3263.600398 | 1.322881463 | 0.012912005 | 0.52 |
| YIR036W-A | 10.01589679 | 0.845460528 | 0.011771285 | 0.52 |
| YJR150C | 147.0716932 | 0.906632553 | 0.011669432 | 0.52 |
| YKL086W | 218.7788586 | 1.231879311 | 0.009605165 | 0.52 |
| YKL124W | 623.7006372 | 0.899307374 | 0.009672611 | 0.52 |
| YLL014W | 750.2093462 | -0.895434062 | 0.012855395 | 0.52 |
| YLR033W | 1049.258607 | -0.762499342 | 0.010346712 | 0.52 |
| YLR263W | 707.0718116 | 1.243323603 | 0.010603179 | 0.52 |
| YLR297W | 487.8704755 | 0.870843752 | 0.01256199 | 0.52 |
| YLR378C | 3906.764345 | -0.784633564 | 0.012349757 | 0.52 |
| YLR416C | 28.8196825 | 1.241551347 | 0.009610914 | 0.52 |
| YMR034C | 377.1167506 | 1.364126613 | 0.010637174 | 0.52 |
| YMR103C | 375.8522499 | 1.366255614 | 0.013113673 | 0.52 |
| YMR105W-A | 43.59841519 | 0.935787412 | 0.011288498 | 0.52 |
| YMR182C | 564.4635256 | 0.940510065 | 0.013034424 | 0.52 |
| YNL016W | 4774.57143 | -0.773141031 | 0.012785934 | 0.52 |
| YNL269W | 50.67416069 | 1.482138548 | 0.009297599 | 0.52 |
| YOR278W | 477.7903224 | -0.861082081 | 0.012298907 | 0.52 |
| YOR286W | 1162.614528 | -0.744341565 | 0.010706068 | 0.52 |
| YOR292C | 645.9873627 | 1.070182444 | 0.009757951 | 0.52 |
| YPL129W | 1819.666295 | -0.823091648 | 0.010699658 | 0.52 |
| YPL230W | 2214.598115 | 1.297680087 | 0.011522157 | 0.52 |
| YPR016C | 1938.157861 | -0.979784576 | 0.012642909 | 0.52 |
| snR191 | 16.29529748 | -0.969543773 | 0.009259423 | 0.52 |
| YLR354C | 11943.28845 | -0.694594736 | 0.013404135 | 0.53 |
| YOL161C | 3.517453047 | 0.769841689 | 0.013476153 | 0.53 |
| YCR027C | 255.2084978 | -0.946984449 | 0.013941747 | 0.53 |
| YDL020C | 3590.678769 | 1.033001623 | 0.013975104 | 0.53 |
| YIL046W | 1246.958862 | 0.933512941 | 0.013777568 | 0.53 |

| | | | | |
|---|---|---|---|---|
| YJL206C | 539.1174838 | 0.882986081 | 0.013958371 | 0.53 |
| YKL112W | 917.1897974 | -0.868946379 | 0.01377576 | 0.53 |
| YPR084W | 689.7868138 | 0.534888662 | 0.013664138 | 0.53 |
| YOL058W | 2645.481694 | 1.31982233 | 0.01433862 | 0.54 |
| YER137C | 65.6651259 | 1.168452237 | 0.01464791 | 0.55 |
| YAR033W | 105.2011719 | -0.880233221 | 0.018014688 | 0.55 |
| YBL091C | 4543.955358 | -0.581574436 | 0.016933289 | 0.55 |
| YBR146W | 1432.145995 | -0.731946911 | 0.018306637 | 0.55 |
| YBR187W | 950.8054053 | -0.569697164 | 0.017044172 | 0.55 |
| YBR233W-A | 253.0169158 | -1.068435137 | 0.015982255 | 0.55 |
| YBR255W | 917.9047892 | 0.802405775 | 0.018737945 | 0.55 |
| YBR265W | 1130.650771 | -0.80650664 | 0.015866046 | 0.55 |
| YDL234C | 2388.462086 | 1.043431713 | 0.018830844 | 0.55 |
| YDR017C | 2063.432291 | 0.569931531 | 0.01699933 | 0.55 |
| YDR018C | 446.8147304 | 1.10768882 | 0.01621674 | 0.55 |
| YDR183C-A | 47.62983085 | -0.757492396 | 0.015640514 | 0.55 |
| YDR421W | 1031.149921 | 0.840683826 | 0.017394428 | 0.55 |
| YDR483W | 4802.755253 | -0.62503789 | 0.017902094 | 0.55 |
| YER054C | 1167.402398 | 1.280512642 | 0.016574088 | 0.55 |
| YER081W | 9451.152045 | 1.226870794 | 0.01870667 | 0.55 |
| YER154W | 1666.340326 | -0.621811694 | 0.016215785 | 0.55 |
| YFL057C | 160.9643278 | 0.831397718 | 0.014881566 | 0.55 |
| YGL059W | 1140.532396 | 0.980817919 | 0.014850221 | 0.55 |
| YGL225W | 1789.038412 | -0.667759982 | 0.015715832 | 0.55 |
| YGR106C | 3546.453004 | -0.721354085 | 0.016986492 | 0.55 |
| YGR108W | 622.4932134 | -1.105252821 | 0.016787788 | 0.55 |
| YGR155W | 6507.68049 | -0.790403555 | 0.017885997 | 0.55 |
| YHR153C | 73.13873436 | -0.909511972 | 0.018112836 | 0.55 |
| YIL035C | 1690.415511 | -0.694047357 | 0.015192744 | 0.55 |
| YIL050W | 1083.890647 | 1.001556995 | 0.015768174 | 0.55 |
| YIR017C | 6.4006803 | 0.806919239 | 0.017208543 | 0.55 |
| YIR041W | 5.637300496 | 0.79550664 | 0.017363807 | 0.55 |
| YKL080W | 5080.565803 | -0.812324993 | 0.015412578 | 0.55 |
| YKL174C | 903.8221117 | 0.580658362 | 0.017868335 | 0.55 |
| YKL188C | 680.4138758 | 0.943959503 | 0.01645854 | 0.55 |
| YKL202W | 21.88607995 | 1.180568659 | 0.016359981 | 0.55 |
| YKR074W | 1111.118511 | -0.820248389 | 0.01633932 | 0.55 |
| YLR116W | 1156.147687 | 0.622727782 | 0.015509471 | 0.55 |
| YLR176C | 916.0927001 | 0.879299674 | 0.01530915 | 0.55 |
| YLR229C | 1318.158779 | -0.741093666 | 0.017459377 | 0.55 |
| YLR372W | 2301.005993 | -0.602769229 | 0.018333484 | 0.55 |
| YML019W | 694.8842248 | -0.758334548 | 0.016463531 | 0.55 |
| YMR079W | 1633.407315 | -0.791362396 | 0.018193321 | 0.55 |

| | | | | |
|---|---|---|---|---|
| YMR125W | 2280.761082 | -0.838147348 | 0.018089108 | 0.55 |
| YNL206C | 942.6169972 | -0.729641769 | 0.01784369 | 0.55 |
| YNL295W | 664.526791 | 0.79353966 | 0.018767202 | 0.55 |
| YNL328C | 68.99382563 | -0.92767471 | 0.015918238 | 0.55 |
| YOL012C | 810.102208 | -0.757629589 | 0.018633573 | 0.55 |
| YOL120C | 9488.699457 | -0.399906516 | 0.016288882 | 0.55 |
| YOR003W | 499.5835225 | 0.876296175 | 0.016790071 | 0.55 |
| YPL164C | 886.5830673 | 0.950786747 | 0.018074411 | 0.55 |
| YPL264C | 197.1729292 | 0.820323292 | 0.018656444 | 0.55 |
| YPR165W | 4589.151851 | -0.784217242 | 0.016104272 | 0.55 |
| snR49 | 77.03202196 | -0.305179704 | 0.018618805 | 0.55 |
| YDL036C | 422.8116952 | -0.856833152 | 0.019443592 | 0.55 |
| YDL054C | 853.6605887 | 1.001092959 | 0.019791029 | 0.55 |
| YEL035C | 12.59177912 | 1.059748952 | 0.019557118 | 0.55 |
| YER091C | 3130.444513 | 1.085068009 | 0.020080245 | 0.55 |
| YER162C | 931.7095211 | 0.889553995 | 0.020287411 | 0.55 |
| YGL125W | 958.6893776 | 1.037652325 | 0.019700169 | 0.55 |
| YHL021C | 7761.479525 | 1.362331699 | 0.01918322 | 0.55 |
| YKL074C | 747.5415477 | 0.686074525 | 0.019998895 | 0.55 |
| YLR002C | 907.6259007 | -0.802739472 | 0.020392809 | 0.55 |
| YLR335W | 2838.169755 | -0.543760133 | 0.019765233 | 0.55 |
| YMR137C | 598.7111524 | 0.855833317 | 0.019551889 | 0.55 |
| YMR181C | 1531.918567 | 0.923118631 | 0.020317136 | 0.55 |
| YNL196C | 204.2397635 | 1.344564854 | 0.020265895 | 0.55 |
| YOL084W | 492.0065357 | 0.96580213 | 0.019960276 | 0.55 |
| YOR004W | 494.9946687 | -0.443549007 | 0.019638541 | 0.55 |
| YOR331C | 254.0815524 | 0.731751587 | 0.019885387 | 0.55 |
| YBR070C | 438.6999546 | -0.73168849 | 0.024840136 | 0.56 |
| YCR005C | 6272.799933 | 0.993993241 | 0.021789205 | 0.56 |
| YCR038C | 411.3006768 | 0.704926963 | 0.022627918 | 0.56 |
| YDL017W | 421.2237274 | 0.686429259 | 0.021528423 | 0.56 |
| YDL079C | 1497.675817 | 0.924283813 | 0.025081238 | 0.56 |
| YDL194W | 1083.690954 | 1.141816032 | 0.022920218 | 0.56 |
| YDL214C | 464.7760152 | 0.937453068 | 0.021039716 | 0.56 |
| YDL226C | 1711.631594 | -0.911783617 | 0.022623895 | 0.56 |
| YDR059C | 828.9203049 | 1.159862293 | 0.021308968 | 0.56 |
| YDR273W | 249.707739 | 0.91344579 | 0.024620641 | 0.56 |
| YDR277C | 2772.925083 | 0.961596639 | 0.022681087 | 0.56 |
| YDR538W | 133.1462476 | -0.776518876 | 0.021993895 | 0.56 |
| YER043C | 7073.679738 | -0.683123287 | 0.024423785 | 0.56 |
| YER096W | 439.4636005 | 1.161931074 | 0.023058811 | 0.56 |
| YFR017C | 3258.918594 | 1.241287211 | 0.023509161 | 0.56 |
| YGR131W | 100.6509022 | 0.77157334 | 0.021727782 | 0.56 |

| | | | | |
|---|---|---|---|---|
| YHL002C-A | 8.009981089 | 1.096776842 | 0.023364238 | 0.56 |
| YHL011C | 1690.318772 | -0.711985687 | 0.023094476 | 0.56 |
| YHR021W-A | 41.37134005 | 0.536998761 | 0.024483559 | 0.56 |
| YHR033W | 1072.035586 | 0.982345023 | 0.020547716 | 0.56 |
| YHR096C | 1194.376628 | 0.713390205 | 0.021899732 | 0.56 |
| YIL118W | 1012.394304 | -0.938433793 | 0.022903707 | 0.56 |
| YIL139C | 280.0724876 | -0.744493937 | 0.021737659 | 0.56 |
| YIR021W-A | 4.66148073 | -1.103932744 | 0.023807353 | 0.56 |
| YJL103C | 874.1846601 | 0.996299403 | 0.02498691 | 0.56 |
| YJL106W | 294.2742704 | 0.906719236 | 0.022950415 | 0.56 |
| YJR025C | 1368.606225 | -1.104826313 | 0.021194483 | 0.56 |
| YJR061W | 732.5988419 | 0.82369247 | 0.022810589 | 0.56 |
| YJR155W | 885.0501737 | 0.850055357 | 0.022198641 | 0.56 |
| YKL081W | 14463.37631 | -0.573749262 | 0.024640278 | 0.56 |
| YKL129C | 2233.401318 | 0.847438035 | 0.023322788 | 0.56 |
| YKL147C | 17.64954721 | 0.894028384 | 0.024326842 | 0.56 |
| YKL184W | 1354.204344 | -0.840328845 | 0.02317577 | 0.56 |
| YKR059W | 1551.844916 | -0.679767203 | 0.020861391 | 0.56 |
| YLR012C | 11.76101632 | 1.555710845 | 0.0212614 | 0.56 |
| YLR195C | 1211.827498 | -0.710749974 | 0.022445292 | 0.56 |
| YLR205C | 412.8411079 | 0.825980819 | 0.023585917 | 0.56 |
| YLR268W | 967.406712 | -0.695892092 | 0.022943745 | 0.56 |
| YLR415C | 24.41748512 | 1.193430261 | 0.023231801 | 0.56 |
| YMR141C | 167.2172498 | 0.910583157 | 0.02486311 | 0.56 |
| YMR202W | 2009.003075 | -0.450347909 | 0.023310016 | 0.56 |
| YNL195C | 538.1405229 | 0.787834617 | 0.024325169 | 0.56 |
| YOL002C | 621.7558878 | -0.990178766 | 0.025103171 | 0.56 |
| YOL119C | 1253.305663 | 1.056747233 | 0.023758472 | 0.56 |
| YOL130W | 1532.382846 | 0.650702282 | 0.023415684 | 0.56 |
| YOR018W | 939.5138966 | 0.691257057 | 0.023107788 | 0.56 |
| YOR164C | 1461.403289 | -0.675983299 | 0.024918111 | 0.56 |
| YOR210W | 598.8398742 | -0.790376888 | 0.022779257 | 0.56 |
| YPL032C | 2772.776631 | -0.796634529 | 0.024957042 | 0.56 |
| YPL150W | 1067.29309 | 0.832316078 | 0.024793129 | 0.56 |
| YPR030W | 2156.768509 | 1.289277662 | 0.025124112 | 0.56 |
| YPR061C | 393.0462601 | 0.96556077 | 0.024324574 | 0.56 |
| YPR092W | 28.34006108 | 0.823242193 | 0.021769911 | 0.56 |
| YPR181C | 4442.448271 | -0.651308102 | 0.024798003 | 0.56 |
| snR3 | 32.48912912 | -0.598279999 | 0.020858205 | 0.56 |
| YDR320W-B | 5.22193975 | 1.002452783 | 0.025461383 | 0.56 |
| YGL015C | 22.18286877 | 0.893610526 | 0.025567637 | 0.56 |
| YLR046C | 345.7378309 | 0.544202379 | 0.0256706 | 0.56 |
| YLR134W | 556.7123022 | 0.880621923 | 0.025862362 | 0.56 |

| | | | | |
|---|---|---|---|---|
| YAR019C | 904.0673159 | 0.828608131 | 0.026257631 | 0.56 |
| YDR302W | 389.0867763 | -0.784084906 | 0.025984607 | 0.56 |
| YJR115W | 567.164948 | 0.529664273 | 0.026270116 | 0.56 |
| YOR161C | 928.1721785 | 0.831646262 | 0.026300081 | 0.56 |
| YPL112C | 806.8455711 | -0.809093745 | 0.026319952 | 0.56 |
| YOR315W | 1158.774212 | -1.071540362 | 0.026472045 | 0.57 |
| YAL023C | 4478.973811 | -0.642237008 | 0.03240686 | 0.57 |
| YAL032C | 524.818116 | 0.577261384 | 0.031661898 | 0.57 |
| YAL034C | 1899.332638 | 0.899659678 | 0.027793554 | 0.57 |
| YBL026W | 682.7010545 | -0.599649817 | 0.031326191 | 0.57 |
| YBR027C | 8.649310737 | 0.538129414 | 0.027926538 | 0.57 |
| YBR088C | 178.1843606 | -0.773222678 | 0.029994667 | 0.57 |
| YBR105C | 2095.906659 | 0.888535542 | 0.026852318 | 0.57 |
| YBR114W | 883.2430403 | 0.824792966 | 0.031855615 | 0.57 |
| YBR191W | 11850.59787 | -0.37336806 | 0.031879087 | 0.57 |
| YBR221C | 5794.723277 | -0.659371827 | 0.028981976 | 0.57 |
| YBR245C | 1580.844401 | -0.556839729 | 0.032648331 | 0.57 |
| YDL083C | 6219.514303 | -0.495450301 | 0.02706537 | 0.57 |
| YDL110C | 2473.385616 | 1.05549812 | 0.030452646 | 0.57 |
| YDL212W | 1994.74877 | -0.734115955 | 0.026989175 | 0.57 |
| YDR003W-A | 67.78719135 | 0.781308702 | 0.027696909 | 0.57 |
| YDR004W | 628.5806399 | 0.579346499 | 0.031172019 | 0.57 |
| YDR007W | 490.6292629 | -0.63086409 | 0.031419419 | 0.57 |
| YDR064W | 9582.184184 | -0.339483591 | 0.031996131 | 0.57 |
| YDR084C | 664.1695601 | -0.608063987 | 0.029184443 | 0.57 |
| YDR321W | 1332.488079 | -0.52187164 | 0.029212663 | 0.57 |
| YDR331W | 571.3069512 | -0.766625609 | 0.032487698 | 0.57 |
| YDR374C | 71.50374083 | 1.04393999 | 0.02711084 | 0.57 |
| YDR382W | 7559.11107 | -0.688966535 | 0.032289224 | 0.57 |
| YDR425W | 1030.779588 | 0.817492898 | 0.032676938 | 0.57 |
| YDR454C | 2624.226849 | -0.626144826 | 0.029199041 | 0.57 |
| YEL002C | 3293.830024 | -0.625024688 | 0.028219831 | 0.57 |
| YER146W | 140.8937845 | -0.671570154 | 0.032761463 | 0.57 |
| YGL036W | 2417.923989 | 0.701968221 | 0.029038958 | 0.57 |
| YGL081W | 335.8828397 | 1.256147837 | 0.031752556 | 0.57 |
| YGL128C | 505.7413235 | 0.756371571 | 0.028586769 | 0.57 |
| YGL147C | 5508.177604 | -0.2931178 | 0.029433306 | 0.57 |
| YGL195W | 6234.838225 | -0.650615046 | 0.028831099 | 0.57 |
| YGL198W | 494.0719347 | 0.96043544 | 0.031986193 | 0.57 |
| YGR034W | 5016.73135 | -0.547425636 | 0.028984656 | 0.57 |
| YGR172C | 884.3306623 | -0.682803184 | 0.031096069 | 0.57 |
| YHL012W | 237.0696554 | 0.588457489 | 0.030672995 | 0.57 |
| YIL099W | 500.7923991 | 1.096675685 | 0.029670341 | 0.57 |

| | | | | |
|---|---|---|---|---|
| YJL141C | 4385.516839 | 1.071897212 | 0.028830459 | 0.57 |
| YJR014W | 1708.927082 | -0.623154276 | 0.032248249 | 0.57 |
| YJR036C | 438.6482321 | 1.041005027 | 0.031124495 | 0.57 |
| YJR118C | 868.4456577 | -0.784694605 | 0.030179623 | 0.57 |
| YKL006W | 2404.725348 | -0.35950964 | 0.02761306 | 0.57 |
| YKR075C | 2465.019565 | 0.739242961 | 0.031587757 | 0.57 |
| YKR085C | 837.2824572 | -0.72505533 | 0.030741547 | 0.57 |
| YLR008C | 578.2897755 | -0.756298321 | 0.031342185 | 0.57 |
| YLR083C | 1368.675892 | -0.852825799 | 0.027484512 | 0.57 |
| YLR142W | 1292.784521 | 0.65684055 | 0.030520926 | 0.57 |
| YLR177W | 3573.62477 | 0.999161118 | 0.029676088 | 0.57 |
| YLR247C | 1785.222725 | 0.683836414 | 0.029625856 | 0.57 |
| YLR301W | 1464.12998 | -0.562880187 | 0.03226899 | 0.57 |
| YLR324W | 1737.806437 | 0.913594904 | 0.031268716 | 0.57 |
| YML023C | 695.7405584 | 0.568101192 | 0.029997941 | 0.57 |
| YML067C | 1114.441541 | -0.563248957 | 0.030664394 | 0.57 |
| YMR104C | 1639.396102 | 1.298098632 | 0.027553727 | 0.57 |
| YNL223W | 533.8185201 | 0.770294597 | 0.032182842 | 0.57 |
| YNL290W | 854.0353401 | -0.709760416 | 0.028504983 | 0.57 |
| YOL097W-A | 14.8332561 | -1.307272457 | 0.03152344 | 0.57 |
| YOR074C | 848.568103 | -0.676010694 | 0.03278531 | 0.57 |
| YOR095C | 614.382855 | -0.72328075 | 0.030817334 | 0.57 |
| YOR165W | 1637.432086 | -0.658773312 | 0.031058499 | 0.57 |
| YOR200W | 9.969440735 | 0.686231072 | 0.029285015 | 0.57 |
| YOR293W | 11411.33478 | -0.438242169 | 0.032286013 | 0.57 |
| YPL038W-A | 14.34188488 | 0.895981829 | 0.027067949 | 0.57 |
| YPL178W | 1119.186631 | -0.546331031 | 0.031761913 | 0.57 |
| YPL202C | 824.3595251 | 0.588441931 | 0.032106486 | 0.57 |
| YPL219W | 632.4863798 | 0.764277639 | 0.028776324 | 0.57 |
| YPL247C | 2827.171119 | 1.057391707 | 0.031347073 | 0.57 |
| YPR059C | 12.62828581 | -0.357240634 | 0.028946618 | 0.57 |
| YPR106W | 760.0851492 | 0.666035137 | 0.02679397 | 0.57 |
| YPR147C | 688.0952487 | -0.676663757 | 0.02794089 | 0.57 |
| YPR151C | 43.40930441 | 0.870128882 | 0.030276095 | 0.57 |
| YPR187W | 1425.65006 | -0.858425242 | 0.030756929 | 0.57 |
| YLL045C | 7512.686429 | -0.579390705 | 0.03287939 | 0.57 |
| YBR222C | 4562.01386 | 1.13693955 | 0.033097313 | 0.57 |
| YDR284C | 1216.7009 | -0.460720445 | 0.033435363 | 0.57 |
| YEL054C | 3213.78471 | -0.494173586 | 0.033157657 | 0.57 |
| YGR234W | 5131.339655 | -0.668947801 | 0.033217823 | 0.57 |
| YLL059C | 81.93358505 | 0.856290264 | 0.033263207 | 0.57 |
| YNL146W | 85.60940575 | 0.803425784 | 0.033592196 | 0.57 |
| YNL311C | 782.0547648 | 0.768585768 | 0.033511074 | 0.57 |

| | | | | |
|---|---|---|---|---|
| YPL185W | 39.28495564 | 1.0034197 | 0.033490523 | 0.57 |
| YDL084W | 7170.004042 | -0.676067054 | 0.033743106 | 0.57 |
| YDL081C | 10855.42483 | -0.551643609 | 0.034124128 | 0.57 |
| YFR035C | 184.0960805 | -0.553377893 | 0.034300815 | 0.57 |
| YHR026W | 2431.689374 | -0.692041128 | 0.034069036 | 0.57 |
| YKL165C | 1154.053703 | -0.711898695 | 0.034169915 | 0.57 |
| YMR024W | 882.2534561 | -0.59840637 | 0.034215755 | 0.57 |
| YPL248C | 793.6864415 | 0.878348808 | 0.033893137 | 0.57 |
| YBL028C | 410.9135571 | -0.619288952 | 0.036627933 | 0.57 |
| YBR189W | 11760.64407 | -0.378442406 | 0.037471247 | 0.57 |
| YBR243C | 997.8241569 | -0.652219288 | 0.037304628 | 0.57 |
| YDL193W | 1212.895326 | 0.654973955 | 0.034933413 | 0.57 |
| YDR085C | 1259.372351 | 0.818323246 | 0.0373925 | 0.57 |
| YDR306C | 1159.558737 | 0.709656027 | 0.037327638 | 0.57 |
| YER036C | 4277.526152 | -0.696696573 | 0.037685342 | 0.57 |
| YER067W | 2559.902963 | 1.306024224 | 0.035697328 | 0.57 |
| YER087C-B | 1311.571506 | -0.570019372 | 0.035410658 | 0.57 |
| YER188C-A | 23.15774642 | 0.582651957 | 0.036104813 | 0.57 |
| YGL104C | 947.9922368 | 0.831703186 | 0.035803034 | 0.57 |
| YGL150C | 2988.328671 | 0.577562418 | 0.037212725 | 0.57 |
| YGR225W | 197.8168855 | 0.89937592 | 0.036024745 | 0.57 |
| YIL021W | 814.8146206 | -0.645507342 | 0.037732594 | 0.57 |
| YIL055C | 666.1215044 | 0.788041532 | 0.037419521 | 0.57 |
| YIR013C | 17.07044727 | 0.893358653 | 0.03745112 | 0.57 |
| YJL074C | 945.199767 | -0.596777625 | 0.037221656 | 0.57 |
| YJL203W | 345.939173 | -0.602083373 | 0.037300972 | 0.57 |
| YJL218W | 146.9624687 | -0.873267668 | 0.035259374 | 0.57 |
| YJR091C | 3359.251305 | 0.831715108 | 0.034509628 | 0.57 |
| YLR037C | 167.7921303 | 0.807268083 | 0.034988611 | 0.57 |
| YLR094C | 1048.349221 | 0.78160222 | 0.035932473 | 0.57 |
| YLR126C | 468.0585451 | -0.952876017 | 0.03639669 | 0.57 |
| YLR189C | 2093.552884 | 0.821469373 | 0.037561951 | 0.57 |
| YML100W | 14497.66976 | 1.314513447 | 0.034498206 | 0.57 |
| YML106W | 2946.051592 | -0.741123292 | 0.037452867 | 0.57 |
| YMR318C | 2412.871237 | -0.591425477 | 0.036894115 | 0.57 |
| YNL125C | 2090.525445 | 0.842671857 | 0.037376825 | 0.57 |
| YNL332W | 25.85198322 | 0.560479291 | 0.036082879 | 0.57 |
| YNR033W | 1803.332991 | 0.777676819 | 0.034788478 | 0.57 |
| YOL091W | 152.6270626 | 0.619348508 | 0.036990393 | 0.57 |
| YOR120W | 2918.156498 | 1.116503824 | 0.036744057 | 0.57 |
| YOR143C | 346.0541178 | -0.732074921 | 0.037151915 | 0.57 |
| YOR190W | 351.1788756 | 0.710510835 | 0.037402515 | 0.57 |
| YPL119C | 673.8655489 | 1.10651065 | 0.035599393 | 0.57 |

| | | | | |
|---|---|---|---|---|
| YPR005C | 426.8209175 | 0.759619031 | 0.035883606 | 0.57 |
| YPR007C | 168.1873056 | 1.052884446 | 0.034865397 | 0.57 |
| YPR064W | 39.20156943 | 0.832050956 | 0.036925375 | 0.57 |
| snR128 | 13.17491061 | -0.21252035 | 0.037378136 | 0.57 |
| YJR101W | 534.1436387 | -0.746226435 | 0.03787264 | 0.57 |
| YBR203W | 994.1971392 | 1.111145943 | 0.038108493 | 0.57 |
| YFR036W-A | 15.15766896 | -0.556468794 | 0.038197941 | 0.57 |
| YOR039W | 1072.032394 | -0.655772108 | 0.038199148 | 0.57 |
| TLC1 | 1417.425959 | -0.589939534 | 0.06038776 | 0.57 |
| YAL010C | 552.0969494 | 0.5243326 | 0.049436364 | 0.57 |
| YAL020C | 521.9507996 | 0.74291898 | 0.053588179 | 0.57 |
| YAL028W | 836.6833612 | 0.784147943 | 0.04046244 | 0.57 |
| YAL037C-B | 133.1022568 | -0.662357917 | 0.053441964 | 0.57 |
| YAL039C | 1453.711163 | 0.770192163 | 0.05304091 | 0.57 |
| YAL040C | 2404.494026 | 0.772895961 | 0.047802257 | 0.57 |
| YAR019W-A | 5.564052438 | 0.113179544 | 0.050162319 | 0.57 |
| YAR020C | 43.79066228 | 0.968286809 | 0.049071431 | 0.57 |
| YBL027W | 10124.8249 | -0.468003521 | 0.055161625 | 0.57 |
| YBL035C | 596.37002 | -0.699639437 | 0.053756359 | 0.57 |
| YBL080C | 486.2802333 | 0.706704451 | 0.05391425 | 0.57 |
| YBL092W | 15431.27049 | -0.286746628 | 0.054431197 | 0.57 |
| YBR048W | 5960.519296 | -0.39410464 | 0.051116641 | 0.57 |
| YBR085C-A | 1789.206511 | 1.080478 | 0.041011233 | 0.57 |
| YBR108W | 1821.037124 | 0.797782022 | 0.042369523 | 0.57 |
| YBR121C | 7616.904588 | -0.495844293 | 0.059203088 | 0.57 |
| YBR159W | 2299.591935 | -0.726094874 | 0.054252056 | 0.57 |
| YBR261C | 744.8069304 | -0.533819938 | 0.056553028 | 0.57 |
| YBR280C | 1033.06933 | 0.892940136 | 0.055629773 | 0.57 |
| YBR283C | 3928.577615 | -0.557435483 | 0.059419872 | 0.57 |
| YCL026C-B | 269.8485152 | 1.290182153 | 0.044403078 | 0.57 |
| YCL037C | 1123.648124 | -0.728702378 | 0.052259263 | 0.57 |
| YCR021C | 10522.50846 | 1.125158092 | 0.052663774 | 0.57 |
| YCR028C-A | 1716.279601 | -0.561994768 | 0.057297164 | 0.57 |
| YDL040C | 3238.410502 | -0.560160399 | 0.055524312 | 0.57 |
| YDL149W | 944.7875532 | 0.920678571 | 0.039457264 | 0.57 |
| YDL164C | 704.8057939 | -0.581712836 | 0.058388577 | 0.57 |
| YDL169C | 664.258116 | 0.902112397 | 0.057981233 | 0.57 |
| YDL179W | 455.9668158 | -0.697458825 | 0.038481089 | 0.57 |
| YDL199C | 559.7723367 | 0.910981026 | 0.040988235 | 0.57 |
| YDL208W | 1913.168285 | -0.640272162 | 0.053507606 | 0.57 |
| YDL219W | 330.3585764 | -0.72335198 | 0.045821664 | 0.57 |
| YDL220C | 416.2502353 | -0.551427195 | 0.058676718 | 0.57 |
| YDL227C | 308.6751335 | -1.053230032 | 0.057797902 | 0.57 |

| | | | | |
|---|---|---|---|---|
| YDL228C | 55.92892077 | -0.66802167 | 0.039044228 | 0.57 |
| YDR014W-A | 36.16776295 | 1.319840929 | 0.053822707 | 0.57 |
| YDR042C | 81.10570396 | 0.978488773 | 0.059959654 | 0.57 |
| YDR043C | 825.8596949 | 0.561934713 | 0.041874323 | 0.57 |
| YDR074W | 10482.83127 | 1.253135166 | 0.044071749 | 0.57 |
| YDR076W | 307.3025521 | 0.728526555 | 0.053079242 | 0.57 |
| YDR096W | 1946.105843 | 0.972221212 | 0.056359 | 0.57 |
| YDR113C | 648.1289584 | -0.63140494 | 0.052103714 | 0.57 |
| YDR164C | 1143.654091 | -0.50544176 | 0.050193504 | 0.57 |
| YDR172W | 3599.759088 | -0.611536807 | 0.052869601 | 0.57 |
| YDR173C | 369.7980333 | 0.669797792 | 0.059421334 | 0.57 |
| YDR223W | 358.2338356 | 0.711886682 | 0.046834641 | 0.57 |
| YDR367W | 606.3521729 | -0.642446769 | 0.059703569 | 0.57 |
| YDR398W | 1282.093858 | -0.5864621 | 0.05382047 | 0.57 |
| YDR412W | 170.5135109 | -0.597071775 | 0.057340963 | 0.57 |
| YDR436W | 1255.862426 | 0.789943276 | 0.046496402 | 0.57 |
| YDR463W | 1963.287278 | 0.749512515 | 0.043918411 | 0.57 |
| YEL009C-A | 8.998175135 | 0.832681559 | 0.038930371 | 0.57 |
| YEL012W | 1045.847779 | 0.919458418 | 0.057846157 | 0.57 |
| YEL020C | 1045.827861 | 0.942714121 | 0.042912959 | 0.57 |
| YEL026W | 1802.083858 | -0.515530806 | 0.044700777 | 0.57 |
| YEL030W | 524.4916329 | 0.79884071 | 0.044504647 | 0.57 |
| YEL053C | 1328.366975 | 0.457497949 | 0.055867706 | 0.57 |
| YER003C | 3520.842344 | -0.447404693 | 0.058730471 | 0.57 |
| YER025W | 3287.005691 | -0.638924534 | 0.056982726 | 0.57 |
| YER044C-A | 102.3682402 | 1.080017647 | 0.053837721 | 0.57 |
| YER066W | 204.0645964 | 0.718222465 | 0.043225354 | 0.57 |
| YER093C | 1449.072568 | 0.60834064 | 0.03960192 | 0.57 |
| YER101C | 634.792482 | 0.757481573 | 0.054382602 | 0.57 |
| YER117W | 4528.496655 | -0.422943198 | 0.048624289 | 0.57 |
| YER131W | 9395.024759 | -0.351741508 | 0.048177413 | 0.57 |
| YER143W | 1597.506506 | 0.774355514 | 0.042185694 | 0.57 |
| YER169W | 1395.485471 | 0.642140814 | 0.051223999 | 0.57 |
| YER183C | 378.9572978 | -0.731682622 | 0.039814234 | 0.57 |
| YFL045C | 5242.345974 | -0.618065594 | 0.050160913 | 0.57 |
| YFL054C | 1511.844206 | 0.886043802 | 0.059938733 | 0.57 |
| YFR015C | 6189.813754 | 0.988208643 | 0.053429566 | 0.57 |
| YFR028C | 1133.072845 | -0.593930483 | 0.050346249 | 0.57 |
| YFR031C | 1205.359626 | 0.556290966 | 0.057561889 | 0.57 |
| YFR031C-A | 6438.897656 | -0.274964948 | 0.055548405 | 0.57 |
| YGL006W | 3158.91505 | 0.727193157 | 0.055063975 | 0.57 |
| YGL010W | 399.2167766 | 0.742612417 | 0.060447193 | 0.57 |
| YGL035C | 1849.649908 | 0.643571129 | 0.04116368 | 0.57 |

| | | | | |
|---|---|---|---|---|
| YGL070C | 462.601344 | -0.700900281 | 0.039936099 | 0.57 |
| YGL076C | 4229.319259 | -0.348313748 | 0.052905736 | 0.57 |
| YGL100W | 1500.808318 | -0.534811139 | 0.041808948 | 0.57 |
| YGL123W | 15975.90289 | -0.568269677 | 0.04410758 | 0.57 |
| YGL133W | 1877.257774 | 0.647135658 | 0.050131474 | 0.57 |
| YGL141W | 1274.03239 | 0.609472507 | 0.048131087 | 0.57 |
| YGL166W | 257.6413584 | 0.723504091 | 0.048058802 | 0.57 |
| YGR001C | 885.4111524 | -0.364292285 | 0.045501005 | 0.57 |
| YGR003W | 648.3625673 | 0.523587345 | 0.053523824 | 0.57 |
| YGR023W | 927.8650533 | 1.087029484 | 0.04054757 | 0.57 |
| YGR038W | 610.0071877 | -0.648645306 | 0.052950502 | 0.57 |
| YGR066C | 223.0827395 | 0.731969327 | 0.050556832 | 0.57 |
| YGR093W | 815.5848449 | 0.652769726 | 0.044174584 | 0.57 |
| YGR097W | 2251.701342 | 0.887398399 | 0.049072069 | 0.57 |
| YGR104C | 540.5492389 | 0.458115729 | 0.055394185 | 0.57 |
| YGR127W | 732.3981957 | 0.567293819 | 0.048186208 | 0.57 |
| YGR148C | 7673.362198 | -0.415973955 | 0.06024028 | 0.57 |
| YGR151C | 2.6142374 | -0.243270518 | 0.043401033 | 0.57 |
| YGR200C | 1330.804216 | -0.758916562 | 0.046100421 | 0.57 |
| YGR214W | 11003.56684 | -0.384079867 | 0.046092113 | 0.57 |
| YGR220C | 677.1113476 | -0.647796287 | 0.058298821 | 0.57 |
| YGR252W | 827.8896718 | -0.541574815 | 0.057627449 | 0.57 |
| YGR261C | 1931.303256 | -0.623026863 | 0.049533107 | 0.57 |
| YGR262C | 876.7108592 | -0.432472475 | 0.057118616 | 0.57 |
| YHL016C | 664.2747126 | 0.717290976 | 0.046211681 | 0.57 |
| YHR007C-A | 61.56889406 | 0.990634959 | 0.040331415 | 0.57 |
| YHR041C | 584.7828437 | -0.762980724 | 0.051366752 | 0.57 |
| YHR068W | 2922.590435 | -0.532069808 | 0.059946351 | 0.57 |
| YHR079C-A | 27.69347479 | 1.672924334 | 0.043014821 | 0.57 |
| YHR080C | 2045.316793 | 0.713698614 | 0.053140852 | 0.57 |
| YHR130C | 24.10330458 | 0.690843007 | 0.039844502 | 0.57 |
| YHR157W | 84.3613017 | 0.871134352 | 0.0415857 | 0.57 |
| YHR160C | 281.5696466 | 0.81353205 | 0.056497324 | 0.57 |
| YHR175W | 775.2689431 | -0.673084653 | 0.048933962 | 0.57 |
| YHR193C | 8396.042296 | -0.708164439 | 0.054346306 | 0.57 |
| YHR202W | 670.4759746 | 0.809374725 | 0.057635525 | 0.57 |
| YIL013C | 704.2538412 | 0.658189919 | 0.049763196 | 0.57 |
| YIL056W | 1630.203256 | 0.745581295 | 0.05911051 | 0.57 |
| YIL086C | 23.45932813 | 0.675237122 | 0.045080613 | 0.57 |
| YIL101C | 1402.735777 | 0.862348132 | 0.039591348 | 0.57 |
| YIL105C | 1531.713028 | 0.70927029 | 0.046192795 | 0.57 |
| YIL107C | 1780.529737 | 0.739881908 | 0.06054226 | 0.57 |
| YIL133C | 6678.70118 | -0.494328968 | 0.050189316 | 0.57 |

| | | | | |
|---|---|---|---|---|
| YIL155C | 2791.794832 | 1.062994529 | 0.057741989 | 0.57 |
| YJL028W | 3.906046468 | 0.879725309 | 0.058743869 | 0.57 |
| YJL032W | 6.67566349 | 0.511740314 | 0.053793761 | 0.57 |
| YJL063C | 928.728548 | -0.713775855 | 0.040467708 | 0.57 |
| YJL089W | 344.4116839 | 1.110524354 | 0.055136616 | 0.57 |
| YJL100W | 1049.188324 | 0.673693902 | 0.058066926 | 0.57 |
| YJL132W | 534.7548177 | 0.681540136 | 0.05555937 | 0.57 |
| YJL142C | 106.3041783 | 0.831880804 | 0.047305763 | 0.57 |
| YJL144W | 785.7166274 | 0.900363552 | 0.05102592 | 0.57 |
| YJL163C | 815.8405692 | 0.850029901 | 0.053210002 | 0.57 |
| YJL185C | 564.2478549 | 0.996376422 | 0.049182329 | 0.57 |
| YJL196C | 1508.45467 | -0.822729997 | 0.057009771 | 0.57 |
| YJR005C-A | 81.403139 | 1.086379497 | 0.042770728 | 0.57 |
| YJR008W | 1161.824512 | 0.967337609 | 0.046266225 | 0.57 |
| YJR024C | 696.9415683 | -0.802935666 | 0.04261848 | 0.57 |
| YJR043C | 619.0166524 | -0.431759615 | 0.059742535 | 0.57 |
| YJR070C | 1785.660344 | -0.508627818 | 0.044028493 | 0.57 |
| YJR076C | 1668.359247 | -0.509101437 | 0.045552959 | 0.57 |
| YKL056C | 11023.03919 | -0.60639 | 0.053412958 | 0.57 |
| YKL093W | 994.1827608 | 0.925611235 | 0.039579388 | 0.57 |
| YKL109W | 7924.096762 | 1.031493675 | 0.055706896 | 0.57 |
| YKL163W | 2089.664663 | 1.104070305 | 0.046481478 | 0.57 |
| YKL183W | 567.8673175 | -0.621567316 | 0.039036685 | 0.57 |
| YKR050W | 1256.621968 | 0.895921153 | 0.055287787 | 0.57 |
| YKR058W | 2130.69981 | 0.826130267 | 0.052955511 | 0.57 |
| YLL013C | 908.878738 | -0.444303208 | 0.057866723 | 0.57 |
| YLL015W | 2174.31995 | 0.771841153 | 0.044348441 | 0.57 |
| YLL030C | 4.099886808 | 0.958224838 | 0.0474445 | 0.57 |
| YLL046C | 28.77526537 | 1.433903401 | 0.05089907 | 0.57 |
| YLL053C | 96.70396409 | 0.685149362 | 0.051428568 | 0.57 |
| YLL058W | 677.8123693 | 0.775599909 | 0.054670503 | 0.57 |
| YLR029C | 12797.37208 | -0.524249235 | 0.045938623 | 0.57 |
| YLR040C | 271.3228626 | -1.521934238 | 0.043357554 | 0.57 |
| YLR108C | 407.9615838 | 0.9264411 | 0.051020024 | 0.57 |
| YLR149C | 1780.729906 | 1.07032028 | 0.049551501 | 0.57 |
| YLR162W-A | 8.884728875 | 0.195577364 | 0.052716799 | 0.57 |
| YLR167W | 28914.13243 | -0.375333338 | 0.053048198 | 0.57 |
| YLR171W | 25.60461968 | 0.61689949 | 0.050547432 | 0.57 |
| YLR173W | 824.1208493 | 0.918326747 | 0.054452618 | 0.57 |
| YLR175W | 4387.69014 | -0.682484235 | 0.052696819 | 0.57 |
| YLR221C | 955.7495307 | -0.621466993 | 0.050088964 | 0.57 |
| YLR252W | 254.6818923 | 0.93369638 | 0.045090012 | 0.57 |
| YLR258W | 9832.7404 | 1.16639414 | 0.043409028 | 0.57 |

| | | | | |
|---|---|---|---|---|
| YLR340W | 21021.60532 | -0.564270129 | 0.052694513 | 0.57 |
| YLR350W | 3571.832696 | 0.737114038 | 0.050691549 | 0.57 |
| YLR358C | 39.08246637 | -0.583712285 | 0.040649864 | 0.57 |
| YLR379W | 289.3293491 | -0.60677086 | 0.044731511 | 0.57 |
| YLR406C | 2407.447144 | -0.479819209 | 0.047096742 | 0.57 |
| YLR413W | 2093.12908 | -0.90907027 | 0.052519693 | 0.57 |
| YML012W | 3525.480868 | -0.603540168 | 0.055584323 | 0.57 |
| YML024W | 8930.152891 | -0.451330045 | 0.038526402 | 0.57 |
| YML026C | 7646.727007 | -0.431695954 | 0.040041118 | 0.57 |
| YML054C | 1101.015099 | 0.437712724 | 0.040848313 | 0.57 |
| YML073C | 8568.253747 | -0.531648278 | 0.059370568 | 0.57 |
| YML100W-A | 6.289539175 | 1.01029749 | 0.057814613 | 0.57 |
| YMR012W | 7425.571889 | -0.530690182 | 0.058548268 | 0.57 |
| YMR031C | 3707.880046 | 0.733015414 | 0.060012946 | 0.57 |
| YMR038C | 1941.251831 | -0.641991957 | 0.047584136 | 0.57 |
| YMR053C | 712.958308 | 0.84636678 | 0.060327517 | 0.57 |
| YMR081C | 3745.685883 | 0.900004045 | 0.043076462 | 0.57 |
| YMR084W | 174.3110976 | 0.748466994 | 0.059054062 | 0.57 |
| YMR096W | 4030.130556 | 1.110589581 | 0.058542737 | 0.57 |
| YMR097C | 670.8866515 | 0.628375202 | 0.043070682 | 0.57 |
| YMR114C | 1303.792032 | 0.96999367 | 0.041170088 | 0.57 |
| YMR116C | 12111.33345 | -0.326177165 | 0.045281114 | 0.57 |
| YMR140W | 1470.388235 | 0.817979265 | 0.042192386 | 0.57 |
| YMR194W | 1484.319594 | -0.546389596 | 0.038405044 | 0.57 |
| YMR235C | 2269.617315 | -0.768646621 | 0.045538828 | 0.57 |
| YMR242C | 8203.831599 | -0.536014216 | 0.043602171 | 0.57 |
| YNL002C | 1677.703778 | -0.639563459 | 0.057722487 | 0.57 |
| YNL012W | 592.4635068 | 0.777529639 | 0.05340135 | 0.57 |
| YNL066W | 1969.929236 | -0.907318372 | 0.047105363 | 0.57 |
| YNL070W | 685.2481589 | -0.619470528 | 0.055415572 | 0.57 |
| YNL110C | 899.4305976 | -0.504061383 | 0.049446628 | 0.57 |
| YNL113W | 221.4093861 | -0.611394405 | 0.054777301 | 0.57 |
| YNL144C | 1767.380134 | 1.051696967 | 0.050141239 | 0.57 |
| YNL147W | 569.2824244 | -0.618842702 | 0.055206478 | 0.57 |
| YNL178W | 12909.44004 | -0.481513193 | 0.053157271 | 0.57 |
| YNL194C | 937.042565 | 0.659930363 | 0.055572348 | 0.57 |
| YNL231C | 1567.48489 | -0.629567593 | 0.042310177 | 0.57 |
| YNL258C | 964.2763945 | -0.639327911 | 0.055936519 | 0.57 |
| YNL274C | 1846.518968 | 1.175193254 | 0.040640452 | 0.57 |
| YNL312W | 897.7476795 | -0.491260296 | 0.048644239 | 0.57 |
| YNR014W | 1455.118637 | 0.68973859 | 0.055614402 | 0.57 |
| YNR016C | 6951.886561 | -0.48706071 | 0.042911792 | 0.57 |
| YNR021W | 3396.726535 | -0.645614761 | 0.060209544 | 0.57 |

| | | | | |
|---|---|---|---|---|
| YNR028W | 918.8142146 | -0.547255304 | 0.060385044 | 0.57 |
| YNR053C | 1005.221474 | -0.970892683 | 0.047239375 | 0.57 |
| YNR061C | 596.8560089 | -0.768155987 | 0.05751079 | 0.57 |
| YNR065C | 953.8955562 | 0.776233307 | 0.051771832 | 0.57 |
| YOL038C-A | 23.65291474 | 0.838168857 | 0.059936461 | 0.57 |
| YOL087C | 2320.349339 | 0.708628225 | 0.059911922 | 0.57 |
| YOL116W | 722.658756 | 0.588039941 | 0.059895913 | 0.57 |
| YOR010C | 455.6940822 | 0.947741398 | 0.051408577 | 0.57 |
| YOR023C | 2117.560133 | 0.782239777 | 0.043819645 | 0.57 |
| YOR069W | 1035.064237 | 0.517135767 | 0.05244852 | 0.57 |
| YOR070C | 1064.02202 | 0.583949877 | 0.056483911 | 0.57 |
| YOR134W | 351.2595047 | 0.920599334 | 0.056139621 | 0.57 |
| YOR160W | 1175.925728 | 0.377674583 | 0.060619414 | 0.57 |
| YOR197W | 2558.031802 | -0.491375504 | 0.041006704 | 0.57 |
| YOR208W | 1224.294967 | 0.969064672 | 0.041120684 | 0.57 |
| YOR209C | 3422.936842 | -0.821970492 | 0.053089519 | 0.57 |
| YOR232W | 2857.566582 | -0.591601347 | 0.057612087 | 0.57 |
| YOR237W | 107.6435638 | 0.774301122 | 0.054767579 | 0.57 |
| YOR239W | 2726.169468 | -0.622449092 | 0.040350028 | 0.57 |
| YOR277C | 6.042620409 | -0.260868767 | 0.058735511 | 0.57 |
| YOR320C | 1070.658594 | -0.628839702 | 0.055549561 | 0.57 |
| YOR326W | 4223.653773 | -0.553363326 | 0.041669426 | 0.57 |
| YOR328W | 830.771978 | 0.947646192 | 0.042902699 | 0.57 |
| YOR361C | 4132.636454 | -0.654453795 | 0.05108419 | 0.57 |
| YOR374W | 6364.239227 | 1.182379721 | 0.043322765 | 0.57 |
| YPL007C | 583.714763 | -0.487736305 | 0.045431447 | 0.57 |
| YPL010W | 2111.463364 | -0.58490981 | 0.058726473 | 0.57 |
| YPL017C | 762.3634934 | 0.743917817 | 0.049620922 | 0.57 |
| YPL027W | 50.11860961 | 0.750644941 | 0.048849113 | 0.57 |
| YPL054W | 345.0757807 | 0.754162234 | 0.049004044 | 0.57 |
| YPL059W | 1008.866988 | -0.556161457 | 0.047432551 | 0.57 |
| YPL066W | 1565.740738 | 0.483629375 | 0.055693314 | 0.57 |
| YPL083C | 495.9825937 | -0.573490872 | 0.044762116 | 0.57 |
| YPL088W | 468.228703 | 0.832640951 | 0.042496764 | 0.57 |
| YPL100W | 911.5930892 | 0.633962069 | 0.057127336 | 0.57 |
| YPL113C | 241.6280316 | 0.696623065 | 0.048119213 | 0.57 |
| YPL167C | 937.9139202 | 0.6926269 | 0.058231305 | 0.57 |
| YPL169C | 1689.790258 | -0.62650762 | 0.046504986 | 0.57 |
| YPL208W | 646.0074609 | -0.645628165 | 0.041588108 | 0.57 |
| YPL241C | 283.7799123 | -0.913741066 | 0.049252909 | 0.57 |
| YPL272C | 344.5733609 | 0.753704547 | 0.039793911 | 0.57 |
| YPR019W | 1285.937535 | -0.480269162 | 0.045195099 | 0.57 |
| YPR052C | 667.2448711 | -0.801956301 | 0.052511331 | 0.57 |

| | | | | |
|---|---|---|---|---|
| YPR150W | 6.435730199 | 1.035900418 | 0.042380757 | 0.57 |
| YPR159C-A | 2.271891006 | 0.731398687 | 0.052303452 | 0.57 |
| YPR166C | 431.6523763 | -0.678822608 | 0.057505746 | 0.57 |
| YPR174C | 411.2973194 | -0.470453866 | 0.044780722 | 0.57 |
| snR56 | 6.041353852 | -0.199804761 | 0.055274412 | 0.57 |
| RUF23 | 8.975541829 | 1.05862138 | 0.086130274 | 0.57 |
| YAL003W | 12374.8526 | -0.35410257 | 0.080936093 | 0.57 |
| YAL016C-A | 61.31300348 | 0.73268998 | 0.097879692 | 0.57 |
| YAL022C | 845.7051212 | -0.616534078 | 0.061628871 | 0.57 |
| YAL042W | 3676.592816 | -0.475017664 | 0.066515063 | 0.57 |
| YAL054C | 1109.295849 | 1.224585356 | 0.065271995 | 0.57 |
| YAL059W | 208.6556716 | -0.59094918 | 0.091262382 | 0.57 |
| YAR008W | 281.6693003 | -0.58932012 | 0.086272467 | 0.57 |
| YAR028W | 513.5910617 | -0.614366648 | 0.092381924 | 0.57 |
| YAR030C | 64.62074774 | 0.5805116 | 0.065297624 | 0.57 |
| YBL018C | 137.4735368 | -0.390908748 | 0.086622991 | 0.57 |
| YBL039C | 1528.083632 | -0.400530472 | 0.098194949 | 0.57 |
| YBL040C | 1584.786506 | -0.519637856 | 0.069790325 | 0.57 |
| YBL057C | 768.8630819 | -0.551613854 | 0.091456535 | 0.57 |
| YBL087C | 3378.344673 | -0.417556396 | 0.097679711 | 0.57 |
| YBR001C | 1318.791969 | 0.725368907 | 0.067333287 | 0.57 |
| YBR006W | 1760.927773 | 0.990687862 | 0.092392187 | 0.57 |
| YBR008C | 906.2949136 | 0.552092845 | 0.069983628 | 0.57 |
| YBR012W-B | 321.6022163 | 1.349126807 | 0.068583542 | 0.57 |
| YBR061C | 630.0720407 | -0.613599546 | 0.079008749 | 0.57 |
| YBR109C | 2942.937927 | -0.484142257 | 0.065947708 | 0.57 |
| YBR113W | 115.7153994 | -0.393561169 | 0.083341916 | 0.57 |
| YBR126C | 9994.323143 | 0.866447808 | 0.071902983 | 0.57 |
| YBR138C | 569.4593536 | 0.690365439 | 0.069643082 | 0.57 |
| YBR152W | 271.4958287 | 0.534592332 | 0.06885049 | 0.57 |
| YBR164C | 1099.977428 | -0.629814933 | 0.090159886 | 0.57 |
| YBR181C | 1794.107741 | -0.280863051 | 0.065137161 | 0.57 |
| YBR191W-A | 25.03964351 | -0.250832655 | 0.081615978 | 0.57 |
| YBR205W | 1374.246383 | -0.481417976 | 0.079448182 | 0.57 |
| YBR206W | 255.7378889 | -0.469157087 | 0.087968225 | 0.57 |
| YBR214W | 5062.049562 | 0.872126007 | 0.082720814 | 0.57 |
| YBR220C | 693.632069 | -0.608487854 | 0.088950456 | 0.57 |
| YBR226C | 66.35832606 | 0.82703048 | 0.071166845 | 0.57 |
| YBR230C | 2523.139558 | 0.979620442 | 0.078372676 | 0.57 |
| YBR249C | 4275.177673 | -0.1755951 | 0.087453721 | 0.57 |
| YBR270C | 464.1113888 | 0.504322815 | 0.09344369 | 0.57 |
| YBR277C | 4.283921599 | 0.196914407 | 0.064199538 | 0.57 |
| YBR279W | 1844.253736 | -0.497045096 | 0.093016506 | 0.57 |

| | | | | |
|---|---|---|---|---|
| YBR288C | 774.8863305 | -0.554120049 | 0.092604755 | 0.57 |
| YBR298C | 1312.084765 | 0.744070455 | 0.079418189 | 0.57 |
| YBR299W | 41.66988027 | 0.591298845 | 0.089674953 | 0.57 |
| YBR300C | 5.482970764 | -0.17046832 | 0.071778807 | 0.57 |
| YCL001W | 1214.350549 | -0.43390836 | 0.097396614 | 0.57 |
| YCL017C | 1588.391961 | -0.430099639 | 0.096679346 | 0.57 |
| YCL024W | 493.5504233 | -0.553574571 | 0.090237827 | 0.57 |
| YCL032W | 909.5801687 | 0.625941282 | 0.066784639 | 0.57 |
| YCL042W | 285.5413665 | 0.974041499 | 0.08104065 | 0.57 |
| YCL051W | 1104.388423 | 0.576239565 | 0.078407057 | 0.57 |
| YCR007C | 318.2333147 | 0.622382528 | 0.077072404 | 0.57 |
| YCR009C | 3474.642891 | -0.571498193 | 0.080720808 | 0.57 |
| YCR014C | 618.1633306 | 0.562007628 | 0.081791921 | 0.57 |
| YCR031C | 11369.4013 | -0.486718468 | 0.071772578 | 0.57 |
| YCR034W | 1456.119271 | -0.502803486 | 0.084447324 | 0.57 |
| YCR035C | 836.2083583 | -0.482250302 | 0.09767727 | 0.57 |
| YCR066W | 426.4825775 | 0.537968895 | 0.095894198 | 0.57 |
| YCR068W | 666.4091136 | 0.713476712 | 0.077055514 | 0.57 |
| YCR079W | 984.6018828 | 0.705248237 | 0.075230966 | 0.57 |
| YCR084C | 3633.028548 | -0.515216999 | 0.077067234 | 0.57 |
| YDL014W | 4306.735257 | -0.543731806 | 0.089123681 | 0.57 |
| YDL026W | 16.56831607 | 0.549377148 | 0.079534213 | 0.57 |
| YDL075W | 5187.384084 | -0.381409893 | 0.083880342 | 0.57 |
| YDL092W | 1065.77792 | -0.615538953 | 0.087648335 | 0.57 |
| YDL113C | 1161.019898 | 0.703986992 | 0.062519912 | 0.57 |
| YDL118W | 21.28319356 | 0.430767585 | 0.094787626 | 0.57 |
| YDL190C | 2381.415921 | 0.493677708 | 0.092691992 | 0.57 |
| YDL192W | 4665.318464 | -0.522199581 | 0.063434347 | 0.57 |
| YDL202W | 808.8072189 | -0.524255551 | 0.092554742 | 0.57 |
| YDL206W | 597.9682559 | 0.49716041 | 0.098289973 | 0.57 |
| YDL213C | 1003.203847 | -0.191524715 | 0.077460602 | 0.57 |
| YDL218W | 253.6505308 | 0.823210353 | 0.090348226 | 0.57 |
| YDL230W | 846.3743425 | 0.70818926 | 0.071175207 | 0.57 |
| YDL239C | 522.0657782 | 0.626828225 | 0.07204859 | 0.57 |
| YDR006C | 2310.266671 | 0.578097809 | 0.091809148 | 0.57 |
| YDR016C | 195.2031569 | -0.61136339 | 0.089925089 | 0.57 |
| YDR025W | 5331.251911 | -0.499437486 | 0.070238362 | 0.57 |
| YDR040C | 132.5886651 | 0.603679138 | 0.084770265 | 0.57 |
| YDR047W | 1762.459456 | -0.628230306 | 0.068704939 | 0.57 |
| YDR053W | 7.370760006 | 0.670406662 | 0.081509355 | 0.57 |
| YDR077W | 88090.2349 | 0.915406521 | 0.091267689 | 0.57 |
| YDR086C | 1629.638329 | -0.672759725 | 0.073965628 | 0.57 |
| YDR107C | 540.536576 | -0.571750735 | 0.074919973 | 0.57 |

| | | | | |
|---|---|---|---|---|
| YDR109C | 536.0726963 | 0.442345355 | 0.078216016 | 0.57 |
| YDR112W | 22.56100124 | -0.937725491 | 0.084904718 | 0.57 |
| YDR171W | 12610.16854 | 0.982320011 | 0.0703396 | 0.57 |
| YDR194C | 2378.698014 | -0.576548042 | 0.081004261 | 0.57 |
| YDR195W | 649.2195359 | 0.457702548 | 0.095385494 | 0.57 |
| YDR209C | 344.9577194 | -1.058402139 | 0.093889923 | 0.57 |
| YDR233C | 7397.691198 | -0.584860638 | 0.083771888 | 0.57 |
| YDR245W | 1164.324708 | -0.722579967 | 0.077660504 | 0.57 |
| YDR280W | 709.9216811 | -0.655992634 | 0.080388637 | 0.57 |
| YDR293C | 6400.531719 | 0.648151571 | 0.097387335 | 0.57 |
| YDR330W | 1850.284876 | 0.716057929 | 0.080870394 | 0.57 |
| YDR342C | 1706.769269 | 0.873417571 | 0.075253992 | 0.57 |
| YDR343C | 868.7911596 | 1.055055062 | 0.082815186 | 0.57 |
| YDR347W | 1409.321423 | -0.39861301 | 0.065908492 | 0.57 |
| YDR358W | 1790.698239 | 0.753163174 | 0.085407306 | 0.57 |
| YDR418W | 2788.265019 | -0.410531445 | 0.097525267 | 0.57 |
| YDR438W | 373.0593006 | 0.617194986 | 0.072773384 | 0.57 |
| YDR447C | 4088.999597 | -0.268451513 | 0.071320928 | 0.57 |
| YDR450W | 8491.535949 | -0.352370608 | 0.078851112 | 0.57 |
| YDR467C | 75.26126934 | 0.677876423 | 0.088873612 | 0.57 |
| YDR481C | 2453.405315 | 0.743205675 | 0.088495198 | 0.57 |
| YDR500C | 6683.941473 | -0.587800863 | 0.07844595 | 0.57 |
| YDR523C | 105.2296251 | 0.412499697 | 0.087357628 | 0.57 |
| YDR524C-A | 2.188681507 | -0.602959427 | 0.079515788 | 0.57 |
| YDR524W-C | 1641.998262 | -0.747253944 | 0.081570135 | 0.57 |
| YDR525W | 4.560372998 | 0.226819262 | 0.087558284 | 0.57 |
| YEL004W | 410.7288824 | -0.685462913 | 0.068331292 | 0.57 |
| YEL011W | 8924.284313 | 1.043512824 | 0.083614823 | 0.57 |
| YEL030C-A | 7.262884205 | 0.949058352 | 0.066498763 | 0.57 |
| YEL034C-A | 2748.462234 | -0.701876516 | 0.077795455 | 0.57 |
| YEL059W | 50.35325287 | 0.525891714 | 0.070974887 | 0.57 |
| YEL060C | 10805.49764 | 1.097361601 | 0.066583365 | 0.57 |
| YER006W | 1977.06385 | -0.268094991 | 0.098166373 | 0.57 |
| YER007C-A | 951.757768 | -0.470163075 | 0.094978238 | 0.57 |
| YER009W | 3909.301197 | -0.5481142 | 0.067742594 | 0.57 |
| YER016W | 838.7636501 | -0.801172117 | 0.06338526 | 0.57 |
| YER017C | 1876.346183 | 0.566654286 | 0.080697719 | 0.57 |
| YER039C | 101.9948572 | 0.580141106 | 0.070785306 | 0.57 |
| YER042W | 653.285425 | 1.105301457 | 0.086910074 | 0.57 |
| YER045C | 814.7735686 | 0.762430142 | 0.065692809 | 0.57 |
| YER055C | 2945.986692 | -0.364766106 | 0.093571396 | 0.57 |
| YER066C-A | 1483.59707 | 1.140419969 | 0.061653966 | 0.57 |
| YER067C-A | 258.7943359 | 1.634227919 | 0.068628991 | 0.57 |

| | | | | |
|---|---|---|---|---|
| YER074W | 4108.232098 | -0.514144253 | 0.072213239 | 0.57 |
| YER086W | 3514.638296 | -0.13416646 | 0.091764468 | 0.57 |
| YER088C-A | 302.6604055 | -0.555368143 | 0.07128808 | 0.57 |
| YER098W | 930.5704049 | 0.675601539 | 0.086254617 | 0.57 |
| YER102W | 2473.647383 | -0.203149241 | 0.097491514 | 0.57 |
| YER105C | 2233.871003 | -0.442214544 | 0.064861495 | 0.57 |
| YER118C | 889.8446464 | -0.400616202 | 0.086282601 | 0.57 |
| YER119C-A | 190.6802462 | -0.753844416 | 0.08250743 | 0.57 |
| YER145C-A | 50.64924556 | -0.613044899 | 0.0789711 | 0.57 |
| YER147C | 529.8378852 | 0.568524296 | 0.096374402 | 0.57 |
| YER150W | 3467.414047 | 0.661096776 | 0.092252221 | 0.57 |
| YER153C | 351.5393272 | 0.81894838 | 0.081666524 | 0.57 |
| YER156C | 1503.86508 | -0.557686046 | 0.097589074 | 0.57 |
| YFL015C | 19.00869044 | 1.118802929 | 0.097115222 | 0.57 |
| YFL020C | 87.69189529 | 0.631843242 | 0.076015624 | 0.57 |
| YFL021W | 926.9442787 | 0.732334975 | 0.082998068 | 0.57 |
| YFL022C | 4340.34362 | -0.526140806 | 0.087509376 | 0.57 |
| YFR032C | 47.36854765 | 0.227879391 | 0.074651534 | 0.57 |
| YFR054C | 20.51319414 | 1.043779047 | 0.083491623 | 0.57 |
| YGL012W | 2257.199778 | -0.511298083 | 0.081492642 | 0.57 |
| YGL017W | 502.0325199 | 0.46357961 | 0.084934559 | 0.57 |
| YGL029W | 383.3919755 | -0.343369431 | 0.092848281 | 0.57 |
| YGL031C | 8332.907395 | -0.326500773 | 0.067051615 | 0.57 |
| YGL040C | 1261.243607 | -0.512596446 | 0.080810599 | 0.57 |
| YGL044C | 659.773802 | 0.396498285 | 0.08694647 | 0.57 |
| YGL045W | 904.8478281 | 0.653525805 | 0.094478724 | 0.57 |
| YGL092W | 1819.13534 | -0.503301744 | 0.089610703 | 0.57 |
| YGL102C | 355.1928609 | -0.388802769 | 0.094599629 | 0.57 |
| YGL137W | 4577.146804 | -0.495438691 | 0.070283899 | 0.57 |
| YGL155W | 739.9393467 | -0.659117694 | 0.078874283 | 0.57 |
| YGL162W | 395.3975497 | 0.29054984 | 0.062749545 | 0.57 |
| YGL173C | 8071.132916 | -0.392117157 | 0.095886749 | 0.57 |
| YGL189C | 20518.43042 | -0.267571493 | 0.07809258 | 0.57 |
| YGL250W | 555.5711852 | 0.701965959 | 0.097561233 | 0.57 |
| YGL262W | 77.76796003 | 1.094714266 | 0.098420458 | 0.57 |
| YGR020C | 3100.566353 | -0.663548885 | 0.074036811 | 0.57 |
| YGR027C | 9077.452786 | -0.508633531 | 0.070911595 | 0.57 |
| YGR053C | 339.3237027 | 0.591499114 | 0.093952717 | 0.57 |
| YGR054W | 2529.613046 | -0.65787361 | 0.065307603 | 0.57 |
| YGR070W | 2198.55855 | 0.907438512 | 0.078183443 | 0.57 |
| YGR082W | 1782.261898 | -0.517349753 | 0.076405262 | 0.57 |
| YGR088W | 3462.726793 | 0.718719319 | 0.067885624 | 0.57 |
| YGR105W | 477.947228 | -0.631163054 | 0.088518576 | 0.57 |

| | | | | |
|---|---|---|---|---|
| YGR123C | 655.6704403 | -0.6693378 | 0.06859762 | 0.57 |
| YGR149W | 1082.04674 | 0.724347922 | 0.090554209 | 0.57 |
| YGR161C | 2432.867838 | 0.747822628 | 0.086875542 | 0.57 |
| YGR187C | 552.0431833 | -0.762925829 | 0.070383463 | 0.57 |
| YGR195W | 892.1333078 | -0.558470068 | 0.069375098 | 0.57 |
| YGR208W | 739.3899853 | -0.329922597 | 0.096444603 | 0.57 |
| YGR290W | 27.11482068 | 0.481442981 | 0.091629017 | 0.57 |
| YHL008C | 1005.082799 | 0.658895636 | 0.08618629 | 0.57 |
| YHL018W | 70.12879735 | 0.645191716 | 0.096072836 | 0.57 |
| YHL030W | 2779.309717 | 0.90664476 | 0.088157969 | 0.57 |
| YHL033C | 4723.149428 | -0.279782632 | 0.08166145 | 0.57 |
| YHL038C | 821.5884324 | 0.539816011 | 0.090514661 | 0.57 |
| YHR010W | 11885.41138 | -0.266825125 | 0.076553316 | 0.57 |
| YHR021C | 5458.86145 | -0.329255445 | 0.073755831 | 0.57 |
| YHR025W | 4807.328396 | -0.251335594 | 0.087254714 | 0.57 |
| YHR052W | 1127.946092 | -0.545352929 | 0.097131693 | 0.57 |
| YHR072W-A | 1183.992345 | -0.661682201 | 0.06360986 | 0.57 |
| YHR087W | 3070.022032 | 0.816165487 | 0.091659178 | 0.57 |
| YHR107C | 1509.6288 | -0.455737144 | 0.073772971 | 0.57 |
| YHR108W | 3358.554888 | -0.492795412 | 0.096050076 | 0.57 |
| YHR127W | 498.1945386 | -0.450401823 | 0.091624782 | 0.57 |
| YHR141C | 3868.981705 | -0.457583779 | 0.096594666 | 0.57 |
| YHR143W-A | 554.7529102 | -0.691182723 | 0.075006409 | 0.57 |
| YHR146W | 3577.393266 | 0.66594841 | 0.094214709 | 0.57 |
| YHR147C | 857.7185719 | -0.522266004 | 0.074712952 | 0.57 |
| YHR164C | 1299.336952 | 0.546833957 | 0.090514192 | 0.57 |
| YHR199C | 2007.264009 | 0.704147105 | 0.083576112 | 0.57 |
| YIL016W | 483.5722155 | -0.556968702 | 0.088768014 | 0.57 |
| YIL018W | 10226.88893 | -0.323252727 | 0.098095927 | 0.57 |
| YIL022W | 2268.335855 | -0.392638 | 0.066868025 | 0.57 |
| YIL024C | 346.2449471 | 1.072351739 | 0.082603123 | 0.57 |
| YIL028W | 59.74955175 | 0.672766333 | 0.080469592 | 0.57 |
| YIL030W-A | 74.85518125 | 0.759081553 | 0.064571951 | 0.57 |
| YIL039W | 1826.424519 | -0.571351957 | 0.071344031 | 0.57 |
| YIL052C | 4784.674042 | -0.368001898 | 0.087905055 | 0.57 |
| YIL067C | 437.2526192 | 0.50854866 | 0.08268838 | 0.57 |
| YIL071C | 381.3119395 | 0.513644063 | 0.061418688 | 0.57 |
| YIL093C | 871.5653388 | -0.553044206 | 0.097539032 | 0.57 |
| YIL096C | 413.8812975 | -0.424176402 | 0.086042941 | 0.57 |
| YIL113W | 384.3761033 | 0.423985962 | 0.098397724 | 0.57 |
| YIL148W | 9852.705743 | -0.408888573 | 0.061249426 | 0.57 |
| YIL159W | 678.524918 | -0.57450666 | 0.064878213 | 0.57 |
| YIL161W | 488.7537738 | -0.664638646 | 0.089079476 | 0.57 |

| | | | | |
|---|---|---|---|---|
| YIR025W | 303.8110514 | 0.485711936 | 0.092786692 | 0.57 |
| YIR043C | 49.74477082 | 0.543091897 | 0.098296393 | 0.57 |
| YJL012C | 2507.923216 | -0.799229919 | 0.087605482 | 0.57 |
| YJL036W | 1275.214013 | 0.636652689 | 0.071692085 | 0.57 |
| YJL056C | 734.4930024 | 0.66492622 | 0.086748252 | 0.57 |
| YJL061W | 1142.158213 | -0.524755249 | 0.088900388 | 0.57 |
| YJL070C | 972.7876692 | 0.697266858 | 0.081075061 | 0.57 |
| YJL080C | 5735.124683 | -0.25478164 | 0.088878884 | 0.57 |
| YJL083W | 610.9975272 | 0.625481309 | 0.071265616 | 0.57 |
| YJL084C | 1615.45903 | 0.655022814 | 0.065181446 | 0.57 |
| YJL099W | 747.8275315 | 0.614024152 | 0.075353517 | 0.57 |
| YJL115W | 931.9259287 | -0.600399045 | 0.064807227 | 0.57 |
| YJL116C | 947.324176 | 0.713376514 | 0.077211636 | 0.57 |
| YJL138C | 1955.737437 | -0.589913654 | 0.065514176 | 0.57 |
| YJL145W | 1081.572718 | -0.663505798 | 0.076507332 | 0.57 |
| YJL155C | 1149.698916 | 0.800580918 | 0.070469494 | 0.57 |
| YJL157C | 1263.741636 | -1.124187267 | 0.09074198 | 0.57 |
| YJL167W | 5562.553131 | -0.566647673 | 0.078539376 | 0.57 |
| YJL177W | 7142.560719 | -0.455921909 | 0.065138248 | 0.57 |
| YJL195C | 5.108398903 | 0.631314983 | 0.08427189 | 0.57 |
| YJL199C | 163.0335414 | 0.484298189 | 0.080977815 | 0.57 |
| YJR007W | 3429.418011 | -0.541324162 | 0.097555455 | 0.57 |
| YJR031C | 1083.680948 | -0.547064286 | 0.081392544 | 0.57 |
| YJR035W | 878.6193365 | 0.684042097 | 0.062119437 | 0.57 |
| YJR039W | 935.4373703 | 0.638484887 | 0.087962742 | 0.57 |
| YJR067C | 196.0036818 | -0.483009523 | 0.0945855 | 0.57 |
| YJR069C | 644.3818806 | -0.525720355 | 0.064390803 | 0.57 |
| YJR072C | 1860.139893 | -0.447623468 | 0.096896627 | 0.57 |
| YJR084W | 669.2500078 | -0.452664005 | 0.09826014 | 0.57 |
| YJR088C | 954.591927 | -0.619443524 | 0.068841707 | 0.57 |
| YJR123W | 26166.00582 | -0.281952089 | 0.086186924 | 0.57 |
| YJR143C | 2572.640819 | -0.461208233 | 0.093808867 | 0.57 |
| YKL033W-A | 1409.695613 | -0.505683692 | 0.079037734 | 0.57 |
| YKL035W | 11294.21789 | 0.726074737 | 0.079875943 | 0.57 |
| YKL058W | 636.5138347 | -0.641557127 | 0.074735253 | 0.57 |
| YKL077W | 3042.957164 | -0.512574155 | 0.077311697 | 0.57 |
| YKL094W | 2161.93051 | 0.554145868 | 0.078974045 | 0.57 |
| YKL095W | 323.4844945 | 0.660379416 | 0.065173073 | 0.57 |
| YKL105C | 1423.737383 | 0.724777778 | 0.077797542 | 0.57 |
| YKL168C | 972.1957599 | 0.574493871 | 0.08441391 | 0.57 |
| YKL192C | 3222.400243 | -0.470134598 | 0.089073688 | 0.57 |
| YKR006C | 1505.723079 | -0.573104903 | 0.082610498 | 0.57 |
| YKR009C | 961.8765586 | 0.835982771 | 0.088592354 | 0.57 |

| | | | | |
|---|---|---|---|---|
| YKR019C | 546.7471303 | 0.660564377 | 0.064206367 | 0.57 |
| YKR021W | 1328.600543 | 0.522892704 | 0.094315469 | 0.57 |
| YKR055W | 476.0228776 | 0.664954594 | 0.067816277 | 0.57 |
| YKR057W | 2516.350053 | -0.639652108 | 0.063444547 | 0.57 |
| YKR070W | 801.1916976 | -0.336409028 | 0.095783809 | 0.57 |
| YKR094C | 5698.150477 | -0.435017734 | 0.069183677 | 0.57 |
| YLL018C | 8408.142457 | -0.469913406 | 0.079154797 | 0.57 |
| YLL028W | 724.5266873 | -0.893050381 | 0.069307343 | 0.57 |
| YLL049W | 333.5985502 | -0.567741074 | 0.085456346 | 0.57 |
| YLL054C | 624.0988471 | -0.604949193 | 0.061227704 | 0.57 |
| YLL056C | 287.6766171 | 0.738092428 | 0.075634311 | 0.57 |
| YLR009W | 1043.475365 | -0.573373303 | 0.095276512 | 0.57 |
| YLR028C | 4846.630925 | -0.325747381 | 0.0746117 | 0.57 |
| YLR031W | 139.756576 | 0.483938184 | 0.089163729 | 0.57 |
| YLR043C | 5741.406917 | -0.479650904 | 0.095637762 | 0.57 |
| YLR056W | 3964.313149 | -0.522266338 | 0.072893988 | 0.57 |
| YLR062C | 194.5995134 | -0.493646753 | 0.080762346 | 0.57 |
| YLR070C | 346.2836033 | 0.758125549 | 0.081898539 | 0.57 |
| YLR073C | 322.9361924 | -0.788930269 | 0.094479378 | 0.57 |
| YLR106C | 6573.73141 | -0.340712359 | 0.073816615 | 0.57 |
| YLR117C | 1230.224284 | 0.604634859 | 0.074957992 | 0.57 |
| YLR121C | 1254.397824 | 0.834555465 | 0.078735064 | 0.57 |
| YLR146C | 511.7510501 | -0.571771852 | 0.071543923 | 0.57 |
| YLR149C-A | 61.1192566 | -0.22790429 | 0.061408833 | 0.57 |
| YLR151C | 483.8723388 | 0.623871835 | 0.064287961 | 0.57 |
| YLR153C | 3246.572959 | -0.537299087 | 0.065514771 | 0.57 |
| YLR164W | 251.3796178 | 0.541332532 | 0.083633305 | 0.57 |
| YLR185W | 4884.991766 | -0.339154436 | 0.064439955 | 0.57 |
| YLR188W | 1233.172467 | -0.492201584 | 0.073406228 | 0.57 |
| YLR208W | 2849.872811 | -0.593121629 | 0.06497891 | 0.57 |
| YLR231C | 929.429053 | -1.118509717 | 0.08688859 | 0.57 |
| YLR240W | 1128.041716 | 0.598450123 | 0.068894987 | 0.57 |
| YLR254C | 554.1871051 | 0.571771342 | 0.082844797 | 0.57 |
| YLR267W | 1321.159087 | 0.549720146 | 0.089376152 | 0.57 |
| YLR272C | 941.972981 | -0.52301078 | 0.069774747 | 0.57 |
| YLR276C | 2051.953288 | -0.672745322 | 0.083968262 | 0.57 |
| YLR285W | 889.692098 | -0.716285903 | 0.06851639 | 0.57 |
| YLR289W | 462.7410856 | 0.68979362 | 0.07110804 | 0.57 |
| YLR308W | 11.06339218 | 0.751657935 | 0.081572235 | 0.57 |
| YLR317W | 2.198595472 | -0.38995554 | 0.075768376 | 0.57 |
| YLR327C | 4824.371416 | 1.087085487 | 0.061880477 | 0.57 |
| YLR331C | 3.488696978 | 0.769523042 | 0.087383355 | 0.57 |
| YLR342W | 12036.59934 | -0.529795775 | 0.069621436 | 0.57 |

| | | | | |
|---|---|---|---|---|
| YLR347C | 1405.641169 | -0.528662591 | 0.067789536 | 0.57 |
| YLR362W | 1083.342817 | 0.822142759 | 0.073483704 | 0.57 |
| YLR366W | 1.593075245 | 0.285767963 | 0.083700795 | 0.57 |
| YLR367W | 2289.066484 | -0.303957462 | 0.097859125 | 0.57 |
| YLR374C | 34.93928504 | 0.651939609 | 0.091756691 | 0.57 |
| YLR388W | 1672.166035 | -0.52406862 | 0.065423573 | 0.57 |
| YLR412W | 993.3823084 | -0.636846858 | 0.094615644 | 0.57 |
| YLR414C | 1706.526088 | 0.641504955 | 0.091720109 | 0.57 |
| YLR423C | 531.1083054 | 0.625978862 | 0.097284919 | 0.57 |
| YLR453C | 293.5520271 | 0.544175842 | 0.086332304 | 0.57 |
| YML008C | 4220.321447 | -0.59377936 | 0.085735644 | 0.57 |
| YML010W | 2999.463402 | -0.53286299 | 0.067595137 | 0.57 |
| YML020W | 864.3038212 | 0.535599996 | 0.080924497 | 0.57 |
| YML022W | 1077.904293 | -0.446632328 | 0.075787096 | 0.57 |
| YML060W | 354.1411795 | -0.335089194 | 0.072304063 | 0.57 |
| YML063W | 16224.08176 | -0.288309153 | 0.067316153 | 0.57 |
| YML064C | 587.9701312 | 0.468885567 | 0.067921603 | 0.57 |
| YML094C-A | 15.34009962 | -0.727540748 | 0.092716649 | 0.57 |
| YML096W | 652.0319638 | -0.564109314 | 0.097513003 | 0.57 |
| YML099W-A | 34.73959572 | 0.665103098 | 0.095958351 | 0.57 |
| YML101C | 335.3248863 | -0.639143073 | 0.075003736 | 0.57 |
| YMR001C | 1386.236499 | -0.627861251 | 0.089978525 | 0.57 |
| YMR001C-A | 16.08451551 | -0.6291735 | 0.075213408 | 0.57 |
| YMR015C | 1137.039851 | -0.983056473 | 0.068016076 | 0.57 |
| YMR030W | 574.6177195 | 0.756953488 | 0.098152234 | 0.57 |
| YMR033W | 1176.392622 | -0.667523388 | 0.081702099 | 0.57 |
| YMR057C | 13.35741525 | 0.12389112 | 0.068477857 | 0.57 |
| YMR059W | 205.0702465 | -0.450975901 | 0.082480883 | 0.57 |
| YMR062C | 1501.153431 | 0.777410359 | 0.085197492 | 0.57 |
| YMR068W | 420.8665046 | 0.647370007 | 0.090550424 | 0.57 |
| YMR091C | 998.5083437 | -0.653206477 | 0.063179313 | 0.57 |
| YMR093W | 713.9051908 | -0.591719346 | 0.090893811 | 0.57 |
| YMR118C | 295.0484477 | 0.639216295 | 0.06144204 | 0.57 |
| YMR120C | 11760.55672 | -0.080377539 | 0.079500778 | 0.57 |
| YMR131C | 2416.896733 | -0.455968995 | 0.095420803 | 0.57 |
| YMR133W | 502.5352221 | 0.666815832 | 0.090274905 | 0.57 |
| YMR142C | 11187.93902 | -0.387981012 | 0.076556672 | 0.57 |
| YMR148W | 862.6307015 | 0.642999638 | 0.095475668 | 0.57 |
| YMR153W | 881.8064044 | -0.455253835 | 0.064381012 | 0.57 |
| YMR158W | 693.5786925 | -0.641439083 | 0.075457694 | 0.57 |
| YMR206W | 572.161077 | 0.7091718 | 0.061241458 | 0.57 |
| YMR213W | 660.1636091 | 0.624496406 | 0.095660513 | 0.57 |
| YMR230W | 6828.386976 | -0.408382523 | 0.092032714 | 0.57 |

| | | | | |
|---|---|---|---|---|
| YMR230W-A | 4.038623613 | 0.525736297 | 0.087272732 | 0.57 |
| YMR237W | 2047.326568 | -0.337777146 | 0.093564394 | 0.57 |
| YMR260C | 3410.062096 | -0.580315992 | 0.07413033 | 0.57 |
| YMR279C | 707.3104958 | 0.551105422 | 0.085925429 | 0.57 |
| YMR298W | 1278.824067 | -0.481303009 | 0.087295064 | 0.57 |
| YMR306W | 518.3037802 | 0.720719907 | 0.098171502 | 0.57 |
| YMR307W | 11748.41017 | -0.529940488 | 0.088179905 | 0.57 |
| YNL067W | 1749.269832 | -0.694072273 | 0.065128185 | 0.57 |
| YNL069C | 13320.02635 | -0.411666207 | 0.071115479 | 0.57 |
| YNL093W | 262.8484398 | 0.73968684 | 0.072724972 | 0.57 |
| YNL126W | 705.9981899 | -0.552370434 | 0.096250096 | 0.57 |
| YNL140C | 6.604161499 | 0.303083758 | 0.072654035 | 0.57 |
| YNL162W | 2350.306083 | -0.525403554 | 0.073188841 | 0.57 |
| YNL165W | 500.9806009 | 0.729212452 | 0.069344864 | 0.57 |
| YNL176C | 1925.34034 | 0.588376896 | 0.095561513 | 0.57 |
| YNL177C | 1301.893277 | -0.436431321 | 0.065583274 | 0.57 |
| YNL184C | 77.22136504 | -0.622395659 | 0.093657949 | 0.57 |
| YNL185C | 474.6914722 | -0.605551949 | 0.066994905 | 0.57 |
| YNL191W | 591.9956262 | 0.646712428 | 0.076308748 | 0.57 |
| YNL216W | 1187.634305 | -0.601710176 | 0.06690326 | 0.57 |
| YNL301C | 1146.542523 | -0.30745946 | 0.083737767 | 0.57 |
| YNL302C | 5878.096109 | -0.306464274 | 0.068565272 | 0.57 |
| YNL313C | 1104.908095 | -0.498026289 | 0.084264774 | 0.57 |
| YNR003C | 554.144982 | -0.599801051 | 0.061390504 | 0.57 |
| YNR034W | 660.948473 | 0.647761786 | 0.069660221 | 0.57 |
| YOL013C | 1216.146773 | 0.453470224 | 0.090982075 | 0.57 |
| YOL013W-B | 4.383672181 | -0.836718736 | 0.072532309 | 0.57 |
| YOL040C | 16073.93418 | -0.391185084 | 0.093282987 | 0.57 |
| YOL061W | 1754.163485 | -0.58256032 | 0.068554644 | 0.57 |
| YOL085C | 30.97884527 | 0.755630594 | 0.072435809 | 0.57 |
| YOL089C | 836.6158144 | 0.563600697 | 0.085653549 | 0.57 |
| YOL097C | 2168.680575 | -0.51326533 | 0.065300478 | 0.57 |
| YOL108C | 360.9401457 | 0.716530622 | 0.089261357 | 0.57 |
| YOL127W | 9841.72384 | -0.371724848 | 0.077257325 | 0.57 |
| YOL146W | 364.144711 | -0.542110755 | 0.091116045 | 0.57 |
| YOL149W | 642.6006688 | -0.627802137 | 0.096438382 | 0.57 |
| YOR019W | 709.5102111 | 0.561448008 | 0.083820496 | 0.57 |
| YOR036W | 1460.876014 | 0.604341494 | 0.096179086 | 0.57 |
| YOR063W | 34774.24031 | -0.163612176 | 0.095811124 | 0.57 |
| YOR072W-A | 14.05662092 | 0.637403428 | 0.09321447 | 0.57 |
| YOR096W | 6750.76705 | -0.439727116 | 0.090741987 | 0.57 |
| YOR102W | 49.45521107 | -0.637032599 | 0.093353777 | 0.57 |
| YOR110W | 712.9241656 | 0.57124991 | 0.075156839 | 0.57 |

| | | | | |
|---|---|---|---|---|
| YOR114W | 311.2740858 | 1.034116962 | 0.067657861 | 0.57 |
| YOR123C | 2187.626634 | -0.406299855 | 0.083884843 | 0.57 |
| YOR133W | 1425.340865 | -0.541524403 | 0.079174142 | 0.57 |
| YOR137C | 579.6398816 | 0.669569875 | 0.093510653 | 0.57 |
| YOR145C | 791.624005 | -0.658333798 | 0.077591605 | 0.57 |
| YOR162C | 795.3084144 | 0.43727349 | 0.066809362 | 0.57 |
| YOR178C | 2730.240932 | 0.940337853 | 0.094084904 | 0.57 |
| YOR182C | 1767.616754 | -0.288258045 | 0.097551396 | 0.57 |
| YOR236W | 448.7599801 | -0.484961281 | 0.065683763 | 0.57 |
| YOR243C | 1479.808571 | -0.539099488 | 0.075074291 | 0.57 |
| YOR253W | 809.4249218 | -0.717030428 | 0.064161237 | 0.57 |
| YOR276W | 906.6874329 | -0.564692132 | 0.081529776 | 0.57 |
| YOR309C | 407.1266153 | -0.473453891 | 0.078182181 | 0.57 |
| YOR312C | 9424.205996 | -0.426503189 | 0.061797143 | 0.57 |
| YOR340C | 775.5259752 | -0.366024622 | 0.063074189 | 0.57 |
| YOR342C | 1114.452608 | -0.654543272 | 0.088043444 | 0.57 |
| YOR369C | 15497.1337 | -0.520413539 | 0.087276069 | 0.57 |
| YOR377W | 997.8606777 | 0.647800265 | 0.06827287 | 0.57 |
| YPL043W | 1057.06526 | -0.530154524 | 0.066419436 | 0.57 |
| YPL050C | 1693.46287 | -0.683616615 | 0.079891247 | 0.57 |
| YPL056C | 172.8020928 | 0.631079901 | 0.071148803 | 0.57 |
| YPL079W | 6102.629794 | -0.399438006 | 0.068163038 | 0.57 |
| YPL119C-A | 58.40963327 | 0.92680938 | 0.064756216 | 0.57 |
| YPL142C | 228.9766843 | -0.554155809 | 0.068586399 | 0.57 |
| YPL151C | 670.1565001 | 0.453377584 | 0.064516035 | 0.57 |
| YPL165C | 810.9716846 | 0.528423452 | 0.091380403 | 0.57 |
| YPL184C | 1810.247002 | -0.704122105 | 0.066640521 | 0.57 |
| YPL237W | 2463.364966 | -0.616378044 | 0.067047345 | 0.57 |
| YPR004C | 1678.278628 | -0.685484611 | 0.097436446 | 0.57 |
| YPR010C | 3009.126793 | -0.478493183 | 0.087577539 | 0.57 |
| YPR012W | 66.76164095 | 0.771353721 | 0.081981607 | 0.57 |
| YPR017C | 239.3606668 | -0.675665594 | 0.071914029 | 0.57 |
| YPR026W | 1343.846593 | 0.786294327 | 0.072666694 | 0.57 |
| YPR036W | 6660.698271 | -0.545949583 | 0.083962593 | 0.57 |
| YPR040W | 901.3829077 | 0.731685898 | 0.094455536 | 0.57 |
| YPR041W | 3343.881383 | -0.618414974 | 0.075580875 | 0.57 |
| YPR044C | 114.0311076 | -0.500796745 | 0.097107294 | 0.57 |
| YPR050C | 1.567602133 | -1.054612914 | 0.090798276 | 0.57 |
| YPR085C | 667.2581144 | 0.517128628 | 0.073620256 | 0.57 |
| YPR102C | 3154.540675 | -0.567619303 | 0.078744362 | 0.57 |
| YPR132W | 5266.120182 | -0.538669222 | 0.087811841 | 0.57 |
| YPR133W-A | 1114.918643 | -0.497487883 | 0.072574082 | 0.57 |
| YPR154W | 2486.651934 | 0.622610667 | 0.096878477 | 0.57 |

| | | | | |
|---|---|---|---|---|
| YPR155C | 883.1515984 | 0.661897341 | 0.08040829 | 0.57 |
| YPR183W | 3494.823853 | -0.463129811 | 0.081292396 | 0.57 |
| snR69 | 9.747852558 | 0.671786518 | 0.069482925 | 0.57 |
| YLR293C | 8683.145693 | -0.47335309 | 0.098936087 | 0.57 |
| YLR352W | 926.4882915 | 0.545884528 | 0.098839461 | 0.57 |
| YMR101C | 112.86694 | 1.163638755 | 0.098734922 | 0.57 |
| YOR247W | 5496.630424 | -0.426827986 | 0.098894728 | 0.57 |
| YJR149W | 627.5970928 | 0.723177218 | 0.099278088 | 0.57 |
| YLL025W | 92.59299699 | 0.603876343 | 0.099494067 | 0.58 |
| YDL048C | 3537.203448 | 0.836954231 | 0.099702849 | 0.58 |
| YDR003W | 807.4785109 | 0.621846154 | 0.099781807 | 0.58 |
| YAR073W | 171.304609 | 1.265403657 | 0.100760244 | 0.58 |
| YBL004W | 2589.356792 | -0.46700338 | 0.100912136 | 0.58 |
| YBR158W | 2964.217702 | -0.897811053 | 0.102334059 | 0.58 |
| YBR201C-A | 201.3285883 | 0.74163912 | 0.100892495 | 0.58 |
| YBR287W | 2396.597397 | 0.554728606 | 0.100367696 | 0.58 |
| YDL003W | 760.5875817 | -0.706790005 | 0.101660021 | 0.58 |
| YDR216W | 2843.606015 | 0.969807284 | 0.100325877 | 0.58 |
| YDR297W | 2184.395638 | -0.464195345 | 0.101849966 | 0.58 |
| YDR484W | 738.7697722 | 0.369772937 | 0.101455663 | 0.58 |
| YDR524C | 904.6436686 | -0.627178885 | 0.102424473 | 0.58 |
| YEL033W | 1403.07415 | -0.69442341 | 0.101127154 | 0.58 |
| YGL159W | 407.2189918 | -0.766223839 | 0.102471218 | 0.58 |
| YGR078C | 839.5108181 | -0.686450551 | 0.101373051 | 0.58 |
| YGR112W | 838.4700607 | 0.747298393 | 0.102244157 | 0.58 |
| YIL040W | 372.5621769 | -0.64320021 | 0.100556283 | 0.58 |
| YJL122W | 643.9700151 | -0.496008612 | 0.102049591 | 0.58 |
| YJR057W | 461.680688 | -0.475579863 | 0.101276638 | 0.58 |
| YKL180W | 12036.25542 | -0.452206109 | 0.101881855 | 0.58 |
| YLR212C | 601.9431586 | -0.547922655 | 0.101518242 | 0.58 |
| YML097C | 708.2199714 | -0.518550673 | 0.10078152 | 0.58 |
| YNR006W | 1506.041195 | 0.508643037 | 0.102330496 | 0.58 |
| YNR064C | 215.5475844 | 0.128402841 | 0.102200825 | 0.58 |
| YOR008W-B | 4.890160442 | 0.125281276 | 0.102128616 | 0.58 |
| YOR045W | 622.8710581 | -0.522895957 | 0.101564231 | 0.58 |
| YOR310C | 3221.373678 | -0.472599199 | 0.101614751 | 0.58 |
| YOR390W | 45.23234325 | -0.968427316 | 0.102297012 | 0.58 |
| YPL090C | 2392.913651 | -0.348817067 | 0.102237312 | 0.58 |
| YPL166W | 366.5892241 | 0.554497101 | 0.101598432 | 0.58 |
| YPR078C | 88.79558885 | 0.47669122 | 0.10236568 | 0.58 |
| YPR142C | 12.52047692 | 0.091236322 | 0.100717592 | 0.58 |
| YDR327W | 614.5550695 | -0.475861116 | 0.102900225 | 0.58 |
| YJL023C | 307.217395 | 0.47091577 | 0.102912371 | 0.58 |

| | | | | |
|---|---|---|---|---|
| YNL010W | 2912.199227 | -0.487692602 | 0.102959458 | 0.58 |
| YNL075W | 678.2006525 | -0.28979401 | 0.102977053 | 0.58 |
| YNL096C | 3035.774305 | -0.361591144 | 0.102685325 | 0.58 |
| YMR284W | 845.7287906 | 0.52579396 | 0.103108332 | 0.58 |
| YCL005W-A | 247.3743167 | -0.50950154 | 0.103378415 | 0.58 |
| YNL307C | 2792.714562 | -0.57833781 | 0.103575583 | 0.58 |
| YAR002C-A | 2600.594621 | -0.488653566 | 0.103793182 | 0.58 |
| YNL021W | 893.5874562 | -0.391166662 | 0.103767694 | 0.58 |
| YBR102C | 889.0122882 | -0.396664911 | 0.104328719 | 0.58 |
| YDR350C | 1056.007335 | 0.54143431 | 0.104408197 | 0.58 |
| YGR009C | 1451.464581 | 0.476754022 | 0.10435318 | 0.58 |
| YLR312C | 417.3618401 | 0.741979901 | 0.104297791 | 0.58 |
| YMR105C | 9001.650714 | 0.965978576 | 0.104488139 | 0.58 |
| YMR201C | 569.2749285 | 0.624822854 | 0.103976614 | 0.58 |
| YPR111W | 665.5579675 | 0.699701074 | 0.104465518 | 0.58 |
| YLR299W | 1242.489768 | 0.656336959 | 0.104824607 | 0.58 |
| YHR013C | 1158.208101 | -0.516557892 | 0.105070009 | 0.58 |
| YJL091C | 524.9582628 | -0.537493552 | 0.105057387 | 0.58 |
| YCL025C | 3652.132376 | 0.629552667 | 0.10550016 | 0.58 |
| YCR036W | 1075.104784 | 0.418062745 | 0.10561214 | 0.58 |
| YER069W | 925.6063303 | 0.764704637 | 0.105670733 | 0.58 |
| YJL188C | 893.4798181 | -0.588487996 | 0.105681363 | 0.58 |
| YLL026W | 27987.51895 | 1.085211824 | 0.105236454 | 0.58 |
| YNL025C | 730.0152463 | 0.488041807 | 0.105598835 | 0.58 |
| YBR247C | 781.1952554 | -0.361354314 | 0.106272453 | 0.58 |
| YHL032C | 2561.780761 | 0.788147212 | 0.106135281 | 0.58 |
| YIL097W | 1123.344615 | 0.703533063 | 0.106200075 | 0.58 |
| YJR136C | 423.3917741 | 0.392551128 | 0.106241987 | 0.58 |
| YKL072W | 554.6822525 | 0.510086979 | 0.105858752 | 0.58 |
| YLR329W | 110.7708514 | 0.768768335 | 0.106011169 | 0.58 |
| YBR259W | 488.2154125 | -0.465778796 | 0.106462255 | 0.58 |
| YDR459C | 457.0405938 | 0.394405567 | 0.107044416 | 0.58 |
| YFR037C | 1666.566511 | -0.525511269 | 0.106865478 | 0.58 |
| YGR238C | 880.0595867 | 0.651936338 | 0.106591964 | 0.58 |
| YHR144C | 383.3281151 | -0.565773218 | 0.106699322 | 0.58 |
| YJL088W | 187.9199165 | 0.888224115 | 0.107071733 | 0.58 |
| YOL100W | 1483.492555 | 0.628105335 | 0.10686883 | 0.58 |
| YOR006C | 758.4019018 | 0.44939466 | 0.107034306 | 0.58 |
| YBR154C | 1262.041062 | -0.650075521 | 0.107399479 | 0.58 |
| YGL226C-A | 628.3645694 | -0.523347317 | 0.10743313 | 0.58 |
| YPR107C | 422.2754774 | 0.442591124 | 0.107378834 | 0.58 |
| YPR056W | 512.7372419 | -0.437606491 | 0.107790518 | 0.58 |
| YDR258C | 9048.121163 | 1.105745374 | 0.108119723 | 0.59 |

| | | | | |
|---|---|---|---|---|
| YAL046C | 313.7043981 | -0.503716751 | 0.108759231 | 0.59 |
| YBL046W | 691.3216368 | 0.446730509 | 0.113200293 | 0.59 |
| YBR032W | 28.5083245 | 0.411207133 | 0.112219615 | 0.59 |
| YBR117C | 1336.758461 | 0.585483981 | 0.112060299 | 0.59 |
| YBR143C | 4538.418741 | -0.584937647 | 0.110533196 | 0.59 |
| YBR182C | 159.913284 | 0.720203636 | 0.111226177 | 0.59 |
| YCL059C | 1691.420964 | -0.335262001 | 0.112910348 | 0.59 |
| YCL073C | 26.5180597 | 0.660828439 | 0.10941613 | 0.59 |
| YCR091W | 1488.499833 | 0.647847498 | 0.112684139 | 0.59 |
| YDL235C | 741.3905909 | -0.574610428 | 0.111464357 | 0.59 |
| YDL238C | 479.1219346 | 0.641183005 | 0.113185905 | 0.59 |
| YDR310C | 1697.719061 | -0.481955841 | 0.110290033 | 0.59 |
| YDR429C | 3118.563913 | -0.474205639 | 0.108996569 | 0.59 |
| YEL039C | 1783.57122 | 0.672373817 | 0.112846804 | 0.59 |
| YER027C | 1561.298312 | -0.444088055 | 0.11328355 | 0.59 |
| YER074W-A | 546.9637993 | -0.46278965 | 0.111780383 | 0.59 |
| YER165C-A | 28.00543952 | -0.536620572 | 0.109495281 | 0.59 |
| YER186C | 678.5275475 | -0.786314668 | 0.109914472 | 0.59 |
| YGL091C | 545.7319922 | 0.389900512 | 0.112723589 | 0.59 |
| YGL095C | 1579.460918 | -0.4622178 | 0.112955999 | 0.59 |
| YGR052W | 642.70953 | -0.034244799 | 0.112227322 | 0.59 |
| YGR130C | 4694.98638 | 0.685252545 | 0.108683026 | 0.59 |
| YGR249W | 613.9099062 | 0.694002614 | 0.112461228 | 0.59 |
| YHL015W | 17106.50707 | -0.184298866 | 0.108756964 | 0.59 |
| YHL022C | 136.8231823 | 1.023985543 | 0.112529212 | 0.59 |
| YHR075C | 548.2435732 | 0.495993291 | 0.11125612 | 0.59 |
| YJR021C | 884.0147888 | 0.530989537 | 0.11165028 | 0.59 |
| YJR030C | 708.7385852 | -0.513828347 | 0.112053452 | 0.59 |
| YJR145C | 6502.704468 | -0.389833978 | 0.111070417 | 0.59 |
| YKL010C | 2290.07323 | 0.42552223 | 0.110536531 | 0.59 |
| YKL156W | 2081.100672 | -0.637928152 | 0.109602769 | 0.59 |
| YKR078W | 765.3811944 | 0.524394101 | 0.113385628 | 0.59 |
| YKR081C | 1261.655674 | -0.400642694 | 0.113008785 | 0.59 |
| YLL002W | 366.0264394 | -0.619169224 | 0.113261055 | 0.59 |
| YLL052C | 77.86079202 | 0.555374501 | 0.109561025 | 0.59 |
| YLR172C | 1976.093187 | -0.459240579 | 0.110924266 | 0.59 |
| YLR330W | 3759.858385 | -0.400009047 | 0.109114716 | 0.59 |
| YLR447C | 5880.017489 | -0.447458025 | 0.112649733 | 0.59 |
| YML078W | 2283.677343 | -0.581991477 | 0.112322802 | 0.59 |
| YML083C | 177.180785 | 0.608683058 | 0.112464849 | 0.59 |
| YML085C | 4155.33263 | -0.354564433 | 0.11201837 | 0.59 |
| YMR270C | 708.6444493 | -0.574843871 | 0.109176273 | 0.59 |
| YMR300C | 1920.32889 | -0.619627486 | 0.113201419 | 0.59 |

| | | | | |
|---|---|---|---|---|
| YMR309C | 2879.75717 | -0.669654492 | 0.109248958 | 0.59 |
| YMR316C-A | 33.48519679 | 0.62519226 | 0.110572346 | 0.59 |
| YNL205C | 11.3162081 | 0.970301676 | 0.109894081 | 0.59 |
| YOL039W | 5643.898973 | -0.37897279 | 0.112480131 | 0.59 |
| YOR031W | 276.3450169 | 0.579611162 | 0.109134284 | 0.59 |
| YOR067C | 1025.694017 | -0.491153048 | 0.109720439 | 0.59 |
| YOR214C | 33.33902204 | 0.466746375 | 0.113230505 | 0.59 |
| YOR251C | 1296.455018 | -0.392495688 | 0.112253488 | 0.59 |
| YOR298W | 171.3347166 | 0.584259613 | 0.11342426 | 0.59 |
| YPL145C | 2994.997329 | -0.596285306 | 0.110705708 | 0.59 |
| YPL226W | 2793.678288 | -0.441702383 | 0.112283163 | 0.59 |
| YPL231W | 11637.98709 | -0.33429267 | 0.109306794 | 0.59 |
| YPL261C | 48.62531076 | 0.574443827 | 0.111787209 | 0.59 |
| YPL263C | 1770.940205 | -0.504344583 | 0.110649299 | 0.59 |
| YHR079C | 1259.061897 | 0.492970468 | 0.113751874 | 0.59 |
| YJL143W | 1167.282332 | -0.495887204 | 0.113677014 | 0.59 |
| YLR285C-A | 41.10839909 | -0.616153016 | 0.113763516 | 0.59 |
| YDR093W | 1759.127414 | -0.461080364 | 0.115248312 | 0.59 |
| YGL190C | 1598.926776 | 0.412054251 | 0.115139534 | 0.59 |
| YGR287C | 287.959068 | 0.671506534 | 0.114553391 | 0.59 |
| YIL152W | 734.4896336 | 0.761183416 | 0.114908301 | 0.59 |
| YJL192C | 2669.267366 | -0.450096409 | 0.114597245 | 0.59 |
| YJR052W | 627.3978725 | 0.456420732 | 0.115196324 | 0.59 |
| YJR063W | 331.564801 | -0.557674744 | 0.11503294 | 0.59 |
| YML016C | 1375.407219 | 0.522194033 | 0.115099379 | 0.59 |
| YMR143W | 7020.519511 | -0.388390867 | 0.114075262 | 0.59 |
| YOR199W | 28.95971662 | -0.431105116 | 0.11430285 | 0.59 |
| YOR314W | 17.17022364 | -0.620349509 | 0.115239201 | 0.59 |
| YOR341W | 3809.897668 | -0.360611407 | 0.114913242 | 0.59 |
| YPL211W | 668.7255746 | -0.405919989 | 0.114835888 | 0.59 |
| YPR170W-B | 216.8408172 | -0.432913053 | 0.114685137 | 0.59 |
| YFR030W | 1464.99151 | 0.737992619 | 0.115541327 | 0.59 |
| YIL043C | 4110.226673 | -0.492738138 | 0.115628946 | 0.59 |
| YLR325C | 5760.035975 | -0.512335204 | 0.115696115 | 0.59 |
| YNL143C | 26.87962177 | -0.914145007 | 0.115671221 | 0.59 |
| YHR156C | 433.8648918 | 0.612861498 | 0.115798106 | 0.59 |
| YFL017C | 717.9144755 | -0.469351148 | 0.116052966 | 0.59 |
| YOL003C | 636.2491343 | -0.487778498 | 0.116050623 | 0.59 |
| YDL029W | 3675.857405 | -0.322010362 | 0.116209999 | 0.59 |
| YGR258C | 1398.526849 | 0.596898999 | 0.116339664 | 0.59 |
| YBR046C | 1727.421703 | 0.947400924 | 0.116676264 | 0.59 |
| YDR386W | 496.5773156 | 0.574744504 | 0.116713262 | 0.59 |
| YGR271C-A | 454.6880839 | -0.279787751 | 0.116752197 | 0.59 |

| | | | | |
|---|---|---|---|---|
| YLR448W | 5493.030487 | -0.387209599 | 0.11671689 | 0.59 |
| YER115C | 529.525795 | 0.53522381 | 0.117123093 | 0.59 |
| YML041C | 413.9344954 | 0.480604629 | 0.117083524 | 0.59 |
| YMR272C | 4343.78123 | -0.617663124 | 0.116987612 | 0.59 |
| YPL121C | 25.34339254 | 1.161833443 | 0.117409479 | 0.59 |
| YIL007C | 541.263808 | 0.56853606 | 0.117786993 | 0.59 |
| YKR066C | 4134.117453 | 0.725431474 | 0.117674095 | 0.59 |
| YOR099W | 2318.668874 | -0.572274292 | 0.117801471 | 0.59 |
| YBR034C | 1111.628509 | -0.583379357 | 0.118260782 | 0.59 |
| YDL170W | 672.20876 | 0.501239081 | 0.118109544 | 0.59 |
| YDR264C | 2392.197288 | -0.510721093 | 0.118473274 | 0.59 |
| YEL042W | 1714.856815 | -0.424746429 | 0.118346975 | 0.59 |
| YGR245C | 1499.530458 | -0.298937011 | 0.118128579 | 0.59 |
| YKR049C | 1528.555977 | 0.690191444 | 0.118533462 | 0.59 |
| YML089C | 19.45677984 | 1.085372541 | 0.118514687 | 0.59 |
| YNL214W | 315.125881 | 0.418912562 | 0.118020719 | 0.59 |
| YJL062W | 1004.118516 | -0.414328389 | 0.118762399 | 0.59 |
| snR17b | 27.76593566 | 0.139646699 | 0.118754637 | 0.59 |
| YGR162W | 4487.780728 | -0.590778385 | 0.119091598 | 0.59 |
| YGR194C | 2007.303758 | 0.742070946 | 0.119207756 | 0.59 |
| YLR435W | 440.7011003 | -0.537713049 | 0.119144859 | 0.59 |
| snR30 | 275.4218461 | -0.165551791 | 0.118945867 | 0.59 |
| YFR052C-A | 3217.393186 | 1.112586579 | 0.119478971 | 0.59 |
| YDL045C | 1046.513314 | 0.375866337 | 0.119651943 | 0.59 |
| YEL051W | 3229.319305 | -0.493788285 | 0.119904406 | 0.59 |
| YPR192W | 327.8980894 | 0.116264049 | 0.120121137 | 0.59 |
| YGR008C | 4582.699628 | 0.849266179 | 0.120433003 | 0.59 |
| YIL009C-A | 425.8589138 | -0.784044355 | 0.120271012 | 0.59 |
| YNL259C | 262.1284302 | -0.540408957 | 0.120427837 | 0.59 |
| YBL049W | 377.5575063 | 0.529804235 | 0.120589664 | 0.60 |
| YBL078C | 809.4078099 | 0.941258379 | 0.121589104 | 0.60 |
| YDL137W | 4443.238326 | -0.482251938 | 0.121615956 | 0.60 |
| YDR433W | 87.14790436 | -0.706268531 | 0.122022764 | 0.60 |
| YER077C | 789.2511196 | 0.38404399 | 0.121349295 | 0.60 |
| YFR029W | 834.7381381 | 0.709591193 | 0.121661059 | 0.60 |
| YJL071W | 590.1206135 | 0.553964609 | 0.121385291 | 0.60 |
| YJL140W | 1600.235301 | -0.467245406 | 0.121732908 | 0.60 |
| YKL062W | 2465.244192 | 0.457962727 | 0.122074078 | 0.60 |
| YLR089C | 5710.768483 | 1.016893664 | 0.121099565 | 0.60 |
| YLR345W | 2235.165084 | 0.627662482 | 0.121809414 | 0.60 |
| YMR290C | 2185.404412 | -0.29925111 | 0.121564504 | 0.60 |
| YNL039W | 1798.324816 | 0.461039233 | 0.122016537 | 0.60 |
| YNL131W | 2877.827714 | -0.477552493 | 0.121007332 | 0.60 |

| | | | | |
|---|---|---|---|---|
| YDR410C | 388.474414 | -0.886307259 | 0.122655583 | 0.60 |
| YGR223C | 1134.541576 | 0.573951097 | 0.12259843 | 0.60 |
| YHR017W | 1256.836076 | 0.581985654 | 0.1224025 | 0.60 |
| YKL009W | 1409.022835 | -0.417679282 | 0.122316989 | 0.60 |
| YLL060C | 550.1729711 | 0.504315729 | 0.122509982 | 0.60 |
| YNL273W | 713.7141054 | -0.441049218 | 0.122318825 | 0.60 |
| YCR061W | 1066.525845 | 0.784327328 | 0.122763059 | 0.60 |
| YNL111C | 246.1352445 | -0.798746542 | 0.122905855 | 0.60 |
| YGL232W | 728.6792639 | -0.543881896 | 0.123036237 | 0.60 |
| YPR123C | 99.99461053 | 0.097000354 | 0.123140341 | 0.60 |
| YOR234C | 2998.326335 | -0.260251256 | 0.12335676 | 0.60 |
| YKR054C | 2620.574744 | -0.394260179 | 0.123578479 | 0.60 |
| YBR244W | 574.3609399 | -0.527893508 | 0.123775808 | 0.60 |
| YER056C-A | 4887.303568 | -0.348799921 | 0.123976031 | 0.60 |
| YML068W | 512.1804017 | 0.41690701 | 0.124047969 | 0.60 |
| YGL138C | 22.53163453 | -0.064115233 | 0.124441683 | 0.60 |
| YJR135W-A | 435.3473743 | -0.50374502 | 0.124784766 | 0.60 |
| YKL171W | 1421.131993 | 0.664055346 | 0.125041993 | 0.60 |
| YLR110C | 106387.3494 | -0.285436637 | 0.125047718 | 0.60 |
| snR63 | 94.54093227 | -0.05952902 | 0.124961652 | 0.60 |
| YBL101C | 1814.259973 | 0.652710325 | 0.125589456 | 0.60 |
| YBR268W | 567.5701433 | -0.520185017 | 0.125309783 | 0.60 |
| YDR461C-A | 326.4764294 | 0.69153148 | 0.12558686 | 0.60 |
| YGR040W | 494.1182396 | -0.710640544 | 0.125446354 | 0.60 |
| YHL030W-A | 22.0345387 | 0.504623593 | 0.125560457 | 0.60 |
| YCR085W | 15.93952821 | -0.300955337 | 0.125785925 | 0.60 |
| YJL158C | 11506.92019 | -0.109282369 | 0.125828281 | 0.60 |
| YOL124C | 564.6964356 | -0.072664268 | 0.125894126 | 0.60 |
| YBL095W | 182.0968065 | 0.562016443 | 0.126502674 | 0.60 |
| YBR091C | 248.9757925 | -0.445974636 | 0.126733161 | 0.60 |
| YCR001W | 33.60447039 | -1.15438442 | 0.126346786 | 0.60 |
| YCR044C | 1029.447304 | -0.532180578 | 0.126846349 | 0.60 |
| YGL210W | 815.3164128 | -0.455491107 | 0.126206634 | 0.60 |
| YJL108C | 258.931357 | 0.570573421 | 0.126149422 | 0.60 |
| YMR266W | 1578.35636 | -0.620421582 | 0.126667044 | 0.60 |
| YOL139C | 5554.572319 | -0.565377449 | 0.126844077 | 0.60 |
| YOR219C | 1764.610148 | 0.422143176 | 0.126923941 | 0.60 |
| YOR223W | 687.6095973 | 0.511715366 | 0.126945923 | 0.60 |
| YPR163C | 5059.100182 | -0.493167983 | 0.126409665 | 0.60 |
| YGR096W | 275.6515308 | 0.57056118 | 0.127380978 | 0.60 |
| YEL070W | 298.2975483 | 0.899504602 | 0.127893194 | 0.61 |
| YFR001W | 1018.480753 | -0.494532269 | 0.127939431 | 0.61 |
| YKR010C | 516.9970849 | -0.568266921 | 0.127779281 | 0.61 |

| | | | | |
|---|---|---|---|---|
| YPL089C | 1858.715154 | 0.644728153 | 0.127854265 | 0.61 |
| YIL142C-A | 7.771319082 | 0.796927567 | 0.128043144 | 0.61 |
| YBL029C-A | 1079.485322 | 0.715920663 | 0.13688602 | 0.61 |
| YBL077W | 139.1714064 | -0.295366667 | 0.132081689 | 0.61 |
| YBR028C | 772.1128092 | -0.698750027 | 0.138122665 | 0.61 |
| YBR031W | 10790.74196 | -0.167537748 | 0.133030365 | 0.61 |
| YBR074W | 1213.234406 | -0.425108077 | 0.139476125 | 0.61 |
| YBR106W | 4289.033597 | -0.436506524 | 0.140018183 | 0.61 |
| YBR139W | 2470.205803 | 0.672062044 | 0.134726493 | 0.61 |
| YBR155W | 833.2201083 | -0.135822854 | 0.132462977 | 0.61 |
| YBR254C | 311.1708692 | -0.581385101 | 0.130801775 | 0.61 |
| YCL075W | 37.39387639 | 1.026992469 | 0.13616457 | 0.61 |
| YCR046C | 1534.125343 | -0.495359569 | 0.134843638 | 0.61 |
| YCR052W | 1334.780982 | -0.470920331 | 0.132550983 | 0.61 |
| YCR094W | 731.6068684 | -0.481255591 | 0.130447848 | 0.61 |
| YDL111C | 757.760422 | -0.510523349 | 0.139945955 | 0.61 |
| YDL236W | 1309.137891 | -0.559305797 | 0.14016699 | 0.61 |
| YDR069C | 1204.898386 | 0.521015749 | 0.132605042 | 0.61 |
| YDR131C | 626.2979697 | 0.420738547 | 0.134120616 | 0.61 |
| YDR170C | 4884.61556 | -0.382812354 | 0.132078845 | 0.61 |
| YDR295C | 798.8722941 | 0.396484291 | 0.133563331 | 0.61 |
| YDR315C | 362.5235632 | -0.513076987 | 0.13143917 | 0.61 |
| YDR337W | 923.5124951 | -0.431873697 | 0.138383137 | 0.61 |
| YDR372C | 1904.188965 | -0.56882085 | 0.139181874 | 0.61 |
| YDR378C | 714.4468599 | -0.419803654 | 0.140040019 | 0.61 |
| YDR387C | 888.6103683 | 0.527386831 | 0.134982293 | 0.61 |
| YDR399W | 3185.764567 | -0.181686883 | 0.134398468 | 0.61 |
| YDR402C | 76.67558535 | 0.955654686 | 0.138762315 | 0.61 |
| YDR448W | 426.123254 | -0.592798135 | 0.129705039 | 0.61 |
| YDR471W | 4845.236011 | -0.387194096 | 0.139543588 | 0.61 |
| YDR542W | 1.676832288 | 0.200249381 | 0.139272022 | 0.61 |
| YEL017C-A | 672.6729484 | -0.601515405 | 0.13342336 | 0.61 |
| YEL049W | 54.33605347 | 0.509848386 | 0.136659863 | 0.61 |
| YER048W-A | 659.0895735 | -0.354445717 | 0.129042167 | 0.61 |
| YER049W | 1494.465582 | -0.470125361 | 0.139530514 | 0.61 |
| YER079C-A | 18.60445932 | 0.385149333 | 0.130653056 | 0.61 |
| YER128W | 281.9899223 | 0.586871817 | 0.130334093 | 0.61 |
| YER152C | 1264.809581 | 0.75109855 | 0.131370486 | 0.61 |
| YFL017W-A | 221.0675394 | -0.551686039 | 0.130714277 | 0.61 |
| YFL056C | 232.3744781 | 0.425660084 | 0.133313813 | 0.61 |
| YFR014C | 1739.196956 | 0.554324611 | 0.134808413 | 0.61 |
| YFR027W | 221.6537022 | 0.593099883 | 0.13489125 | 0.61 |
| YFR032C-A | 2429.038295 | -0.293517124 | 0.131352918 | 0.61 |

| | | | | |
|---|---|---|---|---|
| YFR053C | 31755.34613 | 1.063427552 | 0.138523312 | 0.61 |
| YGL027C | 1397.054052 | -0.41945406 | 0.138586422 | 0.61 |
| YGL077C | 1372.265624 | -0.453928465 | 0.133423272 | 0.61 |
| YGL200C | 3717.842992 | -0.396293486 | 0.138325702 | 0.61 |
| YGL229C | 708.5937653 | 0.54782472 | 0.130273951 | 0.61 |
| YGL234W | 9210.041986 | 0.002190845 | 0.136494394 | 0.61 |
| YGL255W | 1201.748656 | 0.628352851 | 0.139546858 | 0.61 |
| YGR022C | 1.81368062 | 0.833242043 | 0.14017918 | 0.61 |
| YGR032W | 3378.619384 | 0.742865361 | 0.138699867 | 0.61 |
| YGR061C | 5651.499061 | -0.378311935 | 0.136658963 | 0.61 |
| YGR084C | 1041.679379 | -0.42028867 | 0.134358553 | 0.61 |
| YGR099W | 582.6782539 | -0.423839846 | 0.134845309 | 0.61 |
| YGR113W | 643.4472036 | -0.350211737 | 0.132016037 | 0.61 |
| YGR133W | 486.059603 | 0.500672315 | 0.138208945 | 0.61 |
| YGR179C | 503.8137475 | -0.41676797 | 0.13173896 | 0.61 |
| YGR248W | 2366.128573 | 0.555823008 | 0.135519641 | 0.61 |
| YHL009C | 603.1387723 | -0.552843292 | 0.129077774 | 0.61 |
| YHR004C | 921.2706765 | 0.51046604 | 0.138106327 | 0.61 |
| YHR039C | 2261.246675 | -0.563369627 | 0.130741308 | 0.61 |
| YHR084W | 1211.807628 | -0.943844576 | 0.132623316 | 0.61 |
| YHR088W | 1005.281896 | -0.341127154 | 0.139120802 | 0.61 |
| YHR089C | 2056.029582 | -0.46202891 | 0.13833653 | 0.61 |
| YHR133C | 977.6515666 | -0.528657854 | 0.128911463 | 0.61 |
| YHR159W | 783.3955676 | 0.617048022 | 0.133455764 | 0.61 |
| YHR171W | 640.7534835 | 0.68437555 | 0.139854214 | 0.61 |
| YHR175W-A | 167.202498 | -0.389436375 | 0.130267675 | 0.61 |
| YHR195W | 1250.379878 | 0.457213767 | 0.138919748 | 0.61 |
| YIL017C | 1438.735299 | 0.573509938 | 0.138429721 | 0.61 |
| YIL045W | 1955.748951 | 0.647865017 | 0.134271704 | 0.61 |
| YIL114C | 411.0869055 | -0.694472415 | 0.136486604 | 0.61 |
| YIL166C | 439.1240138 | 0.52784445 | 0.131432283 | 0.61 |
| YIR001C | 869.6360964 | -0.438592194 | 0.133037655 | 0.61 |
| YJL011C | 110.249814 | -0.303726708 | 0.137255134 | 0.61 |
| YJL057C | 1203.837592 | 0.561822308 | 0.133166531 | 0.61 |
| YJL075C | 78.82663296 | 0.385423704 | 0.137086281 | 0.61 |
| YJL096W | 663.1647142 | -0.475743692 | 0.137853188 | 0.61 |
| YJL104W | 974.9529404 | -0.444226623 | 0.12961048 | 0.61 |
| YJL107C | 129.4785502 | 0.783867123 | 0.135943155 | 0.61 |
| YJL148W | 1042.119748 | -0.410796033 | 0.129365711 | 0.61 |
| YJR010C-A | 212.1436636 | -0.696805572 | 0.136140757 | 0.61 |
| YJR119C | 341.555846 | 0.416098385 | 0.137064704 | 0.61 |
| YJR139C | 4827.24467 | -0.404865345 | 0.132788079 | 0.61 |
| YKL007W | 2212.362326 | -0.335149635 | 0.132972119 | 0.61 |

| | | | | |
|---|---|---|---|---|
| YKL113C | 707.9229549 | -0.428187199 | 0.13295783 | 0.61 |
| YKL207W | 2006.977142 | -0.485313578 | 0.138478233 | 0.61 |
| YKR017C | 571.9042365 | 0.459713605 | 0.129207042 | 0.61 |
| YKR098C | 1187.222635 | 0.744710421 | 0.135499172 | 0.61 |
| YLL020C | 30.70630028 | 0.581701864 | 0.129168812 | 0.61 |
| YLR021W | 410.9321978 | -0.451426629 | 0.137920217 | 0.61 |
| YLR048W | 13755.73587 | -0.341642263 | 0.129047904 | 0.61 |
| YLR076C | 95.93637703 | -0.17825921 | 0.132433842 | 0.61 |
| YLR080W | 639.2701305 | 0.709508484 | 0.138659951 | 0.61 |
| YLR132C | 337.4595812 | 0.453217893 | 0.136502743 | 0.61 |
| YLR150W | 21689.73235 | -0.321296276 | 0.131284275 | 0.61 |
| YLR227C | 512.1356286 | 0.334946638 | 0.140076263 | 0.61 |
| YLR230W | 287.1818195 | -0.45925883 | 0.137345448 | 0.61 |
| YLR287C-A | 5234.314487 | -0.448050069 | 0.137735675 | 0.61 |
| YLR332W | 1136.948721 | 0.583109313 | 0.130914594 | 0.61 |
| YLR442C | 902.5050218 | 0.451597189 | 0.129505717 | 0.61 |
| YML031C-A | 2.97142876 | 0.481272042 | 0.133966567 | 0.61 |
| YML047W-A | 2.39499993 | 0.00158044 | 0.13926604 | 0.61 |
| YMR009W | 601.1334204 | 0.527243141 | 0.131204004 | 0.61 |
| YMR010W | 733.3907547 | -0.587490444 | 0.133622328 | 0.61 |
| YMR076C | 1386.678863 | -0.405375151 | 0.13875445 | 0.61 |
| YMR280C | 1028.334214 | 0.874523281 | 0.138991973 | 0.61 |
| YNL014W | 784.7565686 | 0.841191294 | 0.134560816 | 0.61 |
| YNL018C | 116.9826504 | 0.885857384 | 0.133194887 | 0.61 |
| YNL064C | 7494.727656 | -0.336759737 | 0.13210153 | 0.61 |
| YNL074C | 2355.063484 | 0.497053684 | 0.136332453 | 0.61 |
| YNL209W | 13780.76239 | -0.354397182 | 0.137718926 | 0.61 |
| YNL285W | 3.926788582 | 0.494512523 | 0.135126138 | 0.61 |
| YNL287W | 4413.325689 | -0.344473871 | 0.136300936 | 0.61 |
| YOL048C | 1220.015423 | 0.503149775 | 0.131226338 | 0.61 |
| YOL077C | 1854.977032 | -0.340000405 | 0.128366692 | 0.61 |
| YOL105C | 760.4552602 | 0.460627502 | 0.138475524 | 0.61 |
| YOL113W | 1261.758159 | 0.78412948 | 0.133597676 | 0.61 |
| YOR044W | 320.4821165 | 0.517151959 | 0.13193587 | 0.61 |
| YOR189W | 408.1511083 | -0.549717448 | 0.137961682 | 0.61 |
| YOR275C | 1566.16532 | 0.372873905 | 0.138608918 | 0.61 |
| YOR332W | 2877.284969 | -0.397932883 | 0.130992956 | 0.61 |
| YOR345C | 1.957434997 | 0.045902242 | 0.131586396 | 0.61 |
| YPL028W | 3100.321144 | -0.448139977 | 0.133744612 | 0.61 |
| YPL052W | 344.0806128 | 0.689374772 | 0.128395671 | 0.61 |
| YPL057C | 1620.500312 | 0.642987657 | 0.138089966 | 0.61 |
| YPL222W | 848.2140163 | 0.771914806 | 0.133404023 | 0.61 |
| YPL244C | 876.1384791 | -0.541142211 | 0.138371598 | 0.61 |

| | | | | |
|---|---|---|---|---|
| snR46 | 23.07413489 | -0.318208473 | 0.136167879 | 0.61 |
| snR80 | 3.471418478 | -0.18236301 | 0.134252072 | 0.61 |
| snR81 | 9.309990788 | -0.11517683 | 0.130386793 | 0.61 |
| YDL052C | 1575.894255 | -0.453317534 | 0.140468266 | 0.61 |
| YDL082W | 7404.050186 | -0.29889912 | 0.140591055 | 0.61 |
| YDR237W | 832.9477874 | -0.485079578 | 0.140436234 | 0.61 |
| YGR128C | 1414.526792 | -0.486084045 | 0.140610818 | 0.61 |
| YJL039C | 2015.018256 | -0.476952689 | 0.140756309 | 0.61 |
| YHL017W | 1099.085857 | 0.551301318 | 0.141309856 | 0.61 |
| YPR072W | 2174.009813 | -0.441383835 | 0.141543196 | 0.61 |
| snR7-L | 6.909111604 | -0.857934241 | 0.141702724 | 0.61 |
| YBR018C | 184.0387102 | 0.458159358 | 0.14312729 | 0.61 |
| YBR084C-A | 8707.666036 | -0.293202359 | 0.143873807 | 0.61 |
| YDL132W | 2917.981172 | 0.45295697 | 0.142046839 | 0.61 |
| YDL142C | 615.3529618 | 0.486921615 | 0.142398938 | 0.61 |
| YDL224C | 1328.137391 | -0.58638652 | 0.142304721 | 0.61 |
| YFL037W | 5301.265498 | -0.337365551 | 0.141971545 | 0.61 |
| YFL051C | 87.84562174 | 0.726585126 | 0.144313824 | 0.61 |
| YGL213C | 596.0286848 | -0.460409883 | 0.143438182 | 0.61 |
| YGR029W | 703.0611046 | -0.481818374 | 0.143844977 | 0.61 |
| YGR201C | 974.6666707 | 0.629687365 | 0.143454444 | 0.61 |
| YHL013C | 503.0447213 | -0.419121979 | 0.143380294 | 0.61 |
| YHR097C | 2968.055756 | 0.617902109 | 0.142441317 | 0.61 |
| YJL101C | 2271.091311 | 0.524672594 | 0.144097672 | 0.61 |
| YJL213W | 560.8105438 | 0.63785381 | 0.144507585 | 0.61 |
| YJR126C | 1827.592541 | 0.413857843 | 0.142964884 | 0.61 |
| YLR394W | 730.9999237 | 0.495615543 | 0.144615572 | 0.61 |
| YMR165C | 1579.930507 | 0.572057491 | 0.142643263 | 0.61 |
| YMR274C | 249.323644 | -0.609905338 | 0.144228743 | 0.61 |
| YNL056W | 768.1896527 | -0.429200498 | 0.142531813 | 0.61 |
| YNL187W | 311.4857358 | 0.46000326 | 0.143501534 | 0.61 |
| YNR029C | 598.402815 | -0.537878636 | 0.142475914 | 0.61 |
| YOR161C-C | 9.014220677 | 0.256841244 | 0.144291331 | 0.61 |
| YOR297C | 445.9780063 | -0.494513193 | 0.142740482 | 0.61 |
| YPL037C | 6009.036088 | -0.562578261 | 0.144141564 | 0.61 |
| YPL081W | 1016.551966 | -0.69718458 | 0.14454609 | 0.61 |
| YPL130W | 24.21083011 | 0.441830978 | 0.144026326 | 0.61 |
| YPL146C | 923.2028372 | -0.372903073 | 0.142445477 | 0.61 |
| YPL175W | 580.2634042 | -0.482146579 | 0.144648542 | 0.61 |
| YPL212C | 1188.64058 | -0.411808619 | 0.142807704 | 0.61 |
| YPR029C | 1930.500091 | -0.389901226 | 0.144566372 | 0.61 |
| YPR079W | 754.8054031 | 0.405470001 | 0.144357915 | 0.61 |
| YBR132C | 1205.603555 | 0.653824109 | 0.145309739 | 0.61 |

| | | | | |
|---|---|---|---|---|
| YBR295W | 678.020174 | 0.621018371 | 0.145334725 | 0.61 |
| YOL052C | 1181.165007 | -0.422113118 | 0.145294322 | 0.61 |
| YOL153C | 1352.172864 | 0.517689491 | 0.145244861 | 0.61 |
| YOR101W | 962.4280426 | -0.594203332 | 0.145173742 | 0.61 |
| YOR204W | 2305.73227 | -0.464852653 | 0.145202562 | 0.61 |
| YJR059W | 3492.702056 | 0.576990274 | 0.145456645 | 0.61 |
| YLR399C | 2272.649731 | 0.477367218 | 0.145600256 | 0.61 |
| YLR135W | 729.3825255 | 0.452245468 | 0.1457425 | 0.61 |
| YAL063C | 272.6184868 | 0.51093919 | 0.151224375 | 0.61 |
| YAR015W | 7474.541729 | -0.11507877 | 0.150880659 | 0.61 |
| YAR023C | 164.0948317 | 0.311921137 | 0.153581143 | 0.61 |
| YBL009W | 764.0807255 | -0.371394617 | 0.155245719 | 0.61 |
| YBL064C | 4712.365868 | 0.862929061 | 0.152567315 | 0.61 |
| YBL107C | 641.7007619 | -0.540630499 | 0.156569419 | 0.61 |
| YBR005W | 1310.131508 | 0.60818948 | 0.150826045 | 0.61 |
| YBR017C | 2405.834993 | -0.454456602 | 0.151973876 | 0.61 |
| YBR079C | 5290.065687 | -0.518744768 | 0.160212617 | 0.61 |
| YBR096W | 835.4279583 | -0.453326551 | 0.161011662 | 0.61 |
| YBR200W | 1077.368125 | -0.389974467 | 0.157909959 | 0.61 |
| YBR212W | 2837.682435 | 0.599695274 | 0.155422917 | 0.61 |
| YBR213W | 228.053134 | -0.3604255 | 0.156172922 | 0.61 |
| YBR238C | 642.2716869 | -0.72626873 | 0.157961028 | 0.61 |
| YBR258C | 250.7365151 | -0.503115798 | 0.157270776 | 0.61 |
| YBR271W | 387.9475512 | -0.461801604 | 0.16050358 | 0.61 |
| YCL038C | 682.2522644 | 0.428865611 | 0.160950274 | 0.61 |
| YCL046W | 22.38513549 | -0.513592318 | 0.158826685 | 0.61 |
| YCL050C | 4005.943189 | -0.247353045 | 0.154914698 | 0.61 |
| YCR022C | 20.80731429 | 0.534299456 | 0.155384462 | 0.61 |
| YDL022W | 5632.112705 | 0.621727796 | 0.147394581 | 0.61 |
| YDL159W-A | 1.037341025 | 0.42663884 | 0.155448843 | 0.61 |
| YDL166C | 434.2604045 | -0.579092376 | 0.14863594 | 0.61 |
| YDL167C | 1031.301724 | -0.711093761 | 0.160197575 | 0.61 |
| YDL195W | 7573.167842 | -0.361746749 | 0.159664513 | 0.61 |
| YDL198C | 1156.426648 | -0.338777872 | 0.147083248 | 0.61 |
| YDL247W | 3.022100866 | -0.251892759 | 0.148895139 | 0.61 |
| YDR012W | 4314.996695 | -0.018065008 | 0.156943963 | 0.61 |
| YDR098C | 1711.271644 | -0.345580651 | 0.155247172 | 0.61 |
| YDR134C | 8310.682574 | 0.663895502 | 0.156370386 | 0.61 |
| YDR144C | 842.4657292 | -0.450449425 | 0.160106949 | 0.61 |
| YDR202C | 884.8533036 | 0.537412935 | 0.146250541 | 0.61 |
| YDR210C-D | 43.7176532 | 1.097711091 | 0.149246414 | 0.61 |
| YDR221W | 936.1837264 | -0.407095553 | 0.154138053 | 0.61 |
| YDR225W | 3245.605018 | -0.731643168 | 0.155460577 | 0.61 |

| | | | | |
|---|---|---|---|---|
| YDR244W | 923.2552739 | 0.439852568 | 0.154296409 | 0.61 |
| YDR360W | 36.19514521 | -0.546146941 | 0.155124347 | 0.61 |
| YDR379W | 1429.073535 | 0.505488724 | 0.156958583 | 0.61 |
| YDR385W | 1851.31364 | -0.518750624 | 0.147868295 | 0.61 |
| YDR390C | 1683.184066 | -0.373522878 | 0.159420989 | 0.61 |
| YDR400W | 652.8362371 | -0.462556128 | 0.150716213 | 0.61 |
| YDR404C | 728.6666751 | -0.528828941 | 0.156223087 | 0.61 |
| YDR519W | 900.2252274 | -0.419717712 | 0.160943516 | 0.61 |
| YDR524C-B | 18745.499 | -0.449512354 | 0.158406392 | 0.61 |
| YEL022W | 2372.854744 | -0.456944768 | 0.157544707 | 0.61 |
| YEL040W | 3165.37713 | -0.788029132 | 0.147678134 | 0.61 |
| YER053C | 3873.936022 | 0.687102848 | 0.161065143 | 0.61 |
| YER076W-A | 76.7457419 | 0.444124339 | 0.14965503 | 0.61 |
| YFL013W-A | 84.24870204 | 0.589677805 | 0.160604812 | 0.61 |
| YFL024C | 1322.826078 | 0.399205845 | 0.15686566 | 0.61 |
| YFR009W | 2637.309837 | -0.462388806 | 0.149596034 | 0.61 |
| YGL022W | 3864.561612 | -0.360822191 | 0.158133663 | 0.61 |
| YGL037C | 14451.53815 | 0.698383217 | 0.147315061 | 0.61 |
| YGL056C | 2031.538502 | 0.601954117 | 0.147294159 | 0.61 |
| YGL062W | 7360.866755 | 1.026402836 | 0.150794014 | 0.61 |
| YGL066W | 1160.200958 | 0.451126858 | 0.156932934 | 0.61 |
| YGL073W | 2411.314838 | 0.503640048 | 0.150312014 | 0.61 |
| YGL079W | 448.9297125 | -0.419943763 | 0.149290895 | 0.61 |
| YGL090W | 485.8897575 | 0.403372981 | 0.150439904 | 0.61 |
| YGL134W | 814.0702966 | 0.559300275 | 0.152450637 | 0.61 |
| YGL135W | 2224.916712 | -0.399842967 | 0.146155379 | 0.61 |
| YGL154C | 461.1131948 | 0.38265571 | 0.159846729 | 0.61 |
| YGL163C | 996.8369865 | 0.526138536 | 0.158752084 | 0.61 |
| YGL170C | 69.87350446 | 0.429970726 | 0.146942632 | 0.61 |
| YGL178W | 2073.150568 | 0.594174158 | 0.160771652 | 0.61 |
| YGL227W | 1564.861142 | 0.602599064 | 0.161332679 | 0.61 |
| YGR036C | 221.213258 | -0.619826647 | 0.157215908 | 0.61 |
| YGR089W | 1598.86435 | 0.415907362 | 0.150140297 | 0.61 |
| YGR159C | 2044.642627 | -0.353364226 | 0.149259231 | 0.61 |
| YGR160W | 19.24237613 | -0.373826055 | 0.154618154 | 0.61 |
| YGR161C-D | 41.89517477 | 1.018427911 | 0.160483922 | 0.61 |
| YGR175C | 3323.313876 | -0.597829238 | 0.158783232 | 0.61 |
| YGR192C | 181337.4741 | -0.243866339 | 0.148532604 | 0.61 |
| YGR243W | 1286.002734 | 0.644964619 | 0.152715074 | 0.61 |
| YHL003C | 1333.653462 | -0.52319886 | 0.159736613 | 0.61 |
| YHR034C | 511.3439371 | 0.506113367 | 0.158848677 | 0.61 |
| YHR083W | 1675.31411 | -0.417211691 | 0.160121497 | 0.61 |
| YHR098C | 2961.484376 | -0.370865515 | 0.152347897 | 0.61 |

| | | | | |
|---|---|---|---|---|
| YHR104W | 3547.736943 | 0.576898058 | 0.152698969 | 0.61 |
| YHR128W | 1328.56215 | -0.377441916 | 0.151902956 | 0.61 |
| YHR139C-A | 36.30751378 | 0.621755754 | 0.157835548 | 0.61 |
| YHR152W | 1107.927047 | -0.387334395 | 0.156442333 | 0.61 |
| YHR181W | 756.8304441 | -0.429366947 | 0.154706226 | 0.61 |
| YIL006W | 374.9591791 | 0.497143842 | 0.147753282 | 0.61 |
| YIL076W | 2351.989968 | -0.452799037 | 0.155936009 | 0.61 |
| YIL078W | 8337.626041 | -0.257317457 | 0.157354316 | 0.61 |
| YIL122W | 600.3807645 | 0.473817317 | 0.147979039 | 0.61 |
| YIL123W | 4293.520238 | -0.421732277 | 0.157388625 | 0.61 |
| YIR010W | 867.0379525 | 0.378441076 | 0.14665494 | 0.61 |
| YIR024C | 366.6225047 | 0.434440134 | 0.150859879 | 0.61 |
| YIR034C | 2228.546885 | 0.66538917 | 0.155355572 | 0.61 |
| YJL009W | 66.31151028 | -0.486854444 | 0.147900158 | 0.61 |
| YJL048C | 1114.208522 | 0.68422364 | 0.147149165 | 0.61 |
| YJL136C | 5415.03448 | -0.127211798 | 0.155049963 | 0.61 |
| YJL147C | 286.6419753 | 0.435651342 | 0.161480621 | 0.61 |
| YJL169W | 25.36216702 | -0.707768158 | 0.152465015 | 0.61 |
| YJR032W | 585.2468444 | -0.488072856 | 0.147676929 | 0.61 |
| YJR097W | 96.20405896 | -0.20619148 | 0.153139497 | 0.61 |
| YKL076C | 34.14810955 | 0.610056256 | 0.157857149 | 0.61 |
| YKL121W | 1073.818565 | 0.448797887 | 0.15198372 | 0.61 |
| YKL143W | 1110.759937 | -0.279714579 | 0.160242702 | 0.61 |
| YKL170W | 81.53735369 | -0.506397414 | 0.151246372 | 0.61 |
| YKL181W | 2705.955695 | -0.56473803 | 0.150580495 | 0.61 |
| YKR067W | 2101.266331 | 0.692315803 | 0.159830025 | 0.61 |
| YLL039C | 7097.91074 | 0.704636331 | 0.149907738 | 0.61 |
| YLR030W | 185.5800298 | 0.298165782 | 0.152452945 | 0.61 |
| YLR044C | 136554.4679 | -0.090245785 | 0.158970909 | 0.61 |
| YLR092W | 690.3329885 | 0.514762418 | 0.157992666 | 0.61 |
| YLR232W | 2.842598345 | -0.588757945 | 0.161148287 | 0.61 |
| YLR249W | 32716.59454 | -0.414678568 | 0.155805457 | 0.61 |
| YLR287C | 617.6178514 | -0.43654113 | 0.150423292 | 0.61 |
| YLR363W-A | 269.4630453 | -0.351764907 | 0.158715192 | 0.61 |
| YLR422W | 2135.341687 | 0.508223658 | 0.153070385 | 0.61 |
| YLR462W | 4.00598291 | 0.368472539 | 0.154496917 | 0.61 |
| YML004C | 3324.450509 | 0.512544649 | 0.160168973 | 0.61 |
| YML035C | 2451.342109 | 0.520750989 | 0.161511903 | 0.61 |
| YML036W | 292.5583071 | -0.585638771 | 0.149391406 | 0.61 |
| YML105C | 1352.843987 | -0.44537919 | 0.152293337 | 0.61 |
| YML119W | 1027.81525 | 0.514936851 | 0.152940232 | 0.61 |
| YMR017W | 57.51133406 | 0.420098831 | 0.147146121 | 0.61 |
| YMR041C | 591.2121868 | 0.425660394 | 0.151933427 | 0.61 |

| | | | | |
|---|---|---|---|---|
| YMR078C | 885.7332334 | -0.45588979 | 0.147880548 | 0.61 |
| YMR158C-A | 22.93483608 | 0.43366881 | 0.154368703 | 0.61 |
| YMR182W-A | 4.590688512 | 0.408446474 | 0.154566692 | 0.61 |
| YMR203W | 3020.852994 | -0.408181087 | 0.157201321 | 0.61 |
| YMR261C | 3042.037261 | 0.563695317 | 0.156911825 | 0.61 |
| YMR294W-A | 229.7802387 | 0.444146308 | 0.155183844 | 0.61 |
| YMR325W | 4.52283349 | 0.939550924 | 0.147267637 | 0.61 |
| YNL103W | 1585.703492 | 0.551629036 | 0.15797405 | 0.61 |
| YNL172W | 1164.569131 | 0.543049786 | 0.152074148 | 0.61 |
| YNL188W | 592.8677499 | -0.755300986 | 0.154567457 | 0.61 |
| YNL189W | 2792.970706 | -0.452590827 | 0.147356097 | 0.61 |
| YNL222W | 196.3405424 | -0.694242165 | 0.157980067 | 0.61 |
| YNL262W | 2257.387419 | -0.215016546 | 0.161456318 | 0.61 |
| YNR007C | 864.4677932 | 0.690243561 | 0.146967745 | 0.61 |
| YNR039C | 749.4014615 | 0.324461951 | 0.154807533 | 0.61 |
| YOL047C | 302.1448912 | 0.164157609 | 0.159005726 | 0.61 |
| YOL080C | 199.8794327 | -0.236679854 | 0.157271969 | 0.61 |
| YOL088C | 665.7794028 | -0.610666933 | 0.158194177 | 0.61 |
| YOL093W | 662.3279101 | -0.561970332 | 0.155872495 | 0.61 |
| YOL103W-B | 37.1755698 | 0.230283049 | 0.158677842 | 0.61 |
| YOL121C | 3681.856963 | -0.452472662 | 0.146199191 | 0.61 |
| YOR008C-A | 9.788861467 | -0.358697579 | 0.153762671 | 0.61 |
| YOR015W | 345.4012917 | 0.510371382 | 0.153530581 | 0.61 |
| YOR020C | 3482.406455 | -0.346591098 | 0.14609991 | 0.61 |
| YOR052C | 2749.089296 | 0.554704053 | 0.158018466 | 0.61 |
| YOR059C | 1230.819833 | 0.456962625 | 0.15754705 | 0.61 |
| YOR091W | 1558.309002 | -0.398284397 | 0.148442963 | 0.61 |
| YOR121C | 64.33732395 | 1.042485405 | 0.15956921 | 0.61 |
| YOR122C | 4814.102226 | -0.417061695 | 0.160301766 | 0.61 |
| YOR138C | 1254.624127 | 0.423832781 | 0.148000422 | 0.61 |
| YOR163W | 1216.735785 | -0.45573268 | 0.153537954 | 0.61 |
| YOR206W | 2506.527701 | -0.514640597 | 0.151818171 | 0.61 |
| YOR283W | 529.1178643 | -0.596450969 | 0.150142337 | 0.61 |
| YOR295W | 256.0382172 | -0.507008075 | 0.15610425 | 0.61 |
| YOR300W | 17.40293291 | 0.559095847 | 0.149035417 | 0.61 |
| YPL039W | 426.2421359 | 0.48916489 | 0.157797244 | 0.61 |
| YPL094C | 2895.773005 | -0.410391501 | 0.160204551 | 0.61 |
| YPL117C | 2245.25569 | -0.407132021 | 0.160199024 | 0.61 |
| YPL123C | 1037.088707 | 0.599280063 | 0.148779645 | 0.61 |
| YPL186C | 1114.795126 | 0.614921353 | 0.160890346 | 0.61 |
| YPL227C | 650.9518323 | -0.488224947 | 0.160442831 | 0.61 |
| YPR046W | 225.601515 | 0.503004968 | 0.155885158 | 0.61 |
| YPR074C | 10075.28978 | -0.36080992 | 0.146836107 | 0.61 |

| | | | | |
|---|---|---|---|---|
| YPR110C | 2091.025042 | -0.393034417 | 0.149758156 | 0.61 |
| YPR118W | 1223.179848 | -0.457974655 | 0.160481432 | 0.61 |
| YPR140W | 900.9897118 | -0.621871335 | 0.155056762 | 0.61 |
| snR34 | 37.83791355 | -0.245811362 | 0.15439535 | 0.61 |
| YCR043C | 438.4206899 | -0.544178798 | 0.162125755 | 0.61 |
| YDL015C | 3069.696513 | -0.368273594 | 0.162113598 | 0.61 |
| YGR181W | 835.8007421 | -0.457472895 | 0.162050353 | 0.61 |
| YIR019C | 622.3080892 | 0.667952347 | 0.161781687 | 0.61 |
| YPL003W | 547.4702758 | 0.434537508 | 0.161750273 | 0.61 |
| YPL255W | 357.750249 | -0.57293209 | 0.161929686 | 0.61 |
| YNL129W | 332.9229067 | -0.468918739 | 0.162237239 | 0.61 |
| YDR124W | 300.8040212 | 0.501875957 | 0.162355102 | 0.61 |
| YOL134C | 23.62819378 | 0.303308109 | 0.16249908 | 0.61 |
| YER110C | 3689.619393 | -0.189005118 | 0.162856892 | 0.61 |
| YMR169C | 812.630576 | 0.422437507 | 0.162876715 | 0.61 |
| YKR077W | 759.0676767 | 0.626616256 | 0.162971053 | 0.61 |
| YJR060W | 664.6110196 | -0.406315182 | 0.163106992 | 0.61 |
| YHL023C | 1253.971871 | 0.415095252 | 0.16354628 | 0.61 |
| YHR203C | 5825.930229 | -0.128497818 | 0.163703861 | 0.61 |
| YKL122C | 1563.613747 | -0.47041348 | 0.163829146 | 0.61 |
| YNL078W | 1260.908163 | -0.660378196 | 0.163672537 | 0.61 |
| YNL135C | 5166.62026 | -0.423222366 | 0.163669683 | 0.61 |
| YNL154C | 2670.283788 | -0.56803137 | 0.16347443 | 0.61 |
| YOR346W | 878.2111125 | -0.577384083 | 0.163817501 | 0.61 |
| YOR221C | 1523.763907 | 0.505385025 | 0.163977607 | 0.61 |
| YNR009W | 344.9832981 | -0.628572921 | 0.164135546 | 0.61 |
| YNL130C | 1326.028586 | -0.451151329 | 0.164681147 | 0.61 |
| YBR056W | 2794.886987 | 0.533981744 | 0.164889923 | 0.61 |
| YBR148W | 178.9686946 | 0.547307715 | 0.165380407 | 0.62 |
| YGL245W | 8715.959502 | -0.310299981 | 0.1655236 | 0.62 |
| YGR051C | 51.37808205 | 0.313942206 | 0.165535311 | 0.62 |
| YNL305C | 2021.497779 | 0.718709675 | 0.165486611 | 0.62 |
| YOR316C-A | 8.072827245 | -0.894156721 | 0.165591118 | 0.62 |
| YAR064W | 14.8431874 | -0.631879142 | 0.165712198 | 0.62 |
| YDR200C | 868.5642679 | 0.361532435 | 0.166202312 | 0.62 |
| YJL093C | 1063.991787 | 0.375820138 | 0.166640036 | 0.62 |
| YBR297W | 465.5986861 | 0.889173464 | 0.166880571 | 0.62 |
| YLR023C | 1130.122098 | 0.421312201 | 0.167039633 | 0.62 |
| YCR090C | 636.078747 | -0.560861108 | 0.16727657 | 0.62 |
| YBL032W | 1731.697141 | -0.36629324 | 0.172085071 | 0.62 |
| YBL042C | 1258.710363 | -0.887503584 | 0.168079267 | 0.62 |
| YBL072C | 3997.128198 | -0.396516911 | 0.169013311 | 0.62 |
| YBR210W | 464.9591183 | -0.534190461 | 0.16847413 | 0.62 |

| | | | | |
|---|---|---|---|---|
| YDL061C | 2048.312712 | -0.290328361 | 0.171602413 | 0.62 |
| YDL064W | 884.0529801 | -0.379721441 | 0.168823861 | 0.62 |
| YDL130W | 4709.707857 | -0.37828997 | 0.172201602 | 0.62 |
| YDL145C | 6294.266606 | -0.336469233 | 0.170706699 | 0.62 |
| YDR075W | 328.511722 | -0.690620799 | 0.170430041 | 0.62 |
| YDR370C | 533.8393634 | -0.530031847 | 0.169540823 | 0.62 |
| YEL001C | 2451.407888 | -0.423195285 | 0.171878149 | 0.62 |
| YFR019W | 2589.385757 | 0.539416308 | 0.170210317 | 0.62 |
| YGL033W | 39.3922468 | 1.040791423 | 0.171213618 | 0.62 |
| YGL156W | 2401.006767 | 0.849104605 | 0.171039252 | 0.62 |
| YGL215W | 3518.182479 | -0.381373334 | 0.171478568 | 0.62 |
| YGR025W | 100.9215658 | -0.407401236 | 0.170035673 | 0.62 |
| YGR264C | 2825.266453 | -0.397366421 | 0.171715557 | 0.62 |
| YIL059C | 89.463829 | 0.205299311 | 0.168571957 | 0.62 |
| YJL014W | 2604.88629 | -0.441178286 | 0.16855135 | 0.62 |
| YJL092W | 1243.946562 | -0.35968349 | 0.170526023 | 0.62 |
| YJL097W | 1713.191405 | -0.424877974 | 0.168923226 | 0.62 |
| YJL121C | 2601.821139 | -0.369528064 | 0.167852892 | 0.62 |
| YJL190C | 9254.873132 | -0.250207822 | 0.170243164 | 0.62 |
| YKL115C | 42.82706794 | -0.53084677 | 0.171920995 | 0.62 |
| YKL119C | 426.0044347 | -0.544274669 | 0.168146012 | 0.62 |
| YKL127W | 1973.346678 | -0.214605844 | 0.171693961 | 0.62 |
| YKL153W | 3671.352718 | -0.474740823 | 0.171154303 | 0.62 |
| YKR045C | 368.478154 | -0.452449544 | 0.170771355 | 0.62 |
| YLR022C | 541.692103 | -0.52407982 | 0.170909776 | 0.62 |
| YLR065C | 843.2361186 | -0.600358239 | 0.171470674 | 0.62 |
| YLR179C | 2935.995773 | -0.233125869 | 0.169989885 | 0.62 |
| YLR225C | 1476.977135 | 0.51916894 | 0.17119677 | 0.62 |
| YLR292C | 1010.798537 | -0.454038978 | 0.168621368 | 0.62 |
| YLR318W | 479.1884801 | 0.376569837 | 0.168514676 | 0.62 |
| YMR129W | 1909.721365 | -0.335156484 | 0.172087598 | 0.62 |
| YMR216C | 1336.629718 | -0.522105898 | 0.168335701 | 0.62 |
| YMR303C | 445.8492587 | 0.181626529 | 0.170671946 | 0.62 |
| YNR031C | 1352.088361 | -0.403243214 | 0.168740349 | 0.62 |
| YNR075C-A | 12.77963038 | -0.201914945 | 0.170229354 | 0.62 |
| YOL007C | 839.2247362 | -0.550958578 | 0.170626004 | 0.62 |
| YOL044W | 473.6470695 | 0.583274332 | 0.170666094 | 0.62 |
| YOL138C | 1055.538589 | 0.386290608 | 0.168223353 | 0.62 |
| YOL155C | 4950.84233 | 0.966695431 | 0.169234731 | 0.62 |
| YOR013W | 86.58484926 | -0.701870376 | 0.168874166 | 0.62 |
| YOR061W | 1662.759146 | -0.442255001 | 0.169550378 | 0.62 |
| YOR077W | 225.6560466 | 0.45956185 | 0.172149358 | 0.62 |
| YOR187W | 4877.755588 | -0.384463642 | 0.169901118 | 0.62 |

| | | | | |
|---|---|---|---|---|
| YOR252W | 480.3993858 | -0.297670243 | 0.167929926 | 0.62 |
| YOR298C-A | 6204.241243 | -0.43795816 | 0.170465977 | 0.62 |
| snR51 | 25.09375178 | 0.360398929 | 0.171977037 | 0.62 |
| YER177W | 16691.55629 | -0.410588612 | 0.172385444 | 0.62 |
| YDL056W | 1411.798857 | 0.462292588 | 0.173020772 | 0.62 |
| YDR395W | 2175.578678 | -0.457260612 | 0.173101055 | 0.62 |
| YJR094W-A | 1689.898779 | -0.307008138 | 0.172989296 | 0.62 |
| YKL096W | 1798.684379 | -0.491145701 | 0.173048146 | 0.62 |
| YKL190W | 813.5201171 | -0.540495348 | 0.172935744 | 0.62 |
| YMR087W | 259.2299237 | 0.446152682 | 0.173164296 | 0.62 |
| YMR233W | 360.9564304 | -0.794350159 | 0.172973226 | 0.62 |
| YNL291C | 628.873147 | -0.443571061 | 0.172777189 | 0.62 |
| YBL084C | 817.2451769 | 0.451970918 | 0.173535165 | 0.62 |
| YDL229W | 7732.532873 | -0.153594371 | 0.173624869 | 0.62 |
| YLL024C | 20764.11389 | 0.115976773 | 0.173685289 | 0.62 |
| YPL144W | 173.5181567 | -0.4384444 | 0.173706639 | 0.62 |
| YAL025C | 983.4578298 | -0.477426654 | 0.176304642 | 0.62 |
| YBR290W | 1203.208406 | 0.410207354 | 0.175860652 | 0.62 |
| YDL023C | 219.5343501 | 0.617682144 | 0.175177458 | 0.62 |
| YDL150W | 525.2641532 | -0.458987089 | 0.176349902 | 0.62 |
| YDL201W | 813.4402377 | -0.462295208 | 0.176457819 | 0.62 |
| YDR114C | 36.82268978 | 0.349513611 | 0.174100924 | 0.62 |
| YDR160W | 879.6911624 | 0.334053829 | 0.176488708 | 0.62 |
| YDR254W | 574.0903535 | 0.38859027 | 0.176570004 | 0.62 |
| YDR326C | 1698.894045 | 0.443647163 | 0.174220783 | 0.62 |
| YDR440W | 548.6066441 | -0.457899351 | 0.175424817 | 0.62 |
| YDR465C | 1088.193625 | -0.282546031 | 0.175704943 | 0.62 |
| YEL053W-A | 186.8810681 | -0.509846236 | 0.176264373 | 0.62 |
| YER078C | 919.3267409 | 0.403697094 | 0.176061113 | 0.62 |
| YGL093W | 1066.84026 | 0.353079888 | 0.17407248 | 0.62 |
| YGL096W | 785.2336716 | 0.5378704 | 0.174227204 | 0.62 |
| YGL258W-A | 56.62039045 | 0.853181193 | 0.175718344 | 0.62 |
| YKL133C | 601.5672696 | 0.508920935 | 0.174327226 | 0.62 |
| YKR064W | 1141.244604 | 0.383840239 | 0.17607415 | 0.62 |
| YLL050C | 6100.834604 | -0.378123356 | 0.176523714 | 0.62 |
| YLR007W | 741.9239686 | 0.509582684 | 0.174671995 | 0.62 |
| YLR206W | 3965.73873 | 0.415318006 | 0.175081652 | 0.62 |
| YLR260W | 983.6383575 | 0.623451244 | 0.17480224 | 0.62 |
| YMR011W | 5664.342762 | 0.498581925 | 0.174914812 | 0.62 |
| YMR146C | 5181.578577 | -0.429686777 | 0.176365897 | 0.62 |
| YNL035C | 765.5856918 | -0.50959141 | 0.176539309 | 0.62 |
| YNL233W | 992.805205 | -0.340325249 | 0.175114587 | 0.62 |
| YOR293C-A | 2.608736587 | -0.559914077 | 0.175203982 | 0.62 |

| | | | | |
|---|---|---|---|---|
| YPL267W | 353.8623989 | -0.390150499 | 0.174566655 | 0.62 |
| YPR049C | 968.2983267 | 0.654346676 | 0.175674495 | 0.62 |
| YCR065W | 756.8632472 | -0.721653712 | 0.176799544 | 0.62 |
| YDL158C | 89.30551169 | -0.426572332 | 0.176792789 | 0.62 |
| YER104W | 241.2969985 | 0.519813152 | 0.17730695 | 0.62 |
| YIL044C | 893.1546159 | -0.489886974 | 0.177327024 | 0.62 |
| YBR068C | 7312.593911 | 0.565564795 | 0.177550712 | 0.62 |
| YDR490C | 1094.352116 | 0.513662992 | 0.177864793 | 0.62 |
| YGL113W | 616.1660547 | 0.444017414 | 0.177757891 | 0.62 |
| YMR191W | 2004.986398 | 0.901847694 | 0.177936673 | 0.62 |
| YPR160W | 12134.47846 | 0.921627546 | 0.1778308 | 0.62 |
| YBR067C | 11273.4348 | 0.370145553 | 0.178155639 | 0.62 |
| YOR080W | 677.2163658 | 0.294475476 | 0.178180465 | 0.62 |
| YJR033C | 1583.049905 | 0.430608908 | 0.178283251 | 0.62 |
| YBR080C | 2198.347068 | -0.386854686 | 0.179728678 | 0.62 |
| YBR128C | 336.3131952 | 0.518848701 | 0.178597265 | 0.62 |
| YBR196C-A | 7.414882286 | 0.177484267 | 0.179786359 | 0.62 |
| YDR057W | 882.7471612 | 0.423582021 | 0.179782923 | 0.62 |
| YDR060W | 3331.110803 | -0.412938025 | 0.179116828 | 0.62 |
| YDR353W | 5059.752159 | -0.397945462 | 0.179203749 | 0.62 |
| YDR391C | 1742.594469 | 0.597586305 | 0.178586244 | 0.62 |
| YGL001C | 1350.851203 | -0.616583401 | 0.179811426 | 0.62 |
| YGL003C | 856.5289623 | 0.377652536 | 0.179885637 | 0.62 |
| YGR154C | 333.0792836 | 0.393223364 | 0.178996157 | 0.62 |
| YIL020C-A | 191.6015626 | -0.515947354 | 0.178684503 | 0.62 |
| YIL158W | 651.6543059 | -0.478068503 | 0.179587436 | 0.62 |
| YLR349W | 23.19539208 | 0.738956968 | 0.17888761 | 0.62 |
| YLR393W | 270.1892054 | 0.55482007 | 0.178944579 | 0.62 |
| YMR049C | 1525.516535 | -0.298477546 | 0.179529151 | 0.62 |
| YNL249C | 367.0811931 | 0.358703492 | 0.179293617 | 0.62 |
| YER020W | 2824.873338 | 0.583505758 | 0.180049131 | 0.62 |
| YDL140C | 8072.034708 | -0.342437973 | 0.180453752 | 0.62 |
| YKL201C | 1146.760044 | 0.514850496 | 0.1806769 | 0.62 |
| YKR072C | 2109.757496 | 0.557044569 | 0.180945223 | 0.62 |
| YNR034W-A | 4345.183948 | 0.697772931 | 0.180616928 | 0.62 |
| YOL142W | 655.5671288 | -0.352494818 | 0.180852909 | 0.62 |
| YPL093W | 1935.182804 | -0.511431884 | 0.180797993 | 0.62 |
| YDL102W | 928.5717968 | -0.290921449 | 0.181322252 | 0.63 |
| YDL237W | 1508.528864 | -0.448599534 | 0.181359494 | 0.63 |
| YOL126C | 3179.903333 | 0.44148082 | 0.181413643 | 0.63 |
| YJR013W | 645.2124988 | -0.376169825 | 0.181798933 | 0.63 |
| YOL049W | 2360.13263 | -0.465554043 | 0.181793163 | 0.63 |
| YCL069W | 36.22888945 | 0.403058728 | 0.18209455 | 0.63 |

| | | | | |
|---|---|---|---|---|
| YER075C | 845.204575 | 0.359944784 | 0.182267825 | 0.63 |
| YNL202W | 289.1619632 | 0.478719712 | 0.182245247 | 0.63 |
| YPL189C-A | 341.9617668 | 0.431044724 | 0.182360628 | 0.63 |
| YPL203W | 1453.358861 | 0.470991706 | 0.182482526 | 0.63 |
| YDR313C | 770.3934735 | 0.510386636 | 0.182623227 | 0.63 |
| YER155C | 4972.789594 | -0.423647656 | 0.182851026 | 0.63 |
| YIL109C | 4372.511551 | -0.324094334 | 0.183163975 | 0.63 |
| YCR015C | 255.0577346 | -0.437971393 | 0.183792576 | 0.63 |
| YER148W | 733.3492834 | -0.589187536 | 0.183723084 | 0.63 |
| YLR120C | 2959.418798 | 0.586603028 | 0.183728539 | 0.63 |
| YML098W | 504.9087228 | -0.487416298 | 0.183723233 | 0.63 |
| YBR147W | 531.1286909 | 0.347890919 | 0.184553202 | 0.63 |
| YDR516C | 6053.645479 | 0.831179081 | 0.18468739 | 0.63 |
| YIL042C | 814.6115743 | 0.567615046 | 0.184615477 | 0.63 |
| YIR014W | 256.3884009 | 0.766400857 | 0.184490739 | 0.63 |
| YLR143W | 501.3258454 | -0.454146902 | 0.184547238 | 0.63 |
| YNR054C | 724.980203 | -0.36411382 | 0.184633677 | 0.63 |
| YMR217W | 5540.397432 | -0.225860638 | 0.184889571 | 0.63 |
| YOR349W | 572.160655 | 0.384786143 | 0.184884029 | 0.63 |
| YJR002W | 1689.001321 | -0.391008242 | 0.185106522 | 0.63 |
| YLR099W-A | 174.4936248 | -0.534298338 | 0.1850951 | 0.63 |
| YDL134C | 1962.281647 | 0.219043535 | 0.185711001 | 0.63 |
| YDR432W | 4551.301879 | -0.387492169 | 0.185368674 | 0.63 |
| YLR441C | 16750.1152 | -0.330740746 | 0.185711936 | 0.63 |
| YMR219W | 3001.660673 | 0.619163341 | 0.185506512 | 0.63 |
| YOR301W | 501.5102259 | -0.491966496 | 0.185602903 | 0.63 |
| YPL082C | 1537.0483 | -0.488143824 | 0.185664512 | 0.63 |
| YBL044W | 3.828780377 | 0.31956433 | 0.186275777 | 0.63 |
| YJL042W | 3416.022292 | 0.555981424 | 0.186176476 | 0.63 |
| YJR075W | 2038.161429 | -0.367259369 | 0.186281651 | 0.63 |
| YOR335C | 3110.388093 | -0.414252903 | 0.186345015 | 0.63 |
| YGL239C | 18.18533761 | -0.597112262 | 0.186647152 | 0.63 |
| YBR153W | 413.971492 | 0.419536546 | 0.186975859 | 0.63 |
| YDR102C | 12.68433027 | -1.126661691 | 0.187053361 | 0.63 |
| YIL065C | 779.4474724 | 0.563152099 | 0.187076167 | 0.63 |
| YIL143C | 1830.368647 | 0.528276356 | 0.187513995 | 0.63 |
| YBR225W | 1153.176358 | 0.497393352 | 0.187673003 | 0.63 |
| YIL126W | 2338.60398 | -0.429042564 | 0.188047037 | 0.63 |
| YPL249C | 1511.229449 | 0.429451342 | 0.188011995 | 0.63 |
| YPL229W | 514.8866138 | 0.564781571 | 0.188205115 | 0.63 |
| YML002W | 853.1962719 | 0.400702778 | 0.188351621 | 0.63 |
| YHL039W | 1028.011858 | -0.465634164 | 0.189430234 | 0.63 |
| YJR045C | 25492.38268 | -0.301148573 | 0.188937756 | 0.63 |

| | | | | |
|---|---|---|---|---|
| YMR251W-A | 5878.547147 | 0.684443908 | 0.188659885 | 0.63 |
| YNL260C | 268.8863879 | -0.433606361 | 0.18885766 | 0.63 |
| YPL070W | 1079.494033 | 0.455590536 | 0.189350357 | 0.63 |
| YPL152W-A | 1.958413515 | -0.307520423 | 0.189195107 | 0.63 |
| YPL273W | 2617.65817 | -0.479862787 | 0.1892204 | 0.63 |
| YPR003C | 637.4553595 | 0.533503002 | 0.18947283 | 0.63 |
| YPR143W | 537.9356283 | -0.285671284 | 0.189052641 | 0.63 |
| snR47 | 5.450594875 | -0.004125508 | 0.189333712 | 0.63 |
| YAL031C | 1097.389337 | 0.423760177 | 0.195693646 | 0.64 |
| YAL067C | 87.22318015 | -0.700861908 | 0.192227729 | 0.64 |
| YBL056W | 3616.602261 | -0.287961214 | 0.195224441 | 0.64 |
| YBR072C-A | 2.769886284 | -0.598714623 | 0.194971864 | 0.64 |
| YCL008C | 1030.586387 | 0.377325148 | 0.190831639 | 0.64 |
| YCR010C | 273.3869127 | 0.497077551 | 0.195891538 | 0.64 |
| YDL045W-A | 694.1101578 | -0.533085799 | 0.193890874 | 0.64 |
| YDL094C | 21.86703582 | 0.798506451 | 0.194849398 | 0.64 |
| YDR001C | 5262.491506 | 0.560642693 | 0.195199176 | 0.64 |
| YDR024W | 7.360211428 | 0.384584316 | 0.190542164 | 0.64 |
| YDR033W | 12974.70679 | -0.441550768 | 0.194346027 | 0.64 |
| YDR062W | 2339.714713 | -0.399563055 | 0.192538515 | 0.64 |
| YDR120C | 1240.636645 | -0.270471059 | 0.193130852 | 0.64 |
| YDR174W | 3004.282379 | -0.416363532 | 0.192256926 | 0.64 |
| YDR226W | 6443.35181 | -0.402097413 | 0.194769446 | 0.64 |
| YDR279W | 575.5796481 | -0.379708675 | 0.194111591 | 0.64 |
| YDR286C | 156.0831168 | -0.589055647 | 0.192070599 | 0.64 |
| YDR352W | 489.6613604 | -0.471519876 | 0.190430558 | 0.64 |
| YDR507C | 1606.54523 | -0.331768983 | 0.191689741 | 0.64 |
| YEL058W | 1683.65634 | -0.380280101 | 0.192713571 | 0.64 |
| YER144C | 1434.76784 | 0.508348989 | 0.194361002 | 0.64 |
| YER147C-A | 275.4373645 | -0.584004427 | 0.193866711 | 0.64 |
| YGL171W | 777.7803509 | -0.318266658 | 0.194602967 | 0.64 |
| YGL230C | 33.65395104 | 0.262956391 | 0.195318978 | 0.64 |
| YGL257C | 1193.728393 | -0.436352268 | 0.192717456 | 0.64 |
| YGR035W-A | 93.40510398 | -0.528731809 | 0.194467877 | 0.64 |
| YGR073C | 7.437514438 | -1.337329883 | 0.190574202 | 0.64 |
| YGR103W | 3122.539574 | -0.112192613 | 0.194820097 | 0.64 |
| YHL028W | 1137.369035 | -0.893706569 | 0.192145526 | 0.64 |
| YHR006W | 1327.030831 | 0.535965669 | 0.195621431 | 0.64 |
| YHR022C-A | 3.937421936 | -0.052304497 | 0.19472352 | 0.64 |
| YHR042W | 4638.852766 | -0.312936306 | 0.19583012 | 0.64 |
| YHR095W | 5.929557488 | -0.270773722 | 0.192000212 | 0.64 |
| YJL002C | 4648.101511 | -0.442249606 | 0.195322403 | 0.64 |
| YJR096W | 2414.849421 | 0.568554605 | 0.191478724 | 0.64 |

| | | | | |
|---|---|---|---|---|
| YJR130C | 1173.075557 | 0.505756705 | 0.190420566 | 0.64 |
| YJR152W | 334.6737416 | 0.332959142 | 0.195468635 | 0.64 |
| YKL036C | 247.9148773 | 0.547215581 | 0.193651581 | 0.64 |
| YKL125W | 592.2613366 | 0.368111775 | 0.192562497 | 0.64 |
| YKL150W | 7424.00365 | 0.783146192 | 0.190587524 | 0.64 |
| YKR026C | 956.8364878 | -0.463771665 | 0.191944669 | 0.64 |
| YLR041W | 27.97790779 | -1.084267777 | 0.191533665 | 0.64 |
| YLR047C | 709.1954371 | 0.323282 | 0.1918719 | 0.64 |
| YML082W | 766.1872079 | -0.420374948 | 0.194659973 | 0.64 |
| YMR036C | 631.5049431 | 0.409304864 | 0.194995652 | 0.64 |
| YMR115W | 1106.490843 | 0.516077975 | 0.19260195 | 0.64 |
| YMR304W | 1821.67097 | 0.508751566 | 0.192963461 | 0.64 |
| YNL226W | 19.21780824 | 0.761143113 | 0.19132678 | 0.64 |
| YNL300W | 1681.600577 | -0.203879262 | 0.191856911 | 0.64 |
| YNL326C | 366.0651518 | -0.376933033 | 0.195290728 | 0.64 |
| YOL090W | 1084.862315 | -0.424455533 | 0.195409049 | 0.64 |
| YOL106W | 14.51135789 | 0.552139751 | 0.189814903 | 0.64 |
| YOR020W-A | 1169.212022 | 0.621466277 | 0.191506667 | 0.64 |
| YOR072W | 29.04315607 | 0.423627917 | 0.19290583 | 0.64 |
| YOR279C | 315.7050157 | -0.540327091 | 0.19033541 | 0.64 |
| YPL074W | 944.5738824 | 0.421427752 | 0.195314612 | 0.64 |
| YPL128C | 628.8955352 | -0.448769709 | 0.195130602 | 0.64 |
| YPL196W | 943.9259177 | 0.541857756 | 0.194518309 | 0.64 |
| YPL243W | 2085.694288 | -0.502794438 | 0.195743646 | 0.64 |
| YPR009W | 346.2208437 | -0.931416305 | 0.193233511 | 0.64 |
| YPR051W | 72.40075259 | -0.636261007 | 0.192073141 | 0.64 |
| YPR131C | 480.7629468 | -0.433578186 | 0.190888836 | 0.64 |
| YPR158W-B | 29.96033516 | 1.082604539 | 0.194501598 | 0.64 |
| YPR185W | 1716.783017 | 0.467815559 | 0.194361616 | 0.64 |
| YDL174C | 2966.298062 | 0.552370795 | 0.196256532 | 0.64 |
| YDR045C | 241.2309895 | -0.529591638 | 0.196159005 | 0.64 |
| YLR075W | 9637.869009 | -0.193666159 | 0.196297044 | 0.64 |
| YNL150W | 83.29807569 | -0.19548324 | 0.196439299 | 0.64 |
| YAL065C | 13.48444795 | -0.488591655 | 0.196701296 | 0.64 |
| YER179W | 293.0579118 | 0.508190427 | 0.196719014 | 0.64 |
| YMR003W | 306.5250482 | -0.279405928 | 0.196751576 | 0.64 |
| YMR123W | 1008.572929 | -0.308790592 | 0.196856621 | 0.64 |
| YBL097W | 438.5923101 | -0.316237441 | 0.197303163 | 0.64 |
| YER116C | 578.3983863 | 0.531285404 | 0.197337735 | 0.64 |
| YGL140C | 1656.222341 | 0.361237808 | 0.197416622 | 0.64 |
| YJL201W | 1030.90288 | 0.336812352 | 0.197212536 | 0.64 |
| YOR220W | 2249.332399 | 0.524678963 | 0.197070125 | 0.64 |
| YGL054C | 1677.23256 | -0.445619128 | 0.197782169 | 0.64 |

| | | | | |
|---|---|---|---|---|
| YGR289C | 671.1201555 | 0.350323173 | 0.197751199 | 0.64 |
| YHR189W | 309.5122663 | 0.513690194 | 0.197946125 | 0.64 |
| YJR038C | 33.2004855 | 0.568152588 | 0.197867891 | 0.64 |
| YMR124W | 1653.834225 | 0.486956628 | 0.198036711 | 0.64 |
| YMR251W | 257.8128007 | 0.568636453 | 0.197985392 | 0.64 |
| YCL033C | 876.4314652 | 0.768066024 | 0.199234993 | 0.64 |
| YCR020C | 245.5464449 | 0.316235439 | 0.199348391 | 0.64 |
| YDR083W | 1132.402572 | -0.409229294 | 0.198792899 | 0.64 |
| YDR121W | 548.7039891 | -0.365939384 | 0.199446317 | 0.64 |
| YDR281C | 42.74964081 | -0.8817596 | 0.198667889 | 0.64 |
| YER023W | 2339.829047 | -0.450361827 | 0.199321017 | 0.64 |
| YGL223C | 914.5737412 | -0.371946718 | 0.199013358 | 0.64 |
| YGR024C | 991.4360363 | -0.343351512 | 0.199015067 | 0.64 |
| YHR201C | 1092.809897 | -0.586539723 | 0.198881931 | 0.64 |
| YIR023C-A | 13.57792364 | -0.429753074 | 0.198434655 | 0.64 |
| YLR053C | 106.3162181 | 0.637775523 | 0.199137982 | 0.64 |
| YLR312W-A | 784.781583 | -0.427441376 | 0.199337053 | 0.64 |
| YNL197C | 1704.51539 | -0.500547085 | 0.19895603 | 0.64 |
| YGL049C | 2260.025541 | 0.331290788 | 0.199605722 | 0.64 |
| YER099C | 790.1735302 | 0.321762883 | 0.199902562 | 0.64 |
| YGR114C | 36.93060097 | 0.059643948 | 0.199898324 | 0.64 |
| YML076C | 1190.303539 | 0.447628953 | 0.199729399 | 0.64 |
| YKL161C | 387.0663817 | 0.214491097 | 0.200353225 | 0.64 |
| YLL027W | 1273.25217 | 0.481628984 | 0.200394719 | 0.64 |
| YLR059C | 1264.300383 | -0.382467208 | 0.200436543 | 0.64 |
| YMR243C | 2759.15272 | -0.391936548 | 0.200584593 | 0.64 |
| YPR091C | 1815.236332 | 0.386499276 | 0.200105795 | 0.64 |
| snR4 | 89.67640266 | 0.001196714 | 0.200562907 | 0.64 |
| YBR052C | 2651.004465 | -0.464370994 | 0.201236535 | 0.64 |
| YDR165W | 1014.540812 | -0.388827156 | 0.201113579 | 0.64 |
| YIR035C | 1381.941223 | -0.490842585 | 0.200912775 | 0.64 |
| YLR241W | 1618.382874 | 0.412657541 | 0.200990036 | 0.64 |
| YNL112W | 1857.470789 | -0.353906463 | 0.200834152 | 0.64 |
| YNL158W | 434.8285056 | 0.336874235 | 0.201211384 | 0.64 |
| YBL030C | 9599.31549 | -0.07372275 | 0.202145157 | 0.64 |
| YGR017W | 1237.157084 | -0.384370599 | 0.201477326 | 0.64 |
| YGR150C | 1213.396537 | -0.371866858 | 0.20158528 | 0.64 |
| YGR240C | 9640.703605 | -0.483629813 | 0.202118564 | 0.64 |
| YGR293C | 7.546648827 | -0.371230506 | 0.202141117 | 0.64 |
| YIL165C | 180.1950386 | 0.325377255 | 0.201886005 | 0.64 |
| YLR255C | 23.50063941 | 0.773771811 | 0.201912586 | 0.64 |
| YPL250W-A | 88.80290187 | -0.577157354 | 0.202088014 | 0.64 |
| YER079W | 1934.057175 | 0.679884973 | 0.202764599 | 0.64 |

| | | | | |
|---|---|---|---|---|
| YKL220C | 368.1832027 | 0.345028039 | 0.202678812 | 0.64 |
| YML126C | 4399.479194 | -0.332027089 | 0.202861813 | 0.64 |
| YDL136W | 915.4579307 | -0.37814792 | 0.203199207 | 0.64 |
| YDR324C | 1092.059972 | -0.409450114 | 0.203205377 | 0.64 |
| YGR174C | 1048.598299 | 0.559010833 | 0.203198723 | 0.64 |
| YDR262W | 1927.726265 | 0.416150574 | 0.203468776 | 0.64 |
| YMR220W | 1203.063636 | -0.421037422 | 0.203491146 | 0.64 |
| YDR426C | 18.4381422 | 0.345516367 | 0.203663093 | 0.64 |
| YMR215W | 1997.531105 | -0.47419394 | 0.203751141 | 0.64 |
| YBR130C | 1160.408881 | -0.497392266 | 0.205697251 | 0.64 |
| YDR340W | 4.60165433 | -0.742939932 | 0.205758483 | 0.64 |
| YER149C | 245.5366344 | -0.337592743 | 0.20530392 | 0.64 |
| YER175C | 1510.549619 | 0.621827013 | 0.20575566 | 0.64 |
| YFL031C-A | 101.8569422 | -0.480215911 | 0.204289367 | 0.64 |
| YGL207W | 4342.654723 | -0.372556591 | 0.204797973 | 0.64 |
| YGR268C | 875.2811265 | 0.366621392 | 0.205834883 | 0.64 |
| YHR123W | 699.5030842 | -0.344492906 | 0.205747976 | 0.64 |
| YIR016W | 1147.104809 | 0.460809297 | 0.205227617 | 0.64 |
| YKL046C | 1814.036225 | -0.339424008 | 0.20422087 | 0.64 |
| YLR404W | 245.17757 | -0.540125775 | 0.204384801 | 0.64 |
| YLR409C | 1307.908763 | -0.407313994 | 0.205803025 | 0.64 |
| YMR099C | 2686.030784 | -0.313221395 | 0.205535375 | 0.64 |
| YMR160W | 946.3755314 | 0.443464771 | 0.204725489 | 0.64 |
| YNL132W | 1681.161762 | -0.184314721 | 0.204612046 | 0.64 |
| YNL248C | 1500.899157 | -0.345942072 | 0.204774003 | 0.64 |
| YPL260W | 2621.937685 | 0.434633935 | 0.205020586 | 0.64 |
| YPR190C | 965.2019513 | -0.329520955 | 0.204658737 | 0.64 |
| YPR202W | 17.57397274 | -0.035271917 | 0.205805386 | 0.64 |
| YHR215W | 110.5268911 | -0.809190448 | 0.205970212 | 0.64 |
| YDL215C | 3694.89417 | 0.741180739 | 0.206510373 | 0.64 |
| YDR496C | 1668.42234 | -0.384724176 | 0.206933743 | 0.64 |
| YFR006W | 3007.667423 | -0.352034457 | 0.206798538 | 0.64 |
| YHR197W | 861.9145076 | -0.372997555 | 0.206806965 | 0.64 |
| YLR209C | 881.8739647 | -0.292978794 | 0.206908475 | 0.64 |
| YLR262C-A | 800.3537696 | -0.498964791 | 0.206699518 | 0.64 |
| YMR291W | 2090.324495 | 0.455168938 | 0.2067833 | 0.64 |
| YAR031W | 151.5423779 | 0.512688089 | 0.207249206 | 0.64 |
| YDR190C | 2078.525809 | -0.473996043 | 0.207177853 | 0.64 |
| YCL007C | 102.0392471 | -0.363652007 | 0.207807367 | 0.64 |
| YNL327W | 5207.566398 | -0.422804617 | 0.207717477 | 0.64 |
| YNL254C | 501.4634557 | 0.46579764 | 0.208432712 | 0.64 |
| YKR092C | 1945.135383 | -0.390040885 | 0.208641153 | 0.64 |
| YOR389W | 203.2493311 | 0.595035619 | 0.208585232 | 0.64 |

| | | | | |
|---|---|---|---|---|
| YDR473C | 858.9200777 | 0.407080237 | 0.208781672 | 0.64 |
| YCL012C | 422.2313505 | 0.654394026 | 0.209452033 | 0.64 |
| YCR057C | 1417.95789 | -0.462347364 | 0.209241229 | 0.64 |
| YDL222C | 906.6701758 | 0.276090364 | 0.209333081 | 0.64 |
| YMR205C | 7427.558626 | -0.508272802 | 0.209074438 | 0.64 |
| YMR293C | 413.7964048 | 0.465393784 | 0.209137579 | 0.64 |
| YNR073C | 195.2436335 | 1.033063171 | 0.209488042 | 0.64 |
| YOR267C | 3236.118357 | 0.457439754 | 0.20891459 | 0.64 |
| YJR015W | 1958.669387 | -0.430378211 | 0.209606653 | 0.64 |
| YAR002W | 1776.649309 | -0.459081491 | 0.210174759 | 0.65 |
| YBL067C | 1161.166699 | 0.3172723 | 0.211253801 | 0.65 |
| YBR219C | 1.059885032 | -0.061400536 | 0.210459399 | 0.65 |
| YDR002W | 3692.839207 | -0.316727736 | 0.211106835 | 0.65 |
| YDR379C-A | 462.1957351 | 0.436678 | 0.210790946 | 0.65 |
| YEL017W | 1022.681916 | -0.270070387 | 0.21117875 | 0.65 |
| YGL175C | 162.4549988 | 0.418697437 | 0.210188369 | 0.65 |
| YGR122C-A | 15.60192201 | -1.025352585 | 0.210225026 | 0.65 |
| YGR152C | 296.8399677 | -0.434392699 | 0.211098051 | 0.65 |
| YJR120W | 18.09619799 | 0.31502674 | 0.210053793 | 0.65 |
| YKL166C | 506.1904776 | -0.52544762 | 0.210887772 | 0.65 |
| YKR003W | 1162.972466 | -0.485274705 | 0.210050781 | 0.65 |
| YKR041W | 32.67705338 | -0.431088103 | 0.211095245 | 0.65 |
| YMR031W-A | 40.58383151 | 0.508027521 | 0.211298286 | 0.65 |
| YOL057W | 2160.171024 | -0.414180854 | 0.210458821 | 0.65 |
| YPR167C | 421.1211 | 0.587730476 | 0.211014135 | 0.65 |
| YCR064C | 12.51518247 | -0.35338216 | 0.212229812 | 0.65 |
| YDR192C | 476.3597916 | 0.373809377 | 0.212376411 | 0.65 |
| YER152W-A | 37.9606133 | 0.534082282 | 0.212297074 | 0.65 |
| YIR026C | 772.7201823 | -0.183872971 | 0.211978811 | 0.65 |
| YKL026C | 613.714189 | 0.497525757 | 0.21234015 | 0.65 |
| YOL076W | 856.9107843 | -0.586344827 | 0.212281962 | 0.65 |
| YOR254C | 3232.98471 | -0.358472198 | 0.211851136 | 0.65 |
| YPL160W | 4849.742262 | -0.218487444 | 0.211937353 | 0.65 |
| YPR200C | 70.27684883 | -0.688446135 | 0.211954345 | 0.65 |
| YGL153W | 279.9517565 | 0.448545236 | 0.213119566 | 0.65 |
| YER107W-A | 76.0626659 | -0.437799375 | 0.213591092 | 0.65 |
| YMR149W | 2739.886665 | -0.440404778 | 0.213500638 | 0.65 |
| YNL115C | 1894.716258 | 0.549298513 | 0.213835683 | 0.65 |
| YGR060W | 3114.502526 | -0.460666903 | 0.214029918 | 0.65 |
| YBR169C | 4813.961053 | 0.684424917 | 0.214508007 | 0.65 |
| YBR171W | 1056.70053 | -0.366640475 | 0.214438619 | 0.65 |
| YDR267C | 660.9899359 | -0.329661073 | 0.215232185 | 0.65 |
| YGL106W | 2381.873892 | -0.439311091 | 0.215403767 | 0.65 |

| | | | | |
|---|---|---|---|---|
| YHR172W | 514.8081565 | -0.391012149 | 0.215037621 | 0.65 |
| YIL029C | 174.946717 | 0.403709063 | 0.21543307 | 0.65 |
| YKL004W | 2100.85667 | -0.416517965 | 0.214994401 | 0.65 |
| YKL012W | 1327.340355 | 0.303710546 | 0.214734361 | 0.65 |
| YKL106C-A | 1.684117879 | -0.591759009 | 0.215296803 | 0.65 |
| YLR431C | 452.0758432 | 0.36367675 | 0.214682375 | 0.65 |
| YMR308C | 3347.564301 | -0.499365303 | 0.215415037 | 0.65 |
| YNL015W | 3387.233502 | 0.919743653 | 0.214792042 | 0.65 |
| YNR047W | 1084.837265 | 0.595745343 | 0.215236785 | 0.65 |
| YIL131C | 677.2016628 | -0.365800838 | 0.215783884 | 0.65 |
| YLR005W | 967.4119109 | 0.348297531 | 0.215731178 | 0.65 |
| YBR136W | 1309.50165 | 0.587757746 | 0.21632372 | 0.65 |
| YDL108W | 550.3870487 | 0.338114684 | 0.216360092 | 0.65 |
| YNL293W | 734.6319527 | 0.466086809 | 0.216222906 | 0.65 |
| YPL220W | 1761.369957 | -0.278904133 | 0.216152973 | 0.65 |
| YDL085W | 757.5796414 | 0.518564536 | 0.216594919 | 0.65 |
| YLR180W | 3585.379162 | -0.143289999 | 0.216597765 | 0.65 |
| YML077W | 412.7293887 | -0.479272689 | 0.216752863 | 0.65 |
| YDL059C | 182.996654 | 0.506745854 | 0.217094743 | 0.65 |
| YGR118W | 4349.282861 | -0.47069999 | 0.217179375 | 0.65 |
| YDR413C | 10.51150758 | -0.375268866 | 0.217514348 | 0.65 |
| YIL144W | 1230.130031 | 0.45496096 | 0.217673603 | 0.65 |
| YJL082W | 1623.02873 | 0.551062027 | 0.217550069 | 0.65 |
| YKL097C | 10.27058523 | 0.924684725 | 0.217605864 | 0.65 |
| YBR282W | 512.5901703 | -0.318329709 | 0.217909793 | 0.65 |
| YNL160W | 19462.39163 | 0.566880713 | 0.217949802 | 0.65 |
| YER087W | 372.1981021 | 0.491961311 | 0.218383983 | 0.65 |
| YML009C | 743.0693468 | -0.313584138 | 0.21826444 | 0.65 |
| YOR098C | 2501.292329 | -0.354238839 | 0.218406567 | 0.65 |
| YKL090W | 556.2185917 | 0.263357705 | 0.218657002 | 0.65 |
| YMR111C | 424.0046487 | 0.344069478 | 0.218765571 | 0.65 |
| YGL158W | 260.9463165 | -0.644157052 | 0.219236396 | 0.65 |
| YOL016C | 3937.741808 | 0.521372191 | 0.219125782 | 0.65 |
| YOL022C | 1058.507084 | -0.296034895 | 0.219236844 | 0.65 |
| YPR124W | 1891.733227 | -0.075389749 | 0.219249822 | 0.65 |
| YLR344W | 3310.960859 | -0.318478299 | 0.219588712 | 0.65 |
| YDR091C | 1728.536947 | -0.403873218 | 0.22014272 | 0.65 |
| YDR236C | 550.9804022 | -0.409463418 | 0.219924798 | 0.65 |
| YHL020C | 961.5750146 | 0.440164836 | 0.220146344 | 0.65 |
| YJL111W | 2828.454028 | -0.427837003 | 0.220507422 | 0.65 |
| YJR089W | 953.5011996 | 0.365198554 | 0.22025679 | 0.65 |
| YKL033W | 666.8103831 | -0.374452179 | 0.219900293 | 0.65 |
| YML069W | 2256.742432 | -0.497788292 | 0.21992969 | 0.65 |

| | | | | |
|---|---|---|---|---|
| YNL334C | 19.40823403 | -0.970208139 | 0.220351937 | 0.65 |
| YOR384W | 1087.084219 | 0.495371255 | 0.220460147 | 0.65 |
| YER039C-A | 49.35745349 | 0.241166141 | 0.220654137 | 0.65 |
| YAR035C-A | 1.313184395 | 0.282118908 | 0.221574448 | 0.65 |
| YBR162C | 7471.337212 | -0.354116239 | 0.222439857 | 0.65 |
| YBR242W | 549.7068913 | -0.384361678 | 0.221085526 | 0.65 |
| YDR183W | 267.9198317 | -0.398974833 | 0.221631306 | 0.65 |
| YDR212W | 2389.189098 | -0.430424381 | 0.22259695 | 0.65 |
| YDR299W | 1150.958917 | -0.43562117 | 0.221264103 | 0.65 |
| YDR368W | 2244.977391 | -0.424088154 | 0.222246217 | 0.65 |
| YDR381W | 4059.585724 | -0.437026261 | 0.223396294 | 0.65 |
| YER005W | 999.8103274 | 0.258649259 | 0.221491359 | 0.65 |
| YER047C | 911.6547256 | 0.408154923 | 0.221898496 | 0.65 |
| YFR025C | 947.4315897 | -0.250894607 | 0.223474653 | 0.65 |
| YGL067W | 999.1477018 | -0.405639959 | 0.223575035 | 0.65 |
| YGL089C | 132.0787924 | 0.377467182 | 0.223492574 | 0.65 |
| YGR121C | 1176.230558 | 0.464002757 | 0.22303193 | 0.65 |
| YGR237C | 1848.787726 | 0.501434197 | 0.223166954 | 0.65 |
| YKL137W | 420.0820122 | -0.395156449 | 0.221684831 | 0.65 |
| YKL151C | 2678.459393 | 0.580075134 | 0.222751708 | 0.65 |
| YLR131C | 1246.543234 | 0.451643399 | 0.221150195 | 0.65 |
| YLR259C | 10469.29817 | -0.232286456 | 0.221986566 | 0.65 |
| YLR264W | 525.6829336 | -0.063622998 | 0.222393201 | 0.65 |
| YLR425W | 1120.132672 | 0.384296174 | 0.223510544 | 0.65 |
| YML025C | 1270.924858 | 0.30062511 | 0.223461455 | 0.65 |
| YML128C | 9230.440998 | 0.605056551 | 0.22295989 | 0.65 |
| YMR225C | 419.9415237 | -0.359919768 | 0.221641956 | 0.65 |
| YNL167C | 1564.476742 | 0.433170958 | 0.222610242 | 0.65 |
| YNL208W | 6496.983749 | 0.466380221 | 0.22327114 | 0.65 |
| YPL106C | 19467.56403 | -0.109193747 | 0.222901213 | 0.65 |
| YPL277C | 206.5316036 | 0.550855476 | 0.221135384 | 0.65 |
| YMR090W | 1879.223335 | 0.597334022 | 0.223695851 | 0.65 |
| YGL117W | 750.1030158 | 0.493466459 | 0.223971768 | 0.65 |
| YNL072W | 456.0809998 | -0.376269961 | 0.223898332 | 0.65 |
| YBR252W | 608.885071 | -0.290775908 | 0.22426141 | 0.65 |
| YIL069C | 2018.584495 | -0.43864266 | 0.224226698 | 0.65 |
| YHR149C | 909.9713222 | -0.497080713 | 0.224637896 | 0.66 |
| YER124C | 2039.69695 | -0.465304874 | 0.224818721 | 0.66 |
| YFL001W | 478.2892235 | -0.380394405 | 0.22518652 | 0.66 |
| YIL027C | 793.9722774 | -0.450438665 | 0.225248698 | 0.66 |
| YML107C | 389.7043417 | 0.343197169 | 0.225082166 | 0.66 |
| YDR230W | 158.0472236 | 0.598127109 | 0.225354897 | 0.66 |
| YCR020C-A | 266.9746924 | -0.346543943 | 0.225473063 | 0.66 |

| | | | | |
|---|---|---|---|---|
| YLR257W | 6468.084537 | 0.538554722 | 0.225628679 | 0.66 |
| YCL035C | 4330.054767 | 0.59169347 | 0.225744818 | 0.66 |
| YBL024W | 2012.558866 | -0.467068828 | 0.22622258 | 0.66 |
| YCL010C | 597.9803619 | 0.434049074 | 0.226301446 | 0.66 |
| YGR257C | 921.5740821 | -0.454369064 | 0.226391391 | 0.66 |
| YOL021C | 1670.215668 | -0.34672828 | 0.226210093 | 0.66 |
| YKR043C | 2568.82772 | -0.033432466 | 0.226606758 | 0.66 |
| YJR110W | 661.1509666 | 0.314208164 | 0.226773084 | 0.66 |
| YOL163W | 57.10674557 | 0.514722208 | 0.226882351 | 0.66 |
| YBR043C | 1241.74171 | 0.512300831 | 0.227885991 | 0.66 |
| YEL046C | 2157.479841 | -0.473601128 | 0.227158395 | 0.66 |
| YGL216W | 962.5040676 | 0.35880779 | 0.227691921 | 0.66 |
| YGR019W | 2682.062979 | 0.662801369 | 0.227246335 | 0.66 |
| YLR392C | 638.9786435 | 0.431137522 | 0.227705793 | 0.66 |
| YNL095C | 702.5523068 | 0.388729422 | 0.227929225 | 0.66 |
| YOL055C | 533.3864453 | 0.385498623 | 0.22789951 | 0.66 |
| YOL128C | 161.4112011 | 0.370629888 | 0.227422882 | 0.66 |
| YPL198W | 414.9903406 | 0.236317879 | 0.227829808 | 0.66 |
| YDR037W | 9604.757078 | -0.050217886 | 0.229540846 | 0.66 |
| YDR104C | 711.7535955 | 0.4319747 | 0.228755281 | 0.66 |
| YDR231C | 627.5516275 | 0.530785124 | 0.229147362 | 0.66 |
| YER187W | 132.2577032 | -1.014912563 | 0.228859521 | 0.66 |
| YGL139W | 1235.197549 | 0.352909703 | 0.228641 | 0.66 |
| YGR013W | 916.8674279 | 0.396990266 | 0.229058481 | 0.66 |
| YHR022C | 158.1795629 | -0.035074283 | 0.229507572 | 0.66 |
| YIR011C | 1166.213259 | 0.367435751 | 0.229380209 | 0.66 |
| YJL007C | 3.350774349 | 0.19700719 | 0.228261694 | 0.66 |
| YJL173C | 671.1948643 | -0.253442589 | 0.22903153 | 0.66 |
| YKL060C | 152874.8661 | -0.077829054 | 0.228551995 | 0.66 |
| YLR311C | 30.19399839 | 0.688282674 | 0.229428002 | 0.66 |
| YOR090C | 1602.791656 | -0.297456902 | 0.228961822 | 0.66 |
| YPL055C | 1072.396721 | 0.442309439 | 0.228970809 | 0.66 |
| YPL266W | 1132.936461 | -0.089835818 | 0.229325273 | 0.66 |
| YEL028W | 113.7795172 | 0.312185611 | 0.230379517 | 0.66 |
| YJL183W | 732.387825 | -0.297856074 | 0.230353644 | 0.66 |
| YJR004C | 1252.603429 | 0.417400229 | 0.230154861 | 0.66 |
| YLR373C | 1362.586121 | 0.360197022 | 0.229961676 | 0.66 |
| YNL061W | 2817.950295 | -0.286586015 | 0.230385224 | 0.66 |
| YOR302W | 66.2485987 | 0.529148184 | 0.230136114 | 0.66 |
| YKL172W | 2863.993194 | -0.393221155 | 0.230703734 | 0.66 |
| YOR136W | 7814.887983 | -0.257515566 | 0.230597008 | 0.66 |
| YPL048W | 7812.579467 | -0.343779283 | 0.230695642 | 0.66 |
| YOL082W | 2476.571718 | 0.376479287 | 0.230881332 | 0.66 |

| | | | | |
|---|---|---|---|---|
| YCL031C | 1450.481764 | -0.487785975 | 0.231503132 | 0.66 |
| YHR040W | 740.6643017 | 0.676693308 | 0.23150654 | 0.66 |
| YOL096C | 679.2678424 | 0.38460905 | 0.231532273 | 0.66 |
| YAR066W | 12.65917974 | -0.631188009 | 0.233834324 | 0.66 |
| YBR269C | 1256.661555 | 0.576450803 | 0.233858937 | 0.66 |
| YDL030W | 676.7024249 | 0.395889761 | 0.233228102 | 0.66 |
| YER028C | 507.0764562 | 0.544244072 | 0.233683724 | 0.66 |
| YFL052W | 300.3969613 | 0.576289159 | 0.234215014 | 0.66 |
| YGL020C | 1129.205349 | -0.377331731 | 0.233056334 | 0.66 |
| YGR094W | 6966.605497 | -0.281561151 | 0.234187639 | 0.66 |
| YGR102C | 796.5538343 | 0.428694716 | 0.233811774 | 0.66 |
| YGR167W | 3020.873713 | -0.326487754 | 0.232936563 | 0.66 |
| YJL129C | 2269.469864 | 0.309341293 | 0.231841251 | 0.66 |
| YJL179W | 307.4969877 | -0.374406255 | 0.234055787 | 0.66 |
| YJR005W | 1677.202028 | 0.35268311 | 0.234411184 | 0.66 |
| YKL191W | 1251.863612 | -0.233635549 | 0.233696547 | 0.66 |
| YLR384C | 2550.492331 | -0.270523189 | 0.233039666 | 0.66 |
| YLR452C | 1814.469836 | -1.030946813 | 0.231887149 | 0.66 |
| YML032C | 1432.723324 | 0.479440339 | 0.232291807 | 0.66 |
| YMR086C-A | 82.59944884 | 0.4511005 | 0.233448856 | 0.66 |
| YMR210W | 1188.233193 | 0.304168545 | 0.234105969 | 0.66 |
| YNL108C | 290.0576123 | -0.399466214 | 0.234325169 | 0.66 |
| YNL173C | 2504.496471 | 0.528840726 | 0.233620475 | 0.66 |
| YNL318C | 22.2967731 | 0.492238264 | 0.233710427 | 0.66 |
| YOL092W | 1398.16665 | -0.331671516 | 0.232019075 | 0.66 |
| YOR092W | 648.6032701 | 0.270582828 | 0.233563457 | 0.66 |
| YOR118W | 1462.615579 | 0.292921175 | 0.232789554 | 0.66 |
| YOR272W | 1204.374523 | -0.276249678 | 0.232393481 | 0.66 |
| YOR363C | 1212.557429 | 0.395471275 | 0.233598956 | 0.66 |
| YPR065W | 1458.659767 | -0.520144076 | 0.234024253 | 0.66 |
| snR33 | 120.9149393 | -0.605070105 | 0.233966301 | 0.66 |
| YER126C | 1138.972805 | -0.354902 | 0.234665641 | 0.66 |
| YKR011C | 1595.285427 | 0.55856822 | 0.234616617 | 0.66 |
| YEL050C | 1046.082243 | -0.348019005 | 0.23494823 | 0.66 |
| YOR159C | 320.0573697 | -0.482263793 | 0.23494579 | 0.66 |
| YGL105W | 8369.251023 | -0.330887785 | 0.235058486 | 0.66 |
| YOR367W | 630.0179766 | -0.50788003 | 0.235409636 | 0.66 |
| YDL013W | 806.2108951 | 0.393692448 | 0.235933074 | 0.66 |
| YDR048C | 13.8983324 | 0.132150766 | 0.235747406 | 0.66 |
| YLR163C | 1121.057782 | 0.423941365 | 0.235725516 | 0.66 |
| YMR268C | 351.0386447 | 0.329812107 | 0.236038035 | 0.66 |
| YOR380W | 392.7551247 | 0.355419047 | 0.236036043 | 0.66 |
| YAL017W | 3309.438838 | 0.5898667 | 0.236413051 | 0.66 |

| | | | | |
|---|---|---|---|---|
| YLR114C | 1321.213644 | -0.375526429 | 0.236348979 | 0.66 |
| YGR145W | 1225.275886 | -0.227000854 | 0.236576929 | 0.66 |
| YLR322W | 7.310944976 | -0.149567422 | 0.236971127 | 0.66 |
| YOL117W | 709.0122211 | 0.551922699 | 0.23680446 | 0.66 |
| YPL163C | 3377.565521 | 0.144607996 | 0.236962267 | 0.66 |
| YJL161W | 501.2537696 | 0.336714214 | 0.237376513 | 0.66 |
| YJL212C | 1268.474729 | 0.639392309 | 0.237243448 | 0.66 |
| YKL214C | 653.6592896 | -0.339185626 | 0.237405167 | 0.66 |
| YCR016W | 907.9551315 | -0.218643254 | 0.238632889 | 0.66 |
| YEL005C | 730.8016186 | 0.299666145 | 0.238281347 | 0.66 |
| YFR026C | 243.3675063 | 0.712135529 | 0.238859859 | 0.66 |
| YHR191C | 210.2719774 | 0.289671448 | 0.238930095 | 0.66 |
| YJL168C | 610.0870071 | -0.468554498 | 0.238891372 | 0.66 |
| YKR035C | 23.50975344 | 0.181406242 | 0.23878006 | 0.66 |
| YKR096W | 1782.315311 | 0.454529476 | 0.23815372 | 0.66 |
| YLR198C | 85.68816582 | -0.367393572 | 0.238465304 | 0.66 |
| YMR234W | 552.7610201 | -0.374464481 | 0.238820874 | 0.66 |
| YDL130W-A | 2765.774643 | 0.799939839 | 0.239366996 | 0.66 |
| YDR341C | 4249.5314 | -0.194710039 | 0.239667935 | 0.66 |
| YDR351W | 1313.626306 | 0.33675275 | 0.239237082 | 0.66 |
| YER142C | 720.8941802 | 0.611654326 | 0.239622469 | 0.66 |
| YGR085C | 1835.900279 | -0.310800121 | 0.239153309 | 0.66 |
| YHR116W | 348.981213 | 0.566372925 | 0.239295635 | 0.66 |
| YOR227W | 3483.244777 | 0.571959871 | 0.239662493 | 0.66 |
| YDL232W | 292.6096052 | -0.458092866 | 0.239844606 | 0.67 |
| YNL038W | 473.5438124 | -0.486043894 | 0.239971396 | 0.67 |
| YBL031W | 539.4437355 | -0.428907441 | 0.240297298 | 0.67 |
| YOL011W | 1428.931177 | 0.297775747 | 0.240198551 | 0.67 |
| YOR144C | 421.6026059 | -0.422705492 | 0.240177763 | 0.67 |
| YDL121C | 1113.601796 | -0.158086359 | 0.240545675 | 0.67 |
| YMR311C | 3644.500712 | 0.517580832 | 0.240527978 | 0.67 |
| YGR087C | 287.4531423 | 0.467281378 | 0.240827561 | 0.67 |
| YIR032C | 113.4974223 | 0.504033307 | 0.241112236 | 0.67 |
| YBR084W | 2864.241089 | -0.129538313 | 0.241546197 | 0.67 |
| YHR002W | 437.310458 | 0.440821163 | 0.241577439 | 0.67 |
| YJL159W | 37900.6137 | 0.707684876 | 0.241276111 | 0.67 |
| YOR108W | 1787.909268 | -0.330830325 | 0.241471261 | 0.67 |
| YCR002C | 1893.656785 | -0.354704033 | 0.241939673 | 0.67 |
| YDL155W | 763.3718322 | -0.421939761 | 0.242364808 | 0.67 |
| YGL061C | 446.8633151 | -0.310059838 | 0.242138098 | 0.67 |
| YJL191W | 879.6997619 | -0.471974273 | 0.242277789 | 0.67 |
| YMR247W-A | 22.89498367 | 0.32701548 | 0.242211768 | 0.67 |
| YAL016W | 2950.826603 | 0.384066151 | 0.242685786 | 0.67 |

| | | | | |
|---|---|---|---|---|
| YJR003C | 586.7430541 | -0.500959974 | 0.242737232 | 0.67 |
| YDR334W | 1700.440041 | 0.340847063 | 0.24289956 | 0.67 |
| YBR016W | 1677.943304 | 0.295610322 | 0.24426719 | 0.67 |
| YBR025C | 6377.054239 | -0.269595427 | 0.243624038 | 0.67 |
| YBR076W | 114.1570412 | 0.261434344 | 0.24420364 | 0.67 |
| YDL187C | 3.68310749 | 0.437882933 | 0.244092967 | 0.67 |
| YFL042C | 1495.188388 | 0.61252436 | 0.244479909 | 0.67 |
| YHR076W | 1124.266555 | -0.367518618 | 0.243553383 | 0.67 |
| YHR092C | 9528.876224 | 0.826013098 | 0.243926654 | 0.67 |
| YJR017C | 953.2894932 | -0.379809337 | 0.244550207 | 0.67 |
| YJR105W | 3472.841224 | -0.237972549 | 0.244020625 | 0.67 |
| YKL154W | 669.8187345 | -0.291024759 | 0.244547372 | 0.67 |
| YLR186W | 830.8214037 | -0.339722208 | 0.243903708 | 0.67 |
| YLR450W | 2180.849651 | 0.480875499 | 0.244639074 | 0.67 |
| YMR082C | 5.748276588 | -0.64567523 | 0.243754506 | 0.67 |
| YMR159C | 22.87110149 | 0.420432624 | 0.244066619 | 0.67 |
| YNR068C | 378.9501574 | 0.498200124 | 0.244627092 | 0.67 |
| YPL143W | 740.5431297 | -0.289667939 | 0.244219566 | 0.67 |
| RNA170 | 10.42132142 | 0.88265195 | 0.247000081 | 0.67 |
| YBL076C | 7280.098127 | -0.120820652 | 0.246569205 | 0.67 |
| YDR008C | 37.32753147 | -0.35993024 | 0.246515305 | 0.67 |
| YDR014W | 486.9454278 | 0.347152207 | 0.246743271 | 0.67 |
| YEL037C | 3155.723536 | -0.314190252 | 0.246830448 | 0.67 |
| YER121W | 184.2483 | 0.207744211 | 0.246646324 | 0.67 |
| YGR039W | 40.79718114 | -0.575117155 | 0.24763609 | 0.67 |
| YGR143W | 1625.66982 | 0.289157214 | 0.246459452 | 0.67 |
| YIR018C-A | 31.48971955 | 0.520169828 | 0.246532908 | 0.67 |
| YJR137C | 1214.839856 | 0.417716578 | 0.245596729 | 0.67 |
| YLR052W | 1071.581443 | 0.206132153 | 0.246253029 | 0.67 |
| YLR060W | 5384.459239 | -0.332653043 | 0.247254516 | 0.67 |
| YLR390W-A | 12426.05045 | 0.440341037 | 0.247423754 | 0.67 |
| YML094W | 513.2141562 | -0.314659297 | 0.245981889 | 0.67 |
| YML111W | 1306.672498 | -0.388459648 | 0.246085445 | 0.67 |
| YML123C | 2594.845401 | -0.770328013 | 0.247253588 | 0.67 |
| YNL151C | 742.0050839 | -0.379136656 | 0.245470821 | 0.67 |
| YNL281W | 1354.281332 | -0.380909726 | 0.246219872 | 0.67 |
| YOL086C | 59663.42444 | -0.242952509 | 0.246499246 | 0.67 |
| YOR284W | 207.2496443 | -0.591624698 | 0.245222865 | 0.67 |
| YOR323C | 2339.246441 | -0.317146021 | 0.245421224 | 0.67 |
| YPL012W | 1836.869245 | -0.271124313 | 0.247470264 | 0.67 |
| YPL131W | 24700.79965 | -0.160065438 | 0.247404828 | 0.67 |
| YPL140C | 588.07308 | 0.322355473 | 0.245867564 | 0.67 |
| YPR042C | 1338.934548 | 0.439108176 | 0.246170623 | 0.67 |

| | | | | |
|---|---|---|---|---|
| YPR077C | 1.201353204 | -0.070919485 | 0.245708707 | 0.67 |
| YPR137W | 608.8911311 | -0.458574872 | 0.247608773 | 0.67 |
| YLR181C | 881.938971 | 0.391540399 | 0.247876901 | 0.67 |
| YBL002W | 4494.23128 | -0.302217524 | 0.253681042 | 0.67 |
| YBL010C | 437.6652565 | -0.29474603 | 0.254297274 | 0.67 |
| YBL093C | 1069.915355 | 0.33176771 | 0.253987282 | 0.67 |
| YBL109W | 16.60491535 | 0.271655626 | 0.252796996 | 0.67 |
| YBR087W | 809.749781 | -0.399284705 | 0.252983845 | 0.67 |
| YCL028W | 2867.256977 | -0.324938412 | 0.253917343 | 0.67 |
| YCL054W | 1476.642208 | -0.267416059 | 0.24963803 | 0.67 |
| YCR073W-A | 1340.952291 | 0.615035302 | 0.252665189 | 0.67 |
| YDL070W | 4583.373904 | 0.496150196 | 0.254096791 | 0.67 |
| YDL141W | 1031.602218 | -0.303674545 | 0.24893413 | 0.67 |
| YDL204W | 5424.611742 | 0.595933943 | 0.250543649 | 0.67 |
| YDR009W | 246.880862 | 0.52763037 | 0.254022365 | 0.67 |
| YDR119W | 1442.902962 | -0.492593745 | 0.251716808 | 0.67 |
| YDR161W | 1294.242267 | -0.313650464 | 0.252343887 | 0.67 |
| YDR247W | 1026.990415 | 0.616789459 | 0.248948505 | 0.67 |
| YDR307W | 749.860008 | -0.376397404 | 0.250067907 | 0.67 |
| YDR338C | 695.8448786 | 0.359147182 | 0.254292138 | 0.67 |
| YDR470C | 683.2878102 | 0.271902273 | 0.254465009 | 0.67 |
| YER182W | 1071.612284 | 0.588399741 | 0.253982472 | 0.67 |
| YFR005C | 788.6190865 | -0.496669757 | 0.252726557 | 0.67 |
| YFR021W | 746.0129726 | 0.287444705 | 0.254612017 | 0.67 |
| YFR051C | 3059.620652 | -0.311453281 | 0.251507262 | 0.67 |
| YGL099W | 1111.20672 | -0.3283287 | 0.254617321 | 0.67 |
| YGL101W | 448.9089719 | -0.375899883 | 0.249461491 | 0.67 |
| YGL120C | 2147.315673 | -0.185949712 | 0.25084693 | 0.67 |
| YGR063C | 116.1850697 | -0.465022564 | 0.252372792 | 0.67 |
| YGR224W | 693.0347479 | 0.252565867 | 0.251513525 | 0.67 |
| YGR227W | 1061.56859 | -0.526405761 | 0.251303806 | 0.67 |
| YGR285C | 10708.64158 | -0.027056264 | 0.249015706 | 0.67 |
| YHR184W | 81.12732022 | -1.028650621 | 0.253790638 | 0.67 |
| YIL060W | 54.27024793 | 0.332510729 | 0.248801839 | 0.67 |
| YIL104C | 354.0212252 | 0.3764955 | 0.252885681 | 0.67 |
| YIL127C | 639.7542142 | -0.197360821 | 0.253442064 | 0.67 |
| YJR019C | 596.1541169 | 0.484620866 | 0.248665598 | 0.67 |
| YJR054W | 539.7820263 | -0.454625397 | 0.251450755 | 0.67 |
| YKL079W | 964.5499269 | 0.295536928 | 0.253682949 | 0.67 |
| YKL096W-A | 39924.89359 | -0.154740009 | 0.254187788 | 0.67 |
| YKL099C | 686.8097655 | -0.313519889 | 0.253953052 | 0.67 |
| YLR017W | 1271.451959 | -0.251277711 | 0.252388912 | 0.67 |
| YLR307C-A | 38.91718273 | 0.030687669 | 0.248917335 | 0.67 |

| | | | | |
|---|---|---|---|---|
| YLR356W | 2015.024039 | 0.581773762 | 0.249315361 | 0.67 |
| YLR418C | 850.5544851 | -0.353972099 | 0.251690213 | 0.67 |
| YLR426W | 420.438408 | -0.371517327 | 0.251481577 | 0.67 |
| YML127W | 993.038483 | -0.350371093 | 0.251000686 | 0.67 |
| YMR172C-A | 11.37396813 | 0.464714588 | 0.252651398 | 0.67 |
| YMR302C | 2126.727676 | 0.508286417 | 0.254132781 | 0.67 |
| YMR319C | 2635.007516 | 0.534554114 | 0.251411448 | 0.67 |
| YNL141W | 1283.346108 | -0.363224141 | 0.253550329 | 0.67 |
| YNL277W-A | 6.085182075 | 0.228940886 | 0.254323842 | 0.67 |
| YOL032W | 1063.876828 | 0.409474316 | 0.249891337 | 0.67 |
| YOL035C | 58.4740954 | 0.509120356 | 0.249290695 | 0.67 |
| YOR012W | 60.32598792 | 0.273538769 | 0.254141093 | 0.67 |
| YOR054C | 904.8796156 | 0.428131073 | 0.250218665 | 0.67 |
| YOR127W | 1893.705383 | 0.357264871 | 0.252948049 | 0.67 |
| YOR207C | 1745.865963 | -0.346638004 | 0.249642066 | 0.67 |
| YOR339C | 48.09399516 | 0.016100139 | 0.252403518 | 0.67 |
| YPL110C | 931.3902179 | 0.436994021 | 0.251384561 | 0.67 |
| YPL250C | 749.2294185 | 0.65039202 | 0.250306884 | 0.67 |
| YPL254W | 446.984746 | -0.533847547 | 0.249661989 | 0.67 |
| YPR184W | 5293.583109 | 0.820256738 | 0.251952964 | 0.67 |
| snR31 | 65.06005548 | -0.405351549 | 0.253431718 | 0.67 |
| snR41 | 28.2332558 | -0.488568743 | 0.253666878 | 0.67 |
| YDR098C-B | 19.81780983 | 0.856267037 | 0.255941986 | 0.67 |
| YDR392W | 654.5457347 | 0.275412641 | 0.255811673 | 0.67 |
| YLL019C | 1900.382246 | 0.602875677 | 0.256084627 | 0.67 |
| YLL021W | 2378.696108 | -0.334684469 | 0.256373147 | 0.67 |
| YML131W | 1116.004425 | 0.773803928 | 0.256160791 | 0.67 |
| YOR173W | 2098.419186 | 0.533274126 | 0.256206496 | 0.67 |
| YPR170W-A | 19.17302177 | -0.293497593 | 0.256321279 | 0.67 |
| YML057C-A | 86.62699701 | 0.440922831 | 0.256584904 | 0.67 |
| YNL303W | 97.83657889 | -0.362352525 | 0.257098491 | 0.67 |
| YMR014W | 1014.309235 | -0.298873549 | 0.257341538 | 0.67 |
| YOR235W | 3.373424746 | 0.234991687 | 0.257264022 | 0.67 |
| YDR424C | 383.6752581 | -0.29965191 | 0.257605473 | 0.67 |
| YNL044W | 495.5352853 | -0.333543177 | 0.257562141 | 0.67 |
| YHR066W | 786.1240577 | -0.155247271 | 0.258000246 | 0.68 |
| YBR241C | 1365.325299 | 0.512991457 | 0.258104173 | 0.68 |
| YNR022C | 907.2227542 | -0.37964699 | 0.258240459 | 0.68 |
| SCR1 | 181.7417356 | 0.428328506 | 0.261560205 | 0.68 |
| YAL007C | 1549.491558 | -0.220089445 | 0.259655993 | 0.68 |
| YAL011W | 648.981818 | 0.459487106 | 0.259608171 | 0.68 |
| YBL066C | 1237.560937 | 0.682201974 | 0.262233035 | 0.68 |
| YBR042C | 577.29725 | -0.360435019 | 0.260471627 | 0.68 |

| | | | | |
|---|---|---|---|---|
| YBR208C | 3562.373289 | 0.685518579 | 0.260819681 | 0.68 |
| YBR211C | 479.4755282 | 0.298441384 | 0.259893542 | 0.68 |
| YCR102W-A | 9.4716423 | 0.439442737 | 0.264214426 | 0.68 |
| YDL016C | 38.72923011 | -0.588402667 | 0.262746046 | 0.68 |
| YDL060W | 1265.050122 | -0.28653161 | 0.261086693 | 0.68 |
| YDL124W | 12433.87204 | 0.573895746 | 0.263577288 | 0.68 |
| YDR056C | 1611.517655 | -0.298086184 | 0.260759396 | 0.68 |
| YDR119W-A | 458.2326718 | 0.178889579 | 0.261982576 | 0.68 |
| YDR181C | 744.5427058 | 0.473677648 | 0.264531897 | 0.68 |
| YDR493W | 345.6632951 | -0.400366645 | 0.262532189 | 0.68 |
| YEL031W | 4279.639928 | -0.240642754 | 0.259757385 | 0.68 |
| YER046W-A | 13.59530817 | 0.246394172 | 0.264534083 | 0.68 |
| YER092W | 378.6245953 | -0.34067731 | 0.259961516 | 0.68 |
| YER114C | 1610.317658 | 0.30656222 | 0.262282653 | 0.68 |
| YFL055W | 265.1999324 | 0.582218579 | 0.260518195 | 0.68 |
| YGL075C | 312.3284424 | 0.361609037 | 0.261568697 | 0.68 |
| YGL108C | 328.3815974 | 0.362572765 | 0.260324449 | 0.68 |
| YGR111W | 1711.371366 | 0.371805939 | 0.260761067 | 0.68 |
| YGR165W | 998.5420975 | -0.399449955 | 0.262840151 | 0.68 |
| YIL031W | 1396.422579 | 0.369620314 | 0.262334879 | 0.68 |
| YIL032C | 13.12625335 | 0.761725637 | 0.263927968 | 0.68 |
| YJL085W | 484.172969 | -0.5019179 | 0.260865289 | 0.68 |
| YJL134W | 603.1655535 | -0.294197006 | 0.262823843 | 0.68 |
| YJR010W | 677.4384961 | 0.250129038 | 0.260428731 | 0.68 |
| YLL062C | 209.8685733 | 0.561814361 | 0.262441869 | 0.68 |
| YLR103C | 462.3096359 | -0.420417896 | 0.258581122 | 0.68 |
| YLR152C | 1272.570191 | 0.441818151 | 0.264505225 | 0.68 |
| YLR264C-A | 117.4733387 | -0.154608132 | 0.262687445 | 0.68 |
| YLR281C | 33.92288358 | -0.177960232 | 0.261192789 | 0.68 |
| YLR338W | 19.58277131 | 0.720529387 | 0.258864291 | 0.68 |
| YNL026W | 973.5528021 | -0.281617517 | 0.263987534 | 0.68 |
| YNL030W | 3372.305745 | -0.393985427 | 0.264343021 | 0.68 |
| YNL054W-B | 451.4424074 | 1.059608966 | 0.262676553 | 0.68 |
| YNL087W | 1914.774752 | -0.315662613 | 0.264427418 | 0.68 |
| YNL109W | 19.36367514 | -0.658383434 | 0.263385181 | 0.68 |
| YNL148C | 367.0903378 | -0.469896511 | 0.261092219 | 0.68 |
| YNL156C | 1413.410846 | 0.357502173 | 0.26229653 | 0.68 |
| YNL237W | 756.7694566 | 0.336475096 | 0.264457864 | 0.68 |
| YNL331C | 347.7391832 | 0.289420811 | 0.263880746 | 0.68 |
| YOL015W | 325.5388955 | 0.279671133 | 0.259095933 | 0.68 |
| YOL041C | 1321.58671 | -0.277686049 | 0.261497817 | 0.68 |
| YOR011W | 696.3198627 | -0.466617859 | 0.261975276 | 0.68 |
| YOR255W | 23.76826284 | 0.255133278 | 0.263461488 | 0.68 |

| | | | | |
|---|---|---|---|---|
| YOR265W | 445.8131326 | -0.294421685 | 0.263864739 | 0.68 |
| YPL005W | 369.5032375 | 0.379084766 | 0.263668972 | 0.68 |
| YPL099C | 334.8201937 | -0.446806193 | 0.259270693 | 0.68 |
| YPL215W | 1126.607522 | -0.363337664 | 0.263603935 | 0.68 |
| YPL265W | 2999.757451 | 0.387659554 | 0.261494015 | 0.68 |
| YPR032W | 901.7856576 | 0.38562892 | 0.264588563 | 0.68 |
| YPR083W | 614.8326597 | 0.588787108 | 0.264508328 | 0.68 |
| YPR139C | 1065.385559 | -0.482161334 | 0.261943631 | 0.68 |
| YPR168W | 170.9858215 | 0.341341156 | 0.264148191 | 0.68 |
| YPR175W | 704.7198488 | -0.291790021 | 0.262956583 | 0.68 |
| snR37 | 153.9250937 | -0.072630659 | 0.264429705 | 0.68 |
| YDR023W | 6264.678536 | -0.265894028 | 0.26530555 | 0.68 |
| YDR300C | 1204.265207 | -0.393396586 | 0.265193465 | 0.68 |
| YHR062C | 561.4502383 | -0.300974606 | 0.265595141 | 0.68 |
| YKR104W | 184.9350151 | 0.538157714 | 0.265639006 | 0.68 |
| YLR098C | 706.1029207 | 0.37063374 | 0.264914156 | 0.68 |
| YMR208W | 1260.14289 | -0.318170739 | 0.265285278 | 0.68 |
| YNL171C | 59.8874962 | 0.691585342 | 0.265387917 | 0.68 |
| YPL036W | 288.1590263 | 0.248863079 | 0.265504339 | 0.68 |
| YGR207C | 1419.844599 | -0.264084604 | 0.265782902 | 0.68 |
| YOL063C | 983.1121387 | 0.319636926 | 0.265879854 | 0.68 |
| YJL050W | 1800.303824 | -0.264982587 | 0.266036219 | 0.68 |
| YGL085W | 385.7388603 | 0.347835941 | 0.266178815 | 0.68 |
| YOL111C | 1821.468977 | -0.27693687 | 0.266268374 | 0.68 |
| YPR164W | 1248.455329 | -0.252100675 | 0.266406164 | 0.68 |
| YER053C-A | 179.222376 | 0.46409885 | 0.266631134 | 0.68 |
| YBR218C | 7147.332039 | 0.543705102 | 0.266847476 | 0.68 |
| YER111C | 1455.121069 | -0.3246241 | 0.267314258 | 0.68 |
| YLR013W | 3.60279815 | 0.819249027 | 0.267412922 | 0.68 |
| YDL159W | 513.5387781 | 0.375490579 | 0.268194496 | 0.68 |
| YER010C | 1015.031336 | -0.336717082 | 0.267898601 | 0.68 |
| YGL201C | 994.5170532 | -0.340371997 | 0.267965191 | 0.68 |
| YHL026C | 386.086378 | 0.518683447 | 0.268315985 | 0.68 |
| YIR005W | 167.3355793 | 0.30235904 | 0.267651528 | 0.68 |
| YKR029C | 884.4269483 | 0.313519573 | 0.268241769 | 0.68 |
| YNL094W | 1768.042353 | 0.344062497 | 0.268295192 | 0.68 |
| YNL292W | 950.1678617 | -0.323989779 | 0.268324505 | 0.68 |
| YOR032W-A | 11.5403846 | -0.350175854 | 0.268453127 | 0.68 |
| YAL002W | 1569.483592 | 0.303047296 | 0.269148689 | 0.68 |
| YIR015W | 351.8671415 | 0.523363283 | 0.269076214 | 0.68 |
| YKL098W | 807.0315194 | 0.362356447 | 0.269054358 | 0.68 |
| YLR298C | 806.4703735 | 0.415863828 | 0.268905932 | 0.68 |
| YIL103W | 637.74726 | -0.372206191 | 0.269522502 | 0.68 |

| | | | | |
|---|---|---|---|---|
| YMR272W-B | 3.438828838 | 0.379139722 | 0.269626577 | 0.68 |
| YNL175C | 613.6716753 | -0.190194197 | 0.269729859 | 0.68 |
| YBR267W | 453.4917639 | -0.43825664 | 0.270141876 | 0.68 |
| YMR242W-A | 1.825788145 | -0.081607534 | 0.270052165 | 0.68 |
| YPL195W | 2059.660784 | -0.269717381 | 0.270068734 | 0.68 |
| YER040W | 900.0166243 | 0.547278256 | 0.270792702 | 0.68 |
| YOR270C | 5846.210473 | -0.409770507 | 0.270889506 | 0.68 |
| YHR007C | 4279.432771 | -0.370710275 | 0.271043217 | 0.68 |
| YNL183C | 1035.511832 | 0.409089096 | 0.271259566 | 0.68 |
| YCR048W | 995.8304266 | 0.441785406 | 0.271669595 | 0.68 |
| YHR174W | 94635.8071 | 0.063322768 | 0.271626846 | 0.68 |
| YMR063W | 168.4962103 | 0.399876588 | 0.271842173 | 0.68 |
| YGR045C | 23.80619598 | 0.421660532 | 0.272075189 | 0.68 |
| YHR190W | 5071.299505 | 0.446235012 | 0.272200062 | 0.68 |
| YMR241W | 1393.972884 | -0.418314074 | 0.272487535 | 0.68 |
| YCR013C | 2806.322738 | -0.195792987 | 0.272834356 | 0.68 |
| YDR510W | 2598.34956 | -0.350741103 | 0.272915448 | 0.68 |
| YDL091C | 1187.448267 | 0.373012244 | 0.273516545 | 0.68 |
| YFL047W | 1142.35423 | -0.554216633 | 0.273556073 | 0.68 |
| YHR161C | 2280.636973 | 0.498927109 | 0.273416686 | 0.68 |
| YLR217W | 16.95655225 | 0.571057883 | 0.27338317 | 0.68 |
| YMR058W | 5542.022572 | -0.749502833 | 0.273401124 | 0.68 |
| YBL020W | 997.2826168 | -0.254074688 | 0.274275082 | 0.68 |
| YDL157C | 952.2291983 | -0.26452686 | 0.274426402 | 0.68 |
| YNL118C | 3319.845542 | 0.284329687 | 0.274400569 | 0.68 |
| YNL218W | 370.2566133 | -0.34802363 | 0.27438037 | 0.68 |
| YNL242W | 1398.740797 | 0.564094021 | 0.274255368 | 0.68 |
| YOR051C | 2007.894821 | -0.385414779 | 0.274051908 | 0.68 |
| YPL200W | 21.68345646 | 0.518665811 | 0.274367711 | 0.68 |
| YPR036W-A | 8058.010483 | 0.584721965 | 0.274600506 | 0.68 |
| YPR082C | 431.92377 | -0.398647294 | 0.274542276 | 0.68 |
| YIR022W | 1186.448374 | -0.346909426 | 0.275038315 | 0.68 |
| YLR003C | 782.5858822 | -0.147458851 | 0.27500024 | 0.68 |
| YOR201C | 630.2633868 | -0.35541001 | 0.275049103 | 0.68 |
| YBR063C | 508.4120941 | 0.289075549 | 0.275813024 | 0.68 |
| YDR071C | 2792.908431 | -0.325385694 | 0.275895822 | 0.68 |
| YEL074W | 35.0602757 | 0.503691378 | 0.275870733 | 0.68 |
| YGL249W | 273.3583474 | 0.235164905 | 0.275465142 | 0.68 |
| YIL092W | 426.021742 | 0.273974885 | 0.275770876 | 0.68 |
| YKR088C | 850.0044261 | -0.296922932 | 0.275549933 | 0.68 |
| YOR325W | 4.631536169 | -0.585467089 | 0.275329123 | 0.68 |
| YPR080W | 5311.40989 | -0.115189799 | 0.275789817 | 0.68 |
| YDR117C | 1080.281382 | -0.295205734 | 0.276411944 | 0.69 |

| | | | | |
|---|---|---|---|---|
| YGL152C | 7.084888852 | 0.277987451 | 0.276310933 | 0.69 |
| YKL031W | 35.11063328 | 0.874968333 | 0.276725803 | 0.69 |
| YLR251W | 493.6892026 | 0.389644549 | 0.276821298 | 0.69 |
| YNL090W | 89.81803356 | -0.478597192 | 0.276748928 | 0.69 |
| YOR009W | 307.6360112 | 0.284447312 | 0.277054412 | 0.69 |
| YDL074C | 1505.723255 | -0.462051329 | 0.277573678 | 0.69 |
| YDR329C | 1759.988143 | 0.515340817 | 0.277586986 | 0.69 |
| YIL009W | 578.8586392 | -0.62596329 | 0.277572224 | 0.69 |
| YNR069C | 436.7652166 | 0.458667361 | 0.277710029 | 0.69 |
| YOR032C | 566.9607469 | 0.353360723 | 0.278012678 | 0.69 |
| YPL126W | 2185.981195 | -0.254835948 | 0.278126573 | 0.69 |
| YIL147C | 2000.977172 | 0.422771101 | 0.278499937 | 0.69 |
| YLR337C | 2433.363455 | 0.423147234 | 0.278416432 | 0.69 |
| YER035W | 1131.143378 | 0.397900167 | 0.278816708 | 0.69 |
| YLR417W | 736.9680806 | 0.332097438 | 0.278833312 | 0.69 |
| YBL098W | 1032.627619 | -0.281047959 | 0.279260743 | 0.69 |
| YGR196C | 1285.850548 | 0.268656856 | 0.279267132 | 0.69 |
| YHR169W | 496.2218428 | -0.331696861 | 0.278979373 | 0.69 |
| YJR129C | 222.0216409 | -0.068700113 | 0.279260905 | 0.69 |
| YLR191W | 472.4930163 | 0.381994104 | 0.279396206 | 0.69 |
| YNL240C | 1001.196601 | 0.471440608 | 0.279747786 | 0.69 |
| YPL256C | 1702.304083 | -0.408998236 | 0.279895704 | 0.69 |
| YDR435C | 760.2512519 | 0.420155986 | 0.280108199 | 0.69 |
| YKL107W | 249.2824796 | 0.634619052 | 0.280440094 | 0.69 |
| YCR081W | 992.4265821 | 0.375030225 | 0.28086066 | 0.69 |
| YMR161W | 724.2845416 | -0.317069463 | 0.280853748 | 0.69 |
| YNL306W | 1534.453742 | -0.339970539 | 0.280758 | 0.69 |
| YCR077C | 3337.529518 | -0.30334246 | 0.281116791 | 0.69 |
| YDL243C | 368.4938618 | 0.623026862 | 0.281278169 | 0.69 |
| YLR078C | 672.0208317 | 0.346333987 | 0.281352922 | 0.69 |
| YMR110C | 3066.365125 | 0.526212958 | 0.281458343 | 0.69 |
| YAL068C | 5.759798843 | 0.577938556 | 0.281823224 | 0.69 |
| YBR060C | 829.7516444 | 0.263394532 | 0.281731301 | 0.69 |
| YNL076W | 1058.446405 | 0.299667642 | 0.281819762 | 0.69 |
| YAL015C | 634.3594781 | 0.464289708 | 0.282269354 | 0.69 |
| YBR050C | 396.3143748 | 0.415518659 | 0.282862881 | 0.69 |
| YDR239C | 796.3576703 | 0.330831526 | 0.282645129 | 0.69 |
| YKL212W | 2578.339284 | -0.279960226 | 0.28283539 | 0.69 |
| YLR027C | 7716.105415 | -0.224200998 | 0.282752457 | 0.69 |
| YMR004W | 1474.539468 | 0.311353262 | 0.282436488 | 0.69 |
| YMR168C | 528.3657431 | -0.37658272 | 0.282754787 | 0.69 |
| YOR167C | 4868.170026 | -0.288407284 | 0.282568707 | 0.69 |
| YOR203W | 129.138005 | 0.463748472 | 0.282658029 | 0.69 |

| | | | | |
|---|---|---|---|---|
| YPL086C | 1089.007189 | -0.173319167 | 0.282928221 | 0.69 |
| YOL072W | 594.5699607 | 0.315364341 | 0.283290419 | 0.69 |
| YDR128W | 2008.877758 | 0.489061311 | 0.283648088 | 0.69 |
| YHR210C | 320.0019552 | -0.571120393 | 0.283601762 | 0.69 |
| YLR207W | 1228.065049 | 0.358983974 | 0.283872331 | 0.69 |
| YBR185C | 732.2746685 | -0.25881309 | 0.284693145 | 0.69 |
| YDR453C | 889.6622872 | 0.392017819 | 0.284548768 | 0.69 |
| YLR214W | 1682.159024 | -0.653891373 | 0.28461099 | 0.69 |
| YLR321C | 729.7163579 | -0.405791609 | 0.285226349 | 0.69 |
| YGR100W | 1479.985263 | 0.244857868 | 0.285705295 | 0.69 |
| snR67 | 6.800863812 | 0.9590742 | 0.285844354 | 0.69 |
| YDL217C | 416.6172424 | -0.321331249 | 0.287268292 | 0.69 |
| YJL010C | 755.184634 | -0.321725916 | 0.286310635 | 0.69 |
| YJL126W | 376.2312005 | 0.302477126 | 0.287348482 | 0.69 |
| YKL205W | 1310.174229 | -0.39300936 | 0.286825971 | 0.69 |
| YLR036C | 360.058471 | -0.46934844 | 0.287187284 | 0.69 |
| YLR165C | 405.6673301 | 0.207556021 | 0.286087199 | 0.69 |
| YLR397C | 907.763381 | -0.300359733 | 0.287295704 | 0.69 |
| YLR459W | 960.9364848 | -0.457674682 | 0.287137971 | 0.69 |
| YMR040W | 194.5584091 | 0.756828923 | 0.286559361 | 0.69 |
| YMR043W | 2143.509763 | 0.351224555 | 0.287033416 | 0.69 |
| YMR186W | 31490.05511 | -0.057608917 | 0.28717857 | 0.69 |
| YMR246W | 3290.570155 | -0.254479932 | 0.286837857 | 0.69 |
| YOL010W | 524.4137956 | -0.402188291 | 0.286874247 | 0.69 |
| YPL127C | 2242.895143 | -0.174853257 | 0.28634467 | 0.69 |
| YDR238C | 4318.542325 | -0.22925195 | 0.287567418 | 0.69 |
| YNL278W | 749.3245241 | 0.396226486 | 0.287503285 | 0.69 |
| YML034C-A | 23.30372078 | 0.462010792 | 0.287785772 | 0.70 |
| YOR146W | 6.962391281 | -0.561784776 | 0.287884118 | 0.70 |
| YDR449C | 532.8954421 | -0.360942797 | 0.288419133 | 0.70 |
| YOR153W | 11636.99259 | -0.590458326 | 0.288398137 | 0.70 |
| YDL191W | 1342.250708 | -0.288144797 | 0.2886358 | 0.70 |
| YBL074C | 232.3789197 | 0.302910194 | 0.289244846 | 0.70 |
| YER031C | 2004.561338 | -0.317356809 | 0.289229195 | 0.70 |
| YFR020W | 385.8178556 | 0.318950944 | 0.289050503 | 0.70 |
| YDR396W | 125.4408117 | -0.383127239 | 0.289796586 | 0.70 |
| YLR244C | 3158.461036 | -0.330398862 | 0.289772962 | 0.70 |
| YMR151W | 69.13225197 | 0.265991235 | 0.289896655 | 0.70 |
| snR60 | 11.58332067 | 0.268499628 | 0.289990348 | 0.70 |
| YAL013W | 1274.881408 | 0.351685233 | 0.291827006 | 0.70 |
| YDR219C | 850.7025719 | 0.304086627 | 0.290854923 | 0.70 |
| YDR303C | 1032.483515 | -0.452018199 | 0.29211896 | 0.70 |
| YER002W | 1431.744689 | -0.280388215 | 0.290973283 | 0.70 |

| | | | | |
|---|---|---|---|---|
| YER084W | 124.9336603 | 0.155876527 | 0.290895334 | 0.70 |
| YHL019W-A | 10.01596588 | 0.451158425 | 0.29161667 | 0.70 |
| YHR019C | 9735.850466 | -0.183236682 | 0.291490025 | 0.70 |
| YHR074W | 2090.525023 | -0.302869751 | 0.29170034 | 0.70 |
| YJL127W-A | 12.80997666 | -0.201761085 | 0.29091805 | 0.70 |
| YJL178C | 1325.547528 | -0.297851779 | 0.290510968 | 0.70 |
| YJR056C | 431.2081935 | 0.344504192 | 0.290239079 | 0.70 |
| YLR236C | 21.91002832 | 0.621960447 | 0.29164169 | 0.70 |
| YLR284C | 336.5089725 | 0.216945727 | 0.290556996 | 0.70 |
| YMR158W-B | 1.160282167 | -0.243254773 | 0.291853291 | 0.70 |
| YOR043W | 1570.483617 | -0.430366269 | 0.292066237 | 0.70 |
| YOR062C | 585.4618256 | -0.037878242 | 0.291606115 | 0.70 |
| YOR066W | 1115.721778 | 0.248479238 | 0.291471957 | 0.70 |
| YOR188W | 1393.160444 | -0.291404714 | 0.292125511 | 0.70 |
| YOR245C | 966.5341375 | 0.397085653 | 0.290348666 | 0.70 |
| YNL155W | 1378.201426 | 0.465682586 | 0.292550054 | 0.70 |
| YJL087C | 474.7532502 | -0.487725523 | 0.293098044 | 0.70 |
| YMR183C | 2217.146681 | -0.426285738 | 0.293437207 | 0.70 |
| snR161 | 8.988952116 | 0.60368728 | 0.293483259 | 0.70 |
| YOL043C | 577.8464057 | 0.278181644 | 0.29371236 | 0.70 |
| YDR344C | 46.73109222 | -0.105849897 | 0.294051744 | 0.70 |
| YDR214W | 2305.632085 | -0.180526719 | 0.294315215 | 0.70 |
| YLR197W | 4160.668892 | -0.427057318 | 0.294230098 | 0.70 |
| YIL160C | 460.4774311 | 0.181676786 | 0.294425297 | 0.70 |
| YMR259C | 1208.426766 | -0.410496432 | 0.294687987 | 0.70 |
| YBR033W | 734.5856264 | 0.503432926 | 0.295830878 | 0.70 |
| YDL050C | 34.4743764 | -0.285079205 | 0.29587807 | 0.70 |
| YDL240W | 734.1749038 | -0.687676814 | 0.295505417 | 0.70 |
| YDR034C-A | 4.06626248 | -0.281898278 | 0.295868029 | 0.70 |
| YDR364C | 596.6987029 | 0.227918699 | 0.295733926 | 0.70 |
| YDR417C | 435.6659374 | -0.523470172 | 0.295720845 | 0.70 |
| YGR185C | 4359.206315 | -0.319362503 | 0.295517058 | 0.70 |
| YJL130C | 21427.36188 | -0.239137315 | 0.295222302 | 0.70 |
| YLR300W | 3505.549383 | -0.541560045 | 0.295946019 | 0.70 |
| YLR369W | 1148.442381 | 0.459297031 | 0.295853143 | 0.70 |
| YNL024C-A | 249.9046551 | -0.38161026 | 0.295258357 | 0.70 |
| YNL137C | 1635.261502 | -0.287775623 | 0.296016209 | 0.70 |
| YLR449W | 2574.095007 | -0.047878858 | 0.296189271 | 0.70 |
| YDR146C | 1456.032171 | -0.35433667 | 0.296394216 | 0.70 |
| YPR162C | 653.0734249 | -0.412266423 | 0.296739724 | 0.70 |
| YER107C | 624.5620743 | -0.419916102 | 0.296959195 | 0.70 |
| YHR060W | 732.5771452 | -0.353403128 | 0.296964556 | 0.70 |
| YDR505C | 1755.081331 | 0.477409702 | 0.297258766 | 0.70 |

| | | | | |
|---|---|---|---|---|
| YML017W | 751.7187367 | -0.31809888 | 0.297470801 | 0.70 |
| YBR231C | 448.4132605 | 0.414114761 | 0.298199467 | 0.70 |
| YDL189W | 1070.015735 | -0.447837114 | 0.29806651 | 0.70 |
| YEL010W | 4.427753031 | 0.356633054 | 0.297863835 | 0.70 |
| YGR169C-A | 750.5850996 | 0.260702183 | 0.298272317 | 0.70 |
| YGR288W | 284.2488249 | 0.328587323 | 0.298092692 | 0.70 |
| YPR022C | 1532.191812 | 0.310228454 | 0.297795714 | 0.70 |
| YBL068W | 791.3128306 | -0.296458406 | 0.299200432 | 0.70 |
| YCL063W | 606.1625178 | -0.370514914 | 0.298582463 | 0.70 |
| YJL109C | 2576.133571 | -0.308075844 | 0.298994656 | 0.70 |
| YJR141W | 396.4509931 | -0.382055532 | 0.299246792 | 0.70 |
| YNL121C | 5779.129433 | -0.267127958 | 0.299020873 | 0.70 |
| YOL094C | 437.8167325 | -0.40418294 | 0.298632638 | 0.70 |
| YOR135C | 405.7747912 | -0.136395988 | 0.299101964 | 0.70 |
| YOR166C | 666.3506234 | -0.317948977 | 0.299159103 | 0.70 |
| YPL159C | 943.1509361 | 0.339227033 | 0.29882533 | 0.70 |
| YKR031C | 1778.007412 | 0.368655634 | 0.299508337 | 0.70 |
| YNR005C | 1.56429782 | 0.10414776 | 0.299496872 | 0.70 |
| YNL277W | 546.4771573 | 0.243681856 | 0.299671617 | 0.70 |
| YDL035C | 1472.135402 | 0.394334545 | 0.300181868 | 0.70 |
| YLR178C | 3271.315417 | 0.444860347 | 0.300083361 | 0.70 |
| YDL018C | 497.6776418 | -0.300027238 | 0.300525587 | 0.71 |
| YBL071C | 215.271228 | 0.381096408 | 0.301596862 | 0.71 |
| YCL016C | 442.1886812 | 0.330979806 | 0.30202299 | 0.71 |
| YCR025C | 44.00460202 | 0.387343831 | 0.300962387 | 0.71 |
| YDR179W-A | 321.5877637 | -0.367258615 | 0.301636007 | 0.71 |
| YFL050C | 563.7473526 | 0.323749997 | 0.301891273 | 0.71 |
| YGL242C | 665.893361 | -0.334095644 | 0.301124683 | 0.71 |
| YGR015C | 438.1021585 | 0.281465411 | 0.301756026 | 0.71 |
| YJL187C | 1084.409994 | -0.287774941 | 0.300886404 | 0.71 |
| YJR047C | 407.0063415 | 0.846708281 | 0.301483339 | 0.71 |
| YLR102C | 476.3043851 | 0.360815664 | 0.301030329 | 0.71 |
| YMR163C | 451.0209425 | 0.295374679 | 0.301923059 | 0.71 |
| YOR089C | 2098.945136 | 0.308771808 | 0.301918742 | 0.71 |
| snR38 | 6.274197792 | 0.306030763 | 0.302002457 | 0.71 |
| YGR271W | 1597.427923 | -0.283627505 | 0.302578286 | 0.71 |
| YKL020C | 2087.802283 | 0.363492088 | 0.302545698 | 0.71 |
| YKL087C | 723.7436146 | 0.429312697 | 0.302585995 | 0.71 |
| YLL057C | 252.3608873 | 0.304913218 | 0.302265756 | 0.71 |
| YLR410W | 2881.535918 | 0.398988786 | 0.302518216 | 0.71 |
| YFL030W | 1073.611575 | 0.819343776 | 0.30298331 | 0.71 |
| YKL019W | 1109.764572 | -0.335452882 | 0.303007799 | 0.71 |
| YLL023C | 2955.471664 | 0.448531801 | 0.302731932 | 0.71 |

| | | | | |
|---|---|---|---|---|
| YOL064C | 847.1293742 | -0.288358603 | 0.303146456 | 0.71 |
| YPL199C | 1017.89655 | -0.363680411 | 0.303123585 | 0.71 |
| YDR488C | 747.4804028 | 0.567954894 | 0.303448222 | 0.71 |
| YLR020C | 866.2619741 | -0.356710861 | 0.303817612 | 0.71 |
| YBR145W | 1370.985162 | 0.379245798 | 0.304532979 | 0.71 |
| YDR129C | 5762.337606 | 0.319345709 | 0.30468676 | 0.71 |
| YDR361C | 744.7914399 | -0.239732466 | 0.30486668 | 0.71 |
| YJL029C | 1512.843835 | 0.259584916 | 0.305025009 | 0.71 |
| YOR183W | 10.65070105 | 0.185667126 | 0.305066621 | 0.71 |
| YOR383C | 13192.95676 | 0.726567753 | 0.304935076 | 0.71 |
| YHR045W | 983.5331682 | -0.457698407 | 0.305272545 | 0.71 |
| YDR188W | 2974.432832 | -0.307731971 | 0.305616605 | 0.71 |
| YDL027C | 1559.088359 | 0.390715087 | 0.306010267 | 0.71 |
| YJL118W | 178.5211681 | -0.344811103 | 0.30630394 | 0.71 |
| YJR135C | 188.8236464 | -0.399705443 | 0.306242277 | 0.71 |
| YML009C-A | 52.04514359 | -0.234572277 | 0.306283843 | 0.71 |
| YOR360C | 883.5535106 | -0.578734244 | 0.306241695 | 0.71 |
| YPL188W | 716.8167679 | 0.37825945 | 0.306023502 | 0.71 |
| YDL053C | 1309.962591 | -0.269147467 | 0.306463178 | 0.71 |
| YGR210C | 1863.079648 | -0.285588701 | 0.306635579 | 0.71 |
| YLR026C | 1116.422754 | -0.336180252 | 0.306622762 | 0.71 |
| YOL162W | 65.72006374 | 0.511316885 | 0.306856235 | 0.71 |
| YPR153W | 81.58409703 | 0.340703054 | 0.306749866 | 0.71 |
| YGR046W | 906.1577966 | 0.312531784 | 0.30701632 | 0.71 |
| YMR195W | 1764.107553 | 0.363973754 | 0.307221233 | 0.71 |
| YOL046C | 5.940874887 | 0.033654716 | 0.307211104 | 0.71 |
| YLR220W | 930.48531 | 0.30656202 | 0.307379796 | 0.71 |
| YDR531W | 969.2146497 | -0.38434329 | 0.307555792 | 0.71 |
| YOR148C | 393.9247418 | -0.361381018 | 0.307699388 | 0.71 |
| YHL019C | 544.6940331 | 0.469624246 | 0.308100833 | 0.71 |
| YBL043W | 324.5838638 | 0.823801594 | 0.30844481 | 0.71 |
| YBR012C | 184.3218002 | 0.821051588 | 0.309907724 | 0.71 |
| YBR188C | 339.0880427 | -0.257816483 | 0.309303626 | 0.71 |
| YDR186C | 2571.215729 | 0.353501237 | 0.309540943 | 0.71 |
| YFL041W-A | 23.17885335 | 0.154629823 | 0.309223093 | 0.71 |
| YGR129W | 435.9932211 | 0.479417337 | 0.309537018 | 0.71 |
| YJR077C | 4794.775519 | -0.059488774 | 0.309919779 | 0.71 |
| YKL209C | 3091.85579 | -1.007034519 | 0.308871849 | 0.71 |
| YLR211C | 375.8938446 | 0.307402812 | 0.308824247 | 0.71 |
| YLR458W | 2.467406493 | 0.194834346 | 0.309758629 | 0.71 |
| YML117W | 3482.921508 | 0.324986061 | 0.309782427 | 0.71 |
| YOL069W | 510.0668666 | -0.308194912 | 0.308799442 | 0.71 |
| YOR150W | 702.6825687 | -0.415597505 | 0.309538243 | 0.71 |

| | | | | |
|---|---|---|---|---|
| YOR271C | 1520.089436 | 0.000887113 | 0.308810447 | 0.71 |
| YPL077C | 211.278667 | 0.291683359 | 0.309591939 | 0.71 |
| YPR145C-A | 45.3897147 | 0.334994034 | 0.309551353 | 0.71 |
| YBR246W | 994.4891912 | -0.287596766 | 0.311685661 | 0.71 |
| YDR408C | 1967.993247 | -0.382542486 | 0.312397034 | 0.71 |
| YDR485C | 1410.914463 | 0.457667994 | 0.31208682 | 0.71 |
| YDR486C | 1425.158924 | 0.350834897 | 0.313054462 | 0.71 |
| YER112W | 1280.704105 | -0.291889524 | 0.313152127 | 0.71 |
| YGR242W | 37.21832003 | -0.505939806 | 0.31256906 | 0.71 |
| YIL077C | 964.8137743 | 0.339842981 | 0.312311719 | 0.71 |
| YJL005W | 4176.860231 | 0.484273126 | 0.313029345 | 0.71 |
| YJL008C | 2286.818099 | -0.300970197 | 0.312543738 | 0.71 |
| YKL042W | 561.3749184 | -0.428558786 | 0.311202006 | 0.71 |
| YKL100C | 2362.710996 | 0.326617976 | 0.313099775 | 0.71 |
| YLR243W | 616.8482711 | -0.209375775 | 0.312472323 | 0.71 |
| YLR306W | 266.7883268 | -0.470453861 | 0.311756659 | 0.71 |
| YMR086W | 2080.250939 | 0.343882692 | 0.313219211 | 0.71 |
| YMR232W | 290.2765579 | -0.099865535 | 0.31145093 | 0.71 |
| YNL080C | 1115.372394 | -0.362473375 | 0.31266251 | 0.71 |
| YOL025W | 668.044846 | 0.272362854 | 0.312736502 | 0.71 |
| YOR186W | 27.95959597 | 0.197427134 | 0.312644597 | 0.71 |
| YPL011C | 726.714861 | 0.286888709 | 0.312050929 | 0.71 |
| YPL210C | 1902.73736 | -0.405569475 | 0.313035194 | 0.71 |
| YBR051W | 12.61851356 | 0.255806916 | 0.313506553 | 0.71 |
| YCL044C | 1238.471005 | 0.468658618 | 0.313951086 | 0.71 |
| YJL027C | 10.13200317 | 0.687060832 | 0.314216236 | 0.71 |
| YKL126W | 2370.590453 | -0.272190592 | 0.314264843 | 0.71 |
| YLR129W | 1609.244371 | -0.448817237 | 0.314250474 | 0.71 |
| YOR078W | 697.5265362 | -0.390302604 | 0.314208065 | 0.71 |
| YOR048C | 2248.629889 | -0.214072926 | 0.314440349 | 0.71 |
| YCR006C | 58.71736424 | -0.677274873 | 0.315470862 | 0.71 |
| YCR083W | 821.7942376 | 0.607757332 | 0.314954748 | 0.71 |
| YDL063C | 706.2977789 | -0.27244807 | 0.315181353 | 0.71 |
| YGL217C | 8.839690893 | -0.515737283 | 0.315032912 | 0.71 |
| YHR179W | 8288.962491 | -0.2041397 | 0.314961697 | 0.71 |
| YIL169C | 2177.35422 | -0.610954545 | 0.315540946 | 0.71 |
| YKL138C | 823.724232 | -0.406236519 | 0.315569482 | 0.71 |
| YOR281C | 1009.23062 | -0.418613388 | 0.315404939 | 0.71 |
| YPR101W | 539.0655902 | 0.396961414 | 0.315287357 | 0.71 |
| YPR152C | 381.9563274 | 0.331481713 | 0.314959126 | 0.71 |
| YBR144C | 8.772859821 | 0.449014114 | 0.316075458 | 0.71 |
| YDR080W | 1103.850689 | 0.434971526 | 0.316440609 | 0.71 |
| YDR414C | 448.7530762 | -0.488543954 | 0.316118624 | 0.71 |

| | | | | |
|---|---|---|---|---|
| YEL006W | 705.1819844 | -0.423733359 | 0.316553695 | 0.71 |
| YIL136W | 7495.331143 | 0.60343931 | 0.31650732 | 0.71 |
| YLR064W | 1412.390781 | 0.359505728 | 0.316050711 | 0.71 |
| YLR077W | 986.4922801 | 0.277818816 | 0.316268031 | 0.71 |
| YNL263C | 1345.46904 | -0.293340625 | 0.316354201 | 0.71 |
| YFR043C | 412.3285976 | 0.290213306 | 0.31698969 | 0.71 |
| YNL082W | 573.887894 | -0.344411913 | 0.317178012 | 0.71 |
| YNR020C | 315.5485889 | -0.36873955 | 0.317114082 | 0.71 |
| YPR013C | 1083.106605 | 0.247841127 | 0.316931134 | 0.71 |
| YPR115W | 2825.752527 | 0.402492212 | 0.317013364 | 0.71 |
| YBR170C | 1144.125576 | 0.311649133 | 0.31787943 | 0.71 |
| YHR063W-A | 174.3401986 | -0.145743522 | 0.317915481 | 0.71 |
| YML050W | 378.8334393 | 0.395905974 | 0.318011253 | 0.71 |
| YMR316C-B | 10.98411933 | 0.301524205 | 0.317897196 | 0.71 |
| YNL101W | 1154.001663 | 0.3676171 | 0.318041137 | 0.71 |
| YMR072W | 3717.086972 | -0.281297934 | 0.318229347 | 0.71 |
| YPL030W | 552.0003609 | -0.37548684 | 0.318263632 | 0.71 |
| YBR199W | 2706.736809 | 0.189573257 | 0.318834943 | 0.72 |
| YDL085C-A | 569.0392384 | -0.519508555 | 0.318582931 | 0.72 |
| YDR228C | 901.0372572 | 0.173024044 | 0.319028043 | 0.72 |
| YDR335W | 3040.69958 | 0.381708838 | 0.318768097 | 0.72 |
| YIL066C | 409.4135175 | 0.569135831 | 0.319145931 | 0.72 |
| YMR224C | 500.4037572 | 0.25497623 | 0.319240859 | 0.72 |
| YNL323W | 681.0384119 | -0.309767126 | 0.319138858 | 0.72 |
| YBR150C | 848.9681448 | 0.389253986 | 0.319557916 | 0.72 |
| YCR075C | 329.3656461 | -0.343288634 | 0.3202436 | 0.72 |
| YER122C | 4064.222569 | -0.371481609 | 0.32020421 | 0.72 |
| YHL034W-A | 585.1049194 | -0.191792882 | 0.320199384 | 0.72 |
| YNL234W | 710.7866789 | 0.212659256 | 0.320176677 | 0.72 |
| YPR024W | 1909.485964 | 0.356512931 | 0.319898847 | 0.72 |
| YJR124C | 485.3455454 | -0.513415086 | 0.320475456 | 0.72 |
| YGL144C | 301.1617021 | 0.350678442 | 0.320785729 | 0.72 |
| YKL179C | 1404.902826 | 0.251357279 | 0.321030334 | 0.72 |
| YER159C | 1569.026083 | -0.43632589 | 0.321864074 | 0.72 |
| YGR221C | 838.7959631 | -0.458252112 | 0.321686078 | 0.72 |
| YJR148W | 6070.611654 | 0.595734643 | 0.321432748 | 0.72 |
| YLR339C | 501.9621213 | -0.390149326 | 0.321579759 | 0.72 |
| YMR229C | 3400.401838 | -0.17942782 | 0.321731504 | 0.72 |
| YMR263W | 652.3046851 | 0.336020122 | 0.32185545 | 0.72 |
| YPL171C | 308.1636844 | 0.851526505 | 0.321442879 | 0.72 |
| YDL095W | 3378.459132 | -0.219929973 | 0.322192094 | 0.72 |
| YGR292W | 46.2187461 | 0.127365036 | 0.322583673 | 0.72 |
| YLR125W | 49.15160341 | 0.228423351 | 0.322532126 | 0.72 |

| | | | | |
|---|---|---|---|---|
| YPL207W | 1844.153383 | 0.476099793 | 0.322629874 | 0.72 |
| YBR058C-A | 528.5056217 | -0.310240134 | 0.323746224 | 0.72 |
| YDR296W | 1076.454873 | -0.301493039 | 0.32366713 | 0.72 |
| YIL171W | 27.73883289 | 0.276261516 | 0.323601254 | 0.72 |
| YKL193C | 1757.604095 | 0.453908503 | 0.323610047 | 0.72 |
| YPR028W | 4646.798863 | -0.336756446 | 0.323824497 | 0.72 |
| YPL134C | 1579.391088 | 0.8165522 | 0.324010238 | 0.72 |
| YBR002C | 607.1395928 | -0.327168591 | 0.324702783 | 0.72 |
| YDL114W | 43.38451628 | 0.425699053 | 0.324805647 | 0.72 |
| YDR167W | 803.2507476 | -0.27680198 | 0.324900753 | 0.72 |
| YFL058W | 7.345596516 | 0.215323054 | 0.324605945 | 0.72 |
| YGL219C | 1273.719314 | 0.424100675 | 0.324731337 | 0.72 |
| YKL176C | 1422.060464 | -0.397788204 | 0.324332475 | 0.72 |
| YMR155W | 707.9678984 | 0.278629423 | 0.324432686 | 0.72 |
| YPL109C | 614.5552418 | 0.328814498 | 0.324965954 | 0.72 |
| YDL112W | 1910.821894 | -0.289510045 | 0.325247324 | 0.72 |
| YNL041C | 1185.16925 | -0.311708491 | 0.325185873 | 0.72 |
| YOR224C | 1759.04057 | -0.231264046 | 0.325311484 | 0.72 |
| YOR351C | 100.6577504 | 0.201033359 | 0.325434937 | 0.72 |
| YGR174W-A | 47.91809201 | 0.177563765 | 0.325808341 | 0.72 |
| YGR048W | 875.7732177 | 0.352517435 | 0.32610192 | 0.72 |
| YOR149C | 790.0054864 | -0.295532796 | 0.326042641 | 0.72 |
| YDR336W | 301.9017741 | 0.263952527 | 0.327271566 | 0.72 |
| YGL103W | 19.59323296 | -0.890085311 | 0.32678267 | 0.72 |
| YLR466C-B | 12.5768575 | -0.014641412 | 0.327236377 | 0.72 |
| YNL253W | 477.8813502 | -0.255062843 | 0.326962217 | 0.72 |
| YOR124C | 2791.881139 | 0.404912575 | 0.327218776 | 0.72 |
| YPL084W | 2075.944666 | 0.230498401 | 0.327124037 | 0.72 |
| YDR242W | 619.518258 | 0.279675456 | 0.327870185 | 0.72 |
| YGL148W | 4268.613078 | -0.127872449 | 0.327669042 | 0.72 |
| YNR063W | 178.9442947 | -0.337458717 | 0.327853796 | 0.72 |
| YOR260W | 2245.68858 | -0.361922936 | 0.327516234 | 0.72 |
| YPR176C | 535.2741493 | -0.334943574 | 0.327727886 | 0.72 |
| YDR374W-A | 251.03932 | -0.537786299 | 0.328055205 | 0.72 |
| YNL071W | 6818.92242 | -0.203511804 | 0.328443134 | 0.72 |
| YOR228C | 505.0466268 | 0.317544698 | 0.32848981 | 0.72 |
| YDL129W | 428.0062351 | 0.332896431 | 0.328908788 | 0.72 |
| YER127W | 556.9285849 | -0.089027019 | 0.328850697 | 0.72 |
| YMR145C | 7155.424672 | -0.185797123 | 0.328900406 | 0.72 |
| YBR289W | 1372.875089 | 0.414886 | 0.329159647 | 0.72 |
| YNL319W | 5.59444275 | -0.417322853 | 0.329173516 | 0.72 |
| YHL034C | 6448.755806 | -0.207193091 | 0.329584969 | 0.72 |
| YJL152W | 449.6754094 | 0.383921816 | 0.329880085 | 0.72 |

| | | | | |
|---|---|---|---|---|
| YKL017C | 627.7815414 | 0.315556632 | 0.329835226 | 0.72 |
| YKL165C-A | 6.96473103 | 0.418914859 | 0.329882274 | 0.72 |
| YNL181W | 1100.958937 | -0.220099775 | 0.329535433 | 0.72 |
| snR18 | 14.34494847 | -0.306142758 | 0.329704846 | 0.72 |
| YJL164C | 1399.537088 | 0.35312334 | 0.330607276 | 0.72 |
| YKL083W | 46.77827339 | -0.502504783 | 0.330602754 | 0.72 |
| YAR027W | 1481.56887 | 0.273828553 | 0.331089795 | 0.72 |
| YBL003C | 4258.595501 | -0.283532369 | 0.331247719 | 0.72 |
| YDR020C | 268.9612697 | -0.435929664 | 0.331185682 | 0.72 |
| YKL071W | 436.0298233 | 0.254969724 | 0.331343524 | 0.72 |
| YLR194C | 2445.708189 | 0.745076668 | 0.330881725 | 0.72 |
| YOR152C | 552.1803625 | 0.48276392 | 0.331057051 | 0.72 |
| YDR462W | 726.8986569 | -0.299085021 | 0.331862845 | 0.72 |
| YKR040C | 4.454389 | -0.167372712 | 0.332000121 | 0.72 |
| YGR286C | 924.5538048 | 0.670399921 | 0.332395802 | 0.73 |
| YKL091C | 918.9800837 | 0.309828713 | 0.332725734 | 0.73 |
| YLL055W | 836.4946306 | 0.432532226 | 0.332814149 | 0.73 |
| YLR437C | 329.593239 | -0.327252025 | 0.33275981 | 0.73 |
| YLR455W | 572.5909218 | -0.339576971 | 0.332561328 | 0.73 |
| YNL011C | 614.0144377 | 0.345298766 | 0.3326571 | 0.73 |
| YBR095C | 539.8071954 | 0.292216409 | 0.333148974 | 0.73 |
| YLR290C | 1190.940137 | 0.337056367 | 0.333101719 | 0.73 |
| YML116W | 562.6593031 | 0.460942916 | 0.333246689 | 0.73 |
| YBR049C | 1277.509775 | -0.37920685 | 0.333397004 | 0.73 |
| YDR475C | 1828.530627 | 0.444181514 | 0.333678489 | 0.73 |
| YFL005W | 2091.331245 | -0.314137468 | 0.333775229 | 0.73 |
| YNL168C | 1117.770385 | -0.21066348 | 0.333688295 | 0.73 |
| YHL027W | 1629.795881 | 0.396960923 | 0.334299288 | 0.73 |
| YNR041C | 337.6551047 | -0.259605271 | 0.334494789 | 0.73 |
| YPL096C-A | 145.3313639 | -0.368802154 | 0.334419781 | 0.73 |
| YNL092W | 338.5129162 | 0.503678186 | 0.3347225 | 0.73 |
| YOL098C | 2415.997601 | -0.269503389 | 0.33469672 | 0.73 |
| YIR031C | 471.7138035 | 0.186797147 | 0.334948 | 0.73 |
| YAL026C-A | 170.5922701 | 0.307130591 | 0.335138848 | 0.73 |
| YAL051W | 1222.388192 | 0.299295079 | 0.335556681 | 0.73 |
| YDL106C | 1181.738765 | 0.35404691 | 0.335558744 | 0.73 |
| YIL142W | 3130.184557 | -0.258373541 | 0.335490196 | 0.73 |
| YOR294W | 771.7312013 | -0.185001034 | 0.335346161 | 0.73 |
| YAR029W | 75.14481749 | 0.255821548 | 0.336918379 | 0.73 |
| YBL103C | 858.4740089 | 0.304070808 | 0.336687972 | 0.73 |
| YDL019C | 2417.237505 | 0.371217668 | 0.337662223 | 0.73 |
| YDL216C | 462.7023205 | 0.263480048 | 0.338218668 | 0.73 |
| YDR157W | 53.23769391 | 0.439114965 | 0.337131778 | 0.73 |

| | | | | |
|---|---|---|---|---|
| YDR198C | 539.1037114 | 0.268500426 | 0.336407059 | 0.73 |
| YDR201W | 235.4443499 | -0.391439056 | 0.338055691 | 0.73 |
| YDR458C | 1309.381626 | 0.276427967 | 0.338156431 | 0.73 |
| YEL055C | 1803.113225 | -0.233398146 | 0.338162301 | 0.73 |
| YEL071W | 3890.356415 | 0.374687717 | 0.337347727 | 0.73 |
| YER133W | 3950.455243 | -0.275148743 | 0.337677064 | 0.73 |
| YER137W-A | 3.086112355 | -0.085199972 | 0.337742284 | 0.73 |
| YGR188C | 503.2494794 | 0.438700343 | 0.336317468 | 0.73 |
| YGR213C | 279.0368426 | 0.380964649 | 0.337166207 | 0.73 |
| YGR229C | 1916.022007 | -0.313086827 | 0.337055815 | 0.73 |
| YHR170W | 1876.524363 | -0.253638279 | 0.336078608 | 0.73 |
| YIR009W | 172.3613335 | 0.269307596 | 0.338002688 | 0.73 |
| YJL041W | 2583.756357 | -0.223615118 | 0.335973305 | 0.73 |
| YMR174C | 615.2184477 | 0.442478287 | 0.337059465 | 0.73 |
| YOR119C | 842.963497 | 0.341679717 | 0.336357369 | 0.73 |
| YOR179C | 265.678804 | -0.46325758 | 0.336893927 | 0.73 |
| YPL172C | 484.663762 | -0.267790001 | 0.33785517 | 0.73 |
| YPL183C | 913.1902093 | -0.35057157 | 0.338167061 | 0.73 |
| YDL008W | 381.8332185 | -0.39169484 | 0.33858135 | 0.73 |
| YPR015C | 179.9077993 | 0.270486663 | 0.33853038 | 0.73 |
| YFL018C | 6022.957872 | -0.196925338 | 0.338799055 | 0.73 |
| YPL152W | 506.4565509 | 0.259268345 | 0.339099971 | 0.73 |
| YAL058W | 573.3781549 | -0.309298502 | 0.340509061 | 0.73 |
| YBR029C | 1375.442278 | -0.258763779 | 0.340079328 | 0.73 |
| YBR275C | 1768.794249 | -0.175453737 | 0.341689805 | 0.73 |
| YDL115C | 894.0269756 | 0.242078333 | 0.341841116 | 0.73 |
| YDR055W | 3654.499745 | 0.506923663 | 0.341768538 | 0.73 |
| YDR248C | 767.0999761 | 0.402643634 | 0.341937097 | 0.73 |
| YEL015W | 1435.206957 | -0.27211156 | 0.340883781 | 0.73 |
| YGR071C | 709.1837053 | 0.244985394 | 0.341877339 | 0.73 |
| YGR274C | 1344.663557 | 0.340641278 | 0.341254193 | 0.73 |
| YIR003W | 1806.796543 | 0.345391594 | 0.341086872 | 0.73 |
| YJL098W | 1058.267184 | -0.32738279 | 0.340445463 | 0.73 |
| YJR064W | 2718.261132 | -0.224488592 | 0.340653898 | 0.73 |
| YJR071W | 37.86358004 | 0.076688125 | 0.341302344 | 0.73 |
| YKL047W | 711.1080868 | -0.352338543 | 0.341483114 | 0.73 |
| YKL078W | 299.5473506 | -0.115449365 | 0.341811245 | 0.73 |
| YKL140W | 1755.744554 | 0.253502992 | 0.341767305 | 0.73 |
| YKL187C | 740.5340162 | 0.369045584 | 0.339835327 | 0.73 |
| YLL008W | 1923.071209 | -0.124930962 | 0.341548341 | 0.73 |
| YNL337W | 0.659062417 | 0.269623306 | 0.339786579 | 0.73 |
| YOR073W-A | 11.62724158 | -0.590281323 | 0.341803157 | 0.73 |
| YOR147W | 1293.879256 | 0.397444666 | 0.341427536 | 0.73 |

| | | | | |
|---|---|---|---|---|
| YOR318C | 25.32067042 | -0.61354109 | 0.341505886 | 0.73 |
| YPR034W | 1306.898874 | -0.346592881 | 0.342248516 | 0.73 |
| YBR195C | 288.8600834 | -0.383429586 | 0.343383895 | 0.73 |
| YDR082W | 550.4664737 | 0.2219038 | 0.343458244 | 0.73 |
| YGR215W | 453.6817551 | -0.31115878 | 0.343145317 | 0.73 |
| YHR112C | 1466.272889 | 0.376413089 | 0.343053085 | 0.73 |
| YIL150C | 295.6540585 | -0.215212994 | 0.343255939 | 0.73 |
| YJL045W | 286.1990705 | 0.400601492 | 0.343717263 | 0.73 |
| YML122C | 47.85383085 | -0.304037028 | 0.34328788 | 0.73 |
| YNL123W | 2060.867856 | -0.301939844 | 0.343690027 | 0.73 |
| YOL135C | 544.3922283 | -0.272269998 | 0.343586389 | 0.73 |
| YEL016C | 826.6367047 | -0.313054986 | 0.34410738 | 0.73 |
| YDL153C | 1719.824839 | -0.146822909 | 0.344819657 | 0.73 |
| YDL231C | 1653.703222 | 0.241938461 | 0.34455114 | 0.73 |
| YKL198C | 1074.404315 | 0.284934492 | 0.344638081 | 0.73 |
| YOR128C | 2990.654925 | -0.297265029 | 0.344711478 | 0.73 |
| YPR198W | 724.9139692 | -0.452899947 | 0.344715606 | 0.73 |
| YBR024W | 889.4330761 | 0.4976298 | 0.345415358 | 0.73 |
| YCR012W | 43096.09714 | -0.020637795 | 0.346249176 | 0.73 |
| YDL088C | 1480.552376 | 0.251299668 | 0.346183205 | 0.73 |
| YDL205C | 1039.950483 | -0.329987897 | 0.346071222 | 0.73 |
| YDR101C | 1247.120699 | -0.378975446 | 0.345538025 | 0.73 |
| YGL237C | 638.8350245 | 0.306351646 | 0.345739512 | 0.73 |
| YIL154C | 2075.807869 | 0.403357731 | 0.345331067 | 0.73 |
| YKL134C | 542.9022847 | 0.270806223 | 0.345146451 | 0.73 |
| YKR090W | 850.9383093 | -0.262343055 | 0.345988081 | 0.73 |
| YPR127W | 1734.545853 | 0.350682743 | 0.345109177 | 0.73 |
| snR48 | 2.926793808 | -0.388736664 | 0.346078555 | 0.73 |
| snR53 | 0.80815136 | -0.15254148 | 0.345990283 | 0.73 |
| YDR210W-B | 26.10421514 | 0.190397162 | 0.346674745 | 0.73 |
| YNL153C | 847.6509295 | -0.346154013 | 0.346794645 | 0.73 |
| YNR043W | 2170.763038 | -0.245032578 | 0.34716136 | 0.73 |
| YPL252C | 462.6644596 | -0.300100258 | 0.347167903 | 0.73 |
| YBR301W | 22.99326035 | 0.513824824 | 0.347706569 | 0.73 |
| YPL279C | 37.92327787 | -0.785993727 | 0.347649943 | 0.73 |
| YDR491C | 102.3610769 | -0.155976503 | 0.347830694 | 0.73 |
| YGL019W | 1587.591155 | -0.215340283 | 0.347974859 | 0.73 |
| YLR113W | 3162.32221 | -0.34867327 | 0.348110452 | 0.73 |
| YNL085W | 3478.261348 | -0.263387405 | 0.348209541 | 0.73 |
| YHR150W | 670.7977171 | 0.431039058 | 0.348688482 | 0.73 |
| YKR091W | 1094.538594 | 0.350213381 | 0.348709851 | 0.73 |
| YML099C | 1070.053648 | 0.386237303 | 0.348526016 | 0.73 |
| YMR130W | 646.8299959 | -0.387958067 | 0.348545753 | 0.73 |

| | | | | |
|---|---|---|---|---|
| YKR086W | 1098.768049 | 0.316571144 | 0.349182457 | 0.73 |
| YLR202C | 55.2829566 | -0.324403204 | 0.348982908 | 0.73 |
| YOL026C | 840.8767019 | 0.255067094 | 0.349137489 | 0.73 |
| YKR001C | 4012.227533 | -0.204377658 | 0.349335397 | 0.73 |
| YPL236C | 570.2921643 | 0.3054679 | 0.349551222 | 0.73 |
| YBL006C | 611.4550235 | -0.419931794 | 0.350521456 | 0.73 |
| YDR197W | 858.0497536 | 0.307005953 | 0.350415892 | 0.73 |
| YGR033C | 1013.438168 | -0.272530234 | 0.349951469 | 0.73 |
| YNL077W | 1205.161408 | 0.441143192 | 0.350017597 | 0.73 |
| YNL182C | 900.9513347 | -0.344448668 | 0.350545935 | 0.73 |
| YPL019C | 1922.931958 | -0.518922955 | 0.350530438 | 0.73 |
| YPL116W | 1071.716343 | 0.284987057 | 0.349807553 | 0.73 |
| YPR054W | 265.2938746 | -0.762838555 | 0.350360422 | 0.73 |
| YLR400W | 49.34542517 | 0.332271278 | 0.350969057 | 0.73 |
| YMR077C | 308.7109529 | 0.250304004 | 0.351135113 | 0.73 |
| YGR018C | 8.799849222 | 0.157362528 | 0.351601675 | 0.73 |
| YGR125W | 1519.849762 | 0.308098921 | 0.351702859 | 0.73 |
| YHR099W | 4737.26123 | -0.184170816 | 0.351760783 | 0.73 |
| YLR403W | 956.7749718 | 0.265676968 | 0.351855052 | 0.73 |
| YOR225W | 54.88254174 | -0.049795113 | 0.351787019 | 0.73 |
| YOR354C | 1688.06875 | -0.252069028 | 0.351833601 | 0.73 |
| YAL037W | 195.7634153 | -0.649164649 | 0.352412734 | 0.73 |
| YBR234C | 4120.481034 | -0.264340414 | 0.353194624 | 0.73 |
| YCR024C-B | 2025.586967 | -0.43484939 | 0.354500522 | 0.73 |
| YDR142C | 534.1730573 | 0.262293134 | 0.356074685 | 0.73 |
| YDR256C | 561.4619469 | 0.363662063 | 0.354610886 | 0.73 |
| YEL018W | 225.091547 | -0.258898976 | 0.356119269 | 0.73 |
| YER136W | 4119.832209 | -0.262450593 | 0.353845925 | 0.73 |
| YGL109W | 79.04302724 | 0.307289862 | 0.355764477 | 0.73 |
| YGR157W | 3036.942102 | -0.25502624 | 0.354626673 | 0.73 |
| YGR280C | 1635.461549 | -0.204290989 | 0.355894028 | 0.73 |
| YHR131C | 966.4131265 | 0.247136787 | 0.353173904 | 0.73 |
| YIL003W | 534.8841649 | 0.322478188 | 0.355168147 | 0.73 |
| YIL145C | 1126.99064 | -0.28873759 | 0.355197879 | 0.73 |
| YJL026W | 4324.794509 | -0.276106511 | 0.355058704 | 0.73 |
| YJL172W | 1932.176519 | -0.274349914 | 0.355357105 | 0.73 |
| YJR112W | 374.389265 | -0.258051503 | 0.353172343 | 0.73 |
| YKL023C-A | 532.0907152 | -0.139181254 | 0.356000094 | 0.73 |
| YKL208W | 299.0743362 | -0.445722562 | 0.353322436 | 0.73 |
| YKR076W | 1274.281879 | 0.306748988 | 0.355013263 | 0.73 |
| YMR095C | 600.2590168 | 0.874547693 | 0.35314279 | 0.73 |
| YMR196W | 3241.463398 | 0.522873482 | 0.355307499 | 0.73 |
| YMR278W | 1574.862371 | 0.341417696 | 0.355477014 | 0.73 |

| | | | | |
|---|---|---|---|---|
| YNL001W | 629.9634099 | -0.324463196 | 0.355909549 | 0.73 |
| YNL034W | 79.93693595 | 0.735306739 | 0.354349648 | 0.73 |
| YNL042W | 593.1711026 | 0.160918604 | 0.355567131 | 0.73 |
| YNL122C | 325.2385534 | -0.331750166 | 0.352482757 | 0.73 |
| YNL133C | 297.07226 | 0.395901203 | 0.353537135 | 0.73 |
| YNL174W | 7.421723425 | -0.187732153 | 0.352511021 | 0.73 |
| YNL330C | 1017.919377 | -0.588910261 | 0.355501632 | 0.73 |
| YNR018W | 1764.18714 | -0.219775685 | 0.354231134 | 0.73 |
| YOL164W-A | 24.89009234 | 0.387170575 | 0.355468594 | 0.73 |
| YOR085W | 1806.530962 | -0.302368798 | 0.355108185 | 0.73 |
| YOR388C | 10.58733181 | 0.213988903 | 0.355806422 | 0.73 |
| YPL162C | 476.9994766 | 0.347527974 | 0.355753777 | 0.73 |
| YPR001W | 533.0217534 | 0.522088059 | 0.35236746 | 0.73 |
| YPR112C | 1587.350101 | -0.193736507 | 0.353786041 | 0.73 |
| YNL284C-B | 190.6577576 | 0.780777533 | 0.35637367 | 0.74 |
| YDR070C | 1811.421734 | 0.40378271 | 0.356756433 | 0.74 |
| YCL048W-A | 89.40578665 | -0.220023835 | 0.357723917 | 0.74 |
| YDL143W | 3248.97051 | -0.278107107 | 0.358485081 | 0.74 |
| YDR204W | 1297.475345 | 0.535166721 | 0.357553834 | 0.74 |
| YGR086C | 9907.562538 | -0.336605624 | 0.358460531 | 0.74 |
| YGR250C | 3005.303955 | 0.419237938 | 0.358433681 | 0.74 |
| YHL050C | 143.7644195 | 0.382428409 | 0.357554863 | 0.74 |
| YHR070C-A | 129.6976848 | -0.225898488 | 0.35837595 | 0.74 |
| YKL021C | 1160.236373 | -0.324899245 | 0.357795098 | 0.74 |
| YML046W | 728.0597624 | -0.465097479 | 0.3581377 | 0.74 |
| YMR207C | 1385.212155 | 0.40188154 | 0.35804921 | 0.74 |
| YMR228W | 316.010038 | -0.286536024 | 0.357651667 | 0.74 |
| YMR317W | 752.1354442 | 0.304477897 | 0.35768467 | 0.74 |
| YOR024W | 4.641545194 | 0.13651144 | 0.358565843 | 0.74 |
| YOR212W | 1199.500024 | -0.891058991 | 0.357633126 | 0.74 |
| YGR064W | 14.24950712 | -0.843468546 | 0.3590771 | 0.74 |
| YLR127C | 955.0961502 | 0.275921936 | 0.359148264 | 0.74 |
| YGR241C | 1428.572307 | 0.287371739 | 0.359419938 | 0.74 |
| SRG1 | 1084.659985 | 0.036338942 | 0.359556395 | 0.74 |
| YCR030C | 2439.320353 | 0.386465946 | 0.359882955 | 0.74 |
| YJL131C | 712.8647158 | 0.343234985 | 0.359936086 | 0.74 |
| YOR086C | 3224.270398 | 0.300245365 | 0.360333019 | 0.74 |
| YPL004C | 14429.78474 | 0.355960471 | 0.360382608 | 0.74 |
| YNL236W | 1176.860728 | -0.283778537 | 0.360634137 | 0.74 |
| YBL048W | 125.8109055 | 0.087118502 | 0.362328603 | 0.74 |
| YBR013C | 207.4427581 | 0.287876522 | 0.361772588 | 0.74 |
| YBR166C | 859.3257193 | -0.307723505 | 0.362252837 | 0.74 |
| YDL138W | 666.9826314 | 0.315851938 | 0.362074392 | 0.74 |

| | | | | |
|---|---|---|---|---|
| YGL082W | 1508.755452 | -0.253885795 | 0.362182042 | 0.74 |
| YGR083C | 2563.158798 | -0.145365754 | 0.361930535 | 0.74 |
| YHR070W | 688.4026548 | -0.178915121 | 0.362435805 | 0.74 |
| YHR117W | 2228.445201 | -0.186691697 | 0.36144418 | 0.74 |
| YIR021W | 261.891268 | -0.318980148 | 0.362140665 | 0.74 |
| YOR353C | 984.6133479 | 0.288765641 | 0.361763043 | 0.74 |
| YPR095C | 1088.661894 | 0.278571805 | 0.36222867 | 0.74 |
| YER148W-A | 0.781507553 | -0.606902338 | 0.363027649 | 0.74 |
| YDL032W | 4.179935405 | 0.199935991 | 0.363232242 | 0.74 |
| YIL120W | 363.836718 | -0.612413312 | 0.363288568 | 0.74 |
| YER178W | 9418.417677 | -0.206703446 | 0.363413336 | 0.74 |
| YNR056C | 522.7352776 | 0.187960792 | 0.363684583 | 0.74 |
| YAL064W-B | 18.20372016 | -0.463994568 | 0.364130241 | 0.74 |
| YLR222C | 1088.378581 | -0.408979943 | 0.364297249 | 0.74 |
| YMR054W | 1855.817154 | -0.298758057 | 0.364187498 | 0.74 |
| YDL025C | 1555.623411 | 0.385958919 | 0.364523175 | 0.74 |
| YDR068W | 1258.879245 | 0.187676028 | 0.364657297 | 0.74 |
| YOL052C-A | 1535.771374 | 0.164906757 | 0.365003272 | 0.74 |
| YLR359W | 9224.21068 | 0.021451543 | 0.365206534 | 0.74 |
| YAL056C-A | 23.07572137 | -0.083490139 | 0.365416874 | 0.74 |
| YBR059C | 2706.219813 | 0.375444749 | 0.365461661 | 0.74 |
| YBR007C | 740.5673067 | -0.343121538 | 0.365873416 | 0.74 |
| YDL009C | 106.1081082 | -0.369744128 | 0.365919166 | 0.74 |
| YOL099C | 11.41692104 | 0.634441914 | 0.365773017 | 0.74 |
| YBR116C | 35.41794033 | 0.027303509 | 0.366314373 | 0.74 |
| YNL321W | 1495.615211 | 0.287117222 | 0.366121637 | 0.74 |
| snR71 | 12.58626851 | 0.153045479 | 0.366274418 | 0.74 |
| YER089C | 1927.936186 | -0.335562951 | 0.3670437 | 0.74 |
| YLL022C | 865.5611491 | -0.169885645 | 0.36714678 | 0.74 |
| LSR1 | 186.1998782 | 0.384493822 | 0.367654829 | 0.74 |
| YIL036W | 2051.229049 | 0.369033471 | 0.367641779 | 0.74 |
| YOR030W | 929.3153858 | 0.278646467 | 0.36783362 | 0.74 |
| YBR160W | 885.0084099 | -0.253660544 | 0.368626806 | 0.74 |
| YCL036W | 402.9926525 | 0.546627569 | 0.368368816 | 0.74 |
| YGR037C | 2287.491924 | -0.298420412 | 0.368359107 | 0.74 |
| YIL002W-A | 495.5817191 | -0.281859129 | 0.368591006 | 0.74 |
| YKL152C | 30648.47356 | -0.225477976 | 0.368750468 | 0.74 |
| YLR066W | 1030.577166 | -0.239787946 | 0.368350386 | 0.74 |
| YPL155C | 690.291389 | 0.365564138 | 0.368166424 | 0.74 |
| YPL239W | 1334.632299 | -0.182102698 | 0.368755675 | 0.74 |
| YCL045C | 3038.105977 | -0.276234282 | 0.368872648 | 0.74 |
| YBL059C-A | 504.0749236 | -0.197604875 | 0.369019423 | 0.74 |
| YDL233W | 547.8994161 | 0.378651174 | 0.369274876 | 0.74 |

| | | | | |
|---|---|---|---|---|
| YDR090C | 473.3492687 | 0.274522852 | 0.369193788 | 0.74 |
| YAR075W | 64.72959559 | 1.00614627 | 0.36957174 | 0.74 |
| YDR115W | 505.144241 | -0.30324253 | 0.370073767 | 0.74 |
| YDR234W | 1972.622196 | 0.433005376 | 0.370139531 | 0.74 |
| YER190W | 15.31140442 | -0.177854462 | 0.370056979 | 0.74 |
| YGL078C | 1925.6458 | -0.222498924 | 0.370070699 | 0.74 |
| YIL164C | 260.9748713 | 0.172460791 | 0.369685701 | 0.74 |
| YNL062C | 805.5436558 | -0.285958715 | 0.369896352 | 0.74 |
| YOR007C | 8060.679187 | -0.252056158 | 0.370516182 | 0.75 |
| YPR189W | 1450.192608 | -0.284217896 | 0.370538439 | 0.75 |
| YFR003C | 1017.431306 | 0.286833458 | 0.370818949 | 0.75 |
| YBR196C-B | 11.09747944 | -0.686837821 | 0.371445672 | 0.75 |
| YEL038W | 943.0798183 | -0.303051381 | 0.371676744 | 0.75 |
| YMR128W | 2322.124433 | 0.119861958 | 0.371415292 | 0.75 |
| YMR136W | 2477.221542 | 0.472950233 | 0.371224098 | 0.75 |
| YNL210W | 106.0752581 | 0.09556448 | 0.37154551 | 0.75 |
| YOL051W | 2569.23447 | 0.423930054 | 0.371583367 | 0.75 |
| YLR314C | 3042.070083 | -0.166292401 | 0.372128673 | 0.75 |
| YOR185C | 2527.056266 | 0.453443504 | 0.372213381 | 0.75 |
| YFL027C | 673.7008422 | -0.77546301 | 0.372824706 | 0.75 |
| YLR090W | 931.4547969 | 0.346352825 | 0.372928846 | 0.75 |
| YCL039W | 1768.904925 | 0.430569932 | 0.373652752 | 0.75 |
| YCL074W | 140.2978941 | 1.512233046 | 0.373632209 | 0.75 |
| YEL063C | 1734.961887 | 0.25092151 | 0.373881557 | 0.75 |
| YGR124W | 7850.205704 | -0.083838901 | 0.373666568 | 0.75 |
| YML039W | 1.131094598 | -0.29565802 | 0.373097248 | 0.75 |
| YMR250W | 7553.858966 | 0.476499324 | 0.373585202 | 0.75 |
| YNL204C | 121.2325338 | 0.268918347 | 0.37365872 | 0.75 |
| YPL041C | 160.4382575 | -0.318533807 | 0.373874797 | 0.75 |
| YHR093W | 35.27687991 | 0.365961537 | 0.374240447 | 0.75 |
| YJR006W | 1054.347239 | -0.295704045 | 0.374361693 | 0.75 |
| YOL017W | 430.0355777 | -0.293238035 | 0.374341092 | 0.75 |
| YPR149W | 11719.12039 | 0.358062622 | 0.374536203 | 0.75 |
| YHR194W | 859.058911 | 0.375712226 | 0.374727633 | 0.75 |
| YBL081W | 976.0547384 | -0.368030615 | 0.375620747 | 0.75 |
| YBR076C-A | 84.52994015 | 0.226004589 | 0.378062053 | 0.75 |
| YBR089C-A | 560.6070076 | -0.126780105 | 0.376765261 | 0.75 |
| YCR072C | 412.6699919 | -0.370852249 | 0.37792387 | 0.75 |
| YCR100C | 233.2545617 | 0.137049121 | 0.377861253 | 0.75 |
| YDR320C-A | 373.904445 | 0.308001019 | 0.378086729 | 0.75 |
| YDR445C | 3.142353253 | -0.518134657 | 0.376476249 | 0.75 |
| YDR504C | 165.3464621 | 0.288997484 | 0.376918172 | 0.75 |
| YER172C | 2101.126018 | -0.200221514 | 0.377999947 | 0.75 |

| | | | | |
|---|---|---|---|---|
| YER181C | 6.5610832 | 0.657865201 | 0.37709221 | 0.75 |
| YFL011W | 70.61598103 | -0.017788169 | 0.377783424 | 0.75 |
| YGL065C | 1014.84416 | -0.21991519 | 0.377265125 | 0.75 |
| YGL136C | 674.1887477 | 0.237122235 | 0.37683496 | 0.75 |
| YGR072W | 838.1301736 | 0.294808483 | 0.375071856 | 0.75 |
| YGR161W-C | 303.4318056 | 0.31406519 | 0.376599598 | 0.75 |
| YGR276C | 1041.106244 | 0.228308442 | 0.376993136 | 0.75 |
| YJL077C | 162.6090211 | 0.237858026 | 0.375567221 | 0.75 |
| YJL211C | 56.87482788 | 0.463466633 | 0.375745793 | 0.75 |
| YJR009C | 33615.59982 | -0.073916902 | 0.377330334 | 0.75 |
| YKL059C | 865.3974506 | 0.203305617 | 0.375718374 | 0.75 |
| YMR190C | 1250.870637 | 0.514046585 | 0.376079557 | 0.75 |
| YNL027W | 1180.449634 | 0.269172145 | 0.375182582 | 0.75 |
| YNL097C | 784.7174827 | 0.360356949 | 0.375411776 | 0.75 |
| YNL200C | 2258.279724 | 0.393279647 | 0.375993161 | 0.75 |
| YOL083W | 724.7720388 | 0.283607649 | 0.375272107 | 0.75 |
| YPL049C | 2306.704815 | -0.385348904 | 0.376672612 | 0.75 |
| YPL092W | 1197.47636 | 0.456832823 | 0.378060693 | 0.75 |
| YPR062W | 2367.986615 | -0.208486425 | 0.376153376 | 0.75 |
| snR61 | 12.13748594 | -0.141407353 | 0.377928101 | 0.75 |
| YDR257C | 688.7119517 | -0.399335991 | 0.37843634 | 0.75 |
| YER167W | 1421.029082 | 0.464335126 | 0.378472713 | 0.75 |
| YIL140W | 1095.766571 | -0.316541657 | 0.378270036 | 0.75 |
| YAL034W-A | 187.40184 | -0.244222323 | 0.379430544 | 0.75 |
| YJL078C | 3272.163203 | 0.501294517 | 0.379115111 | 0.75 |
| YKL008C | 1304.945966 | -0.408886975 | 0.379392367 | 0.75 |
| YKL023W | 1129.71016 | 0.347329067 | 0.379400196 | 0.75 |
| YLR360W | 653.1816733 | -0.261844271 | 0.379458833 | 0.75 |
| YMR214W | 1048.466602 | -0.187944357 | 0.379255406 | 0.75 |
| YPL181W | 1430.008347 | 0.404728258 | 0.379445801 | 0.75 |
| YBR030W | 579.6269936 | -0.254740385 | 0.3798849 | 0.75 |
| YCL052C | 1614.200073 | 0.257305879 | 0.380235773 | 0.75 |
| YCL061C | 1120.832529 | -0.241958433 | 0.380832624 | 0.75 |
| YDR466W | 1549.812513 | 0.278654787 | 0.381098525 | 0.75 |
| YER013W | 829.6032387 | 0.271028678 | 0.381058803 | 0.75 |
| YER033C | 1095.361309 | 0.409607987 | 0.380609949 | 0.75 |
| YHR091C | 596.7962742 | -0.206480563 | 0.380489113 | 0.75 |
| YHR199C-A | 146.5882374 | 0.12726612 | 0.381413276 | 0.75 |
| YKL001C | 376.5252013 | 0.28132205 | 0.380779674 | 0.75 |
| YMR194C-B | 137.2124333 | 0.240670106 | 0.381343269 | 0.75 |
| YNL227C | 372.4616162 | -0.162709158 | 0.380554367 | 0.75 |
| YNL251C | 2085.43933 | 0.203553506 | 0.379634541 | 0.75 |
| YOR285W | 4227.371531 | 0.525265558 | 0.380829208 | 0.75 |

| | | | | |
|---|---|---|---|---|
| YOR330C | 1527.333333 | 0.287363326 | 0.381443686 | 0.75 |
| YOR352W | 729.6147343 | 0.244741216 | 0.380709611 | 0.75 |
| YPL101W | 1017.044188 | -0.223148873 | 0.380758394 | 0.75 |
| snR9 | 75.73439823 | -0.235061409 | 0.381027912 | 0.75 |
| YHR005C | 1534.146963 | -0.88074946 | 0.381793936 | 0.75 |
| YMR271C | 754.1723733 | 0.269991205 | 0.38180359 | 0.75 |
| YOR198C | 6733.130979 | -0.144682618 | 0.381663873 | 0.75 |
| YCR095W-A | 20.07196858 | 0.029361102 | 0.382094751 | 0.75 |
| YBL060W | 624.3164774 | 0.244844789 | 0.382775847 | 0.75 |
| YDL147W | 5209.098548 | 0.318157181 | 0.38395578 | 0.75 |
| YDR207C | 1436.193266 | 0.439061864 | 0.384817293 | 0.75 |
| YDR322W | 1031.616879 | -0.291870698 | 0.382891854 | 0.75 |
| YDR406W-A | 12.68467301 | 0.25939742 | 0.382574386 | 0.75 |
| YFR036W | 228.7837864 | 0.384442312 | 0.384677525 | 0.75 |
| YHR020W | 3665.040866 | -0.12563283 | 0.383254318 | 0.75 |
| YHR064C | 6332.657586 | -0.06171495 | 0.384850074 | 0.75 |
| YJR111C | 500.5453341 | 0.37313168 | 0.384862071 | 0.75 |
| YKR052C | 558.5950233 | 0.266656887 | 0.382513864 | 0.75 |
| YML001W | 1453.0376 | -0.231678406 | 0.384767817 | 0.75 |
| YMR240C | 592.2234425 | 0.236162492 | 0.383246741 | 0.75 |
| YOL075C | 712.6275688 | 0.38812398 | 0.38385025 | 0.75 |
| YOL150C | 3.831416512 | 0.544749126 | 0.383669329 | 0.75 |
| YOR021C | 1473.922183 | -0.147807744 | 0.383065003 | 0.75 |
| YPL029W | 470.8569792 | -0.246698021 | 0.382623438 | 0.75 |
| YPL072W | 410.1680408 | -0.292640069 | 0.384414033 | 0.75 |
| YPL115C | 1185.518438 | -0.299794193 | 0.383575817 | 0.75 |
| YPL124W | 247.3933842 | -0.177310195 | 0.382695132 | 0.75 |
| YPL238C | 56.97812647 | -0.425355461 | 0.384368977 | 0.75 |
| YPR037C | 567.1835844 | -0.25186198 | 0.383318317 | 0.75 |
| YPR089W | 1983.427567 | 0.243482429 | 0.384839657 | 0.75 |
| YPR135W | 846.2041741 | -0.259027076 | 0.384015515 | 0.75 |
| snR74 | 1.214578707 | 0.028657996 | 0.384296786 | 0.75 |
| snR79 | 6.23839996 | 0.16392851 | 0.385020316 | 0.75 |
| YCL040W | 21911.49524 | 0.737099071 | 0.385394099 | 0.75 |
| YPL235W | 1507.98677 | -0.313900297 | 0.385320054 | 0.75 |
| YPR125W | 636.6611003 | -0.252429896 | 0.385475281 | 0.75 |
| YDR411C | 1171.452043 | -0.290181088 | 0.386031825 | 0.75 |
| YCL004W | 588.1019431 | 0.28898593 | 0.38615358 | 0.75 |
| YER044C | 1096.772117 | -0.376111758 | 0.386481984 | 0.75 |
| YIL034C | 3381.997418 | -0.195913844 | 0.386693197 | 0.75 |
| YNL324W | 2.168671185 | 0.308674253 | 0.386726339 | 0.75 |
| YFL002C | 1100.586944 | 0.216511977 | 0.387103129 | 0.75 |
| YLL004W | 381.8327061 | -0.436228837 | 0.387143747 | 0.75 |

| | | | | |
|---|---|---|---|---|
| YLR333C | 2119.808061 | -0.260820529 | 0.387267461 | 0.75 |
| YPL206C | 1410.718758 | -0.292727427 | 0.387314912 | 0.75 |
| snR44 | 201.2193719 | -0.369970488 | 0.387090837 | 0.75 |
| YPL133C | 580.4116075 | 0.297218962 | 0.387494601 | 0.75 |
| YFR010W | 3072.848472 | 0.317888757 | 0.38766668 | 0.75 |
| YJL020C | 4552.045062 | 0.519549629 | 0.387978578 | 0.75 |
| YPL038W | 299.6200602 | -0.543979746 | 0.388198171 | 0.75 |
| YBR120C | 395.0224857 | -0.306575123 | 0.388380891 | 0.75 |
| YCL023C | 8.286348364 | 0.428195841 | 0.388644766 | 0.75 |
| YDR137W | 962.3953612 | 0.154473172 | 0.388620181 | 0.75 |
| YKL217W | 2399.793467 | 0.683385161 | 0.388867396 | 0.75 |
| YLR271W | 587.0390943 | 0.404665405 | 0.38876116 | 0.75 |
| YPR002W | 1192.998192 | 0.69183494 | 0.389005344 | 0.75 |
| YEL044W | 582.0177494 | -0.336670689 | 0.389589521 | 0.75 |
| YEL067C | 149.0124198 | 0.345979906 | 0.38960103 | 0.75 |
| YGR117C | 888.1364455 | -0.379983506 | 0.389543782 | 0.75 |
| YJL033W | 1448.230029 | -0.215631498 | 0.389347362 | 0.75 |
| YBR011C | 18392.14922 | -0.254335951 | 0.389975374 | 0.75 |
| YDL087C | 335.3146294 | 0.323340633 | 0.390534064 | 0.75 |
| YLR343W | 1249.632226 | 0.139937925 | 0.390544382 | 0.75 |
| YJR018W | 57.57815418 | 0.052944369 | 0.390866306 | 0.75 |
| YNR026C | 756.2847135 | -0.320753187 | 0.390936214 | 0.75 |
| YKL194C | 593.5236151 | 0.328003978 | 0.391229095 | 0.75 |
| YIL033C | 4130.82095 | 0.438497769 | 0.391357268 | 0.75 |
| YDR428C | 292.2817321 | -0.392213195 | 0.392065676 | 0.75 |
| YLR123C | 17.47407493 | -0.491385851 | 0.391953266 | 0.75 |
| YOL030W | 6136.814993 | 0.382167689 | 0.39157432 | 0.75 |
| YPR060C | 619.4282416 | -0.213939888 | 0.391710466 | 0.75 |
| YPR172W | 1219.274092 | 0.407341215 | 0.391929092 | 0.75 |
| YPR179C | 773.0608325 | 0.260022473 | 0.392153421 | 0.75 |
| YGL196W | 1262.908167 | 0.520351082 | 0.392423846 | 0.75 |
| YGL256W | 1304.621904 | 0.295287565 | 0.392831652 | 0.75 |
| YKL164C | 7734.898504 | -0.003176429 | 0.392737596 | 0.75 |
| YNL264C | 1013.510724 | -0.224704783 | 0.392781385 | 0.75 |
| YPR021C | 1682.395818 | 0.296276167 | 0.392614607 | 0.75 |
| YDL042C | 548.3117902 | -0.113230265 | 0.393305876 | 0.75 |
| YDR457W | 5588.916733 | -0.180645341 | 0.393219692 | 0.75 |
| YIL046W-A | 7.695578865 | 0.645054149 | 0.393316229 | 0.75 |
| YLL017W | 60.73191178 | 0.226718249 | 0.393007278 | 0.75 |
| YGL214W | 9.34196156 | -0.500018743 | 0.393443272 | 0.75 |
| YGL130W | 854.7206722 | -0.147146421 | 0.393746551 | 0.75 |
| YDR229W | 1110.14149 | 0.240704421 | 0.394022805 | 0.75 |
| YDR249C | 239.973621 | 0.254922431 | 0.394774337 | 0.76 |

| | | | | |
|---|---|---|---|---|
| YHR198C | 872.0543676 | 0.39088494 | 0.394657697 | 0.76 |
| YKR084C | 1262.417524 | -0.293850916 | 0.394696713 | 0.76 |
| YBL061C | 962.4150781 | -0.29528282 | 0.395122576 | 0.76 |
| YGL205W | 247.6080273 | 0.544126066 | 0.395296089 | 0.76 |
| YIL008W | 681.0099622 | -0.230241794 | 0.395404294 | 0.76 |
| YKR028W | 1923.731148 | -0.336405725 | 0.395196787 | 0.76 |
| NME1 | 51.73748825 | 0.217573562 | 0.395728092 | 0.76 |
| YLR058C | 11510.40084 | 0.089902239 | 0.396434873 | 0.76 |
| YOL060C | 2188.941543 | 0.445386043 | 0.396341373 | 0.76 |
| YLL047W | 4.30351014 | 0.4268599 | 0.396676293 | 0.76 |
| YOR246C | 1262.599493 | -0.363251524 | 0.397300017 | 0.76 |
| YMR245W | 80.78367086 | 0.543641042 | 0.397542738 | 0.76 |
| YOR028C | 706.5725488 | -0.213377325 | 0.397854449 | 0.76 |
| YCL021W-A | 277.7575158 | -0.169721939 | 0.398562857 | 0.76 |
| YGL241W | 933.7531382 | -0.229461629 | 0.398522026 | 0.76 |
| YGR284C | 2427.894256 | -0.181581528 | 0.398695241 | 0.76 |
| YDL051W | 1178.81834 | -0.055608654 | 0.399243888 | 0.76 |
| YJL064W | 3.253557565 | -0.70968383 | 0.399282563 | 0.76 |
| YNL116W | 793.4753006 | 0.396131 | 0.399224437 | 0.76 |
| YER004W | 2233.162604 | 0.372073597 | 0.399547234 | 0.76 |
| YOR192C | 247.398385 | 0.529162297 | 0.399563339 | 0.76 |
| YGR267C | 1681.678228 | -0.197658502 | 0.399766203 | 0.76 |
| YGR004W | 331.5972983 | 0.257576495 | 0.400100076 | 0.76 |
| YDL156W | 331.6691228 | -0.272651968 | 0.400295192 | 0.76 |
| YNL207W | 833.7029402 | -0.22131899 | 0.400946958 | 0.76 |
| YDR235W | 670.6497022 | -0.36142226 | 0.401257352 | 0.76 |
| YKR034W | 49.93480154 | 0.069434402 | 0.401279637 | 0.76 |
| YER001W | 1455.504127 | -0.311757572 | 0.401823102 | 0.76 |
| YER171W | 948.7525289 | -0.28074296 | 0.401578879 | 0.76 |
| YLR368W | 767.6389144 | 0.224511345 | 0.401753118 | 0.76 |
| YMR197C | 923.7331716 | 0.238729617 | 0.401851535 | 0.76 |
| YAL034C-B | 9.086766243 | -0.287574635 | 0.401998869 | 0.76 |
| YBL019W | 515.3822708 | 0.248998282 | 0.403140012 | 0.76 |
| YCR032W | 1688.564969 | 0.375892824 | 0.40290373 | 0.76 |
| YDR513W | 3464.363476 | 0.324274093 | 0.402814207 | 0.76 |
| YIL026C | 1582.807889 | -0.241042318 | 0.402532082 | 0.76 |
| YJL208C | 631.1153592 | -0.112762846 | 0.402317864 | 0.76 |
| YJL217W | 2088.445581 | 0.731297775 | 0.402679706 | 0.76 |
| YLL003W | 928.3757705 | 0.177261815 | 0.403033484 | 0.76 |
| YLR079W | 2171.764341 | 0.341545564 | 0.402950547 | 0.76 |
| YLR196W | 1358.156963 | -0.097097777 | 0.402465706 | 0.76 |
| YPL136W | 0.97000745 | 0.132226394 | 0.403148713 | 0.76 |
| YGL005C | 924.1279481 | 0.272367654 | 0.404478831 | 0.76 |

| | | | | |
|---|---|---|---|---|
| YJR058C | 464.9848205 | -0.26753689 | 0.403867915 | 0.76 |
| YLR045C | 1304.494239 | -0.135770529 | 0.404677743 | 0.76 |
| YLR054C | 138.716349 | 0.145111602 | 0.404381189 | 0.76 |
| YLR118C | 939.2371521 | -0.231421171 | 0.404512421 | 0.76 |
| YOR001W | 1523.051345 | -0.309464079 | 0.403991511 | 0.76 |
| YPL258C | 548.5976694 | 0.18722306 | 0.404433894 | 0.76 |
| snR84 | 162.4042194 | -0.346329094 | 0.40458024 | 0.76 |
| YPR138C | 1408.826707 | 0.217147738 | 0.404873032 | 0.76 |
| YNL198C | 23.75937545 | 0.08019825 | 0.405035829 | 0.76 |
| YBR174C | 5.135026563 | 0.264569353 | 0.405475879 | 0.76 |
| YGL034C | 9.87258369 | 0.237566492 | 0.405516227 | 0.76 |
| YKL178C | 114.5867776 | -0.821437239 | 0.405661356 | 0.76 |
| YLL011W | 676.0042616 | -0.400634072 | 0.405635623 | 0.76 |
| YMR056C | 319.9319845 | 0.330322688 | 0.405524895 | 0.76 |
| YNL048W | 1078.322808 | -0.211063386 | 0.405779897 | 0.76 |
| YBR062C | 819.8607412 | 0.325458497 | 0.406339493 | 0.76 |
| YDL114W-A | 24.54000287 | 0.229935158 | 0.406660811 | 0.76 |
| YDL160C | 1362.561057 | -0.362693755 | 0.406300629 | 0.76 |
| YDR031W | 342.8572195 | 0.449422652 | 0.406681507 | 0.76 |
| YLR228C | 906.7831503 | 0.336250741 | 0.406280241 | 0.76 |
| YMR037C | 1272.708272 | -0.3396848 | 0.406619648 | 0.76 |
| YPL174C | 736.6858257 | 0.198376752 | 0.406347937 | 0.76 |
| YJR109C | 3577.314205 | 0.576156098 | 0.406799436 | 0.76 |
| YJL127C | 657.9035431 | 0.227571598 | 0.406925311 | 0.76 |
| YPL204W | 2957.690828 | -0.274144374 | 0.407271864 | 0.76 |
| YJL004C | 587.5114449 | -0.3092072 | 0.407600968 | 0.77 |
| YJL034W | 12765.00653 | -0.099456395 | 0.408215148 | 0.77 |
| YKL082C | 1057.576571 | -0.154011947 | 0.408588918 | 0.77 |
| YBR142W | 1284.846746 | -0.116710028 | 0.409137124 | 0.77 |
| YGR240C-A | 46.68747277 | -0.616738568 | 0.409438323 | 0.77 |
| YAL044C | 2900.555908 | -0.209818661 | 0.413329522 | 0.77 |
| YAR007C | 1627.673812 | -0.219865784 | 0.411739441 | 0.77 |
| YDL103C | 1449.48201 | -0.210446637 | 0.411609805 | 0.77 |
| YDR028C | 3835.45418 | 0.30836025 | 0.413835931 | 0.77 |
| YDR111C | 552.8400061 | -0.374429707 | 0.412141989 | 0.77 |
| YDR169C-A | 13.43403719 | -0.347018149 | 0.410974448 | 0.77 |
| YDR241W | 32.77525977 | 0.06597116 | 0.412004846 | 0.77 |
| YFL036W | 2229.86904 | 0.357818202 | 0.412891532 | 0.77 |
| YGL042C | 343.2123261 | -0.322731318 | 0.412341368 | 0.77 |
| YGL157W | 1718.848717 | -0.221639028 | 0.411780102 | 0.77 |
| YGL160W | 1159.891272 | 0.189883527 | 0.411287302 | 0.77 |
| YGL218W | 11.00510591 | 0.054511077 | 0.410913178 | 0.77 |
| YGR169C | 437.5350343 | -0.282979773 | 0.413925219 | 0.77 |

| | | | | |
|---|---|---|---|---|
| YGR275W | 659.6729657 | -0.260210564 | 0.413614492 | 0.77 |
| YHL046C | 9.174623277 | 0.450988238 | 0.410750804 | 0.77 |
| YHR037W | 3383.544 | 0.272705682 | 0.410369383 | 0.77 |
| YHR155W | 1225.572051 | 0.187644477 | 0.412479561 | 0.77 |
| YIL110W | 498.0390037 | -0.247361477 | 0.411862877 | 0.77 |
| YIR039C | 1029.181658 | 0.594006101 | 0.412756328 | 0.77 |
| YJL120W | 37.94521827 | -0.254392093 | 0.413971923 | 0.77 |
| YJR127C | 3525.789326 | 0.290030701 | 0.413074659 | 0.77 |
| YKL128C | 1025.460861 | -0.279813968 | 0.41255011 | 0.77 |
| YLR437C-A | 26.88165292 | 0.404905926 | 0.411766232 | 0.77 |
| YML055W | 811.1131816 | -0.197401357 | 0.413465904 | 0.77 |
| YML121W | 941.5006143 | 0.257207079 | 0.411509291 | 0.77 |
| YMR144W | 347.0360461 | -0.17556084 | 0.411816199 | 0.77 |
| YMR194C-A | 22.19182237 | -0.414928706 | 0.412912435 | 0.77 |
| YNL008C | 1006.822611 | 0.227021354 | 0.41243902 | 0.77 |
| YNL335W | 13.42162756 | -0.60423789 | 0.410978725 | 0.77 |
| YNR046W | 947.7260883 | -0.181845217 | 0.413455088 | 0.77 |
| YPR018W | 922.6100673 | -0.20402439 | 0.413861618 | 0.77 |
| YPR119W | 682.8278436 | -0.386324479 | 0.412756994 | 0.77 |
| YPR173C | 1842.223717 | -0.266946697 | 0.410927021 | 0.77 |
| YPR182W | 302.4381877 | -0.249951846 | 0.411708126 | 0.77 |
| snR19 | 122.3490857 | 0.009037358 | 0.414134418 | 0.77 |
| snR57 | 17.27289396 | -0.114833943 | 0.414745116 | 0.77 |
| YCR063W | 60.70870613 | -0.288144547 | 0.415010979 | 0.77 |
| YFR048W | 703.4879681 | 0.189882754 | 0.41501588 | 0.77 |
| YDR199W | 75.09684199 | 0.267592121 | 0.416366545 | 0.77 |
| YGL087C | 1504.49763 | 0.42642236 | 0.415614542 | 0.77 |
| YGR075C | 302.7071594 | -0.37666898 | 0.415876115 | 0.77 |
| YIL061C | 381.0352818 | -0.225373737 | 0.415608644 | 0.77 |
| YJL047C | 709.4046862 | 0.255133369 | 0.415710257 | 0.77 |
| YKL068W-A | 172.0547206 | -0.255816207 | 0.416292924 | 0.77 |
| YLR363C | 329.7816062 | 0.135859148 | 0.415818903 | 0.77 |
| YMR221C | 1062.231053 | -0.285228677 | 0.416588013 | 0.77 |
| YOL009C | 380.9976744 | 0.201688212 | 0.416681872 | 0.77 |
| YOR058C | 1144.395841 | 0.235095665 | 0.416018346 | 0.77 |
| YOR073W | 590.4176603 | -0.267521925 | 0.416371432 | 0.77 |
| YOR311C | 1031.860298 | -0.218454091 | 0.416573379 | 0.77 |
| YOR373W | 745.8071516 | -0.227635622 | 0.416137005 | 0.77 |
| YPL096W | 660.7866918 | 0.187303939 | 0.416314628 | 0.77 |
| YEL003W | 487.0478849 | -0.223027228 | 0.417357932 | 0.77 |
| YGR197C | 588.1649926 | 0.382714363 | 0.417304645 | 0.77 |
| YIR020C-B | 5.673146071 | -0.012719918 | 0.417511051 | 0.77 |
| YOL103W | 1490.877623 | -0.198274204 | 0.417417271 | 0.77 |

| | | | | |
|---|---|---|---|---|
| YPL157W | 499.4077603 | -0.241698663 | 0.417412813 | 0.77 |
| YMR315W-A | 6.002010905 | 0.28497734 | 0.417925428 | 0.77 |
| YOL037C | 71.18780393 | 0.492548798 | 0.418006296 | 0.77 |
| YBL015W | 5485.33881 | 0.943575866 | 0.419564192 | 0.77 |
| YJL127C-B | 263.7714489 | -0.37799755 | 0.419441598 | 0.77 |
| YKR039W | 765.789828 | 0.043610634 | 0.419544608 | 0.77 |
| YLR100W | 1453.024103 | -0.166812413 | 0.41913731 | 0.77 |
| YLR316C | 329.0837142 | 0.212745869 | 0.419586264 | 0.77 |
| YLR412C-A | 1.32995427 | -0.383501511 | 0.419328164 | 0.77 |
| YDR240C | 401.4217535 | -0.210444395 | 0.419773006 | 0.77 |
| YMR170C | 596.7981642 | 0.299074132 | 0.419875053 | 0.77 |
| YBL100W-C | 6.925265405 | 0.120691156 | 0.420515157 | 0.77 |
| YFL008W | 1609.672053 | -0.239725914 | 0.420713095 | 0.77 |
| YLR096W | 1948.609356 | 0.323093335 | 0.421004731 | 0.77 |
| YNL252C | 974.4576049 | -0.213221639 | 0.420912599 | 0.77 |
| YOL034W | 735.3581814 | 0.269166789 | 0.421215135 | 0.77 |
| YAR071W | 43.79477313 | -0.613594608 | 0.422596099 | 0.77 |
| YBR192W | 512.8528961 | -0.2391342 | 0.423011872 | 0.77 |
| YDL127W | 422.9534044 | -0.338723756 | 0.421863469 | 0.77 |
| YDL188C | 1068.696328 | -0.300377239 | 0.422932724 | 0.77 |
| YDR276C | 2763.13579 | -0.303264982 | 0.422298428 | 0.77 |
| YEL057C | 119.2283707 | -0.242354421 | 0.422669876 | 0.77 |
| YHR036W | 340.3963184 | -0.364411284 | 0.421998621 | 0.77 |
| YJR027W | 207.2279874 | -0.262679464 | 0.423382419 | 0.77 |
| YKL040C | 858.7842501 | -0.261010625 | 0.423332432 | 0.77 |
| YMR075W | 761.6920474 | 0.272691003 | 0.422169833 | 0.77 |
| YMR320W | 113.0593369 | -0.02730228 | 0.422978827 | 0.77 |
| YNL315C | 1636.297254 | -0.251243651 | 0.423472332 | 0.77 |
| YOR017W | 870.202076 | 0.201020902 | 0.42334978 | 0.77 |
| YOR033C | 730.7503196 | 0.227216903 | 0.422135043 | 0.77 |
| YOR049C | 767.2844506 | 0.188600442 | 0.423437983 | 0.77 |
| YPL201C | 234.758662 | 0.496157901 | 0.4225915 | 0.77 |
| YNR001C | 10779.43573 | 0.614419319 | 0.423619018 | 0.77 |
| YDL057W | 396.4937822 | 0.321665758 | 0.424349452 | 0.77 |
| YDR153C | 1161.100144 | 0.209834885 | 0.424170079 | 0.77 |
| YIL135C | 1336.175722 | 0.290212742 | 0.42418342 | 0.77 |
| YLL034C | 960.4010793 | -0.125781607 | 0.424403274 | 0.77 |
| YLR436C | 2590.336596 | 0.336834468 | 0.42430453 | 0.77 |
| YOR344C | 1985.74596 | -0.43870607 | 0.424795943 | 0.78 |
| YBR228W | 415.0546527 | 0.178006035 | 0.425214764 | 0.78 |
| YLR166C | 1318.264285 | -0.215119698 | 0.425279954 | 0.78 |
| YDR312W | 598.3889984 | -0.182216464 | 0.425636128 | 0.78 |
| YNL013C | 5.629906296 | 0.048871171 | 0.425722858 | 0.78 |

| | | | | |
|---|---|---|---|---|
| YOL019W | 928.0908204 | -0.383301475 | 0.425879309 | 0.78 |
| YGL228W | 2378.82392 | 0.261832184 | 0.426212519 | 0.78 |
| YIL066W-A | 3.308676015 | 0.404782596 | 0.426107177 | 0.78 |
| YDR052C | 1279.720929 | -0.251130947 | 0.426585409 | 0.78 |
| YGL050W | 416.7826537 | -0.411361648 | 0.427107728 | 0.78 |
| YIL108W | 1452.736233 | 0.301907371 | 0.427196163 | 0.78 |
| YBR081C | 1882.249025 | 0.2728155 | 0.427544266 | 0.78 |
| YER185W | 79.91884135 | 0.631369707 | 0.427640779 | 0.78 |
| YIL079C | 348.2789788 | -0.284017741 | 0.427902077 | 0.78 |
| YLR351C | 1673.534289 | -0.214307444 | 0.428054434 | 0.78 |
| YNL099C | 699.7662248 | -0.201131088 | 0.428214915 | 0.78 |
| YNR027W | 549.891736 | -0.1768143 | 0.427404074 | 0.78 |
| YOR215C | 2322.968103 | 0.464463575 | 0.428188181 | 0.78 |
| snR54 | 8.052861878 | 0.344860645 | 0.427773109 | 0.78 |
| YDR156W | 897.9312179 | -0.205142585 | 0.4289316 | 0.78 |
| YFL003C | 165.623347 | 0.195311967 | 0.428893706 | 0.78 |
| YGR198W | 1689.551002 | -0.291283547 | 0.428904668 | 0.78 |
| YMR316W | 898.4662527 | 0.16057626 | 0.428701556 | 0.78 |
| YOR040W | 503.3947844 | 0.406979415 | 0.428903727 | 0.78 |
| YDR502C | 1141.466361 | 0.517910994 | 0.429200431 | 0.78 |
| YAL049C | 1497.895567 | 0.32318969 | 0.430192681 | 0.78 |
| YAR035W | 1245.571751 | 0.562264372 | 0.429899313 | 0.78 |
| YGR266W | 1152.085768 | 0.325258415 | 0.429912113 | 0.78 |
| YHR071W | 3724.97766 | 0.072420617 | 0.429953908 | 0.78 |
| YML021C | 560.3287148 | -0.226801979 | 0.430300949 | 0.78 |
| YOL132W | 124.9358227 | -0.062594366 | 0.430088227 | 0.78 |
| YOL166C | 2.808993483 | 0.871259732 | 0.429758745 | 0.78 |
| YBR021W | 496.9097626 | -0.429071184 | 0.430773967 | 0.78 |
| YGR260W | 2070.398504 | -0.474732462 | 0.430550921 | 0.78 |
| YLR057W | 1092.547525 | 0.245806609 | 0.430747284 | 0.78 |
| YLR370C | 2370.21826 | 0.28324568 | 0.43104178 | 0.78 |
| snR66 | 7.293652703 | -0.147864932 | 0.431017903 | 0.78 |
| YKR046C | 2535.087216 | 0.313911325 | 0.431268466 | 0.78 |
| YBL073W | 6.010181019 | -0.533728904 | 0.431532287 | 0.78 |
| YFL012W-A | 14.51855676 | -0.591367981 | 0.431422603 | 0.78 |
| RPR1 | 36.63381865 | 0.187380591 | 0.432528338 | 0.78 |
| YDL033C | 356.3492838 | 0.320435556 | 0.431972881 | 0.78 |
| YGR144W | 97.38216331 | 0.067058042 | 0.432735834 | 0.78 |
| YJR050W | 528.5017956 | 0.271674817 | 0.432431035 | 0.78 |
| YKR065C | 1072.834618 | -0.19450964 | 0.432259116 | 0.78 |
| YLR105C | 592.2672821 | -0.325674473 | 0.432355929 | 0.78 |
| YLR341W | 104.3766662 | -0.584315347 | 0.432320924 | 0.78 |
| YML086C | 1768.316639 | -0.212005378 | 0.432634009 | 0.78 |

| | | | | |
|---|---|---|---|---|
| YMR262W | 628.7777309 | 0.233058662 | 0.432281964 | 0.78 |
| YCL041C | 119.8839828 | 0.545064969 | 0.433274912 | 0.78 |
| YAL012W | 6420.264953 | -0.060287175 | 0.434284706 | 0.78 |
| YBR162W-A | 580.1318829 | -0.231888493 | 0.434565975 | 0.78 |
| YCR097W-A | 1.60235674 | 0.383461019 | 0.434775155 | 0.78 |
| YDR506C | 448.3958033 | 0.250421086 | 0.435831196 | 0.78 |
| YDR541C | 514.5384389 | -0.48434576 | 0.434882959 | 0.78 |
| YEL064C | 340.0873494 | 0.327027449 | 0.434441616 | 0.78 |
| YER026C | 2306.614486 | -0.243309112 | 0.436182792 | 0.78 |
| YGR270W | 2956.664275 | 0.231185952 | 0.43524222 | 0.78 |
| YHR115C | 896.8338505 | 0.242572331 | 0.435261782 | 0.78 |
| YJL216C | 203.2730397 | 0.018451191 | 0.435374805 | 0.78 |
| YKL022C | 1018.942477 | 0.258249591 | 0.43603934 | 0.78 |
| YLL029W | 3427.507563 | 0.367152158 | 0.435051448 | 0.78 |
| YLR122C | 30.05120318 | 0.454581333 | 0.433904701 | 0.78 |
| YLR200W | 774.013828 | -0.218272342 | 0.434155817 | 0.78 |
| YLR242C | 302.599813 | -0.29694819 | 0.434892842 | 0.78 |
| YLR305C | 2169.127998 | -0.231897559 | 0.435618614 | 0.78 |
| YLR389C | 2863.752537 | 0.268031461 | 0.43455305 | 0.78 |
| YML080W | 296.6855777 | -0.294330656 | 0.433752862 | 0.78 |
| YMR204C | 437.305049 | 0.232680704 | 0.435958728 | 0.78 |
| YMR226C | 4108.834639 | 0.297901537 | 0.435741748 | 0.78 |
| YNL211C | 132.5864716 | -0.377246268 | 0.435274372 | 0.78 |
| YPR035W | 16997.33407 | 0.344906447 | 0.436184264 | 0.78 |
| snR78 | 1.974544426 | 0.970196691 | 0.433943293 | 0.78 |
| YOR217W | 1207.361787 | -0.219902864 | 0.436378521 | 0.78 |
| YGR153W | 193.310946 | 0.315928309 | 0.436801112 | 0.78 |
| YDR422C | 1006.650102 | 0.236575775 | 0.437460196 | 0.78 |
| YDR294C | 2749.492975 | 0.260165117 | 0.438621544 | 0.78 |
| YDR354W | 871.1284672 | -0.062145814 | 0.438712262 | 0.78 |
| YDR479C | 1259.888261 | 0.270926218 | 0.438663401 | 0.78 |
| YDR544C | 2.336721419 | 0.417905912 | 0.438788107 | 0.78 |
| YGR021W | 818.8357092 | 0.347327978 | 0.437998888 | 0.78 |
| YGR069W | 7.64600337 | -0.632200319 | 0.438668128 | 0.78 |
| YHR167W | 161.5282471 | -0.249515346 | 0.438766461 | 0.78 |
| YLR315W | 96.87354306 | 0.240803071 | 0.438585473 | 0.78 |
| YNL243W | 3599.331124 | -0.166045462 | 0.438497267 | 0.78 |
| YOL107W | 336.081692 | -0.325160744 | 0.438543581 | 0.78 |
| YOR394C-A | 4.135778786 | -0.63370794 | 0.438804548 | 0.78 |
| YHR102W | 1800.525655 | 0.282091259 | 0.439536527 | 0.78 |
| YHR109W | 446.2718996 | -0.216759057 | 0.439548176 | 0.78 |
| YHR003C | 820.5090626 | -0.27970886 | 0.439696705 | 0.78 |
| YIL094C | 3348.951104 | -0.205566344 | 0.439950381 | 0.78 |

| | | | | |
|---|---|---|---|---|
| YER051W | 512.8476233 | 0.332721603 | 0.440292618 | 0.78 |
| YMR277W | 1413.809425 | -0.256853441 | 0.440320368 | 0.78 |
| YLL061W | 447.0050952 | 0.34042245 | 0.440494785 | 0.78 |
| YMR295C | 1636.767669 | 0.212769499 | 0.440565176 | 0.78 |
| YBL021C | 336.3286264 | -0.210515765 | 0.442165203 | 0.78 |
| YBR078W | 17879.15973 | -0.245722009 | 0.441920055 | 0.78 |
| YBR196C | 23554.20363 | -0.142756742 | 0.441755902 | 0.78 |
| YBR223C | 483.3688068 | 0.239219025 | 0.441763605 | 0.78 |
| YDR208W | 1907.44225 | 0.206308404 | 0.442418178 | 0.78 |
| YER175W-A | 27.41241786 | 0.10243375 | 0.441049433 | 0.78 |
| YGR081C | 443.1057485 | -0.197773985 | 0.442731688 | 0.78 |
| YHR058C | 391.0666567 | -0.204544644 | 0.442041926 | 0.78 |
| YHR065C | 1260.588152 | -0.198969055 | 0.440846274 | 0.78 |
| YJL146W | 600.4909491 | 0.188499357 | 0.44188678 | 0.78 |
| YKL173W | 1269.160496 | 0.187476327 | 0.44257334 | 0.78 |
| YLR201C | 1199.611196 | 0.273898614 | 0.442460304 | 0.78 |
| YLR380W | 1647.260273 | -0.24758112 | 0.442184221 | 0.78 |
| YLR401C | 1341.53932 | -0.220790083 | 0.4422036 | 0.78 |
| YOR097C | 623.6386998 | 0.207741691 | 0.442664674 | 0.78 |
| YPR053C | 166.5759994 | -0.638484984 | 0.44199671 | 0.78 |
| YPR063C | 1556.762503 | -0.24667637 | 0.441469406 | 0.78 |
| YCR089W | 1832.043276 | 0.316120136 | 0.44292997 | 0.78 |
| YIR017W-A | 3.982442675 | 0.395552122 | 0.442985192 | 0.78 |
| YMR139W | 1446.676083 | 0.269659156 | 0.443552419 | 0.78 |
| YNL063W | 522.2571621 | 0.18734183 | 0.443411233 | 0.78 |
| YNL272C | 1304.645862 | -0.222091287 | 0.443711203 | 0.78 |
| YOL159C-A | 58.15453861 | -0.518625667 | 0.443738975 | 0.78 |
| YOR113W | 1039.201076 | 0.398594957 | 0.443319089 | 0.78 |
| YHR038W | 426.523879 | -0.197685667 | 0.443965153 | 0.78 |
| YPL025C | 15.86269313 | 0.133781132 | 0.444047643 | 0.78 |
| YDL090C | 724.6090506 | -0.29442143 | 0.444531915 | 0.78 |
| YMR236W | 1041.170124 | -0.234867834 | 0.444613387 | 0.78 |
| YNL142W | 1026.084342 | 0.283213142 | 0.444433063 | 0.78 |
| YMR322C | 22.28920933 | 0.376433379 | 0.444911234 | 0.78 |
| YAL061W | 2752.646997 | 0.155074022 | 0.446225263 | 0.78 |
| YDL116W | 1341.682086 | -0.263561622 | 0.445832354 | 0.78 |
| YGL179C | 645.0803373 | 0.444355096 | 0.446205157 | 0.78 |
| YJL053W | 1232.219717 | 0.241334864 | 0.446262964 | 0.78 |
| YJL068C | 1815.839599 | 0.344221288 | 0.446325889 | 0.78 |
| YKL014C | 1967.775598 | -0.180586322 | 0.4463591 | 0.78 |
| YLR112W | 96.84022753 | -0.400590113 | 0.445516094 | 0.78 |
| YMR032W | 652.0530589 | -0.23943561 | 0.446470557 | 0.78 |
| YNL040W | 832.5546055 | 0.275867327 | 0.445541303 | 0.78 |

| | | | | |
|---|---|---|---|---|
| YNL276C | 5.892936072 | -0.172610681 | 0.445159209 | 0.78 |
| YPL033C | 16.58126869 | 0.202160589 | 0.446150738 | 0.78 |
| YPL180W | 1077.43255 | 0.186326178 | 0.446181451 | 0.78 |
| YBR216C | 879.1909045 | 0.216365059 | 0.446793611 | 0.79 |
| YHR057C | 1425.81956 | 0.260939902 | 0.447038339 | 0.79 |
| YPL021W | 14.06563719 | 0.462087901 | 0.447116208 | 0.79 |
| YKL218C | 773.1891416 | 0.284085693 | 0.447413131 | 0.79 |
| YPL168W | 515.986609 | 0.319157258 | 0.447816936 | 0.79 |
| YAL038W | 41041.91188 | 0.077383135 | 0.450144128 | 0.79 |
| YBR041W | 1541.15066 | -0.235191015 | 0.45021632 | 0.79 |
| YDR125C | 356.5508843 | 0.267298841 | 0.449082219 | 0.79 |
| YDR456W | 1671.501856 | 0.241065914 | 0.448772278 | 0.79 |
| YEL029C | 331.9088854 | -0.224039385 | 0.450029627 | 0.79 |
| YFL026W | 6326.346816 | -0.750056291 | 0.44999339 | 0.79 |
| YGR171C | 566.9877726 | -0.251767287 | 0.44979419 | 0.79 |
| YKL146W | 1758.494435 | 0.464356792 | 0.448883299 | 0.79 |
| YKL175W | 2025.203634 | 0.270611037 | 0.449849269 | 0.79 |
| YLL041C | 3315.150059 | 0.488983912 | 0.448323058 | 0.79 |
| YNL047C | 720.3103825 | 0.229592385 | 0.448939279 | 0.79 |
| YNL284C | 1102.406706 | -0.236675096 | 0.449983866 | 0.79 |
| YNR037C | 584.4173485 | -0.19282928 | 0.450131604 | 0.79 |
| YOL073C | 1132.754224 | 0.294648476 | 0.449680522 | 0.79 |
| YOR372C | 1526.129326 | 0.267940348 | 0.44910963 | 0.79 |
| YPL251W | 76.6529722 | -0.349632984 | 0.448994729 | 0.79 |
| YPL278C | 8.613341367 | -0.118338658 | 0.44864021 | 0.79 |
| YPR144C | 1036.348245 | -0.223710085 | 0.449328802 | 0.79 |
| YDR108W | 495.5770673 | 0.216755557 | 0.450511176 | 0.79 |
| YEL041W | 348.1575551 | 0.328362443 | 0.450759793 | 0.79 |
| YLR386W | 1077.114184 | 0.227332114 | 0.450905553 | 0.79 |
| YGL115W | 1145.902181 | -0.205472764 | 0.451372696 | 0.79 |
| YDL037C | 92.66763176 | 0.395616748 | 0.451894021 | 0.79 |
| YJL110C | 684.2542418 | -0.406874451 | 0.452424505 | 0.79 |
| YNL255C | 4025.75008 | -0.174024714 | 0.452165547 | 0.79 |
| YNR013C | 2464.582186 | -0.274354881 | 0.451877682 | 0.79 |
| YPL102C | 13.71806551 | 0.102605942 | 0.452393977 | 0.79 |
| YPL138C | 483.5208101 | 0.236834012 | 0.452286835 | 0.79 |
| YPL221W | 1548.403188 | 0.298924359 | 0.451880114 | 0.79 |
| snR83 | 177.6034874 | -0.292169763 | 0.451997958 | 0.79 |
| YLR157C-B | 0.750636836 | -0.187649803 | 0.452843912 | 0.79 |
| YNL241C | 6779.137631 | 0.570249896 | 0.452898604 | 0.79 |
| YJL072C | 353.6349072 | -0.17365881 | 0.453306911 | 0.79 |
| YOR129C | 585.8837465 | -0.221555064 | 0.453217698 | 0.79 |
| YOR157C | 2318.284571 | -0.178295215 | 0.453506631 | 0.79 |

| | | | | |
|---|---|---|---|---|
| YOR316C | 1558.064909 | -0.319313683 | 0.453739231 | 0.79 |
| YDR205W | 692.1640308 | 0.197057458 | 0.454234691 | 0.79 |
| YEL068C | 304.3910786 | -0.375855813 | 0.454309442 | 0.79 |
| YGR140W | 791.394714 | -0.222867961 | 0.454054916 | 0.79 |
| YOR376W | 25.04102901 | 0.267084052 | 0.454121351 | 0.79 |
| YKR100C | 1038.230895 | 0.211863398 | 0.454789708 | 0.79 |
| YOR002W | 2112.418166 | -0.256654657 | 0.454881169 | 0.79 |
| YPR199C | 796.7118888 | -0.439304883 | 0.454745995 | 0.79 |
| YCR011C | 2808.118139 | 0.267947668 | 0.455183499 | 0.79 |
| YFL063W | 4.881871603 | 0.512556428 | 0.455227022 | 0.79 |
| YNL271C | 3740.617123 | 0.34742263 | 0.455621556 | 0.79 |
| YJR093C | 790.9035242 | 0.161047719 | 0.455964536 | 0.79 |
| YMR016C | 1109.247117 | -0.312107326 | 0.456244043 | 0.79 |
| YDR036C | 1560.530814 | 0.489514693 | 0.456726241 | 0.79 |
| YER173W | 470.1702433 | 0.166738634 | 0.45673516 | 0.79 |
| YHR183W | 10217.98699 | 0.135567117 | 0.457099674 | 0.79 |
| YBL071C-B | 1.645198606 | 0.669731717 | 0.45977933 | 0.79 |
| YBR133C | 1160.137007 | -0.370709126 | 0.458307026 | 0.79 |
| YDR333C | 1359.98522 | -0.25830865 | 0.457782442 | 0.79 |
| YDR439W | 511.333794 | -0.294693864 | 0.458884811 | 0.79 |
| YEL045C | 16.86688846 | 0.69234517 | 0.459494235 | 0.79 |
| YER008C | 2582.281632 | 0.347979037 | 0.459150601 | 0.79 |
| YFL009W | 860.3502379 | 0.178124369 | 0.460162696 | 0.79 |
| YGL009C | 9382.784081 | 0.964683622 | 0.458377642 | 0.79 |
| YGL024W | 7.765530916 | -0.361773582 | 0.458917401 | 0.79 |
| YGL074C | 37.00723578 | -0.14516332 | 0.460142336 | 0.79 |
| YHR046C | 620.1635411 | -0.3756102 | 0.457555679 | 0.79 |
| YIL051C | 4289.573013 | -0.345030562 | 0.458125745 | 0.79 |
| YIL112W | 2132.63932 | 0.413454338 | 0.459008035 | 0.79 |
| YJL166W | 2486.295448 | 0.403785728 | 0.459982211 | 0.79 |
| YJR147W | 698.6444516 | -0.323286024 | 0.458928256 | 0.79 |
| YKL030W | 149.8142583 | 0.152955811 | 0.459876844 | 0.79 |
| YKR089C | 3634.661722 | 0.235759397 | 0.459299004 | 0.79 |
| YLR420W | 2792.570329 | -0.086245304 | 0.458979144 | 0.79 |
| YML007W | 2781.196668 | 0.276870283 | 0.459376543 | 0.79 |
| YML109W | 760.9435844 | 0.383160238 | 0.4580375 | 0.79 |
| YML118W | 395.2746742 | 0.280338989 | 0.458599687 | 0.79 |
| YNL244C | 2167.186944 | -0.251308453 | 0.460101688 | 0.79 |
| YOL164W | 729.2266525 | 0.200364708 | 0.458454281 | 0.79 |
| YOR131C | 796.6694001 | 0.275667142 | 0.457420762 | 0.79 |
| YOR241W | 933.790879 | -0.279861333 | 0.459697237 | 0.79 |
| YAL055W | 312.0534701 | 0.314502842 | 0.460676457 | 0.79 |
| YDR469W | 524.7044195 | -0.275781811 | 0.460613665 | 0.79 |

| | | | | |
|---|---|---|---|---|
| YFL029C | 966.6378114 | 0.35129964 | 0.460723148 | 0.79 |
| YDR149C | 14.02290454 | -0.020875853 | 0.460945948 | 0.79 |
| YJL013C | 787.2498623 | 0.197374463 | 0.461136355 | 0.79 |
| YLR084C | 1027.133398 | -0.393254455 | 0.461042408 | 0.79 |
| YBL016W | 1090.490568 | -0.662704276 | 0.461989466 | 0.79 |
| YBR104W | 350.9392427 | -0.237428385 | 0.4624691 | 0.79 |
| YGL016W | 794.4716524 | -0.346601061 | 0.461901437 | 0.79 |
| YGL084C | 1084.484417 | -0.203462128 | 0.463139543 | 0.79 |
| YGL097W | 1307.008078 | -0.103271637 | 0.462661041 | 0.79 |
| YGL127C | 488.4323072 | 0.210013234 | 0.462839934 | 0.79 |
| YGL233W | 1565.055775 | -0.20722962 | 0.461690604 | 0.79 |
| YGR216C | 759.5937639 | 0.170225854 | 0.462045579 | 0.79 |
| YJR073C | 2980.954697 | 0.487104413 | 0.462322937 | 0.79 |
| YMR002W | 4776.428936 | 0.409513145 | 0.462960764 | 0.79 |
| YMR112C | 344.1352482 | -0.221127333 | 0.462658147 | 0.79 |
| YMR269W | 489.6101925 | -0.182826487 | 0.462193165 | 0.79 |
| YNL289W | 446.6126721 | -0.252149737 | 0.462393384 | 0.79 |
| YOR231W | 881.0751627 | 0.263897848 | 0.462146752 | 0.79 |
| YOR291W | 1661.347699 | 0.207606958 | 0.462299111 | 0.79 |
| YPL249C-A | 77.54661876 | -0.180217536 | 0.463031433 | 0.79 |
| YNL104C | 8329.192681 | 0.729412409 | 0.463427573 | 0.79 |
| YPL161C | 652.0526344 | 0.188856464 | 0.46351432 | 0.79 |
| YGL131C | 1060.332301 | 0.301965018 | 0.464038967 | 0.79 |
| YKL101W | 3903.982421 | 0.062762294 | 0.46410312 | 0.79 |
| YLR248W | 3268.331309 | 0.3069915 | 0.463962951 | 0.79 |
| YLR429W | 3551.790145 | -0.245434058 | 0.464276926 | 0.79 |
| YML062C | 782.2227983 | 0.182194017 | 0.464357053 | 0.79 |
| YNL050C | 614.5421658 | -0.202650959 | 0.464220328 | 0.79 |
| YHR214W | 18.69123236 | 0.284987608 | 0.464559225 | 0.79 |
| YBL094C | 5.123149382 | 0.601957871 | 0.46511169 | 0.79 |
| YDR323C | 314.788297 | 0.131198272 | 0.465043096 | 0.79 |
| YKL142W | 4281.67462 | 0.2806784 | 0.465083506 | 0.79 |
| YDL128W | 1754.772543 | -0.260462876 | 0.465449931 | 0.79 |
| YKL011C | 416.7502341 | 0.178892511 | 0.465450202 | 0.79 |
| YPR204W | 106.9528255 | -0.244786245 | 0.465558891 | 0.79 |
| ICR1 | 346.589135 | 0.339979679 | 0.465777186 | 0.79 |
| YMR299C | 640.5800002 | -0.312004709 | 0.466028649 | 0.79 |
| YKL063C | 828.820505 | -0.239225976 | 0.466291634 | 0.79 |
| YDR135C | 3272.34562 | 0.2689821 | 0.46684773 | 0.79 |
| YKR103W | 526.4204375 | 0.474020073 | 0.466735734 | 0.79 |
| YLR282C | 4.588773458 | 0.357863635 | 0.466536826 | 0.79 |
| YML093W | 1683.648354 | -0.215985192 | 0.466776705 | 0.79 |
| YCL034W | 1930.992154 | 0.232392628 | 0.467138813 | 0.79 |

| | | | | |
|---|---|---|---|---|
| YBR022W | 363.9709831 | -0.304442478 | 0.467921426 | 0.79 |
| YEL047C | 2163.496037 | 0.434273001 | 0.467559391 | 0.79 |
| YER176W | 1918.703807 | -0.110508145 | 0.467805616 | 0.79 |
| YGL202W | 6879.756501 | -0.179527629 | 0.46784943 | 0.79 |
| YOR035C | 1268.519702 | 0.269639965 | 0.467470331 | 0.79 |
| YBL014C | 840.8039898 | 0.272577223 | 0.470568444 | 0.79 |
| YBL047C | 5698.969568 | 0.262742641 | 0.469239917 | 0.79 |
| YBR058C | 1360.304304 | -0.158642649 | 0.469878928 | 0.79 |
| YBR236C | 870.5665803 | -0.317868349 | 0.46873909 | 0.79 |
| YCR024C | 641.2724541 | 0.172895074 | 0.468410684 | 0.79 |
| YGR277C | 611.2932905 | -0.254319223 | 0.469199501 | 0.79 |
| YHL002W | 1516.40559 | 0.250601383 | 0.46963202 | 0.79 |
| YJL154C | 2489.813399 | 0.161859019 | 0.468620472 | 0.79 |
| YJL202C | 29.15207009 | -0.365197671 | 0.470476347 | 0.79 |
| YKR073C | 18.81130727 | -0.208562396 | 0.469334646 | 0.79 |
| YLR001C | 1116.57839 | 0.203795138 | 0.468516805 | 0.79 |
| YLR269C | 113.3738155 | 0.278680379 | 0.470269943 | 0.79 |
| YNL009W | 277.6331838 | 0.184249998 | 0.47003615 | 0.79 |
| YNL020C | 852.0700517 | 0.189434806 | 0.470482723 | 0.79 |
| YNL083W | 670.395816 | 0.253681009 | 0.469341485 | 0.79 |
| YNL106C | 963.8876185 | 0.248783434 | 0.470066105 | 0.79 |
| YOR105W | 21.94253227 | 0.117320973 | 0.470356312 | 0.79 |
| YOR305W | 732.3331193 | -0.305946578 | 0.470078551 | 0.79 |
| YPR108W-A | 1.090591333 | 0.03079532 | 0.469768525 | 0.79 |
| snR58 | 5.413360595 | -1.135157713 | 0.469225673 | 0.79 |
| YNL215W | 1103.384408 | 0.182415409 | 0.470787247 | 0.79 |
| YAL019W-A | 5.164115052 | 0.469258052 | 0.471817452 | 0.80 |
| YAL047W-A | 1.181246705 | -0.280744562 | 0.471789918 | 0.80 |
| YDR130C | 501.1168326 | -0.255419274 | 0.471743334 | 0.80 |
| YGR217W | 881.6554263 | -0.126662502 | 0.470992281 | 0.80 |
| YHR069C | 771.2357851 | -0.226949444 | 0.471727083 | 0.80 |
| YKL057C | 1611.669894 | -0.304456676 | 0.471328796 | 0.80 |
| YKL215C | 1029.488394 | 0.29309489 | 0.471670994 | 0.80 |
| YPL232W | 2081.428239 | -0.289814565 | 0.471434403 | 0.80 |
| YNL079C | 8141.583459 | -0.235320597 | 0.471991391 | 0.80 |
| YBR090C | 79.12993298 | -0.000237486 | 0.472709068 | 0.80 |
| YDL152W | 5.390111654 | -0.881302592 | 0.472777214 | 0.80 |
| YKL104C | 3816.518475 | -0.129868082 | 0.472459865 | 0.80 |
| YOR027W | 11708.63489 | -0.057678187 | 0.472331699 | 0.80 |
| YOR158W | 1525.323441 | -0.196552939 | 0.472563522 | 0.80 |
| YBR112C | 3373.883429 | -0.178433877 | 0.473900721 | 0.80 |
| YBR193C | 556.4922631 | 0.248663821 | 0.473847978 | 0.80 |
| YDR152W | 1737.468978 | -0.245862603 | 0.473450819 | 0.80 |

| | | | | |
|---|---|---|---|---|
| YGR043C | 1676.63942 | -0.022574739 | 0.473294633 | 0.80 |
| YOR303W | 3621.710527 | 0.472952962 | 0.47387487 | 0.80 |
| snR62 | 6.46452847 | 0.325085732 | 0.473837703 | 0.80 |
| YGR158C | 631.124363 | -0.241202928 | 0.474103622 | 0.80 |
| YBL059W | 273.8243146 | -0.171079871 | 0.476422619 | 0.80 |
| YDL043C | 369.3400281 | -0.284981559 | 0.477052461 | 0.80 |
| YDR116C | 1042.960963 | -0.224276676 | 0.47617451 | 0.80 |
| YDR384C | 561.1890906 | -0.323270386 | 0.475603271 | 0.80 |
| YER113C | 743.4950149 | -0.272684068 | 0.474377349 | 0.80 |
| YFL031W | 3441.588262 | -0.286539609 | 0.476593354 | 0.80 |
| YGL258W | 52.00397703 | 0.150771894 | 0.477120887 | 0.80 |
| YGR178C | 3642.360299 | -0.228531791 | 0.474939625 | 0.80 |
| YHL045W | 26.03413018 | 0.588994962 | 0.476311967 | 0.80 |
| YHR001W | 1515.116914 | 0.217091365 | 0.474943591 | 0.80 |
| YHR120W | 606.7255558 | 0.143585996 | 0.476725826 | 0.80 |
| YHR126C | 18.42016663 | 0.093192548 | 0.474908092 | 0.80 |
| YJL112W | 1088.839683 | 0.293335626 | 0.47473942 | 0.80 |
| YJR106W | 683.5325213 | 0.335352755 | 0.47518534 | 0.80 |
| YLR383W | 888.8933966 | -0.091801549 | 0.476153295 | 0.80 |
| YLR406C-A | 2.560257022 | 0.428522115 | 0.47632372 | 0.80 |
| YLR424W | 404.0091344 | 0.165676298 | 0.476414824 | 0.80 |
| YNL138W | 4814.982276 | -0.183403916 | 0.476183003 | 0.80 |
| YNL247W | 4374.768503 | -0.054513364 | 0.47494322 | 0.80 |
| YOL095C | 382.7305205 | 0.175260227 | 0.475087641 | 0.80 |
| YOR156C | 466.9299753 | 0.20876163 | 0.477009213 | 0.80 |
| YOR216C | 1165.7555 | -0.211069422 | 0.477085284 | 0.80 |
| YPL183W-A | 354.0512636 | -0.261742906 | 0.475988519 | 0.80 |
| YPR048W | 424.7867171 | -0.20748826 | 0.476466223 | 0.80 |
| YDR123C | 353.8570592 | 0.319980437 | 0.477477098 | 0.80 |
| YDR141C | 2153.367361 | -0.170366816 | 0.477581445 | 0.80 |
| YMR006C | 894.5004784 | -0.228045174 | 0.477470785 | 0.80 |
| YLR006C | 1262.134466 | 0.380418004 | 0.477843287 | 0.80 |
| YML061C | 1035.299962 | 0.291219225 | 0.477738067 | 0.80 |
| YBR183W | 786.7022482 | 0.182356285 | 0.478246779 | 0.80 |
| YKR020W | 476.7599628 | 0.239491784 | 0.478209908 | 0.80 |
| YLR296W | 4.726955448 | 0.649230452 | 0.478321574 | 0.80 |
| YBR019C | 339.0837633 | 0.119297555 | 0.478957402 | 0.80 |
| YDR019C | 2663.732875 | -0.25483135 | 0.479009906 | 0.80 |
| YPL031C | 1442.118487 | -0.219973439 | 0.478771565 | 0.80 |
| YPR109W | 650.4688515 | 0.173305339 | 0.478740159 | 0.80 |
| YHR050W-A | 204.2549426 | -0.348660236 | 0.479724813 | 0.80 |
| YLR238W | 614.7184945 | 0.170654361 | 0.479900884 | 0.80 |
| YOL029C | 366.4284423 | -0.253737326 | 0.480111813 | 0.80 |

| | | | | |
|---|---|---|---|---|
| snR8 | 32.91757815 | -0.280881979 | 0.480090396 | 0.80 |
| YAR068W | 4.842260677 | 0.192829943 | 0.480533526 | 0.80 |
| YDR169C | 1729.231302 | 0.297628671 | 0.48090731 | 0.80 |
| YPL270W | 1768.066094 | 0.308319127 | 0.480908208 | 0.80 |
| YFL040W | 216.9489399 | 0.113410746 | 0.481775321 | 0.80 |
| YHR056C | 832.3445362 | 0.200715699 | 0.481701432 | 0.80 |
| snR76 | 8.640963018 | -0.284076959 | 0.481532688 | 0.80 |
| YDR220C | 3.958958383 | 0.262277207 | 0.482093945 | 0.80 |
| snR10 | 11.52677515 | -0.076498257 | 0.482538918 | 0.80 |
| YBR255C-A | 311.2384417 | 0.205149352 | 0.483504917 | 0.80 |
| YKL075C | 477.0577168 | 0.214938319 | 0.483295971 | 0.80 |
| YNL225C | 446.0966712 | 0.208994209 | 0.483451405 | 0.80 |
| YGR065C | 698.7091847 | 0.321099205 | 0.484019349 | 0.80 |
| YLR270W | 2664.450215 | 0.29199427 | 0.483792922 | 0.80 |
| YNR051C | 2926.448335 | -0.072577889 | 0.484022917 | 0.80 |
| YPR069C | 2243.3628 | -0.221430655 | 0.483781909 | 0.80 |
| YDR215C | 67.50004526 | 0.524811412 | 0.484673313 | 0.80 |
| YGL111W | 1214.214479 | -0.147471235 | 0.484544746 | 0.80 |
| YGR282C | 11912.38121 | -0.160306084 | 0.484524552 | 0.80 |
| YHR090C | 634.1320105 | 0.236725482 | 0.48505625 | 0.80 |
| YJL058C | 326.9260026 | 0.29241062 | 0.484826806 | 0.80 |
| YKL003C | 652.5154668 | -0.276961742 | 0.484984729 | 0.80 |
| YKL041W | 833.3256523 | 0.201763975 | 0.485351366 | 0.80 |
| YKR004C | 281.3110563 | 0.204984473 | 0.485282371 | 0.80 |
| YNL003C | 715.0467326 | -0.355769822 | 0.485361756 | 0.80 |
| YPR148C | 2535.151148 | -0.185338712 | 0.485509461 | 0.80 |
| YDR118W | 391.9679351 | -0.180653016 | 0.485698776 | 0.80 |
| YDL162C | 2.875843937 | -0.569494768 | 0.485971692 | 0.80 |
| YMR275C | 1521.663704 | 0.331672074 | 0.486273095 | 0.80 |
| YHR016C | 1896.688185 | 0.325494433 | 0.486536195 | 0.80 |
| YNL145W | 1813.203416 | -0.609865251 | 0.486572094 | 0.80 |
| YLR265C | 441.14354 | -0.503930905 | 0.486737206 | 0.80 |
| YGL098W | 531.3017666 | 0.146646618 | 0.487482805 | 0.80 |
| YJR116W | 691.1206507 | 0.299038096 | 0.487680887 | 0.80 |
| YNL105W | 8.716167534 | 0.306872921 | 0.487692622 | 0.80 |
| YNL114C | 62.61749004 | -0.308692652 | 0.487264913 | 0.80 |
| YOR287C | 297.2565187 | -0.062715331 | 0.487362785 | 0.80 |
| YNR023W | 727.2388585 | -0.225368568 | 0.487963894 | 0.80 |
| YJR157W | 1.945110995 | -0.806109615 | 0.488312274 | 0.80 |
| YKL144C | 415.786387 | -0.185040122 | 0.488299932 | 0.80 |
| YMR199W | 1528.60034 | -0.1494707 | 0.48851713 | 0.80 |
| YGL007W | 19.49434054 | 0.052753995 | 0.489177233 | 0.80 |
| YMR305C | 6521.89708 | -0.117228906 | 0.489197106 | 0.80 |

| | | | | |
|---|---|---|---|---|
| YOR104W | 252.8227861 | 0.244973225 | 0.488893635 | 0.80 |
| YPL259C | 962.6624104 | 0.153444741 | 0.489047248 | 0.80 |
| YKR095W-A | 172.6357846 | -0.308933334 | 0.48937461 | 0.80 |
| YDR288W | 555.4180316 | 0.119006558 | 0.490580569 | 0.80 |
| YFR022W | 849.2747048 | 0.10305473 | 0.489852081 | 0.80 |
| YGL110C | 882.4868812 | 0.199622097 | 0.490442389 | 0.80 |
| YHR103W | 1865.030837 | 0.250977988 | 0.490682756 | 0.80 |
| YJL156W-A | 83.97878785 | 0.161325694 | 0.490066825 | 0.80 |
| YLR433C | 1164.793625 | -0.343705242 | 0.489761645 | 0.80 |
| YMR080C | 1907.455726 | -0.219871725 | 0.490523223 | 0.80 |
| YNL049C | 1288.390098 | -0.085557086 | 0.490219423 | 0.80 |
| YPL022W | 1488.077225 | 0.214885076 | 0.490214641 | 0.80 |
| YDL148C | 2147.255757 | -0.150795067 | 0.491566805 | 0.80 |
| YDR127W | 3620.75176 | -0.069447326 | 0.491277951 | 0.80 |
| YGR026W | 4521.168167 | -0.244119985 | 0.491010022 | 0.80 |
| YKL051W | 1194.148337 | 0.106811526 | 0.491327342 | 0.80 |
| YLR086W | 1588.915329 | 0.329881752 | 0.491290998 | 0.80 |
| YOR358W | 499.4768456 | 0.220107972 | 0.491479032 | 0.80 |
| YGR206W | 264.0711736 | -0.330542837 | 0.491741153 | 0.80 |
| YHR059W | 401.0845698 | -0.191662983 | 0.491978259 | 0.81 |
| YBR140C | 4380.796749 | -0.104724769 | 0.492243128 | 0.81 |
| YOL085W-A | 20.71787704 | 0.158798581 | 0.492325422 | 0.81 |
| YDR509W | 13.68760112 | -0.069784746 | 0.493313341 | 0.81 |
| YGL182C | 10.33516089 | 0.712700167 | 0.493126995 | 0.81 |
| YGR077C | 1152.926191 | 0.158422167 | 0.492766417 | 0.81 |
| YGR211W | 3320.437529 | -0.064032721 | 0.493334636 | 0.81 |
| YNL037C | 5393.729596 | -0.124286287 | 0.493250464 | 0.81 |
| YPR067W | 664.4484323 | -0.185387378 | 0.493053227 | 0.81 |
| YPR081C | 985.9306388 | 0.259731864 | 0.493057047 | 0.81 |
| YHL006W-A | 55.86611926 | -0.265897247 | 0.493751946 | 0.81 |
| YMR100W | 811.7962257 | 0.188517708 | 0.494468491 | 0.81 |
| YBR240C | 118.1647641 | 0.086861607 | 0.495102872 | 0.81 |
| YBR296C | 209.9913106 | 0.271815992 | 0.496674039 | 0.81 |
| YDR255C | 870.1597032 | 0.232677585 | 0.496424654 | 0.81 |
| YDR304C | 4614.533404 | -0.175266806 | 0.495073119 | 0.81 |
| YDR377W | 1422.817631 | -0.138699799 | 0.497664097 | 0.81 |
| YFL019C | 43.77288423 | -0.077327304 | 0.495283844 | 0.81 |
| YGL048C | 4170.423196 | -0.150400575 | 0.495895717 | 0.81 |
| YGL252C | 1693.94272 | -0.163064177 | 0.496882268 | 0.81 |
| YGL259W | 86.59077684 | 0.432371007 | 0.497021954 | 0.81 |
| YGL261C | 11.97177178 | 0.700719193 | 0.497253249 | 0.81 |
| YIL068W-A | 5.981971104 | 0.289155998 | 0.495747231 | 0.81 |
| YJR042W | 1389.72183 | -0.273088042 | 0.496102515 | 0.81 |

| | | | | |
|---|---|---|---|---|
| YJR151C | 560.5480031 | 0.274593108 | 0.497346835 | 0.81 |
| YLR371W | 1780.890302 | 0.272237613 | 0.497243977 | 0.81 |
| YML066C | 127.9180195 | -0.634930713 | 0.497526496 | 0.81 |
| YMR276W | 5216.693498 | 0.268882001 | 0.496828734 | 0.81 |
| YMR283C | 413.1862547 | 0.167966745 | 0.496622496 | 0.81 |
| YNL107W | 513.6532475 | 0.214404386 | 0.494824447 | 0.81 |
| YNL149C | 291.2466735 | -0.103206169 | 0.495149262 | 0.81 |
| YNR019W | 1395.465767 | 0.37618559 | 0.495324655 | 0.81 |
| YOL036W | 1888.124512 | 0.322878612 | 0.49635248 | 0.81 |
| YOL065C | 517.0688786 | 0.210677523 | 0.497068672 | 0.81 |
| YOR025W | 737.1839475 | -0.238674611 | 0.495995175 | 0.81 |
| YOR249C | 647.5824875 | 0.110607845 | 0.497561511 | 0.81 |
| YPR025C | 1080.062057 | 0.230323929 | 0.497460579 | 0.81 |
| YNL068C | 1421.244951 | 0.318959934 | 0.498001845 | 0.81 |
| snR70 | 64.50681846 | 0.199371852 | 0.497920633 | 0.81 |
| YGL244W | 2087.284965 | -0.23366337 | 0.498219071 | 0.81 |
| YJR087W | 189.3064917 | -0.298246417 | 0.49837743 | 0.81 |
| YEL014C | 21.68196635 | -0.143632862 | 0.498722158 | 0.81 |
| YKL103C | 3936.941911 | 0.557799146 | 0.499794347 | 0.81 |
| YKR061W | 573.6088479 | 0.067328812 | 0.499716008 | 0.81 |
| YLR261C | 10.17984332 | -0.083200202 | 0.499756806 | 0.81 |
| YMR106C | 546.3577506 | 0.247483581 | 0.499511039 | 0.81 |
| YKL052C | 501.3271381 | -0.186845933 | 0.500106166 | 0.81 |
| YKR008W | 1399.761894 | -0.243278982 | 0.500182473 | 0.81 |
| YBL023C | 762.982123 | -0.164487465 | 0.501090288 | 0.81 |
| YDR389W | 894.8507584 | -0.317047388 | 0.501169638 | 0.81 |
| YFR032C-B | 19.14700684 | -0.119160475 | 0.500938983 | 0.81 |
| YKL027W | 701.3136707 | 0.342521014 | 0.501098691 | 0.81 |
| YLR402W | 27.81687827 | 0.187439243 | 0.501163048 | 0.81 |
| YOR014W | 3387.711726 | 0.259679687 | 0.500507813 | 0.81 |
| YOR046C | 2190.495993 | -0.127916393 | 0.501036354 | 0.81 |
| YPL173W | 902.2046084 | -0.222447694 | 0.501722971 | 0.81 |
| YDL089W | 963.5832446 | 0.211358447 | 0.502145084 | 0.81 |
| YDR369C | 596.1613966 | 0.304236688 | 0.503033861 | 0.81 |
| YER133W-A | 95.44590047 | -0.45232246 | 0.502580167 | 0.81 |
| YJL060W | 1402.078784 | -0.084218288 | 0.502925522 | 0.81 |
| YOR375C | 17984.54886 | 0.747326442 | 0.502878257 | 0.81 |
| YPL062W | 4.463478236 | 0.148942591 | 0.503068123 | 0.81 |
| YAL029C | 2871.619875 | 0.20103843 | 0.503433289 | 0.81 |
| YOR365C | 309.6078398 | -0.006362421 | 0.503627545 | 0.81 |
| YOR391C | 10.43191586 | -0.443271154 | 0.503544029 | 0.81 |
| YPR193C | 57.30247136 | 0.302366188 | 0.503976784 | 0.81 |
| YJR121W | 23130.92602 | 0.091002311 | 0.504117603 | 0.81 |

| | | | | |
|---|---|---|---|---|
| YMR055C | 275.6053073 | -0.23144567 | 0.50450816 | 0.81 |
| YDR029W | 118.6548652 | 0.394568262 | 0.5048232 | 0.81 |
| YER172C-A | 0.861078059 | -0.275768411 | 0.504898798 | 0.81 |
| YMR113W | 461.667001 | 0.153982246 | 0.505032458 | 0.81 |
| YDR049W | 886.770398 | 0.155996882 | 0.505359865 | 0.81 |
| YGR251W | 285.9824856 | -0.251454474 | 0.505873175 | 0.81 |
| YDR476C | 1082.582617 | 0.218913662 | 0.506285587 | 0.81 |
| YFR045W | 698.908615 | 0.277452869 | 0.506249591 | 0.81 |
| YDR126W | 326.9726421 | 0.143446768 | 0.50672536 | 0.81 |
| YBR082C | 6782.616874 | -0.158445533 | 0.506935817 | 0.81 |
| YOR116C | 1548.881592 | -0.234007556 | 0.507149415 | 0.81 |
| YOR268C | 11.87888145 | 0.761685262 | 0.507447114 | 0.81 |
| YOR029W | 6.24277143 | -0.086428732 | 0.507629989 | 0.81 |
| YJR040W | 752.4605261 | -0.235095269 | 0.507835437 | 0.81 |
| YMR020W | 992.3354434 | 0.218302166 | 0.50802823 | 0.81 |
| YCR047C | 558.0855162 | -0.080699958 | 0.508191098 | 0.81 |
| YHR214W-A | 22.9859353 | 0.192000592 | 0.508315816 | 0.81 |
| YJL019W | 605.7582629 | -0.241183651 | 0.508407076 | 0.81 |
| YDL246C | 6.191078768 | 0.391182389 | 0.508701804 | 0.81 |
| YJL207C | 2307.188321 | -0.210173931 | 0.508749033 | 0.81 |
| YLL037W | 13.52665894 | 0.115265314 | 0.508928492 | 0.81 |
| YCR004C | 3730.565062 | -0.265649036 | 0.509205002 | 0.82 |
| YGR116W | 3744.798928 | -0.156516793 | 0.509360366 | 0.82 |
| YML005W | 433.1547412 | -0.162606106 | 0.509463426 | 0.82 |
| YDL100C | 5446.231089 | -0.001330339 | 0.510133482 | 0.82 |
| YDR444W | 770.8185864 | 0.238070451 | 0.510073883 | 0.82 |
| YBL036C | 1223.509384 | -0.204586362 | 0.510654246 | 0.82 |
| YFL061W | 11.58614878 | -0.314146901 | 0.510703828 | 0.82 |
| YPR157W | 448.7836668 | 0.194929474 | 0.510843593 | 0.82 |
| YAL033W | 350.8440347 | -0.2045605 | 0.514383488 | 0.82 |
| YAL048C | 583.0194511 | 0.22652167 | 0.511374241 | 0.82 |
| YAL064C-A | 47.35030167 | -0.524612535 | 0.515010777 | 0.82 |
| YBR003W | 1468.050788 | 0.183795382 | 0.512798581 | 0.82 |
| YBR161W | 517.5371153 | 0.142694821 | 0.513128489 | 0.82 |
| YBR182C-A | 7.795727576 | 0.215783831 | 0.514829405 | 0.82 |
| YBR292C | 15.20409235 | 0.32383307 | 0.513341058 | 0.82 |
| YBR296C-A | 3.00001637 | 0.133396875 | 0.513001791 | 0.82 |
| YDL011C | 7.393723597 | -0.670153613 | 0.512815344 | 0.82 |
| YDL176W | 672.0622412 | 0.194608432 | 0.511102438 | 0.82 |
| YHL014C | 453.0111576 | -0.248160786 | 0.512366327 | 0.82 |
| YHR129C | 426.0946941 | -0.288635905 | 0.514987784 | 0.82 |
| YJL066C | 1041.928086 | 0.384170761 | 0.51318652 | 0.82 |
| YJR048W | 1046.168345 | 0.11014156 | 0.514239266 | 0.82 |

| | | | | |
|---|---|---|---|---|
| YJR107W | 474.7713781 | 0.202700378 | 0.513689694 | 0.82 |
| YJR108W | 40.18860279 | -0.032162512 | 0.511862244 | 0.82 |
| YKR018C | 3562.456922 | 0.415691405 | 0.514815479 | 0.82 |
| YLR072W | 910.1580571 | 0.212720523 | 0.515003846 | 0.82 |
| YML057W | 1931.217961 | 0.20273363 | 0.514107315 | 0.82 |
| YML115C | 1520.660585 | -0.239409166 | 0.514172244 | 0.82 |
| YMR071C | 1073.602658 | -0.122455328 | 0.514021933 | 0.82 |
| YMR141W-A | 54.52612508 | 0.255877121 | 0.514851019 | 0.82 |
| YMR193C-A | 123.8016037 | 0.18076854 | 0.513002906 | 0.82 |
| YNL086W | 191.101467 | 0.197084309 | 0.513964111 | 0.82 |
| YNL201C | 1076.736749 | -0.206424913 | 0.512847598 | 0.82 |
| YNL217W | 1768.453008 | -0.176252234 | 0.514296477 | 0.82 |
| YNL229C | 1349.306513 | -0.153724903 | 0.514170931 | 0.82 |
| YNL294C | 1109.535055 | 0.230328519 | 0.514614735 | 0.82 |
| YNR060W | 291.930447 | 0.119797958 | 0.512320172 | 0.82 |
| YOR381W | 508.0002714 | 0.12855994 | 0.514162571 | 0.82 |
| YPL177C | 1454.555337 | 0.05436808 | 0.513784221 | 0.82 |
| snR42 | 83.98583809 | 0.267533618 | 0.513854912 | 0.82 |
| snR64 | 4.034418541 | 0.196658093 | 0.514278342 | 0.82 |
| YGR090W | 2037.446504 | -0.330448629 | 0.515145452 | 0.82 |
| YFL034C-A | 576.1615287 | -0.440053999 | 0.515898416 | 0.82 |
| YKL167C | 608.6144606 | -0.228115218 | 0.515796356 | 0.82 |
| YHR105W | 228.2746463 | 0.299267411 | 0.516223018 | 0.82 |
| YJL086C | 9.129343542 | -0.172229928 | 0.516123895 | 0.82 |
| YNR074C | 411.2345917 | -0.327945962 | 0.516324482 | 0.82 |
| YOR304C-A | 201.0708852 | -0.317887914 | 0.516606144 | 0.82 |
| YOR385W | 901.596547 | 0.252629149 | 0.516666269 | 0.82 |
| YBR198C | 1683.847961 | -0.20244327 | 0.517463817 | 0.82 |
| YDL049C | 168.7509482 | 0.071803016 | 0.517545651 | 0.82 |
| YOR381W-A | 9.096748957 | -0.035606153 | 0.517324431 | 0.82 |
| YPL042C | 568.6329785 | 0.171583021 | 0.517283644 | 0.82 |
| YPR178W | 455.7725824 | 0.165515342 | 0.517686288 | 0.82 |
| YDR058C | 448.3672811 | 0.230701854 | 0.51793597 | 0.82 |
| YAL035W | 9252.51472 | -0.085505037 | 0.519164673 | 0.82 |
| YGL043W | 1337.950161 | -0.209699268 | 0.519060632 | 0.82 |
| YGL208W | 855.2756461 | 0.368301181 | 0.519165833 | 0.82 |
| YJL151C | 3281.2426 | 0.287466304 | 0.518608354 | 0.82 |
| YJR001W | 1763.85175 | -0.166181476 | 0.518862843 | 0.82 |
| YKL169C | 32.06083426 | -0.409742955 | 0.519219138 | 0.82 |
| YMR132C | 213.3288234 | -0.151621015 | 0.518712151 | 0.82 |
| YOR319W | 365.6759573 | -0.20061959 | 0.519213434 | 0.82 |
| YPR076W | 41.22373968 | 0.427431335 | 0.51839132 | 0.82 |
| YBR180W | 89.72716224 | -0.304048464 | 0.520220413 | 0.82 |

| | | | | |
|---|---|---|---|---|
| YIR018W | 398.3013208 | 0.227885188 | 0.520315771 | 0.82 |
| YLR326W | 402.567239 | 0.446353708 | 0.519905564 | 0.82 |
| YOR296W | 873.0215241 | -0.240015591 | 0.520180539 | 0.82 |
| YPL080C | 11.15479793 | -0.618478433 | 0.520259742 | 0.82 |
| YBR077C | 472.5529414 | -0.313269252 | 0.520456641 | 0.82 |
| YJL031C | 712.299831 | 0.208244199 | 0.520949634 | 0.82 |
| YNL166C | 1009.247872 | -0.117359484 | 0.521003312 | 0.82 |
| YOL018C | 882.7418897 | 0.138728854 | 0.521657134 | 0.82 |
| YFL033C | 2297.741868 | 0.314319323 | 0.521930284 | 0.82 |
| YNL007C | 10095.98848 | -0.146560101 | 0.522236454 | 0.82 |
| YPL098C | 678.1300721 | -0.167150847 | 0.522246083 | 0.82 |
| YPR136C | 12.04995248 | 0.611089625 | 0.522282394 | 0.82 |
| YDL080C | 732.479931 | 0.137094363 | 0.523489718 | 0.82 |
| YDR140W | 525.20568 | -0.163988574 | 0.523186424 | 0.82 |
| YGL047W | 625.4320388 | -0.070431001 | 0.523148198 | 0.82 |
| YML084W | 2.063062303 | 0.131289321 | 0.523563793 | 0.82 |
| YOL118C | 79.69627109 | 0.319060744 | 0.523437108 | 0.82 |
| YOR192C-C | 17.72254902 | 0.030896129 | 0.523262031 | 0.82 |
| YDL022C-A | 6.814086928 | 0.202090924 | 0.523748567 | 0.82 |
| YGL068W | 385.5514446 | -0.156227518 | 0.52382331 | 0.82 |
| YBR098W | 547.1579177 | 0.248345815 | 0.524274048 | 0.82 |
| YJR086W | 331.1910032 | -0.426180266 | 0.524439752 | 0.82 |
| YOL157C | 78.40010831 | 0.208719776 | 0.524458431 | 0.82 |
| YPL001W | 324.2471356 | -0.063696802 | 0.524111066 | 0.82 |
| YJL156C | 869.0736001 | -0.254301936 | 0.52459724 | 0.82 |
| YGR002C | 912.0190954 | 0.168815961 | 0.525159874 | 0.82 |
| YPL047W | 255.7983457 | 0.105545936 | 0.525293829 | 0.82 |
| YCR024C-A | 1033.453894 | -0.304684315 | 0.52543659 | 0.82 |
| YGL253W | 16247.71261 | 0.083230403 | 0.525618019 | 0.82 |
| YER125W | 2425.853452 | -0.189470307 | 0.525993237 | 0.82 |
| YGR098C | 1335.545417 | -0.154113031 | 0.526393802 | 0.82 |
| YIR012W | 1025.35668 | -0.035644302 | 0.526336598 | 0.82 |
| YKR012C | 142.5434976 | 0.004580478 | 0.526455651 | 0.82 |
| YBR107C | 343.2589721 | 0.224445874 | 0.526852935 | 0.82 |
| YDR359C | 1438.122379 | -0.198895725 | 0.526987573 | 0.82 |
| YER088C | 4317.842613 | 0.507928201 | 0.526935193 | 0.82 |
| YOR139C | 5.37072037 | 0.586863846 | 0.526719424 | 0.82 |
| YNR030W | 1357.008714 | -0.167931943 | 0.527339131 | 0.82 |
| YKL064W | 2390.234747 | 0.268340017 | 0.527742207 | 0.82 |
| YMR289W | 967.6114857 | -0.190232146 | 0.527936513 | 0.82 |
| YAL056W | 1700.26388 | 0.209514877 | 0.529798597 | 0.82 |
| YBL029W | 662.4144794 | 0.213503878 | 0.52880111 | 0.82 |
| YCR041W | 16.4079087 | -0.029718682 | 0.529259709 | 0.82 |

| | | | | |
|---|---|---|---|---|
| YDL065C | 1387.029558 | 0.357565319 | 0.530293904 | 0.82 |
| YDL119C | 290.4231748 | 0.201615899 | 0.528454809 | 0.82 |
| YDR349C | 1807.872356 | -0.162563539 | 0.529470066 | 0.82 |
| YFR008W | 307.6442666 | 0.168504121 | 0.530217727 | 0.82 |
| YGL186C | 1156.394032 | -0.20673796 | 0.528630431 | 0.82 |
| YJR099W | 741.8529597 | -0.183644162 | 0.529728455 | 0.82 |
| YKL210W | 8749.432253 | 0.2600537 | 0.529770735 | 0.82 |
| YLR039C | 1333.277955 | 0.191885729 | 0.530185261 | 0.82 |
| YLR133W | 1071.862832 | -0.108645819 | 0.529596648 | 0.82 |
| YLR210W | 386.6683639 | -0.24718407 | 0.528205391 | 0.82 |
| YML038C | 1118.066731 | 0.174216799 | 0.529075954 | 0.82 |
| YMR238W | 935.1001777 | -0.153926553 | 0.529243367 | 0.82 |
| YMR304C-A | 599.6439516 | -0.197860537 | 0.530228127 | 0.82 |
| YOL115W | 646.4767705 | -0.315782602 | 0.529475931 | 0.82 |
| YOR171C | 1509.079417 | 0.249260705 | 0.529417167 | 0.82 |
| YGL194C-A | 142.847988 | -0.626159847 | 0.530491048 | 0.82 |
| YBL100W-B | 175.5232464 | 0.483581153 | 0.530792783 | 0.82 |
| YHR148W | 337.7936716 | -0.063719229 | 0.530839826 | 0.82 |
| YIL014C-A | 253.291496 | -0.38056822 | 0.531119243 | 0.82 |
| YLR407W | 355.7060391 | 0.138333375 | 0.53131047 | 0.82 |
| YDR526C | 4.767823074 | 0.664786853 | 0.531714506 | 0.83 |
| YHR151C | 595.1830514 | 0.288672965 | 0.531730786 | 0.83 |
| YDR151C | 1113.580859 | 0.193145012 | 0.532165421 | 0.83 |
| YIL047C-A | 10.41380776 | -0.203983107 | 0.532357444 | 0.83 |
| YLR227W-B | 558.9698813 | -0.100123324 | 0.532370696 | 0.83 |
| YDR078C | 147.0952169 | 0.219920375 | 0.532502828 | 0.83 |
| YBL083C | 7.154873991 | 0.646013588 | 0.533156507 | 0.83 |
| YBR023C | 1741.008098 | -0.222943135 | 0.533400445 | 0.83 |
| YBR101C | 3697.675555 | -0.130282428 | 0.533577343 | 0.83 |
| YBR202W | 984.15721 | -0.27217477 | 0.533809271 | 0.83 |
| YDR050C | 39610.42901 | -0.262306388 | 0.533863806 | 0.83 |
| YGL221C | 1756.035958 | -0.149415849 | 0.532873834 | 0.83 |
| YHR047C | 1885.311487 | -0.112884933 | 0.533448926 | 0.83 |
| YIL012W | 10.18010829 | -0.040569361 | 0.533754508 | 0.83 |
| YMR135C | 1809.513775 | 0.280151603 | 0.533806501 | 0.83 |
| YOR011W-A | 3.164669454 | 0.229159408 | 0.533403453 | 0.83 |
| YDL135C | 948.2549942 | -0.242143069 | 0.534236428 | 0.83 |
| YDR159W | 1698.043068 | 0.177593338 | 0.534450108 | 0.83 |
| YLR434C | 48.33837913 | -0.106875771 | 0.534245797 | 0.83 |
| YNR045W | 861.3548214 | 0.273969098 | 0.534333079 | 0.83 |
| YGL142C | 1131.213732 | 0.237146417 | 0.534630514 | 0.83 |
| YDL101C | 860.843029 | 0.401685591 | 0.535334785 | 0.83 |
| YDR092W | 1261.21405 | -0.116561116 | 0.534853626 | 0.83 |

| | | | | |
|---|---|---|---|---|
| YDR430C | 1380.899649 | 0.147314828 | 0.535397831 | 0.83 |
| YER151C | 2013.443986 | -0.144147192 | 0.535405831 | 0.83 |
| YJL171C | 1558.602902 | 0.284186308 | 0.535228534 | 0.83 |
| snR45 | 13.49639451 | -0.083351445 | 0.535187743 | 0.83 |
| YLR457C | 335.6948937 | -0.152739453 | 0.535769901 | 0.83 |
| YHR192W | 784.2571123 | -0.163441254 | 0.536258829 | 0.83 |
| YBR004C | 976.5395701 | 0.219533789 | 0.536811181 | 0.83 |
| YGL088W | 0.432313012 | 0.170136794 | 0.53661055 | 0.83 |
| YLR382C | 967.5198513 | 0.208613932 | 0.536717188 | 0.83 |
| YMR257C | 582.2395624 | 0.192102768 | 0.536776309 | 0.83 |
| YBL071W-A | 297.3219208 | 0.253611486 | 0.53710579 | 0.83 |
| YDR301W | 1341.743029 | -0.264841161 | 0.537360783 | 0.83 |
| YIL087C | 960.1717362 | 0.155561762 | 0.537464148 | 0.83 |
| YKL108W | 391.5450461 | -0.205597595 | 0.537519185 | 0.83 |
| YNR066C | 265.0332103 | 0.458095219 | 0.537210029 | 0.83 |
| YLR139C | 783.7450537 | 0.201011048 | 0.537947831 | 0.83 |
| YMR045C | 326.5494461 | 0.468058872 | 0.537928302 | 0.83 |
| YER041W | 647.7988303 | 0.123288644 | 0.538649246 | 0.83 |
| YIR002C | 1205.196083 | 0.188718284 | 0.538891008 | 0.83 |
| YKR047W | 143.8726915 | -0.150297774 | 0.538452501 | 0.83 |
| YKR099W | 800.367483 | -0.199354786 | 0.538719991 | 0.83 |
| YNR001W-A | 2.398336405 | -0.203533669 | 0.538954428 | 0.83 |
| YOL028C | 419.6545084 | -0.230780161 | 0.538517084 | 0.83 |
| YOR154W | 667.1567549 | 0.196360444 | 0.53901148 | 0.83 |
| YPL170W | 622.4061116 | 0.186924659 | 0.538575692 | 0.83 |
| YGL126W | 1063.33427 | -0.104496556 | 0.539382261 | 0.83 |
| YOR321W | 1083.064286 | -0.088683829 | 0.53948271 | 0.83 |
| YHL043W | 209.5139873 | -0.041038449 | 0.539788726 | 0.83 |
| YKL018W | 496.5039326 | -0.179373539 | 0.540443329 | 0.83 |
| YPL120W | 588.9261154 | 0.189445727 | 0.540554565 | 0.83 |
| snR13 | 24.86462889 | 0.104409109 | 0.541118263 | 0.83 |
| YDL160C-A | 95.22130843 | -0.358579348 | 0.542175077 | 0.83 |
| YDR185C | 264.7477216 | 0.234678107 | 0.542157317 | 0.83 |
| YFL028C | 788.0732688 | 0.222459529 | 0.541889971 | 0.83 |
| YGL231C | 1251.65783 | -0.167499717 | 0.542069005 | 0.83 |
| YNR025C | 48.20698063 | 0.116276944 | 0.541716253 | 0.83 |
| snR189 | 9.691874789 | 0.172537596 | 0.541814436 | 0.83 |
| YOL005C | 1554.462517 | -0.254395069 | 0.542533088 | 0.83 |
| YPL197C | 174.3224877 | 0.15409679 | 0.543000745 | 0.83 |
| YER188W | 461.4895098 | -0.43241515 | 0.543351808 | 0.83 |
| YBR165W | 515.6336536 | 0.144917106 | 0.543844619 | 0.83 |
| YHR136C | 132.1966559 | -0.793663227 | 0.543968711 | 0.83 |
| YNL029C | 467.6043522 | 0.228954091 | 0.543963846 | 0.83 |

| | | | | |
|---|---|---|---|---|
| YAR042W | 3250.319067 | 0.324075083 | 0.545281893 | 0.83 |
| YBL099W | 11787.99535 | 0.195026666 | 0.545387923 | 0.83 |
| YDR139C | 266.4093081 | -0.120035483 | 0.545417004 | 0.83 |
| YDR328C | 1849.787592 | -0.166973845 | 0.544717276 | 0.83 |
| YGR294W | 10.95232902 | -0.32456219 | 0.54475369 | 0.83 |
| YKL053C-A | 473.2870725 | -0.195472602 | 0.545332789 | 0.83 |
| YMR064W | 1294.977576 | 0.178775434 | 0.545103458 | 0.83 |
| YOL129W | 2548.890951 | -0.16645502 | 0.545298941 | 0.83 |
| YOR289W | 937.4069291 | 0.242542101 | 0.544294146 | 0.83 |
| YPR120C | 354.4464034 | -0.351849549 | 0.544976311 | 0.83 |
| YPL233W | 252.806959 | -0.182210484 | 0.545954031 | 0.83 |
| YAL014C | 833.2543927 | 0.176359513 | 0.546150559 | 0.83 |
| RUF20 | 86.17514557 | -0.358706192 | 0.547823403 | 0.83 |
| YAL069W | 0.918104495 | 0.345353138 | 0.547696949 | 0.83 |
| YBL051C | 3664.33556 | -0.090079797 | 0.547906951 | 0.83 |
| YDR178W | 2932.392626 | 0.541084003 | 0.547506323 | 0.83 |
| YDR314C | 384.5141242 | 0.333186921 | 0.546982375 | 0.83 |
| YER019C-A | 1962.324665 | -0.127355852 | 0.547609958 | 0.83 |
| YFL041W | 1108.963199 | 0.231803682 | 0.546800477 | 0.83 |
| YFR004W | 2781.166019 | 0.173972029 | 0.547042073 | 0.83 |
| YGR273C | 7.812625483 | 0.523303886 | 0.547650732 | 0.83 |
| YHR094C | 525.1336673 | 0.008506899 | 0.548100595 | 0.83 |
| YJR117W | 3603.291074 | -0.139050406 | 0.547146734 | 0.83 |
| YKR062W | 891.4921763 | -0.163710222 | 0.547521813 | 0.83 |
| YNR048W | 344.510222 | 0.211327805 | 0.547625023 | 0.83 |
| YOL023W | 636.5704075 | 0.197701969 | 0.548020962 | 0.83 |
| YOL071W | 1035.627926 | 0.243733252 | 0.546595966 | 0.83 |
| YOR233W | 853.5809378 | 0.218031293 | 0.548281239 | 0.83 |
| YBR115C | 2897.179739 | 0.403185955 | 0.548831332 | 0.83 |
| YOR313C | 174.7411276 | -0.407533814 | 0.54873988 | 0.83 |
| YPR073C | 916.4024804 | -0.135566925 | 0.548590523 | 0.83 |
| YCR050C | 17.22525728 | 0.293904216 | 0.549969728 | 0.83 |
| YDL039C | 373.0125142 | 0.328352939 | 0.549930856 | 0.83 |
| YEL061C | 1022.169501 | -0.167283541 | 0.549892456 | 0.83 |
| YGL260W | 1.415207671 | 0.030558927 | 0.549926393 | 0.83 |
| YOR109W | 2024.801836 | -0.174027393 | 0.550063806 | 0.83 |
| YOR262W | 658.3672371 | 0.299932152 | 0.54994525 | 0.83 |
| YPR158W | 1022.854173 | 0.364958733 | 0.549668331 | 0.83 |
| YLR279W | 36.37996098 | -0.024339672 | 0.550305102 | 0.83 |
| YAL036C | 1045.610431 | 0.107558499 | 0.550612188 | 0.83 |
| YCL058C | 103.80183 | 0.170674922 | 0.551109784 | 0.83 |
| YDL073W | 971.6916673 | -0.207137303 | 0.551034878 | 0.83 |
| YFR047C | 2554.161875 | -0.303557136 | 0.550940431 | 0.83 |

| | | | | |
|---|---|---|---|---|
| YDR270W | 829.6044968 | 0.214427676 | 0.551562949 | 0.83 |
| YIR027C | 227.1514133 | 0.015138455 | 0.55152507 | 0.83 |
| YMR122C | 59.11544635 | 0.081269285 | 0.551799218 | 0.83 |
| YJL137C | 923.3034038 | 0.657546867 | 0.552055309 | 0.83 |
| YDR537C | 28.61472288 | -0.094129059 | 0.552267495 | 0.83 |
| YBL005W | 1420.668611 | -0.301635826 | 0.553571147 | 0.83 |
| YBR178W | 20.74423611 | -0.00060963 | 0.553749045 | 0.83 |
| YCL027W | 675.5367401 | -0.914023671 | 0.553602476 | 0.83 |
| YDR087C | 817.7593103 | -0.115594575 | 0.554194751 | 0.83 |
| YDR540C | 110.2480797 | -0.264654709 | 0.55421318 | 0.83 |
| YLR405W | 359.0345633 | -0.245698652 | 0.55415101 | 0.83 |
| YMR026C | 666.726482 | -0.202870289 | 0.55379351 | 0.83 |
| YMR273C | 1360.501919 | 0.225192916 | 0.553895969 | 0.83 |
| YMR323W | 206.5265565 | 0.520376959 | 0.552915463 | 0.83 |
| YNL228W | 12.1493579 | 0.013000081 | 0.553470588 | 0.83 |
| YOL001W | 429.8453508 | 0.039766627 | 0.553951712 | 0.83 |
| YOL110W | 433.4687434 | 0.225556637 | 0.553458766 | 0.83 |
| YOR022C | 608.0869392 | 0.269143431 | 0.554105524 | 0.83 |
| YDR492W | 689.8420252 | -0.154358243 | 0.554845676 | 0.84 |
| YER134C | 346.3610386 | -0.323004622 | 0.55468998 | 0.84 |
| YHR048W | 458.0188382 | 0.186695771 | 0.555152818 | 0.84 |
| YJR153W | 431.5896283 | 0.181422269 | 0.554843676 | 0.84 |
| YMR184W | 674.607137 | -0.097222118 | 0.555116885 | 0.84 |
| YPR008W | 1796.059535 | 0.154016082 | 0.554904887 | 0.84 |
| YCL029C | 771.878883 | -0.119161216 | 0.556123881 | 0.84 |
| YCR054C | 374.3747854 | 0.228657997 | 0.556112346 | 0.84 |
| YGR119C | 2276.598641 | -0.165905611 | 0.555826652 | 0.84 |
| YGR170W | 2142.624103 | 0.203108144 | 0.555636457 | 0.84 |
| YHL031C | 431.2428898 | 0.249551625 | 0.556060799 | 0.84 |
| YHR072W | 1833.218722 | -0.214157795 | 0.555943459 | 0.84 |
| YPR197C | 5.599447647 | -0.46819318 | 0.555467618 | 0.84 |
| YJL210W | 543.5019911 | 0.222469484 | 0.556445704 | 0.84 |
| YLR182W | 1421.620302 | 0.17879125 | 0.556472295 | 0.84 |
| YOR324C | 520.2886814 | -0.27766812 | 0.556567487 | 0.84 |
| YDR133C | 1765.716231 | 0.42018512 | 0.557296775 | 0.84 |
| YDR184C | 548.4562077 | -0.160361078 | 0.557289335 | 0.84 |
| YEL052W | 1984.785259 | 0.261731368 | 0.557115085 | 0.84 |
| YGL240W | 171.0354676 | 0.256122742 | 0.557568682 | 0.84 |
| YNL031C | 9825.730059 | -0.197086334 | 0.557540972 | 0.84 |
| YDR512C | 741.4198181 | 0.174459719 | 0.558014356 | 0.84 |
| YGL052W | 83.8653604 | 0.209734952 | 0.55841184 | 0.84 |
| YGR028W | 820.509086 | 0.166359612 | 0.558074566 | 0.84 |
| YJL051W | 1072.541155 | 0.197396639 | 0.558353011 | 0.84 |

| | | | | |
|---|---|---|---|---|
| YMR177W | 436.2483597 | -0.283204599 | 0.558171017 | 0.84 |
| YPL187W | 17.17000227 | 0.00576999 | 0.558220902 | 0.84 |
| YPL246C | 2119.845638 | -0.09817737 | 0.558581008 | 0.84 |
| YMR212C | 1054.541147 | -0.243309174 | 0.558888016 | 0.84 |
| YOR050C | 67.3008437 | -0.393464888 | 0.559039529 | 0.84 |
| YKR025W | 782.5030894 | -0.143831467 | 0.559173794 | 0.84 |
| YDR539W | 882.7410932 | -0.34516733 | 0.559873628 | 0.84 |
| YHR166C | 498.330332 | 0.229062942 | 0.559948438 | 0.84 |
| YJR011C | 218.4017674 | -0.220015251 | 0.559577702 | 0.84 |
| YPR039W | 9.707681236 | 0.49167251 | 0.559820788 | 0.84 |
| YDR061W | 1142.905546 | 0.138548088 | 0.560247898 | 0.84 |
| YGR247W | 484.9641046 | 0.185691137 | 0.560392435 | 0.84 |
| YLR218C | 291.942794 | 0.141941401 | 0.560275992 | 0.84 |
| YLR144C | 964.8658379 | 0.211688646 | 0.560601206 | 0.84 |
| YJR098C | 316.068717 | 0.21087254 | 0.561509302 | 0.84 |
| YKL054C | 4561.844374 | -0.11758936 | 0.562188129 | 0.84 |
| YKL061W | 244.3100746 | 0.169298001 | 0.56216785 | 0.84 |
| YLR010C | 127.8600135 | 0.194315642 | 0.562117305 | 0.84 |
| YLR460C | 187.4228264 | -0.192550308 | 0.562111891 | 0.84 |
| YAL053W | 2097.963607 | 0.464660586 | 0.564139459 | 0.84 |
| YCR053W | 6107.267981 | -0.091448505 | 0.563972972 | 0.84 |
| YDL066W | 3505.644872 | 0.079320114 | 0.563214033 | 0.84 |
| YDL109C | 380.7977413 | 0.23284446 | 0.564726275 | 0.84 |
| YDR376W | 985.9130091 | 0.202234601 | 0.563943107 | 0.84 |
| YER037W | 745.1407721 | 0.069370222 | 0.563633595 | 0.84 |
| YER082C | 1253.635463 | -0.186886751 | 0.564340004 | 0.84 |
| YGL177W | 36.28800868 | 0.497050925 | 0.563082895 | 0.84 |
| YIL168W | 92.19596636 | -0.398959898 | 0.564644917 | 0.84 |
| YJL215C | 44.17616213 | -0.540477266 | 0.564823184 | 0.84 |
| YLL018C-A | 305.6957299 | 0.176524905 | 0.563704883 | 0.84 |
| YLL066W-B | 2.220126499 | 0.187400592 | 0.563268827 | 0.84 |
| YLR170C | 116.6208147 | -0.276970025 | 0.564235837 | 0.84 |
| YNL190W | 6079.357238 | -0.215888397 | 0.56479936 | 0.84 |
| YOL053W | 775.6100204 | 0.245594951 | 0.56480051 | 0.84 |
| YPR058W | 588.0067957 | -0.015737454 | 0.564278754 | 0.84 |
| YPR098C | 1032.023923 | 0.352826737 | 0.563740148 | 0.84 |
| YIL005W | 1687.731259 | -0.162939521 | 0.565222158 | 0.84 |
| YDR521W | 13.62209314 | -0.241239395 | 0.565532637 | 0.84 |
| YDR527W | 984.8264926 | 0.23322905 | 0.565691971 | 0.84 |
| YER129W | 2291.22394 | 0.133624954 | 0.565741983 | 0.84 |
| YJL182C | 1.793622545 | -0.440989442 | 0.565515796 | 0.84 |
| YOL056W | 561.7509431 | -0.177858997 | 0.566203713 | 0.84 |
| YOR083W | 334.3141069 | 0.172510436 | 0.566255205 | 0.84 |

| | | | | |
|---|---|---|---|---|
| YPL149W | 607.7809907 | 0.200641447 | 0.566158931 | 0.84 |
| YDR046C | 2206.66233 | 0.116776234 | 0.567215761 | 0.84 |
| YIL002C | 1259.267656 | 0.148202239 | 0.567259894 | 0.84 |
| YML029W | 1034.498713 | 0.18713266 | 0.567156174 | 0.84 |
| YHR165W-A | 6.420816 | 0.809580012 | 0.567720355 | 0.84 |
| YJL049W | 674.8429005 | 0.129299846 | 0.567525331 | 0.84 |
| YNL280C | 1498.81009 | -0.148154024 | 0.567762593 | 0.84 |
| YGL211W | 636.0371021 | 0.152730575 | 0.568032439 | 0.84 |
| YLR411W | 132.197405 | 0.040494423 | 0.56804691 | 0.84 |
| YAL027W | 394.2818606 | 0.152142321 | 0.568644829 | 0.84 |
| YBL034C | 918.1202298 | -0.139388802 | 0.568816744 | 0.84 |
| YCL009C | 7647.273312 | -0.028973907 | 0.56882276 | 0.84 |
| YBR094W | 1036.817856 | -0.125991132 | 0.569650288 | 0.84 |
| YBR253W | 396.1883397 | -0.200168421 | 0.569573856 | 0.84 |
| YHR008C | 8653.583569 | 0.495892557 | 0.569804215 | 0.84 |
| YLR067C | 640.6258307 | 0.14274187 | 0.569313317 | 0.84 |
| YMR025W | 229.9295307 | 0.1952037 | 0.569726765 | 0.84 |
| YBR056W-A | 43.21007303 | 0.017015041 | 0.570331429 | 0.84 |
| YJR053W | 291.1479634 | -0.129947883 | 0.570936224 | 0.84 |
| YLR234W | 283.065063 | 0.465111949 | 0.570936126 | 0.84 |
| YNR052C | 1154.692673 | -0.180158662 | 0.571040567 | 0.84 |
| YBL013W | 160.0267849 | 0.336874661 | 0.572691086 | 0.84 |
| YBR026C | 2145.982583 | 0.255268305 | 0.571295871 | 0.84 |
| YBR036C | 2605.132077 | 0.196326742 | 0.571796677 | 0.84 |
| YDL211C | 175.1726895 | -0.134632417 | 0.574819489 | 0.84 |
| YEL020W-A | 192.2562057 | -0.2015086 | 0.573627558 | 0.84 |
| YFL048C | 1163.307711 | -0.241812697 | 0.574640268 | 0.84 |
| YHR135C | 3300.939762 | -0.158122665 | 0.573583787 | 0.84 |
| YHR140W | 190.2878906 | -0.274882809 | 0.573011169 | 0.84 |
| YIL048W | 1213.90232 | 0.151700162 | 0.57382269 | 0.84 |
| YIR023W | 1089.743487 | -0.204639377 | 0.57310235 | 0.84 |
| YJL180C | 680.1821358 | -0.111424972 | 0.574478111 | 0.84 |
| YJR154W | 284.604362 | 0.249145946 | 0.574589134 | 0.84 |
| YLL042C | 71.42312394 | 0.517324637 | 0.573046354 | 0.84 |
| YLR250W | 1395.702651 | -0.209246725 | 0.571840352 | 0.84 |
| YLR355C | 9452.460146 | 0.278375624 | 0.574011018 | 0.84 |
| YLR361C | 559.6543534 | 0.141804087 | 0.574363965 | 0.84 |
| YLR456W | 255.5161749 | 0.185859082 | 0.573378009 | 0.84 |
| YML003W | 222.6670877 | 0.142997082 | 0.574475664 | 0.84 |
| YML120C | 4283.544511 | 0.669967446 | 0.571683847 | 0.84 |
| YMR108W | 5770.966467 | 0.093414295 | 0.573201896 | 0.84 |
| YNL084C | 2582.808264 | -0.130958271 | 0.571586991 | 0.84 |
| YNL179C | 130.2526057 | 0.14971063 | 0.57386768 | 0.84 |

| | | | | |
|---|---|---|---|---|
| YNL250W | 994.2437698 | 0.271644123 | 0.573599487 | 0.84 |
| YOL068C | 1498.234041 | -0.196355798 | 0.573930351 | 0.84 |
| YOR037W | 696.733578 | 0.189679085 | 0.574307892 | 0.84 |
| YOR322C | 1313.48176 | 0.164988013 | 0.574842842 | 0.84 |
| YPL040C | 952.0787614 | 0.201065979 | 0.57422316 | 0.84 |
| YPL182C | 43.60254699 | -0.213875215 | 0.571963324 | 0.84 |
| YPR170C | 83.5281119 | -0.338404657 | 0.574550519 | 0.84 |
| YMR189W | 2797.782123 | -0.118831166 | 0.57588953 | 0.84 |
| YHR086W | 910.6374675 | -0.142695661 | 0.576708037 | 0.85 |
| YMR211W | 431.1277672 | 0.111811905 | 0.576907164 | 0.85 |
| YOR071C | 651.6605909 | -0.413971871 | 0.5767947 | 0.85 |
| YCR098C | 141.8503964 | 0.586182158 | 0.577353981 | 0.85 |
| YDR407C | 1886.152895 | -0.162549818 | 0.577763048 | 0.85 |
| YEL062W | 560.1717097 | 0.085537309 | 0.577918106 | 0.85 |
| YER158C | 1028.126375 | 0.097758841 | 0.577245944 | 0.85 |
| YGL060W | 769.6330407 | 0.150992276 | 0.577872904 | 0.85 |
| YGR168C | 490.2041357 | 0.155938166 | 0.577978696 | 0.85 |
| YHL041W | 3.288023996 | 0.292239455 | 0.577470556 | 0.85 |
| YNL299W | 424.5188557 | -0.073524113 | 0.577808308 | 0.85 |
| YBR177C | 2530.709586 | -0.233952491 | 0.57842304 | 0.85 |
| YDR011W | 4201.778813 | -0.185743515 | 0.578461671 | 0.85 |
| YPL190C | 2551.586572 | -0.068464855 | 0.578236125 | 0.85 |
| YML007C-A | 2.984197591 | -0.484531384 | 0.578789021 | 0.85 |
| YBR009C | 3036.286227 | -0.278637301 | 0.578987928 | 0.85 |
| YGR256W | 1242.341744 | 0.026080269 | 0.579379272 | 0.85 |
| YAL042C-A | 8.764234173 | -0.044911145 | 0.579690374 | 0.85 |
| YEL059C-A | 318.8027126 | 0.3867333 | 0.579894884 | 0.85 |
| YBL039W-B | 151.3574507 | 0.210052963 | 0.580992922 | 0.85 |
| YBR235W | 1364.419973 | 0.325651516 | 0.580080362 | 0.85 |
| YDL178W | 1397.749906 | -0.135023474 | 0.580452766 | 0.85 |
| YGL185C | 473.3669461 | 0.202973188 | 0.580655529 | 0.85 |
| YKL055C | 140.4891808 | 0.109541438 | 0.58046506 | 0.85 |
| YKR056W | 1143.695694 | -0.121400422 | 0.580493823 | 0.85 |
| YNL268W | 2190.621688 | -0.246762013 | 0.581121653 | 0.85 |
| YOR202W | 1479.782603 | 0.36945648 | 0.580826357 | 0.85 |
| YOR269W | 302.0905548 | -0.225967233 | 0.580903839 | 0.85 |
| YBL088C | 1866.759984 | 0.23698482 | 0.581374897 | 0.85 |
| YHR119W | 1485.712969 | -0.112895432 | 0.581511829 | 0.85 |
| YBL075C | 2650.065607 | 0.274224747 | 0.58459459 | 0.85 |
| YBR129C | 610.3882792 | -0.26259465 | 0.583336173 | 0.85 |
| YDL165W | 697.5260403 | -0.13468699 | 0.583293969 | 0.85 |
| YDR132C | 453.7495244 | -0.277944421 | 0.584368417 | 0.85 |
| YDR441C | 376.4209723 | 0.153588331 | 0.583138667 | 0.85 |

| | | | | |
|---|---|---|---|---|
| YEL036C | 1490.088365 | -0.127853778 | 0.582192143 | 0.85 |
| YER180C-A | 68.54860649 | 0.125073655 | 0.583204971 | 0.85 |
| YFL025C | 800.9686963 | 0.093063466 | 0.584519481 | 0.85 |
| YFR011C | 642.0102271 | -0.016517639 | 0.582197741 | 0.85 |
| YGL032C | 1976.542723 | -0.474444131 | 0.583100243 | 0.85 |
| YHL005C | 24.92643688 | -0.248298409 | 0.584442215 | 0.85 |
| YHR031C | 752.7001486 | -0.066213379 | 0.582294441 | 0.85 |
| YHR050W | 1110.031456 | -0.002026624 | 0.584016477 | 0.85 |
| YHR082C | 4569.103984 | 0.385636818 | 0.582868223 | 0.85 |
| YHR176W | 373.1735066 | 0.086419717 | 0.584535283 | 0.85 |
| YJR094C | 253.8151399 | -0.459246305 | 0.58364851 | 0.85 |
| YLR004C | 257.8975095 | 0.127507047 | 0.584309493 | 0.85 |
| YML056C | 4012.776938 | -0.149872269 | 0.582397759 | 0.85 |
| YMR138W | 253.6379214 | -0.148478253 | 0.582487312 | 0.85 |
| YMR187C | 357.2988229 | 0.209785856 | 0.583118752 | 0.85 |
| YMR192W | 859.8324449 | -0.100036757 | 0.583653754 | 0.85 |
| YOL147C | 576.9213691 | 0.019065707 | 0.583825665 | 0.85 |
| YOR348C | 282.9627385 | 0.183243893 | 0.583738507 | 0.85 |
| YLR357W | 954.9234442 | -0.192787418 | 0.585015564 | 0.85 |
| YBR257W | 391.5498488 | -0.110878856 | 0.585288526 | 0.85 |
| YPL060W | 360.6791713 | 0.234424801 | 0.585372556 | 0.85 |
| YDR357C | 553.215464 | -0.251527911 | 0.585584926 | 0.85 |
| YDR243C | 512.1645164 | 0.139044329 | 0.586452189 | 0.85 |
| YDR452W | 4294.15743 | 0.168438969 | 0.586304652 | 0.85 |
| YER029C | 871.4448752 | -0.15174565 | 0.586215544 | 0.85 |
| YHR163W | 2218.862273 | 0.070496235 | 0.586505482 | 0.85 |
| YIL134W | 238.1456546 | -0.07459076 | 0.585961436 | 0.85 |
| YOR041C | 37.3273586 | -0.401387088 | 0.585976717 | 0.85 |
| YOR329W-A | 20.44917844 | -0.38045682 | 0.586070528 | 0.85 |
| YDL196W | 15.17669232 | 0.237575568 | 0.587049898 | 0.85 |
| YAL045C | 8.89025023 | -0.433627151 | 0.58732149 | 0.85 |
| YDR246W-A | 23.46436155 | -0.34025036 | 0.587398749 | 0.85 |
| YBR141C | 286.9036203 | -0.160939638 | 0.587836014 | 0.85 |
| YNL006W | 936.4455341 | -0.206396157 | 0.588428611 | 0.85 |
| YGR147C | 943.7636147 | -0.160151906 | 0.589183121 | 0.85 |
| YLR082C | 386.5514271 | -0.275984433 | 0.589158955 | 0.85 |
| YMR200W | 1797.595559 | -0.149058348 | 0.588827174 | 0.85 |
| YNL322C | 3831.740608 | 0.320751736 | 0.588941647 | 0.85 |
| YBL085W | 2143.648048 | 0.1712692 | 0.590998555 | 0.85 |
| YBR020W | 310.6456705 | 0.093256281 | 0.589915647 | 0.85 |
| YDL086W | 1034.431113 | 0.349087866 | 0.590943229 | 0.85 |
| YIL084C | 434.165782 | 0.125380161 | 0.59125822 | 0.85 |
| YJL114W | 3.593194288 | -0.03984393 | 0.591165174 | 0.85 |

| | | | | |
|---|---|---|---|---|
| YKL197C | 902.4571586 | 0.248436703 | 0.589704919 | 0.85 |
| YLR085C | 664.5290334 | 0.131251693 | 0.590351 | 0.85 |
| YMR061W | 1751.530821 | -0.202590491 | 0.590331368 | 0.85 |
| YMR066W | 755.7979974 | 0.118706034 | 0.591008883 | 0.85 |
| YMR166C | 428.7049362 | 0.119252113 | 0.590898811 | 0.85 |
| YMR188C | 1203.74754 | 0.218965885 | 0.589562497 | 0.85 |
| YMR310C | 757.4056331 | -0.250683124 | 0.590262464 | 0.85 |
| YNL220W | 4979.508941 | 0.054093256 | 0.589964755 | 0.85 |
| YOL038W | 4309.883987 | 0.253174346 | 0.590481616 | 0.85 |
| YPR117W | 2597.357181 | 0.207875151 | 0.589952032 | 0.85 |
| YNL232W | 718.0441816 | -0.130262506 | 0.591528588 | 0.85 |
| YCL001W-A | 26.77980078 | -0.533293768 | 0.592037489 | 0.85 |
| YJL022W | 2.660856745 | 0.12052558 | 0.592371812 | 0.85 |
| YOR256C | 1260.844527 | 0.155445162 | 0.592476242 | 0.85 |
| YLR256W | 4392.10169 | -0.175286695 | 0.592907634 | 0.85 |
| YAR047C | 21.66468268 | 0.20117883 | 0.59366106 | 0.85 |
| YIL156W-B | 291.2591429 | 0.170003986 | 0.593626924 | 0.85 |
| YNR032W | 607.7463894 | 0.213194992 | 0.593744544 | 0.85 |
| YFL007W | 4593.396937 | 0.360748125 | 0.593877635 | 0.85 |
| YHR035W | 231.8124454 | 0.322977159 | 0.594076171 | 0.85 |
| YLR141W | 304.974896 | 0.154510875 | 0.594346581 | 0.85 |
| snR190 | 17.01701234 | 0.747599646 | 0.594329262 | 0.85 |
| YDR265W | 518.8074155 | 0.159039638 | 0.595053779 | 0.85 |
| YGR135W | 3309.689244 | -0.122232433 | 0.594964107 | 0.85 |
| YHR071C-A | 39.05421358 | -0.048325464 | 0.594744509 | 0.85 |
| YIL100W | 3.722972412 | 0.400165405 | 0.594618892 | 0.85 |
| YMR253C | 606.2113411 | -0.252545634 | 0.595192747 | 0.85 |
| snR86 | 894.0383097 | 0.042537896 | 0.595266835 | 0.85 |
| YDR317W | 303.4699364 | 0.01944142 | 0.595558754 | 0.85 |
| YNL282W | 259.9576363 | 0.174561405 | 0.59555887 | 0.85 |
| YCR026C | 1309.803177 | 0.162744772 | 0.596133069 | 0.85 |
| YER060W-A | 584.2480367 | -0.063599938 | 0.595995337 | 0.85 |
| YER095W | 1267.382435 | 0.363245677 | 0.595854025 | 0.85 |
| YGL235W | 1.903973228 | 0.156736565 | 0.596223581 | 0.85 |
| snR35 | 41.6109753 | 0.41008249 | 0.596178173 | 0.85 |
| YBL058W | 3576.371774 | 0.168510604 | 0.596588456 | 0.85 |
| YER130C | 1003.132192 | -0.273894921 | 0.597105516 | 0.85 |
| YBR065C | 528.4307138 | 0.119524393 | 0.600237715 | 0.85 |
| YBR073W | 1901.278512 | -0.292053258 | 0.598176684 | 0.85 |
| YBR239C | 441.6078035 | 0.06132006 | 0.59890705 | 0.85 |
| YBR272C | 606.5333168 | 0.141943209 | 0.598549834 | 0.85 |
| YCL065W | 1.135078196 | 0.137889228 | 0.598575355 | 0.85 |
| YCR051W | 1038.832374 | -0.149747508 | 0.600127624 | 0.85 |

| | | | | |
|---|---|---|---|---|
| YDL062W | 21.94852935 | 0.542021301 | 0.599247276 | 0.85 |
| YDR034C | 1497.715383 | 0.099354128 | 0.599790879 | 0.85 |
| YDR044W | 1839.127613 | -0.144849697 | 0.598783123 | 0.85 |
| YFL053W | 516.855343 | 0.744939432 | 0.59916539 | 0.85 |
| YHR085W | 335.931842 | 0.056293 | 0.59795031 | 0.85 |
| YJR083C | 550.0152446 | 0.145420679 | 0.599076534 | 0.85 |
| YKL073W | 1776.826393 | -0.010306275 | 0.599987161 | 0.85 |
| YKR024C | 515.1058632 | -0.225773067 | 0.597737136 | 0.85 |
| YKR102W | 639.1097035 | 0.475769694 | 0.599851987 | 0.85 |
| YMR244C-A | 108.6393324 | 0.181718639 | 0.599249189 | 0.85 |
| YMR314W | 2980.188475 | -0.106776039 | 0.599998042 | 0.85 |
| YNL058C | 1080.922928 | 0.069084427 | 0.598341028 | 0.85 |
| YNL130C-A | 3.921452687 | 0.098978279 | 0.600339556 | 0.85 |
| YNL138W-A | 114.4265169 | -0.256876892 | 0.60022576 | 0.85 |
| YOL154W | 197.7584736 | 0.03258935 | 0.599845693 | 0.85 |
| YOR261C | 3484.142585 | -0.120113022 | 0.599644334 | 0.85 |
| snR50 | 8.290616263 | -0.012527645 | 0.599640889 | 0.85 |
| YHR028W-A | 38.81881491 | -0.324498651 | 0.600721897 | 0.85 |
| YAL031W-A | 36.5218107 | 0.221783616 | 0.600872896 | 0.85 |
| YBR111W-A | 229.152125 | -0.195896973 | 0.603282461 | 0.85 |
| YCR069W | 2249.27644 | 0.133267619 | 0.601874105 | 0.85 |
| YGL063W | 321.9330079 | -0.198946434 | 0.603497905 | 0.85 |
| YGL145W | 1156.588747 | 0.131700918 | 0.601248994 | 0.85 |
| YHR012W | 963.21973 | -0.160598734 | 0.603247507 | 0.85 |
| YHR077C | 2925.324128 | 0.13827941 | 0.602897642 | 0.85 |
| YHR186C | 1649.917336 | -0.118076466 | 0.601380191 | 0.85 |
| YIL064W | 505.7570311 | 0.194647605 | 0.602491047 | 0.85 |
| YJR065C | 3820.877578 | -0.123210227 | 0.602187518 | 0.85 |
| YJR090C | 1653.355955 | 0.11811104 | 0.602103562 | 0.85 |
| YKR032W | 0.979223699 | 0.21778933 | 0.603522242 | 0.85 |
| YLL016W | 1012.667191 | 0.098826102 | 0.601827621 | 0.85 |
| YLR111W | 15.73315875 | 0.201192392 | 0.603129107 | 0.85 |
| YLR245C | 188.3206886 | -0.257912613 | 0.6035732 | 0.85 |
| YLR277C | 732.8482737 | -0.178327197 | 0.604029107 | 0.85 |
| YLR320W | 753.4878726 | 0.144397826 | 0.602794849 | 0.85 |
| YML048W | 4451.705992 | -0.084815357 | 0.603088111 | 0.85 |
| YML124C | 2648.714801 | -0.067253782 | 0.603983682 | 0.85 |
| YMR030W-A | 15.82228062 | 0.10061627 | 0.60401687 | 0.85 |
| YNL325C | 453.4130958 | 0.150029888 | 0.60388231 | 0.85 |
| YNR059W | 692.5145797 | 0.071723433 | 0.60149044 | 0.85 |
| YPL122C | 549.6598792 | -0.150184935 | 0.60345217 | 0.85 |
| YPR104C | 970.815797 | -0.180395568 | 0.603964883 | 0.85 |
| YPR113W | 3675.90658 | -0.227339315 | 0.602959301 | 0.85 |

| | | | | |
|---|---|---|---|---|
| YFL065C | 1.354222162 | -0.046658285 | 0.604377826 | 0.85 |
| YGR059W | 83.10299922 | -0.024575354 | 0.604341031 | 0.85 |
| YDR261C | 1216.328788 | -0.207817747 | 0.605791033 | 0.85 |
| YDR477W | 1886.671826 | 0.14192833 | 0.606122404 | 0.85 |
| YDR511W | 614.1187303 | -0.198799626 | 0.606036841 | 0.85 |
| YER158W-A | 247.6290209 | -0.251627944 | 0.605993927 | 0.85 |
| YFR052W | 2758.803837 | 0.226115986 | 0.604844749 | 0.85 |
| YJL214W | 204.5476536 | -0.382356099 | 0.606148639 | 0.85 |
| YKL092C | 887.6693114 | -0.152188585 | 0.606024884 | 0.85 |
| YLR430W | 3065.658429 | -0.040081187 | 0.605402316 | 0.85 |
| YMR231W | 864.2847521 | -0.126227843 | 0.605835081 | 0.85 |
| YOR084W | 822.8839971 | 0.129615737 | 0.605575204 | 0.85 |
| YPL107W | 480.1557675 | 0.208357408 | 0.606215785 | 0.85 |
| YPR011C | 560.157333 | -0.242591165 | 0.605608214 | 0.85 |
| YDR261C-D | 88.68167414 | -0.436740528 | 0.606682461 | 0.85 |
| YDR514C | 275.4543514 | -0.211874426 | 0.606506086 | 0.85 |
| YOR106W | 543.2030625 | -0.157288456 | 0.606767415 | 0.85 |
| YPR122W | 1503.146832 | -0.62077552 | 0.606389959 | 0.85 |
| YJL176C | 1289.4859 | 0.19521235 | 0.607149341 | 0.85 |
| YKL089W | 415.1913401 | 0.253247721 | 0.60722824 | 0.85 |
| YPL023C | 1340.132366 | -0.108054686 | 0.607111203 | 0.85 |
| YKL088W | 1161.516822 | 0.138355255 | 0.607572696 | 0.85 |
| YCR087W | 24.15940244 | 0.293288226 | 0.607979044 | 0.86 |
| YCR092C | 872.6618833 | 0.138002847 | 0.608274737 | 0.86 |
| YDR437W | 89.57715872 | -0.133273428 | 0.608936195 | 0.86 |
| YGL121C | 888.9061681 | 0.264295742 | 0.609025282 | 0.86 |
| YOR376W-A | 83.93269889 | -0.187456288 | 0.609052047 | 0.86 |
| YBR038W | 1800.515437 | -0.048424728 | 0.611393377 | 0.86 |
| YBR156C | 807.2285578 | 0.106540022 | 0.611081207 | 0.86 |
| YCR099C | 126.2405817 | -0.034572896 | 0.610154078 | 0.86 |
| YDR365W-B | 142.2689351 | -0.196513759 | 0.611464717 | 0.86 |
| YER050C | 1168.128624 | -0.156113409 | 0.610776734 | 0.86 |
| YER106W | 34.98872113 | 0.221031457 | 0.610276974 | 0.86 |
| YGL149W | 44.44871907 | -0.061624132 | 0.61027839 | 0.86 |
| YGL197W | 2610.651509 | 0.23025702 | 0.610753697 | 0.86 |
| YHR030C | 2133.377408 | 0.276640368 | 0.610774974 | 0.86 |
| YIL106W | 893.3105346 | -0.146350366 | 0.609862897 | 0.86 |
| YLR224W | 1026.056679 | 0.095954646 | 0.611583732 | 0.86 |
| YML087C | 330.7889367 | 0.39177524 | 0.610820851 | 0.86 |
| YMR065W | 380.856771 | 0.004538993 | 0.609479065 | 0.86 |
| YNL053W | 544.6626736 | -0.232466637 | 0.611357258 | 0.86 |
| YOL042W | 569.9826336 | 0.140771584 | 0.609640059 | 0.86 |
| YOL066C | 908.4756316 | 0.136126373 | 0.610907214 | 0.86 |

| | | | | |
|---|---|---|---|---|
| YOR226C | 454.5609099 | 0.092151027 | 0.611222828 | 0.86 |
| YPL189W | 119.6862802 | -0.018147005 | 0.61135048 | 0.86 |
| YIL102C-A | 112.0491515 | -0.336873888 | 0.611963786 | 0.86 |
| YML058W-A | 99.58508334 | 0.025388995 | 0.612029194 | 0.86 |
| YLR148W | 974.658438 | 0.173234435 | 0.612260518 | 0.86 |
| YPL044C | 1.74310561 | -0.170171537 | 0.612323032 | 0.86 |
| YJL006C | 336.1143852 | 0.204382788 | 0.612614826 | 0.86 |
| YLR385C | 52.50375543 | 0.39333863 | 0.61278741 | 0.86 |
| YML018C | 644.4081194 | -0.209322409 | 0.613024424 | 0.86 |
| YMR218C | 1133.912405 | 0.162155539 | 0.61308922 | 0.86 |
| YLR025W | 2034.136821 | 0.127231002 | 0.613508583 | 0.86 |
| YOL144W | 766.01586 | -0.044874183 | 0.613549293 | 0.86 |
| YDR289C | 776.5077999 | 0.182093646 | 0.614903858 | 0.86 |
| YEL043W | 2331.741844 | 0.208620433 | 0.614469945 | 0.86 |
| YER064C | 1674.781352 | 0.238825285 | 0.614313166 | 0.86 |
| YFR012W | 37.52047909 | -0.766527626 | 0.614539982 | 0.86 |
| YGL146C | 201.790122 | 0.013652523 | 0.614796589 | 0.86 |
| YJR122W | 745.1766826 | 0.189896172 | 0.6147949 | 0.86 |
| YLR346C | 131.9806369 | 0.200820181 | 0.614584342 | 0.86 |
| YMR209C | 810.1729969 | -0.272054701 | 0.614903905 | 0.86 |
| YGR139W | 13.99976418 | 0.378647486 | 0.615403683 | 0.86 |
| YLR050C | 545.8709868 | -0.092874985 | 0.615532264 | 0.86 |
| YNL298W | 1195.295385 | -0.241720055 | 0.61569762 | 0.86 |
| YDL181W | 1061.423493 | -0.130813677 | 0.616256006 | 0.86 |
| YDR416W | 716.6573712 | 0.156886967 | 0.616457296 | 0.86 |
| YGL011C | 3486.685044 | -0.157016474 | 0.616587154 | 0.86 |
| YKL006C-A | 282.6893408 | -0.157202792 | 0.61647303 | 0.86 |
| YOL024W | 52.91060507 | 0.421046013 | 0.616212688 | 0.86 |
| YEL072W | 207.4067694 | 0.057523154 | 0.616773914 | 0.86 |
| YKL029C | 3291.164423 | 0.267672862 | 0.616978531 | 0.86 |
| YIL015W | 2424.316267 | -0.689318385 | 0.617598675 | 0.86 |
| YLR246W | 470.2308665 | 0.23530092 | 0.617638981 | 0.86 |
| YMR122W-A | 5293.670712 | -0.125002738 | 0.617491323 | 0.86 |
| YMR157C | 556.2957769 | -0.20672744 | 0.617532526 | 0.86 |
| YOR273C | 1760.091485 | 0.288374313 | 0.617680876 | 0.86 |
| YML116W-A | 166.4577303 | 0.21129264 | 0.617847864 | 0.86 |
| YML072C | 6041.463861 | 0.281417807 | 0.618049662 | 0.86 |
| YBR035C | 2483.374886 | 0.219722425 | 0.62037851 | 0.86 |
| YBR083W | 2339.410841 | -0.69691411 | 0.620089387 | 0.86 |
| YGR005C | 1346.509283 | 0.12019124 | 0.619372969 | 0.86 |
| YGR074W | 145.6756142 | -0.289253341 | 0.619169417 | 0.86 |
| YHR168W | 379.0957302 | 0.194195134 | 0.618881871 | 0.86 |
| YHR196W | 929.0161741 | -0.293514771 | 0.620281733 | 0.86 |

| | | | | |
|---|---|---|---|---|
| YKL025C | 1634.326653 | 0.142384941 | 0.619547115 | 0.86 |
| YKL038W | 1738.382543 | 0.343698304 | 0.619653623 | 0.86 |
| YKR053C | 486.0346558 | 0.024343841 | 0.619051145 | 0.86 |
| YLR162W | 2.387178616 | 0.290932637 | 0.618580777 | 0.86 |
| YMR075C-A | 15.14864191 | 0.054925667 | 0.620552913 | 0.86 |
| YNL057W | 19.3681916 | 0.153157975 | 0.62056403 | 0.86 |
| YOL086W-A | 318.8786531 | 0.28904835 | 0.618709716 | 0.86 |
| YPL148C | 142.8618352 | -0.246598511 | 0.619433805 | 0.86 |
| YPR020W | 1129.530709 | 0.051502112 | 0.619634348 | 0.86 |
| YPR033C | 3601.656962 | -0.05243201 | 0.620349439 | 0.86 |
| YPR055W | 1160.769461 | 0.135822099 | 0.620386176 | 0.86 |
| YLR184W | 0.707088895 | 0.244469256 | 0.62092608 | 0.86 |
| YPL091W | 3000.977941 | -0.072508378 | 0.620950956 | 0.86 |
| YDR105C | 933.9584198 | -0.195297391 | 0.621526388 | 0.86 |
| YJL102W | 772.7229577 | 0.56900618 | 0.621728453 | 0.86 |
| YML052W | 1751.911847 | -0.266982673 | 0.621717384 | 0.86 |
| YNL266W | 15.42456766 | 0.234699591 | 0.621574766 | 0.86 |
| YJL038C | 194.4642451 | -0.346354608 | 0.621915419 | 0.86 |
| YDR529C | 3824.487001 | 0.464560085 | 0.622555182 | 0.86 |
| YER048C | 2682.830499 | -0.195460187 | 0.622382107 | 0.86 |
| YLR120W-A | 22.81140064 | 0.529348811 | 0.622514804 | 0.86 |
| YBL091C-A | 508.1746239 | 0.171957954 | 0.623045697 | 0.86 |
| YCR082W | 1363.300313 | 0.083297088 | 0.623504423 | 0.86 |
| YHR214C-B | 235.6005636 | 0.004918836 | 0.623199634 | 0.86 |
| YIL080W | 141.3218365 | 0.074496473 | 0.623498667 | 0.86 |
| YJL123C | 2650.72374 | 0.210910184 | 0.623351095 | 0.86 |
| YEL024W | 1858.219132 | 0.353266635 | 0.623745393 | 0.86 |
| YDR311W | 727.3776002 | 0.080886501 | 0.624482233 | 0.86 |
| YGL181W | 1793.449442 | 0.170253077 | 0.62446682 | 0.86 |
| YIL047C | 1300.508261 | -0.221735974 | 0.624409303 | 0.86 |
| YKR082W | 1772.94275 | 0.186854193 | 0.624102524 | 0.86 |
| YDR170W-A | 128.0923816 | 0.036294377 | 0.625095027 | 0.86 |
| YMR134W | 453.7137928 | -0.104629322 | 0.625283908 | 0.86 |
| YLL012W | 268.5425901 | -0.457175441 | 0.625533706 | 0.86 |
| YCL056C | 401.4808141 | 0.080339396 | 0.626136189 | 0.86 |
| YBL012C | 41.44037077 | 0.231754414 | 0.626687655 | 0.86 |
| YBL082C | 673.6132807 | -0.065102923 | 0.626459671 | 0.86 |
| YFL044C | 776.7355212 | 0.221132447 | 0.62631289 | 0.86 |
| YHR018C | 3547.596925 | 0.311612923 | 0.626585247 | 0.86 |
| YDR253C | 229.9334863 | 0.183059556 | 0.626979841 | 0.86 |
| YNR032C-A | 183.6670278 | -0.144591943 | 0.627346126 | 0.86 |
| YAL041W | 1345.73957 | 0.091978718 | 0.627825613 | 0.86 |
| YJL047C-A | 19.06572924 | 0.362393708 | 0.627835302 | 0.86 |

| | | | | |
|---|---|---|---|---|
| YOL125W | 710.1287675 | -0.16747379 | 0.627835241 | 0.86 |
| YHR056W-A | 0.867587117 | 0.466881247 | 0.628111096 | 0.86 |
| snR39B | 13.58422976 | -0.100300836 | 0.628192775 | 0.86 |
| YDR394W | 3757.628989 | -0.136888942 | 0.628518759 | 0.86 |
| YMR028W | 753.5245359 | 0.114418577 | 0.628649777 | 0.86 |
| YIR007W | 898.4409432 | 0.138897094 | 0.628982496 | 0.86 |
| YMR267W | 919.8795189 | -0.110796634 | 0.629650642 | 0.86 |
| YNR038W | 1115.811097 | -0.19302473 | 0.629922987 | 0.86 |
| YFL059W | 70.79713485 | -0.161574311 | 0.63025082 | 0.87 |
| YDR442W | 17.52253569 | -0.057983022 | 0.630416798 | 0.87 |
| YDL154W | 661.9950458 | -0.151244558 | 0.631988641 | 0.87 |
| YDR501W | 295.5069374 | -0.162749435 | 0.632517447 | 0.87 |
| YGL143C | 536.4770627 | -0.208482691 | 0.631465343 | 0.87 |
| YGR030C | 291.5946813 | -0.007999008 | 0.631202939 | 0.87 |
| YHL001W | 1027.533504 | -0.093600762 | 0.631154651 | 0.87 |
| YIL001W | 471.4395924 | 0.142345157 | 0.632520268 | 0.87 |
| YIL091C | 989.7593991 | -0.232934089 | 0.631441347 | 0.87 |
| YKL034W | 1217.582813 | 0.258452613 | 0.631901885 | 0.87 |
| YLR262C | 283.7623055 | -0.111049948 | 0.631940461 | 0.87 |
| YLR390W | 418.8392228 | 0.178129229 | 0.631305054 | 0.87 |
| YNL169C | 1167.217459 | -0.115683571 | 0.631175631 | 0.87 |
| YOR055W | 14.54954616 | -0.153995211 | 0.632180143 | 0.87 |
| YOR175C | 2021.823765 | -0.102443581 | 0.631674202 | 0.87 |
| YPR097W | 1353.173051 | 0.1641244 | 0.631889636 | 0.87 |
| YPR128C | 787.6455183 | 0.234951999 | 0.632491831 | 0.87 |
| YGR047C | 884.2267464 | 0.13533136 | 0.632928581 | 0.87 |
| YMR092C | 2509.767947 | 0.182361943 | 0.632882354 | 0.87 |
| YDR278C | 0.649229666 | 0.268460863 | 0.633613924 | 0.87 |
| YPL068C | 273.8997414 | -0.027673338 | 0.633888077 | 0.87 |
| YIR033W | 2182.959679 | 0.233133837 | 0.634027078 | 0.87 |
| YEL077C | 280.3488731 | -0.036151078 | 0.634271825 | 0.87 |
| YPL275W | 31.2658679 | 0.096681113 | 0.634828459 | 0.87 |
| YGL069C | 85.63194987 | -0.289779012 | 0.634984193 | 0.87 |
| YBR054W | 4277.398072 | 0.308535978 | 0.635338489 | 0.87 |
| YBR274W | 529.7886878 | -0.143264334 | 0.635501531 | 0.87 |
| YHL029C | 889.4266622 | 0.11931217 | 0.635537987 | 0.87 |
| YLR051C | 783.2029443 | -0.049860969 | 0.635369774 | 0.87 |
| YDR461W | 1830.189511 | -0.551603211 | 0.636041078 | 0.87 |
| YKR087C | 773.1699867 | 0.136607864 | 0.636237432 | 0.87 |
| YAR053W | 9.523462605 | 0.033164098 | 0.636626113 | 0.87 |
| YLR427W | 1359.694674 | -0.091104886 | 0.636879026 | 0.87 |
| YOL160W | 15.67971505 | 0.104757499 | 0.636764754 | 0.87 |
| YKL015W | 1286.742474 | 0.275444315 | 0.63716014 | 0.87 |

| | | | | |
|---|---|---|---|---|
| YOL136C | 1016.324902 | -0.317089789 | 0.637796991 | 0.87 |
| YJR103W | 3899.480126 | 0.410221198 | 0.638134194 | 0.87 |
| YDR305C | 387.9870014 | -0.043942197 | 0.638334413 | 0.87 |
| YNR015W | 1147.190779 | -0.040407241 | 0.638759794 | 0.87 |
| YBR044C | 837.2738234 | -0.176370449 | 0.639186671 | 0.87 |
| YLR304C | 13028.11852 | 0.09844236 | 0.639356095 | 0.87 |
| YGL057C | 252.5789153 | -0.153654652 | 0.639597911 | 0.87 |
| YDL185W | 10910.49268 | -0.077893138 | 0.64018904 | 0.87 |
| YGL222C | 1086.796722 | 0.122515352 | 0.640489579 | 0.87 |
| YHL025W | 1272.013263 | 0.160720177 | 0.640287183 | 0.87 |
| YNL055C | 20028.30046 | 0.464322257 | 0.640232497 | 0.87 |
| YNL119W | 513.2444289 | -0.140391716 | 0.640381053 | 0.87 |
| YNL224C | 704.2866123 | 0.138058495 | 0.639916627 | 0.87 |
| YBR285W | 449.6014678 | -0.108864111 | 0.640897965 | 0.87 |
| YIL088C | 2503.905976 | -0.125359734 | 0.641407099 | 0.87 |
| YJL194W | 154.6134221 | -0.482658067 | 0.640918872 | 0.87 |
| YMR156C | 369.432093 | 0.205003272 | 0.641327386 | 0.87 |
| YMR185W | 803.0326457 | -0.164356426 | 0.640794642 | 0.87 |
| YNL023C | 950.809325 | 0.202523757 | 0.641721579 | 0.87 |
| YNL320W | 323.6388355 | -0.094677814 | 0.641851028 | 0.87 |
| YOL067C | 283.6975055 | 0.163435347 | 0.641453303 | 0.87 |
| YOL159C | 274.8863973 | -0.186150583 | 0.641297066 | 0.87 |
| YOR111W | 390.0644929 | 0.144170022 | 0.641843078 | 0.87 |
| YBR093C | 949.3588178 | -0.17895058 | 0.642561144 | 0.87 |
| YFL062W | 38.28792094 | 0.699333844 | 0.64253819 | 0.87 |
| YBL105C | 2392.516684 | -0.140405414 | 0.643356512 | 0.87 |
| YMR222C | 712.4011542 | 0.157161891 | 0.644402573 | 0.87 |
| YMR265C | 688.687566 | 0.113178661 | 0.644331355 | 0.87 |
| YOR053W | 4.40231234 | 0.032670844 | 0.644380804 | 0.87 |
| YAL047C | 589.7842721 | 0.174850383 | 0.645069305 | 0.87 |
| YLR101C | 65.86852303 | 0.151418814 | 0.645510434 | 0.87 |
| YHR063C | 1432.676855 | -0.055004808 | 0.645820741 | 0.87 |
| YHR207C | 677.6706609 | -0.061773451 | 0.645771713 | 0.87 |
| YBR163W | 719.9181119 | -0.28256829 | 0.646785268 | 0.87 |
| YHR032W | 1019.136565 | -0.148892298 | 0.646616907 | 0.87 |
| YLR104W | 635.3306236 | -0.062379533 | 0.646767537 | 0.87 |
| snR14 | 19.85838705 | -0.013028257 | 0.646699345 | 0.87 |
| YGR186W | 2388.772284 | 0.204543499 | 0.64726012 | 0.87 |
| YMR126C | 404.3389603 | 0.053883374 | 0.647233273 | 0.87 |
| YPL053C | 1363.897708 | -0.161391565 | 0.647044324 | 0.87 |
| YIL128W | 1273.394888 | 0.07661256 | 0.647633474 | 0.88 |
| YMR244W | 46.42719777 | -0.414095432 | 0.647918752 | 0.88 |
| YBR248C | 1552.941582 | 0.107482348 | 0.648285668 | 0.88 |

| | | | | |
|---|---|---|---|---|
| YIL089W | 155.6273305 | 0.096545892 | 0.648489359 | 0.88 |
| YML088W | 784.353378 | -0.08717249 | 0.648554474 | 0.88 |
| YGR230W | 330.0627222 | 0.154814866 | 0.648905134 | 0.88 |
| YGR233C | 1147.708039 | -0.211629031 | 0.648915971 | 0.88 |
| YGR166W | 818.3285643 | 0.183046525 | 0.649383296 | 0.88 |
| YOL059W | 2587.338376 | 0.094100726 | 0.649447568 | 0.88 |
| YPR195C | 54.58354851 | -0.352831423 | 0.64928332 | 0.88 |
| YGR121W-A | 8.749518959 | 0.536186286 | 0.649623671 | 0.88 |
| YJL035C | 163.2101994 | -0.188950272 | 0.649867284 | 0.88 |
| YML037C | 139.9825756 | -0.151090667 | 0.650284375 | 0.88 |
| YNL159C | 572.5140174 | 0.044762404 | 0.650171896 | 0.88 |
| YPL067C | 285.3555196 | 0.071447003 | 0.65013106 | 0.88 |
| snR11 | 261.7635615 | 0.28737907 | 0.650258067 | 0.88 |
| YPL209C | 162.5627938 | 0.261491836 | 0.65071571 | 0.88 |
| YCR108C | 1.912873177 | 0.077973603 | 0.650886869 | 0.88 |
| YGR011W | 95.88242328 | -0.124962642 | 0.651153572 | 0.88 |
| YGL263W | 253.3051631 | 0.381597409 | 0.651661757 | 0.88 |
| YLR387C | 1423.423086 | 0.221596431 | 0.651683535 | 0.88 |
| YPL158C | 1350.795148 | 0.270289649 | 0.651458752 | 0.88 |
| YBR209W | 5.517408049 | -0.132266278 | 0.652405946 | 0.88 |
| YGL192W | 1154.271063 | -0.125127754 | 0.652248107 | 0.88 |
| YKL159C | 432.357669 | -0.187637774 | 0.652376708 | 0.88 |
| YLR097C | 748.1202732 | 0.106946819 | 0.652339978 | 0.88 |
| YPR045C | 1121.270528 | 0.167972651 | 0.652490345 | 0.88 |
| YER068W | 1672.188885 | 0.173626019 | 0.653253751 | 0.88 |
| YHR185C | 7.642313868 | -0.201496972 | 0.65373007 | 0.88 |
| YIL151C | 1650.033227 | 0.109150778 | 0.653505536 | 0.88 |
| YKL024C | 1589.092419 | -0.063943323 | 0.65354894 | 0.88 |
| YLR454W | 3660.437031 | 0.252882727 | 0.652946448 | 0.88 |
| YNL005C | 1578.642517 | -0.110654818 | 0.653213421 | 0.88 |
| YOR370C | 2847.263179 | -0.139606271 | 0.653672465 | 0.88 |
| YGL039W | 719.9626894 | -0.14740526 | 0.655133005 | 0.88 |
| YGL220W | 745.8780305 | -0.162817179 | 0.655908426 | 0.88 |
| YHR061C | 329.4509042 | 0.053140443 | 0.654419507 | 0.88 |
| YIR029W | 388.4137592 | 0.079728788 | 0.656258211 | 0.88 |
| YJL052C-A | 1.688849253 | -0.317099589 | 0.655771567 | 0.88 |
| YJR062C | 945.4060688 | 0.098991646 | 0.655462343 | 0.88 |
| YKR044W | 684.1987114 | 0.10928656 | 0.656411836 | 0.88 |
| YLR187W | 889.7815093 | 0.220292465 | 0.655109635 | 0.88 |
| YLR237W | 1061.722449 | 0.056038195 | 0.65606368 | 0.88 |
| YLR266C | 1095.178781 | -0.002592028 | 0.655453907 | 0.88 |
| YML081C-A | 1028.969006 | 0.469400459 | 0.655695176 | 0.88 |
| YMR285C | 827.213202 | -0.132923057 | 0.656572686 | 0.88 |

| | | | | |
|---|---|---|---|---|
| YNL186W | 1470.462386 | -0.159135549 | 0.656692638 | 0.88 |
| YNL309W | 553.1266537 | -0.089635533 | 0.655741341 | 0.88 |
| YOR026W | 568.5531387 | -0.208451715 | 0.655832928 | 0.88 |
| YOR047C | 1007.814108 | -0.031198594 | 0.655060023 | 0.88 |
| YPL071C | 441.9177148 | -0.206577602 | 0.654895194 | 0.88 |
| YPL097W | 440.8818332 | 0.202664399 | 0.65661433 | 0.88 |
| YPR094W | 154.1332011 | 0.158557733 | 0.655060577 | 0.88 |
| YDL122W | 1760.896516 | -0.145150039 | 0.657092555 | 0.88 |
| YLR019W | 1293.150392 | 0.084484754 | 0.656995937 | 0.88 |
| YNL317W | 817.8553177 | -0.116535611 | 0.657258772 | 0.88 |
| YOR366W | 3.244594662 | -0.968229175 | 0.657253021 | 0.88 |
| YLR093C | 1657.689066 | -0.220101861 | 0.657920053 | 0.88 |
| YHL035C | 3064.413674 | 0.158416131 | 0.658303671 | 0.88 |
| YIL083C | 1169.719343 | 0.217633818 | 0.658387287 | 0.88 |
| YOR174W | 544.9117735 | -0.11075537 | 0.658312689 | 0.88 |
| YNR040W | 422.1360755 | 0.325428709 | 0.658573328 | 0.88 |
| YOL054W | 494.8918395 | 0.163302605 | 0.658698876 | 0.88 |
| YBR167C | 275.1433522 | -0.249192203 | 0.659617208 | 0.88 |
| YDR472W | 675.6541934 | -0.126395217 | 0.660548102 | 0.88 |
| YFL015W-A | 1.143841678 | -0.440352035 | 0.659832989 | 0.88 |
| YJL162C | 484.7743599 | 0.115485972 | 0.660508641 | 0.88 |
| YKR080W | 1412.607481 | 0.105003022 | 0.659955283 | 0.88 |
| YLR136C | 655.8429852 | -0.286497617 | 0.660307493 | 0.88 |
| YLR204W | 619.6469206 | -0.150097629 | 0.660401411 | 0.88 |
| YML014W | 592.0947476 | 0.069825188 | 0.66053455 | 0.88 |
| YMR296C | 3004.03015 | -0.100942086 | 0.659589732 | 0.88 |
| YOR094W | 410.9080628 | -0.168468419 | 0.659052395 | 0.88 |
| YOR177C | 142.866635 | -0.664004468 | 0.659504967 | 0.88 |
| YPL076W | 380.2952376 | 0.013885614 | 0.660525256 | 0.88 |
| snR85 | 18.57666508 | -0.151115976 | 0.659118974 | 0.88 |
| YIL053W | 6767.756759 | -0.027032769 | 0.661305383 | 0.88 |
| YJL030W | 384.5980723 | -0.096464694 | 0.661261513 | 0.88 |
| YML074C | 7957.336564 | -0.14731333 | 0.661155632 | 0.88 |
| YNR067C | 2460.572138 | -0.064355058 | 0.661079871 | 0.88 |
| YBR014C | 1010.158039 | -0.038844432 | 0.661469817 | 0.88 |
| YOL033W | 423.9709748 | -0.121835452 | 0.6617513 | 0.88 |
| YMR127C | 400.3588714 | 0.120144589 | 0.66193071 | 0.88 |
| YJL077W-B | 26.1550574 | 0.364596071 | 0.662569514 | 0.88 |
| YJR034W | 337.4607648 | -0.162557169 | 0.662447401 | 0.88 |
| YML101C-A | 7.327177589 | 0.268814065 | 0.662487025 | 0.88 |
| YPL276W | 15.28614474 | 0.083369927 | 0.662225376 | 0.88 |
| YOR195W | 1326.708475 | -0.041502499 | 0.663114989 | 0.88 |
| YOR314W-A | 0.578252409 | -0.045862289 | 0.663108501 | 0.88 |

| | | | | |
|---|---|---|---|---|
| YCL068C | 4.465179414 | 0.048089747 | 0.664137073 | 0.88 |
| YDL225W | 1750.506735 | -0.125862116 | 0.663986134 | 0.88 |
| YKL068W | 1722.980976 | -0.167977253 | 0.664496951 | 0.88 |
| YOL020W | 976.6345917 | -0.347141509 | 0.664257009 | 0.88 |
| YOR008C | 1383.478149 | -0.22886633 | 0.664457337 | 0.88 |
| YOR264W | 932.8104963 | -0.184336662 | 0.664467358 | 0.88 |
| YPL060C-A | 3.51919416 | -0.016805653 | 0.663819917 | 0.88 |
| YML110C | 4679.227772 | -0.045103058 | 0.664693917 | 0.88 |
| YDR318W | 508.7245087 | -0.186586457 | 0.664995741 | 0.88 |
| YOR064C | 481.1143371 | 0.207712812 | 0.665140862 | 0.88 |
| YCR019W | 382.8804523 | -0.163147842 | 0.665392224 | 0.88 |
| YOR115C | 581.5147795 | -0.122193149 | 0.665519339 | 0.88 |
| YLR035C | 528.4128752 | -0.242390895 | 0.665686123 | 0.88 |
| YDL207W | 1207.359156 | -0.094471734 | 0.666262456 | 0.88 |
| YFR041C | 978.3498425 | -0.102128149 | 0.666232957 | 0.88 |
| YGR191W | 1267.875828 | -0.202799005 | 0.66605006 | 0.88 |
| YLL051C | 1170.125266 | -0.153431772 | 0.665996229 | 0.88 |
| YDR282C | 383.6210181 | 0.055595907 | 0.666529199 | 0.88 |
| YLR199C | 824.459216 | 0.128556554 | 0.667007702 | 0.88 |
| YOR180C | 257.2908648 | 0.001050957 | 0.667024979 | 0.88 |
| YAL037C-A | 1.195205434 | -0.178316146 | 0.668703024 | 0.88 |
| YAR018C | 738.6960091 | 0.110492585 | 0.668766138 | 0.88 |
| YBR103C-A | 1.570441558 | 0.179435718 | 0.669468434 | 0.88 |
| YCL002C | 469.4119103 | -0.17552812 | 0.66974834 | 0.88 |
| YDL168W | 1463.24926 | 0.345218525 | 0.669046217 | 0.88 |
| YDL248W | 224.0356228 | 0.168541388 | 0.671546513 | 0.88 |
| YDR100W | 731.7492012 | -0.235244778 | 0.671048496 | 0.88 |
| YDR138W | 884.1934286 | -0.154955418 | 0.671649301 | 0.88 |
| YDR168W | 2674.002145 | 0.299736359 | 0.672490034 | 0.88 |
| YDR206W | 1789.144398 | -0.176348822 | 0.672960454 | 0.88 |
| YDR224C | 8028.945616 | -0.194262438 | 0.670375366 | 0.88 |
| YDR268W | 586.9385642 | -0.059982963 | 0.668113728 | 0.88 |
| YDR365C | 4849.022737 | 0.184600409 | 0.670941184 | 0.88 |
| YDR532C | 597.6785104 | 0.14650795 | 0.668151277 | 0.88 |
| YEL069C | 125.5249506 | 0.216827737 | 0.672353833 | 0.88 |
| YER021W | 5518.8989 | 0.187892873 | 0.671152697 | 0.88 |
| YFL049W | 806.3019635 | 0.061515792 | 0.667944259 | 0.88 |
| YGL086W | 1388.801297 | 0.165708789 | 0.669046514 | 0.88 |
| YGL114W | 1125.03715 | 0.192231212 | 0.67212092 | 0.88 |
| YGL187C | 1692.180783 | -0.064698423 | 0.669939938 | 0.88 |
| YGR092W | 957.140587 | -0.00254379 | 0.672279348 | 0.88 |
| YGR095C | 969.1047325 | -0.075767762 | 0.672990014 | 0.88 |
| YJL073W | 1063.600671 | -0.021387736 | 0.669689594 | 0.88 |

| | | | | |
|---|---|---|---|---|
| YJL174W | 2482.371966 | -0.08412676 | 0.671930009 | 0.88 |
| YJR055W | 115.4214659 | -0.091772289 | 0.667729211 | 0.88 |
| YJR138W | 1461.069987 | 0.203240636 | 0.672829094 | 0.88 |
| YKL065C | 4326.57925 | 0.149559785 | 0.671580796 | 0.88 |
| YLR071C | 1557.164849 | 0.122810618 | 0.672663869 | 0.88 |
| YMR147W | 251.4438843 | 0.402250766 | 0.672136332 | 0.88 |
| YNL004W | 1614.267096 | -0.126481456 | 0.672476775 | 0.88 |
| YNL180C | 2386.54101 | 0.14584176 | 0.668588969 | 0.88 |
| YNR075W | 186.6264191 | -0.173963248 | 0.670079712 | 0.88 |
| YOL101C | 182.685157 | 0.103347403 | 0.670523573 | 0.88 |
| YOR191W | 2372.954185 | 0.143656676 | 0.669173947 | 0.88 |
| YOR193W | 437.421385 | 0.01063552 | 0.669250915 | 0.88 |
| YOR222W | 1007.50754 | -0.167123667 | 0.670924794 | 0.88 |
| YOR244W | 816.0935801 | -0.150002385 | 0.672125924 | 0.88 |
| YPL026C | 1074.82514 | -0.109434941 | 0.670551212 | 0.88 |
| YPL141C | 1338.030401 | 0.164555299 | 0.668416311 | 0.88 |
| YPL217C | 2396.710824 | 0.0240222 | 0.670293431 | 0.88 |
| YPR088C | 2257.094394 | -0.099301202 | 0.671921326 | 0.88 |
| YNL019C | 10.29601328 | -0.080389493 | 0.673288174 | 0.88 |
| YNL088W | 2851.433881 | 0.264375812 | 0.673223565 | 0.88 |
| YJL076W | 2985.809725 | -0.064312085 | 0.673945454 | 0.88 |
| YMR013C | 348.9768631 | 0.058540239 | 0.673969232 | 0.88 |
| YDR063W | 1183.071975 | -0.150828626 | 0.674107665 | 0.88 |
| YDR158W | 6048.209084 | 0.149395482 | 0.674247553 | 0.88 |
| YIR036C | 938.6857022 | 0.156316875 | 0.674828601 | 0.88 |
| YMR288W | 774.5062547 | 0.136628155 | 0.675057021 | 0.88 |
| YOR075W | 634.1618193 | 0.27455149 | 0.675206875 | 0.88 |
| YGL188C | 26.0866713 | 0.199225612 | 0.675685175 | 0.88 |
| YPL118W | 2044.672532 | -0.112519155 | 0.675941241 | 0.88 |
| YOR076C | 709.8237452 | 0.074959106 | 0.676610484 | 0.88 |
| YMR178W | 1157.445412 | -0.125812054 | 0.676943753 | 0.88 |
| YLR032W | 823.8658356 | -0.077312386 | 0.677607855 | 0.89 |
| YMR286W | 654.5341281 | 0.164345948 | 0.677744062 | 0.89 |
| snR77 | 9.325617253 | 0.492214069 | 0.677836774 | 0.89 |
| YJL025W | 374.9847479 | -0.455472466 | 0.678231864 | 0.89 |
| YML006C | 696.4182875 | -0.148507304 | 0.67829061 | 0.89 |
| YMR019W | 977.1459301 | 0.126349512 | 0.678103543 | 0.89 |
| YML043C | 371.5090553 | -0.313380409 | 0.678440871 | 0.89 |
| YBL090W | 640.7121751 | -0.072320612 | 0.680251257 | 0.89 |
| YCR042C | 1593.772478 | 0.095749295 | 0.678985435 | 0.89 |
| YDR051C | 1050.532227 | 0.075642349 | 0.680078303 | 0.89 |
| YER056C | 2408.22934 | -0.278727952 | 0.679375883 | 0.89 |
| YGL199C | 2.062524245 | 0.901556872 | 0.679962656 | 0.89 |

| | | | | |
|---|---|---|---|---|
| YKL043W | 3024.896012 | -0.106207714 | 0.679212167 | 0.89 |
| YKR036C | 843.7972906 | 0.080337642 | 0.679221452 | 0.89 |
| YLR034C | 1366.422953 | -0.359134406 | 0.680139872 | 0.89 |
| YMR150C | 511.1258272 | -0.148994823 | 0.680249222 | 0.89 |
| YMR172W | 685.1558079 | -0.147848872 | 0.679778334 | 0.89 |
| YPL216W | 730.7653507 | 0.098901694 | 0.679954949 | 0.89 |
| YBL055C | 719.3024768 | -0.137125554 | 0.682466175 | 0.89 |
| YBR190W | 3.930160671 | -0.588001195 | 0.681908707 | 0.89 |
| YDR339C | 1137.925374 | -0.128434723 | 0.682121489 | 0.89 |
| YDR381C-A | 448.1284196 | 0.17183739 | 0.682339611 | 0.89 |
| YDR478W | 183.0152712 | -0.107449887 | 0.681065098 | 0.89 |
| YER038C | 327.709464 | 0.070967372 | 0.680552307 | 0.89 |
| YGL006W-A | 3.943531239 | 0.113856276 | 0.682201767 | 0.89 |
| YGL025C | 784.0082065 | 0.124638619 | 0.6814067 | 0.89 |
| YHL007C | 2278.179389 | -0.097998567 | 0.682268849 | 0.89 |
| YHR187W | 633.7803075 | -0.07649904 | 0.681302087 | 0.89 |
| YIL020C | 394.2399705 | -0.169739433 | 0.681577719 | 0.89 |
| YLR099C | 1018.600053 | 0.26330405 | 0.681821558 | 0.89 |
| YML113W | 567.9041791 | 0.19321972 | 0.681778509 | 0.89 |
| YOR132W | 1718.595531 | 0.192561477 | 0.680863126 | 0.89 |
| YOR238W | 549.7060682 | -0.229521349 | 0.681145871 | 0.89 |
| YPR186C | 328.1330267 | -0.136730991 | 0.682360597 | 0.89 |
| YNL100W | 1262.153408 | 0.181302521 | 0.682611344 | 0.89 |
| YDR187C | 25.29702409 | -0.156900955 | 0.682857584 | 0.89 |
| YIL070C | 1550.574512 | 0.146740019 | 0.682931814 | 0.89 |
| YLR348C | 661.9197696 | 0.277253573 | 0.683047097 | 0.89 |
| YBL089W | 493.6798455 | -0.134000857 | 0.683292284 | 0.89 |
| YIL054W | 8.069700908 | 0.32385839 | 0.683490653 | 0.89 |
| YMR162C | 1643.610674 | 0.149997744 | 0.683428884 | 0.89 |
| YGR058W | 565.2067235 | -0.125558134 | 0.684001292 | 0.89 |
| YGR109W-B | 857.2496031 | 0.166999862 | 0.684066891 | 0.89 |
| YNL286W | 504.2093629 | 0.126027319 | 0.683940769 | 0.89 |
| YPR108W | 2749.924825 | -0.079166871 | 0.683713302 | 0.89 |
| YDR038C | 125.3516014 | 0.131170368 | 0.684768817 | 0.89 |
| YKR048C | 4547.539769 | -0.113737748 | 0.684902601 | 0.89 |
| YDR065W | 277.2481422 | -0.225382991 | 0.685138106 | 0.89 |
| YHR188C | 2845.286709 | -0.066114609 | 0.68564942 | 0.89 |
| YJL150W | 10.09973158 | 0.143335794 | 0.68572187 | 0.89 |
| YKL066W | 1252.501288 | 0.372701477 | 0.685554698 | 0.89 |
| YNL257C | 1662.623362 | 0.127099498 | 0.685609616 | 0.89 |
| YDL242W | 58.94192665 | -0.179328397 | 0.687026413 | 0.89 |
| YDR525W-A | 275.582834 | 0.14898796 | 0.687153385 | 0.89 |
| YFR050C | 3854.503578 | 0.20697983 | 0.686700115 | 0.89 |

| | | | | |
|---|---|---|---|---|
| YHR044C | 186.9132602 | 0.227406715 | 0.686564644 | 0.89 |
| YJR082C | 608.3949667 | 0.085750778 | 0.686257314 | 0.89 |
| YMR198W | 512.8655409 | 0.115756515 | 0.686946083 | 0.89 |
| YOL152W | 484.6917609 | -0.153782028 | 0.686778445 | 0.89 |
| YOR107W | 180.9819764 | -0.158748291 | 0.686356455 | 0.89 |
| YDR041W | 1214.356247 | -0.168362265 | 0.688665606 | 0.89 |
| YDR094W | 4.116771726 | -0.221516103 | 0.68786606 | 0.89 |
| YDR316W | 792.8873428 | -0.101178253 | 0.688454051 | 0.89 |
| YDR373W | 760.4394828 | 0.135282101 | 0.688565454 | 0.89 |
| YDR503C | 368.4756431 | -0.136193431 | 0.688413349 | 0.89 |
| YFL014W | 10835.63862 | 0.125790626 | 0.688943736 | 0.89 |
| YGR254W | 35005.12746 | 0.409143611 | 0.688103176 | 0.89 |
| YLR088W | 986.2933263 | -0.118767948 | 0.688737582 | 0.89 |
| YOL070C | 1742.264613 | 0.243610261 | 0.688865021 | 0.89 |
| YPL154C | 14185.44316 | 0.254198967 | 0.68862339 | 0.89 |
| YPR158C-D | 130.4698158 | -0.195978856 | 0.688270566 | 0.89 |
| snR5 | 32.14065526 | -0.231580414 | 0.687344828 | 0.89 |
| snR75 | 1.574604411 | -0.028463724 | 0.688948251 | 0.89 |
| YPL035C | 100.9354993 | -0.315449752 | 0.689347845 | 0.89 |
| YGR218W | 3998.195199 | -0.087924154 | 0.689727294 | 0.89 |
| YDR072C | 1794.903187 | 0.082678835 | 0.690089729 | 0.89 |
| YNL139C | 1577.578713 | -0.094740568 | 0.690051021 | 0.89 |
| YDR460W | 574.0402083 | 0.079865339 | 0.69073896 | 0.89 |
| YGR041W | 794.2535473 | 0.042333814 | 0.690519397 | 0.89 |
| YHR182W | 713.7981983 | 0.200046004 | 0.691051737 | 0.89 |
| YIR044C | 6.331727114 | -0.120222935 | 0.690918685 | 0.89 |
| YJL135W | 5.319281498 | 0.03641817 | 0.690354384 | 0.89 |
| YLR239C | 753.9388732 | 0.124086233 | 0.690956276 | 0.89 |
| YOR169C | 39.59953477 | -0.172981185 | 0.690733572 | 0.89 |
| YKL048C | 583.1360838 | -0.009372428 | 0.69121578 | 0.89 |
| YKL044W | 194.2159883 | -0.177641459 | 0.691759646 | 0.89 |
| YLR439W | 1150.159115 | 0.300698797 | 0.691574176 | 0.89 |
| YPL156C | 858.4762505 | -0.100053881 | 0.69163525 | 0.89 |
| YPR057W | 656.961666 | -0.178056301 | 0.691825769 | 0.89 |
| YGR253C | 3517.823624 | -0.088682962 | 0.69216069 | 0.89 |
| YKL189W | 702.1789039 | -0.045124661 | 0.692300862 | 0.89 |
| YFR002W | 1555.318988 | -0.149330934 | 0.692681712 | 0.89 |
| YPL193W | 620.6543173 | 0.106779756 | 0.692959064 | 0.89 |
| YKR063C | 686.126014 | 0.124829785 | 0.693350413 | 0.89 |
| YLR069C | 1475.235001 | 0.243418643 | 0.693453358 | 0.89 |
| YCR101C | 105.2670668 | -0.0878344 | 0.693777943 | 0.89 |
| YCR018C | 861.2432726 | -0.188105049 | 0.695872179 | 0.89 |
| YCR023C | 1565.302118 | 0.124447133 | 0.695781631 | 0.89 |

| | | | | |
|---|---|---|---|---|
| YDR218C | 78.04916244 | 0.153636826 | 0.69567292 | 0.89 |
| YDR325W | 750.6903469 | -0.089833655 | 0.695582178 | 0.89 |
| YDR388W | 5322.26629 | -0.070563378 | 0.694891203 | 0.89 |
| YIL038C | 1659.427705 | -0.156409392 | 0.695828192 | 0.89 |
| YLR042C | 282.794653 | -0.152555975 | 0.694306033 | 0.89 |
| YLR049C | 719.3459985 | -0.099654819 | 0.695401534 | 0.89 |
| YLR140W | 51.21550356 | 0.129338055 | 0.694460316 | 0.89 |
| YMR088C | 861.7053416 | -0.11112457 | 0.695295039 | 0.89 |
| YMR227C | 799.7644018 | -0.142214297 | 0.695655407 | 0.89 |
| YOR333C | 3.992636415 | 0.027769781 | 0.695852479 | 0.89 |
| YPR180W | 730.2139766 | 0.121076337 | 0.694992231 | 0.89 |
| YEL019C | 187.0813824 | 0.327223932 | 0.696457422 | 0.89 |
| YMR256C | 1012.905586 | 0.276250495 | 0.696400978 | 0.89 |
| YOR343C | 63.14207102 | 0.077195964 | 0.696482989 | 0.89 |
| YFL034W | 949.1269789 | 0.13488919 | 0.696968313 | 0.89 |
| YDR203W | 15.81601128 | -0.327912591 | 0.697374283 | 0.89 |
| YDR468C | 586.9211769 | -0.198754529 | 0.697454377 | 0.89 |
| YBR230W-A | 211.0787097 | 0.069184842 | 0.697628464 | 0.89 |
| YEL025C | 1231.625482 | 0.078318399 | 0.698149634 | 0.89 |
| YIL075C | 7548.161259 | 0.283638918 | 0.698815766 | 0.89 |
| YJL205C | 132.3580317 | 0.276056983 | 0.698876794 | 0.89 |
| YLR353W | 907.9621406 | -0.072400095 | 0.699085154 | 0.89 |
| YER097W | 28.63993449 | -0.283927159 | 0.700252252 | 0.89 |
| YJR080C | 1270.85858 | -0.018155493 | 0.700343214 | 0.89 |
| YKL132C | 214.3347933 | 0.125383378 | 0.700874122 | 0.89 |
| YGL018C | 470.8004765 | 0.142499072 | 0.701367074 | 0.89 |
| YKL155C | 791.2689341 | 0.151023726 | 0.701282955 | 0.89 |
| YMR247C | 1642.766432 | 0.157703785 | 0.701423342 | 0.89 |
| YCR071C | 461.1120902 | -0.211506452 | 0.702078637 | 0.89 |
| YNL091W | 4224.416881 | -0.026517164 | 0.701845563 | 0.89 |
| YOR141C | 1550.614472 | 0.176437113 | 0.702086817 | 0.89 |
| YOR038C | 676.9008791 | -0.110159079 | 0.702605425 | 0.90 |
| YDL175C | 355.6103486 | -0.221273757 | 0.702946618 | 0.90 |
| YAL005C | 15020.34614 | 0.501384934 | 0.706231091 | 0.90 |
| YAL060W | 4666.556071 | 0.372920902 | 0.705095207 | 0.90 |
| YDL021W | 1151.606846 | 0.225086554 | 0.706432509 | 0.90 |
| YDR162C | 515.4142839 | 0.112370063 | 0.70355488 | 0.90 |
| YDR259C | 452.5952789 | 0.283202305 | 0.704073427 | 0.90 |
| YDR275W | 363.6731352 | -0.007567238 | 0.705288682 | 0.90 |
| YDR298C | 3635.360191 | -0.033975578 | 0.705161738 | 0.90 |
| YEL066W | 476.9220074 | -0.0706935 | 0.706233908 | 0.90 |
| YER012W | 3318.681234 | -0.073200452 | 0.705109246 | 0.90 |
| YER061C | 958.2654695 | -0.135374965 | 0.705809286 | 0.90 |

| | | | | |
|---|---|---|---|---|
| YFR034C | 832.1624053 | -0.098803461 | 0.706708897 | 0.90 |
| YGL236C | 488.164276 | 0.224628727 | 0.704931749 | 0.90 |
| YGR006W | 239.4957069 | -0.124943999 | 0.703851742 | 0.90 |
| YGR231C | 3688.450165 | 0.216744282 | 0.706629097 | 0.90 |
| YHR015W | 104.6987227 | -0.252624093 | 0.705765899 | 0.90 |
| YIL011W | 777.5395764 | 0.103608546 | 0.705059986 | 0.90 |
| YIL058W | 79.78173189 | -0.092827792 | 0.704818563 | 0.90 |
| YKR002W | 1434.178622 | 0.090306828 | 0.703425109 | 0.90 |
| YLR018C | 838.7588705 | -0.134107657 | 0.704885195 | 0.90 |
| YML075C | 1371.306293 | -0.199896102 | 0.706506674 | 0.90 |
| YMR022W | 1239.988654 | 0.11849849 | 0.706384035 | 0.90 |
| YNL245C | 408.0242862 | 0.084423766 | 0.705184915 | 0.90 |
| YNL308C | 3277.042014 | 0.015749785 | 0.706244221 | 0.90 |
| YOL013W-A | 5.152321446 | 0.017849143 | 0.705887665 | 0.90 |
| YOL104C | 616.0912562 | -0.36761928 | 0.703455284 | 0.90 |
| YPL192C | 128.9945052 | 0.041723628 | 0.705064659 | 0.90 |
| YPR116W | 103.378319 | 0.031075951 | 0.705700217 | 0.90 |
| YBR072W | 9138.655064 | 0.276237707 | 0.707868292 | 0.90 |
| YBR251W | 1030.356421 | -0.154776352 | 0.707977158 | 0.90 |
| YDR319C | 485.2377065 | -0.229958265 | 0.707736785 | 0.90 |
| YML103C | 3218.889825 | -0.150318894 | 0.707901903 | 0.90 |
| YOL019W-A | 0.720433626 | 0.109987318 | 0.707675512 | 0.90 |
| YAL009W | 404.8562545 | -0.128464878 | 0.708206969 | 0.90 |
| YOR337W | 623.7332196 | -0.118667159 | 0.708546173 | 0.90 |
| YJL186W | 1797.326564 | -0.052486924 | 0.708897458 | 0.90 |
| YDL133C-A | 72.78703027 | -0.237949242 | 0.709683424 | 0.90 |
| YDL244W | 51.51006224 | 0.242952955 | 0.710636554 | 0.90 |
| YDR498C | 1014.743074 | -0.088340023 | 0.711265841 | 0.90 |
| YFR007W | 462.3779448 | 0.230471321 | 0.709569689 | 0.90 |
| YGR146C-A | 13.60921533 | 0.199813304 | 0.710408392 | 0.90 |
| YGR190C | 89.92851229 | -0.111710778 | 0.711134855 | 0.90 |
| YHR143W | 3407.641358 | -0.089084863 | 0.710159791 | 0.90 |
| YHR205W | 2582.464034 | 0.173905164 | 0.709773257 | 0.90 |
| YJL067W | 49.69168928 | 0.210428609 | 0.711230212 | 0.90 |
| YJL090C | 629.2586035 | 0.05308899 | 0.709878869 | 0.90 |
| YJR078W | 516.4472306 | -0.256139023 | 0.711316061 | 0.90 |
| YKL085W | 9612.778538 | 0.565104488 | 0.709765217 | 0.90 |
| YKR060W | 417.425685 | -0.145423567 | 0.709795288 | 0.90 |
| YKR101W | 347.6961828 | 0.020237524 | 0.71125042 | 0.90 |
| YMR252C | 396.7363367 | 0.132279898 | 0.710570401 | 0.90 |
| YOL148C | 713.6397629 | -0.086663982 | 0.710770534 | 0.90 |
| YOR126C | 423.6858153 | 0.128636825 | 0.711027917 | 0.90 |
| YDL185C-A | 100.139643 | 0.202702225 | 0.71210648 | 0.90 |

| | | | | |
|---|---|---|---|---|
| YDR150W | 4371.450407 | 0.386649356 | 0.712097572 | 0.90 |
| YDR528W | 366.4782046 | -0.346471809 | 0.712003211 | 0.90 |
| YPL046C | 139.4253961 | -0.176145406 | 0.712090742 | 0.90 |
| YCR075W-A | 129.8943621 | 0.184269044 | 0.712586304 | 0.90 |
| YPL078C | 4809.066893 | 0.058032891 | 0.712768792 | 0.90 |
| YDR355C | 4.46006282 | -0.151154952 | 0.713289604 | 0.90 |
| YGL041C-B | 21.05072236 | 0.041757337 | 0.713386613 | 0.90 |
| YGR204C-A | 13.54650687 | -0.027621927 | 0.713146668 | 0.90 |
| YNL157W | 1475.809734 | 0.084800368 | 0.713022547 | 0.90 |
| YOR192C-B | 24.91802453 | -0.119061564 | 0.71349306 | 0.90 |
| YBR086C | 4766.880566 | 0.123805672 | 0.714127448 | 0.90 |
| YHR081W | 846.2639082 | 0.18154031 | 0.714199476 | 0.90 |
| YHL040C | 1041.983492 | -0.185374446 | 0.71481918 | 0.90 |
| YLL032C | 878.0650576 | 0.131566338 | 0.714934775 | 0.90 |
| YEL013W | 1708.200753 | -0.151923508 | 0.71520067 | 0.90 |
| YBR207W | 1493.475156 | -0.178688076 | 0.715608371 | 0.90 |
| YJR104C | 12449.61979 | 0.137459151 | 0.716404683 | 0.90 |
| YNL163C | 1921.119184 | 0.287074699 | 0.716355057 | 0.90 |
| YDR480W | 507.0349407 | -0.064487728 | 0.716834167 | 0.90 |
| YPL014W | 2476.516022 | 0.557457407 | 0.717127468 | 0.90 |
| YIR004W | 1446.338254 | -0.081301759 | 0.717314948 | 0.90 |
| YPL002C | 492.3419627 | 0.07615725 | 0.717668573 | 0.90 |
| YOL122C | 1128.161192 | 0.051054931 | 0.717844379 | 0.90 |
| YJL062W-A | 1034.508423 | 0.051109264 | 0.718240214 | 0.90 |
| YBR123C | 731.156432 | 0.107043884 | 0.718561217 | 0.90 |
| YDR380W | 4921.676121 | -0.025368509 | 0.7186522 | 0.90 |
| YJR044C | 1233.447551 | 0.194124924 | 0.719032238 | 0.90 |
| YOL077W-A | 1163.041858 | 0.015589185 | 0.719074607 | 0.90 |
| YBL069W | 640.7564823 | 0.183732057 | 0.719276668 | 0.90 |
| YER180C | 405.292426 | -0.213960232 | 0.719859298 | 0.90 |
| YMR254C | 26.76419027 | -0.152131697 | 0.71997794 | 0.90 |
| YEL048C | 96.72112286 | -0.351756123 | 0.720180793 | 0.90 |
| YNL329C | 1184.177512 | -0.114520203 | 0.720330324 | 0.90 |
| YDL105W | 423.3140946 | -0.087002307 | 0.720520486 | 0.90 |
| YDR431W | 11.62652329 | -0.324995516 | 0.720802633 | 0.90 |
| YDR533C | 1411.920932 | -0.467801534 | 0.72141476 | 0.90 |
| YGL058W | 714.9977444 | -0.030586695 | 0.721307507 | 0.90 |
| YKL116C | 778.7224786 | -0.09082904 | 0.721163995 | 0.90 |
| YPL064C | 883.912951 | 0.060238502 | 0.721080565 | 0.90 |
| YBR053C | 1693.976608 | 0.174073695 | 0.722013472 | 0.90 |
| YDR166C | 1344.376251 | 0.121381601 | 0.72189266 | 0.90 |
| YGL212W | 286.2266484 | -0.088027674 | 0.722041671 | 0.90 |
| YGR010W | 527.974638 | 0.118999937 | 0.721679873 | 0.90 |

| | | | | |
|---|---|---|---|---|
| YJL054W | 1689.096692 | -0.062346538 | 0.722176673 | 0.90 |
| snR87 | 6.424473744 | -0.058860549 | 0.722512357 | 0.90 |
| YBL070C | 21.83445099 | -0.081932569 | 0.725835454 | 0.90 |
| YBR015C | 2379.48263 | -0.068744811 | 0.723950661 | 0.90 |
| YBR256C | 1957.968457 | 0.114940542 | 0.726603024 | 0.90 |
| YCR086W | 338.9363584 | 0.287101148 | 0.722769405 | 0.90 |
| YDL151C | 4.616503935 | -0.175969679 | 0.724146212 | 0.90 |
| YDL171C | 4109.615185 | 0.324682625 | 0.726523101 | 0.90 |
| YDR189W | 2199.458913 | -0.106694936 | 0.726195804 | 0.90 |
| YFL023W | 930.5339775 | 0.313028089 | 0.725882047 | 0.90 |
| YGL004C | 1031.613443 | -0.17407217 | 0.723706136 | 0.90 |
| YGL014W | 2156.619039 | -0.191779035 | 0.72537463 | 0.90 |
| YGL055W | 3060.50732 | 0.041184352 | 0.726007669 | 0.90 |
| YGL161C | 1505.046156 | 0.085757711 | 0.722979742 | 0.90 |
| YGL238W | 2547.939909 | -0.107186323 | 0.726574285 | 0.90 |
| YHR101C | 345.7524508 | 0.084994252 | 0.726394915 | 0.90 |
| YHR113W | 1875.547688 | -0.057464796 | 0.723487225 | 0.90 |
| YIL156W-A | 6.740450198 | -0.128502733 | 0.726508956 | 0.90 |
| YKL136W | 25.19505527 | -0.301683191 | 0.72579618 | 0.90 |
| YLL036C | 873.0418955 | 0.083976955 | 0.723360675 | 0.90 |
| YLR091W | 340.1364791 | -0.149955018 | 0.725301477 | 0.90 |
| YML058W | 2577.833646 | 0.064446839 | 0.724451052 | 0.90 |
| YMR083W | 8478.832219 | -0.031425568 | 0.726008891 | 0.90 |
| YMR119W-A | 41.58140006 | 0.165426725 | 0.723793588 | 0.90 |
| YNL261W | 688.6478875 | -0.193857632 | 0.723505291 | 0.90 |
| YOL078W | 796.3214789 | 0.146188704 | 0.724551702 | 0.90 |
| YOR034C-A | 45.66152085 | 0.311375726 | 0.724607822 | 0.90 |
| YOR317W | 4625.356852 | 0.245729586 | 0.724808897 | 0.90 |
| YPL024W | 252.5020697 | -0.080533038 | 0.723005713 | 0.90 |
| YPL213W | 340.9140453 | 0.139819487 | 0.725139006 | 0.90 |
| snR73 | 0.982818557 | 0.358749112 | 0.725916096 | 0.90 |
| YDR211W | 2365.993285 | -0.081891034 | 0.726795999 | 0.90 |
| YER119C | 1204.859566 | 0.205740353 | 0.727064565 | 0.90 |
| YNR008W | 899.8873602 | 0.022139425 | 0.726994654 | 0.90 |
| YBR276C | 1528.649976 | -0.084168865 | 0.727316238 | 0.90 |
| YER062C | 1876.234052 | 0.022246131 | 0.727366731 | 0.90 |
| YAL043C | 1560.889418 | 0.167386139 | 0.727604328 | 0.90 |
| YDR110W | 523.1803163 | 0.135517058 | 0.727836685 | 0.90 |
| YFR018C | 1513.157132 | -0.079161163 | 0.728976525 | 0.90 |
| YJR156C | 47.49998049 | 0.227879941 | 0.728893226 | 0.90 |
| YLR145W | 262.3006502 | 0.16291591 | 0.7288246 | 0.90 |
| YPR126C | 82.68269767 | 0.119217857 | 0.728832868 | 0.90 |
| YHR029C | 965.1862947 | 0.285818829 | 0.72964415 | 0.90 |

| | | | | |
|---|---|---|---|---|
| YJL069C | 705.1093909 | -0.231982293 | 0.729791727 | 0.90 |
| YNL238W | 1373.585003 | -0.156448212 | 0.729629579 | 0.90 |
| YOL081W | 5088.832739 | 0.105206674 | 0.729761377 | 0.90 |
| YJR114W | 33.45969567 | 0.01811284 | 0.729942326 | 0.90 |
| YJL079C | 9145.850933 | 0.063979073 | 0.730120754 | 0.90 |
| YBR200W-A | 19.77475914 | -0.147356285 | 0.731015233 | 0.90 |
| YER007W | 271.3998551 | 0.044031054 | 0.73099412 | 0.90 |
| YGL026C | 4379.972897 | 0.313563281 | 0.73093636 | 0.90 |
| YJR022W | 18.26801566 | 0.187409603 | 0.731028469 | 0.90 |
| YKL160W | 1725.224125 | -0.080206063 | 0.731674594 | 0.91 |
| YBR194W | 226.3319567 | 0.114982061 | 0.733489806 | 0.91 |
| YDL071C | 49.14928453 | 0.004409317 | 0.734093117 | 0.91 |
| YEL009C | 21167.59097 | -0.107612394 | 0.734011512 | 0.91 |
| YHR043C | 1297.179518 | 0.026945274 | 0.734128847 | 0.91 |
| YHR193C-A | 1.288094432 | -0.04195785 | 0.734127926 | 0.91 |
| YIL098C | 444.8949209 | -0.150585823 | 0.733511207 | 0.91 |
| YIL157C | 795.6414589 | -0.095928817 | 0.733348454 | 0.91 |
| YJL209W | 626.0794008 | 0.069216771 | 0.73295414 | 0.91 |
| YJR049C | 1397.771565 | 0.184200599 | 0.73315892 | 0.91 |
| YML049C | 1684.51308 | 0.242396898 | 0.733550836 | 0.91 |
| YML114C | 955.0918143 | -0.107027907 | 0.73409299 | 0.91 |
| YNR055C | 1151.83484 | -0.003341964 | 0.732463084 | 0.91 |
| YOR359W | 584.0345426 | -0.333567827 | 0.734029066 | 0.91 |
| YPR070W | 926.8338353 | -0.092836309 | 0.734190943 | 0.91 |
| YCL043C | 17735.82228 | 0.397860976 | 0.734561338 | 0.91 |
| YIL041W | 4311.545468 | -0.044926121 | 0.734803118 | 0.91 |
| YML125C | 2139.072841 | -0.001492165 | 0.734923292 | 0.91 |
| YKL148C | 4538.261302 | 0.637475577 | 0.735759918 | 0.91 |
| YOR194C | 1465.560671 | 0.052106325 | 0.73601702 | 0.91 |
| YJL044C | 1379.126824 | -0.166934228 | 0.736187832 | 0.91 |
| YDL093W | 807.8112275 | -0.005480625 | 0.737121147 | 0.91 |
| YER006C-A | 2.225353622 | 0.260310028 | 0.736997569 | 0.91 |
| YHR051W | 2680.129991 | 0.095868415 | 0.736781661 | 0.91 |
| YHR216W | 3639.691774 | 0.486814304 | 0.736659301 | 0.91 |
| YKL186C | 1093.769583 | 0.107107383 | 0.737124772 | 0.91 |
| YMR152W | 1727.69017 | 0.098397109 | 0.737285361 | 0.91 |
| YKL211C | 1647.230083 | 0.137234713 | 0.73753957 | 0.91 |
| YOR151C | 4196.260493 | -0.119944544 | 0.737654607 | 0.91 |
| YDR363W-A | 1308.889566 | -0.124841743 | 0.738424651 | 0.91 |
| YJR046W | 1243.286799 | 0.00697663 | 0.738241233 | 0.91 |
| YOL123W | 4114.119417 | -0.217949289 | 0.738503004 | 0.91 |
| YPL262W | 5873.810031 | 0.233298002 | 0.738872923 | 0.91 |
| YBL102W | 1291.754797 | -0.085473494 | 0.740554197 | 0.91 |

| | | | | |
|---|---|---|---|---|
| YBR122C | 1316.133892 | -0.04715252 | 0.73983014 | 0.91 |
| YER014W | 690.0676914 | 0.159875058 | 0.740584968 | 0.91 |
| YMR117C | 204.4706004 | 0.173221918 | 0.740556426 | 0.91 |
| YNL081C | 921.3541155 | -0.126709732 | 0.740065515 | 0.91 |
| YOL165C | 22.45477283 | 1.20422905 | 0.740069141 | 0.91 |
| YOR081C | 1945.1418 | 0.074823994 | 0.740254209 | 0.91 |
| YGR219W | 32.06784529 | -0.112068287 | 0.741135294 | 0.91 |
| YML011C | 267.8459983 | 0.077354082 | 0.741359693 | 0.91 |
| YPL009C | 4156.098331 | 0.266070001 | 0.741350139 | 0.91 |
| YPL224C | 949.2587858 | 0.043981115 | 0.741344547 | 0.91 |
| YFR010W-A | 16.18554247 | 0.220178181 | 0.741546306 | 0.91 |
| YDR022C | 413.59294 | 0.043652107 | 0.74220491 | 0.91 |
| YGL094C | 1570.488055 | 0.151017953 | 0.741921343 | 0.91 |
| YGR134W | 745.1524745 | -0.103001629 | 0.741902047 | 0.91 |
| YGR177C | 234.3513182 | -0.458821824 | 0.742290381 | 0.91 |
| YLL007C | 554.0715465 | -0.003670922 | 0.742094384 | 0.91 |
| YHL009W-B | 5.78416984 | 0.180404809 | 0.742450982 | 0.91 |
| YAR014C | 1593.739874 | -0.134404654 | 0.742863734 | 0.91 |
| YML071C | 1055.622411 | 0.099935037 | 0.742918344 | 0.91 |
| YDL200C | 293.2384509 | -0.103608009 | 0.743226677 | 0.91 |
| YGL132W | 16.53764806 | 0.267468246 | 0.743192614 | 0.91 |
| YIL130W | 1279.051362 | 0.023816968 | 0.743350296 | 0.91 |
| YFL034C-B | 921.0599209 | 0.209981521 | 0.743936888 | 0.91 |
| YIL010W | 1043.363564 | -0.05535472 | 0.743549967 | 0.91 |
| YOR266W | 469.4838324 | -0.10696019 | 0.743784387 | 0.91 |
| YOR350C | 702.3901449 | 0.073101201 | 0.743841701 | 0.91 |
| YGR049W | 549.8511767 | -0.024465887 | 0.744569303 | 0.91 |
| YKL141W | 4100.397187 | 0.392476225 | 0.744602564 | 0.91 |
| YLR278C | 1940.968689 | -0.079167986 | 0.744331045 | 0.91 |
| YKL117W | 7112.851481 | 0.122777502 | 0.744834182 | 0.91 |
| YDL010W | 362.0027109 | -0.121585736 | 0.745151577 | 0.91 |
| YBR071W | 990.0527368 | 0.321321635 | 0.745860794 | 0.91 |
| YER168C | 714.524667 | 0.08417724 | 0.745630298 | 0.91 |
| YHR001W-A | 1152.222923 | 0.329843848 | 0.745467195 | 0.91 |
| YKL096C-B | 8.469479675 | 0.116788357 | 0.745778979 | 0.91 |
| YOR060C | 313.4302218 | -0.167807181 | 0.745601369 | 0.91 |
| YDR499W | 868.5785284 | 0.073295895 | 0.74627896 | 0.91 |
| YOR336W | 1255.740018 | 0.131409402 | 0.746177393 | 0.91 |
| YKL070W | 302.0799748 | 0.079184103 | 0.74691542 | 0.91 |
| YGR163W | 840.8307527 | -0.099053782 | 0.747112173 | 0.91 |
| YFL038C | 5600.505944 | -0.068148853 | 0.748177681 | 0.91 |
| YGR222W | 726.7683442 | 0.026167503 | 0.74763984 | 0.91 |
| YKL032C | 3133.578964 | 0.178604791 | 0.748003277 | 0.91 |

| | | | | |
|---|---|---|---|---|
| YKR095W | 4316.14506 | 0.156210061 | 0.74830292 | 0.91 |
| YPR006C | 675.2062996 | 0.324152592 | 0.748056242 | 0.91 |
| YPR068C | 284.1750342 | -0.138689981 | 0.748265027 | 0.91 |
| YCL049C | 935.682406 | 0.211244146 | 0.749050006 | 0.91 |
| YDR345C | 12971.54144 | 0.333882308 | 0.749534283 | 0.91 |
| YDR543C | 0.805732287 | 0.05308792 | 0.74949406 | 0.91 |
| YJL052W | 23370.62399 | -0.064141563 | 0.748703328 | 0.91 |
| YLR216C | 5399.303942 | 0.115766317 | 0.74952769 | 0.91 |
| YMR008C | 3679.477965 | 0.252775935 | 0.748753842 | 0.91 |
| YOR112W | 1049.770792 | -0.096208569 | 0.749606029 | 0.91 |
| YOR327C | 881.0062218 | -0.210152235 | 0.748824498 | 0.91 |
| YPR121W | 333.11981 | -0.497462313 | 0.749511009 | 0.91 |
| YEL020C-B | 34.39724677 | -0.296131594 | 0.749940676 | 0.91 |
| YHR023W | 2927.24121 | 0.281455488 | 0.749875362 | 0.91 |
| YNL316C | 479.8242776 | -0.094289902 | 0.750179789 | 0.91 |
| YOL133W | 270.3290721 | -0.004862087 | 0.750117797 | 0.91 |
| YBR037C | 852.7234834 | 0.131006542 | 0.752374926 | 0.91 |
| YBR291C | 379.9577466 | -0.02750019 | 0.752336763 | 0.91 |
| YCR018C-A | 8.72999001 | -0.413063526 | 0.752199221 | 0.91 |
| YDL076C | 828.6133083 | 0.108472578 | 0.751698769 | 0.91 |
| YDR013W | 271.9851003 | -0.124276198 | 0.75215963 | 0.91 |
| YDR196C | 880.6195121 | 0.081560877 | 0.75141668 | 0.91 |
| YDR266C | 1529.276069 | -0.152733646 | 0.752236836 | 0.91 |
| YDR383C | 354.9421834 | -0.222308497 | 0.75159624 | 0.91 |
| YHR125W | 4.057342841 | 0.118211907 | 0.752578671 | 0.91 |
| YMR173W-A | 260.3952806 | -0.177090168 | 0.751865325 | 0.91 |
| YOL031C | 1089.984363 | 0.237711695 | 0.752654354 | 0.91 |
| YOR357C | 469.3315493 | -0.11504802 | 0.752282444 | 0.91 |
| YBR221W-A | 0.837044541 | -0.127693048 | 0.753082474 | 0.91 |
| YIL117C | 1107.127061 | -0.010285352 | 0.753092815 | 0.91 |
| YHR073W | 1977.66425 | -0.090140059 | 0.753438833 | 0.91 |
| YBR229C | 1792.230472 | -0.04262008 | 0.753844683 | 0.91 |
| YMR321C | 24.07796373 | 0.177412079 | 0.754060307 | 0.91 |
| YHR132C | 1760.575956 | -0.073559811 | 0.754838844 | 0.91 |
| YHR209W | 694.2958547 | 0.085744886 | 0.754492799 | 0.91 |
| YIL068C | 981.7929067 | -0.077927192 | 0.754668228 | 0.91 |
| YOR307C | 1226.108407 | 0.029227536 | 0.7547773 | 0.91 |
| YDL203C | 667.0945544 | -0.076403394 | 0.755292751 | 0.91 |
| YDR535C | 20.65811424 | 0.085071062 | 0.755435139 | 0.91 |
| YGR132C | 2816.110199 | -0.052362039 | 0.755581749 | 0.91 |
| YGL083W | 940.5046339 | -0.081995851 | 0.75580436 | 0.91 |
| YLL038C | 326.4334207 | -0.24734431 | 0.756012142 | 0.92 |
| YLR063W | 280.3060801 | 0.166315042 | 0.75636284 | 0.92 |

| | | | | |
|---|---|---|---|---|
| YBR069C | 2910.073987 | -0.042719846 | 0.756944258 | 0.92 |
| YER046W | 177.0920564 | 0.080302276 | 0.756792161 | 0.92 |
| YER170W | 376.8018239 | 0.142925006 | 0.757533157 | 0.92 |
| YHR178W | 530.4654676 | -0.092788441 | 0.75733822 | 0.92 |
| YKL203C | 2057.606082 | -0.027892051 | 0.757196515 | 0.92 |
| YLR328W | 869.5602715 | -0.159464515 | 0.757418366 | 0.92 |
| YMR029C | 1228.193843 | 0.091065507 | 0.757351933 | 0.92 |
| YNL193W | 690.484771 | -0.133685841 | 0.757041523 | 0.92 |
| YKR023W | 810.8865639 | 0.190422153 | 0.757786164 | 0.92 |
| YPR031W | 763.0479333 | 0.143461164 | 0.757857966 | 0.92 |
| YJL222W | 13.76304146 | -0.551852069 | 0.758365062 | 0.92 |
| YJR023C | 6.716930152 | -0.146383577 | 0.758271998 | 0.92 |
| YLL001W | 1443.400677 | 0.176037118 | 0.758439296 | 0.92 |
| YDL028C | 1026.396259 | 0.116047272 | 0.758782863 | 0.92 |
| YHR182C-A | 130.5541577 | 0.692482245 | 0.758929381 | 0.92 |
| YBR103W | 849.3179685 | 0.070385866 | 0.759269324 | 0.92 |
| YDR362C | 764.8048646 | -0.105071242 | 0.759886033 | 0.92 |
| YGR180C | 8240.520071 | 0.020452444 | 0.759942124 | 0.92 |
| YCR088W | 5938.374856 | -0.082102855 | 0.760889471 | 0.92 |
| YDR401W | 0.227003213 | -0.01762107 | 0.760723456 | 0.92 |
| YGL224C | 544.3672231 | 0.002779894 | 0.76059794 | 0.92 |
| YML059C | 1781.099171 | 0.030158562 | 0.760916677 | 0.92 |
| YDR423C | 1205.3164 | 0.039883882 | 0.761060611 | 0.92 |
| YER161C | 1183.000093 | -0.165797125 | 0.76154905 | 0.92 |
| YML054C-A | 11.9894805 | -0.056790913 | 0.761478671 | 0.92 |
| snR82 | 454.6100305 | 0.118184664 | 0.761470634 | 0.92 |
| YFR046C | 527.2575594 | 0.08891414 | 0.761773069 | 0.92 |
| YMR042W | 272.8014796 | -0.218111052 | 0.762002068 | 0.92 |
| YER032W | 1018.880889 | -0.066903168 | 0.76221905 | 0.92 |
| YDR482C | 359.2978323 | 0.050763996 | 0.762939341 | 0.92 |
| YER138W-A | 1.112909077 | 0.439166866 | 0.762667586 | 0.92 |
| YKR007W | 738.2939867 | -0.098345392 | 0.76293145 | 0.92 |
| YKR022C | 681.4384392 | 0.056466284 | 0.763002758 | 0.92 |
| YLL005C | 615.6111971 | 0.155281387 | 0.763502832 | 0.92 |
| YJR041C | 1150.543526 | -0.083456996 | 0.76367035 | 0.92 |
| YDR227W | 1551.753891 | 0.249048273 | 0.765045978 | 0.92 |
| YLR014C | 353.4875684 | 0.113892807 | 0.765098887 | 0.92 |
| YEL008W | 30.07335462 | -0.476420032 | 0.765682792 | 0.92 |
| YOR364W | 7.913433737 | 0.286558826 | 0.765568898 | 0.92 |
| YDL117W | 1322.092729 | 0.075602756 | 0.765957837 | 0.92 |
| YKL183C-A | 2.603283005 | -0.226756438 | 0.766388217 | 0.92 |
| YOR257W | 743.3095021 | 0.075123 | 0.766371006 | 0.92 |
| YBR233W | 474.558741 | -0.071418794 | 0.766565509 | 0.92 |

| | | | | |
|---|---|---|---|---|
| YDR005C | 1131.574288 | 0.029417816 | 0.766921171 | 0.92 |
| YGR183C | 999.4364709 | 0.261121872 | 0.767345319 | 0.92 |
| YHL006C | 116.1902464 | -0.296119887 | 0.767338377 | 0.92 |
| YOR196C | 1243.855488 | 0.112869481 | 0.767034526 | 0.92 |
| YPR010C-A | 733.8429902 | 0.458497214 | 0.767271556 | 0.92 |
| YPR023C | 1837.667458 | -0.034784148 | 0.767534264 | 0.92 |
| YNR049C | 561.0637018 | 0.091002999 | 0.76792981 | 0.92 |
| YBR260C | 1246.430976 | 0.213649145 | 0.768237541 | 0.92 |
| YPL139C | 637.4930321 | -0.101039748 | 0.768423093 | 0.92 |
| YFL046W | 596.7158239 | -0.082843099 | 0.768865923 | 0.92 |
| YML092C | 2758.585269 | -0.033531287 | 0.769123086 | 0.92 |
| YKR030W | 649.1186189 | 0.078300496 | 0.769413168 | 0.92 |
| YDR346C | 2820.625886 | 0.138098496 | 0.769568926 | 0.92 |
| YER085C | 1.090623589 | 0.220599516 | 0.770462806 | 0.92 |
| YLR396C | 522.7183585 | -0.102388018 | 0.770134486 | 0.92 |
| YOR355W | 1652.733408 | -0.257624787 | 0.770409117 | 0.92 |
| snR43 | 20.09245082 | -0.07490446 | 0.770236621 | 0.92 |
| YBR273C | 1305.311458 | 0.089753672 | 0.771596462 | 0.92 |
| YDR182W | 1361.286881 | -0.086735221 | 0.771707234 | 0.92 |
| YIL023C | 838.1130985 | 0.082353303 | 0.77123475 | 0.92 |
| YIL074C | 1391.44586 | -0.060359799 | 0.771527194 | 0.92 |
| YIR028W | 189.55325 | -0.347995841 | 0.771596205 | 0.92 |
| YLR256W-A | 92.61969338 | 0.063294129 | 0.771128776 | 0.92 |
| YNL230C | 466.9526022 | 0.106416112 | 0.771386069 | 0.92 |
| YOR386W | 897.2436472 | -0.057305646 | 0.770953017 | 0.92 |
| YGR291C | 1.566792632 | 0.067865427 | 0.773237297 | 0.92 |
| YLL033W | 166.5445731 | -0.165694427 | 0.773144656 | 0.92 |
| YDL002C | 564.8870678 | -0.126409249 | 0.774246924 | 0.92 |
| YIL121W | 2357.398488 | -0.453675955 | 0.774032485 | 0.92 |
| YMR282C | 677.7599955 | -0.129699327 | 0.774239945 | 0.92 |
| YBR110W | 1287.287964 | -0.106713717 | 0.774950902 | 0.92 |
| YJL198W | 970.0448734 | -0.046841698 | 0.775149331 | 0.92 |
| YJR144W | 1299.865299 | -0.036754986 | 0.774851254 | 0.92 |
| YLR011W | 284.831779 | -0.11442254 | 0.775143732 | 0.92 |
| YLR432W | 4140.896048 | 0.173234679 | 0.774439177 | 0.92 |
| YOL166W-A | 0.347622843 | 0.069566137 | 0.774918144 | 0.92 |
| YBR204C | 1064.479878 | 0.194470666 | 0.776084337 | 0.93 |
| YJR012C | 286.1622087 | -0.151929626 | 0.776143366 | 0.93 |
| YLR398C | 2110.607765 | -0.038204754 | 0.776342829 | 0.93 |
| YNL265C | 447.0625511 | 0.070938613 | 0.776427491 | 0.93 |
| RUF21 | 43.48350584 | 0.244073146 | 0.776756122 | 0.93 |
| YDL031W | 1680.703139 | 0.10608813 | 0.77757543 | 0.93 |
| YGL188C-A | 18.77666255 | 0.164094665 | 0.777779181 | 0.93 |

| | | | | |
|---|---|---|---|---|
| YBR039W | 6878.817001 | 0.103199888 | 0.778253756 | 0.93 |
| YJL184W | 578.763074 | 0.149441117 | 0.778399232 | 0.93 |
| YLL009C | 581.7236454 | 0.313642944 | 0.77852624 | 0.93 |
| YNL022C | 938.4563852 | -0.024672008 | 0.778240611 | 0.93 |
| YPL214C | 1002.445323 | 0.05667255 | 0.778095345 | 0.93 |
| YCL076W | 24.7210401 | 0.575547683 | 0.7788052 | 0.93 |
| YKL028W | 1511.073698 | -0.165223165 | 0.779009129 | 0.93 |
| YHL044W | 369.968591 | -0.455321482 | 0.779188751 | 0.93 |
| YAL008W | 865.7638687 | 0.040139046 | 0.780272347 | 0.93 |
| YGR142W | 4162.125638 | 0.468624071 | 0.780341266 | 0.93 |
| YLR109W | 26967.32056 | 0.315937563 | 0.780304116 | 0.93 |
| YML015C | 859.349761 | 0.118125135 | 0.779868243 | 0.93 |
| YOR229W | 494.4703852 | -0.078419043 | 0.780420932 | 0.93 |
| YOL079W | 9.64455915 | -0.063031732 | 0.780790981 | 0.93 |
| YMR258C | 733.537512 | -0.013545259 | 0.781230056 | 0.93 |
| YBR179C | 1264.938407 | 0.168187664 | 0.781830735 | 0.93 |
| YDR403W | 327.0898139 | 0.087662781 | 0.781893265 | 0.93 |
| YGL023C | 1541.23519 | 0.138288012 | 0.78156677 | 0.93 |
| YOR368W | 442.6825001 | 0.025239898 | 0.781672463 | 0.93 |
| YAL016C-B | 15.45888685 | 0.352015572 | 0.783253746 | 0.93 |
| YAR009C | 236.1235724 | -0.065013625 | 0.783299725 | 0.93 |
| YBR151W | 3044.846839 | -0.124021152 | 0.783867764 | 0.93 |
| YDL144C | 951.4980585 | -0.031006827 | 0.785637904 | 0.93 |
| YDR136C | 21.36116146 | -0.161993828 | 0.783461906 | 0.93 |
| YDR147W | 507.4257469 | -0.130545885 | 0.785077949 | 0.93 |
| YDR191W | 146.592055 | -0.233514006 | 0.783869914 | 0.93 |
| YER184C | 397.7514223 | 0.009146493 | 0.784203884 | 0.93 |
| YFR012W-A | 2.137477958 | -0.128518995 | 0.785001885 | 0.93 |
| YFR023W | 96.03552642 | -0.306604077 | 0.785181995 | 0.93 |
| YGL013C | 1490.03588 | 0.129903106 | 0.782976923 | 0.93 |
| YGL193C | 193.4216907 | -0.313905403 | 0.785231234 | 0.93 |
| YHR039C-A | 2505.228571 | -0.031829684 | 0.785809988 | 0.93 |
| YIL019W | 543.1334136 | 0.13732873 | 0.785795901 | 0.93 |
| YIL149C | 3262.267006 | 0.21938146 | 0.784970289 | 0.93 |
| YJL133W | 754.0202192 | -0.202681349 | 0.78624108 | 0.93 |
| YJL197W | 1228.673265 | -0.041361477 | 0.786304824 | 0.93 |
| YKL018C-A | 261.9017664 | 0.102954896 | 0.785777697 | 0.93 |
| YKR033C | 0.132511194 | 0.015763102 | 0.783986764 | 0.93 |
| YLR128W | 336.2576961 | -0.191209809 | 0.783068369 | 0.93 |
| YLR336C | 993.7017255 | 0.267857924 | 0.785206078 | 0.93 |
| YLR443W | 472.8026492 | 0.08260403 | 0.786275491 | 0.93 |
| YMR109W | 3069.730753 | -0.064513455 | 0.783242963 | 0.93 |
| YMR176W | 1256.155137 | 0.070989641 | 0.784254392 | 0.93 |

| | | | | |
|---|---|---|---|---|
| YNL067W-B | 0.339190409 | -0.019160574 | 0.786049814 | 0.93 |
| YNL073W | 1409.044777 | 0.027899377 | 0.78388851 | 0.93 |
| YNL333W | 85.09307581 | -0.104658352 | 0.784478803 | 0.93 |
| YOR282W | 14.73409586 | -0.361312796 | 0.785595353 | 0.93 |
| YPL191C | 278.5044925 | 0.091749981 | 0.782827222 | 0.93 |
| YDR194W-A | 0.453289942 | 0.095031852 | 0.786603916 | 0.93 |
| YHR118C | 526.5681517 | -0.05171027 | 0.786868456 | 0.93 |
| YLR147C | 302.8969457 | 0.095747674 | 0.787032124 | 0.93 |
| YPR066W | 302.1036782 | 0.126912557 | 0.787189839 | 0.93 |
| YGR136W | 2019.817506 | 0.141711918 | 0.787829071 | 0.93 |
| YGR295C | 1429.22247 | -0.098134584 | 0.788279741 | 0.93 |
| YJR092W | 2324.987782 | -0.02948625 | 0.787988918 | 0.93 |
| YJR100C | 772.6621126 | 0.038920491 | 0.788104199 | 0.93 |
| YKL106W | 462.66436 | -0.216277033 | 0.787883549 | 0.93 |
| YMR180C | 510.7680392 | -0.095890096 | 0.788291383 | 0.93 |
| YOR218C | 5.229072963 | -0.510300084 | 0.788348524 | 0.93 |
| YPL225W | 2696.542417 | -0.048404831 | 0.7879532 | 0.93 |
| YMR154C | 501.7366665 | 0.015081113 | 0.788572008 | 0.93 |
| YDL099W | 2397.811912 | 0.139035266 | 0.789945262 | 0.93 |
| YDL161W | 1895.847844 | 0.056629086 | 0.789111764 | 0.93 |
| YDR443C | 1163.053889 | 0.172533293 | 0.789538554 | 0.93 |
| YDR515W | 694.9239372 | 0.143547505 | 0.789854356 | 0.93 |
| YIL049W | 383.4643737 | -0.025248444 | 0.790060167 | 0.93 |
| YML053C | 803.1994724 | 0.112332108 | 0.789467397 | 0.93 |
| YMR297W | 8256.874804 | -0.090837106 | 0.790107529 | 0.93 |
| YPL240C | 19482.15572 | 0.521420795 | 0.789455877 | 0.93 |
| YPR105C | 1243.404818 | 0.069353466 | 0.789820171 | 0.93 |
| YBL100C | 649.5552869 | 0.525260065 | 0.790479612 | 0.93 |
| YHR067W | 339.2689826 | -0.087177879 | 0.790806695 | 0.93 |
| YNL239W | 3388.853667 | 0.196239332 | 0.79057103 | 0.93 |
| YOR125C | 859.4168859 | 0.10105972 | 0.790781033 | 0.93 |
| YER030W | 1421.194205 | -0.106191736 | 0.792313545 | 0.93 |
| YER080W | 3267.440935 | 0.16192249 | 0.791618181 | 0.93 |
| YGR137W | 236.2341776 | 0.130455744 | 0.791616307 | 0.93 |
| YGR239C | 410.924928 | -0.031495044 | 0.791743065 | 0.93 |
| YIL134C-A | 3.121789303 | 0.105986836 | 0.791913657 | 0.93 |
| YKR013W | 4305.106637 | 0.125926319 | 0.791894371 | 0.93 |
| YLL010C | 1089.845187 | 0.057448325 | 0.792111707 | 0.93 |
| YLR035C-A | 40.57446476 | 0.737656026 | 0.792339805 | 0.93 |
| YDR222W | 466.2824915 | -0.230156844 | 0.792849971 | 0.93 |
| YDR272W | 1625.208653 | 0.071539905 | 0.793289734 | 0.93 |
| YER058W | 286.3082478 | 0.181194754 | 0.793011207 | 0.93 |
| YGR091W | 441.73971 | 0.0542697 | 0.793265824 | 0.93 |

| | | | | |
|---|---|---|---|---|
| YNL089C | 16.65282477 | -0.050789816 | 0.792849022 | 0.93 |
| YOL141W | 189.2930206 | 0.061865163 | 0.792926174 | 0.93 |
| YHL048C-A | 2.76683096 | 0.375727321 | 0.793614476 | 0.93 |
| YIL115W-A | 23.70554908 | 0.129738124 | 0.793919961 | 0.93 |
| YDR163W | 579.7588355 | -0.080761093 | 0.794411933 | 0.93 |
| YFR033C | 3330.696407 | 0.315541299 | 0.794356955 | 0.93 |
| YAL044W-A | 228.0761832 | -0.132651685 | 0.795008751 | 0.93 |
| YCL011C | 2495.29652 | 0.102267022 | 0.79474452 | 0.93 |
| YDR032C | 4060.720659 | 0.081479958 | 0.795579063 | 0.93 |
| YJR074W | 487.5630441 | 0.27109004 | 0.795632796 | 0.93 |
| YKR079C | 916.529157 | -0.074910178 | 0.795609917 | 0.93 |
| YLR203C | 2343.4384 | -0.04769063 | 0.795538267 | 0.93 |
| YLR222C-A | 13.05158283 | -0.144439516 | 0.795405814 | 0.93 |
| YOR263C | 29.84403306 | 0.043798968 | 0.794818959 | 0.93 |
| YNL199C | 1154.934737 | -0.066759919 | 0.79598837 | 0.93 |
| YPL271W | 1376.293801 | 0.268822365 | 0.79598157 | 0.93 |
| YCR076C | 593.9023385 | 0.094430947 | 0.796748355 | 0.93 |
| YJR128W | 14.84064674 | 0.280414683 | 0.796964688 | 0.93 |
| YDR530C | 1226.853067 | 0.049990111 | 0.797673857 | 0.93 |
| YGR156W | 932.64115 | -0.089003463 | 0.797770593 | 0.93 |
| YNL267W | 1939.373479 | -0.068858794 | 0.798108569 | 0.93 |
| YDR148C | 4531.818706 | 0.38025243 | 0.798273549 | 0.93 |
| YAL064W | 3.131576444 | -0.508281955 | 0.801586311 | 0.93 |
| YAL066W | 0.770254238 | 0.044530056 | 0.802322379 | 0.93 |
| YBR097W | 952.5745358 | -0.070099189 | 0.800086207 | 0.93 |
| YBR119W | 605.7970902 | 0.092585394 | 0.798765677 | 0.93 |
| YBR286W | 17567.33584 | 0.105644906 | 0.798777035 | 0.93 |
| YDL107W | 383.3863117 | 0.005467302 | 0.803148813 | 0.93 |
| YER018C | 488.1395433 | 0.061265606 | 0.801466519 | 0.93 |
| YER091C-A | 0.485272411 | -0.084399539 | 0.799376892 | 0.93 |
| YER166W | 2859.051695 | 0.15940317 | 0.802334127 | 0.93 |
| YGR044C | 1106.237884 | -0.238012206 | 0.80115503 | 0.93 |
| YGR050C | 100.6325721 | -0.237003088 | 0.800936431 | 0.93 |
| YHR134W | 365.9132551 | -0.001632437 | 0.799643336 | 0.93 |
| YIR008C | 729.6399623 | 0.061756054 | 0.801556909 | 0.93 |
| YJR161C | 124.4036715 | 0.093352003 | 0.802514649 | 0.93 |
| YKL130C | 628.0827477 | -0.080662718 | 0.803172562 | 0.93 |
| YKR093W | 2668.023949 | -0.083791458 | 0.80107271 | 0.93 |
| YLL031C | 1740.005527 | -0.040356864 | 0.799727947 | 0.93 |
| YLR055C | 1018.744824 | 0.041155334 | 0.800002063 | 0.93 |
| YLR286C | 14573.14708 | -0.022445754 | 0.802746363 | 0.93 |
| YLR361C-A | 262.7999201 | 0.138141428 | 0.801965018 | 0.93 |
| YML028W | 27162.20889 | 0.046998534 | 0.802761814 | 0.93 |

| | | | | |
|---|---|---|---|---|
| YNL043C | 66.28583242 | -0.143695535 | 0.80248747 | 0.93 |
| YNL152W | 687.0558678 | -0.108308996 | 0.800745699 | 0.93 |
| YNR010W | 212.8810604 | 0.05156638 | 0.799141482 | 0.93 |
| YOL143C | 5652.711754 | -0.061303844 | 0.803106654 | 0.93 |
| YOR042W | 2668.884695 | 0.105657539 | 0.800436574 | 0.93 |
| YOR079C | 215.4299264 | -0.211709561 | 0.80048783 | 0.93 |
| YOR093C | 1072.002934 | -0.09293944 | 0.802253614 | 0.93 |
| YOR299W | 1060.38632 | -0.108838236 | 0.802396966 | 0.93 |
| YOR308C | 716.1626853 | 0.301381672 | 0.803167446 | 0.93 |
| YOR334W | 302.0782196 | 0.054701232 | 0.79929992 | 0.93 |
| YPL069C | 455.5840403 | -0.11219487 | 0.802022008 | 0.93 |
| YPR027C | 342.9171324 | -0.398758037 | 0.802109492 | 0.93 |
| YBR040W | 134.4656267 | 0.057904005 | 0.804530628 | 0.93 |
| YBR134W | 28.00305569 | -0.001218279 | 0.804446847 | 0.93 |
| YCL005W | 658.6999475 | 0.058880849 | 0.804356353 | 0.93 |
| YDL245C | 34.45281204 | 0.251256771 | 0.804143949 | 0.93 |
| YDR283C | 1675.592568 | 0.086425434 | 0.803946784 | 0.93 |
| YJR016C | 6106.441237 | 0.101601898 | 0.804061571 | 0.93 |
| YLL044W | 94.44590477 | -0.54925598 | 0.804598562 | 0.93 |
| YNL024C | 148.4866774 | 0.184147261 | 0.804559198 | 0.93 |
| YNL164C | 278.0976276 | -0.151599737 | 0.803781127 | 0.93 |
| YHR122W | 579.3180386 | 0.334648729 | 0.80514925 | 0.93 |
| YNL103W-A | 104.3621644 | 0.860887084 | 0.804930724 | 0.93 |
| YPL075W | 2226.650529 | -0.030320459 | 0.805084196 | 0.93 |
| YCR093W | 4708.644101 | -0.016908522 | 0.805651485 | 0.93 |
| YHR100C | 468.2098929 | 0.003463939 | 0.805635717 | 0.93 |
| YIR042C | 196.4981605 | -0.025895372 | 0.805434254 | 0.93 |
| YBL050W | 1823.454847 | -0.067798449 | 0.806699296 | 0.93 |
| YNL054W | 1269.220328 | -0.015487874 | 0.806690921 | 0.93 |
| YER123W | 1029.367215 | -0.131655497 | 0.807646763 | 0.93 |
| YER141W | 2454.288383 | -0.064114265 | 0.807666952 | 0.93 |
| YGR122W | 624.9209214 | -0.249150611 | 0.807720336 | 0.93 |
| YGR202C | 1624.722457 | -0.037465955 | 0.807774296 | 0.93 |
| YJR085C | 2130.783694 | -0.034258474 | 0.807837481 | 0.93 |
| YNR062C | 190.9739726 | -0.172830428 | 0.807613881 | 0.93 |
| YOR378W | 266.6242043 | 0.30235426 | 0.807163171 | 0.93 |
| YER164W | 2428.238913 | 0.107253165 | 0.808767495 | 0.93 |
| YMR052W | 219.8800281 | 0.095795778 | 0.808579427 | 0.93 |
| YOR056C | 1339.644393 | -0.049075692 | 0.808756696 | 0.93 |
| YBR172C | 2003.113678 | 0.077008023 | 0.809435107 | 0.93 |
| YDR180W | 980.8547337 | 0.183965865 | 0.809081555 | 0.93 |
| YER052C | 2274.895414 | 0.225326201 | 0.80912638 | 0.93 |
| YGL071W | 1392.981634 | 0.063510199 | 0.809491466 | 0.93 |

| | | | | |
|---|---|---|---|---|
| YMR239C | 477.7892356 | -0.004800539 | 0.809420617 | 0.93 |
| YBL005W-B | 450.3577253 | 0.257761034 | 0.809679721 | 0.93 |
| YLR235C | 18.73304548 | 0.51565128 | 0.810106011 | 0.93 |
| YMR294W | 447.832337 | 0.039053868 | 0.810123155 | 0.93 |
| YML012C-A | 4.439298629 | -0.118961595 | 0.810352419 | 0.93 |
| YNL213C | 587.1355028 | -0.147198369 | 0.810478027 | 0.93 |
| YNL124W | 1308.097286 | 0.170753542 | 0.811012919 | 0.94 |
| YNL314W | 384.9311948 | 0.081483781 | 0.811467959 | 0.94 |
| YOL062C | 1303.409819 | 0.049329649 | 0.811347546 | 0.94 |
| YBR066C | 407.0918605 | -0.364060809 | 0.811649309 | 0.94 |
| YJL219W | 39.81053365 | 0.29936919 | 0.812180506 | 0.94 |
| YPL242C | 2035.672722 | 0.118016266 | 0.812178126 | 0.94 |
| YDL046W | 2942.234491 | -0.115736518 | 0.812860734 | 0.94 |
| YGR281W | 2050.830805 | -0.079939832 | 0.813115155 | 0.94 |
| YKL118W | 29.53687639 | -0.056033195 | 0.812954069 | 0.94 |
| YPR159W | 2743.676738 | -0.06447118 | 0.813068801 | 0.94 |
| YBR281C | 961.503364 | 0.161796192 | 0.814314162 | 0.94 |
| YIL163C | 43.51788641 | -0.079255469 | 0.81387779 | 0.94 |
| YJL193W | 438.7222894 | -0.143894622 | 0.814094211 | 0.94 |
| YJR160C | 11.90922489 | 0.081119305 | 0.814507037 | 0.94 |
| YLL066C | 1.397171575 | 0.374808574 | 0.814547327 | 0.94 |
| YLR146W-A | 192.5995673 | 0.177759965 | 0.814387297 | 0.94 |
| YNR024W | 280.6447987 | -0.042996944 | 0.814799207 | 0.94 |
| YPR145W | 8044.845508 | 0.27435184 | 0.814944042 | 0.94 |
| YPR169W-A | 32.89680766 | 0.096337867 | 0.815238711 | 0.94 |
| HRA1 | 34.22771229 | 0.462679708 | 0.816073286 | 0.94 |
| YCL048W | 92.73705911 | -0.102100663 | 0.816138101 | 0.94 |
| YDL006W | 361.6658456 | -0.115665791 | 0.815840371 | 0.94 |
| YGR265W | 63.64327138 | 0.094591506 | 0.816066761 | 0.94 |
| YJL059W | 446.5285115 | 0.051934905 | 0.816183755 | 0.94 |
| YLR461W | 7.039838185 | -0.017493929 | 0.815821523 | 0.94 |
| YBR250W | 80.81966717 | -0.214901049 | 0.817605084 | 0.94 |
| YDL069C | 126.889784 | 0.085750758 | 0.817840782 | 0.94 |
| YDL241W | 250.4921648 | 0.005970154 | 0.817257832 | 0.94 |
| YDR508C | 2566.963439 | 0.018895522 | 0.816759931 | 0.94 |
| YER038W-A | 0.198141795 | -0.067749313 | 0.817536428 | 0.94 |
| YHR180W | 212.5255364 | -0.247163475 | 0.81706417 | 0.94 |
| YIL115C | 2357.200299 | -0.035703584 | 0.817088847 | 0.94 |
| YKL110C | 846.8769824 | -0.085337312 | 0.817703 | 0.94 |
| YNL052W | 2172.615322 | 0.1425729 | 0.817213366 | 0.94 |
| YOR259C | 3766.746436 | 0.084477925 | 0.817285664 | 0.94 |
| YPR161C | 1370.124428 | 0.106911525 | 0.817988089 | 0.94 |
| YKL114C | 901.3872837 | -0.119349327 | 0.818326707 | 0.94 |

| | | | | |
|---|---|---|---|---|
| YAL026C | 1397.824394 | -0.03651084 | 0.818843242 | 0.94 |
| YBR197C | 458.4082334 | -0.108848738 | 0.819481818 | 0.94 |
| YDL007W | 5356.140616 | 0.139357438 | 0.819637554 | 0.94 |
| YDR375C | 907.9536977 | -0.05155348 | 0.82084157 | 0.94 |
| YFR055W | 975.2716539 | 0.164584025 | 0.821119186 | 0.94 |
| YGL226W | 346.3222245 | -0.03138232 | 0.819477472 | 0.94 |
| YIL014W | 862.4746572 | 0.028110256 | 0.820500122 | 0.94 |
| YJL024C | 668.6768051 | -0.084774575 | 0.820733356 | 0.94 |
| YJL065C | 199.2698224 | -0.091732502 | 0.818725297 | 0.94 |
| YJR112W-A | 356.5088081 | 0.049784515 | 0.821713742 | 0.94 |
| YLR253W | 1177.596684 | -0.09089854 | 0.820821809 | 0.94 |
| YLR451W | 1124.900564 | 0.029403976 | 0.821465554 | 0.94 |
| YML027W | 523.9520569 | -0.102851417 | 0.821699281 | 0.94 |
| YML133C | 9.588165535 | -0.119776496 | 0.820330193 | 0.94 |
| YMR023C | 334.2712538 | -0.092562538 | 0.820256084 | 0.94 |
| YMR167W | 578.0579077 | -0.119187473 | 0.820694693 | 0.94 |
| YMR313C | 988.4819991 | 0.109320462 | 0.821032673 | 0.94 |
| YNL162W-A | 54.12294867 | 0.076387621 | 0.819027906 | 0.94 |
| YOL140W | 502.3006349 | -0.073500918 | 0.821393648 | 0.94 |
| YOR230W | 8762.727513 | 0.214034686 | 0.8216911 | 0.94 |
| YOR250C | 655.6893916 | -0.052988982 | 0.820627824 | 0.94 |
| YPR201W | 177.3808268 | -0.276913646 | 0.819765216 | 0.94 |
| snR17a | 59.62016603 | 0.542977453 | 0.820901924 | 0.94 |
| YIL037C | 352.6358083 | -0.161671172 | 0.822292305 | 0.94 |
| YLL006W-A | 2.814822079 | -0.008362137 | 0.822294348 | 0.94 |
| YOR072W-B | 0.082611248 | 0.01008074 | 0.822431701 | 0.94 |
| YOR371C | 940.0880721 | 0.119446893 | 0.82218125 | 0.94 |
| YBR125C | 746.1231571 | 0.116444829 | 0.826033435 | 0.94 |
| YBR135W | 639.2382675 | 0.128018999 | 0.823716409 | 0.94 |
| YCR037C | 1044.612368 | -0.090798406 | 0.826723925 | 0.94 |
| YDL096C | 9.504570723 | -0.298597295 | 0.825113752 | 0.94 |
| YDR081C | 1262.178022 | 0.040033369 | 0.823311445 | 0.94 |
| YDR103W | 991.2634766 | -0.363819009 | 0.824191806 | 0.94 |
| YDR106W | 46.31624816 | -0.344674637 | 0.823266941 | 0.94 |
| YDR322C-A | 892.3853165 | 0.141556777 | 0.82507089 | 0.94 |
| YDR487C | 2644.625402 | -0.005624756 | 0.82489269 | 0.94 |
| YEL050W-A | 71.35641565 | -0.037273929 | 0.825833247 | 0.94 |
| YGL191W | 2937.202908 | 0.315705242 | 0.826678642 | 0.94 |
| YHR011W | 509.988716 | 0.201370321 | 0.82620133 | 0.94 |
| YIL137C | 2189.193159 | 0.026571245 | 0.825409377 | 0.94 |
| YKL039W | 2424.281879 | -0.051211954 | 0.825052179 | 0.94 |
| YKL120W | 1575.003705 | 0.307582957 | 0.825512369 | 0.94 |
| YKL139W | 677.7577592 | 0.025602187 | 0.824988435 | 0.94 |

| | | | | |
|---|---|---|---|---|
| YML051W | 1749.382621 | -0.10077906 | 0.826529311 | 0.94 |
| YMR060C | 493.2206546 | 0.063573957 | 0.825744792 | 0.94 |
| YNL065W | 531.9531231 | -0.147522699 | 0.823581929 | 0.94 |
| YNL097C-B | 13.11125706 | -0.154030251 | 0.825635857 | 0.94 |
| YNL136W | 1295.54364 | 0.13992256 | 0.825543557 | 0.94 |
| YNL221C | 1802.455751 | -0.054884385 | 0.82446797 | 0.94 |
| YNR036C | 1583.864307 | -0.087298151 | 0.826273037 | 0.94 |
| YOL006C | 1492.750425 | 0.001298568 | 0.826247114 | 0.94 |
| YOL114C | 133.5426332 | -0.028457221 | 0.824708119 | 0.94 |
| YOR329C | 1982.861977 | 0.120366901 | 0.826387688 | 0.94 |
| YPL020C | 1208.520388 | 0.062541118 | 0.824727902 | 0.94 |
| YPL257W | 181.2677801 | 0.113823568 | 0.824482123 | 0.94 |
| YAL021C | 1595.906288 | 0.065363594 | 0.827026029 | 0.94 |
| YGR278W | 442.5587334 | -0.095609378 | 0.827382125 | 0.94 |
| YCR028C | 527.4192161 | 0.038710416 | 0.827802377 | 0.94 |
| YHR114W | 1802.212821 | -0.044792846 | 0.828034141 | 0.94 |
| YJL046W | 632.8219176 | -0.070076355 | 0.82802583 | 0.94 |
| YML129C | 536.8329539 | 0.115375969 | 0.828095171 | 0.94 |
| YDR494W | 1564.77137 | -0.039812619 | 0.828460991 | 0.94 |
| YPL125W | 1168.744838 | 0.10538946 | 0.828460726 | 0.94 |
| YBR085W | 162.0814879 | 0.259954878 | 0.828733411 | 0.94 |
| YDR309C | 2613.4939 | -0.361272856 | 0.828809055 | 0.94 |
| YDL072C | 3778.461007 | 0.084875382 | 0.829424945 | 0.94 |
| YDL197C | 997.8504474 | 0.145999349 | 0.829397673 | 0.94 |
| YBR057C | 866.0815496 | 0.018279057 | 0.830137756 | 0.94 |
| YBR293W | 445.7090525 | -0.530869631 | 0.830131867 | 0.94 |
| YHL042W | 88.3151764 | -0.232474308 | 0.829906921 | 0.94 |
| YIL062C | 1798.738473 | 0.086468377 | 0.829734798 | 0.94 |
| YJR140C | 1086.888351 | -0.004192333 | 0.830225506 | 0.94 |
| YMR179W | 627.9998767 | -0.160000135 | 0.83073982 | 0.94 |
| YBR224W | 27.9935508 | 0.179409602 | 0.831265673 | 0.94 |
| YMR098C | 840.7673515 | -0.121054248 | 0.831269621 | 0.94 |
| YPR191W | 6094.519828 | 0.383823564 | 0.831087961 | 0.94 |
| YGR042W | 764.6829972 | 0.005068483 | 0.83154116 | 0.94 |
| YIL116W | 2678.75766 | 0.074537081 | 0.832021441 | 0.94 |
| YJR132W | 1306.62171 | -0.057906575 | 0.831998991 | 0.94 |
| YOR387C | 76.98828773 | -0.240258138 | 0.832437186 | 0.94 |
| YGL007C-A | 2.085025417 | -0.237635557 | 0.832815705 | 0.94 |
| YGR138C | 1416.124697 | 0.063592349 | 0.833047201 | 0.94 |
| YGL164C | 861.3475827 | 0.010386794 | 0.833458978 | 0.94 |
| YLR275W | 260.6216284 | 0.104477289 | 0.833504951 | 0.94 |
| YDR290W | 8.55083412 | -0.04905344 | 0.834202376 | 0.94 |
| YIL173W | 14.66085042 | -0.450648778 | 0.834355745 | 0.94 |

| | | | | |
|---|---|---|---|---|
| YNL120C | 0.227028385 | -0.015986396 | 0.834256846 | 0.94 |
| YCR017C | 2986.294618 | 0.040574806 | 0.836027756 | 0.94 |
| YCR097W | 1013.079937 | 0.376378053 | 0.836121058 | 0.94 |
| YDR035W | 3632.420651 | 0.192652513 | 0.836363239 | 0.94 |
| YDR088C | 645.3535973 | 0.031628984 | 0.835082177 | 0.94 |
| YDR193W | 12.98139175 | -0.166847485 | 0.834760763 | 0.94 |
| YFL021C-A | 41.5780796 | -0.089401793 | 0.8363373 | 0.94 |
| YFR034W-A | 2.98827912 | 0.184188716 | 0.836380729 | 0.94 |
| YGL051W | 80.44841429 | -0.146796888 | 0.835766433 | 0.94 |
| YGL064C | 383.7385393 | 0.128433658 | 0.83590994 | 0.94 |
| YML081W | 1706.956949 | -0.063513685 | 0.836302906 | 0.94 |
| YOR181W | 1611.589546 | 0.142826014 | 0.834933936 | 0.94 |
| YPL135W | 1241.708085 | 0.15467974 | 0.83487352 | 0.94 |
| YPR134W | 542.0304792 | 0.028087322 | 0.836309643 | 0.94 |
| YDR510C-A | 439.1253795 | -0.273529174 | 0.836757793 | 0.94 |
| YIL156W | 1109.423201 | 0.127104679 | 0.836720589 | 0.94 |
| YGL251C | 608.1790507 | 0.022286843 | 0.837326479 | 0.94 |
| YOR117W | 3352.309698 | 0.087137925 | 0.83807731 | 0.94 |
| YCL047C | 434.4156263 | 0.031293969 | 0.839053841 | 0.94 |
| YDR464W | 2373.668914 | 0.133431785 | 0.839191139 | 0.94 |
| YMR306C-A | 3.882802162 | 0.561633065 | 0.839002387 | 0.94 |
| YGR199W | 855.4590705 | 0.019280548 | 0.839510284 | 0.94 |
| YML079W | 1250.560699 | -0.092811095 | 0.839761944 | 0.94 |
| YDR393W | 653.7265915 | 0.053381397 | 0.840019685 | 0.94 |
| YPR086W | 1622.357285 | 0.024286721 | 0.840588958 | 0.94 |
| YDL131W | 1846.278027 | 0.078993026 | 0.840892919 | 0.94 |
| YIL090W | 898.4488003 | 0.054249868 | 0.841008673 | 0.94 |
| YPL104W | 889.4598617 | 0.124332927 | 0.841089789 | 0.94 |
| YCR095C | 483.8306058 | -0.104449844 | 0.841367215 | 0.95 |
| YKL013C | 2002.473422 | -0.029538983 | 0.841895269 | 0.95 |
| YNL279W | 510.6368292 | -0.217498787 | 0.842009718 | 0.95 |
| YIL170W | 172.5743318 | -0.173898324 | 0.843172197 | 0.95 |
| snR52 | 0.834128967 | -0.031167623 | 0.843392231 | 0.95 |
| YBR175W | 264.1838842 | 0.07437727 | 0.844422797 | 0.95 |
| YGR209C | 5181.82162 | -0.010571568 | 0.84436853 | 0.95 |
| YOR082C | 31.57473045 | -0.027046341 | 0.844413637 | 0.95 |
| YBR131W | 285.0666359 | 0.162787296 | 0.844985834 | 0.95 |
| YDL177C | 218.4776378 | 0.023525888 | 0.845817668 | 0.95 |
| YDR179C | 161.4536724 | -0.00490848 | 0.845809313 | 0.95 |
| YLR419W | 1247.341194 | -0.009922936 | 0.845752975 | 0.95 |
| YMR287C | 795.4088763 | 0.057210352 | 0.845790696 | 0.95 |
| YPR087W | 20.87255534 | -0.229996077 | 0.845950886 | 0.95 |
| YDR415C | 529.3228874 | -0.098919037 | 0.846548476 | 0.95 |

| | | | | |
|---|---|---|---|---|
| YLR444C | 25.68564702 | -0.367002553 | 0.846313376 | 0.95 |
| YOR343W-B | 107.9180972 | -0.330469959 | 0.846208666 | 0.95 |
| YPL013C | 664.0145708 | 0.057305258 | 0.84647334 | 0.95 |
| YML047C | 45.4588878 | -0.207152772 | 0.846715724 | 0.95 |
| YBL108W | 0.561379936 | -0.062881697 | 0.847693318 | 0.95 |
| YFR040W | 3140.971961 | 0.19599397 | 0.84748501 | 0.95 |
| YLR295C | 2976.605613 | 0.063064432 | 0.847599081 | 0.95 |
| YNL256W | 1084.449905 | 0.04702629 | 0.847386921 | 0.95 |
| YAR070C | 0.925768202 | -0.004338267 | 0.847968703 | 0.95 |
| YBR064W | 3.158811985 | -0.588484137 | 0.848048269 | 0.95 |
| YBL007C | 3937.826759 | 0.084166825 | 0.84834849 | 0.95 |
| YER140W | 473.4616837 | 0.005327651 | 0.848220343 | 0.95 |
| YJR079W | 11.62148612 | 0.788604581 | 0.848689605 | 0.95 |
| YDR079C-A | 102.409036 | 0.016164205 | 0.849308465 | 0.95 |
| YBL008W | 564.4162656 | 0.124535875 | 0.850571813 | 0.95 |
| YBR168W | 396.8925406 | -0.077984998 | 0.850340196 | 0.95 |
| YCR104W | 19.10661275 | 0.13198512 | 0.850975655 | 0.95 |
| YDL173W | 1092.267621 | 0.235179107 | 0.850081819 | 0.95 |
| YDR320C | 1271.658982 | -0.041396631 | 0.850077621 | 0.95 |
| YDR366C | 7.819864358 | -0.145055907 | 0.851033686 | 0.95 |
| YER100W | 1553.364761 | -0.028308873 | 0.850383576 | 0.95 |
| YHR032W-A | 60.96537533 | -0.114541791 | 0.850792542 | 0.95 |
| YLL043W | 1838.000712 | -0.066025833 | 0.850548197 | 0.95 |
| YNL235C | 7.928094321 | 0.033537322 | 0.850837282 | 0.95 |
| YDR232W | 2313.644723 | 0.002505627 | 0.851302103 | 0.95 |
| YER093C-A | 366.8172657 | 0.215775364 | 0.851569152 | 0.95 |
| YOR288C | 898.6918954 | -0.076156063 | 0.851434928 | 0.95 |
| YAL068W-A | 0.241654238 | 0.043849207 | 0.851913941 | 0.95 |
| YNR011C | 902.2961481 | 0.114864072 | 0.852107146 | 0.95 |
| YIL124W | 2615.425395 | -0.088368614 | 0.852267898 | 0.95 |
| YLR294C | 1804.172401 | 0.079739176 | 0.852482128 | 0.95 |
| YPR014C | 81.20498688 | -0.019681896 | 0.852591235 | 0.95 |
| YBR176W | 422.8364361 | 0.079916153 | 0.852782755 | 0.95 |
| YCL057W | 3573.183746 | 0.186101212 | 0.853558184 | 0.95 |
| YDL024C | 228.1924469 | -0.131220466 | 0.853969423 | 0.95 |
| YDL041W | 0.97089065 | 0.267792189 | 0.854072558 | 0.95 |
| YGR203W | 222.1887092 | -0.181073839 | 0.853939424 | 0.95 |
| YJL095W | 1317.430087 | 0.120650039 | 0.853481815 | 0.95 |
| YJL204C | 683.8445536 | -0.007003852 | 0.853378218 | 0.95 |
| YJR020W | 8.221696255 | 0.555967474 | 0.853272969 | 0.95 |
| YML070W | 2763.480937 | 0.07559143 | 0.853999719 | 0.95 |
| YOR172W | 824.7713828 | 0.067360969 | 0.853163638 | 0.95 |
| YBL053W | 19.02715806 | -0.112292127 | 0.854404535 | 0.95 |

| | | | | |
|---|---|---|---|---|
| YDL012C | 999.0124699 | 0.116996905 | 0.854599691 | 0.95 |
| YDR260C | 314.9401976 | 0.019424005 | 0.855044315 | 0.95 |
| YDR534C | 224.6689044 | -0.257233113 | 0.855260791 | 0.95 |
| YGR228W | 14.43502374 | 0.192217929 | 0.854719321 | 0.95 |
| YHL048W | 1666.986201 | -0.067560174 | 0.855190412 | 0.95 |
| YLR130C | 1253.100793 | -0.158463426 | 0.854706213 | 0.95 |
| YMR039C | 1898.602952 | 0.094343494 | 0.855294299 | 0.95 |
| YLL035W | 581.4295973 | 0.013416441 | 0.855785666 | 0.95 |
| YMR089C | 2367.558674 | 0.214291896 | 0.855890875 | 0.95 |
| YPL176C | 1288.392675 | 0.086079173 | 0.855691432 | 0.95 |
| YHR204W | 1257.868137 | 0.094668891 | 0.856216176 | 0.95 |
| YDR122W | 2792.779664 | 0.153545946 | 0.856400695 | 0.95 |
| YJL128C | 1506.583064 | -0.040202423 | 0.856545773 | 0.95 |
| YLR274W | 1064.782127 | -0.090429532 | 0.856943078 | 0.95 |
| YBR109W-A | 20.49364363 | 0.064793733 | 0.857341856 | 0.95 |
| YDR034C-D | 94.28631679 | 0.018769778 | 0.857226825 | 0.95 |
| YDL172C | 10.58915184 | -0.020564455 | 0.85772975 | 0.95 |
| YDR026C | 840.5603663 | -0.062151077 | 0.857911716 | 0.95 |
| YPL253C | 481.3229549 | -0.064112485 | 0.857935777 | 0.95 |
| YCR107W | 236.9080811 | -0.055399254 | 0.858535858 | 0.95 |
| YBL096C | 11.76381799 | -0.259140646 | 0.859004212 | 0.95 |
| YHR142W | 1336.189865 | -0.024853411 | 0.858929144 | 0.95 |
| YIL119C | 838.0211886 | -0.262245552 | 0.858802083 | 0.95 |
| YOR016C | 992.9033585 | -0.007505528 | 0.859750293 | 0.95 |
| YLR024C | 1454.81679 | 0.184341572 | 0.860367974 | 0.95 |
| YJL225C | 0.280381926 | 0.019010508 | 0.860908047 | 0.95 |
| YCR008W | 1187.11323 | 0.098263985 | 0.86218055 | 0.95 |
| YDR217C | 884.5738376 | 0.115512757 | 0.862899167 | 0.95 |
| YGL169W | 682.8701947 | 0.089868525 | 0.86285282 | 0.95 |
| YGL247W | 197.7711302 | 0.046843858 | 0.861361667 | 0.95 |
| YGR283C | 529.6205161 | 0.057147686 | 0.861464942 | 0.95 |
| YIL063C | 1081.563362 | 0.031998437 | 0.86286606 | 0.95 |
| YJL055W | 2135.334979 | 0.066352591 | 0.862053231 | 0.95 |
| YKL222C | 519.5767688 | -0.161330896 | 0.862646321 | 0.95 |
| YLR421C | 1264.683888 | 0.036272003 | 0.861153361 | 0.95 |
| YOR034C | 667.8994742 | -0.090400974 | 0.86214587 | 0.95 |
| YPL153C | 363.2997459 | 0.003866115 | 0.862203785 | 0.95 |
| snR32 | 25.20381189 | 0.29802084 | 0.862651218 | 0.95 |
| snR39 | 11.68210812 | -0.016952854 | 0.862349944 | 0.95 |
| YGR079W | 359.5947801 | 0.107635767 | 0.863071431 | 0.95 |
| YPL016W | 1163.813997 | 0.057964259 | 0.863340279 | 0.95 |
| YBR173C | 2339.401983 | 0.130299429 | 0.863682047 | 0.95 |
| YLR223C | 2252.47739 | 0.155643665 | 0.863831185 | 0.95 |

| | | | | |
|---|---|---|---|---|
| YHR124W | 347.8549107 | -0.250794029 | 0.864017449 | 0.95 |
| YDR155C | 17689.29075 | 0.103849292 | 0.864251798 | 0.95 |
| YCR105W | 210.4429239 | 0.093356197 | 0.864589385 | 0.95 |
| YLR410W-B | 66.3723325 | -0.153064169 | 0.864524374 | 0.95 |
| YLR074C | 1095.319455 | 0.173613374 | 0.865369074 | 0.95 |
| YDL104C | 329.3472486 | 0.103595104 | 0.866786502 | 0.95 |
| YDR271C | 28.54979964 | 0.391416071 | 0.866239934 | 0.95 |
| YDR427W | 2923.459081 | 0.018141881 | 0.867316581 | 0.95 |
| YIL125W | 5872.779859 | 0.461638227 | 0.867114847 | 0.95 |
| YLL067C | 0.999643856 | -0.151370824 | 0.867271378 | 0.95 |
| YLR169W | 310.0591441 | 0.03066449 | 0.866362914 | 0.95 |
| YLR190W | 1584.336567 | 0.148963376 | 0.866844227 | 0.95 |
| YLR309C | 2329.969598 | 0.230338954 | 0.865909588 | 0.95 |
| YLR376C | 277.9237935 | 0.048440813 | 0.867302213 | 0.95 |
| YNL046W | 536.2842104 | 0.009046336 | 0.866844148 | 0.95 |
| YPR169W | 1659.207747 | -0.04231406 | 0.865807037 | 0.95 |
| snR59 | 2.356669998 | 0.218092311 | 0.866872784 | 0.95 |
| YFR013W | 1397.161166 | 0.034742389 | 0.868357594 | 0.96 |
| YHR086W-A | 3.107825184 | -0.077644852 | 0.86831781 | 0.96 |
| YCL055W | 639.3184492 | -0.26274308 | 0.868508356 | 0.96 |
| YGR035C | 329.4527926 | -0.237305747 | 0.868765824 | 0.96 |
| YDL058W | 1998.717774 | 0.133409229 | 0.871442116 | 0.96 |
| YDL098C | 246.2120054 | -0.110497636 | 0.871879107 | 0.96 |
| YDR143C | 1247.877225 | 0.015427156 | 0.872551757 | 0.96 |
| YDR497C | 3420.957422 | -0.000470096 | 0.871438335 | 0.96 |
| YDR536W | 315.8429448 | 0.028597933 | 0.869781996 | 0.96 |
| YEL065W | 2278.709245 | -0.278048916 | 0.872035666 | 0.96 |
| YER019W | 1037.358398 | 0.070859846 | 0.87140705 | 0.96 |
| YER071C | 272.1086234 | -0.108271843 | 0.869041528 | 0.96 |
| YFR024C-A | 2497.30533 | 0.082519014 | 0.870501032 | 0.96 |
| YHR111W | 861.1772037 | -0.025013305 | 0.872289288 | 0.96 |
| YIL138C | 1495.862581 | -0.05107838 | 0.871217538 | 0.96 |
| YIR038C | 2847.18512 | -0.025534188 | 0.871997784 | 0.96 |
| YJL160C | 84.43696653 | 0.125682779 | 0.869302901 | 0.96 |
| YKL216W | 7973.975155 | -0.026067933 | 0.871785042 | 0.96 |
| YKL225W | 0.626369128 | 0.023924976 | 0.871710963 | 0.96 |
| YMR027W | 2670.4492 | -0.00527625 | 0.871461695 | 0.96 |
| YNL310C | 605.4678235 | -0.011191432 | 0.872474853 | 0.96 |
| YOL112W | 963.5678133 | -0.053458712 | 0.8714861 | 0.96 |
| YOR087W | 1019.099344 | -0.114866291 | 0.871249021 | 0.96 |
| YOR213C | 459.655666 | -0.065767145 | 0.870943305 | 0.96 |
| YPL045W | 887.7248714 | 0.049843008 | 0.869973113 | 0.96 |
| YPL114W | 1.969765016 | -0.251066133 | 0.870364241 | 0.96 |

| | | | | |
|---|---|---|---|---|
| YPR071W | 302.2135475 | -0.001010788 | 0.872247768 | 0.96 |
| YPR099C | 24.60634511 | -0.431582074 | 0.870550976 | 0.96 |
| tR(ACG)J | 0.2026169 | -0.069184576 | 0.8719556 | 0.96 |
| YPR188C | 534.7061592 | -0.042468615 | 0.872715569 | 0.96 |
| YBR237W | 937.0201345 | -0.001742542 | 0.873402187 | 0.96 |
| YDL126C | 11516.69474 | 0.046002822 | 0.873287842 | 0.96 |
| YGL122C | 2999.105342 | 0.103448166 | 0.873062386 | 0.96 |
| YGL174W | 430.9199773 | -0.019875893 | 0.87359102 | 0.96 |
| YLR137W | 268.6657732 | -0.238563044 | 0.873494439 | 0.96 |
| YBL025W | 212.6114748 | -0.00498445 | 0.874823086 | 0.96 |
| YBR149W | 7039.888751 | 0.055375727 | 0.874447299 | 0.96 |
| YDR099W | 8428.443895 | 0.067880587 | 0.874359404 | 0.96 |
| YDR246W | 916.4165129 | 0.038470223 | 0.874952176 | 0.96 |
| YKL204W | 2110.527656 | -0.041788084 | 0.873980242 | 0.96 |
| YLR087C | 2042.517235 | 0.109247 | 0.874706448 | 0.96 |
| YNL042W-B | 14.4556208 | 0.211743346 | 0.874600685 | 0.96 |
| YNR012W | 597.0846222 | 0.083627836 | 0.874920999 | 0.96 |
| YNR070W | 728.5997471 | 0.061582339 | 0.875098272 | 0.96 |
| YPR096C | 16.87382422 | 0.045458742 | 0.875609881 | 0.96 |
| YDR177W | 1727.041157 | 0.127853919 | 0.876180378 | 0.96 |
| YDR175C | 1042.356621 | -0.034414984 | 0.876588056 | 0.96 |
| YJL113W | 119.706187 | 0.148306381 | 0.876661083 | 0.96 |
| YNL338W | 0.154202725 | 0.012252902 | 0.876839072 | 0.96 |
| YPL268W | 929.1963896 | -0.043063111 | 0.877257548 | 0.96 |
| YER103W | 3269.463849 | 0.22433602 | 0.877698527 | 0.96 |
| YLR307W | 82.37745124 | -0.591223326 | 0.877875837 | 0.96 |
| YAR003W | 588.5199738 | -0.131882566 | 0.879791608 | 0.96 |
| YER057C | 2325.694126 | 0.078250038 | 0.879954584 | 0.96 |
| YOR290C | 3907.230484 | 0.082991009 | 0.879994562 | 0.96 |
| YPL058C | 2629.343095 | -0.128773499 | 0.879571324 | 0.96 |
| YER135C | 23.51565939 | 0.297465935 | 0.880406669 | 0.96 |
| YKL185W | 2577.028628 | 0.173051993 | 0.880554002 | 0.96 |
| YCL014W | 1364.401777 | 0.030644497 | 0.88298598 | 0.96 |
| YCL030C | 8176.65539 | 0.231501692 | 0.882381067 | 0.96 |
| YDL001W | 483.443293 | 0.026465349 | 0.882438277 | 0.96 |
| YDL210W | 142.1653441 | -0.072965456 | 0.882904752 | 0.96 |
| YER132C | 1149.8348 | 0.079578337 | 0.882923707 | 0.96 |
| YFL032W | 146.5638958 | -0.214185749 | 0.881842238 | 0.96 |
| YJR159W | 6.187748904 | 0.146300638 | 0.882047768 | 0.96 |
| YLR016C | 414.1257784 | 0.022154825 | 0.881555376 | 0.96 |
| YLR095C | 1759.144757 | 0.071686538 | 0.881241707 | 0.96 |
| YLR193C | 745.5545212 | 0.094336921 | 0.881233853 | 0.96 |
| YMR102C | 1653.341009 | 0.056178805 | 0.881049524 | 0.96 |

| | | | | |
|---|---|---|---|---|
| YMR264W | 1111.60862 | -0.013659496 | 0.881629279 | 0.96 |
| YNL032W | 670.5607595 | 0.007443096 | 0.881717183 | 0.96 |
| YNL033W | 5.453513622 | 0.285131674 | 0.881815934 | 0.96 |
| YNL146C-A | 29.52790111 | -0.783910496 | 0.882614833 | 0.96 |
| YOR258W | 218.0314211 | -0.097439859 | 0.882290506 | 0.96 |
| YMR281W | 530.4325943 | -0.110107998 | 0.883446813 | 0.96 |
| YNR050C | 3301.317883 | 0.274596372 | 0.883497288 | 0.96 |
| YAL018C | 88.41004538 | -0.36708188 | 0.884469534 | 0.96 |
| YBL063W | 882.9945164 | 0.04103314 | 0.884551276 | 0.96 |
| YBR278W | 245.0159717 | 0.047292212 | 0.884879551 | 0.96 |
| YGR205W | 695.7192252 | -0.120220697 | 0.884808373 | 0.96 |
| YMR073C | 579.5582083 | -0.049230328 | 0.884554122 | 0.96 |
| YMR324C | 2.573506675 | 0.327007285 | 0.884144932 | 0.96 |
| YNL203C | 26.22039237 | 0.031703889 | 0.884957089 | 0.96 |
| YOR392W | 4.708188968 | -0.114672069 | 0.884750798 | 0.96 |
| YPL137C | 1258.265301 | 0.044985544 | 0.885041047 | 0.96 |
| YBL052C | 898.9203912 | 0.034025305 | 0.885688888 | 0.96 |
| YCR067C | 1807.391426 | 0.0214162 | 0.886377671 | 0.96 |
| YDR269C | 1.063098833 | -0.03556043 | 0.885945954 | 0.96 |
| YKL157W | 4702.190162 | 0.224500921 | 0.886230923 | 0.96 |
| YKR042W | 18766.32248 | -0.231124716 | 0.886157133 | 0.96 |
| YLR395C | 1418.063046 | 0.386842793 | 0.886182689 | 0.96 |
| YHR028C | 2661.078301 | 0.025154572 | 0.886667328 | 0.96 |
| YFR044C | 5702.058987 | 0.08004323 | 0.887437954 | 0.96 |
| YGR080W | 1465.883709 | -0.039459628 | 0.887535743 | 0.96 |
| YHR132W-A | 721.2397547 | 0.069492868 | 0.888497426 | 0.96 |
| YAL024C | 799.2694521 | -0.037990711 | 0.889909487 | 0.96 |
| YCR033W | 2347.054478 | 0.059330206 | 0.889296722 | 0.96 |
| YER060W | 750.1023347 | -0.067685953 | 0.889805976 | 0.96 |
| YGL038C | 1572.160144 | -0.038485793 | 0.889745088 | 0.96 |
| YGR173W | 798.4212433 | 0.050837888 | 0.88877463 | 0.96 |
| YHR173C | 88.05682976 | 0.041799205 | 0.890042625 | 0.96 |
| YLR138W | 1645.971988 | 0.136497577 | 0.889411464 | 0.96 |
| YLR283W | 409.1057668 | 0.036422975 | 0.889960145 | 0.96 |
| YML091C | 3874.618724 | 0.462735349 | 0.889417955 | 0.96 |
| YMR067C | 1220.387108 | 0.05625345 | 0.889318888 | 0.96 |
| YML065W | 967.920824 | -0.056621371 | 0.890203603 | 0.96 |
| YAL030W | 729.4057845 | 0.075735121 | 0.891538856 | 0.96 |
| YGL248W | 1428.88377 | -0.14534636 | 0.891580067 | 0.96 |
| YNL051W | 867.7540206 | 0.042149705 | 0.891438472 | 0.96 |
| YNL275W | 352.1195966 | -0.001107829 | 0.891132986 | 0.96 |
| YPL008W | 976.9490792 | -0.055930538 | 0.891261935 | 0.96 |
| YPL065W | 569.9929052 | 0.01194765 | 0.890907799 | 0.96 |

| | | | | |
|---|---|---|---|---|
| YML013W | 1108.615798 | 0.098547397 | 0.891937433 | 0.96 |
| YFL016C | 3333.34381 | 0.190253788 | 0.892458149 | 0.96 |
| YEL007W | 3320.15412 | -0.120524817 | 0.893051313 | 0.96 |
| YER059W | 910.2356733 | -0.02383643 | 0.893566915 | 0.96 |
| YGR056W | 1165.037029 | -0.044912726 | 0.892838792 | 0.96 |
| YGR109C | 138.0694558 | 0.125540662 | 0.893544527 | 0.96 |
| YJR131W | 581.4205166 | 0.04285627 | 0.893323789 | 0.96 |
| YOL158C | 1769.752458 | -0.133836248 | 0.89350922 | 0.96 |
| YFR042W | 519.19235 | 0.164233088 | 0.894223887 | 0.96 |
| YKL138C-A | 204.185803 | -0.022494179 | 0.894136523 | 0.96 |
| snR72 | 8.104041704 | 0.637972293 | 0.89436209 | 0.96 |
| YDR291W | 877.4001219 | -0.04806728 | 0.894757194 | 0.96 |
| YBR126W-B | 156.4452467 | -0.074081633 | 0.895153217 | 0.96 |
| YDR451C | 513.5586606 | -0.052805368 | 0.895097116 | 0.96 |
| snR68 | 10.29817386 | 0.039670523 | 0.895248257 | 0.96 |
| YPR130C | 10.79124991 | 0.150944985 | 0.895826955 | 0.97 |
| YKR106W | 110.7787445 | 0.034344936 | 0.896190728 | 0.97 |
| YPR156C | 1539.255789 | -0.111188397 | 0.896174782 | 0.97 |
| YDL240C-A | 1.077122984 | -0.519155384 | 0.896520602 | 0.97 |
| YDR021W | 377.9993278 | 0.155834058 | 0.896395064 | 0.97 |
| YJR134C | 1708.030399 | -0.010916473 | 0.896869649 | 0.97 |
| YGL002W | 690.1637106 | -0.110962774 | 0.89702912 | 0.97 |
| YNL283C | 1549.072925 | 0.057303178 | 0.897462295 | 0.97 |
| YJL175W | 4.732791699 | -0.460675396 | 0.899580969 | 0.97 |
| YJR142W | 1091.921242 | -0.030790613 | 0.899628759 | 0.97 |
| YLR310C | 3300.526123 | -0.008384822 | 0.899363183 | 0.97 |
| YOR280C | 559.6900755 | 0.020195072 | 0.899235566 | 0.97 |
| YEL032W | 1185.048516 | -0.070266337 | 0.900734354 | 0.97 |
| YGL167C | 2492.369981 | 0.040586587 | 0.900485624 | 0.97 |
| YGR014W | 3080.238337 | -0.025284963 | 0.900178114 | 0.97 |
| YGR016W | 101.0221921 | -0.132675419 | 0.900031414 | 0.97 |
| YMR018W | 66.38637523 | -0.189223559 | 0.90042035 | 0.97 |
| YMR255W | 1118.071772 | 0.007010633 | 0.899887356 | 0.97 |
| YPL228W | 971.5333391 | 0.006214526 | 0.900640212 | 0.97 |
| YDL007C-A | 3.853033258 | 0.446829494 | 0.901271998 | 0.97 |
| YHL010C | 843.3803368 | 0.063914598 | 0.901294971 | 0.97 |
| YER139C | 313.6167797 | -0.026694888 | 0.901572074 | 0.97 |
| YFR038W | 1186.649674 | 0.078404042 | 0.902491346 | 0.97 |
| YJL105W | 247.8507967 | -0.283366557 | 0.902479365 | 0.97 |
| YKL067W | 3440.072656 | 0.165636751 | 0.902134416 | 0.97 |
| YMR290W-A | 11.59226843 | -0.323581857 | 0.902332699 | 0.97 |
| YGL172W | 1277.929348 | -0.052663319 | 0.903139511 | 0.97 |
| YGR176W | 11.59700922 | -0.428936775 | 0.903143459 | 0.97 |

| | | | | |
|---|---|---|---|---|
| YPR100W | 188.3881606 | -0.128005005 | 0.902922277 | 0.97 |
| YDL044C | 428.1351393 | 0.020394895 | 0.903559686 | 0.97 |
| YPR047W | 774.0936392 | 0.02650447 | 0.903753124 | 0.97 |
| YGR012W | 677.5451845 | 0.030153204 | 0.903999546 | 0.97 |
| YNL161W | 1329.930942 | 0.068390341 | 0.904273048 | 0.97 |
| YHR131W-A | 19.588359 | -0.297657699 | 0.904951988 | 0.97 |
| YKR068C | 742.7822083 | 0.050802496 | 0.904879745 | 0.97 |
| YNL134C | 7013.757129 | 0.209031206 | 0.904808838 | 0.97 |
| YPR075C | 1855.120278 | 0.132351459 | 0.904535924 | 0.97 |
| YDL146W | 626.6868048 | -0.038053442 | 0.905902948 | 0.97 |
| YDR067C | 673.9208846 | 0.052198824 | 0.905728171 | 0.97 |
| YDR251W | 2664.249522 | -0.081780873 | 0.90561368 | 0.97 |
| YOR211C | 1075.422511 | -0.105543206 | 0.906009503 | 0.97 |
| YPR093C | 386.3642488 | 0.206904757 | 0.905623203 | 0.97 |
| YOL014W | 428.9922641 | -0.206398256 | 0.906359443 | 0.97 |
| YKL002W | 1601.062288 | -0.051734804 | 0.906807236 | 0.97 |
| YBL104C | 644.4185242 | -0.010560206 | 0.90972664 | 0.97 |
| YFR016C | 3711.04018 | 0.161689686 | 0.907962057 | 0.97 |
| YGL028C | 1677.067671 | -0.137111193 | 0.908526102 | 0.97 |
| YGL041W-A | 345.8516558 | -0.034890851 | 0.909067078 | 0.97 |
| YGL080W | 877.4500948 | -0.045601944 | 0.908626118 | 0.97 |
| YGL206C | 5144.593261 | 0.042008433 | 0.908089177 | 0.97 |
| YGR101W | 624.5372953 | 0.035607502 | 0.908248525 | 0.97 |
| YHL047C | 1687.543568 | -0.125014708 | 0.909324217 | 0.97 |
| YHR177W | 301.5515693 | 0.008508079 | 0.909673087 | 0.97 |
| YIR037W | 2878.72569 | 0.028544174 | 0.909065993 | 0.97 |
| YJR037W | 10.84309032 | 0.216448083 | 0.908664168 | 0.97 |
| YKL053W | 17.58730316 | -0.264838467 | 0.908396518 | 0.97 |
| YKL196C | 2255.642273 | 0.050641195 | 0.909677298 | 0.97 |
| YLR428C | 15.96009415 | -0.182018665 | 0.909143343 | 0.97 |
| YML095C | 255.548255 | -0.073293836 | 0.909301612 | 0.97 |
| YMR005W | 1198.923005 | 0.018996444 | 0.907598693 | 0.97 |
| YMR052C-A | 61.89756679 | 0.056055162 | 0.908876158 | 0.97 |
| YNL219C | 1817.162227 | 0.020055178 | 0.907966789 | 0.97 |
| YPL269W | 614.929882 | -0.065195273 | 0.908209099 | 0.97 |
| YBL062W | 0.924462825 | 0.260379005 | 0.910773265 | 0.97 |
| YBR010W | 4241.31261 | 0.057104586 | 0.910943461 | 0.97 |
| YDL180W | 735.4223348 | 0.030235646 | 0.910471489 | 0.97 |
| YHR165C | 3249.517075 | 0.067974572 | 0.910996252 | 0.97 |
| YIL132C | 143.1249549 | -0.040096815 | 0.911218813 | 0.97 |
| YMR035W | 535.4792987 | 0.15332492 | 0.91112987 | 0.97 |
| YOL137W | 547.8586694 | 0.016019722 | 0.910387699 | 0.97 |
| tR(CCG)L | 0.085991782 | -0.041179339 | 0.910615394 | 0.97 |

| | | | | |
|---|---|---|---|---|
| YDR054C | 1985.938955 | -0.074359966 | 0.911763344 | 0.97 |
| YER090W | 2828.394196 | 0.137834932 | 0.912038239 | 0.97 |
| YHR213W-B | 12.16949824 | 0.46655669 | 0.911765681 | 0.97 |
| YIL129C | 2649.636452 | 0.082668466 | 0.911977645 | 0.97 |
| YOR382W | 2866.519088 | 0.272376828 | 0.911450303 | 0.97 |
| YDR252W | 88.93914678 | -0.186634461 | 0.913908947 | 0.97 |
| YFL012W | 27.05345673 | -0.487814011 | 0.913442213 | 0.97 |
| YFL067W | 57.17560859 | 0.030535763 | 0.913837836 | 0.97 |
| YGL168W | 276.6572721 | 0.002493802 | 0.913763321 | 0.97 |
| YGL203C | 1571.604512 | 0.045899927 | 0.912634864 | 0.97 |
| YIL141W | 35.62085203 | 0.205307727 | 0.913302646 | 0.97 |
| YJL043W | 47.06314645 | -0.049566067 | 0.912310683 | 0.97 |
| YLL006W | 812.6766975 | -0.01036738 | 0.913306724 | 0.97 |
| YMR164C | 711.3389073 | 0.040256529 | 0.912895354 | 0.97 |
| YOL027C | 2880.865523 | 0.093016868 | 0.91376621 | 0.97 |
| YPL006W | 1523.461434 | 0.037738143 | 0.913280362 | 0.97 |
| YPL108W | 159.4425969 | 0.043660738 | 0.913596369 | 0.97 |
| YOL004W | 4094.946984 | 0.069230034 | 0.91414759 | 0.97 |
| YBR186W | 272.8183513 | 0.137059834 | 0.915190529 | 0.97 |
| YER011W | 1426.306813 | -0.250491904 | 0.915226807 | 0.97 |
| YER073W | 2130.104252 | 0.474861048 | 0.915653042 | 0.97 |
| YGR164W | 4.051111594 | -0.073352375 | 0.915815193 | 0.97 |
| YIL025C | 3.736035911 | 0.385102637 | 0.915562196 | 0.97 |
| YJR051W | 1291.045072 | 0.07079249 | 0.915254934 | 0.97 |
| YJR113C | 864.1046061 | 0.010503796 | 0.914778396 | 0.97 |
| YKR037C | 412.1583252 | 0.037183308 | 0.915784483 | 0.97 |
| YLR115W | 1204.082371 | 0.042792782 | 0.914837903 | 0.97 |
| YPL105C | 3260.005512 | 0.030356986 | 0.915283086 | 0.97 |
| YCR020W-B | 152.1239298 | -0.166799253 | 0.915989109 | 0.97 |
| YJL139C | 556.1081958 | 0.079304724 | 0.916250518 | 0.97 |
| YJL181W | 405.181886 | 0.049927851 | 0.916578116 | 0.97 |
| YLR438W | 1335.790008 | 0.141097775 | 0.916737716 | 0.97 |
| YML090W | 156.4475378 | 0.560133903 | 0.916728717 | 0.97 |
| YOL156W | 95.75918972 | -0.166753252 | 0.916559578 | 0.97 |
| YNL098C | 3291.418535 | 0.008464151 | 0.917039318 | 0.97 |
| YAL019W | 1321.957689 | 0.089968948 | 0.918112598 | 0.97 |
| YCR106W | 394.726015 | 0.039273362 | 0.918754383 | 0.97 |
| YDL047W | 1531.758576 | 0.061508038 | 0.918419141 | 0.97 |
| YDR520C | 591.1970741 | -0.009256422 | 0.918944188 | 0.97 |
| YER094C | 2268.208556 | 0.072773291 | 0.917940006 | 0.97 |
| YGR189C | 5490.403188 | 0.25206221 | 0.919363664 | 0.97 |
| YHR009C | 3439.835058 | 0.045848636 | 0.919494598 | 0.97 |
| YHR180W-A | 0.36648756 | -0.009854806 | 0.919023062 | 0.97 |

| | | | | |
|---|---|---|---|---|
| YIL085C | 1498.284283 | -0.062517817 | 0.918464164 | 0.97 |
| YJL125C | 686.7482617 | -0.022989758 | 0.918562837 | 0.97 |
| YJR102C | 338.1234787 | -0.082108482 | 0.918514356 | 0.97 |
| YJR146W | 99.77471099 | 0.096165291 | 0.91909126 | 0.97 |
| YNL128W | 87.91000712 | -0.092657358 | 0.918027134 | 0.97 |
| YOR170W | 97.11015677 | 0.085985562 | 0.919299643 | 0.97 |
| snR40 | 6.538551087 | 0.223085419 | 0.917354319 | 0.97 |
| snR65 | 5.68201134 | 0.036485008 | 0.919423383 | 0.97 |
| YDR522C | 159.2048735 | -0.437695907 | 0.919951167 | 0.97 |
| YDR405W | 1203.736123 | 0.023970117 | 0.920368998 | 0.97 |
| YFL013C | 826.2401076 | 0.039204056 | 0.920766642 | 0.97 |
| YGR246C | 1106.975461 | -0.007594917 | 0.920651928 | 0.97 |
| YML102W | 604.0898534 | -0.003728286 | 0.9208583 | 0.97 |
| YMR121C | 804.8314095 | 0.106841188 | 0.920999334 | 0.97 |
| YHR137W | 2665.655739 | 0.280100056 | 0.921463237 | 0.97 |
| YLL065W | 0.583562943 | -0.00432248 | 0.921337024 | 0.97 |
| YEL018C-A | 0.416927005 | 0.034766299 | 0.921744754 | 0.97 |
| YPR171W | 883.9863961 | -0.002206821 | 0.921764368 | 0.97 |
| tL(GAG)G | 0.599720226 | -0.073247458 | 0.922048379 | 0.97 |
| YGR204W | 10459.91176 | 0.096552795 | 0.9224061 | 0.97 |
| YHR032C-A | 12.55755435 | -0.09606741 | 0.922454932 | 0.97 |
| YGR232W | 824.5178442 | -0.008919904 | 0.923119115 | 0.97 |
| YLR233C | 453.3027113 | -0.020881222 | 0.922834861 | 0.97 |
| YNL336W | 467.9927857 | -0.112521315 | 0.923017522 | 0.97 |
| YEL073C | 122.6223904 | -0.142163324 | 0.923755605 | 0.97 |
| YHR158C | 1689.928736 | 0.077167156 | 0.924082226 | 0.97 |
| YHR208W | 7460.933205 | 0.508748948 | 0.923978246 | 0.97 |
| YJL170C | 341.173457 | -0.243146528 | 0.923946637 | 0.97 |
| YKL102C | 18.56354003 | 0.421244597 | 0.924320747 | 0.97 |
| YMR153C-A | 60.08017136 | -0.046726989 | 0.924354186 | 0.97 |
| YNR044W | 3311.34449 | -0.174103581 | 0.92473336 | 0.97 |
| YOL145C | 2673.103928 | -0.056966939 | 0.924702532 | 0.97 |
| YPR129W | 1545.384555 | 0.033709246 | 0.924995988 | 0.97 |
| YCR102C | 144.1370507 | -0.000206065 | 0.925723627 | 0.97 |
| YDL120W | 712.7094342 | 0.025234197 | 0.925598107 | 0.97 |
| YFR039C | 736.7528646 | 0.019495473 | 0.925648869 | 0.97 |
| tG(CCC)O | 0.202424979 | -0.039794093 | 0.925719108 | 0.97 |
| YCL026C-A | 44.13033287 | -0.117671748 | 0.926371941 | 0.97 |
| YDR406W | 3009.969096 | 0.099091078 | 0.926314146 | 0.97 |
| YJR066W | 2254.068141 | 0.084783755 | 0.926427456 | 0.97 |
| YKR071C | 2094.595777 | 0.166982935 | 0.926067166 | 0.97 |
| YDR363W | 641.2311125 | -0.090877676 | 0.926950644 | 0.97 |
| YKL195W | 3025.520768 | 0.083754568 | 0.926660055 | 0.97 |

| | | | | |
|---|---|---|---|---|
| YPR002C-A | 1.918449357 | -0.131262735 | 0.92689643 | 0.97 |
| YLR334C | 3.854398197 | 0.471957301 | 0.927132103 | 0.97 |
| YPL274W | 1036.210642 | -0.213259713 | 0.927374762 | 0.97 |
| YIL095W | 775.5717487 | -0.035598177 | 0.927652255 | 0.97 |
| YLR213C | 172.0132438 | -0.332003955 | 0.927739476 | 0.97 |
| YPL103C | 590.7071612 | 0.020836049 | 0.927826807 | 0.97 |
| YBR227C | 856.5044097 | -0.001295086 | 0.928811702 | 0.97 |
| YGR141W | 841.9095851 | 0.17606178 | 0.928421054 | 0.97 |
| YJL200C | 1688.489143 | 0.048158324 | 0.928412313 | 0.97 |
| YNL192W | 2879.150646 | 0.028290997 | 0.928143593 | 0.97 |
| YOR140W | 905.6577941 | 0.055072421 | 0.92882458 | 0.97 |
| YPL085W | 4050.180373 | 0.109688524 | 0.928639979 | 0.97 |
| YDL184C | 118.3516335 | 0.177419822 | 0.929040287 | 0.97 |
| YGL254W | 331.7870674 | -0.019655607 | 0.929439557 | 0.97 |
| YDR118W-A | 11.72360647 | -0.304855437 | 0.930722391 | 0.97 |
| YDR455C | 13.39180516 | -0.131549129 | 0.931646022 | 0.97 |
| YGL129C | 1219.243345 | 0.016320637 | 0.931386582 | 0.97 |
| YHR217C | 1.225581631 | 0.263452208 | 0.930445153 | 0.97 |
| YJL003W | 223.2586603 | -0.050631368 | 0.93127684 | 0.97 |
| YJL119C | 7.109202711 | 0.121090675 | 0.931492559 | 0.97 |
| YLR365W | 4.223598425 | -0.252364869 | 0.930277347 | 0.97 |
| YML104C | 923.4299097 | -0.053140118 | 0.931842213 | 0.97 |
| YML130C | 2617.920044 | 0.296910885 | 0.931870473 | 0.97 |
| YOR362C | 3907.49208 | 0.004346256 | 0.930874058 | 0.97 |
| YPL034W | 171.3658521 | -0.139195693 | 0.930687772 | 0.97 |
| YPL073C | 1.14139142 | 0.069921517 | 0.931716241 | 0.97 |
| snR6 | 0.308944371 | 0.066853481 | 0.93141319 | 0.97 |
| YCR003W | 578.0986481 | -0.026104675 | 0.932283918 | 0.97 |
| YLL040C | 5253.851757 | 0.205305154 | 0.932245797 | 0.97 |
| YBL011W | 1645.010666 | 0.002340118 | 0.933351016 | 0.97 |
| YBR111C | 3318.167647 | 0.128812451 | 0.933473005 | 0.97 |
| YDR095C | 5.026425328 | -0.237423528 | 0.93352512 | 0.97 |
| YGR255C | 1753.194799 | -0.021924348 | 0.933072379 | 0.97 |
| YIL029W-A | 8.108002149 | 0.480147846 | 0.933210806 | 0.97 |
| YKL037W | 43.69720214 | -0.09692319 | 0.933063081 | 0.97 |
| YKR038C | 1110.135443 | -0.049640774 | 0.932845073 | 0.97 |
| YIL111W | 1481.52504 | -0.10821278 | 0.933855681 | 0.98 |
| YBR201W | 456.7810734 | 0.073219552 | 0.934661769 | 0.98 |
| YDR089W | 994.7212521 | -0.118033611 | 0.934540926 | 0.98 |
| YFR049W | 788.1728678 | 0.135178394 | 0.934764069 | 0.98 |
| YJL037W | 69.21169324 | -0.527643655 | 0.934671156 | 0.98 |
| YMR223W | 516.9298345 | -0.069114327 | 0.934200222 | 0.98 |
| YPR016W-A | 37.91128357 | -0.299100746 | 0.934265168 | 0.98 |

| | | | | |
|---|---|---|---|---|
| YNL297C | 2104.946506 | -0.004988738 | 0.934949045 | 0.98 |
| YDR213W | 568.9307813 | 0.046785796 | 0.935492036 | 0.98 |
| YKL016C | 3252.389125 | 0.223870977 | 0.935666315 | 0.98 |
| YNL059C | 1845.140766 | -0.049570962 | 0.93528502 | 0.98 |
| YOL109W | 9848.185932 | -0.109800231 | 0.935541706 | 0.98 |
| YMR094W | 306.3491696 | 0.241307136 | 0.9361554 | 0.98 |
| YMR301C | 1249.636815 | 0.053344127 | 0.936271838 | 0.98 |
| YDL182W | 2531.684902 | 0.098830809 | 0.936567571 | 0.98 |
| YLR313C | 272.7990365 | 0.001716021 | 0.936876325 | 0.98 |
| YOL151W | 926.9123911 | 0.151430779 | 0.936832264 | 0.98 |
| YOR057W | 859.8202576 | -0.029573259 | 0.936596988 | 0.98 |
| YIR040C | 0.167590483 | 0.013771792 | 0.937643115 | 0.98 |
| YBR126W-A | 695.852559 | -0.066052768 | 0.939235389 | 0.98 |
| YER087C-A | 36.6678806 | 0.469275384 | 0.93797018 | 0.98 |
| YER109C | 1065.439283 | -0.051162376 | 0.939050628 | 0.98 |
| YFL066C | 2.343434624 | 0.024699512 | 0.939099624 | 0.98 |
| YGL246C | 646.4635665 | 0.020778863 | 0.938205847 | 0.98 |
| YHR206W | 1203.868698 | -0.08466473 | 0.9381268 | 0.98 |
| YJR068W | 1198.277592 | 0.080269635 | 0.938795132 | 0.98 |
| YNR004W | 166.7322391 | -0.066882201 | 0.938947095 | 0.98 |
| YPL205C | 28.51967725 | 0.039036715 | 0.938441529 | 0.98 |
| YPR074W-A | 0.407755058 | 0.002399508 | 0.939096679 | 0.98 |
| YOR168W | 3054.619284 | 0.021141101 | 0.939395413 | 0.98 |
| YPR038W | 18.2580769 | -0.127108137 | 0.940036036 | 0.98 |
| YDR495C | 1040.925179 | 0.006025358 | 0.940402309 | 0.98 |
| YKL135C | 1473.22037 | -0.049864592 | 0.940845251 | 0.98 |
| YLR215C | 433.3667117 | 0.013461324 | 0.94108899 | 0.98 |
| YLR319C | 843.3891881 | 0.006378945 | 0.940986757 | 0.98 |
| snR55 | 9.080532067 | 0.252467851 | 0.940628094 | 0.98 |
| tT(CGU)K | 0.429968112 | -0.002397253 | 0.940947046 | 0.98 |
| YIR030W-A | 17.57088172 | 0.17551554 | 0.942260327 | 0.98 |
| YMR193W | 776.035786 | 0.134851875 | 0.942216183 | 0.98 |
| YOR184W | 6279.813274 | 0.179088796 | 0.942134529 | 0.98 |
| YIL177C | 0.203940714 | -0.013424841 | 0.942537237 | 0.98 |
| YJR125C | 2124.757426 | 0.020940406 | 0.942892253 | 0.98 |
| YOR347C | 1046.083791 | -0.091501591 | 0.942810253 | 0.98 |
| YBL008W-A | 2.748531795 | -0.365895805 | 0.945523269 | 0.98 |
| YBL022C | 3958.243828 | 0.0831789 | 0.943849751 | 0.98 |
| YBR089W | 6.735755394 | 0.023246654 | 0.945906373 | 0.98 |
| YBR215W | 1081.67593 | -0.021492954 | 0.943636032 | 0.98 |
| YCR049C | 7.558405616 | -0.236870016 | 0.945757479 | 0.98 |
| YDR434W | 1180.144828 | -0.027957087 | 0.945575278 | 0.98 |
| YEL023C | 623.1672116 | -0.167613547 | 0.945621266 | 0.98 |

| | | | | |
|---|---|---|---|---|
| YER084W-A | 0.719422121 | -0.091302462 | 0.944103705 | 0.98 |
| YKR051W | 666.9246654 | -0.07647895 | 0.945691298 | 0.98 |
| YLR038C | 1207.825369 | 0.217201842 | 0.944000227 | 0.98 |
| YLR119W | 415.3763546 | -0.048369479 | 0.94411946 | 0.98 |
| YLR364C-A | 0.303554837 | -0.023020544 | 0.944667643 | 0.98 |
| YOR103C | 111.7299261 | 0.097013409 | 0.945217466 | 0.98 |
| YOR274W | 529.1848909 | -0.152968225 | 0.9458542 | 0.98 |
| YPL087W | 1694.761164 | 0.022193627 | 0.945520585 | 0.98 |
| YPL179W | 1527.082249 | 0.047563679 | 0.943575645 | 0.98 |
| snR24 | 8.183960631 | 0.090546014 | 0.944525946 | 0.98 |
| snR36 | 18.91914442 | -0.049945559 | 0.944489278 | 0.98 |
| YKL162C-A | 16.50216676 | -0.126029096 | 0.946209977 | 0.98 |
| YOL008W | 381.7473344 | -0.020931509 | 0.946319057 | 0.98 |
| YDR079W | 403.891688 | -0.051879978 | 0.946474927 | 0.98 |
| YOR304W | 1385.923767 | -0.01628868 | 0.946954169 | 0.98 |
| YDR371W | 631.8329655 | -0.132153278 | 0.947324346 | 0.98 |
| YKL224C | 0.174623486 | 0.018508201 | 0.947361131 | 0.98 |
| RUF22 | 1.126113591 | 0.030247521 | 0.950201347 | 0.98 |
| YAL067W-A | 0.291097411 | -0.010323821 | 0.949067162 | 0.98 |
| YDL005C | 1599.265031 | -0.038880019 | 0.948962125 | 0.98 |
| YDL034W | 29.64111763 | 0.162022258 | 0.950373419 | 0.98 |
| YDL078C | 3642.136284 | 0.220724289 | 0.949803483 | 0.98 |
| YDL221W | 36.90750404 | 0.035146473 | 0.950444402 | 0.98 |
| YDR274C | 62.88697688 | -0.337289877 | 0.950186684 | 0.98 |
| YDR419W | 525.8031882 | -0.042056101 | 0.94988131 | 0.98 |
| YGL107C | 935.2339762 | -0.023945815 | 0.94980598 | 0.98 |
| YGR126W | 190.2789593 | -0.010172481 | 0.949809049 | 0.98 |
| YGR212W | 361.1538773 | -0.038952966 | 0.949585653 | 0.98 |
| YIL030C | 2092.978365 | 0.104129591 | 0.949365566 | 0.98 |
| YKL219W | 500.3272007 | -0.002833158 | 0.948258394 | 0.98 |
| YML031W | 1341.560963 | 0.012061368 | 0.949257587 | 0.98 |
| YML108W | 173.1265371 | 0.044334792 | 0.949445458 | 0.98 |
| YOR065W | 3707.159237 | 0.203842608 | 0.949859697 | 0.98 |
| YPL194W | 451.5443271 | 0.023448403 | 0.949242375 | 0.98 |
| YPR196W | 243.4727794 | -0.144552296 | 0.949681708 | 0.98 |
| YDR145W | 1845.179932 | 0.018511154 | 0.950867968 | 0.98 |
| YDR073W | 640.4950618 | -0.192410639 | 0.951097431 | 0.98 |
| YBL045C | 7514.298009 | 0.337599453 | 0.952057228 | 0.98 |
| YBR262C | 557.8168642 | 0.08864019 | 0.951565847 | 0.98 |
| YDL163W | 0.057163039 | -0.013982486 | 0.951969391 | 0.98 |
| YDL209C | 238.6569979 | 0.008191236 | 0.951823548 | 0.98 |
| YLL063C | 302.5298246 | -0.223554205 | 0.951635894 | 0.98 |
| tR(CCU)J | 0.0941958 | -0.015171956 | 0.951963004 | 0.98 |

| | | | | |
|---|---|---|---|---|
| YGL194C | 571.5770023 | 0.031746193 | 0.952480084 | 0.98 |
| YHR200W | 3347.025252 | 0.104839121 | 0.952917756 | 0.98 |
| YGR055W | 1342.856307 | -0.184536161 | 0.954100371 | 0.98 |
| YMR069W | 79.44065746 | 0.271368754 | 0.954199361 | 0.98 |
| YNR035C | 3358.997637 | 0.059137783 | 0.954930774 | 0.98 |
| tQ(UUG)E2 | 0.083088204 | 0.014662475 | 0.954948171 | 0.98 |
| YGR057C | 395.64537 | -0.011757266 | 0.955123128 | 0.98 |
| YBR232C | 24.08316402 | 0.279236488 | 0.955713715 | 0.98 |
| YHR145C | 0.415497853 | -0.04849064 | 0.95582368 | 0.98 |
| YML112W | 572.6104301 | 0.041257269 | 0.955896063 | 0.98 |
| YMR315W | 4197.096296 | 0.255078426 | 0.955570814 | 0.98 |
| YLR226W | 384.869465 | -0.031034894 | 0.956125535 | 0.98 |
| YFR056C | 41.82065454 | -0.229516145 | 0.956375231 | 0.98 |
| YGR244C | 5502.307672 | 0.19738304 | 0.956725117 | 0.98 |
| YMR135W-A | 18.48988281 | 0.118233208 | 0.956956815 | 0.98 |
| YNR072W | 156.9410464 | 0.247283298 | 0.956826953 | 0.98 |
| YER157W | 974.6953246 | -0.006093985 | 0.957387064 | 0.98 |
| YNR042W | 0.825755426 | -0.338974078 | 0.957461646 | 0.98 |
| YBR124W | 16.74512171 | -0.146384024 | 0.959412207 | 0.98 |
| YBR264C | 359.5103772 | -0.081476517 | 0.958150055 | 0.98 |
| YCL001W-B | 30.32607662 | -0.287377092 | 0.959862172 | 0.98 |
| YDR356W | 1114.540575 | 0.092820998 | 0.960414886 | 0.98 |
| YDR518W | 1466.448011 | 0.149563842 | 0.958417204 | 0.98 |
| YEL056W | 977.5251167 | -0.049545303 | 0.959270072 | 0.98 |
| YER022W | 948.246119 | 0.0258269 | 0.958629196 | 0.98 |
| YFL010C | 2890.139798 | -0.026695107 | 0.96061917 | 0.98 |
| YFL010W-A | 308.9446486 | -0.043098008 | 0.95869987 | 0.98 |
| YGL118C | 2.533670207 | -0.401895103 | 0.959755662 | 0.98 |
| YGR184C | 3421.297226 | 0.040469044 | 0.958205713 | 0.98 |
| YGR263C | 454.4543224 | -0.025372393 | 0.958489905 | 0.98 |
| YHR219W | 0.159331785 | 0.014181743 | 0.95898685 | 0.98 |
| YKL131W | 17.12686382 | -0.338228029 | 0.960110334 | 0.98 |
| YKL149C | 355.2699607 | -0.080405153 | 0.960469098 | 0.98 |
| YLR124W | 42.32116711 | -0.605266405 | 0.95951198 | 0.98 |
| YLR323C | 363.5088652 | 0.111817955 | 0.960343748 | 0.98 |
| YOR155C | 1051.816257 | 0.00641214 | 0.959040256 | 0.98 |
| YOR356W | 1275.917226 | 0.031593942 | 0.960178733 | 0.98 |
| YNL127W | 1075.477959 | 0.053645653 | 0.961336495 | 0.98 |
| YDR097C | 218.1671058 | 0.10777385 | 0.961515137 | 0.98 |
| YHR069C-A | 0.116121444 | 0.013499113 | 0.962423169 | 0.98 |
| YKL049C | 577.3829424 | 0.044799437 | 0.962438099 | 0.98 |
| YKL177W | 2.5279419 | -0.432805372 | 0.962181415 | 0.98 |
| YPL132W | 679.0950944 | 0.001045913 | 0.962436027 | 0.98 |

| | | | | |
|---|---|---|---|---|
| YDR409W | 969.523209 | -0.008329774 | 0.96311764 | 0.98 |
| YDR489W | 358.8647918 | -0.025754517 | 0.962997988 | 0.98 |
| YNL170W | 85.88966089 | 0.270732472 | 0.962985503 | 0.98 |
| YAL063C-A | 8.265236292 | -0.476801613 | 0.963850757 | 0.98 |
| YDR287W | 832.7406735 | 0.016654361 | 0.963526646 | 0.98 |
| YGR107W | 13.44440994 | -0.069547928 | 0.963917928 | 0.98 |
| YNL028W | 6.323808974 | -0.128607997 | 0.96397356 | 0.98 |
| YMR173W | 743.4251563 | 0.125182745 | 0.965105256 | 0.99 |
| YBR157C | 589.2630079 | -0.771768876 | 0.965610159 | 0.99 |
| YLR302C | 0.305761563 | 0.022774083 | 0.965427661 | 0.99 |
| YLR438C-A | 659.2030513 | 0.09842741 | 0.965525861 | 0.99 |
| YBR184W | 226.9887473 | 0.05594864 | 0.965813464 | 0.99 |
| tQ(UUG)B | 0.084186605 | 0.039559704 | 0.966457164 | 0.99 |
| YPL245W | 615.8162164 | -0.037140716 | 0.966780401 | 0.99 |
| YBR137W | 861.5111114 | 0.030300848 | 0.967019246 | 0.99 |
| YDR517W | 2232.436932 | 0.016228055 | 0.967411239 | 0.99 |
| YIL153W | 970.2297401 | 0.038144119 | 0.967656993 | 0.99 |
| YPL015C | 1643.336309 | -0.049569747 | 0.967333787 | 0.99 |
| YPR133C | 1652.526675 | 0.007813834 | 0.967596089 | 0.99 |
| YGR068C | 535.7602898 | -0.037801677 | 0.967811748 | 0.99 |
| YBL113C | 286.5211358 | -0.073916041 | 0.968382455 | 0.99 |
| YDL097C | 4494.471702 | 0.000783302 | 0.96812345 | 0.99 |
| YGR235C | 1048.42614 | 0.061185578 | 0.96825002 | 0.99 |
| YBL054W | 675.0753311 | -0.163232514 | 0.968912839 | 0.99 |
| YNL304W | 349.9176759 | -0.011317837 | 0.968844849 | 0.99 |
| YCL057C-A | 1206.33424 | 0.209988668 | 0.969449713 | 0.99 |
| YNL045W | 3993.621371 | 0.330665345 | 0.969462551 | 0.99 |
| YMR119W | 1094.314263 | -0.014423329 | 0.970452743 | 0.99 |
| YGL021W | 1221.378254 | 0.003049783 | 0.970616582 | 0.99 |
| YLR081W | 221.380283 | -0.039083522 | 0.970785394 | 0.99 |
| YAR050W | 236.1195337 | 0.04840322 | 0.970956483 | 0.99 |
| YGR062C | 462.3300151 | -0.022277193 | 0.971116504 | 0.99 |
| YML030W | 762.1544554 | -0.06525783 | 0.971725336 | 0.99 |
| YOR142W | 3533.236265 | 0.165700371 | 0.971685643 | 0.99 |
| YPR177C | 1.844393858 | 0.019931003 | 0.971672515 | 0.99 |
| YDR066C | 173.6197188 | -0.159603949 | 0.972275119 | 0.99 |
| YDL125C | 2310.617952 | 0.097891625 | 0.973025736 | 0.99 |
| YDR261W-B | 58.90647804 | -0.022176716 | 0.97266845 | 0.99 |
| YGL209W | 1219.680335 | -0.191691941 | 0.972769573 | 0.99 |
| YHR162W | 4274.252821 | 0.065391455 | 0.973135167 | 0.99 |
| YIL102C | 13.70446277 | 0.310592085 | 0.973522203 | 0.99 |
| YKR027W | 720.4128865 | -0.050523146 | 0.973527771 | 0.99 |
| YKR105C | 96.82105819 | -0.089811819 | 0.973798498 | 0.99 |

| | | | | |
|---|---|---|---|---|
| YOR306C | 1215.578289 | 0.063847218 | 0.973776124 | 0.99 |
| YPL051W | 362.5719952 | 0.026829213 | 0.973376309 | 0.99 |
| YCR045C | 398.9252367 | -0.124336345 | 0.974216219 | 0.99 |
| YGR031W | 376.7830976 | -0.030986582 | 0.974434679 | 0.99 |
| tD(GUC)N | 0.865673422 | -0.08351117 | 0.974391965 | 0.99 |
| YKL223W | 0.064032033 | 0.013938421 | 0.974634397 | 0.99 |
| YER163C | 538.8237534 | 0.12755857 | 0.97507796 | 0.99 |
| YIL100C-A | 6.139255363 | 0.194549215 | 0.974983867 | 0.99 |
| YKL221W | 211.2597824 | -0.146693696 | 0.975248694 | 0.99 |
| YLR015W | 562.8104697 | 0.068615808 | 0.975980225 | 0.99 |
| YDR154C | 3531.276985 | 0.024670066 | 0.976346973 | 0.99 |
| YAL001C | 1631.198822 | -0.045675975 | 0.976536874 | 0.99 |
| YPL061W | 6576.454349 | 0.019489358 | 0.977267979 | 0.99 |
| YLR364W | 131.7044529 | -0.046786579 | 0.977872959 | 0.99 |
| YCR060W | 437.7609902 | 0.03762286 | 0.978029087 | 0.99 |
| YBL079W | 2598.015595 | 0.034769742 | 0.979532124 | 0.99 |
| YKL145W | 5482.552156 | 0.049630439 | 0.979540674 | 0.99 |
| YER189W | 0.915658928 | 0.032046759 | 0.980217374 | 0.99 |
| YMR047C | 2343.921258 | -0.004858069 | 0.98055541 | 0.99 |
| YDL186W | 120.8658792 | -0.17710432 | 0.980823533 | 0.99 |
| YBL017C | 3980.2423 | -0.010968909 | 0.982036912 | 0.99 |
| YBR092C | 2232.790112 | -0.175430307 | 0.98345271 | 0.99 |
| YBR217W | 353.7676501 | -0.051105648 | 0.984345743 | 0.99 |
| YFL064C | 3.015892609 | 0.059037165 | 0.984442693 | 0.99 |
| YGL116W | 933.3944118 | 0.026565166 | 0.982399081 | 0.99 |
| YGL151W | 1728.404115 | -0.040955374 | 0.983647309 | 0.99 |
| YGR193C | 2486.614356 | 0.015790991 | 0.984273753 | 0.99 |
| YHL015W-A | 4.315791202 | 0.010674996 | 0.982109411 | 0.99 |
| YHR078W | 1137.159604 | 0.008844072 | 0.982791376 | 0.99 |
| YHR214C-E | 1.813646714 | -0.470042688 | 0.982232069 | 0.99 |
| YIL004C | 332.265247 | -0.034136128 | 0.982399967 | 0.99 |
| YJL136W-A | 0.34890375 | -0.021621896 | 0.983964235 | 0.99 |
| YJR133W | 618.7266674 | 0.132810342 | 0.982154488 | 0.99 |
| YLR381W | 607.7067419 | -0.008655613 | 0.983816707 | 0.99 |
| YLR440C | 889.0207433 | -0.015632616 | 0.98191952 | 0.99 |
| YMR070W | 1232.669923 | 0.181593192 | 0.983390802 | 0.99 |
| YMR074C | 1117.334751 | -0.100941379 | 0.981091473 | 0.99 |
| YNL067W-A | 0.709210999 | -0.061876432 | 0.983544084 | 0.99 |
| YOR130C | 372.7954083 | -0.00482288 | 0.983277939 | 0.99 |
| YOR176W | 2270.643704 | 0.004087608 | 0.981489469 | 0.99 |
| YOR205C | 770.9234478 | -0.063391249 | 0.982648332 | 0.99 |
| YPR103W | 3867.741987 | 0.110830066 | 0.984157585 | 0.99 |
| YPR141C | 623.3342075 | 0.017588299 | 0.984269511 | 0.99 |

| | | | | |
|---|---|---|---|---|
| YBL038W | 1166.095027 | 0.128174227 | 0.985077104 | 0.99 |
| YDL068W | 4.863206233 | -0.085666156 | 0.985131444 | 0.99 |
| YER083C | 1868.758699 | 0.004425162 | 0.985889357 | 0.99 |
| YHR024C | 1377.744705 | 0.008889311 | 0.985796541 | 0.99 |
| YNL212W | 1282.46661 | 0.068518173 | 0.985734613 | 0.99 |
| YIL167W | 108.465152 | -0.130542543 | 0.986059965 | 0.99 |
| YLR375W | 1937.904522 | -0.032241983 | 0.986683325 | 0.99 |
| YKR083C | 244.3380659 | -0.154508959 | 0.987229259 | 0.99 |
| YPR194C | 1121.813284 | 0.071641933 | 0.987856551 | 0.99 |
| YDL067C | 1308.526792 | 0.373218064 | 0.988366462 | 0.99 |
| YLR068W | 534.5964698 | 0.174060993 | 0.988355022 | 0.99 |
| YER174C | 586.1669315 | -0.045252205 | 0.988956082 | 1.00 |
| YGL119W | 689.6648811 | 0.034390729 | 0.98906651 | 1.00 |
| YKL084W | 479.3447424 | 0.079836591 | 0.988662053 | 1.00 |
| YLL048C | 4130.734576 | 0.060320956 | 0.9889842 | 1.00 |
| YGR076C | 617.1306078 | 0.014261813 | 0.989733145 | 1.00 |
| YIR006C | 5095.375019 | 0.04101808 | 0.989589611 | 1.00 |
| YMR326C | 0.406623722 | 0.011421336 | 0.989965716 | 1.00 |
| YJL001W | 1978.350649 | 0.013786997 | 0.990491056 | 1.00 |
| YNL288W | 932.0705303 | -0.01784864 | 0.990435997 | 1.00 |
| YGR182C | 629.4422912 | -0.010410494 | 0.991708395 | 1.00 |
| YDR420W | 1513.909661 | 0.041193715 | 0.991999506 | 1.00 |
| YCR059C | 1130.721478 | 0.015369007 | 0.992393104 | 1.00 |
| YKR016W | 2416.547829 | 0.186782726 | 0.992351378 | 1.00 |
| YCL064C | 3239.199963 | -0.271783806 | 0.992727854 | 1.00 |
| YHL046W-A | 47.85481112 | -0.018859826 | 0.993140133 | 1.00 |
| YHR027C | 5786.339997 | 0.179627377 | 0.993158665 | 1.00 |
| YLR168C | 702.4056037 | 0.215259176 | 0.992901731 | 1.00 |
| YDR176W | 1688.228212 | -0.029923825 | 0.99381686 | 1.00 |
| YNR077C | 0.24124529 | -0.018276524 | 0.993763939 | 1.00 |
| YDR348C | 1537.308452 | 0.006435824 | 0.994283651 | 1.00 |
| YLR408C | 156.1371802 | 0.050150825 | 0.994140263 | 1.00 |
| YDL004W | 2874.441994 | 0.252083737 | 0.994488457 | 1.00 |
| YBL001C | 1539.933777 | -0.127162491 | 0.995571873 | 1.00 |
| YHR211W | 534.044984 | 0.000846621 | 0.995659184 | 1.00 |
| YKR014C | 1917.457601 | -0.045396775 | 0.99571531 | 1.00 |
| YNL017C | 1.615564791 | 0.510948391 | 0.995140199 | 1.00 |
| YOL050C | 16.33482394 | 0.333069564 | 0.995322902 | 1.00 |
| YBL037W | 1072.022099 | -0.021603296 | 0.995938162 | 1.00 |
| YNL296W | 0.17572267 | 0.040241341 | 0.996164822 | 1.00 |
| YBR294W | 590.8243932 | -0.244959388 | 0.997104506 | 1.00 |
| YMR171C | 1518.290743 | -0.141099233 | 0.99697749 | 1.00 |
| YOR379C | 0.107512111 | 0.010861535 | 0.996814524 | 1.00 |

| | | | | |
|---|---|---|---|---|
| YBR298C-A | 3.08929577 | -0.43085114 | 0.997290725 | 1.00 |
| YBL041W | 3024.581572 | 0.011794353 | 0.99797861 | 1.00 |
| YJL117W | 1366.561839 | 0.081648354 | 0.997907323 | 1.00 |
| YMR044W | 903.5982709 | -0.065066244 | 0.997703587 | 1.00 |
| YER034W | 308.6813768 | 0.198813055 | 0.998532771 | 1.00 |
| YGR259C | 0.780911468 | 0.020478478 | 0.998573073 | 1.00 |
| YKL206C | 563.6605725 | 0.001955471 | 0.998314584 | 1.00 |
| YGL053W | 527.1405514 | -0.081988929 | 0.999110097 | 1.00 |
| YMR312W | 719.0027121 | 0.087121889 | 0.999004276 | 1.00 |
| YNL246W | 1195.836863 | -0.017851282 | 0.999110774 | 1.00 |
| YIR030C | 90.29914152 | 0.019566471 | 0.999283321 | 1.00 |
| YAL062W | 2248.194645 | 0.128328249 | NA | NA |
| YDL223C | 1796.784019 | 0.414810997 | NA | NA |
| YER024W | 2782.885664 | 0.287379145 | NA | NA |
| YER065C | 3480.324226 | 0.177826262 | NA | NA |
| YGR067C | 601.4253985 | 0.325309936 | NA | NA |
| YGR236C | 237.8810767 | 0.304212402 | NA | NA |
| YIL057C | 383.2825996 | 0.760655178 | NA | NA |
| YJR095W | 1024.664601 | 0.393432397 | NA | NA |
| YKL065W-A | 60.35011298 | -0.122280721 | NA | NA |
| YKR097W | 1618.765281 | 0.400757392 | NA | NA |
| YLR174W | 1214.876474 | 0.52513636 | NA | NA |
| YLR377C | 564.6863452 | 0.25839914 | NA | NA |
| YML042W | 2686.768376 | 0.959170252 | NA | NA |
| YMR107W | 2022.649302 | 0.153428948 | NA | NA |
| YMR175W | 966.6429644 | 0.107229051 | NA | NA |
| YMR175W-A | 25.9663659 | 0.257879267 | NA | NA |
| YNL036W | 2481.742596 | 0.212882887 | NA | NA |
| YNL117W | 3801.736785 | 0.332444342 | NA | NA |
| YNR002C | 907.6027094 | 0.575924104 | NA | NA |
| YNR071C | 317.5831201 | 0.465128595 | NA | NA |
| YOR100C | 795.4961629 | 0.944397834 | NA | NA |
| YPL223C | 915.9171589 | 0.037548728 | NA | NA |
