## Supplementary material for "A stress response that allows highly mutated eukaryotic cells to survive and proliferate": Table S6

| Gene | baseMean | log2FoldChange | pvalue | padj |
|---|---|---|---|---|
| YER091C | 3130.444513 | 1.697553504 | 4.95E-07 | 0.00 |
| YGL183C | 72.4162606 | 2.594162166 | 4.25E-07 | 0.00 |
| YDR402C | 76.67558535 | 1.907952137 | 4.56E-05 | 0.04 |
| YER095W | 1267.382435 | 1.099239267 | 2.94E-05 | 0.04 |
| YJL153C | 833.1459195 | 2.093060973 | 2.31E-05 | 0.04 |
| YMR108W | 5770.966467 | 1.239939605 | 3.29E-05 | 0.04 |
| YPL111W | 2984.765722 | 1.195561409 | 4.10E-05 | 0.04 |
| YDL218W | 253.6505308 | 1.729799838 | 0.000104 | 0.06 |
| YDR403W | 327.0898139 | 2.138123375 | 7.99E-05 | 0.06 |
| YML034W | 2522.763719 | 0.804491795 | 0.000106895 | 0.06 |
| YPL095C | 828.2325169 | -1.257836988 | 8.55E-05 | 0.06 |
| YPR007C | 168.1873056 | 1.852390362 | 0.000107315 | 0.06 |
| YOL131W | 75.70252965 | 2.034063445 | 0.000122794 | 0.06 |
| YCL074W | 140.2978941 | 2.040583607 | 0.000135876 | 0.06 |
| YPL275W | 31.2658679 | 2.059159929 | 0.000148499 | 0.06 |
| YKL174C | 903.8221117 | 0.568643609 | 0.000190097 | 0.07 |
| YLR012C | 11.76101632 | 2.176856294 | 0.000183981 | 0.07 |
| YMR323W | 206.5265565 | 2.168057136 | 0.000203603 | 0.07 |
| YHL028W | 1137.369035 | -1.531125974 | 0.000255172 | 0.08 |
| YMR096W | 4030.130556 | 2.125874231 | 0.000253722 | 0.08 |
| YMR101C | 112.86694 | 1.930706123 | 0.000271086 | 0.08 |
| YJL102W | 772.7229577 | 1.133794957 | 0.000318178 | 0.09 |
| YAL054C | 1109.295849 | 1.894252647 | 0.000416598 | 0.11 |
| YBR157C | 589.2630079 | -1.817209595 | 0.000403676 | 0.11 |
| YGL184C | 275.5662791 | 1.446245041 | 0.00040622 | 0.11 |
| YGL009C | 9382.784081 | 1.830498008 | 0.000439083 | 0.11 |
| YJL217W | 2088.445581 | 1.558395158 | 0.00047454 | 0.11 |
| YAR035W | 1245.571751 | 2.029183878 | 0.000498217 | 0.12 |
| YJL089W | 344.4116839 | 1.859360828 | 0.000538603 | 0.12 |
| YKL107W | 249.2824796 | 1.943083767 | 0.000527395 | 0.12 |
| YDL085C-A | 569.0392384 | -1.237945731 | 0.000580708 | 0.12 |
| YHR079C-A | 27.69347479 | 1.948086521 | 0.000618501 | 0.13 |
| YDR059C | 828.9203049 | 1.059542933 | 0.000784078 | 0.14 |
| YDR209C | 344.9577194 | -1.190833728 | 0.000766615 | 0.14 |
| YER185W | 79.91884135 | 1.341019629 | 0.000811942 | 0.14 |
| YGR095C | 969.1047325 | 0.996608977 | 0.000834338 | 0.14 |
| YMR094W | 306.3491696 | 1.120294578 | 0.000824252 | 0.14 |
| YNR057C | 196.7239761 | 1.087783311 | 0.00084075 | 0.14 |
| YBL065W | 30.86397396 | 1.605560033 | 0.001008848 | 0.15 |
| YBR006W | 1760.927773 | 1.361066144 | 0.001011242 | 0.15 |

| | | | | |
|---|---|---|---|---|
| YBR222C | 4562.01386 | 1.219292206 | 0.000985014 | 0.15 |
| YCL073C | 26.5180597 | 1.074754129 | 0.000941141 | 0.15 |
| YEL047C | 2163.496037 | 0.928367823 | 0.001036951 | 0.15 |
| YNL104C | 8329.192681 | 1.575545249 | 0.000954514 | 0.15 |
| YOL165C | 22.45477283 | 1.845511821 | 0.001054632 | 0.15 |
| YBR279W | 1844.253736 | -0.630458226 | 0.001161458 | 0.16 |
| YCR010C | 273.3869127 | 1.870949276 | 0.001180825 | 0.16 |
| YER044C-A | 102.3682402 | 1.388934308 | 0.00134744 | 0.17 |
| YKL031W | 35.11063328 | 1.60017511 | 0.001356939 | 0.17 |
| YLL059C | 81.93358505 | 1.031973146 | 0.00132745 | 0.17 |
| YMR095C | 600.2590168 | 1.642836625 | 0.00126745 | 0.17 |
| YNL014W | 784.7565686 | 1.674849301 | 0.001364003 | 0.17 |
| YOL097W-A | 14.8332561 | -1.321815876 | 0.001455903 | 0.18 |
| YOR328W | 830.771978 | 1.332525508 | 0.001459784 | 0.18 |
| YBR047W | 453.8190728 | 1.376918378 | 0.001543343 | 0.18 |
| YIL072W | 537.0665215 | 1.142079532 | 0.001499704 | 0.18 |
| YKR071C | 2094.595777 | 0.889123961 | 0.001541145 | 0.18 |
| YFL062W | 38.28792094 | 1.678496787 | 0.001641908 | 0.18 |
| YHR084W | 1211.807628 | -1.315650968 | 0.001658623 | 0.18 |
| YPL119C | 673.8655489 | 1.33301297 | 0.001688457 | 0.18 |
| YGL255W | 1201.748656 | 1.563969028 | 0.001820433 | 0.19 |
| YGL107C | 935.2339762 | -0.362488979 | 0.00191945 | 0.20 |
| YMR324C | 2.573506675 | 1.614381669 | 0.001925174 | 0.20 |
| YEL064C | 340.0873494 | 0.913581716 | 0.00198793 | 0.20 |
| YJL137C | 923.3034038 | 1.509723926 | 0.001974163 | 0.20 |
| YDR014W-A | 36.16776295 | 1.643629716 | 0.002080317 | 0.20 |
| YLL026W | 27987.51895 | 1.082866972 | 0.002204382 | 0.21 |
| YML072C | 6041.463861 | 0.718370025 | 0.002226912 | 0.21 |
| YFL029C | 966.6378114 | 1.290489316 | 0.002291704 | 0.22 |
| YLR047C | 709.1954371 | 0.474884546 | 0.002483963 | 0.22 |
| YLR099W-A | 174.4936248 | -0.841236614 | 0.00250843 | 0.22 |
| YMR015C | 1137.039851 | -1.03975029 | 0.002518768 | 0.22 |
| YNL326C | 366.0651518 | -0.490386867 | 0.00244024 | 0.22 |
| YPL279C | 37.92327787 | -1.390389919 | 0.002486931 | 0.22 |
| YFR057W | 37.29733796 | 0.882189162 | 0.002606341 | 0.23 |
| YNL103W-A | 104.3621644 | 1.533418976 | 0.00285085 | 0.24 |
| YAR075W | 64.72959559 | 1.055090957 | 0.003672469 | 0.25 |
| YBR045C | 255.946774 | 0.952728965 | 0.003593933 | 0.25 |
| YBR258C | 250.7365151 | -0.627652457 | 0.003315455 | 0.25 |
| YDR309C | 2613.4939 | -1.309555313 | 0.003437317 | 0.25 |
| YER179W | 293.0579118 | 1.262790838 | 0.003505081 | 0.25 |
| YFL047W | 1142.35423 | -0.860044469 | 0.003432587 | 0.25 |
| YHR122W | 579.3180386 | 0.891700109 | 0.003073545 | 0.25 |

| | | | | |
|---|---|---|---|---|
| YJL110C | 684.2542418 | -0.731052593 | 0.003630594 | 0.25 |
| YJR046W | 1243.286799 | 0.627997494 | 0.003006073 | 0.25 |
| YLL015W | 2174.31995 | 0.775658556 | 0.003643888 | 0.25 |
| YLR445W | 32.03943686 | 1.446459266 | 0.003633567 | 0.25 |
| YNL240C | 1001.196601 | 0.862149121 | 0.003165677 | 0.25 |
| YNL241C | 6779.137631 | 1.068842025 | 0.002972391 | 0.25 |
| YOL060C | 2188.941543 | 0.811009337 | 0.003197132 | 0.25 |
| YOR338W | 133.0538926 | 0.986074372 | 0.003557156 | 0.25 |
| YOR378W | 266.6242043 | 1.089227756 | 0.003411345 | 0.25 |
| YPL121C | 25.34339254 | 1.510390174 | 0.003225591 | 0.25 |
| YPL254W | 446.984746 | -0.725666424 | 0.003176956 | 0.25 |
| YPL277C | 206.5316036 | 1.363444255 | 0.003363073 | 0.25 |
| snR58 | 5.413360595 | -1.967571632 | 0.003229186 | 0.25 |
| YOR080W | 677.2163658 | -0.480816424 | 0.003778573 | 0.25 |
| YDL030W | 676.7024249 | 0.583934937 | 0.003944979 | 0.26 |
| YOL119C | 1253.305663 | 1.206932405 | 0.004031466 | 0.26 |
| YNL270C | 240.3573834 | 1.432914956 | 0.004163886 | 0.27 |
| YPR111W | 665.5579675 | 0.885098567 | 0.004162046 | 0.27 |
| YDR228C | 901.0372572 | -0.662672547 | 0.004282387 | 0.27 |
| YIL024C | 346.2449471 | 1.14764933 | 0.004288277 | 0.27 |
| YMR322C | 22.28920933 | 1.734096634 | 0.004245291 | 0.27 |
| YMR034C | 377.1167506 | 1.17135238 | 0.004340011 | 0.27 |
| YDL240W | 734.1749038 | -1.040707898 | 0.004447012 | 0.27 |
| YNL152W | 687.0558678 | -0.480310992 | 0.004447252 | 0.27 |
| YKL119C | 426.0044347 | -0.730978861 | 0.004507583 | 0.27 |
| YOR119C | 842.963497 | 0.687369812 | 0.004713979 | 0.28 |
| YGL262W | 77.76796003 | 1.544164601 | 0.004870796 | 0.29 |
| YLR267W | 1321.159087 | 1.627044259 | 0.004881982 | 0.29 |
| YAL005C | 15020.34614 | 0.836587262 | 0.00500672 | 0.29 |
| YMR118C | 295.0484477 | 1.282499558 | 0.005038965 | 0.29 |
| YOR304W | 1385.923767 | -0.441370909 | 0.005039979 | 0.29 |
| YOR114W | 311.2740858 | 1.013598091 | 0.005140495 | 0.29 |
| YCR027C | 255.2084978 | -0.721152496 | 0.005280147 | 0.29 |
| YGR087C | 287.4531423 | 1.624007754 | 0.005232873 | 0.29 |
| YLR173W | 824.1208493 | 1.037503793 | 0.005299642 | 0.29 |
| YLL058W | 677.8123693 | 0.74010707 | 0.00546133 | 0.30 |
| YDL224C | 1328.137391 | -0.707192837 | 0.005524442 | 0.30 |
| YDL093W | 807.8112275 | 0.66281597 | 0.005586475 | 0.30 |
| YFL030W | 1073.611575 | 1.598746579 | 0.005724762 | 0.30 |
| YJR148W | 6070.611654 | 1.145588148 | 0.005732175 | 0.30 |
| YBR158W | 2964.217702 | -0.972502641 | 0.005866037 | 0.31 |
| YER099C | 790.1735302 | 0.441563407 | 0.006007607 | 0.31 |
| YER145C | 1496.957868 | -0.955137993 | 0.005979606 | 0.31 |

| | | | | |
|---|---|---|---|---|
| YKR005C | 237.9372028 | 0.960666081 | 0.005915884 | 0.31 |
| YNR073C | 195.2436335 | 1.586605739 | 0.005921923 | 0.31 |
| YBR083W | 2339.410841 | -1.107918559 | 0.00613235 | 0.31 |
| YER096W | 439.4636005 | 1.470882252 | 0.006139222 | 0.31 |
| YBR218C | 7147.332039 | 0.759445606 | 0.006317235 | 0.31 |
| YDR526C | 4.767823074 | 1.593506183 | 0.006350562 | 0.31 |
| YAR073W | 171.304609 | 1.213777767 | 0.006481675 | 0.32 |
| YCL075W | 37.39387639 | 1.540407033 | 0.006497255 | 0.32 |
| YPL038W | 299.6200602 | -0.89213289 | 0.006716002 | 0.32 |
| YDL085W | 757.5796414 | 1.595700613 | 0.006866031 | 0.33 |
| YOL161C | 3.517453047 | 1.173139052 | 0.00697338 | 0.33 |
| YOR371C | 940.0880721 | 0.746774621 | 0.006938565 | 0.33 |
| YPR001W | 533.0217534 | 1.493010455 | 0.00703079 | 0.33 |
| YAR020C | 43.79066228 | 1.188177869 | 0.007089856 | 0.33 |
| YGL081W | 335.8828397 | 1.324332857 | 0.007296792 | 0.33 |
| YMR325W | 4.52283349 | 1.360058562 | 0.007250359 | 0.33 |
| YNL242W | 1398.740797 | 0.711727228 | 0.007321126 | 0.33 |
| YPL276W | 15.28614474 | 1.403132473 | 0.007218987 | 0.33 |
| YPL164C | 886.5830673 | 0.810422595 | 0.007419847 | 0.33 |
| YER048W-A | 659.0895735 | -0.364300935 | 0.007862565 | 0.34 |
| YGR269W | 48.40957868 | 0.878757126 | 0.007746785 | 0.34 |
| YJR126C | 1827.592541 | 0.454106189 | 0.00779553 | 0.34 |
| YMR191W | 2004.986398 | 1.314153718 | 0.00787148 | 0.34 |
| YBL015W | 5485.33881 | 1.532798325 | 0.008031994 | 0.35 |
| YML089C | 19.45677984 | 1.280670945 | 0.007980624 | 0.35 |
| YOR154W | 667.1567549 | 0.526525067 | 0.008209473 | 0.35 |
| YPL092W | 1197.47636 | 1.131521026 | 0.008156031 | 0.35 |
| YEL062W | 560.1717097 | -0.619066545 | 0.008265576 | 0.35 |
| YDR235W | 670.6497022 | -0.536199144 | 0.00863872 | 0.36 |
| YDR481C | 2453.405315 | 0.881888854 | 0.008754313 | 0.36 |
| YGL233W | 1565.055775 | -0.440246129 | 0.009158153 | 0.38 |
| YML100W-A | 6.289539175 | 1.371611427 | 0.009215298 | 0.38 |
| YOR367W | 630.0179766 | -0.66972941 | 0.009227129 | 0.38 |
| YDR389W | 894.8507584 | -0.653637773 | 0.00952862 | 0.38 |
| YLL046C | 28.77526537 | 1.421117837 | 0.009563276 | 0.38 |
| YLR326W | 402.567239 | 0.812845304 | 0.009457431 | 0.38 |
| YPL177C | 1454.555337 | -0.607882903 | 0.009506618 | 0.38 |
| YNL012W | 592.4635068 | 0.894258775 | 0.009680403 | 0.38 |
| YMR306W | 518.3037802 | 0.80552297 | 0.009776557 | 0.39 |
| YCR061W | 1066.525845 | 0.878904473 | 0.010004586 | 0.39 |
| YJR061W | 732.5988419 | 0.651013544 | 0.010124798 | 0.39 |
| YNL188W | 592.8677499 | -0.876097565 | 0.010166773 | 0.39 |
| YGL198W | 494.0719347 | 0.744395561 | 0.010483156 | 0.40 |

| | | | | |
|---|---|---|---|---|
| YGR065C | 698.7091847 | 0.896386246 | 0.010485434 | 0.40 |
| YIL161W | 488.7537738 | -0.637892801 | 0.010598359 | 0.40 |
| YLR053C | 106.3162181 | 1.150263499 | 0.010623193 | 0.40 |
| YMR133W | 502.5352221 | 1.012539679 | 0.010537297 | 0.40 |
| YGL062W | 7360.866755 | 1.379566782 | 0.010853824 | 0.41 |
| YBR130C | 1160.408881 | -0.654390152 | 0.01148869 | 0.41 |
| YDL045W-A | 694.1101578 | -0.776452213 | 0.011227719 | 0.41 |
| YDL054C | 853.6605887 | 0.771936687 | 0.012230122 | 0.41 |
| YDL214C | 464.7760152 | 1.292786723 | 0.011575083 | 0.41 |
| YDR158W | 6048.209084 | 0.865739591 | 0.011598574 | 0.41 |
| YDR331W | 571.3069512 | -0.562327428 | 0.011586902 | 0.41 |
| YDR371W | 631.8329655 | -0.728586632 | 0.011527101 | 0.41 |
| YFL053W | 516.855343 | 1.360098114 | 0.012237855 | 0.41 |
| YGL033W | 39.3922468 | 1.270236615 | 0.012015139 | 0.41 |
| YGL106W | 2381.873892 | -0.546548875 | 0.011784423 | 0.41 |
| YGL182C | 10.33516089 | 1.19149421 | 0.012164468 | 0.41 |
| YGL205W | 247.6080273 | 1.06078642 | 0.011798866 | 0.41 |
| YGR108W | 622.4932134 | -0.833613313 | 0.012053657 | 0.41 |
| YKR063C | 686.126014 | 0.448244801 | 0.01195952 | 0.41 |
| YLR255C | 23.50063941 | 1.05377709 | 0.012342937 | 0.41 |
| YLR308W | 11.06339218 | 1.29381497 | 0.012174987 | 0.41 |
| YLR335W | 2838.169755 | 0.473667886 | 0.012005449 | 0.41 |
| YNL125C | 2090.525445 | 0.800538104 | 0.011175497 | 0.41 |
| YNL274C | 1846.518968 | 1.02978827 | 0.012054428 | 0.41 |
| YOR315W | 1158.774212 | -1.163514061 | 0.012305202 | 0.41 |
| YPR002W | 1192.998192 | 1.457991788 | 0.012484078 | 0.41 |
| snR55 | 9.080532067 | 1.415216142 | 0.012446261 | 0.41 |
| YFL027C | 673.7008422 | -0.927575249 | 0.012569256 | 0.41 |
| YBL026W | 682.7010545 | -0.449895975 | 0.012777427 | 0.42 |
| YLR362W | 1083.342817 | 0.861110293 | 0.012819902 | 0.42 |
| YIL150C | 295.6540585 | 0.643868041 | 0.01313178 | 0.43 |
| YDL044C | 428.1351393 | -0.535754937 | 0.01333367 | 0.43 |
| YEL030W | 524.4916329 | 0.720279743 | 0.013267683 | 0.43 |
| YNL081C | 921.3541155 | -0.506278451 | 0.013347088 | 0.43 |
| RNA170 | 10.42132142 | 1.160883772 | 0.013712411 | 0.43 |
| YGL075C | 312.3284424 | 0.524540418 | 0.013653585 | 0.43 |
| YHL022C | 136.8231823 | 1.140530656 | 0.013743326 | 0.43 |
| YMR319C | 2635.007516 | 0.942052478 | 0.013560846 | 0.43 |
| YMR187C | 357.2988229 | 0.68036109 | 0.013886658 | 0.43 |
| YKR050W | 1256.621968 | 0.832218396 | 0.014098109 | 0.44 |
| YPL091W | 3000.977941 | 0.57848178 | 0.014150814 | 0.44 |
| YDL179W | 455.9668158 | -0.538975083 | 0.014384854 | 0.44 |
| YGL124C | 833.4516769 | 0.551189378 | 0.014366602 | 0.44 |

| | | | | |
|---|---|---|---|---|
| YGR096W | 275.6515308 | 0.710255346 | 0.014505874 | 0.44 |
| YJR079W | 11.62148612 | 1.348355512 | 0.014441893 | 0.44 |
| YPL034W | 171.3658521 | -0.646466153 | 0.014381458 | 0.44 |
| YPL134C | 1579.391088 | 1.279009283 | 0.014619161 | 0.44 |
| YLR263W | 707.0718116 | 1.094617388 | 0.014693458 | 0.44 |
| YDL215C | 3694.89417 | 1.19687754 | 0.014875772 | 0.44 |
| YLR089C | 5710.768483 | 1.187753368 | 0.014997805 | 0.45 |
| YDL168W | 1463.24926 | 0.843035285 | 0.015597176 | 0.45 |
| YGL125W | 958.6893776 | 0.883629876 | 0.015233451 | 0.45 |
| YHL037C | 20.29589679 | 1.233215791 | 0.015438015 | 0.45 |
| YHR202W | 670.4759746 | 0.754911088 | 0.015298071 | 0.45 |
| YLL056C | 287.6766171 | 0.957975031 | 0.015383113 | 0.45 |
| YLL062C | 209.8685733 | 0.852592013 | 0.0153228 | 0.45 |
| YNL045W | 3993.621371 | 1.303933419 | 0.01563215 | 0.45 |
| YOL044W | 473.6470695 | 0.75463835 | 0.015514994 | 0.45 |
| YOR392W | 4.708188968 | 1.178255015 | 0.015642308 | 0.45 |
| YDL230W | 846.3743425 | 0.661674122 | 0.01593414 | 0.45 |
| YPL119C-A | 58.40963327 | 1.188274941 | 0.016038047 | 0.45 |
| YBR052C | 2651.004465 | -0.535281088 | 0.016376497 | 0.45 |
| YGL240W | 171.0354676 | 0.577008325 | 0.016409018 | 0.45 |
| YLR015W | 562.8104697 | 0.621252594 | 0.016201951 | 0.45 |
| YLR219W | 2484.562083 | 0.70574097 | 0.016401112 | 0.45 |
| YMR166C | 428.7049362 | -0.403788802 | 0.016155595 | 0.45 |
| YDR421W | 1031.149921 | 0.583911247 | 0.016578122 | 0.46 |
| YER081W | 9451.152045 | 1.278883715 | 0.016781916 | 0.46 |
| YHR208W | 7460.933205 | 1.066727705 | 0.016908495 | 0.46 |
| YJL140W | 1600.235301 | -0.466675493 | 0.016781637 | 0.46 |
| YNL254C | 501.4634557 | 0.721588902 | 0.016886771 | 0.46 |
| YMR161W | 724.2845416 | -0.426333802 | 0.017008576 | 0.46 |
| YHR014W | 138.7802733 | 0.976902551 | 0.017223543 | 0.46 |
| YHR160C | 281.5696466 | 0.897957162 | 0.017193208 | 0.46 |
| YKL209C | 3091.85579 | -1.195538945 | 0.017083144 | 0.46 |
| YJR098C | 316.068717 | 0.631317311 | 0.017362343 | 0.46 |
| YCL027W | 675.5367401 | -1.200465654 | 0.017481231 | 0.46 |
| YLR077W | 986.4922801 | 0.396672421 | 0.01772739 | 0.47 |
| YDR488C | 747.4804028 | 0.757335607 | 0.017801511 | 0.47 |
| YKL170W | 81.53735369 | -0.71832062 | 0.017887083 | 0.47 |
| YNL013C | 5.629906296 | 1.506381069 | 0.017985316 | 0.47 |
| YNL196C | 204.2397635 | 1.232437754 | 0.018436581 | 0.48 |
| YBR046C | 1727.421703 | 0.926254936 | 0.018823024 | 0.48 |
| YBR253W | 396.1883397 | -0.448553395 | 0.019198603 | 0.48 |
| YDL139C | 356.8731646 | 0.514715631 | 0.019447757 | 0.48 |
| YDL194W | 1083.690954 | 0.898787196 | 0.019471728 | 0.48 |

| | | | | |
|---|---|---|---|---|
| YEL070W | 298.2975483 | 1.409319343 | 0.019322638 | 0.48 |
| YFL007W | 4593.396937 | 0.602355102 | 0.018992254 | 0.48 |
| YLR393W | 270.1892054 | 0.731856274 | 0.019267449 | 0.48 |
| YLR415C | 24.41748512 | 1.115942927 | 0.019436656 | 0.48 |
| YLR416C | 28.8196825 | 1.152401334 | 0.019460322 | 0.48 |
| YNL116W | 793.4753006 | 0.805109967 | 0.018817887 | 0.48 |
| YOL106W | 14.51135789 | 1.022181819 | 0.019027814 | 0.48 |
| YPL150W | 1067.29309 | 0.59626981 | 0.018861379 | 0.48 |
| YPR064W | 39.20156943 | 0.964195135 | 0.019549267 | 0.48 |
| YEL040W | 3165.37713 | -1.039110057 | 0.019744479 | 0.48 |
| YBR233W-A | 253.0169158 | -0.765931159 | 0.020039645 | 0.49 |
| YGL082W | 1508.755452 | -0.379763131 | 0.020278502 | 0.49 |
| YGR125W | 1519.849762 | 0.44731388 | 0.020286758 | 0.49 |
| YGR142W | 4162.125638 | 1.014344236 | 0.019903242 | 0.49 |
| YIR010W | 867.0379525 | -0.57474503 | 0.020333594 | 0.49 |
| YJL157C | 1263.741636 | -1.105954296 | 0.020227626 | 0.49 |
| YOR077W | 225.6560466 | 0.625192985 | 0.020321407 | 0.49 |
| YNL165W | 500.9806009 | 0.652509603 | 0.02058039 | 0.49 |
| YKL058W | 636.5138347 | -0.515331362 | 0.02075222 | 0.49 |
| YCR003W | 578.0986481 | -0.692123634 | 0.020884424 | 0.49 |
| YOL088C | 665.7794028 | -0.624597696 | 0.021063243 | 0.50 |
| YOR093C | 1072.002934 | -0.39516783 | 0.021178459 | 0.50 |
| YLR163C | 1121.057782 | 0.502930826 | 0.021298208 | 0.50 |
| YJL127C | 657.9035431 | 0.376384556 | 0.021716697 | 0.50 |
| YJR004C | 1252.603429 | 0.955232058 | 0.021901365 | 0.50 |
| YLR214W | 1682.159024 | -0.885834002 | 0.02170551 | 0.50 |
| YNL258C | 964.2763945 | -0.465791855 | 0.021932119 | 0.50 |
| YNR058W | 282.8560691 | 0.732057635 | 0.022020126 | 0.50 |
| YOR390W | 45.23234325 | -1.015052694 | 0.021988306 | 0.50 |
| YPL066W | 1565.740738 | -0.483889197 | 0.022076639 | 0.50 |
| YPL147W | 471.9776317 | 0.938251832 | 0.022099321 | 0.50 |
| YDR018C | 446.8147304 | 1.037317892 | 0.022547343 | 0.51 |
| YGL114W | 1125.03715 | 0.593345254 | 0.022895463 | 0.51 |
| YJR011C | 218.4017674 | -0.460541359 | 0.022831728 | 0.51 |
| YNL269W | 50.67416069 | 1.182499987 | 0.02294694 | 0.51 |
| YCL076W | 24.7210401 | 1.298110438 | 0.023145394 | 0.51 |
| YKL179C | 1404.902826 | -0.464095532 | 0.023198676 | 0.51 |
| YLR013W | 3.60279815 | 1.26381464 | 0.023151403 | 0.51 |
| YGR188C | 503.2494794 | 0.649033813 | 0.023340733 | 0.52 |
| YCL040W | 21911.49524 | 1.007461626 | 0.023504639 | 0.52 |
| YCL033C | 876.4314652 | 0.967521883 | 0.024057227 | 0.52 |
| YDR063W | 1183.071975 | -0.401068512 | 0.024038 | 0.52 |
| YGL204C | 78.35386912 | -0.901769253 | 0.024177537 | 0.52 |

| | | | | |
|---|---|---|---|---|
| YKL208W | 299.0743362 | -0.569486849 | 0.024164272 | 0.52 |
| YLR189C | 2093.552884 | 0.619635164 | 0.023970648 | 0.52 |
| YMR236W | 1041.170124 | -0.374012796 | 0.023662152 | 0.52 |
| YOR364W | 7.913433737 | 1.163553669 | 0.024040111 | 0.52 |
| YPL007C | 583.714763 | -0.349704158 | 0.024070753 | 0.52 |
| YDR363W-A | 1308.889566 | -0.548104016 | 0.024426602 | 0.52 |
| YOR237W | 107.6435638 | 1.051407704 | 0.024387286 | 0.52 |
| YPL162C | 476.9994766 | 0.631894433 | 0.024505601 | 0.52 |
| YHR155W | 1225.572051 | -0.389229152 | 0.024770612 | 0.52 |
| YCL023C | 8.286348364 | 1.067965089 | 0.02491054 | 0.53 |
| YDR374C | 71.50374083 | 1.250157862 | 0.025287685 | 0.53 |
| YEL009C-A | 8.998175135 | 1.028865377 | 0.024973453 | 0.53 |
| YER042W | 653.285425 | 0.904010032 | 0.025408592 | 0.53 |
| YFR025C | 947.4315897 | 0.392196663 | 0.02538278 | 0.53 |
| YGL165C | 78.38274117 | 0.575612409 | 0.025446676 | 0.53 |
| YLR253W | 1177.596684 | 0.83189906 | 0.025195024 | 0.53 |
| YOL146W | 364.144711 | -0.590226604 | 0.025163835 | 0.53 |
| YCL069W | 36.22888945 | 0.539951503 | 0.025713992 | 0.53 |
| YGL059W | 1140.532396 | 0.736809839 | 0.025686056 | 0.53 |
| YJR054W | 539.7820263 | -0.555807455 | 0.025814362 | 0.53 |
| YGR035C | 329.4527926 | -0.998335819 | 0.026146508 | 0.53 |
| YHR106W | 1096.925363 | 0.762353938 | 0.026182291 | 0.53 |
| YPR049C | 968.2983267 | 0.780183588 | 0.026321512 | 0.53 |
| YCL026C-B | 269.8485152 | 0.902627067 | 0.026698337 | 0.53 |
| YEL030C-A | 7.262884205 | 1.105137188 | 0.026735249 | 0.53 |
| YIL009C-A | 425.8589138 | -0.624010426 | 0.026776403 | 0.53 |
| YMR085W | 377.7619403 | 1.002854527 | 0.026596172 | 0.53 |
| YOL038C-A | 23.65291474 | -0.870252204 | 0.026774126 | 0.53 |
| YGL151W | 1728.404115 | -0.520658732 | 0.027151609 | 0.54 |
| YLR296W | 4.726955448 | 1.220932534 | 0.027176739 | 0.54 |
| YMR182C | 564.4635256 | 0.652096929 | 0.027174196 | 0.54 |
| YOR389W | 203.2493311 | 0.871945886 | 0.027275925 | 0.54 |
| YMR114C | 1303.792032 | 0.932941044 | 0.027472631 | 0.54 |
| YDR258C | 9048.121163 | 0.933719817 | 0.027842901 | 0.55 |
| YAR047C | 21.66468268 | 1.108175366 | 0.028090844 | 0.55 |
| YCR021C | 10522.50846 | 1.292715811 | 0.028012429 | 0.55 |
| YDR518W | 1466.448011 | 0.785400235 | 0.02841299 | 0.55 |
| YBR108W | 1821.037124 | 0.506101318 | 0.029042935 | 0.55 |
| YDR230W | 158.0472236 | 0.828136663 | 0.028727841 | 0.55 |
| YGL091C | 545.7319922 | 0.329987839 | 0.028875024 | 0.55 |
| YGR276C | 1041.106244 | 0.338726186 | 0.028878709 | 0.55 |
| YKL217W | 2399.793467 | 1.288257373 | 0.02903833 | 0.55 |
| YNL023C | 950.809325 | 0.470938707 | 0.028599009 | 0.55 |

| | | | | |
|---|---|---|---|---|
| YPL240C | 19482.15572 | 0.906276124 | 0.028783734 | 0.55 |
| YCL056C | 401.4808141 | -0.399718318 | 0.02932187 | 0.56 |
| YCR060W | 437.7609902 | 0.647728775 | 0.02973186 | 0.56 |
| YDL078C | 3642.136284 | 0.787569486 | 0.029717975 | 0.56 |
| YDR204W | 1297.475345 | 0.895522709 | 0.029793542 | 0.56 |
| YER175C | 1510.549619 | 1.091093525 | 0.029803181 | 0.56 |
| YKR018C | 3562.456922 | 0.789118194 | 0.029737495 | 0.56 |
| YKR070W | 801.1916976 | 0.312798528 | 0.029899977 | 0.56 |
| YBR255W | 917.9047892 | 0.482685992 | 0.030140265 | 0.56 |
| YMR148W | 862.6307015 | 0.573286025 | 0.030086575 | 0.56 |
| YHR086W | 910.6374675 | 0.580950292 | 0.030347179 | 0.56 |
| YML121W | 941.5006143 | 0.421109293 | 0.030292305 | 0.56 |
| YDR407C | 1886.152895 | -0.484961242 | 0.030865612 | 0.56 |
| YGR110W | 407.9203872 | 0.742982558 | 0.030793971 | 0.56 |
| YLR035C-A | 40.57446476 | 0.422309809 | 0.030637506 | 0.56 |
| YMR233W | 360.9564304 | -0.780173956 | 0.03073138 | 0.56 |
| YNL171C | 59.8874962 | 0.736350302 | 0.031075524 | 0.56 |
| YCL058C | 103.80183 | 0.836064947 | 0.031600992 | 0.57 |
| YPL232W | 2081.428239 | -0.481259027 | 0.031799048 | 0.57 |
| YBR204C | 1064.479878 | 0.633768223 | 0.03197777 | 0.58 |
| YBL075C | 2650.065607 | 1.284626785 | 0.032557932 | 0.58 |
| YDL232W | 292.6096052 | -0.660661071 | 0.03259953 | 0.58 |
| YDR310C | 1697.719061 | -0.435097939 | 0.032581103 | 0.58 |
| YPL204W | 2957.690828 | -0.40094183 | 0.032429093 | 0.58 |
| YJL106W | 294.2742704 | 0.758324546 | 0.032801126 | 0.58 |
| YCR005C | 6272.799933 | 1.088037622 | 0.03299994 | 0.58 |
| YJR042W | 1389.72183 | -0.485201457 | 0.033125785 | 0.59 |
| YLL014W | 750.2093462 | -0.546878994 | 0.034037746 | 0.60 |
| YDR239C | 796.3576703 | 0.406487223 | 0.034347797 | 0.60 |
| YBR066C | 407.0918605 | -0.987681749 | 0.034842841 | 0.60 |
| YDR090C | 473.3492687 | 0.486657469 | 0.035465802 | 0.60 |
| YGR063C | 116.1850697 | -0.445666742 | 0.035284674 | 0.60 |
| YGR161C-D | 41.89517477 | 1.040416688 | 0.035182797 | 0.60 |
| YGR201C | 974.6666707 | 1.137608389 | 0.034750456 | 0.60 |
| YIL033C | 4130.82095 | 0.628332361 | 0.034600228 | 0.60 |
| YJR016C | 6106.441237 | 0.814148628 | 0.035283706 | 0.60 |
| YKL111C | 22.34564795 | -0.917933047 | 0.035449482 | 0.60 |
| YLR286C | 14573.14708 | 0.576208028 | 0.035368097 | 0.60 |
| YLR324W | 1737.806437 | 0.665010127 | 0.035263735 | 0.60 |
| YMR170C | 596.7981642 | 0.91365454 | 0.03463233 | 0.60 |
| YPL110C | 931.3902179 | 0.554574821 | 0.034799253 | 0.60 |
| YNL307C | 2792.714562 | -0.507831688 | 0.035877712 | 0.61 |
| YPR121W | 333.11981 | -0.721065997 | 0.035929706 | 0.61 |

| | | | | |
|---|---|---|---|---|
| YBR098W | 547.1579177 | 0.444867587 | 0.036037013 | 0.61 |
| YBR012W-B | 321.6022163 | 0.975033434 | 0.0373384 | 0.61 |
| YBR234C | 4120.481034 | -0.3873432 | 0.03708011 | 0.61 |
| YCR028C | 527.4192161 | 0.363097256 | 0.036603571 | 0.61 |
| YDL101C | 860.843029 | 0.614924022 | 0.036958567 | 0.61 |
| YDL248W | 224.0356228 | 0.890251013 | 0.036872232 | 0.61 |
| YDR201W | 235.4443499 | -0.597102114 | 0.037062884 | 0.61 |
| YDR424C | 383.6752581 | -0.417381297 | 0.036870751 | 0.61 |
| YER015W | 725.609463 | 0.72927912 | 0.038094945 | 0.61 |
| YER017C | 1876.346183 | 0.408703992 | 0.03627777 | 0.61 |
| YER070W | 2200.790509 | -0.879175036 | 0.036974538 | 0.61 |
| YER071C | 272.1086234 | -0.544766652 | 0.038233974 | 0.61 |
| YHR005C | 1534.146963 | -1.090826289 | 0.038377185 | 0.61 |
| YIL013C | 704.2538412 | 0.530408745 | 0.038111595 | 0.61 |
| YIL142C-A | 7.771319082 | 0.815609788 | 0.037918627 | 0.61 |
| YJL045W | 286.1990705 | 1.025540883 | 0.038202731 | 0.61 |
| YJR149W | 627.5970928 | 0.902152488 | 0.036749239 | 0.61 |
| YKL138C | 823.724232 | -0.631388212 | 0.036914747 | 0.61 |
| YLR126C | 468.0585451 | -0.712301174 | 0.037424081 | 0.61 |
| YLR307C-A | 38.91718273 | 1.232717268 | 0.038384094 | 0.61 |
| YNL054W-B | 451.4424074 | 0.935499606 | 0.038053412 | 0.61 |
| YNL227C | 372.4616162 | 0.690117016 | 0.038190917 | 0.61 |
| YOR211C | 1075.422511 | -0.463497946 | 0.038208183 | 0.61 |
| YOR303W | 3621.710527 | 0.998614643 | 0.037191842 | 0.61 |
| YOR354C | 1688.06875 | -0.358195957 | 0.038291762 | 0.61 |
| YKR103W | 526.4204375 | 0.849447951 | 0.039260314 | 0.62 |
| YBL067C | 1161.166699 | 0.316259829 | 0.039646159 | 0.62 |
| YLR133W | 1071.862832 | 0.454351346 | 0.039400343 | 0.62 |
| YML103C | 3218.889825 | -0.556499706 | 0.039636355 | 0.62 |
| YNL034W | 79.93693595 | 0.960147972 | 0.039643784 | 0.62 |
| YPR075C | 1855.120278 | 0.568448047 | 0.039776014 | 0.62 |
| YDL169C | 664.258116 | 1.015759358 | 0.041148826 | 0.62 |
| YDR076W | 307.3025521 | 0.556058319 | 0.041021089 | 0.62 |
| YDR210C-D | 43.7176532 | 1.006816042 | 0.04044245 | 0.62 |
| YDR224C | 8028.945616 | -0.663992637 | 0.040587842 | 0.62 |
| YER085C | 1.090623589 | 0.968056988 | 0.040893857 | 0.62 |
| YJR047C | 407.0063415 | 1.074913512 | 0.040028499 | 0.62 |
| YLR033W | 1049.258607 | -0.387787021 | 0.040858124 | 0.62 |
| YLR216C | 5399.303942 | 0.635417013 | 0.040302746 | 0.62 |
| YLR234W | 283.065063 | 0.721081969 | 0.041081232 | 0.62 |
| YLR452C | 1814.469836 | -0.92245897 | 0.040693649 | 0.62 |
| YMR174C | 615.2184477 | 1.242676257 | 0.041084066 | 0.62 |
| YOL152W | 484.6917609 | 1.036751545 | 0.04108494 | 0.62 |

| | | | | |
|---|---|---|---|---|
| YPR092W | 28.34006108 | 0.724658223 | 0.041102476 | 0.62 |
| snR14 | 19.85838705 | -0.895039598 | 0.040654101 | 0.62 |
| YDL016C | 38.72923011 | -0.558405971 | 0.041269614 | 0.62 |
| YAR031W | 151.5423779 | 0.701055001 | 0.042409433 | 0.63 |
| YCL042W | 285.5413665 | 0.846475319 | 0.042351562 | 0.63 |
| YDR446W | 60.61919013 | 1.044842882 | 0.042205676 | 0.63 |
| YER104W | 241.2969985 | 0.548111625 | 0.041898971 | 0.63 |
| YGR093W | 815.5848449 | 0.494711504 | 0.042233303 | 0.63 |
| YMR103C | 375.8522499 | 0.894063938 | 0.042475016 | 0.63 |
| YPL201C | 234.758662 | 1.170250866 | 0.042156349 | 0.63 |
| YBR028C | 772.1128092 | -0.655373311 | 0.042744992 | 0.63 |
| YGR113W | 643.4472036 | -0.292123345 | 0.042711873 | 0.63 |
| YJR155W | 885.0501737 | 0.993702304 | 0.042890312 | 0.63 |
| YOR391C | 10.43191586 | 1.251007645 | 0.042966839 | 0.63 |
| YLR132C | 337.4595812 | 0.391914903 | 0.04308697 | 0.63 |
| YDL243C | 368.4938618 | 1.023377819 | 0.0437495 | 0.64 |
| YOL058W | 2645.481694 | 0.967484845 | 0.043678091 | 0.64 |
| YOL155C | 4950.84233 | 0.855163091 | 0.043889259 | 0.64 |
| YOR343C | 63.14207102 | 0.951228028 | 0.044160594 | 0.64 |
| YIL099W | 500.7923991 | 1.03033447 | 0.044557661 | 0.65 |
| YDR030C | 444.8820441 | 0.650759468 | 0.044741174 | 0.65 |
| YKL131W | 17.12686382 | -0.934140588 | 0.04493718 | 0.65 |
| YMR056C | 319.9319845 | 0.990627524 | 0.044915437 | 0.65 |
| YKL097C | 10.27058523 | 1.08374086 | 0.045109101 | 0.65 |
| YCR075W-A | 129.8943621 | -0.623190962 | 0.045829116 | 0.65 |
| YDL149W | 944.7875532 | 0.689916333 | 0.04579442 | 0.65 |
| YGL162W | 395.3975497 | -0.985501447 | 0.045734443 | 0.65 |
| YJR129C | 222.0216409 | 0.750311449 | 0.045564662 | 0.65 |
| YKL050C | 526.5764301 | 0.697192004 | 0.045580132 | 0.65 |
| snR78 | 1.974544426 | 0.729072 | 0.045866372 | 0.65 |
| RUF20 | 86.17514557 | -0.64535472 | 0.046170825 | 0.65 |
| YCR098C | 141.8503964 | 1.144797813 | 0.046199734 | 0.65 |
| YFR026C | 243.3675063 | 1.05664142 | 0.046315892 | 0.65 |
| YBR068C | 7312.593911 | 0.806754026 | 0.04679786 | 0.66 |
| YHL032C | 2561.780761 | 0.908467839 | 0.046742107 | 0.66 |
| YNL212W | 1282.46661 | 0.492395736 | 0.047100674 | 0.66 |
| YIR013C | 17.07044727 | 0.939904061 | 0.047312762 | 0.66 |
| YOL134C | 23.62819378 | -0.76071853 | 0.047214982 | 0.66 |
| YGR129W | 435.9932211 | 0.692042116 | 0.047572779 | 0.66 |
| YLR217W | 16.95655225 | 0.881027121 | 0.047478127 | 0.66 |
| YGR022C | 1.81368062 | 0.923385566 | 0.047839967 | 0.66 |
| YOR152C | 552.1803625 | 0.770255495 | 0.047921365 | 0.66 |
| YBL109W | 16.60491535 | 0.875468244 | 0.048143459 | 0.67 |

| | | | | |
|---|---|---|---|---|
| YOR159C | 320.0573697 | -0.52501688 | 0.048362653 | 0.67 |
| YDR392W | 654.5457347 | -0.388311212 | 0.048596286 | 0.67 |
| YGR206W | 264.0711736 | -0.492096549 | 0.048693938 | 0.67 |
| YHR213W-B | 12.16949824 | 0.939958266 | 0.048809006 | 0.67 |
| YOR388C | 10.58733181 | 1.071287894 | 0.048792723 | 0.67 |
| YBR280C | 1033.06933 | 0.876046434 | 0.049030392 | 0.67 |
| YBR297W | 465.5986861 | 0.839400736 | 0.048966417 | 0.67 |
| YPR027C | 342.9171324 | -1.069009629 | 0.049186024 | 0.67 |
| YBR259W | 488.2154125 | -0.34430468 | 0.049633836 | 0.67 |
| YGR149W | 1082.04674 | 0.579629822 | 0.049505697 | 0.67 |
| YGR216C | 759.5937639 | 0.343295223 | 0.049368864 | 0.67 |
| YPR174C | 411.2973194 | 0.528986233 | 0.049649738 | 0.67 |
| YOR186W | 27.95959597 | 0.971517114 | 0.049775506 | 0.67 |
| YBR290W | 1203.208406 | -0.518864029 | 0.050776488 | 0.67 |
| YDL003W | 760.5875817 | -0.729446055 | 0.050418823 | 0.67 |
| YDR229W | 1110.14149 | -0.439839382 | 0.050109211 | 0.67 |
| YDR490C | 1094.352116 | 0.503160737 | 0.050342309 | 0.67 |
| YER178W | 9418.417677 | 0.420037224 | 0.050497336 | 0.67 |
| YGR164W | 4.051111594 | -1.279884102 | 0.050854173 | 0.67 |
| YJR077C | 4794.775519 | 0.827569305 | 0.050657829 | 0.67 |
| YKR027W | 720.4128865 | -0.378447961 | 0.050379727 | 0.67 |
| YKR102W | 639.1097035 | 0.992698404 | 0.050908814 | 0.67 |
| YPL021W | 14.06563719 | 1.037569743 | 0.050669379 | 0.67 |
| YDR137W | 962.3953612 | -0.301702468 | 0.051697703 | 0.67 |
| YDR268W | 586.9385642 | 0.509481584 | 0.051674223 | 0.67 |
| YDR273W | 249.707739 | 0.691818231 | 0.051883605 | 0.67 |
| YJL154C | 2489.813399 | 0.240990821 | 0.05170035 | 0.67 |
| YLR034C | 1366.422953 | -0.58660426 | 0.051414299 | 0.67 |
| YMR001C | 1386.236499 | -0.514985421 | 0.051795396 | 0.67 |
| YPR052C | 667.2448711 | -0.638744901 | 0.051337113 | 0.67 |
| YEL056W | 977.5251167 | -0.345606509 | 0.051998689 | 0.67 |
| YGL047W | 625.4320388 | 0.433277085 | 0.053282704 | 0.68 |
| YGL177W | 36.28800868 | 0.704210266 | 0.053251238 | 0.68 |
| YGR150C | 1213.396537 | -0.380571129 | 0.052647072 | 0.68 |
| YIL166C | 439.1240138 | 0.696487613 | 0.053026164 | 0.68 |
| YJR049C | 1397.771565 | 0.567749609 | 0.05225422 | 0.68 |
| YMR040W | 194.5584091 | 0.837368622 | 0.052537893 | 0.68 |
| YMR065W | 380.856771 | -0.448251622 | 0.05312565 | 0.68 |
| YNL143C | 26.87962177 | -0.862121976 | 0.052390581 | 0.68 |
| YOR126C | 423.6858153 | 0.329256047 | 0.052773275 | 0.68 |
| YPR056W | 512.7372419 | -0.330414771 | 0.052846683 | 0.68 |
| YPR172W | 1219.274092 | 0.626080426 | 0.053139344 | 0.68 |
| YIL002W-A | 495.5817191 | -0.422257238 | 0.053398322 | 0.68 |

| | | | | |
|---|---|---|---|---|
| YKL118W | 29.53687639 | -0.771560624 | 0.053539103 | 0.68 |
| LSR1 | 186.1998782 | 0.923248147 | 0.05400331 | 0.68 |
| YKL147C | 17.64954721 | 0.913229076 | 0.053905108 | 0.68 |
| YLR273C | 457.5126435 | 0.505186614 | 0.053859926 | 0.68 |
| YIL146C | 613.496777 | 0.714968654 | 0.054483162 | 0.68 |
| YLL022C | 865.5611491 | 0.515638364 | 0.054715344 | 0.69 |
| YPR128C | 787.6455183 | 0.404227028 | 0.05490773 | 0.69 |
| YER152C | 1264.809581 | 0.667165247 | 0.055023971 | 0.69 |
| YAL020C | 521.9507996 | 0.533327698 | 0.056595468 | 0.69 |
| YCR004C | 3730.565062 | -0.437346285 | 0.056301888 | 0.69 |
| YDL017W | 421.2237274 | 0.379811411 | 0.056732258 | 0.69 |
| YDL197C | 997.8504474 | 0.510792137 | 0.055441692 | 0.69 |
| YDR103W | 991.2634766 | -0.867855164 | 0.056974665 | 0.69 |
| YDR303C | 1032.483515 | -0.596889969 | 0.057303237 | 0.69 |
| YER155C | 4972.789594 | -0.441273832 | 0.057231299 | 0.69 |
| YGL111W | 1214.214479 | 0.589630463 | 0.05715213 | 0.69 |
| YGR003W | 648.3625673 | 0.350681905 | 0.05641409 | 0.69 |
| YGR039W | 40.79718114 | -0.779326451 | 0.057261538 | 0.69 |
| YGR075C | 302.7071594 | -0.45843064 | 0.056542456 | 0.69 |
| YJR088C | 954.591927 | -0.455846 | 0.057052156 | 0.69 |
| YKL157W | 4702.190162 | 0.63587861 | 0.055983486 | 0.69 |
| YLR028C | 4846.630925 | -0.212380883 | 0.056711027 | 0.69 |
| YLR437C-A | 26.88165292 | 0.743506728 | 0.057158703 | 0.69 |
| YML116W | 562.6593031 | 0.783897625 | 0.05682696 | 0.69 |
| YNR025C | 48.20698063 | -0.79860562 | 0.056249818 | 0.69 |
| YOR346W | 878.2111125 | -0.588772098 | 0.056063155 | 0.69 |
| YPL238C | 56.97812647 | -0.678856834 | 0.055997102 | 0.69 |
| YPR019W | 1285.937535 | -0.304266596 | 0.056532373 | 0.69 |
| YJR090C | 1653.355955 | -0.400236461 | 0.057645542 | 0.69 |
| YER116C | 578.3983863 | 0.605778326 | 0.057780799 | 0.69 |
| YCR068W | 666.4091136 | 0.622707686 | 0.058164467 | 0.69 |
| YDR294C | 2749.492975 | 0.368665202 | 0.058161417 | 0.69 |
| YPL055C | 1072.396721 | 0.419843299 | 0.058055016 | 0.69 |
| YDL189W | 1070.015735 | -0.539950782 | 0.058447075 | 0.69 |
| YPL049C | 2306.704815 | -0.441255825 | 0.058402002 | 0.69 |
| YBR168W | 396.8925406 | -0.475400441 | 0.06047969 | 0.69 |
| YBR244W | 574.3609399 | -0.789059917 | 0.06121225 | 0.69 |
| YDL002C | 564.8870678 | -0.3421225 | 0.059501942 | 0.69 |
| YDR073W | 640.4950618 | -0.670525836 | 0.060754205 | 0.69 |
| YGL115W | 1145.902181 | -0.285306142 | 0.061148342 | 0.69 |
| YGL134W | 814.0702966 | 0.559993458 | 0.060147245 | 0.69 |
| YGL180W | 843.8043135 | 0.550443267 | 0.060825876 | 0.69 |
| YGR225W | 197.8168855 | 0.946777932 | 0.060280679 | 0.69 |

| | | | | |
|---|---|---|---|---|
| YGR273C | 7.812625483 | 0.973981979 | 0.058836293 | 0.69 |
| YHR096C | 1194.376628 | 0.992296228 | 0.06080066 | 0.69 |
| YIL160C | 460.4774311 | 0.97300129 | 0.060979033 | 0.69 |
| YJL212C | 1268.474729 | 0.869620863 | 0.059207472 | 0.69 |
| YKL029C | 3291.164423 | 0.910535167 | 0.061025969 | 0.69 |
| YKL034W | 1217.582813 | 0.573390671 | 0.060026769 | 0.69 |
| YKL197C | 902.4571586 | 0.525594646 | 0.059767991 | 0.69 |
| YLR119W | 415.3763546 | -0.415356337 | 0.060889819 | 0.69 |
| YLR169W | 310.0591441 | -0.439819923 | 0.059309777 | 0.69 |
| YLR226W | 384.869465 | 0.362947777 | 0.05899174 | 0.69 |
| YLR462W | 4.00598291 | 1.0679798 | 0.061153499 | 0.69 |
| YML052W | 1751.911847 | -0.46653632 | 0.058983202 | 0.69 |
| YNL071W | 6818.92242 | 0.435724674 | 0.061188833 | 0.69 |
| YNL239W | 3388.853667 | 0.490275886 | 0.060097197 | 0.69 |
| YNR023W | 727.2388585 | -0.402424268 | 0.058672575 | 0.69 |
| YOR153W | 11636.99259 | -0.875349616 | 0.060812816 | 0.69 |
| YPR059C | 12.62828581 | 0.884131152 | 0.060492424 | 0.69 |
| YBR159W | 2299.591935 | -0.571156466 | 0.061716439 | 0.70 |
| YGR287C | 287.959068 | 0.632395959 | 0.061945991 | 0.70 |
| YJL077W-B | 26.1550574 | 0.841782262 | 0.061639899 | 0.70 |
| YJR056C | 431.2081935 | 0.386740775 | 0.062082715 | 0.70 |
| YKR028W | 1923.731148 | -0.432017409 | 0.062192267 | 0.70 |
| YMR035W | 535.4792987 | 0.569156162 | 0.061913259 | 0.70 |
| YOR214C | 33.33902204 | 0.92133998 | 0.061832463 | 0.70 |
| YPL259C | 962.6624104 | -0.346706936 | 0.06224926 | 0.70 |
| YLL004W | 381.8327061 | -0.548665068 | 0.06237023 | 0.70 |
| YNL147W | 569.2824244 | -0.443881793 | 0.062740885 | 0.70 |
| YAL068C | 5.759798843 | 1.019205154 | 0.064704917 | 0.70 |
| YBR089C-A | 560.6070076 | 0.54451066 | 0.065386804 | 0.70 |
| YBR162W-A | 580.1318829 | -0.58267342 | 0.065011502 | 0.70 |
| YDL006W | 361.6658456 | -0.331382095 | 0.064948477 | 0.70 |
| YDL092W | 1065.77792 | -0.462986186 | 0.063380039 | 0.70 |
| YDR256C | 561.4619469 | 0.75451502 | 0.064712138 | 0.70 |
| YDR410C | 388.474414 | -0.786330927 | 0.065331248 | 0.70 |
| YER087W | 372.1981021 | 0.595572808 | 0.064965626 | 0.70 |
| YFL015C | 19.00869044 | 0.868112955 | 0.065196619 | 0.70 |
| YFL017W-A | 221.0675394 | -0.566217342 | 0.064575104 | 0.70 |
| YGR265W | 63.64327138 | 0.756326233 | 0.065549886 | 0.70 |
| YHR005C-A | 2208.832297 | -0.535347453 | 0.065659505 | 0.70 |
| YJL051W | 1072.541155 | 0.326736718 | 0.06493527 | 0.70 |
| YJL071W | 590.1206135 | 0.471950337 | 0.064373038 | 0.70 |
| YKL105C | 1423.737383 | 0.499750745 | 0.064468119 | 0.70 |
| YKR030W | 649.1186189 | -0.348135341 | 0.064529221 | 0.70 |

| | | | | |
|---|---|---|---|---|
| YKR053C | 486.0346558 | -0.540474515 | 0.063631318 | 0.70 |
| YLR271W | 587.0390943 | 0.741394553 | 0.06383663 | 0.70 |
| YMR075W | 761.6920474 | 0.370604403 | 0.065684097 | 0.70 |
| YNL232W | 718.0441816 | 0.475737648 | 0.063323069 | 0.70 |
| YOR031W | 276.3450169 | 1.010347592 | 0.065285875 | 0.70 |
| YOR142W | 3533.236265 | 0.552540219 | 0.063754202 | 0.70 |
| YPL035C | 100.9354993 | -0.586803558 | 0.065282512 | 0.70 |
| YPR158W-B | 29.96033516 | 0.89170041 | 0.065193898 | 0.70 |
| snR85 | 18.57666508 | -1.005672775 | 0.063432651 | 0.70 |
| YDR157W | 53.23769391 | 0.669032358 | 0.06613686 | 0.70 |
| YHR034C | 511.3439371 | 0.406426142 | 0.066029562 | 0.70 |
| YJL204C | 683.8445536 | -0.432933177 | 0.065826934 | 0.70 |
| YKR098C | 1187.222635 | 0.654083048 | 0.066069313 | 0.70 |
| HRA1 | 34.22771229 | 0.730735401 | 0.067413979 | 0.70 |
| YBL020W | 997.2826168 | 0.421184001 | 0.069637866 | 0.70 |
| YBR085C-A | 1789.206511 | 0.764143027 | 0.06911073 | 0.70 |
| YDR057W | 882.7471612 | 0.419774848 | 0.069743843 | 0.70 |
| YDR370C | 533.8393634 | -0.410216427 | 0.069025548 | 0.70 |
| YDR510C-A | 439.1253795 | -0.621038729 | 0.066655411 | 0.70 |
| YDR524C | 904.6436686 | -0.433710367 | 0.069666993 | 0.70 |
| YER062C | 1876.234052 | -0.663933388 | 0.068427776 | 0.70 |
| YER088C | 4317.842613 | 0.620350431 | 0.066893488 | 0.70 |
| YER180C | 405.292426 | -0.40567738 | 0.067790283 | 0.70 |
| YFL005W | 2091.331245 | -0.416245361 | 0.069689535 | 0.70 |
| YFL028C | 788.0732688 | 0.407972701 | 0.069486645 | 0.70 |
| YGR262C | 876.7108592 | 0.275809897 | 0.067956156 | 0.70 |
| YHL045W | 26.03413018 | 0.952589558 | 0.067427025 | 0.70 |
| YHR147C | 857.7185719 | -0.362715036 | 0.06732665 | 0.70 |
| YHR159W | 783.3955676 | 0.54414852 | 0.069600511 | 0.70 |
| YHR214W-A | 22.9859353 | 0.798344248 | 0.068027374 | 0.70 |
| YIL092W | 426.021742 | 0.309277549 | 0.068615738 | 0.70 |
| YIL095W | 775.5717487 | -0.341724566 | 0.069426768 | 0.70 |
| YIL112W | 2132.63932 | 0.646107401 | 0.066345609 | 0.70 |
| YIL152W | 734.4896336 | 0.67540756 | 0.068737744 | 0.70 |
| YJR159W | 6.187748904 | 0.983812216 | 0.068626306 | 0.70 |
| YKL120W | 1575.003705 | 0.966276982 | 0.067066731 | 0.70 |
| YKR074W | 1111.118511 | -0.440610671 | 0.068425873 | 0.70 |
| YKR096W | 1782.315311 | 0.520216919 | 0.069361041 | 0.70 |
| YKR104W | 184.9350151 | 0.720592663 | 0.067581349 | 0.70 |
| YLR007W | 741.9239686 | 0.542251457 | 0.068231525 | 0.70 |
| YLR268W | 967.406712 | -0.393689382 | 0.069663291 | 0.70 |
| YMR084W | 174.3110976 | 0.822791536 | 0.066822594 | 0.70 |
| YNL111C | 246.1352445 | -0.800159682 | 0.066979506 | 0.70 |

| | | | | |
|---|---|---|---|---|
| YOL166C | 2.808993483 | 0.819975926 | 0.067948814 | 0.70 |
| YOR189W | 408.1511083 | -0.429680247 | 0.067462894 | 0.70 |
| YPL018W | 471.7341201 | 0.660636973 | 0.066392456 | 0.70 |
| YAL060W | 4666.556071 | 0.916855564 | 0.071412761 | 0.71 |
| YDL049C | 168.7509482 | -0.633234119 | 0.071784404 | 0.71 |
| YDL171C | 4109.615185 | 0.584310446 | 0.071997502 | 0.71 |
| YDL235C | 741.3905909 | -0.462711372 | 0.07085905 | 0.71 |
| YDR433W | 87.14790436 | -0.623726957 | 0.07124232 | 0.71 |
| YFL042C | 1495.188388 | 0.73964903 | 0.070549707 | 0.71 |
| YGL042C | 343.2123261 | -0.5405126 | 0.071989942 | 0.71 |
| YGR286C | 924.5538048 | 0.840199982 | 0.071875178 | 0.71 |
| YJL025W | 374.9847479 | -0.735249389 | 0.071398849 | 0.71 |
| YJR073C | 2980.954697 | 0.898921533 | 0.070611487 | 0.71 |
| YKL096W | 1798.684379 | -0.798776461 | 0.070263306 | 0.71 |
| YKR007W | 738.2939867 | -0.379816297 | 0.071470167 | 0.71 |
| YKR073C | 18.81130727 | 0.750067644 | 0.070780537 | 0.71 |
| YML119W | 1027.81525 | 0.415834708 | 0.07016187 | 0.71 |
| YMR201C | 569.2749285 | 0.592103236 | 0.071603769 | 0.71 |
| YOL117W | 709.0122211 | 0.675568562 | 0.070860826 | 0.71 |
| YOR022C | 608.0869392 | 0.532067735 | 0.070998425 | 0.71 |
| YOR028C | 706.5725488 | -0.964249211 | 0.071827916 | 0.71 |
| YPL249C | 1511.229449 | 0.336016941 | 0.072323415 | 0.71 |
| YJR036C | 438.6482321 | 0.652743694 | 0.072576974 | 0.71 |
| YNR066C | 265.0332103 | 0.628573581 | 0.072636525 | 0.71 |
| YJL027C | 10.13200317 | 0.865774075 | 0.07282334 | 0.71 |
| YMR254C | 26.76419027 | -0.742542862 | 0.073228206 | 0.71 |
| YBR167C | 275.1433522 | -0.481151033 | 0.073403852 | 0.71 |
| YHR171W | 640.7534835 | 0.590972194 | 0.07387998 | 0.72 |
| YDR528W | 366.4782046 | -0.902759674 | 0.074303294 | 0.72 |
| YHR139C-A | 36.30751378 | 0.844826815 | 0.074407567 | 0.72 |
| YJR010W | 677.4384961 | -0.631108891 | 0.074332505 | 0.72 |
| YNL222W | 196.3405424 | -0.615714399 | 0.074314642 | 0.72 |
| YEL034C-A | 2748.462234 | -0.652557258 | 0.074541782 | 0.72 |
| YGL048C | 4170.423196 | 0.360943392 | 0.074666459 | 0.72 |
| YGR019W | 2682.062979 | 0.820605725 | 0.075128871 | 0.72 |
| YMR250W | 7553.858966 | 0.938843339 | 0.075438505 | 0.72 |
| YMR271C | 754.1723733 | 0.94428592 | 0.075294571 | 0.72 |
| YOL002C | 621.7558878 | -0.587267161 | 0.07547344 | 0.72 |
| YCR085W | 15.93952821 | -0.910525474 | 0.076003119 | 0.72 |
| YDL043C | 369.3400281 | -0.402838487 | 0.076005692 | 0.72 |
| YDL123W | 682.3334241 | 0.533556954 | 0.076189737 | 0.72 |
| YLL028W | 724.5266873 | -0.646097231 | 0.076307787 | 0.72 |
| YLR191W | 472.4930163 | 0.427259854 | 0.076249678 | 0.72 |

| | | | | |
|---|---|---|---|---|
| YOR212W | 1199.500024 | -0.904207882 | 0.075993979 | 0.72 |
| YOR360C | 883.5535106 | -0.568758917 | 0.075910949 | 0.72 |
| YOR105W | 21.94253227 | -0.784110437 | 0.07657261 | 0.72 |
| YCL057W | 3573.183746 | 0.4327665 | 0.076744028 | 0.72 |
| YCR073C | 984.1918819 | 0.334063559 | 0.076956217 | 0.72 |
| YLR082C | 386.5514271 | -0.427517813 | 0.077062998 | 0.72 |
| YPL178W | 1119.186631 | -0.287957195 | 0.076884952 | 0.72 |
| YMR183C | 2217.146681 | -0.483061834 | 0.077848661 | 0.73 |
| YNL146C-A | 29.52790111 | -0.980425099 | 0.077761475 | 0.73 |
| YPL138C | 483.5208101 | 0.395734631 | 0.07769163 | 0.73 |
| YCL037C | 1123.648124 | -0.924154667 | 0.078085873 | 0.73 |
| YBR174C | 5.135026563 | 0.923121368 | 0.078706745 | 0.73 |
| YDR116C | 1042.960963 | -0.404211516 | 0.078880122 | 0.73 |
| YPL274W | 1036.210642 | 0.800993981 | 0.078585144 | 0.73 |
| YPR114W | 2316.846378 | -0.331017876 | 0.078832321 | 0.73 |
| YPR133W-A | 1114.918643 | -0.613817038 | 0.078766901 | 0.73 |
| YER147C | 529.8378852 | 0.447444628 | 0.079190359 | 0.73 |
| YBR114W | 883.2430403 | 0.465705544 | 0.079434345 | 0.73 |
| YOR076C | 709.8237452 | -0.402926738 | 0.079442547 | 0.73 |
| YBL049W | 377.5575063 | 0.866295212 | 0.079670407 | 0.73 |
| YDR374W-A | 251.03932 | -0.512295734 | 0.079946209 | 0.74 |
| YBL061C | 962.4150781 | -0.440297563 | 0.083179201 | 0.74 |
| YBR179C | 1264.938407 | 0.442683427 | 0.082894816 | 0.74 |
| YBR208C | 3562.373289 | 0.900936485 | 0.083395094 | 0.74 |
| YCR009C | 3474.642891 | -0.389903778 | 0.081318646 | 0.74 |
| YCR018C | 861.2432726 | -0.781068761 | 0.082935153 | 0.74 |
| YDL074C | 1505.723255 | -0.486514709 | 0.082078539 | 0.74 |
| YDL116W | 1341.682086 | -0.356526147 | 0.082428441 | 0.74 |
| YDR220C | 3.958958383 | 0.794719053 | 0.082006611 | 0.74 |
| YEL014C | 21.68196635 | -0.871905477 | 0.083503757 | 0.74 |
| YER069W | 925.6063303 | 0.624611541 | 0.08346325 | 0.74 |
| YER147C-A | 275.4373645 | -0.578949945 | 0.08220748 | 0.74 |
| YGR141W | 841.9095851 | 0.708175767 | 0.08262185 | 0.74 |
| YHL046C | 9.174623277 | 0.951543115 | 0.082870868 | 0.74 |
| YKL063C | 828.820505 | -0.401816747 | 0.082462152 | 0.74 |
| YML115C | 1520.660585 | -0.340707646 | 0.081645054 | 0.74 |
| YMR135C | 1809.513775 | 0.814867944 | 0.081887611 | 0.74 |
| YMR156C | 369.432093 | 0.764820515 | 0.083422434 | 0.74 |
| YMR216C | 1336.629718 | -0.405477817 | 0.082661663 | 0.74 |
| YMR313C | 988.4819991 | 0.406553903 | 0.08151468 | 0.74 |
| YNL145W | 1813.203416 | -0.846446862 | 0.082974053 | 0.74 |
| YNL260C | 268.8863879 | -0.367895165 | 0.082534016 | 0.74 |
| YOR109W | 2024.801836 | -0.340403974 | 0.08274453 | 0.74 |

| | | | | |
|---|---|---|---|---|
| YPL027W | 50.11860961 | 0.81686348 | 0.081874679 | 0.74 |
| YPL250W-A | 88.80290187 | -0.724249649 | 0.081674192 | 0.74 |
| YPR009W | 346.2208437 | -0.78320598 | 0.083030934 | 0.74 |
| YDR463W | 1963.287278 | 0.462555634 | 0.083805729 | 0.74 |
| RUF21 | 43.48350584 | 0.655453914 | 0.103598762 | 0.74 |
| YAR033W | 105.2011719 | -0.384902792 | 0.104797806 | 0.74 |
| YAR050W | 236.1195337 | 0.740890017 | 0.097481248 | 0.74 |
| YBL005W | 1420.668611 | -0.444090955 | 0.097894538 | 0.74 |
| YBL052C | 898.9203912 | 0.352665175 | 0.085718825 | 0.74 |
| YBL100C | 649.5552869 | 0.790500452 | 0.09106831 | 0.74 |
| YBR055C | 925.8039594 | -0.295060338 | 0.087351897 | 0.74 |
| YBR088C | 178.1843606 | -0.538348846 | 0.102412151 | 0.74 |
| YBR105C | 2095.906659 | 0.638977667 | 0.088936119 | 0.74 |
| YBR138C | 569.4593536 | 0.394574724 | 0.102261651 | 0.74 |
| YBR210W | 464.9591183 | -0.462192116 | 0.089369194 | 0.74 |
| YBR257W | 391.5498488 | 0.362503203 | 0.091490041 | 0.74 |
| YCR044C | 1029.447304 | -0.361937504 | 0.104697322 | 0.74 |
| YCR047C | 558.0855162 | 0.505349006 | 0.095058174 | 0.74 |
| YCR073W-A | 1340.952291 | 0.587177506 | 0.085007425 | 0.74 |
| YCR082W | 1363.300313 | -0.559412979 | 0.088711688 | 0.74 |
| YDL062W | 21.94852935 | 0.891311134 | 0.089711014 | 0.74 |
| YDL122W | 1760.896516 | -0.319736756 | 0.104015283 | 0.74 |
| YDL147W | 5209.098548 | 0.341692134 | 0.104104893 | 0.74 |
| YDL161W | 1895.847844 | -0.412077261 | 0.10384533 | 0.74 |
| YDL193W | 1212.895326 | 0.325198498 | 0.107212101 | 0.74 |
| YDL199C | 559.7723367 | 0.653283358 | 0.106475982 | 0.74 |
| YDL226C | 1711.631594 | -0.522674338 | 0.090855217 | 0.74 |
| YDR068W | 1258.879245 | 0.203282653 | 0.105119207 | 0.74 |
| YDR146C | 1456.032171 | -0.539613376 | 0.092279326 | 0.74 |
| YDR171W | 12610.16854 | 0.697486634 | 0.099278738 | 0.74 |
| YDR178W | 2932.392626 | 0.836127558 | 0.102072549 | 0.74 |
| YDR191W | 146.592055 | -0.535366354 | 0.106405863 | 0.74 |
| YDR194C | 2378.698014 | -0.415081488 | 0.098436208 | 0.74 |
| YDR231C | 627.5516275 | 0.653482326 | 0.090729392 | 0.74 |
| YDR232W | 2313.644723 | 0.489801984 | 0.102367754 | 0.74 |
| YDR236C | 550.9804022 | -0.396318262 | 0.102702167 | 0.74 |
| YDR284C | 1216.7009 | -0.215712821 | 0.099331355 | 0.74 |
| YDR301W | 1341.743029 | -0.488686131 | 0.09443166 | 0.74 |
| YDR318W | 508.7245087 | -0.418072236 | 0.091181781 | 0.74 |
| YDR319C | 485.2377065 | -0.461693161 | 0.084485709 | 0.74 |
| YDR329C | 1759.988143 | 0.541434596 | 0.104404116 | 0.74 |
| YDR343C | 868.7911596 | 0.919006258 | 0.107349836 | 0.74 |
| YDR386W | 496.5773156 | 0.450432139 | 0.101075132 | 0.74 |

| | | | | |
|---|---|---|---|---|
| YDR396W | 125.4408117 | -0.523158161 | 0.086303918 | 0.74 |
| YDR502C | 1141.466361 | 0.615457786 | 0.090508512 | 0.74 |
| YDR536W | 315.8429448 | 0.750483442 | 0.106984325 | 0.74 |
| YEL011W | 8924.284313 | 0.916591445 | 0.101164093 | 0.74 |
| YEL013W | 1708.200753 | -0.337506197 | 0.099900941 | 0.74 |
| YEL033W | 1403.07415 | -0.63138478 | 0.10576753 | 0.74 |
| YEL045C | 16.86688846 | 0.78181327 | 0.101178301 | 0.74 |
| YEL060C | 10805.49764 | 0.753378062 | 0.085029577 | 0.74 |
| YER005W | 999.8103274 | 0.250025714 | 0.09670889 | 0.74 |
| YER014W | 690.0676914 | 0.397113602 | 0.089968948 | 0.74 |
| YER030W | 1421.194205 | -0.457953385 | 0.090682521 | 0.74 |
| YER039C | 101.9948572 | 0.581207233 | 0.089911345 | 0.74 |
| YER076W-A | 76.7457419 | -0.467904758 | 0.103416911 | 0.74 |
| YER114C | 1610.317658 | 0.257469132 | 0.105495643 | 0.74 |
| YER119C | 1204.859566 | 0.526212369 | 0.093219607 | 0.74 |
| YER148W | 733.3492834 | -0.444497422 | 0.105392299 | 0.74 |
| YER162C | 931.7095211 | 0.525589565 | 0.102988106 | 0.74 |
| YER163C | 538.8237534 | 0.544728418 | 0.097958045 | 0.74 |
| YFL026W | 6326.346816 | -0.812052036 | 0.099793213 | 0.74 |
| YFR005C | 788.6190865 | -0.437927481 | 0.086146548 | 0.74 |
| YFR034C | 832.1624053 | 0.470522605 | 0.106019128 | 0.74 |
| YGL055W | 3060.50732 | 0.618427756 | 0.103378166 | 0.74 |
| YGL069C | 85.63194987 | -0.663619283 | 0.106265239 | 0.74 |
| YGL077C | 1372.265624 | 0.389017369 | 0.095539615 | 0.74 |
| YGL170C | 69.87350446 | 0.769533658 | 0.103154157 | 0.74 |
| YGL192W | 1154.271063 | -0.706174306 | 0.092349063 | 0.74 |
| YGL193C | 193.4216907 | -0.955416931 | 0.104187842 | 0.74 |
| YGL194C-A | 142.847988 | -0.611449097 | 0.106360061 | 0.74 |
| YGL208W | 855.2756461 | 0.67357549 | 0.106414902 | 0.74 |
| YGL209W | 1219.680335 | -0.637352631 | 0.095365221 | 0.74 |
| YGL241W | 933.7531382 | -0.321157213 | 0.102284441 | 0.74 |
| YGR017W | 1237.157084 | -0.313987225 | 0.106781593 | 0.74 |
| YGR055W | 1342.856307 | -0.591111801 | 0.085820503 | 0.74 |
| YGR060W | 3114.502526 | -0.64577355 | 0.106952814 | 0.74 |
| YGR073C | 7.437514438 | -1.173563271 | 0.09120984 | 0.74 |
| YGR078C | 839.5108181 | -0.493981331 | 0.10729975 | 0.74 |
| YGR112W | 838.4700607 | 0.622431643 | 0.084339781 | 0.74 |
| YGR139W | 13.99976418 | 0.790671751 | 0.099605093 | 0.74 |
| YGR154C | 333.0792836 | 0.50045995 | 0.100557019 | 0.74 |
| YGR179C | 503.8137475 | -0.285311729 | 0.105017573 | 0.74 |
| YGR227W | 1061.56859 | -0.515608242 | 0.087607176 | 0.74 |
| YGR244C | 5502.307672 | 0.623330278 | 0.095522069 | 0.74 |
| YGR246C | 1106.975461 | 0.430625507 | 0.094574618 | 0.74 |

| | | | | |
|---|---|---|---|---|
| YHL016C | 664.2747126 | 0.779882932 | 0.096213476 | 0.74 |
| YHL024W | 910.8922734 | 0.633678425 | 0.087796351 | 0.74 |
| YHR040W | 740.6643017 | 0.646242888 | 0.096721412 | 0.74 |
| YHR050W-A | 204.2549426 | -0.541191751 | 0.089857977 | 0.74 |
| YHR070W | 688.4026548 | 0.539990042 | 0.102841318 | 0.74 |
| YHR086W-A | 3.107825184 | -0.760622485 | 0.09607727 | 0.74 |
| YHR091C | 596.7962742 | -0.273575843 | 0.094091336 | 0.74 |
| YHR149C | 909.9713222 | -0.615212285 | 0.08813232 | 0.74 |
| YHR154W | 1331.473724 | -0.57264488 | 0.093994852 | 0.74 |
| YHR193C | 8396.042296 | -0.496995025 | 0.10361163 | 0.74 |
| YHR201C | 1092.809897 | -0.555166701 | 0.087280147 | 0.74 |
| YHR214W | 18.69123236 | 0.79614798 | 0.098909364 | 0.74 |
| YIL009W | 578.8586392 | -0.622828577 | 0.089706683 | 0.74 |
| YIL029W-A | 8.108002149 | 0.755076421 | 0.090219369 | 0.74 |
| YIL051C | 4289.573013 | -0.611376197 | 0.084890908 | 0.74 |
| YIL066C | 409.4135175 | 0.744478116 | 0.087232572 | 0.74 |
| YIL118W | 1012.394304 | -0.558134889 | 0.09515144 | 0.74 |
| YIL136W | 7495.331143 | 1.012411705 | 0.091840453 | 0.74 |
| YIL155C | 2791.794832 | 0.819781427 | 0.104988215 | 0.74 |
| YJL049W | 674.8429005 | -0.520098101 | 0.090732463 | 0.74 |
| YJL078C | 3272.163203 | 0.517617228 | 0.096451804 | 0.74 |
| YJL105W | 247.8507967 | -0.599324564 | 0.099406446 | 0.74 |
| YJL155C | 1149.698916 | 0.682667001 | 0.085645521 | 0.74 |
| YJL206C | 539.1174838 | 0.498177413 | 0.087560133 | 0.74 |
| YJR008W | 1161.824512 | 0.880021105 | 0.085408202 | 0.74 |
| YJR024C | 696.9415683 | -0.513966139 | 0.101403948 | 0.74 |
| YJR086W | 331.1910032 | -0.700126127 | 0.106275055 | 0.74 |
| YJR087W | 189.3064917 | -0.43394868 | 0.104905359 | 0.74 |
| YJR103W | 3899.480126 | 0.903734206 | 0.103782931 | 0.74 |
| YJR106W | 683.5325213 | 0.42279255 | 0.101487852 | 0.74 |
| YKL093W | 994.1827608 | 0.785504974 | 0.097449584 | 0.74 |
| YKL122C | 1563.613747 | -0.527023246 | 0.094641714 | 0.74 |
| YKL139W | 677.7577592 | -0.367551593 | 0.100226576 | 0.74 |
| YKL148C | 4538.261302 | 0.947432752 | 0.102134433 | 0.74 |
| YKL166C | 506.1904776 | -0.510035125 | 0.090222014 | 0.74 |
| YKL187C | 740.5340162 | 0.745162052 | 0.087958751 | 0.74 |
| YKL204W | 2110.527656 | -0.414479955 | 0.098063314 | 0.74 |
| YKR006C | 1505.723079 | -0.417053903 | 0.094756516 | 0.74 |
| YKR009C | 961.8765586 | 0.752239474 | 0.088916936 | 0.74 |
| YKR019C | 546.7471303 | 0.448741852 | 0.086645638 | 0.74 |
| YKR085C | 837.2824572 | -0.4408282 | 0.092301889 | 0.74 |
| YKR088C | 850.0044261 | 0.348204815 | 0.098965321 | 0.74 |
| YLL023C | 2955.471664 | 0.576297524 | 0.089609117 | 0.74 |

| | | | | |
|---|---|---|---|---|
| YLL057C | 252.3608873 | 0.70218796 | 0.098180754 | 0.74 |
| YLL061W | 447.0050952 | 0.533075873 | 0.106854104 | 0.74 |
| YLR080W | 639.2701305 | 0.675617653 | 0.09649506 | 0.74 |
| YLR113W | 3162.32221 | -0.373918806 | 0.105998669 | 0.74 |
| YLR122C | 30.05120318 | 0.770975371 | 0.098048608 | 0.74 |
| YLR149C | 1780.729906 | 0.763456855 | 0.104350311 | 0.74 |
| YLR162W | 2.387178616 | 0.792794952 | 0.096779577 | 0.74 |
| YLR209C | 881.8739647 | 0.542475137 | 0.084723827 | 0.74 |
| YLR300W | 3505.549383 | -0.659614691 | 0.100635462 | 0.74 |
| YLR355C | 9452.460146 | 0.588086854 | 0.095654083 | 0.74 |
| YLR386W | 1077.114184 | 0.290358371 | 0.100038325 | 0.74 |
| YLR428C | 15.96009415 | -0.765623244 | 0.099415326 | 0.74 |
| YLR442C | 902.5050218 | 0.308526539 | 0.098930287 | 0.74 |
| YLR461W | 7.039838185 | 0.977496577 | 0.086413941 | 0.74 |
| YML036W | 292.5583071 | -0.55891608 | 0.090679094 | 0.74 |
| YML059C | 1781.099171 | -0.407966238 | 0.10404425 | 0.74 |
| YML062C | 782.2227983 | -0.368957336 | 0.091070465 | 0.74 |
| YMR021C | 731.4049333 | 0.380050366 | 0.10596638 | 0.74 |
| YMR069W | 79.44065746 | 0.829939947 | 0.087121446 | 0.74 |
| YMR074C | 1117.334751 | -0.50135753 | 0.087738928 | 0.74 |
| YMR083W | 8478.832219 | 0.490487442 | 0.099787328 | 0.74 |
| YMR097C | 670.8866515 | 0.357315894 | 0.096541556 | 0.74 |
| YMR104C | 1639.396102 | 0.785696897 | 0.089639059 | 0.74 |
| YMR147W | 251.4438843 | 0.689541298 | 0.104296956 | 0.74 |
| YMR301C | 1249.636815 | 0.393257383 | 0.085877001 | 0.74 |
| YMR321C | 24.07796373 | 0.769368463 | 0.09156788 | 0.74 |
| YNL022C | 938.4563852 | 0.454056313 | 0.084861115 | 0.74 |
| YNL066W | 1969.929236 | -0.568435569 | 0.084204294 | 0.74 |
| YNL070W | 685.2481589 | -0.456486738 | 0.09311618 | 0.74 |
| YNL107W | 513.6532475 | 0.284972202 | 0.093809405 | 0.74 |
| YNL138W-A | 114.4265169 | -0.430029389 | 0.093096556 | 0.74 |
| YNL181W | 1100.958937 | 0.340955874 | 0.104311012 | 0.74 |
| YNL191W | 591.9956262 | 0.446077354 | 0.087024217 | 0.74 |
| YNL236W | 1176.860728 | -0.382781155 | 0.08559037 | 0.74 |
| YNL297C | 2104.946506 | -0.341852888 | 0.096620937 | 0.74 |
| YNL305C | 2021.497779 | 0.65701135 | 0.100194763 | 0.74 |
| YNL310C | 605.4678235 | 0.314907122 | 0.104011923 | 0.74 |
| YNL318C | 22.2967731 | 0.833469098 | 0.105897718 | 0.74 |
| YNL334C | 19.40823403 | -0.904600215 | 0.098589442 | 0.74 |
| YNR015W | 1147.190779 | 0.423383787 | 0.088846666 | 0.74 |
| YNR020C | 315.5485889 | -0.430313132 | 0.094550458 | 0.74 |
| YNR065C | 953.8955562 | 0.453355988 | 0.104426941 | 0.74 |
| YOL005C | 1554.462517 | -0.528165692 | 0.092833523 | 0.74 |

| | | | | |
|---|---|---|---|---|
| YOL006C | 1492.750425 | -0.38502057 | 0.092082821 | 0.74 |
| YOL013W-A | 5.152321446 | -1.105597708 | 0.105498067 | 0.74 |
| YOL019W | 928.0908204 | -0.508671704 | 0.091100077 | 0.74 |
| YOL027C | 2880.865523 | 0.493210659 | 0.095782165 | 0.74 |
| YOL075C | 712.6275688 | 0.497504228 | 0.08428935 | 0.74 |
| YOL147C | 576.9213691 | 0.616595622 | 0.091981726 | 0.74 |
| YOR010C | 455.6940822 | 0.819580829 | 0.093073341 | 0.74 |
| YOR011W | 696.3198627 | 0.527976553 | 0.097974318 | 0.74 |
| YOR040W | 503.3947844 | 0.744339084 | 0.089448923 | 0.74 |
| YOR082C | 31.57473045 | -0.799730308 | 0.097452325 | 0.74 |
| YOR120W | 2918.156498 | 0.835928792 | 0.098256279 | 0.74 |
| YOR143C | 346.0541178 | -0.506565471 | 0.107210654 | 0.74 |
| YOR149C | 790.0054864 | 0.354465368 | 0.091837012 | 0.74 |
| YOR209C | 3422.936842 | -0.533927985 | 0.091124554 | 0.74 |
| YOR232W | 2857.566582 | -0.437816123 | 0.088155248 | 0.74 |
| YOR268C | 11.87888145 | 0.919834445 | 0.106792475 | 0.74 |
| YOR274W | 529.1848909 | -0.47596727 | 0.095039124 | 0.74 |
| YOR324C | 520.2886814 | -0.467847942 | 0.087526654 | 0.74 |
| YOR334W | 302.0782196 | -0.374910503 | 0.106998923 | 0.74 |
| YOR348C | 282.9627385 | 0.932553078 | 0.09495222 | 0.74 |
| YOR357C | 469.3315493 | -0.334308023 | 0.095160918 | 0.74 |
| YOR366W | 3.244594662 | -0.887180271 | 0.093464106 | 0.74 |
| YOR375C | 17984.54886 | 0.880149009 | 0.096525801 | 0.74 |
| YPL050C | 1693.46287 | -0.519937156 | 0.093500301 | 0.74 |
| YPL052W | 344.0806128 | 0.483378092 | 0.103443327 | 0.74 |
| YPL061W | 6576.454349 | -0.696638134 | 0.102385207 | 0.74 |
| YPL082C | 1537.0483 | -0.436911459 | 0.101094515 | 0.74 |
| YPL175W | 580.2634042 | -0.363579711 | 0.099040954 | 0.74 |
| YPL185W | 39.28495564 | 0.683649303 | 0.098668305 | 0.74 |
| YPL189W | 119.6862802 | 0.515615218 | 0.096269119 | 0.74 |
| YPL222W | 848.2140163 | 0.806605774 | 0.093448044 | 0.74 |
| YPL248C | 793.6864415 | 0.502197937 | 0.103313988 | 0.74 |
| YPL255W | 357.750249 | -0.543763056 | 0.102457983 | 0.74 |
| YPR006C | 675.2062996 | 0.764754628 | 0.090778113 | 0.74 |
| YPR122W | 1503.146832 | -0.858632091 | 0.097486995 | 0.74 |
| YPR131C | 480.7629468 | -0.355964931 | 0.097474171 | 0.74 |
| snR33 | 120.9149393 | -0.501573359 | 0.100742562 | 0.74 |
| YJR037W | 10.84309032 | 0.701392188 | 0.107519248 | 0.75 |
| YOR351C | 100.6577504 | 0.589276305 | 0.107722412 | 0.75 |
| YDR333C | 1359.98522 | -0.314416665 | 0.107912284 | 0.75 |
| YLR164W | 251.3796178 | 0.785305625 | 0.108410671 | 0.75 |
| YNL170W | 85.88966089 | 0.580880293 | 0.108316113 | 0.75 |
| YNL202W | 289.1619632 | 0.751932112 | 0.108285014 | 0.75 |

| | | | | |
|---|---|---|---|---|
| YPL071C | 441.9177148 | -0.476102497 | 0.108426289 | 0.75 |
| YBR012C | 184.3218002 | 0.871570062 | 0.108586907 | 0.75 |
| YOR092W | 648.6032701 | -0.598210861 | 0.10866438 | 0.75 |
| YDL225W | 1750.506735 | -0.414791282 | 0.108970711 | 0.75 |
| YGR014W | 3080.238337 | -0.741564096 | 0.109036139 | 0.75 |
| YOR192C | 247.398385 | 0.674369723 | 0.109085976 | 0.75 |
| YBL064C | 4712.365868 | 0.694330105 | 0.111187028 | 0.75 |
| YBR033W | 734.5856264 | 0.790231122 | 0.11224646 | 0.75 |
| YBR043C | 1241.74171 | 0.667738727 | 0.112203856 | 0.75 |
| YBR165W | 515.6336536 | -0.340472908 | 0.111719357 | 0.75 |
| YBR182C | 159.913284 | 0.667900118 | 0.111636983 | 0.75 |
| YBR288C | 774.8863305 | -0.361611237 | 0.11019022 | 0.75 |
| YBR293W | 445.7090525 | -0.712602522 | 0.110900584 | 0.75 |
| YDR004W | 628.5806399 | 0.296078451 | 0.111318364 | 0.75 |
| YDR118W-A | 11.72360647 | -0.824096755 | 0.112258747 | 0.75 |
| YDR207C | 1436.193266 | 0.456912172 | 0.112211548 | 0.75 |
| YDR493W | 345.6632951 | -0.462133431 | 0.110642877 | 0.75 |
| YGL054C | 1677.23256 | -0.359963575 | 0.111644068 | 0.75 |
| YGL156W | 2401.006767 | 0.763931614 | 0.110839302 | 0.75 |
| YIL046W | 1246.958862 | 0.505611033 | 0.11088494 | 0.75 |
| YIL119C | 838.0211886 | -0.876925152 | 0.111217987 | 0.75 |
| YIL128W | 1273.394888 | -0.460611332 | 0.110203803 | 0.75 |
| YJL156W-A | 83.97878785 | -0.468502781 | 0.111081419 | 0.75 |
| YLR136C | 655.8429852 | -0.745917353 | 0.111092859 | 0.75 |
| YLR323C | 363.5088652 | 0.417303619 | 0.111328118 | 0.75 |
| YLR394W | 730.9999237 | 0.385726798 | 0.109994851 | 0.75 |
| YML046W | 728.0597624 | -0.436751394 | 0.110082742 | 0.75 |
| YMR306C-A | 3.882802162 | 0.903241449 | 0.111814128 | 0.75 |
| YPL200W | 21.68345646 | 0.873597071 | 0.110738945 | 0.75 |
| YPL265W | 2999.757451 | 0.89794697 | 0.112328139 | 0.75 |
| YOR132W | 1718.595531 | 0.437951786 | 0.112472106 | 0.75 |
| YDL036C | 422.8116952 | -0.390651732 | 0.112636477 | 0.75 |
| YNL018C | 116.9826504 | 0.738820288 | 0.112947334 | 0.75 |
| YOR302W | 66.2485987 | 0.731480607 | 0.113055311 | 0.75 |
| YPL181W | 1430.008347 | 0.527578003 | 0.112972116 | 0.75 |
| YCR023C | 1565.302118 | 0.491435246 | 0.113324248 | 0.75 |
| YOL081W | 5088.832739 | 0.564905647 | 0.113316812 | 0.75 |
| YDL210W | 142.1653441 | 0.699149278 | 0.113846754 | 0.75 |
| YIL027C | 793.9722774 | -0.389239446 | 0.113912158 | 0.75 |
| YKL027W | 701.3136707 | 0.550754797 | 0.113769399 | 0.75 |
| YCR065W | 756.8632472 | -0.587155821 | 0.114345575 | 0.75 |
| YDL107W | 383.3863117 | -0.434394194 | 0.114705592 | 0.75 |
| YER067C-A | 258.7943359 | 0.957397628 | 0.11616808 | 0.75 |

| | | | | |
|---|---|---|---|---|
| YER158W-A | 247.6290209 | -0.594985519 | 0.115530118 | 0.75 |
| YHL021C | 7761.479525 | 0.809122393 | 0.115545039 | 0.75 |
| YHR137W | 2665.655739 | 0.877054493 | 0.115146412 | 0.75 |
| YIR036C | 938.6857022 | 0.426301304 | 0.116112268 | 0.75 |
| YJR096W | 2414.849421 | 0.834415628 | 0.11535711 | 0.75 |
| YKR008W | 1399.761894 | -0.364929373 | 0.114239726 | 0.75 |
| YLR248W | 3268.331309 | 0.470850129 | 0.115525877 | 0.75 |
| YLR389C | 2863.752537 | 0.299832224 | 0.115586525 | 0.75 |
| YLR397C | 907.763381 | 0.426700465 | 0.11568245 | 0.75 |
| YML099W-A | 34.73959572 | 0.452892073 | 0.115562126 | 0.75 |
| YML105C | 1352.843987 | -0.342847971 | 0.11630285 | 0.75 |
| YMR058W | 5542.022572 | -0.703892271 | 0.116149639 | 0.75 |
| YNL009W | 277.6331838 | 0.701339218 | 0.116470247 | 0.75 |
| YNL079C | 8141.583459 | -0.448706472 | 0.115423477 | 0.75 |
| YNR052C | 1154.692673 | -0.3016664 | 0.114651837 | 0.75 |
| YOL076W | 856.9107843 | -0.557730632 | 0.116304824 | 0.75 |
| YOL101C | 182.685157 | 0.60505923 | 0.116448406 | 0.75 |
| YOL109W | 9848.185932 | -0.497128326 | 0.11459137 | 0.75 |
| YOR262W | 658.3672371 | 0.480950889 | 0.116129948 | 0.75 |
| YFR002W | 1555.318988 | -0.422646286 | 0.116706181 | 0.75 |
| YMR132C | 213.3288234 | 0.390610143 | 0.116807952 | 0.75 |
| YDR102C | 12.68433027 | -0.773524657 | 0.117327581 | 0.76 |
| YHR008C | 8653.583569 | 0.675777127 | 0.117339763 | 0.76 |
| YER039C-A | 49.35745349 | 0.590458811 | 0.117637547 | 0.76 |
| YFL060C | 16.02228195 | -0.814477834 | 0.117816943 | 0.76 |
| YFR012W-A | 2.137477958 | 0.560802947 | 0.117812315 | 0.76 |
| YNL267W | 1939.373479 | -0.237316601 | 0.117929297 | 0.76 |
| YPL072W | 410.1680408 | -0.399544294 | 0.117675519 | 0.76 |
| YFR029W | 834.7381381 | 0.494197057 | 0.118151079 | 0.76 |
| YGR153W | 193.310946 | 0.480064631 | 0.118582634 | 0.76 |
| YHR187W | 633.7803075 | 0.299589339 | 0.118634428 | 0.76 |
| YBR169C | 4813.961053 | 0.738459159 | 0.11996776 | 0.76 |
| YBR238C | 642.2716869 | -0.61811559 | 0.119815725 | 0.76 |
| YCR083W | 821.7942376 | 0.640975222 | 0.119679292 | 0.76 |
| YDL155W | 763.3718322 | -0.350119697 | 0.120068311 | 0.76 |
| YDL202W | 808.8072189 | -0.409009051 | 0.120640507 | 0.76 |
| YER078W-A | 4.281084499 | 0.66033385 | 0.120708929 | 0.76 |
| YJL068C | 1815.839599 | 0.451777475 | 0.12076718 | 0.76 |
| YJR010C-A | 212.1436636 | -0.535694949 | 0.120221716 | 0.76 |
| YLR321C | 729.7163579 | -0.437549237 | 0.119292058 | 0.76 |
| YML085C | 4155.33263 | 0.400311131 | 0.120630715 | 0.76 |
| YMR062C | 1501.153431 | 0.628270131 | 0.120199348 | 0.76 |
| YMR158W | 693.5786925 | -0.475874112 | 0.120145516 | 0.76 |

| | | | | |
|---|---|---|---|---|
| YMR270C | 708.6444493 | -0.352753925 | 0.119339831 | 0.76 |
| YNR001C | 10779.43573 | 0.807810345 | 0.12019357 | 0.76 |
| YOR185C | 2527.056266 | 0.637700268 | 0.119976594 | 0.76 |
| YPL037C | 6009.036088 | -0.478571593 | 0.119876963 | 0.76 |
| YPR058W | 588.0067957 | 0.560130981 | 0.12036977 | 0.76 |
| YBR294W | 590.8243932 | -0.536161754 | 0.121105762 | 0.76 |
| YDR199W | 75.09684199 | -0.526406655 | 0.121226194 | 0.76 |
| YIR044C | 6.331727114 | 0.861678252 | 0.121274094 | 0.76 |
| YER113C | 743.4950149 | -0.326359419 | 0.121590393 | 0.76 |
| YFL057C | 160.9643278 | 0.527152176 | 0.121826684 | 0.76 |
| YKL146W | 1758.494435 | 0.654806133 | 0.121619758 | 0.76 |
| YLR331C | 3.488696978 | 0.870110059 | 0.12191671 | 0.76 |
| YOR251C | 1296.455018 | 0.277288936 | 0.121879954 | 0.76 |
| YBR147W | 531.1286909 | 0.863376309 | 0.122650745 | 0.76 |
| YBR148W | 178.9686946 | 0.58688716 | 0.12231532 | 0.76 |
| YER063W | 1431.45469 | -0.37033443 | 0.12253579 | 0.76 |
| YNL015W | 3387.233502 | 0.783375886 | 0.122349447 | 0.76 |
| YNR034W | 660.948473 | 0.443406785 | 0.122657511 | 0.76 |
| YOL163W | 57.10674557 | 0.756111758 | 0.122618599 | 0.76 |
| YJL163C | 815.8405692 | 0.661758236 | 0.122929564 | 0.76 |
| YAL016W | 2950.826603 | 0.351381917 | 0.123603682 | 0.76 |
| YJL219W | 39.81053365 | 0.833204684 | 0.123647892 | 0.76 |
| YMR068W | 420.8665046 | 0.442433491 | 0.12349465 | 0.76 |
| YOR226C | 454.5609099 | 0.705118623 | 0.123883805 | 0.76 |
| YAL041W | 1345.73957 | -0.471746161 | 0.125790039 | 0.77 |
| YBL042C | 1258.710363 | -0.576451975 | 0.126919902 | 0.77 |
| YBR156C | 807.2285578 | -0.294775441 | 0.124931361 | 0.77 |
| YBR276C | 1528.649976 | -0.205198709 | 0.12524365 | 0.77 |
| YCR097W-A | 1.60235674 | 0.42071197 | 0.126929109 | 0.77 |
| YDR010C | 32.06483583 | 0.442387861 | 0.126998576 | 0.77 |
| YER019C-A | 1962.324665 | -0.362556936 | 0.12567847 | 0.77 |
| YER133W-A | 95.44590047 | -0.588656779 | 0.126274524 | 0.77 |
| YFL021W | 926.9442787 | 0.548319753 | 0.126064944 | 0.77 |
| YGL045W | 904.8478281 | 0.615396104 | 0.126881355 | 0.77 |
| YGL168W | 276.6572721 | -0.580687444 | 0.126389965 | 0.77 |
| YHR035W | 231.8124454 | 0.474999358 | 0.127078636 | 0.77 |
| YHR217C | 1.225581631 | 0.624656455 | 0.125712382 | 0.77 |
| YIL003W | 534.8841649 | 0.323115096 | 0.12580912 | 0.77 |
| YJL099W | 747.8275315 | 0.367330327 | 0.12649678 | 0.77 |
| YJL124C | 1576.189687 | -0.327726792 | 0.126439534 | 0.77 |
| YMR180C | 510.7680392 | -0.409152417 | 0.125363671 | 0.77 |
| YNL185C | 474.6914722 | -0.410049691 | 0.126844904 | 0.77 |
| YNL199C | 1154.934737 | -0.184658813 | 0.125937162 | 0.77 |

| | | | | |
|---|---|---|---|---|
| YOL011W | 1428.931177 | -0.534744445 | 0.126166311 | 0.77 |
| YOL111C | 1821.468977 | -0.249544618 | 0.126519913 | 0.77 |
| YOR305W | 732.3331193 | -0.430712263 | 0.124591702 | 0.77 |
| YPL017C | 762.3634934 | 0.483410713 | 0.126048134 | 0.77 |
| YPL196W | 943.9259177 | 0.633833479 | 0.124690465 | 0.77 |
| YPR050C | 1.567602133 | -1.001418541 | 0.126239231 | 0.77 |
| YBR186W | 272.8183513 | 0.666293302 | 0.127652159 | 0.77 |
| YDL102W | 928.5717968 | 0.296115415 | 0.128447748 | 0.77 |
| YGL103W | 19.59323296 | -1.221104137 | 0.127428084 | 0.77 |
| YGR205W | 695.7192252 | -0.596121665 | 0.128091107 | 0.77 |
| YIL032C | 13.12625335 | 0.683598085 | 0.127657087 | 0.77 |
| YJL194W | 154.6134221 | -0.740932781 | 0.128487786 | 0.77 |
| YLL025W | 92.59299699 | 0.620204441 | 0.128482944 | 0.77 |
| YLR108C | 407.9615838 | 0.546859661 | 0.128067525 | 0.77 |
| YLR384C | 2550.492331 | 0.325626112 | 0.128180205 | 0.77 |
| YMR105W-A | 43.59841519 | 0.419492065 | 0.127931738 | 0.77 |
| YNL330C | 1017.919377 | -0.597394136 | 0.128043084 | 0.77 |
| YOR248W | 35.3813631 | -0.607196217 | 0.128746801 | 0.77 |
| snR43 | 20.09245082 | -0.6341462 | 0.129046295 | 0.77 |
| snR56 | 6.041353852 | 0.871441673 | 0.129078342 | 0.77 |
| YMR055C | 275.6053073 | 0.391952154 | 0.129338812 | 0.77 |
| YKL003C | 652.5154668 | -0.417121638 | 0.12959443 | 0.77 |
| YLR430W | 3065.658429 | 0.351231283 | 0.129630584 | 0.77 |
| YEL012W | 1045.847779 | 0.601111211 | 0.130629932 | 0.77 |
| YGL020C | 1129.205349 | -0.375902539 | 0.130401119 | 0.77 |
| YKL169C | 32.06083426 | -0.636068708 | 0.130695816 | 0.77 |
| YLR446W | 527.2109038 | 0.439626735 | 0.130520588 | 0.77 |
| YOR376W | 25.04102901 | 0.812439253 | 0.130911529 | 0.77 |
| YBR282W | 512.5901703 | -0.376036404 | 0.134089038 | 0.77 |
| YCL009C | 7647.273312 | 0.456933556 | 0.131934398 | 0.77 |
| YCR036W | 1075.104784 | -0.325027686 | 0.134677622 | 0.77 |
| YDL068W | 4.863206233 | -0.772494202 | 0.134702751 | 0.77 |
| YDR034C-A | 4.06626248 | -0.745229656 | 0.135195053 | 0.77 |
| YDR334W | 1700.440041 | -0.433256281 | 0.133167229 | 0.77 |
| YDR383C | 354.9421834 | -0.418864606 | 0.133833096 | 0.77 |
| YDR525W | 4.560372998 | 0.745910901 | 0.132337719 | 0.77 |
| YEL018W | 225.091547 | -0.303392754 | 0.134940463 | 0.77 |
| YER101C | 634.792482 | 0.498122645 | 0.134261194 | 0.77 |
| YER135C | 23.51565939 | 0.719065041 | 0.133360871 | 0.77 |
| YFL034C-A | 576.1615287 | -0.704674569 | 0.133507484 | 0.77 |
| YFR017C | 3258.918594 | 0.802080048 | 0.132117664 | 0.77 |
| YGL130W | 854.7206722 | 0.18422537 | 0.133766745 | 0.77 |
| YIL083C | 1169.719343 | 0.579373268 | 0.13152278 | 0.77 |

| | | | | |
|---|---|---|---|---|
| YIL139C | 280.0724876 | -0.290968514 | 0.135538255 | 0.77 |
| YIR021W | 261.891268 | -0.361851601 | 0.132155937 | 0.77 |
| YJL185C | 564.2478549 | 0.621945591 | 0.135544516 | 0.77 |
| YJL214W | 204.5476536 | -0.493557914 | 0.134214454 | 0.77 |
| YKL064W | 2390.234747 | 0.374034204 | 0.132615884 | 0.77 |
| YLR311C | 30.19399839 | 0.795524709 | 0.132627312 | 0.77 |
| YLR351C | 1673.534289 | 0.322331621 | 0.135253635 | 0.77 |
| YLR385C | 52.50375543 | 0.472512484 | 0.133645951 | 0.77 |
| YLR406C-A | 2.560257022 | 0.502986483 | 0.13545901 | 0.77 |
| YMR053C | 712.958308 | 0.482013117 | 0.131420453 | 0.77 |
| YMR106C | 546.3577506 | 0.402659912 | 0.135341264 | 0.77 |
| YMR109W | 3069.730753 | -0.366425688 | 0.132107758 | 0.77 |
| YMR127C | 400.3588714 | 0.399463686 | 0.133890392 | 0.77 |
| YMR165C | 1579.930507 | 0.656870631 | 0.133014308 | 0.77 |
| YNL105W | 8.716167534 | 0.692109612 | 0.135341016 | 0.77 |
| YNL282W | 259.9576363 | 0.386609452 | 0.134207127 | 0.77 |
| YNL283C | 1549.072925 | 0.336625792 | 0.132246489 | 0.77 |
| YNL295W | 664.526791 | 0.37329194 | 0.132049542 | 0.77 |
| YNR006W | 1506.041195 | 0.293797874 | 0.13290876 | 0.77 |
| YOR025W | 737.1839475 | -0.414958089 | 0.13545169 | 0.77 |
| YOR045W | 622.8710581 | -0.478442219 | 0.13188389 | 0.77 |
| YPR061C | 393.0462601 | 0.636502228 | 0.133019996 | 0.77 |
| YDR339C | 1137.925374 | -0.510384524 | 0.136027187 | 0.78 |
| YGR080W | 1465.883709 | -0.343801871 | 0.135889096 | 0.78 |
| YIL064W | 505.7570311 | 0.66190818 | 0.135958695 | 0.78 |
| YLR070C | 346.2836033 | 0.616417337 | 0.136315851 | 0.78 |
| YCL048W | 92.73705911 | 0.751164827 | 0.137011201 | 0.78 |
| YDL127W | 422.9534044 | -0.483352456 | 0.137098799 | 0.78 |
| YEL022W | 2372.854744 | -0.451232784 | 0.137234894 | 0.78 |
| YFL048C | 1163.307711 | -0.413664424 | 0.137020914 | 0.78 |
| YGR251W | 285.9824856 | 0.42531271 | 0.137015037 | 0.78 |
| YIL050W | 1083.890647 | 0.466512287 | 0.136911258 | 0.78 |
| YNR010W | 212.8810604 | -0.400041539 | 0.1371724 | 0.78 |
| YML013W | 1108.615798 | 0.436077836 | 0.137358275 | 0.78 |
| YOR307C | 1226.108407 | -0.378541887 | 0.137535508 | 0.78 |
| YER188C-A | 23.15774642 | 0.492630127 | 0.137863064 | 0.78 |
| YLR262C-A | 800.3537696 | -0.425253086 | 0.137902994 | 0.78 |
| YOR171C | 1509.079417 | 0.368551453 | 0.13774553 | 0.78 |
| YBR145W | 1370.985162 | 0.6802444 | 0.138763679 | 0.78 |
| YBR251W | 1030.356421 | -0.388137551 | 0.138164764 | 0.78 |
| YDL238C | 479.1219346 | 0.49707241 | 0.139584228 | 0.78 |
| YDR096W | 1946.105843 | 0.559882986 | 0.139833657 | 0.78 |
| YDR295C | 798.8722941 | -0.279804831 | 0.139990966 | 0.78 |

| | | | | |
|---|---|---|---|---|
| YFL004W | 2271.622342 | -0.28975988 | 0.140172061 | 0.78 |
| YFR003C | 1017.431306 | -0.429217055 | 0.140338796 | 0.78 |
| YGR194C | 2007.303758 | 0.544240836 | 0.140502787 | 0.78 |
| YGR252W | 827.8896718 | -0.254451366 | 0.14059067 | 0.78 |
| YGR256W | 1242.341744 | 0.886123926 | 0.139688415 | 0.78 |
| YHR059W | 401.0845698 | -0.359446894 | 0.138250681 | 0.78 |
| YIR023C-A | 13.57792364 | -0.988027364 | 0.139494249 | 0.78 |
| YJL211C | 56.87482788 | 0.574788509 | 0.140429352 | 0.78 |
| YKR042W | 18766.32248 | -0.515192961 | 0.139099619 | 0.78 |
| YLR081W | 221.380283 | 0.599948995 | 0.139311832 | 0.78 |
| YLR137W | 268.6657732 | -0.529775279 | 0.139674327 | 0.78 |
| YLR329W | 110.7708514 | 0.581785429 | 0.139014555 | 0.78 |
| YML077W | 412.7293887 | -0.461287022 | 0.138541953 | 0.78 |
| YMR059W | 205.0702465 | 0.280525026 | 0.140119947 | 0.78 |
| YMR125W | 2280.761082 | -0.432399652 | 0.13924705 | 0.78 |
| YNL197C | 1704.51539 | -0.427450838 | 0.138324114 | 0.78 |
| YNL317W | 817.8553177 | -0.228248065 | 0.139036094 | 0.78 |
| YER153C | 351.5393272 | 0.514658854 | 0.14090709 | 0.78 |
| YLR087C | 2042.517235 | 0.384874303 | 0.140807453 | 0.78 |
| YDR086C | 1629.638329 | -0.441023468 | 0.141992795 | 0.78 |
| YGR026W | 4521.168167 | -0.340834561 | 0.141733292 | 0.78 |
| YHR007C-A | 61.56889406 | 0.564003021 | 0.14182964 | 0.78 |
| YKL191W | 1251.863612 | 0.488224674 | 0.141774612 | 0.78 |
| YKR093W | 2668.023949 | -0.632718637 | 0.141122423 | 0.78 |
| YLL012W | 268.5425901 | -0.705175468 | 0.141316154 | 0.78 |
| YMR149W | 2739.886665 | -0.389412019 | 0.141642196 | 0.78 |
| YOR355W | 1652.733408 | -0.571851446 | 0.141853358 | 0.78 |
| YPR016C | 1938.157861 | -0.38259884 | 0.142040104 | 0.78 |
| YGL095C | 1579.460918 | -0.259260445 | 0.142610657 | 0.78 |
| YMR137C | 598.7111524 | 0.414664864 | 0.142431305 | 0.78 |
| YPL210C | 1902.73736 | -0.464375029 | 0.142609876 | 0.78 |
| YDR223W | 358.2338356 | 0.470479143 | 0.142824899 | 0.78 |
| YJL048C | 1114.208522 | 0.587739035 | 0.143109855 | 0.78 |
| YNL176C | 1925.34034 | 0.365451438 | 0.143101314 | 0.78 |
| YBR081C | 1882.249025 | -0.417409838 | 0.143476519 | 0.78 |
| YIL143C | 1830.368647 | 0.470622899 | 0.143836636 | 0.78 |
| YLR218C | 291.942794 | -0.419696792 | 0.14440019 | 0.79 |
| YPR085C | 667.2581144 | 0.301165491 | 0.144805884 | 0.79 |
| YBR164C | 1099.977428 | -0.404296315 | 0.14632016 | 0.79 |
| YCR069W | 2249.27644 | -0.502413094 | 0.145666437 | 0.79 |
| YDR314C | 384.5141242 | 0.522653348 | 0.145565708 | 0.79 |
| YDR349C | 1807.872356 | 0.441111007 | 0.14640409 | 0.79 |
| YJL012C | 2507.923216 | -0.519713669 | 0.146451435 | 0.79 |

| | | | | |
|---|---|---|---|---|
| YKR072C | 2109.757496 | 0.423543004 | 0.146527909 | 0.79 |
| YLR453C | 293.5520271 | 0.333244867 | 0.145804216 | 0.79 |
| YML061C | 1035.299962 | 0.412039128 | 0.145974639 | 0.79 |
| YOL118C | 79.69627109 | 0.735672938 | 0.145634937 | 0.79 |
| YOR121C | 64.33732395 | 0.83280234 | 0.146475373 | 0.79 |
| YPL036W | 288.1590263 | 0.463525197 | 0.146073661 | 0.79 |
| YPL112C | 806.8455711 | -0.345470534 | 0.146201737 | 0.79 |
| YPR150W | 6.435730199 | 0.724503064 | 0.145473164 | 0.79 |
| YDR456W | 1671.501856 | 0.307924039 | 0.147141846 | 0.79 |
| YFL055W | 265.1999324 | 0.657946848 | 0.147244698 | 0.79 |
| YGR021W | 818.8357092 | 0.393845797 | 0.147088047 | 0.79 |
| YLL001W | 1443.400677 | 0.401022649 | 0.14735721 | 0.79 |
| YCL017C | 1588.391961 | 0.35261124 | 0.147509294 | 0.79 |
| YBR265W | 1130.650771 | -0.317977288 | 0.147906678 | 0.79 |
| YNL187W | 311.4857358 | 0.361953652 | 0.147804545 | 0.79 |
| YDR320C-A | 373.904445 | -0.428327826 | 0.148585843 | 0.79 |
| YER073W | 2130.104252 | 0.703946738 | 0.148477288 | 0.79 |
| YLR291C | 1399.72151 | -0.282358767 | 0.148432598 | 0.79 |
| YAL063C | 272.6184868 | 0.521431292 | 0.150326591 | 0.79 |
| YBR212W | 2837.682435 | 0.460479345 | 0.150089915 | 0.79 |
| YCR001W | 33.60447039 | -0.634878715 | 0.149970278 | 0.79 |
| YDR124W | 300.8040212 | 0.539367191 | 0.150837169 | 0.79 |
| YGL196W | 1262.908167 | 0.602782793 | 0.15085421 | 0.79 |
| YGR165W | 998.5420975 | -0.342944676 | 0.14907615 | 0.79 |
| YHR182W | 713.7981983 | 0.417727712 | 0.150008312 | 0.79 |
| YKR083C | 244.3380659 | -0.517166187 | 0.150531709 | 0.79 |
| YLL038C | 326.4334207 | -0.472569931 | 0.150214686 | 0.79 |
| YLL041C | 3315.150059 | 0.712567157 | 0.150681816 | 0.79 |
| YLL055W | 836.4946306 | 0.696849585 | 0.150581863 | 0.79 |
| YLR459W | 960.9364848 | -0.377122777 | 0.150016232 | 0.79 |
| YMR280C | 1028.334214 | 0.662917943 | 0.150524781 | 0.79 |
| YNL041C | 1185.16925 | -0.283121293 | 0.150031828 | 0.79 |
| YNL140C | 6.604161499 | -0.861828107 | 0.149673388 | 0.79 |
| YNL332W | 25.85198322 | 0.807950953 | 0.15037756 | 0.79 |
| YOL028C | 419.6545084 | -0.317072876 | 0.150435676 | 0.79 |
| YOR292C | 645.9873627 | 0.475640204 | 0.150048458 | 0.79 |
| YPR055W | 1160.769461 | -0.393751556 | 0.151167433 | 0.79 |
| YCL032W | 909.5801687 | 0.354431689 | 0.152135241 | 0.79 |
| YDL227C | 308.6751335 | -0.740996818 | 0.152204815 | 0.79 |
| YGL250W | 555.5711852 | 0.507220493 | 0.152317882 | 0.79 |
| YHL009W-B | 5.78416984 | -0.489046266 | 0.152223855 | 0.79 |
| YIL134C-A | 3.121789303 | 0.732414061 | 0.151756992 | 0.79 |
| YKR029C | 884.4269483 | -0.373362977 | 0.15223709 | 0.79 |

| | | | | |
|---|---|---|---|---|
| YLR312C | 417.3618401 | 0.759921761 | 0.152525578 | 0.79 |
| YOR005C | 988.1912317 | 0.415762498 | 0.151909742 | 0.79 |
| YOR021C | 1473.922183 | 0.45329209 | 0.151892283 | 0.79 |
| YPL262W | 5873.810031 | 0.787532215 | 0.152546415 | 0.79 |
| snR161 | 8.988952116 | 0.556755157 | 0.152652289 | 0.79 |
| YGL242C | 665.893361 | -0.494254228 | 0.153002498 | 0.80 |
| YNL101W | 1154.001663 | 0.397871436 | 0.153377736 | 0.80 |
| YKL112W | 917.1897974 | -0.399428062 | 0.153945709 | 0.80 |
| YGL258W | 52.00397703 | 0.618411031 | 0.154127381 | 0.80 |
| YPR012W | 66.76164095 | 0.45975024 | 0.154287505 | 0.80 |
| YLR410W | 2881.535918 | 0.339429318 | 0.154757176 | 0.80 |
| YNL097C | 784.7174827 | 0.401736096 | 0.154795302 | 0.80 |
| YEL069C | 125.5249506 | 0.773224487 | 0.154982328 | 0.80 |
| YDR448W | 426.123254 | -0.419517986 | 0.155204496 | 0.80 |
| YBL048W | 125.8109055 | 0.611487778 | 0.155940183 | 0.80 |
| YFR035C | 184.0960805 | 0.348343231 | 0.155622822 | 0.80 |
| YGL163C | 996.8369865 | 0.374848382 | 0.155842719 | 0.80 |
| YKL057C | 1611.669894 | -0.389308455 | 0.15596292 | 0.80 |
| YML031C-A | 2.97142876 | 0.579846121 | 0.155928741 | 0.80 |
| YNL289W | 446.6126721 | -0.724148017 | 0.155439724 | 0.80 |
| YBR073W | 1901.278512 | -0.663682375 | 0.156418368 | 0.80 |
| YHL030W | 2779.309717 | 0.459545242 | 0.156455033 | 0.80 |
| YJL108C | 258.931357 | 0.472793817 | 0.156533489 | 0.80 |
| YFL054C | 1511.844206 | 0.759687816 | 0.157069323 | 0.80 |
| YPL235W | 1507.98677 | -0.310494222 | 0.157007915 | 0.80 |
| YML109W | 760.9435844 | 0.356651122 | 0.157255142 | 0.80 |
| YBL066C | 1237.560937 | 0.545950655 | 0.158916028 | 0.81 |
| YBR139W | 2470.205803 | 0.479144532 | 0.159545992 | 0.81 |
| YBR163W | 719.9181119 | -0.416944717 | 0.158254138 | 0.81 |
| YCL012C | 422.2313505 | 0.49824732 | 0.159249735 | 0.81 |
| YDL110C | 2473.385616 | 0.555400329 | 0.157808881 | 0.81 |
| YDL118W | 21.28319356 | -0.614312345 | 0.159736817 | 0.81 |
| YDR070C | 1811.421734 | 0.822297953 | 0.159584431 | 0.81 |
| YDR181C | 744.5427058 | 0.458589897 | 0.158107618 | 0.81 |
| YGL206C | 5144.593261 | -0.302063701 | 0.159791124 | 0.81 |
| YJL003W | 223.2586603 | -0.375379476 | 0.15979171 | 0.81 |
| YKL048C | 583.1360838 | -0.518033794 | 0.159702953 | 0.81 |
| YLR207W | 1228.065049 | 0.306868601 | 0.158513199 | 0.81 |
| YML099C | 1070.053648 | 0.45009501 | 0.159220473 | 0.81 |
| YOL162W | 65.72006374 | 0.686441089 | 0.158762818 | 0.81 |
| YOR026W | 568.5531387 | -0.437752112 | 0.158596849 | 0.81 |
| YPR031W | 763.0479333 | 0.402911826 | 0.159488519 | 0.81 |
| YPR057W | 656.961666 | -0.306191012 | 0.159065105 | 0.81 |

| | | | | |
|---|---|---|---|---|
| YAL039C | 1453.711163 | 0.391637039 | 0.161181525 | 0.81 |
| YBR022W | 363.9709831 | -0.468696049 | 0.162248853 | 0.81 |
| YBR120C | 395.0224857 | -0.382132169 | 0.161553927 | 0.81 |
| YBR144C | 8.772859821 | 0.759455941 | 0.161992062 | 0.81 |
| YBR241C | 1365.325299 | 0.69620882 | 0.160282093 | 0.81 |
| YCR015C | 255.0577346 | -0.290362587 | 0.161150903 | 0.81 |
| YDR381W | 4059.585724 | -0.412999337 | 0.161868123 | 0.81 |
| YEL041W | 348.1575551 | 0.519701337 | 0.162158958 | 0.81 |
| YER077C | 789.2511196 | -0.264535447 | 0.161282112 | 0.81 |
| YER171W | 948.7525289 | -0.286997133 | 0.161162656 | 0.81 |
| YFR036W | 228.7837864 | 0.34580191 | 0.160591809 | 0.81 |
| YJR074W | 487.5630441 | 0.533610475 | 0.1618405 | 0.81 |
| YKR055W | 476.0228776 | 0.361869976 | 0.161684646 | 0.81 |
| YLR040C | 271.3228626 | -0.706857326 | 0.162360517 | 0.81 |
| YLR423C | 531.1083054 | 0.466974164 | 0.160218773 | 0.81 |
| YLR438W | 1335.790008 | 0.570710552 | 0.160619375 | 0.81 |
| YMR047C | 2343.921258 | -0.321631559 | 0.160795782 | 0.81 |
| YMR153C-A | 60.08017136 | -0.509863266 | 0.161420545 | 0.81 |
| YPL060W | 360.6791713 | 0.504598136 | 0.160160306 | 0.81 |
| YPL184C | 1810.247002 | -0.417139388 | 0.161534229 | 0.81 |
| YAL055W | 312.0534701 | 0.449380992 | 0.16361356 | 0.81 |
| YDL077C | 1087.970908 | 0.217643315 | 0.16317244 | 0.81 |
| YDR391C | 1742.594469 | 0.564864676 | 0.162946647 | 0.81 |
| YGL011C | 3486.685044 | -0.271932708 | 0.162818679 | 0.81 |
| YLR361C | 559.6543534 | 0.30498953 | 0.163441285 | 0.81 |
| YML100W | 14497.66976 | 0.646152065 | 0.1635372 | 0.81 |
| YNL238W | 1373.585003 | -0.33367255 | 0.162621775 | 0.81 |
| YOR215C | 2322.968103 | 0.560465888 | 0.16327883 | 0.81 |
| YOR329W-A | 20.44917844 | -0.574215852 | 0.163180562 | 0.81 |
| YDL186W | 120.8658792 | 0.553600234 | 0.164234948 | 0.81 |
| YGL117W | 750.1030158 | 0.695744256 | 0.16423053 | 0.81 |
| YNR032C-A | 183.6670278 | -0.359648601 | 0.164496714 | 0.81 |
| YOR327C | 881.0062218 | -0.349872338 | 0.164327163 | 0.81 |
| YPL070W | 1079.494033 | 0.326970471 | 0.164056654 | 0.81 |
| YPR166C | 431.6523763 | -0.38035037 | 0.163838699 | 0.81 |
| snR57 | 17.27289396 | 0.825327275 | 0.164443692 | 0.81 |
| YBR056W-A | 43.21007303 | -0.667744137 | 0.169434594 | 0.81 |
| YBR074W | 1213.234406 | -0.246448166 | 0.166861222 | 0.81 |
| YBR296C | 209.9913106 | 0.780825553 | 0.167726284 | 0.81 |
| YDR034C | 1497.715383 | 0.433211718 | 0.169313598 | 0.81 |
| YDR138W | 884.1934286 | -0.317650328 | 0.168823027 | 0.81 |
| YDR234W | 1972.622196 | 0.530205948 | 0.165660644 | 0.81 |
| YDR299W | 1150.958917 | -0.550936915 | 0.166205821 | 0.81 |

| | | | | |
|---|---|---|---|---|
| YDR351W | 1313.626306 | -0.394287923 | 0.167575254 | 0.81 |
| YDR506C | 448.3958033 | 0.292325623 | 0.165764748 | 0.81 |
| YGL199C | 2.062524245 | 0.68183427 | 0.167430944 | 0.81 |
| YGR066C | 223.0827395 | 0.610401757 | 0.165983829 | 0.81 |
| YGR267C | 1681.678228 | 0.243337288 | 0.168781909 | 0.81 |
| YHR157W | 84.3613017 | 0.446726285 | 0.166820532 | 0.81 |
| YHR164C | 1299.336952 | 0.28880614 | 0.169250313 | 0.81 |
| YIL004C | 332.265247 | -0.257724861 | 0.168914903 | 0.81 |
| YJR020W | 8.221696255 | 0.660846201 | 0.167301792 | 0.81 |
| YKL175W | 2025.203634 | 0.323802384 | 0.167790149 | 0.81 |
| YKR032W | 0.979223699 | 0.510489006 | 0.166884836 | 0.81 |
| YKR049C | 1528.555977 | 0.681781744 | 0.16715706 | 0.81 |
| YML029W | 1034.498713 | 0.276362405 | 0.168866261 | 0.81 |
| YML069W | 2256.742432 | -0.414425858 | 0.168306167 | 0.81 |
| YML111W | 1306.672498 | -0.313712647 | 0.168970432 | 0.81 |
| YMR239C | 477.7892356 | 0.501496144 | 0.169175531 | 0.81 |
| YNL153C | 847.6509295 | -0.459005178 | 0.165436769 | 0.81 |
| YNL172W | 1164.569131 | 0.308467515 | 0.166067897 | 0.81 |
| YNL223W | 533.8185201 | 0.403529709 | 0.166318148 | 0.81 |
| YNL226W | 19.21780824 | 0.557807577 | 0.169465185 | 0.81 |
| YNL245C | 408.0242862 | 0.36410403 | 0.166789973 | 0.81 |
| YNR040W | 422.1360755 | 0.403930317 | 0.168977564 | 0.81 |
| YNR072W | 156.9410464 | 0.680461565 | 0.166347502 | 0.81 |
| YOL141W | 189.2930206 | 0.512660059 | 0.168203649 | 0.81 |
| YOR162C | 795.3084144 | 0.204999387 | 0.167565945 | 0.81 |
| YPL029W | 470.8569792 | -0.273448149 | 0.169361803 | 0.81 |
| YPR038W | 18.2580769 | -0.600069522 | 0.16552465 | 0.81 |
| YPR083W | 614.8326597 | 0.619461257 | 0.169458805 | 0.81 |
| YAL043C | 1560.889418 | 0.323880964 | 0.172387393 | 0.81 |
| YBR007C | 740.5673067 | -0.438247605 | 0.171312082 | 0.81 |
| YBR024W | 889.4330761 | 0.492797664 | 0.169804244 | 0.81 |
| YCR097W | 1013.079937 | 0.657135627 | 0.172059858 | 0.81 |
| YDL114W-A | 24.54000287 | 0.607755101 | 0.172446199 | 0.81 |
| YDR009W | 246.880862 | 0.576219367 | 0.171483187 | 0.81 |
| YDR044W | 1839.127613 | -0.508247029 | 0.171697787 | 0.81 |
| YDR098C-B | 19.81780983 | 0.635975698 | 0.170731536 | 0.81 |
| YDR372C | 1904.188965 | -0.382988122 | 0.172741903 | 0.81 |
| YER124C | 2039.69695 | 0.407827805 | 0.171693981 | 0.81 |
| YER128W | 281.9899223 | 0.361543774 | 0.171511147 | 0.81 |
| YER137C | 65.6651259 | 0.519518016 | 0.171146876 | 0.81 |
| YGL057C | 252.5789153 | -0.255734149 | 0.172708556 | 0.81 |
| YGL153W | 279.9517565 | 0.349513646 | 0.172675083 | 0.81 |
| YGL229C | 708.5937653 | 0.340055646 | 0.16990473 | 0.81 |

| | | | | |
|---|---|---|---|---|
| YHR143W | 3407.641358 | 0.513345997 | 0.17257349 | 0.81 |
| YJL047C | 709.4046862 | 0.331991332 | 0.170764923 | 0.81 |
| YJL075C | 78.82663296 | -0.530627908 | 0.173010384 | 0.81 |
| YLR079W | 2171.764341 | 0.417802434 | 0.172236453 | 0.81 |
| YLR208W | 2849.872811 | -0.330386366 | 0.172817417 | 0.81 |
| YMR088C | 861.7053416 | 0.396898177 | 0.172610727 | 0.81 |
| YNL306W | 1534.453742 | -0.288667044 | 0.171503298 | 0.81 |
| YOL084W | 492.0065357 | 0.627708576 | 0.170305791 | 0.81 |
| YOL115W | 646.4767705 | -0.407629322 | 0.170062618 | 0.81 |
| YOR229W | 494.4703852 | 0.329489819 | 0.170810064 | 0.81 |
| YOR249C | 647.5824875 | -0.316068968 | 0.172693421 | 0.81 |
| YPL004C | 14429.78474 | 0.395024149 | 0.172962106 | 0.81 |
| YBR050C | 396.3143748 | 0.762819547 | 0.17444563 | 0.81 |
| YBR131W | 285.0666359 | 0.352175036 | 0.173946539 | 0.81 |
| YBR221C | 5794.723277 | 0.358867496 | 0.17380631 | 0.81 |
| YDR498C | 1014.743074 | -0.176995233 | 0.174286167 | 0.81 |
| YIL098C | 444.8949209 | -0.40613958 | 0.173762294 | 0.81 |
| YNL059C | 1845.140766 | -0.32464938 | 0.174207324 | 0.81 |
| YOR043W | 1570.483617 | -0.343679026 | 0.17429214 | 0.81 |
| YPL156C | 858.4762505 | -0.627746523 | 0.173972447 | 0.81 |
| YPR091C | 1815.236332 | 0.267146303 | 0.174427043 | 0.81 |
| YPR123C | 99.99461053 | 0.737120559 | 0.17357806 | 0.81 |
| YLR292C | 1010.798537 | -0.469040677 | 0.174576707 | 0.81 |
| YLR102C | 476.3043851 | 0.441343883 | 0.17533135 | 0.82 |
| YBL031W | 539.4437355 | -0.432434712 | 0.181131979 | 0.82 |
| YBL097W | 438.5923101 | -0.409776184 | 0.177279283 | 0.82 |
| YBR230C | 2523.139558 | 0.635108918 | 0.181405854 | 0.82 |
| YDL011C | 7.393723597 | -0.749159808 | 0.179903709 | 0.82 |
| YDL022C-A | 6.814086928 | 0.752671477 | 0.181913902 | 0.82 |
| YDL246C | 6.191078768 | 0.753086688 | 0.179561365 | 0.82 |
| YDR179C | 161.4536724 | 0.277872041 | 0.18194707 | 0.82 |
| YDR225W | 3245.605018 | -0.479470677 | 0.178243491 | 0.82 |
| YDR278C | 0.649229666 | 0.496805727 | 0.179261806 | 0.82 |
| YDR475C | 1828.530627 | 0.492076081 | 0.176343746 | 0.82 |
| YDR522C | 159.2048735 | -0.559503267 | 0.179478918 | 0.82 |
| YEL020C-B | 34.39724677 | -0.510963423 | 0.176948272 | 0.82 |
| YEL024W | 1858.219132 | 0.67819348 | 0.176965278 | 0.82 |
| YER061C | 958.2654695 | 0.379950914 | 0.181942058 | 0.82 |
| YER093C | 1449.072568 | 0.250764611 | 0.178288411 | 0.82 |
| YER129W | 2291.22394 | 0.1826375 | 0.179796 | 0.82 |
| YER148W-A | 0.781507553 | -0.732137133 | 0.181720394 | 0.82 |
| YFL058W | 7.345596516 | 0.759320181 | 0.180361379 | 0.82 |
| YFR038W | 1186.649674 | 0.330882091 | 0.179386129 | 0.82 |

| | | | | |
|---|---|---|---|---|
| YGL027C | 1397.054052 | 0.365053328 | 0.179538865 | 0.82 |
| YGR239C | 410.924928 | 0.390704787 | 0.177090935 | 0.82 |
| YIL015W | 2424.316267 | -0.707988832 | 0.180345804 | 0.82 |
| YIL020C-A | 191.6015626 | -0.390261958 | 0.176754564 | 0.82 |
| YIL049W | 383.4643737 | -0.5163921 | 0.179673132 | 0.82 |
| YIR014W | 256.3884009 | 0.497131347 | 0.175848298 | 0.82 |
| YJL116C | 947.324176 | 0.655271853 | 0.178189464 | 0.82 |
| YJL133W | 754.0202192 | -0.375366934 | 0.181528854 | 0.82 |
| YJR109C | 3577.314205 | 0.660617259 | 0.179846538 | 0.82 |
| YJR118C | 868.4456577 | -0.366345453 | 0.181520138 | 0.82 |
| YJR154W | 284.604362 | 0.573901808 | 0.178797335 | 0.82 |
| YKL035W | 11294.21789 | 0.407306521 | 0.180512694 | 0.82 |
| YKL150W | 7424.00365 | 0.689244213 | 0.176244076 | 0.82 |
| YLL042C | 71.42312394 | 0.498614836 | 0.178858326 | 0.82 |
| YLR042C | 282.794653 | -0.71553143 | 0.178861971 | 0.82 |
| YMR033W | 1176.392622 | -0.388131114 | 0.177954015 | 0.82 |
| YMR124W | 1653.834225 | 0.319893307 | 0.176542779 | 0.82 |
| YMR213W | 660.1636091 | 0.337030362 | 0.181860312 | 0.82 |
| YNL156C | 1413.410846 | 0.300717321 | 0.177558844 | 0.82 |
| YNL174W | 7.421723425 | 0.303685635 | 0.175678449 | 0.82 |
| YNL256W | 1084.449905 | 0.416681061 | 0.180910021 | 0.82 |
| YNR053C | 1005.221474 | -0.542698012 | 0.181169855 | 0.82 |
| YNR061C | 596.8560089 | -0.392426219 | 0.179353108 | 0.82 |
| YOL029C | 366.4284423 | -0.268394659 | 0.179127076 | 0.82 |
| YOL045W | 1132.896142 | 0.220077378 | 0.178891012 | 0.82 |
| YOL139C | 5554.572319 | -0.454923203 | 0.181154695 | 0.82 |
| YOR050C | 67.3008437 | -0.43259516 | 0.181381176 | 0.82 |
| YOR155C | 1051.816257 | 0.311524851 | 0.177019664 | 0.82 |
| YOR221C | 1523.763907 | 0.334701415 | 0.176520206 | 0.82 |
| YPR010C-A | 733.8429902 | 0.727942123 | 0.181523505 | 0.82 |
| YPR029C | 1930.500091 | -0.230912422 | 0.177723276 | 0.82 |
| YPR160W | 12134.47846 | 0.741805216 | 0.180529103 | 0.82 |
| YPR193C | 57.30247136 | 0.635879448 | 0.176498239 | 0.82 |
| YBR085W | 162.0814879 | 0.803900523 | 0.185546961 | 0.82 |
| YBR171W | 1056.70053 | -0.428110144 | 0.183018425 | 0.82 |
| YBR230W-A | 211.0787097 | -0.624073205 | 0.184964759 | 0.82 |
| YBR291C | 379.9577466 | -0.454776225 | 0.184516121 | 0.82 |
| YBR298C-A | 3.08929577 | -0.837680907 | 0.185051319 | 0.82 |
| YBR301W | 22.99326035 | 0.7327016 | 0.1834415 | 0.82 |
| YDL072C | 3778.461007 | 0.247436944 | 0.184816685 | 0.82 |
| YDL114W | 43.38451628 | 0.696548737 | 0.184841359 | 0.82 |
| YDL234C | 2388.462086 | 0.487157806 | 0.183780708 | 0.82 |
| YDL245C | 34.45281204 | 0.697858879 | 0.18240392 | 0.82 |

| | | | | |
|---|---|---|---|---|
| YDR075W | 328.511722 | -0.434695364 | 0.185675531 | 0.82 |
| YDR544C | 2.336721419 | 0.568901418 | 0.184336145 | 0.82 |
| YER046W | 177.0920564 | 0.577519835 | 0.184465549 | 0.82 |
| YGL181W | 1793.449442 | 0.293725164 | 0.185266158 | 0.82 |
| YGL223C | 914.5737412 | -0.240315438 | 0.185125342 | 0.82 |
| YGR294W | 10.95232902 | -0.759450919 | 0.182635464 | 0.82 |
| YHR143W-A | 554.7529102 | -0.403963355 | 0.182462082 | 0.82 |
| YHR216W | 3639.691774 | 0.735537404 | 0.182220561 | 0.82 |
| YJL023C | 307.217395 | 0.324412344 | 0.184200588 | 0.82 |
| YJL173C | 671.1948643 | 0.424376945 | 0.184613024 | 0.82 |
| YJR051W | 1291.045072 | 0.494692755 | 0.184522577 | 0.82 |
| YKL026C | 613.714189 | 0.760592112 | 0.185478284 | 0.82 |
| YKL161C | 387.0663817 | -0.587437157 | 0.184715662 | 0.82 |
| YLR154C | 335.1264893 | -0.262790311 | 0.184873968 | 0.82 |
| YNL163C | 1921.119184 | 0.403199472 | 0.185725589 | 0.82 |
| YOR210W | 598.8398742 | -0.424893329 | 0.183518393 | 0.82 |
| YOR265W | 445.8131326 | -0.329953782 | 0.183044344 | 0.82 |
| YOR298W | 171.3347166 | 0.413089077 | 0.18271015 | 0.82 |
| YPR165W | 4589.151851 | -0.337670969 | 0.184908288 | 0.82 |
| snR45 | 13.49639451 | 0.744086364 | 0.184194268 | 0.82 |
| YLR026C | 1116.422754 | -0.357907527 | 0.185888387 | 0.82 |
| YGL051W | 80.44841429 | 0.454455893 | 0.186184812 | 0.82 |
| YEL055C | 1803.113225 | 0.398977077 | 0.187047503 | 0.82 |
| YFL010C | 2890.139798 | -0.421310861 | 0.186701552 | 0.82 |
| YGR092W | 957.140587 | 0.444236373 | 0.187111128 | 0.82 |
| YGR261C | 1931.303256 | -0.332645152 | 0.187027867 | 0.82 |
| YKL012W | 1327.340355 | 0.203240824 | 0.187048251 | 0.82 |
| YKL049C | 577.3829424 | 0.328869718 | 0.186913151 | 0.82 |
| YMR110C | 3066.365125 | 0.564059564 | 0.186383534 | 0.82 |
| YMR304W | 1821.67097 | 0.334730137 | 0.187317339 | 0.82 |
| YOR062C | 585.4618256 | -0.612296047 | 0.187909804 | 0.82 |
| YBL099W | 11787.99535 | 0.706508112 | 0.189677256 | 0.82 |
| YBR026C | 2145.982583 | 0.405878392 | 0.188766244 | 0.82 |
| YCL055W | 639.3184492 | -0.495738192 | 0.189223466 | 0.82 |
| YDL126C | 11516.69474 | 0.312155462 | 0.190290511 | 0.82 |
| YER112W | 1280.704105 | -0.304157408 | 0.190242792 | 0.82 |
| YGL014W | 2156.619039 | -0.486123853 | 0.190056463 | 0.82 |
| YGL263W | 253.3051631 | 0.684442048 | 0.189328423 | 0.82 |
| YGR119C | 2276.598641 | -0.252699213 | 0.188360835 | 0.82 |
| YHL019C | 544.6940331 | 0.39823173 | 0.18947791 | 0.82 |
| YIL125W | 5872.779859 | 0.671760147 | 0.189927647 | 0.82 |
| YJR055W | 115.4214659 | 0.363274014 | 0.189939209 | 0.82 |
| YKL188C | 680.4138758 | 0.482452656 | 0.189741702 | 0.82 |

| | | | | |
|---|---|---|---|---|
| YLL048C | 4130.734576 | 0.312888173 | 0.190313087 | 0.82 |
| YML071C | 1055.622411 | -0.342037513 | 0.18901036 | 0.82 |
| YMR197C | 923.7331716 | -0.261260838 | 0.189833255 | 0.82 |
| YNR019W | 1395.465767 | 0.422324911 | 0.18952246 | 0.82 |
| YPR023C | 1837.667458 | 0.484064478 | 0.188983466 | 0.82 |
| YPR094W | 154.1332011 | -0.377621301 | 0.190690416 | 0.82 |
| YLR299W | 1242.489768 | 0.407917803 | 0.19090229 | 0.82 |
| YLR400W | 49.34542517 | 0.535183897 | 0.191039843 | 0.82 |
| YGL025C | 784.0082065 | 0.27975228 | 0.191295993 | 0.82 |
| YOL049W | 2360.13263 | -0.351644596 | 0.191296366 | 0.82 |
| YJR012C | 286.1622087 | -0.286988881 | 0.191465262 | 0.82 |
| YBL036C | 1223.509384 | -0.27305657 | 0.192092942 | 0.82 |
| YER051W | 512.8476233 | 0.430128985 | 0.192028948 | 0.82 |
| YKL053W | 17.58730316 | -0.594041494 | 0.192237215 | 0.82 |
| YKL189W | 702.1789039 | -0.437692451 | 0.19267868 | 0.83 |
| YFL010W-A | 308.9446486 | -0.392107031 | 0.192992337 | 0.83 |
| YBR203W | 994.1971392 | 0.641030589 | 0.193433028 | 0.83 |
| YHR194W | 859.058911 | 0.412020875 | 0.193916047 | 0.83 |
| YNL195C | 538.1405229 | 0.697745649 | 0.193826954 | 0.83 |
| YPL167C | 937.9139202 | 0.342086413 | 0.193959451 | 0.83 |
| YMR255W | 1118.071772 | -0.422277619 | 0.194111288 | 0.83 |
| YBR154C | 1262.041062 | -0.402833496 | 0.194889197 | 0.83 |
| YER004W | 2233.162604 | 0.365004267 | 0.195837952 | 0.83 |
| YNL040W | 832.5546055 | 0.430864015 | 0.195889997 | 0.83 |
| YNR063W | 178.9442947 | 0.423578366 | 0.195818221 | 0.83 |
| YDL183C | 401.7373652 | 0.441017213 | 0.196595669 | 0.83 |
| YDR104C | 711.7535955 | 0.355260422 | 0.196375822 | 0.83 |
| YDR128W | 2008.877758 | 0.373813911 | 0.196664597 | 0.83 |
| YHR022C | 158.1795629 | -0.549236386 | 0.196575911 | 0.83 |
| YKL106C-A | 1.684117879 | 0.704657867 | 0.196264119 | 0.83 |
| YDR200C | 868.5642679 | -0.229337516 | 0.197041223 | 0.84 |
| YPL193W | 620.6543173 | 0.262849343 | 0.197088134 | 0.84 |
| YGR084C | 1041.679379 | -0.268840595 | 0.197373497 | 0.84 |
| YER146W | 140.8937845 | -0.449069019 | 0.19794406 | 0.84 |
| YJR114W | 33.45969567 | -0.533781911 | 0.197841921 | 0.84 |
| YKL085W | 9612.778538 | 0.733729206 | 0.197583586 | 0.84 |
| YPR169W-A | 32.89680766 | 0.43214269 | 0.197937052 | 0.84 |
| YBL071C-B | 1.645198606 | 0.64578032 | 0.199967628 | 0.84 |
| YBR201C-A | 201.3285883 | 0.577024107 | 0.200551259 | 0.84 |
| YDR302W | 389.0867763 | -0.359964683 | 0.199225711 | 0.84 |
| YGL109W | 79.04302724 | -0.370137177 | 0.200347578 | 0.84 |
| YGR288W | 284.2488249 | 0.589053459 | 0.200630836 | 0.84 |
| YHR175W-A | 167.202498 | -0.585884353 | 0.200637127 | 0.84 |

| | | | | |
|---|---|---|---|---|
| YLR139C | 783.7450537 | -0.400418717 | 0.200240286 | 0.84 |
| YLR260W | 983.6383575 | 0.397995182 | 0.199388722 | 0.84 |
| YML075C | 1371.306293 | -0.271815524 | 0.20058454 | 0.84 |
| YNL052W | 2172.615322 | 0.705478436 | 0.200576309 | 0.84 |
| YOR012W | 60.32598792 | -0.382269848 | 0.199413379 | 0.84 |
| YOR285W | 4227.371531 | 0.487534673 | 0.200568358 | 0.84 |
| YPL215W | 1126.607522 | -0.294887659 | 0.198979883 | 0.84 |
| YPR093C | 386.3642488 | 0.595950614 | 0.200252 | 0.84 |
| YDR524W-C | 1641.998262 | -0.454963879 | 0.201029145 | 0.84 |
| YIL162W | 3263.600398 | 0.653812387 | 0.200951955 | 0.84 |
| YJR050W | 528.5017956 | 0.358448302 | 0.201191808 | 0.84 |
| YBR111W-A | 229.152125 | -0.367890307 | 0.201610561 | 0.84 |
| YIR039C | 1029.181658 | 0.678527407 | 0.201665636 | 0.84 |
| YJL091C | 524.9582628 | -0.280978323 | 0.201614948 | 0.84 |
| YKR086W | 1098.768049 | 0.293723295 | 0.201773147 | 0.84 |
| YKL163W | 2089.664663 | 0.723917138 | 0.20226266 | 0.84 |
| YAL032C | 524.818116 | -0.25591791 | 0.203854034 | 0.84 |
| YBR102C | 889.0122882 | -0.19914471 | 0.204664167 | 0.84 |
| YBR289W | 1372.875089 | -0.388580936 | 0.206196182 | 0.84 |
| YDL167C | 1031.301724 | -0.530416628 | 0.205790303 | 0.84 |
| YDL200C | 293.2384509 | 0.320820864 | 0.206424751 | 0.84 |
| YDR438W | 373.0593006 | 0.341925334 | 0.206432035 | 0.84 |
| YEL010W | 4.427753031 | 0.67371437 | 0.205870719 | 0.84 |
| YER134C | 346.3610386 | -0.437350285 | 0.204190464 | 0.84 |
| YER140W | 473.4616837 | -0.217543512 | 0.205853606 | 0.84 |
| YGL041W-A | 345.8516558 | -0.339043615 | 0.205870527 | 0.84 |
| YGR090W | 2037.446504 | -0.515779459 | 0.206406161 | 0.84 |
| YHR028W-A | 38.81881491 | 0.44764028 | 0.204360341 | 0.84 |
| YHR031C | 752.7001486 | 0.448692794 | 0.204313264 | 0.84 |
| YHR189W | 309.5122663 | 0.459397822 | 0.202750281 | 0.84 |
| YJL207C | 2307.188321 | -0.386535752 | 0.204487137 | 0.84 |
| YJR161C | 124.4036715 | 0.541510596 | 0.203230185 | 0.84 |
| YKL074C | 747.5415477 | 0.250166166 | 0.202899122 | 0.84 |
| YKR067W | 2101.266331 | 0.456620206 | 0.202800469 | 0.84 |
| YLL007C | 554.0715465 | -0.391470461 | 0.206283738 | 0.84 |
| YLR176C | 916.0927001 | 0.313576808 | 0.206179417 | 0.84 |
| YLR277C | 732.8482737 | -0.29433131 | 0.205017406 | 0.84 |
| YML120C | 4283.544511 | 0.683988593 | 0.203392912 | 0.84 |
| YMR075C-A | 15.14864191 | 0.507687913 | 0.204997935 | 0.84 |
| YMR209C | 810.1729969 | -0.376913638 | 0.204566523 | 0.84 |
| YMR294W-A | 229.7802387 | -0.27660126 | 0.204392925 | 0.84 |
| YNL276C | 5.892936072 | -0.692645849 | 0.203022968 | 0.84 |
| YNR034W-A | 4345.183948 | 0.743171478 | 0.205632957 | 0.84 |

| | | | | |
|---|---|---|---|---|
| YNR069C | 436.7652166 | 0.50637065 | 0.203597024 | 0.84 |
| YOR296W | 873.0215241 | -0.250686071 | 0.2055566 | 0.84 |
| YPR068C | 284.1750342 | -0.228155782 | 0.204477565 | 0.84 |
| YPR088C | 2257.094394 | 0.314988433 | 0.203995184 | 0.84 |
| YPR189W | 1450.192608 | -0.312316464 | 0.203479319 | 0.84 |
| YBL016W | 1090.490568 | -0.678671277 | 0.207013916 | 0.84 |
| YGL108C | 328.3815974 | -0.341137163 | 0.206867694 | 0.84 |
| YLL036C | 873.0418955 | 0.188946842 | 0.207136763 | 0.84 |
| YLR258W | 9832.7404 | 0.593249544 | 0.206995207 | 0.84 |
| YNL284C-B | 190.6577576 | 0.599662953 | 0.206979068 | 0.84 |
| YNL266W | 15.42456766 | 0.479734766 | 0.207311034 | 0.84 |
| YPL270W | 1768.066094 | 0.399979545 | 0.207461052 | 0.84 |
| YDL154W | 661.9950458 | -0.433922869 | 0.208217397 | 0.84 |
| YDR241W | 32.77525977 | -0.537626488 | 0.208486855 | 0.84 |
| YDR263C | 317.4342443 | 0.364661043 | 0.20868875 | 0.84 |
| YEL008W | 30.07335462 | -0.651184176 | 0.209045179 | 0.84 |
| YIL023C | 838.1130985 | 0.331094041 | 0.208492663 | 0.84 |
| YIL151C | 1650.033227 | -0.320055967 | 0.209159447 | 0.84 |
| YJL215C | 44.17616213 | -0.600828452 | 0.209132993 | 0.84 |
| YKL023W | 1129.71016 | 0.429269899 | 0.207929561 | 0.84 |
| YKL213C | 1964.500982 | 0.220758595 | 0.208921626 | 0.84 |
| YMR312W | 719.0027121 | 0.327238962 | 0.20892214 | 0.84 |
| YOL083W | 724.7720388 | 0.535019895 | 0.209131148 | 0.84 |
| YOL123W | 4114.119417 | -0.426979706 | 0.208424196 | 0.84 |
| YPR139C | 1065.385559 | -0.377682472 | 0.207672117 | 0.84 |
| YDR080W | 1103.850689 | 0.359135822 | 0.209793169 | 0.85 |
| YCL045C | 3038.105977 | -0.38460398 | 0.210711331 | 0.85 |
| YDR034W-B | 127.2105401 | 0.691766188 | 0.210599931 | 0.85 |
| YGR070W | 2198.55855 | 0.524177655 | 0.210597632 | 0.85 |
| YKL001C | 376.5252013 | -0.468766588 | 0.210629417 | 0.85 |
| YML014W | 592.0947476 | 0.462639597 | 0.21055519 | 0.85 |
| YJR034W | 337.4607648 | -0.385034339 | 0.211004 | 0.85 |
| YOR008C | 1383.478149 | -0.332572671 | 0.210888402 | 0.85 |
| YMR078C | 885.7332334 | -0.260823203 | 0.211202888 | 0.85 |
| YCR102C | 144.1370507 | 0.524910045 | 0.21250298 | 0.85 |
| YEL016C | 826.6367047 | -0.230620698 | 0.211913381 | 0.85 |
| YFL063W | 4.881871603 | 0.420237186 | 0.211702063 | 0.85 |
| YGL244W | 2087.284965 | -0.332298342 | 0.211859837 | 0.85 |
| YHR094C | 525.1336673 | 0.645188281 | 0.212290176 | 0.85 |
| YMR115W | 1106.490843 | 0.32695406 | 0.212385132 | 0.85 |
| YOR056C | 1339.644393 | 0.300848445 | 0.211922789 | 0.85 |
| YPL207W | 1844.153383 | 0.395734816 | 0.212368258 | 0.85 |
| YPR151C | 43.40930441 | 0.65275064 | 0.211810963 | 0.85 |

| | | | | |
|---|---|---|---|---|
| snR7-L | 6.909111604 | -0.736022296 | 0.212534289 | 0.85 |
| YDR182W | 1361.286881 | -0.239690182 | 0.212728459 | 0.85 |
| YEL074W | 35.0602757 | 0.533722932 | 0.213683651 | 0.85 |
| YGL158W | 260.9463165 | -0.695721978 | 0.213725819 | 0.85 |
| YGR121C | 1176.230558 | 0.393452229 | 0.213441849 | 0.85 |
| YKL190W | 813.5201171 | -0.337547971 | 0.213679093 | 0.85 |
| YLR008C | 578.2897755 | -0.352909316 | 0.213083596 | 0.85 |
| YOR184W | 6279.813274 | 0.536661302 | 0.213794764 | 0.85 |
| YOR325W | 4.631536169 | -0.612635762 | 0.213415317 | 0.85 |
| YBR117C | 1336.758461 | 0.731024533 | 0.214161195 | 0.85 |
| YFR054C | 20.51319414 | 0.520753773 | 0.214191896 | 0.85 |
| YPL171C | 308.1636844 | 0.680667284 | 0.21402523 | 0.85 |
| YDL080C | 732.479931 | -0.233209016 | 0.214531768 | 0.85 |
| YDR035W | 3632.420651 | 0.426493029 | 0.214610075 | 0.85 |
| YPL278C | 8.613341367 | 0.521538748 | 0.215004092 | 0.85 |
| YOR174W | 544.9117735 | 0.261397847 | 0.215269915 | 0.85 |
| YBR254C | 311.1708692 | -0.415003809 | 0.216771559 | 0.85 |
| YEL020C | 1045.827861 | 0.455930319 | 0.216254255 | 0.85 |
| YER142C | 720.8941802 | 0.422084145 | 0.216518881 | 0.85 |
| YER159C | 1569.026083 | -0.442750608 | 0.21587502 | 0.85 |
| YGR257C | 921.5740821 | -0.3068306 | 0.21578391 | 0.85 |
| YJR102C | 338.1234787 | -0.317858111 | 0.216593594 | 0.85 |
| YNL291C | 628.873147 | -0.317368235 | 0.216528794 | 0.85 |
| YOR343W-B | 107.9180972 | -0.594220542 | 0.216698057 | 0.85 |
| YPL106C | 19467.56403 | 0.471971897 | 0.216012321 | 0.85 |
| YBR146W | 1432.145995 | -0.2635574 | 0.217084855 | 0.85 |
| YGL127C | 488.4323072 | -0.373599834 | 0.2172018 | 0.85 |
| YAL051W | 1222.388192 | 0.270839871 | 0.217521186 | 0.85 |
| YGR228W | 14.43502374 | 0.615559241 | 0.217784988 | 0.85 |
| YPR053C | 166.5759994 | -0.473950464 | 0.217840221 | 0.85 |
| YAL049C | 1497.895567 | 0.354507355 | 0.220231912 | 0.85 |
| YBR111C | 3318.167647 | 0.517791483 | 0.219818508 | 0.85 |
| YBR200W | 1077.368125 | -0.22226966 | 0.220722765 | 0.85 |
| YCL041C | 119.8839828 | 0.424057389 | 0.220476498 | 0.85 |
| YDR254W | 574.0903535 | 0.248853786 | 0.220803656 | 0.85 |
| YER028C | 507.0764562 | 0.57923828 | 0.219570055 | 0.85 |
| YER100W | 1553.364761 | -0.284146661 | 0.220430073 | 0.85 |
| YER138W-A | 1.112909077 | 0.628277335 | 0.219708151 | 0.85 |
| YGR224W | 693.0347479 | -0.384146368 | 0.21955744 | 0.85 |
| YHR041C | 584.7828437 | -0.431768487 | 0.219243781 | 0.85 |
| YIR029W | 388.4137592 | 0.605711435 | 0.220657629 | 0.85 |
| YJL005W | 4176.860231 | 0.405909718 | 0.218808262 | 0.85 |
| YJR092W | 2324.987782 | -0.400707836 | 0.219785441 | 0.85 |

| | | | | |
|---|---|---|---|---|
| YLR010C | 127.8600135 | 0.365375098 | 0.220372925 | 0.85 |
| YLR078C | 672.0208317 | 0.264297148 | 0.21935895 | 0.85 |
| YLR188W | 1233.172467 | 0.266357178 | 0.220856881 | 0.85 |
| YLR401C | 1341.53932 | 0.309737046 | 0.220266129 | 0.85 |
| YML005W | 433.1547412 | 0.192591033 | 0.220502669 | 0.85 |
| YML008C | 4220.321447 | -0.429669484 | 0.220376102 | 0.85 |
| YNL142W | 1026.084342 | 0.638571304 | 0.219995767 | 0.85 |
| YNR068C | 378.9501574 | 0.509798886 | 0.220068321 | 0.85 |
| YPL190C | 2551.586572 | 0.319895102 | 0.21801431 | 0.85 |
| snR82 | 454.6100305 | -0.614094983 | 0.219095056 | 0.85 |
| YBL001C | 1539.933777 | -0.405103092 | 0.224245118 | 0.85 |
| YBR023C | 1741.008098 | -0.332013431 | 0.221317193 | 0.85 |
| YDR053W | 7.370760006 | -0.612954469 | 0.223284118 | 0.85 |
| YDR111C | 552.8400061 | 0.546034688 | 0.223230763 | 0.85 |
| YDR113C | 648.1289584 | -0.273457211 | 0.223466217 | 0.85 |
| YDR426C | 18.4381422 | -0.706517086 | 0.222716049 | 0.85 |
| YDR437W | 89.57715872 | 0.416363006 | 0.224134836 | 0.85 |
| YDR501W | 295.5069374 | 0.276872829 | 0.222360079 | 0.85 |
| YDR524C-B | 18745.499 | -0.460325343 | 0.223741986 | 0.85 |
| YDR527W | 984.8264926 | 0.363087469 | 0.222837002 | 0.85 |
| YGR186W | 2388.772284 | 0.323006816 | 0.221954035 | 0.85 |
| YGR222W | 726.7683442 | -0.299140686 | 0.222897129 | 0.85 |
| YIL156W-B | 291.2591429 | -0.296773374 | 0.222932816 | 0.85 |
| YJL070C | 972.7876692 | 0.368061734 | 0.222908532 | 0.85 |
| YJL168C | 610.0870071 | -0.435541522 | 0.222463726 | 0.85 |
| YKL159C | 432.357669 | -0.382748953 | 0.224296265 | 0.85 |
| YLR129W | 1609.244371 | -0.469541927 | 0.223784792 | 0.85 |
| YMR210W | 1188.233193 | -0.287957895 | 0.221829658 | 0.85 |
| YNL055C | 20028.30046 | 0.611212298 | 0.222804158 | 0.85 |
| YOL007C | 839.2247362 | -0.463199024 | 0.222744214 | 0.85 |
| YOL145C | 2673.103928 | -0.333618536 | 0.224032677 | 0.85 |
| YOR023C | 2117.560133 | 0.356573346 | 0.221848557 | 0.85 |
| YOR175C | 2021.823765 | 0.265528585 | 0.222556978 | 0.85 |
| YOR247W | 5496.630424 | -0.535355936 | 0.223956922 | 0.85 |
| YPL173W | 902.2046084 | -0.295276785 | 0.223739428 | 0.85 |
| YBL086C | 1904.650929 | 0.347230717 | 0.225210917 | 0.85 |
| YDL130W-A | 2765.774643 | 0.579501154 | 0.22600171 | 0.85 |
| YDR055W | 3654.499745 | 0.36356253 | 0.225439324 | 0.85 |
| YDR261C-D | 88.68167414 | -0.701670524 | 0.226573061 | 0.85 |
| YEL019C | 187.0813824 | 0.432449459 | 0.226576247 | 0.85 |
| YER048C | 2682.830499 | -0.294413923 | 0.226506664 | 0.85 |
| YER067W | 2559.902963 | 0.688865337 | 0.226186368 | 0.85 |
| YGL142C | 1131.213732 | 0.340506035 | 0.224703529 | 0.85 |

| | | | | |
|---|---|---|---|---|
| YHR182C-A | 130.5541577 | 0.769667231 | 0.226466398 | 0.85 |
| YIR019C | 622.3080892 | 0.453004751 | 0.226057867 | 0.85 |
| YJL156C | 869.0736001 | -0.256885545 | 0.225955805 | 0.85 |
| YKL218C | 773.1891416 | 0.516595798 | 0.225451673 | 0.85 |
| YKR031C | 1778.007412 | -0.369949715 | 0.22482207 | 0.85 |
| YML118W | 395.2746742 | 0.596399767 | 0.225174276 | 0.85 |
| YNL131W | 2877.827714 | -0.295472033 | 0.226521115 | 0.85 |
| YOR245C | 966.5341375 | 0.37931964 | 0.226704777 | 0.85 |
| YPR008W | 1796.059535 | -0.284850426 | 0.224554821 | 0.85 |
| YPR100W | 188.3881606 | -0.388256123 | 0.225384138 | 0.85 |
| YMR019W | 977.1459301 | 0.380871485 | 0.226875584 | 0.85 |
| YBR084W | 2864.241089 | 0.520128264 | 0.232051214 | 0.85 |
| YBR235W | 1364.419973 | 0.397489054 | 0.231935983 | 0.85 |
| YBR248C | 1552.941582 | 0.552444841 | 0.230505591 | 0.85 |
| YDL094C | 21.86703582 | 0.539239442 | 0.230132302 | 0.85 |
| YDR093W | 1759.127414 | -0.267708074 | 0.229741796 | 0.85 |
| YEL053C | 1328.366975 | 0.178282678 | 0.228772755 | 0.85 |
| YEL066W | 476.9220074 | 0.459256898 | 0.231607979 | 0.85 |
| YFL044C | 776.7355212 | 0.400500894 | 0.231746707 | 0.85 |
| YGL089C | 132.0787924 | 0.258401727 | 0.230522332 | 0.85 |
| YGL227W | 1564.861142 | 0.421236076 | 0.23202884 | 0.85 |
| YGL258W-A | 56.62039045 | 0.584720786 | 0.229636465 | 0.85 |
| YGR220C | 677.1113476 | -0.286920379 | 0.230819732 | 0.85 |
| YHL019W-A | 10.01596588 | 0.425029784 | 0.229684402 | 0.85 |
| YHR111W | 861.1772037 | 0.28253078 | 0.227947311 | 0.85 |
| YHR121W | 1103.128777 | -0.250175348 | 0.232042023 | 0.85 |
| YIL054W | 8.069700908 | 0.579280293 | 0.22939856 | 0.85 |
| YIL065C | 779.4474724 | 0.405001622 | 0.228235384 | 0.85 |
| YIL140W | 1095.766571 | -0.453710397 | 0.227384776 | 0.85 |
| YIR030W-A | 17.57088172 | 0.564541639 | 0.229833917 | 0.85 |
| YJL159W | 37900.6137 | 0.423970028 | 0.229857452 | 0.85 |
| YKR003W | 1162.972466 | -0.329662818 | 0.232184523 | 0.85 |
| YLL005C | 615.6111971 | 0.458751942 | 0.227490857 | 0.85 |
| YLR123C | 17.47407493 | 0.631101924 | 0.231186016 | 0.85 |
| YLR220W | 930.48531 | 0.322236749 | 0.228297998 | 0.85 |
| YLR304C | 13028.11852 | 0.553503692 | 0.228453618 | 0.85 |
| YLR334C | 3.854398197 | 0.587111856 | 0.230294311 | 0.85 |
| YLR466C-B | 12.5768575 | 0.467360713 | 0.230853185 | 0.85 |
| YML009C | 743.0693468 | -0.361607787 | 0.23137797 | 0.85 |
| YML127W | 993.038483 | -0.284669543 | 0.228153028 | 0.85 |
| YMR228W | 316.010038 | -0.265141521 | 0.227832351 | 0.85 |
| YNL092W | 338.5129162 | 0.432248109 | 0.230077254 | 0.85 |
| YNL220W | 4979.508941 | 0.519231468 | 0.232095535 | 0.85 |

| | | | | |
|---|---|---|---|---|
| YNL272C | 1304.645862 | -0.233013095 | 0.230876073 | 0.85 |
| YNL280C | 1498.81009 | 0.450775218 | 0.232016453 | 0.85 |
| YNR017W | 1847.028777 | -0.245557125 | 0.230736877 | 0.85 |
| YOR049C | 767.2844506 | -0.491883341 | 0.227582856 | 0.85 |
| YOR072W | 29.04315607 | 0.443949022 | 0.227069058 | 0.85 |
| YOR085W | 1806.530962 | -0.382273775 | 0.228294083 | 0.85 |
| YPL102C | 13.71806551 | 0.465429434 | 0.231851162 | 0.85 |
| YPR107C | 422.2754774 | -0.227476219 | 0.229929242 | 0.85 |
| YIL075C | 7548.161259 | 0.334904354 | 0.232329547 | 0.85 |
| YHR125W | 4.057342841 | 0.516687451 | 0.232703444 | 0.85 |
| YLR265C | 441.14354 | -0.592213399 | 0.232650953 | 0.85 |
| YBR072W | 9138.655064 | 0.765175679 | 0.232982992 | 0.85 |
| YMR138W | 253.6379214 | 0.29203136 | 0.23286136 | 0.85 |
| YAR002W | 1776.649309 | -0.334282639 | 0.23624448 | 0.86 |
| YDL128W | 1754.772543 | -0.243449359 | 0.235614742 | 0.86 |
| YDR298C | 3635.360191 | 0.606691917 | 0.23592588 | 0.86 |
| YER050C | 1168.128624 | -0.382794413 | 0.235136456 | 0.86 |
| YER189W | 0.915658928 | -0.495582857 | 0.238096945 | 0.86 |
| YFL039C | 29152.57267 | 0.225382866 | 0.233998655 | 0.86 |
| YFL040W | 216.9489399 | 0.456751734 | 0.236542112 | 0.86 |
| YFR016C | 3711.04018 | 0.384092111 | 0.237792774 | 0.86 |
| YFR020W | 385.8178556 | 0.281288936 | 0.237925107 | 0.86 |
| YFR049W | 788.1728678 | 0.521091767 | 0.234850818 | 0.86 |
| YGL026C | 4379.972897 | 0.446502173 | 0.237411601 | 0.86 |
| YGL056C | 2031.538502 | 0.370221746 | 0.233673831 | 0.86 |
| YGL152C | 7.084888852 | -0.572796892 | 0.23702813 | 0.86 |
| YGR197C | 588.1649926 | 0.465011138 | 0.238313202 | 0.86 |
| YGR221C | 838.7959631 | -0.343990642 | 0.235098566 | 0.86 |
| YGR293C | 7.546648827 | -0.697357871 | 0.234486633 | 0.86 |
| YHL036W | 545.2189465 | 0.481603016 | 0.238355059 | 0.86 |
| YHR070C-A | 129.6976848 | 0.456485608 | 0.235847554 | 0.86 |
| YJL085W | 484.172969 | -0.382240572 | 0.236831676 | 0.86 |
| YJL132W | 534.7548177 | 0.416485898 | 0.234514414 | 0.86 |
| YJR101W | 534.1436387 | -0.32211769 | 0.234069019 | 0.86 |
| YJR110W | 661.1509666 | 0.200808346 | 0.236868972 | 0.86 |
| YKL141W | 4100.397187 | 0.59758334 | 0.237664584 | 0.86 |
| YKL184W | 1354.204344 | -0.282974472 | 0.237054882 | 0.86 |
| YLR038C | 1207.825369 | 0.627494279 | 0.236628023 | 0.86 |
| YLR380W | 1647.260273 | -0.380119106 | 0.234131654 | 0.86 |
| YLR450W | 2180.849651 | 0.368461594 | 0.236199999 | 0.86 |
| YML078W | 2283.677343 | -0.352756256 | 0.237510662 | 0.86 |
| YML104C | 923.4299097 | 0.25816199 | 0.237653933 | 0.86 |
| YML131W | 1116.004425 | 0.621390469 | 0.23735089 | 0.86 |

| | | | | |
|---|---|---|---|---|
| YMR018W | 66.38637523 | 0.453567637 | 0.238156091 | 0.86 |
| YMR140W | 1470.388235 | 0.365436044 | 0.23417003 | 0.86 |
| YMR225C | 419.9415237 | -0.344506424 | 0.234132543 | 0.86 |
| YNL029C | 467.6043522 | 0.315250375 | 0.237876631 | 0.86 |
| YNL077W | 1205.161408 | 0.440825909 | 0.237816432 | 0.86 |
| YOL026C | 840.8767019 | -0.272167528 | 0.238092105 | 0.86 |
| YOR019W | 709.5102111 | 0.535375319 | 0.236028157 | 0.86 |
| YOR321W | 1083.064286 | 0.307103412 | 0.2351489 | 0.86 |
| YPR120C | 354.4464034 | -0.3344747 | 0.238171007 | 0.86 |
| YPL078C | 4809.066893 | 0.530003334 | 0.238753279 | 0.86 |
| YDL240C-A | 1.077122984 | -0.699528198 | 0.239756292 | 0.86 |
| YDR461W | 1830.189511 | -0.622049711 | 0.239702246 | 0.86 |
| YER074W-A | 546.9637993 | -0.237988398 | 0.239680611 | 0.86 |
| YFL002C | 1100.586944 | 0.181654801 | 0.239466708 | 0.86 |
| YHR210C | 320.0019552 | -0.531339258 | 0.239620276 | 0.86 |
| YJR084W | 669.2500078 | -0.198305173 | 0.239335778 | 0.86 |
| YER040W | 900.0166243 | 0.368721685 | 0.240170752 | 0.86 |
| YHR105W | 228.2746463 | 0.390885982 | 0.24067062 | 0.86 |
| YLL011W | 676.0042616 | -0.402748134 | 0.240638926 | 0.86 |
| YLR290C | 1190.940137 | 0.333582135 | 0.240665319 | 0.86 |
| YOR202W | 1479.782603 | 0.418472551 | 0.240503589 | 0.86 |
| YGR254W | 35005.12746 | 0.666989768 | 0.240985523 | 0.86 |
| YBL043W | 324.5838638 | 0.718511147 | 0.241310312 | 0.86 |
| YDL140C | 8072.034708 | -0.327361353 | 0.241438627 | 0.86 |
| YPL075W | 2226.650529 | -0.353357861 | 0.241682006 | 0.86 |
| YBR017C | 2405.834993 | -0.26167442 | 0.242475237 | 0.86 |
| YBR136W | 1309.50165 | 0.326662809 | 0.24215786 | 0.86 |
| YDR040C | 132.5886651 | 0.481182706 | 0.242733704 | 0.86 |
| YER161C | 1183.000093 | -0.286972359 | 0.242139122 | 0.86 |
| YIR021W-A | 4.66148073 | -0.601492681 | 0.242314827 | 0.86 |
| YNL337W | 0.659062417 | 0.338276789 | 0.242631683 | 0.86 |
| YOL093W | 662.3279101 | -0.365928874 | 0.242654028 | 0.86 |
| YER022W | 948.246119 | 0.172434725 | 0.243576656 | 0.86 |
| YMR281W | 530.4325943 | -0.309794296 | 0.243451996 | 0.86 |
| YMR300C | 1920.32889 | -0.506504757 | 0.243065499 | 0.86 |
| YPL246C | 2119.845638 | 0.244633632 | 0.243270715 | 0.86 |
| YPR081C | 985.9306388 | 0.31122652 | 0.243435911 | 0.86 |
| YGR032W | 3378.619384 | 0.497457707 | 0.244244932 | 0.86 |
| YIR022W | 1186.448374 | -0.271857683 | 0.244570793 | 0.86 |
| YKL013C | 2002.473422 | -0.245510422 | 0.244079441 | 0.86 |
| YLL066W-B | 2.220126499 | 0.421180411 | 0.244445277 | 0.86 |
| YNL214W | 315.125881 | 0.1961389 | 0.244480612 | 0.86 |
| YNL323W | 681.0384119 | -0.237205137 | 0.2445489 | 0.86 |

| | | | | |
|---|---|---|---|---|
| YDR072C | 1794.903187 | -0.403482097 | 0.245041216 | 0.86 |
| YER130C | 1003.132192 | -0.493325994 | 0.24483667 | 0.86 |
| YNL184C | 77.22136504 | -0.437952667 | 0.244972146 | 0.86 |
| YBR246W | 994.4891912 | -0.211735388 | 0.245950765 | 0.86 |
| YDR135C | 3272.34562 | 0.231778465 | 0.246130953 | 0.86 |
| YIR018C-A | 31.48971955 | 0.406120174 | 0.245813859 | 0.86 |
| YLR235C | 18.73304548 | 0.503877936 | 0.246035846 | 0.86 |
| YLR383W | 888.8933966 | 0.280064157 | 0.245872362 | 0.86 |
| YOR073W-A | 11.62724158 | -0.576210562 | 0.246181679 | 0.86 |
| YPR184W | 5293.583109 | 0.634380844 | 0.24556208 | 0.86 |
| YIR037W | 2878.72569 | 0.347549197 | 0.246410629 | 0.86 |
| YNL262W | 2257.387419 | 0.387771008 | 0.246718972 | 0.86 |
| YPR187W | 1425.65006 | -0.414517838 | 0.246772747 | 0.86 |
| YDR366C | 7.819864358 | 0.563856629 | 0.247684614 | 0.86 |
| YDR442W | 17.52253569 | -0.589086001 | 0.247686854 | 0.86 |
| YDR473C | 858.9200777 | 0.240409939 | 0.247929844 | 0.86 |
| YDR510W | 2598.34956 | -0.264601265 | 0.24771674 | 0.86 |
| YGR072W | 838.1301736 | 0.296127809 | 0.248108554 | 0.86 |
| YKL162C | 452.2195831 | 0.349335974 | 0.247299279 | 0.86 |
| YML083C | 177.180785 | 0.530027919 | 0.247522297 | 0.86 |
| YOL066C | 908.4756316 | -0.33702827 | 0.248020655 | 0.86 |
| YCL068C | 4.465179414 | 0.14379906 | 0.24838227 | 0.86 |
| YJL172W | 1932.176519 | 0.433654742 | 0.248740786 | 0.87 |
| YLL030C | 4.099886808 | 0.623292535 | 0.249896815 | 0.87 |
| YBR056W | 2794.886987 | 0.420816353 | 0.250582863 | 0.87 |
| YER137W-A | 3.086112355 | 0.354145125 | 0.250425268 | 0.87 |
| YJR150C | 147.0716932 | 0.607270517 | 0.250438803 | 0.87 |
| YLR146W-A | 192.5995673 | -0.39281696 | 0.250217976 | 0.87 |
| YMR181C | 1531.918567 | 0.360686938 | 0.250660154 | 0.87 |
| YOL150C | 3.831416512 | 0.679175263 | 0.250909272 | 0.87 |
| YDR453C | 889.6622872 | 0.630820657 | 0.251069319 | 0.87 |
| YGR076C | 617.1306078 | -0.281224636 | 0.251387286 | 0.87 |
| YGL104C | 947.9922368 | 0.392224646 | 0.251686069 | 0.87 |
| YDR381C-A | 448.1284196 | 0.441693345 | 0.252709907 | 0.87 |
| YDR461C-A | 326.4764294 | 0.445101377 | 0.252536735 | 0.87 |
| YDR492W | 689.8420252 | 0.433258732 | 0.25250627 | 0.87 |
| YIL090W | 898.4488003 | 0.229645051 | 0.252683476 | 0.87 |
| YOL054W | 494.8918395 | 0.253646901 | 0.252373164 | 0.87 |
| YOL153C | 1352.172864 | 0.495084 | 0.252622271 | 0.87 |
| YPL089C | 1858.715154 | 0.332755174 | 0.252731664 | 0.87 |
| YCR045C | 398.9252367 | -0.322500458 | 0.253031189 | 0.87 |
| YPR148C | 2535.151148 | 0.227664839 | 0.253012904 | 0.87 |
| YGL024W | 7.765530916 | 0.550999274 | 0.253204542 | 0.87 |

| | | | | |
|---|---|---|---|---|
| YCR032W | 1688.564969 | 0.332421989 | 0.253341014 | 0.87 |
| YBR239C | 441.6078035 | -0.268233935 | 0.253627696 | 0.87 |
| YGL140C | 1656.222341 | 0.189253152 | 0.253871645 | 0.87 |
| YHR211W | 534.044984 | 0.495128588 | 0.25403822 | 0.87 |
| YKL024C | 1589.092419 | 0.356684593 | 0.254214766 | 0.87 |
| YOR224C | 1759.04057 | -0.39815738 | 0.254291278 | 0.87 |
| YBL013W | 160.0267849 | 0.372865329 | 0.255335494 | 0.87 |
| YCL030C | 8176.65539 | 0.427975783 | 0.255361012 | 0.87 |
| YDR215C | 67.50004526 | 0.606414103 | 0.255049401 | 0.87 |
| YFR046C | 527.2575594 | 0.267986617 | 0.255212038 | 0.87 |
| YKL160W | 1725.224125 | -0.256806041 | 0.255278403 | 0.87 |
| YLR322W | 7.310944976 | 0.393210744 | 0.255706807 | 0.87 |
| YDL134C | 1962.281647 | 0.118554514 | 0.256271001 | 0.87 |
| YDL164C | 704.8057939 | -0.237358313 | 0.256292911 | 0.87 |
| YIL002C | 1259.267656 | -0.243055572 | 0.256440781 | 0.87 |
| YLR018C | 838.7588705 | -0.24934458 | 0.256096142 | 0.87 |
| YLR228C | 906.7831503 | 0.365026991 | 0.256927023 | 0.87 |
| YPL051W | 362.5719952 | -0.288118518 | 0.256826746 | 0.87 |
| YPL129W | 1819.666295 | -0.256630585 | 0.256721734 | 0.87 |
| YPL228W | 971.5333391 | -0.291661465 | 0.256737168 | 0.87 |
| YPR040W | 901.3829077 | 0.326097818 | 0.256687878 | 0.87 |
| YBL096C | 11.76381799 | -0.711857115 | 0.257098616 | 0.87 |
| YEL067C | 149.0124198 | 0.383525822 | 0.257273133 | 0.87 |
| YMR207C | 1385.212155 | 0.27410471 | 0.257575885 | 0.88 |
| YDL056W | 1411.798857 | -0.342341826 | 0.258142912 | 0.88 |
| YDR046C | 2206.66233 | -0.354926768 | 0.26033273 | 0.88 |
| YDR123C | 353.8570592 | 0.359760861 | 0.260007948 | 0.88 |
| YDR148C | 4531.818706 | 0.559760818 | 0.259867314 | 0.88 |
| YER011W | 1426.306813 | -0.633904931 | 0.26108623 | 0.88 |
| YER165C-A | 28.00543952 | 0.409970608 | 0.259375379 | 0.88 |
| YGL166W | 257.6413584 | 0.309104491 | 0.259253203 | 0.88 |
| YGR054W | 2529.613046 | -0.379132323 | 0.260735572 | 0.88 |
| YGR169C-A | 750.5850996 | -0.310333352 | 0.26042803 | 0.88 |
| YHR033W | 1072.035586 | 0.466127036 | 0.260879324 | 0.88 |
| YHR113W | 1875.547688 | 0.299211436 | 0.261095514 | 0.88 |
| YHR178W | 530.4654676 | 0.380075991 | 0.258295037 | 0.88 |
| YIL012W | 10.18010829 | 0.505094003 | 0.260878108 | 0.88 |
| YIL088C | 2503.905976 | 0.330498452 | 0.257952266 | 0.88 |
| YJR083C | 550.0152446 | -0.289381801 | 0.261031424 | 0.88 |
| YKR035C | 23.50975344 | -0.405239082 | 0.259753755 | 0.88 |
| YLR184W | 0.707088895 | 0.379554542 | 0.259964016 | 0.88 |
| YMR134W | 453.7137928 | 0.331470657 | 0.258893895 | 0.88 |
| YNL065W | 531.9531231 | 0.481804268 | 0.259847944 | 0.88 |

| | | | | |
|---|---|---|---|---|
| YNL115C | 1894.716258 | 0.492564438 | 0.258787238 | 0.88 |
| YOL024W | 52.91060507 | 0.517705949 | 0.259031242 | 0.88 |
| YOL085C | 30.97884527 | 0.528391303 | 0.260373433 | 0.88 |
| YOR094W | 410.9080628 | -0.294120483 | 0.257813494 | 0.88 |
| YOR344C | 1985.74596 | -0.468110999 | 0.260130154 | 0.88 |
| YPL130W | 24.21083011 | 0.554076614 | 0.259784046 | 0.88 |
| YPR037C | 567.1835844 | 0.306464516 | 0.259971206 | 0.88 |
| YBR009C | 3036.286227 | -0.461457567 | 0.262033226 | 0.88 |
| YDR431W | 11.62652329 | -0.54429285 | 0.262125618 | 0.88 |
| YER013W | 829.6032387 | -0.266107912 | 0.262347144 | 0.88 |
| YFR045W | 698.908615 | 0.36276058 | 0.262453846 | 0.88 |
| YGR177C | 234.3513182 | -0.548564219 | 0.261977965 | 0.88 |
| YHR049C-A | 129.3206508 | -0.453856436 | 0.262316036 | 0.88 |
| YJL087C | 474.7532502 | -0.355543592 | 0.262337639 | 0.88 |
| YJR121W | 23130.92602 | 0.599766385 | 0.261942343 | 0.88 |
| YOR158W | 1525.323441 | -0.276293677 | 0.261737931 | 0.88 |
| YBL033C | 1274.437806 | 0.227029654 | 0.264150288 | 0.88 |
| YBR162C | 7471.337212 | -0.386441131 | 0.264586628 | 0.88 |
| YCR048W | 995.8304266 | 0.274142141 | 0.262911581 | 0.88 |
| YDL185W | 10910.49268 | 0.228095556 | 0.263937607 | 0.88 |
| YDR535C | 20.65811424 | 0.498767163 | 0.264282444 | 0.88 |
| YDR542W | 1.676832288 | 0.503167006 | 0.2627218 | 0.88 |
| YGL005C | 924.1279481 | 0.281239849 | 0.263184364 | 0.88 |
| YGR121W-A | 8.749518959 | 0.589173004 | 0.264500824 | 0.88 |
| YKL047W | 711.1080868 | -0.240166032 | 0.264612743 | 0.88 |
| YKR089C | 3634.661722 | -0.317442369 | 0.264570222 | 0.88 |
| YLR101C | 65.86852303 | 0.342348656 | 0.262908178 | 0.88 |
| YNL043C | 66.28583242 | -0.478024844 | 0.264189389 | 0.88 |
| YNL118C | 3319.845542 | -0.209755161 | 0.264575754 | 0.88 |
| YOL100W | 1483.492555 | 0.317475478 | 0.263580661 | 0.88 |
| YOR383C | 13192.95676 | 0.640173684 | 0.264515549 | 0.88 |
| YPL107W | 480.1557675 | 0.333084597 | 0.263191777 | 0.88 |
| YLR229C | 1318.158779 | -0.248829726 | 0.264798952 | 0.88 |
| YKL033W | 666.8103831 | 0.211812691 | 0.265695983 | 0.88 |
| YMR234W | 552.7610201 | -0.214953584 | 0.265696602 | 0.88 |
| YBL057C | 768.8630819 | -0.260780514 | 0.265907153 | 0.88 |
| YHR110W | 1458.33201 | 0.235858799 | 0.266317322 | 0.88 |
| YLR098C | 706.1029207 | 0.297273262 | 0.266343982 | 0.88 |
| YLR232W | 2.842598345 | -0.664835501 | 0.266437097 | 0.88 |
| YDR267C | 660.9899359 | 0.222170723 | 0.266949307 | 0.88 |
| YGL167C | 2492.369981 | -0.324489423 | 0.267173793 | 0.88 |
| YGR025W | 100.9215658 | -0.389714314 | 0.266835671 | 0.88 |
| YGR122C-A | 15.60192201 | -0.554028766 | 0.267287559 | 0.88 |

| | | | | |
|---|---|---|---|---|
| YKR046C | 2535.087216 | 0.363535922 | 0.267248374 | 0.88 |
| YMR158W-B | 1.160282167 | -0.606648712 | 0.266898408 | 0.88 |
| YHR150W | 670.7977171 | 0.386052139 | 0.267470208 | 0.88 |
| YDL042C | 548.3117902 | 0.514664441 | 0.267916891 | 0.88 |
| YLL043W | 1838.000712 | -0.265879287 | 0.267924588 | 0.88 |
| YNR007C | 864.4677932 | 0.475395074 | 0.268117304 | 0.88 |
| YOR193W | 437.421385 | -0.324112897 | 0.268183769 | 0.88 |
| YPL256C | 1702.304083 | -0.42742419 | 0.267780267 | 0.88 |
| YBL024W | 2012.558866 | -0.377572105 | 0.269443622 | 0.88 |
| YBL045C | 7514.298009 | 0.551216257 | 0.269981937 | 0.88 |
| YBR245C | 1580.844401 | -0.214654932 | 0.269242437 | 0.88 |
| YCR042C | 1593.772478 | -0.190410658 | 0.269227417 | 0.88 |
| YDR409W | 969.523209 | -0.265898382 | 0.269821629 | 0.88 |
| YGL169W | 682.8701947 | 0.359708085 | 0.269769395 | 0.88 |
| YHR044C | 186.9132602 | 0.525741666 | 0.26859921 | 0.88 |
| YJL179W | 307.4969877 | -0.361220272 | 0.270178827 | 0.88 |
| YLR036C | 360.058471 | -0.291295413 | 0.27001096 | 0.88 |
| YML039W | 1.131094598 | -0.582209203 | 0.268952407 | 0.88 |
| YNL168C | 1117.770385 | 0.361886636 | 0.2684302 | 0.88 |
| YOL094C | 437.8167325 | -0.343617984 | 0.269371288 | 0.88 |
| YOR317W | 4625.356852 | 0.451579257 | 0.268980716 | 0.88 |
| YPL267W | 353.8623989 | -0.37106372 | 0.269204596 | 0.88 |
| snR191 | 16.29529748 | -0.471006189 | 0.270248154 | 0.88 |
| YJL203W | 345.939173 | -0.212508312 | 0.270600639 | 0.88 |
| YFL017C | 717.9144755 | -0.231298323 | 0.271096969 | 0.88 |
| YGL032C | 1976.542723 | -0.575968911 | 0.271331322 | 0.88 |
| YKR091W | 1094.538594 | -0.52254304 | 0.271335586 | 0.88 |
| YLR056W | 3964.313149 | -0.422382424 | 0.271285801 | 0.88 |
| YLR241W | 1618.382874 | 0.234525356 | 0.270921788 | 0.88 |
| YIL126W | 2338.60398 | -0.356709966 | 0.271472927 | 0.88 |
| YFL025C | 800.9686963 | -0.152437128 | 0.27177038 | 0.88 |
| YPR070W | 926.8338353 | -0.147435971 | 0.271794663 | 0.88 |
| YNL050C | 614.5421658 | -0.219510087 | 0.272014927 | 0.88 |
| YAR030C | 64.62074774 | 0.355957909 | 0.272904814 | 0.88 |
| YBL113C | 286.5211358 | 0.381499352 | 0.273330174 | 0.88 |
| YCL049C | 935.682406 | 0.503720722 | 0.272891352 | 0.88 |
| YDL207W | 1207.359156 | 0.169694633 | 0.272808911 | 0.88 |
| YGR079W | 359.5947801 | 0.434973207 | 0.274079885 | 0.88 |
| YIL076W | 2351.989968 | -0.306221796 | 0.273996239 | 0.88 |
| YIL158W | 651.6543059 | -0.42160422 | 0.27394649 | 0.88 |
| YJL056C | 734.4930024 | 0.364448908 | 0.272513662 | 0.88 |
| YKL106W | 462.66436 | -0.369844407 | 0.2730924 | 0.88 |
| YKR078W | 765.3811944 | 0.280175829 | 0.274038267 | 0.88 |

| | | | | |
|---|---|---|---|---|
| YNL249C | 367.0811931 | 0.218200992 | 0.273541491 | 0.88 |
| YNL298W | 1195.295385 | -0.372029242 | 0.273280673 | 0.88 |
| YOR075W | 634.1618193 | 0.399471226 | 0.273703986 | 0.88 |
| YPL032C | 2772.776631 | -0.328108277 | 0.273049372 | 0.88 |
| YPL065W | 569.9929052 | -0.292611313 | 0.272622255 | 0.88 |
| YHR120W | 606.7255558 | -0.198865222 | 0.274481495 | 0.88 |
| YLR375W | 1937.904522 | 0.215029295 | 0.27452349 | 0.88 |
| YER166W | 2859.051695 | 0.263116584 | 0.275116351 | 0.88 |
| YHR067W | 339.2689826 | 0.414172526 | 0.275237221 | 0.88 |
| YLR396C | 522.7183585 | -0.193528884 | 0.275253105 | 0.88 |
| YOR138C | 1254.624127 | 0.204381822 | 0.274871358 | 0.88 |
| YPL074W | 944.5738824 | 0.285396463 | 0.275238749 | 0.88 |
| YDR136C | 21.36116146 | -0.397403626 | 0.275472285 | 0.88 |
| YBL002W | 4494.23128 | -0.459692153 | 0.276113618 | 0.88 |
| YFL049W | 806.3019635 | -0.207467214 | 0.276246013 | 0.88 |
| YIL047C-A | 10.41380776 | -0.535489732 | 0.275947246 | 0.88 |
| YKL055C | 140.4891808 | 0.315207748 | 0.275985375 | 0.88 |
| YKR105C | 96.82105819 | 0.449350591 | 0.275837767 | 0.88 |
| YBR090C | 79.12993298 | 0.431103924 | 0.276829742 | 0.88 |
| YBR178W | 20.74423611 | 0.511455841 | 0.277799416 | 0.88 |
| YCR089W | 1832.043276 | 0.384968206 | 0.277620957 | 0.88 |
| YDR099W | 8428.443895 | 0.185952756 | 0.277415659 | 0.88 |
| YDR376W | 985.9130091 | 0.254724398 | 0.27773238 | 0.88 |
| YDR529C | 3824.487001 | 0.527614543 | 0.278162737 | 0.88 |
| YGR156W | 932.64115 | -0.309996883 | 0.27755721 | 0.88 |
| YHR016C | 1896.688185 | 0.32470878 | 0.277263444 | 0.88 |
| YJL062W | 1004.118516 | 0.222915736 | 0.276665992 | 0.88 |
| YJR122W | 745.1766826 | 0.306141213 | 0.278076379 | 0.88 |
| YKL042W | 561.3749184 | -0.340119843 | 0.278175308 | 0.88 |
| YMR006C | 894.5004784 | 0.311168927 | 0.278180522 | 0.88 |
| YNL049C | 1288.390098 | 0.254345582 | 0.277109882 | 0.88 |
| YPR153W | 81.58409703 | 0.334423395 | 0.27677291 | 0.88 |
| YDR107C | 540.536576 | 0.252175085 | 0.278325804 | 0.88 |
| YAL064W-B | 18.20372016 | -0.529944461 | 0.278925034 | 0.88 |
| YHR060W | 732.5771452 | -0.270850208 | 0.278798634 | 0.88 |
| YFL056C | 232.3744781 | 0.416555913 | 0.279738603 | 0.89 |
| YDR451C | 513.5586606 | -0.409065696 | 0.279938377 | 0.89 |
| YML038C | 1118.066731 | -0.271948814 | 0.280060818 | 0.89 |
| YDL131W | 1846.278027 | 0.366610205 | 0.280443765 | 0.89 |
| YGR255C | 1753.194799 | -0.275658806 | 0.280580042 | 0.89 |
| YMR044W | 903.5982709 | -0.268918408 | 0.280459991 | 0.89 |
| YKL185W | 2577.028628 | 0.34618805 | 0.280730307 | 0.89 |
| YNL103W | 1585.703492 | 0.308043633 | 0.280871068 | 0.89 |

| | | | | |
|---|---|---|---|---|
| YFL018C | 6022.957872 | 0.331622903 | 0.281225184 | 0.89 |
| YOR017W | 870.202076 | -0.220645608 | 0.281338645 | 0.89 |
| YBR296C-A | 3.00001637 | 0.588892894 | 0.28191951 | 0.89 |
| YDL170W | 672.20876 | 0.347156929 | 0.282070585 | 0.89 |
| YIR035C | 1381.941223 | -0.282419626 | 0.281842586 | 0.89 |
| YML128C | 9230.440998 | 0.648127674 | 0.282060757 | 0.89 |
| YNL038W | 473.5438124 | -0.329199471 | 0.281944402 | 0.89 |
| YNR030W | 1357.008714 | -0.396372049 | 0.282286172 | 0.89 |
| YDR444W | 770.8185864 | 0.265228721 | 0.282609596 | 0.89 |
| YML122C | 47.85383085 | 0.458095928 | 0.28258876 | 0.89 |
| YDR051C | 1050.532227 | 0.182971673 | 0.28330261 | 0.89 |
| YNL011C | 614.0144377 | 0.322001362 | 0.283677435 | 0.89 |
| YBR061C | 630.0720407 | -0.267238158 | 0.283886201 | 0.89 |
| YAL028W | 836.6833612 | 0.371304353 | 0.286187011 | 0.89 |
| YAR070C | 0.925768202 | 0.372049439 | 0.289082125 | 0.89 |
| YBR040W | 134.4656267 | 0.634547835 | 0.287107891 | 0.89 |
| YBR097W | 952.5745358 | -0.221986664 | 0.289010191 | 0.89 |
| YBR198C | 1683.847961 | -0.233313918 | 0.287855457 | 0.89 |
| YBR233W | 474.558741 | -0.342041705 | 0.287474347 | 0.89 |
| YCR077C | 3337.529518 | -0.213248138 | 0.28574128 | 0.89 |
| YDL020C | 3590.678769 | 0.387460486 | 0.287328848 | 0.89 |
| YDR014W | 486.9454278 | 0.223810791 | 0.288759636 | 0.89 |
| YDR216W | 2843.606015 | 0.51176099 | 0.288014352 | 0.89 |
| YDR369C | 596.1613966 | 0.304007665 | 0.289077064 | 0.89 |
| YDR380W | 4921.676121 | 0.626630818 | 0.285839956 | 0.89 |
| YER020W | 2824.873338 | 0.429806495 | 0.28846119 | 0.89 |
| YFL061W | 11.58614878 | 0.497601962 | 0.288270862 | 0.89 |
| YGL092W | 1819.13534 | -0.268129402 | 0.284201694 | 0.89 |
| YGR086C | 9907.562538 | -0.228735207 | 0.284988578 | 0.89 |
| YHR080C | 2045.316793 | 0.281255846 | 0.286020097 | 0.89 |
| YIR043C | 49.74477082 | 0.509219542 | 0.28458845 | 0.89 |
| YJL145W | 1081.572718 | -0.331897473 | 0.285134018 | 0.89 |
| YJL196C | 1508.45467 | -0.372976419 | 0.286744194 | 0.89 |
| YJR153W | 431.5896283 | 0.524022067 | 0.2854872 | 0.89 |
| YKL114C | 901.3872837 | -0.28014849 | 0.289079058 | 0.89 |
| YKL137W | 420.0820122 | -0.313605819 | 0.285083701 | 0.89 |
| YKL186C | 1093.769583 | -0.281394145 | 0.286468666 | 0.89 |
| YLR084C | 1027.133398 | -0.38829256 | 0.286517614 | 0.89 |
| YLR345W | 2235.165084 | 0.355322894 | 0.28588112 | 0.89 |
| YMR010W | 733.3907547 | -0.356823175 | 0.288824864 | 0.89 |
| YMR020W | 992.3354434 | 0.470289003 | 0.288789109 | 0.89 |
| YMR158C-A | 22.93483608 | 0.415951376 | 0.287324509 | 0.89 |
| YMR173W | 743.4251563 | -0.344604047 | 0.287098077 | 0.89 |

| | | | | |
|---|---|---|---|---|
| YNL161W | 1329.930942 | 0.185361465 | 0.28813387 | 0.89 |
| YNL218W | 370.2566133 | 0.284392212 | 0.286465533 | 0.89 |
| YOL125W | 710.1287675 | 0.308468298 | 0.287586856 | 0.89 |
| YOR118W | 1462.615579 | 0.176560469 | 0.289132852 | 0.89 |
| YOR150W | 702.6825687 | -0.298616992 | 0.287420942 | 0.89 |
| YOR230W | 8762.727513 | 0.342367633 | 0.286264088 | 0.89 |
| YOR235W | 3.373424746 | 0.392540588 | 0.288710816 | 0.89 |
| YOR300W | 17.40293291 | 0.384913934 | 0.287232098 | 0.89 |
| YPL203W | 1453.358861 | 0.3546554 | 0.289329458 | 0.89 |
| YDL066W | 3505.644872 | 0.551522208 | 0.289989353 | 0.89 |
| YDR500C | 6683.941473 | -0.383873854 | 0.29047644 | 0.89 |
| YGL256W | 1304.621904 | 0.277654238 | 0.289948364 | 0.89 |
| YNL228W | 12.1493579 | -0.581728628 | 0.290219245 | 0.89 |
| YOL087C | 2320.349339 | 0.275747121 | 0.290058636 | 0.89 |
| YOL159C-A | 58.15453861 | -0.460547366 | 0.290104593 | 0.89 |
| YPR098C | 1032.023923 | 0.416702799 | 0.290230744 | 0.89 |
| tD(GUC)N | 0.865673422 | -0.414709168 | 0.290498361 | 0.89 |
| YBR091C | 248.9757925 | -0.24894225 | 0.290755097 | 0.89 |
| YGR219W | 32.06784529 | -0.371656591 | 0.290918264 | 0.89 |
| YKL171W | 1421.131993 | 0.390738037 | 0.291506483 | 0.89 |
| YKL183C-A | 2.603283005 | 0.385271091 | 0.291968079 | 0.89 |
| YPR035W | 16997.33407 | 0.486639386 | 0.291905598 | 0.89 |
| YGL080W | 877.4500948 | -0.304195018 | 0.292538444 | 0.89 |
| YJL178C | 1325.547528 | -0.197642361 | 0.292681351 | 0.89 |
| YLR240W | 1128.041716 | 0.239781877 | 0.292784573 | 0.89 |
| YPR003C | 637.4553595 | 0.32062026 | 0.292713566 | 0.89 |
| YBR079C | 5290.065687 | -0.339168064 | 0.294290172 | 0.89 |
| YBR286W | 17567.33584 | 0.327421931 | 0.293790603 | 0.89 |
| YDR011W | 4201.778813 | -0.380263399 | 0.294587346 | 0.89 |
| YHL048C-A | 2.76683096 | 0.552674278 | 0.294784289 | 0.89 |
| YHR100C | 468.2098929 | 0.2407621 | 0.294246565 | 0.89 |
| YJL198W | 970.0448734 | 0.373428851 | 0.293776691 | 0.89 |
| YJR131W | 581.4205166 | 0.2936774 | 0.294647299 | 0.89 |
| YLR074C | 1095.319455 | 0.378816126 | 0.294369284 | 0.89 |
| YML067C | 1114.441541 | -0.17608605 | 0.293661538 | 0.89 |
| YNL084C | 2582.808264 | -0.161928791 | 0.294291924 | 0.89 |
| YNR016C | 6951.886561 | 0.379838472 | 0.294614859 | 0.89 |
| YOR033C | 730.7503196 | 0.185957303 | 0.293457877 | 0.89 |
| YOR181W | 1611.589546 | -0.345942869 | 0.294761045 | 0.89 |
| YPR191W | 6094.519828 | 0.563373942 | 0.293291279 | 0.89 |
| YBL034C | 918.1202298 | -0.292683137 | 0.294980708 | 0.89 |
| YBL079W | 2598.015595 | -0.193459519 | 0.295796607 | 0.90 |
| YFR039C | 736.7528646 | -0.284113415 | 0.296098165 | 0.90 |

| | | | | |
|---|---|---|---|---|
| YGR231C | 3688.450165 | 0.32593128 | 0.29651814 | 0.90 |
| YHR190W | 5071.299505 | 0.290798538 | 0.296826408 | 0.90 |
| YIL097W | 1123.344615 | 0.420740677 | 0.296442708 | 0.90 |
| YML110C | 4679.227772 | 0.387220396 | 0.296047142 | 0.90 |
| YOL004W | 4094.946984 | -0.21217155 | 0.296619489 | 0.90 |
| YOL122C | 1128.161192 | 0.390718796 | 0.296489076 | 0.90 |
| YOL143C | 5652.711754 | -0.408992469 | 0.296012085 | 0.90 |
| YOR163W | 1216.735785 | -0.22362444 | 0.296277446 | 0.90 |
| YOR281C | 1009.23062 | -0.299297053 | 0.297006044 | 0.90 |
| YPR077C | 1.201353204 | -0.526905633 | 0.296937807 | 0.90 |
| YOR035C | 1268.519702 | 0.353713059 | 0.297263999 | 0.90 |
| YKL221W | 211.2597824 | 0.509835823 | 0.297510198 | 0.90 |
| YNL154C | 2670.283788 | -0.301903528 | 0.29757311 | 0.90 |
| YNL290W | 854.0353401 | -0.26105536 | 0.298206393 | 0.90 |
| YOR352W | 729.6147343 | -0.229159875 | 0.298089932 | 0.90 |
| YER016W | 838.7636501 | -0.379539793 | 0.298484631 | 0.90 |
| YAR035C-A | 1.313184395 | 0.230377212 | 0.298642943 | 0.90 |
| YOR293C-A | 2.608736587 | 0.355276835 | 0.299272821 | 0.90 |
| YBL095W | 182.0968065 | 0.367362012 | 0.302173657 | 0.90 |
| YBR069C | 2910.073987 | 0.383413953 | 0.301358676 | 0.90 |
| YCR093W | 4708.644101 | -0.299856496 | 0.301591695 | 0.90 |
| YDL177C | 218.4776378 | -0.32057261 | 0.299988541 | 0.90 |
| YDR141C | 2153.367361 | -0.173514732 | 0.29962733 | 0.90 |
| YEL015W | 1435.206957 | -0.183847799 | 0.300076581 | 0.90 |
| YHL004W | 1819.11881 | -0.199609851 | 0.301606285 | 0.90 |
| YIL017C | 1438.735299 | 0.344463058 | 0.301221758 | 0.90 |
| YJL066C | 1041.928086 | 0.479744073 | 0.301611135 | 0.90 |
| YJL067W | 49.69168928 | 0.454076929 | 0.302267692 | 0.90 |
| YJL101C | 2271.091311 | 0.239363212 | 0.302045996 | 0.90 |
| YJR108W | 40.18860279 | -0.417801278 | 0.302124382 | 0.90 |
| YKL165C-A | 6.96473103 | 0.459362333 | 0.30107761 | 0.90 |
| YLL047W | 4.30351014 | 0.589485852 | 0.300319398 | 0.90 |
| YLR152C | 1272.570191 | 0.425457644 | 0.300436656 | 0.90 |
| YLR306W | 266.7883268 | -0.32908199 | 0.301116492 | 0.90 |
| YOL051W | 2569.23447 | 0.268157387 | 0.302169506 | 0.90 |
| YOL103W | 1490.877623 | -0.181795598 | 0.300169655 | 0.90 |
| YOR311C | 1031.860298 | 0.285906634 | 0.301945495 | 0.90 |
| YPR062W | 2367.986615 | 0.214812554 | 0.301721216 | 0.90 |
| YPR066W | 302.1036782 | 0.340313908 | 0.299660477 | 0.90 |
| YKL030W | 149.8142583 | 0.33763242 | 0.30260946 | 0.90 |
| YBL085W | 2143.648048 | -0.294448699 | 0.306588876 | 0.90 |
| YBR010W | 4241.31261 | -0.377654037 | 0.30399229 | 0.90 |
| YBR054W | 4277.398072 | 0.578938763 | 0.307864362 | 0.90 |

| | | | | |
|---|---|---|---|---|
| YBR141C | 286.9036203 | 0.282668642 | 0.304996403 | 0.90 |
| YCL061C | 1120.832529 | -0.276502618 | 0.304900205 | 0.90 |
| YCR017C | 2986.294618 | 0.330320532 | 0.304800317 | 0.90 |
| YCR090C | 636.078747 | -0.348932898 | 0.306994746 | 0.90 |
| YDL244W | 51.51006224 | 0.591219975 | 0.307310792 | 0.90 |
| YDR114C | 36.82268978 | -0.399203853 | 0.303126187 | 0.90 |
| YEL035C | 12.59177912 | 0.460755672 | 0.304365917 | 0.90 |
| YER029C | 871.4448752 | -0.191798694 | 0.307678228 | 0.90 |
| YER079W | 1934.057175 | 0.492232792 | 0.305957238 | 0.90 |
| YFL050C | 563.7473526 | 0.3439617 | 0.304887414 | 0.90 |
| YFR018C | 1513.157132 | 0.30841354 | 0.305637975 | 0.90 |
| YGL001C | 1350.851203 | -0.348633864 | 0.305195198 | 0.90 |
| YGL251C | 608.1790507 | 0.266674588 | 0.306569931 | 0.90 |
| YGL261C | 11.97177178 | 0.576779773 | 0.307738575 | 0.90 |
| YGR057C | 395.64537 | 0.217910013 | 0.304570044 | 0.90 |
| YGR163W | 840.8307527 | 0.230034713 | 0.305021329 | 0.90 |
| YIL019W | 543.1334136 | 0.477568295 | 0.303431142 | 0.90 |
| YIL086C | 23.45932813 | -0.478821034 | 0.305635075 | 0.90 |
| YLR352W | 926.4882915 | 0.225159691 | 0.307548854 | 0.90 |
| YLR363C | 329.7816062 | -0.307900126 | 0.302966755 | 0.90 |
| YML043C | 371.5090553 | -0.481146157 | 0.303652957 | 0.90 |
| YMR285C | 827.213202 | -0.180603831 | 0.306816997 | 0.90 |
| YNL068C | 1421.244951 | 0.269180854 | 0.306066286 | 0.90 |
| YNL324W | 2.168671185 | 0.489156167 | 0.306508958 | 0.90 |
| YNR011C | 902.2961481 | 0.259602738 | 0.30404469 | 0.90 |
| YOL069W | 510.0668666 | 0.251497292 | 0.307604203 | 0.90 |
| YOL108C | 360.9401457 | 0.386234508 | 0.307968905 | 0.90 |
| YOR024W | 4.641545194 | 0.475325718 | 0.305068988 | 0.90 |
| YOR059C | 1230.819833 | 0.239577357 | 0.306990635 | 0.90 |
| YOR227W | 3483.244777 | 0.316112446 | 0.305052112 | 0.90 |
| YOR374W | 6364.239227 | 0.568255682 | 0.307520638 | 0.90 |
| YPL113C | 241.6280316 | 0.329538148 | 0.307573909 | 0.90 |
| YPR026W | 1343.846593 | 0.408202955 | 0.306700206 | 0.90 |
| YPR158W | 1022.854173 | 0.42479629 | 0.304150203 | 0.90 |
| YPR185W | 1716.783017 | 0.233495203 | 0.30615127 | 0.90 |
| YDL159W-A | 1.037341025 | 0.337457502 | 0.309548448 | 0.90 |
| YGR187C | 552.0431833 | -0.38135738 | 0.308981154 | 0.90 |
| YLR388W | 1672.166035 | -0.415078004 | 0.309522395 | 0.90 |
| YML054C-A | 11.9894805 | 0.462651391 | 0.308874701 | 0.90 |
| YMR235C | 2269.617315 | -0.337363537 | 0.309448405 | 0.90 |
| YMR307W | 11748.41017 | -0.405489437 | 0.309007746 | 0.90 |
| YOL104C | 616.0912562 | -0.440111354 | 0.30905843 | 0.90 |
| YOR219C | 1764.610148 | -0.229349454 | 0.309236941 | 0.90 |

| | | | | |
|---|---|---|---|---|
| YEL068C | 304.3910786 | -0.316682545 | 0.310194608 | 0.90 |
| YKL225W | 0.626369128 | 0.274586154 | 0.310337349 | 0.90 |
| YIL060W | 54.27024793 | 0.524861515 | 0.310506303 | 0.90 |
| YPR198W | 724.9139692 | -0.372527867 | 0.310647581 | 0.90 |
| YPR126C | 82.68269767 | -0.478356952 | 0.310986262 | 0.90 |
| YBL069W | 640.7564823 | 0.355806406 | 0.314179916 | 0.90 |
| YCL005W-A | 247.3743167 | -0.248594733 | 0.312688581 | 0.90 |
| YDL141W | 1031.602218 | 0.202160396 | 0.312765662 | 0.90 |
| YDR106W | 46.31624816 | -0.381215015 | 0.312133985 | 0.90 |
| YDR125C | 356.5508843 | 0.401298598 | 0.314262628 | 0.90 |
| YDR219C | 850.7025719 | -0.254270292 | 0.314163521 | 0.90 |
| YDR358W | 1790.698239 | 0.355437369 | 0.314106397 | 0.90 |
| YGR004W | 331.5972983 | 0.2437737 | 0.313162868 | 0.90 |
| YIL093C | 871.5653388 | -0.251699724 | 0.311951721 | 0.90 |
| YIR041W | 5.637300496 | 0.557661299 | 0.312643679 | 0.90 |
| YKL078W | 299.5473506 | 0.513319775 | 0.313173871 | 0.90 |
| YLL016W | 1012.667191 | 0.419879816 | 0.311525329 | 0.90 |
| YLR200W | 774.013828 | -0.237348307 | 0.312610194 | 0.90 |
| YLR390W-A | 12426.05045 | 0.219653627 | 0.313264134 | 0.90 |
| YNL016W | 4774.57143 | -0.247905155 | 0.313461542 | 0.90 |
| YNL122C | 325.2385534 | -0.301621218 | 0.313340243 | 0.90 |
| YNL235C | 7.928094321 | -0.584498866 | 0.312270066 | 0.90 |
| YOL110W | 433.4687434 | 0.444002779 | 0.313249954 | 0.90 |
| YOR255W | 23.76826284 | 0.523261567 | 0.314082569 | 0.90 |
| YPL081W | 1016.551966 | -0.346250751 | 0.314049273 | 0.90 |
| YPR155C | 883.1515984 | 0.321024089 | 0.313457359 | 0.90 |
| YOR160W | 1175.925728 | -0.138569308 | 0.314482855 | 0.90 |
| YMR090W | 1879.223335 | 0.552437464 | 0.314656987 | 0.90 |
| YGL254W | 331.7870674 | 0.318925252 | 0.315123799 | 0.91 |
| YGR181W | 835.8007421 | -0.232872532 | 0.315522285 | 0.91 |
| YJR018W | 57.57815418 | 0.268993788 | 0.315431564 | 0.91 |
| YMR244W | 46.42719777 | 0.493718726 | 0.315302844 | 0.91 |
| YDR183W | 267.9198317 | -0.209588665 | 0.316359036 | 0.91 |
| YGL186C | 1156.394032 | 0.319945248 | 0.316472002 | 0.91 |
| YOL085W-A | 20.71787704 | 0.549222887 | 0.315979177 | 0.91 |
| YPL026C | 1074.82514 | -0.477145434 | 0.316169281 | 0.91 |
| snR31 | 65.06005548 | -0.430905431 | 0.316415835 | 0.91 |
| snR72 | 8.104041704 | 0.100254809 | 0.316085233 | 0.91 |
| YJL029C | 1512.843835 | -0.179942393 | 0.31662263 | 0.91 |
| YBR195C | 288.8600834 | -0.275243233 | 0.317095222 | 0.91 |
| YLR142W | 1292.784521 | 0.519175682 | 0.317192705 | 0.91 |
| YOR339C | 48.09399516 | -0.465966579 | 0.317156112 | 0.91 |
| YIR033W | 2182.959679 | 0.238695008 | 0.318118546 | 0.91 |

| | | | | |
|---|---|---|---|---|
| YKL046C | 1814.036225 | -0.18830707 | 0.318111604 | 0.91 |
| YLR004C | 257.8975095 | 0.312821528 | 0.317954705 | 0.91 |
| YLR342W | 12036.59934 | -0.304361184 | 0.317640577 | 0.91 |
| YOL012C | 810.102208 | -0.220150961 | 0.317629301 | 0.91 |
| YOR048C | 2248.629889 | 0.25654456 | 0.317895894 | 0.91 |
| YBR196C-B | 11.09747944 | -0.575339555 | 0.320676494 | 0.91 |
| YBR274W | 529.7886878 | 0.211509748 | 0.320841993 | 0.91 |
| YDR169C-A | 13.43403719 | -0.446867252 | 0.323131795 | 0.91 |
| YDR491C | 102.3610769 | 0.466674445 | 0.320239506 | 0.91 |
| YDR531W | 969.2146497 | -0.296065838 | 0.323177187 | 0.91 |
| YEL052W | 1984.785259 | 0.310164092 | 0.318942538 | 0.91 |
| YGR266W | 1152.085768 | 0.296131632 | 0.323298607 | 0.91 |
| YHR050W | 1110.031456 | 0.434899156 | 0.322474436 | 0.91 |
| YIL061C | 381.0352818 | 0.244978592 | 0.322954393 | 0.91 |
| YIL165C | 180.1950386 | 0.413476517 | 0.320817288 | 0.91 |
| YIR032C | 113.4974223 | 0.469690732 | 0.321048557 | 0.91 |
| YJL209W | 626.0794008 | -0.214261626 | 0.32197875 | 0.91 |
| YJR080C | 1270.85858 | 0.394527616 | 0.322309856 | 0.91 |
| YJR082C | 608.3949667 | -0.254660859 | 0.321117999 | 0.91 |
| YJR091C | 3359.251305 | 0.259705486 | 0.321656643 | 0.91 |
| YKL044W | 194.2159883 | -0.495762441 | 0.322160462 | 0.91 |
| YLR049C | 719.3459985 | 0.360307692 | 0.323246793 | 0.91 |
| YLR144C | 964.8658379 | 0.267383907 | 0.323021147 | 0.91 |
| YLR261C | 10.17984332 | 0.367539775 | 0.318974163 | 0.91 |
| YLR281C | 33.92288358 | 0.522882303 | 0.319337317 | 0.91 |
| YMR005W | 1198.923005 | -0.26512194 | 0.32002951 | 0.91 |
| YMR081C | 3745.685883 | 0.602695758 | 0.320695463 | 0.91 |
| YMR122W-A | 5293.670712 | -0.288706302 | 0.320688445 | 0.91 |
| YMR135W-A | 18.48988281 | 0.471173804 | 0.321027858 | 0.91 |
| YMR272C | 4343.78123 | -0.251339611 | 0.322755238 | 0.91 |
| YNL231C | 1567.48489 | -0.227313242 | 0.320978272 | 0.91 |
| YNR031C | 1352.088361 | -0.264304964 | 0.320719715 | 0.91 |
| YOR140W | 905.6577941 | 0.253635425 | 0.319857212 | 0.91 |
| YOR266W | 469.4838324 | -0.170058282 | 0.320939639 | 0.91 |
| YOR298C-A | 6204.241243 | -0.213684117 | 0.322142402 | 0.91 |
| YPL125W | 1168.744838 | 0.303130531 | 0.319473015 | 0.91 |
| YPL169C | 1689.790258 | -0.220067 | 0.321844112 | 0.91 |
| YPL268W | 929.1963896 | 0.323851766 | 0.322212693 | 0.91 |
| YPR034W | 1306.898874 | -0.24448444 | 0.322463201 | 0.91 |
| YPR054W | 265.2938746 | -0.419614608 | 0.319686278 | 0.91 |
| YER052C | 2274.895414 | 0.502116984 | 0.323537675 | 0.91 |
| YMR045C | 326.5494461 | 0.411399939 | 0.323676372 | 0.91 |
| YMR275C | 1521.663704 | 0.243950157 | 0.324073883 | 0.91 |

| | | | | |
|---|---|---|---|---|
| YKR023W | 810.8865639 | 0.218815004 | 0.324315764 | 0.91 |
| YDL005C | 1599.265031 | -0.190979399 | 0.324519298 | 0.91 |
| YIR027C | 227.1514133 | 0.503277929 | 0.324904535 | 0.91 |
| YLR259C | 10469.29817 | 0.263631967 | 0.325788959 | 0.91 |
| YNL094W | 1768.042353 | 0.205518215 | 0.326011681 | 0.91 |
| YNR033W | 1803.332991 | 0.241243025 | 0.325715141 | 0.91 |
| YPL039W | 426.2421359 | 0.297737965 | 0.325956236 | 0.91 |
| YDR203W | 15.81601128 | -0.679510452 | 0.326526125 | 0.91 |
| YGL190C | 1598.926776 | -0.275577058 | 0.326409398 | 0.91 |
| YPR199C | 796.7118888 | -0.324182394 | 0.326419089 | 0.91 |
| YNL328C | 68.99382563 | -0.279424315 | 0.327042151 | 0.91 |
| YBL030C | 9599.31549 | 0.572796776 | 0.327760457 | 0.91 |
| YCL057C-A | 1206.33424 | 0.430243803 | 0.328543524 | 0.91 |
| YCR054C | 374.3747854 | 0.282144701 | 0.328269936 | 0.91 |
| YDR251W | 2664.249522 | -0.228593367 | 0.329135988 | 0.91 |
| YER080W | 3267.440935 | 0.305846233 | 0.328942194 | 0.91 |
| YGL052W | 83.8653604 | 0.350816565 | 0.329327665 | 0.91 |
| YGL070C | 462.601344 | -0.257089846 | 0.32749936 | 0.91 |
| YGL144C | 301.1617021 | 0.266363361 | 0.328834948 | 0.91 |
| YGR133W | 486.059603 | 0.211537943 | 0.329575991 | 0.91 |
| YIR023W | 1089.743487 | -0.26993594 | 0.327789475 | 0.91 |
| YKL151C | 2678.459393 | 0.477205964 | 0.329265837 | 0.91 |
| YLR072W | 910.1580571 | 0.211283259 | 0.32854116 | 0.91 |
| YLR353W | 907.9621406 | 0.177122186 | 0.329441692 | 0.91 |
| YMR232W | 290.2765579 | -0.529688788 | 0.329474406 | 0.91 |
| YNL020C | 852.0700517 | -0.296540302 | 0.328515236 | 0.91 |
| YNL124W | 1308.097286 | 0.325270686 | 0.327683122 | 0.91 |
| YOL003C | 636.2491343 | -0.205578191 | 0.328794487 | 0.91 |
| YPL266W | 1132.936461 | 0.47270106 | 0.328188439 | 0.91 |
| YDR478W | 183.0152712 | 0.240747447 | 0.329933731 | 0.91 |
| YGR088W | 3462.726793 | 0.555915089 | 0.329959767 | 0.91 |
| YKR076W | 1274.281879 | 0.485557937 | 0.330088208 | 0.91 |
| YDR445C | 3.142353253 | -0.595697444 | 0.330577413 | 0.91 |
| YHR072W-A | 1183.992345 | -0.353839028 | 0.33057176 | 0.91 |
| YOR283W | 529.1178643 | -0.266427285 | 0.330826266 | 0.91 |
| YDL065C | 1387.029558 | 0.293472854 | 0.331406002 | 0.91 |
| YGR204C-A | 13.54650687 | -0.487525054 | 0.331256918 | 0.91 |
| YHR036W | 340.3963184 | -0.304060053 | 0.331727839 | 0.91 |
| YKL202W | 21.88607995 | 0.347495948 | 0.331799295 | 0.91 |
| YML058W-A | 99.58508334 | -0.552787108 | 0.331758836 | 0.91 |
| YGR174C | 1048.598299 | 0.380290052 | 0.332947077 | 0.91 |
| YHR138C | 1476.126547 | 0.349030854 | 0.332866601 | 0.91 |
| YJR128W | 14.84064674 | 0.427908516 | 0.33280891 | 0.91 |

| | | | | |
|---|---|---|---|---|
| YLR205C | 412.8411079 | 0.418793453 | 0.332694815 | 0.91 |
| YML129C | 536.8329539 | -0.309271319 | 0.332520336 | 0.91 |
| YOR192C-C | 17.72254902 | -0.615986109 | 0.332992785 | 0.91 |
| YPL233W | 252.806959 | 0.205858956 | 0.332287457 | 0.91 |
| YBL044W | 3.828780377 | 0.531032952 | 0.333815427 | 0.92 |
| YBR263W | 3971.725103 | 0.207386737 | 0.333713316 | 0.92 |
| YLR312W-A | 784.781583 | -0.240192762 | 0.33370489 | 0.92 |
| YNL192W | 2879.150646 | -0.414185499 | 0.3335087 | 0.92 |
| YAL026C | 1397.824394 | -0.234177936 | 0.334793824 | 0.92 |
| YAR027W | 1481.56887 | -0.232525666 | 0.335334834 | 0.92 |
| YBR284W | 530.3934343 | 0.373568958 | 0.337606832 | 0.92 |
| YCR011C | 2808.118139 | 0.219508493 | 0.336659014 | 0.92 |
| YDL088C | 1480.552376 | -0.246495101 | 0.336805411 | 0.92 |
| YDR435C | 760.2512519 | 0.301082733 | 0.337601467 | 0.92 |
| YGL079W | 448.9297125 | -0.185659487 | 0.334938592 | 0.92 |
| YGR040W | 494.1182396 | -0.326999969 | 0.334288068 | 0.92 |
| YGR102C | 796.5538343 | 0.335051872 | 0.336156049 | 0.92 |
| YHR151C | 595.1830514 | 0.313283128 | 0.33421587 | 0.92 |
| YHR199C | 2007.264009 | 0.344322961 | 0.337392488 | 0.92 |
| YJL191W | 879.6997619 | -0.449792475 | 0.334711488 | 0.92 |
| YLR246W | 470.2308665 | 0.321251965 | 0.337404138 | 0.92 |
| YLR365W | 4.223598425 | 0.477570812 | 0.336415183 | 0.92 |
| YMR309C | 2879.75717 | -0.324367806 | 0.337494436 | 0.92 |
| YNL051W | 867.7540206 | -0.207801866 | 0.335832927 | 0.92 |
| YNL089C | 16.65282477 | -0.431960772 | 0.33740243 | 0.92 |
| YNL129W | 332.9229067 | -0.358645696 | 0.334535453 | 0.92 |
| YNL206C | 942.6169972 | -0.188607778 | 0.335244985 | 0.92 |
| YOL020W | 976.6345917 | -0.330183705 | 0.337069506 | 0.92 |
| YOR081C | 1945.1418 | -0.268173092 | 0.337172924 | 0.92 |
| YOR135C | 405.7747912 | 0.301831633 | 0.33690117 | 0.92 |
| YPL013C | 664.0145708 | -0.289304924 | 0.336560933 | 0.92 |
| YPL115C | 1185.518438 | -0.257547706 | 0.337544848 | 0.92 |
| YPR032W | 901.7856576 | 0.211450472 | 0.336167484 | 0.92 |
| YPR180W | 730.2139766 | -0.22453026 | 0.337182987 | 0.92 |
| YJL112W | 1088.839683 | 0.269646339 | 0.337790005 | 0.92 |
| YDR036C | 1560.530814 | 0.340364013 | 0.338465491 | 0.92 |
| YBR272C | 606.5333168 | 0.218165137 | 0.339543478 | 0.92 |
| YDR436W | 1255.862426 | 0.296574484 | 0.339203226 | 0.92 |
| YGL093W | 1066.84026 | 0.147082684 | 0.339565293 | 0.92 |
| YJR125C | 2124.757426 | -0.200358377 | 0.338986598 | 0.92 |
| YLR204W | 619.6469206 | -0.278821669 | 0.339605191 | 0.92 |
| YNL157W | 1475.809734 | -0.178680504 | 0.33930654 | 0.92 |
| YOL072W | 594.5699607 | -0.219297158 | 0.340255197 | 0.92 |

| | | | | |
|---|---|---|---|---|
| YBR177C | 2530.709586 | -0.290236707 | 0.34246558 | 0.92 |
| YDR127W | 3620.75176 | 0.366940544 | 0.342549437 | 0.92 |
| YFR023W | 96.03552642 | 0.457415736 | 0.342456671 | 0.92 |
| YHR124W | 347.8549107 | 0.413854611 | 0.342398597 | 0.92 |
| YHR176W | 373.1735066 | -0.26643348 | 0.342141105 | 0.92 |
| YIL059C | 89.463829 | 0.454793615 | 0.342393477 | 0.92 |
| YKL194C | 593.5236151 | 0.229961313 | 0.342434999 | 0.92 |
| YKR056W | 1143.695694 | 0.343974399 | 0.341297782 | 0.92 |
| YLL052C | 77.86079202 | 0.407005386 | 0.341389343 | 0.92 |
| YMR164C | 711.3389073 | 0.223699983 | 0.342125078 | 0.92 |
| YNL205C | 11.3162081 | 0.381069684 | 0.341694191 | 0.92 |
| YOR047C | 1007.814108 | -0.454219003 | 0.340858358 | 0.92 |
| snR66 | 7.293652703 | 0.421110836 | 0.342450291 | 0.92 |
| YDR497C | 3420.957422 | 0.201642299 | 0.342859619 | 0.92 |
| YPR170W-B | 216.8408172 | 0.349022956 | 0.342922511 | 0.92 |
| YJL213W | 560.8105438 | 0.424854466 | 0.343369818 | 0.92 |
| YOL099C | 11.41692104 | -0.427516857 | 0.343295001 | 0.92 |
| YPL023C | 1340.132366 | 0.19914271 | 0.343581781 | 0.92 |
| YDR052C | 1279.720929 | -0.207122099 | 0.344244132 | 0.92 |
| YDR378C | 714.4468599 | -0.176828789 | 0.344127395 | 0.92 |
| YPR204W | 106.9528255 | 0.382077258 | 0.34389121 | 0.92 |
| snR52 | 0.834128967 | -0.363721039 | 0.344313742 | 0.92 |
| YAL024C | 799.2694521 | -0.279021054 | 0.346212517 | 0.92 |
| YDR043C | 825.8596949 | -0.429212599 | 0.346025773 | 0.92 |
| YIL114C | 411.0869055 | -0.315276208 | 0.345733064 | 0.92 |
| YJR057W | 461.680688 | 0.207548371 | 0.344716701 | 0.92 |
| YLR448W | 5493.030487 | -0.423738738 | 0.344916755 | 0.92 |
| YML108W | 173.1265371 | 0.34689422 | 0.346181365 | 0.92 |
| YMR220W | 1203.063636 | -0.186585984 | 0.345891141 | 0.92 |
| YMR268C | 351.0386447 | 0.189077663 | 0.34512125 | 0.92 |
| YNR037C | 584.4173485 | -0.306624387 | 0.344749703 | 0.92 |
| YOL019W-A | 0.720433626 | 0.193715921 | 0.345956457 | 0.92 |
| YOR034C | 667.8994742 | 0.270184131 | 0.34617548 | 0.92 |
| YOR037W | 696.733578 | 0.216488243 | 0.346137228 | 0.92 |
| YOR197W | 2558.031802 | 0.180284065 | 0.345675181 | 0.92 |
| YPR194C | 1121.813284 | 0.481331506 | 0.346474189 | 0.92 |
| YDL100C | 5446.231089 | 0.310119236 | 0.347644805 | 0.92 |
| YDR242W | 619.518258 | 0.252014626 | 0.347915085 | 0.92 |
| YER106W | 34.98872113 | 0.435579111 | 0.347789699 | 0.92 |
| YGL036W | 2417.923989 | 0.198541119 | 0.346963147 | 0.92 |
| YJL088W | 187.9199165 | 0.458648661 | 0.347066713 | 0.92 |
| YML020W | 864.3038212 | -0.237581708 | 0.347521853 | 0.92 |
| YMR031W-A | 40.58383151 | 0.428110804 | 0.346868077 | 0.92 |

| | | | | |
|---|---|---|---|---|
| YOL133W | 270.3290721 | -0.29601216 | 0.34746384 | 0.92 |
| YOR337W | 623.7332196 | 0.249057592 | 0.347447041 | 0.92 |
| YPL142C | 228.9766843 | -0.372967784 | 0.347717899 | 0.92 |
| YAL015C | 634.3594781 | 0.28975127 | 0.359637618 | 0.92 |
| YAL042W | 3676.592816 | -0.163698416 | 0.359071294 | 0.92 |
| YAL047C | 589.7842721 | 0.193363012 | 0.351415739 | 0.92 |
| YAL048C | 583.0194511 | 0.249793077 | 0.358169416 | 0.92 |
| YBL068W | 791.3128306 | 0.290275655 | 0.348317111 | 0.92 |
| YBL084C | 817.2451769 | -0.283117497 | 0.355123011 | 0.92 |
| YBR003W | 1468.050788 | -0.185696694 | 0.354392045 | 0.92 |
| YBR072C-A | 2.769886284 | -0.53367315 | 0.364404508 | 0.92 |
| YBR101C | 3697.675555 | 0.303805229 | 0.35905071 | 0.92 |
| YBR192W | 512.8528961 | -0.180095997 | 0.363663477 | 0.92 |
| YCL031C | 1450.481764 | -0.253227165 | 0.360825798 | 0.92 |
| YCR022C | 20.80731429 | 0.353396227 | 0.352439405 | 0.92 |
| YCR088W | 5938.374856 | -0.249731627 | 0.359488109 | 0.92 |
| YDL025C | 1555.623411 | 0.287798443 | 0.357825234 | 0.92 |
| YDL067C | 1308.526792 | 0.510906205 | 0.355840388 | 0.92 |
| YDL109C | 380.7977413 | 0.276444683 | 0.359767409 | 0.92 |
| YDL121C | 1113.601796 | 0.318104792 | 0.357758241 | 0.92 |
| YDL136W | 915.4579307 | -0.327982815 | 0.358131675 | 0.92 |
| YDL160C | 1362.561057 | -0.287604478 | 0.361979907 | 0.92 |
| YDL166C | 434.2604045 | -0.27599332 | 0.350959731 | 0.92 |
| YDL174C | 2966.298062 | 0.406853506 | 0.358396163 | 0.92 |
| YDL208W | 1913.168285 | -0.301617263 | 0.363733804 | 0.92 |
| YDR152W | 1737.468978 | -0.219314842 | 0.359595543 | 0.92 |
| YDR211W | 2365.993285 | 0.279516631 | 0.35144149 | 0.92 |
| YDR259C | 452.5952789 | 0.387631918 | 0.349420766 | 0.92 |
| YDR266C | 1529.276069 | -0.236768508 | 0.355582223 | 0.92 |
| YDR306C | 1159.558737 | 0.24346502 | 0.353131222 | 0.92 |
| YDR308C | 370.9057891 | -0.253622594 | 0.355032721 | 0.92 |
| YDR315C | 362.5235632 | -0.245999885 | 0.349811013 | 0.92 |
| YDR357C | 553.215464 | -0.317414792 | 0.360638562 | 0.92 |
| YDR523C | 105.2296251 | 0.322808952 | 0.349385876 | 0.92 |
| YEL020W-A | 192.2562057 | -0.234547865 | 0.362637541 | 0.92 |
| YEL072W | 207.4067694 | -0.41947546 | 0.363147152 | 0.92 |
| YER023W | 2339.829047 | -0.24319748 | 0.359276166 | 0.92 |
| YER066C-A | 1483.59707 | 0.543568887 | 0.359669387 | 0.92 |
| YER086W | 3514.638296 | 0.501715301 | 0.350135396 | 0.92 |
| YER087C-A | 36.6678806 | 0.367417145 | 0.364360817 | 0.92 |
| YFR036W-A | 15.15766896 | -0.270740791 | 0.357327084 | 0.92 |
| YFR055W | 975.2716539 | 0.49688458 | 0.358404132 | 0.92 |
| YGL007C-A | 2.085025417 | -0.295046135 | 0.358305532 | 0.92 |

| | | | | |
|---|---|---|---|---|
| YGL017W | 502.0325199 | 0.171318174 | 0.358429438 | 0.92 |
| YGL122C | 2999.105342 | -0.277457634 | 0.357045411 | 0.92 |
| YGL172W | 1277.929348 | -0.233644247 | 0.353703104 | 0.92 |
| YGL211W | 636.0371021 | 0.178705326 | 0.361431234 | 0.92 |
| YGL259W | 86.59077684 | 0.467485868 | 0.348350134 | 0.92 |
| YGR037C | 2287.491924 | -0.288924021 | 0.361610374 | 0.92 |
| YGR052W | 642.70953 | -0.544616789 | 0.35143787 | 0.92 |
| YGR056W | 1165.037029 | -0.262055988 | 0.361646581 | 0.92 |
| YGR127W | 732.3981957 | -0.344147474 | 0.352332249 | 0.92 |
| YGR175C | 3323.313876 | -0.298290628 | 0.36354167 | 0.92 |
| YGR263C | 454.4543224 | 0.198225057 | 0.356192767 | 0.92 |
| YGR271W | 1597.427923 | -0.197201755 | 0.354076116 | 0.92 |
| YHL034C | 6448.755806 | 0.21003605 | 0.361595875 | 0.92 |
| YHR029C | 965.1862947 | 0.364537438 | 0.351065769 | 0.92 |
| YHR056W-A | 0.867587117 | 0.319566709 | 0.350886286 | 0.92 |
| YHR077C | 2925.324128 | -0.19217826 | 0.364116465 | 0.92 |
| YHR092C | 9528.876224 | 0.571999083 | 0.356761703 | 0.92 |
| YHR156C | 433.8648918 | 0.265837313 | 0.358977361 | 0.92 |
| YHR192W | 784.2571123 | -0.204730201 | 0.364118975 | 0.92 |
| YIL040W | 372.5621769 | -0.263176969 | 0.362054069 | 0.92 |
| YIL104C | 354.0212252 | 0.237501485 | 0.35488209 | 0.92 |
| YIL164C | 260.9748713 | 0.382051219 | 0.357093536 | 0.92 |
| YJL020C | 4552.045062 | 0.314844916 | 0.356968466 | 0.92 |
| YJL120W | 37.94521827 | -0.407214793 | 0.355856424 | 0.92 |
| YJL218W | 146.9624687 | 0.444761303 | 0.355177531 | 0.92 |
| YJR078W | 516.4472306 | 0.461975639 | 0.360457122 | 0.92 |
| YJR127C | 3525.789326 | 0.222040126 | 0.358485048 | 0.92 |
| YKL116C | 778.7224786 | -0.133308212 | 0.362790259 | 0.92 |
| YKL129C | 2233.401318 | 0.222275638 | 0.352670512 | 0.92 |
| YKL156W | 2081.100672 | -0.381123561 | 0.353747887 | 0.92 |
| YKR020W | 476.7599628 | -0.226204487 | 0.349376485 | 0.92 |
| YKR084C | 1262.417524 | -0.217143335 | 0.364271723 | 0.92 |
| YLR037C | 167.7921303 | 0.331085419 | 0.355416821 | 0.92 |
| YLR065C | 843.2361186 | -0.295329224 | 0.363725291 | 0.92 |
| YLR115W | 1204.082371 | 0.150841999 | 0.351838161 | 0.92 |
| YLR181C | 881.938971 | 0.196490447 | 0.363731102 | 0.92 |
| YLR199C | 824.459216 | 0.148159126 | 0.361220328 | 0.92 |
| YLR310C | 3300.526123 | -0.251037475 | 0.353611977 | 0.92 |
| YLR314C | 3042.070083 | 0.193290068 | 0.353789992 | 0.92 |
| YLR436C | 2590.336596 | -0.272848744 | 0.350877333 | 0.92 |
| YML026C | 7646.727007 | -0.340516407 | 0.36112036 | 0.92 |
| YML058W | 2577.833646 | -0.301546117 | 0.363632489 | 0.92 |
| YML066C | 127.9180195 | -0.493360576 | 0.355705197 | 0.92 |

| | | | | |
|---|---|---|---|---|
| YML095C | 255.548255 | -0.222550583 | 0.364346119 | 0.92 |
| YMR023C | 334.2712538 | -0.257076461 | 0.354952858 | 0.92 |
| YMR067C | 1220.387108 | -0.192016359 | 0.352580511 | 0.92 |
| YMR190C | 1250.870637 | 0.310415585 | 0.361936178 | 0.92 |
| YMR206W | 572.161077 | 0.523176811 | 0.356464398 | 0.92 |
| YMR243C | 2759.15272 | 0.289151826 | 0.351162588 | 0.92 |
| YMR266W | 1578.35636 | -0.241149234 | 0.356544289 | 0.92 |
| YNL006W | 936.4455341 | 0.254654918 | 0.36401326 | 0.92 |
| YNL093W | 262.8484398 | 0.476789202 | 0.358089079 | 0.92 |
| YNL126W | 705.9981899 | -0.274605024 | 0.357903526 | 0.92 |
| YNL207W | 833.7029402 | 0.289234765 | 0.357471775 | 0.92 |
| YNL288W | 932.0705303 | -0.186544779 | 0.348943673 | 0.92 |
| YOL090W | 1084.862315 | -0.270744006 | 0.360012487 | 0.92 |
| YOR007C | 8060.679187 | 0.267462222 | 0.361672047 | 0.92 |
| YOR190W | 351.1788756 | 0.28396768 | 0.356771133 | 0.92 |
| YOR194C | 1465.560671 | -0.301070547 | 0.358742004 | 0.92 |
| YOR253W | 809.4249218 | -0.263451039 | 0.35460308 | 0.92 |
| YOR286W | 1162.614528 | -0.192697995 | 0.36161386 | 0.92 |
| YOR308C | 716.1626853 | 0.30370984 | 0.36096741 | 0.92 |
| YOR313C | 174.7411276 | -0.51435123 | 0.358363766 | 0.92 |
| YOR356W | 1275.917226 | 0.266370282 | 0.351965593 | 0.92 |
| YPL024W | 252.5020697 | 0.394129895 | 0.35122835 | 0.92 |
| YPL104W | 889.4598617 | 0.268460851 | 0.35639887 | 0.92 |
| YPL105C | 3260.005512 | -0.192985571 | 0.35669731 | 0.92 |
| YPL117C | 2245.25569 | 0.234794766 | 0.364404611 | 0.92 |
| YPL168W | 515.986609 | 0.308250113 | 0.348820563 | 0.92 |
| YPL191C | 278.5044925 | 0.310290796 | 0.349252591 | 0.92 |
| YPL221W | 1548.403188 | -0.278873591 | 0.361616198 | 0.92 |
| YPL247C | 2827.171119 | 0.453806569 | 0.350698737 | 0.92 |
| YPL258C | 548.5976694 | 0.319734823 | 0.355686082 | 0.92 |
| YPR095C | 1088.661894 | -0.189342597 | 0.363526041 | 0.92 |
| YPR144C | 1036.348245 | 0.351695839 | 0.35734699 | 0.92 |
| YPR159C-A | 2.271891006 | 0.439778937 | 0.363242831 | 0.92 |
| YDL204W | 5424.611742 | 0.555236614 | 0.365036775 | 0.92 |
| YDR320W-B | 5.22193975 | 0.431262015 | 0.365049022 | 0.92 |
| YKR092C | 1945.135383 | -0.293851827 | 0.365079664 | 0.92 |
| YOR020W-A | 1169.212022 | 0.351889916 | 0.364794199 | 0.92 |
| YGL218W | 11.00510591 | -0.487386544 | 0.365399923 | 0.92 |
| YMR027W | 2670.4492 | 0.288178865 | 0.365408089 | 0.92 |
| YBR078W | 17879.15973 | -0.328391577 | 0.366651002 | 0.92 |
| YBR126C | 9994.323143 | 0.371377059 | 0.366777983 | 0.92 |
| YBR220C | 693.632069 | 0.30861113 | 0.367215228 | 0.92 |
| YDR074W | 10482.83127 | 0.45197808 | 0.366595756 | 0.92 |

| | | | | |
|---|---|---|---|---|
| YDR140W | 525.20568 | 0.235658633 | 0.367269285 | 0.92 |
| YDR503C | 368.4756431 | 0.236502872 | 0.367349858 | 0.92 |
| YER181C | 6.5610832 | 0.529146113 | 0.366880695 | 0.92 |
| YGL231C | 1251.65783 | -0.174594561 | 0.365845599 | 0.92 |
| YLR121C | 1254.397824 | 0.286073088 | 0.367326465 | 0.92 |
| YMR105C | 9001.650714 | 0.504057493 | 0.36642627 | 0.92 |
| YPL155C | 690.291389 | 0.237182211 | 0.367278351 | 0.92 |
| YPL257W | 181.2677801 | 0.410251905 | 0.366375749 | 0.92 |
| YBR080C | 2198.347068 | -0.218852919 | 0.372107264 | 0.92 |
| YBR221W-A | 0.837044541 | 0.199201241 | 0.371351983 | 0.92 |
| YDR071C | 2792.908431 | -0.222814034 | 0.371736183 | 0.92 |
| YER174C | 586.1669315 | 0.25090466 | 0.371747355 | 0.92 |
| YFR037C | 1666.566511 | -0.208679411 | 0.368261821 | 0.92 |
| YGL074C | 37.00723578 | 0.399651747 | 0.370261697 | 0.92 |
| YGL246C | 646.4635665 | 0.18466393 | 0.368398827 | 0.92 |
| YHR038W | 426.523879 | -0.213154146 | 0.368070404 | 0.92 |
| YHR046C | 620.1635411 | -0.314905071 | 0.371819056 | 0.92 |
| YIL138C | 1495.862581 | -0.274823794 | 0.372073644 | 0.92 |
| YJR048W | 1046.168345 | 0.613388135 | 0.370655816 | 0.92 |
| YJR058C | 464.9848205 | -0.197979485 | 0.371374283 | 0.92 |
| YKL020C | 2087.802283 | -0.26312509 | 0.370041657 | 0.92 |
| YKL066W | 1252.501288 | 0.333007114 | 0.370915904 | 0.92 |
| YKL178C | 114.5867776 | -0.493214047 | 0.370151296 | 0.92 |
| YLR426W | 420.438408 | -0.246176361 | 0.372018685 | 0.92 |
| YMR208W | 1260.14289 | -0.225866671 | 0.370067141 | 0.92 |
| YMR260C | 3410.062096 | -0.26060116 | 0.368145319 | 0.92 |
| YNL102W | 1324.90783 | -0.227169004 | 0.37183104 | 0.92 |
| YNL173C | 2504.496471 | 0.294200157 | 0.370801092 | 0.92 |
| YNL211C | 132.5864716 | -0.297665642 | 0.369157829 | 0.92 |
| YNR021W | 3396.726535 | -0.258103532 | 0.371359745 | 0.92 |
| YOL136C | 1016.324902 | -0.358193236 | 0.36946372 | 0.92 |
| YOR027W | 11708.63489 | 0.316333417 | 0.371142371 | 0.92 |
| YOR064C | 481.1143371 | 0.34008726 | 0.370573597 | 0.92 |
| YOR173W | 2098.419186 | 0.472474307 | 0.369492058 | 0.92 |
| YPL006W | 1523.461434 | 0.277552604 | 0.368598032 | 0.92 |
| YPL011C | 726.714861 | 0.177862532 | 0.370251751 | 0.92 |
| YPL031C | 1442.118487 | 0.244590919 | 0.370629261 | 0.92 |
| YPR065W | 1458.659767 | -0.355467027 | 0.369188552 | 0.92 |
| YPR067W | 664.4484323 | -0.158756538 | 0.369519919 | 0.92 |
| YPR124W | 1891.733227 | 0.492353913 | 0.369794537 | 0.92 |
| snR5 | 32.14065526 | -0.358266719 | 0.37016977 | 0.92 |
| YGR097W | 2251.701342 | 0.267168771 | 0.372302766 | 0.92 |
| YBL006C | 611.4550235 | -0.257567526 | 0.373098724 | 0.92 |

| | | | | |
|---|---|---|---|---|
| YBR096W | 835.4279583 | -0.280794549 | 0.373676418 | 0.92 |
| YBR125C | 746.1231571 | 0.288741798 | 0.374373515 | 0.92 |
| YCR076C | 593.9023385 | 0.297947237 | 0.37398539 | 0.92 |
| YDR530C | 1226.853067 | 0.331312294 | 0.374579518 | 0.92 |
| YER009W | 3909.301197 | -0.267418912 | 0.374942306 | 0.92 |
| YER122C | 4064.222569 | -0.230992434 | 0.373500997 | 0.92 |
| YFL020C | 87.69189529 | 0.429664324 | 0.375293626 | 0.92 |
| YGR151C | 2.6142374 | -0.296835067 | 0.375318103 | 0.92 |
| YHR061C | 329.4509042 | -0.237839326 | 0.373705505 | 0.92 |
| YJL095W | 1317.430087 | 0.163162291 | 0.373942962 | 0.92 |
| YJR005C-A | 81.403139 | 0.501047026 | 0.373609102 | 0.92 |
| YML007C-A | 2.984197591 | -0.455100167 | 0.373905185 | 0.92 |
| YMR063W | 168.4962103 | 0.259709913 | 0.375188613 | 0.92 |
| YMR077C | 308.7109529 | -0.24204055 | 0.373034385 | 0.92 |
| YMR086W | 2080.250939 | 0.196898433 | 0.374762454 | 0.92 |
| YNR009W | 344.9832981 | -0.339531792 | 0.374796051 | 0.92 |
| YNR059W | 692.5145797 | -0.258876713 | 0.373567188 | 0.92 |
| YOL164W | 729.2266525 | 0.309572335 | 0.375282954 | 0.92 |
| YPL096C-A | 145.3313639 | -0.29342591 | 0.373692973 | 0.92 |
| YPR036W-A | 8058.010483 | 0.489055205 | 0.374293775 | 0.92 |
| YAL013W | 1274.881408 | 0.207263373 | 0.380196339 | 0.92 |
| YAL030W | 729.4057845 | 0.218275579 | 0.377958695 | 0.92 |
| YAL056W | 1700.26388 | 0.305342002 | 0.379904406 | 0.92 |
| YBL021C | 336.3286264 | 0.215372482 | 0.379172387 | 0.92 |
| YBR193C | 556.4922631 | 0.20802718 | 0.375744039 | 0.92 |
| YBR215W | 1081.67593 | -0.162552761 | 0.379149259 | 0.92 |
| YCL016C | 442.1886812 | 0.219706604 | 0.380750621 | 0.92 |
| YCR028C-A | 1716.279601 | -0.224449887 | 0.376437266 | 0.92 |
| YDL117W | 1322.092729 | -0.185833429 | 0.376714317 | 0.92 |
| YDL192W | 4665.318464 | -0.259688466 | 0.379856415 | 0.92 |
| YDR126W | 326.9726421 | 0.266785696 | 0.379015343 | 0.92 |
| YDR212W | 2389.189098 | -0.235570483 | 0.378656966 | 0.92 |
| YDR253C | 229.9334863 | 0.234095681 | 0.38032253 | 0.92 |
| YDR332W | 604.027567 | -0.187175274 | 0.378704446 | 0.92 |
| YER136W | 4119.832209 | -0.151797688 | 0.381185441 | 0.92 |
| YFL009W | 860.3502379 | 0.144290159 | 0.37856432 | 0.92 |
| YGL073W | 2411.314838 | 0.203389509 | 0.377779688 | 0.92 |
| YGL086W | 1388.801297 | -0.256200616 | 0.381296856 | 0.92 |
| YGL137W | 4577.146804 | -0.183217132 | 0.380617289 | 0.92 |
| YHR039C-A | 2505.228571 | -0.185329796 | 0.381041666 | 0.92 |
| YIL038C | 1659.427705 | -0.27252877 | 0.380152491 | 0.92 |
| YJL043W | 47.06314645 | 0.414723803 | 0.379960566 | 0.92 |
| YJL141C | 4385.516839 | 0.375710994 | 0.379904423 | 0.92 |

| | | | | |
|---|---|---|---|---|
| YJR039W | 935.4373703 | 0.301711788 | 0.379781576 | 0.92 |
| YKL037W | 43.69720214 | -0.343814606 | 0.378286778 | 0.92 |
| YKL172W | 2863.993194 | -0.305601832 | 0.377407829 | 0.92 |
| YKL215C | 1029.488394 | 0.221731376 | 0.381005643 | 0.92 |
| YLR063W | 280.3060801 | 0.37203283 | 0.380436237 | 0.92 |
| YNL073W | 1409.044777 | -0.256176784 | 0.375798494 | 0.92 |
| YNL087W | 1914.774752 | 0.23716834 | 0.379835536 | 0.92 |
| YNL095C | 702.5523068 | 0.248957308 | 0.376512022 | 0.92 |
| YNL127W | 1075.477959 | 0.26500747 | 0.379400902 | 0.92 |
| YNL134C | 7013.757129 | 0.447138461 | 0.381053283 | 0.92 |
| YNL201C | 1076.736749 | -0.154414946 | 0.379667175 | 0.92 |
| YNL233W | 992.805205 | -0.171166499 | 0.377424099 | 0.92 |
| YPL086C | 1089.007189 | 0.381050397 | 0.37607323 | 0.92 |
| YPL101W | 1017.044188 | 0.23165584 | 0.376353498 | 0.92 |
| YPR022C | 1532.191812 | 0.16713696 | 0.377191497 | 0.92 |
| YPR157W | 448.7836668 | 0.418321377 | 0.377344808 | 0.92 |
| YPR164W | 1248.455329 | 0.175843891 | 0.379814993 | 0.92 |
| snR54 | 8.052861878 | 0.531788195 | 0.376308877 | 0.92 |
| YBR032W | 28.5083245 | 0.36710774 | 0.38181626 | 0.92 |
| YDL024C | 228.1924469 | 0.437345592 | 0.38208037 | 0.92 |
| YFL033C | 2297.741868 | 0.192362453 | 0.382078379 | 0.92 |
| YGR011W | 95.88242328 | 0.207226148 | 0.382122515 | 0.92 |
| YOL008W | 381.7473344 | -0.163147597 | 0.381699696 | 0.92 |
| YOR261C | 3484.142585 | 0.277048423 | 0.382165973 | 0.92 |
| YER068C-A | 10.30512685 | 0.317302266 | 0.382471373 | 0.92 |
| YGL234W | 9210.041986 | 0.51447715 | 0.382451883 | 0.92 |
| YBL029W | 662.4144794 | 0.263609826 | 0.382967214 | 0.93 |
| YLR109W | 26967.32056 | 0.274520146 | 0.383069478 | 0.93 |
| YPL001W | 324.2471356 | -0.271375851 | 0.383083253 | 0.93 |
| YBL070C | 21.83445099 | -0.404584888 | 0.383333605 | 0.93 |
| YER084W-A | 0.719422121 | 0.26155315 | 0.383595053 | 0.93 |
| YER183C | 378.9572978 | -0.1944335 | 0.383820691 | 0.93 |
| YDL222C | 906.6701758 | 0.554300665 | 0.384207764 | 0.93 |
| YOR203W | 129.138005 | 0.291944707 | 0.384112118 | 0.93 |
| YAL038W | 41041.91188 | 0.418280651 | 0.385200107 | 0.93 |
| YCR033W | 2347.054478 | -0.291335799 | 0.384848582 | 0.93 |
| YDR430C | 1380.899649 | -0.116927698 | 0.386009458 | 0.93 |
| YFR034W-A | 2.98827912 | 0.438919258 | 0.386254225 | 0.93 |
| YHL041W | 3.288023996 | 0.426746051 | 0.38586456 | 0.93 |
| YKL109W | 7924.096762 | 0.369527795 | 0.385681806 | 0.93 |
| YLR149C-A | 61.1192566 | 0.508750002 | 0.385551214 | 0.93 |
| YLR412W | 993.3823084 | -0.222088814 | 0.384592277 | 0.93 |
| YMR177W | 436.2483597 | -0.241236857 | 0.386156999 | 0.93 |

| | | | | |
|---|---|---|---|---|
| YMR199W | 1528.60034 | -0.352640142 | 0.385092096 | 0.93 |
| YNL098C | 3291.418535 | -0.188702221 | 0.384929404 | 0.93 |
| YOR297C | 445.9780063 | -0.203090212 | 0.384538002 | 0.93 |
| YPR039W | 9.707681236 | 0.374524188 | 0.385963999 | 0.93 |
| snR62 | 6.46452847 | 0.428173898 | 0.385316255 | 0.93 |
| YMR089C | 2367.558674 | 0.296194646 | 0.386441396 | 0.93 |
| YBR112C | 3373.883429 | -0.241828639 | 0.386730453 | 0.93 |
| YJL096W | 663.1647142 | -0.216836446 | 0.386739654 | 0.93 |
| YDL086W | 1034.431113 | 0.311846102 | 0.387058065 | 0.93 |
| YLR210W | 386.6683639 | -0.217454525 | 0.387104232 | 0.93 |
| YLR444C | 25.68564702 | -0.706946709 | 0.387202899 | 0.93 |
| YAL035W | 9252.51472 | -0.290339606 | 0.387713768 | 0.93 |
| YBL051C | 3664.33556 | -0.332968315 | 0.387696369 | 0.93 |
| YCR071C | 461.1120902 | -0.294635953 | 0.387565694 | 0.93 |
| YCR049C | 7.558405616 | -0.486800942 | 0.388353912 | 0.93 |
| YDR062W | 2339.714713 | -0.213767529 | 0.388575515 | 0.93 |
| YGR192C | 181337.4741 | 0.409705982 | 0.388319603 | 0.93 |
| YIL100W | 3.722972412 | 0.496625325 | 0.388163543 | 0.93 |
| YKL167C | 608.6144606 | -0.221310479 | 0.387904721 | 0.93 |
| YOR275C | 1566.16532 | -0.172240417 | 0.38850026 | 0.93 |
| YPR024W | 1909.485964 | 0.187688217 | 0.388851874 | 0.93 |
| YNL251C | 2085.43933 | -0.394071365 | 0.38950538 | 0.93 |
| YBL078C | 809.4078099 | 0.401993599 | 0.392260592 | 0.93 |
| YBR071W | 990.0527368 | -0.297656613 | 0.390954352 | 0.93 |
| YBR202W | 984.15721 | -0.237357702 | 0.392203016 | 0.93 |
| YBR268W | 567.5701433 | -0.227283556 | 0.393795992 | 0.93 |
| YCR002C | 1893.656785 | -0.213663092 | 0.391677905 | 0.93 |
| YDL045C | 1046.513314 | 0.133017341 | 0.391082676 | 0.93 |
| YDR269C | 1.063098833 | -0.24868669 | 0.391970927 | 0.93 |
| YDR541C | 514.5384389 | -0.446968654 | 0.390009293 | 0.93 |
| YEL059W | 50.35325287 | -0.403118634 | 0.393250872 | 0.93 |
| YER066W | 204.0645964 | 0.388135819 | 0.393402395 | 0.93 |
| YER143W | 1597.506506 | 0.2178371 | 0.391753307 | 0.93 |
| YFR030W | 1464.99151 | 0.312568133 | 0.393116994 | 0.93 |
| YGL121C | 888.9061681 | 0.476273424 | 0.391107581 | 0.93 |
| YGL149W | 44.44871907 | -0.357269516 | 0.390362993 | 0.93 |
| YGL226C-A | 628.3645694 | -0.231741487 | 0.389692486 | 0.93 |
| YGR105W | 477.947228 | -0.23380284 | 0.393659095 | 0.93 |
| YGR155W | 6507.68049 | -0.196041585 | 0.391020519 | 0.93 |
| YGR185C | 4359.206315 | -0.332301109 | 0.391717783 | 0.93 |
| YGR241C | 1428.572307 | -0.20308072 | 0.393504616 | 0.93 |
| YHR162W | 4274.252821 | 0.34230076 | 0.393918607 | 0.93 |
| YHR214C-B | 235.6005636 | -0.536114396 | 0.393966565 | 0.93 |

| | | | | |
|---|---|---|---|---|
| YJL061W | 1142.158213 | -0.166172492 | 0.392188035 | 0.93 |
| YJR038C | 33.2004855 | 0.37107597 | 0.390926601 | 0.93 |
| YKL043W | 3024.896012 | -0.403571423 | 0.39356065 | 0.93 |
| YKR061W | 573.6088479 | -0.393561585 | 0.391640375 | 0.93 |
| YLR238W | 614.7184945 | -0.217965002 | 0.393329902 | 0.93 |
| YLR325C | 5760.035975 | -0.29901088 | 0.392830081 | 0.93 |
| YMR004W | 1474.539468 | -0.193945233 | 0.390803813 | 0.93 |
| YNR026C | 756.2847135 | -0.222827592 | 0.393775472 | 0.93 |
| YOR377W | 997.8606777 | 0.231575084 | 0.392707529 | 0.93 |
| snR51 | 25.09375178 | 0.304847908 | 0.39169692 | 0.93 |
| YIL117C | 1107.127061 | -0.42126057 | 0.394732072 | 0.93 |
| YJR146W | 99.77471099 | 0.435527464 | 0.394590868 | 0.93 |
| YJL039C | 2015.018256 | -0.206957919 | 0.395731909 | 0.93 |
| YJL103C | 874.1846601 | 0.3092505 | 0.395751308 | 0.93 |
| YNL024C-A | 249.9046551 | -0.187277917 | 0.39535798 | 0.93 |
| YOL001W | 429.8453508 | -0.308426743 | 0.395405103 | 0.93 |
| YPL269W | 614.929882 | -0.248005273 | 0.395645907 | 0.93 |
| YPR182W | 302.4381877 | -0.200570855 | 0.395484994 | 0.93 |
| RPR1 | 36.63381865 | -0.386020249 | 0.397056713 | 0.93 |
| YBR132C | 1205.603555 | 0.384073379 | 0.39967665 | 0.93 |
| YCL038C | 682.2522644 | 0.214263794 | 0.397776153 | 0.93 |
| YDL035C | 1472.135402 | 0.216784423 | 0.400332122 | 0.93 |
| YDL162C | 2.875843937 | -0.618545906 | 0.401558426 | 0.93 |
| YDR005C | 1131.574288 | -0.204248854 | 0.399098597 | 0.93 |
| YDR208W | 1907.44225 | -0.234189539 | 0.401110775 | 0.93 |
| YDR422C | 1006.650102 | 0.192396597 | 0.398566983 | 0.93 |
| YEL001C | 2451.407888 | -0.275696639 | 0.400454037 | 0.93 |
| YER007W | 271.3998551 | 0.229145539 | 0.399598661 | 0.93 |
| YER168C | 714.524667 | 0.149795595 | 0.396992358 | 0.93 |
| YFR015C | 6189.813754 | 0.430433493 | 0.396907477 | 0.93 |
| YGL219C | 1273.719314 | 0.26963304 | 0.398081259 | 0.93 |
| YHL023C | 1253.971871 | 0.156360112 | 0.398027293 | 0.93 |
| YHR017W | 1256.836076 | 0.243292969 | 0.400653493 | 0.93 |
| YHR146W | 3577.393266 | 0.242064886 | 0.399206092 | 0.93 |
| YIL137C | 2189.193159 | 0.283446301 | 0.400076457 | 0.93 |
| YJL054W | 1689.096692 | 0.230648885 | 0.39816216 | 0.93 |
| YJL199C | 163.0335414 | -0.212117324 | 0.401100419 | 0.93 |
| YJR111C | 500.5453341 | 0.282220037 | 0.401586164 | 0.93 |
| YJR133W | 618.7266674 | 0.304233731 | 0.399124428 | 0.93 |
| YKL010C | 2290.07323 | 0.138833433 | 0.39698853 | 0.93 |
| YLL021W | 2378.696108 | -0.262387499 | 0.400587844 | 0.93 |
| YLL050C | 6100.834604 | -0.204895757 | 0.399001192 | 0.93 |
| YLR022C | 541.692103 | -0.242079001 | 0.399804909 | 0.93 |

| | | | | |
|---|---|---|---|---|
| YLR274W | 1064.782127 | -0.221484155 | 0.401215977 | 0.93 |
| YLR283W | 409.1057668 | -0.152869249 | 0.401529828 | 0.93 |
| YLR338W | 19.58277131 | 0.325767571 | 0.39899426 | 0.93 |
| YML076C | 1190.303539 | 0.208291737 | 0.397993907 | 0.93 |
| YMR041C | 591.2121868 | 0.342272664 | 0.400730653 | 0.93 |
| YMR214W | 1048.466602 | -0.238541302 | 0.400641238 | 0.93 |
| YNL030W | 3372.305745 | -0.295627494 | 0.396718688 | 0.93 |
| YNL246W | 1195.836863 | 0.137016261 | 0.401279561 | 0.93 |
| YNR045W | 861.3548214 | 0.335261851 | 0.399381642 | 0.93 |
| YPL005W | 369.5032375 | 0.249465395 | 0.399175191 | 0.93 |
| YPL135W | 1241.708085 | 0.480031705 | 0.400657439 | 0.93 |
| YPR197C | 5.599447647 | 0.457620486 | 0.399654107 | 0.93 |
| YLR225C | 1476.977135 | -0.335278712 | 0.402029002 | 0.93 |
| YGL007W | 19.49434054 | -0.377503574 | 0.40249007 | 0.93 |
| YKR066C | 4134.117453 | 0.314534406 | 0.402545747 | 0.93 |
| YBL007C | 3937.826759 | -0.178304491 | 0.402769273 | 0.93 |
| YGL116W | 933.3944118 | 0.222593034 | 0.403535469 | 0.93 |
| YGR190C | 89.92851229 | 0.297773239 | 0.403573805 | 0.93 |
| YMR242W-A | 1.825788145 | 0.472533579 | 0.403248486 | 0.93 |
| YPR044C | 114.0311076 | -0.36514876 | 0.403381583 | 0.93 |
| YJR031C | 1083.680948 | -0.212410257 | 0.403758019 | 0.93 |
| YPR013C | 1083.106605 | -0.291211333 | 0.403918511 | 0.93 |
| YBL028C | 410.9135571 | -0.27525808 | 0.404370895 | 0.93 |
| YGL050W | 416.7826537 | -0.277299063 | 0.404844003 | 0.93 |
| YGR048W | 875.7732177 | -0.196518749 | 0.404429065 | 0.93 |
| YMR176W | 1256.155137 | 0.206297876 | 0.404710165 | 0.93 |
| YOR072W-A | 14.05662092 | 0.328920836 | 0.404585053 | 0.93 |
| YBR191W | 11850.59787 | -0.331085511 | 0.405223345 | 0.93 |
| YOR246C | 1262.599493 | -0.241607625 | 0.405213346 | 0.93 |
| YBR127C | 10020.63379 | 0.194911973 | 0.406548377 | 0.93 |
| YBR180W | 89.72716224 | 0.401354612 | 0.409697958 | 0.93 |
| YDL081C | 10855.42483 | -0.259436115 | 0.409679741 | 0.93 |
| YDL175C | 355.6103486 | 0.280618626 | 0.407758078 | 0.93 |
| YDR003W | 807.4785109 | 0.253618917 | 0.408655028 | 0.93 |
| YDR007W | 490.6292629 | -0.150036907 | 0.407185713 | 0.93 |
| YDR180W | 980.8547337 | 0.235461843 | 0.40938748 | 0.93 |
| YDR353W | 5059.752159 | -0.222353367 | 0.409514207 | 0.93 |
| YDR413C | 10.51150758 | -0.413921936 | 0.4094615 | 0.93 |
| YHR153C | 73.13873436 | -0.290830817 | 0.408159272 | 0.93 |
| YIL021W | 814.8146206 | -0.179580552 | 0.405828884 | 0.93 |
| YIL071W-A | 73.5379669 | -0.239236953 | 0.40570208 | 0.93 |
| YJL107C | 129.4785502 | 0.370216381 | 0.409032437 | 0.93 |
| YJL133C-A | 1683.932792 | 0.395022166 | 0.409343498 | 0.93 |

| | | | | |
|---|---|---|---|---|
| YJL188C | 893.4798181 | -0.263603267 | 0.407789459 | 0.93 |
| YKL038W | 1738.382543 | 0.352254172 | 0.407481682 | 0.93 |
| YKL193C | 1757.604095 | 0.304158057 | 0.408310536 | 0.93 |
| YKR037C | 412.1583252 | 0.142838736 | 0.407918285 | 0.93 |
| YLL017W | 60.73191178 | 0.322051306 | 0.408429721 | 0.93 |
| YLR062C | 194.5995134 | -0.374620173 | 0.408616713 | 0.93 |
| YLR316C | 329.0837142 | 0.158624841 | 0.406891238 | 0.93 |
| YMR057C | 13.35741525 | 0.37160395 | 0.40613797 | 0.93 |
| YMR179W | 627.9998767 | -0.341314978 | 0.406125296 | 0.93 |
| YNR013C | 2464.582186 | -0.214492827 | 0.408977177 | 0.93 |
| YOL018C | 882.7418897 | -0.11524617 | 0.40741682 | 0.93 |
| YOR250C | 655.6893916 | 0.163074238 | 0.406812119 | 0.93 |
| YPL069C | 455.5840403 | -0.236445471 | 0.409042909 | 0.93 |
| YPR079W | 754.8054031 | -0.164353926 | 0.408750624 | 0.93 |
| snR128 | 13.17491061 | 0.656100921 | 0.406674512 | 0.93 |
| YDR016C | 195.2031569 | -0.196739159 | 0.410263308 | 0.94 |
| YEL004W | 410.7288824 | -0.186212662 | 0.410304578 | 0.94 |
| YDR029W | 118.6548652 | 0.302017691 | 0.412924697 | 0.94 |
| YER109C | 1065.439283 | -0.189056263 | 0.412371778 | 0.94 |
| YER157W | 974.6953246 | -0.142121446 | 0.413234191 | 0.94 |
| YFL066C | 2.343434624 | 0.503945066 | 0.413309445 | 0.94 |
| YGL065C | 1014.84416 | -0.154346842 | 0.411864772 | 0.94 |
| YGL100W | 1500.808318 | -0.135864786 | 0.413076582 | 0.94 |
| YGR050C | 100.6325721 | -0.274010629 | 0.413367305 | 0.94 |
| YGR238C | 880.0595867 | 0.222404893 | 0.413404468 | 0.94 |
| YHR207C | 677.6706609 | 0.259907222 | 0.411022099 | 0.94 |
| YJL028W | 3.906046468 | 0.369246445 | 0.412403877 | 0.94 |
| YJL037W | 69.21169324 | 0.384496065 | 0.41292877 | 0.94 |
| YJL097W | 1713.191405 | -0.252044909 | 0.412370471 | 0.94 |
| YLR318W | 479.1884801 | -0.222570704 | 0.41160976 | 0.94 |
| YLR413W | 2093.12908 | -0.400860885 | 0.413127955 | 0.94 |
| YLR417W | 736.9680806 | 0.179272884 | 0.412466141 | 0.94 |
| YMR130W | 646.8299959 | -0.312220799 | 0.412138306 | 0.94 |
| YMR141C | 167.2172498 | 0.273939453 | 0.412774951 | 0.94 |
| YMR311C | 3644.500712 | 0.257604829 | 0.412823396 | 0.94 |
| YOR157C | 2318.284571 | -0.143298822 | 0.411443446 | 0.94 |
| YPL010W | 2111.463364 | -0.189942862 | 0.411516463 | 0.94 |
| YPR158C-D | 130.4698158 | -0.528438331 | 0.410745894 | 0.94 |
| YAL007C | 1549.491558 | -0.261704109 | 0.421010758 | 0.94 |
| YAL011W | 648.981818 | 0.199530009 | 0.420559835 | 0.94 |
| YAL018C | 88.41004538 | 0.419201187 | 0.417163687 | 0.94 |
| YAL066W | 0.770254238 | 0.265138368 | 0.427311365 | 0.94 |
| YAR042W | 3250.319067 | 0.253374877 | 0.424849996 | 0.94 |

| | | | | |
|---|---|---|---|---|
| YBL035C | 596.37002 | -0.256641862 | 0.4211002 | 0.94 |
| YBL071W-A | 297.3219208 | 0.255816596 | 0.413783121 | 0.94 |
| YBR039W | 6878.817001 | 0.418542239 | 0.418600991 | 0.94 |
| YBR143C | 4538.418741 | -0.245375381 | 0.42189448 | 0.94 |
| YBR161W | 517.5371153 | -0.310643761 | 0.416984914 | 0.94 |
| YBR199W | 2706.736809 | -0.100632908 | 0.419169174 | 0.94 |
| YBR211C | 479.4755282 | -0.233666263 | 0.424406564 | 0.94 |
| YCL025C | 3652.132376 | 0.359532071 | 0.422857812 | 0.94 |
| YCL036W | 402.9926525 | 0.346907467 | 0.421206779 | 0.94 |
| YCR012W | 43096.09714 | 0.447615897 | 0.421949271 | 0.94 |
| YCR035C | 836.2083583 | -0.147264394 | 0.42224857 | 0.94 |
| YCR052W | 1334.780982 | -0.176252148 | 0.422422231 | 0.94 |
| YDL059C | 182.996654 | 0.229187253 | 0.42605329 | 0.94 |
| YDL132W | 2917.981172 | 0.148701523 | 0.421151659 | 0.94 |
| YDL182W | 2531.684902 | 0.426928875 | 0.422515289 | 0.94 |
| YDR001C | 5262.491506 | 0.256787008 | 0.426348108 | 0.94 |
| YDR264C | 2392.197288 | -0.172421184 | 0.427260441 | 0.94 |
| YDR347W | 1409.321423 | 0.126788465 | 0.421272033 | 0.94 |
| YDR404C | 728.6666751 | -0.189467141 | 0.423754744 | 0.94 |
| YDR417C | 435.6659374 | -0.425439033 | 0.424220066 | 0.94 |
| YDR420W | 1513.909661 | -0.286797082 | 0.41532831 | 0.94 |
| YDR517W | 2232.436932 | 0.255740979 | 0.421982482 | 0.94 |
| YDR524C-A | 2.188681507 | -0.362245454 | 0.426925 | 0.94 |
| YEL057C | 119.2283707 | -0.467897889 | 0.418353811 | 0.94 |
| YEL077C | 280.3488731 | 0.335774546 | 0.420708622 | 0.94 |
| YER018C | 488.1395433 | 0.36842576 | 0.416809132 | 0.94 |
| YER038C | 327.709464 | 0.277039338 | 0.425059343 | 0.94 |
| YER072W | 2000.095671 | -0.22727238 | 0.426254971 | 0.94 |
| YER092W | 378.6245953 | 0.301566098 | 0.425031841 | 0.94 |
| YFL014W | 10835.63862 | 0.592080788 | 0.422460198 | 0.94 |
| YFL034C-B | 921.0599209 | -0.274942304 | 0.414899201 | 0.94 |
| YFL045C | 5242.345974 | -0.257933534 | 0.422105722 | 0.94 |
| YFL046W | 596.7158239 | -0.170288133 | 0.420502459 | 0.94 |
| YGL146C | 201.790122 | 0.362135801 | 0.424924761 | 0.94 |
| YGR023W | 927.8650533 | 0.374065908 | 0.425125454 | 0.94 |
| YGR028W | 820.509086 | -0.227076263 | 0.423589777 | 0.94 |
| YGR035W-A | 93.40510398 | -0.326338311 | 0.420483279 | 0.94 |
| YGR157W | 3036.942102 | 0.296070447 | 0.42700233 | 0.94 |
| YGR193C | 2486.614356 | 0.1681226 | 0.421642254 | 0.94 |
| YHR032C-A | 12.55755435 | 0.409716974 | 0.421046287 | 0.94 |
| YHR032W-A | 60.96537533 | 0.281086295 | 0.421439212 | 0.94 |
| YHR161C | 2280.636973 | 0.305727112 | 0.422090169 | 0.94 |
| YHR167W | 161.5282471 | -0.288897704 | 0.425807263 | 0.94 |

| | | | | |
|---|---|---|---|---|
| YIL062C | 1798.738473 | -0.17668672 | 0.414912348 | 0.94 |
| YIL085C | 1498.284283 | -0.164523327 | 0.417316427 | 0.94 |
| YIL133C | 6678.70118 | -0.275136172 | 0.427044222 | 0.94 |
| YIL145C | 1126.99064 | 0.276475685 | 0.420556608 | 0.94 |
| YJL167W | 5562.553131 | -0.195467568 | 0.417528404 | 0.94 |
| YJR007W | 3429.418011 | -0.24423517 | 0.42572415 | 0.94 |
| YJR124C | 485.3455454 | -0.313222039 | 0.427188751 | 0.94 |
| YJR132W | 1306.62171 | 0.190178754 | 0.424632984 | 0.94 |
| YKL054C | 4561.844374 | -0.240613439 | 0.418153501 | 0.94 |
| YKL062W | 2465.244192 | -0.366784205 | 0.417851487 | 0.94 |
| YKL067W | 3440.072656 | 0.39384041 | 0.414958812 | 0.94 |
| YKR058W | 2130.69981 | 0.353008601 | 0.419791706 | 0.94 |
| YLL044W | 94.44590477 | -0.016007498 | 0.42546779 | 0.94 |
| YLL067C | 0.999643856 | 0.143638601 | 0.426707994 | 0.94 |
| YLR045C | 1304.494239 | 0.219730322 | 0.418161269 | 0.94 |
| YLR073C | 322.9361924 | -0.350305302 | 0.415916091 | 0.94 |
| YLR192C | 3457.976113 | -0.172920445 | 0.426130657 | 0.94 |
| YLR412C-A | 1.32995427 | -0.409213556 | 0.422830485 | 0.94 |
| YLR431C | 452.0758432 | 0.1762264 | 0.424072407 | 0.94 |
| YLR457C | 335.6948937 | 0.156892617 | 0.422651549 | 0.94 |
| YML019W | 694.8842248 | -0.167106044 | 0.418731687 | 0.94 |
| YML074C | 7957.336564 | -0.260746755 | 0.417728408 | 0.94 |
| YMR052C-A | 61.89756679 | 0.24786441 | 0.419502926 | 0.94 |
| YMR086C-A | 82.59944884 | 0.233643209 | 0.41897351 | 0.94 |
| YMR152W | 1727.69017 | -0.169282972 | 0.418712297 | 0.94 |
| YMR194W | 1484.319594 | -0.234275086 | 0.420981537 | 0.94 |
| YMR251W-A | 5878.547147 | 0.381425651 | 0.423170572 | 0.94 |
| YMR277W | 1413.809425 | -0.194536896 | 0.423005154 | 0.94 |
| YMR286W | 654.5341281 | -0.248096321 | 0.424667311 | 0.94 |
| YMR304C-A | 599.6439516 | -0.340449075 | 0.416137116 | 0.94 |
| YNL003C | 715.0467326 | -0.254395889 | 0.425391538 | 0.94 |
| YNL044W | 495.5352853 | -0.249562825 | 0.423034563 | 0.94 |
| YNL180C | 2386.54101 | 0.310660611 | 0.416625277 | 0.94 |
| YNL327W | 5207.566398 | -0.318566088 | 0.415002055 | 0.94 |
| YNR070W | 728.5997471 | 0.26947217 | 0.420744402 | 0.94 |
| YOL037C | 71.18780393 | 0.327117482 | 0.426744928 | 0.94 |
| YOL114C | 133.5426332 | -0.2907609 | 0.415390977 | 0.94 |
| YOR273C | 1760.091485 | -0.329254859 | 0.422963155 | 0.94 |
| YOR277C | 6.042620409 | -0.248920309 | 0.422137429 | 0.94 |
| YOR301W | 501.5102259 | -0.29543677 | 0.41977417 | 0.94 |
| YOR336W | 1255.740018 | 0.185451474 | 0.414125942 | 0.94 |
| YPL020C | 1208.520388 | -0.131784201 | 0.421693305 | 0.94 |
| YPL033C | 16.58126869 | 0.402572638 | 0.423595566 | 0.94 |

| | | | | |
|---|---|---|---|---|
| YPL209C | 162.5627938 | 0.256327809 | 0.41548131 | 0.94 |
| YPL216W | 730.7653507 | 0.186760862 | 0.423967411 | 0.94 |
| YPR101W | 539.0655902 | -0.303629841 | 0.417311616 | 0.94 |
| snR70 | 64.50681846 | 0.370184964 | 0.417608901 | 0.94 |
| YER044C | 1096.772117 | -0.232467172 | 0.429432024 | 0.94 |
| YGR091W | 441.73971 | 0.219860399 | 0.428604075 | 0.94 |
| YGR212W | 361.1538773 | -0.168469187 | 0.429028711 | 0.94 |
| YGR278W | 442.5587334 | -0.179290651 | 0.427772848 | 0.94 |
| YIL163C | 43.51788641 | -0.318720834 | 0.428564243 | 0.94 |
| YJL002C | 4648.101511 | -0.226186481 | 0.429525967 | 0.94 |
| YJL193W | 438.7222894 | -0.201143222 | 0.428409997 | 0.94 |
| YKL133C | 601.5672696 | 0.297670384 | 0.429154895 | 0.94 |
| YLR363W-A | 269.4630453 | -0.253618972 | 0.42886662 | 0.94 |
| YLR454W | 3660.437031 | 0.210064769 | 0.429329736 | 0.94 |
| YNL083W | 670.395816 | 0.288599837 | 0.428600396 | 0.94 |
| YNL287W | 4413.325689 | -0.123886235 | 0.429017824 | 0.94 |
| YOL036W | 1888.124512 | 0.309578008 | 0.428019732 | 0.94 |
| YPL077C | 211.278667 | -0.198469701 | 0.428575668 | 0.94 |
| YPR069C | 2243.3628 | -0.297317103 | 0.428284647 | 0.94 |
| YPL230W | 2214.598115 | 0.392189296 | 0.429722717 | 0.94 |
| YOR167C | 4868.170026 | -0.273198639 | 0.429958351 | 0.94 |
| YJL100W | 1049.188324 | 0.213104286 | 0.430131931 | 0.94 |
| YOL013W-B | 4.383672181 | -0.459517679 | 0.430453776 | 0.94 |
| YPL197C | 174.3224877 | -0.363656899 | 0.430566945 | 0.94 |
| YPR109W | 650.4688515 | -0.203036149 | 0.43068262 | 0.94 |
| YJL052W | 23370.62399 | 0.474983224 | 0.431342356 | 0.94 |
| YOR314W-A | 0.578252409 | 0.221511653 | 0.431329728 | 0.94 |
| YGR233C | 1147.708039 | -0.255761457 | 0.432014804 | 0.94 |
| YIL006W | 374.9591791 | -0.398047404 | 0.431937427 | 0.94 |
| YMR028W | 753.5245359 | -0.129218788 | 0.431801798 | 0.94 |
| YOL079W | 9.64455915 | -0.451311484 | 0.431834822 | 0.94 |
| YLR005W | 967.4119109 | 0.155324685 | 0.432293073 | 0.94 |
| YGR043C | 1676.63942 | 0.499187795 | 0.432891101 | 0.94 |
| YGR131W | 100.6509022 | 0.302488476 | 0.433004458 | 0.94 |
| YMR139W | 1446.676083 | 0.265869098 | 0.433019086 | 0.94 |
| YCR024C-B | 2025.586967 | -0.238999561 | 0.433674821 | 0.94 |
| YDR414C | 448.7530762 | -0.284416998 | 0.433513813 | 0.94 |
| YKL102C | 18.56354003 | 0.415828609 | 0.433745163 | 0.94 |
| YNL063W | 522.2571621 | -0.165121044 | 0.433404665 | 0.94 |
| YNL067W-A | 0.709210999 | 0.242182553 | 0.433965659 | 0.94 |
| YNR064C | 215.5475844 | 0.299992299 | 0.433850703 | 0.94 |
| YLR257W | 6468.084537 | 0.314859594 | 0.434219321 | 0.94 |
| YMR037C | 1272.708272 | -0.27461483 | 0.434401353 | 0.94 |

| | | | | |
|---|---|---|---|---|
| YER045C | 814.7735686 | 0.251805323 | 0.434716235 | 0.94 |
| YLR206W | 3965.73873 | 0.158622476 | 0.434594492 | 0.94 |
| YDR144C | 842.4657292 | -0.336596957 | 0.435418939 | 0.94 |
| YDR279W | 575.5796481 | 0.182655014 | 0.435669448 | 0.94 |
| YIL036W | 2051.229049 | 0.218505213 | 0.43550429 | 0.94 |
| YIR011C | 1166.213259 | 0.182089423 | 0.435647484 | 0.94 |
| YLR327C | 4824.371416 | 0.495890534 | 0.435011767 | 0.94 |
| YNR056C | 522.7352776 | 0.337928071 | 0.435241559 | 0.94 |
| YPL133C | 580.4116075 | 0.231109364 | 0.435869798 | 0.94 |
| YIL102C | 13.70446277 | 0.446445142 | 0.436544299 | 0.94 |
| YJR040W | 752.4605261 | -0.18881756 | 0.436571106 | 0.94 |
| YKL196C | 2255.642273 | 0.176954549 | 0.436264505 | 0.94 |
| YOL067C | 283.6975055 | 0.229856435 | 0.436594331 | 0.94 |
| YAL047W-A | 1.181246705 | 0.094539626 | 0.437231788 | 0.94 |
| YHR133C | 977.6515666 | -0.286295024 | 0.436998071 | 0.94 |
| YLR364W | 131.7044529 | -0.268384977 | 0.436852397 | 0.94 |
| YPL002C | 492.3419627 | -0.159515948 | 0.437242536 | 0.94 |
| YPL250C | 749.2294185 | 0.350066306 | 0.437352654 | 0.94 |
| YAL045C | 8.89025023 | -0.522013881 | 0.441134687 | 0.94 |
| YBL003C | 4258.595501 | -0.223307687 | 0.443268893 | 0.94 |
| YBL012C | 41.44037077 | -0.342270411 | 0.445437895 | 0.94 |
| YBL104C | 644.4185242 | -0.139673477 | 0.44176345 | 0.94 |
| YCL065W | 1.135078196 | 0.080923874 | 0.442431161 | 0.94 |
| YCR081W | 992.4265821 | 0.156217448 | 0.440728365 | 0.94 |
| YCR108C | 1.912873177 | 0.067593839 | 0.441239784 | 0.94 |
| YDL219W | 330.3585764 | -0.223223083 | 0.442694587 | 0.94 |
| YDR189W | 2199.458913 | -0.178388877 | 0.446045605 | 0.94 |
| YDR194W-A | 0.453289942 | 0.195999246 | 0.445472757 | 0.94 |
| YDR237W | 832.9477874 | -0.189157316 | 0.445473982 | 0.94 |
| YDR307W | 749.860008 | -0.167078215 | 0.442740757 | 0.94 |
| YDR419W | 525.8031882 | 0.213979037 | 0.44187611 | 0.94 |
| YDR468C | 586.9211769 | -0.19251259 | 0.4437111 | 0.94 |
| YDR485C | 1410.914463 | 0.215585427 | 0.445111256 | 0.94 |
| YEL028W | 113.7795172 | 0.217873488 | 0.44040304 | 0.94 |
| YEL061C | 1022.169501 | -0.248504317 | 0.442134766 | 0.94 |
| YER058W | 286.3082478 | -0.201913523 | 0.439983757 | 0.94 |
| YER060W-A | 584.2480367 | 0.263094256 | 0.445581903 | 0.94 |
| YGL010W | 399.2167766 | 0.279898528 | 0.444802334 | 0.94 |
| YGL083W | 940.5046339 | 0.162512388 | 0.441161791 | 0.94 |
| YGL160W | 1159.891272 | -0.145141839 | 0.445817392 | 0.94 |
| YGL195W | 6234.838225 | -0.215128567 | 0.438957522 | 0.94 |
| YGL210W | 815.3164128 | -0.165931312 | 0.442480937 | 0.94 |
| YGR033C | 1013.438168 | 0.14859619 | 0.442656831 | 0.94 |

| | | | | |
|---|---|---|---|---|
| YGR071C | 709.1837053 | -0.180127633 | 0.443310779 | 0.94 |
| YGR146C | 1344.611896 | 0.305156832 | 0.44546635 | 0.94 |
| YGR161W-C | 303.4318056 | -0.303879925 | 0.44425381 | 0.94 |
| YHL044W | 369.968591 | -0.373368431 | 0.442277123 | 0.94 |
| YHL046W-A | 47.85481112 | 0.398565367 | 0.442722693 | 0.94 |
| YIL007C | 541.263808 | 0.229985323 | 0.441119959 | 0.94 |
| YIL028W | 59.74955175 | 0.287978505 | 0.439336564 | 0.94 |
| YIL089W | 155.6273305 | 0.355159025 | 0.440486594 | 0.94 |
| YIR030C | 90.29914152 | 0.368144191 | 0.441707834 | 0.94 |
| YJL004C | 587.5114449 | -0.159250025 | 0.443452472 | 0.94 |
| YJL024C | 668.6768051 | -0.150850795 | 0.445044508 | 0.94 |
| YJL186W | 1797.326564 | 0.171785215 | 0.439028459 | 0.94 |
| YJR001W | 1763.85175 | 0.152425884 | 0.444309037 | 0.94 |
| YKL060C | 152874.8661 | 0.383258587 | 0.439065805 | 0.94 |
| YKL068W-A | 172.0547206 | 0.302058432 | 0.438995613 | 0.94 |
| YKR095W-A | 172.6357846 | -0.261578282 | 0.438892658 | 0.94 |
| YLR020C | 866.2619741 | -0.203097912 | 0.445147781 | 0.94 |
| YLR024C | 1454.81679 | 0.213782167 | 0.445865947 | 0.94 |
| YLR085C | 664.5290334 | 0.131868042 | 0.441678254 | 0.94 |
| YLR094C | 1048.349221 | 0.210237587 | 0.445800161 | 0.94 |
| YLR250W | 1395.702651 | 0.197683568 | 0.443105344 | 0.94 |
| YLR301W | 1464.12998 | -0.201369571 | 0.442167766 | 0.94 |
| YML087C | 330.7889367 | 0.384693097 | 0.441941958 | 0.94 |
| YMR073C | 579.5582083 | 0.225589121 | 0.445902983 | 0.94 |
| YNL253W | 477.8813502 | -0.114897504 | 0.44130312 | 0.94 |
| YNL293W | 734.6319527 | 0.240045865 | 0.444581689 | 0.94 |
| YNR018W | 1764.18714 | 0.281286239 | 0.442303524 | 0.94 |
| YOR279C | 315.7050157 | -0.236136478 | 0.441610799 | 0.94 |
| YOR359W | 584.0345426 | -0.336191324 | 0.443598052 | 0.94 |
| YPL244C | 876.1384791 | -0.206560344 | 0.443493937 | 0.94 |
| YPR041W | 3343.881383 | -0.221849001 | 0.439150346 | 0.94 |
| snR37 | 153.9250937 | 0.485071099 | 0.438761168 | 0.94 |
| snR39 | 11.68210812 | -0.423339236 | 0.438761989 | 0.94 |
| YBR103W | 849.3179685 | 0.176193585 | 0.447477029 | 0.94 |
| YDR250C | 6.490017442 | 0.348899359 | 0.448159932 | 0.94 |
| YDR287W | 832.7406735 | 0.272078522 | 0.449292413 | 0.94 |
| YDR359C | 1438.122379 | 0.193560742 | 0.447137449 | 0.94 |
| YDR515W | 694.9239372 | -0.248630113 | 0.449710136 | 0.94 |
| YFR004W | 2781.166019 | -0.145191647 | 0.449614847 | 0.94 |
| YFR027W | 221.6537022 | 0.19606958 | 0.447979039 | 0.94 |
| YGL221C | 1756.035958 | 0.206983688 | 0.449358592 | 0.94 |
| YGR104C | 540.5492389 | 0.119090485 | 0.448932583 | 0.94 |
| YGR178C | 3642.360299 | 0.169832712 | 0.447106518 | 0.94 |

| | | | | |
|---|---|---|---|---|
| YIL068W-A | 5.981971104 | 0.350259247 | 0.447972551 | 0.94 |
| YJR015W | 1958.669387 | -0.187417784 | 0.448913013 | 0.94 |
| YLR197W | 4160.668892 | -0.251949954 | 0.449639531 | 0.94 |
| YLR256W-A | 92.61969338 | -0.305112336 | 0.449694554 | 0.94 |
| YMR117C | 204.4706004 | 0.229823664 | 0.449015815 | 0.94 |
| YMR293C | 413.7964048 | 0.193525824 | 0.449767618 | 0.94 |
| YNL278W | 749.3245241 | 0.170596271 | 0.447720251 | 0.94 |
| YOR011W-A | 3.164669454 | 0.377140226 | 0.448128826 | 0.94 |
| YOR234C | 2998.326335 | -0.265559467 | 0.449421815 | 0.94 |
| YPL127C | 2242.895143 | 0.228868452 | 0.447008892 | 0.94 |
| YPL143W | 740.5431297 | -0.250637807 | 0.446703934 | 0.94 |
| YPR005C | 426.8209175 | 0.240032377 | 0.447574529 | 0.94 |
| YPR192W | 327.8980894 | -0.374087941 | 0.449633882 | 0.94 |
| YDR352W | 489.6613604 | 0.27436428 | 0.450123278 | 0.94 |
| YHR058C | 391.0666567 | 0.190138568 | 0.450161192 | 0.94 |
| YAL061W | 2752.646997 | -0.416019746 | 0.451610661 | 0.94 |
| YAL063C-A | 8.265236292 | 0.257536136 | 0.452783638 | 0.94 |
| YBL091C | 4543.955358 | -0.121395328 | 0.450845096 | 0.94 |
| YDR255C | 870.1597032 | 0.249977588 | 0.452054698 | 0.94 |
| YER075C | 845.204575 | -0.167649919 | 0.452134162 | 0.94 |
| YGL023C | 1541.23519 | 0.182763593 | 0.451789278 | 0.94 |
| YGL138C | 22.53163453 | -0.491346853 | 0.452288687 | 0.94 |
| YGL159W | 407.2189918 | -0.238235509 | 0.452194376 | 0.94 |
| YGR291C | 1.566792632 | 0.469096494 | 0.452441247 | 0.94 |
| YJL009W | 66.31151028 | -0.234111643 | 0.453202279 | 0.94 |
| YJR093C | 790.9035242 | -0.117260784 | 0.451155291 | 0.94 |
| YKR016W | 2416.547829 | 0.344066534 | 0.453074661 | 0.94 |
| YLL006W-A | 2.814822079 | 0.36806537 | 0.450645444 | 0.94 |
| YLL027W | 1273.25217 | 0.219710639 | 0.453131371 | 0.94 |
| YLR178C | 3271.315417 | 0.39891197 | 0.450962349 | 0.94 |
| YOR299W | 1060.38632 | -0.224627286 | 0.452549223 | 0.94 |
| YPL044C | 1.74310561 | 0.114057205 | 0.451561998 | 0.94 |
| YPL128C | 628.8955352 | -0.198624994 | 0.453267349 | 0.94 |
| YPR076W | 41.22373968 | 0.325084197 | 0.451195246 | 0.94 |
| YPR117W | 2597.357181 | 0.156429508 | 0.452912004 | 0.94 |
| YDR081C | 1262.178022 | -0.174310809 | 0.453955458 | 0.94 |
| YDR202C | 884.8533036 | 0.260084251 | 0.453762635 | 0.94 |
| YER182W | 1071.612284 | 0.297526978 | 0.454092122 | 0.94 |
| YNL312W | 897.7476795 | 0.210577461 | 0.453920408 | 0.94 |
| snR87 | 6.424473744 | 0.402873469 | 0.453583774 | 0.94 |
| YER169W | 1395.485471 | 0.188055565 | 0.454262135 | 0.94 |
| YLR458W | 2.467406493 | 0.284685154 | 0.45462771 | 0.94 |
| YFL016C | 3333.34381 | 0.232988582 | 0.454849077 | 0.94 |

| | | | | |
|---|---|---|---|---|
| YBR049C | 1277.509775 | -0.199108865 | 0.455219603 | 0.94 |
| YER123W | 1029.367215 | -0.164084547 | 0.45512799 | 0.94 |
| ICR1 | 346.589135 | 0.242179741 | 0.46160071 | 0.94 |
| RUF23 | 8.975541829 | 0.384223713 | 0.465860303 | 0.94 |
| YBL008W | 564.4162656 | 0.220072864 | 0.462901851 | 0.94 |
| YBL098W | 1032.627619 | 0.431641527 | 0.459887406 | 0.94 |
| YBR035C | 2483.374886 | 0.181062777 | 0.46044881 | 0.94 |
| YBR213W | 228.053134 | 0.20075147 | 0.461493443 | 0.94 |
| YDL008W | 381.8332185 | -0.218086537 | 0.465941436 | 0.94 |
| YDL061C | 2048.312712 | -0.247035473 | 0.460955292 | 0.94 |
| YDL185C-A | 100.139643 | 0.229421265 | 0.462611817 | 0.94 |
| YDL241W | 250.4921648 | 0.378113755 | 0.45592724 | 0.94 |
| YDR013W | 271.9851003 | -0.215035119 | 0.46569997 | 0.94 |
| YDR031W | 342.8572195 | 0.309150469 | 0.46052647 | 0.94 |
| YDR049W | 886.770398 | 0.107408885 | 0.465683678 | 0.94 |
| YDR069C | 1204.898386 | 0.195477444 | 0.464885872 | 0.94 |
| YDR094W | 4.116771726 | -0.448223056 | 0.461379697 | 0.94 |
| YDR162C | 515.4142839 | -0.154863511 | 0.461355608 | 0.94 |
| YDR174W | 3004.282379 | -0.149319539 | 0.465958018 | 0.94 |
| YDR182W-A | 24.64411227 | 0.214283497 | 0.46072262 | 0.94 |
| YDR365C | 4849.022737 | 0.203306177 | 0.465814719 | 0.94 |
| YEL039C | 1783.57122 | 0.402232832 | 0.464761548 | 0.94 |
| YER032W | 1018.880889 | -0.245311271 | 0.457264623 | 0.94 |
| YER098W | 930.5704049 | 0.22647382 | 0.457369112 | 0.94 |
| YER158C | 1028.126375 | -0.302441887 | 0.461328678 | 0.94 |
| YFL019C | 43.77288423 | 0.401935122 | 0.460822714 | 0.94 |
| YGL150C | 2988.328671 | 0.126319677 | 0.455914997 | 0.94 |
| YGL243W | 434.8625344 | -0.107700631 | 0.463888788 | 0.94 |
| YGR162W | 4487.780728 | -0.239999654 | 0.460549528 | 0.94 |
| YGR202C | 1624.722457 | -0.138702004 | 0.465565406 | 0.94 |
| YGR210C | 1863.079648 | 0.239927275 | 0.461857552 | 0.94 |
| YGR258C | 1398.526849 | 0.179380622 | 0.464135871 | 0.94 |
| YHR022C-A | 3.937421936 | -0.41502085 | 0.458931022 | 0.94 |
| YHR062C | 561.4502383 | 0.259426833 | 0.458482185 | 0.94 |
| YHR139C | 204.3743283 | 0.296977745 | 0.459686472 | 0.94 |
| YIL141W | 35.62085203 | 0.328506165 | 0.463972213 | 0.94 |
| YJL104W | 974.9529404 | -0.189995497 | 0.459835264 | 0.94 |
| YJL183W | 732.387825 | -0.212277035 | 0.458046258 | 0.94 |
| YJR134C | 1708.030399 | -0.195234736 | 0.456378337 | 0.94 |
| YKL084W | 479.3447424 | 0.256824079 | 0.462099719 | 0.94 |
| YKL205W | 1310.174229 | -0.235055923 | 0.463611014 | 0.94 |
| YLR055C | 1018.744824 | -0.23712588 | 0.459920049 | 0.94 |
| YLR112W | 96.84022753 | -0.325905207 | 0.456773035 | 0.94 |

| | | | | |
|---|---|---|---|---|
| YLR145W | 262.3006502 | -0.176473816 | 0.455640729 | 0.94 |
| YLR213C | 172.0132438 | -0.282898841 | 0.464416691 | 0.94 |
| YLR369W | 1148.442381 | 0.247985146 | 0.45907092 | 0.94 |
| YLR402W | 27.81687827 | -0.270295902 | 0.464684051 | 0.94 |
| YLR410W-B | 66.3723325 | 0.261593685 | 0.459228984 | 0.94 |
| YLR460C | 187.4228264 | 0.251591883 | 0.465775483 | 0.94 |
| YML027W | 523.9520569 | -0.337461221 | 0.462860496 | 0.94 |
| YMR112C | 344.1352482 | -0.19803089 | 0.459044967 | 0.94 |
| YMR136W | 2477.221542 | 0.308111919 | 0.461534354 | 0.94 |
| YMR196W | 3241.463398 | 0.339781403 | 0.456922235 | 0.94 |
| YMR215W | 1997.531105 | -0.268408177 | 0.460384703 | 0.94 |
| YMR316W | 898.4662527 | -0.344344936 | 0.462065215 | 0.94 |
| YNL032W | 670.5607595 | -0.162729359 | 0.461270291 | 0.94 |
| YNL053W | 544.6626736 | -0.166287521 | 0.465795827 | 0.94 |
| YNL300W | 1681.600577 | -0.363233438 | 0.459922959 | 0.94 |
| YNR012W | 597.0846222 | 0.248586018 | 0.466014869 | 0.94 |
| YOL130W | 1532.382846 | 0.145927684 | 0.462430988 | 0.94 |
| YOR097C | 623.6386998 | -0.181223691 | 0.463839942 | 0.94 |
| YOR125C | 859.4168859 | 0.25629836 | 0.456847869 | 0.94 |
| YOR134W | 351.2595047 | 0.362654765 | 0.463632589 | 0.94 |
| YOR280C | 559.6900755 | -0.198674426 | 0.462809716 | 0.94 |
| YOR309C | 407.1266153 | -0.39450985 | 0.45758592 | 0.94 |
| YPL166W | 366.5892241 | 0.293249721 | 0.456295602 | 0.94 |
| YPL186C | 1114.795126 | 0.397468757 | 0.463720524 | 0.94 |
| YPL194W | 451.5443271 | 0.156596148 | 0.46056487 | 0.94 |
| YPL217C | 2396.710824 | 0.223571726 | 0.464327503 | 0.94 |
| YPL273W | 2617.65817 | -0.270779261 | 0.458394733 | 0.94 |
| YPR073C | 916.4024804 | 0.195517129 | 0.459534933 | 0.94 |
| YPR141C | 623.3342075 | -0.228679078 | 0.462780631 | 0.94 |
| YPR163C | 5059.100182 | -0.22493434 | 0.461169951 | 0.94 |
| YPR195C | 54.58354851 | -0.358590814 | 0.461226477 | 0.94 |
| snR11 | 261.7635615 | 0.306073474 | 0.458338554 | 0.94 |
| snR79 | 6.23839996 | 0.191367198 | 0.456824113 | 0.94 |
| YOR306C | 1215.578289 | -0.343291151 | 0.466167982 | 0.94 |
| YER053C-A | 179.222376 | -0.350733129 | 0.467146891 | 0.94 |
| YJR003C | 586.7430541 | -0.254413501 | 0.467284152 | 0.94 |
| YLR359W | 9224.21068 | 0.464089418 | 0.467126789 | 0.94 |
| YMR193C-A | 123.8016037 | -0.332689527 | 0.466609623 | 0.94 |
| YOR213C | 459.655666 | 0.181713946 | 0.466809612 | 0.94 |
| YPR048W | 424.7867171 | 0.255671925 | 0.467011677 | 0.94 |
| YDL180W | 735.4223348 | 0.179273022 | 0.46751761 | 0.94 |
| YGR204W | 10459.91176 | 0.210834136 | 0.467617925 | 0.94 |
| YAL010C | 552.0969494 | -0.137600087 | 0.471797969 | 0.94 |

| | | | | |
|---|---|---|---|---|
| YBR128C | 336.3131952 | 0.259003308 | 0.47064104 | 0.94 |
| YBR176W | 422.8364361 | 0.201061282 | 0.470851984 | 0.94 |
| YDL221W | 36.90750404 | 0.292089202 | 0.470500391 | 0.94 |
| YDL242W | 58.94192665 | 0.293801644 | 0.471432846 | 0.94 |
| YDR233C | 7397.691198 | -0.214640649 | 0.469739774 | 0.94 |
| YDR457W | 5588.916733 | 0.176679188 | 0.47214438 | 0.94 |
| YDR464W | 2373.668914 | 0.244531631 | 0.470234074 | 0.94 |
| YEL025C | 1231.625482 | -0.190809998 | 0.471005526 | 0.94 |
| YEL032W | 1185.048516 | -0.146631401 | 0.469202708 | 0.94 |
| YER152W-A | 37.9606133 | 0.26829204 | 0.468792518 | 0.94 |
| YER154W | 1666.340326 | -0.132512852 | 0.471589584 | 0.94 |
| YGL222C | 1086.796722 | -0.369267299 | 0.47019796 | 0.94 |
| YHR076W | 1124.266555 | -0.17112415 | 0.469231046 | 0.94 |
| YIL073C | 530.2219401 | 0.140241289 | 0.471558669 | 0.94 |
| YIL087C | 960.1717362 | 0.338780894 | 0.470295735 | 0.94 |
| YJL080C | 5735.124683 | 0.351533767 | 0.468815171 | 0.94 |
| YKL005C | 1108.015733 | 0.129262987 | 0.470423702 | 0.94 |
| YKL096W-A | 39924.89359 | -0.332363667 | 0.469307084 | 0.94 |
| YKR052C | 558.5950233 | -0.263617593 | 0.469541583 | 0.94 |
| YLL040C | 5253.851757 | 0.268081665 | 0.468366256 | 0.94 |
| YLR006C | 1262.134466 | 0.207482305 | 0.471268089 | 0.94 |
| YLR175W | 4387.69014 | -0.21404055 | 0.472205564 | 0.94 |
| YLR287C-A | 5234.314487 | -0.239245187 | 0.469563643 | 0.94 |
| YMR076C | 1386.678863 | -0.235494873 | 0.472044332 | 0.94 |
| YMR173W-A | 260.3952806 | -0.302402023 | 0.46884036 | 0.94 |
| YNL078W | 1260.908163 | -0.240215319 | 0.468290519 | 0.94 |
| YOR058C | 1144.395841 | -0.194253249 | 0.46896791 | 0.94 |
| YOR099W | 2318.668874 | -0.20878957 | 0.472114291 | 0.94 |
| YOR385W | 901.596547 | -0.296776808 | 0.467990725 | 0.94 |
| YPR115W | 2825.752527 | 0.23942102 | 0.470688563 | 0.94 |
| YAL027W | 394.2818606 | 0.147357827 | 0.476473851 | 0.94 |
| YBR014C | 1010.158039 | 0.16044922 | 0.476257915 | 0.94 |
| YBR029C | 1375.442278 | -0.188771155 | 0.472547933 | 0.94 |
| YBR118W | 10015.00281 | 0.3467835 | 0.474255458 | 0.94 |
| YDR089W | 994.7212521 | -0.208172387 | 0.476384345 | 0.94 |
| YDR221W | 936.1837264 | 0.148836768 | 0.476113857 | 0.94 |
| YER056C | 2408.22934 | 0.268544758 | 0.476226105 | 0.94 |
| YER094C | 2268.208556 | 0.157883635 | 0.474798648 | 0.94 |
| YGR085C | 1835.900279 | -0.263349147 | 0.473737588 | 0.94 |
| YHR009C | 3439.835058 | -0.163045692 | 0.473560815 | 0.94 |
| YHR071W | 3724.97766 | -0.370358517 | 0.475868029 | 0.94 |
| YIL142W | 3130.184557 | -0.175184685 | 0.473555703 | 0.94 |
| YIR016W | 1147.104809 | -0.264273556 | 0.474756122 | 0.94 |

| | | | | |
|---|---|---|---|---|
| YJL079C | 9145.850933 | -0.342761347 | 0.473781067 | 0.94 |
| YKL051W | 1194.148337 | -0.270968267 | 0.473834044 | 0.94 |
| YKR041W | 32.67705338 | -0.273487565 | 0.476116112 | 0.94 |
| YLR099C | 1018.600053 | -0.274471224 | 0.475421785 | 0.94 |
| YLR406C | 2407.447144 | -0.220734523 | 0.473732692 | 0.94 |
| YML107C | 389.7043417 | -0.203122596 | 0.474442933 | 0.94 |
| YMR257C | 582.2395624 | -0.246255526 | 0.476450858 | 0.94 |
| YMR318C | 2412.871237 | -0.205175493 | 0.476510995 | 0.94 |
| YOL053W | 775.6100204 | 0.270693536 | 0.475903894 | 0.94 |
| YOL091W | 152.6270626 | -0.279955486 | 0.472655731 | 0.94 |
| YOL137W | 547.8586694 | 0.131585212 | 0.475618294 | 0.94 |
| YOL154W | 197.7584736 | 0.181344258 | 0.473992615 | 0.94 |
| YOR013W | 86.58484926 | -0.232362265 | 0.475468984 | 0.94 |
| YOR206W | 2506.527701 | -0.234237055 | 0.473515897 | 0.94 |
| YPR142C | 12.52047692 | -0.429489398 | 0.473528434 | 0.94 |
| snR60 | 11.58332067 | 0.137994649 | 0.476219188 | 0.94 |
| YCL054W | 1476.642208 | -0.263174616 | 0.478685989 | 0.94 |
| YCR043C | 438.4206899 | -0.219016702 | 0.479454301 | 0.94 |
| YCR095W-A | 20.07196858 | 0.258245684 | 0.478059426 | 0.94 |
| YDL031W | 1680.703139 | 0.288822158 | 0.476978466 | 0.94 |
| YDL053C | 1309.962591 | 0.1684243 | 0.480232224 | 0.94 |
| YDL111C | 757.760422 | -0.182498846 | 0.477759371 | 0.94 |
| YDL113C | 1161.019898 | 0.203175643 | 0.477550461 | 0.94 |
| YDL178W | 1397.749906 | 0.127125764 | 0.480181695 | 0.94 |
| YDR115W | 505.144241 | -0.158108253 | 0.479226361 | 0.94 |
| YDR150W | 4371.450407 | 0.275772164 | 0.477476611 | 0.94 |
| YDR159W | 1698.043068 | -0.139554556 | 0.480143887 | 0.94 |
| YDR246W | 916.4165129 | -0.134402768 | 0.478876953 | 0.94 |
| YDR469W | 524.7044195 | -0.19491611 | 0.478233425 | 0.94 |
| YFR048W | 703.4879681 | -0.195341958 | 0.476918071 | 0.94 |
| YGL035C | 1849.649908 | -0.282059266 | 0.479928686 | 0.94 |
| YLL019C | 1900.382246 | 0.292202243 | 0.480212774 | 0.94 |
| YNL096C | 3035.774305 | -0.209185598 | 0.479328359 | 0.94 |
| YNL213C | 587.1355028 | -0.287825886 | 0.477671373 | 0.94 |
| YNR041C | 337.6551047 | 0.222769223 | 0.4779642 | 0.94 |
| YOL023W | 636.5704075 | -0.208377681 | 0.477040519 | 0.94 |
| YOR104W | 252.8227861 | 0.185910589 | 0.478181949 | 0.94 |
| YOR151C | 4196.260493 | -0.193053127 | 0.480083394 | 0.94 |
| YPL064C | 883.912951 | -0.119361272 | 0.47892082 | 0.94 |
| YPL087W | 1694.761164 | 0.244198777 | 0.479338796 | 0.94 |
| snR30 | 275.4218461 | 0.448022737 | 0.478895314 | 0.94 |
| YAL025C | 983.4578298 | -0.250600061 | 0.480611133 | 0.94 |
| YNL088W | 2851.433881 | 0.181345621 | 0.480667743 | 0.94 |

| | | | | |
|---|---|---|---|---|
| YOR065W | 3707.159237 | 0.428010136 | 0.480957581 | 0.94 |
| YER187W | 132.2577032 | -0.383000167 | 0.48127675 | 0.95 |
| YOR316C | 1558.064909 | -0.222716439 | 0.481414619 | 0.95 |
| YOR166C | 666.3506234 | -0.125397441 | 0.481750446 | 0.95 |
| YAL065C | 13.48444795 | 0.402887178 | 0.48210154 | 0.95 |
| YDR406W-A | 12.68467301 | -0.424944318 | 0.481934115 | 0.95 |
| YNL301C | 1146.542523 | -0.137629063 | 0.482220842 | 0.95 |
| YHR109W | 446.2718996 | 0.182050574 | 0.482723557 | 0.95 |
| YGL131C | 1060.332301 | 0.229930085 | 0.483438996 | 0.95 |
| YGR042W | 764.6829972 | -0.209968585 | 0.48291194 | 0.95 |
| YHR049W | 2093.457795 | -0.20552864 | 0.483866734 | 0.95 |
| YJL042W | 3416.022292 | 0.184150594 | 0.483099164 | 0.95 |
| YJR032W | 585.2468444 | -0.21635573 | 0.484295545 | 0.95 |
| YKL018W | 496.5039326 | -0.145326013 | 0.484280644 | 0.95 |
| YKL132C | 214.3347933 | 0.198288565 | 0.48375411 | 0.95 |
| YLR264C-A | 117.4733387 | -0.294981842 | 0.484346212 | 0.95 |
| YNL076W | 1058.446405 | -0.146056145 | 0.484310207 | 0.95 |
| YPL096W | 660.7866918 | -0.149320921 | 0.484296284 | 0.95 |
| YPR104C | 970.815797 | -0.153916028 | 0.483213623 | 0.95 |
| YDL165W | 697.5260403 | 0.127697331 | 0.484707701 | 0.95 |
| YNL215W | 1103.384408 | 0.199499024 | 0.484626615 | 0.95 |
| YAL012W | 6420.264953 | -0.246406041 | 0.485196197 | 0.95 |
| YHL050C | 143.7644195 | 0.308971121 | 0.48525577 | 0.95 |
| YML123C | 2594.845401 | -0.287582773 | 0.485282358 | 0.95 |
| YDL012C | 999.0124699 | -0.202040277 | 0.486082235 | 0.95 |
| YNL135C | 5166.62026 | -0.194545211 | 0.486097302 | 0.95 |
| YAR064W | 14.8431874 | 0.280703122 | 0.487007443 | 0.95 |
| YMR203W | 3020.852994 | -0.129019351 | 0.486890175 | 0.95 |
| YMR251W | 257.8128007 | 0.349564188 | 0.48689921 | 0.95 |
| YLR103C | 462.3096359 | -0.20569073 | 0.487218396 | 0.95 |
| YOR290C | 3907.230484 | -0.166311417 | 0.487357312 | 0.95 |
| YDR505C | 1755.081331 | 0.225848417 | 0.488070261 | 0.95 |
| YFL041W-A | 23.17885335 | 0.304854406 | 0.488374795 | 0.95 |
| YGR098C | 1335.545417 | -0.191757291 | 0.487771574 | 0.95 |
| YJR115W | 567.164948 | -0.406570001 | 0.488217685 | 0.95 |
| YKL053C-A | 473.2870725 | -0.206803909 | 0.488023684 | 0.95 |
| YPL152W-A | 1.958413515 | -0.399624336 | 0.488395326 | 0.95 |
| YDL220C | 416.2502353 | 0.277112525 | 0.489031569 | 0.95 |
| YDR214W | 2305.632085 | 0.297351462 | 0.48895043 | 0.95 |
| YLR193C | 745.5545212 | 0.21800678 | 0.488719639 | 0.95 |
| YNR075W | 186.6264191 | 0.298522547 | 0.488833618 | 0.95 |
| YAL037C-A | 1.195205434 | 0.320837302 | 0.493372801 | 0.95 |
| YBL037W | 1072.022099 | -0.149881267 | 0.49439806 | 0.95 |

| | | | | |
|---|---|---|---|---|
| YBL091C-A | 508.1746239 | -0.205512309 | 0.493266658 | 0.95 |
| YBR264C | 359.5103772 | -0.138281065 | 0.494356211 | 0.95 |
| YBR275C | 1768.794249 | 0.191944454 | 0.489823646 | 0.95 |
| YDL034W | 29.64111763 | 0.286297958 | 0.493155239 | 0.95 |
| YDL160C-A | 95.22130843 | -0.327425762 | 0.494313515 | 0.95 |
| YDR153C | 1161.100144 | -0.170880154 | 0.490257598 | 0.95 |
| YER033C | 1095.361309 | 0.248427028 | 0.492637107 | 0.95 |
| YER121W | 184.2483 | -0.351637574 | 0.491497062 | 0.95 |
| YFR032C-B | 19.14700684 | 0.304004636 | 0.493556361 | 0.95 |
| YGL207W | 4342.654723 | -0.160167981 | 0.494346728 | 0.95 |
| YGL238W | 2547.939909 | -0.146744194 | 0.492622643 | 0.95 |
| YGR116W | 3744.798928 | -0.153096805 | 0.493027063 | 0.95 |
| YHL015W-A | 4.315791202 | -0.36729032 | 0.494626856 | 0.95 |
| YHR032W | 1019.136565 | 0.276030867 | 0.491057205 | 0.95 |
| YHR045W | 983.5331682 | -0.220235095 | 0.490662911 | 0.95 |
| YHR173C | 88.05682976 | -0.292914124 | 0.493881256 | 0.95 |
| YHR188C | 2845.286709 | 0.207873319 | 0.493673062 | 0.95 |
| YJL139C | 556.1081958 | 0.245545192 | 0.493507684 | 0.95 |
| YJR062C | 945.4060688 | -0.160744571 | 0.494062984 | 0.95 |
| YJR136C | 423.3917741 | 0.113856895 | 0.491609688 | 0.95 |
| YKL028W | 1511.073698 | -0.179141039 | 0.489415298 | 0.95 |
| YKR022C | 681.4384392 | -0.148582552 | 0.493164592 | 0.95 |
| YLL029W | 3427.507563 | 0.2242393 | 0.494510541 | 0.95 |
| YLR014C | 353.4875684 | 0.193821741 | 0.492256825 | 0.95 |
| YLR016C | 414.1257784 | 0.152080992 | 0.493658262 | 0.95 |
| YLR288C | 449.0939426 | 0.110081032 | 0.491027926 | 0.95 |
| YML035C | 2451.342109 | 0.200510519 | 0.49257656 | 0.95 |
| YMR001C-A | 16.08451551 | -0.168039705 | 0.490160216 | 0.95 |
| YMR048W | 313.0835867 | -0.098410331 | 0.49195922 | 0.95 |
| YMR155W | 707.9678984 | 0.199300898 | 0.492002726 | 0.95 |
| YNL200C | 2258.279724 | 0.338738767 | 0.492544252 | 0.95 |
| YOR030W | 929.3153858 | -0.211429842 | 0.493932668 | 0.95 |
| YOR320C | 1070.658594 | -0.156079359 | 0.493700856 | 0.95 |
| YOR361C | 4132.636454 | -0.186395374 | 0.490503289 | 0.95 |
| YPL080C | 11.15479793 | -0.336677841 | 0.493989717 | 0.95 |
| YPR116W | 103.378319 | 0.292578434 | 0.490420377 | 0.95 |
| YBR126W-A | 695.852559 | -0.298103447 | 0.494782625 | 0.95 |
| SRG1 | 1084.659985 | -0.353368804 | 0.508240817 | 0.95 |
| YAL009W | 404.8562545 | -0.134155924 | 0.513274952 | 0.95 |
| YBL053W | 19.02715806 | 0.085897111 | 0.512381274 | 0.95 |
| YBL063W | 882.9945164 | 0.172342447 | 0.496480698 | 0.95 |
| YBR122C | 1316.133892 | 0.188939539 | 0.506339572 | 0.95 |
| YBR184W | 226.9887473 | 0.263577639 | 0.513230453 | 0.95 |

| | | | | |
|---|---|---|---|---|
| YBR295W | 678.020174 | 0.217014225 | 0.505778116 | 0.95 |
| YBR298C | 1312.084765 | 0.369496013 | 0.497225204 | 0.95 |
| YCL014W | 1364.401777 | -0.188072636 | 0.512645064 | 0.95 |
| YCL021W-A | 277.7575158 | 0.22563017 | 0.510930335 | 0.95 |
| YDL004W | 2874.441994 | 0.352042949 | 0.503382206 | 0.95 |
| YDL009C | 106.1081082 | -0.166580455 | 0.509476166 | 0.95 |
| YDL057W | 396.4937822 | 0.265310398 | 0.500701683 | 0.95 |
| YDL144C | 951.4980585 | 0.177288196 | 0.505845754 | 0.95 |
| YDL191W | 1342.250708 | -0.227732801 | 0.508177639 | 0.95 |
| YDR025W | 5331.251911 | -0.245184364 | 0.510754375 | 0.95 |
| YDR098C | 1711.271644 | 0.135169621 | 0.510002544 | 0.95 |
| YDR240C | 401.4217535 | 0.122910758 | 0.501807036 | 0.95 |
| YDR286C | 156.0831168 | -0.176868897 | 0.51271045 | 0.95 |
| YDR340W | 4.60165433 | -0.439161284 | 0.505581557 | 0.95 |
| YDR350C | 1056.007335 | 0.176449463 | 0.503921129 | 0.95 |
| YDR361C | 744.7914399 | 0.343627831 | 0.503054918 | 0.95 |
| YDR375C | 907.9536977 | 0.121794295 | 0.51208089 | 0.95 |
| YDR387C | 888.6103683 | 0.181819502 | 0.497977248 | 0.95 |
| YDR482C | 359.2978323 | -0.127614276 | 0.505106322 | 0.95 |
| YDR508C | 2566.963439 | -0.29332958 | 0.506668951 | 0.95 |
| YDR540C | 110.2480797 | 0.320102736 | 0.495439684 | 0.95 |
| YEL048C | 96.72112286 | -0.208985024 | 0.511162475 | 0.95 |
| YER056C-A | 4887.303568 | -0.265757995 | 0.501128046 | 0.95 |
| YER131W | 9395.024759 | -0.271265798 | 0.496907896 | 0.95 |
| YER186C | 678.5275475 | -0.200602671 | 0.50634351 | 0.95 |
| YER188W | 461.4895098 | -0.331825137 | 0.508119242 | 0.95 |
| YFL034W | 949.1269789 | -0.119847727 | 0.5118127 | 0.95 |
| YFR028C | 1133.072845 | -0.137796164 | 0.504352251 | 0.95 |
| YFR040W | 3140.971961 | 0.249868228 | 0.51071924 | 0.95 |
| YFR053C | 31755.34613 | 0.395426211 | 0.506519835 | 0.95 |
| YGL012W | 2257.199778 | -0.210495583 | 0.511184051 | 0.95 |
| YGL088W | 0.432313012 | 0.240203494 | 0.500487328 | 0.95 |
| YGL113W | 616.1660547 | 0.157361941 | 0.501736966 | 0.95 |
| YGL157W | 1718.848717 | -0.298350081 | 0.513029645 | 0.95 |
| YGR020C | 3100.566353 | -0.197447306 | 0.512961763 | 0.95 |
| YGR099W | 582.6782539 | 0.179713357 | 0.510050074 | 0.95 |
| YGR122W | 624.9209214 | -0.207042295 | 0.498220585 | 0.95 |
| YGR130C | 4694.98638 | 0.220459875 | 0.511212665 | 0.95 |
| YGR132C | 2816.110199 | 0.183430374 | 0.496546395 | 0.95 |
| YGR217W | 881.6554263 | 0.189151656 | 0.49611661 | 0.95 |
| YHR078W | 1137.159604 | 0.215449 | 0.507934273 | 0.95 |
| YHR131C | 966.4131265 | -0.213488854 | 0.504033318 | 0.95 |
| YHR158C | 1689.928736 | -0.169712539 | 0.502597568 | 0.95 |

| | | | | |
|---|---|---|---|---|
| YHR214C-E | 1.813646714 | 0.179299021 | 0.5122383 | 0.95 |
| YIL010W | 1043.363564 | 0.187460608 | 0.501677364 | 0.95 |
| YIL068C | 981.7929067 | -0.09274302 | 0.511479583 | 0.95 |
| YIL103W | 637.74726 | 0.209184615 | 0.497741871 | 0.95 |
| YIL109C | 4372.511551 | -0.116608638 | 0.511063938 | 0.95 |
| YIL115W-A | 23.70554908 | -0.274331187 | 0.506086823 | 0.95 |
| YIL167W | 108.465152 | 0.222229966 | 0.508073097 | 0.95 |
| YIR009W | 172.3613335 | -0.192430603 | 0.50567676 | 0.95 |
| YJL022W | 2.660856745 | 0.374955004 | 0.508858603 | 0.95 |
| YJL031C | 712.299831 | -0.170496223 | 0.497670979 | 0.95 |
| YJL115W | 931.9259287 | -0.196721407 | 0.504973318 | 0.95 |
| YJL122W | 643.9700151 | -0.250576314 | 0.500355696 | 0.95 |
| YJL162C | 484.7743599 | -0.14338324 | 0.509001701 | 0.95 |
| YJR002W | 1689.001321 | 0.224633775 | 0.495980828 | 0.95 |
| YJR005W | 1677.202028 | -0.151358989 | 0.504999028 | 0.95 |
| YJR025C | 1368.606225 | -0.348186356 | 0.498173382 | 0.95 |
| YJR030C | 708.7385852 | 0.179115183 | 0.507912335 | 0.95 |
| YJR104C | 12449.61979 | 0.192485198 | 0.511885727 | 0.95 |
| YJR156C | 47.49998049 | 0.360915992 | 0.509499129 | 0.95 |
| YKL004W | 2100.85667 | 0.211469367 | 0.502261156 | 0.95 |
| YKL069W | 336.0140703 | -0.164290685 | 0.511697999 | 0.95 |
| YKL086W | 218.7788586 | 0.254651014 | 0.49952219 | 0.95 |
| YKL180W | 12036.25542 | -0.253058227 | 0.50243895 | 0.95 |
| YKL207W | 2006.977142 | -0.154297681 | 0.503650294 | 0.95 |
| YKL211C | 1647.230083 | 0.315519349 | 0.501219212 | 0.95 |
| YKR010C | 516.9970849 | -0.196747499 | 0.503338694 | 0.95 |
| YKR013W | 4305.106637 | -0.252828489 | 0.504982113 | 0.95 |
| YKR047W | 143.8726915 | 0.147758814 | 0.510510714 | 0.95 |
| YKR069W | 646.4230382 | 0.180529358 | 0.5108977 | 0.95 |
| YLR021W | 410.9321978 | -0.129974332 | 0.508431546 | 0.95 |
| YLR023C | 1130.122098 | 0.140676567 | 0.500301633 | 0.95 |
| YLR029C | 12797.37208 | -0.207023886 | 0.50110809 | 0.95 |
| YLR032W | 823.8658356 | 0.174852531 | 0.50446949 | 0.95 |
| YLR051C | 783.2029443 | 0.245352523 | 0.511921799 | 0.95 |
| YLR067C | 640.6258307 | -0.159717531 | 0.501689636 | 0.95 |
| YLR170C | 116.6208147 | -0.20813783 | 0.497688878 | 0.95 |
| YLR194C | 2445.708189 | 0.298550654 | 0.50129635 | 0.95 |
| YLR203C | 2343.4384 | 0.146983426 | 0.510139181 | 0.95 |
| YLR285W | 889.692098 | -0.193624295 | 0.51003976 | 0.95 |
| YLR356W | 2015.024039 | 0.266990747 | 0.507948628 | 0.95 |
| YLR411W | 132.197405 | -0.229322396 | 0.511345824 | 0.95 |
| YML002W | 853.1962719 | 0.155003434 | 0.512888018 | 0.95 |
| YML024W | 8930.152891 | -0.211293318 | 0.505383115 | 0.95 |

| | | | | |
|---|---|---|---|---|
| YML081W | 1706.956949 | -0.208832482 | 0.499119842 | 0.95 |
| YML116W-A | 166.4577303 | -0.193872935 | 0.499959713 | 0.95 |
| YMR052W | 219.8800281 | 0.231426707 | 0.500138829 | 0.95 |
| YMR072W | 3717.086972 | -0.129265939 | 0.503771896 | 0.95 |
| YMR202W | 2009.003075 | -0.189455301 | 0.503219327 | 0.95 |
| YMR302C | 2126.727676 | 0.219593553 | 0.503415157 | 0.95 |
| YMR320W | 113.0593369 | 0.241422084 | 0.502569245 | 0.95 |
| YNL017C | 1.615564791 | 0.326720155 | 0.505415289 | 0.95 |
| YNL039W | 1798.324816 | -0.147565795 | 0.506933108 | 0.95 |
| YNL047C | 720.3103825 | -0.219668877 | 0.498925861 | 0.95 |
| YNL090W | 89.81803356 | -0.220032691 | 0.51014118 | 0.95 |
| YNL162W-A | 54.12294867 | 0.325883445 | 0.508897538 | 0.95 |
| YNL243W | 3599.331124 | -0.122967627 | 0.499732302 | 0.95 |
| YNL259C | 262.1284302 | -0.207021618 | 0.505386645 | 0.95 |
| YNL286W | 504.2093629 | 0.20399215 | 0.502834076 | 0.95 |
| YNL309W | 553.1266537 | 0.159088685 | 0.512671542 | 0.95 |
| YOL031C | 1089.984363 | -0.274831542 | 0.512562641 | 0.95 |
| YOL149W | 642.6006688 | -0.187761129 | 0.509408402 | 0.95 |
| YOL164W-A | 24.89009234 | 0.295837834 | 0.496600462 | 0.95 |
| YOR078W | 697.5265362 | -0.212354801 | 0.508742378 | 0.95 |
| YOR086C | 3224.270398 | 0.138206983 | 0.505755345 | 0.95 |
| YOR284W | 207.2496443 | -0.296906481 | 0.509719175 | 0.95 |
| YPL073C | 1.14139142 | -0.207081097 | 0.506904189 | 0.95 |
| YPL118W | 2044.672532 | -0.169772434 | 0.495285118 | 0.95 |
| YPL160W | 4849.742262 | 0.286742653 | 0.50076833 | 0.95 |
| YPL192C | 128.9945052 | 0.226982907 | 0.500458745 | 0.95 |
| YPL225W | 2696.542417 | -0.144446262 | 0.504425667 | 0.95 |
| YPL241C | 283.7799123 | -0.220993687 | 0.504055079 | 0.95 |
| YPR011C | 560.157333 | 0.243455528 | 0.504628928 | 0.95 |
| YPR014C | 81.20498688 | -0.26310924 | 0.500713093 | 0.95 |
| YPR042C | 1338.934548 | 0.168586251 | 0.511993851 | 0.95 |
| YPR046W | 225.601515 | -0.213067572 | 0.510288226 | 0.95 |
| YPR119W | 682.8278436 | -0.281712022 | 0.510476434 | 0.95 |
| snR19 | 122.3490857 | -0.272217766 | 0.508290939 | 0.95 |
| YBR062C | 819.8607412 | 0.19164217 | 0.514894336 | 0.95 |
| YDR449C | 532.8954421 | -0.224204456 | 0.515309172 | 0.95 |
| YGR007W | 1236.749371 | -0.123129346 | 0.515188202 | 0.95 |
| YGR249W | 613.9099062 | 0.374697964 | 0.514093573 | 0.95 |
| YHR081W | 846.2639082 | 0.197173373 | 0.51555102 | 0.95 |
| YIR026C | 772.7201823 | 0.368159259 | 0.514800752 | 0.95 |
| YJL135W | 5.319281498 | -0.416503929 | 0.51474494 | 0.95 |
| YLR239C | 753.9388732 | 0.164509882 | 0.514138399 | 0.95 |
| YMR258C | 733.537512 | -0.181598708 | 0.513592785 | 0.95 |

| | | | | |
|---|---|---|---|---|
| YMR295C | 1636.767669 | -0.110397894 | 0.515419022 | 0.95 |
| YOR074C | 848.568103 | -0.176718636 | 0.514664568 | 0.95 |
| YOR141C | 1550.614472 | 0.144762538 | 0.513731609 | 0.95 |
| YOR216C | 1165.7555 | -0.215693455 | 0.514360421 | 0.95 |
| YOR244W | 816.0935801 | 0.162772647 | 0.515015848 | 0.95 |
| YPR080W | 5311.40989 | 0.397917361 | 0.515234681 | 0.95 |
| YML073C | 8568.253747 | -0.249497066 | 0.515764562 | 0.95 |
| YCL039W | 1768.904925 | 0.21095949 | 0.515927394 | 0.95 |
| YLR395C | 1418.063046 | 0.340115135 | 0.516087453 | 0.95 |
| YPR102C | 3154.540675 | -0.221925396 | 0.516409706 | 0.95 |
| YAL016C-B | 15.45888685 | -0.269168219 | 0.522572352 | 0.95 |
| YAL026C-A | 170.5922701 | -0.251045586 | 0.524416091 | 0.95 |
| YBL081W | 976.0547384 | -0.292109902 | 0.522721771 | 0.95 |
| YBL101C | 1814.259973 | 0.176396556 | 0.522747412 | 0.95 |
| YBR201W | 456.7810734 | -0.15719747 | 0.525164735 | 0.95 |
| YBR269C | 1256.661555 | 0.249289838 | 0.519278579 | 0.95 |
| YCL004W | 588.1019431 | -0.170776146 | 0.520007565 | 0.95 |
| YCR059C | 1130.721478 | 0.172721195 | 0.523943969 | 0.95 |
| YDL014W | 4306.735257 | -0.224503548 | 0.520038 | 0.95 |
| YDL052C | 1575.894255 | -0.217773446 | 0.52232797 | 0.95 |
| YDL055C | 11798.61054 | -0.15289169 | 0.523466867 | 0.95 |
| YDR163W | 579.7588355 | -0.16242526 | 0.526332356 | 0.95 |
| YDR188W | 2974.432832 | -0.12803524 | 0.521461529 | 0.95 |
| YDR282C | 383.6210181 | 0.213676975 | 0.517979452 | 0.95 |
| YEL026W | 1802.083858 | -0.219047475 | 0.521962535 | 0.95 |
| YEL054C | 3213.78471 | -0.240837952 | 0.517811967 | 0.95 |
| YER144C | 1434.76784 | 0.165404833 | 0.525992285 | 0.95 |
| YFL008W | 1609.672053 | -0.198689629 | 0.527267396 | 0.95 |
| YFL023W | 930.5339775 | 0.224569155 | 0.52260116 | 0.95 |
| YFR021W | 746.0129726 | 0.105068009 | 0.51879363 | 0.95 |
| YGL119W | 689.6648811 | 0.126617691 | 0.517158578 | 0.95 |
| YGL174W | 430.9199773 | -0.138226915 | 0.526711193 | 0.95 |
| YGR290W | 27.11482068 | -0.312323846 | 0.521169935 | 0.95 |
| YHR003C | 820.5090626 | 0.172100465 | 0.523308298 | 0.95 |
| YHR021C | 5458.86145 | -0.24515015 | 0.523893295 | 0.95 |
| YHR073W | 1977.66425 | -0.166230852 | 0.523336805 | 0.95 |
| YHR102W | 1800.525655 | 0.128069081 | 0.522273295 | 0.95 |
| YHR131W-A | 19.588359 | -0.28937318 | 0.522641537 | 0.95 |
| YIL025C | 3.736035911 | 0.34592469 | 0.526405988 | 0.95 |
| YIL034C | 3381.997418 | -0.099913611 | 0.518126114 | 0.95 |
| YIL035C | 1690.415511 | -0.119662278 | 0.525638338 | 0.95 |
| YIL055C | 666.1215044 | 0.217874275 | 0.524834575 | 0.95 |
| YIL106W | 893.3105346 | -0.138139869 | 0.521316465 | 0.95 |

| | | | | |
|---|---|---|---|---|
| YIL116W | 2678.75766 | 0.261916562 | 0.518090568 | 0.95 |
| YIL149C | 3262.267006 | -0.226546251 | 0.522258779 | 0.95 |
| YIR008C | 729.6399623 | -0.119727654 | 0.522490588 | 0.95 |
| YIR017C | 6.4006803 | -0.302261156 | 0.5271387 | 0.95 |
| YJL019W | 605.7582629 | 0.176657655 | 0.52144826 | 0.95 |
| YJL086C | 9.129343542 | -0.280174385 | 0.526955941 | 0.95 |
| YKL015W | 1286.742474 | 0.236752796 | 0.517371209 | 0.95 |
| YKL098W | 807.0315194 | 0.172508629 | 0.517209671 | 0.95 |
| YKL182W | 10979.47839 | -0.147034123 | 0.519243172 | 0.95 |
| YKR094C | 5698.150477 | -0.226902817 | 0.517143025 | 0.95 |
| YKR101W | 347.6961828 | 0.152580596 | 0.520529829 | 0.95 |
| YLL039C | 7097.91074 | 0.249151464 | 0.526050062 | 0.95 |
| YLR002C | 907.6259007 | -0.187215106 | 0.517484756 | 0.95 |
| YLR043C | 5741.406917 | -0.16569864 | 0.523846923 | 0.95 |
| YLR110C | 106387.3494 | -0.259059937 | 0.523878273 | 0.95 |
| YLR223C | 2252.47739 | -0.273032976 | 0.519201781 | 0.95 |
| YLR237W | 1061.722449 | 0.279756023 | 0.527123641 | 0.95 |
| YLR344W | 3310.960859 | -0.221109181 | 0.523187915 | 0.95 |
| YLR346C | 131.9806369 | 0.312673096 | 0.522596667 | 0.95 |
| YLR404W | 245.17757 | -0.175306235 | 0.526011922 | 0.95 |
| YLR437C | 329.593239 | -0.184660022 | 0.521968381 | 0.95 |
| YMR012W | 7425.571889 | -0.150186772 | 0.522697405 | 0.95 |
| YMR066W | 755.7979974 | -0.185897761 | 0.526664697 | 0.95 |
| YMR272W-B | 3.438828838 | -0.069655032 | 0.525588775 | 0.95 |
| YMR287C | 795.4088763 | -0.140900373 | 0.526479041 | 0.95 |
| YNL331C | 347.7391832 | 0.203411568 | 0.524602826 | 0.95 |
| YOL073C | 1132.754224 | 0.226539421 | 0.519372152 | 0.95 |
| YOL113W | 1261.758159 | 0.247974621 | 0.52475236 | 0.95 |
| YOL116W | 722.658756 | 0.154376643 | 0.526165124 | 0.95 |
| YOR032W-A | 11.5403846 | 0.2253627 | 0.526533268 | 0.95 |
| YOR111W | 390.0644929 | 0.15669486 | 0.522459779 | 0.95 |
| YOR228C | 505.0466268 | 0.254864141 | 0.51873493 | 0.95 |
| YOR231W | 881.0751627 | 0.166903802 | 0.526969345 | 0.95 |
| YOR304C-A | 201.0708852 | -0.190128858 | 0.517164491 | 0.95 |
| YPL145C | 2994.997329 | -0.180112075 | 0.520800818 | 0.95 |
| YPR129W | 1545.384555 | 0.159911204 | 0.525590777 | 0.95 |
| YPR173C | 1842.223717 | -0.168194577 | 0.524013607 | 0.95 |
| YPR175W | 704.7198488 | 0.22039515 | 0.519121924 | 0.95 |
| snR18 | 14.34494847 | -0.248642385 | 0.520915299 | 0.95 |
| snR75 | 1.574604411 | 0.322081744 | 0.522862389 | 0.95 |
| YBR093C | 949.3588178 | 0.331339293 | 0.528280847 | 0.95 |
| YDR245W | 1164.324708 | -0.173727703 | 0.528321097 | 0.95 |
| YDR368W | 2244.977391 | 0.197938748 | 0.528528434 | 0.95 |

| | | | | |
|---|---|---|---|---|
| YJL094C | 1236.455255 | 0.148228015 | 0.528315496 | 0.95 |
| YLR035C | 528.4128752 | 0.17146936 | 0.528738962 | 0.95 |
| YLR143W | 501.3258454 | -0.166298301 | 0.52899114 | 0.95 |
| YMR119W | 1094.314263 | -0.164222015 | 0.528114837 | 0.95 |
| YOR139C | 5.37072037 | 0.323495148 | 0.529047097 | 0.95 |
| YOR182C | 1767.616754 | -0.223570195 | 0.527779836 | 0.95 |
| YOR254C | 3232.98471 | -0.203186643 | 0.528470895 | 0.95 |
| YPR084W | 689.7868138 | -0.090148873 | 0.529076773 | 0.95 |
| YBL076C | 7280.098127 | 0.302284396 | 0.53033994 | 0.95 |
| YDR504C | 165.3464621 | -0.182615287 | 0.531688842 | 0.95 |
| YER091C-A | 0.485272411 | -0.209371018 | 0.531440569 | 0.95 |
| YER115C | 529.525795 | 0.172668798 | 0.531091239 | 0.95 |
| YFL015W-A | 1.143841678 | 0.214730119 | 0.532242334 | 0.95 |
| YGR036C | 221.213258 | -0.174865691 | 0.530328429 | 0.95 |
| YHL012W | 237.0696554 | 0.168265799 | 0.529644321 | 0.95 |
| YJR059W | 3492.702056 | -0.192893916 | 0.530701855 | 0.95 |
| YJR152W | 334.6737416 | 0.244071962 | 0.531143106 | 0.95 |
| YLR071C | 1557.164849 | -0.165122928 | 0.531554403 | 0.95 |
| YLR187W | 889.7815093 | 0.124003071 | 0.529692992 | 0.95 |
| YLR378C | 3906.764345 | -0.155897559 | 0.530687817 | 0.95 |
| YMR193W | 776.035786 | 0.225174717 | 0.531215424 | 0.95 |
| YMR194C-B | 137.2124333 | 0.324767173 | 0.532352835 | 0.95 |
| YNL123W | 2060.867856 | -0.203853128 | 0.532266537 | 0.95 |
| YNL194C | 937.042565 | 0.36868825 | 0.52974887 | 0.95 |
| YNL316C | 479.8242776 | -0.144876479 | 0.530715619 | 0.95 |
| YNL319W | 5.59444275 | 0.057789923 | 0.529900609 | 0.95 |
| YOR147W | 1293.879256 | 0.24519047 | 0.530957403 | 0.95 |
| YPL243W | 2085.694288 | -0.176385877 | 0.529932146 | 0.95 |
| YPR136C | 12.04995248 | 0.292974773 | 0.532076718 | 0.95 |
| snR17b | 27.76593566 | 0.388802966 | 0.532334078 | 0.95 |
| YCR072C | 412.6699919 | -0.259118534 | 0.532831499 | 0.95 |
| YDR160W | 879.6911624 | 0.105343461 | 0.532750572 | 0.95 |
| YDL083C | 6219.514303 | -0.143522832 | 0.533588783 | 0.95 |
| YER172C | 2101.126018 | 0.098225824 | 0.533468637 | 0.95 |
| YJR013W | 645.2124988 | -0.109941118 | 0.533516195 | 0.95 |
| YMR061W | 1751.530821 | -0.137905983 | 0.533057844 | 0.95 |
| YOR369C | 15497.1337 | -0.166964101 | 0.533232017 | 0.95 |
| YML079W | 1250.560699 | -0.208430909 | 0.533771751 | 0.95 |
| YBR124W | 16.74512171 | 0.223038754 | 0.53450909 | 0.95 |
| YDR384C | 561.1890906 | 0.421220966 | 0.534535781 | 0.95 |
| YFL038C | 5600.505944 | -0.121373101 | 0.534707421 | 0.95 |
| YHL035C | 3064.413674 | 0.256115519 | 0.534098544 | 0.95 |
| YMR291W | 2090.324495 | -0.327607166 | 0.534774018 | 0.95 |

| | | | | |
|---|---|---|---|---|
| YOR350C | 702.3901449 | 0.173197802 | 0.534460226 | 0.95 |
| YCR091W | 1488.499833 | 0.279583873 | 0.53548589 | 0.95 |
| YER089C | 1927.936186 | -0.227359865 | 0.535633337 | 0.95 |
| YFL051C | 87.84562174 | 0.310794765 | 0.535718664 | 0.95 |
| YOR129C | 585.8837465 | -0.115640684 | 0.535273835 | 0.95 |
| YPL114W | 1.969765016 | 0.316394399 | 0.535755775 | 0.95 |
| YBR094W | 1036.817856 | 0.143102174 | 0.537119112 | 0.95 |
| YCR013C | 2806.322738 | 0.289354005 | 0.536924897 | 0.95 |
| YDR425W | 1030.779588 | 0.168677573 | 0.535994819 | 0.95 |
| YDR516C | 6053.645479 | 0.313474089 | 0.536986803 | 0.95 |
| YLR097C | 748.1202732 | 0.158647743 | 0.536531421 | 0.95 |
| YML094C-A | 15.34009962 | 0.227121075 | 0.53702563 | 0.95 |
| YNL033W | 5.453513622 | 0.360681058 | 0.537107386 | 0.95 |
| YOR169C | 39.59953477 | -0.328751854 | 0.537147318 | 0.95 |
| YOR363C | 1212.557429 | 0.163789429 | 0.537073246 | 0.95 |
| YBR240C | 118.1647641 | 0.273235382 | 0.539162524 | 0.95 |
| YDL146W | 626.6868048 | 0.257784875 | 0.53966141 | 0.95 |
| YDR032C | 4060.720659 | -0.184038425 | 0.540125507 | 0.95 |
| YDR429C | 3118.563913 | -0.150699206 | 0.538707895 | 0.95 |
| YER021W | 5518.8989 | 0.161533965 | 0.539396365 | 0.95 |
| YGR117C | 888.1364455 | -0.15293018 | 0.538185627 | 0.95 |
| YGR118W | 4349.282861 | -0.174536783 | 0.540277551 | 0.95 |
| YHL033C | 4723.149428 | -0.151092286 | 0.539141993 | 0.95 |
| YIL071C | 381.3119395 | 0.122032419 | 0.538533269 | 0.95 |
| YJL077C | 162.6090211 | -0.211186249 | 0.539427611 | 0.95 |
| YJR130C | 1173.075557 | 0.229193729 | 0.539700223 | 0.95 |
| YKL019W | 1109.764572 | -0.129421118 | 0.538640909 | 0.95 |
| YKL220C | 368.1832027 | 0.221796375 | 0.539734737 | 0.95 |
| YKR087C | 773.1699867 | 0.141489873 | 0.538226575 | 0.95 |
| YLR198C | 85.68816582 | -0.288696362 | 0.539314572 | 0.95 |
| YML057W | 1931.217961 | -0.130128014 | 0.540177971 | 0.95 |
| YML094W | 513.2141562 | -0.154318008 | 0.540320828 | 0.95 |
| YML106W | 2946.051592 | -0.193738618 | 0.538775571 | 0.95 |
| YMR054W | 1855.817154 | -0.154715395 | 0.53813587 | 0.95 |
| YNL336W | 467.9927857 | 0.286051094 | 0.539013769 | 0.95 |
| YDR327W | 614.5550695 | -0.124417092 | 0.540917551 | 0.95 |
| YLR162W-A | 8.884728875 | -0.21992413 | 0.54157675 | 0.95 |
| YPL183W-A | 354.0512636 | -0.14463725 | 0.541903627 | 0.95 |
| YIL157C | 795.6414589 | -0.187872595 | 0.542277813 | 0.96 |
| YLR215C | 433.3667117 | -0.119843354 | 0.542732232 | 0.96 |
| YBR058C-A | 528.5056217 | -0.144678697 | 0.543545518 | 0.96 |
| YGL188C-A | 18.77666255 | -0.357421023 | 0.54351845 | 0.96 |
| snR17a | 59.62016603 | 0.368842895 | 0.54356422 | 0.96 |

| | | | | |
|---|---|---|---|---|
| YBR005W | 1310.131508 | -0.255539068 | 0.543787325 | 0.96 |
| YER103W | 3269.463849 | 0.301990608 | 0.544057441 | 0.96 |
| YER176W | 1918.703807 | 0.157665683 | 0.544069923 | 0.96 |
| YDR118W | 391.9679351 | -0.09250354 | 0.544298579 | 0.96 |
| YAL058W | 573.3781549 | 0.156823907 | 0.551136771 | 0.96 |
| YBL056W | 3616.602261 | 0.101296098 | 0.553497973 | 0.96 |
| YBL087C | 3378.344673 | -0.164459627 | 0.552510237 | 0.96 |
| YBR020W | 310.6456705 | 0.236997118 | 0.545556239 | 0.96 |
| YBR041W | 1541.15066 | -0.145974647 | 0.547158518 | 0.96 |
| YBR053C | 1693.976608 | 0.236301231 | 0.554095667 | 0.96 |
| YCL063W | 606.1625178 | -0.187028817 | 0.548665789 | 0.96 |
| YCR007C | 318.2333147 | 0.17433584 | 0.548169319 | 0.96 |
| YDL130W | 4709.707857 | -0.186097883 | 0.548274285 | 0.96 |
| YDL236W | 1309.137891 | -0.163978362 | 0.5504125 | 0.96 |
| YDR028C | 3835.45418 | 0.146604123 | 0.555722758 | 0.96 |
| YDR038C | 125.3516014 | -0.178815108 | 0.554205068 | 0.96 |
| YDR244W | 923.2552739 | 0.12413574 | 0.551406611 | 0.96 |
| YDR275W | 363.6731352 | 0.247968261 | 0.555494527 | 0.96 |
| YDR285W | 510.8247622 | 0.205057259 | 0.54879888 | 0.96 |
| YDR330W | 1850.284876 | 0.200668956 | 0.552900521 | 0.96 |
| YDR489W | 358.8647918 | 0.154796009 | 0.553689932 | 0.96 |
| YER173W | 470.1702433 | -0.162851749 | 0.548834954 | 0.96 |
| YFL032W | 146.5638958 | -0.331935402 | 0.547618029 | 0.96 |
| YFR032C-A | 2429.038295 | -0.273221241 | 0.552210321 | 0.96 |
| YFR042W | 519.19235 | 0.168504137 | 0.54793918 | 0.96 |
| YGL040C | 1261.243607 | 0.193239187 | 0.546907379 | 0.96 |
| YGL090W | 485.8897575 | 0.130917375 | 0.55519611 | 0.96 |
| YGL094C | 1570.488055 | -0.156886598 | 0.553563043 | 0.96 |
| YGL176C | 596.0143813 | 0.113492552 | 0.549548887 | 0.96 |
| YGL215W | 3518.182479 | 0.181378463 | 0.550004697 | 0.96 |
| YGR209C | 5181.82162 | -0.1282434 | 0.553673671 | 0.96 |
| YGR250C | 3005.303955 | 0.192811584 | 0.548644143 | 0.96 |
| YGR275W | 659.6729657 | -0.1467854 | 0.545140682 | 0.96 |
| YHL005C | 24.92643688 | 0.193643906 | 0.546303703 | 0.96 |
| YIL014C-A | 253.291496 | -0.314164032 | 0.546371626 | 0.96 |
| YIL042C | 814.6115743 | 0.224985666 | 0.552514809 | 0.96 |
| YIL159W | 678.524918 | -0.152929414 | 0.549537686 | 0.96 |
| YIL168W | 92.19596636 | 0.215364809 | 0.55451533 | 0.96 |
| YJL072C | 353.6349072 | 0.185997385 | 0.551625565 | 0.96 |
| YJL164C | 1399.537088 | 0.1621993 | 0.553534133 | 0.96 |
| YJL197W | 1228.673265 | 0.109982422 | 0.553308142 | 0.96 |
| YJR063W | 331.564801 | -0.178336702 | 0.555423764 | 0.96 |
| YJR143C | 2572.640819 | 0.244125048 | 0.548827302 | 0.96 |

| | | | | |
|---|---|---|---|---|
| YKL080W | 5080.565803 | -0.143958743 | 0.551620007 | 0.96 |
| YKL210W | 8749.432253 | 0.128601177 | 0.554527955 | 0.96 |
| YLL060C | 550.1729711 | 0.242458769 | 0.547844722 | 0.96 |
| YLR064W | 1412.390781 | -0.14814376 | 0.548189931 | 0.96 |
| YLR104W | 635.3306236 | -0.189616255 | 0.550131551 | 0.96 |
| YLR185W | 4884.991766 | -0.204140207 | 0.545893155 | 0.96 |
| YLR251W | 493.6892026 | -0.204072217 | 0.555332264 | 0.96 |
| YLR264W | 525.6829336 | -0.082759612 | 0.551525741 | 0.96 |
| YLR447C | 5880.017489 | 0.155692695 | 0.549650293 | 0.96 |
| YML050W | 378.8334393 | 0.193401944 | 0.551020677 | 0.96 |
| YML101C | 335.3248863 | -0.167581816 | 0.555433622 | 0.96 |
| YML133C | 9.588165535 | 0.335219963 | 0.555854452 | 0.96 |
| YMR009W | 601.1334204 | 0.161293012 | 0.550396001 | 0.96 |
| YMR242C | 8203.831599 | -0.198715106 | 0.550109075 | 0.96 |
| YMR292W | 429.0322087 | -0.139677291 | 0.552169839 | 0.96 |
| YNL216W | 1187.634305 | -0.136564501 | 0.545410273 | 0.96 |
| YNR043W | 2170.763038 | 0.209616095 | 0.555447877 | 0.96 |
| YOL048C | 1220.015423 | 0.197778478 | 0.55032188 | 0.96 |
| YOL092W | 1398.16665 | 0.229214476 | 0.555882559 | 0.96 |
| YOL102C | 532.3526812 | -0.107049717 | 0.548010091 | 0.96 |
| YOR200W | 9.969440735 | -0.591190072 | 0.546092953 | 0.96 |
| YOR278W | 477.7903224 | -0.138086849 | 0.552997862 | 0.96 |
| YOR282W | 14.73409586 | -0.377879724 | 0.552971921 | 0.96 |
| YOR295W | 256.0382172 | -0.14009072 | 0.555811485 | 0.96 |
| YOR332W | 2877.284969 | -0.121034925 | 0.547736802 | 0.96 |
| YOR384W | 1087.084219 | 0.202994339 | 0.545189933 | 0.96 |
| YPL088W | 468.228703 | 0.22828759 | 0.549584476 | 0.96 |
| YPL137C | 1258.265301 | -0.210638795 | 0.545328114 | 0.96 |
| YPL151C | 670.1565001 | 0.09512109 | 0.553251393 | 0.96 |
| YPL180W | 1077.43255 | 0.097647786 | 0.550313679 | 0.96 |
| YPL224C | 949.2587858 | 0.193493619 | 0.54640053 | 0.96 |
| YPL239W | 1334.632299 | 0.248529115 | 0.548491669 | 0.96 |
| YPL272C | 344.5733609 | 0.173148357 | 0.552836642 | 0.96 |
| YPR030W | 2156.768509 | 0.331627024 | 0.547876032 | 0.96 |
| YPR033C | 3601.656962 | 0.218575925 | 0.550516478 | 0.96 |
| YPR103W | 3867.741987 | -0.143951365 | 0.552843078 | 0.96 |
| YPR156C | 1539.255789 | 0.209965594 | 0.554855148 | 0.96 |
| snR189 | 9.691874789 | 0.327537804 | 0.555001922 | 0.96 |
| YAR023C | 164.0948317 | -0.252482406 | 0.556563515 | 0.96 |
| YDL209C | 238.6569979 | -0.181044654 | 0.556202252 | 0.96 |
| YER083C | 1868.758699 | -0.102208868 | 0.557502896 | 0.96 |
| YHL039W | 1028.011858 | 0.221739755 | 0.556784826 | 0.96 |
| YJL030W | 384.5980723 | -0.136147639 | 0.55712204 | 0.96 |

| | | | | |
|---|---|---|---|---|
| YJL127C-B | 263.7714489 | -0.149155477 | 0.557328914 | 0.96 |
| YML097C | 708.2199714 | -0.140003078 | 0.557384144 | 0.96 |
| YNL026W | 973.5528021 | -0.092022352 | 0.557061785 | 0.96 |
| YNR049C | 561.0637018 | -0.116101103 | 0.55713944 | 0.96 |
| YOR271C | 1520.089436 | 0.348238486 | 0.556042446 | 0.96 |
| YPR099C | 24.60634511 | -0.356148873 | 0.557467937 | 0.96 |
| YLR418C | 850.5544851 | -0.115658659 | 0.557815925 | 0.96 |
| YNL028W | 6.323808974 | 0.174000887 | 0.557794762 | 0.96 |
| YDR538W | 133.1462476 | -0.136783985 | 0.558397182 | 0.96 |
| YGR240C-A | 46.68747277 | -0.296458889 | 0.558425277 | 0.96 |
| YOL103W-B | 37.1755698 | -0.394222088 | 0.558390902 | 0.96 |
| snR76 | 8.640963018 | -0.322958854 | 0.558345405 | 0.96 |
| YLR279W | 36.37996098 | 0.322026342 | 0.558904858 | 0.96 |
| YOL040C | 16073.93418 | -0.218898795 | 0.558909845 | 0.96 |
| YLR135W | 729.3825255 | 0.130802266 | 0.560211272 | 0.96 |
| YLR182W | 1421.620302 | -0.170036667 | 0.560122802 | 0.96 |
| snR9 | 75.73439823 | 0.254284709 | 0.560377635 | 0.96 |
| YDR156W | 897.9312179 | 0.163014976 | 0.56067742 | 0.96 |
| YER054C | 1167.402398 | 0.296082442 | 0.560960487 | 0.96 |
| YCR104W | 19.10661275 | 0.336926897 | 0.56142616 | 0.96 |
| YBR002C | 607.1395928 | -0.169450516 | 0.562357242 | 0.96 |
| YDL133C-A | 72.78703027 | -0.379356199 | 0.561875331 | 0.96 |
| YDR300C | 1204.265207 | -0.160299587 | 0.562715375 | 0.96 |
| YEL051W | 3229.319305 | -0.139994357 | 0.562325477 | 0.96 |
| YGL072C | 15.1439202 | 0.161801361 | 0.562534999 | 0.96 |
| YGR183C | 999.4364709 | 0.317200979 | 0.56244424 | 0.96 |
| YJL121C | 2601.821139 | -0.118073061 | 0.562803004 | 0.96 |
| YKR064W | 1141.244604 | -0.140407786 | 0.562153655 | 0.96 |
| YML004C | 3324.450509 | 0.176655698 | 0.562334092 | 0.96 |
| YJL144W | 785.7166274 | 0.259316748 | 0.563126094 | 0.96 |
| YLR435W | 440.7011003 | -0.198702631 | 0.563420047 | 0.96 |
| TLC1 | 1417.425959 | -0.212436095 | 0.597922542 | 0.96 |
| YAL023C | 4478.973811 | -0.131583612 | 0.600067949 | 0.96 |
| YAL034C-B | 9.086766243 | -0.290958496 | 0.57281806 | 0.96 |
| YAL034W-A | 187.40184 | -0.154732641 | 0.573462852 | 0.96 |
| YAL036C | 1045.610431 | 0.263594648 | 0.593454553 | 0.96 |
| YAR008W | 281.6693003 | -0.155954395 | 0.59084514 | 0.96 |
| YBL005W-B | 450.3577253 | 0.218058827 | 0.592264078 | 0.96 |
| YBL027W | 10124.8249 | -0.145584343 | 0.576592934 | 0.96 |
| YBL059C-A | 504.0749236 | 0.240617623 | 0.592182413 | 0.96 |
| YBL092W | 15431.27049 | -0.134880544 | 0.593314935 | 0.96 |
| YBR016W | 1677.943304 | -0.122983321 | 0.582831332 | 0.96 |
| YBR021W | 496.9097626 | -0.180243445 | 0.572878394 | 0.96 |

| | | | | |
|---|---|---|---|---|
| YBR034C | 1111.628509 | -0.160694738 | 0.595089726 | 0.96 |
| YBR113W | 115.7153994 | -0.159218458 | 0.585747082 | 0.96 |
| YBR116C | 35.41794033 | 0.314654425 | 0.588465194 | 0.96 |
| YBR142W | 1284.846746 | 0.235560289 | 0.589952752 | 0.96 |
| YBR152W | 271.4958287 | 0.10447298 | 0.58659536 | 0.96 |
| YBR197C | 458.4082334 | 0.178596589 | 0.576568022 | 0.96 |
| YBR214W | 5062.049562 | 0.216019863 | 0.58054525 | 0.96 |
| YBR229C | 1792.230472 | 0.098163262 | 0.580765853 | 0.96 |
| YBR277C | 4.283921599 | -0.47872204 | 0.56760154 | 0.96 |
| YCL024W | 493.5504233 | -0.154138595 | 0.598716503 | 0.96 |
| YCL047C | 434.4156263 | 0.183616093 | 0.56874768 | 0.96 |
| YCR008W | 1187.11323 | 0.182105297 | 0.585164894 | 0.96 |
| YCR051W | 1038.832374 | 0.20317541 | 0.574829317 | 0.96 |
| YCR092C | 872.6618833 | -0.124656819 | 0.587038777 | 0.96 |
| YCR105W | 210.4429239 | 0.261937182 | 0.59771579 | 0.96 |
| YCR107W | 236.9080811 | 0.276525713 | 0.575199842 | 0.96 |
| YDL007W | 5356.140616 | 0.141764715 | 0.582282342 | 0.96 |
| YDL050C | 34.4743764 | -0.287957799 | 0.5673304 | 0.96 |
| YDL060W | 1265.050122 | 0.219063592 | 0.596749404 | 0.96 |
| YDL099W | 2397.811912 | -0.138957421 | 0.576344126 | 0.96 |
| YDL108W | 550.3870487 | 0.09647028 | 0.588703082 | 0.96 |
| YDL120W | 712.7094342 | 0.143887525 | 0.57440762 | 0.96 |
| YDL135C | 948.2549942 | -0.134522254 | 0.579331924 | 0.96 |
| YDL145C | 6294.266606 | -0.117278259 | 0.591915673 | 0.96 |
| YDL172C | 10.58915184 | -0.233106371 | 0.594497895 | 0.96 |
| YDL176W | 672.0622412 | 0.12527117 | 0.599246658 | 0.96 |
| YDL187C | 3.68310749 | -0.378178109 | 0.595355622 | 0.96 |
| YDL195W | 7573.167842 | -0.13731303 | 0.591176879 | 0.96 |
| YDL211C | 175.1726895 | -0.207481307 | 0.57652489 | 0.96 |
| YDL212W | 1994.74877 | -0.114340514 | 0.580796359 | 0.96 |
| YDL213C | 1003.203847 | 0.314010762 | 0.591953855 | 0.96 |
| YDL237W | 1508.528864 | 0.163875428 | 0.595958902 | 0.96 |
| YDR024W | 7.360211428 | -0.288626294 | 0.579574567 | 0.96 |
| YDR027C | 1560.996774 | 0.067188115 | 0.582634116 | 0.96 |
| YDR037W | 9604.757078 | 0.353493861 | 0.571136834 | 0.96 |
| YDR065W | 277.2481422 | 0.154955154 | 0.586782768 | 0.96 |
| YDR078C | 147.0952169 | 0.206865086 | 0.582078207 | 0.96 |
| YDR101C | 1247.120699 | -0.17716404 | 0.600669812 | 0.96 |
| YDR110W | 523.1803163 | 0.169260418 | 0.58187427 | 0.96 |
| YDR112W | 22.56100124 | 0.329200595 | 0.57458683 | 0.96 |
| YDR176W | 1688.228212 | -0.128759867 | 0.590461215 | 0.96 |
| YDR177W | 1727.041157 | 0.153513164 | 0.585536814 | 0.96 |
| YDR190C | 2078.525809 | -0.144713752 | 0.576367392 | 0.96 |

| | | | | |
|---|---|---|---|---|
| YDR222W | 466.2824915 | 0.206513269 | 0.579018345 | 0.96 |
| YDR270W | 829.6044968 | -0.166633166 | 0.601101121 | 0.96 |
| YDR322C-A | 892.3853165 | 0.255546871 | 0.587110091 | 0.96 |
| YDR338C | 695.8448786 | 0.1369842 | 0.583855595 | 0.96 |
| YDR345C | 12971.54144 | 0.343565599 | 0.595223163 | 0.96 |
| YDR364C | 596.6987029 | 0.086273608 | 0.60058573 | 0.96 |
| YDR379C-A | 462.1957351 | -0.206729012 | 0.587338523 | 0.96 |
| YDR390C | 1683.184066 | -0.093367958 | 0.600023675 | 0.96 |
| YDR405W | 1203.736123 | -0.110580463 | 0.591252826 | 0.96 |
| YDR471W | 4845.236011 | -0.188626783 | 0.59136113 | 0.96 |
| YEL002C | 3293.830024 | -0.105317477 | 0.568182062 | 0.96 |
| YEL050W-A | 71.35641565 | -0.214113647 | 0.566680359 | 0.96 |
| YER010C | 1015.031336 | -0.113103529 | 0.581546475 | 0.96 |
| YER012W | 3318.681234 | -0.103024419 | 0.592162979 | 0.96 |
| YER059W | 910.2356733 | -0.157038268 | 0.597302788 | 0.96 |
| YER127W | 556.9285849 | 0.298276161 | 0.585797714 | 0.96 |
| YER139C | 313.6167797 | -0.124184463 | 0.594077275 | 0.96 |
| YER151C | 2013.443986 | -0.120436744 | 0.581257586 | 0.96 |
| YER184C | 397.7514223 | 0.153673274 | 0.57771235 | 0.96 |
| YFL041W | 1108.963199 | 0.151501217 | 0.580307074 | 0.96 |
| YFL065C | 1.354222162 | 0.249639152 | 0.592258781 | 0.96 |
| YFR019W | 2589.385757 | 0.122264554 | 0.593959255 | 0.96 |
| YFR024C-A | 2497.30533 | 0.137699439 | 0.570863402 | 0.96 |
| YFR032C | 47.36854765 | 0.141580873 | 0.586150092 | 0.96 |
| YFR047C | 2554.161875 | 0.260501589 | 0.589915366 | 0.96 |
| YGL013C | 1490.03588 | 0.121737243 | 0.587521094 | 0.96 |
| YGL044C | 659.773802 | 0.079263945 | 0.587923108 | 0.96 |
| YGL068W | 385.5514446 | -0.157931484 | 0.570057745 | 0.96 |
| YGL245W | 8715.959502 | 0.214195559 | 0.568427065 | 0.96 |
| YGL252C | 1693.94272 | -0.076455172 | 0.596885599 | 0.96 |
| YGR002C | 912.0190954 | -0.125770607 | 0.584852401 | 0.96 |
| YGR029W | 703.0611046 | -0.12968567 | 0.567916416 | 0.96 |
| YGR044C | 1106.237884 | -0.259684788 | 0.571934842 | 0.96 |
| YGR059W | 83.10299922 | 0.290527061 | 0.580732149 | 0.96 |
| YGR069W | 7.64600337 | -0.275053561 | 0.571745216 | 0.96 |
| YGR107W | 13.44440994 | 0.218757415 | 0.5994894 | 0.96 |
| YGR137W | 236.2341776 | 0.190596363 | 0.573041706 | 0.96 |
| YGR148C | 7673.362198 | -0.184599504 | 0.592734105 | 0.96 |
| YGR172C | 884.3306623 | -0.112344693 | 0.567681154 | 0.96 |
| YGR176W | 11.59700922 | -0.386022903 | 0.591204294 | 0.96 |
| YGR182C | 629.4422912 | 0.193035773 | 0.583929961 | 0.96 |
| YGR237C | 1848.787726 | 0.14513985 | 0.564813149 | 0.96 |
| YGR268C | 875.2811265 | -0.150241997 | 0.587167888 | 0.96 |

| | | | | |
|---|---|---|---|---|
| YGR292W | 46.2187461 | -0.280674691 | 0.583019556 | 0.96 |
| YHL007C | 2278.179389 | -0.153682634 | 0.565513147 | 0.96 |
| YHL010C | 843.3803368 | 0.154585279 | 0.595128692 | 0.96 |
| YHR001W-A | 1152.222923 | 0.279459852 | 0.590720086 | 0.96 |
| YHR013C | 1158.208101 | -0.150984109 | 0.600350354 | 0.96 |
| YHR051W | 2680.129991 | 0.313467992 | 0.593481774 | 0.96 |
| YHR075C | 548.2435732 | 0.124994996 | 0.572981113 | 0.96 |
| YHR082C | 4569.103984 | 0.173179684 | 0.572492329 | 0.96 |
| YHR089C | 2056.029582 | -0.141826912 | 0.5877927 | 0.96 |
| YHR116W | 348.981213 | 0.170225814 | 0.568329318 | 0.96 |
| YHR134W | 365.9132551 | -0.196831326 | 0.581076357 | 0.96 |
| YHR166C | 498.330332 | 0.14929368 | 0.600930717 | 0.96 |
| YIL011W | 777.5395764 | -0.255538068 | 0.597407336 | 0.96 |
| YIL016W | 483.5722155 | -0.146023686 | 0.587387377 | 0.96 |
| YIL022W | 2268.335855 | -0.073045209 | 0.583154675 | 0.96 |
| YIL030W-A | 74.85518125 | 0.135376628 | 0.568462848 | 0.96 |
| YIL044C | 893.1546159 | -0.143255325 | 0.566614217 | 0.96 |
| YIL066W-A | 3.308676015 | -0.227937349 | 0.577620821 | 0.96 |
| YIL069C | 2018.584495 | -0.169172298 | 0.566017413 | 0.96 |
| YIR015W | 351.8671415 | 0.157762871 | 0.590772872 | 0.96 |
| YIR020C-B | 5.673146071 | -0.347259598 | 0.582831902 | 0.96 |
| YIR024C | 366.6225047 | -0.118346946 | 0.582543054 | 0.96 |
| YJL038C | 194.4642451 | 0.228570644 | 0.578775229 | 0.96 |
| YJL064W | 3.253557565 | 0.072800232 | 0.590902922 | 0.96 |
| YJL083W | 610.9975272 | 0.137290207 | 0.582802215 | 0.96 |
| YJL128C | 1506.583064 | -0.106629855 | 0.567720175 | 0.96 |
| YJL147C | 286.6419753 | 0.110329601 | 0.584092611 | 0.96 |
| YJR035W | 878.6193365 | 0.157167018 | 0.566586752 | 0.96 |
| YJR068W | 1198.277592 | 0.155873806 | 0.591172436 | 0.96 |
| YJR069C | 644.3818806 | -0.147124482 | 0.56618919 | 0.96 |
| YJR075W | 2038.161429 | -0.126947419 | 0.584782132 | 0.96 |
| YJR076C | 1668.359247 | -0.090512709 | 0.583738532 | 0.96 |
| YJR089W | 953.5011996 | -0.126090046 | 0.581176033 | 0.96 |
| YJR119C | 341.555846 | -0.186223291 | 0.567676241 | 0.96 |
| YJR144W | 1299.865299 | 0.11407075 | 0.583737843 | 0.96 |
| YKL018C-A | 261.9017664 | 0.112928739 | 0.574577856 | 0.96 |
| YKL032C | 3133.578964 | 0.199060206 | 0.597335055 | 0.96 |
| YKL036C | 247.9148773 | 0.192549195 | 0.597481837 | 0.96 |
| YKL072W | 554.6822525 | 0.111588291 | 0.586347573 | 0.96 |
| YKL108W | 391.5450461 | -0.195243059 | 0.581902263 | 0.96 |
| YKL117W | 7112.851481 | -0.132923714 | 0.593442315 | 0.96 |
| YKL155C | 791.2689341 | 0.125286125 | 0.566763315 | 0.96 |
| YKL192C | 3222.400243 | -0.101049231 | 0.593884228 | 0.96 |

| | | | | |
|---|---|---|---|---|
| YKR038C | 1110.135443 | -0.107577463 | 0.586647664 | 0.96 |
| YKR051W | 666.9246654 | -0.132542551 | 0.598579971 | 0.96 |
| YKR057W | 2516.350053 | -0.105390467 | 0.577043255 | 0.96 |
| YKR062W | 891.4921763 | 0.09489631 | 0.596262893 | 0.96 |
| YLL002W | 366.0264394 | -0.158704348 | 0.568354798 | 0.96 |
| YLL018C-A | 305.6957299 | 0.228693218 | 0.57569721 | 0.96 |
| YLL053C | 96.70396409 | 0.213371712 | 0.569145932 | 0.96 |
| YLL066C | 1.397171575 | 0.266188165 | 0.581945391 | 0.96 |
| YLR068W | 534.5964698 | 0.19599845 | 0.586513711 | 0.96 |
| YLR083C | 1368.675892 | -0.158695541 | 0.594744836 | 0.96 |
| YLR090W | 931.4547969 | 0.177947946 | 0.56410523 | 0.96 |
| YLR128W | 336.2576961 | -0.188154411 | 0.59163505 | 0.96 |
| YLR134W | 556.7123022 | -0.232064304 | 0.57623488 | 0.96 |
| YLR148W | 974.658438 | -0.11376619 | 0.600132694 | 0.96 |
| YLR171W | 25.60461968 | 0.145015205 | 0.600041096 | 0.96 |
| YLR172C | 1976.093187 | 0.230270142 | 0.567311583 | 0.96 |
| YLR269C | 113.3738155 | 0.138955332 | 0.578348933 | 0.96 |
| YLR278C | 1940.968689 | -0.094864267 | 0.597910328 | 0.96 |
| YLR284C | 336.5089725 | 0.167001151 | 0.599548196 | 0.96 |
| YLR295C | 2976.605613 | 0.247341243 | 0.574984414 | 0.96 |
| YLR320W | 753.4878726 | 0.111322326 | 0.585360695 | 0.96 |
| YLR336C | 993.7017255 | 0.202506334 | 0.588658287 | 0.96 |
| YLR340W | 21021.60532 | -0.205841501 | 0.573621464 | 0.96 |
| YLR341W | 104.3766662 | -0.241551995 | 0.565777781 | 0.96 |
| YLR354C | 11943.28845 | 0.184213093 | 0.600914752 | 0.96 |
| YLR371W | 1780.890302 | -0.166221575 | 0.585894892 | 0.96 |
| YLR376C | 277.9237935 | -0.182001167 | 0.572161605 | 0.96 |
| YLR422W | 2135.341687 | 0.125080281 | 0.583376469 | 0.96 |
| YLR440C | 889.0207433 | 0.161572017 | 0.571140364 | 0.96 |
| YML009C-A | 52.04514359 | 0.254762585 | 0.588189666 | 0.96 |
| YML034C-A | 23.30372078 | 0.230835497 | 0.573316824 | 0.96 |
| YML047C | 45.4588878 | 0.226853794 | 0.598077215 | 0.96 |
| YML070W | 2763.480937 | -0.15580009 | 0.595065501 | 0.96 |
| YMR014W | 1014.309235 | -0.164973801 | 0.592501526 | 0.96 |
| YMR017W | 57.51133406 | 0.224351068 | 0.572328436 | 0.96 |
| YMR030W | 574.6177195 | 0.227424626 | 0.578302332 | 0.96 |
| YMR079W | 1633.407315 | -0.156976477 | 0.570435466 | 0.96 |
| YMR159C | 22.87110149 | 0.163549091 | 0.582846241 | 0.96 |
| YMR182W-A | 4.590688512 | 0.090384511 | 0.591047474 | 0.96 |
| YMR192W | 859.8324449 | -0.138170538 | 0.572237295 | 0.96 |
| YMR204C | 437.305049 | -0.169134971 | 0.580652679 | 0.96 |
| YMR219W | 3001.660673 | 0.170407758 | 0.567035355 | 0.96 |
| YMR230W | 6828.386976 | -0.193881463 | 0.588969475 | 0.96 |

| | | | | |
|---|---|---|---|---|
| YMR230W-A | 4.038623613 | 0.246535788 | 0.588926425 | 0.96 |
| YMR247C | 1642.766432 | 0.130875966 | 0.568913765 | 0.96 |
| YMR265C | 688.687566 | 0.154455905 | 0.598006363 | 0.96 |
| YMR274C | 249.323644 | -0.180360095 | 0.569492455 | 0.96 |
| YMR278W | 1574.862371 | 0.164305982 | 0.5944747 | 0.96 |
| YMR279C | 707.3104958 | 0.09946261 | 0.589879644 | 0.96 |
| YMR296C | 3004.03015 | -0.169623066 | 0.591671736 | 0.96 |
| YMR316C-B | 10.98411933 | 0.212686664 | 0.60026061 | 0.96 |
| YNL002C | 1677.703778 | -0.143682538 | 0.567746788 | 0.96 |
| YNL075W | 678.2006525 | 0.335795829 | 0.568210108 | 0.96 |
| YNL086W | 191.101467 | 0.159603584 | 0.576921082 | 0.96 |
| YNL100W | 1262.153408 | 0.217962883 | 0.591078515 | 0.96 |
| YNL106C | 963.8876185 | 0.104689717 | 0.575228835 | 0.96 |
| YNL109W | 19.36367514 | 0.251497629 | 0.576118984 | 0.96 |
| YNL144C | 1767.380134 | 0.29431609 | 0.56975605 | 0.96 |
| YNL148C | 367.0903378 | 0.249831907 | 0.580592908 | 0.96 |
| YNL158W | 434.8285056 | -0.155882551 | 0.592068495 | 0.96 |
| YNL164C | 278.0976276 | -0.163685319 | 0.594496895 | 0.96 |
| YNL219C | 1817.162227 | -0.127432756 | 0.583368897 | 0.96 |
| YNL329C | 1184.177512 | 0.11969752 | 0.571419114 | 0.96 |
| YNR001W-A | 2.398336405 | -0.355680576 | 0.588118018 | 0.96 |
| YNR075C-A | 12.77963038 | -0.321794021 | 0.59827936 | 0.96 |
| YOL013C | 1216.146773 | -0.115188136 | 0.567127193 | 0.96 |
| YOL014W | 428.9922641 | -0.220363022 | 0.599109345 | 0.96 |
| YOL065C | 517.0688786 | 0.13885559 | 0.592632956 | 0.96 |
| YOL078W | 796.3214789 | 0.155314301 | 0.567843544 | 0.96 |
| YOL086C | 59663.42444 | 0.317206691 | 0.588870723 | 0.96 |
| YOL121C | 3681.856963 | -0.157276831 | 0.582397307 | 0.96 |
| YOL135C | 544.3922283 | -0.134378624 | 0.580092969 | 0.96 |
| YOR018W | 939.5138966 | -0.146310059 | 0.588522508 | 0.96 |
| YOR036W | 1460.876014 | -0.183206383 | 0.585079001 | 0.96 |
| YOR053W | 4.40231234 | -0.358357846 | 0.581762081 | 0.96 |
| YOR054C | 904.8796156 | 0.155909225 | 0.600378519 | 0.96 |
| YOR061W | 1662.759146 | -0.126476227 | 0.57116905 | 0.96 |
| YOR066W | 1115.721778 | -0.21611069 | 0.582845189 | 0.96 |
| YOR070C | 1064.02202 | -0.14092502 | 0.569619646 | 0.96 |
| YOR131C | 796.6694001 | 0.143157875 | 0.566904441 | 0.96 |
| YOR137C | 579.6398816 | -0.196558818 | 0.599557653 | 0.96 |
| YOR148C | 393.9247418 | -0.111325162 | 0.590746051 | 0.96 |
| YOR222W | 1007.50754 | 0.213635839 | 0.571453658 | 0.96 |
| YOR238W | 549.7060682 | -0.149267327 | 0.598708897 | 0.96 |
| YOR252W | 480.3993858 | 0.220265275 | 0.569852636 | 0.96 |
| YOR263C | 29.84403306 | -0.211273173 | 0.590328019 | 0.96 |

| | | | | |
|---|---|---|---|---|
| YOR288C | 898.6918954 | -0.171183287 | 0.577678738 | 0.96 |
| YOR312C | 9424.205996 | -0.186369836 | 0.570315772 | 0.96 |
| YOR314W | 17.17022364 | -0.280035133 | 0.574302483 | 0.96 |
| YOR323C | 2339.246441 | 0.199083943 | 0.572471779 | 0.96 |
| YOR331C | 254.0815524 | 0.109270642 | 0.586524215 | 0.96 |
| YPL046C | 139.4253961 | -0.152342409 | 0.598630206 | 0.96 |
| YPL116W | 1071.716343 | -0.141070358 | 0.581875338 | 0.96 |
| YPL144W | 173.5181567 | 0.235717328 | 0.600617383 | 0.96 |
| YPL157W | 499.4077603 | 0.210640781 | 0.60099975 | 0.96 |
| YPL158C | 1350.795148 | 0.195533286 | 0.564612745 | 0.96 |
| YPL165C | 810.9716846 | 0.179688758 | 0.595710292 | 0.96 |
| YPL176C | 1288.392675 | 0.139216404 | 0.583952604 | 0.96 |
| YPL218W | 2797.046919 | -0.10436501 | 0.599979737 | 0.96 |
| YPL236C | 570.2921643 | 0.164199778 | 0.596305581 | 0.96 |
| YPR020W | 1129.530709 | 0.315805404 | 0.586824195 | 0.96 |
| YPR063C | 1556.762503 | -0.118473326 | 0.58813726 | 0.96 |
| YPR072W | 2174.009813 | -0.116967666 | 0.566736464 | 0.96 |
| YPR135W | 846.2041741 | 0.121205971 | 0.600047395 | 0.96 |
| YPR170W-A | 19.17302177 | 0.252453598 | 0.594534374 | 0.96 |
| YPR171W | 883.9863961 | -0.12427825 | 0.573802191 | 0.96 |
| YPR202W | 17.57397274 | 0.314401389 | 0.60062302 | 0.96 |
| snR36 | 18.91914442 | 0.216189132 | 0.593045146 | 0.96 |
| snR4 | 89.67640266 | 0.314429585 | 0.584904024 | 0.96 |
| snR41 | 28.2332558 | -0.207856027 | 0.590224255 | 0.96 |
| snR63 | 94.54093227 | -0.096692084 | 0.585883952 | 0.96 |
| snR71 | 12.58626851 | -0.287937458 | 0.580320643 | 0.96 |
| snR77 | 9.325617253 | 0.21235469 | 0.576905498 | 0.96 |
| YGL145W | 1156.588747 | -0.09865904 | 0.601343111 | 0.96 |
| YBL080C | 486.2802333 | 0.130574804 | 0.601626025 | 0.96 |
| YAL014C | 833.2543927 | -0.155258548 | 0.602489566 | 0.96 |
| YBL107C | 641.7007619 | -0.183207944 | 0.602510344 | 0.96 |
| YBR133C | 1160.137007 | -0.203269284 | 0.603579053 | 0.96 |
| YCL035C | 4330.054767 | 0.216059187 | 0.602099322 | 0.96 |
| YDR297W | 2184.395638 | 0.176310341 | 0.603884987 | 0.96 |
| YEL071W | 3890.356415 | 0.272761302 | 0.603689034 | 0.96 |
| YHR174W | 94635.8071 | 0.380991546 | 0.60226725 | 0.96 |
| YKL095W | 323.4844945 | 0.14569692 | 0.60354764 | 0.96 |
| YLR096W | 1948.609356 | 0.121292092 | 0.602579443 | 0.96 |
| YLR111W | 15.73315875 | 0.245200094 | 0.603097216 | 0.96 |
| YNL019C | 10.29601328 | 0.246442771 | 0.603014933 | 0.96 |
| YOL120C | 9488.699457 | -0.077279908 | 0.604007602 | 0.96 |
| YOR340C | 775.5259752 | -0.145344171 | 0.603771055 | 0.96 |
| YPL076W | 380.2952376 | 0.129434331 | 0.60335689 | 0.96 |

| | | | | |
|---|---|---|---|---|
| snR84 | 162.4042194 | -0.151265234 | 0.60302226 | 0.96 |
| tL(GAG)G | 0.599720226 | 0.17562209 | 0.602352997 | 0.96 |
| YHR117W | 2228.445201 | 0.157261862 | 0.60432518 | 0.96 |
| YOR394C-A | 4.135778786 | -0.488335801 | 0.604178339 | 0.96 |
| YDL090C | 724.6090506 | -0.127403373 | 0.604482016 | 0.96 |
| YBR011C | 18392.14922 | -0.141484234 | 0.605573127 | 0.96 |
| YBR109C | 2942.937927 | -0.084343 | 0.605085957 | 0.96 |
| YBR236C | 870.5665803 | -0.12838757 | 0.606415179 | 0.96 |
| YCL007C | 102.0392471 | -0.164108602 | 0.6060617 | 0.96 |
| YDR155C | 17689.29075 | 0.217366705 | 0.605775813 | 0.96 |
| YDR458C | 1309.381626 | -0.145041122 | 0.60596855 | 0.96 |
| YER008C | 2582.281632 | -0.178102237 | 0.606273725 | 0.96 |
| YER107C | 624.5620743 | -0.150126934 | 0.605281294 | 0.96 |
| YIL153W | 970.2297401 | 0.068512272 | 0.605518181 | 0.96 |
| YLR025W | 2034.136821 | -0.135857513 | 0.605089768 | 0.96 |
| YLR434C | 48.33837913 | -0.269797603 | 0.606401345 | 0.96 |
| YML081C-A | 1028.969006 | 0.272268259 | 0.604982344 | 0.96 |
| YOL086W-A | 318.8786531 | -0.175438481 | 0.605905543 | 0.96 |
| YBR001C | 1318.791969 | 0.163376165 | 0.607137368 | 0.96 |
| YCR084C | 3633.028548 | -0.137081738 | 0.607282414 | 0.96 |
| YDL106C | 1181.738765 | -0.159201899 | 0.60698652 | 0.96 |
| YDR348C | 1537.308452 | 0.071327738 | 0.608300832 | 0.96 |
| YER006C-A | 2.225353622 | 0.164043794 | 0.60772056 | 0.96 |
| YFR043C | 412.3285976 | 0.107639195 | 0.607075708 | 0.96 |
| YGL141W | 1274.03239 | 0.100994648 | 0.607608249 | 0.96 |
| YHR021W-A | 41.37134005 | -0.289223894 | 0.608036071 | 0.96 |
| YKR025W | 782.5030894 | 0.17865547 | 0.608060983 | 0.96 |
| YML102W | 604.0898534 | 0.101620395 | 0.608291118 | 0.96 |
| YOR014W | 3387.711726 | 0.136612452 | 0.607445045 | 0.96 |
| YPL063W | 2649.060944 | 0.110226808 | 0.608049284 | 0.96 |
| YGR198W | 1689.551002 | -0.148426561 | 0.608763401 | 0.96 |
| YAL056C-A | 23.07572137 | 0.211126295 | 0.608967456 | 0.96 |
| YLL031C | 1740.005527 | 0.124309168 | 0.609752238 | 0.96 |
| YOL157C | 78.40010831 | 0.258104587 | 0.609725564 | 0.96 |
| YOR293W | 11411.33478 | -0.178135254 | 0.609541193 | 0.96 |
| YCR031C | 11369.4013 | -0.157057402 | 0.610106123 | 0.96 |
| YNL114C | 62.61749004 | -0.206849183 | 0.610027756 | 0.96 |
| YDR337W | 923.5124951 | -0.102489545 | 0.610313007 | 0.96 |
| YDR355C | 4.46006282 | -0.289003046 | 0.610763248 | 0.96 |
| YLR120W-A | 22.81140064 | -0.235176902 | 0.610857391 | 0.96 |
| YMR299C | 640.5800002 | -0.152444761 | 0.610516481 | 0.96 |
| YNL279W | 510.6368292 | -0.285073157 | 0.61076795 | 0.96 |
| YJL098W | 1058.267184 | -0.165603695 | 0.611345535 | 0.96 |

| | | | | |
|---|---|---|---|---|
| YOL055C | 533.3864453 | 0.157509906 | 0.611358369 | 0.96 |
| YDL010W | 362.0027109 | 0.116691969 | 0.613425555 | 0.96 |
| YDR418W | 2788.265019 | -0.147905036 | 0.613434965 | 0.96 |
| YEL029C | 331.9088854 | 0.200080384 | 0.613421147 | 0.96 |
| YFR010W | 3072.848472 | -0.124739081 | 0.613658864 | 0.96 |
| YIL113W | 384.3761033 | -0.235297266 | 0.612802892 | 0.96 |
| YIL130W | 1279.051362 | -0.2019997 | 0.613437552 | 0.96 |
| YIR038C | 2847.18512 | -0.18402162 | 0.61401195 | 0.96 |
| YJL044C | 1379.126824 | 0.137358845 | 0.613317066 | 0.96 |
| YLR050C | 545.8709868 | 0.205987298 | 0.613494393 | 0.96 |
| YLR060W | 5384.459239 | -0.155181144 | 0.613352218 | 0.96 |
| YNL160W | 19462.39163 | 0.229841285 | 0.613951143 | 0.96 |
| YNL263C | 1345.46904 | -0.124583379 | 0.613968899 | 0.96 |
| YOL047C | 302.1448912 | -0.294884371 | 0.612360329 | 0.96 |
| YJL200C | 1688.489143 | -0.187467836 | 0.614531497 | 0.96 |
| YLR201C | 1199.611196 | 0.140394425 | 0.614814035 | 0.96 |
| YLR276C | 2051.953288 | -0.152481561 | 0.614720908 | 0.96 |
| YJL136C | 5415.03448 | -0.139267724 | 0.615276463 | 0.97 |
| YGL015C | 22.18286877 | 0.205815267 | 0.616921687 | 0.97 |
| YGR041W | 794.2535473 | -0.127340409 | 0.616973088 | 0.97 |
| YIL078W | 8337.626041 | 0.256822628 | 0.616205357 | 0.97 |
| YJR019C | 596.1541169 | 0.144383033 | 0.616830562 | 0.97 |
| YKL014C | 1967.775598 | -0.188924461 | 0.616816002 | 0.97 |
| YKL219W | 500.3272007 | -0.167846423 | 0.615648143 | 0.97 |
| YLR092W | 690.3329885 | 0.160672093 | 0.616371319 | 0.97 |
| YML092C | 2758.585269 | -0.088135791 | 0.616460405 | 0.97 |
| YNL277W-A | 6.085182075 | 0.24201207 | 0.615998938 | 0.97 |
| YOL158C | 1769.752458 | -0.179136648 | 0.616112518 | 0.97 |
| YOR039W | 1072.032394 | -0.102021789 | 0.617161516 | 0.97 |
| YGL071W | 1392.981634 | -0.141981895 | 0.617593109 | 0.97 |
| YIR028W | 189.55325 | 0.22722438 | 0.61772773 | 0.97 |
| YMR143W | 7020.519511 | -0.151281912 | 0.618103087 | 0.97 |
| YLR333C | 2119.808061 | -0.148684927 | 0.618737352 | 0.97 |
| YCR050C | 17.22525728 | -0.344207541 | 0.619035475 | 0.97 |
| YDR047W | 1762.459456 | -0.103159075 | 0.619139293 | 0.97 |
| YDR095C | 5.026425328 | -0.394166964 | 0.620017362 | 0.97 |
| YGL019W | 1587.591155 | 0.080918125 | 0.620239283 | 0.97 |
| YNL237W | 756.7694566 | 0.138641124 | 0.620128314 | 0.97 |
| YPL097W | 440.8818332 | -0.120118901 | 0.620466955 | 0.97 |
| YPL098C | 678.1300721 | -0.099302114 | 0.62035075 | 0.97 |
| YPL227C | 650.9518323 | -0.152930336 | 0.620266932 | 0.97 |
| YPR190C | 965.2019513 | 0.208413766 | 0.620433696 | 0.97 |
| YLR307W | 82.37745124 | -0.304873263 | 0.620964491 | 0.97 |

| | | | | |
|---|---|---|---|---|
| YOR095C | 614.382855 | -0.067743783 | 0.620952506 | 0.97 |
| YAL031W-A | 36.5218107 | -0.209184601 | 0.623976549 | 0.97 |
| YAL033W | 350.8440347 | -0.183716768 | 0.622832674 | 0.97 |
| YCL010C | 597.9803619 | 0.148582394 | 0.623790466 | 0.97 |
| YDL013W | 806.2108951 | 0.132939935 | 0.624865246 | 0.97 |
| YDL217C | 416.6172424 | 0.128656486 | 0.624268115 | 0.97 |
| YDR291W | 877.4001219 | 0.080805196 | 0.624606102 | 0.97 |
| YEL007W | 3320.15412 | -0.161322324 | 0.62494824 | 0.97 |
| YFL003C | 165.623347 | 0.180417031 | 0.622272463 | 0.97 |
| YFR044C | 5702.058987 | 0.166152201 | 0.622985412 | 0.97 |
| YFR052C-A | 3217.393186 | 0.233283505 | 0.624787084 | 0.97 |
| YFR052W | 2758.803837 | 0.155433173 | 0.622354217 | 0.97 |
| YGL194C | 571.5770023 | 0.096683422 | 0.622775249 | 0.97 |
| YHL018W | 70.12879735 | 0.144125981 | 0.624720231 | 0.97 |
| YHR030C | 2133.377408 | -0.239488142 | 0.622797138 | 0.97 |
| YHR200W | 3347.025252 | 0.125139105 | 0.623144783 | 0.97 |
| YIL105C | 1531.713028 | -0.149056006 | 0.622261651 | 0.97 |
| YJL143W | 1167.282332 | -0.10112271 | 0.624883247 | 0.97 |
| YJL169W | 25.36216702 | -0.268378197 | 0.624505728 | 0.97 |
| YJR094W-A | 1689.898779 | -0.118883826 | 0.622489891 | 0.97 |
| YLR009W | 1043.475365 | -0.132814045 | 0.625193865 | 0.97 |
| YMR024W | 882.2534561 | -0.09897408 | 0.625160761 | 0.97 |
| YNL046W | 536.2842104 | 0.14575934 | 0.624948494 | 0.97 |
| YNL210W | 106.0752581 | 0.141164955 | 0.625040034 | 0.97 |
| YNL294C | 1109.535055 | 0.158606492 | 0.624691556 | 0.97 |
| YPL040C | 952.0787614 | 0.133556631 | 0.622121825 | 0.97 |
| YPL054W | 345.0757807 | 0.193357016 | 0.624088265 | 0.97 |
| YBL047C | 5698.969568 | -0.104024713 | 0.625818483 | 0.97 |
| YDR064W | 9582.184184 | -0.112101951 | 0.625439471 | 0.97 |
| YNR047W | 1084.837265 | 0.166625241 | 0.625692713 | 0.97 |
| YOR069W | 1035.064237 | -0.092471626 | 0.625796957 | 0.97 |
| YGR158C | 631.124363 | -0.137468867 | 0.626386345 | 0.97 |
| YDR328C | 1849.787592 | 0.101297995 | 0.627524869 | 0.97 |
| YER074W | 4108.232098 | -0.058102091 | 0.627298047 | 0.97 |
| YER087C-B | 1311.571506 | -0.125352128 | 0.627944576 | 0.97 |
| YJL063C | 928.728548 | -0.118324181 | 0.628075026 | 0.97 |
| YKL022C | 1018.942477 | 0.107673299 | 0.627603622 | 0.97 |
| YLR367W | 2289.066484 | -0.126252494 | 0.627131839 | 0.97 |
| YML060W | 354.1411795 | 0.24557277 | 0.628007514 | 0.97 |
| YMR113W | 461.667001 | -0.128394429 | 0.627731 | 0.97 |
| YNL138W | 4814.982276 | -0.101547816 | 0.62782604 | 0.97 |
| YOL061W | 1754.163485 | -0.115468966 | 0.626826496 | 0.97 |
| YPL109C | 614.5552418 | 0.195029675 | 0.628458282 | 0.97 |

| | | | | |
|---|---|---|---|---|
| YLR168C | 702.4056037 | 0.224326875 | 0.62867824 | 0.97 |
| YOL077C | 1854.977032 | -0.132296918 | 0.628787837 | 0.97 |
| YBR182C-A | 7.795727576 | 0.213948389 | 0.629547913 | 0.97 |
| YDL091C | 1187.448267 | 0.16977762 | 0.629357562 | 0.97 |
| YDR290W | 8.55083412 | -0.199292189 | 0.629505258 | 0.97 |
| YOR079C | 215.4299264 | -0.187581243 | 0.629580751 | 0.97 |
| YOR108W | 1787.909268 | 0.243562034 | 0.629580333 | 0.97 |
| YBR262C | 557.8168642 | 0.215458077 | 0.630094102 | 0.97 |
| YDL129W | 428.0062351 | 0.132334924 | 0.630047981 | 0.97 |
| YDR455C | 13.39180516 | -0.221825884 | 0.630331038 | 0.97 |
| YFL067W | 57.17560859 | 0.165504481 | 0.630329353 | 0.97 |
| YHR004C | 921.2706765 | 0.131299378 | 0.630559982 | 0.97 |
| YLR382C | 967.5198513 | -0.100538433 | 0.630645629 | 0.97 |
| YJL074C | 945.199767 | -0.111607393 | 0.63113168 | 0.97 |
| YJL125C | 686.7482617 | 0.163124411 | 0.631250934 | 0.97 |
| YKL092C | 887.6693114 | -0.121181845 | 0.631059881 | 0.97 |
| YPL187W | 17.17000227 | -0.29435532 | 0.631204765 | 0.97 |
| YBR166C | 859.3257193 | -0.110566604 | 0.632164771 | 0.97 |
| YDL039C | 373.0125142 | -0.260761387 | 0.632535549 | 0.97 |
| YER078C | 919.3267409 | 0.104499138 | 0.632848702 | 0.97 |
| YER141W | 2454.288383 | 0.153379237 | 0.632934967 | 0.97 |
| YGL202W | 6879.756501 | 0.156142444 | 0.631539699 | 0.97 |
| YGR111W | 1711.371366 | 0.132481987 | 0.632860796 | 0.97 |
| YHR095W | 5.929557488 | -0.280078544 | 0.632622135 | 0.97 |
| YHR144C | 383.3281151 | -0.168875384 | 0.632425655 | 0.97 |
| YIL122W | 600.3807645 | -0.208156306 | 0.631985505 | 0.97 |
| YKR015C | 299.9934309 | 0.093249853 | 0.632711097 | 0.97 |
| YLR031W | 139.756576 | 0.185969278 | 0.631953142 | 0.97 |
| YLR100W | 1453.024103 | 0.197492719 | 0.633053009 | 0.97 |
| YBL008W-A | 2.748531795 | -0.359807103 | 0.635535026 | 0.97 |
| YDL079C | 1497.675817 | 0.244587975 | 0.633306686 | 0.97 |
| YDL184C | 118.3516335 | -0.168002641 | 0.634324235 | 0.97 |
| YDR131C | 626.2979697 | -0.114280456 | 0.634964133 | 0.97 |
| YDR406W | 3009.969096 | -0.232876345 | 0.635372339 | 0.97 |
| YER088C-A | 302.6604055 | -0.249957244 | 0.633818405 | 0.97 |
| YFR033C | 3330.696407 | 0.198726146 | 0.634863149 | 0.97 |
| YIL005W | 1687.731259 | -0.097436483 | 0.634318145 | 0.97 |
| YJL084C | 1615.45903 | 0.111950159 | 0.635033121 | 0.97 |
| YKL065C | 4326.57925 | -0.10228186 | 0.635628864 | 0.97 |
| YKL079W | 964.5499269 | 0.079622303 | 0.635373916 | 0.97 |
| YKL125W | 592.2613366 | 0.106837836 | 0.63439317 | 0.97 |
| YML096W | 652.0319638 | -0.110529576 | 0.634042554 | 0.97 |
| YNL058C | 1080.922928 | 0.218691873 | 0.634982467 | 0.97 |

| | | | | |
|---|---|---|---|---|
| YOL070C | 1742.264613 | 0.152413273 | 0.633655461 | 0.97 |
| YOR242C | 110.2984083 | -0.249223018 | 0.635586801 | 0.97 |
| YPL153C | 363.2997459 | 0.136030122 | 0.633623868 | 0.97 |
| YHR063W-A | 174.3401986 | -0.155339376 | 0.636037744 | 0.97 |
| YOL052C-A | 1535.771374 | 0.27700746 | 0.635941472 | 0.97 |
| YGR030C | 291.5946813 | 0.176364226 | 0.636747514 | 0.97 |
| YHR184W | 81.12732022 | -0.263325403 | 0.636518221 | 0.97 |
| YKL045W | 918.6422418 | 0.125784149 | 0.636694495 | 0.97 |
| YMR038C | 1941.251831 | -0.11041829 | 0.636330945 | 0.97 |
| YNL035C | 765.5856918 | 0.121698557 | 0.637055809 | 0.97 |
| YGL037C | 14451.53815 | 0.163421473 | 0.637250518 | 0.97 |
| YLL010C | 1089.845187 | 0.083772402 | 0.637415985 | 0.97 |
| YEL063C | 1734.961887 | 0.140017438 | 0.637780565 | 0.97 |
| YMR082C | 5.748276588 | 0.20938169 | 0.637829465 | 0.97 |
| YAL019W | 1321.957689 | -0.147216532 | 0.641077607 | 0.97 |
| YAL034C | 1899.332638 | 0.175429865 | 0.639701407 | 0.97 |
| YBR107C | 343.2589721 | 0.138114554 | 0.638848804 | 0.97 |
| YCR038C | 411.3006768 | 0.096468558 | 0.639617957 | 0.97 |
| YDR344C | 46.73109222 | 0.266570065 | 0.639765759 | 0.97 |
| YGR144W | 97.38216331 | 0.187542536 | 0.639723065 | 0.97 |
| YGR284C | 2427.894256 | 0.158529067 | 0.640990955 | 0.97 |
| YIL048W | 1213.90232 | -0.089431218 | 0.639337413 | 0.97 |
| YIL123W | 4293.520238 | -0.156504039 | 0.640562774 | 0.97 |
| YIL148W | 9852.705743 | -0.133559305 | 0.641176342 | 0.97 |
| YKL016C | 3252.389125 | 0.202789042 | 0.640397014 | 0.97 |
| YKR045C | 368.478154 | 0.118135974 | 0.640425291 | 0.97 |
| YKR079C | 916.529157 | 0.13273106 | 0.640100519 | 0.97 |
| YLR127C | 955.0961502 | -0.120380418 | 0.639326965 | 0.97 |
| YLR224W | 1026.056679 | -0.139187262 | 0.639770162 | 0.97 |
| YLR272C | 941.972981 | 0.123726045 | 0.640575764 | 0.97 |
| YMR025W | 229.9295307 | 0.173330058 | 0.63916837 | 0.97 |
| YMR308C | 3347.564301 | -0.155211076 | 0.639652561 | 0.97 |
| YOR276W | 906.6874329 | -0.122603513 | 0.641153948 | 0.97 |
| YOR382W | 2866.519088 | 0.38250748 | 0.64115112 | 0.97 |
| YDR472W | 675.6541934 | -0.094266847 | 0.641441453 | 0.97 |
| YGR082W | 1782.261898 | -0.10023361 | 0.641488357 | 0.97 |
| YBR065C | 528.4307138 | 0.139998597 | 0.641747505 | 0.97 |
| YGL155W | 739.9393467 | -0.113183081 | 0.64205373 | 0.97 |
| YGL236C | 488.164276 | 0.14647944 | 0.641921182 | 0.97 |
| YHR132W-A | 721.2397547 | 0.147079548 | 0.642386325 | 0.97 |
| YBL072C | 3997.128198 | -0.14814398 | 0.643246556 | 0.97 |
| YCR106W | 394.726015 | 0.243669177 | 0.643490556 | 0.97 |
| YGL249W | 273.3583474 | -0.233839303 | 0.643539943 | 0.97 |

| | | | | |
|---|---|---|---|---|
| YLR222C-A | 13.05158283 | -0.237861181 | 0.643348735 | 0.97 |
| YMR212C | 1054.541147 | -0.145091895 | 0.643451146 | 0.97 |
| YOL022C | 1058.507084 | 0.212149133 | 0.643047362 | 0.97 |
| YPR010C | 3009.126793 | -0.143343908 | 0.643059749 | 0.97 |
| YPL146C | 923.2028372 | 0.222195596 | 0.644145814 | 0.97 |
| YBL004W | 2589.356792 | -0.15312945 | 0.644768651 | 0.97 |
| YDR210W-B | 26.10421514 | -0.275217265 | 0.645485816 | 0.97 |
| YDR415C | 529.3228874 | -0.13285634 | 0.644702862 | 0.97 |
| YGL096W | 785.2336716 | 0.231614559 | 0.645211846 | 0.97 |
| YGL175C | 162.4549988 | 0.10493498 | 0.64489803 | 0.97 |
| YKL154W | 669.8187345 | -0.140653682 | 0.645546458 | 0.97 |
| YML023C | 695.7405584 | 0.077319985 | 0.644674827 | 0.97 |
| YOR256C | 1260.844527 | -0.11092196 | 0.645080524 | 0.97 |
| YPR146C | 99.73914934 | -0.109962497 | 0.64549992 | 0.97 |
| YHR085W | 335.931842 | 0.212399713 | 0.646042726 | 0.97 |
| YLR017W | 1271.451959 | 0.185127027 | 0.645974416 | 0.97 |
| YOR003W | 499.5835225 | 0.123201547 | 0.645934186 | 0.97 |
| snR10 | 11.52677515 | 0.400119582 | 0.646328122 | 0.97 |
| YCR034W | 1456.119271 | -0.116199355 | 0.646916875 | 0.97 |
| YDR382W | 7559.11107 | -0.113935725 | 0.647052686 | 0.97 |
| YGR274C | 1344.663557 | -0.148264721 | 0.647170055 | 0.97 |
| YNL048W | 1078.322808 | -0.072644267 | 0.647094596 | 0.97 |
| YNL074C | 2355.063484 | -0.12638119 | 0.647112189 | 0.97 |
| YBR086C | 4766.880566 | -0.096467615 | 0.649004826 | 0.97 |
| YBR153W | 413.971492 | 0.098744922 | 0.64930943 | 0.97 |
| YCL028W | 2867.256977 | -0.098876624 | 0.647812345 | 0.97 |
| YDL028C | 1026.396259 | -0.11533132 | 0.649295656 | 0.97 |
| YER046W-A | 13.59530817 | 0.072039916 | 0.648542724 | 0.97 |
| YFL012W-A | 14.51855676 | -0.201697658 | 0.648790345 | 0.97 |
| YGR049W | 549.8511767 | 0.134485513 | 0.648834487 | 0.97 |
| YLR058C | 11510.40084 | 0.376543304 | 0.64881298 | 0.97 |
| YLR125W | 49.15160341 | 0.176969876 | 0.647420015 | 0.97 |
| YMR141W-A | 54.52612508 | 0.244423549 | 0.649144924 | 0.97 |
| YMR298W | 1278.824067 | 0.138567989 | 0.649344219 | 0.97 |
| YNL179C | 130.2526057 | 0.159494096 | 0.647949043 | 0.97 |
| YPL085W | 4050.180373 | -0.135565198 | 0.648830058 | 0.97 |
| YPR170C | 83.5281119 | -0.168697655 | 0.647987427 | 0.97 |
| YOL129W | 2548.890951 | -0.077343549 | 0.649572105 | 0.97 |
| YHR181W | 756.8304441 | -0.140566863 | 0.649790249 | 0.97 |
| YJL146W | 600.4909491 | 0.078475771 | 0.650943715 | 0.98 |
| YAL053W | 2097.963607 | 0.203023117 | 0.653238003 | 0.98 |
| YAR014C | 1593.739874 | 0.116101051 | 0.669124 | 0.98 |
| YBL046W | 691.3216368 | -0.087022336 | 0.672358129 | 0.98 |

| | | | | |
|---|---|---|---|---|
| YBL105C | 2392.516684 | -0.111656114 | 0.658628548 | 0.98 |
| YBR031W | 10790.74196 | 0.25946575 | 0.660107022 | 0.98 |
| YBR151W | 3044.846839 | -0.123248997 | 0.661783996 | 0.98 |
| YBR205W | 1374.246383 | 0.118149879 | 0.656902438 | 0.98 |
| YBR247C | 781.1952554 | 0.227820144 | 0.662125642 | 0.98 |
| YBR285W | 449.6014678 | -0.240527501 | 0.652266232 | 0.98 |
| YBR299W | 41.66988027 | 0.159319566 | 0.663634663 | 0.98 |
| YCR102W-A | 9.4716423 | 0.211769612 | 0.661161774 | 0.98 |
| YDL027C | 1559.088359 | 0.119927653 | 0.670657204 | 0.98 |
| YDL032W | 4.179935405 | 0.164542889 | 0.662579214 | 0.98 |
| YDL033C | 356.3492838 | 0.136869702 | 0.662259816 | 0.98 |
| YDL047W | 1531.758576 | 0.112632728 | 0.658156032 | 0.98 |
| YDL071C | 49.14928453 | 0.17176064 | 0.661856251 | 0.98 |
| YDL082W | 7404.050186 | -0.130150531 | 0.65965792 | 0.98 |
| YDL247W | 3.022100866 | -0.168702591 | 0.658838066 | 0.98 |
| YDR026C | 840.5603663 | 0.079835556 | 0.653842696 | 0.98 |
| YDR082W | 550.4664737 | 0.128747454 | 0.65943404 | 0.98 |
| YDR172W | 3599.759088 | -0.090565752 | 0.662583281 | 0.98 |
| YDR173C | 369.7980333 | 0.130821967 | 0.670283203 | 0.98 |
| YDR293C | 6400.531719 | 0.122366666 | 0.651865443 | 0.98 |
| YDR317W | 303.4699364 | -0.159015839 | 0.655130208 | 0.98 |
| YDR388W | 5322.26629 | 0.07956019 | 0.661536126 | 0.98 |
| YDR399W | 3185.764567 | -0.070831506 | 0.669085499 | 0.98 |
| YDR460W | 574.0402083 | -0.062848656 | 0.670684447 | 0.98 |
| YDR462W | 726.8986569 | -0.100875485 | 0.664381698 | 0.98 |
| YDR487C | 2644.625402 | 0.097739748 | 0.658494094 | 0.98 |
| YDR495C | 1040.925179 | -0.129205568 | 0.654241523 | 0.98 |
| YDR513W | 3464.363476 | 0.195764061 | 0.669474588 | 0.98 |
| YEL009C | 21167.59097 | 0.180236681 | 0.656246984 | 0.98 |
| YER053C | 3873.936022 | 0.230585667 | 0.654624392 | 0.98 |
| YER090W | 2828.394196 | 0.188849328 | 0.66887976 | 0.98 |
| YER097W | 28.63993449 | 0.195188465 | 0.660425199 | 0.98 |
| YER117W | 4528.496655 | -0.067082013 | 0.658211867 | 0.98 |
| YFR001W | 1018.480753 | -0.09972886 | 0.66030882 | 0.98 |
| YFR011C | 642.0102271 | 0.212670092 | 0.6725306 | 0.98 |
| YFR031C | 1205.359626 | 0.075023468 | 0.67314833 | 0.98 |
| YGL038C | 1572.160144 | -0.121521285 | 0.664848101 | 0.98 |
| YGL039W | 719.9626894 | -0.124195572 | 0.667209466 | 0.98 |
| YGL066W | 1160.200958 | 0.086801253 | 0.669468716 | 0.98 |
| YGL191W | 2937.202908 | 0.247981798 | 0.668122875 | 0.98 |
| YGR038W | 610.0071877 | -0.106813811 | 0.657777844 | 0.98 |
| YGR128C | 1414.526792 | -0.15864528 | 0.653781586 | 0.98 |
| YGR166W | 818.3285643 | 0.108367742 | 0.671164882 | 0.98 |

| | | | | |
|---|---|---|---|---|
| YGR215W | 453.6817551 | -0.111814116 | 0.668363777 | 0.98 |
| YHL006W-A | 55.86611926 | -0.148106024 | 0.663084902 | 0.98 |
| YHR007C | 4279.432771 | -0.142257539 | 0.664590286 | 0.98 |
| YHR069C | 771.2357851 | 0.158834268 | 0.66823596 | 0.98 |
| YHR087W | 3070.022032 | 0.252510032 | 0.67224925 | 0.98 |
| YHR108W | 3358.554888 | -0.112680504 | 0.651903794 | 0.98 |
| YHR172W | 514.8081565 | -0.144641507 | 0.66795587 | 0.98 |
| YIL014W | 862.4746572 | -0.118463392 | 0.66409332 | 0.98 |
| YIL043C | 4110.226673 | -0.091501009 | 0.672556294 | 0.98 |
| YIL077C | 964.8137743 | 0.195891991 | 0.659379054 | 0.98 |
| YIL100C-A | 6.139255363 | 0.221253747 | 0.655525051 | 0.98 |
| YIL101C | 1402.735777 | -0.228455373 | 0.670865996 | 0.98 |
| YIL107C | 1780.529737 | 0.126923197 | 0.666412695 | 0.98 |
| YIR036W-A | 10.01589679 | 0.133034637 | 0.671099103 | 0.98 |
| YJL007C | 3.350774349 | -0.433033174 | 0.661270455 | 0.98 |
| YJL026W | 4324.794509 | -0.098964881 | 0.668711598 | 0.98 |
| YJL093C | 1063.991787 | 0.095365992 | 0.652894359 | 0.98 |
| YJL177W | 7142.560719 | -0.12310995 | 0.657994027 | 0.98 |
| YJL182C | 1.793622545 | -0.1869236 | 0.667137459 | 0.98 |
| YJL192C | 2669.267366 | -0.115902252 | 0.651625184 | 0.98 |
| YJR021C | 884.0147888 | 0.111903467 | 0.663980282 | 0.98 |
| YJR135W-A | 435.3473743 | -0.090862419 | 0.662984912 | 0.98 |
| YJR137C | 1214.839856 | 0.117044076 | 0.670300269 | 0.98 |
| YJR140C | 1086.888351 | 0.11822925 | 0.672834833 | 0.98 |
| YKL059C | 865.3974506 | -0.075143337 | 0.672874455 | 0.98 |
| YKL075C | 477.0577168 | 0.106550651 | 0.667092207 | 0.98 |
| YKL103C | 3936.941911 | 0.239744271 | 0.660792762 | 0.98 |
| YKL113C | 707.9229549 | -0.104652715 | 0.673067083 | 0.98 |
| YKL165C | 1154.053703 | -0.117812382 | 0.670185946 | 0.98 |
| YKR034W | 49.93480154 | 0.171185558 | 0.671673038 | 0.98 |
| YLL018C | 8408.142457 | 0.122037597 | 0.669807613 | 0.98 |
| YLL024C | 20764.11389 | 0.300442218 | 0.654378972 | 0.98 |
| YLL035W | 581.4295973 | 0.138864682 | 0.667320331 | 0.98 |
| YLL037W | 13.52665894 | -0.219389001 | 0.672780395 | 0.98 |
| YLL051C | 1170.125266 | 0.109069883 | 0.666603002 | 0.98 |
| YLL054C | 624.0988471 | -0.095848313 | 0.658997636 | 0.98 |
| YLR039C | 1333.277955 | -0.101467742 | 0.653862551 | 0.98 |
| YLR266C | 1095.178781 | -0.136668648 | 0.672219845 | 0.98 |
| YLR357W | 954.9234442 | -0.125940753 | 0.655399703 | 0.98 |
| YLR372W | 2301.005993 | -0.083872207 | 0.666370084 | 0.98 |
| YLR451W | 1124.900564 | 0.076712912 | 0.66743577 | 0.98 |
| YML007W | 2781.196668 | 0.124242217 | 0.653273049 | 0.98 |
| YML047W-A | 2.39499993 | -0.079024465 | 0.657331853 | 0.98 |

| | | | | |
|---|---|---|---|---|
| YML051W | 1749.382621 | -0.143898146 | 0.669426082 | 0.98 |
| YML091C | 3874.618724 | 0.283022108 | 0.658950401 | 0.98 |
| YMR030W-A | 15.82228062 | -0.236804903 | 0.666514037 | 0.98 |
| YMR091C | 998.5083437 | -0.103622974 | 0.663499021 | 0.98 |
| YMR142C | 11187.93902 | -0.085135306 | 0.659008035 | 0.98 |
| YMR171C | 1518.290743 | -0.182076043 | 0.670024047 | 0.98 |
| YMR172C-A | 11.37396813 | 0.141431275 | 0.661100614 | 0.98 |
| YMR178W | 1157.445412 | 0.119997605 | 0.667233121 | 0.98 |
| YMR223W | 516.9298345 | -0.077221933 | 0.667323208 | 0.98 |
| YMR245W | 80.78367086 | 0.162705548 | 0.671377142 | 0.98 |
| YMR253C | 606.2113411 | -0.129553628 | 0.659092326 | 0.98 |
| YMR263W | 652.3046851 | 0.115638185 | 0.666212155 | 0.98 |
| YMR305C | 6521.89708 | -0.180308711 | 0.66545023 | 0.98 |
| YMR314W | 2980.188475 | 0.091427765 | 0.654719724 | 0.98 |
| YNL054W | 1269.220328 | -0.102677004 | 0.6714703 | 0.98 |
| YNL085W | 3478.261348 | -0.119306687 | 0.668233566 | 0.98 |
| YNL091W | 4224.416881 | 0.114947308 | 0.658113461 | 0.98 |
| YNL099C | 699.7662248 | -0.094506763 | 0.659032599 | 0.98 |
| YNL162W | 2350.306083 | -0.120823402 | 0.658000937 | 0.98 |
| YNL189W | 2792.970706 | -0.091836793 | 0.664417215 | 0.98 |
| YNL230C | 466.9526022 | 0.117183059 | 0.67253783 | 0.98 |
| YNL321W | 1495.615211 | 0.098688567 | 0.655122097 | 0.98 |
| YNL335W | 13.42162756 | -0.235784125 | 0.662342437 | 0.98 |
| YNR036C | 1583.864307 | 0.169777059 | 0.659336461 | 0.98 |
| YNR054C | 724.980203 | -0.140992796 | 0.656260666 | 0.98 |
| YOL148C | 713.6397629 | -0.094920623 | 0.655090935 | 0.98 |
| YOL156W | 95.75918972 | 0.188507927 | 0.654399806 | 0.98 |
| YOR006C | 758.4019018 | 0.085056729 | 0.673089076 | 0.98 |
| YOR034C-A | 45.66152085 | 0.141817843 | 0.673240915 | 0.98 |
| YOR051C | 2007.894821 | -0.143496235 | 0.669334022 | 0.98 |
| YOR110W | 712.9241656 | 0.093695954 | 0.66720308 | 0.98 |
| YOR146W | 6.962391281 | -0.185452889 | 0.654740818 | 0.98 |
| YOR192C-B | 24.91802453 | -0.288602507 | 0.658695193 | 0.98 |
| YOR196C | 1243.855488 | 0.124481725 | 0.660820389 | 0.98 |
| YOR358W | 499.4768456 | 0.126478983 | 0.655918401 | 0.98 |
| YOR368W | 442.6825001 | -0.089455836 | 0.662993251 | 0.98 |
| YPL003W | 547.4702758 | -0.145314834 | 0.665022101 | 0.98 |
| YPL059W | 1008.866988 | -0.080065139 | 0.656860689 | 0.98 |
| YPL084W | 2075.944666 | -0.076733702 | 0.655594834 | 0.98 |
| YPL093W | 1935.182804 | -0.138373359 | 0.663413096 | 0.98 |
| YPL103C | 590.7071612 | 0.105420526 | 0.670429884 | 0.98 |
| YPL132W | 679.0950944 | -0.11656652 | 0.660752708 | 0.98 |
| YPL148C | 142.8618352 | 0.17093681 | 0.663693598 | 0.98 |

| | | | | |
|---|---|---|---|---|
| YPL219W | 632.4863798 | 0.110671954 | 0.660876215 | 0.98 |
| YPL234C | 1583.101568 | -0.056872806 | 0.672866401 | 0.98 |
| YPL264C | 197.1729292 | 0.108151953 | 0.653751696 | 0.98 |
| YPR018W | 922.6100673 | -0.122161919 | 0.667204255 | 0.98 |
| YPR045C | 1121.270528 | 0.108714305 | 0.659312931 | 0.98 |
| YPR097W | 1353.173051 | 0.078595966 | 0.668692071 | 0.98 |
| YPR108W | 2749.924825 | -0.090965842 | 0.655777617 | 0.98 |
| YPR113W | 3675.90658 | -0.103081304 | 0.671810607 | 0.98 |
| YPR125W | 636.6611003 | -0.108444451 | 0.655889388 | 0.98 |
| YPR130C | 10.79124991 | 0.166025612 | 0.6547163 | 0.98 |
| YPR137W | 608.8911311 | -0.139142907 | 0.665035769 | 0.98 |
| YPR149W | 11719.12039 | 0.162492222 | 0.669844527 | 0.98 |
| YPR154W | 2486.651934 | 0.098559063 | 0.666150694 | 0.98 |
| snR190 | 17.01701234 | 0.235217729 | 0.657051145 | 0.98 |
| snR3 | 32.48912912 | 0.250211084 | 0.659425358 | 0.98 |
| snR40 | 6.538551087 | -0.247810389 | 0.657532186 | 0.98 |
| YER019W | 1037.358398 | 0.164957705 | 0.673824808 | 0.98 |
| YGR229C | 1916.022007 | -0.107610085 | 0.673842689 | 0.98 |
| YPL122C | 549.6598792 | -0.074953786 | 0.673546203 | 0.98 |
| YBR082C | 6782.616874 | -0.136232639 | 0.674114655 | 0.98 |
| snR53 | 0.80815136 | -0.081522988 | 0.674256166 | 0.98 |
| YDR149C | 14.02290454 | 0.227998464 | 0.674554705 | 0.98 |
| YJL053W | 1232.219717 | -0.103120192 | 0.674805198 | 0.98 |
| YDR227W | 1551.753891 | -0.134284122 | 0.67550482 | 0.98 |
| YEL058W | 1683.65634 | -0.096449945 | 0.676332422 | 0.98 |
| YHL026C | 386.086378 | 0.134349219 | 0.67622196 | 0.98 |
| YKR036C | 843.7972906 | -0.080368677 | 0.676099134 | 0.98 |
| YLR319C | 843.3891881 | -0.067211901 | 0.675274323 | 0.98 |
| YMR129W | 1909.721365 | -0.073983127 | 0.676020467 | 0.98 |
| YMR315W | 4197.096296 | 0.18273124 | 0.676007802 | 0.98 |
| YNL150W | 83.29807569 | 0.207667376 | 0.676189821 | 0.98 |
| YNL308C | 3277.042014 | 0.182265293 | 0.675262097 | 0.98 |
| YPL079W | 6102.629794 | -0.103058445 | 0.675843417 | 0.98 |
| YAL069W | 0.918104495 | 0.044524442 | 0.679415347 | 0.98 |
| YAR015W | 7474.541729 | 0.306761245 | 0.690521775 | 0.98 |
| YAR066W | 12.65917974 | 0.209933464 | 0.68696919 | 0.98 |
| YBL032W | 1731.697141 | -0.087517949 | 0.69732216 | 0.98 |
| YBL039C | 1528.083632 | -0.088921964 | 0.677228721 | 0.98 |
| YBR015C | 2379.48263 | 0.088773802 | 0.694726508 | 0.98 |
| YBR018C | 184.0387102 | 0.165310495 | 0.693775062 | 0.98 |
| YBR057C | 866.0815496 | 0.120664232 | 0.685334719 | 0.98 |
| YBR059C | 2706.219813 | -0.118341502 | 0.686664456 | 0.98 |
| YBR104W | 350.9392427 | 0.251039062 | 0.695461087 | 0.98 |

| | | | | |
|---|---|---|---|---|
| YBR149W | 7039.888751 | 0.104306887 | 0.694442926 | 0.98 |
| YBR187W | 950.8054053 | -0.040285685 | 0.694647966 | 0.98 |
| YBR249C | 4275.177673 | 0.333743371 | 0.687911698 | 0.98 |
| YCL046W | 22.38513549 | -0.128513218 | 0.694967953 | 0.98 |
| YCR079W | 984.6018828 | 0.119425386 | 0.695927553 | 0.98 |
| YDL051W | 1178.81834 | 0.241036763 | 0.697667021 | 0.98 |
| YDL075W | 5187.384084 | -0.068455032 | 0.695891174 | 0.98 |
| YDL150W | 525.2641532 | -0.119280534 | 0.683694097 | 0.98 |
| YDL152W | 5.390111654 | -0.083562383 | 0.690384314 | 0.98 |
| YDL190C | 2381.415921 | 0.072359544 | 0.688982492 | 0.98 |
| YDR041W | 1214.356247 | -0.096211892 | 0.697119033 | 0.98 |
| YDR048C | 13.8983324 | -0.211875916 | 0.686173612 | 0.98 |
| YDR056C | 1611.517655 | 0.091031383 | 0.696851686 | 0.98 |
| YDR083W | 1132.402572 | -0.111652797 | 0.687492493 | 0.98 |
| YDR109C | 536.0726963 | -0.079933897 | 0.685104529 | 0.98 |
| YDR169C | 1729.231302 | -0.125292335 | 0.693171205 | 0.98 |
| YDR196C | 880.6195121 | 0.116594991 | 0.686091955 | 0.98 |
| YDR217C | 884.5738376 | -0.110194156 | 0.690496147 | 0.98 |
| YDR272W | 1625.208653 | 0.130993815 | 0.697016884 | 0.98 |
| YDR365W-B | 142.2689351 | -0.244992205 | 0.67960111 | 0.98 |
| YDR394W | 3757.628989 | -0.079257583 | 0.686354796 | 0.98 |
| YDR441C | 376.4209723 | 0.09113577 | 0.694718518 | 0.98 |
| YDR479C | 1259.888261 | 0.116318195 | 0.678119005 | 0.98 |
| YDR480W | 507.0349407 | -0.058253701 | 0.679952438 | 0.98 |
| YDR520C | 591.1970741 | 0.123613841 | 0.693771484 | 0.98 |
| YDR532C | 597.6785104 | -0.144354063 | 0.686940073 | 0.98 |
| YDR543C | 0.805732287 | 0.020598733 | 0.697330559 | 0.98 |
| YEL003W | 487.0478849 | -0.10333765 | 0.696547621 | 0.98 |
| YER001W | 1455.504127 | -0.143168835 | 0.677528089 | 0.98 |
| YER031C | 2004.561338 | -0.10426414 | 0.681513312 | 0.98 |
| YER084W | 124.9336603 | 0.155307842 | 0.686713457 | 0.98 |
| YFL031C-A | 101.8569422 | 0.130106286 | 0.681011437 | 0.98 |
| YFR010W-A | 16.18554247 | 0.301281126 | 0.679925875 | 0.98 |
| YFR013W | 1397.161166 | -0.101752124 | 0.685681302 | 0.98 |
| YGL022W | 3864.561612 | -0.089537008 | 0.6924993 | 0.98 |
| YGL028C | 1677.067671 | -0.140443626 | 0.685203971 | 0.98 |
| YGL110C | 882.4868812 | -0.084149995 | 0.685885995 | 0.98 |
| YGL123W | 15975.90289 | -0.089081455 | 0.687016682 | 0.98 |
| YGL128C | 505.7413235 | 0.094726273 | 0.683287151 | 0.98 |
| YGL161C | 1505.046156 | -0.058449325 | 0.686293804 | 0.98 |
| YGL188C | 26.0866713 | 0.224547383 | 0.696034024 | 0.98 |
| YGL203C | 1571.604512 | 0.103095926 | 0.688981263 | 0.98 |
| YGL225W | 1789.038412 | -0.087020446 | 0.689116131 | 0.98 |

| | | | | |
|---|---|---|---|---|
| YGL237C | 638.8350245 | 0.126573933 | 0.678850325 | 0.98 |
| YGR006W | 239.4957069 | 0.091928474 | 0.687552802 | 0.98 |
| YGR053C | 339.3237027 | 0.134099369 | 0.685729399 | 0.98 |
| YGR074W | 145.6756142 | -0.096834599 | 0.694824023 | 0.98 |
| YGR100W | 1479.985263 | 0.057902125 | 0.683511691 | 0.98 |
| YGR167W | 3020.873713 | -0.072293373 | 0.682324172 | 0.98 |
| YGR189C | 5490.403188 | -0.161276215 | 0.683784233 | 0.98 |
| YGR199W | 855.4590705 | 0.0976211 | 0.695622224 | 0.98 |
| YGR207C | 1419.844599 | 0.100275571 | 0.67970048 | 0.98 |
| YGR218W | 3998.195199 | 0.052649343 | 0.693604897 | 0.98 |
| YGR234W | 5131.339655 | -0.091209855 | 0.689875846 | 0.98 |
| YGR242W | 37.21832003 | -0.125889383 | 0.687712073 | 0.98 |
| YGR247W | 484.9641046 | 0.132748901 | 0.690465037 | 0.98 |
| YHL002C-A | 8.009981089 | 0.158777095 | 0.685278237 | 0.98 |
| YHL008C | 1005.082799 | -0.175045711 | 0.685413728 | 0.98 |
| YHR037W | 3383.544 | -0.119731727 | 0.684934081 | 0.98 |
| YHR039C | 2261.246675 | -0.115649097 | 0.68529998 | 0.98 |
| YHR052W | 1127.946092 | -0.090066814 | 0.694680049 | 0.98 |
| YHR123W | 699.5030842 | -0.14184769 | 0.691355978 | 0.98 |
| YHR165C | 3249.517075 | -0.140072621 | 0.688601022 | 0.98 |
| YHR198C | 872.0543676 | 0.172193815 | 0.697244886 | 0.98 |
| YHR204W | 1257.868137 | 0.105081028 | 0.681437782 | 0.98 |
| YIL026C | 1582.807889 | -0.107363825 | 0.68645562 | 0.98 |
| YIL046W-A | 7.695578865 | 0.092379001 | 0.683442267 | 0.98 |
| YIL063C | 1081.563362 | 0.151314157 | 0.693652403 | 0.98 |
| YIL080W | 141.3218365 | -0.354169648 | 0.685833585 | 0.98 |
| YIR017W-A | 3.982442675 | 0.111542665 | 0.69378741 | 0.98 |
| YJL006C | 336.1143852 | -0.103929031 | 0.683375149 | 0.98 |
| YJL046W | 632.8219176 | -0.085836201 | 0.684652291 | 0.98 |
| YJL134W | 603.1655535 | 0.151603671 | 0.694621764 | 0.98 |
| YJL161W | 501.2537696 | -0.196314423 | 0.690109835 | 0.98 |
| YJL180C | 680.1821358 | -0.083735614 | 0.686291362 | 0.98 |
| YJL190C | 9254.873132 | -0.114507306 | 0.682994973 | 0.98 |
| YJL222W | 13.76304146 | -0.261452208 | 0.68998824 | 0.98 |
| YKL025C | 1634.326653 | -0.086946807 | 0.679125209 | 0.98 |
| YKL115C | 42.82706794 | -0.125417955 | 0.693154454 | 0.98 |
| YKL143W | 1110.759937 | 0.237288431 | 0.697538332 | 0.98 |
| YKL145W | 5482.552156 | 0.12091292 | 0.690273405 | 0.98 |
| YKL212W | 2578.339284 | -0.081207507 | 0.690980004 | 0.98 |
| YKR043C | 2568.82772 | 0.250930764 | 0.679429568 | 0.98 |
| YKR044W | 684.1987114 | 0.079885876 | 0.686329625 | 0.98 |
| YKR095W | 4316.14506 | -0.09859721 | 0.69100063 | 0.98 |
| YLL008W | 1923.071209 | 0.218015537 | 0.690803528 | 0.98 |

| | | | | |
|---|---|---|---|---|
| YLR048W | 13755.73587 | -0.122372971 | 0.697530367 | 0.98 |
| YLR069C | 1475.235001 | -0.102084639 | 0.685415922 | 0.98 |
| YLR222C | 1088.378581 | -0.155939071 | 0.68650022 | 0.98 |
| YLR231C | 929.429053 | -0.224228455 | 0.68329155 | 0.98 |
| YLR254C | 554.1871051 | -0.118013804 | 0.688308758 | 0.98 |
| YLR297W | 487.8704755 | 0.137368099 | 0.677344985 | 0.98 |
| YLR364C-A | 0.303554837 | -0.054127122 | 0.687421546 | 0.98 |
| YLR407W | 355.7060391 | 0.092754755 | 0.690650703 | 0.98 |
| YLR455W | 572.5909218 | -0.105555902 | 0.67966095 | 0.98 |
| YML032C | 1432.723324 | -0.14810686 | 0.693103611 | 0.98 |
| YML063W | 16224.08176 | -0.093683927 | 0.680709001 | 0.98 |
| YMR031C | 3707.880046 | 0.14287032 | 0.694190285 | 0.98 |
| YMR111C | 424.0046487 | -0.092797269 | 0.690728592 | 0.98 |
| YMR128W | 2322.124433 | 0.237414682 | 0.676685092 | 0.98 |
| YMR188C | 1203.74754 | -0.10482066 | 0.689581457 | 0.98 |
| YMR227C | 799.7644018 | -0.125396283 | 0.682545077 | 0.98 |
| YMR237W | 2047.326568 | 0.051056749 | 0.691187794 | 0.98 |
| YMR247W-A | 22.89498367 | 0.208419305 | 0.684666761 | 0.98 |
| YMR262W | 628.7777309 | 0.126094595 | 0.693036621 | 0.98 |
| YNL069C | 13320.02635 | -0.119366422 | 0.686513642 | 0.98 |
| YNL080C | 1115.372394 | -0.095855838 | 0.694488541 | 0.98 |
| YNL190W | 6079.357238 | 0.101863371 | 0.677668563 | 0.98 |
| YNL198C | 23.75937545 | 0.096507754 | 0.695335866 | 0.98 |
| YNL284C | 1102.406706 | -0.086518705 | 0.685452892 | 0.98 |
| YNL325C | 453.4130958 | 0.105077469 | 0.682691447 | 0.98 |
| YNR074C | 411.2345917 | -0.152726564 | 0.695018709 | 0.98 |
| YOL035C | 58.4740954 | 0.109057299 | 0.688727473 | 0.98 |
| YOL105C | 760.4552602 | 0.097454784 | 0.696271181 | 0.98 |
| YOL142W | 655.5671288 | 0.131434423 | 0.685634783 | 0.98 |
| YOR008C-A | 9.788861467 | 0.231777004 | 0.691038865 | 0.98 |
| YOR008W-B | 4.890160442 | -0.227575183 | 0.696363964 | 0.98 |
| YOR101W | 962.4280426 | -0.0959321 | 0.696596469 | 0.98 |
| YOR180C | 257.2908648 | 0.127667512 | 0.68181396 | 0.98 |
| YOR205C | 770.9234478 | -0.105984545 | 0.69629269 | 0.98 |
| YOR380W | 392.7551247 | 0.09420562 | 0.679371491 | 0.98 |
| YOR387C | 76.98828773 | 0.121113508 | 0.686303477 | 0.98 |
| YPL012W | 1836.869245 | -0.106566696 | 0.692052929 | 0.98 |
| YPL030W | 552.0003609 | -0.133709272 | 0.680140655 | 0.98 |
| YPL260W | 2621.937685 | -0.117285081 | 0.687691685 | 0.98 |
| YPR002C-A | 1.918449357 | -0.069556706 | 0.684036583 | 0.98 |
| YPR143W | 537.9356283 | 0.172055584 | 0.694915022 | 0.98 |
| YPR152C | 381.9563274 | 0.099805549 | 0.67998341 | 0.98 |
| YPR169W | 1659.207747 | 0.116383306 | 0.688340702 | 0.98 |

| | | | | |
|---|---|---|---|---|
| snR61 | 12.13748594 | 0.226253811 | 0.67938774 | 0.98 |
| snR65 | 5.68201134 | -0.211215646 | 0.696932714 | 0.98 |
| YGR147C | 943.7636147 | 0.091046751 | 0.69820125 | 0.98 |
| YML101C-A | 7.327177589 | -0.236938024 | 0.698164858 | 0.98 |
| YNR003C | 554.144982 | -0.103476811 | 0.69812497 | 0.98 |
| YBR170C | 1144.125576 | 0.073335162 | 0.698836242 | 0.98 |
| YEL065W | 2278.709245 | 0.237327016 | 0.698400854 | 0.98 |
| YGL189C | 20518.43042 | -0.139306393 | 0.698552156 | 0.98 |
| YHR068W | 2922.590435 | -0.074239588 | 0.698805109 | 0.98 |
| YFL031W | 3441.588262 | -0.167259882 | 0.699284777 | 0.98 |
| YDL022W | 5632.112705 | 0.158365805 | 0.69970932 | 0.98 |
| YOL056W | 561.7509431 | -0.128885157 | 0.699748071 | 0.98 |
| YAL037W | 195.7634153 | -0.161384892 | 0.707494038 | 0.98 |
| YBL103C | 858.4740089 | -0.113927989 | 0.704622269 | 0.98 |
| YBR256C | 1957.968457 | 0.127640154 | 0.704799187 | 0.98 |
| YDL181W | 1061.423493 | 0.165276156 | 0.705399834 | 0.98 |
| YDR003W-A | 67.78719135 | 0.106109345 | 0.707279216 | 0.98 |
| YDR079C-A | 102.409036 | 0.101968947 | 0.704017004 | 0.98 |
| YDR108W | 495.5770673 | -0.080888024 | 0.707607972 | 0.98 |
| YDR121W | 548.7039891 | -0.074438135 | 0.699993492 | 0.98 |
| YDR213W | 568.9307813 | -0.127209326 | 0.707550241 | 0.98 |
| YDR326C | 1698.894045 | -0.096061726 | 0.707568315 | 0.98 |
| YDR423C | 1205.3164 | -0.130784868 | 0.700538243 | 0.98 |
| YDR499W | 868.5785284 | -0.145241226 | 0.703053015 | 0.98 |
| YER107W-A | 76.0626659 | -0.128014824 | 0.705008953 | 0.98 |
| YGL030W | 7624.667323 | 0.036878935 | 0.702766118 | 0.98 |
| YGL098W | 531.3017666 | 0.103244907 | 0.700707411 | 0.98 |
| YGR062C | 462.3300151 | -0.098543385 | 0.705784159 | 0.98 |
| YGR152C | 296.8399677 | -0.100112997 | 0.706762594 | 0.98 |
| YGR184C | 3421.297226 | 0.067703162 | 0.700145676 | 0.98 |
| YGR195W | 892.1333078 | -0.070789798 | 0.707199517 | 0.98 |
| YIL029C | 174.946717 | 0.152820735 | 0.706849799 | 0.98 |
| YIL115C | 2357.200299 | 0.050502672 | 0.704487915 | 0.98 |
| YIR002C | 1205.196083 | -0.12478546 | 0.706426202 | 0.98 |
| YJL058C | 326.9260026 | 0.095907489 | 0.705033929 | 0.98 |
| YJL142C | 106.3041783 | 0.111859487 | 0.706874283 | 0.98 |
| YJR022W | 18.26801566 | -0.184878918 | 0.702916767 | 0.98 |
| YJR120W | 18.09619799 | -0.205631999 | 0.702930417 | 0.98 |
| YKL009W | 1409.022835 | -0.076587265 | 0.701494327 | 0.98 |
| YKL087C | 723.7436146 | 0.090639828 | 0.705638905 | 0.98 |
| YKL121W | 1073.818565 | 0.092880815 | 0.700821927 | 0.98 |
| YKL124W | 623.7006372 | 0.096708886 | 0.704488374 | 0.98 |
| YKR024C | 515.1058632 | -0.140189544 | 0.703858936 | 0.98 |

| | | | | |
|---|---|---|---|---|
| YKR040C | 4.454389 | 0.293158757 | 0.706585132 | 0.98 |
| YKR080W | 1412.607481 | 0.364593172 | 0.704975725 | 0.98 |
| YKR090W | 850.9383093 | -0.092767999 | 0.704481671 | 0.98 |
| YLR001C | 1116.57839 | 0.079912252 | 0.701275336 | 0.98 |
| YLR059C | 1264.300383 | 0.108550859 | 0.703585932 | 0.98 |
| YLR381W | 607.7067419 | 0.079417867 | 0.702012413 | 0.98 |
| YMR032W | 652.0530589 | -0.105032974 | 0.706695088 | 0.98 |
| YMR222C | 712.4011542 | 0.078326999 | 0.700683468 | 0.98 |
| YMR288W | 774.5062547 | 0.081571622 | 0.705238524 | 0.98 |
| YMR315W-A | 6.002010905 | 0.142193621 | 0.701589312 | 0.98 |
| YNL031C | 9825.730059 | -0.106591555 | 0.702687516 | 0.98 |
| YNL229C | 1349.306513 | -0.053260015 | 0.703696583 | 0.98 |
| YNR032W | 607.7463894 | 0.089399419 | 0.703716919 | 0.98 |
| YOL021C | 1670.215668 | -0.080517175 | 0.70550998 | 0.98 |
| YOL057W | 2160.171024 | -0.100447858 | 0.70165045 | 0.98 |
| YOR016C | 992.9033585 | 0.088193749 | 0.707249775 | 0.98 |
| YOR057W | 859.8202576 | 0.109197918 | 0.706665986 | 0.98 |
| YOR083W | 334.3141069 | -0.109164679 | 0.705522362 | 0.98 |
| YOR096W | 6750.76705 | -0.03240979 | 0.706249313 | 0.98 |
| YOR370C | 2847.263179 | 0.092136408 | 0.704143842 | 0.98 |
| snR69 | 9.747852558 | 0.130050034 | 0.703573202 | 0.98 |
| YCR020C | 245.5464449 | 0.140152018 | 0.708421386 | 0.98 |
| YER093C-A | 366.8172657 | 0.113176675 | 0.708561462 | 0.98 |
| YGR277C | 611.2932905 | -0.095056057 | 0.708553319 | 0.98 |
| YOR310C | 3221.373678 | -0.055829451 | 0.708153592 | 0.98 |
| YOR319W | 365.6759573 | 0.088504509 | 0.708647299 | 0.98 |
| YPL043W | 1057.06526 | -0.110801538 | 0.708058412 | 0.98 |
| YER047C | 911.6547256 | 0.084473592 | 0.708853951 | 0.98 |
| YJR067C | 196.0036818 | 0.095435997 | 0.709002206 | 0.98 |
| YOR233W | 853.5809378 | 0.083560274 | 0.709176791 | 0.98 |
| YDR483W | 4802.755253 | -0.059095759 | 0.709463224 | 0.98 |
| YDR132C | 453.7495244 | 0.108480471 | 0.709677222 | 0.98 |
| YDR006C | 2310.266671 | 0.085521236 | 0.710271756 | 0.98 |
| YOR349W | 572.160655 | 0.077422383 | 0.710157811 | 0.98 |
| YPL245W | 615.8162164 | -0.146379546 | 0.710309025 | 0.98 |
| YNR028W | 918.8142146 | -0.070330564 | 0.710838437 | 0.98 |
| YOL082W | 2476.571718 | -0.118023737 | 0.710705997 | 0.98 |
| YAL021C | 1595.906288 | -0.067712573 | 0.732407449 | 0.98 |
| YAL029C | 2871.619875 | 0.056224097 | 0.73476153 | 0.98 |
| YAL042C-A | 8.764234173 | -0.305984882 | 0.732087089 | 0.98 |
| YAR007C | 1627.673812 | -0.075882325 | 0.725907592 | 0.98 |
| YBL019W | 515.3822708 | 0.096345738 | 0.727797485 | 0.98 |
| YBL022C | 3958.243828 | -0.080337173 | 0.738155884 | 0.98 |

| | | | | |
|---|---|---|---|---|
| YBL029C-A | 1079.485322 | 0.131391007 | 0.734028938 | 0.98 |
| YBL055C | 719.3024768 | 0.090097439 | 0.736825841 | 0.98 |
| YBL071C | 215.271228 | 0.088436237 | 0.733622327 | 0.98 |
| YBL082C | 673.6132807 | 0.113582492 | 0.732020457 | 0.98 |
| YBL093C | 1069.915355 | 0.066463525 | 0.728545885 | 0.98 |
| YBR027C | 8.649310737 | -0.220254077 | 0.716653349 | 0.98 |
| YBR060C | 829.7516444 | -0.069045287 | 0.735847893 | 0.98 |
| YBR076C-A | 84.52994015 | 0.139848017 | 0.718442978 | 0.98 |
| YBR135W | 639.2382675 | -0.124946166 | 0.722519718 | 0.98 |
| YBR140C | 4380.796749 | 0.098461815 | 0.718488989 | 0.98 |
| YBR207W | 1493.475156 | -0.093589652 | 0.730316143 | 0.98 |
| YBR281C | 961.503364 | 0.094479683 | 0.737275782 | 0.98 |
| YCL001W | 1214.350549 | -0.090346477 | 0.719260297 | 0.98 |
| YCL008C | 1030.586387 | -0.09359102 | 0.720505139 | 0.98 |
| YCR006C | 58.71736424 | -0.148536287 | 0.722799152 | 0.98 |
| YCR024C | 641.2724541 | 0.057549146 | 0.72441737 | 0.98 |
| YCR024C-A | 1033.453894 | -0.100453604 | 0.713551891 | 0.98 |
| YCR037C | 1044.612368 | 0.085263483 | 0.727435483 | 0.98 |
| YCR046C | 1534.125343 | -0.078722363 | 0.730410513 | 0.98 |
| YDL015C | 3069.696513 | -0.053765703 | 0.72998714 | 0.98 |
| YDL023C | 219.5343501 | 0.176638397 | 0.721122287 | 0.98 |
| YDL076C | 828.6133083 | 0.063463821 | 0.734641465 | 0.98 |
| YDL097C | 4494.471702 | 0.080387651 | 0.733217264 | 0.98 |
| YDL105W | 423.3140946 | 0.076497418 | 0.737061091 | 0.98 |
| YDL138W | 666.9826314 | 0.079833863 | 0.732383447 | 0.98 |
| YDL142C | 615.3529618 | 0.090023437 | 0.719132479 | 0.98 |
| YDL143W | 3248.97051 | -0.076694136 | 0.726684506 | 0.98 |
| YDL196W | 15.17669232 | 0.156038265 | 0.712699107 | 0.98 |
| YDR042C | 81.10570396 | 0.112451764 | 0.733907807 | 0.98 |
| YDR077W | 88090.2349 | 0.12257708 | 0.714881565 | 0.98 |
| YDR117C | 1080.281382 | -0.062605519 | 0.721947759 | 0.98 |
| YDR161W | 1294.242267 | 0.179599398 | 0.719873197 | 0.98 |
| YDR197W | 858.0497536 | -0.105546686 | 0.722711675 | 0.98 |
| YDR257C | 688.7119517 | 0.131631629 | 0.725799218 | 0.98 |
| YDR280W | 709.9216811 | -0.075462499 | 0.737088102 | 0.98 |
| YDR316W | 792.8873428 | -0.087921653 | 0.730788486 | 0.98 |
| YDR322W | 1031.616879 | -0.093215042 | 0.716913259 | 0.98 |
| YDR336W | 301.9017741 | 0.088617866 | 0.721213306 | 0.98 |
| YDR395W | 2175.578678 | -0.096484803 | 0.734886168 | 0.98 |
| YDR408C | 1967.993247 | 0.158264169 | 0.731228433 | 0.98 |
| YDR432W | 4551.301879 | -0.092971921 | 0.726506629 | 0.98 |
| YDR443C | 1163.053889 | -0.120067206 | 0.721421905 | 0.98 |
| YDR521W | 13.62209314 | -0.19062266 | 0.735957807 | 0.98 |

| | | | | |
|---|---|---|---|---|
| YDR534C | 224.6689044 | -0.200560966 | 0.725457566 | 0.98 |
| YEL006W | 705.1819844 | -0.084846892 | 0.718052232 | 0.98 |
| YEL027W | 5744.186516 | -0.055004342 | 0.735033339 | 0.98 |
| YER006W | 1977.06385 | 0.245952552 | 0.718646754 | 0.98 |
| YER038W-A | 0.198141795 | -0.065537477 | 0.730848074 | 0.98 |
| YER079C-A | 18.60445932 | -0.277858231 | 0.727919577 | 0.98 |
| YER119C-A | 190.6802462 | -0.125885177 | 0.714819215 | 0.98 |
| YFL037W | 5301.265498 | 0.125252716 | 0.738436094 | 0.98 |
| YFR008W | 307.6442666 | -0.096475798 | 0.716252217 | 0.98 |
| YFR050C | 3854.503578 | 0.093004235 | 0.717072361 | 0.98 |
| YGL006W-A | 3.943531239 | -0.121113066 | 0.73709663 | 0.98 |
| YGL102C | 355.1928609 | -0.129170155 | 0.713056689 | 0.98 |
| YGL105W | 8369.251023 | -0.090369122 | 0.732225637 | 0.98 |
| YGL112C | 1298.501075 | 0.060685514 | 0.737317513 | 0.98 |
| YGL136C | 674.1887477 | -0.085549072 | 0.736809573 | 0.98 |
| YGL178W | 2073.150568 | 0.094895353 | 0.718753339 | 0.98 |
| YGL187C | 1692.180783 | 0.222735671 | 0.734231295 | 0.98 |
| YGL253W | 16247.71261 | 0.217552958 | 0.727990003 | 0.98 |
| YGR008C | 4582.699628 | 0.203300282 | 0.718918115 | 0.98 |
| YGR027C | 9077.452786 | -0.063766812 | 0.734993337 | 0.98 |
| YGR031W | 376.7830976 | 0.105186472 | 0.734036247 | 0.98 |
| YGR051C | 51.37808205 | 0.088012147 | 0.73676102 | 0.98 |
| YGR089W | 1598.86435 | -0.085528781 | 0.729125735 | 0.98 |
| YGR103W | 3122.539574 | 0.287797089 | 0.737028016 | 0.98 |
| YGR180C | 8240.520071 | 0.120002349 | 0.728848598 | 0.98 |
| YGR191W | 1267.875828 | 0.111585943 | 0.730089554 | 0.98 |
| YGR200C | 1330.804216 | -0.093253325 | 0.711158872 | 0.98 |
| YGR208W | 739.3899853 | -0.069101968 | 0.729313485 | 0.98 |
| YGR260W | 2070.398504 | 0.177429318 | 0.719113754 | 0.98 |
| YGR295C | 1429.22247 | 0.176487227 | 0.726368234 | 0.98 |
| YHL003C | 1333.653462 | -0.092359512 | 0.713344062 | 0.98 |
| YHL017W | 1099.085857 | 0.09224894 | 0.723839571 | 0.98 |
| YHL029C | 889.4266622 | -0.169061529 | 0.724084123 | 0.98 |
| YHR010W | 11885.41138 | -0.10564382 | 0.721681063 | 0.98 |
| YHR012W | 963.21973 | -0.080402399 | 0.737911784 | 0.98 |
| YHR015W | 104.6987227 | 0.140481903 | 0.722085839 | 0.98 |
| YHR027C | 5786.339997 | 0.147427399 | 0.720381591 | 0.98 |
| YHR047C | 1885.311487 | -0.087091227 | 0.73859507 | 0.98 |
| YHR063C | 1432.676855 | 0.145085368 | 0.735365489 | 0.98 |
| YHR083W | 1675.31411 | 0.085431002 | 0.718655619 | 0.98 |
| YHR104W | 3547.736943 | 0.105509925 | 0.712349724 | 0.98 |
| YHR115C | 896.8338505 | 0.084737011 | 0.723896471 | 0.98 |
| YHR186C | 1649.917336 | -0.072834447 | 0.725536599 | 0.98 |

| | | | | |
|---|---|---|---|---|
| YIL047C | 1300.508261 | -0.089290545 | 0.727081013 | 0.98 |
| YIL070C | 1550.574512 | 0.165187159 | 0.716252608 | 0.98 |
| YIL110W | 498.0390037 | 0.147320242 | 0.730204471 | 0.98 |
| YIR003W | 1806.796543 | 0.092191538 | 0.721005289 | 0.98 |
| YJL010C | 755.184634 | 0.160897923 | 0.732265584 | 0.98 |
| YJL015C | 72.86729941 | 0.040625826 | 0.72653197 | 0.98 |
| YJL032W | 6.67566349 | -0.302745852 | 0.73813083 | 0.98 |
| YJL069C | 705.1093909 | -0.14831985 | 0.727599245 | 0.98 |
| YJL114W | 3.593194288 | 0.247647682 | 0.713101439 | 0.98 |
| YJL136W-A | 0.34890375 | 0.062248746 | 0.724330051 | 0.98 |
| YJL170C | 341.173457 | -0.203743605 | 0.721574167 | 0.98 |
| YJR071W | 37.86358004 | 0.209651255 | 0.71588502 | 0.98 |
| YJR072C | 1860.139893 | 0.122084409 | 0.735555684 | 0.98 |
| YJR147W | 698.6444516 | 0.160374152 | 0.733559122 | 0.98 |
| YKL071W | 436.0298233 | -0.095359154 | 0.71846223 | 0.98 |
| YKL076C | 34.14810955 | 0.136534665 | 0.721096133 | 0.98 |
| YKL100C | 2362.710996 | 0.096407642 | 0.717257203 | 0.98 |
| YKL110C | 846.8769824 | -0.087782803 | 0.723349445 | 0.98 |
| YKL126W | 2370.590453 | -0.070268907 | 0.716761674 | 0.98 |
| YKL198C | 1074.404315 | -0.138752038 | 0.711378366 | 0.98 |
| YKR004C | 281.3110563 | -0.081642311 | 0.734988993 | 0.98 |
| YKR026C | 956.8364878 | -0.100850334 | 0.71282255 | 0.98 |
| YLR105C | 592.2672821 | -0.088345758 | 0.727704223 | 0.98 |
| YLR120C | 2959.418798 | -0.143120366 | 0.737428795 | 0.98 |
| YLR140W | 51.21550356 | -0.09974552 | 0.726188101 | 0.98 |
| YLR177W | 3573.62477 | 0.133172796 | 0.717195682 | 0.98 |
| YLR180W | 3585.379162 | 0.288191635 | 0.711765219 | 0.98 |
| YLR211C | 375.8938446 | 0.111612484 | 0.731483072 | 0.98 |
| YLR233C | 453.3027113 | 0.114761392 | 0.718847593 | 0.98 |
| YLR236C | 21.91002832 | 0.134906296 | 0.732815555 | 0.98 |
| YLR270W | 2664.450215 | 0.142334594 | 0.737749486 | 0.98 |
| YLR328W | 869.5602715 | 0.122637024 | 0.719800426 | 0.98 |
| YLR348C | 661.9197696 | -0.127452493 | 0.729256608 | 0.98 |
| YLR366W | 1.593075245 | 0.105885602 | 0.736147184 | 0.98 |
| YLR374C | 34.93928504 | 0.082307943 | 0.717449739 | 0.98 |
| YLR387C | 1423.423086 | 0.117517761 | 0.729105434 | 0.98 |
| YLR420W | 2792.570329 | 0.161650704 | 0.734798676 | 0.98 |
| YML006C | 696.4182875 | -0.070409809 | 0.738042052 | 0.98 |
| YML025C | 1270.924858 | -0.05534162 | 0.716641238 | 0.98 |
| YML084W | 2.063062303 | 0.254185481 | 0.715183693 | 0.98 |
| YML086C | 1768.316639 | 0.075262706 | 0.727022297 | 0.98 |
| YML093W | 1683.648354 | -0.111734246 | 0.734707392 | 0.98 |
| YML130C | 2617.920044 | 0.153010334 | 0.720307829 | 0.98 |

| | | | | |
|---|---|---|---|---|
| YMR071C | 1073.602658 | 0.042477124 | 0.737114153 | 0.98 |
| YMR163C | 451.0209425 | -0.071704419 | 0.735229201 | 0.98 |
| YMR167W | 578.0579077 | 0.07956478 | 0.72341209 | 0.98 |
| YMR246W | 3290.570155 | 0.181759564 | 0.734036372 | 0.98 |
| YMR294W | 447.832337 | 0.107146638 | 0.734689858 | 0.98 |
| YNL008C | 1006.822611 | 0.09288866 | 0.717102382 | 0.98 |
| YNL027W | 1180.449634 | -0.094436206 | 0.726411936 | 0.98 |
| YNL057W | 19.3681916 | -0.158269499 | 0.718391869 | 0.98 |
| YNL113W | 221.4093861 | -0.070982461 | 0.734149455 | 0.98 |
| YNL247W | 4374.768503 | 0.179686239 | 0.719687429 | 0.98 |
| YNL304W | 349.9176759 | 0.089573794 | 0.716113957 | 0.98 |
| YNL311C | 782.0547648 | 0.087214171 | 0.716985846 | 0.98 |
| YNR005C | 1.56429782 | 0.047456247 | 0.729264205 | 0.98 |
| YNR022C | 907.2227542 | -0.108363624 | 0.720876076 | 0.98 |
| YNR027W | 549.891736 | 0.120099437 | 0.714059606 | 0.98 |
| YNR060W | 291.930447 | 0.081282066 | 0.732966136 | 0.98 |
| YOL034W | 735.3581814 | -0.105675189 | 0.722023317 | 0.98 |
| YOL039W | 5643.898973 | -0.065094213 | 0.737224374 | 0.98 |
| YOL043C | 577.8464057 | -0.122333493 | 0.724626193 | 0.98 |
| YOL126C | 3179.903333 | -0.242692054 | 0.72887342 | 0.98 |
| YOL144W | 766.01586 | 0.15625049 | 0.737978962 | 0.98 |
| YOL151W | 926.9123911 | 0.155059558 | 0.718257991 | 0.98 |
| YOR029W | 6.24277143 | -0.195247636 | 0.722783503 | 0.98 |
| YOR084W | 822.8839971 | -0.108670451 | 0.738027182 | 0.98 |
| YOR127W | 1893.705383 | 0.070973755 | 0.717907359 | 0.98 |
| YOR161C | 928.1721785 | 0.13621968 | 0.733741181 | 0.98 |
| YOR161C-C | 9.014220677 | 0.053841081 | 0.719402931 | 0.98 |
| YOR195W | 1326.708475 | -0.102323631 | 0.722399946 | 0.98 |
| YOR201C | 630.2633868 | 0.084235852 | 0.733826207 | 0.98 |
| YOR217W | 1207.361787 | 0.072100544 | 0.726204519 | 0.98 |
| YOR257W | 743.3095021 | -0.138830488 | 0.714204964 | 0.98 |
| YOR260W | 2245.68858 | -0.120759233 | 0.722951832 | 0.98 |
| YOR264W | 932.8104963 | 0.098668703 | 0.717364419 | 0.98 |
| YOR289W | 937.4069291 | 0.178579242 | 0.728257222 | 0.98 |
| YOR322C | 1313.48176 | 0.073632663 | 0.72035374 | 0.98 |
| YPL108W | 159.4425969 | -0.086576469 | 0.717999305 | 0.98 |
| YPL140C | 588.07308 | -0.10014832 | 0.724013138 | 0.98 |
| YPL141C | 1338.030401 | -0.122355856 | 0.738500252 | 0.98 |
| YPL149W | 607.7809907 | 0.13158577 | 0.730886411 | 0.98 |
| YPL163C | 3377.565521 | 0.165100253 | 0.725267462 | 0.98 |
| YPL174C | 736.6858257 | 0.0601244 | 0.725627963 | 0.98 |
| YPL252C | 462.6644596 | -0.106370615 | 0.723590601 | 0.98 |
| YPR060C | 619.4282416 | -0.106245541 | 0.731694225 | 0.98 |

| | | | | |
|---|---|---|---|---|
| YPR082C | 431.92377 | -0.094229133 | 0.71309428 | 0.98 |
| YPR118W | 1223.179848 | -0.0879697 | 0.727530718 | 0.98 |
| YPR127W | 1734.545853 | 0.140421491 | 0.732267196 | 0.98 |
| YPR134W | 542.0304792 | 0.070424751 | 0.72180473 | 0.98 |
| YPR201W | 177.3808268 | 0.18009463 | 0.722884332 | 0.98 |
| snR35 | 41.6109753 | 0.166332576 | 0.7341539 | 0.98 |
| YFR007W | 462.3779448 | 0.11454936 | 0.738804063 | 0.98 |
| YDR017C | 2063.432291 | 0.04689574 | 0.739144271 | 0.98 |
| YER180C-A | 68.54860649 | -0.092919244 | 0.739244622 | 0.98 |
| YFR051C | 3059.620652 | -0.085006849 | 0.739587636 | 0.98 |
| YGL053W | 527.1405514 | -0.131689193 | 0.739443245 | 0.98 |
| YGL135W | 2224.916712 | -0.044896667 | 0.739475015 | 0.98 |
| YBR183W | 786.7022482 | 0.157927309 | 0.740256108 | 0.98 |
| YHR129C | 426.0946941 | -0.087957537 | 0.740145222 | 0.98 |
| YHR193C-A | 1.288094432 | -0.152241658 | 0.740567569 | 0.98 |
| YPL053C | 1363.897708 | -0.088064133 | 0.740689809 | 0.98 |
| snR8 | 32.91757815 | 0.184256733 | 0.740588796 | 0.98 |
| YDL229W | 7732.532873 | 0.256127162 | 0.740893128 | 0.98 |
| YMR290W-A | 11.59226843 | -0.305067425 | 0.741048259 | 0.98 |
| YDR292C | 1617.849006 | -0.057134597 | 0.741220611 | 0.98 |
| YDR440W | 548.6066441 | 0.115019446 | 0.741418697 | 0.98 |
| YOL046C | 5.940874887 | -0.34720025 | 0.741615189 | 0.98 |
| YDR060W | 3331.110803 | -0.091239043 | 0.742429035 | 0.98 |
| YGL133W | 1877.257774 | 0.064774124 | 0.742394267 | 0.98 |
| YAL031C | 1097.389337 | -0.110587887 | 0.742650173 | 0.98 |
| YEL073C | 122.6223904 | 0.190309021 | 0.742982888 | 0.98 |
| YMR273C | 1360.501919 | -0.106850435 | 0.742948678 | 0.98 |
| YBR008C | 906.2949136 | 0.08003355 | 0.745160239 | 0.98 |
| YDL026W | 16.56831607 | -0.213700416 | 0.744903006 | 0.98 |
| YFL036W | 2229.86904 | -0.109978462 | 0.744674458 | 0.98 |
| YGR259C | 0.780911468 | 0.088043626 | 0.744354942 | 0.98 |
| YKL023C-A | 532.0907152 | 0.140345103 | 0.744901872 | 0.98 |
| YKL068W | 1722.980976 | 0.069971736 | 0.744936135 | 0.98 |
| YLR456W | 255.5161749 | 0.082924953 | 0.744197403 | 0.98 |
| YNL204C | 121.2325338 | 0.097276552 | 0.745071685 | 0.98 |
| YOL068C | 1498.234041 | -0.07489355 | 0.744021055 | 0.98 |
| YOR168W | 3054.619284 | 0.108259027 | 0.744779121 | 0.98 |
| YOR241W | 933.790879 | -0.081389481 | 0.744650562 | 0.98 |
| YPL062W | 4.463478236 | 0.076831968 | 0.744537602 | 0.98 |
| YJR006W | 1054.347239 | 0.084434888 | 0.745513593 | 0.98 |
| YML012W | 3525.480868 | -0.056773292 | 0.74569451 | 0.98 |
| YPL183C | 913.1902093 | 0.120278293 | 0.745659226 | 0.98 |
| YEL036C | 1490.088365 | -0.097788276 | 0.746324406 | 0.98 |

| | | | | |
|---|---|---|---|---|
| YBL010C | 437.6652565 | 0.063519995 | 0.747146542 | 0.98 |
| YDR130C | 501.1168326 | -0.082550498 | 0.747197061 | 0.98 |
| YKL130C | 628.0827477 | -0.080645138 | 0.747186503 | 0.98 |
| YML114C | 955.0918143 | -0.066101219 | 0.747449601 | 0.98 |
| YMR160W | 946.3755314 | 0.079883253 | 0.747372307 | 0.98 |
| YJR099W | 741.8529597 | 0.078131518 | 0.7478642 | 0.98 |
| YHL009C | 603.1387723 | 0.100264807 | 0.749056174 | 0.98 |
| YOL025W | 668.044846 | 0.083580069 | 0.749085688 | 0.98 |
| YBR227C | 856.5044097 | -0.087997577 | 0.749757344 | 0.98 |
| YGL061C | 446.8633151 | 0.118390212 | 0.749760345 | 0.98 |
| YHR048W | 458.0188382 | 0.115939275 | 0.749979557 | 0.98 |
| YBL040C | 1584.786506 | -0.057237513 | 0.750708765 | 0.98 |
| YDL069C | 126.889784 | -0.083624954 | 0.750593753 | 0.98 |
| YOL124C | 564.6964356 | 0.282045773 | 0.750406751 | 0.98 |
| YNL264C | 1013.510724 | 0.136419602 | 0.751076954 | 0.99 |
| YBL106C | 800.8986155 | 0.057785501 | 0.75152145 | 0.99 |
| YBR134W | 28.00305569 | -0.131616733 | 0.752034291 | 0.99 |
| YDR012W | 4314.996695 | 0.247324377 | 0.752001935 | 0.99 |
| YHR128W | 1328.56215 | -0.073149889 | 0.75197978 | 0.99 |
| YMR267W | 919.8795189 | 0.132848162 | 0.751976486 | 0.99 |
| YNR008W | 899.8873602 | -0.078114076 | 0.752277563 | 0.99 |
| YGR168C | 490.2041357 | -0.071631268 | 0.752451693 | 0.99 |
| YGR196C | 1285.850548 | -0.071171815 | 0.752905041 | 0.99 |
| YLR166C | 1318.264285 | -0.052464736 | 0.752953888 | 0.99 |
| YJL111W | 2828.454028 | -0.074052131 | 0.75314802 | 0.99 |
| YIL169C | 2177.35422 | -0.120378637 | 0.753349185 | 0.99 |
| YDL018C | 497.6776418 | 0.120429349 | 0.753791255 | 0.99 |
| YAL017W | 3309.438838 | 0.109108238 | 0.754061604 | 0.99 |
| YIL052C | 4784.674042 | -0.047780088 | 0.754181211 | 0.99 |
| YOL127W | 9841.72384 | -0.021791695 | 0.75450097 | 0.99 |
| YPL253C | 481.3229549 | -0.090020788 | 0.754732874 | 0.99 |
| YAL044C | 2900.555908 | 0.191871419 | 0.763832676 | 0.99 |
| YAR018C | 738.6960091 | -0.078023187 | 0.760174266 | 0.99 |
| YAR029W | 75.14481749 | 0.094916376 | 0.758843771 | 0.99 |
| YBL074C | 232.3789197 | 0.08536711 | 0.763535531 | 0.99 |
| YBR219C | 1.059885032 | -0.042674802 | 0.764019627 | 0.99 |
| YCL034W | 1930.992154 | 0.083438673 | 0.764273155 | 0.99 |
| YCR067C | 1807.391426 | 0.072462124 | 0.75853965 | 0.99 |
| YDL073W | 971.6916673 | -0.081081936 | 0.760862543 | 0.99 |
| YDL158C | 89.30551169 | 0.138380941 | 0.762804671 | 0.99 |
| YDR129C | 5762.337606 | -0.074648565 | 0.759772556 | 0.99 |
| YDR252W | 88.93914678 | 0.106562809 | 0.765112464 | 0.99 |
| YDR377W | 1422.817631 | 0.179899939 | 0.761193085 | 0.99 |

| | | | | |
|---|---|---|---|---|
| YDR398W | 1282.093858 | -0.044386826 | 0.760257824 | 0.99 |
| YDR465C | 1088.193625 | 0.231029652 | 0.764772688 | 0.99 |
| YEL017C-A | 672.6729484 | -0.089266465 | 0.758493436 | 0.99 |
| YER055C | 2945.986692 | -0.048139482 | 0.76414843 | 0.99 |
| YER110C | 3689.619393 | 0.213416804 | 0.75725626 | 0.99 |
| YFL064C | 3.015892609 | 0.105345924 | 0.762135844 | 0.99 |
| YFR006W | 3007.667423 | -0.052739086 | 0.76130634 | 0.99 |
| YGL064C | 383.7385393 | 0.072114995 | 0.764544581 | 0.99 |
| YGL129C | 1219.243345 | 0.080285838 | 0.757813772 | 0.99 |
| YGL147C | 5508.177604 | 0.055638831 | 0.765041595 | 0.99 |
| YGL200C | 3717.842992 | -0.050549638 | 0.763817016 | 0.99 |
| YGR047C | 884.2267464 | -0.066405148 | 0.759339908 | 0.99 |
| YGR077C | 1152.926191 | -0.071740452 | 0.755157262 | 0.99 |
| YGR213C | 279.0368426 | 0.086420369 | 0.760939522 | 0.99 |
| YHL001W | 1027.533504 | -0.050627572 | 0.7603531 | 0.99 |
| YHR118C | 526.5681517 | 0.046814082 | 0.758704826 | 0.99 |
| YHR148W | 337.7936716 | 0.172464934 | 0.761774214 | 0.99 |
| YIL030C | 2092.978365 | 0.090184074 | 0.759051382 | 0.99 |
| YIL031W | 1396.422579 | 0.067371844 | 0.756054368 | 0.99 |
| YIL135C | 1336.175722 | -0.094562484 | 0.758836384 | 0.99 |
| YJL052C-A | 1.688849253 | 0.301295581 | 0.760040588 | 0.99 |
| YJL081C | 1547.994145 | 0.056456181 | 0.763509083 | 0.99 |
| YJL109C | 2576.133571 | -0.096917941 | 0.759282573 | 0.99 |
| YJL148W | 1042.119748 | -0.049656565 | 0.763589104 | 0.99 |
| YJR023C | 6.716930152 | 0.123663578 | 0.761937694 | 0.99 |
| YJR060W | 664.6110196 | 0.057730482 | 0.764420125 | 0.99 |
| YJR117W | 3603.291074 | -0.091330978 | 0.758709376 | 0.99 |
| YKL007W | 2212.362326 | -0.041507685 | 0.762222675 | 0.99 |
| YKL033W-A | 1409.695613 | 0.085079527 | 0.763912107 | 0.99 |
| YKL056C | 11023.03919 | -0.028936446 | 0.759304275 | 0.99 |
| YKL088W | 1161.516822 | -0.107989852 | 0.755449791 | 0.99 |
| YKL216W | 7973.975155 | -0.16459441 | 0.764293913 | 0.99 |
| YLL020C | 30.70630028 | 0.142674726 | 0.763597412 | 0.99 |
| YLR147C | 302.8969457 | -0.068297032 | 0.763175352 | 0.99 |
| YLR302C | 0.305761563 | 0.064792613 | 0.764949139 | 0.99 |
| YLR443W | 472.8026492 | 0.089889152 | 0.757339406 | 0.99 |
| YMR157C | 556.2957769 | -0.083299842 | 0.758878669 | 0.99 |
| YMR218C | 1133.912405 | -0.061858508 | 0.760983271 | 0.99 |
| YMR229C | 3400.401838 | -0.100754899 | 0.764096006 | 0.99 |
| YMR264W | 1111.60862 | 0.077225013 | 0.759850317 | 0.99 |
| YNL021W | 893.5874562 | -0.045896287 | 0.757715245 | 0.99 |
| YNL178W | 12909.44004 | -0.018186538 | 0.757314425 | 0.99 |
| YNL208W | 6496.983749 | 0.085869638 | 0.76505491 | 0.99 |

| | | | | |
|---|---|---|---|---|
| YNL265C | 447.0625511 | 0.062108302 | 0.760703892 | 0.99 |
| YOL132W | 124.9358227 | -0.200419505 | 0.763511178 | 0.99 |
| YOR041C | 37.3273586 | -0.105732483 | 0.756332533 | 0.99 |
| YOR102W | 49.45521107 | -0.091720618 | 0.763818475 | 0.99 |
| YOR176W | 2270.643704 | -0.099577993 | 0.764022416 | 0.99 |
| YOR208W | 1224.294967 | 0.095754333 | 0.763930118 | 0.99 |
| YPR047W | 774.0936392 | 0.075386987 | 0.75808343 | 0.99 |
| YPR105C | 1243.404818 | -0.064819719 | 0.757759435 | 0.99 |
| YPR132W | 5266.120182 | 0.002637262 | 0.758304294 | 0.99 |
| snR39B | 13.58422976 | -0.160123802 | 0.758244126 | 0.99 |
| snR42 | 83.98583809 | 0.182716911 | 0.764784224 | 0.99 |
| snR86 | 894.0383097 | 0.222151964 | 0.757947046 | 0.99 |
| YDR512C | 741.4198181 | -0.082841542 | 0.765877065 | 0.99 |
| YHL006C | 116.1902464 | -0.119989389 | 0.76576536 | 0.99 |
| YOR272W | 1204.374523 | 0.176914794 | 0.765612809 | 0.99 |
| YPL048W | 7812.579467 | 0.06981281 | 0.765585869 | 0.99 |
| YCR053W | 6107.267981 | 0.149620809 | 0.766657757 | 0.99 |
| YGR174W-A | 47.91809201 | 0.14613203 | 0.766087338 | 0.99 |
| YGR211W | 3320.437529 | 0.178372379 | 0.766623687 | 0.99 |
| YKL142W | 4281.67462 | 0.108456089 | 0.766800359 | 0.99 |
| YMR064W | 1294.977576 | -0.084117778 | 0.766433434 | 0.99 |
| YNR038W | 1115.811097 | -0.098579664 | 0.766473195 | 0.99 |
| YBR044C | 837.2738234 | -0.076628179 | 0.767065485 | 0.99 |
| YCL050C | 4005.943189 | 0.204575216 | 0.768660964 | 0.99 |
| YDL021W | 1151.606846 | 0.18670863 | 0.7686254 | 0.99 |
| YDL104C | 329.3472486 | 0.086180093 | 0.767605805 | 0.99 |
| YDR119W-A | 458.2326718 | 0.242587622 | 0.768397348 | 0.99 |
| YGL043W | 1337.950161 | -0.070739526 | 0.767759906 | 0.99 |
| YKL134C | 542.9022847 | 0.062365473 | 0.767526771 | 0.99 |
| YKR021W | 1328.600543 | 0.056892875 | 0.767744102 | 0.99 |
| YMR122C | 59.11544635 | 0.101184268 | 0.768853855 | 0.99 |
| YNL042W | 593.1711026 | -0.135480948 | 0.767907193 | 0.99 |
| YNR067C | 2460.572138 | 0.111673048 | 0.768605357 | 0.99 |
| YOL089C | 836.6158144 | 0.059017817 | 0.768097082 | 0.99 |
| YOR124C | 2791.881139 | 0.075239837 | 0.768725743 | 0.99 |
| YIL171W | 27.73883289 | 0.136517494 | 0.76939004 | 0.99 |
| YLR114C | 1321.213644 | -0.064740809 | 0.769371544 | 0.99 |
| NME1 | 51.73748825 | 0.193173005 | 0.773556266 | 0.99 |
| YAR003W | 588.5199738 | 0.09137389 | 0.779562981 | 0.99 |
| YBL025W | 212.6114748 | 0.090422913 | 0.784030975 | 0.99 |
| YBL083C | 7.154873991 | 0.151281938 | 0.782638334 | 0.99 |
| YBR048W | 5960.519296 | -0.075300768 | 0.770237821 | 0.99 |
| YBR058C | 1360.304304 | 0.042085442 | 0.779797481 | 0.99 |

| | | | | |
|---|---|---|---|---|
| YBR063C | 508.4120941 | -0.075454681 | 0.7835357 | 0.99 |
| YBR087W | 809.749781 | -0.066680431 | 0.774194628 | 0.99 |
| YBR095C | 539.8071954 | 0.056160701 | 0.784319965 | 0.99 |
| YBR224W | 27.9935508 | 0.135622966 | 0.773997092 | 0.99 |
| YCL002C | 469.4119103 | 0.082164049 | 0.784226581 | 0.99 |
| YDL019C | 2417.237505 | -0.070406244 | 0.784049398 | 0.99 |
| YDL124W | 12433.87204 | 0.100061434 | 0.777986421 | 0.99 |
| YDL233W | 547.8994161 | -0.074788242 | 0.773368415 | 0.99 |
| YDR198C | 539.1037114 | 0.061184556 | 0.77641221 | 0.99 |
| YDR320C | 1271.658982 | 0.055509258 | 0.772606382 | 0.99 |
| YDR427W | 2923.459081 | -0.057383259 | 0.783005931 | 0.99 |
| YDR476C | 1082.582617 | 0.073911939 | 0.782625051 | 0.99 |
| YDR539W | 882.7410932 | -0.125820053 | 0.784092018 | 0.99 |
| YEL005C | 730.8016186 | -0.110809508 | 0.78023009 | 0.99 |
| YEL038W | 943.0798183 | 0.080930309 | 0.773558692 | 0.99 |
| YER111C | 1455.121069 | -0.081254066 | 0.769596031 | 0.99 |
| YER145C-A | 50.64924556 | -0.08652018 | 0.772980972 | 0.99 |
| YGL029W | 383.3919755 | -0.022893408 | 0.777049984 | 0.99 |
| YGL031C | 8332.907395 | -0.014053322 | 0.783417129 | 0.99 |
| YGL049C | 2260.025541 | 0.042064342 | 0.782447105 | 0.99 |
| YGL060W | 769.6330407 | 0.059806811 | 0.779162472 | 0.99 |
| YGL078C | 1925.6458 | 0.14308484 | 0.78411306 | 0.99 |
| YGL099W | 1111.20672 | 0.127697993 | 0.77444999 | 0.99 |
| YGL154C | 461.1131948 | 0.058448457 | 0.78098005 | 0.99 |
| YGL179C | 645.0803373 | -0.153567102 | 0.779261829 | 0.99 |
| YGR009C | 1451.464581 | -0.058491401 | 0.776619923 | 0.99 |
| YGR034W | 5016.73135 | -0.00920913 | 0.784223418 | 0.99 |
| YGR140W | 791.394714 | 0.084033168 | 0.783517182 | 0.99 |
| YGR143W | 1625.66982 | -0.105720042 | 0.77089913 | 0.99 |
| YGR160W | 19.24237613 | -0.073457167 | 0.771046678 | 0.99 |
| YGR203W | 222.1887092 | -0.11155742 | 0.781891982 | 0.99 |
| YGR214W | 11003.56684 | -0.040795215 | 0.776384694 | 0.99 |
| YGR283C | 529.6205161 | 0.08373547 | 0.777265251 | 0.99 |
| YHR101C | 345.7524508 | 0.060876883 | 0.775219924 | 0.99 |
| YHR140W | 190.2878906 | -0.116798027 | 0.780920935 | 0.99 |
| YHR179W | 8288.962491 | 0.123920333 | 0.775555796 | 0.99 |
| YHR191C | 210.2719774 | -0.058207308 | 0.774341143 | 0.99 |
| YIL037C | 352.6358083 | 0.101298224 | 0.773820133 | 0.99 |
| YIL096C | 413.8812975 | 0.167297353 | 0.773028504 | 0.99 |
| YIL124W | 2615.425395 | -0.081697073 | 0.771310215 | 0.99 |
| YIR018W | 398.3013208 | 0.104292112 | 0.773199788 | 0.99 |
| YJL065C | 199.2698224 | -0.064315263 | 0.770907465 | 0.99 |
| YJL195C | 5.108398903 | 0.132152155 | 0.781792634 | 0.99 |

| | | | | |
|---|---|---|---|---|
| YJR065C | 3820.877578 | 0.05502537 | 0.771491884 | 0.99 |
| YJR070C | 1785.660344 | 0.006339404 | 0.780443232 | 0.99 |
| YKL002W | 1601.062288 | 0.06795012 | 0.774000936 | 0.99 |
| YKL008C | 1304.945966 | 0.135066109 | 0.783784819 | 0.99 |
| YKL123W | 31.46802918 | -0.158153254 | 0.7778634 | 0.99 |
| YKL203C | 2057.606082 | 0.060362024 | 0.777165241 | 0.99 |
| YKR001C | 4012.227533 | 0.077394439 | 0.777415079 | 0.99 |
| YLL045C | 7512.686429 | -0.014026487 | 0.772697019 | 0.99 |
| YLR057W | 1092.547525 | -0.076535706 | 0.781612503 | 0.99 |
| YLR093C | 1657.689066 | 0.082261112 | 0.772364624 | 0.99 |
| YLR107W | 1035.729202 | -0.098199642 | 0.777946731 | 0.99 |
| YLR141W | 304.974896 | -0.084987879 | 0.783216415 | 0.99 |
| YLR196W | 1358.156963 | -0.055497358 | 0.782567232 | 0.99 |
| YLR360W | 653.1816733 | -0.053268248 | 0.770056536 | 0.99 |
| YLR390W | 418.8392228 | -0.064514251 | 0.772595028 | 0.99 |
| YLR429W | 3551.790145 | -0.106713902 | 0.782383306 | 0.99 |
| YLR438C-A | 659.2030513 | 0.098945656 | 0.775422589 | 0.99 |
| YML010W | 2999.463402 | -0.055116358 | 0.776479777 | 0.99 |
| YMR008C | 3679.477965 | 0.106407857 | 0.778403177 | 0.99 |
| YMR144W | 347.0360461 | 0.116432317 | 0.781176175 | 0.99 |
| YMR244C-A | 108.6393324 | -0.08612351 | 0.772514324 | 0.99 |
| YMR283C | 413.1862547 | 0.054631462 | 0.77489636 | 0.99 |
| YMR310C | 757.4056331 | -0.111596325 | 0.775715694 | 0.99 |
| YMR317W | 752.1354442 | 0.082376711 | 0.778205028 | 0.99 |
| YMR326C | 0.406623722 | 0.017971201 | 0.774923741 | 0.99 |
| YNL005C | 1578.642517 | 0.062705925 | 0.774805412 | 0.99 |
| YNL067W | 1749.269832 | -0.018840717 | 0.769876086 | 0.99 |
| YNL097C-B | 13.11125706 | 0.079908042 | 0.77576999 | 0.99 |
| YNL333W | 85.09307581 | 0.140595133 | 0.783460797 | 0.99 |
| YNR035C | 3358.997637 | -0.061691328 | 0.770024379 | 0.99 |
| YOL059W | 2587.338376 | 0.098676023 | 0.771461576 | 0.99 |
| YOR046C | 2190.495993 | -0.072751108 | 0.770390786 | 0.99 |
| YOR145C | 791.624005 | -0.042061169 | 0.780175017 | 0.99 |
| YOR225W | 54.88254174 | 0.194005038 | 0.780906964 | 0.99 |
| YOR326W | 4223.653773 | -0.048077112 | 0.773727585 | 0.99 |
| YOR365C | 309.6078398 | -0.175377181 | 0.772139195 | 0.99 |
| YOR372C | 1526.129326 | -0.077026778 | 0.770621511 | 0.99 |
| YOR373W | 745.8071516 | 0.08697249 | 0.782942185 | 0.99 |
| YOR376W-A | 83.93269889 | -0.092707339 | 0.780017995 | 0.99 |
| YPL123C | 1037.088707 | 0.116211472 | 0.783844498 | 0.99 |
| YPL124W | 247.3933842 | -0.062016074 | 0.783801607 | 0.99 |
| YPL152W | 506.4565509 | -0.055529932 | 0.784144269 | 0.99 |
| YPL154C | 14185.44316 | 0.128753462 | 0.782755629 | 0.99 |

| | | | | |
|---|---|---|---|---|
| YPL188W | 716.8167679 | 0.076009298 | 0.778352281 | 0.99 |
| YPL212C | 1188.64058 | 0.16831944 | 0.77407742 | 0.99 |
| YPL271W | 1376.293801 | 0.107875495 | 0.782817536 | 0.99 |
| YPR071W | 302.2135475 | 0.091743519 | 0.783366348 | 0.99 |
| YPR168W | 170.9858215 | 0.086250076 | 0.780054332 | 0.99 |
| snR83 | 177.6034874 | -0.142578688 | 0.77777694 | 0.99 |
| YDR484W | 738.7697722 | 0.03773563 | 0.784540358 | 0.99 |
| YCR064C | 12.51518247 | -0.124987416 | 0.784854462 | 0.99 |
| YDR170C | 4884.61556 | -0.054459148 | 0.784849277 | 0.99 |
| YJL174W | 2482.371966 | -0.04429725 | 0.785005377 | 0.99 |
| YKR106W | 110.7787445 | 0.105261893 | 0.785175503 | 0.99 |
| YOL064C | 847.1293742 | 0.123705799 | 0.785382929 | 0.99 |
| YGR123C | 655.6704403 | -0.066182245 | 0.785729948 | 0.99 |
| YDR459C | 457.0405938 | -0.053183338 | 0.786082481 | 0.99 |
| YER060W | 750.1023347 | -0.100315768 | 0.786404189 | 0.99 |
| YOR179C | 265.678804 | -0.075510937 | 0.786358286 | 0.99 |
| YDL095W | 3378.459132 | -0.065429829 | 0.787266395 | 0.99 |
| YJR094C | 253.8151399 | 0.085893593 | 0.787478065 | 0.99 |
| YJR112W-A | 356.5088081 | 0.069019898 | 0.787936584 | 0.99 |
| YLR011W | 284.831779 | -0.063563321 | 0.787130703 | 0.99 |
| YLR368W | 767.6389144 | 0.063510383 | 0.787991165 | 0.99 |
| YNL167C | 1564.476742 | -0.084678681 | 0.787869804 | 0.99 |
| YOR381W | 508.0002714 | 0.105709774 | 0.787870679 | 0.99 |
| YPL057C | 1620.500312 | -0.099246939 | 0.787273974 | 0.99 |
| YBR270C | 464.1113888 | 0.05661533 | 0.788237463 | 0.99 |
| YAL016C-A | 61.31300348 | -0.119101379 | 0.790831293 | 0.99 |
| YAL037C-B | 133.1022568 | 0.173703723 | 0.790007016 | 0.99 |
| YAR068W | 4.842260677 | 0.190247957 | 0.789239952 | 0.99 |
| YBR217W | 353.7676501 | -0.061778667 | 0.790790375 | 0.99 |
| YCR025C | 44.00460202 | 0.095382629 | 0.788949175 | 0.99 |
| YCR041W | 16.4079087 | -0.119666359 | 0.789934998 | 0.99 |
| YDL148C | 2147.255757 | -0.078719495 | 0.790179968 | 0.99 |
| YFR031C-A | 6438.897656 | 0.015447415 | 0.790751869 | 0.99 |
| YGL008C | 26806.32689 | 0.033584587 | 0.790801241 | 0.99 |
| YHR093W | 35.27687991 | 0.066106939 | 0.790752953 | 0.99 |
| YHR141C | 3868.981705 | -0.032600929 | 0.789565248 | 0.99 |
| YNL062C | 805.5436558 | 0.150344252 | 0.790476826 | 0.99 |
| YNL130C-A | 3.921452687 | -0.154050637 | 0.789315334 | 0.99 |
| YNL268W | 2190.621688 | -0.063210437 | 0.789798296 | 0.99 |
| YOR170W | 97.11015677 | 0.08504081 | 0.790819519 | 0.99 |
| YPL220W | 1761.369957 | 0.166754576 | 0.788866564 | 0.99 |
| YER002W | 1431.744689 | 0.107238248 | 0.791535383 | 0.99 |
| YER170W | 376.8018239 | -0.05261724 | 0.791661116 | 0.99 |

| | | | | |
|---|---|---|---|---|
| YHR114W | 1802.212821 | 0.042840452 | 0.791699637 | 0.99 |
| YNL209W | 13780.76239 | 0.164039591 | 0.79118934 | 0.99 |
| snR59 | 2.356669998 | 0.143688633 | 0.791629133 | 0.99 |
| RUF22 | 1.126113591 | 0.054129511 | 0.808133795 | 0.99 |
| SCR1 | 181.7417356 | 0.201791083 | 0.813700648 | 0.99 |
| YAL001C | 1631.198822 | -0.057453928 | 0.816642339 | 0.99 |
| YAL068W-A | 0.241654238 | 0.021959617 | 0.81903171 | 0.99 |
| YBL014C | 840.8039898 | 0.068900399 | 0.797774439 | 0.99 |
| YBR030W | 579.6269936 | 0.092931501 | 0.804558089 | 0.99 |
| YBR051W | 12.61851356 | 0.076777466 | 0.80918757 | 0.99 |
| YBR077C | 472.5529414 | -0.054854827 | 0.81075946 | 0.99 |
| YBR126W-B | 156.4452467 | 0.075908907 | 0.816839676 | 0.99 |
| YBR181C | 1794.107741 | 0.055050182 | 0.820833383 | 0.99 |
| YBR189W | 11760.64407 | 0.013681776 | 0.796334633 | 0.99 |
| YBR194W | 226.3319567 | 0.063854946 | 0.823380672 | 0.99 |
| YBR209W | 5.517408049 | -0.166080691 | 0.808318545 | 0.99 |
| YBR225W | 1153.176358 | -0.084848836 | 0.820879771 | 0.99 |
| YCL043C | 17735.82228 | 0.098557249 | 0.807743308 | 0.99 |
| YCL048W-A | 89.40578665 | 0.101645632 | 0.814868134 | 0.99 |
| YCR086W | 338.9363584 | 0.0992673 | 0.797323409 | 0.99 |
| YDL007C-A | 3.853033258 | 0.039159095 | 0.801611915 | 0.99 |
| YDL029W | 3675.857405 | -0.028460214 | 0.806219029 | 0.99 |
| YDL041W | 0.97089065 | -0.125998195 | 0.806551879 | 0.99 |
| YDL089W | 963.5832446 | 0.08334616 | 0.801145727 | 0.99 |
| YDL133W | 835.1418998 | -0.046481939 | 0.82412454 | 0.99 |
| YDL153C | 1719.824839 | 0.154596035 | 0.80414268 | 0.99 |
| YDL157C | 952.2291983 | -0.047429477 | 0.804404324 | 0.99 |
| YDL188C | 1068.696328 | -0.070099018 | 0.803731997 | 0.99 |
| YDL206W | 597.9682559 | -0.074788679 | 0.822015676 | 0.99 |
| YDR008C | 37.32753147 | 0.075276307 | 0.81865601 | 0.99 |
| YDR020C | 268.9612697 | 0.075928982 | 0.819383285 | 0.99 |
| YDR088C | 645.3535973 | -0.065440561 | 0.822804926 | 0.99 |
| YDR091C | 1728.536947 | -0.045725647 | 0.81879837 | 0.99 |
| YDR097C | 218.1671058 | 0.368731666 | 0.803750465 | 0.99 |
| YDR100W | 731.7492012 | -0.077057001 | 0.82167605 | 0.99 |
| YDR133C | 1765.716231 | 0.147579144 | 0.797129796 | 0.99 |
| YDR139C | 266.4093081 | -0.049132285 | 0.802611517 | 0.99 |
| YDR142C | 534.1730573 | 0.064941616 | 0.797138358 | 0.99 |
| YDR165W | 1014.540812 | 0.129865704 | 0.817319933 | 0.99 |
| YDR166C | 1344.376251 | 0.056729372 | 0.793974467 | 0.99 |
| YDR168W | 2674.002145 | 0.101832084 | 0.799536398 | 0.99 |
| YDR184C | 548.4562077 | 0.100085584 | 0.8205214 | 0.99 |
| YDR271C | 28.54979964 | -0.138693089 | 0.815608698 | 0.99 |

| | | | | |
|---|---|---|---|---|
| YDR288W | 555.4180316 | -0.069158398 | 0.819135358 | 0.99 |
| YDR304C | 4614.533404 | -0.052153868 | 0.800470987 | 0.99 |
| YDR305C | 387.9870014 | 0.085241575 | 0.823794038 | 0.99 |
| YDR335W | 3040.69958 | 0.059249834 | 0.806657962 | 0.99 |
| YDR341C | 4249.5314 | 0.156847177 | 0.818571247 | 0.99 |
| YDR362C | 764.8048646 | -0.045920169 | 0.802278861 | 0.99 |
| YDR411C | 1171.452043 | -0.066040365 | 0.822239935 | 0.99 |
| YDR434W | 1180.144828 | -0.06902648 | 0.821129711 | 0.99 |
| YDR454C | 2624.226849 | -0.029597293 | 0.818790289 | 0.99 |
| YDR467C | 75.26126934 | 0.037928667 | 0.821981271 | 0.99 |
| YDR507C | 1606.54523 | -0.067642247 | 0.805473508 | 0.99 |
| YDR511W | 614.1187303 | -0.064015909 | 0.81565768 | 0.99 |
| YEL049W | 54.33605347 | 0.087390721 | 0.799709444 | 0.99 |
| YER035W | 1131.143378 | -0.116721554 | 0.802568885 | 0.99 |
| YER068W | 1672.188885 | -0.077308057 | 0.803123615 | 0.99 |
| YER120W | 4329.489139 | -0.030481819 | 0.824103246 | 0.99 |
| YER126C | 1138.972805 | -0.054849685 | 0.81365543 | 0.99 |
| YER175W-A | 27.41241786 | 0.057853317 | 0.817930301 | 0.99 |
| YFL052W | 300.3969613 | 0.163270638 | 0.816675216 | 0.99 |
| YGL003C | 856.5289623 | 0.052232073 | 0.802341466 | 0.99 |
| YGL041C-B | 21.05072236 | -0.002712557 | 0.792087585 | 0.99 |
| YGL173C | 8071.132916 | -0.040788714 | 0.800224201 | 0.99 |
| YGL213C | 596.0286848 | -0.046051079 | 0.807688809 | 0.99 |
| YGL217C | 8.839690893 | -0.105349652 | 0.794716039 | 0.99 |
| YGL235W | 1.903973228 | 0.123441813 | 0.818211901 | 0.99 |
| YGR015C | 438.1021585 | -0.048343333 | 0.808257135 | 0.99 |
| YGR101W | 624.5372953 | 0.049864564 | 0.801583421 | 0.99 |
| YGR120C | 376.5121986 | -0.02454847 | 0.817842656 | 0.99 |
| YGR134W | 745.1524745 | -0.057872537 | 0.817429174 | 0.99 |
| YGR170W | 2142.624103 | 0.046451827 | 0.823859431 | 0.99 |
| YGR223C | 1134.541576 | 0.060173736 | 0.824165769 | 0.99 |
| YGR240C | 9640.703605 | 0.112921228 | 0.796200764 | 0.99 |
| YGR245C | 1499.530458 | 0.176979753 | 0.821658306 | 0.99 |
| YGR279C | 11712.04252 | -0.048617646 | 0.808810098 | 0.99 |
| YHL025W | 1272.013263 | -0.0822393 | 0.822845814 | 0.99 |
| YHL034W-A | 585.1049194 | -0.080620817 | 0.813852705 | 0.99 |
| YHL043W | 209.5139873 | -0.115109096 | 0.819283147 | 0.99 |
| YHR006W | 1327.030831 | 0.079318013 | 0.811058878 | 0.99 |
| YHR043C | 1297.179518 | -0.045377421 | 0.806879991 | 0.99 |
| YHR090C | 634.1320105 | 0.071989465 | 0.822064525 | 0.99 |
| YHR098C | 2961.484376 | -0.056234625 | 0.793011268 | 0.99 |
| YHR130C | 24.10330458 | -0.136844868 | 0.818006885 | 0.99 |
| YHR163W | 2218.862273 | 0.174756269 | 0.801857275 | 0.99 |

| | | | | |
|---|---|---|---|---|
| YHR180W | 212.5255364 | -0.121913095 | 0.806509169 | 0.99 |
| YHR195W | 1250.379878 | -0.099749516 | 0.813553407 | 0.99 |
| YIL018W | 10226.88893 | 0.028131096 | 0.815878344 | 0.99 |
| YIL039W | 1826.424519 | -0.051082105 | 0.796949975 | 0.99 |
| YIL058W | 79.78173189 | 0.044351305 | 0.821590631 | 0.99 |
| YIL091C | 989.7593991 | -0.092191676 | 0.819546401 | 0.99 |
| YIL102C-A | 112.0491515 | -0.07248845 | 0.805040896 | 0.99 |
| YIL121W | 2357.398488 | 0.066759415 | 0.797976003 | 0.99 |
| YIL134W | 238.1456546 | 0.119321782 | 0.803172479 | 0.99 |
| YIL156W-A | 6.740450198 | -0.238126023 | 0.824153516 | 0.99 |
| YIL173W | 14.66085042 | -0.246028386 | 0.793109801 | 0.99 |
| YJL016W | 1543.218497 | -0.085592142 | 0.82206842 | 0.99 |
| YJL036W | 1275.214013 | 0.063173901 | 0.817438443 | 0.99 |
| YJL047C-A | 19.06572924 | 0.109672072 | 0.798066493 | 0.99 |
| YJL050W | 1800.303824 | 0.134302903 | 0.803241239 | 0.99 |
| YJL062W-A | 1034.508423 | -0.068465534 | 0.810401262 | 0.99 |
| YJL076W | 2985.809725 | -0.057228905 | 0.814705108 | 0.99 |
| YJL113W | 119.706187 | 0.088238876 | 0.814674859 | 0.99 |
| YJL117W | 1366.561839 | -0.045186066 | 0.815474538 | 0.99 |
| YJL123C | 2650.72374 | 0.065895461 | 0.792644809 | 0.99 |
| YJL126W | 376.2312005 | 0.046390835 | 0.810740102 | 0.99 |
| YJL129C | 2269.469864 | 0.032073629 | 0.818849481 | 0.99 |
| YJL166W | 2486.295448 | 0.139687493 | 0.806306716 | 0.99 |
| YJL201W | 1030.90288 | -0.061228647 | 0.792533623 | 0.99 |
| YJL210W | 543.5019911 | 0.07528691 | 0.821019176 | 0.99 |
| YJR053W | 291.1479634 | 0.061974777 | 0.822000883 | 0.99 |
| YJR066W | 2254.068141 | 0.060467406 | 0.80492317 | 0.99 |
| YJR100C | 772.6621126 | -0.049153897 | 0.813465901 | 0.99 |
| YJR107W | 474.7713781 | 0.066648053 | 0.795621267 | 0.99 |
| YJR145C | 6502.704468 | 0.003780836 | 0.81157308 | 0.99 |
| YKL021C | 1160.236373 | 0.109016998 | 0.822499085 | 0.99 |
| YKL039W | 2424.281879 | 0.049354114 | 0.813121034 | 0.99 |
| YKL040C | 858.7842501 | -0.044556798 | 0.819895856 | 0.99 |
| YKL081W | 14463.37631 | 0.001763033 | 0.814038943 | 0.99 |
| YKL091C | 918.9800837 | 0.101810339 | 0.813254052 | 0.99 |
| YKL104C | 3816.518475 | -0.058668961 | 0.80910547 | 0.99 |
| YKL152C | 30648.47356 | 0.141144238 | 0.819065275 | 0.99 |
| YKL201C | 1146.760044 | -0.079912025 | 0.8061168 | 0.99 |
| YKL206C | 563.6605725 | 0.063852026 | 0.818109583 | 0.99 |
| YKR077W | 759.0676767 | -0.121902804 | 0.796720004 | 0.99 |
| YKR100C | 1038.230895 | -0.060300321 | 0.80621365 | 0.99 |
| YLL013C | 908.878738 | 0.091162816 | 0.811762898 | 0.99 |
| YLL032C | 878.0650576 | 0.050008561 | 0.811293988 | 0.99 |

| | | | | |
|---|---|---|---|---|
| YLL065W | 0.583562943 | 0.005222787 | 0.810015121 | 0.99 |
| YLR046C | 345.7378309 | 0.028875883 | 0.814285593 | 0.99 |
| YLR054C | 138.716349 | 0.098272714 | 0.815820772 | 0.99 |
| YLR153C | 3246.572959 | 0.113993545 | 0.803827903 | 0.99 |
| YLR221C | 955.7495307 | -0.020680089 | 0.822682985 | 0.99 |
| YLR256W | 4392.10169 | 0.077716079 | 0.801207341 | 0.99 |
| YLR275W | 260.6216284 | 0.073639859 | 0.793456626 | 0.99 |
| YLR293C | 8683.145693 | -0.040873844 | 0.81795866 | 0.99 |
| YLR330W | 3759.858385 | 0.044233035 | 0.816895257 | 0.99 |
| YLR337C | 2433.363455 | -0.063667801 | 0.796741858 | 0.99 |
| YLR339C | 501.9621213 | -0.089905183 | 0.80673892 | 0.99 |
| YLR349W | 23.19539208 | -0.150413248 | 0.795321308 | 0.99 |
| YLR361C-A | 262.7999201 | -0.051279047 | 0.818537998 | 0.99 |
| YLR398C | 2110.607765 | -0.054522852 | 0.799759807 | 0.99 |
| YLR403W | 956.7749718 | -0.083244257 | 0.815221745 | 0.99 |
| YLR439W | 1150.159115 | 0.07701085 | 0.804633736 | 0.99 |
| YML003W | 222.6670877 | -0.07918136 | 0.817831859 | 0.99 |
| YML012C-A | 4.439298629 | 0.173991035 | 0.79719355 | 0.99 |
| YML030W | 762.1544554 | 0.093169969 | 0.817861017 | 0.99 |
| YML049C | 1684.51308 | -0.079784028 | 0.793376085 | 0.99 |
| YML055W | 811.1131816 | 0.064286067 | 0.824021782 | 0.99 |
| YML088W | 784.353378 | 0.083433975 | 0.798804201 | 0.99 |
| YML112W | 572.6104301 | -0.057188277 | 0.803203314 | 0.99 |
| YML125C | 2139.072841 | 0.126982254 | 0.818622832 | 0.99 |
| YMR026C | 666.726482 | 0.06017266 | 0.794501575 | 0.99 |
| YMR093W | 713.9051908 | -0.057518842 | 0.797997203 | 0.99 |
| YMR098C | 840.7673515 | 0.063546567 | 0.806268909 | 0.99 |
| YMR123W | 1008.572929 | 0.145520755 | 0.795613818 | 0.99 |
| YMR145C | 7155.424672 | -0.015226127 | 0.820039317 | 0.99 |
| YMR153W | 881.8064044 | 0.05575265 | 0.793266231 | 0.99 |
| YMR184W | 674.607137 | 0.068457644 | 0.806781724 | 0.99 |
| YMR186W | 31490.05511 | 0.196790926 | 0.796342455 | 0.99 |
| YMR194C-A | 22.19182237 | 0.128969928 | 0.793382224 | 0.99 |
| YMR241W | 1393.972884 | 0.115056531 | 0.797923888 | 0.99 |
| YMR256C | 1012.905586 | 0.153775261 | 0.80234721 | 0.99 |
| YMR261C | 3042.037261 | 0.069200105 | 0.802779032 | 0.99 |
| YMR276W | 5216.693498 | -0.053730528 | 0.796426392 | 0.99 |
| YMR289W | 967.6114857 | 0.0591104 | 0.797067527 | 0.99 |
| YNL064C | 7494.727656 | 0.185628562 | 0.794807409 | 0.99 |
| YNL072W | 456.0809998 | -0.051155458 | 0.807334514 | 0.99 |
| YNL108C | 290.0576123 | -0.055268443 | 0.822134501 | 0.99 |
| YNL121C | 5779.129433 | -0.048530295 | 0.814044381 | 0.99 |
| YNL203C | 26.22039237 | 0.07508796 | 0.815019217 | 0.99 |

| | | | | |
|---|---|---|---|---|
| YNL224C | 704.2866123 | -0.045061565 | 0.816960737 | 0.99 |
| YNL234W | 710.7866789 | 0.09346552 | 0.813350941 | 0.99 |
| YNL244C | 2167.186944 | -0.047214415 | 0.820057271 | 0.99 |
| YNL292W | 950.1678617 | 0.086213721 | 0.813165634 | 0.99 |
| YNL299W | 424.5188557 | 0.105549366 | 0.798630939 | 0.99 |
| YNL313C | 1104.908095 | -0.048894591 | 0.815979345 | 0.99 |
| YNR004W | 166.7322391 | 0.069094691 | 0.81657745 | 0.99 |
| YNR029C | 598.402815 | -0.056192951 | 0.796361433 | 0.99 |
| YNR039C | 749.4014615 | 0.03332095 | 0.807189373 | 0.99 |
| YNR044W | 3311.34449 | -0.21480515 | 0.805754256 | 0.99 |
| YNR055C | 1151.83484 | -0.070212782 | 0.800449581 | 0.99 |
| YOL017W | 430.0355777 | -0.053083052 | 0.817756187 | 0.99 |
| YOL030W | 6136.814993 | -0.073697735 | 0.814341156 | 0.99 |
| YOL071W | 1035.627926 | -0.103967887 | 0.799044837 | 0.99 |
| YOL107W | 336.081692 | -0.075537654 | 0.822379462 | 0.99 |
| YOL112W | 963.5678133 | -0.060244652 | 0.796275576 | 0.99 |
| YOL138C | 1055.538589 | -0.061206976 | 0.816363339 | 0.99 |
| YOL166W-A | 0.347622843 | 0.020026928 | 0.796143374 | 0.99 |
| YOR009W | 307.6360112 | -0.153413762 | 0.794544289 | 0.99 |
| YOR067C | 1025.694017 | 0.062855392 | 0.823379337 | 0.99 |
| YOR089C | 2098.945136 | -0.078978305 | 0.80188381 | 0.99 |
| YOR091W | 1558.309002 | -0.04342657 | 0.823584386 | 0.99 |
| YOR112W | 1049.770792 | -0.044905362 | 0.805094434 | 0.99 |
| YOR113W | 1039.201076 | 0.09747105 | 0.805403928 | 0.99 |
| YOR177C | 142.866635 | 0.113670708 | 0.792916059 | 0.99 |
| YOR236W | 448.7599801 | -0.039721897 | 0.821342879 | 0.99 |
| YOR267C | 3236.118357 | 0.051437845 | 0.822392689 | 0.99 |
| YOR291W | 1661.347699 | -0.045257907 | 0.820800567 | 0.99 |
| YOR330C | 1527.333333 | 0.044297109 | 0.820082044 | 0.99 |
| YOR342C | 1114.452608 | -0.066811551 | 0.792045276 | 0.99 |
| YPL015C | 1643.336309 | 0.059497672 | 0.820698295 | 0.99 |
| YPL022W | 1488.077225 | 0.060123866 | 0.804718771 | 0.99 |
| YPL038W-A | 14.34188488 | -0.150617495 | 0.801613396 | 0.99 |
| YPL094C | 2895.773005 | 0.067613205 | 0.823931123 | 0.99 |
| YPL100W | 911.5930892 | -0.070503771 | 0.811530478 | 0.99 |
| YPL231W | 11637.98709 | -0.056948664 | 0.800110907 | 0.99 |
| YPL242C | 2035.672722 | -0.064813162 | 0.809793 | 0.99 |
| YPR017C | 239.3606668 | -0.043182329 | 0.817026877 | 0.99 |
| YPR051W | 72.40075259 | -0.061513583 | 0.807716913 | 0.99 |
| YPR074C | 10075.28978 | -0.056579086 | 0.797893422 | 0.99 |
| YPR086W | 1622.357285 | 0.064674837 | 0.80800137 | 0.99 |
| YPR138C | 1408.826707 | -0.109662916 | 0.806416849 | 0.99 |
| snR24 | 8.183960631 | -0.041001141 | 0.823978938 | 0.99 |

| | | | | |
|---|---|---|---|---|
| tR(ACG)J | 0.2026169 | -0.064756253 | 0.821831489 | 0.99 |
| YAL022C | 845.7051212 | 0.08393309 | 0.824808515 | 0.99 |
| YAR019C | 904.0673159 | 0.04710599 | 0.834574304 | 0.99 |
| YBL062W | 0.924462825 | -0.092794261 | 0.827861937 | 0.99 |
| YBR110W | 1287.287964 | -0.050276504 | 0.825691788 | 0.99 |
| YBR150C | 848.9681448 | -0.056640386 | 0.828629339 | 0.99 |
| YBR175W | 264.1838842 | -0.069514366 | 0.83660844 | 0.99 |
| YDL001W | 483.443293 | -0.044347217 | 0.830263509 | 0.99 |
| YDL063C | 706.2977789 | 0.153508975 | 0.83086162 | 0.99 |
| YDL151C | 4.616503935 | -0.073746188 | 0.829256933 | 0.99 |
| YDL198C | 1156.426648 | 0.122266757 | 0.825974537 | 0.99 |
| YDL203C | 667.0945544 | 0.034062754 | 0.832361498 | 0.99 |
| YDR019C | 2663.732875 | 0.181449722 | 0.831984761 | 0.99 |
| YDR087C | 817.7593103 | 0.126227503 | 0.831200638 | 0.99 |
| YDR143C | 1247.877225 | -0.058661858 | 0.825293553 | 0.99 |
| YDR175C | 1042.356621 | -0.054413515 | 0.832424169 | 0.99 |
| YDR186C | 2571.215729 | 0.056119699 | 0.82964316 | 0.99 |
| YDR218C | 78.04916244 | 0.101391784 | 0.835480042 | 0.99 |
| YDR274C | 62.88697688 | 0.037294099 | 0.837879464 | 0.99 |
| YDR289C | 776.5077999 | 0.060227398 | 0.837593681 | 0.99 |
| YDR296W | 1076.454873 | -0.04446548 | 0.834969448 | 0.99 |
| YEL042W | 1714.856815 | 0.132269327 | 0.826843864 | 0.99 |
| YEL050C | 1046.082243 | -0.039670933 | 0.830683353 | 0.99 |
| YER025W | 3287.005691 | 0.089662108 | 0.829078989 | 0.99 |
| YER082C | 1253.635463 | -0.06102748 | 0.826709313 | 0.99 |
| YER165W | 11478.83612 | -0.027909727 | 0.83727737 | 0.99 |
| YER190W | 15.31140442 | 0.160289472 | 0.829625676 | 0.99 |
| YFL001W | 478.2892235 | -0.043688057 | 0.835567522 | 0.99 |
| YFR041C | 978.3498425 | -0.041904538 | 0.832628015 | 0.99 |
| YGL002W | 690.1637106 | -0.053521514 | 0.827555656 | 0.99 |
| YGL006W | 3158.91505 | 0.052899473 | 0.831089066 | 0.99 |
| YGL248W | 1428.88377 | -0.090670358 | 0.838381277 | 0.99 |
| YGR005C | 1346.509283 | -0.03953995 | 0.833293805 | 0.99 |
| YGR253C | 3517.823624 | -0.034689968 | 0.832948951 | 0.99 |
| YHL030W-A | 22.0345387 | 0.001351005 | 0.830339599 | 0.99 |
| YHL042W | 88.3151764 | 0.072191455 | 0.83603041 | 0.99 |
| YHR024C | 1377.744705 | -0.035565946 | 0.834114967 | 0.99 |
| YHR103W | 1865.030837 | 0.051552217 | 0.82857391 | 0.99 |
| YHR136C | 132.1966559 | -0.143082533 | 0.831361931 | 0.99 |
| YIL053W | 6767.756759 | -0.047409633 | 0.831620911 | 0.99 |
| YIL084C | 434.165782 | -0.06417208 | 0.834269052 | 0.99 |
| YIR007W | 898.4409432 | 0.043703673 | 0.832778678 | 0.99 |
| YIR031C | 471.7138035 | -0.110342292 | 0.835229506 | 0.99 |

| | | | | |
|---|---|---|---|---|
| YJL011C | 110.249814 | -0.00115872 | 0.829318923 | 0.99 |
| YJL034W | 12765.00653 | 0.115517982 | 0.838898886 | 0.99 |
| YJL057C | 1203.837592 | 0.071240138 | 0.830640264 | 0.99 |
| YJL205C | 132.3580317 | 0.070965035 | 0.829029576 | 0.99 |
| YJR009C | 33615.59982 | 0.175693757 | 0.838393064 | 0.99 |
| YJR014W | 1708.927082 | -0.030861442 | 0.835608885 | 0.99 |
| YJR041C | 1150.543526 | -0.077551987 | 0.836566574 | 0.99 |
| YJR097W | 96.20405896 | 0.147534251 | 0.835659617 | 0.99 |
| YJR141W | 396.4509931 | -0.041355502 | 0.833853846 | 0.99 |
| YKL006W | 2404.725348 | 0.095427984 | 0.833285404 | 0.99 |
| YKL090W | 556.2185917 | -0.08756646 | 0.836446798 | 0.99 |
| YKL128C | 1025.460861 | -0.05618487 | 0.836012784 | 0.99 |
| YKR048C | 4547.539769 | 0.060118466 | 0.837450081 | 0.99 |
| YLL006W | 812.6766975 | -0.078342479 | 0.826018203 | 0.99 |
| YLL009C | 581.7236454 | -0.071223952 | 0.832410264 | 0.99 |
| YLL049W | 333.5985502 | 0.087568146 | 0.838701564 | 0.99 |
| YLR116W | 1156.147687 | 0.027986114 | 0.830533342 | 0.99 |
| YLR183C | 714.9768804 | 0.009597011 | 0.836662686 | 0.99 |
| YLR227C | 512.1356286 | -0.049999215 | 0.836410109 | 0.99 |
| YLR294C | 1804.172401 | 0.0692414 | 0.834781485 | 0.99 |
| YLR313C | 272.7990365 | -0.069219618 | 0.830230928 | 0.99 |
| YLR399C | 2272.649731 | 0.04046844 | 0.838994124 | 0.99 |
| YLR405W | 359.0345633 | -0.061465829 | 0.826577527 | 0.99 |
| YLR408C | 156.1371802 | -0.044195259 | 0.835033646 | 0.99 |
| YLR409C | 1307.908763 | -0.065876063 | 0.832411743 | 0.99 |
| YMR119W-A | 41.58140006 | -0.100463952 | 0.838069523 | 0.99 |
| YMR185W | 803.0326457 | 0.071513194 | 0.834630936 | 0.99 |
| YMR231W | 864.2847521 | 0.046579258 | 0.832629814 | 0.99 |
| YMR297W | 8256.874804 | -0.051338958 | 0.834479522 | 0.99 |
| YNL137C | 1635.261502 | -0.038476096 | 0.835133635 | 0.99 |
| YNL151C | 742.0050839 | -0.035172293 | 0.836587221 | 0.99 |
| YNL183C | 1035.511832 | -0.057172934 | 0.829790564 | 0.99 |
| YNL261W | 688.6478875 | -0.064667889 | 0.833086386 | 0.99 |
| YNL314W | 384.9311948 | 0.05156995 | 0.827429474 | 0.99 |
| YNR046W | 947.7260883 | -0.058453227 | 0.835008679 | 0.99 |
| YOL096C | 679.2678424 | -0.057650507 | 0.824501261 | 0.99 |
| YOR071C | 651.6605909 | -0.096252056 | 0.826288761 | 0.99 |
| YOR178C | 2730.240932 | 0.145660772 | 0.835297071 | 0.99 |
| YOR183W | 10.65070105 | -0.145549392 | 0.833651507 | 0.99 |
| YOR187W | 4877.755588 | -0.037468053 | 0.826773629 | 0.99 |
| YOR335C | 3110.388093 | 0.091064762 | 0.838367994 | 0.99 |
| YOR347C | 1046.083791 | -0.08080368 | 0.832502902 | 0.99 |
| YOR386W | 897.2436472 | 0.098122058 | 0.825330188 | 0.99 |

| | | | | |
|---|---|---|---|---|
| YPL014W | 2476.516022 | 0.14185143 | 0.824971898 | 0.99 |
| YPL016W | 1163.813997 | 0.050815012 | 0.824758277 | 0.99 |
| YPL025C | 15.86269313 | -0.028921411 | 0.832750326 | 0.99 |
| YPL182C | 43.60254699 | 0.069974123 | 0.838791158 | 0.99 |
| YPL202C | 824.3595251 | -0.056818971 | 0.832490793 | 0.99 |
| YPL251W | 76.6529722 | -0.10793572 | 0.828614532 | 0.99 |
| YPR074W-A | 0.407755058 | 0.019463133 | 0.83777298 | 0.99 |
| YPR078C | 88.79558885 | 0.071832747 | 0.83661431 | 0.99 |
| YPR087W | 20.87255534 | 0.020720981 | 0.837644406 | 0.99 |
| YPR112C | 1587.350101 | -0.032346634 | 0.832308086 | 0.99 |
| YPR140W | 900.9897118 | -0.057125915 | 0.839096839 | 0.99 |
| YPR159W | 2743.676738 | 0.065284786 | 0.833916362 | 0.99 |
| tT(CGU)K | 0.429968112 | 0.006212424 | 0.83757676 | 0.99 |
| YMR003W | 306.5250482 | 0.097362405 | 0.839271314 | 0.99 |
| YBR129C | 610.3882792 | -0.056274806 | 0.839483958 | 0.99 |
| YAL046C | 313.7043981 | -0.025910585 | 0.840721652 | 0.99 |
| YBL102W | 1291.754797 | 0.090960433 | 0.840151884 | 0.99 |
| YBR070C | 438.6999546 | 0.137281298 | 0.839863841 | 0.99 |
| YER043C | 7073.679738 | -0.005563363 | 0.840654064 | 0.99 |
| YFR012W | 37.52047909 | 0.014703132 | 0.841009573 | 0.99 |
| YGL214W | 9.34196156 | -0.12165599 | 0.840130455 | 0.99 |
| YMR205C | 7427.558626 | -0.051904515 | 0.840253156 | 0.99 |
| YNL303W | 97.83657889 | -0.129598454 | 0.840254235 | 0.99 |
| YOR144C | 421.6026059 | 0.055081828 | 0.840826055 | 0.99 |
| YPR179C | 773.0608325 | -0.048046641 | 0.840954089 | 0.99 |
| YDR367W | 606.3521729 | -0.03218675 | 0.841319927 | 0.99 |
| YNR014W | 1455.118637 | 0.069900896 | 0.841240104 | 0.99 |
| YAL002W | 1569.483592 | 0.032052483 | 0.842523406 | 0.99 |
| YAL064C-A | 47.35030167 | -0.135871651 | 0.842088461 | 0.99 |
| YBR160W | 885.0084099 | -0.038397449 | 0.843160137 | 0.99 |
| YDR067C | 673.9208846 | 0.04428698 | 0.84217533 | 0.99 |
| YGR264C | 2825.266453 | -0.024509673 | 0.842880468 | 0.99 |
| YHL048W | 1666.986201 | -0.092805255 | 0.842370258 | 0.99 |
| YJL008C | 2286.818099 | 0.078144627 | 0.843106169 | 0.99 |
| YJL176C | 1289.4859 | -0.060592649 | 0.843008704 | 0.99 |
| YKL214C | 653.6592896 | 0.050345969 | 0.843377333 | 0.99 |
| YMR198W | 512.8655409 | -0.046513555 | 0.842178006 | 0.99 |
| YOL016C | 3937.741808 | 0.108796981 | 0.842840052 | 0.99 |
| YPR176C | 535.2741493 | -0.038834038 | 0.842705827 | 0.99 |
| YPR188C | 534.7061592 | -0.040107357 | 0.843335067 | 0.99 |
| YPL261C | 48.62531076 | -0.091193218 | 0.843668448 | 0.99 |
| YLR066W | 1030.577166 | -0.029224834 | 0.843883404 | 0.99 |
| YHR071C-A | 39.05421358 | -0.093601442 | 0.844137857 | 0.99 |

| | | | | |
|---|---|---|---|---|
| YAR028W | 513.5910617 | -0.038733692 | 0.848346851 | 0.99 |
| YBL011W | 1645.010666 | -0.041500882 | 0.852155128 | 0.99 |
| YBL060W | 624.3164774 | 0.038991822 | 0.847899631 | 0.99 |
| YBL094C | 5.123149382 | 0.123956652 | 0.854525079 | 0.99 |
| YBL100W-B | 175.5232464 | -0.11749008 | 0.856641026 | 0.99 |
| YBR037C | 852.7234834 | 0.053329741 | 0.859038592 | 0.99 |
| YBR064W | 3.158811985 | -0.24005443 | 0.851839547 | 0.99 |
| YCL005W | 658.6999475 | -0.027883506 | 0.854262027 | 0.99 |
| YCL011C | 2495.29652 | 0.054778694 | 0.850619394 | 0.99 |
| YCL029C | 771.878883 | 0.040342902 | 0.856989634 | 0.99 |
| YCR030C | 2439.320353 | 0.047251661 | 0.855822848 | 0.99 |
| YDL046W | 2942.234491 | -0.043651137 | 0.851904012 | 0.99 |
| YDL070W | 4583.373904 | 0.058482478 | 0.854527921 | 0.99 |
| YDL112W | 1910.821894 | -0.053869773 | 0.854616686 | 0.99 |
| YDL216C | 462.7023205 | 0.046698627 | 0.850851731 | 0.99 |
| YDR022C | 413.59294 | 0.072257058 | 0.856657056 | 0.99 |
| YDR050C | 39610.42901 | -0.037867812 | 0.846802642 | 0.99 |
| YDR058C | 448.3672811 | -0.063417935 | 0.849277215 | 0.99 |
| YDR151C | 1113.580859 | 0.038504846 | 0.853876506 | 0.99 |
| YDR193W | 12.98139175 | 0.118953195 | 0.857972526 | 0.99 |
| YDR206W | 1789.144398 | -0.043727087 | 0.859723257 | 0.99 |
| YDR238C | 4318.542325 | 0.050571926 | 0.850180681 | 0.99 |
| YDR246W-A | 23.46436155 | 0.085407888 | 0.858884411 | 0.99 |
| YDR262W | 1927.726265 | 0.059144818 | 0.855206359 | 0.99 |
| YDR312W | 598.3889984 | -0.028038538 | 0.859719689 | 0.99 |
| YDR321W | 1332.488079 | 0.202149342 | 0.846111964 | 0.99 |
| YDR325W | 750.6903469 | -0.047834983 | 0.859447231 | 0.99 |
| YDR393W | 653.7265915 | 0.034445958 | 0.856523918 | 0.99 |
| YDR412W | 170.5135109 | -0.026367219 | 0.853054428 | 0.99 |
| YER105C | 2233.871003 | 0.032560986 | 0.859525268 | 0.99 |
| YFL012W | 27.05345673 | 0.061543408 | 0.852911269 | 0.99 |
| YFL024C | 1322.826078 | 0.033744126 | 0.849296743 | 0.99 |
| YGL021W | 1221.378254 | 0.041949582 | 0.850410898 | 0.99 |
| YGL139W | 1235.197549 | -0.041430943 | 0.853181154 | 0.99 |
| YGL212W | 286.2266484 | -0.035829963 | 0.859722361 | 0.99 |
| YGL216W | 962.5040676 | 0.026223566 | 0.858196773 | 0.99 |
| YGL224C | 544.3672231 | -0.03604544 | 0.858808728 | 0.99 |
| YGL226W | 346.3222245 | 0.047456155 | 0.855077262 | 0.99 |
| YGL230C | 33.65395104 | 0.082758872 | 0.84877439 | 0.99 |
| YGL239C | 18.18533761 | -0.04866765 | 0.860044049 | 0.99 |
| YGR010W | 527.974638 | 0.050582419 | 0.854162866 | 0.99 |
| YGR046W | 906.1577966 | 0.061005814 | 0.853209318 | 0.99 |
| YGR094W | 6966.605497 | 0.121175547 | 0.854443742 | 0.99 |

| | | | | |
|---|---|---|---|---|
| YGR159C | 2044.642627 | -0.039772058 | 0.8554477 | 0.99 |
| YGR161C | 2432.867838 | -0.087925723 | 0.846492126 | 0.99 |
| YHL011C | 1690.318772 | 0.147973889 | 0.85947595 | 0.99 |
| YHL014C | 453.0111576 | 0.053517174 | 0.858452836 | 0.99 |
| YHR028C | 2661.078301 | 0.046309667 | 0.855821515 | 0.99 |
| YHR142W | 1336.189865 | 0.078472796 | 0.856361774 | 0.99 |
| YHR219W | 0.159331785 | 0.019790496 | 0.859680674 | 0.99 |
| YIL074C | 1391.44586 | 0.086504813 | 0.854936933 | 0.99 |
| YIL079C | 348.2789788 | -0.100230077 | 0.852603177 | 0.99 |
| YIL147C | 2000.977172 | 0.038996274 | 0.854209767 | 0.99 |
| YJL118W | 178.5211681 | -0.070076756 | 0.849981285 | 0.99 |
| YJL127W-A | 12.80997666 | -0.043243629 | 0.856746046 | 0.99 |
| YJL171C | 1558.602902 | -0.045081405 | 0.847741797 | 0.99 |
| YJL225C | 0.280381926 | 0.027823207 | 0.856167136 | 0.99 |
| YJR113C | 864.1046061 | 0.059347587 | 0.855523095 | 0.99 |
| YKL017C | 627.7815414 | 0.036647065 | 0.855841619 | 0.99 |
| YKL136W | 25.19505527 | -0.090222503 | 0.84634943 | 0.99 |
| YKL149C | 355.2699607 | -0.048533264 | 0.856777775 | 0.99 |
| YKR081C | 1261.655674 | -0.004426671 | 0.848861675 | 0.99 |
| YLR030W | 185.5800298 | 0.036160771 | 0.846933991 | 0.99 |
| YLR106C | 6573.73141 | -0.021319552 | 0.854431126 | 0.99 |
| YLR124W | 42.32116711 | 0.04230769 | 0.857391135 | 0.99 |
| YLR130C | 1253.100793 | 0.049251129 | 0.851837748 | 0.99 |
| YLR186W | 830.8214037 | -0.031431613 | 0.847951717 | 0.99 |
| YLR285C-A | 41.10839909 | -0.016879653 | 0.848288803 | 0.99 |
| YLR332W | 1136.948721 | -0.075880221 | 0.859880096 | 0.99 |
| YLR358C | 39.08246637 | 0.037545326 | 0.853613174 | 0.99 |
| YML113W | 567.9041791 | -0.077193491 | 0.859983773 | 0.99 |
| YML126C | 4399.479194 | 0.099028249 | 0.848120263 | 0.99 |
| YMR016C | 1109.247117 | -0.070641844 | 0.854814524 | 0.99 |
| YMR029C | 1228.193843 | -0.048959651 | 0.850115141 | 0.99 |
| YMR070W | 1232.669923 | -0.09274609 | 0.856455918 | 0.99 |
| YMR195W | 1764.107553 | -0.098633221 | 0.856099292 | 0.99 |
| YMR224C | 500.4037572 | 0.030660414 | 0.852685077 | 0.99 |
| YMR284W | 845.7287906 | 0.041429451 | 0.854934241 | 0.99 |
| YNL004W | 1614.267096 | -0.035154632 | 0.850543741 | 0.99 |
| YNL136W | 1295.54364 | -0.060067372 | 0.8586329 | 0.99 |
| YNL141W | 1283.346108 | 0.010485114 | 0.85168906 | 0.99 |
| YNL302C | 5878.096109 | 0.04234117 | 0.851579525 | 0.99 |
| YNR048W | 344.510222 | 0.041793895 | 0.851282621 | 0.99 |
| YOL077W-A | 1163.041858 | -0.071549165 | 0.857204645 | 0.99 |
| YOL098C | 2415.997601 | -0.039080554 | 0.850390681 | 0.99 |
| YOR002W | 2112.418166 | 0.039416263 | 0.845329417 | 0.99 |

| | | | | |
|---|---|---|---|---|
| YOR004W | 494.9946687 | 0.047095389 | 0.857452603 | 0.99 |
| YOR020C | 3482.406455 | -0.023476395 | 0.859614401 | 0.99 |
| YOR156C | 466.9299753 | 0.032781033 | 0.848430273 | 0.99 |
| YOR220W | 2249.332399 | 0.062393369 | 0.852084601 | 0.99 |
| YOR239W | 2726.169468 | -0.027312145 | 0.847694813 | 0.99 |
| YOR345C | 1.957434997 | 0.113103317 | 0.859910462 | 0.99 |
| YPL019C | 1922.931958 | -0.043460773 | 0.85880391 | 0.99 |
| YPR016W-A | 37.91128357 | -0.121055248 | 0.849855444 | 0.99 |
| YPR025C | 1080.062057 | 0.049009497 | 0.846012556 | 0.99 |
| snR13 | 24.86462889 | 0.09788409 | 0.850145869 | 0.99 |
| snR48 | 2.926793808 | -0.092294564 | 0.850162744 | 0.99 |
| snR49 | 77.03202196 | 0.186004321 | 0.848493847 | 0.99 |
| YDR452W | 4294.15743 | -0.050141362 | 0.860840027 | 0.99 |
| YHR145C | 0.415497853 | -0.03337707 | 0.861612578 | 0.99 |
| YIR006C | 5095.375019 | -0.03322954 | 0.861818276 | 0.99 |
| YJL014W | 2604.88629 | -0.0214785 | 0.861462572 | 0.99 |
| YJL202C | 29.15207009 | -0.081860826 | 0.860314115 | 0.99 |
| YJR064W | 2718.261132 | -0.025166491 | 0.861548615 | 0.99 |
| YKR059W | 1551.844916 | -0.009931145 | 0.860972295 | 0.99 |
| YML022W | 1077.904293 | 0.02087074 | 0.861622828 | 0.99 |
| YNL271C | 3740.617123 | 0.036291894 | 0.861778181 | 0.99 |
| YOL128C | 161.4112011 | 0.035632308 | 0.861339448 | 0.99 |
| YOR060C | 313.4302218 | -0.049835855 | 0.860599337 | 0.99 |
| YOR130C | 372.7954083 | 0.087507344 | 0.862026035 | 0.99 |
| YPL090C | 2392.913651 | 0.079485858 | 0.862140273 | 0.99 |
| YAR071W | 43.79477313 | -0.046768416 | 0.862821416 | 0.99 |
| YBR019C | 339.0837633 | 0.045057375 | 0.862641259 | 0.99 |
| YER177W | 16691.55629 | -0.036220256 | 0.862382947 | 0.99 |
| YMR217W | 5540.397432 | 0.178947357 | 0.862788707 | 0.99 |
| YJL175W | 4.732791699 | -0.031640945 | 0.863234121 | 0.99 |
| YML028W | 27162.20889 | -0.024029306 | 0.863530729 | 0.99 |
| YLR289W | 462.7410856 | 0.04603328 | 0.863700537 | 0.99 |
| YCL026C-A | 44.13033287 | -0.08877816 | 0.864273066 | 0.99 |
| YER064C | 1674.781352 | -0.064414583 | 0.864131071 | 0.99 |
| YGR235C | 1048.42614 | 0.066121251 | 0.864749638 | 0.99 |
| YLR414C | 1706.526088 | -0.066904644 | 0.864928943 | 0.99 |
| YEL031W | 4279.639928 | -0.038243392 | 0.865442811 | 0.99 |
| YBR038W | 1800.515437 | 0.051584687 | 0.865603809 | 0.99 |
| YJL151C | 3281.2426 | -0.072248679 | 0.86586889 | 0.99 |
| YBL039W-B | 151.3574507 | 0.05155846 | 0.867454105 | 0.99 |
| YBR076W | 114.1570412 | -0.099956874 | 0.869473821 | 0.99 |
| YBR172C | 2003.113678 | -0.057099134 | 0.868790541 | 0.99 |
| YBR196C | 23554.20363 | 0.074599529 | 0.868725758 | 0.99 |

| | | | | |
|---|---|---|---|---|
| YBR300C | 5.482970764 | 0.156992321 | 0.869519424 | 0.99 |
| YDR066C | 173.6197188 | 0.040348876 | 0.867513475 | 0.99 |
| YDR379W | 1429.073535 | -0.049448396 | 0.867682239 | 0.99 |
| YDR477W | 1886.671826 | -0.026335362 | 0.868803918 | 0.99 |
| YER041W | 647.7988303 | -0.05738876 | 0.869294604 | 0.99 |
| YGR138C | 1416.124697 | 0.111211351 | 0.868065501 | 0.99 |
| YGR169C | 437.5350343 | 0.064439773 | 0.867210203 | 0.99 |
| YIL041W | 4311.545468 | -0.028574147 | 0.867976123 | 0.99 |
| YJL184W | 578.763074 | 0.04147814 | 0.866585677 | 0.99 |
| YKL181W | 2705.955695 | -0.0238796 | 0.866523347 | 0.99 |
| YKL224C | 0.174623486 | 0.022229079 | 0.868649441 | 0.99 |
| YLR167W | 28914.13243 | 0.037894778 | 0.869373608 | 0.99 |
| YLR230W | 287.1818195 | -0.028250235 | 0.867123104 | 0.99 |
| YLR262C | 283.7623055 | 0.058083608 | 0.86757412 | 0.99 |
| YOR116C | 1548.881592 | -0.046357567 | 0.869497826 | 0.99 |
| YOR123C | 2187.626634 | -0.022428204 | 0.869506384 | 0.99 |
| YPL047W | 255.7983457 | -0.052036538 | 0.869617406 | 0.99 |
| YPR028W | 4646.798863 | -0.028324881 | 0.868503257 | 0.99 |
| YDL096C | 9.504570723 | -0.169838929 | 0.870197569 | 0.99 |
| YJL216C | 203.2730397 | -0.100998586 | 0.870424608 | 0.99 |
| YLR202C | 55.2829566 | -0.062481371 | 0.870324281 | 0.99 |
| YMR042W | 272.8014796 | -0.055151779 | 0.870425871 | 0.99 |
| YPL198W | 414.9903406 | 0.146328366 | 0.870339125 | 0.99 |
| YAL008W | 865.7638687 | 0.065866665 | 0.874373614 | 0.99 |
| YBL041W | 3024.581572 | -0.018244726 | 0.885623736 | 0.99 |
| YBL054W | 675.0753311 | -0.086343021 | 0.884062573 | 0.99 |
| YBL059W | 273.8243146 | -0.040034342 | 0.879749125 | 0.99 |
| YBL090W | 640.7121751 | -0.02244678 | 0.873452693 | 0.99 |
| YBR119W | 605.7970902 | -0.0510063 | 0.883221396 | 0.99 |
| YBR121C | 7616.904588 | 0.120166049 | 0.875945902 | 0.99 |
| YBR123C | 731.156432 | 0.028425035 | 0.881719012 | 0.99 |
| YBR191W-A | 25.03964351 | 0.472669404 | 0.882371509 | 0.99 |
| YBR196C-A | 7.414882286 | 0.113879073 | 0.878847646 | 0.99 |
| YBR231C | 448.4132605 | 0.043097723 | 0.887250258 | 0.99 |
| YBR252W | 608.885071 | 0.00937439 | 0.880025775 | 0.99 |
| YCR087W | 24.15940244 | 0.06259936 | 0.887392932 | 0.99 |
| YDR033W | 12974.70679 | -0.0066806 | 0.876128555 | 0.99 |
| YDR034C-D | 94.28631679 | -0.077722245 | 0.88346061 | 0.99 |
| YDR045C | 241.2309895 | -0.013936404 | 0.874869419 | 0.99 |
| YDR061W | 1142.905546 | -0.030073125 | 0.87512789 | 0.99 |
| YDR084C | 664.1695601 | 0.059031185 | 0.872799995 | 0.99 |
| YDR119W | 1442.902962 | -0.052137188 | 0.877112369 | 0.99 |
| YDR134C | 8310.682574 | 0.045678555 | 0.885530006 | 0.99 |

| | | | | |
|---|---|---|---|---|
| YDR164C | 1143.654091 | 0.032370756 | 0.87839216 | 0.99 |
| YDR247W | 1026.990415 | 0.076351285 | 0.878216092 | 0.99 |
| YDR276C | 2763.13579 | -0.031190306 | 0.884348855 | 0.99 |
| YDR277C | 2772.925083 | 0.05871998 | 0.873665716 | 0.99 |
| YDR283C | 1675.592568 | -0.042106876 | 0.885363277 | 0.99 |
| YDR323C | 314.788297 | -0.032031513 | 0.884308334 | 0.99 |
| YDR356W | 1114.540575 | 0.044143464 | 0.873270944 | 0.99 |
| YEL037C | 3155.723536 | -0.019784831 | 0.880962629 | 0.99 |
| YER026C | 2306.614486 | -0.017558209 | 0.880152131 | 0.99 |
| YER049W | 1494.465582 | 0.087833287 | 0.88367755 | 0.99 |
| YER132C | 1149.8348 | -0.052225509 | 0.886376216 | 0.99 |
| YER133W | 3950.455243 | -0.026926729 | 0.871796571 | 0.99 |
| YFL011W | 70.61598103 | -0.11393203 | 0.87757169 | 0.99 |
| YFR056C | 41.82065454 | 0.054456874 | 0.872857361 | 0.99 |
| YGL034C | 9.87258369 | 0.069794496 | 0.878978018 | 0.99 |
| YGL120C | 2147.315673 | 0.144524892 | 0.882759062 | 0.99 |
| YGR045C | 23.80619598 | 0.026534572 | 0.885661703 | 0.99 |
| YGR136W | 2019.817506 | 0.039561612 | 0.887166538 | 0.99 |
| YGR146C-A | 13.60921533 | -0.098966189 | 0.87151111 | 0.99 |
| YGR171C | 566.9877726 | -0.030574015 | 0.871798354 | 0.99 |
| YHL040C | 1041.983492 | -0.021086096 | 0.880523402 | 0.99 |
| YHR056C | 832.3445362 | 0.026877323 | 0.882304371 | 0.99 |
| YHR057C | 1425.81956 | -0.035747093 | 0.886579554 | 0.99 |
| YHR066W | 786.1240577 | 0.013493154 | 0.884097158 | 0.99 |
| YHR088W | 1005.281896 | -0.000770633 | 0.87414641 | 0.99 |
| YHR112C | 1466.272889 | 0.047997268 | 0.873172602 | 0.99 |
| YHR152W | 1107.927047 | 0.050262707 | 0.873651939 | 0.99 |
| YIL129C | 2649.636452 | 0.029159229 | 0.887238645 | 0.99 |
| YIL131C | 677.2016628 | 0.071896183 | 0.873097961 | 0.99 |
| YIL144W | 1230.130031 | 0.034386838 | 0.879472834 | 0.99 |
| YIR001C | 869.6360964 | 0.032565593 | 0.885154182 | 0.99 |
| YJL055W | 2135.334979 | -0.033331442 | 0.880019152 | 0.99 |
| YJL073W | 1063.600671 | 0.04914892 | 0.87881435 | 0.99 |
| YJL082W | 1623.02873 | 0.056002225 | 0.880739419 | 0.99 |
| YJR027W | 207.2279874 | 0.013065012 | 0.878598727 | 0.99 |
| YJR043C | 619.0166524 | -0.008135219 | 0.881487894 | 0.99 |
| YJR112W | 374.389265 | 0.055817988 | 0.882180823 | 0.99 |
| YJR123W | 26166.00582 | 0.049497844 | 0.886274072 | 0.99 |
| YKL099C | 686.8097655 | -0.01197323 | 0.881637827 | 0.99 |
| YKL153W | 3671.352718 | -0.081389694 | 0.873753878 | 0.99 |
| YKL164C | 7734.898504 | 0.040546273 | 0.885360548 | 0.99 |
| YKL176C | 1422.060464 | -0.043715402 | 0.874055832 | 0.99 |
| YKR012C | 142.5434976 | -0.067970233 | 0.877769238 | 0.99 |

| | | | | |
|---|---|---|---|---|
| YKR017C | 571.9042365 | 0.034229116 | 0.887320401 | 0.99 |
| YKR033C | 0.132511194 | 0.019024685 | 0.873062277 | 0.99 |
| YLL034C | 960.4010793 | -0.023019426 | 0.87238803 | 0.99 |
| YLR076C | 95.93637703 | 0.112491445 | 0.873226015 | 0.99 |
| YLR086W | 1588.915329 | 0.033465648 | 0.886351174 | 0.99 |
| YLR088W | 986.2933263 | 0.029546233 | 0.878286124 | 0.99 |
| YLR151C | 483.8723388 | -0.041382365 | 0.882813261 | 0.99 |
| YLR190W | 1584.336567 | 0.062766799 | 0.874381637 | 0.99 |
| YLR370C | 2370.21826 | -0.037255567 | 0.881924833 | 0.99 |
| YML017W | 751.7187367 | -0.03619619 | 0.881943442 | 0.99 |
| YML041C | 413.9344954 | 0.030143181 | 0.885155644 | 0.99 |
| YML090W | 156.4475378 | 0.12912664 | 0.880414882 | 0.99 |
| YMR049C | 1525.516535 | -0.006822447 | 0.875993774 | 0.99 |
| YMR100W | 811.7962257 | -0.069089224 | 0.881417345 | 0.99 |
| YMR162C | 1643.610674 | 0.029363454 | 0.879837447 | 0.99 |
| YMR169C | 812.630576 | -0.089681804 | 0.875849085 | 0.99 |
| YMR269W | 489.6101925 | -0.017636786 | 0.884136335 | 0.99 |
| YMR316C-A | 33.48519679 | 0.013563474 | 0.885164963 | 0.99 |
| YNL025C | 730.0152463 | 0.025665678 | 0.883350997 | 0.99 |
| YNL042W-B | 14.4556208 | 0.171540362 | 0.876246038 | 0.99 |
| YNL130C | 1326.028586 | -0.024532509 | 0.885471395 | 0.99 |
| YNL133C | 297.07226 | 0.04734712 | 0.871788685 | 0.99 |
| YNL169C | 1167.217459 | 0.025370151 | 0.876294736 | 0.99 |
| YNL225C | 446.0966712 | -0.028708379 | 0.881978729 | 0.99 |
| YNL255C | 4025.75008 | -0.026094943 | 0.877228833 | 0.99 |
| YNL273W | 713.7141054 | -0.032305954 | 0.885205326 | 0.99 |
| YNR042W | 0.825755426 | -0.397507652 | 0.873310579 | 0.99 |
| YOL009C | 380.9976744 | -0.042751219 | 0.882454896 | 0.99 |
| YOL033W | 423.9709748 | -0.031492461 | 0.875939776 | 0.99 |
| YOL038W | 4309.883987 | 0.043669594 | 0.882383006 | 0.99 |
| YOL042W | 569.9826336 | 0.038116297 | 0.88731664 | 0.99 |
| YOR032C | 566.9607469 | 0.042313616 | 0.887169167 | 0.99 |
| YOR073W | 590.4176603 | 0.048719803 | 0.874981447 | 0.99 |
| YOR107W | 180.9819764 | 0.011351118 | 0.880350632 | 0.99 |
| YOR165W | 1637.432086 | -0.018008072 | 0.8796484 | 0.99 |
| YOR198C | 6733.130979 | -0.015692887 | 0.875885215 | 0.99 |
| YOR269W | 302.0905548 | -0.038085433 | 0.876333787 | 0.99 |
| YOR362C | 3907.49208 | -0.025875102 | 0.880709652 | 0.99 |
| YPL136W | 0.97000745 | 0.079173387 | 0.881016229 | 0.99 |
| YPL159C | 943.1509361 | -0.056304015 | 0.877412731 | 0.99 |
| YPL179W | 1527.082249 | 0.038310648 | 0.883748936 | 0.99 |
| YPR196W | 243.4727794 | 0.030148785 | 0.885293285 | 0.99 |
| snR32 | 25.20381189 | -0.041633139 | 0.874090038 | 0.99 |

| | | | | |
|---|---|---|---|---|
| snR38 | 6.274197792 | -0.048690936 | 0.873156081 | 0.99 |
| snR44 | 201.2193719 | -0.020625353 | 0.886088937 | 0.99 |
| snR64 | 4.034418541 | -0.062428315 | 0.881199932 | 0.99 |
| tQ(UUG)E2 | 0.083088204 | 0.018837575 | 0.883633208 | 0.99 |
| tR(CCG)L | 0.085991782 | -0.037076141 | 0.888004223 | 0.99 |
| YAL019W-A | 5.164115052 | -0.083662132 | 0.889479625 | 0.99 |
| YBR036C | 2605.132077 | 0.042805554 | 0.889485595 | 0.99 |
| YCL052C | 1614.200073 | -0.041061727 | 0.889394858 | 0.99 |
| YDL119C | 290.4231748 | -0.038454563 | 0.889726486 | 0.99 |
| YDR170W-A | 128.0923816 | 0.089920206 | 0.889331878 | 0.99 |
| YHR065C | 1260.588152 | -0.021174363 | 0.889419619 | 0.99 |
| YKL041W | 833.3256523 | -0.040603514 | 0.890071185 | 0.99 |
| YKL162C-A | 16.50216676 | 0.057979109 | 0.88865277 | 0.99 |
| YLR421C | 1264.683888 | -0.023717536 | 0.889684505 | 0.99 |
| YMR013C | 348.9768631 | -0.05333252 | 0.889960583 | 0.99 |
| YOR259C | 3766.746436 | -0.026860289 | 0.888255338 | 0.99 |
| YPL056C | 172.8020928 | -0.12150723 | 0.888485999 | 0.99 |
| YPR162C | 653.0734249 | -0.037279444 | 0.889998095 | 0.99 |
| YAR053W | 9.523462605 | 0.026080985 | 0.892679893 | 0.99 |
| YBR155W | 833.2201083 | 0.173752817 | 0.892857639 | 0.99 |
| YDR195W | 649.2195359 | 0.017307588 | 0.89284453 | 0.99 |
| YDR346C | 2820.625886 | 0.042046673 | 0.892310375 | 0.99 |
| YEL017W | 1022.681916 | 0.128132364 | 0.892149137 | 0.99 |
| YFL013W-A | 84.24870204 | -0.064730679 | 0.89074426 | 0.99 |
| YFR022W | 849.2747048 | 0.016479309 | 0.891498418 | 0.99 |
| YHL038C | 821.5884324 | 0.022733801 | 0.892990182 | 0.99 |
| YHR135C | 3300.939762 | -0.028972987 | 0.892532484 | 0.99 |
| YJL138C | 1955.737437 | -0.013249906 | 0.891313475 | 0.99 |
| YKR054C | 2620.574744 | 0.039543196 | 0.891623768 | 0.99 |
| YLR019W | 1293.150392 | -0.037609222 | 0.891705 | 0.99 |
| YLR150W | 21689.73235 | 0.107601125 | 0.891110774 | 0.99 |
| YLR309C | 2329.969598 | 0.061149472 | 0.892476465 | 0.99 |
| YNL338W | 0.154202725 | 0.019000942 | 0.89251301 | 0.99 |
| YOR318C | 25.32067042 | -0.051659132 | 0.89231507 | 0.99 |
| YPL263C | 1770.940205 | -0.019431451 | 0.891908153 | 0.99 |
| YNR024W | 280.6447987 | -0.025544099 | 0.893640214 | 0.99 |
| snR50 | 8.290616263 | -0.028496482 | 0.893500025 | 0.99 |
| YDL115C | 894.0269756 | -0.02847953 | 0.893986367 | 1.00 |
| YHR196W | 929.0161741 | -0.064333846 | 0.89415976 | 1.00 |
| YNL296W | 0.17572267 | 0.018983681 | 0.894148991 | 1.00 |
| YLR287C | 617.6178514 | 0.101959745 | 0.894378139 | 1.00 |
| YDR187C | 25.29702409 | 0.072041823 | 0.894763299 | 1.00 |
| YLR317W | 2.198595472 | 0.12622082 | 0.894708283 | 1.00 |

| | | | | |
|---|---|---|---|---|
| YPR089W | 1983.427567 | -0.03052473 | 0.894912662 | 1.00 |
| YIL008W | 681.0099622 | -0.034828734 | 0.895259054 | 1.00 |
| YLR243W | 616.8482711 | 0.074980277 | 0.895335333 | 1.00 |
| YBR292C | 15.20409235 | -0.119968248 | 0.895499948 | 1.00 |
| YBR185C | 732.2746685 | 0.045631245 | 0.898962488 | 1.00 |
| YBR261C | 744.8069304 | 0.00217164 | 0.896668384 | 1.00 |
| YBR271W | 387.9475512 | 0.050968604 | 0.899358115 | 1.00 |
| YCL064C | 3239.199963 | 0.154970495 | 0.897779143 | 1.00 |
| YDR145W | 1845.179932 | -0.035742 | 0.897815755 | 1.00 |
| YDR185C | 264.7477216 | 0.056997044 | 0.896392198 | 1.00 |
| YDR226W | 6443.35181 | -0.006748779 | 0.896843681 | 1.00 |
| YDR447C | 4088.999597 | 0.106575152 | 0.895967419 | 1.00 |
| YGL220W | 745.8780305 | 0.037637124 | 0.898526467 | 1.00 |
| YGR106C | 3546.453004 | 0.060127906 | 0.896231888 | 1.00 |
| YHR023W | 2927.24121 | -0.056578914 | 0.897630928 | 1.00 |
| YHR026W | 2431.689374 | -0.002032338 | 0.897429638 | 1.00 |
| YHR127W | 498.1945386 | -0.010617613 | 0.896168653 | 1.00 |
| YHR170W | 1876.524363 | 0.097990537 | 0.8959402 | 1.00 |
| YHR199C-A | 146.5882374 | -0.073992234 | 0.899022177 | 1.00 |
| YIL120W | 363.836718 | 0.038037714 | 0.897316789 | 1.00 |
| YJR138W | 1461.069987 | 0.027174143 | 0.898618868 | 1.00 |
| YKL127W | 1973.346678 | 0.131223481 | 0.8988632 | 1.00 |
| YKR002W | 1434.178622 | 0.023323834 | 0.898836862 | 1.00 |
| YNL110C | 899.4305976 | 0.047439496 | 0.898218921 | 1.00 |
| YNL132W | 1681.161762 | 0.095364886 | 0.899217068 | 1.00 |
| YOL010W | 524.4137956 | -0.006086671 | 0.896786433 | 1.00 |
| YOR164C | 1461.403289 | -0.006611823 | 0.899267242 | 1.00 |
| YPL120W | 588.9261154 | -0.03821032 | 0.898665329 | 1.00 |
| YPL213W | 340.9140453 | -0.028935539 | 0.898973996 | 1.00 |
| YDR313C | 770.3934735 | 0.030087814 | 0.899870423 | 1.00 |
| YAR019W-A | 5.564052438 | 0.026676641 | 0.900229949 | 1.00 |
| YBL077W | 139.1714064 | -0.042347605 | 0.901540102 | 1.00 |
| YBR025C | 6377.054239 | 0.044118986 | 0.901559298 | 1.00 |
| YDR260C | 314.9401976 | -0.035870234 | 0.900278546 | 1.00 |
| YGR126W | 190.2789593 | -0.045369605 | 0.901511893 | 1.00 |
| YHL002W | 1516.40559 | -0.047684974 | 0.901203966 | 1.00 |
| YHR020W | 3665.040866 | 0.098335777 | 0.900452281 | 1.00 |
| YHR072W | 1833.218722 | -0.040678748 | 0.90104988 | 1.00 |
| YHR126C | 18.42016663 | -0.079008864 | 0.901008918 | 1.00 |
| YJR116W | 691.1206507 | -0.036755067 | 0.900919248 | 1.00 |
| YPL131W | 24700.79965 | 0.119592173 | 0.901368314 | 1.00 |
| YGR064W | 14.24950712 | -0.029279326 | 0.901914368 | 1.00 |
| YLR392C | 638.9786435 | 0.03534341 | 0.901779635 | 1.00 |

| | | | | |
|---|---|---|---|---|
| YGR016W | 101.0221921 | -0.02885233 | 0.90259555 | 1.00 |
| YDR023W | 6264.678536 | 0.071759246 | 0.90302804 | 1.00 |
| YGR173W | 798.4212433 | -0.011801699 | 0.902876964 | 1.00 |
| YDR261W-B | 58.90647804 | 0.026479177 | 0.903523386 | 1.00 |
| YOR218C | 5.229072963 | -0.135340975 | 0.903350871 | 1.00 |
| YPL214C | 1002.445323 | 0.028774942 | 0.903541937 | 1.00 |
| YML016C | 1375.407219 | 0.021153963 | 0.903784605 | 1.00 |
| YAL003W | 12374.8526 | 0.133854581 | 0.956096877 | 1.00 |
| YAL044W-A | 228.0761832 | -0.048585105 | 0.939321677 | 1.00 |
| YAL059W | 208.6556716 | 0.021104477 | 0.939477466 | 1.00 |
| YAR009C | 236.1235724 | -0.002444498 | 0.944670498 | 1.00 |
| YBL009W | 764.0807255 | 0.046802015 | 0.943813433 | 1.00 |
| YBL017C | 3980.2423 | -0.030941314 | 0.935231399 | 1.00 |
| YBL058W | 3576.371774 | 0.019786748 | 0.935501627 | 1.00 |
| YBL073W | 6.010181019 | 0.008479212 | 0.953003449 | 1.00 |
| YBL100W-C | 6.925265405 | -0.132942893 | 0.923207652 | 1.00 |
| YBL108W | 0.561379936 | -0.052885944 | 0.931610819 | 1.00 |
| YBR004C | 976.5395701 | 0.028169435 | 0.934514272 | 1.00 |
| YBR042C | 577.29725 | -0.004156908 | 0.948029289 | 1.00 |
| YBR067C | 11273.4348 | -0.040877254 | 0.958740836 | 1.00 |
| YBR089W | 6.735755394 | -0.021877079 | 0.90547838 | 1.00 |
| YBR092C | 2232.790112 | 0.050623139 | 0.91233077 | 1.00 |
| YBR103C-A | 1.570441558 | -0.001492549 | 0.922353754 | 1.00 |
| YBR137W | 861.5111114 | 0.052368317 | 0.953927787 | 1.00 |
| YBR173C | 2339.401983 | -0.019096042 | 0.909666716 | 1.00 |
| YBR188C | 339.0880427 | 0.02090696 | 0.949816441 | 1.00 |
| YBR190W | 3.930160671 | -0.161194235 | 0.944756727 | 1.00 |
| YBR200W-A | 19.77475914 | -0.079084836 | 0.920295934 | 1.00 |
| YBR206W | 255.7378889 | -0.027797745 | 0.924928938 | 1.00 |
| YBR216C | 879.1909045 | -0.017896501 | 0.932002174 | 1.00 |
| YBR223C | 483.3688068 | 0.024960802 | 0.907929691 | 1.00 |
| YBR228W | 415.0546527 | 0.011192608 | 0.959193995 | 1.00 |
| YBR232C | 24.08316402 | 0.051572407 | 0.948090494 | 1.00 |
| YBR237W | 937.0201345 | 0.014180523 | 0.953071393 | 1.00 |
| YBR242W | 549.7068913 | 0.034531096 | 0.916093691 | 1.00 |
| YBR250W | 80.81966717 | -0.008211428 | 0.932396054 | 1.00 |
| YBR255C-A | 311.2384417 | 0.023384638 | 0.924414551 | 1.00 |
| YBR273C | 1305.311458 | 0.023207942 | 0.916552545 | 1.00 |
| YBR278W | 245.0159717 | -0.037613601 | 0.934207642 | 1.00 |
| YBR283C | 3928.577615 | 0.008126464 | 0.927039179 | 1.00 |
| YCL001W-A | 26.77980078 | 0.035930897 | 0.908705613 | 1.00 |
| YCL001W-B | 30.32607662 | -0.048566045 | 0.95842515 | 1.00 |
| YCL044C | 1238.471005 | -0.030972615 | 0.940366221 | 1.00 |

| | | | | |
|---|---|---|---|---|
| YCL051W | 1104.388423 | -0.027640783 | 0.946159068 | 1.00 |
| YCL059C | 1691.420964 | 0.113473356 | 0.924591841 | 1.00 |
| YCR014C | 618.1633306 | 0.008347644 | 0.921861815 | 1.00 |
| YCR016W | 907.9551315 | 0.093974114 | 0.932487654 | 1.00 |
| YCR020W-B | 152.1239298 | -0.022477679 | 0.928739747 | 1.00 |
| YCR026C | 1309.803177 | -0.030366939 | 0.946963952 | 1.00 |
| YCR075C | 329.3656461 | -0.011701214 | 0.938718073 | 1.00 |
| YCR094W | 731.6068684 | -0.011769826 | 0.935044415 | 1.00 |
| YCR099C | 126.2405817 | 0.021157503 | 0.919082965 | 1.00 |
| YCR100C | 233.2545617 | -0.00439047 | 0.948729063 | 1.00 |
| YCR101C | 105.2670668 | -0.066339566 | 0.940286294 | 1.00 |
| YDL037C | 92.66763176 | 0.064484196 | 0.914961867 | 1.00 |
| YDL040C | 3238.410502 | 0.00036333 | 0.954327446 | 1.00 |
| YDL048C | 3537.203448 | 0.048753278 | 0.905499505 | 1.00 |
| YDL058W | 1998.717774 | 0.016487071 | 0.940043992 | 1.00 |
| YDL064W | 884.0529801 | 0.061354317 | 0.905326975 | 1.00 |
| YDL084W | 7170.004042 | 0.014538427 | 0.920288836 | 1.00 |
| YDL098C | 246.2120054 | -0.027601097 | 0.922923255 | 1.00 |
| YDL103C | 1449.48201 | -0.016155847 | 0.910023402 | 1.00 |
| YDL125C | 2310.617952 | 0.040843826 | 0.935095342 | 1.00 |
| YDL163W | 0.057163039 | -0.00999052 | 0.958353975 | 1.00 |
| YDL173W | 1092.267621 | 0.077328501 | 0.91082547 | 1.00 |
| YDL205C | 1039.950483 | -0.025532008 | 0.938784528 | 1.00 |
| YDL228C | 55.92892077 | -0.086669508 | 0.932359132 | 1.00 |
| YDL231C | 1653.703222 | -0.0025875 | 0.95593188 | 1.00 |
| YDL239C | 522.0657782 | -0.037309669 | 0.946160504 | 1.00 |
| YDR002W | 3692.839207 | 0.108152118 | 0.921729126 | 1.00 |
| YDR021W | 377.9993278 | 0.043560814 | 0.958110192 | 1.00 |
| YDR054C | 1985.938955 | 0.020335285 | 0.947727441 | 1.00 |
| YDR085C | 1259.372351 | 0.023887097 | 0.91253365 | 1.00 |
| YDR092W | 1261.21405 | -0.003489832 | 0.955870401 | 1.00 |
| YDR105C | 933.9584198 | -0.012974134 | 0.958819646 | 1.00 |
| YDR120C | 1240.636645 | 0.10940635 | 0.918632182 | 1.00 |
| YDR179W-A | 321.5877637 | 0.043357851 | 0.917039913 | 1.00 |
| YDR192C | 476.3597916 | -0.025262809 | 0.934894434 | 1.00 |
| YDR248C | 767.0999761 | -0.020784116 | 0.931103503 | 1.00 |
| YDR249C | 239.973621 | -0.043826079 | 0.916902488 | 1.00 |
| YDR261C | 1216.328788 | -0.021739856 | 0.933385238 | 1.00 |
| YDR265W | 518.8074155 | 0.01777952 | 0.939541236 | 1.00 |
| YDR324C | 1092.059972 | -0.002915317 | 0.927561874 | 1.00 |
| YDR342C | 1706.769269 | 0.010724713 | 0.927457671 | 1.00 |
| YDR354W | 871.1284672 | 0.0949801 | 0.930833075 | 1.00 |
| YDR360W | 36.19514521 | -0.027245102 | 0.906616399 | 1.00 |

| | | | | |
|---|---|---|---|---|
| YDR363W | 641.2311125 | -0.023284625 | 0.937283721 | 1.00 |
| YDR373W | 760.4394828 | -0.010041156 | 0.942362303 | 1.00 |
| YDR385W | 1851.31364 | -0.003597259 | 0.925187175 | 1.00 |
| YDR400W | 652.8362371 | 0.026231014 | 0.954387074 | 1.00 |
| YDR401W | 0.227003213 | -0.008997121 | 0.91516004 | 1.00 |
| YDR439W | 511.333794 | -0.017853374 | 0.949852003 | 1.00 |
| YDR470C | 683.2878102 | -0.017183 | 0.93448662 | 1.00 |
| YDR494W | 1564.77137 | -0.01740298 | 0.929238175 | 1.00 |
| YDR496C | 1668.42234 | 0.068787414 | 0.95164269 | 1.00 |
| YDR509W | 13.68760112 | -0.107739684 | 0.948598792 | 1.00 |
| YDR519W | 900.2252274 | 0.019872334 | 0.940294895 | 1.00 |
| YDR537C | 28.61472288 | -0.017813554 | 0.918277398 | 1.00 |
| YEL018C-A | 0.416927005 | 0.016479628 | 0.937805475 | 1.00 |
| YEL023C | 623.1672116 | -0.041242214 | 0.924532355 | 1.00 |
| YER034W | 308.6813768 | 0.056167469 | 0.909057369 | 1.00 |
| YER036C | 4277.526152 | 0.068630585 | 0.9323373 | 1.00 |
| YER037W | 745.1407721 | -0.024123277 | 0.943678048 | 1.00 |
| YER057C | 2325.694126 | 0.022366297 | 0.948935251 | 1.00 |
| YER076C | 386.5202214 | 0.003904941 | 0.950538054 | 1.00 |
| YER102W | 2473.647383 | 0.073376843 | 0.911963007 | 1.00 |
| YER118C | 889.8446464 | 0.077137139 | 0.950452599 | 1.00 |
| YER125W | 2425.853452 | 0.01798318 | 0.935760941 | 1.00 |
| YER149C | 245.5366344 | 0.084385558 | 0.926962768 | 1.00 |
| YER150W | 3467.414047 | 0.030734317 | 0.956072931 | 1.00 |
| YER164W | 2428.238913 | -0.017181919 | 0.946627056 | 1.00 |
| YER172C-A | 0.861078059 | -0.219943711 | 0.959412829 | 1.00 |
| YFL013C | 826.2401076 | -0.026586965 | 0.904376962 | 1.00 |
| YFL022C | 4340.34362 | 0.005687622 | 0.915355442 | 1.00 |
| YFL059W | 70.79713485 | -0.045264505 | 0.913197933 | 1.00 |
| YFR009W | 2637.309837 | 0.036729915 | 0.951360564 | 1.00 |
| YFR014C | 1739.196956 | 0.023843027 | 0.942340776 | 1.00 |
| YGL004C | 1031.613443 | 0.017543093 | 0.949137513 | 1.00 |
| YGL016W | 794.4716524 | 0.015491083 | 0.952374596 | 1.00 |
| YGL018C | 470.8004765 | -0.024472116 | 0.920381779 | 1.00 |
| YGL067W | 999.1477018 | -0.020959618 | 0.917856416 | 1.00 |
| YGL076C | 4229.319259 | 0.115590446 | 0.941344385 | 1.00 |
| YGL084C | 1084.484417 | -0.005922749 | 0.959102728 | 1.00 |
| YGL101W | 448.9089719 | 0.043241551 | 0.956473843 | 1.00 |
| YGL143C | 536.4770627 | -0.021258463 | 0.936465879 | 1.00 |
| YGL148W | 4268.613078 | 0.108039843 | 0.917592155 | 1.00 |
| YGL164C | 861.3475827 | -0.017025209 | 0.954769957 | 1.00 |
| YGL171W | 777.7803509 | 0.092969783 | 0.946998001 | 1.00 |
| YGL185C | 473.3669461 | -0.017340838 | 0.953811809 | 1.00 |

| | | | | |
|---|---|---|---|---|
| YGL197W | 2610.651509 | -0.033217273 | 0.939604368 | 1.00 |
| YGL201C | 994.5170532 | -0.008221387 | 0.950805732 | 1.00 |
| YGL228W | 2378.82392 | -0.01933943 | 0.949326807 | 1.00 |
| YGL232W | 728.6792639 | 0.005570649 | 0.944764254 | 1.00 |
| YGL257C | 1193.728393 | -0.010955506 | 0.934059138 | 1.00 |
| YGL260W | 1.415207671 | -0.052407901 | 0.909775181 | 1.00 |
| YGR001C | 885.4111524 | 0.12860592 | 0.911180114 | 1.00 |
| YGR013W | 916.8674279 | 0.019680247 | 0.912883325 | 1.00 |
| YGR018C | 8.799849222 | -0.122934505 | 0.940846906 | 1.00 |
| YGR024C | 991.4360363 | -0.00688359 | 0.947247555 | 1.00 |
| YGR058W | 565.2067235 | -0.022449227 | 0.913991701 | 1.00 |
| YGR061C | 5651.499061 | 0.109180299 | 0.919189208 | 1.00 |
| YGR081C | 443.1057485 | -0.026584685 | 0.908918397 | 1.00 |
| YGR083C | 2563.158798 | 0.005915512 | 0.939080067 | 1.00 |
| YGR109C | 138.0694558 | 0.020885688 | 0.935615218 | 1.00 |
| YGR232W | 824.5178442 | -0.002673824 | 0.954802099 | 1.00 |
| YGR243W | 1286.002734 | 0.088480577 | 0.955799809 | 1.00 |
| YGR248W | 2366.128573 | 0.035518235 | 0.945865594 | 1.00 |
| YGR270W | 2956.664275 | -0.022946432 | 0.917103792 | 1.00 |
| YGR281W | 2050.830805 | -0.019889953 | 0.930982503 | 1.00 |
| YGR285C | 10708.64158 | 0.121786489 | 0.921482825 | 1.00 |
| YGR289C | 671.1201555 | -0.029361712 | 0.934346204 | 1.00 |
| YHL015W | 17106.50707 | 0.038446142 | 0.905724762 | 1.00 |
| YHL027W | 1629.795881 | -0.028969951 | 0.955484392 | 1.00 |
| YHL047C | 1687.543568 | -0.004252234 | 0.944241505 | 1.00 |
| YHR002W | 437.310458 | 0.021097421 | 0.946791136 | 1.00 |
| YHR011W | 509.988716 | -0.025426738 | 0.937876411 | 1.00 |
| YHR025W | 4807.328396 | 0.096460516 | 0.922808124 | 1.00 |
| YHR042W | 4638.852766 | -0.013443184 | 0.916450874 | 1.00 |
| YHR074W | 2090.525023 | -0.014670385 | 0.929643302 | 1.00 |
| YHR079C | 1259.061897 | 0.012415745 | 0.933258226 | 1.00 |
| YHR099W | 4737.26123 | 0.043793049 | 0.922203747 | 1.00 |
| YHR107C | 1509.6288 | 0.046187387 | 0.951622479 | 1.00 |
| YHR119W | 1485.712969 | 0.027073283 | 0.910234086 | 1.00 |
| YHR132C | 1760.575956 | 0.029361655 | 0.944436033 | 1.00 |
| YHR169W | 496.2218428 | -0.000122147 | 0.947429024 | 1.00 |
| YHR177W | 301.5515693 | -0.05072137 | 0.904650795 | 1.00 |
| YHR180W-A | 0.36648756 | -0.037505204 | 0.947517672 | 1.00 |
| YHR185C | 7.642313868 | -0.002239478 | 0.948715174 | 1.00 |
| YHR209W | 694.2958547 | -0.017442589 | 0.948190921 | 1.00 |
| YHR215W | 110.5268911 | 0.046792612 | 0.921135388 | 1.00 |
| YIL001W | 471.4395924 | -0.045553826 | 0.916445606 | 1.00 |
| YIL020C | 394.2399705 | 0.036360587 | 0.94584771 | 1.00 |

| | | | | |
|---|---|---|---|---|
| YIL045W | 1955.748951 | 0.053998026 | 0.911692983 | 1.00 |
| YIL056W | 1630.203256 | 0.018718254 | 0.908136826 | 1.00 |
| YIL108W | 1452.736233 | -0.033469277 | 0.955286609 | 1.00 |
| YIL111W | 1481.52504 | 0.0279865 | 0.933212592 | 1.00 |
| YIL127C | 639.7542142 | 0.076713786 | 0.946984242 | 1.00 |
| YIL132C | 143.1249549 | -0.042407453 | 0.937537093 | 1.00 |
| YIL170W | 172.5743318 | -0.008242525 | 0.920450045 | 1.00 |
| YIL177C | 0.203940714 | -0.011193549 | 0.945541941 | 1.00 |
| YIR004W | 1446.338254 | -0.012684423 | 0.94114399 | 1.00 |
| YIR012W | 1025.35668 | 0.038405802 | 0.942452066 | 1.00 |
| YIR025W | 303.8110514 | 0.01947926 | 0.904772752 | 1.00 |
| YIR034C | 2228.546885 | -0.022219815 | 0.94666949 | 1.00 |
| YIR040C | 0.167590483 | 0.041432527 | 0.955159317 | 1.00 |
| YIR042C | 196.4981605 | -0.001055307 | 0.945113082 | 1.00 |
| YJL001W | 1978.350649 | 0.017237116 | 0.94325963 | 1.00 |
| YJL013C | 787.2498623 | 0.008860515 | 0.94804392 | 1.00 |
| YJL035C | 163.2101994 | -0.028361734 | 0.930936084 | 1.00 |
| YJL041W | 2583.756357 | 0.017556347 | 0.953208753 | 1.00 |
| YJL059W | 446.5285115 | -0.030950056 | 0.908380379 | 1.00 |
| YJL060W | 1402.078784 | -0.010229606 | 0.914152905 | 1.00 |
| YJL090C | 629.2586035 | -0.021626026 | 0.957527138 | 1.00 |
| YJL119C | 7.109202711 | 0.006823445 | 0.934244223 | 1.00 |
| YJL130C | 21427.36188 | -0.032818156 | 0.933873097 | 1.00 |
| YJL131C | 712.8647158 | 0.029498558 | 0.916342883 | 1.00 |
| YJL149W | 1033.010855 | -0.028661746 | 0.931960336 | 1.00 |
| YJL150W | 10.09973158 | 0.021860591 | 0.951847302 | 1.00 |
| YJL158C | 11506.92019 | -0.001079899 | 0.913166069 | 1.00 |
| YJL160C | 84.43696653 | 0.045832596 | 0.944487402 | 1.00 |
| YJL187C | 1084.409994 | -0.012228088 | 0.94812821 | 1.00 |
| YJL208C | 631.1153592 | 0.089504103 | 0.95752662 | 1.00 |
| YJR033C | 1583.049905 | 0.014069891 | 0.926234961 | 1.00 |
| YJR045C | 25492.38268 | 0.072632825 | 0.957083547 | 1.00 |
| YJR052W | 627.3978725 | 0.01716187 | 0.914367062 | 1.00 |
| YJR105W | 3472.841224 | 0.073692339 | 0.951190145 | 1.00 |
| YJR151C | 560.5480031 | 0.0088297 | 0.927615612 | 1.00 |
| YJR160C | 11.90922489 | -0.105751492 | 0.94799294 | 1.00 |
| YKL006C-A | 282.6893408 | 0.024461665 | 0.950678 | 1.00 |
| YKL052C | 501.3271381 | -0.021804134 | 0.906179706 | 1.00 |
| YKL061W | 244.3100746 | -0.029381555 | 0.923170687 | 1.00 |
| YKL070W | 302.0799748 | 0.015298446 | 0.936351555 | 1.00 |
| YKL077W | 3042.957164 | -0.012374657 | 0.910573365 | 1.00 |
| YKL082C | 1057.576571 | 0.056730401 | 0.951313896 | 1.00 |
| YKL083W | 46.77827339 | -0.022992046 | 0.957594125 | 1.00 |

| | | | | |
|---|---|---|---|---|
| YKL089W | 415.1913401 | -0.040868663 | 0.915785568 | 1.00 |
| YKL096C-B | 8.469479675 | 0.145947168 | 0.957429827 | 1.00 |
| YKL101W | 3903.982421 | -0.020539581 | 0.95898284 | 1.00 |
| YKL135C | 1473.22037 | -0.020858147 | 0.914046698 | 1.00 |
| YKL138C-A | 204.185803 | -0.017625153 | 0.92302775 | 1.00 |
| YKL140W | 1755.744554 | -0.015566083 | 0.930073226 | 1.00 |
| YKL177W | 2.5279419 | -0.017214332 | 0.959047137 | 1.00 |
| YKL183W | 567.8673175 | 0.06855209 | 0.927778803 | 1.00 |
| YKL195W | 3025.520768 | -0.012795277 | 0.923042292 | 1.00 |
| YKL223W | 0.064032033 | 0.017966338 | 0.916924002 | 1.00 |
| YKR014C | 1917.457601 | 0.037289891 | 0.917494371 | 1.00 |
| YKR039W | 765.789828 | -0.009215139 | 0.947591151 | 1.00 |
| YKR060W | 417.425685 | 0.04982325 | 0.906386739 | 1.00 |
| YKR068C | 742.7822083 | 0.019774619 | 0.938345887 | 1.00 |
| YKR082W | 1772.94275 | -0.023945233 | 0.940998451 | 1.00 |
| YKR099W | 800.367483 | -0.026655941 | 0.941756036 | 1.00 |
| YLL033W | 166.5445731 | -0.01337264 | 0.959435355 | 1.00 |
| YLR003C | 782.5858822 | 0.029761051 | 0.954443988 | 1.00 |
| YLR041W | 27.97790779 | -0.078548062 | 0.910641145 | 1.00 |
| YLR044C | 136554.4679 | 0.095802664 | 0.943511071 | 1.00 |
| YLR052W | 1071.581443 | -0.013344745 | 0.916096105 | 1.00 |
| YLR075W | 9637.869009 | 0.148850691 | 0.914463608 | 1.00 |
| YLR091W | 340.1364791 | 0.008748592 | 0.957581324 | 1.00 |
| YLR117C | 1230.224284 | 0.017403813 | 0.914014562 | 1.00 |
| YLR146C | 511.7510501 | -0.009555901 | 0.91269715 | 1.00 |
| YLR157C-B | 0.750636836 | -0.141181757 | 0.953116263 | 1.00 |
| YLR179C | 2935.995773 | 0.097605508 | 0.94326316 | 1.00 |
| YLR195C | 1211.827498 | 0.004712865 | 0.948672399 | 1.00 |
| YLR227W-B | 558.9698813 | -0.039170794 | 0.958403995 | 1.00 |
| YLR242C | 302.599813 | -0.021158236 | 0.913685514 | 1.00 |
| YLR245C | 188.3206886 | 0.054833985 | 0.910324737 | 1.00 |
| YLR247C | 1785.222725 | -0.029202643 | 0.944590251 | 1.00 |
| YLR252W | 254.6818923 | 0.009957307 | 0.952476099 | 1.00 |
| YLR298C | 806.4703735 | 0.011068101 | 0.956801344 | 1.00 |
| YLR305C | 2169.127998 | -0.032701984 | 0.905338483 | 1.00 |
| YLR315W | 96.87354306 | 0.031162828 | 0.925974672 | 1.00 |
| YLR342W-A | 51.47657195 | -0.059696863 | 0.927943407 | 1.00 |
| YLR350W | 3571.832696 | -0.031265941 | 0.945070336 | 1.00 |
| YLR419W | 1247.341194 | -0.024428969 | 0.941309851 | 1.00 |
| YLR424W | 404.0091344 | 0.007442764 | 0.955380487 | 1.00 |
| YLR432W | 4140.896048 | 0.134402129 | 0.91514925 | 1.00 |
| YLR433C | 1164.793625 | 0.030178272 | 0.918145956 | 1.00 |
| YLR441C | 16750.1152 | 0.030864677 | 0.916042442 | 1.00 |

| | | | | |
|---|---|---|---|---|
| YML011C | 267.8459983 | 0.020643618 | 0.913927276 | 1.00 |
| YML015C | 859.349761 | -0.028480671 | 0.906351382 | 1.00 |
| YML056C | 4012.776938 | 0.073350431 | 0.939960369 | 1.00 |
| YML065W | 967.920824 | 0.012330604 | 0.944487493 | 1.00 |
| YML080W | 296.6855777 | 0.071328079 | 0.913026505 | 1.00 |
| YML082W | 766.1872079 | -0.02118765 | 0.908497163 | 1.00 |
| YML098W | 504.9087228 | 0.032669077 | 0.940683097 | 1.00 |
| YML117W | 3482.921508 | -0.022331584 | 0.951067056 | 1.00 |
| YMR007W | 90.19118303 | -0.040595771 | 0.928624033 | 1.00 |
| YMR011W | 5664.342762 | 0.138836041 | 0.939384063 | 1.00 |
| YMR022W | 1239.988654 | -0.010240842 | 0.956612271 | 1.00 |
| YMR036C | 631.5049431 | -0.02871909 | 0.936479575 | 1.00 |
| YMR043W | 2143.509763 | -0.054889336 | 0.920800928 | 1.00 |
| YMR080C | 1907.455726 | 0.018304543 | 0.95782171 | 1.00 |
| YMR092C | 2509.767947 | 0.033946585 | 0.907952717 | 1.00 |
| YMR102C | 1653.341009 | 0.066372548 | 0.918053378 | 1.00 |
| YMR116C | 12111.33345 | 0.20356357 | 0.958945123 | 1.00 |
| YMR120C | 11760.55672 | 0.182774659 | 0.922383992 | 1.00 |
| YMR126C | 404.3389603 | -0.041137193 | 0.911611853 | 1.00 |
| YMR146C | 5181.578577 | -0.008502901 | 0.921443129 | 1.00 |
| YMR150C | 511.1258272 | 0.02798259 | 0.917039298 | 1.00 |
| YMR168C | 528.3657431 | -0.024930628 | 0.906490677 | 1.00 |
| YMR189W | 2797.782123 | 0.061849139 | 0.929038817 | 1.00 |
| YMR200W | 1797.595559 | -0.013929488 | 0.937114485 | 1.00 |
| YMR221C | 1062.231053 | -0.011452036 | 0.948582917 | 1.00 |
| YMR240C | 592.2234425 | 0.008241673 | 0.941905231 | 1.00 |
| YMR252C | 396.7363367 | 0.026928446 | 0.922347837 | 1.00 |
| YMR282C | 677.7599955 | -0.021259046 | 0.929338159 | 1.00 |
| YMR290C | 2185.404412 | 0.082999211 | 0.957127954 | 1.00 |
| YNL001W | 629.9634099 | -0.008920029 | 0.924727326 | 1.00 |
| YNL007C | 10095.98848 | 0.036046343 | 0.938275231 | 1.00 |
| YNL010W | 2912.199227 | 0.01442783 | 0.920599073 | 1.00 |
| YNL056W | 768.1896527 | 0.03541064 | 0.909226555 | 1.00 |
| YNL061W | 2817.950295 | 0.076919037 | 0.929930471 | 1.00 |
| YNL067W-B | 0.339190409 | -0.004044151 | 0.909832246 | 1.00 |
| YNL082W | 573.887894 | -0.01897273 | 0.934298069 | 1.00 |
| YNL119W | 513.2444289 | 0.030861965 | 0.948864753 | 1.00 |
| YNL128W | 87.91000712 | -0.057492647 | 0.921406299 | 1.00 |
| YNL139C | 1577.578713 | 0.011286656 | 0.948048917 | 1.00 |
| YNL159C | 572.5140174 | -0.032885206 | 0.927533982 | 1.00 |
| YNL175C | 613.6716753 | 0.106036844 | 0.959459376 | 1.00 |
| YNL177C | 1301.893277 | -0.00552889 | 0.953510874 | 1.00 |
| YNL186W | 1470.462386 | 0.01854816 | 0.925927466 | 1.00 |

| | | | | |
|---|---|---|---|---|
| YNL193W | 690.484771 | -0.034396601 | 0.934203728 | 1.00 |
| YNL217W | 1768.453008 | -0.01340982 | 0.919969923 | 1.00 |
| YNL248C | 1500.899157 | 0.032957564 | 0.936235322 | 1.00 |
| YNL252C | 974.4576049 | -0.00963048 | 0.92580842 | 1.00 |
| YNL275W | 352.1195966 | -0.025454964 | 0.93327493 | 1.00 |
| YNL277W | 546.4771573 | -0.058755308 | 0.943001744 | 1.00 |
| YNL285W | 3.926788582 | -0.101892899 | 0.95659848 | 1.00 |
| YNL315C | 1636.297254 | -0.011355853 | 0.933429042 | 1.00 |
| YNR062C | 190.9739726 | 0.023637857 | 0.916448475 | 1.00 |
| YOL041C | 1321.58671 | 0.056827134 | 0.938258128 | 1.00 |
| YOL050C | 16.33482394 | -0.029343121 | 0.954804002 | 1.00 |
| YOL052C | 1181.165007 | 0.029574985 | 0.906306911 | 1.00 |
| YOL062C | 1303.409819 | -0.011349981 | 0.942789239 | 1.00 |
| YOL097C | 2168.680575 | 0.057303699 | 0.945825039 | 1.00 |
| YOL159C | 274.8863973 | -0.0115728 | 0.952482802 | 1.00 |
| YOL160W | 15.67971505 | -0.008861494 | 0.948188067 | 1.00 |
| YOR001W | 1523.051345 | 0.026354844 | 0.928010836 | 1.00 |
| YOR038C | 676.9008791 | 0.010081506 | 0.941690347 | 1.00 |
| YOR055W | 14.54954616 | 0.042027751 | 0.938706839 | 1.00 |
| YOR087W | 1019.099344 | -0.030136016 | 0.909575665 | 1.00 |
| YOR090C | 1602.791656 | -0.018714122 | 0.905299308 | 1.00 |
| YOR103C | 111.7299261 | -0.002218306 | 0.934292337 | 1.00 |
| YOR128C | 2990.654925 | 0.027446293 | 0.949168913 | 1.00 |
| YOR133W | 1425.340865 | 0.003146405 | 0.93729757 | 1.00 |
| YOR136W | 7814.887983 | 0.058354 | 0.953127909 | 1.00 |
| YOR188W | 1393.160444 | 0.017877839 | 0.932767083 | 1.00 |
| YOR191W | 2372.954185 | -0.034358078 | 0.915438172 | 1.00 |
| YOR204W | 2305.73227 | 0.040417892 | 0.959188612 | 1.00 |
| YOR223W | 687.6095973 | 0.00895167 | 0.949519625 | 1.00 |
| YOR243C | 1479.808571 | 0.02263119 | 0.956102981 | 1.00 |
| YOR258W | 218.0314211 | -0.031026152 | 0.917098568 | 1.00 |
| YOR316C-A | 8.072827245 | 0.093978818 | 0.946511207 | 1.00 |
| YOR333C | 3.992636415 | -0.071790675 | 0.955249366 | 1.00 |
| YOR379C | 0.107512111 | -0.010713225 | 0.923759424 | 1.00 |
| YOR381W-A | 9.096748957 | -0.134326381 | 0.931430571 | 1.00 |
| YPL008W | 976.9490792 | 0.014533658 | 0.928469524 | 1.00 |
| YPL041C | 160.4382575 | -0.043800657 | 0.910619785 | 1.00 |
| YPL042C | 568.6329785 | -0.024913873 | 0.952065254 | 1.00 |
| YPL045W | 887.7248714 | 0.017111551 | 0.936669107 | 1.00 |
| YPL060C-A | 3.51919416 | 0.072560849 | 0.959062528 | 1.00 |
| YPL067C | 285.3555196 | -0.007054893 | 0.951795652 | 1.00 |
| YPL099C | 334.8201937 | -0.019796615 | 0.917093259 | 1.00 |
| YPL170W | 622.4061116 | 0.019096722 | 0.927229101 | 1.00 |

| | | | | |
|---|---|---|---|---|
| YPL172C | 484.663762 | 0.028850993 | 0.908790623 | 1.00 |
| YPL189C-A | 341.9617668 | 0.021465706 | 0.942181134 | 1.00 |
| YPL195W | 2059.660784 | -0.015442837 | 0.913272151 | 1.00 |
| YPL208W | 646.0074609 | 0.00245966 | 0.926421348 | 1.00 |
| YPL226W | 2793.678288 | 0.004906612 | 0.906808746 | 1.00 |
| YPL237W | 2463.364966 | 0.059061357 | 0.949252067 | 1.00 |
| YPR004C | 1678.278628 | 0.05693305 | 0.922683223 | 1.00 |
| YPR036W | 6660.698271 | 0.048618269 | 0.90787347 | 1.00 |
| YPR106W | 760.0851492 | 0.014040087 | 0.915316948 | 1.00 |
| YPR108W-A | 1.090591333 | -0.054949508 | 0.943312856 | 1.00 |
| YPR133C | 1652.526675 | -0.010458931 | 0.933623648 | 1.00 |
| YPR145C-A | 45.3897147 | -0.032351577 | 0.946852002 | 1.00 |
| YPR145W | 8044.845508 | 0.051226708 | 0.916998079 | 1.00 |
| YPR161C | 1370.124428 | -0.022648256 | 0.937676003 | 1.00 |
| YPR178W | 455.7725824 | -0.01839015 | 0.947530265 | 1.00 |
| YPR181C | 4442.448271 | 0.035317884 | 0.948453177 | 1.00 |
| YPR183W | 3494.823853 | 0.040558709 | 0.942665682 | 1.00 |
| YPR186C | 328.1330267 | -0.025454012 | 0.926461182 | 1.00 |
| snR34 | 37.83791355 | 0.187192895 | 0.956388632 | 1.00 |
| snR46 | 23.07413489 | 0.032024754 | 0.910745384 | 1.00 |
| snR67 | 6.800863812 | 0.087252302 | 0.927922167 | 1.00 |
| snR73 | 0.982818557 | -0.014272361 | 0.922888206 | 1.00 |
| snR74 | 1.214578707 | 0.196868382 | 0.929452168 | 1.00 |
| snR80 | 3.471418478 | 0.143845349 | 0.957971743 | 1.00 |
| tR(CCU)J | 0.0941958 | -0.009916254 | 0.958353139 | 1.00 |
| YDL137W | 4443.238326 | 0.005127966 | 0.960170247 | 1.00 |
| YAL064W | 3.131576444 | -0.095075664 | 0.983633247 | 1.00 |
| YAL067C | 87.22318015 | -0.029181242 | 0.982109329 | 1.00 |
| YAL067W-A | 0.291097411 | -0.032096492 | 0.98424787 | 1.00 |
| YAR002C-A | 2600.594621 | 0.02321539 | 0.971591799 | 1.00 |
| YBL023C | 762.982123 | -0.000662582 | 0.981336405 | 1.00 |
| YBL038W | 1166.095027 | 0.010038768 | 0.980670942 | 1.00 |
| YBL050W | 1823.454847 | 0.014098728 | 0.968964989 | 1.00 |
| YBL088C | 1866.759984 | 0.001516325 | 0.981714546 | 1.00 |
| YBL089W | 493.6798455 | 0.004995194 | 0.98520199 | 1.00 |
| YBR013C | 207.4427581 | -0.017780497 | 0.98311577 | 1.00 |
| YBR084C-A | 8707.666036 | 0.045356992 | 0.963709532 | 1.00 |
| YBR115C | 2897.179739 | 0.012206797 | 0.984991947 | 1.00 |
| YBR243C | 997.8241569 | 0.020610107 | 0.976925999 | 1.00 |
| YBR260C | 1246.430976 | 0.013702659 | 0.963063219 | 1.00 |
| YBR267W | 453.4917639 | 0.022989204 | 0.983270678 | 1.00 |
| YCR019W | 382.8804523 | -0.007377244 | 0.96764415 | 1.00 |
| YCR020C-A | 266.9746924 | 0.025781636 | 0.982725247 | 1.00 |

| | | | | |
|---|---|---|---|---|
| YCR057C | 1417.95789 | 0.031638159 | 0.972226301 | 1.00 |
| YCR063W | 60.70870613 | 0.012107007 | 0.960755584 | 1.00 |
| YDL087C | 335.3146294 | -0.014410794 | 0.964458948 | 1.00 |
| YDL156W | 331.6691228 | 0.014511126 | 0.974309909 | 1.00 |
| YDR122W | 2792.779664 | -0.007010394 | 0.967877639 | 1.00 |
| YDR147W | 507.4257469 | -0.003069687 | 0.980960907 | 1.00 |
| YDR154C | 3531.276985 | 0.004681433 | 0.980776473 | 1.00 |
| YDR167W | 803.2507476 | 0.01679737 | 0.971690523 | 1.00 |
| YDR183C-A | 47.62983085 | -0.008134585 | 0.976093408 | 1.00 |
| YDR205W | 692.1640308 | -0.002155957 | 0.983872006 | 1.00 |
| YDR243C | 512.1645164 | -0.009329924 | 0.975939321 | 1.00 |
| YDR428C | 292.2817321 | 0.042388044 | 0.964435636 | 1.00 |
| YDR450W | 8491.535949 | 0.073370723 | 0.980015879 | 1.00 |
| YDR486C | 1425.158924 | 0.002389037 | 0.985031029 | 1.00 |
| YDR514C | 275.4543514 | -0.004331893 | 0.971976094 | 1.00 |
| YDR525W-A | 275.582834 | 0.040511219 | 0.964852055 | 1.00 |
| YDR533C | 1411.920932 | -0.015464667 | 0.968290453 | 1.00 |
| YEL046C | 2157.479841 | 0.072706075 | 0.969039657 | 1.00 |
| YEL053W-A | 186.8810681 | -0.111555124 | 0.963157136 | 1.00 |
| YEL059C-A | 318.8027126 | -0.010014408 | 0.967120725 | 1.00 |
| YER003C | 3520.842344 | 0.049217533 | 0.9764287 | 1.00 |
| YER027C | 1561.298312 | 0.00491228 | 0.983373565 | 1.00 |
| YER156C | 1503.86508 | 0.039209252 | 0.980941923 | 1.00 |
| YFL021C-A | 41.5780796 | -0.050846323 | 0.965194059 | 1.00 |
| YGL063W | 321.9330079 | 0.02161315 | 0.966120815 | 1.00 |
| YGL085W | 385.7388603 | 0.007501481 | 0.960978326 | 1.00 |
| YGL087C | 1504.49763 | 0.006481368 | 0.965753575 | 1.00 |
| YGL097W | 1307.008078 | 0.066184019 | 0.966038776 | 1.00 |
| YGL118C | 2.533670207 | -0.163965133 | 0.979693949 | 1.00 |
| YGL126W | 1063.33427 | 0.012293677 | 0.978657793 | 1.00 |
| YGL132W | 16.53764806 | -0.03678252 | 0.970476693 | 1.00 |
| YGL247W | 197.7711302 | 0.012738693 | 0.975053543 | 1.00 |
| YGR012W | 677.5451845 | 0.012158794 | 0.965238647 | 1.00 |
| YGR068C | 535.7602898 | -0.000507556 | 0.984410628 | 1.00 |
| YGR114C | 36.93060097 | -0.229663753 | 0.985374116 | 1.00 |
| YGR124W | 7850.205704 | 0.066792676 | 0.975520503 | 1.00 |
| YGR135W | 3309.689244 | 0.023001853 | 0.980796596 | 1.00 |
| YGR230W | 330.0627222 | -0.007118227 | 0.975509577 | 1.00 |
| YGR282C | 11912.38121 | 0.017484515 | 0.983652831 | 1.00 |
| YHL013C | 503.0447213 | 0.049814019 | 0.984399211 | 1.00 |
| YHL020C | 961.5750146 | -0.011986886 | 0.979741528 | 1.00 |
| YHL031C | 431.2428898 | 0.007987615 | 0.978439897 | 1.00 |
| YHR018C | 3547.596925 | 0.074542674 | 0.966369304 | 1.00 |

| | | | | |
|---|---|---|---|---|
| YHR019C | 9735.850466 | 0.046423453 | 0.978874183 | 1.00 |
| YHR097C | 2968.055756 | -0.026921666 | 0.979735009 | 1.00 |
| YHR183W | 10217.98699 | 0.033237134 | 0.966054256 | 1.00 |
| YHR197W | 861.9145076 | 0.046747126 | 0.9652147 | 1.00 |
| YHR205W | 2582.464034 | 0.001878125 | 0.982869939 | 1.00 |
| YIL067C | 437.2526192 | 0.003307633 | 0.96355002 | 1.00 |
| YIL094C | 3348.951104 | 0.007341286 | 0.963656491 | 1.00 |
| YIL154C | 2075.807869 | 0.006350629 | 0.974126217 | 1.00 |
| YIL156W | 1109.423201 | 0.006214843 | 0.966062217 | 1.00 |
| YIR005W | 167.3355793 | -0.011203806 | 0.983151469 | 1.00 |
| YJL033W | 1448.230029 | 0.062236229 | 0.968217782 | 1.00 |
| YJL092W | 1243.946562 | 0.019002414 | 0.967758979 | 1.00 |
| YJL165C | 2120.172359 | -0.021727287 | 0.969595723 | 1.00 |
| YJR017C | 953.2894932 | 0.006788946 | 0.977272052 | 1.00 |
| YJR135C | 188.8236464 | 0.014730717 | 0.975867284 | 1.00 |
| YJR139C | 4827.24467 | 0.054828149 | 0.984800508 | 1.00 |
| YJR142W | 1091.921242 | -0.001682958 | 0.983024042 | 1.00 |
| YKL011C | 416.7502341 | 0.005464835 | 0.972159863 | 1.00 |
| YKL094W | 2161.93051 | -0.000396945 | 0.97814164 | 1.00 |
| YKL144C | 415.786387 | 0.034739258 | 0.975200718 | 1.00 |
| YKL168C | 972.1957599 | -0.010583125 | 0.983182205 | 1.00 |
| YKL222C | 519.5767688 | 0.002443609 | 0.969976923 | 1.00 |
| YKR011C | 1595.285427 | 0.001660634 | 0.965504218 | 1.00 |
| YKR065C | 1072.834618 | 0.009000099 | 0.975495007 | 1.00 |
| YKR075C | 2465.019565 | 0.016131038 | 0.973256572 | 1.00 |
| YLL003W | 928.3757705 | -0.01219289 | 0.968677108 | 1.00 |
| YLR027C | 7716.105415 | 0.033354882 | 0.974283091 | 1.00 |
| YLR095C | 1759.144757 | -0.012247161 | 0.963267359 | 1.00 |
| YLR118C | 939.2371521 | -0.000965021 | 0.971192342 | 1.00 |
| YLR131C | 1246.543234 | -0.017044692 | 0.983654893 | 1.00 |
| YLR212C | 601.9431586 | 0.044323669 | 0.982490517 | 1.00 |
| YLR249W | 32716.59454 | 0.118970853 | 0.973531457 | 1.00 |
| YLR282C | 4.588773458 | 0.036916968 | 0.969582378 | 1.00 |
| YLR343W | 1249.632226 | -0.090012782 | 0.97432698 | 1.00 |
| YLR347C | 1405.641169 | 0.013989437 | 0.9701917 | 1.00 |
| YLR373C | 1362.586121 | 0.00306226 | 0.968701476 | 1.00 |
| YLR449W | 2574.095007 | 0.081822048 | 0.969258549 | 1.00 |
| YML021C | 560.3287148 | -0.005336902 | 0.972105969 | 1.00 |
| YML031W | 1341.560963 | 0.011632016 | 0.965407777 | 1.00 |
| YML037C | 139.9825756 | 0.006893798 | 0.979801853 | 1.00 |
| YML048W | 4451.705992 | 0.001648278 | 0.970781726 | 1.00 |
| YML054C | 1101.015099 | -0.006108994 | 0.964366753 | 1.00 |
| YML057C-A | 86.62699701 | -0.016599617 | 0.965727362 | 1.00 |

| | | | | |
|---|---|---|---|---|
| YML064C | 587.9701312 | 0.000511338 | 0.972158418 | 1.00 |
| YML068W | 512.1804017 | 0.002715668 | 0.970080496 | 1.00 |
| YML124C | 2648.714801 | 0.030875687 | 0.975808245 | 1.00 |
| YMR002W | 4776.428936 | 0.024199783 | 0.962294243 | 1.00 |
| YMR039C | 1898.602952 | 0.003823735 | 0.983311107 | 1.00 |
| YMR121C | 804.8314095 | 0.011146405 | 0.976640965 | 1.00 |
| YMR131C | 2416.896733 | 0.048046585 | 0.983859391 | 1.00 |
| YMR154C | 501.7366665 | 0.00277445 | 0.975919657 | 1.00 |
| YMR172W | 685.1558079 | -0.011518076 | 0.970733677 | 1.00 |
| YMR211W | 431.1277672 | 0.004621279 | 0.972875761 | 1.00 |
| YMR226C | 4108.834639 | 0.005915009 | 0.980910225 | 1.00 |
| YMR238W | 935.1001777 | 0.049445083 | 0.974508872 | 1.00 |
| YMR303C | 445.8492587 | -0.031058541 | 0.972207055 | 1.00 |
| YNL024C | 148.4866774 | -0.055339214 | 0.971892825 | 1.00 |
| YNL037C | 5393.729596 | 0.054720374 | 0.974907804 | 1.00 |
| YNL112W | 1857.470789 | 0.179634008 | 0.962897254 | 1.00 |
| YNL120C | 0.227028385 | 0.015196243 | 0.965369544 | 1.00 |
| YNL155W | 1378.201426 | 0.022452595 | 0.964598643 | 1.00 |
| YNL182C | 900.9513347 | 0.039880862 | 0.963912718 | 1.00 |
| YNL221C | 1802.455751 | 0.003690723 | 0.963363906 | 1.00 |
| YNL250W | 994.2437698 | 0.000973186 | 0.982849087 | 1.00 |
| YNL281W | 1354.281332 | 0.025697926 | 0.970533669 | 1.00 |
| YNR050C | 3301.317883 | 0.024972255 | 0.977992643 | 1.00 |
| YNR077C | 0.24124529 | -0.010651764 | 0.965411056 | 1.00 |
| YOL032W | 1063.876828 | 0.012105187 | 0.963560733 | 1.00 |
| YOL063C | 983.1121387 | 0.002445919 | 0.969989756 | 1.00 |
| YOL080C | 199.8794327 | 0.069891183 | 0.977949404 | 1.00 |
| YOL095C | 382.7305205 | -0.01762794 | 0.982049815 | 1.00 |
| YOR015W | 345.4012917 | -0.021833249 | 0.981097204 | 1.00 |
| YOR042W | 2668.884695 | 0.014387262 | 0.962024342 | 1.00 |
| YOR044W | 320.4821165 | -0.024629309 | 0.976498809 | 1.00 |
| YOR063W | 34774.24031 | 0.124248963 | 0.973283951 | 1.00 |
| YOR098C | 2501.292329 | -0.00558306 | 0.974718184 | 1.00 |
| YOR106W | 543.2030625 | -0.008669731 | 0.970423314 | 1.00 |
| YOR122C | 4814.102226 | 0.011584936 | 0.981028295 | 1.00 |
| YOR199W | 28.95971662 | 0.026600118 | 0.963868342 | 1.00 |
| YOR270C | 5846.210473 | 0.004639049 | 0.981131272 | 1.00 |
| YOR287C | 297.2565187 | 0.040225676 | 0.963823959 | 1.00 |
| YOR294W | 771.7312013 | 0.078211639 | 0.963566988 | 1.00 |
| YOR341W | 3809.897668 | 0.055873625 | 0.96751789 | 1.00 |
| YPL028W | 3100.321144 | 0.041975634 | 0.980814188 | 1.00 |
| YPL068C | 273.8997414 | 0.000361602 | 0.98477036 | 1.00 |
| YPL161C | 652.0526344 | 0.004930042 | 0.972261988 | 1.00 |

| | | | | |
|---|---|---|---|---|
| YPL199C | 1017.89655 | 0.012381656 | 0.979242169 | 1.00 |
| YPL206C | 1410.718758 | 0.01842211 | 0.978033153 | 1.00 |
| YPL211W | 668.7255746 | 0.16417872 | 0.984994028 | 1.00 |
| YPL229W | 514.8866138 | 0.005142693 | 0.965350432 | 1.00 |
| YPL249C-A | 77.54661876 | -0.048126886 | 0.980025999 | 1.00 |
| YPR015C | 179.9077993 | -0.0097215 | 0.985470111 | 1.00 |
| YPR021C | 1682.395818 | -0.025715444 | 0.977944394 | 1.00 |
| YPR096C | 16.87382422 | -0.015436056 | 0.979100558 | 1.00 |
| YPR110C | 2091.025042 | 0.04194126 | 0.969046239 | 1.00 |
| YPR147C | 688.0952487 | 0.001693778 | 0.98140955 | 1.00 |
| YPR167C | 421.1211 | -0.001766197 | 0.979022755 | 1.00 |
| YPR200C | 70.27684883 | 0.012572512 | 0.978785026 | 1.00 |
| snR47 | 5.450594875 | -0.099520493 | 0.970197343 | 1.00 |
| snR6 | 0.308944371 | -0.010777098 | 0.96804077 | 1.00 |
| snR68 | 10.29817386 | -0.018675833 | 0.971284324 | 1.00 |
| snR81 | 9.309990788 | 0.03819088 | 0.976064273 | 1.00 |
| tG(CCC)O | 0.202424979 | -0.009612957 | 0.975280533 | 1.00 |
| YJL152W | 449.6754094 | -0.021495712 | 0.985627584 | 1.00 |
| YOR072W-B | 0.082611248 | 0.01638723 | 0.98614529 | 1.00 |
| YOR353C | 984.6133479 | -0.009661798 | 0.985914754 | 1.00 |
| tQ(UUG)B | 0.084186605 | 0.015608443 | 0.986093423 | 1.00 |
| YCR095C | 483.8306058 | -0.001661775 | 0.987136239 | 1.00 |
| YGL058W | 714.9977444 | 0.012465604 | 0.986998651 | 1.00 |
| YHR001W | 1515.116914 | -0.004488196 | 0.987056578 | 1.00 |
| YJL181W | 405.181886 | -0.009162016 | 0.987763768 | 1.00 |
| YLR244C | 3158.461036 | 0.025886299 | 0.987405005 | 1.00 |
| YNL149C | 291.2466735 | 0.02847686 | 0.987894223 | 1.00 |
| YOR115C | 581.5147795 | 0.00485893 | 0.987904238 | 1.00 |
| YOR172W | 824.7713828 | 0.001810718 | 0.987362972 | 1.00 |
| YPR177C | 1.844393858 | 0.095920047 | 0.987600666 | 1.00 |
| YBR109W-A | 20.49364363 | -0.007647301 | 0.988220473 | 1.00 |
| YHR206W | 1203.868698 | -0.006512158 | 0.988340702 | 1.00 |
| YJR044C | 1233.447551 | -0.002150184 | 0.988769523 | 1.00 |
| YLR427W | 1359.694674 | 0.001091614 | 0.988632543 | 1.00 |
| YAL040C | 2404.494026 | -0.041817259 | 0.989117827 | 1.00 |
| YGR280C | 1635.461549 | 0.052768023 | 0.989130569 | 1.00 |
| YBL018C | 137.4735368 | 0.104247688 | 0.989359585 | 1.00 |
| YMR151W | 69.13225197 | -0.025633971 | 0.989504027 | 1.00 |
| YDR466W | 1549.812513 | -0.00740475 | 0.990255565 | 1.00 |
| YJR085C | 2130.783694 | 0.01292639 | 0.99015145 | 1.00 |
| YLR379W | 289.3293491 | 0.032360589 | 0.989848675 | 1.00 |
| YOR052C | 2749.089296 | -0.005596261 | 0.990081288 | 1.00 |
| YBR226C | 66.35832606 | -0.017116373 | 0.990703862 | 1.00 |

| | | | | |
|---|---|---|---|---|
| YEL044W | 582.0177494 | 0.001801539 | 0.990513244 | 1.00 |
| YGR271C-A | 454.6880839 | 0.087758229 | 0.990720469 | 1.00 |
| YBR287W | 2396.597397 | -0.00659826 | 0.992584657 | 1.00 |
| YCR066W | 426.4825775 | -0.012252839 | 0.991927143 | 1.00 |
| YDL159W | 513.5387781 | -0.000405841 | 0.991921875 | 1.00 |
| YDL201W | 813.4402377 | 0.035486886 | 0.99226678 | 1.00 |
| YDR311W | 727.3776002 | -0.005570606 | 0.99269984 | 1.00 |
| YEL043W | 2331.741844 | -0.006405577 | 0.992274244 | 1.00 |
| YGR145W | 1225.275886 | 0.084415513 | 0.992992 | 1.00 |
| YHR069C-A | 0.116121444 | -0.009045017 | 0.993015577 | 1.00 |
| YHR203C | 5825.930229 | 0.116474534 | 0.992717583 | 1.00 |
| YML001W | 1453.0376 | 0.014565566 | 0.992175874 | 1.00 |
| YML053C | 803.1994724 | -0.0037085 | 0.99140277 | 1.00 |
| YMR099C | 2686.030784 | 0.009592537 | 0.992861568 | 1.00 |
| YNL257C | 1662.623362 | -0.000515919 | 0.992079642 | 1.00 |
| YPL083C | 495.9825937 | 0.008455641 | 0.991525838 | 1.00 |
| YDR079W | 403.891688 | 0.01387618 | 0.993550163 | 1.00 |
| YKL173W | 1269.160496 | -0.003089726 | 0.993989738 | 1.00 |
| YML018C | 644.4081194 | 0.006876392 | 0.994069875 | 1.00 |
| YDR281C | 42.74964081 | -0.040070822 | 0.995732759 | 1.00 |
| YDR416W | 716.6573712 | -0.000210791 | 0.996858479 | 1.00 |
| YER007C-A | 951.757768 | 0.049596945 | 0.995663304 | 1.00 |
| YER167W | 1421.029082 | -0.007527025 | 0.996670848 | 1.00 |
| YGR109W-B | 857.2496031 | -0.122843115 | 0.994776683 | 1.00 |
| YHR168W | 379.0957302 | 0.001190769 | 0.996037726 | 1.00 |
| YHR175W | 775.2689431 | 0.02603923 | 0.995074668 | 1.00 |
| YJR157W | 1.945110995 | -0.271895462 | 0.996410036 | 1.00 |
| YKL073W | 1776.826393 | 0.017354963 | 0.995671271 | 1.00 |
| YLR165C | 405.6673301 | -0.020224382 | 0.99545731 | 1.00 |
| YMR087W | 259.2299237 | -0.010550959 | 0.995957432 | 1.00 |
| YNL146W | 85.60940575 | -0.013744091 | 0.995270075 | 1.00 |
| YNL322C | 3831.740608 | -0.001822781 | 0.996289643 | 1.00 |
| YOL015W | 325.5388955 | -0.029369081 | 0.996677141 | 1.00 |
| YOR117W | 3352.309698 | 0.00932497 | 0.99562635 | 1.00 |
| YPL058C | 2629.343095 | -0.017020534 | 0.99473117 | 1.00 |
| YPL139C | 637.4930321 | 0.001276859 | 0.996810515 | 1.00 |
| YBR106W | 4289.033597 | 0.11456641 | 0.998175171 | 1.00 |
| YCR018C-A | 8.72999001 | 0.028486323 | 0.998629279 | 1.00 |
| YHR064C | 6332.657586 | 0.058871967 | 0.999416314 | 1.00 |
| YHR165W-A | 6.420816 | -0.022151389 | 0.999622616 | 1.00 |
| YLL063C | 302.5298246 | -0.016142264 | 0.99856847 | 1.00 |
| YLR138W | 1645.971988 | 0.001309738 | 0.999115461 | 1.00 |
| YLR425W | 1120.132672 | -0.005909687 | 0.997613484 | 1.00 |

| | | | | |
|---|---|---|---|---|
| YMR060C | 493.2206546 | 0.005223068 | 0.999813711 | 1.00 |
| YMR259C | 1208.426766 | 0.001742436 | 0.999836511 | 1.00 |
| YNL166C | 1009.247872 | 0.022754088 | 0.99951489 | 1.00 |
| YNL320W | 323.6388355 | 0.015820795 | 0.99819747 | 1.00 |
| YNR051C | 2926.448335 | 0.042909625 | 0.998033837 | 1.00 |
| YOL140W | 502.3006349 | 0.004281221 | 0.998883751 | 1.00 |
| YOR207C | 1745.865963 | 0.015153287 | 0.998265531 | 1.00 |
| YOR329C | 1982.861977 | -0.006794479 | 0.998490496 | 1.00 |
| YPL009C | 4156.098331 | 0.021509471 | 0.999913771 | 1.00 |
| YPL126W | 2185.981195 | 0.04690219 | 0.999826829 | 1.00 |
| YPL205C | 28.51967725 | 0.001767934 | 0.997997248 | 1.00 |
| YAL062W | 2248.194645 | 1.57241574 | NA | NA |
| YDL223C | 1796.784019 | 0.923761952 | NA | NA |
| YER024W | 2782.885664 | 1.283866874 | NA | NA |
| YER065C | 3480.324226 | 0.801244933 | NA | NA |
| YGR067C | 601.4253985 | 0.772224307 | NA | NA |
| YGR236C | 237.8810767 | 1.099443492 | NA | NA |
| YIL057C | 383.2825996 | 1.910864868 | NA | NA |
| YJR095W | 1024.664601 | 1.497045046 | NA | NA |
| YKL065W-A | 60.35011298 | 0.962962127 | NA | NA |
| YKR097W | 1618.765281 | 1.522974944 | NA | NA |
| YLR174W | 1214.876474 | 1.597183536 | NA | NA |
| YLR377C | 564.6863452 | 1.046239335 | NA | NA |
| YML042W | 2686.768376 | 1.311794212 | NA | NA |
| YMR107W | 2022.649302 | 1.5107923 | NA | NA |
| YMR175W | 966.6429644 | 1.751311277 | NA | NA |
| YMR175W-A | 25.9663659 | 1.596706792 | NA | NA |
| YNL036W | 2481.742596 | 0.708075624 | NA | NA |
| YNL117W | 3801.736785 | 1.41311495 | NA | NA |
| YNR002C | 907.6027094 | 1.321863331 | NA | NA |
| YNR071C | 317.5831201 | 1.347593887 | NA | NA |
| YOR100C | 795.4961629 | 2.243177687 | NA | NA |
| YPL223C | 915.9171589 | 1.454483093 | NA | NA |
