## Supplementary material for "A stress response that allows highly mutated eukaryotic cells to survive and proliferate": Table S7

| Gene | baseMean | log2FoldChange | pvalue | padj | qvalue |
|---|---|---|---|---|---|
| YLL026W | 27987.51895 | 1.926467331 | 2.71E-12 | 0.00 | 0.00 |
| YAL005C | 15020.34614 | 1.566275424 | 1.24E-11 | 0.00 | 0.00 |
| YPL177C | 1454.555337 | -1.063555635 | 2.58E-09 | 0.00 | 0.00 |
| YML034W | 2522.763719 | 0.949805977 | 3.77E-09 | 0.00 | 0.00 |
| YDR258C | 9048.121163 | 1.893738959 | 2.70E-08 | 0.00 | 0.00 |
| YNL199C | 1154.934737 | -0.49608732 | 4.28E-08 | 0.00 | 0.00 |
| YBL042C | 1258.710363 | -1.598651303 | 5.54E-08 | 0.00 | 0.00 |
| YEL047C | 2163.496037 | 1.122754286 | 2.94E-07 | 0.00 | 0.00 |
| YFL062W | 38.28792094 | 2.512412299 | 3.43E-07 | 0.00 | 0.00 |
| YGL183C | 72.4162606 | 2.271368683 | 3.12E-07 | 0.00 | 0.00 |
| YNL172W | 1164.569131 | 0.884266383 | 4.37E-07 | 0.00 | 0.00 |
| YJL110C | 684.2542418 | -0.966201645 | 5.47E-07 | 0.00 | 0.00 |
| YER048W-A | 659.0895735 | -0.511147409 | 8.53E-07 | 0.00 | 0.00 |
| YNL242W | 1398.740797 | 1.020750236 | 8.21E-07 | 0.00 | 0.00 |
| YFL033C | 2297.741868 | 0.830548887 | 1.77E-06 | 0.00 | 0.00 |
| YLL058W | 677.8123693 | 0.971955065 | 2.89E-06 | 0.00 | 0.00 |
| YCL074W | 140.2978941 | 2.415585924 | 5.72E-06 | 0.00 | 0.00 |
| YKL111C | 22.34564795 | -1.240245135 | 6.60E-06 | 0.00 | 0.00 |
| YNR057C | 196.7239761 | 1.185523487 | 6.76E-06 | 0.00 | 0.00 |
| YBL026W | 682.7010545 | -0.618814072 | 9.37E-06 | 0.00 | 0.00 |
| YDL179W | 455.9668158 | -0.753752669 | 9.34E-06 | 0.00 | 0.00 |
| YBR012W-B | 321.6022163 | 1.955469364 | 1.25E-05 | 0.00 | 0.00 |
| YBR108W | 1821.037124 | 0.78993117 | 1.22E-05 | 0.00 | 0.00 |
| YNL034W | 79.93693595 | 1.758013352 | 1.36E-05 | 0.00 | 0.00 |
| YNR065C | 953.8955562 | 0.990893445 | 1.38E-05 | 0.00 | 0.00 |
| YPL111W | 2984.765722 | 0.993539409 | 1.34E-05 | 0.00 | 0.00 |
| YGR035C | 329.4527926 | -1.675271401 | 1.66E-05 | 0.00 | 0.00 |
| YNL152W | 687.0558678 | -0.544697147 | 1.90E-05 | 0.00 | 0.00 |
| YPL178W | 1119.186631 | -0.541225745 | 2.03E-05 | 0.00 | 0.00 |
| YPL249C | 1511.229449 | 0.615281305 | 2.02E-05 | 0.00 | 0.00 |
| YPL240C | 19482.15572 | 1.356152923 | 2.17E-05 | 0.00 | 0.00 |
| YHL037C | 20.29589679 | 2.010766991 | 2.44E-05 | 0.00 | 0.00 |
| YJL102W | 772.7229577 | 1.047163435 | 2.46E-05 | 0.00 | 0.00 |
| YCL073C | 26.5180597 | 1.942151322 | 2.76E-05 | 0.01 | 0.00 |
| YGR108W | 622.4932134 | -1.110638612 | 2.85E-05 | 0.01 | 0.00 |
| YHL022C | 136.8231823 | 1.659949467 | 2.82E-05 | 0.01 | 0.00 |
| YKL058W | 636.5138347 | -0.728418794 | 3.21E-05 | 0.01 | 0.00 |
| YBR218C | 7147.332039 | 0.904751071 | 3.55E-05 | 0.01 | 0.00 |
| YCR021C | 10522.50846 | 1.962492656 | 3.54E-05 | 0.01 | 0.00 |
| YKL174C | 903.8221117 | 0.483820209 | 3.49E-05 | 0.01 | 0.00 |

| | | | | | |
|---|---|---|---|---|---|
| YHR164C | 1299.336952 | 0.690734256 | 3.73E-05 | 0.01 | 0.00 |
| YOR192C | 247.398385 | 1.488306573 | 3.88E-05 | 0.01 | 0.00 |
| YHR216W | 3639.691774 | 1.870591776 | 3.99E-05 | 0.01 | 0.00 |
| YNL012W | 592.4635068 | 1.147280683 | 4.59E-05 | 0.01 | 0.00 |
| YPR016C | 1938.157861 | -0.864945951 | 4.60E-05 | 0.01 | 0.00 |
| YGL184C | 275.5662791 | 1.372117739 | 5.05E-05 | 0.01 | 0.00 |
| YHL036W | 545.2189465 | 1.411425327 | 4.95E-05 | 0.01 | 0.00 |
| YMR180C | 510.7680392 | -0.826438766 | 5.09E-05 | 0.01 | 0.00 |
| YDL030W | 676.7024249 | 0.633109341 | 5.21E-05 | 0.01 | 0.00 |
| YCL030C | 8176.65539 | 1.167775706 | 6.18E-05 | 0.01 | 0.01 |
| YPR007C | 168.1873056 | 1.623162757 | 6.58E-05 | 0.01 | 0.01 |
| YBR083W | 2339.410841 | -1.28133125 | 7.22E-05 | 0.01 | 0.01 |
| YBR258C | 250.7365151 | -0.628195969 | 7.42E-05 | 0.01 | 0.01 |
| YIL067C | 437.2526192 | 0.58043815 | 6.95E-05 | 0.01 | 0.01 |
| YKL050C | 526.5764301 | 1.171414525 | 7.07E-05 | 0.01 | 0.01 |
| YNL188W | 592.8677499 | -1.070207151 | 7.41E-05 | 0.01 | 0.01 |
| YOL165C | 22.45477283 | 2.170319776 | 7.02E-05 | 0.01 | 0.01 |
| YBR045C | 255.946774 | 1.047423255 | 8.05E-05 | 0.01 | 0.01 |
| YBR158W | 2964.217702 | -1.115165289 | 8.19E-05 | 0.01 | 0.01 |
| YDL227C | 308.6751335 | -1.880090479 | 8.12E-05 | 0.01 | 0.01 |
| YDR402C | 76.67558535 | 1.60005062 | 8.09E-05 | 0.01 | 0.01 |
| YER044C-A | 102.3682402 | 1.408392241 | 8.32E-05 | 0.01 | 0.01 |
| YGL093W | 1066.84026 | 0.465328124 | 8.37E-05 | 0.01 | 0.01 |
| YPL095C | 828.2325169 | -0.990307597 | 7.80E-05 | 0.01 | 0.01 |
| YAR075W | 64.72959559 | 1.69268317 | 8.86E-05 | 0.01 | 0.01 |
| YOR114W | 311.2740858 | 1.134268415 | 9.17E-05 | 0.01 | 0.01 |
| YIL072W | 537.0665215 | 1.139010861 | 9.54E-05 | 0.01 | 0.01 |
| YPL110C | 931.3902179 | 0.807711039 | 0.000100746 | 0.01 | 0.01 |
| YBR279W | 1844.253736 | -0.581699968 | 0.000104325 | 0.01 | 0.01 |
| YHR029C | 965.1862947 | 1.166948935 | 0.000107155 | 0.01 | 0.01 |
| YPL150W | 1067.29309 | 0.771843944 | 0.00011349 | 0.01 | 0.01 |
| YMR236W | 1041.170124 | -0.487984426 | 0.000116315 | 0.01 | 0.01 |
| YHR182C-A | 130.5541577 | 1.887245541 | 0.000119218 | 0.01 | 0.01 |
| YMR207C | 1385.212155 | 0.737692428 | 0.000119869 | 0.01 | 0.01 |
| YHR122W | 579.3180386 | 0.900763354 | 0.000123234 | 0.01 | 0.01 |
| YER069W | 925.6063303 | 1.142825607 | 0.000129354 | 0.01 | 0.01 |
| YDR365C | 4849.022737 | 0.884728155 | 0.000133284 | 0.01 | 0.01 |
| YOL102C | 532.3526812 | -0.564313479 | 0.000135091 | 0.01 | 0.01 |
| YDL114W | 43.38451628 | 2.01520932 | 0.000142543 | 0.01 | 0.01 |
| YER095W | 1267.382435 | 0.778204274 | 0.000143827 | 0.01 | 0.01 |
| YNL171C | 59.8874962 | 1.02554676 | 0.000139957 | 0.01 | 0.01 |
| YPL038W | 299.6200602 | -1.004870799 | 0.000141468 | 0.01 | 0.01 |
| YDR396W | 125.4408117 | -0.916111229 | 0.000146193 | 0.01 | 0.01 |

| | | | | | |
|---|---|---|---|---|---|
| YHL023C | 1253.971871 | 0.54487747 | 0.000151354 | 0.01 | 0.01 |
| YOR338W | 133.0538926 | 1.041650735 | 0.000150636 | 0.01 | 0.01 |
| YJR126C | 1827.592541 | 0.494857351 | 0.000158077 | 0.01 | 0.01 |
| YBR016W | 1677.943304 | -0.615358982 | 0.000167521 | 0.01 | 0.01 |
| YDR171W | 12610.16854 | 1.272732257 | 0.000168672 | 0.01 | 0.01 |
| YHR104W | 3547.736943 | 0.837515651 | 0.000165834 | 0.01 | 0.01 |
| YCL026C-B | 269.8485152 | 1.226750675 | 0.000171729 | 0.01 | 0.01 |
| YBR047W | 453.8190728 | 1.299220408 | 0.000178081 | 0.01 | 0.01 |
| YGR063C | 116.1850697 | -0.609597959 | 0.000183845 | 0.01 | 0.01 |
| YGR139W | 13.99976418 | 1.609239218 | 0.000180149 | 0.01 | 0.01 |
| YLR216C | 5399.303942 | 0.884235123 | 0.000182493 | 0.01 | 0.01 |
| YER042W | 653.285425 | 1.204408679 | 0.000188111 | 0.01 | 0.01 |
| YGL140C | 1656.222341 | 0.481135049 | 0.000186298 | 0.01 | 0.01 |
| YGR142W | 4162.125638 | 1.303703681 | 0.000193071 | 0.01 | 0.01 |
| YLR073C | 322.9361924 | -1.226582172 | 0.000195019 | 0.01 | 0.01 |
| YOR214C | 33.33902204 | 1.968442104 | 0.000192904 | 0.01 | 0.01 |
| YDR498C | 1014.743074 | -0.368431076 | 0.000198996 | 0.01 | 0.01 |
| YBR148W | 178.9686946 | 1.203215706 | 0.000203189 | 0.01 | 0.01 |
| YOR120W | 2918.156498 | 1.527879608 | 0.000203285 | 0.01 | 0.01 |
| YFR057W | 37.29733796 | 1.410527057 | 0.00023175 | 0.01 | 0.01 |
| YGR113W | 643.4472036 | -0.40272998 | 0.000229718 | 0.01 | 0.01 |
| YGL262W | 77.76796003 | 2.056493203 | 0.000250805 | 0.02 | 0.01 |
| YNL274C | 1846.518968 | 1.187179141 | 0.000254463 | 0.02 | 0.01 |
| YOR121C | 64.33732395 | 1.753697422 | 0.000252351 | 0.02 | 0.01 |
| YPL251W | 76.6529722 | -0.988076336 | 0.000250392 | 0.02 | 0.01 |
| YNR073C | 195.2436335 | 1.911718264 | 0.000258897 | 0.02 | 0.01 |
| YIL024C | 346.2449471 | 1.214705777 | 0.000261544 | 0.02 | 0.01 |
| YBR265W | 1130.650771 | -0.628382846 | 0.000282344 | 0.02 | 0.01 |
| YCR081W | 992.4265821 | 0.57983867 | 0.000282421 | 0.02 | 0.01 |
| YEL030W | 524.4916329 | 0.838901972 | 0.000276753 | 0.02 | 0.01 |
| YER070W | 2200.790509 | -1.286857566 | 0.000283332 | 0.02 | 0.01 |
| YGL081W | 335.8828397 | 1.577332071 | 0.000277454 | 0.02 | 0.01 |
| YJL124C | 1576.189687 | -0.624231485 | 0.000285547 | 0.02 | 0.01 |
| YMR325W | 4.52283349 | 2.30623515 | 0.000272915 | 0.02 | 0.01 |
| YDR075W | 328.511722 | -0.944890221 | 0.000291191 | 0.02 | 0.01 |
| YLL015W | 2174.31995 | 0.760570939 | 0.00028928 | 0.02 | 0.01 |
| YNL270C | 240.3573834 | 1.570013486 | 0.000297672 | 0.02 | 0.01 |
| YJL157C | 1263.741636 | -1.501406056 | 0.000301784 | 0.02 | 0.01 |
| YMR272C | 4343.78123 | -0.73360743 | 0.000309899 | 0.02 | 0.01 |
| YKL129C | 2233.401318 | 0.6979955 | 0.000320221 | 0.02 | 0.01 |
| YAR047C | 21.66468268 | 1.566039236 | 0.000323224 | 0.02 | 0.01 |
| YBR259W | 488.2154125 | -0.475504406 | 0.000331467 | 0.02 | 0.01 |
| YDL077C | 1087.970908 | 0.433702067 | 0.000338128 | 0.02 | 0.01 |

| | | | | | |
|---|---|---|---|---|---|
| YNR069C | 436.7652166 | 1.185927423 | 0.000344458 | 0.02 | 0.01 |
| YAR073W | 171.304609 | 1.765947565 | 0.00035016 | 0.02 | 0.01 |
| YHR112C | 1466.272889 | 0.869043936 | 0.000349005 | 0.02 | 0.01 |
| YFR025C | 947.4315897 | 0.475872854 | 0.00035646 | 0.02 | 0.01 |
| YER081W | 9451.152045 | 1.750334991 | 0.000365377 | 0.02 | 0.01 |
| YAL063C | 272.6184868 | 1.093892776 | 0.000380948 | 0.02 | 0.01 |
| YGL028C | 1677.067671 | -0.93130041 | 0.000377909 | 0.02 | 0.01 |
| YHL028W | 1137.369035 | -1.182654303 | 0.000378748 | 0.02 | 0.01 |
| YHL030W | 2779.309717 | 0.928632736 | 0.000378235 | 0.02 | 0.01 |
| YIL020C-A | 191.6015626 | -0.766740038 | 0.000390274 | 0.02 | 0.01 |
| YJR046W | 1243.286799 | 0.576124979 | 0.000404248 | 0.02 | 0.01 |
| YOL058W | 2645.481694 | 1.489910787 | 0.000431041 | 0.02 | 0.01 |
| YLR273C | 457.5126435 | 0.739370781 | 0.000438799 | 0.02 | 0.01 |
| YNR068C | 378.9501574 | 1.240507723 | 0.000438301 | 0.02 | 0.01 |
| YBR222C | 4562.01386 | 1.029836985 | 0.000451032 | 0.02 | 0.01 |
| YDL171C | 4109.615185 | 0.907716209 | 0.000467462 | 0.02 | 0.01 |
| YDR174W | 3004.282379 | -0.593575729 | 0.000464344 | 0.02 | 0.01 |
| YLR187W | 889.7815093 | 0.549336723 | 0.000461524 | 0.02 | 0.01 |
| YJR154W | 284.604362 | 1.267671022 | 0.000473015 | 0.02 | 0.01 |
| YNR058W | 282.8560691 | 0.916908769 | 0.000476571 | 0.02 | 0.01 |
| YDL224C | 1328.137391 | -0.683768895 | 0.000485911 | 0.02 | 0.01 |
| YFL053W | 516.855343 | 1.733910601 | 0.000492154 | 0.02 | 0.01 |
| YBL033C | 1274.437806 | 0.546552573 | 0.000499948 | 0.02 | 0.01 |
| YOR106W | 543.2030625 | -0.457034123 | 0.000503651 | 0.02 | 0.01 |
| YBR233W-A | 253.0169158 | -0.92125422 | 0.000514053 | 0.02 | 0.01 |
| snR58 | 5.413360595 | -1.512237722 | 0.000515024 | 0.02 | 0.01 |
| YDR158W | 6048.209084 | 0.931172398 | 0.000522663 | 0.02 | 0.01 |
| YDR389W | 894.8507584 | -0.671369958 | 0.000532045 | 0.02 | 0.01 |
| YLR033W | 1049.258607 | -0.502983519 | 0.000554565 | 0.02 | 0.02 |
| YML100W | 14497.66976 | 1.285555951 | 0.00055621 | 0.02 | 0.02 |
| YHR159W | 783.3955676 | 0.814001662 | 0.000600281 | 0.02 | 0.02 |
| YHR202W | 670.4759746 | 0.85000175 | 0.000598496 | 0.02 | 0.02 |
| YIL153W | 970.2297401 | 0.335825992 | 0.000608609 | 0.02 | 0.02 |
| YDL043C | 369.3400281 | -0.606607079 | 0.000629299 | 0.03 | 0.02 |
| YCR061W | 1066.525845 | 0.93824286 | 0.000636068 | 0.03 | 0.02 |
| YBR255W | 917.9047892 | 0.587713311 | 0.000668614 | 0.03 | 0.02 |
| YOL131W | 75.70252965 | 1.520806966 | 0.000669019 | 0.03 | 0.02 |
| YJR106W | 683.5325213 | 0.686892399 | 0.000676866 | 0.03 | 0.02 |
| YDR331W | 571.3069512 | -0.582825165 | 0.00068537 | 0.03 | 0.02 |
| YGL107C | 935.2339762 | -0.294731488 | 0.000732824 | 0.03 | 0.02 |
| YER145C | 1496.957868 | -0.948665942 | 0.000747928 | 0.03 | 0.02 |
| YPL121C | 25.34339254 | 1.519584707 | 0.000747955 | 0.03 | 0.02 |
| YBL065W | 30.86397396 | 1.488882798 | 0.000762619 | 0.03 | 0.02 |

| | | | | | |
|---|---|---|---|---|---|
| YLR214W | 1682.159024 | -1.037072608 | 0.000763244 | 0.03 | 0.02 |
| YNR066C | 265.0332103 | 0.93693748 | 0.000778549 | 0.03 | 0.02 |
| YJL071W | 590.1206135 | 0.672041372 | 0.000813254 | 0.03 | 0.02 |
| YCR073C | 984.1918819 | 0.489852746 | 0.000870722 | 0.03 | 0.02 |
| YOR028C | 706.5725488 | -1.565868783 | 0.000870001 | 0.03 | 0.02 |
| YAR031W | 151.5423779 | 0.941568666 | 0.000886463 | 0.03 | 0.02 |
| YHL021C | 7761.479525 | 1.375465164 | 0.000884843 | 0.03 | 0.02 |
| YHR079C-A | 27.69347479 | 1.657650364 | 0.000902068 | 0.03 | 0.02 |
| YHR182W | 713.7981983 | 0.782061428 | 0.000901068 | 0.03 | 0.02 |
| YPL254W | 446.984746 | -0.622100709 | 0.000896903 | 0.03 | 0.02 |
| YDR225W | 3245.605018 | -1.117553204 | 0.000915895 | 0.03 | 0.02 |
| YLR087C | 2042.517235 | 0.683899877 | 0.000914122 | 0.03 | 0.02 |
| YMR015C | 1137.039851 | -0.909680705 | 0.000935184 | 0.03 | 0.02 |
| YKL097C | 10.27058523 | 1.763684078 | 0.000941377 | 0.03 | 0.02 |
| YBR297W | 465.5986861 | 1.148368326 | 0.000969063 | 0.03 | 0.02 |
| YER093C | 1449.072568 | 0.48709003 | 0.000963202 | 0.03 | 0.02 |
| YGR086C | 9907.562538 | -0.560571101 | 0.000972222 | 0.03 | 0.02 |
| YMR104C | 1639.396102 | 1.236168582 | 0.000964504 | 0.03 | 0.02 |
| YPL018W | 471.7341201 | 0.983265413 | 0.000962176 | 0.03 | 0.02 |
| YBR157C | 589.2630079 | -1.515964892 | 0.000980367 | 0.03 | 0.02 |
| YLR263W | 707.0718116 | 1.334451178 | 0.00098745 | 0.03 | 0.02 |
| YGR017W | 1237.157084 | -0.493017366 | 0.001003827 | 0.03 | 0.02 |
| YCL075W | 37.39387639 | 1.964487979 | 0.001024398 | 0.03 | 0.02 |
| YJL159W | 37900.6137 | 0.922570336 | 0.001024542 | 0.03 | 0.02 |
| YPR111W | 665.5579675 | 0.808443434 | 0.00103663 | 0.03 | 0.02 |
| YJL156W-A | 83.97878785 | -0.733266097 | 0.00105429 | 0.04 | 0.02 |
| YPL049C | 2306.704815 | -0.586069418 | 0.001080825 | 0.04 | 0.02 |
| YMR108W | 5770.966467 | 0.765925014 | 0.001089348 | 0.04 | 0.02 |
| YHR106W | 1096.925363 | 0.895384097 | 0.001128699 | 0.04 | 0.02 |
| YOR292C | 645.9873627 | 0.861107372 | 0.001159748 | 0.04 | 0.03 |
| YLR416C | 28.8196825 | 1.477295668 | 0.001175616 | 0.04 | 0.03 |
| YPR117W | 2597.357181 | 0.532521688 | 0.001171983 | 0.04 | 0.03 |
| YDR224C | 8028.945616 | -0.873191809 | 0.001186559 | 0.04 | 0.03 |
| YDR488C | 747.4804028 | 0.82518073 | 0.001207425 | 0.04 | 0.03 |
| YGL180W | 843.8043135 | 0.761006334 | 0.001204821 | 0.04 | 0.03 |
| YMR101C | 112.86694 | 1.496466052 | 0.001200788 | 0.04 | 0.03 |
| YNL326C | 366.0651518 | -0.395286964 | 0.001195229 | 0.04 | 0.03 |
| YER029C | 871.4448752 | -0.464130385 | 0.001277245 | 0.04 | 0.03 |
| YMR103C | 375.8522499 | 1.181205832 | 0.001285742 | 0.04 | 0.03 |
| YER091C | 3130.444513 | 0.844810495 | 0.001324214 | 0.04 | 0.03 |
| YJR079W | 11.62148612 | 1.671552467 | 0.00131275 | 0.04 | 0.03 |
| YOR375C | 17984.54886 | 1.480545056 | 0.001321219 | 0.04 | 0.03 |
| YDL092W | 1065.77792 | -0.63072958 | 0.001345534 | 0.04 | 0.03 |

| | | | | | |
|---|---|---|---|---|---|
| YMR317W | 752.1354442 | 0.76416813 | 0.001335648 | 0.04 | 0.03 |
| YPR120C | 354.4464034 | -0.700017867 | 0.001346013 | 0.04 | 0.03 |
| YDR421W | 1031.149921 | 0.612415002 | 0.00136594 | 0.04 | 0.03 |
| YNR052C | 1154.692673 | -0.472213304 | 0.001366957 | 0.04 | 0.03 |
| YGL049C | 2260.025541 | 0.377092127 | 0.00137606 | 0.04 | 0.03 |
| YPL193W | 620.6543173 | -0.511188418 | 0.00138946 | 0.04 | 0.03 |
| YDL093W | 807.8112275 | 0.592721064 | 0.001429038 | 0.04 | 0.03 |
| YJL004C | 587.5114449 | -0.526427708 | 0.001449598 | 0.04 | 0.03 |
| YLR259C | 10469.29817 | 0.617664682 | 0.001447407 | 0.04 | 0.03 |
| YNR053C | 1005.221474 | -1.084697598 | 0.001473018 | 0.04 | 0.03 |
| YJL153C | 833.1459195 | 1.31517713 | 0.001495695 | 0.04 | 0.03 |
| YDL248W | 224.0356228 | 1.096046155 | 0.001509019 | 0.04 | 0.03 |
| YGL047W | 625.4320388 | 0.540979432 | 0.001520334 | 0.04 | 0.03 |
| YBR284W | 530.3934343 | 1.076443174 | 0.001532828 | 0.04 | 0.03 |
| YBR250W | 80.81966717 | 1.283912462 | 0.001556011 | 0.04 | 0.03 |
| YIL009C-A | 425.8589138 | -0.696899467 | 0.001563325 | 0.04 | 0.03 |
| YJL089W | 344.4116839 | 1.505682514 | 0.001557816 | 0.04 | 0.03 |
| YPR138C | 1408.826707 | 0.90719911 | 0.001565592 | 0.04 | 0.03 |
| YLR415C | 24.41748512 | 1.294463061 | 0.001578012 | 0.04 | 0.03 |
| YKR023W | 810.8865639 | 0.532401054 | 0.001590295 | 0.04 | 0.03 |
| YBR061C | 630.0720407 | -0.649046658 | 0.001631517 | 0.05 | 0.03 |
| YEL034C-A | 2748.462234 | -0.942695996 | 0.001625849 | 0.05 | 0.03 |
| YBR244W | 574.3609399 | -1.25016605 | 0.001640789 | 0.05 | 0.03 |
| YNL040W | 832.5546055 | 0.83017837 | 0.001651381 | 0.05 | 0.03 |
| YDR209C | 344.9577194 | -0.895398444 | 0.00167645 | 0.05 | 0.03 |
| YOR142W | 3533.236265 | 0.721521339 | 0.001694042 | 0.05 | 0.03 |
| YPL021W | 14.06563719 | 1.83592335 | 0.001684322 | 0.05 | 0.03 |
| YPR160W | 12134.47846 | 1.464517303 | 0.001696808 | 0.05 | 0.03 |
| YFL027C | 673.7008422 | -0.925442472 | 0.001727483 | 0.05 | 0.03 |
| YER185W | 79.91884135 | 1.019496375 | 0.001758211 | 0.05 | 0.03 |
| YCL057W | 3573.183746 | 0.586553143 | 0.001776897 | 0.05 | 0.03 |
| YBR114W | 883.2430403 | 0.650107237 | 0.001845032 | 0.05 | 0.03 |
| YBR130C | 1160.408881 | -0.628835289 | 0.001827814 | 0.05 | 0.03 |
| YGR078C | 839.5108181 | -0.784019374 | 0.001856125 | 0.05 | 0.03 |
| YKL213C | 1964.500982 | 0.425662421 | 0.001842679 | 0.05 | 0.03 |
| YLR133W | 1071.862832 | 0.530259934 | 0.001812616 | 0.05 | 0.03 |
| YLR326W | 402.567239 | 0.770169615 | 0.001797603 | 0.05 | 0.03 |
| YMR055C | 275.6053073 | -0.63342452 | 0.001818207 | 0.05 | 0.03 |
| YMR075W | 761.6920474 | 0.484657384 | 0.001848325 | 0.05 | 0.03 |
| YMR321C | 24.07796373 | 1.411278403 | 0.001849488 | 0.05 | 0.03 |
| YOL019W | 928.0908204 | -0.733342762 | 0.001799993 | 0.05 | 0.03 |
| YBR006W | 1760.927773 | 1.001900781 | 0.001891548 | 0.05 | 0.03 |
| YMR048W | 313.0835867 | -0.337755485 | 0.001904289 | 0.05 | 0.03 |

| | | | | | |
|---|---|---|---|---|---|
| YDL240W | 734.1749038 | -0.889920799 | 0.001943153 | 0.05 | 0.03 |
| YDR218C | 78.04916244 | 1.755792508 | 0.001939656 | 0.05 | 0.03 |
| YDR152W | 1737.468978 | -0.602936307 | 0.001958316 | 0.05 | 0.03 |
| YOR367W | 630.0179766 | -0.622342168 | 0.001957822 | 0.05 | 0.03 |
| YBR169C | 4813.961053 | 1.246827447 | 0.001990776 | 0.05 | 0.03 |
| YMR306W | 518.3037802 | 0.785152253 | 0.001994157 | 0.05 | 0.03 |
| YFL063W | 4.881871603 | 2.005638026 | 0.002013153 | 0.05 | 0.03 |
| YOR153W | 11636.99259 | -1.279897751 | 0.002012482 | 0.05 | 0.03 |
| YGR258C | 1398.526849 | 0.600670779 | 0.002052246 | 0.05 | 0.03 |
| YOR027W | 11708.63489 | 0.819091002 | 0.00205894 | 0.05 | 0.03 |
| YPL185W | 39.28495564 | 1.086720292 | 0.002063974 | 0.05 | 0.03 |
| YMR096W | 4030.130556 | 1.474082662 | 0.0021272 | 0.05 | 0.03 |
| YNL054W-B | 451.4424074 | 1.3091148 | 0.002135475 | 0.05 | 0.03 |
| YDL139C | 356.8731646 | 0.518163382 | 0.002162795 | 0.05 | 0.03 |
| YBR085C-A | 1789.206511 | 1.016423766 | 0.002204229 | 0.05 | 0.03 |
| YLR035C-A | 40.57446476 | 1.300159758 | 0.002205639 | 0.05 | 0.03 |
| YPL027W | 50.11860961 | 1.32363893 | 0.00220869 | 0.05 | 0.03 |
| YDR059C | 828.9203049 | 0.749212437 | 0.002226377 | 0.05 | 0.04 |
| YKL209C | 3091.85579 | -1.351854211 | 0.002244201 | 0.05 | 0.04 |
| YKL031W | 35.11063328 | 1.321810866 | 0.002284281 | 0.05 | 0.04 |
| YDR386W | 496.5773156 | 0.663030185 | 0.002343225 | 0.06 | 0.04 |
| YGL059W | 1140.532396 | 0.79671357 | 0.002354647 | 0.06 | 0.04 |
| YCL069W | 36.22888945 | 1.054766466 | 0.002397108 | 0.06 | 0.04 |
| YBR054W | 4277.398072 | 1.41257051 | 0.002462718 | 0.06 | 0.04 |
| YCL009C | 7647.273312 | 0.714695128 | 0.002446854 | 0.06 | 0.04 |
| YGL229C | 708.5937653 | 0.594761211 | 0.002461632 | 0.06 | 0.04 |
| YJR060W | 664.6110196 | -0.432647971 | 0.002422822 | 0.06 | 0.04 |
| YOR005C | 988.1912317 | 0.711762116 | 0.002452485 | 0.06 | 0.04 |
| YPL129W | 1819.666295 | -0.552610468 | 0.002451442 | 0.06 | 0.04 |
| YJR098C | 316.068717 | 0.629974584 | 0.002488916 | 0.06 | 0.04 |
| YKR103W | 526.4204375 | 1.073718534 | 0.002496839 | 0.06 | 0.04 |
| YLR452C | 1814.469836 | -1.079637213 | 0.00250293 | 0.06 | 0.04 |
| YDR055W | 3654.499745 | 0.697976619 | 0.002548065 | 0.06 | 0.04 |
| YHR171W | 640.7534835 | 0.792509401 | 0.002550818 | 0.06 | 0.04 |
| YJL051W | 1072.541155 | 0.411375751 | 0.002578938 | 0.06 | 0.04 |
| YJR109C | 3577.314205 | 1.371822069 | 0.002588526 | 0.06 | 0.04 |
| YGR188C | 503.2494794 | 0.692035725 | 0.002608194 | 0.06 | 0.04 |
| YLL046C | 28.77526537 | 1.469754649 | 0.002611261 | 0.06 | 0.04 |
| YGR257C | 921.5740821 | -0.593199008 | 0.002668151 | 0.06 | 0.04 |
| YMR053C | 712.958308 | 0.772800118 | 0.002683622 | 0.06 | 0.04 |
| YKL010C | 2290.07323 | 0.380063493 | 0.002693389 | 0.06 | 0.04 |
| LSR1 | 186.1998782 | 1.184397587 | 0.002703892 | 0.06 | 0.04 |
| YDR294C | 2749.492975 | 0.449171665 | 0.002735376 | 0.06 | 0.04 |

| | | | | | |
|---|---|---|---|---|---|
| YFR020W | 385.8178556 | 0.565625245 | 0.0027484 | 0.06 | 0.04 |
| YDR490C | 1094.352116 | 0.604565723 | 0.002761631 | 0.06 | 0.04 |
| YFL015C | 19.00869044 | 1.112201119 | 0.002771791 | 0.06 | 0.04 |
| YPR064W | 39.20156943 | 1.09064699 | 0.002806948 | 0.06 | 0.04 |
| YOL119C | 1253.305663 | 1.025343954 | 0.002921762 | 0.06 | 0.04 |
| YNL103W-A | 104.3621644 | 1.262599745 | 0.002940008 | 0.06 | 0.04 |
| YAR020C | 43.79066228 | 1.115248125 | 0.002966002 | 0.06 | 0.04 |
| YBR154C | 1262.041062 | -0.777598501 | 0.002986187 | 0.06 | 0.04 |
| YFR028C | 1133.072845 | -0.484215025 | 0.002989764 | 0.06 | 0.04 |
| YIR030C | 90.29914152 | 1.406192072 | 0.002971608 | 0.06 | 0.04 |
| YML089C | 19.45677984 | 1.365911028 | 0.003034649 | 0.06 | 0.04 |
| YDL190C | 2381.415921 | 0.430107037 | 0.003052181 | 0.06 | 0.04 |
| YML105C | 1352.843987 | -0.511873542 | 0.003063264 | 0.06 | 0.04 |
| YOR315W | 1158.774212 | -1.163502795 | 0.003081932 | 0.06 | 0.04 |
| YBR046C | 1727.421703 | 0.916696441 | 0.003129718 | 0.06 | 0.04 |
| YCR048W | 995.8304266 | 0.561670797 | 0.003132308 | 0.06 | 0.04 |
| YGL198W | 494.0719347 | 0.648834339 | 0.003133104 | 0.06 | 0.04 |
| YMR161W | 724.2845416 | -0.404821458 | 0.003122673 | 0.06 | 0.04 |
| RNA170 | 10.42132142 | 1.232861067 | 0.003186078 | 0.07 | 0.04 |
| YOR359W | 584.0345426 | -0.896952152 | 0.003181309 | 0.07 | 0.04 |
| YKR054C | 2620.574744 | 0.556961433 | 0.003210833 | 0.07 | 0.04 |
| YCR065W | 756.8632472 | -0.851466932 | 0.003242778 | 0.07 | 0.04 |
| YOL060C | 2188.941543 | 0.630536374 | 0.003240943 | 0.07 | 0.04 |
| YLL062C | 209.8685733 | 0.845556993 | 0.003291141 | 0.07 | 0.04 |
| YLR362W | 1083.342817 | 0.816790976 | 0.003300906 | 0.07 | 0.04 |
| YLR268W | 967.406712 | -0.504882502 | 0.003343787 | 0.07 | 0.04 |
| YOR107W | 180.9819764 | -0.655335852 | 0.003343102 | 0.07 | 0.04 |
| YPL264C | 197.1729292 | 0.62113573 | 0.003325941 | 0.07 | 0.04 |
| YBR264C | 359.5103772 | -0.46303943 | 0.003357987 | 0.07 | 0.04 |
| YGL124C | 833.4516769 | 0.515390552 | 0.003383619 | 0.07 | 0.04 |
| HRA1 | 34.22771229 | 0.907938814 | 0.003415154 | 0.07 | 0.04 |
| YOL161C | 3.517453047 | 1.470578383 | 0.003426432 | 0.07 | 0.04 |
| YDR135C | 3272.34562 | 0.453455369 | 0.003475902 | 0.07 | 0.04 |
| YKR104W | 184.9350151 | 0.980841605 | 0.003472603 | 0.07 | 0.04 |
| YER067C-A | 258.7943359 | 1.177536245 | 0.003512303 | 0.07 | 0.05 |
| YGR007W | 1236.749371 | -0.460834578 | 0.00352149 | 0.07 | 0.05 |
| YGR120C | 376.5121986 | -0.259221173 | 0.003510769 | 0.07 | 0.05 |
| YGR250C | 3005.303955 | 0.733970858 | 0.003525989 | 0.07 | 0.05 |
| YKR070W | 801.1916976 | 0.318440444 | 0.0035421 | 0.07 | 0.05 |
| YMR306C-A | 3.882802162 | 1.56569658 | 0.003560779 | 0.07 | 0.05 |
| YGL141W | 1274.03239 | 0.453248469 | 0.003585038 | 0.07 | 0.05 |
| YCR044C | 1029.447304 | -0.508438034 | 0.003630163 | 0.07 | 0.05 |
| YGR252W | 827.8896718 | -0.385653653 | 0.003635518 | 0.07 | 0.05 |

| | | | | | |
|---|---|---|---|---|---|
| YDR018C | 446.8147304 | 1.103158361 | 0.003657489 | 0.07 | 0.05 |
| YGR221C | 838.7959631 | -0.638305469 | 0.00366502 | 0.07 | 0.05 |
| YDL194W | 1083.690954 | 0.891222293 | 0.003750507 | 0.07 | 0.05 |
| YKR005C | 237.9372028 | 0.823762509 | 0.003731362 | 0.07 | 0.05 |
| YLR461W | 7.039838185 | 1.639921424 | 0.00374777 | 0.07 | 0.05 |
| YNL318C | 22.2967731 | 1.777649236 | 0.003794543 | 0.07 | 0.05 |
| YNL206C | 942.6169972 | -0.441397183 | 0.003810104 | 0.07 | 0.05 |
| YKR098C | 1187.222635 | 0.831193852 | 0.00382332 | 0.07 | 0.05 |
| YPR185W | 1716.783017 | 0.51517146 | 0.003834759 | 0.07 | 0.05 |
| YML072C | 6041.463861 | 0.526024153 | 0.00384882 | 0.07 | 0.05 |
| YDL017W | 421.2237274 | 0.445313775 | 0.003875776 | 0.07 | 0.05 |
| YEL060C | 10805.49764 | 1.006134457 | 0.003946626 | 0.07 | 0.05 |
| YJR110W | 661.1509666 | 0.377987746 | 0.00394075 | 0.07 | 0.05 |
| YPL259C | 962.6624104 | -0.409551359 | 0.003976355 | 0.07 | 0.05 |
| YPR121W | 333.11981 | -0.763072463 | 0.003969131 | 0.07 | 0.05 |
| YOL024W | 52.91060507 | 1.669705596 | 0.004007138 | 0.07 | 0.05 |
| YNL125C | 2090.525445 | 0.723445583 | 0.004051248 | 0.07 | 0.05 |
| YLR018C | 838.7588705 | -0.486393189 | 0.004103107 | 0.07 | 0.05 |
| YJR036C | 438.6482321 | 0.837927454 | 0.004115201 | 0.07 | 0.05 |
| YGL209W | 1219.680335 | -0.860009199 | 0.004205634 | 0.08 | 0.05 |
| YDL016C | 38.72923011 | -0.595410784 | 0.00427649 | 0.08 | 0.05 |
| YDL234C | 2388.462086 | 0.834147796 | 0.004341296 | 0.08 | 0.05 |
| YDR481C | 2453.405315 | 0.749000022 | 0.004343901 | 0.08 | 0.05 |
| YDL127W | 422.9534044 | -0.724435567 | 0.004474949 | 0.08 | 0.05 |
| YJL027C | 10.13200317 | 1.183239899 | 0.004468573 | 0.08 | 0.05 |
| YKL105C | 1423.737383 | 0.607334545 | 0.004452318 | 0.08 | 0.05 |
| YNL288W | 932.0705303 | -0.43519959 | 0.004467686 | 0.08 | 0.05 |
| YOL146W | 364.144711 | -0.590978001 | 0.004577134 | 0.08 | 0.05 |
| YPL092W | 1197.47636 | 0.996687313 | 0.004576212 | 0.08 | 0.05 |
| YFR023W | 96.03552642 | 1.611871515 | 0.004595595 | 0.08 | 0.05 |
| YLR383W | 888.8933966 | 0.525996069 | 0.004615952 | 0.08 | 0.05 |
| YML052W | 1751.911847 | -0.542521782 | 0.00461721 | 0.08 | 0.05 |
| YLR412W | 993.3823084 | -0.576837768 | 0.004634465 | 0.08 | 0.05 |
| YOR101W | 962.4280426 | -0.680152372 | 0.004684085 | 0.08 | 0.05 |
| YER148W | 733.3492834 | -0.633370695 | 0.004727532 | 0.08 | 0.05 |
| YIL066C | 409.4135175 | 1.046582032 | 0.004713905 | 0.08 | 0.05 |
| YPL134C | 1579.391088 | 1.221251008 | 0.004717741 | 0.08 | 0.05 |
| YEL010W | 4.427753031 | 1.677861489 | 0.004751986 | 0.08 | 0.05 |
| YGL150C | 2988.328671 | 0.382928037 | 0.004852587 | 0.08 | 0.05 |
| YGR075C | 302.7071594 | -0.5200217 | 0.004814438 | 0.08 | 0.05 |
| YHL050C | 143.7644195 | 1.27411911 | 0.00480302 | 0.08 | 0.05 |
| YNL104C | 8329.192681 | 1.083056375 | 0.004832095 | 0.08 | 0.05 |
| YPR056W | 512.7372419 | -0.366719677 | 0.004841773 | 0.08 | 0.05 |

| | | | | | |
|---|---|---|---|---|---|
| YPR202W | 17.57397274 | 1.557972772 | 0.004847277 | 0.08 | 0.05 |
| YDR446W | 60.61919013 | 1.43019063 | 0.004890824 | 0.08 | 0.05 |
| YDR333C | 1359.98522 | -0.4239889 | 0.004976542 | 0.08 | 0.05 |
| YGR225W | 197.8168855 | 1.223337639 | 0.004946347 | 0.08 | 0.05 |
| YHR208W | 7460.933205 | 1.017986495 | 0.004966342 | 0.08 | 0.05 |
| YJL065C | 199.2698224 | -0.434422432 | 0.00497472 | 0.08 | 0.05 |
| YJL133W | 754.0202192 | -0.606342848 | 0.004951455 | 0.08 | 0.05 |
| YMR062C | 1501.153431 | 0.911864411 | 0.004988049 | 0.08 | 0.05 |
| YPR053C | 166.5759994 | -0.800137939 | 0.004959808 | 0.08 | 0.05 |
| YDL128W | 1754.772543 | -0.445941115 | 0.005112134 | 0.08 | 0.06 |
| YAL018C | 88.41004538 | 1.830567982 | 0.0052241 | 0.09 | 0.06 |
| YBR098W | 547.1579177 | 0.459124851 | 0.005213631 | 0.09 | 0.06 |
| YMR072W | 3717.086972 | -0.45000793 | 0.005192998 | 0.09 | 0.06 |
| YNL222W | 196.3405424 | -0.778424405 | 0.005223112 | 0.09 | 0.06 |
| YHR015W | 104.6987227 | 1.554855512 | 0.005256326 | 0.09 | 0.06 |
| YPL232W | 2081.428239 | -0.484969922 | 0.005272666 | 0.09 | 0.06 |
| YGL114W | 1125.03715 | 0.567599342 | 0.005318087 | 0.09 | 0.06 |
| YPR167C | 421.1211 | 0.762130649 | 0.005324683 | 0.09 | 0.06 |
| YLL061W | 447.0050952 | 0.756536447 | 0.005375264 | 0.09 | 0.06 |
| YLL011W | 676.0042616 | -0.774194696 | 0.005388778 | 0.09 | 0.06 |
| YBR012C | 184.3218002 | 1.359381354 | 0.00544231 | 0.09 | 0.06 |
| YGR282C | 11912.38121 | 0.461259049 | 0.005429604 | 0.09 | 0.06 |
| YNL078W | 1260.908163 | -0.762773532 | 0.005425815 | 0.09 | 0.06 |
| YER067W | 2559.902963 | 1.28210135 | 0.005477579 | 0.09 | 0.06 |
| YJL137C | 923.3034038 | 1.139595847 | 0.00548691 | 0.09 | 0.06 |
| YNL138W-A | 114.4265169 | -0.548602042 | 0.005552028 | 0.09 | 0.06 |
| YJL127C | 657.9035431 | 0.348589898 | 0.005595258 | 0.09 | 0.06 |
| YOL104C | 616.0912562 | 1.001351903 | 0.005601795 | 0.09 | 0.06 |
| YHR079C | 1259.061897 | 0.438985421 | 0.005748431 | 0.09 | 0.06 |
| YHR207C | 677.6706609 | 0.654790692 | 0.005785438 | 0.09 | 0.06 |
| YPR081C | 985.9306388 | 0.581609244 | 0.005786688 | 0.09 | 0.06 |
| YBR052C | 2651.004465 | -0.484959541 | 0.005858211 | 0.09 | 0.06 |
| YDR057W | 882.7471612 | 0.498268098 | 0.005879755 | 0.09 | 0.06 |
| YBL064C | 4712.365868 | 0.927688249 | 0.005932782 | 0.09 | 0.06 |
| YGL243W | 434.8625344 | -0.311377162 | 0.005943187 | 0.09 | 0.06 |
| YHR138C | 1476.126547 | 0.79224728 | 0.005907815 | 0.09 | 0.06 |
| YML100W-A | 6.289539175 | 1.276458128 | 0.005940258 | 0.09 | 0.06 |
| YHL016C | 664.2747126 | 1.232649914 | 0.005975565 | 0.09 | 0.06 |
| YER119C-A | 190.6802462 | -0.688063677 | 0.006013123 | 0.09 | 0.06 |
| YHL019C | 544.6940331 | 0.666343206 | 0.006023304 | 0.09 | 0.06 |
| YNL239W | 3388.853667 | 0.547419784 | 0.006019005 | 0.09 | 0.06 |
| YFL028C | 788.0732688 | 0.471547793 | 0.006059547 | 0.09 | 0.06 |
| YKR008W | 1399.761894 | -0.492126256 | 0.006053489 | 0.09 | 0.06 |

| | | | | | |
|---|---|---|---|---|---|
| YBR170C | 1144.125576 | 0.386879242 | 0.006093497 | 0.09 | 0.06 |
| YHL024W | 910.8922734 | 0.860677693 | 0.006094165 | 0.09 | 0.06 |
| YPL279C | 37.92327787 | -0.920254381 | 0.006119047 | 0.09 | 0.06 |
| YCL040W | 21911.49524 | 0.951764667 | 0.006226628 | 0.09 | 0.06 |
| YLR276C | 2051.953288 | -0.831862693 | 0.006234597 | 0.09 | 0.06 |
| YPR194C | 1121.813284 | 1.342557262 | 0.006245802 | 0.09 | 0.06 |
| YGL033W | 39.3922468 | 1.152841371 | 0.006290249 | 0.09 | 0.06 |
| YHR043C | 1297.179518 | 0.600206773 | 0.006314048 | 0.09 | 0.06 |
| YLR053C | 106.3162181 | 1.029833717 | 0.006308087 | 0.09 | 0.06 |
| YHR035W | 231.8124454 | 0.68102607 | 0.0063446 | 0.09 | 0.06 |
| YMR275C | 1521.663704 | 0.531029238 | 0.006360129 | 0.09 | 0.06 |
| YDR035W | 3632.420651 | 0.735110117 | 0.00639449 | 0.09 | 0.06 |
| YDR074W | 10482.83127 | 1.040718972 | 0.006482034 | 0.10 | 0.06 |
| YMR065W | 380.856771 | -0.480136027 | 0.006532075 | 0.10 | 0.06 |
| YHR156C | 433.8648918 | 0.640498242 | 0.006556335 | 0.10 | 0.06 |
| YOR342C | 1114.452608 | -0.606354713 | 0.00657089 | 0.10 | 0.06 |
| YMR034C | 377.1167506 | 0.886608466 | 0.006606848 | 0.10 | 0.06 |
| YKL085W | 9612.778538 | 1.256433463 | 0.006665254 | 0.10 | 0.06 |
| YMR270C | 708.6444493 | -0.479616093 | 0.006679594 | 0.10 | 0.06 |
| YEL035C | 12.59177912 | 1.514158048 | 0.00673549 | 0.10 | 0.06 |
| YLR321C | 729.7163579 | -0.605343057 | 0.006730176 | 0.10 | 0.06 |
| YJR051W | 1291.045072 | 0.800731967 | 0.006766518 | 0.10 | 0.06 |
| YKL188C | 680.4138758 | 0.833616521 | 0.006821578 | 0.10 | 0.07 |
| YLL014W | 750.2093462 | -0.55361075 | 0.006932432 | 0.10 | 0.07 |
| YML114C | 955.0918143 | -0.415463319 | 0.006934991 | 0.10 | 0.07 |
| YPR052C | 667.2448711 | -0.725986604 | 0.006910388 | 0.10 | 0.07 |
| YDR369C | 596.1613966 | 0.618516677 | 0.007053343 | 0.10 | 0.07 |
| YBR136W | 1309.50165 | 0.605436665 | 0.007103162 | 0.10 | 0.07 |
| YBL015W | 5485.33881 | 1.329226651 | 0.007196773 | 0.10 | 0.07 |
| YCR027C | 255.2084978 | -0.532722025 | 0.007164719 | 0.10 | 0.07 |
| YDR524C | 904.6436686 | -0.488046628 | 0.007186838 | 0.10 | 0.07 |
| YEL033W | 1403.07415 | -0.858360634 | 0.007204535 | 0.10 | 0.07 |
| YJL025W | 374.9847479 | -0.869176851 | 0.007180014 | 0.10 | 0.07 |
| YLR189C | 2093.552884 | 0.57300325 | 0.007354153 | 0.10 | 0.07 |
| YNL269W | 50.67416069 | 1.408074493 | 0.007370433 | 0.10 | 0.07 |
| YOR376W | 25.04102901 | 1.398166877 | 0.007472223 | 0.10 | 0.07 |
| YOL097W-A | 14.8332561 | -0.878911732 | 0.007496113 | 0.11 | 0.07 |
| YLR080W | 639.2701305 | 0.886935993 | 0.007542697 | 0.11 | 0.07 |
| YML019W | 694.8842248 | -0.448721012 | 0.00761521 | 0.11 | 0.07 |
| YFR053C | 31755.34613 | 1.156187231 | 0.007654615 | 0.11 | 0.07 |
| YGL009C | 9382.784081 | 1.130985295 | 0.007669721 | 0.11 | 0.07 |
| YJR077C | 4794.775519 | 0.897693764 | 0.007718258 | 0.11 | 0.07 |
| YHL045W | 26.03413018 | 1.329574738 | 0.007744613 | 0.11 | 0.07 |

| | | | | | |
|---|---|---|---|---|---|
| YDR544C | 2.336721419 | 1.712455916 | 0.007787615 | 0.11 | 0.07 |
| YLL059C | 81.93358505 | 0.679880117 | 0.007795514 | 0.11 | 0.07 |
| YLR308W | 11.06339218 | 1.445223246 | 0.007851872 | 0.11 | 0.07 |
| YDR387C | 888.6103683 | 0.572336738 | 0.007937354 | 0.11 | 0.07 |
| YGL175C | 162.4549988 | 0.495426676 | 0.007943211 | 0.11 | 0.07 |
| YDR246W | 916.4165129 | -0.385337186 | 0.008008578 | 0.11 | 0.07 |
| YDR502C | 1141.466361 | 0.752393524 | 0.008083029 | 0.11 | 0.07 |
| YMR187C | 357.2988229 | 0.578028947 | 0.008076223 | 0.11 | 0.07 |
| YDR014W-A | 36.16776295 | 1.24900903 | 0.008106267 | 0.11 | 0.07 |
| YMR323W | 206.5265565 | 1.262878768 | 0.0082178 | 0.11 | 0.07 |
| YBR088C | 178.1843606 | -0.755946857 | 0.008261721 | 0.11 | 0.07 |
| YKL064W | 2390.234747 | 0.51224224 | 0.008257155 | 0.11 | 0.07 |
| YEL009C-A | 8.998175135 | 1.107260903 | 0.008317458 | 0.11 | 0.07 |
| YER129W | 2291.22394 | 0.277271934 | 0.00835118 | 0.11 | 0.07 |
| YIR039C | 1029.181658 | 1.241381471 | 0.008360665 | 0.11 | 0.07 |
| YLR258W | 9832.7404 | 0.991963368 | 0.008338018 | 0.11 | 0.07 |
| YHR084W | 1211.807628 | -0.897832502 | 0.008410852 | 0.11 | 0.07 |
| YCR032W | 1688.564969 | 0.624164952 | 0.008444771 | 0.11 | 0.07 |
| YDR374C | 71.50374083 | 1.372699447 | 0.008456751 | 0.11 | 0.07 |
| YLR291C | 1399.72151 | -0.404222729 | 0.008483164 | 0.11 | 0.07 |
| YGL258W-A | 56.62039045 | 1.170178666 | 0.008526625 | 0.11 | 0.07 |
| YLR137W | 268.6657732 | -0.740160466 | 0.008544205 | 0.11 | 0.07 |
| YJL108C | 258.931357 | 0.735542104 | 0.00856972 | 0.11 | 0.07 |
| YER175C | 1510.549619 | 1.100792636 | 0.008620813 | 0.11 | 0.07 |
| YPL066W | 1565.740738 | -0.423185645 | 0.008703068 | 0.11 | 0.08 |
| YHR213W-B | 12.16949824 | 1.255966199 | 0.008867268 | 0.12 | 0.08 |
| YOR303W | 3621.710527 | 1.080425061 | 0.008860365 | 0.12 | 0.08 |
| YML119W | 1027.81525 | 0.466692432 | 0.008960584 | 0.12 | 0.08 |
| YNL114C | 62.61749004 | -0.878703834 | 0.008990869 | 0.12 | 0.08 |
| YGR237C | 1848.787726 | 0.511017695 | 0.009054889 | 0.12 | 0.08 |
| YBL113C | 286.5211358 | 0.768770552 | 0.009101526 | 0.12 | 0.08 |
| YBR208C | 3562.373289 | 1.298699269 | 0.009110485 | 0.12 | 0.08 |
| YDL074C | 1505.723255 | -0.566738909 | 0.009167748 | 0.12 | 0.08 |
| YHR075C | 548.2435732 | 0.46087216 | 0.009179154 | 0.12 | 0.08 |
| YMR216C | 1336.629718 | -0.462924793 | 0.009292487 | 0.12 | 0.08 |
| YJL156C | 869.0736001 | -0.424003067 | 0.0093339 | 0.12 | 0.08 |
| YMR094W | 306.3491696 | 0.687479737 | 0.009390519 | 0.12 | 0.08 |
| YPL164C | 886.5830673 | 0.612900742 | 0.009385403 | 0.12 | 0.08 |
| YCL058C | 103.80183 | 0.824102099 | 0.009437502 | 0.12 | 0.08 |
| YPL235W | 1507.98677 | -0.447319326 | 0.009425779 | 0.12 | 0.08 |
| YJL133C-A | 1683.932792 | 1.055081591 | 0.009474558 | 0.12 | 0.08 |
| YGR097W | 2251.701342 | 0.630402659 | 0.009717355 | 0.12 | 0.08 |
| YKR034W | 49.93480154 | 1.725790886 | 0.009766847 | 0.12 | 0.08 |

| | | | | | |
|---|---|---|---|---|---|
| YIL002W-A | 495.5817191 | -0.442302755 | 0.009843137 | 0.12 | 0.08 |
| YDR263C | 317.4342443 | 0.649923907 | 0.009880018 | 0.12 | 0.08 |
| YFR030W | 1464.99151 | 0.788692799 | 0.009913473 | 0.12 | 0.08 |
| YIL051C | 4289.573013 | -0.77355519 | 0.00994364 | 0.13 | 0.08 |
| YFL018C | 6022.957872 | 0.593510008 | 0.010016115 | 0.13 | 0.08 |
| YJR057W | 461.680688 | -0.414841946 | 0.010002824 | 0.13 | 0.08 |
| YBR288C | 774.8863305 | -0.46092664 | 0.010111792 | 0.13 | 0.08 |
| YDR343C | 868.7911596 | 1.229099621 | 0.010106244 | 0.13 | 0.08 |
| YOL151W | 926.9123911 | -0.711799638 | 0.010112771 | 0.13 | 0.08 |
| YHR014W | 138.7802733 | 0.889827193 | 0.010169093 | 0.13 | 0.08 |
| YHR016C | 1896.688185 | 0.588181412 | 0.010206196 | 0.13 | 0.08 |
| YDL155W | 763.3718322 | -0.442314465 | 0.01028884 | 0.13 | 0.08 |
| YHL042W | 88.3151764 | 0.783714121 | 0.010256355 | 0.13 | 0.08 |
| YHR139C-A | 36.30751378 | 1.002523128 | 0.010320924 | 0.13 | 0.08 |
| YKL215C | 1029.488394 | 0.51527667 | 0.010314692 | 0.13 | 0.08 |
| YNL015W | 3387.233502 | 1.079828046 | 0.010320314 | 0.13 | 0.08 |
| YPR091C | 1815.236332 | 0.388489213 | 0.010352258 | 0.13 | 0.08 |
| YOR206W | 2506.527701 | -0.775289379 | 0.010398493 | 0.13 | 0.08 |
| YEL011W | 8924.284313 | 1.110043981 | 0.010462339 | 0.13 | 0.08 |
| YBR055C | 925.8039594 | -0.336235459 | 0.010486211 | 0.13 | 0.08 |
| YGL041W-A | 345.8516558 | -0.539068252 | 0.01054718 | 0.13 | 0.08 |
| YFR054C | 20.51319414 | 1.057943118 | 0.010613229 | 0.13 | 0.09 |
| YLR386W | 1077.114184 | 0.346794055 | 0.010675656 | 0.13 | 0.09 |
| YPR009W | 346.2208437 | -0.999457594 | 0.010659061 | 0.13 | 0.09 |
| YBR165W | 515.6336536 | -0.418750885 | 0.010838032 | 0.13 | 0.09 |
| YER158W-A | 247.6290209 | -0.726423414 | 0.011199214 | 0.14 | 0.09 |
| YCL031C | 1450.481764 | -0.569705001 | 0.011313952 | 0.14 | 0.09 |
| YCR023C | 1565.302118 | 0.623293006 | 0.011264917 | 0.14 | 0.09 |
| YDL116W | 1341.682086 | -0.399042458 | 0.011311763 | 0.14 | 0.09 |
| YHR008C | 8653.583569 | 0.853154276 | 0.011282007 | 0.14 | 0.09 |
| YGL082W | 1508.755452 | -0.319290972 | 0.011380617 | 0.14 | 0.09 |
| YHL031C | 431.2428898 | 0.404420678 | 0.011367684 | 0.14 | 0.09 |
| YOL075C | 712.6275688 | 0.580888862 | 0.011401129 | 0.14 | 0.09 |
| YNR017W | 1847.028777 | -0.431404057 | 0.011448784 | 0.14 | 0.09 |
| YGL032C | 1976.542723 | -1.489212222 | 0.011601904 | 0.14 | 0.09 |
| YGL194C-A | 142.847988 | -0.738060176 | 0.011853252 | 0.14 | 0.09 |
| YHR120W | 606.7255558 | 0.343039336 | 0.011876557 | 0.14 | 0.09 |
| YHR125W | 4.057342841 | 1.488229779 | 0.011876193 | 0.14 | 0.09 |
| YML130C | 2617.920044 | 0.821530984 | 0.011855271 | 0.14 | 0.09 |
| YDL167C | 1031.301724 | -0.890402223 | 0.011925301 | 0.14 | 0.09 |
| YER166W | 2859.051695 | 0.480718089 | 0.011921456 | 0.14 | 0.09 |
| YBR101C | 3697.675555 | 0.633302986 | 0.011951024 | 0.14 | 0.09 |
| YBR267W | 453.4917639 | -0.65169976 | 0.012074301 | 0.14 | 0.09 |

| | | | | | |
|---|---|---|---|---|---|
| YER066C-A | 1483.59707 | 1.105949502 | 0.012063629 | 0.14 | 0.09 |
| YLR162W | 2.387178616 | 1.542903935 | 0.012062152 | 0.14 | 0.09 |
| YPR187W | 1425.65006 | -0.805681299 | 0.012163748 | 0.14 | 0.09 |
| YJR091C | 3359.251305 | 0.527317916 | 0.012221767 | 0.14 | 0.09 |
| YLR241W | 1618.382874 | 0.416370755 | 0.012238428 | 0.14 | 0.09 |
| YOL045W | 1132.896142 | 0.319523709 | 0.012221447 | 0.14 | 0.09 |
| YNL258C | 964.2763945 | -0.390814539 | 0.012375683 | 0.14 | 0.09 |
| YPR024W | 1909.485964 | 0.422562552 | 0.01235599 | 0.14 | 0.09 |
| YIL015W | 2424.316267 | -1.42349949 | 0.012450548 | 0.14 | 0.09 |
| YIL118W | 1012.394304 | -0.690587557 | 0.012511905 | 0.14 | 0.09 |
| YIL167W | 108.465152 | 0.693519798 | 0.012473997 | 0.14 | 0.09 |
| YMR324C | 2.573506675 | 1.460818692 | 0.012518013 | 0.14 | 0.09 |
| YCL023C | 8.286348364 | 1.210860353 | 0.012574728 | 0.14 | 0.09 |
| YBR066C | 407.0918605 | -1.014099146 | 0.01272975 | 0.14 | 0.10 |
| YCL048W | 92.73705911 | 1.268721441 | 0.012729764 | 0.14 | 0.10 |
| YGL204C | 78.35386912 | -0.783719868 | 0.012707073 | 0.14 | 0.10 |
| YKL194C | 593.5236151 | 0.462545919 | 0.012773685 | 0.14 | 0.10 |
| YLR013W | 3.60279815 | 1.460747985 | 0.012711644 | 0.14 | 0.10 |
| YNL311C | 782.0547648 | 0.531570729 | 0.012687205 | 0.14 | 0.10 |
| YPR032W | 901.7856576 | 0.435344058 | 0.012771335 | 0.14 | 0.10 |
| YCL025C | 3652.132376 | 1.001789292 | 0.012906249 | 0.15 | 0.10 |
| YDL054C | 853.6605887 | 0.598134287 | 0.0129303 | 0.15 | 0.10 |
| YGL178W | 2073.150568 | 0.564683923 | 0.012923865 | 0.15 | 0.10 |
| YHR126C | 18.42016663 | 1.430002449 | 0.013072638 | 0.15 | 0.10 |
| YKL170W | 81.53735369 | -0.587766649 | 0.013104528 | 0.15 | 0.10 |
| YNL201C | 1076.736749 | -0.331541565 | 0.013111397 | 0.15 | 0.10 |
| YPL070W | 1079.494033 | 0.4570123 | 0.013131517 | 0.15 | 0.10 |
| YDR473C | 858.9200777 | 0.400645574 | 0.013215386 | 0.15 | 0.10 |
| YKL103C | 3936.941911 | 0.959781846 | 0.013291245 | 0.15 | 0.10 |
| YBL100C | 649.5552869 | 0.960603264 | 0.013415783 | 0.15 | 0.10 |
| YJL219W | 39.81053365 | 1.247597916 | 0.013388733 | 0.15 | 0.10 |
| YOR203W | 129.138005 | 0.795765392 | 0.013394649 | 0.15 | 0.10 |
| YPL119C | 673.8655489 | 0.836949189 | 0.013466758 | 0.15 | 0.10 |
| YER146W | 140.8937845 | -0.716494376 | 0.013637027 | 0.15 | 0.10 |
| YGR035W-A | 93.40510398 | -0.743035061 | 0.013614475 | 0.15 | 0.10 |
| YOL087C | 2320.349339 | 0.514207716 | 0.013591897 | 0.15 | 0.10 |
| YOR123C | 2187.626634 | -0.285023017 | 0.013614674 | 0.15 | 0.10 |
| YOR321W | 1083.064286 | 0.492720492 | 0.013583218 | 0.15 | 0.10 |
| YNL253W | 477.8813502 | -0.283675348 | 0.013730477 | 0.15 | 0.10 |
| YGL261C | 11.97177178 | 1.325045491 | 0.013754153 | 0.15 | 0.10 |
| YMR293C | 413.7964048 | 0.494352587 | 0.013862941 | 0.15 | 0.10 |
| YER103W | 3269.463849 | 1.03373885 | 0.013964592 | 0.15 | 0.10 |
| YGR195W | 892.1333078 | -0.428216257 | 0.013964061 | 0.15 | 0.10 |

| | | | | | |
|---|---|---|---|---|---|
| YCL042W | 285.5413665 | 0.875563327 | 0.014027038 | 0.15 | 0.10 |
| YNL145W | 1813.203416 | -1.426971621 | 0.014034748 | 0.15 | 0.10 |
| YBR253W | 396.1883397 | -0.362855781 | 0.014155148 | 0.15 | 0.10 |
| YCR035C | 836.2083583 | -0.354341426 | 0.014301276 | 0.15 | 0.10 |
| YDL185W | 10910.49268 | 0.378344156 | 0.014251136 | 0.15 | 0.10 |
| YIR041W | 5.637300496 | 1.356493455 | 0.014254216 | 0.15 | 0.10 |
| YJR049C | 1397.771565 | 0.561435162 | 0.01425741 | 0.15 | 0.10 |
| YJR137C | 1214.839856 | 0.578752227 | 0.014241482 | 0.15 | 0.10 |
| YKL166C | 506.1904776 | -0.593649133 | 0.01421655 | 0.15 | 0.10 |
| YKR095W-A | 172.6357846 | -0.699999129 | 0.014290307 | 0.15 | 0.10 |
| YLR089C | 5710.768483 | 0.998326189 | 0.01431557 | 0.15 | 0.10 |
| YOR212W | 1199.500024 | -1.158991635 | 0.014074162 | 0.15 | 0.10 |
| YOR355W | 1652.733408 | -0.744073639 | 0.01410354 | 0.15 | 0.10 |
| snR55 | 9.080532067 | 1.332927045 | 0.014215029 | 0.15 | 0.10 |
| YER124C | 2039.69695 | -0.557599601 | 0.014375575 | 0.15 | 0.10 |
| YLL038C | 326.4334207 | -0.631321843 | 0.014382001 | 0.15 | 0.10 |
| YKR102W | 639.1097035 | 1.153562581 | 0.014504856 | 0.15 | 0.10 |
| YCL076W | 24.7210401 | 1.497310841 | 0.014585009 | 0.15 | 0.10 |
| YHR190W | 5071.299505 | 0.520962995 | 0.014605658 | 0.15 | 0.10 |
| YML115C | 1520.660585 | -0.367467216 | 0.014601242 | 0.15 | 0.10 |
| YJR149W | 627.5970928 | 0.918171852 | 0.014655159 | 0.15 | 0.10 |
| YPL112C | 806.8455711 | -0.461450127 | 0.014648766 | 0.15 | 0.10 |
| YOR371C | 940.0880721 | 0.522037621 | 0.014756994 | 0.15 | 0.10 |
| YDR083W | 1132.402572 | -0.69220514 | 0.014951955 | 0.16 | 0.10 |
| YGL062W | 7360.866755 | 1.117581382 | 0.014948213 | 0.16 | 0.10 |
| YLR077W | 986.4922801 | 0.312042278 | 0.014924044 | 0.16 | 0.10 |
| YEL030C-A | 7.262884205 | 1.027403119 | 0.015047818 | 0.16 | 0.10 |
| YLR152C | 1272.570191 | 0.816573292 | 0.015035583 | 0.16 | 0.10 |
| YLR240W | 1128.041716 | 0.440069524 | 0.015013991 | 0.16 | 0.10 |
| YJL037W | 69.21169324 | 1.311675099 | 0.015089083 | 0.16 | 0.10 |
| YKR035C | 23.50975344 | -0.699383639 | 0.015180729 | 0.16 | 0.10 |
| YJL160C | 84.43696653 | 0.908808197 | 0.015336882 | 0.16 | 0.10 |
| YGR287C | 287.959068 | 0.63872113 | 0.015382467 | 0.16 | 0.10 |
| YNL147W | 569.2824244 | -0.45719549 | 0.015419362 | 0.16 | 0.10 |
| YEL020C-B | 34.39724677 | -0.699858659 | 0.015448556 | 0.16 | 0.10 |
| YFR019W | 2589.385757 | 0.457397542 | 0.015520684 | 0.16 | 0.10 |
| YHL038C | 821.5884324 | 0.417986477 | 0.015526625 | 0.16 | 0.10 |
| YNL328C | 68.99382563 | -0.541734403 | 0.015701739 | 0.16 | 0.11 |
| YFL055W | 265.1999324 | 0.969499706 | 0.015826261 | 0.16 | 0.11 |
| YER180C | 405.292426 | -0.402074087 | 0.015872484 | 0.16 | 0.11 |
| YGR112W | 838.4700607 | 0.680550819 | 0.015950231 | 0.16 | 0.11 |
| YKL163W | 2089.664663 | 1.304372428 | 0.016050252 | 0.16 | 0.11 |
| YHR018C | 3547.596925 | 1.173117236 | 0.016093177 | 0.16 | 0.11 |

| | | | | | |
|---|---|---|---|---|---|
| YNL081C | 921.3541155 | -0.379798451 | 0.016190759 | 0.16 | 0.11 |
| YBL024W | 2012.558866 | -0.676800024 | 0.016446542 | 0.17 | 0.11 |
| YBR145W | 1370.985162 | 0.876162833 | 0.01645965 | 0.17 | 0.11 |
| YFR031C | 1205.359626 | 0.346078705 | 0.016479216 | 0.17 | 0.11 |
| YOR159C | 320.0573697 | -0.495934101 | 0.016510738 | 0.17 | 0.11 |
| YNR001C | 10779.43573 | 1.01624517 | 0.01654449 | 0.17 | 0.11 |
| YFR005C | 788.6190865 | -0.47356157 | 0.016728329 | 0.17 | 0.11 |
| YLR413W | 2093.12908 | -1.044646477 | 0.01673065 | 0.17 | 0.11 |
| YNL049C | 1288.390098 | 0.428687049 | 0.016759172 | 0.17 | 0.11 |
| YOL143C | 5652.711754 | -0.772756816 | 0.016736479 | 0.17 | 0.11 |
| YJR078W | 516.4472306 | 1.371259031 | 0.016828086 | 0.17 | 0.11 |
| YPL171C | 308.1636844 | 1.245371104 | 0.016985231 | 0.17 | 0.11 |
| YEL077C | 280.3488731 | 0.942729516 | 0.017085921 | 0.17 | 0.11 |
| YGR165W | 998.5420975 | -0.448861489 | 0.017058273 | 0.17 | 0.11 |
| YJL165C | 2120.172359 | 0.569259611 | 0.017126261 | 0.17 | 0.11 |
| YOR283W | 529.1178643 | -0.550178793 | 0.017103584 | 0.17 | 0.11 |
| YBR115C | 2897.179739 | 0.685478632 | 0.017168474 | 0.17 | 0.11 |
| YOL111C | 1821.468977 | -0.30098693 | 0.017187968 | 0.17 | 0.11 |
| YER153C | 351.5393272 | 0.685877031 | 0.017225299 | 0.17 | 0.11 |
| YNL018C | 116.9826504 | 1.019735067 | 0.017291857 | 0.17 | 0.11 |
| YDR378C | 714.4468599 | -0.356968458 | 0.017480367 | 0.17 | 0.11 |
| YGR138C | 1416.124697 | 1.123366631 | 0.017517406 | 0.17 | 0.11 |
| YIL027C | 793.9722774 | -0.471668132 | 0.017556636 | 0.17 | 0.11 |
| YPL250C | 749.2294185 | 0.89599053 | 0.017533022 | 0.17 | 0.11 |
| YDR076W | 307.3025521 | 0.50879432 | 0.017647505 | 0.17 | 0.11 |
| YER030W | 1421.194205 | -0.512244976 | 0.017627627 | 0.17 | 0.11 |
| YNL317W | 817.8553177 | -0.275883657 | 0.017643481 | 0.17 | 0.11 |
| YPL147W | 471.9776317 | 0.787556632 | 0.017687501 | 0.17 | 0.11 |
| YKL184W | 1354.204344 | -0.438094337 | 0.017833404 | 0.17 | 0.11 |
| YOR138C | 1254.624127 | 0.343375106 | 0.017836953 | 0.17 | 0.11 |
| YPL033C | 16.58126869 | 1.281828785 | 0.017802385 | 0.17 | 0.11 |
| YAL068C | 5.759798843 | 1.286945422 | 0.01788119 | 0.17 | 0.11 |
| YGL259W | 86.59077684 | 1.117934656 | 0.017924509 | 0.17 | 0.11 |
| YGR217W | 881.6554263 | 0.521782287 | 0.017954515 | 0.17 | 0.11 |
| YKR074W | 1111.118511 | -0.453393183 | 0.017969528 | 0.17 | 0.11 |
| YGR070W | 2198.55855 | 0.833850394 | 0.018077803 | 0.17 | 0.11 |
| YIR014W | 256.3884009 | 0.66548092 | 0.018074412 | 0.17 | 0.11 |
| YML083C | 177.180785 | 0.985777893 | 0.018236293 | 0.17 | 0.12 |
| YOR278W | 477.7903224 | -0.441198453 | 0.018241212 | 0.17 | 0.12 |
| YHL034C | 6448.755806 | 0.400663899 | 0.018383381 | 0.17 | 0.12 |
| YOR360C | 883.5535106 | -0.588402781 | 0.018376477 | 0.17 | 0.12 |
| YCR098C | 141.8503964 | 1.264779102 | 0.018418576 | 0.18 | 0.12 |
| YDR404C | 728.6666751 | -0.447781893 | 0.018472534 | 0.18 | 0.12 |

| | | | | | |
|---|---|---|---|---|---|
| YHR080C | 2045.316793 | 0.509442722 | 0.018453973 | 0.18 | 0.12 |
| YBR246W | 994.4891912 | -0.330396608 | 0.018651186 | 0.18 | 0.12 |
| YOL117W | 709.0122211 | 0.701732079 | 0.018707917 | 0.18 | 0.12 |
| YKR083C | 244.3380659 | -0.69073558 | 0.018777347 | 0.18 | 0.12 |
| YFL026W | 6326.346816 | -1.285360456 | 0.018813647 | 0.18 | 0.12 |
| YDL085C-A | 569.0392384 | -0.667380919 | 0.018892839 | 0.18 | 0.12 |
| YMR105C | 9001.650714 | 1.008290578 | 0.018890413 | 0.18 | 0.12 |
| YBR024W | 889.4330761 | 0.650675842 | 0.018920607 | 0.18 | 0.12 |
| YER053C | 3873.936022 | 0.922785057 | 0.018949483 | 0.18 | 0.12 |
| YHR105W | 228.2746463 | 0.622939839 | 0.019028322 | 0.18 | 0.12 |
| YKL172W | 2863.993194 | -0.784131141 | 0.019030883 | 0.18 | 0.12 |
| YPL130W | 24.21083011 | 1.583104859 | 0.019078936 | 0.18 | 0.12 |
| YDR016C | 195.2031569 | -0.473361931 | 0.01914407 | 0.18 | 0.12 |
| YDR299W | 1150.958917 | -0.876783508 | 0.019142749 | 0.18 | 0.12 |
| YNR034W-A | 4345.183948 | 1.140281593 | 0.019297227 | 0.18 | 0.12 |
| YGL177W | 36.28800868 | 0.660736751 | 0.019363375 | 0.18 | 0.12 |
| YNL146C-A | 29.52790111 | -1.122122472 | 0.019377933 | 0.18 | 0.12 |
| YMR095C | 600.2590168 | 0.981981044 | 0.019420271 | 0.18 | 0.12 |
| YKL096W | 1798.684379 | -0.971095775 | 0.019459031 | 0.18 | 0.12 |
| YPR146C | 99.73914934 | -0.501805876 | 0.019572864 | 0.18 | 0.12 |
| YLR358C | 39.08246637 | -0.475487402 | 0.019695913 | 0.18 | 0.12 |
| YMR304W | 1821.67097 | 0.465701263 | 0.019690803 | 0.18 | 0.12 |
| YDR030C | 444.8820441 | 0.599674251 | 0.019832008 | 0.18 | 0.12 |
| YHL048C-A | 2.76683096 | 1.629591898 | 0.019778491 | 0.18 | 0.12 |
| YJL171C | 1558.602902 | 0.536002316 | 0.019828402 | 0.18 | 0.12 |
| YJR014W | 1708.927082 | -0.350666652 | 0.019923406 | 0.18 | 0.12 |
| YPL248C | 793.6864415 | 0.579585014 | 0.019912763 | 0.18 | 0.12 |
| YEL056W | 977.5251167 | -0.316703813 | 0.0199554 | 0.18 | 0.12 |
| YCL027W | 675.5367401 | -0.980986246 | 0.020014011 | 0.18 | 0.12 |
| YKL150W | 7424.00365 | 0.97939682 | 0.020052154 | 0.18 | 0.12 |
| YER098W | 930.5704049 | 0.587334612 | 0.020131741 | 0.18 | 0.12 |
| YHR005C | 1534.146963 | -1.141923854 | 0.020136493 | 0.18 | 0.12 |
| YNR018W | 1764.18714 | 0.648496746 | 0.020111359 | 0.18 | 0.12 |
| YHR044C | 186.9132602 | 0.920302615 | 0.020208152 | 0.18 | 0.12 |
| YHR093W | 35.27687991 | 0.792174882 | 0.020427135 | 0.18 | 0.12 |
| YJL141C | 4385.516839 | 0.808506665 | 0.020363957 | 0.18 | 0.12 |
| YJR090C | 1653.355955 | -0.372350161 | 0.020409121 | 0.18 | 0.12 |
| YNR009W | 344.9832981 | -0.7211359 | 0.020296257 | 0.18 | 0.12 |
| YOR268C | 11.87888145 | 1.20175838 | 0.020414679 | 0.18 | 0.12 |
| YPR204W | 106.9528255 | 1.029234501 | 0.020346229 | 0.18 | 0.12 |
| YCL033C | 876.4314652 | 0.784847078 | 0.020662881 | 0.18 | 0.12 |
| YDL100C | 5446.231089 | 0.564141432 | 0.020679807 | 0.18 | 0.12 |
| YFR052C-A | 3217.393186 | 1.075905369 | 0.020653823 | 0.18 | 0.12 |

| | | | | | |
|---|---|---|---|---|---|
| YDL036C | 422.8116952 | -0.444079127 | 0.020758635 | 0.19 | 0.12 |
| YJR010C-A | 212.1436636 | -0.65773597 | 0.020843236 | 0.19 | 0.12 |
| YLR219W | 2484.562083 | 0.535384299 | 0.020840688 | 0.19 | 0.12 |
| YJL099W | 747.8275315 | 0.436463809 | 0.020977801 | 0.19 | 0.12 |
| YKL202W | 21.88607995 | 0.627744503 | 0.021002119 | 0.19 | 0.12 |
| YLR353W | 907.9621406 | 0.327511948 | 0.020988406 | 0.19 | 0.12 |
| YML120C | 4283.544511 | 1.000245733 | 0.021007231 | 0.19 | 0.12 |
| YMR191W | 2004.986398 | 0.915844998 | 0.02091656 | 0.19 | 0.12 |
| YDR116C | 1042.960963 | -0.418099441 | 0.021050585 | 0.19 | 0.12 |
| YGL156W | 2401.006767 | 0.919436324 | 0.021076311 | 0.19 | 0.12 |
| YHR034C | 511.3439371 | 0.392639424 | 0.021124997 | 0.19 | 0.12 |
| YDR538W | 133.1462476 | -0.44239366 | 0.021181405 | 0.19 | 0.12 |
| YDR235W | 670.6497022 | -0.360112201 | 0.02141213 | 0.19 | 0.12 |
| YFR036W-A | 15.15766896 | -0.706379454 | 0.021772526 | 0.19 | 0.13 |
| YKR038C | 1110.135443 | -0.347012476 | 0.02174401 | 0.19 | 0.13 |
| YLR311C | 30.19399839 | 1.083947736 | 0.021767357 | 0.19 | 0.13 |
| YNL241C | 6779.137631 | 0.639962468 | 0.021763473 | 0.19 | 0.13 |
| YOL026C | 840.8767019 | -0.409850619 | 0.021826867 | 0.19 | 0.13 |
| YBR295W | 678.020174 | 0.638917574 | 0.021953749 | 0.19 | 0.13 |
| YHL018W | 70.12879735 | 0.575475147 | 0.021961474 | 0.19 | 0.13 |
| YOR022C | 608.0869392 | 0.532826422 | 0.021965407 | 0.19 | 0.13 |
| YOR072W | 29.04315607 | 0.739869675 | 0.021896371 | 0.19 | 0.13 |
| YGL196W | 1262.908167 | 0.77985392 | 0.022037267 | 0.19 | 0.13 |
| YDL235C | 741.3905909 | -0.466194031 | 0.02207657 | 0.19 | 0.13 |
| YBR299W | 41.66988027 | 0.726626839 | 0.022140006 | 0.19 | 0.13 |
| YFL034C-A | 576.1615287 | -0.926918206 | 0.022191505 | 0.19 | 0.13 |
| YPR012W | 66.76164095 | 0.583273761 | 0.022164268 | 0.19 | 0.13 |
| YEL014C | 21.68196635 | -0.957151207 | 0.022258598 | 0.19 | 0.13 |
| YGL115W | 1145.902181 | -0.267902042 | 0.022308531 | 0.19 | 0.13 |
| YLL039C | 7097.91074 | 0.643440741 | 0.022280401 | 0.19 | 0.13 |
| YCL005W | 658.6999475 | -0.263875045 | 0.02240041 | 0.19 | 0.13 |
| YHR166C | 498.330332 | 0.524885151 | 0.022371164 | 0.19 | 0.13 |
| YPR126C | 82.68269767 | -0.722735444 | 0.022399487 | 0.19 | 0.13 |
| YDR114C | 36.82268978 | -0.599382 | 0.022760321 | 0.19 | 0.13 |
| YER145C-A | 50.64924556 | -0.784851142 | 0.022627571 | 0.19 | 0.13 |
| YIL139C | 280.0724876 | -0.343121143 | 0.02278629 | 0.19 | 0.13 |
| YJL088W | 187.9199165 | 1.138372266 | 0.022810931 | 0.19 | 0.13 |
| YKL216W | 7973.975155 | -1.003459852 | 0.022769378 | 0.19 | 0.13 |
| YKL221W | 211.2597824 | 1.074405603 | 0.022794745 | 0.19 | 0.13 |
| YKR032W | 0.979223699 | 1.121120645 | 0.022619751 | 0.19 | 0.13 |
| YMR040W | 194.5584091 | 0.786437471 | 0.022788603 | 0.19 | 0.13 |
| YOR202W | 1479.782603 | 0.661653851 | 0.022742337 | 0.19 | 0.13 |
| YPL216W | 730.7653507 | 0.429287938 | 0.022676217 | 0.19 | 0.13 |

| | | | | | |
|---|---|---|---|---|---|
| YPL241C | 283.7799123 | -0.624802419 | 0.022643293 | 0.19 | 0.13 |
| YPR030W | 2156.768509 | 1.117205909 | 0.02259806 | 0.19 | 0.13 |
| YFL007W | 4593.396937 | 0.454496291 | 0.023005729 | 0.19 | 0.13 |
| YGL125W | 958.6893776 | 0.653066824 | 0.023016181 | 0.19 | 0.13 |
| RUF20 | 86.17514557 | -0.579866414 | 0.023161978 | 0.19 | 0.13 |
| YHR160C | 281.5696466 | 0.705711957 | 0.02323241 | 0.19 | 0.13 |
| YLR300W | 3505.549383 | -0.772605192 | 0.023225342 | 0.19 | 0.13 |
| YMR233W | 360.9564304 | -0.64421273 | 0.02325656 | 0.19 | 0.13 |
| YLR002C | 907.6259007 | -0.601681542 | 0.023301986 | 0.19 | 0.13 |
| YHR086W | 910.6374675 | 0.475419008 | 0.023420708 | 0.19 | 0.13 |
| YLR462W | 4.00598291 | 1.270685443 | 0.023463574 | 0.19 | 0.13 |
| YOR211C | 1075.422511 | -0.390399808 | 0.023457193 | 0.19 | 0.13 |
| YOR190W | 351.1788756 | 0.608234966 | 0.023524835 | 0.20 | 0.13 |
| YNR007C | 864.4677932 | 0.790912949 | 0.023689346 | 0.20 | 0.13 |
| YAR009C | 236.1235724 | 0.777987051 | 0.023783728 | 0.20 | 0.13 |
| YER161C | 1183.000093 | -0.427675248 | 0.023764024 | 0.20 | 0.13 |
| YDR413C | 10.51150758 | -0.933234927 | 0.023870516 | 0.20 | 0.13 |
| YBR034C | 1111.628509 | -0.703554617 | 0.023912788 | 0.20 | 0.13 |
| YAL059W | 208.6556716 | -0.643973353 | 0.024078272 | 0.20 | 0.13 |
| YLL005C | 615.6111971 | 0.744986433 | 0.024070961 | 0.20 | 0.13 |
| YNR033W | 1803.332991 | 0.435854673 | 0.024165312 | 0.20 | 0.13 |
| YPR128C | 787.6455183 | 0.36322927 | 0.024222919 | 0.20 | 0.13 |
| YER017C | 1876.346183 | 0.336793948 | 0.024426513 | 0.20 | 0.13 |
| YGL026C | 4379.972897 | 0.65084002 | 0.02438173 | 0.20 | 0.13 |
| YJR024C | 696.9415683 | -0.574663695 | 0.024400165 | 0.20 | 0.13 |
| YLR285C-A | 41.10839909 | -0.763433153 | 0.02433527 | 0.20 | 0.13 |
| YMR033W | 1176.392622 | -0.524510875 | 0.024416867 | 0.20 | 0.13 |
| YPR158W | 1022.854173 | 0.762698961 | 0.024341495 | 0.20 | 0.13 |
| YEL040W | 3165.37713 | -0.920372846 | 0.024621252 | 0.20 | 0.13 |
| YGR076C | 617.1306078 | -0.454564007 | 0.024568285 | 0.20 | 0.13 |
| YOL106W | 14.51135789 | 0.877504366 | 0.024577071 | 0.20 | 0.13 |
| YOR316C-A | 8.072827245 | -0.975901567 | 0.024617779 | 0.20 | 0.13 |
| YOR255W | 23.76826284 | 1.544457686 | 0.024715793 | 0.20 | 0.13 |
| YJR054W | 539.7820263 | -0.424475312 | 0.024775108 | 0.20 | 0.13 |
| YMR071C | 1073.602658 | -0.205018442 | 0.024854502 | 0.20 | 0.13 |
| YDL003W | 760.5875817 | -0.673967943 | 0.024951922 | 0.20 | 0.13 |
| YKL068W | 1722.980976 | -0.410309501 | 0.024949583 | 0.20 | 0.13 |
| YBL109W | 16.60491535 | 1.037684856 | 0.025024711 | 0.20 | 0.13 |
| YNL142W | 1026.084342 | 1.213509471 | 0.025003186 | 0.20 | 0.13 |
| YER188C-A | 23.15774642 | 0.865319301 | 0.0253846 | 0.20 | 0.13 |
| YHR005C-A | 2208.832297 | -0.521296591 | 0.025268827 | 0.20 | 0.13 |
| YJL197W | 1228.673265 | 0.324415229 | 0.02537011 | 0.20 | 0.13 |
| YLR022C | 541.692103 | -0.516998613 | 0.025331317 | 0.20 | 0.13 |

| | | | | | |
|---|---|---|---|---|---|
| YLR086W | 1588.915329 | 0.44671582 | 0.02526805 | 0.20 | 0.13 |
| YMR201C | 569.2749285 | 0.586323844 | 0.025219689 | 0.20 | 0.13 |
| YNL186W | 1470.462386 | -0.405308537 | 0.025155599 | 0.20 | 0.13 |
| YOR092W | 648.6032701 | -0.653816576 | 0.025335617 | 0.20 | 0.13 |
| YOR095C | 614.382855 | -0.705799043 | 0.025177355 | 0.20 | 0.13 |
| YPR158W-B | 29.96033516 | 1.029859993 | 0.025239325 | 0.20 | 0.13 |
| YDR063W | 1183.071975 | -0.306007518 | 0.025641793 | 0.20 | 0.13 |
| YLR445W | 32.03943686 | 0.935714601 | 0.025679797 | 0.20 | 0.13 |
| YGL095C | 1579.460918 | -0.301995697 | 0.025779624 | 0.20 | 0.13 |
| YJL154C | 2489.813399 | 0.21012408 | 0.025811129 | 0.20 | 0.13 |
| YBL045C | 7514.298009 | 0.898908668 | 0.025916149 | 0.20 | 0.13 |
| YJR161C | 124.4036715 | 0.765597912 | 0.025961752 | 0.20 | 0.14 |
| YIR010W | 867.0379525 | -0.425752386 | 0.026018701 | 0.20 | 0.14 |
| YOR328W | 830.771978 | 0.756534605 | 0.026042061 | 0.20 | 0.14 |
| YDR122W | 2792.779664 | 0.394338988 | 0.026199856 | 0.21 | 0.14 |
| YIL013C | 704.2538412 | 0.452061198 | 0.026244151 | 0.21 | 0.14 |
| YMR182C | 564.4635256 | 0.523827645 | 0.02620644 | 0.21 | 0.14 |
| YNL197C | 1704.51539 | -0.499719334 | 0.026235974 | 0.21 | 0.14 |
| YPL091W | 3000.977941 | 0.402218235 | 0.026301986 | 0.21 | 0.14 |
| YBR174C | 5.135026563 | 1.070377795 | 0.026340281 | 0.21 | 0.14 |
| YPR002W | 1192.998192 | 1.094394412 | 0.026503785 | 0.21 | 0.14 |
| YGR125W | 1519.849762 | 0.330180283 | 0.026563856 | 0.21 | 0.14 |
| YEL018W | 225.091547 | -0.3356745 | 0.026622872 | 0.21 | 0.14 |
| YNL066W | 1969.929236 | -0.574868962 | 0.026688912 | 0.21 | 0.14 |
| YBR198C | 1683.847961 | -0.376178992 | 0.026806149 | 0.21 | 0.14 |
| YIL054W | 8.069700908 | 1.132170141 | 0.027033171 | 0.21 | 0.14 |
| YML002W | 853.1962719 | 0.436149711 | 0.027071516 | 0.21 | 0.14 |
| YPL017C | 762.3634934 | 0.569349133 | 0.027088724 | 0.21 | 0.14 |
| YHR007C-A | 61.56889406 | 0.814433057 | 0.027166282 | 0.21 | 0.14 |
| YNL092W | 338.5129162 | 0.63808981 | 0.027156867 | 0.21 | 0.14 |
| YIL033C | 4130.82095 | 0.50370309 | 0.027201692 | 0.21 | 0.14 |
| YDL215C | 3694.89417 | 0.893175925 | 0.027252946 | 0.21 | 0.14 |
| YKL148C | 4538.261302 | 1.083592605 | 0.027493272 | 0.21 | 0.14 |
| YOR145C | 791.624005 | -0.587026529 | 0.02746644 | 0.21 | 0.14 |
| snR190 | 17.01701234 | 1.081353504 | 0.027555924 | 0.21 | 0.14 |
| YBR301W | 22.99326035 | 1.083372716 | 0.027942601 | 0.21 | 0.14 |
| YCR072C | 412.6699919 | -0.800235878 | 0.028076703 | 0.21 | 0.14 |
| YDR475C | 1828.530627 | 0.654941073 | 0.027917556 | 0.21 | 0.14 |
| YER054C | 1167.402398 | 0.958619233 | 0.027964764 | 0.21 | 0.14 |
| YER096W | 439.4636005 | 0.988284161 | 0.027987792 | 0.21 | 0.14 |
| YFR016C | 3711.04018 | 0.567328551 | 0.027986435 | 0.21 | 0.14 |
| YGL163C | 996.8369865 | 0.464904049 | 0.027788 | 0.21 | 0.14 |
| YGR219W | 32.06784529 | -0.604665435 | 0.028030902 | 0.21 | 0.14 |

| | | | | | |
|---|---|---|---|---|---|
| YJL077W-B | 26.1550574 | 0.842412944 | 0.028035699 | 0.21 | 0.14 |
| YLR070C | 346.2836033 | 0.773038909 | 0.027858083 | 0.21 | 0.14 |
| YMR058W | 5542.022572 | -0.850198634 | 0.027759506 | 0.21 | 0.14 |
| YOR135C | 405.7747912 | 0.501606906 | 0.028096801 | 0.21 | 0.14 |
| YJR035W | 878.6193365 | 0.517818478 | 0.028158647 | 0.21 | 0.14 |
| YDL185C-A | 100.139643 | 0.524444808 | 0.028250186 | 0.21 | 0.14 |
| YFR017C | 3258.918594 | 0.983478254 | 0.028265663 | 0.21 | 0.14 |
| YBR203W | 994.1971392 | 0.868779616 | 0.028308756 | 0.21 | 0.14 |
| YLR297W | 487.8704755 | 0.708359161 | 0.02837263 | 0.21 | 0.14 |
| YDL055C | 11798.61054 | -0.732182119 | 0.028415078 | 0.21 | 0.14 |
| YJL038C | 194.4642451 | 0.859971535 | 0.028529945 | 0.21 | 0.14 |
| YLR126C | 468.0585451 | -0.606032939 | 0.028533233 | 0.21 | 0.14 |
| YLR459W | 960.9364848 | -0.440004924 | 0.028522668 | 0.21 | 0.14 |
| YAL060W | 4666.556071 | 0.921126287 | 0.028612678 | 0.21 | 0.14 |
| YHR183W | 10217.98699 | 1.022544755 | 0.028615582 | 0.21 | 0.14 |
| YLR008C | 578.2897755 | -0.531980116 | 0.028688335 | 0.21 | 0.14 |
| YEL041W | 348.1575551 | 0.649937076 | 0.028743107 | 0.21 | 0.14 |
| YDL208W | 1913.168285 | -0.685756953 | 0.02879376 | 0.21 | 0.14 |
| YIL099W | 500.7923991 | 0.986758555 | 0.028833663 | 0.21 | 0.14 |
| YMR261C | 3042.037261 | 0.450841491 | 0.028893187 | 0.21 | 0.14 |
| YGR003W | 648.3625673 | 0.312023328 | 0.02893202 | 0.21 | 0.14 |
| YOR134W | 351.2595047 | 0.905014516 | 0.02901928 | 0.21 | 0.14 |
| YML109W | 760.9435844 | 0.431860629 | 0.029122692 | 0.22 | 0.14 |
| YDR261W-B | 58.90647804 | 0.979359886 | 0.029508938 | 0.22 | 0.14 |
| YER014W | 690.0676914 | 0.388661479 | 0.029517415 | 0.22 | 0.14 |
| YKL063C | 828.820505 | -0.396116271 | 0.029607597 | 0.22 | 0.14 |
| YPR015C | 179.9077993 | 0.803992052 | 0.029579353 | 0.22 | 0.14 |
| YER101C | 634.792482 | 0.576049438 | 0.029674831 | 0.22 | 0.14 |
| YCL055W | 639.3184492 | -0.686827839 | 0.029724474 | 0.22 | 0.14 |
| YNR061C | 596.8560089 | -0.528600856 | 0.029815276 | 0.22 | 0.14 |
| YDR250C | 6.490017442 | 0.945124263 | 0.029891098 | 0.22 | 0.14 |
| YKL018W | 496.5039326 | -0.354980058 | 0.029870906 | 0.22 | 0.14 |
| YER140W | 473.4616837 | -0.279562344 | 0.029926194 | 0.22 | 0.14 |
| YEL020W-A | 192.2562057 | -0.465161203 | 0.029977641 | 0.22 | 0.14 |
| YLR162W-A | 8.884728875 | 1.313651474 | 0.030120093 | 0.22 | 0.14 |
| YMR009W | 601.1334204 | 0.461238524 | 0.030101773 | 0.22 | 0.14 |
| YGL199C | 2.062524245 | 0.976818795 | 0.030198531 | 0.22 | 0.15 |
| YAL002W | 1569.483592 | 0.31337048 | 0.030329959 | 0.22 | 0.15 |
| YJR052W | 627.3978725 | 0.360804348 | 0.030356378 | 0.22 | 0.15 |
| YDL111C | 757.760422 | -0.468173138 | 0.030454533 | 0.22 | 0.15 |
| YDR374W-A | 251.03932 | -0.512246163 | 0.030661085 | 0.22 | 0.15 |
| YDR506C | 448.3958033 | 0.350158857 | 0.030711617 | 0.22 | 0.15 |
| YGL015C | 22.18286877 | 0.908746571 | 0.030801218 | 0.22 | 0.15 |

| | | | | | |
|---|---|---|---|---|---|
| YGR032W | 3378.619384 | 0.78756214 | 0.030699527 | 0.22 | 0.15 |
| YHR136C | 132.1966559 | -1.144336123 | 0.0304827 | 0.22 | 0.15 |
| YJR015W | 1958.669387 | -0.427570629 | 0.030589513 | 0.22 | 0.15 |
| YJR039W | 935.4373703 | 0.625393544 | 0.030641821 | 0.22 | 0.15 |
| YKL157W | 4702.190162 | 0.559877387 | 0.030612706 | 0.22 | 0.15 |
| YLR457C | 335.6948937 | 0.320883102 | 0.030742279 | 0.22 | 0.15 |
| YNL334C | 19.40823403 | -1.0035196 | 0.030568251 | 0.22 | 0.15 |
| YOL020W | 976.6345917 | -0.589800586 | 0.030627949 | 0.22 | 0.15 |
| YPL168W | 515.986609 | 0.577184557 | 0.030778635 | 0.22 | 0.15 |
| YBR242W | 549.7068913 | -0.391552406 | 0.030963217 | 0.22 | 0.15 |
| YGR059W | 83.10299922 | 1.277464149 | 0.030968941 | 0.22 | 0.15 |
| YGR153W | 193.310946 | 0.530430823 | 0.030925953 | 0.22 | 0.15 |
| YGR200C | 1330.804216 | -0.532810748 | 0.031008437 | 0.22 | 0.15 |
| YML034C-A | 23.30372078 | 0.738128377 | 0.0310192 | 0.22 | 0.15 |
| YOR237W | 107.6435638 | 0.920332207 | 0.03088776 | 0.22 | 0.15 |
| YGL071W | 1392.981634 | -0.447052361 | 0.031113124 | 0.22 | 0.15 |
| YOR166C | 666.3506234 | -0.287978345 | 0.031118981 | 0.22 | 0.15 |
| YER147C-A | 275.4373645 | -0.579676178 | 0.031164997 | 0.22 | 0.15 |
| YPL082C | 1537.0483 | -0.43587807 | 0.031242841 | 0.22 | 0.15 |
| YPR131C | 480.7629468 | -0.36409665 | 0.031266582 | 0.22 | 0.15 |
| YGL106W | 2381.873892 | -0.364997827 | 0.031325775 | 0.22 | 0.15 |
| YGL251C | 608.1790507 | 0.45477201 | 0.031380809 | 0.22 | 0.15 |
| YKL087C | 723.7436146 | 0.399331003 | 0.031449607 | 0.22 | 0.15 |
| YDR320W-B | 5.22193975 | 0.925042371 | 0.031539047 | 0.22 | 0.15 |
| YKL211C | 1647.230083 | 0.735171493 | 0.03154952 | 0.22 | 0.15 |
| YML085C | 4155.33263 | 0.411387127 | 0.031703857 | 0.22 | 0.15 |
| YIL121W | 2357.398488 | -1.011884798 | 0.031762614 | 0.22 | 0.15 |
| YKL042W | 561.3749184 | -0.546436992 | 0.031778144 | 0.22 | 0.15 |
| YLL042C | 71.42312394 | 0.654045525 | 0.031861774 | 0.22 | 0.15 |
| YCL037C | 1123.648124 | -0.886387983 | 0.031972874 | 0.22 | 0.15 |
| YCR083W | 821.7942376 | 0.722731632 | 0.032121621 | 0.22 | 0.15 |
| YGR155W | 6507.68049 | -0.423897471 | 0.032126313 | 0.22 | 0.15 |
| YGR206W | 264.0711736 | -0.41349281 | 0.032129223 | 0.22 | 0.15 |
| YKL156W | 2081.100672 | -0.785903785 | 0.032109467 | 0.22 | 0.15 |
| YGL117W | 750.1030158 | 0.918762403 | 0.03221107 | 0.22 | 0.15 |
| YGL091C | 545.7319922 | 0.246507416 | 0.032310623 | 0.22 | 0.15 |
| YJL107C | 129.4785502 | 0.832071272 | 0.032297128 | 0.22 | 0.15 |
| YER073W | 2130.104252 | 0.864869968 | 0.032437841 | 0.22 | 0.15 |
| YNL307C | 2792.714562 | -0.405828765 | 0.032431705 | 0.22 | 0.15 |
| YOL164W-A | 24.89009234 | 0.882789868 | 0.032449176 | 0.22 | 0.15 |
| YFL042C | 1495.188388 | 0.708372946 | 0.032593521 | 0.22 | 0.15 |
| YLR329W | 110.7708514 | 0.699158368 | 0.032509621 | 0.22 | 0.15 |
| YMR135W-A | 18.48988281 | 0.896772679 | 0.032545772 | 0.22 | 0.15 |

| | | | | | |
|---|---|---|---|---|---|
| YPR034W | 1306.898874 | -0.41549971 | 0.032562424 | 0.22 | 0.15 |
| YGR077C | 1152.926191 | -0.344603683 | 0.03263862 | 0.22 | 0.15 |
| YBR184W | 226.9887473 | 0.736447934 | 0.032884599 | 0.23 | 0.15 |
| YIL169C | 2177.35422 | -0.705335851 | 0.032838648 | 0.23 | 0.15 |
| YOR269W | 302.0905548 | -0.330646814 | 0.032902231 | 0.23 | 0.15 |
| YPL093W | 1935.182804 | -0.747763974 | 0.032931557 | 0.23 | 0.15 |
| YOR232W | 2857.566582 | -0.437434392 | 0.033112698 | 0.23 | 0.15 |
| YDR153C | 1161.100144 | -0.394116589 | 0.033179489 | 0.23 | 0.15 |
| YGR149W | 1082.04674 | 0.491554201 | 0.033310825 | 0.23 | 0.15 |
| YGR273C | 7.812625483 | 1.251436633 | 0.033315358 | 0.23 | 0.15 |
| YJL140W | 1600.235301 | -0.317988898 | 0.033302664 | 0.23 | 0.15 |
| YAL065C | 13.48444795 | 1.017820288 | 0.033370696 | 0.23 | 0.15 |
| YDR204W | 1297.475345 | 0.702227707 | 0.033424347 | 0.23 | 0.15 |
| YOL091W | 152.6270626 | 0.581480467 | 0.033408523 | 0.23 | 0.15 |
| YGL182C | 10.33516089 | 0.868218322 | 0.033532339 | 0.23 | 0.15 |
| YIL102C-A | 112.0491515 | -0.526064212 | 0.033543991 | 0.23 | 0.15 |
| YIL125W | 5872.779859 | 0.967684516 | 0.03350822 | 0.23 | 0.15 |
| YPL210C | 1902.73736 | -0.546724123 | 0.033635079 | 0.23 | 0.15 |
| YGR293C | 7.546648827 | -1.121448308 | 0.033809689 | 0.23 | 0.15 |
| YLR430W | 3065.658429 | 0.387228723 | 0.033795188 | 0.23 | 0.15 |
| YMR008C | 3679.477965 | 0.56097832 | 0.033786541 | 0.23 | 0.15 |
| YNL212W | 1282.46661 | 0.408005832 | 0.034125276 | 0.23 | 0.15 |
| YEL053W-A | 186.8810681 | -0.917626924 | 0.03425941 | 0.23 | 0.15 |
| YER176W | 1918.703807 | 0.412926253 | 0.034389266 | 0.23 | 0.15 |
| YGL233W | 1565.055775 | -0.269824013 | 0.034285901 | 0.23 | 0.15 |
| YGR022C | 1.81368062 | 1.05446004 | 0.034355142 | 0.23 | 0.15 |
| YGR134W | 745.1524745 | 0.394622982 | 0.034251416 | 0.23 | 0.15 |
| YPL267W | 353.8623989 | -0.606530959 | 0.034340177 | 0.23 | 0.15 |
| YHR151C | 595.1830514 | 0.55301653 | 0.034624727 | 0.23 | 0.15 |
| YMR137C | 598.7111524 | 0.471866937 | 0.034869672 | 0.23 | 0.15 |
| YBR280C | 1033.06933 | 0.758833889 | 0.035176316 | 0.23 | 0.15 |
| YDL049C | 168.7509482 | -0.581663939 | 0.035229297 | 0.23 | 0.15 |
| YIL032C | 13.12625335 | 1.264514384 | 0.035191898 | 0.23 | 0.15 |
| YJL129C | 2269.469864 | 0.279849036 | 0.035199925 | 0.23 | 0.15 |
| YNL111C | 246.1352445 | -0.766412015 | 0.035171022 | 0.23 | 0.15 |
| YOR094W | 410.9080628 | -0.437723726 | 0.035026492 | 0.23 | 0.15 |
| YOR364W | 7.913433737 | 0.938582921 | 0.035050011 | 0.23 | 0.15 |
| YBL028C | 410.9135571 | -0.708110066 | 0.035355057 | 0.23 | 0.15 |
| YDR391C | 1742.594469 | 0.696301393 | 0.035430776 | 0.23 | 0.15 |
| YGL179C | 645.0803373 | 0.965499283 | 0.035388805 | 0.23 | 0.15 |
| YKL047W | 711.1080868 | -0.350508364 | 0.035440122 | 0.23 | 0.15 |
| YPR075C | 1855.120278 | 0.45436807 | 0.035351014 | 0.23 | 0.15 |
| YDR376W | 985.9130091 | 0.384189289 | 0.035651591 | 0.23 | 0.15 |

| | | | | | |
|---|---|---|---|---|---|
| YEL015W | 1435.206957 | -0.287379606 | 0.03564643 | 0.23 | 0.15 |
| YOR011W-A | 3.164669454 | 1.361450714 | 0.035593212 | 0.23 | 0.15 |
| YOR086C | 3224.270398 | 0.342747762 | 0.035612811 | 0.23 | 0.15 |
| YDR194C | 2378.698014 | -0.412811769 | 0.035828471 | 0.24 | 0.16 |
| YPL047W | 255.7983457 | -0.391011145 | 0.035848264 | 0.24 | 0.16 |
| YBR234C | 4120.481034 | -0.304504665 | 0.03591533 | 0.24 | 0.16 |
| YIL158W | 651.6543059 | -0.723637533 | 0.036028556 | 0.24 | 0.16 |
| YDL226C | 1711.631594 | -0.524735203 | 0.036113027 | 0.24 | 0.16 |
| YLR146C | 511.7510501 | -0.393527587 | 0.036112849 | 0.24 | 0.16 |
| YGR073C | 7.437514438 | -0.842276219 | 0.036245498 | 0.24 | 0.16 |
| YOR034C-A | 45.66152085 | 0.659870887 | 0.036381349 | 0.24 | 0.16 |
| YDL002C | 564.8870678 | -0.291620377 | 0.036506431 | 0.24 | 0.16 |
| YDR327W | 614.5550695 | -0.333285659 | 0.036481586 | 0.24 | 0.16 |
| YGL192W | 1154.271063 | -0.698985451 | 0.036434639 | 0.24 | 0.16 |
| YEL070W | 298.2975483 | 1.066218916 | 0.03668674 | 0.24 | 0.16 |
| YGR162W | 4487.780728 | -0.578503216 | 0.036696531 | 0.24 | 0.16 |
| YGR184C | 3421.297226 | 0.283659479 | 0.036716088 | 0.24 | 0.16 |
| YHL046C | 9.174623277 | 1.087805291 | 0.036612221 | 0.24 | 0.16 |
| YKL009W | 1409.022835 | -0.809671643 | 0.036578877 | 0.24 | 0.16 |
| YDR303C | 1032.483515 | -0.514048099 | 0.036780991 | 0.24 | 0.16 |
| YOR199W | 28.95971662 | 0.835910316 | 0.03691516 | 0.24 | 0.16 |
| YBR146W | 1432.145995 | -0.351014169 | 0.037050118 | 0.24 | 0.16 |
| YDR516C | 6053.645479 | 0.768750369 | 0.037129886 | 0.24 | 0.16 |
| YHR099W | 4737.26123 | 0.552597132 | 0.037264042 | 0.24 | 0.16 |
| YJL122W | 643.9700151 | -0.824216455 | 0.037237135 | 0.24 | 0.16 |
| YLL066C | 1.397171575 | 1.367932228 | 0.037167493 | 0.24 | 0.16 |
| YNL196C | 204.2397635 | 0.924187922 | 0.03720771 | 0.24 | 0.16 |
| YOL081W | 5088.832739 | 0.601512322 | 0.03737587 | 0.24 | 0.16 |
| YPL078C | 4809.066893 | 0.70533701 | 0.03739775 | 0.24 | 0.16 |
| YPL119C-A | 58.40963327 | 0.854173996 | 0.037420226 | 0.24 | 0.16 |
| YPL167C | 937.9139202 | 0.440414542 | 0.037323648 | 0.24 | 0.16 |
| YPL192C | 128.9945052 | -0.647699871 | 0.037085475 | 0.24 | 0.16 |
| snR7-L | 6.909111604 | -0.989659321 | 0.037333651 | 0.24 | 0.16 |
| YDL150W | 525.2641532 | -0.527822299 | 0.037639957 | 0.24 | 0.16 |
| YAL015C | 634.3594781 | 0.507013083 | 0.037871485 | 0.24 | 0.16 |
| YBR079C | 5290.065687 | -0.564221107 | 0.037724503 | 0.24 | 0.16 |
| YBR239C | 441.6078035 | -0.360109187 | 0.037806791 | 0.24 | 0.16 |
| YGL231C | 1251.65783 | -0.318490247 | 0.037839815 | 0.24 | 0.16 |
| YHR028W-A | 38.81881491 | 0.58574394 | 0.037871697 | 0.24 | 0.16 |
| YKL118W | 29.53687639 | -0.537225145 | 0.037872726 | 0.24 | 0.16 |
| YJL164C | 1399.537088 | 0.460263904 | 0.038015102 | 0.24 | 0.16 |
| YHR114W | 1802.212821 | 0.262163847 | 0.03823969 | 0.24 | 0.16 |
| YKL027W | 701.3136707 | 0.588287204 | 0.038265707 | 0.24 | 0.16 |

| | | | | | |
|---|---|---|---|---|---|
| YDR073W | 640.4950618 | -0.582484867 | 0.038623364 | 0.24 | 0.16 |
| YOL093W | 662.3279101 | -0.520921294 | 0.038534392 | 0.24 | 0.16 |
| YOL115W | 646.4767705 | -0.481268031 | 0.038511726 | 0.24 | 0.16 |
| YOL123W | 4114.119417 | -0.586334883 | 0.038471537 | 0.24 | 0.16 |
| YOR072W-A | 14.05662092 | 0.808729852 | 0.03860748 | 0.24 | 0.16 |
| YPL043W | 1057.06526 | -0.657968519 | 0.038568074 | 0.24 | 0.16 |
| YMR001C | 1386.236499 | -0.415191688 | 0.03880732 | 0.24 | 0.16 |
| YPL247C | 2827.171119 | 0.854541644 | 0.038921706 | 0.24 | 0.16 |
| YOR354C | 1688.06875 | -0.273451128 | 0.03901319 | 0.24 | 0.16 |
| YDR245W | 1164.324708 | -0.511286093 | 0.039160621 | 0.25 | 0.16 |
| YIL046W-A | 7.695578865 | 1.280244595 | 0.039249676 | 0.25 | 0.16 |
| YLL056C | 287.6766171 | 0.65804838 | 0.039270402 | 0.25 | 0.16 |
| YOL010W | 524.4137956 | -0.63900155 | 0.039314956 | 0.25 | 0.16 |
| YKL122C | 1563.613747 | -0.529896435 | 0.039401193 | 0.25 | 0.16 |
| YDL168W | 1463.24926 | 0.554608411 | 0.039596228 | 0.25 | 0.16 |
| YHR177W | 301.5515693 | 0.560499808 | 0.039741761 | 0.25 | 0.16 |
| YIL009W | 578.8586392 | -0.605234041 | 0.039796069 | 0.25 | 0.16 |
| YHR155W | 1225.572051 | 0.269142456 | 0.039844036 | 0.25 | 0.16 |
| YCR043C | 438.4206899 | -0.54155613 | 0.040153743 | 0.25 | 0.16 |
| YER107C | 624.5620743 | -0.514083997 | 0.040180217 | 0.25 | 0.16 |
| YGR194C | 2007.303758 | 0.594709858 | 0.04008878 | 0.25 | 0.16 |
| YHR047C | 1885.311487 | 0.593528708 | 0.040112201 | 0.25 | 0.16 |
| YKL217W | 2399.793467 | 1.073240997 | 0.040070805 | 0.25 | 0.16 |
| YLR248W | 3268.331309 | 0.483918834 | 0.039937127 | 0.25 | 0.16 |
| YOR025W | 737.1839475 | -0.44373214 | 0.040069657 | 0.25 | 0.16 |
| YPL215W | 1126.607522 | -0.376277146 | 0.040095794 | 0.25 | 0.16 |
| YNL271C | 3740.617123 | 0.371456546 | 0.040275311 | 0.25 | 0.16 |
| YFL016C | 3333.34381 | 0.479356426 | 0.040369317 | 0.25 | 0.16 |
| YKL035W | 11294.21789 | 0.479531451 | 0.040347891 | 0.25 | 0.16 |
| YLR224W | 1026.056679 | -0.453403014 | 0.040591114 | 0.25 | 0.16 |
| YMR091C | 998.5083437 | -0.450062609 | 0.040600635 | 0.25 | 0.16 |
| YPR041W | 3343.881383 | -0.541974439 | 0.040532367 | 0.25 | 0.16 |
| YML043C | 371.5090553 | -0.846697451 | 0.040703221 | 0.25 | 0.16 |
| YMR043W | 2143.509763 | -0.488470576 | 0.040800063 | 0.25 | 0.16 |
| YOR297C | 445.9780063 | -0.374656633 | 0.040848518 | 0.25 | 0.16 |
| YER074W-A | 546.9637993 | -0.325261171 | 0.040918052 | 0.25 | 0.17 |
| YJR148W | 6070.611654 | 0.663299669 | 0.041012248 | 0.25 | 0.17 |
| YNL216W | 1187.634305 | -0.372248775 | 0.041074837 | 0.25 | 0.17 |
| YFL030W | 1073.611575 | 0.982565592 | 0.041209211 | 0.25 | 0.17 |
| YOR045W | 622.8710581 | -0.529767165 | 0.041195133 | 0.25 | 0.17 |
| YDR241W | 32.77525977 | -0.65801247 | 0.041368498 | 0.25 | 0.17 |
| YHR017W | 1256.836076 | 0.472899656 | 0.041430602 | 0.25 | 0.17 |
| YIL022W | 2268.335855 | -0.218682298 | 0.041401477 | 0.25 | 0.17 |

| | | | | | |
|---|---|---|---|---|---|
| YLR176C | 916.0927001 | 0.402565762 | 0.041336269 | 0.25 | 0.17 |
| YBL100W-B | 175.5232464 | 0.906953477 | 0.041731921 | 0.25 | 0.17 |
| YGL133W | 1877.257774 | 0.357174663 | 0.041632053 | 0.25 | 0.17 |
| YJL101C | 2271.091311 | 0.37135115 | 0.041615871 | 0.25 | 0.17 |
| YJL144W | 785.7166274 | 0.783216303 | 0.041620667 | 0.25 | 0.17 |
| YJR042W | 1389.72183 | -0.35736157 | 0.041739102 | 0.25 | 0.17 |
| YKL112W | 917.1897974 | -0.451911191 | 0.04171917 | 0.25 | 0.17 |
| YMR133W | 502.5352221 | 0.65774853 | 0.041629036 | 0.25 | 0.17 |
| YBL099W | 11787.99535 | 0.852863062 | 0.041822073 | 0.25 | 0.17 |
| YGR019W | 2682.062979 | 0.743168163 | 0.041856795 | 0.25 | 0.17 |
| YDR410C | 388.474414 | -0.788968214 | 0.041917167 | 0.25 | 0.17 |
| YLR234W | 283.065063 | 0.573428915 | 0.041974963 | 0.25 | 0.17 |
| YCL026C-A | 44.13033287 | 0.746892899 | 0.042016967 | 0.25 | 0.17 |
| YDR102C | 12.68433027 | -0.80610687 | 0.042152074 | 0.25 | 0.17 |
| YLR215C | 433.3667117 | -0.307344335 | 0.04212793 | 0.25 | 0.17 |
| YLR221C | 955.7495307 | -0.506390204 | 0.042090277 | 0.25 | 0.17 |
| YJL113W | 119.706187 | 0.639513331 | 0.04228401 | 0.25 | 0.17 |
| YBR251W | 1030.356421 | -0.42851692 | 0.042361627 | 0.25 | 0.17 |
| YOR282W | 14.73409586 | -0.756958686 | 0.042482998 | 0.25 | 0.17 |
| YDL149W | 944.7875532 | 0.562948372 | 0.042528226 | 0.25 | 0.17 |
| YJR155W | 885.0501737 | 0.883049028 | 0.042600433 | 0.25 | 0.17 |
| YPL204W | 2957.690828 | -0.289042775 | 0.042598032 | 0.25 | 0.17 |
| YDR001C | 5262.491506 | 0.516586963 | 0.042846097 | 0.25 | 0.17 |
| YFL017C | 717.9144755 | -0.337196511 | 0.042840767 | 0.25 | 0.17 |
| YGR227W | 1061.56859 | -0.475815101 | 0.042732301 | 0.25 | 0.17 |
| YLR324W | 1737.806437 | 0.493646464 | 0.042808838 | 0.25 | 0.17 |
| YKL119C | 426.0044347 | -0.406839327 | 0.042983657 | 0.25 | 0.17 |
| snR17a | 59.62016603 | 0.92500983 | 0.043086017 | 0.25 | 0.17 |
| YGL227W | 1564.861142 | 0.570107747 | 0.043361866 | 0.26 | 0.17 |
| YJR008W | 1161.824512 | 0.844318594 | 0.043287457 | 0.26 | 0.17 |
| YKL059C | 865.3974506 | -0.264574231 | 0.043328437 | 0.26 | 0.17 |
| YKR057W | 2516.350053 | -0.748308324 | 0.043284533 | 0.26 | 0.17 |
| snR17b | 27.76593566 | 1.087340006 | 0.043237753 | 0.26 | 0.17 |
| YDR228C | 901.0372572 | -0.357449808 | 0.043468071 | 0.26 | 0.17 |
| YLR177W | 3573.62477 | 0.627916673 | 0.043512817 | 0.26 | 0.17 |
| YBL020W | 997.2826168 | 0.361684962 | 0.043612643 | 0.26 | 0.17 |
| YOR208W | 1224.294967 | 0.592520665 | 0.043589828 | 0.26 | 0.17 |
| YDR253C | 229.9334863 | 0.435838596 | 0.043777458 | 0.26 | 0.17 |
| YGL104C | 947.9922368 | 0.557556232 | 0.043790705 | 0.26 | 0.17 |
| YGL127C | 488.4323072 | -0.472612566 | 0.04382457 | 0.26 | 0.17 |
| YOR281C | 1009.23062 | -0.471010799 | 0.043771964 | 0.26 | 0.17 |
| YGR187C | 552.0431833 | -0.666791011 | 0.043939988 | 0.26 | 0.17 |
| YBR204C | 1064.479878 | 0.454415789 | 0.044210994 | 0.26 | 0.17 |

| | | | | | |
|---|---|---|---|---|---|
| YEL074W | 35.0602757 | 0.996275696 | 0.044176442 | 0.26 | 0.17 |
| YPL063W | 2649.060944 | -0.276782139 | 0.04416149 | 0.26 | 0.17 |
| YOR264W | 932.8104963 | -0.424409591 | 0.044335038 | 0.26 | 0.17 |
| YER152C | 1264.809581 | 0.557712602 | 0.044404776 | 0.26 | 0.17 |
| YDR309C | 2613.4939 | -0.760228774 | 0.044468885 | 0.26 | 0.17 |
| YIR030W-A | 17.57088172 | 0.988470692 | 0.044633189 | 0.26 | 0.17 |
| YLR037C | 167.7921303 | 0.634469703 | 0.044618695 | 0.26 | 0.17 |
| YLR173W | 824.1208493 | 0.597277244 | 0.044702449 | 0.26 | 0.17 |
| YLR260W | 983.6383575 | 0.486337202 | 0.044782846 | 0.26 | 0.17 |
| YMR281W | 530.4325943 | -0.419607376 | 0.044753122 | 0.26 | 0.17 |
| YDL078C | 3642.136284 | 0.563682031 | 0.044866647 | 0.26 | 0.17 |
| YKR085C | 837.2824572 | -0.415409617 | 0.044850812 | 0.26 | 0.17 |
| YDR181C | 744.5427058 | 0.520548407 | 0.045005388 | 0.26 | 0.17 |
| YGL258W | 52.00397703 | 0.805348869 | 0.044969994 | 0.26 | 0.17 |
| YNL002C | 1677.703778 | -0.638089703 | 0.04507572 | 0.26 | 0.17 |
| YGR254W | 35005.12746 | 0.86547966 | 0.045485444 | 0.26 | 0.17 |
| YNL153C | 847.6509295 | -0.543413751 | 0.045463283 | 0.26 | 0.17 |
| YNL154C | 2670.283788 | -0.459624494 | 0.045428314 | 0.26 | 0.17 |
| YOL139C | 5554.572319 | -0.582510471 | 0.045452398 | 0.26 | 0.17 |
| YNL141W | 1283.346108 | -0.821091728 | 0.045573813 | 0.26 | 0.17 |
| YJR070C | 1785.660344 | -0.756485694 | 0.045882849 | 0.26 | 0.17 |
| YML046W | 728.0597624 | -0.422653194 | 0.04588891 | 0.26 | 0.17 |
| YER106W | 34.98872113 | 0.978291161 | 0.045969927 | 0.26 | 0.17 |
| YNL071W | 6818.92242 | 0.347211568 | 0.045937711 | 0.26 | 0.17 |
| YJR121W | 23130.92602 | 0.880837019 | 0.046042298 | 0.26 | 0.17 |
| YDR127W | 3620.75176 | 0.616019164 | 0.04609833 | 0.26 | 0.17 |
| YHR179W | 8288.962491 | 0.517951471 | 0.046324844 | 0.26 | 0.17 |
| YLR207W | 1228.065049 | 0.338570453 | 0.046328872 | 0.26 | 0.17 |
| YNL306W | 1534.453742 | -0.33057152 | 0.046285606 | 0.26 | 0.17 |
| YBR138C | 569.4593536 | 0.369704954 | 0.046424142 | 0.26 | 0.17 |
| YDR014W | 486.9454278 | 0.331159565 | 0.046421035 | 0.26 | 0.17 |
| YIR021W | 261.891268 | -0.366958645 | 0.046582486 | 0.26 | 0.17 |
| YLR285W | 889.692098 | -0.537751637 | 0.046648655 | 0.26 | 0.17 |
| YOL162W | 65.72006374 | 0.902811204 | 0.046615819 | 0.26 | 0.17 |
| YGR244C | 5502.307672 | 0.585246169 | 0.046791057 | 0.26 | 0.17 |
| YGR286C | 924.5538048 | 0.743874645 | 0.046864643 | 0.26 | 0.17 |
| YJL106W | 294.2742704 | 0.570540719 | 0.046864595 | 0.26 | 0.17 |
| YOR357C | 469.3315493 | -0.313649133 | 0.046773 | 0.26 | 0.17 |
| YDR052C | 1279.720929 | -0.329813312 | 0.046963619 | 0.26 | 0.17 |
| YJR140C | 1086.888351 | 0.450136126 | 0.047073652 | 0.26 | 0.18 |
| YHR214W | 18.69123236 | 0.89672413 | 0.047126025 | 0.27 | 0.18 |
| YDR370C | 533.8393634 | -0.342016134 | 0.04717713 | 0.27 | 0.18 |
| YEL012W | 1045.847779 | 0.630015933 | 0.047539061 | 0.27 | 0.18 |

| | | | | | |
|---|---|---|---|---|---|
| YPL060C-A | 3.51919416 | 1.0661384 | 0.047590758 | 0.27 | 0.18 |
| YMR093W | 713.9051908 | -0.613510834 | 0.04765236 | 0.27 | 0.18 |
| YAR050W | 236.1195337 | 0.803661759 | 0.047905146 | 0.27 | 0.18 |
| YCR004C | 3730.565062 | -0.360080598 | 0.047838075 | 0.27 | 0.18 |
| YDR077W | 88090.2349 | 0.663951442 | 0.047782393 | 0.27 | 0.18 |
| YDR496C | 1668.42234 | -0.648601935 | 0.047897798 | 0.27 | 0.18 |
| YGR052W | 642.70953 | -0.969786735 | 0.047826773 | 0.27 | 0.18 |
| YOR309C | 407.1266153 | -0.849348786 | 0.04780398 | 0.27 | 0.18 |
| YJL063C | 928.728548 | -0.445674981 | 0.047960425 | 0.27 | 0.18 |
| snR38 | 6.274197792 | 1.183991736 | 0.048176324 | 0.27 | 0.18 |
| YGL006W | 3158.91505 | 0.423950626 | 0.048288453 | 0.27 | 0.18 |
| YDR461W | 1830.189511 | -1.191892885 | 0.04841025 | 0.27 | 0.18 |
| YJL012C | 2507.923216 | -0.607791838 | 0.048435375 | 0.27 | 0.18 |
| YKR060W | 417.425685 | -0.607048001 | 0.048436716 | 0.27 | 0.18 |
| YKL086W | 218.7788586 | 0.663535251 | 0.048488607 | 0.27 | 0.18 |
| YCR011C | 2808.118139 | 0.35382187 | 0.048661859 | 0.27 | 0.18 |
| YOR285W | 4227.371531 | 0.595892885 | 0.048646413 | 0.27 | 0.18 |
| YPL122C | 549.6598792 | -0.267273454 | 0.048701477 | 0.27 | 0.18 |
| YDL110C | 2473.385616 | 0.603375604 | 0.048836708 | 0.27 | 0.18 |
| YDL243C | 368.4938618 | 0.83761171 | 0.048931767 | 0.27 | 0.18 |
| YMR245W | 80.78367086 | 0.634614496 | 0.049035549 | 0.27 | 0.18 |
| YDR293C | 6400.531719 | 0.454470189 | 0.049082748 | 0.27 | 0.18 |
| YDL239C | 522.0657782 | 0.407144896 | 0.049184124 | 0.27 | 0.18 |
| YBR092C | 2232.790112 | -0.905774685 | 0.049473561 | 0.27 | 0.18 |
| YER186C | 678.5275475 | -0.491690391 | 0.049485039 | 0.27 | 0.18 |
| YLR120W-A | 22.81140064 | 0.751494278 | 0.049262116 | 0.27 | 0.18 |
| YLR130C | 1253.100793 | -0.444826792 | 0.049459054 | 0.27 | 0.18 |
| YLR299W | 1242.489768 | 0.489180397 | 0.049334308 | 0.27 | 0.18 |
| YPL037C | 6009.036088 | -0.499286167 | 0.049336466 | 0.27 | 0.18 |
| YPL189W | 119.6862802 | 0.478468356 | 0.049440312 | 0.27 | 0.18 |
| YML029W | 1034.498713 | 0.306087518 | 0.049558423 | 0.27 | 0.18 |
| YKL208W | 299.0743362 | -0.384011395 | 0.049708045 | 0.27 | 0.18 |
| YDR500C | 6683.941473 | -0.64475414 | 0.049781772 | 0.27 | 0.18 |
| YGL170C | 69.87350446 | 0.938226988 | 0.049860705 | 0.27 | 0.18 |
| YOR077W | 225.6560466 | 0.413251748 | 0.049843142 | 0.27 | 0.18 |
| YCR005C | 6272.799933 | 0.808468926 | 0.050079734 | 0.27 | 0.18 |
| YEL045C | 16.86688846 | 0.772552583 | 0.050068826 | 0.27 | 0.18 |
| YKR071C | 2094.595777 | 0.424644526 | 0.050065972 | 0.27 | 0.18 |
| YDR320C-A | 373.904445 | -0.445088421 | 0.050190459 | 0.27 | 0.18 |
| YJL060W | 1402.078784 | 0.420857568 | 0.050218425 | 0.27 | 0.18 |
| snR54 | 8.052861878 | 1.113078136 | 0.0502051 | 0.27 | 0.18 |
| YHR118C | 526.5681517 | 0.224237994 | 0.050375901 | 0.27 | 0.18 |
| YNL205C | 11.3162081 | 0.893499483 | 0.050372532 | 0.27 | 0.18 |

| | | | | | |
|---|---|---|---|---|---|
| YDL124W | 12433.87204 | 0.581581901 | 0.05056098 | 0.27 | 0.18 |
| YDL126C | 11516.69474 | 0.356125736 | 0.050603826 | 0.27 | 0.18 |
| YJL174W | 2482.371966 | 0.272133462 | 0.050505226 | 0.27 | 0.18 |
| YLR054C | 138.716349 | 0.915402015 | 0.050541965 | 0.27 | 0.18 |
| YNR063W | 178.9442947 | 0.518747064 | 0.050591175 | 0.27 | 0.18 |
| YPR150W | 6.435730199 | 0.940073826 | 0.050737975 | 0.27 | 0.18 |
| YER128W | 281.9899223 | 0.397240004 | 0.051096905 | 0.27 | 0.18 |
| YJL105W | 247.8507967 | -0.559239539 | 0.051059678 | 0.27 | 0.18 |
| YMR078C | 885.7332334 | -0.310966024 | 0.051048411 | 0.27 | 0.18 |
| YDL021W | 1151.606846 | 0.983216355 | 0.051247029 | 0.28 | 0.18 |
| YMR279C | 707.3104958 | 0.31018009 | 0.051274177 | 0.28 | 0.18 |
| YFR047C | 2554.161875 | 0.766198439 | 0.051519676 | 0.28 | 0.18 |
| YKL147C | 17.64954721 | 0.935138951 | 0.051490773 | 0.28 | 0.18 |
| YOR390W | 45.23234325 | -0.652507309 | 0.051570195 | 0.28 | 0.18 |
| YKL072W | 554.6822525 | 0.319930746 | 0.052188713 | 0.28 | 0.18 |
| YNL041C | 1185.16925 | -0.294385709 | 0.052218641 | 0.28 | 0.18 |
| YNL053W | 544.6626736 | -0.32751597 | 0.052358144 | 0.28 | 0.18 |
| YAL025C | 983.4578298 | -0.711789328 | 0.052463399 | 0.28 | 0.18 |
| YDL066W | 3505.644872 | 0.857091628 | 0.052489896 | 0.28 | 0.18 |
| YNL220W | 4979.508941 | 0.652700146 | 0.052418161 | 0.28 | 0.18 |
| YDL166C | 434.2604045 | -0.504084752 | 0.052550988 | 0.28 | 0.18 |
| YER068C-A | 10.30512685 | 0.994327792 | 0.052788124 | 0.28 | 0.19 |
| YBR206W | 255.7378889 | -0.389247946 | 0.052851428 | 0.28 | 0.19 |
| YDL094C | 21.86703582 | 0.772529317 | 0.052901155 | 0.28 | 0.19 |
| YOL105C | 760.4552602 | 0.576552003 | 0.053100843 | 0.28 | 0.19 |
| YLR454W | 3660.437031 | 0.414279014 | 0.05320572 | 0.28 | 0.19 |
| YPL243W | 2085.694288 | -0.465272605 | 0.053243272 | 0.28 | 0.19 |
| YPR154W | 2486.651934 | 0.375072796 | 0.053198376 | 0.28 | 0.19 |
| YLR121C | 1254.397824 | 0.498374571 | 0.053290333 | 0.28 | 0.19 |
| YBL054W | 675.0753311 | -0.708786373 | 0.053377031 | 0.28 | 0.19 |
| YGR090W | 2037.446504 | -0.611183905 | 0.053513185 | 0.28 | 0.19 |
| YBL036C | 1223.509384 | -0.319163854 | 0.053910309 | 0.28 | 0.19 |
| YDL161W | 1895.847844 | -0.379382354 | 0.054022163 | 0.28 | 0.19 |
| YEL024W | 1858.219132 | 0.800723234 | 0.054123112 | 0.28 | 0.19 |
| YFL017W-A | 221.0675394 | -0.457277139 | 0.054246558 | 0.28 | 0.19 |
| YGR123C | 655.6704403 | -0.738192144 | 0.054162001 | 0.28 | 0.19 |
| YHR140W | 190.2878906 | 0.769795857 | 0.053855269 | 0.28 | 0.19 |
| YKL067W | 3440.072656 | 0.747940142 | 0.053824176 | 0.28 | 0.19 |
| YKL203C | 2057.606082 | 0.334887348 | 0.053714301 | 0.28 | 0.19 |
| YML075C | 1371.306293 | -0.308883264 | 0.053860598 | 0.28 | 0.19 |
| YMR235C | 2269.617315 | -0.567045118 | 0.053774776 | 0.28 | 0.19 |
| YMR315W | 4197.096296 | 0.632440685 | 0.05412602 | 0.28 | 0.19 |
| YOR137C | 579.6398816 | 0.52672026 | 0.054040968 | 0.28 | 0.19 |

| | | | | | |
|---|---|---|---|---|---|
| YOR361C | 4132.636454 | -0.486355561 | 0.054223407 | 0.28 | 0.19 |
| YPL184C | 1810.247002 | -0.463131596 | 0.054248637 | 0.28 | 0.19 |
| YPL006W | 1523.461434 | 0.478710979 | 0.054365678 | 0.28 | 0.19 |
| YKL161C | 387.0663817 | -0.660604054 | 0.05461248 | 0.28 | 0.19 |
| YLL041C | 3315.150059 | 0.799926646 | 0.054573989 | 0.28 | 0.19 |
| YMR162C | 1643.610674 | 0.399945955 | 0.054620221 | 0.28 | 0.19 |
| YIL136W | 7495.331143 | 1.013480826 | 0.054672571 | 0.28 | 0.19 |
| YPL096C-A | 145.3313639 | -0.489684258 | 0.054789707 | 0.28 | 0.19 |
| YER179W | 293.0579118 | 0.67102334 | 0.054938249 | 0.28 | 0.19 |
| YPR058W | 588.0067957 | 0.537475478 | 0.054899438 | 0.28 | 0.19 |
| YBR271W | 387.9475512 | -0.398264222 | 0.055158576 | 0.28 | 0.19 |
| YDR310C | 1697.719061 | -0.297838149 | 0.055220174 | 0.28 | 0.19 |
| YDR436W | 1255.862426 | 0.482473179 | 0.055156791 | 0.28 | 0.19 |
| YGL202W | 6879.756501 | 0.462054345 | 0.055269605 | 0.28 | 0.19 |
| YKR026C | 956.8364878 | -0.48650075 | 0.055305451 | 0.28 | 0.19 |
| YMR266W | 1578.35636 | -0.389178348 | 0.055162074 | 0.28 | 0.19 |
| YNL014W | 784.7565686 | 0.806671121 | 0.055291779 | 0.28 | 0.19 |
| YOR307C | 1226.108407 | -0.379944474 | 0.055260432 | 0.28 | 0.19 |
| RUF23 | 8.975541829 | 0.949636055 | 0.055906852 | 0.28 | 0.19 |
| YAR002W | 1776.649309 | -0.434703087 | 0.056000602 | 0.28 | 0.19 |
| YDL101C | 860.843029 | 0.438442445 | 0.055863325 | 0.28 | 0.19 |
| YDR101C | 1247.120699 | -0.718460533 | 0.055879395 | 0.28 | 0.19 |
| YGL042C | 343.2123261 | -0.437572906 | 0.055887921 | 0.28 | 0.19 |
| YGL144C | 301.1617021 | 0.432101956 | 0.055856512 | 0.28 | 0.19 |
| YGR270W | 2956.664275 | 0.277053957 | 0.055460881 | 0.28 | 0.19 |
| YHR082C | 4569.103984 | 0.476346636 | 0.055385189 | 0.28 | 0.19 |
| YHR092C | 9528.876224 | 0.895287722 | 0.056012732 | 0.28 | 0.19 |
| YIL168W | 92.19596636 | 0.576608007 | 0.056019451 | 0.28 | 0.19 |
| YJL034W | 12765.00653 | 0.675464553 | 0.055827504 | 0.28 | 0.19 |
| YJR018W | 57.57815418 | -0.54259429 | 0.055723342 | 0.28 | 0.19 |
| YLR328W | 869.5602715 | -0.482052605 | 0.055936703 | 0.28 | 0.19 |
| YMR166C | 428.7049362 | -0.24180459 | 0.055750104 | 0.28 | 0.19 |
| YMR196W | 3241.463398 | 0.724970679 | 0.056024661 | 0.28 | 0.19 |
| YPR067W | 664.4484323 | -0.259887082 | 0.0555099 | 0.28 | 0.19 |
| YJL103C | 874.1846601 | 0.588261902 | 0.056297242 | 0.29 | 0.19 |
| YBL016W | 1090.490568 | -1.032288576 | 0.056552765 | 0.29 | 0.19 |
| YDR239C | 796.3576703 | 0.283419565 | 0.056652356 | 0.29 | 0.19 |
| YKR042W | 18766.32248 | -0.529087891 | 0.056577325 | 0.29 | 0.19 |
| YLR265C | 441.14354 | -0.951255105 | 0.056649544 | 0.29 | 0.19 |
| YLR393W | 270.1892054 | 0.464526993 | 0.056512099 | 0.29 | 0.19 |
| YMR209C | 810.1729969 | -0.447734817 | 0.05660399 | 0.29 | 0.19 |
| YGL037C | 14451.53815 | 0.510404261 | 0.056944335 | 0.29 | 0.19 |
| YJR150C | 147.0716932 | 0.98369033 | 0.057020868 | 0.29 | 0.19 |

| | | | | | |
|---|---|---|---|---|---|
| YFL051C | 87.84562174 | 0.904782101 | 0.057328823 | 0.29 | 0.19 |
| YHR037W | 3383.544 | 0.443442757 | 0.057257006 | 0.29 | 0.19 |
| YJL115W | 931.9259287 | -0.485744178 | 0.057173899 | 0.29 | 0.19 |
| YKL107W | 249.2824796 | 0.848095753 | 0.057168959 | 0.29 | 0.19 |
| YLR092W | 690.3329885 | 0.554280633 | 0.057250069 | 0.29 | 0.19 |
| YOR098C | 2501.292329 | -0.354560232 | 0.057302537 | 0.29 | 0.19 |
| YOL088C | 665.7794028 | -0.400450144 | 0.05750099 | 0.29 | 0.19 |
| YDL130W-A | 2765.774643 | 0.778898232 | 0.05759118 | 0.29 | 0.19 |
| YNL325C | 453.4130958 | 0.424175119 | 0.057753049 | 0.29 | 0.19 |
| YPR170C | 83.5281119 | -0.439573207 | 0.057714549 | 0.29 | 0.19 |
| YPL155C | 690.291389 | 0.397477325 | 0.057862403 | 0.29 | 0.19 |
| YBR043C | 1241.74171 | 0.671267607 | 0.05795655 | 0.29 | 0.19 |
| YHR158C | 1689.928736 | 0.382710483 | 0.0581233 | 0.29 | 0.19 |
| YDR487C | 2644.625402 | 0.307766181 | 0.05832088 | 0.29 | 0.19 |
| YLR023C | 1130.122098 | 0.320047401 | 0.058435441 | 0.29 | 0.19 |
| YML027W | 523.9520569 | -0.72026787 | 0.058440405 | 0.29 | 0.19 |
| YNL323W | 681.0384119 | -0.298353679 | 0.058459946 | 0.29 | 0.19 |
| YPR047W | 774.0936392 | 0.330530059 | 0.058501187 | 0.29 | 0.19 |
| YLR425W | 1120.132672 | 0.316703031 | 0.058812247 | 0.29 | 0.19 |
| YGR267C | 1681.678228 | 0.252521715 | 0.058883325 | 0.29 | 0.19 |
| YBR076W | 114.1570412 | 0.883478908 | 0.059021725 | 0.29 | 0.19 |
| YER052C | 2274.895414 | 0.822000974 | 0.058936644 | 0.29 | 0.19 |
| YHR143W-A | 554.7529102 | -0.480838711 | 0.058974618 | 0.29 | 0.19 |
| YJL209W | 626.0794008 | -0.308937959 | 0.059064411 | 0.29 | 0.19 |
| YCR028C-A | 1716.279601 | -0.425055286 | 0.059370885 | 0.29 | 0.19 |
| YJL095W | 1317.430087 | 0.268852435 | 0.059370621 | 0.29 | 0.19 |
| YPL050C | 1693.46287 | -0.463405366 | 0.059648926 | 0.30 | 0.19 |
| YKR092C | 1945.135383 | -0.590107379 | 0.05983584 | 0.30 | 0.20 |
| YMR322C | 22.28920933 | 1.067089713 | 0.059844573 | 0.30 | 0.20 |
| YGR054W | 2529.613046 | -0.527419656 | 0.059990549 | 0.30 | 0.20 |
| YMR194W | 1484.319594 | -0.630094343 | 0.060024533 | 0.30 | 0.20 |
| YNL007C | 10095.98848 | 0.442586754 | 0.05993688 | 0.30 | 0.20 |
| YLR289W | 462.7410856 | 0.478552817 | 0.060144767 | 0.30 | 0.20 |
| YLR453C | 293.5520271 | 0.336299344 | 0.060180883 | 0.30 | 0.20 |
| YDR298C | 3635.360191 | 0.778049208 | 0.060392063 | 0.30 | 0.20 |
| YGR213C | 279.0368426 | 0.56282365 | 0.060383878 | 0.30 | 0.20 |
| YIR044C | 6.331727114 | 0.996961423 | 0.060480882 | 0.30 | 0.20 |
| YDL152W | 5.390111654 | -1.05244099 | 0.06066341 | 0.30 | 0.20 |
| YNL260C | 268.8863879 | -0.296536944 | 0.060638267 | 0.30 | 0.20 |
| YDR080W | 1103.850689 | 0.416755437 | 0.060909789 | 0.30 | 0.20 |
| YBR033W | 734.5856264 | 0.821192825 | 0.061137829 | 0.30 | 0.20 |
| YPL014W | 2476.516022 | 0.849322608 | 0.061218047 | 0.30 | 0.20 |
| YBL019W | 515.3822708 | 0.424516732 | 0.061321862 | 0.30 | 0.20 |

| | | | | | |
|---|---|---|---|---|---|
| YBL098W | 1032.627619 | 1.054316007 | 0.0614076 | 0.30 | 0.20 |
| YPR177C | 1.844393858 | 1.08060742 | 0.061433021 | 0.30 | 0.20 |
| YLR229C | 1318.158779 | -0.339653998 | 0.061508011 | 0.30 | 0.20 |
| YKL218C | 773.1891416 | 0.674924176 | 0.06160554 | 0.30 | 0.20 |
| YNR070W | 728.5997471 | 0.523060644 | 0.06156657 | 0.30 | 0.20 |
| YBR126C | 9994.323143 | 0.597242293 | 0.06186184 | 0.30 | 0.20 |
| YLR190W | 1584.336567 | 0.496680665 | 0.062219037 | 0.30 | 0.20 |
| YLR091W | 340.1364791 | -0.299070138 | 0.062288579 | 0.30 | 0.20 |
| YJL185C | 564.2478549 | 0.635756703 | 0.062452567 | 0.30 | 0.20 |
| YOR227W | 3483.244777 | 0.469081804 | 0.062483696 | 0.30 | 0.20 |
| YPR122W | 1503.146832 | -0.81519222 | 0.062393011 | 0.30 | 0.20 |
| YDR451C | 513.5586606 | -0.538976562 | 0.062615943 | 0.30 | 0.20 |
| YER083C | 1868.758699 | -0.253963372 | 0.062875242 | 0.30 | 0.20 |
| YER123W | 1029.367215 | -0.311470731 | 0.062914026 | 0.30 | 0.20 |
| YIR027C | 227.1514133 | 1.088309266 | 0.062903026 | 0.30 | 0.20 |
| YLL066W-B | 2.220126499 | 1.143123306 | 0.062862096 | 0.30 | 0.20 |
| YPL265W | 2999.757451 | 0.90648469 | 0.063085717 | 0.30 | 0.20 |
| YGL260W | 1.415207671 | 1.376942039 | 0.063170622 | 0.30 | 0.20 |
| YGR204W | 10459.91176 | 0.415762353 | 0.063182192 | 0.30 | 0.20 |
| YOL068C | 1498.234041 | -0.334174587 | 0.063293825 | 0.30 | 0.20 |
| YLR314C | 3042.070083 | 0.287657392 | 0.063349477 | 0.30 | 0.20 |
| YIL116W | 2678.75766 | 0.586463953 | 0.063411041 | 0.30 | 0.20 |
| YDR128W | 2008.877758 | 0.432525192 | 0.06362257 | 0.31 | 0.20 |
| YDR358W | 1790.698239 | 0.511055034 | 0.063669139 | 0.31 | 0.20 |
| YLR247C | 1785.222725 | 0.328793413 | 0.063543184 | 0.31 | 0.20 |
| YLR422W | 2135.341687 | 0.354032396 | 0.063624517 | 0.31 | 0.20 |
| YDR266C | 1529.276069 | -0.362492902 | 0.063974375 | 0.31 | 0.20 |
| YKR106W | 110.7787445 | 0.949380413 | 0.063945681 | 0.31 | 0.20 |
| YKL066W | 1252.501288 | 0.675262068 | 0.064093319 | 0.31 | 0.20 |
| YMR228W | 316.010038 | -0.323291073 | 0.064106635 | 0.31 | 0.20 |
| YLR284C | 336.5089725 | -0.546722606 | 0.064165098 | 0.31 | 0.20 |
| YNL093W | 262.8484398 | 0.826274866 | 0.064244238 | 0.31 | 0.20 |
| YOR104W | 252.8227861 | 0.382033525 | 0.064483261 | 0.31 | 0.20 |
| YAL016C-B | 15.45888685 | 0.743407807 | 0.064726893 | 0.31 | 0.20 |
| YBR201C-A | 201.3285883 | 0.73295652 | 0.06465818 | 0.31 | 0.20 |
| YER063W | 1431.45469 | -0.352038574 | 0.064745506 | 0.31 | 0.20 |
| YJR093C | 790.9035242 | -0.216345054 | 0.064799645 | 0.31 | 0.20 |
| YPR133W-A | 1114.918643 | -0.521491126 | 0.064879951 | 0.31 | 0.20 |
| YBR001C | 1318.791969 | 0.48642797 | 0.065529319 | 0.31 | 0.20 |
| YDL050C | 34.4743764 | -0.730150706 | 0.065035915 | 0.31 | 0.20 |
| YFR026C | 243.3675063 | 0.89904028 | 0.065261713 | 0.31 | 0.20 |
| YGR220C | 677.1113476 | -0.355029391 | 0.065156186 | 0.31 | 0.20 |
| YHR004C | 921.2706765 | 0.408241779 | 0.065121094 | 0.31 | 0.20 |

| | | | | | |
|---|---|---|---|---|---|
| YJR038C | 33.2004855 | 0.721355932 | 0.065585803 | 0.31 | 0.20 |
| YKL182W | 10979.47839 | -0.349328027 | 0.065231841 | 0.31 | 0.20 |
| YKL197C | 902.4571586 | 0.403479402 | 0.065225189 | 0.31 | 0.20 |
| YLR009W | 1043.475365 | -0.613597871 | 0.06536416 | 0.31 | 0.20 |
| YLR154C | 335.1264893 | -0.28078155 | 0.065589129 | 0.31 | 0.20 |
| YNL016W | 4774.57143 | -0.370490407 | 0.065592685 | 0.31 | 0.20 |
| YNL226W | 19.21780824 | 0.594829363 | 0.06556109 | 0.31 | 0.20 |
| YNL254C | 501.4634557 | 0.436620461 | 0.065123062 | 0.31 | 0.20 |
| YNL255C | 4025.75008 | -0.499935658 | 0.065401899 | 0.31 | 0.20 |
| YOR062C | 585.4618256 | -0.707729266 | 0.065439612 | 0.31 | 0.20 |
| YKL139W | 677.7577592 | -0.313633205 | 0.065731867 | 0.31 | 0.20 |
| YDL136W | 915.4579307 | -0.63883257 | 0.065826621 | 0.31 | 0.20 |
| YLR222C | 1088.378581 | -0.621693823 | 0.065917533 | 0.31 | 0.20 |
| YDR148C | 4531.818706 | 0.741630571 | 0.066084523 | 0.31 | 0.20 |
| YOR189W | 408.1511083 | -0.342116731 | 0.066207052 | 0.31 | 0.20 |
| YNL101W | 1154.001663 | 0.405525594 | 0.066367941 | 0.31 | 0.21 |
| YDL045W-A | 694.1101578 | -0.438344514 | 0.0665366 | 0.31 | 0.21 |
| YML116W | 562.6593031 | 0.634397854 | 0.066508078 | 0.31 | 0.21 |
| YDR339C | 1137.925374 | -0.531298965 | 0.066693346 | 0.31 | 0.21 |
| YEL064C | 340.0873494 | 0.421415941 | 0.066690196 | 0.31 | 0.21 |
| YHR102W | 1800.525655 | 0.294460815 | 0.066692395 | 0.31 | 0.21 |
| YMR300C | 1920.32889 | -0.711225156 | 0.066742729 | 0.31 | 0.21 |
| YDL005C | 1599.265031 | -0.272291256 | 0.066795104 | 0.31 | 0.21 |
| YBR068C | 7312.593911 | 0.602483774 | 0.066974609 | 0.31 | 0.21 |
| YOR110W | 712.9241656 | 0.324531892 | 0.06699573 | 0.31 | 0.21 |
| YGL155W | 739.9393467 | -0.360361239 | 0.067123224 | 0.31 | 0.21 |
| YDL219W | 330.3585764 | -0.547586673 | 0.067378856 | 0.31 | 0.21 |
| YDR280W | 709.9216811 | -0.460132898 | 0.067415926 | 0.31 | 0.21 |
| YHR028C | 2661.078301 | 0.367837291 | 0.067293879 | 0.31 | 0.21 |
| YOL030W | 6136.814993 | 0.442762671 | 0.067432413 | 0.31 | 0.21 |
| YNR008W | 899.8873602 | 0.427942127 | 0.067500789 | 0.31 | 0.21 |
| YOR024W | 4.641545194 | 1.100917669 | 0.067584881 | 0.31 | 0.21 |
| YOR026W | 568.5531387 | -0.4459619 | 0.067539658 | 0.31 | 0.21 |
| YBR235W | 1364.419973 | 0.493178348 | 0.067998441 | 0.31 | 0.21 |
| YDR119W-A | 458.2326718 | 0.971123265 | 0.067831807 | 0.31 | 0.21 |
| YJR011C | 218.4017674 | -0.287826375 | 0.067750331 | 0.31 | 0.21 |
| YKL204W | 2110.527656 | -0.354823788 | 0.068051512 | 0.31 | 0.21 |
| YLR038C | 1207.825369 | 0.770156157 | 0.068001728 | 0.31 | 0.21 |
| YLR175W | 4387.69014 | -0.500341989 | 0.067988475 | 0.31 | 0.21 |
| YOR374W | 6364.239227 | 0.774534956 | 0.068006016 | 0.31 | 0.21 |
| snR67 | 6.800863812 | 0.884890709 | 0.068049641 | 0.31 | 0.21 |
| YBL030C | 9599.31549 | 0.808972846 | 0.068234774 | 0.31 | 0.21 |
| YDL020C | 3590.678769 | 0.566637278 | 0.068416983 | 0.31 | 0.21 |

| | | | | | |
|---|---|---|---|---|---|
| YDR178W | 2932.392626 | 0.760233659 | 0.068711948 | 0.31 | 0.21 |
| YDR367W | 606.3521729 | -0.379632953 | 0.068203768 | 0.31 | 0.21 |
| YDR462W | 726.8986569 | -0.357196047 | 0.068703363 | 0.31 | 0.21 |
| YIL021W | 814.8146206 | -0.332362763 | 0.068573539 | 0.31 | 0.21 |
| YJR130C | 1173.075557 | 0.55674914 | 0.068516526 | 0.31 | 0.21 |
| YKR028W | 1923.731148 | -0.320586623 | 0.068670224 | 0.31 | 0.21 |
| YLR403W | 956.7749718 | -0.392767463 | 0.068419856 | 0.31 | 0.21 |
| YMR045C | 326.5494461 | 0.679168121 | 0.06874095 | 0.31 | 0.21 |
| YOR003W | 499.5835225 | 0.412378974 | 0.068621049 | 0.31 | 0.21 |
| YPR003C | 637.4553595 | 0.446096728 | 0.068300119 | 0.31 | 0.21 |
| YPR010C-A | 733.8429902 | 0.856961475 | 0.068393992 | 0.31 | 0.21 |
| YLR325C | 5760.035975 | -0.633269132 | 0.068891301 | 0.31 | 0.21 |
| YMR183C | 2217.146681 | -0.397677096 | 0.068869369 | 0.31 | 0.21 |
| YNL026W | 973.5528021 | -0.223915564 | 0.068804656 | 0.31 | 0.21 |
| YCR068W | 666.4091136 | 0.473701035 | 0.069022579 | 0.31 | 0.21 |
| YDR449C | 532.8954421 | -0.689826813 | 0.069696181 | 0.31 | 0.21 |
| YER120W | 4329.489139 | -0.264708879 | 0.069784576 | 0.31 | 0.21 |
| YGL165C | 78.38274117 | 0.355453189 | 0.069815651 | 0.31 | 0.21 |
| YJL033W | 1448.230029 | -0.630006512 | 0.069804868 | 0.31 | 0.21 |
| YLR099W-A | 174.4936248 | -0.381600593 | 0.069377432 | 0.31 | 0.21 |
| YLR120C | 2959.418798 | 0.693685637 | 0.069752233 | 0.31 | 0.21 |
| YLR390W-A | 12426.05045 | 0.305468488 | 0.069192302 | 0.31 | 0.21 |
| YML113W | 567.9041791 | -0.482105581 | 0.069626177 | 0.31 | 0.21 |
| YMR234W | 552.7610201 | -0.268526216 | 0.069835582 | 0.31 | 0.21 |
| YNL112W | 1857.470789 | -0.769226789 | 0.069421746 | 0.31 | 0.21 |
| YOL002C | 621.7558878 | -0.481079094 | 0.069768127 | 0.31 | 0.21 |
| YOL044W | 473.6470695 | 0.437482495 | 0.069466107 | 0.31 | 0.21 |
| YOL076W | 856.9107843 | -0.509650838 | 0.069265427 | 0.31 | 0.21 |
| YOR305W | 732.3331193 | -0.402226164 | 0.069815548 | 0.31 | 0.21 |
| YPR112C | 1587.350101 | -0.639639665 | 0.069607939 | 0.31 | 0.21 |
| YPR157W | 448.7836668 | 0.807688992 | 0.069825338 | 0.31 | 0.21 |
| YPR182W | 302.4381877 | -0.337045124 | 0.069438471 | 0.31 | 0.21 |
| YFL024C | 1322.826078 | -0.346506143 | 0.070019437 | 0.31 | 0.21 |
| YDR390C | 1683.184066 | -0.25278253 | 0.070088329 | 0.31 | 0.21 |
| YER159C | 1569.026083 | -0.534733064 | 0.070585245 | 0.32 | 0.21 |
| YNL174W | 7.421723425 | -0.991597848 | 0.070613219 | 0.32 | 0.21 |
| YCR073W-A | 1340.952291 | 0.468865727 | 0.070784031 | 0.32 | 0.21 |
| YBR162W-A | 580.1318829 | -0.450569228 | 0.07084866 | 0.32 | 0.21 |
| YOR119C | 842.963497 | 0.340121035 | 0.07096018 | 0.32 | 0.21 |
| YPR001W | 533.0217534 | 0.814286522 | 0.070912883 | 0.32 | 0.21 |
| YNL038W | 473.5438124 | -0.446371166 | 0.071105825 | 0.32 | 0.21 |
| YEL032W | 1185.048516 | -0.271952966 | 0.071374883 | 0.32 | 0.21 |
| YER112W | 1280.704105 | -0.328785309 | 0.071335849 | 0.32 | 0.21 |

| | | | | | |
|---|---|---|---|---|---|
| YKL089W | 415.1913401 | 0.451100689 | 0.071346323 | 0.32 | 0.21 |
| YLR352W | 926.4882915 | 0.312376043 | 0.071322786 | 0.32 | 0.21 |
| YGR101W | 624.5372953 | -0.26900923 | 0.071487473 | 0.32 | 0.21 |
| YGR146C | 1344.611896 | 0.601759494 | 0.071486939 | 0.32 | 0.21 |
| YGR092W | 957.140587 | 0.485206643 | 0.071538099 | 0.32 | 0.21 |
| YLR380W | 1647.260273 | -0.474252824 | 0.071634944 | 0.32 | 0.21 |
| YDL147W | 5209.098548 | 0.287932855 | 0.071954501 | 0.32 | 0.21 |
| YLL001W | 1443.400677 | 0.388673773 | 0.071923625 | 0.32 | 0.21 |
| YLR139C | 783.7450537 | -0.429916661 | 0.071904039 | 0.32 | 0.21 |
| YOL137W | 547.8586694 | 0.258089487 | 0.072074226 | 0.32 | 0.21 |
| YGR141W | 841.9095851 | 0.593676904 | 0.072303282 | 0.32 | 0.21 |
| YHR157W | 84.3613017 | 0.465511373 | 0.072479329 | 0.32 | 0.21 |
| YKL136W | 25.19505527 | -0.487332591 | 0.072507731 | 0.32 | 0.21 |
| YER116C | 578.3983863 | 0.451987886 | 0.072617557 | 0.32 | 0.21 |
| YNL127W | 1075.477959 | 0.430881209 | 0.072594456 | 0.32 | 0.21 |
| YJL082W | 1623.02873 | 0.509540629 | 0.072690982 | 0.32 | 0.21 |
| YDL072C | 3778.461007 | 0.251514465 | 0.072804335 | 0.32 | 0.21 |
| YBR010W | 4241.31261 | -0.60290528 | 0.072934767 | 0.32 | 0.21 |
| YDR505C | 1755.081331 | 0.475533664 | 0.073019619 | 0.32 | 0.21 |
| YFR001W | 1018.480753 | -0.424134962 | 0.073061607 | 0.32 | 0.21 |
| YCR028C | 527.4192161 | 0.240219136 | 0.073166112 | 0.32 | 0.21 |
| YGL100W | 1500.808318 | -0.233068211 | 0.073435885 | 0.32 | 0.21 |
| YJR063W | 331.564801 | -0.658275207 | 0.073418683 | 0.32 | 0.21 |
| YBL057C | 768.8630819 | -0.336465723 | 0.073797656 | 0.32 | 0.21 |
| YBL101C | 1814.259973 | 0.414115717 | 0.073600323 | 0.32 | 0.21 |
| YBR139W | 2470.205803 | 0.471774008 | 0.073794927 | 0.32 | 0.21 |
| YDL198C | 1156.426648 | 0.524349554 | 0.073874388 | 0.32 | 0.21 |
| YDR072C | 1794.903187 | -0.491364708 | 0.073924111 | 0.32 | 0.21 |
| YDR429C | 3118.563913 | -0.439355188 | 0.073841701 | 0.32 | 0.21 |
| YGR161C | 2432.867838 | 0.728210722 | 0.073955981 | 0.32 | 0.21 |
| YLR312C | 417.3618401 | 0.818315397 | 0.073937286 | 0.32 | 0.21 |
| YLR361C | 559.6543534 | 0.307612451 | 0.073607559 | 0.32 | 0.21 |
| YMR148W | 862.6307015 | 0.36434204 | 0.073609731 | 0.32 | 0.21 |
| YGL211W | 636.0371021 | -0.273721158 | 0.074201287 | 0.32 | 0.21 |
| YOL149W | 642.6006688 | -0.439340015 | 0.074240984 | 0.32 | 0.21 |
| YHR149C | 909.9713222 | -0.49761054 | 0.074326334 | 0.32 | 0.21 |
| YJL191W | 879.6997619 | -0.783964892 | 0.074417039 | 0.32 | 0.21 |
| YBR143C | 4538.418741 | -0.497046823 | 0.074532717 | 0.32 | 0.21 |
| YMR105W-A | 43.59841519 | 0.373220051 | 0.074495246 | 0.32 | 0.21 |
| YOR302W | 66.2485987 | 0.754065863 | 0.07458779 | 0.32 | 0.21 |
| YBR188C | 339.0880427 | -0.283477638 | 0.074996355 | 0.32 | 0.21 |
| YDR398W | 1282.093858 | -0.537668008 | 0.074920246 | 0.32 | 0.21 |
| YLR217W | 16.95655225 | 0.631760482 | 0.074958636 | 0.32 | 0.21 |

| | | | | | |
|---|---|---|---|---|---|
| YCR057C | 1417.95789 | -0.532406265 | 0.07514243 | 0.32 | 0.21 |
| YDR223W | 358.2338356 | 0.481152069 | 0.075229204 | 0.32 | 0.21 |
| YER087W | 372.1981021 | 0.447187491 | 0.07519072 | 0.32 | 0.21 |
| YLR197W | 4160.668892 | -0.645515612 | 0.07537174 | 0.33 | 0.21 |
| YNL231C | 1567.48489 | -0.332689324 | 0.075489591 | 0.33 | 0.22 |
| YDL022C-A | 6.814086928 | 0.887266607 | 0.07557523 | 0.33 | 0.22 |
| YDL189W | 1070.015735 | -0.384799292 | 0.075924118 | 0.33 | 0.22 |
| YBR275C | 1768.794249 | 0.384218662 | 0.076008686 | 0.33 | 0.22 |
| YAR035W | 1245.571751 | 0.763888322 | 0.076557553 | 0.33 | 0.22 |
| YDR156W | 897.9312179 | -0.390370787 | 0.076564332 | 0.33 | 0.22 |
| YGR129W | 435.9932211 | 0.491196466 | 0.076569992 | 0.33 | 0.22 |
| YMR280C | 1028.334214 | 0.690061028 | 0.07653664 | 0.33 | 0.22 |
| YBL081W | 976.0547384 | -0.61888139 | 0.077297969 | 0.33 | 0.22 |
| YDL109C | 380.7977413 | 0.435024156 | 0.077234058 | 0.33 | 0.22 |
| YDR121W | 548.7039891 | -0.284411848 | 0.077188366 | 0.33 | 0.22 |
| YGR065C | 698.7091847 | 0.4863087 | 0.077057479 | 0.33 | 0.22 |
| YHL017W | 1099.085857 | 0.41154739 | 0.077176092 | 0.33 | 0.22 |
| YJL194W | 154.6134221 | -0.741994636 | 0.07686822 | 0.33 | 0.22 |
| YLR149C | 1780.729906 | 0.664246778 | 0.07710197 | 0.33 | 0.22 |
| YLR385C | 52.50375543 | 0.439041819 | 0.077289378 | 0.33 | 0.22 |
| YMR260C | 3410.062096 | -0.451807565 | 0.077192838 | 0.33 | 0.22 |
| YOL166C | 2.808993483 | 1.004289164 | 0.077075784 | 0.33 | 0.22 |
| YOR274W | 529.1848909 | -0.388800251 | 0.076927125 | 0.33 | 0.22 |
| YGR029W | 703.0611046 | -0.323669792 | 0.077355534 | 0.33 | 0.22 |
| YDL081C | 10855.42483 | -0.594013258 | 0.077724804 | 0.33 | 0.22 |
| YDL023C | 219.5343501 | 0.549708729 | 0.077839475 | 0.33 | 0.22 |
| YNL131W | 2877.827714 | -0.34641288 | 0.077984249 | 0.33 | 0.22 |
| YKR069W | 646.4230382 | 0.42674404 | 0.078158102 | 0.33 | 0.22 |
| YNL332W | 25.85198322 | 1.000019496 | 0.078289619 | 0.33 | 0.22 |
| YHR193C | 8396.042296 | -0.434897178 | 0.078352311 | 0.33 | 0.22 |
| YGL203C | 1571.604512 | 0.376217665 | 0.078476311 | 0.33 | 0.22 |
| YPR139C | 1065.385559 | -0.421154131 | 0.078486767 | 0.33 | 0.22 |
| YDR412W | 170.5135109 | -0.449943294 | 0.078749753 | 0.33 | 0.22 |
| YKL076C | 34.14810955 | 0.630060535 | 0.078781589 | 0.33 | 0.22 |
| YOL109W | 9848.185932 | -0.462326939 | 0.078714123 | 0.33 | 0.22 |
| YHR056W-A | 0.867587117 | 0.830167646 | 0.078915898 | 0.33 | 0.22 |
| YBR210W | 464.9591183 | -0.375470108 | 0.079114041 | 0.33 | 0.22 |
| YBR028C | 772.1128092 | -0.439398346 | 0.07944567 | 0.33 | 0.22 |
| YMR309C | 2879.75717 | -0.489997707 | 0.079428386 | 0.33 | 0.22 |
| YMR310C | 757.4056331 | -0.521652684 | 0.079413381 | 0.33 | 0.22 |
| YER154W | 1666.340326 | -0.26084226 | 0.079956453 | 0.34 | 0.22 |
| YKR007W | 738.2939867 | -0.285689613 | 0.079924752 | 0.34 | 0.22 |
| YLR246W | 470.2308665 | 0.486238316 | 0.079866319 | 0.34 | 0.22 |

| | | | | | |
|---|---|---|---|---|---|
| YLR388W | 1672.166035 | -0.653290633 | 0.079831769 | 0.34 | 0.22 |
| YMR250W | 7553.858966 | 0.746962481 | 0.079866719 | 0.34 | 0.22 |
| YOR336W | 1255.740018 | 0.317274685 | 0.079722793 | 0.34 | 0.22 |
| YLR424W | 404.0091344 | 0.24066633 | 0.080029545 | 0.34 | 0.22 |
| YMR020W | 992.3354434 | 0.654465994 | 0.080463262 | 0.34 | 0.22 |
| YOR148C | 393.9247418 | -0.290562385 | 0.08045607 | 0.34 | 0.22 |
| YOR327C | 881.0062218 | -0.354949824 | 0.080445499 | 0.34 | 0.22 |
| YGR074W | 145.6756142 | -0.351404447 | 0.080591838 | 0.34 | 0.22 |
| YBR039W | 6878.817001 | 0.738250821 | 0.081240599 | 0.34 | 0.22 |
| YCR097W-A | 1.60235674 | 0.629247298 | 0.080933168 | 0.34 | 0.22 |
| YDL048C | 3537.203448 | 0.677610312 | 0.081096128 | 0.34 | 0.22 |
| YIL093C | 871.5653388 | -0.346636767 | 0.08112706 | 0.34 | 0.22 |
| YIR029W | 388.4137592 | 0.789338997 | 0.08109499 | 0.34 | 0.22 |
| YJR096W | 2414.849421 | 0.786369728 | 0.081027355 | 0.34 | 0.22 |
| YKL214C | 653.6592896 | -0.285597694 | 0.081114779 | 0.34 | 0.22 |
| YML069W | 2256.742432 | -0.425919528 | 0.080933977 | 0.34 | 0.22 |
| YOR004W | 494.9946687 | -0.679353467 | 0.081163082 | 0.34 | 0.22 |
| YPR063C | 1556.762503 | -0.321644578 | 0.081209697 | 0.34 | 0.22 |
| YNL275W | 352.1195966 | -0.291368455 | 0.081428879 | 0.34 | 0.22 |
| YPR155C | 883.1515984 | 0.445165452 | 0.081428275 | 0.34 | 0.22 |
| YNL108C | 290.0576123 | -0.393975241 | 0.081516166 | 0.34 | 0.22 |
| YOR304W | 1385.923767 | -0.206882406 | 0.081601133 | 0.34 | 0.22 |
| YDR160W | 879.6911624 | 0.230541419 | 0.081740928 | 0.34 | 0.22 |
| YGR060W | 3114.502526 | -0.619993616 | 0.081819633 | 0.34 | 0.22 |
| YGR238C | 880.0595867 | 0.383550562 | 0.081731886 | 0.34 | 0.22 |
| YPL106C | 19467.56403 | 0.501864487 | 0.081841127 | 0.34 | 0.22 |
| YER178W | 9418.417677 | 0.283956169 | 0.081901319 | 0.34 | 0.22 |
| YMR061W | 1751.530821 | -0.304929197 | 0.082087565 | 0.34 | 0.22 |
| YMR038C | 1941.251831 | -0.377630965 | 0.082142513 | 0.34 | 0.22 |
| YIR013C | 17.07044727 | 0.771930638 | 0.082238674 | 0.34 | 0.22 |
| YDR172W | 3599.759088 | -0.327746605 | 0.082482912 | 0.34 | 0.22 |
| YOL016C | 3937.741808 | 0.611225499 | 0.082512905 | 0.34 | 0.22 |
| YDR105C | 933.9584198 | -0.283858503 | 0.082737408 | 0.34 | 0.23 |
| YEL049W | 54.33605347 | 0.603607992 | 0.082965099 | 0.34 | 0.23 |
| YIL107C | 1780.529737 | 0.431830039 | 0.082944404 | 0.34 | 0.23 |
| YKL074C | 747.5415477 | 0.264953987 | 0.083046479 | 0.34 | 0.23 |
| YNL017C | 1.615564791 | 1.181602574 | 0.083250384 | 0.34 | 0.23 |
| YCR097W | 1013.079937 | 0.707214757 | 0.083480127 | 0.34 | 0.23 |
| YMR319C | 2635.007516 | 0.528167101 | 0.083553258 | 0.34 | 0.23 |
| YJL070C | 972.7876692 | 0.409221011 | 0.083732922 | 0.34 | 0.23 |
| YPR191W | 6094.519828 | 0.756126045 | 0.083847072 | 0.34 | 0.23 |
| YJR118C | 868.4456577 | -0.388774187 | 0.083910999 | 0.34 | 0.23 |
| YDR457W | 5588.916733 | 0.334731441 | 0.084053959 | 0.34 | 0.23 |

| | | | | | |
|---|---|---|---|---|---|
| YPL011C | 726.714861 | -0.275399858 | 0.084321102 | 0.34 | 0.23 |
| YEL016C | 826.6367047 | -0.242412175 | 0.08449448 | 0.35 | 0.23 |
| YOR392W | 4.708188968 | 0.88177478 | 0.084515791 | 0.35 | 0.23 |
| YJL169W | 25.36216702 | -0.630033929 | 0.084677601 | 0.35 | 0.23 |
| YJL206C | 539.1174838 | 0.398426172 | 0.084711774 | 0.35 | 0.23 |
| YDR215C | 67.50004526 | 0.854802661 | 0.084820918 | 0.35 | 0.23 |
| YDL014W | 4306.735257 | -0.668189134 | 0.084979529 | 0.35 | 0.23 |
| YOR078W | 697.5265362 | -0.452179228 | 0.085133702 | 0.35 | 0.23 |
| YCR006C | 58.71736424 | -0.593035058 | 0.085246603 | 0.35 | 0.23 |
| YER015W | 725.609463 | 0.476730556 | 0.085302882 | 0.35 | 0.23 |
| YIR015W | 351.8671415 | 0.403779637 | 0.085363154 | 0.35 | 0.23 |
| YOR130C | 372.7954083 | 0.591570469 | 0.085444069 | 0.35 | 0.23 |
| YGR150C | 1213.396537 | -0.255820349 | 0.085688122 | 0.35 | 0.23 |
| YLR331C | 3.488696978 | 0.860937477 | 0.085691872 | 0.35 | 0.23 |
| YBR049C | 1277.509775 | -0.357095539 | 0.085779438 | 0.35 | 0.23 |
| YHR117W | 2228.445201 | 0.373871376 | 0.086094074 | 0.35 | 0.23 |
| YLR264W | 525.6829336 | -0.618541321 | 0.086087063 | 0.35 | 0.23 |
| YPR002C-A | 1.918449357 | 1.182967391 | 0.08623092 | 0.35 | 0.23 |
| YOL100W | 1483.492555 | 0.39491871 | 0.08630591 | 0.35 | 0.23 |
| YER025W | 3287.005691 | -0.388965651 | 0.086564314 | 0.35 | 0.23 |
| YFL005W | 2091.331245 | -0.310019299 | 0.086665624 | 0.35 | 0.23 |
| YFL039C | 29152.57267 | 0.243042032 | 0.086403242 | 0.35 | 0.23 |
| YMR188C | 1203.74754 | -0.373442238 | 0.086744277 | 0.35 | 0.23 |
| YMR290W-A | 11.59226843 | -0.819160125 | 0.086760922 | 0.35 | 0.23 |
| YNL144C | 1767.380134 | 0.705795422 | 0.086721192 | 0.35 | 0.23 |
| YNL165W | 500.9806009 | 0.371381561 | 0.086547124 | 0.35 | 0.23 |
| YPL256C | 1702.304083 | -0.506318419 | 0.086668667 | 0.35 | 0.23 |
| YDL065C | 1387.029558 | 0.395293404 | 0.087535454 | 0.35 | 0.23 |
| YFL058W | 7.345596516 | 1.085496958 | 0.087608401 | 0.35 | 0.23 |
| YFR049W | 788.1728678 | 0.600308361 | 0.087617309 | 0.35 | 0.23 |
| YHR001W-A | 1152.222923 | 0.729846943 | 0.087690341 | 0.35 | 0.23 |
| YMR292W | 429.0322087 | -0.436454328 | 0.0876276 | 0.35 | 0.23 |
| YOL021C | 1670.215668 | -0.296037968 | 0.087734568 | 0.35 | 0.23 |
| YJL213W | 560.8105438 | 0.648969663 | 0.087836862 | 0.35 | 0.23 |
| YOR324C | 520.2886814 | -0.355634763 | 0.087875044 | 0.35 | 0.23 |
| snR57 | 17.27289396 | 0.909600694 | 0.087907531 | 0.35 | 0.23 |
| YPL262W | 5873.810031 | 0.761250621 | 0.087999345 | 0.35 | 0.23 |
| YDL123W | 682.3334241 | 0.400953579 | 0.088238493 | 0.35 | 0.23 |
| YHL026C | 386.086378 | 0.482200471 | 0.088198907 | 0.35 | 0.23 |
| YIL004C | 332.265247 | -0.245006359 | 0.088348352 | 0.35 | 0.23 |
| YOR091W | 1558.309002 | -0.455220394 | 0.08841036 | 0.35 | 0.23 |
| YHR124W | 347.8549107 | 0.636512156 | 0.088487096 | 0.35 | 0.23 |
| YML087C | 330.7889367 | 0.756230601 | 0.088670867 | 0.35 | 0.23 |

| | | | | | |
|---|---|---|---|---|---|
| YKL140W | 1755.744554 | 0.224733329 | 0.088751553 | 0.35 | 0.23 |
| YBR120C | 395.0224857 | -0.372180372 | 0.088806838 | 0.35 | 0.23 |
| YGR177C | 234.3513182 | -0.661567866 | 0.088902739 | 0.35 | 0.23 |
| YIL150C | 295.6540585 | 0.345420474 | 0.089023009 | 0.35 | 0.23 |
| YBL088C | 1866.759984 | 0.349296607 | 0.089110411 | 0.35 | 0.23 |
| YER135C | 23.51565939 | 0.666672422 | 0.089187751 | 0.35 | 0.23 |
| YNL080C | 1115.372394 | -0.347374273 | 0.089209963 | 0.35 | 0.23 |
| YBR256C | 1957.968457 | 0.439781919 | 0.089287658 | 0.35 | 0.23 |
| YBL091C | 4543.955358 | -0.222156151 | 0.08945194 | 0.35 | 0.23 |
| YOL077C | 1854.977032 | -0.629496877 | 0.089421756 | 0.35 | 0.23 |
| YOL122C | 1128.161192 | 0.507675847 | 0.089475862 | 0.35 | 0.23 |
| YJL180C | 680.1821358 | -0.294867392 | 0.089601875 | 0.35 | 0.23 |
| YJL061W | 1142.158213 | -0.253043481 | 0.089670762 | 0.35 | 0.23 |
| YAL047C | 589.7842721 | 0.268912744 | 0.090373568 | 0.36 | 0.24 |
| YCL038C | 682.2522644 | 0.335405675 | 0.090261531 | 0.36 | 0.24 |
| YER004W | 2233.162604 | 0.35709335 | 0.090636675 | 0.36 | 0.24 |
| YGL205W | 247.6080273 | 0.590390808 | 0.090479394 | 0.36 | 0.24 |
| YJL094C | 1236.455255 | 0.333684909 | 0.090598593 | 0.36 | 0.24 |
| YNL284C-B | 190.6577576 | 0.819713992 | 0.090192523 | 0.36 | 0.24 |
| YNL330C | 1017.919377 | -0.556118021 | 0.090571101 | 0.36 | 0.24 |
| YOR304C-A | 201.0708852 | -0.425125036 | 0.0906175 | 0.36 | 0.24 |
| YOR346W | 878.2111125 | -0.40294148 | 0.090492409 | 0.36 | 0.24 |
| YPR054W | 265.2938746 | -0.553018592 | 0.090142495 | 0.36 | 0.24 |
| YPR076W | 41.22373968 | 0.57347858 | 0.090646524 | 0.36 | 0.24 |
| YPR039W | 9.707681236 | 0.722365532 | 0.090831789 | 0.36 | 0.24 |
| YBR009C | 3036.286227 | -0.647621581 | 0.091011162 | 0.36 | 0.24 |
| YKR050W | 1256.621968 | 0.455009472 | 0.090952834 | 0.36 | 0.24 |
| YNL077W | 1205.161408 | 0.501252994 | 0.091102095 | 0.36 | 0.24 |
| YOR185C | 2527.056266 | 0.540733095 | 0.091111065 | 0.36 | 0.24 |
| YPR166C | 431.6523763 | -0.375057081 | 0.091122543 | 0.36 | 0.24 |
| YBR214W | 5062.049562 | 0.513695072 | 0.091297882 | 0.36 | 0.24 |
| YGR179C | 503.8137475 | -0.222923681 | 0.09128998 | 0.36 | 0.24 |
| YDR268W | 586.9385642 | 0.341976464 | 0.091607412 | 0.36 | 0.24 |
| YEL039C | 1783.57122 | 0.733329437 | 0.091660829 | 0.36 | 0.24 |
| YNL055C | 20028.30046 | 0.65298841 | 0.091406523 | 0.36 | 0.24 |
| YOL145C | 2673.103928 | -0.365251684 | 0.091497944 | 0.36 | 0.24 |
| YOR001W | 1523.051345 | -0.327293202 | 0.091520439 | 0.36 | 0.24 |
| YPR017C | 239.3606668 | -0.375457713 | 0.09156329 | 0.36 | 0.24 |
| YMR203W | 3020.852994 | -0.258309817 | 0.09179817 | 0.36 | 0.24 |
| YLR024C | 1454.81679 | 0.381657748 | 0.09198005 | 0.36 | 0.24 |
| YPR027C | 342.9171324 | -0.856927525 | 0.092077829 | 0.36 | 0.24 |
| YHR186C | 1649.917336 | 0.266783012 | 0.092233863 | 0.36 | 0.24 |
| YKR094C | 5698.150477 | -0.692638166 | 0.092230878 | 0.36 | 0.24 |

| | | | | | |
|---|---|---|---|---|---|
| YMR114C | 1303.792032 | 0.561659536 | 0.092387906 | 0.36 | 0.24 |
| YNL032W | 670.5607595 | -0.280836971 | 0.092359111 | 0.36 | 0.24 |
| YDR203W | 15.81601128 | -0.618390316 | 0.092469616 | 0.36 | 0.24 |
| YMR014W | 1014.309235 | -0.505101629 | 0.092586036 | 0.36 | 0.24 |
| YPL005W | 369.5032375 | 0.407381998 | 0.092595879 | 0.36 | 0.24 |
| YGL162W | 395.3975497 | -0.675137993 | 0.092818972 | 0.36 | 0.24 |
| YBR176W | 422.8364361 | 0.356574811 | 0.092875144 | 0.36 | 0.24 |
| YER051W | 512.8476233 | 0.452047551 | 0.093069732 | 0.36 | 0.24 |
| YER144C | 1434.76784 | 0.359625068 | 0.093104373 | 0.36 | 0.24 |
| ICR1 | 346.589135 | 0.444724799 | 0.093349075 | 0.36 | 0.24 |
| YBR178W | 20.74423611 | 0.636625438 | 0.093453157 | 0.36 | 0.24 |
| YBR230C | 2523.139558 | 0.638719391 | 0.093443164 | 0.36 | 0.24 |
| YER078W-A | 4.281084499 | 0.743549533 | 0.093310079 | 0.36 | 0.24 |
| YOL064C | 847.1293742 | 0.493143819 | 0.093445679 | 0.36 | 0.24 |
| YLR111W | 15.73315875 | 0.731740946 | 0.0936204 | 0.36 | 0.24 |
| YOR368W | 442.6825001 | -0.255952488 | 0.093672095 | 0.36 | 0.24 |
| YAR019C | 904.0673159 | 0.385378307 | 0.093867779 | 0.36 | 0.24 |
| snR33 | 120.9149393 | -0.409213876 | 0.093993405 | 0.36 | 0.24 |
| YGR064W | 14.24950712 | -0.640344537 | 0.094173566 | 0.36 | 0.24 |
| YGL131C | 1060.332301 | 0.46737074 | 0.094274013 | 0.36 | 0.24 |
| YBR111C | 3318.167647 | 0.54735856 | 0.094531236 | 0.36 | 0.24 |
| YFL050C | 563.7473526 | 0.466919012 | 0.094433698 | 0.36 | 0.24 |
| YMR140W | 1470.388235 | 0.409850672 | 0.094529796 | 0.36 | 0.24 |
| YGL036W | 2417.923989 | 0.279340891 | 0.094592222 | 0.36 | 0.24 |
| YBR087W | 809.749781 | -0.372238945 | 0.094864588 | 0.36 | 0.24 |
| YHR027C | 5786.339997 | 0.486056085 | 0.094885738 | 0.36 | 0.24 |
| YNR029C | 598.402815 | -0.300191725 | 0.094761221 | 0.36 | 0.24 |
| YOL113W | 1261.758159 | 0.545843972 | 0.094809979 | 0.36 | 0.24 |
| YOR193W | 437.421385 | -0.379329928 | 0.094808643 | 0.36 | 0.24 |
| YPL142C | 228.9766843 | -0.643182244 | 0.094936237 | 0.36 | 0.24 |
| YBL066C | 1237.560937 | 0.529504853 | 0.095448405 | 0.36 | 0.24 |
| YHL009W-B | 5.78416984 | 0.92876191 | 0.095326691 | 0.36 | 0.24 |
| YJL068C | 1815.839599 | 0.369271368 | 0.095320497 | 0.36 | 0.24 |
| YKR024C | 515.1058632 | -0.488746579 | 0.095204997 | 0.36 | 0.24 |
| YOR040W | 503.3947844 | 0.586363389 | 0.095423367 | 0.36 | 0.24 |
| YPL032C | 2772.776631 | -0.381208308 | 0.095337652 | 0.36 | 0.24 |
| YNL045W | 3993.621371 | 0.72050567 | 0.095617308 | 0.36 | 0.24 |
| YML059C | 1781.099171 | -0.314644611 | 0.095739546 | 0.36 | 0.24 |
| YOR143C | 346.0541178 | -0.414000001 | 0.095720758 | 0.36 | 0.24 |
| YJL118W | 178.5211681 | -0.507035542 | 0.095887348 | 0.36 | 0.24 |
| YMR025W | 229.9295307 | 0.501173085 | 0.096000166 | 0.36 | 0.24 |
| YDR086C | 1629.638329 | -0.410108873 | 0.096228645 | 0.36 | 0.24 |
| YHR111W | 861.1772037 | 0.297105471 | 0.09624398 | 0.36 | 0.24 |

| | | | | | |
|---|---|---|---|---|---|
| YLR047C | 709.1954371 | 0.198922369 | 0.096338045 | 0.36 | 0.24 |
| YPR192W | 327.8980894 | -0.599853799 | 0.096757193 | 0.37 | 0.24 |
| YER152W-A | 37.9606133 | 0.607040916 | 0.09698307 | 0.37 | 0.24 |
| YDL230W | 846.3743425 | 0.34756303 | 0.097385238 | 0.37 | 0.24 |
| YFR012W | 37.52047909 | 0.863284786 | 0.097307142 | 0.37 | 0.24 |
| YLR195C | 1211.827498 | -0.324011205 | 0.097385466 | 0.37 | 0.24 |
| YCR102C | 144.1370507 | 0.567381051 | 0.097555974 | 0.37 | 0.24 |
| YFR018C | 1513.157132 | 0.398094782 | 0.097524951 | 0.37 | 0.24 |
| YKL141W | 4100.397187 | 0.671584926 | 0.097612579 | 0.37 | 0.24 |
| YPL064C | 883.912951 | -0.213701407 | 0.097637386 | 0.37 | 0.24 |
| YDR009W | 246.880862 | 0.577194033 | 0.098088025 | 0.37 | 0.24 |
| YDR113C | 648.1289584 | -0.286343147 | 0.098096803 | 0.37 | 0.24 |
| YHR189W | 309.5122663 | 0.492438986 | 0.098090568 | 0.37 | 0.24 |
| YLL028W | 724.5266873 | -0.482380215 | 0.09818513 | 0.37 | 0.24 |
| YHL048W | 1666.986201 | 0.641184762 | 0.098311449 | 0.37 | 0.24 |
| YHR204W | 1257.868137 | 0.37187926 | 0.098311317 | 0.37 | 0.24 |
| YEL054C | 3213.78471 | -0.654529268 | 0.098382905 | 0.37 | 0.24 |
| YKL109W | 7924.096762 | 0.539381143 | 0.098440931 | 0.37 | 0.24 |
| YGR005C | 1346.509283 | -0.236404195 | 0.098654428 | 0.37 | 0.24 |
| YJR103W | 3899.480126 | 0.759822231 | 0.098721951 | 0.37 | 0.24 |
| YBR112C | 3373.883429 | -0.359144136 | 0.098908457 | 0.37 | 0.24 |
| YFL034W | 949.1269789 | 0.231836401 | 0.098970135 | 0.37 | 0.24 |
| YKL083W | 46.77827339 | -0.472229432 | 0.098951502 | 0.37 | 0.24 |
| YPR163C | 5059.100182 | -0.495990124 | 0.098927856 | 0.37 | 0.24 |
| YNL170W | 85.88966089 | 0.477557753 | 0.099065845 | 0.37 | 0.24 |
| YOR171C | 1509.079417 | 0.315026802 | 0.099295208 | 0.37 | 0.24 |
| YEL062W | 560.1717097 | -0.293133191 | 0.099448447 | 0.37 | 0.24 |
| YIL028W | 59.74955175 | 0.51289509 | 0.099480207 | 0.37 | 0.24 |
| YMR049C | 1525.516535 | -0.572367773 | 0.099567517 | 0.37 | 0.24 |
| YHR194W | 859.058911 | 0.416729334 | 0.099629285 | 0.37 | 0.24 |
| YJR160C | 11.90922489 | 0.744034178 | 0.100078059 | 0.37 | 0.25 |
| YDR010C | 32.06483583 | 0.41007817 | 0.100337462 | 0.37 | 0.25 |
| YGL193C | 193.4216907 | -0.909652183 | 0.100368015 | 0.37 | 0.25 |
| YLL040C | 5253.851757 | 0.498192851 | 0.100332247 | 0.37 | 0.25 |
| YNL128W | 87.91000712 | 0.664953376 | 0.100250474 | 0.37 | 0.25 |
| YIL162W | 3263.600398 | 0.726064441 | 0.100457411 | 0.37 | 0.25 |
| YLR140W | 51.21550356 | -0.326901585 | 0.100666633 | 0.37 | 0.25 |
| YML093W | 1683.648354 | -0.500542081 | 0.100736523 | 0.37 | 0.25 |
| YPR137W | 608.8911311 | -0.476669784 | 0.100800243 | 0.37 | 0.25 |
| YGR128C | 1414.526792 | -0.607884672 | 0.101161568 | 0.37 | 0.25 |
| YLL006W-A | 2.814822079 | 1.033893318 | 0.10109692 | 0.37 | 0.25 |
| YLR198C | 85.68816582 | -0.67671312 | 0.101171058 | 0.37 | 0.25 |
| YDL245C | 34.45281204 | 0.892427255 | 0.101275394 | 0.37 | 0.25 |

| | | | | | |
|---|---|---|---|---|---|
| YLR264C-A | 117.4733387 | -0.708909997 | 0.101509424 | 0.37 | 0.25 |
| YDL193W | 1212.895326 | 0.25153054 | 0.101650472 | 0.37 | 0.25 |
| YMR131C | 2416.896733 | -0.533360352 | 0.101599447 | 0.37 | 0.25 |
| YER163C | 538.8237534 | 0.421655101 | 0.102070319 | 0.38 | 0.25 |
| YLL055W | 836.4946306 | 0.663406324 | 0.102202622 | 0.38 | 0.25 |
| YBL002W | 4494.23128 | -0.681834229 | 0.102423951 | 0.38 | 0.25 |
| YFL060C | 16.02228195 | -0.637758203 | 0.102429292 | 0.38 | 0.25 |
| YIL134C-A | 3.121789303 | 0.843262814 | 0.102559546 | 0.38 | 0.25 |
| YJR141W | 396.4509931 | -0.317369624 | 0.102666154 | 0.38 | 0.25 |
| YKL132C | 214.3347933 | 0.371355207 | 0.102898193 | 0.38 | 0.25 |
| YPR065W | 1458.659767 | -0.5437506 | 0.102999194 | 0.38 | 0.25 |
| YJR047C | 407.0063415 | 0.660728561 | 0.103069042 | 0.38 | 0.25 |
| YOR049C | 767.2844506 | -0.543943556 | 0.103119545 | 0.38 | 0.25 |
| YBR164C | 1099.977428 | -0.368954775 | 0.103249116 | 0.38 | 0.25 |
| YPL199C | 1017.89655 | -0.438955951 | 0.103264451 | 0.38 | 0.25 |
| YGL110C | 882.4868812 | 0.253903431 | 0.103358239 | 0.38 | 0.25 |
| YGL054C | 1677.23256 | -0.292161437 | 0.10348444 | 0.38 | 0.25 |
| YOR267C | 3236.118357 | 0.371667038 | 0.103429505 | 0.38 | 0.25 |
| YBR074W | 1213.234406 | -0.219540918 | 0.10362957 | 0.38 | 0.25 |
| YLR183C | 714.9768804 | -0.417723831 | 0.103707588 | 0.38 | 0.25 |
| YER133W-A | 95.44590047 | -0.513114921 | 0.103888435 | 0.38 | 0.25 |
| YLL010C | 1089.845187 | -0.23387127 | 0.103840779 | 0.38 | 0.25 |
| YML107C | 389.7043417 | -0.356406453 | 0.103984142 | 0.38 | 0.25 |
| YOL163W | 57.10674557 | 0.72444127 | 0.103992893 | 0.38 | 0.25 |
| YML017W | 751.7187367 | 0.283168056 | 0.104184774 | 0.38 | 0.25 |
| RUF21 | 43.48350584 | 0.527309396 | 0.104385722 | 0.38 | 0.25 |
| YBL037W | 1072.022099 | -0.271443787 | 0.104678214 | 0.38 | 0.25 |
| YBR038W | 1800.515437 | 0.310985027 | 0.104794845 | 0.38 | 0.25 |
| YBR131W | 285.0666359 | 0.322395305 | 0.104934175 | 0.38 | 0.25 |
| YDR363W-A | 1308.889566 | -0.311247649 | 0.104981614 | 0.38 | 0.25 |
| YIL048W | 1213.90232 | 0.2366781 | 0.104817555 | 0.38 | 0.25 |
| YJR087W | 189.3064917 | -0.324784586 | 0.104913167 | 0.38 | 0.25 |
| YKL003C | 652.5154668 | -0.357232603 | 0.104984021 | 0.38 | 0.25 |
| YOL003C | 636.2491343 | -0.267586777 | 0.104814653 | 0.38 | 0.25 |
| YOR377W | 997.8606777 | 0.356962708 | 0.104482851 | 0.38 | 0.25 |
| YPR042C | 1338.934548 | 0.331909418 | 0.104567683 | 0.38 | 0.25 |
| YPR049C | 968.2983267 | 0.451196539 | 0.104965912 | 0.38 | 0.25 |
| YNL151C | 742.0050839 | -0.479268869 | 0.105201291 | 0.38 | 0.25 |
| YCL047C | 434.4156263 | 0.392277711 | 0.105493321 | 0.38 | 0.25 |
| YDR091C | 1728.536947 | -0.529059662 | 0.105460673 | 0.38 | 0.25 |
| YOR222W | 1007.50754 | 0.497794529 | 0.10554388 | 0.38 | 0.25 |
| YDR027C | 1560.996774 | 0.156089774 | 0.105682515 | 0.38 | 0.25 |
| YCL041C | 119.8839828 | 0.430163082 | 0.106117866 | 0.38 | 0.25 |

| | | | | | |
|---|---|---|---|---|---|
| YEL013W | 1708.200753 | -0.252812843 | 0.105954469 | 0.38 | 0.25 |
| YLR028C | 4846.630925 | -0.137534462 | 0.106083886 | 0.38 | 0.25 |
| YNL327W | 5207.566398 | -0.53405651 | 0.106065771 | 0.38 | 0.25 |
| YDR096W | 1946.105843 | 0.489044927 | 0.106245587 | 0.38 | 0.25 |
| YHL019W-A | 10.01596588 | 0.628431831 | 0.106256465 | 0.38 | 0.25 |
| YMR219W | 3001.660673 | 0.412748012 | 0.106358147 | 0.38 | 0.25 |
| YML077W | 412.7293887 | -0.39828987 | 0.10650612 | 0.38 | 0.25 |
| YGR008C | 4582.699628 | 0.683131797 | 0.106697647 | 0.38 | 0.25 |
| YGR119C | 2276.598641 | -0.239192781 | 0.106640851 | 0.38 | 0.25 |
| YDR414C | 448.7530762 | -0.470163426 | 0.106794771 | 0.38 | 0.25 |
| YMR313C | 988.4819991 | 0.29171468 | 0.106895537 | 0.38 | 0.25 |
| YPL250W-A | 88.80290187 | -0.52403584 | 0.106962631 | 0.38 | 0.25 |
| YKL034W | 1217.582813 | 0.389768948 | 0.107279244 | 0.38 | 0.25 |
| YDR236C | 550.9804022 | -0.303091106 | 0.107546009 | 0.38 | 0.25 |
| YOR007C | 8060.679187 | 0.360393502 | 0.107601357 | 0.38 | 0.25 |
| YDL015C | 3069.696513 | -0.216258696 | 0.107776614 | 0.39 | 0.25 |
| YOR039W | 1072.032394 | -0.282030502 | 0.108078045 | 0.39 | 0.26 |
| YBL097W | 438.5923101 | 0.406439847 | 0.10842137 | 0.39 | 0.26 |
| YER097W | 28.63993449 | 0.578245226 | 0.108650057 | 0.39 | 0.26 |
| YFL004W | 2271.622342 | -0.244068806 | 0.108335071 | 0.39 | 0.26 |
| YGL002W | 690.1637106 | -0.32498712 | 0.108728064 | 0.39 | 0.26 |
| YKR016W | 2416.547829 | 0.57014108 | 0.108526389 | 0.39 | 0.26 |
| YML074C | 7957.336564 | -0.468008915 | 0.108558903 | 0.39 | 0.26 |
| YMR081C | 3745.685883 | 0.772583945 | 0.108357428 | 0.39 | 0.26 |
| YMR268C | 351.0386447 | -0.26016951 | 0.108320582 | 0.39 | 0.26 |
| YNL160W | 19462.39163 | 0.596200713 | 0.108742107 | 0.39 | 0.26 |
| YOL095C | 382.7305205 | 0.306214563 | 0.108671132 | 0.39 | 0.26 |
| YJR027W | 207.2279874 | 0.658091836 | 0.108884876 | 0.39 | 0.26 |
| YNL070W | 685.2481589 | -0.33570243 | 0.108948409 | 0.39 | 0.26 |
| YDL086W | 1034.431113 | 0.43258408 | 0.109059132 | 0.39 | 0.26 |
| YBR129C | 610.3882792 | -0.372942403 | 0.109416907 | 0.39 | 0.26 |
| YOL062C | 1303.409819 | -0.258658339 | 0.109476226 | 0.39 | 0.26 |
| YGL142C | 1131.213732 | 0.357583461 | 0.109722577 | 0.39 | 0.26 |
| YDL210W | 142.1653441 | 0.631850336 | 0.10987266 | 0.39 | 0.26 |
| YKR006C | 1505.723079 | -0.310260346 | 0.109885296 | 0.39 | 0.26 |
| YLL057C | 252.3608873 | 0.570964743 | 0.109902044 | 0.39 | 0.26 |
| YKL175W | 2025.203634 | 0.288873992 | 0.110272458 | 0.39 | 0.26 |
| YLR062C | 194.5995134 | -0.633345055 | 0.110113068 | 0.39 | 0.26 |
| YNL050C | 614.5421658 | -0.243873997 | 0.110245476 | 0.39 | 0.26 |
| YOL006C | 1492.750425 | -0.276858092 | 0.110275349 | 0.39 | 0.26 |
| YOL012C | 810.102208 | -0.292863191 | 0.110099397 | 0.39 | 0.26 |
| YPR184W | 5293.583109 | 0.754618763 | 0.11000946 | 0.39 | 0.26 |
| snR24 | 8.183960631 | 1.027160773 | 0.110652349 | 0.39 | 0.26 |

| | | | | | |
|---|---|---|---|---|---|
| YER088C | 4317.842613 | 0.431566072 | 0.110855359 | 0.39 | 0.26 |
| YJR005C-A | 81.403139 | 0.750430292 | 0.11086124 | 0.39 | 0.26 |
| YDR125C | 356.5508843 | 0.535776326 | 0.111036336 | 0.39 | 0.26 |
| YBR200W-A | 19.77475914 | 1.187782338 | 0.111187038 | 0.39 | 0.26 |
| YIL113W | 384.3761033 | -0.606645382 | 0.111306105 | 0.39 | 0.26 |
| YIL133C | 6678.70118 | -0.626809668 | 0.11133471 | 0.39 | 0.26 |
| YOL130W | 1532.382846 | 0.2546118 | 0.111529296 | 0.39 | 0.26 |
| YLL002W | 366.0264394 | -0.365284095 | 0.111698098 | 0.39 | 0.26 |
| YLR122C | 30.05120318 | 0.64285563 | 0.111647075 | 0.39 | 0.26 |
| YMR153C-A | 60.08017136 | -0.397961581 | 0.1117418 | 0.39 | 0.26 |
| YDR165W | 1014.540812 | -0.504798389 | 0.112033974 | 0.39 | 0.26 |
| YAL028W | 836.6833612 | 0.454432149 | 0.113027387 | 0.39 | 0.26 |
| YAL067C | 87.22318015 | 0.905623238 | 0.113238099 | 0.39 | 0.26 |
| YDL148C | 2147.255757 | -0.539668435 | 0.11309603 | 0.39 | 0.26 |
| YDL222C | 906.6701758 | -1.002267429 | 0.113053819 | 0.39 | 0.26 |
| YER131W | 9395.024759 | -0.664751976 | 0.11316284 | 0.39 | 0.26 |
| YHR076W | 1124.266555 | 0.321561882 | 0.113193339 | 0.39 | 0.26 |
| YKL123W | 31.46802918 | 0.419652922 | 0.112933964 | 0.39 | 0.26 |
| YKR018C | 3562.456922 | 0.446315696 | 0.113254495 | 0.39 | 0.26 |
| YLR252W | 254.6818923 | 0.503375925 | 0.112916256 | 0.39 | 0.26 |
| YMR158C-A | 22.93483608 | 0.532482132 | 0.113392425 | 0.39 | 0.26 |
| YPR010C | 3009.126793 | -0.530877356 | 0.11340392 | 0.39 | 0.26 |
| YPR026W | 1343.846593 | 0.537280785 | 0.113422637 | 0.39 | 0.26 |
| YPR072W | 2174.009813 | -0.263350734 | 0.113114449 | 0.39 | 0.26 |
| YPR156C | 1539.255789 | 0.443643154 | 0.113659336 | 0.39 | 0.26 |
| YER016W | 838.7636501 | -0.473209836 | 0.113813828 | 0.40 | 0.26 |
| YCR042C | 1593.772478 | 0.205932884 | 0.114180543 | 0.40 | 0.26 |
| YDR124W | 300.8040212 | 0.500029801 | 0.114268397 | 0.40 | 0.26 |
| YNL072W | 456.0809998 | -0.280906991 | 0.114373976 | 0.40 | 0.26 |
| YIR028W | 189.55325 | 0.913073437 | 0.114655944 | 0.40 | 0.26 |
| YGR117C | 888.1364455 | -0.304590799 | 0.114733078 | 0.40 | 0.26 |
| YDR292C | 1617.849006 | -0.276449589 | 0.114805617 | 0.40 | 0.26 |
| YPL181W | 1430.008347 | 0.416680458 | 0.114952801 | 0.40 | 0.26 |
| YBR191W | 11850.59787 | -0.664274929 | 0.115059104 | 0.40 | 0.26 |
| YLR369W | 1148.442381 | 0.42815746 | 0.115330137 | 0.40 | 0.26 |
| YDR201W | 235.4443499 | -0.354230399 | 0.115766955 | 0.40 | 0.26 |
| YLR287C-A | 5234.314487 | -0.458538462 | 0.115537163 | 0.40 | 0.26 |
| YMR145C | 7155.424672 | 0.544786868 | 0.115748424 | 0.40 | 0.26 |
| YOR008C | 1383.478149 | -0.326283143 | 0.115687048 | 0.40 | 0.26 |
| YPL097W | 440.8818332 | 0.315473368 | 0.115667268 | 0.40 | 0.26 |
| YGR164W | 4.051111594 | -0.75267559 | 0.115985784 | 0.40 | 0.26 |
| YMR284W | 845.7287906 | 0.34168085 | 0.116051861 | 0.40 | 0.26 |
| YIL159W | 678.524918 | -0.319139531 | 0.116235462 | 0.40 | 0.26 |

| | | | | | |
|---|---|---|---|---|---|
| YML026C | 7646.727007 | -0.608862331 | 0.116258279 | 0.40 | 0.26 |
| YKL120W | 1575.003705 | 0.697136706 | 0.116353338 | 0.40 | 0.26 |
| YPL034W | 171.3658521 | -0.317898626 | 0.116553881 | 0.40 | 0.26 |
| YGL010W | 399.2167766 | 0.466962275 | 0.116889548 | 0.40 | 0.26 |
| YLR417W | 736.9680806 | 0.269610851 | 0.116786478 | 0.40 | 0.26 |
| YNL089C | 16.65282477 | -0.65568318 | 0.116891634 | 0.40 | 0.26 |
| YAL054C | 1109.295849 | 0.6194417 | 0.117174474 | 0.40 | 0.27 |
| YIL069C | 2018.584495 | -0.571737773 | 0.117236569 | 0.40 | 0.27 |
| YMR085W | 377.7619403 | 0.588614468 | 0.117095301 | 0.40 | 0.27 |
| YMR242C | 8203.831599 | -0.593170162 | 0.117208382 | 0.40 | 0.27 |
| YBL079W | 2598.015595 | -0.220038906 | 0.117333285 | 0.40 | 0.27 |
| YKR081C | 1261.655674 | -0.525219823 | 0.117386152 | 0.40 | 0.27 |
| YGR159C | 2044.642627 | -0.711167509 | 0.117640201 | 0.40 | 0.27 |
| YNL305C | 2021.497779 | 0.478059426 | 0.117678121 | 0.40 | 0.27 |
| YDR405W | 1203.736123 | -0.266533777 | 0.117905234 | 0.40 | 0.27 |
| YKL151C | 2678.459393 | 0.648285883 | 0.117899621 | 0.40 | 0.27 |
| YMR147W | 251.4438843 | 0.543467015 | 0.118176694 | 0.40 | 0.27 |
| YOR373W | 745.8071516 | -0.3706117 | 0.118132934 | 0.40 | 0.27 |
| YDR115W | 505.144241 | -0.295067155 | 0.118354299 | 0.40 | 0.27 |
| YBR089W | 6.735755394 | -0.853034035 | 0.118829112 | 0.40 | 0.27 |
| YDR465C | 1088.193625 | -0.572184017 | 0.119421793 | 0.40 | 0.27 |
| YER085C | 1.090623589 | 0.727969354 | 0.118791356 | 0.40 | 0.27 |
| YER093C-A | 366.8172657 | 0.340328583 | 0.118752042 | 0.40 | 0.27 |
| YER104W | 241.2969985 | 0.324818988 | 0.119503393 | 0.40 | 0.27 |
| YER142C | 720.8941802 | 0.399629561 | 0.119100998 | 0.40 | 0.27 |
| YFL064C | 3.015892609 | 1.139952284 | 0.119008583 | 0.40 | 0.27 |
| YIL075C | 7548.161259 | 0.329749474 | 0.119137246 | 0.40 | 0.27 |
| YJL163C | 815.8405692 | 0.547186377 | 0.119071262 | 0.40 | 0.27 |
| YKL186C | 1093.769583 | -0.328003146 | 0.119267459 | 0.40 | 0.27 |
| YLR435W | 440.7011003 | -0.541672171 | 0.119202343 | 0.40 | 0.27 |
| YOR180C | 257.2908648 | -0.499089349 | 0.119003078 | 0.40 | 0.27 |
| YOR311C | 1031.860298 | 0.331723281 | 0.119375192 | 0.40 | 0.27 |
| YPR029C | 1930.500091 | -0.205250427 | 0.119443026 | 0.40 | 0.27 |
| YDR261C | 1216.328788 | 0.480864982 | 0.119729244 | 0.40 | 0.27 |
| YGL252C | 1693.94272 | -0.175192779 | 0.119647696 | 0.40 | 0.27 |
| YJL086C | 9.129343542 | 0.694663939 | 0.11986097 | 0.40 | 0.27 |
| YLR255C | 23.50063941 | 0.522616411 | 0.119855173 | 0.40 | 0.27 |
| snR84 | 162.4042194 | -0.353447593 | 0.119780637 | 0.40 | 0.27 |
| YBR147W | 531.1286909 | 0.743623882 | 0.120359504 | 0.40 | 0.27 |
| YBR268W | 567.5701433 | -0.338118459 | 0.120257749 | 0.40 | 0.27 |
| YFL040W | 216.9489399 | 0.545727765 | 0.120099035 | 0.40 | 0.27 |
| YGR161C-D | 41.89517477 | 0.683810087 | 0.120373473 | 0.40 | 0.27 |
| YIL076W | 2351.989968 | -0.3729781 | 0.120247337 | 0.40 | 0.27 |

| | | | | | |
|---|---|---|---|---|---|
| YMR069W | 79.44065746 | 0.683955468 | 0.120337572 | 0.40 | 0.27 |
| YOR369C | 15497.1337 | -0.522884265 | 0.120364967 | 0.40 | 0.27 |
| YOR146W | 6.962391281 | -0.718964875 | 0.120635405 | 0.41 | 0.27 |
| YBR171W | 1056.70053 | -0.423963706 | 0.120769768 | 0.41 | 0.27 |
| YEL019C | 187.0813824 | 0.448382429 | 0.120850704 | 0.41 | 0.27 |
| YIL089W | 155.6273305 | 0.650154946 | 0.120854023 | 0.41 | 0.27 |
| YIL119C | 838.0211886 | -0.756533973 | 0.12096017 | 0.41 | 0.27 |
| YOR263C | 29.84403306 | -0.468875819 | 0.120971428 | 0.41 | 0.27 |
| YOL007C | 839.2247362 | -0.470094135 | 0.121061413 | 0.41 | 0.27 |
| YLR046C | 345.7378309 | -0.206839093 | 0.121136119 | 0.41 | 0.27 |
| YIL040W | 372.5621769 | -0.358402359 | 0.121395713 | 0.41 | 0.27 |
| YNL301C | 1146.542523 | -0.570197078 | 0.121426639 | 0.41 | 0.27 |
| YFL048C | 1163.307711 | -0.340420859 | 0.121716728 | 0.41 | 0.27 |
| YHR132C | 1760.575956 | 0.333215803 | 0.121758141 | 0.41 | 0.27 |
| YOR043W | 1570.483617 | -0.297350485 | 0.121754356 | 0.41 | 0.27 |
| YDR060W | 3331.110803 | -0.46072627 | 0.122156554 | 0.41 | 0.27 |
| YNL250W | 994.2437698 | 0.368603465 | 0.122188857 | 0.41 | 0.27 |
| YPR093C | 386.3642488 | 0.608427662 | 0.122274701 | 0.41 | 0.27 |
| YBR293W | 445.7090525 | -0.585236571 | 0.122354906 | 0.41 | 0.27 |
| YDR444W | 770.8185864 | 0.300856144 | 0.122477401 | 0.41 | 0.27 |
| YOR126C | 423.6858153 | 0.199377075 | 0.122416957 | 0.41 | 0.27 |
| YCR017C | 2986.294618 | 0.392557627 | 0.122556883 | 0.41 | 0.27 |
| YBR202W | 984.15721 | -0.332758913 | 0.122902279 | 0.41 | 0.27 |
| YER165W | 11478.83612 | -0.355880419 | 0.123027 | 0.41 | 0.27 |
| YJL045W | 286.1990705 | 0.687569534 | 0.122955221 | 0.41 | 0.27 |
| YOR124C | 2791.881139 | 0.322623415 | 0.12299679 | 0.41 | 0.27 |
| YHR162W | 4274.252821 | 0.469098394 | 0.123206576 | 0.41 | 0.27 |
| YKL054C | 4561.844374 | -0.356225432 | 0.123149198 | 0.41 | 0.27 |
| YLR389C | 2863.752537 | 0.226875045 | 0.123308903 | 0.41 | 0.27 |
| YDR200C | 868.5642679 | 0.205962405 | 0.123613221 | 0.41 | 0.27 |
| YPL104W | 889.4598617 | 0.353484397 | 0.12380408 | 0.41 | 0.27 |
| YDL202W | 808.8072189 | -0.321139129 | 0.124038103 | 0.41 | 0.27 |
| YKL176C | 1422.060464 | -0.296619445 | 0.12416739 | 0.41 | 0.27 |
| snR56 | 6.041353852 | 0.875470726 | 0.124206447 | 0.41 | 0.27 |
| YLR113W | 3162.32221 | -0.277326522 | 0.124329158 | 0.41 | 0.27 |
| YBL006C | 611.4550235 | -0.36295299 | 0.124733043 | 0.41 | 0.27 |
| YML061C | 1035.299962 | 0.343731288 | 0.124825556 | 0.41 | 0.27 |
| YBL039C | 1528.083632 | -0.549700436 | 0.125019593 | 0.41 | 0.27 |
| YDR047W | 1762.459456 | -0.273232359 | 0.124981119 | 0.41 | 0.27 |
| YCR009C | 3474.642891 | -0.269228552 | 0.125740778 | 0.41 | 0.27 |
| YGL263W | 253.3051631 | 0.831960897 | 0.125591392 | 0.41 | 0.27 |
| YIL146C | 613.496777 | 0.456799883 | 0.125769735 | 0.41 | 0.27 |
| YJL090C | 629.2586035 | -0.248514957 | 0.12572053 | 0.41 | 0.27 |

| | | | | | |
|---|---|---|---|---|---|
| YKR053C | 486.0346558 | -0.336688437 | 0.125534966 | 0.41 | 0.27 |
| YNL243W | 3599.331124 | 0.22644846 | 0.125758469 | 0.41 | 0.27 |
| YPL173W | 902.2046084 | -0.298719392 | 0.125640573 | 0.41 | 0.27 |
| YPR168W | 170.9858215 | 0.375378556 | 0.125896087 | 0.41 | 0.27 |
| YNL175C | 613.6716753 | -0.552067442 | 0.125963877 | 0.41 | 0.27 |
| YDL024C | 228.1924469 | 0.69656818 | 0.126127228 | 0.41 | 0.27 |
| YDR103W | 991.2634766 | -0.575674325 | 0.126511467 | 0.41 | 0.27 |
| YHR051W | 2680.129991 | 0.683236652 | 0.126596515 | 0.41 | 0.27 |
| YJL114W | 3.593194288 | 0.820932545 | 0.126625956 | 0.41 | 0.27 |
| YOR323C | 2339.246441 | 0.382367404 | 0.126575688 | 0.41 | 0.27 |
| YHR217C | 1.225581631 | 0.859050879 | 0.126882603 | 0.42 | 0.27 |
| YGR122W | 624.9209214 | -0.35956335 | 0.127216904 | 0.42 | 0.27 |
| YHL012W | 237.0696554 | 0.455603883 | 0.12717548 | 0.42 | 0.27 |
| YEL065W | 2278.709245 | -0.715170005 | 0.127380346 | 0.42 | 0.28 |
| YDR529C | 3824.487001 | 0.621917363 | 0.127457164 | 0.42 | 0.28 |
| YGR284C | 2427.894256 | 0.356825436 | 0.127572321 | 0.42 | 0.28 |
| YJR153W | 431.5896283 | 0.741346804 | 0.128047491 | 0.42 | 0.28 |
| YLR039C | 1333.277955 | 0.250772105 | 0.128136726 | 0.42 | 0.28 |
| YMR247C | 1642.766432 | 0.281200578 | 0.128143532 | 0.42 | 0.28 |
| YNL058C | 1080.922928 | 0.519520328 | 0.128360473 | 0.42 | 0.28 |
| YDR247W | 1026.990415 | 0.524590199 | 0.128620638 | 0.42 | 0.28 |
| YFL047W | 1142.35423 | -0.340033987 | 0.128831053 | 0.42 | 0.28 |
| YJR003C | 586.7430541 | -0.434279717 | 0.128991916 | 0.42 | 0.28 |
| YLR405W | 359.0345633 | -0.369602862 | 0.129016911 | 0.42 | 0.28 |
| YGL166W | 257.6413584 | 0.322192956 | 0.129170348 | 0.42 | 0.28 |
| YBR260C | 1246.430976 | 0.334117047 | 0.129373046 | 0.42 | 0.28 |
| YOL072W | 594.5699607 | -0.264539569 | 0.129348147 | 0.42 | 0.28 |
| YOR178C | 2730.240932 | 0.753546439 | 0.129291027 | 0.42 | 0.28 |
| YER190W | 15.31140442 | 0.77716133 | 0.129453238 | 0.42 | 0.28 |
| YDL044C | 428.1351393 | -0.250363416 | 0.129728276 | 0.42 | 0.28 |
| YLR307W | 82.37745124 | 0.925892261 | 0.130167965 | 0.42 | 0.28 |
| YDL201W | 813.4402377 | -0.422229185 | 0.130510919 | 0.42 | 0.28 |
| YER019C-A | 1962.324665 | -0.278639763 | 0.1306229 | 0.42 | 0.28 |
| YLR144C | 964.8658379 | 0.324281716 | 0.130559518 | 0.42 | 0.28 |
| YMR247W-A | 22.89498367 | 0.716391815 | 0.130687549 | 0.42 | 0.28 |
| YMR277W | 1413.809425 | -0.280446887 | 0.130670717 | 0.42 | 0.28 |
| YOL136C | 1016.324902 | -0.48443393 | 0.130418536 | 0.42 | 0.28 |
| snR45 | 13.49639451 | 0.756942137 | 0.130338559 | 0.42 | 0.28 |
| YJL132W | 534.7548177 | 0.446354609 | 0.130984684 | 0.42 | 0.28 |
| YLR142W | 1292.784521 | 0.697909854 | 0.131750411 | 0.43 | 0.28 |
| YBR281C | 961.503364 | 0.319714391 | 0.132116311 | 0.43 | 0.28 |
| YPL123C | 1037.088707 | 0.48420642 | 0.132098601 | 0.43 | 0.28 |
| YLR334C | 3.854398197 | 0.820335969 | 0.132231237 | 0.43 | 0.28 |

| | | | | | |
|---|---|---|---|---|---|
| YDR332W | 604.027567 | 0.230690108 | 0.132406117 | 0.43 | 0.28 |
| YGL012W | 2257.199778 | -0.534848313 | 0.132566988 | 0.43 | 0.28 |
| YLR406C | 2407.447144 | -0.599458134 | 0.132603862 | 0.43 | 0.28 |
| YBL049W | 377.5575063 | 0.639321084 | 0.133070408 | 0.43 | 0.28 |
| YBR032W | 28.5083245 | 0.657881648 | 0.133151734 | 0.43 | 0.28 |
| YLR135W | 729.3825255 | 0.274925935 | 0.133387161 | 0.43 | 0.28 |
| YDR284C | 1216.7009 | -0.148642769 | 0.133750914 | 0.43 | 0.28 |
| snR40 | 6.538551087 | 0.77246924 | 0.133753043 | 0.43 | 0.28 |
| YJR065C | 3820.877578 | 0.204583017 | 0.133965893 | 0.43 | 0.28 |
| YGL149W | 44.44871907 | -0.451124833 | 0.134067163 | 0.43 | 0.28 |
| YDL132W | 2917.981172 | 0.212171092 | 0.134291266 | 0.43 | 0.28 |
| YDR137W | 962.3953612 | -0.175215816 | 0.13439968 | 0.43 | 0.28 |
| YMR259C | 1208.426766 | -0.368748046 | 0.134347974 | 0.43 | 0.28 |
| YJR094W-A | 1689.898779 | -0.497916163 | 0.134496321 | 0.43 | 0.28 |
| YEL007W | 3320.15412 | -0.364404819 | 0.134595313 | 0.43 | 0.28 |
| YJL047C | 709.4046862 | 0.285630022 | 0.134692959 | 0.43 | 0.28 |
| YJL217W | 2088.445581 | 0.493840272 | 0.134877251 | 0.43 | 0.28 |
| YLR312W-A | 784.781583 | -0.29919815 | 0.134964353 | 0.43 | 0.28 |
| YKL016C | 3252.389125 | 0.50284208 | 0.135260007 | 0.43 | 0.29 |
| YOR081C | 1945.1418 | -0.320712833 | 0.135430122 | 0.43 | 0.29 |
| YKL220C | 368.1832027 | 0.532781988 | 0.135530811 | 0.43 | 0.29 |
| YBL068W | 791.3128306 | -0.328139116 | 0.135624628 | 0.43 | 0.29 |
| YJL212C | 1268.474729 | 0.593414642 | 0.135770612 | 0.43 | 0.29 |
| YNL336W | 467.9927857 | 0.556366588 | 0.135765551 | 0.43 | 0.29 |
| YCR082W | 1363.300313 | -0.402481598 | 0.136071359 | 0.43 | 0.29 |
| YJR059W | 3492.702056 | 0.34271312 | 0.136142854 | 0.43 | 0.29 |
| YMR124W | 1653.834225 | 0.276896109 | 0.136190806 | 0.43 | 0.29 |
| YPL012W | 1836.869245 | -0.503089861 | 0.136153794 | 0.43 | 0.29 |
| YPL045W | 887.7248714 | 0.236358806 | 0.136174159 | 0.43 | 0.29 |
| YJR061W | 732.5988419 | 0.29413147 | 0.136342798 | 0.43 | 0.29 |
| YPR016W-A | 37.91128357 | -0.509129017 | 0.136326791 | 0.43 | 0.29 |
| YIL155C | 2791.794832 | 0.591227303 | 0.136455733 | 0.43 | 0.29 |
| YAL048C | 583.0194511 | 0.324384742 | 0.137090231 | 0.43 | 0.29 |
| YBL106C | 800.8986155 | 0.330395322 | 0.137446862 | 0.43 | 0.29 |
| YCR034W | 1456.119271 | -0.556680888 | 0.137356005 | 0.43 | 0.29 |
| YDR025W | 5331.251911 | -0.569213376 | 0.137385457 | 0.43 | 0.29 |
| YGR042W | 764.6829972 | -0.35082079 | 0.137271427 | 0.43 | 0.29 |
| YLR354C | 11943.28845 | 0.351183219 | 0.137376849 | 0.43 | 0.29 |
| YML124C | 2648.714801 | 0.411374277 | 0.136960033 | 0.43 | 0.29 |
| YMR112C | 344.1352482 | -0.319266087 | 0.137341965 | 0.43 | 0.29 |
| YOR230W | 8762.727513 | 0.360488182 | 0.13737331 | 0.43 | 0.29 |
| YPL211W | 668.7255746 | -0.588964208 | 0.137287402 | 0.43 | 0.29 |
| YDR526C | 4.767823074 | 0.558678621 | 0.137539599 | 0.43 | 0.29 |

| | | | | | |
|---|---|---|---|---|---|
| YJL203W | 345.939173 | -0.219332323 | 0.137741952 | 0.43 | 0.29 |
| YBR056W | 2794.886987 | 0.412765805 | 0.13808815 | 0.43 | 0.29 |
| YCL068C | 4.465179414 | 0.331992845 | 0.138372244 | 0.43 | 0.29 |
| YDL159W-A | 1.037341025 | 0.739036039 | 0.138283977 | 0.43 | 0.29 |
| YJL028W | 3.906046468 | 0.735310649 | 0.138535167 | 0.43 | 0.29 |
| YJL127C-B | 263.7714489 | -0.294658003 | 0.138450148 | 0.43 | 0.29 |
| YJL155C | 1149.698916 | 0.469266735 | 0.138596323 | 0.43 | 0.29 |
| YKL026C | 613.714189 | 0.675456252 | 0.138378968 | 0.43 | 0.29 |
| YMR017W | 57.51133406 | 0.572123579 | 0.138519384 | 0.43 | 0.29 |
| YNL177C | 1301.893277 | -0.17710418 | 0.138407077 | 0.43 | 0.29 |
| YOR235W | 3.373424746 | 0.752564553 | 0.13846173 | 0.43 | 0.29 |
| YPR006C | 675.2062996 | 0.51945837 | 0.138563403 | 0.43 | 0.29 |
| YKL201C | 1146.760044 | 0.373294007 | 0.138667107 | 0.43 | 0.29 |
| YDL001W | 483.443293 | -0.227684027 | 0.139129527 | 0.44 | 0.29 |
| YHR214W-A | 22.9859353 | 0.660287286 | 0.139192865 | 0.44 | 0.29 |
| YOL079W | 9.64455915 | -0.715161154 | 0.139177248 | 0.44 | 0.29 |
| YPL221W | 1548.403188 | 0.367455958 | 0.139045862 | 0.44 | 0.29 |
| YBR172C | 2003.113678 | -0.339603477 | 0.139567217 | 0.44 | 0.29 |
| YER181C | 6.5610832 | 0.754030695 | 0.139439469 | 0.44 | 0.29 |
| YGR265W | 63.64327138 | 0.489006534 | 0.139555569 | 0.44 | 0.29 |
| YBR021W | 496.9097626 | -0.353904971 | 0.139998198 | 0.44 | 0.29 |
| YIL036W | 2051.229049 | 0.328267535 | 0.140020408 | 0.44 | 0.29 |
| YDL026W | 16.56831607 | 0.612101229 | 0.140202924 | 0.44 | 0.29 |
| YDR335W | 3040.69958 | 0.297127392 | 0.140345026 | 0.44 | 0.29 |
| YPR073C | 916.4024804 | 0.287926918 | 0.140334118 | 0.44 | 0.29 |
| snR44 | 201.2193719 | -0.591392266 | 0.140412691 | 0.44 | 0.29 |
| NME1 | 51.73748825 | 0.658964152 | 0.140544447 | 0.44 | 0.29 |
| YJL166W | 2486.295448 | 0.613038458 | 0.141062152 | 0.44 | 0.29 |
| YBL005W | 1420.668611 | -0.304386441 | 0.141274141 | 0.44 | 0.29 |
| YDR394W | 3757.628989 | -0.240099004 | 0.141520095 | 0.44 | 0.29 |
| YNL236W | 1176.860728 | -0.248317597 | 0.141695411 | 0.44 | 0.29 |
| YPL139C | 637.4930321 | -0.189484869 | 0.141653676 | 0.44 | 0.29 |
| YGL011C | 3486.685044 | -0.226411055 | 0.14181067 | 0.44 | 0.29 |
| YBR298C | 1312.084765 | 0.628851663 | 0.141891937 | 0.44 | 0.29 |
| YIL045W | 1955.748951 | 0.681605331 | 0.141985633 | 0.44 | 0.29 |
| YDR041W | 1214.356247 | -0.303001377 | 0.142423154 | 0.44 | 0.29 |
| YGL013C | 1490.03588 | -0.261883845 | 0.142355195 | 0.44 | 0.29 |
| YHR030C | 2133.377408 | 0.685496804 | 0.142408788 | 0.44 | 0.29 |
| YKL106W | 462.66436 | -0.375239186 | 0.142232002 | 0.44 | 0.29 |
| YPL089C | 1858.715154 | 0.34812456 | 0.14246319 | 0.44 | 0.29 |
| YGR295C | 1429.22247 | 0.552528615 | 0.142613404 | 0.44 | 0.29 |
| YLR004C | 257.8975095 | 0.381068499 | 0.142564328 | 0.44 | 0.29 |
| YCR020W-B | 152.1239298 | -0.46801533 | 0.142866968 | 0.44 | 0.29 |

| | | | | | |
|---|---|---|---|---|---|
| YDR212W | 2389.189098 | -0.325252136 | 0.142784489 | 0.44 | 0.29 |
| YDR248C | 767.0999761 | 0.352549433 | 0.142897459 | 0.44 | 0.29 |
| YIL141W | 35.62085203 | 0.519669099 | 0.143025466 | 0.44 | 0.29 |
| YER141W | 2454.288383 | 0.36095407 | 0.143213488 | 0.44 | 0.29 |
| YPR159W | 2743.676738 | 0.360945453 | 0.143216082 | 0.44 | 0.29 |
| YER002W | 1431.744689 | -0.338797135 | 0.143326682 | 0.44 | 0.29 |
| YIL071C | 381.3119395 | 0.228676342 | 0.143541196 | 0.44 | 0.29 |
| YJL170C | 341.173457 | -0.857832368 | 0.143644116 | 0.44 | 0.29 |
| YKR015C | 299.9934309 | 0.271391503 | 0.143794348 | 0.44 | 0.29 |
| YOR127W | 1893.705383 | 0.250746501 | 0.143930817 | 0.44 | 0.29 |
| YDL058W | 1998.717774 | 0.261743782 | 0.144127194 | 0.44 | 0.29 |
| YKL144C | 415.786387 | -0.394267691 | 0.14417427 | 0.44 | 0.29 |
| YMR031W-A | 40.58383151 | 0.532149597 | 0.144475103 | 0.44 | 0.29 |
| YNL106C | 963.8876185 | 0.216882344 | 0.144438926 | 0.44 | 0.29 |
| YCR007C | 318.2333147 | 0.367037459 | 0.14468318 | 0.44 | 0.29 |
| YGL208W | 855.2756461 | 0.494134369 | 0.144901765 | 0.44 | 0.29 |
| YJL009W | 66.31151028 | -0.374687313 | 0.145010747 | 0.44 | 0.29 |
| YMR159C | 22.87110149 | 0.35667078 | 0.145004934 | 0.44 | 0.29 |
| YPR083W | 614.8326597 | 0.538895455 | 0.144806759 | 0.44 | 0.29 |
| YBR286W | 17567.33584 | -0.377719575 | 0.145111933 | 0.44 | 0.29 |
| YLR404W | 245.17757 | -0.312486932 | 0.145196674 | 0.44 | 0.29 |
| YNL011C | 614.0144377 | 0.346757549 | 0.145253298 | 0.44 | 0.29 |
| YDR520C | 591.1970741 | 0.404539792 | 0.145374397 | 0.44 | 0.29 |
| YGL246C | 646.4635665 | -0.227507247 | 0.145422391 | 0.44 | 0.29 |
| YPL234C | 1583.101568 | -0.186451606 | 0.145931791 | 0.45 | 0.29 |
| YIR032C | 113.4974223 | 0.581041392 | 0.146842451 | 0.45 | 0.30 |
| YJR138W | 1461.069987 | 0.349832033 | 0.146843662 | 0.45 | 0.30 |
| YDL004W | 2874.441994 | 0.568941777 | 0.146993837 | 0.45 | 0.30 |
| YDR448W | 426.123254 | -0.32429532 | 0.146942872 | 0.45 | 0.30 |
| YJL075C | 78.82663296 | -0.368421504 | 0.147052412 | 0.45 | 0.30 |
| YBR072W | 9138.655064 | 0.703638583 | 0.147182081 | 0.45 | 0.30 |
| YNL061W | 2817.950295 | -0.433864984 | 0.147301897 | 0.45 | 0.30 |
| YDL067C | 1308.526792 | 0.65136954 | 0.147482203 | 0.45 | 0.30 |
| YIL058W | 79.78173189 | -0.40982138 | 0.147539685 | 0.45 | 0.30 |
| YDR471W | 4845.236011 | -0.547034781 | 0.147666034 | 0.45 | 0.30 |
| YBR226C | 66.35832606 | 0.420461793 | 0.147965452 | 0.45 | 0.30 |
| YDR190C | 2078.525809 | -0.315613519 | 0.148008877 | 0.45 | 0.30 |
| YGL118C | 2.533670207 | 1.053152241 | 0.147985922 | 0.45 | 0.30 |
| YBR168W | 396.8925406 | -0.282831955 | 0.148344027 | 0.45 | 0.30 |
| snR9 | 75.73439823 | 0.475715854 | 0.148343125 | 0.45 | 0.30 |
| YML131W | 1116.004425 | 0.583448277 | 0.148667768 | 0.45 | 0.30 |
| YKL205W | 1310.174229 | -0.372264712 | 0.14882674 | 0.45 | 0.30 |
| YGR205W | 695.7192252 | -0.459031423 | 0.149032354 | 0.45 | 0.30 |

| | | | | | |
|---|---|---|---|---|---|
| YMR230W | 6828.386976 | -0.513483989 | 0.14910924 | 0.45 | 0.30 |
| YBL032W | 1731.697141 | -0.349087798 | 0.149310249 | 0.45 | 0.30 |
| YJR104C | 12449.61979 | -0.322715158 | 0.149427095 | 0.45 | 0.30 |
| YAR071W | 43.79477313 | -0.671684124 | 0.151256819 | 0.45 | 0.30 |
| YCR104W | 19.10661275 | 0.679540814 | 0.15119023 | 0.45 | 0.30 |
| YDL071C | 49.14928453 | 0.431850179 | 0.151117165 | 0.45 | 0.30 |
| YDR081C | 1262.178022 | -0.251730065 | 0.150256217 | 0.45 | 0.30 |
| YER056C-A | 4887.303568 | -0.549995611 | 0.15068544 | 0.45 | 0.30 |
| YER182W | 1071.612284 | 0.443178017 | 0.150063281 | 0.45 | 0.30 |
| YER183C | 378.9572978 | -0.25866448 | 0.15067358 | 0.45 | 0.30 |
| YGL066W | 1160.200958 | 0.253584837 | 0.150409485 | 0.45 | 0.30 |
| YGR121W-A | 8.749518959 | 0.663327999 | 0.150041739 | 0.45 | 0.30 |
| YHR163W | 2218.862273 | 0.599147584 | 0.150857124 | 0.45 | 0.30 |
| YIL166C | 439.1240138 | 0.422021377 | 0.151026855 | 0.45 | 0.30 |
| YLL004W | 381.8327061 | -0.322723781 | 0.15115501 | 0.45 | 0.30 |
| YLR279W | 36.37996098 | 0.632565868 | 0.151271671 | 0.45 | 0.30 |
| YLR292C | 1010.798537 | -0.401930502 | 0.151087228 | 0.45 | 0.30 |
| YML053C | 803.1994724 | -0.342274164 | 0.150281613 | 0.45 | 0.30 |
| YML110C | 4679.227772 | 0.387885339 | 0.150386625 | 0.45 | 0.30 |
| YMR074C | 1117.334751 | -0.341833902 | 0.151114043 | 0.45 | 0.30 |
| YMR251W-A | 5878.547147 | 0.508922088 | 0.150835518 | 0.45 | 0.30 |
| YNL062C | 805.5436558 | -0.50291597 | 0.15014301 | 0.45 | 0.30 |
| YOR029W | 6.24277143 | 0.761406656 | 0.151330741 | 0.45 | 0.30 |
| YOR132W | 1718.595531 | 0.310276578 | 0.149978567 | 0.45 | 0.30 |
| YOR340C | 775.5259752 | -0.531432443 | 0.151385978 | 0.45 | 0.30 |
| YPL169C | 1689.790258 | -0.252857244 | 0.151335375 | 0.45 | 0.30 |
| YPL207W | 1844.153383 | 0.366729067 | 0.1502282 | 0.45 | 0.30 |
| YPL252C | 462.6644596 | -0.39150266 | 0.151350195 | 0.45 | 0.30 |
| YGL240W | 171.0354676 | 0.266925036 | 0.151528808 | 0.45 | 0.30 |
| YGR181W | 835.8007421 | -0.26318525 | 0.151463768 | 0.45 | 0.30 |
| YBR263W | 3971.725103 | 0.231375471 | 0.151703755 | 0.45 | 0.30 |
| YKL036C | 247.9148773 | 0.405226378 | 0.151673621 | 0.45 | 0.30 |
| YLR026C | 1116.422754 | -0.306179173 | 0.151758716 | 0.45 | 0.30 |
| YNR019W | 1395.465767 | 0.356953647 | 0.152066322 | 0.45 | 0.30 |
| YNR028W | 918.8142146 | -0.220040642 | 0.152102482 | 0.45 | 0.30 |
| YKL021C | 1160.236373 | -0.406330595 | 0.152186119 | 0.45 | 0.30 |
| YGR240C-A | 46.68747277 | -0.553905166 | 0.152389719 | 0.45 | 0.30 |
| YOR378W | 266.6242043 | 0.426344059 | 0.152333583 | 0.45 | 0.30 |
| YJL148W | 1042.119748 | -0.551870649 | 0.152489014 | 0.45 | 0.30 |
| YCR012W | 43096.09714 | 0.570520154 | 0.153059634 | 0.45 | 0.30 |
| YKL190W | 813.5201171 | -0.312487995 | 0.153130554 | 0.45 | 0.30 |
| YLR065C | 843.2361186 | -0.399003447 | 0.153155831 | 0.45 | 0.30 |
| YLR069C | 1475.235001 | 0.310735173 | 0.153059444 | 0.45 | 0.30 |

| | | | | | |
|---|---|---|---|---|---|
| YLR330W | 3759.858385 | -0.185802115 | 0.15289601 | 0.45 | 0.30 |
| YDL207W | 1207.359156 | -0.168889258 | 0.153408129 | 0.45 | 0.30 |
| YJL100W | 1049.188324 | 0.313596035 | 0.153485569 | 0.45 | 0.30 |
| YPL007C | 583.714763 | -0.16620052 | 0.153440211 | 0.45 | 0.30 |
| YDR143C | 1247.877225 | -0.252953439 | 0.153687608 | 0.45 | 0.30 |
| YGR087C | 287.4531423 | 0.728070055 | 0.153618604 | 0.45 | 0.30 |
| YHR109W | 446.2718996 | 0.290573179 | 0.153777475 | 0.46 | 0.30 |
| YAR070C | 0.925768202 | 0.772599304 | 0.154165589 | 0.46 | 0.30 |
| YHR103W | 1865.030837 | 0.292118963 | 0.154156839 | 0.46 | 0.30 |
| YJL126W | 376.2312005 | 0.199629755 | 0.154034379 | 0.46 | 0.30 |
| YNL090W | 89.81803356 | -0.449239673 | 0.154535271 | 0.46 | 0.30 |
| YBR014C | 1010.158039 | 0.233287234 | 0.154688136 | 0.46 | 0.30 |
| YJL096W | 663.1647142 | -0.293279944 | 0.154797497 | 0.46 | 0.30 |
| YNL023C | 950.809325 | 0.237774591 | 0.155124496 | 0.46 | 0.30 |
| YNL223W | 533.8185201 | 0.322194067 | 0.155056426 | 0.46 | 0.30 |
| YHR031C | 752.7001486 | 0.400035348 | 0.15583148 | 0.46 | 0.30 |
| YLR012C | 11.76101632 | 0.581202197 | 0.155857229 | 0.46 | 0.30 |
| snR70 | 64.50681846 | 0.539853711 | 0.15597088 | 0.46 | 0.30 |
| YIL173W | 14.66085042 | 0.702111527 | 0.156060727 | 0.46 | 0.30 |
| YDR504C | 165.3464621 | 0.298283929 | 0.15647335 | 0.46 | 0.30 |
| YFL011W | 70.61598103 | 0.825041622 | 0.156423888 | 0.46 | 0.30 |
| YPL187W | 17.17000227 | 0.882660037 | 0.156421102 | 0.46 | 0.30 |
| YDR302W | 389.0867763 | -0.317111846 | 0.156697784 | 0.46 | 0.30 |
| YEL020C | 1045.827861 | 0.419710643 | 0.156548498 | 0.46 | 0.30 |
| YMR211W | 431.1277672 | -0.200831952 | 0.156760176 | 0.46 | 0.30 |
| YNL163C | 1921.119184 | 0.341445022 | 0.156637319 | 0.46 | 0.30 |
| YDL070W | 4583.373904 | 0.370846948 | 0.156932335 | 0.46 | 0.30 |
| YDL063C | 706.2977789 | -0.539345732 | 0.157729432 | 0.46 | 0.31 |
| YGR172C | 884.3306623 | -0.242038848 | 0.15813432 | 0.46 | 0.31 |
| YJL190C | 9254.873132 | -0.700282255 | 0.158154103 | 0.46 | 0.31 |
| YJR020W | 8.221696255 | 0.540134657 | 0.15824285 | 0.46 | 0.31 |
| YLR034C | 1366.422953 | -0.32433513 | 0.158069244 | 0.46 | 0.31 |
| YML103C | 3218.889825 | -0.29187241 | 0.15804459 | 0.46 | 0.31 |
| YNL184C | 77.22136504 | -0.418519555 | 0.157890925 | 0.46 | 0.31 |
| YOL051W | 2569.23447 | 0.28772485 | 0.158022226 | 0.46 | 0.31 |
| YOR236W | 448.7599801 | -0.434443212 | 0.157678591 | 0.46 | 0.31 |
| YOR239W | 2726.169468 | -0.295128149 | 0.158296293 | 0.46 | 0.31 |
| YPL094C | 2895.773005 | -0.261025281 | 0.158179376 | 0.46 | 0.31 |
| YDL164C | 704.8057939 | -0.225337472 | 0.158863675 | 0.46 | 0.31 |
| YDL180W | 735.4223348 | 0.279632329 | 0.158661494 | 0.46 | 0.31 |
| YDR403W | 327.0898139 | 0.600492059 | 0.158729119 | 0.46 | 0.31 |
| YHL032C | 2561.780761 | 0.517545041 | 0.158448074 | 0.46 | 0.31 |
| YHR100C | 468.2098929 | 0.243250676 | 0.159009125 | 0.46 | 0.31 |

| | | | | | |
|---|---|---|---|---|---|
| YJL020C | 4552.045062 | 0.384838279 | 0.159004166 | 0.46 | 0.31 |
| YKR067W | 2101.266331 | 0.385699569 | 0.158980588 | 0.46 | 0.31 |
| YOR234C | 2998.326335 | -0.540150248 | 0.158533406 | 0.46 | 0.31 |
| YPL270W | 1768.066094 | 0.356524552 | 0.158532396 | 0.46 | 0.31 |
| YPR031W | 763.0479333 | 0.322583042 | 0.158641913 | 0.46 | 0.31 |
| YOR155C | 1051.816257 | 0.246264623 | 0.159397029 | 0.46 | 0.31 |
| YBL013W | 160.0267849 | 0.364643664 | 0.159865935 | 0.46 | 0.31 |
| YIR017C | 6.4006803 | 0.616529715 | 0.159898345 | 0.46 | 0.31 |
| YNL248C | 1500.899157 | -0.555441386 | 0.159825113 | 0.46 | 0.31 |
| SCR1 | 181.7417356 | 0.737770841 | 0.160145877 | 0.46 | 0.31 |
| YEL068C | 304.3910786 | -0.363474448 | 0.160084075 | 0.46 | 0.31 |
| YER006C-A | 2.225353622 | 0.805909487 | 0.160301401 | 0.46 | 0.31 |
| YGL188C | 26.0866713 | 0.689352823 | 0.160392173 | 0.46 | 0.31 |
| YHR153C | 73.13873436 | -0.417335833 | 0.160218956 | 0.46 | 0.31 |
| YJL050W | 1800.303824 | -0.421120287 | 0.160328765 | 0.46 | 0.31 |
| YOR224C | 1759.04057 | -0.439189489 | 0.160526176 | 0.46 | 0.31 |
| YDR120C | 1240.636645 | -0.411642183 | 0.160913274 | 0.46 | 0.31 |
| YDR438W | 373.0593006 | 0.303287774 | 0.16122415 | 0.46 | 0.31 |
| YER132C | 1149.8348 | 0.331152139 | 0.161230888 | 0.46 | 0.31 |
| YER156C | 1503.86508 | -0.384846682 | 0.161060729 | 0.46 | 0.31 |
| YIL114C | 411.0869055 | -0.393192143 | 0.161347683 | 0.46 | 0.31 |
| YLR410W | 2881.535918 | 0.25842815 | 0.161184264 | 0.46 | 0.31 |
| YMR308C | 3347.564301 | -0.358697625 | 0.161226213 | 0.46 | 0.31 |
| YNR075W | 186.6264191 | 0.537756561 | 0.161288096 | 0.46 | 0.31 |
| YPL255W | 357.750249 | -0.358804519 | 0.161641707 | 0.47 | 0.31 |
| YCR071C | 461.1120902 | -0.392130101 | 0.162248143 | 0.47 | 0.31 |
| YDL011C | 7.393723597 | -0.623742455 | 0.161788979 | 0.47 | 0.31 |
| YDR319C | 485.2377065 | -0.29103654 | 0.162427949 | 0.47 | 0.31 |
| YGR294W | 10.95232902 | -0.683611719 | 0.162190735 | 0.47 | 0.31 |
| YJR089W | 953.5011996 | 0.226284714 | 0.162031276 | 0.47 | 0.31 |
| YKL008C | 1304.945966 | -0.420246292 | 0.162383804 | 0.47 | 0.31 |
| YLR426W | 420.438408 | -0.294043997 | 0.162257997 | 0.47 | 0.31 |
| YMR223W | 516.9298345 | -0.180261446 | 0.162331908 | 0.47 | 0.31 |
| YNR064C | 215.5475844 | 0.565035982 | 0.162387172 | 0.47 | 0.31 |
| YOR059C | 1230.819833 | 0.25807263 | 0.162472729 | 0.47 | 0.31 |
| YPL126W | 2185.981195 | -0.468739138 | 0.161961835 | 0.47 | 0.31 |
| YNR003C | 554.144982 | -0.450809133 | 0.162805368 | 0.47 | 0.31 |
| YJR086W | 331.1910032 | -0.715543187 | 0.163227414 | 0.47 | 0.31 |
| YOR152C | 552.1803625 | 0.421655607 | 0.163145485 | 0.47 | 0.31 |
| YPL186C | 1114.795126 | 0.642828759 | 0.163329914 | 0.47 | 0.31 |
| YPR057W | 656.961666 | -0.230309067 | 0.163344642 | 0.47 | 0.31 |
| YNL116W | 793.4753006 | 0.373988835 | 0.163429482 | 0.47 | 0.31 |
| YBR200W | 1077.368125 | -0.196880058 | 0.163592473 | 0.47 | 0.31 |

| | | | | | |
|---|---|---|---|---|---|
| YDL228C | 55.92892077 | -0.736883329 | 0.163534563 | 0.47 | 0.31 |
| YCL054W | 1476.642208 | -0.516918982 | 0.16382599 | 0.47 | 0.31 |
| YGR118W | 4349.282861 | -0.435387797 | 0.163958074 | 0.47 | 0.31 |
| YHR052W | 1127.946092 | -0.449924643 | 0.163934861 | 0.47 | 0.31 |
| YOR173W | 2098.419186 | 0.596624871 | 0.16377173 | 0.47 | 0.31 |
| YLR442C | 902.5050218 | 0.199579809 | 0.164039211 | 0.47 | 0.31 |
| YIL161W | 488.7537738 | -0.261465337 | 0.164164479 | 0.47 | 0.31 |
| YDL083C | 6219.514303 | -0.497453732 | 0.164290741 | 0.47 | 0.31 |
| YBR026C | 2145.982583 | 0.327522335 | 0.164598506 | 0.47 | 0.31 |
| YBR091C | 248.9757925 | -0.267663712 | 0.164668599 | 0.47 | 0.31 |
| YIL147C | 2000.977172 | 0.306662647 | 0.164986566 | 0.47 | 0.31 |
| YDR210C-D | 43.7176532 | 0.609998507 | 0.165064628 | 0.47 | 0.31 |
| YBR103C-A | 1.570441558 | 0.993397592 | 0.165177511 | 0.47 | 0.31 |
| YJL005W | 4176.860231 | 0.368469352 | 0.165234803 | 0.47 | 0.31 |
| YBR254C | 311.1708692 | -0.364832146 | 0.165606489 | 0.47 | 0.31 |
| YCL057C-A | 1206.33424 | 0.470053431 | 0.16558087 | 0.47 | 0.31 |
| YGR100W | 1479.985263 | 0.152288513 | 0.165485096 | 0.47 | 0.31 |
| YOR205C | 770.9234478 | -0.274624643 | 0.165425224 | 0.47 | 0.31 |
| YDR523C | 105.2296251 | 0.584101197 | 0.165749965 | 0.47 | 0.31 |
| YNL219C | 1817.162227 | -0.254390497 | 0.165779208 | 0.47 | 0.31 |
| YDR364C | 596.6987029 | -0.194001776 | 0.166088023 | 0.47 | 0.31 |
| YLR448W | 5493.030487 | -0.540875411 | 0.165998122 | 0.47 | 0.31 |
| YOR177C | 142.866635 | 0.644926559 | 0.166079772 | 0.47 | 0.31 |
| YDR104C | 711.7535955 | 0.302593374 | 0.166521379 | 0.47 | 0.31 |
| YIL002C | 1259.267656 | 0.219041263 | 0.1666701 | 0.47 | 0.31 |
| YML096W | 652.0319638 | -0.259253565 | 0.166593905 | 0.47 | 0.31 |
| YOR329W-A | 20.44917844 | -0.413234862 | 0.16660929 | 0.47 | 0.31 |
| YPL143W | 740.5431297 | -0.485886522 | 0.166498735 | 0.47 | 0.31 |
| YPR102C | 3154.540675 | -0.465229421 | 0.166833649 | 0.47 | 0.31 |
| YDL199C | 559.7723367 | 0.444415647 | 0.167259609 | 0.47 | 0.31 |
| YDR425W | 1030.779588 | 0.30268011 | 0.167188085 | 0.47 | 0.31 |
| YGR024C | 991.4360363 | -0.175211692 | 0.167260253 | 0.47 | 0.31 |
| YJR007W | 3429.418011 | -0.374237628 | 0.167013013 | 0.47 | 0.31 |
| YPR110C | 2091.025042 | -0.440848943 | 0.167223207 | 0.47 | 0.31 |
| YLR129W | 1609.244371 | -0.421206818 | 0.167615208 | 0.47 | 0.31 |
| YMR301C | 1249.636815 | 0.244449336 | 0.167920274 | 0.47 | 0.31 |
| YLR199C | 824.459216 | 0.171135372 | 0.168078394 | 0.47 | 0.31 |
| YPL105C | 3260.005512 | -0.22235974 | 0.168100224 | 0.47 | 0.31 |
| YCL051W | 1104.388423 | 0.274764848 | 0.168690156 | 0.47 | 0.31 |
| YHR040W | 740.6643017 | 0.426491097 | 0.168625733 | 0.47 | 0.31 |
| YHR041C | 584.7828437 | -0.373119799 | 0.168621991 | 0.47 | 0.31 |
| YLR327C | 4824.371416 | 0.601432115 | 0.168717277 | 0.47 | 0.31 |
| YBR291C | 379.9577466 | -0.371903022 | 0.168919861 | 0.47 | 0.31 |

| | | | | | |
|---|---|---|---|---|---|
| YPR133C | 1652.526675 | -0.235690911 | 0.168934773 | 0.47 | 0.31 |
| YMR117C | 204.4706004 | 0.323328787 | 0.169238569 | 0.47 | 0.31 |
| YMR243C | 2759.15272 | -0.302763493 | 0.169186593 | 0.47 | 0.31 |
| YNL240C | 1001.196601 | 0.307357041 | 0.16923332 | 0.47 | 0.31 |
| YBR058C | 1360.304304 | -0.152404892 | 0.169338936 | 0.47 | 0.31 |
| YJL058C | 326.9260026 | 0.276212032 | 0.169397016 | 0.47 | 0.31 |
| YNL098C | 3291.418535 | -0.23516077 | 0.169586517 | 0.47 | 0.31 |
| YAL031W-A | 36.5218107 | 0.357762172 | 0.16987701 | 0.47 | 0.31 |
| YDL122W | 1760.896516 | -0.204129756 | 0.169962171 | 0.47 | 0.31 |
| YNL052W | 2172.615322 | 0.616660379 | 0.170082243 | 0.47 | 0.31 |
| SRG1 | 1084.659985 | -0.594059591 | 0.170314827 | 0.47 | 0.31 |
| YDR524W-C | 1641.998262 | -0.421445598 | 0.170876112 | 0.47 | 0.31 |
| YIL129C | 2649.636452 | 0.240279008 | 0.170865051 | 0.47 | 0.31 |
| YIR031C | 471.7138035 | 0.445294927 | 0.170618145 | 0.47 | 0.31 |
| YLR363W-A | 269.4630453 | -0.423332863 | 0.170864191 | 0.47 | 0.31 |
| YLR409C | 1307.908763 | -0.45177763 | 0.170885157 | 0.47 | 0.31 |
| YMR290C | 2185.404412 | -0.566539159 | 0.170775219 | 0.47 | 0.31 |
| YOL001W | 429.8453508 | -0.38227763 | 0.170468707 | 0.47 | 0.31 |
| YOR301W | 501.5102259 | -0.404489348 | 0.170505864 | 0.47 | 0.31 |
| YOR343C | 63.14207102 | 0.573319869 | 0.170370626 | 0.47 | 0.31 |
| YDR435C | 760.2512519 | 0.323782686 | 0.171114777 | 0.48 | 0.31 |
| YPR199C | 796.7118888 | -0.356732548 | 0.171054099 | 0.48 | 0.31 |
| YLR021W | 410.9321978 | -0.214283763 | 0.171255182 | 0.48 | 0.31 |
| YDR542W | 1.676832288 | 0.731484236 | 0.171455686 | 0.48 | 0.31 |
| YLR225C | 1476.977135 | 0.434913051 | 0.171732974 | 0.48 | 0.31 |
| YLR131C | 1246.543234 | 0.256114484 | 0.17213253 | 0.48 | 0.32 |
| YPL042C | 568.6329785 | -0.345108832 | 0.172140359 | 0.48 | 0.32 |
| YPL146C | 923.2028372 | -0.376996105 | 0.172118982 | 0.48 | 0.32 |
| YDL119C | 290.4231748 | 0.266326309 | 0.172547232 | 0.48 | 0.32 |
| YLR213C | 172.0132438 | -0.37392519 | 0.172612481 | 0.48 | 0.32 |
| YMR030W | 574.6177195 | 0.442534693 | 0.172955347 | 0.48 | 0.32 |
| YDL046W | 2942.234491 | -0.299769245 | 0.173298556 | 0.48 | 0.32 |
| YEL071W | 3890.356415 | 0.577517237 | 0.173263665 | 0.48 | 0.32 |
| YNL244C | 2167.186944 | -0.340941059 | 0.173502669 | 0.48 | 0.32 |
| YNR038W | 1115.811097 | -0.333229323 | 0.173504533 | 0.48 | 0.32 |
| YMR129W | 1909.721365 | -0.182713059 | 0.174311099 | 0.48 | 0.32 |
| snR85 | 18.57666508 | -0.578660626 | 0.174351858 | 0.48 | 0.32 |
| YGL207W | 4342.654723 | -0.259569456 | 0.17462165 | 0.48 | 0.32 |
| YGR066C | 223.0827395 | 0.484329882 | 0.174736812 | 0.48 | 0.32 |
| YKL023W | 1129.71016 | 0.366348086 | 0.174694366 | 0.48 | 0.32 |
| YOR031W | 276.3450169 | 0.643529713 | 0.17452619 | 0.48 | 0.32 |
| YNL283C | 1549.072925 | 0.240642431 | 0.174872482 | 0.48 | 0.32 |
| YHR161C | 2280.636973 | 0.437477489 | 0.175155617 | 0.48 | 0.32 |

| | | | | | |
|---|---|---|---|---|---|
| YEL038W | 943.0798183 | -0.250996541 | 0.175404359 | 0.48 | 0.32 |
| YLR446W | 527.2109038 | 0.318369592 | 0.175343771 | 0.48 | 0.32 |
| YPL081W | 1016.551966 | -0.44672626 | 0.176003284 | 0.48 | 0.32 |
| YDR308C | 370.9057891 | -0.30010546 | 0.176587747 | 0.49 | 0.32 |
| YJL069C | 705.1093909 | -0.489580921 | 0.176885583 | 0.49 | 0.32 |
| YAR008W | 281.6693003 | -0.369603367 | 0.176974148 | 0.49 | 0.32 |
| YBR019C | 339.0837633 | -0.509453918 | 0.177152397 | 0.49 | 0.32 |
| YNL110C | 899.4305976 | -0.438643182 | 0.177254684 | 0.49 | 0.32 |
| YOL080C | 199.8794327 | -0.510335289 | 0.177464592 | 0.49 | 0.32 |
| YBR140C | 4380.796749 | 0.282297466 | 0.177817432 | 0.49 | 0.32 |
| YBR151W | 3044.846839 | -0.300495441 | 0.178283066 | 0.49 | 0.32 |
| YGL020C | 1129.205349 | -0.267929767 | 0.178264901 | 0.49 | 0.32 |
| YJL168C | 610.0870071 | -0.377075242 | 0.178322132 | 0.49 | 0.32 |
| YLR378C | 3906.764345 | -0.286209422 | 0.178048104 | 0.49 | 0.32 |
| YNL214W | 315.125881 | 0.17082029 | 0.178298739 | 0.49 | 0.32 |
| YMR006C | 894.5004784 | -0.298691477 | 0.178414083 | 0.49 | 0.32 |
| YDL027C | 1559.088359 | 0.311827462 | 0.178747444 | 0.49 | 0.32 |
| YDR042C | 81.10570396 | 0.572315891 | 0.178801125 | 0.49 | 0.32 |
| YGL119W | 689.6648811 | 0.197260509 | 0.178800454 | 0.49 | 0.32 |
| YIL156W | 1109.423201 | 0.27819324 | 0.17874628 | 0.49 | 0.32 |
| YLR423C | 531.1083054 | 0.347947935 | 0.178826455 | 0.49 | 0.32 |
| YBR296C-A | 3.00001637 | 0.806286221 | 0.179189949 | 0.49 | 0.32 |
| YDL022W | 5632.112705 | 0.409654106 | 0.179100461 | 0.49 | 0.32 |
| YER167W | 1421.029082 | 0.352339885 | 0.179340281 | 0.49 | 0.32 |
| YFL052W | 300.3969613 | 0.636564558 | 0.179154173 | 0.49 | 0.32 |
| YJR016C | 6106.441237 | 0.410891767 | 0.179067958 | 0.49 | 0.32 |
| YNL113W | 221.4093861 | -0.435192245 | 0.179212449 | 0.49 | 0.32 |
| YNR054C | 724.980203 | -0.431570104 | 0.179357246 | 0.49 | 0.32 |
| YGR148C | 7673.362198 | -0.499815401 | 0.179472712 | 0.49 | 0.32 |
| YER057C | 2325.694126 | 0.256762035 | 0.179551284 | 0.49 | 0.32 |
| YMR181C | 1531.918567 | 0.342616458 | 0.179887196 | 0.49 | 0.32 |
| YPL118W | 2044.672532 | -0.268400996 | 0.17995078 | 0.49 | 0.32 |
| YAL035W | 9252.51472 | -0.42661782 | 0.180506206 | 0.49 | 0.32 |
| YBR013C | 207.4427581 | 0.404332926 | 0.180584256 | 0.49 | 0.32 |
| YPR061C | 393.0462601 | 0.454433359 | 0.180595042 | 0.49 | 0.32 |
| YDR202C | 884.8533036 | 0.364056728 | 0.181002619 | 0.49 | 0.32 |
| YGR122C-A | 15.60192201 | -0.522761506 | 0.181232776 | 0.49 | 0.32 |
| YKR095W | 4316.14506 | 0.244523204 | 0.181176707 | 0.49 | 0.32 |
| YNL047C | 720.3103825 | -0.332674474 | 0.18120569 | 0.49 | 0.32 |
| YNL298W | 1195.295385 | -0.345591761 | 0.180918805 | 0.49 | 0.32 |
| YOR131C | 796.6694001 | 0.25360667 | 0.181225861 | 0.49 | 0.32 |
| YOR244W | 816.0935801 | -0.260823578 | 0.180993951 | 0.49 | 0.32 |
| YGL070C | 462.601344 | -0.291755021 | 0.181366205 | 0.49 | 0.32 |

| | | | | | |
|---|---|---|---|---|---|
| YPL040C | 952.0787614 | 0.298718652 | 0.181632441 | 0.49 | 0.32 |
| YKL102C | 18.56354003 | 0.777383827 | 0.181762474 | 0.49 | 0.32 |
| YBR058C-A | 528.5056217 | -0.268192395 | 0.182223312 | 0.49 | 0.33 |
| YHR185C | 7.642313868 | 0.831371688 | 0.182448322 | 0.49 | 0.33 |
| YHR199C | 2007.264009 | 0.380264793 | 0.182379189 | 0.49 | 0.33 |
| YNL043C | 66.28583242 | -0.415549541 | 0.182520784 | 0.49 | 0.33 |
| YHR197W | 861.9145076 | -0.382156919 | 0.182606704 | 0.49 | 0.33 |
| YJR131W | 581.4205166 | 0.294274048 | 0.182964292 | 0.49 | 0.33 |
| YIL092W | 426.021742 | 0.172871999 | 0.183206296 | 0.49 | 0.33 |
| YDR231C | 627.5516275 | 0.388692343 | 0.183339229 | 0.49 | 0.33 |
| YDR349C | 1807.872356 | 0.321838984 | 0.183476044 | 0.49 | 0.33 |
| YKL180W | 12036.25542 | -0.480832431 | 0.183744251 | 0.49 | 0.33 |
| YCR038C | 411.3006768 | 0.262454887 | 0.183908919 | 0.49 | 0.33 |
| YGR145W | 1225.275886 | -0.470872762 | 0.183917505 | 0.49 | 0.33 |
| YJL145W | 1081.572718 | -0.340532483 | 0.184029981 | 0.49 | 0.33 |
| YPR174C | 411.2973194 | 0.28379072 | 0.184063112 | 0.49 | 0.33 |
| YDR029W | 118.6548652 | 0.386432984 | 0.18417626 | 0.49 | 0.33 |
| YGR084C | 1041.679379 | -0.215312276 | 0.184490813 | 0.49 | 0.33 |
| YOL052C | 1181.165007 | -0.208634879 | 0.184464598 | 0.49 | 0.33 |
| YOL101C | 182.685157 | 0.421943979 | 0.184464428 | 0.49 | 0.33 |
| snR35 | 41.6109753 | 0.493001036 | 0.184498546 | 0.49 | 0.33 |
| YDL238C | 479.1219346 | 0.347399962 | 0.185010941 | 0.50 | 0.33 |
| YLR082C | 386.5514271 | -0.248017619 | 0.184968177 | 0.50 | 0.33 |
| YNL293W | 734.6319527 | 0.329839358 | 0.184993152 | 0.50 | 0.33 |
| YEL008W | 30.07335462 | -0.562079657 | 0.185223555 | 0.50 | 0.33 |
| YGL213C | 596.0286848 | -0.231100855 | 0.185288029 | 0.50 | 0.33 |
| YLR212C | 601.9431586 | -0.374926951 | 0.185211113 | 0.50 | 0.33 |
| YHR038W | 426.523879 | 0.272631388 | 0.185485197 | 0.50 | 0.33 |
| YLR218C | 291.942794 | -0.306764969 | 0.185517245 | 0.50 | 0.33 |
| YDR324C | 1092.059972 | -0.470695842 | 0.185687699 | 0.50 | 0.33 |
| YPL176C | 1288.392675 | 0.268515264 | 0.185693433 | 0.50 | 0.33 |
| YCL005W-A | 247.3743167 | -0.247238429 | 0.185783486 | 0.50 | 0.33 |
| YBL087C | 3378.344673 | -0.454514412 | 0.185919576 | 0.50 | 0.33 |
| YCR037C | 1044.612368 | -0.238050675 | 0.186267545 | 0.50 | 0.33 |
| YOR128C | 2990.654925 | -0.440039883 | 0.186642662 | 0.50 | 0.33 |
| YDR141C | 2153.367361 | 0.17327246 | 0.186764412 | 0.50 | 0.33 |
| YGL210W | 815.3164128 | -0.225652215 | 0.186808844 | 0.50 | 0.33 |
| YDR229W | 1110.14149 | -0.230354385 | 0.186960011 | 0.50 | 0.33 |
| YKR003W | 1162.972466 | -0.283608808 | 0.187053559 | 0.50 | 0.33 |
| YGL176C | 596.0143813 | 0.211421689 | 0.187257895 | 0.50 | 0.33 |
| YEL053C | 1328.366975 | 0.149932734 | 0.187562967 | 0.50 | 0.33 |
| YFL023W | 930.5339775 | 0.381950971 | 0.187681105 | 0.50 | 0.33 |
| YJL150W | 10.09973158 | 0.626741573 | 0.187739192 | 0.50 | 0.33 |

| | | | | | |
|---|---|---|---|---|---|
| YKL056C | 11023.03919 | -0.376785236 | 0.18797785 | 0.50 | 0.33 |
| YOL061W | 1754.163485 | -0.282810469 | 0.187934124 | 0.50 | 0.33 |
| YDR150W | 4371.450407 | 0.44389317 | 0.188164688 | 0.50 | 0.33 |
| YNL030W | 3372.305745 | -0.438560456 | 0.188332308 | 0.50 | 0.33 |
| YNL182C | 900.9513347 | -0.467140689 | 0.188352028 | 0.50 | 0.33 |
| YDR524C-B | 18745.499 | -0.429098427 | 0.188435097 | 0.50 | 0.33 |
| YBL008W | 564.4162656 | 0.313817889 | 0.188733103 | 0.50 | 0.33 |
| YOR149C | 790.0054864 | 0.216091433 | 0.188895026 | 0.50 | 0.33 |
| YDR326C | 1698.894045 | 0.237628094 | 0.189053687 | 0.50 | 0.33 |
| YKR063C | 686.126014 | 0.180794465 | 0.189218405 | 0.50 | 0.33 |
| YLR344W | 3310.960859 | -0.474058283 | 0.189867726 | 0.50 | 0.33 |
| YDR291W | 877.4001219 | 0.170832674 | 0.190039153 | 0.50 | 0.33 |
| YIR024C | 366.6225047 | 0.213810471 | 0.189971574 | 0.50 | 0.33 |
| YBR109W-A | 20.49364363 | 0.501895176 | 0.190408378 | 0.50 | 0.33 |
| YCL035C | 4330.054767 | 0.424116503 | 0.190581604 | 0.50 | 0.33 |
| YHL034W-A | 585.1049194 | 0.252716063 | 0.190847843 | 0.50 | 0.33 |
| YJL109C | 2576.133571 | -0.370431789 | 0.190857276 | 0.50 | 0.33 |
| YJL178C | 1325.547528 | -0.195355588 | 0.190559753 | 0.50 | 0.33 |
| YMR146C | 5181.578577 | -0.284080017 | 0.190625287 | 0.50 | 0.33 |
| YNL278W | 749.3245241 | 0.238715707 | 0.190648116 | 0.50 | 0.33 |
| YOL029C | 366.4284423 | -0.193281575 | 0.190849536 | 0.50 | 0.33 |
| YBR132C | 1205.603555 | 0.477377553 | 0.191093121 | 0.50 | 0.33 |
| YML104C | 923.4299097 | 0.226664992 | 0.191102375 | 0.50 | 0.33 |
| YOL040C | 16073.93418 | -0.494014345 | 0.191260887 | 0.50 | 0.33 |
| YPL162C | 476.9994766 | 0.285650723 | 0.191202313 | 0.50 | 0.33 |
| YCR086W | 338.9363584 | 0.323068767 | 0.192019621 | 0.50 | 0.33 |
| YDR214W | 2305.632085 | 0.439666982 | 0.192030358 | 0.50 | 0.33 |
| YMR088C | 861.7053416 | 0.298342635 | 0.191887836 | 0.50 | 0.33 |
| YOR294W | 771.7312013 | -0.430194793 | 0.191854347 | 0.50 | 0.33 |
| YGL090W | 485.8897575 | 0.234146554 | 0.192205569 | 0.50 | 0.33 |
| YEL044W | 582.0177494 | -0.244844905 | 0.19228879 | 0.50 | 0.33 |
| YCL012C | 422.2313505 | 0.350541796 | 0.192623107 | 0.50 | 0.33 |
| YOL135C | 544.3922283 | -0.2533654 | 0.192898072 | 0.51 | 0.33 |
| YDR430C | 1380.899649 | 0.133952343 | 0.193155008 | 0.51 | 0.33 |
| YER082C | 1253.635463 | -0.329373478 | 0.193386073 | 0.51 | 0.33 |
| YLR343W | 1249.632226 | 0.490932187 | 0.193349342 | 0.51 | 0.33 |
| YOL094C | 437.8167325 | -0.326737567 | 0.193339647 | 0.51 | 0.33 |
| YOR044W | 320.4821165 | 0.294538036 | 0.19324917 | 0.51 | 0.33 |
| YOR050C | 67.3008437 | -0.316548187 | 0.193328857 | 0.51 | 0.33 |
| YGR158C | 631.124363 | -0.349513003 | 0.193519937 | 0.51 | 0.33 |
| YAL020C | 521.9507996 | 0.279355181 | 0.193673375 | 0.51 | 0.33 |
| YNL168C | 1117.770385 | 0.322623318 | 0.19373532 | 0.51 | 0.33 |
| YOL133W | 270.3290721 | -0.330921272 | 0.19388414 | 0.51 | 0.33 |

| | | | | | |
|---|---|---|---|---|---|
| YOR093C | 1072.002934 | -0.167678991 | 0.193961477 | 0.51 | 0.33 |
| YPR135W | 846.2041741 | 0.231139928 | 0.193944178 | 0.51 | 0.33 |
| YCL043C | 17735.82228 | 0.364169792 | 0.194408235 | 0.51 | 0.33 |
| YGL080W | 877.4500948 | -0.301206693 | 0.194130475 | 0.51 | 0.33 |
| YGR082W | 1782.261898 | -0.254023729 | 0.194243932 | 0.51 | 0.33 |
| YLR277C | 732.8482737 | -0.231213902 | 0.19438843 | 0.51 | 0.33 |
| YOR293W | 11411.33478 | -0.49879214 | 0.19437517 | 0.51 | 0.33 |
| YDR314C | 384.5141242 | 0.369364673 | 0.19452294 | 0.51 | 0.33 |
| YEL026W | 1802.083858 | -0.48562161 | 0.194730147 | 0.51 | 0.33 |
| YFR002W | 1555.318988 | -0.270150622 | 0.194594489 | 0.51 | 0.33 |
| YML013W | 1108.615798 | 0.296379648 | 0.194734166 | 0.51 | 0.33 |
| YBR238C | 642.2716869 | -0.397737024 | 0.194928183 | 0.51 | 0.33 |
| YDL131W | 1846.278027 | 0.34636324 | 0.195053807 | 0.51 | 0.33 |
| YDL170W | 672.20876 | 0.345105216 | 0.195040634 | 0.51 | 0.33 |
| YKL049C | 577.3829424 | 0.247762006 | 0.195112637 | 0.51 | 0.33 |
| YPL203W | 1453.358861 | 0.331489983 | 0.195185022 | 0.51 | 0.33 |
| YBR017C | 2405.834993 | -0.225974864 | 0.195265549 | 0.51 | 0.33 |
| YFR045W | 698.908615 | 0.33175648 | 0.195695877 | 0.51 | 0.33 |
| YIR023W | 1089.743487 | -0.273258845 | 0.195444469 | 0.51 | 0.33 |
| YML058W | 2577.833646 | -0.356020597 | 0.195589374 | 0.51 | 0.33 |
| YMR168C | 528.3657431 | -0.218214172 | 0.195733828 | 0.51 | 0.33 |
| YNR046W | 947.7260883 | -0.421931165 | 0.195508141 | 0.51 | 0.33 |
| YGL190C | 1598.926776 | -0.274704485 | 0.196232493 | 0.51 | 0.33 |
| YIL049W | 383.4643737 | -0.40027839 | 0.196148033 | 0.51 | 0.33 |
| YML098W | 504.9087228 | -0.287966074 | 0.196224086 | 0.51 | 0.33 |
| YPR095C | 1088.661894 | 0.202776432 | 0.196010368 | 0.51 | 0.33 |
| YKL210W | 8749.432253 | 0.20987694 | 0.196383041 | 0.51 | 0.33 |
| YMR007W | 90.19118303 | 0.457216026 | 0.196409191 | 0.51 | 0.33 |
| YER009W | 3909.301197 | -0.358454834 | 0.197074687 | 0.51 | 0.34 |
| YBL069W | 640.7564823 | 0.360930693 | 0.197353378 | 0.51 | 0.34 |
| YGR283C | 529.6205161 | -0.288029491 | 0.197451703 | 0.51 | 0.34 |
| YHR001W | 1515.116914 | 0.231955315 | 0.197808663 | 0.51 | 0.34 |
| YPL069C | 455.5840403 | -0.28448642 | 0.19775633 | 0.51 | 0.34 |
| YGR152C | 296.8399677 | -0.390384389 | 0.197978656 | 0.51 | 0.34 |
| YGL230C | 33.65395104 | 0.578361409 | 0.198098004 | 0.51 | 0.34 |
| YDR351W | 1313.626306 | -0.285348616 | 0.198378486 | 0.51 | 0.34 |
| YER184C | 397.7514223 | 0.299577936 | 0.198317764 | 0.51 | 0.34 |
| YDL191W | 1342.250708 | -0.434356377 | 0.198855451 | 0.51 | 0.34 |
| YDL244W | 51.51006224 | 0.706224214 | 0.198758468 | 0.51 | 0.34 |
| YDR463W | 1963.287278 | 0.264270718 | 0.198750343 | 0.51 | 0.34 |
| YGR137W | 236.2341776 | 0.321172462 | 0.199055395 | 0.51 | 0.34 |
| YHR050W-A | 204.2549426 | -0.316651653 | 0.198958231 | 0.51 | 0.34 |
| YKL078W | 299.5473506 | -0.507969049 | 0.198792317 | 0.51 | 0.34 |

| | | | | | |
|---|---|---|---|---|---|
| YLR333C | 2119.808061 | -0.468404025 | 0.199106448 | 0.51 | 0.34 |
| YOR226C | 454.5609099 | 0.486185081 | 0.198943521 | 0.51 | 0.34 |
| YPL269W | 614.929882 | -0.281375968 | 0.199400549 | 0.51 | 0.34 |
| YIL025C | 3.736035911 | 0.71102751 | 0.19960749 | 0.51 | 0.34 |
| YDL029W | 3675.857405 | 0.134351137 | 0.199717167 | 0.51 | 0.34 |
| YOR352W | 729.6147343 | -0.222593391 | 0.199816461 | 0.51 | 0.34 |
| YDR011W | 4201.778813 | -0.372883211 | 0.2002483 | 0.51 | 0.34 |
| YJR127C | 3525.789326 | 0.249684472 | 0.200180659 | 0.51 | 0.34 |
| YOL142W | 655.5671288 | -0.249294876 | 0.20031654 | 0.51 | 0.34 |
| YBR085W | 162.0814879 | 0.634691792 | 0.200730515 | 0.51 | 0.34 |
| YER134C | 346.3610386 | -0.374403981 | 0.200685169 | 0.51 | 0.34 |
| YHR137W | 2665.655739 | 0.592824404 | 0.20070497 | 0.51 | 0.34 |
| YOR238W | 549.7060682 | -0.279962311 | 0.200461544 | 0.51 | 0.34 |
| YOR253W | 809.4249218 | -0.308237253 | 0.200650153 | 0.51 | 0.34 |
| YBR072C-A | 2.769886284 | -0.716064507 | 0.200843558 | 0.51 | 0.34 |
| YLR045C | 1304.494239 | 0.269073801 | 0.201010797 | 0.51 | 0.34 |
| YOR260W | 2245.68858 | -0.364084405 | 0.201024653 | 0.51 | 0.34 |
| YBL027W | 10124.8249 | -0.450859562 | 0.201261237 | 0.51 | 0.34 |
| YER171W | 948.7525289 | -0.197990017 | 0.20151797 | 0.51 | 0.34 |
| YAL053W | 2097.963607 | 0.457103911 | 0.201942066 | 0.51 | 0.34 |
| YCR045C | 398.9252367 | -0.272098977 | 0.202438626 | 0.51 | 0.34 |
| YGR036C | 221.213258 | -0.284821145 | 0.201773738 | 0.51 | 0.34 |
| YJL188C | 893.4798181 | -0.420544018 | 0.202531943 | 0.51 | 0.34 |
| YKR010C | 516.9970849 | -0.291080363 | 0.202477775 | 0.51 | 0.34 |
| YLL024C | 20764.11389 | 0.552464444 | 0.202231679 | 0.51 | 0.34 |
| YML073C | 8568.253747 | -0.471268715 | 0.201825062 | 0.51 | 0.34 |
| YNL130C-A | 3.921452687 | 0.638124793 | 0.202173279 | 0.51 | 0.34 |
| YOR139C | 5.37072037 | 0.579880912 | 0.202460708 | 0.51 | 0.34 |
| YPL101W | 1017.044188 | -0.244483105 | 0.202285254 | 0.51 | 0.34 |
| YPL194W | 451.5443271 | 0.215418149 | 0.201682428 | 0.51 | 0.34 |
| YPL230W | 2214.598115 | 0.559754784 | 0.202510682 | 0.51 | 0.34 |
| YDR098C-B | 19.81780983 | 0.535430813 | 0.202708938 | 0.51 | 0.34 |
| YNR037C | 584.4173485 | -0.340279006 | 0.202875444 | 0.51 | 0.34 |
| YLR271W | 587.0390943 | 0.401570614 | 0.203076895 | 0.51 | 0.34 |
| YOR272W | 1204.374523 | -0.405081758 | 0.203104608 | 0.51 | 0.34 |
| YAL066W | 0.770254238 | 0.608116463 | 0.203433744 | 0.51 | 0.34 |
| YBR278W | 245.0159717 | 0.338344382 | 0.203542812 | 0.51 | 0.34 |
| YLR124W | 42.32116711 | -0.640229923 | 0.20323723 | 0.51 | 0.34 |
| YMR154C | 501.7366665 | 0.260158056 | 0.203484469 | 0.51 | 0.34 |
| YOR061W | 1662.759146 | -0.234101036 | 0.203433651 | 0.51 | 0.34 |
| YMR251W | 257.8128007 | 0.504600581 | 0.203968905 | 0.51 | 0.34 |
| YLL048C | 4130.734576 | 0.237442044 | 0.204307784 | 0.52 | 0.34 |
| YDL125C | 2310.617952 | 0.32022727 | 0.204532305 | 0.52 | 0.34 |

| | | | | | |
|---|---|---|---|---|---|
| YHR188C | 2845.286709 | 0.299485985 | 0.204795942 | 0.52 | 0.34 |
| YLR083C | 1368.675892 | -0.375965907 | 0.204860835 | 0.52 | 0.34 |
| YML127W | 993.038483 | -0.234410141 | 0.20476179 | 0.52 | 0.34 |
| YML080W | 296.6855777 | -0.390177686 | 0.205132696 | 0.52 | 0.34 |
| YDR379W | 1429.073535 | 0.244662892 | 0.20556198 | 0.52 | 0.34 |
| YGR201C | 974.6666707 | 0.535952352 | 0.20557893 | 0.52 | 0.34 |
| YIL011W | 777.5395764 | -0.640037281 | 0.205454977 | 0.52 | 0.34 |
| YPL158C | 1350.795148 | -0.358863578 | 0.205607637 | 0.52 | 0.34 |
| YPR099C | 24.60634511 | -0.375369601 | 0.20534182 | 0.52 | 0.34 |
| YGL132W | 16.53764806 | 0.620154825 | 0.205769086 | 0.52 | 0.34 |
| YLR450W | 2180.849651 | 0.316779186 | 0.205907976 | 0.52 | 0.34 |
| YOR136W | 7814.887983 | 0.31459005 | 0.206023939 | 0.52 | 0.34 |
| YOL049W | 2360.13263 | -0.267859872 | 0.206180038 | 0.52 | 0.34 |
| YGL121C | 888.9061681 | 0.623225716 | 0.206321647 | 0.52 | 0.34 |
| YGR051C | 51.37808205 | 0.355519027 | 0.206628818 | 0.52 | 0.34 |
| YGR271W | 1597.427923 | 0.217593226 | 0.206703443 | 0.52 | 0.34 |
| YIL091C | 989.7593991 | -0.412117705 | 0.206662934 | 0.52 | 0.34 |
| YDR497C | 3420.957422 | 0.206472981 | 0.20687675 | 0.52 | 0.34 |
| YDR119W | 1442.902962 | -0.387264069 | 0.207552982 | 0.52 | 0.34 |
| YDR131C | 626.2979697 | 0.213960441 | 0.207539757 | 0.52 | 0.34 |
| YER113C | 743.4950149 | -0.201181457 | 0.20778228 | 0.52 | 0.34 |
| YJL011C | 110.249814 | -0.404310465 | 0.20767448 | 0.52 | 0.34 |
| YJL222W | 13.76304146 | 0.537606737 | 0.207815732 | 0.52 | 0.34 |
| YMR244C-A | 108.6393324 | 0.278815307 | 0.207503613 | 0.52 | 0.34 |
| YOR320C | 1070.658594 | -0.227669704 | 0.207441046 | 0.52 | 0.34 |
| YOR325W | 4.631536169 | -0.561517156 | 0.207766824 | 0.52 | 0.34 |
| YER122C | 4064.222569 | -0.254357126 | 0.207902247 | 0.52 | 0.34 |
| YCR054C | 374.3747854 | 0.289152678 | 0.208634157 | 0.52 | 0.34 |
| YDL146W | 626.6868048 | 0.417622731 | 0.208564215 | 0.52 | 0.34 |
| YDR177W | 1727.041157 | 0.267894577 | 0.208586682 | 0.52 | 0.34 |
| YER143W | 1597.506506 | 0.24251958 | 0.208817285 | 0.52 | 0.34 |
| YER172C-A | 0.861078059 | -0.724207586 | 0.20816222 | 0.52 | 0.34 |
| YGR045C | 23.80619598 | 0.437279339 | 0.208500329 | 0.52 | 0.34 |
| YGR280C | 1635.461549 | -0.409906133 | 0.208231135 | 0.52 | 0.34 |
| YKL143W | 1110.759937 | -0.393401432 | 0.208889768 | 0.52 | 0.34 |
| YLR029C | 12797.37208 | -0.46367115 | 0.208930175 | 0.52 | 0.34 |
| YML112W | 572.6104301 | -0.221859259 | 0.20844401 | 0.52 | 0.34 |
| YMR044W | 903.5982709 | -0.247617741 | 0.208088452 | 0.52 | 0.34 |
| YOR248W | 35.3813631 | -0.350475897 | 0.208730822 | 0.52 | 0.34 |
| YBR159W | 2299.591935 | -0.306545726 | 0.209465358 | 0.52 | 0.34 |
| YER048C | 2682.830499 | -0.244864932 | 0.209539586 | 0.52 | 0.34 |
| YGR103W | 3122.539574 | -0.482113872 | 0.209164182 | 0.52 | 0.34 |
| YGR241C | 1428.572307 | -0.225392138 | 0.209548783 | 0.52 | 0.34 |

| | | | | | |
|---|---|---|---|---|---|
| YLR340W | 21021.60532 | -0.431741916 | 0.209543963 | 0.52 | 0.34 |
| YNL200C | 2258.279724 | 0.517397331 | 0.209278398 | 0.52 | 0.34 |
| YOL126C | 3179.903333 | -0.561195413 | 0.209117264 | 0.52 | 0.34 |
| YER162C | 931.7095211 | 0.313695589 | 0.209705829 | 0.52 | 0.34 |
| YIL065C | 779.4474724 | 0.322354138 | 0.209801741 | 0.52 | 0.34 |
| YPL153C | 363.2997459 | 0.287376907 | 0.210182023 | 0.52 | 0.34 |
| YGL089C | 132.0787924 | 0.208161135 | 0.210697196 | 0.52 | 0.34 |
| YLR196W | 1358.156963 | -0.392429961 | 0.210635301 | 0.52 | 0.34 |
| YML118W | 395.2746742 | 0.519759986 | 0.210719167 | 0.52 | 0.34 |
| YMR047C | 2343.921258 | -0.219514421 | 0.210421717 | 0.52 | 0.34 |
| YNL229C | 1349.306513 | -0.138274121 | 0.210483698 | 0.52 | 0.34 |
| YPL079W | 6102.629794 | -0.467438171 | 0.210415638 | 0.52 | 0.34 |
| YLR165C | 405.6673301 | -0.320162843 | 0.210945381 | 0.52 | 0.34 |
| YER079W | 1934.057175 | 0.4625156 | 0.211191592 | 0.52 | 0.34 |
| YLL045C | 7512.686429 | -0.45250722 | 0.211550265 | 0.52 | 0.34 |
| YML099C | 1070.053648 | 0.321136758 | 0.211511827 | 0.52 | 0.34 |
| YOR167C | 4868.170026 | -0.410049444 | 0.211666409 | 0.52 | 0.34 |
| YOR113W | 1039.201076 | 0.393133451 | 0.212162997 | 0.52 | 0.35 |
| YBR224W | 27.9935508 | 0.463203116 | 0.212411193 | 0.52 | 0.35 |
| YDR064W | 9582.184184 | -0.476450647 | 0.212808101 | 0.52 | 0.35 |
| YDR528W | 366.4782046 | -0.532160169 | 0.212602068 | 0.52 | 0.35 |
| YHR072W-A | 1183.992345 | -0.365652261 | 0.212822949 | 0.52 | 0.35 |
| YIL063C | 1081.563362 | 0.337273959 | 0.212776137 | 0.52 | 0.35 |
| YJR156C | 47.49998049 | 0.621781874 | 0.212894276 | 0.52 | 0.35 |
| YPL030W | 552.0003609 | -0.306797113 | 0.212727861 | 0.52 | 0.35 |
| YKL138C | 823.724232 | -0.291105012 | 0.213078542 | 0.52 | 0.35 |
| YKR045C | 368.478154 | -0.244577258 | 0.213290043 | 0.52 | 0.35 |
| YLR382C | 967.5198513 | 0.20351825 | 0.213222463 | 0.52 | 0.35 |
| YMR316C-B | 10.98411933 | 0.484438209 | 0.213193698 | 0.52 | 0.35 |
| YDL181W | 1061.423493 | 0.483668606 | 0.213399313 | 0.52 | 0.35 |
| YNL068C | 1421.244951 | 0.266249243 | 0.213637519 | 0.52 | 0.35 |
| YBL043W | 324.5838638 | 0.616103289 | 0.213839407 | 0.52 | 0.35 |
| YDR045C | 241.2309895 | -0.298848301 | 0.213974214 | 0.52 | 0.35 |
| YGR049W | 549.8511767 | 0.28745443 | 0.213877032 | 0.52 | 0.35 |
| YMR220W | 1203.063636 | -0.188839337 | 0.214003918 | 0.52 | 0.35 |
| YDL069C | 126.889784 | 0.277044452 | 0.214158348 | 0.52 | 0.35 |
| YPL013C | 664.0145708 | -0.311422854 | 0.214545793 | 0.52 | 0.35 |
| YJR080C | 1270.85858 | 0.382221838 | 0.214984209 | 0.53 | 0.35 |
| YDL232W | 292.6096052 | -0.304792086 | 0.215403758 | 0.53 | 0.35 |
| YOR065W | 3707.159237 | 0.558052312 | 0.215409405 | 0.53 | 0.35 |
| YHR078W | 1137.159604 | 0.336296134 | 0.215644123 | 0.53 | 0.35 |
| YPL222W | 848.2140163 | 0.497172752 | 0.215563771 | 0.53 | 0.35 |
| YBR196C | 23554.20363 | 0.330856599 | 0.215854719 | 0.53 | 0.35 |

| | | | | | |
|---|---|---|---|---|---|
| YDR013W | 271.9851003 | -0.293293281 | 0.215814808 | 0.53 | 0.35 |
| YML094W | 513.2141562 | -0.247544615 | 0.215967831 | 0.53 | 0.35 |
| YNL249C | 367.0811931 | 0.193872447 | 0.216118895 | 0.53 | 0.35 |
| YOR341W | 3809.897668 | -0.390468323 | 0.216075495 | 0.53 | 0.35 |
| YDR179C | 161.4536724 | 0.196019866 | 0.216491006 | 0.53 | 0.35 |
| YDR522C | 159.2048735 | -0.396884971 | 0.216461173 | 0.53 | 0.35 |
| YKR019C | 546.7471303 | 0.251362288 | 0.216925328 | 0.53 | 0.35 |
| YLR366W | 1.593075245 | 0.677616147 | 0.216987208 | 0.53 | 0.35 |
| YER147C | 529.8378852 | 0.242784939 | 0.217188227 | 0.53 | 0.35 |
| YJR033C | 1583.049905 | 0.213292647 | 0.217483013 | 0.53 | 0.35 |
| YMR090W | 1879.223335 | 0.532423412 | 0.217433049 | 0.53 | 0.35 |
| YOR317W | 4625.356852 | 0.379423989 | 0.21739532 | 0.53 | 0.35 |
| YPL257W | 181.2677801 | 0.500542799 | 0.21728215 | 0.53 | 0.35 |
| YGR176W | 11.59700922 | -0.611736162 | 0.217705397 | 0.53 | 0.35 |
| YMR271C | 754.1723733 | 0.550220512 | 0.217856771 | 0.53 | 0.35 |
| YHR110W | 1458.33201 | 0.19916857 | 0.217979058 | 0.53 | 0.35 |
| YLR052W | 1071.581443 | -0.104413996 | 0.218072281 | 0.53 | 0.35 |
| YAL051W | 1222.388192 | 0.21023584 | 0.218537913 | 0.53 | 0.35 |
| YDR183W | 267.9198317 | -0.202914049 | 0.218403291 | 0.53 | 0.35 |
| YFR011C | 642.0102271 | 0.425764187 | 0.218461768 | 0.53 | 0.35 |
| YLR372W | 2301.005993 | -0.433460509 | 0.218260698 | 0.53 | 0.35 |
| YNR032W | 607.7463894 | 0.224858877 | 0.218300993 | 0.53 | 0.35 |
| YAL017W | 3309.438838 | 0.359707228 | 0.218664656 | 0.53 | 0.35 |
| YMR082C | 5.748276588 | 0.824639441 | 0.219340477 | 0.53 | 0.35 |
| YPL137C | 1258.265301 | -0.329089923 | 0.219387519 | 0.53 | 0.35 |
| YAL014C | 833.2543927 | -0.29020862 | 0.219817754 | 0.53 | 0.35 |
| YFL054C | 1511.844206 | 0.546059587 | 0.220526212 | 0.53 | 0.35 |
| YGL075C | 312.3284424 | 0.199610164 | 0.220013039 | 0.53 | 0.35 |
| YIR025W | 303.8110514 | 0.197168505 | 0.220117233 | 0.53 | 0.35 |
| YJL077C | 162.6090211 | 0.313132626 | 0.220737671 | 0.53 | 0.35 |
| YKR056W | 1143.695694 | -0.332983806 | 0.220305174 | 0.53 | 0.35 |
| YLR015W | 562.8104697 | 0.247194672 | 0.220066323 | 0.53 | 0.35 |
| YLR208W | 2849.872811 | -0.234264808 | 0.219845453 | 0.53 | 0.35 |
| YMR092C | 2509.767947 | 0.230498721 | 0.220394808 | 0.53 | 0.35 |
| YOL121C | 3681.856963 | -0.419454023 | 0.219636902 | 0.53 | 0.35 |
| YOL134C | 23.62819378 | -0.329676269 | 0.220801968 | 0.53 | 0.35 |
| YOR020W-A | 1169.212022 | 0.358380101 | 0.220206743 | 0.53 | 0.35 |
| YOR041C | 37.3273586 | -0.412974159 | 0.219715347 | 0.53 | 0.35 |
| YPL059W | 1008.866988 | -0.191751373 | 0.219539496 | 0.53 | 0.35 |
| YPR153W | 81.58409703 | 0.305991797 | 0.22061475 | 0.53 | 0.35 |
| YPR193C | 57.30247136 | 0.480310246 | 0.22077145 | 0.53 | 0.35 |
| snR128 | 13.17491061 | 0.836232085 | 0.220759657 | 0.53 | 0.35 |
| YLR323C | 363.5088652 | 0.246908462 | 0.2211389 | 0.53 | 0.35 |

| | | | | | |
|---|---|---|---|---|---|
| YPL065W | 569.9929052 | -0.259462472 | 0.221765285 | 0.53 | 0.35 |
| YDL057W | 396.4937822 | 0.392482302 | 0.22214182 | 0.53 | 0.35 |
| YGL019W | 1587.591155 | -0.146435783 | 0.222069233 | 0.53 | 0.35 |
| YDL082W | 7404.050186 | -0.440596004 | 0.222636549 | 0.53 | 0.35 |
| YGR156W | 932.64115 | -0.264825543 | 0.222610523 | 0.53 | 0.35 |
| YML091C | 3874.618724 | 0.590470174 | 0.222538868 | 0.53 | 0.35 |
| YPR109W | 650.4688515 | -0.242241477 | 0.222881341 | 0.53 | 0.35 |
| YGR261C | 1931.303256 | -0.23650574 | 0.223109719 | 0.53 | 0.35 |
| YMR089C | 2367.558674 | 0.323670876 | 0.223187759 | 0.53 | 0.35 |
| YBL004W | 2589.356792 | -0.346628833 | 0.223794902 | 0.53 | 0.35 |
| YGL255W | 1201.748656 | 0.450913713 | 0.22376002 | 0.53 | 0.35 |
| YKR049C | 1528.555977 | 0.484040552 | 0.223354155 | 0.53 | 0.35 |
| YMR153W | 881.8064044 | -0.165173358 | 0.223819395 | 0.53 | 0.35 |
| YMR186W | 31490.05511 | 0.467250383 | 0.223741657 | 0.53 | 0.35 |
| YOL005C | 1554.462517 | -0.301900156 | 0.22346794 | 0.53 | 0.35 |
| YOR056C | 1339.644393 | -0.226098339 | 0.223696906 | 0.53 | 0.35 |
| YLR098C | 706.1029207 | 0.260646396 | 0.224118589 | 0.53 | 0.35 |
| YLR194C | 2445.708189 | 0.453506375 | 0.224204174 | 0.53 | 0.35 |
| YML128C | 9230.440998 | 0.595003092 | 0.224297485 | 0.53 | 0.35 |
| YOR150W | 702.6825687 | -0.280914607 | 0.2241565 | 0.53 | 0.35 |
| YPL067C | 285.3555196 | 0.414773501 | 0.224278894 | 0.53 | 0.35 |
| YIR011C | 1166.213259 | -0.238804922 | 0.22464912 | 0.54 | 0.35 |
| YDR285W | 510.8247622 | 0.355392201 | 0.22515059 | 0.54 | 0.35 |
| YMR176W | 1256.155137 | 0.245950719 | 0.225126383 | 0.54 | 0.35 |
| YNL181W | 1100.958937 | 0.194266768 | 0.225274971 | 0.54 | 0.35 |
| YGL134W | 814.0702966 | 0.279638938 | 0.225535621 | 0.54 | 0.35 |
| YML006C | 696.4182875 | -0.186151863 | 0.225477834 | 0.54 | 0.35 |
| YIL102C | 13.70446277 | 0.582203907 | 0.22616315 | 0.54 | 0.36 |
| YIL148W | 9852.705743 | -0.442086057 | 0.226101723 | 0.54 | 0.36 |
| YDR192C | 476.3597916 | -0.220677786 | 0.226808092 | 0.54 | 0.36 |
| YER119C | 1204.859566 | 0.295359693 | 0.226465924 | 0.54 | 0.36 |
| YJR005W | 1677.202028 | -0.209626652 | 0.226801659 | 0.54 | 0.36 |
| YKL198C | 1074.404315 | -0.315685377 | 0.226703904 | 0.54 | 0.36 |
| YLR006C | 1262.134466 | 0.279466777 | 0.226414797 | 0.54 | 0.36 |
| YLR427W | 1359.694674 | -0.099418512 | 0.226866919 | 0.54 | 0.36 |
| YNR021W | 3396.726535 | -0.303977066 | 0.226470883 | 0.54 | 0.36 |
| YOR284W | 207.2496443 | -0.516648248 | 0.226824218 | 0.54 | 0.36 |
| YBR073W | 1901.278512 | -0.486147889 | 0.227196945 | 0.54 | 0.36 |
| YGL050W | 416.7826537 | -0.305937167 | 0.227115511 | 0.54 | 0.36 |
| YDR130C | 501.1168326 | -0.245743566 | 0.227457449 | 0.54 | 0.36 |
| YPL152W-A | 1.958413515 | -0.604587914 | 0.227758565 | 0.54 | 0.36 |
| YDL019C | 2417.237505 | 0.227652761 | 0.227861891 | 0.54 | 0.36 |
| YGL224C | 544.3672231 | 0.337127518 | 0.22797161 | 0.54 | 0.36 |

| | | | | | |
|---|---|---|---|---|---|
| YNL157W | 1475.809734 | -0.172421294 | 0.228336359 | 0.54 | 0.36 |
| YNR013C | 2464.582186 | -0.241981657 | 0.228531055 | 0.54 | 0.36 |
| YPL228W | 971.5333391 | -0.243217627 | 0.228678785 | 0.54 | 0.36 |
| YGR136W | 2019.817506 | 0.242987163 | 0.228872478 | 0.54 | 0.36 |
| YJL052W | 23370.62399 | 0.551097981 | 0.22889284 | 0.54 | 0.36 |
| YHR170W | 1876.524363 | -0.432548849 | 0.229066627 | 0.54 | 0.36 |
| YNL284C | 1102.406706 | -0.214688744 | 0.229152936 | 0.54 | 0.36 |
| YJL016W | 1543.218497 | 0.242868204 | 0.22930886 | 0.54 | 0.36 |
| YER078C | 919.3267409 | 0.207957106 | 0.229490496 | 0.54 | 0.36 |
| YHR089C | 2056.029582 | -0.388631758 | 0.22956452 | 0.54 | 0.36 |
| YDL096C | 9.504570723 | -0.53478313 | 0.230380134 | 0.54 | 0.36 |
| YDL130W | 4709.707857 | -0.406972036 | 0.2304246 | 0.54 | 0.36 |
| YJR151C | 560.5480031 | 0.387560414 | 0.230232203 | 0.54 | 0.36 |
| YMR173W-A | 260.3952806 | -0.397142364 | 0.230160985 | 0.54 | 0.36 |
| YNL155W | 1378.201426 | 0.269515269 | 0.230376885 | 0.54 | 0.36 |
| YNL262W | 2257.387419 | 0.316391874 | 0.230223001 | 0.54 | 0.36 |
| YOR389W | 203.2493311 | 0.377863743 | 0.230097783 | 0.54 | 0.36 |
| YNL246W | 1195.836863 | -0.153355146 | 0.230799848 | 0.54 | 0.36 |
| YIL096C | 413.8812975 | -0.342960019 | 0.230969309 | 0.54 | 0.36 |
| YNL162W | 2350.306083 | -0.440677579 | 0.231273521 | 0.54 | 0.36 |
| YLR134W | 556.7123022 | 0.33285319 | 0.231514364 | 0.54 | 0.36 |
| YGR055W | 1342.856307 | -0.324922955 | 0.231658847 | 0.54 | 0.36 |
| YAL058W | 573.3781549 | 0.244278826 | 0.231762254 | 0.54 | 0.36 |
| YOL041C | 1321.58671 | -0.425958159 | 0.232052548 | 0.54 | 0.36 |
| YDR259C | 452.5952789 | 0.3980289 | 0.232188688 | 0.54 | 0.36 |
| YKL177W | 2.5279419 | 0.747932702 | 0.232353417 | 0.54 | 0.36 |
| YDL196W | 15.17669232 | 0.428891931 | 0.23263352 | 0.54 | 0.36 |
| YJR152W | 334.6737416 | 0.423415006 | 0.232610372 | 0.54 | 0.36 |
| YOR163W | 1216.735785 | -0.199909477 | 0.23260035 | 0.54 | 0.36 |
| YNR044W | 3311.34449 | -0.612273822 | 0.232733865 | 0.54 | 0.36 |
| YBR292C | 15.20409235 | 0.566514388 | 0.233212698 | 0.55 | 0.36 |
| YNR023W | 727.2388585 | -0.19216064 | 0.233306637 | 0.55 | 0.36 |
| YKL057C | 1611.669894 | -0.246985839 | 0.233748967 | 0.55 | 0.36 |
| YJL142C | 106.3041783 | 0.396995617 | 0.234012508 | 0.55 | 0.36 |
| YGL003C | 856.5289623 | -0.241201186 | 0.234238585 | 0.55 | 0.36 |
| YFL045C | 5242.345974 | -0.355355314 | 0.234535599 | 0.55 | 0.36 |
| YCR003W | 578.0986481 | -0.281618089 | 0.23511571 | 0.55 | 0.36 |
| YDR482C | 359.2978323 | -0.170265931 | 0.234984014 | 0.55 | 0.36 |
| YKR068C | 742.7822083 | -0.205985423 | 0.23513274 | 0.55 | 0.36 |
| YML008C | 4220.321447 | -0.34141182 | 0.234890566 | 0.55 | 0.36 |
| YNL091W | 4224.416881 | 0.237617792 | 0.235060937 | 0.55 | 0.36 |
| YOR251C | 1296.455018 | -0.158646797 | 0.234747933 | 0.55 | 0.36 |
| snR62 | 6.46452847 | 0.540071624 | 0.235277627 | 0.55 | 0.36 |

| | | | | | |
|---|---|---|---|---|---|
| YGR271C-A | 454.6880839 | -0.394656997 | 0.235473104 | 0.55 | 0.36 |
| YMR151W | 69.13225197 | 0.289249694 | 0.235474863 | 0.55 | 0.36 |
| YNL126W | 705.9981899 | -0.268120182 | 0.235695298 | 0.55 | 0.36 |
| YNL161W | 1329.930942 | 0.159464306 | 0.235763093 | 0.55 | 0.36 |
| YBR193C | 556.4922631 | 0.212869698 | 0.236450087 | 0.55 | 0.36 |
| YDR406W | 3009.969096 | -0.530336002 | 0.236357093 | 0.55 | 0.36 |
| YEL048C | 96.72112286 | -0.362857974 | 0.236461622 | 0.55 | 0.36 |
| YER126C | 1138.972805 | -0.409501831 | 0.236597185 | 0.55 | 0.36 |
| YGL006W-A | 3.943531239 | 0.630822066 | 0.236546655 | 0.55 | 0.36 |
| YGR216C | 759.5937639 | 0.159137503 | 0.236261617 | 0.55 | 0.36 |
| YLR178C | 3271.315417 | 0.49751883 | 0.236447648 | 0.55 | 0.36 |
| YDR416W | 716.6573712 | 0.200757176 | 0.236699731 | 0.55 | 0.36 |
| YGR004W | 331.5972983 | 0.225299398 | 0.23690113 | 0.55 | 0.36 |
| YJL002C | 4648.101511 | -0.273369102 | 0.237021548 | 0.55 | 0.36 |
| YOR286W | 1162.614528 | -0.201053161 | 0.23730207 | 0.55 | 0.36 |
| YER066W | 204.0645964 | 0.472304377 | 0.237419834 | 0.55 | 0.36 |
| YOL039W | 5643.898973 | -0.413241896 | 0.237508799 | 0.55 | 0.36 |
| YOL118C | 79.69627109 | 0.504934858 | 0.237592361 | 0.55 | 0.36 |
| YML122C | 47.85383085 | 0.452475567 | 0.238183517 | 0.55 | 0.36 |
| YPL053C | 1363.897708 | -0.320230535 | 0.238111043 | 0.55 | 0.36 |
| YOR246C | 1262.599493 | -0.275390219 | 0.23837277 | 0.55 | 0.36 |
| YML018C | 644.4081194 | -0.344180901 | 0.23868121 | 0.55 | 0.36 |
| YMR170C | 596.7981642 | 0.394559446 | 0.238927486 | 0.55 | 0.36 |
| YNL024C-A | 249.9046551 | -0.212192637 | 0.238875771 | 0.55 | 0.36 |
| YCR046C | 1534.125343 | -0.236759783 | 0.239381137 | 0.55 | 0.36 |
| YGR222W | 726.7683442 | -0.220586219 | 0.239351945 | 0.55 | 0.36 |
| YLR016C | 414.1257784 | -0.204386812 | 0.239315377 | 0.55 | 0.36 |
| YDR233C | 7397.691198 | -0.296677976 | 0.239562305 | 0.55 | 0.36 |
| YGR041W | 794.2535473 | -0.212326531 | 0.239877815 | 0.55 | 0.36 |
| YIL055C | 666.1215044 | 0.364508157 | 0.23981676 | 0.55 | 0.36 |
| YJL196C | 1508.45467 | -0.353127893 | 0.239862683 | 0.55 | 0.36 |
| YLL033W | 166.5445731 | -0.191838024 | 0.239664916 | 0.55 | 0.36 |
| YLR373C | 1362.586121 | 0.182363255 | 0.239898444 | 0.55 | 0.36 |
| YIL097W | 1123.344615 | 0.374466835 | 0.239985721 | 0.55 | 0.36 |
| YHR070W | 688.4026548 | 0.309604392 | 0.240269942 | 0.55 | 0.36 |
| YNL021W | 893.5874562 | 0.147120739 | 0.240225683 | 0.55 | 0.36 |
| YBR144C | 8.772859821 | 0.530309315 | 0.240655226 | 0.55 | 0.36 |
| YDL186W | 120.8658792 | 0.390166722 | 0.240572767 | 0.55 | 0.36 |
| YIR017W-A | 3.982442675 | 0.706081608 | 0.24077455 | 0.55 | 0.36 |
| YMR158W | 693.5786925 | -0.278963852 | 0.240608142 | 0.55 | 0.36 |
| YOL112W | 963.5678133 | 0.234214832 | 0.240790573 | 0.55 | 0.36 |
| YPL010W | 2111.463364 | -0.221205595 | 0.240762698 | 0.55 | 0.36 |
| YKL195W | 3025.520768 | 0.189235676 | 0.241340878 | 0.55 | 0.37 |

| | | | | | |
|---|---|---|---|---|---|
| YML024W | 8930.152891 | -0.404182754 | 0.241328775 | 0.55 | 0.37 |
| YPR085C | 667.2581144 | 0.187461117 | 0.241263125 | 0.55 | 0.37 |
| YMR190C | 1250.870637 | 0.328905782 | 0.241548497 | 0.55 | 0.37 |
| YEL003W | 487.0478849 | -0.305806003 | 0.241711546 | 0.55 | 0.37 |
| YHR098C | 2961.484376 | 0.24698294 | 0.241748011 | 0.55 | 0.37 |
| YPL188W | 716.8167679 | 0.252062755 | 0.241830841 | 0.55 | 0.37 |
| YKL060C | 152874.8661 | 0.436918934 | 0.242028469 | 0.55 | 0.37 |
| YGR081C | 443.1057485 | -0.413451642 | 0.242195896 | 0.55 | 0.37 |
| YIL035C | 1690.415511 | -0.180101082 | 0.242199756 | 0.55 | 0.37 |
| YCR077C | 3337.529518 | -0.180117563 | 0.242304194 | 0.55 | 0.37 |
| YIR043C | 49.74477082 | 0.582960528 | 0.242383678 | 0.55 | 0.37 |
| YGL122C | 2999.105342 | -0.29696364 | 0.242579941 | 0.55 | 0.37 |
| YGL123W | 15975.90289 | -0.384358275 | 0.242823558 | 0.55 | 0.37 |
| YBR187W | 950.8054053 | -0.389536264 | 0.243061237 | 0.55 | 0.37 |
| YNL235C | 7.928094321 | -0.565528798 | 0.243012687 | 0.55 | 0.37 |
| YPL077C | 211.278667 | 0.21287399 | 0.243172882 | 0.55 | 0.37 |
| YAR068W | 4.842260677 | 0.748933089 | 0.243500687 | 0.55 | 0.37 |
| YGR279C | 11712.04252 | 0.230529966 | 0.243443703 | 0.55 | 0.37 |
| YHR091C | 596.7962742 | 0.149107277 | 0.243492634 | 0.55 | 0.37 |
| YFR032C-A | 2429.038295 | -0.454554691 | 0.244172265 | 0.56 | 0.37 |
| YIL094C | 3348.951104 | 0.250948298 | 0.24415618 | 0.56 | 0.37 |
| YOR011W | 696.3198627 | 0.29922882 | 0.244183872 | 0.56 | 0.37 |
| YPL128C | 628.8955352 | -0.226352794 | 0.244121228 | 0.56 | 0.37 |
| YGL158W | 260.9463165 | -0.71251124 | 0.244698819 | 0.56 | 0.37 |
| YIL128W | 1273.394888 | -0.257848817 | 0.245041727 | 0.56 | 0.37 |
| YJL195C | 5.108398903 | 0.495346727 | 0.245161961 | 0.56 | 0.37 |
| YML111W | 1306.672498 | -0.200321484 | 0.245242823 | 0.56 | 0.37 |
| YDR062W | 2339.714713 | -0.231467122 | 0.245482193 | 0.56 | 0.37 |
| YDR510W | 2598.34956 | -0.206716088 | 0.245518864 | 0.56 | 0.37 |
| YMR143W | 7020.519511 | -0.415957781 | 0.245503361 | 0.56 | 0.37 |
| YCR084C | 3633.028548 | -0.27073663 | 0.24567365 | 0.56 | 0.37 |
| YMR026C | 666.726482 | -0.201260548 | 0.245759672 | 0.56 | 0.37 |
| YHR176W | 373.1735066 | 0.243539937 | 0.246060654 | 0.56 | 0.37 |
| YNL042W | 593.1711026 | -0.333274487 | 0.246209712 | 0.56 | 0.37 |
| YMR307W | 11748.41017 | -0.363812565 | 0.246855537 | 0.56 | 0.37 |
| YPR092W | 28.34006108 | 0.343167682 | 0.246883094 | 0.56 | 0.37 |
| YDR377W | 1422.817631 | 0.471710637 | 0.247272032 | 0.56 | 0.37 |
| YDL064W | 884.0529801 | -0.266389481 | 0.247611657 | 0.56 | 0.37 |
| YER077C | 789.2511196 | 0.160359185 | 0.247815487 | 0.56 | 0.37 |
| YGR170W | 2142.624103 | 0.21594116 | 0.248057868 | 0.56 | 0.37 |
| YIL046W | 1246.958862 | 0.288723607 | 0.247730386 | 0.56 | 0.37 |
| YIL086C | 23.45932813 | 0.38733956 | 0.248030664 | 0.56 | 0.37 |
| YIL095W | 775.5717487 | -0.162943695 | 0.248113387 | 0.56 | 0.37 |

| | | | | | |
|---|---|---|---|---|---|
| YJR037W | 10.84309032 | 0.436801682 | 0.248141922 | 0.56 | 0.37 |
| YPL108W | 159.4425969 | -0.254517947 | 0.248063815 | 0.56 | 0.37 |
| YIR033W | 2182.959679 | 0.217918219 | 0.248513029 | 0.56 | 0.37 |
| YLR185W | 4884.991766 | -0.419716089 | 0.248665312 | 0.56 | 0.37 |
| YBL031W | 539.4437355 | -0.290213213 | 0.248901767 | 0.56 | 0.37 |
| YML009C | 743.0693468 | -0.284279025 | 0.249143199 | 0.56 | 0.37 |
| YDL138W | 666.9826314 | 0.230766438 | 0.249630198 | 0.56 | 0.37 |
| YEL009C | 21167.59097 | 0.374204375 | 0.249520075 | 0.56 | 0.37 |
| YNL300W | 1681.600577 | -0.483373465 | 0.249642131 | 0.56 | 0.37 |
| YOR266W | 469.4838324 | -0.151064717 | 0.249431691 | 0.56 | 0.37 |
| YHL002C-A | 8.009981089 | 0.451098184 | 0.24983976 | 0.56 | 0.37 |
| YPL133C | 580.4116075 | 0.27567014 | 0.249827413 | 0.56 | 0.37 |
| YNR034W | 660.948473 | 0.249392164 | 0.249982818 | 0.56 | 0.37 |
| YIL103W | 637.74726 | -0.253816313 | 0.250289459 | 0.56 | 0.37 |
| YLR108C | 407.9615838 | 0.31400077 | 0.25031483 | 0.56 | 0.37 |
| snR83 | 177.6034874 | -0.462326734 | 0.250203924 | 0.56 | 0.37 |
| YKL019W | 1109.764572 | -0.195100985 | 0.250434162 | 0.56 | 0.37 |
| YDR089W | 994.7212521 | -0.252312874 | 0.250623961 | 0.56 | 0.37 |
| YDR273W | 249.707739 | 0.324003024 | 0.250616104 | 0.56 | 0.37 |
| YGL099W | 1111.20672 | -0.301984924 | 0.250739171 | 0.56 | 0.37 |
| YDR007W | 490.6292629 | -0.162673827 | 0.251164637 | 0.56 | 0.37 |
| YHR023W | 2927.24121 | 0.318366126 | 0.251180123 | 0.56 | 0.37 |
| YHR215W | 110.5268911 | -0.573154896 | 0.251086744 | 0.56 | 0.37 |
| YBR029C | 1375.442278 | -0.253483519 | 0.251483178 | 0.56 | 0.37 |
| YML097C | 708.2199714 | -0.21976068 | 0.251493593 | 0.56 | 0.37 |
| YPL249C-A | 77.54661876 | -0.475928306 | 0.251476046 | 0.56 | 0.37 |
| YCR022C | 20.80731429 | 0.385984088 | 0.252228742 | 0.56 | 0.37 |
| YGL045W | 904.8478281 | 0.385290559 | 0.252182555 | 0.56 | 0.37 |
| YLR295C | 2976.605613 | 0.38827854 | 0.252100417 | 0.56 | 0.37 |
| YLR449W | 2574.095007 | -0.472668234 | 0.252905868 | 0.57 | 0.37 |
| YFR036W | 228.7837864 | 0.218689702 | 0.253655498 | 0.57 | 0.37 |
| YML133C | 9.588165535 | 0.576733472 | 0.253838193 | 0.57 | 0.37 |
| YMR084W | 174.3110976 | 0.419879936 | 0.253761436 | 0.57 | 0.37 |
| YNL096C | 3035.774305 | -0.356752638 | 0.25378969 | 0.57 | 0.37 |
| YNL290W | 854.0353401 | -0.229066913 | 0.253216908 | 0.57 | 0.37 |
| YOL089C | 836.6158144 | 0.204225386 | 0.253768432 | 0.57 | 0.37 |
| YOR074C | 848.568103 | -0.279871119 | 0.253511479 | 0.57 | 0.37 |
| YPR020W | 1129.530709 | 0.472952068 | 0.253779563 | 0.57 | 0.37 |
| YPR070W | 926.8338353 | -0.114135585 | 0.253414483 | 0.57 | 0.37 |
| YDL034W | 29.64111763 | 0.368685995 | 0.25415562 | 0.57 | 0.37 |
| YPL191C | 278.5044925 | 0.297514383 | 0.254151506 | 0.57 | 0.37 |
| YPR159C-A | 2.271891006 | 0.511885384 | 0.25403985 | 0.57 | 0.37 |
| YNL044W | 495.5352853 | -0.308790031 | 0.254257072 | 0.57 | 0.37 |

| | | | | | |
|---|---|---|---|---|---|
| YCR019W | 382.8804523 | -0.174531921 | 0.25491448 | 0.57 | 0.37 |
| YDR470C | 683.2878102 | -0.163859167 | 0.255013492 | 0.57 | 0.37 |
| YML031C-A | 2.97142876 | 0.528870737 | 0.254784956 | 0.57 | 0.37 |
| YMR155W | 707.9678984 | 0.264667615 | 0.254991823 | 0.57 | 0.37 |
| YOL013W-B | 4.383672181 | 0.818646013 | 0.254984666 | 0.57 | 0.37 |
| YKR058W | 2130.69981 | 0.411170928 | 0.255271503 | 0.57 | 0.37 |
| YJR045C | 25492.38268 | 0.365761247 | 0.255489558 | 0.57 | 0.37 |
| YML090W | 156.4475378 | 0.569010914 | 0.255545161 | 0.57 | 0.37 |
| YER138W-A | 1.112909077 | 0.448471021 | 0.256027951 | 0.57 | 0.37 |
| YGR259C | 0.780911468 | 0.47446325 | 0.255998001 | 0.57 | 0.37 |
| YJL087C | 474.7532502 | -0.273574284 | 0.2559852 | 0.57 | 0.37 |
| YLR266C | 1095.178781 | -0.269870994 | 0.255808902 | 0.57 | 0.37 |
| YDL102W | 928.5717968 | 0.172433926 | 0.256283642 | 0.57 | 0.38 |
| YJR083C | 550.0152446 | -0.225709979 | 0.256320295 | 0.57 | 0.38 |
| YOR109W | 2024.801836 | -0.166735618 | 0.256393219 | 0.57 | 0.38 |
| YER071C | 272.1086234 | -0.236056202 | 0.256484957 | 0.57 | 0.38 |
| YPR142C | 12.52047692 | -0.428175725 | 0.256593611 | 0.57 | 0.38 |
| YJL062W | 1004.118516 | 0.180485 | 0.256748527 | 0.57 | 0.38 |
| YBR276C | 1528.649976 | -0.114133706 | 0.257032196 | 0.57 | 0.38 |
| YOL055C | 533.3864453 | 0.288523176 | 0.257154494 | 0.57 | 0.38 |
| YIL152W | 734.4896336 | 0.321597703 | 0.257502278 | 0.57 | 0.38 |
| YDL007C-A | 3.853033258 | 0.905875656 | 0.258348107 | 0.57 | 0.38 |
| YKL124W | 623.7006372 | 0.247488701 | 0.258221139 | 0.57 | 0.38 |
| YNL115C | 1894.716258 | 0.389472761 | 0.258335864 | 0.57 | 0.38 |
| YIR022W | 1186.448374 | -0.211624551 | 0.258792588 | 0.57 | 0.38 |
| YJL048C | 1114.208522 | 0.356536639 | 0.258686178 | 0.57 | 0.38 |
| YLR257W | 6468.084537 | 0.374364521 | 0.258773484 | 0.57 | 0.38 |
| YBL035C | 596.37002 | -0.276227002 | 0.259808181 | 0.57 | 0.38 |
| YCL064C | 3239.199963 | -0.788674672 | 0.260377804 | 0.57 | 0.38 |
| YDL006W | 361.6658456 | -0.153802271 | 0.260259803 | 0.57 | 0.38 |
| YDL035C | 1472.135402 | 0.232751743 | 0.260112577 | 0.57 | 0.38 |
| YEL022W | 2372.854744 | -0.259580392 | 0.260382322 | 0.57 | 0.38 |
| YER049W | 1494.465582 | -0.302546134 | 0.259985284 | 0.57 | 0.38 |
| YER074W | 4108.232098 | -0.349453041 | 0.259692716 | 0.57 | 0.38 |
| YGL112C | 1298.501075 | -0.136448376 | 0.259898856 | 0.57 | 0.38 |
| YHR063C | 1432.676855 | 0.356832189 | 0.259249997 | 0.57 | 0.38 |
| YJL003W | 223.2586603 | -0.240574992 | 0.260287475 | 0.57 | 0.38 |
| YJR001W | 1763.85175 | 0.172042541 | 0.259351996 | 0.57 | 0.38 |
| YJR022W | 18.26801566 | -0.476466134 | 0.260100826 | 0.57 | 0.38 |
| YNR010W | 212.8810604 | -0.232709068 | 0.260306031 | 0.57 | 0.38 |
| YOL096C | 679.2678424 | 0.234306312 | 0.259119549 | 0.57 | 0.38 |
| YPL085W | 4050.180373 | 0.253836237 | 0.260395132 | 0.57 | 0.38 |
| YPR147C | 688.0952487 | -0.185764596 | 0.26001089 | 0.57 | 0.38 |

| | | | | | |
|---|---|---|---|---|---|
| YML055W | 811.1131816 | 0.206160964 | 0.260493257 | 0.57 | 0.38 |
| YGR169C-A | 750.5850996 | -0.239548048 | 0.260943588 | 0.57 | 0.38 |
| YLR357W | 954.9234442 | -0.240025829 | 0.26094147 | 0.57 | 0.38 |
| YOR184W | 6279.813274 | 0.385555356 | 0.261294484 | 0.57 | 0.38 |
| YBL075C | 2650.065607 | 0.519248843 | 0.261675176 | 0.57 | 0.38 |
| YBR048W | 5960.519296 | -0.451177114 | 0.2619684 | 0.57 | 0.38 |
| YBR162C | 7471.337212 | -0.345151607 | 0.261885511 | 0.57 | 0.38 |
| YDL160C-A | 95.22130843 | -0.553078607 | 0.261704864 | 0.57 | 0.38 |
| YDR208W | 1907.44225 | -0.235622345 | 0.261779262 | 0.57 | 0.38 |
| YML081C-A | 1028.969006 | 0.434237298 | 0.261548015 | 0.57 | 0.38 |
| YPR094W | 154.1332011 | 0.25822982 | 0.261847017 | 0.57 | 0.38 |
| YIR036W-A | 10.01589679 | 0.356896103 | 0.262219642 | 0.57 | 0.38 |
| YMR303C | 445.8492587 | -0.429898041 | 0.262209348 | 0.57 | 0.38 |
| YHR119W | 1485.712969 | 0.224485423 | 0.262423907 | 0.57 | 0.38 |
| YOL015W | 325.5388955 | 0.382494604 | 0.262374617 | 0.57 | 0.38 |
| YBR217W | 353.7676501 | -0.202491342 | 0.263576392 | 0.57 | 0.38 |
| YBR248C | 1552.941582 | 0.424708252 | 0.263586909 | 0.57 | 0.38 |
| YDR191W | 146.592055 | -0.286466587 | 0.263499395 | 0.57 | 0.38 |
| YGL120C | 2147.315673 | -0.409879491 | 0.263592312 | 0.57 | 0.38 |
| YKL164C | 7734.898504 | -0.382553313 | 0.263289146 | 0.57 | 0.38 |
| YKL178C | 114.5867776 | 0.570435471 | 0.26347055 | 0.57 | 0.38 |
| YNR011C | 902.2961481 | 0.22324862 | 0.26347466 | 0.57 | 0.38 |
| YNL130C | 1326.028586 | -0.320112293 | 0.26386576 | 0.57 | 0.38 |
| YJL116C | 947.324176 | 0.517933597 | 0.2640528 | 0.57 | 0.38 |
| YBL089W | 493.6798455 | 0.203852443 | 0.264622581 | 0.58 | 0.38 |
| YDR288W | 555.4180316 | -0.238832203 | 0.264625316 | 0.58 | 0.38 |
| YLR209C | 881.8739647 | 0.276461738 | 0.264574959 | 0.58 | 0.38 |
| YAR053W | 9.523462605 | 0.452180007 | 0.26583293 | 0.58 | 0.38 |
| YDL233W | 547.8994161 | 0.239492207 | 0.265787983 | 0.58 | 0.38 |
| YDR185C | 264.7477216 | 0.343004612 | 0.265611453 | 0.58 | 0.38 |
| YFR007W | 462.3779448 | 0.300099477 | 0.265709634 | 0.58 | 0.38 |
| YFR044C | 5702.058987 | 0.256144336 | 0.265265877 | 0.58 | 0.38 |
| YIL088C | 2503.905976 | 0.251946426 | 0.265733075 | 0.58 | 0.38 |
| YJL139C | 556.1081958 | 0.323347023 | 0.26551341 | 0.58 | 0.38 |
| YMR244W | 46.42719777 | 0.54150784 | 0.26581104 | 0.58 | 0.38 |
| YJR071W | 37.86358004 | -0.440950697 | 0.265942476 | 0.58 | 0.38 |
| YMR122W-A | 5293.670712 | -0.270151347 | 0.266106996 | 0.58 | 0.38 |
| YDR234W | 1972.622196 | 0.341802402 | 0.266591112 | 0.58 | 0.38 |
| YJL128C | 1506.583064 | -0.157263472 | 0.266666682 | 0.58 | 0.38 |
| YAL013W | 1274.881408 | -0.214833603 | 0.26756335 | 0.58 | 0.38 |
| YDR433W | 87.14790436 | -0.282648099 | 0.267580039 | 0.58 | 0.38 |
| YFL031C-A | 101.8569422 | 0.298993568 | 0.267680004 | 0.58 | 0.38 |
| YJL207C | 2307.188321 | -0.265634396 | 0.26751482 | 0.58 | 0.38 |

| | | | | | |
|---|---|---|---|---|---|
| YKR073C | 18.81130727 | 0.348012473 | 0.267650596 | 0.58 | 0.38 |
| YMR285C | 827.213202 | -0.148865225 | 0.26721446 | 0.58 | 0.38 |
| YNL059C | 1845.140766 | -0.206646058 | 0.267735799 | 0.58 | 0.38 |
| YNL146W | 85.60940575 | 0.253722319 | 0.266952183 | 0.58 | 0.38 |
| YOR312C | 9424.205996 | -0.399476253 | 0.267527307 | 0.58 | 0.38 |
| YOR350C | 702.3901449 | 0.251626274 | 0.267574721 | 0.58 | 0.38 |
| YPL238C | 56.97812647 | -0.29067725 | 0.26693722 | 0.58 | 0.38 |
| YGR034W | 5016.73135 | -0.387154116 | 0.26783289 | 0.58 | 0.38 |
| YHR214C-B | 235.6005636 | 0.541488312 | 0.268022061 | 0.58 | 0.38 |
| YIL038C | 1659.427705 | -0.275373574 | 0.267962794 | 0.58 | 0.38 |
| YHR024C | 1377.744705 | 0.160399201 | 0.268207434 | 0.58 | 0.38 |
| YMR109W | 3069.730753 | -0.203380026 | 0.268415522 | 0.58 | 0.38 |
| YNL176C | 1925.34034 | 0.215495653 | 0.268451557 | 0.58 | 0.38 |
| YIL066W-A | 3.308676015 | 0.660370235 | 0.268815463 | 0.58 | 0.38 |
| YJR002W | 1689.001321 | -0.251223205 | 0.26918369 | 0.58 | 0.38 |
| YOR047C | 1007.814108 | -0.413892309 | 0.26921672 | 0.58 | 0.38 |
| YOR310C | 3221.373678 | -0.367944777 | 0.269089344 | 0.58 | 0.38 |
| YGL189C | 20518.43042 | -0.395151673 | 0.269394443 | 0.58 | 0.38 |
| YMR240C | 592.2234425 | 0.268042198 | 0.269595786 | 0.58 | 0.38 |
| YBL084C | 817.2451769 | -0.261317758 | 0.270007882 | 0.58 | 0.38 |
| YCR050C | 17.22525728 | 0.380181637 | 0.269913089 | 0.58 | 0.38 |
| YGR183C | 999.4364709 | 0.466542773 | 0.270074787 | 0.58 | 0.38 |
| YIL110W | 498.0390037 | -0.295840466 | 0.269760497 | 0.58 | 0.38 |
| YOR181W | 1611.589546 | -0.284641258 | 0.269983791 | 0.58 | 0.38 |
| YFR015C | 6189.813754 | 0.38370444 | 0.270457614 | 0.58 | 0.38 |
| YGL035C | 1849.649908 | -0.33739143 | 0.270443406 | 0.58 | 0.38 |
| YGL096W | 785.2336716 | 0.498409133 | 0.270306443 | 0.58 | 0.38 |
| YGL029W | 383.3919755 | -0.35398728 | 0.270996011 | 0.58 | 0.38 |
| YJR124C | 485.3455454 | -0.334878713 | 0.270839165 | 0.58 | 0.38 |
| YOR158W | 1525.323441 | -0.215867243 | 0.271028558 | 0.58 | 0.38 |
| YOR172W | 824.7713828 | 0.177159239 | 0.271008439 | 0.58 | 0.38 |
| YAL056C-A | 23.07572137 | 0.334145036 | 0.271145646 | 0.58 | 0.38 |
| YBR060C | 829.7516444 | -0.160137953 | 0.27142908 | 0.58 | 0.38 |
| YOR035C | 1268.519702 | 0.299389859 | 0.27158925 | 0.58 | 0.38 |
| YDR312W | 598.3889984 | -0.308756082 | 0.272140578 | 0.58 | 0.38 |
| YDR514C | 275.4543514 | -0.279680881 | 0.272148277 | 0.58 | 0.38 |
| YIL061C | 381.0352818 | -0.201421103 | 0.272255842 | 0.58 | 0.38 |
| YIL047C | 1300.508261 | -0.212173998 | 0.272534751 | 0.58 | 0.38 |
| YLL047W | 4.30351014 | 0.599242092 | 0.272492641 | 0.58 | 0.38 |
| YLR437C-A | 26.88165292 | 0.313149674 | 0.272453091 | 0.58 | 0.38 |
| YKR066C | 4134.117453 | 0.322988 | 0.272813151 | 0.58 | 0.38 |
| YBR050C | 396.3143748 | 0.513469492 | 0.273257281 | 0.58 | 0.38 |
| YCR067C | 1807.391426 | 0.199120089 | 0.273143239 | 0.58 | 0.38 |

| | | | | | |
|---|---|---|---|---|---|
| YEL028W | 113.7795172 | -0.291030283 | 0.273197306 | 0.58 | 0.38 |
| YEL059C-A | 318.8027126 | 0.327169745 | 0.273771473 | 0.58 | 0.38 |
| YJL015C | 72.86729941 | 0.275340616 | 0.273618375 | 0.58 | 0.38 |
| YJL211C | 56.87482788 | 0.352337162 | 0.273725304 | 0.58 | 0.38 |
| YKL017C | 627.7815414 | 0.210284169 | 0.272974371 | 0.58 | 0.38 |
| YNL102W | 1324.90783 | -0.212787745 | 0.273525407 | 0.58 | 0.38 |
| YOR076C | 709.8237452 | -0.194376307 | 0.273674958 | 0.58 | 0.38 |
| YPR051W | 72.40075259 | -0.371163486 | 0.273507468 | 0.58 | 0.38 |
| YKL024C | 1589.092419 | 0.263301559 | 0.274232368 | 0.58 | 0.39 |
| YDR230W | 158.0472236 | 0.306790393 | 0.274385972 | 0.58 | 0.39 |
| YMR027W | 2670.4492 | 0.262460759 | 0.27471251 | 0.58 | 0.39 |
| YMR218C | 1133.912405 | 0.16731548 | 0.274733446 | 0.58 | 0.39 |
| YER087C-B | 1311.571506 | -0.287933645 | 0.275150745 | 0.58 | 0.39 |
| YJL179W | 307.4969877 | -0.297891218 | 0.27553737 | 0.58 | 0.39 |
| YBR180W | 89.72716224 | 0.512858649 | 0.275772152 | 0.58 | 0.39 |
| YDR099W | 8428.443895 | 0.141195346 | 0.275794598 | 0.58 | 0.39 |
| YFR033C | 3330.696407 | 0.437342352 | 0.27599296 | 0.58 | 0.39 |
| YKL073W | 1776.826393 | 0.360781151 | 0.276144754 | 0.58 | 0.39 |
| snR19 | 122.3490857 | 0.617200958 | 0.276068469 | 0.58 | 0.39 |
| YER072W | 2000.095671 | -0.286814211 | 0.276435511 | 0.59 | 0.39 |
| YCR030C | 2439.320353 | 0.221615506 | 0.276637979 | 0.59 | 0.39 |
| YKL101W | 3903.982421 | 0.386674017 | 0.277003608 | 0.59 | 0.39 |
| YKL183W | 567.8673175 | -0.26047877 | 0.277026808 | 0.59 | 0.39 |
| YPL227C | 650.9518323 | -0.314056541 | 0.276905767 | 0.59 | 0.39 |
| YIL034C | 3381.997418 | -0.129920283 | 0.277162209 | 0.59 | 0.39 |
| YDR301W | 1341.743029 | -0.242265718 | 0.277335252 | 0.59 | 0.39 |
| YBL072C | 3997.128198 | -0.398622862 | 0.277502843 | 0.59 | 0.39 |
| YDR182W | 1361.286881 | -0.160017638 | 0.277701831 | 0.59 | 0.39 |
| YCR105W | 210.4429239 | 0.535400897 | 0.278227844 | 0.59 | 0.39 |
| YHL046W-A | 47.85481112 | 0.555995986 | 0.278428053 | 0.59 | 0.39 |
| YKL005C | 1108.015733 | 0.155575555 | 0.278529729 | 0.59 | 0.39 |
| YLL025W | 92.59299699 | 0.356720687 | 0.278349489 | 0.59 | 0.39 |
| YLR040C | 271.3228626 | -0.417235219 | 0.278510836 | 0.59 | 0.39 |
| YNL169C | 1167.217459 | -0.128245666 | 0.278308309 | 0.59 | 0.39 |
| YPL242C | 2035.672722 | 0.212778031 | 0.278352474 | 0.59 | 0.39 |
| YLR200W | 774.013828 | -0.200761992 | 0.278787961 | 0.59 | 0.39 |
| YLR262C-A | 800.3537696 | -0.255059438 | 0.278855162 | 0.59 | 0.39 |
| YDR321W | 1332.488079 | -0.358210476 | 0.27971497 | 0.59 | 0.39 |
| YDR345C | 12971.54144 | 0.588217288 | 0.27973045 | 0.59 | 0.39 |
| YHR130C | 24.10330458 | 0.419285447 | 0.279982442 | 0.59 | 0.39 |
| YIL020C | 394.2399705 | -0.306463437 | 0.279522044 | 0.59 | 0.39 |
| YJR013W | 645.2124988 | -0.144421651 | 0.279890634 | 0.59 | 0.39 |
| YJR040W | 752.4605261 | 0.21529502 | 0.280140773 | 0.59 | 0.39 |

| | | | | | |
|---|---|---|---|---|---|
| YLR455W | 572.5909218 | -0.218307815 | 0.279793268 | 0.59 | 0.39 |
| YNL035C | 765.5856918 | -0.211736426 | 0.280076791 | 0.59 | 0.39 |
| YPR164W | 1248.455329 | 0.167324284 | 0.280219577 | 0.59 | 0.39 |
| YPR028W | 4646.798863 | -0.244317044 | 0.280350024 | 0.59 | 0.39 |
| YPL154C | 14185.44316 | 0.338421716 | 0.280674132 | 0.59 | 0.39 |
| YJL066C | 1041.928086 | 0.380790646 | 0.280804708 | 0.59 | 0.39 |
| YBR113W | 115.7153994 | -0.2624945 | 0.281005749 | 0.59 | 0.39 |
| YBR294W | 590.8243932 | -0.281557014 | 0.281228497 | 0.59 | 0.39 |
| YML099W-A | 34.73959572 | 0.233939053 | 0.28115575 | 0.59 | 0.39 |
| YFL020C | 87.69189529 | 0.492789777 | 0.281347141 | 0.59 | 0.39 |
| YGL067W | 999.1477018 | 0.22774112 | 0.281544792 | 0.59 | 0.39 |
| YMR278W | 1574.862371 | 0.256719859 | 0.281642733 | 0.59 | 0.39 |
| YER099C | 790.1735302 | 0.130944265 | 0.28180979 | 0.59 | 0.39 |
| YDR382W | 7559.11107 | -0.321567245 | 0.282173545 | 0.59 | 0.39 |
| YFL065C | 1.354222162 | 0.759769006 | 0.282187952 | 0.59 | 0.39 |
| YLL023C | 2955.471664 | 0.271460565 | 0.282097638 | 0.59 | 0.39 |
| YOR020C | 3482.406455 | 0.285141239 | 0.281960573 | 0.59 | 0.39 |
| YDL062W | 21.94852935 | 0.435639741 | 0.282297306 | 0.59 | 0.39 |
| YMR141C | 167.2172498 | 0.290944539 | 0.282384428 | 0.59 | 0.39 |
| YOL019W-A | 0.720433626 | 0.496032326 | 0.282512394 | 0.59 | 0.39 |
| YDL089W | 963.5832446 | 0.307277851 | 0.283217873 | 0.59 | 0.39 |
| YER117W | 4528.496655 | -0.3609784 | 0.283131146 | 0.59 | 0.39 |
| YGR116W | 3744.798928 | -0.193885581 | 0.283302003 | 0.59 | 0.39 |
| YGR215W | 453.6817551 | -0.270701221 | 0.283050436 | 0.59 | 0.39 |
| YML058W-A | 99.58508334 | -0.509119802 | 0.283111557 | 0.59 | 0.39 |
| YMR005W | 1198.923005 | -0.231414669 | 0.283023164 | 0.59 | 0.39 |
| YNL178W | 12909.44004 | -0.370225455 | 0.283204302 | 0.59 | 0.39 |
| YOR122C | 4814.102226 | -0.253535794 | 0.28295822 | 0.59 | 0.39 |
| YDR311W | 727.3776002 | -0.212261059 | 0.283416074 | 0.59 | 0.39 |
| YCL059C | 1691.420964 | -0.360005974 | 0.284093654 | 0.59 | 0.39 |
| YCR036W | 1075.104784 | -0.184223217 | 0.284631296 | 0.59 | 0.39 |
| YDR071C | 2792.908431 | -0.224447215 | 0.28426558 | 0.59 | 0.39 |
| YEL066W | 476.9220074 | 0.307200308 | 0.284020221 | 0.59 | 0.39 |
| YFR042W | 519.19235 | 0.223637176 | 0.283918444 | 0.59 | 0.39 |
| YGL041C-B | 21.05072236 | -0.35645541 | 0.284530882 | 0.59 | 0.39 |
| YGR255C | 1753.194799 | 0.213758607 | 0.284536104 | 0.59 | 0.39 |
| YIL029W-A | 8.108002149 | 0.420290661 | 0.283980085 | 0.59 | 0.39 |
| YJL081C | 1547.994145 | -0.136982133 | 0.284049767 | 0.59 | 0.39 |
| YLR109W | 26967.32056 | 0.252516514 | 0.284689821 | 0.59 | 0.39 |
| YLR145W | 262.3006502 | 0.205936256 | 0.283823866 | 0.59 | 0.39 |
| YLR294C | 1804.172401 | 0.354851197 | 0.284539897 | 0.59 | 0.39 |
| YMR231W | 864.2847521 | 0.183527429 | 0.284225037 | 0.59 | 0.39 |
| YER028C | 507.0764562 | 0.420829724 | 0.285014214 | 0.59 | 0.39 |

| | | | | | |
|---|---|---|---|---|---|
| YLR117C | 1230.224284 | 0.230730777 | 0.285028036 | 0.59 | 0.39 |
| YOR182C | 1767.616754 | -0.345105587 | 0.285490937 | 0.59 | 0.39 |
| YAL034C-B | 9.086766243 | -0.445832045 | 0.285748369 | 0.59 | 0.39 |
| YBR191W-A | 25.03964351 | -0.245342385 | 0.285766684 | 0.59 | 0.39 |
| YER038C | 327.709464 | 0.303358258 | 0.285864195 | 0.59 | 0.39 |
| YJL136C | 5415.03448 | -0.357570363 | 0.285850478 | 0.59 | 0.39 |
| YHR097C | 2968.055756 | 0.39782596 | 0.286354279 | 0.59 | 0.39 |
| YAR042W | 3250.319067 | 0.270969878 | 0.286668504 | 0.59 | 0.39 |
| YER050C | 1168.128624 | -0.281821306 | 0.286936073 | 0.59 | 0.39 |
| YGR240C | 9640.703605 | -0.293047292 | 0.287342199 | 0.59 | 0.39 |
| YGL014W | 2156.619039 | -0.316140594 | 0.287550428 | 0.59 | 0.39 |
| YGL030W | 7624.667323 | -0.307726538 | 0.288168732 | 0.59 | 0.39 |
| YGL077C | 1372.265624 | -0.180435084 | 0.288021864 | 0.59 | 0.39 |
| YLR249W | 32716.59454 | -0.406170824 | 0.287670596 | 0.59 | 0.39 |
| YMR002W | 4776.428936 | 0.23533474 | 0.287921289 | 0.59 | 0.39 |
| YMR010W | 733.3907547 | -0.272741412 | 0.288114878 | 0.59 | 0.39 |
| YOL110W | 433.4687434 | 0.385415349 | 0.287897719 | 0.59 | 0.39 |
| YPR145W | 8044.845508 | 0.475418355 | 0.287542682 | 0.59 | 0.39 |
| YPR201W | 177.3808268 | 0.53266064 | 0.287718042 | 0.59 | 0.39 |
| YGR105W | 477.947228 | -0.242817681 | 0.288588789 | 0.59 | 0.39 |
| YGR186W | 2388.772284 | 0.21747065 | 0.288336965 | 0.59 | 0.39 |
| YHR152W | 1107.927047 | 0.208628757 | 0.28847925 | 0.59 | 0.39 |
| YOL120C | 9488.699457 | -0.370056001 | 0.288547691 | 0.59 | 0.39 |
| YMR125W | 2280.761082 | -0.23100772 | 0.288741657 | 0.59 | 0.39 |
| YAR028W | 513.5910617 | 0.212399481 | 0.289571146 | 0.59 | 0.39 |
| YCL010C | 597.9803619 | 0.256412181 | 0.289131623 | 0.59 | 0.39 |
| YDR087C | 817.7593103 | -0.349138091 | 0.289626748 | 0.59 | 0.39 |
| YDR329C | 1759.988143 | 0.268754035 | 0.289006311 | 0.59 | 0.39 |
| YER149C | 245.5366344 | -0.334191532 | 0.289554554 | 0.59 | 0.39 |
| YLR064W | 1412.390781 | 0.200429809 | 0.289619351 | 0.59 | 0.39 |
| YLR269C | 113.3738155 | 0.197201416 | 0.289055362 | 0.59 | 0.39 |
| YMR164C | 711.3389073 | 0.194050273 | 0.289180594 | 0.59 | 0.39 |
| YNR062C | 190.9739726 | 0.354987133 | 0.289324035 | 0.59 | 0.39 |
| YKL037W | 43.69720214 | -0.331892856 | 0.28973643 | 0.59 | 0.39 |
| YHR022C | 158.1795629 | -0.350528885 | 0.289848161 | 0.59 | 0.39 |
| YKL179C | 1404.902826 | -0.165588865 | 0.290121429 | 0.59 | 0.39 |
| YLR336C | 993.7017255 | 0.30632408 | 0.290007739 | 0.59 | 0.39 |
| YLR398C | 2110.607765 | 0.175955817 | 0.290160071 | 0.59 | 0.39 |
| YGL044C | 659.773802 | 0.120636679 | 0.290531973 | 0.59 | 0.39 |
| YGL237C | 638.8350245 | -0.286356473 | 0.290519122 | 0.59 | 0.39 |
| YIL016W | 483.5722155 | -0.273817702 | 0.290622414 | 0.59 | 0.39 |
| YDR058C | 448.3672811 | 0.302844217 | 0.291039523 | 0.59 | 0.39 |
| YHL025W | 1272.013263 | 0.283964566 | 0.290882664 | 0.59 | 0.39 |

| | | | | | |
|---|---|---|---|---|---|
| YHR032W | 1019.136565 | 0.345116041 | 0.290974858 | 0.59 | 0.39 |
| YPL196W | 943.9259177 | 0.336002382 | 0.2907739 | 0.59 | 0.39 |
| YDL165W | 697.5260403 | -0.14385001 | 0.291245629 | 0.59 | 0.39 |
| YOL038C-A | 23.65291474 | 0.270926926 | 0.29118968 | 0.59 | 0.39 |
| YER094C | 2268.208556 | 0.175725226 | 0.29135749 | 0.59 | 0.39 |
| YCR013C | 2806.322738 | 0.404883487 | 0.291754999 | 0.60 | 0.39 |
| YPL226W | 2793.678288 | -0.316706622 | 0.291609048 | 0.60 | 0.39 |
| snR75 | 1.574604411 | 0.745713041 | 0.291764843 | 0.60 | 0.39 |
| YDR286C | 156.0831168 | -0.244094521 | 0.292272558 | 0.60 | 0.39 |
| YFL067W | 57.17560859 | 0.773394129 | 0.292095745 | 0.60 | 0.39 |
| YGL078C | 1925.6458 | -0.32243504 | 0.291952091 | 0.60 | 0.39 |
| YIL130W | 1279.051362 | -0.309961372 | 0.292185245 | 0.60 | 0.39 |
| YNL107W | 513.6532475 | 0.134881116 | 0.292005914 | 0.60 | 0.39 |
| YDL052C | 1575.894255 | -0.310338518 | 0.29281105 | 0.60 | 0.39 |
| YER006W | 1977.06385 | -0.32706492 | 0.29280308 | 0.60 | 0.39 |
| YML070W | 2763.480937 | 0.256950513 | 0.292675725 | 0.60 | 0.39 |
| YMR052W | 219.8800281 | 0.271917246 | 0.292490428 | 0.60 | 0.39 |
| YNL324W | 2.168671185 | 0.626068206 | 0.292652185 | 0.60 | 0.39 |
| YDR338C | 695.8448786 | 0.20843434 | 0.29302109 | 0.60 | 0.39 |
| YDR510C-A | 439.1253795 | -0.256929086 | 0.293162432 | 0.60 | 0.39 |
| YER036C | 4277.526152 | -0.28666202 | 0.293451319 | 0.60 | 0.39 |
| YGL191W | 2937.202908 | 0.431233489 | 0.293411388 | 0.60 | 0.39 |
| YKR099W | 800.367483 | -0.276034458 | 0.293506699 | 0.60 | 0.39 |
| YBR152W | 271.4958287 | 0.158785789 | 0.294154015 | 0.60 | 0.39 |
| YIL062C | 1798.738473 | -0.185488279 | 0.294167559 | 0.60 | 0.39 |
| YJR101W | 534.1436387 | -0.222259605 | 0.293907124 | 0.60 | 0.39 |
| YOL038W | 4309.883987 | 0.195181023 | 0.294018501 | 0.60 | 0.39 |
| YHR209W | 694.2958547 | 0.382851217 | 0.29463811 | 0.60 | 0.39 |
| YBR287W | 2396.597397 | 0.225453243 | 0.295379682 | 0.60 | 0.40 |
| YDR269C | 1.063098833 | -0.46063639 | 0.295389061 | 0.60 | 0.40 |
| YEL072W | 207.4067694 | 0.420538917 | 0.295673704 | 0.60 | 0.40 |
| YFL046W | 596.7158239 | 0.174503544 | 0.296027793 | 0.60 | 0.40 |
| YJL200C | 1688.489143 | 0.355984824 | 0.295765233 | 0.60 | 0.40 |
| YJR048W | 1046.168345 | 0.6426373 | 0.295966876 | 0.60 | 0.40 |
| YKL193C | 1757.604095 | 0.289605565 | 0.295349279 | 0.60 | 0.40 |
| YLR180W | 3585.379162 | -0.406098224 | 0.295940896 | 0.60 | 0.40 |
| YLR345W | 2235.165084 | 0.267861938 | 0.296203714 | 0.60 | 0.40 |
| YMR035W | 535.4792987 | 0.244965303 | 0.295032414 | 0.60 | 0.40 |
| YMR206W | 572.161077 | 0.525994008 | 0.295299242 | 0.60 | 0.40 |
| YNL202W | 289.1619632 | 0.429413672 | 0.295605948 | 0.60 | 0.40 |
| YOR084W | 822.8839971 | -0.241653921 | 0.296112907 | 0.60 | 0.40 |
| YPL052W | 344.0806128 | 0.234239811 | 0.295953852 | 0.60 | 0.40 |
| YAL032C | 524.818116 | -0.160411095 | 0.296700577 | 0.60 | 0.40 |

| | | | | | |
|---|---|---|---|---|---|
| YBR090C | 79.12993298 | 0.319621228 | 0.296793608 | 0.60 | 0.40 |
| YDR501W | 295.5069374 | 0.185158786 | 0.296750781 | 0.60 | 0.40 |
| YNR026C | 756.2847135 | -0.211257967 | 0.296711945 | 0.60 | 0.40 |
| YPR118W | 1223.179848 | -0.345398405 | 0.296866933 | 0.60 | 0.40 |
| YBR077C | 472.5529414 | -0.203768151 | 0.297160525 | 0.60 | 0.40 |
| YHR012W | 963.21973 | 0.228553556 | 0.297712199 | 0.60 | 0.40 |
| YDR443C | 1163.053889 | 0.254281118 | 0.298121038 | 0.60 | 0.40 |
| YGL147C | 5508.177604 | -0.350560876 | 0.298118071 | 0.60 | 0.40 |
| YNL232W | 718.0441816 | -0.20436249 | 0.297984 | 0.60 | 0.40 |
| YKL028W | 1511.073698 | -0.214076143 | 0.298385308 | 0.60 | 0.40 |
| YPL087W | 1694.761164 | 0.259141105 | 0.298495462 | 0.60 | 0.40 |
| YIL106W | 893.3105346 | -0.17038138 | 0.298878605 | 0.60 | 0.40 |
| YMR302C | 2126.727676 | 0.271852746 | 0.299099687 | 0.60 | 0.40 |
| YPL201C | 234.758662 | 0.444124669 | 0.299121167 | 0.60 | 0.40 |
| YPR044C | 114.0311076 | -0.359980592 | 0.299082763 | 0.60 | 0.40 |
| YGL073W | 2411.314838 | 0.189867411 | 0.299283181 | 0.60 | 0.40 |
| YLR307C-A | 38.91718273 | 0.831709326 | 0.299683727 | 0.60 | 0.40 |
| YLR414C | 1706.526088 | 0.292981062 | 0.299670042 | 0.60 | 0.40 |
| YMR130W | 646.8299959 | -0.320188391 | 0.299603311 | 0.60 | 0.40 |
| YOL066C | 908.4756316 | -0.239738868 | 0.299554939 | 0.60 | 0.40 |
| YDR189W | 2199.458913 | -0.189608588 | 0.299965174 | 0.60 | 0.40 |
| YDL080C | 732.479931 | -0.145963356 | 0.300335538 | 0.60 | 0.40 |
| YOR165W | 1637.432086 | -0.188983553 | 0.300311681 | 0.60 | 0.40 |
| YDR242W | 619.518258 | 0.229092354 | 0.300713576 | 0.60 | 0.40 |
| YOR298W | 171.3347166 | 0.2669944 | 0.300667499 | 0.60 | 0.40 |
| YJL147C | 286.6419753 | 0.16451386 | 0.300885333 | 0.60 | 0.40 |
| YIL068C | 981.7929067 | -0.112186447 | 0.301015138 | 0.60 | 0.40 |
| YDR518W | 1466.448011 | 0.281608987 | 0.301112402 | 0.60 | 0.40 |
| YHR087W | 3070.022032 | 0.425581588 | 0.301408095 | 0.60 | 0.40 |
| YAL016W | 2950.826603 | 0.177353183 | 0.301672348 | 0.60 | 0.40 |
| YDR408C | 1967.993247 | -0.32243953 | 0.301966152 | 0.60 | 0.40 |
| YOL078W | 796.3214789 | 0.234924502 | 0.302124966 | 0.60 | 0.40 |
| YNL022C | 938.4563852 | 0.211852821 | 0.302271658 | 0.60 | 0.40 |
| YDR464W | 2373.668914 | 0.2737782 | 0.302435359 | 0.60 | 0.40 |
| YGR130C | 4694.98638 | 0.271036697 | 0.30270473 | 0.60 | 0.40 |
| YOL127W | 9841.72384 | -0.364213969 | 0.302712678 | 0.60 | 0.40 |
| YBL059W | 273.8243146 | 0.277374663 | 0.303153979 | 0.60 | 0.40 |
| YPL029W | 470.8569792 | -0.152276304 | 0.303241176 | 0.60 | 0.40 |
| YML117W | 3482.921508 | 0.188435513 | 0.303825564 | 0.61 | 0.40 |
| YDR418W | 2788.265019 | -0.358507727 | 0.304075164 | 0.61 | 0.40 |
| YPL055C | 1072.396721 | 0.171968234 | 0.304180237 | 0.61 | 0.40 |
| YDL169C | 664.258116 | 0.380180974 | 0.304476665 | 0.61 | 0.40 |
| YLR205C | 412.8411079 | 0.353602624 | 0.304598153 | 0.61 | 0.40 |

| | | | | | |
|---|---|---|---|---|---|
| YPL236C | 570.2921643 | 0.260270593 | 0.304550712 | 0.61 | 0.40 |
| YDL241W | 250.4921648 | 0.438913138 | 0.304911435 | 0.61 | 0.40 |
| YEL001C | 2451.407888 | -0.284924984 | 0.304908744 | 0.61 | 0.40 |
| YEL063C | 1734.961887 | 0.266176944 | 0.304992145 | 0.61 | 0.40 |
| YDR347W | 1409.321423 | -0.116156412 | 0.305418486 | 0.61 | 0.40 |
| YMR221C | 1062.231053 | -0.227244029 | 0.30534483 | 0.61 | 0.40 |
| YOR174W | 544.9117735 | -0.164369334 | 0.305322874 | 0.61 | 0.40 |
| YPR033C | 3601.656962 | 0.269156566 | 0.305400994 | 0.61 | 0.40 |
| YCR049C | 7.558405616 | -0.370759056 | 0.305656066 | 0.61 | 0.40 |
| YHR165W-A | 6.420816 | 0.464766576 | 0.305704685 | 0.61 | 0.40 |
| YPL166W | 366.5892241 | 0.331233511 | 0.305670598 | 0.61 | 0.40 |
| YGL187C | 1692.180783 | 0.480293488 | 0.306064661 | 0.61 | 0.40 |
| YOL018C | 882.7418897 | 0.108558678 | 0.306076663 | 0.61 | 0.40 |
| YGL172W | 1277.929348 | -0.203004807 | 0.306176971 | 0.61 | 0.40 |
| YDL117W | 1322.092729 | -0.16065618 | 0.306868243 | 0.61 | 0.40 |
| YDL204W | 5424.611742 | 0.528145012 | 0.306841963 | 0.61 | 0.40 |
| YDR070C | 1811.421734 | 0.539056573 | 0.30640573 | 0.61 | 0.40 |
| YER018C | 488.1395433 | 0.400466212 | 0.306419876 | 0.61 | 0.40 |
| YGL072C | 15.1439202 | 0.276439564 | 0.306810648 | 0.61 | 0.40 |
| YIL156W-A | 6.740450198 | -0.450221173 | 0.30668381 | 0.61 | 0.40 |
| YOR079C | 215.4299264 | -0.34823555 | 0.306800183 | 0.61 | 0.40 |
| YDL214C | 464.7760152 | 0.429071346 | 0.307032574 | 0.61 | 0.40 |
| YPL046C | 139.4253961 | -0.233553339 | 0.307198818 | 0.61 | 0.40 |
| YDL061C | 2048.312712 | -0.329520911 | 0.30739179 | 0.61 | 0.40 |
| YGR230W | 330.0627222 | -0.18498441 | 0.307331655 | 0.61 | 0.40 |
| YHR011W | 509.988716 | 0.331988335 | 0.307514065 | 0.61 | 0.40 |
| YBR186W | 272.8183513 | 0.386388935 | 0.307956003 | 0.61 | 0.40 |
| YCL063W | 606.1625178 | -0.248611729 | 0.308039234 | 0.61 | 0.40 |
| YDR278C | 0.649229666 | 0.367074775 | 0.3080296 | 0.61 | 0.40 |
| YHR165C | 3249.517075 | 0.27350829 | 0.307979174 | 0.61 | 0.40 |
| YOR241W | 933.790879 | -0.208472248 | 0.308065931 | 0.61 | 0.40 |
| YBL067C | 1161.166699 | 0.119289261 | 0.308450442 | 0.61 | 0.40 |
| YDL085W | 757.5796414 | 0.475215269 | 0.308544395 | 0.61 | 0.40 |
| YDR297W | 2184.395638 | -0.219564932 | 0.308320124 | 0.61 | 0.40 |
| YER007W | 271.3998551 | 0.223300601 | 0.308727216 | 0.61 | 0.40 |
| YGL079W | 448.9297125 | -0.150130046 | 0.308633292 | 0.61 | 0.40 |
| YOR319W | 365.6759573 | -0.173598306 | 0.308699121 | 0.61 | 0.40 |
| YGL055W | 3060.50732 | -0.283812064 | 0.309075729 | 0.61 | 0.40 |
| YNL013C | 5.629906296 | 0.515062189 | 0.309086099 | 0.61 | 0.40 |
| YBR166C | 859.3257193 | -0.1963361 | 0.310042221 | 0.61 | 0.40 |
| YFR037C | 1666.566511 | -0.195905302 | 0.310022175 | 0.61 | 0.40 |
| YHR006W | 1327.030831 | 0.32311501 | 0.31023888 | 0.61 | 0.40 |
| YHR196W | 929.0161741 | -0.352588983 | 0.30996285 | 0.61 | 0.40 |

| | | | | | |
|---|---|---|---|---|---|
| YIL060W | 54.27024793 | 0.462823257 | 0.310174469 | 0.61 | 0.40 |
| YMR237W | 2047.326568 | 0.098487024 | 0.310095978 | 0.61 | 0.40 |
| YOR085W | 1806.530962 | -0.256319435 | 0.310166301 | 0.61 | 0.40 |
| YOR179C | 265.678804 | -0.269275511 | 0.310347394 | 0.61 | 0.40 |
| YDL114W-A | 24.54000287 | 0.380198277 | 0.310549723 | 0.61 | 0.40 |
| YOL099C | 11.41692104 | 0.365179513 | 0.310465495 | 0.61 | 0.40 |
| YLR100W | 1453.024103 | 0.302729957 | 0.310831832 | 0.61 | 0.40 |
| YNL039W | 1798.324816 | 0.163412493 | 0.310996458 | 0.61 | 0.40 |
| YPL182C | 43.60254699 | 0.258107339 | 0.31098985 | 0.61 | 0.40 |
| YNL268W | 2190.621688 | -0.19679861 | 0.311123808 | 0.61 | 0.40 |
| YNL143C | 26.87962177 | -0.306505421 | 0.311222327 | 0.61 | 0.40 |
| YMR111C | 424.0046487 | 0.166874015 | 0.311695668 | 0.61 | 0.40 |
| YMR272W-B | 3.438828838 | 0.514144251 | 0.311442083 | 0.61 | 0.40 |
| YNR072W | 156.9410464 | 0.420119031 | 0.311648548 | 0.61 | 0.40 |
| YOL154W | 197.7584736 | -0.225868161 | 0.311611909 | 0.61 | 0.40 |
| YDL103C | 1449.48201 | 0.190734935 | 0.312002224 | 0.61 | 0.40 |
| YDR256C | 561.4619469 | 0.339615378 | 0.31193741 | 0.61 | 0.40 |
| YMR037C | 1272.708272 | -0.276207522 | 0.312060517 | 0.61 | 0.40 |
| YBL086C | 1904.650929 | 0.218591801 | 0.312167016 | 0.61 | 0.40 |
| YIL052C | 4784.674042 | -0.342764699 | 0.31242315 | 0.61 | 0.40 |
| YLR375W | 1937.904522 | -0.158575045 | 0.312390216 | 0.61 | 0.40 |
| YBL092W | 15431.27049 | -0.347908192 | 0.312628549 | 0.61 | 0.40 |
| snR68 | 10.29817386 | 0.507181896 | 0.312654611 | 0.61 | 0.40 |
| YDL059C | 182.996654 | 0.224344937 | 0.31286544 | 0.61 | 0.40 |
| YGR027C | 9077.452786 | -0.339121245 | 0.313095537 | 0.61 | 0.40 |
| YOR300W | 17.40293291 | 0.323988774 | 0.313094111 | 0.61 | 0.40 |
| YDR385W | 1851.31364 | -0.317607889 | 0.313222747 | 0.61 | 0.40 |
| YPR021C | 1682.395818 | 0.229377547 | 0.313380087 | 0.61 | 0.40 |
| YJL186W | 1797.326564 | 0.173121799 | 0.31369543 | 0.61 | 0.40 |
| YKL062W | 2465.244192 | -0.363003793 | 0.313687583 | 0.61 | 0.40 |
| YBR272C | 606.5333168 | 0.175970285 | 0.314018347 | 0.61 | 0.40 |
| YLR007W | 741.9239686 | 0.227890436 | 0.314504287 | 0.61 | 0.40 |
| YGR085C | 1835.900279 | -0.319499687 | 0.314906236 | 0.61 | 0.40 |
| YKL192C | 3222.400243 | -0.159453126 | 0.314863302 | 0.61 | 0.40 |
| YML049C | 1684.51308 | 0.23957932 | 0.314760252 | 0.61 | 0.40 |
| YIR037W | 2878.72569 | 0.221016461 | 0.31512974 | 0.61 | 0.40 |
| YFR012W-A | 2.137477958 | 0.483792118 | 0.315537611 | 0.61 | 0.41 |
| YNL185C | 474.6914722 | -0.202639945 | 0.315749267 | 0.61 | 0.41 |
| YPL026C | 1074.82514 | -0.364590628 | 0.315755068 | 0.61 | 0.41 |
| YPL271W | 1376.293801 | 0.374940306 | 0.315573513 | 0.61 | 0.41 |
| YPR140W | 900.9897118 | -0.244250486 | 0.315874573 | 0.61 | 0.41 |
| YOR196C | 1243.855488 | 0.209058714 | 0.316194566 | 0.61 | 0.41 |
| YPR145C-A | 45.3897147 | 0.440948278 | 0.316327171 | 0.61 | 0.41 |

| | | | | | |
|---|---|---|---|---|---|
| YLR411W | 132.197405 | 0.244680084 | 0.316726071 | 0.61 | 0.41 |
| YNR060W | 291.930447 | 0.232599454 | 0.316773763 | 0.61 | 0.41 |
| YHR056C | 832.3445362 | 0.176489506 | 0.316972314 | 0.61 | 0.41 |
| YMR173W | 743.4251563 | 0.282239584 | 0.316893951 | 0.61 | 0.41 |
| YPR008W | 1796.059535 | -0.179890891 | 0.317146176 | 0.61 | 0.41 |
| YBR277C | 4.283921599 | -0.643538522 | 0.317379197 | 0.61 | 0.41 |
| YDR535C | 20.65811424 | 0.37602206 | 0.317542071 | 0.61 | 0.41 |
| YGL069C | 85.63194987 | -0.295936162 | 0.317400872 | 0.61 | 0.41 |
| YMR119W-A | 41.58140006 | 0.31981961 | 0.317474961 | 0.61 | 0.41 |
| YHR025W | 4807.328396 | 0.439939488 | 0.317721527 | 0.61 | 0.41 |
| YGR276C | 1041.106244 | 0.11690528 | 0.31822194 | 0.61 | 0.41 |
| YKL095W | 323.4844945 | 0.232695576 | 0.318227484 | 0.61 | 0.41 |
| YOR295W | 256.0382172 | -0.189834232 | 0.318113528 | 0.61 | 0.41 |
| YOR366W | 3.244594662 | -0.533753181 | 0.317911485 | 0.61 | 0.41 |
| YOR386W | 897.2436472 | 0.36857709 | 0.31795714 | 0.61 | 0.41 |
| YNL005C | 1578.642517 | 0.158783425 | 0.318363757 | 0.61 | 0.41 |
| YFL044C | 776.7355212 | 0.254757539 | 0.318583516 | 0.61 | 0.41 |
| YBL003C | 4258.595501 | -0.314681418 | 0.319103527 | 0.62 | 0.41 |
| YBL100W-C | 6.925265405 | 0.426673821 | 0.319347286 | 0.62 | 0.41 |
| YGR185C | 4359.206315 | -0.339673525 | 0.318978349 | 0.62 | 0.41 |
| YHR192W | 784.2571123 | 0.188172605 | 0.319170165 | 0.62 | 0.41 |
| YIL087C | 960.1717362 | 0.408312264 | 0.319005711 | 0.62 | 0.41 |
| YLR136C | 655.8429852 | -0.414043582 | 0.319191142 | 0.62 | 0.41 |
| YNL067W | 1749.269832 | -0.306065137 | 0.319311056 | 0.62 | 0.41 |
| YCR014C | 618.1633306 | 0.204894009 | 0.319585548 | 0.62 | 0.41 |
| YHL008C | 1005.082799 | 0.291971955 | 0.320102612 | 0.62 | 0.41 |
| YHR071W | 3724.97766 | 0.39979522 | 0.320079186 | 0.62 | 0.41 |
| YLR188W | 1233.172467 | 0.168000894 | 0.320191716 | 0.62 | 0.41 |
| YOL159C | 274.8863973 | 0.326931471 | 0.320076022 | 0.62 | 0.41 |
| YKL146W | 1758.494435 | 0.334408748 | 0.320332509 | 0.62 | 0.41 |
| YAL044C | 2900.555908 | -0.430632037 | 0.320835923 | 0.62 | 0.41 |
| YJR053W | 291.1479634 | 0.209870592 | 0.320809693 | 0.62 | 0.41 |
| YLR003C | 782.5858822 | -0.31707297 | 0.320635855 | 0.62 | 0.41 |
| YPR014C | 81.20498688 | -0.313060511 | 0.320771275 | 0.62 | 0.41 |
| YCR031C | 11369.4013 | -0.329040313 | 0.321248714 | 0.62 | 0.41 |
| YGR175C | 3323.313876 | -0.274312521 | 0.320996874 | 0.62 | 0.41 |
| YIL059C | 89.463829 | 0.448595714 | 0.321163975 | 0.62 | 0.41 |
| YKL004W | 2100.85667 | -0.238291011 | 0.321151702 | 0.62 | 0.41 |
| YBR030W | 579.6269936 | -0.226143937 | 0.321586857 | 0.62 | 0.41 |
| YHL004W | 1819.11881 | -0.143415751 | 0.321832386 | 0.62 | 0.41 |
| YLR103C | 462.3096359 | -0.21752447 | 0.321663803 | 0.62 | 0.41 |
| YMR019W | 977.1459301 | 0.251227834 | 0.321583179 | 0.62 | 0.41 |
| YOR385W | 901.596547 | 0.352772942 | 0.321361065 | 0.62 | 0.41 |

| | | | | | |
|---|---|---|---|---|---|
| snR59 | 2.356669998 | 0.739321244 | 0.321788133 | 0.62 | 0.41 |
| YNL302C | 5878.096109 | -0.328398175 | 0.322194486 | 0.62 | 0.41 |
| YKL181W | 2705.955695 | -0.290409837 | 0.322489479 | 0.62 | 0.41 |
| YOR256C | 1260.844527 | 0.17835402 | 0.32249574 | 0.62 | 0.41 |
| YDR495C | 1040.925179 | 0.230256453 | 0.322720458 | 0.62 | 0.41 |
| YGL023C | 1541.23519 | 0.185255607 | 0.323052453 | 0.62 | 0.41 |
| YDR340W | 4.60165433 | -0.475614744 | 0.323647699 | 0.62 | 0.41 |
| YLR050C | 545.8709868 | 0.297683786 | 0.323465181 | 0.62 | 0.41 |
| YLR148W | 974.658438 | 0.170235902 | 0.323566376 | 0.62 | 0.41 |
| YDR237W | 832.9477874 | -0.191992629 | 0.324766962 | 0.62 | 0.41 |
| YDR469W | 524.7044195 | -0.220464358 | 0.324786819 | 0.62 | 0.41 |
| YHR059W | 401.0845698 | -0.187812612 | 0.323926934 | 0.62 | 0.41 |
| YIL126W | 2338.60398 | -0.243060572 | 0.324910054 | 0.62 | 0.41 |
| YIR034C | 2228.546885 | 0.393208133 | 0.324258948 | 0.62 | 0.41 |
| YLR017W | 1271.451959 | -0.249016209 | 0.324925741 | 0.62 | 0.41 |
| YNL065W | 531.9531231 | -0.41221045 | 0.324332529 | 0.62 | 0.41 |
| YOL153C | 1352.172864 | 0.345392663 | 0.324861652 | 0.62 | 0.41 |
| YOL164W | 729.2266525 | 0.280162243 | 0.324677997 | 0.62 | 0.41 |
| YOR144C | 421.6026059 | -0.204921777 | 0.323899695 | 0.62 | 0.41 |
| YPL035C | 100.9354993 | -0.241216276 | 0.32471551 | 0.62 | 0.41 |
| YPR025C | 1080.062057 | -0.227481737 | 0.324152957 | 0.62 | 0.41 |
| snR78 | 1.974544426 | 0.320731368 | 0.324641691 | 0.62 | 0.41 |
| YDR395W | 2175.578678 | -0.259486157 | 0.325027144 | 0.62 | 0.41 |
| YMR172W | 685.1558079 | -0.195514319 | 0.325297383 | 0.62 | 0.41 |
| YAL055W | 312.0534701 | 0.246528277 | 0.326207948 | 0.62 | 0.41 |
| YGR151C | 2.6142374 | -0.486274122 | 0.326203995 | 0.62 | 0.41 |
| YGR198W | 1689.551002 | -0.227005736 | 0.32616952 | 0.62 | 0.41 |
| YGR192C | 181337.4741 | 0.348412223 | 0.327242292 | 0.62 | 0.41 |
| YIL037C | 352.6358083 | 0.398878295 | 0.327167437 | 0.62 | 0.41 |
| YHR050W | 1110.031456 | 0.359089222 | 0.327720968 | 0.62 | 0.41 |
| YCL002C | 469.4119103 | -0.218476109 | 0.328293913 | 0.62 | 0.41 |
| YHR058C | 391.0666567 | 0.190076519 | 0.328173524 | 0.62 | 0.41 |
| YJL177W | 7142.560719 | -0.335970412 | 0.328421833 | 0.62 | 0.41 |
| YPL107W | 480.1557675 | 0.225305157 | 0.328362066 | 0.62 | 0.41 |
| YPR048W | 424.7867171 | -0.24755629 | 0.327980064 | 0.62 | 0.41 |
| snR51 | 25.09375178 | 0.357542691 | 0.328372923 | 0.62 | 0.41 |
| YDR036C | 1560.530814 | 0.27321951 | 0.328650143 | 0.62 | 0.41 |
| YDL079C | 1497.675817 | 0.441378062 | 0.329021827 | 0.62 | 0.41 |
| YFR040W | 3140.971961 | 0.30194704 | 0.330052308 | 0.62 | 0.41 |
| YJL125C | 686.7482617 | -0.253614671 | 0.330094641 | 0.62 | 0.41 |
| YBR216C | 879.1909045 | 0.156984945 | 0.330299096 | 0.63 | 0.41 |
| YGL092W | 1819.13534 | -0.181641073 | 0.330642801 | 0.63 | 0.41 |
| YGR069W | 7.64600337 | -0.439719644 | 0.330842279 | 0.63 | 0.41 |

| | | | | | |
|---|---|---|---|---|---|
| YDR407C | 1886.152895 | -0.162764941 | 0.331199037 | 0.63 | 0.41 |
| YGR023W | 927.8650533 | 0.373912647 | 0.331432846 | 0.63 | 0.41 |
| YIL017C | 1438.735299 | 0.252726037 | 0.331572288 | 0.63 | 0.41 |
| YKR009C | 961.8765586 | 0.336968675 | 0.331215398 | 0.63 | 0.41 |
| YLR210W | 386.6683639 | -0.185792906 | 0.331499675 | 0.63 | 0.41 |
| YPL060W | 360.6791713 | 0.27405614 | 0.331327072 | 0.63 | 0.41 |
| YAL037W | 195.7634153 | 0.302341014 | 0.331746949 | 0.63 | 0.41 |
| YJR133W | 618.7266674 | 0.265906306 | 0.331832369 | 0.63 | 0.41 |
| YFL003C | 165.623347 | 0.339672347 | 0.332055878 | 0.63 | 0.41 |
| YIL064W | 505.7570311 | 0.358207302 | 0.332299613 | 0.63 | 0.41 |
| YKR025W | 782.5030894 | -0.230396433 | 0.332746943 | 0.63 | 0.41 |
| YLL036C | 873.0418955 | -0.112111409 | 0.332590121 | 0.63 | 0.41 |
| YOR245C | 966.5341375 | 0.237272882 | 0.332679076 | 0.63 | 0.41 |
| YER148W-A | 0.781507553 | -0.475350371 | 0.333133408 | 0.63 | 0.41 |
| YGL039W | 719.9626894 | 0.292699248 | 0.33321484 | 0.63 | 0.41 |
| YGL188C-A | 18.77666255 | 0.556064924 | 0.333162233 | 0.63 | 0.41 |
| YIL030W-A | 74.85518125 | 0.194931569 | 0.333270303 | 0.63 | 0.41 |
| YJR067C | 196.0036818 | -0.160984477 | 0.333282855 | 0.63 | 0.41 |
| YJL001W | 1978.350649 | -0.15052697 | 0.333536303 | 0.63 | 0.41 |
| YML101C | 335.3248863 | -0.226046485 | 0.333650927 | 0.63 | 0.41 |
| YDR507C | 1606.54523 | -0.292656959 | 0.333850209 | 0.63 | 0.41 |
| YGR124W | 7850.205704 | -0.391277924 | 0.33413463 | 0.63 | 0.41 |
| YLR395C | 1418.063046 | 0.409954518 | 0.334120112 | 0.63 | 0.41 |
| YMR142C | 11187.93902 | -0.309006507 | 0.334013383 | 0.63 | 0.41 |
| YMR149W | 2739.886665 | -0.196915989 | 0.334175278 | 0.63 | 0.41 |
| YDR184C | 548.4562077 | -0.308127669 | 0.334948673 | 0.63 | 0.42 |
| YML086C | 1768.316639 | -0.146786906 | 0.334794523 | 0.63 | 0.42 |
| YNL074C | 2355.063484 | 0.18977307 | 0.334728956 | 0.63 | 0.42 |
| YOR259C | 3766.746436 | -0.218204825 | 0.334900047 | 0.63 | 0.42 |
| YCL049C | 935.682406 | 0.359057647 | 0.335163277 | 0.63 | 0.42 |
| YDL088C | 1480.552376 | -0.190041595 | 0.335084818 | 0.63 | 0.42 |
| YGR214W | 11003.56684 | -0.332501734 | 0.33546934 | 0.63 | 0.42 |
| YKL038W | 1738.382543 | 0.352324557 | 0.33558292 | 0.63 | 0.42 |
| YDR254W | 574.0903535 | 0.151048843 | 0.335726343 | 0.63 | 0.42 |
| YAR029W | 75.14481749 | 0.313043397 | 0.335857227 | 0.63 | 0.42 |
| YCR108C | 1.912873177 | 0.225451419 | 0.33602736 | 0.63 | 0.42 |
| YBR195C | 288.8600834 | -0.200344382 | 0.336144902 | 0.63 | 0.42 |
| YPL113C | 241.6280316 | 0.244672669 | 0.336366024 | 0.63 | 0.42 |
| YMR075C-A | 15.14864191 | 0.327836978 | 0.336551753 | 0.63 | 0.42 |
| YER037W | 745.1407721 | 0.457486392 | 0.33669372 | 0.63 | 0.42 |
| YLR114C | 1321.213644 | 0.185346134 | 0.336759552 | 0.63 | 0.42 |
| YBR247C | 781.1952554 | -0.273347641 | 0.337671205 | 0.63 | 0.42 |
| YDL236W | 1309.137891 | -0.249821158 | 0.337888979 | 0.63 | 0.42 |

| | | | | | |
|---|---|---|---|---|---|
| YDR450W | 8491.535949 | -0.357661019 | 0.337106011 | 0.63 | 0.42 |
| YJL079C | 9145.850933 | 0.372013153 | 0.337344268 | 0.63 | 0.42 |
| YMR032W | 652.0530589 | -0.213552397 | 0.337803308 | 0.63 | 0.42 |
| YMR273C | 1360.501919 | 0.208662105 | 0.337493231 | 0.63 | 0.42 |
| YNL263C | 1345.46904 | -0.212872608 | 0.337850204 | 0.63 | 0.42 |
| YOL052C-A | 1535.771374 | -0.656206818 | 0.337521931 | 0.63 | 0.42 |
| YOR289W | 937.4069291 | 0.392645549 | 0.337430169 | 0.63 | 0.42 |
| YPL099C | 334.8201937 | -0.2100952 | 0.33716753 | 0.63 | 0.42 |
| YPR127W | 1734.545853 | 0.30149393 | 0.337860703 | 0.63 | 0.42 |
| YBL001C | 1539.933777 | -0.267821636 | 0.338046703 | 0.63 | 0.42 |
| YNL331C | 347.7391832 | 0.270460991 | 0.338086409 | 0.63 | 0.42 |
| YGR020C | 3100.566353 | -0.24637268 | 0.338184803 | 0.63 | 0.42 |
| YJL104W | 974.9529404 | -0.202169981 | 0.338343666 | 0.63 | 0.42 |
| YHR173C | 88.05682976 | -0.297278188 | 0.338664845 | 0.63 | 0.42 |
| YJR088C | 954.591927 | -0.177933392 | 0.338645339 | 0.63 | 0.42 |
| YHR168W | 379.0957302 | 0.172100283 | 0.33900429 | 0.63 | 0.42 |
| YDR182W-A | 24.64411227 | 0.217449597 | 0.3391031 | 0.63 | 0.42 |
| YBR236C | 870.5665803 | -0.191046721 | 0.33958988 | 0.63 | 0.42 |
| YCR051W | 1038.832374 | -0.245645952 | 0.339810611 | 0.63 | 0.42 |
| YLR304C | 13028.11852 | 0.344684104 | 0.33992484 | 0.63 | 0.42 |
| YDR417C | 435.6659374 | -0.399994636 | 0.340208685 | 0.63 | 0.42 |
| YLR149C-A | 61.1192566 | -0.444890718 | 0.340622263 | 0.63 | 0.42 |
| YDL075W | 5187.384084 | -0.316623519 | 0.340759259 | 0.63 | 0.42 |
| YER059W | 910.2356733 | -0.217710665 | 0.341059735 | 0.63 | 0.42 |
| YPL183W-A | 354.0512636 | -0.186350957 | 0.340973653 | 0.63 | 0.42 |
| YOR293C-A | 2.608736587 | -0.570388131 | 0.341466903 | 0.63 | 0.42 |
| YGL063W | 321.9330079 | -0.273712143 | 0.341880249 | 0.63 | 0.42 |
| YGR033C | 1013.438168 | -0.137842555 | 0.341947947 | 0.63 | 0.42 |
| YHL006C | 116.1902464 | -0.299972056 | 0.342015531 | 0.63 | 0.42 |
| YKL011C | 416.7502341 | 0.222849536 | 0.342184359 | 0.63 | 0.42 |
| YKL113C | 707.9229549 | -0.221206105 | 0.342211617 | 0.63 | 0.42 |
| YPR179C | 773.0608325 | 0.164704623 | 0.342338593 | 0.63 | 0.42 |
| YGL171W | 777.7803509 | -0.29703966 | 0.342752419 | 0.63 | 0.42 |
| YPL258C | 548.5976694 | 0.288496048 | 0.342946549 | 0.63 | 0.42 |
| YAL001C | 1631.198822 | 0.170408556 | 0.343462294 | 0.63 | 0.42 |
| YDR216W | 2843.606015 | 0.376864678 | 0.343286609 | 0.63 | 0.42 |
| YMR042W | 272.8014796 | -0.300143037 | 0.343364755 | 0.63 | 0.42 |
| YPL163C | 3377.565521 | 0.369431253 | 0.343413835 | 0.63 | 0.42 |
| YNL119W | 513.2444289 | -0.25163896 | 0.343812015 | 0.64 | 0.42 |
| YFR046C | 527.2575594 | 0.17173956 | 0.344507727 | 0.64 | 0.42 |
| YMR056C | 319.9319845 | 0.356750713 | 0.344358359 | 0.64 | 0.42 |
| YOL085C | 30.97884527 | 0.414079912 | 0.344463402 | 0.64 | 0.42 |
| YAL041W | 1345.73957 | -0.224780343 | 0.344729984 | 0.64 | 0.42 |

| | | | | | |
|---|---|---|---|---|---|
| YAL064W-B | 18.20372016 | 0.46237086 | 0.345451794 | 0.64 | 0.42 |
| YOL063C | 983.1121387 | 0.160537206 | 0.345528014 | 0.64 | 0.42 |
| YBR240C | 118.1647641 | 0.391741466 | 0.345638798 | 0.64 | 0.42 |
| YOR215C | 2322.968103 | 0.283311151 | 0.345785408 | 0.64 | 0.42 |
| YDR028C | 3835.45418 | 0.189030121 | 0.346009356 | 0.64 | 0.42 |
| YOR096W | 6750.76705 | -0.294180099 | 0.346119901 | 0.64 | 0.42 |
| YJL111W | 2828.454028 | -0.235725649 | 0.346231336 | 0.64 | 0.42 |
| YGR056W | 1165.037029 | -0.205927873 | 0.346412834 | 0.64 | 0.42 |
| YLR286C | 14573.14708 | -0.206347902 | 0.346472325 | 0.64 | 0.42 |
| YCL014W | 1364.401777 | 0.213138521 | 0.346604175 | 0.64 | 0.42 |
| YBL040C | 1584.786506 | -0.189425429 | 0.346882474 | 0.64 | 0.42 |
| YMR210W | 1188.233193 | -0.170857814 | 0.34731954 | 0.64 | 0.42 |
| YOR343W-B | 107.9180972 | -0.35426369 | 0.347306706 | 0.64 | 0.42 |
| YPL174C | 736.6858257 | 0.134121232 | 0.34723156 | 0.64 | 0.42 |
| YBR125C | 746.1231571 | 0.242651853 | 0.347782184 | 0.64 | 0.42 |
| YLL021W | 2378.696108 | -0.23166105 | 0.347596967 | 0.64 | 0.42 |
| YNL046W | 536.2842104 | -0.227910121 | 0.347520728 | 0.64 | 0.42 |
| YOR370C | 2847.263179 | -0.177991817 | 0.347700165 | 0.64 | 0.42 |
| YAL056W | 1700.26388 | 0.266815976 | 0.348123318 | 0.64 | 0.42 |
| YDL153C | 1719.824839 | -0.267906167 | 0.348191858 | 0.64 | 0.42 |
| YMR141W-A | 54.52612508 | 0.482882279 | 0.348239765 | 0.64 | 0.42 |
| YKR004C | 281.3110563 | 0.169435357 | 0.348535735 | 0.64 | 0.42 |
| YLR407W | 355.7060391 | -0.188567117 | 0.348590051 | 0.64 | 0.42 |
| YMR255W | 1118.071772 | -0.24823844 | 0.348798114 | 0.64 | 0.42 |
| YDR381C-A | 448.1284196 | 0.28524899 | 0.349010873 | 0.64 | 0.42 |
| YER088C-A | 302.6604055 | -0.342693978 | 0.34893268 | 0.64 | 0.42 |
| YNL291C | 628.873147 | -0.178262034 | 0.34911528 | 0.64 | 0.42 |
| YFL066C | 2.343434624 | 0.71593823 | 0.349474859 | 0.64 | 0.42 |
| YOR194C | 1465.560671 | -0.248042992 | 0.349638721 | 0.64 | 0.42 |
| YGR025W | 100.9215658 | -0.251077388 | 0.349774083 | 0.64 | 0.42 |
| YIL137C | 2189.193159 | 0.241817263 | 0.349883544 | 0.64 | 0.42 |
| YGL068W | 385.5514446 | -0.240415915 | 0.350515992 | 0.64 | 0.42 |
| YDL183C | 401.7373652 | 0.248265024 | 0.350920575 | 0.64 | 0.42 |
| YJL175W | 4.732791699 | -0.468479609 | 0.351070641 | 0.64 | 0.42 |
| snR3 | 32.48912912 | 0.382606433 | 0.351015633 | 0.64 | 0.42 |
| YLR364W | 131.7044529 | 0.278385129 | 0.351432288 | 0.64 | 0.42 |
| YNR024W | 280.6447987 | -0.203898449 | 0.351615541 | 0.64 | 0.42 |
| YJL216C | 203.2730397 | -0.316040296 | 0.352206329 | 0.64 | 0.42 |
| YFL036W | 2229.86904 | 0.244884052 | 0.352611405 | 0.64 | 0.43 |
| YMR021C | 731.4049333 | 0.163750652 | 0.352528107 | 0.64 | 0.43 |
| YDR210W-B | 26.10421514 | 0.359534427 | 0.352925998 | 0.64 | 0.43 |
| YDR020C | 268.9612697 | -0.263334187 | 0.353480093 | 0.64 | 0.43 |
| YDR315C | 362.5235632 | -0.190124284 | 0.353649595 | 0.64 | 0.43 |

| | | | | | |
|---|---|---|---|---|---|
| YEL050C | 1046.082243 | -0.156968442 | 0.353754514 | 0.64 | 0.43 |
| YHR116W | 348.981213 | 0.21493533 | 0.353339985 | 0.64 | 0.43 |
| YLR056W | 3964.313149 | -0.267553159 | 0.353781858 | 0.64 | 0.43 |
| YOR279C | 315.7050157 | -0.219964028 | 0.353278725 | 0.64 | 0.43 |
| snR39 | 11.68210812 | 0.660764861 | 0.353603488 | 0.64 | 0.43 |
| YDR217C | 884.5738376 | 0.196961448 | 0.354216356 | 0.64 | 0.43 |
| YMR126C | 404.3389603 | -0.199499473 | 0.354214035 | 0.64 | 0.43 |
| YBR189W | 11760.64407 | -0.33679082 | 0.354447313 | 0.64 | 0.43 |
| YKL121W | 1073.818565 | 0.197391331 | 0.354685903 | 0.64 | 0.43 |
| YDR061W | 1142.905546 | 0.12563815 | 0.355293933 | 0.65 | 0.43 |
| YBR196C-B | 11.09747944 | -0.389851444 | 0.355476157 | 0.65 | 0.43 |
| YOR306C | 1215.578289 | 0.412802379 | 0.355626227 | 0.65 | 0.43 |
| YDR183C-A | 47.62983085 | -0.312260042 | 0.355904957 | 0.65 | 0.43 |
| YGR229C | 1916.022007 | -0.206841703 | 0.356117827 | 0.65 | 0.43 |
| YKR072C | 2109.757496 | 0.208222366 | 0.356175266 | 0.65 | 0.43 |
| YOL008W | 381.7473344 | -0.135674927 | 0.356142287 | 0.65 | 0.43 |
| YFL001W | 478.2892235 | -0.20348132 | 0.356551995 | 0.65 | 0.43 |
| YGL226C-A | 628.3645694 | -0.199520091 | 0.357129834 | 0.65 | 0.43 |
| YIL014W | 862.4746572 | 0.18942142 | 0.357480955 | 0.65 | 0.43 |
| YJR113C | 864.1046061 | -0.201458228 | 0.357409751 | 0.65 | 0.43 |
| YHR154W | 1331.473724 | -0.223277295 | 0.357597046 | 0.65 | 0.43 |
| YMR135C | 1809.513775 | 0.350870947 | 0.357716116 | 0.65 | 0.43 |
| YHL020C | 961.5750146 | -0.211478742 | 0.358141147 | 0.65 | 0.43 |
| YIL164C | 260.9748713 | 0.30911211 | 0.358098838 | 0.65 | 0.43 |
| YOR276W | 906.6874329 | -0.263253703 | 0.358336274 | 0.65 | 0.43 |
| YPR171W | 883.9863961 | -0.152213181 | 0.358296117 | 0.65 | 0.43 |
| YIL144W | 1230.130031 | 0.199061606 | 0.358647643 | 0.65 | 0.43 |
| YGR011W | 95.88242328 | 0.158027022 | 0.358856855 | 0.65 | 0.43 |
| YBR167C | 275.1433522 | -0.189286555 | 0.358969904 | 0.65 | 0.43 |
| YCR106W | 394.726015 | 0.455812624 | 0.359566671 | 0.65 | 0.43 |
| YGR249W | 613.9099062 | 0.454871797 | 0.359344357 | 0.65 | 0.43 |
| YJL138C | 1955.737437 | -0.187153133 | 0.359731354 | 0.65 | 0.43 |
| YLR036C | 360.058471 | -0.184621252 | 0.359801096 | 0.65 | 0.43 |
| YLR048W | 13755.73587 | -0.322946167 | 0.359760879 | 0.65 | 0.43 |
| YLR143W | 501.3258454 | -0.19003429 | 0.359485602 | 0.65 | 0.43 |
| YOR363C | 1212.557429 | 0.206710513 | 0.35984454 | 0.65 | 0.43 |
| YDR533C | 1411.920932 | -0.455173219 | 0.360083687 | 0.65 | 0.43 |
| YGL088W | 0.432313012 | 0.421966378 | 0.360029045 | 0.65 | 0.43 |
| YBR182C | 159.913284 | 0.317712802 | 0.360281079 | 0.65 | 0.43 |
| YKL187C | 740.5340162 | 0.308080749 | 0.360533341 | 0.65 | 0.43 |
| YOR348C | 282.9627385 | 0.475201667 | 0.360611109 | 0.65 | 0.43 |
| YDR424C | 383.6752581 | -0.136458883 | 0.360831885 | 0.65 | 0.43 |
| snR77 | 9.325617253 | 0.425570151 | 0.360833293 | 0.65 | 0.43 |

| | | | | | |
|---|---|---|---|---|---|
| YLR227W-B | 558.9698813 | 0.36473821 | 0.361008981 | 0.65 | 0.43 |
| YAL008W | 865.7638687 | 0.261589012 | 0.361659442 | 0.65 | 0.43 |
| YDR108W | 495.5770673 | 0.143965073 | 0.361330696 | 0.65 | 0.43 |
| YHR210C | 320.0019552 | 0.38799113 | 0.361643436 | 0.65 | 0.43 |
| YIL165C | 180.1950386 | 0.320051681 | 0.361543836 | 0.65 | 0.43 |
| YOL013W-A | 5.152321446 | -0.378865118 | 0.361455626 | 0.65 | 0.43 |
| YNL173C | 2504.496471 | 0.215793767 | 0.361773297 | 0.65 | 0.43 |
| YER114C | 1610.317658 | 0.111238233 | 0.362278797 | 0.65 | 0.43 |
| YGL222C | 1086.796722 | -0.382921129 | 0.362257849 | 0.65 | 0.43 |
| YHR073W | 1977.66425 | 0.190978905 | 0.362377247 | 0.65 | 0.43 |
| YCR052W | 1334.780982 | -0.155408985 | 0.362845987 | 0.65 | 0.43 |
| YGR107W | 13.44440994 | 0.387850672 | 0.362762154 | 0.65 | 0.43 |
| YKL093W | 994.1827608 | 0.344965681 | 0.362703455 | 0.65 | 0.43 |
| YPR071W | 302.2135475 | 0.291731512 | 0.362648447 | 0.65 | 0.43 |
| YCL039W | 1768.904925 | 0.234149175 | 0.363152788 | 0.65 | 0.43 |
| YLR116W | 1156.147687 | -0.128743392 | 0.363218307 | 0.65 | 0.43 |
| YMR004W | 1474.539468 | 0.152908839 | 0.363032806 | 0.65 | 0.43 |
| YPR023C | 1837.667458 | 0.265214268 | 0.363279755 | 0.65 | 0.43 |
| YBL083C | 7.154873991 | -0.379051268 | 0.364476105 | 0.65 | 0.43 |
| YBR229C | 1792.230472 | 0.122586512 | 0.364629456 | 0.65 | 0.43 |
| YDR447C | 4088.999597 | -0.299252684 | 0.364548644 | 0.65 | 0.43 |
| YER068W | 1672.188885 | -0.204877422 | 0.364675469 | 0.65 | 0.43 |
| YGL057C | 252.5789153 | -0.125458779 | 0.36453757 | 0.65 | 0.43 |
| YJR032W | 585.2468444 | -0.244104137 | 0.36449796 | 0.65 | 0.43 |
| YLR390W | 418.8392228 | -0.143999522 | 0.363847757 | 0.65 | 0.43 |
| YML088W | 784.353378 | 0.219297293 | 0.364350025 | 0.65 | 0.43 |
| YMR287C | 795.4088763 | 0.153774799 | 0.364264297 | 0.65 | 0.43 |
| YNL295W | 664.526791 | 0.172199983 | 0.363875357 | 0.65 | 0.43 |
| YOR169C | 39.59953477 | -0.356517921 | 0.364021733 | 0.65 | 0.43 |
| YOR358W | 499.4768456 | 0.198319752 | 0.364588449 | 0.65 | 0.43 |
| YPL237W | 2463.364966 | -0.207237175 | 0.364824252 | 0.65 | 0.43 |
| YNL337W | 0.659062417 | 0.288625407 | 0.364972662 | 0.65 | 0.43 |
| YHL027W | 1629.795881 | 0.255185986 | 0.365184978 | 0.65 | 0.43 |
| YMR229C | 3400.401838 | -0.268722285 | 0.365760902 | 0.65 | 0.43 |
| YBR231C | 448.4132605 | 0.214587474 | 0.36631376 | 0.65 | 0.43 |
| YDR085C | 1259.372351 | 0.296017802 | 0.366038692 | 0.65 | 0.43 |
| YLR228C | 906.7831503 | 0.228045311 | 0.366181888 | 0.65 | 0.43 |
| YNL037C | 5393.729596 | 0.246491209 | 0.366333774 | 0.65 | 0.43 |
| YNL333W | 85.09307581 | 0.390006811 | 0.366527395 | 0.65 | 0.43 |
| YOR051C | 2007.894821 | -0.366715796 | 0.36654686 | 0.65 | 0.43 |
| YCL021W-A | 277.7575158 | 0.321140674 | 0.367316047 | 0.65 | 0.43 |
| YCR102W-A | 9.4716423 | 0.387066961 | 0.367672328 | 0.65 | 0.43 |
| YDL221W | 36.90750404 | 0.351795151 | 0.367903851 | 0.65 | 0.43 |

| | | | | | |
|---|---|---|---|---|---|
| YER164W | 2428.238913 | 0.142426175 | 0.367230892 | 0.65 | 0.43 |
| YGL135W | 2224.916712 | -0.271219622 | 0.367461978 | 0.65 | 0.43 |
| YHR066W | 786.1240577 | -0.332302601 | 0.367497205 | 0.65 | 0.43 |
| YJL091C | 524.9582628 | -0.146033456 | 0.367436613 | 0.65 | 0.43 |
| YLR093C | 1657.689066 | -0.20021906 | 0.367720033 | 0.65 | 0.43 |
| YPL202C | 824.3595251 | -0.153525955 | 0.367911809 | 0.65 | 0.43 |
| snR61 | 12.13748594 | 0.552015271 | 0.367907344 | 0.65 | 0.43 |
| YDR187C | 25.29702409 | 0.319720439 | 0.369017953 | 0.66 | 0.43 |
| YDR485C | 1410.914463 | 0.195488456 | 0.36897025 | 0.66 | 0.43 |
| YDL218W | 253.6505308 | 0.270179737 | 0.369285633 | 0.66 | 0.43 |
| YLR408C | 156.1371802 | 0.169628559 | 0.369343571 | 0.66 | 0.43 |
| YOR383C | 13192.95676 | 0.364945754 | 0.369478461 | 0.66 | 0.43 |
| YLR309C | 2329.969598 | 0.235839546 | 0.369600751 | 0.66 | 0.43 |
| YFL019C | 43.77288423 | 0.476920556 | 0.369904016 | 0.66 | 0.43 |
| YJL097W | 1713.191405 | -0.233898234 | 0.370110505 | 0.66 | 0.43 |
| YML036W | 292.5583071 | -0.220052863 | 0.369876114 | 0.66 | 0.43 |
| YNL006W | 936.4455341 | 0.199177699 | 0.370048546 | 0.66 | 0.43 |
| YNL048W | 1078.322808 | -0.111011106 | 0.369971485 | 0.66 | 0.43 |
| YNL276C | 5.892936072 | 0.612458324 | 0.370259354 | 0.66 | 0.43 |
| YBR076C-A | 84.52994015 | 0.360623084 | 0.370637844 | 0.66 | 0.43 |
| YBR207W | 1493.475156 | -0.221810565 | 0.370543442 | 0.66 | 0.43 |
| YGL103W | 19.59323296 | -0.371234177 | 0.370741129 | 0.66 | 0.43 |
| YLR281C | 33.92288358 | 0.399884011 | 0.370987225 | 0.66 | 0.43 |
| YDL197C | 997.8504474 | 0.186319926 | 0.371305144 | 0.66 | 0.43 |
| YGR114C | 36.93060097 | -0.506849313 | 0.371221211 | 0.66 | 0.43 |
| YGR245C | 1499.530458 | -0.28672138 | 0.371265047 | 0.66 | 0.43 |
| YBL010C | 437.6652565 | -0.125170878 | 0.371669904 | 0.66 | 0.43 |
| YHR101C | 345.7524508 | 0.156819868 | 0.37183388 | 0.66 | 0.43 |
| YKL032C | 3133.578964 | -0.296773783 | 0.371692151 | 0.66 | 0.43 |
| YOR125C | 859.4168859 | 0.23582169 | 0.37180763 | 0.66 | 0.43 |
| YKL137W | 420.0820122 | -0.213688748 | 0.372237991 | 0.66 | 0.43 |
| YOR308C | 716.1626853 | 0.223850938 | 0.372136054 | 0.66 | 0.43 |
| YPL140C | 588.07308 | 0.17867599 | 0.372138525 | 0.66 | 0.43 |
| YGR131W | 100.6509022 | 0.279804181 | 0.372557117 | 0.66 | 0.43 |
| YJL143W | 1167.282332 | -0.162358513 | 0.372536484 | 0.66 | 0.43 |
| YML057W | 1931.217961 | -0.146744895 | 0.372589841 | 0.66 | 0.43 |
| YDL051W | 1178.81834 | -0.288854228 | 0.372877888 | 0.66 | 0.43 |
| YDR144C | 842.4657292 | -0.295915814 | 0.372938839 | 0.66 | 0.43 |
| YNL031C | 9825.730059 | -0.249005139 | 0.373003907 | 0.66 | 0.43 |
| YFL022C | 4340.34362 | -0.234119318 | 0.373331515 | 0.66 | 0.43 |
| YGL225W | 1789.038412 | -0.26768434 | 0.373409374 | 0.66 | 0.43 |
| YPR170W-A | 19.17302177 | 0.33200752 | 0.373435465 | 0.66 | 0.43 |
| YIL073C | 530.2219401 | 0.140548455 | 0.374709232 | 0.66 | 0.44 |

| | | | | | |
|---|---|---|---|---|---|
| YOR243C | 1479.808571 | -0.270535615 | 0.374934151 | 0.66 | 0.44 |
| YJL205C | 132.3580317 | 0.23066506 | 0.375370612 | 0.66 | 0.44 |
| YBR031W | 10790.74196 | 0.40517907 | 0.376235963 | 0.66 | 0.44 |
| YDR155C | 17689.29075 | 0.260369452 | 0.376262778 | 0.66 | 0.44 |
| YDR350C | 1056.007335 | 0.186080945 | 0.376359422 | 0.66 | 0.44 |
| YGL139W | 1235.197549 | -0.138942118 | 0.376504438 | 0.66 | 0.44 |
| YKL116C | 778.7224786 | -0.097785734 | 0.376485703 | 0.66 | 0.44 |
| YMR217W | 5540.397432 | -0.3142703 | 0.376213118 | 0.66 | 0.44 |
| YGL219C | 1273.719314 | 0.221733388 | 0.377098902 | 0.66 | 0.44 |
| YNL238W | 1373.585003 | -0.15956964 | 0.377064271 | 0.66 | 0.44 |
| YLR431C | 452.0758432 | 0.152702635 | 0.377563748 | 0.66 | 0.44 |
| YJR129C | 222.0216409 | 0.26656897 | 0.377884811 | 0.66 | 0.44 |
| YJR074W | 487.5630441 | 0.257103401 | 0.378042221 | 0.66 | 0.44 |
| YGR133W | 486.059603 | 0.142109023 | 0.378557582 | 0.66 | 0.44 |
| YER105C | 2233.871003 | -0.10865382 | 0.378669439 | 0.66 | 0.44 |
| YGR013W | 916.8674279 | 0.164528047 | 0.3789569 | 0.66 | 0.44 |
| YNR047W | 1084.837265 | 0.269687004 | 0.378884433 | 0.66 | 0.44 |
| YDR240C | 401.4217535 | -0.119728665 | 0.379225774 | 0.66 | 0.44 |
| YDR322C-A | 892.3853165 | 0.335339955 | 0.379266205 | 0.66 | 0.44 |
| YBR142W | 1284.846746 | -0.252397325 | 0.379632598 | 0.66 | 0.44 |
| YBR150C | 848.9681448 | 0.181970111 | 0.379598331 | 0.66 | 0.44 |
| YDR043C | 825.8596949 | -0.315982854 | 0.379901881 | 0.66 | 0.44 |
| YDR265W | 518.8074155 | -0.172214893 | 0.380095255 | 0.66 | 0.44 |
| YHR042W | 4638.852766 | 0.185124413 | 0.380126377 | 0.66 | 0.44 |
| YMR256C | 1012.905586 | 0.393623677 | 0.380987789 | 0.67 | 0.44 |
| YPL183C | 913.1902093 | -0.224562013 | 0.38173961 | 0.67 | 0.44 |
| YKL048C | 583.1360838 | -0.259232187 | 0.381863965 | 0.67 | 0.44 |
| YER043C | 7073.679738 | -0.225296444 | 0.382337223 | 0.67 | 0.44 |
| YGL105W | 8369.251023 | -0.239090517 | 0.38256657 | 0.67 | 0.44 |
| YHR074W | 2090.525023 | 0.165546512 | 0.382600869 | 0.67 | 0.44 |
| YML001W | 1453.0376 | -0.138594418 | 0.38249949 | 0.67 | 0.44 |
| YJL184W | 578.763074 | 0.157164008 | 0.382855259 | 0.67 | 0.44 |
| YLR341W | 104.3766662 | -0.261749323 | 0.382796799 | 0.67 | 0.44 |
| YMR063W | 168.4962103 | 0.202573499 | 0.383030968 | 0.67 | 0.44 |
| YPR096C | 16.87382422 | 0.353621685 | 0.383213568 | 0.67 | 0.44 |
| YIL050W | 1083.890647 | 0.201787939 | 0.383546249 | 0.67 | 0.44 |
| YBR027C | 8.649310737 | 0.311139929 | 0.383883008 | 0.67 | 0.44 |
| YIL029C | 174.946717 | 0.274375207 | 0.384311055 | 0.67 | 0.44 |
| YJR116W | 691.1206507 | 0.208807765 | 0.384240132 | 0.67 | 0.44 |
| YKL014C | 1967.775598 | -0.265572117 | 0.384245649 | 0.67 | 0.44 |
| YMR169C | 812.630576 | -0.50213971 | 0.383927256 | 0.67 | 0.44 |
| YNR006W | 1506.041195 | 0.128876925 | 0.383716401 | 0.67 | 0.44 |
| YOR013W | 86.58484926 | -0.21230042 | 0.384027318 | 0.67 | 0.44 |

| | | | | | |
|---|---|---|---|---|---|
| YNL088W | 2851.433881 | 0.169629091 | 0.384485622 | 0.67 | 0.44 |
| YML126C | 4399.479194 | 0.260093527 | 0.384724245 | 0.67 | 0.44 |
| YIR019C | 622.3080892 | 0.275219079 | 0.384922371 | 0.67 | 0.44 |
| YNR004W | 166.7322391 | 0.1898308 | 0.385005712 | 0.67 | 0.44 |
| YKR086W | 1098.768049 | 0.154621041 | 0.38513724 | 0.67 | 0.44 |
| YDR354W | 871.1284672 | 0.392754153 | 0.385729513 | 0.67 | 0.44 |
| YDR517W | 2232.436932 | 0.213309198 | 0.385771751 | 0.67 | 0.44 |
| YGR132C | 2816.110199 | 0.168595792 | 0.385775165 | 0.67 | 0.44 |
| YNL069C | 13320.02635 | -0.296897215 | 0.385643961 | 0.67 | 0.44 |
| YPR040W | 901.3829077 | 0.18949341 | 0.385765508 | 0.67 | 0.44 |
| YAL061W | 2752.646997 | -0.47536322 | 0.386351108 | 0.67 | 0.44 |
| YPL244C | 876.1384791 | -0.184870053 | 0.386377783 | 0.67 | 0.44 |
| YAL019W-A | 5.164115052 | 0.455700731 | 0.386594807 | 0.67 | 0.44 |
| YPR046W | 225.601515 | -0.21748255 | 0.386535319 | 0.67 | 0.44 |
| YIL079C | 348.2789788 | -0.300205462 | 0.387081921 | 0.67 | 0.44 |
| YJL074C | 945.199767 | -0.171350472 | 0.386897565 | 0.67 | 0.44 |
| YOR209C | 3422.936842 | -0.204339749 | 0.387056945 | 0.67 | 0.44 |
| YPR143W | 537.9356283 | -0.20122102 | 0.387063607 | 0.67 | 0.44 |
| YDR112W | 22.56100124 | -0.309937986 | 0.387806268 | 0.67 | 0.44 |
| YIL163C | 43.51788641 | 0.276343267 | 0.387635573 | 0.67 | 0.44 |
| YLR088W | 986.2933263 | 0.127219473 | 0.387646529 | 0.67 | 0.44 |
| YMR189W | 2797.782123 | -0.372604837 | 0.38749998 | 0.67 | 0.44 |
| YMR286W | 654.5341281 | -0.212815321 | 0.387823851 | 0.67 | 0.44 |
| YKL207W | 2006.977142 | -0.167132877 | 0.38814182 | 0.67 | 0.44 |
| YML121W | 941.5006143 | 0.125128501 | 0.388103854 | 0.67 | 0.44 |
| YLR441C | 16750.1152 | -0.285762257 | 0.388367947 | 0.67 | 0.44 |
| YBL078C | 809.4078099 | 0.277333209 | 0.388521135 | 0.67 | 0.44 |
| YPR082C | 431.92377 | -0.188968302 | 0.388726187 | 0.67 | 0.44 |
| YDR031W | 342.8572195 | 0.25636132 | 0.38936901 | 0.67 | 0.44 |
| YHR107C | 1509.6288 | 0.2045463 | 0.389307796 | 0.67 | 0.44 |
| YLL030C | 4.099886808 | 0.317653956 | 0.389382382 | 0.67 | 0.44 |
| YBR063C | 508.4120941 | -0.161714798 | 0.389650242 | 0.67 | 0.44 |
| YPL272C | 344.5733609 | 0.251056926 | 0.389710425 | 0.67 | 0.44 |
| YOL077W-A | 1163.041858 | 0.323540294 | 0.389992417 | 0.67 | 0.44 |
| YBR233W | 474.558741 | -0.216770377 | 0.390714205 | 0.67 | 0.44 |
| YDL225W | 1750.506735 | -0.172080737 | 0.39068298 | 0.67 | 0.44 |
| YIR008C | 729.6399623 | -0.124084155 | 0.390539491 | 0.67 | 0.44 |
| YMR295C | 1636.767669 | 0.111336335 | 0.391066949 | 0.67 | 0.44 |
| YBR005W | 1310.131508 | 0.287429835 | 0.391522611 | 0.67 | 0.44 |
| YGL113W | 616.1660547 | 0.166570116 | 0.391703649 | 0.67 | 0.44 |
| YLR101C | 65.86852303 | 0.2015945 | 0.391657878 | 0.67 | 0.44 |
| YPL068C | 273.8997414 | -0.303859232 | 0.3916615 | 0.67 | 0.44 |
| YPR132W | 5266.120182 | -0.245644471 | 0.391517277 | 0.67 | 0.44 |

| | | | | | |
|---|---|---|---|---|---|
| snR4 | 89.67640266 | 0.439116711 | 0.391579957 | 0.67 | 0.44 |
| YJL013C | 787.2498623 | 0.119495393 | 0.392064249 | 0.67 | 0.44 |
| YKL055C | 140.4891808 | -0.219178412 | 0.39206169 | 0.67 | 0.44 |
| YKL108W | 391.5450461 | -0.237887156 | 0.391943326 | 0.67 | 0.44 |
| YBR241C | 1365.325299 | 0.322213218 | 0.392191164 | 0.67 | 0.44 |
| YDR196C | 880.6195121 | -0.201107049 | 0.39257845 | 0.67 | 0.45 |
| YJR017C | 953.2894932 | -0.165553159 | 0.393058624 | 0.67 | 0.45 |
| YPR119W | 682.8278436 | -0.284623434 | 0.392958923 | 0.67 | 0.45 |
| YBL011W | 1645.010666 | -0.147708813 | 0.393531138 | 0.67 | 0.45 |
| YDR090C | 473.3492687 | 0.149648259 | 0.393739493 | 0.67 | 0.45 |
| YDR531W | 969.2146497 | -0.203706112 | 0.39363443 | 0.67 | 0.45 |
| YJR145C | 6502.704468 | -0.280644656 | 0.393565665 | 0.67 | 0.45 |
| YJR146W | 99.77471099 | 0.386635064 | 0.393723543 | 0.67 | 0.45 |
| YFR034C | 832.1624053 | 0.195919486 | 0.393949078 | 0.67 | 0.45 |
| YLL008W | 1923.071209 | -0.263262768 | 0.394021784 | 0.67 | 0.45 |
| YAL038W | 41041.91188 | 0.307917681 | 0.394259976 | 0.68 | 0.45 |
| YDR334W | 1700.440041 | -0.19141792 | 0.394510128 | 0.68 | 0.45 |
| YJR114W | 33.45969567 | -0.277679415 | 0.394522588 | 0.68 | 0.45 |
| YML106W | 2946.051592 | -0.228351279 | 0.394808179 | 0.68 | 0.45 |
| YDR188W | 2974.432832 | -0.135473152 | 0.395027852 | 0.68 | 0.45 |
| YNL207W | 833.7029402 | -0.19616939 | 0.395020181 | 0.68 | 0.45 |
| YLR090W | 931.4547969 | 0.209466964 | 0.395146335 | 0.68 | 0.45 |
| YDR168W | 2674.002145 | 0.226679844 | 0.395413063 | 0.68 | 0.45 |
| YOR014W | 3387.711726 | 0.179874781 | 0.395780445 | 0.68 | 0.45 |
| YDR300C | 1204.265207 | -0.194751944 | 0.396263148 | 0.68 | 0.45 |
| YKR011C | 1595.285427 | 0.218885649 | 0.396042848 | 0.68 | 0.45 |
| YOR326W | 4223.653773 | 0.204995944 | 0.396116 | 0.68 | 0.45 |
| YPL109C | 614.5552418 | 0.292187968 | 0.396147076 | 0.68 | 0.45 |
| YPR196W | 243.4727794 | 0.236922894 | 0.396348375 | 0.68 | 0.45 |
| YEL055C | 1803.113225 | 0.206886745 | 0.396835528 | 0.68 | 0.45 |
| YLR351C | 1673.534289 | 0.139036286 | 0.397087376 | 0.68 | 0.45 |
| YPL175W | 580.2634042 | -0.136014901 | 0.397251751 | 0.68 | 0.45 |
| YDR460W | 574.0402083 | -0.095308127 | 0.397903237 | 0.68 | 0.45 |
| YOL124C | 564.6964356 | -0.271422921 | 0.398009698 | 0.68 | 0.45 |
| YHL044W | 369.968591 | 0.34888937 | 0.398174019 | 0.68 | 0.45 |
| snR191 | 16.29529748 | 0.407495286 | 0.39845191 | 0.68 | 0.45 |
| YDL013W | 806.2108951 | 0.191535625 | 0.398816357 | 0.68 | 0.45 |
| YMR174C | 615.2184477 | 0.354498853 | 0.398886121 | 0.68 | 0.45 |
| YNL277W-A | 6.085182075 | 0.542631386 | 0.398699238 | 0.68 | 0.45 |
| YBR282W | 512.5901703 | -0.163729127 | 0.399199984 | 0.68 | 0.45 |
| YGR256W | 1242.341744 | 0.533197303 | 0.399077131 | 0.68 | 0.45 |
| YKL071W | 436.0298233 | -0.159868198 | 0.399156762 | 0.68 | 0.45 |
| YIL044C | 893.1546159 | -0.16899423 | 0.399366433 | 0.68 | 0.45 |

| | | | | | |
|---|---|---|---|---|---|
| YDR069C | 1204.898386 | 0.179459477 | 0.39950524 | 0.68 | 0.45 |
| YPL145C | 2994.997329 | -0.191709928 | 0.399792533 | 0.68 | 0.45 |
| YPL212C | 1188.64058 | -0.273086475 | 0.400239848 | 0.68 | 0.45 |
| YFR029W | 834.7381381 | 0.203062228 | 0.400462735 | 0.68 | 0.45 |
| YMR199W | 1528.60034 | -0.25167321 | 0.400389346 | 0.68 | 0.45 |
| YML037C | 139.9825756 | 0.192270872 | 0.400592235 | 0.68 | 0.45 |
| YOR066W | 1115.721778 | -0.227446167 | 0.400944534 | 0.68 | 0.45 |
| YDR524C-A | 2.188681507 | -0.269163753 | 0.401405944 | 0.68 | 0.45 |
| YOR333C | 3.992636415 | 0.439442799 | 0.401402208 | 0.68 | 0.45 |
| YNR030W | 1357.008714 | -0.251765274 | 0.401548403 | 0.68 | 0.45 |
| YDR493W | 345.6632951 | -0.183186326 | 0.402164341 | 0.68 | 0.45 |
| YKL169C | 32.06083426 | -0.249037134 | 0.402160516 | 0.68 | 0.45 |
| YLR296W | 4.726955448 | 0.312721436 | 0.402107606 | 0.68 | 0.45 |
| YLR346C | 131.9806369 | -0.454120844 | 0.401970874 | 0.68 | 0.45 |
| YDR147W | 507.4257469 | -0.206088956 | 0.402516365 | 0.68 | 0.45 |
| YJR066W | 2254.068141 | 0.172070128 | 0.402513903 | 0.68 | 0.45 |
| YKR091W | 1094.538594 | 0.359587431 | 0.402482695 | 0.68 | 0.45 |
| YML064C | 587.9701312 | 0.106261432 | 0.40258629 | 0.68 | 0.45 |
| YBL082C | 673.6132807 | 0.197441371 | 0.402918157 | 0.68 | 0.45 |
| YKL041W | 833.3256523 | 0.160345218 | 0.40282868 | 0.68 | 0.45 |
| YMR070W | 1232.669923 | -0.259502221 | 0.402712046 | 0.68 | 0.45 |
| YAR033W | 105.2011719 | -0.150527954 | 0.403890769 | 0.68 | 0.45 |
| YCR020C-A | 266.9746924 | 0.206776012 | 0.403392525 | 0.68 | 0.45 |
| YDR257C | 688.7119517 | -0.217143335 | 0.40329536 | 0.68 | 0.45 |
| YDR283C | 1675.592568 | 0.15090217 | 0.403851506 | 0.68 | 0.45 |
| YEL002C | 3293.830024 | -0.126905333 | 0.403884598 | 0.68 | 0.45 |
| YGR040W | 494.1182396 | -0.20257059 | 0.403817627 | 0.68 | 0.45 |
| YGR268C | 875.2811265 | -0.179035717 | 0.403567182 | 0.68 | 0.45 |
| YJR012C | 286.1622087 | -0.140671232 | 0.403374063 | 0.68 | 0.45 |
| YLR270W | 2664.450215 | 0.25056463 | 0.404191622 | 0.68 | 0.45 |
| YPR141C | 623.3342075 | 0.205598794 | 0.40421771 | 0.68 | 0.45 |
| YGL256W | 1304.621904 | 0.172334927 | 0.40443458 | 0.68 | 0.45 |
| YGR044C | 1106.237884 | -0.426792999 | 0.404443695 | 0.68 | 0.45 |
| YDL212W | 1994.74877 | -0.169706925 | 0.404719804 | 0.68 | 0.45 |
| YLR244C | 3158.461036 | -0.210712716 | 0.404697515 | 0.68 | 0.45 |
| YBR181C | 1794.107741 | -0.27335151 | 0.405016018 | 0.68 | 0.45 |
| YKR076W | 1274.281879 | 0.332529219 | 0.405004727 | 0.68 | 0.45 |
| YEL050W-A | 71.35641565 | -0.229095623 | 0.405365046 | 0.68 | 0.45 |
| YGR014W | 3080.238337 | -0.28891947 | 0.405495148 | 0.68 | 0.45 |
| YGR180C | 8240.520071 | 0.195735039 | 0.405644825 | 0.68 | 0.45 |
| YDR066C | 173.6197188 | 0.217991522 | 0.405993438 | 0.68 | 0.45 |
| YIR007W | 898.4409432 | 0.147405025 | 0.406149851 | 0.68 | 0.45 |
| YCL007C | 102.0392471 | 0.301023143 | 0.406608365 | 0.68 | 0.45 |

| | | | | | |
|---|---|---|---|---|---|
| YDR046C | 2206.66233 | 0.217130968 | 0.406512064 | 0.68 | 0.45 |
| YDR534C | 224.6689044 | -0.369323854 | 0.406934052 | 0.68 | 0.45 |
| YGL101W | 448.9089719 | -0.212436162 | 0.406881823 | 0.68 | 0.45 |
| YNL079C | 8141.583459 | -0.18762603 | 0.406873688 | 0.68 | 0.45 |
| YOR207C | 1745.865963 | -0.187942529 | 0.407118414 | 0.68 | 0.45 |
| YGL001C | 1350.851203 | -0.234709864 | 0.407820139 | 0.68 | 0.45 |
| YGR154C | 333.0792836 | 0.195013395 | 0.407963892 | 0.68 | 0.45 |
| YHR003C | 820.5090626 | 0.173276118 | 0.407553149 | 0.68 | 0.45 |
| YKR020W | 476.7599628 | -0.16215903 | 0.40783294 | 0.68 | 0.45 |
| YLR071C | 1557.164849 | -0.164925188 | 0.408009096 | 0.68 | 0.45 |
| YMR296C | 3004.03015 | -0.211657389 | 0.408029972 | 0.68 | 0.45 |
| YOL141W | 189.2930206 | 0.249416215 | 0.407620902 | 0.68 | 0.45 |
| YBR133C | 1160.137007 | -0.246824193 | 0.408206941 | 0.68 | 0.45 |
| YLR290C | 1190.940137 | 0.175858165 | 0.408605463 | 0.68 | 0.45 |
| YPL116W | 1071.716343 | -0.160486825 | 0.408675318 | 0.68 | 0.45 |
| YKL006W | 2404.725348 | -0.232932373 | 0.409134261 | 0.68 | 0.45 |
| YMR160W | 946.3755314 | 0.181366532 | 0.409171847 | 0.68 | 0.45 |
| YER086W | 3514.638296 | 0.385300195 | 0.409433582 | 0.69 | 0.45 |
| YBL021C | 336.3286264 | -0.149239687 | 0.409606544 | 0.69 | 0.45 |
| YPL071C | 441.9177148 | -0.188698262 | 0.409969205 | 0.69 | 0.45 |
| YGL017W | 502.0325199 | 0.11664841 | 0.410220244 | 0.69 | 0.45 |
| YLR317W | 2.198595472 | 0.604738618 | 0.41053049 | 0.69 | 0.45 |
| YKL099C | 686.8097655 | -0.253978505 | 0.410662209 | 0.69 | 0.45 |
| YBR002C | 607.1395928 | -0.189564214 | 0.410816704 | 0.69 | 0.45 |
| YKL149C | 355.2699607 | -0.17001956 | 0.411074638 | 0.69 | 0.45 |
| YFR031C-A | 6438.897656 | -0.25271967 | 0.411552194 | 0.69 | 0.45 |
| YGR262C | 876.7108592 | -0.091771604 | 0.41148898 | 0.69 | 0.45 |
| YPL054W | 345.0757807 | 0.293359264 | 0.411568325 | 0.69 | 0.45 |
| YFR027W | 221.6537022 | 0.16334134 | 0.411804135 | 0.69 | 0.45 |
| YNL137C | 1635.261502 | -0.122673403 | 0.411775716 | 0.69 | 0.45 |
| YHL010C | 843.3803368 | 0.197670561 | 0.412067379 | 0.69 | 0.45 |
| YLR227C | 512.1356286 | 0.115108173 | 0.412154271 | 0.69 | 0.45 |
| YIR038C | 2847.18512 | -0.281910228 | 0.412340975 | 0.69 | 0.45 |
| YOR262W | 658.3672371 | 0.192995838 | 0.412576397 | 0.69 | 0.45 |
| YGR096W | 275.6515308 | 0.175080004 | 0.413038031 | 0.69 | 0.45 |
| YNL075W | 678.2006525 | -0.240221126 | 0.412941042 | 0.69 | 0.45 |
| YNL132W | 1681.161762 | -0.251221644 | 0.412908466 | 0.69 | 0.45 |
| YOL032W | 1063.876828 | 0.220263222 | 0.413002429 | 0.69 | 0.45 |
| YKL117W | 7112.851481 | -0.173102471 | 0.413253211 | 0.69 | 0.45 |
| YNL135C | 5166.62026 | -0.194009674 | 0.41316002 | 0.69 | 0.45 |
| YAL049C | 1497.895567 | 0.176914621 | 0.413552032 | 0.69 | 0.45 |
| YGL215W | 3518.182479 | 0.195675845 | 0.413703082 | 0.69 | 0.45 |
| YPL120W | 588.9261154 | -0.178902929 | 0.413899653 | 0.69 | 0.45 |

| | | | | | |
|---|---|---|---|---|---|
| YJR125C | 2124.757426 | -0.138058347 | 0.41416992 | 0.69 | 0.45 |
| YKL039W | 2424.281879 | 0.130075118 | 0.414163844 | 0.69 | 0.45 |
| YOR218C | 5.229072963 | 0.440633412 | 0.414258097 | 0.69 | 0.45 |
| YNL140C | 6.604161499 | -0.387996101 | 0.414369947 | 0.69 | 0.45 |
| YDR049W | 886.770398 | 0.09226192 | 0.414558468 | 0.69 | 0.45 |
| YDR161W | 1294.242267 | -0.209459348 | 0.41472078 | 0.69 | 0.45 |
| YJL024C | 668.6768051 | -0.124979635 | 0.414911688 | 0.69 | 0.46 |
| YEL037C | 3155.723536 | -0.157218197 | 0.415583972 | 0.69 | 0.46 |
| YLR138W | 1645.971988 | 0.205656509 | 0.415570248 | 0.69 | 0.46 |
| YAL010C | 552.0969494 | 0.113045191 | 0.415761558 | 0.69 | 0.46 |
| YDL033C | 356.3492838 | -0.22750858 | 0.415813291 | 0.69 | 0.46 |
| YGR189C | 5490.403188 | 0.342811658 | 0.416294221 | 0.69 | 0.46 |
| YLR399C | 2272.649731 | 0.143688743 | 0.416226004 | 0.69 | 0.46 |
| YAL069W | 0.918104495 | 0.398374308 | 0.416972318 | 0.69 | 0.46 |
| YCR107W | 236.9080811 | 0.390041242 | 0.416983787 | 0.69 | 0.46 |
| YHR095W | 5.929557488 | 0.563346877 | 0.416941914 | 0.69 | 0.46 |
| YNL282W | 259.9576363 | 0.161758049 | 0.41679888 | 0.69 | 0.46 |
| YLR104W | 635.3306236 | -0.206662091 | 0.417123746 | 0.69 | 0.46 |
| YLR193C | 745.5545212 | 0.191958937 | 0.417248702 | 0.69 | 0.46 |
| YBR156C | 807.2285578 | -0.117397698 | 0.417763473 | 0.69 | 0.46 |
| YOR339C | 48.09399516 | 0.280918088 | 0.418044294 | 0.69 | 0.46 |
| YMR097C | 670.8866515 | 0.129785252 | 0.41820256 | 0.69 | 0.46 |
| YDL112W | 1910.821894 | -0.213892657 | 0.418562467 | 0.69 | 0.46 |
| YFL057C | 160.9643278 | 0.210434226 | 0.418503673 | 0.69 | 0.46 |
| YKL006C-A | 282.6893408 | -0.179597984 | 0.418808948 | 0.69 | 0.46 |
| YOR048C | 2248.629889 | 0.164062876 | 0.418663181 | 0.69 | 0.46 |
| YOR052C | 2749.089296 | 0.237229016 | 0.418399136 | 0.69 | 0.46 |
| YPR151C | 43.40930441 | 0.384030895 | 0.418849845 | 0.69 | 0.46 |
| RPR1 | 36.63381865 | 0.498755348 | 0.419164165 | 0.69 | 0.46 |
| YBR223C | 483.3688068 | 0.185989548 | 0.419614721 | 0.69 | 0.46 |
| YKR021W | 1328.600543 | 0.130238547 | 0.419540206 | 0.69 | 0.46 |
| YOR291W | 1661.347699 | 0.115145305 | 0.419563214 | 0.69 | 0.46 |
| YJL044C | 1379.126824 | -0.168045787 | 0.419728526 | 0.69 | 0.46 |
| YHR002W | 437.310458 | 0.24684055 | 0.419883325 | 0.69 | 0.46 |
| YFL002C | 1100.586944 | 0.09483936 | 0.420080022 | 0.69 | 0.46 |
| YIL001W | 471.4395924 | 0.225572008 | 0.420332803 | 0.69 | 0.46 |
| YNL001W | 629.9634099 | -0.171426666 | 0.420561998 | 0.69 | 0.46 |
| YIL070C | 1550.574512 | 0.254664567 | 0.421593771 | 0.69 | 0.46 |
| YIR023C-A | 13.57792364 | -0.310159055 | 0.42154975 | 0.69 | 0.46 |
| YLR060W | 5384.459239 | -0.251104362 | 0.421683341 | 0.69 | 0.46 |
| YCL036W | 402.9926525 | 0.3150858 | 0.421974538 | 0.69 | 0.46 |
| YPR055W | 1160.769461 | -0.172305518 | 0.421954813 | 0.69 | 0.46 |
| YCR079W | 984.6018828 | 0.197302748 | 0.422679261 | 0.69 | 0.46 |

| | | | | | |
|---|---|---|---|---|---|
| YER056C | 2408.22934 | -0.209862231 | 0.422580998 | 0.69 | 0.46 |
| YLR192C | 3457.976113 | -0.137698303 | 0.4227031 | 0.69 | 0.46 |
| YMR115W | 1106.490843 | 0.157262521 | 0.422405734 | 0.69 | 0.46 |
| YDR048C | 13.8983324 | 0.372738001 | 0.42372669 | 0.70 | 0.46 |
| YHL043W | 209.5139873 | 0.246705893 | 0.423816935 | 0.70 | 0.46 |
| YPR188C | 534.7061592 | -0.148611225 | 0.423835636 | 0.70 | 0.46 |
| snR64 | 4.034418541 | 0.434576644 | 0.423539678 | 0.70 | 0.46 |
| YMR076C | 1386.678863 | -0.205360407 | 0.42426311 | 0.70 | 0.46 |
| YOR252W | 480.3993858 | -0.168874679 | 0.424307361 | 0.70 | 0.46 |
| YML084W | 2.063062303 | 0.466185494 | 0.424912115 | 0.70 | 0.46 |
| YPR198W | 724.9139692 | -0.230899913 | 0.425022452 | 0.70 | 0.46 |
| YBR123C | 731.156432 | 0.118297878 | 0.425278864 | 0.70 | 0.46 |
| YOL152W | 484.6917609 | 0.342843418 | 0.425319145 | 0.70 | 0.46 |
| YBL076C | 7280.098127 | 0.290114956 | 0.425568144 | 0.70 | 0.46 |
| YGL025C | 784.0082065 | -0.138963909 | 0.425674073 | 0.70 | 0.46 |
| YDR372C | 1904.188965 | -0.169855969 | 0.426557011 | 0.70 | 0.46 |
| YGR224W | 693.0347479 | -0.189588694 | 0.426453293 | 0.70 | 0.46 |
| YLR014C | 353.4875684 | 0.184539064 | 0.426436785 | 0.70 | 0.46 |
| YLR256W-A | 92.61969338 | 0.255069932 | 0.426355132 | 0.70 | 0.46 |
| YLR181C | 881.938971 | 0.129997173 | 0.426710839 | 0.70 | 0.46 |
| YOL046C | 5.940874887 | -0.360479292 | 0.427157548 | 0.70 | 0.46 |
| YJL214W | 204.5476536 | -0.193740898 | 0.427292349 | 0.70 | 0.46 |
| YLR342W-A | 51.47657195 | 0.244338941 | 0.427770672 | 0.70 | 0.46 |
| YDR305C | 387.9870014 | 0.22456701 | 0.428326715 | 0.70 | 0.46 |
| YKR093W | 2668.023949 | -0.256457592 | 0.428438375 | 0.70 | 0.46 |
| YDR034W-B | 127.2105401 | 0.37833571 | 0.428835976 | 0.70 | 0.46 |
| YJL198W | 970.0448734 | -0.218144057 | 0.428749002 | 0.70 | 0.46 |
| YOR242C | 110.2984083 | 0.231176635 | 0.42902732 | 0.70 | 0.46 |
| snR39B | 13.58422976 | -0.357524531 | 0.429405206 | 0.70 | 0.46 |
| YPL019C | 1922.931958 | -0.287636466 | 0.429553367 | 0.70 | 0.46 |
| YMR282C | 677.7599955 | -0.139276671 | 0.429752405 | 0.70 | 0.46 |
| YMR083W | 8478.832219 | 0.180438231 | 0.429925999 | 0.70 | 0.46 |
| YEL029C | 331.9088854 | 0.238996718 | 0.430293049 | 0.70 | 0.46 |
| YGR231C | 3688.450165 | 0.182097394 | 0.430976709 | 0.70 | 0.46 |
| YKR048C | 4547.539769 | -0.169549176 | 0.43093258 | 0.70 | 0.46 |
| YMR113W | 461.667001 | -0.161498717 | 0.431024504 | 0.70 | 0.46 |
| YPL263C | 1770.940205 | -0.182682278 | 0.430797573 | 0.70 | 0.46 |
| YPR050C | 1.567602133 | -0.362916454 | 0.430847551 | 0.70 | 0.46 |
| YNL335W | 13.42162756 | -0.305527856 | 0.431225225 | 0.70 | 0.46 |
| YCL065W | 1.135078196 | 0.211657992 | 0.431828849 | 0.70 | 0.46 |
| YAL027W | 394.2818606 | -0.130081503 | 0.432069461 | 0.70 | 0.46 |
| YKL131W | 17.12686382 | -0.215924221 | 0.432202725 | 0.70 | 0.46 |
| YHR198C | 872.0543676 | 0.256990536 | 0.432362017 | 0.70 | 0.46 |

| | | | | | |
|---|---|---|---|---|---|
| YOR334W | 302.0782196 | -0.135974886 | 0.432470631 | 0.70 | 0.46 |
| YBR025C | 6377.054239 | -0.30429051 | 0.432818671 | 0.70 | 0.46 |
| YFL008W | 1609.672053 | -0.195782376 | 0.432721085 | 0.70 | 0.46 |
| YLR147C | 302.8969457 | -0.133403537 | 0.432778766 | 0.70 | 0.46 |
| YGL034C | 9.87258369 | 0.362792889 | 0.433332636 | 0.70 | 0.47 |
| YLR253W | 1177.596684 | -0.25720811 | 0.433532881 | 0.70 | 0.47 |
| YHR057C | 1425.81956 | 0.161721947 | 0.433663802 | 0.70 | 0.47 |
| YKL162C | 452.2195831 | 0.176991477 | 0.433809394 | 0.70 | 0.47 |
| YDR219C | 850.7025719 | -0.156652751 | 0.434006537 | 0.70 | 0.47 |
| YJR023C | 6.716930152 | -0.372569255 | 0.434212547 | 0.70 | 0.47 |
| YKL142W | 4281.67462 | 0.218495051 | 0.434463437 | 0.70 | 0.47 |
| YLR458W | 2.467406493 | 0.392912274 | 0.434431679 | 0.70 | 0.47 |
| YBR270C | 464.1113888 | 0.174671573 | 0.435022253 | 0.70 | 0.47 |
| YCR064C | 12.51518247 | -0.244782173 | 0.434959682 | 0.70 | 0.47 |
| YDL108W | 550.3870487 | 0.106277293 | 0.434962028 | 0.70 | 0.47 |
| YDR118W-A | 11.72360647 | -0.271779115 | 0.435107822 | 0.70 | 0.47 |
| YMR177W | 436.2483597 | -0.165196213 | 0.434966831 | 0.70 | 0.47 |
| YMR023C | 334.2712538 | -0.165841374 | 0.435327323 | 0.71 | 0.47 |
| YCL032W | 909.5801687 | 0.144135757 | 0.436071489 | 0.71 | 0.47 |
| YDR543C | 0.805732287 | 0.468879337 | 0.435682833 | 0.71 | 0.47 |
| YHR128W | 1328.56215 | -0.266705653 | 0.436016125 | 0.71 | 0.47 |
| YJL039C | 2015.018256 | -0.13958321 | 0.435857636 | 0.71 | 0.47 |
| YMR118C | 295.0484477 | 0.248561097 | 0.43605413 | 0.71 | 0.47 |
| YJR006W | 1054.347239 | -0.148062219 | 0.436282098 | 0.71 | 0.47 |
| YDL037C | 92.66763176 | -0.355360417 | 0.437135334 | 0.71 | 0.47 |
| YDR051C | 1050.532227 | -0.10765012 | 0.437183627 | 0.71 | 0.47 |
| YER005W | 999.8103274 | 0.088336792 | 0.437556461 | 0.71 | 0.47 |
| YHR113W | 1875.547688 | 0.15437675 | 0.436929333 | 0.71 | 0.47 |
| YHR127W | 498.1945386 | -0.234963624 | 0.43761317 | 0.71 | 0.47 |
| YHR146W | 3577.393266 | 0.156195789 | 0.436723673 | 0.71 | 0.47 |
| YJL073W | 1063.600671 | 0.217746017 | 0.437612389 | 0.71 | 0.47 |
| YLL009C | 581.7236454 | 0.308759168 | 0.437675019 | 0.71 | 0.47 |
| YNL230C | 466.9526022 | -0.186789567 | 0.437126763 | 0.71 | 0.47 |
| YNL309W | 553.1266537 | -0.151966806 | 0.437246222 | 0.71 | 0.47 |
| YNL320W | 323.6388355 | 0.207348914 | 0.437711737 | 0.71 | 0.47 |
| YOL017W | 430.0355777 | -0.138192482 | 0.436565354 | 0.71 | 0.47 |
| YOR221C | 1523.763907 | 0.144538391 | 0.436795735 | 0.71 | 0.47 |
| YBR194W | 226.3319567 | 0.194229924 | 0.438238168 | 0.71 | 0.47 |
| YLR067C | 640.6258307 | -0.136563305 | 0.438275156 | 0.71 | 0.47 |
| YGL168W | 276.6572721 | -0.225073468 | 0.439201988 | 0.71 | 0.47 |
| YKL225W | 0.626369128 | 0.266076031 | 0.43909556 | 0.71 | 0.47 |
| YNL139C | 1577.578713 | 0.113302832 | 0.439304157 | 0.71 | 0.47 |
| YOL097C | 2168.680575 | -0.233271202 | 0.43918414 | 0.71 | 0.47 |

| | | | | | |
|---|---|---|---|---|---|
| YPR060C | 619.4282416 | -0.256898265 | 0.439433314 | 0.71 | 0.47 |
| YLR011W | 284.831779 | -0.149243845 | 0.439949623 | 0.71 | 0.47 |
| YJR069C | 644.3818806 | -0.218002142 | 0.440166803 | 0.71 | 0.47 |
| YGL024W | 7.765530916 | 0.398234644 | 0.440370782 | 0.71 | 0.47 |
| YJR112W | 374.389265 | -0.165691873 | 0.440623544 | 0.71 | 0.47 |
| YKL030W | 149.8142583 | -0.35903251 | 0.440647572 | 0.71 | 0.47 |
| YLR055C | 1018.744824 | 0.197689328 | 0.441097905 | 0.71 | 0.47 |
| YMR192W | 859.8324449 | 0.162685691 | 0.441092444 | 0.71 | 0.47 |
| YHR150W | 670.7977171 | 0.208301044 | 0.441382651 | 0.71 | 0.47 |
| YER090W | 2828.394196 | 0.236060923 | 0.441719455 | 0.71 | 0.47 |
| YEL036C | 1490.088365 | -0.204621418 | 0.442108197 | 0.71 | 0.47 |
| YLL006W | 812.6766975 | 0.245981593 | 0.442032717 | 0.71 | 0.47 |
| YPL245W | 615.8162164 | -0.256816789 | 0.441977228 | 0.71 | 0.47 |
| YHL013C | 503.0447213 | -0.17581153 | 0.442251533 | 0.71 | 0.47 |
| YGR191W | 1267.875828 | -0.165992247 | 0.442622699 | 0.71 | 0.47 |
| YMR316C-A | 33.48519679 | 0.335463245 | 0.44263536 | 0.71 | 0.47 |
| YBR290W | 1203.208406 | -0.16224409 | 0.443298341 | 0.71 | 0.47 |
| YCR018C | 861.2432726 | 0.282123086 | 0.443453908 | 0.71 | 0.47 |
| YDL182W | 2531.684902 | -0.260317831 | 0.442938574 | 0.71 | 0.47 |
| YIR026C | 772.7201823 | -0.278934299 | 0.443081407 | 0.71 | 0.47 |
| YJL215C | 44.17616213 | -0.250913058 | 0.443177168 | 0.71 | 0.47 |
| YPL220W | 1761.369957 | -0.272589562 | 0.443443053 | 0.71 | 0.47 |
| YER092W | 378.6245953 | -0.216477725 | 0.443971874 | 0.71 | 0.47 |
| YJL098W | 1058.267184 | -0.181913432 | 0.44377497 | 0.71 | 0.47 |
| YPL273W | 2617.65817 | -0.265320416 | 0.443812726 | 0.71 | 0.47 |
| YPR186C | 328.1330267 | -0.157402895 | 0.444002544 | 0.71 | 0.47 |
| YAL034C | 1899.332638 | 0.282773545 | 0.444360813 | 0.71 | 0.47 |
| YPL144W | 173.5181567 | -0.217607826 | 0.444327279 | 0.71 | 0.47 |
| YIL143C | 1830.368647 | 0.18471735 | 0.445355412 | 0.71 | 0.47 |
| YMR262W | 628.7777309 | 0.199997198 | 0.445298021 | 0.71 | 0.47 |
| YDR249C | 239.973621 | 0.18744295 | 0.446034429 | 0.71 | 0.47 |
| YGR098C | 1335.545417 | 0.171709355 | 0.445822881 | 0.71 | 0.47 |
| YGR232W | 824.5178442 | 0.144127174 | 0.445716597 | 0.71 | 0.47 |
| YLR356W | 2015.024039 | 0.230816272 | 0.445969993 | 0.71 | 0.47 |
| YOL037C | 71.18780393 | 0.263549181 | 0.445902898 | 0.71 | 0.47 |
| YOR273C | 1760.091485 | 0.291426459 | 0.44550541 | 0.71 | 0.47 |
| YPR005C | 426.8209175 | 0.191328937 | 0.44618436 | 0.71 | 0.47 |
| YDR328C | 1849.787592 | -0.117754032 | 0.446802476 | 0.71 | 0.47 |
| YHR135C | 3300.939762 | -0.119330284 | 0.446640002 | 0.71 | 0.47 |
| YIL140W | 1095.766571 | -0.218747965 | 0.44669977 | 0.71 | 0.47 |
| YIR036C | 938.6857022 | 0.151765165 | 0.446828639 | 0.71 | 0.47 |
| YDR136C | 21.36116146 | -0.2130552 | 0.447323563 | 0.71 | 0.47 |
| YBL062W | 0.924462825 | 0.415849019 | 0.448255038 | 0.71 | 0.47 |

| | | | | | |
|---|---|---|---|---|---|
| YDR017C | 2063.432291 | 0.092220066 | 0.448201956 | 0.71 | 0.47 |
| YGL031C | 8332.907395 | -0.257285438 | 0.44823229 | 0.71 | 0.47 |
| YKR002W | 1434.178622 | 0.126914013 | 0.447862108 | 0.71 | 0.47 |
| YOL144W | 766.01586 | -0.236438366 | 0.448175743 | 0.71 | 0.47 |
| YOR060C | 313.4302218 | -0.161884838 | 0.447892366 | 0.71 | 0.47 |
| YDL113C | 1161.019898 | 0.165922409 | 0.448776536 | 0.71 | 0.47 |
| YDR359C | 1438.122379 | 0.162951556 | 0.448583197 | 0.71 | 0.47 |
| YLL019C | 1900.382246 | 0.246709141 | 0.448785549 | 0.71 | 0.47 |
| YPL197C | 174.3224877 | -0.279187948 | 0.448808478 | 0.71 | 0.47 |
| YFL061W | 11.58614878 | 0.372919802 | 0.448957046 | 0.71 | 0.47 |
| YML063W | 16224.08176 | -0.236307659 | 0.449089105 | 0.71 | 0.47 |
| YBR221C | 5794.723277 | 0.153770703 | 0.449992628 | 0.71 | 0.47 |
| YDL209C | 238.6569979 | -0.195635052 | 0.450003939 | 0.71 | 0.47 |
| YDR353W | 5059.752159 | -0.165293346 | 0.449775155 | 0.71 | 0.47 |
| YJL007C | 3.350774349 | -0.359060049 | 0.450067507 | 0.71 | 0.47 |
| YKL081W | 14463.37631 | -0.231180521 | 0.449590135 | 0.71 | 0.47 |
| YLL034C | 960.4010793 | -0.267082589 | 0.44972957 | 0.71 | 0.47 |
| YGL174W | 430.9199773 | 0.132498655 | 0.450354932 | 0.71 | 0.47 |
| YDR117C | 1080.281382 | -0.103563026 | 0.450504187 | 0.71 | 0.47 |
| YBR219C | 1.059885032 | -0.585906179 | 0.450965173 | 0.71 | 0.47 |
| YDR356W | 1114.540575 | 0.167347805 | 0.451064678 | 0.71 | 0.47 |
| YHL033C | 4723.149428 | -0.212885582 | 0.450688411 | 0.71 | 0.47 |
| YHR115C | 896.8338505 | 0.160713268 | 0.451168295 | 0.71 | 0.47 |
| YIL127C | 639.7542142 | -0.216197422 | 0.45105865 | 0.71 | 0.47 |
| YOL103W-B | 37.1755698 | 0.271206551 | 0.450813029 | 0.71 | 0.47 |
| YER061C | 958.2654695 | 0.167980393 | 0.451506664 | 0.71 | 0.47 |
| YJR073C | 2980.954697 | 0.263208648 | 0.451875886 | 0.71 | 0.47 |
| YPL115C | 1185.518438 | -0.151193647 | 0.451777594 | 0.71 | 0.47 |
| YKR078W | 765.3811944 | 0.147794989 | 0.452400762 | 0.72 | 0.47 |
| YNL097C-B | 13.11125706 | -0.34984299 | 0.45242435 | 0.72 | 0.47 |
| YBL048W | 125.8109055 | 0.25999649 | 0.452615612 | 0.72 | 0.47 |
| YBL108W | 0.561379936 | 0.514890876 | 0.453053621 | 0.72 | 0.47 |
| YEL027W | 5744.186516 | -0.125271956 | 0.453176749 | 0.72 | 0.47 |
| YGL185C | 473.3669461 | 0.188501428 | 0.453209035 | 0.72 | 0.47 |
| YML047W-A | 2.39499993 | 0.388218283 | 0.452927367 | 0.72 | 0.47 |
| YMR184W | 674.607137 | 0.142223944 | 0.453119884 | 0.72 | 0.47 |
| YNL292W | 950.1678617 | -0.165557014 | 0.453082839 | 0.72 | 0.47 |
| YDR337W | 923.5124951 | -0.123739776 | 0.453516951 | 0.72 | 0.47 |
| YJL042W | 3416.022292 | 0.156964734 | 0.45361965 | 0.72 | 0.47 |
| YJR030C | 708.7385852 | 0.165720064 | 0.453519072 | 0.72 | 0.47 |
| YHR187W | 633.7803075 | 0.111050403 | 0.453804984 | 0.72 | 0.47 |
| YPL090C | 2392.913651 | -0.197179748 | 0.454004796 | 0.72 | 0.47 |
| YCR041W | 16.4079087 | 0.28865791 | 0.454707933 | 0.72 | 0.47 |

| | | | | | |
|---|---|---|---|---|---|
| YJL202C | 29.15207009 | -0.213249499 | 0.454742801 | 0.72 | 0.47 |
| YMR195W | 1764.107553 | -0.38998095 | 0.455247565 | 0.72 | 0.47 |
| YOR033C | 730.7503196 | 0.102821747 | 0.455175122 | 0.72 | 0.47 |
| YDR503C | 368.4756431 | -0.153144671 | 0.456461647 | 0.72 | 0.47 |
| YLR267W | 1321.159087 | 0.274307879 | 0.456383772 | 0.72 | 0.47 |
| YMR215W | 1997.531105 | -0.223254633 | 0.456595777 | 0.72 | 0.47 |
| YPR113W | 3675.90658 | -0.150873742 | 0.457165978 | 0.72 | 0.48 |
| YLR043C | 5741.406917 | -0.171601686 | 0.45750399 | 0.72 | 0.48 |
| YHR211W | 534.044984 | 0.281643028 | 0.45789144 | 0.72 | 0.48 |
| YJR111C | 500.5453341 | 0.197305558 | 0.458161567 | 0.72 | 0.48 |
| YNL134C | 7013.757129 | 0.278574143 | 0.458164208 | 0.72 | 0.48 |
| YEL061C | 1022.169501 | -0.185544501 | 0.458341828 | 0.72 | 0.48 |
| YPL044C | 1.74310561 | -0.507487052 | 0.459036147 | 0.72 | 0.48 |
| YHR013C | 1158.208101 | -0.18541971 | 0.45916793 | 0.72 | 0.48 |
| YGL094C | 1570.488055 | 0.153911974 | 0.459465339 | 0.72 | 0.48 |
| YMR110C | 3066.365125 | 0.233122739 | 0.45970886 | 0.72 | 0.48 |
| YIL003W | 534.8841649 | 0.116746249 | 0.459991388 | 0.72 | 0.48 |
| YER062C | 1876.234052 | -0.224093296 | 0.460162065 | 0.72 | 0.48 |
| YDR111C | 552.8400061 | -0.242277782 | 0.460311143 | 0.72 | 0.48 |
| YCR060W | 437.7609902 | 0.164232352 | 0.460854522 | 0.72 | 0.48 |
| YLR365W | 4.223598425 | 0.307154761 | 0.460829648 | 0.72 | 0.48 |
| YDL060W | 1265.050122 | -0.192585398 | 0.462772157 | 0.72 | 0.48 |
| YJR043C | 619.0166524 | -0.223781594 | 0.462442693 | 0.72 | 0.48 |
| YKR059W | 1551.844916 | -0.207082456 | 0.462684659 | 0.72 | 0.48 |
| snR66 | 7.293652703 | 0.455791842 | 0.462575941 | 0.72 | 0.48 |
| YOR162C | 795.3084144 | 0.082626287 | 0.463277291 | 0.73 | 0.48 |
| YPL023C | 1340.132366 | -0.114170724 | 0.463555993 | 0.73 | 0.48 |
| YPL004C | 14429.78474 | 0.154901077 | 0.463962483 | 0.73 | 0.48 |
| YLR322W | 7.310944976 | -0.359295206 | 0.465062462 | 0.73 | 0.48 |
| YOR265W | 445.8131326 | -0.139461975 | 0.465041827 | 0.73 | 0.48 |
| YDL039C | 373.0125142 | -0.311678727 | 0.465703564 | 0.73 | 0.48 |
| YHL030W-A | 22.0345387 | 0.190062948 | 0.465412497 | 0.73 | 0.48 |
| YHR144C | 383.3281151 | -0.218976402 | 0.465582214 | 0.73 | 0.48 |
| YLL050C | 6100.834604 | -0.151481272 | 0.465710297 | 0.73 | 0.48 |
| YDR162C | 515.4142839 | -0.119614496 | 0.466193667 | 0.73 | 0.48 |
| YGR030C | 291.5946813 | -0.176890835 | 0.466515168 | 0.73 | 0.48 |
| YBR004C | 976.5395701 | 0.191707891 | 0.466806973 | 0.73 | 0.48 |
| YBR035C | 2483.374886 | 0.126110262 | 0.46687438 | 0.73 | 0.48 |
| YMR258C | 733.537512 | -0.154720435 | 0.467304923 | 0.73 | 0.48 |
| YDR034C-D | 94.28631679 | 0.2548326 | 0.467711185 | 0.73 | 0.48 |
| YER033C | 1095.361309 | 0.224446381 | 0.467755882 | 0.73 | 0.48 |
| YNL084C | 2582.808264 | -0.086714739 | 0.467601444 | 0.73 | 0.48 |
| YBR249C | 4275.177673 | 0.421441459 | 0.468394181 | 0.73 | 0.48 |

| | | | | | |
|---|---|---|---|---|---|
| YER079C-A | 18.60445932 | -0.353554013 | 0.468168064 | 0.73 | 0.48 |
| YGR178C | 3642.360299 | 0.126386855 | 0.468315965 | 0.73 | 0.48 |
| YNL057W | 19.3681916 | 0.368557777 | 0.468456883 | 0.73 | 0.48 |
| YBR300C | 5.482970764 | 0.423223697 | 0.468593884 | 0.73 | 0.48 |
| YLR440C | 889.0207433 | 0.170029417 | 0.46891843 | 0.73 | 0.48 |
| YIL008W | 681.0099622 | -0.213657413 | 0.469313603 | 0.73 | 0.48 |
| YIL053W | 6767.756759 | -0.311359837 | 0.46922269 | 0.73 | 0.48 |
| YDR441C | 376.4209723 | -0.130522046 | 0.469528011 | 0.73 | 0.48 |
| YIL030C | 2092.978365 | 0.169388014 | 0.469640953 | 0.73 | 0.48 |
| YGR289C | 671.1201555 | 0.284009571 | 0.470067742 | 0.73 | 0.48 |
| YLR370C | 2370.21826 | 0.140302274 | 0.47010873 | 0.73 | 0.48 |
| YMR194C-B | 137.2124333 | -0.35980953 | 0.470158858 | 0.73 | 0.48 |
| YGL043W | 1337.950161 | -0.148851339 | 0.470500405 | 0.73 | 0.48 |
| YHR077C | 2925.324128 | 0.11414088 | 0.470588714 | 0.73 | 0.48 |
| YDR316W | 792.8873428 | 0.161582325 | 0.470817583 | 0.73 | 0.48 |
| YJL014W | 2604.88629 | -0.166939024 | 0.470721698 | 0.73 | 0.48 |
| YOL056W | 561.7509431 | -0.220973517 | 0.471040287 | 0.73 | 0.48 |
| YPL157W | 499.4077603 | -0.20559417 | 0.471054145 | 0.73 | 0.48 |
| YLL067C | 0.999643856 | 0.378613551 | 0.471273295 | 0.73 | 0.48 |
| YLR001C | 1116.57839 | 0.121656192 | 0.472058513 | 0.73 | 0.48 |
| YOR067C | 1025.694017 | 0.141483871 | 0.471961145 | 0.73 | 0.48 |
| YKL125W | 592.2613366 | -0.165470941 | 0.472888985 | 0.73 | 0.48 |
| YPR114W | 2316.846378 | -0.097697216 | 0.473439683 | 0.73 | 0.49 |
| YDL206W | 597.9682559 | 0.145079678 | 0.473852939 | 0.73 | 0.49 |
| YDL042C | 548.3117902 | 0.287261888 | 0.47401369 | 0.73 | 0.49 |
| YBR255C-A | 311.2384417 | -0.203973063 | 0.47433891 | 0.74 | 0.49 |
| YMR031C | 3707.880046 | 0.199657209 | 0.474537789 | 0.74 | 0.49 |
| YNL190W | 6079.357238 | 0.149931875 | 0.474532511 | 0.74 | 0.49 |
| YBL022C | 3958.243828 | 0.155529522 | 0.474910909 | 0.74 | 0.49 |
| YGR248W | 2366.128573 | 0.324518282 | 0.474900882 | 0.74 | 0.49 |
| YNL198C | 23.75937545 | -0.361365074 | 0.474788073 | 0.74 | 0.49 |
| YPR066W | 302.1036782 | 0.181181585 | 0.475203529 | 0.74 | 0.49 |
| YKR040C | 4.454389 | -0.391383634 | 0.475490391 | 0.74 | 0.49 |
| YGR083C | 2563.158798 | -0.183971668 | 0.475786599 | 0.74 | 0.49 |
| YLR293C | 8683.145693 | -0.187011587 | 0.475806979 | 0.74 | 0.49 |
| YIR042C | 196.4981605 | 0.417472584 | 0.476056177 | 0.74 | 0.49 |
| YBR192W | 512.8528961 | -0.106010532 | 0.476359708 | 0.74 | 0.49 |
| YBR211C | 479.4755282 | -0.166043093 | 0.476612489 | 0.74 | 0.49 |
| YLR444C | 25.68564702 | -0.303974034 | 0.476605886 | 0.74 | 0.49 |
| YML051W | 1749.382621 | -0.189860254 | 0.476493377 | 0.74 | 0.49 |
| YCR088W | 5938.374856 | -0.158878162 | 0.477240283 | 0.74 | 0.49 |
| YDR371W | 631.8329655 | -0.157224525 | 0.476981669 | 0.74 | 0.49 |
| YJL119C | 7.109202711 | 0.336430816 | 0.477075644 | 0.74 | 0.49 |

| | | | | | |
|---|---|---|---|---|---|
| YKL015W | 1286.742474 | 0.21064364 | 0.477275512 | 0.74 | 0.49 |
| YDR532C | 597.6785104 | 0.208493594 | 0.477439573 | 0.74 | 0.49 |
| YPL100W | 911.5930892 | 0.144310494 | 0.477638849 | 0.74 | 0.49 |
| YAL033W | 350.8440347 | -0.218283609 | 0.477858633 | 0.74 | 0.49 |
| YFR055W | 975.2716539 | 0.343065228 | 0.477878241 | 0.74 | 0.49 |
| YDR206W | 1789.144398 | -0.120583834 | 0.478103124 | 0.74 | 0.49 |
| YDR262W | 1927.726265 | 0.226477099 | 0.480135065 | 0.74 | 0.49 |
| YFL029C | 966.6378114 | 0.197090334 | 0.479713518 | 0.74 | 0.49 |
| YMR150C | 511.1258272 | 0.126751325 | 0.480098297 | 0.74 | 0.49 |
| YOR133W | 1425.340865 | -0.176683805 | 0.480038265 | 0.74 | 0.49 |
| YOR187W | 4877.755588 | 0.149920482 | 0.47980649 | 0.74 | 0.49 |
| YPL038W-A | 14.34188488 | 0.239348334 | 0.480051116 | 0.74 | 0.49 |
| YPL219W | 632.4863798 | 0.149967683 | 0.47952266 | 0.74 | 0.49 |
| YGL145W | 1156.588747 | -0.102277849 | 0.480284606 | 0.74 | 0.49 |
| YCL017C | 1588.391961 | 0.131768471 | 0.480564653 | 0.74 | 0.49 |
| YAL003W | 12374.8526 | -0.253918608 | 0.481462931 | 0.74 | 0.49 |
| YAL011W | 648.981818 | 0.131078822 | 0.481484413 | 0.74 | 0.49 |
| YGL076C | 4229.319259 | -0.225538021 | 0.481147372 | 0.74 | 0.49 |
| YJL134W | 603.1655535 | 0.190775471 | 0.481052755 | 0.74 | 0.49 |
| YLR102C | 476.3043851 | 0.170435907 | 0.481580983 | 0.74 | 0.49 |
| YLR451W | 1124.900564 | -0.1050072 | 0.481661902 | 0.74 | 0.49 |
| YNL082W | 573.887894 | -0.139284688 | 0.481381214 | 0.74 | 0.49 |
| YNL316C | 479.8242776 | -0.126319133 | 0.481346012 | 0.74 | 0.49 |
| YHR184W | 81.12732022 | 0.355492826 | 0.481795941 | 0.74 | 0.49 |
| YAR035C-A | 1.313184395 | 0.346618821 | 0.482347288 | 0.74 | 0.49 |
| YBR084C-A | 8707.666036 | -0.24586678 | 0.482394116 | 0.74 | 0.49 |
| YDL104C | 329.3472486 | 0.156119964 | 0.482210612 | 0.74 | 0.49 |
| YFR043C | 412.3285976 | 0.111065931 | 0.482359543 | 0.74 | 0.49 |
| YLR057W | 1092.547525 | 0.131980172 | 0.482191063 | 0.74 | 0.49 |
| YPR013C | 1083.106605 | -0.189092276 | 0.482528647 | 0.74 | 0.49 |
| YKL045W | 918.6422418 | -0.116531234 | 0.482741797 | 0.74 | 0.49 |
| YNL189W | 2792.970706 | -0.131071438 | 0.482875673 | 0.74 | 0.49 |
| YGL253W | 16247.71261 | 0.338511635 | 0.483161972 | 0.74 | 0.49 |
| YAR030C | 64.62074774 | 0.198680596 | 0.483707111 | 0.74 | 0.49 |
| YDL047W | 1531.758576 | -0.128952141 | 0.483913566 | 0.74 | 0.49 |
| YKL171W | 1421.131993 | 0.207848717 | 0.484230421 | 0.74 | 0.49 |
| YLR105C | 592.2672821 | -0.129132245 | 0.484208641 | 0.74 | 0.49 |
| YLR128W | 336.2576961 | -0.195019362 | 0.484140763 | 0.74 | 0.49 |
| YMR165C | 1579.930507 | 0.23822847 | 0.484046357 | 0.74 | 0.49 |
| YHR200W | 3347.025252 | 0.131656778 | 0.48437953 | 0.74 | 0.49 |
| YDL216C | 462.7023205 | 0.184451141 | 0.485192376 | 0.74 | 0.49 |
| YDR361C | 744.7914399 | -0.211939922 | 0.485283891 | 0.74 | 0.49 |
| YER001W | 1455.504127 | -0.177799387 | 0.48529095 | 0.74 | 0.49 |

| | | | | | |
|---|---|---|---|---|---|
| YLR184W | 0.707088895 | 0.274419767 | 0.485423951 | 0.74 | 0.49 |
| YOR228C | 505.0466268 | 0.210773897 | 0.485462245 | 0.74 | 0.49 |
| YOR249C | 647.5824875 | -0.124979469 | 0.485484423 | 0.74 | 0.49 |
| YGR264C | 2825.266453 | -0.203766931 | 0.485753429 | 0.74 | 0.49 |
| YOL157C | 78.40010831 | 0.299961827 | 0.486380217 | 0.74 | 0.49 |
| YPL074W | 944.5738824 | 0.141957254 | 0.486681163 | 0.74 | 0.49 |
| YHR147C | 857.7185719 | -0.104930065 | 0.486841861 | 0.74 | 0.49 |
| YLR320W | 753.4878726 | 0.11567033 | 0.48701571 | 0.74 | 0.49 |
| YFL012W | 27.05345673 | 0.400248141 | 0.487433611 | 0.74 | 0.49 |
| YGR182C | 629.4422912 | 0.205023622 | 0.487616991 | 0.74 | 0.49 |
| YLR032W | 823.8658356 | 0.145311919 | 0.488469976 | 0.74 | 0.49 |
| YFR006W | 3007.667423 | 0.104314658 | 0.488791738 | 0.74 | 0.49 |
| YLR238W | 614.7184945 | -0.136184017 | 0.489219615 | 0.75 | 0.49 |
| YNL251C | 2085.43933 | -0.244538297 | 0.489512379 | 0.75 | 0.49 |
| YOR380W | 392.7551247 | 0.136951668 | 0.4898428 | 0.75 | 0.49 |
| YOL033W | 423.9709748 | 0.125441338 | 0.490107305 | 0.75 | 0.49 |
| YHR021C | 5458.86145 | -0.214901775 | 0.490499904 | 0.75 | 0.49 |
| YDR008C | 37.32753147 | 0.149390452 | 0.490628205 | 0.75 | 0.49 |
| YFL056C | 232.3744781 | 0.218796196 | 0.490897667 | 0.75 | 0.49 |
| YJL072C | 353.6349072 | 0.166390185 | 0.490873558 | 0.75 | 0.49 |
| YGL102C | 355.1928609 | -0.229742647 | 0.49129029 | 0.75 | 0.49 |
| YEL073C | 122.6223904 | 0.499835109 | 0.491528917 | 0.75 | 0.49 |
| YNL191W | 591.9956262 | 0.128678091 | 0.491743318 | 0.75 | 0.49 |
| YBL005W-B | 450.3577253 | 0.274049752 | 0.491969264 | 0.75 | 0.49 |
| YMR003W | 306.5250482 | -0.162707145 | 0.492004481 | 0.75 | 0.49 |
| YBR245C | 1580.844401 | -0.097420845 | 0.492183759 | 0.75 | 0.49 |
| YGR037C | 2287.491924 | -0.173474082 | 0.492261965 | 0.75 | 0.49 |
| YBL047C | 5698.969568 | 0.110504153 | 0.492497065 | 0.75 | 0.49 |
| YDL240C-A | 1.077122984 | -0.309804363 | 0.492672975 | 0.75 | 0.49 |
| YHR201C | 1092.809897 | -0.161442329 | 0.492825013 | 0.75 | 0.49 |
| YKR001C | 4012.227533 | 0.129535755 | 0.492665999 | 0.75 | 0.49 |
| snR14 | 19.85838705 | -0.269080915 | 0.492898783 | 0.75 | 0.49 |
| YDR423C | 1205.3164 | -0.173658573 | 0.493440097 | 0.75 | 0.49 |
| YNL287W | 4413.325689 | 0.08746165 | 0.493686466 | 0.75 | 0.49 |
| YPR195C | 54.58354851 | 0.304373911 | 0.49361512 | 0.75 | 0.49 |
| YBR230W-A | 211.0787097 | -0.379257615 | 0.494567145 | 0.75 | 0.50 |
| YER136W | 4119.832209 | -0.090952502 | 0.494465385 | 0.75 | 0.50 |
| YMR099C | 2686.030784 | 0.104171621 | 0.495029632 | 0.75 | 0.50 |
| YOL048C | 1220.015423 | 0.179826991 | 0.495068634 | 0.75 | 0.50 |
| YPR172W | 1219.274092 | 0.159511868 | 0.494878099 | 0.75 | 0.50 |
| YEL023C | 623.1672116 | 0.151242015 | 0.495391363 | 0.75 | 0.50 |
| YLR283W | 409.1057668 | -0.096176821 | 0.49549295 | 0.75 | 0.50 |
| YLR363C | 329.7816062 | -0.162024085 | 0.495794994 | 0.75 | 0.50 |

| | | | | | |
|---|---|---|---|---|---|
| YJR072C | 1860.139893 | -0.128739817 | 0.496498394 | 0.75 | 0.50 |
| YKR100C | 1038.230895 | 0.11956678 | 0.496319347 | 0.75 | 0.50 |
| YPR115W | 2825.752527 | 0.185105639 | 0.496487429 | 0.75 | 0.50 |
| YDR194W-A | 0.453289942 | 0.203287484 | 0.496722784 | 0.75 | 0.50 |
| YKL033W-A | 1409.695613 | 0.131271165 | 0.496925901 | 0.75 | 0.50 |
| YGR169C | 437.5350343 | -0.172221045 | 0.49711908 | 0.75 | 0.50 |
| YDL192W | 4665.318464 | -0.160321314 | 0.497676596 | 0.75 | 0.50 |
| YGR147C | 943.7636147 | -0.112459441 | 0.497541707 | 0.75 | 0.50 |
| YHR048W | 458.0188382 | 0.23255941 | 0.497585908 | 0.75 | 0.50 |
| YHR132W-A | 721.2397547 | 0.160257201 | 0.497412341 | 0.75 | 0.50 |
| YDL200C | 293.2384509 | 0.128651913 | 0.498182234 | 0.75 | 0.50 |
| YEL046C | 2157.479841 | -0.187060163 | 0.498066679 | 0.75 | 0.50 |
| YLR287C | 617.6178514 | -0.180245862 | 0.498150049 | 0.75 | 0.50 |
| YGL027C | 1397.054052 | 0.149350556 | 0.498428467 | 0.75 | 0.50 |
| YGL087C | 1504.49763 | 0.158593729 | 0.498744703 | 0.75 | 0.50 |
| YGR095C | 969.1047325 | -0.171619808 | 0.498893127 | 0.75 | 0.50 |
| YLR460C | 187.4228264 | 0.172517122 | 0.498895543 | 0.75 | 0.50 |
| YPR148C | 2535.151148 | -0.102599498 | 0.498773165 | 0.75 | 0.50 |
| YLR123C | 17.47407493 | 0.323055984 | 0.499021449 | 0.75 | 0.50 |
| YIL112W | 2132.63932 | 0.172509687 | 0.499192264 | 0.75 | 0.50 |
| YCL034W | 1930.992154 | -0.157142589 | 0.499556518 | 0.75 | 0.50 |
| YLR118C | 939.2371521 | 0.118800791 | 0.499575844 | 0.75 | 0.50 |
| YPR100W | 188.3881606 | -0.170740912 | 0.499374283 | 0.75 | 0.50 |
| YJR019C | 596.1541169 | 0.147549206 | 0.499861601 | 0.75 | 0.50 |
| YDR211W | 2365.993285 | 0.161528532 | 0.500228599 | 0.75 | 0.50 |
| YER102W | 2473.647383 | -0.226896537 | 0.500228135 | 0.75 | 0.50 |
| YPL080C | 11.15479793 | -0.149911244 | 0.500291634 | 0.75 | 0.50 |
| YAR003W | 588.5199738 | 0.177838509 | 0.500529817 | 0.75 | 0.50 |
| YNL315C | 1636.297254 | -0.135519519 | 0.500484621 | 0.75 | 0.50 |
| YHR131C | 966.4131265 | 0.157110281 | 0.500898197 | 0.75 | 0.50 |
| YIL019W | 543.1334136 | -0.23934571 | 0.500884633 | 0.75 | 0.50 |
| YFR051C | 3059.620652 | -0.162109493 | 0.501062557 | 0.75 | 0.50 |
| YBR274W | 529.7886878 | -0.108541449 | 0.501663539 | 0.75 | 0.50 |
| YCL056C | 401.4808141 | -0.092280802 | 0.50153479 | 0.75 | 0.50 |
| YER020W | 2824.873338 | 0.19709649 | 0.501457675 | 0.75 | 0.50 |
| YGL126W | 1063.33427 | 0.253274048 | 0.501520732 | 0.75 | 0.50 |
| snR79 | 6.23839996 | 0.318984411 | 0.50168727 | 0.75 | 0.50 |
| YBR177C | 2530.709586 | -0.168801251 | 0.50203956 | 0.75 | 0.50 |
| YOR210W | 598.8398742 | -0.156957854 | 0.502234283 | 0.75 | 0.50 |
| YDL158C | 89.30551169 | 0.216801556 | 0.502757602 | 0.75 | 0.50 |
| YDR342C | 1706.769269 | 0.275965692 | 0.502557245 | 0.75 | 0.50 |
| YKL153W | 3671.352718 | -0.253902179 | 0.5027846 | 0.75 | 0.50 |
| YPL218W | 2797.046919 | -0.114930582 | 0.502871146 | 0.75 | 0.50 |

| | | | | | |
|---|---|---|---|---|---|
| YJR159W | 6.187748904 | 0.294744832 | 0.503625013 | 0.75 | 0.50 |
| YDR142C | 534.1730573 | -0.162822148 | 0.503884686 | 0.75 | 0.50 |
| YLR058C | 11510.40084 | -0.239545924 | 0.504000616 | 0.75 | 0.50 |
| YDR078C | 147.0952169 | 0.19615302 | 0.504590139 | 0.75 | 0.50 |
| YDR348C | 1537.308452 | 0.071132383 | 0.504662056 | 0.75 | 0.50 |
| YIL047C-A | 10.41380776 | -0.253022519 | 0.504428036 | 0.75 | 0.50 |
| YLR433C | 1164.793625 | -0.163275027 | 0.504593501 | 0.75 | 0.50 |
| snR69 | 9.747852558 | 0.257100619 | 0.504406582 | 0.75 | 0.50 |
| YDR366C | 7.819864358 | 0.34439949 | 0.504805748 | 0.75 | 0.50 |
| YDR480W | 507.0349407 | -0.072280017 | 0.505036847 | 0.75 | 0.50 |
| YNR056C | 522.7352776 | 0.269467817 | 0.505151668 | 0.75 | 0.50 |
| YOL059W | 2587.338376 | 0.177490942 | 0.505414684 | 0.75 | 0.50 |
| YOL160W | 15.67971505 | 0.226558562 | 0.505485182 | 0.75 | 0.50 |
| YPL020C | 1208.520388 | -0.083230341 | 0.505484168 | 0.75 | 0.50 |
| YPL061W | 6576.454349 | -0.186834495 | 0.505367871 | 0.75 | 0.50 |
| YHL041W | 3.288023996 | 0.349783367 | 0.505934856 | 0.75 | 0.50 |
| YJL006C | 336.1143852 | -0.142009065 | 0.506016752 | 0.75 | 0.50 |
| YLR151C | 483.8723388 | 0.130405166 | 0.506267812 | 0.75 | 0.50 |
| YMR067C | 1220.387108 | -0.106979752 | 0.506228624 | 0.75 | 0.50 |
| YMR213W | 660.1636091 | 0.127366558 | 0.506161049 | 0.75 | 0.50 |
| YDR322W | 1031.616879 | -0.162565676 | 0.506403351 | 0.75 | 0.50 |
| YDR521W | 13.62209314 | -0.294534172 | 0.506949719 | 0.76 | 0.50 |
| YLL012W | 268.5425901 | -0.222178207 | 0.506898828 | 0.76 | 0.50 |
| YMR119W | 1094.314263 | 0.139482127 | 0.506923924 | 0.76 | 0.50 |
| YPL072W | 410.1680408 | -0.123513535 | 0.507041885 | 0.76 | 0.50 |
| YOL022C | 1058.507084 | -0.167728155 | 0.507238223 | 0.76 | 0.50 |
| YNL217W | 1768.453008 | 0.257242527 | 0.507381312 | 0.76 | 0.50 |
| YLR206W | 3965.73873 | 0.104909531 | 0.507546183 | 0.76 | 0.50 |
| YBL080C | 486.2802333 | 0.126670813 | 0.507694331 | 0.76 | 0.50 |
| YER089C | 1927.936186 | -0.188094123 | 0.508154474 | 0.76 | 0.50 |
| YGL085W | 385.7388603 | 0.117591496 | 0.508322434 | 0.76 | 0.50 |
| YMR178W | 1157.445412 | 0.13392002 | 0.508960571 | 0.76 | 0.50 |
| YKL160W | 1725.224125 | -0.115127839 | 0.509145286 | 0.76 | 0.50 |
| YAL039C | 1453.711163 | 0.131060984 | 0.509629204 | 0.76 | 0.50 |
| YDL068W | 4.863206233 | -0.251564826 | 0.509862528 | 0.76 | 0.50 |
| YER008C | 2582.281632 | 0.1656711 | 0.509642828 | 0.76 | 0.50 |
| YNL179C | 130.2526057 | -0.250697178 | 0.509863326 | 0.76 | 0.50 |
| YDR019C | 2663.732875 | -0.282802363 | 0.510366257 | 0.76 | 0.50 |
| YFL037W | 5301.265498 | 0.160833764 | 0.510453326 | 0.76 | 0.50 |
| YGL040C | 1261.243607 | 0.160391515 | 0.510477848 | 0.76 | 0.50 |
| YGL244W | 2087.284965 | -0.136253445 | 0.510295135 | 0.76 | 0.50 |
| YMR185W | 803.0326457 | -0.159787757 | 0.510285252 | 0.76 | 0.50 |
| YJL130C | 21427.36188 | 0.295862642 | 0.511198078 | 0.76 | 0.50 |

| | | | | | |
|---|---|---|---|---|---|
| YML076C | 1190.303539 | 0.127440839 | 0.511288474 | 0.76 | 0.50 |
| YER110C | 3689.619393 | -0.186357168 | 0.511570607 | 0.76 | 0.50 |
| YJL161W | 501.2537696 | 0.240648298 | 0.511481757 | 0.76 | 0.50 |
| YOR391C | 10.43191586 | 0.348317301 | 0.51172177 | 0.76 | 0.50 |
| YDR006C | 2310.266671 | -0.150631794 | 0.512097796 | 0.76 | 0.50 |
| YOR116C | 1548.881592 | -0.140339403 | 0.512025834 | 0.76 | 0.50 |
| YDR095C | 5.026425328 | -0.238092589 | 0.512885592 | 0.76 | 0.50 |
| YGR009C | 1451.464581 | 0.096017426 | 0.513513224 | 0.76 | 0.50 |
| YHR191C | 210.2719774 | 0.085250998 | 0.513494028 | 0.76 | 0.50 |
| YLL003W | 928.3757705 | -0.085728625 | 0.513318957 | 0.76 | 0.50 |
| YLR338W | 19.58277131 | 0.176250775 | 0.513483421 | 0.76 | 0.50 |
| YOL028C | 419.6545084 | -0.106819815 | 0.513281761 | 0.76 | 0.50 |
| YOL071W | 1035.627926 | 0.192582451 | 0.51334251 | 0.76 | 0.50 |
| YOR097C | 623.6386998 | 0.114640661 | 0.512913851 | 0.76 | 0.50 |
| YPL151C | 670.1565001 | 0.081388158 | 0.513135992 | 0.76 | 0.50 |
| YLR099C | 1018.600053 | 0.20765296 | 0.514184101 | 0.76 | 0.50 |
| YLR387C | 1423.423086 | 0.169602619 | 0.514099238 | 0.76 | 0.50 |
| YOR195W | 1326.708475 | 0.182747587 | 0.514040799 | 0.76 | 0.50 |
| YNL159C | 572.5140174 | -0.143810826 | 0.514505197 | 0.76 | 0.50 |
| YLR097C | 748.1202732 | 0.125348583 | 0.514778319 | 0.76 | 0.50 |
| YIL083C | 1169.719343 | 0.187434728 | 0.515460948 | 0.76 | 0.50 |
| YNL149C | 291.2466735 | -0.182198866 | 0.515417344 | 0.76 | 0.50 |
| YLR168C | 702.4056037 | 0.239562172 | 0.515716306 | 0.76 | 0.50 |
| YMR079W | 1633.407315 | -0.146006134 | 0.51578196 | 0.76 | 0.50 |
| YBR094W | 1036.817856 | -0.111365016 | 0.516308774 | 0.76 | 0.50 |
| YCR018C-A | 8.72999001 | 0.291449522 | 0.517097002 | 0.76 | 0.50 |
| YGR290W | 27.11482068 | 0.269679649 | 0.517077043 | 0.76 | 0.50 |
| YLR233C | 453.3027113 | 0.182139179 | 0.517548979 | 0.76 | 0.50 |
| YLR251W | 493.6892026 | 0.1766901 | 0.517530772 | 0.76 | 0.50 |
| YLR171W | 25.60461968 | 0.141581877 | 0.518095205 | 0.76 | 0.50 |
| YIL018W | 10226.88893 | -0.178843863 | 0.51864639 | 0.76 | 0.50 |
| YPR123C | 99.99461053 | 0.317169835 | 0.518823998 | 0.76 | 0.50 |
| YNL286W | 504.2093629 | 0.156901653 | 0.519045897 | 0.76 | 0.50 |
| YFR013W | 1397.161166 | -0.123054344 | 0.519321599 | 0.76 | 0.51 |
| YLL029W | 3427.507563 | 0.158461062 | 0.519717374 | 0.76 | 0.51 |
| YOR381W | 508.0002714 | 0.236397382 | 0.519635839 | 0.76 | 0.51 |
| YCR063W | 60.70870613 | -0.132280557 | 0.519839506 | 0.76 | 0.51 |
| YGR088W | 3462.726793 | 0.313313473 | 0.520007702 | 0.76 | 0.51 |
| YKR012C | 142.5434976 | 0.194551052 | 0.520302445 | 0.76 | 0.51 |
| YDR456W | 1671.501856 | 0.103132288 | 0.520559237 | 0.77 | 0.51 |
| YGR288W | 284.2488249 | 0.233081105 | 0.520606632 | 0.77 | 0.51 |
| YDL246C | 6.191078768 | 0.295568463 | 0.521336189 | 0.77 | 0.51 |
| YOR197W | 2558.031802 | 0.095644542 | 0.521516449 | 0.77 | 0.51 |

| | | | | | |
|---|---|---|---|---|---|
| YIL142C-A | 7.771319082 | 0.168993959 | 0.52185917 | 0.77 | 0.51 |
| YDL211C | 175.1726895 | -0.16971727 | 0.522913424 | 0.77 | 0.51 |
| YDR003W-A | 67.78719135 | 0.144315678 | 0.522954079 | 0.77 | 0.51 |
| YDR243C | 512.1645164 | -0.10761964 | 0.523022151 | 0.77 | 0.51 |
| YGL097W | 1307.008078 | -0.222922045 | 0.522312506 | 0.77 | 0.51 |
| YKR084C | 1262.417524 | -0.120897152 | 0.522450218 | 0.77 | 0.51 |
| YMR276W | 5216.693498 | -0.116147921 | 0.522910214 | 0.77 | 0.51 |
| YOR099W | 2318.668874 | -0.146248497 | 0.522615316 | 0.77 | 0.51 |
| YPR078C | 88.79558885 | 0.267784357 | 0.522908418 | 0.77 | 0.51 |
| YBR111W-A | 229.152125 | -0.149038317 | 0.523478543 | 0.77 | 0.51 |
| YDR271C | 28.54979964 | 0.224850906 | 0.523371697 | 0.77 | 0.51 |
| YDR467C | 75.26126934 | 0.127566612 | 0.523536114 | 0.77 | 0.51 |
| YMR120C | 11760.55672 | -0.232915888 | 0.523541883 | 0.77 | 0.51 |
| YCR095C | 483.8306058 | 0.125277808 | 0.524285416 | 0.77 | 0.51 |
| YKL154W | 669.8187345 | -0.183814587 | 0.524407705 | 0.77 | 0.51 |
| YLL065W | 0.583562943 | 0.386505675 | 0.524591697 | 0.77 | 0.51 |
| YMR012W | 7425.571889 | -0.110627247 | 0.524587717 | 0.77 | 0.51 |
| YMR318C | 2412.871237 | -0.139123522 | 0.524637571 | 0.77 | 0.51 |
| YNL211C | 132.5864716 | -0.161296919 | 0.524042398 | 0.77 | 0.51 |
| YPR144C | 1036.348245 | -0.188907214 | 0.524209792 | 0.77 | 0.51 |
| YDL115C | 894.0269756 | 0.096591451 | 0.525026887 | 0.77 | 0.51 |
| YIL098C | 444.8949209 | -0.151212457 | 0.525130749 | 0.77 | 0.51 |
| YDR255C | 870.1597032 | 0.16797429 | 0.525358801 | 0.77 | 0.51 |
| YDR093W | 1759.127414 | 0.126152756 | 0.526036665 | 0.77 | 0.51 |
| YJL010C | 755.184634 | -0.175141575 | 0.525928199 | 0.77 | 0.51 |
| YLR274W | 1064.782127 | -0.127990139 | 0.526074439 | 0.77 | 0.51 |
| YMR156C | 369.432093 | 0.213774541 | 0.525674855 | 0.77 | 0.51 |
| YNL313C | 1104.908095 | -0.147591707 | 0.5259549 | 0.77 | 0.51 |
| YPR129W | 1545.384555 | 0.117372995 | 0.526384963 | 0.77 | 0.51 |
| YER044C | 1096.772117 | -0.161565878 | 0.527187714 | 0.77 | 0.51 |
| snR8 | 32.91757815 | 0.277430315 | 0.527509899 | 0.77 | 0.51 |
| YDR306C | 1159.558737 | 0.121246981 | 0.527710897 | 0.77 | 0.51 |
| YDL154W | 661.9950458 | -0.173549624 | 0.528155333 | 0.77 | 0.51 |
| YPL165C | 810.9716846 | 0.180138344 | 0.528355629 | 0.77 | 0.51 |
| YCR016W | 907.9551315 | -0.173119496 | 0.52849182 | 0.77 | 0.51 |
| YBR044C | 837.2738234 | 0.133061523 | 0.52888951 | 0.77 | 0.51 |
| YJR123W | 26166.00582 | -0.201379138 | 0.529116364 | 0.77 | 0.51 |
| YJL035C | 163.2101994 | -0.176798211 | 0.529651519 | 0.77 | 0.51 |
| snR46 | 23.07413489 | -0.210412676 | 0.529709973 | 0.77 | 0.51 |
| YPL125W | 1168.744838 | 0.157056149 | 0.530181847 | 0.77 | 0.51 |
| YGR242W | 37.21832003 | -0.167488469 | 0.530498919 | 0.77 | 0.51 |
| YKL061W | 244.3100746 | 0.168970907 | 0.530481154 | 0.77 | 0.51 |
| YDR146C | 1456.032171 | -0.140047143 | 0.53090769 | 0.77 | 0.51 |

| | | | | | |
|---|---|---|---|---|---|
| YML067C | 1114.441541 | -0.079807113 | 0.531109089 | 0.77 | 0.51 |
| YCR090C | 636.078747 | -0.163505657 | 0.531666523 | 0.77 | 0.51 |
| YJL085W | 484.172969 | -0.143442801 | 0.531825741 | 0.77 | 0.51 |
| YKR064W | 1141.244604 | 0.104765939 | 0.531964089 | 0.77 | 0.51 |
| YOL083W | 724.7720388 | 0.201794769 | 0.532037889 | 0.77 | 0.51 |
| YNL100W | 1262.153408 | 0.187734361 | 0.53221291 | 0.77 | 0.51 |
| YGR208W | 739.3899853 | -0.174643743 | 0.532462504 | 0.77 | 0.51 |
| YDL141W | 1031.602218 | 0.097061965 | 0.532704149 | 0.77 | 0.51 |
| YLR432W | 4140.896048 | 0.335038626 | 0.532867932 | 0.77 | 0.51 |
| YMR132C | 213.3288234 | 0.120289089 | 0.532947148 | 0.77 | 0.51 |
| YOR156C | 466.9299753 | -0.122964816 | 0.532712669 | 0.77 | 0.51 |
| YLR429W | 3551.790145 | -0.239904132 | 0.53312349 | 0.77 | 0.51 |
| YHR022C-A | 3.937421936 | -0.32843806 | 0.533295332 | 0.77 | 0.51 |
| YDL095W | 3378.459132 | -0.131044762 | 0.533486272 | 0.77 | 0.51 |
| YOL147C | 576.9213691 | 0.187056634 | 0.533535499 | 0.77 | 0.51 |
| YBL056W | 3616.602261 | 0.079451375 | 0.533802995 | 0.77 | 0.51 |
| YBR199W | 2706.736809 | 0.058052343 | 0.533841277 | 0.77 | 0.51 |
| YCL052C | 1614.200073 | 0.123071059 | 0.534320776 | 0.77 | 0.51 |
| YGL223C | 914.5737412 | -0.084252294 | 0.534158879 | 0.77 | 0.51 |
| YOR157C | 2318.284571 | 0.092782185 | 0.53431035 | 0.77 | 0.51 |
| YBR163W | 719.9181119 | -0.134909386 | 0.534897968 | 0.77 | 0.51 |
| YIL014C-A | 253.291496 | 0.378069229 | 0.535637443 | 0.78 | 0.51 |
| YIL134W | 238.1456546 | 0.242728936 | 0.535862773 | 0.78 | 0.51 |
| YGR062C | 462.3300151 | -0.131457993 | 0.536402835 | 0.78 | 0.51 |
| YDR134C | 8310.682574 | -0.230593778 | 0.536668381 | 0.78 | 0.51 |
| YER026C | 2306.614486 | -0.165501759 | 0.536785273 | 0.78 | 0.51 |
| YBL007C | 3937.826759 | 0.104031577 | 0.537157778 | 0.78 | 0.51 |
| YMR036C | 631.5049431 | -0.12750346 | 0.537222727 | 0.78 | 0.51 |
| YML101C-A | 7.327177589 | -0.247555111 | 0.537468425 | 0.78 | 0.51 |
| YNL321W | 1495.615211 | 0.108553513 | 0.538205934 | 0.78 | 0.51 |
| YGL116W | 933.3944118 | 0.12903973 | 0.538336362 | 0.78 | 0.51 |
| YKR101W | 347.6961828 | 0.119363116 | 0.539152834 | 0.78 | 0.51 |
| YDR170C | 4884.61556 | 0.115722084 | 0.540140192 | 0.78 | 0.52 |
| YDR260C | 314.9401976 | -0.123552248 | 0.540299345 | 0.78 | 0.52 |
| YDR453C | 889.6622872 | 0.261440972 | 0.540200859 | 0.78 | 0.52 |
| YGR121C | 1176.230558 | 0.150212803 | 0.540062865 | 0.78 | 0.52 |
| YLR049C | 719.3459985 | -0.173716473 | 0.540460761 | 0.78 | 0.52 |
| YBL046W | 691.3216368 | 0.092202238 | 0.540902287 | 0.78 | 0.52 |
| YOR384W | 1087.084219 | 0.170505803 | 0.540922359 | 0.78 | 0.52 |
| YNL164C | 278.0976276 | -0.138581004 | 0.541056475 | 0.78 | 0.52 |
| YOR288C | 898.6918954 | -0.150740778 | 0.54196251 | 0.78 | 0.52 |
| YBL061C | 962.4150781 | -0.116455075 | 0.542255029 | 0.78 | 0.52 |
| YAR018C | 738.6960091 | 0.107922403 | 0.543027318 | 0.78 | 0.52 |

| | | | | | |
|---|---|---|---|---|---|
| YDR175C | 1042.356621 | -0.133949692 | 0.543091956 | 0.78 | 0.52 |
| YMR001C-A | 16.08451551 | -0.157617058 | 0.543298077 | 0.78 | 0.52 |
| YOR069W | 1035.064237 | 0.083726445 | 0.543387334 | 0.78 | 0.52 |
| YER084W-A | 0.719422121 | 0.415935687 | 0.543643906 | 0.78 | 0.52 |
| YGR161W-C | 303.4318056 | -0.222099027 | 0.543950502 | 0.78 | 0.52 |
| YLR371W | 1780.890302 | 0.141000921 | 0.5439904 | 0.78 | 0.52 |
| YNL019C | 10.29601328 | 0.273852556 | 0.543835876 | 0.78 | 0.52 |
| YOR223W | 687.6095973 | 0.131372186 | 0.54400586 | 0.78 | 0.52 |
| YGL136C | 674.1887477 | -0.1138957 | 0.544158994 | 0.78 | 0.52 |
| YHR020W | 3665.040866 | 0.263576737 | 0.544276715 | 0.78 | 0.52 |
| YLR204W | 619.6469206 | -0.142979676 | 0.544877591 | 0.78 | 0.52 |
| YLR349W | 23.19539208 | 0.217272473 | 0.545042741 | 0.78 | 0.52 |
| YIR020C-B | 5.673146071 | -0.283125767 | 0.54589958 | 0.78 | 0.52 |
| YPL209C | 162.5627938 | 0.147756615 | 0.546230501 | 0.78 | 0.52 |
| YER053C-A | 179.222376 | -0.304460231 | 0.547044414 | 0.79 | 0.52 |
| YJL032W | 6.67566349 | -0.317804391 | 0.546930905 | 0.79 | 0.52 |
| YCR053W | 6107.267981 | 0.201186048 | 0.547193564 | 0.79 | 0.52 |
| TLC1 | 1417.425959 | -0.182462739 | 0.548019117 | 0.79 | 0.52 |
| YDR513W | 3464.363476 | 0.195970599 | 0.54786514 | 0.79 | 0.52 |
| YGR277C | 611.2932905 | -0.127498885 | 0.547689762 | 0.79 | 0.52 |
| YNR012W | 597.0846222 | -0.170512662 | 0.547946852 | 0.79 | 0.52 |
| YBR196C-A | 7.414882286 | 0.334891485 | 0.548556151 | 0.79 | 0.52 |
| YGL058W | 714.9977444 | 0.157367696 | 0.548472549 | 0.79 | 0.52 |
| YMR116C | 12111.33345 | -0.170931192 | 0.548550027 | 0.79 | 0.52 |
| YPL200W | 21.68345646 | 0.283738685 | 0.548695228 | 0.79 | 0.52 |
| YAL016C-A | 61.31300348 | 0.174431928 | 0.549376018 | 0.79 | 0.52 |
| YBL063W | 882.9945164 | 0.119614171 | 0.549495392 | 0.79 | 0.52 |
| YCR099C | 126.2405817 | 0.252696397 | 0.549625994 | 0.79 | 0.52 |
| YER045C | 814.7735686 | 0.158520414 | 0.549154681 | 0.79 | 0.52 |
| YGL056C | 2031.538502 | 0.137867937 | 0.549560533 | 0.79 | 0.52 |
| YBR080C | 2198.347068 | -0.113149536 | 0.550145439 | 0.79 | 0.52 |
| YDR126W | 326.9726421 | 0.142032302 | 0.550442317 | 0.79 | 0.52 |
| YOL036W | 1888.124512 | 0.192853322 | 0.550408265 | 0.79 | 0.52 |
| YAL029C | 2871.619875 | 0.079536333 | 0.550677198 | 0.79 | 0.52 |
| YBL029W | 662.4144794 | 0.14188737 | 0.550759043 | 0.79 | 0.52 |
| YAL030W | 729.4057845 | 0.10704187 | 0.551130733 | 0.79 | 0.52 |
| YDR318W | 508.7245087 | -0.108631558 | 0.552761633 | 0.79 | 0.52 |
| YGR144W | 97.38216331 | -0.241445448 | 0.551922448 | 0.79 | 0.52 |
| YLR355C | 9452.460146 | 0.1673814 | 0.552786357 | 0.79 | 0.52 |
| YML022W | 1077.904293 | -0.132039251 | 0.552503606 | 0.79 | 0.52 |
| YNL158W | 434.8285056 | -0.132201925 | 0.55230149 | 0.79 | 0.52 |
| YOL086C | 59663.42444 | 0.249365944 | 0.552318287 | 0.79 | 0.52 |
| YOR115C | 581.5147795 | -0.113813725 | 0.552542164 | 0.79 | 0.52 |

| | | | | | |
|---|---|---|---|---|---|
| YOR219C | 1764.610148 | 0.095451999 | 0.552200081 | 0.79 | 0.52 |
| YOR231W | 881.0751627 | 0.125687943 | 0.551420523 | 0.79 | 0.52 |
| YPL036W | 288.1590263 | 0.157860952 | 0.552113892 | 0.79 | 0.52 |
| YPL117C | 2245.25569 | -0.108559645 | 0.552631056 | 0.79 | 0.52 |
| YPR097W | 1353.173051 | 0.084055459 | 0.552415119 | 0.79 | 0.52 |
| YPR169W-A | 32.89680766 | 0.14924577 | 0.551628041 | 0.79 | 0.52 |
| YHR139C | 204.3743283 | 0.250135298 | 0.553008677 | 0.79 | 0.52 |
| YJL062W-A | 1034.508423 | 0.158462203 | 0.554645077 | 0.79 | 0.52 |
| YJL067W | 49.69168928 | 0.214157886 | 0.554417859 | 0.79 | 0.52 |
| YKL162C-A | 16.50216676 | 0.243714967 | 0.554563531 | 0.79 | 0.52 |
| YER188W | 461.4895098 | 0.312789202 | 0.554918785 | 0.79 | 0.52 |
| YGR212W | 361.1538773 | 0.097746525 | 0.555464031 | 0.79 | 0.52 |
| YJL023C | 307.217395 | 0.104729568 | 0.554993953 | 0.79 | 0.52 |
| YJR102C | 338.1234787 | 0.120772525 | 0.555531772 | 0.79 | 0.52 |
| YKL002W | 1601.062288 | -0.105457043 | 0.555414941 | 0.79 | 0.52 |
| YMR157C | 556.2957769 | -0.133271333 | 0.555097766 | 0.79 | 0.52 |
| YOL004W | 4094.946984 | 0.093246844 | 0.555572306 | 0.79 | 0.52 |
| YCR024C-B | 2025.586967 | -0.125708183 | 0.55661951 | 0.79 | 0.52 |
| YHR061C | 329.4509042 | -0.116964332 | 0.557398635 | 0.79 | 0.52 |
| YNL122C | 325.2385534 | -0.138855619 | 0.557752813 | 0.79 | 0.52 |
| snR73 | 0.982818557 | 0.502095087 | 0.557710799 | 0.79 | 0.52 |
| YIR003W | 1806.796543 | 0.116385977 | 0.557879866 | 0.79 | 0.52 |
| YBR103W | 849.3179685 | 0.10317758 | 0.558413869 | 0.79 | 0.52 |
| YGL022W | 3864.561612 | 0.141435164 | 0.558368217 | 0.79 | 0.52 |
| YOR299W | 1060.38632 | 0.148956985 | 0.558335655 | 0.79 | 0.52 |
| YMR263W | 652.3046851 | -0.147543352 | 0.558556779 | 0.79 | 0.52 |
| YOR191W | 2372.954185 | 0.110262692 | 0.558768203 | 0.79 | 0.52 |
| YML031W | 1341.560963 | -0.114024666 | 0.559014548 | 0.79 | 0.52 |
| YNL056W | 768.1896527 | -0.088019481 | 0.558965282 | 0.79 | 0.52 |
| YDR388W | 5322.26629 | 0.078017349 | 0.55936582 | 0.79 | 0.52 |
| YOL057W | 2160.171024 | -0.130038059 | 0.56042757 | 0.79 | 0.52 |
| YGL250W | 555.5711852 | 0.144617026 | 0.560665222 | 0.79 | 0.52 |
| YNL227C | 372.4616162 | 0.158251279 | 0.560683526 | 0.79 | 0.52 |
| YOR365C | 309.6078398 | -0.251458576 | 0.561062497 | 0.79 | 0.53 |
| YDR145W | 1845.179932 | -0.129687965 | 0.561430678 | 0.79 | 0.53 |
| YGR278W | 442.5587334 | -0.099322173 | 0.561628842 | 0.79 | 0.53 |
| YJR115W | 567.164948 | -0.312994305 | 0.561351247 | 0.79 | 0.53 |
| YMR257C | 582.2395624 | 0.153569664 | 0.561528167 | 0.79 | 0.53 |
| YOL009C | 380.9976744 | 0.106689622 | 0.561679707 | 0.79 | 0.53 |
| YGL051W | 80.44841429 | 0.154307072 | 0.561980953 | 0.80 | 0.53 |
| YBR059C | 2706.219813 | 0.119200385 | 0.563110911 | 0.80 | 0.53 |
| YER137C | 65.6651259 | 0.14382307 | 0.562900888 | 0.80 | 0.53 |
| YGL200C | 3717.842992 | 0.099470128 | 0.56288175 | 0.80 | 0.53 |

| | | | | | |
|---|---|---|---|---|---|
| YMR066W | 755.7979974 | 0.124708088 | 0.563179585 | 0.80 | 0.53 |
| YNR045W | 861.3548214 | 0.187085872 | 0.563182994 | 0.80 | 0.53 |
| YBL051C | 3664.33556 | -0.190477656 | 0.563858677 | 0.80 | 0.53 |
| YFL031W | 3441.588262 | 0.203247812 | 0.564048239 | 0.80 | 0.53 |
| YHR067W | 339.2689826 | 0.173530981 | 0.563936643 | 0.80 | 0.53 |
| YPL149W | 607.7809907 | -0.159490647 | 0.564290087 | 0.80 | 0.53 |
| YPR176C | 535.2741493 | -0.105029444 | 0.564233415 | 0.80 | 0.53 |
| YDR133C | 1765.716231 | -0.196681505 | 0.564583311 | 0.80 | 0.53 |
| YLR169W | 310.0591441 | -0.103069785 | 0.565468844 | 0.80 | 0.53 |
| YLR392C | 638.9786435 | 0.156021313 | 0.565429932 | 0.80 | 0.53 |
| YDR458C | 1309.381626 | -0.126195366 | 0.565826098 | 0.80 | 0.53 |
| YDL009C | 106.1081082 | -0.111925524 | 0.566358623 | 0.80 | 0.53 |
| YJR134C | 1708.030399 | -0.119563173 | 0.566576092 | 0.80 | 0.53 |
| YOR017W | 870.202076 | -0.089584914 | 0.567101205 | 0.80 | 0.53 |
| YOR036W | 1460.876014 | 0.134962074 | 0.567424559 | 0.80 | 0.53 |
| YHR094C | 525.1336673 | -0.266487106 | 0.567700962 | 0.80 | 0.53 |
| snR87 | 6.424473744 | 0.282410543 | 0.56759056 | 0.80 | 0.53 |
| YJR105W | 3472.841224 | 0.195035615 | 0.568003304 | 0.80 | 0.53 |
| YGR281W | 2050.830805 | -0.088753855 | 0.568787321 | 0.80 | 0.53 |
| YHR088W | 1005.281896 | -0.142106193 | 0.56861094 | 0.80 | 0.53 |
| YOR012W | 60.32598792 | -0.12800362 | 0.568544763 | 0.80 | 0.53 |
| YPL008W | 976.9490792 | 0.09927251 | 0.568723466 | 0.80 | 0.53 |
| YEL067C | 149.0124198 | 0.140972259 | 0.569218148 | 0.80 | 0.53 |
| YGR050C | 100.6325721 | -0.149030142 | 0.569041091 | 0.80 | 0.53 |
| YLR439W | 1150.159115 | 0.140970988 | 0.569160325 | 0.80 | 0.53 |
| YDR003W | 807.4785109 | 0.127118763 | 0.569449566 | 0.80 | 0.53 |
| YJR119C | 341.555846 | -0.134417293 | 0.569551109 | 0.80 | 0.53 |
| YLR211C | 375.8938446 | 0.144546169 | 0.56975078 | 0.80 | 0.53 |
| YCR091W | 1488.499833 | 0.200702944 | 0.570406465 | 0.80 | 0.53 |
| YGR234W | 5131.339655 | -0.152002095 | 0.570525556 | 0.80 | 0.53 |
| YOR362C | 3907.49208 | -0.089856393 | 0.570460269 | 0.80 | 0.53 |
| YOL082W | 2476.571718 | 0.122253965 | 0.570825468 | 0.80 | 0.53 |
| YGL007C-A | 2.085025417 | 0.251123845 | 0.571138518 | 0.80 | 0.53 |
| YDL032W | 4.179935405 | -0.289855169 | 0.571450776 | 0.80 | 0.53 |
| YDL173W | 1092.267621 | 0.209710292 | 0.571426679 | 0.80 | 0.53 |
| YDR246W-A | 23.46436155 | -0.201376241 | 0.57160758 | 0.80 | 0.53 |
| YPL132W | 679.0950944 | 0.123684375 | 0.57273732 | 0.80 | 0.53 |
| YIR009W | 172.3613335 | -0.130974709 | 0.573210336 | 0.80 | 0.53 |
| YJR135C | 188.8236464 | -0.116417583 | 0.573142926 | 0.80 | 0.53 |
| YPL189C-A | 341.9617668 | 0.182969585 | 0.573448656 | 0.80 | 0.53 |
| YMR077C | 308.7109529 | -0.120870031 | 0.573604456 | 0.80 | 0.53 |
| YOR037W | 696.733578 | 0.097908666 | 0.573926196 | 0.80 | 0.53 |
| YER023W | 2339.829047 | -0.121133208 | 0.574433504 | 0.80 | 0.53 |

| | | | | | |
|---|---|---|---|---|---|
| YPL103C | 590.7071612 | 0.109431853 | 0.574387884 | 0.80 | 0.53 |
| YBR227C | 856.5044097 | 0.134292752 | 0.57487052 | 0.80 | 0.53 |
| YER169W | 1395.485471 | 0.113124261 | 0.575102313 | 0.80 | 0.53 |
| YLR332W | 1136.948721 | 0.143711942 | 0.575091913 | 0.80 | 0.53 |
| YDR176W | 1688.228212 | -0.102279843 | 0.576199596 | 0.81 | 0.53 |
| YGR135W | 3309.689244 | -0.10936396 | 0.575861217 | 0.81 | 0.53 |
| YKL167C | 608.6144606 | -0.114680294 | 0.576038011 | 0.81 | 0.53 |
| YLR106C | 6573.73141 | 0.211516337 | 0.576280665 | 0.81 | 0.53 |
| YNL136W | 1295.54364 | 0.159308851 | 0.576311441 | 0.81 | 0.53 |
| YNL183C | 1035.511832 | 0.104956638 | 0.57612558 | 0.81 | 0.53 |
| YPL048W | 7812.579467 | -0.094412913 | 0.576327597 | 0.81 | 0.53 |
| YDR276C | 2763.13579 | -0.12496451 | 0.57659498 | 0.81 | 0.53 |
| YER151C | 2013.443986 | -0.091634036 | 0.576530458 | 0.81 | 0.53 |
| YAL026C | 1397.824394 | 0.110811142 | 0.577008139 | 0.81 | 0.53 |
| YBR126W-B | 156.4452467 | 0.182747365 | 0.577530277 | 0.81 | 0.53 |
| YGL007W | 19.49434054 | 0.211466925 | 0.577495917 | 0.81 | 0.53 |
| YIL142W | 3130.184557 | -0.101296144 | 0.577252278 | 0.81 | 0.53 |
| YNL267W | 1939.373479 | -0.06334781 | 0.577394877 | 0.81 | 0.53 |
| YLR084C | 1027.133398 | -0.145406193 | 0.57769922 | 0.81 | 0.53 |
| YLR262C | 283.7623055 | 0.118437067 | 0.577989243 | 0.81 | 0.53 |
| YBR051W | 12.61851356 | 0.2809227 | 0.579258602 | 0.81 | 0.53 |
| YDR213W | 568.9307813 | 0.168955137 | 0.579036261 | 0.81 | 0.53 |
| YDR420W | 1513.909661 | -0.145577048 | 0.578382036 | 0.81 | 0.53 |
| YGL048C | 4170.423196 | 0.086221272 | 0.578740741 | 0.81 | 0.53 |
| YHR141C | 3868.981705 | -0.159509049 | 0.579493613 | 0.81 | 0.53 |
| YLR166C | 1318.264285 | -0.071249338 | 0.57930475 | 0.81 | 0.53 |
| YLR306W | 266.7883268 | -0.139223691 | 0.579029795 | 0.81 | 0.53 |
| YMR024W | 882.2534561 | -0.086037077 | 0.578752196 | 0.81 | 0.53 |
| YNL252C | 974.4576049 | -0.13585437 | 0.578790998 | 0.81 | 0.53 |
| YNR051C | 2926.448335 | -0.14690202 | 0.578838028 | 0.81 | 0.53 |
| YOR220W | 2249.332399 | 0.151863496 | 0.579480389 | 0.81 | 0.53 |
| YGL161C | 1505.046156 | -0.062582429 | 0.580605884 | 0.81 | 0.53 |
| YGR266W | 1152.085768 | 0.12239248 | 0.580069173 | 0.81 | 0.53 |
| YLR447C | 5880.017489 | 0.109631446 | 0.580172405 | 0.81 | 0.53 |
| YMR205C | 7427.558626 | -0.157371175 | 0.580685921 | 0.81 | 0.53 |
| YOL011W | 1428.931177 | -0.148520846 | 0.580623501 | 0.81 | 0.53 |
| YOR164C | 1461.403289 | -0.098418413 | 0.580401069 | 0.81 | 0.53 |
| snR13 | 24.86462889 | -0.223995746 | 0.580344911 | 0.81 | 0.53 |
| YGR197C | 588.1649926 | 0.170790918 | 0.581038298 | 0.81 | 0.53 |
| YIL131C | 677.2016628 | -0.152931966 | 0.581181568 | 0.81 | 0.53 |
| YPR181C | 4442.448271 | -0.094550884 | 0.581089212 | 0.81 | 0.53 |
| snR76 | 8.640963018 | -0.176940775 | 0.581422868 | 0.81 | 0.53 |
| YLR420W | 2792.570329 | -0.184193119 | 0.582012987 | 0.81 | 0.53 |

| | | | | | |
|---|---|---|---|---|---|
| YGL245W | 8715.959502 | 0.157445412 | 0.582252378 | 0.81 | 0.53 |
| YJL158C | 11506.92019 | 0.231003509 | 0.582362251 | 0.81 | 0.53 |
| YGR160W | 19.24237613 | -0.204678741 | 0.582582826 | 0.81 | 0.53 |
| YLR027C | 7716.105415 | -0.105314173 | 0.583335317 | 0.81 | 0.53 |
| YDR100W | 731.7492012 | -0.193688267 | 0.584114644 | 0.81 | 0.54 |
| YOL053W | 775.6100204 | 0.152653769 | 0.584372239 | 0.81 | 0.54 |
| YLR182W | 1421.620302 | -0.121594466 | 0.584540814 | 0.81 | 0.54 |
| YGR209C | 5181.82162 | -0.100675291 | 0.584766856 | 0.81 | 0.54 |
| YLR254C | 554.1871051 | -0.124439228 | 0.584883966 | 0.81 | 0.54 |
| YJL036W | 1275.214013 | 0.111451498 | 0.585040442 | 0.81 | 0.54 |
| YAL042C-A | 8.764234173 | 0.230108366 | 0.585439727 | 0.81 | 0.54 |
| YFR034W-A | 2.98827912 | 0.248703078 | 0.585491667 | 0.81 | 0.54 |
| YGL148W | 4268.613078 | 0.218014881 | 0.585306474 | 0.81 | 0.54 |
| YMR274C | 249.323644 | -0.138137998 | 0.586098767 | 0.81 | 0.54 |
| YPL268W | 929.1963896 | -0.157462907 | 0.586007833 | 0.81 | 0.54 |
| YBR062C | 819.8607412 | 0.115797394 | 0.586690265 | 0.81 | 0.54 |
| YDR159W | 1698.043068 | -0.080615504 | 0.586874151 | 0.81 | 0.54 |
| YLR153C | 3246.572959 | -0.094223989 | 0.58676178 | 0.81 | 0.54 |
| YMR039C | 1898.602952 | -0.142612462 | 0.58690071 | 0.81 | 0.54 |
| YMR102C | 1653.341009 | -0.202782828 | 0.58641023 | 0.81 | 0.54 |
| YOR090C | 1602.791656 | 0.065183888 | 0.586360978 | 0.81 | 0.54 |
| YNL237W | 756.7694566 | -0.145056144 | 0.587546145 | 0.81 | 0.54 |
| YNL195C | 538.1405229 | 0.284910863 | 0.587920793 | 0.81 | 0.54 |
| YPR101W | 539.0655902 | 0.162363828 | 0.588200785 | 0.81 | 0.54 |
| YDR267C | 660.9899359 | -0.075645442 | 0.58881837 | 0.81 | 0.54 |
| YMR080C | 1907.455726 | -0.10748637 | 0.588819811 | 0.81 | 0.54 |
| YOL092W | 1398.16665 | -0.128524573 | 0.588787196 | 0.81 | 0.54 |
| YML035C | 2451.342109 | 0.121596905 | 0.589002131 | 0.81 | 0.54 |
| YOR186W | 27.95959597 | 0.19480796 | 0.589730729 | 0.81 | 0.54 |
| YLR167W | 28914.13243 | -0.149651627 | 0.590486379 | 0.81 | 0.54 |
| YGL052W | 83.8653604 | 0.156162643 | 0.591004699 | 0.81 | 0.54 |
| YGL226W | 346.3222245 | -0.097197984 | 0.591016883 | 0.81 | 0.54 |
| YMR136W | 2477.221542 | 0.177221124 | 0.590988076 | 0.81 | 0.54 |
| YLR220W | 930.48531 | 0.106711084 | 0.591482353 | 0.82 | 0.54 |
| YOR271C | 1520.089436 | 0.253244977 | 0.591712338 | 0.82 | 0.54 |
| YHR064C | 6332.657586 | 0.196590738 | 0.591935631 | 0.82 | 0.54 |
| YOR087W | 1019.099344 | 0.113610785 | 0.592102019 | 0.82 | 0.54 |
| YBL060W | 624.3164774 | 0.097606535 | 0.593496588 | 0.82 | 0.54 |
| YCR069W | 2249.27644 | -0.150701512 | 0.593243782 | 0.82 | 0.54 |
| YDR173C | 369.7980333 | 0.127292961 | 0.593535124 | 0.82 | 0.54 |
| YER040W | 900.0166243 | 0.125657139 | 0.593951501 | 0.82 | 0.54 |
| YGL218W | 11.00510591 | -0.220546141 | 0.594434854 | 0.82 | 0.54 |
| YIR016W | 1147.104809 | 0.141998291 | 0.593201858 | 0.82 | 0.54 |

| | | | | | |
|---|---|---|---|---|---|
| YJL059W | 446.5285115 | 0.117520098 | 0.592699177 | 0.82 | 0.54 |
| YJR009C | 33615.59982 | 0.259576093 | 0.593472329 | 0.82 | 0.54 |
| YKL165C-A | 6.96473103 | 0.173467188 | 0.59445506 | 0.82 | 0.54 |
| YML095C | 255.548255 | -0.098156295 | 0.594071501 | 0.82 | 0.54 |
| YNL289W | 446.6126721 | -0.18923397 | 0.593128915 | 0.82 | 0.54 |
| YNL303W | 97.83657889 | -0.192339066 | 0.593762758 | 0.82 | 0.54 |
| YNR039C | 749.4014615 | -0.071464116 | 0.594407372 | 0.82 | 0.54 |
| YOR290C | 3907.230484 | 0.098514551 | 0.594185096 | 0.82 | 0.54 |
| YPR035W | 16997.33407 | 0.197980637 | 0.593743999 | 0.82 | 0.54 |
| YKL114C | 901.3872837 | -0.111388643 | 0.595239001 | 0.82 | 0.54 |
| YDL143W | 3248.97051 | -0.1093738 | 0.595537419 | 0.82 | 0.54 |
| YML039W | 1.131094598 | -0.189285568 | 0.595692794 | 0.82 | 0.54 |
| YNL004W | 1614.267096 | 0.080853383 | 0.59603544 | 0.82 | 0.54 |
| YPR130C | 10.79124991 | -0.224912518 | 0.596061517 | 0.82 | 0.54 |
| snR82 | 454.6100305 | -0.301206428 | 0.596087604 | 0.82 | 0.54 |
| YKR052C | 558.5950233 | -0.162201895 | 0.596614355 | 0.82 | 0.54 |
| YOR141C | 1550.614472 | 0.090848951 | 0.596643544 | 0.82 | 0.54 |
| YDR169C | 1729.231302 | -0.127159999 | 0.597223552 | 0.82 | 0.54 |
| YLR466C-B | 12.5768575 | 0.289168855 | 0.597116734 | 0.82 | 0.54 |
| YHL035C | 3064.413674 | 0.186396142 | 0.597616046 | 0.82 | 0.54 |
| tR(ACG)J | 0.2026169 | -0.142094046 | 0.597654188 | 0.82 | 0.54 |
| YDL205C | 1039.950483 | 0.184877655 | 0.598221415 | 0.82 | 0.54 |
| YPL152W | 506.4565509 | -0.084574134 | 0.598231411 | 0.82 | 0.54 |
| YCL050C | 4005.943189 | 0.229798628 | 0.598453866 | 0.82 | 0.54 |
| YNL064C | 7494.727656 | 0.256368602 | 0.598567837 | 0.82 | 0.54 |
| YBL071C-B | 1.645198606 | 0.167139999 | 0.598740594 | 0.82 | 0.54 |
| YBR137W | 861.5111114 | 0.161460109 | 0.598855245 | 0.82 | 0.54 |
| YFR032C-B | 19.14700684 | 0.196718382 | 0.599269517 | 0.82 | 0.54 |
| YIL115C | 2357.200299 | 0.054188073 | 0.599468642 | 0.82 | 0.54 |
| YLR230W | 287.1818195 | -0.109651015 | 0.599464402 | 0.82 | 0.54 |
| YOR382W | 2866.519088 | -0.274269469 | 0.60011676 | 0.82 | 0.54 |
| YAL040C | 2404.494026 | 0.144751717 | 0.600362932 | 0.82 | 0.54 |
| YLR085C | 664.5290334 | -0.071311194 | 0.60058235 | 0.82 | 0.54 |
| YDL213C | 1003.203847 | -0.132179142 | 0.600947807 | 0.82 | 0.54 |
| YPR134W | 542.0304792 | -0.083813458 | 0.600968095 | 0.82 | 0.54 |
| YNL265C | 447.0625511 | 0.07654366 | 0.60141045 | 0.82 | 0.54 |
| YML011C | 267.8459983 | -0.082225584 | 0.601600712 | 0.82 | 0.54 |
| YNL105W | 8.716167534 | 0.18286212 | 0.602363269 | 0.82 | 0.54 |
| YPL246C | 2119.845638 | -0.078164281 | 0.602294686 | 0.82 | 0.54 |
| YKL115C | 42.82706794 | -0.106065535 | 0.602823095 | 0.82 | 0.54 |
| YIL039W | 1826.424519 | 0.117805088 | 0.603002451 | 0.82 | 0.54 |
| YGR012W | 677.5451845 | -0.088349206 | 0.603362086 | 0.82 | 0.54 |
| YIL007C | 541.263808 | 0.115779815 | 0.603437432 | 0.82 | 0.54 |

| | | | | | |
|---|---|---|---|---|---|
| YHL002W | 1516.40559 | -0.128761762 | 0.60357758 | 0.82 | 0.54 |
| YKL080W | 5080.565803 | -0.099156688 | 0.604089139 | 0.82 | 0.54 |
| YIL117C | 1107.127061 | 0.309559631 | 0.60448186 | 0.82 | 0.54 |
| YER172C | 2101.126018 | 0.064067252 | 0.604991279 | 0.82 | 0.54 |
| YGR010W | 527.974638 | 0.109094896 | 0.605207675 | 0.82 | 0.54 |
| YML082W | 766.1872079 | -0.088230294 | 0.605146197 | 0.82 | 0.54 |
| YML094C-A | 15.34009962 | -0.136809289 | 0.605375845 | 0.82 | 0.54 |
| YNL033W | 5.453513622 | 0.263308963 | 0.605834627 | 0.82 | 0.54 |
| YGL216W | 962.5040676 | 0.075439526 | 0.606232291 | 0.82 | 0.54 |
| YNL180C | 2386.54101 | -0.192236963 | 0.60644976 | 0.82 | 0.54 |
| YDR106W | 46.31624816 | -0.145437965 | 0.606788346 | 0.82 | 0.55 |
| YDR527W | 984.8264926 | -0.141869684 | 0.606746694 | 0.82 | 0.55 |
| YEL006W | 705.1819844 | -0.091184268 | 0.607578585 | 0.83 | 0.55 |
| YER165C-A | 28.00543952 | 0.153007828 | 0.607634941 | 0.83 | 0.55 |
| YML078W | 2283.677343 | -0.123975293 | 0.60752089 | 0.83 | 0.55 |
| snR42 | 83.98583809 | 0.237065833 | 0.607809118 | 0.83 | 0.55 |
| YMR326C | 0.406623722 | 0.194449197 | 0.607973852 | 0.83 | 0.55 |
| YDR244W | 923.2552739 | 0.079796238 | 0.608320798 | 0.83 | 0.55 |
| YML003W | 222.6670877 | 0.128246094 | 0.608408402 | 0.83 | 0.55 |
| YOR192C-C | 17.72254902 | -0.270088747 | 0.608421298 | 0.83 | 0.55 |
| YHL009C | 603.1387723 | -0.110681339 | 0.608945842 | 0.83 | 0.55 |
| YNL166C | 1009.247872 | 0.159168774 | 0.608959755 | 0.83 | 0.55 |
| YBL059C-A | 504.0749236 | 0.163276184 | 0.609288238 | 0.83 | 0.55 |
| YGL221C | 1756.035958 | 0.105513517 | 0.609760527 | 0.83 | 0.55 |
| YDL028C | 1026.396259 | 0.098526793 | 0.609912441 | 0.83 | 0.55 |
| YBR252W | 608.885071 | -0.140429682 | 0.610372202 | 0.83 | 0.55 |
| YFR009W | 2637.309837 | -0.107692769 | 0.610269405 | 0.83 | 0.55 |
| YLR278C | 1940.968689 | -0.06695539 | 0.610599408 | 0.83 | 0.55 |
| YNL029C | 467.6043522 | 0.106378672 | 0.610790503 | 0.83 | 0.55 |
| YNL099C | 699.7662248 | -0.090097421 | 0.61100678 | 0.83 | 0.55 |
| YFR021W | 746.0129726 | -0.066818073 | 0.611150466 | 0.83 | 0.55 |
| YMR212C | 1054.541147 | -0.119223297 | 0.611418107 | 0.83 | 0.55 |
| YAL064C-A | 47.35030167 | 0.257881359 | 0.612099842 | 0.83 | 0.55 |
| YAR002C-A | 2600.594621 | -0.105213944 | 0.612411787 | 0.83 | 0.55 |
| YCR001W | 33.60447039 | -0.156385514 | 0.612555453 | 0.83 | 0.55 |
| YGL169W | 682.8701947 | -0.133293243 | 0.613605493 | 0.83 | 0.55 |
| YGL197W | 2610.651509 | 0.122058029 | 0.61166407 | 0.83 | 0.55 |
| YGR246C | 1106.975461 | 0.09818081 | 0.612482403 | 0.83 | 0.55 |
| YJR092W | 2324.987782 | 0.148300293 | 0.61326568 | 0.83 | 0.55 |
| YKR022C | 681.4384392 | -0.086402315 | 0.613227529 | 0.83 | 0.55 |
| YLR079W | 2171.764341 | -0.134373171 | 0.613135273 | 0.83 | 0.55 |
| YLR235C | 18.73304548 | 0.171922662 | 0.613195481 | 0.83 | 0.55 |
| YLR261C | 10.17984332 | -0.224493058 | 0.612379687 | 0.83 | 0.55 |

| | | | | | |
|---|---|---|---|---|---|
| YLR275W | 260.6216284 | -0.118717486 | 0.612830194 | 0.83 | 0.55 |
| YLR400W | 49.34542517 | 0.176912792 | 0.613390157 | 0.83 | 0.55 |
| YML071C | 1055.622411 | -0.10427338 | 0.613510958 | 0.83 | 0.55 |
| YML123C | 2594.845401 | -0.164632826 | 0.612840423 | 0.83 | 0.55 |
| YNL118C | 3319.845542 | -0.073264717 | 0.613628654 | 0.83 | 0.55 |
| YPL098C | 678.1300721 | -0.083071454 | 0.613658727 | 0.83 | 0.55 |
| YJL057C | 1203.837592 | 0.170371494 | 0.614156612 | 0.83 | 0.55 |
| YGL186C | 1156.394032 | 0.135792067 | 0.614429777 | 0.83 | 0.55 |
| YCL028W | 2867.256977 | -0.087743889 | 0.614981594 | 0.83 | 0.55 |
| YDR084C | 664.1695601 | 0.098639433 | 0.615010735 | 0.83 | 0.55 |
| YOR042W | 2668.884695 | -0.125284753 | 0.615051931 | 0.83 | 0.55 |
| YPL277C | 206.5316036 | 0.142530993 | 0.615072288 | 0.83 | 0.55 |
| YGL083W | 940.5046339 | -0.081343525 | 0.6152023 | 0.83 | 0.55 |
| YDL142C | 615.3529618 | 0.097902598 | 0.615388145 | 0.83 | 0.55 |
| YFL013W-A | 84.24870204 | 0.097659457 | 0.615710858 | 0.83 | 0.55 |
| YML048W | 4451.705992 | 0.095053785 | 0.615738191 | 0.83 | 0.55 |
| YDR375C | 907.9536977 | -0.070899783 | 0.615920553 | 0.83 | 0.55 |
| YDR478W | 183.0152712 | -0.093540115 | 0.616678239 | 0.83 | 0.55 |
| YJR132W | 1306.62171 | 0.097066255 | 0.616617802 | 0.83 | 0.55 |
| YKL069W | 336.0140703 | -0.099239199 | 0.616065002 | 0.83 | 0.55 |
| YKR051W | 666.9246654 | 0.097834842 | 0.616437795 | 0.83 | 0.55 |
| YOR071C | 651.6605909 | 0.15875253 | 0.616598965 | 0.83 | 0.55 |
| YPR098C | 1032.023923 | 0.139540573 | 0.616393923 | 0.83 | 0.55 |
| YGL167C | 2492.369981 | 0.119055478 | 0.61731956 | 0.83 | 0.55 |
| YGR218W | 3998.195199 | 0.052158187 | 0.617311173 | 0.83 | 0.55 |
| YJL049W | 674.8429005 | -0.121828055 | 0.616986832 | 0.83 | 0.55 |
| YPL225W | 2696.542417 | 0.09041881 | 0.617283408 | 0.83 | 0.55 |
| YBR070C | 438.6999546 | -0.113808711 | 0.617664984 | 0.83 | 0.55 |
| YDR466W | 1549.812513 | 0.079790737 | 0.617959472 | 0.83 | 0.55 |
| YNL087W | 1914.774752 | 0.111347645 | 0.617854833 | 0.83 | 0.55 |
| YDR222W | 466.2824915 | -0.19929617 | 0.618519933 | 0.83 | 0.55 |
| YEL005C | 730.8016186 | -0.142064291 | 0.618448989 | 0.83 | 0.55 |
| YMR022W | 1239.988654 | -0.083933262 | 0.61854226 | 0.83 | 0.55 |
| YJR128W | 14.84064674 | 0.208525586 | 0.619055996 | 0.83 | 0.55 |
| YML025C | 1270.924858 | 0.056985647 | 0.618960232 | 0.83 | 0.55 |
| YML062C | 782.2227983 | -0.084910261 | 0.618917592 | 0.83 | 0.55 |
| YOR021C | 1473.922183 | 0.127428558 | 0.618846706 | 0.83 | 0.55 |
| YGR071C | 709.1837053 | -0.0885502 | 0.619346584 | 0.83 | 0.55 |
| YBR042C | 577.29725 | 0.100724201 | 0.619738311 | 0.83 | 0.55 |
| YDR281C | 42.74964081 | -0.261014494 | 0.619872362 | 0.83 | 0.55 |
| YLR302C | 0.305761563 | 0.180996929 | 0.620088251 | 0.83 | 0.55 |
| YPR019W | 1285.937535 | -0.056261574 | 0.620904078 | 0.83 | 0.55 |
| YDR021W | 377.9993278 | -0.168345521 | 0.621636165 | 0.83 | 0.55 |

| | | | | | |
|---|---|---|---|---|---|
| YFR024C-A | 2497.30533 | 0.08743582 | 0.621742272 | 0.83 | 0.55 |
| YJR058C | 464.9848205 | 0.091023578 | 0.621392453 | 0.83 | 0.55 |
| YOR117W | 3352.309698 | 0.093037147 | 0.62166425 | 0.83 | 0.55 |
| YOL159C-A | 58.15453861 | 0.198428649 | 0.62190168 | 0.83 | 0.55 |
| YGR058W | 565.2067235 | -0.083641442 | 0.622363792 | 0.83 | 0.55 |
| YMR139W | 1446.676083 | 0.121913879 | 0.622422632 | 0.83 | 0.55 |
| YGL242C | 665.893361 | -0.113451728 | 0.622794702 | 0.83 | 0.55 |
| YKL096W-A | 39924.89359 | -0.159069432 | 0.622677469 | 0.83 | 0.55 |
| YDR499W | 868.5785284 | 0.133793706 | 0.623763112 | 0.83 | 0.55 |
| YKL091C | 918.9800837 | 0.162450735 | 0.623302903 | 0.83 | 0.55 |
| YKR089C | 3634.661722 | 0.106298634 | 0.623480624 | 0.83 | 0.55 |
| YML016C | 1375.407219 | 0.093172068 | 0.623334611 | 0.83 | 0.55 |
| YMR179W | 627.9998767 | -0.150371056 | 0.623717241 | 0.83 | 0.55 |
| YOL138C | 1055.538589 | 0.076862804 | 0.623098994 | 0.83 | 0.55 |
| snR30 | 275.4218461 | 0.287169711 | 0.623641808 | 0.83 | 0.55 |
| YIL104C | 354.0212252 | 0.094842455 | 0.623981361 | 0.83 | 0.55 |
| YDL106C | 1181.738765 | -0.121493657 | 0.624598133 | 0.83 | 0.55 |
| YNL024C | 148.4866774 | 0.253052236 | 0.624774679 | 0.83 | 0.55 |
| YER012W | 3318.681234 | 0.081136294 | 0.625313343 | 0.83 | 0.55 |
| YOR176W | 2270.643704 | -0.140997134 | 0.625646404 | 0.83 | 0.55 |
| YDR479C | 1259.888261 | 0.10969751 | 0.626744319 | 0.83 | 0.55 |
| YER100W | 1553.364761 | -0.088368006 | 0.626329759 | 0.83 | 0.55 |
| YIL085C | 1498.284283 | -0.075193432 | 0.626510136 | 0.83 | 0.55 |
| YJL131C | 712.8647158 | 0.107591643 | 0.62664285 | 0.83 | 0.55 |
| YLR110C | 106387.3494 | -0.148210151 | 0.626676585 | 0.83 | 0.55 |
| YOR198C | 6733.130979 | -0.108787571 | 0.626647039 | 0.83 | 0.55 |
| YGR269W | 48.40957868 | 0.093340813 | 0.626991891 | 0.83 | 0.55 |
| YBR107C | 343.2589721 | 0.115954172 | 0.6273124 | 0.83 | 0.55 |
| YML066C | 127.9180195 | -0.178824014 | 0.62731008 | 0.83 | 0.55 |
| YHR019C | 9735.850466 | 0.190452499 | 0.627800827 | 0.84 | 0.55 |
| YNL138W | 4814.982276 | -0.084492786 | 0.62789072 | 0.84 | 0.55 |
| YHL003C | 1333.653462 | -0.127706917 | 0.628214796 | 0.84 | 0.55 |
| YMR232W | 290.2765579 | -0.174388041 | 0.62851086 | 0.84 | 0.55 |
| YDR012W | 4314.996695 | 0.315087305 | 0.629270485 | 0.84 | 0.55 |
| YHR133C | 977.6515666 | -0.125199286 | 0.629414081 | 0.84 | 0.55 |
| YNL245C | 408.0242862 | -0.104626604 | 0.629331789 | 0.84 | 0.55 |
| YNL310C | 605.4678235 | 0.069963442 | 0.629215139 | 0.84 | 0.55 |
| YNL322C | 3831.740608 | 0.124984512 | 0.628831791 | 0.84 | 0.55 |
| YOR233W | 853.5809378 | 0.103620272 | 0.629061789 | 0.84 | 0.55 |
| YPL206C | 1410.718758 | 0.131310609 | 0.629816858 | 0.84 | 0.55 |
| YGR126W | 190.2789593 | 0.117045487 | 0.630044425 | 0.84 | 0.55 |
| YBR213W | 228.053134 | -0.08591394 | 0.630712465 | 0.84 | 0.55 |
| YDR054C | 1985.938955 | -0.109399912 | 0.631058817 | 0.84 | 0.55 |

| | | | | | |
|---|---|---|---|---|---|
| YDR110W | 523.1803163 | 0.119719292 | 0.631926385 | 0.84 | 0.55 |
| YDR289C | 776.5077999 | 0.110105704 | 0.630331599 | 0.84 | 0.55 |
| YER076W-A | 76.7457419 | -0.106228686 | 0.632417454 | 0.84 | 0.55 |
| YFR032C | 47.36854765 | 0.359679879 | 0.631632996 | 0.84 | 0.55 |
| YGL008C | 26806.32689 | 0.34232297 | 0.631539555 | 0.84 | 0.55 |
| YHR032C-A | 12.55755435 | 0.215990485 | 0.631614688 | 0.84 | 0.55 |
| YIL122W | 600.3807645 | -0.154559983 | 0.632310916 | 0.84 | 0.55 |
| YJR010W | 677.4384961 | -0.135449515 | 0.632466857 | 0.84 | 0.55 |
| YKL206C | 563.6605725 | -0.097295246 | 0.631744325 | 0.84 | 0.55 |
| YLR348C | 661.9197696 | -0.181599809 | 0.631974864 | 0.84 | 0.55 |
| YLR350W | 3571.832696 | -0.130008159 | 0.631330001 | 0.84 | 0.55 |
| YLR397C | 907.763381 | 0.109598645 | 0.630552469 | 0.84 | 0.55 |
| YMR057C | 13.35741525 | 0.208807131 | 0.632122203 | 0.84 | 0.55 |
| YMR222C | 712.4011542 | -0.074234644 | 0.632043487 | 0.84 | 0.55 |
| YOR064C | 481.1143371 | 0.1407344 | 0.630869771 | 0.84 | 0.55 |
| YPL058C | 2629.343095 | 0.143735887 | 0.632200412 | 0.84 | 0.55 |
| YDR362C | 764.8048646 | 0.07035019 | 0.632882339 | 0.84 | 0.55 |
| YDR454C | 2624.226849 | -0.104019989 | 0.633227341 | 0.84 | 0.55 |
| YJR056C | 431.2081935 | -0.080385633 | 0.633271221 | 0.84 | 0.55 |
| snR63 | 94.54093227 | -0.036227434 | 0.633984885 | 0.84 | 0.55 |
| YKR029C | 884.4269483 | -0.099034447 | 0.634526959 | 0.84 | 0.55 |
| YPL022W | 1488.077225 | -0.110818544 | 0.634573377 | 0.84 | 0.55 |
| YFL041W | 1108.963199 | 0.099994287 | 0.634912724 | 0.84 | 0.55 |
| YDL056W | 1411.798857 | -0.113811362 | 0.635088408 | 0.84 | 0.55 |
| YKL020C | 2087.802283 | -0.109624754 | 0.635294735 | 0.84 | 0.55 |
| YPR036W-A | 8058.010483 | 0.177482404 | 0.63585558 | 0.84 | 0.55 |
| YBL073W | 6.010181019 | 0.32396917 | 0.635991041 | 0.84 | 0.55 |
| YAL021C | 1595.906288 | -0.072344183 | 0.636835193 | 0.84 | 0.55 |
| YDR381W | 4059.585724 | -0.109963171 | 0.636795213 | 0.84 | 0.55 |
| YER031C | 2004.561338 | 0.132096306 | 0.63686539 | 0.84 | 0.55 |
| YHR174W | 94635.8071 | 0.262447324 | 0.636693275 | 0.84 | 0.55 |
| YIL074C | 1391.44586 | 0.155288113 | 0.636546979 | 0.84 | 0.55 |
| YER007C-A | 951.757768 | -0.087442925 | 0.637750036 | 0.84 | 0.56 |
| YFL059W | 70.79713485 | 0.272036568 | 0.637824066 | 0.84 | 0.56 |
| YAL042W | 3676.592816 | -0.06214603 | 0.638318799 | 0.84 | 0.56 |
| YER087C-A | 36.6678806 | 0.161481585 | 0.638362887 | 0.84 | 0.56 |
| YHR071C-A | 39.05421358 | 0.163614259 | 0.638239549 | 0.84 | 0.56 |
| YML015C | 859.349761 | -0.123703386 | 0.638176791 | 0.84 | 0.56 |
| YLR282C | 4.588773458 | 0.24547415 | 0.638909693 | 0.84 | 0.56 |
| YDR411C | 1171.452043 | -0.122499448 | 0.639291955 | 0.84 | 0.56 |
| YGL109W | 79.04302724 | 0.095818994 | 0.639399544 | 0.84 | 0.56 |
| YNL150W | 83.29807569 | -0.110113498 | 0.639324556 | 0.84 | 0.56 |
| YBL008W-A | 2.748531795 | -0.160783328 | 0.639701898 | 0.84 | 0.56 |

| | | | | | |
|---|---|---|---|---|---|
| YMR253C | 606.2113411 | 0.122971632 | 0.639911935 | 0.84 | 0.56 |
| YGL086W | 1388.801297 | 0.107218985 | 0.640395242 | 0.84 | 0.56 |
| YKL222C | 519.5767688 | 0.12091515 | 0.640431196 | 0.84 | 0.56 |
| YMR320W | 113.0593369 | 0.180361495 | 0.640262254 | 0.84 | 0.56 |
| YDL217C | 416.6172424 | -0.082474275 | 0.640613036 | 0.84 | 0.56 |
| YBR197C | 458.4082334 | -0.119449783 | 0.640780034 | 0.84 | 0.56 |
| YGR243W | 1286.002734 | -0.312544661 | 0.641365049 | 0.84 | 0.56 |
| YNR040W | 422.1360755 | 0.101298537 | 0.641245947 | 0.84 | 0.56 |
| YBR232C | 24.08316402 | 0.16649955 | 0.641601303 | 0.84 | 0.56 |
| YOL108C | 360.9401457 | 0.136853229 | 0.641636364 | 0.84 | 0.56 |
| YBL094C | 5.123149382 | 0.23120506 | 0.641948644 | 0.84 | 0.56 |
| YER032W | 1018.880889 | -0.114147065 | 0.642469348 | 0.84 | 0.56 |
| YFL034C-B | 921.0599209 | -0.127212876 | 0.642640207 | 0.84 | 0.56 |
| YOR314W-A | 0.578252409 | 0.170390711 | 0.642590972 | 0.84 | 0.56 |
| YBL023C | 762.982123 | 0.091069386 | 0.643290183 | 0.84 | 0.56 |
| YDL091C | 1187.448267 | 0.132047969 | 0.64309895 | 0.84 | 0.56 |
| YGR099W | 582.6782539 | 0.104964996 | 0.643320876 | 0.84 | 0.56 |
| YJR034W | 337.4607648 | -0.113329251 | 0.643082457 | 0.84 | 0.56 |
| YKL104C | 3816.518475 | 0.167818476 | 0.64338711 | 0.84 | 0.56 |
| YMR238W | 935.1001777 | 0.18162428 | 0.643640675 | 0.84 | 0.56 |
| YPR165W | 4589.151851 | -0.077786085 | 0.643806006 | 0.84 | 0.56 |
| YBR093C | 949.3588178 | -0.232254903 | 0.644353415 | 0.84 | 0.56 |
| YCL001W-A | 26.77980078 | 0.202195464 | 0.64475525 | 0.84 | 0.56 |
| YGR167W | 3020.873713 | -0.06633605 | 0.644660899 | 0.84 | 0.56 |
| YHL015W | 17106.50707 | -0.123998443 | 0.644699601 | 0.84 | 0.56 |
| YHR068W | 2922.590435 | 0.199130347 | 0.644820374 | 0.84 | 0.56 |
| YOL166W-A | 0.347622843 | 0.159529412 | 0.644297574 | 0.84 | 0.56 |
| YBR078W | 17879.15973 | -0.132462632 | 0.645079522 | 0.84 | 0.56 |
| snR60 | 11.58332067 | 0.18472248 | 0.645155313 | 0.84 | 0.56 |
| YKR037C | 412.1583252 | -0.060516825 | 0.646161047 | 0.84 | 0.56 |
| YML021C | 560.3287148 | 0.093590698 | 0.646428022 | 0.85 | 0.56 |
| YAL064W | 3.131576444 | 0.330952599 | 0.647325067 | 0.85 | 0.56 |
| YBR096W | 835.4279583 | -0.100570705 | 0.647017842 | 0.85 | 0.56 |
| YDR092W | 1261.21405 | -0.076446367 | 0.647382524 | 0.85 | 0.56 |
| YDR330W | 1850.284876 | 0.105422415 | 0.646746638 | 0.85 | 0.56 |
| YGR039W | 40.79718114 | -0.11120778 | 0.646962929 | 0.85 | 0.56 |
| YGR199W | 855.4590705 | 0.093019808 | 0.647142642 | 0.85 | 0.56 |
| YDR022C | 413.59294 | 0.144132257 | 0.648383922 | 0.85 | 0.56 |
| YGL151W | 1728.404115 | -0.080936472 | 0.648473848 | 0.85 | 0.56 |
| YJL008C | 2286.818099 | -0.091995294 | 0.648980646 | 0.85 | 0.56 |
| YLR186W | 830.8214037 | -0.106624337 | 0.648804121 | 0.85 | 0.56 |
| YMR299C | 640.5800002 | -0.102399083 | 0.648739497 | 0.85 | 0.56 |
| YOL014W | 428.9922641 | -0.200707926 | 0.648223579 | 0.85 | 0.56 |

| | | | | | |
|---|---|---|---|---|---|
| YOR103C | 111.7299261 | 0.108501403 | 0.648200358 | 0.85 | 0.56 |
| YOR108W | 1787.909268 | -0.124415576 | 0.64834861 | 0.85 | 0.56 |
| YPR106W | 760.0851492 | -0.093507042 | 0.648868848 | 0.85 | 0.56 |
| YNL210W | 106.0752581 | -0.140577777 | 0.649144681 | 0.85 | 0.56 |
| YNL156C | 1413.410846 | 0.072954973 | 0.64957306 | 0.85 | 0.56 |
| YOR313C | 174.7411276 | 0.325189031 | 0.649593952 | 0.85 | 0.56 |
| YDL073W | 971.6916673 | 0.09901749 | 0.649748659 | 0.85 | 0.56 |
| YMR122C | 59.11544635 | 0.14113228 | 0.649975079 | 0.85 | 0.56 |
| YKR105C | 96.82105819 | -0.170034166 | 0.650346367 | 0.85 | 0.56 |
| YDL188C | 1068.696328 | -0.093057191 | 0.650738339 | 0.85 | 0.56 |
| YDR123C | 353.8570592 | -0.133545513 | 0.651305988 | 0.85 | 0.56 |
| YNR036C | 1583.864307 | -0.127824357 | 0.651319042 | 0.85 | 0.56 |
| YDR468C | 586.9211769 | -0.085990824 | 0.651662611 | 0.85 | 0.56 |
| YDR193W | 12.98139175 | 0.204878272 | 0.652072876 | 0.85 | 0.56 |
| YBL107C | 641.7007619 | -0.124397584 | 0.653126664 | 0.85 | 0.56 |
| YBR298C-A | 3.08929577 | -0.11146017 | 0.653881839 | 0.85 | 0.56 |
| YGR173W | 798.4212433 | -0.119036675 | 0.653827989 | 0.85 | 0.56 |
| YIR006C | 5095.375019 | 0.066335645 | 0.654133889 | 0.85 | 0.56 |
| YKL075C | 477.0577168 | 0.084364539 | 0.653720892 | 0.85 | 0.56 |
| YLR222C-A | 13.05158283 | -0.155925339 | 0.654095435 | 0.85 | 0.56 |
| YMR208W | 1260.14289 | 0.102026002 | 0.65401653 | 0.85 | 0.56 |
| YDR287W | 832.7406735 | 0.116532743 | 0.654767019 | 0.85 | 0.56 |
| YLR368W | 767.6389144 | 0.099158674 | 0.655131509 | 0.85 | 0.56 |
| YOL027C | 2880.865523 | 0.097687754 | 0.655160613 | 0.85 | 0.56 |
| YBR036C | 2605.132077 | 0.091415311 | 0.656819132 | 0.85 | 0.56 |
| YIL071W-A | 73.5379669 | 0.072544624 | 0.656474331 | 0.85 | 0.56 |
| YIL108W | 1452.736233 | 0.132382123 | 0.656692227 | 0.85 | 0.56 |
| YJR107W | 474.7713781 | 0.120014398 | 0.656739689 | 0.85 | 0.56 |
| YKL082C | 1057.576571 | -0.113491741 | 0.656238178 | 0.85 | 0.56 |
| YNL280C | 1498.81009 | 0.136615695 | 0.65637045 | 0.85 | 0.56 |
| YOL085W-A | 20.71787704 | 0.21405015 | 0.656715647 | 0.85 | 0.56 |
| YPL062W | 4.463478236 | 0.275672317 | 0.656708494 | 0.85 | 0.56 |
| YPL233W | 252.806959 | 0.073762692 | 0.655898565 | 0.85 | 0.56 |
| YBR011C | 18392.14922 | -0.108182583 | 0.657641039 | 0.85 | 0.56 |
| YDR431W | 11.62652329 | 0.202866741 | 0.657471641 | 0.85 | 0.56 |
| YIL031W | 1396.422579 | 0.082274738 | 0.657722606 | 0.85 | 0.56 |
| YDL174C | 2966.298062 | 0.145367526 | 0.65816999 | 0.85 | 0.56 |
| YER055C | 2945.986692 | -0.100429767 | 0.65870006 | 0.85 | 0.56 |
| YFL013C | 826.2401076 | 0.085439031 | 0.658585646 | 0.85 | 0.56 |
| YGR072W | 838.1301736 | 0.080999214 | 0.658506014 | 0.85 | 0.56 |
| YKL155C | 791.2689341 | 0.073896154 | 0.658170558 | 0.85 | 0.56 |
| YMR018W | 66.38637523 | 0.153298641 | 0.658696089 | 0.85 | 0.56 |
| YNL294C | 1109.535055 | 0.115168549 | 0.658529734 | 0.85 | 0.56 |

| | | | | | |
|---|---|---|---|---|---|
| YBR121C | 7616.904588 | -0.100368181 | 0.659020447 | 0.85 | 0.56 |
| YLR107W | 1035.729202 | 0.080427466 | 0.659066967 | 0.85 | 0.56 |
| YPL170W | 622.4061116 | -0.070191053 | 0.659253346 | 0.85 | 0.56 |
| YBR285W | 449.6014678 | 0.203817478 | 0.659610064 | 0.85 | 0.56 |
| YDL162C | 2.875843937 | -0.087086927 | 0.65955367 | 0.85 | 0.56 |
| YBR126W-A | 695.852559 | -0.16342528 | 0.660432211 | 0.85 | 0.56 |
| YDL184C | 118.3516335 | -0.138379292 | 0.66054835 | 0.85 | 0.56 |
| YDR179W-A | 321.5877637 | -0.092356972 | 0.660114663 | 0.85 | 0.56 |
| YML116W-A | 166.4577303 | 0.088963752 | 0.660384054 | 0.85 | 0.56 |
| YNL218W | 370.2566133 | 0.098738273 | 0.660309799 | 0.85 | 0.56 |
| YAR007C | 1627.673812 | -0.083025616 | 0.661013124 | 0.85 | 0.56 |
| YNL003C | 715.0467326 | -0.109328083 | 0.660893325 | 0.85 | 0.56 |
| YOL035C | 58.4740954 | 0.116548756 | 0.661022084 | 0.85 | 0.56 |
| YFR052W | 2758.803837 | 0.094717246 | 0.661284113 | 0.85 | 0.56 |
| YJL029C | 1512.843835 | -0.060648958 | 0.662134413 | 0.85 | 0.56 |
| YLR072W | 910.1580571 | 0.072235826 | 0.662132861 | 0.85 | 0.56 |
| YML092C | 2758.585269 | -0.06317444 | 0.661970103 | 0.85 | 0.56 |
| snR47 | 5.450594875 | 0.272705883 | 0.662250784 | 0.85 | 0.56 |
| YBR225W | 1153.176358 | 0.106467161 | 0.662988453 | 0.85 | 0.56 |
| YIL109C | 4372.511551 | -0.058657749 | 0.662820726 | 0.85 | 0.56 |
| YOR112W | 1049.770792 | -0.059960454 | 0.663021913 | 0.85 | 0.56 |
| YJL078C | 3272.163203 | 0.102821903 | 0.663194564 | 0.85 | 0.56 |
| YBL044W | 3.828780377 | -0.219114624 | 0.663485211 | 0.85 | 0.56 |
| YPR103W | 3867.741987 | -0.092007842 | 0.663655428 | 0.85 | 0.56 |
| YHR199C-A | 146.5882374 | 0.172838558 | 0.663896224 | 0.85 | 0.56 |
| YBR106W | 4289.033597 | -0.158570935 | 0.664093557 | 0.85 | 0.56 |
| YOR332W | 2877.284969 | -0.071981934 | 0.664387078 | 0.85 | 0.56 |
| YPR178W | 455.7725824 | 0.07190396 | 0.664400572 | 0.85 | 0.56 |
| YOR381W-A | 9.096748957 | 0.219735282 | 0.664628842 | 0.85 | 0.56 |
| YJL183W | 732.387825 | -0.084779888 | 0.66484083 | 0.85 | 0.56 |
| YDR040C | 132.5886651 | 0.121764084 | 0.665045899 | 0.85 | 0.56 |
| YDR525W | 4.560372998 | -0.243734096 | 0.66539576 | 0.85 | 0.56 |
| YHR142W | 1336.189865 | 0.136150908 | 0.665428229 | 0.85 | 0.56 |
| YNL083W | 670.395816 | 0.115856836 | 0.665455158 | 0.85 | 0.56 |
| YEL052W | 1984.785259 | 0.103731214 | 0.665787988 | 0.85 | 0.56 |
| YLR245C | 188.3206886 | 0.119920957 | 0.665679242 | 0.85 | 0.56 |
| YBL071C | 215.271228 | -0.127056461 | 0.666059648 | 0.85 | 0.56 |
| YER170W | 376.8018239 | 0.077142484 | 0.666580754 | 0.86 | 0.57 |
| YGR046W | 906.1577966 | -0.137165126 | 0.666481724 | 0.86 | 0.57 |
| YPL261C | 48.62531076 | 0.103337216 | 0.66661591 | 0.86 | 0.57 |
| YBL039W-B | 151.3574507 | 0.104266835 | 0.667047802 | 0.86 | 0.57 |
| YDR277C | 2772.925083 | 0.148961511 | 0.667154427 | 0.86 | 0.57 |
| YDR346C | 2820.625886 | 0.096891493 | 0.667339346 | 0.86 | 0.57 |

| | | | | | |
|---|---|---|---|---|---|
| YLR059C | 1264.300383 | 0.089110262 | 0.667436463 | 0.86 | 0.57 |
| YBL074C | 232.3789197 | 0.104429251 | 0.667816116 | 0.86 | 0.57 |
| YDL176W | 672.0622412 | 0.082075022 | 0.668065183 | 0.86 | 0.57 |
| YGR204C-A | 13.54650687 | 0.19918303 | 0.668358298 | 0.86 | 0.57 |
| YNL095C | 702.5523068 | 0.096519014 | 0.668345882 | 0.86 | 0.57 |
| YNL272C | 1304.645862 | -0.063196886 | 0.668109054 | 0.86 | 0.57 |
| YNR027W | 549.891736 | 0.106467005 | 0.667966627 | 0.86 | 0.57 |
| YOR073W | 590.4176603 | -0.103195136 | 0.668298152 | 0.86 | 0.57 |
| YDR399W | 3185.764567 | -0.031633473 | 0.669243696 | 0.86 | 0.57 |
| YGL212W | 286.2266484 | -0.078245378 | 0.66974401 | 0.86 | 0.57 |
| YPL229W | 514.8866138 | -0.129167883 | 0.670607126 | 0.86 | 0.57 |
| YER021W | 5518.8989 | 0.080541927 | 0.670796001 | 0.86 | 0.57 |
| YER064C | 1674.781352 | -0.120829771 | 0.670986809 | 0.86 | 0.57 |
| YBR183W | 786.7022482 | 0.14934296 | 0.671581574 | 0.86 | 0.57 |
| YHL039W | 1028.011858 | -0.096981353 | 0.671533813 | 0.86 | 0.57 |
| YDR426C | 18.4381422 | -0.167997107 | 0.672864565 | 0.86 | 0.57 |
| YDR477W | 1886.671826 | 0.05589215 | 0.672072177 | 0.86 | 0.57 |
| YGL016W | 794.4716524 | -0.123773238 | 0.672725951 | 0.86 | 0.57 |
| YGR080W | 1465.883709 | 0.075849409 | 0.672484982 | 0.86 | 0.57 |
| YHL001W | 1027.533504 | -0.101720473 | 0.672265638 | 0.86 | 0.57 |
| YJL218W | 146.9624687 | -0.150401502 | 0.673152 | 0.86 | 0.57 |
| YMR064W | 1294.977576 | -0.087065674 | 0.673031607 | 0.86 | 0.57 |
| YMR193W | 776.035786 | 0.114145286 | 0.673154642 | 0.86 | 0.57 |
| YOL086W-A | 318.8786531 | 0.1440653 | 0.673042683 | 0.86 | 0.57 |
| YOR330C | 1527.333333 | -0.070557093 | 0.67227011 | 0.86 | 0.57 |
| YBL025W | 212.6114748 | 0.113420172 | 0.673612718 | 0.86 | 0.57 |
| YJR025C | 1368.606225 | 0.267867524 | 0.673643731 | 0.86 | 0.57 |
| YLR132C | 337.4595812 | 0.058656413 | 0.673716931 | 0.86 | 0.57 |
| YDR065W | 277.2481422 | -0.08839663 | 0.674849015 | 0.86 | 0.57 |
| YDR220C | 3.958958383 | -0.266669028 | 0.674784621 | 0.86 | 0.57 |
| YDR296W | 1076.454873 | -0.087762815 | 0.674508754 | 0.86 | 0.57 |
| YMR059W | 205.0702465 | -0.058439489 | 0.67473648 | 0.86 | 0.57 |
| YNL076W | 1058.446405 | -0.065979166 | 0.675431544 | 0.86 | 0.57 |
| YLL007C | 554.0715465 | -0.099655722 | 0.675955555 | 0.86 | 0.57 |
| YOR008W-B | 4.890160442 | -0.201903737 | 0.676043665 | 0.86 | 0.57 |
| snR18 | 14.34494847 | 0.316828182 | 0.676037824 | 0.86 | 0.57 |
| YLR172C | 1976.093187 | -0.103106739 | 0.676276632 | 0.86 | 0.57 |
| YOR111W | 390.0644929 | 0.074063096 | 0.676703525 | 0.86 | 0.57 |
| YLR250W | 1395.702651 | -0.082079883 | 0.67698565 | 0.86 | 0.57 |
| YOL067C | 283.6975055 | 0.095641804 | 0.677016022 | 0.86 | 0.57 |
| YNL042W-B | 14.4556208 | 0.27253227 | 0.677404117 | 0.86 | 0.57 |
| YNL123W | 2060.867856 | -0.100445973 | 0.677607076 | 0.86 | 0.57 |
| YAL026C-A | 170.5922701 | -0.121002184 | 0.678510947 | 0.86 | 0.57 |

| | | | | | |
|---|---|---|---|---|---|
| YLR347C | 1405.641169 | 0.07400364 | 0.678525537 | 0.86 | 0.57 |
| YJR076C | 1668.359247 | 0.062483787 | 0.678707438 | 0.86 | 0.57 |
| YHR180W | 212.5255364 | 0.141278437 | 0.679030553 | 0.86 | 0.57 |
| YBL050W | 1823.454847 | -0.065551196 | 0.681256132 | 0.86 | 0.57 |
| YBR095C | 539.8071954 | -0.071376826 | 0.679645049 | 0.86 | 0.57 |
| YBR104W | 350.9392427 | -0.10904439 | 0.67984179 | 0.86 | 0.57 |
| YCR010C | 273.3869127 | 0.095236956 | 0.680793654 | 0.86 | 0.57 |
| YDR227W | 1551.753891 | 0.11580747 | 0.679943861 | 0.86 | 0.57 |
| YDR274C | 62.88697688 | 0.144939884 | 0.679481495 | 0.86 | 0.57 |
| YDR279W | 575.5796481 | -0.06459889 | 0.680289312 | 0.86 | 0.57 |
| YEL057C | 119.2283707 | 0.236714534 | 0.681434633 | 0.86 | 0.57 |
| YFL010C | 2890.139798 | -0.107648094 | 0.68142768 | 0.86 | 0.57 |
| YGR026W | 4521.168167 | -0.076062595 | 0.680233447 | 0.86 | 0.57 |
| YKL051W | 1194.148337 | -0.121643769 | 0.681698897 | 0.86 | 0.57 |
| YLL020C | 30.70630028 | 0.162250733 | 0.681023021 | 0.86 | 0.57 |
| YLR242C | 302.599813 | -0.077331598 | 0.681656949 | 0.86 | 0.57 |
| YMR060C | 493.2206546 | -0.072039807 | 0.680835855 | 0.86 | 0.57 |
| YMR200W | 1797.595559 | 0.079893408 | 0.680075698 | 0.86 | 0.57 |
| YMR267W | 919.8795189 | -0.104253318 | 0.680800291 | 0.86 | 0.57 |
| YNL097C | 784.7174827 | 0.084056579 | 0.680994775 | 0.86 | 0.57 |
| YNR020C | 315.5485889 | -0.071642005 | 0.681576191 | 0.86 | 0.57 |
| YPR200C | 70.27684883 | -0.147558468 | 0.68076892 | 0.86 | 0.57 |
| YBR003W | 1468.050788 | 0.062193655 | 0.681871535 | 0.86 | 0.57 |
| YGL241W | 933.7531382 | -0.056583435 | 0.682077177 | 0.86 | 0.57 |
| YOR247W | 5496.630424 | -0.093150554 | 0.682300889 | 0.86 | 0.57 |
| YBR057C | 866.0815496 | -0.102907272 | 0.682917156 | 0.86 | 0.57 |
| YDR295C | 798.8722941 | 0.057296939 | 0.682946233 | 0.86 | 0.57 |
| YGL159W | 407.2189918 | -0.099280506 | 0.683263966 | 0.86 | 0.57 |
| YAL068W-A | 0.241654238 | 0.146115896 | 0.683898818 | 0.86 | 0.57 |
| YDR056C | 1611.517655 | 0.068007953 | 0.683924699 | 0.86 | 0.57 |
| YKL077W | 3042.957164 | -0.06336122 | 0.684033403 | 0.86 | 0.57 |
| YGL153W | 279.9517565 | 0.073816114 | 0.684742019 | 0.86 | 0.57 |
| YMR068W | 420.8665046 | 0.080957079 | 0.684675696 | 0.86 | 0.57 |
| YGR202C | 1624.722457 | 0.061759941 | 0.685337464 | 0.87 | 0.57 |
| YOR006C | 758.4019018 | -0.077202002 | 0.685277073 | 0.87 | 0.57 |
| YOL140W | 502.3006349 | 0.171009409 | 0.685478326 | 0.87 | 0.57 |
| YOL128C | 161.4112011 | -0.100579711 | 0.685742096 | 0.87 | 0.57 |
| YBR081C | 1882.249025 | 0.085065727 | 0.685879507 | 0.87 | 0.57 |
| YDR373W | 760.4394828 | 0.061342082 | 0.686427139 | 0.87 | 0.57 |
| snR37 | 153.9250937 | 0.244618437 | 0.686519054 | 0.87 | 0.57 |
| YCR033W | 2347.054478 | -0.099206281 | 0.686955197 | 0.87 | 0.57 |
| YHR175W | 775.2689431 | 0.114467676 | 0.687080315 | 0.87 | 0.57 |
| YNL067W-A | 0.709210999 | 0.23273524 | 0.686936104 | 0.87 | 0.57 |

| | | | | | |
|---|---|---|---|---|---|
| YGR174W-A | 47.91809201 | 0.208311678 | 0.687435584 | 0.87 | 0.57 |
| YNL209W | 13780.76239 | -0.123660886 | 0.687459825 | 0.87 | 0.57 |
| YAL031C | 1097.389337 | 0.090250025 | 0.688301312 | 0.87 | 0.57 |
| YCR100C | 233.2545617 | 0.169005435 | 0.688097636 | 0.87 | 0.57 |
| YGL239C | 18.18533761 | 0.153962982 | 0.688089734 | 0.87 | 0.57 |
| YGR193C | 2486.614356 | 0.062337796 | 0.688177438 | 0.87 | 0.57 |
| YOR016C | 992.9033585 | 0.069805637 | 0.688183339 | 0.87 | 0.57 |
| YPR183W | 3494.823853 | -0.074237334 | 0.687962496 | 0.87 | 0.57 |
| YDL018C | 497.6776418 | 0.118389385 | 0.688629877 | 0.87 | 0.57 |
| YMR312W | 719.0027121 | -0.083334286 | 0.689082883 | 0.87 | 0.57 |
| YBL055C | 719.3024768 | -0.075207756 | 0.689350788 | 0.87 | 0.57 |
| YER155C | 4972.789594 | -0.06451644 | 0.689849042 | 0.87 | 0.57 |
| YIL120W | 363.836718 | 0.116963881 | 0.690270066 | 0.87 | 0.57 |
| YEL042W | 1714.856815 | -0.086259083 | 0.690583569 | 0.87 | 0.57 |
| YOL034W | 735.3581814 | 0.083184591 | 0.691161683 | 0.87 | 0.57 |
| YDR491C | 102.3610769 | 0.160709391 | 0.691499692 | 0.87 | 0.57 |
| YNL264C | 1013.510724 | 0.118667772 | 0.691332549 | 0.87 | 0.57 |
| YPL224C | 949.2587858 | 0.095683692 | 0.691596308 | 0.87 | 0.57 |
| YDR163W | 579.7588355 | 0.078130507 | 0.692261991 | 0.87 | 0.57 |
| YFL041W-A | 23.17885335 | 0.129322895 | 0.692419725 | 0.87 | 0.57 |
| YHR169W | 496.2218428 | -0.105706073 | 0.69246109 | 0.87 | 0.57 |
| YLR066W | 1030.577166 | -0.071903092 | 0.693344563 | 0.87 | 0.58 |
| YDR336W | 301.9017741 | -0.089732767 | 0.693924069 | 0.87 | 0.58 |
| YHR060W | 732.5771452 | 0.084603552 | 0.693846876 | 0.87 | 0.58 |
| YNR075C-A | 12.77963038 | -0.131555337 | 0.693861898 | 0.87 | 0.58 |
| YOR188W | 1393.160444 | -0.070527025 | 0.693578834 | 0.87 | 0.58 |
| YLR401C | 1341.53932 | -0.07294519 | 0.694194691 | 0.87 | 0.58 |
| YGR223C | 1134.541576 | 0.09275653 | 0.694578894 | 0.87 | 0.58 |
| YJL225C | 0.280381926 | 0.120945035 | 0.694812302 | 0.87 | 0.58 |
| YJR041C | 1150.543526 | -0.1062165 | 0.694693818 | 0.87 | 0.58 |
| YPR069C | 2243.3628 | -0.12807596 | 0.695066696 | 0.87 | 0.58 |
| YBR067C | 11273.4348 | -0.125457045 | 0.69604385 | 0.87 | 0.58 |
| YEL069C | 125.5249506 | 0.174291388 | 0.696133279 | 0.87 | 0.58 |
| YJL030W | 384.5980723 | -0.074962242 | 0.696088812 | 0.87 | 0.58 |
| YJL123C | 2650.72374 | -0.076679704 | 0.695694933 | 0.87 | 0.58 |
| YOR105W | 21.94253227 | -0.114155669 | 0.695787368 | 0.87 | 0.58 |
| YJR082C | 608.3949667 | -0.082501485 | 0.696302882 | 0.87 | 0.58 |
| YOL132W | 124.9358227 | 0.176649244 | 0.696510057 | 0.87 | 0.58 |
| YMR194C-A | 22.19182237 | -0.128868478 | 0.696647642 | 0.87 | 0.58 |
| YBL018C | 137.4735368 | -0.081116256 | 0.69683034 | 0.87 | 0.58 |
| YPR045C | 1121.270528 | 0.073128061 | 0.697021154 | 0.87 | 0.58 |
| YDR325W | 750.6903469 | 0.09019468 | 0.697968379 | 0.87 | 0.58 |
| YLR042C | 282.794653 | 0.26803404 | 0.698289367 | 0.87 | 0.58 |

| | | | | | |
|---|---|---|---|---|---|
| YPR125W | 636.6611003 | -0.07503967 | 0.698624942 | 0.87 | 0.58 |
| YAL045C | 8.89025023 | -0.207250362 | 0.699209442 | 0.87 | 0.58 |
| YNL247W | 4374.768503 | -0.114097726 | 0.699116939 | 0.87 | 0.58 |
| YOR394C-A | 4.135778786 | -0.106810158 | 0.698986658 | 0.87 | 0.58 |
| YBR155W | 833.2201083 | -0.071337372 | 0.69957795 | 0.87 | 0.58 |
| YJL149W | 1033.010855 | 0.060538148 | 0.699783728 | 0.87 | 0.58 |
| YKL096C-B | 8.469479675 | 0.362206851 | 0.699973365 | 0.87 | 0.58 |
| YNL279W | 510.6368292 | 0.169514852 | 0.700020014 | 0.87 | 0.58 |
| YLR428C | 15.96009415 | -0.141289244 | 0.700162371 | 0.87 | 0.58 |
| YIR002C | 1205.196083 | 0.087634036 | 0.700329709 | 0.87 | 0.58 |
| YKR065C | 1072.834618 | -0.069744522 | 0.700509561 | 0.87 | 0.58 |
| YGL220W | 745.8780305 | -0.07629832 | 0.700818237 | 0.87 | 0.58 |
| YLR226W | 384.869465 | 0.055932938 | 0.70086384 | 0.87 | 0.58 |
| YBR020W | 310.6456705 | -0.149181023 | 0.701310546 | 0.87 | 0.58 |
| YOR388C | 10.58733181 | 0.226919539 | 0.701309979 | 0.87 | 0.58 |
| YLR402W | 27.81687827 | -0.094122966 | 0.70196117 | 0.87 | 0.58 |
| YER075C | 845.204575 | -0.064037682 | 0.703523506 | 0.88 | 0.58 |
| YGL129C | 1219.243345 | -0.071782724 | 0.703289877 | 0.88 | 0.58 |
| YKL133C | 601.5672696 | 0.096768998 | 0.703151441 | 0.88 | 0.58 |
| YOR225W | 54.88254174 | 0.237427436 | 0.703025577 | 0.88 | 0.58 |
| YOR322C | 1313.48176 | -0.067782228 | 0.703430314 | 0.88 | 0.58 |
| YPL138C | 483.5208101 | 0.061875203 | 0.703699724 | 0.88 | 0.58 |
| YGR239C | 410.924928 | 0.090925155 | 0.703940241 | 0.88 | 0.58 |
| YLR310C | 3300.526123 | 0.083663087 | 0.70440504 | 0.88 | 0.58 |
| YBL029C-A | 1079.485322 | 0.119940062 | 0.70524231 | 0.88 | 0.58 |
| YKL025C | 1634.326653 | -0.061895361 | 0.705064015 | 0.88 | 0.58 |
| YKR080W | 1412.607481 | 0.31441408 | 0.705251047 | 0.88 | 0.58 |
| YLR020C | 866.2619741 | 0.097291328 | 0.705168537 | 0.88 | 0.58 |
| YLR051C | 783.2029443 | -0.103369678 | 0.705417864 | 0.88 | 0.58 |
| YBR097W | 952.5745358 | -0.059337678 | 0.707231546 | 0.88 | 0.58 |
| YDR169C-A | 13.43403719 | -0.133948205 | 0.706340535 | 0.88 | 0.58 |
| YGR061C | 5651.499061 | -0.07512882 | 0.707019334 | 0.88 | 0.58 |
| YIL124W | 2615.425395 | -0.097900497 | 0.707364365 | 0.88 | 0.58 |
| YIR012W | 1025.35668 | -0.085534606 | 0.706887452 | 0.88 | 0.58 |
| YKL098W | 807.0315194 | -0.089132571 | 0.706969408 | 0.88 | 0.58 |
| YMR087W | 259.2299237 | -0.090113122 | 0.706532284 | 0.88 | 0.58 |
| YOR147W | 1293.879256 | 0.105421613 | 0.707249849 | 0.88 | 0.58 |
| YPL086C | 1089.007189 | -0.09225549 | 0.706941777 | 0.88 | 0.58 |
| YDL137W | 4443.238326 | 0.07610923 | 0.707610731 | 0.88 | 0.58 |
| YDR139C | 266.4093081 | 0.082208994 | 0.708072778 | 0.88 | 0.58 |
| YDR445C | 3.142353253 | -0.086596624 | 0.707741205 | 0.88 | 0.58 |
| YJL084C | 1615.45903 | 0.073082199 | 0.708139996 | 0.88 | 0.58 |
| YOR009W | 307.6360112 | -0.139302785 | 0.708249112 | 0.88 | 0.58 |

| | | | | | |
|---|---|---|---|---|---|
| YPR108W | 2749.924825 | -0.063244714 | 0.70813554 | 0.88 | 0.58 |
| YMR127C | 400.3588714 | -0.084518964 | 0.708802258 | 0.88 | 0.58 |
| YIL115W-A | 23.70554908 | -0.09341339 | 0.709014374 | 0.88 | 0.58 |
| YBL017C | 3980.2423 | 0.090761485 | 0.70938784 | 0.88 | 0.58 |
| YIL154C | 2075.807869 | 0.08040372 | 0.709289623 | 0.88 | 0.58 |
| YJL053W | 1232.219717 | 0.065570611 | 0.710139418 | 0.88 | 0.58 |
| YDR383C | 354.9421834 | -0.079502384 | 0.710537179 | 0.88 | 0.58 |
| YCL045C | 3038.105977 | -0.077380975 | 0.710709721 | 0.88 | 0.58 |
| YEL004W | 410.7288824 | -0.062322087 | 0.711041001 | 0.88 | 0.58 |
| YGL061C | 446.8633151 | -0.084683833 | 0.7114055 | 0.88 | 0.58 |
| YLR068W | 534.5964698 | -0.110873262 | 0.711401364 | 0.88 | 0.58 |
| YDR270W | 829.6044968 | 0.092363677 | 0.712130471 | 0.88 | 0.58 |
| YKL224C | 0.174623486 | 0.113740683 | 0.712154851 | 0.88 | 0.58 |
| YLR150W | 21689.73235 | -0.099748538 | 0.711809321 | 0.88 | 0.58 |
| YMR016C | 1109.247117 | -0.093144602 | 0.711959233 | 0.88 | 0.58 |
| YMR304C-A | 599.6439516 | -0.109196223 | 0.711727472 | 0.88 | 0.58 |
| YLR406C-A | 2.560257022 | 0.133788477 | 0.712409821 | 0.88 | 0.58 |
| YDR427W | 2923.459081 | 0.076994014 | 0.712706408 | 0.88 | 0.58 |
| YGL005C | 924.1279481 | 0.06647471 | 0.712844777 | 0.88 | 0.58 |
| YJL204C | 683.8445536 | -0.067549392 | 0.713246337 | 0.88 | 0.58 |
| YER174C | 586.1669315 | -0.08962074 | 0.713446562 | 0.88 | 0.58 |
| YBL093C | 1069.915355 | -0.057542262 | 0.713981223 | 0.88 | 0.58 |
| YBR262C | 557.8168642 | 0.118522252 | 0.714950894 | 0.88 | 0.58 |
| YDL107W | 383.3863117 | -0.079445241 | 0.714420873 | 0.88 | 0.58 |
| YDL242W | 58.94192665 | 0.119461163 | 0.714237313 | 0.88 | 0.58 |
| YDR195W | 649.2195359 | 0.056275546 | 0.714939709 | 0.88 | 0.58 |
| YDR205W | 692.1640308 | 0.059159684 | 0.714777414 | 0.88 | 0.58 |
| YER107W-A | 76.0626659 | -0.101241688 | 0.715051213 | 0.88 | 0.58 |
| YHR205W | 2582.464034 | -0.095756235 | 0.713652479 | 0.88 | 0.58 |
| YMR224C | 500.4037572 | 0.050860458 | 0.714488411 | 0.88 | 0.58 |
| YOR168W | 3054.619284 | -0.084806694 | 0.715098925 | 0.88 | 0.58 |
| YOR213C | 459.655666 | -0.068826788 | 0.714304765 | 0.88 | 0.58 |
| YPL253C | 481.3229549 | -0.075357964 | 0.714751744 | 0.88 | 0.58 |
| YMR289W | 967.6114857 | 0.062105745 | 0.715310622 | 0.88 | 0.58 |
| YGR089W | 1598.86435 | 0.058623379 | 0.715623557 | 0.88 | 0.58 |
| YAL037C-B | 133.1022568 | -0.102966802 | 0.716611712 | 0.88 | 0.58 |
| YDL076C | 828.6133083 | -0.052786101 | 0.716241338 | 0.88 | 0.58 |
| YMR172C-A | 11.37396813 | -0.147438355 | 0.716645221 | 0.88 | 0.58 |
| YOR058C | 1144.395841 | 0.068184679 | 0.716383958 | 0.88 | 0.58 |
| YPL159C | 943.1509361 | -0.105820876 | 0.716245215 | 0.88 | 0.58 |
| YBR022W | 363.9709831 | -0.087711562 | 0.717138096 | 0.88 | 0.58 |
| YBR205W | 1374.246383 | 0.071299473 | 0.717064459 | 0.88 | 0.58 |
| YIL157C | 795.6414589 | -0.091194805 | 0.717057737 | 0.88 | 0.58 |

| | | | | | |
|---|---|---|---|---|---|
| YDR197W | 858.0497536 | 0.076407281 | 0.718261185 | 0.88 | 0.58 |
| YGL160W | 1159.891272 | -0.053479232 | 0.71770001 | 0.88 | 0.58 |
| YHR032W-A | 60.96537533 | -0.128991794 | 0.718198709 | 0.88 | 0.58 |
| YJR031C | 1083.680948 | -0.057557016 | 0.718149183 | 0.88 | 0.58 |
| YLR201C | 1199.611196 | 0.07143063 | 0.717920475 | 0.88 | 0.58 |
| YLR305C | 2169.127998 | 0.088388743 | 0.71823835 | 0.88 | 0.58 |
| YPL141C | 1338.030401 | -0.093927961 | 0.717472777 | 0.88 | 0.58 |
| YDR409W | 969.523209 | -0.064596688 | 0.718703214 | 0.88 | 0.58 |
| YLL031C | 1740.005527 | 0.068971723 | 0.718681263 | 0.88 | 0.58 |
| YOR018W | 939.5138966 | 0.057188974 | 0.718659383 | 0.88 | 0.58 |
| YGL214W | 9.34196156 | -0.129407192 | 0.719179048 | 0.88 | 0.58 |
| YBR296C | 209.9913106 | 0.121029949 | 0.71958235 | 0.88 | 0.58 |
| YML057C-A | 86.62699701 | 0.068342121 | 0.720541258 | 0.88 | 0.58 |
| YPL131W | 24700.79965 | -0.103211447 | 0.720929579 | 0.88 | 0.58 |
| YFL021W | 926.9442787 | 0.077266568 | 0.72131856 | 0.88 | 0.58 |
| YHL040C | 1041.983492 | -0.10730969 | 0.721522872 | 0.88 | 0.58 |
| YOL043C | 577.8464057 | 0.080146378 | 0.721525791 | 0.88 | 0.58 |
| YPL024W | 252.5020697 | -0.104961369 | 0.721800345 | 0.88 | 0.58 |
| YER013W | 829.6032387 | -0.066709716 | 0.722293014 | 0.88 | 0.58 |
| YKR096W | 1782.315311 | 0.070016141 | 0.722205967 | 0.88 | 0.58 |
| YBR040W | 134.4656267 | -0.206314948 | 0.722652022 | 0.89 | 0.58 |
| YPL073C | 1.14139142 | -0.128813235 | 0.723082992 | 0.89 | 0.59 |
| YCR066W | 426.4825775 | 0.064235733 | 0.723366221 | 0.89 | 0.59 |
| YER127W | 556.9285849 | -0.128548775 | 0.723602185 | 0.89 | 0.59 |
| YOR075W | 634.1618193 | 0.092745748 | 0.723754248 | 0.89 | 0.59 |
| YIL171W | 27.73883289 | 0.165570925 | 0.72391621 | 0.89 | 0.59 |
| YLR005W | 967.4119109 | 0.0504707 | 0.724363975 | 0.89 | 0.59 |
| YJL055W | 2135.334979 | 0.065924551 | 0.724582666 | 0.89 | 0.59 |
| YEL017C-A | 672.6729484 | 0.127248646 | 0.725145604 | 0.89 | 0.59 |
| YJL152W | 449.6754094 | 0.080335382 | 0.72548825 | 0.89 | 0.59 |
| YJL210W | 543.5019911 | -0.101356416 | 0.725737762 | 0.89 | 0.59 |
| YNL129W | 332.9229067 | 0.178341204 | 0.725583646 | 0.89 | 0.59 |
| YOR140W | 905.6577941 | 0.072780429 | 0.725791837 | 0.89 | 0.59 |
| YLR364C-A | 0.303554837 | -0.058609352 | 0.726069199 | 0.89 | 0.59 |
| YDR290W | 8.55083412 | -0.153463394 | 0.726639459 | 0.89 | 0.59 |
| YGL254W | 331.7870674 | 0.086097366 | 0.726396156 | 0.89 | 0.59 |
| YPL056C | 172.8020928 | -0.13591723 | 0.726660162 | 0.89 | 0.59 |
| YBR023C | 1741.008098 | 0.087388797 | 0.72739168 | 0.89 | 0.59 |
| YBR135W | 639.2382675 | 0.123379716 | 0.727684548 | 0.89 | 0.59 |
| YJL080C | 5735.124683 | 0.15523142 | 0.72739687 | 0.89 | 0.59 |
| YJL127W-A | 12.80997666 | 0.139250991 | 0.727393191 | 0.89 | 0.59 |
| YLR337C | 2433.363455 | -0.062418143 | 0.72790117 | 0.89 | 0.59 |
| YLR360W | 653.1816733 | -0.047026532 | 0.727521241 | 0.89 | 0.59 |

| | | | | | |
|---|---|---|---|---|---|
| YOL023W | 636.5704075 | 0.080516606 | 0.727853504 | 0.89 | 0.59 |
| YDL105W | 423.3140946 | -0.06475935 | 0.728695722 | 0.89 | 0.59 |
| YDR232W | 2313.644723 | 0.081895908 | 0.729263332 | 0.89 | 0.59 |
| YBR124W | 16.74512171 | -0.131996741 | 0.730779063 | 0.89 | 0.59 |
| YDR368W | 2244.977391 | -0.080889782 | 0.730933327 | 0.89 | 0.59 |
| YER039C | 101.9948572 | 0.075344062 | 0.731101764 | 0.89 | 0.59 |
| YGL004C | 1031.613443 | -0.078825066 | 0.730156827 | 0.89 | 0.59 |
| YGL206C | 5144.593261 | 0.056810028 | 0.730503342 | 0.89 | 0.59 |
| YGR018C | 8.799849222 | -0.183024289 | 0.729900656 | 0.89 | 0.59 |
| YIL005W | 1687.731259 | 0.05660344 | 0.731189363 | 0.89 | 0.59 |
| YJL047C-A | 19.06572924 | -0.115089982 | 0.729744761 | 0.89 | 0.59 |
| YKR039W | 765.789828 | 0.192375968 | 0.72966769 | 0.89 | 0.59 |
| YLR239C | 753.9388732 | 0.065072014 | 0.730209329 | 0.89 | 0.59 |
| YLR256W | 4392.10169 | 0.097402585 | 0.730640424 | 0.89 | 0.59 |
| YMR225C | 419.9415237 | 0.103890754 | 0.730080793 | 0.89 | 0.59 |
| YNR032C-A | 183.6670278 | -0.066356602 | 0.730781843 | 0.89 | 0.59 |
| YPR104C | 970.815797 | -0.053803944 | 0.731181781 | 0.89 | 0.59 |
| YBL012C | 41.44037077 | -0.114142022 | 0.73252111 | 0.89 | 0.59 |
| YCR008W | 1187.11323 | -0.092622071 | 0.732425052 | 0.89 | 0.59 |
| YDR360W | 36.19514521 | 0.135113812 | 0.732374546 | 0.89 | 0.59 |
| YMR029C | 1228.193843 | 0.068562462 | 0.732615679 | 0.89 | 0.59 |
| YOL047C | 302.1448912 | 0.207125014 | 0.732056688 | 0.89 | 0.59 |
| YOR183W | 10.65070105 | -0.152615795 | 0.732547297 | 0.89 | 0.59 |
| YLR288C | 449.0939426 | -0.050173943 | 0.733015766 | 0.89 | 0.59 |
| YLR384C | 2550.492331 | 0.061982674 | 0.732954477 | 0.89 | 0.59 |
| snR32 | 25.20381189 | 0.216759306 | 0.733345907 | 0.89 | 0.59 |
| YBR179C | 1264.938407 | 0.06544186 | 0.733857783 | 0.89 | 0.59 |
| YOL073C | 1132.754224 | 0.081567557 | 0.734243574 | 0.89 | 0.59 |
| snR52 | 0.834128967 | 0.497616439 | 0.734163628 | 0.89 | 0.59 |
| tT(CGU)K | 0.429968112 | 0.084936918 | 0.733997105 | 0.89 | 0.59 |
| YDR422C | 1006.650102 | 0.059460607 | 0.735708009 | 0.89 | 0.59 |
| YDL247W | 3.022100866 | 0.389117202 | 0.73656426 | 0.89 | 0.59 |
| YMR239C | 477.7892356 | -0.094227342 | 0.736453522 | 0.89 | 0.59 |
| YOL025W | 668.044846 | 0.074334967 | 0.736308882 | 0.89 | 0.59 |
| YDR199W | 75.09684199 | -0.077677089 | 0.737583921 | 0.89 | 0.59 |
| YJR147W | 698.6444516 | -0.090496879 | 0.737639174 | 0.89 | 0.59 |
| YKR017C | 571.9042365 | 0.083524895 | 0.737399519 | 0.89 | 0.59 |
| YLL032C | 878.0650576 | 0.058645121 | 0.737494882 | 0.89 | 0.59 |
| YOR298C-A | 6204.241243 | -0.052462822 | 0.737093035 | 0.89 | 0.59 |
| YOR337W | 623.7332196 | 0.075276087 | 0.736988672 | 0.89 | 0.59 |
| YPR004C | 1678.278628 | -0.081903818 | 0.737716601 | 0.89 | 0.59 |
| YPR116W | 103.378319 | 0.118903736 | 0.737527322 | 0.89 | 0.59 |
| YNL085W | 3478.261348 | 0.082454049 | 0.738006816 | 0.89 | 0.59 |

| | | | | | |
|---|---|---|---|---|---|
| YDR461C-A | 326.4764294 | 0.075384163 | 0.739155917 | 0.89 | 0.59 |
| YEL025C | 1231.625482 | 0.066367222 | 0.739452924 | 0.89 | 0.59 |
| YFL012W-A | 14.51855676 | -0.123986889 | 0.739750889 | 0.89 | 0.59 |
| YGR106C | 3546.453004 | 0.082372813 | 0.739620431 | 0.89 | 0.59 |
| YHR145C | 0.415497853 | 0.244693086 | 0.73921942 | 0.89 | 0.59 |
| YIL026C | 1582.807889 | -0.068578185 | 0.738882554 | 0.89 | 0.59 |
| YKL007W | 2212.362326 | -0.035966767 | 0.739580264 | 0.89 | 0.59 |
| YKL088W | 1161.516822 | -0.087665804 | 0.739166584 | 0.89 | 0.59 |
| YKR043C | 2568.82772 | 0.15917127 | 0.739022254 | 0.89 | 0.59 |
| YLR301W | 1464.12998 | -0.038576883 | 0.738394851 | 0.89 | 0.59 |
| YNR049C | 561.0637018 | -0.051600516 | 0.739358052 | 0.89 | 0.59 |
| YLL043W | 1838.000712 | 0.065464799 | 0.740132044 | 0.89 | 0.59 |
| YLL016W | 1012.667191 | 0.108920073 | 0.740408517 | 0.89 | 0.59 |
| YHR178W | 530.4654676 | 0.091969383 | 0.740717536 | 0.89 | 0.59 |
| YBL105C | 2392.516684 | -0.060595007 | 0.741035839 | 0.89 | 0.59 |
| YDR304C | 4614.533404 | 0.067224042 | 0.741184681 | 0.89 | 0.59 |
| YPL160W | 4849.742262 | 0.125091478 | 0.741095429 | 0.89 | 0.59 |
| YLR191W | 472.4930163 | 0.055998142 | 0.741614157 | 0.89 | 0.59 |
| YMR121C | 804.8314095 | -0.089720579 | 0.74196257 | 0.89 | 0.59 |
| YDL151C | 4.616503935 | -0.098702246 | 0.742917601 | 0.89 | 0.59 |
| YGR068C | 535.7602898 | -0.053938744 | 0.74272452 | 0.89 | 0.59 |
| YHR086W-A | 3.107825184 | 0.258182525 | 0.742359863 | 0.89 | 0.59 |
| YJR108W | 40.18860279 | -0.09649453 | 0.743178608 | 0.89 | 0.59 |
| YKL018C-A | 261.9017664 | 0.049525047 | 0.742243205 | 0.89 | 0.59 |
| YML010W | 2999.463402 | -0.055347758 | 0.743075928 | 0.89 | 0.59 |
| YMR052C-A | 61.89756679 | 0.076786281 | 0.742963984 | 0.89 | 0.59 |
| YNL297C | 2104.946506 | -0.049940152 | 0.743090422 | 0.89 | 0.59 |
| YDR508C | 2566.963439 | -0.09558665 | 0.743363548 | 0.89 | 0.59 |
| YGR015C | 438.1021585 | 0.044933594 | 0.743457005 | 0.89 | 0.59 |
| YDR352W | 489.6613604 | 0.106389398 | 0.744117198 | 0.90 | 0.59 |
| YJR044C | 1233.447551 | 0.060161228 | 0.744541856 | 0.90 | 0.59 |
| YJL092W | 1243.946562 | 0.0830734 | 0.744708188 | 0.90 | 0.59 |
| YBR128C | 336.3131952 | 0.078502232 | 0.745726239 | 0.90 | 0.59 |
| YHR134W | 365.9132551 | 0.091705463 | 0.745425558 | 0.90 | 0.59 |
| YKR041W | 32.67705338 | -0.052278812 | 0.74604277 | 0.90 | 0.59 |
| YLR243W | 616.8482711 | -0.067527183 | 0.745545275 | 0.90 | 0.59 |
| YNL103W | 1585.703492 | 0.064558025 | 0.745857869 | 0.90 | 0.59 |
| YOR192C-B | 24.91802453 | 0.176524847 | 0.745884951 | 0.90 | 0.59 |
| YOR356W | 1275.917226 | 0.067799339 | 0.746140293 | 0.90 | 0.59 |
| YFR038W | 1186.649674 | -0.066455885 | 0.746673372 | 0.90 | 0.59 |
| YML007W | 2781.196668 | -0.08044625 | 0.746716077 | 0.90 | 0.59 |
| YNL281W | 1354.281332 | 0.137104569 | 0.746370291 | 0.90 | 0.59 |
| YOL031C | 1089.984363 | -0.09617538 | 0.746744383 | 0.90 | 0.59 |

| | | | | | |
|---|---|---|---|---|---|
| snR41 | 28.2332558 | -0.066881293 | 0.746971165 | 0.90 | 0.59 |
| YOR053W | 4.40231234 | 0.209968151 | 0.747302793 | 0.90 | 0.59 |
| YDR539W | 882.7410932 | 0.139348008 | 0.747682118 | 0.90 | 0.59 |
| YGR057C | 395.64537 | 0.051774101 | 0.747792604 | 0.90 | 0.59 |
| YBR273C | 1305.311458 | -0.051397686 | 0.748971131 | 0.90 | 0.59 |
| YER019W | 1037.358398 | 0.110628288 | 0.748965319 | 0.90 | 0.59 |
| YGL108C | 328.3815974 | -0.072619216 | 0.74892562 | 0.90 | 0.59 |
| YGR253C | 3517.823624 | -0.050262852 | 0.748973496 | 0.90 | 0.59 |
| YNR005C | 1.56429782 | 0.108310168 | 0.748709813 | 0.90 | 0.59 |
| YPL217C | 2396.710824 | -0.066336895 | 0.748297051 | 0.90 | 0.59 |
| snR161 | 8.988952116 | 0.084982125 | 0.748998745 | 0.90 | 0.59 |
| YBR221W-A | 0.837044541 | 0.132384376 | 0.749785074 | 0.90 | 0.59 |
| YDL041W | 0.97089065 | -0.273624719 | 0.749582179 | 0.90 | 0.59 |
| YJL151C | 3281.2426 | 0.081499323 | 0.749635692 | 0.90 | 0.59 |
| YKL130C | 628.0827477 | -0.072456316 | 0.74969989 | 0.90 | 0.59 |
| YKR031C | 1778.007412 | -0.080202272 | 0.749886895 | 0.90 | 0.59 |
| YPR170W-B | 216.8408172 | -0.045489107 | 0.74937304 | 0.90 | 0.59 |
| YGR093W | 815.5848449 | 0.04980694 | 0.750090472 | 0.90 | 0.59 |
| YFR014C | 1739.196956 | -0.110476907 | 0.751100839 | 0.90 | 0.59 |
| YJL181W | 405.181886 | 0.056916649 | 0.751110078 | 0.90 | 0.59 |
| YJL201W | 1030.90288 | 0.043961882 | 0.751107581 | 0.90 | 0.59 |
| YKL068W-A | 172.0547206 | 0.107686442 | 0.750528463 | 0.90 | 0.59 |
| YKL173W | 1269.160496 | 0.036973198 | 0.750284107 | 0.90 | 0.59 |
| YLL018C | 8408.142457 | 0.068629394 | 0.751081232 | 0.90 | 0.59 |
| YOL156W | 95.75918972 | 0.170026254 | 0.750814472 | 0.90 | 0.59 |
| YPR169W | 1659.207747 | -0.06977699 | 0.751266205 | 0.90 | 0.59 |
| YDR434W | 1180.144828 | -0.082057505 | 0.752025316 | 0.90 | 0.59 |
| YDR440W | 548.6066441 | -0.057404431 | 0.752478216 | 0.90 | 0.59 |
| YGL257C | 1193.728393 | 0.07269102 | 0.751717406 | 0.90 | 0.59 |
| YGR110W | 407.9203872 | 0.055904597 | 0.752443386 | 0.90 | 0.59 |
| YKR027W | 720.4128865 | -0.044282626 | 0.752534582 | 0.90 | 0.59 |
| YLR010C | 127.8600135 | -0.083084359 | 0.752233397 | 0.90 | 0.59 |
| YMR134W | 453.7137928 | 0.077801485 | 0.752529358 | 0.90 | 0.59 |
| YPL075W | 2226.650529 | 0.073002631 | 0.751960759 | 0.90 | 0.59 |
| YPR059C | 12.62828581 | 0.122429111 | 0.751725199 | 0.90 | 0.59 |
| YOL114C | 133.5426332 | -0.097515937 | 0.752683575 | 0.90 | 0.59 |
| tD(GUC)N | 0.865673422 | 0.247669977 | 0.752911119 | 0.90 | 0.59 |
| YPL239W | 1334.632299 | 0.107540947 | 0.753814253 | 0.90 | 0.59 |
| YOR331C | 254.0815524 | 0.034929202 | 0.754003218 | 0.90 | 0.59 |
| YIR004W | 1446.338254 | -0.050777602 | 0.754373297 | 0.90 | 0.59 |
| YGR174C | 1048.598299 | 0.077497555 | 0.755055871 | 0.90 | 0.59 |
| YIL100W | 3.722972412 | 0.185027071 | 0.755161883 | 0.90 | 0.59 |
| YFL015W-A | 1.143841678 | -0.135783702 | 0.755563315 | 0.90 | 0.59 |

| | | | | | |
|---|---|---|---|---|---|
| YPL156C | 858.4762505 | -0.11405252 | 0.75543443 | 0.90 | 0.59 |
| YDR067C | 673.9208846 | 0.049587646 | 0.756064868 | 0.90 | 0.59 |
| YDR406W-A | 12.68467301 | -0.210838956 | 0.756135723 | 0.90 | 0.59 |
| YHR219W | 0.159331785 | 0.083088649 | 0.755927743 | 0.90 | 0.59 |
| YMR128W | 2322.124433 | -0.094354182 | 0.756353385 | 0.90 | 0.59 |
| YOL107W | 336.081692 | -0.078409489 | 0.7567233 | 0.90 | 0.59 |
| YBR116C | 35.41794033 | -0.165129343 | 0.756862379 | 0.90 | 0.59 |
| YHR090C | 634.1320105 | -0.08752577 | 0.757150142 | 0.90 | 0.60 |
| YIL068W-A | 5.981971104 | -0.204694345 | 0.757614338 | 0.90 | 0.60 |
| YIL080W | 141.3218365 | 0.17310522 | 0.757499447 | 0.90 | 0.60 |
| YPR018W | 922.6100673 | 0.085920069 | 0.757455765 | 0.90 | 0.60 |
| YER022W | 948.246119 | 0.034683888 | 0.757899783 | 0.90 | 0.60 |
| YBR160W | 885.0084099 | -0.05648655 | 0.758268689 | 0.90 | 0.60 |
| YJR112W-A | 356.5088081 | 0.060302555 | 0.758357827 | 0.90 | 0.60 |
| YFL009W | 860.3502379 | 0.037456382 | 0.760038306 | 0.90 | 0.60 |
| YHR131W-A | 19.588359 | 0.123184903 | 0.759945408 | 0.90 | 0.60 |
| YLR025W | 2034.136821 | -0.069679963 | 0.760155656 | 0.90 | 0.60 |
| YBR134W | 28.00305569 | 0.175298176 | 0.760341945 | 0.90 | 0.60 |
| YER118C | 889.8446464 | -0.057676834 | 0.760557714 | 0.90 | 0.60 |
| YBL038W | 1166.095027 | -0.096393655 | 0.761050485 | 0.90 | 0.60 |
| YDL187C | 3.68310749 | 0.220116908 | 0.761138148 | 0.90 | 0.60 |
| YMR269W | 489.6101925 | -0.066543682 | 0.760987051 | 0.90 | 0.60 |
| YPR107C | 422.2754774 | 0.041749672 | 0.760956201 | 0.90 | 0.60 |
| YBR053C | 1693.976608 | 0.073795537 | 0.761643983 | 0.90 | 0.60 |
| YGR048W | 875.7732177 | -0.058398434 | 0.761971681 | 0.90 | 0.60 |
| YJL136W-A | 0.34890375 | 0.1306456 | 0.761980307 | 0.90 | 0.60 |
| YOL116W | 722.658756 | -0.077505848 | 0.762061023 | 0.90 | 0.60 |
| YOR154W | 667.1567549 | 0.045059678 | 0.761778518 | 0.90 | 0.60 |
| YBR127C | 10020.63379 | -0.04396113 | 0.762768021 | 0.90 | 0.60 |
| YPL260W | 2621.937685 | 0.06090777 | 0.762709825 | 0.90 | 0.60 |
| YBR008C | 906.2949136 | 0.061326012 | 0.763511503 | 0.90 | 0.60 |
| YMR291W | 2090.324495 | -0.136444721 | 0.763571068 | 0.90 | 0.60 |
| YIL105C | 1531.713028 | -0.081151697 | 0.763772448 | 0.90 | 0.60 |
| YGR260W | 2070.398504 | 0.131942199 | 0.764273166 | 0.90 | 0.60 |
| snR86 | 894.0383097 | 0.19174333 | 0.76450895 | 0.90 | 0.60 |
| YJR100C | 772.6621126 | -0.049524866 | 0.764942631 | 0.91 | 0.60 |
| YFR035C | 184.0960805 | -0.04257166 | 0.765742723 | 0.91 | 0.60 |
| YPL266W | 1132.936461 | -0.073830537 | 0.765951779 | 0.91 | 0.60 |
| YHR214C-E | 1.813646714 | 0.339985648 | 0.766721396 | 0.91 | 0.60 |
| YDL025C | 1555.623411 | 0.065127061 | 0.767435347 | 0.91 | 0.60 |
| YJR097W | 96.20405896 | -0.069532341 | 0.767510519 | 0.91 | 0.60 |
| YKL185W | 2577.028628 | 0.077899812 | 0.767240469 | 0.91 | 0.60 |
| YLR164W | 251.3796178 | 0.085872229 | 0.767634344 | 0.91 | 0.60 |

| | | | | | |
|---|---|---|---|---|---|
| YBR089C-A | 560.6070076 | 0.072872193 | 0.768534919 | 0.91 | 0.60 |
| YCL024W | 493.5504233 | -0.052917784 | 0.76841775 | 0.91 | 0.60 |
| YDR154C | 3531.276985 | 0.076853886 | 0.768690268 | 0.91 | 0.60 |
| YFR008W | 307.6442666 | -0.065943485 | 0.768619123 | 0.91 | 0.60 |
| YLR031W | 139.756576 | 0.072155869 | 0.76880457 | 0.91 | 0.60 |
| YCL046W | 22.38513549 | -0.074356037 | 0.769413736 | 0.91 | 0.60 |
| YMR226C | 4108.834639 | 0.053233183 | 0.769410253 | 0.91 | 0.60 |
| YNL329C | 1184.177512 | -0.047323517 | 0.769183528 | 0.91 | 0.60 |
| YDR380W | 4921.676121 | 0.139299634 | 0.770654626 | 0.91 | 0.60 |
| YDR439W | 511.333794 | 0.057067437 | 0.771128804 | 0.91 | 0.60 |
| YER109C | 1065.439283 | -0.04952625 | 0.770879378 | 0.91 | 0.60 |
| YGR251W | 285.9824856 | -0.064789374 | 0.771173592 | 0.91 | 0.60 |
| YJL208C | 631.1153592 | -0.07009042 | 0.770832471 | 0.91 | 0.60 |
| YPL096W | 660.7866918 | 0.043586023 | 0.771018919 | 0.91 | 0.60 |
| YFL049W | 806.3019635 | -0.042904939 | 0.772041169 | 0.91 | 0.60 |
| YGL236C | 488.164276 | 0.08030543 | 0.771957676 | 0.91 | 0.60 |
| YAL063C-A | 8.265236292 | 0.172983375 | 0.772812338 | 0.91 | 0.60 |
| YER180C-A | 68.54860649 | -0.062235639 | 0.773221059 | 0.91 | 0.60 |
| YMR086C-A | 82.59944884 | 0.060143793 | 0.773312822 | 0.91 | 0.60 |
| YBR215W | 1081.67593 | 0.040857597 | 0.773907164 | 0.91 | 0.60 |
| YDR323C | 314.788297 | 0.035303133 | 0.773574564 | 0.91 | 0.60 |
| YLR203C | 2343.4384 | 0.04878636 | 0.773825979 | 0.91 | 0.60 |
| YNR055C | 1151.83484 | -0.057394866 | 0.773870493 | 0.91 | 0.60 |
| YKL135C | 1473.22037 | -0.044865112 | 0.774322185 | 0.91 | 0.60 |
| YCR075W-A | 129.8943621 | 0.077609889 | 0.774615089 | 0.91 | 0.60 |
| YLR381W | 607.7067419 | -0.046512384 | 0.774856649 | 0.91 | 0.60 |
| YKL001C | 376.5252013 | 0.097559617 | 0.775254756 | 0.91 | 0.60 |
| YOR082C | 31.57473045 | 0.102729677 | 0.775618083 | 0.91 | 0.60 |
| YER038W-A | 0.198141795 | -0.035072562 | 0.77618211 | 0.91 | 0.60 |
| YLR436C | 2590.336596 | 0.060979778 | 0.776213553 | 0.91 | 0.60 |
| YJR157W | 1.945110995 | 0.30794503 | 0.77648889 | 0.91 | 0.60 |
| YOL148C | 713.6397629 | 0.049435158 | 0.77669826 | 0.91 | 0.60 |
| YLR232W | 2.842598345 | -0.006574003 | 0.776913071 | 0.91 | 0.60 |
| YNL054W | 1269.220328 | 0.057440665 | 0.777260388 | 0.91 | 0.60 |
| YDL159W | 513.5387781 | 0.05861129 | 0.777674576 | 0.91 | 0.60 |
| YML054C-A | 11.9894805 | -0.152944705 | 0.777949185 | 0.91 | 0.60 |
| YOR296W | 873.0215241 | 0.04244143 | 0.777812884 | 0.91 | 0.60 |
| YIR035C | 1381.941223 | -0.051775475 | 0.778930528 | 0.91 | 0.60 |
| YLR359W | 9224.21068 | 0.177470437 | 0.778677971 | 0.91 | 0.60 |
| snR49 | 77.03202196 | 0.228448133 | 0.778818003 | 0.91 | 0.60 |
| snR53 | 0.80815136 | -0.022527549 | 0.778694872 | 0.91 | 0.60 |
| YCR059C | 1130.721478 | 0.056341127 | 0.77970706 | 0.91 | 0.60 |
| YDL084W | 7170.004042 | -0.043805491 | 0.779916728 | 0.91 | 0.60 |

| | | | | | |
|---|---|---|---|---|---|
| YDR207C | 1436.193266 | -0.072312626 | 0.779554121 | 0.91 | 0.60 |
| YKR033C | 0.132511194 | 0.081851394 | 0.779954754 | 0.91 | 0.60 |
| YOR217W | 1207.361787 | 0.046094324 | 0.779589721 | 0.91 | 0.60 |
| YBL058W | 3576.371774 | -0.050580674 | 0.780648928 | 0.91 | 0.60 |
| YMR316W | 898.4662527 | 0.161900948 | 0.780511556 | 0.91 | 0.60 |
| YJR050W | 528.5017956 | 0.05353354 | 0.780974279 | 0.91 | 0.60 |
| YPL102C | 13.71806551 | 0.040395868 | 0.781234854 | 0.91 | 0.60 |
| YIL084C | 434.165782 | -0.065014963 | 0.782062517 | 0.92 | 0.61 |
| YCR024C | 641.2724541 | -0.038215608 | 0.782733768 | 0.92 | 0.61 |
| YDL229W | 7732.532873 | -0.092417724 | 0.782738423 | 0.92 | 0.61 |
| YDR251W | 2664.249522 | -0.049921123 | 0.782808465 | 0.92 | 0.61 |
| YNL133C | 297.07226 | 0.053918859 | 0.782695033 | 0.92 | 0.61 |
| YDR320C | 1271.658982 | 0.040759165 | 0.783259241 | 0.92 | 0.61 |
| YGR109C | 138.0694558 | 0.110354678 | 0.783219164 | 0.92 | 0.61 |
| YBR071W | 990.0527368 | -0.081252959 | 0.784061937 | 0.92 | 0.61 |
| YIL145C | 1126.99064 | 0.081569189 | 0.784593935 | 0.92 | 0.61 |
| YDR034C | 1497.715383 | 0.060714599 | 0.784892896 | 0.92 | 0.61 |
| YGR091W | 441.73971 | 0.05793021 | 0.785110184 | 0.92 | 0.61 |
| YJR120W | 18.09619799 | 0.071685794 | 0.785534883 | 0.92 | 0.61 |
| YEL059W | 50.35325287 | -0.114308663 | 0.786021581 | 0.92 | 0.61 |
| YHR172W | 514.8081565 | 0.123375754 | 0.78601678 | 0.92 | 0.61 |
| YML125C | 2139.072841 | -0.068072472 | 0.786107481 | 0.92 | 0.61 |
| YER041W | 647.7988303 | 0.0559016 | 0.786322164 | 0.92 | 0.61 |
| YDR023W | 6264.678536 | -0.052533888 | 0.786992421 | 0.92 | 0.61 |
| YKL159C | 432.357669 | 0.073327267 | 0.787067424 | 0.92 | 0.61 |
| snR34 | 37.83791355 | 0.251334477 | 0.78699571 | 0.92 | 0.61 |
| YKL100C | 2362.710996 | -0.06599322 | 0.78746676 | 0.92 | 0.61 |
| YML081W | 1706.956949 | 0.0692696 | 0.787414382 | 0.92 | 0.61 |
| YCR025C | 44.00460202 | -0.118418977 | 0.788044241 | 0.92 | 0.61 |
| YDR109C | 536.0726963 | 0.034607202 | 0.788777528 | 0.92 | 0.61 |
| YER139C | 313.6167797 | -0.050927168 | 0.788857063 | 0.92 | 0.61 |
| YFL038C | 5600.505944 | 0.043449767 | 0.78891994 | 0.92 | 0.61 |
| YGL194C | 571.5770023 | -0.046024288 | 0.788379855 | 0.92 | 0.61 |
| YIL111W | 1481.52504 | 0.068240272 | 0.788806278 | 0.92 | 0.61 |
| YPL275W | 31.2658679 | -0.35156351 | 0.788625811 | 0.92 | 0.61 |
| snR189 | 9.691874789 | -0.118136577 | 0.788940472 | 0.92 | 0.61 |
| YCL001W-B | 30.32607662 | 0.112331416 | 0.789315101 | 0.92 | 0.61 |
| YDL007W | 5356.140616 | 0.048164909 | 0.78939496 | 0.92 | 0.61 |
| YLL044W | 94.44590477 | -0.340957478 | 0.78954257 | 0.92 | 0.61 |
| YMR242W-A | 1.825788145 | 0.182073294 | 0.789903798 | 0.92 | 0.61 |
| YNL266W | 15.42456766 | -0.127452834 | 0.789986007 | 0.92 | 0.61 |
| YNL067W-B | 0.339190409 | 0.102708562 | 0.790424773 | 0.92 | 0.61 |
| YLR418C | 850.5544851 | -0.038391016 | 0.791028105 | 0.92 | 0.61 |

| | | | | | |
|---|---|---|---|---|---|
| YPL195W | 2059.660784 | 0.037463326 | 0.790920876 | 0.92 | 0.61 |
| YPR086W | 1622.357285 | 0.054012373 | 0.79104888 | 0.92 | 0.61 |
| YAR019W-A | 5.564052438 | 0.181574771 | 0.791773426 | 0.92 | 0.61 |
| YBR149W | 7039.888751 | 0.047345718 | 0.791932096 | 0.92 | 0.61 |
| YDL145C | 6294.266606 | 0.064811059 | 0.792367261 | 0.92 | 0.61 |
| YDR221W | 936.1837264 | -0.037436946 | 0.792230396 | 0.92 | 0.61 |
| YER003C | 3520.842344 | -0.047583593 | 0.792589109 | 0.92 | 0.61 |
| YER125W | 2425.853452 | -0.040059483 | 0.792233444 | 0.92 | 0.61 |
| YNL273W | 713.7141054 | -0.041624783 | 0.792495406 | 0.92 | 0.61 |
| YBL085W | 2143.648048 | 0.055204252 | 0.793718476 | 0.92 | 0.61 |
| YCR076C | 593.9023385 | 0.061145536 | 0.794188167 | 0.92 | 0.61 |
| YCR092C | 872.6618833 | 0.04261656 | 0.793902083 | 0.92 | 0.61 |
| YDR384C | 561.1890906 | -0.054552033 | 0.793997611 | 0.92 | 0.61 |
| YDR432W | 4551.301879 | 0.084731703 | 0.794281752 | 0.92 | 0.61 |
| YDR536W | 315.8429448 | 0.081292326 | 0.794094051 | 0.92 | 0.61 |
| YGL064C | 383.7385393 | -0.04825052 | 0.792878576 | 0.92 | 0.61 |
| YJL026W | 4324.794509 | 0.057823232 | 0.794331566 | 0.92 | 0.61 |
| YML129C | 536.8329539 | -0.070566106 | 0.794392897 | 0.92 | 0.61 |
| YMR171C | 1518.290743 | 0.103489066 | 0.793014917 | 0.92 | 0.61 |
| YNL208W | 6496.983749 | 0.048985847 | 0.793561992 | 0.92 | 0.61 |
| YOR070C | 1064.02202 | -0.052538789 | 0.793636764 | 0.92 | 0.61 |
| YHR026W | 2431.689374 | 0.069898579 | 0.795135871 | 0.92 | 0.61 |
| YNL094W | 1768.042353 | -0.049572394 | 0.795096328 | 0.92 | 0.61 |
| YNR031C | 1352.088361 | 0.073902036 | 0.794715667 | 0.92 | 0.61 |
| YOR254C | 3232.98471 | -0.047961703 | 0.794882177 | 0.92 | 0.61 |
| YPL076W | 380.2952376 | -0.062982615 | 0.794814717 | 0.92 | 0.61 |
| YBR228W | 415.0546527 | 0.055018777 | 0.795534364 | 0.92 | 0.61 |
| YGR047C | 884.2267464 | -0.043636111 | 0.795632922 | 0.92 | 0.61 |
| YML005W | 433.1547412 | 0.032148569 | 0.795639676 | 0.92 | 0.61 |
| YAL036C | 1045.610431 | 0.10869887 | 0.795965564 | 0.92 | 0.61 |
| YJR122W | 745.1766826 | 0.053204056 | 0.796264916 | 0.92 | 0.61 |
| YNL148C | 367.0903378 | 0.106910453 | 0.796704667 | 0.92 | 0.61 |
| YPR022C | 1532.191812 | 0.036203223 | 0.796576742 | 0.92 | 0.61 |
| YAL009W | 404.8562545 | 0.043625319 | 0.797724356 | 0.92 | 0.61 |
| YHR148W | 337.7936716 | -0.070203929 | 0.797852187 | 0.92 | 0.61 |
| YNL215W | 1103.384408 | 0.049679121 | 0.79758261 | 0.92 | 0.61 |
| YPL180W | 1077.43255 | -0.035008055 | 0.797819841 | 0.92 | 0.61 |
| YPR037C | 567.1835844 | 0.051996252 | 0.79765663 | 0.92 | 0.61 |
| YJL041W | 2583.756357 | -0.034617492 | 0.798127477 | 0.92 | 0.61 |
| YNR014W | 1455.118637 | 0.084958019 | 0.798348956 | 0.92 | 0.61 |
| YML108W | 173.1265371 | -0.069850289 | 0.79895974 | 0.92 | 0.61 |
| tQ(UUG)E2 | 0.083088204 | 0.067251982 | 0.799064498 | 0.92 | 0.61 |
| YML023C | 695.7405584 | -0.04632537 | 0.799227187 | 0.92 | 0.61 |

| | | | | | |
|---|---|---|---|---|---|
| YGL195W | 6234.838225 | -0.032272509 | 0.800071039 | 0.92 | 0.61 |
| snR5 | 32.14065526 | -0.081827971 | 0.800084411 | 0.92 | 0.61 |
| YDR166C | 1344.376251 | 0.043211369 | 0.800304688 | 0.92 | 0.61 |
| YJL056C | 734.4930024 | 0.057191512 | 0.80061717 | 0.92 | 0.61 |
| YFR010W | 3072.848472 | 0.047857095 | 0.800792396 | 0.92 | 0.61 |
| YGR285C | 10708.64158 | 0.132034367 | 0.801058505 | 0.92 | 0.61 |
| YHL015W-A | 4.315791202 | -0.077618886 | 0.801293349 | 0.92 | 0.61 |
| YLR170C | 116.6208147 | 0.069563417 | 0.801484563 | 0.92 | 0.61 |
| YML030W | 762.1544554 | -0.090542972 | 0.801391406 | 0.92 | 0.61 |
| YOR002W | 2112.418166 | -0.040089547 | 0.800951469 | 0.92 | 0.61 |
| YPR036W | 6660.698271 | 0.057111564 | 0.801839673 | 0.92 | 0.61 |
| YER046W | 177.0920564 | -0.113198959 | 0.802868123 | 0.92 | 0.61 |
| YFL032W | 146.5638958 | 0.07255268 | 0.802726773 | 0.92 | 0.61 |
| YGR111W | 1711.371366 | 0.04725633 | 0.803083388 | 0.93 | 0.61 |
| YCR002C | 1893.656785 | 0.058137866 | 0.803501747 | 0.93 | 0.61 |
| YOL065C | 517.0688786 | -0.055004748 | 0.803488141 | 0.93 | 0.61 |
| YAL012W | 6420.264953 | -0.06164392 | 0.803926289 | 0.93 | 0.61 |
| YBR117C | 1336.758461 | 0.106546229 | 0.804295734 | 0.93 | 0.61 |
| YGR002C | 912.0190954 | -0.044651463 | 0.804406777 | 0.93 | 0.61 |
| YLL063C | 302.5298246 | -0.084614629 | 0.804194473 | 0.93 | 0.61 |
| YLR335W | 2838.169755 | 0.041547989 | 0.804110238 | 0.93 | 0.61 |
| YBR269C | 1256.661555 | 0.055224765 | 0.8051246 | 0.93 | 0.61 |
| YML009C-A | 52.04514359 | -0.070562734 | 0.805043199 | 0.93 | 0.61 |
| YNL124W | 1308.097286 | 0.061600487 | 0.804940508 | 0.93 | 0.61 |
| snR36 | 18.91914442 | -0.105559855 | 0.805107819 | 0.93 | 0.61 |
| YER130C | 1003.132192 | 0.139959604 | 0.806269945 | 0.93 | 0.61 |
| YGL157W | 1718.848717 | -0.082351951 | 0.80591014 | 0.93 | 0.61 |
| YJL120W | 37.94521827 | 0.081575136 | 0.805572177 | 0.93 | 0.61 |
| YKL134C | 542.9022847 | -0.050624272 | 0.806191784 | 0.93 | 0.61 |
| YMR202W | 2009.003075 | 0.229358916 | 0.805612056 | 0.93 | 0.61 |
| YPL009C | 4156.098331 | 0.055976325 | 0.806133675 | 0.93 | 0.61 |
| YPR162C | 653.0734249 | 0.06495927 | 0.806252999 | 0.93 | 0.61 |
| YDR540C | 110.2480797 | -0.11833493 | 0.806703464 | 0.93 | 0.61 |
| YML020W | 864.3038212 | -0.050825696 | 0.806996756 | 0.93 | 0.61 |
| YNL314W | 384.9311948 | -0.050542138 | 0.80697026 | 0.93 | 0.61 |
| YOR351C | 100.6577504 | 0.057219679 | 0.807456932 | 0.93 | 0.61 |
| YNL194C | 937.042565 | 0.071689663 | 0.807980093 | 0.93 | 0.61 |
| YCR095W-A | 20.07196858 | 0.063902302 | 0.808240379 | 0.93 | 0.61 |
| YDR180W | 980.8547337 | 0.054348209 | 0.808322155 | 0.93 | 0.61 |
| YNL203C | 26.22039237 | -0.074316662 | 0.808414676 | 0.93 | 0.61 |
| YDR050C | 39610.42901 | 0.081373655 | 0.808812286 | 0.93 | 0.61 |
| YDR149C | 14.02290454 | -0.119913772 | 0.808919484 | 0.93 | 0.61 |
| YGR104C | 540.5492389 | -0.034127069 | 0.809769759 | 0.93 | 0.61 |

| | | | | | |
|---|---|---|---|---|---|
| YER027C | 1561.298312 | -0.036455778 | 0.810002956 | 0.93 | 0.61 |
| YAL022C | 845.7051212 | -0.040406213 | 0.810487864 | 0.93 | 0.61 |
| YOR057W | 859.8202576 | -0.055447104 | 0.810610984 | 0.93 | 0.61 |
| YKL012W | 1327.340355 | 0.026253844 | 0.810815621 | 0.93 | 0.61 |
| YNL228W | 12.1493579 | -0.098900431 | 0.811210672 | 0.93 | 0.61 |
| YNL259C | 262.1284302 | -0.049547683 | 0.811406353 | 0.93 | 0.61 |
| YJL135W | 5.319281498 | -0.089924434 | 0.811724619 | 0.93 | 0.61 |
| YLL022C | 865.5611491 | 0.054808644 | 0.811913423 | 0.93 | 0.61 |
| YLR396C | 522.7183585 | -0.031223798 | 0.81186382 | 0.93 | 0.61 |
| YBR209W | 5.517408049 | -0.086578564 | 0.812244309 | 0.93 | 0.61 |
| YDR400W | 652.8362371 | -0.039731786 | 0.812432189 | 0.93 | 0.61 |
| YDR494W | 1564.77137 | -0.04375882 | 0.812489126 | 0.93 | 0.61 |
| YNR042W | 0.825755426 | -0.152164136 | 0.812455434 | 0.93 | 0.61 |
| YHR033W | 1072.035586 | -0.136203434 | 0.813213113 | 0.93 | 0.61 |
| YOR229W | 494.4703852 | 0.041593227 | 0.813268879 | 0.93 | 0.61 |
| YAL034W-A | 187.40184 | -0.029889112 | 0.814459908 | 0.93 | 0.61 |
| YBR283C | 3928.577615 | -0.029595617 | 0.814928269 | 0.93 | 0.61 |
| YGR168C | 490.2041357 | 0.038797785 | 0.814874895 | 0.93 | 0.61 |
| YHL029C | 889.4266622 | 0.079167535 | 0.814949791 | 0.93 | 0.61 |
| YMR041C | 591.2121868 | -0.136868831 | 0.815009492 | 0.93 | 0.61 |
| YOR019W | 709.5102111 | 0.060970439 | 0.81447198 | 0.93 | 0.61 |
| YNL338W | 0.154202725 | 0.057053613 | 0.81515712 | 0.93 | 0.61 |
| YPL088W | 468.228703 | -0.107905312 | 0.815707822 | 0.93 | 0.62 |
| YOR318C | 25.32067042 | 0.10134175 | 0.81593046 | 0.93 | 0.62 |
| YDR455C | 13.39180516 | 0.101925846 | 0.816262041 | 0.93 | 0.62 |
| YGR102C | 796.5538343 | 0.049801974 | 0.816363128 | 0.93 | 0.62 |
| YDR170W-A | 128.0923816 | -0.119243305 | 0.817229334 | 0.93 | 0.62 |
| YER011W | 1426.306813 | -0.116508421 | 0.817326969 | 0.93 | 0.62 |
| YHR065C | 1260.588152 | 0.09838362 | 0.817096374 | 0.93 | 0.62 |
| YDL121C | 1113.601796 | 0.074039353 | 0.818044675 | 0.93 | 0.62 |
| YLR434C | 48.33837913 | 0.106044718 | 0.818072249 | 0.93 | 0.62 |
| YML050W | 378.8334393 | 0.051712184 | 0.818268345 | 0.93 | 0.62 |
| YOR129C | 585.8837465 | 0.033795465 | 0.818096959 | 0.93 | 0.62 |
| YOR344C | 1985.74596 | -0.047915942 | 0.818132032 | 0.93 | 0.62 |
| YDR088C | 645.3535973 | -0.053188532 | 0.818858867 | 0.93 | 0.62 |
| YGR207C | 1419.844599 | 0.04316707 | 0.818905517 | 0.93 | 0.62 |
| YKL106C-A | 1.684117879 | 0.101736359 | 0.818542463 | 0.93 | 0.62 |
| YMR144W | 347.0360461 | -0.053700966 | 0.818779534 | 0.93 | 0.62 |
| YDR537C | 28.61472288 | -0.070730947 | 0.819422675 | 0.93 | 0.62 |
| YCL001W | 1214.350549 | -0.02865042 | 0.820492572 | 0.93 | 0.62 |
| YLR096W | 1948.609356 | -0.048188596 | 0.82054903 | 0.93 | 0.62 |
| YNR059W | 692.5145797 | -0.049880309 | 0.820388838 | 0.93 | 0.62 |
| YPL001W | 324.2471356 | 0.096912165 | 0.820888676 | 0.93 | 0.62 |

| | | | | | |
|---|---|---|---|---|---|
| YBR220C | 693.632069 | -0.042680031 | 0.821714698 | 0.93 | 0.62 |
| YER060W-A | 584.2480367 | 0.077840666 | 0.821531933 | 0.93 | 0.62 |
| YGR038W | 610.0071877 | -0.027658882 | 0.821349509 | 0.93 | 0.62 |
| YML102W | 604.0898534 | -0.031899155 | 0.821720578 | 0.93 | 0.62 |
| YBR056W-A | 43.21007303 | -0.061451453 | 0.823404089 | 0.93 | 0.62 |
| YDR118W | 391.9679351 | -0.023983777 | 0.82238998 | 0.93 | 0.62 |
| YDR452W | 4294.15743 | -0.060994021 | 0.822870021 | 0.93 | 0.62 |
| YER010C | 1015.031336 | -0.035997214 | 0.823303746 | 0.93 | 0.62 |
| YFR048W | 703.4879681 | -0.047471793 | 0.823389945 | 0.93 | 0.62 |
| YGR146C-A | 13.60921533 | 0.12052671 | 0.822732541 | 0.93 | 0.62 |
| YMR138W | 253.6379214 | -0.041379664 | 0.822942114 | 0.93 | 0.62 |
| YNL063W | 522.2571621 | -0.036916443 | 0.82338353 | 0.93 | 0.62 |
| YNL162W-A | 54.12294867 | 0.113866233 | 0.82241408 | 0.93 | 0.62 |
| YPL127C | 2242.895143 | 0.056370385 | 0.823211641 | 0.93 | 0.62 |
| YPL214C | 1002.445323 | 0.039501822 | 0.82361202 | 0.93 | 0.62 |
| YPL274W | 1036.210642 | 0.074919488 | 0.822010674 | 0.93 | 0.62 |
| YPR149W | 11719.12039 | -0.085467627 | 0.823690939 | 0.93 | 0.62 |
| YDR132C | 453.7495244 | 0.053821098 | 0.824494694 | 0.93 | 0.62 |
| YER158C | 1028.126375 | -0.067208739 | 0.824380909 | 0.93 | 0.62 |
| YKR061W | 573.6088479 | 0.098246318 | 0.824765665 | 0.93 | 0.62 |
| YLR339C | 501.9621213 | -0.070704693 | 0.824634164 | 0.93 | 0.62 |
| YAL007C | 1549.491558 | 0.117029194 | 0.825751894 | 0.93 | 0.62 |
| YDL172C | 10.58915184 | 0.158639354 | 0.825585507 | 0.93 | 0.62 |
| YDR264C | 2392.197288 | -0.031300821 | 0.825324833 | 0.93 | 0.62 |
| YDR392W | 654.5457347 | -0.034654214 | 0.825059993 | 0.93 | 0.62 |
| YHL047C | 1687.543568 | 0.095018353 | 0.825660548 | 0.93 | 0.62 |
| YNR048W | 344.510222 | -0.040866915 | 0.825658625 | 0.93 | 0.62 |
| YBR007C | 740.5673067 | -0.039972258 | 0.827105922 | 0.94 | 0.62 |
| YBR212W | 2837.682435 | -0.076206959 | 0.827223635 | 0.94 | 0.62 |
| YHR081W | 846.2639082 | -0.053411008 | 0.826797092 | 0.94 | 0.62 |
| YNL319W | 5.59444275 | 0.127680021 | 0.827123774 | 0.94 | 0.62 |
| YMR252C | 396.7363367 | 0.044588353 | 0.827653129 | 0.94 | 0.62 |
| YOR161C | 928.1721785 | -0.11181889 | 0.8279898 | 0.94 | 0.62 |
| YPL028W | 3100.321144 | 0.102540847 | 0.828459987 | 0.94 | 0.62 |
| YLR298C | 806.4703735 | 0.036689237 | 0.828974819 | 0.94 | 0.62 |
| YAL023C | 4478.973811 | -0.022771042 | 0.829311714 | 0.94 | 0.62 |
| YLR081W | 221.380283 | 0.067210023 | 0.829254933 | 0.94 | 0.62 |
| YBR161W | 517.5371153 | 0.063031326 | 0.830167946 | 0.94 | 0.62 |
| YDR053W | 7.370760006 | 0.081877423 | 0.830189016 | 0.94 | 0.62 |
| YDR459C | 457.0405938 | -0.031852978 | 0.830929465 | 0.94 | 0.62 |
| YJL046W | 632.8219176 | 0.035966733 | 0.830744019 | 0.94 | 0.62 |
| YKL183C-A | 2.603283005 | -0.200338786 | 0.829682504 | 0.94 | 0.62 |
| YKR014C | 1917.457601 | 0.046628207 | 0.830408245 | 0.94 | 0.62 |

| | | | | | |
|---|---|---|---|---|---|
| YLR095C | 1759.144757 | -0.033307639 | 0.829898337 | 0.94 | 0.62 |
| YMR246W | 3290.570155 | -0.047841465 | 0.8304472 | 0.94 | 0.62 |
| YNR022C | 907.2227542 | -0.04728948 | 0.829798638 | 0.94 | 0.62 |
| YOR151C | 4196.260493 | -0.041481739 | 0.83060685 | 0.94 | 0.62 |
| YOR216C | 1165.7555 | -0.05137476 | 0.830841164 | 0.94 | 0.62 |
| YKL219W | 500.3272007 | -0.060209592 | 0.831329811 | 0.94 | 0.62 |
| YDR098C | 1711.271644 | 0.036304076 | 0.831592548 | 0.94 | 0.62 |
| YIL138C | 1495.862581 | -0.052003708 | 0.831631193 | 0.94 | 0.62 |
| YBR289W | 1372.875089 | -0.052244016 | 0.832226539 | 0.94 | 0.62 |
| YKR055W | 476.0228776 | 0.030879951 | 0.832649031 | 0.94 | 0.62 |
| YLR163C | 1121.057782 | 0.030547902 | 0.832663811 | 0.94 | 0.62 |
| YNL257C | 1662.623362 | -0.030348467 | 0.832334984 | 0.94 | 0.62 |
| YPL002C | 492.3419627 | -0.035126127 | 0.832737757 | 0.94 | 0.62 |
| YJL182C | 1.793622545 | 0.186091367 | 0.833422899 | 0.94 | 0.62 |
| YKR036C | 843.7972906 | -0.030912349 | 0.834372229 | 0.94 | 0.62 |
| YPR088C | 2257.094394 | 0.038789397 | 0.834412511 | 0.94 | 0.62 |
| tR(CCG)L | 0.085991782 | -0.04214211 | 0.8342358 | 0.94 | 0.62 |
| YLL018C-A | 305.6957299 | -0.075496659 | 0.83460393 | 0.94 | 0.62 |
| YLR361C-A | 262.7999201 | 0.044023072 | 0.83486308 | 0.94 | 0.62 |
| YFR039C | 736.7528646 | 0.05314643 | 0.835321609 | 0.94 | 0.62 |
| YKR062W | 891.4921763 | -0.028987422 | 0.835439875 | 0.94 | 0.62 |
| YBR119W | 605.7970902 | 0.065954904 | 0.835881283 | 0.94 | 0.62 |
| YLL017W | 60.73191178 | 0.059496378 | 0.836329043 | 0.94 | 0.62 |
| YER035W | 1131.143378 | 0.049300855 | 0.836692707 | 0.94 | 0.62 |
| YDR034C-A | 4.06626248 | 0.063788835 | 0.837200554 | 0.94 | 0.62 |
| YFR010W-A | 16.18554247 | 0.094128693 | 0.83809676 | 0.94 | 0.62 |
| YJL146W | 600.4909491 | 0.026276939 | 0.837954344 | 0.94 | 0.62 |
| YLR127C | 955.0961502 | 0.035305788 | 0.837776525 | 0.94 | 0.62 |
| YNL020C | 852.0700517 | 0.045907972 | 0.837891561 | 0.94 | 0.62 |
| YGR021W | 818.8357092 | 0.035992185 | 0.838261599 | 0.94 | 0.62 |
| YLR394W | 730.9999237 | 0.027907983 | 0.838792023 | 0.94 | 0.62 |
| YER076C | 386.5202214 | -0.039764714 | 0.839294421 | 0.94 | 0.62 |
| YLR179C | 2935.995773 | 0.136711214 | 0.839678482 | 0.94 | 0.62 |
| YDR238C | 4318.542325 | 0.040347999 | 0.839864797 | 0.94 | 0.62 |
| YGL128C | 505.7413235 | -0.055485983 | 0.840651661 | 0.94 | 0.62 |
| YDR138W | 884.1934286 | -0.031833999 | 0.840948921 | 0.94 | 0.62 |
| YIL132C | 143.1249549 | 0.068337593 | 0.841078967 | 0.94 | 0.62 |
| YGR166W | 818.3285643 | -0.046283053 | 0.841315865 | 0.94 | 0.62 |
| YDL237W | 1508.528864 | 0.047263234 | 0.842192307 | 0.94 | 0.62 |
| YDR261C-D | 88.68167414 | 0.104300533 | 0.84204318 | 0.94 | 0.62 |
| YHR129C | 426.0946941 | 0.04846263 | 0.841750075 | 0.94 | 0.62 |
| YMR204C | 437.305049 | 0.042918956 | 0.841986517 | 0.94 | 0.62 |
| YMR288W | 774.5062547 | 0.034495301 | 0.842144514 | 0.94 | 0.62 |

| | | | | | |
|---|---|---|---|---|---|
| YPL084W | 2075.944666 | -0.027695975 | 0.842015847 | 0.94 | 0.62 |
| YDR038C | 125.3516014 | 0.043394315 | 0.8428957 | 0.94 | 0.62 |
| YDR486C | 1425.158924 | -0.044986724 | 0.842478303 | 0.94 | 0.62 |
| YPR011C | 560.157333 | 0.058522038 | 0.842761029 | 0.94 | 0.62 |
| tL(GAG)G | 0.599720226 | 0.112374961 | 0.842782125 | 0.94 | 0.62 |
| YGL084C | 1084.484417 | -0.026329702 | 0.843479913 | 0.94 | 0.62 |
| YHR046C | 620.1635411 | 0.081396153 | 0.844048729 | 0.94 | 0.62 |
| YOR170W | 97.11015677 | 0.049407956 | 0.84395136 | 0.94 | 0.62 |
| YOR347C | 1046.083791 | 0.070231626 | 0.843780731 | 0.94 | 0.62 |
| YOR353C | 984.6133479 | -0.054957024 | 0.843819176 | 0.94 | 0.62 |
| YHL005C | 24.92643688 | 0.042538942 | 0.844818807 | 0.94 | 0.62 |
| YKR046C | 2535.087216 | 0.043022412 | 0.844834667 | 0.94 | 0.62 |
| YMR106C | 546.3577506 | 0.03438772 | 0.844855966 | 0.94 | 0.62 |
| YOL069W | 510.0668666 | -0.033088445 | 0.844964402 | 0.94 | 0.62 |
| YBR175W | 264.1838842 | -0.050985625 | 0.845377264 | 0.94 | 0.62 |
| YCL061C | 1120.832529 | 0.048564909 | 0.845439875 | 0.94 | 0.62 |
| YDR107C | 540.536576 | -0.029229023 | 0.845623197 | 0.94 | 0.62 |
| YLR443W | 472.8026492 | -0.039924686 | 0.845695642 | 0.94 | 0.62 |
| YKL033W | 666.8103831 | 0.032325869 | 0.845933586 | 0.94 | 0.62 |
| YAL037C-A | 1.195205434 | 0.228183508 | 0.846128773 | 0.94 | 0.62 |
| YGL164C | 861.3475827 | 0.034373706 | 0.846499078 | 0.95 | 0.62 |
| YDL090C | 724.6090506 | -0.037319677 | 0.847146656 | 0.95 | 0.62 |
| YHR010W | 11885.41138 | -0.003228926 | 0.847265583 | 0.95 | 0.62 |
| YBR015C | 2379.48263 | -0.028502217 | 0.848634362 | 0.95 | 0.63 |
| YDR419W | 525.8031882 | 0.043036538 | 0.848498339 | 0.95 | 0.63 |
| YGL234W | 9210.041986 | 0.144224727 | 0.848613007 | 0.95 | 0.63 |
| YGR163W | 840.8307527 | 0.033655705 | 0.848435421 | 0.95 | 0.63 |
| YLL060C | 550.1729711 | 0.045540494 | 0.848474123 | 0.95 | 0.63 |
| YLR316C | 329.0837142 | 0.026829143 | 0.84814292 | 0.95 | 0.63 |
| YNL225C | 446.0966712 | -0.0301781 | 0.848705646 | 0.95 | 0.63 |
| YOR046C | 2190.495993 | -0.036314615 | 0.848099273 | 0.95 | 0.63 |
| YIL151C | 1650.033227 | 0.03665324 | 0.84892557 | 0.95 | 0.63 |
| YAR023C | 164.0948317 | -0.070173067 | 0.849662977 | 0.95 | 0.63 |
| YDR167W | 803.2507476 | -0.027618172 | 0.849743594 | 0.95 | 0.63 |
| YMR198W | 512.8655409 | 0.030869253 | 0.849376652 | 0.95 | 0.63 |
| YNL296W | 0.17572267 | 0.030291598 | 0.849341989 | 0.95 | 0.63 |
| YOL070C | 1742.264613 | 0.039776395 | 0.849753293 | 0.95 | 0.63 |
| YKR079C | 916.529157 | -0.039292131 | 0.849947055 | 0.95 | 0.63 |
| YFR004W | 2781.166019 | -0.029081657 | 0.850465476 | 0.95 | 0.63 |
| YDL118W | 21.28319356 | 0.048377109 | 0.850710337 | 0.95 | 0.63 |
| YDR044W | 1839.127613 | -0.038619538 | 0.851131953 | 0.95 | 0.63 |
| YOR261C | 3484.142585 | 0.040049769 | 0.851460841 | 0.95 | 0.63 |
| snR80 | 3.471418478 | 0.118318281 | 0.851680452 | 0.95 | 0.63 |

| | | | | | |
|---|---|---|---|---|---|
| YAL044W-A | 228.0761832 | 0.0435445 | 0.85340768 | 0.95 | 0.63 |
| YDL099W | 2397.811912 | -0.040017624 | 0.85357669 | 0.95 | 0.63 |
| YDR344C | 46.73109222 | -0.056161812 | 0.853565008 | 0.95 | 0.63 |
| YER034W | 308.6813768 | 0.052101186 | 0.852558461 | 0.95 | 0.63 |
| YER058W | 286.3082478 | -0.038291905 | 0.853267257 | 0.95 | 0.63 |
| YGR210C | 1863.079648 | 0.057335702 | 0.852859125 | 0.95 | 0.63 |
| YHR039C-A | 2505.228571 | 0.036855413 | 0.853591722 | 0.95 | 0.63 |
| YIR018C-A | 31.48971955 | 0.012957946 | 0.853480012 | 0.95 | 0.63 |
| YJR136C | 423.3917741 | -0.0285892 | 0.852095727 | 0.95 | 0.63 |
| YMR167W | 578.0579077 | -0.031308264 | 0.85354565 | 0.95 | 0.63 |
| YNL213C | 587.1355028 | 0.086162084 | 0.853333303 | 0.95 | 0.63 |
| YOR250C | 655.6893916 | 0.029109546 | 0.852299376 | 0.95 | 0.63 |
| YPL114W | 1.969765016 | 0.040804598 | 0.853532798 | 0.95 | 0.63 |
| YGL154C | 461.1131948 | -0.046459355 | 0.854295173 | 0.95 | 0.63 |
| YLR063W | 280.3060801 | -0.076064034 | 0.854273857 | 0.95 | 0.63 |
| YNL304W | 349.9176759 | -0.033251602 | 0.854147481 | 0.95 | 0.63 |
| YOR376W-A | 83.93269889 | 0.063060691 | 0.853864665 | 0.95 | 0.63 |
| YKL126W | 2370.590453 | 0.035911936 | 0.854715252 | 0.95 | 0.63 |
| YPR089W | 1983.427567 | -0.030295646 | 0.85458253 | 0.95 | 0.63 |
| YGR228W | 14.43502374 | -0.147831349 | 0.855095708 | 0.95 | 0.63 |
| YGR053C | 339.3237027 | -0.070387272 | 0.85536177 | 0.95 | 0.63 |
| YBR153W | 413.971492 | 0.027543633 | 0.856240818 | 0.95 | 0.63 |
| YCR026C | 1309.803177 | -0.048212436 | 0.856451923 | 0.95 | 0.63 |
| YDL177C | 218.4776378 | -0.046232784 | 0.856671381 | 0.95 | 0.63 |
| YDR068W | 1258.879245 | 0.016110702 | 0.856255416 | 0.95 | 0.63 |
| YFL021C-A | 41.5780796 | 0.082639417 | 0.856488719 | 0.95 | 0.63 |
| YJR021C | 884.0147888 | -0.059156182 | 0.855689684 | 0.95 | 0.63 |
| YJR099W | 741.8529597 | 0.031065244 | 0.855564989 | 0.95 | 0.63 |
| YKL052C | 501.3271381 | 0.03946657 | 0.856189505 | 0.95 | 0.63 |
| YOR010C | 455.6940822 | -0.131753814 | 0.85670763 | 0.95 | 0.63 |
| YBL053W | 19.02715806 | -0.181482196 | 0.856986596 | 0.95 | 0.63 |
| YBR084W | 2864.241089 | 0.073742243 | 0.857327183 | 0.95 | 0.63 |
| YDR519W | 900.2252274 | -0.032288427 | 0.857565448 | 0.95 | 0.63 |
| snR31 | 65.06005548 | 0.001594054 | 0.857598922 | 0.95 | 0.63 |
| YBR185C | 732.2746685 | 0.038062995 | 0.860425701 | 0.95 | 0.63 |
| YCL004W | 588.1019431 | -0.037316163 | 0.860790431 | 0.95 | 0.63 |
| YCR020C | 245.5464449 | -0.097935374 | 0.858649754 | 0.95 | 0.63 |
| YDR401W | 0.227003213 | 0.061320004 | 0.859113725 | 0.95 | 0.63 |
| YDR511W | 614.1187303 | 0.05692104 | 0.858495634 | 0.95 | 0.63 |
| YER133W | 3950.455243 | 0.033528672 | 0.860553056 | 0.95 | 0.63 |
| YGR016W | 101.0221921 | -0.037362035 | 0.860442255 | 0.95 | 0.63 |
| YGR247W | 484.9641046 | -0.054923297 | 0.859276858 | 0.95 | 0.63 |
| YHR063W-A | 174.3401986 | -0.031330064 | 0.859384222 | 0.95 | 0.63 |

| | | | | | |
|---|---|---|---|---|---|
| YIL090W | 898.4488003 | 0.027920585 | 0.859686484 | 0.95 | 0.63 |
| YJL121C | 2601.821139 | -0.025956322 | 0.860457727 | 0.95 | 0.63 |
| YKL223W | 0.064032033 | 0.046316131 | 0.859502467 | 0.95 | 0.63 |
| YLR374C | 34.93928504 | 0.032436974 | 0.860794129 | 0.95 | 0.63 |
| YML012W | 3525.480868 | -0.020516841 | 0.860699624 | 0.95 | 0.63 |
| YMR182W-A | 4.590688512 | 0.045382811 | 0.860535434 | 0.95 | 0.63 |
| YMR297W | 8256.874804 | -0.042258501 | 0.859206491 | 0.95 | 0.63 |
| YNR043W | 2170.763038 | -0.037795248 | 0.858959726 | 0.95 | 0.63 |
| YPR136C | 12.04995248 | 0.043289371 | 0.860435252 | 0.95 | 0.63 |
| YBR261C | 744.8069304 | -0.013654133 | 0.861602758 | 0.95 | 0.63 |
| YHR009C | 3439.835058 | 0.028427663 | 0.861706373 | 0.95 | 0.63 |
| YMR011W | 5664.342762 | -0.077720063 | 0.861467603 | 0.95 | 0.63 |
| YNL009W | 277.6331838 | -0.102390688 | 0.861282835 | 0.95 | 0.63 |
| YHR083W | 1675.31411 | -0.026525807 | 0.862028608 | 0.95 | 0.63 |
| YIL078W | 8337.626041 | 0.084307334 | 0.862049468 | 0.95 | 0.63 |
| YGR031W | 376.7830976 | -0.046300022 | 0.862207664 | 0.95 | 0.63 |
| YEL043W | 2331.741844 | -0.040652625 | 0.862461564 | 0.95 | 0.63 |
| YPL172C | 484.663762 | 0.03262874 | 0.862521163 | 0.95 | 0.63 |
| YDR282C | 383.6210181 | 0.037764671 | 0.862724128 | 0.95 | 0.63 |
| YGR006W | 239.4957069 | -0.031250779 | 0.863041704 | 0.95 | 0.63 |
| YEL017W | 1022.681916 | -0.007101489 | 0.865052598 | 0.95 | 0.63 |
| YGL248W | 1428.88377 | -0.063942558 | 0.865557413 | 0.95 | 0.63 |
| YGR203W | 222.1887092 | -0.048177785 | 0.86417902 | 0.95 | 0.63 |
| YHR085W | 335.931842 | 0.086846637 | 0.865841025 | 0.95 | 0.63 |
| YJR094C | 253.8151399 | -0.067652608 | 0.864865114 | 0.95 | 0.63 |
| YKL165C | 1154.053703 | -0.008782659 | 0.864463614 | 0.95 | 0.63 |
| YKR013W | 4305.106637 | 0.07351195 | 0.865449641 | 0.95 | 0.63 |
| YKR075C | 2465.019565 | -0.122117185 | 0.865880296 | 0.95 | 0.63 |
| YLR237W | 1061.722449 | 0.037002112 | 0.865151106 | 0.95 | 0.63 |
| YMR028W | 753.5245359 | 0.020140435 | 0.865689281 | 0.95 | 0.63 |
| YPL016W | 1163.813997 | 0.040276948 | 0.864413157 | 0.95 | 0.63 |
| YPR108W-A | 1.090591333 | 0.25199916 | 0.865043902 | 0.95 | 0.63 |
| snR72 | 8.104041704 | 0.032236882 | 0.865471511 | 0.95 | 0.63 |
| YLR367W | 2289.066484 | 0.067946915 | 0.866126501 | 0.95 | 0.63 |
| YFL014W | 10835.63862 | -0.145571833 | 0.866804495 | 0.95 | 0.63 |
| YDR252W | 88.93914678 | 0.054518531 | 0.86697299 | 0.95 | 0.63 |
| YDR186C | 2571.215729 | 0.04069161 | 0.867212383 | 0.95 | 0.63 |
| YKL084W | 479.3447424 | 0.03421606 | 0.867823921 | 0.95 | 0.63 |
| YDR037W | 9604.757078 | 0.107997475 | 0.868353239 | 0.95 | 0.63 |
| YIL006W | 374.9591791 | 0.053410668 | 0.86840282 | 0.95 | 0.63 |
| YAR014C | 1593.739874 | -0.03440273 | 0.86959764 | 0.95 | 0.63 |
| YER168C | 714.524667 | -0.022478742 | 0.869584179 | 0.95 | 0.63 |
| YLR438C-A | 659.2030513 | 0.039214713 | 0.871561014 | 0.96 | 0.63 |

| | | | | | |
|---|---|---|---|---|---|
| YER115C | 529.525795 | -0.051645997 | 0.872445396 | 0.96 | 0.63 |
| YKL189W | 702.1789039 | -0.045553374 | 0.87238828 | 0.96 | 0.63 |
| YLR125W | 49.15160341 | -0.067074187 | 0.872393924 | 0.96 | 0.63 |
| YLR231C | 929.429053 | 0.106423305 | 0.872094839 | 0.96 | 0.63 |
| YOR277C | 6.042620409 | 0.051428663 | 0.872580834 | 0.96 | 0.63 |
| YHL014C | 453.0111576 | 0.041547648 | 0.872879365 | 0.96 | 0.63 |
| YNL086W | 191.101467 | -0.045518313 | 0.872946012 | 0.96 | 0.63 |
| YNL204C | 121.2325338 | -0.067162411 | 0.873312144 | 0.96 | 0.63 |
| YML068W | 512.1804017 | 0.022625681 | 0.873963382 | 0.96 | 0.63 |
| YBL091C-A | 508.1746239 | 0.032181565 | 0.87438982 | 0.96 | 0.63 |
| YBL104C | 644.4185242 | 0.024094413 | 0.875178342 | 0.96 | 0.63 |
| YDL134C | 1962.281647 | 0.011511637 | 0.874934421 | 0.96 | 0.63 |
| YGR028W | 820.509086 | 0.032717424 | 0.874686196 | 0.96 | 0.63 |
| YGR171C | 566.9877726 | -0.022405446 | 0.87511323 | 0.96 | 0.63 |
| YHR181W | 756.8304441 | 0.081595019 | 0.874306007 | 0.96 | 0.63 |
| YKL092C | 887.6693114 | 0.036906711 | 0.874879769 | 0.96 | 0.63 |
| YMR123W | 1008.572929 | -0.024601452 | 0.875410693 | 0.96 | 0.63 |
| YPR197C | 5.599447647 | 0.031189544 | 0.875290388 | 0.96 | 0.63 |
| YPR077C | 1.201353204 | 0.074900025 | 0.875562437 | 0.96 | 0.63 |
| YBL077W | 139.1714064 | -0.028737956 | 0.878518056 | 0.96 | 0.63 |
| YBL096C | 11.76381799 | -0.02597298 | 0.876887449 | 0.96 | 0.63 |
| YBR122C | 1316.133892 | -0.034964451 | 0.87736769 | 0.96 | 0.63 |
| YCR075C | 329.3656461 | 0.028255597 | 0.879721936 | 0.96 | 0.63 |
| YDR363W | 641.2311125 | 0.034946027 | 0.876284298 | 0.96 | 0.63 |
| YFL025C | 800.9686963 | -0.015577313 | 0.879414103 | 0.96 | 0.63 |
| YGL065C | 1014.84416 | -0.01927318 | 0.878660553 | 0.96 | 0.63 |
| YHR069C | 771.2357851 | -0.024482502 | 0.877883923 | 0.96 | 0.63 |
| YHR121W | 1103.128777 | 0.04112567 | 0.878629402 | 0.96 | 0.63 |
| YIL149C | 3262.267006 | -0.046096436 | 0.879248891 | 0.96 | 0.63 |
| YJL162C | 484.7743599 | -0.026194055 | 0.878588757 | 0.96 | 0.63 |
| YJL187C | 1084.409994 | 0.04462924 | 0.876825951 | 0.96 | 0.63 |
| YKL023C-A | 532.0907152 | 0.054063024 | 0.8794263 | 0.96 | 0.63 |
| YKL145W | 5482.552156 | 0.029510129 | 0.878319845 | 0.96 | 0.63 |
| YLL027W | 1273.25217 | 0.024048529 | 0.879407238 | 0.96 | 0.63 |
| YLR157C-B | 0.750636836 | 0.064138689 | 0.877106123 | 0.96 | 0.63 |
| YMR100W | 811.7962257 | 0.033585264 | 0.877453616 | 0.96 | 0.63 |
| YMR294W | 447.832337 | -0.042642673 | 0.876788923 | 0.96 | 0.63 |
| YNL120C | 0.227028385 | -0.058524454 | 0.877265454 | 0.96 | 0.63 |
| YNL221C | 1802.455751 | 0.035629865 | 0.877879228 | 0.96 | 0.63 |
| YNL312W | 897.7476795 | 0.046051034 | 0.879462946 | 0.96 | 0.63 |
| YNR025C | 48.20698063 | -0.047240287 | 0.878223824 | 0.96 | 0.63 |
| YNR077C | 0.24124529 | -0.047267888 | 0.879687785 | 0.96 | 0.63 |
| YOL103W | 1490.877623 | 0.024709619 | 0.876515325 | 0.96 | 0.63 |

| | | | | | |
|---|---|---|---|---|---|
| YOR379C | 0.107512111 | -0.039539103 | 0.87727292 | 0.96 | 0.63 |
| YPL124W | 247.3933842 | 0.048857002 | 0.87646058 | 0.96 | 0.63 |
| YPR180W | 730.2139766 | -0.030444544 | 0.878835282 | 0.96 | 0.63 |
| YBL034C | 918.1202298 | 0.040675584 | 0.88064191 | 0.96 | 0.63 |
| YBL071W-A | 297.3219208 | -0.043231661 | 0.880074521 | 0.96 | 0.63 |
| YGL235W | 1.903973228 | 0.000321431 | 0.88021538 | 0.96 | 0.63 |
| YGR196C | 1285.850548 | -0.027170854 | 0.880262067 | 0.96 | 0.63 |
| YIL041W | 4311.545468 | -0.028789986 | 0.880459472 | 0.96 | 0.63 |
| YIL077C | 964.8137743 | 0.03947169 | 0.880572207 | 0.96 | 0.63 |
| YGR109W-B | 857.2496031 | 0.124088688 | 0.881185536 | 0.96 | 0.63 |
| YCL011C | 2495.29652 | -0.027677585 | 0.881428805 | 0.96 | 0.63 |
| YDR024W | 7.360211428 | 0.139331676 | 0.881597746 | 0.96 | 0.63 |
| YIR040C | 0.167590483 | 0.059415105 | 0.881721489 | 0.96 | 0.63 |
| YHR123W | 699.5030842 | 0.003705978 | 0.882033439 | 0.96 | 0.63 |
| YDR151C | 1113.580859 | 0.023748842 | 0.882751411 | 0.96 | 0.63 |
| YDR530C | 1226.853067 | 0.032253111 | 0.882524157 | 0.96 | 0.63 |
| YKL090W | 556.2185917 | -0.047357585 | 0.882782906 | 0.96 | 0.63 |
| YKL127W | 1973.346678 | -0.000546263 | 0.882704891 | 0.96 | 0.63 |
| YER091C-A | 0.485272411 | 0.282850648 | 0.883621558 | 0.96 | 0.63 |
| YOR118W | 1462.615579 | 0.016151602 | 0.883660639 | 0.96 | 0.63 |
| YGL232W | 728.6792639 | 0.061740107 | 0.883871228 | 0.96 | 0.63 |
| YGR043C | 1676.63942 | 0.033688215 | 0.884175304 | 0.96 | 0.63 |
| YJR004C | 1252.603429 | 0.013123811 | 0.884507339 | 0.96 | 0.63 |
| YOR257W | 743.3095021 | 0.038783373 | 0.884477086 | 0.96 | 0.63 |
| YCL029C | 771.878883 | 0.027392548 | 0.884728941 | 0.96 | 0.63 |
| YCL008C | 1030.586387 | 0.019046366 | 0.885752099 | 0.96 | 0.63 |
| YDL203C | 667.0945544 | -0.017257386 | 0.885214711 | 0.96 | 0.63 |
| YGL053W | 527.1405514 | 0.047821896 | 0.885836293 | 0.96 | 0.63 |
| YHR036W | 340.3963184 | -0.019975147 | 0.885492017 | 0.96 | 0.63 |
| YJL064W | 3.253557565 | -0.073849465 | 0.885031075 | 0.96 | 0.63 |
| YKL046C | 1814.036225 | -0.01531045 | 0.885503024 | 0.96 | 0.63 |
| YKL128C | 1025.460861 | -0.028352604 | 0.885696121 | 0.96 | 0.63 |
| YLR421C | 1264.683888 | -0.025639036 | 0.88601726 | 0.96 | 0.63 |
| YPR074W-A | 0.407755058 | -0.076019402 | 0.886871168 | 0.96 | 0.63 |
| YBR069C | 2910.073987 | -0.02267502 | 0.887152121 | 0.96 | 0.63 |
| YLL053C | 96.70396409 | -0.088032662 | 0.887384786 | 0.96 | 0.63 |
| YKL138C-A | 204.185803 | -0.027034388 | 0.887789097 | 0.96 | 0.63 |
| YMR073C | 579.5582083 | -0.033888195 | 0.887992406 | 0.96 | 0.63 |
| YBR190W | 3.930160671 | -0.042229592 | 0.888660718 | 0.96 | 0.63 |
| YGL238W | 2547.939909 | 0.024509513 | 0.888445733 | 0.96 | 0.63 |
| YJR135W-A | 435.3473743 | -0.015762711 | 0.888605876 | 0.96 | 0.63 |
| YLR094C | 1048.349221 | 0.018804833 | 0.888256066 | 0.96 | 0.63 |
| YML028W | 27162.20889 | -0.018568292 | 0.888919027 | 0.96 | 0.63 |

| | | | | | |
|---|---|---|---|---|---|
| YDR541C | 514.5384389 | -0.022770349 | 0.88965062 | 0.96 | 0.63 |
| YKL044W | 194.2159883 | 0.164469644 | 0.88979287 | 0.96 | 0.63 |
| YOL098C | 2415.997601 | 0.031093022 | 0.889622316 | 0.96 | 0.63 |
| YPR068C | 284.1750342 | -0.018699596 | 0.88972105 | 0.96 | 0.63 |
| YBR065C | 528.4307138 | -0.048608842 | 0.889945887 | 0.96 | 0.63 |
| YAR064W | 14.8431874 | -0.020911807 | 0.89039105 | 0.96 | 0.64 |
| YDR483W | 4802.755253 | -0.013322239 | 0.891126629 | 0.96 | 0.64 |
| YER121W | 184.2483 | 0.030660117 | 0.891050001 | 0.96 | 0.64 |
| YHR049C-A | 129.3206508 | -0.001860854 | 0.891296011 | 0.96 | 0.64 |
| YHR195W | 1250.379878 | 0.024134653 | 0.890916513 | 0.96 | 0.64 |
| YLR074C | 1095.319455 | -0.038979831 | 0.891389162 | 0.96 | 0.64 |
| YPL205C | 28.51967725 | -0.041896068 | 0.890837379 | 0.96 | 0.64 |
| snR65 | 5.68201134 | -0.007725492 | 0.891351867 | 0.96 | 0.64 |
| YLR438W | 1335.790008 | 0.019938567 | 0.891966858 | 0.96 | 0.64 |
| YFR041C | 978.3498425 | 0.039797708 | 0.892519095 | 0.96 | 0.64 |
| YKL065C | 4326.57925 | 0.019263674 | 0.892449875 | 0.96 | 0.64 |
| YML054C | 1101.015099 | 0.07657414 | 0.892346133 | 0.96 | 0.64 |
| YHR021W-A | 41.37134005 | 0.012233939 | 0.893098923 | 0.96 | 0.64 |
| YHR175W-A | 167.202498 | -0.005003215 | 0.893541084 | 0.96 | 0.64 |
| YGL217C | 8.839690893 | 0.112368231 | 0.893769835 | 0.96 | 0.64 |
| YCR024C-A | 1033.453894 | -0.006456917 | 0.894927664 | 0.96 | 0.64 |
| YCR047C | 558.0855162 | -0.020696193 | 0.894901179 | 0.96 | 0.64 |
| YDR317W | 303.4699364 | 0.029993343 | 0.89476222 | 0.96 | 0.64 |
| YER039C-A | 49.35745349 | -0.0714158 | 0.89420574 | 0.96 | 0.64 |
| YER080W | 3267.440935 | 0.023036082 | 0.89429671 | 0.96 | 0.64 |
| YJL054W | 1689.096692 | 0.02900638 | 0.894529785 | 0.96 | 0.64 |
| YJL199C | 163.0335414 | -0.033608126 | 0.894868061 | 0.96 | 0.64 |
| YLL035W | 581.4295973 | 0.038805 | 0.895414546 | 0.96 | 0.64 |
| YOL125W | 710.1287675 | 0.029940026 | 0.895467623 | 0.96 | 0.64 |
| YDL157C | 952.2291983 | -0.033703346 | 0.896373123 | 0.96 | 0.64 |
| YHR180W-A | 0.36648756 | 0.072826465 | 0.896047435 | 0.96 | 0.64 |
| YJL031C | 712.299831 | 0.02352519 | 0.896179452 | 0.96 | 0.64 |
| YLR141W | 304.974896 | -0.034331474 | 0.896226805 | 0.96 | 0.64 |
| YOR201C | 630.2633868 | -0.018414273 | 0.896111814 | 0.96 | 0.64 |
| YCL016C | 442.1886812 | 0.018851387 | 0.897361497 | 0.96 | 0.64 |
| YDR129C | 5762.337606 | 0.020886826 | 0.897133467 | 0.96 | 0.64 |
| YGR127W | 732.3981957 | 0.024799651 | 0.897404843 | 0.96 | 0.64 |
| YIL101C | 1402.735777 | -0.086069311 | 0.897392011 | 0.96 | 0.64 |
| YIL160C | 460.4774311 | -0.086004615 | 0.897609713 | 0.96 | 0.64 |
| YIR018W | 398.3013208 | 0.029166565 | 0.897480347 | 0.96 | 0.64 |
| YMR214W | 1048.466602 | -0.014181123 | 0.897455716 | 0.96 | 0.64 |
| YHR193C-A | 1.288094432 | 0.127966605 | 0.897908076 | 0.96 | 0.64 |
| YOR032W-A | 11.5403846 | 0.089300774 | 0.897934725 | 0.96 | 0.64 |

| | | | | | |
|---|---|---|---|---|---|
| YDR198C | 539.1037114 | 0.020456328 | 0.89863466 | 0.96 | 0.64 |
| YGL074C | 37.00723578 | 0.062920686 | 0.8983671 | 0.96 | 0.64 |
| YIL123W | 4293.520238 | 0.0659897 | 0.898613221 | 0.96 | 0.64 |
| YMR294W-A | 229.7802387 | -0.024304396 | 0.898630423 | 0.96 | 0.64 |
| YPR175W | 704.7198488 | 0.046039558 | 0.899144518 | 0.96 | 0.64 |
| YLR019W | 1293.150392 | -0.024805462 | 0.899483915 | 0.96 | 0.64 |
| YOR270C | 5846.210473 | 0.033092124 | 0.899525367 | 0.96 | 0.64 |
| YCR087W | 24.15940244 | -0.055809467 | 0.900484867 | 0.96 | 0.64 |
| YGL060W | 769.6330407 | -0.027786236 | 0.900509985 | 0.96 | 0.64 |
| YOR280C | 559.6900755 | -0.030277459 | 0.900707595 | 0.96 | 0.64 |
| YBL103C | 858.4740089 | 0.0216257 | 0.901036806 | 0.96 | 0.64 |
| YBR173C | 2339.401983 | -0.022388621 | 0.900996621 | 0.96 | 0.64 |
| YBR082C | 6782.616874 | 0.041153368 | 0.902071878 | 0.96 | 0.64 |
| YDR082W | 550.4664737 | -0.039593558 | 0.901904148 | 0.96 | 0.64 |
| YGR291C | 1.566792632 | -0.137830612 | 0.901325309 | 0.96 | 0.64 |
| YJL022W | 2.660856745 | -0.02858258 | 0.902095495 | 0.96 | 0.64 |
| YLR419W | 1247.341194 | 0.030439878 | 0.901961811 | 0.96 | 0.64 |
| YMR152W | 1727.69017 | -0.022745647 | 0.901915189 | 0.96 | 0.64 |
| YPL003W | 547.4702758 | 0.021203856 | 0.901571141 | 0.96 | 0.64 |
| YNR074C | 411.2345917 | -0.036914943 | 0.902740004 | 0.97 | 0.64 |
| YGR275W | 659.6729657 | -0.02077211 | 0.903025779 | 0.97 | 0.64 |
| YOR314W | 17.17022364 | 0.0734045 | 0.903302337 | 0.97 | 0.64 |
| YGL021W | 1221.378254 | -0.021675456 | 0.903640182 | 0.97 | 0.64 |
| YGL130W | 854.7206722 | 0.011928846 | 0.90352058 | 0.97 | 0.64 |
| YNL308C | 3277.042014 | 0.044279136 | 0.904049439 | 0.97 | 0.64 |
| YNR001W-A | 2.398336405 | 0.123214158 | 0.9046669 | 0.97 | 0.64 |
| YBR041W | 1541.15066 | -0.018787927 | 0.905544635 | 0.97 | 0.64 |
| YGL098W | 531.3017666 | -0.032512461 | 0.90557263 | 0.97 | 0.64 |
| YIR001C | 869.6360964 | -0.016757773 | 0.905073497 | 0.97 | 0.64 |
| YMR298W | 1278.824067 | -0.018388018 | 0.90492148 | 0.97 | 0.64 |
| YNL299W | 424.5188557 | -0.021425166 | 0.905495717 | 0.97 | 0.64 |
| YNR015W | 1147.190779 | 0.024660606 | 0.905400857 | 0.97 | 0.64 |
| YFR022W | 849.2747048 | 0.025891551 | 0.905761293 | 0.97 | 0.64 |
| YER060W | 750.1023347 | 0.041667995 | 0.905939933 | 0.97 | 0.64 |
| YNL025C | 730.0152463 | 0.014759604 | 0.90618196 | 0.97 | 0.64 |
| YKL168C | 972.1957599 | 0.01658222 | 0.906856493 | 0.97 | 0.64 |
| YIL043C | 4110.226673 | -0.017713344 | 0.907042935 | 0.97 | 0.64 |
| YJL172W | 1932.176519 | 0.030398544 | 0.907211961 | 0.97 | 0.64 |
| YHR096C | 1194.376628 | -0.088696081 | 0.907660931 | 0.97 | 0.64 |
| YHR167W | 161.5282471 | -0.005254899 | 0.907785201 | 0.97 | 0.64 |
| YML004C | 3324.450509 | 0.012677167 | 0.907923568 | 0.97 | 0.64 |
| YJL192C | 2669.267366 | 0.043623395 | 0.908534499 | 0.97 | 0.64 |
| YDL040C | 3238.410502 | 0.036580475 | 0.908847984 | 0.97 | 0.64 |

| | | | | | |
|---|---|---|---|---|---|
| YGR157W | 3036.942102 | 0.049414789 | 0.909260155 | 0.97 | 0.64 |
| YKL022C | 1018.942477 | 0.018183627 | 0.909034179 | 0.97 | 0.64 |
| YPR173C | 1842.223717 | -0.015365478 | 0.909213932 | 0.97 | 0.64 |
| YLR412C-A | 1.32995427 | 0.215689144 | 0.909589049 | 0.97 | 0.64 |
| YMR311C | 3644.500712 | 0.010897696 | 0.909752483 | 0.97 | 0.64 |
| YPR087W | 20.87255534 | 0.023429943 | 0.90974889 | 0.97 | 0.64 |
| YDL045C | 1046.513314 | 0.010792153 | 0.910011381 | 0.97 | 0.64 |
| YCR101C | 105.2670668 | 0.059241237 | 0.910491124 | 0.97 | 0.64 |
| YNL121C | 5779.129433 | -0.016178725 | 0.910612723 | 0.97 | 0.64 |
| YBL102W | 1291.754797 | 0.028631066 | 0.912283886 | 0.97 | 0.64 |
| YBR102C | 889.0122882 | 0.016441116 | 0.912220666 | 0.97 | 0.64 |
| YDL156W | 331.6691228 | -0.022583113 | 0.9118706 | 0.97 | 0.64 |
| YER187W | 132.2577032 | 0.096884547 | 0.911404307 | 0.97 | 0.64 |
| YLL037W | 13.52665894 | -0.057709912 | 0.912252733 | 0.97 | 0.64 |
| YLR319C | 843.3891881 | -0.013437878 | 0.911500414 | 0.97 | 0.64 |
| YMR158W-B | 1.160282167 | -0.07276874 | 0.911759145 | 0.97 | 0.64 |
| YOL084W | 492.0065357 | 0.036496357 | 0.911555758 | 0.97 | 0.64 |
| YPL135W | 1241.708085 | -0.03248223 | 0.912240431 | 0.97 | 0.64 |
| snR71 | 12.58626851 | 0.056132794 | 0.911582073 | 0.97 | 0.64 |
| YDL140C | 8072.034708 | 0.043699458 | 0.9133321 | 0.97 | 0.64 |
| YGR274C | 1344.663557 | 0.020780361 | 0.913294334 | 0.97 | 0.64 |
| YHR143W | 3407.641358 | 0.034706163 | 0.913403462 | 0.97 | 0.64 |
| YKL043W | 3024.896012 | 0.135891942 | 0.913185891 | 0.97 | 0.64 |
| YLR236C | 21.91002832 | 0.03524306 | 0.912926708 | 0.97 | 0.64 |
| YLR379W | 289.3293491 | -0.005809769 | 0.91367492 | 0.97 | 0.64 |
| YAR066W | 12.65917974 | -0.002246834 | 0.914288235 | 0.97 | 0.64 |
| YNL261W | 688.6478875 | -0.022096446 | 0.91439896 | 0.97 | 0.64 |
| YCR015C | 255.0577346 | -0.01137058 | 0.915215287 | 0.97 | 0.64 |
| YDR079W | 403.891688 | 0.026867327 | 0.914951381 | 0.97 | 0.64 |
| YDR428C | 292.2817321 | -0.027204661 | 0.915318489 | 0.97 | 0.64 |
| YDR437W | 89.57715872 | -0.024446435 | 0.915749047 | 0.97 | 0.64 |
| YER047C | 911.6547256 | 0.016447927 | 0.91481009 | 0.97 | 0.64 |
| YER111C | 1455.121069 | 0.027101169 | 0.915546904 | 0.97 | 0.64 |
| YGL138C | 22.53163453 | 0.015905062 | 0.915798271 | 0.97 | 0.64 |
| YOR008C-A | 9.788861467 | 0.081707108 | 0.915947837 | 0.97 | 0.64 |
| YOR034C | 667.8994742 | 0.028451939 | 0.915664436 | 0.97 | 0.64 |
| YPL015C | 1643.336309 | -0.022075722 | 0.915818725 | 0.97 | 0.64 |
| YGR143W | 1625.66982 | -0.030434292 | 0.91613971 | 0.97 | 0.64 |
| YBL014C | 840.8039898 | -0.0370868 | 0.917251543 | 0.97 | 0.64 |
| YER150W | 3467.414047 | 0.014142579 | 0.916899136 | 0.97 | 0.64 |
| YER173W | 470.1702433 | 0.017847106 | 0.916765366 | 0.97 | 0.64 |
| YLR313C | 272.7990365 | 0.03556066 | 0.917272708 | 0.97 | 0.64 |
| YMR264W | 1111.60862 | -0.018038805 | 0.917170513 | 0.97 | 0.64 |

| | | | | | |
|---|---|---|---|---|---|
| snR6 | 0.308944371 | 0.020677083 | 0.917152905 | 0.97 | 0.64 |
| YDR026C | 840.5603663 | -0.01380527 | 0.91808283 | 0.97 | 0.64 |
| YGL152C | 7.084888852 | 0.028747054 | 0.918063831 | 0.97 | 0.64 |
| YGR235C | 1048.42614 | 0.022995164 | 0.917870566 | 0.97 | 0.64 |
| YKR047W | 143.8726915 | -0.017758986 | 0.91783171 | 0.97 | 0.64 |
| YOR102W | 49.45521107 | -0.009452284 | 0.917504827 | 0.97 | 0.64 |
| YDR313C | 770.3934735 | 0.014381163 | 0.918420451 | 0.97 | 0.64 |
| YDR525W-A | 275.582834 | 0.026881747 | 0.91882382 | 0.97 | 0.64 |
| YLR410W-B | 66.3723325 | 0.009341286 | 0.918675418 | 0.97 | 0.64 |
| YOR160W | 1175.925728 | -0.011833934 | 0.918714238 | 0.97 | 0.64 |
| YOR200W | 9.969440735 | -0.034038475 | 0.918841296 | 0.97 | 0.64 |
| YBR237W | 937.0201345 | 0.02089413 | 0.919315512 | 0.97 | 0.64 |
| YML060W | 354.1411795 | 0.07129323 | 0.919746246 | 0.97 | 0.64 |
| YLR456W | 255.5161749 | 0.012583952 | 0.919935988 | 0.97 | 0.64 |
| YDL008W | 381.8332185 | -0.016508609 | 0.921242838 | 0.97 | 0.64 |
| YDR515W | 694.9239372 | -0.024585954 | 0.921820327 | 0.97 | 0.64 |
| YER157W | 974.6953246 | 0.013202136 | 0.92207479 | 0.97 | 0.64 |
| YJL052C-A | 1.688849253 | 0.198795056 | 0.920877196 | 0.97 | 0.64 |
| YKL013C | 2002.473422 | -0.016223275 | 0.922095369 | 0.97 | 0.64 |
| YKL029C | 3291.164423 | 0.023809269 | 0.922155743 | 0.97 | 0.64 |
| YLL051C | 1170.125266 | -0.016930044 | 0.921683765 | 0.97 | 0.64 |
| YML056C | 4012.776938 | 0.014538284 | 0.92142727 | 0.97 | 0.64 |
| YMR086W | 2080.250939 | 0.013723518 | 0.921451978 | 0.97 | 0.64 |
| YOL050C | 16.33482394 | -0.039568431 | 0.921039928 | 0.97 | 0.64 |
| YOR030W | 929.3153858 | 0.016695034 | 0.920484079 | 0.97 | 0.64 |
| YOR372C | 1526.129326 | 0.017967135 | 0.921937582 | 0.97 | 0.64 |
| YPR079W | 754.8054031 | -0.01748751 | 0.922087177 | 0.97 | 0.64 |
| YIL170W | 172.5743318 | -0.032562688 | 0.922581877 | 0.97 | 0.64 |
| YLR146W-A | 192.5995673 | -0.034789931 | 0.922717399 | 0.97 | 0.64 |
| YLR202C | 55.2829566 | -0.025603596 | 0.922568792 | 0.97 | 0.64 |
| YDL012C | 999.0124699 | -0.023797662 | 0.923289802 | 0.97 | 0.64 |
| YDL231C | 1653.703222 | -0.030240057 | 0.923156539 | 0.97 | 0.64 |
| YOL150C | 3.831416512 | -0.189459082 | 0.923131091 | 0.97 | 0.64 |
| YGR263C | 454.4543224 | 0.013704824 | 0.923481693 | 0.97 | 0.64 |
| YDL097C | 4494.471702 | -0.017622731 | 0.923674085 | 0.97 | 0.64 |
| YGR292W | 46.2187461 | 0.033129256 | 0.923892288 | 0.97 | 0.64 |
| YHR049W | 2093.457795 | 0.104294322 | 0.924309313 | 0.97 | 0.64 |
| YIR005W | 167.3355793 | 0.009393033 | 0.924412377 | 0.97 | 0.64 |
| YOR258W | 218.0314211 | 0.01956236 | 0.924425567 | 0.97 | 0.64 |
| YJL193W | 438.7222894 | -0.017314023 | 0.925185684 | 0.97 | 0.64 |
| YLR119W | 415.3763546 | -0.016545903 | 0.925058175 | 0.97 | 0.64 |
| YJR062C | 945.4060688 | 0.014737914 | 0.925534301 | 0.97 | 0.64 |
| YLR030W | 185.5800298 | -0.068029119 | 0.925743326 | 0.97 | 0.64 |

| | | | | | |
|---|---|---|---|---|---|
| YOL090W | 1084.862315 | 0.041234999 | 0.925640629 | 0.97 | 0.64 |
| YKL053W | 17.58730316 | -0.029861786 | 0.926049355 | 0.97 | 0.64 |
| YPL278C | 8.613341367 | -0.049037817 | 0.926097199 | 0.97 | 0.64 |
| YBL095W | 182.0968065 | -0.044119707 | 0.927197805 | 0.97 | 0.64 |
| YBR243C | 997.8241569 | 0.002615096 | 0.927344934 | 0.97 | 0.64 |
| YGL201C | 994.5170532 | -0.006156365 | 0.927157722 | 0.97 | 0.64 |
| YJR055W | 115.4214659 | -0.012419454 | 0.927066632 | 0.97 | 0.64 |
| YMR315W-A | 6.002010905 | -0.047818186 | 0.927051004 | 0.97 | 0.64 |
| YDR157W | 53.23769391 | -0.11158345 | 0.92788023 | 0.97 | 0.64 |
| YDR509W | 13.68760112 | -0.048037462 | 0.927823601 | 0.97 | 0.64 |
| YOR175C | 2021.823765 | 0.017528305 | 0.927587661 | 0.97 | 0.64 |
| YER177W | 16691.55629 | 0.030043704 | 0.928523114 | 0.97 | 0.64 |
| YOR329C | 1982.861977 | 0.020032367 | 0.928542757 | 0.97 | 0.64 |
| YJL093C | 1063.991787 | 0.00950403 | 0.928842144 | 0.97 | 0.64 |
| YOR287C | 297.2565187 | -0.001995974 | 0.928857537 | 0.97 | 0.64 |
| YPR161C | 1370.124428 | 0.016472203 | 0.929406232 | 0.97 | 0.64 |
| YPL025C | 15.86269313 | -0.00453852 | 0.929734965 | 0.97 | 0.64 |
| YJR144W | 1299.865299 | -0.011509319 | 0.929950128 | 0.97 | 0.64 |
| YOR335C | 3110.388093 | 0.000508509 | 0.930432809 | 0.97 | 0.64 |
| YAL067W-A | 0.291097411 | 0.062245507 | 0.930844795 | 0.97 | 0.64 |
| YDR002W | 3692.839207 | 7.71E-05 | 0.930805225 | 0.97 | 0.64 |
| YOL158C | 1769.752458 | 0.063212982 | 0.930959842 | 0.97 | 0.64 |
| YDR379C-A | 462.1957351 | 0.006362329 | 0.931433408 | 0.97 | 0.64 |
| YHR206W | 1203.868698 | -0.021174287 | 0.931340103 | 0.97 | 0.64 |
| YBR201W | 456.7810734 | 0.021534966 | 0.93197215 | 0.97 | 0.64 |
| YEL051W | 3229.319305 | -0.006376615 | 0.93179027 | 0.97 | 0.64 |
| YFR003C | 1017.431306 | -0.02889201 | 0.932007265 | 0.97 | 0.64 |
| YDR484W | 738.7697722 | 0.007762909 | 0.932232134 | 0.97 | 0.64 |
| YLL013C | 908.878738 | -0.008211234 | 0.932366176 | 0.97 | 0.64 |
| YIL012W | 10.18010829 | 0.075228013 | 0.93304734 | 0.98 | 0.64 |
| YAL043C | 1560.889418 | -0.016695446 | 0.933338083 | 0.98 | 0.64 |
| YGR079W | 359.5947801 | -0.055445224 | 0.933354977 | 0.98 | 0.64 |
| YPR152C | 381.9563274 | -0.019135082 | 0.93375603 | 0.98 | 0.64 |
| YJR139C | 4827.24467 | 0.008802964 | 0.934158151 | 0.98 | 0.64 |
| YDR415C | 529.3228874 | -0.013284268 | 0.934471641 | 0.98 | 0.64 |
| YIL042C | 814.6115743 | 0.005019461 | 0.934321858 | 0.98 | 0.64 |
| YDR512C | 741.4198181 | -0.022171891 | 0.93486169 | 0.98 | 0.64 |
| YOR275C | 1566.16532 | -0.015409316 | 0.934820591 | 0.98 | 0.64 |
| YBL070C | 21.83445099 | 0.024165187 | 0.935716526 | 0.98 | 0.65 |
| YDL175C | 355.6103486 | 0.025679072 | 0.935592517 | 0.98 | 0.65 |
| YNL028W | 6.323808974 | -0.008691865 | 0.935892458 | 0.98 | 0.65 |
| YDR005C | 1131.574288 | -0.017045722 | 0.936071034 | 0.98 | 0.65 |
| YDR033W | 12974.70679 | 0.01853132 | 0.936557383 | 0.98 | 0.65 |

| | | | | | |
|---|---|---|---|---|---|
| YBR182C-A | 7.795727576 | 0.037783187 | 0.936969776 | 0.98 | 0.65 |
| snR10 | 11.52677515 | 0.128605918 | 0.937474923 | 0.98 | 0.65 |
| YGR001C | 885.4111524 | 0.026836995 | 0.938266296 | 0.98 | 0.65 |
| YHL007C | 2278.179389 | -0.009570417 | 0.938447937 | 0.98 | 0.65 |
| YMR197C | 923.7331716 | -0.015543282 | 0.938456289 | 0.98 | 0.65 |
| YGL173C | 8071.132916 | -0.005674709 | 0.938687895 | 0.98 | 0.65 |
| YPL190C | 2551.586572 | -0.014075495 | 0.94011914 | 0.98 | 0.65 |
| YDL163W | 0.057163039 | -0.008380916 | 0.940672108 | 0.98 | 0.65 |
| YKL152C | 30648.47356 | -0.036090112 | 0.940658387 | 0.98 | 0.65 |
| tR(CCU)J | 0.0941958 | -0.013492084 | 0.940671742 | 0.98 | 0.65 |
| YKL079W | 964.5499269 | 0.007986111 | 0.941261161 | 0.98 | 0.65 |
| YBR086C | 4766.880566 | 0.011997987 | 0.941531599 | 0.98 | 0.65 |
| YOL042W | 569.9826336 | -0.019964679 | 0.941731227 | 0.98 | 0.65 |
| YEL031W | 4279.639928 | 0.012030821 | 0.941984157 | 0.98 | 0.65 |
| YMR241W | 1393.972884 | 0.002095369 | 0.942221473 | 0.98 | 0.65 |
| YML065W | 967.920824 | -0.015735241 | 0.942629006 | 0.98 | 0.65 |
| YOR080W | 677.2163658 | -0.010095016 | 0.942811944 | 0.98 | 0.65 |
| YOR032C | 566.9607469 | -0.062389691 | 0.943651764 | 0.98 | 0.65 |
| YAL024C | 799.2694521 | -0.0111657 | 0.947644914 | 0.98 | 0.65 |
| YDL087C | 335.3146294 | -0.02257814 | 0.944006985 | 0.98 | 0.65 |
| YDL133C-A | 72.78703027 | -0.043008975 | 0.946912494 | 0.98 | 0.65 |
| YDR492W | 689.8420252 | -0.016670506 | 0.944726975 | 0.98 | 0.65 |
| YGL018C | 470.8004765 | 0.01088912 | 0.944759532 | 0.98 | 0.65 |
| YGL146C | 201.790122 | -0.032629227 | 0.946235415 | 0.98 | 0.65 |
| YHL006W-A | 55.86611926 | 0.05513414 | 0.946540543 | 0.98 | 0.65 |
| YHR007C | 4279.432771 | 0.031637026 | 0.946978705 | 0.98 | 0.65 |
| YHR039C | 2261.246675 | 0.039989642 | 0.944824032 | 0.98 | 0.65 |
| YIL010W | 1043.363564 | 0.011548346 | 0.946260318 | 0.98 | 0.65 |
| YIL023C | 838.1130985 | 0.009007742 | 0.947561705 | 0.98 | 0.65 |
| YIL056W | 1630.203256 | 0.003139125 | 0.945058976 | 0.98 | 0.65 |
| YJL019W | 605.7582629 | -0.010030493 | 0.945820766 | 0.98 | 0.65 |
| YJR117W | 3603.291074 | -0.014829786 | 0.946959195 | 0.98 | 0.65 |
| YKL094W | 2161.93051 | 0.004020044 | 0.947435452 | 0.98 | 0.65 |
| YMR314W | 2980.188475 | 0.01041076 | 0.944237664 | 0.98 | 0.65 |
| YNL008C | 1006.822611 | 0.006838773 | 0.947077617 | 0.98 | 0.65 |
| YNL010W | 2912.199227 | -0.001332997 | 0.945202666 | 0.98 | 0.65 |
| YNL193W | 690.484771 | 0.022712724 | 0.947318938 | 0.98 | 0.65 |
| YNL285W | 3.926788582 | -0.02847764 | 0.947768872 | 0.98 | 0.65 |
| YNR067C | 2460.572138 | 0.037208565 | 0.946005744 | 0.98 | 0.65 |
| YOL054W | 494.8918395 | 0.008968684 | 0.947400095 | 0.98 | 0.65 |
| YOR387C | 76.98828773 | 0.015948425 | 0.944662239 | 0.98 | 0.65 |
| YPL208W | 646.0074609 | 0.006765821 | 0.947504022 | 0.98 | 0.65 |
| YPR074C | 10075.28978 | 0.014464791 | 0.946283278 | 0.98 | 0.65 |

| | | | | | |
|---|---|---|---|---|---|
| YPR124W | 1891.733227 | 0.098622984 | 0.945809404 | 0.98 | 0.65 |
| YOR038C | 676.9008791 | -0.008220318 | 0.947978383 | 0.98 | 0.65 |
| YAR027W | 1481.56887 | -0.016534636 | 0.948158256 | 0.98 | 0.65 |
| YNL224C | 704.2866123 | -0.010388295 | 0.948380813 | 0.98 | 0.65 |
| YBL041W | 3024.581572 | -0.007490953 | 0.949338832 | 0.98 | 0.65 |
| YDR164C | 1143.654091 | 0.013450981 | 0.949122913 | 0.98 | 0.65 |
| YGL111W | 1214.214479 | 0.021878666 | 0.94895356 | 0.98 | 0.65 |
| YJL117W | 1366.561839 | 0.01564281 | 0.949386749 | 0.98 | 0.65 |
| YNR050C | 3301.317883 | 0.039679088 | 0.948933665 | 0.98 | 0.65 |
| YPL231W | 11637.98709 | 0.057845101 | 0.949395924 | 0.98 | 0.65 |
| YMR030W-A | 15.82228062 | -0.026650597 | 0.949656219 | 0.98 | 0.65 |
| tQ(UUG)B | 0.084186605 | 0.019760976 | 0.94998853 | 0.98 | 0.65 |
| YGL137W | 4577.146804 | 0.02012295 | 0.950530183 | 0.98 | 0.65 |
| YDL010W | 362.0027109 | -0.011728119 | 0.951697421 | 0.98 | 0.65 |
| YDL053C | 1309.962591 | 0.014936419 | 0.952145611 | 0.98 | 0.65 |
| YDL133W | 835.1418998 | -0.013591579 | 0.951280254 | 0.98 | 0.65 |
| YDR472W | 675.6541934 | 0.011799926 | 0.952020923 | 0.98 | 0.65 |
| YJL076W | 2985.809725 | -0.007980524 | 0.951590189 | 0.98 | 0.65 |
| YKR030W | 649.1186189 | -0.008694482 | 0.950924558 | 0.98 | 0.65 |
| YMR254C | 26.76419027 | 0.023581438 | 0.951258817 | 0.98 | 0.65 |
| YOR015W | 345.4012917 | 0.003419227 | 0.95166313 | 0.98 | 0.65 |
| YOR349W | 572.160655 | -0.018115759 | 0.951932256 | 0.98 | 0.65 |
| YPL148C | 142.8618352 | 0.026861962 | 0.951232108 | 0.98 | 0.65 |
| YDL031W | 1680.703139 | 0.028243857 | 0.952301816 | 0.98 | 0.65 |
| YJR075W | 2038.161429 | 0.024239263 | 0.953701854 | 0.98 | 0.65 |
| YJR085C | 2130.783694 | 0.011617424 | 0.953655385 | 0.98 | 0.65 |
| YOR055W | 14.54954616 | -0.015454057 | 0.953690448 | 0.98 | 0.65 |
| YML041C | 413.9344954 | -0.018893417 | 0.9540364 | 0.98 | 0.65 |
| YNL027W | 1180.449634 | -0.015233233 | 0.955040142 | 0.98 | 0.65 |
| YNR041C | 337.6551047 | -0.000880715 | 0.955978632 | 0.99 | 0.65 |
| YDR476C | 1082.582617 | 0.005931771 | 0.956771925 | 0.99 | 0.65 |
| YGR094W | 6966.605497 | 0.017402011 | 0.956215289 | 0.99 | 0.65 |
| YLR078C | 672.0208317 | 0.005611393 | 0.956452844 | 0.99 | 0.65 |
| YNL187W | 311.4857358 | 0.00258595 | 0.956773246 | 0.99 | 0.65 |
| YNL256W | 1084.449905 | -0.006445535 | 0.956659317 | 0.99 | 0.65 |
| YAL019W | 1321.957689 | -0.004839279 | 0.95752984 | 0.99 | 0.65 |
| YLR437C | 329.593239 | 0.003138961 | 0.957542878 | 0.99 | 0.65 |
| YBR257W | 391.5498488 | -0.006550378 | 0.961140661 | 0.99 | 0.65 |
| YDL129W | 428.0062351 | 0.007145543 | 0.960708192 | 0.99 | 0.65 |
| YDL160C | 1362.561057 | 0.027241876 | 0.959997614 | 0.99 | 0.65 |
| YDR226W | 6443.35181 | 0.029909826 | 0.95980077 | 0.99 | 0.65 |
| YER046W-A | 13.59530817 | -0.014852025 | 0.961064347 | 0.99 | 0.65 |
| YER137W-A | 3.086112355 | -0.153796998 | 0.959792041 | 0.99 | 0.65 |

| | | | | | |
|---|---|---|---|---|---|
| YER175W-A | 27.41241786 | -0.005034966 | 0.958746881 | 0.99 | 0.65 |
| YFR050C | 3854.503578 | 0.005908823 | 0.959639794 | 0.99 | 0.65 |
| YJL043W | 47.06314645 | 0.016157701 | 0.959299797 | 0.99 | 0.65 |
| YJR142W | 1091.921242 | 0.006177725 | 0.960115208 | 0.99 | 0.65 |
| YKL040C | 858.7842501 | -0.005763676 | 0.959849678 | 0.99 | 0.65 |
| YKR090W | 850.9383093 | 0.000455753 | 0.960625351 | 0.99 | 0.65 |
| YLL052C | 77.86079202 | -0.000126994 | 0.959918151 | 0.99 | 0.65 |
| YLR035C | 528.4128752 | 0.012129931 | 0.961089948 | 0.99 | 0.65 |
| YLR318W | 479.1884801 | -0.014715207 | 0.958634752 | 0.99 | 0.65 |
| YML032C | 1432.723324 | -0.023070454 | 0.960501349 | 0.99 | 0.65 |
| YMR265C | 688.687566 | -0.017227368 | 0.958425698 | 0.99 | 0.65 |
| YMR283C | 413.1862547 | 0.005619078 | 0.961145815 | 0.99 | 0.65 |
| YNL277W | 546.4771573 | 0.006488457 | 0.95901824 | 0.99 | 0.65 |
| YOR063W | 34774.24031 | 0.094535142 | 0.958803837 | 0.99 | 0.65 |
| YPL161C | 652.0526344 | -0.008686229 | 0.958148053 | 0.99 | 0.65 |
| YPR084W | 689.7868138 | 0.004076361 | 0.958060841 | 0.99 | 0.65 |
| YLR112W | 96.84022753 | 0.036753133 | 0.961454637 | 0.99 | 0.65 |
| YGL143C | 536.4770627 | -0.004859347 | 0.961855934 | 0.99 | 0.65 |
| YMR098C | 840.7673515 | -0.013365123 | 0.961797109 | 0.99 | 0.65 |
| YDL178W | 1397.749906 | -0.004871353 | 0.963239932 | 0.99 | 0.65 |
| YIL177C | 0.203940714 | 0.023793686 | 0.962910578 | 0.99 | 0.65 |
| YLR075W | 9637.869009 | 0.049951845 | 0.962991325 | 0.99 | 0.65 |
| YNL073W | 1409.044777 | -0.01038335 | 0.963299898 | 0.99 | 0.65 |
| YNL109W | 19.36367514 | -0.002031416 | 0.963195831 | 0.99 | 0.65 |
| YNR016C | 6951.886561 | 0.025260397 | 0.962767667 | 0.99 | 0.65 |
| YPR038W | 18.2580769 | 0.010824115 | 0.962568829 | 0.99 | 0.65 |
| YCR089W | 1832.043276 | -0.038357798 | 0.963603475 | 0.99 | 0.65 |
| YCR094W | 731.6068684 | 0.017901085 | 0.964671578 | 0.99 | 0.65 |
| YDR140W | 525.20568 | -0.005587708 | 0.965311976 | 0.99 | 0.65 |
| YEL018C-A | 0.416927005 | -0.056492349 | 0.965259101 | 0.99 | 0.65 |
| YHR070C-A | 129.6976848 | 0.014271732 | 0.965030248 | 0.99 | 0.65 |
| YKR087C | 773.1699867 | 0.006014145 | 0.96492408 | 0.99 | 0.65 |
| YKR088C | 850.0044261 | 0.009236325 | 0.964993767 | 0.99 | 0.65 |
| YPL276W | 15.28614474 | -0.113318788 | 0.965264857 | 0.99 | 0.65 |
| YPL136W | 0.97000745 | 0.04454518 | 0.965877387 | 0.99 | 0.65 |
| tG(CCC)O | 0.202424979 | -0.007117411 | 0.965967333 | 0.99 | 0.65 |
| YGR211W | 3320.437529 | 0.00335841 | 0.966134425 | 0.99 | 0.65 |
| YER189W | 0.915658928 | 0.119687226 | 0.966332313 | 0.99 | 0.65 |
| YKL053C-A | 473.2870725 | -0.001657532 | 0.966674513 | 0.99 | 0.65 |
| YKL070W | 302.0799748 | 0.006189873 | 0.966680046 | 0.99 | 0.65 |
| YIL135C | 1336.175722 | 0.005316411 | 0.967001225 | 0.99 | 0.65 |
| YDL098C | 246.2120054 | 0.006960539 | 0.967850492 | 0.99 | 0.65 |
| YDL144C | 951.4980585 | -0.007084229 | 0.968303242 | 0.99 | 0.65 |

| | | | | | |
|---|---|---|---|---|---|
| YHR203C | 5825.930229 | 0.059269393 | 0.968247107 | 0.99 | 0.65 |
| YJR068W | 1198.277592 | 0.00990493 | 0.968389266 | 0.99 | 0.65 |
| YOL129W | 2548.890951 | -0.005653464 | 0.968013792 | 0.99 | 0.65 |
| YDR442W | 17.52253569 | 0.047655077 | 0.968541655 | 0.99 | 0.65 |
| YMR305C | 6521.89708 | 0.043547322 | 0.968700128 | 0.99 | 0.65 |
| YDR275W | 363.6731352 | -0.0265685 | 0.969277007 | 0.99 | 0.65 |
| YIL100C-A | 6.139255363 | -0.01459468 | 0.969041273 | 0.99 | 0.65 |
| YMR227C | 799.7644018 | 0.013692932 | 0.969470311 | 0.99 | 0.65 |
| YOL013C | 1216.146773 | -0.009482038 | 0.969475138 | 0.99 | 0.65 |
| YBL052C | 898.9203912 | 0.003833371 | 0.969764874 | 0.99 | 0.65 |
| YHR072W | 1833.218722 | -0.001207626 | 0.969887546 | 0.99 | 0.65 |
| YJR064W | 2718.261132 | 0.035325263 | 0.969939029 | 0.99 | 0.65 |
| YAL047W-A | 1.181246705 | -0.066389278 | 0.971301212 | 0.99 | 0.65 |
| YBR105C | 2095.906659 | -0.011344821 | 0.972920347 | 0.99 | 0.65 |
| YCL048W-A | 89.40578665 | 0.117436976 | 0.971482336 | 0.99 | 0.65 |
| YDR272W | 1625.208653 | -0.004007893 | 0.972856935 | 0.99 | 0.65 |
| YDR357C | 553.215464 | 0.01075468 | 0.971973276 | 0.99 | 0.65 |
| YGL249W | 273.3583474 | -0.014362588 | 0.972372475 | 0.99 | 0.65 |
| YGR190C | 89.92851229 | -0.00372224 | 0.972009374 | 0.99 | 0.65 |
| YHR108W | 3358.554888 | 0.033182323 | 0.970293955 | 0.99 | 0.65 |
| YIL156W-B | 291.2591429 | -0.009746762 | 0.972569922 | 0.99 | 0.65 |
| YJR084W | 669.2500078 | -0.001602878 | 0.972038504 | 0.99 | 0.65 |
| YJR143C | 2572.640819 | 0.036597646 | 0.972350131 | 0.99 | 0.65 |
| YKL110C | 846.8769824 | 0.012252494 | 0.97050422 | 0.99 | 0.65 |
| YKL191W | 1251.863612 | 0.029983815 | 0.970804976 | 0.99 | 0.65 |
| YMR230W-A | 4.038623613 | -0.058429405 | 0.971304677 | 0.99 | 0.65 |
| YNL192W | 2879.150646 | 0.023596697 | 0.973017184 | 0.99 | 0.65 |
| YNL233W | 992.805205 | 0.012690864 | 0.972249194 | 0.99 | 0.65 |
| YNR035C | 3358.997637 | -0.009427409 | 0.970267157 | 0.99 | 0.65 |
| YOR204W | 2305.73227 | 0.019463917 | 0.970639782 | 0.99 | 0.65 |
| YPL057C | 1620.500312 | -0.02486076 | 0.971964536 | 0.99 | 0.65 |
| YPR189W | 1450.192608 | 0.015878497 | 0.972983961 | 0.99 | 0.65 |
| YML014W | 592.0947476 | -0.008233576 | 0.97381011 | 0.99 | 0.65 |
| YDR079C-A | 102.409036 | 0.005588689 | 0.974379741 | 0.99 | 0.65 |
| YPR105C | 1243.404818 | -0.006318296 | 0.974450667 | 0.99 | 0.65 |
| YPL051W | 362.5719952 | -0.003478042 | 0.974956665 | 0.99 | 0.65 |
| YPL041C | 160.4382575 | 0.02215412 | 0.975455104 | 0.99 | 0.65 |
| YOR023C | 2117.560133 | -0.022309257 | 0.975994306 | 0.99 | 0.65 |
| YOR073W-A | 11.62724158 | -0.013009411 | 0.975957141 | 0.99 | 0.65 |
| YPL031C | 1442.118487 | 0.006455905 | 0.97596648 | 0.99 | 0.65 |
| YDR355C | 4.46006282 | -0.067019077 | 0.976203191 | 0.99 | 0.66 |
| YLR272C | 941.972981 | 0.007116533 | 0.977409723 | 0.99 | 0.66 |
| snR74 | 1.214578707 | 0.205089318 | 0.977284927 | 0.99 | 0.66 |

| | | | | | |
|---|---|---|---|---|---|
| YHR045W | 983.5331682 | 0.007713251 | 0.977803136 | 0.99 | 0.66 |
| YDL135C | 948.2549942 | -0.002603829 | 0.978259057 | 0.99 | 0.66 |
| YPL083C | 495.9825937 | 0.009866824 | 0.978465978 | 0.99 | 0.66 |
| snR81 | 9.309990788 | 0.115952346 | 0.978375658 | 0.99 | 0.66 |
| YPL039W | 426.2421359 | -0.020079068 | 0.97878242 | 0.99 | 0.66 |
| YDR365W-B | 142.2689351 | -0.076343168 | 0.979217778 | 0.99 | 0.66 |
| YLR223C | 2252.47739 | -0.015242575 | 0.979120832 | 0.99 | 0.66 |
| YOR316C | 1558.064909 | 0.013356743 | 0.979678548 | 0.99 | 0.66 |
| YER084W | 124.9336603 | 0.05901309 | 0.980362796 | 0.99 | 0.66 |
| YLR315W | 96.87354306 | -0.014077705 | 0.980655893 | 0.99 | 0.66 |
| YDR489W | 358.8647918 | -0.005041031 | 0.980958475 | 0.99 | 0.66 |
| YGL038C | 1572.160144 | -0.001783575 | 0.98113375 | 0.99 | 0.66 |
| YMR163C | 451.0209425 | 0.002263443 | 0.98122347 | 0.99 | 0.66 |
| YLR342W | 12036.59934 | 0.031045076 | 0.981948206 | 0.99 | 0.66 |
| snR11 | 261.7635615 | -0.019565891 | 0.981830723 | 0.99 | 0.66 |
| YAL046C | 313.7043981 | 0.033235609 | 0.982107015 | 0.99 | 0.66 |
| YBR141C | 286.9036203 | -0.001453667 | 0.982853687 | 1.00 | 0.66 |
| YEL058W | 1683.65634 | 0.013692105 | 0.983154105 | 1.00 | 0.66 |
| YDR032C | 4060.720659 | -0.000317733 | 0.983864289 | 1.00 | 0.66 |
| YDR307W | 749.860008 | 0.008234721 | 0.983812892 | 1.00 | 0.66 |
| YOR161C-C | 9.014220677 | -0.08067675 | 0.98386329 | 1.00 | 0.66 |
| YPR080W | 5311.40989 | 0.063614968 | 0.983553482 | 1.00 | 0.66 |
| snR50 | 8.290616263 | -0.009304688 | 0.98413767 | 1.00 | 0.66 |
| YGL228W | 2378.82392 | -0.0023423 | 0.984488109 | 1.00 | 0.66 |
| YOR083W | 334.3141069 | -0.007177631 | 0.984471424 | 1.00 | 0.66 |
| YBL090W | 640.7121751 | 0.012770268 | 0.985010713 | 1.00 | 0.66 |
| YDL120W | 712.7094342 | 0.00377004 | 0.986084589 | 1.00 | 0.66 |
| YDR097C | 218.1671058 | -0.359282453 | 0.985648593 | 1.00 | 0.66 |
| YDR393W | 653.7265915 | -0.003978058 | 0.985469178 | 1.00 | 0.66 |
| YJL112W | 1088.839683 | -0.008096238 | 0.985338419 | 1.00 | 0.66 |
| YJL176C | 1289.4859 | -0.004960957 | 0.985931511 | 1.00 | 0.66 |
| YLR076C | 95.93637703 | 0.115151645 | 0.985698672 | 1.00 | 0.66 |
| YNL051W | 867.7540206 | -0.003859386 | 0.985559867 | 1.00 | 0.66 |
| YNL167C | 1564.476742 | -0.002374246 | 0.985879927 | 1.00 | 0.66 |
| YOL155C | 4950.84233 | -0.024617576 | 0.986629596 | 1.00 | 0.66 |
| YOR072W-B | 0.082611248 | -0.005753941 | 0.986428279 | 1.00 | 0.66 |
| YOR089C | 2098.945136 | -0.01176774 | 0.986633634 | 1.00 | 0.66 |
| YPR190C | 965.2019513 | 0.021398007 | 0.986584345 | 1.00 | 0.66 |
| YBR018C | 184.0387102 | -0.016537464 | 0.98738106 | 1.00 | 0.66 |
| YBR110W | 1287.287964 | 0.003962738 | 0.987392568 | 1.00 | 0.66 |
| YLR041W | 27.97790779 | -0.006109474 | 0.987104653 | 1.00 | 0.66 |
| YOR054C | 904.8796156 | -0.015046471 | 0.987641007 | 1.00 | 0.66 |
| YDL195W | 7573.167842 | 0.009305645 | 0.987828553 | 1.00 | 0.66 |

| | | | | | |
|---|---|---|---|---|---|
| YDR004W | 628.5806399 | -0.003699216 | 0.988240495 | 1.00 | 0.66 |
| snR43 | 20.09245082 | 0.026097875 | 0.988217232 | 1.00 | 0.66 |
| YLR044C | 136554.4679 | 0.043050412 | 0.98876953 | 1.00 | 0.66 |
| YBR109C | 2942.937927 | 0.003019286 | 0.989463412 | 1.00 | 0.66 |
| YKR082W | 1772.94275 | -0.004425064 | 0.989591536 | 1.00 | 0.66 |
| YLR115W | 1204.082371 | -0.001287153 | 0.989722355 | 1.00 | 0.66 |
| YPL198W | 414.9903406 | 0.196938068 | 0.989239508 | 1.00 | 0.66 |
| YPL213W | 340.9140453 | -0.004694072 | 0.989330185 | 1.00 | 0.66 |
| YCL044C | 1238.471005 | -0.008665353 | 0.989934303 | 1.00 | 0.66 |
| snR48 | 2.926793808 | 0.034150064 | 0.990059683 | 1.00 | 0.66 |
| YDL220C | 416.2502353 | 0.030484014 | 0.990215086 | 1.00 | 0.66 |
| YHR062C | 561.4502383 | 0.01468121 | 0.990699164 | 1.00 | 0.66 |
| YML012C-A | 4.439298629 | 0.026441874 | 0.990961746 | 1.00 | 0.66 |
| YIR021W-A | 4.66148073 | 0.065304448 | 0.991522 | 1.00 | 0.66 |
| YPL179W | 1527.082249 | 0.007129537 | 0.991513686 | 1.00 | 0.66 |
| YPR062W | 2367.986615 | 0.003430868 | 0.991935236 | 1.00 | 0.66 |
| RUF22 | 1.126113591 | 0.222657511 | 0.992791987 | 1.00 | 0.66 |
| YBL009W | 764.0807255 | 0.026151927 | 0.993493614 | 1.00 | 0.66 |
| YCR093W | 4708.644101 | 0.010206719 | 0.993064352 | 1.00 | 0.66 |
| YDR094W | 4.116771726 | 0.048407665 | 0.99356285 | 1.00 | 0.66 |
| YJL083W | 610.9975272 | -0.006348062 | 0.993167194 | 1.00 | 0.66 |
| YJL173C | 671.1948643 | 0.010595399 | 0.992719247 | 1.00 | 0.66 |
| YKL196C | 2255.642273 | 0.00042228 | 0.993298657 | 1.00 | 0.66 |
| YKR044W | 684.1987114 | -0.001823515 | 0.993220243 | 1.00 | 0.66 |
| YMR193C-A | 123.8016037 | 0.030457833 | 0.993109818 | 1.00 | 0.66 |
| YNL234W | 710.7866789 | -0.00727347 | 0.992813308 | 1.00 | 0.66 |
| YOR345C | 1.957434997 | 0.000532824 | 0.993651684 | 1.00 | 0.66 |
| YLR376C | 277.9237935 | -0.005559596 | 0.993820558 | 1.00 | 0.66 |
| YLL054C | 624.0988471 | 0.005714626 | 0.994711117 | 1.00 | 0.66 |
| YGL247W | 197.7711302 | 0.002905581 | 0.994950024 | 1.00 | 0.66 |
| YAR015W | 7474.541729 | 0.060056606 | 0.995683687 | 1.00 | 0.66 |
| YGR233C | 1147.708039 | 0.006935966 | 0.995616768 | 1.00 | 0.66 |
| YHR069C-A | 0.116121444 | 0.007443768 | 0.995568058 | 1.00 | 0.66 |
| YML079W | 1250.560699 | -0.003630766 | 0.995704688 | 1.00 | 0.66 |
| YGL181W | 1793.449442 | -0.002837262 | 0.996044856 | 1.00 | 0.66 |
| YBR118W | 10015.00281 | 0.053854712 | 0.996580344 | 1.00 | 0.66 |
| YDR341C | 4249.5314 | 0.037628398 | 0.996608568 | 1.00 | 0.66 |
| YHL011C | 1690.318772 | 0.042172711 | 0.996601423 | 1.00 | 0.66 |
| YCR085W | 15.93952821 | 0.118822827 | 0.996977932 | 1.00 | 0.66 |
| YJL167W | 5562.553131 | 0.014312625 | 0.997407233 | 1.00 | 0.66 |
| YFL010W-A | 308.9446486 | -0.002724391 | 0.99785592 | 1.00 | 0.66 |
| YFR056C | 41.82065454 | -0.001124362 | 0.997936804 | 1.00 | 0.66 |
| YGR140W | 791.394714 | 0.009421857 | 0.997756506 | 1.00 | 0.66 |

| | | | | | |
|---|---|---|---|---|---|
| YKL212W | 2578.339284 | 0.01511822 | 0.998228401 | 1.00 | 0.66 |
| YML047C | 45.4588878 | 0.075776348 | 0.998125936 | 1.00 | 0.66 |
| YBR037C | 852.7234834 | -0.003536088 | 0.999418859 | 1.00 | 0.66 |
| YKR077W | 759.0676767 | -0.007772277 | 0.999108947 | 1.00 | 0.66 |
| YLL049W | 333.5985502 | 0.021262787 | 0.999085385 | 1.00 | 0.66 |
| YML007C-A | 2.984197591 | 0.106424133 | 0.999091695 | 1.00 | 0.66 |
| YML038C | 1118.066731 | -0.000162043 | 0.999696324 | 1.00 | 0.66 |
| YMR013C | 348.9768631 | 0.001807018 | 0.998735696 | 1.00 | 0.66 |
| YMR054W | 1855.817154 | 0.010702924 | 0.999384722 | 1.00 | 0.66 |
| YPR158C-D | 130.4698158 | 0.013980718 | 0.999598551 | 1.00 | 0.66 |
| YBR064W | 3.158811985 | -0.030937094 | 0.999970694 | 1.00 | 0.66 |
| YAL062W | 2248.194645 | 0.611860536 | NA | NA | |
| YDL223C | 1796.784019 | -0.056664028 | NA | NA | |
| YER024W | 2782.885664 | 0.889343779 | NA | NA | |
| YER065C | 3480.324226 | 0.914735156 | NA | NA | |
| YGR067C | 601.4253985 | 0.737691154 | NA | NA | |
| YGR236C | 237.8810767 | -0.346544511 | NA | NA | |
| YIL057C | 383.2825996 | 0.622963751 | NA | NA | |
| YJR095W | 1024.664601 | 1.250008364 | NA | NA | |
| YKL065W-A | 60.35011298 | -0.466318293 | NA | NA | |
| YKR097W | 1618.765281 | 0.794475196 | NA | NA | |
| YLR174W | 1214.876474 | 0.855633868 | NA | NA | |
| YLR377C | 564.6863452 | 0.913514065 | NA | NA | |
| YML042W | 2686.768376 | 1.307495286 | NA | NA | |
| YMR107W | 2022.649302 | -0.038188305 | NA | NA | |
| YMR175W | 966.6429644 | 0.202209217 | NA | NA | |
| YMR175W-A | 25.9663659 | 0.419946709 | NA | NA | |
| YNL036W | 2481.742596 | 0.862184365 | NA | NA | |
| YNL117W | 3801.736785 | 1.200161834 | NA | NA | |
| YNR002C | 907.6027094 | 0.348712174 | NA | NA | |
| YNR071C | 317.5831201 | 1.284013589 | NA | NA | |
| YOR100C | 795.4961629 | 1.53629758 | NA | NA | |
| YPL223C | 915.9171589 | -0.518833208 | NA | NA | |
